# Adaptation of *Pseudomonas aeruginosa* to the lung allograft environment in cystic fibrosis lung transplant recipients

**DOI:** 10.64898/2026.07.06.736721

**Authors:** Thilo Köhler, Léna Falconnet, Alexandre Luscher, Marie Graindorge Beaume, Marc Chanson, Gilbert Greub, Angela Koutsokera, Gregory Berra, Paola M. Soccal, Christian van Delden, the Swiss Transplant Cohort Study (STCS)

## Abstract

Lung transplantation (LT) is the ultimate treatment option for patients suffering from end stage cystic fibrosis (CF). Most LT-patients, colonized pre-LT by *Pseudomonas aeruginosa* witness colonization of their non-CF allograft within a few days or weeks post-LT, thereby compromising graft and life expectancy. How *P. aeruginosa* isolates adapted for years to the specific CF lung environment efficiently colonize and survive in the non-CF allograft environment remains unclear. To address this question, we collected sequential isolates from CF LT-recipients and non-CF LT-recipients and performed phenotypic and genetic analyses of pairs of early and late isolates from LT-patients. We found evidence for mutations compatible with a switch from biofilm to planktonic lifestyle as well as loss of mucoid phenotypes. Hypermutators, characteristic of chronic CF-adapted isolates, were also found in four LT-patients. Their persistence in the non-CF allograft environment suggests a continuous seeding from the sinuses. Our results suggest that in CF LT-recipients efficient colonisation by *P. aeruginosa* of the allograft implies both adaptation and continuous seeding from the sinuses to the lower respiratory tract.

## Introduction

Cystic fibrosis (CF) is an autosomal genetic disease, resulting from mutations in the cystic fibrosis transmembrane conductance regulator (CFTR), which affects multiple organs. Chronic lung infections by *Pseudomonas aeruginosa*, facilitated by thick mucus production and accumulation due to the CFTR dysfunction, remain life-threatening conditions for CF-patients. Although life expectancy of CF-patients has increased over the past 20 years, chronic bacterial colonization and sustained inflammatory responses leading to progressive lung dysfunction remain the major hurdles for therapeutic intervention. The recent development of CFTR modulators has significantly improved the outcome of CF-patients, however not all mutations can be corrected (1) and lung transplantation (LT) remains the ultimate therapy for CF-patients with deteriorating lung function. Unfortunately, chronic lung allograft dysfunction (CLAD) is a major risk, occurring in 50% (2) of LT-recipients after 5 years and in 75% after 10 years (3). The CFTR dysfunction creates a particular environment in the CF-lung to which *P. aeruginosa* progressively adapts over years. This genetic adaptation is facilitated by a hypermutator genotype, characteristic of *P. aeruginosa* isolates recovered from CF-patients (4). Although the allograft is a non-CF environment, colonization by *P. aeruginosa* is observed in most CF LT-recipients colonized by *P. aeruginosa* prior to transplantation. Earlier studies have shown that the genotypes of the *P. aeruginosa* isolates pre-LT correspond to those found after LT (5). This colonization likely occurs by seeding from the sinuses, which remain a CF-environment and are connected to the upper-respiratory tract (6). This has been further supported by a high correlation between pre-transplant sinus cultures and post-transplant bronchoalveolar lavage (BAL) cultures for *P. aeruginosa*, *Staphylococcus aureus* and *Burkholderia* spp. (7). Allograft colonization by *P. aeruginosa* is a risk factor for CLAD and contributes to the lower life expectancy of LT-patients in general, when compared with other solid organ transplantations. However, the question of whether the isolates from the CF-sinus environment adapt to the non-CF allograft environment has not been clearly investigated. We have previously shown that allograft colonization occurs within a few days after LT (8). Our observation also concurred with previous reports showing reduced microbiome richness and diversity following LT in CF-patients. Here we extended this study including CF LT-recipients, as well as non-CF LT-recipients as a control group. The data are in favour of a continuous allograft seeding by CF-adapted *P. aeruginosa* from the sinuses or the upper respiratory tract, with further adaptation of some isolates to the non-CF environment.

## Results

Patients underwent bilateral lung transplantation at the University Hospital Lausanne (CHUV) and were followed thereafter either by the pulmonologist team of the CHUV or the Geneva University Hospitals (HUG). None of the patients had received CFTR modulator therapy before or after LT. Only patients colonized subsequently by *P. aeruginosa* were retained for this study. Forty-two *P. aeruginosa* isolates from seven CF-patients (D, P, H, O, M, G and L) and eight isolates from three non-CF control LT-recipients diagnosed with chronic obstructive pulmonary disease (A), non-CF bronchiectasis (B) or interstitial lung disease (C), were obtained via the local bacteriology laboratories or re-isolated from frozen respiratory samples (Table 1). All patients in this study were colonized pre-LT by *P. aeruginosa*. CF-patient G underwent two LTs (the first on day 0 and the second on day 991 after the first LT). Allograft colonization by *P. aeruginosa* occurred within a few days after LT (data not shown), but at the latest within the first 12 months after LT-surgery.

**Table 1.** Sample and isolate characteristics.

| Patient<sup>1</sup> | Isolate name<sup>2</sup> | Sample type<sup>3</sup> | Mucoid | Days post-LT | <sup>4</sup>DLST | <sup>5</sup>Mutator |
| --- | --- | --- | --- | --- | --- | --- |
| D (CF) | **D0** | ASP | yes | 0 | 18-34 | yes (*mutL* fs) |
|  | D1M | ASP | yes | 1 | 18-34 | yes |
|  | D1NM | ASP | no | 1 | 18-34 | yes |
|  | **D350** | ASP | yes | 350 | 18-34 | yes (*mutL* fs) |
| P (CF) | **P0** | ASP | no | 0 | 08-37 | no |
|  | P183.1M | BAL | yes | 183 | 08-37 | yes |
|  | P183.1NM | BAL | no | 183 | 08-37 | yes |
|  | P183.2 | ASP | no | 183 | 08-37 | no |
|  | P344 | BAL | no | 344 | 08-37 | no |
|  | P610.1 | ASP | no | 610 | 08-37 | no |
|  | P610.2 | ASP | yes | 610 | 08-37 | no |
|  | **P932** | BAL | yes | 932 | 08-37 | yes (*mutS* T493P) |
| H (CF) | H0M | ASP | yes | 0 | 15-188 | no |
|  | **H0NM** | ASP | no | 0 | 15-188 | no |
|  | H4 | ASP | no | 4 | 8-33 | no |
|  | H334 | EXP | no | 334 | 15-188 | no |
|  | **H727** | ASP | no | 727 | 15-188 | no |
| O (CF) | O-30 (1.2) | SIN | no | -32 | 55-58 | yes |
|  | O-30 (1.7) | SIN | no | -32 | 55-58 | no |
|  | **O-13** | ASP | no | -13 | 55-58 | yes (*mutS* fs) |
|  | O29 (8.1) | ASP | no | 29 | 55-180 | yes |
|  | O29 (9.1) | BAL | no | 29 | 55-58 | no |
|  | O29 (9.2) | BAL | no | 29 | 55-180 | yes |
|  | **O49** | ASP | no | 49 | 55-58 | yes (*mutS* fs, *mutL*) |
| M (CF) | **M9** | SIN | no | 9 | 10-191 | yes (*ung*) |
|  | **M351.1** | BAL | no | 351 | 10-191 | yes (*ung*) |
|  | M351.2 | ASP | no | 351 | 10-191 | yes (ND) |
| G (CF) | G-33 | EXP | yes | -33 | 16-04 | no |
|  | **G159** | BAL | no | 159 | 08-37 | no |
|  | G1022.1M | ASP | yes | 1022 | 08-37 | ND |
|  | G1022.1 | ASP | no | 1022 | 08-37 | ND |
|  | G1022.2 | ASP | no | 1022 | 08-37 | no |
|  | G1080.1M | BAL | yes | 1080 | 08-37 | ND |
|  | G1080.1 | BAL | no | 1080 | 08-37 | ND |
|  | G1080.2 | BAL | no | 1080 | 08-37 | no |
|  | G1080.3 | ASP | no | 1080 | 08-37 | ND |
|  | G1080.4 | ASP | no | 1080 | 08-37 | ND |
|  | G1395.1 | BAL | no | 1395 | 08-37 | no |
|  | **G1395.2** | ASP | no | 1395 | 08-37 | no |
| L (CF) | L-879 | NA | yes | -879 | 7-9 | no |
|  | L-388 | EXP | no | -388 | 7-9 | no |
|  | L235 | BAL | yes | 235 | 7-9 | no |
| A (COPD) | A92.1 | BAL | no | 92 | 18-34 | no |
|  | A92.2 | ASP | no | 92 | 18-34 | no |
|  | A755 | BAL | no | 755 | deletion | no |
| B (BR) | B185 | ASP | no | 185 | 8-37 | no |
|  | B1094 | EXP | no | 1094 | 8-37 | no |
|  | B1132 | SIN | no | 1132 | 8-37 | no |
| C (ILD) | C19 | BAL | no | 19 | 48-40 | no |
|  | C45 | ASP | yes | 45 | 25-11 | no |

<sup>1</sup>COPD, chronic obstructive pulmonary disease; BR, non-CF bronchiectasis; ILD, Interstitial lung disease
<sup>2</sup>Genomes of isolates indicated in bold were submitted to WGS; M, mucoid; NM, non-mucoid
<sup>3</sup>ASP, tracheal aspirate; BAL, bronchoalveolar lavage; EXP, expectoration; SIN, sinus swab; NA, not available
<sup>4</sup>DLST, double locus sequence type
<sup>5</sup>ND, not determined

### Genotyping

We first established whether the isolates in the allograft belonged to one or several genotypes using double locus sequence typing (DLST) (9). Out of the seven CF-patients, five (D, P, M, G, L) were colonized post-LT by a single genotype, while two CF (H, O) and one control patient (C) harboured two different genotypes (Table 1). Whether these additional genotypes were also present pre-LT could not be established retrospectively. In CF-patient H, one isolate recovered 4 days post-LT (H4) showed a different genotype (08–33) than the other isolates (15–188), while in CF-patient O, the genotype isolated from the sinus pre-LT (55–58) was also present in all other subsequent BAL and bronchial aspirate (ASP) samples. For this patient, an additional genotype (55–180) was isolated from both BAL and ASP on day 29 post-LT. In CF-patient M, the same genotype (10–191) was recovered from a sinus swab (day 9 post-LT) as well as from BAL and ASP samples (day 351 post-LT). These data are in favour of a colonization of the allograft from the sinuses, presenting a CF-environment, as suggested previously (6). The DLST type 08-37, which corresponds to the international lineage ST111 (9), was the most prominent genotype, isolated from two CF LT-recipients and one control (Table 1).

### General phenotyping

During chronic colonization of the CF-lung, *P. aeruginosa* adapts to the particular conditions of the CF-lung environment (10, 11). These adaptations involve biofilm formation capacities, acquisition of antibiotic resistance, as well as loss of motility, LPS O-side chain synthesis and mutations in the LasRI quorum sensing (QS) system (12). To assess a potential evolution of the CF-adapted isolates to the non-CF allograft, we further characterized the isolates with respect to growth, motility, virulence factor production (mucoidity, cytotoxicity, pyocyanin and rhamnolipid production) as well as biofilm formation and hypermutator phenotypes.

#### Growth

We first tested growth in various media using PAO1 as a reference strain. Isolates from CF-patients reached lower OD_600_ values compared to PAO1 (Fig. 1). In contrast growth of isolates from control patients A, B and C reached OD_600_ values after 24 h incubation comparable to those of PAO1 in minimal media (except isolate C45). Only isolates from CF-patient O did not grow in M9 minimal medium but showed improved growth when supplemented with casamino acids (CAA), reflecting an auxotrophic phenotype frequently encountered in CF-isolates. Iron supplementation showed only a marginal effect on growth and mainly in the iron-limited M9-CAA medium.

**Figure 1.**
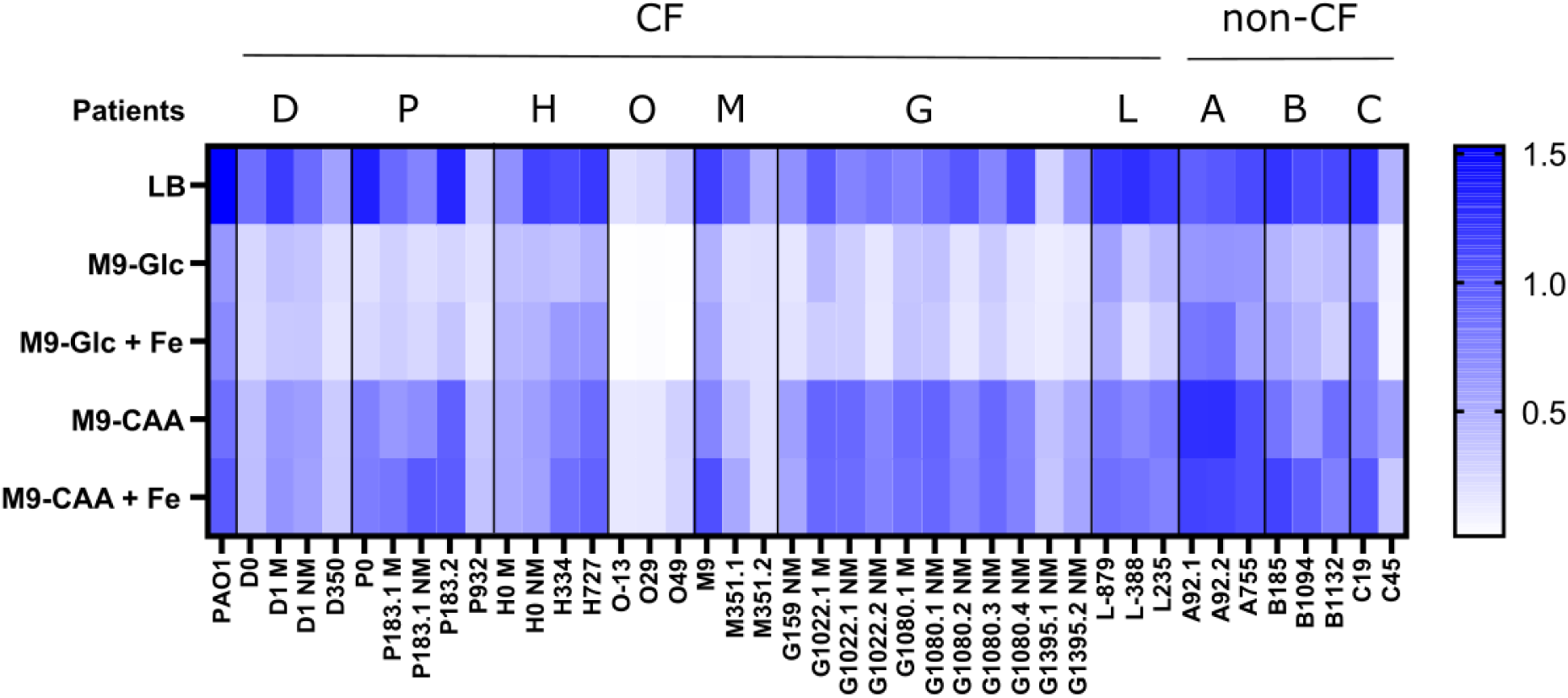
Growth of clinical isolates in rich and defined media with and without Fe-supplementation. Growth was monitored in a plate reader at 37°C under static conditions during 24 h. Values are the average of OD_600_ measurements from technical triplicates after deduction of blank values at 24 h. Glc glucose (0.2% w/v); CAA, casamino acids (0.5% w/v as sole C-source); Fe, FeCl_2_ (10 μM final concentration).

There was no overall trend for growth improvement during LT-colonization under our *in vitro* conditions. Only the late isolates from CF-patients O and H showed higher OD_600_ values in all growth media, when compared to the corresponding initial isolate of the same genotype (Fig. 1), supporting adaptation to the more nutrient restricted non-CF allograft environment.

#### Antibiotic susceptibility

All isolates showed different susceptibility profiles for the three antibiotics tested (ceftazidime, ciprofloxacin, gentamicin). Isolates with gentamicin-resistance were observed in two out of seven CF-patients, but not in isolates from control patients. With the exception of one control (C), all other patients harboured at least one ceftazidime-resistant isolate. Four of the seven CF-patients carried at least one ciprofloxacin resistant isolate, while all isolates from control patients were susceptible (Table 2). The ciprofloxacin resistant isolates carried a T83I substitution in GyrA, known to confer high-level quinolone resistance. Ceftazidime resistance could not be associated to a single genetic modification but resulted from various combinations of mutations identified by WGS in *ampC*, *ampDh3*, *ampD*, *mpl* (AmpC up-regulation) and in *mexR* (MexAB-OprM overexpression) (Table S1). Antibiotic resistance did not increase over time in the tested isolates, suggesting a possible co-existence of susceptible and resistant subpopulations and/or rapid and transient emergence of resistance secondary to a specific antibiotic treatment.

**Table 2.** Antibiotic susceptibilities of *P. aeruginosa* LT-isolates.

|  | MIC (µg/ml) |  |  |
| --- | --- | --- | --- |
|  | CAZ | CIP | GEN |
| PAO1 | 4 | 0,25 | 2 |
| **D0** | 2 | > 4 | 1 |
| **D1M** | 8 | 2 | 1 |
| **D350** | 2 | > 4 | 0,5 |
| P0 | 4 | > 4 | 4 |
| **P183.1NM** | 8 | 4 | 4 |
| **P183.2** | 8 | 4 | 4 |
| **P932** | 16 | > 4 | 0,5 |
| H0M | 4 | > 4 | 16 |
| H0NM | 32 | 8 | 8 |
| H334 | 64 | 4 | 8 |
| H727 | 16 | > 4 | 4 |
| **O-13** | > 32 | > 4 | 16 |
| **O29** | 64 | 2 | >16 |
| **O49** | 4 | > 4 | 1 |
| **M9** | 8 | 1 | >16 |
| **M351.1** | 8 | 0,125 | 4 |
| **M351.2** | 8 | 0,125 | 4 |
| G-33 | 1 | 0,06 | 2 |
| G159 | 32 | 0,125 | 4 |
| G1022.1 | 4 | 0,25 | >16 |
| G1080.3 | 32 | 0,25 | 16 |
| G1395.2 | 16 | 0,25 | >16 |
| L-879 | 1 | 2 | 4 |
| L-388 | 16 | >4 | 1 |
| L235 | <0.5 | >4 | 4 |
| A92.1 | 8 | 0,125 | 4 |
| A92.2 | 4 | 0,06 | 4 |
| A755 | 8 | 0,125 | 4 |
| B185 | 16 | 0,25 | 4 |
| B1094 | 8 | 0,06 | 4 |
| B1132 | 8 | 0,125 | 4 |
| C19 | 4 | <0,06 | 2 |
| C45 | 1 | <0,06 | 4 |

Mutator isolates are shown in bold

#### Motility

CF-isolates are frequently non-motile since they lose the capacity to synthesize flagella (13). While isolates from the CF-patients were either non-motile or partially swimming proficient (< 50% of PAO1), the control isolates showed at least 50% of the swimming capacity of PAO1 (except isolate C45) (Fig. 2). The vast majority of isolates, whether from CF LT-recipients or controls, showed no or drastically reduced swarming motility compared to PAO1 (Fig. 2). Only isolates H4, H755 and C19 were partially swarming proficient (20-50% of PAO1). WGS revealed an identical nonsense mutation in *fliC* (FliC Y36*), coding for the a-type flagellin protein of isolates D0, D350, O-13, O49, M9 and M351.1 (Table S1). This mutation should abrogate flagella synthesis resulting likely in loss of swimming and swarming motilities in these isolates (Fig. 2). There was no indication for recovery of swimming or swarming motility during allograft colonization.

**Figure 2.**
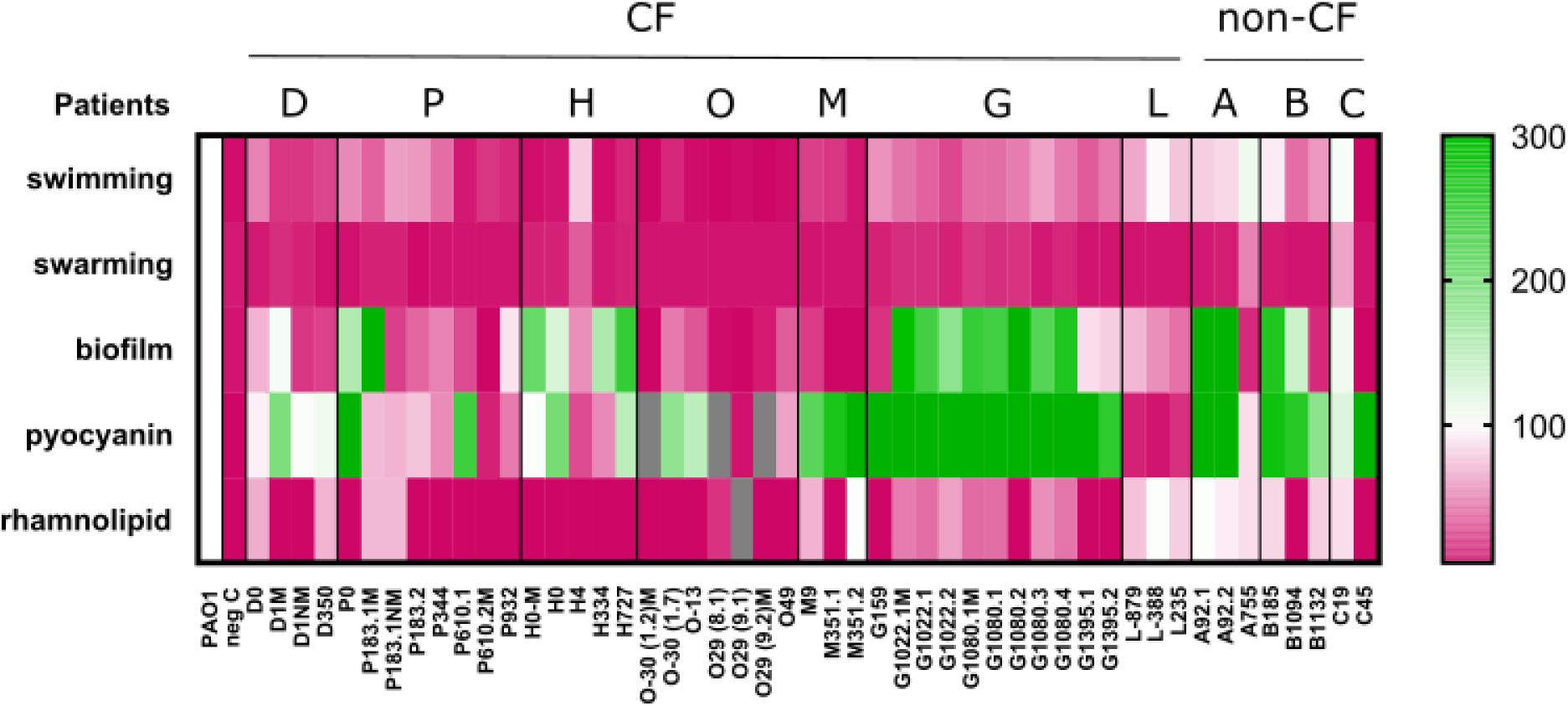
Phenotype heat map of *P. aeruginosa* isolates. Values are given with respect to PAO1 set to 100% (white). Grey boxes indicate isolates that did not grow in the medium used for the assay. Negative control strains (neg C) were PAO1 mutants with defined genetic deletions in *fliC* (swimming and swarming), *pilA* (biofilm), *pqsR* (pyocyanin) or *rhlA* (rhamnolipid).

#### Biofilm formation

Isolates from patients O and M showed only marginal biofilm formation, whereas those from patient H showed constant biofilm formation, except for isolate H4, which belongs to a different genotype (Table 1). For all other isolates, both from CF and non-CF control patients, there was a general trend for decreased biofilm formation capacity over time, which would reflect a transition of isolates from the biofilm to planktonic lifestyle during colonization of the non-CF allograft (Fig. 2 and Fig. S1).

#### Virulence factor production

We next evaluated the production of quorum-sensing (QS)-regulated virulence factors. Pyocyanin is a redox-active secondary metabolite, whose expression is regulated by the Rhl and Pqs QS systems. With the exception of all isolates from CF-patient L and one isolate from CF-patients P and O, all other isolates, both from CF and control patients, produced at least 30% of PAO1 pyocyanin levels. Isolates from CF-patients M and G produced at least twice the amount of pyocyanin compared to PAO1. Three isolates from CF-patient O did not grow in the defined medium used for pyocyanin production, suggesting that these isolates were unable to grow on alanine or glycerol as C-sources (Fig. 1). Two of them belonged to a different genotype. Isolates from the three non-CF control patients produced pyocyanin at similar or higher levels compared to PAO1 (Fig. 2). There was no obvious trend towards either increased or decreased pyocyanin production over time.

Rhamnolipid is a secondary metabolite involved in nutrient acquisition, biofilm architecture and as surfactant for swarming motility (14). Biosynthesis is mainly controlled by the Rhl QS system. Rhamnolipid production was impaired in all CF-isolates, without any trend of recovery during allograft colonization. Indeed, all sequenced early and late isolates harbour missense mutations in one or several rhamnolipid biosynthesis genes (*rhlB*, *rhlC, rhlG*), but not in the main QS regulator gene *rhlR* or the homoserine-lactone (HSL) synthetase gene *rhlI* (Table S1). This could explain the absence of swarming motility in otherwise swimming proficient isolates, while maintaining expression of other Rhl QS-controlled phenotypes (Fig. 2).

#### Cytotoxicity

Cytotoxicity of the isolates was assessed on the respiratory airway epithelial cell line Calu-3 and the isogenic *cftr* knock down cell line, cultivated on Transwell filters at the air-liquid interface (15, 16). Early and late isolates from five CF LT-recipients were evaluated for their ability to affect the transmembrane electrical resistance (TEER), an indicator of epithelial cell integrity (Fig. 3). While the cytotoxic PAO1 control strain decreased TEER to approximately 50 Ω x cm^2^, indicating disruption of the epithelial barrier, all LT-isolates from both CF LT-recipients and controls showed TEER values identical or even higher than the uninfected control well at 24 h post-infection. This indicates that the LT isolates were non-cytotoxic even for the *cftr*-cell line. Except for isolates P0 and P932, there was no difference between early and late isolates, suggesting that colonization of a non-CF environment does not select for increased cytotoxicity.

**Figure 3.**
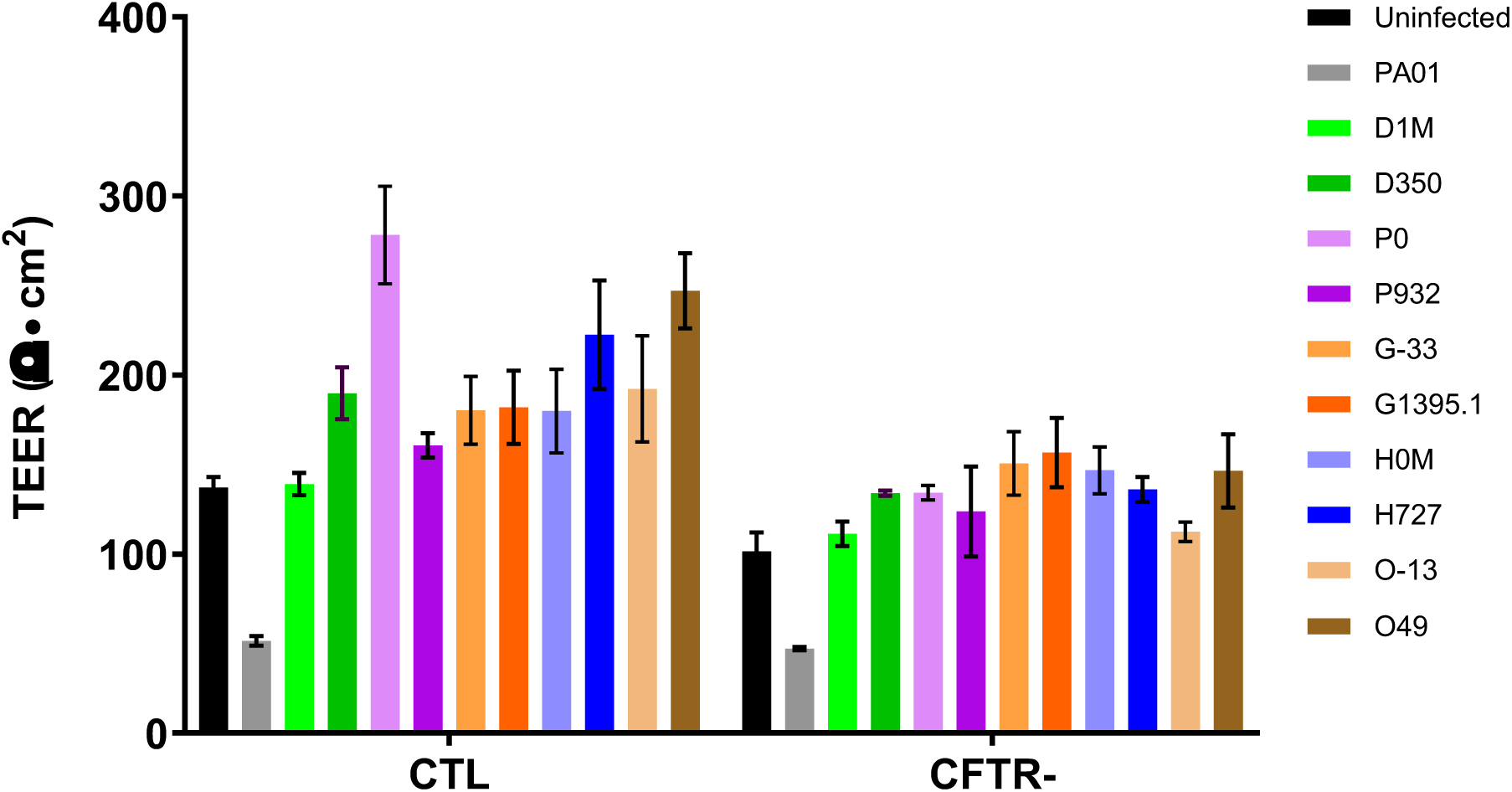
Transepithelial electrical resistance (TEER) determination on Calu-3 (CTL) and *cftr*-epithelial cells cultured in air-liquid interface on Transwell filters and infected with *P. aeruginosa*. TEER was determined after 24 h incubation in a 5% CO_2_ atmosphere. Values are the average and mean of three independent experiments.

Altogether, these observations suggest evolution during allograft colonization towards improved growth and reduced biofilm formation capacity, but also phenotypic stability with respect to other virulence-related traits of the initial allograft isolates (Table S2).

##### Mutator phenotype

The DNA mismatch repair deficient mutants displaying a hypermutator phenotype are a hallmark of *P. aeruginosa* isolates from chronic infections (17). They are present in 30-60% of CF-patients and rarely (<1%) in non-CF-patients (18, 19). In our collection, twelve of thirty-six isolates tested (30%) displayed a hypermutator phenotype, defined as a more than 20-fold higher mutation frequency compared to the PAO1 reference strain. These twelve isolates were recovered from three (D, P, O) of the seven (43%) CF-patients (Tables 1 and 3). The three isolates of the CF-patient M also showed an increased mutation rate of 10-fold compared to PAO1 (data not shown) and are referred to below, together with hypermutators, as mutator strains.

**Table 3.**
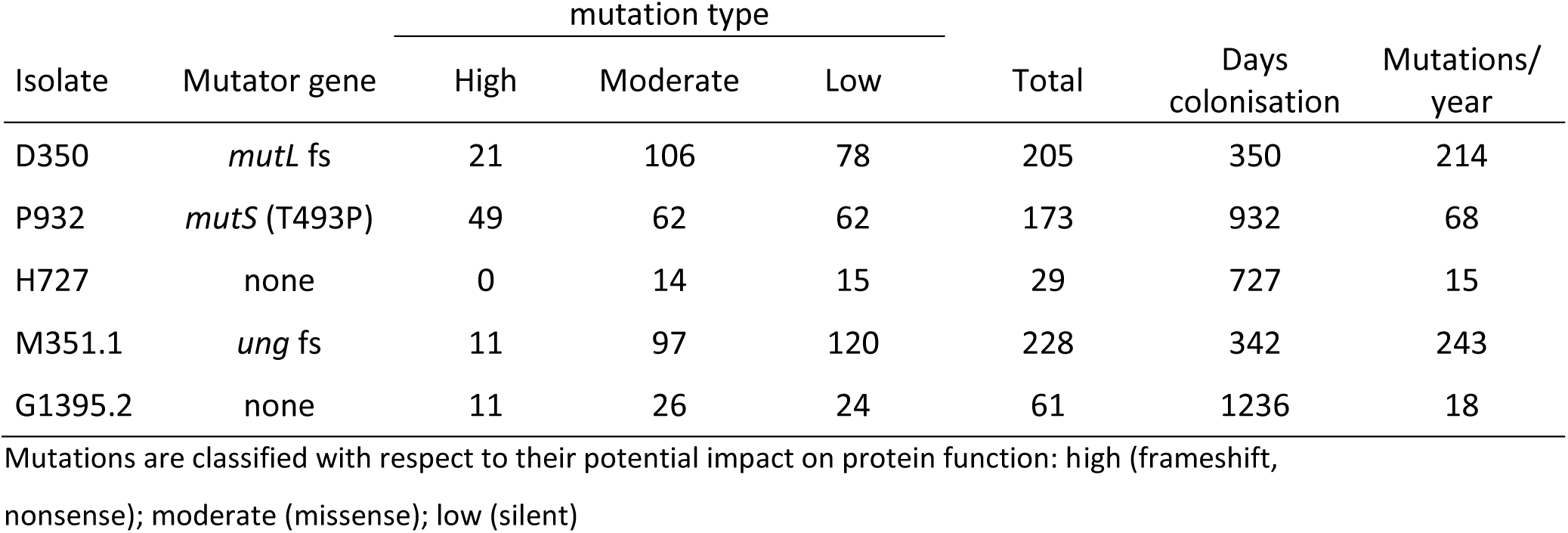
Theoretical mutation rate during allograft colonization.

Mutator and non-mutator isolates could be detected within the same sample (sinus swab (SIN) O-30) or in different samples from the same day (P183.1 from BAL and P183.2 from ASP) (Table 1). Hence, mutators were present in all sampling sites (SIN, ASP, BAL), but without any significant association (i.e. BAL vs ASP; Fisher exact test, p= 0.56). As expected, no mutator phenotype was observed among isolates from the three non-CF patients control A, B and C (Table 1).

##### Genetic evolution

To follow a possible genetic evolution of early isolates within the non-CF allograft environment, we further submitted isogenic pairs of early and late isolates from six CF-patients to WGS (Table 1 and Table S1). All contigs were compared to the genome of the Liverpool epidemic strain LESB58 (20), which showed the highest coverage with the contigs of the twelve sequenced isolates. We compared the number of mutations between the six isogenic pairs of early and late isolates and searched for loci that were altered in at least two lineages. We could identify 16 novel mutations with high impact (frame shift (fs), nonsense) and 21 with moderate impact (codon insertion/deletion, missense) occurring in 30 different loci (Table 4). Mutations emerged preferentially in pyoverdin (*pvdJ*, *pvdI*, *pvdL*) and alginate (*mucA*, *algG*, *algA*) biosynthesis genes, as well as in loci involved in antibiotic resistance (*oprD*, *fusA1*, *ampR, pirR*), cell metabolism (*treA*, *rpoN*, *mntH*, *purL*, *aphA*), biofilm formation (*hsbR*, *bifA, cupE6*) and in the QS regulator gene *lasR*.

**Table 4.** Genes mutated in late vs early *P. aeruginosa* isolates in at least 2 lineages.

| Locus | Mutation type<br>(n mutations) | Lineages<br>(n) | Function, annotation | Isolate<br>(n mutations) |
| --- | --- | --- | --- | --- |
| *hsbR* | fs | 3 | biofilm switch | D350/P932/O49 |
| PALES_18641 | codon insertion | 3 | hypothetical protein (PA3203) | D350/P932/G1395.2 |
| *pirR* | fs | 3 | TCS response regulator | D350/P932/O49 |
| *purL* | missense | 3 | purine synthesis | D350/M351.1/P932 |
| *rpoN* | missense | 3 | sigma factor | O49/D350/M351.1 |
| *algG* | fs (1)/missense (2) | 3 | alginate biosynthesis | D350/G1395.2/O49 |
| *algA* | missense | 2 | alginate biosynthesis | D350/P932 |
| *mucA* | fs/missense | 2 | alginate biosynthesis | G1395.2/O49 |
| *pvdI* | fs/missense | 2 | pyoverdinin synthesis | M351.1(1)/ H727(1) |
| *pvdL* | missense | 2 | pyoverdinin synthesis | O49/M351.1 |
| *pvdJ* | missense | 2 | pyoverdinin synthesis | H727 (3)/O49 (1) |
| *ampR* | missense | 2 | ampC beta-lactamase regulation | D350/M351.1 |
| *amaB* | fs/missense | 2 | amino acid aldehyde DH | O49/M351.1 |
| *treA* | codon insertion | 2 | trehalase (hyperosmotic response) | D350/G1395.2 |
| *apaH* | missense | 2 | purine metabolism | D350/O49 |
| *bifA* | fs/missense | 2 | neg. regulator single species biofilm | G1395.2/O49 |
| *fusA1* | missense | 2 | aminoglycoside resistance | O49 (3)/G1395.2 (1) |
| *lasR* | missense | 2 | las QS regulator | O49/H727 |
| *oprD* | nonsense | 2 | carbapenem uptake | P932/O49 |
| PALES_00411 | missense | 2 | transposase | D350/P932 |
| PALES_03041 | fs | 2 | phospholipase related (PA0308) | P932/G1395.2 |
| PALES_29291 | fs | 2 | vgrG3 typeVI secretion | P932/G1395.2 |
| PALES_33191 | fs | 2 | putative H<sup>+</sup>/gluconate symporter | O49/D350 |
| PALES_34431 | nonsense/missense | 2 | aldehyde DH (PA1880) | M351.1/O49 |
| PALES_40261 | fs | 2 | MFS transporter | P932/O49 |
| PALES_45331 | fs | 2 | manganese transporter MntH | P932/O49 |
| PALES_50391 | fs | 2 | fimbrial adhesin CupE6 | O49/M351.1 |
| *pys2* | fs/missense | 2 | pyocin S2 | D350 (1)/M351.1 (1) |
| PALES_55371 | fs/missense | 2 | lpxH, lipid A (LPS) biosynthesis | P932/O49 |
| *potB* | missense | 2 | polyamine transporter | P932/O49 |

fs, frameshift

Interestingly, fs mutations in *hsbR* occurred in late isolates of three different lineages (patients D, P, O) (Table 4). HsbR is a response regulator involved in the switch between biofilm and planktonic life styles (21, 22). Mutations occurred at position G509 located in the kinase domain (aa position 443-563) of HsbR, leaving intact the phosphatase domain, which acts on HsbA (21). Dephosphorylated HsbA releases the anti-sigma factor FlgM from FliA, which in turn activates transcription of flagella genes (23) and potentially affects c-di-GMP as well as pyocyanin levels (24, 25). Missense mutations in *purL*, involved in purine biosynthesis, occurred also in three different lineages (D, P, M). PurL is required for inosine and GTP biosynthesis, which could indirectly affect c-di-GMP levels (26). These data suggest a possible trend towards decreased c-di-GMP levels and a “switch” from biofilm towards planktonic lifestyle during allograft colonization.

Mucoidy is another phenotype associated with chronic colonization in the CF-environment. Mutations were identified in *mucA* in all sequenced isolates with the exception of the non-mucoid isolates H727, M9 and M351.1 (Table S1). Interestingly, late isolates G1395.2 and O49 were non-mucoid. Further analysis of alginate biosynthesis genes revealed additional mutations in *algG* (fs in G1395.2) and *alg44* and *algQ* in isolate O49, all involved in alginate biosynthesis or transport. These mutations could have emerged during allograft colonization as an adaptation to the non-CF environment, leading to mucoid reversion.

As expected, the majority (74%) of mutations occurred in mutator isolates D350, P932, O49 and M351.1. WGS confirmed the presence of *mutS* and/or *mutL* alterations in the hypermutator strains as well as a fs mutation in the uracil DNA glycosylase gene *ung* in isolates M9 and M351.1, which showed intermediate mutation frequencies (27) (Table 1 and Fig. S1). When comparing the twelve sequenced isolates individually to strain LESB58, we found a significantly higher proportion of high and moderate impact mutations and a lower proportion of low impact mutations in the hypermutator compared to the non-hypermutator isolates (Fig. 4 and Table S1). Mutations with high or moderate impact represented 50-70 % of all mapped mutations, irrespective of the mutator phenotype.

**Figure 4.**
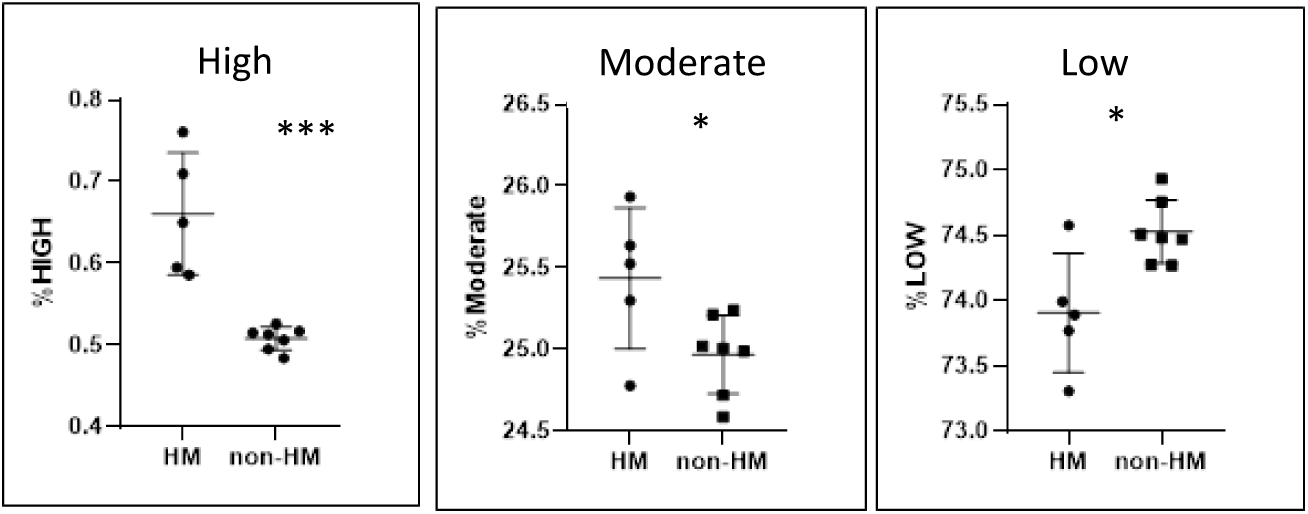
Mutation analysis of five hypermutator (HM) versus seven non-hypermutator (non-HM) isolates. The number of mutations with high (frameshift, nonsense), moderate (codon insertion/deletion, missense) and low (silent) impact are determined with respect to strain LESB58 and expressed as percentage of the total number of mutations for each isolate (Table S1). Student t-test; ***, p ≤ 0.005; *, p ≤ 0.05.

This suggests that frameshift and nonsense mutations as well as missense mutations with potential loss or gain of function consequences occurred preferentially in hypermutator isolates, indicative of a positive selection pressure.

To determine the genetic evolution of the initial post-LT isolate in CF-patients, we calculated the number of mutations accumulated during *de-novo* colonization of the allograft, based on the hypothesis that late isolates evolved from an isogenic ancestor. We excluded the strain couple O-13/O49, since the early isolate was obtained 13 days pre-LT (O-13). In all five other sequenced pairs of isolates from CF patients, we divided the total number of new mutations (SNPs and InDels), identified by WGS between the early and late isolates, by the corresponding days of allograft colonization. We obtained values of 15 and 18 mutations/year for the non-mutator isolates (AVG ± STD: 16.5 ± 1.5) and 68, 214, 243 (AVG ± STD: 175 ± 94) for mutator isolates (Table 3). During adaptation to the CF-lung, isolates were reported to accumulate a median of 3 SNPs/year (range: 0.5 to 14) for non-mutators and 48 SNPs/year (range: 2 to > 350) for hypermutators (28). Hence, our results are within the upper range of the reported mutation frequencies of isolates during chronic lung colonization in CF patients, while maintaining a similar 10-fold higher “in patient” mutation frequency for mutator compared to non-mutator isolates. Thus, we observed no reversion to lower mutation frequencies in isolates recovered from the non-CF allograft.

In summary, our data support the continuous seeding of CF-adapted *P. aeruginosa* isolates from the sinuses of LT-recipients to their non-CF allograft and provide evidence for further adaptation of specific phenotypic traits to the non-CF environment of the allograft.

## Discussion

The aim of this study was to assess whether *P. aeruginosa* isolates that colonized and adapted to the specific lung environment in CF-patients for years further adapt to the novel non-CF allograft environment. To this end, we collected longitudinal isolates both from CF LT-recipients and non-CF control patients and analysed their phenotypic and genotypic characteristics.

In agreement with previous studies, post-LT isolates showed the same genotype as the pre-LT isolates, except for one patient (5). Furthermore, sinus isolates from LT-recipients showed the same genotype as those from BAL and ASP samples, supporting colonization of the allograft from the sinus (29).

As expected for CF-isolates, the majority showed impaired motility, partial loss of QS-dependent phenotypes and reduced cytotoxicity. All isolates were swarming deficient, resulting likely from genetic alterations identified by WGS in flagella and/or rhamnolipid biosynthesis genes. Specific virulence traits like rhamnolipid or pyocyanin production, both controlled by the QS-systems, were genotype specific and did not show any phenotypic reversion during allograft colonization. All strains tested were less cytotoxic than PAO1 on epithelial cells, but without any significant differences between early and late isogenic isolates, suggesting that isolates do not seem to revert to increased virulence or cytotoxicity during allograft colonization.

The most striking evidence for adaptations to the allograft is the general trend, although not statistically significant, for decreased biofilm formation capacities during allograft colonization and loss of mucoid phenotype due to secondary mutations in initially mucoid isolates. This could be explained by the observation that mutations affecting flagella and secondary metabolite synthesis were located in structural or biosynthetic genes, while those leading to biofilm and mucoid phenotypes affect regulatory circuits controlling c-di-GMP or alginate production, which are more likely to be genetically “reversible”.

Several non-exclusive scenarios, depicted in Figure 5, may explain our observations: (i) CF-adapted LT-isolates evolve and acquire novel genetic mutations, which provide a fitness advantage in the non-CF-lung environment. This is supported by highly specific mutations identified in the biofilm regulator HsbR and secondary mutations in alginate biosynthesis genes of mucoid isolates in several lineages, (ii) the nutrient-restricted non-CF lung environment counter-selects auxotrophic CF-adapted isolates. Indeed, only four isolates (<10% of all isolates tested) from one CF-patient (14%) were unable to grow in defined media. This proportion is higher in CF-patients, where up to 40% of isolates from up to 86% of adult CF-patients were auxotrophic mutants (30) (iii) CF-adapted isolates from the sinus survive in a non-CF environment, without further adaptation. Since isolates continuously evolve in the CF-environment of the sinuses, where hypermutators are maintained, their higher phenotypic diversity increases the likelihood of variants compatible with the novel allograft environment explaining their presence among early and late isolates (Fig. 5).

**Figure 5.**
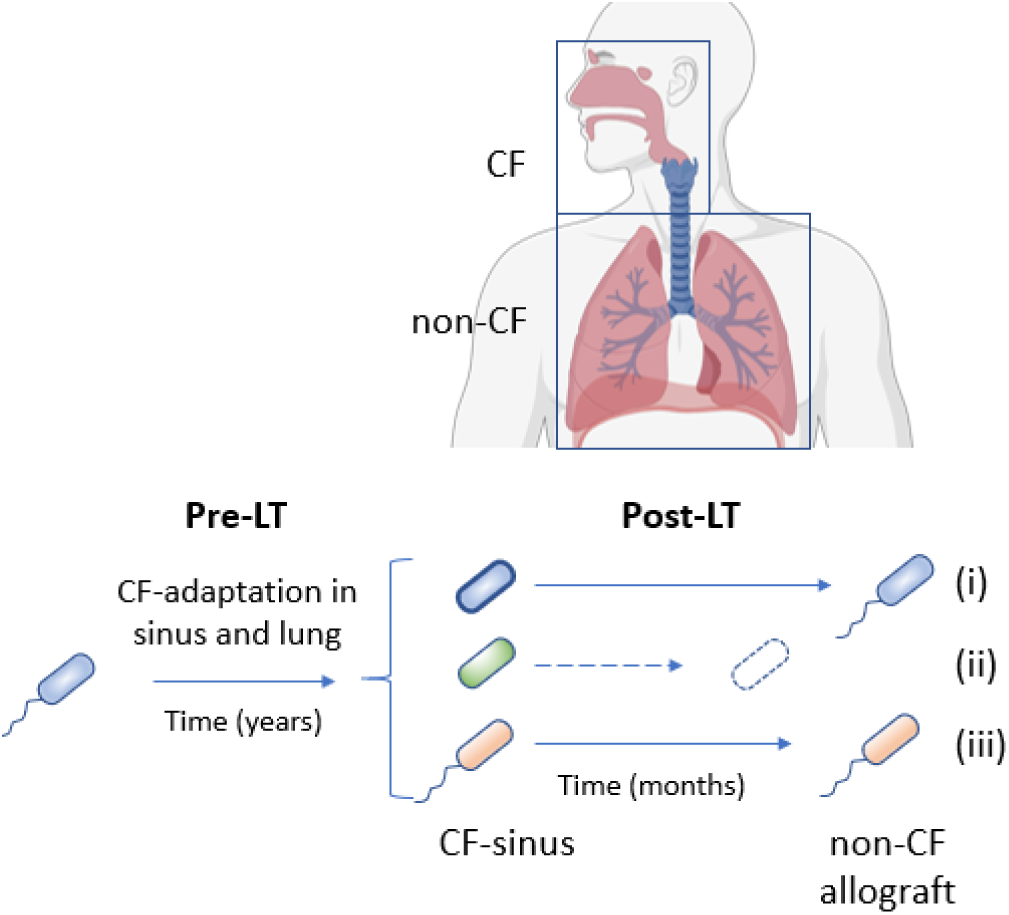
Possible fates of *P. aeruginosa* in the non-CF allograft. (i) adaptation of a CF-adapted variant to a non-CF environment (i.e. loss of biofilm or mucoid phenotype); (ii) extinction of CF-adapted variant due to fitness cost in the non-CF-allograft environment (auxotrophic phenotype); (iii) CF-adapted variant with allograft compatible phenotype. For details see discussion.

The calculated novel mutations acquired per year by the allograft isolates was in the upper range of mutation frequencies observed for both hyper and non-hypermutators during chronic colonization (28). Mutator genotypes were observed in early and/or late isolates in four of the seven CF LT-recipients, suggesting that mutator isolates are likely descending from sinuses or the upper respiratory tract where a CF-environment is maintained.

Colonization of the allograft was reported to occur between 15 to 19 days post-LT. This early colonization very likely occurs from the sinuses or upper trachea or nasopharynx, which remain colonized by this organism in CF-patients (31). Since the colonization of the allograft is associated with CLAD and allograft dysfunction, preventing the seeding from the sinuses could bring a clinical benefit. Several strategies, including pre-LT (32) and/or post-LT sinus surgery with and without sinus lavage have been evaluated in prospective and retrospective studies (33–35). While there was no significant benefit of this intervention with respect to overall survival and frequency of CLAD, one single center study found a significant decrease in allograft infection in the sinus resected LT-patients (33). Given the invasiveness of sinus surgery most centers refrain from this procedure. Our data now further support the continuous seeding from the sinus reservoir of CF-adapted *P. aeruginosa* isolates, reinforcing the potential benefit of preventive strategies. As the prevention of CLAD would bring a major clinical benefit on allograft survival, new preventive strategies, potentially including alternative approaches to classic antibiotics such as phage inhalations should be investigated in clinical trials.

## Materials and methods

### Sample and isolate collection

During routine surveillance bronchoscopies, sequential bronchial aspirates (ASP) and bronchoalveolar lavages (BAL) were collected. ASPs consisted of respiratory secretions aspirated below the bronchial suture representing heterogeneous sampling from both lungs. BAL fluid was obtained by aspiration following instillation of approximately 150 ml of NaCl 0.9% through the bronchoscope wedged in a subsegmental bronchus, representing a more homogeneous sample from a distal respiratory zone. One ml aliquots of native ASP and BAL samples were frozen and stored at − 80 °C. Patients were screened for the presence of *P. aeruginosa* either in ASPs, BALs, sputum or sinus samples. *P. aeruginosa* was isolated by the routine diagnostic microbiology laboratories of the Lausanne University Hospital (CHUV) and the University Hospitals Geneva (HUG), Switzerland. *P. aeruginosa* isolates were sequentially collected during a period of up to 5 years post-LT. Isolates were stored as glycerol stocks at −80 °C.

### Genotyping

For colony PCR amplification, a single colony was resuspended in 0.9% NaCl and heated at 95°C for 10 min. PCR amplification of each locus (ms172 or ms217) was carried out in separate reactions in a total of 30 µl, containing 3 µl of DNA suspension, 5U/µL of TAQ DNA polymerase (Sigma), 1X Taq reaction buffer, 1.5 µL of DMSO, 1.8 µL (10 µM) of each primer, 2.4 µl (2.5 mM) of dNTP. Cycling conditions were 2 min of initial denaturation at 95°C, 30 cycles consisting of 20s at 95°C, 30s at 60°C (ms172) or 64°C (ms217), 45s at 72°C and a final extension for 4 min at 72°C. The sizes of the DNA fragments were respectively 400 bp (ms172) and 350 bp (ms217). The sequences of the forward and reverse primers used were as follows: ms172.forward (5’-3’) GGATTCTCTCGCACGAGGT; ms172.reverse (5’-3’) TACGTGACCTGACGTTGGTG; ms217.forward (5’-3’) TTCTGGCTGTCGCGACTGAT and ms217.reverse (5’-3’) GAACAGCGTCTTTTCCTCGC. The PCR product was sequenced and analysed by mapping the DNA sequences to the DLST database (www.DLST.org) (9).

### Growth measurements

Bacterial suspensions for growth measurements were prepared by diluting overnight cultures grown in LB broth into 0.9% NaCl to an OD_600_ of 1.0. LB medium or M9 salts minimal medium supplemented with 2 mM MgSO_4_ and a carbon source (0.2% glucose) were inoculated with bacterial suspensions (final OD_600_= 0.2). Casamino acids (CAA) medium (M9 salts medium supplemented with 0.5% casamino acids) was used for iron-limited growth with or without FeCl_2_ supplementation at 20 µM final concentration. Growth (OD_600_) at 37°C was monitored under static conditions in 96 well microtiter plates using a plate reader (BioTek Synergy H1). Experiments were performed in duplicates.

### Mucoidity

Mucoidity of isolates, resulting from alginate production, was observed visually after 24-48 h of incubation at 37°C on LB agar plates and scored as mucoid (shiny, mucoid morphology) or non-mucoid (smooth colony phenotype).

### Motility assays

Swimming motility was measured as follows: a colony from a fresh LB plate was stab-inoculated into a 0.3% LB-agar plate and incubated at 37°C for 18 h. Swarming motility was determined by inoculating a single colony from a fresh LB plate, onto a 0.5 % agar plate medium (1X M8 medium supplemented with 0.2 % glucose and 0.05 % glutamate) and incubated during 24h at 37°C. The diameter of the swimming/swarming area was measured and compared to the one of the PAO1 strain, set to 100%.

### Pyocyanin production

A colony from a LB agar plate was grown in GA-C medium (6g/L DL-alanine, 1% glycerol, 0.1 g/L K2HPO_4_, 2 g/L MgSO_4_, 0.018 g/L FeSO_4_) during 18-20 h at 37°C (36). The optical density of the culture was measured at 590 nm. Pyocyanin was extracted from 0.75 ml of culture supernatant with chloroform and extracted again from the organic phase with 0.2 N HCl. Absorbance was measured in the aqueous phase at 520 nm. The ratios of OD520/OD590 were determined and compared to those of PAO1 set to 100%.

### Rhamnolipid production

Five µL of an overnight culture in LB medium were spotted on a rhamnolipid agar medium plate composed of M8 salts 1X, 0.2% glucose, 2mM MgSO4, 0.0005% methylene blue, 0.1% glutamate, 0.02% cetyltrimethylammonium bromide and solidified with agar (1,6% final concentration) (14, 37). Plates were incubated overnight at 37°C, followed by 24h at room temperature and 3-5 hours at 4°C. Rhamnolipid production is indicated by a blue-violet halo forming around the colony. The diameter of this halo, corresponding to a precipitate of rhamnolipids, was measured in mm and compared to the one obtained with PAO1.

### Microtiter plate biofilm assay

After overnight culture in LB medium (37°C), bacteria were grown in M63 medium supplemented with 0.05% casamino acids, 0.4% arginine, 1 µg/ml vitamin B12, 1 µg/ml vitamin B1, 1 mM MgSO_4_ (20 µL overnight culture in 180 µL of supplemented M63 medium) in a 96 well plate. The plate was incubated during 24h at 37°C in static conditions and the absorbance was measured at 590 nm and biofilms were stained with crystal violet (38). Optical density of cultures was measured at 550 nm in a plate reader (BioTek Synergy H1). All experiments were performed in biological duplicates with PAO1 as reference strain for comparison.

### Cytotoxicity

Cytotoxicity was assessed by adding *P. aeruginosa* isolates to a monolayer of Calu-3 (HTB-55TM, ATCC, Manassas, VA, United States) control cells or isogenic Calu-3 *cftr* knock-down cells as described (15, 16). Briefly, overnight LB-grown bacteria were resuspended in saline buffer (NaCl 0.9%, HEPES 10 mM, CaCl_2_ 1.2 mM) and adjusted by serial dilution in saline buffer to approximately 10^5^ CFU/ml. Ten µl of this dilution (approx. inoculum of 10^3^±10^1^ CFUs, MOI of 0.002) were added apically to the Transwell filters containing 5 × 10^5^ cells. Ten µl of saline buffer was added to uninfected control cells. Strain PAO1 was used as a positive cytotoxic control strain. Filters were incubated at 37°C with 5% CO_2_ for 24h (16). At this point, bacterial growth was determined by adding apically 200 µl of saline buffer to the Transwell filter. A 100 µl aliquot was removed apically and diluted 10-fold to perform plate counts on LB agar plates. Transepithelial electrical resistance was measured as before (16) using chopstick electrodes and a voltohm-meter (EVOM, World Precision Instruments, Inc).

### Whole genome sequencing

Genomic DNA from 12 isolates (first and last isolate from CF-patients D, M, O, G, H and P) was extracted using the Qiagen tissue extraction kit according to the manufacturer’s protocol. Whole genome sequencing was performed by NGMicrobes at the University of Birmingham, UK. Contigs were aligned on the genome (NC_011770.1) of the reference isolate LESB58 (20), which showed maximal coverage for the majority of contigs.

### Minimal inhibitory concentration assays (MIC)

MIC determinations were performed according to CLSI guidelines by two-fold serial dilutions in Mueller-Hinton broth medium as described previously (39).

### Mutation frequency measurement

One bacterial colony was resuspended in 20 ml of Mueller-Hinton Broth (MHB) and grown at 37°C overnight. Bacterial cells were then centrifuged at 3’000 rpm for 5 min and resuspended in 1 ml of MHB. A 100-µl sample from this suspension as well as from successive dilutions was plated onto LB plates, with and without rifampicin (300 µg/ml final concentration). In all cases, the isolates were originally susceptible to such concentrations of rifampicin. Colony counting was performed in plates containing 100 to 300 colonies after 24-36 h of incubation. To avoid mutation jackpot (the recovery, by chance, of a vast number of mutants), all experiments were performed at least in triplicate and the mean value was recorded. As previously described (17), one strain was considered a mutator when the corresponding mutation frequencies on rifampicin (300 µg/ml) were at least 20-fold higher than those observed for the PAO1 reference strain.

## Data availability

The datasets generated and/or analysed during the current study are available in the Zenodo repository (https://doi.org/10.5281/zenodo.17779534).

## Acknowledgements

We would like to thank Prof. John-David Aubert as well as Dr. Eric Bernasconi (Service of pneumology, CHUV) for organizing the sample collection and providing access to their LT-patient cohorts. We are grateful to all medical staff members involved in patient follow-up and sample collection at both institutions. We also thank Dr. Juliette Simonin for preparing Calu-3 cell cultures.

## Author contributions

Conception and design, C.vD., T.K.; Investigation, L.F., A.L., M.G-B., Resources, G.B., G.G., M.C., A.K., P.M.S.; Analysis and interpretation, T.K., C.vD., A.L.; Writing-original draft, T.K., C.vD. All authors have reviewed the manuscript and approved the submitted version.

## Declarations

### Ethical approval

Authorization to use all clinical samples for research purposes were obtained from the local ethical committee of the University Hospitals of Geneva (authorization CER 07–301), and the patients provided their written informed consent. All methods and experimental protocols were carried out in accordance with the ethical principles from the Helsinki declaration for medical research involving human participants. The Swiss Transplant Cohort Study is registered under ClinicalTrials.gov number: NCT 01204944.

---

**Figure S1.**
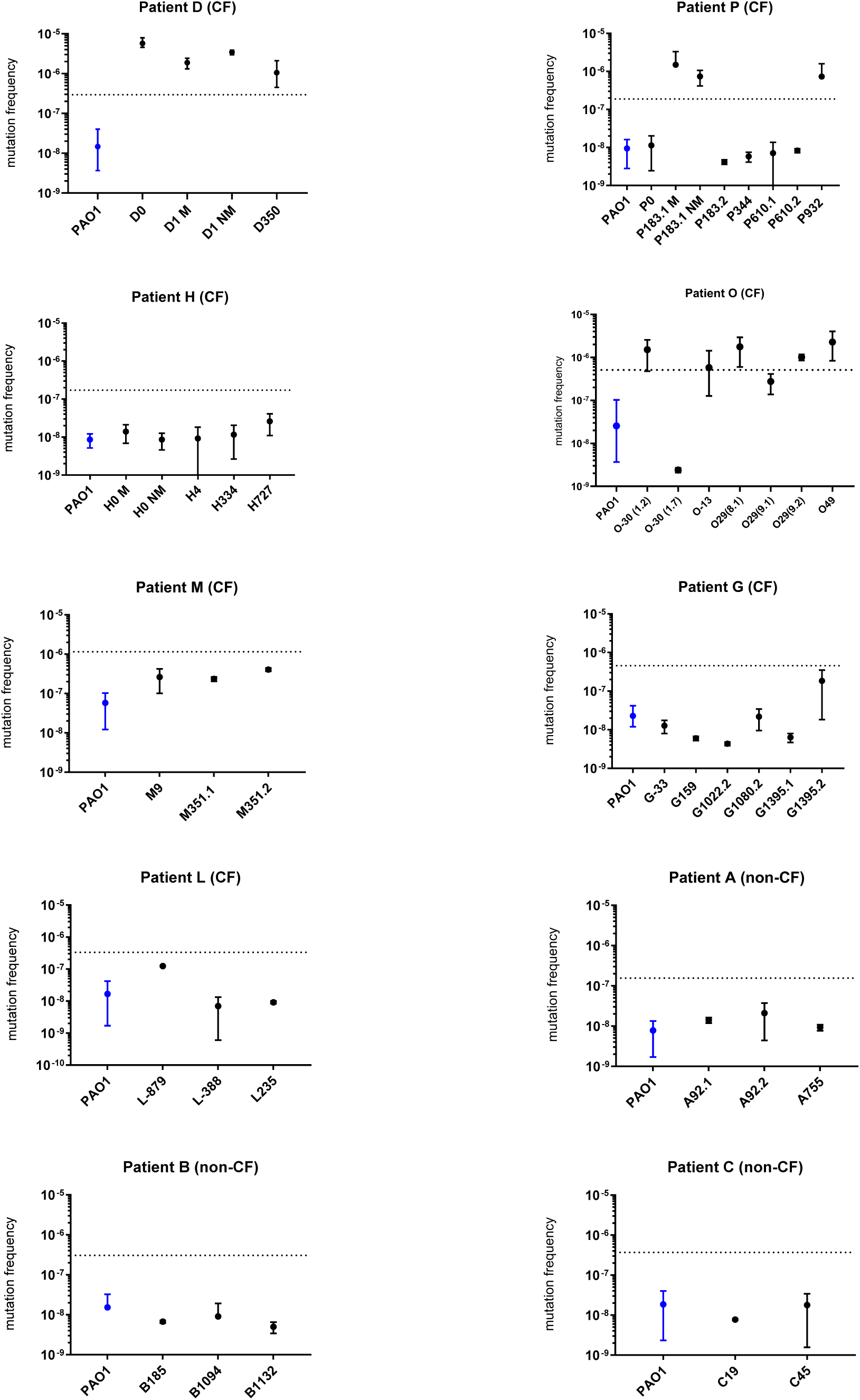
Mutation frequencies of LT-isolates. The dotted line indicates the threshold of a 20-fold increased mutation frequency for classification as a hypermutator

**Figure S2.**
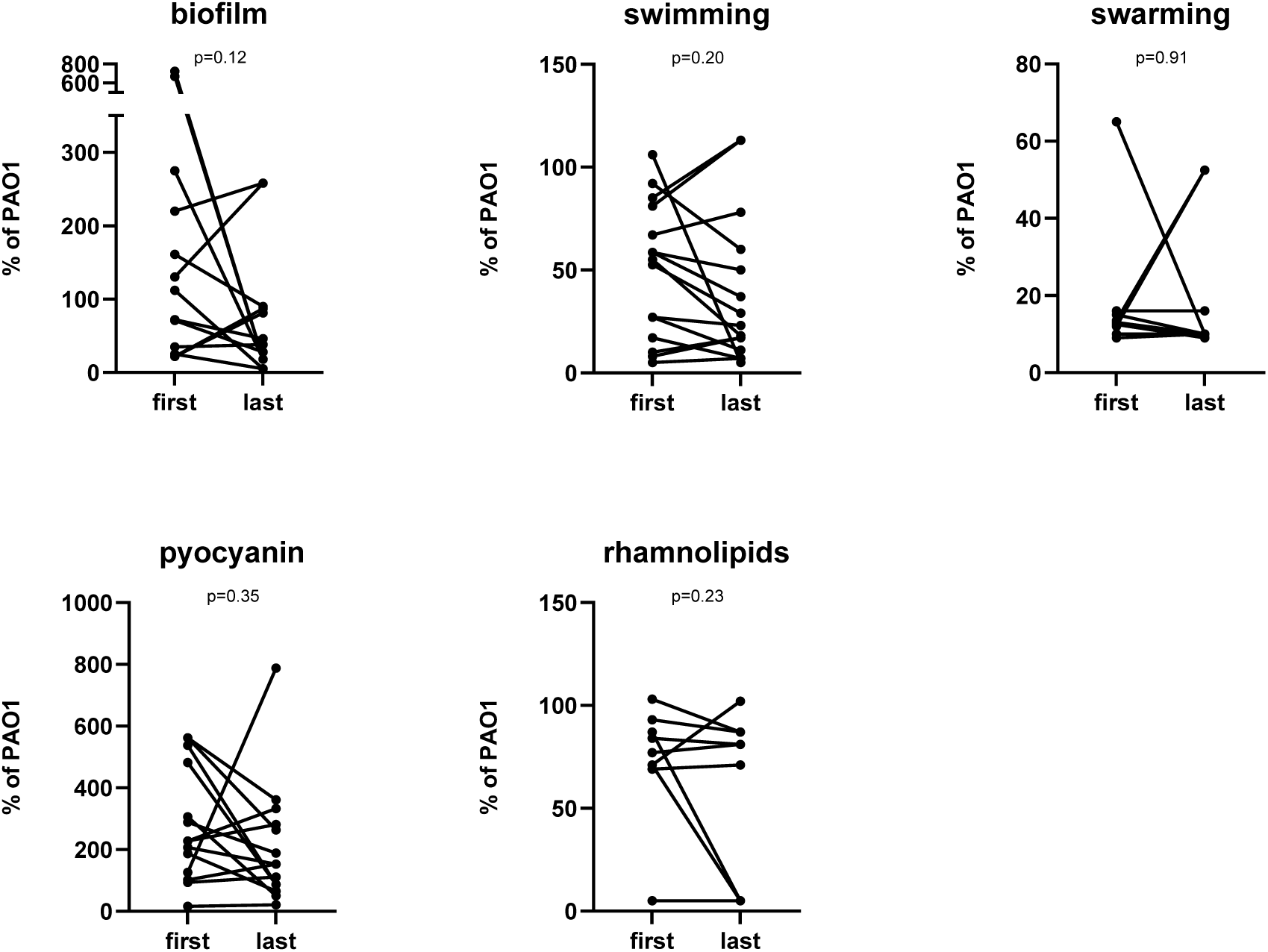
Phenotype evolution between first and last isolate of seven CF and three non-CF LT-patients A paired t-test (GraphPad prism, version 10.6.1) was used to analyse differences between the first and the last isolate available from each of the ten LT-patients. p-values are indicated above each graph

