## Supplemental Table 1 for "Adaptation of *Pseudomonas aeruginosa* to the lung allograft environment in cystic fibrosis lung transplant recipients"

### Table S1. Variant call mapping on isolate LESB58

| Chromosome | Position | Ref_Allele_ | Alt_Alle | Average_ | train | Homoz | Het_ca | Mutation_Type | Mutation_ | Mutation_ | Codon_Su | AminoAcid | Locus_Tag | D0 24845_D0 | D350 24846_D3 | P0 24847_P0 | P932 24848_P9 | G159 24849_G1 | G1395.2 24850_G1 | O-13 24851_O-1 | O49 24852_O4 | H0-NM 24853_H0- | H727 24854_H7 | M9 24855_M9 | M351.1 24856_M3511 |
|---|---|---|---|---|---|---|---|---|---|---|---|---|---|---|---|---|---|---|---|---|---|---|---|---|---|
| NC_011770 | 2677426 | C | T | 48 | 0 | 6 | 0 | NON_SYNONYMOU | MODERAT | MISSENSE | Gac/Aac | D106N | aat | | | | | | | T | T | T | T | T | T |
| NC_011770 | 1506161 | A | G | 75 | 0 | 8 | 0 | NON_SYNONYMOU | MODERAT | MISSENSE | Acg/Gcg | T289A | accA | G | G | | | | | G | G | G | G | G | G |
| NC_011770 | 5775170 | G | A | 70 | 0 | 1 | 0 | NON_SYNONYMOU | MODERAT | MISSENSE | atG/atA | M121I | accB | | | | | | | | | A | | | |
| NC_011770 | 5775163 | A | G | 68 | 0 | 2 | 0 | NON_SYNONYMOU | MODERAT | MISSENSE | gAa/gGa | E119G | accB | | | G | G | | | | | | | | |
| NC_011770 | 5775785 | G | A | 61 | 0 | 1 | 0 | NON_SYNONYMOU | MODERAT | MISSENSE | Ggc/Agc | G164S | accC | | A | | | | | | | | | | |
| NC_011770 | 5775473 | T | C | 54 | 0 | 2 | 0 | NON_SYNONYMOU | MODERAT | MISSENSE | Tac/Cac | Y60H | accC | | | | | | | | | | | C | C |
| NC_011770 | 5775894 | C | T | 74 | 0 | 12 | 0 | NON_SYNONYMOU | MODERAT | MISSENSE | cCg/cTg | P200L | accC | T | T | T | T | T | T | T | T | T | T | T | T |
| NC_011770 | 5775986 | T | C | 84 | 0 | 1 | 0 | NON_SYNONYMOU | MODERAT | MISSENSE | Tcc/Ccc | S231P | accC | | | | | | | C | | | | | |
| NC_011770 | 5776220 | A | G | 82 | 0 | 12 | 0 | NON_SYNONYMOU | MODERAT | MISSENSE | Atc/Gtc | I309V | accC | G | G | G | G | G | G | G | G | G | G | G | G |
| NC_011770 | 2101973 | A | T | 78 | 0 | 2 | 0 | NON_SYNONYMOU | MODERAT | MISSENSE | Atg/Ttg | M122L | accD | | | T | T | | | | | | | | |
| NC_011770 | 2101892 | T | C | 87 | 0 | 1 | 0 | NON_SYNONYMOU | MODERAT | MISSENSE | Tac/Cac | Y95H | accD | | | | | | | | C | | | | |
| NC_011770 | 5972757 | G | A | 90 | 0 | 1 | 0 | NON_SYNONYMOU | MODERAT | MISSENSE | Gcc/Acc | A643T | aceE | | | | | | | A | | | | | |
| NC_011770 | 5973862 | A | G | 27 | 0 | 2 | 0 | NON_SYNONYMOU | MODERAT | MISSENSE | gAa/gGa | E80G | aceF | | | | | | | | | | | G | G |
| NC_011770 | 5973957 | G | A | 31 | 0 | 2 | 0 | NON_SYNONYMOU | MODERAT | MISSENSE | Gcc/Acc | A112T | aceF | A | A | | | | | | | | | | |
| NC_011770 | 5974257 | G | T | 60 | 0 | 2 | 0 | NON_SYNONYMOU | MODERAT | MISSENSE | Gcc/Tcc | A212S | aceF | | | | | | | | | | | T | T |
| NC_011770 | 5974958 | G | C | 78 | 0 | 2 | 0 | NON_SYNONYMOU | MODERAT | MISSENSE | gaG/gaC | E445D | aceF | | | | | | | | | C | C | | |
| NC_011770 | 4216904 | G | GC | 67 | 0 | 1 | 0 | FRAME_SHIFT | HIGH | | ggc/Cggc | G320R? | aceK | | | | | | | | GC | | | | |
| NC_011770 | 4216023 | A | G | 73 | 0 | 1 | 0 | NON_SYNONYMOU | MODERAT | MISSENSE | cAg/cGg | Q24R | aceK | | | | | | | | G | | | | |
| NC_011770 | 4165711 | C | A | 81 | 0 | 2 | 0 | NON_SYNONYMOU | MODERAT | MISSENSE | gaC/gaA | D196E | acnA | | | | | | | | | | | A | A |
| NC_011770 | 4166312 | C | A | 72 | 0 | 2 | 0 | NON_SYNONYMOU | MODERAT | MISSENSE | Ctc/Atc | L397I | acnA | | | | | | | | | | | A | A |
| NC_011770 | 4167419 | C | A | 68 | 0 | 10 | 0 | NON_SYNONYMOU | MODERAT | MISSENSE | Caa/Aaa | Q766K | acnA | A | A | A | A | A | A | A | A | | | A | A |
| NC_011770 | 4167455 | G | C | 74 | 0 | 2 | 0 | NON_SYNONYMOU | MODERAT | MISSENSE | Gag/Cag | E778Q | acnA | | | | | | | | | | | C | C |
| NC_011770 | 4167278 | C | T | 75 | 0 | 1 | 0 | NON_SYNONYMOU | MODERAT | MISSENSE | Ccg/Tcg | P719S | acnA | | | | | | | | T | | | | |
| NC_011770 | 3930939 | C | G | 74 | 0 | 2 | 0 | NON_SYNONYMOU | MODERAT | MISSENSE | Ctc/Gtc | L168V | acnB | G | G | | | | | | | | | | |
| NC_011770 | 3931128 | T | C | 75 | 0 | 2 | 0 | NON_SYNONYMOU | MODERAT | MISSENSE | Ttc/Ctc | F231L | acnB | | | | | | | C | C | | | | |
| NC_011770 | 3932136 | A | T | 71 | 0 | 2 | 0 | NON_SYNONYMOU | MODERAT | MISSENSE | Acc/Tcc | T567S | acnB | | | | | | | | | T | T | | |
| NC_011770 | 847612 | A | G | 83 | 0 | 1 | 0 | NON_SYNONYMOU | MODERAT | MISSENSE | gTg/gCg | V308A | acoB | | | | | | | | G | | | | |
| NC_011770 | 852061 | C | T | 89 | 0 | 2 | 0 | NON_SYNONYMOU | MODERAT | MISSENSE | aCc/aTc | T110I | acoR | | | | | | | T | T | | | | |
| NC_011770 | 852546 | C | G | 83 | 0 | 12 | 0 | NON_SYNONYMOU | MODERAT | MISSENSE | Ctg/Gtg | L272V | acoR | G | G | G | G | G | G | G | G | G | G | G | G |
| NC_011770 | 853437 | G | A | 69 | 0 | 2 | 0 | NON_SYNONYMOU | MODERAT | MISSENSE | Gcc/Acc | A569T | acoR | | | | | | | | | | | A | A |
| NC_011770 | 1991392 | C | T | 70 | 0 | 2 | 0 | NON_SYNONYMOU | MODERAT | MISSENSE | Ccc/Tcc | P155S | acpD | | | | | | | | | T | T | | |
| NC_011770 | 4867045 | T | C | 71 | 0 | 1 | 0 | NON_SYNONYMOU | MODERAT | MISSENSE | Atc/Gtc | I559V | acsA | | C | | | | | | | | | | |
| NC_011770 | 4868568 | G | A | 98 | 0 | 2 | 0 | NON_SYNONYMOU | MODERAT | MISSENSE | cCg/cTg | P51L | acsA | A | A | | | | | | | | | | |
| NC_011770 | 5649459 | A | C | 77 | 0 | 12 | 0 | NON_SYNONYMOU | MODERAT | MISSENSE | gaA/gaC | E36D | acsB | C | C | C | C | C | C | C | C | C | C | C | C |
| NC_011770 | 5649469 | G | A | 80 | 0 | 2 | 0 | NON_SYNONYMOU | MODERAT | MISSENSE | Ggc/Agc | G40S | acsB | | | | | | | | | A | A | | |
| NC_011770 | 5649895 | G | A | 86 | 0 | 2 | 0 | NON_SYNONYMOU | MODERAT | MISSENSE | Gtg/Atg | V182M | acsB | | | | | | | | | | | A | A |
| NC_011770 | 5650097 | C | T | 83 | 0 | 4 | 0 | NON_SYNONYMOU | MODERAT | MISSENSE | gCg/gTg | A249V | acsB | | | T | T | T | T | | | | | | |
| NC_011770 | 5650354 | G | A | 76 | 0 | 2 | 0 | NON_SYNONYMOU | MODERAT | MISSENSE | Gac/Aac | D335N | acsB | | | | | A | A | | | | | | |
| NC_011770 | 1980565 | A | G | 79 | 0 | 1 | 0 | NON_SYNONYMOU | MODERAT | MISSENSE | tAc/tGc | Y301C | actP | | | | | | | G | | | | | |
| NC_011770 | 3542914 | G | T | 72 | 0 | 2 | 0 | NON_SYNONYMOU | MODERAT | MISSENSE | agC/agA | S82R | ada | | | | | | | T | T | | | | |
| NC_011770 | 3543093 | C | T | 60 | 0 | 2 | 0 | NON_SYNONYMOU | MODERAT | MISSENSE | Gcg/Acg | A23T | ada | | | | | | | T | T | | | | |
| NC_011770 | 3543146 | A | G | 67 | 0 | 2 | 0 | NON_SYNONYMOU | MODERAT | MISSENSE | cTc/cCc | L5P | ada | | | | | | | G | G | | | | |
| NC_011770 | 6443739 | G | A | 85 | 0 | 1 | 0 | NON_SYNONYMOU | MODERAT | MISSENSE | Ggc/Agc | G236S | adhA | | | | | | | | | | | | A |
| NC_011770 | 6443599 | A | G | 93 | 0 | 1 | 0 | NON_SYNONYMOU | MODERAT | MISSENSE | tAc/tGc | Y189C | adhA | | | | | | | G | | | | | |
| NC_011770 | 1515528 | C | T | 66 | 0 | 1 | 0 | NON_SYNONYMOU | MODERAT | MISSENSE | gCg/gTg | A71V | adhC | | | | | | | T | | | | | |
| NC_011770 | 1515630 | A | G | 94 | 0 | 1 | 0 | NON_SYNONYMOU | MODERAT | MISSENSE | aAc/aGc | N105S | adhC | | | | | | | | G | | | | |
| NC_011770 | 1410365 | T | C | 77 | 0 | 4 | 0 | NON_SYNONYMOU | MODERAT | MISSENSE | gTt/gCt | V148A | adk | | | C | C | C | C | | | | | | |
| NC_011770 | 4168900 | C | G | 81 | 0 | 2 | 0 | NON_SYNONYMOU | MODERAT | MISSENSE | gaC/gaG | D224E | aer | | | | | | | | | | | G | G |
| NC_011770 | 4169153 | G | A | 77 | 0 | 1 | 0 | NON_SYNONYMOU | MODERAT | MISSENSE | Gcc/Acc | A309T | aer | | A | | | | | | | | | | |
| NC_011770 | 4169240 | G | A | 70 | 0 | 1 | 0 | NON_SYNONYMOU | MODERAT | MISSENSE | Ggg/Agg | G338R | aer | | A | | | | | | | | | | |
| NC_011770 | 199337 | T | A | 80 | 0 | 2 | 0 | NON_SYNONYMOU | MODERAT | MISSENSE | cAa/cTa | Q378L | aer2 | | | | | | | A | A | | | | |
| NC_011770 | 199345 | G | C | 79 | 0 | 2 | 0 | NON_SYNONYMOU | MODERAT | MISSENSE | gaC/gaG | D375E | aer2 | | | | | | | C | C | | | | |
| NC_011770 | 199794 | T | A | 83 | 0 | 2 | 0 | NON_SYNONYMOU | MODERAT | MISSENSE | Atg/Ttg | M226L | aer2 | | | | | | | A | A | | | | |
| NC_011770 | 323071 | A | G | 85 | 0 | 1 | 0 | NON_SYNONYMOU | MODERAT | MISSENSE | Ttc/Ctc | F257L | aguB | | | | | | | G | | | | | |
| NC_011770 | 324666 | ATG | A | 52 | 0 | 12 | 0 | FRAME_SHIFT+STA | HIGH | | atg/ | M1 | aguR | A | A | A | A | A | A | A | A | A | A | A | A |
| NC_011770 | 324416 | G | T | 56 | 0 | 2 | 0 | NON_SYNONYMOU | MODERAT | MISSENSE | gaC/gaA | D84E | aguR | T | T | | | | | | | | | | |
| NC_011770 | 324652 | G | A | 52 | 0 | 12 | 0 | NON_SYNONYMOU | MODERAT | MISSENSE | Ccc/Tcc | P6S | aguR | A | A | A | A | A | A | A | A | A | A | A | A |
| NC_011770 | 157851 | A | G | 78 | 0 | 1 | 0 | NON_SYNONYMOU | MODERAT | MISSENSE | gAt/gGt | D28G | ahpF | | | | | | | G | | | | | |
| NC_011770 | 158660 | A | G | 90 | 0 | 6 | 0 | NON_SYNONYMOU | MODERAT | MISSENSE | Acc/Gcc | T298A | ahpF | | | G | G | G | G | G | G | | | | |
| NC_011770 | 158682 | A | G | 88 | 0 | 6 | 0 | NON_SYNONYMOU | MODERAT | MISSENSE | cAg/cGg | Q305R | ahpF | | | G | G | G | G | G | G | | | | |

| | | | | | | | | | | | | | | | | | | | | | | | | | |
|---|---|---|---|---|---|---|---|---|---|---|---|---|---|---|---|---|---|---|---|---|---|---|---|---|---|
| NC_011770 | 159236 | A | T | 69 | 0 | 2 | 0 | NON_SYNONYMOU | MODERAT | MISSENSE | Acg/Tcg | T490S | ahpF | T | T | | | | | | | | | | |
| NC_011770 | 4304990 | G | C | 77 | 0 | 4 | 0 | NON_SYNONYMOU | MODERAT | MISSENSE | Gcg/Ccg | A124P | alc | C | C | | | | | | | | | C | C |
| NC_011770 | 4305048 | C | A | 82 | 0 | 2 | 0 | NON_SYNONYMOU | MODERAT | MISSENSE | cCc/cAc | P143H | alc | | | | | | | A | A | | | | |
| NC_011770 | 4305135 | A | G | 75 | 0 | 12 | 0 | NON_SYNONYMOU | MODERAT | MISSENSE | aAc/aGc | N172S | alc | G | G | G | G | G | G | G | G | G | G | G | G |
| NC_011770 | 1615016 | C | A | 52 | 0 | 2 | 0 | NON_SYNONYMOU | MODERAT | MISSENSE | Gcc/Tcc | A362S | alg44 | | | | | | | A | A | | | | |
| NC_011770 | 1615176 | G | T | 56 | 0 | 2 | 0 | NON_SYNONYMOU | MODERAT | MISSENSE | aaC/aaA | N308K | alg44 | | | T | T | | | | | | | | |
| NC_011770 | 1614955 | G | A | 48 | 0 | 1 | 0 | NON_SYNONYMOU | MODERAT | MISSENSE | gCc/gTc | A382V | alg44 | | | | | | | | A | | | | |
| NC_011770 | 1617114 | C | T | 87 | 0 | 2 | 0 | NON_SYNONYMOU | MODERAT | MISSENSE | Gcg/Acg | A185T | alg8 | | | | | | | T | T | | | | |
| NC_011770 | 1603211 | C | T | 56 | 0 | 1 | 0 | NON_SYNONYMOU | MODERAT | MISSENSE | Gcc/Acc | A238T | algA | | T | | | | | | | | | | |
| NC_011770 | 1603202 | C | T | 58 | 0 | 1 | 0 | NON_SYNONYMOU | MODERAT | MISSENSE | Gaa/Aaa | E241K | algA | | | | T | | | | | | | | |
| NC_011770 | 6510384 | C | T | 85 | 0 | 1 | 0 | NON_SYNONYMOU | MODERAT | MISSENSE | Cgc/Tgc | R373C | algB | | | | | | | | | | | T | |
| NC_011770 | 6509670 | G | A | 90 | 0 | 1 | 0 | NON_SYNONYMOU | MODERAT | MISSENSE | Gcc/Acc | A135T | algB | | | | | | | A | | | | | |
| NC_011770 | 6509757 | G | A | 88 | 0 | 2 | 0 | NON_SYNONYMOU | MODERAT | MISSENSE | Gta/Ata | V164I | algB | | | A | A | | | | | | | | |
| NC_011770 | 6328831 | C | T | 48 | 0 | 1 | 0 | NON_SYNONYMOU | MODERAT | MISSENSE | gCc/gTc | A168V | algC | | | | | | | | | | | | T |
| NC_011770 | 6329266 | A | G | 76 | 0 | 2 | 0 | NON_SYNONYMOU | MODERAT | MISSENSE | aAg/aGg | K313R | algC | G | G | | | | | | | | | | |
| NC_011770 | 1613392 | C | T | 72 | 0 | 4 | 0 | NON_SYNONYMOU | MODERAT | MISSENSE | gGc/gAc | G34D | algE | | | T | T | T | T | | | | | | |
| NC_011770 | 1613393 | C | T | 73 | 0 | 4 | 0 | NON_SYNONYMOU | MODERAT | MISSENSE | Ggc/Agc | G34S | algE | | | T | T | T | T | | | | | | |
| NC_011770 | 1604529 | C | CG | 48 | 0 | 2 | 0 | FRAME_SHIFT | HIGH | | ccc/cccC | P80P? | algF | | | | CG | | | CG | | | | | |
| NC_011770 | 1611497 | AC | A | 75 | 0 | 1 | 0 | FRAME_SHIFT | HIGH | | gtg/ | V168 | algG | | | | | | A | | | | | | |
| NC_011770 | 1611824 | A | G | 70 | 0 | 1 | 0 | NON_SYNONYMOU | MODERAT | MISSENSE | cTg/cCg | L59P | algG | | G | | | | | | | | | | |
| NC_011770 | 1610883 | C | T | 75 | 0 | 1 | 0 | NON_SYNONYMOU | MODERAT | MISSENSE | Gtc/Atc | V373I | algG | | | T | | | | | | | | | |
| NC_011770 | 1611828 | C | T | 72 | 0 | 3 | 0 | NON_SYNONYMOU | MODERAT | MISSENSE | Ggc/Agc | G58S | algG | T | T | | | | | | T | | | | |
| NC_011770 | 1606973 | C | T | 88 | 0 | 1 | 0 | NON_SYNONYMOU | MODERAT | MISSENSE | Gac/Aac | D208N | algI | | | T | | | | | | | | | |
| NC_011770 | 1607495 | TCAG | T | 76 | 0 | 1 | 0 | CODON_DELETION | MODERATE | | ctg/ | L33 | algI | | | | T | | | | | | | | |
| NC_011770 | 1605903 | C | A | 81 | 0 | 2 | 0 | NON_SYNONYMOU | MODERAT | MISSENSE | Gcc/Tcc | A39S | algJ | | | | | | | A | A | | | | |
| NC_011770 | 1605768 | C | T | 78 | 0 | 1 | 0 | NON_SYNONYMOU | MODERAT | MISSENSE | Ggc/Agc | G84S | algJ | | T | | | | | | | | | | |
| NC_011770 | 1614246 | G | A | 75 | 0 | 12 | 0 | NON_SYNONYMOU | MODERAT | MISSENSE | gCc/gTc | A224V | algK | A | A | A | A | A | A | A | A | A | A | A | A |
| NC_011770 | 1614677 | C | A | 59 | 0 | 2 | 0 | NON_SYNONYMOU | MODERAT | MISSENSE | caG/caT | Q80H | algK | | | | | A | A | | | | | | |
| NC_011770 | 1614683 | G | C | 57 | 0 | 4 | 0 | NON_SYNONYMOU | MODERAT | MISSENSE | agC/agG | S78R | algK | | | C | C | C | C | | | | | | |
| NC_011770 | 1608558 | T | C | 85 | 0 | 12 | 0 | NON_SYNONYMOU | MODERAT | MISSENSE | Acc/Gcc | T124A | algL | C | C | C | C | C | C | C | C | C | C | C | C |
| NC_011770 | 1608791 | T | C | 72 | 0 | 12 | 0 | NON_SYNONYMOU | MODERAT | MISSENSE | aAc/aGc | N46S | algL | C | C | C | C | C | C | C | C | C | C | C | C |
| NC_011770 | 1608828 | G | A | 71 | 0 | 2 | 0 | NON_SYNONYMOU | MODERAT | MISSENSE | Ccc/Tcc | P34S | algL | | | | | | | | | A | A | | |
| NC_011770 | 1607939 | G | C | 73 | 0 | 1 | 0 | NON_SYNONYMOU | MODERAT | MISSENSE | gCc/gGc | A330G | algL | | | | C | | | | | | | | |
| NC_011770 | 6251455 | G | A | 42 | 0 | 1 | 0 | NON_SYNONYMOU | MODERAT | MISSENSE | gCt/gTt | A192V | algP | | | | | A | | | | | | | |
| NC_011770 | 6251099 | C | T | 62 | 0 | 2 | 0 | NON_SYNONYMOU | MODERAT | MISSENSE | Gcc/Acc | A311T | algP | | | | | | | | | T | T | | |
| NC_011770 | 6251104 | G | A | 58 | 0 | 4 | 0 | NON_SYNONYMOU | MODERAT | MISSENSE | gCc/gTc | A309V | algP | | | A | A | A | A | | | | | | |
| NC_011770 | 6251198 | G | T | 30 | 0 | 6 | 0 | NON_SYNONYMOU | MODERAT | MISSENSE | Ccc/Acc | P278T | algP | | | T | T | T | T | | | A | A | | |
| NC_011770 | 6251206 | A | G | 31 | 0 | 6 | 0 | NON_SYNONYMOU | MODERAT | MISSENSE | gTg/gCg | V275A | algP | | | G | G | G | G | | | | | G | G |
| NC_011770 | 6251294 | T | C | 31 | 0 | 4 | 0 | NON_SYNONYMOU | MODERAT | MISSENSE | Acg/Gcg | T246A | algP | | | C | C | C | C | | | | | | |
| NC_011770 | 6251372 | G | A | 43 | 0 | 2 | 0 | NON_SYNONYMOU | MODERAT | MISSENSE | Ccg/Tcg | P220S | algP | A | A | | | | | | | | | | |
| NC_011770 | 6251401 | G | A | 48 | 0 | 4 | 0 | NON_SYNONYMOU | MODERAT | MISSENSE | gCc/gTc | A210V | algP | A | A | | | | | | | | | A | A |
| NC_011770 | 6251468 | T | C | 39 | 0 | 4 | 0 | NON_SYNONYMOU | MODERAT | MISSENSE | Acg/Gcg | T188A | algP | | | C | C | C | C | | | | | | |
| NC_011770 | 6253085 | G | A | 85 | 0 | 1 | 0 | NON_SYNONYMOU | MODERAT | MISSENSE | Cgc/Tgc | R82C | algQ | | | | | | | | A | | | | |
| NC_011770 | 5313545 | G | A | 62 | 0 | 1 | 0 | START_LOST | HIGH | MISSENSE | atG/atA | M1I | algW | | | A | | | | | | | | | |
| NC_011770 | 5314711 | G | A | 57 | 0 | 2 | 0 | SPLICE_SITE_REGI | LOW | SILENT | tGa/tAa | *390 | algW | | | | | | | | | | | A | A |
| NC_011770 | 5313942 | G | A | 31 | 0 | 1 | 0 | NON_SYNONYMOU | MODERAT | MISSENSE | Gcc/Acc | A134T | algW | | | | A | | | | | | | | |
| NC_011770 | 5314698 | A | G | 56 | 0 | 12 | 0 | NON_SYNONYMOU | MODERAT | MISSENSE | Aac/Gac | N386D | algW | G | G | G | G | G | G | G | G | G | G | G | G |
| NC_011770 | 1608948 | A | G | 73 | 0 | 2 | 0 | NON_SYNONYMOU | MODERAT | MISSENSE | Tcg/Ccg | S470P | algX | | | | | | | G | G | | | | |
| NC_011770 | 1610262 | A | C | 56 | 0 | 12 | 0 | NON_SYNONYMOU | MODERAT | MISSENSE | Tcg/Gcg | S32A | algX | C | C | C | C | C | C | C | C | C | C | C | C |
| NC_011770 | 1609062 | C | T | 82 | 0 | 1 | 0 | NON_SYNONYMOU | MODERAT | MISSENSE | Gac/Aac | D432N | algX | | | | T | | | | | | | | |
| NC_011770 | 6260277 | C | A | 65 | 0 | 1 | 0 | NON_SYNONYMOU | MODERAT | MISSENSE | gGc/gTc | G8V | algZ | | | | | A | | | | | | | |
| NC_011770 | 4032861 | A | G | 66 | 0 | 2 | 0 | SPLICE_SITE_REGI | LOW | SILENT | taA/taG | *298 | alkA | | | | | | | | | G | G | | |
| NC_011770 | 4032466 | G | A | 70 | 0 | 2 | 0 | NON_SYNONYMOU | MODERAT | MISSENSE | Ggc/Agc | G167S | alkA | A | A | | | | | | | | | | |
| NC_011770 | 4032664 | A | ACCG | 62 | 0 | 1 | 0 | CODON_CHANGE_ | MODERATE | | gaa/gCCGa | E235AE | alkA | | | | | ACCG | | | | | | | |
| NC_011770 | 4032838 | A | C | 66 | 0 | 4 | 0 | NON_SYNONYMOU | MODERAT | MISSENSE | Aac/Cac | N291H | alkA | | | | | | | | | C | C | C | C |
| NC_011770 | 1896932 | T | C | 80 | 0 | 6 | 0 | NON_SYNONYMOU | MODERAT | MISSENSE | cTa/cCa | L200P | alkB | C | C | C | C | C | C | | | | | | |
| NC_011770 | 2919477 | G | A | 70 | 0 | 2 | 0 | NON_SYNONYMOU | MODERAT | MISSENSE | Gtc/Atc | V108I | alkB1 | | | | | | | | | A | A | | |
| NC_011770 | 5865365 | C | T | 62 | 0 | 2 | 0 | NON_SYNONYMOU | MODERAT | MISSENSE | cGc/cAc | R355H | alr | | | | | | | | | T | T | | |
| NC_011770 | 5865450 | C | G | 60 | 0 | 2 | 0 | NON_SYNONYMOU | MODERAT | MISSENSE | Ggg/Cgg | G327R | alr | | | | | | | | | | | G | G |
| NC_011770 | 5865546 | T | G | 63 | 0 | 2 | 0 | NON_SYNONYMOU | MODERAT | MISSENSE | Acc/Ccc | T295P | alr | | | | | | | | | | | G | G |
| NC_011770 | 5865639 | C | T | 63 | 0 | 2 | 0 | NON_SYNONYMOU | MODERAT | MISSENSE | Gcg/Acg | A264T | alr | | | | | | | | | | | T | T |
| NC_011770 | 5865767 | G | A | 71 | 0 | 2 | 0 | NON_SYNONYMOU | MODERAT | MISSENSE | gCc/gTc | A221V | alr | | | | | | | | | | | A | A |

| | | | | | | | | | | | | | | | | | | | | | | | | | |
|---|---|---|---|---|---|---|---|---|---|---|---|---|---|---|---|---|---|---|---|---|---|---|---|---|---|
| NC_011770 | 5866022 | G | A | 85 | 0 | 2 | 0 | NON_SYNONYMOU | MODERAT | MISSENSE | gCc/gTc | A136V | alr | | | | | | | | | | | A | A |
| NC_011770 | 5866134 | C | T | 87 | 0 | 2 | 0 | NON_SYNONYMOU | MODERAT | MISSENSE | Gtg/Atg | V99M | alr | | | | | | | | | | | T | T |
| NC_011770 | 5866407 | C | T | 80 | 0 | 4 | 0 | NON_SYNONYMOU | MODERAT | MISSENSE | Gtc/Atc | V8I | alr | | | T | T | T | T | | | | | | |
| NC_011770 | 6567804 | G | A | 80 | 0 | 2 | 0 | NON_SYNONYMOU | MODERAT | MISSENSE | gCt/gTt | A372V | amiA | | | | | | | | | A | A | | |
| NC_011770 | 6568127 | C | A | 88 | 0 | 4 | 0 | NON_SYNONYMOU | MODERAT | MISSENSE | gaG/gaT | E264D | amiA | A | A | | | | | | | G | G | | |
| NC_011770 | 6568467 | G | A | 78 | 0 | 2 | 0 | NON_SYNONYMOU | MODERAT | MISSENSE | aCg/aTg | T151M | amiA | | | | | | | | | | | A | A |
| NC_011770 | 5884071 | T | A | 85 | 0 | 4 | 0 | NON_SYNONYMOU | MODERAT | MISSENSE | aAc/aTc | N458I | amiB | | | A | A | A | A | | | | | | |
| NC_011770 | 5884411 | G | A | 70 | 0 | 1 | 0 | NON_SYNONYMOU | MODERAT | MISSENSE | Cgt/Tgt | R345C | amiB | | | | | | | A | | | | | |
| NC_011770 | 1827892 | C | T | 67 | 0 | 4 | 0 | NON_SYNONYMOU | MODERAT | MISSENSE | Ccc/Tcc | P263S | amiC | | | T | T | T | T | | | | | | |
| NC_011770 | 1828019 | G | A | 74 | 0 | 2 | 0 | NON_SYNONYMOU | MODERAT | MISSENSE | aGc/aAc | S305N | amiC | | | | | | | | | | | A | A |
| NC_011770 | 1825662 | A | G | 82 | 0 | 1 | 0 | NON_SYNONYMOU | MODERAT | MISSENSE | aAc/aGc | N274S | amiE | | | | | | | G | | | | | |
| NC_011770 | 1825719 | G | A | 76 | 0 | 2 | 0 | NON_SYNONYMOU | MODERAT | MISSENSE | gGc/gAc | G293D | amiE | | | | | | | | | | | A | A |
| NC_011770 | 1828746 | G | A | 86 | 0 | 4 | 0 | NON_SYNONYMOU | MODERAT | MISSENSE | Gac/Aac | D163N | amiR | | | | | | | A | A | A | A | | |
| NC_011770 | 1828777 | A | G | 82 | 0 | 12 | 0 | NON_SYNONYMOU | MODERAT | MISSENSE | cAg/cGg | Q173R | amiR | G | G | G | G | G | G | G | G | G | G | G | G |
| NC_011770 | 1086662 | C | T | 87 | 0 | 1 | 0 | NON_SYNONYMOU | MODERAT | MISSENSE | Cgc/Tgc | R254C | amn | | | | | | | T | | | | | |
| NC_011770 | 940404 | G | A | 71 | 0 | 2 | 0 | NON_SYNONYMOU | MODERAT | MISSENSE | cCc/cTc | | ampC | A | A | | | | | | | | | | |
| NC_011770 | 940549 | A | G | 84 | 0 | 1 | 0 | NON_SYNONYMOU | MODERAT | MISSENSE | Tac/Cac | Y226H | ampC | | | | | | | G | | | | | |
| NC_011770 | 940989 | C | T | 78 | 0 | 2 | 0 | NON_SYNONYMOU | MODERAT | MISSENSE | cGg/cAg | R79Q | ampC | T | T | | | | | | | | | | |
| NC_011770 | 941206 | G | A | 69 | 0 | 2 | 0 | NON_SYNONYMOU | MODERAT | MISSENSE | Ccc/Tcc | P7S | ampC | | | | | | | A | A | | | | |
| NC_011770 | 940698 | A | C | 84 | 0 | 4 | 0 | NON_SYNONYMOU | MODERAT | MISSENSE | cTc/cGc | L176R | ampC (150 PAO1) | | | | | | | | | C | C | C | C |
| NC_011770 | 940159 | C | T | 53 | 0 | 1 | 0 | NON_SYNONYMOU | MODERAT | MISSENSE | Gtc/Atc | V356I | ampC (330 PAO1) | | | | | | | T | | | | | |
| NC_011770 | 5398707 | G | A | 80 | 0 | 1 | 0 | NON_SYNONYMOU | MODERAT | MISSENSE | aCg/aTg | T124M | ampD | | | | | | | | | | | | A |
| NC_011770 | 5398531 | C | A | 87 | 0 | 2 | 0 | NON_SYNONYMOU | MODERAT | MISSENSE | Gat/Tat | D183Y | ampD | | | | | | | | | A | A | | |
| NC_011770 | 5398554 | A | G | 83 | 0 | 12 | 0 | NON_SYNONYMOU | MODERAT | MISSENSE | tTg/tCg | L175S | ampD | G | G | G | G | G | G | G | G | G | G | G | G |
| NC_011770 | 5398635 | G | C | 81 | 0 | 6 | 0 | NON_SYNONYMOU | MODERAT | MISSENSE | gCg/gGg | A148G | ampD | C | C | | | | | C | C | | | C | C |
| NC_011770 | 5398710 | T | C | 79 | 0 | 1 | 0 | NON_SYNONYMOU | MODERAT | MISSENSE | gAc/gGc | D123G | ampD | | | | | | | C | | | | | |
| NC_011770 | 5398854 | A | G | 73 | 0 | 12 | 0 | NON_SYNONYMOU | MODERAT | MISSENSE | tTc/tCc | F75S | ampD | G | G | G | G | G | G | G | G | G | G | G | G |
| NC_011770 | 5399046 | C | A | 66 | 0 | 2 | 0 | NON_SYNONYMOU | MODERAT | MISSENSE | cGt/cTt | R11L | ampD | | | | | | | | | A | A | | |
| NC_011770 | 4982508 | C | T | 48 | 0 | 8 | 0 | NON_SYNONYMOU | MODERAT | MISSENSE | Gcc/Acc | A219T | ampDh3 | T | T | T | T | T | T | | | T | T | | |
| NC_011770 | 4983026 | T | C | 78 | 0 | 1 | 0 | NON_SYNONYMOU | MODERAT | MISSENSE | cAc/cGc | H46R | ampDh3 | | | | | | | C | | | | | |
| NC_011770 | 941713 | A | C | 66 | 0 | 2 | 0 | NON_SYNONYMOU | MODERAT | MISSENSE | gAg/gCg | E114A | ampR | | | | | | | C | C | | | | |
| NC_011770 | 942220 | G | A | 62 | 0 | 6 | 0 | NON_SYNONYMOU | MODERAT | MISSENSE | gGg/gAg | G283E | ampR | | | A | A | A | A | A | A | | | | |
| NC_011770 | 942235 | T | G | 36 | 4 | 2 | 0 | NON_SYNONYMOU | MODERAT | MISSENSE | aTg/aGg | M288R | ampR | | | . | . | . | . | G | G | | | | |
| NC_011770 | 941644 | A | G | 67 | 0 | 1 | 0 | NON_SYNONYMOU | MODERAT | MISSENSE | cAc/cGc | H91R | ampR | | G | | | | | | | | | | |
| NC_011770 | 941583 | C | T | 68 | 0 | 1 | 0 | NON_SYNONYMOU | MODERAT | MISSENSE | Ccg/Tcg | P71S | ampR | | | | | | | | | | | | T |
| NC_011770 | 6288080 | C | T | 84 | 0 | 4 | 0 | NON_SYNONYMOU | MODERAT | MISSENSE | Gcc/Acc | A95T | amtB | | | T | T | T | T | | | | | | |
| NC_011770 | 695070 | G | T | 71 | 0 | 4 | 0 | NON_SYNONYMOU | MODERAT | MISSENSE | Ctg/Atg | L305M | anmK | | | | | | | T | T | T | T | | |
| NC_011770 | 695130 | T | C | 73 | 0 | 10 | 0 | NON_SYNONYMOU | MODERAT | MISSENSE | Aaa/Gaa | K285E | anmK | C | C | C | C | C | C | C | C | C | C | | |
| NC_011770 | 695253 | A | G | 71 | 0 | 4 | 0 | NON_SYNONYMOU | MODERAT | MISSENSE | Tcc/Ccc | S244P | anmK | | | | | | | G | G | G | G | | |
| NC_011770 | 695343 | T | C | 77 | 0 | 4 | 0 | NON_SYNONYMOU | MODERAT | MISSENSE | Atg/Gtg | M214V | anmK | | | | | | | C | C | C | C | | |
| NC_011770 | 695433 | C | T | 75 | 0 | 1 | 0 | NON_SYNONYMOU | MODERAT | MISSENSE | Ggc/Agc | G184S | anmK | | | | | | | T | | | | | |
| NC_011770 | 695522 | T | C | 72 | 0 | 4 | 0 | NON_SYNONYMOU | MODERAT | MISSENSE | gAt/gGt | D154G | anmK | | | | | | | C | C | C | C | | |
| NC_011770 | 695723 | T | C | 76 | 0 | 2 | 0 | NON_SYNONYMOU | MODERAT | MISSENSE | gAa/gGa | E87G | anmK | | | | | | | | | C | C | | |
| NC_011770 | 695910 | C | T | 81 | 0 | 2 | 0 | NON_SYNONYMOU | MODERAT | MISSENSE | Gat/Aat | D25N | anmK | | | | | | | | | T | T | | |
| NC_011770 | 4186895 | G | A | 64 | 0 | 1 | 0 | NON_SYNONYMOU | MODERAT | MISSENSE | cGc/cAc | R175H | anr | | | | | | | | | | | A | |
| NC_011770 | 3363112 | C | T | 63 | 0 | 12 | 0 | NON_SYNONYMOU | MODERAT | MISSENSE | Gtc/Atc | V112I | ansA | T | T | T | T | T | T | T | T | T | T | T | T |
| NC_011770 | 4252317 | G | C | 80 | 0 | 2 | 0 | NON_SYNONYMOU | MODERAT | MISSENSE | Gaa/Caa | E85Q | ansB | | | | | | | C | C | | | | |
| NC_011770 | 2998406 | C | T | 89 | 0 | 2 | 0 | NON_SYNONYMOU | MODERAT | MISSENSE | Gcc/Acc | A262T | antA | T | T | | | | | | | | | | |
| NC_011770 | 2999053 | T | C | 80 | 0 | 2 | 0 | NON_SYNONYMOU | MODERAT | MISSENSE | gAg/gGg | E46G | antA | | | | | | | C | C | | | | |
| NC_011770 | 2999176 | C | T | 74 | 0 | 4 | 0 | NON_SYNONYMOU | MODERAT | MISSENSE | cGc/cAc | R5H | antA | | | T | T | T | T | | | | | | |
| NC_011770 | 2997490 | T | G | 75 | 0 | 2 | 0 | NON_SYNONYMOU | MODERAT | MISSENSE | gaA/gaC | E103D | antB | | | | | | | | | | | G | G |
| NC_011770 | 2996321 | T | C | 73 | 0 | 2 | 0 | NON_SYNONYMOU | MODERAT | MISSENSE | cAc/cGc | H323R | antC | C | C | | | | | | | | | | |
| NC_011770 | 2996372 | A | G | 71 | 0 | 1 | 0 | NON_SYNONYMOU | MODERAT | MISSENSE | cTg/cCg | L306P | antC | | | | | | | G | | | | | |
| NC_011770 | 2996505 | C | T | 71 | 0 | 2 | 0 | NON_SYNONYMOU | MODERAT | MISSENSE | Gcc/Acc | A262T | antC | | | | | | | | | | | T | T |
| NC_011770 | 2996540 | G | T | 70 | 0 | 4 | 0 | NON_SYNONYMOU | MODERAT | MISSENSE | gCc/gAc | A250D | antC | | | T | T | T | T | | | | | | |
| NC_011770 | 2997099 | C | G | 69 | 0 | 4 | 0 | NON_SYNONYMOU | MODERAT | MISSENSE | Gag/Cag | E64Q | antC | | | G | G | G | G | | | | | | |
| NC_011770 | 4862281 | T | C | 85 | 0 | 12 | 0 | NON_SYNONYMOU | MODERAT | MISSENSE | Atg/Gtg | M166V | aotP | C | C | C | C | C | C | C | C | C | C | C | C |
| NC_011770 | 641631 | T | C | 78 | 0 | 1 | 0 | NON_SYNONYMOU | MODERAT | MISSENSE | gAc/gGc | D13G | apaG | | | | | | | C | | | | | |
| NC_011770 | 640583 | A | G | 77 | 0 | 1 | 0 | NON_SYNONYMOU | MODERAT | MISSENSE | Tgc/Cgc | C236R | apaH (purine metabolism | | **G** | | | | | | | | | | |
| NC_011770 | 640601 | C | T | 78 | 0 | 1 | 0 | NON_SYNONYMOU | MODERAT | MISSENSE | Gcg/Acg | A230T | apaH (purine metabolism) | | | | | | | | **T** | | | | |
| NC_011770 | 641203 | T | C | 85 | 0 | 1 | 0 | NON_SYNONYMOU | MODERAT | MISSENSE | aAg/aGg | K29R | apaH (purine metabolism) | | | | | | | | **C** | | | | |
| NC_011770 | 926908 | T | G | 52 | 0 | 8 | 0 | NON_SYNONYMOU | MODERAT | MISSENSE | Tca/Gca | S13A | aph | | | G | G | G | G | G | G | G | G | | |

| | | | | | | | | | | | | | | | | | | | | | | | | | |
|---|---|---|---|---|---|---|---|---|---|---|---|---|---|---|---|---|---|---|---|---|---|---|---|---|---|
| NC_011770 | 926993 | C | T | 61 | 0 | 4 | 0 | NON_SYNONYMOU | MODERAT | MISSENSE | gCt/gTt | A41V | aph | T | T | | | | | | | | | T | T |
| NC_011770 | 926996 | A | C | 59 | 0 | 12 | 0 | NON_SYNONYMOU | MODERAT | MISSENSE | gAg/gCg | E42A | aph | C | C | C | C | C | C | C | C | C | C | C | C |
| NC_011770 | 927022 | C | A | 62 | 0 | 6 | 0 | NON_SYNONYMOU | MODERAT | MISSENSE | Cag/Aag | Q51K | aph | | | A | A | A | A | | | A | A | | |
| NC_011770 | 926894 | T | TGCC | 52 | 0 | 9 | 0 | CODON_CHANGE_ | MODERATE | | cag/cCGCa | Q10PQ | aph | | TGCC | TGCC | | TGCC | TGCC | | TGCC | TGCC | TGCC | TGCC | TGCC |
| NC_011770 | 4418989 | C | A | 73 | 0 | 2 | 0 | NON_SYNONYMOU | MODERAT | MISSENSE | cGg/cTg | R224L | aphA | | | | | | | | | A | A | | |
| NC_011770 | 4419173 | C | T | 67 | 0 | 1 | 0 | NON_SYNONYMOU | MODERAT | MISSENSE | Ggt/Agt | G163S | aphA | | | | | | | T | | | | | |
| NC_011770 | 4419476 | G | T | 53 | 0 | 12 | 0 | NON_SYNONYMOU | MODERAT | MISSENSE | Ctc/Atc | L62I | aphA | T | T | T | T | T | T | T | T | T | T | T | T |
| NC_011770 | 4419535 | G | A | 57 | 0 | 2 | 0 | NON_SYNONYMOU | MODERAT | MISSENSE | tCg/tTg | S42L | aphA | | | | | | | | | A | A | | |
| NC_011770 | 4484334 | C | T | 70 | 0 | 2 | 0 | NON_SYNONYMOU | MODERAT | MISSENSE | aGc/aAc | S346N | aprA | | | | | T | T | | | | | | |
| NC_011770 | 4484659 | C | T | 74 | 0 | 2 | 0 | NON_SYNONYMOU | MODERAT | MISSENSE | Gcc/Acc | A238T | aprA | | | | | | | | | | | T | T |
| NC_011770 | 4485034 | C | A | 84 | 0 | 12 | 0 | NON_SYNONYMOU | MODERAT | MISSENSE | Gcg/Tcg | A113S | aprA | A | A | A | A | A | A | A | A | A | A | A | A |
| NC_011770 | 4489204 | C | T | 79 | 0 | 1 | 0 | NON_SYNONYMOU | MODERAT | MISSENSE | Ggc/Agc | G351S | aprD | T | | | | | | | | | | | |
| NC_011770 | 4488703 | T | C | 92 | 0 | 10 | 0 | NON_SYNONYMOU | MODERAT | MISSENSE | Agc/Ggc | S518G | aprD | C | C | C | C | C | C | C | C | C | C | | |
| NC_011770 | 4488864 | A | G | 77 | 0 | 2 | 0 | NON_SYNONYMOU | MODERAT | MISSENSE | gTc/gCc | V464A | aprD | | | | | | | | | | | G | G |
| NC_011770 | 4489344 | T | G | 78 | 0 | 2 | 0 | NON_SYNONYMOU | MODERAT | MISSENSE | gAt/gCt | D304A | aprD | | | | | | | | | G | G | | |
| NC_011770 | 4487653 | G | A | 81 | 0 | 1 | 0 | NON_SYNONYMOU | MODERAT | MISSENSE | gCg/gTg | A275V | aprE | | | | | | | A | | | | | |
| NC_011770 | 4485734 | T | G | 87 | 0 | 10 | 0 | NON_SYNONYMOU | MODERAT | MISSENSE | Aat/Cat | N481H | aprF | G | G | G | G | G | G | G | G | | | G | G |
| NC_011770 | 4485743 | C | T | 83 | 0 | 10 | 0 | NON_SYNONYMOU | MODERAT | MISSENSE | Ggc/Agc | G478S | aprF | T | T | T | T | T | T | T | T | | | T | T |
| NC_011770 | 4485830 | T | G | 85 | 0 | 2 | 0 | NON_SYNONYMOU | MODERAT | MISSENSE | Acg/Ccg | T449P | aprF | | | | | | | | | G | G | | |
| NC_011770 | 4485832 | T | G | 86 | 0 | 2 | 0 | NON_SYNONYMOU | MODERAT | MISSENSE | gAg/gCg | E448A | aprF | | | | | | | | | G | G | | |
| NC_011770 | 4187667 | G | C | 74 | 0 | 2 | 0 | NON_SYNONYMOU | MODERAT | MISSENSE | gGc/gCc | G159A | apt | | | | | | | C | C | | | | |
| NC_011770 | 1022597 | C | T | 71 | 2 | 4 | 0 | NON_SYNONYMOU | MODERAT | MISSENSE | gGc/gAc | G125D | aqpZ | | | T | T | T | T | . | . | | | | |
| NC_011770 | 6155853 | G | A | 82 | 0 | 1 | 0 | STOP_GAINED | HIGH | NONSENSE | tGg/tAg | W138* | arcD | | | | | | | | A | | | | |
| NC_011770 | 6155945 | T | C | 84 | 0 | 2 | 0 | NON_SYNONYMOU | MODERAT | MISSENSE | Ttc/Ctc | F169L | arcD | C | C | | | | | | | | | | |
| NC_011770 | 692348 | G | A | 76 | 0 | 2 | 0 | NON_SYNONYMOU | MODERAT | MISSENSE | gGg/gAg | G39E | argC | | | | | | | | | | | A | A |
| NC_011770 | 692420 | T | C | 74 | 0 | 12 | 0 | NON_SYNONYMOU | MODERAT | MISSENSE | gTg/gCg | V63A | argC | C | C | C | C | C | C | C | C | C | C | C | C |
| NC_011770 | 692906 | G | T | 73 | 0 | 4 | 0 | NON_SYNONYMOU | MODERAT | MISSENSE | gGc/gTc | G225V | argC | | | T | T | T | T | | | | | | |
| NC_011770 | 692910 | C | G | 76 | 0 | 6 | 0 | NON_SYNONYMOU | MODERAT | MISSENSE | gaC/gaG | D226E | argC | | | | | | | G | G | G | G | G | G |
| NC_011770 | 692288 | G | A | 70 | 0 | 1 | 0 | NON_SYNONYMOU | MODERAT | MISSENSE | cGc/cAc | R19H | argC | | | | | | | | **A** | | | | |
| NC_011770 | 4859941 | T | G | 78 | 0 | 4 | 0 | NON_SYNONYMOU | MODERAT | MISSENSE | aAg/aCg | K180T | argD | | | | | | | G | G | G | G | | |
| NC_011770 | 6196882 | G | A | 69 | 0 | 2 | 0 | NON_SYNONYMOU | MODERAT | MISSENSE | gCc/gTc | A222V | argE | | | | | | | | | | | A | A |
| NC_011770 | 6196885 | T | C | 66 | 0 | 10 | 0 | NON_SYNONYMOU | MODERAT | MISSENSE | cAg/cGg | Q221R | argE | C | C | C | C | C | C | C | C | C | C | | |
| NC_011770 | 6197069 | C | T | 60 | 0 | 2 | 0 | NON_SYNONYMOU | MODERAT | MISSENSE | Gcc/Acc | A160T | argE | | | | | | | | | | | T | T |
| NC_011770 | 6197317 | G | C | 72 | 0 | 2 | 0 | NON_SYNONYMOU | MODERAT | MISSENSE | aCc/aGc | T77S | argE | | | | | | | C | C | | | | |
| NC_011770 | 6260646 | G | A | 77 | 0 | 2 | 0 | NON_SYNONYMOU | MODERAT | MISSENSE | atG/atA | M41I | argH | | | | | | | | | A | A | | |
| NC_011770 | 6260716 | A | G | 72 | 0 | 10 | 0 | NON_SYNONYMOU | MODERAT | MISSENSE | Atc/Gtc | I65V | argH | G | G | G | G | G | G | G | G | G | G | | |
| NC_011770 | 6260755 | G | A | 71 | 0 | 2 | 0 | NON_SYNONYMOU | MODERAT | MISSENSE | Ggc/Agc | G78S | argH | A | A | | | | | | | | | | |
| NC_011770 | 6261067 | G | A | 72 | 0 | 2 | 0 | NON_SYNONYMOU | MODERAT | MISSENSE | Ggc/Agc | G182S | argH | | | | | | | | | | | A | A |
| NC_011770 | 6261259 | G | A | 75 | 0 | 4 | 0 | NON_SYNONYMOU | MODERAT | MISSENSE | Gcc/Acc | A246T | argH | | | | | | | | | A | A | A | A |
| NC_011770 | 6261529 | A | G | 62 | 0 | 10 | 0 | NON_SYNONYMOU | MODERAT | MISSENSE | Acc/Gcc | T336A | argH | G | G | G | G | G | G | | | G | G | G | G |
| NC_011770 | 6261545 | A | G | 62 | 0 | 10 | 0 | NON_SYNONYMOU | MODERAT | MISSENSE | cAc/cGc | H341R | argH | G | G | G | G | G | G | G | G | | | G | G |
| NC_011770 | 6261835 | G | A | 61 | 0 | 2 | 0 | NON_SYNONYMOU | MODERAT | MISSENSE | Gcc/Acc | A438T | argH | | | | | | | A | A | | | | |
| NC_011770 | 6261886 | G | A | 58 | 0 | 4 | 0 | NON_SYNONYMOU | MODERAT | MISSENSE | Gcc/Acc | A455T | argH | | | A | A | A | A | | | | | | |
| NC_011770 | 5266650 | G | T | 94 | 0 | 2 | 0 | NON_SYNONYMOU | MODERAT | MISSENSE | Ctg/Atg | L261M | argJ | | | | | | | | | | | T | T |
| NC_011770 | 5267006 | C | T | 74 | 0 | 4 | 0 | NON_SYNONYMOU | MODERAT | MISSENSE | cGc/cAc | R142H | argJ | T | T | | | | | | | T | T | | |
| NC_011770 | 6024855 | G | A | 61 | 0 | 6 | 0 | NON_SYNONYMOU | MODERAT | MISSENSE | Gcc/Acc | A72T | argS | A | A | A | A | A | A | | | | | | |
| NC_011770 | 6025188 | A | G | 72 | 0 | 12 | 0 | NON_SYNONYMOU | MODERAT | MISSENSE | Atc/Gtc | I183V | argS | G | G | G | G | G | G | G | G | G | G | G | G |
| NC_011770 | 6025528 | A | G | 87 | 0 | 2 | 0 | NON_SYNONYMOU | MODERAT | MISSENSE | aAg/aGg | K296R | argS | G | G | | | | | | | | | | |
| NC_011770 | 6025549 | T | C | 82 | 0 | 1 | 0 | NON_SYNONYMOU | MODERAT | MISSENSE | cTc/cCc | L303P | argS | | | | | | | C | | | | | |
| NC_011770 | 6026188 | T | C | 74 | 0 | 1 | 0 | NON_SYNONYMOU | MODERAT | MISSENSE | cTg/cCg | L516P | argS | | | | | | | C | | | | | |
| NC_011770 | 1595555 | C | T | 70 | 0 | 2 | 0 | NON_SYNONYMOU | MODERAT | MISSENSE | cGg/cAg | R502Q | arnT | | | | | | | T | T | | | | |
| NC_011770 | 1595723 | C | T | 74 | 0 | 2 | 0 | NON_SYNONYMOU | MODERAT | MISSENSE | cGc/cAc | R446H | arnT | T | T | | | | | | | | | | |
| NC_011770 | 1595733 | T | C | 74 | 0 | 6 | 0 | NON_SYNONYMOU | MODERAT | MISSENSE | Acc/Gcc | T443A | arnT | | | C | C | C | C | | | C | C | | |
| NC_011770 | 1596261 | A | C | 72 | 0 | 8 | 0 | NON_SYNONYMOU | MODERAT | MISSENSE | Tcc/Gcc | S267A | arnT | | | C | C | C | C | | | C | C | C | C |
| NC_011770 | 1596264 | C | T | 71 | 0 | 6 | 0 | NON_SYNONYMOU | MODERAT | MISSENSE | Gtc/Atc | V266I | arnT | | | T | T | T | T | | | T | T | | |
| NC_011770 | 1596594 | G | C | 79 | 0 | 12 | 0 | NON_SYNONYMOU | MODERAT | MISSENSE | Cgg/Ggg | R156G | arnT | C | C | C | C | C | C | C | C | C | C | C | C |
| NC_011770 | 1597039 | A | C | 71 | 0 | 2 | 0 | NON_SYNONYMOU | MODERAT | MISSENSE | tgT/tgG | C7W | arnT | | | | | | | | | | | C | C |
| NC_011770 | 6009723 | G | A | 80 | 0 | 1 | 0 | NON_SYNONYMOU | MODERAT | MISSENSE | cCg/cTg | P30L | aroB | | | | | | | | A | | | | |
| NC_011770 | 6008937 | G | A | 81 | 0 | 2 | 0 | NON_SYNONYMOU | MODERAT | MISSENSE | aCc/aTc | T292I | aroB | | | | | | | | | A | A | | |
| NC_011770 | 6009108 | A | G | 64 | 0 | 12 | 0 | NON_SYNONYMOU | MODERAT | MISSENSE | tTg/tCg | L235S | aroB | G | G | G | G | G | G | G | G | G | G | G | G |
| NC_011770 | 6009136 | G | A | 65 | 0 | 2 | 0 | NON_SYNONYMOU | MODERAT | MISSENSE | Cgc/Tgc | R226C | aroB | | | | | | | | | A | A | | |
| NC_011770 | 4036390 | C | A | 81 | 0 | 2 | 0 | NON_SYNONYMOU | MODERAT | MISSENSE | Gcc/Tcc | A307S | aroC | | | | | | | | | A | A | | |

| | | | | | | | | | | | | | | | | | | | | | | | | | |
|---|---|---|---|---|---|---|---|---|---|---|---|---|---|---|---|---|---|---|---|---|---|---|---|---|---|
| NC_011770 | 4036797 | C | T | 81 | 0 | 2 | 0 | NON_SYNONYMOU | MODERAT | MISSENSE | aGc/aAc | S171N | aroC | | | | | | | | | T | T | | |
| NC_011770 | 4036807 | T | C | 82 | 0 | 1 | 0 | NON_SYNONYMOU | MODERAT | MISSENSE | Agc/Ggc | S168G | aroC | | | | | | | C | | | | | |
| NC_011770 | 4037158 | C | T | 68 | 0 | 1 | 0 | NON_SYNONYMOU | MODERAT | MISSENSE | Ggc/Agc | G51S | aroC | | | | | | | | | | | | T |
| NC_011770 | 27225 | G | C | 81 | 0 | 2 | 0 | NON_SYNONYMOU | MODERAT | MISSENSE | gaG/gaC | E176D | aroE | | | | | | | C | C | | | | |
| NC_011770 | 2236367 | C | T | 63 | 0 | 4 | 0 | NON_SYNONYMOU | MODERAT | MISSENSE | Ccg/Tcg | P123S | aroP1 | | | T | T | T | T | | | | | | |
| NC_011770 | 4890145 | C | A | 82 | 0 | 2 | 0 | NON_SYNONYMOU | MODERAT | MISSENSE | Gtg/Ttg | V347L | aroP2 | | | | | | | | | | | A | A |
| NC_011770 | 4890424 | C | T | 77 | 0 | 2 | 0 | NON_SYNONYMOU | MODERAT | MISSENSE | Gtc/Atc | V254I | aroP2 | | | | | | | | | T | T | | |
| NC_011770 | 270609 | T | G | 86 | 0 | 2 | 0 | NON_SYNONYMOU | MODERAT | MISSENSE | gAc/gCc | D34A | aroQ2 | | | | | | | | | G | G | | |
| NC_011770 | 3336101 | C | T | 71 | 0 | 2 | 0 | NON_SYNONYMOU | MODERAT | MISSENSE | Gtc/Atc | V405I | arsB | T | T | | | | | | | | | | |
| NC_011770 | 3336545 | C | T | 68 | 0 | 4 | 0 | NON_SYNONYMOU | MODERAT | MISSENSE | Gca/Aca | A257T | arsB | | | T | T | T | T | | | | | | |
| NC_011770 | 3336554 | C | T | 67 | 0 | 6 | 0 | NON_SYNONYMOU | MODERAT | MISSENSE | Gcc/Acc | A254T | arsB | | | T | T | T | T | T | T | | | | |
| NC_011770 | 3336749 | T | A | 64 | 0 | 2 | 0 | NON_SYNONYMOU | MODERAT | MISSENSE | Acc/Tcc | T189S | arsB | | | | | | | | | A | A | | |
| NC_011770 | 3337250 | G | T | 73 | 0 | 4 | 0 | NON_SYNONYMOU | MODERAT | MISSENSE | Ctc/Atc | L22I | arsB | | | T | T | T | T | | | | | | |
| NC_011770 | 3335656 | G | C | 94 | 0 | 2 | 0 | NON_SYNONYMOU | MODERAT | MISSENSE | Cgg/Ggg | R115G | arsC | | | | | | | | | C | C | | |
| NC_011770 | 3337327 | C | T | 70 | 0 | 4 | 0 | SPLICE_SITE_REGIC | LOW | SILENT | taG/taA | *117 | arsR | | | T | T | T | T | | | | | | |
| NC_011770 | 3337458 | C | T | 65 | 0 | 4 | 0 | NON_SYNONYMOU | MODERAT | MISSENSE | Gcg/Acg | A74T | arsR | | | T | T | T | T | | | | | | |
| NC_011770 | 4852888 | T | C | 71 | 0 | 1 | 0 | NON_SYNONYMOU | MODERAT | MISSENSE | aAg/aGg | K318R | aruE | | | | | | | C | | | | | |
| NC_011770 | 4853327 | G | C | 76 | 0 | 2 | 0 | NON_SYNONYMOU | MODERAT | MISSENSE | Cag/Gag | Q172E | aruE | | | | | | | C | C | | | | |
| NC_011770 | 4857014 | C | G | 78 | 0 | 2 | 0 | NON_SYNONYMOU | MODERAT | MISSENSE | gGt/gCt | G340A | aruG | | | | | | | | | | | G | G |
| NC_011770 | 4857218 | T | C | 76 | 0 | 2 | 0 | NON_SYNONYMOU | MODERAT | MISSENSE | aAc/aGc | N272S | aruG | | | | | | | | | C | C | | |
| NC_011770 | 5919206 | G | T | 98 | 0 | 4 | 0 | NON_SYNONYMOU | MODERAT | MISSENSE | gaC/gaA | D227E | aruH | | | T | T | T | T | | | | | | |
| NC_011770 | 5919303 | A | G | 103 | 0 | 4 | 0 | NON_SYNONYMOU | MODERAT | MISSENSE | aTg/aCg | M195T | aruH | | | G | G | G | G | | | | | | |
| NC_011770 | 5919636 | C | T | 97 | 0 | 4 | 0 | NON_SYNONYMOU | MODERAT | MISSENSE | cGg/cAg | R84Q | aruH | | | T | T | T | T | | | | | | |
| NC_011770 | 5919730 | C | A | 95 | 0 | 4 | 0 | NON_SYNONYMOU | MODERAT | MISSENSE | Gcg/Tcg | A53S | aruH | | | A | A | A | A | | | | | | |
| NC_011770 | 5919752 | A | C | 95 | 0 | 4 | 0 | NON_SYNONYMOU | MODERAT | MISSENSE | gaT/gaG | D45E | aruH | C | C | | | | | | | C | C | | |
| NC_011770 | 5921588 | T | C | 86 | 0 | 4 | 0 | START_LOST | HIGH | MISSENSE | Atg/Gtg | M1V | aruI | | | C | C | C | C | | | | | | |
| NC_011770 | 5919992 | T | C | 99 | 0 | 10 | 0 | NON_SYNONYMOU | MODERAT | MISSENSE | Aac/Gac | N533D | aruI | C | C | C | C | C | C | C | C | C | C | | |
| NC_011770 | 5920467 | C | A | 71 | 0 | 2 | 0 | NON_SYNONYMOU | MODERAT | MISSENSE | gaG/gaT | E374D | aruI | | | | | | | A | A | | | | |
| NC_011770 | 5920534 | G | A | 78 | 0 | 4 | 0 | NON_SYNONYMOU | MODERAT | MISSENSE | gCg/gTg | A352V | aruI | | | A | A | A | A | | | | | | |
| NC_011770 | 5920759 | G | A | 75 | 0 | 2 | 0 | NON_SYNONYMOU | MODERAT | MISSENSE | gCc/gTc | A277V | aruI | | | | | | | | | | | A | A |
| NC_011770 | 5920783 | A | T | 73 | 0 | 2 | 0 | NON_SYNONYMOU | MODERAT | MISSENSE | cTg/cAg | L269Q | aruI | | | | | | | T | T | | | | |
| NC_011770 | 5920851 | A | C | 71 | 0 | 4 | 0 | NON_SYNONYMOU | MODERAT | MISSENSE | gaT/gaG | D246E | aruI | | | C | C | C | C | | | | | | |
| NC_011770 | 5920939 | G | C | 80 | 0 | 4 | 0 | NON_SYNONYMOU | MODERAT | MISSENSE | aCc/aGc | T217S | aruI | | | C | C | C | C | | | | | | |
| NC_011770 | 5921533 | G | A | 81 | 0 | 2 | 0 | NON_SYNONYMOU | MODERAT | MISSENSE | aCg/aTg | T19M | aruI | | | | | | | A | A | | | | |
| NC_011770 | 2095228 | C | T | 75 | 0 | 2 | 0 | NON_SYNONYMOU | MODERAT | MISSENSE | aCc/aTc | T215I | asd | | | | | | | T | T | | | | |
| NC_011770 | 5912766 | T | G | 66 | 0 | 4 | 0 | NON_SYNONYMOU | MODERAT | MISSENSE | Aag/Cag | K97Q | aspP | | | G | G | G | G | | | | | | |
| NC_011770 | 4788170 | G | C | 63 | 0 | 4 | 0 | NON_SYNONYMOU | MODERAT | MISSENSE | Ccg/Gcg | P588A | aspS | | | | | | | C | C | C | C | | |
| NC_011770 | 4788788 | C | T | 78 | 0 | 2 | 0 | NON_SYNONYMOU | MODERAT | MISSENSE | Gtg/Atg | V382M | aspS | | | | | | | | | T | T | | |
| NC_011770 | 4788837 | T | G | 76 | 0 | 4 | 0 | NON_SYNONYMOU | MODERAT | MISSENSE | gaA/gaC | E365D | aspS | | | G | G | G | G | | | | | | |
| NC_011770 | 4855863 | C | T | 64 | 0 | 2 | 0 | NON_SYNONYMOU | MODERAT | MISSENSE | Gcg/Acg | A369T | astD | | | | | | | | | | | T | T |
| NC_011770 | 4856406 | G | C | 60 | 0 | 6 | 0 | NON_SYNONYMOU | MODERAT | MISSENSE | Cag/Gag | Q188E | astD | | | C | C | C | C | | | | | C | C |
| NC_011770 | 4856624 | C | T | 58 | 0 | 2 | 0 | NON_SYNONYMOU | MODERAT | MISSENSE | cGc/cAc | R115H | astD | T | T | | | | | | | | | | |
| NC_011770 | 4856717 | A | G | 59 | 0 | 2 | 0 | NON_SYNONYMOU | MODERAT | MISSENSE | gTg/gCg | V84A | astD | | | | | | | | | | | G | G |
| NC_011770 | 4856853 | T | C | 63 | 0 | 2 | 0 | NON_SYNONYMOU | MODERAT | MISSENSE | Acc/Gcc | T39A | astD | | | | | | | | | | | C | C |
| NC_011770 | 4856855 | G | A | 63 | 0 | 4 | 0 | NON_SYNONYMOU | MODERAT | MISSENSE | gCc/gTc | A38V | astD | | | A | A | A | A | | | | | | |
| NC_011770 | 3677990 | C | T | 45 | 0 | 2 | 0 | NON_SYNONYMOU | MODERAT | MISSENSE | Gcg/Acg | A256T | atoB | | | | | T | T | | | | | | |
| NC_011770 | 3678245 | A | T | 55 | 0 | 8 | 0 | NON_SYNONYMOU | MODERAT | MISSENSE | Tcc/Acc | S171T | atoB | T | T | T | T | T | T | T | T | | | | |
| NC_011770 | 6587996 | C | T | 70 | 0 | 2 | 0 | NON_SYNONYMOU | MODERAT | MISSENSE | Ggc/Agc | G504S | atpA | T | T | | | | | | | | | | |
| NC_011770 | 6586707 | G | A | 99 | 0 | 1 | 0 | NON_SYNONYMOU | MODERAT | MISSENSE | Cgc/Tgc | R105C | atpD | | | | | | | A | | | | | |
| NC_011770 | 6589759 | T | C | 75 | 0 | 1 | 0 | NON_SYNONYMOU | MODERAT | MISSENSE | aAg/aGg | K101R | atpH | | | | | | | C | | | | | |
| NC_011770 | 6591909 | T | C | 82 | 0 | 2 | 0 | NON_SYNONYMOU | MODERAT | MISSENSE | Acg/Gcg | T89A | atpI | | | | | | | | | | | C | C |
| NC_011770 | 206761 | C | G | 80 | 0 | 2 | 0 | NON_SYNONYMOU | MODERAT | MISSENSE | Gtg/Ctg | V535L | atsA | | | | | | | | | | | G | G |
| NC_011770 | 207124 | T | G | 82 | 0 | 12 | 0 | NON_SYNONYMOU | MODERAT | MISSENSE | Atc/Ctc | I414L | atsA | G | G | G | G | G | G | G | G | G | G | G | G |
| NC_011770 | 207469 | C | T | 73 | 0 | 2 | 0 | NON_SYNONYMOU | MODERAT | MISSENSE | Gac/Aac | D299N | atsA | | | | | | | T | T | | | | |
| NC_011770 | 207579 | G | C | 65 | 0 | 2 | 0 | NON_SYNONYMOU | MODERAT | MISSENSE | aCc/aGc | T262S | atsA | | | | | | | | | C | C | | |
| NC_011770 | 207711 | C | T | 64 | 0 | 2 | 0 | NON_SYNONYMOU | MODERAT | MISSENSE | cGg/cAg | R218Q | atsA | | | | | | | | | T | T | | |
| NC_011770 | 207794 | T | G | 60 | 0 | 2 | 0 | NON_SYNONYMOU | MODERAT | MISSENSE | caA/caC | Q190H | atsA | | | | | | | | | G | G | | |
| NC_011770 | 208123 | C | T | 54 | 0 | 2 | 0 | NON_SYNONYMOU | MODERAT | MISSENSE | Gaa/Aaa | E81K | atsA | | | | | | | T | T | | | | |
| NC_011770 | 2355777 | G | A | 64 | 0 | 12 | 0 | NON_SYNONYMOU | MODERAT | MISSENSE | Cac/Tac | H512Y | atuA | A | A | A | A | A | A | A | A | A | A | A | A |
| NC_011770 | 2355899 | T | C | 69 | 0 | 4 | 0 | NON_SYNONYMOU | MODERAT | MISSENSE | cAc/cGc | H471R | atuA | | | | | | | C | C | C | C | | |
| NC_011770 | 2356259 | T | C | 65 | 0 | 2 | 0 | NON_SYNONYMOU | MODERAT | MISSENSE | gAg/gGg | E351G | atuA | C | C | | | | | | | | | | |
| NC_011770 | 2356448 | A | G | 68 | 0 | 2 | 0 | NON_SYNONYMOU | MODERAT | MISSENSE | gTc/gCc | V288A | atuA | G | G | | | | | | | | | | |

| | | | | | | | | | | | | | | | | | | | | | | | | | |
|---|---|---|---|---|---|---|---|---|---|---|---|---|---|---|---|---|---|---|---|---|---|---|---|---|---|
| NC_011770 | 2356829 | C | T | 79 | 0 | 2 | 0 | NON_SYNONYMOU | MODERAT | MISSENSE | gGc/gAc | G161D | atuA | | | | | | | | | T | T | | |
| NC_011770 | 2356932 | C | A | 73 | 0 | 4 | 0 | NON_SYNONYMOU | MODERAT | MISSENSE | Gcc/Tcc | A127S | atuA | | | A | A | A | A | | | | | | |
| NC_011770 | 2354646 | A | G | 83 | 0 | 4 | 0 | NON_SYNONYMOU | MODERAT | MISSENSE | Ttc/Ctc | F265L | atuB | | | G | G | G | G | | | | | | |
| NC_011770 | 2353134 | G | C | 73 | 0 | 12 | 0 | NON_SYNONYMOU | MODERAT | MISSENSE | gaC/gaG | D475E | atuC | C | C | C | C | C | C | C | C | C | C | C | C |
| NC_011770 | 2352543 | T | G | 65 | 0 | 2 | 0 | NON_SYNONYMOU | MODERAT | MISSENSE | Acc/Ccc | T95P | atuD | G | G | | | | | | | | | | |
| NC_011770 | 2351325 | G | A | 64 | 0 | 2 | 0 | NON_SYNONYMOU | MODERAT | MISSENSE | gCg/gTg | A107V | atuE | | | | | | | A | A | | | | |
| NC_011770 | 2351463 | C | T | 67 | 0 | 4 | 0 | NON_SYNONYMOU | MODERAT | MISSENSE | gGc/gAc | G61D | atuE | | | | | | | T | T | T | T | | |
| NC_011770 | 2351605 | C | T | 73 | 0 | 6 | 0 | NON_SYNONYMOU | MODERAT | MISSENSE | Gtc/Atc | V14I | atuE | | | T | T | T | T | | | | | T | T |
| NC_011770 | 2349051 | A | G | 68 | 0 | 4 | 0 | NON_SYNONYMOU | MODERAT | MISSENSE | gTc/gCc | V585A | atuF | | | | | | | G | G | | | G | G |
| NC_011770 | 2349153 | C | T | 71 | 0 | 4 | 0 | NON_SYNONYMOU | MODERAT | MISSENSE | cGc/cAc | R551H | atuF | | | T | T | T | T | | | | | | |
| NC_011770 | 2349189 | C | T | 75 | 0 | 2 | 0 | NON_SYNONYMOU | MODERAT | MISSENSE | gGa/gAa | G539E | atuF | T | T | | | | | | | | | | |
| NC_011770 | 2349961 | C | T | 70 | 0 | 1 | 0 | NON_SYNONYMOU | MODERAT | MISSENSE | Gag/Aag | E282K | atuF | | | | | | | T | | | | | |
| NC_011770 | 2350262 | G | C | 83 | 0 | 8 | 0 | NON_SYNONYMOU | MODERAT | MISSENSE | gaC/gaG | D181E | atuF | C | C | C | C | C | C | | | | | C | C |
| NC_011770 | 2350612 | T | G | 71 | 0 | 12 | 0 | NON_SYNONYMOU | MODERAT | MISSENSE | Atc/Ctc | I65L | atuF | G | G | G | G | G | G | G | G | G | G | G | G |
| NC_011770 | 2350685 | C | G | 71 | 0 | 10 | 0 | NON_SYNONYMOU | MODERAT | MISSENSE | gaG/gaC | E40D | atuF | G | G | G | G | G | G | G | G | G | G | | |
| NC_011770 | 2346038 | C | T | 70 | 0 | 2 | 0 | NON_SYNONYMOU | MODERAT | MISSENSE | Gtc/Atc | V582I | atuH | T | T | | | | | | | | | | |
| NC_011770 | 2346302 | C | T | 75 | 0 | 1 | 0 | NON_SYNONYMOU | MODERAT | MISSENSE | Ggg/Agg | G494R | atuH | | T | | | | | | | | | | |
| NC_011770 | 2357828 | G | T | 86 | 0 | 2 | 0 | NON_SYNONYMOU | MODERAT | MISSENSE | Gcc/Tcc | A80S | atuR | T | T | | | | | | | | | | |
| NC_011770 | 2358179 | G | A | 90 | 0 | 6 | 0 | NON_SYNONYMOU | MODERAT | MISSENSE | Gac/Aac | D197N | atuR | A | A | A | A | A | A | | | | | | |
| NC_011770 | 3721815 | T | C | 46 | 0 | 12 | 0 | NON_SYNONYMOU | MODERAT | MISSENSE | Atg/Gtg | M277V | bacA | C | C | C | C | C | C | C | C | C | C | C | C |
| NC_011770 | 3722027 | T | A | 53 | 0 | 12 | 0 | NON_SYNONYMOU | MODERAT | MISSENSE | aAg/aTg | K206M | bacA | A | A | A | A | A | A | A | A | A | A | A | A |
| NC_011770 | 3722028 | T | C | 54 | 0 | 12 | 0 | NON_SYNONYMOU | MODERAT | MISSENSE | Aag/Gag | K206E | bacA | C | C | C | C | C | C | C | C | C | C | C | C |
| NC_011770 | 3722163 | C | T | 54 | 0 | 12 | 0 | NON_SYNONYMOU | MODERAT | MISSENSE | Gtc/Atc | V161I | bacA | T | T | T | T | T | T | T | T | T | T | T | T |
| NC_011770 | 3722234 | T | C | 56 | 0 | 12 | 0 | NON_SYNONYMOU | MODERAT | MISSENSE | cAc/cGc | H137R | bacA | C | C | C | C | C | C | C | C | C | C | C | C |
| NC_011770 | 4754893 | G | A | 53 | 0 | 4 | 0 | NON_SYNONYMOU | MODERAT | MISSENSE | Gac/Aac | D11N | bcp | A | A | | | | | | | | | A | A |
| NC_011770 | 4755331 | G | A | 51 | 0 | 6 | 0 | NON_SYNONYMOU | MODERAT | MISSENSE | Gcc/Acc | A157T | bcp | | | A | A | A | A | | | A | A | | |
| NC_011770 | 3675168 | G | C | 72 | 0 | 2 | 0 | NON_SYNONYMOU | MODERAT | MISSENSE | gGc/gCc | G68A | bdhA | C | C | | | | | | | | | | |
| NC_011770 | 3675207 | T | C | 70 | 0 | 2 | 0 | NON_SYNONYMOU | MODERAT | MISSENSE | gTc/gCc | V81A | bdhA | | | | | | | C | C | | | | |
| NC_011770 | 3675311 | T | C | 86 | 0 | 1 | 0 | NON_SYNONYMOU | MODERAT | MISSENSE | Ttc/Ctc | F116L | bdhA | | | | | | | C | | | | | |
| NC_011770 | 3675558 | C | T | 74 | 0 | 8 | 0 | NON_SYNONYMOU | MODERAT | MISSENSE | aCc/aTc | T198I | bdhA | | | T | T | T | T | T | T | | | T | T |
| NC_011770 | 3675712 | G | A | 66 | 0 | 2 | 0 | NON_SYNONYMOU | MODERAT | MISSENSE | atG/atA | M249I | bdhA | A | A | | | | | | | | | | |
| NC_011770 | 2995569 | T | C | 73 | 0 | 4 | 0 | NON_SYNONYMOU | MODERAT | MISSENSE | gTc/gCc | V24A | benD | C | C | | | | | | | | | C | C |
| NC_011770 | 6384364 | G | A | 73 | 0 | 1 | 0 | NON_SYNONYMOU | MODERAT | MISSENSE | Cgc/Tgc | R211C | betA | | A | | | | | | | | | | |
| NC_011770 | 6383364 | G | A | 55 | 0 | 4 | 0 | NON_SYNONYMOU | MODERAT | MISSENSE | gCg/gTg | A544V | betA | | | A | A | A | A | | | | | | |
| NC_011770 | 6383631 | C | T | 68 | 0 | 2 | 0 | NON_SYNONYMOU | MODERAT | MISSENSE | aGc/aAc | S455N | betA | | | | | | | | | | | T | T |
| NC_011770 | 6384282 | G | A | 83 | 0 | 2 | 0 | NON_SYNONYMOU | MODERAT | MISSENSE | tCc/tTc | S238F | betA | | | | | | | | | | | A | A |
| NC_011770 | 6385471 | G | T | 70 | 0 | 10 | 0 | NON_SYNONYMOU | MODERAT | MISSENSE | Cgc/Agc | R378S | betB | T | T | T | T | T | T | T | T | | | T | T |
| NC_011770 | 6385508 | T | A | 74 | 0 | 10 | 0 | NON_SYNONYMOU | MODERAT | MISSENSE | aaA/aaT | K365N | betB | A | A | A | A | A | A | A | A | | | A | A |
| NC_011770 | 6385687 | C | T | 74 | 0 | 2 | 0 | NON_SYNONYMOU | MODERAT | MISSENSE | Gta/Ata | V306I | betB | T | T | | | | | | | | | | |
| NC_011770 | 33009 | C | T | 76 | 0 | 2 | 0 | NON_SYNONYMOU | MODERAT | MISSENSE | cGc/cAc | R497H | betC | T | T | | | | | | | | | | |
| NC_011770 | 33088 | T | C | 81 | 0 | 2 | 0 | NON_SYNONYMOU | MODERAT | MISSENSE | Agc/Ggc | S471G | betC | | | | | | | | | C | C | | |
| NC_011770 | 33784 | C | T | 84 | 0 | 2 | 0 | NON_SYNONYMOU | MODERAT | MISSENSE | Gac/Aac | D239N | betC | | | | | | | T | T | | | | |
| NC_011770 | 34223 | G | C | 70 | 0 | 2 | 0 | NON_SYNONYMOU | MODERAT | MISSENSE | aaC/aaG | N92K | betC | | | | | | | | | C | C | | |
| NC_011770 | 34416 | A | G | 78 | 0 | 4 | 0 | NON_SYNONYMOU | MODERAT | MISSENSE | gTg/gCg | V28A | betC | | | | | | | G | G | G | G | | |
| NC_011770 | 6388532 | C | T | 85 | 0 | 1 | 0 | NON_SYNONYMOU | MODERAT | MISSENSE | gCc/gTc | A315V | betT1 | | | | | | | T | | | | | |
| NC_011770 | 1631535 | C | T | 71 | 0 | 4 | 0 | NON_SYNONYMOU | MODERAT | MISSENSE | Gcc/Acc | A42T | bfrB | | | T | T | T | T | | | | | | |
| NC_011770 | 4000430 | G | A | 87 | 0 | 2 | 0 | NON_SYNONYMOU | MODERAT | MISSENSE | Gtg/Atg | V250M | bglX | | | | | | | | | | | A | A |
| NC_011770 | 4001192 | G | A | 74 | 0 | 2 | 0 | NON_SYNONYMOU | MODERAT | MISSENSE | Gtg/Atg | V504M | bglX | | | | | | | A | A | | | | |
| NC_011770 | 4001453 | A | G | 63 | 0 | 1 | 0 | NON_SYNONYMOU | MODERAT | MISSENSE | Agc/Ggc | S591G | bglX | | | | | | | | G | | | | |
| NC_011770 | 460678 | C | T | 88 | 0 | 2 | 0 | NON_SYNONYMOU | MODERAT | MISSENSE | Gac/Aac | D434N | bioA | T | T | | | | | | | | | | |
| NC_011770 | 461074 | C | T | 87 | 0 | 4 | 0 | NON_SYNONYMOU | MODERAT | MISSENSE | Gtc/Atc | V302I | bioA | | | T | T | T | T | | | | | | |
| NC_011770 | 461320 | C | T | 93 | 0 | 4 | 0 | NON_SYNONYMOU | MODERAT | MISSENSE | Gtc/Atc | V220I | bioA | | | T | T | T | T | | | | | | |
| NC_011770 | 551971 | C | G | 96 | 0 | 2 | 0 | NON_SYNONYMOU | MODERAT | MISSENSE | Ctc/Gtc | L15V | bioB | | | | | | | | | G | G | | |
| NC_011770 | 552172 | A | T | 92 | 0 | 4 | 0 | NON_SYNONYMOU | MODERAT | MISSENSE | Atg/Ttg | M82L | bioB | | | | | | | | | T | T | T | T |
| NC_011770 | 556345 | T | C | 75 | 0 | 12 | 0 | NON_SYNONYMOU | MODERAT | MISSENSE | gTc/gCc | V171A | bioD | C | C | C | C | C | C | C | C | C | C | C | C |
| NC_011770 | 553460 | C | T | 82 | 0 | 2 | 0 | NON_SYNONYMOU | MODERAT | MISSENSE | aCg/aTg | T123M | bioF | T | T | | | | | | | | | | |
| NC_011770 | 554071 | C | T | 75 | 0 | 1 | 0 | NON_SYNONYMOU | MODERAT | MISSENSE | Cgc/Tgc | R327C | bioF | | | | | | | | T | | | | |
| NC_011770 | 704944 | C | G | 85 | 0 | 4 | 0 | NON_SYNONYMOU | MODERAT | MISSENSE | gCt/gGt | A92G | birA | | | G | G | G | G | | | | | | |
| NC_011770 | 705151 | A | G | 67 | 0 | 2 | 0 | NON_SYNONYMOU | MODERAT | MISSENSE | gAc/gGc | D161G | birA | | | | | | | G | G | | | | |
| NC_011770 | 705273 | G | A | 72 | 0 | 2 | 0 | NON_SYNONYMOU | MODERAT | MISSENSE | Gtc/Atc | V202I | birA | | | | | | | A | A | | | | |
| NC_011770 | 705097 | G | A | 70 | 0 | 1 | 0 | NON_SYNONYMOU | MODERAT | MISSENSE | aGc/aAc | S143N | birA | | | | | | A | | | | | | |
| NC_011770 | 3369201 | C | T | 68 | 0 | 1 | 0 | NON_SYNONYMOU | MODERAT | MISSENSE | gGc/gAc | G216D | bkdA2 | | | | | | | T | | | | | |

| | | | | | | | | | | | | | | | | | | | | | | | | | |
|---|---|---|---|---|---|---|---|---|---|---|---|---|---|---|---|---|---|---|---|---|---|---|---|---|---|
| NC_011770 | 3367740 | G | A | 71 | 0 | 2 | 0 | NON_SYNONYMOU | MODERAT | MISSENSE | gCg/gTg | A352V | bkdB | | | | | | | | | A | A | | |
| NC_011770 | 3368098 | C | T | 82 | 0 | 2 | 0 | NON_SYNONYMOU | MODERAT | MISSENSE | Gtc/Atc | V233I | bkdB | | | | | | | T | T | | | | |
| NC_011770 | 3368232 | C | G | 68 | 0 | 2 | 0 | NON_SYNONYMOU | MODERAT | MISSENSE | gGc/gCc | G188A | bkdB | | | | | | | G | G | | | | |
| NC_011770 | 3368422 | C | A | 54 | 0 | 2 | 0 | NON_SYNONYMOU | MODERAT | MISSENSE | Gac/Tac | D125Y | bkdB | | | | | | | | | | | A | A |
| NC_011770 | 3368542 | A | G | 54 | 0 | 12 | 0 | NON_SYNONYMOU | MODERAT | MISSENSE | Ttc/Ctc | F85L | bkdB | G | G | G | G | G | G | G | G | G | G | G | G |
| NC_011770 | 3368506 | GCGCGGCC | G | 53 | 0 | 1 | 0 | CODON_DELETION | MODERATE | | gcgacgccg | ATPAA92 | bkdB | | | | | | | | | | G | | |
| NC_011770 | 6085849 | C | G | 74 | 0 | 2 | 0 | NON_SYNONYMOU | MODERAT | MISSENSE | Gtg/Ctg | V118L | blc | G | G | | | | | | | | | | |
| NC_011770 | 6085935 | T | G | 76 | 0 | 2 | 0 | NON_SYNONYMOU | MODERAT | MISSENSE | gAg/gCg | E89A | blc | G | G | | | | | | | | | | |
| NC_011770 | 6086014 | G | T | 78 | 0 | 2 | 0 | NON_SYNONYMOU | MODERAT | MISSENSE | Cat/Aat | H63N | blc | | | | | | | | | | | T | T |
| NC_011770 | 4135274 | T | C | 90 | 0 | 2 | 0 | NON_SYNONYMOU | MODERAT | MISSENSE | Atc/Gtc | I123V | braB | | | | | | | C | C | | | | |
| NC_011770 | 4135594 | G | A | 80 | 0 | 2 | 0 | NON_SYNONYMOU | MODERAT | MISSENSE | aCg/aTg | T16M | braB | | | | | | | | | | | A | A |
| NC_011770 | 4134466 | T | C | 28 | 0 | 1 | 0 | NON_SYNONYMOU | MODERAT | MISSENSE | aAc/aGc | N392S | braB | | | | | | | | C | | | | |
| NC_011770 | 4685372 | G | A | 78 | 0 | 4 | 0 | NON_SYNONYMOU | MODERAT | MISSENSE | Gcc/Acc | A335T | braC | | | A | A | A | A | | | | | | |
| NC_011770 | 4686305 | C | G | 91 | 0 | 2 | 0 | NON_SYNONYMOU | MODERAT | MISSENSE | gCc/gGc | A185G | braD | | | | | G | G | | | | | | |
| NC_011770 | 3713508 | A | G | 61 | 0 | 2 | 0 | NON_SYNONYMOU | MODERAT | MISSENSE | Atc/Gtc | I255V | braZ | | | | | | | | | | | G | G |
| NC_011770 | 3713554 | T | C | 64 | 0 | 1 | 0 | NON_SYNONYMOU | MODERAT | MISSENSE | gTg/gCg | V270A | braZ | | | | | | | C | | | | | |
| NC_011770 | 3713926 | A | G | 66 | 0 | 2 | 0 | NON_SYNONYMOU | MODERAT | MISSENSE | gAc/gGc | D394G | braZ | G | G | | | | | | | | | | |
| NC_011770 | 5343698 | G | A | 61 | 0 | 1 | 0 | NON_SYNONYMOU | MODERAT | MISSENSE | Cgg/Tgg | R34W | cafA | | | | | | | | | | | | A |
| NC_011770 | 5342551 | G | A | 84 | 0 | 2 | 0 | NON_SYNONYMOU | MODERAT | MISSENSE | aCc/aTc | T416I | cafA | | | | | | | | | | | A | A |
| NC_011770 | 5343379 | T | A | 68 | 0 | 12 | 0 | NON_SYNONYMOU | MODERAT | MISSENSE | gAc/gTc | D140V | cafA | A | A | A | A | A | A | A | A | A | A | A | A |
| NC_011770 | 1948952 | C | T | 53 | 0 | 2 | 0 | NON_SYNONYMOU | MODERAT | MISSENSE | Gtc/Atc | V69I | capB | | | | | | | | | | | T | T |
| NC_011770 | 5678052 | G | T | 86 | 0 | 2 | 0 | NON_SYNONYMOU | MODERAT | MISSENSE | agC/agA | S226R | carA | | | | | | | | | | | T | T |
| NC_011770 | 5674093 | G | A | 68 | 0 | 4 | 0 | NON_SYNONYMOU | MODERAT | MISSENSE | aCc/aTc | T940I | carB | | | A | A | A | A | | | | | | |
| NC_011770 | 5674276 | A | G | 72 | 0 | 12 | 0 | NON_SYNONYMOU | MODERAT | MISSENSE | aTc/aCc | I879T | carB | G | G | G | G | G | G | G | G | G | G | G | G |
| NC_011770 | 5075733 | C | T | 80 | 0 | 2 | 0 | NON_SYNONYMOU | MODERAT | MISSENSE | gCc/gTc | A91V | cat | | | | | | | T | T | | | | |
| NC_011770 | 5075750 | A | G | 78 | 0 | 2 | 0 | NON_SYNONYMOU | MODERAT | MISSENSE | Aac/Gac | N97D | cat | | | | | | | | | G | G | | |
| NC_011770 | 5076017 | A | C | 92 | 0 | 12 | 0 | NON_SYNONYMOU | MODERAT | MISSENSE | Atg/Ctg | M186L | cat | C | C | C | C | C | C | C | C | C | C | C | C |
| NC_011770 | 5076026 | A | G | 97 | 0 | 6 | 0 | NON_SYNONYMOU | MODERAT | MISSENSE | Act/Gct | T189A | cat | G | G | | | | | | | G | G | G | G |
| NC_011770 | 5076074 | A | T | 86 | 0 | 4 | 0 | NON_SYNONYMOU | MODERAT | MISSENSE | Acg/Tcg | T205S | cat | | | T | T | T | T | | | | | | |
| NC_011770 | 3003911 | G | T | 60 | 0 | 6 | 0 | NON_SYNONYMOU | MODERAT | MISSENSE | Gcc/Tcc | A292S | catA | T | T | | | | | | | T | T | T | T |
| NC_011770 | 3001647 | G | C | 54 | 0 | 12 | 0 | NON_SYNONYMOU | MODERAT | MISSENSE | aGc/aCc | S33T | catB | C | C | C | C | C | C | C | C | C | C | C | C |
| NC_011770 | 3002013 | A | G | 67 | 0 | 2 | 0 | NON_SYNONYMOU | MODERAT | MISSENSE | gAg/gGg | E155G | catB | G | G | | | | | | | | | | |
| NC_011770 | 3002594 | C | A | 72 | 0 | 2 | 0 | NON_SYNONYMOU | MODERAT | MISSENSE | Cac/Aac | H349N | catB | | | | | | | | | | | A | A |
| NC_011770 | 3002666 | T | C | 68 | 0 | 12 | 0 | NON_SYNONYMOU | MODERAT | MISSENSE | Tgc/Cgc | C373R | catB | C | C | C | C | C | C | C | C | C | C | C | C |
| NC_011770 | 3000682 | G | C | 86 | 0 | 2 | 0 | NON_SYNONYMOU | MODERAT | MISSENSE | gCg/gGg | A236G | catR | | | | | | | | | | | C | C |
| NC_011770 | 3000875 | T | G | 83 | 0 | 10 | 0 | NON_SYNONYMOU | MODERAT | MISSENSE | Acg/Ccg | T172P | catR | G | G | G | G | G | G | G | G | | | G | G |
| NC_011770 | 4907337 | G | A | 78 | 0 | 2 | 0 | NON_SYNONYMOU | MODERAT | MISSENSE | Gcc/Acc | A258T | cbpD | | | | | | | | | | | A | A |
| NC_011770 | 4907338 | C | A | 78 | 0 | 2 | 0 | NON_SYNONYMOU | MODERAT | MISSENSE | gCc/gAc | A258D | cbpD | | | | | | | | | | | A | A |
| NC_011770 | 4907406 | A | G | 78 | 0 | 6 | 0 | NON_SYNONYMOU | MODERAT | MISSENSE | Atc/Gtc | I281V | cbpD | | | | | | | G | G | G | G | G | G |
| NC_011770 | 4907499 | G | A | 86 | 0 | 4 | 0 | NON_SYNONYMOU | MODERAT | MISSENSE | Gtc/Atc | V312I | cbpD | | | A | A | A | A | | | | | | |
| NC_011770 | 5638323 | C | T | 100 | 0 | 2 | 0 | NON_SYNONYMOU | MODERAT | MISSENSE | aCc/aTc | T165I | cbrA | | | | | | | T | T | | | | |
| NC_011770 | 5638794 | T | C | 88 | 0 | 1 | 0 | NON_SYNONYMOU | MODERAT | MISSENSE | cTg/cCg | L322P | cbrA | | | | | | | C | | | | | |
| NC_011770 | 5641843 | C | T | 67 | 0 | 2 | 0 | NON_SYNONYMOU | MODERAT | MISSENSE | Ccg/Tcg | P346S | cbrB | T | T | | | | | | | | | | |
| NC_011770 | 633114 | A | G | 72 | 0 | 12 | 0 | NON_SYNONYMOU | MODERAT | MISSENSE | Tgt/Cgt | C394R | cca | G | G | G | G | G | G | G | G | G | G | G | G |
| NC_011770 | 633372 | C | T | 72 | 0 | 1 | 0 | NON_SYNONYMOU | MODERAT | MISSENSE | Gcc/Acc | A308T | cca | | | | | | | T | | | | | |
| NC_011770 | 633471 | T | C | 66 | 0 | 12 | 0 | NON_SYNONYMOU | MODERAT | MISSENSE | Act/Gct | T275A | cca | C | C | C | C | C | C | C | C | C | C | C | C |
| NC_011770 | 633822 | C | A | 56 | 0 | 2 | 0 | NON_SYNONYMOU | MODERAT | MISSENSE | Gcc/Tcc | A158S | cca | A | A | | | | | | | | | | |
| NC_011770 | 633840 | C | T | 56 | 0 | 4 | 0 | NON_SYNONYMOU | MODERAT | MISSENSE | Gtc/Atc | V152I | cca | | | | | | | T | T | T | T | | |
| NC_011770 | 4349986 | C | CCCG | 66 | 0 | 2 | 0 | CODON_INSERTIOI | MODERATE | | ggc/gCGG | G230AG | ccmA | CCCG | CCCG | | | | | | | | | | |
| NC_011770 | 4350062 | C | T | 69 | 0 | 8 | 0 | NON_SYNONYMOU | MODERAT | MISSENSE | Gtg/Atg | V205M | ccmA | | | T | T | T | T | | | T | T | T | T |
| NC_011770 | 4350131 | C | T | 73 | 0 | 6 | 0 | NON_SYNONYMOU | MODERAT | MISSENSE | Gcg/Acg | A182T | ccmA | | | T | T | T | T | | | | | T | T |
| NC_011770 | 4350266 | C | T | 76 | 0 | 2 | 0 | NON_SYNONYMOU | MODERAT | MISSENSE | Gtt/Att | V137I | ccmA | | | | | | | | | T | T | | |
| NC_011770 | 4350544 | G | A | 51 | 0 | 12 | 0 | NON_SYNONYMOU | MODERAT | MISSENSE | gCg/gTg | A44V | ccmA | A | A | A | A | A | A | A | A | A | A | A | A |
| NC_011770 | 4348471 | C | T | 91 | 0 | 4 | 0 | NON_SYNONYMOU | MODERAT | MISSENSE | Gcc/Acc | A238T | ccmC | | | T | T | T | T | | | | | | |
| NC_011770 | 4348558 | G | T | 86 | 0 | 12 | 0 | NON_SYNONYMOU | MODERAT | MISSENSE | Ctc/Atc | L209I | ccmC | T | T | T | T | T | T | T | T | T | T | T | T |
| NC_011770 | 4348205 | A | G | 82 | 0 | 6 | 0 | NON_SYNONYMOU | MODERAT | MISSENSE | gTg/gCg | V17A | ccmE | | | G | G | G | G | | | G | G | | |
| NC_011770 | 4346694 | T | C | 64 | 0 | 10 | 0 | NON_SYNONYMOU | MODERAT | MISSENSE | Atg/Gtg | M358V | ccmF | C | C | C | C | C | C | C | C | | | C | C |
| NC_011770 | 4347317 | A | G | 46 | 0 | 2 | 0 | NON_SYNONYMOU | MODERAT | MISSENSE | cTg/cCg | L150P | ccmF | G | G | | | | | | | | | | |
| NC_011770 | 4345225 | T | G | 72 | 0 | 10 | 0 | NON_SYNONYMOU | MODERAT | MISSENSE | Atg/Ctg | M9L | ccmH | G | G | G | G | G | G | G | G | | | G | G |
| NC_011770 | 4345233 | G | A | 72 | 0 | 6 | 0 | NON_SYNONYMOU | MODERAT | MISSENSE | gCc/gTc | A6V | ccmH | A | A | | | | | A | A | | | A | A |
| NC_011770 | 1489552 | A | T | 80 | 0 | 4 | 0 | NON_SYNONYMOU | MODERAT | MISSENSE | gAa/gTa | E50V | cdsA | | | T | T | T | T | | | | | | |
| NC_011770 | 1490005 | C | T | 74 | 0 | 4 | 0 | NON_SYNONYMOU | MODERAT | MISSENSE | gCg/gTg | A201V | cdsA | | | T | T | T | T | | | | | | |

| | | | | | | | | | | | | | | | | | | | | | | | | | |
|---|---|---|---|---|---|---|---|---|---|---|---|---|---|---|---|---|---|---|---|---|---|---|---|---|---|
| NC_011770 | 1490010 | C | T | 77 | 0 | 2 | 0 | NON_SYNONYMOU | MODERAT | MISSENSE | Ctc/Ttc | L203F | cdsA | | | | | | | T | T | | | | |
| NC_011770 | 3948757 | G | A | 73 | 0 | 2 | 0 | NON_SYNONYMOU | MODERAT | MISSENSE | Cgc/Tgc | R2C | cfrX | A | A | | | | | | | | | | |
| NC_011770 | 3312765 | T | TG | 67 | 0 | 1 | 0 | FRAME_SHIFT | HIGH | | ggc/ggGc | G107G? | chiC | | | | | | | TG | | | | | |
| NC_011770 | 3312953 | A | G | 76 | 0 | 12 | 0 | NON_SYNONYMOU | MODERAT | MISSENSE | Acc/Gcc | T168A | chiC | G | G | G | G | G | G | G | G | G | G | G | G |
| NC_011770 | 449874 | C | T | 69 | 0 | 2 | 0 | NON_SYNONYMOU | MODERAT | MISSENSE | Ccg/Tcg | P670S | chpA | | | | | | | T | T | | | | |
| NC_011770 | 450242 | G | C | 70 | 0 | 8 | 0 | NON_SYNONYMOU | MODERAT | MISSENSE | agG/agC | R792S | chpA | | | C | C | C | C | | | C | C | C | C |
| NC_011770 | 450762 | G | A | 66 | 0 | 1 | 0 | NON_SYNONYMOU | MODERAT | MISSENSE | Gcc/Acc | A966T | chpA | | | | | | | A | | | | | |
| NC_011770 | 451879 | A | T | 81 | 0 | 10 | 0 | NON_SYNONYMOU | MODERAT | MISSENSE | tAc/tTc | Y1338F | chpA | | | T | T | T | T | T | T | T | T | T | T |
| NC_011770 | 452266 | C | T | 82 | 0 | 2 | 0 | NON_SYNONYMOU | MODERAT | MISSENSE | cCc/cTc | P1467L | chpA | | | | | | | | | | | T | T |
| NC_011770 | 452295 | G | T | 80 | 0 | 10 | 0 | NON_SYNONYMOU | MODERAT | MISSENSE | Gcg/Tcg | A1477S | chpA | | | T | T | T | T | T | T | T | T | T | T |
| NC_011770 | 452322 | C | G | 82 | 0 | 8 | 0 | NON_SYNONYMOU | MODERAT | MISSENSE | Ctg/Gtg | L1486V | chpA | | | G | G | G | G | | | G | G | G | G |
| NC_011770 | 452382 | A | G | 84 | 0 | 12 | 0 | NON_SYNONYMOU | MODERAT | MISSENSE | Acc/Gcc | T1506A | chpA | G | G | G | G | G | G | G | G | G | G | G | G |
| NC_011770 | 452510 | G | C | 80 | 0 | 2 | 0 | NON_SYNONYMOU | MODERAT | MISSENSE | atG/atC | M1548I | chpA | | | | | | | | | | | C | C |
| NC_011770 | 452992 | G | A | 69 | 0 | 2 | 0 | NON_SYNONYMOU | MODERAT | MISSENSE | cGg/cAg | R1709Q | chpA | | | | | | | | | | | A | A |
| NC_011770 | 453091 | A | T | 58 | 0 | 2 | 0 | NON_SYNONYMOU | MODERAT | MISSENSE | cAa/cTa | Q1742L | chpA | | | | | | | | | | | T | T |
| NC_011770 | 453244 | A | G | 39 | 0 | 2 | 0 | NON_SYNONYMOU | MODERAT | MISSENSE | gAc/gGc | D1793G | chpA | G | G | | | | | | | | | | |
| NC_011770 | 453253 | G | A | 36 | 0 | 8 | 0 | NON_SYNONYMOU | MODERAT | MISSENSE | gGc/gAc | G1796D | chpA | A | A | A | A | A | A | | | | | A | A |
| NC_011770 | 450250 | A | G | 66 | 0 | 1 | 0 | NON_SYNONYMOU | MODERAT | MISSENSE | gAa/gGa | E795G | chpA | | | | | | | | **G** | | | | |
| NC_011770 | 452919 | G | A | 71 | 0 | 1 | 0 | NON_SYNONYMOU | MODERAT | MISSENSE | Gag/Aag | E1685K | chpA | | | | | | | | | | | | **A** |
| NC_011770 | 455291 | C | T | 84 | 0 | 6 | 0 | NON_SYNONYMOU | MODERAT | MISSENSE | gCc/gTc | A5V | chpB | | | T | T | T | T | | | T | T | | |
| NC_011770 | 455321 | C | G | 87 | 0 | 2 | 0 | NON_SYNONYMOU | MODERAT | MISSENSE | aCc/aGc | T15S | chpB | G | G | | | | | | | | | | |
| NC_011770 | 456001 | C | T | 92 | 0 | 6 | 0 | NON_SYNONYMOU | MODERAT | MISSENSE | Ccc/Tcc | P242S | chpB | T | T | | | | | T | T | T | T | | |
| NC_011770 | 456568 | C | G | 72 | 0 | 6 | 0 | NON_SYNONYMOU | MODERAT | MISSENSE | cCg/cGg | P88R | chpC | | | G | G | G | G | G | G | | | | |
| NC_011770 | 456756 | T | C | 70 | 0 | 12 | 0 | NON_SYNONYMOU | MODERAT | MISSENSE | Tgg/Cgg | W151R | chpC | C | C | C | C | C | C | C | C | C | C | C | C |
| NC_011770 | 457324 | C | A | 76 | 0 | 2 | 0 | NON_SYNONYMOU | MODERAT | MISSENSE | Ctg/Atg | L169M | chpD | | | | | | | A | A | | | | |
| NC_011770 | 457364 | A | G | 76 | 0 | 2 | 0 | NON_SYNONYMOU | MODERAT | MISSENSE | gAg/gGg | E182G | chpD | | | | | | | | | | | G | G |
| NC_011770 | 2702288 | T | A | 13 | 8 | 2 | 0 | NON_SYNONYMOU | MODERAT | MISSENSE | ttA/ttT | L227F | cI | | | . | . | A | A | . | . | . | . | . | . |
| NC_011770 | 2702293 | G | T | 13 | 8 | 2 | 0 | NON_SYNONYMOU | MODERAT | MISSENSE | Cct/Act | P226T | cI | | | . | . | T | T | . | . | . | . | . | . |
| NC_011770 | 2702351 | A | C | 14 | 8 | 2 | 0 | NON_SYNONYMOU | MODERAT | MISSENSE | caT/caG | H206Q | cI | | | . | . | C | C | . | . | . | . | . | . |
| NC_011770 | 2702653 | A | C | 6 | 10 | 2 | 0 | NON_SYNONYMOU | MODERAT | MISSENSE | Tcg/Gcg | S106A | cI | C | C | . | . | . | . | . | . | . | . | . | . |
| NC_011770 | 1130732 | C | T | 67 | 0 | 1 | 0 | NON_SYNONYMOU | MODERAT | MISSENSE | Ccg/Tcg | P269S | cioA | | | | | | | T | | | | | |
| NC_011770 | 6503264 | C | G | 80 | 0 | 2 | 0 | NON_SYNONYMOU | MODERAT | MISSENSE | atG/atC | M22I | citA | | | | | | | | | | | G | G |
| NC_011770 | 2673928 | C | T | 100 | 0 | 2 | 0 | NON_SYNONYMOU | MODERAT | MISSENSE | aCc/aTc | T104I | clpA | | | | | | | | | | | T | T |
| NC_011770 | 2674528 | G | A | 86 | 0 | 2 | 0 | NON_SYNONYMOU | MODERAT | MISSENSE | gGg/gAg | G304E | clpA | | | | | | | | | | | A | A |
| NC_011770 | 2673915 | A | G | 103 | 0 | 1 | 0 | NON_SYNONYMOU | MODERAT | MISSENSE | Aag/Gag | K100E | clpA | | | | | | | | G | | | | |
| NC_011770 | 5422564 | A | C | 67 | 0 | 12 | 0 | NON_SYNONYMOU | MODERAT | MISSENSE | Ttg/Gtg | L846V | clpB | C | C | C | C | C | C | C | C | C | C | C | C |
| NC_011770 | 6408699 | G | T | 65 | 0 | 2 | 0 | NON_SYNONYMOU | MODERAT | MISSENSE | Ctg/Atg | L207M | cls | | | | | | | T | T | | | | |
| NC_011770 | 3949761 | T | C | 73 | 0 | 1 | 0 | NON_SYNONYMOU | MODERAT | MISSENSE | tAc/tGc | Y16C | cmaX | | | | | | | | C | | | | |
| NC_011770 | 2052450 | T | G | 73 | 0 | 4 | 0 | NON_SYNONYMOU | MODERAT | MISSENSE | cTc/cGc | L123R | cmk | | | G | G | G | G | | | | | | |
| NC_011770 | 6326142 | C | A | 79 | 0 | 2 | 0 | NON_SYNONYMOU | MODERAT | MISSENSE | cCc/cAc | P243H | coaC | | | | | | | | | | | A | A |
| NC_011770 | 6326588 | G | A | 76 | 0 | 6 | 0 | NON_SYNONYMOU | MODERAT | MISSENSE | Gtc/Atc | V392I | coaC | | | A | A | A | A | | | | | A | A |
| NC_011770 | 400701 | C | G | 82 | 0 | 4 | 0 | NON_SYNONYMOU | MODERAT | MISSENSE | aGc/aCc | S46T | coaD | | | G | G | G | G | | | | | | |
| NC_011770 | 5409222 | G | A | 85 | 0 | 1 | 0 | NON_SYNONYMOU | MODERAT | MISSENSE | Gcc/Acc | A160T | coaE | | | | | | | | A | | | | |
| NC_011770 | 5409125 | G | C | 76 | 0 | 2 | 0 | NON_SYNONYMOU | MODERAT | MISSENSE | caG/caC | Q127H | coaE | C | C | | | | | | | | | | |
| NC_011770 | 5409239 | C | A | 82 | 0 | 2 | 0 | NON_SYNONYMOU | MODERAT | MISSENSE | gaC/gaA | D165E | coaE | A | A | | | | | | | | | | |
| NC_011770 | 3945149 | G | A | 75 | 0 | 4 | 0 | NON_SYNONYMOU | MODERAT | MISSENSE | Gtc/Atc | V34I | cobA | | | A | A | A | A | | | | | | |
| NC_011770 | 3945779 | A | C | 52 | 0 | 4 | 0 | NON_SYNONYMOU | MODERAT | MISSENSE | Agc/Cgc | S244R | cobA | | | C | C | C | C | | | | | | |
| NC_011770 | 4455719 | A | G | 52 | 0 | 12 | 0 | NON_SYNONYMOU | MODERAT | MISSENSE | Tac/Cac | Y293H | cobB | G | G | G | G | G | G | G | G | G | G | G | G |
| NC_011770 | 4455796 | G | A | 59 | 0 | 2 | 0 | NON_SYNONYMOU | MODERAT | MISSENSE | gCg/gTg | A267V | cobB | | | | | | | | | | | A | A |
| NC_011770 | 4455887 | C | T | 60 | 0 | 2 | 0 | NON_SYNONYMOU | MODERAT | MISSENSE | Ggc/Agc | G237S | cobB | | | | | | | | | | | T | T |
| NC_011770 | 4455967 | C | A | 54 | 0 | 2 | 0 | NON_SYNONYMOU | MODERAT | MISSENSE | cGg/cTg | R210L | cobB | A | A | | | | | | | | | | |
| NC_011770 | 4455997 | G | A | 49 | 0 | 4 | 0 | NON_SYNONYMOU | MODERAT | MISSENSE | gCg/gTg | A200V | cobB | A | A | | | | | | | | | A | A |
| NC_011770 | 4456148 | G | A | 46 | 0 | 2 | 0 | NON_SYNONYMOU | MODERAT | MISSENSE | Cgc/Tgc | R150C | cobB | A | A | | | | | | | | | | |
| NC_011770 | 4453005 | C | T | 69 | 0 | 2 | 0 | NON_SYNONYMOU | MODERAT | MISSENSE | aGt/aAt | S233N | cobC | | | | | | | | | T | T | | |
| NC_011770 | 4453456 | A | T | 25 | 0 | 8 | 0 | NON_SYNONYMOU | MODERAT | MISSENSE | Ttg/Atg | L83M | cobC | | | T | T | T | T | T | T | | | T | T |
| NC_011770 | 4453600 | G | A | 41 | 0 | 6 | 0 | NON_SYNONYMOU | MODERAT | MISSENSE | Ccg/Tcg | P35S | cobC | | | A | A | A | A | A | A | | | | |
| NC_011770 | 4453972 | T | C | 61 | 0 | 4 | 0 | NON_SYNONYMOU | MODERAT | MISSENSE | gAc/gGc | D221G | cobD | C | C | | | | | | | C | C | | |
| NC_011770 | 4454270 | G | A | 66 | 0 | 2 | 0 | NON_SYNONYMOU | MODERAT | MISSENSE | Cgt/Tgt | R122C | cobD | | | | | | | | | A | A | | |
| NC_011770 | 4454411 | T | C | 58 | 0 | 10 | 0 | NON_SYNONYMOU | MODERAT | MISSENSE | Acg/Gcg | T75A | cobD | C | C | C | C | C | C | C | C | C | C | | |
| NC_011770 | 4454491 | C | G | 57 | 0 | 6 | 0 | NON_SYNONYMOU | MODERAT | MISSENSE | gGc/gCc | G48A | cobD | G | G | G | G | G | G | | | | | | |
| NC_011770 | 4454602 | A | G | 64 | 0 | 4 | 0 | NON_SYNONYMOU | MODERAT | MISSENSE | gTg/gCg | V11A | cobD | | | | | | | G | G | G | G | | |
| NC_011770 | 2335591 | G | A | 56 | 0 | 2 | 0 | NON_SYNONYMOU | MODERAT | MISSENSE | gGc/gAc | G57D | cobH | A | A | | | | | | | | | | |

| | | | | | | | | | | | | | | | | | | | | | | | | | |
|---|---|---|---|---|---|---|---|---|---|---|---|---|---|---|---|---|---|---|---|---|---|---|---|---|---|
| NC_011770 | 2335620 | G | A | 61 | 0 | 4 | 0 | NON_SYNONYMOU | MODERAT | MISSENSE | Gcc/Acc | A67T | cobH | | | | | | | | | A | A | A | A |
| NC_011770 | 2337231 | A | G | 66 | 0 | 12 | 0 | NON_SYNONYMOU | MODERAT | MISSENSE | Acc/Gcc | T148A | cobJ | G | G | G | G | G | G | G | G | G | G | G | G |
| NC_011770 | 2337307 | A | G | 62 | 0 | 4 | 0 | NON_SYNONYMOU | MODERAT | MISSENSE | gAc/gGc | D173G | cobJ | | | G | G | G | G | | | | | | |
| NC_011770 | 2337343 | A | C | 61 | 0 | 4 | 0 | NON_SYNONYMOU | MODERAT | MISSENSE | gAg/gCg | E185A | cobJ | | | C | C | C | C | | | | | | |
| NC_011770 | 2337600 | G | A | 63 | 0 | 10 | 0 | NON_SYNONYMOU | MODERAT | MISSENSE | Gcc/Acc | A271T | cobJ | | | A | A | A | A | A | A | A | A | A | A |
| NC_011770 | 2337894 | C | T | 79 | 0 | 1 | 0 | NON_SYNONYMOU | MODERAT | MISSENSE | Cgg/Tgg | R369W | cobJ | | | | | | | T | | | | | |
| NC_011770 | 2338110 | G | C | 66 | 0 | 6 | 0 | NON_SYNONYMOU | MODERAT | MISSENSE | Gtg/Ctg | V441L | cobJ | | | C | C | C | C | | | | | C | C |
| NC_011770 | 2338404 | C | T | 74 | 0 | 2 | 0 | NON_SYNONYMOU | MODERAT | MISSENSE | Cgc/Tgc | R539C | cobJ | | | | | | | T | T | | | | |
| NC_011770 | 2330648 | GC | G | 57 | 0 | 2 | 0 | FRAME_SHIFT | HIGH | | gcg/ | A215 | cobK | | | | | | | | | G | G | | |
| NC_011770 | 2330808 | C | T | 54 | 0 | 4 | 0 | NON_SYNONYMOU | MODERAT | MISSENSE | Gag/Aag | E162K | cobK | | | T | T | T | T | | | | | | |
| NC_011770 | 2330883 | G | A | 63 | 0 | 2 | 0 | NON_SYNONYMOU | MODERAT | MISSENSE | Cat/Tat | H137Y | cobK | | | | | | | | | | | A | A |
| NC_011770 | 2331059 | T | C | 65 | 0 | 4 | 0 | NON_SYNONYMOU | MODERAT | MISSENSE | cAg/cGg | Q78R | cobK | C | C | | | | | | | | | C | C |
| NC_011770 | 2331200 | A | G | 60 | 0 | 2 | 0 | NON_SYNONYMOU | MODERAT | MISSENSE | cTg/cCg | L31P | cobK | G | G | | | | | | | | | | |
| NC_011770 | 2333270 | C | T | 59 | 0 | 1 | 0 | STOP_GAINED | HIGH | NONSENSE | tGg/tAg | W120* | cobL | | | | | | | | T | | | | |
| NC_011770 | 2332425 | C | T | 63 | 0 | 2 | 0 | NON_SYNONYMOU | MODERAT | MISSENSE | Gaa/Aaa | E402K | cobL | T | T | | | | | | | | | | |
| NC_011770 | 2332883 | C | T | 58 | 0 | 4 | 0 | NON_SYNONYMOU | MODERAT | MISSENSE | cGc/cAc | R249H | cobL | T | T | | | | | | | | | T | T |
| NC_011770 | 2333076 | G | T | 51 | 0 | 2 | 0 | NON_SYNONYMOU | MODERAT | MISSENSE | Ctc/Atc | L185I | cobL | | | | | | | | | | | T | T |
| NC_011770 | 2333527 | T | G | 62 | 0 | 4 | 0 | NON_SYNONYMOU | MODERAT | MISSENSE | caA/caC | Q34H | cobL | | | G | G | G | G | | | | | | |
| NC_011770 | 2291776 | C | T | 58 | 0 | 2 | 0 | NON_SYNONYMOU | MODERAT | MISSENSE | Ccg/Tcg | P329S | cobN | | | | | | | | | T | T | | |
| NC_011770 | 2292005 | C | T | 51 | 0 | 2 | 0 | NON_SYNONYMOU | MODERAT | MISSENSE | gCg/gTg | A405V | cobN | | | | | | | T | T | | | | |
| NC_011770 | 2293144 | C | A | 67 | 0 | 2 | 0 | NON_SYNONYMOU | MODERAT | MISSENSE | Cgc/Agc | R785S | cobN | A | A | | | | | | | | | | |
| NC_011770 | 2293174 | A | G | 58 | 0 | 8 | 0 | NON_SYNONYMOU | MODERAT | MISSENSE | Acc/Gcc | T795A | cobN | | | G | G | G | G | G | G | G | G | | |
| NC_011770 | 2293801 | A | T | 51 | 0 | 4 | 0 | NON_SYNONYMOU | MODERAT | MISSENSE | Acg/Tcg | T1004S | cobN | | | T | T | T | T | | | | | | |
| NC_011770 | 3765307 | GGAC | G | 59 | 0 | 4 | 0 | CODON_CHANGE_ | MODERATE | | gcgtcc/gcc | AS1224A | cobN | | | G | G | G | G | | | | | | |
| NC_011770 | 3765548 | G | T | 69 | 0 | 10 | 0 | NON_SYNONYMOU | MODERAT | MISSENSE | gCg/gAg | A1145E | cobN | T | T | T | T | T | T | T | T | T | T | | |
| NC_011770 | 3766020 | C | T | 80 | 0 | 4 | 0 | NON_SYNONYMOU | MODERAT | MISSENSE | Ggg/Agg | G988R | cobN | | | T | T | T | T | | | | | | |
| NC_011770 | 3766074 | T | C | 84 | 0 | 4 | 0 | NON_SYNONYMOU | MODERAT | MISSENSE | Acc/Gcc | T970A | cobN | | | C | C | C | C | | | | | | |
| NC_011770 | 3767453 | T | A | 66 | 0 | 10 | 0 | NON_SYNONYMOU | MODERAT | MISSENSE | tAc/tTc | Y510F | cobN | A | A | A | A | A | A | A | A | A | A | | |
| NC_011770 | 3767505 | C | T | 63 | 0 | 2 | 0 | NON_SYNONYMOU | MODERAT | MISSENSE | Ggc/Agc | G493S | cobN | | | | | | | | | T | T | | |
| NC_011770 | 3767715 | A | G | 68 | 0 | 2 | 0 | NON_SYNONYMOU | MODERAT | MISSENSE | Tat/Cat | Y423H | cobN | | | | | | | | | | | G | G |
| NC_011770 | 3768051 | A | T | 67 | 0 | 12 | 0 | NON_SYNONYMOU | MODERAT | MISSENSE | Ttg/Atg | L311M | cobN | T | T | T | T | T | T | T | T | T | T | T | T |
| NC_011770 | 3768086 | G | A | 64 | 0 | 6 | 0 | NON_SYNONYMOU | MODERAT | MISSENSE | gCg/gTg | A299V | cobN | | | A | A | A | A | | | | | A | A |
| NC_011770 | 3768096 | G | A | 67 | 0 | 2 | 0 | NON_SYNONYMOU | MODERAT | MISSENSE | Ccg/Tcg | P296S | cobN | | | | | | | | | A | A | | |
| NC_011770 | 3768458 | C | A | 62 | 0 | 2 | 0 | NON_SYNONYMOU | MODERAT | MISSENSE | gGc/gTc | G175V | cobN | A | A | | | | | | | | | | |
| NC_011770 | 3768537 | T | G | 60 | 0 | 12 | 0 | NON_SYNONYMOU | MODERAT | MISSENSE | Acc/Ccc | T149P | cobN | G | G | G | G | G | G | G | G | G | G | G | G |
| NC_011770 | 3768720 | C | G | 56 | 0 | 2 | 0 | NON_SYNONYMOU | MODERAT | MISSENSE | Gag/Cag | E88Q | cobN | | | | | | | | | G | G | | |
| NC_011770 | 3768771 | T | C | 54 | 0 | 2 | 0 | NON_SYNONYMOU | MODERAT | MISSENSE | Aac/Gac | N71D | cobN | | | | | | | C | C | | | | |
| NC_011770 | 3768783 | T | G | 56 | 0 | 12 | 0 | NON_SYNONYMOU | MODERAT | MISSENSE | Atg/Ctg | M67L | cobN | G | G | G | G | G | G | G | G | G | G | G | G |
| NC_011770 | 3768921 | G | A | 63 | 0 | 4 | 0 | NON_SYNONYMOU | MODERAT | MISSENSE | Ccg/Tcg | P21S | cobN | | | A | A | A | A | | | | | | |
| NC_011770 | 4456716 | C | T | 73 | 0 | 4 | 0 | NON_SYNONYMOU | MODERAT | MISSENSE | Gtc/Atc | V175I | cobO | T | T | | | | | T | T | | | | |
| NC_011770 | 4456868 | T | G | 80 | 0 | 12 | 0 | NON_SYNONYMOU | MODERAT | MISSENSE | gAc/gCc | D124A | cobO | G | G | G | G | G | G | G | G | G | G | G | G |
| NC_011770 | 4457175 | T | C | 81 | 0 | 10 | 0 | NON_SYNONYMOU | MODERAT | MISSENSE | Att/Gtt | I22V | cobO | C | C | C | C | C | C | C | C | C | C | | |
| NC_011770 | 4451492 | A | G | 74 | 0 | 2 | 0 | NON_SYNONYMOU | MODERAT | MISSENSE | gTg/gCg | V408A | cobQ | | | | | | | G | G | | | | |
| NC_011770 | 4451623 | C | G | 70 | 0 | 2 | 0 | NON_SYNONYMOU | MODERAT | MISSENSE | ttG/ttC | L364F | cobQ | G | G | | | | | | | | | | |
| NC_011770 | 4451844 | A | G | 61 | 0 | 2 | 0 | NON_SYNONYMOU | MODERAT | MISSENSE | Tcg/Ccg | S291P | cobQ | G | G | | | | | | | | | | |
| NC_011770 | 4452134 | C | T | 78 | 0 | 8 | 0 | NON_SYNONYMOU | MODERAT | MISSENSE | aGg/aAg | R194K | cobQ | T | T | T | T | T | T | | | T | T | | |
| NC_011770 | 4448426 | T | C | 73 | 0 | 2 | 0 | NON_SYNONYMOU | MODERAT | MISSENSE | Acc/Gcc | T223A | cobS | | | | | | | C | C | | | | |
| NC_011770 | 4448545 | C | A | 78 | 0 | 12 | 0 | NON_SYNONYMOU | MODERAT | MISSENSE | gGg/gTg | G183V | cobS | A | A | A | A | A | A | A | A | A | A | A | A |
| NC_011770 | 4448563 | G | C | 81 | 0 | 2 | 0 | NON_SYNONYMOU | MODERAT | MISSENSE | gCg/gGg | A177G | cobS | | | | | | | C | C | | | | |
| NC_011770 | 4448711 | A | C | 81 | 0 | 12 | 0 | NON_SYNONYMOU | MODERAT | MISSENSE | Tcc/Gcc | S128A | cobS | C | C | C | C | C | C | C | C | C | C | C | C |
| NC_011770 | 4448819 | C | T | 89 | 0 | 4 | 0 | NON_SYNONYMOU | MODERAT | MISSENSE | Gtc/Atc | V92I | cobS | T | T | | | | | | | T | T | | |
| NC_011770 | 4448932 | A | C | 88 | 0 | 6 | 0 | NON_SYNONYMOU | MODERAT | MISSENSE | cTg/cGg | L54R | cobS | C | C | | | | | C | C | C | C | | |
| NC_011770 | 4449726 | C | T | 71 | 0 | 1 | 0 | NON_SYNONYMOU | MODERAT | MISSENSE | Gcg/Acg | A336T | cobT | | | | | | | T | | | | | |
| NC_011770 | 4449734 | A | G | 68 | 0 | 8 | 0 | NON_SYNONYMOU | MODERAT | MISSENSE | gTg/gCg | V333A | cobT | G | G | | | | | G | G | G | G | G | G |
| NC_011770 | 4449948 | C | T | 67 | 0 | 2 | 0 | NON_SYNONYMOU | MODERAT | MISSENSE | Gtc/Atc | V262I | cobT | T | T | | | | | | | | | | |
| NC_011770 | 4450083 | T | C | 66 | 0 | 4 | 0 | NON_SYNONYMOU | MODERAT | MISSENSE | Atc/Gtc | I217V | cobT | | | C | C | C | C | | | | | | |
| NC_011770 | 4450092 | C | T | 65 | 0 | 4 | 0 | NON_SYNONYMOU | MODERAT | MISSENSE | Gcc/Acc | A214T | cobT | T | T | | | | | | | T | T | | |
| NC_011770 | 4450148 | G | A | 65 | 0 | 2 | 0 | NON_SYNONYMOU | MODERAT | MISSENSE | gCg/gTg | A195V | cobT | | | | | | | | | A | A | | |
| NC_011770 | 4450265 | C | T | 67 | 0 | 2 | 0 | NON_SYNONYMOU | MODERAT | MISSENSE | cGc/cAc | R156H | cobT | | | | | | | T | T | | | | |
| NC_011770 | 4450311 | G | A | 66 | 0 | 4 | 0 | NON_SYNONYMOU | MODERAT | MISSENSE | Ccg/Tcg | P141S | cobT | | | A | A | A | A | | | | | | |
| NC_011770 | 4450383 | A | C | 67 | 0 | 4 | 0 | NON_SYNONYMOU | MODERAT | MISSENSE | Ttc/Gtc | F117V | cobT | C | C | | | | | | | C | C | | |
| NC_011770 | 4450509 | T | C | 68 | 0 | 4 | 0 | NON_SYNONYMOU | MODERAT | MISSENSE | Atc/Gtc | I75V | cobT | C | C | | | | | | | C | C | | |

| | | | | | | | | | | | | | | | | | | | | | | | | | |
|---|---|---|---|---|---|---|---|---|---|---|---|---|---|---|---|---|---|---|---|---|---|---|---|---|---|
| NC_011770 | 4450557 | T | G | 68 | 0 | 4 | 0 | NON_SYNONYMOU | MODERAT | MISSENSE | Atc/Ctc | I59L | cobT | G | G | | | | | | | G | G | | |
| NC_011770 | 4450798 | A | G | 69 | 0 | 12 | 0 | NON_SYNONYMOU | MODERAT | MISSENSE | gTc/gCc | V151A | cobU | G | G | G | G | G | G | G | G | G | G | G | G |
| NC_011770 | 4450912 | T | G | 62 | 0 | 2 | 0 | NON_SYNONYMOU | MODERAT | MISSENSE | gAg/gCg | E113A | cobU | G | G | | | | | | | | | | |
| NC_011770 | 4450957 | T | A | 60 | 0 | 2 | 0 | NON_SYNONYMOU | MODERAT | MISSENSE | cAg/cTg | Q98L | cobU | A | A | | | | | | | | | | |
| NC_011770 | 4451027 | C | G | 57 | 0 | 2 | 0 | NON_SYNONYMOU | MODERAT | MISSENSE | Gcc/Ccc | A75P | cobU | G | G | | | | | | | | | | |
| NC_011770 | 483141 | C | G | 82 | 0 | 12 | 0 | NON_SYNONYMOU | MODERAT | MISSENSE | aGc/aCc | S420T | codA | G | G | G | G | G | G | G | G | G | G | G | G |
| NC_011770 | 485071 | G | T | 68 | 0 | 2 | 0 | NON_SYNONYMOU | MODERAT | MISSENSE | cCg/cAg | P190Q | codB | T | T | | | | | | | | | | |
| NC_011770 | 130295 | G | A | 90 | 0 | 2 | 0 | NON_SYNONYMOU | MODERAT | MISSENSE | Ggc/Agc | G252S | colII | | | | | | | A | A | | | | |
| NC_011770 | 5428099 | G | A | 77 | 0 | 1 | 0 | NON_SYNONYMOU | MODERAT | MISSENSE | Ggt/Agt | G338S | comL | | | | | | | | | | | | A |
| NC_011770 | 2435309 | C | G | 68 | 0 | 2 | 0 | NON_SYNONYMOU | MODERAT | MISSENSE | cGa/cCa | R443P | copS | | | | | | | G | G | | | | |
| NC_011770 | 2435658 | C | G | 66 | 0 | 4 | 0 | NON_SYNONYMOU | MODERAT | MISSENSE | Gaa/Caa | E327Q | copS | | | G | G | G | G | | | | | | |
| NC_011770 | 2435678 | C | T | 66 | 0 | 4 | 0 | NON_SYNONYMOU | MODERAT | MISSENSE | cGc/cAc | R320H | copS | | | T | T | T | T | | | | | | |
| NC_011770 | 6270506 | T | C | 78 | 0 | 4 | 0 | NON_SYNONYMOU | MODERAT | MISSENSE | Atc/Gtc | I31V | corA | | | C | C | C | C | | | | | | |
| NC_011770 | 6270544 | T | C | 78 | 0 | 4 | 0 | NON_SYNONYMOU | MODERAT | MISSENSE | gAc/gGc | D18G | corA | | | | | | | | | C | C | C | C |
| NC_011770 | 126654 | G | A | 90 | 0 | 2 | 0 | NON_SYNONYMOU | MODERAT | MISSENSE | Gtc/Atc | V139I | coxB | A | A | | | | | | | | | | |
| NC_011770 | 2459343 | C | T | 76 | 0 | 1 | 0 | NON_SYNONYMOU | MODERAT | MISSENSE | Gcg/Acg | A387T | cpg2 | | | | | | | T | | | | | |
| NC_011770 | 2459472 | T | C | 65 | 0 | 2 | 0 | NON_SYNONYMOU | MODERAT | MISSENSE | Atc/Gtc | I344V | cpg2 | | | | | | | | | C | C | | |
| NC_011770 | 2460026 | C | T | 65 | 0 | 2 | 0 | NON_SYNONYMOU | MODERAT | MISSENSE | gGc/gAc | G159D | cpg2 | | | | | | | | | T | T | | |
| NC_011770 | 2460063 | C | T | 58 | 0 | 8 | 0 | NON_SYNONYMOU | MODERAT | MISSENSE | Gtg/Atg | V147M | cpg2 | | | T | T | T | T | | | T | T | T | T |
| NC_011770 | 2460194 | C | T | 55 | 0 | 2 | 0 | NON_SYNONYMOU | MODERAT | MISSENSE | aGc/aAc | S103N | cpg2 | T | T | | | | | | | | | | |
| NC_011770 | 2460227 | T | G | 51 | 0 | 2 | 0 | NON_SYNONYMOU | MODERAT | MISSENSE | gAc/gCc | D92A | cpg2 | | | | | | | | | G | G | | |
| NC_011770 | 2460369 | C | T | 39 | 0 | 2 | 0 | NON_SYNONYMOU | MODERAT | MISSENSE | Gac/Aac | D45N | cpg2 | | | | | | | | | T | T | | |
| NC_011770 | 2460377 | G | A | 38 | 0 | 4 | 0 | NON_SYNONYMOU | MODERAT | MISSENSE | gCc/gTc | A42V | cpg2 | | | A | A | A | A | | | | | | |
| NC_011770 | 2460441 | G | C | 32 | 0 | 12 | 0 | NON_SYNONYMOU | MODERAT | MISSENSE | Ccc/Gcc | P21A | cpg2 | C | C | C | C | C | C | C | C | C | C | C | C |
| NC_011770 | 2460468 | A | G | 35 | 0 | 8 | 0 | NON_SYNONYMOU | MODERAT | MISSENSE | Ttc/Ctc | F12L | cpg2 | | | G | G | G | G | | | G | G | G | G |
| NC_011770 | 2460492 | T | G | 36 | 0 | 10 | 0 | NON_SYNONYMOU | MODERAT | MISSENSE | Agc/Cgc | S4R | cpg2 | | | G | G | G | G | G | G | G | G | G | G |
| NC_011770 | 2460500 | A | C | 38 | 0 | 2 | 0 | NON_SYNONYMOU | MODERAT | MISSENSE | gTg/gGg | V1G | cpg2 | C | C | | | | | | | | | | |
| NC_011770 | 515913 | G | A | 69 | 0 | 10 | 0 | NON_SYNONYMOU | MODERAT | MISSENSE | gGg/gAg | G128E | creB | | | A | A | A | A | A | A | A | A | A | A |
| NC_011770 | 515978 | C | T | 74 | 0 | 2 | 0 | NON_SYNONYMOU | MODERAT | MISSENSE | Ccg/Tcg | P150S | creB | | | | | | | | | | | T | T |
| NC_011770 | 515573 | G | A | 84 | 0 | 1 | 0 | NON_SYNONYMOU | MODERAT | MISSENSE | Gac/Aac | D15N | creB | | | | | A | | | | | | | |
| NC_011770 | 516862 | C | T | 100 | 0 | 2 | 0 | STOP_GAINED | HIGH | NONSENSE | Cag/Tag | Q215* | creC | | | | | | | | | | | T | T |
| NC_011770 | 516689 | G | C | 91 | 0 | 2 | 0 | NON_SYNONYMOU | MODERAT | MISSENSE | gGc/gCc | G157A | creC | | | | | | | | | C | C | | |
| NC_011770 | 517266 | G | T | 83 | 0 | 4 | 0 | NON_SYNONYMOU | MODERAT | MISSENSE | caG/caT | Q349H | creC | | | T | T | T | T | | | | | | |
| NC_011770 | 517293 | A | T | 85 | 0 | 2 | 0 | NON_SYNONYMOU | MODERAT | MISSENSE | gaA/gaT | E358D | creC | T | T | | | | | | | | | | |
| NC_011770 | 517409 | T | C | 84 | 0 | 6 | 0 | NON_SYNONYMOU | MODERAT | MISSENSE | gTc/gCc | V397A | creC | C | C | | | | | C | C | C | C | | |
| NC_011770 | 517526 | A | T | 83 | 0 | 2 | 0 | NON_SYNONYMOU | MODERAT | MISSENSE | aAg/aTg | K436M | creC | | | | | | | | | | | T | T |
| NC_011770 | 517904 | G | T | 93 | 0 | 4 | 0 | NON_SYNONYMOU | MODERAT | MISSENSE | cGc/cTc | R53L | creD | | | T | T | T | T | | | | | | |
| NC_011770 | 518749 | G | A | 98 | 0 | 4 | 0 | NON_SYNONYMOU | MODERAT | MISSENSE | Gtc/Atc | V335I | creD | A | A | | | | | | | | | A | A |
| NC_011770 | 518926 | G | A | 83 | 0 | 2 | 0 | NON_SYNONYMOU | MODERAT | MISSENSE | Gtc/Atc | V394I | creD | A | A | | | | | | | | | | |
| NC_011770 | 518927 | T | C | 82 | 0 | 8 | 0 | NON_SYNONYMOU | MODERAT | MISSENSE | gTc/gCc | V394A | creD | | | C | C | C | C | C | C | C | C | | |
| NC_011770 | 519025 | G | A | 68 | 0 | 4 | 0 | NON_SYNONYMOU | MODERAT | MISSENSE | Gcg/Acg | A427T | creD | | | A | A | A | A | | | | | | |
| NC_011770 | 519079 | T | C | 67 | 0 | 2 | 0 | NON_SYNONYMOU | MODERAT | MISSENSE | Ttc/Ctc | F445L | creD | C | C | | | | | | | | | | |
| NC_011770 | 519098 | G | A | 68 | 0 | 2 | 0 | NON_SYNONYMOU | MODERAT | MISSENSE | aGg/aAg | R451K | creD | A | A | | | | | | | | | | |
| NC_011770 | 1992821 | C | T | 78 | 0 | 4 | 0 | NON_SYNONYMOU | MODERAT | MISSENSE | Gtc/Atc | V34I | csaA | | | T | T | T | T | | | | | | |
| NC_011770 | 3862121 | G | A | 68 | 0 | 1 | 0 | STOP_GAINED | HIGH | NONSENSE | Cag/Tag | Q237* | cti | | | | | | | A | | | | | |
| NC_011770 | 3860987 | A | G | 85 | 0 | 2 | 0 | NON_SYNONYMOU | MODERAT | MISSENSE | Ttc/Ctc | F615L | cti | | | | | | | G | G | | | | |
| NC_011770 | 3861234 | A | C | 82 | 0 | 12 | 0 | NON_SYNONYMOU | MODERAT | MISSENSE | caT/caG | H532Q | cti | C | C | C | C | C | C | C | C | C | C | C | C |
| NC_011770 | 3861248 | C | T | 82 | 0 | 2 | 0 | NON_SYNONYMOU | MODERAT | MISSENSE | Gtc/Atc | V528I | cti | | | | | | | | | T | T | | |
| NC_011770 | 3861500 | T | G | 74 | 0 | 8 | 0 | NON_SYNONYMOU | MODERAT | MISSENSE | Atc/Ctc | I444L | cti | | | G | G | G | G | G | G | | | G | G |
| NC_011770 | 3861844 | G | A | 67 | 0 | 2 | 0 | NON_SYNONYMOU | MODERAT | MISSENSE | gCc/gTc | A329V | cti | | | | | | | | | A | A | | |
| NC_011770 | 3862520 | G | A | 65 | 0 | 8 | 0 | NON_SYNONYMOU | MODERAT | MISSENSE | Ccg/Tcg | P104S | cti | A | A | A | A | A | A | A | A | | | | |
| NC_011770 | 3860780 | C | T | 83 | 0 | 1 | 0 | NON_SYNONYMOU | MODERAT | MISSENSE | Gtg/Atg | V684M | cti | | | | | | | | | | | T | |
| NC_011770 | 3529007 | T | C | 84 | 0 | 2 | 0 | NON_SYNONYMOU | MODERAT | MISSENSE | Agc/Ggc | S125G | cupA2 | | | | | | | | | | | C | C |
| NC_011770 | 3529171 | G | A | 79 | 0 | 4 | 0 | NON_SYNONYMOU | MODERAT | MISSENSE | gCg/gTg | A70V | cupA2 | | | A | A | A | A | | | | | | |
| NC_011770 | 3526080 | C | G | 73 | 0 | 4 | 0 | NON_SYNONYMOU | MODERAT | MISSENSE | gGt/gCt | G857A | cupA3 | | | G | G | G | G | | | | | | |
| NC_011770 | 3526249 | C | T | 78 | 0 | 4 | 0 | NON_SYNONYMOU | MODERAT | MISSENSE | Ggc/Agc | G801S | cupA3 | | | T | T | T | T | | | | | | |
| NC_011770 | 3527572 | G | T | 75 | 0 | 4 | 0 | NON_SYNONYMOU | MODERAT | MISSENSE | Ctg/Atg | L360M | cupA3 | | | T | T | T | T | | | | | | |
| NC_011770 | 3527956 | T | C | 74 | 0 | 4 | 0 | NON_SYNONYMOU | MODERAT | MISSENSE | Agc/Ggc | S232G | cupA3 | | | C | C | C | C | | | | | | |
| NC_011770 | 3528273 | T | C | 82 | 0 | 2 | 0 | NON_SYNONYMOU | MODERAT | MISSENSE | gAt/gGt | D126G | cupA3 | | | | | | | | | C | C | | |
| NC_011770 | 3525143 | T | C | 74 | 0 | 4 | 0 | NON_SYNONYMOU | MODERAT | MISSENSE | Agc/Ggc | S298G | cupA4 | | | | | | | | | C | C | C | C |
| NC_011770 | 3525167 | A | G | 72 | 0 | 8 | 0 | NON_SYNONYMOU | MODERAT | MISSENSE | Tcc/Ccc | S290P | cupA4 | G | G | G | G | G | G | G | G | | | | |
| NC_011770 | 3525406 | A | G | 71 | 0 | 2 | 0 | NON_SYNONYMOU | MODERAT | MISSENSE | aTc/aCc | I210T | cupA4 | | | | | | | G | G | | | | |

| | | | | | | | | | | | | | | | | | | | | | | | | | |
|---|---|---|---|---|---|---|---|---|---|---|---|---|---|---|---|---|---|---|---|---|---|---|---|---|---|
| NC_011770 | 3525535 | C | G | 73 | 0 | 12 | 0 | NON_SYNONYMOU | MODERAT | MISSENSE | aGc/aCc | S167T | cupA4 | G | G | G | G | G | G | G | G | G | G | G | G |
| NC_011770 | 3525947 | C | T | 68 | 0 | 1 | 0 | NON_SYNONYMOU | MODERAT | MISSENSE | Gtc/Atc | V30I | cupA4 | | | | | | | T | | | | | |
| NC_011770 | 3524062 | G | A | 66 | 0 | 2 | 0 | NON_SYNONYMOU | MODERAT | MISSENSE | Cgg/Tgg | R208W | cupA5 | | | | | | | A | A | | | | |
| NC_011770 | 968764 | T | C | 73 | 0 | 4 | 0 | NON_SYNONYMOU | MODERAT | MISSENSE | gTg/gCg | V25A | cupB1 | | | | | | | C | C | | | C | C |
| NC_011770 | 968994 | A | C | 78 | 0 | 4 | 0 | NON_SYNONYMOU | MODERAT | MISSENSE | Acg/Ccg | T102P | cupB1 | | | C | C | C | C | | | | | | |
| NC_011770 | 968995 | C | A | 77 | 0 | 4 | 0 | NON_SYNONYMOU | MODERAT | MISSENSE | aCg/aAg | T102K | cupB1 | | | A | A | A | A | | | | | | |
| NC_011770 | 969145 | C | T | 70 | 0 | 2 | 0 | NON_SYNONYMOU | MODERAT | MISSENSE | aCc/aTc | T152I | cupB1 | | | | | | | | | | | T | T |
| NC_011770 | 969211 | T | C | 64 | 0 | 4 | 0 | NON_SYNONYMOU | MODERAT | MISSENSE | aTc/aCc | I174T | cupB1 | | | C | C | C | C | | | | | | |
| NC_011770 | 969358 | T | C | 70 | 0 | 8 | 0 | NON_SYNONYMOU | MODERAT | MISSENSE | gTc/gCc | V23A | cupB2 | | | C | C | C | C | C | C | | | C | C |
| NC_011770 | 970354 | T | C | 68 | 0 | 12 | 0 | NON_SYNONYMOU | MODERAT | MISSENSE | gTg/gCg | V1A | cupB3 | C | C | C | C | C | C | C | C | C | C | C | C |
| NC_011770 | 970380 | T | G | 66 | 0 | 6 | 0 | NON_SYNONYMOU | MODERAT | MISSENSE | Ttg/Gtg | L10V | cupB3 | G | G | G | G | G | G | | | | | | |
| NC_011770 | 972604 | A | G | 70 | 0 | 12 | 0 | NON_SYNONYMOU | MODERAT | MISSENSE | aAg/aGg | K751R | cupB3 | G | G | G | G | G | G | G | G | G | G | G | G |
| NC_011770 | 973058 | G | A | 60 | 0 | 2 | 0 | NON_SYNONYMOU | MODERAT | MISSENSE | Gcc/Acc | A59T | cupB4 | | | | | | | | | A | A | | |
| NC_011770 | 973322 | G | T | 50 | 0 | 2 | 0 | NON_SYNONYMOU | MODERAT | MISSENSE | Gcc/Tcc | A147S | cupB4 | | | | | | | | | T | T | | |
| NC_011770 | 973395 | C | T | 45 | 0 | 2 | 0 | NON_SYNONYMOU | MODERAT | MISSENSE | gCg/gTg | A171V | cupB4 | | | | | | | T | T | | | | |
| NC_011770 | 973508 | C | G | 48 | 0 | 2 | 0 | NON_SYNONYMOU | MODERAT | MISSENSE | Ctt/Gtt | L209V | cupB4 | | | | | | | G | G | | | | |
| NC_011770 | 976702 | CA | C | 40 | 0 | 1 | 0 | FRAME_SHIFT | HIGH | | aac/ | N998 | cupB5 | | | | | | C | | | | | | |
| NC_011770 | 973778 | A | T | 69 | 0 | 4 | 0 | NON_SYNONYMOU | MODERAT | MISSENSE | Agt/Tgt | S23C | cupB5 | | | | | | | | | T | T | T | T |
| NC_011770 | 973827 | C | T | 69 | 0 | 6 | 0 | NON_SYNONYMOU | MODERAT | MISSENSE | gCc/gTc | A39V | cupB5 | | | T | T | T | T | T | T | | | | |
| NC_011770 | 973833 | C | T | 71 | 0 | 2 | 0 | NON_SYNONYMOU | MODERAT | MISSENSE | gCc/gTc | A41V | cupB5 | | | | | | | | | | | T | T |
| NC_011770 | 974162 | A | G | 83 | 0 | 2 | 0 | NON_SYNONYMOU | MODERAT | MISSENSE | Aat/Gat | N151D | cupB5 | | | G | G | | | | | | | | |
| NC_011770 | 974272 | C | G | 79 | 0 | 6 | 0 | NON_SYNONYMOU | MODERAT | MISSENSE | caC/caG | H187Q | cupB5 | | | G | G | G | G | G | G | | | | |
| NC_011770 | 974534 | A | T | 81 | 0 | 4 | 0 | NON_SYNONYMOU | MODERAT | MISSENSE | Aac/Tac | N275Y | cupB5 | | | | | | | T | T | T | T | | |
| NC_011770 | 974549 | T | A | 74 | 0 | 4 | 0 | NON_SYNONYMOU | MODERAT | MISSENSE | Tcc/Acc | S280T | cupB5 | | | | | | | A | A | A | A | | |
| NC_011770 | 974665 | C | A | 70 | 0 | 2 | 0 | NON_SYNONYMOU | MODERAT | MISSENSE | aaC/aaA | N318K | cupB5 | A | A | | | | | | | | | | |
| NC_011770 | 974850 | G | A | 63 | 0 | 1 | 0 | NON_SYNONYMOU | MODERAT | MISSENSE | cGg/cAg | R380Q | cupB5 | | | | | | | A | | | | | |
| NC_011770 | 975639 | C | G | 47 | 0 | 2 | 0 | NON_SYNONYMOU | MODERAT | MISSENSE | gCt/gGt | A643G | cupB5 | | | | | | | | | | | G | G |
| NC_011770 | 976113 | T | C | 38 | 0 | 2 | 0 | NON_SYNONYMOU | MODERAT | MISSENSE | aTg/aCg | M801T | cupB5 | | | | | | | | | | | C | C |
| NC_011770 | 976239 | A | G | 41 | 0 | 4 | 0 | NON_SYNONYMOU | MODERAT | MISSENSE | cAc/cGc | H843R | cupB5 | | | G | G | G | G | | | | | | |
| NC_011770 | 976240 | C | A | 41 | 0 | 4 | 0 | NON_SYNONYMOU | MODERAT | MISSENSE | caC/caA | H843Q | cupB5 | | | A | A | A | A | | | | | | |
| NC_011770 | 976343 | T | G | 44 | 0 | 2 | 0 | NON_SYNONYMOU | MODERAT | MISSENSE | Ttc/Gtc | F878V | cupB5 | | | | | | | | | | | G | G |
| NC_011770 | 976455 | G | A | 51 | 0 | 2 | 0 | NON_SYNONYMOU | MODERAT | MISSENSE | aGc/aAc | S915N | cupB5 | | | | | | | | | | | A | A |
| NC_011770 | 976460 | G | A | 50 | 0 | 2 | 0 | NON_SYNONYMOU | MODERAT | MISSENSE | Ggc/Agc | G917S | cupB5 | | | | | | | | | | | A | A |
| NC_011770 | 976601 | A | G | 53 | 0 | 2 | 0 | NON_SYNONYMOU | MODERAT | MISSENSE | Aag/Gag | K964E | cupB5 | | | | | | | | | | | G | G |
| NC_011770 | 976686 | G | A | 42 | 0 | 2 | 0 | NON_SYNONYMOU | MODERAT | MISSENSE | cGt/cAt | R992H | cupB5 | | | | | | | | | | | A | A |
| NC_011770 | 975275 | A | G | 69 | 0 | 1 | 0 | NON_SYNONYMOU | MODERAT | MISSENSE | Acg/Gcg | T522A | cupB5 | G | | | | | | | | | | | |
| NC_011770 | 976922 | C | A | 37 | 0 | 2 | 0 | NON_SYNONYMOU | MODERAT | MISSENSE | Cag/Aag | Q19K | cupB6 | | | | | | | | | | | A | A |
| NC_011770 | 976990 | C | G | 35 | 0 | 2 | 0 | NON_SYNONYMOU | MODERAT | MISSENSE | gaC/gaG | D41E | cupB6 | | | | | | | | | | | G | G |
| NC_011770 | 976991 | A | G | 35 | 0 | 2 | 0 | NON_SYNONYMOU | MODERAT | MISSENSE | Atc/Gtc | I42V | cupB6 | | | | | | | | | | | G | G |
| NC_011770 | 977021 | C | A | 33 | 0 | 2 | 0 | NON_SYNONYMOU | MODERAT | MISSENSE | Cag/Aag | Q52K | cupB6 | | | | | | | | | | | A | A |
| NC_011770 | 977027 | A | T | 32 | 0 | 2 | 0 | NON_SYNONYMOU | MODERAT | MISSENSE | Atg/Ttg | M54L | cupB6 | | | | | | | | | | | T | T |
| NC_011770 | 977088 | C | T | 24 | 0 | 4 | 0 | NON_SYNONYMOU | MODERAT | MISSENSE | gCc/gTc | A74V | cupB6 | | | T | T | T | T | | | | | | |
| NC_011770 | 977112 | G | A | 22 | 0 | 2 | 0 | NON_SYNONYMOU | MODERAT | MISSENSE | aGc/aAc | S82N | cupB6 | | | | | | | | | | | A | A |
| NC_011770 | 977153 | C | A | 17 | 0 | 2 | 0 | NON_SYNONYMOU | MODERAT | MISSENSE | Ctc/Atc | L96I | cupB6 | | | | | | | | | | | A | A |
| NC_011770 | 977294 | A | G | 19 | 0 | 12 | 0 | NON_SYNONYMOU | MODERAT | MISSENSE | Acg/Gcg | T143A | cupB6 | G | G | G | G | G | G | G | G | G | G | G | G |
| NC_011770 | 977549 | G | A | 31 | 0 | 2 | 0 | NON_SYNONYMOU | MODERAT | MISSENSE | Gcc/Acc | A228T | cupB6 | | | | | | | A | A | | | | |
| NC_011770 | 977559 | C | G | 33 | 0 | 2 | 0 | NON_SYNONYMOU | MODERAT | MISSENSE | gCt/gGt | A231G | cupB6 | | | | | | | | | G | G | | |
| NC_011770 | 977699 | G | A | 45 | 0 | 2 | 0 | NON_SYNONYMOU | MODERAT | MISSENSE | Gat/Aat | D278N | cupB6 | A | A | | | | | | | | | | |
| NC_011770 | 4773483 | C | T | 83 | 0 | 6 | 0 | NON_SYNONYMOU | MODERAT | MISSENSE | Gac/Aac | D189N | cupC1 | T | T | | | | | T | T | | | T | T |
| NC_011770 | 4773488 | C | A | 84 | 0 | 2 | 0 | NON_SYNONYMOU | MODERAT | MISSENSE | aGt/aTt | S187I | cupC1 | | | | | | | A | A | | | | |
| NC_011770 | 4773501 | G | C | 82 | 0 | 2 | 0 | NON_SYNONYMOU | MODERAT | MISSENSE | Cct/Gct | P183A | cupC1 | | | | | | | C | C | | | | |
| NC_011770 | 4773506 | A | G | 81 | 0 | 2 | 0 | NON_SYNONYMOU | MODERAT | MISSENSE | gTa/gCa | V181A | cupC1 | | | | | | | G | G | | | | |
| NC_011770 | 4773510 | A | C | 78 | 0 | 2 | 0 | NON_SYNONYMOU | MODERAT | MISSENSE | Tcc/Gcc | S180A | cupC1 | | | | | | | C | C | | | | |
| NC_011770 | 4773544 | C | A | 85 | 0 | 2 | 0 | NON_SYNONYMOU | MODERAT | MISSENSE | aaG/aaT | K168N | cupC1 | | | | | | | A | A | | | | |
| NC_011770 | 4773588 | G | C | 90 | 0 | 2 | 0 | NON_SYNONYMOU | MODERAT | MISSENSE | Caa/Gaa | Q154E | cupC1 | | | | | | | C | C | | | | |
| NC_011770 | 4773591 | C | T | 93 | 0 | 2 | 0 | NON_SYNONYMOU | MODERAT | MISSENSE | Gta/Ata | V153I | cupC1 | | | | | | | T | T | | | | |
| NC_011770 | 4773592 | C | A | 91 | 0 | 2 | 0 | NON_SYNONYMOU | MODERAT | MISSENSE | aaG/aaT | K152N | cupC1 | | | | | | | A | A | | | | |
| NC_011770 | 4773600 | C | T | 93 | 0 | 2 | 0 | NON_SYNONYMOU | MODERAT | MISSENSE | Gat/Aat | D150N | cupC1 | | | | | | | T | T | | | | |
| NC_011770 | 4773612 | C | T | 93 | 0 | 2 | 0 | NON_SYNONYMOU | MODERAT | MISSENSE | Gat/Aat | D146N | cupC1 | | | | | | | T | T | | | | |
| NC_011770 | 4773747 | T | C | 87 | 0 | 2 | 0 | NON_SYNONYMOU | MODERAT | MISSENSE | Aac/Gac | N101D | cupC1 | | | | | | | C | C | | | | |
| NC_011770 | 4773878 | T | C | 87 | 0 | 2 | 0 | NON_SYNONYMOU | MODERAT | MISSENSE | aAc/aGc | N57S | cupC1 | | | | | | | C | C | | | | |
| NC_011770 | 4774041 | T | C | 82 | 0 | 2 | 0 | NON_SYNONYMOU | MODERAT | MISSENSE | Agc/Ggc | S3G | cupC1 | | | | | C | C | | | | | | |

| | | | | | | | | | | | | | | | | | | | | | | | | | |
|---|---|---|---|---|---|---|---|---|---|---|---|---|---|---|---|---|---|---|---|---|---|---|---|---|---|
| NC_011770 | 4772742 | C | T | 79 | 0 | 2 | 0 | NON_SYNONYMOU | MODERAT | MISSENSE | Gtc/Atc | V217I | cupC2 | | | | | | | T | T | | | | |
| NC_011770 | 4772760 | A | G | 77 | 0 | 2 | 0 | NON_SYNONYMOU | MODERAT | MISSENSE | Tcc/Ccc | S211P | cupC2 | | | | | | | G | G | | | | |
| NC_011770 | 4772775 | T | C | 79 | 0 | 2 | 0 | NON_SYNONYMOU | MODERAT | MISSENSE | Atc/Gtc | I206V | cupC2 | | | | | | | C | C | | | | |
| NC_011770 | 4772828 | T | G | 88 | 0 | 2 | 0 | NON_SYNONYMOU | MODERAT | MISSENSE | tAc/tCc | Y188S | cupC2 | | | | | | | G | G | | | | |
| NC_011770 | 4772855 | C | T | 82 | 0 | 2 | 0 | NON_SYNONYMOU | MODERAT | MISSENSE | aGc/aAc | S179N | cupC2 | | | | | | | T | T | | | | |
| NC_011770 | 4772858 | G | A | 82 | 0 | 2 | 0 | NON_SYNONYMOU | MODERAT | MISSENSE | gCc/gTc | A178V | cupC2 | | | A | A | | | | | | | | |
| NC_011770 | 4772860 | C | T | 82 | 0 | 10 | 0 | NON_SYNONYMOU | MODERAT | MISSENSE | ttG/ttC | L177F | cupC2 | | | T | T | T | T | G | G | T | T | T | T |
| NC_011770 | 4772933 | T | C | 78 | 0 | 2 | 0 | NON_SYNONYMOU | MODERAT | MISSENSE | aAc/aGc | N153S | cupC2 | | | | | | | | | C | C | | |
| NC_011770 | 4773235 | C | A | 86 | 0 | 2 | 0 | NON_SYNONYMOU | MODERAT | MISSENSE | aaG/aaT | K52N | cupC2 | | | | | | | A | A | | | | |
| NC_011770 | 4771128 | T | G | 75 | 0 | 4 | 0 | NON_SYNONYMOU | MODERAT | MISSENSE | gaA/gaC | E483D | cupC3 | | | G | G | G | G | | | | | | |
| NC_011770 | 4771313 | A | G | 84 | 0 | 6 | 0 | NON_SYNONYMOU | MODERAT | MISSENSE | Ttt/Ctt | F422L | cupC3 | | | G | G | G | G | G | G | | | | |
| NC_011770 | 4771985 | G | A | 102 | 0 | 1 | 0 | NON_SYNONYMOU | MODERAT | MISSENSE | Cgc/Tgc | R198C | cupC3 | | | | | | | A | | | | | |
| NC_011770 | 4772162 | T | C | 91 | 0 | 2 | 0 | NON_SYNONYMOU | MODERAT | MISSENSE | Atc/Gtc | I139V | cupC3 | | | | | | | C | C | | | | |
| NC_011770 | 4772218 | T | G | 84 | 0 | 2 | 0 | NON_SYNONYMOU | MODERAT | MISSENSE | gAt/gCt | D120A | cupC3 | | | | | | | G | G | | | | |
| NC_011770 | 4772280 | T | C | 86 | 0 | 2 | 0 | NON_SYNONYMOU | MODERAT | MISSENSE | atA/atG | I99M | cupC3 | | | | | | | C | C | | | | |
| NC_011770 | 4772282 | T | C | 86 | 0 | 8 | 0 | NON_SYNONYMOU | MODERAT | MISSENSE | Ata/Gta | I99V | cupC3 | C | C | | | | | C | C | C | C | C | C |
| NC_011770 | 4772288 | C | T | 84 | 0 | 2 | 0 | NON_SYNONYMOU | MODERAT | MISSENSE | Gtc/Atc | V97I | cupC3 | | | | | | | T | T | | | | |
| NC_011770 | 4772315 | G | A | 82 | 0 | 2 | 0 | NON_SYNONYMOU | MODERAT | MISSENSE | Ccc/Tcc | P88S | cupC3 | | | | | | | A | A | | | | |
| NC_011770 | 4772449 | T | C | 82 | 0 | 2 | 0 | NON_SYNONYMOU | MODERAT | MISSENSE | aAc/aGc | N43S | cupC3 | C | C | | | | | | | | | | |
| NC_011770 | 4772513 | T | C | 83 | 0 | 2 | 0 | NON_SYNONYMOU | MODERAT | MISSENSE | Atg/Gtg | M22V | cupC3 | | | | | | | C | C | | | | |
| NC_011770 | 4772543 | A | C | 79 | 0 | 2 | 0 | NON_SYNONYMOU | MODERAT | MISSENSE | Ttg/Gtg | L12V | cupC3 | | | | | | | C | C | | | | |
| NC_011770 | 4772572 | G | A | 77 | 0 | 2 | 0 | NON_SYNONYMOU | MODERAT | MISSENSE | gCc/gTc | A2V | cupC3 | | | | | | | A | A | | | | |
| NC_011770 | 6272959 | C | A | 89 | 0 | 4 | 0 | NON_SYNONYMOU | MODERAT | MISSENSE | cCg/cAg | P221Q | cyaA | | | | | | | | | A | A | A | A |
| NC_011770 | 6274000 | G | A | 89 | 0 | 2 | 0 | NON_SYNONYMOU | MODERAT | MISSENSE | cGg/cAg | R568Q | cyaA | A | A | | | | | | | | | | |
| NC_011770 | 6274555 | C | T | 84 | 0 | 12 | 0 | NON_SYNONYMOU | MODERAT | MISSENSE | cCg/cTg | P753L | cyaA | T | T | T | T | T | T | T | T | T | T | T | T |
| NC_011770 | 6274735 | A | G | 93 | 0 | 12 | 0 | NON_SYNONYMOU | MODERAT | MISSENSE | cAc/cGc | H813R | cyaA | G | G | G | G | G | G | G | G | G | G | G | G |
| NC_011770 | 6274770 | G | T | 91 | 0 | 6 | 0 | NON_SYNONYMOU | MODERAT | MISSENSE | Gcg/Tcg | A825S | cyaA | T | T | T | T | T | T | | | | | | |
| NC_011770 | 6274809 | G | A | 89 | 0 | 1 | 0 | NON_SYNONYMOU | MODERAT | MISSENSE | Ggc/Agc | G838S | cyaA | | | | | | | A | | | | | |
| NC_011770 | 6274815 | G | T | 91 | 0 | 6 | 0 | NON_SYNONYMOU | MODERAT | MISSENSE | Gtg/Ttg | V840L | cyaA | T | T | T | T | T | T | | | | | | |
| NC_011770 | 6275065 | C | T | 86 | 0 | 2 | 0 | NON_SYNONYMOU | MODERAT | MISSENSE | gCc/gTc | A923V | cyaA | | | | | | | T | T | | | | |
| NC_011770 | 6276702 | G | GC | 65 | 0 | 1 | 0 | FRAME_SHIFT | HIGH | | ggc/ggGc | G103G? | cyaY | GC | | | | | | | | | | | |
| NC_011770 | 6276719 | T | C | 66 | 0 | 1 | 0 | NON_SYNONYMOU | MODERAT | MISSENSE | Acc/Gcc | T98A | cyaY | | | | | | | C | | | | | |
| NC_011770 | 6276974 | C | T | 75 | 0 | 4 | 0 | NON_SYNONYMOU | MODERAT | MISSENSE | Gtc/Atc | V13I | cyaY | | | | | | | | | T | T | T | T |
| NC_011770 | 4343585 | G | C | 70 | 0 | 1 | 0 | NON_SYNONYMOU | MODERAT | MISSENSE | Cag/Gag | Q401E | cycH | | | | | | | C | | | | | |
| NC_011770 | 4344223 | C | T | 75 | 0 | 1 | 0 | NON_SYNONYMOU | MODERAT | MISSENSE | gGc/gAc | G188D | cycH | | | | | | | | T | | | | |
| NC_011770 | 3617788 | C | T | 64 | 0 | 2 | 0 | NON_SYNONYMOU | MODERAT | MISSENSE | Gtg/Atg | V253M | cynR | T | T | | | | | | | | | | |
| NC_011770 | 3618088 | C | G | 79 | 0 | 2 | 0 | NON_SYNONYMOU | MODERAT | MISSENSE | Gtg/Ctg | V153L | cynR | G | G | | | | | | | | | | |
| NC_011770 | 3619003 | G | A | 60 | 0 | 10 | 0 | NON_SYNONYMOU | MODERAT | MISSENSE | caG/caC | Q115H | cynT | A | A | C | C | C | C | C | C | | | A | A |
| NC_011770 | 3619206 | A | G | 56 | 0 | 2 | 0 | NON_SYNONYMOU | MODERAT | MISSENSE | gAc/gGc | D183G | cynT | | | | | | | | | | | G | G |
| NC_011770 | 3619299 | A | C | 57 | 0 | 2 | 0 | NON_SYNONYMOU | MODERAT | MISSENSE | gAa/gCa | E214A | cynT | C | C | | | | | | | | | | |
| NC_011770 | 4275120 | G | A | 81 | 0 | 2 | 0 | NON_SYNONYMOU | MODERAT | MISSENSE | Ccg/Tcg | P159S | cyoA | | | | | | | | | | | A | A |
| NC_011770 | 4272299 | G | A | 94 | 0 | 2 | 0 | NON_SYNONYMOU | MODERAT | MISSENSE | gCc/gTc | A105V | cyoC | A | A | | | | | | | | | | |
| NC_011770 | 4271802 | A | G | 98 | 0 | 2 | 0 | NON_SYNONYMOU | MODERAT | MISSENSE | gTg/gCg | V61A | cyoD | | | | | | | | | | | G | G |
| NC_011770 | 4271563 | TC | T | 77 | 0 | 1 | 0 | FRAME_SHIFT | HIGH | | gga/ | G25 | cyoE | | | | | | | T | | | | | |
| NC_011770 | 307105 | G | T | 79 | 0 | 2 | 0 | NON_SYNONYMOU | MODERAT | MISSENSE | gCg/gAg | A264E | cysA | T | T | | | | | | | | | | |
| NC_011770 | 5310601 | C | T | 70 | 0 | 2 | 0 | NON_SYNONYMOU | MODERAT | MISSENSE | Gag/Aag | E212K | cysD | | | | | | | | | | | T | T |
| NC_011770 | 1257620 | A | T | 92 | 0 | 10 | 0 | NON_SYNONYMOU | MODERAT | MISSENSE | Acc/Tcc | T173S | cysE | T | T | T | T | T | T | | | T | T | T | T |
| NC_011770 | 1257623 | G | A | 97 | 0 | 1 | 0 | NON_SYNONYMOU | MODERAT | MISSENSE | Gag/Aag | E174K | cysE | | | | | | | A | | | | | |
| NC_011770 | 1257458 | A | G | 90 | 0 | 1 | 0 | NON_SYNONYMOU | MODERAT | MISSENSE | Acc/Gcc | T119A | cysE | | | | | | | | G | | | | |
| NC_011770 | 2685003 | T | C | 63 | 0 | 6 | 0 | NON_SYNONYMOU | MODERAT | MISSENSE | gTc/gCc | V74A | cysG | C | C | | | | | | | C | C | C | C |
| NC_011770 | 2685054 | G | A | 58 | 0 | 4 | 0 | NON_SYNONYMOU | MODERAT | MISSENSE | cGa/cAa | R91Q | cysG | | | A | A | A | A | | | | | | |
| NC_011770 | 3971707 | TC | T | 42 | 0 | 1 | 0 | FRAME_SHIFT | HIGH | | cag/ | Q43 | cysH | | | | T | | | | | | | | |
| NC_011770 | 3972139 | A | G | 65 | 0 | 1 | 0 | NON_SYNONYMOU | MODERAT | MISSENSE | Acg/Gcg | T185A | cysH | | | | | | | | G | | | | |
| NC_011770 | 3871861 | G | C | 48 | 0 | 2 | 0 | NON_SYNONYMOU | MODERAT | MISSENSE | Gag/Cag | E196Q | cysI | | | | | | | C | C | | | | |
| NC_011770 | 3872173 | G | A | 66 | 0 | 6 | 0 | NON_SYNONYMOU | MODERAT | MISSENSE | Gcc/Acc | A300T | cysI | | | A | A | A | A | | | A | A | | |
| NC_011770 | 3872346 | G | C | 63 | 0 | 2 | 0 | NON_SYNONYMOU | MODERAT | MISSENSE | gaG/gaC | E357D | cysI | | | | | | | C | C | | | | |
| NC_011770 | 4817541 | G | A | 83 | 0 | 1 | 0 | NON_SYNONYMOU | MODERAT | MISSENSE | Ccg/Tcg | P48S | cysM | | | | A | | | | | | | | |
| NC_011770 | 5308469 | C | A | 60 | 0 | 2 | 0 | NON_SYNONYMOU | MODERAT | MISSENSE | Gcg/Tcg | A613S | cysN | | | | | | | | | | | A | A |
| NC_011770 | 5308592 | A | G | 62 | 0 | 2 | 0 | NON_SYNONYMOU | MODERAT | MISSENSE | Tcg/Ccg | S572P | cysN | G | G | | | | | | | | | | |
| NC_011770 | 5308633 | A | G | 69 | 0 | 1 | 0 | NON_SYNONYMOU | MODERAT | MISSENSE | cTg/cCg | L558P | cysN | | | | | | | G | | | | | |
| NC_011770 | 5308729 | G | A | 69 | 0 | 1 | 0 | NON_SYNONYMOU | MODERAT | MISSENSE | aCc/aTc | T526I | cysN | | | | | | | A | | | | | |
| NC_011770 | 5308904 | C | T | 76 | 0 | 2 | 0 | NON_SYNONYMOU | MODERAT | MISSENSE | Gta/Ata | V468I | cysN | | | | | | | | | T | T | | |

| | | | | | | | | | | | | | | | | | | | | | | | | | |
|---|---|---|---|---|---|---|---|---|---|---|---|---|---|---|---|---|---|---|---|---|---|---|---|---|---|
| NC_011770 | 5308967 | C | T | 77 | 0 | 2 | 0 | NON_SYNONYMOU | MODERAT | MISSENSE | Ggt/Agt | G447S | cysN | | | | | | | | | T | T | | |
| NC_011770 | 5308997 | A | G | 77 | 0 | 6 | 0 | NON_SYNONYMOU | MODERAT | MISSENSE | Tcg/Ccg | S437P | cysN | | | G | G | G | G | | | | | G | G |
| NC_011770 | 5309009 | T | C | 78 | 0 | 2 | 0 | NON_SYNONYMOU | MODERAT | MISSENSE | Atc/Gtc | I433V | cysN | | | | | | | | | | | C | C |
| NC_011770 | 5309794 | C | T | 85 | 0 | 10 | 0 | NON_SYNONYMOU | MODERAT | MISSENSE | gGc/gAc | G171D | cysN | T | T | T | T | T | T | | | T | T | T | T |
| NC_011770 | 4332426 | G | A | 46 | 0 | 12 | 0 | NON_SYNONYMOU | MODERAT | MISSENSE | aGg/aAg | R306K | cysP | A | A | A | A | A | A | A | A | A | A | A | A |
| NC_011770 | 6164056 | C | G | 41 | 0 | 2 | 0 | NON_SYNONYMOU | MODERAT | MISSENSE | Gag/Cag | E260Q | cysQ | | | | | | | G | G | | | | |
| NC_011770 | 6164410 | C | T | 68 | 0 | 2 | 0 | NON_SYNONYMOU | MODERAT | MISSENSE | Ggc/Agc | G142S | cysQ | | | | | | | T | T | | | | |
| NC_011770 | 6164602 | G | A | 79 | 0 | 4 | 0 | NON_SYNONYMOU | MODERAT | MISSENSE | Cac/Tac | H78Y | cysQ | | | A | A | A | A | | | | | | |
| NC_011770 | 6164701 | A | C | 91 | 0 | 10 | 0 | NON_SYNONYMOU | MODERAT | MISSENSE | Tcc/Gcc | S45A | cysQ | C | C | C | C | C | C | C | C | C | C | | |
| NC_011770 | 6164748 | G | A | 92 | 0 | 2 | 0 | NON_SYNONYMOU | MODERAT | MISSENSE | gCg/gTg | A29V | cysQ | A | A | | | | | | | | | | |
| NC_011770 | 3922020 | T | C | 51 | 0 | 6 | 0 | NON_SYNONYMOU | MODERAT | MISSENSE | cAg/cGg | Q439R | cysS | | | | | | | C | C | C | C | C | C |
| NC_011770 | 3922332 | G | A | 66 | 0 | 4 | 0 | NON_SYNONYMOU | MODERAT | MISSENSE | gCg/gTg | A335V | cysS | | | A | A | A | A | | | | | | |
| NC_011770 | 3922956 | G | A | 82 | 0 | 6 | 0 | NON_SYNONYMOU | MODERAT | MISSENSE | aCc/aTc | T127I | cysS | | | A | A | A | A | | | | | A | A |
| NC_011770 | 309032 | A | G | 80 | 0 | 2 | 0 | NON_SYNONYMOU | MODERAT | MISSENSE | gTg/gCg | V189A | cysT | | | | | | | G | G | | | | |
| NC_011770 | 308372 | A | G | 76 | 0 | 2 | 0 | NON_SYNONYMOU | MODERAT | MISSENSE | Tac/Cac | Y133H | cysW | | | | | G | G | | | | | | |
| NC_011770 | 2988236 | A | G | 70 | 0 | 2 | 0 | NON_SYNONYMOU | MODERAT | MISSENSE | aAg/aGg | K33R | czcA | | | | | | | | | G | G | | |
| NC_011770 | 2989922 | G | A | 67 | 0 | 4 | 0 | NON_SYNONYMOU | MODERAT | MISSENSE | aGg/aAg | R595K | czcA | | | A | A | A | A | | | | | | |
| NC_011770 | 2988436 | G | A | 61 | 0 | 1 | 0 | NON_SYNONYMOU | MODERAT | MISSENSE | Ggc/Agc | G100S | czcA | | | | | | | | | | | | A |
| NC_011770 | 2986686 | G | C | 72 | 0 | 2 | 0 | NON_SYNONYMOU | MODERAT | MISSENSE | Gtg/Ctg | V9L | czcB | | | | | | | | | | | C | C |
| NC_011770 | 2986758 | G | T | 69 | 0 | 2 | 0 | NON_SYNONYMOU | MODERAT | MISSENSE | Gcc/Tcc | A33S | czcB | | | | | | | | | T | T | | |
| NC_011770 | 2985992 | G | T | 76 | 0 | 2 | 0 | NON_SYNONYMOU | MODERAT | MISSENSE | Gcc/Tcc | A224S | czcC | | | | | | | | | | | T | T |
| NC_011770 | 6308808 | G | T | 86 | 0 | 2 | 0 | NON_SYNONYMOU | MODERAT | MISSENSE | gCg/gAg | A208E | dadA | | | | | | | | | | | T | T |
| NC_011770 | 6306687 | G | T | 65 | 0 | 4 | 0 | NON_SYNONYMOU | MODERAT | MISSENSE | Ctg/Atg | L343M | dadX | | | T | T | T | T | | | | | | |
| NC_011770 | 6306841 | A | C | 52 | 0 | 12 | 0 | NON_SYNONYMOU | MODERAT | MISSENSE | aaT/aaG | N291K | dadX | C | C | C | C | C | C | C | C | C | C | C | C |
| NC_011770 | 6307436 | T | A | 55 | 0 | 2 | 0 | NON_SYNONYMOU | MODERAT | MISSENSE | cAc/cTc | H93L | dadX | A | A | | | | | | | | | | |
| NC_011770 | 5679019 | C | T | 87 | 0 | 4 | 0 | NON_SYNONYMOU | MODERAT | MISSENSE | Gct/Act | A234T | dapB | | | T | T | T | T | | | | | | |
| NC_011770 | 5679529 | C | T | 79 | 0 | 2 | 0 | NON_SYNONYMOU | MODERAT | MISSENSE | Ggc/Agc | G64S | dapB | | | | | | | T | T | | | | |
| NC_011770 | 4588908 | G | A | 71 | 0 | 2 | 0 | NON_SYNONYMOU | MODERAT | MISSENSE | Gag/Aag | E128K | dapE | | | | | | | | | | | A | A |
| NC_011770 | 4589197 | C | T | 79 | 0 | 2 | 0 | NON_SYNONYMOU | MODERAT | MISSENSE | gCg/gTg | A224V | dapE | T | T | | | | | | | | | | |
| NC_011770 | 4589211 | G | A | 79 | 0 | 2 | 0 | NON_SYNONYMOU | MODERAT | MISSENSE | Gac/Aac | D229N | dapE | | | | | | | A | A | | | | |
| NC_011770 | 4589500 | A | G | 76 | 0 | 2 | 0 | NON_SYNONYMOU | MODERAT | MISSENSE | cAg/cGg | Q325R | dapE | G | G | | | | | | | | | | |
| NC_011770 | 6278795 | A | G | 87 | 0 | 6 | 0 | NON_SYNONYMOU | MODERAT | MISSENSE | Atc/Gtc | I42V | dapF | G | G | | | | | | | G | G | G | G |
| NC_011770 | 505963 | T | A | 86 | 0 | 8 | 0 | NON_SYNONYMOU | MODERAT | MISSENSE | Agc/Tgc | S245C | dbpA | A | A | | | | | A | A | A | A | A | A |
| NC_011770 | 506443 | C | T | 70 | 0 | 2 | 0 | NON_SYNONYMOU | MODERAT | MISSENSE | Gcc/Acc | A85T | dbpA | | | | | | | | | T | T | | |
| NC_011770 | 5276534 | C | T | 76 | 0 | 4 | 0 | NON_SYNONYMOU | MODERAT | MISSENSE | Gaa/Aaa | E272K | ddl | | | | | | | T | T | T | T | | |
| NC_011770 | 5276618 | T | G | 73 | 0 | 12 | 0 | NON_SYNONYMOU | MODERAT | MISSENSE | Agc/Cgc | S244R | ddl | G | G | G | G | G | G | G | G | G | G | G | G |
| NC_011770 | 5276839 | A | T | 69 | 0 | 12 | 0 | NON_SYNONYMOU | MODERAT | MISSENSE | gTc/gAc | V170D | ddl | T | T | T | T | T | T | T | T | T | T | T | T |
| NC_011770 | 5277200 | T | C | 74 | 0 | 12 | 0 | NON_SYNONYMOU | MODERAT | MISSENSE | Agg/Ggg | R50G | ddl | C | C | C | C | C | C | C | C | C | C | C | C |
| NC_011770 | 791316 | G | A | 78 | 0 | 6 | 0 | NON_SYNONYMOU | MODERAT | MISSENSE | cGc/cAc | R57H | ddlA | | | A | A | A | A | | | A | A | | |
| NC_011770 | 791756 | G | C | 74 | 0 | 2 | 0 | NON_SYNONYMOU | MODERAT | MISSENSE | Gag/Cag | E204Q | ddlA | | | | | | | | | C | C | | |
| NC_011770 | 791847 | G | A | 71 | 0 | 2 | 0 | NON_SYNONYMOU | MODERAT | MISSENSE | gGt/gAt | G234D | ddlA | | | | | | | | | A | A | | |
| NC_011770 | 5817211 | A | G | 85 | 0 | 1 | 0 | NON_SYNONYMOU | MODERAT | MISSENSE | Tgg/Cgg | W68R | desB | | | | | | | | G | | | | |
| NC_011770 | 5816305 | C | T | 71 | 0 | 2 | 0 | NON_SYNONYMOU | MODERAT | MISSENSE | Ggc/Agc | G370S | desB | | | | | | | | | T | T | | |
| NC_011770 | 5816532 | A | G | 82 | 0 | 12 | 0 | NON_SYNONYMOU | MODERAT | MISSENSE | cTg/cCg | L294P | desB | G | G | G | G | G | G | G | G | G | G | G | G |
| NC_011770 | 5816665 | C | T | 83 | 0 | 12 | 0 | NON_SYNONYMOU | MODERAT | MISSENSE | Gcc/Acc | A250T | desB | T | T | T | T | T | T | T | T | T | T | T | T |
| NC_011770 | 5817318 | A | T | 88 | 0 | 2 | 0 | NON_SYNONYMOU | MODERAT | MISSENSE | cTg/cAg | L32Q | desB | | | | | | | | | T | T | | |
| NC_011770 | 5817321 | C | T | 83 | 0 | 4 | 0 | NON_SYNONYMOU | MODERAT | MISSENSE | cGc/cAc | R31H | desB | T | T | | | | | | | | | T | T |
| NC_011770 | 5819296 | G | A | 57 | 0 | 2 | 0 | SPLICE_SITE_REGIO | LOW | SILENT | tGa/tAa | *210 | desT | | | | | | | A | A | | | | |
| NC_011770 | 5819293 | G | A | 57 | 0 | 2 | 0 | NON_SYNONYMOU | MODERAT | MISSENSE | gGc/gAc | G209D | desT | | | | | | | A | A | | | | |
| NC_011770 | 4629278 | C | A | 69 | 0 | 2 | 0 | NON_SYNONYMOU | MODERAT | MISSENSE | caG/caT | Q466H | dgt | | | | | | | A | A | | | | |
| NC_011770 | 4629952 | G | T | 79 | 0 | 4 | 0 | NON_SYNONYMOU | MODERAT | MISSENSE | Cgc/Agc | R242S | dgt | T | T | | | | | | | T | T | | |
| NC_011770 | 4630225 | T | C | 77 | 0 | 4 | 0 | NON_SYNONYMOU | MODERAT | MISSENSE | Ata/Gta | I151V | dgt | | | C | C | C | C | | | | | | |
| NC_011770 | 4630377 | G | A | 71 | 0 | 10 | 0 | NON_SYNONYMOU | MODERAT | MISSENSE | gCc/gTc | A100V | dgt | A | A | A | A | A | A | A | A | A | A | | |
| NC_011770 | 489594 | C | G | 80 | 0 | 8 | 0 | NON_SYNONYMOU | MODERAT | MISSENSE | Ggg/Cgg | G135R | dht | G | G | | | | | G | G | G | G | G | G |
| NC_011770 | 488634 | G | A | 73 | 0 | 1 | 0 | NON_SYNONYMOU | MODERAT | MISSENSE | Cgc/Tgc | R455C | dht | | | | | | | | A | | | | |
| NC_011770 | 4829638 | T | C | 68 | 2 | 2 | 0 | NON_SYNONYMOU | MODERAT | MISSENSE | gAg/gGg | E268G | dinB | | | | | | | C | C | | | . | . |
| NC_011770 | 4717425 | A | T | 87 | 0 | 2 | 0 | NON_SYNONYMOU | MODERAT | MISSENSE | gaT/gaA | D158E | dinG | T | T | | | | | | | | | | |
| NC_011770 | 4717433 | T | A | 89 | 0 | 4 | 0 | NON_SYNONYMOU | MODERAT | MISSENSE | Aac/Tac | N156Y | dinG | | | | | | | A | A | A | A | | |
| NC_011770 | 4717774 | G | A | 81 | 0 | 4 | 0 | NON_SYNONYMOU | MODERAT | MISSENSE | gCg/gTg | A42V | dinG | | | A | A | A | A | | | | | | |
| NC_011770 | 5772761 | A | G | 75 | 0 | 4 | 0 | NON_SYNONYMOU | MODERAT | MISSENSE | Aag/Gag | K114E | dipZ | G | G | | | | | G | G | | | | |
| NC_011770 | 5772946 | C | G | 82 | 0 | 2 | 0 | NON_SYNONYMOU | MODERAT | MISSENSE | agC/agG | S175R | dipZ | | | | | | | | | | | G | G |
| NC_011770 | 830085 | A | C | 81 | 0 | 12 | 0 | NON_SYNONYMOU | MODERAT | MISSENSE | gAg/gCg | E168A | dkgB | C | C | C | C | C | C | C | C | C | C | C | C |

| | | | | | | | | | | | | | | | | | | | | | | | | | |
|---|---|---|---|---|---|---|---|---|---|---|---|---|---|---|---|---|---|---|---|---|---|---|---|---|---|
| NC_011770 | 1294 | G | T | 99 | 0 | 1 | 0 | NON_SYNONYMOU | MODERAT | MISSENSE | cGc/cTc | R271L | dnaA | | | | | | | T | | | | | |
| NC_011770 | 5867204 | T | C | 87 | 0 | 1 | 0 | NON_SYNONYMOU | MODERAT | MISSENSE | Acc/Gcc | T230A | dnaB | | | | | | | C | | | | | |
| NC_011770 | 5125052 | A | G | 41 | 0 | 6 | 0 | NON_SYNONYMOU | MODERAT | MISSENSE | Atc/Gtc | I71V | dnaE2 | | | G | G | G | G | G | G | | | | |
| NC_011770 | 5125248 | G | A | 60 | 0 | 2 | 0 | NON_SYNONYMOU | MODERAT | MISSENSE | gGg/gAg | G136E | dnaE2 | | | | | | | A | A | | | | |
| NC_011770 | 5125382 | C | T | 56 | 0 | 4 | 0 | NON_SYNONYMOU | MODERAT | MISSENSE | Ccc/Tcc | P181S | dnaE2 | | | T | T | T | T | | | | | | |
| NC_011770 | 5125473 | G | T | 63 | 0 | 4 | 0 | NON_SYNONYMOU | MODERAT | MISSENSE | aGc/aTc | S211I | dnaE2 | | | T | T | T | T | | | | | | |
| NC_011770 | 5125536 | A | T | 66 | 0 | 2 | 0 | NON_SYNONYMOU | MODERAT | MISSENSE | cAg/cTg | Q232L | dnaE2 | | | | | | | T | T | | | | |
| NC_011770 | 5125743 | A | G | 71 | 0 | 2 | 0 | NON_SYNONYMOU | MODERAT | MISSENSE | gAg/gGg | E301G | dnaE2 | | | | | | | | | | | G | G |
| NC_011770 | 5125829 | G | A | 79 | 0 | 6 | 0 | NON_SYNONYMOU | MODERAT | MISSENSE | Gaa/Aaa | E330K | dnaE2 | A | A | | | | | | | A | A | A | A |
| NC_011770 | 5126087 | G | A | 78 | 0 | 2 | 0 | NON_SYNONYMOU | MODERAT | MISSENSE | Gtg/Atg | V416M | dnaE2 | | | | | | | A | A | | | | |
| NC_011770 | 5126250 | A | G | 87 | 0 | 4 | 0 | NON_SYNONYMOU | MODERAT | MISSENSE | gAc/gGc | D470G | dnaE2 | | | G | G | G | G | | | | | | |
| NC_011770 | 5126420 | A | G | 76 | 0 | 12 | 0 | NON_SYNONYMOU | MODERAT | MISSENSE | Atc/Gtc | I527V | dnaE2 | G | G | G | G | G | G | G | G | G | G | G | G |
| NC_011770 | 5126753 | G | A | 73 | 0 | 6 | 0 | NON_SYNONYMOU | MODERAT | MISSENSE | Gaa/Aaa | E638K | dnaE2 | | | | | | | A | A | A | A | A | A |
| NC_011770 | 5126822 | G | A | 63 | 0 | 2 | 0 | NON_SYNONYMOU | MODERAT | MISSENSE | Gcc/Acc | A661T | dnaE2 | | | | | | | A | A | | | | |
| NC_011770 | 5126884 | T | G | 67 | 0 | 2 | 0 | NON_SYNONYMOU | MODERAT | MISSENSE | caT/caG | H681Q | dnaE2 | G | G | | | | | | | | | | |
| NC_011770 | 5126935 | G | C | 67 | 0 | 6 | 0 | NON_SYNONYMOU | MODERAT | MISSENSE | aaG/aaC | K698N | dnaE2 | | | C | C | C | C | | | C | C | | |
| NC_011770 | 5127155 | G | A | 67 | 0 | 2 | 0 | NON_SYNONYMOU | MODERAT | MISSENSE | Gcc/Acc | A772T | dnaE2 | | | | | | | | | | | A | A |
| NC_011770 | 5127212 | T | C | 68 | 0 | 8 | 0 | NON_SYNONYMOU | MODERAT | MISSENSE | Tcg/Ccg | S791P | dnaE2 | | | C | C | C | C | | | C | C | C | C |
| NC_011770 | 5127311 | G | C | 71 | 0 | 10 | 0 | NON_SYNONYMOU | MODERAT | MISSENSE | Gag/Cag | E824Q | dnaE2 | | | C | C | C | C | C | C | C | C | C | C |
| NC_011770 | 5127483 | C | T | 65 | 0 | 10 | 0 | NON_SYNONYMOU | MODERAT | MISSENSE | gCc/gTc | A881V | dnaE2 | | | T | T | T | T | T | T | T | T | T | T |
| NC_011770 | 5127495 | G | C | 65 | 0 | 4 | 0 | NON_SYNONYMOU | MODERAT | MISSENSE | aGc/aCc | S885T | dnaE2 | | | | | | | | | C | C | C | C |
| NC_011770 | 5127611 | C | T | 62 | 0 | 4 | 0 | NON_SYNONYMOU | MODERAT | MISSENSE | Cgc/Tgc | R924C | dnaE2 | | | | | | | | | T | T | T | T |
| NC_011770 | 5127683 | G | A | 65 | 0 | 4 | 0 | NON_SYNONYMOU | MODERAT | MISSENSE | Gtc/Atc | V948I | dnaE2 | | | | | | | | | A | A | A | A |
| NC_011770 | 5127752 | G | C | 71 | 0 | 4 | 0 | NON_SYNONYMOU | MODERAT | MISSENSE | Gtc/Ctc | V971L | dnaE2 | | | | | | | | | C | C | C | C |
| NC_011770 | 627885 | T | C | 79 | 0 | 8 | 0 | NON_SYNONYMOU | MODERAT | MISSENSE | Acg/Gcg | T515A | dnaG | C | C | | | | | C | C | C | C | C | C |
| NC_011770 | 627992 | C | A | 84 | 0 | 2 | 0 | NON_SYNONYMOU | MODERAT | MISSENSE | gGc/gTc | G479V | dnaG | | | | | | | A | A | | | | |
| NC_011770 | 628028 | A | G | 75 | 0 | 12 | 0 | NON_SYNONYMOU | MODERAT | MISSENSE | cTc/cCc | L467P | dnaG | G | G | G | G | G | G | G | G | G | G | G | G |
| NC_011770 | 628059 | G | A | 77 | 0 | 12 | 0 | NON_SYNONYMOU | MODERAT | MISSENSE | Ctc/Ttc | L457F | dnaG | A | A | A | A | A | A | A | A | A | A | A | A |
| NC_011770 | 628079 | G | A | 76 | 0 | 6 | 0 | NON_SYNONYMOU | MODERAT | MISSENSE | gCc/gTc | A450V | dnaG | | | | | | | A | A | A | A | A | A |
| NC_011770 | 628080 | C | T | 78 | 0 | 2 | 0 | NON_SYNONYMOU | MODERAT | MISSENSE | Gcc/Acc | A450T | dnaG | T | T | | | | | | | | | | |
| NC_011770 | 628583 | C | T | 74 | 0 | 2 | 0 | NON_SYNONYMOU | MODERAT | MISSENSE | cGc/cAc | R282H | dnaG | T | T | | | | | | | | | | |
| NC_011770 | 628706 | A | G | 76 | 0 | 1 | 0 | NON_SYNONYMOU | MODERAT | MISSENSE | gTc/gCc | V241A | dnaG | | | | | | | G | | | | | |
| NC_011770 | 628895 | T | G | 72 | 0 | 2 | 0 | NON_SYNONYMOU | MODERAT | MISSENSE | aAc/aCc | N178T | dnaG | | | | | | | | | G | G | | |
| NC_011770 | 629190 | G | T | 74 | 0 | 4 | 0 | NON_SYNONYMOU | MODERAT | MISSENSE | Ctg/Atg | L80M | dnaG | | | T | T | T | T | | | | | | |
| NC_011770 | 5681172 | C | T | 56 | 0 | 12 | 0 | NON_SYNONYMOU | MODERAT | MISSENSE | aGc/aAc | S589N | dnaK | T | T | T | T | T | T | T | T | T | T | T | T |
| NC_011770 | 5681358 | C | T | 62 | 0 | 1 | 0 | NON_SYNONYMOU | MODERAT | MISSENSE | cGc/cAc | R527H | dnaK | | | | | | | T | | | | | |
| NC_011770 | 2303 | C | T | 92 | 0 | 1 | 0 | NON_SYNONYMOU | MODERAT | MISSENSE | cCg/cTg | P83L | dnaN | | | | | | | T | | | | | |
| NC_011770 | 2587 | G | A | 101 | 0 | 2 | 0 | NON_SYNONYMOU | MODERAT | MISSENSE | Gcc/Acc | A178T | dnaN | | | | | | | | | | | A | A |
| NC_011770 | 4197933 | T | C | 81 | 0 | 1 | 0 | NON_SYNONYMOU | MODERAT | MISSENSE | Acc/Gcc | T104A | dnaX | C | | | | | | | | | | | |
| NC_011770 | 4196431 | G | T | 35 | 0 | 2 | 0 | NON_SYNONYMOU | MODERAT | MISSENSE | caC/caA | H604Q | dnaX | | | | | | | | | | | T | T |
| NC_011770 | 4196438 | C | T | 31 | 0 | 8 | 0 | NON_SYNONYMOU | MODERAT | MISSENSE | gGc/gAc | G602D | dnaX | T | T | T | T | T | T | | | | | T | T |
| NC_011770 | 4196468 | C | G | 35 | 0 | 4 | 0 | NON_SYNONYMOU | MODERAT | MISSENSE | gGt/gCt | G592A | dnaX | | | | | | | G | G | | | G | G |
| NC_011770 | 4196744 | G | T | 32 | 0 | 2 | 0 | NON_SYNONYMOU | MODERAT | MISSENSE | gCg/gAg | A500E | dnaX | | | | | | | | | T | T | | |
| NC_011770 | 4196760 | T | C | 31 | 0 | 10 | 0 | NON_SYNONYMOU | MODERAT | MISSENSE | Atc/Gtc | I495V | dnaX | C | C | C | C | C | C | C | C | | | C | C |
| NC_011770 | 4196949 | T | C | 36 | 0 | 4 | 0 | NON_SYNONYMOU | MODERAT | MISSENSE | Acg/Gcg | T432A | dnaX | | | | | | | | | C | C | C | C |
| NC_011770 | 4196969 | A | G | 32 | 0 | 10 | 0 | NON_SYNONYMOU | MODERAT | MISSENSE | gTg/gCg | V425A | dnaX | | | G | G | G | G | G | G | G | G | G | G |
| NC_011770 | 4197003 | C | T | 35 | 0 | 4 | 0 | NON_SYNONYMOU | MODERAT | MISSENSE | Gct/Act | A414T | dnaX | | | T | T | T | T | | | | | | |
| NC_011770 | 4197048 | T | C | 39 | 0 | 4 | 0 | NON_SYNONYMOU | MODERAT | MISSENSE | Acc/Gcc | T399A | dnaX | | | | | | | | | C | C | C | C |
| NC_011770 | 4197146 | G | A | 49 | 0 | 4 | 0 | NON_SYNONYMOU | MODERAT | MISSENSE | cCg/gTg | A366V | dnaX | | | | | | | A | A | A | A | | |
| NC_011770 | 578320 | C | T | 81 | 0 | 2 | 0 | NON_SYNONYMOU | MODERAT | MISSENSE | cGc/cAc | R209H | dnr | | | | | | | T | T | | | | |
| NC_011770 | 6517075 | C | T | 85 | 0 | 1 | 0 | NON_SYNONYMOU | MODERAT | MISSENSE | Gac/Aac | D140N | dsbA | | | | | | | | | | | | T |
| NC_011770 | 589292 | A | G | 84 | 0 | 2 | 0 | NON_SYNONYMOU | MODERAT | MISSENSE | gTc/gCc | V157A | dsbB | | | | | | | | | | | G | G |
| NC_011770 | 589551 | C | T | 69 | 0 | 2 | 0 | NON_SYNONYMOU | MODERAT | MISSENSE | Ggc/Agc | G71S | dsbB | | | | | | | | | | | T | T |
| NC_011770 | 1347457 | A | G | 73 | 0 | 1 | 0 | NON_SYNONYMOU | MODERAT | MISSENSE | Atc/Gtc | I148V | dsbC | | G | | | | | | | | | | |
| NC_011770 | 3034226 | T | C | 78 | 0 | 12 | 0 | NON_SYNONYMOU | MODERAT | MISSENSE | gTg/gCg | V190A | dsbG | C | C | C | C | C | C | C | C | C | C | C | C |
| NC_011770 | 3034310 | C | A | 78 | 0 | 12 | 0 | NON_SYNONYMOU | MODERAT | MISSENSE | gCg/gAg | A218E | dsbG | A | A | A | A | A | A | A | A | A | A | A | A |
| NC_011770 | 6253837 | A | G | 90 | 0 | 2 | 0 | NON_SYNONYMOU | MODERAT | MISSENSE | gTa/gCa | V88A | dsbH | G | G | | | | | | | | | | |
| NC_011770 | 6253928 | T | C | 84 | 0 | 4 | 0 | NON_SYNONYMOU | MODERAT | MISSENSE | Atc/Gtc | I58V | dsbH | C | C | | | | | | | | | C | C |
| NC_011770 | 1833358 | C | T | 81 | 0 | 12 | 0 | NON_SYNONYMOU | MODERAT | MISSENSE | Gcg/Acg | A434T | dsdA | T | T | T | T | T | T | T | T | T | T | T | T |
| NC_011770 | 1833981 | C | T | 62 | 0 | 2 | 0 | NON_SYNONYMOU | MODERAT | MISSENSE | cGc/cAc | R226H | dsdA | | | | | | | | | T | T | | |
| NC_011770 | 1834243 | C | T | 76 | 0 | 2 | 0 | NON_SYNONYMOU | MODERAT | MISSENSE | Ggc/Agc | G139S | dsdA | | | | | T | T | | | | | | |
| NC_011770 | 1834368 | G | A | 69 | 0 | 2 | 0 | NON_SYNONYMOU | MODERAT | MISSENSE | gCc/gTc | A97V | dsdA | | | | | | | | | A | A | | |

| | | | | | | | | | | | | | | | | | | | | | | | | | |
|---|---|---|---|---|---|---|---|---|---|---|---|---|---|---|---|---|---|---|---|---|---|---|---|---|---|
| NC_011770 | 1834549 | T | C | 70 | 0 | 12 | 0 | NON_SYNONYMOU | MODERAT | MISSENSE | Acc/Gcc | T37A | dsdA | C | C | C | C | C | C | C | C | C | C | C | C |
| NC_011770 | 1834623 | A | G | 76 | 0 | 12 | 0 | NON_SYNONYMOU | MODERAT | MISSENSE | gTc/gCc | V12A | dsdA | G | G | G | G | G | G | G | G | G | G | G | G |
| NC_011770 | 1834642 | A | G | 83 | 0 | 2 | 0 | NON_SYNONYMOU | MODERAT | MISSENSE | Tcc/Ccc | S6P | dsdA | | | | | | | | | | | G | G |
| NC_011770 | 1490444 | A | G | 72 | 0 | 2 | 0 | NON_SYNONYMOU | MODERAT | MISSENSE | Atc/Gtc | I77V | dxr | | | | | | | | | | | G | G |
| NC_011770 | 1009696 | G | A | 62 | 0 | 1 | 0 | NON_SYNONYMOU | MODERAT | MISSENSE | Ccg/Tcg | P475S | dxs | | | | | | | A | | | | | |
| NC_011770 | 2015547 | A | G | 85 | 0 | 2 | 0 | NON_SYNONYMOU | MODERAT | MISSENSE | gAa/gGa | E7G | edd | | | | | | | | | | | G | G |
| NC_011770 | 2016789 | G | A | 75 | 0 | 4 | 0 | NON_SYNONYMOU | MODERAT | MISSENSE | cGc/cAc | R421H | edd | | | A | A | A | A | | | | | | |
| NC_011770 | 2015907 | C | T | 66 | 0 | 1 | 0 | NON_SYNONYMOU | MODERAT | MISSENSE | gCc/gTc | A127V | edd | | | | | | | | T | | | | |
| NC_011770 | 2390607 | A | G | 52 | 0 | 2 | 0 | NON_SYNONYMOU | MODERAT | MISSENSE | gAg/gGg | E181G | efp | G | G | | | | | | | | | | |
| NC_011770 | 2492578 | C | T | 70 | 0 | 2 | 0 | NON_SYNONYMOU | MODERAT | MISSENSE | Gcc/Acc | A58T | endA | | | | | | | T | T | | | | |
| NC_011770 | 2492256 | C | T | 70 | 0 | 1 | 0 | NON_SYNONYMOU | MODERAT | MISSENSE | gGg/gAg | G165E | endA | | | | | | | | T | | | | |
| NC_011770 | 1272003 | T | C | 74 | 0 | 12 | 0 | NON_SYNONYMOU | MODERAT | MISSENSE | gTg/gCg | V167A | engA | C | C | C | C | C | C | C | C | C | C | C | C |
| NC_011770 | 6519005 | C | T | 67 | 0 | 2 | 0 | NON_SYNONYMOU | MODERAT | MISSENSE | gCc/gTc | A73V | engB | | | | | | | | | T | T | | |
| NC_011770 | 6519358 | A | G | 49 | 0 | 10 | 0 | NON_SYNONYMOU | MODERAT | MISSENSE | Atc/Gtc | I191V | engB | | | G | G | G | G | G | G | G | G | G | G |
| NC_011770 | 6187706 | T | C | 61 | 0 | 2 | 0 | NON_SYNONYMOU | MODERAT | MISSENSE | Atc/Gtc | I260V | envZ | | | | | | | | | | | C | C |
| NC_011770 | 6187718 | C | T | 64 | 0 | 2 | 0 | NON_SYNONYMOU | MODERAT | MISSENSE | Gtc/Atc | V256I | envZ | | | | | | | | | T | T | | |
| NC_011770 | 6188122 | C | A | 51 | 0 | 12 | 0 | NON_SYNONYMOU | MODERAT | MISSENSE | gGg/gTg | G121V | envZ | A | A | A | A | A | A | A | A | A | A | A | A |
| NC_011770 | 5028089 | C | T | 81 | 0 | 4 | 0 | NON_SYNONYMOU | MODERAT | MISSENSE | Gta/Ata | V157I | era | | | T | T | T | T | | | | | | |
| NC_011770 | 6089042 | C | T | 70 | 0 | 2 | 0 | NON_SYNONYMOU | MODERAT | MISSENSE | Ggc/Agc | G624S | estA | | | | | | | T | T | | | | |
| NC_011770 | 6089286 | C | A | 72 | 0 | 2 | 0 | NON_SYNONYMOU | MODERAT | MISSENSE | aaG/aaT | K542N | estA | A | A | | | | | | | | | | |
| NC_011770 | 6089311 | G | C | 69 | 0 | 8 | 0 | NON_SYNONYMOU | MODERAT | MISSENSE | aCc/aGc | T534S | estA | C | C | C | C | C | C | | | C | C | | |
| NC_011770 | 6089560 | T | C | 75 | 0 | 12 | 0 | NON_SYNONYMOU | MODERAT | MISSENSE | cAc/cGc | H451R | estA | C | C | C | C | C | C | C | C | C | C | C | C |
| NC_011770 | 6089774 | GGCC | G | 65 | 0 | 1 | 0 | CODON_DELETION | MODERATE | | ggc/ | G379 | estA | | | | | | | G | | | | | |
| NC_011770 | 6090470 | C | G | 83 | 0 | 12 | 0 | NON_SYNONYMOU | MODERAT | MISSENSE | Gtg/Ctg | V148L | estA | G | G | G | G | G | G | G | G | G | G | G | G |
| NC_011770 | 3950762 | C | T | 50 | 0 | 2 | 0 | NON_SYNONYMOU | MODERAT | MISSENSE | cGc/cAc | R280H | estX | | | | | | | | | T | T | | |
| NC_011770 | 3950806 | G | T | 48 | 0 | 2 | 0 | NON_SYNONYMOU | MODERAT | MISSENSE | ttC/ttA | F265L | estX | T | T | | | | | | | | | | |
| NC_011770 | 3950904 | C | T | 48 | 0 | 2 | 0 | NON_SYNONYMOU | MODERAT | MISSENSE | Gac/Aac | D233N | estX | | | | | | | | | | | T | T |
| NC_011770 | 3950946 | A | G | 45 | 0 | 12 | 0 | NON_SYNONYMOU | MODERAT | MISSENSE | Tgt/Cgt | C219R | estX | G | G | G | G | G | G | G | G | G | G | G | G |
| NC_011770 | 3950964 | G | C | 44 | 0 | 4 | 0 | NON_SYNONYMOU | MODERAT | MISSENSE | Ctg/Gtg | L213V | estX | | | | | | | C | C | | | C | C |
| NC_011770 | 1031821 | C | T | 55 | 0 | 1 | 0 | NON_SYNONYMOU | MODERAT | MISSENSE | Gcc/Acc | A113T | eutB | | | | | | | | T | | | | |
| NC_011770 | 3697154 | G | C | 84 | 0 | 2 | 0 | NON_SYNONYMOU | MODERAT | MISSENSE | Gtg/Ctg | V163L | exaA | | | | | | | | | | | C | C |
| NC_011770 | 3697763 | C | G | 69 | 0 | 12 | 0 | NON_SYNONYMOU | MODERAT | MISSENSE | Cac/Gac | H366D | exaA | G | G | G | G | G | G | G | G | G | G | G | G |
| NC_011770 | 3698201 | G | C | 55 | 0 | 2 | 0 | NON_SYNONYMOU | MODERAT | MISSENSE | Gtg/Ctg | V512L | exaA | | | | | | | | | C | C | | |
| NC_011770 | 3698057 | G | A | 59 | 0 | 1 | 0 | NON_SYNONYMOU | MODERAT | MISSENSE | Gcc/Acc | A464T | exaA | | | | | | | | A | | | | |
| NC_011770 | 3698279 | A | G | 55 | 0 | 1 | 0 | NON_SYNONYMOU | MODERAT | MISSENSE | Acc/Gcc | T538A | exaA | | | | | | | | G | | | | |
| NC_011770 | 3696339 | G | A | 72 | 0 | 2 | 0 | NON_SYNONYMOU | MODERAT | MISSENSE | gCc/gTc | A6V | exaB | | | | | | | | | | | A | A |
| NC_011770 | 227328 | G | A | 83 | 0 | 2 | 0 | NON_SYNONYMOU | MODERAT | MISSENSE | Gcc/Acc | A93T | exbB1 | | | | | | | | | A | A | | |
| NC_011770 | 5089314 | A | G | 64 | 0 | 8 | 0 | NON_SYNONYMOU | MODERAT | MISSENSE | gTc/gCc | V306A | exbB2 | G | G | | | | | G | G | G | G | G | G |
| NC_011770 | 5089326 | G | A | 64 | 0 | 10 | 0 | NON_SYNONYMOU | MODERAT | MISSENSE | gCc/gTc | A302V | exbB2 | A | A | C | C | C | C | A | A | A | A | | |
| NC_011770 | 5090090 | C | G | 68 | 0 | 8 | 0 | NON_SYNONYMOU | MODERAT | MISSENSE | gaG/gaC | E47D | exbB2 | G | G | | | | | G | G | G | G | G | G |
| NC_011770 | 228121 | G | A | 86 | 0 | 2 | 0 | NON_SYNONYMOU | MODERAT | MISSENSE | Gct/Act | A117T | exbD1 | A | A | | | | | | | | | | |
| NC_011770 | 1225833 | G | A | 91 | 0 | 4 | 0 | NON_SYNONYMOU | MODERAT | MISSENSE | aGc/aAc | S121N | exoS | | | A | A | A | A | | | | | | |
| NC_011770 | 1226631 | A | G | 78 | 0 | 2 | 0 | NON_SYNONYMOU | MODERAT | MISSENSE | gAa/gGa | E387G | exoS | G | G | | | | | | | | | | |
| NC_011770 | 58995 | T | C | 81 | 0 | 12 | 0 | NON_SYNONYMOU | MODERAT | MISSENSE | gTg/gCg | V135A | exoT | C | C | C | C | C | C | C | C | C | C | C | C |
| NC_011770 | 59129 | G | C | 92 | 0 | 2 | 0 | NON_SYNONYMOU | MODERAT | MISSENSE | Ggg/Cgg | G180R | exoT | | | | | | | C | C | | | | |
| NC_011770 | 59238 | C | T | 103 | 0 | 12 | 0 | NON_SYNONYMOU | MODERAT | MISSENSE | gCa/gTa | A216V | exoT | T | T | T | T | T | T | T | T | T | T | T | T |
| NC_011770 | 59496 | G | A | 89 | 0 | 6 | 0 | NON_SYNONYMOU | MODERAT | MISSENSE | cGg/cAg | R302Q | exoT | | | | | | | A | A | A | A | A | A |
| NC_011770 | 59645 | G | A | 86 | 0 | 12 | 0 | NON_SYNONYMOU | MODERAT | MISSENSE | Ggc/Agc | G352S | exoT | A | A | A | A | A | A | A | A | A | A | A | A |
| NC_011770 | 59670 | A | T | 86 | 0 | 6 | 0 | NON_SYNONYMOU | MODERAT | MISSENSE | cAg/cTg | Q360L | exoT | T | T | T | T | T | T | | | | | | |
| NC_011770 | 3461963 | TC | T | 66 | 0 | 1 | 0 | FRAME_SHIFT | HIGH | | cgg/ | R314 | exoY | | | | | | | T | | | | | |
| NC_011770 | 3461043 | C | T | 67 | 0 | 2 | 0 | NON_SYNONYMOU | MODERAT | MISSENSE | Cat/Tat | H6Y | exoY | | | | | | | | | T | T | | |
| NC_011770 | 3461091 | C | T | 68 | 0 | 2 | 0 | NON_SYNONYMOU | MODERAT | MISSENSE | Cgc/Tgc | R22C | exoY | | | | | | | | | | | T | T |
| NC_011770 | 3461154 | C | T | 69 | 0 | 2 | 0 | NON_SYNONYMOU | MODERAT | MISSENSE | Ccc/Tcc | P43S | exoY | | | | | | | T | T | | | | |
| NC_011770 | 3461355 | G | A | 63 | 0 | 4 | 0 | NON_SYNONYMOU | MODERAT | MISSENSE | Gac/Aac | D110N | exoY | | | | | | | A | A | A | A | | |
| NC_011770 | 3461568 | G | C | 68 | 0 | 2 | 0 | NON_SYNONYMOU | MODERAT | MISSENSE | Gag/Cag | E181Q | exoY | | | | | | | C | C | | | | |
| NC_011770 | 3461604 | C | T | 65 | 0 | 4 | 0 | NON_SYNONYMOU | MODERAT | MISSENSE | Cgg/Tgg | R193W | exoY | | | T | T | T | T | | | | | | |
| NC_011770 | 3461985 | G | C | 66 | 0 | 6 | 0 | NON_SYNONYMOU | MODERAT | MISSENSE | Gtc/Ctc | V320L | exoY | C | C | | | | | C | C | C | C | | |
| NC_011770 | 3462100 | A | C | 63 | 0 | 2 | 0 | NON_SYNONYMOU | MODERAT | MISSENSE | gAc/gCc | D358A | exoY | | | | | | | | | | | C | C |
| NC_011770 | 3462130 | A | G | 63 | 0 | 2 | 0 | NON_SYNONYMOU | MODERAT | MISSENSE | cAg/cGg | Q368R | exoY | G | G | | | | | | | | | | |
| NC_011770 | 4010929 | G | A | 64 | 0 | 1 | 0 | NON_SYNONYMOU | MODERAT | MISSENSE | gCc/gTc | A4V | exsA | | | | | | | | A | | | | |
| NC_011770 | 4011272 | C | T | 57 | 0 | 2 | 0 | NON_SYNONYMOU | MODERAT | MISSENSE | Gcc/Acc | A127T | exsB | | | | | | | | | T | T | | |
| NC_011770 | 4011280 | T | C | 56 | 0 | 4 | 0 | NON_SYNONYMOU | MODERAT | MISSENSE | aAt/aGt | N124S | exsB | | | C | C | C | C | | | | | | |

| | | | | | | | | | | | | | | | | | | | | | | | | | |
|---|---|---|---|---|---|---|---|---|---|---|---|---|---|---|---|---|---|---|---|---|---|---|---|---|---|
| NC_011770 | 4011337 | T | C | 56 | 0 | 6 | 0 | NON_SYNONYMOU | MODERAT | MISSENSE | cAg/cGg | Q105R | exsB | | | C | C | C | C | | | | | C | C |
| NC_011770 | 4011383 | G | A | 54 | 0 | 2 | 0 | NON_SYNONYMOU | MODERAT | MISSENSE | Ctt/Ttt | L90F | exsB | | | A | A | | | | | | | | |
| NC_011770 | 4011412 | T | C | 59 | 0 | 2 | 0 | NON_SYNONYMOU | MODERAT | MISSENSE | gAg/gGg | E80G | exsB | C | C | | | | | | | | | | |
| NC_011770 | 4011520 | C | T | 66 | 0 | 4 | 0 | NON_SYNONYMOU | MODERAT | MISSENSE | cGc/cAc | R44H | exsB | | | T | T | T | T | | | | | | |
| NC_011770 | 4009693 | A | C | 66 | 0 | 4 | 0 | NON_SYNONYMOU | MODERAT | MISSENSE | Tcc/Gcc | S105A | exsD | | | | | | | | | C | C | C | C |
| NC_011770 | 4115922 | G | C | 75 | 0 | 10 | 0 | NON_SYNONYMOU | MODERAT | MISSENSE | gaG/gaC | E211D | fabB | C | C | C | C | C | C | C | C | | | C | C |
| NC_011770 | 2268235 | G | A | 83 | 0 | 6 | 0 | NON_SYNONYMOU | MODERAT | MISSENSE | cGc/cAc | R56H | fabD | A | A | A | A | A | A | | | | | | |
| NC_011770 | 2268750 | T | G | 76 | 0 | 8 | 0 | NON_SYNONYMOU | MODERAT | MISSENSE | Tcg/Gcg | S228A | fabD | G | G | | | | | G | G | G | G | G | G |
| NC_011770 | 2268868 | A | G | 84 | 0 | 4 | 0 | NON_SYNONYMOU | MODERAT | MISSENSE | gAa/gGa | E267G | fabD | | | G | G | G | G | | | | | | |
| NC_011770 | 4219848 | C | T | 79 | 0 | 6 | 0 | NON_SYNONYMOU | MODERAT | MISSENSE | gCc/gTc | A57V | fabF2 | | | T | T | T | T | | | T | T | | |
| NC_011770 | 4219851 | A | G | 82 | 0 | 4 | 0 | NON_SYNONYMOU | MODERAT | MISSENSE | cAg/cGg | Q58R | fabF2 | G | G | | | | | G | G | | | | |
| NC_011770 | 4220151 | G | A | 84 | 0 | 4 | 0 | NON_SYNONYMOU | MODERAT | MISSENSE | cGc/cAc | R158H | fabF2 | A | A | | | | | A | A | | | | |
| NC_011770 | 4220498 | G | T | 63 | 0 | 4 | 0 | NON_SYNONYMOU | MODERAT | MISSENSE | Gcc/Tcc | A274S | fabF2 | | | T | T | T | T | | | | | | |
| NC_011770 | 4220678 | A | G | 58 | 0 | 4 | 0 | NON_SYNONYMOU | MODERAT | MISSENSE | Acc/Gcc | T334A | fabF2 | | | G | G | G | G | | | | | | |
| NC_011770 | 4220858 | G | A | 53 | 0 | 2 | 0 | NON_SYNONYMOU | MODERAT | MISSENSE | Gcc/Acc | A394T | fabF2 | | | | | | | | | | | A | A |
| NC_011770 | 4220937 | G | A | 41 | 0 | 4 | 0 | NON_SYNONYMOU | MODERAT | MISSENSE | cGc/cAc | R420H | fabF2 | | | A | A | A | A | | | | | | |
| NC_011770 | 2269186 | G | A | 83 | 0 | 2 | 0 | NON_SYNONYMOU | MODERAT | MISSENSE | Ggc/Agc | G53S | fabG | | | | | | | | | | | A | A |
| NC_011770 | 5252965 | G | A | 77 | 0 | 1 | 0 | NON_SYNONYMOU | MODERAT | MISSENSE | atG/atA | M72I | fabG | | | | | | | | | | | | A |
| NC_011770 | 5252882 | G | A | 72 | 0 | 2 | 0 | NON_SYNONYMOU | MODERAT | MISSENSE | Gcc/Acc | A45T | fabG | A | A | | | | | | | | | | |
| NC_011770 | 5253050 | G | A | 77 | 0 | 1 | 0 | NON_SYNONYMOU | MODERAT | MISSENSE | Gtc/Atc | V101I | fabG | | | | | | | A | | | | | |
| NC_011770 | 1859449 | C | T | 77 | 0 | 2 | 0 | NON_SYNONYMOU | MODERAT | MISSENSE | Gac/Aac | D234N | fabH2 | | | | | | | T | T | | | | |
| NC_011770 | 2221054 | A | G | 86 | 0 | 1 | 0 | NON_SYNONYMOU | MODERAT | MISSENSE | Acc/Gcc | T104A | fadA | | | | | | | G | | | | | |
| NC_011770 | 2221289 | A | G | 98 | 0 | 1 | 0 | NON_SYNONYMOU | MODERAT | MISSENSE | aAg/aGg | K182R | fadA | | | | | | | G | | | | | |
| NC_011770 | 2221417 | G | C | 90 | 0 | 6 | 0 | NON_SYNONYMOU | MODERAT | MISSENSE | Gag/Cag | E225Q | fadA | | | C | C | C | C | C | C | | | | |
| NC_011770 | 2221915 | A | G | 71 | 0 | 2 | 0 | NON_SYNONYMOU | MODERAT | MISSENSE | Atc/Gtc | I391V | fadA | | | | | | | G | G | | | | |
| NC_011770 | 2221307 | A | G | 97 | 0 | 1 | 0 | NON_SYNONYMOU | MODERAT | MISSENSE | aAg/aGg | K188R | fadA | | | | | | | | G | | | | |
| NC_011770 | 2220074 | T | C | 89 | 0 | 1 | 0 | NON_SYNONYMOU | MODERAT | MISSENSE | gTg/gCg | V503A | fadB | | | | | | | C | | | | | |
| NC_011770 | 2219755 | G | A | 83 | 0 | 3 | 0 | NON_SYNONYMOU | MODERAT | MISSENSE | Gtc/Atc | V397I | fadB | | | | | | | | | A | A | | A |
| NC_011770 | 2429602 | C | T | 55 | 0 | 4 | 0 | NON_SYNONYMOU | MODERAT | MISSENSE | Gcc/Acc | A771T | fadE | | | | | | | | | T | T | T | T |
| NC_011770 | 2123740 | A | G | 63 | 0 | 4 | 0 | NON_SYNONYMOU | MODERAT | MISSENSE | gAc/gGc | D454G | fadH1 | | | G | G | G | G | | | | | | |
| NC_011770 | 2124036 | G | A | 66 | 0 | 4 | 0 | NON_SYNONYMOU | MODERAT | MISSENSE | Gtc/Atc | V553I | fadH1 | | | A | A | A | A | | | | | | |
| NC_011770 | 5737848 | A | AG | 41 | 0 | 7 | 0 | FRAME_SHIFT | HIGH | | gcg/gGcg | A679G? | fadH2 | | AG | AG | | AG | AG | AG | | AG | | AG | |
| NC_011770 | 5737488 | G | A | 64 | 0 | 4 | 0 | NON_SYNONYMOU | MODERAT | MISSENSE | cGg/cAg | R562Q | fadH2 | | | A | A | A | A | | | | | | |
| NC_011770 | 5737665 | T | A | 51 | 0 | 12 | 0 | NON_SYNONYMOU | MODERAT | MISSENSE | gTg/gAg | V621E | fadH2 | A | A | A | A | A | A | A | A | A | A | A | A |
| NC_011770 | 5737738 | A | C | 42 | 0 | 12 | 0 | NON_SYNONYMOU | MODERAT | MISSENSE | agA/agC | R645S | fadH2 | C | C | C | C | C | C | C | C | C | C | C | C |
| NC_011770 | 3668498 | A | T | 60 | 0 | 12 | 0 | NON_SYNONYMOU | MODERAT | MISSENSE | cAg/cTg | Q35L | fahA | T | T | T | T | T | T | T | T | T | T | T | T |
| NC_011770 | 5728875 | T | C | 78 | 0 | 12 | 0 | NON_SYNONYMOU | MODERAT | MISSENSE | Ata/Gta | I162V | fdhE | C | C | C | C | C | C | C | C | C | C | C | C |
| NC_011770 | 5731302 | C | T | 85 | 0 | 2 | 0 | NON_SYNONYMOU | MODERAT | MISSENSE | cGc/cAc | R756H | fdnG | | | | | | | | | | | T | T |
| NC_011770 | 5732413 | G | A | 83 | 0 | 2 | 0 | NON_SYNONYMOU | MODERAT | MISSENSE | Cgc/Tgc | R386C | fdnG | A | A | | | | | | | | | | |
| NC_011770 | 5730537 | T | G | 69 | 0 | 6 | 0 | NON_SYNONYMOU | MODERAT | MISSENSE | gaA/gaC | E194D | fdnH | | | G | G | G | G | | | G | G | | |
| NC_011770 | 1522914 | T | C | 54 | 0 | 8 | 0 | NON_SYNONYMOU | MODERAT | MISSENSE | Atc/Gtc | I78V | fdxA | C | C | C | C | C | C | | | C | C | | |
| NC_011770 | 1165840 | A | G | 79 | 0 | 2 | 0 | NON_SYNONYMOU | MODERAT | MISSENSE | aTc/aCc | I541T | fecA | G | G | | | | | | | | | | |
| NC_011770 | 1166437 | G | A | 82 | 0 | 4 | 0 | NON_SYNONYMOU | MODERAT | MISSENSE | gCg/gTg | A342V | fecA | | | A | A | A | A | | | | | | |
| NC_011770 | 1166447 | C | T | 81 | 0 | 8 | 0 | NON_SYNONYMOU | MODERAT | MISSENSE | Gcg/Acg | A339T | fecA | | | T | T | T | T | | | T | T | T | T |
| NC_011770 | 1167178 | G | A | 66 | 0 | 4 | 0 | NON_SYNONYMOU | MODERAT | MISSENSE | gCc/gTc | A95V | fecA | | | A | A | A | A | | | | | | |
| NC_011770 | 5220475 | C | A | 59 | 0 | 12 | 0 | STOP_GAINED | HIGH | NONSENSE | Gag/Tag | E165* | feoB | A | A | A | A | A | A | A | A | A | A | A | A |
| NC_011770 | 5218969 | C | T | 87 | 0 | 1 | 0 | NON_SYNONYMOU | MODERAT | MISSENSE | Gtc/Atc | V667I | feoB | | | | | | | | | | | T | |
| NC_011770 | 5218777 | G | C | 85 | 0 | 2 | 0 | NON_SYNONYMOU | MODERAT | MISSENSE | Ctc/Gtc | L731V | feoB | | | | | | | | | | | C | C |
| NC_011770 | 5219287 | C | T | 93 | 0 | 4 | 0 | NON_SYNONYMOU | MODERAT | MISSENSE | Gtg/Atg | V561M | feoB | | | T | T | T | T | | | | | | |
| NC_011770 | 5220276 | T | C | 77 | 0 | 4 | 0 | NON_SYNONYMOU | MODERAT | MISSENSE | cAg/cGg | Q231R | feoB | C | C | | | | | C | C | | | | |
| NC_011770 | 5220412 | T | C | 75 | 0 | 12 | 0 | NON_SYNONYMOU | MODERAT | MISSENSE | Agc/Ggc | S186G | feoB | C | C | C | C | C | C | C | C | C | C | C | C |
| NC_011770 | 5220466 | C | G | 59 | 0 | 12 | 0 | NON_SYNONYMOU | MODERAT | MISSENSE | Gtc/Ctc | V168L | feoB | G | G | G | G | G | G | G | G | G | G | G | G |
| NC_011770 | 5220473 | C | G | 59 | 0 | 12 | 0 | NON_SYNONYMOU | MODERAT | MISSENSE | gaG/gaC | E165D | feoB | G | G | G | G | G | G | G | G | G | G | G | G |
| NC_011770 | 5220474 | T | G | 59 | 0 | 12 | 0 | NON_SYNONYMOU | MODERAT | MISSENSE | gAg/gCg | E165A | feoB | G | G | G | G | G | G | G | G | G | G | G | G |
| NC_011770 | 5218978 | T | C | 88 | 0 | 1 | 0 | NON_SYNONYMOU | MODERAT | MISSENSE | Agc/Ggc | S664G | feoB | | | | | | | | C | | | | |
| NC_011770 | 838083 | T | TG | 70 | 0 | 1 | 0 | FRAME_SHIFT | HIGH | | cag/cCag | Q199P? | fepB | | | | | | | TG | | | | | |
| NC_011770 | 838053 | C | T | 73 | 0 | 10 | 0 | NON_SYNONYMOU | MODERAT | MISSENSE | cGg/cAg | R209Q | fepB | T | T | T | T | T | T | T | T | T | T | | |
| NC_011770 | 838171 | T | C | 75 | 0 | 12 | 0 | NON_SYNONYMOU | MODERAT | MISSENSE | Acg/Gcg | T170A | fepB | C | C | C | C | C | C | C | C | C | C | C | C |
| NC_011770 | 838811 | T | C | 95 | 0 | 12 | 0 | NON_SYNONYMOU | MODERAT | MISSENSE | Tat/Cat | Y3H | fepC | C | C | C | C | C | C | C | C | C | C | C | C |
| NC_011770 | 839538 | C | T | 86 | 0 | 2 | 0 | NON_SYNONYMOU | MODERAT | MISSENSE | gCc/gTc | A245V | fepC | | | | | | | T | T | | | | |
| NC_011770 | 837045 | C | T | 61 | 0 | 2 | 0 | NON_SYNONYMOU | MODERAT | MISSENSE | cGg/cAg | R233Q | fepD | | | | | | | | | | | T | T |
| NC_011770 | 837415 | C | T | 66 | 0 | 12 | 0 | NON_SYNONYMOU | MODERAT | MISSENSE | Gtc/Atc | V110I | fepD | T | T | T | T | T | T | T | T | T | T | T | T |

| | | | | | | | | | | | | | | | | | | | | | | | | | |
|---|---|---|---|---|---|---|---|---|---|---|---|---|---|---|---|---|---|---|---|---|---|---|---|---|---|
| NC_011770 | 837583 | C | T | 70 | 0 | 1 | 0 | NON_SYNONYMOU | MODERAT | MISSENSE | Gat/Aat | D54N | fepD | | | | | | | T | | | | | |
| NC_011770 | 835691 | C | CT | 77 | 0 | 10 | 0 | FRAME_SHIFT+SPL | HIGH | | tga/tgaA | *344*? | fepG | CT | CT | CT | CT | CT | CT | CT | CT | CT | CT | | |
| NC_011770 | 835964 | G | T | 77 | 0 | 10 | 0 | NON_SYNONYMOU | MODERAT | MISSENSE | Ctg/Atg | L254M | fepG | T | T | T | T | T | T | T | T | T | T | | |
| NC_011770 | 836093 | A | C | 69 | 0 | 12 | 0 | NON_SYNONYMOU | MODERAT | MISSENSE | Tcg/Gcg | S211A | fepG | C | C | C | C | C | C | C | C | C | C | C | C |
| NC_011770 | 4198466 | C | T | 77 | 0 | 8 | 0 | INTRAGENIC | MODIFIER | | | | ffs | T | T | T | T | T | T | | | | | T | T |
| NC_011770 | 4198540 | G | A | 76 | 0 | 4 | 0 | INTRAGENIC | MODIFIER | | | | ffs | | | A | A | A | A | | | | | | |
| NC_011770 | 4198903 | G | A | 70 | 0 | 2 | 0 | INTRAGENIC | MODIFIER | | | | ffs | | | | | | | | | A | A | | |
| NC_011770 | 4199124 | G | A | 64 | 0 | 4 | 0 | INTRAGENIC | MODIFIER | | | | ffs | | | A | A | A | A | | | | | | |
| NC_011770 | 4199148 | G | A | 62 | 0 | 2 | 0 | INTRAGENIC | MODIFIER | | | | ffs | | | | | | | | | A | A | | |
| NC_011770 | 4199174 | G | C | 62 | 0 | 2 | 0 | INTRAGENIC | MODIFIER | | | | ffs | C | C | | | | | | | | | | |
| NC_011770 | 4199195 | C | T | 77 | 0 | 2 | 0 | INTRAGENIC | MODIFIER | | | | ffs | | | | | | | | | T | T | | |
| NC_011770 | 4199200 | G | A | 80 | 0 | 2 | 0 | INTRAGENIC | MODIFIER | | | | ffs | A | A | | | | | | | | | | |
| NC_011770 | 4199391 | A | G | 68 | 0 | 2 | 0 | INTRAGENIC | MODIFIER | | | | ffs | | | | | | | | | G | G | | |
| NC_011770 | 4664874 | CGA | C | 51 | 0 | 6 | 0 | FRAME_SHIFT | HIGH | | ctc/ | L1620 | fgtA | C | C | | | | | C | C | | | C | C |
| NC_011770 | 4664879 | A | AGG | 50 | 0 | 6 | 0 | FRAME_SHIFT | HIGH | | gtg/gCCtg | V1619A? | fgtA | AGG | AGG | | | | | AGG | AGG | | | AGG | AGG |
| NC_011770 | 4664775 | C | A | 35 | 6 | 2 | 0 | NON_SYNONYMOU | MODERAT | MISSENSE | Gcg/Tcg | A1654S | fgtA | . | . | | | | | . | . | A | A | . | . |
| NC_011770 | 4664785 | C | G | 54 | 0 | 2 | 0 | NON_SYNONYMOU | MODERAT | MISSENSE | gaG/gaC | E1650D | fgtA | G | G | | | | | | | | | | |
| NC_011770 | 4664790 | G | C | 54 | 0 | 6 | 0 | NON_SYNONYMOU | MODERAT | MISSENSE | Ctg/Gtg | L1649V | fgtA | C | C | | | | | C | C | | | C | C |
| NC_011770 | 4664797 | C | G | 54 | 0 | 6 | 0 | NON_SYNONYMOU | MODERAT | MISSENSE | agG/agC | R1646S | fgtA | G | G | | | | | G | G | | | G | G |
| NC_011770 | 4664801 | G | A | 54 | 0 | 6 | 0 | NON_SYNONYMOU | MODERAT | MISSENSE | tCc/tTc | S1645F | fgtA | A | A | | | | | A | A | | | A | A |
| NC_011770 | 4664831 | C | T | 46 | 0 | 6 | 0 | NON_SYNONYMOU | MODERAT | MISSENSE | aGt/aAt | S1635N | fgtA | T | T | | | | | T | T | | | T | T |
| NC_011770 | 4664832 | T | C | 45 | 0 | 6 | 0 | NON_SYNONYMOU | MODERAT | MISSENSE | Agt/Ggt | S1635G | fgtA | C | C | | | | | C | C | | | C | C |
| NC_011770 | 4664837 | C | T | 46 | 0 | 6 | 0 | NON_SYNONYMOU | MODERAT | MISSENSE | cGc/cAc | R1633H | fgtA | T | T | | | | | T | T | | | T | T |
| NC_011770 | 4664846 | A | C | 46 | 0 | 6 | 0 | NON_SYNONYMOU | MODERAT | MISSENSE | cTt/cGt | L1630R | fgtA | C | C | | | | | C | C | | | C | C |
| NC_011770 | 4664862 | T | C | 52 | 0 | 6 | 0 | NON_SYNONYMOU | MODERAT | MISSENSE | Atc/Gtc | I1625V | fgtA | C | C | | | | | C | C | | | C | C |
| NC_011770 | 4664924 | C | A | 54 | 0 | 6 | 0 | NON_SYNONYMOU | MODERAT | MISSENSE | cGc/cTc | R1604L | fgtA | A | A | | | | | A | A | | | A | A |
| NC_011770 | 4664937 | A | C | 49 | 0 | 8 | 0 | NON_SYNONYMOU | MODERAT | MISSENSE | Ttg/Gtg | L1600V | fgtA | C | C | | | | | C | C | G | G | C | C |
| NC_011770 | 4665419 | C | T | 37 | 6 | 2 | 0 | NON_SYNONYMOU | MODERAT | MISSENSE | cGg/cAg | R1439Q | fgtA | . | . | | | | | . | . | T | T | . | . |
| NC_011770 | 4665783 | C | T | 33 | 6 | 2 | 0 | NON_SYNONYMOU | MODERAT | MISSENSE | Gtg/Atg | V1318M | fgtA | . | . | | | | | . | . | T | T | . | . |
| NC_011770 | 4666142 | C | T | 27 | 6 | 6 | 0 | NON_SYNONYMOU | MODERAT | MISSENSE | cGc/cAc | R1198H | fgtA | . | . | T | T | T | T | . | . | T | T | . | . |
| NC_011770 | 4667274 | G | A | 37 | 6 | 6 | 0 | NON_SYNONYMOU | MODERAT | MISSENSE | Ctc/Ttc | L821F | fgtA | . | . | A | A | A | A | . | . | A | A | . | . |
| NC_011770 | 4669159 | C | A | 31 | 6 | 2 | 0 | NON_SYNONYMOU | MODERAT | MISSENSE | gaG/gaT | E192D | fgtA | . | . | | | | | . | . | A | A | . | . |
| NC_011770 | 4669313 | C | T | 26 | 6 | 4 | 0 | NON_SYNONYMOU | MODERAT | MISSENSE | cGc/cAc | R141H | fgtA | . | . | T | T | T | T | . | . | | | . | . |
| NC_011770 | 4669394 | G | C | 24 | 6 | 6 | 0 | NON_SYNONYMOU | MODERAT | MISSENSE | tCc/tGc | S114C | fgtA | . | . | C | C | C | C | . | . | C | C | . | . |
| NC_011770 | 4669406 | G | A | 25 | 6 | 4 | 0 | NON_SYNONYMOU | MODERAT | MISSENSE | gCt/gTt | A110V | fgtA | . | . | A | A | A | A | . | . | | | . | . |
| NC_011770 | 4669556 | A | G | 23 | 6 | 4 | 0 | NON_SYNONYMOU | MODERAT | MISSENSE | gTc/gCc | V60A | fgtA | . | . | G | G | G | G | . | . | | | . | . |
| NC_011770 | 4668666 | T | C | 37 | 6 | 1 | 0 | NON_SYNONYMOU | MODERAT | MISSENSE | Agc/Ggc | S357G | fgtA | . | . | | C | | | . | . | | | . | . |
| NC_011770 | 2623506 | G | T | 76 | 0 | 6 | 0 | NON_SYNONYMOU | MODERAT | MISSENSE | gaG/gaT | E149D | fhp | | | | | | | T | T | T | T | T | T |
| NC_011770 | 2624162 | G | A | 70 | 0 | 2 | 0 | NON_SYNONYMOU | MODERAT | MISSENSE | cGc/cAc | R368H | fhp | | | | | | | | | | | A | A |
| NC_011770 | 5432844 | C | G | 63 | 2 | 8 | 0 | NON_SYNONYMOU | MODERAT | MISSENSE | Cag/Gag | Q41E | fimT | | | G | G | G | G | G | G | . | . | G | G |
| NC_011770 | 5433000 | G | A | 63 | 2 | 2 | 0 | NON_SYNONYMOU | MODERAT | MISSENSE | Gaa/Aaa | E93K | fimT | A | A | | | | | | | . | . | | |
| NC_011770 | 5433046 | G | T | 65 | 2 | 4 | 0 | NON_SYNONYMOU | MODERAT | MISSENSE | cGc/cTc | R108L | fimT | | | T | T | T | T | | | . | . | | |
| NC_011770 | 2097413 | T | C | 64 | 0 | 12 | 0 | NON_SYNONYMOU | MODERAT | MISSENSE | Tcg/Ccg | S146P | fimV | C | C | C | C | C | C | C | C | C | C | C | C |
| NC_011770 | 2097492 | A | G | 67 | 0 | 12 | 0 | NON_SYNONYMOU | MODERAT | MISSENSE | gAc/gGc | D172G | fimV | G | G | G | G | G | G | G | G | G | G | G | G |
| NC_011770 | 2097679 | G | C | 74 | 0 | 2 | 0 | NON_SYNONYMOU | MODERAT | MISSENSE | caG/caC | Q234H | fimV | | | | | | | | | C | C | | |
| NC_011770 | 2097771 | C | T | 68 | 0 | 2 | 0 | NON_SYNONYMOU | MODERAT | MISSENSE | gCt/gTt | A265V | fimV | | | | | | | | | | | T | T |
| NC_011770 | 2097920 | C | A | 66 | 0 | 6 | 0 | NON_SYNONYMOU | MODERAT | MISSENSE | Ctc/Atc | L315I | fimV | A | A | | | | | A | A | | | A | A |
| NC_011770 | 2098362 | G | A | 68 | 0 | 12 | 0 | NON_SYNONYMOU | MODERAT | MISSENSE | aGc/aAc | S462N | fimV | A | A | A | A | A | A | A | A | A | A | A | A |
| NC_011770 | 2098484 | G | A | 75 | 0 | 12 | 0 | NON_SYNONYMOU | MODERAT | MISSENSE | Gcc/Acc | A503T | fimV | A | A | A | A | A | A | A | A | A | A | A | A |
| NC_011770 | 2098575 | C | T | 72 | 0 | 8 | 0 | NON_SYNONYMOU | MODERAT | MISSENSE | gCt/gTt | A533V | fimV | T | T | | | | | T | T | T | T | T | T |
| NC_011770 | 2099025 | A | G | 66 | 0 | 2 | 0 | NON_SYNONYMOU | MODERAT | MISSENSE | gAc/gGc | D683G | fimV | | | | | | | | | G | G | | |
| NC_011770 | 2099111 | G | C | 62 | 0 | 4 | 0 | NON_SYNONYMOU | MODERAT | MISSENSE | Gcg/Ccg | A712P | fimV | C | C | | | | | | | | | C | C |
| NC_011770 | 5898214 | CCG | C | 71 | 0 | 1 | 0 | FRAME_SHIFT | HIGH | | cgc/ | R132 | fimX | | | | | | | | C | | | | |
| NC_011770 | 5898099 | G | A | 74 | 0 | 2 | 0 | NON_SYNONYMOU | MODERAT | MISSENSE | aGc/aAc | S91N | fimX | A | A | | | | | | | | | | |
| NC_011770 | 5781237 | A | G | 83 | 0 | 1 | 0 | NON_SYNONYMOU | MODERAT | MISSENSE | Acc/Gcc | T27A | fis | | | | | | | | G | | | | |
| NC_011770 | 5781270 | T | G | 81 | 0 | 12 | 0 | NON_SYNONYMOU | MODERAT | MISSENSE | Tac/Gac | Y38D | fis | G | G | G | G | G | G | G | G | G | G | G | G |
| NC_011770 | 523465 | G | GTA | 23 | 5 | 3 | 0 | FRAME_SHIFT | HIGH | | atc/atTAc | I419I? | fiuA | . | GTA | . | . | . | . | | | | | GTA | GTA |
| NC_011770 | 523468 | ACT | A | 23 | 5 | 3 | 0 | FRAME_SHIFT | HIGH | | agt/ | S418 | fiuA | . | A | . | . | . | . | | | | | A | A |
| NC_011770 | 522308 | C | T | 52 | 0 | 2 | 0 | SPLICE_SITE_REGIC | LOW | SILENT | tGa/tAa | *805 | fiuA | T | T | | | | | | | | | | |
| NC_011770 | 522585 | C | T | 85 | 0 | 10 | 0 | NON_SYNONYMOU | MODERAT | MISSENSE | Gcg/Acg | A713T | fiuA | | | T | T | T | T | T | T | T | T | T | T |
| NC_011770 | 522687 | C | T | 81 | 0 | 2 | 0 | NON_SYNONYMOU | MODERAT | MISSENSE | Gcc/Acc | A679T | fiuA | T | T | | | | | | | | | | |
| NC_011770 | 522699 | A | C | 82 | 0 | 8 | 0 | NON_SYNONYMOU | MODERAT | MISSENSE | Tcc/Gcc | S675A | fiuA | C | C | C | C | C | C | | | | | C | C |

| | | | | | | | | | | | | | | | | | | | | | | | | | |
|---|---|---|---|---|---|---|---|---|---|---|---|---|---|---|---|---|---|---|---|---|---|---|---|---|---|
| NC_011770 | 522708 | C | T | 77 | 0 | 8 | 0 | NON_SYNONYMOU | MODERAT | MISSENSE | Gtg/Atg | V672M | fiuA | T | T | T | T | T | T | | | | | T | T |
| NC_011770 | 522894 | G | T | 39 | 2 | 6 | 0 | NON_SYNONYMOU | MODERAT | MISSENSE | Ctg/Atg | L610M | fiuA | . | . | T | T | T | T | | | | | T | T |
| NC_011770 | 522959 | C | G | 20 | 4 | 4 | 0 | NON_SYNONYMOU | MODERAT | MISSENSE | aGc/aCc | S588T | fiuA | . | . | G | G | . | . | | | | | G | G |
| NC_011770 | 522987 | G | T | 64 | 0 | 8 | 0 | NON_SYNONYMOU | MODERAT | MISSENSE | Ctt/Att | L579I | fiuA | T | T | T | T | T | T | | | | | T | T |
| NC_011770 | 522999 | G | C | 73 | 0 | 8 | 0 | NON_SYNONYMOU | MODERAT | MISSENSE | Ctg/Gtg | L575V | fiuA | C | C | C | C | C | C | | | | | C | C |
| NC_011770 | 523073 | T | C | 66 | 0 | 8 | 0 | NON_SYNONYMOU | MODERAT | MISSENSE | aAc/aGc | N550S | fiuA | C | C | C | C | C | C | | | | | C | C |
| NC_011770 | 523074 | T | C | 66 | 0 | 8 | 0 | NON_SYNONYMOU | MODERAT | MISSENSE | Aac/Gac | N550D | fiuA | C | C | C | C | C | C | | | | | C | C |
| NC_011770 | 523077 | G | T | 66 | 0 | 8 | 0 | NON_SYNONYMOU | MODERAT | MISSENSE | Cag/Aag | Q549K | fiuA | T | T | T | T | T | T | | | | | T | T |
| NC_011770 | 523098 | C | T | 67 | 0 | 8 | 0 | NON_SYNONYMOU | MODERAT | MISSENSE | Gcc/Acc | A542T | fiuA | T | T | T | T | T | T | | | | | T | T |
| NC_011770 | 523193 | C | T | 32 | 4 | 4 | 0 | NON_SYNONYMOU | MODERAT | MISSENSE | cGa/cAa | R510Q | fiuA | T | T | . | . | . | . | | | | | T | T |
| NC_011770 | 523205 | A | T | 33 | 4 | 4 | 0 | NON_SYNONYMOU | MODERAT | MISSENSE | tTc/tAc | F506Y | fiuA | T | T | . | . | . | . | | | | | T | T |
| NC_011770 | 523211 | C | T | 32 | 4 | 4 | 0 | NON_SYNONYMOU | MODERAT | MISSENSE | aGt/aAt | S504N | fiuA | T | T | . | . | . | . | | | | | T | T |
| NC_011770 | 523212 | T | C | 32 | 4 | 4 | 0 | NON_SYNONYMOU | MODERAT | MISSENSE | Agt/Ggt | S504G | fiuA | C | C | . | . | . | . | | | | | C | C |
| NC_011770 | 523221 | A | G | 30 | 4 | 4 | 0 | NON_SYNONYMOU | MODERAT | MISSENSE | Tcg/Ccg | S501P | fiuA | G | G | . | . | . | . | | | | | G | G |
| NC_011770 | 523256 | T | A | 38 | 3 | 1 | 0 | NON_SYNONYMOU | MODERAT | MISSENSE | aAc/aTc | N489I | fiuA | | | . | A | . | . | | | | | | |
| NC_011770 | 523260 | C | T | 38 | 3 | 4 | 0 | NON_SYNONYMOU | MODERAT | MISSENSE | Gtc/Atc | V488I | fiuA | T | T | . | | . | . | | | | | T | T |
| NC_011770 | 523265 | G | C | 38 | 3 | 1 | 0 | NON_SYNONYMOU | MODERAT | MISSENSE | aCc/aGc | T486S | fiuA | | | . | C | . | . | | | | | | |
| NC_011770 | 523276 | ACCG | A | 27 | 4 | 4 | 0 | CODON_CHANGE_ | MODERATE | | accggt/act | TG481T | fiuA | A | A | . | . | . | . | | | | | A | A |
| NC_011770 | 523286 | T | C | 27 | 4 | 4 | 0 | NON_SYNONYMOU | MODERAT | MISSENSE | aAc/aGc | N479S | fiuA | C | C | . | . | . | . | | | | | C | C |
| NC_011770 | 523287 | T | C | 27 | 4 | 4 | 0 | NON_SYNONYMOU | MODERAT | MISSENSE | Aac/Gac | N479D | fiuA | C | C | . | . | . | . | | | | | C | C |
| NC_011770 | 523289 | T | A | 27 | 4 | 4 | 0 | NON_SYNONYMOU | MODERAT | MISSENSE | tAc/tTc | Y478F | fiuA | A | A | . | . | . | . | | | | | A | A |
| NC_011770 | 523303 | G | T | 27 | 4 | 4 | 0 | NON_SYNONYMOU | MODERAT | MISSENSE | gaC/gaA | D473E | fiuA | T | T | . | . | . | . | | | | | T | T |
| NC_011770 | 523307 | G | T | 27 | 4 | 4 | 0 | NON_SYNONYMOU | MODERAT | MISSENSE | aCc/aAc | T472N | fiuA | T | T | . | . | . | . | | | | | T | T |
| NC_011770 | 523310 | C | G | 26 | 4 | 4 | 0 | NON_SYNONYMOU | MODERAT | MISSENSE | cGc/cCc | R471P | fiuA | G | G | . | . | . | . | | | | | G | G |
| NC_011770 | 523311 | G | T | 27 | 4 | 4 | 0 | NON_SYNONYMOU | MODERAT | MISSENSE | Cgc/Agc | R471S | fiuA | T | T | . | . | . | . | | | | | T | T |
| NC_011770 | 523327 | G | C | 27 | 4 | 4 | 0 | NON_SYNONYMOU | MODERAT | MISSENSE | atC/atG | I465M | fiuA | C | C | . | . | . | . | | | | | C | C |
| NC_011770 | 523329 | T | G | 27 | 4 | 4 | 0 | NON_SYNONYMOU | MODERAT | MISSENSE | Atc/Ctc | I465L | fiuA | G | G | . | . | . | . | | | | | G | G |
| NC_011770 | 523363 | C | G | 30 | 4 | 4 | 0 | NON_SYNONYMOU | MODERAT | MISSENSE | gaG/gaC | E453D | fiuA | G | G | . | . | . | . | | | | | G | G |
| NC_011770 | 523407 | T | C | 38 | 0 | 8 | 0 | NON_SYNONYMOU | MODERAT | MISSENSE | Atc/Gtc | I439V | fiuA | C | C | C | C | C | C | | | | | C | C |
| NC_011770 | 523427 | G | C | 22 | 5 | 3 | 0 | NON_SYNONYMOU | MODERAT | MISSENSE | aCg/aGg | T432R | fiuA | . | C | . | . | . | . | | | | | C | C |
| NC_011770 | 523429 | C | G | 22 | 5 | 3 | 0 | NON_SYNONYMOU | MODERAT | MISSENSE | atG/atC | M431I | fiuA | . | G | . | . | . | . | | | | | G | G |
| NC_011770 | 523430 | A | G | 22 | 5 | 3 | 0 | NON_SYNONYMOU | MODERAT | MISSENSE | aTg/aCg | M431T | fiuA | . | G | . | . | . | . | | | | | G | G |
| NC_011770 | 523451 | G | T | 21 | 5 | 3 | 0 | NON_SYNONYMOU | MODERAT | MISSENSE | gCc/gAc | A424D | fiuA | . | T | . | . | . | . | | | | | T | T |
| NC_011770 | 523463 | T | G | 22 | 5 | 3 | 0 | NON_SYNONYMOU | MODERAT | MISSENSE | gAt/gCt | D420A | fiuA | . | G | . | . | . | . | | | | | G | G |
| NC_011770 | 523476 | T | C | 23 | 5 | 3 | 0 | NON_SYNONYMOU | MODERAT | MISSENSE | Acc/Gcc | T416A | fiuA | . | C | . | . | . | . | | | | | C | C |
| NC_011770 | 523488 | T | C | 24 | 5 | 3 | 0 | NON_SYNONYMOU | MODERAT | MISSENSE | Acg/Gcg | T412A | fiuA | . | C | . | . | . | . | | | | | C | C |
| NC_011770 | 523566 | A | C | 84 | 0 | 8 | 0 | NON_SYNONYMOU | MODERAT | MISSENSE | Tcc/Gcc | S386A | fiuA | C | C | C | C | C | C | | | | | C | C |
| NC_011770 | 523598 | A | T | 84 | 0 | 8 | 0 | NON_SYNONYMOU | MODERAT | MISSENSE | tTc/tAc | F375Y | fiuA | T | T | T | T | T | T | | | | | T | T |
| NC_011770 | 523600 | G | T | 84 | 0 | 8 | 0 | NON_SYNONYMOU | MODERAT | MISSENSE | gaC/gaA | D374E | fiuA | T | T | T | T | T | T | | | | | T | T |
| NC_011770 | 523638 | T | C | 74 | 0 | 8 | 0 | NON_SYNONYMOU | MODERAT | MISSENSE | Att/Gtt | I362V | fiuA | C | C | C | C | C | C | | | | | C | C |
| NC_011770 | 523694 | G | A | 64 | 0 | 8 | 0 | NON_SYNONYMOU | MODERAT | MISSENSE | gCc/gTc | A343V | fiuA | A | A | A | A | A | A | | | | | A | A |
| NC_011770 | 523695 | C | T | 64 | 0 | 8 | 0 | NON_SYNONYMOU | MODERAT | MISSENSE | Gcc/Acc | A343T | fiuA | T | T | T | T | T | T | | | | | T | T |
| NC_011770 | 523697 | G | C | 64 | 0 | 8 | 0 | NON_SYNONYMOU | MODERAT | MISSENSE | gCc/gGc | A342G | fiuA | C | C | C | C | C | C | | | | | C | C |
| NC_011770 | 523722 | T | C | 80 | 0 | 8 | 0 | NON_SYNONYMOU | MODERAT | MISSENSE | Agc/Ggc | S334G | fiuA | C | C | C | C | C | C | | | | | C | C |
| NC_011770 | 523839 | G | C | 81 | 0 | 10 | 0 | NON_SYNONYMOU | MODERAT | MISSENSE | Ctc/Gtc | L295V | fiuA | C | C | C | C | C | C | | | C | C | C | C |
| NC_011770 | 523869 | T | G | 80 | 0 | 10 | 0 | NON_SYNONYMOU | MODERAT | MISSENSE | Atc/Ctc | I285L | fiuA | G | G | G | G | G | G | | | G | G | G | G |
| NC_011770 | 523917 | T | C | 81 | 0 | 10 | 0 | NON_SYNONYMOU | MODERAT | MISSENSE | Atc/Gtc | I269V | fiuA | C | C | C | C | C | C | | | C | C | C | C |
| NC_011770 | 523929 | G | C | 74 | 0 | 10 | 0 | NON_SYNONYMOU | MODERAT | MISSENSE | Ctg/Gtg | L265V | fiuA | C | C | C | C | C | C | | | C | C | C | C |
| NC_011770 | 523944 | T | C | 74 | 0 | 10 | 0 | NON_SYNONYMOU | MODERAT | MISSENSE | Atc/Gtc | I260V | fiuA | C | C | C | C | C | C | | | C | C | C | C |
| NC_011770 | 523946 | C | T | 73 | 0 | 10 | 0 | NON_SYNONYMOU | MODERAT | MISSENSE | gGc/gAc | G259D | fiuA | T | T | T | T | T | T | | | T | T | T | T |
| NC_011770 | 523947 | C | T | 72 | 0 | 10 | 0 | NON_SYNONYMOU | MODERAT | MISSENSE | Ggc/Agc | G259S | fiuA | T | T | T | T | T | T | | | T | T | T | T |
| NC_011770 | 524025 | G | A | 78 | 0 | 8 | 0 | NON_SYNONYMOU | MODERAT | MISSENSE | Ctc/Ttc | L233F | fiuA | A | A | A | A | A | A | | | | | A | A |
| NC_011770 | 524034 | G | C | 75 | 0 | 8 | 0 | NON_SYNONYMOU | MODERAT | MISSENSE | Ctc/Gtc | L230V | fiuA | C | C | C | C | C | C | | | | | C | C |
| NC_011770 | 524075 | G | A | 64 | 0 | 8 | 0 | NON_SYNONYMOU | MODERAT | MISSENSE | tCa/tTa | S216L | fiuA | A | A | A | A | A | A | | | | | A | A |
| NC_011770 | 524076 | A | C | 64 | 0 | 8 | 0 | NON_SYNONYMOU | MODERAT | MISSENSE | Tca/Gca | S216A | fiuA | C | C | C | C | C | C | | | | | C | C |
| NC_011770 | 524081 | G | A | 62 | 0 | 8 | 0 | NON_SYNONYMOU | MODERAT | MISSENSE | aCa/aTa | T214I | fiuA | A | A | A | A | A | A | | | | | A | A |
| NC_011770 | 524082 | T | C | 63 | 0 | 8 | 0 | NON_SYNONYMOU | MODERAT | MISSENSE | Aca/Gca | T214A | fiuA | C | C | C | C | C | C | | | | | C | C |
| NC_011770 | 524151 | A | T | 64 | 0 | 8 | 0 | NON_SYNONYMOU | MODERAT | MISSENSE | Tcc/Acc | S191T | fiuA | T | T | T | T | T | T | | | | | T | T |
| NC_011770 | 5456147 | G | A | 73 | 0 | 1 | 0 | NON_SYNONYMOU | MODERAT | MISSENSE | gCc/gTc | A80V | fklB | | | | | | | | | | | | A |
| NC_011770 | 5456174 | G | A | 76 | 0 | 2 | 0 | NON_SYNONYMOU | MODERAT | MISSENSE | gCc/gTc | A71V | fklB | | | | | | | A | A | | | | |
| NC_011770 | 4658565 | T | C | 65 | 0 | 1 | 0 | NON_SYNONYMOU | MODERAT | MISSENSE | cAa/cGa | Q399R | fleQ | | | | | | | C | | | | | |
| NC_011770 | 4655799 | GGGCACC | G | 85 | 0 | 2 | 0 | CODON_CHANGE_ | MODERATE | | ccggtgccc/ | PVP386P | fleR | | | | | | | | | | | G | G |

| | | | | | | | | | | | | | | | | | | | | | | | | | |
|---|---|---|---|---|---|---|---|---|---|---|---|---|---|---|---|---|---|---|---|---|---|---|---|---|---|
| NC_011770 | 4656643 | G | A | 70 | 0 | 2 | 0 | NON_SYNONYMOU | MODERAT | MISSENSE | gCg/gTg | A107V | fleR | | | | | | | | | | | A | A |
| NC_011770 | 4656980 | A | G | 74 | 0 | 12 | 0 | NON_SYNONYMOU | MODERAT | MISSENSE | gTg/gCg | V399A | fleS | G | G | G | G | G | G | G | G | G | G | G | G |
| NC_011770 | 4657250 | G | A | 74 | 0 | 6 | 0 | NON_SYNONYMOU | MODERAT | MISSENSE | cCc/cTc | P309L | fleS | | | A | A | A | A | | | A | A | | |
| NC_011770 | 4658160 | T | G | 69 | 0 | 6 | 0 | NON_SYNONYMOU | MODERAT | MISSENSE | Aac/Cac | N6H | fleS | | | G | G | G | G | | | G | G | | |
| NC_011770 | 1840449 | G | A | 83 | 0 | 2 | 0 | NON_SYNONYMOU | MODERAT | MISSENSE | Ccg/Tcg | P145S | flgA | | | | | | | | | A | A | | |
| NC_011770 | 4682657 | C | T | 76 | 0 | 1 | 0 | START_LOST | HIGH | MISSENSE | atG/atA | M1I | flgC | | | | | | | T | | | | | |
| NC_011770 | 4682436 | T | C | 79 | 0 | 1 | 0 | NON_SYNONYMOU | MODERAT | MISSENSE | cAg/cGg | Q75R | flgC | | | | | | | C | | | | | |
| NC_011770 | 4681817 | G | C | 44 | 0 | 2 | 0 | NON_SYNONYMOU | MODERAT | MISSENSE | aaC/aaG | N130K | flgD | C | C | | | | | | | | | | |
| NC_011770 | 4680277 | A | C | 74 | 0 | 4 | 0 | NON_SYNONYMOU | MODERAT | MISSENSE | Tcg/Gcg | S397A | flgE | | | C | C | C | C | | | | | | |
| NC_011770 | 4678835 | T | C | 73 | 0 | 1 | 0 | NON_SYNONYMOU | MODERAT | MISSENSE | Acc/Gcc | T77A | flgG | | | | | | | C | | | | | |
| NC_011770 | 4677719 | C | T | 75 | 0 | 1 | 0 | NON_SYNONYMOU | MODERAT | MISSENSE | Ggc/Agc | G172S | flgH | | | | | | | | | | | T | |
| NC_011770 | 4675714 | C | T | 74 | 0 | 2 | 0 | NON_SYNONYMOU | MODERAT | MISSENSE | aGg/aAg | R231K | flgJ | | | | | | | T | T | | | | |
| NC_011770 | 4675895 | C | T | 65 | 0 | 4 | 0 | NON_SYNONYMOU | MODERAT | MISSENSE | Gcc/Acc | A171T | flgJ | | | | | | | | | T | T | T | T |
| NC_011770 | 4676057 | C | T | 76 | 0 | 4 | 0 | NON_SYNONYMOU | MODERAT | MISSENSE | Ggc/Agc | G117S | flgJ | T | T | | | | | T | T | | | | |
| NC_011770 | 4673168 | A | T | 72 | 0 | 6 | 0 | NON_SYNONYMOU | MODERAT | MISSENSE | Tcg/Acg | S673T | flgK | T | T | | | | | T | T | | | T | T |
| NC_011770 | 4673201 | T | C | 81 | 0 | 6 | 0 | NON_SYNONYMOU | MODERAT | MISSENSE | Agc/Ggc | S662G | flgK | C | C | | | | | C | C | | | C | C |
| NC_011770 | 4673307 | G | C | 64 | 0 | 6 | 0 | NON_SYNONYMOU | MODERAT | MISSENSE | gaC/gaG | D626E | flgK | C | C | | | | | C | C | | | C | C |
| NC_011770 | 4673311 | G | C | 65 | 0 | 6 | 0 | NON_SYNONYMOU | MODERAT | MISSENSE | gCc/gGc | A625G | flgK | C | C | | | | | C | C | | | C | C |
| NC_011770 | 4673312 | C | T | 65 | 0 | 6 | 0 | NON_SYNONYMOU | MODERAT | MISSENSE | Gcc/Acc | A625T | flgK | T | T | | | | | T | T | | | T | T |
| NC_011770 | 4673323 | G | A | 77 | 0 | 6 | 0 | NON_SYNONYMOU | MODERAT | MISSENSE | gCg/gTg | A621V | flgK | A | A | | | | | A | A | | | A | A |
| NC_011770 | 4673370 | G | C | 77 | 0 | 6 | 0 | NON_SYNONYMOU | MODERAT | MISSENSE | gaC/gaG | D605E | flgK | C | C | | | | | C | C | | | C | C |
| NC_011770 | 4673392 | C | T | 72 | 0 | 6 | 0 | NON_SYNONYMOU | MODERAT | MISSENSE | aGc/aAc | S598N | flgK | T | T | | | | | T | T | | | T | T |
| NC_011770 | 4673395 | T | C | 73 | 0 | 2 | 0 | NON_SYNONYMOU | MODERAT | MISSENSE | gAc/gGc | D597G | flgK | | | | | | | | | C | C | | |
| NC_011770 | 4673430 | A | C | 71 | 0 | 6 | 0 | NON_SYNONYMOU | MODERAT | MISSENSE | aaT/aaG | N585K | flgK | C | C | | | | | C | C | | | C | C |
| NC_011770 | 4673431 | T | A | 71 | 0 | 6 | 0 | NON_SYNONYMOU | MODERAT | MISSENSE | aAt/aTt | N585I | flgK | A | A | | | | | A | A | | | A | A |
| NC_011770 | 4673432 | T | G | 72 | 0 | 6 | 0 | NON_SYNONYMOU | MODERAT | MISSENSE | Aat/Cat | N585H | flgK | G | G | | | | | G | G | | | G | G |
| NC_011770 | 4673459 | C | A | 43 | 6 | 2 | 0 | NON_SYNONYMOU | MODERAT | MISSENSE | Gcg/Tcg | A576S | flgK | . | . | | | | | . | . | A | A | . | . |
| NC_011770 | 4673596 | T | G | 69 | 0 | 6 | 0 | NON_SYNONYMOU | MODERAT | MISSENSE | aAc/aCc | N530T | flgK | G | G | | | | | G | G | | | G | G |
| NC_011770 | 4673615 | G | T | 66 | 0 | 6 | 0 | NON_SYNONYMOU | MODERAT | MISSENSE | Ccc/Acc | P524T | flgK | T | T | | | | | T | T | | | T | T |
| NC_011770 | 4673669 | T | A | 66 | 0 | 6 | 0 | NON_SYNONYMOU | MODERAT | MISSENSE | Acc/Tcc | T506S | flgK | A | A | | | | | A | A | | | A | A |
| NC_011770 | 4673672 | T | G | 69 | 0 | 6 | 0 | NON_SYNONYMOU | MODERAT | MISSENSE | Atg/Ctg | M505L | flgK | G | G | | | | | G | G | | | G | G |
| NC_011770 | 4673699 | C | G | 69 | 0 | 6 | 0 | NON_SYNONYMOU | MODERAT | MISSENSE | Gtc/Ctc | V496L | flgK | G | G | | | | | G | G | | | G | G |
| NC_011770 | 4673773 | T | C | 49 | 4 | 6 | 0 | NON_SYNONYMOU | MODERAT | MISSENSE | gAc/gGc | D471G | flgK | . | . | C | C | C | C | | | C | C | . | . |
| NC_011770 | 4673879 | T | C | 79 | 0 | 6 | 0 | NON_SYNONYMOU | MODERAT | MISSENSE | Acg/Gcg | T436A | flgK | C | C | | | | | C | C | | | C | C |
| NC_011770 | 4673889 | G | C | 79 | 0 | 6 | 0 | NON_SYNONYMOU | MODERAT | MISSENSE | agC/agG | S432R | flgK | C | C | | | | | C | C | | | C | C |
| NC_011770 | 4673890 | C | G | 78 | 0 | 6 | 0 | NON_SYNONYMOU | MODERAT | MISSENSE | aGc/aCc | S432T | flgK | G | G | | | | | G | G | | | G | G |
| NC_011770 | 4673891 | T | C | 77 | 0 | 6 | 0 | NON_SYNONYMOU | MODERAT | MISSENSE | Agc/Ggc | S432G | flgK | C | C | | | | | C | C | | | C | C |
| NC_011770 | 4673957 | C | G | 85 | 0 | 6 | 0 | NON_SYNONYMOU | MODERAT | MISSENSE | Gtg/Ctg | V410L | flgK | G | G | | | | | G | G | | | G | G |
| NC_011770 | 4673959 | C | T | 86 | 0 | 6 | 0 | NON_SYNONYMOU | MODERAT | MISSENSE | aGc/aAc | S409N | flgK | T | T | | | | | T | T | | | T | T |
| NC_011770 | 4674070 | T | C | 92 | 0 | 4 | 0 | NON_SYNONYMOU | MODERAT | MISSENSE | aAt/aGt | N372S | flgK | C | C | | | | | | | | | C | C |
| NC_011770 | 4674272 | C | T | 76 | 0 | 6 | 0 | NON_SYNONYMOU | MODERAT | MISSENSE | Gtc/Atc | V305I | flgK | T | T | | | | | T | T | | | T | T |
| NC_011770 | 4674284 | A | G | 43 | 5 | 1 | 0 | NON_SYNONYMOU | MODERAT | MISSENSE | Ttc/Ctc | F301L | flgK | . | . | | | | | . | . | | | G | . |
| NC_011770 | 4674554 | C | T | 80 | 0 | 6 | 0 | NON_SYNONYMOU | MODERAT | MISSENSE | Gtc/Atc | V211I | flgK | T | T | | | | | T | T | | | T | T |
| NC_011770 | 4674596 | G | T | 82 | 0 | 6 | 0 | NON_SYNONYMOU | MODERAT | MISSENSE | Ctg/Atg | L197M | flgK | T | T | | | | | T | T | | | T | T |
| NC_011770 | 4674651 | C | G | 65 | 0 | 6 | 0 | NON_SYNONYMOU | MODERAT | MISSENSE | gaG/gaC | E178D | flgK | G | G | | | | | G | G | | | G | G |
| NC_011770 | 4674652 | T | C | 63 | 0 | 6 | 0 | NON_SYNONYMOU | MODERAT | MISSENSE | gAg/gGg | E178G | flgK | C | C | | | | | C | C | | | C | C |
| NC_011770 | 4674653 | C | T | 64 | 0 | 6 | 0 | NON_SYNONYMOU | MODERAT | MISSENSE | Gag/Aag | E178K | flgK | T | T | | | | | T | T | | | T | T |
| NC_011770 | 4674666 | G | C | 64 | 0 | 6 | 0 | NON_SYNONYMOU | MODERAT | MISSENSE | agC/agG | S173R | flgK | C | C | | | | | C | C | | | C | C |
| NC_011770 | 4674667 | C | G | 63 | 0 | 6 | 0 | NON_SYNONYMOU | MODERAT | MISSENSE | aGc/aCc | S173T | flgK | G | G | | | | | G | G | | | G | G |
| NC_011770 | 4674668 | T | A | 62 | 0 | 6 | 0 | NON_SYNONYMOU | MODERAT | MISSENSE | Agc/Tgc | S173C | flgK | A | A | | | | | A | A | | | A | A |
| NC_011770 | 4674674 | T | G | 61 | 0 | 6 | 0 | NON_SYNONYMOU | MODERAT | MISSENSE | Aat/Cat | N171H | flgK | G | G | | | | | G | G | | | G | G |
| NC_011770 | 4674689 | T | C | 61 | 0 | 6 | 0 | NON_SYNONYMOU | MODERAT | MISSENSE | Acc/Gcc | T166A | flgK | C | C | | | | | C | C | | | C | C |
| NC_011770 | 4674764 | T | C | 76 | 0 | 6 | 0 | NON_SYNONYMOU | MODERAT | MISSENSE | Agc/Ggc | S141G | flgK | C | C | | | | | C | C | | | C | C |
| NC_011770 | 4674773 | G | C | 74 | 0 | 6 | 0 | NON_SYNONYMOU | MODERAT | MISSENSE | Cag/Gag | Q138E | flgK | C | C | | | | | C | C | | | C | C |
| NC_011770 | 4674819 | C | G | 42 | 0 | 6 | 0 | NON_SYNONYMOU | MODERAT | MISSENSE | caG/caC | Q122H | flgK | G | G | | | | | G | G | | | G | G |
| NC_011770 | 4674821 | G | T | 42 | 0 | 6 | 0 | NON_SYNONYMOU | MODERAT | MISSENSE | Cag/Aag | Q122K | flgK | T | T | | | | | T | T | | | T | T |
| NC_011770 | 4674830 | T | C | 43 | 0 | 6 | 0 | NON_SYNONYMOU | MODERAT | MISSENSE | Acc/Gcc | T119A | flgK | C | C | | | | | C | C | | | C | C |
| NC_011770 | 4674842 | A | C | 41 | 0 | 6 | 0 | NON_SYNONYMOU | MODERAT | MISSENSE | Tcc/Gcc | S115A | flgK | C | C | | | | | C | C | | | C | C |
| NC_011770 | 4674880 | T | G | 44 | 0 | 6 | 0 | NON_SYNONYMOU | MODERAT | MISSENSE | aAc/aCc | N102T | flgK | G | G | | | | | G | G | | | G | G |
| NC_011770 | 4674884 | C | T | 43 | 0 | 6 | 0 | NON_SYNONYMOU | MODERAT | MISSENSE | Gat/Aat | D101N | flgK | T | T | | | | | T | T | | | T | T |
| NC_011770 | 4674887 | C | A | 43 | 0 | 6 | 0 | NON_SYNONYMOU | MODERAT | MISSENSE | Gca/Tca | A100S | flgK | A | A | | | | | A | A | | | A | A |
| NC_011770 | 4674895 | G | C | 44 | 0 | 6 | 0 | NON_SYNONYMOU | MODERAT | MISSENSE | tCg/tGg | S97W | flgK | C | C | | | | | C | C | | | C | C |

| | | | | | | | | | | | | | | | | | | | | | | | | | |
|---|---|---|---|---|---|---|---|---|---|---|---|---|---|---|---|---|---|---|---|---|---|---|---|---|---|
| NC_011770 | 4674896 | A | C | 42 | 0 | 6 | 0 | NON_SYNONYMOU | MODERAT | MISSENSE | Tcg/Gcg | S97A | flgK | C | C | | | | | C | C | | | C | C |
| NC_011770 | 4674899 | T | C | 43 | 0 | 6 | 0 | NON_SYNONYMOU | MODERAT | MISSENSE | Aac/Gac | N96D | flgK | C | C | | | | | C | C | | | C | C |
| NC_011770 | 4674906 | G | C | 42 | 0 | 6 | 0 | NON_SYNONYMOU | MODERAT | MISSENSE | gaC/gaG | D93E | flgK | C | C | | | | | C | C | | | C | C |
| NC_011770 | 4674917 | C | T | 42 | 0 | 6 | 0 | NON_SYNONYMOU | MODERAT | MISSENSE | Ggc/Agc | G90S | flgK | T | T | | | | | T | T | | | T | T |
| NC_011770 | 4674919 | A | C | 42 | 0 | 6 | 0 | NON_SYNONYMOU | MODERAT | MISSENSE | cTg/cGg | L89R | flgK | C | C | | | | | C | C | | | C | C |
| NC_011770 | 4674928 | T | C | 43 | 0 | 6 | 0 | NON_SYNONYMOU | MODERAT | MISSENSE | aAc/aGc | N86S | flgK | C | C | | | | | C | C | | | C | C |
| NC_011770 | 4675052 | T | C | 76 | 0 | 6 | 0 | NON_SYNONYMOU | MODERAT | MISSENSE | Atc/Gtc | I45V | flgK | C | C | | | | | C | C | | | C | C |
| NC_011770 | 4675054 | C | T | 77 | 0 | 6 | 0 | NON_SYNONYMOU | MODERAT | MISSENSE | cGg/cAg | R44Q | flgK | T | T | | | | | T | T | | | T | T |
| NC_011770 | 4675124 | C | T | 79 | 0 | 6 | 0 | NON_SYNONYMOU | MODERAT | MISSENSE | Gtc/Atc | V21I | flgK | T | T | | | | | T | T | | | T | T |
| NC_011770 | 4674709 | T | C | 84 | 0 | 1 | 0 | NON_SYNONYMOU | MODERAT | MISSENSE | aAc/aGc | N159S | flgK | | | | | | | | C | | | | |
| NC_011770 | 4671917 | G | C | 50 | 5 | 1 | 0 | NON_SYNONYMOU | MODERAT | MISSENSE | atC/atG | I397M | flgL | . | . | | | | | . | . | | | C | . |
| NC_011770 | 4671925 | C | A | 50 | 5 | 1 | 0 | NON_SYNONYMOU | MODERAT | MISSENSE | Gcc/Tcc | A395S | flgL | . | . | | | | | . | . | | | A | . |
| NC_011770 | 4671928 | T | C | 49 | 5 | 1 | 0 | NON_SYNONYMOU | MODERAT | MISSENSE | Acc/Gcc | T394A | flgL | . | . | | | | | . | . | | | C | . |
| NC_011770 | 4671939 | G | T | 50 | 5 | 1 | 0 | NON_SYNONYMOU | MODERAT | MISSENSE | aCc/aAc | T390N | flgL | . | . | | | | | . | . | | | T | . |
| NC_011770 | 4672506 | G | A | 42 | 6 | 2 | 0 | NON_SYNONYMOU | MODERAT | MISSENSE | cCg/cTg | P201L | flgL | . | . | | | | | . | . | A | A | . | . |
| NC_011770 | 4672518 | T | C | 41 | 6 | 2 | 0 | NON_SYNONYMOU | MODERAT | MISSENSE | gAc/gGc | D197G | flgL | . | . | | | | | . | . | C | C | . | . |
| NC_011770 | 4672813 | T | C | 41 | 5 | 1 | 0 | NON_SYNONYMOU | MODERAT | MISSENSE | Aac/Gac | N99D | flgL | . | . | | | | | C | . | | | . | . |
| NC_011770 | 4672824 | A | C | 42 | 5 | 1 | 0 | NON_SYNONYMOU | MODERAT | MISSENSE | gTg/gGg | V95G | flgL | . | . | | | | | C | . | | | . | . |
| NC_011770 | 4672844 | A | C | 43 | 5 | 1 | 0 | NON_SYNONYMOU | MODERAT | MISSENSE | caT/caG | H88Q | flgL | . | . | | | | | C | . | | | . | . |
| NC_011770 | 4672849 | T | G | 43 | 5 | 1 | 0 | NON_SYNONYMOU | MODERAT | MISSENSE | Atc/Ctc | I87L | flgL | . | . | | | | | G | . | | | . | . |
| NC_011770 | 4672854 | G | T | 40 | 5 | 1 | 0 | NON_SYNONYMOU | MODERAT | MISSENSE | aCg/aAg | T85K | flgL | . | . | | | | | T | . | | | . | . |
| NC_011770 | 4672866 | T | C | 41 | 5 | 1 | 0 | NON_SYNONYMOU | MODERAT | MISSENSE | aAt/aGt | N81S | flgL | . | . | | | | | C | . | | | . | . |
| NC_011770 | 4672886 | C | G | 40 | 5 | 1 | 0 | NON_SYNONYMOU | MODERAT | MISSENSE | caG/caC | Q74H | flgL | . | . | | | | | G | . | | | . | . |
| NC_011770 | 4672887 | T | C | 40 | 5 | 1 | 0 | NON_SYNONYMOU | MODERAT | MISSENSE | cAg/cGg | Q74R | flgL | . | . | | | | | C | . | | | . | . |
| NC_011770 | 4672888 | G | T | 40 | 5 | 1 | 0 | NON_SYNONYMOU | MODERAT | MISSENSE | Cag/Aag | Q74K | flgL | . | . | | | | | T | . | | | . | . |
| NC_011770 | 4673002 | G | T | 47 | 0 | 6 | 0 | NON_SYNONYMOU | MODERAT | MISSENSE | Ctg/Atg | L36M | flgL | T | T | | | | | T | T | | | T | T |
| NC_011770 | 1839605 | C | T | 69 | 0 | 6 | 0 | NON_SYNONYMOU | MODERAT | MISSENSE | aGc/aAc | S19N | flgN | | | T | T | T | T | T | T | | | | |
| NC_011770 | 4372513 | C | T | 75 | 0 | 2 | 0 | NON_SYNONYMOU | MODERAT | MISSENSE | Gtc/Atc | V26I | flhA | | | | | | | | | | | T | T |
| NC_011770 | 4369750 | ACGC | A | 83 | 0 | 1 | 0 | CODON_CHANGE_ | MODERATE | | ggcgta/gta | GV210V | flhF | A | | | | | | | | | | | |
| NC_011770 | 4369136 | T | G | 91 | 0 | 2 | 0 | NON_SYNONYMOU | MODERAT | MISSENSE | Atg/Ctg | M416L | flhF | | | | | | | G | G | | | | |
| NC_011770 | 4369784 | T | G | 75 | 0 | 2 | 0 | NON_SYNONYMOU | MODERAT | MISSENSE | Acc/Ccc | T200P | flhF | | | | | | | | | G | G | | |
| NC_011770 | 4370131 | G | T | 64 | 0 | 8 | 0 | NON_SYNONYMOU | MODERAT | MISSENSE | aCc/aAc | T84N | flhF | T | T | T | T | T | T | | | | | T | T |
| NC_011770 | 4370044 | G | A | 75 | 0 | 1 | 0 | NON_SYNONYMOU | MODERAT | MISSENSE | cCc/cTc | P113L | flhF | | | | | | | | A | | | | |
| NC_011770 | 4367628 | A | C | 93 | 0 | 4 | 0 | NON_SYNONYMOU | MODERAT | MISSENSE | Ttg/Gtg | L163V | fliA | C | C | | | | | | | C | C | | |
| NC_011770 | 4664182 | G | C | 50 | 0 | 6 | 0 | STOP_GAINED | HIGH | NONSENSE | taC/taG | Y36* | fliC | C | C | | | | | C | C | | | C | C |
| NC_011770 | 4662846 | C | A | 62 | 0 | 6 | 0 | NON_SYNONYMOU | MODERAT | MISSENSE | Gcg/Tcg | A482S | fliC | A | A | | | | | A | A | | | A | A |
| NC_011770 | 4662912 | A | T | 59 | 0 | 6 | 0 | NON_SYNONYMOU | MODERAT | MISSENSE | Tcg/Acg | S460T | fliC | T | T | | | | | T | T | | | T | T |
| NC_011770 | 4662917 | G | T | 56 | 0 | 6 | 0 | NON_SYNONYMOU | MODERAT | MISSENSE | gCg/gAg | A458E | fliC | T | T | | | | | T | T | | | T | T |
| NC_011770 | 4662918 | C | T | 55 | 0 | 6 | 0 | NON_SYNONYMOU | MODERAT | MISSENSE | Gcg/Acg | A458T | fliC | T | T | | | | | T | T | | | T | T |
| NC_011770 | 4662999 | C | T | 52 | 0 | 6 | 0 | NON_SYNONYMOU | MODERAT | MISSENSE | Gac/Aac | D431N | fliC | T | T | | | | | T | T | | | T | T |
| NC_011770 | 4663009 | C | G | 51 | 0 | 6 | 0 | NON_SYNONYMOU | MODERAT | MISSENSE | aaG/aaC | K427N | fliC | G | G | | | | | G | G | | | G | G |
| NC_011770 | 4663011 | T | C | 51 | 0 | 6 | 0 | NON_SYNONYMOU | MODERAT | MISSENSE | Aag/Gag | K427E | fliC | C | C | | | | | C | C | | | C | C |
| NC_011770 | 4663242 | T | C | 40 | 6 | 2 | 0 | NON_SYNONYMOU | MODERAT | MISSENSE | Atc/Gtc | I350V | fliC | . | . | | | | | . | . | C | C | . | . |
| NC_011770 | 4663857 | T | C | 58 | 0 | 6 | 0 | NON_SYNONYMOU | MODERAT | MISSENSE | Acc/Gcc | T145A | fliC | C | C | | | | | C | C | | | C | C |
| NC_011770 | 4663859 | G | A | 58 | 0 | 6 | 0 | NON_SYNONYMOU | MODERAT | MISSENSE | aCc/aTc | T144I | fliC | A | A | | | | | A | A | | | A | A |
| NC_011770 | 4663860 | T | C | 57 | 0 | 6 | 0 | NON_SYNONYMOU | MODERAT | MISSENSE | Acc/Gcc | T144A | fliC | C | C | | | | | C | C | | | C | C |
| NC_011770 | 4663966 | G | C | 38 | 4 | 2 | 0 | NON_SYNONYMOU | MODERAT | MISSENSE | gaC/gaG | D108E | fliC | . | . | | | | | . | C | | | . | C |
| NC_011770 | 4663971 | C | A | 39 | 3 | 3 | 0 | NON_SYNONYMOU | MODERAT | MISSENSE | Gcc/Tcc | A107S | fliC | A | A | | | | | . | . | | | . | A |
| NC_011770 | 4664004 | C | A | 53 | 0 | 6 | 0 | NON_SYNONYMOU | MODERAT | MISSENSE | Gcc/Tcc | A96S | fliC | A | A | | | | | A | A | | | A | A |
| NC_011770 | 4664014 | G | C | 55 | 0 | 6 | 0 | NON_SYNONYMOU | MODERAT | MISSENSE | atC/atG | I92M | fliC | C | C | | | | | C | C | | | C | C |
| NC_011770 | 4664093 | C | T | 49 | 0 | 6 | 0 | NON_SYNONYMOU | MODERAT | MISSENSE | cGc/cAc | R66H | fliC | T | T | | | | | T | T | | | T | T |
| NC_011770 | 4664094 | G | T | 49 | 0 | 6 | 0 | NON_SYNONYMOU | MODERAT | MISSENSE | Cgc/Agc | R66S | fliC | T | T | | | | | T | T | | | T | T |
| NC_011770 | 4664114 | C | T | 49 | 0 | 6 | 0 | NON_SYNONYMOU | MODERAT | MISSENSE | aGc/aAc | S59N | fliC | T | T | | | | | T | T | | | T | T |
| NC_011770 | 4664118 | T | C | 50 | 0 | 6 | 0 | NON_SYNONYMOU | MODERAT | MISSENSE | Atc/Gtc | I58V | fliC | C | C | | | | | C | C | | | C | C |
| NC_011770 | 4664123 | T | C | 50 | 0 | 6 | 0 | NON_SYNONYMOU | MODERAT | MISSENSE | aAc/aGc | N56S | fliC | C | C | | | | | C | C | | | C | C |
| NC_011770 | 4664127 | A | T | 50 | 0 | 6 | 0 | NON_SYNONYMOU | MODERAT | MISSENSE | Tcc/Acc | S55T | fliC | T | T | | | | | T | T | | | T | T |
| NC_011770 | 4664139 | A | C | 52 | 0 | 6 | 0 | NON_SYNONYMOU | MODERAT | MISSENSE | Tcc/Gcc | S51A | fliC | C | C | | | | | C | C | | | C | C |
| NC_011770 | 4664183 | T | G | 50 | 0 | 6 | 0 | NON_SYNONYMOU | MODERAT | MISSENSE | tAc/tCc | Y36S | fliC | G | G | | | | | G | G | | | G | G |
| NC_011770 | 4664193 | T | A | 50 | 0 | 6 | 0 | NON_SYNONYMOU | MODERAT | MISSENSE | Acc/Tcc | T33S | fliC | A | A | | | | | A | A | | | A | A |
| NC_011770 | 4661196 | C | G | 44 | 4 | 2 | 0 | NON_SYNONYMOU | MODERAT | MISSENSE | gaG/gaC | E364D | fliD | . | . | | | | | . | . | | | G | G |
| NC_011770 | 4661438 | C | T | 69 | 0 | 6 | 0 | NON_SYNONYMOU | MODERAT | MISSENSE | Gtc/Atc | V284I | fliD | T | T | | | | | T | T | | | T | T |
| NC_011770 | 4655194 | A | G | 84 | 0 | 10 | 0 | NON_SYNONYMOU | MODERAT | MISSENSE | gTg/gCg | V34A | fliE | G | G | G | G | G | G | | | G | G | G | G |

| | | | | | | | | | | | | | | | | | | | | | | | | | |
|---|---|---|---|---|---|---|---|---|---|---|---|---|---|---|---|---|---|---|---|---|---|---|---|---|---|
| NC_011770 | 4655212 | A | G | 84 | 0 | 2 | 0 | NON_SYNONYMOU | MODERAT | MISSENSE | gTt/gCt | V28A | fliE | | | | | | | | | | | G | G |
| NC_011770 | 4653528 | A | G | 77 | 0 | 12 | 0 | NON_SYNONYMOU | MODERAT | MISSENSE | gTc/gCc | V472A | fliF | G | G | G | G | G | G | G | G | G | G | G | G |
| NC_011770 | 4654425 | A | T | 86 | 0 | 2 | 0 | NON_SYNONYMOU | MODERAT | MISSENSE | gTg/gAg | V173E | fliF | | | | | | | T | T | | | | |
| NC_011770 | 4652245 | T | C | 75 | 0 | 1 | 0 | NON_SYNONYMOU | MODERAT | MISSENSE | aAg/aGg | K299R | fliG | C | | | | | | | | | | | |
| NC_011770 | 4652746 | G | A | 87 | 0 | 1 | 0 | NON_SYNONYMOU | MODERAT | MISSENSE | cCg/cTg | P132L | fliG | | | | | | | A | | | | | |
| NC_011770 | 4651348 | A | G | 62 | 0 | 12 | 0 | NON_SYNONYMOU | MODERAT | MISSENSE | Ttc/Ctc | F259L | fliH | G | G | G | G | G | G | G | G | G | G | G | G |
| NC_011770 | 4651977 | G | A | 56 | 0 | 4 | 0 | NON_SYNONYMOU | MODERAT | MISSENSE | gCc/gTc | A49V | fliH | | | A | A | A | A | | | | | | |
| NC_011770 | 4652094 | C | T | 65 | 0 | 12 | 0 | NON_SYNONYMOU | MODERAT | MISSENSE | aGt/aAt | S10N | fliH | T | T | T | T | T | T | T | T | T | T | T | T |
| NC_011770 | 4650063 | G | A | 79 | 0 | 2 | 0 | NON_SYNONYMOU | MODERAT | MISSENSE | Cgc/Tgc | R422C | fliI | | | | | | | | | A | A | | |
| NC_011770 | 4649713 | G | A | 82 | 0 | 2 | 0 | NON_SYNONYMOU | MODERAT | MISSENSE | gCc/gTc | A82V | fliJ | | | | | | | | | | | A | A |
| NC_011770 | 4379529 | G | A | 96 | 0 | 2 | 0 | NON_SYNONYMOU | MODERAT | MISSENSE | Cac/Tac | H275Y | fliM | A | A | | | | | | | | | | |
| NC_011770 | 4378886 | A | G | 83 | 0 | 1 | 0 | NON_SYNONYMOU | MODERAT | MISSENSE | cTg/cCg | L156P | fliN | | | | | | | G | | | | | |
| NC_011770 | 4379211 | G | A | 86 | 0 | 2 | 0 | NON_SYNONYMOU | MODERAT | MISSENSE | Ccg/Tcg | P48S | fliN | | | | | | | | | | | A | A |
| NC_011770 | 4378588 | T | G | 84 | 0 | 2 | 0 | NON_SYNONYMOU | MODERAT | MISSENSE | cAg/cCg | Q97P | fliO | | | | | | | | | | | G | G |
| NC_011770 | 4377951 | C | T | 72 | 0 | 2 | 0 | NON_SYNONYMOU | MODERAT | MISSENSE | Gtc/Atc | V160I | fliP | | | | | | | | | T | T | | |
| NC_011770 | 4377396 | A | G | 72 | 0 | 1 | 0 | NON_SYNONYMOU | MODERAT | MISSENSE | aTg/aCg | M74T | fliQ | | | | | | | G | | | | | |
| NC_011770 | 4376779 | G | A | 74 | 0 | 2 | 0 | NON_SYNONYMOU | MODERAT | MISSENSE | gCg/gTg | A190V | fliR | | | | | | | | | | | A | A |
| NC_011770 | 4377185 | C | T | 79 | 0 | 2 | 0 | NON_SYNONYMOU | MODERAT | MISSENSE | Gtg/Atg | V55M | fliR | | | | | | | | | T | T | | |
| NC_011770 | 4660530 | G | C | 46 | 5 | 1 | 0 | NON_SYNONYMOU | MODERAT | MISSENSE | gCg/gGg | A71G | fliS | . | . | | | | | C | . | | | . | . |
| NC_011770 | 4660549 | T | C | 60 | 0 | 6 | 0 | NON_SYNONYMOU | MODERAT | MISSENSE | Atc/Gtc | I65V | fliS | C | C | | | | | C | C | | | C | C |
| NC_011770 | 4660554 | C | G | 62 | 0 | 6 | 0 | NON_SYNONYMOU | MODERAT | MISSENSE | gGc/gCc | G63A | fliS | G | G | | | | | G | G | | | G | G |
| NC_011770 | 5170705 | T | C | 52 | 0 | 12 | 0 | NON_SYNONYMOU | MODERAT | MISSENSE | Ttc/Ctc | F7L | flp | C | C | C | C | C | C | C | C | C | C | C | C |
| NC_011770 | 5170720 | C | G | 52 | 0 | 4 | 0 | NON_SYNONYMOU | MODERAT | MISSENSE | Ctt/Gtt | L12V | flp | | | G | G | G | G | | | | | | |
| NC_011770 | 5170870 | A | G | 51 | 0 | 12 | 0 | NON_SYNONYMOU | MODERAT | MISSENSE | Acc/Gcc | T62A | flp | G | G | G | G | G | G | G | G | G | G | G | G |
| NC_011770 | 5170871 | C | A | 50 | 0 | 12 | 0 | NON_SYNONYMOU | MODERAT | MISSENSE | aCc/aAc | T62N | flp | A | A | A | A | A | A | A | A | A | A | A | A |
| NC_011770 | 20141 | A | C | 79 | 0 | 6 | 0 | NON_SYNONYMOU | MODERAT | MISSENSE | Ttg/Gtg | L287V | fmt | | | | | | | C | C | C | C | C | C |
| NC_011770 | 20239 | C | T | 85 | 0 | 2 | 0 | NON_SYNONYMOU | MODERAT | MISSENSE | gGg/gAg | G254E | fmt | | | | | | | | | | | T | T |
| NC_011770 | 20414 | T | C | 87 | 0 | 4 | 0 | NON_SYNONYMOU | MODERAT | MISSENSE | Atc/Gtc | I196V | fmt | | | C | C | C | C | | | | | | |
| NC_011770 | 20459 | T | C | 85 | 0 | 12 | 0 | NON_SYNONYMOU | MODERAT | MISSENSE | Atc/Gtc | I181V | fmt | C | C | C | C | C | C | C | C | C | C | C | C |
| NC_011770 | 20839 | T | C | 91 | 0 | 12 | 0 | NON_SYNONYMOU | MODERAT | MISSENSE | aAc/aGc | N54S | fmt | C | C | C | C | C | C | C | C | C | C | C | C |
| NC_011770 | 20954 | C | A | 80 | 0 | 2 | 0 | NON_SYNONYMOU | MODERAT | MISSENSE | Gcc/Tcc | A16S | fmt | | | | | | | | | | | A | A |
| NC_011770 | 387256 | G | C | 74 | 0 | 2 | 0 | NON_SYNONYMOU | MODERAT | MISSENSE | gGc/gCc | G70A | folA | | | | | | | | | C | C | | |
| NC_011770 | 387502 | T | C | 63 | 0 | 10 | 0 | NON_SYNONYMOU | MODERAT | MISSENSE | gTc/gCc | V152A | folA | | | C | C | C | C | C | C | C | C | C | C |
| NC_011770 | 2102749 | G | A | 87 | 0 | 2 | 0 | NON_SYNONYMOU | MODERAT | MISSENSE | Gtc/Atc | V91I | folC | | | | | | | A | A | | | | |
| NC_011770 | 3921272 | G | A | 42 | 0 | 2 | 0 | NON_SYNONYMOU | MODERAT | MISSENSE | Gac/Aac | D78N | folD | | | | | | | | | | | A | A |
| NC_011770 | 5644722 | G | A | 79 | 0 | 1 | 0 | NON_SYNONYMOU | MODERAT | MISSENSE | Gcg/Acg | A60T | folK | | | | | | | | A | | | | |
| NC_011770 | 5644947 | G | A | 84 | 0 | 2 | 0 | NON_SYNONYMOU | MODERAT | MISSENSE | Gac/Aac | D135N | folK | A | A | | | | | | | | | | |
| NC_011770 | 5645001 | T | C | 85 | 0 | 2 | 0 | NON_SYNONYMOU | MODERAT | MISSENSE | Ttc/Ctc | F153L | folK | | | | | | | C | C | | | | |
| NC_011770 | 5669070 | T | C | 91 | 0 | 8 | 0 | NON_SYNONYMOU | MODERAT | MISSENSE | Aat/Gat | N177D | folP | C | C | | | | | C | C | C | C | C | C |
| NC_011770 | 5669432 | C | A | 82 | 0 | 2 | 0 | NON_SYNONYMOU | MODERAT | MISSENSE | cGg/cTg | R56L | folP | | | | | | | A | A | | | | |
| NC_011770 | 3043511 | C | T | 65 | 0 | 1 | 0 | NON_SYNONYMOU | MODERAT | MISSENSE | gCc/gTc | A35V | foxA | | | | | | | T | | | | | |
| NC_011770 | 3044380 | G | A | 69 | 0 | 10 | 0 | NON_SYNONYMOU | MODERAT | MISSENSE | Gtc/Atc | V325I | foxA | A | A | A | A | A | A | A | A | | | A | A |
| NC_011770 | 3045049 | G | A | 75 | 0 | 12 | 0 | NON_SYNONYMOU | MODERAT | MISSENSE | Gtc/Atc | V548I | foxA | A | A | A | A | A | A | A | A | A | A | A | A |
| NC_011770 | 3045054 | G | C | 74 | 0 | 12 | 0 | NON_SYNONYMOU | MODERAT | MISSENSE | gaG/gaC | E549D | foxA | C | C | C | C | C | C | C | C | C | C | C | C |
| NC_011770 | 3041854 | GCGCGGC( | G | 94 | 0 | 2 | 0 | CODON_DELETION | MODERATE | | gcggcggac, | AAD39 | foxI | | | | | G | G | | | | | | |
| NC_011770 | 3042360 | C | G | 89 | 0 | 2 | 0 | NON_SYNONYMOU | MODERAT | MISSENSE | gCc/gGc | A28G | foxR | | | | | | | | | G | G | | |
| NC_011770 | 3042460 | A | C | 90 | 0 | 2 | 0 | NON_SYNONYMOU | MODERAT | MISSENSE | caA/caC | Q61H | foxR | | | | | | | | | C | C | | |
| NC_011770 | 3043095 | C | G | 63 | 0 | 2 | 0 | NON_SYNONYMOU | MODERAT | MISSENSE | gCc/gGc | A273G | foxR | | | | | | | | | G | G | | |
| NC_011770 | 3043178 | G | A | 65 | 0 | 2 | 0 | NON_SYNONYMOU | MODERAT | MISSENSE | Gac/Aac | D301N | foxR | | | | | | | | | A | A | | |
| NC_011770 | 5159986 | A | G | 66 | 0 | 2 | 0 | NON_SYNONYMOU | MODERAT | MISSENSE | aAg/aGg | K113R | fppA | | | | | | | | | G | G | | |
| NC_011770 | 5160011 | G | T | 66 | 0 | 2 | 0 | NON_SYNONYMOU | MODERAT | MISSENSE | aaG/aaT | K121N | fppA | | | | | | | | | | | T | T |
| NC_011770 | 5160046 | C | T | 66 | 0 | 2 | 0 | NON_SYNONYMOU | MODERAT | MISSENSE | gCa/gTa | A133V | fppA | T | T | | | | | | | | | | |
| NC_011770 | 5160087 | A | G | 65 | 0 | 2 | 0 | NON_SYNONYMOU | MODERAT | MISSENSE | Agc/Ggc | S147G | fppA | | | | | | | | | G | G | | |
| NC_011770 | 5160118 | G | A | 68 | 0 | 2 | 0 | NON_SYNONYMOU | MODERAT | MISSENSE | aGt/aAt | S157N | fppA | | | | | | | | | A | A | | |
| NC_011770 | 767131 | T | G | 84 | 0 | 2 | 0 | NON_SYNONYMOU | MODERAT | MISSENSE | Tcc/Gcc | S281A | fptA | | | | | | | | | G | G | | |
| NC_011770 | 767345 | C | T | 78 | 0 | 6 | 0 | NON_SYNONYMOU | MODERAT | MISSENSE | aCg/aTg | T352M | fptA | T | T | T | T | T | T | | | | | | |
| NC_011770 | 3177448 | G | C | 12 | 8 | 4 | 0 | NON_SYNONYMOU | MODERAT | MISSENSE | aGc/aCc | S234T | fpvA | . | . | C | C | C | C | . | . | . | . | . | . |
| NC_011770 | 3182669 | G | C | 13 | 8 | 4 | 0 | NON_SYNONYMOU | MODERAT | MISSENSE | aGc/aCc | S234T | fpvA2 | . | . | C | C | C | C | . | . | . | . | . | . |
| NC_011770 | 3187890 | G | C | 10 | 8 | 4 | 0 | NON_SYNONYMOU | MODERAT | MISSENSE | aGc/aCc | S234T | fpvA3 | . | . | C | C | C | C | . | . | . | . | . | . |
| NC_011770 | 826962 | T | C | 85 | 0 | 10 | 0 | NON_SYNONYMOU | MODERAT | MISSENSE | aAc/aGc | N760S | fpvB | C | C | C | C | C | C | C | C | C | C | | |
| NC_011770 | 827043 | T | C | 79 | 0 | 2 | 0 | NON_SYNONYMOU | MODERAT | MISSENSE | cAg/cGg | Q733R | fpvB | | | | | | | | | | | C | C |
| NC_011770 | 827230 | T | C | 68 | 0 | 2 | 0 | NON_SYNONYMOU | MODERAT | MISSENSE | Agc/Ggc | S671G | fpvB | | | | | | | | | | | C | C |

| | | | | | | | | | | | | | | | | | | | | | | | | | |
|---|---|---|---|---|---|---|---|---|---|---|---|---|---|---|---|---|---|---|---|---|---|---|---|---|---|
| NC_011770 | 827787 | A | G | 77 | 0 | 12 | 0 | NON_SYNONYMOU | MODERAT | MISSENSE | cTc/cCc | L485P | fpvB | G | G | G | G | G | G | G | G | G | G | G | G |
| NC_011770 | 1592089 | T | G | 73 | 0 | 12 | 0 | NON_SYNONYMOU | MODERAT | MISSENSE | Tgg/Ggg | W246G | fruA | G | G | G | G | G | G | G | G | G | G | G | G |
| NC_011770 | 1592203 | A | T | 69 | 0 | 4 | 0 | NON_SYNONYMOU | MODERAT | MISSENSE | Acg/Tcg | T284S | fruA | | | T | T | T | T | | | | | | |
| NC_011770 | 1592910 | C | G | 79 | 0 | 2 | 0 | NON_SYNONYMOU | MODERAT | MISSENSE | ttC/ttG | F519L | fruA | | | G | G | | | | | | | | |
| NC_011770 | 1590762 | G | T | 63 | 0 | 2 | 0 | NON_SYNONYMOU | MODERAT | MISSENSE | Gcg/Tcg | A119S | fruK | | | | | | | T | T | | | | |
| NC_011770 | 1590966 | G | C | 72 | 0 | 2 | 0 | NON_SYNONYMOU | MODERAT | MISSENSE | Gtc/Ctc | V187L | fruK | | | | | | | | | C | C | | |
| NC_011770 | 1591012 | C | T | 72 | 0 | 12 | 0 | NON_SYNONYMOU | MODERAT | MISSENSE | gCc/gTc | A202V | fruK | T | T | T | T | T | T | T | T | T | T | T | T |
| NC_011770 | 1591046 | G | T | 72 | 0 | 6 | 0 | NON_SYNONYMOU | MODERAT | MISSENSE | caG/caT | Q213H | fruK | | | T | T | T | T | | | | | T | T |
| NC_011770 | 1591075 | A | G | 69 | 0 | 2 | 0 | NON_SYNONYMOU | MODERAT | MISSENSE | cAg/cGg | Q223R | fruK | G | G | | | | | | | | | | |
| NC_011770 | 1586257 | G | A | 78 | 0 | 2 | 0 | NON_SYNONYMOU | MODERAT | MISSENSE | Cac/Tac | H319Y | fruR | | | | | | | | | A | A | | |
| NC_011770 | 1586275 | C | T | 75 | 0 | 2 | 0 | NON_SYNONYMOU | MODERAT | MISSENSE | Gac/Aac | D313N | fruR | | | | | | | | | T | T | | |
| NC_011770 | 412708 | A | C | 99 | 0 | 4 | 0 | NON_SYNONYMOU | MODERAT | MISSENSE | cAc/cCc | H26P | ftsE | | | C | C | C | C | | | | | | |
| NC_011770 | 412884 | T | C | 91 | 0 | 2 | 0 | NON_SYNONYMOU | MODERAT | MISSENSE | Ttc/Ctc | F85L | ftsE | | | | | | | | | C | C | | |
| NC_011770 | 5671008 | G | A | 70 | 0 | 1 | 0 | NON_SYNONYMOU | MODERAT | MISSENSE | Cgc/Tgc | R174C | ftsH | | | | | | | | A | | | | |
| NC_011770 | 5287867 | C | T | 86 | 0 | 2 | 0 | NON_SYNONYMOU | MODERAT | MISSENSE | gGc/gAc | G63D | ftsI | | T | | | | | T | | | | | |
| NC_011770 | 5287259 | G | A | 88 | 0 | 1 | 0 | NON_SYNONYMOU | MODERAT | MISSENSE | Ccc/Tcc | P266S | ftsI | | | | A | | | | | | | | |
| NC_011770 | 5286545 | A | G | 81 | 0 | 12 | 0 | NON_SYNONYMOU | MODERAT | MISSENSE | Tgc/Cgc | C504R | ftsI | G | G | G | G | G | G | G | G | G | G | G | G |
| NC_011770 | 5287330 | T | C | 89 | 0 | 1 | 0 | NON_SYNONYMOU | MODERAT | MISSENSE | aAc/aGc | N242S | ftsI | | | | | | | | | | | | |
| NC_011770 | 5671956 | C | A | 70 | 0 | 2 | 0 | NON_SYNONYMOU | MODERAT | MISSENSE | Gcc/Tcc | A132S | ftsJ | | | | | | | | | A | A | | |
| NC_011770 | 5672236 | C | A | 83 | 0 | 2 | 0 | NON_SYNONYMOU | MODERAT | MISSENSE | gaG/gaT | E38D | ftsJ | | | | | | | | | | | A | A |
| NC_011770 | 2680386 | A | G | 74 | 0 | 2 | 0 | NON_SYNONYMOU | MODERAT | MISSENSE | aAc/aGc | N477S | ftsK | G | G | | | | | | | | | | |
| NC_011770 | 5275712 | C | T | 78 | 0 | 1 | 0 | NON_SYNONYMOU | MODERAT | MISSENSE | Gtc/Atc | V225I | ftsQ | | | | | | | | | | | T | |
| NC_011770 | 411535 | A | G | 38 | 0 | 6 | 0 | NON_SYNONYMOU | MODERAT | MISSENSE | Agt/Ggt | S91G | ftsY | G | G | G | G | G | G | | | | | | |
| NC_011770 | 411577 | G | C | 41 | 0 | 2 | 0 | NON_SYNONYMOU | MODERAT | MISSENSE | Gag/Cag | E105Q | ftsY | | | | | | | | | | | C | C |
| NC_011770 | 411613 | GCCGAGCC | G | 41 | 0 | 1 | 0 | CODON_CHANGE_ | MODERATE | | gagcccgag | EPEPV128V | ftsY | | | | | | | | | | | | **G** |
| NC_011770 | 411503 | A | AGCC | 36 | 0 | 5 | 0 | CODON_INSERTIOI | MODERATE | | gcg/CCGgc | A82PA | ftsY | AGCC | AGCC | | AGCC | AGCC | AGCC | | | | | | |
| NC_011770 | 411357 | GCAGCCTG | G | 46 | 0 | 6 | 0 | CODON_CHANGE_ | MODERATE | | gccgatcagc | ADQPV38V | ftsY | G | G | | | | G | G | | G | | G | |
| NC_011770 | 5273165 | C | T | 81 | 0 | 1 | 0 | NON_SYNONYMOU | MODERAT | MISSENSE | cGc/cAc | R344H | ftsZ | | | | | | | T | | | | | |
| NC_011770 | 5273190 | T | C | 86 | 0 | 10 | 0 | NON_SYNONYMOU | MODERAT | MISSENSE | Acc/Gcc | T336A | ftsZ | | | C | C | C | C | C | C | C | C | C | C |
| NC_011770 | 5333169 | G | A | 88 | 0 | 2 | 0 | NON_SYNONYMOU | MODERAT | MISSENSE | Cgc/Tgc | R153C | fumC | | | | | | | | | A | A | | |
| NC_011770 | 5333279 | C | A | 97 | 0 | 2 | 0 | NON_SYNONYMOU | MODERAT | MISSENSE | cGt/cTt | R116L | fumC | A | A | | | | | | | | | | |
| NC_011770 | 4904516 | G | T | 54 | 0 | 2 | 0 | NON_SYNONYMOU | MODERAT | MISSENSE | Ggc/Tgc | G81C | fumC2 | | | | | | | T | T | | | | |
| NC_011770 | 722567 | A | G | 95 | 0 | 2 | 0 | NON_SYNONYMOU | MODERAT | MISSENSE | gAc/gGc | D213G | fusA1 | | | | | G | G | | | | | | |
| NC_011770 | 723370 | G | A | 79 | 0 | 1 | 0 | NON_SYNONYMOU | MODERAT | MISSENSE | Gcc/Acc | A481T | fusA1 | | | | | | | A | | | | | |
| NC_011770 | 723692 | G | A | 82 | 0 | 12 | 0 | NON_SYNONYMOU | MODERAT | MISSENSE | gGc/gAc | G588D | fusA1 | A | A | A | A | A | A | A | A | A | A | A | A |
| NC_011770 | 723782 | C | T | 84 | 0 | 2 | 0 | NON_SYNONYMOU | MODERAT | MISSENSE | cCg/cTg | P618L | fusA1 | T | T | | | | | | | | | | |
| NC_011770 | 723871 | A | G | 74 | 0 | 1 | 0 | NON_SYNONYMOU | MODERAT | MISSENSE | Atg/Gtg | M648V | fusA1 | | | | | | | G | | | | | |
| NC_011770 | 723967 | C | T | 73 | 0 | 1 | 0 | NON_SYNONYMOU | MODERAT | MISSENSE | Cgc/Tgc | R680C | fusA1 | | | | | | | T | | | | | |
| NC_011770 | 723700 | T | G | 84 | 0 | 1 | 0 | NON_SYNONYMOU | MODERAT | MISSENSE | Tcc/Gcc | S591A | fusA1 | | | | | | G | | | | | | |
| NC_011770 | 721991 | C | T | 92 | 0 | 1 | 0 | NON_SYNONYMOU | MODERAT | MISSENSE | gCg/gTg | A21V | fusA1 | | | | | | | | | T | | | |
| NC_011770 | 722167 | G | A | 81 | 0 | 1 | 0 | NON_SYNONYMOU | MODERAT | MISSENSE | Gac/Aac | D80N | fusA1 | | | | | | | | **A** | | | | |
| NC_011770 | 722518 | G | A | 91 | 0 | 1 | 0 | NON_SYNONYMOU | MODERAT | MISSENSE | Gac/Aac | D197N | fusA1 | | | | | | | | **A** | | | | |
| NC_011770 | 723296 | C | T | 86 | 0 | 1 | 0 | NON_SYNONYMOU | MODERAT | MISSENSE | aCc/aTc | T456I | fusA1 | | | | | | | | T | | | | |
| NC_011770 | 3591246 | C | T | 60 | 0 | 2 | 0 | NON_SYNONYMOU | MODERAT | MISSENSE | Gtc/Atc | V697I | fusA2 | | | | | | | T | T | | | | |
| NC_011770 | 3591532 | T | G | 73 | 0 | 4 | 0 | NON_SYNONYMOU | MODERAT | MISSENSE | caA/caC | Q601H | fusA2 | | | G | G | G | G | | | | | | |
| NC_011770 | 3591900 | C | T | 48 | 2 | 2 | 0 | NON_SYNONYMOU | MODERAT | MISSENSE | Ggc/Agc | G479S | fusA2 | | | | | . | . | T | T | | | | |
| NC_011770 | 3592043 | C | T | 59 | 2 | 10 | 0 | NON_SYNONYMOU | MODERAT | MISSENSE | aGg/aAg | R431K | fusA2 | T | T | T | T | . | . | T | T | T | T | T | T |
| NC_011770 | 3592745 | G | T | 79 | 0 | 2 | 0 | NON_SYNONYMOU | MODERAT | MISSENSE | gCc/gAc | A197D | fusA2 | | | | | | | | | | | T | T |
| NC_011770 | 3592809 | C | A | 79 | 0 | 4 | 0 | NON_SYNONYMOU | MODERAT | MISSENSE | Gcg/Tcg | A176S | fusA2 | A | A | | | | | | | A | A | | |
| NC_011770 | 295149 | G | C | 75 | 0 | 4 | 0 | NON_SYNONYMOU | MODERAT | MISSENSE | Gtc/Ctc | V328L | gabT | | | C | C | C | C | | | | | | |
| NC_011770 | 4824309 | G | A | 71 | 0 | 2 | 0 | NON_SYNONYMOU | MODERAT | MISSENSE | Gac/Aac | D558N | gacS | | | | | | | | | | | A | A |
| NC_011770 | 4825212 | C | G | 87 | 0 | 2 | 0 | NON_SYNONYMOU | MODERAT | MISSENSE | Cac/Gac | H859D | gacS | | | | | | | | | | | G | G |
| NC_011770 | 3650006 | G | A | 62 | 0 | 1 | 0 | STOP_GAINED | HIGH | NONSENSE | Cag/Tag | Q239* | galU | | | | | A | | | | | | | |
| NC_011770 | 3650012 | G | A | 63 | 0 | 1 | 0 | STOP_GAINED | HIGH | NONSENSE | Cag/Tag | Q237* | galU | | | | | | A | | | | | | |
| NC_011770 | 2014579 | C | A | 84 | 0 | 4 | 0 | NON_SYNONYMOU | MODERAT | MISSENSE | Ggc/Tgc | G274C | gapA | | | A | A | A | A | | | | | | |
| NC_011770 | 2014644 | C | T | 89 | 0 | 4 | 0 | NON_SYNONYMOU | MODERAT | MISSENSE | cGg/cAg | R252Q | gapA | | | T | T | T | T | | | | | | |
| NC_011770 | 2015009 | G | T | 85 | 0 | 2 | 0 | NON_SYNONYMOU | MODERAT | MISSENSE | gaC/gaA | D130E | gapA | T | T | | | | | | | | | | |
| NC_011770 | 2015232 | C | T | 85 | 0 | 2 | 0 | NON_SYNONYMOU | MODERAT | MISSENSE | cGc/cAc | R56H | gapA | T | T | | | | | | | | | | |
| NC_011770 | 2015325 | T | C | 90 | 0 | 2 | 0 | NON_SYNONYMOU | MODERAT | MISSENSE | cAc/cGc | H25R | gapA | | | | | | | | | | | C | C |
| NC_011770 | 5347999 | G | A | 63 | 0 | 12 | 0 | NON_SYNONYMOU | MODERAT | MISSENSE | Gcc/Acc | A110T | gatA | A | A | A | A | A | A | A | A | A | A | A | A |
| NC_011770 | 5348091 | C | G | 70 | 0 | 8 | 0 | NON_SYNONYMOU | MODERAT | MISSENSE | aaC/aaG | N140K | gatA | G | G | | | | | G | G | G | G | G | G |
| NC_011770 | 5348503 | G | T | 85 | 0 | 2 | 0 | NON_SYNONYMOU | MODERAT | MISSENSE | Gtg/Ttg | V278L | gatA | | | | | | | T | T | | | | |

| | | | | | | | | | | | | | | | | | | | | | | | | | |
|---|---|---|---|---|---|---|---|---|---|---|---|---|---|---|---|---|---|---|---|---|---|---|---|---|---|
| NC_011770 | 5349573 | C | G | 80 | 0 | 6 | 0 | NON_SYNONYMOU | MODERAT | MISSENSE | aCc/aGc | T114S | gatB | | | G | G | G | G | G | G | | | | |
| NC_011770 | 5349848 | G | A | 77 | 0 | 1 | 0 | NON_SYNONYMOU | MODERAT | MISSENSE | Gcc/Acc | A206T | gatB | | | | | | | A | | | | | |
| NC_011770 | 5350394 | C | A | 51 | 0 | 8 | 0 | NON_SYNONYMOU | MODERAT | MISSENSE | Ctc/Atc | L388I | gatB | A | A | A | A | A | A | A | A | | | | |
| NC_011770 | 5347655 | A | C | 48 | 0 | 8 | 0 | STOP_LOST+SPLICE | HIGH | MISSENSE | tAa/tCa | *96S | gatC | C | C | | | | | C | C | C | C | C | C |
| NC_011770 | 2135314 | T | C | 72 | 0 | 6 | 0 | NON_SYNONYMOU | MODERAT | MISSENSE | gTc/gCc | V22A | gbt | C | C | | | | | C | C | | | C | C |
| NC_011770 | 2135682 | A | G | 77 | 0 | 2 | 0 | NON_SYNONYMOU | MODERAT | MISSENSE | Acc/Gcc | T145A | gbt | | | | | | | | | G | G | | |
| NC_011770 | 2136085 | T | C | 76 | 0 | 12 | 0 | NON_SYNONYMOU | MODERAT | MISSENSE | cTc/cCc | L279P | gbt | C | C | C | C | C | C | C | C | C | C | C | C |
| NC_011770 | 4405906 | G | A | 43 | 0 | 2 | 0 | NON_SYNONYMOU | MODERAT | MISSENSE | Ggc/Agc | G79S | gbuA | | | | | | | | | | | A | A |
| NC_011770 | 4406122 | G | A | 58 | 0 | 2 | 0 | NON_SYNONYMOU | MODERAT | MISSENSE | Gtc/Atc | V151I | gbuA | | | | | | | | | | | A | A |
| NC_011770 | 4406512 | A | G | 71 | 0 | 4 | 0 | NON_SYNONYMOU | MODERAT | MISSENSE | Atc/Gtc | I281V | gbuA | | | | | | | G | G | | | G | G |
| NC_011770 | 4404730 | C | T | 84 | 0 | 1 | 0 | NON_SYNONYMOU | MODERAT | MISSENSE | Gcc/Acc | A246T | gbuR | | | | | | | T | | | | | |
| NC_011770 | 4404903 | C | T | 81 | 0 | 2 | 0 | NON_SYNONYMOU | MODERAT | MISSENSE | aGa/aAa | R188K | gbuR | | | | | | | | | | | T | T |
| NC_011770 | 4405215 | C | T | 72 | 0 | 8 | 0 | NON_SYNONYMOU | MODERAT | MISSENSE | gGa/gAa | G84E | gbuR | T | T | T | T | T | T | | | T | T | | |
| NC_011770 | 3323203 | G | A | 75 | 0 | 2 | 0 | NON_SYNONYMOU | MODERAT | MISSENSE | cGc/cAc | R6H | gcd | | | | | | | A | A | | | | |
| NC_011770 | 3323534 | C | T | 77 | 0 | 4 | 0 | NON_SYNONYMOU | MODERAT | MISSENSE | gaC/gaG | D116E | gcd | | | | | | | T | T | G | G | | |
| NC_011770 | 3324060 | A | G | 63 | 0 | 12 | 0 | NON_SYNONYMOU | MODERAT | MISSENSE | Acg/Gcg | T292A | gcd | G | G | G | G | G | G | G | G | G | G | G | G |
| NC_011770 | 3324061 | C | A | 64 | 0 | 2 | 0 | NON_SYNONYMOU | MODERAT | MISSENSE | aCg/aAg | T292K | gcd | | | | | | | | | A | A | | |
| NC_011770 | 3324235 | G | A | 55 | 0 | 1 | 0 | NON_SYNONYMOU | MODERAT | MISSENSE | gGc/gAc | G350D | gcd | | | | | | | A | | | | | |
| NC_011770 | 3325023 | T | G | 72 | 0 | 2 | 0 | NON_SYNONYMOU | MODERAT | MISSENSE | Tcg/Gcg | S613A | gcd | | | | | | | | | G | G | | |
| NC_011770 | 3325480 | C | T | 46 | 0 | 2 | 0 | NON_SYNONYMOU | MODERAT | MISSENSE | cCg/cTg | P765L | gcd | | | T | T | | | | | | | | |
| NC_011770 | 495348 | A | G | 68 | 0 | 2 | 0 | NON_SYNONYMOU | MODERAT | MISSENSE | gTg/gCg | V274A | gcdH | G | G | | | | | | | | | | |
| NC_011770 | 496123 | C | T | 77 | 0 | 1 | 0 | NON_SYNONYMOU | MODERAT | MISSENSE | Gac/Aac | D16N | gcdH | | | | | | | | **T** | | | | |
| NC_011770 | 4320945 | T | C | 51 | 0 | 1 | 0 | NON_SYNONYMOU | MODERAT | MISSENSE | cTg/cCg | L257P | gcl | | | | | | | C | | | | | |
| NC_011770 | 4321791 | G | A | 78 | 0 | 2 | 0 | NON_SYNONYMOU | MODERAT | MISSENSE | cGc/cAc | R539H | gcl | A | A | | | | | | | | | | |
| NC_011770 | 630824 | G | A | 72 | 0 | 2 | 0 | NON_SYNONYMOU | MODERAT | MISSENSE | cGg/cAg | R127Q | gcp | A | A | | | | | | | | | | |
| NC_011770 | 630916 | G | A | 69 | 0 | 6 | 0 | NON_SYNONYMOU | MODERAT | MISSENSE | Gtg/Atg | V158M | gcp | | | A | A | A | A | | | A | A | | |
| NC_011770 | 631010 | T | G | 60 | 0 | 2 | 0 | NON_SYNONYMOU | MODERAT | MISSENSE | cTc/cGc | L189R | gcp | | | | | | | | | | | G | G |
| NC_011770 | 3087235 | C | T | 71 | 0 | 2 | 0 | NON_SYNONYMOU | MODERAT | MISSENSE | Ctc/Ttc | L44F | gcvH2 | | | | | | | | | | | T | T |
| NC_011770 | 6205268 | G | A | 68 | 0 | 1 | 0 | STOP_GAINED | HIGH | NONSENSE | Cag/Tag | Q573* | gcvP1 | | | | | | | A | | | | | |
| NC_011770 | 6204214 | T | C | 68 | 0 | 4 | 0 | NON_SYNONYMOU | MODERAT | MISSENSE | tAt/tGt | Y924C | gcvP1 | | | C | C | C | C | | | | | | |
| NC_011770 | 6204215 | A | G | 68 | 0 | 12 | 0 | NON_SYNONYMOU | MODERAT | MISSENSE | Tat/Cat | Y924H | gcvP1 | G | G | G | G | G | G | G | G | G | G | G | G |
| NC_011770 | 6205109 | A | G | 64 | 0 | 4 | 0 | NON_SYNONYMOU | MODERAT | MISSENSE | Tgc/Cgc | C626R | gcvP1 | | | | | | | G | G | | | G | G |
| NC_011770 | 6205556 | C | T | 77 | 0 | 2 | 0 | NON_SYNONYMOU | MODERAT | MISSENSE | Gtg/Atg | V477M | gcvP1 | | | T | T | | | | | | | | |
| NC_011770 | 6205744 | C | T | 72 | 0 | 2 | 0 | NON_SYNONYMOU | MODERAT | MISSENSE | cGc/cAc | R414H | gcvP1 | T | T | | | | | | | | | | |
| NC_011770 | 6205793 | G | A | 69 | 0 | 2 | 0 | NON_SYNONYMOU | MODERAT | MISSENSE | Cgt/Tgt | R398C | gcvP1 | A | A | | | | | | | | | | |
| NC_011770 | 6205859 | C | T | 71 | 0 | 4 | 0 | NON_SYNONYMOU | MODERAT | MISSENSE | Gaa/Aaa | E376K | gcvP1 | | | T | T | T | T | | | | | | |
| NC_011770 | 6205867 | G | A | 71 | 0 | 2 | 0 | NON_SYNONYMOU | MODERAT | MISSENSE | gCg/gTg | A373V | gcvP1 | A | A | | | | | | | | | | |
| NC_011770 | 6206693 | A | G | 65 | 0 | 10 | 0 | NON_SYNONYMOU | MODERAT | MISSENSE | Tcg/Ccg | S98P | gcvP1 | G | G | G | G | G | G | G | G | | | G | G |
| NC_011770 | 6206800 | G | A | 70 | 0 | 2 | 0 | NON_SYNONYMOU | MODERAT | MISSENSE | gCc/gTc | A62V | gcvP1 | | | | | | | A | A | | | | |
| NC_011770 | 6206962 | T | G | 61 | 0 | 10 | 0 | NON_SYNONYMOU | MODERAT | MISSENSE | gAc/gCc | D8A | gcvP1 | | | G | G | G | G | G | G | G | G | G | G |
| NC_011770 | 3088124 | G | A | 67 | 0 | 2 | 0 | NON_SYNONYMOU | MODERAT | MISSENSE | Gcc/Acc | A209T | gcvP2 | | | | | | | | | A | A | | |
| NC_011770 | 3088539 | T | A | 78 | 0 | 12 | 0 | NON_SYNONYMOU | MODERAT | MISSENSE | aTc/aAc | I347N | gcvP2 | A | A | A | A | A | A | A | A | A | A | A | A |
| NC_011770 | 3088868 | G | A | 79 | 0 | 4 | 0 | NON_SYNONYMOU | MODERAT | MISSENSE | Gcc/Acc | A457T | gcvP2 | | | | | | | A | A | A | A | | |
| NC_011770 | 3090152 | C | T | 51 | 0 | 2 | 0 | NON_SYNONYMOU | MODERAT | MISSENSE | Cgc/Tgc | R885C | gcvP2 | | | | | | | | | T | T | | |
| NC_011770 | 3089426 | A | G | 86 | 0 | 1 | 0 | NON_SYNONYMOU | MODERAT | MISSENSE | Agc/Ggc | S643G | gcvP2 | | | | | | | | G | | | | |
| NC_011770 | 6207857 | C | A | 62 | 0 | 2 | 0 | NON_SYNONYMOU | MODERAT | MISSENSE | Gcc/Tcc | A274S | gcvT | | | | | | | A | A | | | | |
| NC_011770 | 3093903 | C | T | 38 | 0 | 4 | 0 | NON_SYNONYMOU | MODERAT | MISSENSE | gCc/gTc | A203V | gcvT2 | | | T | T | T | T | | | | | | |
| NC_011770 | 5472883 | G | A | 82 | 0 | 4 | 0 | NON_SYNONYMOU | MODERAT | MISSENSE | gCc/gTc | A335V | gdhA | | | A | A | A | A | | | | | | |
| NC_011770 | 5473415 | C | T | 79 | 0 | 2 | 0 | NON_SYNONYMOU | MODERAT | MISSENSE | Gcc/Acc | A158T | gdhA | | | | | | | | | | | T | T |
| NC_011770 | 5473602 | G | T | 78 | 0 | 4 | 0 | NON_SYNONYMOU | MODERAT | MISSENSE | ttC/ttA | F95L | gdhA | | | T | T | T | T | | | | | | |
| NC_011770 | 2154269 | A | G | 92 | 0 | 4 | 0 | NON_SYNONYMOU | MODERAT | MISSENSE | aAg/aGg | K176R | gdhB | | | G | G | G | G | | | | | | |
| NC_011770 | 2154340 | C | T | 89 | 0 | 2 | 0 | NON_SYNONYMOU | MODERAT | MISSENSE | Cgc/Tgc | R200C | gdhB | | | T | T | | | | | | | | |
| NC_011770 | 2154467 | G | C | 84 | 0 | 2 | 0 | NON_SYNONYMOU | MODERAT | MISSENSE | aGc/aCc | S242T | gdhB | | | | | | | C | C | | | | |
| NC_011770 | 2154694 | G | C | 82 | 0 | 2 | 0 | NON_SYNONYMOU | MODERAT | MISSENSE | Ggc/Cgc | G318R | gdhB | C | C | | | | | | | | | | |
| NC_011770 | 2155018 | C | T | 63 | 0 | 1 | 0 | NON_SYNONYMOU | MODERAT | MISSENSE | Ccg/Tcg | P426S | gdhB | | | | | | | T | | | | | |
| NC_011770 | 2156525 | G | A | 74 | 0 | 1 | 0 | NON_SYNONYMOU | MODERAT | MISSENSE | gGc/gAc | G928D | gdhB | | | | | | | A | | | | | |
| NC_011770 | 2157455 | G | A | 77 | 0 | 12 | 0 | NON_SYNONYMOU | MODERAT | MISSENSE | aGg/aAg | R1238K | gdhB | A | A | A | A | A | A | A | A | A | A | A | A |
| NC_011770 | 2157670 | A | G | 72 | 0 | 1 | 0 | NON_SYNONYMOU | MODERAT | MISSENSE | Acc/Gcc | T1310A | gdhB | | | | | | | G | | | | | |
| NC_011770 | 2158048 | A | G | 71 | 0 | 2 | 0 | NON_SYNONYMOU | MODERAT | MISSENSE | Atc/Gtc | I1436V | gdhB | | | | | | | G | G | | | | |
| NC_011770 | 2154218 | A | G | 96 | 0 | 1 | 0 | NON_SYNONYMOU | MODERAT | MISSENSE | tAc/tGc | Y159C | gdhB | | | | | | | | G | | | | |
| NC_011770 | 4250312 | C | T | 47 | 0 | 2 | 0 | NON_SYNONYMOU | MODERAT | MISSENSE | Cct/Tct | P3S | ggt | | | | | | | | | | | T | T |
| NC_011770 | 4251090 | T | C | 77 | 0 | 1 | 0 | NON_SYNONYMOU | MODERAT | MISSENSE | cTc/cCc | L262P | ggt | | | | | | | C | | | | | |

| | | | | | | | | | | | | | | | | | | | | | | | | | |
|---|---|---|---|---|---|---|---|---|---|---|---|---|---|---|---|---|---|---|---|---|---|---|---|---|---|
| NC_011770 | 4251288 | T | A | 61 | 0 | 2 | 0 | NON_SYNONYMOU | MODERAT | MISSENSE | gTg/gAg | V328E | ggt | A | A | | | | | | | | | | |
| NC_011770 | 4251530 | G | A | 59 | 0 | 1 | 0 | NON_SYNONYMOU | MODERAT | MISSENSE | Gcc/Acc | A409T | ggt | | | | | | | A | | | | | |
| NC_011770 | 4251594 | G | T | 63 | 0 | 1 | 0 | NON_SYNONYMOU | MODERAT | MISSENSE | gGc/gTc | G430V | ggt | | | | | | | T | | | | | |
| NC_011770 | 6594693 | G | A | 71 | 0 | 1 | 0 | NON_SYNONYMOU | MODERAT | MISSENSE | Cgc/Tgc | R624C | gidA | | | | | | | A | | | | | |
| NC_011770 | 6594891 | C | A | 68 | 0 | 10 | 0 | NON_SYNONYMOU | MODERAT | MISSENSE | Gcc/Tcc | A558S | gidA | A | A | A | A | A | A | A | A | A | A | | |
| NC_011770 | 6595139 | C | T | 79 | 0 | 2 | 0 | NON_SYNONYMOU | MODERAT | MISSENSE | cGc/cAc | R475H | gidA | | | | | | | | | T | T | | |
| NC_011770 | 6595814 | C | A | 78 | 0 | 2 | 0 | NON_SYNONYMOU | MODERAT | MISSENSE | cGg/cTg | R250L | gidA | | | | | | | | | | | A | A |
| NC_011770 | 535047 | C | T | 83 | 0 | 1 | 0 | NON_SYNONYMOU | MODERAT | MISSENSE | gGc/gAc | G529D | glcB | | | | | | | T | | | | | |
| NC_011770 | 535527 | G | C | 75 | 0 | 2 | 0 | NON_SYNONYMOU | MODERAT | MISSENSE | aCc/aGc | T369S | glcB | | | C | C | | | | | | | | |
| NC_011770 | 6360070 | A | G | 85 | 0 | 12 | 0 | NON_SYNONYMOU | MODERAT | MISSENSE | Aac/Gac | N73D | glcC | G | G | G | G | G | G | G | G | G | G | G | G |
| NC_011770 | 6360086 | T | C | 85 | 0 | 6 | 0 | NON_SYNONYMOU | MODERAT | MISSENSE | gTc/gCc | V78A | glcC | C | C | C | C | C | C | | | | | | |
| NC_011770 | 6360259 | A | G | 110 | 0 | 6 | 0 | NON_SYNONYMOU | MODERAT | MISSENSE | Acc/Gcc | T136A | glcC | G | G | G | G | G | G | | | | | | |
| NC_011770 | 6358190 | T | C | 65 | 0 | 4 | 0 | NON_SYNONYMOU | MODERAT | MISSENSE | cAc/cGc | H487R | glcD | | | C | C | C | C | | | | | | |
| NC_011770 | 6359301 | A | G | 88 | 0 | 10 | 0 | NON_SYNONYMOU | MODERAT | MISSENSE | Tcg/Ccg | S117P | glcD | G | G | G | G | G | G | G | G | G | G | | |
| NC_011770 | 6359559 | C | G | 93 | 0 | 2 | 0 | NON_SYNONYMOU | MODERAT | MISSENSE | Gac/Cac | D31H | glcD | G | G | | | | | | | | | | |
| NC_011770 | 6357163 | G | A | 81 | 0 | 2 | 0 | NON_SYNONYMOU | MODERAT | MISSENSE | Ccg/Tcg | P330S | glcE | A | A | | | | | | | | | | |
| NC_011770 | 6357229 | C | T | 78 | 0 | 4 | 0 | NON_SYNONYMOU | MODERAT | MISSENSE | Gcc/Acc | A308T | glcE | | | T | T | T | T | | | | | | |
| NC_011770 | 6357232 | C | T | 76 | 0 | 6 | 0 | NON_SYNONYMOU | MODERAT | MISSENSE | Gcc/Acc | A307T | glcE | T | T | | | | | T | T | T | T | | |
| NC_011770 | 6357419 | C | A | 76 | 0 | 2 | 0 | NON_SYNONYMOU | MODERAT | MISSENSE | gaG/gaT | E244D | glcE | A | A | | | | | | | | | | |
| NC_011770 | 6357567 | G | A | 82 | 0 | 2 | 0 | NON_SYNONYMOU | MODERAT | MISSENSE | aCg/aTg | T195M | glcE | A | A | | | | | | | | | | |
| NC_011770 | 6355846 | C | T | 91 | 0 | 4 | 0 | NON_SYNONYMOU | MODERAT | MISSENSE | Gga/Aga | G406R | glcF | | | T | T | T | T | | | | | | |
| NC_011770 | 6355888 | G | A | 89 | 0 | 4 | 0 | NON_SYNONYMOU | MODERAT | MISSENSE | Ccg/Tcg | P392S | glcF | | | A | A | A | A | | | | | | |
| NC_011770 | 6355938 | A | G | 95 | 0 | 10 | 0 | NON_SYNONYMOU | MODERAT | MISSENSE | gTc/gCc | V375A | glcF | | | G | G | G | G | G | G | G | G | G | G |
| NC_011770 | 6355995 | C | T | 95 | 0 | 2 | 0 | NON_SYNONYMOU | MODERAT | MISSENSE | cGg/cAg | R356Q | glcF | T | T | | | | | | | | | | |
| NC_011770 | 6356097 | G | A | 93 | 0 | 4 | 0 | NON_SYNONYMOU | MODERAT | MISSENSE | gCg/gTg | A322V | glcF | | | A | A | A | A | | | | | | |
| NC_011770 | 6356185 | A | T | 92 | 0 | 2 | 0 | NON_SYNONYMOU | MODERAT | MISSENSE | Tcc/Acc | S293T | glcF | | | | | | | | | | | T | T |
| NC_011770 | 6356244 | C | T | 92 | 0 | 4 | 0 | NON_SYNONYMOU | MODERAT | MISSENSE | cGc/cAc | R273H | glcF | | | | | | | | | T | T | T | T |
| NC_011770 | 6356329 | C | T | 92 | 0 | 2 | 0 | NON_SYNONYMOU | MODERAT | MISSENSE | Gtg/Atg | V245M | glcF | T | T | | | | | | | | | | |
| NC_011770 | 6356658 | A | G | 66 | 0 | 12 | 0 | NON_SYNONYMOU | MODERAT | MISSENSE | aTt/aCt | I135T | glcF | G | G | G | G | G | G | G | G | G | G | G | G |
| NC_011770 | 6356659 | T | C | 65 | 0 | 8 | 0 | NON_SYNONYMOU | MODERAT | MISSENSE | Att/Gtt | I135V | glcF | C | C | C | C | C | C | C | C | | | | |
| NC_011770 | 6357004 | G | T | 81 | 0 | 2 | 0 | NON_SYNONYMOU | MODERAT | MISSENSE | Cgc/Agc | R20S | glcF | | | | | | | | | | | T | T |
| NC_011770 | 3483548 | A | G | 87 | 0 | 12 | 0 | NON_SYNONYMOU | MODERAT | MISSENSE | Acc/Gcc | T309A | glgA | G | G | G | G | G | G | G | G | G | G | G | G |
| NC_011770 | 3483630 | A | G | 84 | 0 | 1 | 0 | NON_SYNONYMOU | MODERAT | MISSENSE | cAg/cGg | Q336R | glgA | | | | | | | G | | | | | |
| NC_011770 | 3483545 | C | T | 88 | 0 | 1 | 0 | NON_SYNONYMOU | MODERAT | MISSENSE | Cgc/Tgc | R308C | glgA | | | | | | | | | | | T | |
| NC_011770 | 3499203 | C | T | 92 | 0 | 1 | 0 | STOP_GAINED | HIGH | NONSENSE | tgG/tgA | W633* | glgB | | | | | | | T | | | | | |
| NC_011770 | 3500369 | G | A | 77 | 0 | 4 | 0 | NON_SYNONYMOU | MODERAT | MISSENSE | Cgc/Tgc | R245C | glgB | | | A | A | A | A | | | | | | |
| NC_011770 | 3500402 | G | A | 74 | 0 | 2 | 0 | NON_SYNONYMOU | MODERAT | MISSENSE | Cgc/Tgc | R234C | glgB | | | | | | | | | | | A | A |
| NC_011770 | 3500840 | G | T | 62 | 0 | 2 | 0 | NON_SYNONYMOU | MODERAT | MISSENSE | Cgc/Agc | R88S | glgB | T | T | | | | | | | | | | |
| NC_011770 | 3511914 | C | A | 80 | 0 | 2 | 0 | NON_SYNONYMOU | MODERAT | MISSENSE | gGg/gTg | G745V | glgP | | | | | | | | | | | A | A |
| NC_011770 | 3513023 | C | T | 92 | 0 | 2 | 0 | NON_SYNONYMOU | MODERAT | MISSENSE | atG/atA | M375I | glgP | T | T | | | | | | | | | | |
| NC_011770 | 3513139 | C | T | 94 | 0 | 1 | 0 | NON_SYNONYMOU | MODERAT | MISSENSE | Gtg/Atg | V337M | glgP | | | | | | | T | | | | | |
| NC_011770 | 2018387 | G | A | 73 | 0 | 2 | 0 | NON_SYNONYMOU | MODERAT | MISSENSE | Gcc/Acc | A311T | glk | | | | | | | | | | | A | A |
| NC_011770 | 5667589 | C | T | 66 | 0 | 6 | 0 | NON_SYNONYMOU | MODERAT | MISSENSE | aGc/aAc | S381N | glmM | T | T | | | | | T | T | | | T | T |
| NC_011770 | 6582344 | C | T | 78 | 0 | 6 | 0 | NON_SYNONYMOU | MODERAT | MISSENSE | Gcc/Acc | A253T | glmR | T | T | | | | | T | T | T | T | | |
| NC_011770 | 6582392 | G | A | 81 | 0 | 8 | 0 | NON_SYNONYMOU | MODERAT | MISSENSE | Cct/Tct | P237S | glmR | A | A | A | A | A | A | | | A | A | | |
| NC_011770 | 6582572 | G | T | 81 | 0 | 12 | 0 | NON_SYNONYMOU | MODERAT | MISSENSE | Ctc/Atc | L177I | glmR | T | T | T | T | T | T | T | T | T | T | T | T |
| NC_011770 | 6580892 | C | T | 81 | 0 | 1 | 0 | NON_SYNONYMOU | MODERAT | MISSENSE | Ggc/Agc | G474S | glmS | | | | | | | | T | | | | |
| NC_011770 | 6582029 | G | C | 83 | 0 | 2 | 0 | NON_SYNONYMOU | MODERAT | MISSENSE | Ctg/Gtg | L95V | glmS | | | | | | | C | C | | | | |
| NC_011770 | 6584193 | T | C | 90 | 0 | 12 | 0 | NON_SYNONYMOU | MODERAT | MISSENSE | cAc/cGc | H288R | glmU | C | C | C | C | C | C | C | C | C | C | C | C |
| NC_011770 | 6584545 | C | A | 94 | 0 | 2 | 0 | NON_SYNONYMOU | MODERAT | MISSENSE | Gcg/Tcg | A171S | glmU | | | | | | | | | A | A | | |
| NC_011770 | 1483663 | C | T | 93 | 0 | 2 | 0 | NON_SYNONYMOU | MODERAT | MISSENSE | gGc/gAc | G175D | glnD | | | | | | | T | T | | | | |
| NC_011770 | 1483672 | C | T | 93 | 0 | 4 | 0 | NON_SYNONYMOU | MODERAT | MISSENSE | cGa/cAa | R172Q | glnD | | | T | T | T | T | | | | | | |
| NC_011770 | 1483972 | T | C | 80 | 0 | 2 | 0 | NON_SYNONYMOU | MODERAT | MISSENSE | gAc/gGc | D72G | glnD | | | | | | | | | C | C | | |
| NC_011770 | 1484045 | C | T | 77 | 0 | 2 | 0 | NON_SYNONYMOU | MODERAT | MISSENSE | Gac/Aac | D48N | glnD | | | | | | | | | T | T | | |
| NC_011770 | 5968336 | C | G | 82 | 0 | 4 | 0 | NON_SYNONYMOU | MODERAT | MISSENSE | caG/caC | Q738H | glnE | | | G | G | G | G | | | | | | |
| NC_011770 | 5968809 | T | C | 69 | 0 | 2 | 0 | NON_SYNONYMOU | MODERAT | MISSENSE | Acc/Gcc | T581A | glnE | | | | | | | | | | | C | C |
| NC_011770 | 5969900 | A | T | 79 | 0 | 12 | 0 | NON_SYNONYMOU | MODERAT | MISSENSE | cTg/cAg | L217Q | glnE | T | T | T | T | T | T | T | T | T | T | T | T |
| NC_011770 | 5970298 | C | G | 66 | 0 | 2 | 0 | NON_SYNONYMOU | MODERAT | MISSENSE | caG/caC | Q84H | glnE | G | G | | | | | | | | | | |
| NC_011770 | 3923506 | T | C | 70 | 0 | 10 | 0 | NON_SYNONYMOU | MODERAT | MISSENSE | Acc/Gcc | T504A | glnS | C | C | C | C | C | C | | | C | C | C | C |
| NC_011770 | 3924505 | C | T | 72 | 0 | 2 | 0 | NON_SYNONYMOU | MODERAT | MISSENSE | Gcc/Acc | A171T | glnS | | | | | | | | | | | T | T |
| NC_011770 | 3924525 | G | A | 75 | 0 | 1 | 0 | NON_SYNONYMOU | MODERAT | MISSENSE | gCc/gTc | A164V | glnS | | | | | | | A | | | | | |
| NC_011770 | 3924664 | C | T | 71 | 0 | 4 | 0 | NON_SYNONYMOU | MODERAT | MISSENSE | Gcc/Acc | A118T | glnS | | | T | T | T | T | | | | | | |

| | | | | | | | | | | | | | | | | | | | | | | | | | |
|---|---|---|---|---|---|---|---|---|---|---|---|---|---|---|---|---|---|---|---|---|---|---|---|---|---|
| NC_011770 | 3924679 | T | C | 70 | 0 | 4 | 0 | NON_SYNONYMOU | MODERAT | MISSENSE | Atc/Gtc | I113V | glnS | | | | | | | C | C | C | C | | |
| NC_011770 | 5072701 | C | T | 84 | 0 | 2 | 0 | NON_SYNONYMOU | MODERAT | MISSENSE | cGc/cAc | R98H | gloA2 | T | T | | | | | | | | | | |
| NC_011770 | 5072755 | T | C | 80 | 0 | 2 | 0 | NON_SYNONYMOU | MODERAT | MISSENSE | gAg/gGg | E80G | gloA2 | C | C | | | | | | | | | | |
| NC_011770 | 5072794 | G | C | 86 | 0 | 4 | 0 | NON_SYNONYMOU | MODERAT | MISSENSE | aCc/aGc | T67S | gloA2 | | | | | | | | | C | C | C | C |
| NC_011770 | 5072801 | C | T | 82 | 0 | 4 | 0 | NON_SYNONYMOU | MODERAT | MISSENSE | Ggc/Agc | G65S | gloA2 | | | | | | | | | T | T | T | T |
| NC_011770 | 5072846 | G | A | 77 | 0 | 2 | 0 | NON_SYNONYMOU | MODERAT | MISSENSE | Cgc/Tgc | R50C | gloA2 | A | A | | | | | | | | | | |
| NC_011770 | 5072886 | T | G | 78 | 0 | 2 | 0 | NON_SYNONYMOU | MODERAT | MISSENSE | gaA/gaC | E36D | gloA2 | G | G | | | | | | | | | | |
| NC_011770 | 5072920 | T | C | 82 | 0 | 6 | 0 | NON_SYNONYMOU | MODERAT | MISSENSE | gAc/gGc | D25G | gloA2 | | | C | C | C | C | C | C | | | | |
| NC_011770 | 5072924 | G | T | 82 | 0 | 4 | 0 | NON_SYNONYMOU | MODERAT | MISSENSE | Ctg/Atg | L24M | gloA2 | | | | | | | | | T | T | T | T |
| NC_011770 | 1562554 | G | A | 67 | 0 | 2 | 0 | NON_SYNONYMOU | MODERAT | MISSENSE | Cgc/Tgc | R437C | glpD | | | | | | | | | A | A | | |
| NC_011770 | 1562734 | T | C | 56 | 0 | 2 | 0 | NON_SYNONYMOU | MODERAT | MISSENSE | Acg/Gcg | T377A | glpD | | | | | | | | | C | C | | |
| NC_011770 | 1562740 | C | T | 56 | 0 | 4 | 0 | NON_SYNONYMOU | MODERAT | MISSENSE | Gcg/Acg | A375T | glpD | | | T | T | T | T | | | | | | |
| NC_011770 | 1563042 | C | T | 77 | 0 | 4 | 0 | NON_SYNONYMOU | MODERAT | MISSENSE | cGt/cAt | R274H | glpD | | | T | T | T | T | | | | | | |
| NC_011770 | 1567342 | T | C | 44 | 0 | 4 | 0 | NON_SYNONYMOU | MODERAT | MISSENSE | Atc/Gtc | I53V | glpF | C | C | | | | | | | C | C | | |
| NC_011770 | 1561975 | G | A | 86 | 0 | 2 | 0 | NON_SYNONYMOU | MODERAT | MISSENSE | Ctc/Ttc | L73F | glpM | | | | | | | | | A | A | | |
| NC_011770 | 383840 | G | C | 65 | 0 | 4 | 0 | NON_SYNONYMOU | MODERAT | MISSENSE | gCc/gGc | A266G | glpQ | | | | | | | | | C | C | C | C |
| NC_011770 | 383841 | C | T | 65 | 0 | 4 | 0 | NON_SYNONYMOU | MODERAT | MISSENSE | Gcc/Acc | A266T | glpQ | | | | | | | | | T | T | T | T |
| NC_011770 | 384258 | C | T | 69 | 0 | 2 | 0 | NON_SYNONYMOU | MODERAT | MISSENSE | Gtc/Atc | V127I | glpQ | T | T | | | | | | | | | | |
| NC_011770 | 6230181 | T | G | 69 | 0 | 2 | 0 | STOP_LOST+SPLICE | HIGH | MISSENSE | Tga/Gga | *449G | glpT | | | | | | | G | G | | | | |
| NC_011770 | 4148234 | T | C | 86 | 0 | 2 | 0 | NON_SYNONYMOU | MODERAT | MISSENSE | Tcc/Ccc | S149P | gltA | | | C | C | | | | | | | | |
| NC_011770 | 6004970 | G | A | 99 | 0 | 2 | 0 | NON_SYNONYMOU | MODERAT | MISSENSE | Ccc/Tcc | P619S | gltB | | | | | | | | | A | A | | |
| NC_011770 | 6005168 | T | C | 93 | 0 | 1 | 0 | NON_SYNONYMOU | MODERAT | MISSENSE | Acc/Gcc | T553A | gltB | | | | | | | C | | | | | |
| NC_011770 | 2036057 | C | T | 77 | 0 | 2 | 0 | NON_SYNONYMOU | MODERAT | MISSENSE | Ggc/Agc | G204S | gltS | T | T | | | | | | | | | | |
| NC_011770 | 2078764 | A | G | 87 | 0 | 12 | 0 | NON_SYNONYMOU | MODERAT | MISSENSE | Aag/Gag | K161E | gltX | G | G | G | G | G | G | G | G | G | G | G | G |
| NC_011770 | 2078801 | A | C | 85 | 0 | 2 | 0 | NON_SYNONYMOU | MODERAT | MISSENSE | gAt/gCt | D173A | gltX | | | | | | | | | C | C | | |
| NC_011770 | 2078978 | G | A | 82 | 0 | 4 | 0 | NON_SYNONYMOU | MODERAT | MISSENSE | gGc/gAc | G232D | gltX | | | A | A | A | A | | | | | | |
| NC_011770 | 5497126 | A | G | 78 | 0 | 6 | 0 | SPLICE_SITE_REGIC | LOW | SILENT | taA/taG | *418 | glyA | G | G | | | | | G | G | G | G | | |
| NC_011770 | 5495901 | G | A | 70 | 0 | 2 | 0 | NON_SYNONYMOU | MODERAT | MISSENSE | cGc/cAc | R10H | glyA | A | A | | | | | | | | | | |
| NC_011770 | 6430840 | G | A | 80 | 0 | 2 | 0 | NON_SYNONYMOU | MODERAT | MISSENSE | atG/atA | M298I | glyA1 | | | A | A | | | | | | | | |
| NC_011770 | 6430857 | G | C | 74 | 0 | 2 | 0 | NON_SYNONYMOU | MODERAT | MISSENSE | gGg/gCg | G304A | glyA1 | | | | | | | C | C | | | | |
| NC_011770 | 3090787 | A | C | 68 | 0 | 6 | 0 | NON_SYNONYMOU | MODERAT | MISSENSE | gAc/gCc | D89A | glyA2 | | | C | C | C | C | C | C | | | | |
| NC_011770 | 3091494 | A | C | 63 | 0 | 2 | 0 | NON_SYNONYMOU | MODERAT | MISSENSE | Aag/Cag | K325Q | glyA2 | | | | | C | C | | | | | | |
| NC_011770 | 3091495 | A | G | 62 | 0 | 2 | 0 | NON_SYNONYMOU | MODERAT | MISSENSE | aAg/aGg | K325R | glyA2 | | | | | G | G | | | | | | |
| NC_011770 | 3091496 | G | C | 63 | 0 | 2 | 0 | NON_SYNONYMOU | MODERAT | MISSENSE | aaG/aaC | K325N | glyA2 | | | | | C | C | | | | | | |
| NC_011770 | 3091516 | C | A | 58 | 0 | 2 | 0 | NON_SYNONYMOU | MODERAT | MISSENSE | gCc/gAc | A332D | glyA2 | | | | | A | A | | | | | | |
| NC_011770 | 3091731 | G | A | 78 | 0 | 2 | 0 | NON_SYNONYMOU | MODERAT | MISSENSE | Ggc/Agc | G404S | glyA2 | A | A | | | | | | | | | | |
| NC_011770 | 3091732 | G | A | 77 | 0 | 2 | 0 | NON_SYNONYMOU | MODERAT | MISSENSE | gGc/gAc | G404D | glyA2 | A | A | | | | | | | | | | |
| NC_011770 | 3091251 | G | T | 60 | 0 | 1 | 0 | NON_SYNONYMOU | MODERAT | MISSENSE | Gcc/Tcc | A244S | glyA2 | | | | | | | | | | | T | |
| NC_011770 | 11333 | A | C | 86 | 0 | 10 | 0 | NON_SYNONYMOU | MODERAT | MISSENSE | Tcg/Gcg | S382A | glyS | | | C | C | C | C | C | C | C | C | C | C |
| NC_011770 | 12130 | T | C | 78 | 0 | 2 | 0 | NON_SYNONYMOU | MODERAT | MISSENSE | cAg/cGg | Q116R | glyS | | | | | | | | | C | C | | |
| NC_011770 | 12165 | C | A | 79 | 0 | 2 | 0 | NON_SYNONYMOU | MODERAT | MISSENSE | caG/caT | Q104H | glyS | | | | | | | A | A | | | | |
| NC_011770 | 12356 | T | C | 82 | 0 | 2 | 0 | NON_SYNONYMOU | MODERAT | MISSENSE | Agc/Ggc | S41G | glyS | C | C | | | | | | | | | | |
| NC_011770 | 12269 | C | T | 80 | 0 | 1 | 0 | NON_SYNONYMOU | MODERAT | MISSENSE | Gtc/Atc | V70I | glyS | | T | | | | | | | | | | |
| NC_011770 | 6479115 | G | A | 100 | 0 | 1 | 0 | INTRAGENIC | MODIFIER | | | | gmd | | | | | | | A | | | | | |
| NC_011770 | 6479155 | G | A | 96 | 0 | 4 | 0 | INTRAGENIC | MODIFIER | | | | gmd | | | A | A | A | A | | | | | | |
| NC_011770 | 6479427 | C | T | 105 | 0 | 2 | 0 | INTRAGENIC | MODIFIER | | | | gmd | | | | | | | | | | | T | T |
| NC_011770 | 6341404 | G | A | 84 | 0 | 4 | 0 | NON_SYNONYMOU | MODERAT | MISSENSE | Gag/Aag | E87K | gmk | | | A | A | A | A | | | | | | |
| NC_011770 | 3647870 | G | T | 64 | 0 | 4 | 0 | INTRAGENIC | MODIFIER | | | | gor | | | T | T | T | T | | | | | | |
| NC_011770 | 3647888 | A | G | 67 | 0 | 4 | 0 | INTRAGENIC | MODIFIER | | | | gor | | | G | G | G | G | | | | | | |
| NC_011770 | 3647933 | A | G | 71 | 0 | 4 | 0 | INTRAGENIC | MODIFIER | | | | gor | | | G | G | G | G | | | | | | |
| NC_011770 | 3648101 | C | T | 72 | 0 | 2 | 0 | INTRAGENIC | MODIFIER | | | | gor | T | T | | | | | | | | | | |
| NC_011770 | 3648116 | G | A | 70 | 0 | 2 | 0 | INTRAGENIC | MODIFIER | | | | gor | | | | | | | A | A | | | | |
| NC_011770 | 3648143 | G | A | 71 | 0 | 6 | 0 | INTRAGENIC | MODIFIER | | | | gor | A | A | A | A | A | A | | | | | | |
| NC_011770 | 3648146 | A | G | 73 | 0 | 6 | 0 | INTRAGENIC | MODIFIER | | | | gor | G | G | G | G | G | G | | | | | | |
| NC_011770 | 3648147 | T | G | 74 | 0 | 2 | 0 | INTRAGENIC | MODIFIER | | | | gor | | | | | | | | | | | G | G |
| NC_011770 | 3648166 | A | G | 71 | 0 | 6 | 0 | INTRAGENIC | MODIFIER | | | | gor | G | G | G | G | G | G | | | | | | |
| NC_011770 | 3648167 | T | C | 71 | 0 | 6 | 0 | INTRAGENIC | MODIFIER | | | | gor | C | C | C | C | C | C | | | | | | |
| NC_011770 | 3648173 | T | C | 72 | 0 | 8 | 0 | INTRAGENIC | MODIFIER | | | | gor | C | C | C | C | C | C | | | | | C | C |
| NC_011770 | 3648185 | G | A | 73 | 0 | 2 | 0 | INTRAGENIC | MODIFIER | | | | gor | | | | | | | | | A | A | | |
| NC_011770 | 3648230 | G | A | 72 | 0 | 4 | 0 | INTRAGENIC | MODIFIER | | | | gor | | | A | A | A | A | | | | | | |
| NC_011770 | 3648233 | A | G | 73 | 0 | 8 | 0 | INTRAGENIC | MODIFIER | | | | gor | G | G | G | G | G | G | | | | | G | G |
| NC_011770 | 3648245 | G | T | 75 | 0 | 2 | 0 | INTRAGENIC | MODIFIER | | | | gor | | | | | | | | | T | T | | |

| | | | | | | | | | | | | | | | | | | | | | | | | | |
|---|---|---|---|---|---|---|---|---|---|---|---|---|---|---|---|---|---|---|---|---|---|---|---|---|---|
| NC_011770 | 3648254 | C | T | 73 | 0 | 2 | 0 | INTRAGENIC | MODIFIER | | | | gor | | | | | | | | | T | T | | |
| NC_011770 | 3648269 | G | A | 80 | 0 | 4 | 0 | INTRAGENIC | MODIFIER | | | | gor | | | A | A | A | A | | | | | | |
| NC_011770 | 3648311 | A | T | 75 | 0 | 4 | 0 | INTRAGENIC | MODIFIER | | | | gor | | | T | T | T | T | | | | | | |
| NC_011770 | 3648312 | G | C | 77 | 0 | 4 | 0 | INTRAGENIC | MODIFIER | | | | gor | | | C | C | C | C | | | | | | |
| NC_011770 | 3648314 | T | G | 76 | 0 | 4 | 0 | INTRAGENIC | MODIFIER | | | | gor | | | G | G | G | G | | | | | | |
| NC_011770 | 3648335 | G | A | 78 | 0 | 4 | 0 | INTRAGENIC | MODIFIER | | | | gor | | | A | A | A | A | | | | | | |
| NC_011770 | 3648362 | G | A | 76 | 0 | 4 | 0 | INTRAGENIC | MODIFIER | | | | gor | | | A | A | A | A | | | | | | |
| NC_011770 | 3648378 | G | GC | 76 | 0 | 12 | 0 | INTRAGENIC | MODIFIER | | | | gor | GC | GC | GC | GC | GC | GC | GC | GC | GC | GC | GC | GC |
| NC_011770 | 3648445 | G | A | 73 | 0 | 2 | 0 | INTRAGENIC | MODIFIER | | | | gor | | | | | | | | | A | A | | |
| NC_011770 | 3648557 | G | A | 69 | 0 | 4 | 0 | INTRAGENIC | MODIFIER | | | | gor | | | A | A | A | A | | | | | | |
| NC_011770 | 3648595 | C | T | 65 | 0 | 2 | 0 | INTRAGENIC | MODIFIER | | | | gor | | | | | | | | | T | T | | |
| NC_011770 | 3648727 | A | G | 69 | 0 | 10 | 0 | INTRAGENIC | MODIFIER | | | | gor | G | G | G | G | G | G | | | G | G | G | G |
| NC_011770 | 3649087 | G | A | 37 | 2 | 2 | 0 | INTRAGENIC | MODIFIER | | | | gor | | | . | . | | | A | A | | | | |
| NC_011770 | 3648711 | G | A | 68 | 0 | 1 | 0 | INTRAGENIC | MODIFIER | | | | gor | A | | | | | | | | | | | |
| NC_011770 | 4109300 | G | C | 53 | 0 | 12 | 0 | NON_SYNONYMOU | MODERAT | MISSENSE | aCa/aGa | T45R | gpsA | C | C | C | C | C | C | C | C | C | C | C | C |
| NC_011770 | 317379 | C | T | 77 | 0 | 1 | 0 | STOP_GAINED | HIGH | NONSENSE | Cag/Tag | Q266* | gpuA | | | | T | | | | | | | | |
| NC_011770 | 315786 | A | T | 73 | 0 | 2 | 0 | NON_SYNONYMOU | MODERAT | MISSENSE | gAc/gTc | D215V | gpuP | T | T | | | | | | | | | | |
| NC_011770 | 316308 | T | C | 66 | 0 | 1 | 0 | NON_SYNONYMOU | MODERAT | MISSENSE | cTg/cCg | L389P | gpuP | | | | | | | C | | | | | |
| NC_011770 | 318184 | G | A | 77 | 0 | 8 | 0 | NON_SYNONYMOU | MODERAT | MISSENSE | Gcg/Acg | A198T | gpuR | A | A | | | | | A | A | A | A | A | A |
| NC_011770 | 2382123 | G | T | 79 | 0 | 2 | 0 | NON_SYNONYMOU | MODERAT | MISSENSE | gCg/gAg | A167E | greB | | | | | | | | | T | T | | |
| NC_011770 | 2382616 | G | T | 84 | 0 | 2 | 0 | NON_SYNONYMOU | MODERAT | MISSENSE | Cgc/Agc | R3S | greB | | | | | T | T | | | | | | |
| NC_011770 | 5249249 | T | C | 72 | 0 | 1 | 0 | NON_SYNONYMOU | MODERAT | MISSENSE | aAg/aGg | K528R | groEL | | | | | | | C | | | | | |
| NC_011770 | 5250146 | T | C | 68 | 0 | 1 | 0 | NON_SYNONYMOU | MODERAT | MISSENSE | aAc/aGc | N229S | groEL | | | | | | | C | | | | | |
| NC_011770 | 5683191 | A | G | 72 | 0 | 12 | 0 | NON_SYNONYMOU | MODERAT | MISSENSE | Tac/Cac | Y133H | grpE | G | G | G | G | G | G | G | G | G | G | G | G |
| NC_011770 | 6111823 | T | A | 61 | 0 | 2 | 0 | NON_SYNONYMOU | MODERAT | MISSENSE | Atc/Ttc | I32F | grx | A | A | | | | | | | | | | |
| NC_011770 | 6192704 | C | G | 58 | 0 | 12 | 0 | NON_SYNONYMOU | MODERAT | MISSENSE | gCc/gGc | A83G | gshA | G | G | G | G | G | G | G | G | G | G | G | G |
| NC_011770 | 6192829 | C | A | 67 | 0 | 10 | 0 | NON_SYNONYMOU | MODERAT | MISSENSE | Ctg/Atg | L125M | gshA | A | A | A | A | A | A | A | A | A | A | | |
| NC_011770 | 6192931 | G | A | 72 | 0 | 2 | 0 | NON_SYNONYMOU | MODERAT | MISSENSE | Gag/Aag | E159K | gshA | | | | | | | | | A | A | | |
| NC_011770 | 6193657 | C | A | 78 | 0 | 6 | 0 | NON_SYNONYMOU | MODERAT | MISSENSE | Cag/Aag | Q401K | gshA | | | A | A | A | A | A | A | | | | |
| NC_011770 | 3039576 | T | G | 51 | 0 | 8 | 0 | NON_SYNONYMOU | MODERAT | MISSENSE | Acc/Ccc | T350P | gtdA | G | G | G | G | G | G | | | | | G | G |
| NC_011770 | 3039728 | C | T | 65 | 0 | 2 | 0 | NON_SYNONYMOU | MODERAT | MISSENSE | gGc/gAc | G299D | gtdA | T | T | | | | | | | | | | |
| NC_011770 | 3039809 | C | T | 64 | 0 | 2 | 0 | NON_SYNONYMOU | MODERAT | MISSENSE | gGc/gAc | G272D | gtdA | | | | | | | | | | | T | T |
| NC_011770 | 3039938 | G | A | 56 | 0 | 2 | 0 | NON_SYNONYMOU | MODERAT | MISSENSE | gCc/gTc | A229V | gtdA | A | A | | | | | | | | | | |
| NC_011770 | 3040076 | T | C | 66 | 0 | 4 | 0 | NON_SYNONYMOU | MODERAT | MISSENSE | gAt/gGt | D183G | gtdA | C | C | | | | | | | | | C | C |
| NC_011770 | 3040080 | G | A | 66 | 0 | 4 | 0 | NON_SYNONYMOU | MODERAT | MISSENSE | Cat/Tat | H182Y | gtdA | A | A | | | | | | | | | A | A |
| NC_011770 | 3040082 | T | C | 65 | 0 | 4 | 0 | NON_SYNONYMOU | MODERAT | MISSENSE | aAc/aGc | N181S | gtdA | C | C | | | | | | | | | C | C |
| NC_011770 | 3040170 | C | G | 73 | 0 | 4 | 0 | NON_SYNONYMOU | MODERAT | MISSENSE | Gcc/Ccc | A152P | gtdA | G | G | | | | | | | | | G | G |
| NC_011770 | 3040209 | T | G | 73 | 0 | 2 | 0 | NON_SYNONYMOU | MODERAT | MISSENSE | Atc/Ctc | I139L | gtdA | | | | | | | G | G | | | | |
| NC_011770 | 3040266 | T | C | 69 | 0 | 12 | 0 | NON_SYNONYMOU | MODERAT | MISSENSE | Agc/Ggc | S120G | gtdA | C | C | C | C | C | C | C | C | C | C | C | C |
| NC_011770 | 3040374 | C | A | 66 | 0 | 6 | 0 | NON_SYNONYMOU | MODERAT | MISSENSE | Gcg/Tcg | A84S | gtdA | | | A | A | A | A | A | A | | | | |
| NC_011770 | 1304908 | A | T | 48 | 0 | 2 | 0 | NON_SYNONYMOU | MODERAT | MISSENSE | gaA/gaT | E491D | guaA | | | | | | | T | T | | | | |
| NC_011770 | 1304900 | C | T | 51 | 0 | 1 | 0 | NON_SYNONYMOU | MODERAT | MISSENSE | Ccc/Tcc | P489S | guaA | | | | | | | | T | | | | |
| NC_011770 | 2043878 | T | C | 84 | 0 | 6 | 0 | NON_SYNONYMOU | MODERAT | MISSENSE | aTc/aCc | I83T | gyrA | | | | | C | C | | | C | C | C | C |
| NC_011770 | 2045236 | A | G | 85 | 0 | 1 | 0 | NON_SYNONYMOU | MODERAT | MISSENSE | Acc/Gcc | T536A | gyrA | | | | | | | G | | | | | |
| NC_011770 | 2045642 | T | C | 88 | 0 | 2 | 0 | NON_SYNONYMOU | MODERAT | MISSENSE | gTc/gCc | V671A | gyrA | | | | | | | | | | | C | C |
| NC_011770 | 2043889 | G | A | 93 | 0 | 1 | 0 | NON_SYNONYMOU | MODERAT | MISSENSE | Gac/Aac | D87N | gyrA | | | | | | | | | A | | | |
| NC_011770 | 2043781 | G | T | 86 | 0 | 1 | 0 | NON_SYNONYMOU | MODERAT | MISSENSE | Gcc/Tcc | A51S | gyrA | | | | | | | | T | | | | |
| NC_011770 | 5692 | T | C | 102 | 0 | 1 | 0 | NON_SYNONYMOU | MODERAT | MISSENSE | aTc/aCc | I473T | gyrB | | | | | | | C | | | | | |
| NC_011770 | 4705 | A | G | 88 | 0 | 1 | 0 | NON_SYNONYMOU | MODERAT | MISSENSE | gAa/gGa | E144G | **gyrB** | | | | | | | | G | | | | |
| NC_011770 | 5679 | G | T | 101 | 0 | 1 | 0 | NON_SYNONYMOU | MODERAT | MISSENSE | Gtc/Ttc | V469F | gyrB | | | | | | | | | | T | | |
| NC_011770 | 1779952 | T | C | 73 | 0 | 12 | 0 | NON_SYNONYMOU | MODERAT | MISSENSE | Tgg/Cgg | W231R | hasD | C | C | C | C | C | C | C | C | C | C | C | C |
| NC_011770 | 1780062 | A | T | 73 | 0 | 2 | 0 | NON_SYNONYMOU | MODERAT | MISSENSE | gaA/gaT | E267D | hasD | | | | | | | T | T | | | | |
| NC_011770 | 1780458 | G | C | 72 | 0 | 6 | 0 | NON_SYNONYMOU | MODERAT | MISSENSE | caG/caC | Q399H | hasD | | | C | C | C | C | | | C | C | | |
| NC_011770 | 1780812 | G | C | 69 | 0 | 10 | 0 | NON_SYNONYMOU | MODERAT | MISSENSE | caG/caC | Q517H | hasD | C | C | C | C | C | C | | | C | C | C | C |
| NC_011770 | 1781382 | C | G | 83 | 0 | 2 | 0 | NON_SYNONYMOU | MODERAT | MISSENSE | Ctg/Gtg | L112V | hasE | | | | | | | G | G | | | | |
| NC_011770 | 1775706 | AG | A | 63 | 0 | 1 | 0 | FRAME_SHIFT | HIGH | | gcg/ | A16 | hasR | | | | A | | | | | | | | |
| NC_011770 | 1775787 | A | G | 65 | 0 | 2 | 0 | NON_SYNONYMOU | MODERAT | MISSENSE | gAc/gGc | D41G | hasR | | | | | | | | | G | G | | |
| NC_011770 | 1775954 | G | T | 63 | 0 | 4 | 0 | NON_SYNONYMOU | MODERAT | MISSENSE | Gtg/Ttg | V97L | hasR | | | | | | | T | T | T | T | | |
| NC_011770 | 1776155 | G | A | 68 | 0 | 2 | 0 | NON_SYNONYMOU | MODERAT | MISSENSE | Gag/Aag | E164K | hasR | A | A | | | | | | | | | | |
| NC_011770 | 1776187 | A | G | 69 | 0 | 8 | 0 | NON_SYNONYMOU | MODERAT | MISSENSE | atA/atG | I174M | hasR | G | G | | | | | G | G | G | G | G | G |
| NC_011770 | 1776894 | C | T | 76 | 0 | 2 | 0 | NON_SYNONYMOU | MODERAT | MISSENSE | gCg/gTg | A410V | hasR | | | | | | | | | T | T | | |
| NC_011770 | 1777250 | C | T | 78 | 0 | 2 | 0 | NON_SYNONYMOU | MODERAT | MISSENSE | Ctc/Ttc | L529F | hasR | | | | | | | | | T | T | | |

| | | | | | | | | | | | | | | | | | | | | | | | | | |
|---|---|---|---|---|---|---|---|---|---|---|---|---|---|---|---|---|---|---|---|---|---|---|---|---|---|
| NC_011770 | 1777308 | A | G | 73 | 0 | 1 | 0 | NON_SYNONYMOU | MODERAT | MISSENSE | tAc/tGc | Y548C | hasR | | | | | | | G | | | | | |
| NC_011770 | 1778246 | A | G | 79 | 0 | 2 | 0 | NON_SYNONYMOU | MODERAT | MISSENSE | Acc/Gcc | T861A | hasR | | | | | | | | | G | G | | |
| NC_011770 | 1776629 | G | A | 64 | 0 | 1 | 0 | NON_SYNONYMOU | MODERAT | MISSENSE | Gac/Aac | D322N | hasR | | A | | | | | | | | | | |
| NC_011770 | 3459653 | G | A | 53 | 0 | 2 | 0 | NON_SYNONYMOU | MODERAT | MISSENSE | Cca/Tca | P104S | hcnA | | | | | | | A | A | | | | |
| NC_011770 | 3458649 | C | T | 64 | 0 | 2 | 0 | NON_SYNONYMOU | MODERAT | MISSENSE | Gcc/Acc | A335T | hcnB | | | | | T | T | | | | | | |
| NC_011770 | 3458655 | G | A | 65 | 0 | 2 | 0 | NON_SYNONYMOU | MODERAT | MISSENSE | Cgc/Tgc | R333C | hcnB | A | A | | | | | | | | | | |
| NC_011770 | 3458962 | A | C | 55 | 0 | 12 | 0 | NON_SYNONYMOU | MODERAT | MISSENSE | gaT/gaG | D230E | hcnB | C | C | C | C | C | C | C | C | C | C | C | C |
| NC_011770 | 3459350 | C | T | 65 | 0 | 2 | 0 | NON_SYNONYMOU | MODERAT | MISSENSE | cGg/cAg | R101Q | hcnB | | | | | | | T | T | | | | |
| NC_011770 | 3459377 | C | T | 65 | 0 | 2 | 0 | NON_SYNONYMOU | MODERAT | MISSENSE | cGc/cAc | R92H | hcnB | T | T | | | | | | | | | | |
| NC_011770 | 3459624 | C | T | 47 | 0 | 2 | 0 | NON_SYNONYMOU | MODERAT | MISSENSE | Ggc/Agc | G10S | hcnB | | | T | T | | | | | | | | |
| NC_011770 | 3457267 | C | T | 59 | 0 | 2 | 0 | NON_SYNONYMOU | MODERAT | MISSENSE | Gag/Aag | E330K | hcnC | | | | | | | | | T | T | | |
| NC_011770 | 3457527 | G | A | 66 | 0 | 2 | 0 | NON_SYNONYMOU | MODERAT | MISSENSE | gCc/gTc | A243V | hcnC | | | | | | | | | A | A | | |
| NC_011770 | 3457629 | G | A | 71 | 0 | 4 | 0 | NON_SYNONYMOU | MODERAT | MISSENSE | cCg/cTg | P209L | hcnC | | | A | A | A | A | | | | | | |
| NC_011770 | 3457876 | T | C | 74 | 0 | 1 | 0 | NON_SYNONYMOU | MODERAT | MISSENSE | Acc/Gcc | T127A | hcnC | | C | | | | | | | | | | |
| NC_011770 | 3458067 | A | G | 72 | 0 | 1 | 0 | NON_SYNONYMOU | MODERAT | MISSENSE | gTg/gCg | V63A | hcnC | | | | | | | | G | | | | |
| NC_011770 | 4308306 | C | G | 54 | 0 | 1 | 0 | NON_SYNONYMOU | MODERAT | MISSENSE | aCc/aGc | T60S | hcpA | | | | | | | | G | | | | |
| NC_011770 | 6240707 | T | C | 76 | 0 | 1 | 0 | NON_SYNONYMOU | MODERAT | MISSENSE | tAc/tGc | Y283C | hemB | | | | C | | | | | | | | |
| NC_011770 | 6240815 | G | A | 80 | 0 | 2 | 0 | NON_SYNONYMOU | MODERAT | MISSENSE | gCc/gTc | A247V | hemB | | | | | | | | | A | A | | |
| NC_011770 | 6257801 | C | T | 72 | 0 | 4 | 0 | NON_SYNONYMOU | MODERAT | MISSENSE | aGt/aAt | S188N | hemC | | | | | | | T | T | T | T | | |
| NC_011770 | 6257932 | C | A | 71 | 0 | 4 | 0 | NON_SYNONYMOU | MODERAT | MISSENSE | caG/caT | Q144H | hemC | | | A | A | A | A | | | | | | |
| NC_011770 | 6000060 | T | C | 68 | 0 | 6 | 0 | NON_SYNONYMOU | MODERAT | MISSENSE | Acc/Gcc | T226A | hemE | C | C | | | | | C | C | C | C | | |
| NC_011770 | 6000140 | T | C | 75 | 0 | 2 | 0 | NON_SYNONYMOU | MODERAT | MISSENSE | cAc/cGc | H199R | hemE | C | C | | | | | | | | | | |
| NC_011770 | 6000207 | G | T | 77 | 0 | 6 | 0 | NON_SYNONYMOU | MODERAT | MISSENSE | Cag/Aag | Q177K | hemE | T | T | | | | | T | T | T | T | | |
| NC_011770 | 25733 | A | G | 69 | 0 | 12 | 0 | NON_SYNONYMOU | MODERAT | MISSENSE | cAt/cGt | H4R | hemF | G | G | G | G | G | G | G | G | G | G | G | G |
| NC_011770 | 5557693 | G | C | 68 | 0 | 2 | 0 | NON_SYNONYMOU | MODERAT | MISSENSE | Ccg/Gcg | P339A | hemH | | | | | | | | | | | C | C |
| NC_011770 | 5557715 | C | G | 70 | 0 | 2 | 0 | NON_SYNONYMOU | MODERAT | MISSENSE | gaG/gaC | E331D | hemH | | | | | | | | | | | G | G |
| NC_011770 | 5558358 | C | T | 79 | 0 | 2 | 0 | NON_SYNONYMOU | MODERAT | MISSENSE | cGg/cAg | R117Q | hemH | | | | | | | | | | | T | T |
| NC_011770 | 5566805 | A | G | 90 | 0 | 8 | 0 | NON_SYNONYMOU | MODERAT | MISSENSE | gTa/gCa | V275A | hemK | G | G | G | G | G | G | | | | | G | G |
| NC_011770 | 5567471 | C | A | 87 | 0 | 4 | 0 | NON_SYNONYMOU | MODERAT | MISSENSE | cGg/cTg | R53L | hemK | | | | | | | A | A | | | A | A |
| NC_011770 | 1893798 | A | G | 81 | 0 | 2 | 0 | NON_SYNONYMOU | MODERAT | MISSENSE | Tgg/Cgg | W647R | hepA | | | | | | | G | G | | | | |
| NC_011770 | 1895451 | A | G | 75 | 0 | 8 | 0 | NON_SYNONYMOU | MODERAT | MISSENSE | Tgc/Cgc | C96R | hepA | G | G | G | G | G | G | G | G | | | | |
| NC_011770 | 5877625 | A | G | 71 | 0 | 1 | 0 | NON_SYNONYMOU | MODERAT | MISSENSE | gTg/gCg | V166A | hflC | | | | | | | G | | | | | |
| NC_011770 | 5879240 | ACCG | A | 79 | 0 | 1 | 0 | CODON_CHANGE_ | MODERATE | | ggcggt/ggt | GG27G | hflK | | | | | | | | | | A | | |
| NC_011770 | 5879137 | C | T | 89 | 0 | 1 | 0 | NON_SYNONYMOU | MODERAT | MISSENSE | Ggc/Agc | G63S | hflK | | | | | | | | | | | | T |
| NC_011770 | 5314882 | G | A | 69 | 0 | 2 | 0 | NON_SYNONYMOU | MODERAT | MISSENSE | aCc/aTc | T310I | hisC1 | | | | | | | A | A | | | | |
| NC_011770 | 5314909 | T | C | 72 | 0 | 12 | 0 | NON_SYNONYMOU | MODERAT | MISSENSE | cAg/cGg | Q301R | hisC1 | C | C | C | C | C | C | C | C | C | C | C | C |
| NC_011770 | 5315779 | C | T | 74 | 0 | 12 | 0 | NON_SYNONYMOU | MODERAT | MISSENSE | gGc/gAc | G11D | hisC1 | T | T | T | T | T | T | T | T | T | T | T | T |
| NC_011770 | 2048846 | G | A | 67 | 0 | 2 | 0 | NON_SYNONYMOU | MODERAT | MISSENSE | Gcg/Acg | A36T | hisC2 | A | A | | | | | | | | | | |
| NC_011770 | 2049690 | G | A | 65 | 0 | 4 | 0 | NON_SYNONYMOU | MODERAT | MISSENSE | cGc/cAc | R317H | hisC2 | | | | | | | | | A | A | A | A |
| NC_011770 | 5316655 | A | T | 101 | 0 | 2 | 0 | NON_SYNONYMOU | MODERAT | MISSENSE | Tcc/Acc | S161T | hisD | | | | | | | | | T | T | | |
| NC_011770 | 5317770 | A | G | 88 | 0 | 4 | 0 | NON_SYNONYMOU | MODERAT | MISSENSE | Tcc/Ccc | S43P | hisG | | | G | G | G | G | | | | | | |
| NC_011770 | 5317835 | G | C | 85 | 0 | 4 | 0 | NON_SYNONYMOU | MODERAT | MISSENSE | gCg/gGg | A21G | hisG | | | C | C | C | C | | | | | | |
| NC_011770 | 2314331 | A | G | 94 | 0 | 1 | 0 | NON_SYNONYMOU | MODERAT | MISSENSE | Tgg/Cgg | W219R | hisM | | | | | | | G | | | | | |
| NC_011770 | 2314540 | C | T | 88 | 0 | 2 | 0 | NON_SYNONYMOU | MODERAT | MISSENSE | cGc/cAc | R149H | hisM | | | | | | | | | T | T | | |
| NC_011770 | 2314823 | G | T | 91 | 0 | 2 | 0 | NON_SYNONYMOU | MODERAT | MISSENSE | Ctg/Atg | L55M | hisM | | | | | | | | | | | T | T |
| NC_011770 | 2313561 | G | C | 78 | 0 | 2 | 0 | NON_SYNONYMOU | MODERAT | MISSENSE | Caa/Gaa | Q231E | hisP | | | | | | | | | | | C | C |
| NC_011770 | 2313593 | A | G | 77 | 0 | 2 | 0 | NON_SYNONYMOU | MODERAT | MISSENSE | gTg/gCg | V220A | hisP | | | | | | | G | G | | | | |
| NC_011770 | 2313866 | C | T | 83 | 0 | 2 | 0 | NON_SYNONYMOU | MODERAT | MISSENSE | cGc/cAc | R129H | hisP | | | | | | | | | | | T | T |
| NC_011770 | 2313992 | C | T | 90 | 0 | 4 | 0 | NON_SYNONYMOU | MODERAT | MISSENSE | aGg/aAg | R87K | hisP | T | T | | | | | | | T | T | | |
| NC_011770 | 2314035 | C | T | 88 | 0 | 2 | 0 | NON_SYNONYMOU | MODERAT | MISSENSE | Gcc/Acc | A73T | hisP | | | | | | | | | | | T | T |
| NC_011770 | 1269601 | A | C | 90 | 0 | 10 | 0 | NON_SYNONYMOU | MODERAT | MISSENSE | gAa/gCa | E427A | hisS | | | C | C | C | C | C | C | C | C | C | C |
| NC_011770 | 1268356 | A | G | 84 | 0 | 1 | 0 | NON_SYNONYMOU | MODERAT | MISSENSE | aAc/aGc | N12S | hisS | | G | | | | | | | | | | |
| NC_011770 | 5875894 | G | A | 86 | 0 | 4 | 0 | NON_SYNONYMOU | MODERAT | MISSENSE | gCg/gTg | A348V | hisZ | A | A | | | | | | | A | A | | |
| NC_011770 | 5875991 | T | A | 83 | 0 | 2 | 0 | NON_SYNONYMOU | MODERAT | MISSENSE | Acc/Tcc | T316S | hisZ | | | | | | | | | | | A | A |
| NC_011770 | 5593040 | A | G | 74 | 0 | 10 | 0 | INTRAGENIC | MODIFIER | | | | hitA | G | G | G | G | G | G | G | G | | | G | G |
| NC_011770 | 5593070 | A | G | 72 | 0 | 4 | 0 | INTRAGENIC | MODIFIER | | | | hitA | | | | | | | | | G | G | G | G |
| NC_011770 | 5593202 | T | C | 63 | 0 | 12 | 0 | INTRAGENIC | MODIFIER | | | | hitA | C | C | C | C | C | C | C | C | C | C | C | C |
| NC_011770 | 5593367 | A | C | 54 | 0 | 4 | 0 | INTRAGENIC | MODIFIER | | | | hitA | | | | | | | C | C | | | C | C |
| NC_011770 | 5593447 | A | G | 45 | 0 | 1 | 0 | INTRAGENIC | MODIFIER | | | | hitA | | | | | | | G | | | | | |
| NC_011770 | 5593472 | G | A | 47 | 0 | 2 | 0 | INTRAGENIC | MODIFIER | | | | hitA | | | | | | | A | A | | | | |
| NC_011770 | 5593818 | C | A | 55 | 0 | 2 | 0 | INTRAGENIC | MODIFIER | | | | hitA | | | | | | | A | A | | | | |
| NC_011770 | 5594058 | T | C | 53 | 0 | 12 | 0 | INTRAGENIC | MODIFIER | | | | hitA | C | C | C | C | C | C | C | C | C | C | C | C |

| | | | | | | | | | | | | | | | | | | | | | | | | | |
|---|---|---|---|---|---|---|---|---|---|---|---|---|---|---|---|---|---|---|---|---|---|---|---|---|---|
| NC_011770 | 5594094 | C | T | 60 | 0 | 6 | 0 | INTRAGENIC | MODIFIER | | | | hitA | T | T | T | T | T | T | | | | | | |
| NC_011770 | 5594102 | A | G | 58 | 0 | 12 | 0 | INTRAGENIC | MODIFIER | | | | hitA | G | G | G | G | G | G | G | G | G | G | G | G |
| NC_011770 | 5594106 | G | T | 59 | 0 | 2 | 0 | INTRAGENIC | MODIFIER | | | | hitA | | | | | | | T | T | | | | |
| NC_011770 | 5594147 | CT | C | 60 | 0 | 10 | 0 | INTRAGENIC | MODIFIER | | | | hitA | C | C | C | C | C | C | C | C | | | C | C |
| NC_011770 | 5594158 | G | A | 60 | 0 | 2 | 0 | INTRAGENIC | MODIFIER | | | | hitA | | | | | | | | | | | A | A |
| NC_011770 | 3667732 | G | A | 65 | 0 | 2 | 0 | NON_SYNONYMOU | MODERAT | MISSENSE | gGc/gAc | G214D | hmgA | A | A | | | | | | | | | | |
| NC_011770 | 5619333 | T | C | 86 | 0 | 1 | 0 | NON_SYNONYMOU | MODERAT | MISSENSE | Acc/Gcc | T176A | hmuV | | | | | | | C | | | | | |
| NC_011770 | 1065126 | C | T | 46 | 0 | 2 | 0 | NON_SYNONYMOU | MODERAT | MISSENSE | aGc/aAc | S299N | holA | T | T | | | | | | | | | | |
| NC_011770 | 1065958 | C | T | 53 | 0 | 2 | 0 | NON_SYNONYMOU | MODERAT | MISSENSE | Gtc/Atc | V22I | holA | | | | | | | | | T | T | | |
| NC_011770 | 2274258 | G | A | 72 | 0 | 2 | 0 | NON_SYNONYMOU | MODERAT | MISSENSE | Gcc/Acc | A53T | holB | | | | | | | | | | | A | A |
| NC_011770 | 2274886 | A | G | 78 | 0 | 2 | 0 | NON_SYNONYMOU | MODERAT | MISSENSE | cAc/cGc | H262R | holB | | | | | | | | | | | G | G |
| NC_011770 | 1239451 | C | T | 66 | 0 | 12 | 0 | NON_SYNONYMOU | MODERAT | MISSENSE | cGg/cAg | R34Q | holC | T | T | T | T | T | T | T | T | T | T | T | T |
| NC_011770 | 1348024 | G | A | 73 | 0 | 4 | 0 | NON_SYNONYMOU | MODERAT | MISSENSE | Gaa/Aaa | E23K | hom | | | | | | | A | A | A | A | | |
| NC_011770 | 1348364 | T | C | 76 | 0 | 1 | 0 | NON_SYNONYMOU | MODERAT | MISSENSE | gTg/gCg | V136A | hom | | | | | | | | C | | | | |
| NC_011770 | 963880 | C | CG | 75 | 0 | 1 | 0 | FRAME_SHIFT | HIGH | | ccc/cccC | P231P? | hpaA | | | | | | | | CG | | | | |
| NC_011770 | 963135 | T | C | 78 | 0 | 12 | 0 | NON_SYNONYMOU | MODERAT | MISSENSE | gAg/gGg | E480G | hpaA | C | C | C | C | C | C | C | C | C | C | C | C |
| NC_011770 | 963589 | A | G | 85 | 0 | 1 | 0 | NON_SYNONYMOU | MODERAT | MISSENSE | Tcc/Ccc | S329P | hpaA | | | | | | | G | | | | | |
| NC_011770 | 964030 | C | T | 65 | 0 | 2 | 0 | NON_SYNONYMOU | MODERAT | MISSENSE | Gtc/Atc | V182I | hpaA | | | | | | | | | T | T | | |
| NC_011770 | 962784 | C | T | 78 | 0 | 2 | 0 | NON_SYNONYMOU | MODERAT | MISSENSE | Gcc/Acc | A67T | hpaC | | | | | | | T | T | | | | |
| NC_011770 | 922288 | C | T | 96 | 0 | 1 | 0 | NON_SYNONYMOU | MODERAT | MISSENSE | Ggc/Agc | G122S | hpcB | | | | | | | T | | | | | |
| NC_011770 | 922330 | C | T | 97 | 0 | 2 | 0 | NON_SYNONYMOU | MODERAT | MISSENSE | Gcg/Acg | A108T | hpcB | | | | | | | | | T | T | | |
| NC_011770 | 924138 | C | T | 70 | 0 | 2 | 0 | NON_SYNONYMOU | MODERAT | MISSENSE | Gcg/Acg | A36T | hpcC | | | | | | | | | | | T | T |
| NC_011770 | 924218 | C | T | 79 | 0 | 2 | 0 | NON_SYNONYMOU | MODERAT | MISSENSE | cGc/cAc | R9H | hpcC | | | | | | | | | T | T | | |
| NC_011770 | 921551 | C | T | 92 | 0 | 4 | 0 | NON_SYNONYMOU | MODERAT | MISSENSE | aGg/aAg | R56K | hpcD | T | T | | | | | T | T | | | | |
| NC_011770 | 921668 | T | C | 87 | 0 | 2 | 0 | NON_SYNONYMOU | MODERAT | MISSENSE | gAc/gGc | D17G | hpcD | C | C | | | | | | | | | | |
| NC_011770 | 4891873 | C | T | 76 | 0 | 2 | 0 | NON_SYNONYMOU | MODERAT | MISSENSE | Gat/Aat | D228N | hpd | | | | | | | | | | | T | T |
| NC_011770 | 4892187 | C | G | 75 | 0 | 12 | 0 | NON_SYNONYMOU | MODERAT | MISSENSE | tGc/tCc | C123S | hpd | G | G | G | G | G | G | G | G | G | G | G | G |
| NC_011770 | 4892412 | T | C | 65 | 0 | 1 | 0 | NON_SYNONYMOU | MODERAT | MISSENSE | aAg/aGg | K48R | hpd | | | | | | | C | | | | | |
| NC_011770 | 5528411 | G | A | 71 | 0 | 12 | 0 | NON_SYNONYMOU | MODERAT | MISSENSE | cGc/cAc | R26H | hprA | A | A | A | A | A | A | A | A | A | A | A | A |
| NC_011770 | 5528507 | C | T | 66 | 0 | 2 | 0 | NON_SYNONYMOU | MODERAT | MISSENSE | gCg/gTg | A58V | hprA | | | | | | | T | T | | | | |
| NC_011770 | 5528725 | A | G | 69 | 0 | 8 | 0 | NON_SYNONYMOU | MODERAT | MISSENSE | Agc/Ggc | S131G | hprA | G | G | | | | | G | G | G | G | G | G |
| NC_011770 | 1261858 | G | C | 80 | 0 | 10 | 0 | NON_SYNONYMOU | MODERAT | MISSENSE | Gag/Cag | E252Q | hscA | C | C | C | C | C | C | C | C | C | C | | |
| NC_011770 | 1262794 | G | A | 91 | 0 | 2 | 0 | NON_SYNONYMOU | MODERAT | MISSENSE | Gcg/Acg | A564T | hscA | | | | | | | | | | | A | A |
| NC_011770 | 6182403 | C | CG | 70 | 0 | 1 | 0 | FRAME_SHIFT | HIGH | | ccc/cccC | P180P? | hslO | | | | | | | CG | | | | | |
| NC_011770 | 6182579 | T | C | 74 | 0 | 2 | 0 | NON_SYNONYMOU | MODERAT | MISSENSE | aAg/aGg | K122R | hslO | | | | | | | C | C | | | | |
| NC_011770 | 2411401 | TCG | T | 52 | 0 | 1 | 0 | FRAME_SHIFT | HIGH | | cccgac/ | PD253 | htpX | | | | | | | T | | | | | |
| NC_011770 | 2411567 | C | CGAA | 56 | 0 | 1 | 0 | CODON_INSERTIOI | MODERATE | | ttc/ttcTTC | F198FF | htpX | | | CGAA | | | | | | | | | |
| NC_011770 | 6066806 | T | C | 69 | 0 | 12 | 0 | NON_SYNONYMOU | MODERAT | MISSENSE | cAc/cGc | H262R | hutG | C | C | C | C | C | C | C | C | C | C | C | C |
| NC_011770 | 6067521 | G | A | 94 | 0 | 2 | 0 | NON_SYNONYMOU | MODERAT | MISSENSE | Cgc/Tgc | R24C | hutG | A | A | | | | | | | | | | |
| NC_011770 | 6067774 | T | C | 88 | 0 | 8 | 0 | NON_SYNONYMOU | MODERAT | MISSENSE | Acg/Gcg | T340A | hutI | C | C | C | C | C | C | | | | | C | C |
| NC_011770 | 6068511 | G | C | 74 | 0 | 4 | 0 | NON_SYNONYMOU | MODERAT | MISSENSE | gCc/gGc | A94G | hutI | | | C | C | C | C | | | | | | |
| NC_011770 | 6068523 | A | T | 75 | 0 | 12 | 0 | NON_SYNONYMOU | MODERAT | MISSENSE | gTg/gAg | V90E | hutI | T | T | T | T | T | T | T | T | T | T | T | T |
| NC_011770 | 6068631 | C | T | 83 | 0 | 4 | 0 | NON_SYNONYMOU | MODERAT | MISSENSE | gGt/gAt | G54D | hutI | T | T | | | | | | | | | T | T |
| NC_011770 | 6068658 | C | T | 79 | 0 | 2 | 0 | NON_SYNONYMOU | MODERAT | MISSENSE | gGc/gAc | G45D | hutI | | | | | | | | | | | T | T |
| NC_011770 | 6068731 | C | G | 83 | 0 | 4 | 0 | NON_SYNONYMOU | MODERAT | MISSENSE | Gtc/Ctc | V21L | hutI | | | G | G | G | G | | | | | | |
| NC_011770 | 6077929 | T | C | 73 | 0 | 1 | 0 | NON_SYNONYMOU | MODERAT | MISSENSE | cAg/cGg | Q500R | hutU | | | | C | | | | | | | | |
| NC_011770 | 6077947 | ACGC | A | 73 | 0 | 1 | 0 | CODON_CHANGE_ | MODERATE | | ggcgta/gta | GV493V | hutU | | | | | | | | A | | | | |
| NC_011770 | 6077889 | C | G | 76 | 0 | 2 | 0 | NON_SYNONYMOU | MODERAT | MISSENSE | gaG/gaC | E513D | hutU | G | G | | | | | | | | | | |
| NC_011770 | 2672521 | A | G | 43 | 0 | 2 | 0 | NON_SYNONYMOU | MODERAT | MISSENSE | gAc/gGc | D376G | icd | | | G | G | | | | | | | | |
| NC_011770 | 2672199 | A | G | 71 | 0 | 1 | 0 | NON_SYNONYMOU | MODERAT | MISSENSE | Aag/Gag | K269E | icd | | | | | | | | G | | | | |
| NC_011770 | 5225505 | C | G | 62 | 0 | 6 | 0 | NON_SYNONYMOU | MODERAT | MISSENSE | Gcg/Ccg | A149P | iciA | G | G | | | | | | | G | G | G | G |
| NC_011770 | 5022369 | G | T | 51 | 0 | 4 | 0 | NON_SYNONYMOU | MODERAT | MISSENSE | Gac/Tac | D31Y | icp | | | T | T | T | T | | | | | | |
| NC_011770 | 5022371 | C | A | 50 | 0 | 4 | 0 | NON_SYNONYMOU | MODERAT | MISSENSE | gaC/gaA | D31E | icp | | | | | | | A | A | A | A | | |
| NC_011770 | 5022448 | C | G | 42 | 0 | 4 | 0 | NON_SYNONYMOU | MODERAT | MISSENSE | aCc/aGc | T57S | icp | | | G | G | G | G | | | | | | |
| NC_011770 | 2668972 | T | C | 68 | 0 | 2 | 0 | NON_SYNONYMOU | MODERAT | MISSENSE | Aag/Gag | K689E | idh | C | C | | | | | | | | | | |
| NC_011770 | 2669690 | G | C | 68 | 0 | 12 | 0 | NON_SYNONYMOU | MODERAT | MISSENSE | gaC/gaG | D449E | idh | C | C | C | C | C | C | C | C | C | C | C | C |
| NC_011770 | 2670715 | C | G | 73 | 0 | 4 | 0 | NON_SYNONYMOU | MODERAT | MISSENSE | Gtt/Ctt | V108L | idh | | | | | | | G | G | | | G | G |
| NC_011770 | 2670822 | G | A | 69 | 0 | 12 | 0 | NON_SYNONYMOU | MODERAT | MISSENSE | gCc/gTc | A72V | idh | A | A | A | A | A | A | A | A | A | A | A | A |
| NC_011770 | 2503659 | A | C | 75 | 0 | 12 | 0 | NON_SYNONYMOU | MODERAT | MISSENSE | caA/caC | Q41H | ihfA | C | C | T | T | T | T | T | T | C | C | C | C |
| NC_011770 | 5444762 | C | T | 84 | 0 | 1 | 0 | NON_SYNONYMOU | MODERAT | MISSENSE | Gtc/Atc | V52I | ileS | | | | | | T | | | | | | |
| NC_011770 | 5442349 | A | G | 89 | 0 | 8 | 0 | NON_SYNONYMOU | MODERAT | MISSENSE | gTc/gCc | V856A | ileS | G | G | | | | | G | G | G | G | G | G |
| NC_011770 | 5442368 | C | T | 99 | 0 | 2 | 0 | NON_SYNONYMOU | MODERAT | MISSENSE | Gcg/Acg | A850T | ileS | | | | | | | T | T | | | | |

| | | | | | | | | | | | | | | | | | | | | | | | | | |
|---|---|---|---|---|---|---|---|---|---|---|---|---|---|---|---|---|---|---|---|---|---|---|---|---|---|
| NC_011770 | 5443332 | C | G | 86 | 0 | 2 | 0 | NON_SYNONYMOU | MODERAT | MISSENSE | gaG/gaC | E528D | ileS | G | G | | | | | | | | | | |
| NC_011770 | 5443347 | G | C | 79 | 0 | 6 | 0 | NON_SYNONYMOU | MODERAT | MISSENSE | gaC/gaG | D523E | ileS | C | C | | | | | | | C | C | C | C |
| NC_011770 | 4265439 | G | A | 65 | 0 | 6 | 0 | NON_SYNONYMOU | MODERAT | MISSENSE | aCc/aTc | T390I | ilvA2 | | | A | A | A | A | | | A | A | | |
| NC_011770 | 390827 | T | C | 93 | 0 | 1 | 0 | NON_SYNONYMOU | MODERAT | MISSENSE | gAc/gGc | D270G | ilvD | | | | | | | | C | | | | |
| NC_011770 | 5243531 | C | T | 79 | 0 | 4 | 0 | NON_SYNONYMOU | MODERAT | MISSENSE | Gtc/Atc | V95I | inaA | T | T | | | | | | | | | T | T |
| NC_011770 | 5663216 | G | A | 80 | 0 | 8 | 0 | NON_SYNONYMOU | MODERAT | MISSENSE | gCg/gTg | A187V | infB | | | A | A | A | A | A | A | A | A | | |
| NC_011770 | 5663279 | A | G | 77 | 0 | 2 | 0 | NON_SYNONYMOU | MODERAT | MISSENSE | gTt/gCt | V166A | infB | | | | | | | | | | | G | G |
| NC_011770 | 5573001 | A | C | 74 | 0 | 2 | 0 | NON_SYNONYMOU | MODERAT | MISSENSE | aAg/aCg | K155T | ipk | C | C | | | | | | | | | | |
| NC_011770 | 5573042 | G | T | 72 | 0 | 6 | 0 | NON_SYNONYMOU | MODERAT | MISSENSE | Gtg/Ttg | V169L | ipk | T | T | T | T | T | T | | | | | | |
| NC_011770 | 5573313 | A | G | 70 | 0 | 4 | 0 | NON_SYNONYMOU | MODERAT | MISSENSE | gAc/gGc | D259G | ipk | | | | | | | G | G | G | G | | |
| NC_011770 | 5573348 | G | A | 73 | 0 | 6 | 0 | NON_SYNONYMOU | MODERAT | MISSENSE | Gtc/Atc | V271I | ipk | | | | | | | A | A | A | A | A | A |
| NC_011770 | 5813275 | G | A | 52 | 0 | 2 | 0 | NON_SYNONYMOU | MODERAT | MISSENSE | Gtc/Atc | V188I | irlR | | | A | A | | | | | | | | |
| NC_011770 | 5813287 | G | T | 49 | 0 | 2 | 0 | NON_SYNONYMOU | MODERAT | MISSENSE | Gcg/Tcg | A192S | irlR | | | | | | | | | T | T | | |
| NC_011770 | 1258945 | T | C | 90 | 0 | 12 | 0 | NON_SYNONYMOU | MODERAT | MISSENSE | Tcg/Ccg | S134P | iscS | C | C | C | C | C | C | C | C | C | C | C | C |
| NC_011770 | 1259381 | C | T | 94 | 0 | 1 | 0 | NON_SYNONYMOU | MODERAT | MISSENSE | gCg/gTg | A279V | iscS | | | | | | | | T | | | | |
| NC_011770 | 1011508 | A | C | 76 | 0 | 6 | 0 | NON_SYNONYMOU | MODERAT | MISSENSE | Tcg/Gcg | S203A | ispA | | | C | C | C | C | | | | | C | C |
| NC_011770 | 1011532 | C | T | 75 | 0 | 1 | 0 | NON_SYNONYMOU | MODERAT | MISSENSE | Gcc/Acc | A195T | ispA | | T | | | | | | | | | | |
| NC_011770 | 1512179 | G | A | 71 | 0 | 2 | 0 | NON_SYNONYMOU | MODERAT | MISSENSE | Gac/Aac | D62N | ispD | | | | | | | | | | | A | A |
| NC_011770 | 1512246 | C | T | 65 | 0 | 4 | 0 | NON_SYNONYMOU | MODERAT | MISSENSE | gCc/gTc | A84V | ispD | | | T | T | T | T | | | | | | |
| NC_011770 | 1267858 | T | C | 75 | 0 | 1 | 0 | NON_SYNONYMOU | MODERAT | MISSENSE | gTg/gCg | V224A | ispG | | | | | | | C | | | | | |
| NC_011770 | 5440871 | C | T | 67 | 0 | 2 | 0 | NON_SYNONYMOU | MODERAT | MISSENSE | Gac/Aac | D65N | ispH | | | | | | | | | | | T | T |
| NC_011770 | 739380 | G | A | 81 | 0 | 2 | 0 | NON_SYNONYMOU | MODERAT | MISSENSE | Gaa/Aaa | E2K | katA | | | | | | | | | | | A | A |
| NC_011770 | 739385 | G | C | 80 | 0 | 1 | 0 | NON_SYNONYMOU | MODERAT | MISSENSE | gaG/gaC | E3D | katA | | | | | | | | | | | C | |
| NC_011770 | 5505492 | G | A | 81 | 0 | 1 | 0 | NON_SYNONYMOU | MODERAT | MISSENSE | gCc/gTc | A457V | katB | | A | | | | | | | | | | |
| NC_011770 | 5505999 | A | G | 84 | 0 | 2 | 0 | NON_SYNONYMOU | MODERAT | MISSENSE | gTg/gCg | V288A | katB | | | | | | | | | | | G | G |
| NC_011770 | 5506489 | G | T | 80 | 0 | 12 | 0 | NON_SYNONYMOU | MODERAT | MISSENSE | Cac/Aac | H125N | katB | T | T | T | T | T | T | T | T | T | T | T | T |
| NC_011770 | 3508720 | C | T | 76 | 0 | 1 | 0 | NON_SYNONYMOU | MODERAT | MISSENSE | gGc/gAc | G613D | katE | | | | | | | | T | | | | |
| NC_011770 | 3465691 | T | C | 67 | 0 | 8 | 0 | NON_SYNONYMOU | MODERAT | MISSENSE | Acc/Gcc | T276A | katN | C | C | C | C | C | C | | | C | C | | |
| NC_011770 | 3465787 | G | T | 81 | 0 | 4 | 0 | NON_SYNONYMOU | MODERAT | MISSENSE | Ctg/Atg | L244M | katN | T | T | | | | | | | T | T | | |
| NC_011770 | 3465885 | G | A | 79 | 0 | 4 | 0 | NON_SYNONYMOU | MODERAT | MISSENSE | gCc/gTc | A211V | katN | | | A | A | A | A | | | | | | |
| NC_011770 | 3466509 | A | G | 71 | 0 | 2 | 0 | NON_SYNONYMOU | MODERAT | MISSENSE | gTa/gCa | V3A | katN | G | G | | | | | | | | | | |
| NC_011770 | 4091189 | C | T | 76 | 0 | 8 | 0 | NON_SYNONYMOU | MODERAT | MISSENSE | Gtc/Atc | V318I | kdpA | T | T | | | | | T | T | T | T | T | T |
| NC_011770 | 4090040 | G | A | 58 | 0 | 6 | 0 | NON_SYNONYMOU | MODERAT | MISSENSE | gCt/gTt | A132V | kdpB | A | A | | | | | A | A | | | A | A |
| NC_011770 | 4090044 | C | T | 57 | 0 | 2 | 0 | NON_SYNONYMOU | MODERAT | MISSENSE | Gtc/Atc | V131I | kdpB | T | T | | | | | | | | | | |
| NC_011770 | 4089531 | C | T | 62 | 0 | 1 | 0 | NON_SYNONYMOU | MODERAT | MISSENSE | Ggt/Agt | G302S | kdpB | | | | | | | | | | | T | |
| NC_011770 | 4087774 | C | T | 86 | 0 | 2 | 0 | NON_SYNONYMOU | MODERAT | MISSENSE | Gcg/Acg | A179T | kdpC | | | | | | | | | | | T | T |
| NC_011770 | 4088125 | C | T | 88 | 0 | 2 | 0 | NON_SYNONYMOU | MODERAT | MISSENSE | Gac/Aac | D62N | kdpC | | | | | | | | | T | T | | |
| NC_011770 | 4088260 | T | G | 82 | 0 | 2 | 0 | NON_SYNONYMOU | MODERAT | MISSENSE | Atg/Ctg | M17L | kdpC | | | | | | | | | | | G | G |
| NC_011770 | 4085030 | G | A | 80 | 0 | 4 | 0 | NON_SYNONYMOU | MODERAT | MISSENSE | Ccg/Tcg | P874S | kdpD | | | A | A | A | A | | | | | | |
| NC_011770 | 4087340 | G | C | 79 | 0 | 4 | 0 | NON_SYNONYMOU | MODERAT | MISSENSE | Ctg/Gtg | L104V | kdpD | | | C | C | C | C | | | | | | |
| NC_011770 | 4087351 | G | A | 75 | 0 | 2 | 0 | NON_SYNONYMOU | MODERAT | MISSENSE | cCg/cTg | P100L | kdpD | | | | | | | | | | | A | A |
| NC_011770 | 4086077 | GGGC | G | 70 | 0 | 1 | 0 | CODON_DELETION | MODERATE | | gcc/ | A524 | kdpD | | | | | | | | G | | | | |
| NC_011770 | 4084750 | T | C | 81 | 0 | 2 | 0 | NON_SYNONYMOU | MODERAT | MISSENSE | Aac/Gac | N48D | kdpE | C | C | | | | | | | | | | |
| NC_011770 | 2257395 | C | A | 79 | 0 | 8 | 0 | NON_SYNONYMOU | MODERAT | MISSENSE | aCc/aAc | T234N | kdsB | A | A | A | A | A | A | | | | | A | A |
| NC_011770 | 4532015 | G | A | 83 | 0 | 2 | 0 | STOP_GAINED | HIGH | NONSENSE | Cag/Tag | Q113* | kefB | | | | | | | | | | | A | A |
| NC_011770 | 4531813 | C | T | 73 | 0 | 4 | 0 | NON_SYNONYMOU | MODERAT | MISSENSE | gGc/gAc | G180D | kefB | | | | | | | | | T | T | T | T |
| NC_011770 | 4531818 | T | G | 73 | 0 | 12 | 0 | NON_SYNONYMOU | MODERAT | MISSENSE | gaA/gaC | E178D | kefB | G | G | G | G | G | G | G | G | G | G | G | G |
| NC_011770 | 4532051 | G | C | 85 | 0 | 2 | 0 | NON_SYNONYMOU | MODERAT | MISSENSE | Ctg/Gtg | L101V | kefB | C | C | | | | | | | | | | |
| NC_011770 | 641758 | C | T | 83 | 0 | 2 | 0 | NON_SYNONYMOU | MODERAT | MISSENSE | Gcc/Acc | A252T | ksgA | T | T | | | | | | | | | | |
| NC_011770 | 641929 | C | T | 78 | 0 | 2 | 0 | NON_SYNONYMOU | MODERAT | MISSENSE | Gcc/Acc | A195T | ksgA | | | | | | | T | T | | | | |
| NC_011770 | 642136 | G | A | 86 | 0 | 6 | 0 | NON_SYNONYMOU | MODERAT | MISSENSE | Cac/Tac | H126Y | ksgA | | | A | A | A | A | A | A | | | | |
| NC_011770 | 642226 | C | T | 91 | 0 | 2 | 0 | NON_SYNONYMOU | MODERAT | MISSENSE | Gcc/Acc | A96T | ksgA | | | | | | | | | | | T | T |
| NC_011770 | 3574861 | A | G | 80 | 0 | 2 | 0 | NON_SYNONYMOU | MODERAT | MISSENSE | aAc/aGc | N16S | kynB | | | | | | | G | G | | | | |
| NC_011770 | 3575037 | G | A | 81 | 0 | 6 | 0 | NON_SYNONYMOU | MODERAT | MISSENSE | Gtc/Atc | V75I | kynB | A | A | | | | | | | A | A | A | A |
| NC_011770 | 3575074 | G | A | 79 | 0 | 2 | 0 | NON_SYNONYMOU | MODERAT | MISSENSE | gGc/gAc | G87D | kynB | A | A | | | | | | | | | | |
| NC_011770 | 3575227 | G | T | 76 | 0 | 2 | 0 | NON_SYNONYMOU | MODERAT | MISSENSE | cGc/cTc | R138L | kynB | | | | | | | T | T | | | | |
| NC_011770 | 3575266 | A | C | 73 | 0 | 2 | 0 | NON_SYNONYMOU | MODERAT | MISSENSE | gAc/gCc | D151A | kynB | | | | | | | | | | | C | C |
| NC_011770 | 3575310 | C | T | 77 | 0 | 2 | 0 | NON_SYNONYMOU | MODERAT | MISSENSE | Cgc/Tgc | R166C | kynB | | | | | | | T | T | | | | |
| NC_011770 | 1080646 | C | A | 84 | 0 | 2 | 0 | NON_SYNONYMOU | MODERAT | MISSENSE | Ctt/Att | L10I | ladS | | | | | | | | | | | A | A |
| NC_011770 | 1081867 | T | C | 88 | 0 | 12 | 0 | NON_SYNONYMOU | MODERAT | MISSENSE | Tgc/Cgc | C417R | ladS | C | C | C | C | C | C | C | C | C | C | C | C |
| NC_011770 | 1082566 | C | A | 78 | 0 | 4 | 0 | NON_SYNONYMOU | MODERAT | MISSENSE | Ctg/Atg | L650M | ladS | | | A | A | A | A | | | | | | |
| NC_011770 | 3834700 | G | T | 93 | 0 | 2 | 0 | NON_SYNONYMOU | MODERAT | MISSENSE | Ctc/Atc | L230I | lasA | | | | | | | | | T | T | | |

| | | | | | | | | | | | | | | | | | | | | | | | | | |
|---|---|---|---|---|---|---|---|---|---|---|---|---|---|---|---|---|---|---|---|---|---|---|---|---|---|
| NC_011770 | 3835056 | A | G | 96 | 0 | 6 | 0 | NON_SYNONYMOU | MODERAT | MISSENSE | gTc/gCc | V111A | lasA | G | G | | | | | G | G | G | G | | |
| NC_011770 | 3835150 | C | A | 89 | 0 | 6 | 0 | NON_SYNONYMOU | MODERAT | MISSENSE | Gtg/Ttg | V80L | lasA | A | A | | | | | A | A | A | A | | |
| NC_011770 | 3835354 | G | A | 83 | 0 | 6 | 0 | NON_SYNONYMOU | MODERAT | MISSENSE | Ccg/Tcg | P12S | lasA | A | A | | | | | A | A | A | A | | |
| NC_011770 | 1367741 | A | G | 83 | 0 | 6 | 0 | NON_SYNONYMOU | MODERAT | MISSENSE | Agc/Ggc | S241G | lasB | | | G | G | G | G | G | G | | | | |
| NC_011770 | 1368399 | G | C | 85 | 0 | 2 | 0 | NON_SYNONYMOU | MODERAT | MISSENSE | aGc/aCc | S460T | lasB | | | | | | | | | | | C | C |
| NC_011770 | 4394177 | G | A | 85 | 0 | 2 | 0 | STOP_GAINED | HIGH | NONSENSE | Cag/Tag | Q186* | lasR | | | | | A | A | | | | | | |
| NC_011770 | 4394057 | C | T | 72 | 0 | 1 | 0 | NON_SYNONYMOU | MODERAT | MISSENSE | Gta/Ata | V226I | lasR | | | | | | | | | | T | | |
| NC_011770 | 4394510 | T | G | 84 | 0 | 1 | 0 | NON_SYNONYMOU | MODERAT | MISSENSE | Acg/Ccg | T75P | lasR | | | G | | | | | | | | | |
| NC_011770 | 4394072 | C | T | 76 | 0 | 1 | 0 | NON_SYNONYMOU | MODERAT | MISSENSE | Gtg/Atg | V221M | lasR | | | | | | | T | | | | | |
| NC_011770 | 4394155 | G | A | 81 | 0 | 2 | 0 | NON_SYNONYMOU | MODERAT | MISSENSE | aCc/aTc | **T193I** | lasR | | | | | | | | | | | A | A |
| NC_011770 | 4394062 | C | T | 74 | 0 | 1 | 0 | NON_SYNONYMOU | MODERAT | MISSENSE | cGc/cAc | **R224H** | lasR | | | | | | | | T | | | | |
| NC_011770 | 1767997 | C | T | 82 | 0 | 1 | 0 | STOP_GAINED | HIGH | NONSENSE | Cag/Tag | Q257* | ldh | | | | | | | | T | | | | |
| NC_011770 | 1767580 | A | T | 66 | 0 | 2 | 0 | NON_SYNONYMOU | MODERAT | MISSENSE | Agc/Tgc | S118C | ldh | | | | | | | T | T | | | | |
| NC_011770 | 4825440 | G | C | 93 | 0 | 4 | 0 | NON_SYNONYMOU | MODERAT | MISSENSE | aGc/aCc | S8T | ldhA | C | C | | | | | | | C | C | | |
| NC_011770 | 4825760 | A | G | 84 | 0 | 4 | 0 | NON_SYNONYMOU | MODERAT | MISSENSE | Acg/Gcg | T115A | ldhA | | | G | G | G | G | | | | | | |
| NC_011770 | 4825959 | T | C | 85 | 0 | 2 | 0 | NON_SYNONYMOU | MODERAT | MISSENSE | aTc/aCc | I181T | ldhA | | | | | | | C | C | | | | |
| NC_011770 | 1829470 | C | T | 79 | 0 | 1 | 0 | NON_SYNONYMOU | MODERAT | MISSENSE | Ggc/Agc | G115S | lecB | | | | | | | T | | | | | |
| NC_011770 | 5031262 | G | A | 87 | 0 | 1 | 0 | NON_SYNONYMOU | MODERAT | MISSENSE | Ccg/Tcg | P394S | lepA | | | | | | | A | | | | | |
| NC_011770 | 5032011 | T | C | 72 | 0 | 2 | 0 | NON_SYNONYMOU | MODERAT | MISSENSE | gAg/gGg | E144G | lepA | C | C | | | | | | | | | | |
| NC_011770 | 1279092 | T | C | 74 | 0 | 1 | 0 | NON_SYNONYMOU | MODERAT | MISSENSE | Acc/Gcc | T251A | leuA | | | | | | | C | | | | | |
| NC_011770 | 2093973 | C | A | 71 | 0 | 2 | 0 | NON_SYNONYMOU | MODERAT | MISSENSE | Cgc/Agc | R181S | leuB | | | | | | | A | A | | | | |
| NC_011770 | 2094364 | G | A | 61 | 0 | 2 | 0 | NON_SYNONYMOU | MODERAT | MISSENSE | aGc/aAc | S311N | leuB | | | | | | | | | A | A | | |
| NC_011770 | 2094373 | C | T | 58 | 0 | 2 | 0 | NON_SYNONYMOU | MODERAT | MISSENSE | aCc/aTc | T314I | leuB | | | | | | | | | T | T | | |
| NC_011770 | 2090500 | G | A | 89 | 0 | 2 | 0 | NON_SYNONYMOU | MODERAT | MISSENSE | Gac/Aac | D12N | leuC | | | | | | | | | A | A | | |
| NC_011770 | 1067248 | C | T | 85 | 0 | 6 | 0 | NON_SYNONYMOU | MODERAT | MISSENSE | aGc/aAc | S708N | leuS | T | T | T | T | T | T | | | | | | |
| NC_011770 | 1067255 | T | C | 82 | 0 | 12 | 0 | NON_SYNONYMOU | MODERAT | MISSENSE | Acc/Gcc | T706A | leuS | C | C | C | C | C | C | C | C | C | C | C | C |
| NC_011770 | 1067284 | G | A | 81 | 0 | 2 | 0 | NON_SYNONYMOU | MODERAT | MISSENSE | gCc/gTc | A696V | leuS | | | | | | | A | A | | | | |
| NC_011770 | 1067791 | C | T | 64 | 0 | 2 | 0 | NON_SYNONYMOU | MODERAT | MISSENSE | aGg/aAg | R527K | leuS | T | T | | | | | | | | | | |
| NC_011770 | 1067911 | T | C | 65 | 0 | 6 | 0 | NON_SYNONYMOU | MODERAT | MISSENSE | aAc/aGc | N487S | leuS | C | C | C | C | C | C | | | | | | |
| NC_011770 | 1068260 | T | C | 80 | 0 | 6 | 0 | NON_SYNONYMOU | MODERAT | MISSENSE | Aac/Gac | N371D | leuS | C | C | C | C | C | C | | | | | | |
| NC_011770 | 4200643 | T | C | 80 | 0 | 2 | 0 | NON_SYNONYMOU | MODERAT | MISSENSE | Acg/Gcg | T522A | ligA | C | C | | | | | | | | | | |
| NC_011770 | 4201670 | C | A | 80 | 0 | 2 | 0 | NON_SYNONYMOU | MODERAT | MISSENSE | gaG/gaT | E179D | ligA | A | A | | | | | | | | | | |
| NC_011770 | 4201827 | G | A | 75 | 0 | 1 | 0 | NON_SYNONYMOU | MODERAT | MISSENSE | cCg/cTg | P127L | ligA | | | | | | | A | | | | | |
| NC_011770 | 4202059 | C | T | 70 | 0 | 2 | 0 | NON_SYNONYMOU | MODERAT | MISSENSE | Gcc/Acc | A50T | ligA | T | T | | | | | | | | | | |
| NC_011770 | 3518798 | T | TC | 71 | 0 | 1 | 0 | FRAME_SHIFT | HIGH | | ggg/gggG | G209G? | ligD | | | | | | | TC | | | | | |
| NC_011770 | 3516959 | T | C | 76 | 0 | 12 | 0 | NON_SYNONYMOU | MODERAT | MISSENSE | Acc/Gcc | T823A | ligD | C | C | C | C | C | C | C | C | C | C | C | C |
| NC_011770 | 3517541 | G | C | 71 | 0 | 4 | 0 | NON_SYNONYMOU | MODERAT | MISSENSE | Ccg/Gcg | P629A | ligD | | | C | C | C | C | | | | | | |
| NC_011770 | 3517553 | C | G | 69 | 0 | 12 | 0 | NON_SYNONYMOU | MODERAT | MISSENSE | Gcc/Ccc | A625P | ligD | G | G | G | G | G | G | G | G | G | G | G | G |
| NC_011770 | 3517711 | A | T | 61 | 0 | 2 | 0 | NON_SYNONYMOU | MODERAT | MISSENSE | tTc/tAc | F572Y | ligD | T | T | | | | | | | | | | |
| NC_011770 | 3517814 | T | C | 63 | 0 | 12 | 0 | NON_SYNONYMOU | MODERAT | MISSENSE | Agc/Ggc | S538G | ligD | C | C | C | C | C | C | C | C | C | C | C | C |
| NC_011770 | 3518846 | T | C | 74 | 0 | 10 | 0 | NON_SYNONYMOU | MODERAT | MISSENSE | Acc/Gcc | T194A | ligD | C | C | C | C | C | C | C | C | | | C | C |
| NC_011770 | 3518969 | G | A | 84 | 0 | 2 | 0 | NON_SYNONYMOU | MODERAT | MISSENSE | Ccc/Tcc | P153S | ligD | | | | | | | | | | | A | A |
| NC_011770 | 3519122 | C | A | 81 | 0 | 2 | 0 | NON_SYNONYMOU | MODERAT | MISSENSE | Gcc/Tcc | A102S | ligD | | | | | | | | | | | A | A |
| NC_011770 | 3519131 | A | G | 83 | 0 | 12 | 0 | NON_SYNONYMOU | MODERAT | MISSENSE | Tac/Cac | Y99H | ligD | G | G | G | G | G | G | G | G | G | G | G | G |
| NC_011770 | 2381110 | C | G | 42 | 0 | 2 | 0 | NON_SYNONYMOU | MODERAT | MISSENSE | gGt/gCt | G167A | ligT | | | | | | | G | G | | | | |
| NC_011770 | 2381197 | A | G | 35 | 0 | 6 | 0 | NON_SYNONYMOU | MODERAT | MISSENSE | cTa/cCa | L138P | ligT | | | G | G | G | G | G | G | | | | |
| NC_011770 | 2381203 | G | A | 34 | 0 | 2 | 0 | NON_SYNONYMOU | MODERAT | MISSENSE | gCc/gTc | A136V | ligT | | | | | | | A | A | | | | |
| NC_011770 | 2381221 | A | G | 34 | 0 | 10 | 0 | NON_SYNONYMOU | MODERAT | MISSENSE | gTc/gCc | V130A | ligT | | | G | G | G | G | G | G | G | G | G | G |
| NC_011770 | 2381228 | T | G | 35 | 0 | 12 | 0 | NON_SYNONYMOU | MODERAT | MISSENSE | Agc/Cgc | S128R | ligT | G | G | G | G | G | G | G | G | G | G | G | G |
| NC_011770 | 2381230 | G | A | 35 | 0 | 2 | 0 | NON_SYNONYMOU | MODERAT | MISSENSE | gCt/gTt | A127V | ligT | A | A | | | | | | | | | | |
| NC_011770 | 2381246 | G | C | 35 | 0 | 12 | 0 | NON_SYNONYMOU | MODERAT | MISSENSE | Ccc/Gcc | P122A | ligT | C | C | C | C | C | C | C | C | C | C | C | C |
| NC_011770 | 2381326 | C | T | 36 | 0 | 2 | 0 | NON_SYNONYMOU | MODERAT | MISSENSE | gGc/gAc | G95D | ligT | | | | | | | | | | | T | T |
| NC_011770 | 2381456 | G | A | 45 | 0 | 2 | 0 | NON_SYNONYMOU | MODERAT | MISSENSE | Ccg/Tcg | P52S | ligT | | | | | | | A | A | | | | |
| NC_011770 | 2381524 | T | C | 44 | 0 | 10 | 0 | NON_SYNONYMOU | MODERAT | MISSENSE | cAg/cGg | Q29R | ligT | C | C | C | C | C | C | C | C | C | C | | |
| NC_011770 | 2381527 | T | C | 43 | 0 | 2 | 0 | NON_SYNONYMOU | MODERAT | MISSENSE | gAc/gGc | D28G | ligT | C | C | | | | | | | | | | |
| NC_011770 | 2381561 | G | A | 48 | 0 | 2 | 0 | NON_SYNONYMOU | MODERAT | MISSENSE | Ccc/Tcc | P17S | ligT | | | | | | | | | | | A | A |
| NC_011770 | 2381596 | G | A | 49 | 0 | 8 | 0 | NON_SYNONYMOU | MODERAT | MISSENSE | tCt/tTt | S5F | ligT | | | A | A | A | A | A | A | A | A | | |
| NC_011770 | 2380256 | T | C | 59 | 0 | 4 | 0 | NON_SYNONYMOU | MODERAT | MISSENSE | Atc/Gtc | I204V | lipA | | | C | C | C | C | | | | | | |
| NC_011770 | 2380340 | T | C | 57 | 0 | 2 | 0 | NON_SYNONYMOU | MODERAT | MISSENSE | Acc/Gcc | T176A | lipA | C | C | | | | | | | | | | |
| NC_011770 | 2380400 | T | C | 65 | 0 | 6 | 0 | NON_SYNONYMOU | MODERAT | MISSENSE | Atc/Gtc | I156V | lipA | C | C | | | | | C | C | C | C | | |
| NC_011770 | 2380403 | C | T | 65 | 0 | 1 | 0 | NON_SYNONYMOU | MODERAT | MISSENSE | Gca/Aca | A155T | lipA | | | | | | | T | | | | | |
| NC_011770 | 2380672 | C | T | 77 | 0 | 2 | 0 | NON_SYNONYMOU | MODERAT | MISSENSE | gGt/gAt | G65D | lipA | | | T | T | | | | | | | | |

| | | | | | | | | | | | | | | | | | | | | | | | | | |
|---|---|---|---|---|---|---|---|---|---|---|---|---|---|---|---|---|---|---|---|---|---|---|---|---|---|
| NC_011770 | 5734694 | C | A | 81 | 0 | 2 | 0 | NON_SYNONYMOU | MODERAT | MISSENSE | Gcc/Tcc | A251S | lipC | A | A | | | | | | | | | | |
| NC_011770 | 2378980 | T | C | 57 | 0 | 2 | 0 | NON_SYNONYMOU | MODERAT | MISSENSE | Acc/Gcc | T249A | lipH | | | | | | | | | | | C | C |
| NC_011770 | 1059800 | A | G | 80 | 0 | 1 | 0 | NON_SYNONYMOU | MODERAT | MISSENSE | Acc/Gcc | T38A | lis | | | | G | | | | | | | | |
| NC_011770 | 3659544 | G | A | 83 | 0 | 2 | 0 | NON_SYNONYMOU | MODERAT | MISSENSE | Gcc/Acc | A26T | liuA | | | | | | | | | | | A | A |
| NC_011770 | 3660003 | G | C | 78 | 0 | 8 | 0 | NON_SYNONYMOU | MODERAT | MISSENSE | Gcg/Ccg | A179P | liuA | C | C | C | C | C | C | | | C | C | | |
| NC_011770 | 3661913 | G | T | 82 | 0 | 2 | 0 | NON_SYNONYMOU | MODERAT | MISSENSE | Gcc/Tcc | A387S | liuB | | | | | | | | | | | T | T |
| NC_011770 | 3662768 | G | C | 49 | 0 | 12 | 0 | NON_SYNONYMOU | MODERAT | MISSENSE | gaG/gaC | E131D | liuC | C | C | C | C | C | C | C | C | C | C | C | C |
| NC_011770 | 3662778 | T | C | 54 | 0 | 1 | 0 | NON_SYNONYMOU | MODERAT | MISSENSE | Tgc/Cgc | C135R | liuC | | C | | | | | | | | | | |
| NC_011770 | 3663183 | A | G | 51 | 0 | 4 | 0 | NON_SYNONYMOU | MODERAT | MISSENSE | tAc/tGc | Y5C | liuD | | | G | G | G | G | | | | | | |
| NC_011770 | 3663200 | A | C | 50 | 0 | 12 | 0 | NON_SYNONYMOU | MODERAT | MISSENSE | Atc/Ctc | I11L | liuD | C | C | C | C | C | C | C | C | C | C | C | C |
| NC_011770 | 3663329 | G | A | 45 | 0 | 4 | 0 | NON_SYNONYMOU | MODERAT | MISSENSE | Gcc/Acc | A54T | liuD | A | A | | | | | | | A | A | | |
| NC_011770 | 3663335 | G | A | 46 | 0 | 2 | 0 | NON_SYNONYMOU | MODERAT | MISSENSE | Gac/Aac | D56N | liuD | | | | | | | A | A | | | | |
| NC_011770 | 3663357 | C | T | 46 | 0 | 4 | 0 | NON_SYNONYMOU | MODERAT | MISSENSE | gCc/gTc | A63V | liuD | | | | | | | T | T | | | T | T |
| NC_011770 | 3663380 | G | A | 48 | 0 | 12 | 0 | NON_SYNONYMOU | MODERAT | MISSENSE | Ggg/Agg | G71R | liuD | A | A | A | A | A | A | A | A | A | A | A | A |
| NC_011770 | 3663482 | A | G | 53 | 0 | 12 | 0 | NON_SYNONYMOU | MODERAT | MISSENSE | Agc/Ggc | S105G | liuD | G | G | G | G | G | G | G | G | G | G | G | G |
| NC_011770 | 3663608 | G | C | 54 | 0 | 12 | 0 | NON_SYNONYMOU | MODERAT | MISSENSE | Gcc/Ccc | A147P | liuD | C | C | C | C | C | C | C | C | C | C | C | C |
| NC_011770 | 3663609 | C | T | 54 | 0 | 12 | 0 | NON_SYNONYMOU | MODERAT | MISSENSE | gCc/gTc | A147V | liuD | T | T | T | T | T | T | T | T | T | T | T | T |
| NC_011770 | 3663637 | G | C | 55 | 0 | 12 | 0 | NON_SYNONYMOU | MODERAT | MISSENSE | agG/agC | R156S | liuD | C | C | C | C | C | C | C | C | C | C | C | C |
| NC_011770 | 3663705 | A | G | 62 | 0 | 6 | 0 | NON_SYNONYMOU | MODERAT | MISSENSE | cAc/cGc | H179R | liuD | | | G | G | G | G | | | | | G | G |
| NC_011770 | 3663836 | C | A | 59 | 0 | 12 | 0 | NON_SYNONYMOU | MODERAT | MISSENSE | Cgc/Agc | R223S | liuD | A | A | A | A | A | A | A | A | A | A | A | A |
| NC_011770 | 3664527 | A | G | 56 | 0 | 2 | 0 | NON_SYNONYMOU | MODERAT | MISSENSE | gAc/gGc | D453G | liuD | | | | | | | G | G | | | | |
| NC_011770 | 3664682 | G | T | 57 | 0 | 10 | 0 | NON_SYNONYMOU | MODERAT | MISSENSE | Gcc/Tcc | A505S | liuD | T | T | T | T | T | T | T | T | T | T | | |
| NC_011770 | 3664967 | C | T | 72 | 0 | 2 | 0 | NON_SYNONYMOU | MODERAT | MISSENSE | Ccc/Tcc | P600S | liuD | T | T | | | | | | | | | | |
| NC_011770 | 3665510 | A | G | 80 | 0 | 6 | 0 | NON_SYNONYMOU | MODERAT | MISSENSE | aAg/aGg | K118R | liuE | | | G | G | G | G | | | | | G | G |
| NC_011770 | 3665792 | A | G | 72 | 0 | 1 | 0 | NON_SYNONYMOU | MODERAT | MISSENSE | tAc/tGc | Y212C | liuE | | | | | | | | G | | | | |
| NC_011770 | 3659023 | T | C | 69 | 0 | 2 | 0 | NON_SYNONYMOU | MODERAT | MISSENSE | aTc/aCc | I3T | liuR | | | | | | | | | | | C | C |
| NC_011770 | 3659399 | C | A | 72 | 0 | 2 | 0 | NON_SYNONYMOU | MODERAT | MISSENSE | aaC/aaA | N128K | liuR | | | | | | | | | A | A | | |
| NC_011770 | 3659405 | G | T | 70 | 0 | 12 | 0 | NON_SYNONYMOU | MODERAT | MISSENSE | caG/caT | Q130H | liuR | T | T | T | T | T | T | T | T | T | T | A | A |
| NC_011770 | 3659407 | C | T | 71 | 0 | 2 | 0 | NON_SYNONYMOU | MODERAT | MISSENSE | aCa/aTa | T131I | liuR | | | | | | | | | | | T | T |
| NC_011770 | 3659413 | A | C | 71 | 0 | 2 | 0 | NON_SYNONYMOU | MODERAT | MISSENSE | gAa/gCa | E133A | liuR | | | | | | | | | | | C | C |
| NC_011770 | 4686807 | G | A | 95 | 0 | 12 | 0 | NON_SYNONYMOU | MODERAT | MISSENSE | Gcc/Acc | A46T | livM | A | A | A | A | A | A | A | A | A | A | A | A |
| NC_011770 | 4686954 | A | G | 88 | 0 | 12 | 0 | NON_SYNONYMOU | MODERAT | MISSENSE | Atc/Gtc | I95V | livM | G | G | G | G | G | G | G | G | G | G | G | G |
| NC_011770 | 4687767 | G | A | 87 | 0 | 1 | 0 | NON_SYNONYMOU | MODERAT | MISSENSE | Gtg/Atg | V366M | livM | | | | | | | | A | | | | |
| NC_011770 | 3210179 | T | C | 71 | 0 | 2 | 0 | NON_SYNONYMOU | MODERAT | MISSENSE | aTc/aCc | I4T | lldA | C | C | | | | | | | | | | |
| NC_011770 | 3211273 | C | T | 57 | 0 | 4 | 0 | NON_SYNONYMOU | MODERAT | MISSENSE | Cgc/Tgc | R369C | lldA | | | T | T | T | T | | | | | | |
| NC_011770 | 3210778 | G | A | 79 | 0 | 1 | 0 | NON_SYNONYMOU | MODERAT | MISSENSE | Ggt/Agt | G204S | lldA | | | A | | | | | | | | | |
| NC_011770 | 5691245 | G | A | 93 | 0 | 4 | 0 | NON_SYNONYMOU | MODERAT | MISSENSE | Gac/Aac | D66N | lldD | | | A | A | A | A | | | | | | |
| NC_011770 | 5691281 | G | A | 92 | 0 | 2 | 0 | NON_SYNONYMOU | MODERAT | MISSENSE | Gtc/Atc | V78I | lldD | | | | | | | | | | | A | A |
| NC_011770 | 5690080 | A | G | 78 | 0 | 12 | 0 | NON_SYNONYMOU | MODERAT | MISSENSE | Agc/Ggc | S290G | lldP | G | G | G | G | G | G | G | G | G | G | G | G |
| NC_011770 | 5690389 | T | C | 92 | 0 | 6 | 0 | NON_SYNONYMOU | MODERAT | MISSENSE | Ttc/Ctc | F393L | lldP | | | | | | | C | C | C | C | C | C |
| NC_011770 | 5690407 | G | A | 88 | 0 | 6 | 0 | NON_SYNONYMOU | MODERAT | MISSENSE | Gtg/Atg | V399M | lldP | | | | | | | A | A | A | A | A | A |
| NC_011770 | 5690408 | T | C | 86 | 0 | 12 | 0 | NON_SYNONYMOU | MODERAT | MISSENSE | gTg/gCg | V399A | lldP | C | C | C | C | C | C | C | C | C | C | C | C |
| NC_011770 | 5690453 | A | G | 89 | 0 | 2 | 0 | NON_SYNONYMOU | MODERAT | MISSENSE | aAg/aGg | K414R | lldP | | | | | | | | | G | G | | |
| NC_011770 | 1071266 | C | G | 78 | 0 | 10 | 0 | NON_SYNONYMOU | MODERAT | MISSENSE | Gag/Cag | E376Q | lnt | G | G | G | G | G | G | | | G | G | G | G |
| NC_011770 | 1071470 | T | C | 70 | 0 | 12 | 0 | NON_SYNONYMOU | MODERAT | MISSENSE | Acc/Gcc | T308A | lnt | C | C | C | C | C | C | C | C | C | C | C | C |
| NC_011770 | 1072124 | G | A | 82 | 0 | 4 | 0 | NON_SYNONYMOU | MODERAT | MISSENSE | Ctc/Ttc | L90F | lnt | | | | | | | A | A | | | A | A |
| NC_011770 | 1071416 | C | T | 72 | 0 | 1 | 0 | NON_SYNONYMOU | MODERAT | MISSENSE | Ggc/Agc | G326S | lnt | | T | | | | | | | | | | |
| NC_011770 | 5572420 | A | T | 85 | 0 | 2 | 0 | NON_SYNONYMOU | MODERAT | MISSENSE | Acc/Tcc | T168S | lolB | | | | | | | | | T | T | | |
| NC_011770 | 5753678 | C | T | 76 | 0 | 1 | 0 | NON_SYNONYMOU | MODERAT | MISSENSE | Gcc/Acc | A183T | lpd3 | | | | | | | | | | | | T |
| NC_011770 | 5752918 | G | A | 81 | 0 | 1 | 0 | NON_SYNONYMOU | MODERAT | MISSENSE | gCg/gTg | A436V | lpd3 | | | | | | | | A | | | | |
| NC_011770 | 5752957 | A | G | 84 | 0 | 4 | 0 | NON_SYNONYMOU | MODERAT | MISSENSE | aTc/aCc | I423T | lpd3 | | | G | G | G | G | | | | | | |
| NC_011770 | 5752987 | T | C | 89 | 0 | 1 | 0 | NON_SYNONYMOU | MODERAT | MISSENSE | cAc/cGc | H413R | lpd3 | | | | | | | C | | | | | |
| NC_011770 | 5753710 | G | A | 77 | 0 | 12 | 0 | NON_SYNONYMOU | MODERAT | MISSENSE | gCc/gTc | A172V | lpd3 | A | A | A | A | A | A | A | A | A | A | A | A |
| NC_011770 | 5753845 | T | G | 76 | 0 | 12 | 0 | NON_SYNONYMOU | MODERAT | MISSENSE | gAc/gCc | D127A | lpd3 | G | G | G | G | G | G | G | G | G | G | G | G |
| NC_011770 | 5753852 | T | C | 78 | 0 | 12 | 0 | NON_SYNONYMOU | MODERAT | MISSENSE | Agc/Ggc | S125G | lpd3 | C | C | C | C | C | C | C | C | C | C | C | C |
| NC_011770 | 4138370 | C | T | 68 | 0 | 4 | 0 | NON_SYNONYMOU | MODERAT | MISSENSE | gGc/gAc | G467D | lpdG | | | T | T | T | T | | | | | | |
| NC_011770 | 3366607 | G | A | 75 | 0 | 1 | 0 | NON_SYNONYMOU | MODERAT | MISSENSE | Cgc/Tgc | R300C | lpdV | | | | | | | A | | | | | |
| NC_011770 | 3366727 | T | C | 75 | 0 | 12 | 0 | NON_SYNONYMOU | MODERAT | MISSENSE | Aag/Gag | K260E | lpdV | C | C | C | C | C | C | C | C | C | C | C | C |
| NC_011770 | 3366816 | T | C | 77 | 0 | 2 | 0 | NON_SYNONYMOU | MODERAT | MISSENSE | aAg/aGg | K230R | lpdV | | | | | | | C | C | | | | |
| NC_011770 | 3366958 | T | C | 68 | 0 | 12 | 0 | NON_SYNONYMOU | MODERAT | MISSENSE | Atc/Gtc | I183V | lpdV | C | C | C | C | C | C | C | C | C | C | C | C |
| NC_011770 | 3367297 | C | T | 67 | 0 | 2 | 0 | NON_SYNONYMOU | MODERAT | MISSENSE | Ggc/Agc | G70S | lpdV | T | T | | | | | | | | | | |
| NC_011770 | 7270 | T | C | 76 | 0 | 12 | 0 | NON_SYNONYMOU | MODERAT | MISSENSE | Agc/Ggc | S171G | lptA | C | C | C | C | C | C | C | C | C | C | C | C |

| | | | | | | | | | | | | | | | | | | | | | | | | | |
|---|---|---|---|---|---|---|---|---|---|---|---|---|---|---|---|---|---|---|---|---|---|---|---|---|---|
| NC_011770 | 1498639 | C | G | 78 | 0 | 8 | 0 | NON_SYNONYMOU | MODERAT | MISSENSE | Cag/Gag | Q173E | lpxB | G | G | G | G | G | G | | | | | G | G |
| NC_011770 | 1498696 | C | G | 76 | 0 | 2 | 0 | NON_SYNONYMOU | MODERAT | MISSENSE | Ctg/Gtg | L192V | lpxB | G | G | | | | | | | | | | |
| NC_011770 | 1495814 | T | C | 88 | 0 | 2 | 0 | NON_SYNONYMOU | MODERAT | MISSENSE | Tcc/Ccc | S6P | lpxD | C | C | | | | | | | | | | |
| NC_011770 | 1495856 | G | C | 77 | 0 | 2 | 0 | NON_SYNONYMOU | MODERAT | MISSENSE | Gtc/Ctc | V20L | lpxD | | | | | | | | | C | C | | |
| NC_011770 | 1496015 | G | T | 67 | 0 | 4 | 0 | NON_SYNONYMOU | MODERAT | MISSENSE | Gcc/Tcc | A73S | lpxD | | | T | T | T | T | | | | | | |
| NC_011770 | 2255486 | G | A | 76 | 0 | 2 | 0 | NON_SYNONYMOU | MODERAT | MISSENSE | Gag/Aag | E5K | lpxK | A | A | | | | | | | | | | |
| NC_011770 | 5384085 | G | A | 75 | 0 | 2 | 0 | NON_SYNONYMOU | MODERAT | MISSENSE | Gcc/Acc | A207T | lpxO1 | | | | | | | | | | | A | A |
| NC_011770 | 4811878 | T | C | 48 | 0 | 1 | 0 | NON_SYNONYMOU | MODERAT | MISSENSE | cTg/cCg | L234P | lpxO2 | | | | | | | | C | | | | |
| NC_011770 | 6429008 | A | AT | 77 | 0 | 2 | 0 | FRAME_SHIFT+STA | HIGH | | atg/aAtg | M1N? | ltaA | | | | | | | | | | | AT | AT |
| NC_011770 | 6428195 | A | G | 83 | 0 | 12 | 0 | NON_SYNONYMOU | MODERAT | MISSENSE | gTc/gCc | V272A | ltaA | G | G | G | G | G | G | G | G | G | G | G | G |
| NC_011770 | 6429000 | G | T | 75 | 0 | 2 | 0 | NON_SYNONYMOU | MODERAT | MISSENSE | Cac/Aac | H4N | ltaA | T | T | | | | | | | | | | |
| NC_011770 | 4847650 | C | T | 78 | 0 | 2 | 0 | NON_SYNONYMOU | MODERAT | MISSENSE | Gtg/Atg | V384M | lysC | | | | | | | | | | | T | T |
| NC_011770 | 4848514 | T | C | 87 | 0 | 1 | 0 | NON_SYNONYMOU | MODERAT | MISSENSE | Acc/Gcc | T96A | lysC | | | | | | | C | | | | | |
| NC_011770 | 3670134 | G | A | 61 | 0 | 4 | 0 | NON_SYNONYMOU | MODERAT | MISSENSE | Gac/Aac | D149N | maiA | | | | | | | A | A | | | A | A |
| NC_011770 | 4033236 | C | G | 77 | 0 | 2 | 0 | NON_SYNONYMOU | MODERAT | MISSENSE | Gtg/Ctg | V187L | masA | | | | | | | | | G | G | | |
| NC_011770 | 4033556 | A | G | 74 | 0 | 8 | 0 | NON_SYNONYMOU | MODERAT | MISSENSE | gTg/gCg | V80A | masA | | | G | G | G | G | G | G | G | G | | |
| NC_011770 | 4033680 | G | A | 80 | 0 | 2 | 0 | NON_SYNONYMOU | MODERAT | MISSENSE | Cac/Tac | H39Y | masA | | | | | | | A | A | | | | |
| NC_011770 | 4812160 | C | T | 60 | 0 | 10 | 0 | SPLICE_SITE_REGIC | LOW | SILENT | tGa/tAa | *277 | mazG | T | T | T | T | T | T | T | T | | | T | T |
| NC_011770 | 4812475 | G | C | 63 | 0 | 1 | 0 | NON_SYNONYMOU | MODERAT | MISSENSE | cCg/cGg | P172R | mazG | | C | | | | | | | | | | |
| NC_011770 | 4812389 | T | C | 56 | 0 | 10 | 0 | NON_SYNONYMOU | MODERAT | MISSENSE | Agc/Ggc | S201G | mazG | C | C | C | C | C | C | C | C | C | C | | |
| NC_011770 | 229586 | C | G | 87 | 0 | 2 | 0 | NON_SYNONYMOU | MODERAT | MISSENSE | Cgc/Ggc | R38G | mdcA | G | G | | | | | | | | | | |
| NC_011770 | 230690 | T | C | 71 | 0 | 12 | 0 | NON_SYNONYMOU | MODERAT | MISSENSE | Tcc/Ccc | S406P | mdcA | C | C | C | C | C | C | C | C | C | C | C | C |
| NC_011770 | 230982 | A | T | 88 | 0 | 6 | 0 | NON_SYNONYMOU | MODERAT | MISSENSE | cAg/cTg | Q503L | mdcA | | | T | T | T | T | T | T | | | | |
| NC_011770 | 229890 | C | T | 84 | 0 | 1 | 0 | NON_SYNONYMOU | MODERAT | MISSENSE | gCg/gTg | A139V | mdcA | | | | | | | | | | | | T |
| NC_011770 | 232217 | C | T | 63 | 0 | 12 | 0 | NON_SYNONYMOU | MODERAT | MISSENSE | Ccc/Tcc | P66S | mdcC | T | T | T | T | T | T | T | T | T | T | T | T |
| NC_011770 | 232298 | C | G | 78 | 0 | 2 | 0 | NON_SYNONYMOU | MODERAT | MISSENSE | Ctg/Gtg | L93V | mdcC | | | | | | | | | G | G | | |
| NC_011770 | 233094 | A | G | 73 | 0 | 10 | 0 | NON_SYNONYMOU | MODERAT | MISSENSE | Agc/Ggc | S261G | mdcD | G | G | G | G | G | G | | | G | G | G | G |
| NC_011770 | 233430 | C | T | 76 | 0 | 2 | 0 | NON_SYNONYMOU | MODERAT | MISSENSE | cCa/cTa | P86L | mdcE | | | | | | | T | T | | | | |
| NC_011770 | 233451 | C | T | 81 | 0 | 2 | 0 | NON_SYNONYMOU | MODERAT | MISSENSE | gCc/gTc | A93V | mdcE | | | | | | | T | T | | | | |
| NC_011770 | 233739 | C | T | 87 | 0 | 2 | 0 | NON_SYNONYMOU | MODERAT | MISSENSE | gCg/gTg | A189V | mdcE | | | | | | | T | T | | | | |
| NC_011770 | 5831080 | G | C | 70 | 0 | 2 | 0 | NON_SYNONYMOU | MODERAT | MISSENSE | aCc/aGc | T525S | mdlC | | | | | | | | | | | C | C |
| NC_011770 | 5831194 | A | G | 69 | 0 | 12 | 0 | NON_SYNONYMOU | MODERAT | MISSENSE | gTc/gCc | V487A | mdlC | G | G | G | G | G | G | G | G | G | G | G | G |
| NC_011770 | 5832109 | G | A | 60 | 0 | 2 | 0 | NON_SYNONYMOU | MODERAT | MISSENSE | gCc/gTc | A182V | mdlC | | | | | | | | | A | A | | |
| NC_011770 | 5832148 | A | G | 65 | 0 | 12 | 0 | NON_SYNONYMOU | MODERAT | MISSENSE | gTc/gCc | V169A | mdlC | G | G | G | G | G | G | G | G | G | G | G | G |
| NC_011770 | 5832217 | G | A | 71 | 0 | 2 | 0 | NON_SYNONYMOU | MODERAT | MISSENSE | gCc/gTc | A146V | mdlC | A | A | | | | | | | | | | |
| NC_011770 | 6050281 | A | G | 108 | 0 | 1 | 0 | NON_SYNONYMOU | MODERAT | MISSENSE | Tgg/Cgg | W418R | mdoH | | | | | | | | G | | | | |
| NC_011770 | 2882203 | AGC | A | 1 | 10 | 2 | 0 | FRAME_SHIFT | HIGH | | ctgctg/ | LL469 | merA | . | . | . | . | . | . | A | A | . | . | . | . |
| NC_011770 | 2882307 | C | CG | 7 | 8 | 4 | 0 | FRAME_SHIFT | HIGH | | ctg/ctCg | L435L? | merA | CG | CG | . | . | . | . | CG | CG | . | . | . | . |
| NC_011770 | 2881979 | G | A | 9 | 8 | 2 | 0 | NON_SYNONYMOU | MODERAT | MISSENSE | Ctc/Ttc | L545F | merA | | | . | . | . | . | A | A | . | . | . | . |
| NC_011770 | 2882181 | C | G | 1 | 10 | 2 | 0 | NON_SYNONYMOU | MODERAT | MISSENSE | agG/agC | R477S | merA | . | . | . | . | . | . | G | G | . | . | . | . |
| NC_011770 | 2882225 | T | G | 11 | 8 | 4 | 0 | NON_SYNONYMOU | MODERAT | MISSENSE | Atc/Ctc | I463L | merA | G | G | . | . | . | . | G | G | . | . | . | . |
| NC_011770 | 2882230 | T | G | 10 | 8 | 4 | 0 | NON_SYNONYMOU | MODERAT | MISSENSE | gAc/gCc | D461A | merA | G | G | . | . | . | . | G | G | . | . | . | . |
| NC_011770 | 2882232 | G | T | 10 | 8 | 2 | 0 | NON_SYNONYMOU | MODERAT | MISSENSE | caC/caA | H460Q | merA | T | T | . | . | . | . | | | . | . | . | . |
| NC_011770 | 2882254 | T | A | 12 | 8 | 2 | 0 | NON_SYNONYMOU | MODERAT | MISSENSE | tAc/tTc | Y453F | merA | A | A | . | . | . | . | | | . | . | . | . |
| NC_011770 | 2882312 | C | T | 6 | 8 | 4 | 0 | NON_SYNONYMOU | MODERAT | MISSENSE | Gac/Aac | D434N | merA | T | T | . | . | . | . | T | T | . | . | . | . |
| NC_011770 | 2882317 | G | A | 7 | 8 | 2 | 0 | NON_SYNONYMOU | MODERAT | MISSENSE | gCg/gTg | A432V | merA | A | A | . | . | . | . | | | . | . | . | . |
| NC_011770 | 2882318 | C | T | 7 | 8 | 4 | 0 | NON_SYNONYMOU | MODERAT | MISSENSE | Gcg/Acg | A432T | merA | T | T | . | . | . | . | T | T | . | . | . | . |
| NC_011770 | 2882322 | G | C | 7 | 8 | 4 | 0 | NON_SYNONYMOU | MODERAT | MISSENSE | gaC/gaG | D430E | merA | C | C | . | . | . | . | C | C | . | . | . | . |
| NC_011770 | 2882353 | G | T | 14 | 8 | 2 | 0 | NON_SYNONYMOU | MODERAT | MISSENSE | aCg/aAg | T420K | merA | T | T | . | . | . | . | | | . | . | . | . |
| NC_011770 | 2882393 | C | T | 14 | 8 | 2 | 0 | NON_SYNONYMOU | MODERAT | MISSENSE | Gac/Aac | D407N | merA | T | T | . | . | . | . | | | . | . | . | . |
| NC_011770 | 2882543 | T | G | 0 | 11 | 1 | 0 | NON_SYNONYMOU | MODERAT | MISSENSE | Aag/Cag | K357Q | merA | . | . | . | . | . | . | G | . | . | . | . | . |
| NC_011770 | 2882702 | T | A | 10 | 8 | 2 | 0 | NON_SYNONYMOU | MODERAT | MISSENSE | Acg/Tcg | T304S | merA | A | A | . | . | . | . | | | . | . | . | . |
| NC_011770 | 2882704 | C | T | 10 | 8 | 4 | 0 | NON_SYNONYMOU | MODERAT | MISSENSE | aGc/aAc | S303N | merA | T | T | . | . | . | . | T | T | . | . | . | . |
| NC_011770 | 2882705 | T | G | 9 | 8 | 2 | 0 | NON_SYNONYMOU | MODERAT | MISSENSE | Agc/Cgc | S303R | merA | | | . | . | . | . | G | G | . | . | . | . |
| NC_011770 | 2882724 | C | G | 8 | 8 | 2 | 0 | NON_SYNONYMOU | MODERAT | MISSENSE | caG/caC | Q296H | merA | G | G | . | . | . | . | | | . | . | . | . |
| NC_011770 | 2882725 | T | C | 8 | 8 | 2 | 0 | NON_SYNONYMOU | MODERAT | MISSENSE | cAg/cGg | Q296R | merA | C | C | . | . | . | . | | | . | . | . | . |
| NC_011770 | 2882726 | G | T | 8 | 8 | 2 | 0 | NON_SYNONYMOU | MODERAT | MISSENSE | Cag/Aag | Q296K | merA | | | . | . | . | . | T | T | . | . | . | . |
| NC_011770 | 2882897 | T | C | 11 | 8 | 2 | 0 | NON_SYNONYMOU | MODERAT | MISSENSE | Att/Gtt | I239V | merA | | | . | . | . | . | C | C | . | . | . | . |
| NC_011770 | 2883013 | G | T | 6 | 10 | 2 | 0 | NON_SYNONYMOU | MODERAT | MISSENSE | gCa/gAa | A200E | merA | T | T | . | . | . | . | . | . | . | . | . | . |
| NC_011770 | 2883048 | C | G | 6 | 10 | 2 | 0 | NON_SYNONYMOU | MODERAT | MISSENSE | gaG/gaC | E188D | merA | G | G | . | . | . | . | . | . | . | . | . | . |
| NC_011770 | 2883094 | G | A | 3 | 10 | 2 | 0 | NON_SYNONYMOU | MODERAT | MISSENSE | gCt/gTt | A173V | merA | A | A | . | . | . | . | . | . | . | . | . | . |
| NC_011770 | 2883095 | C | G | 3 | 10 | 2 | 0 | NON_SYNONYMOU | MODERAT | MISSENSE | Gct/Cct | A173P | merA | G | G | . | . | . | . | . | . | . | . | . | . |

| | | | | | | | | | | | | | | | | | | | | | | | | | |
|---|---|---|---|---|---|---|---|---|---|---|---|---|---|---|---|---|---|---|---|---|---|---|---|---|---|
| NC_011770 | 2883101 | C | G | 3 | 10 | 2 | 0 | NON_SYNONYMOU | MODERAT | MISSENSE | Gcg/Ccg | A171P | merA | G | G | . | . | . | . | . | . | . | . | . | . |
| NC_011770 | 2883104 | C | T | 3 | 10 | 2 | 0 | NON_SYNONYMOU | MODERAT | MISSENSE | Gca/Aca | A170T | merA | T | T | . | . | . | . | . | . | . | . | . | . |
| NC_011770 | 2883118 | G | C | 3 | 10 | 2 | 0 | NON_SYNONYMOU | MODERAT | MISSENSE | gCt/gGt | A165G | merA | C | C | . | . | . | . | . | . | . | . | . | . |
| NC_011770 | 2883148 | A | G | 5 | 10 | 2 | 0 | NON_SYNONYMOU | MODERAT | MISSENSE | gTc/gCc | V155A | merA | G | G | . | . | . | . | . | . | . | . | . | . |
| NC_011770 | 2883150 | G | C | 5 | 10 | 2 | 0 | NON_SYNONYMOU | MODERAT | MISSENSE | atC/atG | I154M | merA | C | C | . | . | . | . | . | . | . | . | . | . |
| NC_011770 | 2883152 | T | C | 5 | 10 | 2 | 0 | NON_SYNONYMOU | MODERAT | MISSENSE | Atc/Gtc | I154V | merA | C | C | . | . | . | . | . | . | . | . | . | . |
| NC_011770 | 2883217 | G | A | 11 | 8 | 4 | 0 | NON_SYNONYMOU | MODERAT | MISSENSE | aCc/aTc | T132I | merA | A | A | . | . | . | . | A | A | . | . | . | . |
| NC_011770 | 2883694 | C | A | 3 | 10 | 2 | 0 | NON_SYNONYMOU | MODERAT | MISSENSE | gaG/gaT | E72D | merP | . | . | . | . | . | . | A | A | . | . | . | . |
| NC_011770 | 2884116 | T | G | 4 | 10 | 2 | 0 | NON_SYNONYMOU | MODERAT | MISSENSE | Atc/Ctc | I53L | merT | . | . | . | . | . | . | G | G | . | . | . | . |
| NC_011770 | 3761511 | G | A | 89 | 0 | 1 | 0 | STOP_GAINED | HIGH | NONSENSE | Cag/Tag | Q115* | metE | | | | | | | A | | | | | |
| NC_011770 | 3761793 | C | CT | 78 | 0 | 1 | 0 | FRAME_SHIFT | HIGH | | aaa/aaaA | K20K? | metE | | | | | | | CT | | | | | |
| NC_011770 | 3760738 | C | G | 96 | 0 | 6 | 0 | NON_SYNONYMOU | MODERAT | MISSENSE | gaG/gaC | E372D | metE | G | G | | | | | T | T | T | T | | |
| NC_011770 | 3761330 | C | G | 91 | 0 | 8 | 0 | NON_SYNONYMOU | MODERAT | MISSENSE | gGc/gCc | G175A | metE | | | G | G | G | G | | | G | G | G | G |
| NC_011770 | 3761841 | G | A | 78 | 0 | 1 | 0 | NON_SYNONYMOU | MODERAT | MISSENSE | Cac/Tac | H5Y | metE | | | | | | | | | | | | A |
| NC_011770 | 475571 | A | G | 92 | 0 | 1 | 0 | NON_SYNONYMOU | MODERAT | MISSENSE | cTg/cCg | L80P | metF | | | | | | | G | | | | | |
| NC_011770 | 1660970 | C | G | 73 | 0 | 10 | 0 | NON_SYNONYMOU | MODERAT | MISSENSE | Gag/Cag | E659Q | metG | | | G | G | G | G | G | G | G | G | G | G |
| NC_011770 | 1661398 | C | T | 68 | 0 | 4 | 0 | NON_SYNONYMOU | MODERAT | MISSENSE | cGg/cAg | R516Q | metG | T | T | | | | | T | T | | | | |
| NC_011770 | 1661510 | C | T | 74 | 0 | 1 | 0 | NON_SYNONYMOU | MODERAT | MISSENSE | Ggc/Agc | G479S | metG | | | | | | | T | | | | | |
| NC_011770 | 1662092 | T | C | 81 | 0 | 2 | 0 | NON_SYNONYMOU | MODERAT | MISSENSE | Agt/Ggt | S285G | metG | C | C | | | | | | | | | | |
| NC_011770 | 1662140 | C | T | 77 | 0 | 8 | 0 | NON_SYNONYMOU | MODERAT | MISSENSE | Gcg/Acg | A269T | metG | | | T | T | T | T | | | T | T | T | T |
| NC_011770 | 3867818 | G | A | 78 | 0 | 1 | 0 | STOP_GAINED | HIGH | NONSENSE | tgG/tgA | W1110* | metH | | | | | | | A | | | | | |
| NC_011770 | 3864946 | T | C | 68 | 0 | 1 | 0 | NON_SYNONYMOU | MODERAT | MISSENSE | gTg/gCg | V153A | metH | | | | | | | C | | | | | |
| NC_011770 | 3866163 | C | A | 79 | 0 | 12 | 0 | NON_SYNONYMOU | MODERAT | MISSENSE | Cac/Aac | H559N | metH | A | A | A | A | A | A | A | A | A | A | A | A |
| NC_011770 | 3867499 | G | A | 76 | 0 | 2 | 0 | NON_SYNONYMOU | MODERAT | MISSENSE | cGc/cAc | R1004H | metH | | | | | | | | | A | A | | |
| NC_011770 | 3867534 | G | A | 85 | 0 | 2 | 0 | NON_SYNONYMOU | MODERAT | MISSENSE | Gag/Aag | E1016K | metH | | | | | | | A | A | | | | |
| NC_011770 | 3867708 | G | A | 86 | 0 | 1 | 0 | NON_SYNONYMOU | MODERAT | MISSENSE | Ggg/Agg | G1074R | metH | | | | | | | A | | | | | |
| NC_011770 | 3867987 | G | A | 74 | 0 | 2 | 0 | NON_SYNONYMOU | MODERAT | MISSENSE | Ggc/Agc | G1167S | metH | | | | | | | A | A | | | | |
| NC_011770 | 3867267 | G | A | 85 | 0 | 1 | 0 | NON_SYNONYMOU | MODERAT | MISSENSE | Ggc/Agc | G927S | metH | | | | | | | | | | | A | |
| NC_011770 | 596215 | C | T | 84 | 0 | 2 | 0 | NON_SYNONYMOU | MODERAT | MISSENSE | Gtc/Atc | V322I | metK | | | | | | | | | | | T | T |
| NC_011770 | 596220 | C | G | 82 | 0 | 2 | 0 | NON_SYNONYMOU | MODERAT | MISSENSE | gGc/gCc | G320A | metK | G | G | | | | | | | | | | |
| NC_011770 | 596710 | C | A | 93 | 0 | 2 | 0 | NON_SYNONYMOU | MODERAT | MISSENSE | Ggc/Tgc | G157C | metK | A | A | | | | | | | | | | |
| NC_011770 | 6531555 | C | T | 75 | 0 | 1 | 0 | NON_SYNONYMOU | MODERAT | MISSENSE | Cat/Tat | H39Y | metN | | | | | | | T | | | | | |
| NC_011770 | 1560029 | C | T | 80 | 0 | 6 | 0 | NON_SYNONYMOU | MODERAT | MISSENSE | Ggt/Agt | G178S | metR | T | T | | | | | T | T | | | T | T |
| NC_011770 | 1560110 | T | C | 78 | 0 | 6 | 0 | NON_SYNONYMOU | MODERAT | MISSENSE | Atc/Gtc | I151V | metR | C | C | | | | | C | C | | | C | C |
| NC_011770 | 2893701 | T | G | 22 | 8 | 4 | 0 | NON_SYNONYMOU | MODERAT | MISSENSE | gaT/gaG | D129E | metR | G | G | G | G | . | . | . | . | . | . | . | . |
| NC_011770 | 424482 | CT | C | 86 | 0 | 1 | 0 | FRAME_SHIFT | HIGH | | aag/ | K343 | metX | | | | C | | | | | | | | |
| NC_011770 | 424909 | G | A | 82 | 0 | 12 | 0 | NON_SYNONYMOU | MODERAT | MISSENSE | aCt/aTt | T201I | metX | A | A | A | A | A | A | A | A | A | A | A | A |
| NC_011770 | 425051 | C | T | 74 | 0 | 4 | 0 | NON_SYNONYMOU | MODERAT | MISSENSE | Gtc/Atc | V154I | metX | | | T | T | T | T | | | | | | |
| NC_011770 | 425113 | T | C | 77 | 0 | 1 | 0 | NON_SYNONYMOU | MODERAT | MISSENSE | gAc/gGc | D133G | metX | | | | | | | C | | | | | |
| NC_011770 | 425171 | T | G | 81 | 0 | 4 | 0 | NON_SYNONYMOU | MODERAT | MISSENSE | Atc/Ctc | I114L | metX | | | G | G | G | G | | | | | | |
| NC_011770 | 5991319 | T | TG | 73 | 0 | 1 | 0 | FRAME_SHIFT | HIGH | | ggc/ggGc | G334G? | metY | | | | | | | | TG | | | | |
| NC_011770 | 5990916 | A | G | 54 | 0 | 2 | 0 | NON_SYNONYMOU | MODERAT | MISSENSE | Atc/Gtc | I198V | metY | G | G | | | | | | | | | | |
| NC_011770 | 5990989 | A | G | 61 | 0 | 2 | 0 | NON_SYNONYMOU | MODERAT | MISSENSE | aAa/aGa | K222R | metY | G | G | | | | | | | | | | |
| NC_011770 | 5991087 | A | G | 63 | 0 | 12 | 0 | NON_SYNONYMOU | MODERAT | MISSENSE | Acc/Gcc | T255A | metY | G | G | G | G | G | G | G | G | G | G | G | G |
| NC_011770 | 5991400 | A | G | 75 | 0 | 4 | 0 | NON_SYNONYMOU | MODERAT | MISSENSE | aAg/aGg | K359R | metY | | | G | G | G | G | | | | | | |
| NC_011770 | 2107692 | T | C | 46 | 0 | 12 | 0 | NON_SYNONYMOU | MODERAT | MISSENSE | cTg/cCg | L301P | metZ | C | C | C | C | C | C | C | C | C | C | C | C |
| NC_011770 | 2107772 | T | C | 46 | 0 | 12 | 0 | NON_SYNONYMOU | MODERAT | MISSENSE | Tgc/Cgc | C328R | metZ | C | C | C | C | C | C | C | C | C | C | C | C |
| NC_011770 | 469593 | GC | G | 86 | 0 | 1 | 0 | FRAME_SHIFT | HIGH | | gcg/ | A888 | **mexB** | | | | | | | | **G** | | | | |
| NC_011770 | 469795 | GC | G | 80 | 0 | 1 | 0 | FRAME_SHIFT | HIGH | | ggc/ | G955 | mexB | | | | | | | G | | | | | |
| NC_011770 | 467703 | T | C | 98 | 0 | 2 | 0 | NON_SYNONYMOU | MODERAT | MISSENSE | Tcc/Ccc | S258P | mexB | | | C | C | | | | | | | | |
| NC_011770 | 468946 | G | T | 87 | 0 | 12 | 0 | NON_SYNONYMOU | MODERAT | MISSENSE | cGg/cTg | R672L | mexB | T | T | T | T | T | T | T | T | T | T | T | T |
| NC_011770 | 468126 | G | A | 92 | 0 | 1 | 0 | NON_SYNONYMOU | MODERAT | MISSENSE | Gtg/Atg | V399M | mexB | | | | | | | | | | | | **A** |
| NC_011770 | 5489389 | A | G | 65 | 2 | 10 | 0 | NON_SYNONYMOU | MODERAT | MISSENSE | Tct/Cct | S383P | mexC | G | G | G | G | G | G | G | G | . | . | G | G |
| NC_011770 | 5489548 | C | A | 64 | 2 | 4 | 0 | NON_SYNONYMOU | MODERAT | MISSENSE | Gcg/Tcg | A330S | mexC | A | A | | | | | | | . | . | A | A |
| NC_011770 | 5489607 | C | T | 68 | 2 | 10 | 0 | NON_SYNONYMOU | MODERAT | MISSENSE | cGc/cAc | R310H | mexC | T | T | T | T | T | T | T | T | . | . | T | T |
| NC_011770 | 5490309 | T | C | 80 | 2 | 10 | 0 | NON_SYNONYMOU | MODERAT | MISSENSE | cAg/cGg | Q76R | mexC | C | C | C | C | C | C | C | C | . | . | C | C |
| NC_011770 | 5488113 | G | A | 66 | 2 | 1 | 0 | NON_SYNONYMOU | MODERAT | MISSENSE | gCc/gTc | A411V | mexD | | | | | | | | | . | . | A | |
| NC_011770 | 5486467 | T | G | 92 | 2 | 2 | 0 | NON_SYNONYMOU | MODERAT | MISSENSE | Atc/Ctc | I960L | mexD | G | G | | | | | | | . | . | | |
| NC_011770 | 5486470 | C | A | 90 | 2 | 2 | 0 | NON_SYNONYMOU | MODERAT | MISSENSE | Gcc/Tcc | A959S | mexD | | | | | | | A | A | . | . | | |
| NC_011770 | 5486812 | C | A | 81 | 2 | 2 | 0 | NON_SYNONYMOU | MODERAT | MISSENSE | Gcg/Tcg | A845S | mexD | A | A | | | | | | | . | . | | |
| NC_011770 | 5488576 | G | C | 91 | 2 | 4 | 0 | NON_SYNONYMOU | MODERAT | MISSENSE | Cag/Gag | Q257E | mexD | C | C | | | | | C | C | . | . | | |
| NC_011770 | 3019848 | A | AG | 69 | 0 | 1 | 0 | FRAME_SHIFT | HIGH | | ctc/cCtc | L13P? | mexE | | | | | | | | AG | | | | |

| | | | | | | | | | | | | | | | | | | | | | | | | | |
|---|---|---|---|---|---|---|---|---|---|---|---|---|---|---|---|---|---|---|---|---|---|---|---|---|---|
| NC_011770 | 3018973 | T | A | 64 | 0 | 1 | 0 | NON_SYNONYMOU | MODERAT | MISSENSE | Atc/Ttc | I305F | mexE | | | | | | | A | | | | | |
| NC_011770 | 3019033 | T | C | 57 | 0 | 1 | 0 | NON_SYNONYMOU | MODERAT | MISSENSE | Acc/Gcc | T285A | mexE | | | | | | | C | | | | | |
| NC_011770 | 3019042 | C | T | 61 | 0 | 2 | 0 | NON_SYNONYMOU | MODERAT | MISSENSE | Ggc/Agc | G282S | mexE | T | T | | | | | | | | | | |
| NC_011770 | 3019653 | A | G | 67 | 0 | 1 | 0 | NON_SYNONYMOU | MODERAT | MISSENSE | gTg/gCg | V78A | mexE | | | | | | | G | | | | | |
| NC_011770 | 3019732 | A | G | 68 | 0 | 1 | 0 | NON_SYNONYMOU | MODERAT | MISSENSE | Tgg/Cgg | W52R | mexE | | | | | | | | G | | | | |
| NC_011770 | 3015701 | T | C | 86 | 0 | 12 | 0 | INTRAGENIC | MODIFIER | | | | mexF | C | C | C | C | C | C | C | C | C | C | C | C |
| NC_011770 | 3015950 | G | A | 91 | 0 | 2 | 0 | INTRAGENIC | MODIFIER | | | | mexF | | | | | | | | | | | A | A |
| NC_011770 | 3016225 | A | G | 94 | 0 | 4 | 0 | INTRAGENIC | MODIFIER | | | | mexF | G | G | | | | | G | G | | | | |
| NC_011770 | 3016283 | A | G | 94 | 0 | 2 | 0 | INTRAGENIC | MODIFIER | | | | mexF | | | | | | | | | | | G | G |
| NC_011770 | 3016445 | C | T | 85 | 0 | 6 | 0 | INTRAGENIC | MODIFIER | | | | mexF | T | T | T | T | T | T | | | | | | |
| NC_011770 | 3016658 | C | A | 80 | 0 | 2 | 0 | INTRAGENIC | MODIFIER | | | | mexF | | | | | | | A | A | | | | |
| NC_011770 | 3016778 | C | T | 81 | 0 | 4 | 0 | INTRAGENIC | MODIFIER | | | | mexF | T | T | | | | | | | T | T | | |
| NC_011770 | 3016827 | G | T | 80 | 0 | 2 | 0 | INTRAGENIC | MODIFIER | | | | mexF | T | T | | | | | | | | | | |
| NC_011770 | 3016835 | A | C | 84 | 0 | 4 | 0 | INTRAGENIC | MODIFIER | | | | mexF | | | C | C | C | C | | | | | | |
| NC_011770 | 3016952 | T | C | 77 | 0 | 4 | 0 | INTRAGENIC | MODIFIER | | | | mexF | | | | | | | C | C | | | C | C |
| NC_011770 | 3016979 | G | A | 80 | 0 | 4 | 0 | INTRAGENIC | MODIFIER | | | | mexF | A | A | | | | | | | A | A | | |
| NC_011770 | 3017303 | C | T | 79 | 0 | 2 | 0 | INTRAGENIC | MODIFIER | | | | mexF | | | | | | | T | T | | | | |
| NC_011770 | 3017414 | C | T | 85 | 0 | 2 | 0 | INTRAGENIC | MODIFIER | | | | mexF | | | | | | | | | T | T | | |
| NC_011770 | 3017549 | T | C | 86 | 0 | 12 | 0 | INTRAGENIC | MODIFIER | | | | mexF | C | C | C | C | C | C | C | C | C | C | C | C |
| NC_011770 | 3017627 | G | A | 87 | 0 | 6 | 0 | INTRAGENIC | MODIFIER | | | | mexF | A | A | A | A | A | A | | | | | | |
| NC_011770 | 3017654 | T | C | 89 | 0 | 2 | 0 | INTRAGENIC | MODIFIER | | | | mexF | | | | | | | C | C | | | | |
| NC_011770 | 3017884 | A | C | 72 | 0 | 12 | 0 | INTRAGENIC | MODIFIER | | | | mexF | C | C | C | C | C | C | C | C | C | C | C | C |
| NC_011770 | 3017930 | A | G | 72 | 0 | 4 | 0 | INTRAGENIC | MODIFIER | | | | mexF | | | | | | | G | G | | | G | G |
| NC_011770 | 3017960 | G | A | 72 | 0 | 2 | 0 | INTRAGENIC | MODIFIER | | | | mexF | A | A | | | | | | | | | | |
| NC_011770 | 3018044 | G | A | 76 | 0 | 4 | 0 | INTRAGENIC | MODIFIER | | | | mexF | | | | | | | A | A | | | A | A |
| NC_011770 | 3018071 | G | A | 78 | 0 | 4 | 0 | INTRAGENIC | MODIFIER | | | | mexF | | | | | | | A | A | | | A | A |
| NC_011770 | 3018158 | G | A | 87 | 0 | 2 | 0 | INTRAGENIC | MODIFIER | | | | mexF | | | | | | | | | | | A | A |
| NC_011770 | 3018287 | G | A | 87 | 0 | 2 | 0 | INTRAGENIC | MODIFIER | | | | mexF | A | A | | | | | | | | | | |
| NC_011770 | 3018551 | A | G | 75 | 0 | 2 | 0 | INTRAGENIC | MODIFIER | | | | mexF | | | | | | | G | G | | | | |
| NC_011770 | 3018563 | C | T | 76 | 0 | 2 | 0 | INTRAGENIC | MODIFIER | | | | mexF | | | | | | | | | T | T | | |
| NC_011770 | 3018283 | C | T | 84 | 0 | 1 | 0 | INTRAGENIC | MODIFIER | | | | mexF | | T | | | | | | | | | | |
| NC_011770 | 3017395 | C | T | 87 | 0 | 1 | 0 | INTRAGENIC | MODIFIER | | | | mexF | | | | | | | | T | | | | |
| NC_011770 | 785988 | C | T | 82 | 0 | 2 | 0 | NON_SYNONYMOU | MODERAT | MISSENSE | Gcg/Acg | A233T | mexH | | | | | T | T | | | | | | |
| NC_011770 | 782942 | A | G | 88 | 0 | 1 | 0 | NON_SYNONYMOU | MODERAT | MISSENSE | cTg/cCg | L873P | mexI | | | | | | | G | | | | | |
| NC_011770 | 783215 | G | T | 84 | 0 | 6 | 0 | NON_SYNONYMOU | MODERAT | MISSENSE | gCg/gAg | A782E | mexI | | | T | T | T | T | | | T | T | | |
| NC_011770 | 465421 | T | TG | 68 | 0 | 1 | 0 | FRAME_SHIFT | HIGH | | cgc/cgcC | R23R? | mexR | | | | | | | | | TG | | | |
| NC_011770 | 465096 | A | G | 68 | 0 | 2 | 0 | NON_SYNONYMOU | MODERAT | MISSENSE | gTg/gCg | V132A | mexR | G | G | | | | | | | | | | |
| NC_011770 | 465114 | A | T | 63 | 0 | 6 | 0 | NON_SYNONYMOU | MODERAT | MISSENSE | gTg/gAg | V126E | mexR | T | T | T | T | T | T | | | | | | |
| NC_011770 | 465244 | A | G | 66 | 0 | 12 | 0 | NON_SYNONYMOU | MODERAT | MISSENSE | Tgc/Cgc | C83R | mexR | G | G | G | G | G | G | G | G | G | G | G | G |
| NC_011770 | 3020047 | A | C | 73 | 0 | 2 | 0 | INTRAGENIC | MODIFIER | | | | mexT | | | | | | | | | | | C | C |
| NC_011770 | 3020054 | C | G | 71 | 0 | 2 | 0 | INTRAGENIC | MODIFIER | | | | mexT | | | | | | | | | | | G | G |
| NC_011770 | 3020063 | T | C | 70 | 0 | 12 | 0 | INTRAGENIC | MODIFIER | | | | mexT | C | C | C | C | C | C | C | C | C | C | C | C |
| NC_011770 | 3020074 | T | C | 71 | 0 | 2 | 0 | INTRAGENIC | MODIFIER | | | | mexT | | | | | | | | | | | C | C |
| NC_011770 | 3020077 | C | T | 71 | 0 | 8 | 0 | INTRAGENIC | MODIFIER | | | | mexT | T | T | | | | | T | T | T | T | T | T |
| NC_011770 | 3020173 | G | C | 69 | 0 | 2 | 0 | INTRAGENIC | MODIFIER | | | | mexT | | | | | | | | | | | C | C |
| NC_011770 | 3020224 | G | C | 69 | 0 | 2 | 0 | INTRAGENIC | MODIFIER | | | | mexT | | | | | | | C | C | | | | |
| NC_011770 | 3020374 | C | T | 61 | 0 | 2 | 0 | INTRAGENIC | MODIFIER | | | | mexT | | | | | | | | | T | T | | |
| NC_011770 | 3020425 | T | G | 63 | 0 | 6 | 0 | INTRAGENIC | MODIFIER | | | | mexT | G | G | | | | | G | G | G | G | | |
| NC_011770 | 3020458 | A | G | 63 | 0 | 2 | 0 | INTRAGENIC | MODIFIER | | | | mexT | | | | | | | | | G | G | | |
| NC_011770 | 3020527 | G | A | 60 | 0 | 2 | 0 | INTRAGENIC | MODIFIER | | | | mexT | | | | | | | | | | | A | A |
| NC_011770 | 3020659 | C | T | 57 | 0 | 2 | 0 | INTRAGENIC | MODIFIER | | | | mexT | | | | | | | T | T | | | | |
| NC_011770 | 3020688 | A | G | 56 | 0 | 2 | 0 | INTRAGENIC | MODIFIER | | | | mexT | | | | | | | G | G | | | | |
| NC_011770 | 3020716 | G | T | 54 | 0 | 4 | 0 | INTRAGENIC | MODIFIER | | | | mexT | | | T | T | T | T | | | | | | |
| NC_011770 | 3020767 | A | G | 54 | 0 | 2 | 0 | INTRAGENIC | MODIFIER | | | | mexT | | | | | | | | | | | G | G |
| NC_011770 | 3020788 | A | G | 52 | 0 | 2 | 0 | INTRAGENIC | MODIFIER | | | | mexT | | | | | | | | | | | G | G |
| NC_011770 | 3020839 | T | C | 51 | 0 | 6 | 0 | INTRAGENIC | MODIFIER | | | | mexT | C | C | | | | | | | C | C | C | C |
| NC_011770 | 3020860 | G | A | 50 | 0 | 2 | 0 | INTRAGENIC | MODIFIER | | | | mexT | | | | | | | | | | | A | A |
| NC_011770 | 3020908 | G | A | 48 | 0 | 2 | 0 | INTRAGENIC | MODIFIER | | | | mexT | | | | | | | | | | | A | A |
| NC_011770 | 3020974 | A | G | 45 | 0 | 8 | 0 | INTRAGENIC | MODIFIER | | | | mexT | G | G | G | G | G | G | | | G | G | | |
| NC_011770 | 3021133 | C | T | 34 | 0 | 10 | 0 | INTRAGENIC | MODIFIER | | | | mexT | | | T | T | T | T | T | T | T | T | T | T |
| NC_011770 | 2232537 | T | C | 71 | 0 | 2 | 0 | NON_SYNONYMOU | MODERAT | MISSENSE | gAt/gGt | D586G | mfd | | | C | C | | | | | | | | |
| NC_011770 | 2233198 | C | T | 50 | 0 | 2 | 0 | NON_SYNONYMOU | MODERAT | MISSENSE | Gcc/Acc | A366T | mfd | | | | | | | | | T | T | | |

| | | | | | | | | | | | | | | | | | | | | | | | | | |
|---|---|---|---|---|---|---|---|---|---|---|---|---|---|---|---|---|---|---|---|---|---|---|---|---|---|
| NC_011770 | 2234259 | C | G | 69 | 0 | 2 | 0 | NON_SYNONYMOU | MODERAT | MISSENSE | aGc/aCc | S12T | mfd | | | | | | | | | G | G | | |
| NC_011770 | 5749113 | T | C | 80 | 0 | 12 | 0 | NON_SYNONYMOU | MODERAT | MISSENSE | Acc/Gcc | T504A | mgtA | C | C | C | C | C | C | C | C | C | C | C | C |
| NC_011770 | 5749941 | G | A | 72 | 0 | 2 | 0 | NON_SYNONYMOU | MODERAT | MISSENSE | Cgg/Tgg | R228W | mgtA | | | | | | | A | A | | | | |
| NC_011770 | 4841855 | T | C | 87 | 0 | 1 | 0 | NON_SYNONYMOU | MODERAT | MISSENSE | cAg/cGg | Q166R | mgtE | | | | | | | C | | | | | |
| NC_011770 | 4842189 | C | T | 74 | 0 | 2 | 0 | NON_SYNONYMOU | MODERAT | MISSENSE | Gta/Ata | V55I | mgtE | | | | | | | | | | | T | T |
| NC_011770 | 4841075 | G | A | 46 | 0 | 1 | 0 | NON_SYNONYMOU | MODERAT | MISSENSE | gCc/gTc | A426V | mgtE | | | | | | | | A | | | | |
| NC_011770 | 5881660 | T | A | 88 | 0 | 2 | 0 | NON_SYNONYMOU | MODERAT | MISSENSE | cAg/cTg | Q133L | miaA | | | | | | | | | | | A | A |
| NC_011770 | 5881793 | T | C | 79 | 0 | 12 | 0 | NON_SYNONYMOU | MODERAT | MISSENSE | Aag/Gag | K89E | miaA | C | C | C | C | C | C | C | C | C | C | C | C |
| NC_011770 | 5881982 | C | A | 81 | 0 | 2 | 0 | NON_SYNONYMOU | MODERAT | MISSENSE | Gcc/Tcc | A26S | miaA | | | | | | | | | A | A | | |
| NC_011770 | 1971655 | G | A | 64 | 0 | 2 | 0 | NON_SYNONYMOU | MODERAT | MISSENSE | gCc/gTc | A135V | minC | A | A | | | | | | | | | | |
| NC_011770 | 5311768 | G | A | 34 | 0 | 4 | 0 | NON_SYNONYMOU | MODERAT | MISSENSE | cGc/cAc | R64H | mltB1 | | | A | A | A | A | | | | | | |
| NC_011770 | 5312377 | A | G | 47 | 0 | 12 | 0 | NON_SYNONYMOU | MODERAT | MISSENSE | aAc/aGc | N267S | mltB1 | G | G | G | G | G | G | G | G | G | G | G | G |
| NC_011770 | 3897798 | G | GC | 72 | 0 | 1 | 0 | FRAME_SHIFT | HIGH | | cag/cCag | Q321P? | mltD | | | | | | | | GC | | | | |
| NC_011770 | 3896931 | T | C | 58 | 0 | 12 | 0 | NON_SYNONYMOU | MODERAT | MISSENSE | gTc/gCc | V30A | mltD | C | C | C | C | C | C | C | C | C | C | C | C |
| NC_011770 | 3898238 | C | A | 77 | 0 | 12 | 0 | NON_SYNONYMOU | MODERAT | MISSENSE | Ccc/Acc | P466T | mltD | A | A | A | A | A | A | A | A | A | A | A | A |
| NC_011770 | 1578478 | G | A | 79 | 0 | 2 | 0 | NON_SYNONYMOU | MODERAT | MISSENSE | Gtc/Atc | V44I | mmsA | | | | | | | | | | | A | A |
| NC_011770 | 1579972 | A | G | 77 | 0 | 2 | 0 | NON_SYNONYMOU | MODERAT | MISSENSE | Agc/Ggc | S39G | mmsB | | | | | | | | | G | G | | |
| NC_011770 | 1580305 | A | G | 73 | 0 | 2 | 0 | NON_SYNONYMOU | MODERAT | MISSENSE | Atc/Gtc | I150V | mmsB | | | | | | | G | G | | | | |
| NC_011770 | 1580524 | C | T | 87 | 0 | 2 | 0 | NON_SYNONYMOU | MODERAT | MISSENSE | Ccc/Tcc | P223S | mmsB | | | | | | | | | | | T | T |
| NC_011770 | 1580706 | G | C | 87 | 0 | 2 | 0 | NON_SYNONYMOU | MODERAT | MISSENSE | gaG/gaC | E283D | mmsB | | | | | | | | | | | C | C |
| NC_011770 | 1577956 | C | T | 40 | 0 | 2 | 0 | NON_SYNONYMOU | MODERAT | MISSENSE | Gcc/Acc | A88T | mmsR | | | | | | | | | | | T | T |
| NC_011770 | 1577957 | C | G | 42 | 0 | 4 | 0 | NON_SYNONYMOU | MODERAT | MISSENSE | gaG/gaC | E87D | mmsR | | | | | | | G | G | G | G | | |
| NC_011770 | 2667126 | A | G | 72 | 0 | 2 | 0 | NON_SYNONYMOU | MODERAT | MISSENSE | gTg/gCg | V362A | mnmA | | | | | | | G | G | | | | |
| NC_011770 | 2667228 | G | A | 71 | 0 | 6 | 0 | NON_SYNONYMOU | MODERAT | MISSENSE | aCc/aTc | T328I | mnmA | | | A | A | A | A | | | A | A | | |
| NC_011770 | 2667241 | C | T | 76 | 0 | 4 | 0 | NON_SYNONYMOU | MODERAT | MISSENSE | Gtc/Atc | V324I | mnmA | | | | | | | T | T | | | T | T |
| NC_011770 | 2667528 | A | G | 86 | 0 | 10 | 0 | NON_SYNONYMOU | MODERAT | MISSENSE | aTc/aCc | I228T | mnmA | | | G | G | G | G | G | G | G | G | G | G |
| NC_011770 | 1730729 | A | G | 61 | 0 | 4 | 0 | NON_SYNONYMOU | MODERAT | MISSENSE | gTc/gCc | V576A | mnmC | G | G | | | | | G | G | | | | |
| NC_011770 | 1731458 | C | A | 62 | 0 | 2 | 0 | NON_SYNONYMOU | MODERAT | MISSENSE | cGc/cTc | R333L | mnmC | | | | | | | A | A | | | | |
| NC_011770 | 1732101 | C | T | 71 | 0 | 2 | 0 | NON_SYNONYMOU | MODERAT | MISSENSE | Gag/Aag | E119K | mnmC | T | T | | | | | | | | | | |
| NC_011770 | 1202270 | G | C | 89 | 0 | 12 | 0 | NON_SYNONYMOU | MODERAT | MISSENSE | Ggg/Cgg | G107R | moaA | C | C | C | C | C | C | C | C | C | C | C | C |
| NC_011770 | 4317603 | C | T | 82 | 0 | 4 | 0 | NON_SYNONYMOU | MODERAT | MISSENSE | Gac/Aac | D175N | moaA2 | T | T | | | | | T | T | | | | |
| NC_011770 | 2201247 | T | A | 71 | 0 | 12 | 0 | NON_SYNONYMOU | MODERAT | MISSENSE | cTg/cAg | L32Q | moaB2 | A | A | A | A | A | A | A | A | A | A | A | A |
| NC_011770 | 2200576 | C | T | 60 | 0 | 2 | 0 | NON_SYNONYMOU | MODERAT | MISSENSE | Gag/Aag | E171K | mobA | | | | | | | T | T | | | | |
| NC_011770 | 2200761 | C | T | 60 | 0 | 1 | 0 | NON_SYNONYMOU | MODERAT | MISSENSE | cGc/cAc | R109H | mobA | | | | | | | T | | | | | |
| NC_011770 | 2200936 | C | T | 62 | 0 | 2 | 0 | NON_SYNONYMOU | MODERAT | MISSENSE | Gac/Aac | D51N | mobA | | | | | | | T | T | | | | |
| NC_011770 | 2200986 | C | T | 56 | 0 | 4 | 0 | NON_SYNONYMOU | MODERAT | MISSENSE | gGc/gAc | G34D | mobA | | | T | T | T | T | | | | | | |
| NC_011770 | 3845166 | C | T | 54 | 0 | 2 | 0 | NON_SYNONYMOU | MODERAT | MISSENSE | Ccg/Tcg | P84S | modA | | | | | | | | | | | T | T |
| NC_011770 | 3845638 | T | C | 71 | 0 | 4 | 0 | NON_SYNONYMOU | MODERAT | MISSENSE | gTc/gCc | V241A | modA | C | C | | | | | | | C | C | | |
| NC_011770 | 3846578 | G | A | 69 | 0 | 6 | 0 | NON_SYNONYMOU | MODERAT | MISSENSE | cGc/cAc | R69H | modC | A | A | A | A | A | A | | | | | | |
| NC_011770 | 3846575 | G | A | 68 | 0 | 1 | 0 | NON_SYNONYMOU | MODERAT | MISSENSE | cGc/cAc | R68H | modC | | | | A | | | | | | | | |
| NC_011770 | 1151218 | A | C | 67 | 0 | 4 | 0 | NON_SYNONYMOU | MODERAT | MISSENSE | cAt/cCt | H56P | moeA1 | | | C | C | C | C | | | | | | |
| NC_011770 | 1151478 | C | T | 61 | 0 | 2 | 0 | NON_SYNONYMOU | MODERAT | MISSENSE | Cgc/Tgc | R143C | moeA1 | T | T | | | | | | | | | | |
| NC_011770 | 2201709 | G | T | 78 | 0 | 2 | 0 | NON_SYNONYMOU | MODERAT | MISSENSE | caG/caT | Q7H | moeA2 | | | | | | | T | T | | | | |
| NC_011770 | 2201959 | G | A | 71 | 0 | 8 | 0 | NON_SYNONYMOU | MODERAT | MISSENSE | Gag/Aag | E91K | moeA2 | A | A | A | A | A | A | | | A | A | | |
| NC_011770 | 2201975 | G | A | 74 | 0 | 2 | 0 | NON_SYNONYMOU | MODERAT | MISSENSE | gGt/gAt | G96D | moeA2 | | | | | | | | | A | A | | |
| NC_011770 | 2202095 | C | T | 65 | 0 | 8 | 0 | NON_SYNONYMOU | MODERAT | MISSENSE | gCg/gTg | A136V | moeA2 | T | T | T | T | T | T | | | T | T | | |
| NC_011770 | 2202719 | C | T | 58 | 0 | 2 | 0 | NON_SYNONYMOU | MODERAT | MISSENSE | gCg/gTg | A344V | moeA2 | | | | | | | | | | | T | T |
| NC_011770 | 5566401 | G | A | 69 | 0 | 2 | 0 | NON_SYNONYMOU | MODERAT | MISSENSE | gCg/gTg | A135V | moeB | | | | | | | | | | | A | A |
| NC_011770 | 5566477 | T | C | 78 | 0 | 2 | 0 | NON_SYNONYMOU | MODERAT | MISSENSE | Aag/Gag | K110E | moeB | C | C | | | | | | | | | | |
| NC_011770 | 5566617 | G | A | 88 | 0 | 2 | 0 | NON_SYNONYMOU | MODERAT | MISSENSE | aCg/aTg | T63M | moeB | | | | | | | | | A | A | | |
| NC_011770 | 5494253 | C | G | 89 | 0 | 1 | 0 | NON_SYNONYMOU | MODERAT | MISSENSE | agC/agG | S917R | morA | | | | | | | | | | | | G |
| NC_011770 | 5492704 | G | A | 83 | 0 | 1 | 0 | NON_SYNONYMOU | MODERAT | MISSENSE | gGc/gAc | G401D | morA | | | | | | | | A | | | | |
| NC_011770 | 5494908 | G | A | 95 | 0 | 1 | 0 | NON_SYNONYMOU | MODERAT | MISSENSE | Ggc/Agc | G1136S | morA | | | | | | | | A | | | | |
| NC_011770 | 5491794 | A | G | 59 | 2 | 10 | 0 | NON_SYNONYMOU | MODERAT | MISSENSE | Aac/Gac | N98D | morA | G | G | G | G | G | G | G | G | . | . | G | G |
| NC_011770 | 5491795 | A | G | 59 | 2 | 10 | 0 | NON_SYNONYMOU | MODERAT | MISSENSE | aAc/aGc | N98S | morA | G | G | G | G | G | G | G | G | . | . | G | G |
| NC_011770 | 5491864 | C | T | 58 | 2 | 2 | 0 | NON_SYNONYMOU | MODERAT | MISSENSE | cCg/cTg | P121L | morA | | | | | | | T | T | . | . | | |
| NC_011770 | 5491873 | G | A | 57 | 2 | 10 | 0 | NON_SYNONYMOU | MODERAT | MISSENSE | gGc/gAc | G124D | morA | A | A | A | A | A | A | A | A | . | . | A | A |
| NC_011770 | 5493174 | G | A | 79 | 0 | 2 | 0 | NON_SYNONYMOU | MODERAT | MISSENSE | Gaa/Aaa | E558K | morA | | | | | | | | | | | A | A |
| NC_011770 | 5494981 | A | T | 92 | 0 | 2 | 0 | NON_SYNONYMOU | MODERAT | MISSENSE | gAc/gTc | D1160V | morA | T | T | | | | | | | | | | |
| NC_011770 | 5495487 | G | A | 91 | 0 | 2 | 0 | NON_SYNONYMOU | MODERAT | MISSENSE | Gtg/Atg | V1329M | morA | | | A | A | | | | | | | | |
| NC_011770 | 2307027 | C | T | 53 | 0 | 2 | 0 | NON_SYNONYMOU | MODERAT | MISSENSE | cGc/cAc | R262H | morB | | | | | | | T | T | | | | |
| NC_011770 | 2307162 | A | T | 62 | 0 | 2 | 0 | NON_SYNONYMOU | MODERAT | MISSENSE | aTc/aAc | I217N | morB | | | | | | | | | | | T | T |

| | | | | | | | | | | | | | | | | | | | | | | | | | |
|---|---|---|---|---|---|---|---|---|---|---|---|---|---|---|---|---|---|---|---|---|---|---|---|---|---|
| NC_011770 | 2307803 | G | T | 88 | 0 | 2 | 0 | NON_SYNONYMOU | MODERAT | MISSENSE | aaC/aaA | N3K | morB | | | | | | | | | | | T | T |
| NC_011770 | 5891499 | G | T | 84 | 0 | 2 | 0 | NON_SYNONYMOU | MODERAT | MISSENSE | Ctg/Atg | L237M | motA | T | T | | | | | | | | | | |
| NC_011770 | 4362348 | G | A | 79 | 0 | 10 | 0 | NON_SYNONYMOU | MODERAT | MISSENSE | aCc/aTc | T10I | motC | | | A | A | A | A | A | A | A | A | A | A |
| NC_011770 | 4360876 | T | C | 48 | 0 | 12 | 0 | NON_SYNONYMOU | MODERAT | MISSENSE | Atc/Gtc | I250V | motD | C | C | C | C | C | C | C | C | C | C | C | C |
| NC_011770 | 1037925 | A | AC | 72 | 1 | 1 | 0 | FRAME_SHIFT | HIGH | | atg/Catg | M38H? | mpl | | | | . | AC | | | | | | | |
| NC_011770 | 1037931 | C | T | 55 | 2 | 1 | 0 | NON_SYNONYMOU | MODERAT | MISSENSE | cCc/cTc | P37L | mpl | . | T | | . | | | | | | | | |
| NC_011770 | 1038255 | T | G | 73 | 1 | 2 | 0 | NON_SYNONYMOU | MODERAT | MISSENSE | gTt/gGt | V145G | mpl | | | | . | G | G | | | | | | |
| NC_011770 | 1038564 | AGGC | A | 77 | 1 | 2 | 0 | CODON_DELETION | MODERATE | | gcg/ | A249 | mpl | | | | . | | | | | | | A | A |
| NC_011770 | 1038656 | C | T | 79 | 1 | 1 | 0 | NON_SYNONYMOU | MODERAT | MISSENSE | Cac/Tac | H279Y | mpl | | | T | . | | | | | | | | |
| NC_011770 | 1038710 | A | G | 70 | 1 | 4 | 0 | NON_SYNONYMOU | MODERAT | MISSENSE | Atg/Gtg | M297V | mpl | | | | . | | | G | G | G | G | | |
| NC_011770 | 1038729 | C | T | 67 | 1 | 2 | 0 | NON_SYNONYMOU | MODERAT | MISSENSE | gCg/gTg | A303V | mpl | | | | . | | | | | T | T | | |
| NC_011770 | 1039053 | A | C | 76 | 0 | 6 | 0 | NON_SYNONYMOU | MODERAT | MISSENSE | gAt/gCt | D411A | mpl | | | C | C | C | C | | | | | C | C |
| NC_011770 | 5544216 | G | A | 62 | 0 | 4 | 0 | NON_SYNONYMOU | MODERAT | MISSENSE | cGc/cAc | R205H | mqoB | A | A | | | | | | | | | A | A |
| NC_011770 | 5288847 | C | T | 79 | 0 | 4 | 0 | NON_SYNONYMOU | MODERAT | MISSENSE | Gcc/Acc | A146T | mraW | T | T | | | | | | | | | T | T |
| NC_011770 | 5288994 | T | C | 72 | 0 | 12 | 0 | NON_SYNONYMOU | MODERAT | MISSENSE | Atc/Gtc | I97V | mraW | C | C | C | C | C | C | C | C | C | C | C | C |
| NC_011770 | 5611972 | A | T | 71 | 0 | 4 | 0 | NON_SYNONYMOU | MODERAT | MISSENSE | aTg/aAg | M715K | mrcB | | | T | T | T | T | | | | | | |
| NC_011770 | 5612296 | T | C | 70 | 0 | 4 | 0 | NON_SYNONYMOU | MODERAT | MISSENSE | aAg/aGg | K607R | mrcB | | | C | C | C | C | | | | | | |
| NC_011770 | 5613058 | T | A | 83 | 0 | 6 | 0 | NON_SYNONYMOU | MODERAT | MISSENSE | cAg/cTg | Q353L | mrcB | | | A | A | A | A | | | A | A | | |
| NC_011770 | 5614043 | T | C | 50 | 0 | 4 | 0 | NON_SYNONYMOU | MODERAT | MISSENSE | Agc/Ggc | S25G | mrcB | C | C | | | | | | | | | C | C |
| NC_011770 | 5345024 | T | C | 76 | 0 | 12 | 0 | NON_SYNONYMOU | MODERAT | MISSENSE | cAc/cGc | H330R | mreC | C | C | C | C | C | C | C | C | C | C | C | C |
| NC_011770 | 5345070 | G | C | 71 | 0 | 2 | 0 | NON_SYNONYMOU | MODERAT | MISSENSE | Cca/Gca | P315A | mreC | | | | | | | | | C | C | | |
| NC_011770 | 5345138 | G | A | 68 | 0 | 2 | 0 | NON_SYNONYMOU | MODERAT | MISSENSE | gCc/gTc | A292V | mreC | | | | | | | | | | | A | A |
| NC_011770 | 5948965 | C | T | 81 | 0 | 2 | 0 | NON_SYNONYMOU | MODERAT | MISSENSE | Gtt/Att | V419I | msbA | | | | | | | | | | | T | T |
| NC_011770 | 5948977 | G | T | 82 | 0 | 4 | 0 | NON_SYNONYMOU | MODERAT | MISSENSE | Ctc/Atc | L415I | msbA | | | | | | | T | T | | | T | T |
| NC_011770 | 5949085 | T | C | 84 | 0 | 4 | 0 | NON_SYNONYMOU | MODERAT | MISSENSE | Atc/Gtc | I379V | msbA | | | | | | | C | C | | | C | C |
| NC_011770 | 5949183 | G | C | 83 | 0 | 4 | 0 | NON_SYNONYMOU | MODERAT | MISSENSE | aCc/aGc | T346S | msbA | | | | | | | C | C | | | C | C |
| NC_011770 | 5949480 | A | T | 83 | 0 | 2 | 0 | NON_SYNONYMOU | MODERAT | MISSENSE | cTc/cAc | L247H | msbA | | | | | | | | | | | T | T |
| NC_011770 | 5507470 | G | A | 64 | 0 | 2 | 0 | NON_SYNONYMOU | MODERAT | MISSENSE | Gcc/Acc | A28T | mscL | A | A | | | | | | | | | | |
| NC_011770 | 5978527 | C | T | 74 | 0 | 2 | 0 | NON_SYNONYMOU | MODERAT | MISSENSE | cCg/cTg | P17L | msrA | | | | | | | | | | | T | T |
| NC_011770 | 5978946 | T | A | 71 | 0 | 10 | 0 | NON_SYNONYMOU | MODERAT | MISSENSE | Ttg/Atg | L157M | msrA | A | A | A | A | A | A | A | A | | | A | A |
| NC_011770 | 3241616 | G | C | 71 | 0 | 2 | 0 | NON_SYNONYMOU | MODERAT | MISSENSE | Gtc/Ctc | V129L | msuD | C | C | | | | | | | | | | |
| NC_011770 | 3259424 | T | C | 63 | 0 | 1 | 0 | NON_SYNONYMOU | MODERAT | MISSENSE | cAg/cGg | Q27R | mtlD | | | | | | | C | | | | | |
| NC_011770 | 3264263 | T | C | 67 | 0 | 1 | 0 | NON_SYNONYMOU | MODERAT | MISSENSE | cAc/cGc | H214R | mtlR | | | | | | | C | | | | | |
| NC_011770 | 3264791 | T | C | 44 | 0 | 1 | 0 | NON_SYNONYMOU | MODERAT | MISSENSE | cAc/cGc | H38R | mtlR | | | | | | | | C | | | | |
| NC_011770 | 3256552 | G | T | 84 | 0 | 2 | 0 | NON_SYNONYMOU | MODERAT | MISSENSE | Cgt/Agt | R494S | mtlY | | | | | | | | | T | T | | |
| NC_011770 | 3256632 | C | T | 77 | 0 | 4 | 0 | NON_SYNONYMOU | MODERAT | MISSENSE | cGg/cAg | R467Q | mtlY | | | T | T | T | T | | | | | | |
| NC_011770 | 3256785 | T | C | 66 | 0 | 4 | 0 | NON_SYNONYMOU | MODERAT | MISSENSE | aAc/aGc | N416S | mtlY | | | C | C | C | C | | | | | | |
| NC_011770 | 3256813 | C | T | 63 | 0 | 10 | 0 | NON_SYNONYMOU | MODERAT | MISSENSE | Gtc/Atc | V407I | mtlY | | | T | T | T | T | T | T | T | T | T | T |
| NC_011770 | 3256954 | G | A | 70 | 0 | 4 | 0 | NON_SYNONYMOU | MODERAT | MISSENSE | Cac/Tac | H360Y | mtlY | | | A | A | A | A | | | | | | |
| NC_011770 | 3257290 | A | C | 59 | 0 | 8 | 0 | NON_SYNONYMOU | MODERAT | MISSENSE | Tgc/Ggc | C248G | mtlY | C | C | C | C | C | C | | | | | C | C |
| NC_011770 | 3257415 | A | G | 56 | 0 | 1 | 0 | NON_SYNONYMOU | MODERAT | MISSENSE | cTg/cCg | L206P | mtlY | | | | | | | G | | | | | |
| NC_011770 | 3258022 | C | T | 55 | 0 | 2 | 0 | NON_SYNONYMOU | MODERAT | MISSENSE | Gag/Aag | E4K | mtlY | | | | | | | T | T | | | | |
| NC_011770 | 3256131 | T | G | 74 | 0 | 2 | 0 | NON_SYNONYMOU | MODERAT | MISSENSE | Acg/Ccg | T118P | mtlZ | | | | | | | | | | | G | G |
| NC_011770 | 3256171 | G | C | 69 | 0 | 4 | 0 | NON_SYNONYMOU | MODERAT | MISSENSE | gaC/gaG | D104E | mtlZ | | | C | C | C | C | | | | | | |
| NC_011770 | 3256302 | T | C | 80 | 0 | 4 | 0 | NON_SYNONYMOU | MODERAT | MISSENSE | Acg/Gcg | T61A | mtlZ | | | C | C | C | C | | | | | | |
| NC_011770 | 3256407 | A | G | 83 | 0 | 1 | 0 | NON_SYNONYMOU | MODERAT | MISSENSE | Ttc/Ctc | F26L | mtlZ | | G | | | | | | | | | | |
| NC_011770 | 6451286 | C | T | 96 | 0 | 2 | 0 | NON_SYNONYMOU | MODERAT | MISSENSE | aGg/aAg | R317K | mtr | | | | | | | | | T | T | | |
| NC_011770 | 6451733 | C | G | 82 | 0 | 2 | 0 | NON_SYNONYMOU | MODERAT | MISSENSE | gGc/gCc | G168A | mtr | | | | | | | G | G | | | | |
| NC_011770 | 6451788 | G | T | 84 | 0 | 4 | 0 | NON_SYNONYMOU | MODERAT | MISSENSE | Cgc/Agc | R150S | mtr | | | T | T | T | T | | | | | | |
| NC_011770 | 2375697 | C | T | 56 | 0 | 4 | 0 | NON_SYNONYMOU | MODERAT | MISSENSE | Ggc/Agc | G262S | mttC | | | T | T | T | T | | | | | | |
| NC_011770 | 2375745 | C | A | 58 | 0 | 4 | 0 | NON_SYNONYMOU | MODERAT | MISSENSE | Gca/Tca | A246S | mttC | A | A | | | | | | | | | A | A |
| NC_011770 | 2376111 | C | T | 72 | 0 | 1 | 0 | NON_SYNONYMOU | MODERAT | MISSENSE | Gag/Aag | E124K | mttC | | | | | | | T | | | | | |
| NC_011770 | 2376321 | G | A | 66 | 0 | 2 | 0 | NON_SYNONYMOU | MODERAT | MISSENSE | Ctc/Ttc | L54F | mttC | | | | | | | | | | | A | A |
| NC_011770 | 5035620 | GC | G | 72 | 0 | 4 | 0 | FRAME_SHIFT | HIGH | | gcg/ | A144 | mucA | G | G | | | | G | | GCC | | | | |
| NC_011770 | 5036032 | G | A | 96 | 0 | 1 | 0 | STOP_GAINED | HIGH | NONSENSE | Cag/Tag | Q7* | mucA | | | | | | | | A | | | | |
| NC_011770 | 5035616 | C | CG | 74 | 0 | 1 | 0 | FRAME_SHIFT | HIGH | | ccg/ccCg | P145P? | mucA | | | | | | | | | CG | | | |
| NC_011770 | 5035485 | G | A | 76 | 0 | 1 | 0 | NON_SYNONYMOU | MODERAT | MISSENSE | gCc/gTc | A189V | mucA | | | | A | | | | | | | | |
| NC_011770 | 5035029 | G | GT | 73 | 0 | 2 | 0 | FRAME_SHIFT | HIGH | | gac/gaAc | D143E? | mucB | | | GT | GT | | | | | | | | |
| NC_011770 | 5034827 | C | T | 71 | 0 | 12 | 0 | NON_SYNONYMOU | MODERAT | MISSENSE | Gcc/Acc | A211T | mucB | T | T | T | T | T | T | T | T | T | T | T | T |
| NC_011770 | 5032695 | T | C | 72 | 0 | 12 | 0 | NON_SYNONYMOU | MODERAT | MISSENSE | Atc/Gtc | I441V | mucD | C | C | C | C | C | C | C | C | C | C | C | C |
| NC_011770 | 5033343 | C | G | 66 | 0 | 12 | 0 | NON_SYNONYMOU | MODERAT | MISSENSE | Gag/Cag | E225Q | mucD | G | G | G | G | G | G | G | G | G | G | G | G |
| NC_011770 | 5033607 | T | C | 69 | 0 | 6 | 0 | NON_SYNONYMOU | MODERAT | MISSENSE | Atc/Gtc | I137V | mucD | C | C | | | | | | | C | C | C | C |

| | | | | | | | | | | | | | | | | | | | | | | | | | |
|---|---|---|---|---|---|---|---|---|---|---|---|---|---|---|---|---|---|---|---|---|---|---|---|---|---|
| NC_011770 | 5033738 | TCGCGCTG | T | 66 | 0 | 1 | 0 | CODON_CHANGE_ | MODERATE | | ggccagcag | GQQRE89I | mucD | | | | | | | T | | | | | |
| NC_011770 | 4742410 | G | A | 83 | 0 | 1 | 0 | NON_SYNONYMOU | MODERAT | MISSENSE | gCc/gTc | A311V | mucK | | A | | | | | | | | | | |
| NC_011770 | 4742773 | G | A | 70 | 0 | 1 | 0 | NON_SYNONYMOU | MODERAT | MISSENSE | gCc/gTc | A190V | mucK | | | | | | | | | | | A | |
| NC_011770 | 4742003 | G | T | 77 | 0 | 2 | 0 | NON_SYNONYMOU | MODERAT | MISSENSE | Cgc/Agc | R447S | mucK | | | | | | | | | | | T | T |
| NC_011770 | 4742020 | A | C | 83 | 0 | 12 | 0 | NON_SYNONYMOU | MODERAT | MISSENSE | gTg/gGg | V441G | mucK | C | C | C | C | C | C | C | C | C | C | C | C |
| NC_011770 | 4742815 | G | A | 71 | 0 | 2 | 0 | NON_SYNONYMOU | MODERAT | MISSENSE | aCc/aTc | T176I | mucK | A | A | | | | | | | | | | |
| NC_011770 | 4742816 | T | C | 69 | 0 | 6 | 0 | NON_SYNONYMOU | MODERAT | MISSENSE | Acc/Gcc | T176A | mucK | C | C | | | | | | | C | C | C | C |
| NC_011770 | 4742920 | T | C | 85 | 0 | 10 | 0 | NON_SYNONYMOU | MODERAT | MISSENSE | aAg/aGg | K141R | mucK | C | C | C | C | C | C | | | C | C | C | C |
| NC_011770 | 5318182 | T | C | 97 | 0 | 1 | 0 | NON_SYNONYMOU | MODERAT | MISSENSE | aAc/aGc | N333S | murA | | | | | | | | C | | | | |
| NC_011770 | 2257983 | C | T | 71 | 0 | 2 | 0 | NON_SYNONYMOU | MODERAT | MISSENSE | Cgc/Tgc | R22C | murB | | | | | | | | | T | T | | |
| NC_011770 | 2258059 | T | C | 73 | 0 | 2 | 0 | NON_SYNONYMOU | MODERAT | MISSENSE | tTg/tCg | L47S | murB | | | | | C | C | | | | | | |
| NC_011770 | 2258664 | G | A | 57 | 0 | 6 | 0 | NON_SYNONYMOU | MODERAT | MISSENSE | Gcg/Acg | A249T | murB | A | A | | | | | | | A | A | A | A |
| NC_011770 | 2258860 | A | G | 54 | 0 | 8 | 0 | NON_SYNONYMOU | MODERAT | MISSENSE | cAg/cGg | Q314R | murB | G | G | | | | | G | G | G | G | G | G |
| NC_011770 | 5278387 | C | A | 66 | 0 | 2 | 0 | NON_SYNONYMOU | MODERAT | MISSENSE | Gcc/Tcc | A134S | murC | | | | | | | | | A | A | | |
| NC_011770 | 5281240 | C | T | 83 | 0 | 2 | 0 | NON_SYNONYMOU | MODERAT | MISSENSE | gGg/gAg | G383E | murD | | | | | | | | | | | T | T |
| NC_011770 | 5281478 | G | A | 74 | 0 | 12 | 0 | NON_SYNONYMOU | MODERAT | MISSENSE | Ccc/Tcc | P304S | murD | A | A | A | A | A | A | A | A | A | A | A | A |
| NC_011770 | 5285045 | T | C | 74 | 0 | 4 | 0 | NON_SYNONYMOU | MODERAT | MISSENSE | gAc/gGc | D424G | murE | | | C | C | C | C | | | | | | |
| NC_011770 | 5285084 | G | A | 75 | 0 | 2 | 0 | NON_SYNONYMOU | MODERAT | MISSENSE | gCg/gTg | A411V | murE | | | | | | | | | | | A | A |
| NC_011770 | 5283937 | T | C | 85 | 0 | 4 | 0 | NON_SYNONYMOU | MODERAT | MISSENSE | gAt/gGt | D308G | murF | | | | | | | C | C | C | C | | |
| NC_011770 | 5284579 | G | A | 79 | 0 | 2 | 0 | NON_SYNONYMOU | MODERAT | MISSENSE | gCg/gTg | A94V | murF | | | | | | | A | A | | | | |
| NC_011770 | 5278939 | G | A | 68 | 0 | 2 | 0 | NON_SYNONYMOU | MODERAT | MISSENSE | cCg/cTg | P305L | murG | | | | | | | A | A | | | | |
| NC_011770 | 5565315 | T | C | 69 | 0 | 2 | 0 | NON_SYNONYMOU | MODERAT | MISSENSE | Agg/Ggg | R248G | murI | C | C | | | | | | | | | | |
| NC_011770 | 5565329 | G | T | 69 | 0 | 2 | 0 | NON_SYNONYMOU | MODERAT | MISSENSE | cCg/cAg | P243Q | murI | | | | | | | | | T | T | | |
| NC_011770 | 5882168 | A | AGGTCG | 77 | 0 | 1 | 0 | FRAME_SHIFT | HIGH | | acc/accCG | T616TR? | mutL | | | | | | | | AGGTCG | | | | |
| NC_011770 | 5882977 | G | GC | 73 | 0 | 2 | 0 | FRAME_SHIFT | HIGH | | gcg/gGcg | A347G? | mutL | GC | GC | | | | | | | | | | |
| NC_011770 | 5882776 | C | T | 65 | 0 | 4 | 0 | NON_SYNONYMOU | MODERAT | MISSENSE | gGt/gAt | G414D | mutL | | | T | T | T | T | | | | | | |
| NC_011770 | 5882878 | G | A | 69 | 0 | 2 | 0 | NON_SYNONYMOU | MODERAT | MISSENSE | gCg/gTg | A380V | mutL | | | | | | | A | A | | | | |
| NC_011770 | 5883056 | C | T | 81 | 0 | 4 | 0 | NON_SYNONYMOU | MODERAT | MISSENSE | Gtc/Atc | V321I | mutL | | | T | T | T | T | | | | | | |
| NC_011770 | 395621 | C | G | 77 | 0 | 4 | 0 | NON_SYNONYMOU | MODERAT | MISSENSE | aCc/aGc | T251S | mutM | | | G | G | G | G | | | | | | |
| NC_011770 | 1525498 | GCCGATCT | G | 72 | 0 | 1 | 0 | FRAME_SHIFT | HIGH | | gagcagatc | EQIG91 | mutS | | | | | | | G | | | | | |
| NC_011770 | 1523697 | C | T | 80 | 0 | 1 | 0 | NON_SYNONYMOU | MODERAT | MISSENSE | Ggc/Agc | G695S | mutS | | | | | | | T | | | | | |
| NC_011770 | 1525221 | A | C | 64 | 0 | 8 | 0 | NON_SYNONYMOU | MODERAT | MISSENSE | Tcg/Gcg | S187A | mutS | C | C | | | | | C | C | C | C | C | C |
| NC_011770 | 1524303 | T | G | 75 | 0 | 1 | 0 | NON_SYNONYMOU | MODERAT | MISSENSE | Acc/Ccc | T493P | mutS | | | | G | | | | | | | | |
| NC_011770 | 6131412 | G | A | 68 | 0 | 2 | 0 | NON_SYNONYMOU | MODERAT | MISSENSE | Ggc/Agc | G311S | mutY | A | A | | | | | | | | | | |
| NC_011770 | 6131541 | G | A | 57 | 0 | 6 | 0 | NON_SYNONYMOU | MODERAT | MISSENSE | Gcc/Acc | A354T | mutY | | | A | A | A | A | | | | | A | A |
| NC_011770 | 4761049 | G | A | 71 | 0 | 1 | 0 | NON_SYNONYMOU | MODERAT | MISSENSE | Ggc/Agc | G262S | mvfR | | | A | | | | | | | | | |
| NC_011770 | 4760298 | G | A | 82 | 0 | 1 | 0 | NON_SYNONYMOU | MODERAT | MISSENSE | atG/atA | M11I | mvfR | | | | | | | | | | | | A |
| NC_011770 | 4759358 | C | G | 86 | 0 | 2 | 0 | NON_SYNONYMOU | MODERAT | MISSENSE | Gtg/Ctg | V54L | nadA | | | | | | | G | G | | | | |
| NC_011770 | 5037519 | C | T | 89 | 0 | 1 | 0 | NON_SYNONYMOU | MODERAT | MISSENSE | Cgg/Tgg | R150W | nadB | | | | | | | T | | | | | |
| NC_011770 | 5037670 | C | G | 85 | 0 | 8 | 0 | NON_SYNONYMOU | MODERAT | MISSENSE | aCc/aGc | T200S | nadB | G | G | G | G | G | G | | | G | G | | |
| NC_011770 | 5401817 | A | G | 76 | 0 | 12 | 0 | NON_SYNONYMOU | MODERAT | MISSENSE | Atc/Gtc | I17V | nadC | G | G | G | G | G | G | G | G | G | G | G | G |
| NC_011770 | 5402156 | G | A | 81 | 0 | 2 | 0 | NON_SYNONYMOU | MODERAT | MISSENSE | Gcg/Acg | A130T | nadC | | | | | | | | | | | A | A |
| NC_011770 | 5402324 | G | C | 78 | 0 | 2 | 0 | NON_SYNONYMOU | MODERAT | MISSENSE | Gaa/Caa | E186Q | nadC | C | C | | | | | | | | | | |
| NC_011770 | 5402513 | G | A | 71 | 0 | 10 | 0 | NON_SYNONYMOU | MODERAT | MISSENSE | Ggc/Agc | G249S | nadC | A | A | A | A | A | A | | | A | A | A | A |
| NC_011770 | 1051051 | A | G | 79 | 0 | 4 | 0 | NON_SYNONYMOU | MODERAT | MISSENSE | cAg/cGg | Q78R | nadD | | | G | G | G | G | | | | | | |
| NC_011770 | 5852782 | C | T | 62 | 0 | 4 | 0 | NON_SYNONYMOU | MODERAT | MISSENSE | Ccg/Tcg | P249S | nadE | | | T | T | T | T | | | | | | |
| NC_011770 | 4574894 | C | T | 86 | 0 | 2 | 0 | NON_SYNONYMOU | MODERAT | MISSENSE | gCg/gTg | A580V | napA | | | | | | | | | T | T | | |
| NC_011770 | 4575170 | G | A | 81 | 0 | 1 | 0 | NON_SYNONYMOU | MODERAT | MISSENSE | cGc/cAc | R672H | napA | | | | | | | A | | | | | |
| NC_011770 | 4575251 | C | T | 72 | 0 | 10 | 0 | NON_SYNONYMOU | MODERAT | MISSENSE | gCc/gTc | A699V | napA | T | T | T | T | T | T | | | T | T | T | T |
| NC_011770 | 4575632 | T | C | 44 | 0 | 1 | 0 | NON_SYNONYMOU | MODERAT | MISSENSE | gTt/gCt | V826A | napA | | | | | | | C | | | | | |
| NC_011770 | 4576156 | C | A | 45 | 0 | 12 | 0 | NON_SYNONYMOU | MODERAT | MISSENSE | aaC/aaA | N162K | napB | A | A | A | A | A | A | A | A | A | A | A | A |
| NC_011770 | 4576240 | A | G | 42 | 0 | 12 | 0 | NON_SYNONYMOU | MODERAT | MISSENSE | aAc/aGc | N23S | napC | G | G | G | G | G | G | G | G | G | G | G | G |
| NC_011770 | 4572605 | G | C | 70 | 0 | 12 | 0 | NON_SYNONYMOU | MODERAT | MISSENSE | Ggc/Cgc | G82R | napF | C | C | C | C | C | C | C | C | C | C | C | C |
| NC_011770 | 4572720 | C | G | 69 | 0 | 2 | 0 | NON_SYNONYMOU | MODERAT | MISSENSE | gCc/gGc | A120G | napF | G | G | | | | | | | | | | |
| NC_011770 | 4572749 | G | A | 70 | 0 | 4 | 0 | NON_SYNONYMOU | MODERAT | MISSENSE | Gac/Aac | D130N | napF | | | A | A | A | A | | | | | | |
| NC_011770 | 1194331 | A | T | 73 | 0 | 2 | 0 | NON_SYNONYMOU | MODERAT | MISSENSE | aAc/aTc | N27I | narG | | | | | | | | | T | T | | |
| NC_011770 | 1194682 | T | C | 84 | 0 | 10 | 0 | NON_SYNONYMOU | MODERAT | MISSENSE | gTc/gCc | V144A | narG | C | C | C | C | C | C | C | C | | | C | C |
| NC_011770 | 1196518 | C | T | 92 | 0 | 4 | 0 | NON_SYNONYMOU | MODERAT | MISSENSE | aCc/aTc | T756I | narG | | | T | T | T | T | | | | | | |
| NC_011770 | 1196898 | G | A | 58 | 0 | 2 | 0 | NON_SYNONYMOU | MODERAT | MISSENSE | Gtg/Atg | V883M | narG | | | | | | | | | | | A | A |
| NC_011770 | 1199207 | C | G | 81 | 0 | 4 | 0 | NON_SYNONYMOU | MODERAT | MISSENSE | Cag/Gag | Q387E | narH | | | G | G | G | G | | | | | | |
| NC_011770 | 1199579 | G | A | 79 | 0 | 12 | 0 | NON_SYNONYMOU | MODERAT | MISSENSE | Gtc/Atc | V511I | narH | A | A | A | A | A | A | A | A | A | A | A | A |
| NC_011770 | 1200589 | G | A | 91 | 0 | 12 | 0 | NON_SYNONYMOU | MODERAT | MISSENSE | aGc/aAc | S84N | narI | A | A | A | A | A | A | A | A | A | A | A | A |

| | | | | | | | | | | | | | | | | | | | | | | | | | |
|---|---|---|---|---|---|---|---|---|---|---|---|---|---|---|---|---|---|---|---|---|---|---|---|---|---|
| NC_011770 | 1193816 | C | A | 61 | 0 | 2 | 0 | NON_SYNONYMOU | MODERAT | MISSENSE | aaC/aaA | N349K | narK2 | | | | | | | A | A | | | | |
| NC_011770 | 1193920 | C | T | 61 | 0 | 2 | 0 | NON_SYNONYMOU | MODERAT | MISSENSE | gCc/gTc | A384V | narK2 | | | | | | | T | T | | | | |
| NC_011770 | 1193941 | A | G | 60 | 0 | 4 | 0 | NON_SYNONYMOU | MODERAT | MISSENSE | cAg/cGg | Q391R | narK2 | | | G | G | G | G | | | | | | |
| NC_011770 | 1189039 | C | T | 80 | 0 | 2 | 0 | NON_SYNONYMOU | MODERAT | MISSENSE | Gcc/Acc | A129T | narL | | | | | | | T | T | | | | |
| NC_011770 | 3936390 | G | A | 77 | 0 | 1 | 0 | STOP_GAINED | HIGH | NONSENSE | tgG/tgA | W28* | nasA | | | | | | | | | | | A | |
| NC_011770 | 3936614 | C | A | 77 | 0 | 4 | 0 | NON_SYNONYMOU | MODERAT | MISSENSE | gCg/gAg | A103E | nasA | | | | | | | A | A | A | A | | |
| NC_011770 | 3937195 | T | A | 83 | 0 | 2 | 0 | NON_SYNONYMOU | MODERAT | MISSENSE | Tcc/Acc | S297T | nasA | A | A | | | | | | | | | | |
| NC_011770 | 3933408 | A | G | 69 | 0 | 6 | 0 | NON_SYNONYMOU | MODERAT | MISSENSE | Acc/Gcc | T62A | nasS | G | G | G | G | G | G | | | | | | |
| NC_011770 | 3934299 | T | C | 84 | 0 | 4 | 0 | NON_SYNONYMOU | MODERAT | MISSENSE | Tgc/Cgc | C359R | nasS | | | C | C | C | C | | | | | | |
| NC_011770 | 3934426 | A | C | 81 | 0 | 2 | 0 | NON_SYNONYMOU | MODERAT | MISSENSE | gAc/gCc | D401A | nasS | | | | | | | C | C | | | | |
| NC_011770 | 3934429 | T | C | 82 | 0 | 2 | 0 | NON_SYNONYMOU | MODERAT | MISSENSE | cTc/cCc | L402P | nasS | | | | | | | C | C | | | | |
| NC_011770 | 3934586 | G | A | 83 | 0 | 2 | 0 | NON_SYNONYMOU | MODERAT | MISSENSE | cGt/cAt | R46H | nasT | | | | | | | | | | | A | A |
| NC_011770 | 3934831 | A | G | 78 | 0 | 10 | 0 | NON_SYNONYMOU | MODERAT | MISSENSE | Act/Gct | T128A | nasT | G | G | G | G | G | G | G | G | G | G | | |
| NC_011770 | 1263873 | C | A | 75 | 0 | 2 | 0 | NON_SYNONYMOU | MODERAT | MISSENSE | Ctg/Atg | L32M | ndk | | | | | | | | | | | A | A |
| NC_011770 | 5491248 | G | GC | 60 | 2 | 1 | 0 | FRAME_SHIFT | HIGH | | gcg/gcCg | A185A? | nfxB | | | GC | | | | | | . | . | | |
| NC_011770 | 3888525 | A | G | 80 | 0 | 4 | 0 | NON_SYNONYMOU | MODERAT | MISSENSE | Acc/Gcc | T129A | nhaB | | | G | G | G | G | | | | | | |
| NC_011770 | 3939691 | A | C | 73 | 0 | 2 | 0 | NON_SYNONYMOU | MODERAT | MISSENSE | gAc/gCc | D79A | nirB | | | | | | | | | | | C | C |
| NC_011770 | 3939789 | G | C | 74 | 0 | 2 | 0 | NON_SYNONYMOU | MODERAT | MISSENSE | Gtc/Ctc | V112L | nirB | | | | | | | | | | | C | C |
| NC_011770 | 3939963 | G | C | 73 | 0 | 2 | 0 | NON_SYNONYMOU | MODERAT | MISSENSE | Gac/Cac | D170H | nirB | C | C | | | | | | | | | | |
| NC_011770 | 3941332 | T | C | 56 | 0 | 1 | 0 | NON_SYNONYMOU | MODERAT | MISSENSE | aTg/aCg | M626T | nirB | | | | | | | C | | | | | |
| NC_011770 | 3941880 | C | T | 60 | 0 | 2 | 0 | NON_SYNONYMOU | MODERAT | MISSENSE | Ccg/Tcg | P809S | nirB | | | | | | | T | T | | | | |
| NC_011770 | 570032 | AG | A | 63 | 0 | 1 | 0 | FRAME_SHIFT | HIGH | | ccc/ | P95 | nirC | | | | | | | | A | | | | |
| NC_011770 | 570287 | C | T | 56 | 0 | 2 | 0 | NON_SYNONYMOU | MODERAT | MISSENSE | Gcc/Acc | A11T | nirC | | | | | | | | | | | T | T |
| NC_011770 | 3941980 | T | G | 56 | 0 | 8 | 0 | NON_SYNONYMOU | MODERAT | MISSENSE | Tcc/Gcc | S8A | nirD | G | G | G | G | G | G | | | | | G | G |
| NC_011770 | 569928 | C | T | 57 | 0 | 2 | 0 | NON_SYNONYMOU | MODERAT | MISSENSE | Gcc/Acc | A12T | nirF | T | T | | | | | | | | | | |
| NC_011770 | 566394 | A | G | 71 | 0 | 12 | 0 | NON_SYNONYMOU | MODERAT | MISSENSE | gTg/gCg | V163A | nirJ | G | G | G | G | G | G | G | G | G | G | G | G |
| NC_011770 | 566826 | C | T | 65 | 0 | 2 | 0 | NON_SYNONYMOU | MODERAT | MISSENSE | cGc/cAc | R19H | nirJ | | | | | | | | | | | T | T |
| NC_011770 | 570438 | G | A | 65 | 0 | 2 | 0 | NON_SYNONYMOU | MODERAT | MISSENSE | gCg/gTg | A64V | nirM | | | | | | | | | | | A | A |
| NC_011770 | 563736 | A | G | 70 | 0 | 2 | 0 | NON_SYNONYMOU | MODERAT | MISSENSE | gTc/gCc | V381A | nirN | | | | | | | | | G | G | | |
| NC_011770 | 563962 | C | T | 76 | 0 | 2 | 0 | NON_SYNONYMOU | MODERAT | MISSENSE | Gac/Aac | D306N | nirN | T | T | | | | | | | | | | |
| NC_011770 | 564031 | T | C | 80 | 0 | 2 | 0 | NON_SYNONYMOU | MODERAT | MISSENSE | Aac/Gac | N283D | nirN | | | | | | | | | C | C | | |
| NC_011770 | 564570 | G | C | 69 | 0 | 10 | 0 | NON_SYNONYMOU | MODERAT | MISSENSE | aCc/aGc | T103S | nirN | C | C | C | C | C | C | C | C | | | C | C |
| NC_011770 | 572631 | A | G | 93 | 0 | 1 | 0 | NON_SYNONYMOU | MODERAT | MISSENSE | Acc/Gcc | T11A | nirQ | | | | | | | G | | | | | |
| NC_011770 | 573159 | A | G | 80 | 0 | 2 | 0 | NON_SYNONYMOU | MODERAT | MISSENSE | Acc/Gcc | T187A | nirQ | G | G | | | | | | | | | | |
| NC_011770 | 571095 | G | A | 82 | 0 | 2 | 0 | NON_SYNONYMOU | MODERAT | MISSENSE | Ccg/Tcg | P431S | nirS | A | A | | | | | | | | | | |
| NC_011770 | 571362 | C | T | 93 | 0 | 4 | 0 | NON_SYNONYMOU | MODERAT | MISSENSE | Gtc/Atc | V342I | nirS | | | T | T | T | T | | | | | | |
| NC_011770 | 572363 | A | G | 103 | 0 | 10 | 0 | NON_SYNONYMOU | MODERAT | MISSENSE | gTg/gCg | V8A | nirS | G | G | G | G | G | G | | | G | G | G | G |
| NC_011770 | 571646 | A | G | 97 | 0 | 1 | 0 | NON_SYNONYMOU | MODERAT | MISSENSE | aTc/aCc | I247T | nirS | | G | | | | | | | | | | |
| NC_011770 | 575223 | A | G | 88 | 0 | 12 | 0 | NON_SYNONYMOU | MODERAT | MISSENSE | Atg/Gtg | M162V | norB | G | G | G | G | G | G | G | G | G | G | G | G |
| NC_011770 | 1792814 | C | T | 74 | 0 | 12 | 0 | NON_SYNONYMOU | MODERAT | MISSENSE | gGg/gAg | G425E | nosD | T | T | T | T | T | T | T | T | T | T | T | T |
| NC_011770 | 1793078 | C | T | 85 | 0 | 2 | 0 | NON_SYNONYMOU | MODERAT | MISSENSE | aGc/aAc | S337N | nosD | | | | | | | | | T | T | | |
| NC_011770 | 1793485 | C | G | 88 | 0 | 4 | 0 | NON_SYNONYMOU | MODERAT | MISSENSE | gaG/gaC | E201D | nosD | | | | | | | | | G | G | G | G |
| NC_011770 | 1793733 | C | G | 71 | 0 | 2 | 0 | NON_SYNONYMOU | MODERAT | MISSENSE | Gtg/Ctg | V119L | nosD | | | | | | | | | | | G | G |
| NC_011770 | 1794066 | G | T | 77 | 0 | 2 | 0 | NON_SYNONYMOU | MODERAT | MISSENSE | Ctg/Atg | L8M | nosD | | | | | | | | | T | T | | |
| NC_011770 | 1792986 | C | T | 84 | 0 | 1 | 0 | NON_SYNONYMOU | MODERAT | MISSENSE | Gcc/Acc | A368T | nosD | | | | | | | | | | | | T |
| NC_011770 | 1790535 | C | A | 60 | 0 | 4 | 0 | NON_SYNONYMOU | MODERAT | MISSENSE | cGg/cTg | R174L | nosL | | | | | | | A | A | A | A | | |
| NC_011770 | 1796251 | C | T | 89 | 0 | 2 | 0 | NON_SYNONYMOU | MODERAT | MISSENSE | aGg/aAg | R648K | nosR | | | | | | | | | T | T | | |
| NC_011770 | 1796371 | A | G | 93 | 0 | 4 | 0 | NON_SYNONYMOU | MODERAT | MISSENSE | gTg/gCg | V608A | nosR | | | | | | | | | G | G | G | G |
| NC_011770 | 1796792 | T | A | 100 | 0 | 2 | 0 | NON_SYNONYMOU | MODERAT | MISSENSE | Agc/Tgc | S468C | nosR | | | | | | | A | A | | | | |
| NC_011770 | 1796930 | C | T | 93 | 0 | 4 | 0 | NON_SYNONYMOU | MODERAT | MISSENSE | Gtc/Atc | V422I | nosR | T | T | | | | | T | T | | | | |
| NC_011770 | 1798055 | C | G | 74 | 0 | 2 | 0 | NON_SYNONYMOU | MODERAT | MISSENSE | Gag/Cag | E47Q | nosR | G | G | | | | | | | | | | |
| NC_011770 | 1798071 | A | C | 76 | 0 | 12 | 0 | NON_SYNONYMOU | MODERAT | MISSENSE | caT/caG | H41Q | nosR | C | C | C | C | C | C | C | C | C | C | C | C |
| NC_011770 | 1798095 | C | A | 74 | 0 | 2 | 0 | NON_SYNONYMOU | MODERAT | MISSENSE | gaG/gaT | E33D | nosR | | | | | | | A | A | | | | |
| NC_011770 | 1798097 | C | T | 72 | 0 | 4 | 0 | NON_SYNONYMOU | MODERAT | MISSENSE | Gag/Aag | E33K | nosR | | | T | T | T | T | | | | | | |
| NC_011770 | 1798174 | G | T | 78 | 0 | 2 | 0 | NON_SYNONYMOU | MODERAT | MISSENSE | tCc/tAc | S7Y | nosR | | | | | T | T | | | | | | |
| NC_011770 | 1796771 | C | T | 97 | 0 | 1 | 0 | NON_SYNONYMOU | MODERAT | MISSENSE | Gtc/Atc | V475I | nosR | T | | | | | | | | | | | |
| NC_011770 | 1791101 | G | T | 52 | 0 | 2 | 0 | NON_SYNONYMOU | MODERAT | MISSENSE | Cac/Aac | H267N | nosY | | | | | | | | | T | T | | |
| NC_011770 | 1791395 | T | C | 51 | 0 | 2 | 0 | NON_SYNONYMOU | MODERAT | MISSENSE | Agc/Ggc | S169G | nosY | | | | | | | | | C | C | | |
| NC_011770 | 1791514 | G | A | 47 | 0 | 4 | 0 | NON_SYNONYMOU | MODERAT | MISSENSE | gCg/gTg | A129V | nosY | A | A | | | | | | | A | A | | |
| NC_011770 | 1791779 | A | C | 41 | 0 | 12 | 0 | NON_SYNONYMOU | MODERAT | MISSENSE | Tcc/Gcc | S41A | nosY | C | C | C | C | C | C | C | C | C | C | C | C |
| NC_011770 | 1795853 | C | T | 83 | 0 | 4 | 0 | NON_SYNONYMOU | MODERAT | MISSENSE | Ggc/Agc | G51S | nosZ | | | T | T | T | T | | | | | | |
| NC_011770 | 2237835 | C | T | 75 | 0 | 4 | 0 | NON_SYNONYMOU | MODERAT | MISSENSE | gCg/gTg | A56V | nqrA | | | T | T | T | T | | | | | | |

| | | | | | | | | | | | | | | | | | | | | | | | | | |
|---|---|---|---|---|---|---|---|---|---|---|---|---|---|---|---|---|---|---|---|---|---|---|---|---|---|
| NC_011770 | 2238299 | T | G | 72 | 0 | 2 | 0 | NON_SYNONYMOU | MODERAT | MISSENSE | Tcc/Gcc | S211A | nqrA | | | | | | | G | G | | | | |
| NC_011770 | 2239496 | A | C | 58 | 0 | 2 | 0 | NON_SYNONYMOU | MODERAT | MISSENSE | Acc/Ccc | T163P | nqrB | C | C | | | | | | | | | | |
| NC_011770 | 2240736 | A | C | 78 | 0 | 2 | 0 | NON_SYNONYMOU | MODERAT | MISSENSE | Acc/Ccc | T175P | nqrC | | | | | | | | | C | C | | |
| NC_011770 | 2241651 | C | G | 81 | 0 | 2 | 0 | NON_SYNONYMOU | MODERAT | MISSENSE | tCc/tGc | S219C | nqrD | | | | | | | G | G | | | | |
| NC_011770 | 2242469 | G | A | 81 | 0 | 2 | 0 | NON_SYNONYMOU | MODERAT | MISSENSE | Gcg/Acg | A61T | nqrF | | | | | | | A | A | | | | |
| NC_011770 | 2242844 | T | C | 82 | 0 | 8 | 0 | NON_SYNONYMOU | MODERAT | MISSENSE | Tcg/Ccg | S186P | nqrF | | | C | C | C | C | C | C | C | C | | |
| NC_011770 | 2242949 | A | C | 75 | 0 | 2 | 0 | NON_SYNONYMOU | MODERAT | MISSENSE | Aag/Cag | K221Q | nqrF | | | | | | | C | C | | | | |
| NC_011770 | 4597897 | G | A | 94 | 0 | 2 | 0 | NON_SYNONYMOU | MODERAT | MISSENSE | gGc/gAc | G28D | nrdB | | | | | | | | | | | A | A |
| NC_011770 | 6526917 | C | T | 87 | 0 | 1 | 0 | NON_SYNONYMOU | MODERAT | MISSENSE | Ggc/Agc | G98S | nrdJa | | | | | | | | | | | T | |
| NC_011770 | 6525828 | T | C | 83 | 0 | 2 | 0 | NON_SYNONYMOU | MODERAT | MISSENSE | Agg/Ggg | R461G | nrdJa | | | | | | | | | | | C | C |
| NC_011770 | 6526806 | T | G | 100 | 0 | 2 | 0 | NON_SYNONYMOU | MODERAT | MISSENSE | Aac/Cac | N135H | nrdJa | | | | | | | G | G | | | | |
| NC_011770 | 6526969 | G | T | 92 | 0 | 4 | 0 | NON_SYNONYMOU | MODERAT | MISSENSE | gaC/gaA | D80E | nrdJa | T | T | | | | | | | | | T | T |
| NC_011770 | 6527198 | G | A | 92 | 0 | 2 | 0 | NON_SYNONYMOU | MODERAT | MISSENSE | aCg/aTg | T4M | nrdJa | | | | | | | | | | | A | A |
| NC_011770 | 3884872 | A | AC | 93 | 0 | 1 | 0 | FRAME_SHIFT | HIGH | | gtc/gGtc | V169G? | nudC | | | | | | | AC | | | | | |
| NC_011770 | 3885373 | A | G | 84 | 0 | 12 | 0 | NON_SYNONYMOU | MODERAT | MISSENSE | gTc/gCc | V2A | nudC | G | G | G | G | G | G | G | G | G | G | G | G |
| NC_011770 | 6164961 | C | T | 90 | 0 | 2 | 0 | NON_SYNONYMOU | MODERAT | MISSENSE | Ggc/Agc | G146S | nudE | T | T | | | | | | | | | | |
| NC_011770 | 6165377 | A | G | 85 | 0 | 2 | 0 | NON_SYNONYMOU | MODERAT | MISSENSE | gTa/gCa | V7A | nudE | G | G | | | | | | | | | | |
| NC_011770 | 162246 | T | C | 44 | 0 | 2 | 0 | NON_SYNONYMOU | MODERAT | MISSENSE | Tgc/Cgc | C8R | nuh | | | | | | | | | | | C | C |
| NC_011770 | 162601 | G | C | 53 | 0 | 6 | 0 | NON_SYNONYMOU | MODERAT | MISSENSE | aGc/aCc | S126T | nuh | | | C | C | C | C | | | | | C | C |
| NC_011770 | 162976 | A | G | 72 | 0 | 6 | 0 | NON_SYNONYMOU | MODERAT | MISSENSE | cAg/cGg | Q251R | nuh | | | G | G | G | G | | | | | G | G |
| NC_011770 | 163251 | A | G | 76 | 0 | 2 | 0 | NON_SYNONYMOU | MODERAT | MISSENSE | Acc/Gcc | T343A | nuh | | | | | | | | | | | G | G |
| NC_011770 | 2651552 | G | A | 66 | 0 | 1 | 0 | NON_SYNONYMOU | MODERAT | MISSENSE | Cgc/Tgc | R135C | nuoF | | | | | | | A | | | | | |
| NC_011770 | 2648073 | C | T | 72 | 0 | 1 | 0 | NON_SYNONYMOU | MODERAT | MISSENSE | Gcc/Acc | A802T | nuoG | | | | | | | T | | | | | |
| NC_011770 | 2648723 | C | T | 80 | 0 | 12 | 0 | NON_SYNONYMOU | MODERAT | MISSENSE | aGg/aAg | R585K | nuoG | T | T | T | T | T | T | T | T | T | T | T | T |
| NC_011770 | 2649180 | C | T | 68 | 0 | 2 | 0 | NON_SYNONYMOU | MODERAT | MISSENSE | Gcg/Acg | A433T | nuoG | | | | | | | | | T | T | | |
| NC_011770 | 2649161 | A | G | 70 | 0 | 1 | 0 | NON_SYNONYMOU | MODERAT | MISSENSE | gTg/gCg | V439A | nuoG | | | | | | | | G | | | | |
| NC_011770 | 2643781 | C | T | 47 | 0 | 4 | 0 | NON_SYNONYMOU | MODERAT | MISSENSE | Gtc/Atc | V522I | nuoL | | | T | T | T | T | | | | | | |
| NC_011770 | 2643810 | G | C | 45 | 0 | 4 | 0 | NON_SYNONYMOU | MODERAT | MISSENSE | gCc/gGc | A512G | nuoL | | | C | C | C | C | | | | | | |
| NC_011770 | 2644279 | C | T | 51 | 0 | 4 | 0 | NON_SYNONYMOU | MODERAT | MISSENSE | Gtc/Atc | V356I | nuoL | | | T | T | T | T | | | | | | |
| NC_011770 | 2643753 | C | T | 46 | 0 | 1 | 0 | NON_SYNONYMOU | MODERAT | MISSENSE | gGt/gAt | G531D | nuoL | | | | | | | | T | | | | |
| NC_011770 | 2642428 | T | C | 80 | 0 | 2 | 0 | NON_SYNONYMOU | MODERAT | MISSENSE | Atc/Gtc | I348V | nuoM | | | | | | | | | | | C | C |
| NC_011770 | 2642779 | C | T | 75 | 0 | 2 | 0 | NON_SYNONYMOU | MODERAT | MISSENSE | Gtc/Atc | V231I | nuoM | | | | | | | | | | | T | T |
| NC_011770 | 5664539 | G | A | 83 | 0 | 1 | 0 | NON_SYNONYMOU | MODERAT | MISSENSE | cCg/cTg | P249L | nusA | | | | | | | A | | | | | |
| NC_011770 | 1003779 | C | T | 83 | 0 | 2 | 0 | NON_SYNONYMOU | MODERAT | MISSENSE | Ccc/Tcc | P41S | nusB | | | | | | | T | T | | | | |
| NC_011770 | 1003809 | A | G | 83 | 0 | 2 | 0 | NON_SYNONYMOU | MODERAT | MISSENSE | Acc/Gcc | T51A | nusB | | | | | | | | | G | G | | |
| NC_011770 | 1003916 | C | A | 91 | 0 | 4 | 0 | NON_SYNONYMOU | MODERAT | MISSENSE | gaC/gaA | D86E | nusB | | | A | A | A | A | | | | | | |
| NC_011770 | 709857 | G | A | 78 | 0 | 2 | 0 | SPLICE_SITE_REGIO | LOW | SILENT | taG/taA | *178 | nusG | A | A | | | | | | | | | | |
| NC_011770 | 709621 | A | G | 75 | 0 | 2 | 0 | NON_SYNONYMOU | MODERAT | MISSENSE | Acc/Gcc | T100A | nusG | | | | | | | G | G | | | | |
| NC_011770 | 709783 | A | ATCC | 78 | 0 | 1 | 0 | CODON_CHANGE_ | MODERATE | | caa/cTCCa | Q155LQ | nusG | | ATCC | | | | | | | | | | |
| NC_011770 | 4769658 | T | C | 72 | 0 | 12 | 0 | NON_SYNONYMOU | MODERAT | MISSENSE | tTc/tCc | F68S | ogt | C | C | C | C | C | C | C | C | C | C | C | C |
| NC_011770 | 5686459 | C | T | 38 | 0 | 1 | 0 | NON_SYNONYMOU | MODERAT | MISSENSE | Ctc/Ttc | L153F | omlA | | | T | | | | | | | | | |
| NC_011770 | 5686306 | G | A | 52 | 0 | 1 | 0 | NON_SYNONYMOU | MODERAT | MISSENSE | Gcc/Acc | A102T | omlA | | | | | | | A | | | | | |
| NC_011770 | 5686415 | A | C | 41 | 0 | 6 | 0 | NON_SYNONYMOU | MODERAT | MISSENSE | gAg/gCg | E138A | omlA | | | C | C | C | C | C | C | | | | |
| NC_011770 | 2581841 | G | A | 80 | 0 | 4 | 0 | NON_SYNONYMOU | MODERAT | MISSENSE | Ctc/Ttc | L397F | opdB | | | A | A | A | A | | | | | | |
| NC_011770 | 2581913 | T | G | 74 | 0 | 12 | 0 | NON_SYNONYMOU | MODERAT | MISSENSE | Acc/Ccc | T373P | opdB | G | G | G | G | G | G | G | G | G | G | G | G |
| NC_011770 | 2582357 | C | A | 76 | 0 | 2 | 0 | NON_SYNONYMOU | MODERAT | MISSENSE | Gcg/Tcg | A225S | opdB | A | A | | | | | | | | | | |
| NC_011770 | 2582824 | G | A | 59 | 0 | 2 | 0 | NON_SYNONYMOU | MODERAT | MISSENSE | gCc/gTc | A69V | opdB | A | A | | | | | | | | | | |
| NC_011770 | 2582864 | C | T | 53 | 0 | 2 | 0 | NON_SYNONYMOU | MODERAT | MISSENSE | Ggg/Agg | G56R | opdB | | | | | | | T | T | | | | |
| NC_011770 | 183745 | G | T | 93 | 0 | 2 | 0 | NON_SYNONYMOU | MODERAT | MISSENSE | Gcg/Tcg | A108S | opdC | | | | | | | T | T | | | | |
| NC_011770 | 183866 | A | G | 82 | 0 | 2 | 0 | NON_SYNONYMOU | MODERAT | MISSENSE | aAc/aGc | N148S | opdC | G | G | | | | | | | | | | |
| NC_011770 | 183929 | C | T | 88 | 0 | 2 | 0 | NON_SYNONYMOU | MODERAT | MISSENSE | aCc/aTc | T169I | opdC | | | | | | | | | T | T | | |
| NC_011770 | 5373826 | G | A | 60 | 0 | 1 | 0 | NON_SYNONYMOU | MODERAT | MISSENSE | Gac/Aac | D410N | opdD | | | | | | | | A | | | | |
| NC_011770 | 5372692 | C | T | 74 | 0 | 4 | 0 | NON_SYNONYMOU | MODERAT | MISSENSE | Ccg/Tcg | P32S | opdD | | | T | T | T | T | | | | | | |
| NC_011770 | 5043806 | C | T | 72 | 0 | 2 | 0 | NON_SYNONYMOU | MODERAT | MISSENSE | tCg/tTg | S14L | opdH | | | | | | | | | | | T | T |
| NC_011770 | 5043850 | A | G | 71 | 0 | 12 | 0 | NON_SYNONYMOU | MODERAT | MISSENSE | Agc/Ggc | S29G | opdH | G | G | G | G | G | G | G | G | G | G | G | G |
| NC_011770 | 5044943 | T | C | 76 | 0 | 2 | 0 | NON_SYNONYMOU | MODERAT | MISSENSE | cTg/cCg | L393P | opdH | | | | | | | | | | | C | C |
| NC_011770 | 5827215 | G | C | 73 | 0 | 6 | 0 | NON_SYNONYMOU | MODERAT | MISSENSE | Ctc/Gtc | L276V | opdK | C | C | | | | | | | C | C | C | C |
| NC_011770 | 5827689 | T | C | 76 | 0 | 2 | 0 | NON_SYNONYMOU | MODERAT | MISSENSE | Aac/Gac | N118D | opdK | C | C | | | | | | | | | | |
| NC_011770 | 5827936 | C | A | 76 | 0 | 12 | 0 | NON_SYNONYMOU | MODERAT | MISSENSE | gaG/gaT | E35D | opdK | A | A | A | A | A | A | A | A | A | A | A | A |
| NC_011770 | 3548036 | G | A | 68 | 0 | 2 | 0 | NON_SYNONYMOU | MODERAT | MISSENSE | Gcc/Acc | A107T | opdO | | | | | | | | | | | A | A |
| NC_011770 | 3548385 | G | A | 74 | 0 | 2 | 0 | NON_SYNONYMOU | MODERAT | MISSENSE | aGg/aAg | R223K | opdO | | | | | | | A | A | | | | |
| NC_011770 | 3548503 | A | C | 67 | 0 | 2 | 0 | NON_SYNONYMOU | MODERAT | MISSENSE | gaA/gaC | E262D | opdO | | | | | | | C | C | | | | |

| | | | | | | | | | | | | | | | | | | | | | | | | | |
|---|---|---|---|---|---|---|---|---|---|---|---|---|---|---|---|---|---|---|---|---|---|---|---|---|---|
| NC_011770 | 3548771 | G | A | 85 | 0 | 4 | 0 | NON_SYNONYMOU | MODERAT | MISSENSE | Ggc/Agc | G352S | opdO | | | | | | | A | A | A | A | | |
| NC_011770 | 3548936 | A | G | 96 | 0 | 4 | 0 | NON_SYNONYMOU | MODERAT | MISSENSE | Acg/Gcg | T407A | opdO | | | | | | | G | G | G | G | | |
| NC_011770 | 3004925 | G | A | 74 | 0 | 2 | 0 | NON_SYNONYMOU | MODERAT | MISSENSE | Gtg/Atg | V73M | opdT | | | | | | | | | A | A | | |
| NC_011770 | 3005445 | C | T | 79 | 0 | 2 | 0 | NON_SYNONYMOU | MODERAT | MISSENSE | gCc/gTc | A246V | opdT | | | | | | | | | T | T | | |
| NC_011770 | 3005889 | A | G | 77 | 0 | 1 | 0 | NON_SYNONYMOU | MODERAT | MISSENSE | aAc/aGc | N394S | opdT | | | | | | | G | | | | | |
| NC_011770 | 781069 | C | T | 65 | 0 | 2 | 0 | NON_SYNONYMOU | MODERAT | MISSENSE | Gac/Aac | D469N | opmD | | | | | | | | | | | T | T |
| NC_011770 | 781114 | C | T | 63 | 0 | 2 | 0 | NON_SYNONYMOU | MODERAT | MISSENSE | Gtc/Atc | V454I | opmD | | | | | | | T | T | | | | |
| NC_011770 | 781320 | T | C | 67 | 0 | 2 | 0 | NON_SYNONYMOU | MODERAT | MISSENSE | gAc/gGc | D385G | opmD | C | C | | | | | | | | | | |
| NC_011770 | 781627 | C | T | 61 | 0 | 4 | 0 | NON_SYNONYMOU | MODERAT | MISSENSE | Gtc/Atc | V283I | opmD | | | T | T | T | T | | | | | | |
| NC_011770 | 3198795 | GT | G | 66 | 0 | 1 | 0 | FRAME_SHIFT | HIGH | | aac/ | N342 | opmQ | | | | G | | | | | | | | |
| NC_011770 | 3198968 | C | T | 76 | 0 | 8 | 0 | NON_SYNONYMOU | MODERAT | MISSENSE | Ggc/Agc | G285S | opmQ | T | T | | | | | T | T | T | T | T | T |
| NC_011770 | 3199241 | T | A | 77 | 0 | 4 | 0 | NON_SYNONYMOU | MODERAT | MISSENSE | Agg/Tgg | R194W | opmQ | | | A | A | A | A | | | | | | |
| NC_011770 | 3199343 | C | A | 80 | 0 | 2 | 0 | NON_SYNONYMOU | MODERAT | MISSENSE | Gcc/Tcc | A160S | opmQ | | | | | | | | | A | A | | |
| NC_011770 | 3199378 | C | T | 78 | 0 | 2 | 0 | NON_SYNONYMOU | MODERAT | MISSENSE | cGg/cAg | R148Q | opmQ | | | | | | | | | | | T | T |
| NC_011770 | 3199483 | T | C | 76 | 0 | 2 | 0 | NON_SYNONYMOU | MODERAT | MISSENSE | gAc/gGc | D113G | opmQ | | | | | | | | | | | C | C |
| NC_011770 | 3199556 | C | T | 68 | 0 | 2 | 0 | NON_SYNONYMOU | MODERAT | MISSENSE | Gtg/Atg | V89M | opmQ | | | | | | | T | T | | | | |
| NC_011770 | 3199723 | G | C | 60 | 0 | 10 | 0 | NON_SYNONYMOU | MODERAT | MISSENSE | cCt/cGt | P33R | opmQ | C | C | C | C | C | C | C | C | | | C | C |
| NC_011770 | 3199724 | G | T | 58 | 0 | 10 | 0 | NON_SYNONYMOU | MODERAT | MISSENSE | Cct/Act | P33T | opmQ | T | T | T | T | T | T | T | T | | | T | T |
| NC_011770 | 3198800 | C | T | 66 | 0 | 1 | 0 | NON_SYNONYMOU | MODERAT | MISSENSE | Gcc/Acc | A341T | opmQ | | | | | | | | T | | | | |
| NC_011770 | 2025643 | A | C | 83 | 0 | 2 | 0 | NON_SYNONYMOU | MODERAT | MISSENSE | cAc/cCc | H30P | oprB | | | | | | | C | C | | | | |
| NC_011770 | 1280815 | C | T | 61 | 0 | 2 | 0 | NON_SYNONYMOU | MODERAT | MISSENSE | aCt/aTt | T7I | oprC | T | T | | | | | | | | | | |
| NC_011770 | 1280970 | T | C | 69 | 0 | 12 | 0 | NON_SYNONYMOU | MODERAT | MISSENSE | Tac/Cac | Y59H | oprC | C | C | C | C | C | C | C | C | C | C | C | C |
| NC_011770 | 1281696 | G | C | 91 | 0 | 2 | 0 | NON_SYNONYMOU | MODERAT | MISSENSE | Gtc/Ctc | V301L | oprC | | | | | | | | | C | C | | |
| NC_011770 | 1281997 | G | A | 90 | 0 | 2 | 0 | NON_SYNONYMOU | MODERAT | MISSENSE | aGc/aAc | S401N | oprC | A | A | | | | | | | | | | |
| NC_011770 | 1282394 | C | A | 87 | 0 | 2 | 0 | NON_SYNONYMOU | MODERAT | MISSENSE | gaC/gaA | D533E | oprC | | | | | | | A | A | | | | |
| NC_011770 | 1282828 | T | C | 70 | 0 | 10 | 0 | NON_SYNONYMOU | MODERAT | MISSENSE | gTc/gCc | V678A | oprC | C | C | C | C | C | C | C | C | C | C | | |
| NC_011770 | 1281870 | A | G | 88 | 0 | 1 | 0 | NON_SYNONYMOU | MODERAT | MISSENSE | Acc/Gcc | T359A | oprC | | | | | | | | G | | | | |
| NC_011770 | 4794321 | G | A | 79 | 0 | 1 | 0 | STOP_GAINED | HIGH | NONSENSE | tgG/tgA | W415* | oprD | | | | | | | | A | | | | |
| NC_011770 | 4794320 | G | A | 79 | 0 | 1 | 0 | STOP_GAINED | HIGH | NONSENSE | tGg/tAg | W415* | oprD | | | | A | | | | | | | | |
| NC_011770 | 4793907 | G | A | 82 | 0 | 4 | 0 | STOP_GAINED | HIGH | NONSENSE | tgG/tgA | W277* | oprD | A | A | | | | | | | | | A | A |
| NC_011770 | 4793203 | A | G | 86 | 0 | 4 | 0 | NON_SYNONYMOU | MODERAT | MISSENSE | Aac/Gac | N43D | oprD | | | G | G | G | G | | | | | | |
| NC_011770 | 4793245 | G | A | 73 | 0 | 4 | 0 | NON_SYNONYMOU | MODERAT | MISSENSE | Gaa/Aaa | E57K | oprD | | | A | A | A | A | | | | | | |
| NC_011770 | 4793246 | A | G | 74 | 0 | 4 | 0 | NON_SYNONYMOU | MODERAT | MISSENSE | gAa/gGa | E57G | oprD | | | G | G | G | G | | | | | | |
| NC_011770 | 4793247 | A | C | 73 | 0 | 4 | 0 | NON_SYNONYMOU | MODERAT | MISSENSE | gaA/gaC | E57D | oprD | | | C | C | C | C | | | | | | |
| NC_011770 | 4793455 | G | C | 85 | 0 | 4 | 0 | NON_SYNONYMOU | MODERAT | MISSENSE | Gtg/Ctg | V127L | oprD | | | C | C | C | C | | | | | | |
| NC_011770 | 4793629 | G | C | 73 | 0 | 4 | 0 | NON_SYNONYMOU | MODERAT | MISSENSE | Gag/Cag | E185Q | oprD | | | C | C | C | C | | | | | | |
| NC_011770 | 4793632 | C | G | 70 | 0 | 4 | 0 | NON_SYNONYMOU | MODERAT | MISSENSE | Ccg/Gcg | P186A | oprD | | | G | G | G | G | | | | | | |
| NC_011770 | 4793633 | C | G | 72 | 0 | 4 | 0 | NON_SYNONYMOU | MODERAT | MISSENSE | cCg/cGg | P186R | oprD | | | G | G | G | G | | | | | | |
| NC_011770 | 4793641 | G | A | 73 | 0 | 4 | 0 | NON_SYNONYMOU | MODERAT | MISSENSE | Gtc/Atc | V189I | oprD | | | A | A | A | A | | | | | | |
| NC_011770 | 4793642 | T | C | 73 | 0 | 4 | 0 | NON_SYNONYMOU | MODERAT | MISSENSE | gTc/gCc | V189A | oprD | | | C | C | C | C | | | | | | |
| NC_011770 | 4793875 | T | G | 84 | 0 | 4 | 0 | NON_SYNONYMOU | MODERAT | MISSENSE | Tcc/Gcc | S267A | oprD | | | G | G | G | G | | | | | | |
| NC_011770 | 4793902 | A | G | 82 | 0 | 4 | 0 | NON_SYNONYMOU | MODERAT | MISSENSE | Acc/Gcc | T276A | oprD | | | G | G | G | G | | | | | | |
| NC_011770 | 4793908 | T | C | 82 | 0 | 1 | 0 | NON_SYNONYMOU | MODERAT | MISSENSE | Tcc/Ccc | S278P | oprD | | | | | | | C | | | | | |
| NC_011770 | 4793909 | C | T | 83 | 0 | 1 | 0 | NON_SYNONYMOU | MODERAT | MISSENSE | tCc/tTc | S278F | oprD | | | | | | | T | | | | | |
| NC_011770 | 4794005 | G | A | 73 | 0 | 4 | 0 | NON_SYNONYMOU | MODERAT | MISSENSE | gGc/gAc | G310D | oprD | | | A | A | A | A | | | | | | |
| NC_011770 | 4794010 | G | C | 74 | 0 | 4 | 0 | NON_SYNONYMOU | MODERAT | MISSENSE | Ggt/Cgt | G312R | oprD | | | C | C | C | C | | | | | | |
| NC_011770 | 4794020 | C | G | 73 | 0 | 4 | 0 | NON_SYNONYMOU | MODERAT | MISSENSE | gCc/gGc | A315G | oprD | | | G | G | G | G | | | | | | |
| NC_011770 | 4794115 | C | A | 78 | 0 | 4 | 0 | NON_SYNONYMOU | MODERAT | MISSENSE | Ctg/Atg | L347M | oprD | | | A | A | A | A | | | | | | |
| NC_011770 | 4794140 | T | C | 76 | 0 | 1 | 0 | NON_SYNONYMOU | MODERAT | MISSENSE | cTg/cCg | L355P | oprD | | | | | | | C | | | | | |
| NC_011770 | 4794151 | C | G | 76 | 0 | 4 | 0 | NON_SYNONYMOU | MODERAT | MISSENSE | Ctg/Gtg | L359V | oprD | | | G | G | G | G | | | | | | |
| NC_011770 | 4794277 | T | G | 75 | 0 | 4 | 0 | NON_SYNONYMOU | MODERAT | MISSENSE | Tcc/Gcc | S401A | oprD | | | G | G | G | G | | | | | | |
| NC_011770 | 4794340 | C | G | 71 | 0 | 4 | 0 | NON_SYNONYMOU | MODERAT | MISSENSE | Cag/Gag | Q422E | oprD | | | G | G | G | G | | | | | | |
| NC_011770 | 5485408 | T | C | 47 | 2 | 4 | 0 | NON_SYNONYMOU | MODERAT | MISSENSE | cAg/cGg | Q267R | oprJ | | | C | C | C | C | | | . | . | | |
| NC_011770 | 5485852 | G | A | 69 | 2 | 1 | 0 | NON_SYNONYMOU | MODERAT | MISSENSE | gCc/gTc | A119V | oprJ | | | | | | | A | | . | . | | |
| NC_011770 | 5486003 | T | C | 76 | 2 | 10 | 0 | NON_SYNONYMOU | MODERAT | MISSENSE | Atg/Gtg | M69V | oprJ | C | C | C | C | C | C | C | C | . | . | C | C |
| NC_011770 | 5486005 | T | C | 76 | 2 | 4 | 0 | NON_SYNONYMOU | MODERAT | MISSENSE | gAc/gGc | D68G | oprJ | C | C | | | | | | | . | . | C | C |
| NC_011770 | 471308 | A | G | 78 | 0 | 2 | 0 | NON_SYNONYMOU | MODERAT | MISSENSE | gAg/gGg | E412G | oprM | | | | | | | | | G | G | | |
| NC_011770 | 3014267 | C | T | 64 | 0 | 1 | 0 | NON_SYNONYMOU | MODERAT | MISSENSE | Gcc/Acc | A390T | oprN | | | | | | | T | | | | | |
| NC_011770 | 3015352 | A | G | 80 | 0 | 1 | 0 | NON_SYNONYMOU | MODERAT | MISSENSE | gTg/gCg | V28A | oprN | | | | | | | G | | | | | |
| NC_011770 | 3015398 | A | G | 83 | 0 | 2 | 0 | NON_SYNONYMOU | MODERAT | MISSENSE | Tcc/Ccc | S13P | oprN | | | | | | | | | | | G | G |
| NC_011770 | 1928676 | C | T | 72 | 0 | 1 | 0 | STOP_GAINED | HIGH | NONSENSE | Cag/Tag | Q60* | oprO | | | | | | | | T | | | | |
| NC_011770 | 1929387 | C | T | 81 | 0 | 1 | 0 | NON_SYNONYMOU | MODERAT | MISSENSE | Ccg/Tcg | P297S | oprO | | | | | | | T | | | | | |

| | | | | | | | | | | | | | | | | | | | | | | | | | |
|---|---|---|---|---|---|---|---|---|---|---|---|---|---|---|---|---|---|---|---|---|---|---|---|---|---|
| NC_011770 | 1930566 | C | T | 74 | 0 | 10 | 0 | NON_SYNONYMOU | MODERAT | MISSENSE | gCc/gTc | A98V | oprP | T | T | T | T | T | T | T | T | T | T | | |
| NC_011770 | 1685295 | A | AGT | 10 | 10 | 2 | 0 | FRAME_SHIFT | HIGH | | aga/agTGa | R95S? | orf2 | . | . | . | . | AGT | AGT | . | . | . | . | . | . |
| NC_011770 | 1685305 | CAG | C | 10 | 10 | 2 | 0 | FRAME_SHIFT | HIGH | | ccaggt/ | PG98 | orf2 | . | . | . | . | C | C | . | . | . | . | . | . |
| NC_011770 | 1685047 | C | T | 18 | 10 | 2 | 0 | NON_SYNONYMOU | MODERAT | MISSENSE | gCc/gTc | A12V | orf2 | . | . | . | . | T | T | . | . | . | . | . | . |
| NC_011770 | 1685079 | A | G | 20 | 10 | 2 | 0 | NON_SYNONYMOU | MODERAT | MISSENSE | Acc/Gcc | T23A | orf2 | . | . | . | . | G | G | . | . | . | . | . | . |
| NC_011770 | 1685118 | A | G | 20 | 10 | 2 | 0 | NON_SYNONYMOU | MODERAT | MISSENSE | Aac/Gac | N36D | orf2 | . | . | . | . | G | G | . | . | . | . | . | . |
| NC_011770 | 1685155 | C | A | 20 | 10 | 2 | 0 | NON_SYNONYMOU | MODERAT | MISSENSE | aCc/aAc | T48N | orf2 | . | . | . | . | A | A | . | . | . | . | . | . |
| NC_011770 | 1685196 | A | G | 20 | 10 | 2 | 0 | NON_SYNONYMOU | MODERAT | MISSENSE | Aac/Gac | N62D | orf2 | . | . | . | . | G | G | . | . | . | . | . | . |
| NC_011770 | 1685240 | A | C | 20 | 10 | 2 | 0 | NON_SYNONYMOU | MODERAT | MISSENSE | gaA/gaC | E76D | orf2 | . | . | . | . | C | C | . | . | . | . | . | . |
| NC_011770 | 1685282 | G | C | 10 | 10 | 2 | 0 | NON_SYNONYMOU | MODERAT | MISSENSE | ttG/ttC | L90F | orf2 | . | . | . | . | C | C | . | . | . | . | . | . |
| NC_011770 | 1685283 | C | A | 10 | 10 | 2 | 0 | NON_SYNONYMOU | MODERAT | MISSENSE | Cag/Aag | Q91K | orf2 | . | . | . | . | A | A | . | . | . | . | . | . |
| NC_011770 | 1685293 | T | A | 10 | 10 | 2 | 0 | NON_SYNONYMOU | MODERAT | MISSENSE | gTc/gAc | V94D | orf2 | . | . | . | . | A | A | . | . | . | . | . | . |
| NC_011770 | 1685316 | G | A | 0 | 10 | 2 | 0 | NON_SYNONYMOU | MODERAT | MISSENSE | Gaa/Aaa | E102K | orf2 | . | . | . | . | A | A | . | . | . | . | . | . |
| NC_011770 | 1685323 | C | A | 18 | 10 | 2 | 0 | NON_SYNONYMOU | MODERAT | MISSENSE | aCc/aAc | T104N | orf2 | . | . | . | . | A | A | . | . | . | . | . | . |
| NC_011770 | 1685359 | T | C | 19 | 10 | 2 | 0 | NON_SYNONYMOU | MODERAT | MISSENSE | gTg/gCg | V116A | orf2 | . | . | . | . | C | C | . | . | . | . | . | . |
| NC_011770 | 1693747 | C | A | 16 | 10 | 2 | 0 | NON_SYNONYMOU | MODERAT | MISSENSE | aaC/aaA | N69K | orf20 | . | . | . | . | A | A | . | . | . | . | . | . |
| NC_011770 | 1693774 | G | T | 17 | 10 | 2 | 0 | NON_SYNONYMOU | MODERAT | MISSENSE | caG/caT | Q78H | orf20 | . | . | . | . | T | T | . | . | . | . | . | . |
| NC_011770 | 1694105 | T | C | 20 | 10 | 2 | 0 | NON_SYNONYMOU | MODERAT | MISSENSE | aTc/aCc | I48T | orf21 | . | . | . | . | C | C | . | . | . | . | . | . |
| NC_011770 | 1694222 | T | C | 22 | 10 | 2 | 0 | NON_SYNONYMOU | MODERAT | MISSENSE | tTg/tCg | L87S | orf21 | . | . | . | . | C | C | . | . | . | . | . | . |
| NC_011770 | 1694828 | A | C | 21 | 10 | 2 | 0 | NON_SYNONYMOU | MODERAT | MISSENSE | Aag/Cag | K71Q | orf22 | . | . | . | . | C | C | . | . | . | . | . | . |
| NC_011770 | 1694960 | T | G | 19 | 10 | 2 | 0 | NON_SYNONYMOU | MODERAT | MISSENSE | Tcc/Gcc | S115A | orf22 | . | . | . | . | G | G | . | . | . | . | . | . |
| NC_011770 | 1695005 | A | G | 20 | 10 | 2 | 0 | NON_SYNONYMOU | MODERAT | MISSENSE | Agt/Ggt | S130G | orf22 | . | . | . | . | G | G | . | . | . | . | . | . |
| NC_011770 | 2739102 | GCT | G | 9 | 10 | 2 | 0 | FRAME_SHIFT | HIGH | | ctg/ | L103 | orf32 | . | . | . | . | G | G | . | . | . | . | . | . |
| NC_011770 | 2739108 | C | CGT | 0 | 10 | 1 | 0 | FRAME_SHIFT | HIGH | | cag/GTcag | Q105V? | orf32 | . | . | . | . | . | CGT | . | . | . | . | . | . |
| NC_011770 | 2739017 | A | G | 10 | 10 | 2 | 0 | NON_SYNONYMOU | MODERAT | MISSENSE | gAt/gGt | D74G | orf32 | . | . | . | . | G | G | . | . | . | . | . | . |
| NC_011770 | 2739049 | C | A | 10 | 10 | 2 | 0 | NON_SYNONYMOU | MODERAT | MISSENSE | Ctg/Atg | L85M | orf32 | . | . | . | . | A | A | . | . | . | . | . | . |
| NC_011770 | 2739056 | A | G | 12 | 10 | 2 | 0 | NON_SYNONYMOU | MODERAT | MISSENSE | cAg/cGg | Q87R | orf32 | . | . | . | . | G | G | . | . | . | . | . | . |
| NC_011770 | 2739080 | G | A | 10 | 10 | 2 | 0 | NON_SYNONYMOU | MODERAT | MISSENSE | cGa/cAa | R95Q | orf32 | . | . | . | . | A | A | . | . | . | . | . | . |
| NC_011770 | 2739089 | C | G | 10 | 10 | 2 | 0 | NON_SYNONYMOU | MODERAT | MISSENSE | gCg/gGg | A98G | orf32 | . | . | . | . | G | G | . | . | . | . | . | . |
| NC_011770 | 2739144 | T | G | 13 | 10 | 2 | 0 | NON_SYNONYMOU | MODERAT | MISSENSE | gaT/gaG | D116E | orf32 | . | . | . | . | G | G | . | . | . | . | . | . |
| NC_011770 | 2739535 | C | CG | 14 | 10 | 2 | 0 | FRAME_SHIFT+STA | HIGH | | atg/Gatg | M1D? | orf34 | . | . | . | . | CG | CG | . | . | . | . | . | . |
| NC_011770 | 2739653 | G | C | 12 | 10 | 2 | 0 | NON_SYNONYMOU | MODERAT | MISSENSE | Gag/Cag | E40Q | orf34 | . | . | . | . | C | C | . | . | . | . | . | . |
| NC_011770 | 2739731 | T | G | 10 | 10 | 2 | 0 | NON_SYNONYMOU | MODERAT | MISSENSE | Tcg/Gcg | S66A | orf34 | . | . | . | . | G | G | . | . | . | . | . | . |
| NC_011770 | 1712055 | CAA | C | 13 | 8 | 4 | 0 | FRAME_SHIFT | HIGH | | gcaacg/ | AT8 | orf46e | . | . | . | . | C | C | . | . | C | C | . | . |
| NC_011770 | 1712059 | G | GTC | 13 | 6 | 4 | 0 | FRAME_SHIFT | HIGH | | aac/TCaac | N10S? | orf46e | | | . | . | GTC | GTC | . | . | GTC | GTC | . | . |
| NC_011770 | 1712045 | A | T | 12 | 8 | 4 | 0 | NON_SYNONYMOU | MODERAT | MISSENSE | Atc/Ttc | I5F | orf46e | . | . | . | . | T | T | . | . | T | T | . | . |
| NC_011770 | 1712106 | A | G | 37 | 6 | 4 | 0 | NON_SYNONYMOU | MODERAT | MISSENSE | aAc/aGc | N25S | orf46e | | | . | . | G | G | . | . | G | G | . | . |
| NC_011770 | 1712142 | A | T | 37 | 6 | 2 | 0 | NON_SYNONYMOU | MODERAT | MISSENSE | aAg/aTg | K37M | orf46e | | | . | . | T | T | . | . | | | . | . |
| NC_011770 | 1712186 | G | A | 37 | 6 | 2 | 0 | NON_SYNONYMOU | MODERAT | MISSENSE | Gtg/Atg | V52M | orf46e | A | A | . | . | | | . | . | | | . | . |
| NC_011770 | 1712225 | A | G | 41 | 7 | 1 | 0 | NON_SYNONYMOU | MODERAT | MISSENSE | Acc/Gcc | T65A | orf46e | G | . | . | . | | | . | . | | | . | . |
| NC_011770 | 1712226 | C | A | 40 | 7 | 1 | 0 | NON_SYNONYMOU | MODERAT | MISSENSE | aCc/aAc | T65N | orf46e | A | . | . | . | | | . | . | | | . | . |
| NC_011770 | 1712231 | A | G | 40 | 7 | 2 | 0 | NON_SYNONYMOU | MODERAT | MISSENSE | Agc/Ggc | S67G | orf46e | | . | . | . | | | . | . | G | G | . | . |
| NC_011770 | 1712232 | G | C | 40 | 7 | 2 | 0 | NON_SYNONYMOU | MODERAT | MISSENSE | aGc/aCc | S67T | orf46e | | . | . | . | | | . | . | C | C | . | . |
| NC_011770 | 1712238 | A | T | 42 | 7 | 3 | 0 | NON_SYNONYMOU | MODERAT | MISSENSE | tAc/tTc | Y69F | orf46e | T | . | . | . | T | T | . | . | | | . | . |
| NC_011770 | 1712246 | A | G | 40 | 7 | 1 | 0 | NON_SYNONYMOU | MODERAT | MISSENSE | Att/Gtt | I72V | orf46e | G | . | . | . | | | . | . | | | . | . |
| NC_011770 | 1712252 | A | T | 40 | 7 | 3 | 0 | NON_SYNONYMOU | MODERAT | MISSENSE | Agc/Tgc | S74C | orf46e | T | . | . | . | T | T | . | . | | | . | . |
| NC_011770 | 1712253 | G | C | 40 | 7 | 3 | 0 | NON_SYNONYMOU | MODERAT | MISSENSE | aGc/aCc | S74T | orf46e | C | . | . | . | C | C | . | . | | | . | . |
| NC_011770 | 1712285 | C | G | 46 | 6 | 4 | 0 | NON_SYNONYMOU | MODERAT | MISSENSE | Ctt/Gtt | L85V | orf46e | G | G | . | . | G | G | . | . | | | . | . |
| NC_011770 | 1712324 | A | C | 46 | 6 | 2 | 0 | NON_SYNONYMOU | MODERAT | MISSENSE | Atc/Ctc | I98L | orf46e | C | C | . | . | | | . | . | | | . | . |
| NC_011770 | 1712343 | A | T | 49 | 6 | 2 | 0 | NON_SYNONYMOU | MODERAT | MISSENSE | tAc/tTc | Y104F | orf46e | T | T | . | . | | | . | . | | | . | . |
| NC_011770 | 1712358 | C | T | 48 | 6 | 2 | 0 | NON_SYNONYMOU | MODERAT | MISSENSE | gCt/gTt | A109V | orf46e | T | T | . | . | | | . | . | | | . | . |
| NC_011770 | 1712379 | AGGC | A | 27 | 7 | 3 | 0 | CODON_DELETION | MODERATE | | gcg/ | A117 | orf46e | A | A | . | . | . | A | . | . | | | . | . |
| NC_011770 | 1712389 | G | GGCC | 41 | 8 | 2 | 0 | CODON_INSERTIOI | MODERATE | | tgg/GCCtg | W120AW | orf46e | . | . | . | . | GGCC | GGCC | . | . | | | . | . |
| NC_011770 | 1712402 | T | A | 44 | 6 | 6 | 0 | NON_SYNONYMOU | MODERAT | MISSENSE | Ttg/Atg | L124M | orf46e | A | A | . | . | A | A | . | . | A | A | . | . |
| NC_011770 | 1712441 | A | G | 51 | 6 | 2 | 0 | NON_SYNONYMOU | MODERAT | MISSENSE | Atc/Gtc | I137V | orf46e | G | G | . | . | | | . | . | | | . | . |
| NC_011770 | 1712517 | A | T | 45 | 6 | 2 | 0 | NON_SYNONYMOU | MODERAT | MISSENSE | tAt/tTt | Y162F | orf46e | T | T | . | . | | | . | . | | | . | . |
| NC_011770 | 1712840 | A | T | 48 | 6 | 2 | 0 | NON_SYNONYMOU | MODERAT | MISSENSE | Aac/Tac | N270Y | orf46e | T | T | . | . | | | . | . | | | . | . |
| NC_011770 | 1712865 | T | G | 46 | 6 | 2 | 0 | NON_SYNONYMOU | MODERAT | MISSENSE | tTc/tGc | F278C | orf46e | G | G | . | . | | | . | . | | | . | . |
| NC_011770 | 1712866 | C | G | 46 | 6 | 2 | 0 | NON_SYNONYMOU | MODERAT | MISSENSE | ttC/ttG | F278L | orf46e | G | G | . | . | | | . | . | | | . | . |
| NC_011770 | 1713913 | G | A | 52 | 6 | 2 | 0 | SPLICE_SITE_REGIC | LOW | SILENT | tGa/tAa | *308 | orf48 | A | A | . | . | | | . | . | | | . | . |
| NC_011770 | 1713016 | A | G | 45 | 7 | 1 | 0 | NON_SYNONYMOU | MODERAT | MISSENSE | aAc/aGc | N9S | orf48 | . | G | . | . | | | . | . | | | . | . |
| NC_011770 | 1713141 | G | A | 50 | 6 | 2 | 0 | NON_SYNONYMOU | MODERAT | MISSENSE | Gga/Aga | G51R | orf48 | | | . | . | | | . | . | A | A | . | . |

| | | | | | | | | | | | | | | | | | | | | | | | | | |
|---|---|---|---|---|---|---|---|---|---|---|---|---|---|---|---|---|---|---|---|---|---|---|---|---|---|
| NC_011770 | 1713229 | A | C | 51 | 6 | 2 | 0 | NON_SYNONYMOU | MODERAT | MISSENSE | aAc/aCc | N80T | orf48 | C | C | . | . | | | . | . | | | . | . |
| NC_011770 | 1713244 | A | C | 47 | 6 | 2 | 0 | NON_SYNONYMOU | MODERAT | MISSENSE | aAg/aCg | K85T | orf48 | C | C | . | . | | | . | . | | | . | . |
| NC_011770 | 1713245 | G | T | 47 | 6 | 2 | 0 | NON_SYNONYMOU | MODERAT | MISSENSE | aaG/aaT | K85N | orf48 | T | T | . | . | | | . | . | | | . | . |
| NC_011770 | 1713256 | T | A | 50 | 6 | 2 | 0 | NON_SYNONYMOU | MODERAT | MISSENSE | gTc/gAc | V89D | orf48 | A | A | . | . | | | . | . | | | . | . |
| NC_011770 | 1713271 | G | C | 52 | 6 | 2 | 0 | NON_SYNONYMOU | MODERAT | MISSENSE | gGt/gCt | G94A | orf48 | C | C | . | . | | | . | . | | | . | . |
| NC_011770 | 1713360 | A | T | 46 | 7 | 1 | 0 | NON_SYNONYMOU | MODERAT | MISSENSE | Atg/Ttg | M124L | orf48 | . | T | . | . | | | . | . | | | . | . |
| NC_011770 | 1713363 | A | G | 45 | 7 | 1 | 0 | NON_SYNONYMOU | MODERAT | MISSENSE | Att/Gtt | I125V | orf48 | . | G | . | . | | | . | . | | | . | . |
| NC_011770 | 1713554 | C | G | 47 | 6 | 2 | 0 | NON_SYNONYMOU | MODERAT | MISSENSE | gaC/gaG | D188E | orf48 | G | G | . | . | | | . | . | | | . | . |
| NC_011770 | 1713623 | A | C | 51 | 6 | 2 | 0 | NON_SYNONYMOU | MODERAT | MISSENSE | agA/agC | R211S | orf48 | C | C | . | . | | | . | . | | | . | . |
| NC_011770 | 1713688 | C | A | 53 | 6 | 2 | 0 | NON_SYNONYMOU | MODERAT | MISSENSE | aCc/aAc | T233N | orf48 | A | A | . | . | | | . | . | | | . | . |
| NC_011770 | 1713690 | A | G | 52 | 6 | 2 | 0 | NON_SYNONYMOU | MODERAT | MISSENSE | Act/Gct | T234A | orf48 | G | G | . | . | | | . | . | | | . | . |
| NC_011770 | 1713691 | C | T | 53 | 6 | 2 | 0 | NON_SYNONYMOU | MODERAT | MISSENSE | aCt/aTt | T234I | orf48 | T | T | . | . | | | . | . | | | . | . |
| NC_011770 | 1714570 | C | A | 45 | 8 | 2 | 0 | NON_SYNONYMOU | MODERAT | MISSENSE | gaC/gaA | D207E | orf49 | . | . | . | . | A | A | . | . | | | . | . |
| NC_011770 | 1715107 | T | G | 46 | 8 | 2 | 0 | NON_SYNONYMOU | MODERAT | MISSENSE | atT/atG | I386M | orf49 | . | . | . | . | G | G | . | . | | | . | . |
| NC_011770 | 1715183 | C | G | 46 | 8 | 2 | 0 | NON_SYNONYMOU | MODERAT | MISSENSE | Cgg/Ggg | R412G | orf49 | . | . | . | . | G | G | . | . | | | . | . |
| NC_011770 | 1715526 | C | G | 50 | 8 | 2 | 0 | NON_SYNONYMOU | MODERAT | MISSENSE | aCc/aGc | T526S | orf49 | . | . | . | . | G | G | . | . | | | . | . |
| NC_011770 | 1686213 | C | T | 21 | 10 | 2 | 0 | NON_SYNONYMOU | MODERAT | MISSENSE | gCg/gTg | A70V | orf6 | . | . | . | . | T | T | . | . | . | . | . | . |
| NC_011770 | 1686308 | A | C | 22 | 10 | 2 | 0 | NON_SYNONYMOU | MODERAT | MISSENSE | Atg/Ctg | M102L | orf6 | . | . | . | . | C | C | . | . | . | . | . | . |
| NC_011770 | 1686957 | C | T | 23 | 10 | 2 | 0 | NON_SYNONYMOU | MODERAT | MISSENSE | aCc/aTc | T318I | orf6 | . | . | . | . | T | T | . | . | . | . | . | . |
| NC_011770 | 1686968 | A | G | 22 | 10 | 2 | 0 | NON_SYNONYMOU | MODERAT | MISSENSE | Att/Gtt | I322V | orf6 | . | . | . | . | G | G | . | . | . | . | . | . |
| NC_011770 | 1687402 | G | T | 18 | 10 | 2 | 0 | NON_SYNONYMOU | MODERAT | MISSENSE | gaG/gaT | E466D | orf6 | . | . | . | . | T | T | . | . | . | . | . | . |
| NC_011770 | 1687850 | A | G | 12 | 10 | 2 | 0 | NON_SYNONYMOU | MODERAT | MISSENSE | Atg/Gtg | M616V | orf6 | . | . | . | . | G | G | . | . | . | . | . | . |
| NC_011770 | 1687892 | A | G | 12 | 10 | 2 | 0 | NON_SYNONYMOU | MODERAT | MISSENSE | Acc/Gcc | T630A | orf6 | . | . | . | . | G | G | . | . | . | . | . | . |
| NC_011770 | 1687895 | G | A | 11 | 10 | 2 | 0 | NON_SYNONYMOU | MODERAT | MISSENSE | Gcc/Acc | A631T | orf6 | . | . | . | . | A | A | . | . | . | . | . | . |
| NC_011770 | 1687992 | A | T | 13 | 10 | 2 | 0 | NON_SYNONYMOU | MODERAT | MISSENSE | cAa/cTa | Q663L | orf6 | . | . | . | . | T | T | . | . | . | . | . | . |
| NC_011770 | 1688120 | G | A | 13 | 10 | 2 | 0 | NON_SYNONYMOU | MODERAT | MISSENSE | aGc/aAc | S17N | orf7 | . | . | . | . | A | A | . | . | . | . | . | . |
| NC_011770 | 1688549 | A | T | 15 | 10 | 2 | 0 | NON_SYNONYMOU | MODERAT | MISSENSE | tAc/tTc | Y160F | orf7 | . | . | . | . | T | T | . | . | . | . | . | . |
| NC_011770 | 1688838 | G | A | 19 | 10 | 2 | 0 | NON_SYNONYMOU | MODERAT | MISSENSE | Gtg/Atg | V1M | orf8 | . | . | . | . | A | A | . | . | . | . | . | . |
| NC_011770 | 1689067 | G | A | 20 | 10 | 2 | 0 | NON_SYNONYMOU | MODERAT | MISSENSE | cGg/cAg | R77Q | orf8 | . | . | . | . | A | A | . | . | . | . | . | . |
| NC_011770 | 1689072 | A | G | 20 | 10 | 2 | 0 | NON_SYNONYMOU | MODERAT | MISSENSE | Atc/Gtc | I79V | orf8 | . | . | . | . | G | G | . | . | . | . | . | . |
| NC_011770 | 1689180 | T | G | 22 | 10 | 2 | 0 | NON_SYNONYMOU | MODERAT | MISSENSE | Tcg/Gcg | S115A | orf8 | . | . | . | . | G | G | . | . | . | . | . | . |
| NC_011770 | 1689225 | C | A | 21 | 10 | 2 | 0 | NON_SYNONYMOU | MODERAT | MISSENSE | Ccc/Acc | P130T | orf8 | . | . | . | . | A | A | . | . | . | . | . | . |
| NC_011770 | 2713525 | T | A | 7 | 10 | 2 | 0 | NON_SYNONYMOU | MODERAT | MISSENSE | Tgc/Agc | C13S | orf91 | A | A | . | . | . | . | . | . | . | . | . | . |
| NC_011770 | 2713553 | A | G | 6 | 10 | 2 | 0 | NON_SYNONYMOU | MODERAT | MISSENSE | aAa/aGa | K22R | orf91 | G | G | . | . | . | . | . | . | . | . | . | . |
| NC_011770 | 2713574 | C | G | 7 | 10 | 2 | 0 | NON_SYNONYMOU | MODERAT | MISSENSE | tCc/tGc | S29C | orf91 | G | G | . | . | . | . | . | . | . | . | . | . |
| NC_011770 | 2713714 | G | A | 5 | 10 | 2 | 0 | NON_SYNONYMOU | MODERAT | MISSENSE | Ggc/Agc | G76S | orf91 | A | A | . | . | . | . | . | . | . | . | . | . |
| NC_011770 | 2713717 | A | C | 5 | 10 | 2 | 0 | NON_SYNONYMOU | MODERAT | MISSENSE | Aca/Cca | T77P | orf91 | C | C | . | . | . | . | . | . | . | . | . | . |
| NC_011770 | 2713726 | A | G | 5 | 10 | 2 | 0 | NON_SYNONYMOU | MODERAT | MISSENSE | Aaa/Gaa | K80E | orf91 | G | G | . | . | . | . | . | . | . | . | . | . |
| NC_011770 | 2713919 | C | A | 17 | 8 | 4 | 0 | NON_SYNONYMOU | MODERAT | MISSENSE | aCg/aAg | T21K | orf93 | A | A | . | . | A | A | . | . | . | . | . | . |
| NC_011770 | 2713971 | T | G | 16 | 8 | 4 | 0 | NON_SYNONYMOU | MODERAT | MISSENSE | gaT/gaG | D38E | orf93 | G | G | . | . | G | G | . | . | . | . | . | . |
| NC_011770 | 2713980 | A | C | 4 | 10 | 2 | 0 | NON_SYNONYMOU | MODERAT | MISSENSE | gaA/gaC | E41D | orf93 | C | C | . | . | . | . | . | . | . | . | . | . |
| NC_011770 | 2713985 | C | T | 4 | 10 | 2 | 0 | NON_SYNONYMOU | MODERAT | MISSENSE | gCg/gTg | A43V | orf93 | T | T | . | . | . | . | . | . | . | . | . | . |
| NC_011770 | 2714171 | C | CG | 25 | 8 | 2 | 0 | FRAME_SHIFT | HIGH | | ggc/ggGc | G52G? | orf94 | | | . | . | CG | CG | . | . | . | . | . | . |
| NC_011770 | 2714222 | C | A | 24 | 8 | 2 | 0 | STOP_GAINED | HIGH | NONSENSE | tgC/tgA | C68* | orf94 | A | A | . | . | | | . | . | . | . | . | . |
| NC_011770 | 2714227 | C | A | 24 | 8 | 2 | 0 | STOP_GAINED | HIGH | NONSENSE | tCg/tAg | S70* | orf94 | A | A | . | . | | | . | . | . | . | . | . |
| NC_011770 | 2714325 | T | C | 17 | 10 | 2 | 0 | STOP_LOST+SPLICE | HIGH | MISSENSE | Tga/Cga | *103R | orf94 | . | . | . | . | C | C | . | . | . | . | . | . |
| NC_011770 | 2714050 | C | G | 19 | 8 | 2 | 0 | NON_SYNONYMOU | MODERAT | MISSENSE | aCc/aGc | T11S | orf94 | | | . | . | G | G | . | . | . | . | . | . |
| NC_011770 | 2714061 | T | G | 19 | 8 | 4 | 0 | NON_SYNONYMOU | MODERAT | MISSENSE | Tct/Gct | S15A | orf94 | G | G | . | . | G | G | . | . | . | . | . | . |
| NC_011770 | 2714176 | T | A | 25 | 8 | 2 | 0 | NON_SYNONYMOU | MODERAT | MISSENSE | tTt/tAt | F53Y | orf94 | A | A | . | . | | | . | . | . | . | . | . |
| NC_011770 | 2714181 | C | G | 26 | 8 | 2 | 0 | NON_SYNONYMOU | MODERAT | MISSENSE | Cat/Gat | H55D | orf94 | G | G | . | . | | | . | . | . | . | . | . |
| NC_011770 | 2714182 | A | T | 25 | 8 | 2 | 0 | NON_SYNONYMOU | MODERAT | MISSENSE | cAt/cTt | H55L | orf94 | T | T | . | . | | | . | . | . | . | . | . |
| NC_011770 | 2714191 | C | T | 25 | 8 | 2 | 0 | NON_SYNONYMOU | MODERAT | MISSENSE | aCg/aTg | T58M | orf94 | T | T | . | . | | | . | . | . | . | . | . |
| NC_011770 | 2714212 | T | C | 24 | 8 | 2 | 0 | NON_SYNONYMOU | MODERAT | MISSENSE | cTg/cCg | L65P | orf94 | C | C | . | . | | | . | . | . | . | . | . |
| NC_011770 | 2714215 | G | A | 24 | 8 | 2 | 0 | NON_SYNONYMOU | MODERAT | MISSENSE | aGt/aAt | S66N | orf94 | A | A | . | . | | | . | . | . | . | . | . |
| NC_011770 | 2714216 | T | G | 24 | 8 | 2 | 0 | NON_SYNONYMOU | MODERAT | MISSENSE | agT/agG | S66R | orf94 | G | G | . | . | | | . | . | . | . | . | . |
| NC_011770 | 2714218 | G | C | 24 | 8 | 2 | 0 | NON_SYNONYMOU | MODERAT | MISSENSE | cGc/cCc | R67P | orf94 | C | C | . | . | | | . | . | . | . | . | . |
| NC_011770 | 2714233 | A | G | 25 | 8 | 2 | 0 | NON_SYNONYMOU | MODERAT | MISSENSE | aAc/aGc | N72S | orf94 | G | G | . | . | | | . | . | . | . | . | . |
| NC_011770 | 2714242 | T | G | 25 | 8 | 2 | 0 | NON_SYNONYMOU | MODERAT | MISSENSE | gTa/gGa | V75G | orf94 | G | G | . | . | | | . | . | . | . | . | . |
| NC_011770 | 2714265 | T | C | 19 | 10 | 2 | 0 | NON_SYNONYMOU | MODERAT | MISSENSE | Tat/Cat | Y83H | orf94 | . | . | . | . | C | C | . | . | . | . | . | . |
| NC_011770 | 644945 | T | C | 80 | 0 | 8 | 0 | NON_SYNONYMOU | MODERAT | MISSENSE | Acg/Gcg | T910A | ostA | C | C | C | C | C | C | | | | | C | C |
| NC_011770 | 644996 | T | C | 80 | 0 | 2 | 0 | NON_SYNONYMOU | MODERAT | MISSENSE | Atc/Gtc | I893V | ostA | | | | | | | C | C | | | | |
| NC_011770 | 645013 | T | G | 64 | 0 | 2 | 0 | NON_SYNONYMOU | MODERAT | MISSENSE | gAa/gCa | E887A | ostA | | | | | | | G | G | | | | |

| | | | | | | | | | | | | | | | | | | | | | | | | | |
|---|---|---|---|---|---|---|---|---|---|---|---|---|---|---|---|---|---|---|---|---|---|---|---|---|---|
| NC_011770 | 645016 | T | G | 64 | 0 | 6 | 0 | NON_SYNONYMOU | MODERAT | MISSENSE | aAt/aCt | N886T | ostA | | | G | G | G | G | | | C | C | | |
| NC_011770 | 645021 | G | T | 63 | 0 | 4 | 0 | NON_SYNONYMOU | MODERAT | MISSENSE | aaC/aaA | N884K | ostA | | | | | | | T | T | T | T | | |
| NC_011770 | 645022 | T | A | 64 | 0 | 8 | 0 | NON_SYNONYMOU | MODERAT | MISSENSE | aAc/aTc | N884I | ostA | | | A | A | A | A | G | G | A | A | | |
| NC_011770 | 645023 | T | C | 63 | 0 | 6 | 0 | NON_SYNONYMOU | MODERAT | MISSENSE | Aac/Gac | N884D | ostA | | | C | C | C | C | | | C | C | | |
| NC_011770 | 645026 | G | A | 69 | 0 | 2 | 0 | NON_SYNONYMOU | MODERAT | MISSENSE | Ctc/Ttc | L883F | ostA | | | | | | | | | A | A | | |
| NC_011770 | 645179 | G | T | 63 | 0 | 2 | 0 | NON_SYNONYMOU | MODERAT | MISSENSE | Ctg/Atg | L832M | ostA | | | | | | | T | T | | | | |
| NC_011770 | 645221 | T | C | 42 | 0 | 12 | 0 | NON_SYNONYMOU | MODERAT | MISSENSE | Aac/Gac | N818D | ostA | C | C | C | C | C | C | C | C | C | C | C | C |
| NC_011770 | 645234 | C | T | 38 | 0 | 6 | 0 | NON_SYNONYMOU | MODERAT | MISSENSE | gaG/gaC | E813D | ostA | | | T | T | T | T | | | G | G | | |
| NC_011770 | 645235 | T | A | 38 | 0 | 8 | 0 | NON_SYNONYMOU | MODERAT | MISSENSE | gAg/gTg | E813V | ostA | A | A | | | | | A | A | A | A | A | A |
| NC_011770 | 645236 | C | T | 38 | 0 | 6 | 0 | NON_SYNONYMOU | MODERAT | MISSENSE | Gag/Aag | E813K | ostA | T | T | | | | | T | T | | | T | T |
| NC_011770 | 645238 | A | T | 39 | 0 | 4 | 0 | NON_SYNONYMOU | MODERAT | MISSENSE | tTc/tAc | F812Y | ostA | T | T | | | | | | | | | T | T |
| NC_011770 | 645251 | A | C | 40 | 0 | 4 | 0 | NON_SYNONYMOU | MODERAT | MISSENSE | Tct/Gct | S808A | ostA | C | C | | | | | | | | | C | C |
| NC_011770 | 645253 | C | G | 41 | 0 | 4 | 0 | NON_SYNONYMOU | MODERAT | MISSENSE | aGc/aCc | S807T | ostA | | | G | G | G | G | | | | | | |
| NC_011770 | 645254 | T | C | 41 | 0 | 4 | 0 | NON_SYNONYMOU | MODERAT | MISSENSE | Agc/Ggc | S807G | ostA | C | C | | | | | | | | | C | C |
| NC_011770 | 645269 | A | C | 32 | 4 | 8 | 0 | NON_SYNONYMOU | MODERAT | MISSENSE | Tcc/Gcc | S802A | ostA | C | C | . | . | . | . | C | C | C | C | C | C |
| NC_011770 | 645382 | A | T | 39 | 4 | 2 | 0 | NON_SYNONYMOU | MODERAT | MISSENSE | tTt/tAt | F764Y | ostA | | | . | . | . | . | | | T | T | | |
| NC_011770 | 645395 | C | G | 36 | 4 | 2 | 0 | NON_SYNONYMOU | MODERAT | MISSENSE | Gtc/Ctc | V760L | ostA | | | . | . | . | . | | | G | G | | |
| NC_011770 | 645396 | T | C | 37 | 4 | 8 | 0 | NON_SYNONYMOU | MODERAT | MISSENSE | agA/agT | R759S | ostA | C | C | . | . | . | . | C | C | A | A | C | C |
| NC_011770 | 645700 | C | T | 34 | 4 | 2 | 0 | NON_SYNONYMOU | MODERAT | MISSENSE | aGc/aAc | S658N | ostA | | | . | . | . | . | | | T | T | | |
| NC_011770 | 645738 | C | G | 12 | 8 | 4 | 0 | NON_SYNONYMOU | MODERAT | MISSENSE | gaG/gaC | E645D | ostA | G | G | . | . | . | . | . | . | . | . | G | G |
| NC_011770 | 645810 | T | A | 7 | 8 | 4 | 0 | NON_SYNONYMOU | MODERAT | MISSENSE | ttA/ttT | L621F | ostA | A | A | . | . | . | . | . | . | . | . | A | A |
| NC_011770 | 645958 | T | A | 31 | 2 | 10 | 0 | NON_SYNONYMOU | MODERAT | MISSENSE | tAc/tTc | Y572F | ostA | A | A | A | A | A | A | A | A | . | . | A | A |
| NC_011770 | 645991 | C | T | 34 | 2 | 4 | 0 | NON_SYNONYMOU | MODERAT | MISSENSE | cGc/cAc | R561H | ostA | T | T | | | | | | | . | . | T | T |
| NC_011770 | 646028 | C | T | 12 | 6 | 4 | 0 | NON_SYNONYMOU | MODERAT | MISSENSE | Gta/Ata | V549I | ostA | T | T | . | . | . | . | . | . | | | T | T |
| NC_011770 | 646058 | G | T | 4 | 8 | 4 | 0 | NON_SYNONYMOU | MODERAT | MISSENSE | Caa/Aaa | Q539K | ostA | T | T | . | . | . | . | . | . | . | . | T | T |
| NC_011770 | 646086 | C | A | 4 | 8 | 4 | 0 | NON_SYNONYMOU | MODERAT | MISSENSE | gaG/gaT | E529D | ostA | A | A | . | . | . | . | . | . | . | . | A | A |
| NC_011770 | 646087 | T | A | 4 | 8 | 4 | 0 | NON_SYNONYMOU | MODERAT | MISSENSE | gAg/gTg | E529V | ostA | A | A | . | . | . | . | . | . | . | . | A | A |
| NC_011770 | 646112 | T | C | 17 | 8 | 4 | 0 | NON_SYNONYMOU | MODERAT | MISSENSE | Att/Gtt | I521V | ostA | C | C | . | . | . | . | . | . | . | . | C | C |
| NC_011770 | 646163 | A | G | 13 | 8 | 4 | 0 | NON_SYNONYMOU | MODERAT | MISSENSE | Ttc/Ctc | F504L | ostA | G | G | . | . | . | . | . | . | . | . | G | G |
| NC_011770 | 646169 | C | A | 17 | 6 | 2 | 0 | NON_SYNONYMOU | MODERAT | MISSENSE | Gtg/Ttg | V502L | ostA | | | . | . | . | . | . | . | A | A | | |
| NC_011770 | 646208 | T | C | 8 | 8 | 4 | 0 | NON_SYNONYMOU | MODERAT | MISSENSE | Atc/Gtc | I489V | ostA | . | . | . | . | . | . | C | C | C | C | . | . |
| NC_011770 | 646221 | C | G | 8 | 8 | 4 | 0 | NON_SYNONYMOU | MODERAT | MISSENSE | gaG/gaC | E484D | ostA | . | . | . | . | . | . | G | G | G | G | . | . |
| NC_011770 | 646223 | C | T | 8 | 8 | 2 | 0 | NON_SYNONYMOU | MODERAT | MISSENSE | Gag/Aag | E484K | ostA | . | . | . | . | . | . | T | T | | | . | . |
| NC_011770 | 646325 | C | G | 15 | 8 | 4 | 0 | NON_SYNONYMOU | MODERAT | MISSENSE | Gtc/Ctc | V450L | ostA | G | G | . | . | . | . | . | . | . | . | G | G |
| NC_011770 | 646355 | T | A | 43 | 4 | 4 | 0 | NON_SYNONYMOU | MODERAT | MISSENSE | Act/Tct | T440S | ostA | A | A | . | . | . | . | | | | | A | A |
| NC_011770 | 646497 | G | T | 87 | 0 | 2 | 0 | NON_SYNONYMOU | MODERAT | MISSENSE | gaC/gaA | D392E | ostA | | | | | | | T | T | | | | |
| NC_011770 | 646521 | G | C | 84 | 0 | 4 | 0 | NON_SYNONYMOU | MODERAT | MISSENSE | aaC/aaG | N384K | ostA | | | C | C | C | C | | | | | | |
| NC_011770 | 646522 | T | G | 84 | 0 | 4 | 0 | NON_SYNONYMOU | MODERAT | MISSENSE | aAc/aCc | N384T | ostA | | | G | G | G | G | | | | | | |
| NC_011770 | 646528 | G | A | 85 | 0 | 4 | 0 | NON_SYNONYMOU | MODERAT | MISSENSE | aCc/aTc | T382I | ostA | | | A | A | A | A | | | | | | |
| NC_011770 | 647443 | C | G | 63 | 0 | 10 | 0 | NON_SYNONYMOU | MODERAT | MISSENSE | aGc/aCc | S77T | ostA | G | G | G | G | G | G | | | G | G | G | G |
| NC_011770 | 646529 | T | C | 85 | 0 | 1 | 0 | NON_SYNONYMOU | MODERAT | MISSENSE | Acc/Gcc | T382A | ostA | | | | | | | | C | | | | |
| NC_011770 | 645088 | C | T | 73 | 0 | 1 | 0 | NON_SYNONYMOU | MODERAT | MISSENSE | aGc/aAc | S862N | ostA | | | | | | | | | | | | **T** |
| NC_011770 | 3969518 | A | G | 60 | 0 | 2 | 0 | NON_SYNONYMOU | MODERAT | MISSENSE | Atg/Gtg | M30V | pabB | | | | | | | G | G | | | | |
| NC_011770 | 3969917 | G | A | 67 | 0 | 12 | 0 | NON_SYNONYMOU | MODERAT | MISSENSE | Gcg/Acg | A163T | pabB | A | A | A | A | A | A | A | A | A | A | A | A |
| NC_011770 | 3970652 | A | G | 64 | 0 | 2 | 0 | NON_SYNONYMOU | MODERAT | MISSENSE | Aac/Gac | N408D | pabB | G | G | | | | | | | | | | |
| NC_011770 | 3970638 | C | T | 63 | 0 | 1 | 0 | NON_SYNONYMOU | MODERAT | MISSENSE | aCc/aTc | T403I | pabB | | | | | | | | | | | | T |
| NC_011770 | 2271755 | G | A | 77 | 0 | 4 | 0 | NON_SYNONYMOU | MODERAT | MISSENSE | Gcc/Acc | A60T | pabC | | | | | | | | | A | A | A | A |
| NC_011770 | 2271908 | G | A | 81 | 0 | 4 | 0 | NON_SYNONYMOU | MODERAT | MISSENSE | Gcc/Acc | A111T | pabC | | | A | A | A | A | | | | | | |
| NC_011770 | 5564605 | T | C | 63 | 0 | 1 | 0 | STOP_LOST+SPLICE | HIGH | MISSENSE | tAg/tGg | *174W | pagL | | | | | | | | C | | | | |
| NC_011770 | 5565023 | G | A | 72 | 0 | 1 | 0 | STOP_GAINED | HIGH | NONSENSE | Cag/Tag | Q35* | pagL | | | | | | | A | | | | | |
| NC_011770 | 5564740 | C | T | 65 | 0 | 12 | 0 | NON_SYNONYMOU | MODERAT | MISSENSE | gGc/gAc | G129D | pagL | T | T | T | T | T | T | T | T | T | T | T | T |
| NC_011770 | 4127163 | A | G | 86 | 0 | 2 | 0 | NON_SYNONYMOU | MODERAT | MISSENSE | cAg/cGg | Q16R | panB | G | G | | | | | | | | | | |
| NC_011770 | 5645554 | G | T | 80 | 0 | 12 | 0 | NON_SYNONYMOU | MODERAT | MISSENSE | caG/caT | Q91H | panB | T | T | T | T | T | T | T | T | T | T | T | T |
| NC_011770 | 5645570 | C | T | 80 | 0 | 1 | 0 | NON_SYNONYMOU | MODERAT | MISSENSE | Ctt/Ttt | L97F | panB | | | | | | | T | | | | | |
| NC_011770 | 5646287 | A | C | 68 | 0 | 8 | 0 | NON_SYNONYMOU | MODERAT | MISSENSE | aAa/aCa | K70T | panC | C | C | C | C | C | C | | | C | C | | |
| NC_011770 | 5646372 | G | A | 73 | 0 | 4 | 0 | NON_SYNONYMOU | MODERAT | MISSENSE | atG/atA | M98I | panC | | | A | A | A | A | | | | | | |
| NC_011770 | 5646604 | C | A | 75 | 0 | 2 | 0 | NON_SYNONYMOU | MODERAT | MISSENSE | Ccg/Acg | P176T | panC | A | A | | | | | | | | | | |
| NC_011770 | 5646658 | A | G | 69 | 0 | 10 | 0 | NON_SYNONYMOU | MODERAT | MISSENSE | Aac/Gac | N194D | panC | G | G | G | G | G | G | G | G | G | G | | |
| NC_011770 | 5646872 | T | C | 73 | 0 | 8 | 0 | NON_SYNONYMOU | MODERAT | MISSENSE | gTc/gCc | V265A | panC | | | C | C | C | C | C | C | | | C | C |
| NC_011770 | 5646916 | G | A | 76 | 0 | 2 | 0 | NON_SYNONYMOU | MODERAT | MISSENSE | Gtc/Atc | V280I | panC | | | | | | | A | A | | | | |
| NC_011770 | 5647132 | G | A | 78 | 0 | 2 | 0 | NON_SYNONYMOU | MODERAT | MISSENSE | Gga/Aga | G37R | panD | A | A | | | | | | | | | | |
| NC_011770 | 5261059 | G | A | 64 | 0 | 1 | 0 | NON_SYNONYMOU | MODERAT | MISSENSE | Ggc/Agc | G106S | panE | | | | | | | A | | | | | |

| | | | | | | | | | | | | | | | | | | | | | | | | | |
|---|---|---|---|---|---|---|---|---|---|---|---|---|---|---|---|---|---|---|---|---|---|---|---|---|---|
| NC_011770 | 5261204 | G | A | 63 | 0 | 2 | 0 | NON_SYNONYMOU | MODERAT | MISSENSE | aGc/aAc | S154N | panE | | | | | | | | | A | A | | |
| NC_011770 | 5261221 | G | A | 65 | 0 | 4 | 0 | NON_SYNONYMOU | MODERAT | MISSENSE | Gcg/Acg | A160T | panE | | | | | | | | | A | A | A | A |
| NC_011770 | 5261591 | G | A | 57 | 0 | 2 | 0 | NON_SYNONYMOU | MODERAT | MISSENSE | cGc/cAc | R283H | panE | | | | | | | | | A | A | | |
| NC_011770 | 3950007 | C | A | 67 | 0 | 2 | 0 | NON_SYNONYMOU | MODERAT | MISSENSE | aaG/aaT | K159N | panE2 | A | A | | | | | | | | | | |
| NC_011770 | 3950230 | T | C | 68 | 0 | 4 | 0 | NON_SYNONYMOU | MODERAT | MISSENSE | aAc/aGc | N85S | panE2 | | | C | C | C | C | | | | | | |
| NC_011770 | 5904886 | C | A | 72 | 0 | 2 | 0 | NON_SYNONYMOU | MODERAT | MISSENSE | Gtg/Ttg | V646L | parC | | | | | | | | | | | A | A |
| NC_011770 | 5906776 | A | C | 90 | 0 | 2 | 0 | NON_SYNONYMOU | MODERAT | MISSENSE | Tcg/Gcg | S16A | parC | | | | | | | | | C | C | | |
| NC_011770 | 5908683 | G | A | 82 | 0 | 1 | 0 | NON_SYNONYMOU | MODERAT | MISSENSE | Ccg/Tcg | P524S | parE | | | | | | | | A | | | | |
| NC_011770 | 5909829 | T | C | 68 | 0 | 12 | 0 | NON_SYNONYMOU | MODERAT | MISSENSE | Aac/Gac | N142D | parE | C | C | C | C | C | C | C | C | C | C | C | C |
| NC_011770 | 4745366 | T | C | 98 | 0 | 1 | 0 | NON_SYNONYMOU | MODERAT | MISSENSE | cAg/cGg | Q352R | pauA | | C | | | | | | | | | | |
| NC_011770 | 4744476 | A | G | 60 | 0 | 4 | 0 | NON_SYNONYMOU | MODERAT | MISSENSE | Ttc/Ctc | F649L | pauA | | | G | G | G | G | | | | | | |
| NC_011770 | 4745414 | A | G | 98 | 0 | 2 | 0 | NON_SYNONYMOU | MODERAT | MISSENSE | gTc/gCc | V336A | pauA | | | | | | | | | G | G | | |
| NC_011770 | 4745475 | C | G | 99 | 0 | 2 | 0 | NON_SYNONYMOU | MODERAT | MISSENSE | Gtg/Ctg | V316L | pauA | G | G | | | | | | | | | | |
| NC_011770 | 4745555 | C | T | 90 | 0 | 2 | 0 | NON_SYNONYMOU | MODERAT | MISSENSE | gGc/gAc | G289D | pauA | T | T | | | | | | | | | | |
| NC_011770 | 4745577 | T | C | 84 | 0 | 2 | 0 | NON_SYNONYMOU | MODERAT | MISSENSE | Aag/Gag | K282E | pauA | | | | | | | | | C | C | | |
| NC_011770 | 4745839 | G | C | 91 | 0 | 10 | 0 | NON_SYNONYMOU | MODERAT | MISSENSE | tgC/tgG | C194W | pauA | C | C | C | C | C | C | C | C | C | C | | |
| NC_011770 | 4745841 | A | G | 91 | 0 | 12 | 0 | NON_SYNONYMOU | MODERAT | MISSENSE | Tgc/Cgc | C194R | pauA | G | G | G | G | G | G | G | G | G | G | G | G |
| NC_011770 | 4745919 | C | A | 88 | 0 | 2 | 0 | NON_SYNONYMOU | MODERAT | MISSENSE | Gta/Tta | V168L | pauA | A | A | | | | | | | | | | |
| NC_011770 | 4746012 | A | G | 81 | 0 | 8 | 0 | NON_SYNONYMOU | MODERAT | MISSENSE | Tac/Cac | Y137H | pauA | G | G | G | G | G | G | G | G | | | | |
| NC_011770 | 4746170 | G | A | 88 | 0 | 8 | 0 | NON_SYNONYMOU | MODERAT | MISSENSE | gCg/gTg | A84V | pauA | A | A | A | A | A | A | A | A | | | | |
| NC_011770 | 1052379 | T | C | 67 | 0 | 2 | 0 | NON_SYNONYMOU | MODERAT | MISSENSE | gTg/gCg | V17A | pbpA | C | C | | | | | | | | | | |
| NC_011770 | 1053635 | C | T | 72 | 0 | 1 | 0 | NON_SYNONYMOU | MODERAT | MISSENSE | Cgc/Tgc | R436C | pbpA | | | | | | | | T | | | | |
| NC_011770 | 3342262 | C | T | 73 | 0 | 12 | 0 | NON_SYNONYMOU | MODERAT | MISSENSE | Gcg/Acg | A104T | pbpC | T | T | G | G | G | G | G | G | G | G | G | G |
| NC_011770 | 3342540 | A | G | 82 | 0 | 1 | 0 | NON_SYNONYMOU | MODERAT | MISSENSE | aTc/aCc | I11T | pbpC | | | | | | | G | | | | | |
| NC_011770 | 4888564 | G | A | 92 | 0 | 1 | 0 | STOP_GAINED | HIGH | NONSENSE | tgG/tgA | W275* | pbpG | | | | | | | | A | | | | |
| NC_011770 | 4888488 | G | A | 96 | 0 | 2 | 0 | NON_SYNONYMOU | MODERAT | MISSENSE | aGc/aAc | S250N | pbpG | | | | | | | A | A | | | | |
| NC_011770 | 252557 | T | C | 86 | 0 | 10 | 0 | NON_SYNONYMOU | MODERAT | MISSENSE | aTg/aCg | M15T | pcaB | | | C | C | C | C | C | C | C | C | C | C |
| NC_011770 | 253021 | C | T | 91 | 0 | 2 | 0 | NON_SYNONYMOU | MODERAT | MISSENSE | Ctc/Ttc | L170F | pcaB | | | | | | | | | | | T | T |
| NC_011770 | 253528 | G | T | 82 | 0 | 2 | 0 | NON_SYNONYMOU | MODERAT | MISSENSE | Gcc/Tcc | A339S | pcaB | | | | | | | | | | | T | T |
| NC_011770 | 253875 | A | C | 81 | 0 | 12 | 0 | NON_SYNONYMOU | MODERAT | MISSENSE | ttA/ttC | L454F | pcaB | C | C | C | C | C | C | C | C | C | C | C | C |
| NC_011770 | 255106 | C | G | 93 | 0 | 2 | 0 | NON_SYNONYMOU | MODERAT | MISSENSE | Cgg/Ggg | R133G | pcaC | | | | | | | | | G | G | | |
| NC_011770 | 254419 | G | A | 98 | 0 | 2 | 0 | NON_SYNONYMOU | MODERAT | MISSENSE | cGc/cAc | R171H | pcaD | | | | | | | A | A | | | | |
| NC_011770 | 254473 | C | T | 95 | 0 | 2 | 0 | NON_SYNONYMOU | MODERAT | MISSENSE | gCt/gTt | A189V | pcaD | | | | | T | T | | | | | | |
| NC_011770 | 254652 | G | A | 94 | 0 | 2 | 0 | NON_SYNONYMOU | MODERAT | MISSENSE | Ggc/Agc | G249S | pcaD | | | | | | | | | A | A | | |
| NC_011770 | 250574 | A | G | 91 | 0 | 6 | 0 | NON_SYNONYMOU | MODERAT | MISSENSE | aAg/aGg | K246R | pcaF | | | G | G | G | G | | | G | G | | |
| NC_011770 | 174461 | A | G | 84 | 0 | 6 | 0 | NON_SYNONYMOU | MODERAT | MISSENSE | gAc/gGc | D44G | pcaG | | | G | G | G | G | G | G | | | | |
| NC_011770 | 174467 | C | G | 87 | 0 | 2 | 0 | NON_SYNONYMOU | MODERAT | MISSENSE | cCg/cGg | P46R | pcaG | | | | | | | G | G | | | | |
| NC_011770 | 174719 | C | T | 82 | 0 | 2 | 0 | NON_SYNONYMOU | MODERAT | MISSENSE | tCg/tTg | S130L | pcaG | | | | | | | | | | | T | T |
| NC_011770 | 174778 | A | G | 80 | 0 | 2 | 0 | NON_SYNONYMOU | MODERAT | MISSENSE | Acc/Gcc | T150A | pcaG | | | | | | | | | | | G | G |
| NC_011770 | 174851 | G | A | 76 | 0 | 6 | 0 | NON_SYNONYMOU | MODERAT | MISSENSE | cGg/cAg | R174Q | pcaG | A | A | | | | | A | A | A | A | | |
| NC_011770 | 174866 | A | G | 80 | 0 | 1 | 0 | NON_SYNONYMOU | MODERAT | MISSENSE | gAc/gGc | D179G | pcaG | | | | | | | G | | | | | |
| NC_011770 | 174096 | A | G | 83 | 0 | 12 | 0 | NON_SYNONYMOU | MODERAT | MISSENSE | Atc/Gtc | I166V | pcaH | G | G | G | G | G | G | G | G | G | G | G | G |
| NC_011770 | 174195 | C | T | 83 | 0 | 2 | 0 | NON_SYNONYMOU | MODERAT | MISSENSE | Ccg/Tcg | P199S | pcaH | T | T | | | | | | | | | | |
| NC_011770 | 174199 | G | A | 81 | 0 | 12 | 0 | NON_SYNONYMOU | MODERAT | MISSENSE | gGg/gAg | G200E | pcaH | A | A | A | A | A | A | A | A | A | A | A | A |
| NC_011770 | 173117 | T | C | 82 | 0 | 6 | 0 | NON_SYNONYMOU | MODERAT | MISSENSE | cTg/cCg | L160P | pcaQ | C | C | | | | | | | C | C | C | C |
| NC_011770 | 173209 | G | C | 81 | 0 | 2 | 0 | NON_SYNONYMOU | MODERAT | MISSENSE | Gcg/Ccg | A191P | pcaQ | | | | | | | | | C | C | | |
| NC_011770 | 746749 | C | T | 62 | 0 | 2 | 0 | NON_SYNONYMOU | MODERAT | MISSENSE | Ggc/Agc | G408S | pchA | T | T | | | | | | | | | | |
| NC_011770 | 747261 | A | G | 77 | 0 | 12 | 0 | NON_SYNONYMOU | MODERAT | MISSENSE | cTg/cCg | L237P | pchA | G | G | G | G | G | G | G | G | G | G | G | G |
| NC_011770 | 747414 | T | C | 79 | 0 | 12 | 0 | NON_SYNONYMOU | MODERAT | MISSENSE | gAg/gGg | E186G | pchA | C | C | C | C | C | C | C | C | C | C | C | C |
| NC_011770 | 747751 | T | A | 83 | 0 | 4 | 0 | NON_SYNONYMOU | MODERAT | MISSENSE | Agc/Tgc | S74C | pchA | A | A | | | | | | | A | A | | |
| NC_011770 | 747889 | G | T | 94 | 0 | 2 | 0 | NON_SYNONYMOU | MODERAT | MISSENSE | Ctc/Atc | L28I | pchA | T | T | | | | | | | | | | |
| NC_011770 | 748378 | C | T | 91 | 0 | 2 | 0 | NON_SYNONYMOU | MODERAT | MISSENSE | cGc/cAc | R217H | pchC | | | | | | | | | | | T | T |
| NC_011770 | 748480 | A | G | 88 | 0 | 12 | 0 | NON_SYNONYMOU | MODERAT | MISSENSE | gTc/gCc | V183A | pchC | G | G | G | G | G | G | G | G | G | G | G | G |
| NC_011770 | 748711 | A | G | 89 | 0 | 12 | 0 | NON_SYNONYMOU | MODERAT | MISSENSE | gTc/gCc | V106A | pchC | G | G | G | G | G | G | G | G | G | G | G | G |
| NC_011770 | 748823 | A | C | 88 | 0 | 12 | 0 | NON_SYNONYMOU | MODERAT | MISSENSE | Tcc/Gcc | S69A | pchC | C | C | C | C | C | C | C | C | C | C | C | C |
| NC_011770 | 748849 | G | T | 89 | 0 | 2 | 0 | NON_SYNONYMOU | MODERAT | MISSENSE | gCg/gAg | A60E | pchC | | | | | | | | | T | T | | |
| NC_011770 | 748981 | C | T | 87 | 0 | 6 | 0 | NON_SYNONYMOU | MODERAT | MISSENSE | cGc/cAc | R16H | pchC | | | T | T | T | T | T | T | | | | |
| NC_011770 | 749539 | C | T | 77 | 0 | 2 | 0 | NON_SYNONYMOU | MODERAT | MISSENSE | Gac/Aac | D377N | pchD | T | T | | | | | | | | | | |
| NC_011770 | 749718 | C | T | 76 | 0 | 1 | 0 | NON_SYNONYMOU | MODERAT | MISSENSE | gGc/gAc | G317D | pchD | | | | | | | T | | | | | |
| NC_011770 | 749763 | C | T | 70 | 0 | 2 | 0 | NON_SYNONYMOU | MODERAT | MISSENSE | aGc/aAc | S302N | pchD | T | T | | | | | | | | | | |
| NC_011770 | 750189 | C | T | 56 | 0 | 6 | 0 | NON_SYNONYMOU | MODERAT | MISSENSE | cGg/cAg | R160Q | pchD | | | T | T | T | T | T | T | | | | |
| NC_011770 | 750544 | C | T | 40 | 0 | 2 | 0 | NON_SYNONYMOU | MODERAT | MISSENSE | Gcc/Acc | A42T | pchD | T | T | | | | | | | | | | |

| | | | | | | | | | | | | | | | | | | | | | | | | | |
|---|---|---|---|---|---|---|---|---|---|---|---|---|---|---|---|---|---|---|---|---|---|---|---|---|---|
| NC_011770 | 750564 | T | C | 40 | 0 | 4 | 0 | NON_SYNONYMOU | MODERAT | MISSENSE | aAc/aGc | N35S | pchD | C | C | | | | | | | | | C | C |
| NC_011770 | 752269 | G | A | 50 | 0 | 6 | 0 | NON_SYNONYMOU | MODERAT | MISSENSE | Gcg/Acg | A97T | pchE | A | A | A | A | A | A | | | | | | |
| NC_011770 | 752279 | A | C | 52 | 0 | 4 | 0 | NON_SYNONYMOU | MODERAT | MISSENSE | gAt/gCt | D100A | pchE | | | C | C | C | C | | | | | | |
| NC_011770 | 752389 | A | G | 56 | 0 | 6 | 0 | NON_SYNONYMOU | MODERAT | MISSENSE | Acg/Gcg | T137A | pchE | | | | | | | G | G | G | G | G | G |
| NC_011770 | 752398 | A | G | 59 | 0 | 12 | 0 | NON_SYNONYMOU | MODERAT | MISSENSE | Atc/Gtc | I140V | pchE | G | G | G | G | G | G | G | G | G | G | G | G |
| NC_011770 | 752425 | A | G | 58 | 0 | 12 | 0 | NON_SYNONYMOU | MODERAT | MISSENSE | Acg/Gcg | T149A | pchE | G | G | G | G | G | G | G | G | G | G | G | G |
| NC_011770 | 752470 | C | T | 59 | 0 | 4 | 0 | NON_SYNONYMOU | MODERAT | MISSENSE | Ctc/Ttc | L164F | pchE | | | T | T | T | T | | | | | | |
| NC_011770 | 752509 | T | C | 62 | 0 | 4 | 0 | NON_SYNONYMOU | MODERAT | MISSENSE | Tcc/Ccc | S177P | pchE | | | C | C | C | C | | | | | | |
| NC_011770 | 752804 | G | A | 55 | 0 | 6 | 0 | NON_SYNONYMOU | MODERAT | MISSENSE | cGc/cAc | R275H | pchE | A | A | A | A | A | A | | | | | | |
| NC_011770 | 753287 | A | G | 63 | 0 | 6 | 0 | NON_SYNONYMOU | MODERAT | MISSENSE | cAg/cGg | Q436R | pchE | | | | | | | G | G | G | G | G | G |
| NC_011770 | 753557 | G | A | 75 | 0 | 12 | 0 | NON_SYNONYMOU | MODERAT | MISSENSE | gGg/gAg | G526E | pchE | A | A | A | A | A | A | A | A | A | A | A | A |
| NC_011770 | 753901 | A | G | 77 | 0 | 12 | 0 | NON_SYNONYMOU | MODERAT | MISSENSE | Aca/Gca | T641A | pchE | G | G | G | G | G | G | G | G | G | G | G | G |
| NC_011770 | 754684 | A | G | 75 | 0 | 6 | 0 | NON_SYNONYMOU | MODERAT | MISSENSE | Agc/Ggc | S902G | pchE | | | G | G | G | G | | | | | G | G |
| NC_011770 | 755600 | A | C | 68 | 0 | 4 | 0 | NON_SYNONYMOU | MODERAT | MISSENSE | gAg/gCg | E1207A | pchE | | | C | C | C | C | | | | | | |
| NC_011770 | 755666 | C | T | 64 | 0 | 12 | 0 | NON_SYNONYMOU | MODERAT | MISSENSE | gCg/gTg | A1229V | pchE | T | T | T | T | T | T | T | T | T | T | T | T |
| NC_011770 | 755812 | A | C | 64 | 0 | 10 | 0 | NON_SYNONYMOU | MODERAT | MISSENSE | Atc/Ctc | I1278L | pchE | C | C | C | C | C | C | | | C | C | C | C |
| NC_011770 | 752559 | GCGCGACT | G | 62 | 0 | 1 | 0 | CODON_CHANGE_ | MODERATE | | caggcgctac | QALRDWR | pchE | | | | | G | | | | | | | |
| NC_011770 | 756601 | C | T | 68 | 0 | 6 | 0 | NON_SYNONYMOU | MODERAT | MISSENSE | gCc/gTc | A103V | pchF | | | T | T | T | T | | | | | T | T |
| NC_011770 | 756658 | C | T | 80 | 0 | 2 | 0 | NON_SYNONYMOU | MODERAT | MISSENSE | cCg/cTg | P122L | pchF | | | | | | | | | | | T | T |
| NC_011770 | 757755 | A | C | 72 | 0 | 12 | 0 | NON_SYNONYMOU | MODERAT | MISSENSE | Aag/Cag | K488Q | pchF | C | C | C | C | C | C | C | C | C | C | C | C |
| NC_011770 | 757784 | G | T | 66 | 0 | 12 | 0 | NON_SYNONYMOU | MODERAT | MISSENSE | gaG/gaT | E497D | pchF | T | T | T | T | T | T | T | T | T | T | T | T |
| NC_011770 | 757888 | T | C | 66 | 0 | 2 | 0 | NON_SYNONYMOU | MODERAT | MISSENSE | gTg/gCg | V532A | pchF | C | C | | | | | | | | | | |
| NC_011770 | 757942 | G | C | 64 | 0 | 2 | 0 | NON_SYNONYMOU | MODERAT | MISSENSE | aGc/aCc | S550T | pchF | C | C | | | | | | | | | | |
| NC_011770 | 759232 | C | T | 72 | 0 | 1 | 0 | NON_SYNONYMOU | MODERAT | MISSENSE | gCc/gTc | A980V | pchF | | | | | | | T | | | | | |
| NC_011770 | 759531 | G | A | 65 | 0 | 4 | 0 | NON_SYNONYMOU | MODERAT | MISSENSE | Gcc/Acc | A1080T | pchF | | | A | A | A | A | | | | | | |
| NC_011770 | 759813 | A | G | 67 | 0 | 2 | 0 | NON_SYNONYMOU | MODERAT | MISSENSE | Acg/Gcg | T1174A | pchF | G | G | | | | | | | | | | |
| NC_011770 | 759853 | T | C | 68 | 0 | 6 | 0 | NON_SYNONYMOU | MODERAT | MISSENSE | gTg/gCg | V1187A | pchF | C | C | C | C | C | C | | | | | | |
| NC_011770 | 760164 | T | C | 64 | 0 | 10 | 0 | NON_SYNONYMOU | MODERAT | MISSENSE | Tgc/Cgc | C1291R | pchF | C | C | C | C | C | C | C | C | | | C | C |
| NC_011770 | 760464 | G | A | 61 | 0 | 2 | 0 | NON_SYNONYMOU | MODERAT | MISSENSE | Ggc/Agc | G1391S | pchF | | | | | | | A | A | | | | |
| NC_011770 | 760760 | C | A | 64 | 0 | 6 | 0 | NON_SYNONYMOU | MODERAT | MISSENSE | gaC/gaA | D1489E | pchF | | | A | A | A | A | A | A | | | | |
| NC_011770 | 760789 | C | T | 63 | 0 | 4 | 0 | NON_SYNONYMOU | MODERAT | MISSENSE | gCg/gTg | A1499V | pchF | | | T | T | T | T | | | | | | |
| NC_011770 | 760810 | C | A | 65 | 0 | 2 | 0 | NON_SYNONYMOU | MODERAT | MISSENSE | cCg/cAg | P1506Q | pchF | | | | | | | A | A | | | | |
| NC_011770 | 761073 | C | G | 74 | 0 | 2 | 0 | NON_SYNONYMOU | MODERAT | MISSENSE | Cgc/Ggc | R1594G | pchF | | | | | | | G | G | | | | |
| NC_011770 | 761308 | A | T | 69 | 0 | 4 | 0 | NON_SYNONYMOU | MODERAT | MISSENSE | cAg/cTg | Q1672L | pchF | | | | | | | T | T | | | T | T |
| NC_011770 | 761329 | C | T | 69 | 0 | 2 | 0 | NON_SYNONYMOU | MODERAT | MISSENSE | gCg/gTg | A1679V | pchF | | | | | | | | | | | T | T |
| NC_011770 | 761350 | T | G | 68 | 0 | 10 | 0 | NON_SYNONYMOU | MODERAT | MISSENSE | gTc/gGc | V1686G | pchF | G | G | G | G | G | G | G | G | | | G | G |
| NC_011770 | 761359 | C | T | 69 | 0 | 2 | 0 | NON_SYNONYMOU | MODERAT | MISSENSE | gCg/gTg | A1689V | pchF | | | | | | | | | | | T | T |
| NC_011770 | 761379 | C | G | 71 | 0 | 2 | 0 | NON_SYNONYMOU | MODERAT | MISSENSE | Ctg/Gtg | L1696V | pchF | | | | | | | G | G | | | | |
| NC_011770 | 761443 | T | C | 69 | 0 | 8 | 0 | NON_SYNONYMOU | MODERAT | MISSENSE | gTc/gCc | V1717A | pchF | | | C | C | C | C | C | C | | | C | C |
| NC_011770 | 761479 | G | A | 76 | 0 | 2 | 0 | NON_SYNONYMOU | MODERAT | MISSENSE | tGt/tAt | C1729Y | pchF | | | | | | | | | A | A | | |
| NC_011770 | 761506 | A | C | 71 | 0 | 6 | 0 | NON_SYNONYMOU | MODERAT | MISSENSE | gAg/gCg | E1738A | pchF | C | C | C | C | C | C | | | | | | |
| NC_011770 | 761676 | C | A | 76 | 0 | 2 | 0 | NON_SYNONYMOU | MODERAT | MISSENSE | Ctg/Atg | L1795M | pchF | | | | | | | | | | | A | A |
| NC_011770 | 761686 | C | T | 77 | 0 | 6 | 0 | NON_SYNONYMOU | MODERAT | MISSENSE | tCc/tTc | S1798F | pchF | T | T | | | | | T | T | | | T | T |
| NC_011770 | 761706 | G | A | 79 | 0 | 2 | 0 | NON_SYNONYMOU | MODERAT | MISSENSE | Gag/Aag | E1805K | pchF | | | | | | | A | A | | | | |
| NC_011770 | 759874 | C | T | 67 | 0 | 1 | 0 | NON_SYNONYMOU | MODERAT | MISSENSE | aCc/aTc | T1194I | pchF | | | | | | | | | | | | **T** |
| NC_011770 | 762132 | G | A | 64 | 0 | 2 | 0 | NON_SYNONYMOU | MODERAT | MISSENSE | cGc/cAc | R138H | pchG | | | | | | | | | | | A | A |
| NC_011770 | 762323 | C | T | 62 | 0 | 4 | 0 | NON_SYNONYMOU | MODERAT | MISSENSE | Ctc/Ttc | L202F | pchG | | | T | T | T | T | | | | | | |
| NC_011770 | 762324 | T | A | 62 | 0 | 10 | 0 | NON_SYNONYMOU | MODERAT | MISSENSE | cTc/cAc | L202H | pchG | A | A | A | A | A | A | A | A | | | A | A |
| NC_011770 | 762338 | C | A | 62 | 0 | 2 | 0 | NON_SYNONYMOU | MODERAT | MISSENSE | Ctg/Atg | L207M | pchG | A | A | | | | | | | | | | |
| NC_011770 | 762570 | A | G | 66 | 0 | 1 | 0 | NON_SYNONYMOU | MODERAT | MISSENSE | cAc/cGc | H284R | pchG | | | | | | | G | | | | | |
| NC_011770 | 762620 | G | A | 62 | 0 | 8 | 0 | NON_SYNONYMOU | MODERAT | MISSENSE | Gac/Aac | D301N | pchG | | | A | A | A | A | A | A | | | A | A |
| NC_011770 | 6293273 | C | T | 105 | 0 | 2 | 0 | NON_SYNONYMOU | MODERAT | MISSENSE | Gtg/Atg | V302M | pchP | T | T | | | | | | | | | | |
| NC_011770 | 6293425 | G | A | 98 | 0 | 4 | 0 | NON_SYNONYMOU | MODERAT | MISSENSE | aCc/aTc | T251I | pchP | | | A | A | A | A | | | | | | |
| NC_011770 | 6293698 | T | C | 94 | 0 | 2 | 0 | NON_SYNONYMOU | MODERAT | MISSENSE | gAc/gGc | D160G | pchP | | | | | | | C | C | | | | |
| NC_011770 | 6180941 | T | C | 56 | 0 | 1 | 0 | NON_SYNONYMOU | MODERAT | MISSENSE | Acc/Gcc | T332A | pckA | C | | | | | | | | | | | |
| NC_011770 | 6181466 | T | C | 73 | 0 | 1 | 0 | NON_SYNONYMOU | MODERAT | MISSENSE | Acc/Gcc | T157A | pckA | | | | | | | | C | | | | |
| NC_011770 | 6180579 | G | C | 66 | 0 | 10 | 0 | NON_SYNONYMOU | MODERAT | MISSENSE | gaC/gaG | D452E | pckA | | | C | C | C | C | C | C | C | C | C | C |
| NC_011770 | 6181655 | T | C | 70 | 0 | 12 | 0 | NON_SYNONYMOU | MODERAT | MISSENSE | Att/Gtt | I94V | pckA | C | C | C | C | C | C | C | C | C | C | C | C |
| NC_011770 | 1520197 | C | G | 72 | 0 | 4 | 0 | NON_SYNONYMOU | MODERAT | MISSENSE | gaC/gaG | D162E | pcm | | | G | G | G | G | | | | | | |
| NC_011770 | 5643500 | C | T | 82 | 0 | 1 | 0 | NON_SYNONYMOU | MODERAT | MISSENSE | gCc/gTc | A119V | pcnB | | | | | | | | | | | | T |
| NC_011770 | 3601684 | A | G | 89 | 0 | 1 | 0 | NON_SYNONYMOU | MODERAT | MISSENSE | Atg/Gtg | M290V | pcoA | | | | | | | G | | | | | |
| NC_011770 | 3601983 | C | A | 97 | 0 | 2 | 0 | NON_SYNONYMOU | MODERAT | MISSENSE | agC/agA | S389R | pcoA | | | | | | | A | A | | | | |

| | | | | | | | | | | | | | | | | | | | | | | | | | |
|---|---|---|---|---|---|---|---|---|---|---|---|---|---|---|---|---|---|---|---|---|---|---|---|---|---|
| NC_011770 | 3602116 | A | G | 101 | 0 | 2 | 0 | NON_SYNONYMOU | MODERAT | MISSENSE | Acc/Gcc | T434A | pcoA | | | | | | | | | | | G | G |
| NC_011770 | 3602566 | G | A | 95 | 0 | 1 | 0 | NON_SYNONYMOU | MODERAT | MISSENSE | Gcc/Acc | A584T | pcoA | | | | | | | A | | | | | |
| NC_011770 | 4584536 | A | G | 69 | 0 | 2 | 0 | NON_SYNONYMOU | MODERAT | MISSENSE | Acc/Gcc | T163A | pcpS | | | | | | | G | G | | | | |
| NC_011770 | 4016842 | G | A | 82 | 0 | 2 | 0 | STOP_GAINED | HIGH | NONSENSE | Cag/Tag | Q658* | pcrD | | | | | | | | | | | A | A |
| NC_011770 | 4017390 | C | T | 75 | 0 | 2 | 0 | NON_SYNONYMOU | MODERAT | MISSENSE | aGc/aAc | S475N | pcrD | | | | | | | | | | | T | T |
| NC_011770 | 4017840 | A | G | 58 | 0 | 10 | 0 | NON_SYNONYMOU | MODERAT | MISSENSE | cTc/cCc | L325P | pcrD | G | G | G | G | G | G | G | G | G | G | | |
| NC_011770 | 4017841 | G | C | 58 | 0 | 10 | 0 | NON_SYNONYMOU | MODERAT | MISSENSE | Ctc/Gtc | L325V | pcrD | C | C | C | C | C | C | C | C | C | C | | |
| NC_011770 | 4016111 | G | A | 75 | 0 | 2 | 0 | NON_SYNONYMOU | MODERAT | MISSENSE | Ctt/Ttt | L42F | pcrG | | | | | | | A | A | | | | |
| NC_011770 | 4016230 | C | T | 70 | 0 | 4 | 0 | NON_SYNONYMOU | MODERAT | MISSENSE | gGc/gAc | G2D | pcrG | | | T | T | T | T | | | | | | |
| NC_011770 | 4016591 | G | A | 82 | 0 | 2 | 0 | NON_SYNONYMOU | MODERAT | MISSENSE | Cat/Tat | H36Y | pcrR | A | A | | | | | | | | | | |
| NC_011770 | 4015075 | C | A | 85 | 0 | 2 | 0 | NON_SYNONYMOU | MODERAT | MISSENSE | aGc/aTc | S285I | pcrV | | | | | | | | | | | A | A |
| NC_011770 | 4015244 | G | A | 86 | 0 | 2 | 0 | NON_SYNONYMOU | MODERAT | MISSENSE | Ccc/Tcc | P229S | pcrV | | | | | | | A | A | | | | |
| NC_011770 | 1213228 | C | T | 92 | 0 | 12 | 0 | NON_SYNONYMOU | MODERAT | MISSENSE | aCc/aTc | T27I | pcs | T | T | T | T | T | T | T | T | T | T | T | T |
| NC_011770 | 1213627 | C | T | 76 | 0 | 2 | 0 | NON_SYNONYMOU | MODERAT | MISSENSE | gCc/gTc | A160V | pcs | | | | | | | | | T | T | | |
| NC_011770 | 5171157 | C | G | 70 | 0 | 12 | 0 | NON_SYNONYMOU | MODERAT | MISSENSE | aCc/aGc | T24S | pctC | G | G | G | G | G | G | G | G | G | G | G | G |
| NC_011770 | 5171321 | G | A | 76 | 0 | 2 | 0 | NON_SYNONYMOU | MODERAT | MISSENSE | Gac/Aac | D79N | pctC | | | | | | | A | A | | | | |
| NC_011770 | 5171417 | G | A | 75 | 0 | 12 | 0 | NON_SYNONYMOU | MODERAT | MISSENSE | Gcc/Acc | A111T | pctC | A | A | A | A | A | A | A | A | A | A | A | A |
| NC_011770 | 5171493 | T | C | 75 | 0 | 1 | 0 | NON_SYNONYMOU | MODERAT | MISSENSE | cTg/cCg | L136P | pctC | | | | | | | C | | | | | |
| NC_011770 | 5171810 | C | A | 82 | 0 | 4 | 0 | NON_SYNONYMOU | MODERAT | MISSENSE | Cgc/Agc | R242S | pctC | | | A | A | A | A | | | | | | |
| NC_011770 | 5172665 | G | A | 89 | 0 | 2 | 0 | NON_SYNONYMOU | MODERAT | MISSENSE | Gtc/Atc | V527I | pctC | | | | | | | | | | | A | A |
| NC_011770 | 5172961 | G | C | 42 | 0 | 2 | 0 | NON_SYNONYMOU | MODERAT | MISSENSE | caG/caC | Q625H | pctC | | | | | | | | | | | C | C |
| NC_011770 | 642896 | C | T | 83 | 0 | 6 | 0 | NON_SYNONYMOU | MODERAT | MISSENSE | Gtc/Atc | V240I | pdxA | T | T | | | | | T | T | | | T | T |
| NC_011770 | 643394 | C | T | 49 | 0 | 4 | 0 | NON_SYNONYMOU | MODERAT | MISSENSE | Gta/Ata | V74I | pdxA | | | T | T | T | T | | | | | | |
| NC_011770 | 643597 | G | A | 32 | 0 | 2 | 0 | NON_SYNONYMOU | MODERAT | MISSENSE | gCg/gTg | A6V | pdxA | A | A | | | | | | | | | | |
| NC_011770 | 3440379 | C | T | 78 | 0 | 4 | 0 | NON_SYNONYMOU | MODERAT | MISSENSE | cGc/cAc | R335H | pdxA | | | T | T | T | T | | | | | | |
| NC_011770 | 3440928 | A | C | 74 | 0 | 2 | 0 | NON_SYNONYMOU | MODERAT | MISSENSE | tTc/tGc | F152C | pdxA | C | C | | | | | | | | | | |
| NC_011770 | 3440929 | A | C | 75 | 0 | 2 | 0 | NON_SYNONYMOU | MODERAT | MISSENSE | Ttc/Gtc | F152V | pdxA | C | C | | | | | | | | | | |
| NC_011770 | 3441157 | T | C | 75 | 0 | 2 | 0 | NON_SYNONYMOU | MODERAT | MISSENSE | Acg/Gcg | T76A | pdxA | | | | | | | | | C | C | | |
| NC_011770 | 3441277 | T | G | 80 | 0 | 6 | 0 | NON_SYNONYMOU | MODERAT | MISSENSE | Aag/Cag | K36Q | pdxA | G | G | | | | | | | G | G | G | G |
| NC_011770 | 4217750 | T | C | 51 | 0 | 4 | 0 | NON_SYNONYMOU | MODERAT | MISSENSE | gAg/gGg | E367G | pdxB | | | C | C | C | C | | | | | | |
| NC_011770 | 4217753 | A | G | 51 | 0 | 12 | 0 | NON_SYNONYMOU | MODERAT | MISSENSE | gTg/gCg | V366A | pdxB | G | G | G | G | G | G | G | G | G | G | G | G |
| NC_011770 | 4217756 | C | T | 50 | 0 | 4 | 0 | NON_SYNONYMOU | MODERAT | MISSENSE | cGg/cAg | R365Q | pdxB | | | T | T | T | T | | | | | | |
| NC_011770 | 4217840 | G | T | 53 | 0 | 4 | 0 | NON_SYNONYMOU | MODERAT | MISSENSE | aCc/aAc | T337N | pdxB | | | T | T | T | T | | | | | | |
| NC_011770 | 4217930 | A | G | 61 | 0 | 12 | 0 | NON_SYNONYMOU | MODERAT | MISSENSE | gTc/gCc | V307A | pdxB | G | G | G | G | G | G | G | G | G | G | G | G |
| NC_011770 | 4218110 | C | T | 60 | 0 | 4 | 0 | NON_SYNONYMOU | MODERAT | MISSENSE | cGc/cAc | R247H | pdxB | | | | | | | | | T | T | T | T |
| NC_011770 | 4218314 | C | T | 54 | 0 | 2 | 0 | NON_SYNONYMOU | MODERAT | MISSENSE | cGc/cAc | R179H | pdxB | | | | | | | | | T | T | | |
| NC_011770 | 4218401 | T | C | 59 | 0 | 2 | 0 | NON_SYNONYMOU | MODERAT | MISSENSE | cAg/cGg | Q150R | pdxB | | | | | | | | | C | C | | |
| NC_011770 | 4710336 | G | C | 54 | 0 | 2 | 0 | NON_SYNONYMOU | MODERAT | MISSENSE | tCg/tGg | S209W | pdxH | C | C | | | | | | | | | | |
| NC_011770 | 4710337 | A | G | 54 | 0 | 2 | 0 | NON_SYNONYMOU | MODERAT | MISSENSE | Tcg/Ccg | S209P | pdxH | G | G | | | | | | | | | | |
| NC_011770 | 4710485 | T | G | 59 | 0 | 8 | 0 | NON_SYNONYMOU | MODERAT | MISSENSE | gaA/gaC | E159D | pdxH | G | G | G | G | G | G | | | | | G | G |
| NC_011770 | 4710651 | G | A | 61 | 0 | 2 | 0 | NON_SYNONYMOU | MODERAT | MISSENSE | gCg/gTg | A104V | pdxH | | | | | | | A | A | | | | |
| NC_011770 | 4710792 | G | A | 67 | 0 | 4 | 0 | NON_SYNONYMOU | MODERAT | MISSENSE | gCc/gTc | A57V | pdxH | | | A | A | A | A | | | | | | |
| NC_011770 | 5026150 | T | C | 74 | 0 | 4 | 0 | NON_SYNONYMOU | MODERAT | MISSENSE | Acc/Gcc | T246A | pdxJ | | | C | C | C | C | | | | | | |
| NC_011770 | 5026372 | T | C | 81 | 0 | 12 | 0 | NON_SYNONYMOU | MODERAT | MISSENSE | Acc/Gcc | T172A | pdxJ | C | C | C | C | C | C | C | C | C | C | C | C |
| NC_011770 | 5026678 | T | C | 82 | 0 | 1 | 0 | NON_SYNONYMOU | MODERAT | MISSENSE | Acg/Gcg | T70A | pdxJ | | | | | | | C | | | | | |
| NC_011770 | 6544858 | G | A | 84 | 0 | 10 | 0 | NON_SYNONYMOU | MODERAT | MISSENSE | cGa/cAa | R51Q | pdxY | A | A | A | A | A | A | A | A | | | A | A |
| NC_011770 | 6545187 | A | G | 81 | 0 | 6 | 0 | NON_SYNONYMOU | MODERAT | MISSENSE | Aac/Gac | N161D | pdxY | | | G | G | G | G | | | G | G | | |
| NC_011770 | 6545280 | G | T | 74 | 0 | 4 | 0 | NON_SYNONYMOU | MODERAT | MISSENSE | Gcc/Tcc | A192S | pdxY | T | T | | | | | T | T | | | | |
| NC_011770 | 6545323 | C | T | 71 | 0 | 4 | 0 | NON_SYNONYMOU | MODERAT | MISSENSE | gCc/gTc | A206V | pdxY | | | T | T | T | T | | | | | | |
| NC_011770 | 6545415 | G | C | 72 | 0 | 12 | 0 | NON_SYNONYMOU | MODERAT | MISSENSE | Gtc/Ctc | V237L | pdxY | C | C | C | C | C | C | C | C | C | C | C | C |
| NC_011770 | 2162708 | T | TG | 48 | 0 | 3 | 0 | FRAME_SHIFT | HIGH | | ggc/ggGc | G474G? | pelA | TG | | | TG | | | TG | | | | | |
| NC_011770 | 2163487 | GCCGTACT/ | G | 76 | 0 | 1 | 0 | FRAME_SHIFT | HIGH | | tactacggct | YYGFR734 | pelA | | | | | | | | | G | | | |
| NC_011770 | 2162712 | GC | G | 36 | 3 | 1 | 0 | FRAME_SHIFT | HIGH | | ggc/ | G474 | pelA | . | G | | . | | | . | | | | | |
| NC_011770 | 2161347 | T | C | 65 | 0 | 6 | 0 | NON_SYNONYMOU | MODERAT | MISSENSE | aTc/aCc | I19T | pelA | C | C | C | C | C | C | | | | | | |
| NC_011770 | 2161364 | T | C | 60 | 0 | 6 | 0 | NON_SYNONYMOU | MODERAT | MISSENSE | Tgc/Cgc | C25R | pelA | C | C | C | C | C | C | | | | | | |
| NC_011770 | 2161425 | A | C | 59 | 0 | 6 | 0 | NON_SYNONYMOU | MODERAT | MISSENSE | gAt/gCt | D45A | pelA | C | C | C | C | C | C | | | | | | |
| NC_011770 | 2161449 | C | T | 59 | 0 | 2 | 0 | NON_SYNONYMOU | MODERAT | MISSENSE | gCg/gTg | A53V | pelA | | | | | | | | | T | T | | |
| NC_011770 | 2161638 | G | C | 73 | 0 | 2 | 0 | NON_SYNONYMOU | MODERAT | MISSENSE | cGg/cCg | R116P | pelA | | | | | | | | | C | C | | |
| NC_011770 | 2161712 | C | T | 68 | 0 | 4 | 0 | NON_SYNONYMOU | MODERAT | MISSENSE | Cac/Tac | H141Y | pelA | T | T | | | | | | | T | T | | |
| NC_011770 | 2161920 | G | A | 56 | 0 | 4 | 0 | NON_SYNONYMOU | MODERAT | MISSENSE | gGc/gAc | G210D | pelA | | | A | A | A | A | | | | | | |
| NC_011770 | 2162002 | C | G | 62 | 0 | 4 | 0 | NON_SYNONYMOU | MODERAT | MISSENSE | gaC/gaG | D237E | pelA | | | | | | | | | G | G | G | G |
| NC_011770 | 2162105 | A | G | 63 | 0 | 8 | 0 | NON_SYNONYMOU | MODERAT | MISSENSE | Acg/Gcg | T272A | pelA | | | G | G | G | G | G | G | | | G | G |

| | | | | | | | | | | | | | | | | | | | | | | | | | |
|---|---|---|---|---|---|---|---|---|---|---|---|---|---|---|---|---|---|---|---|---|---|---|---|---|---|
| NC_011770 | 2162576 | T | G | 52 | 0 | 2 | 0 | NON_SYNONYMOU | MODERAT | MISSENSE | Tcc/Gcc | S429A | pelA | | | | | | | G | G | | | | |
| NC_011770 | 2162603 | G | A | 51 | 0 | 10 | 0 | NON_SYNONYMOU | MODERAT | MISSENSE | Gtc/Atc | V438I | pelA | A | A | A | A | A | A | | | A | A | A | A |
| NC_011770 | 2162640 | C | G | 50 | 0 | 4 | 0 | NON_SYNONYMOU | MODERAT | MISSENSE | gCg/gGg | A450G | pelA | | | G | G | G | G | | | | | | |
| NC_011770 | 2162648 | G | A | 50 | 0 | 12 | 0 | NON_SYNONYMOU | MODERAT | MISSENSE | Gcc/Acc | A453T | pelA | A | A | A | A | A | A | A | A | A | A | A | A |
| NC_011770 | 2163645 | A | G | 98 | 0 | 2 | 0 | NON_SYNONYMOU | MODERAT | MISSENSE | cAt/cGt | H785R | pelA | | | | | | | | | | | G | G |
| NC_011770 | 2163801 | G | C | 80 | 0 | 2 | 0 | NON_SYNONYMOU | MODERAT | MISSENSE | gGc/gCc | G837A | pelA | | | | | | | | | | | C | C |
| NC_011770 | 2163905 | C | T | 77 | 0 | 2 | 0 | NON_SYNONYMOU | MODERAT | MISSENSE | Cgg/Tgg | R872W | pelA | | | | | | | | | T | T | | |
| NC_011770 | 2164149 | C | A | 70 | 0 | 4 | 0 | NON_SYNONYMOU | MODERAT | MISSENSE | Cag/Aag | Q12K | pelB | | | A | A | A | A | | | | | | |
| NC_011770 | 2164749 | C | T | 50 | 0 | 4 | 0 | NON_SYNONYMOU | MODERAT | MISSENSE | Ccg/Tcg | P212S | pelB | T | T | | | | | | | | | T | T |
| NC_011770 | 2164824 | T | G | 57 | 0 | 2 | 0 | NON_SYNONYMOU | MODERAT | MISSENSE | Ttg/Gtg | L237V | pelB | | | | | | | G | G | | | | |
| NC_011770 | 2164830 | C | G | 60 | 0 | 2 | 0 | NON_SYNONYMOU | MODERAT | MISSENSE | Ctg/Gtg | L239V | pelB | | | | | | | G | G | | | | |
| NC_011770 | 2164876 | T | C | 59 | 0 | 10 | 0 | NON_SYNONYMOU | MODERAT | MISSENSE | gTc/gCc | V254A | pelB | C | C | C | C | C | C | C | C | | | C | C |
| NC_011770 | 2164887 | G | C | 58 | 0 | 2 | 0 | NON_SYNONYMOU | MODERAT | MISSENSE | Gag/Cag | E258Q | pelB | | | | | | | C | C | | | | |
| NC_011770 | 2165022 | A | G | 59 | 0 | 10 | 0 | NON_SYNONYMOU | MODERAT | MISSENSE | Atc/Gtc | I303V | pelB | G | G | G | G | G | G | G | G | | | G | G |
| NC_011770 | 2165047 | A | G | 57 | 0 | 8 | 0 | NON_SYNONYMOU | MODERAT | MISSENSE | cAg/cGg | Q311R | pelB | G | G | G | G | G | G | | | | | G | G |
| NC_011770 | 2165079 | G | A | 53 | 0 | 10 | 0 | NON_SYNONYMOU | MODERAT | MISSENSE | Gcg/Acg | A322T | pelB | A | A | A | A | A | A | A | A | | | A | A |
| NC_011770 | 2165091 | A | G | 56 | 0 | 10 | 0 | NON_SYNONYMOU | MODERAT | MISSENSE | Aat/Gat | N326D | pelB | G | G | G | G | G | G | G | G | | | G | G |
| NC_011770 | 2165131 | G | A | 54 | 0 | 8 | 0 | NON_SYNONYMOU | MODERAT | MISSENSE | cGg/cAg | R339Q | pelB | A | A | A | A | A | A | | | | | A | A |
| NC_011770 | 2165142 | T | C | 56 | 0 | 8 | 0 | NON_SYNONYMOU | MODERAT | MISSENSE | Ttt/Ctt | F343L | pelB | C | C | C | C | C | C | | | | | C | C |
| NC_011770 | 2165144 | T | G | 56 | 0 | 8 | 0 | NON_SYNONYMOU | MODERAT | MISSENSE | ttT/ttG | F343L | pelB | G | G | G | G | G | G | | | | | G | G |
| NC_011770 | 2165461 | C | T | 70 | 0 | 2 | 0 | NON_SYNONYMOU | MODERAT | MISSENSE | gCc/gTc | A449V | pelB | | | | | T | T | | | | | | |
| NC_011770 | 2166246 | C | A | 87 | 0 | 2 | 0 | NON_SYNONYMOU | MODERAT | MISSENSE | Cag/Aag | Q711K | pelB | A | A | | | | | | | | | | |
| NC_011770 | 2166471 | C | T | 81 | 0 | 2 | 0 | NON_SYNONYMOU | MODERAT | MISSENSE | Ccg/Tcg | P786S | pelB | | | | | | | | | T | T | | |
| NC_011770 | 2166697 | G | T | 84 | 0 | 2 | 0 | NON_SYNONYMOU | MODERAT | MISSENSE | cGc/cTc | R861L | pelB | | | | | | | T | T | | | | |
| NC_011770 | 2166927 | G | T | 87 | 0 | 4 | 0 | NON_SYNONYMOU | MODERAT | MISSENSE | Gcc/Tcc | A938S | pelB | | | T | T | T | T | | | | | | |
| NC_011770 | 2167023 | G | T | 83 | 0 | 8 | 0 | NON_SYNONYMOU | MODERAT | MISSENSE | Gtg/Ttg | V970L | pelB | | | T | T | T | T | | | T | T | T | T |
| NC_011770 | 2167341 | G | C | 83 | 0 | 2 | 0 | NON_SYNONYMOU | MODERAT | MISSENSE | Gtg/Ctg | V1076L | pelB | | | | | | | C | C | | | | |
| NC_011770 | 2167555 | G | A | 68 | 0 | 2 | 0 | NON_SYNONYMOU | MODERAT | MISSENSE | gGc/gAc | G1147D | pelB | | | | | A | A | | | | | | |
| NC_011770 | 2168786 | G | A | 72 | 0 | 4 | 0 | NON_SYNONYMOU | MODERAT | MISSENSE | Ggc/Agc | G176S | pelD | | | A | A | A | A | | | | | | |
| NC_011770 | 2168791 | G | T | 74 | 0 | 8 | 0 | NON_SYNONYMOU | MODERAT | MISSENSE | gaG/gaT | E177D | pelD | T | T | | | | | T | T | T | T | T | T |
| NC_011770 | 2168906 | C | T | 81 | 0 | 2 | 0 | NON_SYNONYMOU | MODERAT | MISSENSE | Ccc/Tcc | P216S | pelD | | | | | | | | | T | T | | |
| NC_011770 | 2170185 | C | T | 53 | 0 | 2 | 0 | NON_SYNONYMOU | MODERAT | MISSENSE | Ctc/Ttc | L194F | pelE | T | T | | | | | | | | | | |
| NC_011770 | 2170221 | G | T | 54 | 0 | 2 | 0 | NON_SYNONYMOU | MODERAT | MISSENSE | Ggc/Tgc | G206C | pelE | | | | | | | | | T | T | | |
| NC_011770 | 2170347 | A | G | 67 | 0 | 6 | 0 | NON_SYNONYMOU | MODERAT | MISSENSE | Aca/Gca | T248A | pelE | | | | | | | G | G | G | G | G | G |
| NC_011770 | 2170417 | G | A | 70 | 0 | 2 | 0 | NON_SYNONYMOU | MODERAT | MISSENSE | gGg/gAg | G271E | pelE | | | | | | | | | A | A | | |
| NC_011770 | 2170420 | A | G | 69 | 0 | 8 | 0 | NON_SYNONYMOU | MODERAT | MISSENSE | cAc/cGc | H272R | pelE | G | G | | | | | G | G | G | G | G | G |
| NC_011770 | 2170622 | G | C | 68 | 0 | 4 | 0 | NON_SYNONYMOU | MODERAT | MISSENSE | Gtc/Ctc | V11L | pelF | | | C | C | C | C | | | | | | |
| NC_011770 | 2170923 | C | T | 80 | 0 | 2 | 0 | NON_SYNONYMOU | MODERAT | MISSENSE | cCg/cTg | P111L | pelF | T | T | | | | | | | | | | |
| NC_011770 | 2171117 | G | A | 77 | 0 | 2 | 0 | NON_SYNONYMOU | MODERAT | MISSENSE | Gcg/Acg | A176T | pelF | A | A | | | | | | | | | | |
| NC_011770 | 2171277 | A | G | 85 | 0 | 4 | 0 | NON_SYNONYMOU | MODERAT | MISSENSE | aAc/aGc | N229S | pelF | | | G | G | G | G | | | | | | |
| NC_011770 | 2171624 | T | G | 87 | 0 | 12 | 0 | NON_SYNONYMOU | MODERAT | MISSENSE | Tca/Gca | S345A | pelF | G | G | G | G | G | G | G | G | G | G | G | G |
| NC_011770 | 1240194 | A | T | 95 | 0 | 2 | 0 | NON_SYNONYMOU | MODERAT | MISSENSE | cTg/cAg | L281Q | pepA | | | | | | | | | T | T | | |
| NC_011770 | 1240710 | G | A | 95 | 0 | 4 | 0 | NON_SYNONYMOU | MODERAT | MISSENSE | gCg/gTg | A109V | pepA | | | | | | | A | A | A | A | | |
| NC_011770 | 1240956 | T | C | 94 | 0 | 2 | 0 | NON_SYNONYMOU | MODERAT | MISSENSE | aAg/aGg | K27R | pepA | | | | | | | C | C | | | | |
| NC_011770 | 2132351 | G | A | 86 | 0 | 12 | 0 | NON_SYNONYMOU | MODERAT | MISSENSE | gGa/gAa | G69E | pepN | A | A | A | A | A | A | A | A | A | A | A | A |
| NC_011770 | 2132491 | T | C | 92 | 0 | 1 | 0 | NON_SYNONYMOU | MODERAT | MISSENSE | Tac/Cac | Y116H | pepN | | | | | | | C | | | | | |
| NC_011770 | 2133395 | A | G | 83 | 0 | 1 | 0 | NON_SYNONYMOU | MODERAT | MISSENSE | cAc/cGc | H417R | pepN | | | | | | | G | | | | | |
| NC_011770 | 2133680 | T | C | 80 | 0 | 6 | 0 | NON_SYNONYMOU | MODERAT | MISSENSE | gTg/gCg | V512A | pepN | | | C | C | C | C | C | C | | | | |
| NC_011770 | 6218016 | G | A | 82 | 0 | 1 | 0 | NON_SYNONYMOU | MODERAT | MISSENSE | Cac/Tac | H309Y | pepP | | | | | | | | | | | | A |
| NC_011770 | 6217758 | A | T | 82 | 0 | 4 | 0 | NON_SYNONYMOU | MODERAT | MISSENSE | Tcg/Acg | S395T | pepP | T | T | | | | | | | T | T | | |
| NC_011770 | 6217802 | G | A | 83 | 0 | 2 | 0 | NON_SYNONYMOU | MODERAT | MISSENSE | gCg/gTg | A380V | pepP | | | | | | | | | | | A | A |
| NC_011770 | 6217992 | T | C | 81 | 0 | 1 | 0 | NON_SYNONYMOU | MODERAT | MISSENSE | Acc/Gcc | T317A | pepP | | | | | | | C | | | | | |
| NC_011770 | 6218638 | C | A | 86 | 0 | 4 | 0 | NON_SYNONYMOU | MODERAT | MISSENSE | caG/caT | Q101H | pepP | A | A | | | | | | | | | A | A |
| NC_011770 | 2594782 | T | C | 59 | 0 | 12 | 0 | NON_SYNONYMOU | MODERAT | MISSENSE | Agc/Ggc | S605G | pfeA | C | C | C | C | C | C | C | C | C | C | C | C |
| NC_011770 | 2595409 | C | T | 55 | 0 | 2 | 0 | NON_SYNONYMOU | MODERAT | MISSENSE | Gag/Aag | E396K | pfeA | | | | | | | | | | | T | T |
| NC_011770 | 2598765 | AT | A | 80 | 0 | 2 | 0 | FRAME_SHIFT | HIGH | | aat/ | N62 | pfeR | | | | | | | | | A | A | | |
| NC_011770 | 2598916 | C | T | 83 | 0 | 2 | 0 | STOP_GAINED | HIGH | NONSENSE | tGg/tAg | W12* | pfeR | | | | | | | | | T | T | | |
| NC_011770 | 2598109 | G | A | 81 | 0 | 4 | 0 | NON_SYNONYMOU | MODERAT | MISSENSE | gCg/gTg | A281V | pfeR | | | A | A | A | A | | | | | | |
| NC_011770 | 2598346 | G | A | 79 | 0 | 4 | 0 | NON_SYNONYMOU | MODERAT | MISSENSE | tCg/tTg | S202L | pfeR | | | A | A | A | A | | | | | | |
| NC_011770 | 2598526 | G | A | 86 | 0 | 2 | 0 | NON_SYNONYMOU | MODERAT | MISSENSE | tCg/tTg | S142L | pfeR | | | | | | | | | A | A | | |
| NC_011770 | 2598655 | C | T | 78 | 0 | 2 | 0 | NON_SYNONYMOU | MODERAT | MISSENSE | cGg/cAg | R99Q | pfeR | | | | | | | | | T | T | | |
| NC_011770 | 2598727 | A | T | 82 | 0 | 2 | 0 | NON_SYNONYMOU | MODERAT | MISSENSE | aTc/aAc | I75N | pfeR | | | | | | | | | T | T | | |

| | | | | | | | | | | | | | | | | | | | | | | | | | |
|---|---|---|---|---|---|---|---|---|---|---|---|---|---|---|---|---|---|---|---|---|---|---|---|---|---|
| NC_011770 | 2598799 | C | A | 83 | 0 | 2 | 0 | NON_SYNONYMOU | MODERAT | MISSENSE | gGc/gTc | G51V | pfeR | | | | | | | | | A | A | | |
| NC_011770 | 2598856 | G | T | 83 | 0 | 2 | 0 | NON_SYNONYMOU | MODERAT | MISSENSE | cCa/cAa | P32Q | pfeR | | | | | | | | | T | T | | |
| NC_011770 | 2598857 | G | T | 84 | 0 | 2 | 0 | NON_SYNONYMOU | MODERAT | MISSENSE | Cca/Aca | P32T | pfeR | | | | | | | | | T | T | | |
| NC_011770 | 2598898 | A | G | 82 | 0 | 2 | 0 | NON_SYNONYMOU | MODERAT | MISSENSE | cTc/cCc | L18P | pfeR | | | | | | | | | G | G | | |
| NC_011770 | 2598935 | T | G | 85 | 0 | 2 | 0 | NON_SYNONYMOU | MODERAT | MISSENSE | Agt/Cgt | S6R | pfeR | | | | | | | | | G | G | | |
| NC_011770 | 2596953 | C | T | 80 | 0 | 4 | 0 | NON_SYNONYMOU | MODERAT | MISSENSE | Gac/Aac | D361N | pfeS | | | | | | | T | T | T | T | | |
| NC_011770 | 2597095 | A | C | 73 | 0 | 6 | 0 | NON_SYNONYMOU | MODERAT | MISSENSE | gaT/gaG | D313E | pfeS | | | C | C | C | C | C | C | | | | |
| NC_011770 | 2597643 | C | G | 71 | 0 | 2 | 0 | NON_SYNONYMOU | MODERAT | MISSENSE | Gga/Cga | G131R | pfeS | G | G | | | | | | | | | | |
| NC_011770 | 2597742 | C | T | 78 | 0 | 2 | 0 | NON_SYNONYMOU | MODERAT | MISSENSE | Gcc/Acc | A98T | pfeS | | | | | | | | | | | T | T |
| NC_011770 | 2597775 | G | A | 74 | 0 | 2 | 0 | NON_SYNONYMOU | MODERAT | MISSENSE | Cac/Tac | H87Y | pfeS | | | | | | | A | A | | | | |
| NC_011770 | 2597854 | G | T | 74 | 0 | 4 | 0 | NON_SYNONYMOU | MODERAT | MISSENSE | agC/agA | S60R | pfeS | | | T | T | T | T | | | | | | |
| NC_011770 | 2597900 | C | T | 71 | 0 | 6 | 0 | NON_SYNONYMOU | MODERAT | MISSENSE | aGc/aAc | S45N | pfeS | | | T | T | T | T | | | T | T | | |
| NC_011770 | 2597901 | T | G | 71 | 0 | 6 | 0 | NON_SYNONYMOU | MODERAT | MISSENSE | Agc/Cgc | S45R | pfeS | | | G | G | G | G | | | G | G | | |
| NC_011770 | 393698 | C | G | 72 | 0 | 2 | 0 | NON_SYNONYMOU | MODERAT | MISSENSE | gGt/gCt | G25A | pfpI | | | | | | | G | G | | | | |
| NC_011770 | 603834 | T | C | 79 | 0 | 1 | 0 | NON_SYNONYMOU | MODERAT | MISSENSE | cTg/cCg | L91P | pgk | | | | | | | C | | | | | |
| NC_011770 | 603993 | C | T | 69 | 0 | 2 | 0 | NON_SYNONYMOU | MODERAT | MISSENSE | gCc/gTc | A144V | pgk | | | T | T | | | | | | | | |
| NC_011770 | 604068 | C | T | 64 | 0 | 2 | 0 | NON_SYNONYMOU | MODERAT | MISSENSE | gCc/gTc | A169V | pgk | | | | | T | T | | | | | | |
| NC_011770 | 604098 | A | G | 63 | 0 | 10 | 0 | NON_SYNONYMOU | MODERAT | MISSENSE | gAc/gGc | D179G | pgk | G | G | G | G | G | G | G | G | G | G | | |
| NC_011770 | 604706 | G | A | 73 | 0 | 1 | 0 | NON_SYNONYMOU | MODERAT | MISSENSE | Gag/Aag | E382K | pgk | | A | | | | | | | | | | |
| NC_011770 | 2030713 | G | C | 66 | 0 | 2 | 0 | NON_SYNONYMOU | MODERAT | MISSENSE | Gtc/Ctc | V74L | pgl | | | | | | | | | C | C | | |
| NC_011770 | 2030957 | A | G | 76 | 0 | 4 | 0 | NON_SYNONYMOU | MODERAT | MISSENSE | aAc/aGc | N155S | pgl | | | G | G | G | G | | | | | | |
| NC_011770 | 2031068 | T | C | 87 | 0 | 12 | 0 | NON_SYNONYMOU | MODERAT | MISSENSE | gTg/gCg | V192A | pgl | C | C | C | C | C | C | C | C | C | C | C | C |
| NC_011770 | 6112963 | G | C | 75 | 0 | 2 | 0 | NON_SYNONYMOU | MODERAT | MISSENSE | gGc/gCc | G101A | pgm | | | | | | | | | C | C | | |
| NC_011770 | 6113401 | A | G | 74 | 0 | 2 | 0 | NON_SYNONYMOU | MODERAT | MISSENSE | gAa/gGa | E247G | pgm | | | | | | | | | G | G | | |
| NC_011770 | 6031150 | G | A | 76 | 0 | 2 | 0 | NON_SYNONYMOU | MODERAT | MISSENSE | atG/atA | M369I | phaC1 | A | A | | | | | | | | | | |
| NC_011770 | 6031659 | A | G | 58 | 0 | 8 | 0 | NON_SYNONYMOU | MODERAT | MISSENSE | aAc/aGc | N539S | phaC1 | G | G | G | G | G | G | | | G | G | | |
| NC_011770 | 6033146 | G | T | 110 | 0 | 4 | 0 | NON_SYNONYMOU | MODERAT | MISSENSE | aGc/aTc | S37I | phaC2 | | | T | T | T | T | | | | | | |
| NC_011770 | 6033527 | A | G | 99 | 0 | 2 | 0 | NON_SYNONYMOU | MODERAT | MISSENSE | gAa/gGa | E164G | phaC2 | | | | | | | | | | | G | G |
| NC_011770 | 6034169 | A | G | 91 | 0 | 2 | 0 | NON_SYNONYMOU | MODERAT | MISSENSE | aAc/aGc | N378S | phaC2 | | | | | | | | | G | G | | |
| NC_011770 | 6034642 | A | G | 83 | 0 | 4 | 0 | NON_SYNONYMOU | MODERAT | MISSENSE | Aac/Gac | N536D | phaC2 | | | G | G | G | G | | | | | | |
| NC_011770 | 6032695 | A | G | 100 | 0 | 10 | 0 | NON_SYNONYMOU | MODERAT | MISSENSE | Atc/Gtc | I274V | phaD | G | G | G | G | G | G | | | G | G | G | G |
| NC_011770 | 1902160 | C | T | 78 | 0 | 8 | 0 | NON_SYNONYMOU | MODERAT | MISSENSE | atG/atA | M117I | phaJ1 | T | T | | | | | T | T | T | T | T | T |
| NC_011770 | 5711123 | G | A | 41 | 0 | 2 | 0 | NON_SYNONYMOU | MODERAT | MISSENSE | cGc/cAc | R134H | phaJ3 | A | A | | | | | | | | | | |
| NC_011770 | 5711174 | G | A | 45 | 0 | 6 | 0 | NON_SYNONYMOU | MODERAT | MISSENSE | tGc/tAc | C151Y | phaJ3 | | | A | A | A | A | | | | | A | A |
| NC_011770 | 5711242 | G | A | 42 | 0 | 2 | 0 | NON_SYNONYMOU | MODERAT | MISSENSE | Gtc/Atc | V174I | phaJ3 | A | A | | | | | | | | | | |
| NC_011770 | 2047668 | G | A | 81 | 0 | 2 | 0 | NON_SYNONYMOU | MODERAT | MISSENSE | Gcc/Acc | A32T | pheA | | | A | A | | | | | | | | |
| NC_011770 | 2048319 | C | G | 57 | 0 | 12 | 0 | NON_SYNONYMOU | MODERAT | MISSENSE | Cac/Gac | H249D | pheA | G | G | G | G | G | G | G | G | G | G | G | G |
| NC_011770 | 1669287 | C | T | 77 | 0 | 2 | 0 | NON_SYNONYMOU | MODERAT | MISSENSE | cCg/cTg | P2L | pheC | | | | | | | T | T | | | | |
| NC_011770 | 1669622 | G | A | 82 | 0 | 2 | 0 | NON_SYNONYMOU | MODERAT | MISSENSE | Gcg/Acg | A114T | pheC | | | | | | | | | | | A | A |
| NC_011770 | 1669773 | C | T | 78 | 0 | 4 | 0 | NON_SYNONYMOU | MODERAT | MISSENSE | gCg/gTg | A164V | pheC | | | T | T | T | T | | | | | | |
| NC_011770 | 1670021 | A | G | 85 | 0 | 2 | 0 | NON_SYNONYMOU | MODERAT | MISSENSE | Agc/Ggc | S247G | pheC | G | G | | | | | | | | | | |
| NC_011770 | 2500456 | G | A | 85 | 0 | 12 | 0 | NON_SYNONYMOU | MODERAT | MISSENSE | cGg/cAg | R118Q | pheS | A | A | A | A | A | A | A | A | A | A | A | A |
| NC_011770 | 2501380 | G | A | 77 | 0 | 2 | 0 | NON_SYNONYMOU | MODERAT | MISSENSE | Gtc/Atc | V76I | pheT | | | | | | | | | | | A | A |
| NC_011770 | 2501774 | C | A | 73 | 0 | 4 | 0 | NON_SYNONYMOU | MODERAT | MISSENSE | gCc/gAc | A207D | pheT | A | A | | | | | A | A | | | | |
| NC_011770 | 2502046 | A | G | 75 | 0 | 12 | 0 | NON_SYNONYMOU | MODERAT | MISSENSE | Acc/Gcc | T298A | pheT | G | G | G | G | G | G | G | G | G | G | G | G |
| NC_011770 | 2502941 | A | G | 73 | 0 | 4 | 0 | NON_SYNONYMOU | MODERAT | MISSENSE | aAg/aGg | K596R | pheT | G | G | | | | | | | G | G | | |
| NC_011770 | 2503110 | A | C | 62 | 0 | 4 | 0 | NON_SYNONYMOU | MODERAT | MISSENSE | gaA/gaC | E652D | pheT | C | C | | | | | C | C | | | | |
| NC_011770 | 2503183 | C | T | 70 | 0 | 12 | 0 | NON_SYNONYMOU | MODERAT | MISSENSE | Ctc/Ttc | L677F | pheT | T | T | T | T | T | T | T | T | T | T | T | T |
| NC_011770 | 4885557 | A | T | 77 | 0 | 2 | 0 | NON_SYNONYMOU | MODERAT | MISSENSE | Agc/Tgc | S193C | phhA | | | | | | | T | T | | | | |
| NC_011770 | 4885749 | C | T | 73 | 0 | 1 | 0 | NON_SYNONYMOU | MODERAT | MISSENSE | Ccg/Tcg | P257S | phhA | | | | | | | | | | | | T |
| NC_011770 | 4886813 | G | A | 78 | 0 | 6 | 0 | NON_SYNONYMOU | MODERAT | MISSENSE | cGc/cAc | R196H | phhC | | | A | A | A | A | A | A | | | | |
| NC_011770 | 4886947 | G | A | 79 | 0 | 2 | 0 | NON_SYNONYMOU | MODERAT | MISSENSE | Gtc/Atc | V241I | phhC | | | | | | | | | | | A | A |
| NC_011770 | 4887166 | A | G | 71 | 0 | 12 | 0 | NON_SYNONYMOU | MODERAT | MISSENSE | Agc/Ggc | S314G | phhC | G | G | G | G | G | G | G | G | G | G | G | G |
| NC_011770 | 4883372 | TC | T | 58 | 0 | 2 | 0 | FRAME_SHIFT | HIGH | | gga/ | G442 | phhR | | | | | | | | | T | T | | |
| NC_011770 | 4883149 | C | T | 69 | 0 | 4 | 0 | NON_SYNONYMOU | MODERAT | MISSENSE | Ggc/Agc | G517S | phhR | T | T | | | | | | | | | T | T |
| NC_011770 | 4762008 | C | T | 81 | 0 | 2 | 0 | NON_SYNONYMOU | MODERAT | MISSENSE | aGc/aAc | S461N | phnA | | | | | | | | | T | T | | |
| NC_011770 | 4762713 | G | A | 74 | 0 | 2 | 0 | NON_SYNONYMOU | MODERAT | MISSENSE | gCt/gTt | A226V | phnA | | | | | | | | | A | A | | |
| NC_011770 | 4762956 | A | G | 73 | 0 | 8 | 0 | NON_SYNONYMOU | MODERAT | MISSENSE | gTg/gCg | V145A | phnA | G | G | G | G | G | G | | | G | G | | |
| NC_011770 | 4762987 | G | A | 77 | 0 | 2 | 0 | NON_SYNONYMOU | MODERAT | MISSENSE | Ctc/Ttc | L135F | phnA | A | A | | | | | | | | | | |
| NC_011770 | 4762482 | T | C | 78 | 0 | 1 | 0 | NON_SYNONYMOU | MODERAT | MISSENSE | gAc/gGc | D303G | phnA | | | | | | | | C | | | | |
| NC_011770 | 1802089 | G | A | 81 | 0 | 2 | 0 | NON_SYNONYMOU | MODERAT | MISSENSE | aGg/aAg | R24K | phnC | A | A | | | | | | | | | | |
| NC_011770 | 1802451 | T | C | 69 | 0 | 4 | 0 | NON_SYNONYMOU | MODERAT | MISSENSE | Ttc/Ctc | F145L | phnC | | | | | | | C | C | | | C | C |

| | | | | | | | | | | | | | | | | | | | | | | | | | |
|---|---|---|---|---|---|---|---|---|---|---|---|---|---|---|---|---|---|---|---|---|---|---|---|---|---|
| NC_011770 | 1802764 | A | G | 67 | 0 | 1 | 0 | NON_SYNONYMOU | MODERAT | MISSENSE | tAc/tGc | Y249C | phnC | | | | | | | | G | | | | |
| NC_011770 | 1805762 | G | C | 74 | 0 | 2 | 0 | NON_SYNONYMOU | MODERAT | MISSENSE | gGt/gCt | G79A | phnG | C | C | | | | | | | | | | |
| NC_011770 | 1806418 | C | A | 84 | 0 | 2 | 0 | NON_SYNONYMOU | MODERAT | MISSENSE | gCc/gAc | A145D | phnH | A | A | | | | | | | | | | |
| NC_011770 | 1806480 | G | A | 84 | 0 | 2 | 0 | NON_SYNONYMOU | MODERAT | MISSENSE | Gcc/Acc | A166T | phnH | | | | | | | | | A | A | | |
| NC_011770 | 1807365 | C | T | 76 | 0 | 12 | 0 | NON_SYNONYMOU | MODERAT | MISSENSE | gCc/gTc | A251V | phnI | T | T | T | T | T | T | T | T | T | T | T | T |
| NC_011770 | 1807415 | C | T | 79 | 0 | 2 | 0 | NON_SYNONYMOU | MODERAT | MISSENSE | Ccg/Tcg | P268S | phnI | T | T | | | | | | | | | | |
| NC_011770 | 1808140 | C | T | 69 | 0 | 2 | 0 | NON_SYNONYMOU | MODERAT | MISSENSE | Cct/Tct | P144S | phnJ | | | | | | | T | T | | | | |
| NC_011770 | 1808149 | A | G | 67 | 0 | 12 | 0 | NON_SYNONYMOU | MODERAT | MISSENSE | Acg/Gcg | T147A | phnJ | G | G | G | G | G | G | G | G | G | G | G | G |
| NC_011770 | 1808176 | G | A | 71 | 0 | 2 | 0 | NON_SYNONYMOU | MODERAT | MISSENSE | Gac/Aac | D156N | phnJ | | | | | | | | | | | A | A |
| NC_011770 | 1808330 | G | A | 75 | 0 | 4 | 0 | NON_SYNONYMOU | MODERAT | MISSENSE | aGc/aAc | S207N | phnJ | A | A | | | | | | | | | A | A |
| NC_011770 | 1808566 | T | C | 66 | 0 | 2 | 0 | NON_SYNONYMOU | MODERAT | MISSENSE | Tgc/Cgc | C286R | phnJ | C | C | | | | | | | | | | |
| NC_011770 | 1808623 | A | G | 60 | 0 | 2 | 0 | NON_SYNONYMOU | MODERAT | MISSENSE | aAc/aGc | N11S | phnK | G | G | | | | | | | | | | |
| NC_011770 | 1808942 | G | A | 79 | 0 | 2 | 0 | NON_SYNONYMOU | MODERAT | MISSENSE | atG/atA | M117I | phnK | | | | | | | | | A | A | | |
| NC_011770 | 1810299 | TG | T | 50 | 0 | 1 | 0 | FRAME_SHIFT | HIGH | | ggc/ | G48 | phnM | | | | | | | T | | | | | |
| NC_011770 | 1810212 | G | A | 45 | 0 | 1 | 0 | NON_SYNONYMOU | MODERAT | MISSENSE | cGg/cAg | R17Q | phnM | | | | | | | A | | | | | |
| NC_011770 | 1810271 | C | T | 49 | 0 | 2 | 0 | NON_SYNONYMOU | MODERAT | MISSENSE | Cgc/Tgc | R37C | phnM | T | T | | | | | | | | | | |
| NC_011770 | 1810569 | G | A | 72 | 0 | 2 | 0 | NON_SYNONYMOU | MODERAT | MISSENSE | cGc/cAc | R136H | phnM | A | A | | | | | | | | | | |
| NC_011770 | 1810805 | G | T | 81 | 0 | 2 | 0 | NON_SYNONYMOU | MODERAT | MISSENSE | Gcc/Tcc | A215S | phnM | | | | | | | | | T | T | | |
| NC_011770 | 1812240 | T | G | 86 | 0 | 12 | 0 | NON_SYNONYMOU | MODERAT | MISSENSE | Tcc/Gcc | S123A | phnP | G | G | G | G | G | G | G | G | G | G | G | G |
| NC_011770 | 1812279 | G | T | 85 | 0 | 2 | 0 | NON_SYNONYMOU | MODERAT | MISSENSE | Gcg/Tcg | A136S | phnP | | | | | | | T | T | | | | |
| NC_011770 | 1812466 | G | A | 83 | 0 | 2 | 0 | NON_SYNONYMOU | MODERAT | MISSENSE | cGg/cAg | R198Q | phnP | | | | | | | | | | | A | A |
| NC_011770 | 1812515 | A | C | 89 | 0 | 6 | 0 | NON_SYNONYMOU | MODERAT | MISSENSE | gaA/gaC | E214D | phnP | | | C | C | C | C | | | C | C | | |
| NC_011770 | 4283309 | C | CG | 66 | 0 | 1 | 0 | FRAME_SHIFT | HIGH | | ccc/cccC | P13P? | phnW | | | | CG | | | | | | | | |
| NC_011770 | 4282300 | C | G | 63 | 0 | 12 | 0 | NON_SYNONYMOU | MODERAT | MISSENSE | gGg/gCg | G350A | phnW | G | G | G | G | G | G | G | G | G | G | G | G |
| NC_011770 | 4282727 | C | G | 80 | 0 | 2 | 0 | NON_SYNONYMOU | MODERAT | MISSENSE | Gcc/Ccc | A208P | phnW | | | | | | | | | G | G | | |
| NC_011770 | 4281879 | C | G | 65 | 0 | 12 | 0 | NON_SYNONYMOU | MODERAT | MISSENSE | Gtg/Ctg | V107L | phnX | G | G | G | G | G | G | G | G | G | G | G | G |
| NC_011770 | 1912511 | C | T | 64 | 0 | 12 | 0 | NON_SYNONYMOU | MODERAT | MISSENSE | Gac/Aac | D265N | phoA | T | T | T | T | T | T | T | T | T | T | T | T |
| NC_011770 | 1912492 | G | A | 65 | 0 | 1 | 0 | NON_SYNONYMOU | MODERAT | MISSENSE | gCc/gTc | A271V | phoA | | | | | | | | | A | | | |
| NC_011770 | 4571198 | C | A | 84 | 0 | 2 | 0 | NON_SYNONYMOU | MODERAT | MISSENSE | Gcg/Tcg | A29S | phoP | | | | | | | A | A | | | | |
| NC_011770 | 6363661 | C | G | 89 | 0 | 12 | 0 | NON_SYNONYMOU | MODERAT | MISSENSE | caC/caG | H58Q | phoR | G | G | G | G | G | G | G | G | G | G | G | G |
| NC_011770 | 6364362 | C | T | 90 | 0 | 6 | 0 | NON_SYNONYMOU | MODERAT | MISSENSE | gCc/gTc | A292V | phoR | | | T | T | T | T | T | T | | | | |
| NC_011770 | 6364536 | G | T | 90 | 0 | 2 | 0 | NON_SYNONYMOU | MODERAT | MISSENSE | cGc/cTc | R350L | phoR | | | | | | | | | T | T | | |
| NC_011770 | 5563447 | G | GC | 71 | 0 | 1 | 0 | FRAME_SHIFT | HIGH | | ggc/Cggc | G141R? | phr | | | | GC | | | | | | | | |
| NC_011770 | 5563129 | G | C | 74 | 0 | 4 | 0 | NON_SYNONYMOU | MODERAT | MISSENSE | Gtg/Ctg | V33L | phr | | | C | C | C | C | | | | | | |
| NC_011770 | 5563388 | G | A | 73 | 0 | 4 | 0 | NON_SYNONYMOU | MODERAT | MISSENSE | gGc/gAc | G119D | phr | A | A | | | | | | | | | A | A |
| NC_011770 | 5563972 | T | C | 68 | 0 | 12 | 0 | NON_SYNONYMOU | MODERAT | MISSENSE | Tgc/Cgc | C314R | phr | C | C | C | C | C | C | C | C | C | C | C | C |
| NC_011770 | 5564233 | G | T | 67 | 0 | 4 | 0 | NON_SYNONYMOU | MODERAT | MISSENSE | Gcg/Tcg | A401S | phr | | | T | T | T | T | | | | | | |
| NC_011770 | 5624824 | G | A | 70 | 0 | 1 | 0 | NON_SYNONYMOU | MODERAT | MISSENSE | Gcg/Acg | A596T | phuR | | | A | | | | | | | | | |
| NC_011770 | 5623834 | G | A | 77 | 0 | 1 | 0 | NON_SYNONYMOU | MODERAT | MISSENSE | Gtc/Atc | V266I | phuR | | | | A | | | | | | | | |
| NC_011770 | 5623067 | G | A | 15 | 0 | 2 | 0 | NON_SYNONYMOU | MODERAT | MISSENSE | cGc/cAc | R10H | phuR | | | | | A | A | | | | | | |
| NC_011770 | 5623803 | C | G | 73 | 0 | 2 | 0 | NON_SYNONYMOU | MODERAT | MISSENSE | ttC/ttG | F255L | phuR | G | G | | | | | | | | | | |
| NC_011770 | 5623810 | G | A | 74 | 0 | 2 | 0 | NON_SYNONYMOU | MODERAT | MISSENSE | Gtc/Atc | V258I | phuR | | | | | | | | | | | A | A |
| NC_011770 | 5623873 | G | A | 82 | 0 | 1 | 0 | NON_SYNONYMOU | MODERAT | MISSENSE | Gac/Aac | D279N | phuR | | | | | | | A | | | | | |
| NC_011770 | 5623997 | C | A | 76 | 0 | 6 | 0 | NON_SYNONYMOU | MODERAT | MISSENSE | cCg/cAg | P320Q | phuR | | | A | A | A | A | A | A | | | | |
| NC_011770 | 5624071 | T | C | 71 | 0 | 1 | 0 | NON_SYNONYMOU | MODERAT | MISSENSE | Tac/Cac | Y345H | phuR | | | | | | | C | | | | | |
| NC_011770 | 5624464 | G | A | 76 | 0 | 2 | 0 | NON_SYNONYMOU | MODERAT | MISSENSE | Gtc/Atc | V476I | phuR | A | A | | | | | | | | | | |
| NC_011770 | 5624627 | A | G | 75 | 0 | 1 | 0 | NON_SYNONYMOU | MODERAT | MISSENSE | cAc/cGc | H530R | phuR | | | | | | | G | | | | | |
| NC_011770 | 5621216 | T | A | 55 | 0 | 12 | 0 | NON_SYNONYMOU | MODERAT | MISSENSE | gaA/gaT | E190D | phuT | A | A | A | A | A | A | A | A | A | A | A | A |
| NC_011770 | 5621298 | C | T | 63 | 0 | 2 | 0 | NON_SYNONYMOU | MODERAT | MISSENSE | aGt/aAt | S163N | phuT | | | | | | | | | | | T | T |
| NC_011770 | 5621724 | G | C | 70 | 0 | 2 | 0 | NON_SYNONYMOU | MODERAT | MISSENSE | gCc/gGc | A21G | phuT | C | C | | | | | | | | | | |
| NC_011770 | 779284 | A | G | 82 | 0 | 4 | 0 | NON_SYNONYMOU | MODERAT | MISSENSE | Tac/Cac | Y6H | phzA1 | | | | | | | | | G | G | G | G |
| NC_011770 | 778960 | C | G | 61 | 0 | 5 | 0 | NON_SYNONYMOU | MODERAT | MISSENSE | Gag/Cag | E114Q | phzA1 | G | G | | | G | | | | | | G | G |
| NC_011770 | 778863 | C | T | 69 | 0 | 1 | 0 | NON_SYNONYMOU | MODERAT | MISSENSE | cGt/cAt | R146H | phzA1 | | | | | | | | | T | | | |
| NC_011770 | 778322 | C | G | 79 | 0 | 1 | 0 | NON_SYNONYMOU | MODERAT | MISSENSE | Gaa/Caa | E154Q | phzB1 | | | | | G | | | | | | | |
| NC_011770 | 776959 | A | G | 81 | 0 | 12 | 0 | NON_SYNONYMOU | MODERAT | MISSENSE | Tcg/Ccg | S29P | phzD1 | G | G | G | G | G | G | G | G | G | G | G | G |
| NC_011770 | 3795644 | A | G | 86 | 0 | 12 | 0 | NON_SYNONYMOU | MODERAT | MISSENSE | Tcg/Ccg | S29P | phzD2 | G | G | G | G | G | G | G | G | G | G | G | G |
| NC_011770 | 775729 | T | C | 79 | 0 | 2 | 0 | NON_SYNONYMOU | MODERAT | MISSENSE | gAc/gGc | D232G | phzE1 | | | | | C | C | | | | | | |
| NC_011770 | 776111 | T | G | 67 | 0 | 10 | 0 | NON_SYNONYMOU | MODERAT | MISSENSE | Atg/Ctg | M105L | phzE1 | | | G | G | G | G | G | G | G | G | G | G |
| NC_011770 | 3794414 | T | C | 78 | 0 | 2 | 0 | NON_SYNONYMOU | MODERAT | MISSENSE | gAc/gGc | D232G | phzE2 | | | | | C | C | | | | | | |
| NC_011770 | 3794796 | T | G | 66 | 0 | 10 | 0 | NON_SYNONYMOU | MODERAT | MISSENSE | Atg/Ctg | M105L | phzE2 | | | G | G | G | G | G | G | G | G | G | G |
| NC_011770 | 773746 | T | C | 71 | 0 | 4 | 0 | NON_SYNONYMOU | MODERAT | MISSENSE | Acg/Gcg | T261A | phzF1 | | | C | C | C | C | | | | | | |
| NC_011770 | 3792431 | T | C | 67 | 0 | 4 | 0 | NON_SYNONYMOU | MODERAT | MISSENSE | Acg/Gcg | T261A | phzF2 | | | C | C | C | C | | | | | | |

| | | | | | | | | | | | | | | | | | | | | | | | | | |
|---|---|---|---|---|---|---|---|---|---|---|---|---|---|---|---|---|---|---|---|---|---|---|---|---|---|
| NC_011770 | 3797007 | C | G | 78 | 0 | 1 | 0 | NON_SYNONYMOU | MODERAT | MISSENSE | Gaa/Caa | E154Q | phzG | | | | | G | | | | | | | |
| NC_011770 | 67618 | G | A | 82 | 0 | 1 | 0 | STOP_GAINED | HIGH | NONSENSE | tGg/tAg | W504* | phzH | | | | | | | | | | | A | |
| NC_011770 | 66861 | G | T | 51 | 0 | 8 | 0 | NON_SYNONYMOU | MODERAT | MISSENSE | Gcg/Tcg | A252S | phzH | | | T | T | T | T | | | T | T | T | T |
| NC_011770 | 67171 | T | C | 64 | 0 | 2 | 0 | NON_SYNONYMOU | MODERAT | MISSENSE | tTc/tCc | F355S | phzH | C | C | | | | | | | | | | |
| NC_011770 | 780424 | C | T | 71 | 0 | 2 | 0 | NON_SYNONYMOU | MODERAT | MISSENSE | Cgg/Tgg | R144W | phzM | | | | | | | | | T | T | | |
| NC_011770 | 780734 | A | G | 73 | 0 | 2 | 0 | NON_SYNONYMOU | MODERAT | MISSENSE | gAc/gGc | D247G | phzM | | | | | | | G | G | | | | |
| NC_011770 | 772017 | T | C | 77 | 0 | 10 | 0 | NON_SYNONYMOU | MODERAT | MISSENSE | Aac/Gac | N256D | phzS | C | C | C | C | C | C | C | C | | | C | C |
| NC_011770 | 772322 | A | T | 78 | 0 | 12 | 0 | NON_SYNONYMOU | MODERAT | MISSENSE | cTg/cAg | L154Q | phzS | T | T | T | T | T | T | T | T | T | T | T | T |
| NC_011770 | 5404537 | T | G | 17 | 8 | 2 | 0 | NON_SYNONYMOU | MODERAT | MISSENSE | Att/Ctt | I63L | pilA | | | . | . | . | . | G | G | . | . | . | . |
| NC_011770 | 5404540 | C | T | 17 | 8 | 2 | 0 | NON_SYNONYMOU | MODERAT | MISSENSE | Gag/Aag | E62K | pilA | | | . | . | . | . | T | T | . | . | . | . |
| NC_011770 | 5404588 | C | T | 54 | 4 | 6 | 0 | NON_SYNONYMOU | MODERAT | MISSENSE | Gtc/Atc | V46I | pilA | | | T | T | T | T | T | T | . | . | . | . |
| NC_011770 | 5404719 | T | G | 67 | 0 | 2 | 0 | NON_SYNONYMOU | MODERAT | MISSENSE | aAa/aCa | K2T | pilA | | | | | | | | | G | G | | |
| NC_011770 | 5405720 | ACGGG | A | 87 | 0 | 2 | 0 | FRAME_SHIFT | HIGH | | gggcgg/ | GR257 | pilB | | | | | | | | | | | A | A |
| NC_011770 | 5405023 | A | T | 65 | 2 | 2 | 0 | NON_SYNONYMOU | MODERAT | MISSENSE | Acc/Tcc | T24S | pilB | T | T | | | | | | | . | . | | |
| NC_011770 | 5405030 | G | T | 66 | 2 | 2 | 0 | NON_SYNONYMOU | MODERAT | MISSENSE | cGc/cTc | R26L | pilB | | | | | | | | | . | . | T | T |
| NC_011770 | 5405041 | G | A | 67 | 2 | 2 | 0 | NON_SYNONYMOU | MODERAT | MISSENSE | Gtg/Atg | V30M | pilB | | | | | | | | | . | . | A | A |
| NC_011770 | 5405042 | T | C | 67 | 2 | 2 | 0 | NON_SYNONYMOU | MODERAT | MISSENSE | gTg/gCg | V30A | pilB | | | | | | | | | . | . | C | C |
| NC_011770 | 5405050 | G | C | 62 | 2 | 2 | 0 | NON_SYNONYMOU | MODERAT | MISSENSE | Gcg/Ccg | A33P | pilB | | | | | | | | | . | . | C | C |
| NC_011770 | 5405051 | C | A | 62 | 2 | 2 | 0 | NON_SYNONYMOU | MODERAT | MISSENSE | gCg/gAg | A33E | pilB | | | | | | | | | . | . | A | A |
| NC_011770 | 5405063 | A | T | 63 | 2 | 2 | 0 | NON_SYNONYMOU | MODERAT | MISSENSE | cAg/cTg | Q37L | pilB | | | | | | | | | . | . | T | T |
| NC_011770 | 5405182 | A | T | 78 | 2 | 2 | 0 | NON_SYNONYMOU | MODERAT | MISSENSE | Att/Ttt | I77F | pilB | | | | | | | | | . | . | T | T |
| NC_011770 | 5405188 | A | C | 65 | 2 | 1 | 0 | NON_SYNONYMOU | MODERAT | MISSENSE | Aag/Cag | K79Q | pilB | | | | | | | | | . | . | C | . |
| NC_011770 | 5405189 | A | G | 64 | 2 | 2 | 0 | NON_SYNONYMOU | MODERAT | MISSENSE | aAg/aGg | K79R | pilB | | | | | | | | | . | . | G | G |
| NC_011770 | 5405195 | T | C | 63 | 2 | 2 | 0 | NON_SYNONYMOU | MODERAT | MISSENSE | gTc/gCc | V81A | pilB | | | | | | | | | . | . | C | C |
| NC_011770 | 5405206 | C | A | 62 | 3 | 1 | 0 | NON_SYNONYMOU | MODERAT | MISSENSE | Cgc/Agc | R85S | pilB | | | | | | | | | . | . | A | . |
| NC_011770 | 5405207 | G | A | 63 | 3 | 1 | 0 | NON_SYNONYMOU | MODERAT | MISSENSE | cGc/cAc | R85H | pilB | | | | | | | | | . | . | A | . |
| NC_011770 | 5405365 | G | C | 69 | 2 | 2 | 0 | NON_SYNONYMOU | MODERAT | MISSENSE | Gta/Cta | V138L | pilB | | | | | | | | | . | . | C | C |
| NC_011770 | 5405376 | G | C | 70 | 2 | 2 | 0 | NON_SYNONYMOU | MODERAT | MISSENSE | gaG/gaC | E141D | pilB | | | | | | | | | . | . | C | C |
| NC_011770 | 5405542 | C | A | 78 | 2 | 2 | 0 | NON_SYNONYMOU | MODERAT | MISSENSE | Cgc/Agc | R197S | pilB | | | | | | | | | . | . | A | A |
| NC_011770 | 5405543 | G | A | 78 | 1 | 2 | 0 | NON_SYNONYMOU | MODERAT | MISSENSE | cGc/cAc | R197H | pilB | | | | | | | | | | . | A | A |
| NC_011770 | 5405585 | C | T | 84 | 0 | 4 | 0 | NON_SYNONYMOU | MODERAT | MISSENSE | aCc/aTc | T211I | pilB | | | | | | | | | T | T | T | T |
| NC_011770 | 5405750 | C | A | 84 | 0 | 8 | 0 | NON_SYNONYMOU | MODERAT | MISSENSE | aCc/aAc | T266N | pilB | A | A | A | A | A | A | A | A | | | | |
| NC_011770 | 5406354 | C | G | 80 | 0 | 2 | 0 | NON_SYNONYMOU | MODERAT | MISSENSE | gaC/gaG | D467E | pilB | | | | | | | | | G | G | | |
| NC_011770 | 5406396 | G | T | 83 | 0 | 2 | 0 | NON_SYNONYMOU | MODERAT | MISSENSE | gaG/gaT | E481D | pilB | | | | | | | | | T | T | | |
| NC_011770 | 5406398 | T | G | 83 | 0 | 2 | 0 | NON_SYNONYMOU | MODERAT | MISSENSE | cTg/cGg | L482R | pilB | | | | | | | | | G | G | | |
| NC_011770 | 5406438 | A | C | 79 | 0 | 4 | 0 | NON_SYNONYMOU | MODERAT | MISSENSE | gaA/gaC | E495D | pilB | | | | | | | | | C | C | C | C |
| NC_011770 | 5406439 | A | C | 78 | 0 | 4 | 0 | NON_SYNONYMOU | MODERAT | MISSENSE | Aac/Cac | N496H | pilB | | | | | | | | | C | C | C | C |
| NC_011770 | 5406448 | G | A | 78 | 0 | 4 | 0 | NON_SYNONYMOU | MODERAT | MISSENSE | Ggc/Agc | G499S | pilB | | | | | | | | | A | A | A | A |
| NC_011770 | 5406449 | G | A | 79 | 0 | 4 | 0 | NON_SYNONYMOU | MODERAT | MISSENSE | gGc/gAc | G499D | pilB | | | | | | | | | A | A | A | A |
| NC_011770 | 5406499 | G | T | 82 | 0 | 6 | 0 | NON_SYNONYMOU | MODERAT | MISSENSE | Gcc/Tcc | A516S | pilB | | | T | T | T | T | T | T | | | | |
| NC_011770 | 5407872 | C | G | 60 | 0 | 3 | 0 | CODON_INSERTIOI | MODERATE | | taa/GGAta | *406G* | pilC | | | | | | | | | G | G | | CGGA |
| NC_011770 | 5406691 | G | A | 82 | 0 | 4 | 0 | NON_SYNONYMOU | MODERAT | MISSENSE | Gta/Ata | V12I | pilC | | | | | | | | | A | A | A | A |
| NC_011770 | 5406718 | A | G | 80 | 0 | 2 | 0 | NON_SYNONYMOU | MODERAT | MISSENSE | Acc/Gcc | T21A | pilC | | | | | | | | | | | G | G |
| NC_011770 | 5406719 | C | G | 79 | 0 | 2 | 0 | NON_SYNONYMOU | MODERAT | MISSENSE | aCc/aGc | T21S | pilC | | | | | | | | | G | G | | |
| NC_011770 | 5406739 | T | G | 72 | 0 | 4 | 0 | NON_SYNONYMOU | MODERAT | MISSENSE | Tcc/Gcc | S28A | pilC | | | | | | | | | G | G | A | A |
| NC_011770 | 5406742 | A | G | 72 | 0 | 4 | 0 | NON_SYNONYMOU | MODERAT | MISSENSE | Agc/Ggc | S29G | pilC | | | | | | | | | G | G | G | G |
| NC_011770 | 5406744 | C | G | 71 | 0 | 4 | 0 | NON_SYNONYMOU | MODERAT | MISSENSE | agC/agG | S29R | pilC | | | | | | | | | G | G | G | G |
| NC_011770 | 5406755 | C | T | 71 | 0 | 4 | 0 | NON_SYNONYMOU | MODERAT | MISSENSE | aCc/aTc | T33I | pilC | | | | | | | | | T | T | T | T |
| NC_011770 | 5406771 | A | T | 72 | 0 | 2 | 0 | NON_SYNONYMOU | MODERAT | MISSENSE | caA/caT | Q38H | pilC | | | | | | | | | | | T | T |
| NC_011770 | 5406775 | C | T | 74 | 0 | 1 | 0 | NON_SYNONYMOU | MODERAT | MISSENSE | Cgc/Tgc | R40C | pilC | | | | | | | T | | | | | |
| NC_011770 | 5406791 | C | A | 73 | 0 | 4 | 0 | NON_SYNONYMOU | MODERAT | MISSENSE | aCg/aAg | T45K | pilC | | | | | | | | | A | A | A | A |
| NC_011770 | 5406796 | G | C | 74 | 0 | 4 | 0 | NON_SYNONYMOU | MODERAT | MISSENSE | Gtc/Ctc | V47L | pilC | | | | | | | | | C | C | C | C |
| NC_011770 | 5406820 | T | A | 79 | 0 | 2 | 0 | NON_SYNONYMOU | MODERAT | MISSENSE | Tcc/Acc | S55T | pilC | | | | | | | | | A | A | | |
| NC_011770 | 5406844 | A | G | 78 | 0 | 2 | 0 | NON_SYNONYMOU | MODERAT | MISSENSE | Atc/Gtc | I63V | pilC | | | | | | | | | | | G | G |
| NC_011770 | 5406846 | C | T | 76 | 0 | 4 | 0 | NON_SYNONYMOU | MODERAT | MISSENSE | atC/atG | I63M | pilC | | | | | | | | | T | T | G | G |
| NC_011770 | 5406883 | T | G | 81 | 0 | 2 | 0 | NON_SYNONYMOU | MODERAT | MISSENSE | Tcc/Gcc | S76A | pilC | | | | | | | | | | | G | G |
| NC_011770 | 5406896 | C | G | 79 | 0 | 4 | 0 | NON_SYNONYMOU | MODERAT | MISSENSE | gCc/gGc | A80G | pilC | | | | | | | | | G | G | G | G |
| NC_011770 | 5406934 | A | G | 78 | 0 | 4 | 0 | NON_SYNONYMOU | MODERAT | MISSENSE | Agc/Ggc | S93G | pilC | | | | | | | | | G | G | G | G |
| NC_011770 | 5406948 | C | A | 80 | 0 | 2 | 0 | NON_SYNONYMOU | MODERAT | MISSENSE | gaC/gaA | D97E | pilC | | | | | | | | | A | A | | |
| NC_011770 | 5406975 | G | T | 78 | 0 | 4 | 0 | NON_SYNONYMOU | MODERAT | MISSENSE | gaG/gaT | E106D | pilC | | | | | | | | | T | T | T | T |
| NC_011770 | 5407291 | A | C | 66 | 4 | 4 | 0 | NON_SYNONYMOU | MODERAT | MISSENSE | Atg/Ctg | M212L | pilC | C | C | | | | | C | C | . | . | . | . |
| NC_011770 | 5407292 | T | A | 65 | 4 | 4 | 0 | NON_SYNONYMOU | MODERAT | MISSENSE | aTg/aAg | M212K | pilC | A | A | | | | | A | A | . | . | . | . |

| | | | | | | | | | | | | | | | | | | | | | | | | | |
|---|---|---|---|---|---|---|---|---|---|---|---|---|---|---|---|---|---|---|---|---|---|---|---|---|---|
| NC_011770 | 5407293 | G | A | 65 | 4 | 4 | 0 | NON_SYNONYMOU | MODERAT | MISSENSE | atG/atA | M212I | pilC | A | A | | | | | A | A | . | . | . | . |
| NC_011770 | 5407610 | C | G | 62 | 0 | 2 | 0 | NON_SYNONYMOU | MODERAT | MISSENSE | aCc/aGc | T318S | pilC | | | | | | | | | G | G | | |
| NC_011770 | 5407645 | A | C | 64 | 0 | 4 | 0 | NON_SYNONYMOU | MODERAT | MISSENSE | Aac/Cac | N330H | pilC | | | C | C | C | C | | | | | | |
| NC_011770 | 5407646 | A | G | 63 | 0 | 2 | 0 | NON_SYNONYMOU | MODERAT | MISSENSE | aAc/aGc | N330S | pilC | | | | | | | | | | | G | G |
| NC_011770 | 5407648 | A | G | 65 | 0 | 4 | 0 | NON_SYNONYMOU | MODERAT | MISSENSE | Atc/Gtc | I331V | pilC | | | | | | | | | G | G | G | G |
| NC_011770 | 5407658 | G | A | 61 | 0 | 2 | 0 | NON_SYNONYMOU | MODERAT | MISSENSE | aGc/aAc | S334N | pilC | | | | | | | | | | | A | A |
| NC_011770 | 5407699 | G | T | 56 | 2 | 2 | 0 | NON_SYNONYMOU | MODERAT | MISSENSE | Gcg/Tcg | A348S | pilC | | | | | | | | | . | . | T | T |
| NC_011770 | 5407710 | C | G | 57 | 2 | 2 | 0 | NON_SYNONYMOU | MODERAT | MISSENSE | gaC/gaG | D351E | pilC | | | | | | | | | . | . | G | G |
| NC_011770 | 5407717 | G | A | 58 | 2 | 2 | 0 | NON_SYNONYMOU | MODERAT | MISSENSE | Gcc/Acc | A354T | pilC | | | | | | | | | . | . | A | A |
| NC_011770 | 5407718 | C | G | 58 | 2 | 2 | 0 | NON_SYNONYMOU | MODERAT | MISSENSE | gCc/gGc | A354G | pilC | | | | | | | | | . | . | G | G |
| NC_011770 | 5407729 | G | A | 57 | 2 | 2 | 0 | NON_SYNONYMOU | MODERAT | MISSENSE | Ggc/Agc | G358S | pilC | | | | | | | | | . | . | A | A |
| NC_011770 | 5407733 | T | A | 57 | 2 | 2 | 0 | NON_SYNONYMOU | MODERAT | MISSENSE | tTc/tAc | F359Y | pilC | | | | | | | | | . | . | A | A |
| NC_011770 | 5407741 | C | G | 57 | 2 | 2 | 0 | NON_SYNONYMOU | MODERAT | MISSENSE | Cag/Gag | Q362E | pilC | | | | | | | | | . | . | G | G |
| NC_011770 | 5407774 | G | A | 60 | 2 | 2 | 0 | NON_SYNONYMOU | MODERAT | MISSENSE | Gcc/Acc | A373T | pilC | | | | | | | | | . | . | A | A |
| NC_011770 | 5407831 | A | G | 59 | 0 | 2 | 0 | NON_SYNONYMOU | MODERAT | MISSENSE | Atc/Gtc | I392V | pilC | | | | | | | | | | | G | G |
| NC_011770 | 5407833 | C | G | 60 | 0 | 2 | 0 | NON_SYNONYMOU | MODERAT | MISSENSE | atC/atG | I392M | pilC | | | | | | | | | | | G | G |
| NC_011770 | 5407864 | A | T | 59 | 0 | 2 | 0 | NON_SYNONYMOU | MODERAT | MISSENSE | Aac/Tac | N403Y | pilC | | | | | | | | | T | T | | |
| NC_011770 | 5407865 | A | C | 59 | 0 | 2 | 0 | NON_SYNONYMOU | MODERAT | MISSENSE | aAc/aCc | N403T | pilC | | | | | | | | | C | C | | |
| NC_011770 | 5407875 | A | AAG | 60 | 0 | 4 | 0 | FRAME_SHIFT+STA | HIGH | | atg/aGAtg | M1R? | pilD | | | | | | | | | AAG | AAG | ACTG | ACTG |
| NC_011770 | 5408716 | C | T | 60 | 0 | 1 | 0 | NON_SYNONYMOU | MODERAT | MISSENSE | Cgg/Tgg | R281W | pilD | | | | | | | | | | | T | |
| NC_011770 | 5407879 | A | G | 48 | 2 | 2 | 0 | NON_SYNONYMOU | MODERAT | MISSENSE | Acc/Gcc | T2A | pilD | | | | | | | | | G | G | . | . |
| NC_011770 | 5407880 | C | A | 58 | 0 | 2 | 0 | NON_SYNONYMOU | MODERAT | MISSENSE | aCc/aAc | T2N | pilD | | | | | | | | | A | A | | |
| NC_011770 | 5407927 | A | G | 63 | 0 | 6 | 0 | NON_SYNONYMOU | MODERAT | MISSENSE | Acc/Gcc | T18A | pilD | | | | | | | G | G | G | G | G | G |
| NC_011770 | 5408163 | G | C | 58 | 0 | 4 | 0 | NON_SYNONYMOU | MODERAT | MISSENSE | aaG/aaC | K96N | pilD | | | C | C | C | C | | | | | | |
| NC_011770 | 5408281 | C | T | 66 | 0 | 2 | 0 | NON_SYNONYMOU | MODERAT | MISSENSE | Ctc/Ttc | L136F | pilD | | | | | | | | | T | T | | |
| NC_011770 | 5408431 | G | A | 64 | 0 | 4 | 0 | NON_SYNONYMOU | MODERAT | MISSENSE | Gcg/Acg | A186T | pilD | | | A | A | A | A | | | | | | |
| NC_011770 | 5439771 | AG | A | 66 | 2 | 1 | 0 | FRAME_SHIFT | HIGH | | ggg/ | G43 | pilE | | | | A | | | | | . | . | | |
| NC_011770 | 444252 | G | A | 58 | 0 | 1 | 0 | START_LOST | HIGH | MISSENSE | atG/atA | M1I | pilI | | | | | | | A | | | | | |
| NC_011770 | 445814 | C | T | 73 | 2 | 8 | 0 | NON_SYNONYMOU | MODERAT | MISSENSE | gCc/gTc | A315V | pilJ | | | T | T | T | T | . | . | T | T | T | T |
| NC_011770 | 447398 | T | C | 93 | 0 | 4 | 0 | NON_SYNONYMOU | MODERAT | MISSENSE | gTg/gCg | V140A | pilK | | | | | | | | | C | C | C | C |
| NC_011770 | 447414 | A | C | 91 | 0 | 2 | 0 | NON_SYNONYMOU | MODERAT | MISSENSE | gaA/gaC | E145D | pilK | | | | | | | C | C | | | | |
| NC_011770 | 6014730 | G | A | 68 | 0 | 1 | 0 | STOP_GAINED | HIGH | NONSENSE | Cag/Tag | Q221* | pilM | | | | | | | | | | | A | |
| NC_011770 | 6014646 | G | A | 67 | 0 | 1 | 0 | STOP_GAINED | HIGH | NONSENSE | Cag/Tag | Q249* | pilM | | | | | | | A | | | | | |
| NC_011770 | 6014607 | G | A | 65 | 0 | 2 | 0 | NON_SYNONYMOU | MODERAT | MISSENSE | Cgt/Tgt | R262C | pilM | | | | | | | | | A | A | | |
| NC_011770 | 6013966 | T | C | 88 | 0 | 2 | 0 | NON_SYNONYMOU | MODERAT | MISSENSE | Acc/Gcc | T121A | pilN | | | | | | | | | C | C | | |
| NC_011770 | 6014103 | T | C | 82 | 0 | 2 | 0 | NON_SYNONYMOU | MODERAT | MISSENSE | aAg/aGg | K75R | pilN | | | | | | | | | C | C | | |
| NC_011770 | 6014188 | T | C | 70 | 2 | 4 | 0 | NON_SYNONYMOU | MODERAT | MISSENSE | Atc/Gtc | I47V | pilN | | | C | C | C | C | | | . | . | | |
| NC_011770 | 6013180 | C | T | 78 | 0 | 2 | 0 | NON_SYNONYMOU | MODERAT | MISSENSE | aGc/aAc | S185N | pilO | | | | | | | | | T | T | | |
| NC_011770 | 6013183 | C | T | 77 | 0 | 2 | 0 | NON_SYNONYMOU | MODERAT | MISSENSE | gGc/gAc | G184D | pilO | | | | | | | | | T | T | | |
| NC_011770 | 6013190 | C | A | 75 | 0 | 2 | 0 | NON_SYNONYMOU | MODERAT | MISSENSE | Gcg/Tcg | A182S | pilO | | | | | | | | | A | A | | |
| NC_011770 | 6013192 | A | G | 76 | 0 | 2 | 0 | NON_SYNONYMOU | MODERAT | MISSENSE | gTc/gCc | V181A | pilO | | | | | | | | | G | G | | |
| NC_011770 | 6013325 | C | T | 82 | 0 | 2 | 0 | NON_SYNONYMOU | MODERAT | MISSENSE | Gcc/Acc | A137T | pilO | | | | | | | | | T | T | | |
| NC_011770 | 6013517 | A | C | 86 | 0 | 2 | 0 | NON_SYNONYMOU | MODERAT | MISSENSE | Tcc/Gcc | S73A | pilO | | | | | | | | | C | C | | |
| NC_011770 | 6013535 | T | A | 84 | 0 | 2 | 0 | NON_SYNONYMOU | MODERAT | MISSENSE | Acg/Tcg | T67S | pilO | | | | | | | | | A | A | | |
| NC_011770 | 6013544 | C | G | 86 | 0 | 2 | 0 | NON_SYNONYMOU | MODERAT | MISSENSE | Gaa/Caa | E64Q | pilO | | | | | | | | | G | G | | |
| NC_011770 | 6013568 | C | T | 86 | 0 | 2 | 0 | NON_SYNONYMOU | MODERAT | MISSENSE | Gct/Act | A56T | pilO | | | | | | | | | T | T | | |
| NC_011770 | 6013574 | T | A | 86 | 0 | 2 | 0 | NON_SYNONYMOU | MODERAT | MISSENSE | Atg/Ttg | M54L | pilO | | | | | | | | | A | A | | |
| NC_011770 | 6013580 | T | G | 84 | 0 | 2 | 0 | NON_SYNONYMOU | MODERAT | MISSENSE | Agt/Cgt | S52R | pilO | | | | | | | | | G | G | | |
| NC_011770 | 6013691 | T | C | 92 | 0 | 2 | 0 | NON_SYNONYMOU | MODERAT | MISSENSE | Atc/Gtc | I15V | pilO | | | | | | | | | C | C | | |
| NC_011770 | 6012821 | T | C | 65 | 0 | 1 | 0 | NON_SYNONYMOU | MODERAT | MISSENSE | aAc/aGc | N98S | pilP | | | | | | | | C | | | | |
| NC_011770 | 6012867 | T | C | 70 | 0 | 2 | 0 | NON_SYNONYMOU | MODERAT | MISSENSE | Atc/Gtc | I83V | pilP | | | | | | | | | C | C | | |
| NC_011770 | 6010474 | TA | T | 84 | 0 | 2 | 0 | FRAME_SHIFT | HIGH | | gat/ | D687 | pilQ | | | | | | | | | T | T | | |
| NC_011770 | 6010804 | G | A | 74 | 0 | 4 | 0 | STOP_GAINED | HIGH | NONSENSE | Cag/Tag | Q578* | pilQ | | | | | | | | | A | A | A | A |
| NC_011770 | 6010461 | A | G | 81 | 0 | 4 | 0 | NON_SYNONYMOU | MODERAT | MISSENSE | gTg/gCg | V692A | pilQ | | | | | | | | | G | G | C | C |
| NC_011770 | 6010462 | C | T | 80 | 0 | 4 | 0 | NON_SYNONYMOU | MODERAT | MISSENSE | Gtg/Atg | V692M | pilQ | | | | | | | | | T | T | G | G |
| NC_011770 | 6010468 | A | T | 82 | 0 | 2 | 0 | NON_SYNONYMOU | MODERAT | MISSENSE | Tcg/Acg | S690T | pilQ | | | | | | | | | | | T | T |
| NC_011770 | 6010508 | G | T | 85 | 0 | 2 | 0 | NON_SYNONYMOU | MODERAT | MISSENSE | gaC/gaA | D676E | pilQ | | | | | | | | | | | T | T |
| NC_011770 | 6010529 | G | C | 85 | 0 | 2 | 0 | NON_SYNONYMOU | MODERAT | MISSENSE | gaC/gaG | D669E | pilQ | | | | | | | | | | | C | C |
| NC_011770 | 6010537 | A | C | 82 | 0 | 2 | 0 | NON_SYNONYMOU | MODERAT | MISSENSE | Tcg/Gcg | S667A | pilQ | | | | | | | | | C | C | | |
| NC_011770 | 6010541 | G | T | 84 | 0 | 2 | 0 | NON_SYNONYMOU | MODERAT | MISSENSE | agC/agA | S665R | pilQ | | | | | | | | | T | T | | |
| NC_011770 | 6010542 | C | G | 86 | 0 | 2 | 0 | NON_SYNONYMOU | MODERAT | MISSENSE | aGc/aCc | S665T | pilQ | | | | | | | | | G | G | | |
| NC_011770 | 6010548 | G | T | 87 | 0 | 2 | 0 | NON_SYNONYMOU | MODERAT | MISSENSE | aCc/aAc | T663N | pilQ | | | | | | | | | | | T | T |

| | | | | | | | | | | | | | | | | | | | | | | | | | |
|---|---|---|---|---|---|---|---|---|---|---|---|---|---|---|---|---|---|---|---|---|---|---|---|---|---|
| NC_011770 | 6010549 | T | C | 86 | 0 | 2 | 0 | NON_SYNONYMOU | MODERAT | MISSENSE | Acc/Gcc | T663A | pilQ | | | | | | | | | | | C | C |
| NC_011770 | 6010650 | C | T | 64 | 2 | 2 | 0 | NON_SYNONYMOU | MODERAT | MISSENSE | cGc/cAc | R629H | pilQ | | | | | | | | | T | T | . | . |
| NC_011770 | 6010651 | G | T | 64 | 2 | 2 | 0 | NON_SYNONYMOU | MODERAT | MISSENSE | Cgc/Agc | R629S | pilQ | | | | | | | | | T | T | . | . |
| NC_011770 | 6010652 | A | C | 64 | 2 | 2 | 0 | NON_SYNONYMOU | MODERAT | MISSENSE | gaT/gaG | D628E | pilQ | | | | | | | | | C | C | . | . |
| NC_011770 | 6010654 | C | G | 66 | 2 | 2 | 0 | NON_SYNONYMOU | MODERAT | MISSENSE | Gat/Cat | D628H | pilQ | | | | | | | | | G | G | . | . |
| NC_011770 | 6010798 | C | T | 77 | 0 | 2 | 0 | NON_SYNONYMOU | MODERAT | MISSENSE | Gta/Ata | V580I | pilQ | | | | | | | | | T | T | | |
| NC_011770 | 6010803 | T | G | 75 | 0 | 4 | 0 | NON_SYNONYMOU | MODERAT | MISSENSE | cAg/cCg | Q578P | pilQ | | | | | | | | | G | G | G | G |
| NC_011770 | 6010862 | G | C | 79 | 0 | 2 | 0 | NON_SYNONYMOU | MODERAT | MISSENSE | atC/atG | I558M | pilQ | | | | | | | | | C | C | | |
| NC_011770 | 6010864 | T | C | 78 | 0 | 2 | 0 | NON_SYNONYMOU | MODERAT | MISSENSE | Atc/Gtc | I558V | pilQ | | | | | | | | | C | C | | |
| NC_011770 | 6010969 | T | G | 76 | 2 | 2 | 0 | NON_SYNONYMOU | MODERAT | MISSENSE | Atc/Ctc | I523L | pilQ | | | | | | | | | . | . | G | G |
| NC_011770 | 6011022 | T | C | 75 | 2 | 2 | 0 | NON_SYNONYMOU | MODERAT | MISSENSE | aAc/aGc | N505S | pilQ | | | | | | | | | . | . | C | C |
| NC_011770 | 6011080 | C | T | 83 | 0 | 2 | 0 | NON_SYNONYMOU | MODERAT | MISSENSE | Gac/Aac | D486N | pilQ | | | | | | | | | T | T | | |
| NC_011770 | 6011092 | C | T | 82 | 0 | 2 | 0 | NON_SYNONYMOU | MODERAT | MISSENSE | Ggc/Agc | G482S | pilQ | | | | | | | | | T | T | | |
| NC_011770 | 6011093 | T | G | 78 | 0 | 10 | 0 | NON_SYNONYMOU | MODERAT | MISSENSE | agA/agC | R481S | pilQ | | | G | G | G | G | G | G | G | G | G | G |
| NC_011770 | 6011094 | C | T | 79 | 0 | 8 | 0 | NON_SYNONYMOU | MODERAT | MISSENSE | aGa/aAa | R481K | pilQ | | | T | T | T | T | T | T | T | T | | |
| NC_011770 | 6011105 | C | A | 83 | 0 | 2 | 0 | NON_SYNONYMOU | MODERAT | MISSENSE | aaG/aaT | K477N | pilQ | | | | | | | | | A | A | | |
| NC_011770 | 6011626 | T | C | 92 | 0 | 1 | 0 | NON_SYNONYMOU | MODERAT | MISSENSE | Acc/Gcc | T304A | pilQ | | | | | | | C | | | | | |
| NC_011770 | 6011739 | G | A | 98 | 0 | 2 | 0 | NON_SYNONYMOU | MODERAT | MISSENSE | aCg/aTg | T266M | pilQ | | | | | | | | | A | A | | |
| NC_011770 | 6012133 | T | C | 88 | 0 | 4 | 0 | NON_SYNONYMOU | MODERAT | MISSENSE | Acc/Gcc | T135A | pilQ | | | | | | | | | C | C | C | C |
| NC_011770 | 6012485 | G | T | 72 | 0 | 2 | 0 | NON_SYNONYMOU | MODERAT | MISSENSE | ttC/ttA | F17L | pilQ | T | T | | | | | | | | | | |
| NC_011770 | 5431018 | G | A | 85 | 0 | 1 | 0 | NON_SYNONYMOU | MODERAT | MISSENSE | Gta/Ata | V303I | pilR | | | | | | | | | | | A | |
| NC_011770 | 5431065 | T | A | 82 | 0 | 10 | 0 | NON_SYNONYMOU | MODERAT | MISSENSE | gaT/gaA | D318E | pilR | A | A | A | A | A | A | A | A | | | A | A |
| NC_011770 | 5431255 | C | T | 80 | 0 | 2 | 0 | NON_SYNONYMOU | MODERAT | MISSENSE | Ccg/Tcg | P382S | pilR | | | | | | | | | | | T | T |
| NC_011770 | 5429319 | T | C | 84 | 0 | 12 | 0 | NON_SYNONYMOU | MODERAT | MISSENSE | aTg/aCg | M272T | pilS | C | C | C | C | C | C | C | C | C | C | C | C |
| NC_011770 | 5429708 | G | C | 85 | 0 | 8 | 0 | NON_SYNONYMOU | MODERAT | MISSENSE | Gaa/Caa | E402Q | pilS | | | C | C | C | C | C | C | C | C | | |
| NC_011770 | 5429888 | G | A | 79 | 0 | 6 | 0 | NON_SYNONYMOU | MODERAT | MISSENSE | Gtc/Atc | V462I | pilS | | | A | A | A | A | A | A | | | | |
| NC_011770 | 430536 | C | T | 93 | 0 | 1 | 0 | NON_SYNONYMOU | MODERAT | MISSENSE | Ccg/Tcg | P76S | pilT | | | | | | | T | | | | | |
| NC_011770 | 431644 | C | T | 82 | 0 | 1 | 0 | NON_SYNONYMOU | MODERAT | MISSENSE | cCg/cTg | P41L | pilU | | | | | | | | T | | | | |
| NC_011770 | 5433838 | G | A | 92 | 2 | 1 | 0 | START_LOST | HIGH | MISSENSE | atG/atA | M1I | pilV | | | | | | | | A | . | . | | |
| NC_011770 | 5434371 | A | G | 72 | 2 | 1 | 0 | NON_SYNONYMOU | MODERAT | MISSENSE | tAc/tGc | Y179C | pilV | | | | G | | | | | . | . | | |
| NC_011770 | 5433858 | G | A | 88 | 2 | 2 | 0 | NON_SYNONYMOU | MODERAT | MISSENSE | aGg/aAg | R8K | pilV | A | A | | | | | | | . | . | | |
| NC_011770 | 5433860 | T | C | 87 | 2 | 2 | 0 | NON_SYNONYMOU | MODERAT | MISSENSE | Tcg/Ccg | S9P | pilV | C | C | | | | | | | . | . | | |
| NC_011770 | 5433866 | C | T | 90 | 2 | 2 | 0 | NON_SYNONYMOU | MODERAT | MISSENSE | Cac/Tac | H11Y | pilV | T | T | | | | | | | . | . | | |
| NC_011770 | 5433908 | C | G | 96 | 0 | 2 | 0 | NON_SYNONYMOU | MODERAT | MISSENSE | Ctg/Gtg | L25V | pilV | | | | | | | | | G | G | | |
| NC_011770 | 5433938 | A | G | 91 | 0 | 2 | 0 | NON_SYNONYMOU | MODERAT | MISSENSE | Atc/Gtc | I35V | pilV | | | | | | | | | G | G | | |
| NC_011770 | 5435446 | C | G | 77 | 2 | 2 | 0 | NON_SYNONYMOU | MODERAT | MISSENSE | aCg/aGg | T79R | pilX | G | G | | | | | | | . | . | | |
| NC_011770 | 5435448 | G | C | 74 | 2 | 4 | 0 | NON_SYNONYMOU | MODERAT | MISSENSE | Gcc/Ccc | A80P | pilX | C | C | | | | | A | A | . | . | | |
| NC_011770 | 5435584 | G | C | 83 | 2 | 6 | 0 | NON_SYNONYMOU | MODERAT | MISSENSE | aGc/aCc | S125T | pilX | C | C | C | C | C | C | | | . | . | | |
| NC_011770 | 5436513 | C | A | 77 | 2 | 2 | 0 | STOP_GAINED | HIGH | NONSENSE | tCg/tAg | S235* | pilY1 | | | | | | | A | A | . | . | | |
| NC_011770 | 5435820 | T | C | 84 | 2 | 8 | 0 | NON_SYNONYMOU | MODERAT | MISSENSE | gTa/gCa | V4A | pilY1 | C | C | C | C | C | C | C | C | . | . | | |
| NC_011770 | 5435825 | C | A | 86 | 2 | 4 | 0 | NON_SYNONYMOU | MODERAT | MISSENSE | Cac/Aac | H6N | pilY1 | | | A | A | A | A | | | . | . | | |
| NC_011770 | 5435849 | G | A | 83 | 2 | 2 | 0 | NON_SYNONYMOU | MODERAT | MISSENSE | Gca/Aca | A14T | pilY1 | | | | | | | A | A | . | . | | |
| NC_011770 | 5436035 | G | A | 65 | 4 | 2 | 0 | NON_SYNONYMOU | MODERAT | MISSENSE | Gtc/Atc | V76I | pilY1 | A | A | | | | | . | . | . | . | | |
| NC_011770 | 5436057 | A | T | 70 | 2 | 2 | 0 | NON_SYNONYMOU | MODERAT | MISSENSE | tAc/tTc | Y83F | pilY1 | | | | | | | T | T | . | . | | |
| NC_011770 | 5436072 | T | A | 70 | 2 | 2 | 0 | NON_SYNONYMOU | MODERAT | MISSENSE | tTc/tAc | F88Y | pilY1 | | | | | | | A | A | . | . | | |
| NC_011770 | 5436146 | T | C | 75 | 2 | 2 | 0 | NON_SYNONYMOU | MODERAT | MISSENSE | Tcc/Ccc | S113P | pilY1 | | | | | | | C | C | . | . | | |
| NC_011770 | 5436507 | G | A | 77 | 2 | 2 | 0 | NON_SYNONYMOU | MODERAT | MISSENSE | aGt/aAt | S233N | pilY1 | | | | | | | A | A | . | . | | |
| NC_011770 | 5436519 | T | A | 75 | 2 | 2 | 0 | NON_SYNONYMOU | MODERAT | MISSENSE | cTg/cAg | L237Q | pilY1 | | | | | | | A | A | . | . | | |
| NC_011770 | 5436620 | G | A | 76 | 2 | 2 | 0 | NON_SYNONYMOU | MODERAT | MISSENSE | Gga/Aga | G271R | pilY1 | | | | | | | A | A | . | . | | |
| NC_011770 | 5436621 | G | A | 76 | 2 | 2 | 0 | NON_SYNONYMOU | MODERAT | MISSENSE | gGa/gAa | G271E | pilY1 | | | | | | | A | A | . | . | | |
| NC_011770 | 5436719 | A | G | 84 | 2 | 2 | 0 | NON_SYNONYMOU | MODERAT | MISSENSE | Acc/Gcc | T304A | pilY1 | | | | | | | | | . | . | G | G |
| NC_011770 | 5436722 | T | G | 82 | 2 | 2 | 0 | NON_SYNONYMOU | MODERAT | MISSENSE | Tcg/Gcg | S305A | pilY1 | | | | | | | G | G | . | . | | |
| NC_011770 | 5436873 | C | G | 89 | 2 | 2 | 0 | NON_SYNONYMOU | MODERAT | MISSENSE | gCt/gGt | A355G | pilY1 | | | | | | | G | G | . | . | | |
| NC_011770 | 5436875 | G | A | 89 | 2 | 2 | 0 | NON_SYNONYMOU | MODERAT | MISSENSE | Gcg/Acg | A356T | pilY1 | | | | | | | A | A | . | . | | |
| NC_011770 | 5437086 | A | G | 73 | 2 | 2 | 0 | NON_SYNONYMOU | MODERAT | MISSENSE | aAg/aGg | K426R | pilY1 | | | | | | | G | G | . | . | | |
| NC_011770 | 5437104 | A | T | 73 | 2 | 2 | 0 | NON_SYNONYMOU | MODERAT | MISSENSE | tAc/tTc | Y432F | pilY1 | | | | | | | T | T | . | . | | |
| NC_011770 | 5437118 | G | A | 64 | 2 | 2 | 0 | NON_SYNONYMOU | MODERAT | MISSENSE | Gac/Aac | D437N | pilY1 | | | | | | | A | A | . | . | | |
| NC_011770 | 5437123 | A | T | 63 | 2 | 2 | 0 | NON_SYNONYMOU | MODERAT | MISSENSE | gaA/gaT | E438D | pilY1 | | | | | | | T | T | . | . | | |
| NC_011770 | 5437128 | C | T | 64 | 2 | 2 | 0 | NON_SYNONYMOU | MODERAT | MISSENSE | gCc/gTc | A440V | pilY1 | | | | | | | T | T | . | . | | |
| NC_011770 | 5437136 | C | A | 60 | 2 | 2 | 0 | NON_SYNONYMOU | MODERAT | MISSENSE | Cgc/Agc | R443S | pilY1 | | | | | | | A | A | . | . | | |
| NC_011770 | 5437137 | G | A | 61 | 2 | 2 | 0 | NON_SYNONYMOU | MODERAT | MISSENSE | cGc/cAc | R443H | pilY1 | | | | | | | A | A | . | . | | |
| NC_011770 | 5437152 | A | G | 60 | 2 | 2 | 0 | NON_SYNONYMOU | MODERAT | MISSENSE | aAc/aGc | N448S | pilY1 | | | | | | | G | G | . | . | | |

| | | | | | | | | | | | | | | | | | | | | | | | | | |
|---|---|---|---|---|---|---|---|---|---|---|---|---|---|---|---|---|---|---|---|---|---|---|---|---|---|
| NC_011770 | 5437165 | CCAT | C | 61 | 2 | 2 | 0 | CODON_DELETION | MODERATE | | cat/ | H453 | pilY1 | | | | | | | C | C | . | . | | |
| NC_011770 | 5437170 | G | GCGC | 62 | 2 | 2 | 0 | CODON_INSERTIOI | MODERATE | | aac/GCCaa | N455AN | pilY1 | | | | | | | GCGC | GCGC | . | . | | |
| NC_011770 | 5437589 | C | A | 74 | 2 | 2 | 0 | NON_SYNONYMOU | MODERAT | MISSENSE | Cag/Aag | Q594K | pilY1 | | | | | | | A | A | . | . | | |
| NC_011770 | 5439362 | T | G | 81 | 2 | 2 | 0 | NON_SYNONYMOU | MODERAT | MISSENSE | gaT/gaG | D20E | pilY2 | | | | | | | G | G | . | . | | |
| NC_011770 | 5439445 | A | G | 76 | 2 | 10 | 0 | NON_SYNONYMOU | MODERAT | MISSENSE | cAc/cGc | H48R | pilY2 | G | G | G | G | G | G | G | G | . | . | G | G |
| NC_011770 | 5439558 | A | G | 79 | 2 | 2 | 0 | NON_SYNONYMOU | MODERAT | MISSENSE | Atc/Gtc | I86V | pilY2 | | | | | | | | | . | . | G | G |
| NC_011770 | 4818885 | CG | C | 63 | 0 | 2 | 0 | FRAME_SHIFT | HIGH | | ccc/ | P425 | pirA | | | | | | | C | C | | | | |
| NC_011770 | 4819478 | G | A | 73 | 0 | 2 | 0 | NON_SYNONYMOU | MODERAT | MISSENSE | gCg/gTg | A228V | pirA | A | A | | | | | | | | | | |
| NC_011770 | 821184 | A | G | 59 | 0 | 1 | 0 | NON_SYNONYMOU | MODERAT | MISSENSE | cTg/cCg | L197P | piv | | | | | | | G | | | | | |
| NC_011770 | 821271 | C | T | 63 | 0 | 1 | 0 | NON_SYNONYMOU | MODERAT | MISSENSE | cGt/cAt | R168H | piv | | | | | | | T | | | | | |
| NC_011770 | 821355 | G | A | 57 | 0 | 1 | 0 | NON_SYNONYMOU | MODERAT | MISSENSE | tCg/tTg | S140L | piv | | | | | | | | | | | A | |
| NC_011770 | 27681 | G | A | 66 | 0 | 4 | 0 | NON_SYNONYMOU | MODERAT | MISSENSE | Gcc/Acc | A17T | plcB | | | A | A | A | A | | | | | | |
| NC_011770 | 27820 | C | T | 64 | 0 | 2 | 0 | NON_SYNONYMOU | MODERAT | MISSENSE | gCg/gTg | A63V | plcB | | | | | | | T | T | | | | |
| NC_011770 | 28494 | T | C | 92 | 0 | 12 | 0 | NON_SYNONYMOU | MODERAT | MISSENSE | Ttc/Ctc | F288L | plcB | C | C | C | C | C | C | C | C | C | C | C | C |
| NC_011770 | 4917508 | G | GC | 87 | 0 | 2 | 0 | FRAME_SHIFT | HIGH | | agc/Cagc | S193Q? | plcH | GC | GC | | | | | | | | | | |
| NC_011770 | 4917052 | A | G | 75 | 0 | 4 | 0 | NON_SYNONYMOU | MODERAT | MISSENSE | Atc/Gtc | I39V | plcH | | | | | | | G | G | | | G | G |
| NC_011770 | 4917448 | C | T | 89 | 0 | 2 | 0 | NON_SYNONYMOU | MODERAT | MISSENSE | Cac/Tac | H171Y | plcH | | | | | | | T | T | | | | |
| NC_011770 | 4917849 | C | A | 88 | 0 | 2 | 0 | NON_SYNONYMOU | MODERAT | MISSENSE | aaC/aaA | N304K | plcH | | | | | | | | | A | A | | |
| NC_011770 | 4917951 | G | C | 86 | 0 | 2 | 0 | NON_SYNONYMOU | MODERAT | MISSENSE | gaG/gaC | E338D | plcH | | | | | | | C | C | | | | |
| NC_011770 | 4918106 | C | T | 95 | 0 | 2 | 0 | NON_SYNONYMOU | MODERAT | MISSENSE | gCa/gTa | A390V | plcH | | | | | | | | | T | T | | |
| NC_011770 | 4918199 | T | A | 94 | 0 | 2 | 0 | NON_SYNONYMOU | MODERAT | MISSENSE | tTc/tAc | F421Y | plcH | | | | | | | | | | | A | A |
| NC_011770 | 4918580 | T | C | 88 | 0 | 12 | 0 | NON_SYNONYMOU | MODERAT | MISSENSE | cTg/cCg | L548P | plcH | C | C | C | C | C | C | C | C | C | C | C | C |
| NC_011770 | 4919053 | C | T | 73 | 0 | 4 | 0 | NON_SYNONYMOU | MODERAT | MISSENSE | Ctc/Ttc | L706F | plcH | | | | | | | T | T | T | T | | |
| NC_011770 | 1880993 | T | TG | 68 | 0 | 1 | 0 | FRAME_SHIFT | HIGH | | gcc/gGcc | A306G? | plcN | | | | | | | TG | | | | | |
| NC_011770 | 1880381 | C | G | 77 | 0 | 2 | 0 | NON_SYNONYMOU | MODERAT | MISSENSE | Ccc/Gcc | P100A | plcN | | | G | G | | | | | | | | |
| NC_011770 | 1880528 | A | G | 79 | 0 | 12 | 0 | NON_SYNONYMOU | MODERAT | MISSENSE | Aag/Gag | K149E | plcN | G | G | G | G | G | G | G | G | G | G | G | G |
| NC_011770 | 1881687 | G | A | 73 | 0 | 8 | 0 | NON_SYNONYMOU | MODERAT | MISSENSE | aGc/aAc | S535N | plcN | A | A | A | A | A | A | | | | | A | A |
| NC_011770 | 1881012 | C | T | 76 | 0 | 1 | 0 | NON_SYNONYMOU | MODERAT | MISSENSE | tCc/tTc | S310F | plcN | | | | | | | | | | | | T |
| NC_011770 | 4919220 | C | T | 53 | 0 | 6 | 0 | NON_SYNONYMOU | MODERAT | MISSENSE | Ctt/Ttt | L25F | plcR | T | T | | | | | T | T | | | T | T |
| NC_011770 | 4919365 | T | A | 51 | 0 | 2 | 0 | NON_SYNONYMOU | MODERAT | MISSENSE | tTc/tAc | F73Y | plcR | | | | | | | | | A | A | | |
| NC_011770 | 4919518 | C | T | 60 | 0 | 4 | 0 | NON_SYNONYMOU | MODERAT | MISSENSE | gCc/gTc | A124V | plcR | | | T | T | T | T | | | | | | |
| NC_011770 | 4919592 | G | C | 57 | 0 | 8 | 0 | NON_SYNONYMOU | MODERAT | MISSENSE | Gcg/Ccg | A149P | plcR | C | C | | | | | C | C | C | C | C | C |
| NC_011770 | 4919677 | A | G | 63 | 0 | 6 | 0 | NON_SYNONYMOU | MODERAT | MISSENSE | gAc/gGc | D177G | plcR | G | G | | | | | | | G | G | G | G |
| NC_011770 | 7917 | C | T | 97 | 0 | 4 | 0 | NON_SYNONYMOU | MODERAT | MISSENSE | Gcc/Acc | A138T | PLES_00051 | | | T | T | T | T | | | | | | |
| NC_011770 | 7986 | G | A | 99 | 0 | 2 | 0 | NON_SYNONYMOU | MODERAT | MISSENSE | Cac/Tac | H115Y | PLES_00051 | | | | | | | A | A | | | | |
| NC_011770 | 7989 | C | T | 96 | 0 | 4 | 0 | NON_SYNONYMOU | MODERAT | MISSENSE | Gag/Aag | E114K | PLES_00051 | | | T | T | T | T | | | | | | |
| NC_011770 | 8029 | G | T | 92 | 0 | 2 | 0 | NON_SYNONYMOU | MODERAT | MISSENSE | gaC/gaA | D100E | PLES_00051 | | | | | | | T | T | | | | |
| NC_011770 | 8202 | G | T | 94 | 0 | 6 | 0 | NON_SYNONYMOU | MODERAT | MISSENSE | Cag/Aag | Q43K | PLES_00051 | | | T | T | T | T | T | T | | | | |
| NC_011770 | 8256 | G | A | 91 | 0 | 2 | 0 | NON_SYNONYMOU | MODERAT | MISSENSE | Ctc/Ttc | L25F | PLES_00051 | A | A | | | | | | | | | | |
| NC_011770 | 8705 | C | T | 82 | 0 | 6 | 0 | NON_SYNONYMOU | MODERAT | MISSENSE | gCt/gTt | A16V | PLES_00061 | | | T | T | T | T | T | T | | | | |
| NC_011770 | 9085 | G | T | 82 | 0 | 2 | 0 | NON_SYNONYMOU | MODERAT | MISSENSE | Gcc/Tcc | A143S | PLES_00061 | | | | | | | T | T | | | | |
| NC_011770 | 9152 | C | A | 86 | 0 | 10 | 0 | NON_SYNONYMOU | MODERAT | MISSENSE | cCg/cAg | P165Q | PLES_00061 | A | A | A | A | A | A | A | A | | | A | A |
| NC_011770 | 9697 | G | A | 60 | 0 | 2 | 0 | NON_SYNONYMOU | MODERAT | MISSENSE | Gaa/Aaa | E347K | PLES_00061 | | | | | | | | | A | A | | |
| NC_011770 | 9752 | A | G | 53 | 0 | 2 | 0 | NON_SYNONYMOU | MODERAT | MISSENSE | gAc/gGc | D365G | PLES_00061 | | | G | G | | | | | | | | |
| NC_011770 | 14481 | C | G | 93 | 0 | 1 | 0 | NON_SYNONYMOU | MODERAT | MISSENSE | Ccg/Gcg | P87A | PLES_00101 | | | | | | | G | | | | | |
| NC_011770 | 15326 | T | G | 89 | 0 | 10 | 0 | NON_SYNONYMOU | MODERAT | MISSENSE | gaT/gaG | D44E | PLES_00111 | G | G | G | G | G | G | C | C | G | G | | |
| NC_011770 | 16972 | A | G | 86 | 0 | 2 | 0 | NON_SYNONYMOU | MODERAT | MISSENSE | cTg/cCg | L78P | PLES_00141 | G | G | | | | | | | | | | |
| NC_011770 | 18882 | C | T | 76 | 0 | 2 | 0 | NON_SYNONYMOU | MODERAT | MISSENSE | Ggc/Agc | G393S | PLES_00161 | | | | | | | T | T | | | | |
| NC_011770 | 19346 | A | T | 73 | 0 | 4 | 0 | NON_SYNONYMOU | MODERAT | MISSENSE | cTg/cAg | L238Q | PLES_00161 | T | T | | | | | | | T | T | | |
| NC_011770 | 19476 | T | C | 74 | 0 | 6 | 0 | NON_SYNONYMOU | MODERAT | MISSENSE | Aag/Gag | K195E | PLES_00161 | | | C | C | C | C | | | | | C | C |
| NC_011770 | 19883 | G | A | 78 | 0 | 4 | 0 | NON_SYNONYMOU | MODERAT | MISSENSE | gCg/gTg | A59V | PLES_00161 | | | A | A | A | A | | | | | | |
| NC_011770 | 22426 | C | T | 91 | 0 | 1 | 0 | NON_SYNONYMOU | MODERAT | MISSENSE | cCg/cTg | P243L | PLES_00191 | | | | | | | T | | | | | |
| NC_011770 | 22991 | G | A | 71 | 0 | 2 | 0 | NON_SYNONYMOU | MODERAT | MISSENSE | Ggc/Agc | G45S | PLES_00201 | | | | | | | | | A | A | | |
| NC_011770 | 23104 | G | C | 65 | 0 | 12 | 0 | NON_SYNONYMOU | MODERAT | MISSENSE | caG/caC | Q82H | PLES_00201 | C | C | C | C | C | C | C | C | C | C | C | C |
| NC_011770 | 23186 | G | A | 58 | 0 | 2 | 0 | NON_SYNONYMOU | MODERAT | MISSENSE | Gct/Act | A110T | PLES_00201 | | | | | | | A | A | | | | |
| NC_011770 | 23360 | G | A | 57 | 0 | 2 | 0 | NON_SYNONYMOU | MODERAT | MISSENSE | Gct/Act | A168T | PLES_00201 | | | | | | | A | A | | | | |
| NC_011770 | 23361 | C | T | 56 | 0 | 2 | 0 | NON_SYNONYMOU | MODERAT | MISSENSE | gCt/gTt | A168V | PLES_00201 | | | | | | | T | T | | | | |
| NC_011770 | 23451 | G | A | 57 | 0 | 2 | 0 | NON_SYNONYMOU | MODERAT | MISSENSE | gGt/gAt | G198D | PLES_00201 | | | | | | | A | A | | | | |
| NC_011770 | 23772 | C | T | 53 | 0 | 2 | 0 | NON_SYNONYMOU | MODERAT | MISSENSE | gCg/gTg | A305V | PLES_00201 | | | | | | | T | T | | | | |
| NC_011770 | 28625 | C | T | 95 | 0 | 2 | 0 | NON_SYNONYMOU | MODERAT | MISSENSE | gCc/gTc | A9V | PLES_00261 | | | | | | | T | T | | | | |
| NC_011770 | 28677 | G | T | 102 | 0 | 2 | 0 | NON_SYNONYMOU | MODERAT | MISSENSE | tgG/tgT | W26C | PLES_00261 | T | T | | | | | | | | | | |
| NC_011770 | 28957 | C | T | 92 | 0 | 4 | 0 | NON_SYNONYMOU | MODERAT | MISSENSE | Ctc/Ttc | L120F | PLES_00261 | | | T | T | T | T | | | | | | |

| | | | | | | | | | | | | | | | | | | | | | | | | | |
|---|---|---|---|---|---|---|---|---|---|---|---|---|---|---|---|---|---|---|---|---|---|---|---|---|---|
| NC_011770 | 29027 | G | A | 101 | 0 | 2 | 0 | NON_SYNONYMOU | MODERAT | MISSENSE | aGc/aAc | S143N | PLES_00261 | A | A | | | | | | | | | | |
| NC_011770 | 29385 | T | G | 96 | 0 | 2 | 0 | NON_SYNONYMOU | MODERAT | MISSENSE | gaT/gaG | D262E | PLES_00261 | | | | | | | G | G | | | | |
| NC_011770 | 29542 | G | A | 98 | 0 | 2 | 0 | NON_SYNONYMOU | MODERAT | MISSENSE | Ggc/Agc | G315S | PLES_00261 | | | | | | | A | A | | | | |
| NC_011770 | 29543 | G | A | 97 | 0 | 2 | 0 | NON_SYNONYMOU | MODERAT | MISSENSE | gGc/gAc | G315D | PLES_00261 | | | | | | | A | A | | | | |
| NC_011770 | 29749 | A | G | 94 | 0 | 4 | 0 | NON_SYNONYMOU | MODERAT | MISSENSE | Aag/Gag | K384E | PLES_00261 | | | | | | | G | G | | | G | G |
| NC_011770 | 29765 | C | T | 95 | 0 | 1 | 0 | NON_SYNONYMOU | MODERAT | MISSENSE | cCc/cTc | P389L | PLES_00261 | | | | | | | | | | | | T |
| NC_011770 | 29937 | A | G | 78 | 0 | 6 | 0 | NON_SYNONYMOU | MODERAT | MISSENSE | Acc/Gcc | T19A | PLES_00271 | G | G | | | | | G | G | | | G | G |
| NC_011770 | 29988 | T | C | 72 | 0 | 2 | 0 | NON_SYNONYMOU | MODERAT | MISSENSE | Tct/Cct | S36P | PLES_00271 | | | | | | | C | C | | | | |
| NC_011770 | 29994 | T | C | 73 | 0 | 2 | 0 | NON_SYNONYMOU | MODERAT | MISSENSE | Tcg/Ccg | S38P | PLES_00271 | | | | | | | C | C | | | | |
| NC_011770 | 30009 | G | C | 77 | 0 | 2 | 0 | NON_SYNONYMOU | MODERAT | MISSENSE | Gtt/Ctt | V43L | PLES_00271 | C | C | | | | | | | | | | |
| NC_011770 | 30049 | T | C | 75 | 0 | 4 | 0 | NON_SYNONYMOU | MODERAT | MISSENSE | cTg/cCg | L56P | PLES_00271 | | | | | | | | | C | C | C | C |
| NC_011770 | 30073 | C | T | 84 | 0 | 2 | 0 | NON_SYNONYMOU | MODERAT | MISSENSE | gCa/gTa | A64V | PLES_00271 | | | | | | | T | T | | | | |
| NC_011770 | 30769 | C | T | 90 | 0 | 6 | 0 | NON_SYNONYMOU | MODERAT | MISSENSE | cGc/cAc | R425H | PLES_00281 | | | | | | | T | T | T | T | T | T |
| NC_011770 | 31183 | T | A | 90 | 0 | 2 | 0 | NON_SYNONYMOU | MODERAT | MISSENSE | cAg/cTg | Q287L | PLES_00281 | | | | | | | A | A | | | | |
| NC_011770 | 31288 | T | G | 88 | 0 | 2 | 0 | NON_SYNONYMOU | MODERAT | MISSENSE | gAc/gCc | D252A | PLES_00281 | | | | | G | G | | | | | | |
| NC_011770 | 31321 | C | T | 86 | 0 | 10 | 0 | NON_SYNONYMOU | MODERAT | MISSENSE | aGt/aAt | S241N | PLES_00281 | T | T | T | T | T | T | T | T | | | T | T |
| NC_011770 | 31400 | T | A | 94 | 0 | 2 | 0 | NON_SYNONYMOU | MODERAT | MISSENSE | Acg/Tcg | T215S | PLES_00281 | A | A | | | | | | | | | | |
| NC_011770 | 31543 | T | C | 85 | 0 | 2 | 0 | NON_SYNONYMOU | MODERAT | MISSENSE | aAc/aGc | N167S | PLES_00281 | | | | | | | | | | | C | C |
| NC_011770 | 31687 | A | G | 82 | 0 | 2 | 0 | NON_SYNONYMOU | MODERAT | MISSENSE | gTc/gCc | V119A | PLES_00281 | | | | | | | | | | | G | G |
| NC_011770 | 32047 | T | C | 87 | 0 | 12 | 0 | SPLICE_SITE_REGIC | LOW | SILENT | taA/taG | *308 | PLES_00291 | C | C | C | C | C | C | C | C | C | C | C | C |
| NC_011770 | 32126 | T | C | 88 | 0 | 6 | 0 | NON_SYNONYMOU | MODERAT | MISSENSE | cAg/cGg | Q282R | PLES_00291 | C | C | | | | | | | C | C | C | C |
| NC_011770 | 32162 | G | T | 86 | 0 | 2 | 0 | NON_SYNONYMOU | MODERAT | MISSENSE | gCc/gAc | A270D | PLES_00291 | | | | | | | | | | | T | T |
| NC_011770 | 32588 | C | T | 85 | 0 | 4 | 0 | NON_SYNONYMOU | MODERAT | MISSENSE | cGc/cAc | R128H | PLES_00291 | | | T | T | T | T | | | | | | |
| NC_011770 | 32657 | C | G | 88 | 0 | 2 | 0 | NON_SYNONYMOU | MODERAT | MISSENSE | gGt/gCt | G105A | PLES_00291 | G | G | | | | | | | | | | |
| NC_011770 | 34614 | G | T | 74 | 0 | 4 | 0 | STOP_GAINED | HIGH | NONSENSE | Gga/Tga | G2* | PLES_00311 | T | T | | | | | T | T | | | | |
| NC_011770 | 34618 | A | G | 69 | 0 | 2 | 0 | NON_SYNONYMOU | MODERAT | MISSENSE | gAa/gGa | E3G | PLES_00311 | G | G | | | | | | | | | | |
| NC_011770 | 34621 | G | A | 70 | 0 | 6 | 0 | NON_SYNONYMOU | MODERAT | MISSENSE | cGc/cAc | R4H | PLES_00311 | A | A | | | | | A | A | | | A | A |
| NC_011770 | 34633 | T | C | 67 | 0 | 10 | 0 | NON_SYNONYMOU | MODERAT | MISSENSE | cTg/cCg | L8P | PLES_00311 | C | C | C | C | C | C | C | C | | | C | C |
| NC_011770 | 35993 | T | C | 87 | 0 | 12 | 0 | NON_SYNONYMOU | MODERAT | MISSENSE | Acc/Gcc | T89A | PLES_00321 | C | C | C | C | C | C | C | C | C | C | C | C |
| NC_011770 | 36103 | T | G | 81 | 0 | 2 | 0 | NON_SYNONYMOU | MODERAT | MISSENSE | gAg/gCg | E52A | PLES_00321 | G | G | | | | | | | | | | |
| NC_011770 | 40607 | G | A | 71 | 0 | 1 | 0 | NON_SYNONYMOU | MODERAT | MISSENSE | gGc/gAc | G11D | PLES_00381 | | | | | | | A | | | | | |
| NC_011770 | 41290 | A | C | 66 | 0 | 12 | 0 | NON_SYNONYMOU | MODERAT | MISSENSE | tAc/tCc | Y64S | PLES_00391 | C | C | C | C | C | C | C | C | C | C | C | C |
| NC_011770 | 41296 | A | G | 65 | 0 | 2 | 0 | NON_SYNONYMOU | MODERAT | MISSENSE | gAc/gGc | D66G | PLES_00391 | | | | | | | | | G | G | | |
| NC_011770 | 41586 | G | A | 103 | 0 | 2 | 0 | NON_SYNONYMOU | MODERAT | MISSENSE | Gcg/Acg | A163T | PLES_00391 | | | | | | | | | A | A | | |
| NC_011770 | 42237 | T | C | 105 | 0 | 12 | 0 | NON_SYNONYMOU | MODERAT | MISSENSE | Ttc/Ctc | F380L | PLES_00391 | C | C | C | C | C | C | C | C | C | C | C | C |
| NC_011770 | 42293 | C | G | 105 | 0 | 12 | 0 | NON_SYNONYMOU | MODERAT | MISSENSE | caC/caG | H398Q | PLES_00391 | G | G | G | G | G | G | G | G | G | G | G | G |
| NC_011770 | 42306 | A | C | 110 | 0 | 2 | 0 | NON_SYNONYMOU | MODERAT | MISSENSE | Aat/Cat | N403H | PLES_00391 | | | | | | | | | | | C | C |
| NC_011770 | 42372 | C | T | 111 | 0 | 2 | 0 | NON_SYNONYMOU | MODERAT | MISSENSE | Cgc/Tgc | R425C | PLES_00391 | | | | | | | T | T | | | | |
| NC_011770 | 43054 | A | G | 46 | 0 | 4 | 0 | NON_SYNONYMOU | MODERAT | MISSENSE | Atc/Gtc | I52V | PLES_00401 | G | G | | | | | | | | | G | G |
| NC_011770 | 43079 | G | A | 40 | 0 | 2 | 0 | NON_SYNONYMOU | MODERAT | MISSENSE | gGc/gAc | G60D | PLES_00401 | | | | | | | | | A | A | | |
| NC_011770 | 43705 | A | C | 73 | 0 | 4 | 0 | NON_SYNONYMOU | MODERAT | MISSENSE | Aag/Cag | K269Q | PLES_00401 | C | C | | | | | | | C | C | | |
| NC_011770 | 43706 | A | G | 73 | 0 | 4 | 0 | NON_SYNONYMOU | MODERAT | MISSENSE | aAg/aGg | K269R | PLES_00401 | G | G | | | | | | | G | G | | |
| NC_011770 | 43919 | C | T | 56 | 0 | 4 | 0 | NON_SYNONYMOU | MODERAT | MISSENSE | gCg/gTg | A340V | PLES_00401 | T | T | | | | | | | T | T | | |
| NC_011770 | 44132 | G | C | 64 | 0 | 6 | 0 | NON_SYNONYMOU | MODERAT | MISSENSE | gGc/gCc | G411A | PLES_00401 | | | | | | | C | C | C | C | C | C |
| NC_011770 | 44143 | G | A | 70 | 0 | 2 | 0 | NON_SYNONYMOU | MODERAT | MISSENSE | Gac/Aac | D415N | PLES_00401 | | | | | | | | | | | A | A |
| NC_011770 | 45064 | A | G | 73 | 0 | 4 | 0 | NON_SYNONYMOU | MODERAT | MISSENSE | Acg/Gcg | T722A | PLES_00401 | | | G | G | G | G | | | | | | |
| NC_011770 | 45065 | C | T | 72 | 0 | 4 | 0 | NON_SYNONYMOU | MODERAT | MISSENSE | aCg/aTg | T722M | PLES_00401 | | | T | T | T | T | | | | | | |
| NC_011770 | 45553 | A | C | 50 | 0 | 4 | 0 | NON_SYNONYMOU | MODERAT | MISSENSE | Aag/Cag | K885Q | PLES_00401 | | | | | | | C | C | C | C | | |
| NC_011770 | 45626 | T | C | 46 | 0 | 4 | 0 | NON_SYNONYMOU | MODERAT | MISSENSE | gTa/gCa | V909A | PLES_00401 | | | C | C | C | C | | | | | | |
| NC_011770 | 45715 | G | A | 65 | 0 | 4 | 0 | NON_SYNONYMOU | MODERAT | MISSENSE | Gtc/Atc | V939I | PLES_00401 | | | A | A | A | A | | | | | | |
| NC_011770 | 45736 | G | T | 68 | 0 | 4 | 0 | NON_SYNONYMOU | MODERAT | MISSENSE | Gac/Tac | D946Y | PLES_00401 | | | T | T | T | T | | | | | | |
| NC_011770 | 45790 | A | G | 71 | 0 | 4 | 0 | NON_SYNONYMOU | MODERAT | MISSENSE | Acc/Gcc | T964A | PLES_00401 | | | G | G | G | G | | | | | | |
| NC_011770 | 45887 | C | T | 64 | 0 | 4 | 0 | NON_SYNONYMOU | MODERAT | MISSENSE | gCg/gTg | A996V | PLES_00401 | | | T | T | T | T | | | | | | |
| NC_011770 | 45986 | A | C | 59 | 0 | 6 | 0 | NON_SYNONYMOU | MODERAT | MISSENSE | gAc/gCc | D1029A | PLES_00401 | C | C | | | | | C | C | C | C | | |
| NC_011770 | 46031 | C | G | 61 | 0 | 8 | 0 | NON_SYNONYMOU | MODERAT | MISSENSE | gCc/gGc | A1044G | PLES_00401 | | | G | G | G | G | | | G | G | G | G |
| NC_011770 | 46048 | A | G | 62 | 0 | 8 | 0 | NON_SYNONYMOU | MODERAT | MISSENSE | Acc/Gcc | T1050A | PLES_00401 | G | G | | | | | G | G | G | G | G | G |
| NC_011770 | 46237 | G | A | 55 | 0 | 10 | 0 | NON_SYNONYMOU | MODERAT | MISSENSE | Gac/Aac | D1113N | PLES_00401 | A | A | A | A | A | A | | | A | A | A | A |
| NC_011770 | 46250 | C | G | 56 | 0 | 10 | 0 | NON_SYNONYMOU | MODERAT | MISSENSE | aCc/aGc | T1117S | PLES_00401 | G | G | G | G | G | G | | | G | G | G | G |
| NC_011770 | 46256 | T | C | 58 | 0 | 8 | 0 | NON_SYNONYMOU | MODERAT | MISSENSE | gTc/gCc | V1119A | PLES_00401 | C | C | C | C | C | C | | | C | C | | |
| NC_011770 | 46537 | A | G | 59 | 0 | 4 | 0 | NON_SYNONYMOU | MODERAT | MISSENSE | Atc/Gtc | I1213V | PLES_00401 | | | | | | | | | G | G | G | G |
| NC_011770 | 46895 | G | A | 56 | 0 | 2 | 0 | NON_SYNONYMOU | MODERAT | MISSENSE | gGc/gAc | G1332D | PLES_00401 | | | | | | | | | A | A | | |
| NC_011770 | 46905 | A | C | 58 | 0 | 10 | 0 | NON_SYNONYMOU | MODERAT | MISSENSE | gaA/gaC | E1335D | PLES_00401 | | | C | C | C | C | C | C | C | C | C | C |

| | | | | | | | | | | | | | | | | | | | | | | | | | |
|---|---|---|---|---|---|---|---|---|---|---|---|---|---|---|---|---|---|---|---|---|---|---|---|---|---|
| NC_011770 | 46945 | C | T | 58 | 0 | 4 | 0 | NON_SYNONYMOU | MODERAT | MISSENSE | Cgg/Tgg | R1349W | PLES_00401 | | | T | T | T | T | | | | | | |
| NC_011770 | 47018 | T | C | 58 | 0 | 6 | 0 | NON_SYNONYMOU | MODERAT | MISSENSE | gTc/gCc | V1373A | PLES_00401 | | | | | | | C | C | C | C | C | C |
| NC_011770 | 47224 | A | G | 56 | 0 | 2 | 0 | NON_SYNONYMOU | MODERAT | MISSENSE | Acc/Gcc | T1442A | PLES_00401 | | | | | | | | | G | G | | |
| NC_011770 | 47327 | C | G | 55 | 0 | 10 | 0 | NON_SYNONYMOU | MODERAT | MISSENSE | aCc/aGc | T1476S | PLES_00401 | | | G | G | G | G | G | G | G | G | G | G |
| NC_011770 | 47329 | A | G | 54 | 0 | 10 | 0 | NON_SYNONYMOU | MODERAT | MISSENSE | Acc/Gcc | T1477A | PLES_00401 | | | G | G | G | G | G | G | G | G | G | G |
| NC_011770 | 47378 | A | G | 55 | 0 | 4 | 0 | NON_SYNONYMOU | MODERAT | MISSENSE | gAc/gGc | D1493G | PLES_00401 | | | | | | | G | G | | | G | G |
| NC_011770 | 47396 | T | G | 53 | 0 | 6 | 0 | NON_SYNONYMOU | MODERAT | MISSENSE | gTc/gGc | V1499G | PLES_00401 | G | G | | | | | G | G | | | G | G |
| NC_011770 | 47516 | G | A | 50 | 0 | 2 | 0 | NON_SYNONYMOU | MODERAT | MISSENSE | cGg/cAg | R1539Q | PLES_00401 | | | | | | | | | | | A | A |
| NC_011770 | 47758 | G | A | 42 | 0 | 2 | 0 | NON_SYNONYMOU | MODERAT | MISSENSE | Ggc/Agc | G1620S | PLES_00401 | | | A | A | | | | | | | | |
| NC_011770 | 48053 | G | A | 48 | 0 | 2 | 0 | NON_SYNONYMOU | MODERAT | MISSENSE | gGc/gAc | G1718D | PLES_00401 | | | | | | | A | A | | | | |
| NC_011770 | 48212 | G | C | 70 | 0 | 12 | 0 | NON_SYNONYMOU | MODERAT | MISSENSE | aGc/aCc | S1771T | PLES_00401 | C | C | C | C | C | C | C | C | C | C | C | C |
| NC_011770 | 48399 | G | C | 80 | 0 | 4 | 0 | NON_SYNONYMOU | MODERAT | MISSENSE | ttG/ttC | L1833F | PLES_00401 | | | | | | | C | C | | | C | C |
| NC_011770 | 48408 | G | C | 80 | 0 | 8 | 0 | NON_SYNONYMOU | MODERAT | MISSENSE | gaG/gaC | E1836D | PLES_00401 | C | C | | | | | C | C | C | C | C | C |
| NC_011770 | 48496 | G | T | 90 | 0 | 2 | 0 | NON_SYNONYMOU | MODERAT | MISSENSE | Gcc/Tcc | A1866S | PLES_00401 | | | | | | | | | | | T | T |
| NC_011770 | 49288 | A | G | 56 | 0 | 12 | 0 | NON_SYNONYMOU | MODERAT | MISSENSE | Agc/Ggc | S2130G | PLES_00401 | G | G | G | G | G | G | G | G | G | G | G | G |
| NC_011770 | 49670 | G | T | 55 | 0 | 2 | 0 | NON_SYNONYMOU | MODERAT | MISSENSE | gGc/gTc | G2257V | PLES_00401 | | | | | | | | | | | T | T |
| NC_011770 | 49676 | G | C | 56 | 0 | 2 | 0 | NON_SYNONYMOU | MODERAT | MISSENSE | gGc/gCc | G2259A | PLES_00401 | | | | | | | C | C | | | | |
| NC_011770 | 49757 | T | C | 59 | 0 | 8 | 0 | NON_SYNONYMOU | MODERAT | MISSENSE | gTc/gCc | V2286A | PLES_00401 | C | C | C | C | C | C | | | C | C | | |
| NC_011770 | 49843 | G | A | 64 | 0 | 4 | 0 | NON_SYNONYMOU | MODERAT | MISSENSE | Ggc/Agc | G2315S | PLES_00401 | | | A | A | A | A | | | | | | |
| NC_011770 | 50006 | A | C | 83 | 0 | 4 | 0 | NON_SYNONYMOU | MODERAT | MISSENSE | gAt/gCt | D2369A | PLES_00401 | C | C | | | | | | | C | C | | |
| NC_011770 | 50158 | A | C | 86 | 0 | 6 | 0 | NON_SYNONYMOU | MODERAT | MISSENSE | Aag/Cag | K2420Q | PLES_00401 | | | C | C | C | C | | | | | C | C |
| NC_011770 | 50166 | G | C | 87 | 0 | 6 | 0 | NON_SYNONYMOU | MODERAT | MISSENSE | caG/caC | Q2422H | PLES_00401 | | | C | C | C | C | | | | | C | C |
| NC_011770 | 50207 | T | C | 146 | 0 | 12 | 0 | NON_SYNONYMOU | MODERAT | MISSENSE | gTg/gCg | V2436A | PLES_00401 | C | C | C | C | C | C | C | C | C | C | C | C |
| NC_011770 | 50367 | T | G | 132 | 0 | 2 | 0 | NON_SYNONYMOU | MODERAT | MISSENSE | caT/caG | H2489Q | PLES_00401 | | | | | | | G | G | | | | |
| NC_011770 | 50752 | A | G | 140 | 0 | 12 | 0 | NON_SYNONYMOU | MODERAT | MISSENSE | Atc/Gtc | I2618V | PLES_00401 | G | G | G | G | G | G | G | G | G | G | G | G |
| NC_011770 | 50783 | A | G | 145 | 0 | 4 | 0 | NON_SYNONYMOU | MODERAT | MISSENSE | aAc/aGc | N2628S | PLES_00401 | | | | | | | G | G | | | G | G |
| NC_011770 | 50987 | C | T | 147 | 0 | 2 | 0 | NON_SYNONYMOU | MODERAT | MISSENSE | aCc/aTc | T2696I | PLES_00401 | T | T | | | | | | | | | | |
| NC_011770 | 51101 | A | G | 139 | 0 | 2 | 0 | NON_SYNONYMOU | MODERAT | MISSENSE | gAc/gGc | D2734G | PLES_00401 | | | | | | | | | | | G | G |
| NC_011770 | 51369 | G | A | 146 | 0 | 4 | 0 | NON_SYNONYMOU | MODERAT | MISSENSE | gaG/gaC | E2823D | PLES_00401 | | | | | | | | | A | A | C | C |
| NC_011770 | 51473 | G | C | 122 | 0 | 2 | 0 | NON_SYNONYMOU | MODERAT | MISSENSE | aGc/aCc | S2858T | PLES_00401 | C | C | | | | | | | | | | |
| NC_011770 | 51574 | G | A | 119 | 0 | 2 | 0 | NON_SYNONYMOU | MODERAT | MISSENSE | Gcg/Acg | A2892T | PLES_00401 | | | | | | | A | A | | | | |
| NC_011770 | 51647 | T | G | 109 | 0 | 4 | 0 | NON_SYNONYMOU | MODERAT | MISSENSE | aTg/aGg | M2916R | PLES_00401 | G | G | | | | | | | | | G | G |
| NC_011770 | 52442 | T | C | 93 | 2 | 6 | 0 | NON_SYNONYMOU | MODERAT | MISSENSE | gTg/gCg | V3181A | PLES_00401 | C | C | C | C | C | C | | | . | . | | |
| NC_011770 | 52640 | C | T | 92 | 2 | 8 | 0 | NON_SYNONYMOU | MODERAT | MISSENSE | gCg/gTg | A3247V | PLES_00401 | T | T | T | T | T | T | T | T | . | . | | |
| NC_011770 | 52945 | T | A | 72 | 2 | 4 | 0 | NON_SYNONYMOU | MODERAT | MISSENSE | Tcg/Acg | S3349T | PLES_00401 | | | A | A | A | A | | | . | . | | |
| NC_011770 | 52958 | T | C | 71 | 2 | 2 | 0 | NON_SYNONYMOU | MODERAT | MISSENSE | cTc/cCc | L3353P | PLES_00401 | | | | | C | C | | | . | . | | |
| NC_011770 | 53176 | G | T | 29 | 6 | 2 | 0 | NON_SYNONYMOU | MODERAT | MISSENSE | Gcc/Tcc | A3426S | PLES_00401 | T | T | | | | | . | . | . | . | . | . |
| NC_011770 | 53186 | G | A | 29 | 6 | 2 | 0 | NON_SYNONYMOU | MODERAT | MISSENSE | gGg/gAg | G3429E | PLES_00401 | A | A | | | | | . | . | . | . | . | . |
| NC_011770 | 45310 | G | A | 77 | 0 | 1 | 0 | NON_SYNONYMOU | MODERAT | MISSENSE | Ggc/Agc | G804S | PLES_00401 | | | | **A** | | | | | | | | |
| NC_011770 | 54907 | C | T | 102 | 0 | 3 | 0 | STOP_GAINED | HIGH | NONSENSE | Cag/Tag | Q207* | PLES_00411 | | | | T | T | T | | | | | | |
| NC_011770 | 54375 | AT | A | 134 | 2 | 2 | 0 | FRAME_SHIFT | HIGH | | tgc/ | C31 | PLES_00411 | | | | | | | A | A | | | . | . |
| NC_011770 | 54615 | CGA | C | 136 | 2 | 2 | 0 | FRAME_SHIFT | HIGH | | gat/ | D110 | PLES_00411 | | | | | | | C | C | | | . | . |
| NC_011770 | 54635 | G | A | 135 | 2 | 2 | 0 | STOP_GAINED | HIGH | NONSENSE | tGg/tAg | W116* | PLES_00411 | | | | | | | A | A | | | . | . |
| NC_011770 | 54325 | C | T | 126 | 2 | 2 | 0 | NON_SYNONYMOU | MODERAT | MISSENSE | Ctc/Ttc | L13F | PLES_00411 | | | | | | | T | T | | | . | . |
| NC_011770 | 54400 | C | T | 143 | 2 | 2 | 0 | NON_SYNONYMOU | MODERAT | MISSENSE | Cat/Tat | H38Y | PLES_00411 | | | | | | | T | T | | | . | . |
| NC_011770 | 54427 | C | A | 146 | 2 | 2 | 0 | NON_SYNONYMOU | MODERAT | MISSENSE | Ctg/Atg | L47M | PLES_00411 | | | | | | | A | A | | | . | . |
| NC_011770 | 54442 | C | G | 145 | 2 | 2 | 0 | NON_SYNONYMOU | MODERAT | MISSENSE | Caa/Gaa | Q52E | PLES_00411 | | | | | | | G | G | | | . | . |
| NC_011770 | 54497 | G | A | 144 | 2 | 2 | 0 | NON_SYNONYMOU | MODERAT | MISSENSE | gGc/gAc | G70D | PLES_00411 | | | | | | | A | A | | | . | . |
| NC_011770 | 54637 | T | C | 134 | 2 | 2 | 0 | NON_SYNONYMOU | MODERAT | MISSENSE | Tca/Cca | S117P | PLES_00411 | | | | | | | C | C | | | . | . |
| NC_011770 | 54785 | G | A | 112 | 4 | 2 | 0 | NON_SYNONYMOU | MODERAT | MISSENSE | aGg/aAg | R166K | PLES_00411 | A | A | | | | | . | . | | | . | . |
| NC_011770 | 54839 | T | G | 116 | 2 | 2 | 0 | NON_SYNONYMOU | MODERAT | MISSENSE | gTg/gGg | V184G | PLES_00411 | | | | | | | G | G | | | . | . |
| NC_011770 | 54848 | T | G | 113 | 2 | 2 | 0 | NON_SYNONYMOU | MODERAT | MISSENSE | aTc/aGc | I187S | PLES_00411 | | | | | | | G | G | | | . | . |
| NC_011770 | 54908 | A | T | 101 | 0 | 1 | 0 | NON_SYNONYMOU | MODERAT | MISSENSE | cAg/cTg | Q207L | PLES_00411 | | T | | | | | | | | | | |
| NC_011770 | 54299 | C | T | 123 | 2 | 1 | 0 | NON_SYNONYMOU | MODERAT | MISSENSE | gCc/gTc | A4V | PLES_00411 | | | | | | | | **T** | | | . | . |
| NC_011770 | 54911 | A | G | 99 | 0 | 9 | 0 | NON_SYNONYMOU | MODERAT | MISSENSE | gAa/gGa | E208G | PLES_00411 | | G | | G | G | G | G | . | G | G | G | G |
| NC_011770 | 55286 | A | G | 21 | 8 | 4 | 0 | NON_SYNONYMOU | MODERAT | MISSENSE | aAg/aGg | K27R | PLES_00421 | . | . | . | . | . | . | G | G | . | . | G | G |
| NC_011770 | 55297 | G | A | 21 | 8 | 4 | 0 | NON_SYNONYMOU | MODERAT | MISSENSE | Gca/Aca | A31T | PLES_00421 | . | . | . | . | . | . | A | A | . | . | A | A |
| NC_011770 | 56402 | C | T | 96 | 0 | 2 | 0 | NON_SYNONYMOU | MODERAT | MISSENSE | gCg/gTg | A15V | PLES_00431 | | | | | | | | | | | T | T |
| NC_011770 | 57324 | G | C | 93 | 0 | 4 | 0 | NON_SYNONYMOU | MODERAT | MISSENSE | Ctg/Gtg | L369V | PLES_00441 | | | C | C | C | C | | | | | | |
| NC_011770 | 57326 | G | A | 92 | 0 | 2 | 0 | NON_SYNONYMOU | MODERAT | MISSENSE | gCc/gTc | A368V | PLES_00441 | | | | | | | A | A | | | | |
| NC_011770 | 57339 | T | G | 89 | 0 | 10 | 0 | NON_SYNONYMOU | MODERAT | MISSENSE | Atg/Ctg | M364L | PLES_00441 | | | G | G | G | G | G | G | G | G | G | G |
| NC_011770 | 58349 | C | T | 62 | 0 | 2 | 0 | NON_SYNONYMOU | MODERAT | MISSENSE | cGg/cAg | R27Q | PLES_00441 | | | | | | | T | T | | | | |

| | | | | | | | | | | | | | | | | | | | | | | | | | |
|---|---|---|---|---|---|---|---|---|---|---|---|---|---|---|---|---|---|---|---|---|---|---|---|---|---|
| NC_011770 | 58352 | A | G | 61 | 0 | 2 | 0 | NON_SYNONYMOU | MODERAT | MISSENSE | gTg/gCg | V26A | PLES_00441 | | | | | | | G | G | | | | |
| NC_011770 | 58356 | C | T | 64 | 0 | 2 | 0 | NON_SYNONYMOU | MODERAT | MISSENSE | Gcc/Acc | A25T | PLES_00441 | | | | | | | T | T | | | | |
| NC_011770 | 58373 | A | T | 67 | 0 | 2 | 0 | NON_SYNONYMOU | MODERAT | MISSENSE | cTt/cAt | L19H | PLES_00441 | | | | | | | T | T | | | | |
| NC_011770 | 58377 | C | G | 66 | 0 | 2 | 0 | NON_SYNONYMOU | MODERAT | MISSENSE | Gct/Cct | A18P | PLES_00441 | | | | | | | G | G | | | | |
| NC_011770 | 60482 | A | G | 45 | 0 | 4 | 0 | NON_SYNONYMOU | MODERAT | MISSENSE | Aca/Gca | T8A | PLES_00461 | | | | | | | G | G | | | G | G |
| NC_011770 | 62234 | T | C | 68 | 0 | 6 | 0 | NON_SYNONYMOU | MODERAT | MISSENSE | Acc/Gcc | T120A | PLES_00491 | | | C | C | C | C | | | | | C | C |
| NC_011770 | 62402 | G | C | 74 | 0 | 12 | 0 | NON_SYNONYMOU | MODERAT | MISSENSE | Cag/Gag | Q64E | PLES_00491 | C | C | C | C | C | C | C | C | C | C | C | C |
| NC_011770 | 63389 | T | G | 76 | 0 | 12 | 0 | NON_SYNONYMOU | MODERAT | MISSENSE | ttA/ttC | L382F | PLES_00501 | G | G | G | G | G | G | G | G | G | G | G | G |
| NC_011770 | 64209 | C | G | 54 | 0 | 4 | 0 | NON_SYNONYMOU | MODERAT | MISSENSE | aGc/aCc | S109T | PLES_00501 | | | G | G | G | G | | | | | | |
| NC_011770 | 64495 | T | C | 46 | 0 | 4 | 0 | NON_SYNONYMOU | MODERAT | MISSENSE | Acc/Gcc | T14A | PLES_00501 | | | | | | | C | C | C | C | | |
| NC_011770 | 64521 | C | G | 39 | 0 | 2 | 0 | NON_SYNONYMOU | MODERAT | MISSENSE | cGt/cCt | R5P | PLES_00501 | | | | | | | G | G | | | | |
| NC_011770 | 68400 | C | T | 73 | 0 | 2 | 0 | NON_SYNONYMOU | MODERAT | MISSENSE | Gtg/Atg | V8M | PLES_00531 | | | | | | | | | T | T | | |
| NC_011770 | 68656 | G | C | 73 | 0 | 4 | 0 | INTRAGENIC | MODIFIER | | | | PLES_00541 | | | C | C | C | C | | | | | | |
| NC_011770 | 68714 | A | G | 75 | 0 | 6 | 0 | INTRAGENIC | MODIFIER | | | | PLES_00541 | G | G | | | | | G | G | G | G | | |
| NC_011770 | 68722 | C | T | 72 | 0 | 2 | 0 | INTRAGENIC | MODIFIER | | | | PLES_00541 | T | T | | | | | | | | | | |
| NC_011770 | 68825 | A | G | 73 | 0 | 4 | 0 | INTRAGENIC | MODIFIER | | | | PLES_00541 | | | | | | | G | G | G | G | | |
| NC_011770 | 68959 | A | G | 71 | 0 | 12 | 0 | INTRAGENIC | MODIFIER | | | | PLES_00551 | G | G | G | G | G | G | G | G | G | G | G | G |
| NC_011770 | 68961 | T | C | 72 | 0 | 8 | 0 | INTRAGENIC | MODIFIER | | | | PLES_00551 | C | C | C | C | C | C | C | C | | | | |
| NC_011770 | 69022 | C | T | 75 | 0 | 6 | 0 | INTRAGENIC | MODIFIER | | | | PLES_00551 | T | T | T | T | T | T | | | | | | |
| NC_011770 | 69799 | G | C | 82 | 0 | 2 | 0 | NON_SYNONYMOU | MODERAT | MISSENSE | gCg/gGg | A304G | PLES_00571 | | | | | | | | | C | C | | |
| NC_011770 | 70084 | G | A | 78 | 0 | 4 | 0 | NON_SYNONYMOU | MODERAT | MISSENSE | aCc/aTc | T209I | PLES_00571 | | | A | A | A | A | | | | | | |
| NC_011770 | 71848 | C | T | 90 | 0 | 6 | 0 | NON_SYNONYMOU | MODERAT | MISSENSE | Cgc/Tgc | R28C | PLES_00591 | T | T | | | | | T | T | T | T | | |
| NC_011770 | 72226 | G | T | 73 | 0 | 12 | 0 | NON_SYNONYMOU | MODERAT | MISSENSE | Gcc/Tcc | A154S | PLES_00591 | T | T | T | T | T | T | T | T | T | T | T | T |
| NC_011770 | 72382 | C | T | 82 | 0 | 6 | 0 | NON_SYNONYMOU | MODERAT | MISSENSE | Ctc/Ttc | L206F | PLES_00591 | T | T | | | | | T | T | T | T | | |
| NC_011770 | 72395 | A | G | 81 | 0 | 4 | 0 | NON_SYNONYMOU | MODERAT | MISSENSE | cAg/cGg | Q210R | PLES_00591 | | | G | G | G | G | | | | | | |
| NC_011770 | 72400 | T | C | 81 | 0 | 12 | 0 | NON_SYNONYMOU | MODERAT | MISSENSE | Tgc/Cgc | C212R | PLES_00591 | C | C | C | C | C | C | C | C | C | C | C | C |
| NC_011770 | 72419 | G | A | 80 | 0 | 6 | 0 | NON_SYNONYMOU | MODERAT | MISSENSE | gGt/gAt | G218D | PLES_00591 | | | A | A | A | A | | | | | A | A |
| NC_011770 | 73148 | G | A | 90 | 0 | 6 | 0 | NON_SYNONYMOU | MODERAT | MISSENSE | Gcc/Acc | A10T | PLES_00611 | A | A | A | A | A | A | | | | | | |
| NC_011770 | 73491 | C | T | 74 | 0 | 2 | 0 | NON_SYNONYMOU | MODERAT | MISSENSE | Gcg/Acg | A105T | PLES_00621 | | | | | | | | | T | T | | |
| NC_011770 | 75352 | C | G | 80 | 0 | 2 | 0 | NON_SYNONYMOU | MODERAT | MISSENSE | Caa/Gaa | Q329E | PLES_00641 | | | | | | | | | G | G | | |
| NC_011770 | 75427 | G | T | 77 | 0 | 2 | 0 | NON_SYNONYMOU | MODERAT | MISSENSE | Gcc/Tcc | A354S | PLES_00641 | | | | | | | | | | | T | T |
| NC_011770 | 75652 | GC | G | 71 | 0 | 1 | 0 | FRAME_SHIFT | HIGH | | ggg/ | G278 | PLES_00651 | | | | | | | G | | | | | |
| NC_011770 | 76201 | C | T | 45 | 0 | 4 | 0 | NON_SYNONYMOU | MODERAT | MISSENSE | Gtc/Atc | V96I | PLES_00651 | | | T | T | T | T | | | | | | |
| NC_011770 | 76206 | T | C | 43 | 0 | 2 | 0 | NON_SYNONYMOU | MODERAT | MISSENSE | cAg/cGg | Q94R | PLES_00651 | | | | | | | C | C | | | | |
| NC_011770 | 77259 | T | C | 72 | 0 | 2 | 0 | NON_SYNONYMOU | MODERAT | MISSENSE | aAg/aGg | K154R | PLES_00671 | | | | | | | | | | | C | C |
| NC_011770 | 79930 | A | C | 64 | 0 | 2 | 0 | NON_SYNONYMOU | MODERAT | MISSENSE | gAc/gCc | D31A | PLES_00691 | | | | | | | | | | | C | C |
| NC_011770 | 81002 | G | C | 79 | 0 | 6 | 0 | NON_SYNONYMOU | MODERAT | MISSENSE | gGc/gCc | G267A | PLES_00701 | C | C | C | C | C | C | | | | | | |
| NC_011770 | 81681 | C | T | 79 | 0 | 2 | 0 | NON_SYNONYMOU | MODERAT | MISSENSE | aGc/aAc | S242N | PLES_00711 | | | | | | | | | T | T | | |
| NC_011770 | 83716 | G | A | 74 | 0 | 2 | 0 | NON_SYNONYMOU | MODERAT | MISSENSE | gCg/gTg | A155V | PLES_00721 | | | | | | | | | | | A | A |
| NC_011770 | 84151 | G | C | 71 | 0 | 2 | 0 | NON_SYNONYMOU | MODERAT | MISSENSE | gCc/gGc | A10G | PLES_00721 | | | | | C | C | | | | | | |
| NC_011770 | 84067 | A | G | 62 | 0 | 1 | 0 | NON_SYNONYMOU | MODERAT | MISSENSE | gTc/gCc | V38A | PLES_00721 | G | | | | | | | | | | | |
| NC_011770 | 84261 | C | T | 85 | 0 | 2 | 0 | NON_SYNONYMOU | MODERAT | MISSENSE | Gcc/Acc | A371T | PLES_00731 | | | T | T | | | | | | | | |
| NC_011770 | 84606 | C | T | 91 | 0 | 2 | 0 | NON_SYNONYMOU | MODERAT | MISSENSE | Gag/Aag | E256K | PLES_00731 | | | | | T | T | | | | | | |
| NC_011770 | 84856 | T | A | 83 | 0 | 2 | 0 | NON_SYNONYMOU | MODERAT | MISSENSE | gaA/gaT | E172D | PLES_00731 | | | | | | | A | A | | | | |
| NC_011770 | 84888 | C | T | 78 | 0 | 1 | 0 | NON_SYNONYMOU | MODERAT | MISSENSE | Ggc/Agc | G162S | PLES_00731 | | | | | | | T | | | | | |
| NC_011770 | 84794 | A | G | 87 | 0 | 1 | 0 | NON_SYNONYMOU | MODERAT | MISSENSE | cTg/cCg | L193P | PLES_00731 | tagS1 | | | | | | | **G** | | | | |
| NC_011770 | 85603 | G | A | 69 | 0 | 2 | 0 | NON_SYNONYMOU | MODERAT | MISSENSE | gCg/gTg | A163V | PLES_00741 | | | | | | | | | | | A | A |
| NC_011770 | 85607 | G | T | 68 | 0 | 2 | 0 | NON_SYNONYMOU | MODERAT | MISSENSE | Ctg/Atg | L162M | PLES_00741 | | | | | | | | | | | T | T |
| NC_011770 | 85739 | C | T | 69 | 0 | 2 | 0 | NON_SYNONYMOU | MODERAT | MISSENSE | Gcg/Acg | A118T | PLES_00741 | | | | | | | | | | | T | T |
| NC_011770 | 89384 | T | C | 96 | 0 | 10 | 0 | NON_SYNONYMOU | MODERAT | MISSENSE | Acg/Gcg | T180A | PLES_00761 | C | C | C | C | C | C | C | C | | | C | C |
| NC_011770 | 89822 | C | A | 85 | 0 | 4 | 0 | NON_SYNONYMOU | MODERAT | MISSENSE | Gcg/Tcg | A34S | PLES_00761 | | | A | A | A | A | | | | | | |
| NC_011770 | 90283 | C | T | 75 | 0 | 4 | 0 | NON_SYNONYMOU | MODERAT | MISSENSE | gGc/gAc | G110D | PLES_00771 | | | T | T | T | T | | | | | | |
| NC_011770 | 90419 | C | CG | 72 | 0 | 1 | 0 | FRAME_SHIFT | HIGH | | ccc/cccC | P64P? | PLES_00771 | (hypothetica | CG | | | | | | | | | | |
| NC_011770 | 90897 | C | G | 68 | 0 | 6 | 0 | NON_SYNONYMOU | MODERAT | MISSENSE | aGc/aCc | S1006T | PLES_00781 | | | G | G | G | G | G | G | | | | |
| NC_011770 | 90898 | T | C | 66 | 0 | 6 | 0 | NON_SYNONYMOU | MODERAT | MISSENSE | Agc/Ggc | S1006G | PLES_00781 | | | C | C | C | C | C | C | | | | |
| NC_011770 | 90997 | T | C | 70 | 0 | 6 | 0 | NON_SYNONYMOU | MODERAT | MISSENSE | Acg/Gcg | T973A | PLES_00781 | | | C | C | C | C | C | C | | | | |
| NC_011770 | 91164 | G | C | 76 | 0 | 12 | 0 | NON_SYNONYMOU | MODERAT | MISSENSE | aCc/aGc | T917S | PLES_00781 | C | C | C | C | C | C | C | C | C | C | C | C |
| NC_011770 | 91633 | T | C | 64 | 0 | 10 | 0 | NON_SYNONYMOU | MODERAT | MISSENSE | Acc/Gcc | T761A | PLES_00781 | C | C | C | C | C | C | C | C | | | C | C |
| NC_011770 | 92146 | T | C | 77 | 0 | 10 | 0 | NON_SYNONYMOU | MODERAT | MISSENSE | Atg/Gtg | M590V | PLES_00781 | C | C | C | C | C | C | C | C | | | C | C |
| NC_011770 | 93015 | C | T | 99 | 0 | 1 | 0 | NON_SYNONYMOU | MODERAT | MISSENSE | gGc/gAc | G300D | PLES_00781 | | | | | | | T | | | | | |
| NC_011770 | 94199 | T | C | 81 | 0 | 12 | 0 | NON_SYNONYMOU | MODERAT | MISSENSE | aAc/aGc | N429S | PLES_00791 | C | C | C | C | C | C | C | C | C | C | C | C |
| NC_011770 | 94208 | C | G | 81 | 0 | 8 | 0 | NON_SYNONYMOU | MODERAT | MISSENSE | aGc/aCc | S426T | PLES_00791 | | | G | G | G | G | G | G | G | G | | |

| | | | | | | | | | | | | | | | | | | | | | | | | | |
|---|---|---|---|---|---|---|---|---|---|---|---|---|---|---|---|---|---|---|---|---|---|---|---|---|---|
| NC_011770 | 97024 | C | A | 81 | 0 | 2 | 0 | NON_SYNONYMOU | MODERAT | MISSENSE | gaG/gaT | E94D | PLES_00811 | | | | | | | | | | | A | A |
| NC_011770 | 97915 | A | G | 35 | 0 | 1 | 0 | NON_SYNONYMOU | MODERAT | MISSENSE | cTg/cCg | L308P | PLES_00821 | | | | | | | G | | | | | |
| NC_011770 | 99986 | GC | G | 74 | 0 | 1 | 0 | FRAME_SHIFT | HIGH | | ggc/ | G261 | PLES_00831 | G | | | | | | | | | | | |
| NC_011770 | 99795 | A | T | 83 | 0 | 2 | 0 | NON_SYNONYMOU | MODERAT | MISSENSE | gaA/gaT | E197D | PLES_00831 | T | T | | | | | | | | | | |
| NC_011770 | 99803 | A | T | 81 | 0 | 2 | 0 | NON_SYNONYMOU | MODERAT | MISSENSE | cAg/cTg | Q200L | PLES_00831 | | | | | | | | | | | T | T |
| NC_011770 | 99938 | G | A | 73 | 0 | 2 | 0 | NON_SYNONYMOU | MODERAT | MISSENSE | gGc/gAc | G245D | PLES_00831 | | | | | | | | | | | A | A |
| NC_011770 | 99943 | C | T | 73 | 0 | 2 | 0 | NON_SYNONYMOU | MODERAT | MISSENSE | Ccc/Tcc | P247S | PLES_00831 | | | | | | | | | | | T | T |
| NC_011770 | 99968 | C | T | 71 | 0 | 2 | 0 | NON_SYNONYMOU | MODERAT | MISSENSE | gCc/gTc | A255V | PLES_00831 | | | | | | | | | | | T | T |
| NC_011770 | 99970 | G | A | 73 | 0 | 2 | 0 | NON_SYNONYMOU | MODERAT | MISSENSE | Gcc/Acc | A256T | PLES_00831 | A | A | | | | | | | | | | |
| NC_011770 | 99983 | G | T | 75 | 0 | 2 | 0 | NON_SYNONYMOU | MODERAT | MISSENSE | gGg/gTg | G260V | PLES_00831 | | | | | | | | | | | T | T |
| NC_011770 | 100123 | G | A | 72 | 0 | 2 | 0 | NON_SYNONYMOU | MODERAT | MISSENSE | Gtg/Atg | V307M | PLES_00831 | | | | | | | | | A | A | | |
| NC_011770 | 103780 | G | T | 83 | 0 | 2 | 0 | NON_SYNONYMOU | MODERAT | MISSENSE | Gtc/Ttc | V232F | PLES_00871 | | | | | | | T | T | | | | |
| NC_011770 | 106375 | C | G | 78 | 0 | 2 | 0 | NON_SYNONYMOU | MODERAT | MISSENSE | tCg/tGg | S38W | PLES_00901 | | | | | | | G | G | | | | |
| NC_011770 | 108478 | C | A | 93 | 0 | 6 | 0 | NON_SYNONYMOU | MODERAT | MISSENSE | Cag/Aag | Q393K | PLES_00911 | | | A | A | A | A | | | A | A | | |
| NC_011770 | 108766 | T | G | 74 | 0 | 6 | 0 | NON_SYNONYMOU | MODERAT | MISSENSE | Tcc/Gcc | S489A | PLES_00911 | | | G | G | G | G | | | G | G | | |
| NC_011770 | 108799 | C | T | 77 | 0 | 2 | 0 | NON_SYNONYMOU | MODERAT | MISSENSE | Cgc/Tgc | R500C | PLES_00911 | | | | | | | | | | | T | T |
| NC_011770 | 108887 | G | A | 78 | 0 | 2 | 0 | NON_SYNONYMOU | MODERAT | MISSENSE | gGc/gAc | G529D | PLES_00911 | | | | | | | | | A | A | | |
| NC_011770 | 109097 | G | A | 73 | 0 | 4 | 0 | NON_SYNONYMOU | MODERAT | MISSENSE | cGg/cAg | R599Q | PLES_00911 | A | A | | | | | A | A | | | | |
| NC_011770 | 109189 | A | G | 71 | 0 | 1 | 0 | NON_SYNONYMOU | MODERAT | MISSENSE | Agc/Ggc | S630G | PLES_00911 | | | | | | | G | | | | | |
| NC_011770 | 109498 | G | T | 80 | 0 | 2 | 0 | NON_SYNONYMOU | MODERAT | MISSENSE | Gtg/Ttg | V733L | PLES_00911 | | | | | | | | | T | T | | |
| NC_011770 | 109742 | G | A | 77 | 0 | 2 | 0 | NON_SYNONYMOU | MODERAT | MISSENSE | cGc/cAc | R814H | PLES_00911 | | | | | | | | | | | A | A |
| NC_011770 | 109821 | C | G | 78 | 0 | 2 | 0 | NON_SYNONYMOU | MODERAT | MISSENSE | gaC/gaG | D840E | PLES_00911 | | | | | | | | | G | G | | |
| NC_011770 | 110246 | G | A | 81 | 0 | 2 | 0 | NON_SYNONYMOU | MODERAT | MISSENSE | Ggc/Agc | G64S | PLES_00921 | A | A | | | | | | | | | | |
| NC_011770 | 110426 | G | A | 85 | 0 | 2 | 0 | NON_SYNONYMOU | MODERAT | MISSENSE | Gac/Aac | D124N | PLES_00921 | | | | | | | A | A | | | | |
| NC_011770 | 110625 | G | C | 82 | 0 | 4 | 0 | NON_SYNONYMOU | MODERAT | MISSENSE | gGt/gCt | G190A | PLES_00921 | | | C | C | C | C | | | | | | |
| NC_011770 | 111983 | G | A | 92 | 0 | 2 | 0 | NON_SYNONYMOU | MODERAT | MISSENSE | Ggc/Agc | G643S | PLES_00921 | | | | | | | A | A | | | | |
| NC_011770 | 112230 | C | T | 95 | 0 | 2 | 0 | NON_SYNONYMOU | MODERAT | MISSENSE | cGg/cAg | R53Q | PLES_00931 | T | T | | | | | | | | | | |
| NC_011770 | 112796 | G | A | 82 | 0 | 6 | 0 | NON_SYNONYMOU | MODERAT | MISSENSE | gCc/gTc | A294V | PLES_00941 | | | A | A | A | A | | | | | A | A |
| NC_011770 | 112886 | G | A | 71 | 0 | 2 | 0 | NON_SYNONYMOU | MODERAT | MISSENSE | gCg/gTg | A264V | PLES_00941 | | | | | | | | | A | A | | |
| NC_011770 | 114903 | A | G | 91 | 0 | 2 | 0 | NON_SYNONYMOU | MODERAT | MISSENSE | cAg/cGg | Q175R | PLES_00961 | | | | | | | | | G | G | | |
| NC_011770 | 114906 | G | A | 92 | 0 | 2 | 0 | NON_SYNONYMOU | MODERAT | MISSENSE | aGc/aAc | S176N | PLES_00961 | | | | | | | | | A | A | | |
| NC_011770 | 115351 | G | T | 84 | 0 | 2 | 0 | NON_SYNONYMOU | MODERAT | MISSENSE | gaG/gaT | E324D | PLES_00961 | | | | | | | | | T | T | | |
| NC_011770 | 116525 | A | G | 76 | 0 | 1 | 0 | NON_SYNONYMOU | MODERAT | MISSENSE | Acc/Gcc | T716A | PLES_00961 | | | | | | | G | | | | | |
| NC_011770 | 115654 | C | G | 67 | 0 | 1 | 0 | NON_SYNONYMOU | MODERAT | MISSENSE | atC/atG | I425M | PLES_00961 | | | | | | | | **G** | | | | |
| NC_011770 | 115612 | G | T | 64 | 0 | 1 | 0 | NON_SYNONYMOU | MODERAT | MISSENSE | atG/atT | M411I | PLES_00961 (vgr type VI secretion) | | | | | | | | **T** | | | | |
| NC_011770 | 117012 | G | A | 75 | 0 | 2 | 0 | NON_SYNONYMOU | MODERAT | MISSENSE | cGt/cAt | R127H | PLES_00971 | | | | | | | A | A | | | | |
| NC_011770 | 117022 | T | C | 74 | 0 | 2 | 0 | NON_SYNONYMOU | MODERAT | MISSENSE | aTc/aCc | I4T | PLES_00981 | | | | | | | C | C | | | | |
| NC_011770 | 117025 | T | C | 76 | 0 | 2 | 0 | NON_SYNONYMOU | MODERAT | MISSENSE | tTt/tCt | F5S | PLES_00981 | | | | | | | C | C | | | | |
| NC_011770 | 117246 | A | G | 69 | 0 | 2 | 0 | NON_SYNONYMOU | MODERAT | MISSENSE | Acg/Gcg | T79A | PLES_00981 | | | | | | | G | G | | | | |
| NC_011770 | 117285 | T | C | 67 | 0 | 2 | 0 | NON_SYNONYMOU | MODERAT | MISSENSE | Ttt/Ctt | F92L | PLES_00981 | | | | | | | C | C | | | | |
| NC_011770 | 117432 | A | G | 70 | 0 | 8 | 0 | NON_SYNONYMOU | MODERAT | MISSENSE | Acg/Gcg | T141A | PLES_00981 | | | G | G | G | G | G | G | | | G | G |
| NC_011770 | 117504 | A | C | 72 | 0 | 4 | 0 | NON_SYNONYMOU | MODERAT | MISSENSE | Agc/Cgc | S165R | PLES_00981 | | | | | | | C | C | C | C | | |
| NC_011770 | 118086 | A | G | 71 | 0 | 2 | 0 | NON_SYNONYMOU | MODERAT | MISSENSE | Atc/Gtc | I359V | PLES_00981 | | | | | | | G | G | | | | |
| NC_011770 | 118177 | T | C | 70 | 0 | 10 | 0 | NON_SYNONYMOU | MODERAT | MISSENSE | gTa/gCa | V389A | PLES_00981 | C | C | C | C | C | C | | | C | C | C | C |
| NC_011770 | 118218 | T | C | 73 | 0 | 4 | 0 | NON_SYNONYMOU | MODERAT | MISSENSE | gTc/gCc | V4A | PLES_00991 | | | | | | | C | C | | | C | C |
| NC_011770 | 118990 | C | G | 35 | 0 | 12 | 0 | NON_SYNONYMOU | MODERAT | MISSENSE | gaC/gaG | D261E | PLES_00991 | G | G | G | G | G | G | G | G | G | G | G | G |
| NC_011770 | 119012 | G | T | 37 | 0 | 8 | 0 | NON_SYNONYMOU | MODERAT | MISSENSE | Gcg/Tcg | A269S | PLES_00991 | T | T | | | | | T | T | T | T | T | T |
| NC_011770 | 119018 | G | A | 37 | 0 | 8 | 0 | NON_SYNONYMOU | MODERAT | MISSENSE | Gcc/Acc | A271T | PLES_00991 | A | A | | | | | A | A | A | A | A | A |
| NC_011770 | 119255 | C | G | 39 | 8 | 4 | 0 | NON_SYNONYMOU | MODERAT | MISSENSE | Cag/Gag | Q4E | PLES_01001 | . | . | G | G | G | G | . | . | . | . | . | . |
| NC_011770 | 119471 | C | T | 58 | 0 | 8 | 0 | NON_SYNONYMOU | MODERAT | MISSENSE | Cac/Tac | H76Y | PLES_01001 | T | T | | | | | T | T | T | T | T | T |
| NC_011770 | 119475 | T | A | 59 | 0 | 8 | 0 | NON_SYNONYMOU | MODERAT | MISSENSE | tTc/tAc | F77Y | PLES_01001 | A | A | | | | | A | A | A | A | A | A |
| NC_011770 | 119481 | A | C | 58 | 0 | 8 | 0 | NON_SYNONYMOU | MODERAT | MISSENSE | aAg/aCg | K79T | PLES_01001 | C | C | | | | | C | C | C | C | C | C |
| NC_011770 | 119482 | G | C | 59 | 0 | 8 | 0 | NON_SYNONYMOU | MODERAT | MISSENSE | aaG/aaC | K79N | PLES_01001 | C | C | | | | | C | C | C | C | C | C |
| NC_011770 | 119498 | G | C | 60 | 0 | 8 | 0 | NON_SYNONYMOU | MODERAT | MISSENSE | Gct/Cct | A85P | PLES_01001 | C | C | | | | | C | C | C | C | C | C |
| NC_011770 | 119525 | A | G | 61 | 0 | 8 | 0 | NON_SYNONYMOU | MODERAT | MISSENSE | Atc/Gtc | I94V | PLES_01001 | G | G | | | | | G | G | G | G | G | G |
| NC_011770 | 119539 | T | G | 55 | 2 | 6 | 0 | NON_SYNONYMOU | MODERAT | MISSENSE | aaT/aaG | N98K | PLES_01001 | . | G | | | | | G | G | G | . | G | G |
| NC_011770 | 119541 | C | T | 55 | 1 | 7 | 0 | NON_SYNONYMOU | MODERAT | MISSENSE | aCc/aTc | T99I | PLES_01001 | T | T | | | | | T | T | T | . | T | T |
| NC_011770 | 119542 | C | CAAG | 51 | 4 | 2 | 0 | CODON_INSERTIOI | MODERATE | | ggc/AAGgg | G100KG | PLES_01001 | . | . | | | | | CAAG | . | . | . | . | CAAG |
| NC_011770 | 119573 | G | A | 63 | 0 | 8 | 0 | NON_SYNONYMOU | MODERAT | MISSENSE | Gac/Aac | D110N | PLES_01001 | A | A | | | | | A | A | A | A | A | A |
| NC_011770 | 119582 | A | T | 62 | 0 | 8 | 0 | NON_SYNONYMOU | MODERAT | MISSENSE | Atc/Ttc | I113F | PLES_01001 | T | T | | | | | T | T | T | T | T | T |
| NC_011770 | 119588 | G | A | 63 | 0 | 8 | 0 | NON_SYNONYMOU | MODERAT | MISSENSE | Ggc/Agc | G115S | PLES_01001 | A | A | | | | | A | A | A | A | A | A |
| NC_011770 | 119591 | C | A | 63 | 0 | 8 | 0 | NON_SYNONYMOU | MODERAT | MISSENSE | Ctg/Atg | L116M | PLES_01001 | A | A | | | | | A | A | A | A | A | A |

| | | | | | | | | | | | | | | | | | | | | | | | | | |
|---|---|---|---|---|---|---|---|---|---|---|---|---|---|---|---|---|---|---|---|---|---|---|---|---|---|
| NC_011770 | 119592 | T | A | 62 | 0 | 8 | 0 | NON_SYNONYMOU | MODERAT | MISSENSE | cTg/cAg | L116Q | PLES_01001 | A | A | | | | | A | A | A | A | A | A |
| NC_011770 | 119921 | G | A | 48 | 8 | 4 | 0 | NON_SYNONYMOU | MODERAT | MISSENSE | Gtc/Atc | V226I | PLES_01001 | . | . | A | A | A | A | . | . | . | . | . | . |
| NC_011770 | 119532 | GTAC | G | 64 | 0 | 5 | 0 | CODON_CHANGE_ | MODERATE | | agtacc/agc | ST96S | PLES_01001 | G | G | | | | | | | | G | G | G |
| NC_011770 | 121662 | G | A | 54 | 0 | 4 | 0 | NON_SYNONYMOU | MODERAT | MISSENSE | Gcg/Acg | A180T | PLES_01021 | | | A | A | A | A | | | | | | |
| NC_011770 | 122015 | C | A | 72 | 0 | 2 | 0 | NON_SYNONYMOU | MODERAT | MISSENSE | gaC/gaA | D297E | PLES_01021 | | | | | | | A | A | | | | |
| NC_011770 | 122044 | C | T | 70 | 0 | 8 | 0 | NON_SYNONYMOU | MODERAT | MISSENSE | gCc/gTc | A307V | PLES_01021 | T | T | T | T | T | T | T | T | | | | |
| NC_011770 | 122090 | A | C | 69 | 0 | 8 | 0 | NON_SYNONYMOU | MODERAT | MISSENSE | gaA/gaC | E322D | PLES_01021 | C | C | C | C | C | C | C | C | | | | |
| NC_011770 | 122091 | G | A | 68 | 0 | 8 | 0 | NON_SYNONYMOU | MODERAT | MISSENSE | Gac/Aac | D323N | PLES_01021 | A | A | A | A | A | A | A | A | | | | |
| NC_011770 | 122122 | G | C | 75 | 0 | 2 | 0 | NON_SYNONYMOU | MODERAT | MISSENSE | aGc/aCc | S333T | PLES_01021 | | | | | | | | | C | C | | |
| NC_011770 | 122132 | G | C | 73 | 0 | 2 | 0 | NON_SYNONYMOU | MODERAT | MISSENSE | gaG/gaC | E336D | PLES_01021 | | | | | | | | | C | C | | |
| NC_011770 | 122137 | A | C | 70 | 0 | 2 | 0 | NON_SYNONYMOU | MODERAT | MISSENSE | gAg/gCg | E338A | PLES_01021 | | | | | | | | | C | C | | |
| NC_011770 | 122145 | T | A | 70 | 0 | 2 | 0 | NON_SYNONYMOU | MODERAT | MISSENSE | Tcc/Acc | S341T | PLES_01021 | | | | | | | | | A | A | | |
| NC_011770 | 122163 | A | C | 72 | 0 | 2 | 0 | NON_SYNONYMOU | MODERAT | MISSENSE | Aac/Cac | N347H | PLES_01021 | | | | | | | | | C | C | | |
| NC_011770 | 122224 | G | A | 68 | 0 | 4 | 0 | NON_SYNONYMOU | MODERAT | MISSENSE | cGc/cAc | R367H | PLES_01021 | | | | | | | A | A | A | A | | |
| NC_011770 | 123796 | T | C | 80 | 0 | 12 | 0 | NON_SYNONYMOU | MODERAT | MISSENSE | cTg/cCg | L42P | PLES_01041 | C | C | C | C | C | C | C | C | C | C | C | C |
| NC_011770 | 125413 | G | C | 81 | 0 | 2 | 0 | NON_SYNONYMOU | MODERAT | MISSENSE | Ggg/Cgg | G12R | PLES_01051 | | | | | | | C | C | | | | |
| NC_011770 | 125416 | T | C | 79 | 0 | 10 | 0 | NON_SYNONYMOU | MODERAT | MISSENSE | Ttc/Ctc | F13L | PLES_01051 | C | C | C | C | C | C | C | C | C | C | | |
| NC_011770 | 125449 | C | T | 89 | 0 | 6 | 0 | NON_SYNONYMOU | MODERAT | MISSENSE | Cac/Tac | H24Y | PLES_01051 | | | T | T | T | T | | | T | T | | |
| NC_011770 | 125464 | C | T | 87 | 0 | 8 | 0 | NON_SYNONYMOU | MODERAT | MISSENSE | Cgg/Tgg | R29W | PLES_01051 | T | T | T | T | T | T | | | T | T | | |
| NC_011770 | 125473 | C | T | 85 | 0 | 8 | 0 | NON_SYNONYMOU | MODERAT | MISSENSE | Cgc/Tgc | R32C | PLES_01051 | T | T | T | T | T | T | | | T | T | | |
| NC_011770 | 125485 | A | G | 84 | 0 | 10 | 0 | NON_SYNONYMOU | MODERAT | MISSENSE | Atg/Gtg | M36V | PLES_01051 | G | G | G | G | G | G | G | G | G | G | | |
| NC_011770 | 125539 | T | C | 89 | 0 | 2 | 0 | NON_SYNONYMOU | MODERAT | MISSENSE | Ttc/Ctc | F54L | PLES_01051 | | | | | | | C | C | | | | |
| NC_011770 | 125542 | G | A | 90 | 0 | 2 | 0 | NON_SYNONYMOU | MODERAT | MISSENSE | Gtc/Atc | V55I | PLES_01051 | | | | | | | A | A | | | | |
| NC_011770 | 125722 | G | A | 90 | 0 | 2 | 0 | NON_SYNONYMOU | MODERAT | MISSENSE | Gag/Aag | E115K | PLES_01051 | A | A | | | | | | | | | | |
| NC_011770 | 125872 | G | T | 82 | 0 | 2 | 0 | NON_SYNONYMOU | MODERAT | MISSENSE | Gtc/Ttc | V165F | PLES_01051 | | | | | | | T | T | | | | |
| NC_011770 | 130506 | G | A | 96 | 0 | 4 | 0 | NON_SYNONYMOU | MODERAT | MISSENSE | gCg/gTg | A50V | PLES_01101 | A | A | | | | | A | A | | | | |
| NC_011770 | 130958 | C | G | 71 | 0 | 6 | 0 | NON_SYNONYMOU | MODERAT | MISSENSE | Cag/Gag | Q97E | PLES_01111 | G | G | | | | | | | G | G | G | G |
| NC_011770 | 131109 | T | C | 70 | 0 | 2 | 0 | NON_SYNONYMOU | MODERAT | MISSENSE | cTg/cCg | L147P | PLES_01111 | | | | | | | | | | | C | C |
| NC_011770 | 131288 | G | C | 78 | 0 | 10 | 0 | NON_SYNONYMOU | MODERAT | MISSENSE | Gcg/Ccg | A207P | PLES_01111 | C | C | C | C | C | C | | | C | C | C | C |
| NC_011770 | 131461 | C | T | 92 | 0 | 2 | 0 | NON_SYNONYMOU | MODERAT | MISSENSE | cCt/cTt | P8L | PLES_01121 | T | T | | | | | | | | | | |
| NC_011770 | 131560 | A | G | 86 | 0 | 4 | 0 | NON_SYNONYMOU | MODERAT | MISSENSE | aAt/aGt | N41S | PLES_01121 | | | G | G | G | G | | | | | | |
| NC_011770 | 131866 | C | T | 88 | 0 | 6 | 0 | NON_SYNONYMOU | MODERAT | MISSENSE | gCc/gTc | A143V | PLES_01121 | | | T | T | T | T | T | T | | | | |
| NC_011770 | 131878 | A | C | 83 | 0 | 2 | 0 | NON_SYNONYMOU | MODERAT | MISSENSE | aAg/aCg | K147T | PLES_01121 | C | C | | | | | | | | | | |
| NC_011770 | 131902 | T | A | 79 | 0 | 2 | 0 | NON_SYNONYMOU | MODERAT | MISSENSE | cTg/cAg | L155Q | PLES_01121 | | | | | | | | | | | A | A |
| NC_011770 | 132121 | G | A | 82 | 0 | 4 | 0 | NON_SYNONYMOU | MODERAT | MISSENSE | Gtg/Atg | V14M | PLES_01131 | | | A | A | A | A | | | | | | |
| NC_011770 | 132649 | G | A | 73 | 0 | 6 | 0 | NON_SYNONYMOU | MODERAT | MISSENSE | Gcc/Acc | A190T | PLES_01131 | | | A | A | A | A | | | | | A | A |
| NC_011770 | 133067 | C | A | 72 | 0 | 2 | 0 | NON_SYNONYMOU | MODERAT | MISSENSE | cCg/cAg | P329Q | PLES_01131 | | | | | | | | | | | A | A |
| NC_011770 | 134036 | T | C | 96 | 0 | 2 | 0 | NON_SYNONYMOU | MODERAT | MISSENSE | Ttc/Ctc | F286L | PLES_01141 | | | | | | | | | C | C | | |
| NC_011770 | 134463 | A | G | 79 | 0 | 12 | 0 | NON_SYNONYMOU | MODERAT | MISSENSE | Atc/Gtc | I115V | PLES_01151 | G | G | G | G | G | G | G | G | G | G | G | G |
| NC_011770 | 135437 | G | C | 95 | 0 | 12 | 0 | NON_SYNONYMOU | MODERAT | MISSENSE | Gtc/Ctc | V20L | PLES_01171 | C | C | C | C | C | C | C | C | C | C | C | C |
| NC_011770 | 135650 | C | T | 94 | 0 | 4 | 0 | NON_SYNONYMOU | MODERAT | MISSENSE | Ccg/Tcg | P91S | PLES_01171 | | | T | T | T | T | | | | | | |
| NC_011770 | 137177 | G | T | 67 | 0 | 8 | 0 | NON_SYNONYMOU | MODERAT | MISSENSE | caG/caT | Q105H | PLES_01191 | | | T | T | T | T | | | T | T | T | T |
| NC_011770 | 138668 | A | C | 88 | 0 | 12 | 0 | NON_SYNONYMOU | MODERAT | MISSENSE | cAg/cCg | Q330P | PLES_01201 | C | C | C | C | C | C | C | C | C | C | C | C |
| NC_011770 | 138688 | A | G | 90 | 0 | 2 | 0 | NON_SYNONYMOU | MODERAT | MISSENSE | Atc/Gtc | I337V | PLES_01201 | G | G | | | | | | | | | | |
| NC_011770 | 140418 | C | T | 85 | 0 | 4 | 0 | NON_SYNONYMOU | MODERAT | MISSENSE | gCc/gTc | A185V | PLES_01221 | | | T | T | T | T | | | | | | |
| NC_011770 | 141337 | C | G | 65 | 0 | 2 | 0 | NON_SYNONYMOU | MODERAT | MISSENSE | Ggg/Cgg | G265R | PLES_01241 | | | | | | | | | | | G | G |
| NC_011770 | 141691 | T | G | 65 | 0 | 2 | 0 | NON_SYNONYMOU | MODERAT | MISSENSE | Aat/Cat | N147H | PLES_01241 | | | | | | | G | G | | | | |
| NC_011770 | 142039 | A | C | 74 | 0 | 2 | 0 | NON_SYNONYMOU | MODERAT | MISSENSE | Tcg/Gcg | S31A | PLES_01241 | | | | | | | | | | | C | C |
| NC_011770 | 142086 | C | T | 77 | 0 | 12 | 0 | NON_SYNONYMOU | MODERAT | MISSENSE | gGc/gAc | G15D | PLES_01241 | T | T | T | T | T | T | T | T | T | T | T | T |
| NC_011770 | 142448 | C | T | 84 | 0 | 2 | 0 | NON_SYNONYMOU | MODERAT | MISSENSE | aGg/aAg | R88K | PLES_01251 | T | T | | | | | | | | | | |
| NC_011770 | 142668 | C | T | 75 | 0 | 4 | 0 | NON_SYNONYMOU | MODERAT | MISSENSE | Gcc/Acc | A15T | PLES_01251 | T | T | | | | | T | T | | | | |
| NC_011770 | 142738 | G | A | 78 | 0 | 2 | 0 | NON_SYNONYMOU | MODERAT | MISSENSE | aCc/aTc | T66I | PLES_01261 | | | | | | | A | A | | | | |
| NC_011770 | 144204 | G | A | 89 | 0 | 4 | 0 | NON_SYNONYMOU | MODERAT | MISSENSE | cGg/cAg | R125Q | PLES_01281 | | | | | | | A | A | | | A | A |
| NC_011770 | 147768 | C | T | 88 | 0 | 4 | 0 | NON_SYNONYMOU | MODERAT | MISSENSE | aGg/aAg | R50K | PLES_01311 | | | T | T | T | T | | | | | | |
| NC_011770 | 147892 | T | C | 89 | 0 | 2 | 0 | NON_SYNONYMOU | MODERAT | MISSENSE | Aac/Gac | N9D | PLES_01311 | C | C | | | | | | | | | | |
| NC_011770 | 148795 | G | A | 94 | 0 | 2 | 0 | NON_SYNONYMOU | MODERAT | MISSENSE | aCc/aTc | T280I | PLES_01331 | | | | | | | A | A | | | | |
| NC_011770 | 150684 | G | A | 72 | 0 | 1 | 0 | SPLICE_SITE_REGIO | LOW | SILENT | tGa/tAa | *306 | PLES_01341 | | | | | | | | | | | | A |
| NC_011770 | 150174 | A | G | 83 | 0 | 4 | 0 | NON_SYNONYMOU | MODERAT | MISSENSE | cAg/cGg | Q136R | PLES_01341 | | | G | G | G | G | | | | | | |
| NC_011770 | 150803 | C | T | 77 | 0 | 2 | 0 | NON_SYNONYMOU | MODERAT | MISSENSE | gGg/gAg | G456E | PLES_01351 | T | T | | | | | | | | | | |
| NC_011770 | 150809 | G | A | 77 | 0 | 2 | 0 | NON_SYNONYMOU | MODERAT | MISSENSE | gCg/gTg | A454V | PLES_01351 | | | | | | | | | A | A | | |
| NC_011770 | 150864 | C | G | 78 | 0 | 12 | 0 | NON_SYNONYMOU | MODERAT | MISSENSE | Gtg/Ctg | V436L | PLES_01351 | G | G | G | G | G | G | G | G | G | G | G | G |
| NC_011770 | 150915 | T | G | 77 | 0 | 12 | 0 | NON_SYNONYMOU | MODERAT | MISSENSE | Agc/Cgc | S419R | PLES_01351 | G | G | G | G | G | G | G | G | G | G | G | G |

| | | | | | | | | | | | | | | | | | | | | | | | | | |
|---|---|---|---|---|---|---|---|---|---|---|---|---|---|---|---|---|---|---|---|---|---|---|---|---|---|
| NC_011770 | 150990 | C | T | 81 | 0 | 2 | 0 | NON_SYNONYMOU | MODERAT | MISSENSE | Gtc/Atc | V394I | PLES_01351 | | | | | | | T | T | | | | |
| NC_011770 | 151959 | T | C | 79 | 0 | 2 | 0 | NON_SYNONYMOU | MODERAT | MISSENSE | Acc/Gcc | T71A | PLES_01351 | | | | | | | | | | | C | C |
| NC_011770 | 152144 | T | G | 89 | 0 | 2 | 0 | NON_SYNONYMOU | MODERAT | MISSENSE | aAc/aCc | N9T | PLES_01351 | | | | | | | | | | | G | G |
| NC_011770 | 150824 | C | T | 79 | 0 | 1 | 0 | NON_SYNONYMOU | MODERAT | MISSENSE | gGg/gAg | G449E | PLES_01351 | | | | | | | | | | | T | |
| NC_011770 | 152697 | T | G | 77 | 0 | 4 | 0 | STOP_LOST+SPLICE | HIGH | MISSENSE | Taa/Gaa | *47E | PLES_01361 | | | G | G | G | G | | | | | | |
| NC_011770 | 152637 | G | T | 72 | 0 | 4 | 0 | NON_SYNONYMOU | MODERAT | MISSENSE | Gat/Tat | D27Y | PLES_01361 | | | T | T | T | T | | | | | | |
| NC_011770 | 152686 | A | T | 77 | 0 | 2 | 0 | NON_SYNONYMOU | MODERAT | MISSENSE | cAt/cTt | H43L | PLES_01361 | | | | | | | | | | | T | T |
| NC_011770 | 152689 | A | C | 77 | 0 | 2 | 0 | NON_SYNONYMOU | MODERAT | MISSENSE | cAt/cCt | H44P | PLES_01361 | | | | | | | | | | | C | C |
| NC_011770 | 152690 | T | G | 78 | 0 | 4 | 0 | NON_SYNONYMOU | MODERAT | MISSENSE | caT/caG | H44Q | PLES_01361 | | | G | G | G | G | | | | | | |
| NC_011770 | 153292 | T | A | 94 | 0 | 2 | 0 | NON_SYNONYMOU | MODERAT | MISSENSE | Tcc/Acc | S5T | PLES_01371 | | | | | | | | | | | A | A |
| NC_011770 | 153305 | C | T | 89 | 0 | 2 | 0 | NON_SYNONYMOU | MODERAT | MISSENSE | cCg/cTg | P9L | PLES_01371 | T | T | | | | | | | | | | |
| NC_011770 | 153481 | A | G | 84 | 0 | 4 | 0 | NON_SYNONYMOU | MODERAT | MISSENSE | Atc/Gtc | I68V | PLES_01371 | | | G | G | G | G | | | | | | |
| NC_011770 | 153559 | C | T | 92 | 0 | 1 | 0 | NON_SYNONYMOU | MODERAT | MISSENSE | Cat/Tat | H94Y | PLES_01371 | | | | | | | T | | | | | |
| NC_011770 | 154040 | C | T | 87 | 0 | 4 | 0 | NON_SYNONYMOU | MODERAT | MISSENSE | gCt/gTt | A254V | PLES_01371 | T | T | | | | | T | T | | | | |
| NC_011770 | 154114 | A | G | 80 | 0 | 6 | 0 | NON_SYNONYMOU | MODERAT | MISSENSE | Acc/Gcc | T279A | PLES_01371 | G | G | | | | | G | G | | | G | G |
| NC_011770 | 154256 | C | G | 80 | 0 | 12 | 0 | NON_SYNONYMOU | MODERAT | MISSENSE | gCc/gGc | A326G | PLES_01371 | G | G | G | G | G | G | G | G | G | G | G | G |
| NC_011770 | 154474 | G | A | 75 | 0 | 10 | 0 | NON_SYNONYMOU | MODERAT | MISSENSE | Gcc/Acc | A399T | PLES_01371 | A | A | A | A | A | A | | | A | A | A | A |
| NC_011770 | 154861 | C | T | 86 | 0 | 2 | 0 | NON_SYNONYMOU | MODERAT | MISSENSE | tCc/tTc | S4F | PLES_01381 | | | | | | | | | | | T | T |
| NC_011770 | 155298 | A | G | 81 | 0 | 1 | 0 | NON_SYNONYMOU | MODERAT | MISSENSE | Acc/Gcc | T150A | PLES_01381 | | | | | | | G | | | | | |
| NC_011770 | 155445 | C | A | 92 | 0 | 2 | 0 | NON_SYNONYMOU | MODERAT | MISSENSE | Cac/Aac | H199N | PLES_01381 | | | | | | | A | A | | | | |
| NC_011770 | 155826 | A | T | 90 | 0 | 4 | 0 | NON_SYNONYMOU | MODERAT | MISSENSE | Acc/Tcc | T326S | PLES_01381 | T | T | | | | | | | | | T | T |
| NC_011770 | 156163 | C | T | 82 | 0 | 12 | 0 | NON_SYNONYMOU | MODERAT | MISSENSE | gCg/gTg | A65V | PLES_01391 | T | T | T | T | T | T | T | T | T | T | T | T |
| NC_011770 | 159480 | C | G | 71 | 0 | 4 | 0 | NON_SYNONYMOU | MODERAT | MISSENSE | gaG/gaC | E277D | PLES_01421 | G | G | | | | | | | | | G | G |
| NC_011770 | 161075 | A | C | 84 | 0 | 4 | 0 | NON_SYNONYMOU | MODERAT | MISSENSE | gAa/gCa | E103A | PLES_01431 | | | | | | | C | C | C | C | | |
| NC_011770 | 161428 | C | G | 84 | 0 | 2 | 0 | NON_SYNONYMOU | MODERAT | MISSENSE | Ctg/Gtg | L221V | PLES_01431 | | | | | | | G | G | | | | |
| NC_011770 | 162023 | C | T | 54 | 0 | 2 | 0 | NON_SYNONYMOU | MODERAT | MISSENSE | gCc/gTc | A419V | PLES_01431 | | | | | | | | | T | T | | |
| NC_011770 | 162089 | A | C | 51 | 0 | 4 | 0 | NON_SYNONYMOU | MODERAT | MISSENSE | gAg/gCg | E441A | PLES_01431 | C | C | | | | | | | C | C | | |
| NC_011770 | 162090 | G | T | 51 | 0 | 4 | 0 | NON_SYNONYMOU | MODERAT | MISSENSE | gaG/gaT | E441D | PLES_01431 | T | T | | | | | | | T | T | | |
| NC_011770 | 163364 | G | T | 60 | 0 | 6 | 0 | NON_SYNONYMOU | MODERAT | MISSENSE | Cta/Ata | L190I | PLES_01451 | | | T | T | T | T | | | | | T | T |
| NC_011770 | 163366 | G | C | 61 | 0 | 6 | 0 | NON_SYNONYMOU | MODERAT | MISSENSE | aCg/aGg | T189R | PLES_01451 | | | C | C | C | C | | | | | C | C |
| NC_011770 | 163369 | G | C | 61 | 0 | 6 | 0 | NON_SYNONYMOU | MODERAT | MISSENSE | gCa/gGa | A188G | PLES_01451 | | | C | C | C | C | | | | | C | C |
| NC_011770 | 163379 | C | A | 66 | 0 | 4 | 0 | NON_SYNONYMOU | MODERAT | MISSENSE | Gcg/Tcg | A185S | PLES_01451 | | | A | A | A | A | | | | | | |
| NC_011770 | 163466 | T | C | 67 | 0 | 12 | 0 | NON_SYNONYMOU | MODERAT | MISSENSE | Acc/Gcc | T156A | PLES_01451 | C | C | C | C | C | C | C | C | C | C | C | C |
| NC_011770 | 163917 | C | G | 77 | 0 | 2 | 0 | NON_SYNONYMOU | MODERAT | MISSENSE | aaG/aaC | K5N | PLES_01451 | | | | | | | | | | | G | G |
| NC_011770 | 163927 | T | C | 77 | 0 | 2 | 0 | NON_SYNONYMOU | MODERAT | MISSENSE | aAc/aGc | N2S | PLES_01451 | | | | | | | | | | | C | C |
| NC_011770 | 164195 | T | C | 76 | 0 | 2 | 0 | NON_SYNONYMOU | MODERAT | MISSENSE | cAc/cGc | H125R | PLES_01461 | | | | | | | | | | | C | C |
| NC_011770 | 164199 | C | T | 78 | 0 | 1 | 0 | NON_SYNONYMOU | MODERAT | MISSENSE | Gag/Aag | E124K | PLES_01461 | | | | | | | T | | | | | |
| NC_011770 | 164238 | C | T | 82 | 0 | 6 | 0 | NON_SYNONYMOU | MODERAT | MISSENSE | Gcc/Acc | A111T | PLES_01461 | T | T | T | T | T | T | | | | | | |
| NC_011770 | 164472 | C | G | 82 | 0 | 2 | 0 | NON_SYNONYMOU | MODERAT | MISSENSE | Gtc/Ctc | V33L | PLES_01461 | | | | | | | | | | | G | G |
| NC_011770 | 164504 | G | A | 81 | 0 | 2 | 0 | NON_SYNONYMOU | MODERAT | MISSENSE | aCc/aTc | T22I | PLES_01461 | | | | | | | | | | | A | A |
| NC_011770 | 166025 | C | T | 89 | 0 | 4 | 0 | NON_SYNONYMOU | MODERAT | MISSENSE | Cgt/Tgt | R17C | PLES_01481 | | | T | T | T | T | | | | | | |
| NC_011770 | 166529 | T | G | 84 | 0 | 2 | 0 | NON_SYNONYMOU | MODERAT | MISSENSE | Tcg/Gcg | S185A | PLES_01481 | | | | | | | | | | | G | G |
| NC_011770 | 167060 | A | G | 96 | 0 | 2 | 0 | NON_SYNONYMOU | MODERAT | MISSENSE | aAc/aGc | N6S | PLES_01491 | | | | | | | G | G | | | | |
| NC_011770 | 167717 | A | G | 80 | 0 | 1 | 0 | NON_SYNONYMOU | MODERAT | MISSENSE | gAc/gGc | D225G | PLES_01491 | | | | | | | G | | | | | |
| NC_011770 | 168768 | G | T | 79 | 0 | 2 | 0 | NON_SYNONYMOU | MODERAT | MISSENSE | gaG/gaT | E12D | PLES_01511 | | | | | | | | | | | T | T |
| NC_011770 | 168838 | C | T | 81 | 0 | 2 | 0 | NON_SYNONYMOU | MODERAT | MISSENSE | Cgg/Tgg | R36W | PLES_01511 | T | T | | | | | | | | | | |
| NC_011770 | 169264 | G | A | 66 | 0 | 6 | 0 | NON_SYNONYMOU | MODERAT | MISSENSE | Gcg/Acg | A178T | PLES_01511 | | | A | A | A | A | | | | | A | A |
| NC_011770 | 169346 | C | A | 65 | 0 | 2 | 0 | NON_SYNONYMOU | MODERAT | MISSENSE | gCg/gAg | A205E | PLES_01511 | | | | | | | | | | | A | A |
| NC_011770 | 169457 | A | G | 57 | 0 | 6 | 0 | NON_SYNONYMOU | MODERAT | MISSENSE | aAg/aGg | K242R | PLES_01511 | | | G | G | G | G | G | G | | | | |
| NC_011770 | 169637 | G | A | 58 | 0 | 2 | 0 | NON_SYNONYMOU | MODERAT | MISSENSE | aGc/aAc | S302N | PLES_01511 | | | | | A | A | | | | | | |
| NC_011770 | 169889 | C | T | 44 | 0 | 2 | 0 | NON_SYNONYMOU | MODERAT | MISSENSE | cCg/cTg | P5L | PLES_01521 | T | T | | | | | | | | | | |
| NC_011770 | 169952 | T | C | 52 | 0 | 6 | 0 | NON_SYNONYMOU | MODERAT | MISSENSE | gTg/gCg | V26A | PLES_01521 | | | C | C | C | C | | | C | C | | |
| NC_011770 | 170108 | T | A | 54 | 0 | 12 | 0 | NON_SYNONYMOU | MODERAT | MISSENSE | gTc/gAc | V78D | PLES_01521 | A | A | A | A | A | A | A | A | A | A | A | A |
| NC_011770 | 170305 | C | G | 64 | 0 | 2 | 0 | NON_SYNONYMOU | MODERAT | MISSENSE | Cta/Gta | L144V | PLES_01521 | | | | | | | | | | | G | G |
| NC_011770 | 170313 | A | T | 66 | 0 | 6 | 0 | NON_SYNONYMOU | MODERAT | MISSENSE | gaA/gaT | E146D | PLES_01521 | | | T | T | T | T | | | T | T | | |
| NC_011770 | 170681 | G | A | 80 | 0 | 4 | 0 | NON_SYNONYMOU | MODERAT | MISSENSE | cGc/cAc | R269H | PLES_01521 | A | A | | | | | A | A | | | | |
| NC_011770 | 170980 | A | G | 79 | 0 | 12 | 0 | NON_SYNONYMOU | MODERAT | MISSENSE | Agc/Ggc | S369G | PLES_01521 | G | G | G | G | G | G | G | G | G | G | G | G |
| NC_011770 | 171179 | G | A | 85 | 0 | 2 | 0 | NON_SYNONYMOU | MODERAT | MISSENSE | gGc/gAc | G435D | PLES_01521 | | | | | | | | | A | A | | |
| NC_011770 | 171442 | G | A | 74 | 0 | 1 | 0 | NON_SYNONYMOU | MODERAT | MISSENSE | Ggc/Agc | G523S | PLES_01521 | | | | | | | A | | | | | |
| NC_011770 | 171664 | G | C | 77 | 0 | 4 | 0 | NON_SYNONYMOU | MODERAT | MISSENSE | Gag/Cag | E597Q | PLES_01521 | | | C | C | C | C | | | | | | |
| NC_011770 | 172240 | G | C | 72 | 0 | 2 | 0 | NON_SYNONYMOU | MODERAT | MISSENSE | Gtg/Ctg | V789L | PLES_01521 | | | | | | | | | | | C | C |
| NC_011770 | 176210 | G | A | 83 | 0 | 6 | 0 | NON_SYNONYMOU | MODERAT | MISSENSE | Ggc/Agc | G26S | PLES_01571 | | | A | A | A | A | | | | | A | A |

| | | | | | | | | | | | | | | | | | | | | | | | | | |
|---|---|---|---|---|---|---|---|---|---|---|---|---|---|---|---|---|---|---|---|---|---|---|---|---|---|
| NC_011770 | 177252 | C | T | 80 | 0 | 1 | 0 | NON_SYNONYMOU | MODERAT | MISSENSE | gCc/gTc | A373V | PLES_01571 triA | | | | | | | | | | | | **T** |
| NC_011770 | 177320 | C | T | 79 | 0 | 2 | 0 | NON_SYNONYMOU | MODERAT | MISSENSE | gCc/gTc | A13V | PLES_01581 | | | | | | | | | T | T | | |
| NC_011770 | 177353 | A | C | 77 | 0 | 12 | 0 | NON_SYNONYMOU | MODERAT | MISSENSE | cAg/cCg | Q24P | PLES_01581 | C | C | C | C | C | C | C | C | C | C | C | C |
| NC_011770 | 178120 | G | A | 88 | 0 | 4 | 0 | NON_SYNONYMOU | MODERAT | MISSENSE | Gtg/Atg | V280M | PLES_01581 | | | A | A | A | A | | | | | | |
| NC_011770 | 177385 | A | G | 82 | 0 | 1 | 0 | NON_SYNONYMOU | MODERAT | MISSENSE | Acc/Gcc | T35A | PLES_01581 triB | | G | | | | | | | | | | |
| NC_011770 | 179368 | G | A | 89 | 0 | 4 | 0 | NON_SYNONYMOU | MODERAT | MISSENSE | cGg/cAg | R340Q | PLES_01591 | | | A | A | A | A | | | | | | |
| NC_011770 | 182669 | C | T | 77 | 0 | 2 | 0 | NON_SYNONYMOU | MODERAT | MISSENSE | gCt/gTt | A7V | PLES_01611 | | | | | | | | | | | T | T |
| NC_011770 | 183176 | C | T | 74 | 0 | 2 | 0 | NON_SYNONYMOU | MODERAT | MISSENSE | Cca/Tca | P21S | PLES_01621 | | | | | | | T | T | | | | |
| NC_011770 | 185529 | C | T | 46 | 0 | 2 | 0 | NON_SYNONYMOU | MODERAT | MISSENSE | cGc/cAc | R19H | PLES_01641 | T | T | | | | | | | | | | |
| NC_011770 | 186035 | G | C | 70 | 0 | 4 | 0 | NON_SYNONYMOU | MODERAT | MISSENSE | gGc/gCc | G125A | PLES_01651 | | | C | C | C | C | | | | | | |
| NC_011770 | 188121 | T | C | 67 | 0 | 10 | 0 | NON_SYNONYMOU | MODERAT | MISSENSE | Tcg/Ccg | S58P | PLES_01661 | C | C | C | C | C | C | C | C | | | C | C |
| NC_011770 | 190129 | A | G | 45 | 0 | 2 | 0 | NON_SYNONYMOU | MODERAT | MISSENSE | Acc/Gcc | T368A | PLES_01671 | G | G | | | | | | | | | | |
| NC_011770 | 190186 | C | G | 51 | 0 | 4 | 0 | NON_SYNONYMOU | MODERAT | MISSENSE | Ccg/Gcg | P387A | PLES_01671 | G | G | | | | | | | G | G | | |
| NC_011770 | 191176 | C | T | 76 | 0 | 2 | 0 | NON_SYNONYMOU | MODERAT | MISSENSE | aCc/aTc | T217I | PLES_01681 | T | T | | | | | | | | | | |
| NC_011770 | 191621 | A | C | 80 | 0 | 12 | 0 | NON_SYNONYMOU | MODERAT | MISSENSE | Atc/Ctc | I142L | PLES_01691 | C | C | C | C | C | C | C | C | C | C | C | C |
| NC_011770 | 191675 | G | A | 76 | 0 | 2 | 0 | NON_SYNONYMOU | MODERAT | MISSENSE | Gcc/Acc | A160T | PLES_01691 | | | | | | | | | | | A | A |
| NC_011770 | 191769 | T | C | 71 | 0 | 6 | 0 | NON_SYNONYMOU | MODERAT | MISSENSE | cTg/cCg | L191P | PLES_01691 | | | | | | | C | C | C | C | C | C |
| NC_011770 | 192855 | G | T | 90 | 0 | 4 | 0 | NON_SYNONYMOU | MODERAT | MISSENSE | gCc/gAc | A52D | PLES_01711 | | | | | | | T | T | | | T | T |
| NC_011770 | 194084 | C | T | 86 | 0 | 2 | 0 | NON_SYNONYMOU | MODERAT | MISSENSE | Gcg/Acg | A499T | PLES_01731 | | | | | | | | | T | T | | |
| NC_011770 | 194186 | C | T | 87 | 0 | 6 | 0 | NON_SYNONYMOU | MODERAT | MISSENSE | Gcc/Acc | A465T | PLES_01731 | T | T | | | | | T | T | T | T | | |
| NC_011770 | 194738 | G | T | 88 | 0 | 4 | 0 | NON_SYNONYMOU | MODERAT | MISSENSE | Cag/Aag | Q281K | PLES_01731 | | | T | T | T | T | | | | | | |
| NC_011770 | 194804 | C | T | 89 | 0 | 2 | 0 | NON_SYNONYMOU | MODERAT | MISSENSE | Gcc/Acc | A259T | PLES_01731 | T | T | | | | | | | | | | |
| NC_011770 | 195157 | G | A | 78 | 0 | 4 | 0 | NON_SYNONYMOU | MODERAT | MISSENSE | cCg/cTg | P141L | PLES_01731 | | | A | A | A | A | | | | | | |
| NC_011770 | 195228 | C | G | 73 | 0 | 10 | 0 | NON_SYNONYMOU | MODERAT | MISSENSE | caG/caC | Q117H | PLES_01731 | G | G | G | G | G | G | G | G | | | G | G |
| NC_011770 | 196422 | T | C | 86 | 0 | 2 | 0 | NON_SYNONYMOU | MODERAT | MISSENSE | Atc/Gtc | I157V | PLES_01741 | | | | | | | | | C | C | | |
| NC_011770 | 196833 | C | T | 97 | 0 | 2 | 0 | NON_SYNONYMOU | MODERAT | MISSENSE | Gag/Aag | E20K | PLES_01741 | | | | | | | | | | | T | T |
| NC_011770 | 197131 | C | T | 82 | 0 | 1 | 0 | NON_SYNONYMOU | MODERAT | MISSENSE | Gtc/Atc | V128I | PLES_01751 | | | | | | | T | | | | | |
| NC_011770 | 197410 | C | T | 78 | 0 | 2 | 0 | NON_SYNONYMOU | MODERAT | MISSENSE | Ggc/Agc | G35S | PLES_01751 | | | | | | | T | T | | | | |
| NC_011770 | 197550 | A | G | 72 | 0 | 2 | 0 | NON_SYNONYMOU | MODERAT | MISSENSE | Tcc/Ccc | S271P | PLES_01761 | | | | | | | G | G | | | | |
| NC_011770 | 197879 | T | A | 70 | 0 | 2 | 0 | NON_SYNONYMOU | MODERAT | MISSENSE | cAg/cTg | Q161L | PLES_01761 | | | | | | | A | A | | | | |
| NC_011770 | 201425 | A | G | 87 | 0 | 1 | 0 | NON_SYNONYMOU | MODERAT | MISSENSE | cTg/cCg | L491P | PLES_01791 | | | | | | | G | | | | | |
| NC_011770 | 201555 | T | C | 76 | 0 | 6 | 0 | NON_SYNONYMOU | MODERAT | MISSENSE | Acg/Gcg | T448A | PLES_01791 | C | C | | | | | C | C | | | C | C |
| NC_011770 | 202206 | G | T | 59 | 0 | 6 | 0 | NON_SYNONYMOU | MODERAT | MISSENSE | Ccg/Acg | P231T | PLES_01791 | T | T | | | | | | | T | T | T | T |
| NC_011770 | 202290 | C | T | 57 | 0 | 2 | 0 | NON_SYNONYMOU | MODERAT | MISSENSE | Gcg/Acg | A203T | PLES_01791 | | | | | | | | | T | T | | |
| NC_011770 | 202304 | G | T | 60 | 0 | 4 | 0 | NON_SYNONYMOU | MODERAT | MISSENSE | cCg/cAg | P198Q | PLES_01791 | | | T | T | T | T | | | | | | |
| NC_011770 | 202572 | T | G | 71 | 0 | 6 | 0 | NON_SYNONYMOU | MODERAT | MISSENSE | Agt/Cgt | S109R | PLES_01791 | G | G | G | G | G | G | | | | | | |
| NC_011770 | 205175 | G | A | 82 | 0 | 2 | 0 | NON_SYNONYMOU | MODERAT | MISSENSE | Ccg/Tcg | P204S | PLES_01821 | | | | | | | | | | | A | A |
| NC_011770 | 208461 | T | G | 44 | 0 | 8 | 0 | NON_SYNONYMOU | MODERAT | MISSENSE | Atc/Ctc | I277L | PLES_01851 | | | G | G | G | G | | | G | G | G | G |
| NC_011770 | 208595 | C | T | 50 | 0 | 2 | 0 | NON_SYNONYMOU | MODERAT | MISSENSE | gGc/gAc | G232D | PLES_01851 | | | | | | | | | T | T | | |
| NC_011770 | 209277 | C | T | 80 | 0 | 12 | 0 | NON_SYNONYMOU | MODERAT | MISSENSE | Gcc/Acc | A5T | PLES_01851 | T | T | T | T | T | T | T | T | T | T | T | T |
| NC_011770 | 209308 | G | A | 82 | 0 | 2 | 0 | NON_SYNONYMOU | MODERAT | MISSENSE | gCc/gTc | A532V | PLES_01861 | | | | | | | | | | | A | A |
| NC_011770 | 209369 | G | A | 86 | 0 | 2 | 0 | NON_SYNONYMOU | MODERAT | MISSENSE | Ctc/Ttc | L512F | PLES_01861 | A | A | | | | | | | | | | |
| NC_011770 | 209849 | G | T | 69 | 0 | 4 | 0 | NON_SYNONYMOU | MODERAT | MISSENSE | Ctc/Atc | L352I | PLES_01861 | | | T | T | T | T | | | | | | |
| NC_011770 | 210623 | C | T | 62 | 0 | 4 | 0 | NON_SYNONYMOU | MODERAT | MISSENSE | Ggc/Agc | G94S | PLES_01861 | | | T | T | T | T | | | | | | |
| NC_011770 | 210397 | A | G | 69 | 0 | 1 | 0 | NON_SYNONYMOU | MODERAT | MISSENSE | gTg/gCg | V169A | PLES_01861 | | | | | | | | G | | | | |
| NC_011770 | 211388 | G | A | 71 | 0 | 2 | 0 | NON_SYNONYMOU | MODERAT | MISSENSE | Gcg/Acg | A74T | PLES_01871 | A | A | | | | | | | | | | |
| NC_011770 | 212852 | GC | G | 68 | 0 | 1 | 0 | FRAME_SHIFT | HIGH | | cac/ | H71 | PLES_01881 | | | | G | | | | | | | | |
| NC_011770 | 212789 | C | T | 72 | 0 | 2 | 0 | NON_SYNONYMOU | MODERAT | MISSENSE | Cgc/Tgc | R48C | PLES_01881 | | | | | | | | | | | T | T |
| NC_011770 | 212800 | C | G | 72 | 0 | 12 | 0 | NON_SYNONYMOU | MODERAT | MISSENSE | caC/caG | H51Q | PLES_01881 | G | G | G | G | G | G | G | G | G | G | G | G |
| NC_011770 | 212976 | C | T | 76 | 0 | 2 | 0 | NON_SYNONYMOU | MODERAT | MISSENSE | aCc/aTc | T110I | PLES_01881 | | | | | | | | | | | T | T |
| NC_011770 | 214533 | C | T | 69 | 0 | 2 | 0 | NON_SYNONYMOU | MODERAT | MISSENSE | aGc/aAc | S402N | PLES_01901 | T | T | | | | | | | | | | |
| NC_011770 | 214554 | C | T | 71 | 0 | 6 | 0 | NON_SYNONYMOU | MODERAT | MISSENSE | cGt/cAt | R395H | PLES_01901 | | | T | T | T | T | | | T | T | | |
| NC_011770 | 214915 | T | A | 78 | 2 | 2 | 0 | NON_SYNONYMOU | MODERAT | MISSENSE | Acc/Tcc | T275S | PLES_01901 | | | | | | | A | A | . | . | | |
| NC_011770 | 214957 | C | T | 77 | 2 | 4 | 0 | NON_SYNONYMOU | MODERAT | MISSENSE | Ggc/Agc | G261S | PLES_01901 | | | T | T | T | T | | | . | . | | |
| NC_011770 | 215095 | A | G | 80 | 2 | 10 | 0 | NON_SYNONYMOU | MODERAT | MISSENSE | Tcg/Ccg | S215P | PLES_01901 | G | G | G | G | G | G | G | G | . | . | G | G |
| NC_011770 | 215657 | A | C | 63 | 2 | 4 | 0 | NON_SYNONYMOU | MODERAT | MISSENSE | gaT/gaG | D27E | PLES_01901 | | | C | C | C | C | | | . | . | | |
| NC_011770 | 217105 | G | C | 64 | 0 | 2 | 0 | NON_SYNONYMOU | MODERAT | MISSENSE | Ctg/Gtg | L183V | PLES_01921 | | | | | | | | | C | C | | |
| NC_011770 | 217249 | C | A | 63 | 0 | 4 | 0 | NON_SYNONYMOU | MODERAT | MISSENSE | Gcc/Tcc | A135S | PLES_01921 | | | A | A | A | A | | | | | | |
| NC_011770 | 218007 | C | T | 60 | 0 | 4 | 0 | NON_SYNONYMOU | MODERAT | MISSENSE | Cac/Tac | H3Y | PLES_01931 | | | T | T | T | T | | | | | | |
| NC_011770 | 218041 | C | A | 58 | 0 | 2 | 0 | NON_SYNONYMOU | MODERAT | MISSENSE | cCc/cAc | P14H | PLES_01931 | | | | | | | | | A | A | | |
| NC_011770 | 218148 | A | G | 62 | 0 | 4 | 0 | NON_SYNONYMOU | MODERAT | MISSENSE | Agc/Ggc | S50G | PLES_01931 | | | G | G | G | G | | | | | | |
| NC_011770 | 218154 | C | G | 59 | 0 | 12 | 0 | NON_SYNONYMOU | MODERAT | MISSENSE | Ctc/Gtc | L52V | PLES_01931 | G | G | G | G | G | G | G | G | G | G | G | G |

| | | | | | | | | | | | | | | | | | | | | | | | | | |
|---|---|---|---|---|---|---|---|---|---|---|---|---|---|---|---|---|---|---|---|---|---|---|---|---|---|
| NC_011770 | 218155 | T | C | 60 | 0 | 4 | 0 | NON_SYNONYMOU | MODERAT | MISSENSE | cTc/cCc | L52P | PLES_01931 | | | C | C | C | C | | | | | | |
| NC_011770 | 218161 | A | C | 64 | 0 | 4 | 0 | NON_SYNONYMOU | MODERAT | MISSENSE | gAc/gCc | D54A | PLES_01931 | | | C | C | C | C | | | | | | |
| NC_011770 | 218577 | G | A | 82 | 0 | 4 | 0 | NON_SYNONYMOU | MODERAT | MISSENSE | Gcc/Acc | A193T | PLES_01931 | | | A | A | A | A | | | | | | |
| NC_011770 | 218614 | G | T | 81 | 0 | 4 | 0 | NON_SYNONYMOU | MODERAT | MISSENSE | gGc/gTc | G205V | PLES_01931 | | | T | T | T | T | | | | | | |
| NC_011770 | 218863 | G | C | 83 | 0 | 8 | 0 | NON_SYNONYMOU | MODERAT | MISSENSE | gGc/gCc | G288A | PLES_01931 | | | C | C | C | C | C | C | | | C | C |
| NC_011770 | 219364 | C | G | 84 | 0 | 4 | 0 | NON_SYNONYMOU | MODERAT | MISSENSE | aCc/aGc | T455S | PLES_01931 | | | G | G | G | G | | | | | | |
| NC_011770 | 220459 | A | G | 99 | 0 | 2 | 0 | NON_SYNONYMOU | MODERAT | MISSENSE | Atc/Gtc | I16V | PLES_01941 | G | G | | | | | | | | | | |
| NC_011770 | 228376 | A | C | 81 | 0 | 10 | 0 | NON_SYNONYMOU | MODERAT | MISSENSE | gaT/gaG | D64E | PLES_02011 | C | C | C | C | C | C | C | C | | | C | C |
| NC_011770 | 228909 | A | G | 91 | 0 | 1 | 0 | NON_SYNONYMOU | MODERAT | MISSENSE | Acc/Gcc | T43A | PLES_02021 | | | | | | | G | | | | | |
| NC_011770 | 229158 | A | T | 81 | 0 | 2 | 0 | NON_SYNONYMOU | MODERAT | MISSENSE | Agc/Tgc | S126C | PLES_02021 | | | | | | | | | | | T | T |
| NC_011770 | 229180 | A | T | 82 | 0 | 2 | 0 | NON_SYNONYMOU | MODERAT | MISSENSE | aAc/aTc | N133I | PLES_02021 | | | | | T | T | | | | | | |
| NC_011770 | 231664 | G | T | 78 | 0 | 2 | 0 | NON_SYNONYMOU | MODERAT | MISSENSE | Ggc/Tgc | G176C | PLES_02041 | T | T | | | | | | | | | | |
| NC_011770 | 231772 | G | A | 71 | 0 | 4 | 0 | NON_SYNONYMOU | MODERAT | MISSENSE | Gtc/Atc | V212I | PLES_02041 | | | | | | | A | A | | | A | A |
| NC_011770 | 231824 | T | C | 69 | 0 | 6 | 0 | NON_SYNONYMOU | MODERAT | MISSENSE | gTg/gCg | V229A | PLES_02041 | C | C | | | | | C | C | | | C | C |
| NC_011770 | 231673 | C | T | 75 | 0 | 1 | 0 | NON_SYNONYMOU | MODERAT | MISSENSE | Cgc/Tgc | R179C | PLES_02041 | | | | | | | | **T** | | | | |
| NC_011770 | 234264 | T | G | 81 | 0 | 2 | 0 | NON_SYNONYMOU | MODERAT | MISSENSE | tTg/tGg | L90W | PLES_02081 | | | | | G | G | | | | | | |
| NC_011770 | 234390 | C | G | 91 | 0 | 2 | 0 | NON_SYNONYMOU | MODERAT | MISSENSE | gCc/gGc | A132G | PLES_02081 | | | | | | | | | G | G | | |
| NC_011770 | 234523 | G | C | 90 | 0 | 2 | 0 | NON_SYNONYMOU | MODERAT | MISSENSE | caG/caC | Q176H | PLES_02081 | | | | | | | | | C | C | | |
| NC_011770 | 234668 | A | G | 84 | 0 | 6 | 0 | NON_SYNONYMOU | MODERAT | MISSENSE | Atc/Gtc | I225V | PLES_02081 | G | G | | | | | G | G | | | G | G |
| NC_011770 | 234387 | T | C | 87 | 0 | 1 | 0 | NON_SYNONYMOU | MODERAT | MISSENSE | cTg/cCg | L131P | PLES_02081 | C | | | | | | | | | | | |
| NC_011770 | 234839 | G | A | 79 | 0 | 2 | 0 | NON_SYNONYMOU | MODERAT | MISSENSE | cGc/cAc | R52H | PLES_02091 | | | | | | | | | A | A | | |
| NC_011770 | 236044 | C | G | 57 | 0 | 6 | 0 | NON_SYNONYMOU | MODERAT | MISSENSE | cCg/cGg | P124R | PLES_02101 | G | G | | | | | G | G | | | G | G |
| NC_011770 | 238822 | T | C | 64 | 0 | 12 | 0 | NON_SYNONYMOU | MODERAT | MISSENSE | Atc/Gtc | I260V | PLES_02141 | C | C | C | C | C | C | C | C | C | C | C | C |
| NC_011770 | 239697 | A | G | 70 | 0 | 2 | 0 | NON_SYNONYMOU | MODERAT | MISSENSE | Tcg/Ccg | S37P | PLES_02151 | | | | | | | G | G | | | | |
| NC_011770 | 240388 | G | A | 83 | 0 | 8 | 0 | NON_SYNONYMOU | MODERAT | MISSENSE | Gcc/Acc | A147T | PLES_02161 | | | A | A | A | A | A | A | A | A | | |
| NC_011770 | 240577 | A | G | 86 | 0 | 12 | 0 | NON_SYNONYMOU | MODERAT | MISSENSE | Atg/Gtg | M210V | PLES_02161 | G | G | G | G | G | G | G | G | G | G | G | G |
| NC_011770 | 240651 | G | T | 87 | 0 | 2 | 0 | NON_SYNONYMOU | MODERAT | MISSENSE | gaG/gaT | E234D | PLES_02161 | | | | | | | | | T | T | | |
| NC_011770 | 240704 | G | A | 91 | 0 | 10 | 0 | NON_SYNONYMOU | MODERAT | MISSENSE | cGg/cAg | R252Q | PLES_02161 | A | A | A | A | A | A | A | A | A | A | | |
| NC_011770 | 241112 | A | G | 98 | 0 | 2 | 0 | NON_SYNONYMOU | MODERAT | MISSENSE | gAc/gGc | D388G | PLES_02161 | | | | | | | | | G | G | | |
| NC_011770 | 241113 | C | A | 98 | 0 | 2 | 0 | NON_SYNONYMOU | MODERAT | MISSENSE | gaC/gaA | D388E | PLES_02161 | | | | | | | | | A | A | | |
| NC_011770 | 243113 | A | G | 73 | 0 | 1 | 0 | NON_SYNONYMOU | MODERAT | MISSENSE | aAc/aGc | N51S | PLES_02181 | | | | | | | G | | | | | |
| NC_011770 | 244139 | C | T | 76 | 0 | 2 | 0 | NON_SYNONYMOU | MODERAT | MISSENSE | gCg/gTg | A393V | PLES_02181 | T | T | | | | | | | | | | |
| NC_011770 | 244528 | G | T | 83 | 0 | 4 | 0 | NON_SYNONYMOU | MODERAT | MISSENSE | caG/caT | Q7H | PLES_02191 | | | T | T | T | T | | | | | | |
| NC_011770 | 245786 | G | T | 75 | 0 | 12 | 0 | NON_SYNONYMOU | MODERAT | MISSENSE | Ctc/Atc | L247I | PLES_02201 | T | T | T | T | T | T | T | T | T | T | T | T |
| NC_011770 | 246158 | G | T | 92 | 0 | 2 | 0 | NON_SYNONYMOU | MODERAT | MISSENSE | Cgc/Agc | R123S | PLES_02201 | T | T | | | | | | | | | | |
| NC_011770 | 246239 | C | T | 90 | 0 | 1 | 0 | NON_SYNONYMOU | MODERAT | MISSENSE | Gaa/Aaa | E96K | PLES_02201 | | | | | | | | | | | T | |
| NC_011770 | 246007 | C | T | 88 | 0 | 1 | 0 | NON_SYNONYMOU | MODERAT | MISSENSE | cGc/cAc | R173H | PLES_02201 ergB | | T | | | | | | | | | | |
| NC_011770 | 246923 | G | T | 70 | 0 | 2 | 0 | NON_SYNONYMOU | MODERAT | MISSENSE | gaC/gaA | D148E | PLES_02211 | | | | | | | | | T | T | | |
| NC_011770 | 246948 | T | C | 62 | 0 | 1 | 0 | NON_SYNONYMOU | MODERAT | MISSENSE | cAc/cGc | H140R | PLES_02211 | | | | | | | | **C** | | | | |
| NC_011770 | 247583 | G | C | 78 | 0 | 8 | 0 | NON_SYNONYMOU | MODERAT | MISSENSE | gaG/gaC | E21D | PLES_02221 | C | C | C | C | C | C | | | C | C | | |
| NC_011770 | 249028 | C | T | 74 | 0 | 2 | 0 | NON_SYNONYMOU | MODERAT | MISSENSE | gCc/gTc | A273V | PLES_02231 | T | T | | | | | | | | | | |
| NC_011770 | 256219 | C | G | 91 | 0 | 12 | 0 | NON_SYNONYMOU | MODERAT | MISSENSE | Cag/Gag | Q312E | PLES_02301 | G | G | G | G | G | G | G | G | G | G | G | G |
| NC_011770 | 256641 | G | A | 86 | 0 | 12 | 0 | NON_SYNONYMOU | MODERAT | MISSENSE | gGc/gAc | G75D | PLES_02311 | A | A | A | A | A | A | A | A | A | A | A | A |
| NC_011770 | 257004 | A | G | 85 | 0 | 2 | 0 | NON_SYNONYMOU | MODERAT | MISSENSE | cAt/cGt | H196R | PLES_02311 | | | | | | | | | | | G | G |
| NC_011770 | 257091 | T | C | 82 | 0 | 2 | 0 | NON_SYNONYMOU | MODERAT | MISSENSE | gTc/gCc | V225A | PLES_02311 | | | | | | | | | | | C | C |
| NC_011770 | 260177 | C | T | 84 | 0 | 2 | 0 | NON_SYNONYMOU | MODERAT | MISSENSE | Ccc/Tcc | P211S | PLES_02331 | | | | | | | | | T | T | | |
| NC_011770 | 260603 | A | G | 78 | 0 | 12 | 0 | NON_SYNONYMOU | MODERAT | MISSENSE | Acc/Gcc | T12A | PLES_02341 | G | G | G | G | G | G | G | G | G | G | G | G |
| NC_011770 | 261122 | T | C | 79 | 0 | 12 | 0 | NON_SYNONYMOU | MODERAT | MISSENSE | Tac/Cac | Y185H | PLES_02341 | C | C | C | C | C | C | C | C | C | C | C | C |
| NC_011770 | 261197 | G | A | 77 | 0 | 6 | 0 | NON_SYNONYMOU | MODERAT | MISSENSE | Gtc/Atc | V210I | PLES_02341 | | | | | | | A | A | A | A | A | A |
| NC_011770 | 261435 | T | A | 68 | 0 | 8 | 0 | NON_SYNONYMOU | MODERAT | MISSENSE | gTg/gAg | V289E | PLES_02341 | | | A | A | A | A | | | A | A | A | A |
| NC_011770 | 261449 | G | A | 66 | 0 | 10 | 0 | NON_SYNONYMOU | MODERAT | MISSENSE | Gca/Aca | A294T | PLES_02341 | | | A | A | A | A | A | A | A | A | A | A |
| NC_011770 | 261695 | C | A | 79 | 0 | 4 | 0 | NON_SYNONYMOU | MODERAT | MISSENSE | gaC/gaA | D20E | PLES_02351 | | | A | A | A | A | | | | | | |
| NC_011770 | 261975 | A | C | 76 | 0 | 2 | 0 | NON_SYNONYMOU | MODERAT | MISSENSE | Agc/Cgc | S114R | PLES_02351 | | | | | | | | | | | C | C |
| NC_011770 | 262311 | G | C | 74 | 0 | 4 | 0 | NON_SYNONYMOU | MODERAT | MISSENSE | Gcg/Ccg | A226P | PLES_02351 | | | C | C | C | C | | | | | | |
| NC_011770 | 262395 | A | G | 74 | 0 | 6 | 0 | NON_SYNONYMOU | MODERAT | MISSENSE | Acg/Gcg | T254A | PLES_02351 | G | G | | | | | G | G | | | G | G |
| NC_011770 | 262821 | G | C | 85 | 0 | 6 | 0 | NON_SYNONYMOU | MODERAT | MISSENSE | aGc/aCc | S74T | PLES_02361 | | | C | C | C | C | | | C | C | | |
| NC_011770 | 262991 | C | T | 89 | 0 | 2 | 0 | NON_SYNONYMOU | MODERAT | MISSENSE | Ctc/Ttc | L131F | PLES_02361 | T | T | | | | | | | | | | |
| NC_011770 | 263414 | G | T | 93 | 0 | 2 | 0 | NON_SYNONYMOU | MODERAT | MISSENSE | Gcg/Tcg | A272S | PLES_02361 | | | | | | | | | T | T | | |
| NC_011770 | 263967 | GGC | G | 78 | 0 | 1 | 0 | FRAME_SHIFT | HIGH | | gcc/ | A268 | PLES_02371 | | | | | | | | G | | | | |
| NC_011770 | 264112 | C | G | 78 | 0 | 2 | 0 | NON_SYNONYMOU | MODERAT | MISSENSE | aGc/aCc | S220T | PLES_02371 | | | | | | | | | G | G | | |
| NC_011770 | 264192 | C | G | 68 | 0 | 4 | 0 | NON_SYNONYMOU | MODERAT | MISSENSE | atG/atC | M193I | PLES_02371 | | | G | G | G | G | | | | | | |
| NC_011770 | 264445 | A | G | 68 | 0 | 2 | 0 | NON_SYNONYMOU | MODERAT | MISSENSE | aTc/aCc | I109T | PLES_02371 | | | | | | | | | G | G | | |

| | | | | | | | | | | | | | | | | | | | | | | | | | |
|---|---|---|---|---|---|---|---|---|---|---|---|---|---|---|---|---|---|---|---|---|---|---|---|---|---|
| NC_011770 | 264553 | C | T | 61 | 0 | 2 | 0 | NON_SYNONYMOU | MODERAT | MISSENSE | cGc/cAc | R73H | PLES_02371 | T | T | | | | | | | | | | |
| NC_011770 | 264590 | C | T | 66 | 0 | 2 | 0 | NON_SYNONYMOU | MODERAT | MISSENSE | Gcc/Acc | A61T | PLES_02371 | T | T | | | | | | | | | | |
| NC_011770 | 264595 | C | T | 65 | 0 | 2 | 0 | NON_SYNONYMOU | MODERAT | MISSENSE | gGc/gAc | G59D | PLES_02371 | T | T | | | | | | | | | | |
| NC_011770 | 264643 | G | A | 66 | 0 | 2 | 0 | NON_SYNONYMOU | MODERAT | MISSENSE | gCg/gTg | A43V | PLES_02371 | A | A | | | | | | | | | | |
| NC_011770 | 264763 | T | G | 67 | 0 | 4 | 0 | NON_SYNONYMOU | MODERAT | MISSENSE | cAc/cCc | H3P | PLES_02371 | | | G | G | G | G | | | | | | |
| NC_011770 | 264678 | TTGCCCGG | T | 67 | 0 | 5 | 0 | CODON_CHANGE_ | MODERATE | | cacgtccacg | HVHAGQ2 | PLES_02371 | | | T | | | | T | T | | | T | T |
| NC_011770 | 265273 | T | A | 53 | 0 | 4 | 0 | NON_SYNONYMOU | MODERAT | MISSENSE | Acc/Tcc | T297S | PLES_02381 | | | | | | | A | A | | | A | A |
| NC_011770 | 265465 | GCTC | G | 66 | 0 | 12 | 0 | CODON_DELETION | MODERATE | | gag/ | E232 | PLES_02381 | G | G | G | G | G | G | G | G | G | G | G | G |
| NC_011770 | 265612 | C | T | 66 | 0 | 2 | 0 | NON_SYNONYMOU | MODERAT | MISSENSE | Ggc/Agc | G184S | PLES_02381 | | | | | | | | | T | T | | |
| NC_011770 | 266867 | G | A | 65 | 0 | 8 | 0 | NON_SYNONYMOU | MODERAT | MISSENSE | gGc/gAc | G54D | PLES_02391 | | | A | A | A | A | | | A | A | A | A |
| NC_011770 | 266930 | G | A | 70 | 0 | 4 | 0 | NON_SYNONYMOU | MODERAT | MISSENSE | gGg/gAg | G75E | PLES_02391 | | | | | | | | | A | A | A | A |
| NC_011770 | 267211 | A | G | 78 | 0 | 1 | 0 | NON_SYNONYMOU | MODERAT | MISSENSE | Acc/Gcc | T169A | PLES_02391 | | | | | | | G | | | | | |
| NC_011770 | 267518 | A | G | 76 | 0 | 4 | 0 | NON_SYNONYMOU | MODERAT | MISSENSE | cAc/cGc | H271R | PLES_02391 | G | G | | | | | G | G | | | | |
| NC_011770 | 267628 | G | A | 82 | 0 | 8 | 0 | NON_SYNONYMOU | MODERAT | MISSENSE | Ggc/Agc | G308S | PLES_02391 | | | A | A | A | A | | | A | A | A | A |
| NC_011770 | 267808 | T | C | 74 | 0 | 12 | 0 | NON_SYNONYMOU | MODERAT | MISSENSE | Tgc/Cgc | C368R | PLES_02391 | C | C | C | C | C | C | C | C | C | C | C | C |
| NC_011770 | 268319 | A | G | 100 | 0 | 2 | 0 | NON_SYNONYMOU | MODERAT | MISSENSE | gAg/gGg | E538G | PLES_02391 | G | G | | | | | | | | | | |
| NC_011770 | 268523 | G | A | 91 | 0 | 8 | 0 | NON_SYNONYMOU | MODERAT | MISSENSE | gGc/gAc | G606D | PLES_02391 | | | A | A | A | A | A | A | | | A | A |
| NC_011770 | 268725 | G | C | 87 | 0 | 2 | 0 | NON_SYNONYMOU | MODERAT | MISSENSE | Gtc/Ctc | V8L | PLES_02401 | | | | | | | | | | | C | C |
| NC_011770 | 269462 | T | C | 93 | 0 | 12 | 0 | NON_SYNONYMOU | MODERAT | MISSENSE | Acg/Gcg | T269A | PLES_02411 | C | C | C | C | C | C | C | C | C | C | C | C |
| NC_011770 | 269893 | T | C | 86 | 0 | 2 | 0 | NON_SYNONYMOU | MODERAT | MISSENSE | gAc/gGc | D125G | PLES_02411 | | | | | | | | | | | C | C |
| NC_011770 | 270840 | T | C | 87 | 0 | 2 | 0 | NON_SYNONYMOU | MODERAT | MISSENSE | Acg/Gcg | T498A | PLES_02431 | | | | | | | | | C | C | | |
| NC_011770 | 271266 | C | T | 87 | 0 | 4 | 0 | NON_SYNONYMOU | MODERAT | MISSENSE | Ggc/Agc | G356S | PLES_02431 | | | T | T | T | T | | | | | | |
| NC_011770 | 271835 | T | C | 79 | 0 | 12 | 0 | NON_SYNONYMOU | MODERAT | MISSENSE | cAc/cGc | H166R | PLES_02431 | C | C | C | C | C | C | C | C | C | C | C | C |
| NC_011770 | 271998 | T | C | 69 | 0 | 2 | 0 | NON_SYNONYMOU | MODERAT | MISSENSE | Atc/Gtc | I112V | PLES_02431 | | | | | | | | | C | C | | |
| NC_011770 | 272204 | T | A | 72 | 0 | 8 | 0 | NON_SYNONYMOU | MODERAT | MISSENSE | tAc/tTc | Y43F | PLES_02431 | A | A | | | | | A | A | A | A | A | A |
| NC_011770 | 272054 | A | G | 76 | 0 | 1 | 0 | NON_SYNONYMOU | MODERAT | MISSENSE | cTt/cCt | L93P | PLES_02431 | | | | | | | | **G** | | | | |
| NC_011770 | 273881 | T | C | 64 | 0 | 2 | 0 | NON_SYNONYMOU | MODERAT | MISSENSE | aTc/aCc | I5T | PLES_02451 | | | | | | | | | C | C | | |
| NC_011770 | 274145 | G | A | 76 | 0 | 2 | 0 | NON_SYNONYMOU | MODERAT | MISSENSE | cGc/cAc | R93H | PLES_02451 | A | A | | | | | | | | | | |
| NC_011770 | 274168 | C | G | 71 | 0 | 6 | 0 | NON_SYNONYMOU | MODERAT | MISSENSE | Ctc/Gtc | L101V | PLES_02451 | | | G | G | G | G | G | G | | | | |
| NC_011770 | 274297 | G | A | 72 | 0 | 2 | 0 | NON_SYNONYMOU | MODERAT | MISSENSE | Ggc/Agc | G144S | PLES_02451 | | | | | | | | | | | A | A |
| NC_011770 | 274312 | G | A | 72 | 0 | 2 | 0 | NON_SYNONYMOU | MODERAT | MISSENSE | Gct/Act | A149T | PLES_02451 | | | | | | | | | | | A | A |
| NC_011770 | 274528 | A | G | 82 | 0 | 4 | 0 | NON_SYNONYMOU | MODERAT | MISSENSE | Agc/Ggc | S221G | PLES_02451 | | | | | | | | | G | G | G | G |
| NC_011770 | 274715 | C | T | 88 | 0 | 2 | 0 | NON_SYNONYMOU | MODERAT | MISSENSE | gCt/gTt | A283V | PLES_02451 | | | | | | | | | T | T | | |
| NC_011770 | 274762 | C | A | 83 | 0 | 12 | 0 | NON_SYNONYMOU | MODERAT | MISSENSE | cGg/cTg | R139L | PLES_02461 | A | A | A | A | A | A | A | A | A | A | A | A |
| NC_011770 | 274931 | C | T | 82 | 0 | 8 | 0 | NON_SYNONYMOU | MODERAT | MISSENSE | Gcg/Acg | A83T | PLES_02461 | | | T | T | T | T | T | T | T | T | | |
| NC_011770 | 275027 | C | T | 87 | 0 | 2 | 0 | NON_SYNONYMOU | MODERAT | MISSENSE | Gat/Aat | D51N | PLES_02461 | | | | | | | | | | | T | T |
| NC_011770 | 275164 | A | G | 77 | 0 | 10 | 0 | NON_SYNONYMOU | MODERAT | MISSENSE | cTg/cCg | L5P | PLES_02461 | | | G | G | G | G | G | G | G | G | G | G |
| NC_011770 | 275629 | C | G | 87 | 0 | 4 | 0 | NON_SYNONYMOU | MODERAT | MISSENSE | Gcg/Ccg | A21P | PLES_02471 | | | | | | | | | G | G | G | G |
| NC_011770 | 275953 | TTC | T | 40 | 0 | 12 | 0 | FRAME_SHIFT | HIGH | | gaa/ | E181 | PLES_02481 | T | T | T | T | T | T | T | T | T | T | T | T |
| NC_011770 | 275945 | T | G | 77 | 0 | 12 | 0 | NON_SYNONYMOU | MODERAT | MISSENSE | aAc/aCc | N184T | PLES_02481 | G | G | G | G | G | G | G | G | G | G | G | G |
| NC_011770 | 275976 | T | C | 57 | 0 | 12 | 0 | NON_SYNONYMOU | MODERAT | MISSENSE | Aag/Gag | K174E | PLES_02481 | C | C | C | C | C | C | C | C | C | C | C | C |
| NC_011770 | 275977 | G | C | 57 | 0 | 12 | 0 | NON_SYNONYMOU | MODERAT | MISSENSE | aaC/aaG | N173K | PLES_02481 | C | C | C | C | C | C | C | C | C | C | C | C |
| NC_011770 | 275983 | T | G | 58 | 0 | 12 | 0 | NON_SYNONYMOU | MODERAT | MISSENSE | aaA/aaC | K171N | PLES_02481 | G | G | G | G | G | G | G | G | G | G | G | G |
| NC_011770 | 276050 | T | G | 66 | 0 | 12 | 0 | NON_SYNONYMOU | MODERAT | MISSENSE | gAc/gCc | D149A | PLES_02481 | G | G | G | G | G | G | G | G | G | G | G | G |
| NC_011770 | 276057 | C | T | 68 | 0 | 1 | 0 | NON_SYNONYMOU | MODERAT | MISSENSE | Ggt/Agt | G147S | PLES_02481 | | | | | | | T | | | | | |
| NC_011770 | 276069 | C | A | 65 | 0 | 12 | 0 | NON_SYNONYMOU | MODERAT | MISSENSE | Gcc/Tcc | A143S | PLES_02481 | A | A | A | A | A | A | A | A | A | A | A | A |
| NC_011770 | 276078 | C | T | 56 | 0 | 2 | 0 | NON_SYNONYMOU | MODERAT | MISSENSE | Gaa/Aaa | E140K | PLES_02481 | T | T | | | | | | | | | | |
| NC_011770 | 276080 | C | T | 54 | 0 | 12 | 0 | NON_SYNONYMOU | MODERAT | MISSENSE | gGc/gAc | G139D | PLES_02481 | T | T | T | T | T | T | T | T | T | T | T | T |
| NC_011770 | 276090 | T | C | 53 | 0 | 12 | 0 | NON_SYNONYMOU | MODERAT | MISSENSE | Acc/Gcc | T136A | PLES_02481 | C | C | C | C | C | C | C | C | C | C | C | C |
| NC_011770 | 276121 | G | C | 59 | 0 | 12 | 0 | NON_SYNONYMOU | MODERAT | MISSENSE | caC/caG | H125Q | PLES_02481 | C | C | C | C | C | C | C | C | C | C | C | C |
| NC_011770 | 276151 | G | C | 38 | 0 | 12 | 0 | NON_SYNONYMOU | MODERAT | MISSENSE | agC/agG | S115R | PLES_02481 | C | C | C | C | C | C | C | C | C | C | C | C |
| NC_011770 | 276152 | C | G | 38 | 0 | 12 | 0 | NON_SYNONYMOU | MODERAT | MISSENSE | aGc/aCc | S115T | PLES_02481 | G | G | G | G | G | G | G | G | G | G | G | G |
| NC_011770 | 276153 | T | C | 38 | 0 | 12 | 0 | NON_SYNONYMOU | MODERAT | MISSENSE | Agc/Ggc | S115G | PLES_02481 | C | C | C | C | C | C | C | C | C | C | C | C |
| NC_011770 | 276223 | C | A | 45 | 0 | 12 | 0 | NON_SYNONYMOU | MODERAT | MISSENSE | gaG/gaT | E91D | PLES_02481 | A | A | A | A | A | A | A | A | A | A | A | A |
| NC_011770 | 276236 | A | T | 43 | 0 | 12 | 0 | NON_SYNONYMOU | MODERAT | MISSENSE | gTc/gAc | V87D | PLES_02481 | T | T | T | T | T | T | T | T | T | T | T | T |
| NC_011770 | 276260 | C | G | 50 | 0 | 12 | 0 | NON_SYNONYMOU | MODERAT | MISSENSE | aGc/aCc | S79T | PLES_02481 | G | G | G | G | G | G | G | G | G | G | G | G |
| NC_011770 | 276261 | T | C | 50 | 0 | 6 | 0 | NON_SYNONYMOU | MODERAT | MISSENSE | Agc/Ggc | S79G | PLES_02481 | C | C | | | | | | | C | C | C | C |
| NC_011770 | 276274 | G | C | 45 | 0 | 12 | 0 | NON_SYNONYMOU | MODERAT | MISSENSE | agC/agG | S74R | PLES_02481 | C | C | C | C | C | C | C | C | C | C | C | C |
| NC_011770 | 276275 | C | T | 45 | 0 | 12 | 0 | NON_SYNONYMOU | MODERAT | MISSENSE | aGc/aAc | S74N | PLES_02481 | T | T | T | T | T | T | T | T | T | T | T | T |
| NC_011770 | 276302 | C | G | 48 | 0 | 12 | 0 | NON_SYNONYMOU | MODERAT | MISSENSE | cGg/cCg | R65P | PLES_02481 | G | G | G | G | G | G | G | G | G | G | G | G |
| NC_011770 | 276350 | C | G | 41 | 0 | 12 | 0 | NON_SYNONYMOU | MODERAT | MISSENSE | aGc/aCc | S49T | PLES_02481 | G | G | G | G | G | G | G | G | G | G | G | G |
| NC_011770 | 276359 | T | A | 38 | 0 | 12 | 0 | NON_SYNONYMOU | MODERAT | MISSENSE | cAg/cTg | Q46L | PLES_02481 | A | A | A | A | A | A | A | A | A | A | A | A |

| | | | | | | | | | | | | | | | | | | | | | | | | | |
|---|---|---|---|---|---|---|---|---|---|---|---|---|---|---|---|---|---|---|---|---|---|---|---|---|---|
| NC_011770 | 276368 | C | G | 39 | 0 | 12 | 0 | NON_SYNONYMOU | MODERAT | MISSENSE | gGc/gCc | G43A | PLES_02481 | G | G | G | G | G | G | G | G | G | G | G | G |
| NC_011770 | 276389 | G | A | 29 | 0 | 6 | 0 | NON_SYNONYMOU | MODERAT | MISSENSE | gCa/gTa | A36V | PLES_02481 | | | A | A | A | A | A | A | | | | |
| NC_011770 | 276393 | T | C | 28 | 0 | 12 | 0 | NON_SYNONYMOU | MODERAT | MISSENSE | Acg/Gcg | T35A | PLES_02481 | C | C | C | C | C | C | C | C | C | C | C | C |
| NC_011770 | 276428 | A | G | 21 | 0 | 12 | 0 | NON_SYNONYMOU | MODERAT | MISSENSE | gTc/gCc | V23A | PLES_02481 | G | G | G | G | G | G | G | G | G | G | G | G |
| NC_011770 | 276429 | C | T | 20 | 0 | 12 | 0 | NON_SYNONYMOU | MODERAT | MISSENSE | Gtc/Atc | V23I | PLES_02481 | T | T | T | T | T | T | T | T | T | T | T | T |
| NC_011770 | 276430 | G | C | 20 | 0 | 12 | 0 | NON_SYNONYMOU | MODERAT | MISSENSE | atC/atG | I22M | PLES_02481 | C | C | C | C | C | C | C | C | C | C | C | C |
| NC_011770 | 276434 | C | G | 20 | 0 | 12 | 0 | NON_SYNONYMOU | MODERAT | MISSENSE | aGc/aCc | S21T | PLES_02481 | G | G | G | G | G | G | G | G | G | G | G | G |
| NC_011770 | 276435 | T | C | 20 | 0 | 12 | 0 | NON_SYNONYMOU | MODERAT | MISSENSE | Agc/Ggc | S21G | PLES_02481 | C | C | C | C | C | C | C | C | C | C | C | C |
| NC_011770 | 276455 | T | G | 27 | 0 | 12 | 0 | NON_SYNONYMOU | MODERAT | MISSENSE | gAc/gCc | D14A | PLES_02481 | G | G | G | G | G | G | G | G | G | G | G | G |
| NC_011770 | 276917 | A | G | 54 | 0 | 6 | 0 | NON_SYNONYMOU | MODERAT | MISSENSE | gTc/gCc | V45A | PLES_02491 | | | G | G | G | G | G | G | | | | |
| NC_011770 | 276928 | T | G | 55 | 0 | 6 | 0 | NON_SYNONYMOU | MODERAT | MISSENSE | gaA/gaC | E41D | PLES_02491 | | | G | G | G | G | G | G | | | | |
| NC_011770 | 276971 | G | A | 56 | 0 | 6 | 0 | NON_SYNONYMOU | MODERAT | MISSENSE | aCc/aTc | T27I | PLES_02491 | | | A | A | A | A | A | A | | | | |
| NC_011770 | 277031 | G | A | 63 | 0 | 6 | 0 | NON_SYNONYMOU | MODERAT | MISSENSE | cCc/cTc | P7L | PLES_02491 | | | A | A | A | A | A | A | | | | |
| NC_011770 | 277047 | C | T | 61 | 0 | 6 | 0 | NON_SYNONYMOU | MODERAT | MISSENSE | Ggc/Agc | G2S | PLES_02491 | | | T | T | T | T | T | T | | | | |
| NC_011770 | 277149 | C | G | 76 | 0 | 2 | 0 | NON_SYNONYMOU | MODERAT | MISSENSE | aGg/aCg | R157T | PLES_02501 | G | G | | | | | | | | | | |
| NC_011770 | 277177 | A | G | 76 | 0 | 12 | 0 | NON_SYNONYMOU | MODERAT | MISSENSE | Tgg/Cgg | W148R | PLES_02501 | G | G | G | G | G | G | G | G | G | G | G | G |
| NC_011770 | 277194 | A | G | 78 | 0 | 6 | 0 | NON_SYNONYMOU | MODERAT | MISSENSE | gTc/gCc | V142A | PLES_02501 | | | G | G | G | G | G | G | | | | |
| NC_011770 | 277207 | C | T | 80 | 0 | 2 | 0 | NON_SYNONYMOU | MODERAT | MISSENSE | Gcg/Acg | A138T | PLES_02501 | | | | | | | | | | | T | T |
| NC_011770 | 277225 | A | C | 74 | 0 | 2 | 0 | NON_SYNONYMOU | MODERAT | MISSENSE | Tcc/Gcc | S132A | PLES_02501 | C | C | | | | | | | | | | |
| NC_011770 | 277227 | C | A | 78 | 0 | 2 | 0 | NON_SYNONYMOU | MODERAT | MISSENSE | gGc/gTc | G131V | PLES_02501 | | | | | | | | | | | A | A |
| NC_011770 | 277300 | C | T | 80 | 0 | 2 | 0 | NON_SYNONYMOU | MODERAT | MISSENSE | Gcg/Acg | A107T | PLES_02501 | T | T | | | | | | | | | | |
| NC_011770 | 277868 | T | G | 78 | 0 | 8 | 0 | NON_SYNONYMOU | MODERAT | MISSENSE | Aag/Cag | K428Q | PLES_02511 | | | G | G | G | G | G | G | G | G | | |
| NC_011770 | 278854 | A | G | 66 | 0 | 2 | 0 | NON_SYNONYMOU | MODERAT | MISSENSE | aTg/aCg | M99T | PLES_02511 | G | G | | | | | | | | | | |
| NC_011770 | 278883 | T | A | 66 | 0 | 2 | 0 | NON_SYNONYMOU | MODERAT | MISSENSE | gaA/gaT | E89D | PLES_02511 | | | | | | | | | | | A | A |
| NC_011770 | 280399 | C | G | 98 | 0 | 2 | 0 | NON_SYNONYMOU | MODERAT | MISSENSE | Gcg/Ccg | A223P | PLES_02531 | | | | | | | G | G | | | | |
| NC_011770 | 280471 | C | T | 97 | 0 | 2 | 0 | NON_SYNONYMOU | MODERAT | MISSENSE | Ggc/Agc | G199S | PLES_02531 | | | | | | | T | T | | | | |
| NC_011770 | 280816 | G | C | 95 | 0 | 2 | 0 | NON_SYNONYMOU | MODERAT | MISSENSE | Cat/Gat | H84D | PLES_02531 | | | | | | | | | | | C | C |
| NC_011770 | 280885 | C | T | 85 | 0 | 2 | 0 | NON_SYNONYMOU | MODERAT | MISSENSE | Gac/Aac | D61N | PLES_02531 | | | | | | | | | T | T | | |
| NC_011770 | 282253 | T | G | 53 | 4 | 8 | 0 | NON_SYNONYMOU | MODERAT | MISSENSE | gAt/gCt | D27A | PLES_02541 | G | G | G | G | G | G | . | . | G | G | . | . |
| NC_011770 | 283424 | T | C | 16 | 7 | 1 | 0 | NON_SYNONYMOU | MODERAT | MISSENSE | atA/atG | I175M | PLES_02551 | | | . | . | . | C | . | . | | | . | . |
| NC_011770 | 283426 | T | C | 16 | 7 | 1 | 0 | NON_SYNONYMOU | MODERAT | MISSENSE | Ata/Gta | I175V | PLES_02551 | | | . | . | . | C | . | . | | | . | . |
| NC_011770 | 283463 | T | A | 52 | 4 | 4 | 0 | NON_SYNONYMOU | MODERAT | MISSENSE | agA/agT | R162S | PLES_02551 | | | A | A | A | A | . | . | | | . | . |
| NC_011770 | 283527 | A | G | 58 | 4 | 4 | 0 | NON_SYNONYMOU | MODERAT | MISSENSE | gTc/gCc | V141A | PLES_02551 | | | G | G | G | G | . | . | | | . | . |
| NC_011770 | 283767 | C | T | 52 | 4 | 4 | 0 | NON_SYNONYMOU | MODERAT | MISSENSE | aGg/aAg | R61K | PLES_02551 | | | T | T | T | T | . | . | | | . | . |
| NC_011770 | 283814 | C | T | 49 | 4 | 4 | 0 | NON_SYNONYMOU | MODERAT | MISSENSE | atG/atA | M45I | PLES_02551 | | | T | T | T | T | . | . | | | . | . |
| NC_011770 | 283815 | A | G | 49 | 4 | 4 | 0 | NON_SYNONYMOU | MODERAT | MISSENSE | aTg/aCg | M45T | PLES_02551 | | | G | G | G | G | . | . | | | . | . |
| NC_011770 | 283816 | T | C | 49 | 4 | 4 | 0 | NON_SYNONYMOU | MODERAT | MISSENSE | Atg/Gtg | M45V | PLES_02551 | | | C | C | C | C | . | . | | | . | . |
| NC_011770 | 283882 | C | T | 55 | 4 | 4 | 0 | NON_SYNONYMOU | MODERAT | MISSENSE | Gta/Ata | V23I | PLES_02551 | | | T | T | T | T | . | . | | | . | . |
| NC_011770 | 283920 | T | G | 53 | 4 | 4 | 0 | NON_SYNONYMOU | MODERAT | MISSENSE | aAa/aCa | K10T | PLES_02551 | | | G | G | G | G | . | . | | | . | . |
| NC_011770 | 283927 | C | G | 52 | 4 | 4 | 0 | NON_SYNONYMOU | MODERAT | MISSENSE | Gct/Cct | A8P | PLES_02551 | | | G | G | G | G | . | . | | | . | . |
| NC_011770 | 284835 | C | T | 50 | 4 | 2 | 0 | NON_SYNONYMOU | MODERAT | MISSENSE | Gcc/Acc | A474T | PLES_02561 | T | T | | | | | . | . | | | . | . |
| NC_011770 | 285897 | C | T | 51 | 4 | 4 | 0 | NON_SYNONYMOU | MODERAT | MISSENSE | Gtc/Atc | V120I | PLES_02561 | | | T | T | T | T | . | . | | | . | . |
| NC_011770 | 285933 | C | T | 53 | 4 | 2 | 0 | NON_SYNONYMOU | MODERAT | MISSENSE | Gcc/Acc | A108T | PLES_02561 | T | T | | | | | . | . | | | . | . |
| NC_011770 | 285956 | A | G | 51 | 4 | 6 | 0 | NON_SYNONYMOU | MODERAT | MISSENSE | gTc/gCc | V100A | PLES_02561 | G | G | G | G | G | G | . | . | | | . | . |
| NC_011770 | 286378 | C | T | 50 | 4 | 2 | 0 | NON_SYNONYMOU | MODERAT | MISSENSE | cGg/cAg | R124Q | PLES_02571 | | | | | | | . | . | T | T | . | . |
| NC_011770 | 287246 | C | T | 58 | 4 | 8 | 0 | NON_SYNONYMOU | MODERAT | MISSENSE | Ggt/Agt | G856S | PLES_02581 | T | T | T | T | T | T | . | . | T | T | . | . |
| NC_011770 | 288098 | T | C | 54 | 0 | 10 | 0 | NON_SYNONYMOU | MODERAT | MISSENSE | Aag/Gag | K572E | PLES_02581 | C | C | C | C | C | C | C | C | | | C | C |
| NC_011770 | 288785 | T | G | 79 | 0 | 2 | 0 | NON_SYNONYMOU | MODERAT | MISSENSE | Agc/Cgc | S343R | PLES_02581 | G | G | | | | | | | | | | |
| NC_011770 | 288923 | T | C | 77 | 0 | 2 | 0 | NON_SYNONYMOU | MODERAT | MISSENSE | Atc/Gtc | I297V | PLES_02581 | C | C | | | | | | | | | | |
| NC_011770 | 288940 | C | T | 77 | 0 | 1 | 0 | NON_SYNONYMOU | MODERAT | MISSENSE | gGc/gAc | G291D | PLES_02581 | | | | | | | T | | | | | |
| NC_011770 | 288994 | G | C | 80 | 0 | 6 | 0 | NON_SYNONYMOU | MODERAT | MISSENSE | aCc/aGc | T273S | PLES_02581 | C | C | | | | | C | C | C | C | | |
| NC_011770 | 289060 | C | T | 74 | 0 | 2 | 0 | NON_SYNONYMOU | MODERAT | MISSENSE | gGc/gAc | G251D | PLES_02581 | | | | | | | T | T | | | | |
| NC_011770 | 289663 | C | T | 65 | 0 | 2 | 0 | NON_SYNONYMOU | MODERAT | MISSENSE | aGg/aAg | R50K | PLES_02581 | | | | | | | T | T | | | | |
| NC_011770 | 289702 | T | C | 62 | 0 | 2 | 0 | NON_SYNONYMOU | MODERAT | MISSENSE | aAc/aGc | N37S | PLES_02581 | | | | | | | C | C | | | | |
| NC_011770 | 289764 | T | G | 62 | 0 | 2 | 0 | NON_SYNONYMOU | MODERAT | MISSENSE | aaA/aaC | K16N | PLES_02581 | | | | | | | G | G | | | | |
| NC_011770 | 292161 | T | C | 60 | 0 | 2 | 0 | NON_SYNONYMOU | MODERAT | MISSENSE | aAg/aGg | K96R | PLES_02601 | | | | | | | C | C | | | | |
| NC_011770 | 292227 | C | T | 66 | 0 | 2 | 0 | NON_SYNONYMOU | MODERAT | MISSENSE | gGc/gAc | G74D | PLES_02601 | | | | | | | T | T | | | | |
| NC_011770 | 292252 | C | T | 66 | 0 | 10 | 0 | NON_SYNONYMOU | MODERAT | MISSENSE | Gca/Aca | A66T | PLES_02601 | T | T | T | T | T | T | T | T | | | T | T |
| NC_011770 | 296128 | C | A | 70 | 0 | 4 | 0 | NON_SYNONYMOU | MODERAT | MISSENSE | Ctg/Atg | L119M | PLES_02631 | | | A | A | A | A | | | | | | |
| NC_011770 | 296233 | A | G | 65 | 0 | 12 | 0 | NON_SYNONYMOU | MODERAT | MISSENSE | Acg/Gcg | T154A | PLES_02631 | G | G | G | G | G | G | G | G | G | G | G | G |
| NC_011770 | 296333 | A | C | 70 | 0 | 2 | 0 | NON_SYNONYMOU | MODERAT | MISSENSE | gAt/gCt | D187A | PLES_02631 | C | C | | | | | | | | | | |
| NC_011770 | 296825 | G | A | 80 | 0 | 2 | 0 | NON_SYNONYMOU | MODERAT | MISSENSE | gGc/gAc | G351D | PLES_02631 | A | A | | | | | | | | | | |

| | | | | | | | | | | | | | | | | | | | | | | | | | |
|---|---|---|---|---|---|---|---|---|---|---|---|---|---|---|---|---|---|---|---|---|---|---|---|---|---|
| NC_011770 | 296902 | C | A | 77 | 0 | 2 | 0 | NON_SYNONYMOU | MODERAT | MISSENSE | Ctg/Atg | L377M | PLES_02631 | | | | | | | | | A | A | | |
| NC_011770 | 297419 | C | G | 80 | 0 | 2 | 0 | NON_SYNONYMOU | MODERAT | MISSENSE | Ggg/Cgg | G383R | PLES_02641 | | | | | | | G | G | | | | |
| NC_011770 | 297736 | T | C | 81 | 0 | 12 | 0 | NON_SYNONYMOU | MODERAT | MISSENSE | cAc/cGc | H277R | PLES_02641 | C | C | C | C | C | C | C | C | C | C | C | C |
| NC_011770 | 297925 | G | T | 77 | 0 | 2 | 0 | NON_SYNONYMOU | MODERAT | MISSENSE | cCg/cAg | P214Q | PLES_02641 | T | T | | | | | | | | | | |
| NC_011770 | 298027 | T | C | 72 | 0 | 4 | 0 | NON_SYNONYMOU | MODERAT | MISSENSE | aAc/aGc | N180S | PLES_02641 | | | C | C | C | C | | | | | | |
| NC_011770 | 298385 | C | T | 48 | 0 | 6 | 0 | NON_SYNONYMOU | MODERAT | MISSENSE | Gcc/Acc | A61T | PLES_02641 | | | T | T | T | T | T | T | | | | |
| NC_011770 | 298995 | G | C | 69 | 0 | 6 | 0 | NON_SYNONYMOU | MODERAT | MISSENSE | Gcg/Ccg | A102P | PLES_02651 | | | C | C | C | C | C | C | | | | |
| NC_011770 | 299181 | C | T | 63 | 0 | 2 | 0 | NON_SYNONYMOU | MODERAT | MISSENSE | gCg/gTg | A14V | PLES_02661 | | | | | | | T | T | | | | |
| NC_011770 | 299208 | T | C | 65 | 0 | 4 | 0 | NON_SYNONYMOU | MODERAT | MISSENSE | gTt/gCt | V23A | PLES_02661 | | | C | C | C | C | | | | | | |
| NC_011770 | 299232 | A | G | 66 | 0 | 4 | 0 | NON_SYNONYMOU | MODERAT | MISSENSE | cAc/cGc | H31R | PLES_02661 | | | G | G | G | G | | | | | | |
| NC_011770 | 299241 | T | C | 70 | 0 | 12 | 0 | NON_SYNONYMOU | MODERAT | MISSENSE | gTc/gCc | V34A | PLES_02661 | C | C | C | C | C | C | C | C | C | C | C | C |
| NC_011770 | 299252 | A | T | 68 | 0 | 2 | 0 | NON_SYNONYMOU | MODERAT | MISSENSE | Att/Ttt | I38F | PLES_02661 | | | | | | | | | T | T | | |
| NC_011770 | 299982 | G | A | 77 | 0 | 4 | 0 | NON_SYNONYMOU | MODERAT | MISSENSE | Ccc/Tcc | P272S | PLES_02681 | | | A | A | A | A | | | | | | |
| NC_011770 | 299999 | T | C | 79 | 0 | 4 | 0 | NON_SYNONYMOU | MODERAT | MISSENSE | cAc/cGc | H266R | PLES_02681 | | | C | C | C | C | | | | | | |
| NC_011770 | 300697 | G | C | 103 | 0 | 4 | 0 | NON_SYNONYMOU | MODERAT | MISSENSE | caC/caG | H33Q | PLES_02681 | | | C | C | C | C | | | | | | |
| NC_011770 | 300870 | C | T | 83 | 0 | 4 | 0 | NON_SYNONYMOU | MODERAT | MISSENSE | gGc/gAc | G397D | PLES_02691 | | | T | T | T | T | | | | | | |
| NC_011770 | 300873 | T | C | 84 | 0 | 12 | 0 | NON_SYNONYMOU | MODERAT | MISSENSE | cAa/cGa | Q396R | PLES_02691 | C | C | C | C | C | C | C | C | C | C | C | C |
| NC_011770 | 300877 | G | C | 82 | 0 | 12 | 0 | NON_SYNONYMOU | MODERAT | MISSENSE | Ccg/Gcg | P395A | PLES_02691 | C | C | C | C | C | C | C | C | C | C | C | C |
| NC_011770 | 300888 | T | G | 86 | 0 | 12 | 0 | NON_SYNONYMOU | MODERAT | MISSENSE | gAc/gCc | D391A | PLES_02691 | G | G | G | G | G | G | G | G | G | G | G | G |
| NC_011770 | 300895 | G | T | 94 | 0 | 2 | 0 | NON_SYNONYMOU | MODERAT | MISSENSE | Ctg/Atg | L389M | PLES_02691 | | | | | | | | | T | T | | |
| NC_011770 | 300936 | A | G | 85 | 0 | 2 | 0 | NON_SYNONYMOU | MODERAT | MISSENSE | cTg/cCg | L375P | PLES_02691 | | | | | G | G | | | | | | |
| NC_011770 | 300955 | T | C | 84 | 0 | 4 | 0 | NON_SYNONYMOU | MODERAT | MISSENSE | Acg/Gcg | T369A | PLES_02691 | | | C | C | C | C | | | | | | |
| NC_011770 | 300958 | C | T | 83 | 0 | 4 | 0 | NON_SYNONYMOU | MODERAT | MISSENSE | Gcc/Acc | A368T | PLES_02691 | | | T | T | T | T | | | | | | |
| NC_011770 | 301015 | T | C | 84 | 0 | 4 | 0 | NON_SYNONYMOU | MODERAT | MISSENSE | Atc/Gtc | I349V | PLES_02691 | | | C | C | C | C | | | | | | |
| NC_011770 | 301212 | G | A | 76 | 0 | 2 | 0 | NON_SYNONYMOU | MODERAT | MISSENSE | gCc/gTc | A283V | PLES_02691 | | | | | A | A | | | | | | |
| NC_011770 | 301479 | G | T | 77 | 0 | 4 | 0 | NON_SYNONYMOU | MODERAT | MISSENSE | gCc/gAc | A194D | PLES_02691 | T | T | | | | | T | T | | | | |
| NC_011770 | 301539 | G | A | 67 | 0 | 12 | 0 | NON_SYNONYMOU | MODERAT | MISSENSE | cCc/cTc | P174L | PLES_02691 | A | A | A | A | A | A | A | A | A | A | A | A |
| NC_011770 | 301693 | C | T | 61 | 0 | 2 | 0 | NON_SYNONYMOU | MODERAT | MISSENSE | Gcc/Acc | A123T | PLES_02691 | T | T | | | | | | | | | | |
| NC_011770 | 301840 | C | A | 68 | 0 | 2 | 0 | NON_SYNONYMOU | MODERAT | MISSENSE | Gtc/Ttc | V74F | PLES_02691 | | | | | | | | | | | A | A |
| NC_011770 | 301917 | T | G | 65 | 0 | 4 | 0 | NON_SYNONYMOU | MODERAT | MISSENSE | gAc/gCc | D48A | PLES_02691 | G | G | | | | | | | | | G | G |
| NC_011770 | 302038 | C | A | 50 | 0 | 6 | 0 | NON_SYNONYMOU | MODERAT | MISSENSE | Ggc/Tgc | G8C | PLES_02691 | | | A | A | A | A | | | | | A | A |
| NC_011770 | 302499 | A | G | 66 | 0 | 10 | 0 | NON_SYNONYMOU | MODERAT | MISSENSE | aTt/aCt | I165T | PLES_02701 | G | G | G | G | G | G | | | G | G | G | G |
| NC_011770 | 302589 | T | C | 65 | 0 | 10 | 0 | NON_SYNONYMOU | MODERAT | MISSENSE | gAc/gGc | D135G | PLES_02701 | C | C | C | C | C | C | | | C | C | C | C |
| NC_011770 | 302704 | C | T | 70 | 0 | 2 | 0 | NON_SYNONYMOU | MODERAT | MISSENSE | Gcg/Acg | A97T | PLES_02701 | | | | | | | | | | | T | T |
| NC_011770 | 302739 | C | T | 68 | 0 | 10 | 0 | NON_SYNONYMOU | MODERAT | MISSENSE | gGc/gAc | G85D | PLES_02701 | T | T | T | T | T | T | | | T | T | T | T |
| NC_011770 | 302923 | T | C | 76 | 0 | 10 | 0 | NON_SYNONYMOU | MODERAT | MISSENSE | Acc/Gcc | T24A | PLES_02701 | C | C | C | C | C | C | | | C | C | C | C |
| NC_011770 | 302940 | G | A | 78 | 0 | 10 | 0 | NON_SYNONYMOU | MODERAT | MISSENSE | gCg/gTg | A18V | PLES_02701 | A | A | A | A | A | A | | | A | A | A | A |
| NC_011770 | 302973 | A | G | 80 | 0 | 10 | 0 | NON_SYNONYMOU | MODERAT | MISSENSE | tTc/tCc | F7S | PLES_02701 | G | G | G | G | G | G | | | G | G | G | G |
| NC_011770 | 302974 | A | G | 80 | 0 | 10 | 0 | NON_SYNONYMOU | MODERAT | MISSENSE | Ttc/Ctc | F7L | PLES_02701 | G | G | G | G | G | G | | | G | G | G | G |
| NC_011770 | 302988 | G | A | 82 | 0 | 10 | 0 | NON_SYNONYMOU | MODERAT | MISSENSE | cCc/cTc | P2L | PLES_02701 | A | A | A | A | A | A | | | A | A | A | A |
| NC_011770 | 302695 | C | T | 74 | 0 | 1 | 0 | NON_SYNONYMOU | MODERAT | MISSENSE | Ggc/Agc | G100S | PLES_02701 | HP | T | | | | | | | | | | |
| NC_011770 | 305169 | G | A | 71 | 0 | 2 | 0 | NON_SYNONYMOU | MODERAT | MISSENSE | aCc/aTc | T3I | PLES_02731 | A | A | | | | | | | | | | |
| NC_011770 | 305653 | C | T | 71 | 0 | 4 | 0 | NON_SYNONYMOU | MODERAT | MISSENSE | Gtt/Att | V150I | PLES_02741 | | | T | T | T | T | | | | | | |
| NC_011770 | 305871 | G | A | 63 | 0 | 2 | 0 | NON_SYNONYMOU | MODERAT | MISSENSE | gCc/gTc | A77V | PLES_02741 | | | | | | | A | A | | | | |
| NC_011770 | 306057 | G | C | 63 | 0 | 2 | 0 | NON_SYNONYMOU | MODERAT | MISSENSE | aCc/aGc | T15S | PLES_02741 | | | | | | | | | | | C | C |
| NC_011770 | 306453 | T | C | 67 | 0 | 10 | 0 | NON_SYNONYMOU | MODERAT | MISSENSE | gTc/gCc | V87A | PLES_02751 | C | C | C | C | C | C | C | C | C | C | | |
| NC_011770 | 306524 | C | T | 65 | 0 | 4 | 0 | NON_SYNONYMOU | MODERAT | MISSENSE | Cgc/Tgc | R111C | PLES_02751 | | | T | T | T | T | | | | | | |
| NC_011770 | 306662 | A | G | 64 | 0 | 4 | 0 | NON_SYNONYMOU | MODERAT | MISSENSE | Agg/Ggg | R157G | PLES_02751 | | | G | G | G | G | | | | | | |
| NC_011770 | 306680 | G | T | 67 | 0 | 2 | 0 | NON_SYNONYMOU | MODERAT | MISSENSE | Gcg/Tcg | A163S | PLES_02751 | T | T | | | | | | | | | | |
| NC_011770 | 306687 | T | C | 63 | 0 | 4 | 0 | NON_SYNONYMOU | MODERAT | MISSENSE | gTc/gCc | V165A | PLES_02751 | | | C | C | C | C | | | | | | |
| NC_011770 | 306851 | C | A | 71 | 0 | 2 | 0 | NON_SYNONYMOU | MODERAT | MISSENSE | Cgc/Agc | R220S | PLES_02751 | | | | | | | | | | | A | A |
| NC_011770 | 311297 | C | A | 72 | 0 | 4 | 0 | NON_SYNONYMOU | MODERAT | MISSENSE | Gcg/Tcg | A756S | PLES_02811 | | | | | | | | | A | A | A | A |
| NC_011770 | 312021 | A | C | 93 | 0 | 2 | 0 | NON_SYNONYMOU | MODERAT | MISSENSE | gaT/gaG | D514E | PLES_02811 | | | | | | | C | C | | | | |
| NC_011770 | 312664 | A | G | 87 | 0 | 1 | 0 | NON_SYNONYMOU | MODERAT | MISSENSE | gTg/gCg | V300A | PLES_02811 | | | | | G | | | | | | | |
| NC_011770 | 324885 | C | T | 57 | 0 | 4 | 0 | NON_SYNONYMOU | MODERAT | MISSENSE | Gcc/Acc | A303T | PLES_02911 | | | T | T | T | T | | | | | | |
| NC_011770 | 325229 | C | G | 72 | 0 | 2 | 0 | NON_SYNONYMOU | MODERAT | MISSENSE | gGt/gCt | G188A | PLES_02911 | G | G | | | | | | | | | | |
| NC_011770 | 325431 | C | T | 70 | 0 | 2 | 0 | NON_SYNONYMOU | MODERAT | MISSENSE | Gtc/Atc | V121I | PLES_02911 | T | T | | | | | | | | | | |
| NC_011770 | 326727 | G | A | 89 | 1 | 1 | 0 | NON_SYNONYMOU | MODERAT | MISSENSE | Cac/Tac | H233Y | PLES_02921 | | **A** | | | | | | . | | | | |
| NC_011770 | 327333 | C | A | 81 | 0 | 1 | 0 | NON_SYNONYMOU | MODERAT | MISSENSE | Gca/Tca | A31S | PLES_02921 | | **A** | | | | | | | | | | |
| NC_011770 | 327372 | T | G | 79 | 0 | 1 | 0 | NON_SYNONYMOU | MODERAT | MISSENSE | Aag/Cag | K18Q | PLES_02921 | | **G** | | | | | | | | | | |
| NC_011770 | 327380 | T | A | 74 | 0 | 1 | 0 | NON_SYNONYMOU | MODERAT | MISSENSE | gAg/gTg | E15V | PLES_02921 | | **A** | | | | | | | | | | |
| NC_011770 | 327381 | C | G | 74 | 0 | 1 | 0 | NON_SYNONYMOU | MODERAT | MISSENSE | Gag/Cag | E15Q | PLES_02921 | | **G** | | | | | | | | | | |

| | | | | | | | | | | | | | | | | | | | | | | | | | |
|---|---|---|---|---|---|---|---|---|---|---|---|---|---|---|---|---|---|---|---|---|---|---|---|---|---|
| NC_011770 | 327410 | T | A | 76 | 0 | 1 | 0 | NON_SYNONYMOU | MODERAT | MISSENSE | cAg/cTg | Q5L | PLES_02921 | | **A** | | | | | | | | | | |
| NC_011770 | 326703 | C | T | 95 | 1 | 1 | 0 | NON_SYNONYMOU | MODERAT | MISSENSE | Gtc/Atc | V241I | PLES_02921 spuI Glutam | | **T** | | | | | | . | | | | |
| NC_011770 | 337581 | T | C | 66 | 0 | 2 | 0 | NON_SYNONYMOU | MODERAT | MISSENSE | aAg/aGg | K693R | PLES_03011 | | | | | | | C | C | | | | |
| NC_011770 | 337672 | C | T | 67 | 0 | 10 | 0 | NON_SYNONYMOU | MODERAT | MISSENSE | Ggc/Agc | G663S | PLES_03011 | | | T | T | T | T | T | T | T | T | T | T |
| NC_011770 | 337705 | C | T | 69 | 0 | 6 | 0 | NON_SYNONYMOU | MODERAT | MISSENSE | Gcc/Acc | A652T | PLES_03011 | | | T | T | T | T | T | T | | | | |
| NC_011770 | 337776 | C | G | 75 | 0 | 6 | 0 | NON_SYNONYMOU | MODERAT | MISSENSE | gGc/gCc | G628A | PLES_03011 | G | G | | | | | | | G | G | G | G |
| NC_011770 | 337923 | T | A | 74 | 0 | 2 | 0 | NON_SYNONYMOU | MODERAT | MISSENSE | tAc/tTc | Y579F | PLES_03011 | A | A | | | | | | | | | | |
| NC_011770 | 337931 | G | C | 73 | 0 | 6 | 0 | NON_SYNONYMOU | MODERAT | MISSENSE | aaC/aaG | N576K | PLES_03011 | C | C | C | C | C | C | | | | | | |
| NC_011770 | 338164 | C | T | 69 | 0 | 2 | 0 | NON_SYNONYMOU | MODERAT | MISSENSE | Gcc/Acc | A499T | PLES_03011 | | | | | | | T | T | | | | |
| NC_011770 | 338635 | T | C | 86 | 0 | 8 | 0 | NON_SYNONYMOU | MODERAT | MISSENSE | Aaa/Gaa | K342E | PLES_03011 | C | C | | | | | C | C | C | C | C | C |
| NC_011770 | 338686 | C | T | 84 | 0 | 4 | 0 | NON_SYNONYMOU | MODERAT | MISSENSE | Gtc/Atc | V325I | PLES_03011 | | | T | T | T | T | | | | | | |
| NC_011770 | 339195 | C | T | 82 | 0 | 4 | 0 | NON_SYNONYMOU | MODERAT | MISSENSE | gGc/gAc | G155D | PLES_03011 | | | T | T | T | T | | | | | | |
| NC_011770 | 339220 | C | T | 76 | 0 | 2 | 0 | NON_SYNONYMOU | MODERAT | MISSENSE | Gcg/Acg | A147T | PLES_03011 | T | T | | | | | | | | | | |
| NC_011770 | 339553 | C | G | 71 | 0 | 4 | 0 | NON_SYNONYMOU | MODERAT | MISSENSE | Gtg/Ctg | V36L | PLES_03011 | | | | | | | | | G | G | G | G |
| NC_011770 | 340276 | C | T | 75 | 0 | 4 | 0 | NON_SYNONYMOU | MODERAT | MISSENSE | Cgc/Tgc | R170C | PLES_03021 | | | | | | | | | T | T | T | T |
| NC_011770 | 341418 | T | C | 52 | 0 | 2 | 0 | NON_SYNONYMOU | MODERAT | MISSENSE | gAa/gGa | E201G | PLES_03031 | | | | | | | C | C | | | | |
| NC_011770 | 341809 | C | G | 66 | 0 | 4 | 0 | NON_SYNONYMOU | MODERAT | MISSENSE | Gcg/Ccg | A71P | PLES_03031 | | | G | G | G | G | | | | | | |
| NC_011770 | 341970 | C | T | 72 | 0 | 2 | 0 | NON_SYNONYMOU | MODERAT | MISSENSE | cGc/cAc | R17H | PLES_03031 | | | | | | | T | T | | | | |
| NC_011770 | 341994 | C | T | 71 | 0 | 4 | 0 | NON_SYNONYMOU | MODERAT | MISSENSE | cGt/cAt | R9H | PLES_03031 | | | | | | | | | T | T | T | T |
| NC_011770 | 342547 | GC | G | 76 | 0 | 2 | 0 | FRAME_SHIFT | HIGH | PA0308 | ggc/ | G177 | PLES_03041 | | | | G | | G | | | | | | |
| NC_011770 | 342477 | G | A | 73 | 0 | 12 | 0 | NON_SYNONYMOU | MODERAT | MISSENSE | Gaa/Aaa | E154K | PLES_03041 | A | A | A | A | A | A | A | A | A | A | A | A |
| NC_011770 | 343061 | T | G | 79 | 0 | 12 | 0 | NON_SYNONYMOU | MODERAT | MISSENSE | cAc/cCc | H250P | PLES_03051 | G | G | G | G | G | G | G | G | G | G | G | G |
| NC_011770 | 343599 | C | T | 83 | 0 | 2 | 0 | NON_SYNONYMOU | MODERAT | MISSENSE | Gcc/Acc | A71T | PLES_03051 | | | | | | | T | T | | | | |
| NC_011770 | 343996 | C | T | 97 | 0 | 2 | 0 | NON_SYNONYMOU | MODERAT | MISSENSE | Cac/Tac | H47Y | PLES_03061 | | | | | | | | | | | T | T |
| NC_011770 | 344030 | C | T | 92 | 0 | 4 | 0 | NON_SYNONYMOU | MODERAT | MISSENSE | aCc/aTc | T58I | PLES_03061 | | | T | T | T | T | | | | | | |
| NC_011770 | 344087 | G | C | 89 | 0 | 10 | 0 | NON_SYNONYMOU | MODERAT | MISSENSE | gGg/gCg | G77A | PLES_03061 | C | C | C | C | C | C | C | C | | | C | C |
| NC_011770 | 344594 | T | C | 74 | 0 | 4 | 0 | NON_SYNONYMOU | MODERAT | MISSENSE | aAg/aGg | K180R | PLES_03071 | | | C | C | C | C | | | | | | |
| NC_011770 | 344791 | G | T | 63 | 0 | 2 | 0 | NON_SYNONYMOU | MODERAT | MISSENSE | gaC/gaA | D114E | PLES_03071 | | | | | | | | | | | T | T |
| NC_011770 | 345044 | T | C | 45 | 0 | 1 | 0 | NON_SYNONYMOU | MODERAT | MISSENSE | gAc/gGc | D30G | PLES_03071 HP | | | | | | | | C | | | | |
| NC_011770 | 345780 | C | A | 76 | 0 | 2 | 0 | NON_SYNONYMOU | MODERAT | MISSENSE | gCc/gAc | A128D | PLES_03081 | | | | | | | A | A | | | | |
| NC_011770 | 346929 | CG | C | 75 | 0 | 1 | 0 | FRAME_SHIFT | HIGH | | ccg/ | P167 | PLES_03101 | | | | | | | C | | | | | |
| NC_011770 | 347014 | C | T | 66 | 0 | 12 | 0 | NON_SYNONYMOU | MODERAT | MISSENSE | aGg/aAg | R139K | PLES_03101 | T | T | T | T | T | T | T | T | T | T | T | T |
| NC_011770 | 347933 | C | T | 62 | 0 | 1 | 0 | NON_SYNONYMOU | MODERAT | MISSENSE | cCg/cTg | P71L | PLES_03111 | | | | T | | | | | | | | |
| NC_011770 | 349947 | C | T | 74 | 0 | 12 | 0 | NON_SYNONYMOU | MODERAT | MISSENSE | Ctc/Ttc | L100F | PLES_03131 | T | T | T | T | T | T | T | T | T | T | T | T |
| NC_011770 | 349980 | C | T | 78 | 0 | 2 | 0 | NON_SYNONYMOU | MODERAT | MISSENSE | Ccg/Tcg | P111S | PLES_03131 | T | T | | | | | | | | | | |
| NC_011770 | 350265 | G | C | 84 | 0 | 2 | 0 | NON_SYNONYMOU | MODERAT | MISSENSE | Gtg/Ctg | V206L | PLES_03131 | | | | | | | C | C | | | | |
| NC_011770 | 351718 | T | G | 68 | 0 | 2 | 0 | NON_SYNONYMOU | MODERAT | MISSENSE | Tcc/Gcc | S195A | PLES_03141 | G | G | | | | | | | | | | |
| NC_011770 | 352210 | G | A | 76 | 0 | 4 | 0 | NON_SYNONYMOU | MODERAT | MISSENSE | aGg/aAg | R104K | PLES_03151 | | | A | A | A | A | | | | | | |
| NC_011770 | 354603 | C | T | 76 | 0 | 2 | 0 | NON_SYNONYMOU | MODERAT | MISSENSE | Ggc/Agc | G416S | PLES_03181 | | | | | | | | | | | T | T |
| NC_011770 | 355395 | C | A | 72 | 0 | 2 | 0 | NON_SYNONYMOU | MODERAT | MISSENSE | Ggc/Tgc | G152C | PLES_03181 | A | A | | | | | | | | | | |
| NC_011770 | 355835 | C | T | 63 | 0 | 4 | 0 | NON_SYNONYMOU | MODERAT | MISSENSE | cGc/cAc | R5H | PLES_03181 | | | T | T | T | T | | | | | | |
| NC_011770 | 355839 | C | G | 63 | 0 | 4 | 0 | NON_SYNONYMOU | MODERAT | MISSENSE | Gaa/Caa | E4Q | PLES_03181 | | | G | G | G | G | | | | | | |
| NC_011770 | 356270 | T | C | 48 | 0 | 8 | 0 | SPLICE_SITE_REGIC | LOW | SILENT | tAa/tGa | *348 | PLES_03191 | C | C | C | C | C | C | | | C | C | | |
| NC_011770 | 356406 | A | G | 44 | 0 | 2 | 0 | NON_SYNONYMOU | MODERAT | MISSENSE | Ttc/Ctc | F303L | PLES_03191 | G | G | | | | | | | | | | |
| NC_011770 | 357746 | A | G | 79 | 0 | 4 | 0 | NON_SYNONYMOU | MODERAT | MISSENSE | Ttc/Ctc | F135L | PLES_03201 | | | G | G | G | G | | | | | | |
| NC_011770 | 357851 | T | G | 88 | 0 | 2 | 0 | NON_SYNONYMOU | MODERAT | MISSENSE | Acc/Ccc | T100P | PLES_03201 | G | G | | | | | | | | | | |
| NC_011770 | 357988 | G | A | 90 | 0 | 2 | 0 | NON_SYNONYMOU | MODERAT | MISSENSE | gCg/gTg | A54V | PLES_03201 | | | | | | | | | A | A | | |
| NC_011770 | 358161 | A | C | 82 | 0 | 12 | 0 | NON_SYNONYMOU | MODERAT | MISSENSE | gTc/gGc | V305G | PLES_03211 | C | C | C | C | C | C | C | C | C | C | C | C |
| NC_011770 | 358173 | C | G | 83 | 0 | 4 | 0 | NON_SYNONYMOU | MODERAT | MISSENSE | cGg/cCg | R301P | PLES_03211 | | | G | G | G | G | | | | | | |
| NC_011770 | 358804 | G | T | 82 | 0 | 2 | 0 | NON_SYNONYMOU | MODERAT | MISSENSE | Cag/Aag | Q91K | PLES_03211 | | | | | | | | | | | T | T |
| NC_011770 | 359510 | G | C | 81 | 0 | 2 | 0 | NON_SYNONYMOU | MODERAT | MISSENSE | aCc/aGc | T204S | PLES_03221 | C | C | | | | | | | | | | |
| NC_011770 | 360908 | G | A | 58 | 0 | 2 | 0 | NON_SYNONYMOU | MODERAT | MISSENSE | cGc/cAc | R154H | PLES_03231 | | | | | | | A | A | | | | |
| NC_011770 | 360802 | C | T | 73 | 0 | 1 | 0 | NON_SYNONYMOU | MODERAT | MISSENSE | Cgg/Tgg | R119W | PLES_03231 | | | | | | | | | | | T | |
| NC_011770 | 361566 | C | T | 83 | 0 | 4 | 0 | NON_SYNONYMOU | MODERAT | MISSENSE | Gcg/Acg | A611T | PLES_03241 | | | T | T | T | T | | | | | | |
| NC_011770 | 361608 | T | G | 82 | 0 | 4 | 0 | NON_SYNONYMOU | MODERAT | MISSENSE | Atg/Ctg | M597L | PLES_03241 | | | G | G | G | G | | | | | | |
| NC_011770 | 361616 | A | C | 82 | 0 | 2 | 0 | NON_SYNONYMOU | MODERAT | MISSENSE | gTc/gGc | V594G | PLES_03241 | C | C | | | | | | | | | | |
| NC_011770 | 361850 | T | C | 86 | 0 | 8 | 0 | NON_SYNONYMOU | MODERAT | MISSENSE | cAc/cGc | H516R | PLES_03241 | | | C | C | C | C | | | C | C | C | C |
| NC_011770 | 362241 | C | T | 70 | 0 | 8 | 0 | NON_SYNONYMOU | MODERAT | MISSENSE | Gat/Aat | D386N | PLES_03241 | T | T | T | T | T | T | T | T | | | | |
| NC_011770 | 362534 | G | T | 85 | 0 | 4 | 0 | NON_SYNONYMOU | MODERAT | MISSENSE | aCc/aAc | T288N | PLES_03241 | | | T | T | T | T | | | | | | |
| NC_011770 | 362813 | C | T | 82 | 0 | 4 | 0 | NON_SYNONYMOU | MODERAT | MISSENSE | aGc/aAc | S195N | PLES_03241 | | | | | | | | | T | T | T | T |
| NC_011770 | 362898 | C | T | 87 | 0 | 4 | 0 | NON_SYNONYMOU | MODERAT | MISSENSE | Gcc/Acc | A167T | PLES_03241 | | | | | | | | | T | T | T | T |
| NC_011770 | 363062 | C | T | 90 | 0 | 1 | 0 | NON_SYNONYMOU | MODERAT | MISSENSE | gGc/gAc | G112D | PLES_03241 | | | | | | | T | | | | | |

| | | | | | | | | | | | | | | | | | | | | | | | | | |
|---|---|---|---|---|---|---|---|---|---|---|---|---|---|---|---|---|---|---|---|---|---|---|---|---|---|
| NC_011770 | 362454 | T | C | 79 | 0 | 1 | 0 | NON_SYNONYMOU | MODERAT | MISSENSE | Acc/Gcc | T315A | PLES_03241 | | C | | | | | | | | | | |
| NC_011770 | 363849 | T | G | 67 | 2 | 1 | 0 | NON_SYNONYMOU | MODERAT | MISSENSE | cTg/cGg | L55R | PLES_03251 | . | . | | | | | G | | | | | |
| NC_011770 | 367485 | C | A | 77 | 0 | 4 | 0 | NON_SYNONYMOU | MODERAT | MISSENSE | Ctc/Atc | L5I | PLES_03291 | | | A | A | A | A | | | | | | |
| NC_011770 | 367593 | G | A | 71 | 0 | 4 | 0 | NON_SYNONYMOU | MODERAT | MISSENSE | Gtc/Atc | V41I | PLES_03291 | | | | | | | | | A | A | A | A |
| NC_011770 | 367684 | C | T | 74 | 0 | 12 | 0 | NON_SYNONYMOU | MODERAT | MISSENSE | gCg/gTg | A71V | PLES_03291 | T | T | T | T | T | T | T | T | T | T | T | T |
| NC_011770 | 367803 | G | A | 77 | 0 | 2 | 0 | NON_SYNONYMOU | MODERAT | MISSENSE | Gcc/Acc | A111T | PLES_03291 | | | | | | | A | A | | | | |
| NC_011770 | 367845 | A | G | 84 | 0 | 4 | 0 | NON_SYNONYMOU | MODERAT | MISSENSE | Acc/Gcc | T125A | PLES_03291 | | | G | G | G | G | | | | | | |
| NC_011770 | 368144 | C | T | 72 | 0 | 1 | 0 | NON_SYNONYMOU | MODERAT | MISSENSE | Ctt/Ttt | L52F | PLES_03301 | | | | | | | T | | | | | |
| NC_011770 | 368264 | A | G | 73 | 0 | 4 | 0 | NON_SYNONYMOU | MODERAT | MISSENSE | Aca/Gca | T92A | PLES_03301 | | | G | G | G | G | | | | | | |
| NC_011770 | 368487 | T | C | 81 | 0 | 2 | 0 | NON_SYNONYMOU | MODERAT | MISSENSE | aTt/aCt | I166T | PLES_03301 | C | C | | | | | | | | | | |
| NC_011770 | 368567 | C | T | 80 | 0 | 8 | 0 | NON_SYNONYMOU | MODERAT | MISSENSE | Cgc/Tgc | R193C | PLES_03301 | | | T | T | T | T | | | T | T | T | T |
| NC_011770 | 368679 | T | C | 87 | 0 | 6 | 0 | NON_SYNONYMOU | MODERAT | MISSENSE | gTg/gCg | V230A | PLES_03301 | C | C | | | | | | | C | C | C | C |
| NC_011770 | 368685 | C | T | 85 | 0 | 2 | 0 | NON_SYNONYMOU | MODERAT | MISSENSE | cCg/cTg | P232L | PLES_03301 | T | T | | | | | | | | | | |
| NC_011770 | 368808 | C | T | 84 | 0 | 2 | 0 | NON_SYNONYMOU | MODERAT | MISSENSE | gCg/gTg | A273V | PLES_03301 | T | T | | | | | | | | | | |
| NC_011770 | 368828 | G | A | 80 | 0 | 2 | 0 | NON_SYNONYMOU | MODERAT | MISSENSE | Gcg/Acg | A280T | PLES_03301 | | | | | | | | | | | A | A |
| NC_011770 | 369730 | TC | T | 88 | 0 | 2 | 0 | FRAME_SHIFT | HIGH | | gac/ | D390 | PLES_03311 | T | T | | | | | | | | | | |
| NC_011770 | 369733 | C | G | 80 | 2 | 4 | 0 | NON_SYNONYMOU | MODERAT | MISSENSE | gGg/gCg | G389A | PLES_03311 | . | . | G | G | G | G | | | | | | |
| NC_011770 | 369767 | A | G | 90 | 0 | 4 | 0 | NON_SYNONYMOU | MODERAT | MISSENSE | Ttc/Ctc | F378L | PLES_03311 | G | G | | | | | G | G | | | | |
| NC_011770 | 370325 | C | A | 89 | 0 | 6 | 0 | NON_SYNONYMOU | MODERAT | MISSENSE | Gcc/Tcc | A192S | PLES_03311 | A | A | A | A | A | A | | | | | | |
| NC_011770 | 370363 | T | C | 86 | 0 | 6 | 0 | NON_SYNONYMOU | MODERAT | MISSENSE | aAt/aGt | N179S | PLES_03311 | C | C | | | | | | | C | C | C | C |
| NC_011770 | 370384 | T | C | 82 | 0 | 6 | 0 | NON_SYNONYMOU | MODERAT | MISSENSE | aAg/aGg | K172R | PLES_03311 | C | C | C | C | C | C | | | | | | |
| NC_011770 | 370418 | A | T | 83 | 0 | 6 | 0 | NON_SYNONYMOU | MODERAT | MISSENSE | Ttc/Atc | F161I | PLES_03311 | T | T | | | | | | | G | G | G | G |
| NC_011770 | 370786 | G | A | 68 | 0 | 12 | 0 | NON_SYNONYMOU | MODERAT | MISSENSE | gCg/gTg | A38V | PLES_03311 | A | A | A | A | A | A | A | A | A | A | A | A |
| NC_011770 | 371097 | C | T | 68 | 0 | 1 | 0 | NON_SYNONYMOU | MODERAT | MISSENSE | Gac/Aac | D183N | PLES_03321 | | | | | | | T | | | | | |
| NC_011770 | 374955 | G | A | 76 | 0 | 6 | 0 | NON_SYNONYMOU | MODERAT | MISSENSE | aGc/aAc | S101N | PLES_03351 | | | | | | | A | A | A | A | A | A |
| NC_011770 | 375151 | G | T | 79 | 0 | 4 | 0 | NON_SYNONYMOU | MODERAT | MISSENSE | gaG/gaT | E166D | PLES_03351 | | | T | T | T | T | | | | | | |
| NC_011770 | 376419 | A | AC | 64 | 0 | 1 | 0 | FRAME_SHIFT | HIGH | | gta/gGta | V42G? | PLES_03361 | | | | | | | | AC | | | | |
| NC_011770 | 377328 | G | A | 88 | 0 | 4 | 0 | NON_SYNONYMOU | MODERAT | MISSENSE | Gcc/Acc | A222T | PLES_03371 | | | A | A | A | A | | | | | | |
| NC_011770 | 379544 | C | T | 87 | 0 | 2 | 0 | NON_SYNONYMOU | MODERAT | MISSENSE | Gtc/Atc | V187I | PLES_03401 | T | T | | | | | | | | | | |
| NC_011770 | 379627 | C | T | 88 | 0 | 2 | 0 | NON_SYNONYMOU | MODERAT | MISSENSE | cGc/cAc | R159H | PLES_03401 | | | | | | | | | | | T | T |
| NC_011770 | 380948 | C | T | 75 | 0 | 2 | 0 | NON_SYNONYMOU | MODERAT | MISSENSE | cGg/cAg | R190Q | PLES_03411 | T | T | | | | | | | | | | |
| NC_011770 | 380990 | A | G | 73 | 0 | 1 | 0 | NON_SYNONYMOU | MODERAT | MISSENSE | cTg/cCg | L176P | PLES_03411 | | | | | | | G | | | | | |
| NC_011770 | 382234 | G | A | 75 | 0 | 1 | 0 | STOP_GAINED | HIGH | NONSENSE | Cag/Tag | Q221* | PLES_03421 | | A | | | | | | | | | | |
| NC_011770 | 381898 | C | T | 72 | 0 | 6 | 0 | NON_SYNONYMOU | MODERAT | MISSENSE | Ggc/Agc | G333S | PLES_03421 | T | T | T | T | T | T | | | | | | |
| NC_011770 | 382027 | G | C | 74 | 0 | 4 | 0 | NON_SYNONYMOU | MODERAT | MISSENSE | Cac/Gac | H290D | PLES_03421 | | | | | | | | | C | C | C | C |
| NC_011770 | 382060 | G | A | 77 | 0 | 4 | 0 | NON_SYNONYMOU | MODERAT | MISSENSE | Ctc/Ttc | L279F | PLES_03421 | | | | | | | | | A | A | A | A |
| NC_011770 | 382438 | C | T | 52 | 0 | 2 | 0 | NON_SYNONYMOU | MODERAT | MISSENSE | Gcc/Acc | A153T | PLES_03421 | | | | | | | T | T | | | | |
| NC_011770 | 382461 | C | T | 58 | 0 | 2 | 0 | NON_SYNONYMOU | MODERAT | MISSENSE | gGc/gAc | G145D | PLES_03421 | | | | | | | T | T | | | | |
| NC_011770 | 382516 | A | G | 60 | 0 | 12 | 0 | NON_SYNONYMOU | MODERAT | MISSENSE | Ttc/Ctc | F127L | PLES_03421 | G | G | G | G | G | G | G | G | G | G | G | G |
| NC_011770 | 382795 | C | G | 72 | 0 | 4 | 0 | NON_SYNONYMOU | MODERAT | MISSENSE | Gtt/Ctt | V34L | PLES_03421 | | | | | | | | | G | G | G | G |
| NC_011770 | 381679 | C | T | 74 | 0 | 1 | 0 | NON_SYNONYMOU | MODERAT | MISSENSE | Ggc/Agc | G406S | PLES_03421 | | | | | | | | | | | | **T** |
| NC_011770 | 383300 | T | C | 85 | 0 | 1 | 0 | NON_SYNONYMOU | MODERAT | MISSENSE | gTc/gCc | V72A | PLES_03431 | | | | | | | C | | | | | |
| NC_011770 | 383317 | A | G | 87 | 0 | 4 | 0 | NON_SYNONYMOU | MODERAT | MISSENSE | Atg/Gtg | M78V | PLES_03431 | | | | | | | | | G | G | G | G |
| NC_011770 | 384949 | A | C | 70 | 0 | 4 | 0 | NON_SYNONYMOU | MODERAT | MISSENSE | cAg/cCg | Q35P | PLES_03451 | | | | | | | | | C | C | C | C |
| NC_011770 | 385287 | C | A | 83 | 0 | 2 | 0 | NON_SYNONYMOU | MODERAT | MISSENSE | Ctc/Atc | L148I | PLES_03451 | | | | | | | A | A | | | | |
| NC_011770 | 385334 | G | T | 88 | 0 | 12 | 0 | NON_SYNONYMOU | MODERAT | MISSENSE | agG/agT | R163S | PLES_03451 | T | T | T | T | T | T | T | T | T | T | T | T |
| NC_011770 | 386070 | CG | C | 76 | 0 | 1 | 0 | FRAME_SHIFT | HIGH | | ccg/ | P45 | PLES_03461 | | | | | | | C | | | | | |
| NC_011770 | 385972 | G | T | 84 | 0 | 12 | 0 | NON_SYNONYMOU | MODERAT | MISSENSE | tgG/tgT | W12C | PLES_03461 | T | T | T | T | T | T | T | T | T | T | T | T |
| NC_011770 | 385995 | T | C | 82 | 0 | 12 | 0 | NON_SYNONYMOU | MODERAT | MISSENSE | gTc/gCc | V20A | PLES_03461 | C | C | C | C | C | C | C | C | C | C | C | C |
| NC_011770 | 386514 | A | G | 70 | 0 | 12 | 0 | NON_SYNONYMOU | MODERAT | MISSENSE | aAg/aGg | K193R | PLES_03461 | G | G | G | G | G | G | G | G | G | G | G | G |
| NC_011770 | 386519 | A | G | 63 | 0 | 4 | 0 | NON_SYNONYMOU | MODERAT | MISSENSE | Atc/Gtc | I195V | PLES_03461 | | | | | | | G | G | | | G | G |
| NC_011770 | 386613 | G | A | 68 | 0 | 2 | 0 | NON_SYNONYMOU | MODERAT | MISSENSE | aGc/aAc | S226N | PLES_03461 | | | | | | | | | | | A | A |
| NC_011770 | 386741 | G | A | 73 | 0 | 2 | 0 | NON_SYNONYMOU | MODERAT | MISSENSE | Gtg/Atg | V269M | PLES_03461 | A | A | | | | | | | | | | |
| NC_011770 | 386757 | C | T | 74 | 0 | 2 | 0 | NON_SYNONYMOU | MODERAT | MISSENSE | gCc/gTc | A274V | PLES_03461 | T | T | | | | | | | | | | |
| NC_011770 | 388823 | T | C | 92 | 0 | 10 | 0 | NON_SYNONYMOU | MODERAT | MISSENSE | Ata/Gta | I249V | PLES_03491 | | | C | C | C | C | C | C | C | C | C | C |
| NC_011770 | 392501 | C | T | 85 | 0 | 2 | 0 | NON_SYNONYMOU | MODERAT | MISSENSE | cGg/cAg | R220Q | PLES_03511 | T | T | | | | | | | | | | |
| NC_011770 | 396234 | C | T | 79 | 0 | 4 | 0 | NON_SYNONYMOU | MODERAT | MISSENSE | cCc/cTc | P159L | PLES_03551 | | | | | | | | | T | T | T | T |
| NC_011770 | 397076 | G | A | 80 | 0 | 4 | 0 | NON_SYNONYMOU | MODERAT | MISSENSE | Gta/Ata | V15I | PLES_03571 | | | | | | | | | A | A | A | A |
| NC_011770 | 397107 | T | C | 79 | 0 | 4 | 0 | NON_SYNONYMOU | MODERAT | MISSENSE | gTg/gCg | V25A | PLES_03571 | | | | | | | | | C | C | C | C |
| NC_011770 | 397119 | C | G | 79 | 0 | 4 | 0 | NON_SYNONYMOU | MODERAT | MISSENSE | gCc/gGc | A29G | PLES_03571 | | | | | | | | | G | G | G | G |
| NC_011770 | 397122 | A | C | 79 | 0 | 4 | 0 | NON_SYNONYMOU | MODERAT | MISSENSE | aAg/aCg | K30T | PLES_03571 | | | | | | | | | C | C | C | C |
| NC_011770 | 397518 | G | A | 79 | 0 | 2 | 0 | NON_SYNONYMOU | MODERAT | MISSENSE | aGc/aAc | S162N | PLES_03571 | | | | | | | A | A | | | | |

| | | | | | | | | | | | | | | | | | | | | | | | | | |
|---|---|---|---|---|---|---|---|---|---|---|---|---|---|---|---|---|---|---|---|---|---|---|---|---|---|
| NC_011770 | 397604 | G | A | 74 | 0 | 2 | 0 | NON_SYNONYMOU | MODERAT | MISSENSE | Gcc/Acc | A191T | PLES_03571 | | | | | | | A | A | | | | |
| NC_011770 | 397651 | C | G | 77 | 0 | 6 | 0 | NON_SYNONYMOU | MODERAT | MISSENSE | caC/caG | H206Q | PLES_03571 | G | G | G | G | G | G | | | | | | |
| NC_011770 | 397901 | A | T | 71 | 0 | 12 | 0 | NON_SYNONYMOU | MODERAT | MISSENSE | Acc/Tcc | T290S | PLES_03571 | T | T | T | T | T | T | T | T | T | T | T | T |
| NC_011770 | 397929 | G | A | 74 | 0 | 2 | 0 | NON_SYNONYMOU | MODERAT | MISSENSE | gGc/gAc | G299D | PLES_03571 | | | | | | | A | A | | | | |
| NC_011770 | 397942 | G | T | 75 | 0 | 4 | 0 | NON_SYNONYMOU | MODERAT | MISSENSE | gaG/gaT | E303D | PLES_03571 | | | T | T | T | T | | | | | | |
| NC_011770 | 398633 | C | T | 59 | 0 | 4 | 0 | NON_SYNONYMOU | MODERAT | MISSENSE | gGt/gAt | G409D | PLES_03581 | T | T | | | | | T | T | | | | |
| NC_011770 | 399345 | C | T | 74 | 0 | 4 | 0 | NON_SYNONYMOU | MODERAT | MISSENSE | Gta/Ata | V172I | PLES_03581 | | | | | | | | | T | T | T | T |
| NC_011770 | 399765 | C | T | 66 | 0 | 4 | 0 | NON_SYNONYMOU | MODERAT | MISSENSE | Gac/Aac | D32N | PLES_03581 | | | T | T | T | T | | | | | | |
| NC_011770 | 401152 | G | A | 72 | 0 | 12 | 0 | NON_SYNONYMOU | MODERAT | MISSENSE | gCg/gTg | A478V | PLES_03611 | A | A | A | A | A | A | A | A | A | A | A | A |
| NC_011770 | 401156 | T | C | 73 | 0 | 10 | 0 | NON_SYNONYMOU | MODERAT | MISSENSE | Acc/Gcc | T477A | PLES_03611 | C | C | C | C | C | C | | | C | C | C | C |
| NC_011770 | 401271 | G | T | 81 | 0 | 4 | 0 | NON_SYNONYMOU | MODERAT | MISSENSE | agC/agA | S438R | PLES_03611 | | | T | T | T | T | | | | | | |
| NC_011770 | 401963 | T | C | 78 | 0 | 1 | 0 | NON_SYNONYMOU | MODERAT | MISSENSE | Acc/Gcc | T208A | PLES_03611 | | | | | | | C | | | | | |
| NC_011770 | 403290 | A | AG | 92 | 0 | 2 | 0 | FRAME_SHIFT | HIGH | | ccc/cccC | P461P? | PLES_03631 | AG | | | | | | | AG | | | | |
| NC_011770 | 403652 | T | C | 87 | 0 | 1 | 0 | NON_SYNONYMOU | MODERAT | MISSENSE | aAc/aGc | N341S | PLES_03631 | | | | | | | C | | | | | |
| NC_011770 | 403829 | A | G | 81 | 0 | 1 | 0 | NON_SYNONYMOU | MODERAT | MISSENSE | gTg/gCg | V282A | PLES_03631 | | | | | | | G | | | | | |
| NC_011770 | 404327 | A | G | 89 | 0 | 1 | 0 | NON_SYNONYMOU | MODERAT | MISSENSE | gTg/gCg | V116A | PLES_03631 | | | | | | | | **G** | | | | |
| NC_011770 | 406250 | G | T | 80 | 0 | 4 | 0 | NON_SYNONYMOU | MODERAT | MISSENSE | agC/agA | S272R | PLES_03651 | T | T | | | | | | | | | T | T |
| NC_011770 | 407143 | G | A | 62 | 0 | 2 | 0 | NON_SYNONYMOU | MODERAT | MISSENSE | Cgc/Tgc | R84C | PLES_03661 | A | A | | | | | | | | | | |
| NC_011770 | 407259 | A | G | 61 | 0 | 10 | 0 | NON_SYNONYMOU | MODERAT | MISSENSE | gTc/gCc | V45A | PLES_03661 | G | G | G | G | G | G | | | G | G | G | G |
| NC_011770 | 407301 | G | A | 58 | 0 | 2 | 0 | NON_SYNONYMOU | MODERAT | MISSENSE | gCc/gTc | A31V | PLES_03661 | | | | | | | | | | | A | A |
| NC_011770 | 407732 | C | T | 64 | 0 | 2 | 0 | NON_SYNONYMOU | MODERAT | MISSENSE | cGg/cAg | R179Q | PLES_03671 | | | | | | | | | | | T | T |
| NC_011770 | 407811 | C | T | 65 | 0 | 2 | 0 | NON_SYNONYMOU | MODERAT | MISSENSE | Ggc/Agc | G153S | PLES_03671 | | | T | T | | | | | | | | |
| NC_011770 | 407898 | C | T | 58 | 0 | 2 | 0 | NON_SYNONYMOU | MODERAT | MISSENSE | Gct/Act | A124T | PLES_03671 | T | T | | | | | | | | | | |
| NC_011770 | 408877 | T | C | 57 | 0 | 2 | 0 | NON_SYNONYMOU | MODERAT | MISSENSE | cAc/cGc | H293R | PLES_03681 | | | | | | | | | C | C | | |
| NC_011770 | 408919 | A | G | 69 | 0 | 10 | 0 | NON_SYNONYMOU | MODERAT | MISSENSE | gTc/gCc | V279A | PLES_03681 | G | G | G | G | G | G | | | G | G | G | G |
| NC_011770 | 409070 | T | G | 71 | 0 | 4 | 0 | NON_SYNONYMOU | MODERAT | MISSENSE | Acg/Ccg | T229P | PLES_03681 | | | | | | | | | G | G | G | G |
| NC_011770 | 409076 | C | T | 71 | 0 | 4 | 0 | NON_SYNONYMOU | MODERAT | MISSENSE | Gtt/Att | V227I | PLES_03681 | | | | | | | | | T | T | T | T |
| NC_011770 | 409664 | G | T | 72 | 0 | 8 | 0 | NON_SYNONYMOU | MODERAT | MISSENSE | Ccc/Acc | P31T | PLES_03681 | | | T | T | T | T | | | T | T | T | T |
| NC_011770 | 411072 | G | A | 28 | 0 | 2 | 0 | NON_SYNONYMOU | MODERAT | MISSENSE | Ctt/Ttt | L25F | PLES_03691 | | | | | | | A | A | | | | |
| NC_011770 | 415349 | C | T | 69 | 0 | 2 | 0 | NON_SYNONYMOU | MODERAT | MISSENSE | Cgc/Tgc | R3C | PLES_03741 | T | T | | | | | | | | | | |
| NC_011770 | 415530 | G | A | 67 | 0 | 12 | 0 | NON_SYNONYMOU | MODERAT | MISSENSE | gGt/gAt | G63D | PLES_03741 | A | A | A | A | A | A | A | A | A | A | A | A |
| NC_011770 | 415745 | G | A | 73 | 0 | 2 | 0 | NON_SYNONYMOU | MODERAT | MISSENSE | Gtg/Atg | V135M | PLES_03741 | | | | | | | | | A | A | | |
| NC_011770 | 419608 | C | T | 71 | 0 | 2 | 0 | NON_SYNONYMOU | MODERAT | MISSENSE | Ccg/Tcg | P122S | PLES_03801 | | | | | | | T | T | | | | |
| NC_011770 | 419674 | G | A | 75 | 0 | 2 | 0 | NON_SYNONYMOU | MODERAT | MISSENSE | Ggc/Agc | G144S | PLES_03801 | | | | | | | | | | | A | A |
| NC_011770 | 419942 | G | T | 91 | 0 | 2 | 0 | NON_SYNONYMOU | MODERAT | MISSENSE | gGt/gTt | G233V | PLES_03801 | | | T | T | | | | | | | | |
| NC_011770 | 420054 | C | A | 100 | 0 | 4 | 0 | NON_SYNONYMOU | MODERAT | MISSENSE | gaC/gaA | D270E | PLES_03801 | | | A | A | A | A | | | | | | |
| NC_011770 | 420242 | A | G | 102 | 0 | 4 | 0 | NON_SYNONYMOU | MODERAT | MISSENSE | gAc/gGc | D333G | PLES_03801 | | | G | G | G | G | | | | | | |
| NC_011770 | 421391 | C | T | 77 | 0 | 2 | 0 | NON_SYNONYMOU | MODERAT | MISSENSE | cGc/cAc | R364H | PLES_03831 | T | T | | | | | | | | | | |
| NC_011770 | 421402 | G | C | 77 | 0 | 4 | 0 | NON_SYNONYMOU | MODERAT | MISSENSE | gaC/gaG | D360E | PLES_03831 | | | C | C | C | C | | | | | | |
| NC_011770 | 421727 | C | T | 89 | 0 | 2 | 0 | NON_SYNONYMOU | MODERAT | MISSENSE | cGc/cAc | R252H | PLES_03831 | | | | | | | | | | | T | T |
| NC_011770 | 421778 | T | G | 84 | 0 | 2 | 0 | NON_SYNONYMOU | MODERAT | MISSENSE | gAg/gCg | E235A | PLES_03831 | | | | | | | | | G | G | | |
| NC_011770 | 421890 | A | G | 77 | 0 | 1 | 0 | NON_SYNONYMOU | MODERAT | MISSENSE | Tgg/Cgg | W198R | PLES_03831 | | | | | | | G | | | | | |
| NC_011770 | 422373 | G | T | 35 | 0 | 2 | 0 | NON_SYNONYMOU | MODERAT | MISSENSE | Ccg/Acg | P37T | PLES_03831 | | | | | | | | | T | T | | |
| NC_011770 | 422436 | C | T | 19 | 0 | 2 | 0 | NON_SYNONYMOU | MODERAT | MISSENSE | Gtg/Atg | V16M | PLES_03831 | | | | | | | | | T | T | | |
| NC_011770 | 423394 | G | C | 78 | 0 | 4 | 0 | NON_SYNONYMOU | MODERAT | MISSENSE | gCc/gGc | A73G | PLES_03851 | C | C | | | | | | | C | C | | |
| NC_011770 | 423395 | C | T | 76 | 0 | 4 | 0 | NON_SYNONYMOU | MODERAT | MISSENSE | Gcc/Acc | A73T | PLES_03851 | T | T | | | | | | | T | T | | |
| NC_011770 | 423511 | A | G | 81 | 0 | 1 | 0 | NON_SYNONYMOU | MODERAT | MISSENSE | gTg/gCg | V34A | PLES_03851 | | | | | | | | G | | | | |
| NC_011770 | 424035 | C | A | 88 | 0 | 12 | 0 | NON_SYNONYMOU | MODERAT | MISSENSE | tGc/tTc | C110F | PLES_03861 | A | A | A | A | A | A | A | A | A | A | A | A |
| NC_011770 | 425714 | C | T | 74 | 0 | 2 | 0 | NON_SYNONYMOU | MODERAT | MISSENSE | gGg/gAg | G618E | PLES_03881 | T | T | | | | | | | | | | |
| NC_011770 | 425724 | C | T | 74 | 0 | 1 | 0 | NON_SYNONYMOU | MODERAT | MISSENSE | Ggg/Agg | G615R | PLES_03881 | | | | | | | T | | | | | |
| NC_011770 | 427823 | G | T | 48 | 0 | 2 | 0 | NON_SYNONYMOU | MODERAT | MISSENSE | Cat/Aat | H8N | PLES_03891 | | | | | | | | | T | T | | |
| NC_011770 | 429715 | C | T | 76 | 0 | 4 | 0 | NON_SYNONYMOU | MODERAT | MISSENSE | Gtc/Atc | V128I | PLES_03921 | | | T | T | T | T | | | | | | |
| NC_011770 | 433988 | T | C | 77 | 0 | 2 | 0 | NON_SYNONYMOU | MODERAT | MISSENSE | gTg/gCg | V86A | PLES_03961 | | | | | | | C | C | | | | |
| NC_011770 | 434473 | A | T | 74 | 0 | 4 | 0 | NON_SYNONYMOU | MODERAT | MISSENSE | tAc/tTc | Y32F | PLES_03971 | | | | | | | | | T | T | T | T |
| NC_011770 | 434679 | T | A | 54 | 0 | 12 | 0 | NON_SYNONYMOU | MODERAT | MISSENSE | Tcc/Acc | S101T | PLES_03971 | A | A | A | A | A | A | A | A | A | A | A | A |
| NC_011770 | 435980 | A | G | 81 | 0 | 4 | 0 | NON_SYNONYMOU | MODERAT | MISSENSE | Acc/Gcc | T78A | PLES_03981 | | | | | | | | | G | G | G | G |
| NC_011770 | 436837 | C | A | 69 | 0 | 4 | 0 | NON_SYNONYMOU | MODERAT | MISSENSE | gaC/gaA | D363E | PLES_03981 | | | A | A | A | A | | | | | | |
| NC_011770 | 436847 | G | A | 71 | 0 | 2 | 0 | NON_SYNONYMOU | MODERAT | MISSENSE | Ggt/Agt | G367S | PLES_03981 | A | A | | | | | | | | | | |
| NC_011770 | 441594 | C | G | 75 | 0 | 4 | 0 | NON_SYNONYMOU | MODERAT | MISSENSE | gaG/gaC | E166D | PLES_04041 | | | G | G | G | G | | | | | | |
| NC_011770 | 441850 | G | T | 66 | 0 | 2 | 0 | NON_SYNONYMOU | MODERAT | MISSENSE | aCc/aAc | T81N | PLES_04041 | | | | | | | | | | | T | T |
| NC_011770 | 441923 | A | G | 54 | 0 | 6 | 0 | NON_SYNONYMOU | MODERAT | MISSENSE | Ttc/Ctc | F57L | PLES_04041 | | | G | G | G | G | | | G | G | | |
| NC_011770 | 441992 | C | T | 46 | 0 | 1 | 0 | NON_SYNONYMOU | MODERAT | MISSENSE | Gct/Act | A34T | PLES_04041 | | | | | | | T | | | | | |

| | | | | | | | | | | | | | | | | | | | | | | | | | |
|---|---|---|---|---|---|---|---|---|---|---|---|---|---|---|---|---|---|---|---|---|---|---|---|---|---|
| NC_011770 | 442004 | A | G | 45 | 0 | 1 | 0 | NON_SYNONYMOU | MODERAT | MISSENSE | Tcc/Ccc | S30P | PLES_04041 tonB3 | | | | | | | | **G** | | | | |
| NC_011770 | 458361 | G | C | 72 | 0 | 8 | 0 | NON_SYNONYMOU | MODERAT | MISSENSE | aCg/aGg | T455R | PLES_04161 | | | C | C | C | C | | | C | C | C | C |
| NC_011770 | 458698 | T | C | 68 | 0 | 4 | 0 | NON_SYNONYMOU | MODERAT | MISSENSE | Acc/Gcc | T343A | PLES_04161 | | | C | C | C | C | | | | | | |
| NC_011770 | 458748 | G | T | 74 | 0 | 2 | 0 | NON_SYNONYMOU | MODERAT | MISSENSE | cCc/cAc | P326H | PLES_04161 | T | T | | | | | | | | | | |
| NC_011770 | 458803 | C | T | 66 | 0 | 2 | 0 | NON_SYNONYMOU | MODERAT | MISSENSE | Gtc/Atc | V308I | PLES_04161 | | | | | | | T | T | | | | |
| NC_011770 | 458867 | G | C | 69 | 0 | 4 | 0 | NON_SYNONYMOU | MODERAT | MISSENSE | gaC/gaG | D286E | PLES_04161 | C | C | | | | | | | C | C | | |
| NC_011770 | 458950 | C | T | 69 | 0 | 1 | 0 | NON_SYNONYMOU | MODERAT | MISSENSE | Gac/Aac | D259N | PLES_04161 | | | | | | | T | | | | | |
| NC_011770 | 458953 | C | T | 70 | 0 | 2 | 0 | NON_SYNONYMOU | MODERAT | MISSENSE | Gac/Aac | D258N | PLES_04161 | | | | | | | | | | | T | T |
| NC_011770 | 459183 | C | T | 63 | 0 | 4 | 0 | NON_SYNONYMOU | MODERAT | MISSENSE | cGc/cAc | R181H | PLES_04161 | | | T | T | T | T | | | | | | |
| NC_011770 | 459220 | C | T | 62 | 0 | 4 | 0 | NON_SYNONYMOU | MODERAT | MISSENSE | Gac/Aac | D169N | PLES_04161 | | | T | T | T | T | | | | | | |
| NC_011770 | 459364 | T | C | 57 | 0 | 4 | 0 | NON_SYNONYMOU | MODERAT | MISSENSE | Aac/Gac | N121D | PLES_04161 | | | C | C | C | C | | | | | | |
| NC_011770 | 459415 | G | A | 62 | 0 | 4 | 0 | NON_SYNONYMOU | MODERAT | MISSENSE | Ctc/Ttc | L104F | PLES_04161 | | | A | A | A | A | | | | | | |
| NC_011770 | 459423 | T | C | 59 | 0 | 4 | 0 | NON_SYNONYMOU | MODERAT | MISSENSE | cAg/cGg | Q101R | PLES_04161 | | | C | C | C | C | | | | | | |
| NC_011770 | 459433 | G | T | 58 | 1 | 4 | 0 | NON_SYNONYMOU | MODERAT | MISSENSE | Ccc/Acc | P98T | PLES_04161 | | | T | T | T | T | . | | | | | |
| NC_011770 | 459276 | G | A | 63 | 0 | 1 | 0 | NON_SYNONYMOU | MODERAT | MISSENSE | gCc/gTc | A150V | PLES_04161 | | | | | | | | | | | | **A** |
| NC_011770 | 460306 | T | C | 86 | 0 | 2 | 0 | NON_SYNONYMOU | MODERAT | MISSENSE | aAg/aGg | K59R | PLES_04171 | | | | | | | C | C | | | | |
| NC_011770 | 460418 | C | T | 79 | 0 | 4 | 0 | NON_SYNONYMOU | MODERAT | MISSENSE | Gcc/Acc | A22T | PLES_04171 | | | T | T | T | T | | | | | | |
| NC_011770 | 460445 | A | C | 77 | 0 | 4 | 0 | NON_SYNONYMOU | MODERAT | MISSENSE | Tcc/Gcc | S13A | PLES_04171 | | | C | C | C | C | | | | | | |
| NC_011770 | 459925 | A | G | 81 | 0 | 1 | 0 | NON_SYNONYMOU | MODERAT | MISSENSE | cTg/cCg | L186P | PLES_04171 | | | | G | | | | | | | | |
| NC_011770 | 463553 | T | C | 69 | 0 | 2 | 0 | NON_SYNONYMOU | MODERAT | MISSENSE | aTg/aCg | M468T | PLES_04191 | | | | | | | | | C | C | | |
| NC_011770 | 464050 | A | C | 81 | 0 | 12 | 0 | NON_SYNONYMOU | MODERAT | MISSENSE | Acg/Ccg | T77P | PLES_04201 | C | C | C | C | C | C | C | C | C | C | C | C |
| NC_011770 | 464060 | C | T | 84 | 0 | 2 | 0 | NON_SYNONYMOU | MODERAT | MISSENSE | gCc/gTc | A80V | PLES_04201 | | | | | | | | | | | T | T |
| NC_011770 | 471968 | C | T | 64 | 0 | 6 | 0 | NON_SYNONYMOU | MODERAT | MISSENSE | Ggc/Agc | G527S | PLES_04261 | | | T | T | T | T | | | T | T | | |
| NC_011770 | 472132 | G | A | 69 | 0 | 2 | 0 | NON_SYNONYMOU | MODERAT | MISSENSE | cCc/cTc | P472L | PLES_04261 | | | | | | | A | A | | | | |
| NC_011770 | 472999 | T | G | 82 | 0 | 12 | 0 | NON_SYNONYMOU | MODERAT | MISSENSE | cAc/cCc | H183P | PLES_04261 | G | G | G | G | G | G | G | G | G | G | G | G |
| NC_011770 | 473054 | T | G | 81 | 0 | 2 | 0 | NON_SYNONYMOU | MODERAT | MISSENSE | Atg/Ctg | M165L | PLES_04261 | | | G | G | | | | | | | | |
| NC_011770 | 473113 | C | T | 78 | 0 | 1 | 0 | NON_SYNONYMOU | MODERAT | MISSENSE | gGg/gAg | G145E | PLES_04261 | | | | | | | T | | | | | |
| NC_011770 | 473348 | C | T | 62 | 0 | 1 | 0 | NON_SYNONYMOU | MODERAT | MISSENSE | Ggt/Agt | G67S | PLES_04261 | | | | | T | | | | | | | |
| NC_011770 | 472232 | C | T | 65 | 0 | 1 | 0 | NON_SYNONYMOU | MODERAT | MISSENSE | Gct/Act | A439T | PLES_04261 | | T | | | | | | | | | | |
| NC_011770 | 474149 | G | A | 69 | 0 | 2 | 0 | NON_SYNONYMOU | MODERAT | MISSENSE | tCc/tTc | S240F | PLES_04271 | A | A | | | | | | | | | | |
| NC_011770 | 474592 | G | C | 99 | 0 | 2 | 0 | NON_SYNONYMOU | MODERAT | MISSENSE | gaC/gaG | D92E | PLES_04271 | | | | | | | | | | | C | C |
| NC_011770 | 476267 | G | T | 53 | 0 | 8 | 0 | NON_SYNONYMOU | MODERAT | MISSENSE | gCg/gAg | A47E | PLES_04291 | T | T | | | | | T | T | T | T | T | T |
| NC_011770 | 476322 | T | C | 47 | 0 | 2 | 0 | NON_SYNONYMOU | MODERAT | MISSENSE | Acc/Gcc | T29A | PLES_04291 | | | | | | | | | C | C | | |
| NC_011770 | 476369 | C | T | 44 | 0 | 2 | 0 | NON_SYNONYMOU | MODERAT | MISSENSE | cGc/cAc | R13H | PLES_04291 | | | | | | | | | T | T | | |
| NC_011770 | 478179 | T | A | 58 | 0 | 4 | 0 | NON_SYNONYMOU | MODERAT | MISSENSE | aTc/aAc | I12N | PLES_04311 | | | | | | | A | A | | | A | A |
| NC_011770 | 478370 | T | G | 42 | 0 | 2 | 0 | NON_SYNONYMOU | MODERAT | MISSENSE | Tcc/Gcc | S76A | PLES_04311 | G | G | | | | | | | | | | |
| NC_011770 | 478386 | C | T | 43 | 0 | 2 | 0 | NON_SYNONYMOU | MODERAT | MISSENSE | gCa/gTa | A81V | PLES_04311 | | | | | | | | | | | T | T |
| NC_011770 | 478391 | G | A | 41 | 0 | 2 | 0 | NON_SYNONYMOU | MODERAT | MISSENSE | Gtc/Atc | V83I | PLES_04311 | A | A | | | | | | | | | | |
| NC_011770 | 478838 | C | G | 59 | 0 | 4 | 0 | NON_SYNONYMOU | MODERAT | MISSENSE | gCc/gGc | A45G | PLES_04321 | | | | | | | G | G | | | G | G |
| NC_011770 | 479353 | G | C | 80 | 0 | 12 | 0 | NON_SYNONYMOU | MODERAT | MISSENSE | Gtg/Ctg | V217L | PLES_04321 | C | C | C | C | C | C | C | C | C | C | C | C |
| NC_011770 | 479974 | C | A | 84 | 0 | 4 | 0 | NON_SYNONYMOU | MODERAT | MISSENSE | Ccc/Acc | P424T | PLES_04321 | | | A | A | A | A | | | | | | |
| NC_011770 | 480101 | T | C | 88 | 0 | 2 | 0 | NON_SYNONYMOU | MODERAT | MISSENSE | gTg/gCg | V466A | PLES_04321 | | | | | | | | | C | C | | |
| NC_011770 | 480113 | A | G | 83 | 2 | 8 | 0 | NON_SYNONYMOU | MODERAT | MISSENSE | cAg/cGg | Q470R | PLES_04321 | | | G | G | G | G | G | G | . | . | G | G |
| NC_011770 | 480121 | A | C | 87 | 0 | 2 | 0 | NON_SYNONYMOU | MODERAT | MISSENSE | Acc/Ccc | T473P | PLES_04321 | | | | | | | | | C | C | | |
| NC_011770 | 480163 | C | G | 84 | 0 | 6 | 0 | NON_SYNONYMOU | MODERAT | MISSENSE | Ctc/Gtc | L487V | PLES_04321 | | | | | | | G | G | G | G | G | G |
| NC_011770 | 480731 | C | A | 83 | 0 | 4 | 0 | NON_SYNONYMOU | MODERAT | MISSENSE | aCg/aAg | T676K | PLES_04321 | | | A | A | A | A | | | | | | |
| NC_011770 | 479380 | G | A | 83 | 0 | 1 | 0 | NON_SYNONYMOU | MODERAT | MISSENSE | Ggc/Agc | G226S | PLES_04321 | | | | | | | | **A** | | | | |
| NC_011770 | 481103 | G | A | 76 | 0 | 10 | 0 | NON_SYNONYMOU | MODERAT | MISSENSE | Ggc/Agc | G66S | PLES_04331 | | | A | A | A | A | A | A | A | A | A | A |
| NC_011770 | 481298 | A | T | 82 | 0 | 6 | 0 | NON_SYNONYMOU | MODERAT | MISSENSE | Atc/Ttc | I131F | PLES_04331 | | | T | T | T | T | | | T | T | | |
| NC_011770 | 481499 | T | G | 79 | 0 | 4 | 0 | NON_SYNONYMOU | MODERAT | MISSENSE | Tgg/Ggg | W198G | PLES_04331 | | | | | | | G | G | | | G | G |
| NC_011770 | 481557 | C | T | 87 | 0 | 6 | 0 | NON_SYNONYMOU | MODERAT | MISSENSE | gCg/gTg | A217V | PLES_04331 | | | T | T | T | T | | | T | T | | |
| NC_011770 | 481795 | G | C | 74 | 0 | 12 | 0 | NON_SYNONYMOU | MODERAT | MISSENSE | gaG/gaC | E296D | PLES_04331 | C | C | C | C | C | C | C | C | C | C | C | C |
| NC_011770 | 486165 | C | G | 75 | 0 | 12 | 0 | NON_SYNONYMOU | MODERAT | MISSENSE | Gag/Cag | E312Q | PLES_04371 | G | G | G | G | G | G | G | G | G | G | G | G |
| NC_011770 | 486432 | C | T | 77 | 0 | 2 | 0 | NON_SYNONYMOU | MODERAT | MISSENSE | Ggc/Agc | G223S | PLES_04371 | | | | | | | | | | | T | T |
| NC_011770 | 486600 | T | C | 74 | 0 | 4 | 0 | NON_SYNONYMOU | MODERAT | MISSENSE | Acc/Gcc | T167A | PLES_04371 | | | C | C | C | C | | | | | | |
| NC_011770 | 486883 | C | A | 82 | 0 | 4 | 0 | NON_SYNONYMOU | MODERAT | MISSENSE | caG/caT | Q72H | PLES_04371 | | | A | A | A | A | | | | | | |
| NC_011770 | 487946 | T | C | 84 | 0 | 8 | 0 | NON_SYNONYMOU | MODERAT | MISSENSE | Acc/Gcc | T173A | PLES_04381 | C | C | | | | | C | C | C | C | C | C |
| NC_011770 | 488077 | T | C | 78 | 0 | 8 | 0 | NON_SYNONYMOU | MODERAT | MISSENSE | gAc/gGc | D129G | PLES_04381 | C | C | | | | | C | C | C | C | C | C |
| NC_011770 | 487760 | C | T | 84 | 0 | 1 | 0 | NON_SYNONYMOU | MODERAT | MISSENSE | Ggc/Agc | G235S | PLES_04381 | T | | | | | | | | | | | |
| NC_011770 | 490626 | A | T | 76 | 0 | 2 | 0 | NON_SYNONYMOU | MODERAT | MISSENSE | agA/agT | R5S | PLES_04401 | | | | | | | | | T | T | | |
| NC_011770 | 492049 | C | T | 69 | 0 | 2 | 0 | NON_SYNONYMOU | MODERAT | MISSENSE | Ctc/Ttc | L480F | PLES_04401 | T | T | | | | | | | | | | |
| NC_011770 | 492059 | G | T | 66 | 0 | 2 | 0 | NON_SYNONYMOU | MODERAT | MISSENSE | cGc/cTc | R483L | PLES_04401 | | | | | | | | | | | T | T |

| | | | | | | | | | | | | | | | | | | | | | | | | | |
|---|---|---|---|---|---|---|---|---|---|---|---|---|---|---|---|---|---|---|---|---|---|---|---|---|---|
| NC_011770 | 492171 | T | C | 72 | 0 | 12 | 0 | NON_SYNONYMOU | MODERAT | MISSENSE | gTc/gCc | V4A | PLES_04411 | C | C | C | C | C | C | C | C | C | C | C | C |
| NC_011770 | 492539 | G | A | 68 | 0 | 4 | 0 | NON_SYNONYMOU | MODERAT | MISSENSE | Gcg/Acg | A127T | PLES_04411 | | | A | A | A | A | | | | | | |
| NC_011770 | 493682 | C | T | 25 | 0 | 2 | 0 | NON_SYNONYMOU | MODERAT | MISSENSE | Gcc/Acc | A396T | PLES_04421 | | | | | | | T | T | | | | |
| NC_011770 | 494053 | A | G | 67 | 0 | 1 | 0 | NON_SYNONYMOU | MODERAT | MISSENSE | cTg/cCg | L272P | PLES_04421 | | | | | | | G | | | | | |
| NC_011770 | 494374 | C | T | 87 | 0 | 12 | 0 | NON_SYNONYMOU | MODERAT | MISSENSE | aGc/aAc | S165N | PLES_04421 | T | T | T | T | T | T | T | T | T | T | T | T |
| NC_011770 | 494393 | G | T | 87 | 0 | 2 | 0 | NON_SYNONYMOU | MODERAT | MISSENSE | Ctc/Atc | L159I | PLES_04421 | | | | | | | T | T | | | | |
| NC_011770 | 498278 | T | C | 76 | 0 | 6 | 0 | NON_SYNONYMOU | MODERAT | MISSENSE | cAg/cGg | Q414R | PLES_04461 | C | C | C | C | C | C | | | | | | |
| NC_011770 | 498302 | C | T | 71 | 0 | 2 | 0 | NON_SYNONYMOU | MODERAT | MISSENSE | cGc/cAc | R406H | PLES_04461 | T | T | | | | | | | | | | |
| NC_011770 | 498329 | C | T | 70 | 0 | 4 | 0 | NON_SYNONYMOU | MODERAT | MISSENSE | gGc/gAc | G397D | PLES_04461 | | | T | T | T | T | | | | | | |
| NC_011770 | 498399 | C | G | 71 | 0 | 6 | 0 | NON_SYNONYMOU | MODERAT | MISSENSE | Gaa/Caa | E374Q | PLES_04461 | | | G | G | G | G | | | | | G | G |
| NC_011770 | 498583 | A | C | 88 | 0 | 12 | 0 | NON_SYNONYMOU | MODERAT | MISSENSE | gaT/gaG | D312E | PLES_04461 | C | C | C | C | C | C | C | C | C | C | C | C |
| NC_011770 | 499223 | C | G | 85 | 0 | 12 | 0 | NON_SYNONYMOU | MODERAT | MISSENSE | aGc/aCc | S99T | PLES_04461 | G | G | G | G | G | G | G | G | G | G | G | G |
| NC_011770 | 499341 | T | C | 85 | 0 | 6 | 0 | NON_SYNONYMOU | MODERAT | MISSENSE | Acg/Gcg | T60A | PLES_04461 | C | C | C | C | C | C | | | | | | |
| NC_011770 | 499422 | G | A | 76 | 0 | 6 | 0 | NON_SYNONYMOU | MODERAT | MISSENSE | Ctc/Ttc | L33F | PLES_04461 | A | A | A | A | A | A | | | | | | |
| NC_011770 | 499463 | C | T | 66 | 0 | 12 | 0 | NON_SYNONYMOU | MODERAT | MISSENSE | cGc/cAc | R19H | PLES_04461 | T | T | T | T | T | T | T | T | T | T | T | T |
| NC_011770 | 499493 | T | A | 65 | 0 | 2 | 0 | NON_SYNONYMOU | MODERAT | MISSENSE | gAc/gTc | D9V | PLES_04461 | A | A | | | | | | | | | | |
| NC_011770 | 499908 | C | T | 54 | 0 | 4 | 0 | NON_SYNONYMOU | MODERAT | MISSENSE | cCg/cTg | P4L | PLES_04471 | | | T | T | T | T | | | | | | |
| NC_011770 | 500329 | C | A | 64 | 0 | 12 | 0 | NON_SYNONYMOU | MODERAT | MISSENSE | gaC/gaA | D144E | PLES_04471 | A | A | A | A | A | A | A | A | A | A | A | A |
| NC_011770 | 500403 | C | T | 67 | 0 | 12 | 0 | NON_SYNONYMOU | MODERAT | MISSENSE | gCc/gTc | A169V | PLES_04471 | T | T | T | T | T | T | T | T | T | T | T | T |
| NC_011770 | 500522 | A | G | 72 | 0 | 12 | 0 | NON_SYNONYMOU | MODERAT | MISSENSE | Agc/Ggc | S209G | PLES_04471 | G | G | G | G | G | G | G | G | G | G | G | G |
| NC_011770 | 500705 | G | T | 79 | 0 | 2 | 0 | NON_SYNONYMOU | MODERAT | MISSENSE | Gcg/Tcg | A270S | PLES_04471 | T | T | | | | | | | | | | |
| NC_011770 | 501105 | T | A | 83 | 0 | 6 | 0 | NON_SYNONYMOU | MODERAT | MISSENSE | cTg/cAg | L403Q | PLES_04471 | A | A | A | A | A | A | | | | | | |
| NC_011770 | 502008 | T | A | 100 | 0 | 6 | 0 | NON_SYNONYMOU | MODERAT | MISSENSE | gaT/gaA | D261E | PLES_04481 | A | A | A | A | A | A | | | | | | |
| NC_011770 | 502439 | T | C | 67 | 0 | 4 | 0 | NON_SYNONYMOU | MODERAT | MISSENSE | Aat/Gat | N110D | PLES_04491 | | | C | C | C | C | | | | | | |
| NC_011770 | 502607 | C | T | 59 | 0 | 2 | 0 | NON_SYNONYMOU | MODERAT | MISSENSE | Gag/Aag | E54K | PLES_04491 | T | T | | | | | | | | | | |
| NC_011770 | 502673 | A | G | 55 | 0 | 6 | 0 | NON_SYNONYMOU | MODERAT | MISSENSE | Ttc/Ctc | F32L | PLES_04491 | G | G | | | | | G | G | G | G | | |
| NC_011770 | 502711 | C | T | 58 | 0 | 4 | 0 | NON_SYNONYMOU | MODERAT | MISSENSE | gGt/gAt | G19D | PLES_04491 | | | T | T | T | T | | | | | | |
| NC_011770 | 503688 | C | A | 96 | 0 | 12 | 0 | NON_SYNONYMOU | MODERAT | MISSENSE | gGc/gTc | G457V | PLES_04501 | A | A | A | A | A | A | A | A | A | A | A | A |
| NC_011770 | 503848 | C | A | 101 | 0 | 2 | 0 | NON_SYNONYMOU | MODERAT | MISSENSE | Gcc/Tcc | A404S | PLES_04501 | A | A | | | | | | | | | | |
| NC_011770 | 507605 | A | G | 85 | 0 | 12 | 0 | NON_SYNONYMOU | MODERAT | MISSENSE | aAg/aGg | K68R | PLES_04531 | G | G | G | G | G | G | G | G | G | G | G | G |
| NC_011770 | 507722 | CCGG | C | 87 | 0 | 2 | 0 | CODON_DELETION | MODERATE | | ggc/ | G109 | PLES_04531 | | | C | C | | | | | | | | |
| NC_011770 | 507907 | A | G | 86 | 0 | 2 | 0 | NON_SYNONYMOU | MODERAT | MISSENSE | Agc/Ggc | S169G | PLES_04531 | | | | | | | | | G | G | | |
| NC_011770 | 508344 | G | A | 81 | 0 | 2 | 0 | NON_SYNONYMOU | MODERAT | MISSENSE | Cgc/Tgc | R463C | PLES_04541 | | | | | | | | | A | A | | |
| NC_011770 | 508949 | C | T | 80 | 0 | 2 | 0 | NON_SYNONYMOU | MODERAT | MISSENSE | aGc/aAc | S261N | PLES_04541 | T | T | | | | | | | | | | |
| NC_011770 | 510823 | C | G | 81 | 0 | 4 | 0 | NON_SYNONYMOU | MODERAT | MISSENSE | aaC/aaG | N158K | PLES_04551 | | | G | G | G | G | | | | | | |
| NC_011770 | 511370 | G | A | 80 | 0 | 6 | 0 | NON_SYNONYMOU | MODERAT | MISSENSE | Gtc/Atc | V341I | PLES_04551 | A | A | A | A | A | A | | | | | | |
| NC_011770 | 511800 | A | G | 82 | 0 | 10 | 0 | NON_SYNONYMOU | MODERAT | MISSENSE | aAg/aGg | K484R | PLES_04551 | G | G | G | G | G | G | | | G | G | G | G |
| NC_011770 | 511875 | A | T | 84 | 0 | 4 | 0 | NON_SYNONYMOU | MODERAT | MISSENSE | cAc/cTc | H509L | PLES_04551 | | | T | T | T | T | | | | | | |
| NC_011770 | 512765 | G | A | 74 | 0 | 2 | 0 | NON_SYNONYMOU | MODERAT | MISSENSE | Gcc/Acc | A806T | PLES_04551 | | | | | | | | | A | A | | |
| NC_011770 | 512798 | G | A | 76 | 0 | 2 | 0 | NON_SYNONYMOU | MODERAT | MISSENSE | Ggc/Agc | G817S | PLES_04551 | | | | | | | | | | | A | A |
| NC_011770 | 512817 | A | G | 77 | 0 | 4 | 0 | NON_SYNONYMOU | MODERAT | MISSENSE | aAg/aGg | K823R | PLES_04551 | | | | | | | G | G | | | G | G |
| NC_011770 | 512894 | G | A | 80 | 0 | 6 | 0 | NON_SYNONYMOU | MODERAT | MISSENSE | Gcc/Acc | A849T | PLES_04551 | A | A | | | | | A | A | | | A | A |
| NC_011770 | 514566 | C | T | 76 | 0 | 2 | 0 | NON_SYNONYMOU | MODERAT | MISSENSE | Ggc/Agc | G77S | PLES_04571 | | | | | | | T | T | | | | |
| NC_011770 | 514656 | C | T | 83 | 0 | 2 | 0 | NON_SYNONYMOU | MODERAT | MISSENSE | Ggc/Agc | G47S | PLES_04571 | | | | | | | T | T | | | | |
| NC_011770 | 515232 | A | G | 91 | 0 | 6 | 0 | NON_SYNONYMOU | MODERAT | MISSENSE | aAc/aGc | N164S | PLES_04581 | G | G | G | G | G | G | | | | | | |
| NC_011770 | 519781 | C | T | 75 | 0 | 2 | 0 | NON_SYNONYMOU | MODERAT | MISSENSE | Ggc/Agc | G148S | PLES_04631 | T | T | | | | | | | | | | |
| NC_011770 | 519804 | C | G | 72 | 0 | 6 | 0 | NON_SYNONYMOU | MODERAT | MISSENSE | gGg/gCg | G140A | PLES_04631 | G | G | | | | | | | G | G | G | G |
| NC_011770 | 519835 | T | C | 72 | 0 | 2 | 0 | NON_SYNONYMOU | MODERAT | MISSENSE | Acc/Gcc | T130A | PLES_04631 | C | C | | | | | | | | | | |
| NC_011770 | 519880 | A | T | 72 | 0 | 2 | 0 | NON_SYNONYMOU | MODERAT | MISSENSE | Tcc/Acc | S115T | PLES_04631 | T | T | | | | | | | | | | |
| NC_011770 | 520608 | T | C | 77 | 0 | 8 | 0 | NON_SYNONYMOU | MODERAT | MISSENSE | cAg/cGg | Q195R | PLES_04641 | C | C | C | C | C | C | C | C | | | | |
| NC_011770 | 520615 | C | T | 77 | 0 | 4 | 0 | NON_SYNONYMOU | MODERAT | MISSENSE | Gcc/Acc | A193T | PLES_04641 | | | T | T | T | T | | | | | | |
| NC_011770 | 520820 | G | C | 81 | 0 | 8 | 0 | NON_SYNONYMOU | MODERAT | MISSENSE | gaC/gaG | D124E | PLES_04641 | C | C | | | | | C | C | C | C | C | C |
| NC_011770 | 521158 | A | G | 96 | 0 | 2 | 0 | NON_SYNONYMOU | MODERAT | MISSENSE | Ttc/Ctc | F12L | PLES_04641 | G | G | | | | | | | | | | |
| NC_011770 | 521314 | CG | CGG | 82 | 0 | 2 | 0 | FRAME_SHIFT | HIGH | | ggc/ | G15 | PLES_04651 | CGG | C | | | | | | | | | | |
| NC_011770 | 521706 | C | T | 90 | 0 | 2 | 0 | NON_SYNONYMOU | MODERAT | MISSENSE | Cac/Tac | H144Y | PLES_04651 | T | T | | | | | | | | | | |
| NC_011770 | 524970 | T | C | 65 | 0 | 6 | 0 | NON_SYNONYMOU | MODERAT | MISSENSE | cAg/cGg | Q276R | PLES_04671 | C | C | | | | | C | C | C | C | | |
| NC_011770 | 525051 | G | T | 71 | 0 | 2 | 0 | NON_SYNONYMOU | MODERAT | MISSENSE | gCa/gAa | A249E | PLES_04671 | | | | | | | | | T | T | | |
| NC_011770 | 525600 | G | A | 77 | 0 | 2 | 0 | NON_SYNONYMOU | MODERAT | MISSENSE | gCc/gTc | A66V | PLES_04671 | | | | | | | | | | | A | A |
| NC_011770 | 526733 | T | C | 78 | 0 | 6 | 0 | NON_SYNONYMOU | MODERAT | MISSENSE | gTc/gCc | V85A | PLES_04691 | | | C | C | C | C | C | C | | | | |
| NC_011770 | 526930 | G | A | 75 | 0 | 6 | 0 | NON_SYNONYMOU | MODERAT | MISSENSE | Gtc/Atc | V151I | PLES_04691 | | | A | A | A | A | A | A | | | | |
| NC_011770 | 528109 | A | G | 84 | 0 | 2 | 0 | NON_SYNONYMOU | MODERAT | MISSENSE | gAg/gGg | E96G | PLES_04711 | G | G | | | | | | | | | | |
| NC_011770 | 528472 | C | T | 74 | 0 | 2 | 0 | NON_SYNONYMOU | MODERAT | MISSENSE | cGc/cAc | R561H | PLES_04721 | | | | | | | | | | | T | T |

| | | | | | | | | | | | | | | | | | | | | | | | | | |
|---|---|---|---|---|---|---|---|---|---|---|---|---|---|---|---|---|---|---|---|---|---|---|---|---|---|
| NC_011770 | 528641 | C | G | 74 | 0 | 10 | 0 | NON_SYNONYMOU | MODERAT | MISSENSE | Gga/Cga | G505R | PLES_04721 | G | G | G | G | G | G | G | G | | | G | G |
| NC_011770 | 528931 | A | G | 93 | 0 | 1 | 0 | NON_SYNONYMOU | MODERAT | MISSENSE | gTg/gCg | V408A | PLES_04721 | | | | | | | G | | | | | |
| NC_011770 | 529693 | A | G | 84 | 0 | 1 | 0 | NON_SYNONYMOU | MODERAT | MISSENSE | cTg/cCg | L154P | PLES_04721 | | | | | | | G | | | | | |
| NC_011770 | 528454 | G | A | 71 | 0 | 1 | 0 | NON_SYNONYMOU | MODERAT | MISSENSE | gCg/gTg | A567V | PLES_04721 | PA0476 | A | | | | | | | | | | |
| NC_011770 | 530551 | C | T | 76 | 0 | 2 | 0 | NON_SYNONYMOU | MODERAT | MISSENSE | Gcg/Acg | A293T | PLES_04731 | | | | | | | | | | | T | T |
| NC_011770 | 530648 | G | T | 75 | 0 | 10 | 0 | NON_SYNONYMOU | MODERAT | MISSENSE | gaC/gaA | D260E | PLES_04731 | T | T | T | T | T | T | T | T | | | T | T |
| NC_011770 | 531141 | T | A | 89 | 0 | 2 | 0 | NON_SYNONYMOU | MODERAT | MISSENSE | cAg/cTg | Q96L | PLES_04731 | A | A | | | | | | | | | | |
| NC_011770 | 531911 | C | T | 92 | 0 | 2 | 0 | NON_SYNONYMOU | MODERAT | MISSENSE | Cgc/Tgc | R133C | PLES_04741 | T | T | | | | | | | | | | |
| NC_011770 | 532246 | T | C | 75 | 0 | 12 | 0 | NON_SYNONYMOU | MODERAT | MISSENSE | Agg/Ggg | R259G | PLES_04751 | C | C | C | C | C | C | C | C | C | C | C | C |
| NC_011770 | 532405 | A | G | 78 | 0 | 12 | 0 | NON_SYNONYMOU | MODERAT | MISSENSE | Tcg/Ccg | S206P | PLES_04751 | G | G | G | G | G | G | G | G | G | G | G | G |
| NC_011770 | 532773 | C | T | 74 | 0 | 6 | 0 | NON_SYNONYMOU | MODERAT | MISSENSE | gGc/gAc | G83D | PLES_04751 | T | T | T | T | T | T | | | | | | |
| NC_011770 | 533166 | C | T | 93 | 0 | 2 | 0 | NON_SYNONYMOU | MODERAT | MISSENSE | Ctc/Ttc | L15F | PLES_04761 | | | T | T | | | | | | | | |
| NC_011770 | 533209 | C | T | 92 | 0 | 2 | 0 | NON_SYNONYMOU | MODERAT | MISSENSE | gCg/gTg | A29V | PLES_04761 | | | | | | | | | T | T | | |
| NC_011770 | 533376 | T | C | 91 | 0 | 10 | 0 | NON_SYNONYMOU | MODERAT | MISSENSE | Ttc/Ctc | F85L | PLES_04761 | C | C | C | C | C | C | C | C | | | C | C |
| NC_011770 | 533653 | T | A | 79 | 0 | 4 | 0 | NON_SYNONYMOU | MODERAT | MISSENSE | aTg/aAg | M177K | PLES_04761 | | | A | A | A | A | | | | | | |
| NC_011770 | 533811 | A | G | 83 | 0 | 10 | 0 | NON_SYNONYMOU | MODERAT | MISSENSE | Acg/Gcg | T230A | PLES_04761 | G | G | G | G | G | G | G | G | | | G | G |
| NC_011770 | 533844 | T | C | 84 | 0 | 10 | 0 | NON_SYNONYMOU | MODERAT | MISSENSE | Tcg/Ccg | S241P | PLES_04761 | C | C | C | C | C | C | C | C | | | C | C |
| NC_011770 | 533917 | C | A | 89 | 0 | 10 | 0 | NON_SYNONYMOU | MODERAT | MISSENSE | gCc/gAc | A265D | PLES_04761 | A | A | A | A | A | A | A | A | | | A | A |
| NC_011770 | 534407 | A | G | 64 | 0 | 4 | 0 | STOP_LOST+SPLICE | HIGH | MISSENSE | tgA/tgG | *148W | PLES_04771 | | | G | G | G | G | | | | | | |
| NC_011770 | 534172 | A | G | 78 | 0 | 2 | 0 | NON_SYNONYMOU | MODERAT | MISSENSE | cAc/cGc | H70R | PLES_04771 | | | | | | | G | G | | | | |
| NC_011770 | 534234 | G | A | 77 | 0 | 2 | 0 | NON_SYNONYMOU | MODERAT | MISSENSE | Gag/Aag | E91K | PLES_04771 | | | | | | | | | | | A | A |
| NC_011770 | 534343 | G | A | 69 | 0 | 2 | 0 | NON_SYNONYMOU | MODERAT | MISSENSE | cGt/cAt | R127H | PLES_04771 | | | | | | | | | | | A | A |
| NC_011770 | 539068 | G | T | 94 | 0 | 2 | 0 | NON_SYNONYMOU | MODERAT | MISSENSE | agC/agA | S149R | PLES_04811 | | | | | | | | | T | T | | |
| NC_011770 | 539918 | C | G | 73 | 0 | 2 | 0 | NON_SYNONYMOU | MODERAT | MISSENSE | gaG/gaC | E258D | PLES_04821 | | | | | | | G | G | | | | |
| NC_011770 | 540508 | C | A | 82 | 0 | 2 | 0 | NON_SYNONYMOU | MODERAT | MISSENSE | Gcg/Tcg | A62S | PLES_04821 | A | A | | | | | | | | | | |
| NC_011770 | 540865 | A | G | 94 | 0 | 12 | 0 | NON_SYNONYMOU | MODERAT | MISSENSE | aTc/aCc | I216T | PLES_04831 | G | G | G | G | G | G | G | G | G | G | G | G |
| NC_011770 | 541100 | C | A | 84 | 0 | 2 | 0 | NON_SYNONYMOU | MODERAT | MISSENSE | Gcc/Tcc | A138S | PLES_04831 | A | A | | | | | | | | | | |
| NC_011770 | 541683 | A | G | 74 | 0 | 2 | 0 | NON_SYNONYMOU | MODERAT | MISSENSE | Ttc/Ctc | F73L | PLES_04841 | | | | | | | G | G | | | | |
| NC_011770 | 542020 | T | C | 72 | 0 | 4 | 0 | NON_SYNONYMOU | MODERAT | MISSENSE | cAc/cGc | H216R | PLES_04851 | | | | | | | C | C | | | C | C |
| NC_011770 | 542021 | G | A | 73 | 0 | 2 | 0 | NON_SYNONYMOU | MODERAT | MISSENSE | Cac/Tac | H216Y | PLES_04851 | | | | | | | A | A | | | | |
| NC_011770 | 542027 | T | C | 75 | 0 | 4 | 0 | NON_SYNONYMOU | MODERAT | MISSENSE | Acc/Gcc | T214A | PLES_04851 | | | | | | | C | C | | | C | C |
| NC_011770 | 542195 | A | T | 68 | 0 | 2 | 0 | NON_SYNONYMOU | MODERAT | MISSENSE | Tgc/Agc | C158S | PLES_04851 | | | | | | | T | T | | | | |
| NC_011770 | 542197 | T | C | 68 | 0 | 10 | 0 | NON_SYNONYMOU | MODERAT | MISSENSE | cAc/cGc | H157R | PLES_04851 | C | C | C | C | C | C | C | C | | | C | C |
| NC_011770 | 542209 | T | C | 69 | 0 | 2 | 0 | NON_SYNONYMOU | MODERAT | MISSENSE | gAg/gGg | E153G | PLES_04851 | | | | | | | C | C | | | | |
| NC_011770 | 542354 | C | T | 68 | 0 | 2 | 0 | NON_SYNONYMOU | MODERAT | MISSENSE | Gag/Aag | E105K | PLES_04851 | | | | | | | T | T | | | | |
| NC_011770 | 542998 | C | T | 83 | 0 | 4 | 0 | NON_SYNONYMOU | MODERAT | MISSENSE | gGc/gAc | G34D | PLES_04861 | | | | | | | T | T | | | T | T |
| NC_011770 | 543943 | T | C | 86 | 0 | 12 | 0 | NON_SYNONYMOU | MODERAT | MISSENSE | Agc/Ggc | S46G | PLES_04871 | C | C | C | C | C | C | C | C | C | C | C | C |
| NC_011770 | 545275 | C | T | 99 | 0 | 2 | 0 | NON_SYNONYMOU | MODERAT | MISSENSE | aCc/aTc | T82I | PLES_04891 | | | | | | | T | T | | | | |
| NC_011770 | 545616 | T | G | 86 | 0 | 10 | 0 | NON_SYNONYMOU | MODERAT | MISSENSE | gaT/gaG | D109E | PLES_04901 | G | G | G | G | G | G | | | G | G | G | G |
| NC_011770 | 546275 | A | T | 79 | 0 | 2 | 0 | NON_SYNONYMOU | MODERAT | MISSENSE | gAg/gTg | E329V | PLES_04901 | | | | | | | T | T | | | | |
| NC_011770 | 546295 | G | A | 78 | 0 | 2 | 0 | NON_SYNONYMOU | MODERAT | MISSENSE | Gcg/Acg | A336T | PLES_04901 | A | A | | | | | | | | | | |
| NC_011770 | 546644 | C | G | 80 | 0 | 2 | 0 | NON_SYNONYMOU | MODERAT | MISSENSE | aCc/aGc | T452S | PLES_04901 | | | | | | | G | G | | | | |
| NC_011770 | 546731 | G | T | 82 | 0 | 12 | 0 | NON_SYNONYMOU | MODERAT | MISSENSE | Gcc/Tcc | A22S | PLES_04911 | T | T | T | T | T | T | T | T | T | T | T | T |
| NC_011770 | 546903 | T | C | 81 | 0 | 4 | 0 | NON_SYNONYMOU | MODERAT | MISSENSE | gTc/gCc | V79A | PLES_04911 | | | | | | | C | C | C | C | | |
| NC_011770 | 547071 | G | A | 83 | 0 | 12 | 0 | NON_SYNONYMOU | MODERAT | MISSENSE | cGg/cAg | R135Q | PLES_04911 | A | A | A | A | A | A | A | A | A | A | A | A |
| NC_011770 | 547400 | T | C | 94 | 0 | 12 | 0 | NON_SYNONYMOU | MODERAT | MISSENSE | Tgg/Cgg | W245R | PLES_04911 | C | C | C | C | C | C | C | C | C | C | C | C |
| NC_011770 | 547506 | G | A | 93 | 0 | 2 | 0 | NON_SYNONYMOU | MODERAT | MISSENSE | gGc/gAc | G280D | PLES_04911 | | | | | | | | | A | A | | |
| NC_011770 | 547788 | G | A | 79 | 0 | 2 | 0 | NON_SYNONYMOU | MODERAT | MISSENSE | Gcg/Acg | A85T | PLES_04921 | | | | | | | | | A | A | | |
| NC_011770 | 547941 | G | A | 78 | 0 | 2 | 0 | NON_SYNONYMOU | MODERAT | MISSENSE | Gcg/Acg | A136T | PLES_04921 | A | A | | | | | | | | | | |
| NC_011770 | 548148 | A | T | 79 | 0 | 2 | 0 | NON_SYNONYMOU | MODERAT | MISSENSE | Acc/Tcc | T205S | PLES_04921 | T | T | | | | | | | | | | |
| NC_011770 | 548419 | G | C | 76 | 0 | 4 | 0 | NON_SYNONYMOU | MODERAT | MISSENSE | gGg/gCg | G295A | PLES_04921 | C | C | | | | | | | | | C | C |
| NC_011770 | 548430 | G | C | 75 | 0 | 2 | 0 | NON_SYNONYMOU | MODERAT | MISSENSE | Gcc/Ccc | A299P | PLES_04921 | C | C | | | | | | | | | | |
| NC_011770 | 548583 | C | T | 74 | 0 | 2 | 0 | NON_SYNONYMOU | MODERAT | MISSENSE | Gcg/Acg | A327T | PLES_04931 | | | | | | | | | T | T | | |
| NC_011770 | 549755 | C | T | 94 | 0 | 2 | 0 | NON_SYNONYMOU | MODERAT | MISSENSE | Gcc/Acc | A298T | PLES_04941 | | | | | | | T | T | | | | |
| NC_011770 | 554357 | G | A | 62 | 0 | 2 | 0 | NON_SYNONYMOU | MODERAT | MISSENSE | Gac/Aac | D23N | PLES_04981 | | | | | | | A | A | | | | |
| NC_011770 | 554871 | T | C | 59 | 0 | 2 | 0 | NON_SYNONYMOU | MODERAT | MISSENSE | cTg/cCg | L194P | PLES_04981 | C | C | | | | | | | | | | |
| NC_011770 | 555100 | C | T | 60 | 0 | 2 | 0 | NON_SYNONYMOU | MODERAT | MISSENSE | gCc/gTc | A32V | PLES_04991 | | | | | | | T | T | | | | |
| NC_011770 | 555592 | G | A | 72 | 0 | 2 | 0 | NON_SYNONYMOU | MODERAT | MISSENSE | gGc/gAc | G196D | PLES_04991 | | | | | | | | | | | A | A |
| NC_011770 | 556666 | A | G | 80 | 0 | 4 | 0 | NON_SYNONYMOU | MODERAT | MISSENSE | aAc/aGc | N13S | PLES_05011 | | | G | G | G | G | | | | | | |
| NC_011770 | 556741 | C | T | 78 | 0 | 2 | 0 | NON_SYNONYMOU | MODERAT | MISSENSE | tCg/tTg | S38L | PLES_05011 | | | | | | | | | | | T | T |
| NC_011770 | 556743 | G | C | 76 | 0 | 6 | 0 | NON_SYNONYMOU | MODERAT | MISSENSE | Gcc/Ccc | A39P | PLES_05011 | | | C | C | C | C | C | C | | | | |
| NC_011770 | 556990 | C | T | 76 | 0 | 4 | 0 | NON_SYNONYMOU | MODERAT | MISSENSE | aGc/aAc | S47N | PLES_05021 | | | | | | | T | T | | | T | T |

| | | | | | | | | | | | | | | | | | | | | | | | | | |
|---|---|---|---|---|---|---|---|---|---|---|---|---|---|---|---|---|---|---|---|---|---|---|---|---|---|
| NC_011770 | 557008 | A | C | 77 | 0 | 2 | 0 | NON_SYNONYMOU | MODERAT | MISSENSE | aTg/aGg | M41R | PLES_05021 | C | C | | | | | | | | | | |
| NC_011770 | 557081 | C | T | 82 | 0 | 12 | 0 | NON_SYNONYMOU | MODERAT | MISSENSE | Gct/Act | A17T | PLES_05021 | T | T | T | T | T | T | T | T | T | T | T | T |
| NC_011770 | 558930 | T | C | 73 | 0 | 12 | 0 | NON_SYNONYMOU | MODERAT | MISSENSE | Tac/Cac | Y578H | PLES_05031 | C | C | C | C | C | C | C | C | C | C | C | C |
| NC_011770 | 558958 | G | A | 71 | 0 | 12 | 0 | NON_SYNONYMOU | MODERAT | MISSENSE | aGc/aAc | S587N | PLES_05031 | A | A | A | A | A | A | A | A | A | A | A | A |
| NC_011770 | 559932 | A | C | 103 | 0 | 2 | 0 | NON_SYNONYMOU | MODERAT | MISSENSE | gAa/gCa | E239A | PLES_05041 | | | | | | | C | C | | | | |
| NC_011770 | 559933 | A | C | 101 | 0 | 2 | 0 | NON_SYNONYMOU | MODERAT | MISSENSE | gaA/gaC | E239D | PLES_05041 | | | | | | | C | C | | | | |
| NC_011770 | 559937 | A | G | 104 | 0 | 2 | 0 | NON_SYNONYMOU | MODERAT | MISSENSE | Aac/Gac | N241D | PLES_05041 | | | | | | | G | G | | | | |
| NC_011770 | 559939 | C | G | 104 | 0 | 2 | 0 | NON_SYNONYMOU | MODERAT | MISSENSE | aaC/aaG | N241K | PLES_05041 | | | | | | | G | G | | | | |
| NC_011770 | 559941 | T | C | 102 | 0 | 2 | 0 | NON_SYNONYMOU | MODERAT | MISSENSE | gTg/gCg | V242A | PLES_05041 | | | | | | | C | C | | | | |
| NC_011770 | 559947 | C | T | 99 | 0 | 2 | 0 | NON_SYNONYMOU | MODERAT | MISSENSE | gCg/gTg | A244V | PLES_05041 | | | | | | | T | T | | | | |
| NC_011770 | 560219 | C | A | 93 | 0 | 2 | 0 | NON_SYNONYMOU | MODERAT | MISSENSE | Ccg/Acg | P335T | PLES_05041 | A | A | | | | | | | | | | |
| NC_011770 | 561583 | G | A | 69 | 0 | 4 | 0 | NON_SYNONYMOU | MODERAT | MISSENSE | Gcc/Acc | A100T | PLES_05051 | A | A | | | | | | | | | A | A |
| NC_011770 | 561734 | A | G | 80 | 0 | 4 | 0 | NON_SYNONYMOU | MODERAT | MISSENSE | aAg/aGg | K150R | PLES_05051 | | | G | G | G | G | | | | | | |
| NC_011770 | 561761 | T | C | 80 | 0 | 4 | 0 | NON_SYNONYMOU | MODERAT | MISSENSE | aTc/aCc | I159T | PLES_05051 | | | C | C | C | C | | | | | | |
| NC_011770 | 562426 | G | T | 77 | 0 | 2 | 0 | NON_SYNONYMOU | MODERAT | MISSENSE | Gcg/Tcg | A381S | PLES_05051 | | | | | | | | | T | T | | |
| NC_011770 | 562693 | G | A | 57 | 0 | 4 | 0 | NON_SYNONYMOU | MODERAT | MISSENSE | Gac/Aac | D470N | PLES_05051 | | | A | A | A | A | | | | | | |
| NC_011770 | 565026 | C | A | 76 | 0 | 2 | 0 | NON_SYNONYMOU | MODERAT | MISSENSE | Gcc/Tcc | A228S | PLES_05071 | A | A | | | | | | | | | | |
| NC_011770 | 565095 | T | C | 72 | 0 | 2 | 0 | NON_SYNONYMOU | MODERAT | MISSENSE | Agc/Ggc | S205G | PLES_05071 | C | C | | | | | | | | | | |
| NC_011770 | 565644 | G | C | 55 | 0 | 2 | 0 | NON_SYNONYMOU | MODERAT | MISSENSE | Ctg/Gtg | L22V | PLES_05071 | | | | | | | C | C | | | | |
| NC_011770 | 567562 | C | T | 70 | 0 | 2 | 0 | NON_SYNONYMOU | MODERAT | MISSENSE | Gcg/Acg | A83T | PLES_05101 | T | T | | | | | | | | | | |
| NC_011770 | 567726 | G | T | 64 | 0 | 2 | 0 | NON_SYNONYMOU | MODERAT | MISSENSE | gCc/gAc | A28D | PLES_05101 | T | T | | | | | | | | | | |
| NC_011770 | 567751 | A | G | 64 | 0 | 12 | 0 | NON_SYNONYMOU | MODERAT | MISSENSE | Tcg/Ccg | S20P | PLES_05101 | G | G | G | G | G | G | G | G | G | G | G | G |
| NC_011770 | 568712 | T | G | 65 | 0 | 4 | 0 | NON_SYNONYMOU | MODERAT | MISSENSE | gaA/gaC | E21D | PLES_05121 | G | G | | | | | | | G | G | | |
| NC_011770 | 568713 | T | G | 64 | 0 | 2 | 0 | NON_SYNONYMOU | MODERAT | MISSENSE | gAa/gCa | E21A | PLES_05121 | | | | | | | | | G | G | | |
| NC_011770 | 573865 | A | C | 89 | 0 | 4 | 0 | NON_SYNONYMOU | MODERAT | MISSENSE | Atg/Ctg | M150L | PLES_05181 | | | | | | | | | C | C | C | C |
| NC_011770 | 576781 | G | C | 63 | 0 | 4 | 0 | NON_SYNONYMOU | MODERAT | MISSENSE | Gcg/Ccg | A215P | PLES_05221 | C | C | | | | | C | C | | | | |
| NC_011770 | 576862 | G | A | 67 | 0 | 4 | 0 | NON_SYNONYMOU | MODERAT | MISSENSE | Gtc/Atc | V242I | PLES_05221 | | | A | A | A | A | | | | | | |
| NC_011770 | 577004 | A | G | 80 | 0 | 2 | 0 | NON_SYNONYMOU | MODERAT | MISSENSE | tAc/tGc | Y289C | PLES_05221 | G | G | | | | | | | | | | |
| NC_011770 | 577702 | G | T | 79 | 0 | 4 | 0 | NON_SYNONYMOU | MODERAT | MISSENSE | Ggc/Tgc | G522C | PLES_05221 | | | T | T | T | T | | | | | | |
| NC_011770 | 577991 | G | A | 85 | 0 | 4 | 0 | NON_SYNONYMOU | MODERAT | MISSENSE | gCt/gTt | A63V | PLES_05231 | | | A | A | A | A | | | | | | |
| NC_011770 | 578087 | C | T | 77 | 0 | 2 | 0 | NON_SYNONYMOU | MODERAT | MISSENSE | cGc/cAc | R31H | PLES_05231 | | | | | | | T | T | | | | |
| NC_011770 | 579304 | T | C | 87 | 0 | 12 | 0 | NON_SYNONYMOU | MODERAT | MISSENSE | Atc/Gtc | I292V | PLES_05251 | C | C | C | C | C | C | C | C | C | C | C | C |
| NC_011770 | 580748 | G | A | 87 | 0 | 2 | 0 | NON_SYNONYMOU | MODERAT | MISSENSE | gGg/gAg | G157E | PLES_05261 | | | | | | | | | | | A | A |
| NC_011770 | 580753 | G | C | 87 | 0 | 4 | 0 | NON_SYNONYMOU | MODERAT | MISSENSE | Gag/Cag | E159Q | PLES_05261 | C | C | | | | | C | C | | | | |
| NC_011770 | 580688 | A | G | 88 | 0 | 1 | 0 | NON_SYNONYMOU | MODERAT | MISSENSE | cAg/cGg | Q137R | PLES_05261 | | | | | | | | G | | | | |
| NC_011770 | 581422 | G | T | 100 | 0 | 2 | 0 | NON_SYNONYMOU | MODERAT | MISSENSE | Ggc/Tgc | G73C | PLES_05271 | T | T | | | | | | | | | | |
| NC_011770 | 582883 | A | G | 76 | 0 | 1 | 0 | NON_SYNONYMOU | MODERAT | MISSENSE | aAc/aGc | N166S | PLES_05281 | | | | | | | G | | | | | |
| NC_011770 | 583823 | C | T | 58 | 0 | 1 | 0 | STOP_GAINED | HIGH | NONSENSE | Cag/Tag | Q33* | PLES_05301 | | | | | | | | | | | | T |
| NC_011770 | 585215 | T | TA | 73 | 0 | 4 | 0 | FRAME_SHIFT | HIGH | | ggg/gAgg | G494E? | PLES_05301 | | | TA | TA | TA | TA | | | | | | |
| NC_011770 | 584075 | CTGA | C | 61 | 0 | 1 | 0 | CODON_CHANGE_ | MODERATE | | ctgacc/ccc | LT117P | PLES_05301 | | | | | | | C | | | | | |
| NC_011770 | 584487 | C | T | 83 | 0 | 2 | 0 | NON_SYNONYMOU | MODERAT | MISSENSE | cCg/cTg | P254L | PLES_05301 | | | | | T | T | | | | | | |
| NC_011770 | 584577 | A | G | 90 | 0 | 4 | 0 | NON_SYNONYMOU | MODERAT | MISSENSE | tAc/tGc | Y284C | PLES_05301 | G | G | | | | | G | G | | | | |
| NC_011770 | 584595 | C | T | 85 | 0 | 8 | 0 | NON_SYNONYMOU | MODERAT | MISSENSE | gCg/gTg | A290V | PLES_05301 | T | T | T | T | T | T | T | T | | | | |
| NC_011770 | 585396 | C | CG | 68 | 0 | 1 | 0 | FRAME_SHIFT | HIGH | | ctt/Gctt | L52A? | PLES_05311 | | | | | | | | CG | | | | |
| NC_011770 | 585358 | T | C | 64 | 0 | 2 | 0 | NON_SYNONYMOU | MODERAT | MISSENSE | Tgt/Cgt | C37R | PLES_05311 | | | | | | | | | | | C | C |
| NC_011770 | 585427 | GCCC | G | 67 | 0 | 2 | 0 | CODON_CHANGE_ | MODERATE | | gcccct/gct | AP60A | PLES_05311 | | | | | | | G | G | | | | |
| NC_011770 | 585863 | C | G | 80 | 0 | 2 | 0 | NON_SYNONYMOU | MODERAT | MISSENSE | gaC/gaG | D103E | PLES_05321 | | | | | | | | | | | G | G |
| NC_011770 | 586117 | A | G | 84 | 0 | 2 | 0 | NON_SYNONYMOU | MODERAT | MISSENSE | aAt/aGt | N188S | PLES_05321 | | | | | | | | | | | G | G |
| NC_011770 | 586620 | A | C | 78 | 0 | 2 | 0 | NON_SYNONYMOU | MODERAT | MISSENSE | Atc/Ctc | I356L | PLES_05321 | C | C | | | | | | | | | | |
| NC_011770 | 586699 | C | T | 83 | 0 | 2 | 0 | NON_SYNONYMOU | MODERAT | MISSENSE | gCg/gTg | A382V | PLES_05321 | | | | | | | T | T | | | | |
| NC_011770 | 586099 | C | T | 85 | 0 | 1 | 0 | NON_SYNONYMOU | MODERAT | MISSENSE | cCc/cTc | P182L | PLES_05321 | | | | | | | | T | | | | |
| NC_011770 | 587586 | T | C | 7 | 2 | 4 | 0 | NON_SYNONYMOU | MODERAT | MISSENSE | Agc/Ggc | S323G | PLES_05341 | | . | | | | | C | C | . | | C | C |
| NC_011770 | 588064 | G | T | 74 | 0 | 8 | 0 | NON_SYNONYMOU | MODERAT | MISSENSE | gaC/gaA | D163E | PLES_05341 | T | T | T | T | T | T | | | T | T | | |
| NC_011770 | 588161 | T | C | 83 | 0 | 8 | 0 | NON_SYNONYMOU | MODERAT | MISSENSE | cAg/cGg | Q131R | PLES_05341 | C | C | C | C | C | C | | | C | C | | |
| NC_011770 | 590555 | T | C | 68 | 0 | 2 | 0 | NON_SYNONYMOU | MODERAT | MISSENSE | gTc/gCc | V189A | PLES_05371 | C | C | | | | | | | | | | |
| NC_011770 | 590561 | C | T | 69 | 0 | 2 | 0 | NON_SYNONYMOU | MODERAT | MISSENSE | gCg/gTg | A191V | PLES_05371 | T | T | | | | | | | | | | |
| NC_011770 | 592376 | G | A | 77 | 0 | 4 | 0 | NON_SYNONYMOU | MODERAT | MISSENSE | aCc/aTc | T28I | PLES_05401 | | | A | A | A | A | | | | | | |
| NC_011770 | 592377 | T | C | 76 | 0 | 6 | 0 | NON_SYNONYMOU | MODERAT | MISSENSE | Acc/Gcc | T28A | PLES_05401 | | | | | | | C | C | C | C | C | C |
| NC_011770 | 592380 | T | C | 75 | 0 | 6 | 0 | NON_SYNONYMOU | MODERAT | MISSENSE | Acc/Gcc | T27A | PLES_05401 | | | | | | | C | C | C | C | C | C |
| NC_011770 | 592410 | A | C | 74 | 0 | 6 | 0 | NON_SYNONYMOU | MODERAT | MISSENSE | Tcc/Gcc | S17A | PLES_05401 | | | | | | | C | C | C | C | C | C |
| NC_011770 | 592805 | G | GC | 79 | 0 | 1 | 0 | FRAME_SHIFT | HIGH | | gcg/Cgcg | A36R? | PLES_05411 | | | | | | | | GC | | | | |
| NC_011770 | 592807 | C | A | 63 | 0 | 4 | 0 | NON_SYNONYMOU | MODERAT | MISSENSE | Ccc/Acc | P34T | PLES_05411 | | | A | A | A | A | | | | | | |

| | | | | | | | | | | | | | | | | | | | | | | | | | |
|---|---|---|---|---|---|---|---|---|---|---|---|---|---|---|---|---|---|---|---|---|---|---|---|---|---|
| NC_011770 | 592820 | T | C | 79 | 0 | 12 | 0 | NON_SYNONYMOU | MODERAT | MISSENSE | gTc/gCc | V38A | PLES_05411 | C | C | C | C | C | C | C | C | C | C | C | C |
| NC_011770 | 592897 | T | A | 75 | 0 | 4 | 0 | NON_SYNONYMOU | MODERAT | MISSENSE | Tcc/Acc | S64T | PLES_05411 | | | A | A | A | A | | | | | | |
| NC_011770 | 592936 | C | T | 78 | 0 | 2 | 0 | NON_SYNONYMOU | MODERAT | MISSENSE | Cgg/Tgg | R77W | PLES_05411 | | | | | | | | | T | T | | |
| NC_011770 | 593014 | G | A | 72 | 0 | 2 | 0 | NON_SYNONYMOU | MODERAT | MISSENSE | Gcc/Acc | A103T | PLES_05411 | A | A | | | | | | | | | | |
| NC_011770 | 593152 | T | C | 66 | 0 | 6 | 0 | NON_SYNONYMOU | MODERAT | MISSENSE | Tcg/Ccg | S149P | PLES_05411 | C | C | C | C | C | C | | | | | | |
| NC_011770 | 593188 | G | A | 66 | 0 | 2 | 0 | NON_SYNONYMOU | MODERAT | MISSENSE | Ggc/Agc | G161S | PLES_05411 | A | A | | | | | | | | | | |
| NC_011770 | 593245 | G | A | 70 | 0 | 2 | 0 | NON_SYNONYMOU | MODERAT | MISSENSE | Gcg/Acg | A180T | PLES_05411 | A | A | | | | | | | | | | |
| NC_011770 | 593273 | G | A | 75 | 0 | 2 | 0 | NON_SYNONYMOU | MODERAT | MISSENSE | gGg/gAg | G189E | PLES_05411 | A | A | | | | | | | | | | |
| NC_011770 | 593639 | T | C | 73 | 0 | 2 | 0 | NON_SYNONYMOU | MODERAT | MISSENSE | gTg/gCg | V311A | PLES_05411 | C | C | | | | | | | | | | |
| NC_011770 | 593954 | C | T | 76 | 0 | 12 | 0 | NON_SYNONYMOU | MODERAT | MISSENSE | gGa/gAa | G157E | PLES_05421 | T | T | T | T | T | T | T | T | T | T | T | T |
| NC_011770 | 593978 | T | C | 78 | 0 | 2 | 0 | NON_SYNONYMOU | MODERAT | MISSENSE | cAg/cGg | Q149R | PLES_05421 | C | C | | | | | | | | | | |
| NC_011770 | 594093 | C | T | 79 | 0 | 2 | 0 | NON_SYNONYMOU | MODERAT | MISSENSE | Ggc/Agc | G111S | PLES_05421 | | | | | | | T | T | | | | |
| NC_011770 | 594120 | G | T | 78 | 0 | 12 | 0 | NON_SYNONYMOU | MODERAT | MISSENSE | Ccc/Acc | P102T | PLES_05421 | T | T | T | T | T | T | T | T | T | T | T | T |
| NC_011770 | 594152 | G | A | 77 | 0 | 6 | 0 | NON_SYNONYMOU | MODERAT | MISSENSE | gCg/gTg | A91V | PLES_05421 | | | A | A | A | A | A | A | | | | |
| NC_011770 | 594755 | C | T | 67 | 0 | 6 | 0 | NON_SYNONYMOU | MODERAT | MISSENSE | gCg/gTg | A43V | PLES_05431 | | | T | T | T | T | T | T | | | | |
| NC_011770 | 594772 | T | A | 67 | 0 | 6 | 0 | NON_SYNONYMOU | MODERAT | MISSENSE | Tcg/Acg | S49T | PLES_05431 | | | A | A | A | A | A | A | | | | |
| NC_011770 | 595045 | A | G | 79 | 0 | 6 | 0 | NON_SYNONYMOU | MODERAT | MISSENSE | Atc/Gtc | I140V | PLES_05431 | | | G | G | G | G | G | G | | | | |
| NC_011770 | 595249 | A | G | 84 | 0 | 2 | 0 | NON_SYNONYMOU | MODERAT | MISSENSE | Acg/Gcg | T208A | PLES_05431 | | | | | | | | | | | G | G |
| NC_011770 | 595288 | C | T | 87 | 0 | 2 | 0 | NON_SYNONYMOU | MODERAT | MISSENSE | Cac/Tac | H221Y | PLES_05431 | T | T | | | | | | | | | | |
| NC_011770 | 595289 | A | G | 85 | 0 | 2 | 0 | NON_SYNONYMOU | MODERAT | MISSENSE | cAc/cGc | H221R | PLES_05431 | | | | | | | | | | | G | G |
| NC_011770 | 595628 | C | T | 89 | 0 | 2 | 0 | NON_SYNONYMOU | MODERAT | MISSENSE | gCg/gTg | A334V | PLES_05431 | T | T | | | | | | | | | | |
| NC_011770 | 595774 | C | A | 75 | 0 | 2 | 0 | NON_SYNONYMOU | MODERAT | MISSENSE | Cag/Aag | Q383K | PLES_05431 | A | A | | | | | | | | | | |
| NC_011770 | 597930 | C | T | 89 | 0 | 2 | 0 | NON_SYNONYMOU | MODERAT | MISSENSE | cGc/cAc | R89H | PLES_05451 | | | | | | | T | T | | | | |
| NC_011770 | 601004 | GA | G | 72 | 0 | 1 | 0 | FRAME_SHIFT | HIGH | | gga/ | G189 | PLES_05471 | | | | | | | G | | | | | |
| NC_011770 | 600500 | C | T | 56 | 0 | 2 | 0 | NON_SYNONYMOU | MODERAT | MISSENSE | gCg/gTg | A21V | PLES_05471 | | | | | | | | | | | T | T |
| NC_011770 | 600884 | G | A | 68 | 0 | 2 | 0 | NON_SYNONYMOU | MODERAT | MISSENSE | cGg/cAg | R149Q | PLES_05471 | A | A | | | | | | | | | | |
| NC_011770 | 600971 | A | G | 71 | 0 | 2 | 0 | NON_SYNONYMOU | MODERAT | MISSENSE | gAc/gGc | D178G | PLES_05471 | | | | | | | | | | | G | G |
| NC_011770 | 601249 | G | A | 86 | 0 | 2 | 0 | NON_SYNONYMOU | MODERAT | MISSENSE | Gca/Aca | A271T | PLES_05471 | A | A | | | | | | | | | | |
| NC_011770 | 601592 | G | T | 88 | 0 | 2 | 0 | NON_SYNONYMOU | MODERAT | MISSENSE | gaC/gaA | D230E | PLES_05481 | T | T | | | | | | | | | | |
| NC_011770 | 601714 | C | T | 82 | 0 | 2 | 0 | NON_SYNONYMOU | MODERAT | MISSENSE | Gcc/Acc | A190T | PLES_05481 | | | | | | | T | T | | | | |
| NC_011770 | 601872 | C | T | 80 | 0 | 4 | 0 | NON_SYNONYMOU | MODERAT | MISSENSE | aGg/aAg | R137K | PLES_05481 | | | | | | | | | T | T | T | T |
| NC_011770 | 601873 | T | C | 80 | 0 | 10 | 0 | NON_SYNONYMOU | MODERAT | MISSENSE | Agg/Ggg | R137G | PLES_05481 | C | C | C | C | C | C | | | C | C | C | C |
| NC_011770 | 602130 | T | G | 80 | 0 | 2 | 0 | NON_SYNONYMOU | MODERAT | MISSENSE | gAc/gCc | D51A | PLES_05481 | | | | | | | G | G | | | | |
| NC_011770 | 602239 | A | G | 78 | 0 | 12 | 0 | NON_SYNONYMOU | MODERAT | MISSENSE | Tac/Cac | Y15H | PLES_05481 | G | G | G | G | G | G | G | G | G | G | G | G |
| NC_011770 | 601650 | C | T | 83 | 0 | 1 | 0 | NON_SYNONYMOU | MODERAT | MISSENSE | cGc/cAc | R211H | PLES_05481 | | | | | | | | T | | | | |
| NC_011770 | 605183 | T | C | 70 | 0 | 4 | 0 | NON_SYNONYMOU | MODERAT | MISSENSE | Ttc/Ctc | F9L | PLES_05521 | | | C | C | C | C | | | | | | |
| NC_011770 | 605192 | T | C | 66 | 0 | 4 | 0 | NON_SYNONYMOU | MODERAT | MISSENSE | Ttc/Ctc | F12L | PLES_05521 | | | C | C | C | C | | | | | | |
| NC_011770 | 605226 | C | G | 70 | 0 | 4 | 0 | NON_SYNONYMOU | MODERAT | MISSENSE | gCt/gGt | A23G | PLES_05521 | | | G | G | G | G | | | | | | |
| NC_011770 | 606982 | A | G | 79 | 0 | 2 | 0 | NON_SYNONYMOU | MODERAT | MISSENSE | cAa/cGa | Q78R | PLES_05541 | | | | | | | | | G | G | | |
| NC_011770 | 607123 | C | T | 82 | 0 | 12 | 0 | NON_SYNONYMOU | MODERAT | MISSENSE | aCg/aTg | T125M | PLES_05541 | T | T | T | T | T | T | T | T | T | T | T | T |
| NC_011770 | 607213 | C | T | 76 | 0 | 2 | 0 | NON_SYNONYMOU | MODERAT | MISSENSE | gCg/gTg | A155V | PLES_05541 | | | | | | | | | | | T | T |
| NC_011770 | 607318 | G | T | 81 | 0 | 2 | 0 | NON_SYNONYMOU | MODERAT | MISSENSE | caC/caA | H191Q | PLES_05551 | | | | | | | T | T | | | | |
| NC_011770 | 607365 | G | T | 81 | 0 | 2 | 0 | NON_SYNONYMOU | MODERAT | MISSENSE | Cag/Aag | Q176K | PLES_05551 | | | | | | | T | T | | | | |
| NC_011770 | 607421 | G | T | 79 | 0 | 2 | 0 | NON_SYNONYMOU | MODERAT | MISSENSE | cCg/cAg | P157Q | PLES_05551 | | | | | | | T | T | | | | |
| NC_011770 | 607679 | C | T | 84 | 0 | 1 | 0 | NON_SYNONYMOU | MODERAT | MISSENSE | cGc/cAc | R71H | PLES_05551 | | | | | | | T | | | | | |
| NC_011770 | 607686 | C | T | 76 | 0 | 2 | 0 | NON_SYNONYMOU | MODERAT | MISSENSE | Gtc/Atc | V69I | PLES_05551 | | | | | | | T | T | | | | |
| NC_011770 | 607835 | G | A | 80 | 0 | 2 | 0 | NON_SYNONYMOU | MODERAT | MISSENSE | cCa/cTa | P19L | PLES_05551 | | | | | | | A | A | | | | |
| NC_011770 | 607373 | G | A | 80 | 0 | 1 | 0 | NON_SYNONYMOU | MODERAT | MISSENSE | cCg/cTg | P173L | PLES_05551 | | | | | | | | | | | A | |
| NC_011770 | 608443 | G | T | 90 | 0 | 2 | 0 | NON_SYNONYMOU | MODERAT | MISSENSE | ttG/ttT | L185F | PLES_05561 | | | | | | | T | T | | | | |
| NC_011770 | 608601 | G | A | 82 | 0 | 2 | 0 | NON_SYNONYMOU | MODERAT | MISSENSE | aGg/aAg | R238K | PLES_05561 | | | | | | | A | A | | | | |
| NC_011770 | 608904 | T | C | 62 | 0 | 6 | 0 | NON_SYNONYMOU | MODERAT | MISSENSE | Acc/Gcc | T310A | PLES_05571 | C | C | | | | | C | C | | | C | C |
| NC_011770 | 609044 | C | T | 67 | 0 | 6 | 0 | NON_SYNONYMOU | MODERAT | MISSENSE | aGc/aAc | S263N | PLES_05571 | T | T | | | | | T | T | | | T | T |
| NC_011770 | 609167 | A | G | 68 | 0 | 6 | 0 | NON_SYNONYMOU | MODERAT | MISSENSE | gTg/gCg | V222A | PLES_05571 | G | G | | | | | G | G | | | G | G |
| NC_011770 | 609407 | T | C | 72 | 0 | 6 | 0 | NON_SYNONYMOU | MODERAT | MISSENSE | cAc/cGc | H142R | PLES_05571 | C | C | | | | | C | C | | | C | C |
| NC_011770 | 609416 | G | T | 71 | 0 | 4 | 0 | NON_SYNONYMOU | MODERAT | MISSENSE | gCc/gAc | A139D | PLES_05571 | T | T | | | | | T | T | | | | |
| NC_011770 | 609949 | C | G | 81 | 0 | 2 | 0 | NON_SYNONYMOU | MODERAT | MISSENSE | atC/atG | I2M | PLES_05581 | G | G | | | | | | | | | | |
| NC_011770 | 609977 | G | A | 87 | 0 | 2 | 0 | NON_SYNONYMOU | MODERAT | MISSENSE | Gca/Aca | A12T | PLES_05581 | A | A | | | | | | | | | | |
| NC_011770 | 610181 | G | A | 92 | 0 | 2 | 0 | NON_SYNONYMOU | MODERAT | MISSENSE | Gcc/Acc | A80T | PLES_05581 | | | | | | | A | A | | | | |
| NC_011770 | 610457 | C | T | 71 | 0 | 4 | 0 | NON_SYNONYMOU | MODERAT | MISSENSE | Gcg/Acg | A227T | PLES_05591 | T | T | | | | | | | | | T | T |
| NC_011770 | 610550 | C | G | 76 | 0 | 8 | 0 | NON_SYNONYMOU | MODERAT | MISSENSE | Gag/Cag | E196Q | PLES_05591 | G | G | G | G | G | G | G | G | | | | |
| NC_011770 | 611806 | G | A | 89 | 0 | 10 | 0 | NON_SYNONYMOU | MODERAT | MISSENSE | Ccc/Tcc | P301S | PLES_05611 | A | A | A | A | A | A | A | A | A | A | | |
| NC_011770 | 612537 | C | G | 84 | 0 | 2 | 0 | NON_SYNONYMOU | MODERAT | MISSENSE | gGt/gCt | G57A | PLES_05611 | | | | | | | | | G | G | | |

| | | | | | | | | | | | | | | | | | | | | | | | | | |
|---|---|---|---|---|---|---|---|---|---|---|---|---|---|---|---|---|---|---|---|---|---|---|---|---|---|
| NC_011770 | 612162 | T | C | 94 | 0 | 1 | 0 | NON_SYNONYMOU | MODERAT | MISSENSE | gAc/gGc | D182G | PLES_05611 | C | | | | | | | | | | | |
| NC_011770 | 613292 | C | T | 94 | 0 | 4 | 0 | NON_SYNONYMOU | MODERAT | MISSENSE | Gtg/Atg | V126M | PLES_05631 | | | T | T | T | T | | | | | | |
| NC_011770 | 613406 | C | T | 90 | 0 | 6 | 0 | NON_SYNONYMOU | MODERAT | MISSENSE | Gcg/Acg | A88T | PLES_05631 | | | T | T | T | T | | | | | T | T |
| NC_011770 | 613465 | G | C | 94 | 0 | 6 | 0 | NON_SYNONYMOU | MODERAT | MISSENSE | gCc/gGc | A68G | PLES_05631 | | | C | C | C | C | | | | | C | C |
| NC_011770 | 614775 | G | T | 87 | 0 | 2 | 0 | NON_SYNONYMOU | MODERAT | MISSENSE | Ctg/Atg | L65M | PLES_05661 | | | | | | | | | T | T | | |
| NC_011770 | 614882 | C | T | 83 | 0 | 2 | 0 | NON_SYNONYMOU | MODERAT | MISSENSE | cGc/cAc | R29H | PLES_05661 | | | | | | | | | | | T | T |
| NC_011770 | 614919 | G | T | 84 | 0 | 2 | 0 | NON_SYNONYMOU | MODERAT | MISSENSE | Cgc/Agc | R17S | PLES_05661 | | | | | | | | | | | T | T |
| NC_011770 | 615404 | C | T | 91 | 0 | 2 | 0 | NON_SYNONYMOU | MODERAT | MISSENSE | Gcc/Acc | A177T | PLES_05681 | T | T | | | | | | | | | | |
| NC_011770 | 615428 | C | T | 89 | 0 | 2 | 0 | NON_SYNONYMOU | MODERAT | MISSENSE | Gat/Aat | D169N | PLES_05681 | | | | | | | | | | | T | T |
| NC_011770 | 615445 | T | A | 91 | 0 | 2 | 0 | NON_SYNONYMOU | MODERAT | MISSENSE | gAc/gTc | D163V | PLES_05681 | A | A | | | | | | | | | | |
| NC_011770 | 615566 | T | C | 91 | 0 | 8 | 0 | NON_SYNONYMOU | MODERAT | MISSENSE | Aag/Gag | K123E | PLES_05681 | C | C | | | | | C | C | C | C | C | C |
| NC_011770 | 615800 | T | C | 101 | 0 | 6 | 0 | NON_SYNONYMOU | MODERAT | MISSENSE | Acg/Gcg | T45A | PLES_05681 | | | | | | | C | C | C | C | C | C |
| NC_011770 | 615808 | C | T | 101 | 0 | 2 | 0 | NON_SYNONYMOU | MODERAT | MISSENSE | cGt/cAt | R42H | PLES_05681 | | | | | | | | | | | T | T |
| NC_011770 | 615832 | T | C | 106 | 0 | 8 | 0 | NON_SYNONYMOU | MODERAT | MISSENSE | cAg/cGg | Q34R | PLES_05681 | C | C | | | | | C | C | C | C | C | C |
| NC_011770 | 615852 | A | C | 110 | 0 | 6 | 0 | NON_SYNONYMOU | MODERAT | MISSENSE | atT/atG | I27M | PLES_05681 | | | | | | | C | C | C | C | C | C |
| NC_011770 | 616166 | A | G | 83 | 0 | 2 | 0 | NON_SYNONYMOU | MODERAT | MISSENSE | cTt/cCt | L910P | PLES_05691 | G | G | | | | | | | | | | |
| NC_011770 | 618027 | G | A | 102 | 0 | 2 | 0 | NON_SYNONYMOU | MODERAT | MISSENSE | Cgg/Tgg | R290W | PLES_05691 | | | | | | | | | | | A | A |
| NC_011770 | 618047 | C | G | 106 | 0 | 2 | 0 | NON_SYNONYMOU | MODERAT | MISSENSE | aGg/aCg | R283T | PLES_05691 | | | | | | | | | G | G | | |
| NC_011770 | 618122 | C | T | 104 | 0 | 2 | 0 | NON_SYNONYMOU | MODERAT | MISSENSE | aGc/aAc | S258N | PLES_05691 | | | | | | | | | T | T | | |
| NC_011770 | 618327 | C | T | 92 | 0 | 4 | 0 | NON_SYNONYMOU | MODERAT | MISSENSE | Gcc/Acc | A190T | PLES_05691 | T | T | | | | | | | | | T | T |
| NC_011770 | 618447 | T | C | 95 | 0 | 4 | 0 | NON_SYNONYMOU | MODERAT | MISSENSE | Acc/Gcc | T150A | PLES_05691 | C | C | | | | | | | | | C | C |
| NC_011770 | 619429 | T | G | 95 | 0 | 10 | 0 | STOP_LOST+SPLICE | HIGH | MISSENSE | tAg/tCg | *124S | PLES_05701 | G | G | G | G | G | G | | | G | G | G | G |
| NC_011770 | 619465 | G | C | 82 | 0 | 8 | 0 | STOP_GAINED | HIGH | NONSENSE | tCa/tGa | S112* | PLES_05701 | C | C | C | C | C | C | | | C | C | | |
| NC_011770 | 619708 | C | T | 97 | 0 | 4 | 0 | STOP_GAINED | HIGH | NONSENSE | tGg/tAg | W31* | PLES_05701 | | | T | T | T | T | | | | | | |
| NC_011770 | 619456 | G | C | 83 | 0 | 8 | 0 | NON_SYNONYMOU | MODERAT | MISSENSE | aCc/aGc | T115S | PLES_05701 | C | C | C | C | C | C | | | C | C | | |
| NC_011770 | 619457 | T | G | 82 | 0 | 8 | 0 | NON_SYNONYMOU | MODERAT | MISSENSE | Acc/Ccc | T115P | PLES_05701 | G | G | G | G | G | G | | | G | G | | |
| NC_011770 | 619468 | A | G | 82 | 0 | 8 | 0 | NON_SYNONYMOU | MODERAT | MISSENSE | gTg/gCg | V111A | PLES_05701 | G | G | G | G | G | G | | | G | G | | |
| NC_011770 | 619471 | G | A | 82 | 0 | 8 | 0 | NON_SYNONYMOU | MODERAT | MISSENSE | tCg/tTg | S110L | PLES_05701 | A | A | A | A | A | A | | | A | A | | |
| NC_011770 | 619501 | G | A | 85 | 0 | 8 | 0 | NON_SYNONYMOU | MODERAT | MISSENSE | aCc/aTc | T100I | PLES_05701 | A | A | A | A | A | A | | | A | A | | |
| NC_011770 | 620086 | T | C | 91 | 0 | 6 | 0 | NON_SYNONYMOU | MODERAT | MISSENSE | Atc/Gtc | I310V | PLES_05711 | C | C | C | C | C | C | | | | | | |
| NC_011770 | 620180 | C | G | 95 | 0 | 2 | 0 | NON_SYNONYMOU | MODERAT | MISSENSE | aaG/aaC | K278N | PLES_05711 | G | G | | | | | | | | | | |
| NC_011770 | 620193 | T | G | 95 | 0 | 2 | 0 | NON_SYNONYMOU | MODERAT | MISSENSE | gAa/gCa | E274A | PLES_05711 | G | G | | | | | | | | | | |
| NC_011770 | 620208 | A | T | 94 | 0 | 4 | 0 | NON_SYNONYMOU | MODERAT | MISSENSE | cTg/cAg | L269Q | PLES_05711 | | | T | T | T | T | | | | | | |
| NC_011770 | 620213 | C | G | 96 | 0 | 2 | 0 | NON_SYNONYMOU | MODERAT | MISSENSE | aaG/aaC | K267N | PLES_05711 | G | G | | | | | | | | | | |
| NC_011770 | 620229 | C | G | 93 | 0 | 2 | 0 | NON_SYNONYMOU | MODERAT | MISSENSE | gGg/gCg | G262A | PLES_05711 | G | G | | | | | | | | | | |
| NC_011770 | 620230 | C | T | 93 | 0 | 2 | 0 | NON_SYNONYMOU | MODERAT | MISSENSE | Ggg/Agg | G262R | PLES_05711 | T | T | | | | | | | | | | |
| NC_011770 | 620233 | C | T | 93 | 0 | 2 | 0 | NON_SYNONYMOU | MODERAT | MISSENSE | Ggc/Agc | G261S | PLES_05711 | T | T | | | | | | | | | | |
| NC_011770 | 620681 | C | A | 94 | 0 | 10 | 0 | NON_SYNONYMOU | MODERAT | MISSENSE | caG/caT | Q111H | PLES_05711 | A | A | A | A | A | A | | | A | A | A | A |
| NC_011770 | 620880 | T | C | 87 | 0 | 12 | 0 | NON_SYNONYMOU | MODERAT | MISSENSE | gAg/gGg | E45G | PLES_05711 | C | C | C | C | C | C | C | C | C | C | C | C |
| NC_011770 | 619912 | G | A | 95 | 0 | 1 | 0 | NON_SYNONYMOU | MODERAT | MISSENSE | Cgt/Tgt | R368C | PLES_05711 | | | | | | | | A | | | | |
| NC_011770 | 622110 | T | A | 72 | 0 | 4 | 0 | NON_SYNONYMOU | MODERAT | MISSENSE | cAg/cTg | Q1095L | PLES_05721 | | | | | | | | | A | A | A | A |
| NC_011770 | 622884 | C | T | 85 | 0 | 4 | 0 | NON_SYNONYMOU | MODERAT | MISSENSE | aGt/aAt | S837N | PLES_05721 | | | | | | | | | T | T | T | T |
| NC_011770 | 623388 | G | A | 80 | 0 | 4 | 0 | NON_SYNONYMOU | MODERAT | MISSENSE | gCc/gTc | A669V | PLES_05721 | | | A | A | A | A | | | | | | |
| NC_011770 | 623659 | G | T | 85 | 0 | 4 | 0 | NON_SYNONYMOU | MODERAT | MISSENSE | Cgc/Agc | R579S | PLES_05721 | | | T | T | T | T | | | | | | |
| NC_011770 | 623791 | C | T | 72 | 0 | 2 | 0 | NON_SYNONYMOU | MODERAT | MISSENSE | Gac/Aac | D535N | PLES_05721 | T | T | | | | | | | | | | |
| NC_011770 | 624501 | T | C | 72 | 0 | 12 | 0 | NON_SYNONYMOU | MODERAT | MISSENSE | aAt/aGt | N298S | PLES_05721 | C | C | C | C | C | C | C | C | C | C | C | C |
| NC_011770 | 624994 | G | T | 70 | 0 | 4 | 0 | NON_SYNONYMOU | MODERAT | MISSENSE | Ccg/Acg | P134T | PLES_05721 | | | | | | | | | T | T | T | T |
| NC_011770 | 625161 | A | G | 72 | 0 | 6 | 0 | NON_SYNONYMOU | MODERAT | MISSENSE | cTg/cCg | L78P | PLES_05721 | G | G | | | | | | | G | G | G | G |
| NC_011770 | 625192 | C | T | 70 | 0 | 6 | 0 | NON_SYNONYMOU | MODERAT | MISSENSE | Ggc/Agc | G68S | PLES_05721 | T | T | | | | | | | T | T | T | T |
| NC_011770 | 625195 | C | T | 70 | 0 | 4 | 0 | NON_SYNONYMOU | MODERAT | MISSENSE | Gtc/Atc | V67I | PLES_05721 | | | T | T | T | T | | | | | | |
| NC_011770 | 625239 | A | T | 68 | 0 | 4 | 0 | NON_SYNONYMOU | MODERAT | MISSENSE | tTc/tAc | F52Y | PLES_05721 | | | T | T | T | T | | | | | | |
| NC_011770 | 625332 | C | T | 60 | 0 | 6 | 0 | NON_SYNONYMOU | MODERAT | MISSENSE | gGc/gAc | G21D | PLES_05721 | T | T | | | | | | | T | T | T | T |
| NC_011770 | 629719 | C | T | 77 | 0 | 2 | 0 | NON_SYNONYMOU | MODERAT | MISSENSE | Gct/Act | A81T | PLES_05751 | | | | | | | | | | | T | T |
| NC_011770 | 631664 | A | G | 57 | 0 | 1 | 0 | NON_SYNONYMOU | MODERAT | MISSENSE | cTg/cCg | L152P | PLES_05781 | | | | | | | G | | | | | |
| NC_011770 | 632631 | G | C | 78 | 0 | 12 | 0 | NON_SYNONYMOU | MODERAT | MISSENSE | gGt/gCt | G29A | PLES_05801 | C | C | C | C | C | C | C | C | C | C | C | C |
| NC_011770 | 632648 | C | T | 80 | 0 | 2 | 0 | NON_SYNONYMOU | MODERAT | MISSENSE | Cgt/Tgt | R35C | PLES_05801 | | | | | | | T | T | | | | |
| NC_011770 | 632987 | G | A | 79 | 0 | 8 | 0 | NON_SYNONYMOU | MODERAT | MISSENSE | Gcg/Acg | A148T | PLES_05801 | | | A | A | A | A | | | A | A | A | A |
| NC_011770 | 634356 | T | A | 64 | 0 | 4 | 0 | NON_SYNONYMOU | MODERAT | MISSENSE | cAg/cTg | Q163L | PLES_05821 | | | | | | | A | A | | | A | A |
| NC_011770 | 634636 | C | T | 73 | 0 | 4 | 0 | NON_SYNONYMOU | MODERAT | MISSENSE | Gcc/Acc | A70T | PLES_05821 | | | T | T | T | T | | | | | | |
| NC_011770 | 634755 | C | T | 65 | 0 | 4 | 0 | NON_SYNONYMOU | MODERAT | MISSENSE | aGc/aAc | S30N | PLES_05821 | | | | | | | T | T | | | T | T |
| NC_011770 | 635460 | A | T | 85 | 0 | 12 | 0 | NON_SYNONYMOU | MODERAT | MISSENSE | gaT/gaA | D344E | PLES_05831 | T | T | T | T | T | T | T | T | T | T | T | T |
| NC_011770 | 635644 | T | C | 96 | 0 | 1 | 0 | NON_SYNONYMOU | MODERAT | MISSENSE | cAc/cGc | H283R | PLES_05831 | | | | C | | | | | | | | |

| | | | | | | | | | | | | | | | | | | | | | | | | | |
|---|---|---|---|---|---|---|---|---|---|---|---|---|---|---|---|---|---|---|---|---|---|---|---|---|---|
| NC_011770 | 639197 | A | G | 85 | 0 | 2 | 0 | NON_SYNONYMOU | MODERAT | MISSENSE | Tac/Cac | Y196H | PLES_05851 | | | | | | | G | G | | | | |
| NC_011770 | 648258 | C | G | 69 | 0 | 4 | 0 | NON_SYNONYMOU | MODERAT | MISSENSE | Cag/Gag | Q154E | PLES_05931 | | | G | G | G | G | | | | | | |
| NC_011770 | 648330 | C | A | 68 | 0 | 10 | 0 | NON_SYNONYMOU | MODERAT | MISSENSE | Cag/Aag | Q178K | PLES_05931 | A | A | A | A | A | A | A | A | A | A | | |
| NC_011770 | 648547 | A | G | 69 | 0 | 6 | 0 | NON_SYNONYMOU | MODERAT | MISSENSE | aAt/aGt | N250S | PLES_05931 | | | G | G | G | G | | | | | G | G |
| NC_011770 | 648567 | A | G | 66 | 0 | 2 | 0 | NON_SYNONYMOU | MODERAT | MISSENSE | Acc/Gcc | T257A | PLES_05931 | G | G | | | | | | | | | | |
| NC_011770 | 649013 | A | T | 70 | 0 | 12 | 0 | NON_SYNONYMOU | MODERAT | MISSENSE | Acg/Tcg | T68S | PLES_05941 | T | T | T | T | T | T | T | T | T | T | T | T |
| NC_011770 | 649247 | G | A | 70 | 0 | 2 | 0 | NON_SYNONYMOU | MODERAT | MISSENSE | Ggc/Agc | G146S | PLES_05941 | | | A | A | | | | | | | | |
| NC_011770 | 649409 | T | C | 70 | 0 | 4 | 0 | NON_SYNONYMOU | MODERAT | MISSENSE | Tat/Cat | Y200H | PLES_05941 | C | C | | | | | C | C | | | | |
| NC_011770 | 649597 | G | A | 69 | 0 | 2 | 0 | NON_SYNONYMOU | MODERAT | MISSENSE | cGc/cAc | R37H | PLES_05951 | | | | | | | | | A | A | | |
| NC_011770 | 649661 | C | A | 68 | 0 | 2 | 0 | NON_SYNONYMOU | MODERAT | MISSENSE | gaC/gaA | D58E | PLES_05951 | | | | | | | | | | | A | A |
| NC_011770 | 649718 | G | A | 73 | 0 | 2 | 0 | NON_SYNONYMOU | MODERAT | MISSENSE | atG/atA | M77I | PLES_05951 | | | | | | | | | A | A | | |
| NC_011770 | 649722 | A | C | 70 | 0 | 2 | 0 | NON_SYNONYMOU | MODERAT | MISSENSE | Atc/Ctc | I79L | PLES_05951 | C | C | | | | | | | | | | |
| NC_011770 | 649989 | C | T | 70 | 0 | 12 | 0 | NON_SYNONYMOU | MODERAT | MISSENSE | Ccg/Tcg | P168S | PLES_05951 | T | T | T | T | T | T | T | T | T | T | T | T |
| NC_011770 | 651182 | CG | C | 59 | 0 | 1 | 0 | FRAME_SHIFT | HIGH | | ccc/ | P42 | PLES_05961 | | | | C | | | | | | | | |
| NC_011770 | 650338 | T | G | 63 | 0 | 2 | 0 | NON_SYNONYMOU | MODERAT | MISSENSE | gAc/gCc | D324A | PLES_05961 | | | | | | | | | | | G | G |
| NC_011770 | 650692 | G | A | 63 | 0 | 2 | 0 | NON_SYNONYMOU | MODERAT | MISSENSE | gCg/gTg | A206V | PLES_05961 | A | A | | | | | | | | | | |
| NC_011770 | 650771 | G | C | 55 | 0 | 12 | 0 | NON_SYNONYMOU | MODERAT | MISSENSE | Caa/Gaa | Q180E | PLES_05961 | C | C | C | C | C | C | C | C | C | C | C | C |
| NC_011770 | 650887 | G | C | 54 | 0 | 2 | 0 | NON_SYNONYMOU | MODERAT | MISSENSE | cCt/cGt | P141R | PLES_05961 | C | C | | | | | | | | | | |
| NC_011770 | 651190 | G | T | 57 | 0 | 2 | 0 | NON_SYNONYMOU | MODERAT | MISSENSE | cCg/cAg | P40Q | PLES_05961 | | | | | | | T | T | | | | |
| NC_011770 | 651222 | A | T | 59 | 0 | 10 | 0 | NON_SYNONYMOU | MODERAT | MISSENSE | caT/caA | H29Q | PLES_05961 | T | T | T | T | T | T | | | T | T | T | T |
| NC_011770 | 651274 | G | A | 58 | 0 | 6 | 0 | NON_SYNONYMOU | MODERAT | MISSENSE | gCc/gTc | A12V | PLES_05961 | | | A | A | A | A | | | A | A | | |
| NC_011770 | 653044 | C | T | 106 | 0 | 1 | 0 | STOP_GAINED | HIGH | NONSENSE | Cag/Tag | Q529* | PLES_05971 | | | | | | | | T | | | | |
| NC_011770 | 651488 | T | G | 57 | 0 | 6 | 0 | NON_SYNONYMOU | MODERAT | MISSENSE | tTc/tGc | F10C | PLES_05971 | | | G | G | G | G | | | G | G | | |
| NC_011770 | 651977 | G | T | 59 | 0 | 12 | 0 | NON_SYNONYMOU | MODERAT | MISSENSE | gGg/gTg | G173V | PLES_05971 | T | T | T | T | T | T | T | T | T | T | T | T |
| NC_011770 | 651979 | G | T | 58 | 0 | 12 | 0 | NON_SYNONYMOU | MODERAT | MISSENSE | Gcg/Tcg | A174S | PLES_05971 | T | T | T | T | T | T | T | T | T | T | T | T |
| NC_011770 | 652356 | T | G | 79 | 0 | 4 | 0 | NON_SYNONYMOU | MODERAT | MISSENSE | caT/caG | H299Q | PLES_05971 | | | G | G | G | G | | | | | | |
| NC_011770 | 652691 | C | T | 83 | 0 | 4 | 0 | NON_SYNONYMOU | MODERAT | MISSENSE | gCc/gTc | A411V | PLES_05971 | | | T | T | T | T | | | | | | |
| NC_011770 | 653845 | G | A | 103 | 0 | 4 | 0 | NON_SYNONYMOU | MODERAT | MISSENSE | Ggc/Agc | G796S | PLES_05971 | A | A | | | | | | | A | A | | |
| NC_011770 | 654712 | C | T | 77 | 0 | 2 | 0 | NON_SYNONYMOU | MODERAT | MISSENSE | gCg/gTg | A18V | PLES_05991 | T | T | | | | | | | | | | |
| NC_011770 | 657897 | G | A | 81 | 0 | 6 | 0 | NON_SYNONYMOU | MODERAT | MISSENSE | cGg/cAg | R269Q | PLES_06011 | | | A | A | A | A | | | A | A | | |
| NC_011770 | 658283 | G | A | 79 | 0 | 8 | 0 | NON_SYNONYMOU | MODERAT | MISSENSE | Gcc/Acc | A11T | PLES_06021 | | | A | A | A | A | | | A | A | A | A |
| NC_011770 | 658505 | G | T | 78 | 0 | 2 | 0 | NON_SYNONYMOU | MODERAT | MISSENSE | Gcc/Tcc | A85S | PLES_06021 | | | | | | | T | T | | | | |
| NC_011770 | 658653 | C | T | 85 | 0 | 1 | 0 | NON_SYNONYMOU | MODERAT | MISSENSE | cCc/cTc | P134L | PLES_06021 | | | | | | | T | | | | | |
| NC_011770 | 658736 | A | G | 88 | 0 | 2 | 0 | NON_SYNONYMOU | MODERAT | MISSENSE | Agc/Ggc | S162G | PLES_06021 | | | | | | | | | G | G | | |
| NC_011770 | 658829 | T | C | 84 | 0 | 1 | 0 | NON_SYNONYMOU | MODERAT | MISSENSE | Tac/Cac | Y193H | PLES_06021 | | | | | | | C | | | | | |
| NC_011770 | 658877 | G | A | 86 | 0 | 2 | 0 | NON_SYNONYMOU | MODERAT | MISSENSE | Gtc/Atc | V209I | PLES_06021 | A | A | | | | | | | | | | |
| NC_011770 | 660432 | C | G | 76 | 0 | 8 | 0 | NON_SYNONYMOU | MODERAT | MISSENSE | gCc/gGc | A276G | PLES_06031 | G | G | | | | | G | G | G | G | G | G |
| NC_011770 | 661357 | C | T | 81 | 0 | 2 | 0 | NON_SYNONYMOU | MODERAT | MISSENSE | Ctc/Ttc | L42F | PLES_06051 | T | T | | | | | | | | | | |
| NC_011770 | 661426 | C | T | 78 | 0 | 1 | 0 | NON_SYNONYMOU | MODERAT | MISSENSE | Cgt/Tgt | R65C | PLES_06051 | | | | | | | T | | | | | |
| NC_011770 | 665930 | A | C | 88 | 0 | 2 | 0 | NON_SYNONYMOU | MODERAT | MISSENSE | gAg/gCg | E41A | PLES_06101 | C | C | | | | | | | | | | |
| NC_011770 | 666023 | A | C | 85 | 0 | 2 | 0 | NON_SYNONYMOU | MODERAT | MISSENSE | aAc/aCc | N72T | PLES_06101 | | | | | | | | | C | C | | |
| NC_011770 | 666161 | T | C | 72 | 0 | 6 | 0 | NON_SYNONYMOU | MODERAT | MISSENSE | gTg/gCg | V118A | PLES_06101 | | | C | C | C | C | | | C | C | | |
| NC_011770 | 665898 | CCTGATCG( | C | 90 | 0 | 1 | 0 | CODON_CHANGE_ | MODERATE | | atgctgatcg | MLIAS34S | PLES_06101 | | | | | | **C** | | | | | | |
| NC_011770 | 666606 | CTG | C | 57 | 0 | 2 | 0 | FRAME_SHIFT | HIGH | | ctg/ | L26 | PLES_06111 | C | C | | | | | | | | | | |
| NC_011770 | 666573 | C | G | 57 | 0 | 2 | 0 | NON_SYNONYMOU | MODERAT | MISSENSE | Ccc/Gcc | P15A | PLES_06111 | | | | | | | | | G | G | | |
| NC_011770 | 666576 | G | T | 59 | 0 | 2 | 0 | NON_SYNONYMOU | MODERAT | MISSENSE | Ggg/Tgg | G16W | PLES_06111 | T | T | | | | | | | | | | |
| NC_011770 | 666934 | T | G | 59 | 0 | 2 | 0 | NON_SYNONYMOU | MODERAT | MISSENSE | gTt/gGt | V135G | PLES_06111 | | | | | G | G | | | | | | |
| NC_011770 | 667078 | C | T | 55 | 6 | 4 | 0 | NON_SYNONYMOU | MODERAT | MISSENSE | gCg/gTg | A26V | PLES_06121 | . | . | T | T | T | T | | | . | . | . | . |
| NC_011770 | 667228 | C | G | 56 | 6 | 4 | 0 | NON_SYNONYMOU | MODERAT | MISSENSE | gCc/gGc | A76G | PLES_06121 | . | . | G | G | G | G | | | . | . | . | . |
| NC_011770 | 667302 | C | G | 54 | 6 | 4 | 0 | NON_SYNONYMOU | MODERAT | MISSENSE | Cag/Gag | Q101E | PLES_06121 | . | . | G | G | G | G | | | . | . | . | . |
| NC_011770 | 667709 | A | G | 61 | 6 | 4 | 0 | NON_SYNONYMOU | MODERAT | MISSENSE | Ata/Gta | I66V | PLES_06131 | . | . | G | G | G | G | | | . | . | . | . |
| NC_011770 | 668619 | C | G | 54 | 6 | 6 | 0 | NON_SYNONYMOU | MODERAT | MISSENSE | Cag/Gag | Q25E | PLES_06151 | . | . | G | G | G | G | G | G | . | . | . | . |
| NC_011770 | 668785 | C | T | 54 | 6 | 4 | 0 | NON_SYNONYMOU | MODERAT | MISSENSE | gCc/gTc | A80V | PLES_06151 | . | . | T | T | T | T | | | . | . | . | . |
| NC_011770 | 669523 | A | C | 65 | 6 | 4 | 0 | NON_SYNONYMOU | MODERAT | MISSENSE | Aac/Cac | N33H | PLES_06161 | . | . | C | C | C | C | | | . | . | . | . |
| NC_011770 | 669742 | C | A | 54 | 6 | 4 | 0 | NON_SYNONYMOU | MODERAT | MISSENSE | Ctt/Att | L106I | PLES_06161 | . | . | A | A | A | A | | | . | . | . | . |
| NC_011770 | 669743 | T | C | 54 | 6 | 4 | 0 | NON_SYNONYMOU | MODERAT | MISSENSE | cTt/cCt | L106P | PLES_06161 | . | . | C | C | C | C | | | . | . | . | . |
| NC_011770 | 669890 | T | A | 60 | 4 | 2 | 0 | NON_SYNONYMOU | MODERAT | MISSENSE | gTa/gAa | V155E | PLES_06161 | A | A | | | | | | | . | . | . | . |
| NC_011770 | 669896 | G | T | 59 | 4 | 2 | 0 | NON_SYNONYMOU | MODERAT | MISSENSE | cGg/cTg | R157L | PLES_06161 | T | T | | | | | | | . | . | . | . |
| NC_011770 | 669898 | T | G | 58 | 4 | 2 | 0 | NON_SYNONYMOU | MODERAT | MISSENSE | Tcc/Gcc | S158A | PLES_06161 | G | G | | | | | | | . | . | . | . |
| NC_011770 | 669916 | G | A | 59 | 4 | 4 | 0 | NON_SYNONYMOU | MODERAT | MISSENSE | Gtg/Atg | V164M | PLES_06161 | A | A | | | | | C | C | . | . | . | . |
| NC_011770 | 669917 | T | C | 58 | 4 | 2 | 0 | NON_SYNONYMOU | MODERAT | MISSENSE | gTg/gCg | V164A | PLES_06161 | C | C | | | | | | | . | . | . | . |
| NC_011770 | 669920 | C | T | 58 | 4 | 2 | 0 | NON_SYNONYMOU | MODERAT | MISSENSE | aCc/aTc | T165I | PLES_06161 | T | T | | | | | | | . | . | . | . |

| | | | | | | | | | | | | | | | | | | | | | | | | | |
|---|---|---|---|---|---|---|---|---|---|---|---|---|---|---|---|---|---|---|---|---|---|---|---|---|---|
| NC_011770 | 669943 | T | A | 57 | 4 | 2 | 0 | NON_SYNONYMOU | MODERAT | MISSENSE | Ttc/Atc | F173I | PLES_06161 | A | A | | | | | | | . | . | . | . |
| NC_011770 | 669944 | T | G | 56 | 4 | 2 | 0 | NON_SYNONYMOU | MODERAT | MISSENSE | tTc/tGc | F173C | PLES_06161 | G | G | | | | | | | . | . | . | . |
| NC_011770 | 670265 | C | G | 48 | 4 | 2 | 0 | NON_SYNONYMOU | MODERAT | MISSENSE | Cag/Gag | Q102E | PLES_06171 | G | G | | | | | | | . | . | . | . |
| NC_011770 | 670272 | C | T | 46 | 4 | 2 | 0 | NON_SYNONYMOU | MODERAT | MISSENSE | gCc/gTc | A104V | PLES_06171 | | | | | | | T | T | . | . | . | . |
| NC_011770 | 670295 | T | G | 49 | 4 | 2 | 0 | NON_SYNONYMOU | MODERAT | MISSENSE | Tcc/Gcc | S112A | PLES_06171 | G | G | | | | | | | . | . | . | . |
| NC_011770 | 670314 | A | T | 48 | 4 | 2 | 0 | NON_SYNONYMOU | MODERAT | MISSENSE | tAc/tTc | Y118F | PLES_06171 | T | T | | | | | | | . | . | . | . |
| NC_011770 | 670409 | A | G | 44 | 4 | 2 | 0 | NON_SYNONYMOU | MODERAT | MISSENSE | Aac/Gac | N150D | PLES_06171 | G | G | | | | | | | . | . | . | . |
| NC_011770 | 670418 | A | G | 45 | 5 | 1 | 0 | NON_SYNONYMOU | MODERAT | MISSENSE | Atc/Gtc | I153V | PLES_06171 | G | . | | | | | | | . | . | . | . |
| NC_011770 | 670499 | C | G | 41 | 6 | 6 | 0 | NON_SYNONYMOU | MODERAT | MISSENSE | Ctc/Gtc | L180V | PLES_06171 | . | . | G | G | G | G | G | G | . | . | . | . |
| NC_011770 | 670623 | C | G | 39 | 6 | 6 | 0 | NON_SYNONYMOU | MODERAT | MISSENSE | aCc/aGc | T221S | PLES_06171 | . | . | G | G | G | G | G | G | . | . | . | . |
| NC_011770 | 671055 | A | C | 42 | 6 | 4 | 0 | NON_SYNONYMOU | MODERAT | MISSENSE | aAc/aCc | N365T | PLES_06171 | . | . | C | C | C | C | | | . | . | . | . |
| NC_011770 | 671058 | G | C | 42 | 6 | 4 | 0 | NON_SYNONYMOU | MODERAT | MISSENSE | aGt/aCt | S366T | PLES_06171 | . | . | C | C | C | C | | | . | . | . | . |
| NC_011770 | 671249 | A | G | 26 | 6 | 4 | 0 | NON_SYNONYMOU | MODERAT | MISSENSE | Atc/Gtc | I430V | PLES_06171 | . | . | G | G | G | G | | | . | . | . | . |
| NC_011770 | 671250 | T | C | 24 | 6 | 4 | 0 | NON_SYNONYMOU | MODERAT | MISSENSE | aTc/aCc | I430T | PLES_06171 | . | . | C | C | C | C | | | . | . | . | . |
| NC_011770 | 671257 | G | C | 25 | 6 | 4 | 0 | NON_SYNONYMOU | MODERAT | MISSENSE | gaG/gaC | E432D | PLES_06171 | . | . | C | C | C | C | | | . | . | . | . |
| NC_011770 | 671270 | C | A | 25 | 6 | 4 | 0 | NON_SYNONYMOU | MODERAT | MISSENSE | Cag/Aag | Q437K | PLES_06171 | . | . | A | A | A | A | | | . | . | . | . |
| NC_011770 | 671963 | G | C | 4 | 8 | 4 | 0 | NON_SYNONYMOU | MODERAT | MISSENSE | Gtc/Ctc | V668L | PLES_06171 | . | . | C | C | C | C | . | . | . | . | . | . |
| NC_011770 | 671985 | A | C | 5 | 8 | 4 | 0 | NON_SYNONYMOU | MODERAT | MISSENSE | aAc/aCc | N675T | PLES_06171 | . | . | C | C | C | C | . | . | . | . | . | . |
| NC_011770 | 669967 | GACCAAT | G | 58 | 4 | 1 | 0 | CODON_CHANGE_ | MODERATE | | accaatact/ | TNT3T | PLES_06171 | | **G** | | | | | | | . | . | . | . |
| NC_011770 | 672861 | A | G | 60 | 6 | 6 | 0 | NON_SYNONYMOU | MODERAT | MISSENSE | Atc/Gtc | I102V | PLES_06181 | . | . | G | G | G | G | G | G | . | . | . | . |
| NC_011770 | 673148 | C | G | 59 | 6 | 4 | 0 | NON_SYNONYMOU | MODERAT | MISSENSE | aaC/aaG | N197K | PLES_06181 | . | . | G | G | G | G | | | . | . | . | . |
| NC_011770 | 673242 | G | A | 63 | 6 | 4 | 0 | NON_SYNONYMOU | MODERAT | MISSENSE | Gtc/Atc | V229I | PLES_06181 | . | . | A | A | A | A | | | . | . | . | . |
| NC_011770 | 676738 | TG | T | 47 | 6 | 1 | 0 | FRAME_SHIFT | HIGH | | ggg/ | G661 | PLES_06211 | . | . | | T | | | | | . | . | . | . |
| NC_011770 | 674871 | G | T | 62 | 6 | 6 | 0 | NON_SYNONYMOU | MODERAT | MISSENSE | Gcc/Tcc | A37S | PLES_06211 | . | . | T | T | T | T | T | T | . | . | . | . |
| NC_011770 | 675118 | C | T | 59 | 6 | 2 | 0 | NON_SYNONYMOU | MODERAT | MISSENSE | gCc/gTc | A119V | PLES_06211 | . | . | T | T | | | | | . | . | . | . |
| NC_011770 | 675421 | T | C | 37 | 6 | 6 | 0 | NON_SYNONYMOU | MODERAT | MISSENSE | gTt/gCt | V220A | PLES_06211 | . | . | C | C | C | C | C | C | . | . | . | . |
| NC_011770 | 675669 | C | T | 20 | 6 | 2 | 0 | NON_SYNONYMOU | MODERAT | MISSENSE | Cgg/Tgg | R303W | PLES_06211 | . | . | | | | | T | T | . | . | . | . |
| NC_011770 | 675841 | T | A | 32 | 6 | 4 | 0 | NON_SYNONYMOU | MODERAT | MISSENSE | gTg/gAg | V360E | PLES_06211 | . | . | A | A | A | A | | | . | . | . | . |
| NC_011770 | 676225 | C | G | 48 | 6 | 4 | 0 | NON_SYNONYMOU | MODERAT | MISSENSE | aCc/aGc | T488S | PLES_06211 | . | . | G | G | G | G | | | . | . | . | . |
| NC_011770 | 676410 | C | G | 59 | 6 | 2 | 0 | NON_SYNONYMOU | MODERAT | MISSENSE | Ctg/Gtg | L550V | PLES_06211 | . | . | | | | | G | G | . | . | . | . |
| NC_011770 | 676590 | G | A | 55 | 6 | 6 | 0 | NON_SYNONYMOU | MODERAT | MISSENSE | Gtt/Att | V610I | PLES_06211 | . | . | A | A | A | A | A | A | . | . | . | . |
| NC_011770 | 676783 | C | A | 48 | 6 | 2 | 0 | NON_SYNONYMOU | MODERAT | MISSENSE | gCc/gAc | A674D | PLES_06211 | . | . | | | | | A | A | . | . | . | . |
| NC_011770 | 676806 | G | C | 47 | 6 | 6 | 0 | NON_SYNONYMOU | MODERAT | MISSENSE | Gcc/Ccc | A682P | PLES_06211 | . | . | C | C | C | C | C | C | . | . | . | . |
| NC_011770 | 676810 | T | C | 50 | 6 | 2 | 0 | NON_SYNONYMOU | MODERAT | MISSENSE | gTa/gCa | V683A | PLES_06211 | . | . | | | | | C | C | . | . | . | . |
| NC_011770 | 676822 | G | A | 48 | 6 | 6 | 0 | NON_SYNONYMOU | MODERAT | MISSENSE | gGa/gAa | G687E | PLES_06211 | . | . | A | A | A | A | A | A | . | . | . | . |
| NC_011770 | 676837 | C | T | 50 | 6 | 4 | 0 | NON_SYNONYMOU | MODERAT | MISSENSE | gCg/gTg | A692V | PLES_06211 | . | . | T | T | T | T | | | . | . | . | . |
| NC_011770 | 676933 | A | G | 54 | 6 | 2 | 0 | NON_SYNONYMOU | MODERAT | MISSENSE | cAa/cGa | Q724R | PLES_06211 | . | . | | | | | G | G | . | . | . | . |
| NC_011770 | 677139 | G | T | 60 | 6 | 2 | 0 | NON_SYNONYMOU | MODERAT | MISSENSE | Gcg/Tcg | A44S | PLES_06221 | . | . | | | | | T | T | . | . | . | . |
| NC_011770 | 677415 | C | T | 57 | 6 | 2 | 0 | NON_SYNONYMOU | MODERAT | MISSENSE | Ccg/Tcg | P136S | PLES_06221 | . | . | | | | | T | T | . | . | . | . |
| NC_011770 | 677428 | C | T | 58 | 6 | 2 | 0 | NON_SYNONYMOU | MODERAT | MISSENSE | gCa/gTa | A140V | PLES_06221 | . | . | | | | | T | T | . | . | . | . |
| NC_011770 | 677556 | G | A | 57 | 6 | 2 | 0 | NON_SYNONYMOU | MODERAT | MISSENSE | Gtg/Atg | V183M | PLES_06221 | . | . | | | | | A | A | . | . | . | . |
| NC_011770 | 678244 | A | C | 49 | 6 | 6 | 0 | NON_SYNONYMOU | MODERAT | MISSENSE | Atc/Ctc | I42L | PLES_06241 | . | . | C | C | C | C | C | C | . | . | . | . |
| NC_011770 | 678727 | T | G | 45 | 6 | 2 | 0 | NON_SYNONYMOU | MODERAT | MISSENSE | Ttc/Gtc | F203V | PLES_06241 | . | . | | | | | G | G | . | . | . | . |
| NC_011770 | 678751 | A | G | 47 | 6 | 2 | 0 | NON_SYNONYMOU | MODERAT | MISSENSE | Acg/Gcg | T211A | PLES_06241 | . | . | | | | | G | G | . | . | . | . |
| NC_011770 | 678865 | A | G | 51 | 6 | 6 | 0 | NON_SYNONYMOU | MODERAT | MISSENSE | Acc/Gcc | T249A | PLES_06241 | . | . | G | G | G | G | G | G | . | . | . | . |
| NC_011770 | 679213 | T | C | 66 | 0 | 12 | 0 | NON_SYNONYMOU | MODERAT | MISSENSE | gTg/gCg | V24A | PLES_06251 | C | C | C | C | C | C | C | C | C | C | C | C |
| NC_011770 | 679365 | G | A | 60 | 0 | 10 | 0 | NON_SYNONYMOU | MODERAT | MISSENSE | Ggt/Agt | G75S | PLES_06251 | | | A | A | A | A | A | A | A | A | A | A |
| NC_011770 | 679759 | C | T | 71 | 0 | 4 | 0 | NON_SYNONYMOU | MODERAT | MISSENSE | gCg/gTg | A206V | PLES_06251 | | | | | | | T | T | T | T | | |
| NC_011770 | 679980 | G | C | 76 | 0 | 4 | 0 | NON_SYNONYMOU | MODERAT | MISSENSE | aGg/aCg | R71T | PLES_06261 | | | | | | | C | C | C | C | | |
| NC_011770 | 679995 | T | C | 76 | 0 | 4 | 0 | NON_SYNONYMOU | MODERAT | MISSENSE | gTc/gCc | V76A | PLES_06261 | C | C | | | | | | | | | C | C |
| NC_011770 | 680081 | G | A | 71 | 0 | 4 | 0 | NON_SYNONYMOU | MODERAT | MISSENSE | Gtg/Atg | V105M | PLES_06261 | | | A | A | A | A | | | | | | |
| NC_011770 | 680082 | T | C | 70 | 0 | 4 | 0 | NON_SYNONYMOU | MODERAT | MISSENSE | gTg/gCg | V105A | PLES_06261 | | | C | C | C | C | | | | | | |
| NC_011770 | 680266 | G | A | 68 | 0 | 2 | 0 | NON_SYNONYMOU | MODERAT | MISSENSE | Ggc/Agc | G47S | PLES_06271 | | | | | | | | | | | A | A |
| NC_011770 | 680278 | A | C | 70 | 0 | 2 | 0 | NON_SYNONYMOU | MODERAT | MISSENSE | Aag/Cag | K51Q | PLES_06271 | | | | | | | | | | | C | C |
| NC_011770 | 684506 | G | A | 95 | 0 | 2 | 0 | NON_SYNONYMOU | MODERAT | MISSENSE | Gcc/Acc | A103T | PLES_06321 | A | A | | | | | | | | | | |
| NC_011770 | 686848 | G | A | 90 | 0 | 2 | 0 | NON_SYNONYMOU | MODERAT | MISSENSE | gCg/gTg | A10V | PLES_06351 | | | | | | | | | | | A | A |
| NC_011770 | 687736 | A | C | 81 | 0 | 10 | 0 | NON_SYNONYMOU | MODERAT | MISSENSE | aAg/aCg | K261T | PLES_06361 | | | C | C | C | C | C | C | C | C | C | C |
| NC_011770 | 688222 | T | A | 72 | 0 | 2 | 0 | NON_SYNONYMOU | MODERAT | MISSENSE | cTt/cAt | L423H | PLES_06361 | | | | | | | | | A | A | | |
| NC_011770 | 688328 | G | T | 75 | 0 | 4 | 0 | NON_SYNONYMOU | MODERAT | MISSENSE | gaG/gaT | E458D | PLES_06361 | | | | | | | T | T | T | T | | |
| NC_011770 | 688347 | T | G | 72 | 0 | 4 | 0 | NON_SYNONYMOU | MODERAT | MISSENSE | Tcg/Gcg | S465A | PLES_06361 | | | | | | | G | G | G | G | | |
| NC_011770 | 688489 | A | G | 71 | 0 | 10 | 0 | NON_SYNONYMOU | MODERAT | MISSENSE | gTc/gCc | V263A | PLES_06371 | | | G | G | G | G | G | G | G | G | G | G |
| NC_011770 | 688762 | T | C | 71 | 0 | 2 | 0 | NON_SYNONYMOU | MODERAT | MISSENSE | gAg/gGg | E172G | PLES_06371 | C | C | | | | | | | | | | |

| | | | | | | | | | | | | | | | | | | | | | | | | | |
|---|---|---|---|---|---|---|---|---|---|---|---|---|---|---|---|---|---|---|---|---|---|---|---|---|---|
| NC_011770 | 688897 | A | G | 76 | 0 | 4 | 0 | NON_SYNONYMOU | MODERAT | MISSENSE | gTg/gCg | V127A | PLES_06371 | | | | | | | | | G | G | G | G |
| NC_011770 | 689922 | G | T | 44 | 0 | 4 | 0 | NON_SYNONYMOU | MODERAT | MISSENSE | gaC/gaA | D164E | PLES_06381 | T | T | | | | | T | T | | | | |
| NC_011770 | 689975 | T | A | 44 | 0 | 4 | 0 | NON_SYNONYMOU | MODERAT | MISSENSE | Acc/Tcc | T147S | PLES_06381 | A | A | | | | | A | A | | | | |
| NC_011770 | 689978 | C | T | 45 | 0 | 2 | 0 | NON_SYNONYMOU | MODERAT | MISSENSE | Gcc/Acc | A146T | PLES_06381 | | | | | | | | | T | T | | |
| NC_011770 | 690038 | C | T | 47 | 0 | 2 | 0 | NON_SYNONYMOU | MODERAT | MISSENSE | Gtc/Atc | V126I | PLES_06381 | | | | | | | T | T | | | | |
| NC_011770 | 690539 | C | T | 69 | 0 | 2 | 0 | NON_SYNONYMOU | MODERAT | MISSENSE | cGc/cAc | R322H | PLES_06391 | | | | | | | | | | | T | T |
| NC_011770 | 690561 | G | T | 72 | 0 | 2 | 0 | NON_SYNONYMOU | MODERAT | MISSENSE | Cgc/Agc | R315S | PLES_06391 | | | | | | | T | T | | | | |
| NC_011770 | 690581 | G | T | 72 | 0 | 2 | 0 | NON_SYNONYMOU | MODERAT | MISSENSE | gCc/gAc | A308D | PLES_06391 | | | | | | | T | T | | | | |
| NC_011770 | 691636 | T | G | 86 | 0 | 2 | 0 | NON_SYNONYMOU | MODERAT | MISSENSE | Aag/Cag | K103Q | PLES_06401 | | | | | | | | | | | G | G |
| NC_011770 | 691681 | C | T | 87 | 0 | 1 | 0 | NON_SYNONYMOU | MODERAT | MISSENSE | Gtg/Atg | V88M | PLES_06401 HP | | | | | | | | **T** | | | | |
| NC_011770 | 693958 | A | G | 74 | 0 | 1 | 0 | NON_SYNONYMOU | MODERAT | MISSENSE | cAg/cGg | Q231R | PLES_06421 | | | | | | | G | | | | | |
| NC_011770 | 696053 | T | C | 90 | 0 | 2 | 0 | NON_SYNONYMOU | MODERAT | MISSENSE | Acc/Gcc | T426A | PLES_06461 | | | | | | | | | | | C | C |
| NC_011770 | 696734 | C | T | 90 | 0 | 2 | 0 | NON_SYNONYMOU | MODERAT | MISSENSE | Gtc/Atc | V199I | PLES_06461 | | | | | | | | | | | T | T |
| NC_011770 | 696864 | G | T | 92 | 0 | 2 | 0 | NON_SYNONYMOU | MODERAT | MISSENSE | gaC/gaA | D155E | PLES_06461 | | | | | | | T | T | | | | |
| NC_011770 | 697072 | G | C | 79 | 0 | 2 | 0 | NON_SYNONYMOU | MODERAT | MISSENSE | aCc/aGc | T86S | PLES_06461 | | | | | | | | | | | C | C |
| NC_011770 | 697105 | G | A | 81 | 0 | 2 | 0 | NON_SYNONYMOU | MODERAT | MISSENSE | aCc/aTc | T75I | PLES_06461 | | | | | | | | | | | A | A |
| NC_011770 | 697127 | T | C | 79 | 0 | 8 | 0 | NON_SYNONYMOU | MODERAT | MISSENSE | Aca/Gca | T68A | PLES_06461 | C | C | C | C | C | C | | | | | C | C |
| NC_011770 | 697138 | T | C | 79 | 0 | 4 | 0 | NON_SYNONYMOU | MODERAT | MISSENSE | gAc/gGc | D64G | PLES_06461 | | | C | C | C | C | | | | | C | C |
| NC_011770 | 703403 | A | G | 111 | 0 | 1 | 0 | INTRAGENIC | MODIFIER | | | | PLES_06475 | | | | | | | G | | | | | |
| NC_011770 | 703550 | G | A | 100 | 0 | 1 | 0 | INTRAGENIC | MODIFIER | | | | PLES_06475 | | | | | | | | A | | | | |
| NC_011770 | 703956 | C | T | 96 | 0 | 1 | 0 | INTRAGENIC | MODIFIER | | | | PLES_06475 | | | | | | | | T | | | | |
| NC_011770 | 704478 | C | T | 91 | 0 | 1 | 0 | INTRAGENIC | MODIFIER | | | | PLES_06476 | | | | | | | | T | | | | |
| NC_011770 | 705665 | G | T | 67 | 0 | 2 | 0 | NON_SYNONYMOU | MODERAT | MISSENSE | Gcg/Tcg | A21S | PLES_06491 | | | | | | | T | T | | | | |
| NC_011770 | 706289 | G | T | 76 | 0 | 2 | 0 | NON_SYNONYMOU | MODERAT | MISSENSE | Ggc/Tgc | G229C | PLES_06491 | | | | | | | T | T | | | | |
| NC_011770 | 706535 | C | T | 78 | 0 | 2 | 0 | NON_SYNONYMOU | MODERAT | MISSENSE | tCg/tTg | S59L | PLES_06501 | | | | | | | T | T | | | | |
| NC_011770 | 706555 | G | T | 80 | 0 | 4 | 0 | NON_SYNONYMOU | MODERAT | MISSENSE | Gtg/Ttg | V66L | PLES_06501 | | | T | T | T | T | | | | | | |
| NC_011770 | 706594 | G | A | 81 | 0 | 2 | 0 | NON_SYNONYMOU | MODERAT | MISSENSE | Gcc/Acc | A79T | PLES_06501 | | | | | | | | | A | A | | |
| NC_011770 | 706628 | C | T | 80 | 0 | 2 | 0 | NON_SYNONYMOU | MODERAT | MISSENSE | gCg/gTg | A90V | PLES_06501 | | | | | | | | | T | T | | |
| NC_011770 | 706646 | A | G | 81 | 0 | 4 | 0 | NON_SYNONYMOU | MODERAT | MISSENSE | cAg/cGg | Q96R | PLES_06501 | | | G | G | G | G | | | | | | |
| NC_011770 | 706877 | C | T | 77 | 0 | 10 | 0 | NON_SYNONYMOU | MODERAT | MISSENSE | gCc/gTc | A173V | PLES_06501 | T | T | T | T | T | T | T | T | | | T | T |
| NC_011770 | 706902 | G | C | 83 | 0 | 2 | 0 | NON_SYNONYMOU | MODERAT | MISSENSE | gaG/gaC | E181D | PLES_06501 | | | | | | | | | C | C | | |
| NC_011770 | 763301 | T | A | 69 | 0 | 2 | 0 | NON_SYNONYMOU | MODERAT | MISSENSE | cTg/cAg | L179Q | PLES_07041 | | | | | | | A | A | | | | |
| NC_011770 | 763417 | G | T | 56 | 0 | 6 | 0 | NON_SYNONYMOU | MODERAT | MISSENSE | Gcg/Tcg | A218S | PLES_07041 | | | T | T | T | T | | | | | T | T |
| NC_011770 | 763544 | C | T | 56 | 0 | 2 | 0 | NON_SYNONYMOU | MODERAT | MISSENSE | gCc/gTc | A260V | PLES_07041 | T | T | | | | | | | | | | |
| NC_011770 | 763571 | T | C | 53 | 0 | 10 | 0 | NON_SYNONYMOU | MODERAT | MISSENSE | gTg/gCg | V269A | PLES_07041 | C | C | C | C | C | C | C | C | | | C | C |
| NC_011770 | 763715 | C | T | 48 | 0 | 4 | 0 | NON_SYNONYMOU | MODERAT | MISSENSE | gCc/gTc | A317V | PLES_07041 | | | T | T | T | T | | | | | | |
| NC_011770 | 763727 | C | T | 49 | 0 | 2 | 0 | NON_SYNONYMOU | MODERAT | MISSENSE | tCg/tTg | S321L | PLES_07041 | | | | | | | | | | | T | T |
| NC_011770 | 763927 | A | G | 72 | 0 | 2 | 0 | NON_SYNONYMOU | MODERAT | MISSENSE | Att/Gtt | I388V | PLES_07041 | G | G | | | | | | | | | | |
| NC_011770 | 764281 | A | G | 83 | 0 | 2 | 0 | NON_SYNONYMOU | MODERAT | MISSENSE | Aac/Gac | N506D | PLES_07041 | G | G | | | | | | | | | | |
| NC_011770 | 764779 | C | T | 69 | 0 | 2 | 0 | NON_SYNONYMOU | MODERAT | MISSENSE | cCg/cTg | P102L | PLES_07051 | | | | | | | T | T | | | | |
| NC_011770 | 764952 | C | G | 69 | 0 | 8 | 0 | NON_SYNONYMOU | MODERAT | MISSENSE | Ctt/Gtt | L160V | PLES_07051 | G | G | G | G | G | G | G | G | | | | |
| NC_011770 | 764964 | T | G | 69 | 0 | 8 | 0 | NON_SYNONYMOU | MODERAT | MISSENSE | Ttc/Gtc | F164V | PLES_07051 | G | G | G | G | G | G | G | G | | | | |
| NC_011770 | 765164 | A | T | 67 | 0 | 10 | 0 | NON_SYNONYMOU | MODERAT | MISSENSE | gaA/gaT | E230D | PLES_07051 | T | T | T | T | T | T | T | T | | | T | T |
| NC_011770 | 765455 | C | G | 64 | 0 | 2 | 0 | NON_SYNONYMOU | MODERAT | MISSENSE | agC/agG | S327R | PLES_07051 | G | G | | | | | | | | | | |
| NC_011770 | 765613 | T | C | 61 | 0 | 8 | 0 | NON_SYNONYMOU | MODERAT | MISSENSE | gTc/gCc | V380A | PLES_07051 | C | C | C | C | C | C | C | C | | | | |
| NC_011770 | 765624 | C | T | 64 | 0 | 8 | 0 | NON_SYNONYMOU | MODERAT | MISSENSE | Cgt/Tgt | R384C | PLES_07051 | T | T | T | T | T | T | T | T | | | | |
| NC_011770 | 765682 | G | A | 63 | 0 | 2 | 0 | NON_SYNONYMOU | MODERAT | MISSENSE | aGt/aAt | S403N | PLES_07051 | | | | | | | | | | | A | A |
| NC_011770 | 765918 | A | G | 65 | 0 | 6 | 0 | NON_SYNONYMOU | MODERAT | MISSENSE | Acg/Gcg | T482A | PLES_07051 | | | G | G | G | G | G | G | | | | |
| NC_011770 | 766083 | G | A | 66 | 0 | 6 | 0 | NON_SYNONYMOU | MODERAT | MISSENSE | Gag/Aag | E537K | PLES_07051 | | | A | A | A | A | A | A | | | | |
| NC_011770 | 766108 | T | G | 64 | 0 | 10 | 0 | NON_SYNONYMOU | MODERAT | MISSENSE | cTt/cGt | L545R | PLES_07051 | G | G | G | G | G | G | G | G | | | G | G |
| NC_011770 | 766186 | T | C | 66 | 0 | 8 | 0 | NON_SYNONYMOU | MODERAT | MISSENSE | gTc/gCc | V571A | PLES_07051 | C | C | C | C | C | C | C | C | | | | |
| NC_011770 | 768594 | T | C | 54 | 0 | 12 | 0 | NON_SYNONYMOU | MODERAT | MISSENSE | Ttc/Ctc | F48L | PLES_07071 | C | C | C | C | C | C | C | C | C | C | C | C |
| NC_011770 | 768642 | C | G | 59 | 0 | 2 | 0 | NON_SYNONYMOU | MODERAT | MISSENSE | Ctg/Gtg | L64V | PLES_07071 | G | G | | | | | | | | | | |
| NC_011770 | 768652 | G | A | 60 | 0 | 2 | 0 | NON_SYNONYMOU | MODERAT | MISSENSE | gGt/gAt | G67D | PLES_07071 | A | A | | | | | | | | | | |
| NC_011770 | 768693 | C | G | 65 | 0 | 12 | 0 | NON_SYNONYMOU | MODERAT | MISSENSE | Cta/Gta | L81V | PLES_07071 | G | G | G | G | G | G | G | G | G | G | G | G |
| NC_011770 | 768694 | T | C | 68 | 0 | 2 | 0 | NON_SYNONYMOU | MODERAT | MISSENSE | cTa/cCa | L81P | PLES_07071 | C | C | | | | | | | | | | |
| NC_011770 | 768466 | C | T | 59 | 0 | 1 | 0 | NON_SYNONYMOU | MODERAT | MISSENSE | tCg/tTg | S5L | PLES_07071 | | | | | | | | | | | | T |
| NC_011770 | 769058 | T | C | 61 | 0 | 6 | 0 | START_LOST | HIGH | MISSENSE | aTg/aCg | M1T | PLES_07081 | C | C | | | | | C | C | C | C | | |
| NC_011770 | 769409 | C | T | 49 | 0 | 2 | 0 | NON_SYNONYMOU | MODERAT | MISSENSE | cCg/cTg | P118L | PLES_07081 | | | | | | | T | T | | | | |
| NC_011770 | 769562 | G | A | 45 | 0 | 8 | 0 | NON_SYNONYMOU | MODERAT | MISSENSE | cGg/cAg | R169Q | PLES_07081 | | | A | A | A | A | | | A | A | A | A |
| NC_011770 | 769621 | G | A | 45 | 0 | 2 | 0 | NON_SYNONYMOU | MODERAT | MISSENSE | Gcc/Acc | A189T | PLES_07081 | | | | | | | A | A | | | | |
| NC_011770 | 769631 | G | A | 45 | 0 | 2 | 0 | NON_SYNONYMOU | MODERAT | MISSENSE | gGc/gAc | G192D | PLES_07081 | A | A | | | | | | | | | | |

| | | | | | | | | | | | | | | | | | | | | | | | | | |
|---|---|---|---|---|---|---|---|---|---|---|---|---|---|---|---|---|---|---|---|---|---|---|---|---|---|
| NC_011770 | 769925 | C | T | 33 | 0 | 2 | 0 | NON_SYNONYMOU | MODERAT | MISSENSE | gCg/gTg | A290V | PLES_07081 | | | | | | | | | T | T | | |
| NC_011770 | 770137 | A | T | 18 | 0 | 2 | 0 | NON_SYNONYMOU | MODERAT | MISSENSE | Acc/Tcc | T361S | PLES_07081 | | | | | | | T | T | | | | |
| NC_011770 | 770173 | G | A | 8 | 2 | 2 | 0 | NON_SYNONYMOU | MODERAT | MISSENSE | Gcc/Acc | A373T | PLES_07081 | . | . | | | | | | | | | A | A |
| NC_011770 | 770747 | A | G | 58 | 0 | 8 | 0 | NON_SYNONYMOU | MODERAT | MISSENSE | Acg/Gcg | T172A | PLES_07091 | G | G | | | | | G | G | G | G | G | G |
| NC_011770 | 770762 | A | G | 63 | 0 | 2 | 0 | NON_SYNONYMOU | MODERAT | MISSENSE | Atg/Gtg | M177V | PLES_07091 | | | | | | | | | G | G | | |
| NC_011770 | 770837 | G | A | 66 | 0 | 4 | 0 | NON_SYNONYMOU | MODERAT | MISSENSE | Gca/Aca | A202T | PLES_07091 | | | A | A | A | A | | | | | | |
| NC_011770 | 771101 | G | T | 60 | 0 | 2 | 0 | NON_SYNONYMOU | MODERAT | MISSENSE | Gtg/Ttg | V290L | PLES_07091 | | | | | | | T | T | | | | |
| NC_011770 | 788666 | C | T | 61 | 0 | 2 | 0 | NON_SYNONYMOU | MODERAT | MISSENSE | Gcc/Acc | A97T | PLES_07231 | | | | | | | T | T | | | | |
| NC_011770 | 789045 | A | C | 71 | 0 | 12 | 0 | NON_SYNONYMOU | MODERAT | MISSENSE | Tcc/Gcc | S282A | PLES_07241 | C | C | C | C | C | C | C | C | C | C | C | C |
| NC_011770 | 789057 | G | C | 73 | 0 | 10 | 0 | NON_SYNONYMOU | MODERAT | MISSENSE | Ctg/Gtg | L278V | PLES_07241 | C | C | C | C | C | C | C | C | | | C | C |
| NC_011770 | 789506 | C | T | 65 | 0 | 2 | 0 | NON_SYNONYMOU | MODERAT | MISSENSE | cGc/cAc | R128H | PLES_07241 | T | T | | | | | | | | | | |
| NC_011770 | 789636 | C | T | 60 | 0 | 6 | 0 | NON_SYNONYMOU | MODERAT | MISSENSE | Gcc/Acc | A85T | PLES_07241 | | | T | T | T | T | T | T | | | | |
| NC_011770 | 789779 | A | C | 61 | 0 | 12 | 0 | NON_SYNONYMOU | MODERAT | MISSENSE | cTc/cGc | L37R | PLES_07241 | C | C | C | C | C | C | C | C | C | C | C | C |
| NC_011770 | 789818 | T | C | 63 | 0 | 2 | 0 | NON_SYNONYMOU | MODERAT | MISSENSE | cAg/cGg | Q24R | PLES_07241 | C | C | | | | | | | | | | |
| NC_011770 | 790296 | C | T | 68 | 0 | 4 | 0 | NON_SYNONYMOU | MODERAT | MISSENSE | Ccg/Tcg | P101S | PLES_07251 | T | T | | | | | T | T | | | | |
| NC_011770 | 790303 | T | C | 67 | 0 | 4 | 0 | NON_SYNONYMOU | MODERAT | MISSENSE | gTg/gCg | V103A | PLES_07251 | | | | | | | | | C | C | C | C |
| NC_011770 | 790313 | G | C | 70 | 0 | 4 | 0 | NON_SYNONYMOU | MODERAT | MISSENSE | aaG/aaC | K106N | PLES_07251 | | | | | | | | | C | C | C | C |
| NC_011770 | 790458 | T | A | 66 | 0 | 6 | 0 | NON_SYNONYMOU | MODERAT | MISSENSE | Tgt/Agt | C155S | PLES_07251 | | | A | A | A | A | | | A | A | | |
| NC_011770 | 790776 | C | T | 69 | 0 | 1 | 0 | NON_SYNONYMOU | MODERAT | MISSENSE | Ccg/Tcg | P261S | PLES_07251 HP | | | | | | | | **T** | | | | |
| NC_011770 | 792242 | G | A | 73 | 0 | 2 | 0 | NON_SYNONYMOU | MODERAT | MISSENSE | aCc/aTc | T279I | PLES_07271 | | | | | | | | | | | A | A |
| NC_011770 | 792900 | T | C | 83 | 0 | 2 | 0 | NON_SYNONYMOU | MODERAT | MISSENSE | Acc/Gcc | T60A | PLES_07271 | | | | | | | C | C | | | | |
| NC_011770 | 793416 | C | A | 79 | 0 | 4 | 0 | NON_SYNONYMOU | MODERAT | MISSENSE | Gcg/Tcg | A551S | PLES_07281 | | | A | A | A | A | | | | | | |
| NC_011770 | 793424 | G | A | 78 | 0 | 6 | 0 | NON_SYNONYMOU | MODERAT | MISSENSE | gCc/gTc | A548V | PLES_07281 | A | A | | | | | | | A | A | A | A |
| NC_011770 | 793439 | A | T | 77 | 0 | 2 | 0 | NON_SYNONYMOU | MODERAT | MISSENSE | cTg/cAg | L543Q | PLES_07281 | T | T | | | | | | | | | | |
| NC_011770 | 793708 | C | T | 85 | 0 | 1 | 0 | NON_SYNONYMOU | MODERAT | MISSENSE | atG/atA | M453I | PLES_07281 | | | | | | | T | | | | | |
| NC_011770 | 794043 | T | G | 81 | 0 | 2 | 0 | NON_SYNONYMOU | MODERAT | MISSENSE | Atc/Ctc | I342L | PLES_07281 | | | | | | | | | G | G | | |
| NC_011770 | 795447 | C | T | 90 | 0 | 1 | 0 | NON_SYNONYMOU | MODERAT | MISSENSE | Gag/Aag | E454K | PLES_07291 | | | | | | | T | | | | | |
| NC_011770 | 796734 | C | G | 48 | 0 | 2 | 0 | NON_SYNONYMOU | MODERAT | MISSENSE | Gag/Cag | E25Q | PLES_07291 | | | | | | | | | G | G | | |
| NC_011770 | 798004 | A | AC | 68 | 0 | 2 | 0 | FRAME_SHIFT | HIGH | | cag/cCag | Q320P? | PLES_07301 | AC | AC | | | | | | | | | | |
| NC_011770 | 798033 | C | T | 72 | 0 | 2 | 0 | NON_SYNONYMOU | MODERAT | MISSENSE | Cgc/Tgc | R328C | PLES_07301 | | | | | | | T | T | | | | |
| NC_011770 | 799173 | T | C | 76 | 0 | 12 | 0 | NON_SYNONYMOU | MODERAT | MISSENSE | Tac/Cac | Y708H | PLES_07301 | C | C | C | C | C | C | C | C | C | C | C | C |
| NC_011770 | 798438 | C | T | 66 | 0 | 1 | 0 | NON_SYNONYMOU | MODERAT | MISSENSE | Cgc/Tgc | R463C | PLES_07301 | | | | | | | | | | | T | |
| NC_011770 | 797826 | G | C | 65 | 0 | 1 | 0 | NON_SYNONYMOU | MODERAT | MISSENSE | Ggc/Cgc | G259R | PLES_07301 bfiS | | | | | | **C** | | | | | | |
| NC_011770 | 799449 | G | T | 83 | 0 | 2 | 0 | NON_SYNONYMOU | MODERAT | MISSENSE | cGg/cTg | R40L | PLES_07311 | T | T | | | | | | | | | | |
| NC_011770 | 799551 | T | C | 76 | 0 | 12 | 0 | NON_SYNONYMOU | MODERAT | MISSENSE | gTg/gCg | V74A | PLES_07311 | C | C | C | C | C | C | C | C | C | C | C | C |
| NC_011770 | 799725 | G | A | 73 | 0 | 2 | 0 | NON_SYNONYMOU | MODERAT | MISSENSE | cGc/cAc | R132H | PLES_07311 | A | A | | | | | | | | | | |
| NC_011770 | 799808 | C | T | 80 | 0 | 10 | 0 | NON_SYNONYMOU | MODERAT | MISSENSE | Cat/Tat | H160Y | PLES_07311 | T | T | T | T | T | T | T | T | | | T | T |
| NC_011770 | 800864 | C | T | 85 | 0 | 4 | 0 | NON_SYNONYMOU | MODERAT | MISSENSE | gCg/gTg | A245V | PLES_07321 | | | | | | | T | T | T | T | | |
| NC_011770 | 801088 | C | A | 100 | 0 | 4 | 0 | NON_SYNONYMOU | MODERAT | MISSENSE | Cat/Aat | H15N | PLES_07331 | | | A | A | A | A | | | | | | |
| NC_011770 | 801393 | C | G | 109 | 0 | 2 | 0 | NON_SYNONYMOU | MODERAT | MISSENSE | agC/agG | S116R | PLES_07331 | | | | | | | | | | | G | G |
| NC_011770 | 801527 | T | C | 103 | 0 | 1 | 0 | NON_SYNONYMOU | MODERAT | MISSENSE | cTt/cCt | L161P | PLES_07331 | | | | | | | C | | | | | |
| NC_011770 | 801599 | A | G | 109 | 0 | 1 | 0 | NON_SYNONYMOU | MODERAT | MISSENSE | tAc/tGc | Y185C | PLES_07331 | | | | | | | G | | | | | |
| NC_011770 | 802784 | C | A | 69 | 0 | 6 | 0 | NON_SYNONYMOU | MODERAT | MISSENSE | Cgc/Agc | R115S | PLES_07351 | | | A | A | A | A | A | A | | | | |
| NC_011770 | 802793 | T | A | 70 | 0 | 12 | 0 | NON_SYNONYMOU | MODERAT | MISSENSE | Ttg/Atg | L118M | PLES_07351 | A | A | A | A | A | A | A | A | A | A | A | A |
| NC_011770 | 806208 | A | G | 84 | 0 | 1 | 0 | NON_SYNONYMOU | MODERAT | MISSENSE | Ttc/Ctc | F296L | PLES_07381 | | | | | | | G | | | | | |
| NC_011770 | 806958 | C | T | 68 | 0 | 2 | 0 | NON_SYNONYMOU | MODERAT | MISSENSE | Ggc/Agc | G46S | PLES_07381 | | | | | | | | | T | T | | |
| NC_011770 | 806994 | C | T | 66 | 0 | 2 | 0 | NON_SYNONYMOU | MODERAT | MISSENSE | Ggc/Agc | G34S | PLES_07381 | | | | | | | | | T | T | | |
| NC_011770 | 807462 | G | A | 73 | 0 | 2 | 0 | NON_SYNONYMOU | MODERAT | MISSENSE | cCg/cTg | P179L | PLES_07391 | | | | | | | | | | | A | A |
| NC_011770 | 807922 | C | T | 62 | 0 | 6 | 0 | NON_SYNONYMOU | MODERAT | MISSENSE | Gcc/Acc | A26T | PLES_07391 | T | T | | | | | T | T | T | T | | |
| NC_011770 | 808711 | G | T | 79 | 0 | 12 | 0 | NON_SYNONYMOU | MODERAT | MISSENSE | gCg/gAg | A227E | PLES_07401 | T | T | T | T | T | T | T | T | T | T | T | T |
| NC_011770 | 809236 | C | T | 82 | 0 | 2 | 0 | NON_SYNONYMOU | MODERAT | MISSENSE | aGc/aAc | S52N | PLES_07401 | T | T | | | | | | | | | | |
| NC_011770 | 810023 | T | G | 69 | 0 | 12 | 0 | NON_SYNONYMOU | MODERAT | MISSENSE | Agc/Cgc | S275R | PLES_07411 | G | G | G | G | G | G | G | G | G | G | G | G |
| NC_011770 | 811031 | C | G | 83 | 0 | 12 | 0 | NON_SYNONYMOU | MODERAT | MISSENSE | gaG/gaC | E212D | PLES_07421 | G | G | G | G | G | G | G | G | G | G | G | G |
| NC_011770 | 812696 | G | T | 68 | 0 | 2 | 0 | NON_SYNONYMOU | MODERAT | MISSENSE | Gcg/Tcg | A242S | PLES_07431 | | | | | | | | | T | T | | |
| NC_011770 | 813378 | G | A | 63 | 0 | 12 | 0 | NON_SYNONYMOU | MODERAT | MISSENSE | Gtc/Atc | V120I | PLES_07441 | A | A | A | A | A | A | A | A | A | A | A | A |
| NC_011770 | 813497 | C | T | 61 | 0 | 2 | 0 | NON_SYNONYMOU | MODERAT | MISSENSE | Ggc/Agc | G212S | PLES_07451 | | | | | | | | | | | T | **T** |
| NC_011770 | 813498 | G | C | 60 | 0 | 6 | 0 | NON_SYNONYMOU | MODERAT | MISSENSE | gaC/gaG | D211E | PLES_07451 | | | C | C | C | C | | | | | C | C |
| NC_011770 | 814198 | C | T | 63 | 0 | 4 | 0 | NON_SYNONYMOU | MODERAT | MISSENSE | cGc/cAc | R227H | PLES_07461 | | | T | T | T | T | | | | | | |
| NC_011770 | 814231 | C | T | 66 | 0 | 4 | 0 | NON_SYNONYMOU | MODERAT | MISSENSE | cGc/cAc | R216H | PLES_07461 | | | T | T | T | T | | | | | | |
| NC_011770 | 814318 | C | T | 60 | 0 | 10 | 0 | NON_SYNONYMOU | MODERAT | MISSENSE | cGg/cAg | R187Q | PLES_07461 | T | T | T | T | T | T | T | T | T | T | | |
| NC_011770 | 816643 | A | G | 87 | 0 | 6 | 0 | NON_SYNONYMOU | MODERAT | MISSENSE | cAc/cGc | H489R | PLES_07471 | | | G | G | G | G | G | G | | | | |
| NC_011770 | 816788 | C | G | 83 | 0 | 2 | 0 | NON_SYNONYMOU | MODERAT | MISSENSE | aaC/aaG | N537K | PLES_07471 | | | | | | | G | G | | | | |

| | | | | | | | | | | | | | | | | | | | | | | | | | |
|---|---|---|---|---|---|---|---|---|---|---|---|---|---|---|---|---|---|---|---|---|---|---|---|---|---|
| NC_011770 | 816078 | C | T | 86 | 0 | 1 | 0 | NON_SYNONYMOU | MODERAT | MISSENSE | Ccg/Tcg | P301S | PLES_07471 | | | | T | | | | | | | | |
| NC_011770 | 816996 | G | C | 82 | 0 | 12 | 0 | NON_SYNONYMOU | MODERAT | MISSENSE | atC/atG | I388M | PLES_07481 | C | C | C | C | C | C | C | C | C | C | C | C |
| NC_011770 | 816998 | T | C | 82 | 0 | 12 | 0 | NON_SYNONYMOU | MODERAT | MISSENSE | Atc/Gtc | I388V | PLES_07481 | C | C | C | C | C | C | C | C | C | C | C | C |
| NC_011770 | 817277 | C | G | 62 | 0 | 12 | 0 | NON_SYNONYMOU | MODERAT | MISSENSE | Gag/Cag | E295Q | PLES_07481 | G | G | G | G | G | G | G | G | G | G | G | G |
| NC_011770 | 817391 | G | C | 56 | 0 | 12 | 0 | NON_SYNONYMOU | MODERAT | MISSENSE | Ctc/Gtc | L257V | PLES_07481 | C | C | C | C | C | C | C | C | C | C | C | C |
| NC_011770 | 817619 | G | A | 48 | 0 | 12 | 0 | NON_SYNONYMOU | MODERAT | MISSENSE | Ctc/Ttc | L181F | PLES_07481 | A | A | A | A | A | A | A | A | A | A | A | A |
| NC_011770 | 817787 | G | C | 44 | 0 | 4 | 0 | NON_SYNONYMOU | MODERAT | MISSENSE | Cag/Gag | Q125E | PLES_07481 | | | C | C | C | C | | | | | | |
| NC_011770 | 817583 | C | T | 53 | 0 | 1 | 0 | NON_SYNONYMOU | MODERAT | MISSENSE | Gac/Aac | D193N | PLES_07481 | | | | T | | | | | | | | |
| NC_011770 | 818485 | C | T | 49 | 0 | 2 | 0 | NON_SYNONYMOU | MODERAT | MISSENSE | Gag/Aag | E219K | PLES_07491 | | | | | | | | | | | T | T |
| NC_011770 | 818494 | A | T | 48 | 0 | 12 | 0 | NON_SYNONYMOU | MODERAT | MISSENSE | Tcg/Acg | S216T | PLES_07491 | T | T | T | T | T | T | T | T | T | T | T | T |
| NC_011770 | 818562 | G | A | 59 | 0 | 2 | 0 | NON_SYNONYMOU | MODERAT | MISSENSE | aCc/aTc | T193I | PLES_07491 | | | | | | | | | | | A | A |
| NC_011770 | 819273 | C | T | 81 | 0 | 8 | 0 | NON_SYNONYMOU | MODERAT | MISSENSE | tGc/tAc | C123Y | PLES_07501 | T | T | T | T | T | T | T | T | | | | |
| NC_011770 | 819373 | C | T | 75 | 0 | 4 | 0 | NON_SYNONYMOU | MODERAT | MISSENSE | Gag/Aag | E90K | PLES_07501 | | | | | | | | | T | T | T | T |
| NC_011770 | 819376 | C | T | 75 | 0 | 4 | 0 | NON_SYNONYMOU | MODERAT | MISSENSE | Ggc/Agc | G89S | PLES_07501 | | | T | T | T | T | | | | | | |
| NC_011770 | 819550 | C | T | 76 | 0 | 2 | 0 | NON_SYNONYMOU | MODERAT | MISSENSE | Ggc/Agc | G31S | PLES_07501 | | | | | | | | | | | T | T |
| NC_011770 | 819597 | T | C | 79 | 0 | 2 | 0 | NON_SYNONYMOU | MODERAT | MISSENSE | cAc/cGc | H15R | PLES_07501 | | | | | | | | | | | C | C |
| NC_011770 | 819630 | G | A | 76 | 0 | 2 | 0 | NON_SYNONYMOU | MODERAT | MISSENSE | aCc/aTc | T4I | PLES_07501 | | | | | | | | | | | A | A |
| NC_011770 | 822355 | C | T | 64 | 0 | 2 | 0 | NON_SYNONYMOU | MODERAT | MISSENSE | Gcc/Acc | A293T | PLES_07531 | | | | | | | | | | | T | T |
| NC_011770 | 822600 | T | C | 60 | 0 | 10 | 0 | NON_SYNONYMOU | MODERAT | MISSENSE | gAg/gGg | E211G | PLES_07531 | C | C | C | C | C | C | C | C | C | C | | |
| NC_011770 | 823158 | G | A | 85 | 0 | 2 | 0 | NON_SYNONYMOU | MODERAT | MISSENSE | aCg/aTg | T25M | PLES_07531 | | | | | | | | | | | A | A |
| NC_011770 | 822786 | C | T | 80 | 0 | 1 | 0 | NON_SYNONYMOU | MODERAT | MISSENSE | cGc/cAc | R149H | PLES_07531 | | | | | | | | | | | T | |
| NC_011770 | 823680 | T | C | 75 | 0 | 10 | 0 | NON_SYNONYMOU | MODERAT | MISSENSE | Tac/Cac | Y104H | PLES_07541 | C | C | C | C | C | C | C | C | C | C | | |
| NC_011770 | 825168 | C | T | 74 | 0 | 1 | 0 | NON_SYNONYMOU | MODERAT | MISSENSE | gGc/gAc | G7D | PLES_07561 | | | | | | | T | | | | | |
| NC_011770 | 825402 | G | A | 65 | 0 | 2 | 0 | NON_SYNONYMOU | MODERAT | MISSENSE | gCg/gTg | A285V | PLES_07571 | | | | | | | | | | | A | A |
| NC_011770 | 825643 | T | C | 71 | 0 | 10 | 0 | NON_SYNONYMOU | MODERAT | MISSENSE | Acg/Gcg | T205A | PLES_07571 | C | C | C | C | C | C | C | C | C | C | | |
| NC_011770 | 825906 | C | T | 70 | 0 | 10 | 0 | NON_SYNONYMOU | MODERAT | MISSENSE | cGc/cAc | R117H | PLES_07571 | T | T | T | T | T | T | T | T | T | T | | |
| NC_011770 | 826033 | A | C | 71 | 0 | 12 | 0 | NON_SYNONYMOU | MODERAT | MISSENSE | Tat/Gat | Y75D | PLES_07571 | C | C | C | C | C | C | C | C | C | C | C | C |
| NC_011770 | 826035 | G | A | 70 | 0 | 10 | 0 | NON_SYNONYMOU | MODERAT | MISSENSE | gCc/gTc | A74V | PLES_07571 | A | A | A | A | A | A | A | A | A | A | | |
| NC_011770 | 826238 | C | A | 79 | 0 | 2 | 0 | NON_SYNONYMOU | MODERAT | MISSENSE | ttG/ttT | L6F | PLES_07571 | | | | | | | | | | | A | A |
| NC_011770 | 830745 | C | G | 98 | 0 | 2 | 0 | NON_SYNONYMOU | MODERAT | MISSENSE | aGc/aCc | S61T | PLES_07611 | | | | | | | | | | | G | G |
| NC_011770 | 831544 | C | T | 63 | 0 | 2 | 0 | NON_SYNONYMOU | MODERAT | MISSENSE | Cgc/Tgc | R163C | PLES_07621 | | | | | | | | | | | T | T |
| NC_011770 | 832420 | A | G | 61 | 0 | 12 | 0 | NON_SYNONYMOU | MODERAT | MISSENSE | Acc/Gcc | T455A | PLES_07621 | G | G | G | G | G | G | G | G | G | G | G | G |
| NC_011770 | 832537 | G | A | 55 | 0 | 10 | 0 | NON_SYNONYMOU | MODERAT | MISSENSE | Gcg/Acg | A494T | PLES_07621 | A | A | A | A | A | A | A | A | A | A | | |
| NC_011770 | 832866 | T | C | 39 | 0 | 2 | 0 | NON_SYNONYMOU | MODERAT | MISSENSE | Agc/Ggc | S9G | PLES_07631 | | | | | | | | | | | C | C |
| NC_011770 | 833209 | A | C | 47 | 0 | 12 | 0 | NON_SYNONYMOU | MODERAT | MISSENSE | Ttc/Gtc | F485V | PLES_07641 | C | C | C | C | C | C | C | C | C | C | C | C |
| NC_011770 | 833758 | A | G | 60 | 0 | 1 | 0 | NON_SYNONYMOU | MODERAT | MISSENSE | Tcc/Ccc | S302P | PLES_07641 | | | | | | | G | | | | | |
| NC_011770 | 834229 | T | C | 91 | 0 | 12 | 0 | NON_SYNONYMOU | MODERAT | MISSENSE | Acg/Gcg | T145A | PLES_07641 | C | C | C | C | C | C | C | C | C | C | C | C |
| NC_011770 | 839731 | T | C | 84 | 0 | 2 | 0 | NON_SYNONYMOU | MODERAT | MISSENSE | Acc/Gcc | T223A | PLES_07701 | | | | | | | C | C | | | | |
| NC_011770 | 840753 | T | C | 63 | 0 | 12 | 0 | NON_SYNONYMOU | MODERAT | MISSENSE | cTc/cCc | L34P | PLES_07711 | C | C | C | C | C | C | C | C | C | C | C | C |
| NC_011770 | 840776 | C | A | 66 | 0 | 2 | 0 | NON_SYNONYMOU | MODERAT | MISSENSE | Ctc/Atc | L42I | PLES_07711 | | | | | | | | | A | A | | |
| NC_011770 | 840849 | C | T | 67 | 0 | 2 | 0 | NON_SYNONYMOU | MODERAT | MISSENSE | aCc/aTc | T66I | PLES_07711 | T | T | | | | | | | | | | |
| NC_011770 | 841467 | A | G | 72 | 0 | 12 | 0 | NON_SYNONYMOU | MODERAT | MISSENSE | aAc/aGc | N272S | PLES_07711 | G | G | G | G | G | G | G | G | G | G | G | G |
| NC_011770 | 841712 | A | G | 83 | 0 | 4 | 0 | NON_SYNONYMOU | MODERAT | MISSENSE | Atc/Gtc | I354V | PLES_07711 | G | G | | | | | | | G | G | | |
| NC_011770 | 842171 | C | G | 87 | 0 | 2 | 0 | NON_SYNONYMOU | MODERAT | MISSENSE | Ctg/Gtg | L507V | PLES_07711 | | | | | | | | | G | G | | |
| NC_011770 | 842250 | A | C | 81 | 0 | 12 | 0 | NON_SYNONYMOU | MODERAT | MISSENSE | gAg/gCg | E533A | PLES_07711 | C | C | C | C | C | C | C | C | C | C | C | C |
| NC_011770 | 842459 | G | T | 78 | 0 | 2 | 0 | NON_SYNONYMOU | MODERAT | MISSENSE | Gtc/Ttc | V603F | PLES_07711 | T | T | | | | | | | | | | |
| NC_011770 | 843690 | A | G | 77 | 0 | 6 | 0 | NON_SYNONYMOU | MODERAT | MISSENSE | Acg/Gcg | T296A | PLES_07721 | G | G | | | | | G | G | | | G | G |
| NC_011770 | 844861 | C | T | 69 | 0 | 1 | 0 | NON_SYNONYMOU | MODERAT | MISSENSE | Ccc/Tcc | P105S | PLES_07731 | | | | | | | | **T** | | | | |
| NC_011770 | 845804 | T | A | 77 | 0 | 6 | 0 | NON_SYNONYMOU | MODERAT | MISSENSE | Agc/Tgc | S197C | PLES_07741 | | | A | A | A | A | | | A | A | | |
| NC_011770 | 846115 | C | T | 75 | 0 | 2 | 0 | NON_SYNONYMOU | MODERAT | MISSENSE | gGt/gAt | G93D | PLES_07741 | | | | | | | T | T | | | | |
| NC_011770 | 846179 | C | G | 79 | 0 | 2 | 0 | NON_SYNONYMOU | MODERAT | MISSENSE | Ggc/Cgc | G72R | PLES_07741 | | | | | | | | | | | G | G |
| NC_011770 | 848665 | T | C | 75 | 0 | 12 | 0 | NON_SYNONYMOU | MODERAT | MISSENSE | Atc/Gtc | I293V | PLES_07771 | C | C | C | C | C | C | C | C | C | C | C | C |
| NC_011770 | 849058 | A | C | 60 | 0 | 12 | 0 | NON_SYNONYMOU | MODERAT | MISSENSE | Tcg/Gcg | S162A | PLES_07771 | C | C | C | C | C | C | C | C | C | C | C | C |
| NC_011770 | 850016 | C | T | 73 | 0 | 6 | 0 | NON_SYNONYMOU | MODERAT | MISSENSE | Gtg/Atg | V197M | PLES_07781 | T | T | T | T | T | T | | | | | | |
| NC_011770 | 850522 | C | T | 77 | 0 | 2 | 0 | NON_SYNONYMOU | MODERAT | MISSENSE | gGg/gAg | G28E | PLES_07781 | | | | | | | T | T | | | | |
| NC_011770 | 850540 | C | T | 75 | 0 | 1 | 0 | NON_SYNONYMOU | MODERAT | MISSENSE | cGc/cAc | R22H | PLES_07781 | | | | | | | T | | | | | |
| NC_011770 | 850734 | C | T | 77 | 0 | 1 | 0 | NON_SYNONYMOU | MODERAT | MISSENSE | cGc/cAc | R223H | PLES_07791 | | | | | | | T | | | | | |
| NC_011770 | 854565 | C | T | 74 | 0 | 2 | 0 | NON_SYNONYMOU | MODERAT | MISSENSE | Gag/Aag | E247K | PLES_07821 | | | | | | | | | | | T | T |
| NC_011770 | 856006 | A | C | 112 | 0 | 12 | 0 | NON_SYNONYMOU | MODERAT | MISSENSE | gTc/gGc | V270G | PLES_07831 | C | C | C | C | C | C | C | C | C | C | C | C |
| NC_011770 | 856805 | G | A | 110 | 0 | 2 | 0 | NON_SYNONYMOU | MODERAT | MISSENSE | Ctc/Ttc | L4F | PLES_07831 | | | | | | | A | A | | | | |
| NC_011770 | 857070 | A | G | 132 | 0 | 1 | 0 | NON_SYNONYMOU | MODERAT | MISSENSE | cTg/cCg | L635P | PLES_07841 | | | | | | | G | | | | | |
| NC_011770 | 859402 | C | T | 106 | 0 | 2 | 0 | NON_SYNONYMOU | MODERAT | MISSENSE | Gtc/Atc | V280I | PLES_07851 | | | | | | | T | T | | | | |

| | | | | | | | | | | | | | | | | | | | | | | | | | |
|---|---|---|---|---|---|---|---|---|---|---|---|---|---|---|---|---|---|---|---|---|---|---|---|---|---|
| NC_011770 | 859752 | C | T | 104 | 0 | 10 | 0 | NON_SYNONYMOU | MODERAT | MISSENSE | cGa/cAa | R163Q | PLES_07851 | T | T | T | T | T | T | T | T | T | T | | |
| NC_011770 | 860052 | C | T | 93 | 0 | 8 | 0 | NON_SYNONYMOU | MODERAT | MISSENSE | aGc/aAc | S63N | PLES_07851 | T | T | T | T | T | T | | | T | T | | |
| NC_011770 | 860136 | A | G | 97 | 0 | 12 | 0 | NON_SYNONYMOU | MODERAT | MISSENSE | gTg/gCg | V35A | PLES_07851 | G | G | G | G | G | G | G | G | G | G | G | G |
| NC_011770 | 860386 | T | G | 67 | 0 | 12 | 0 | NON_SYNONYMOU | MODERAT | MISSENSE | Aaa/Caa | K84Q | PLES_07861 | G | G | G | G | G | G | G | G | G | G | G | G |
| NC_011770 | 862020 | G | T | 71 | 0 | 8 | 0 | NON_SYNONYMOU | MODERAT | MISSENSE | Caa/Aaa | Q240K | PLES_07871 | T | T | T | T | T | T | | | T | T | | |
| NC_011770 | 862717 | G | T | 55 | 0 | 8 | 0 | NON_SYNONYMOU | MODERAT | MISSENSE | caC/caA | H7Q | PLES_07871 | T | T | T | T | T | T | | | T | T | | |
| NC_011770 | 862727 | A | G | 54 | 0 | 8 | 0 | NON_SYNONYMOU | MODERAT | MISSENSE | cTc/cCc | L4P | PLES_07871 | G | G | G | G | G | G | | | G | G | | |
| NC_011770 | 867114 | C | T | 1 | 9 | 1 | 0 | NON_SYNONYMOU | MODERAT | MISSENSE | Gct/Act | A381T | PLES_07931 | . | . | . | . | | | . | . | . | . | . | **T** |
| NC_011770 | 867180 | C | T | 17 | 10 | 2 | 0 | NON_SYNONYMOU | MODERAT | MISSENSE | Gac/Aac | D359N | PLES_07931 | . | . | . | . | T | T | . | . | . | . | . | . |
| NC_011770 | 867447 | C | A | 6 | 10 | 2 | 0 | NON_SYNONYMOU | MODERAT | MISSENSE | Gcc/Tcc | A270S | PLES_07931 | . | . | . | . | . | . | . | . | . | . | A | A |
| NC_011770 | 867705 | A | G | 3 | 10 | 2 | 0 | NON_SYNONYMOU | MODERAT | MISSENSE | Tac/Cac | Y184H | PLES_07931 | . | . | . | . | G | G | . | . | . | . | . | . |
| NC_011770 | 867708 | G | T | 3 | 10 | 2 | 0 | NON_SYNONYMOU | MODERAT | MISSENSE | Ccc/Acc | P183T | PLES_07931 | . | . | . | . | T | T | . | . | . | . | . | . |
| NC_011770 | 867710 | C | T | 3 | 10 | 2 | 0 | NON_SYNONYMOU | MODERAT | MISSENSE | cGa/cAa | R182Q | PLES_07931 | . | . | . | . | T | T | . | . | . | . | . | . |
| NC_011770 | 867722 | C | T | 4 | 10 | 2 | 0 | NON_SYNONYMOU | MODERAT | MISSENSE | aGt/aAt | S178N | PLES_07931 | . | . | . | . | T | T | . | . | . | . | . | . |
| NC_011770 | 867738 | G | C | 4 | 10 | 2 | 0 | NON_SYNONYMOU | MODERAT | MISSENSE | Ctg/Gtg | L173V | PLES_07931 | . | . | . | . | C | C | . | . | . | . | . | . |
| NC_011770 | 867785 | C | T | 6 | 10 | 2 | 0 | NON_SYNONYMOU | MODERAT | MISSENSE | gGt/gAt | G157D | PLES_07931 | . | . | . | . | T | T | . | . | . | . | . | . |
| NC_011770 | 867807 | G | C | 9 | 10 | 2 | 0 | NON_SYNONYMOU | MODERAT | MISSENSE | Cag/Gag | Q150E | PLES_07931 | . | . | . | . | C | C | . | . | . | . | . | . |
| NC_011770 | 867828 | G | T | 9 | 10 | 2 | 0 | NON_SYNONYMOU | MODERAT | MISSENSE | Ctc/Atc | L143I | PLES_07931 | . | . | . | . | T | T | . | . | . | . | . | . |
| NC_011770 | 867831 | C | T | 9 | 10 | 2 | 0 | NON_SYNONYMOU | MODERAT | MISSENSE | Gac/Aac | D142N | PLES_07931 | . | . | . | . | T | T | . | . | . | . | . | . |
| NC_011770 | 867851 | A | T | 11 | 10 | 2 | 0 | NON_SYNONYMOU | MODERAT | MISSENSE | cTc/cAc | L135H | PLES_07931 | . | . | . | . | T | T | . | . | . | . | . | . |
| NC_011770 | 867852 | G | A | 11 | 10 | 2 | 0 | NON_SYNONYMOU | MODERAT | MISSENSE | Ctc/Ttc | L135F | PLES_07931 | . | . | . | . | A | A | . | . | . | . | . | . |
| NC_011770 | 867879 | G | C | 11 | 10 | 2 | 0 | NON_SYNONYMOU | MODERAT | MISSENSE | Ctc/Gtc | L126V | PLES_07931 | . | . | . | . | C | C | . | . | . | . | . | . |
| NC_011770 | 867896 | A | T | 11 | 10 | 2 | 0 | NON_SYNONYMOU | MODERAT | MISSENSE | aTt/aAt | I120N | PLES_07931 | . | . | . | . | T | T | . | . | . | . | . | . |
| NC_011770 | 867897 | T | G | 11 | 10 | 2 | 0 | NON_SYNONYMOU | MODERAT | MISSENSE | Att/Ctt | I120L | PLES_07931 | . | . | . | . | G | G | . | . | . | . | . | . |
| NC_011770 | 867902 | C | T | 11 | 10 | 2 | 0 | NON_SYNONYMOU | MODERAT | MISSENSE | gGa/gAa | G118E | PLES_07931 | . | . | . | . | T | T | . | . | . | . | . | . |
| NC_011770 | 867908 | C | G | 11 | 10 | 2 | 0 | NON_SYNONYMOU | MODERAT | MISSENSE | gGt/gCt | G116A | PLES_07931 | . | . | . | . | G | G | . | . | . | . | . | . |
| NC_011770 | 867917 | T | C | 12 | 10 | 2 | 0 | NON_SYNONYMOU | MODERAT | MISSENSE | gAc/gGc | D113G | PLES_07931 | . | . | . | . | C | C | . | . | . | . | . | . |
| NC_011770 | 873136 | C | A | 2 | 10 | 2 | 0 | NON_SYNONYMOU | MODERAT | MISSENSE | aaC/aaA | N10K | PLES_07981 | . | . | . | . | A | A | . | . | . | . | . | . |
| NC_011770 | 874368 | T | G | 15 | 10 | 2 | 0 | NON_SYNONYMOU | MODERAT | MISSENSE | Tct/Gct | S54A | PLES_08001 | . | . | . | . | G | G | . | . | . | . | . | . |
| NC_011770 | 874465 | G | A | 13 | 10 | 2 | 0 | NON_SYNONYMOU | MODERAT | MISSENSE | cGg/cAg | R86Q | PLES_08001 | . | . | . | . | A | A | . | . | . | . | . | . |
| NC_011770 | 874509 | T | C | 14 | 10 | 2 | 0 | NON_SYNONYMOU | MODERAT | MISSENSE | Tca/Cca | S101P | PLES_08001 | . | . | . | . | C | C | . | . | . | . | . | . |
| NC_011770 | 874559 | A | T | 14 | 10 | 2 | 0 | NON_SYNONYMOU | MODERAT | MISSENSE | gaA/gaT | E117D | PLES_08001 | . | . | . | . | T | T | . | . | . | . | . | . |
| NC_011770 | 874581 | G | A | 16 | 10 | 2 | 0 | NON_SYNONYMOU | MODERAT | MISSENSE | Gcc/Acc | A125T | PLES_08001 | . | . | . | . | A | A | . | . | . | . | . | . |
| NC_011770 | 874748 | C | A | 17 | 10 | 2 | 0 | NON_SYNONYMOU | MODERAT | MISSENSE | gaC/gaA | D180E | PLES_08001 | . | . | . | . | A | A | . | . | . | . | . | . |
| NC_011770 | 874816 | A | T | 18 | 10 | 2 | 0 | NON_SYNONYMOU | MODERAT | MISSENSE | tAc/tTc | Y203F | PLES_08001 | . | . | . | . | T | T | . | . | . | . | . | . |
| NC_011770 | 874827 | G | T | 17 | 10 | 2 | 0 | NON_SYNONYMOU | MODERAT | MISSENSE | Gca/Tca | A207S | PLES_08001 | . | . | . | . | T | T | . | . | . | . | . | . |
| NC_011770 | 878149 | G | A | 0 | 11 | 1 | 0 | NON_SYNONYMOU | MODERAT | MISSENSE | Gtg/Atg | V355M | PLES_08031 | . | . | . | . | A | . | . | . | . | . | . | . |
| NC_011770 | 878153 | T | A | 0 | 11 | 1 | 0 | NON_SYNONYMOU | MODERAT | MISSENSE | gTg/gAg | V356E | PLES_08031 | . | . | . | . | A | . | . | . | . | . | . | . |
| NC_011770 | 878230 | C | G | 2 | 10 | 2 | 0 | NON_SYNONYMOU | MODERAT | MISSENSE | Cag/Gag | Q382E | PLES_08031 | . | . | . | . | G | G | . | . | . | . | . | . |
| NC_011770 | 879731 | A | T | 12 | 10 | 2 | 0 | NON_SYNONYMOU | MODERAT | MISSENSE | cAg/cTg | Q61L | PLES_08051 | . | . | . | . | T | T | . | . | . | . | . | . |
| NC_011770 | 880195 | G | GA | 12 | 10 | 1 | 0 | FRAME_SHIFT | HIGH | | gag/gAag | E79E? | PLES_08061 | . | . | . | . | | GA | . | . | . | . | . | . |
| NC_011770 | 880158 | G | A | 12 | 10 | 2 | 0 | NON_SYNONYMOU | MODERAT | MISSENSE | Gtc/Atc | V70I | PLES_08061 | . | . | . | . | A | A | . | . | . | . | . | . |
| NC_011770 | 881742 | CAT | C | 26 | 6 | 4 | 0 | FRAME_SHIFT | HIGH | | atc/ | I21 | PLES_08091 | . | . | . | . | C | C | . | . | C | C | | |
| NC_011770 | 881749 | A | G | 26 | 6 | 4 | 0 | NON_SYNONYMOU | MODERAT | MISSENSE | Att/Gtt | I23V | PLES_08091 | . | . | . | . | G | G | . | . | G | G | | |
| NC_011770 | 881754 | C | A | 27 | 6 | 4 | 0 | NON_SYNONYMOU | MODERAT | MISSENSE | gaC/gaA | D24E | PLES_08091 | . | . | . | . | A | A | . | . | A | A | | |
| NC_011770 | 881791 | A | T | 30 | 6 | 2 | 0 | NON_SYNONYMOU | MODERAT | MISSENSE | Acg/Tcg | T37S | PLES_08091 | . | . | . | . | T | T | . | . | | | | |
| NC_011770 | 881846 | TCAA | T | 31 | 6 | 2 | 0 | CODON_CHANGE_ | MODERATE | | ctcaag/ctg | LK55L | PLES_08091 | . | . | . | . | T | T | . | . | | | | |
| NC_011770 | 881854 | C | CGAT | 33 | 6 | 2 | 0 | CODON_CHANGE_ | MODERATE | | ctg/cGATt[illegible] | L58RL | PLES_08091 | . | . | . | . | CGAT | CGAT | . | . | | | | |
| NC_011770 | 881914 | A | G | 35 | 6 | 2 | 0 | NON_SYNONYMOU | MODERAT | MISSENSE | Acg/Gcg | T78A | PLES_08091 | . | . | . | . | | | . | . | G | G | | |
| NC_011770 | 881917 | C | G | 34 | 6 | 2 | 0 | NON_SYNONYMOU | MODERAT | MISSENSE | Ccg/Gcg | P79A | PLES_08091 | . | . | . | . | | | . | . | | | G | G |
| NC_011770 | 881920 | G | A | 34 | 6 | 4 | 0 | NON_SYNONYMOU | MODERAT | MISSENSE | Gcc/Acc | A80T | PLES_08091 | . | . | . | . | A | A | . | . | | | A | A |
| NC_011770 | 881921 | C | T | 33 | 6 | 2 | 0 | NON_SYNONYMOU | MODERAT | MISSENSE | gCc/gTc | A80V | PLES_08091 | . | . | . | . | | | . | . | T | T | | |
| NC_011770 | 881929 | A | C | 34 | 6 | 2 | 0 | NON_SYNONYMOU | MODERAT | MISSENSE | Aag/Cag | K83Q | PLES_08091 | . | . | . | . | | | . | . | C | C | | |
| NC_011770 | 881930 | A | G | 33 | 6 | 2 | 0 | NON_SYNONYMOU | MODERAT | MISSENSE | aAg/aGg | K83R | PLES_08091 | . | . | . | . | | | . | . | G | G | | |
| NC_011770 | 881931 | G | C | 34 | 6 | 2 | 0 | NON_SYNONYMOU | MODERAT | MISSENSE | aaG/aaC | K83N | PLES_08091 | . | . | . | . | | | . | . | C | C | | |
| NC_011770 | 881944 | C | A | 34 | 6 | 2 | 0 | NON_SYNONYMOU | MODERAT | MISSENSE | Cag/Aag | Q88K | PLES_08091 | . | . | . | . | | | . | . | | | A | A |
| NC_011770 | 882002 | C | A | 28 | 6 | 6 | 0 | NON_SYNONYMOU | MODERAT | MISSENSE | gCc/gAc | A107D | PLES_08091 | . | . | . | . | A | A | . | . | A | A | A | A |
| NC_011770 | 882016 | T | C | 30 | 6 | 2 | 0 | NON_SYNONYMOU | MODERAT | MISSENSE | Tat/Cat | Y112H | PLES_08091 | . | . | . | . | C | C | . | . | | | | |
| NC_011770 | 882055 | T | C | 28 | 6 | 6 | 0 | NON_SYNONYMOU | MODERAT | MISSENSE | Tac/Cac | Y125H | PLES_08091 | . | . | . | . | C | C | . | . | C | C | C | C |
| NC_011770 | 882148 | G | T | 25 | 6 | 2 | 0 | NON_SYNONYMOU | MODERAT | MISSENSE | Gcg/Tcg | A156S | PLES_08091 | . | . | . | . | | | . | . | | | T | T |
| NC_011770 | 882152 | G | C | 25 | 6 | 2 | 0 | NON_SYNONYMOU | MODERAT | MISSENSE | gGg/gCg | G157A | PLES_08091 | . | . | . | . | | | . | . | | | C | C |
| NC_011770 | 882158 | G | T | 26 | 6 | 2 | 0 | NON_SYNONYMOU | MODERAT | MISSENSE | tGg/tTg | W159L | PLES_08091 | . | . | . | . | T | T | . | . | | | | |

| | | | | | | | | | | | | | | | | | | | | | | | | | |
|---|---|---|---|---|---|---|---|---|---|---|---|---|---|---|---|---|---|---|---|---|---|---|---|---|---|
| NC_011770 | 882159 | G | C | 27 | 6 | 2 | 0 | NON_SYNONYMOU | MODERAT | MISSENSE | tgG/tgC | W159C | PLES_08091 | . | . | . | . | C | C | . | . | | | | |
| NC_011770 | 882163 | A | C | 26 | 6 | 4 | 0 | NON_SYNONYMOU | MODERAT | MISSENSE | Aag/Cag | K161Q | PLES_08091 | . | . | . | . | C | C | . | . | C | C | | |
| NC_011770 | 882164 | A | G | 26 | 6 | 2 | 0 | NON_SYNONYMOU | MODERAT | MISSENSE | aAg/aGg | K161R | PLES_08091 | . | . | . | . | | | . | . | G | G | | |
| NC_011770 | 882183 | C | T | 13 | 6 | 6 | 0 | NON_SYNONYMOU | MODERAT | MISSENSE | gaC/gaA | D167E | PLES_08091 | . | . | . | . | T | T | . | . | T | T | A | A |
| NC_011770 | 882229 | G | A | 12 | 8 | 2 | 0 | NON_SYNONYMOU | MODERAT | MISSENSE | Ggc/Agc | G183S | PLES_08091 | . | . | . | . | . | . | . | . | | | A | A |
| NC_011770 | 882249 | G | T | 7 | 10 | 2 | 0 | NON_SYNONYMOU | MODERAT | MISSENSE | gaG/gaT | E189D | PLES_08091 | . | . | . | . | . | . | . | . | . | . | T | T |
| NC_011770 | 882257 | T | C | 7 | 10 | 2 | 0 | NON_SYNONYMOU | MODERAT | MISSENSE | gTg/gCg | V192A | PLES_08091 | . | . | . | . | . | . | . | . | . | . | C | C |
| NC_011770 | 882283 | G | T | 28 | 8 | 2 | 0 | NON_SYNONYMOU | MODERAT | MISSENSE | Gcg/Tcg | A201S | PLES_08091 | . | . | . | . | T | T | . | . | . | . | | |
| NC_011770 | 882573 | C | G | 29 | 8 | 4 | 0 | NON_SYNONYMOU | MODERAT | MISSENSE | gCt/gGt | A13G | PLES_08101 | . | . | . | . | G | G | . | . | . | . | G | G |
| NC_011770 | 882690 | G | T | 19 | 6 | 6 | 0 | NON_SYNONYMOU | MODERAT | MISSENSE | aGc/aTc | S52I | PLES_08101 | . | . | . | . | T | T | . | . | T | T | A | A |
| NC_011770 | 882702 | C | T | 18 | 6 | 2 | 0 | NON_SYNONYMOU | MODERAT | MISSENSE | gCt/gTt | A56V | PLES_08101 | . | . | . | . | | | . | . | | | T | T |
| NC_011770 | 882707 | A | G | 18 | 6 | 4 | 0 | NON_SYNONYMOU | MODERAT | MISSENSE | Acc/Gcc | T58A | PLES_08101 | . | . | . | . | G | G | . | . | G | G | | |
| NC_011770 | 882713 | G | T | 18 | 6 | 2 | 0 | NON_SYNONYMOU | MODERAT | MISSENSE | Gca/Tca | A60S | PLES_08101 | . | . | . | . | | | . | . | T | T | | |
| NC_011770 | 882716 | A | C | 18 | 6 | 6 | 0 | NON_SYNONYMOU | MODERAT | MISSENSE | Atg/Ctg | M61L | PLES_08101 | . | . | . | . | C | C | . | . | C | C | C | C |
| NC_011770 | 882827 | A | G | 20 | 6 | 6 | 0 | NON_SYNONYMOU | MODERAT | MISSENSE | Aca/Gca | T98A | PLES_08101 | . | . | . | . | G | G | . | . | G | G | G | G |
| NC_011770 | 882831 | G | A | 19 | 6 | 6 | 0 | NON_SYNONYMOU | MODERAT | MISSENSE | gGc/gAc | G99D | PLES_08101 | . | . | . | . | A | A | . | . | A | A | A | A |
| NC_011770 | 882843 | A | C | 20 | 6 | 2 | 0 | NON_SYNONYMOU | MODERAT | MISSENSE | cAg/cCg | Q103P | PLES_08101 | . | . | . | . | | | . | . | C | C | | |
| NC_011770 | 882857 | A | T | 18 | 6 | 6 | 0 | NON_SYNONYMOU | MODERAT | MISSENSE | Acc/Tcc | T108S | PLES_08101 | . | . | . | . | T | T | . | . | T | T | T | T |
| NC_011770 | 882869 | G | C | 20 | 6 | 2 | 0 | NON_SYNONYMOU | MODERAT | MISSENSE | Gat/Cat | D112H | PLES_08101 | . | . | . | . | | | . | . | C | C | | |
| NC_011770 | 882878 | A | G | 20 | 6 | 2 | 0 | NON_SYNONYMOU | MODERAT | MISSENSE | Aga/Gga | R115G | PLES_08101 | . | . | . | . | | | . | . | G | G | | |
| NC_011770 | 882956 | A | G | 16 | 6 | 4 | 0 | NON_SYNONYMOU | MODERAT | MISSENSE | Aaa/Gaa | K141E | PLES_08101 | . | . | . | . | G | G | . | . | | | G | G |
| NC_011770 | 882968 | A | G | 13 | 7 | 1 | 0 | NON_SYNONYMOU | MODERAT | MISSENSE | Aag/Gag | K145E | PLES_08101 | . | . | . | . | | | . | . | . | G | | |
| NC_011770 | 882981 | G | C | 11 | 8 | 2 | 0 | NON_SYNONYMOU | MODERAT | MISSENSE | gGc/gCc | G149A | PLES_08101 | . | . | . | . | C | C | . | . | . | . | | |
| NC_011770 | 883291 | T | C | 11 | 9 | 1 | 0 | NON_SYNONYMOU | MODERAT | MISSENSE | Tac/Cac | Y97H | PLES_08111 | . | . | . | . | C | . | . | . | . | . | | |
| NC_011770 | 883676 | A | C | 7 | 10 | 2 | 0 | NON_SYNONYMOU | MODERAT | MISSENSE | gAc/gCc | D225A | PLES_08111 | . | . | . | . | . | . | . | . | . | . | C | C |
| NC_011770 | 883695 | T | G | 7 | 10 | 2 | 0 | NON_SYNONYMOU | MODERAT | MISSENSE | gaT/gaG | D231E | PLES_08111 | . | . | . | . | . | . | . | . | . | . | G | G |
| NC_011770 | 886267 | C | T | 15 | 10 | 2 | 0 | NON_SYNONYMOU | MODERAT | MISSENSE | gCa/gTa | A611V | PLES_08131 | . | . | . | . | . | . | . | . | . | . | T | T |
| NC_011770 | 890432 | C | G | 2 | 10 | 2 | 0 | NON_SYNONYMOU | MODERAT | MISSENSE | Cct/Gct | P18A | PLES_08161 | . | . | . | . | . | . | . | . | . | . | G | G |
| NC_011770 | 890465 | G | A | 0 | 10 | 2 | 0 | NON_SYNONYMOU | MODERAT | MISSENSE | Gta/Ata | V29I | PLES_08161 | . | . | . | . | . | . | . | . | . | . | A | A |
| NC_011770 | 890594 | G | A | 3 | 10 | 2 | 0 | NON_SYNONYMOU | MODERAT | MISSENSE | Gat/Aat | D72N | PLES_08161 | . | . | . | . | . | . | . | . | . | . | A | A |
| NC_011770 | 890610 | C | T | 3 | 10 | 2 | 0 | NON_SYNONYMOU | MODERAT | MISSENSE | gCt/gTt | A77V | PLES_08161 | . | . | . | . | . | . | . | . | . | . | T | T |
| NC_011770 | 890654 | C | G | 5 | 9 | 2 | 0 | NON_SYNONYMOU | MODERAT | MISSENSE | Ccc/Gcc | P92A | PLES_08161 | . | . | . | . | . | | . | . | . | . | G | G |
| NC_011770 | 890973 | T | C | 20 | 8 | 2 | 0 | NON_SYNONYMOU | MODERAT | MISSENSE | Tgc/Cgc | C29R | PLES_08171 | . | . | . | . | | | . | . | . | . | C | C |
| NC_011770 | 891064 | C | G | 17 | 8 | 2 | 0 | NON_SYNONYMOU | MODERAT | MISSENSE | gCg/gGg | A59G | PLES_08171 | . | . | . | . | G | G | . | . | . | . | | |
| NC_011770 | 891377 | T | G | 16 | 8 | 2 | 0 | NON_SYNONYMOU | MODERAT | MISSENSE | gaT/gaG | D163E | PLES_08171 | . | . | . | . | G | G | . | . | . | . | | |
| NC_011770 | 891429 | G | A | 16 | 8 | 2 | 0 | NON_SYNONYMOU | MODERAT | MISSENSE | Gat/Aat | D181N | PLES_08171 | . | . | . | . | | | . | . | . | . | A | A |
| NC_011770 | 891487 | C | G | 15 | 8 | 2 | 0 | NON_SYNONYMOU | MODERAT | MISSENSE | gCg/gGg | A200G | PLES_08171 | . | . | . | . | | | . | . | . | . | G | G |
| NC_011770 | 891522 | T | A | 14 | 8 | 2 | 0 | NON_SYNONYMOU | MODERAT | MISSENSE | Tcg/Acg | S10T | PLES_08181 | . | . | . | . | | | . | . | . | . | A | A |
| NC_011770 | 891540 | T | C | 14 | 8 | 2 | 0 | NON_SYNONYMOU | MODERAT | MISSENSE | Tcg/Ccg | S16P | PLES_08181 | . | . | . | . | | | . | . | . | . | C | C |
| NC_011770 | 891577 | T | C | 15 | 8 | 4 | 0 | NON_SYNONYMOU | MODERAT | MISSENSE | gTc/gCc | V28A | PLES_08181 | . | . | . | . | C | C | . | . | . | . | C | C |
| NC_011770 | 891913 | C | T | 21 | 10 | 2 | 0 | NON_SYNONYMOU | MODERAT | MISSENSE | Cat/Tat | H75Y | PLES_08191 | . | . | . | . | . | . | . | . | . | . | T | T |
| NC_011770 | 891990 | C | A | 21 | 10 | 2 | 0 | NON_SYNONYMOU | MODERAT | MISSENSE | caC/caA | H100Q | PLES_08191 | . | . | . | . | . | . | . | . | . | . | A | A |
| NC_011770 | 891992 | T | C | 21 | 10 | 2 | 0 | NON_SYNONYMOU | MODERAT | MISSENSE | aTc/aCc | I101T | PLES_08191 | . | . | . | . | . | . | . | . | . | . | C | C |
| NC_011770 | 892059 | G | T | 18 | 10 | 2 | 0 | NON_SYNONYMOU | MODERAT | MISSENSE | gaG/gaT | E123D | PLES_08191 | . | . | . | . | . | . | . | . | . | . | T | T |
| NC_011770 | 892121 | C | T | 17 | 10 | 2 | 0 | NON_SYNONYMOU | MODERAT | MISSENSE | aCc/aTc | T144I | PLES_08191 | . | . | . | . | . | . | . | . | . | . | T | T |
| NC_011770 | 892393 | T | G | 18 | 9 | 3 | 0 | NON_SYNONYMOU | MODERAT | MISSENSE | Tca/Gca | S235A | PLES_08191 | . | . | . | . | . | G | . | . | . | . | G | G |
| NC_011770 | 892643 | C | T | 25 | 8 | 2 | 0 | NON_SYNONYMOU | MODERAT | MISSENSE | aCg/aTg | T40M | PLES_08201 | . | . | . | . | | | . | . | . | . | T | T |
| NC_011770 | 892741 | A | G | 24 | 8 | 2 | 0 | NON_SYNONYMOU | MODERAT | MISSENSE | Aca/Gca | T73A | PLES_08201 | . | . | . | . | | | . | . | . | . | G | G |
| NC_011770 | 892799 | G | A | 22 | 8 | 2 | 0 | NON_SYNONYMOU | MODERAT | MISSENSE | gGc/gAc | G92D | PLES_08201 | . | . | . | . | | | . | . | . | . | A | A |
| NC_011770 | 892868 | C | T | 19 | 8 | 2 | 0 | NON_SYNONYMOU | MODERAT | MISSENSE | gCt/gTt | A115V | PLES_08201 | . | . | . | . | | | . | . | . | . | T | T |
| NC_011770 | 892883 | G | A | 18 | 8 | 2 | 0 | NON_SYNONYMOU | MODERAT | MISSENSE | gGg/gAg | G120E | PLES_08201 | . | . | . | . | | | . | . | . | . | A | A |
| NC_011770 | 892969 | C | A | 17 | 8 | 2 | 0 | NON_SYNONYMOU | MODERAT | MISSENSE | Ctt/Att | L149I | PLES_08201 | . | . | . | . | | | . | . | . | . | A | A |
| NC_011770 | 892980 | T | G | 17 | 8 | 1 | 0 | NON_SYNONYMOU | MODERAT | MISSENSE | gaT/gaG | D152E | PLES_08201 | . | . | . | . | . | G | . | . | . | . | | |
| NC_011770 | 893027 | A | C | 18 | 8 | 2 | 0 | NON_SYNONYMOU | MODERAT | MISSENSE | gAt/gCt | D168A | PLES_08201 | . | . | . | . | | | . | . | . | . | C | C |
| NC_011770 | 893120 | G | A | 18 | 8 | 2 | 0 | NON_SYNONYMOU | MODERAT | MISSENSE | gGg/gAg | G199E | PLES_08201 | . | . | . | . | | | . | . | . | . | A | A |
| NC_011770 | 893141 | T | A | 18 | 8 | 2 | 0 | NON_SYNONYMOU | MODERAT | MISSENSE | cTg/cAg | L206Q | PLES_08201 | . | . | . | . | | | . | . | . | . | A | A |
| NC_011770 | 893152 | A | G | 17 | 8 | 2 | 0 | NON_SYNONYMOU | MODERAT | MISSENSE | Act/Gct | T210A | PLES_08201 | . | . | . | . | | | . | . | . | . | G | G |
| NC_011770 | 893938 | T | TAGCCA | 10 | 8 | 1 | 0 | FRAME_SHIFT | HIGH | | gct/gcCAA | A104AK? | PLES_08211 | . | . | . | . | | | . | . | . | . | . | TAGCCA |
| NC_011770 | 893945 | GCTTTC | G | 12 | 8 | 2 | 0 | FRAME_SHIFT | HIGH | | ctttcc/ | LS106 | PLES_08211 | . | . | . | . | | | . | . | . | . | G | G |
| NC_011770 | 893743 | A | G | 24 | 8 | 2 | 0 | NON_SYNONYMOU | MODERAT | MISSENSE | aAg/aGg | K38R | PLES_08211 | . | . | . | . | G | G | . | . | . | . | | |
| NC_011770 | 893937 | A | G | 9 | 8 | 2 | 0 | NON_SYNONYMOU | MODERAT | MISSENSE | Ata/Gta | I103V | PLES_08211 | . | . | . | . | | | . | . | . | . | G | G |
| NC_011770 | 894130 | T | A | 13 | 8 | 2 | 0 | NON_SYNONYMOU | MODERAT | MISSENSE | tTc/tAc | F167Y | PLES_08211 | . | . | . | . | A | A | . | . | . | . | | |

| | | | | | | | | | | | | | | | | | | | | | | | | | |
|---|---|---|---|---|---|---|---|---|---|---|---|---|---|---|---|---|---|---|---|---|---|---|---|---|---|
| NC_011770 | 894154 | G | C | 13 | 8 | 4 | 0 | NON_SYNONYMOU | MODERAT | MISSENSE | aGc/aCc | S175T | PLES_08211 | . | . | . | . | C | C | . | . | . | . | C | C |
| NC_011770 | 894561 | A | G | 14 | 5 | 2 | 0 | NON_SYNONYMOU | MODERAT | MISSENSE | Atc/Gtc | I311V | PLES_08211 | | . | . | . | | | | | . | . | G | G |
| NC_011770 | 894601 | C | T | 14 | 8 | 2 | 0 | NON_SYNONYMOU | MODERAT | MISSENSE | gCg/gTg | A324V | PLES_08211 | . | . | . | . | | | . | . | . | . | T | T |
| NC_011770 | 894645 | T | G | 21 | 8 | 2 | 0 | NON_SYNONYMOU | MODERAT | MISSENSE | Tcc/Gcc | S339A | PLES_08211 | . | . | . | . | | | . | . | . | . | G | G |
| NC_011770 | 894993 | A | G | 29 | 8 | 4 | 0 | NON_SYNONYMOU | MODERAT | MISSENSE | Aat/Gat | N455D | PLES_08211 | . | . | . | . | G | G | . | . | . | . | G | G |
| NC_011770 | 894995 | T | A | 29 | 8 | 4 | 0 | NON_SYNONYMOU | MODERAT | MISSENSE | aaT/aaA | N455K | PLES_08211 | . | . | . | . | A | A | . | . | . | . | A | A |
| NC_011770 | 895044 | T | G | 29 | 8 | 4 | 0 | NON_SYNONYMOU | MODERAT | MISSENSE | Tcg/Gcg | S472A | PLES_08211 | . | . | . | . | G | G | . | . | . | . | G | G |
| NC_011770 | 895045 | C | G | 28 | 8 | 4 | 0 | NON_SYNONYMOU | MODERAT | MISSENSE | tCg/tGg | S472W | PLES_08211 | . | . | . | . | G | G | . | . | . | . | G | G |
| NC_011770 | 895060 | C | A | 30 | 8 | 4 | 0 | NON_SYNONYMOU | MODERAT | MISSENSE | aCt/aAt | T477N | PLES_08211 | . | . | . | . | A | A | . | . | . | . | A | A |
| NC_011770 | 895172 | A | T | 30 | 8 | 4 | 0 | NON_SYNONYMOU | MODERAT | MISSENSE | gaA/gaT | E514D | PLES_08211 | . | . | . | . | T | T | . | . | . | . | T | T |
| NC_011770 | 895311 | G | T | 32 | 8 | 2 | 0 | NON_SYNONYMOU | MODERAT | MISSENSE | Gcg/Tcg | A561S | PLES_08211 | . | . | . | . | T | T | . | . | . | . | | |
| NC_011770 | 895939 | A | C | 26 | 8 | 2 | 0 | NON_SYNONYMOU | MODERAT | MISSENSE | gAg/gCg | E770A | PLES_08211 | . | . | . | . | C | C | . | . | . | . | | |
| NC_011770 | 895975 | C | T | 26 | 8 | 4 | 0 | NON_SYNONYMOU | MODERAT | MISSENSE | cCg/cTg | P782L | PLES_08211 | . | . | . | . | T | T | . | . | . | . | T | T |
| NC_011770 | 897261 | A | T | 34 | 8 | 2 | 0 | NON_SYNONYMOU | MODERAT | MISSENSE | tAc/tTc | Y62F | PLES_08221 | . | . | . | . | | | . | . | . | . | T | T |
| NC_011770 | 897338 | G | A | 31 | 8 | 2 | 0 | NON_SYNONYMOU | MODERAT | MISSENSE | Gtc/Atc | V88I | PLES_08221 | . | . | . | . | | | . | . | . | . | A | A |
| NC_011770 | 897587 | A | G | 19 | 8 | 4 | 0 | NON_SYNONYMOU | MODERAT | MISSENSE | Aga/Gga | R171G | PLES_08221 | . | . | . | . | G | G | . | . | . | . | G | G |
| NC_011770 | 897588 | G | T | 19 | 8 | 4 | 0 | NON_SYNONYMOU | MODERAT | MISSENSE | aGa/aTa | R171I | PLES_08221 | . | . | . | . | T | T | . | . | . | . | T | T |
| NC_011770 | 897589 | A | C | 19 | 8 | 4 | 0 | NON_SYNONYMOU | MODERAT | MISSENSE | agA/agC | R171S | PLES_08221 | . | . | . | . | C | C | . | . | . | . | C | C |
| NC_011770 | 897593 | G | A | 19 | 8 | 4 | 0 | NON_SYNONYMOU | MODERAT | MISSENSE | Gac/Aac | D173N | PLES_08221 | . | . | . | . | A | A | . | . | . | . | A | A |
| NC_011770 | 897778 | C | G | 18 | 8 | 2 | 0 | NON_SYNONYMOU | MODERAT | MISSENSE | gaC/gaG | D234E | PLES_08221 | . | . | . | . | | | . | . | . | . | G | G |
| NC_011770 | 898026 | G | C | 11 | 10 | 2 | 0 | NON_SYNONYMOU | MODERAT | MISSENSE | gGa/gCa | G317A | PLES_08221 | . | . | . | . | . | . | . | . | . | . | C | C |
| NC_011770 | 898086 | C | A | 12 | 10 | 2 | 0 | NON_SYNONYMOU | MODERAT | MISSENSE | gCa/gAa | A337E | PLES_08221 | . | . | . | . | . | . | . | . | . | . | A | A |
| NC_011770 | 898200 | A | C | 13 | 10 | 2 | 0 | NON_SYNONYMOU | MODERAT | MISSENSE | gAg/gCg | E375A | PLES_08221 | . | . | . | . | . | . | . | . | . | . | C | C |
| NC_011770 | 898214 | G | T | 13 | 10 | 2 | 0 | NON_SYNONYMOU | MODERAT | MISSENSE | Gca/Tca | A380S | PLES_08221 | . | . | . | . | . | . | . | . | . | . | T | T |
| NC_011770 | 898281 | G | A | 12 | 9 | 2 | 0 | NON_SYNONYMOU | MODERAT | MISSENSE | aGc/aAc | S402N | PLES_08221 | . | . | . | . | . | . | . | . | . | . | A | A |
| NC_011770 | 898282 | C | A | 12 | 9 | 2 | 0 | NON_SYNONYMOU | MODERAT | MISSENSE | agC/agA | S402R | PLES_08221 | . | . | . | . | | . | . | . | . | . | A | A |
| NC_011770 | 898322 | A | C | 13 | 8 | 2 | 0 | NON_SYNONYMOU | MODERAT | MISSENSE | Atc/Ctc | I416L | PLES_08221 | . | . | . | . | | | . | . | . | . | C | C |
| NC_011770 | 898358 | G | T | 16 | 8 | 2 | 0 | NON_SYNONYMOU | MODERAT | MISSENSE | Gta/Tta | V428L | PLES_08221 | . | . | . | . | | | . | . | . | . | T | T |
| NC_011770 | 898379 | T | C | 17 | 8 | 4 | 0 | NON_SYNONYMOU | MODERAT | MISSENSE | gTg/gCg | V2A | PLES_08231 | . | . | . | . | C | C | . | . | . | . | C | C |
| NC_011770 | 898520 | A | C | 17 | 8 | 4 | 0 | NON_SYNONYMOU | MODERAT | MISSENSE | gAt/gCt | D49A | PLES_08231 | . | . | . | . | C | C | . | . | . | . | C | C |
| NC_011770 | 898555 | G | T | 13 | 8 | 2 | 0 | NON_SYNONYMOU | MODERAT | MISSENSE | Gcg/Tcg | A61S | PLES_08231 | . | . | . | . | | | . | . | . | . | T | T |
| NC_011770 | 898570 | G | A | 13 | 8 | 2 | 0 | NON_SYNONYMOU | MODERAT | MISSENSE | Gtg/Atg | V66M | PLES_08231 | . | . | . | . | A | A | . | . | . | . | | |
| NC_011770 | 898607 | C | A | 10 | 8 | 2 | 0 | NON_SYNONYMOU | MODERAT | MISSENSE | gCc/gAc | A78D | PLES_08231 | . | . | . | . | | | . | . | . | . | A | A |
| NC_011770 | 899365 | G | A | 3 | 8 | 2 | 0 | NON_SYNONYMOU | MODERAT | MISSENSE | Ggc/Agc | G9S | PLES_08251 | . | . | . | . | | | . | . | . | . | A | A |
| NC_011770 | 901611 | C | A | 16 | 8 | 4 | 0 | STOP_GAINED | HIGH | NONSENSE | tCg/tAg | S97* | PLES_08271 | . | . | . | . | A | A | . | . | . | . | A | A |
| NC_011770 | 901601 | A | G | 16 | 8 | 4 | 0 | NON_SYNONYMOU | MODERAT | MISSENSE | Act/Gct | T94A | PLES_08271 | . | . | . | . | G | G | . | . | . | . | G | G |
| NC_011770 | 901610 | T | G | 17 | 8 | 4 | 0 | NON_SYNONYMOU | MODERAT | MISSENSE | Tcg/Gcg | S97A | PLES_08271 | . | . | . | . | G | G | . | . | . | . | G | G |
| NC_011770 | 901751 | A | G | 22 | 8 | 4 | 0 | NON_SYNONYMOU | MODERAT | MISSENSE | Acg/Gcg | T144A | PLES_08271 | . | . | . | . | G | G | . | . | . | . | G | G |
| NC_011770 | 901821 | C | G | 26 | 8 | 2 | 0 | NON_SYNONYMOU | MODERAT | MISSENSE | aCt/aGt | T167S | PLES_08271 | . | . | . | . | G | G | . | . | . | . | | |
| NC_011770 | 901679 | C | T | 21 | 8 | 1 | 0 | NON_SYNONYMOU | MODERAT | MISSENSE | Ctt/Ttt | L120F | PLES_08271 | . | . | . | . | | | . | . | . | . | | T |
| NC_011770 | 902289 | GTC | G | 7 | 8 | 2 | 0 | FRAME_SHIFT | HIGH | | tcg/ | S84 | PLES_08281 | . | . | . | . | | | . | . | . | . | G | G |
| NC_011770 | 902295 | G | GCA | 4 | 9 | 3 | 0 | FRAME_SHIFT | HIGH | | ata/CAata | I86Q? | PLES_08281 | . | . | . | . | GCA | . | . | . | . | . | GCA | GCA |
| NC_011770 | 902599 | TA | T | 20 | 8 | 2 | 0 | FRAME_SHIFT | HIGH | | gcg/ | A184 | PLES_08281 | . | . | . | . | | | . | . | . | . | T | T |
| NC_011770 | 902182 | G | A | 23 | 8 | 2 | 0 | NON_SYNONYMOU | MODERAT | MISSENSE | Gcg/Acg | A48T | PLES_08281 | . | . | . | . | | | . | . | . | . | A | A |
| NC_011770 | 902320 | T | A | 4 | 9 | 3 | 0 | NON_SYNONYMOU | MODERAT | MISSENSE | Tcc/Acc | S94T | PLES_08281 | . | . | . | . | A | . | . | . | . | . | A | A |
| NC_011770 | 902321 | C | G | 4 | 9 | 3 | 0 | NON_SYNONYMOU | MODERAT | MISSENSE | tCc/tGc | S94C | PLES_08281 | . | . | . | . | G | . | . | . | . | . | G | G |
| NC_011770 | 902341 | T | A | 9 | 8 | 2 | 0 | NON_SYNONYMOU | MODERAT | MISSENSE | Tcc/Acc | S101T | PLES_08281 | . | . | . | . | A | A | . | . | . | . | | |
| NC_011770 | 902407 | T | A | 14 | 8 | 2 | 0 | NON_SYNONYMOU | MODERAT | MISSENSE | Tcg/Acg | S123T | PLES_08281 | . | . | . | . | A | A | . | . | . | . | | |
| NC_011770 | 902408 | C | G | 15 | 8 | 2 | 0 | NON_SYNONYMOU | MODERAT | MISSENSE | tCg/tGg | S123W | PLES_08281 | . | . | . | . | G | G | . | . | . | . | | |
| NC_011770 | 902422 | C | A | 20 | 8 | 2 | 0 | NON_SYNONYMOU | MODERAT | MISSENSE | Ctc/Atc | L128I | PLES_08281 | . | . | . | . | | | . | . | . | . | A | A |
| NC_011770 | 902489 | A | T | 25 | 8 | 2 | 0 | NON_SYNONYMOU | MODERAT | MISSENSE | cAg/cTg | Q150L | PLES_08281 | . | . | . | . | | | . | . | . | . | T | T |
| NC_011770 | 902530 | A | G | 14 | 8 | 4 | 0 | NON_SYNONYMOU | MODERAT | MISSENSE | Agc/Ggc | S164G | PLES_08281 | . | . | . | . | G | G | . | . | . | . | G | G |
| NC_011770 | 902538 | T | G | 14 | 8 | 2 | 0 | NON_SYNONYMOU | MODERAT | MISSENSE | gaT/gaG | D166E | PLES_08281 | . | . | . | . | G | G | . | . | . | . | | |
| NC_011770 | 902539 | G | T | 13 | 8 | 4 | 0 | NON_SYNONYMOU | MODERAT | MISSENSE | Ggc/Tgc | G167C | PLES_08281 | . | . | . | . | T | T | . | . | . | . | T | T |
| NC_011770 | 902540 | G | C | 14 | 8 | 4 | 0 | NON_SYNONYMOU | MODERAT | MISSENSE | gGc/gCc | G167A | PLES_08281 | . | . | . | . | C | C | . | . | . | . | C | C |
| NC_011770 | 902557 | G | A | 15 | 8 | 2 | 0 | NON_SYNONYMOU | MODERAT | MISSENSE | Ggc/Agc | G173S | PLES_08281 | . | . | . | . | | | . | . | . | . | A | A |
| NC_011770 | 902558 | G | A | 14 | 8 | 2 | 0 | NON_SYNONYMOU | MODERAT | MISSENSE | gGc/gAc | G173D | PLES_08281 | . | . | . | . | | | . | . | . | . | A | A |
| NC_011770 | 902576 | C | T | 15 | 8 | 2 | 0 | NON_SYNONYMOU | MODERAT | MISSENSE | gCg/gTg | A179V | PLES_08281 | . | . | . | . | | | . | . | . | . | T | T |
| NC_011770 | 902590 | G | A | 18 | 8 | 2 | 0 | NON_SYNONYMOU | MODERAT | MISSENSE | Gcg/Acg | A184T | PLES_08281 | . | . | . | . | | | . | . | . | . | A | A |
| NC_011770 | 902645 | C | A | 14 | 8 | 2 | 0 | NON_SYNONYMOU | MODERAT | MISSENSE | gCc/gAc | A15D | PLES_08291 | . | . | . | . | | | . | . | . | . | A | A |
| NC_011770 | 902647 | G | C | 14 | 8 | 2 | 0 | NON_SYNONYMOU | MODERAT | MISSENSE | Gcg/Ccg | A16P | PLES_08291 | . | . | . | . | | | . | . | . | . | C | C |
| NC_011770 | 902654 | A | C | 14 | 8 | 2 | 0 | NON_SYNONYMOU | MODERAT | MISSENSE | gAg/gCg | E18A | PLES_08291 | . | . | . | . | | | . | . | . | . | C | C |

| | | | | | | | | | | | | | | | | | | | | | | | | | |
|---|---|---|---|---|---|---|---|---|---|---|---|---|---|---|---|---|---|---|---|---|---|---|---|---|---|
| NC_011770 | 902683 | A | G | 14 | 8 | 2 | 0 | NON_SYNONYMOU | MODERAT | MISSENSE | Agc/Ggc | S28G | PLES_08291 | . | . | . | . | | | . | . | . | . | G | G |
| NC_011770 | 902686 | A | G | 15 | 8 | 2 | 0 | NON_SYNONYMOU | MODERAT | MISSENSE | Acc/Gcc | T29A | PLES_08291 | . | . | . | . | G | G | . | . | . | . | | |
| NC_011770 | 903600 | G | A | 5 | 10 | 2 | 0 | STOP_GAINED | HIGH | NONSENSE | tgG/tgA | W57* | PLES_08301 | . | . | . | . | A | A | . | . | . | . | . | . |
| NC_011770 | 903623 | GTGCC | G | 0 | 11 | 1 | 0 | FRAME_SHIFT | HIGH | | cgtgcc/ | RA65 | PLES_08301 | . | . | . | . | . | G | . | . | . | . | . | . |
| NC_011770 | 903577 | G | A | 5 | 10 | 2 | 0 | NON_SYNONYMOU | MODERAT | MISSENSE | Gct/Act | A50T | PLES_08301 | . | . | . | . | A | A | . | . | . | . | . | . |
| NC_011770 | 903599 | G | T | 5 | 10 | 2 | 0 | NON_SYNONYMOU | MODERAT | MISSENSE | tGg/tTg | W57L | PLES_08301 | . | . | . | . | T | T | . | . | . | . | . | . |
| NC_011770 | 903614 | C | T | 0 | 11 | 1 | 0 | NON_SYNONYMOU | MODERAT | MISSENSE | gCa/gTa | A62V | PLES_08301 | . | . | . | . | . | T | . | . | . | . | . | . |
| NC_011770 | 903617 | T | C | 0 | 11 | 1 | 0 | NON_SYNONYMOU | MODERAT | MISSENSE | gTg/gCg | V63A | PLES_08301 | . | . | . | . | . | C | . | . | . | . | . | . |
| NC_011770 | 903622 | C | A | 0 | 11 | 1 | 0 | NON_SYNONYMOU | MODERAT | MISSENSE | Cgt/Agt | R65S | PLES_08301 | . | . | . | . | . | A | . | . | . | . | . | . |
| NC_011770 | 903644 | G | C | 11 | 10 | 2 | 0 | NON_SYNONYMOU | MODERAT | MISSENSE | aGc/aCc | S72T | PLES_08301 | . | . | . | . | C | C | . | . | . | . | . | . |
| NC_011770 | 903665 | A | C | 11 | 10 | 2 | 0 | NON_SYNONYMOU | MODERAT | MISSENSE | gAt/gCt | D79A | PLES_08301 | . | . | . | . | C | C | . | . | . | . | . | . |
| NC_011770 | 903711 | T | G | 12 | 10 | 2 | 0 | NON_SYNONYMOU | MODERAT | MISSENSE | gaT/gaG | D94E | PLES_08301 | . | . | . | . | G | G | . | . | . | . | . | . |
| NC_011770 | 903712 | A | G | 12 | 10 | 2 | 0 | NON_SYNONYMOU | MODERAT | MISSENSE | Acc/Gcc | T95A | PLES_08301 | . | . | . | . | G | G | . | . | . | . | . | . |
| NC_011770 | 903784 | G | A | 13 | 10 | 2 | 0 | NON_SYNONYMOU | MODERAT | MISSENSE | gGa/gAa | G16E | PLES_08311 | . | . | . | . | A | A | . | . | . | . | . | . |
| NC_011770 | 903817 | T | C | 14 | 10 | 2 | 0 | NON_SYNONYMOU | MODERAT | MISSENSE | gTg/gCg | V27A | PLES_08311 | . | . | . | . | C | C | . | . | . | . | . | . |
| NC_011770 | 903894 | G | A | 21 | 8 | 2 | 0 | NON_SYNONYMOU | MODERAT | MISSENSE | Gtg/Atg | V53M | PLES_08311 | . | . | . | . | | | . | . | . | . | A | A |
| NC_011770 | 903895 | T | C | 21 | 8 | 2 | 0 | NON_SYNONYMOU | MODERAT | MISSENSE | gTg/gCg | V53A | PLES_08311 | . | . | . | . | | | . | . | . | . | C | C |
| NC_011770 | 903913 | C | A | 19 | 8 | 2 | 0 | NON_SYNONYMOU | MODERAT | MISSENSE | gCc/gAc | A59D | PLES_08311 | . | . | . | . | | | . | . | . | . | A | A |
| NC_011770 | 903918 | G | A | 18 | 8 | 4 | 0 | NON_SYNONYMOU | MODERAT | MISSENSE | Gtc/Atc | V61I | PLES_08311 | . | . | . | . | A | A | . | . | . | . | A | A |
| NC_011770 | 903960 | G | A | 15 | 10 | 2 | 0 | NON_SYNONYMOU | MODERAT | MISSENSE | Ggt/Agt | G75S | PLES_08311 | . | . | . | . | A | A | . | . | . | . | . | . |
| NC_011770 | 904161 | G | A | 12 | 9 | 1 | 0 | NON_SYNONYMOU | MODERAT | MISSENSE | Gat/Aat | D38N | PLES_08321 | . | . | . | . | | | . | . | A | . | . | . |
| NC_011770 | 904163 | T | G | 12 | 9 | 1 | 0 | NON_SYNONYMOU | MODERAT | MISSENSE | gaT/gaG | D38E | PLES_08321 | . | . | . | . | | | . | . | G | . | . | . |
| NC_011770 | 904227 | T | G | 0 | 10 | 2 | 0 | NON_SYNONYMOU | MODERAT | MISSENSE | Tct/Gct | S60A | PLES_08321 | . | . | . | . | G | G | . | . | . | . | . | . |
| NC_011770 | 904122 | G | A | 13 | 9 | 1 | 0 | NON_SYNONYMOU | MODERAT | MISSENSE | Gac/Aac | D25N | PLES_08321 | . | . | . | . | | A | . | . | | . | . | . |
| NC_011770 | 904143 | G | A | 13 | 9 | 1 | 0 | NON_SYNONYMOU | MODERAT | MISSENSE | Gct/Act | A32T | PLES_08321 | . | . | . | . | | A | . | . | | . | . | . |
| NC_011770 | 907959 | G | C | 85 | 0 | 2 | 0 | STOP_GAINED | HIGH | NONSENSE | taC/taG | Y212* | PLES_08341 | | | | | | | | | | | C | C |
| NC_011770 | 907519 | C | T | 93 | 0 | 2 | 0 | NON_SYNONYMOU | MODERAT | MISSENSE | cGc/cAc | R359H | PLES_08341 | | | | | | | | | | | T | T |
| NC_011770 | 908371 | C | T | 60 | 0 | 8 | 0 | NON_SYNONYMOU | MODERAT | MISSENSE | gGc/gAc | G75D | PLES_08341 | T | T | T | T | T | T | | | T | T | | |
| NC_011770 | 908978 | G | A | 70 | 0 | 2 | 0 | NON_SYNONYMOU | MODERAT | MISSENSE | aGg/aAg | R7K | PLES_08351 | | | | | | | | | | | A | A |
| NC_011770 | 909001 | G | C | 70 | 0 | 10 | 0 | NON_SYNONYMOU | MODERAT | MISSENSE | Gga/Cga | G15R | PLES_08351 | C | C | C | C | C | C | | | C | C | C | C |
| NC_011770 | 909154 | T | G | 73 | 0 | 10 | 0 | NON_SYNONYMOU | MODERAT | MISSENSE | Ttc/Gtc | F66V | PLES_08351 | G | G | G | G | G | G | | | G | G | G | G |
| NC_011770 | 909254 | T | C | 77 | 0 | 10 | 0 | NON_SYNONYMOU | MODERAT | MISSENSE | gTg/gCg | V99A | PLES_08351 | C | C | C | C | C | C | | | C | C | C | C |
| NC_011770 | 909386 | C | T | 78 | 0 | 1 | 0 | NON_SYNONYMOU | MODERAT | MISSENSE | gCa/gTa | A143V | PLES_08351 | T | | | | | | | | | | | |
| NC_011770 | 909592 | C | T | 72 | 0 | 1 | 0 | NON_SYNONYMOU | MODERAT | MISSENSE | Cgg/Tgg | R212W | PLES_08351 | | | | | | | | **T** | | | | |
| NC_011770 | 910461 | C | T | 56 | 0 | 1 | 0 | NON_SYNONYMOU | MODERAT | MISSENSE | gCg/gTg | A70V | PLES_08361 | | | | | | | | **T** | | | | |
| NC_011770 | 913271 | C | T | 77 | 0 | 4 | 0 | NON_SYNONYMOU | MODERAT | MISSENSE | Gcc/Acc | A285T | PLES_08391 | T | T | | | | | | | T | T | | |
| NC_011770 | 913154 | A | G | 66 | 0 | 1 | 0 | NON_SYNONYMOU | MODERAT | MISSENSE | Tgg/Cgg | W324R | PLES_08391 | G | | | | | | | | | | | |
| NC_011770 | 913493 | C | T | 79 | 0 | 1 | 0 | NON_SYNONYMOU | MODERAT | MISSENSE | Gaa/Aaa | E211K | PLES_08391 | | | T | | | | | | | | | |
| NC_011770 | 914271 | G | A | 76 | 0 | 2 | 0 | NON_SYNONYMOU | MODERAT | MISSENSE | Ccg/Tcg | P558S | PLES_08401 | | | | | | | A | A | | | | |
| NC_011770 | 916276 | C | T | 77 | 0 | 6 | 0 | NON_SYNONYMOU | MODERAT | MISSENSE | Cac/Tac | H77Y | PLES_08411 | | | T | T | T | T | | | | | T | T |
| NC_011770 | 916453 | G | A | 77 | 0 | 2 | 0 | NON_SYNONYMOU | MODERAT | MISSENSE | Gcg/Acg | A136T | PLES_08411 | | | A | A | | | | | | | | |
| NC_011770 | 916958 | G | A | 74 | 0 | 6 | 0 | NON_SYNONYMOU | MODERAT | MISSENSE | gGc/gAc | G304D | PLES_08411 | | | A | A | A | A | A | A | | | | |
| NC_011770 | 917439 | C | G | 76 | 0 | 1 | 0 | NON_SYNONYMOU | MODERAT | MISSENSE | tgC/tgG | C464W | PLES_08411 | | | | | | | | G | | | | |
| NC_011770 | 918862 | T | C | 92 | 0 | 1 | 0 | NON_SYNONYMOU | MODERAT | MISSENSE | gAc/gGc | D78G | PLES_08431 | | | | | | | C | | | | | |
| NC_011770 | 918902 | G | C | 93 | 0 | 2 | 0 | NON_SYNONYMOU | MODERAT | MISSENSE | Ccc/Gcc | P65A | PLES_08431 | | | | | | | | | | | C | C |
| NC_011770 | 920454 | A | G | 75 | 0 | 1 | 0 | NON_SYNONYMOU | MODERAT | MISSENSE | gTg/gCg | V260A | PLES_08451 | | | | | | | G | | | | | |
| NC_011770 | 920590 | T | C | 71 | 0 | 4 | 0 | NON_SYNONYMOU | MODERAT | MISSENSE | Acg/Gcg | T215A | PLES_08451 | | | C | C | C | C | | | | | | |
| NC_011770 | 924529 | A | G | 85 | 0 | 1 | 0 | NON_SYNONYMOU | MODERAT | MISSENSE | gTt/gCt | V164A | PLES_08491 | | | | | | | | G | | | | |
| NC_011770 | 924586 | A | G | 86 | 0 | 1 | 0 | NON_SYNONYMOU | MODERAT | MISSENSE | gTg/gCg | V145A | PLES_08491 | | | | | | | | G | | | | |
| NC_011770 | 926036 | C | T | 86 | 0 | 2 | 0 | NON_SYNONYMOU | MODERAT | MISSENSE | gCg/gTg | A43V | PLES_08511 | T | T | | | | | | | | | | |
| NC_011770 | 926143 | A | G | 84 | 0 | 1 | 0 | NON_SYNONYMOU | MODERAT | MISSENSE | Acc/Gcc | T79A | PLES_08511 | | | | | | | G | | | | | |
| NC_011770 | 926380 | C | T | 67 | 0 | 10 | 0 | NON_SYNONYMOU | MODERAT | MISSENSE | Ccc/Tcc | P158S | PLES_08511 | | | T | T | T | T | T | T | T | T | T | T |
| NC_011770 | 926395 | A | G | 73 | 0 | 10 | 0 | NON_SYNONYMOU | MODERAT | MISSENSE | Acg/Gcg | T163A | PLES_08511 | | | G | G | G | G | G | G | G | G | G | G |
| NC_011770 | 926396 | C | T | 73 | 0 | 2 | 0 | NON_SYNONYMOU | MODERAT | MISSENSE | aCg/aTg | T163M | PLES_08511 | T | T | | | | | | | | | | |
| NC_011770 | 926045 | A | G | 87 | 0 | 1 | 0 | NON_SYNONYMOU | MODERAT | MISSENSE | cAc/cGc | H46R | PLES_08511 | | | | | | | | G | | | | |
| NC_011770 | 927828 | TCGCCGAA | T | 93 | 0 | 1 | 0 | FRAME_SHIFT | HIGH | | gtgctttcgg | VLSA177 | PLES_08531 | | | | | | | | T | | | | |
| NC_011770 | 928314 | C | T | 78 | 0 | 4 | 0 | NON_SYNONYMOU | MODERAT | MISSENSE | Gct/Act | A19T | PLES_08531 | T | T | | | | | T | T | | | | |
| NC_011770 | 928568 | A | G | 91 | 0 | 2 | 0 | NON_SYNONYMOU | MODERAT | MISSENSE | Tac/Cac | Y689H | PLES_08541 | | | | | | | | | G | G | | |
| NC_011770 | 928885 | T | C | 81 | 0 | 6 | 0 | NON_SYNONYMOU | MODERAT | MISSENSE | cAg/cGg | Q583R | PLES_08541 | | | | | | | C | C | C | C | C | C |
| NC_011770 | 929434 | A | T | 76 | 0 | 6 | 0 | NON_SYNONYMOU | MODERAT | MISSENSE | gTg/gAg | V400E | PLES_08541 | T | T | T | T | T | T | | | | | | |
| NC_011770 | 930811 | T | C | 90 | 0 | 8 | 0 | NON_SYNONYMOU | MODERAT | MISSENSE | cAg/cGg | Q146R | PLES_08551 | | | C | C | C | C | C | C | | | C | C |
| NC_011770 | 930925 | C | T | 81 | 0 | 4 | 0 | NON_SYNONYMOU | MODERAT | MISSENSE | cGg/cAg | R108Q | PLES_08551 | | | | | | | T | T | | | T | T |

| | | | | | | | | | | | | | | | | | | | | | | | | | |
|---|---|---|---|---|---|---|---|---|---|---|---|---|---|---|---|---|---|---|---|---|---|---|---|---|---|
| NC_011770 | 931016 | C | T | 80 | 0 | 6 | 0 | NON_SYNONYMOU | MODERAT | MISSENSE | Gcc/Acc | A78T | PLES_08551 | T | T | T | T | T | T | | | | | | |
| NC_011770 | 931035 | G | T | 77 | 0 | 2 | 0 | NON_SYNONYMOU | MODERAT | MISSENSE | gaC/gaA | D71E | PLES_08551 | | | | | | | | | T | T | | |
| NC_011770 | 931220 | C | T | 77 | 0 | 2 | 0 | NON_SYNONYMOU | MODERAT | MISSENSE | Gcc/Acc | A10T | PLES_08551 | T | T | | | | | | | | | | |
| NC_011770 | 932239 | T | C | 87 | 0 | 2 | 0 | NON_SYNONYMOU | MODERAT | MISSENSE | aAg/aGg | K171R | PLES_08561 | | | | | | | C | C | | | | |
| NC_011770 | 933630 | A | C | 67 | 0 | 2 | 0 | NON_SYNONYMOU | MODERAT | MISSENSE | aAg/aCg | K154T | PLES_08571 | | | | | | | | | C | C | | |
| NC_011770 | 934337 | A | G | 73 | 0 | 2 | 0 | NON_SYNONYMOU | MODERAT | MISSENSE | Acc/Gcc | T169A | PLES_08581 | | | | | G | G | | | | | | |
| NC_011770 | 934529 | G | A | 87 | 0 | 1 | 0 | NON_SYNONYMOU | MODERAT | MISSENSE | Gag/Aag | E233K | PLES_08581 | | | | | | | A | | | | | |
| NC_011770 | 934554 | G | A | 87 | 0 | 6 | 0 | NON_SYNONYMOU | MODERAT | MISSENSE | cGg/cAg | R241Q | PLES_08581 | A | A | A | A | A | A | | | | | | |
| NC_011770 | 934680 | C | T | 86 | 0 | 6 | 0 | NON_SYNONYMOU | MODERAT | MISSENSE | gCc/gTc | A283V | PLES_08581 | | | T | T | T | T | | | T | T | | |
| NC_011770 | 934745 | A | T | 81 | 0 | 2 | 0 | NON_SYNONYMOU | MODERAT | MISSENSE | Agc/Tgc | S305C | PLES_08581 | | | | | | | | | T | T | | |
| NC_011770 | 934913 | G | A | 89 | 0 | 4 | 0 | NON_SYNONYMOU | MODERAT | MISSENSE | Ggc/Agc | G361S | PLES_08581 | | | A | A | A | A | | | | | | |
| NC_011770 | 934944 | C | T | 90 | 0 | 8 | 0 | NON_SYNONYMOU | MODERAT | MISSENSE | gCg/gTg | A371V | PLES_08581 | | | T | T | T | T | T | T | T | T | | |
| NC_011770 | 935472 | A | G | 68 | 0 | 2 | 0 | NON_SYNONYMOU | MODERAT | MISSENSE | Ata/Gta | I108V | PLES_08591 | | | | | | | G | G | | | | |
| NC_011770 | 938292 | G | A | 60 | 0 | 2 | 0 | NON_SYNONYMOU | MODERAT | MISSENSE | Gtc/Atc | V1048I | PLES_08591 | | | | | | | | | | | A | A |
| NC_011770 | 938317 | A | G | 62 | 0 | 12 | 0 | NON_SYNONYMOU | MODERAT | MISSENSE | cAt/cGt | H1056R | PLES_08591 | G | G | G | G | G | G | G | G | G | G | G | G |
| NC_011770 | 938412 | C | T | 69 | 0 | 1 | 0 | NON_SYNONYMOU | MODERAT | MISSENSE | Ccg/Tcg | P1088S | PLES_08591 | | | | | | | T | | | | | |
| NC_011770 | 938478 | G | A | 75 | 0 | 2 | 0 | NON_SYNONYMOU | MODERAT | MISSENSE | Gcg/Acg | A1110T | PLES_08591 | | | | | | | A | A | | | | |
| NC_011770 | 939229 | G | A | 66 | 0 | 2 | 0 | NON_SYNONYMOU | MODERAT | MISSENSE | aGc/aAc | S1360N | PLES_08591 | | | | | | | A | A | | | | |
| NC_011770 | 939243 | C | T | 65 | 0 | 4 | 0 | NON_SYNONYMOU | MODERAT | MISSENSE | Cgt/Tgt | R1365C | PLES_08591 | | | T | T | T | T | | | | | | |
| NC_011770 | 939262 | A | G | 68 | 0 | 1 | 0 | NON_SYNONYMOU | MODERAT | MISSENSE | cAg/cGg | Q1371R | PLES_08591 | | | | | | | G | | | | | |
| NC_011770 | 939876 | G | T | 21 | 0 | 2 | 0 | NON_SYNONYMOU | MODERAT | MISSENSE | Cgc/Agc | R31S | PLES_08601 | | | | | | | | | | | T | T |
| NC_011770 | 939951 | G | A | 16 | 0 | 2 | 0 | NON_SYNONYMOU | MODERAT | MISSENSE | Ccc/Tcc | P6S | PLES_08601 | | | | | | | | | A | A | | |
| NC_011770 | 942321 | T | C | 33 | 4 | 2 | 0 | NON_SYNONYMOU | MODERAT | MISSENSE | Aaa/Gaa | K61E | PLES_08631 | | | . | . | . | . | C | C | | | | |
| NC_011770 | 942398 | C | T | 35 | 4 | 2 | 0 | NON_SYNONYMOU | MODERAT | MISSENSE | cGc/cAc | R35H | PLES_08631 | | | . | . | . | . | T | T | | | | |
| NC_011770 | 942856 | C | CCG | 59 | 1 | 1 | 0 | FRAME_SHIFT | HIGH | | ggc/gCGgc | G404A? | PLES_08641 | | | | | | | . | CCG | | | | |
| NC_011770 | 942862 | GA | G | 60 | 1 | 1 | 0 | FRAME_SHIFT | HIGH | | tcc/ | S402 | PLES_08641 | | | | | | | G | . | | | | |
| NC_011770 | 942868 | GC | G | 71 | 0 | 2 | 0 | FRAME_SHIFT | HIGH | | gcc/ | A400 | PLES_08641 | | | | | | | G | G | | | | |
| NC_011770 | 942828 | A | C | 69 | 0 | 2 | 0 | NON_SYNONYMOU | MODERAT | MISSENSE | aaT/aaG | N413K | PLES_08641 | | | | | | | C | C | | | | |
| NC_011770 | 942838 | C | T | 69 | 0 | 4 | 0 | NON_SYNONYMOU | MODERAT | MISSENSE | gGa/gAa | G410E | PLES_08641 | | | | | | | | | T | T | T | T |
| NC_011770 | 942866 | T | C | 69 | 0 | 2 | 0 | NON_SYNONYMOU | MODERAT | MISSENSE | Aag/Gag | K401E | PLES_08641 | | | | | | | C | C | | | | |
| NC_011770 | 942911 | C | T | 70 | 0 | 4 | 0 | NON_SYNONYMOU | MODERAT | MISSENSE | Gcc/Acc | A386T | PLES_08641 | | | T | T | T | T | | | | | | |
| NC_011770 | 942968 | G | C | 80 | 0 | 12 | 0 | NON_SYNONYMOU | MODERAT | MISSENSE | Cta/Gta | L367V | PLES_08641 | C | C | C | C | C | C | C | C | C | C | C | C |
| NC_011770 | 943429 | T | G | 90 | 0 | 2 | 0 | NON_SYNONYMOU | MODERAT | MISSENSE | cAg/cCg | Q213P | PLES_08641 | | | | | | | G | G | | | | |
| NC_011770 | 943431 | G | C | 91 | 0 | 4 | 0 | NON_SYNONYMOU | MODERAT | MISSENSE | gaC/gaG | D212E | PLES_08641 | | | | | | | | | C | C | C | C |
| NC_011770 | 943519 | C | T | 92 | 0 | 4 | 0 | NON_SYNONYMOU | MODERAT | MISSENSE | cGc/cAc | R183H | PLES_08641 | | | | | | | | | T | T | T | T |
| NC_011770 | 943535 | G | A | 92 | 0 | 4 | 0 | NON_SYNONYMOU | MODERAT | MISSENSE | Ccc/Tcc | P178S | PLES_08641 | | | | | | | | | A | A | A | A |
| NC_011770 | 943634 | G | C | 93 | 0 | 6 | 0 | NON_SYNONYMOU | MODERAT | MISSENSE | Ctc/Gtc | L145V | PLES_08641 | | | C | C | C | C | | | C | C | | |
| NC_011770 | 943729 | C | T | 91 | 0 | 4 | 0 | NON_SYNONYMOU | MODERAT | MISSENSE | cGc/cAc | R113H | PLES_08641 | | | | | | | | | T | T | T | T |
| NC_011770 | 943895 | T | C | 80 | 0 | 4 | 0 | NON_SYNONYMOU | MODERAT | MISSENSE | Agc/Ggc | S58G | PLES_08641 | | | | | | | | | C | C | C | C |
| NC_011770 | 943949 | C | A | 74 | 0 | 4 | 0 | NON_SYNONYMOU | MODERAT | MISSENSE | Gcc/Tcc | A40S | PLES_08641 | | | | | | | | | A | A | A | A |
| NC_011770 | 943954 | C | T | 74 | 0 | 4 | 0 | NON_SYNONYMOU | MODERAT | MISSENSE | aGc/aAc | S38N | PLES_08641 | | | | | | | | | T | T | T | T |
| NC_011770 | 943975 | C | T | 75 | 0 | 4 | 0 | NON_SYNONYMOU | MODERAT | MISSENSE | aGa/aAa | R31K | PLES_08641 | | | | | | | | | T | T | T | T |
| NC_011770 | 944489 | C | T | 82 | 0 | 2 | 0 | NON_SYNONYMOU | MODERAT | MISSENSE | Ccg/Tcg | P34S | PLES_08651 | T | T | | | | | | | | | | |
| NC_011770 | 944514 | C | T | 78 | 0 | 1 | 0 | NON_SYNONYMOU | MODERAT | MISSENSE | gCg/gTg | A42V | PLES_08651 | | | | | | | T | | | | | |
| NC_011770 | 945050 | A | G | 83 | 0 | 12 | 0 | NON_SYNONYMOU | MODERAT | MISSENSE | gAc/gGc | D40G | PLES_08661 | G | G | G | G | G | G | G | G | G | G | G | G |
| NC_011770 | 945491 | C | T | 75 | 0 | 4 | 0 | NON_SYNONYMOU | MODERAT | MISSENSE | gCc/gTc | A187V | PLES_08661 | | | T | T | T | T | | | | | | |
| NC_011770 | 945989 | A | AC | 70 | 0 | 1 | 0 | FRAME_SHIFT | HIGH | | ctg/cCtg | L81P? | PLES_08671 | | | | AC | | | | | | | | |
| NC_011770 | 945902 | A | G | 70 | 0 | 1 | 0 | NON_SYNONYMOU | MODERAT | MISSENSE | Agc/Ggc | S50G | PLES_08671 | | | | | | | G | | | | | |
| NC_011770 | 946017 | T | G | 50 | 2 | 4 | 0 | NON_SYNONYMOU | MODERAT | MISSENSE | gTc/gGc | V88G | PLES_08671 | | | G | G | G | G | | | | | . | . |
| NC_011770 | 946040 | C | T | 51 | 2 | 4 | 0 | NON_SYNONYMOU | MODERAT | MISSENSE | Cgg/Tgg | R96W | PLES_08671 | | | | | | | T | T | T | T | . | . |
| NC_011770 | 947470 | G | A | 83 | 0 | 2 | 0 | NON_SYNONYMOU | MODERAT | MISSENSE | aCc/aTc | T155I | PLES_08691 | | | | | | | A | A | | | | |
| NC_011770 | 947708 | C | T | 84 | 0 | 2 | 0 | NON_SYNONYMOU | MODERAT | MISSENSE | Gcg/Acg | A76T | PLES_08691 | | | | | | | | | | | T | T |
| NC_011770 | 948740 | G | A | 88 | 0 | 2 | 0 | NON_SYNONYMOU | MODERAT | MISSENSE | Cat/Tat | H210Y | PLES_08701 | | | | | | | | | | | A | A |
| NC_011770 | 948878 | A | G | 81 | 0 | 10 | 0 | NON_SYNONYMOU | MODERAT | MISSENSE | Ttc/Ctc | F164L | PLES_08701 | G | G | G | G | G | G | | | G | G | G | G |
| NC_011770 | 948998 | T | C | 77 | 0 | 2 | 0 | NON_SYNONYMOU | MODERAT | MISSENSE | Agc/Ggc | S124G | PLES_08701 | | | | | | | | | | | C | C |
| NC_011770 | 949009 | T | G | 78 | 0 | 10 | 0 | NON_SYNONYMOU | MODERAT | MISSENSE | aAg/aCg | K120T | PLES_08701 | G | G | G | G | G | G | | | G | G | G | G |
| NC_011770 | 949235 | T | C | 94 | 0 | 1 | 0 | NON_SYNONYMOU | MODERAT | MISSENSE | Atg/Gtg | M45V | PLES_08701 | | | | | | | C | | | | | |
| NC_011770 | 949307 | G | C | 85 | 0 | 12 | 0 | NON_SYNONYMOU | MODERAT | MISSENSE | Ccc/Gcc | P21A | PLES_08701 | C | C | C | C | C | C | C | C | C | C | C | C |
| NC_011770 | 948179 | C | T | 93 | 0 | 1 | 0 | NON_SYNONYMOU | MODERAT | MISSENSE | Ggc/Agc | G397S | PLES_08701 | | | | | | | | T | | | | |
| NC_011770 | 948575 | T | C | 98 | 0 | 1 | 0 | NON_SYNONYMOU | MODERAT | MISSENSE | Aac/Gac | N265D | PLES_08701 | | | | | | | | C | | | | |
| NC_011770 | 949392 | A | G | 89 | 0 | 12 | 0 | NON_SYNONYMOU | MODERAT | MISSENSE | cTg/cCg | L238P | PLES_08711 | G | G | G | G | G | G | G | G | G | G | G | G |
| NC_011770 | 949632 | G | A | 83 | 0 | 2 | 0 | NON_SYNONYMOU | MODERAT | MISSENSE | gCc/gTc | A158V | PLES_08711 | | | | | | | | | | | A | A |

| | | | | | | | | | | | | | | | | | | | | | | | | | |
|---|---|---|---|---|---|---|---|---|---|---|---|---|---|---|---|---|---|---|---|---|---|---|---|---|---|
| NC_011770 | 950614 | A | C | 81 | 0 | 2 | 0 | NON_SYNONYMOU | MODERAT | MISSENSE | Acc/Ccc | T52P | PLES_08721 | | | | | | | C | C | | | | |
| NC_011770 | 951897 | G | A | 76 | 0 | 2 | 0 | STOP_GAINED | HIGH | NONSENSE | Cag/Tag | Q169* | PLES_08731 | | | | | | | | | | | A | A |
| NC_011770 | 951107 | T | C | 75 | 0 | 12 | 0 | NON_SYNONYMOU | MODERAT | MISSENSE | cAg/cGg | Q432R | PLES_08731 | C | C | C | C | C | C | C | C | C | C | C | C |
| NC_011770 | 951638 | C | A | 83 | 0 | 2 | 0 | NON_SYNONYMOU | MODERAT | MISSENSE | cGc/cTc | R255L | PLES_08731 | | | | | | | A | A | | | | |
| NC_011770 | 953135 | G | T | 69 | 0 | 2 | 0 | NON_SYNONYMOU | MODERAT | MISSENSE | gaC/gaA | D242E | PLES_08741 | | | | | | | | | T | T | | |
| NC_011770 | 954807 | G | A | 69 | 0 | 2 | 0 | NON_SYNONYMOU | MODERAT | MISSENSE | Cgc/Tgc | R313C | PLES_08761 | A | A | | | | | | | | | | |
| NC_011770 | 955982 | C | T | 78 | 0 | 2 | 0 | NON_SYNONYMOU | MODERAT | MISSENSE | Ggc/Agc | G340S | PLES_08771 | | | | | | | T | T | | | | |
| NC_011770 | 956379 | T | G | 71 | 0 | 2 | 0 | NON_SYNONYMOU | MODERAT | MISSENSE | gaA/gaC | E207D | PLES_08771 | | | | | | | G | G | | | | |
| NC_011770 | 956492 | A | G | 71 | 0 | 12 | 0 | NON_SYNONYMOU | MODERAT | MISSENSE | Tgt/Cgt | C170R | PLES_08771 | G | G | G | G | G | G | G | G | G | G | G | G |
| NC_011770 | 956504 | C | T | 69 | 0 | 8 | 0 | NON_SYNONYMOU | MODERAT | MISSENSE | Gcc/Acc | A166T | PLES_08771 | T | T | T | T | T | T | | | T | T | | |
| NC_011770 | 956848 | C | T | 62 | 0 | 2 | 0 | NON_SYNONYMOU | MODERAT | MISSENSE | gGc/gAc | G51D | PLES_08771 | | | | | T | T | | | | | | |
| NC_011770 | 956111 | A | G | 84 | 0 | 1 | 0 | NON_SYNONYMOU | MODERAT | MISSENSE | Tgt/Cgt | C297R | PLES_08771 | | | | | | | | G | | | | |
| NC_011770 | 959215 | C | G | 61 | 0 | 4 | 0 | NON_SYNONYMOU | MODERAT | MISSENSE | gCc/gGc | A96G | PLES_08801 | | | G | G | G | G | | | | | | |
| NC_011770 | 960946 | C | G | 39 | 8 | 4 | 0 | NON_SYNONYMOU | MODERAT | MISSENSE | gaG/gaC | E172D | PLES_08821 | . | . | G | G | G | G | . | . | . | . | . | . |
| NC_011770 | 961371 | C | T | 41 | 7 | 4 | 0 | NON_SYNONYMOU | MODERAT | MISSENSE | Gtg/Atg | V31M | PLES_08821 | . | . | T | T | T | T | . | . | . | . | . | . |
| NC_011770 | 964845 | G | A | 51 | 0 | 2 | 0 | NON_SYNONYMOU | MODERAT | MISSENSE | cCc/cTc | P62L | PLES_08861 | | | | | | | A | A | | | | |
| NC_011770 | 965240 | C | CT | 67 | 0 | 12 | 0 | FRAME_SHIFT+SPL | HIGH | | tga/tgaA | *254*? | PLES_08871 | CT | CT | CT | CT | CT | CT | CT | CT | CT | CT | CT | CT |
| NC_011770 | 965715 | C | A | 54 | 0 | 2 | 0 | NON_SYNONYMOU | MODERAT | MISSENSE | caG/caT | Q96H | PLES_08871 | | | | | | | | | A | A | | |
| NC_011770 | 965804 | G | A | 68 | 0 | 2 | 0 | NON_SYNONYMOU | MODERAT | MISSENSE | Cac/Tac | H67Y | PLES_08871 | A | A | | | | | | | | | | |
| NC_011770 | 966160 | C | T | 88 | 0 | 6 | 0 | NON_SYNONYMOU | MODERAT | MISSENSE | Gtc/Atc | V409I | PLES_08881 | T | T | | | | | | | T | T | T | T |
| NC_011770 | 966766 | C | T | 77 | 0 | 1 | 0 | NON_SYNONYMOU | MODERAT | MISSENSE | Gac/Aac | D207N | PLES_08881 | | | | | | | T | | | | | |
| NC_011770 | 966793 | T | C | 71 | 0 | 6 | 0 | NON_SYNONYMOU | MODERAT | MISSENSE | Acc/Gcc | T198A | PLES_08881 | C | C | | | | | C | C | | | C | C |
| NC_011770 | 966795 | G | T | 71 | 0 | 6 | 0 | NON_SYNONYMOU | MODERAT | MISSENSE | gCc/gAc | A197D | PLES_08881 | T | T | | | | | T | T | | | T | T |
| NC_011770 | 967290 | T | A | 73 | 0 | 4 | 0 | NON_SYNONYMOU | MODERAT | MISSENSE | cAg/cTg | Q32L | PLES_08881 | | | A | A | A | A | | | | | | |
| NC_011770 | 967345 | C | T | 75 | 0 | 6 | 0 | NON_SYNONYMOU | MODERAT | MISSENSE | Gcg/Acg | A14T | PLES_08881 | | | T | T | T | T | T | T | | | | |
| NC_011770 | 966306 | G | A | 88 | 0 | 1 | 0 | NON_SYNONYMOU | MODERAT | MISSENSE | gCg/gTg | A360V | PLES_08881 | | | | | | | | | | | | A |
| NC_011770 | 967661 | G | T | 76 | 0 | 4 | 0 | NON_SYNONYMOU | MODERAT | MISSENSE | gCc/gAc | A186D | PLES_08891 | T | T | | | | | | | | | T | T |
| NC_011770 | 967677 | T | C | 78 | 0 | 4 | 0 | NON_SYNONYMOU | MODERAT | MISSENSE | Atc/Gtc | I181V | PLES_08891 | C | C | | | | | | | | | C | C |
| NC_011770 | 967760 | C | T | 80 | 0 | 4 | 0 | NON_SYNONYMOU | MODERAT | MISSENSE | gGc/gAc | G153D | PLES_08891 | T | T | | | | | | | | | T | T |
| NC_011770 | 967806 | C | T | 82 | 0 | 4 | 0 | NON_SYNONYMOU | MODERAT | MISSENSE | Gag/Aag | E138K | PLES_08891 | T | T | | | | | | | | | T | T |
| NC_011770 | 967842 | C | A | 77 | 0 | 4 | 0 | NON_SYNONYMOU | MODERAT | MISSENSE | Gcc/Tcc | A126S | PLES_08891 | A | A | | | | | | | | | A | A |
| NC_011770 | 967866 | C | T | 75 | 0 | 4 | 0 | NON_SYNONYMOU | MODERAT | MISSENSE | Gcg/Acg | A118T | PLES_08891 | T | T | | | | | | | | | T | T |
| NC_011770 | 967958 | T | C | 72 | 0 | 2 | 0 | NON_SYNONYMOU | MODERAT | MISSENSE | aAg/aGg | K87R | PLES_08891 | | | | | | | | | | | C | C |
| NC_011770 | 968112 | C | G | 76 | 0 | 2 | 0 | NON_SYNONYMOU | MODERAT | MISSENSE | Gaa/Caa | E36Q | PLES_08891 | | | | | | | | | | | G | G |
| NC_011770 | 980079 | G | GACC | 86 | 0 | 6 | 0 | CODON_INSERTIOI | MODERATE | | taa/taaGG | *992*G | PLES_08981 | | | GACC | GACC | GACC | GACC | | | GACC | GACC | | |
| NC_011770 | 981510 | A | C | 69 | 0 | 2 | 0 | NON_SYNONYMOU | MODERAT | MISSENSE | Tcc/Gcc | S516A | PLES_08981 | | | | | | | | | C | C | | |
| NC_011770 | 981577 | G | C | 68 | 0 | 2 | 0 | NON_SYNONYMOU | MODERAT | MISSENSE | gaC/gaG | D493E | PLES_08981 | | | | | | | | | C | C | | |
| NC_011770 | 981584 | T | C | 68 | 0 | 2 | 0 | NON_SYNONYMOU | MODERAT | MISSENSE | cAc/cGc | H491R | PLES_08981 | | | | | | | | | C | C | | |
| NC_011770 | 981591 | G | T | 70 | 0 | 2 | 0 | NON_SYNONYMOU | MODERAT | MISSENSE | Cgt/Agt | R489S | PLES_08981 | | | | | | | | | | | T | T |
| NC_011770 | 981731 | T | C | 73 | 0 | 2 | 0 | NON_SYNONYMOU | MODERAT | MISSENSE | gAc/gGc | D442G | PLES_08981 | | | | | | | | | C | C | | |
| NC_011770 | 981734 | T | C | 75 | 0 | 2 | 0 | NON_SYNONYMOU | MODERAT | MISSENSE | cAg/cGg | Q441R | PLES_08981 | | | | | | | | | C | C | | |
| NC_011770 | 981776 | G | C | 76 | 0 | 2 | 0 | NON_SYNONYMOU | MODERAT | MISSENSE | tCc/tGc | S427C | PLES_08981 | | | | | | | | | C | C | | |
| NC_011770 | 982080 | G | T | 76 | 0 | 2 | 0 | NON_SYNONYMOU | MODERAT | MISSENSE | Cct/Act | P326T | PLES_08981 | T | T | | | | | | | | | | |
| NC_011770 | 982316 | G | A | 73 | 0 | 2 | 0 | NON_SYNONYMOU | MODERAT | MISSENSE | gCg/gTg | A247V | PLES_08981 | | | | | | | | | A | A | | |
| NC_011770 | 982329 | C | A | 73 | 0 | 2 | 0 | NON_SYNONYMOU | MODERAT | MISSENSE | Gtg/Ttg | V243L | PLES_08981 | | | | | | | | | A | A | | |
| NC_011770 | 982330 | C | T | 73 | 0 | 2 | 0 | NON_SYNONYMOU | MODERAT | MISSENSE | atG/atA | M242I | PLES_08981 | | | | | | | | | T | T | | |
| NC_011770 | 982469 | G | C | 62 | 0 | 2 | 0 | NON_SYNONYMOU | MODERAT | MISSENSE | tCc/tGc | S196C | PLES_08981 | | | | | | | | | C | C | | |
| NC_011770 | 982522 | A | C | 60 | 0 | 2 | 0 | NON_SYNONYMOU | MODERAT | MISSENSE | aaT/aaG | N178K | PLES_08981 | | | | | | | | | C | C | | |
| NC_011770 | 982524 | T | C | 58 | 0 | 2 | 0 | NON_SYNONYMOU | MODERAT | MISSENSE | Aat/Gat | N178D | PLES_08981 | | | | | | | | | C | C | | |
| NC_011770 | 982544 | C | T | 54 | 0 | 2 | 0 | NON_SYNONYMOU | MODERAT | MISSENSE | aGc/aAc | S171N | PLES_08981 | | | | | | | | | T | T | | |
| NC_011770 | 982545 | T | C | 54 | 0 | 4 | 0 | NON_SYNONYMOU | MODERAT | MISSENSE | Agc/Ggc | S171G | PLES_08981 | | | | | | | | | C | C | C | C |
| NC_011770 | 982556 | G | C | 53 | 0 | 2 | 0 | NON_SYNONYMOU | MODERAT | MISSENSE | gCc/gGc | A167G | PLES_08981 | | | | | | | | | C | C | | |
| NC_011770 | 982557 | C | T | 54 | 0 | 2 | 0 | NON_SYNONYMOU | MODERAT | MISSENSE | Gcc/Acc | A167T | PLES_08981 | | | | | | | | | T | T | | |
| NC_011770 | 982568 | C | T | 55 | 0 | 2 | 0 | NON_SYNONYMOU | MODERAT | MISSENSE | aGt/aAt | S163N | PLES_08981 | | | | | | | | | T | T | | |
| NC_011770 | 982685 | G | A | 47 | 0 | 2 | 0 | NON_SYNONYMOU | MODERAT | MISSENSE | gCg/gTg | A124V | PLES_08981 | | | | | | | | | A | A | | |
| NC_011770 | 982697 | A | G | 45 | 0 | 2 | 0 | NON_SYNONYMOU | MODERAT | MISSENSE | gTc/gCc | V120A | PLES_08981 | G | G | | | | | | | | | | |
| NC_011770 | 982729 | C | A | 43 | 0 | 4 | 0 | NON_SYNONYMOU | MODERAT | MISSENSE | caG/caT | Q109H | PLES_08981 | | | A | A | A | A | | | | | | |
| NC_011770 | 982746 | A | C | 42 | 0 | 2 | 0 | NON_SYNONYMOU | MODERAT | MISSENSE | Ttg/Gtg | L104V | PLES_08981 | | | | | | | | | C | C | | |
| NC_011770 | 982770 | T | C | 38 | 0 | 6 | 0 | NON_SYNONYMOU | MODERAT | MISSENSE | Atc/Gtc | I96V | PLES_08981 | | | C | C | C | C | | | | | C | C |
| NC_011770 | 982775 | C | T | 38 | 0 | 2 | 0 | NON_SYNONYMOU | MODERAT | MISSENSE | cGg/cAg | R94Q | PLES_08981 | | | | | | | | | T | T | | |
| NC_011770 | 982788 | C | T | 37 | 0 | 4 | 0 | NON_SYNONYMOU | MODERAT | MISSENSE | Gac/Aac | D90N | PLES_08981 | | | T | T | T | T | | | | | | |
| NC_011770 | 982889 | A | C | 23 | 0 | 2 | 0 | NON_SYNONYMOU | MODERAT | MISSENSE | cTg/cGg | L56R | PLES_08981 | | | | | | | | | C | C | | |

| | | | | | | | | | | | | | | | | | | | | | | | | | |
|---|---|---|---|---|---|---|---|---|---|---|---|---|---|---|---|---|---|---|---|---|---|---|---|---|---|
| NC_011770 | 982891 | C | G | 23 | 0 | 4 | 0 | NON_SYNONYMOU | MODERAT | MISSENSE | gaG/gaC | E55D | PLES_08981 | | | | | | | G | G | | | T | T |
| NC_011770 | 982897 | C | G | 22 | 0 | 2 | 0 | NON_SYNONYMOU | MODERAT | MISSENSE | caG/caC | Q53H | PLES_08981 | | | | | | | G | G | | | | |
| NC_011770 | 982898 | T | C | 23 | 0 | 12 | 0 | NON_SYNONYMOU | MODERAT | MISSENSE | cAg/cGg | Q53R | PLES_08981 | C | C | C | C | C | C | C | C | C | C | C | C |
| NC_011770 | 983493 | A | G | 55 | 0 | 4 | 0 | NON_SYNONYMOU | MODERAT | MISSENSE | gTc/gCc | V117A | PLES_09001 | | | G | G | G | G | | | | | | |
| NC_011770 | 983505 | A | G | 55 | 0 | 2 | 0 | NON_SYNONYMOU | MODERAT | MISSENSE | gTc/gCc | V113A | PLES_09001 | | | | | | | | | G | G | | |
| NC_011770 | 983584 | T | C | 51 | 0 | 4 | 0 | NON_SYNONYMOU | MODERAT | MISSENSE | Acc/Gcc | T87A | PLES_09001 | C | C | | | | | | | C | C | | |
| NC_011770 | 983763 | G | A | 50 | 0 | 4 | 0 | NON_SYNONYMOU | MODERAT | MISSENSE | tCg/tTg | S27L | PLES_09001 | A | A | | | | | | | A | A | | |
| NC_011770 | 983796 | C | G | 52 | 0 | 2 | 0 | NON_SYNONYMOU | MODERAT | MISSENSE | gGc/gCc | G16A | PLES_09001 | | | | | | | | | G | G | | |
| NC_011770 | 983990 | C | A | 29 | 2 | 8 | 0 | INTRAGENIC | MODIFIER | | | | PLES_09011 | | | A | A | A | A | . | . | A | A | A | A |
| NC_011770 | 984001 | A | G | 38 | 0 | 10 | 0 | INTRAGENIC | MODIFIER | | | | PLES_09011 | | | G | G | G | G | G | G | G | G | G | G |
| NC_011770 | 984028 | C | G | 40 | 0 | 10 | 0 | INTRAGENIC | MODIFIER | | | | PLES_09011 | | | G | G | G | G | G | G | G | G | G | G |
| NC_011770 | 984034 | G | A | 44 | 0 | 2 | 0 | INTRAGENIC | MODIFIER | | | | PLES_09011 | A | A | | | | | | | | | | |
| NC_011770 | 984050 | C | T | 40 | 0 | 12 | 0 | INTRAGENIC | MODIFIER | | | | PLES_09011 | T | T | T | T | T | T | T | T | T | T | T | T |
| NC_011770 | 984054 | T | C | 41 | 0 | 12 | 0 | INTRAGENIC | MODIFIER | | | | PLES_09011 | C | C | C | C | C | C | C | C | C | C | C | C |
| NC_011770 | 984076 | C | T | 42 | 0 | 12 | 0 | INTRAGENIC | MODIFIER | | | | PLES_09011 | T | T | T | T | T | T | T | T | T | T | T | T |
| NC_011770 | 984082 | A | G | 41 | 0 | 12 | 0 | INTRAGENIC | MODIFIER | | | | PLES_09011 | G | G | G | G | G | G | G | G | G | G | G | G |
| NC_011770 | 984086 | C | T | 44 | 0 | 12 | 0 | INTRAGENIC | MODIFIER | | | | PLES_09011 | T | T | T | T | T | T | T | T | T | T | T | T |
| NC_011770 | 984091 | T | C | 43 | 0 | 12 | 0 | INTRAGENIC | MODIFIER | | | | PLES_09011 | C | C | C | C | C | C | C | C | C | C | C | C |
| NC_011770 | 984100 | C | T | 43 | 0 | 12 | 0 | INTRAGENIC | MODIFIER | | | | PLES_09011 | T | T | T | T | T | T | T | T | T | T | T | T |
| NC_011770 | 984173 | A | G | 49 | 0 | 10 | 0 | INTRAGENIC | MODIFIER | | | | PLES_09011 | | | G | G | G | G | G | G | G | G | G | G |
| NC_011770 | 984176 | G | C | 50 | 0 | 10 | 0 | INTRAGENIC | MODIFIER | | | | PLES_09011 | | | C | C | C | C | C | C | C | C | C | C |
| NC_011770 | 984203 | G | A | 51 | 0 | 10 | 0 | INTRAGENIC | MODIFIER | | | | PLES_09011 | | | A | A | A | A | A | A | A | A | A | A |
| NC_011770 | 984249 | A | G | 52 | 0 | 10 | 0 | INTRAGENIC | MODIFIER | | | | PLES_09011 | | | G | G | G | G | G | G | G | G | G | G |
| NC_011770 | 984468 | C | T | 76 | 0 | 6 | 0 | NON_SYNONYMOU | MODERAT | MISSENSE | Gcg/Acg | A157T | PLES_09021 | | | | | | | T | T | T | T | T | T |
| NC_011770 | 984598 | C | A | 78 | 0 | 4 | 0 | NON_SYNONYMOU | MODERAT | MISSENSE | caG/caT | Q113H | PLES_09021 | | | | | | | | | A | A | A | A |
| NC_011770 | 984675 | T | C | 90 | 0 | 4 | 0 | NON_SYNONYMOU | MODERAT | MISSENSE | Aag/Gag | K88E | PLES_09021 | | | | | | | | | C | C | C | C |
| NC_011770 | 984686 | C | T | 89 | 0 | 2 | 0 | NON_SYNONYMOU | MODERAT | MISSENSE | cGc/cAc | R84H | PLES_09021 | | | | | | | | | T | T | | |
| NC_011770 | 984725 | C | T | 92 | 0 | 2 | 0 | NON_SYNONYMOU | MODERAT | MISSENSE | aGc/aAc | S71N | PLES_09021 | | | | | | | | | T | T | | |
| NC_011770 | 984905 | A | G | 80 | 0 | 6 | 0 | NON_SYNONYMOU | MODERAT | MISSENSE | cTg/cCg | L11P | PLES_09021 | | | G | G | G | G | | | G | G | | |
| NC_011770 | 984933 | A | G | 86 | 0 | 6 | 0 | NON_SYNONYMOU | MODERAT | MISSENSE | Tct/Cct | S2P | PLES_09021 | | | G | G | G | G | | | G | G | | |
| NC_011770 | 985373 | G | A | 72 | 0 | 6 | 0 | NON_SYNONYMOU | MODERAT | MISSENSE | aGc/aAc | S70N | PLES_09031 | | | A | A | A | A | | | A | A | | |
| NC_011770 | 985432 | G | A | 79 | 0 | 2 | 0 | NON_SYNONYMOU | MODERAT | MISSENSE | Gtc/Atc | V90I | PLES_09031 | | | | | | | | | A | A | | |
| NC_011770 | 986132 | C | T | 52 | 0 | 2 | 0 | NON_SYNONYMOU | MODERAT | MISSENSE | gCg/gTg | A323V | PLES_09031 | T | T | | | | | | | | | | |
| NC_011770 | 986161 | C | T | 52 | 0 | 2 | 0 | NON_SYNONYMOU | MODERAT | MISSENSE | Ccc/Tcc | P333S | PLES_09031 | | | | | | | | | T | T | | |
| NC_011770 | 985642 | G | A | 77 | 0 | 1 | 0 | NON_SYNONYMOU | MODERAT | MISSENSE | Gtg/Atg | V160M | PLES_09031 | | | | | | | | | | | | A |
| NC_011770 | 986827 | A | T | 62 | 0 | 2 | 0 | NON_SYNONYMOU | MODERAT | MISSENSE | aAc/aTc | N3I | PLES_09041 | | | | | | | T | T | | | | |
| NC_011770 | 987222 | G | A | 82 | 0 | 4 | 0 | NON_SYNONYMOU | MODERAT | MISSENSE | Ggc/Agc | G135S | PLES_09041 | | | A | A | A | A | | | | | | |
| NC_011770 | 987354 | A | G | 84 | 0 | 10 | 0 | NON_SYNONYMOU | MODERAT | MISSENSE | Acc/Gcc | T179A | PLES_09041 | G | G | G | G | G | G | | | G | G | G | G |
| NC_011770 | 987417 | A | G | 81 | 0 | 12 | 0 | NON_SYNONYMOU | MODERAT | MISSENSE | Aac/Gac | N200D | PLES_09041 | G | G | G | G | G | G | G | G | G | G | G | G |
| NC_011770 | 987495 | A | G | 85 | 0 | 10 | 0 | NON_SYNONYMOU | MODERAT | MISSENSE | Acg/Gcg | T226A | PLES_09041 | | | G | G | G | G | G | G | G | G | G | G |
| NC_011770 | 987552 | A | G | 90 | 0 | 2 | 0 | NON_SYNONYMOU | MODERAT | MISSENSE | Atc/Gtc | I245V | PLES_09041 | | | | | | | | | | | G | G |
| NC_011770 | 988069 | T | C | 96 | 0 | 10 | 0 | NON_SYNONYMOU | MODERAT | MISSENSE | gTc/gCc | V417A | PLES_09041 | | | C | C | C | C | C | C | C | C | C | C |
| NC_011770 | 988204 | A | G | 94 | 0 | 2 | 0 | NON_SYNONYMOU | MODERAT | MISSENSE | aAt/aGt | N462S | PLES_09041 | | | | | | | G | G | | | | |
| NC_011770 | 988279 | G | A | 92 | 0 | 4 | 0 | NON_SYNONYMOU | MODERAT | MISSENSE | aGg/aAg | R487K | PLES_09041 | | | A | A | A | A | | | | | | |
| NC_011770 | 986983 | C | T | 73 | 0 | 1 | 0 | NON_SYNONYMOU | MODERAT | MISSENSE | gCg/gTg | A55V | PLES_09041 | | | | | | | | T | | | | |
| NC_011770 | 988637 | A | AG | 83 | 0 | 1 | 0 | FRAME_SHIFT | HIGH | | ggc/ggGc | G120G? | PLES_09051 | | AG | | | | | | | | | | |
| NC_011770 | 988524 | CG | C | 81 | 0 | 3 | 0 | FRAME_SHIFT | HIGH | | ggg/ | G82 | PLES_09051 | | | | | | | C | | | | TG | TG |
| NC_011770 | 988364 | C | A | 91 | 0 | 4 | 0 | NON_SYNONYMOU | MODERAT | MISSENSE | gCc/gAc | A27D | PLES_09051 | | | A | A | A | A | | | | | | |
| NC_011770 | 988600 | A | C | 80 | 0 | 6 | 0 | NON_SYNONYMOU | MODERAT | MISSENSE | Agc/Cgc | S106R | PLES_09051 | | | C | C | C | C | | | | | C | C |
| NC_011770 | 988613 | T | C | 79 | 0 | 6 | 0 | NON_SYNONYMOU | MODERAT | MISSENSE | cTc/cCc | L110P | PLES_09051 | | | C | C | C | C | | | | | C | C |
| NC_011770 | 988652 | T | C | 81 | 0 | 8 | 0 | NON_SYNONYMOU | MODERAT | MISSENSE | gTc/gCc | V123A | PLES_09051 | C | C | C | C | C | C | | | | | C | C |
| NC_011770 | 988727 | A | G | 77 | 0 | 12 | 0 | NON_SYNONYMOU | MODERAT | MISSENSE | aAc/aGc | N148S | PLES_09051 | G | G | G | G | G | G | G | G | G | G | G | G |
| NC_011770 | 988844 | G | A | 79 | 0 | 2 | 0 | NON_SYNONYMOU | MODERAT | MISSENSE | cGc/cAc | R187H | PLES_09051 | A | A | | | | | | | | | | |
| NC_011770 | 988859 | C | T | 83 | 0 | 2 | 0 | NON_SYNONYMOU | MODERAT | MISSENSE | gCc/gTc | A192V | PLES_09051 | T | T | | | | | | | | | | |
| NC_011770 | 989407 | A | C | 68 | 0 | 2 | 0 | NON_SYNONYMOU | MODERAT | MISSENSE | gaA/gaC | E128D | PLES_09061 | | | | | | | | | | | C | C |
| NC_011770 | 989408 | G | C | 71 | 0 | 2 | 0 | NON_SYNONYMOU | MODERAT | MISSENSE | Gag/Cag | E129Q | PLES_09061 | | | | | | | | | | | C | C |
| NC_011770 | 989682 | C | T | 73 | 0 | 2 | 0 | NON_SYNONYMOU | MODERAT | MISSENSE | tCc/tTc | S220F | PLES_09061 | | | | | | | | | T | T | | |
| NC_011770 | 991135 | A | G | 97 | 0 | 2 | 0 | NON_SYNONYMOU | MODERAT | MISSENSE | Atc/Gtc | I6V | PLES_09081 | | | | | | | | | G | G | | |
| NC_011770 | 991240 | C | T | 86 | 0 | 2 | 0 | NON_SYNONYMOU | MODERAT | MISSENSE | Cgc/Tgc | R41C | PLES_09081 | | | | | | | T | T | | | | |
| NC_011770 | 991270 | G | A | 88 | 0 | 2 | 0 | NON_SYNONYMOU | MODERAT | MISSENSE | Gta/Ata | V51I | PLES_09081 | A | A | | | | | | | | | | |
| NC_011770 | 991304 | C | T | 81 | 0 | 2 | 0 | NON_SYNONYMOU | MODERAT | MISSENSE | gCg/gTg | A62V | PLES_09081 | | | | | | | | | | | T | T |
| NC_011770 | 992286 | C | G | 75 | 0 | 10 | 0 | NON_SYNONYMOU | MODERAT | MISSENSE | Gcg/Ccg | A41P | PLES_09091 | G | G | G | G | G | G | G | G | G | G | | |

| | | | | | | | | | | | | | | | | | | | | | | | | | |
|---|---|---|---|---|---|---|---|---|---|---|---|---|---|---|---|---|---|---|---|---|---|---|---|---|---|
| NC_011770 | 992325 | A | G | 74 | 0 | 2 | 0 | NON_SYNONYMOU | MODERAT | MISSENSE | Tgg/Cgg | W28R | PLES_09091 | | | | | | | G | G | | | | |
| NC_011770 | 992387 | G | A | 78 | 0 | 2 | 0 | NON_SYNONYMOU | MODERAT | MISSENSE | aCc/aTc | T7I | PLES_09091 | | | | | A | A | | | | | | |
| NC_011770 | 993634 | A | G | 35 | 0 | 1 | 0 | NON_SYNONYMOU | MODERAT | MISSENSE | cTg/cCg | L74P | PLES_09111 | | | | | | | G | | | | | |
| NC_011770 | 993988 | C | T | 72 | 0 | 4 | 0 | NON_SYNONYMOU | MODERAT | MISSENSE | Gac/Aac | D384N | PLES_09121 | | | T | T | T | T | | | | | | |
| NC_011770 | 995038 | C | T | 69 | 0 | 2 | 0 | NON_SYNONYMOU | MODERAT | MISSENSE | Gcg/Acg | A34T | PLES_09121 | | | | | | | | | | | T | T |
| NC_011770 | 995672 | C | T | 60 | 0 | 2 | 0 | NON_SYNONYMOU | MODERAT | MISSENSE | gGc/gAc | G58D | PLES_09131 | | | | | | | | | T | T | | |
| NC_011770 | 996759 | T | C | 55 | 0 | 2 | 0 | NON_SYNONYMOU | MODERAT | MISSENSE | gTc/gCc | V41A | PLES_09151 | | | | | | | | | | | C | C |
| NC_011770 | 996770 | T | A | 55 | 0 | 12 | 0 | NON_SYNONYMOU | MODERAT | MISSENSE | Tgg/Agg | W45R | PLES_09151 | A | A | A | A | A | A | A | A | A | A | A | A |
| NC_011770 | 996710 | C | T | 59 | 0 | 1 | 0 | NON_SYNONYMOU | MODERAT | MISSENSE | Cgc/Tgc | R25C | PLES_09151 | | | | | | | | T | | | | |
| NC_011770 | 998041 | T | G | 74 | 0 | 4 | 0 | NON_SYNONYMOU | MODERAT | MISSENSE | gaA/gaC | E155D | PLES_09171 | G | G | | | | | | | G | G | | |
| NC_011770 | 998399 | G | T | 60 | 0 | 6 | 0 | NON_SYNONYMOU | MODERAT | MISSENSE | gCg/gAg | A36E | PLES_09171 | T | T | | | | | T | T | T | T | | |
| NC_011770 | 998671 | G | A | 80 | 0 | 2 | 0 | NON_SYNONYMOU | MODERAT | MISSENSE | gCc/gTc | A140V | PLES_09181 | A | A | | | | | | | | | | |
| NC_011770 | 998905 | C | T | 77 | 0 | 6 | 0 | NON_SYNONYMOU | MODERAT | MISSENSE | aGc/aAc | S62N | PLES_09181 | T | T | T | T | T | T | | | | | | |
| NC_011770 | 1005660 | T | C | 78 | 0 | 2 | 0 | NON_SYNONYMOU | MODERAT | MISSENSE | Ttc/Ctc | F7L | PLES_09271 | | | | | | | | | | | C | C |
| NC_011770 | 1005705 | T | G | 73 | 0 | 6 | 0 | NON_SYNONYMOU | MODERAT | MISSENSE | Tcc/Gcc | S22A | PLES_09271 | | | G | G | G | G | | | G | G | | |
| NC_011770 | 1006443 | T | C | 76 | 0 | 4 | 0 | NON_SYNONYMOU | MODERAT | MISSENSE | cTg/cCg | L18P | PLES_09281 | | | C | C | C | C | | | | | | |
| NC_011770 | 1006601 | G | A | 78 | 0 | 4 | 0 | NON_SYNONYMOU | MODERAT | MISSENSE | Gtc/Atc | V71I | PLES_09281 | A | A | | | | | | | | | A | A |
| NC_011770 | 1006617 | A | T | 83 | 0 | 4 | 0 | NON_SYNONYMOU | MODERAT | MISSENSE | gAa/gTa | E76V | PLES_09281 | | | T | T | T | T | | | | | | |
| NC_011770 | 1007872 | A | C | 76 | 0 | 10 | 0 | NON_SYNONYMOU | MODERAT | MISSENSE | ttA/ttC | L31F | PLES_09301 | | | C | C | C | C | C | C | C | C | C | C |
| NC_011770 | 1008032 | A | G | 77 | 0 | 2 | 0 | NON_SYNONYMOU | MODERAT | MISSENSE | Acc/Gcc | T85A | PLES_09301 | | | | | | | G | G | | | | |
| NC_011770 | 1012616 | G | A | 79 | 0 | 2 | 0 | NON_SYNONYMOU | MODERAT | MISSENSE | Gat/Aat | D36N | PLES_09351 | | | | | | | A | A | | | | |
| NC_011770 | 1012620 | T | C | 77 | 0 | 2 | 0 | NON_SYNONYMOU | MODERAT | MISSENSE | aTc/aCc | I37T | PLES_09351 | | | | | | | | | | | C | C |
| NC_011770 | 1013351 | T | G | 58 | 0 | 2 | 0 | NON_SYNONYMOU | MODERAT | MISSENSE | Ttg/Gtg | L281V | PLES_09351 | | | | | | | | | | | G | G |
| NC_011770 | 1014055 | C | A | 54 | 0 | 4 | 0 | NON_SYNONYMOU | MODERAT | MISSENSE | Gcc/Tcc | A289S | PLES_09361 | | | A | A | A | A | | | | | | |
| NC_011770 | 1014174 | G | A | 63 | 0 | 1 | 0 | NON_SYNONYMOU | MODERAT | MISSENSE | tCg/tTg | S249L | PLES_09361 | | | | | | | A | | | | | |
| NC_011770 | 1014279 | C | T | 67 | 0 | 4 | 0 | NON_SYNONYMOU | MODERAT | MISSENSE | cGa/cAa | R214Q | PLES_09361 | | | T | T | T | T | | | | | | |
| NC_011770 | 1014446 | G | C | 72 | 0 | 12 | 0 | NON_SYNONYMOU | MODERAT | MISSENSE | tgC/tgG | C158W | PLES_09361 | C | C | C | C | C | C | C | C | C | C | C | C |
| NC_011770 | 1014708 | G | A | 70 | 0 | 4 | 0 | NON_SYNONYMOU | MODERAT | MISSENSE | aCg/aTg | T71M | PLES_09361 | | | | | | | | | A | A | A | A |
| NC_011770 | 1014864 | A | G | 76 | 0 | 2 | 0 | NON_SYNONYMOU | MODERAT | MISSENSE | aTc/aCc | I19T | PLES_09361 | | | | | | | G | G | | | | |
| NC_011770 | 1014904 | C | T | 75 | 0 | 2 | 0 | NON_SYNONYMOU | MODERAT | MISSENSE | Gta/Ata | V6I | PLES_09361 | | | | | | | T | T | | | | |
| NC_011770 | 1014906 | T | C | 74 | 0 | 2 | 0 | NON_SYNONYMOU | MODERAT | MISSENSE | aAc/aGc | N5S | PLES_09361 | | | | | | | C | C | | | | |
| NC_011770 | 1014910 | G | C | 75 | 0 | 2 | 0 | NON_SYNONYMOU | MODERAT | MISSENSE | Cgt/Ggt | R4G | PLES_09361 | | | | | | | C | C | | | | |
| NC_011770 | 1014971 | A | G | 81 | 0 | 2 | 0 | NON_SYNONYMOU | MODERAT | MISSENSE | Ttc/Ctc | F585L | PLES_09371 | | | | | | | G | G | | | | |
| NC_011770 | 1015024 | C | G | 82 | 0 | 2 | 0 | NON_SYNONYMOU | MODERAT | MISSENSE | gGg/gCg | G567A | PLES_09371 | | | | | | | G | G | | | | |
| NC_011770 | 1015637 | G | A | 79 | 0 | 2 | 0 | NON_SYNONYMOU | MODERAT | MISSENSE | Cac/Tac | H363Y | PLES_09371 | A | A | | | | | | | | | | |
| NC_011770 | 1015967 | C | T | 70 | 0 | 2 | 0 | NON_SYNONYMOU | MODERAT | MISSENSE | Ggc/Agc | G253S | PLES_09371 | | | | | | | | | T | T | | |
| NC_011770 | 1016046 | T | G | 67 | 0 | 4 | 0 | NON_SYNONYMOU | MODERAT | MISSENSE | gaA/gaC | E226D | PLES_09371 | | | | | | | | | G | G | G | G |
| NC_011770 | 1016083 | C | T | 70 | 0 | 4 | 0 | NON_SYNONYMOU | MODERAT | MISSENSE | cGc/cAc | R214H | PLES_09371 | | | T | T | T | T | | | | | | |
| NC_011770 | 1016303 | G | A | 71 | 0 | 2 | 0 | NON_SYNONYMOU | MODERAT | MISSENSE | Ccg/Tcg | P141S | PLES_09371 | A | A | | | | | | | | | | |
| NC_011770 | 1016476 | A | G | 66 | 0 | 2 | 0 | NON_SYNONYMOU | MODERAT | MISSENSE | gTc/gCc | V83A | PLES_09371 | G | G | | | | | | | | | | |
| NC_011770 | 1016558 | C | T | 69 | 0 | 2 | 0 | NON_SYNONYMOU | MODERAT | MISSENSE | Gac/Aac | D56N | PLES_09371 | | | | | | | | | | | T | T |
| NC_011770 | 1016759 | C | T | 69 | 0 | 2 | 0 | NON_SYNONYMOU | MODERAT | MISSENSE | Gtc/Atc | V237I | PLES_09381 | | | | | | | | | T | T | | |
| NC_011770 | 1016854 | A | G | 66 | 0 | 1 | 0 | NON_SYNONYMOU | MODERAT | MISSENSE | gTc/gCc | V205A | PLES_09381 | | | | | | | G | | | | | |
| NC_011770 | 1017295 | G | T | 71 | 0 | 2 | 0 | NON_SYNONYMOU | MODERAT | MISSENSE | cCc/cAc | P58H | PLES_09381 | | | | | | | | | T | T | | |
| NC_011770 | 1017389 | C | T | 69 | 0 | 2 | 0 | NON_SYNONYMOU | MODERAT | MISSENSE | Gta/Ata | V27I | PLES_09381 | | | | | | | T | T | | | | |
| NC_011770 | 1017475 | G | T | 59 | 0 | 8 | 0 | NON_SYNONYMOU | MODERAT | MISSENSE | Ccc/Acc | P306T | PLES_09391 | T | T | | | | | T | T | T | T | T | T |
| NC_011770 | 1017529 | C | A | 58 | 0 | 2 | 0 | NON_SYNONYMOU | MODERAT | MISSENSE | Gtg/Ttg | V288L | PLES_09391 | | | | | | | A | A | | | | |
| NC_011770 | 1017549 | C | T | 56 | 0 | 2 | 0 | NON_SYNONYMOU | MODERAT | MISSENSE | aGc/aAc | S281N | PLES_09391 | | | | | | | T | T | | | | |
| NC_011770 | 1018131 | C | T | 69 | 0 | 2 | 0 | NON_SYNONYMOU | MODERAT | MISSENSE | gGa/gAa | G87E | PLES_09391 | | | | | | | T | T | | | | |
| NC_011770 | 1019875 | C | CG | 74 | 0 | 2 | 0 | FRAME_SHIFT | HIGH | | ccg/ccCg | P407P? | PLES_09401 | CG | CG | | | | | | | | | | |
| NC_011770 | 1019018 | C | T | 69 | 0 | 2 | 0 | NON_SYNONYMOU | MODERAT | MISSENSE | cGg/cAg | R693Q | PLES_09401 | T | T | | | | | | | | | | |
| NC_011770 | 1019194 | C | G | 68 | 0 | 2 | 0 | NON_SYNONYMOU | MODERAT | MISSENSE | caG/caC | Q634H | PLES_09401 | | | | | | | G | G | | | | |
| NC_011770 | 1019465 | C | T | 62 | 0 | 1 | 0 | NON_SYNONYMOU | MODERAT | MISSENSE | cGc/cAc | R544H | PLES_09401 | | | | | | | T | | | | | |
| NC_011770 | 1019654 | T | C | 60 | 0 | 4 | 0 | NON_SYNONYMOU | MODERAT | MISSENSE | gAa/gGa | E481G | PLES_09401 | C | C | | | | | | | | | C | C |
| NC_011770 | 1019912 | C | T | 75 | 0 | 4 | 0 | NON_SYNONYMOU | MODERAT | MISSENSE | cGc/cAc | R395H | PLES_09401 | T | T | | | | | | | | | T | T |
| NC_011770 | 1020071 | G | A | 87 | 0 | 2 | 0 | NON_SYNONYMOU | MODERAT | MISSENSE | gCc/gTc | A342V | PLES_09401 | | | | | | | A | A | | | | |
| NC_011770 | 1020482 | A | G | 75 | 0 | 8 | 0 | NON_SYNONYMOU | MODERAT | MISSENSE | gTc/gCc | V205A | PLES_09401 | | | G | G | G | G | G | G | G | G | | |
| NC_011770 | 1020987 | G | C | 95 | 0 | 2 | 0 | NON_SYNONYMOU | MODERAT | MISSENSE | Ctg/Gtg | L37V | PLES_09401 | | | | | | | C | C | | | | |
| NC_011770 | 1023553 | C | A | 73 | 0 | 1 | 0 | NON_SYNONYMOU | MODERAT | MISSENSE | Gac/Tac | D12Y | PLES_09431 | | | | | | | | A | | | | |
| NC_011770 | 1023934 | G | A | 89 | 0 | 1 | 0 | NON_SYNONYMOU | MODERAT | MISSENSE | Gac/Aac | D52N | PLES_09441 | | | | | | | A | | | | | |
| NC_011770 | 1024112 | G | A | 80 | 0 | 1 | 0 | NON_SYNONYMOU | MODERAT | MISSENSE | cGa/cAa | R111Q | PLES_09441 | | | | | | | A | | | | | |
| NC_011770 | 1026090 | C | T | 77 | 0 | 1 | 0 | NON_SYNONYMOU | MODERAT | MISSENSE | gGc/gAc | G202D | PLES_09471 | | | | | | | | T | | | | |

| | | | | | | | | | | | | | | | | | | | | | | | | | |
|---|---|---|---|---|---|---|---|---|---|---|---|---|---|---|---|---|---|---|---|---|---|---|---|---|---|
| NC_011770 | 1027670 | A | G | 66 | 0 | 2 | 0 | STOP_LOST+SPLICE | HIGH | MISSENSE | Tag/Cag | *74Q | PLES_09481 | | | | | | | G | G | | | | |
| NC_011770 | 1027723 | T | C | 61 | 0 | 2 | 0 | NON_SYNONYMOU | MODERAT | MISSENSE | gAc/gGc | D56G | PLES_09481 | | | | | | | C | C | | | | |
| NC_011770 | 1027807 | C | T | 65 | 0 | 2 | 0 | NON_SYNONYMOU | MODERAT | MISSENSE | gGg/gAg | G28E | PLES_09481 | | | | | | | T | T | | | | |
| NC_011770 | 1028381 | A | G | 68 | 0 | 12 | 0 | NON_SYNONYMOU | MODERAT | MISSENSE | Tcg/Ccg | S293P | PLES_09491 | G | G | G | G | G | G | G | G | G | G | G | G |
| NC_011770 | 1028387 | C | T | 71 | 0 | 2 | 0 | NON_SYNONYMOU | MODERAT | MISSENSE | Gaa/Aaa | E291K | PLES_09491 | | | | | | | T | T | | | | |
| NC_011770 | 1028708 | G | A | 83 | 0 | 2 | 0 | NON_SYNONYMOU | MODERAT | MISSENSE | Cac/Tac | H184Y | PLES_09491 | | | | | | | | | A | A | | |
| NC_011770 | 1028869 | C | A | 88 | 0 | 2 | 0 | NON_SYNONYMOU | MODERAT | MISSENSE | cGt/cTt | R130L | PLES_09491 | | | | | | | A | A | | | | |
| NC_011770 | 1028959 | A | G | 78 | 0 | 12 | 0 | NON_SYNONYMOU | MODERAT | MISSENSE | gTc/gCc | V100A | PLES_09491 | G | G | G | G | G | G | G | G | G | G | G | G |
| NC_011770 | 1029196 | C | T | 80 | 0 | 2 | 0 | NON_SYNONYMOU | MODERAT | MISSENSE | gGc/gAc | G21D | PLES_09491 | | | | | | | | | T | T | | |
| NC_011770 | 1030400 | A | G | 72 | 0 | 1 | 0 | NON_SYNONYMOU | MODERAT | MISSENSE | cTg/cCg | L118P | PLES_09511 | | | | | | | | G | | | | |
| NC_011770 | 1032580 | AC | A | 62 | 0 | 1 | 0 | FRAME_SHIFT | HIGH | | gtg/ | V362 | PLES_09531 | | | | | | | A | | | | | |
| NC_011770 | 1033662 | C | T | 72 | 0 | 1 | 0 | START_LOST | HIGH | MISSENSE | atG/atA | M1I | PLES_09531 | | | | | | | T | | | | | |
| NC_011770 | 1033094 | C | G | 75 | 0 | 2 | 0 | NON_SYNONYMOU | MODERAT | MISSENSE | Gtt/Ctt | V191L | PLES_09531 | | | | | | | G | G | | | | |
| NC_011770 | 1035346 | C | G | 79 | 0 | 2 | 0 | NON_SYNONYMOU | MODERAT | MISSENSE | Gcc/Ccc | A17P | PLES_09541 | | | | | | | G | G | | | | |
| NC_011770 | 1036592 | C | A | 94 | 0 | 2 | 0 | NON_SYNONYMOU | MODERAT | MISSENSE | cCg/cAg | P287Q | PLES_09551 | | | | | | | | | A | A | | |
| NC_011770 | 1036682 | G | C | 86 | 0 | 12 | 0 | NON_SYNONYMOU | MODERAT | MISSENSE | gGc/gCc | G317A | PLES_09551 | C | C | C | C | C | C | C | C | C | C | C | C |
| NC_011770 | 1036749 | G | T | 89 | 0 | 2 | 0 | NON_SYNONYMOU | MODERAT | MISSENSE | gaG/gaT | E339D | PLES_09551 | | | | | | | | | | | T | T |
| NC_011770 | 1037168 | A | G | 89 | 0 | 12 | 0 | NON_SYNONYMOU | MODERAT | MISSENSE | gAg/gGg | E479G | PLES_09551 | G | G | G | G | G | G | G | G | G | G | G | G |
| NC_011770 | 1037479 | C | G | 62 | 0 | 2 | 0 | NON_SYNONYMOU | MODERAT | MISSENSE | Ccc/Gcc | P583A | PLES_09551 | | | | | | | | | G | G | | |
| NC_011770 | 1037559 | C | A | 67 | 1 | 2 | 0 | NON_SYNONYMOU | MODERAT | MISSENSE | gaC/gaA | D609E | PLES_09551 | | | | . | | | | | | | A | A |
| NC_011770 | 1039612 | T | C | 75 | 0 | 2 | 0 | NON_SYNONYMOU | MODERAT | MISSENSE | Ttc/Ctc | F143L | PLES_09571 | | | | | | | | | | | C | C |
| NC_011770 | 1039847 | A | G | 77 | 0 | 6 | 0 | NON_SYNONYMOU | MODERAT | MISSENSE | Agc/Ggc | S11G | PLES_09581 | G | G | | | | | G | G | G | G | | |
| NC_011770 | 1040075 | A | G | 71 | 0 | 6 | 0 | NON_SYNONYMOU | MODERAT | MISSENSE | Atc/Gtc | I87V | PLES_09581 | G | G | | | | | G | G | G | G | | |
| NC_011770 | 1040840 | G | A | 84 | 0 | 4 | 0 | NON_SYNONYMOU | MODERAT | MISSENSE | tCc/tTc | S567F | PLES_09601 | A | A | | | | | | | A | A | | |
| NC_011770 | 1041774 | G | T | 80 | 0 | 6 | 0 | NON_SYNONYMOU | MODERAT | MISSENSE | Cat/Aat | H256N | PLES_09601 | | | T | T | T | T | | | | | T | T |
| NC_011770 | 1042350 | C | T | 62 | 0 | 2 | 0 | NON_SYNONYMOU | MODERAT | MISSENSE | Gcc/Acc | A64T | PLES_09601 | T | T | | | | | | | | | | |
| NC_011770 | 1043736 | T | A | 69 | 0 | 2 | 0 | NON_SYNONYMOU | MODERAT | MISSENSE | Tcc/Acc | S136T | PLES_09621 | | | | | | | | | A | A | | |
| NC_011770 | 1043635 | C | T | 61 | 0 | 1 | 0 | NON_SYNONYMOU | MODERAT | MISSENSE | gCc/gTc | A102V | PLES_09621 | | | | | | | | T | | | | |
| NC_011770 | 1044549 | T | C | 85 | 0 | 1 | 0 | NON_SYNONYMOU | MODERAT | MISSENSE | Tac/Cac | Y27H | PLES_09641 | | | | | | | C | | | | | |
| NC_011770 | 1044843 | G | A | 82 | 0 | 2 | 0 | NON_SYNONYMOU | MODERAT | MISSENSE | Gcc/Acc | A125T | PLES_09641 | | | A | A | | | | | | | | |
| NC_011770 | 1045208 | C | T | 74 | 0 | 4 | 0 | NON_SYNONYMOU | MODERAT | MISSENSE | aGc/aAc | S394N | PLES_09651 | | | | | | | | | T | T | T | T |
| NC_011770 | 1046483 | T | C | 81 | 0 | 8 | 0 | NON_SYNONYMOU | MODERAT | MISSENSE | Acc/Gcc | T208A | PLES_09661 | | | C | C | C | C | C | C | | | C | C |
| NC_011770 | 1046741 | T | C | 77 | 0 | 4 | 0 | NON_SYNONYMOU | MODERAT | MISSENSE | Acg/Gcg | T122A | PLES_09661 | | | C | C | C | C | | | | | | |
| NC_011770 | 1046925 | C | A | 68 | 0 | 2 | 0 | NON_SYNONYMOU | MODERAT | MISSENSE | gaG/gaT | E60D | PLES_09661 | A | A | | | | | | | | | | |
| NC_011770 | 1047570 | A | G | 67 | 0 | 10 | 0 | NON_SYNONYMOU | MODERAT | MISSENSE | Aag/Gag | K135E | PLES_09671 | G | G | G | G | G | G | | | G | G | G | G |
| NC_011770 | 1048127 | C | T | 57 | 0 | 2 | 0 | NON_SYNONYMOU | MODERAT | MISSENSE | gGc/gAc | G377D | PLES_09681 | | | | | | | T | T | | | | |
| NC_011770 | 1048254 | C | T | 58 | 0 | 10 | 0 | NON_SYNONYMOU | MODERAT | MISSENSE | Gtg/Atg | V335M | PLES_09681 | T | T | T | T | T | T | T | T | | | T | T |
| NC_011770 | 1048346 | C | T | 63 | 0 | 2 | 0 | NON_SYNONYMOU | MODERAT | MISSENSE | gGc/gAc | G304D | PLES_09681 | | | | | | | T | T | | | | |
| NC_011770 | 1048955 | C | T | 66 | 0 | 4 | 0 | NON_SYNONYMOU | MODERAT | MISSENSE | cGg/cAg | R101Q | PLES_09681 | | | | | | | | | T | T | T | T |
| NC_011770 | 1049091 | T | C | 68 | 0 | 4 | 0 | NON_SYNONYMOU | MODERAT | MISSENSE | Acc/Gcc | T56A | PLES_09681 | | | C | C | C | C | | | | | | |
| NC_011770 | 1049142 | G | A | 73 | 0 | 4 | 0 | NON_SYNONYMOU | MODERAT | MISSENSE | Cgc/Tgc | R39C | PLES_09681 | | | A | A | A | A | | | | | | |
| NC_011770 | 1049199 | A | G | 73 | 0 | 4 | 0 | NON_SYNONYMOU | MODERAT | MISSENSE | Ttc/Ctc | F20L | PLES_09681 | | | G | G | G | G | | | | | | |
| NC_011770 | 1048104 | G | A | 59 | 0 | 1 | 0 | NON_SYNONYMOU | MODERAT | MISSENSE | Cac/Tac | H385Y | PLES_09681 | | | | | | | | | | | | A |
| NC_011770 | 1048503 | C | T | 61 | 0 | 1 | 0 | NON_SYNONYMOU | MODERAT | MISSENSE | Ggg/Agg | G252R | PLES_09681 | | | | T | | | | | | | | |
| NC_011770 | 1051751 | T | C | 86 | 0 | 1 | 0 | NON_SYNONYMOU | MODERAT | MISSENSE | gTg/gCg | V89A | PLES_09711 | | | | | | | C | | | | | |
| NC_011770 | 1057199 | CA | C | 78 | 0 | 4 | 0 | FRAME_SHIFT | HIGH | | aac/ | N259 | PLES_09761 | C | C | C | C | | | | | | | | |
| NC_011770 | 1057102 | G | A | 75 | 0 | 6 | 0 | NON_SYNONYMOU | MODERAT | MISSENSE | Gcg/Acg | A224T | PLES_09761 | A | A | A | A | A | A | | | | | | |
| NC_011770 | 1057089 | CGCT | C | 80 | 0 | 5 | 0 | CODON_DELETION | MODERATE | | gct/ | A220 | PLES_09761 | | C | C | C | C | C | | | | | | |
| NC_011770 | 1056928 | G | A | 85 | 0 | 1 | 0 | NON_SYNONYMOU | MODERAT | MISSENSE | Gtc/Atc | V166I | PLES_09761 | | | | | | | | | | | A | |
| NC_011770 | 1058914 | G | A | 74 | 0 | 4 | 0 | NON_SYNONYMOU | MODERAT | MISSENSE | Gtc/Atc | V53I | PLES_09781 | A | A | | | | | | | | | A | A |
| NC_011770 | 1061214 | A | G | 71 | 0 | 1 | 0 | NON_SYNONYMOU | MODERAT | MISSENSE | cTg/cCg | L166P | PLES_09811 | | | | | | | G | | | | | |
| NC_011770 | 1061274 | C | T | 80 | 0 | 8 | 0 | NON_SYNONYMOU | MODERAT | MISSENSE | cGg/cAg | R146Q | PLES_09811 | T | T | | | | | T | T | T | T | T | T |
| NC_011770 | 1061842 | A | G | 70 | 0 | 6 | 0 | NON_SYNONYMOU | MODERAT | MISSENSE | Acc/Gcc | T9A | PLES_09821 | G | G | | | | | | | G | G | G | G |
| NC_011770 | 1061932 | A | G | 77 | 0 | 2 | 0 | NON_SYNONYMOU | MODERAT | MISSENSE | Acc/Gcc | T39A | PLES_09821 | G | G | | | | | | | | | | |
| NC_011770 | 1062005 | C | T | 71 | 0 | 2 | 0 | NON_SYNONYMOU | MODERAT | MISSENSE | gCc/gTc | A63V | PLES_09821 | | | | | | | | | | | T | T |
| NC_011770 | 1062007 | C | T | 71 | 0 | 2 | 0 | NON_SYNONYMOU | MODERAT | MISSENSE | Ccg/Tcg | P64S | PLES_09821 | T | T | | | | | | | | | | |
| NC_011770 | 1062037 | C | T | 69 | 0 | 1 | 0 | NON_SYNONYMOU | MODERAT | MISSENSE | Cgc/Tgc | R74C | PLES_09821 | | | | | | | T | | | | | |
| NC_011770 | 1062088 | A | G | 68 | 0 | 2 | 0 | NON_SYNONYMOU | MODERAT | MISSENSE | Agc/Ggc | S91G | PLES_09821 | | | | | | | G | G | | | | |
| NC_011770 | 1062172 | G | T | 60 | 0 | 2 | 0 | NON_SYNONYMOU | MODERAT | MISSENSE | Gta/Tta | V119L | PLES_09821 | | | | | | | T | T | | | | |
| NC_011770 | 1062469 | G | A | 57 | 0 | 4 | 0 | NON_SYNONYMOU | MODERAT | MISSENSE | Gag/Aag | E218K | PLES_09821 | | | A | A | A | A | | | | | | |
| NC_011770 | 1062593 | A | G | 61 | 0 | 2 | 0 | NON_SYNONYMOU | MODERAT | MISSENSE | cAc/cGc | H259R | PLES_09821 | G | G | | | | | | | | | | |
| NC_011770 | 1062631 | A | G | 61 | 0 | 8 | 0 | NON_SYNONYMOU | MODERAT | MISSENSE | Agc/Ggc | S272G | PLES_09821 | G | G | G | G | G | G | | | G | G | | |

| | | | | | | | | | | | | | | | | | | | | | | | | | |
|---|---|---|---|---|---|---|---|---|---|---|---|---|---|---|---|---|---|---|---|---|---|---|---|---|---|
| NC_011770 | 1062664 | G | A | 60 | 0 | 6 | 0 | NON_SYNONYMOU | MODERAT | MISSENSE | Gcc/Acc | A283T | PLES_09821 | | | A | A | A | A | | | A | A | | |
| NC_011770 | 1063362 | T | C | 65 | 0 | 2 | 0 | NON_SYNONYMOU | MODERAT | MISSENSE | aAc/aGc | N268S | PLES_09831 | | | | | | | | | | | C | C |
| NC_011770 | 1063701 | T | C | 67 | 0 | 4 | 0 | NON_SYNONYMOU | MODERAT | MISSENSE | gAc/gGc | D155G | PLES_09831 | | | C | C | C | C | | | | | | |
| NC_011770 | 1064013 | A | G | 73 | 0 | 12 | 0 | NON_SYNONYMOU | MODERAT | MISSENSE | gTg/gCg | V51A | PLES_09831 | G | G | G | G | G | G | G | G | G | G | G | G |
| NC_011770 | 1064025 | T | A | 75 | 0 | 12 | 0 | NON_SYNONYMOU | MODERAT | MISSENSE | cAg/cTg | Q47L | PLES_09831 | A | A | A | A | A | A | A | A | A | A | A | A |
| NC_011770 | 1064341 | T | C | 41 | 0 | 10 | 0 | INTRAGENIC | MODIFIER | | | | PLES_09841 | C | C | C | C | C | C | | | C | C | C | C |
| NC_011770 | 1064374 | C | T | 41 | 0 | 2 | 0 | INTRAGENIC | MODIFIER | | | | PLES_09841 | | | | | | | T | T | | | | |
| NC_011770 | 1064397 | G | T | 42 | 0 | 6 | 0 | INTRAGENIC | MODIFIER | | | | PLES_09841 | T | T | | | | | | | T | T | T | T |
| NC_011770 | 1064508 | T | C | 36 | 0 | 6 | 0 | INTRAGENIC | MODIFIER | | | | PLES_09841 | C | C | | | | | | | C | C | C | C |
| NC_011770 | 1066246 | C | T | 65 | 0 | 2 | 0 | NON_SYNONYMOU | MODERAT | MISSENSE | cGc/cAc | R146H | PLES_09871 | T | T | | | | | | | | | | |
| NC_011770 | 1069965 | G | A | 77 | 0 | 2 | 0 | NON_SYNONYMOU | MODERAT | MISSENSE | gGc/gAc | G142D | PLES_09891 | | | | | | | A | A | | | | |
| NC_011770 | 1069801 | A | ACCCCG | 66 | 0 | 1 | 0 | CODON_CHANGE_ | MODERATE | | caa/cCCCG | Q94PRPQ | PLES_09891 HP | | ACCCCGGCCG | | | | | | | | | | |
| NC_011770 | 1070220 | G | C | 73 | 0 | 6 | 0 | NON_SYNONYMOU | MODERAT | MISSENSE | gGg/gCg | G59A | PLES_09901 | | | C | C | C | C | | | | | C | C |
| NC_011770 | 1073693 | A | G | 78 | 0 | 12 | 0 | NON_SYNONYMOU | MODERAT | MISSENSE | cTc/cCc | L68P | PLES_09931 | G | G | G | G | G | G | G | G | G | G | G | G |
| NC_011770 | 1073949 | T | C | 75 | 0 | 12 | 0 | NON_SYNONYMOU | MODERAT | MISSENSE | aAc/aGc | N320S | PLES_09941 | C | C | C | C | C | C | C | C | C | C | C | C |
| NC_011770 | 1076804 | C | T | 76 | 0 | 1 | 0 | NON_SYNONYMOU | MODERAT | MISSENSE | cCg/cTg | P87L | PLES_09961 | | | | | | | | T | | | | |
| NC_011770 | 1083623 | C | T | 84 | 0 | 12 | 0 | NON_SYNONYMOU | MODERAT | MISSENSE | gCc/gTc | A171V | PLES_10021 | T | T | T | T | T | T | T | T | T | T | T | T |
| NC_011770 | 1083915 | G | A | 83 | 0 | 4 | 0 | NON_SYNONYMOU | MODERAT | MISSENSE | Gcc/Acc | A54T | PLES_10031 | A | A | | | | | | | A | A | | |
| NC_011770 | 1084864 | C | G | 76 | 0 | 2 | 0 | NON_SYNONYMOU | MODERAT | MISSENSE | gCc/gGc | A370G | PLES_10031 | | | | | | | G | G | | | | |
| NC_011770 | 1085104 | G | T | 70 | 0 | 12 | 0 | NON_SYNONYMOU | MODERAT | MISSENSE | gGg/gTg | G450V | PLES_10031 | T | T | T | T | T | T | T | T | T | T | T | T |
| NC_011770 | 1085143 | G | A | 75 | 0 | 2 | 0 | NON_SYNONYMOU | MODERAT | MISSENSE | gGc/gAc | G463D | PLES_10031 | A | A | | | | | | | | | | |
| NC_011770 | 1085400 | A | G | 66 | 0 | 4 | 0 | NON_SYNONYMOU | MODERAT | MISSENSE | Acc/Gcc | T549A | PLES_10031 | | | | | | | G | G | G | G | | |
| NC_011770 | 1085796 | A | G | 77 | 0 | 6 | 0 | NON_SYNONYMOU | MODERAT | MISSENSE | cAc/cGc | H127R | PLES_10041 | | | G | G | G | G | | | G | G | | |
| NC_011770 | 1088108 | T | C | 77 | 0 | 2 | 0 | NON_SYNONYMOU | MODERAT | MISSENSE | gAg/gGg | E352G | PLES_10061 | | | | | | | C | C | | | | |
| NC_011770 | 1088964 | G | A | 57 | 0 | 4 | 0 | NON_SYNONYMOU | MODERAT | MISSENSE | Ccg/Tcg | P67S | PLES_10061 | | | A | A | A | A | | | | | | |
| NC_011770 | 1088973 | C | A | 55 | 0 | 6 | 0 | NON_SYNONYMOU | MODERAT | MISSENSE | Gtc/Ttc | V64F | PLES_10061 | | | A | A | A | A | | | A | A | | |
| NC_011770 | 1089415 | T | G | 54 | 0 | 10 | 0 | NON_SYNONYMOU | MODERAT | MISSENSE | Tta/Gta | L39V | PLES_10071 | | | G | G | G | G | G | G | G | G | G | G |
| NC_011770 | 1089568 | A | G | 55 | 0 | 12 | 0 | NON_SYNONYMOU | MODERAT | MISSENSE | Aag/Gag | K90E | PLES_10071 | G | G | G | G | G | G | G | G | G | G | G | G |
| NC_011770 | 1090950 | A | G | 77 | 0 | 1 | 0 | NON_SYNONYMOU | MODERAT | MISSENSE | gTg/gCg | V123A | PLES_10101 | | | | G | | | | | | | | |
| NC_011770 | 1091709 | G | A | 72 | 0 | 6 | 0 | NON_SYNONYMOU | MODERAT | MISSENSE | aCg/aTg | T259M | PLES_10111 | | | A | A | A | A | | | A | A | | |
| NC_011770 | 1091724 | G | T | 71 | 0 | 6 | 0 | NON_SYNONYMOU | MODERAT | MISSENSE | cCt/cAt | P254H | PLES_10111 | | | T | T | T | T | | | T | T | | |
| NC_011770 | 1091839 | T | C | 66 | 0 | 6 | 0 | NON_SYNONYMOU | MODERAT | MISSENSE | Acc/Gcc | T216A | PLES_10111 | | | C | C | C | C | | | C | C | | |
| NC_011770 | 1091845 | C | T | 67 | 0 | 2 | 0 | NON_SYNONYMOU | MODERAT | MISSENSE | Ggc/Agc | G214S | PLES_10111 | | | | | | | | | | | T | T |
| NC_011770 | 1091847 | G | A | 67 | 0 | 2 | 0 | NON_SYNONYMOU | MODERAT | MISSENSE | gCc/gTc | A213V | PLES_10111 | | | | | | | A | A | | | | |
| NC_011770 | 1091880 | G | A | 74 | 0 | 2 | 0 | NON_SYNONYMOU | MODERAT | MISSENSE | tCc/tTc | S202F | PLES_10111 | A | A | | | | | | | | | | |
| NC_011770 | 1091900 | C | T | 69 | 0 | 6 | 0 | NON_SYNONYMOU | MODERAT | MISSENSE | atG/atA | M195I | PLES_10111 | | | T | T | T | T | | | T | T | | |
| NC_011770 | 1091902 | T | C | 69 | 0 | 6 | 0 | NON_SYNONYMOU | MODERAT | MISSENSE | Atg/Gtg | M195V | PLES_10111 | | | C | C | C | C | | | C | C | | |
| NC_011770 | 1091968 | A | G | 73 | 0 | 2 | 0 | NON_SYNONYMOU | MODERAT | MISSENSE | Tcc/Ccc | S173P | PLES_10111 | | | | | | | | | | | G | G |
| NC_011770 | 1091989 | C | T | 75 | 0 | 2 | 0 | NON_SYNONYMOU | MODERAT | MISSENSE | Gcg/Acg | A166T | PLES_10111 | T | T | | | | | | | | | | |
| NC_011770 | 1092091 | G | A | 78 | 0 | 6 | 0 | NON_SYNONYMOU | MODERAT | MISSENSE | Cgg/Tgg | R132W | PLES_10111 | | | A | A | A | A | | | A | A | | |
| NC_011770 | 1092145 | C | T | 77 | 0 | 6 | 0 | NON_SYNONYMOU | MODERAT | MISSENSE | Gcc/Acc | A114T | PLES_10111 | | | T | T | T | T | | | T | T | | |
| NC_011770 | 1092175 | C | T | 80 | 0 | 6 | 0 | NON_SYNONYMOU | MODERAT | MISSENSE | Gcg/Acg | A104T | PLES_10111 | | | T | T | T | T | | | T | T | | |
| NC_011770 | 1092214 | A | G | 77 | 0 | 6 | 0 | NON_SYNONYMOU | MODERAT | MISSENSE | Tcc/Ccc | S91P | PLES_10111 | | | G | G | G | G | | | G | G | | |
| NC_011770 | 1092219 | T | G | 81 | 0 | 6 | 0 | NON_SYNONYMOU | MODERAT | MISSENSE | aAc/aCc | N89T | PLES_10111 | | | G | G | G | G | | | G | G | | |
| NC_011770 | 1092234 | G | T | 79 | 0 | 6 | 0 | NON_SYNONYMOU | MODERAT | MISSENSE | cCc/cAc | P84H | PLES_10111 | | | T | T | T | T | | | T | T | | |
| NC_011770 | 1092393 | A | G | 74 | 0 | 10 | 0 | NON_SYNONYMOU | MODERAT | MISSENSE | gTc/gCc | V31A | PLES_10111 | G | G | G | G | G | G | | | G | G | G | G |
| NC_011770 | 1091937 | C | T | 73 | 0 | 1 | 0 | NON_SYNONYMOU | MODERAT | MISSENSE | cGg/cAg | R183Q | PLES_10111 | | | | | | | | | | | T | |
| NC_011770 | 1093336 | A | G | 56 | 0 | 1 | 0 | START_LOST | HIGH | MISSENSE | aTg/aCg | M1T | PLES_10121 | | | | | | | | G | | | | |
| NC_011770 | 1093228 | T | C | 63 | 0 | 12 | 0 | NON_SYNONYMOU | MODERAT | MISSENSE | gAt/gGt | D37G | PLES_10121 | C | C | C | C | C | C | C | C | C | C | C | C |
| NC_011770 | 1094560 | G | C | 60 | 0 | 4 | 0 | NON_SYNONYMOU | MODERAT | MISSENSE | Gcg/Ccg | A193P | PLES_10141 | C | C | | | | | | | C | C | | |
| NC_011770 | 1094581 | A | G | 61 | 0 | 1 | 0 | NON_SYNONYMOU | MODERAT | MISSENSE | Atc/Gtc | I200V | PLES_10141 | | | | | | | G | | | | | |
| NC_011770 | 1095196 | A | T | 81 | 0 | 8 | 0 | NON_SYNONYMOU | MODERAT | MISSENSE | Aac/Tac | N405Y | PLES_10141 | T | T | T | T | T | T | T | T | | | | |
| NC_011770 | 1095406 | C | A | 76 | 0 | 10 | 0 | NON_SYNONYMOU | MODERAT | MISSENSE | Cct/Act | P475T | PLES_10141 | A | A | A | A | A | A | A | A | A | A | | |
| NC_011770 | 1095647 | A | T | 82 | 0 | 2 | 0 | NON_SYNONYMOU | MODERAT | MISSENSE | cAg/cTg | Q555L | PLES_10141 | | | | | | | | | | | T | T |
| NC_011770 | 1095911 | G | A | 88 | 0 | 2 | 0 | NON_SYNONYMOU | MODERAT | MISSENSE | cGg/cAg | R643Q | PLES_10141 | | | | | | | | | | | A | A |
| NC_011770 | 1096159 | C | T | 89 | 0 | 1 | 0 | NON_SYNONYMOU | MODERAT | MISSENSE | Ccg/Tcg | P726S | PLES_10141 | | | | | | | T | | | | | |
| NC_011770 | 1096199 | T | C | 75 | 0 | 1 | 0 | NON_SYNONYMOU | MODERAT | MISSENSE | gTc/gCc | V739A | PLES_10141 | | | | | | | C | | | | | |
| NC_011770 | 1096409 | A | G | 85 | 0 | 8 | 0 | NON_SYNONYMOU | MODERAT | MISSENSE | aAg/aGg | K809R | PLES_10141 | G | G | G | G | G | G | | | | | G | G |
| NC_011770 | 1096641 | G | A | 78 | 0 | 10 | 0 | NON_SYNONYMOU | MODERAT | MISSENSE | gCc/gTc | A93V | PLES_10151 | A | A | A | A | A | A | A | A | | | A | A |
| NC_011770 | 1096959 | C | T | 85 | 0 | 2 | 0 | NON_SYNONYMOU | MODERAT | MISSENSE | Gac/Aac | D240N | PLES_10161 | | | | | | | T | T | | | | |
| NC_011770 | 1097305 | C | A | 80 | 0 | 8 | 0 | NON_SYNONYMOU | MODERAT | MISSENSE | caG/caT | Q124H | PLES_10161 | A | A | A | A | A | A | A | A | | | | |
| NC_011770 | 1098733 | A | AG | 76 | 0 | 1 | 0 | FRAME_SHIFT | HIGH | | ctg/cCtg | L58P? | PLES_10171 | | AG | | | | | | | | | | |

| | | | | | | | | | | | | | | | | | | | | | | | | | |
|---|---|---|---|---|---|---|---|---|---|---|---|---|---|---|---|---|---|---|---|---|---|---|---|---|---|
| NC_011770 | 1098002 | T | C | 78 | 0 | 2 | 0 | NON_SYNONYMOU | MODERAT | MISSENSE | Atc/Gtc | I302V | PLES_10171 | | | | | | | | | | | C | C |
| NC_011770 | 1098004 | G | C | 79 | 0 | 2 | 0 | NON_SYNONYMOU | MODERAT | MISSENSE | gCc/gGc | A301G | PLES_10171 | | | | | | | | | C | C | | |
| NC_011770 | 1099388 | A | G | 75 | 0 | 10 | 0 | NON_SYNONYMOU | MODERAT | MISSENSE | cAc/cGc | H141R | PLES_10181 | G | G | G | G | G | G | G | G | | | G | G |
| NC_011770 | 1099408 | A | G | 78 | 0 | 10 | 0 | NON_SYNONYMOU | MODERAT | MISSENSE | Agt/Ggt | S148G | PLES_10181 | G | G | G | G | G | G | G | G | | | G | G |
| NC_011770 | 1099616 | T | G | 75 | 0 | 2 | 0 | NON_SYNONYMOU | MODERAT | MISSENSE | gTg/gGg | V217G | PLES_10181 | G | G | | | | | | | | | | |
| NC_011770 | 1099829 | G | A | 63 | 0 | 2 | 0 | NON_SYNONYMOU | MODERAT | MISSENSE | gCc/gTc | A139V | PLES_10191 | A | A | | | | | | | | | | |
| NC_011770 | 1099952 | C | T | 65 | 0 | 2 | 0 | NON_SYNONYMOU | MODERAT | MISSENSE | cGc/cAc | R98H | PLES_10191 | T | T | | | | | | | | | | |
| NC_011770 | 1099953 | G | C | 67 | 0 | 2 | 0 | NON_SYNONYMOU | MODERAT | MISSENSE | Cgc/Ggc | R98G | PLES_10191 | C | C | | | | | | | | | | |
| NC_011770 | 1100106 | C | T | 67 | 0 | 2 | 0 | NON_SYNONYMOU | MODERAT | MISSENSE | Gta/Ata | V47I | PLES_10191 | | | | | | | | | T | T | | |
| NC_011770 | 1100397 | T | C | 69 | 0 | 4 | 0 | NON_SYNONYMOU | MODERAT | MISSENSE | Acg/Gcg | T191A | PLES_10201 | | | | | | | C | C | | | C | C |
| NC_011770 | 1100406 | C | G | 69 | 0 | 2 | 0 | NON_SYNONYMOU | MODERAT | MISSENSE | Gcc/Ccc | A188P | PLES_10201 | | | | | G | G | | | | | | |
| NC_011770 | 1102704 | C | T | 88 | 0 | 2 | 0 | NON_SYNONYMOU | MODERAT | MISSENSE | Gta/Ata | V61I | PLES_10221 | | | | | | | | | | | T | T |
| NC_011770 | 1102731 | CGGCGGC( | C | 84 | 0 | 2 | 0 | CODON_DELETION | MODERATE | | tggcgcgccg | WRAA48 | PLES_10221 | | | | | | | C | C | | | | |
| NC_011770 | 1102736 | G | A | 70 | 0 | 2 | 0 | NON_SYNONYMOU | MODERAT | MISSENSE | gCc/gTc | A50V | PLES_10221 | | | | | | | | | | | A | A |
| NC_011770 | 1102806 | G | C | 85 | 0 | 2 | 0 | NON_SYNONYMOU | MODERAT | MISSENSE | Cac/Gac | H27D | PLES_10221 | | | | | | | | | | | C | C |
| NC_011770 | 1102854 | A | G | 97 | 0 | 2 | 0 | NON_SYNONYMOU | MODERAT | MISSENSE | Tcc/Ccc | S11P | PLES_10221 | | | | | | | | | | | G | G |
| NC_011770 | 1102875 | C | G | 84 | 0 | 2 | 0 | NON_SYNONYMOU | MODERAT | MISSENSE | Gat/Cat | D4H | PLES_10221 | | | | | | | | | | | G | G |
| NC_011770 | 1103293 | T | C | 74 | 0 | 2 | 0 | NON_SYNONYMOU | MODERAT | MISSENSE | Tcg/Ccg | S57P | PLES_10231 | | | | | | | | | | | C | C |
| NC_011770 | 1103581 | A | C | 69 | 0 | 4 | 0 | NON_SYNONYMOU | MODERAT | MISSENSE | Acc/Ccc | T153P | PLES_10231 | | | | | | | C | C | C | C | | |
| NC_011770 | 1104375 | A | C | 81 | 0 | 2 | 0 | NON_SYNONYMOU | MODERAT | MISSENSE | gAc/gCc | D10A | PLES_10251 | | | | | | | C | C | | | | |
| NC_011770 | 1104621 | C | G | 81 | 0 | 2 | 0 | NON_SYNONYMOU | MODERAT | MISSENSE | gCc/gGc | A92G | PLES_10251 | | | | | | | | | | | G | G |
| NC_011770 | 1104626 | G | A | 77 | 0 | 2 | 0 | NON_SYNONYMOU | MODERAT | MISSENSE | Gtc/Atc | V94I | PLES_10251 | A | A | | | | | | | | | | |
| NC_011770 | 1104771 | A | G | 75 | 0 | 1 | 0 | NON_SYNONYMOU | MODERAT | MISSENSE | aAc/aGc | N142S | PLES_10251 | | | | | | | G | | | | | |
| NC_011770 | 1105041 | C | T | 74 | 0 | 8 | 0 | NON_SYNONYMOU | MODERAT | MISSENSE | gCc/gTc | A232V | PLES_10251 | T | T | | | | | T | T | T | T | T | T |
| NC_011770 | 1105079 | C | T | 71 | 0 | 2 | 0 | NON_SYNONYMOU | MODERAT | MISSENSE | Cgc/Tgc | R245C | PLES_10251 | | | | | | | | | | | T | T |
| NC_011770 | 1105238 | C | A | 72 | 0 | 2 | 0 | NON_SYNONYMOU | MODERAT | MISSENSE | Ctg/Atg | L298M | PLES_10251 | | | | | | | | | A | A | | |
| NC_011770 | 1104933 | T | C | 76 | 0 | 1 | 0 | NON_SYNONYMOU | MODERAT | MISSENSE | cTg/cCg | L196P | PLES_10251 | | | | | | | | C | | | | |
| NC_011770 | 1105991 | T | C | 74 | 0 | 4 | 0 | NON_SYNONYMOU | MODERAT | MISSENSE | Acc/Gcc | T302A | PLES_10261 | | | | | | | C | C | | | C | C |
| NC_011770 | 1106012 | G | A | 69 | 0 | 4 | 0 | NON_SYNONYMOU | MODERAT | MISSENSE | Ctc/Ttc | L295F | PLES_10261 | | | | | | | A | A | | | A | A |
| NC_011770 | 1106167 | C | T | 68 | 0 | 4 | 0 | NON_SYNONYMOU | MODERAT | MISSENSE | cGg/cAg | R243Q | PLES_10261 | | | | | | | T | T | | | T | T |
| NC_011770 | 1106341 | T | C | 66 | 0 | 4 | 0 | NON_SYNONYMOU | MODERAT | MISSENSE | gAc/gGc | D185G | PLES_10261 | | | | | | | C | C | | | C | C |
| NC_011770 | 1106387 | T | C | 65 | 0 | 6 | 0 | NON_SYNONYMOU | MODERAT | MISSENSE | Atc/Gtc | I170V | PLES_10261 | C | C | | | | | C | C | | | C | C |
| NC_011770 | 1106423 | C | T | 66 | 0 | 4 | 0 | NON_SYNONYMOU | MODERAT | MISSENSE | Gac/Aac | D158N | PLES_10261 | | | T | T | T | T | | | | | | |
| NC_011770 | 1106804 | C | T | 56 | 0 | 4 | 0 | NON_SYNONYMOU | MODERAT | MISSENSE | Gtc/Atc | V31I | PLES_10261 | | | T | T | T | T | | | | | | |
| NC_011770 | 1108001 | A | G | 67 | 0 | 4 | 0 | NON_SYNONYMOU | MODERAT | MISSENSE | tTa/tCa | L40S | PLES_10281 | G | G | | | | | | | G | G | | |
| NC_011770 | 1108013 | C | T | 60 | 0 | 4 | 0 | NON_SYNONYMOU | MODERAT | MISSENSE | aGt/aAt | S36N | PLES_10281 | T | T | | | | | | | T | T | | |
| NC_011770 | 1113425 | G | T | 75 | 0 | 8 | 0 | NON_SYNONYMOU | MODERAT | MISSENSE | cGg/cTg | R43L | PLES_10311 | T | T | | | | | T | T | T | T | T | T |
| NC_011770 | 1114156 | C | T | 79 | 0 | 2 | 0 | NON_SYNONYMOU | MODERAT | MISSENSE | gCg/gTg | A81V | PLES_10321 | T | T | | | | | | | | | | |
| NC_011770 | 1114284 | G | A | 80 | 0 | 2 | 0 | NON_SYNONYMOU | MODERAT | MISSENSE | Gcc/Acc | A124T | PLES_10321 | | | | | | | | | | | A | A |
| NC_011770 | 1114411 | A | G | 79 | 0 | 2 | 0 | NON_SYNONYMOU | MODERAT | MISSENSE | gAc/gGc | D166G | PLES_10321 | G | G | | | | | | | | | | |
| NC_011770 | 1114474 | C | T | 76 | 0 | 2 | 0 | NON_SYNONYMOU | MODERAT | MISSENSE | gCc/gTc | A187V | PLES_10321 | | | | | | | | | | | T | T |
| NC_011770 | 1115010 | C | A | 89 | 0 | 6 | 0 | NON_SYNONYMOU | MODERAT | MISSENSE | cCc/cAc | P172H | PLES_10331 | A | A | | | | | A | A | | | A | A |
| NC_011770 | 1115382 | C | T | 86 | 0 | 2 | 0 | NON_SYNONYMOU | MODERAT | MISSENSE | cCg/cTg | P296L | PLES_10331 | | | | | | | | | | | T | T |
| NC_011770 | 1115700 | A | G | 74 | 0 | 6 | 0 | NON_SYNONYMOU | MODERAT | MISSENSE | cAc/cGc | H402R | PLES_10331 | | | G | G | G | G | | | G | G | | |
| NC_011770 | 1115790 | G | C | 84 | 0 | 2 | 0 | NON_SYNONYMOU | MODERAT | MISSENSE | gGc/gCc | G432A | PLES_10331 | | | | | | | C | C | | | | |
| NC_011770 | 1117189 | G | A | 78 | 0 | 2 | 0 | NON_SYNONYMOU | MODERAT | MISSENSE | Gag/Aag | E67K | PLES_10351 | | | | | | | A | A | | | | |
| NC_011770 | 1118477 | C | T | 72 | 0 | 2 | 0 | NON_SYNONYMOU | MODERAT | MISSENSE | gCc/gTc | A39V | PLES_10371 | | | | | | | T | T | | | | |
| NC_011770 | 1119051 | C | G | 55 | 0 | 2 | 0 | NON_SYNONYMOU | MODERAT | MISSENSE | gaC/gaG | D230E | PLES_10371 | | | | | | | | | | | G | G |
| NC_011770 | 1119230 | C | T | 65 | 0 | 1 | 0 | NON_SYNONYMOU | MODERAT | MISSENSE | gCc/gTc | A290V | PLES_10371 | | | | | | | | | | | | T |
| NC_011770 | 1119556 | C | T | 69 | 0 | 1 | 0 | STOP_GAINED | HIGH | NONSENSE | Cag/Tag | Q24* | PLES_10381 | | | | | | | | | | | | T |
| NC_011770 | 1119863 | T | C | 64 | 0 | 8 | 0 | NON_SYNONYMOU | MODERAT | MISSENSE | gTc/gCc | V126A | PLES_10381 | C | C | C | C | C | C | | | C | C | | |
| NC_011770 | 1120849 | C | A | 79 | 0 | 6 | 0 | NON_SYNONYMOU | MODERAT | MISSENSE | gaC/gaA | D95E | PLES_10391 | | | A | A | A | A | | | | | A | A |
| NC_011770 | 1120899 | C | G | 80 | 0 | 4 | 0 | NON_SYNONYMOU | MODERAT | MISSENSE | aCc/aGc | T112S | PLES_10391 | | | | | | | G | G | | | G | G |
| NC_011770 | 1120959 | T | C | 81 | 0 | 2 | 0 | NON_SYNONYMOU | MODERAT | MISSENSE | gTc/gCc | V132A | PLES_10391 | | | | | | | | | | | C | C |
| NC_011770 | 1121307 | C | T | 85 | 0 | 2 | 0 | NON_SYNONYMOU | MODERAT | MISSENSE | gCg/gTg | A248V | PLES_10391 | T | T | | | | | | | | | | |
| NC_011770 | 1125568 | C | T | 91 | 0 | 1 | 0 | NON_SYNONYMOU | MODERAT | MISSENSE | gGc/gAc | G602D | PLES_10431 | | | | | | | T | | | | | |
| NC_011770 | 1125642 | G | C | 90 | 0 | 12 | 0 | NON_SYNONYMOU | MODERAT | MISSENSE | gaC/gaG | D577E | PLES_10431 | C | C | C | C | C | C | C | C | C | C | C | C |
| NC_011770 | 1126178 | T | G | 75 | 0 | 2 | 0 | NON_SYNONYMOU | MODERAT | MISSENSE | Acc/Ccc | T399P | PLES_10431 | | | | | | | | | | | G | G |
| NC_011770 | 1126519 | C | T | 83 | 0 | 2 | 0 | NON_SYNONYMOU | MODERAT | MISSENSE | gGc/gAc | G285D | PLES_10431 | | | | | | | | | T | T | | |
| NC_011770 | 1126591 | A | T | 88 | 0 | 2 | 0 | NON_SYNONYMOU | MODERAT | MISSENSE | cTg/cAg | L261Q | PLES_10431 | | | T | T | | | | | | | | |
| NC_011770 | 1126702 | A | G | 82 | 0 | 8 | 0 | NON_SYNONYMOU | MODERAT | MISSENSE | gTg/gCg | V224A | PLES_10431 | | | G | G | G | G | G | G | G | G | | |
| NC_011770 | 1126715 | C | G | 78 | 0 | 8 | 0 | NON_SYNONYMOU | MODERAT | MISSENSE | Gcg/Ccg | A220P | PLES_10431 | | | G | G | G | G | G | G | G | G | | |

| | | | | | | | | | | | | | | | | | | | | | | | | | |
|---|---|---|---|---|---|---|---|---|---|---|---|---|---|---|---|---|---|---|---|---|---|---|---|---|---|
| NC_011770 | 1126768 | A | G | 76 | 0 | 1 | 0 | NON_SYNONYMOU | MODERAT | MISSENSE | gTg/gCg | V202A | PLES_10431 | | | | | | | | G | | | | |
| NC_011770 | 1127840 | C | T | 54 | 0 | 10 | 0 | NON_SYNONYMOU | MODERAT | MISSENSE | Gtc/Atc | V233I | PLES_10441 | T | T | T | T | T | T | | | T | T | T | T |
| NC_011770 | 1127864 | C | T | 55 | 0 | 2 | 0 | NON_SYNONYMOU | MODERAT | MISSENSE | Gcc/Acc | A225T | PLES_10441 | | | | | | | T | T | | | | |
| NC_011770 | 1127903 | T | C | 50 | 0 | 12 | 0 | NON_SYNONYMOU | MODERAT | MISSENSE | Atc/Gtc | I212V | PLES_10441 | C | C | C | C | C | C | C | C | C | C | C | C |
| NC_011770 | 1128859 | C | T | 55 | 0 | 4 | 0 | NON_SYNONYMOU | MODERAT | MISSENSE | cGc/cAc | R211H | PLES_10451 | | | T | T | T | T | | | | | | |
| NC_011770 | 1129418 | G | T | 57 | 0 | 6 | 0 | NON_SYNONYMOU | MODERAT | MISSENSE | Ctc/Atc | L25I | PLES_10451 | | | T | T | T | T | T | T | | | | |
| NC_011770 | 1129435 | T | C | 55 | 0 | 4 | 0 | NON_SYNONYMOU | MODERAT | MISSENSE | aAc/aGc | N19S | PLES_10451 | | | C | C | C | C | | | | | | |
| NC_011770 | 1132467 | G | C | 80 | 0 | 6 | 0 | NON_SYNONYMOU | MODERAT | MISSENSE | Gcg/Ccg | A17P | PLES_10491 | C | C | C | C | C | C | | | | | | |
| NC_011770 | 1133642 | C | CG | 68 | 0 | 2 | 0 | FRAME_SHIFT | HIGH | | cgc/cCgc | R6P? | PLES_10501 | | | | | | | | | | | CG | CG |
| NC_011770 | 1132890 | C | T | 86 | 0 | 8 | 0 | NON_SYNONYMOU | MODERAT | MISSENSE | Gtc/Atc | V257I | PLES_10501 | T | T | T | T | T | T | | | T | T | | |
| NC_011770 | 1132959 | T | C | 83 | 0 | 2 | 0 | NON_SYNONYMOU | MODERAT | MISSENSE | Agc/Ggc | S234G | PLES_10501 | C | C | | | | | | | | | | |
| NC_011770 | 1133079 | T | C | 80 | 0 | 2 | 0 | NON_SYNONYMOU | MODERAT | MISSENSE | Agc/Ggc | S194G | PLES_10501 | | | | | | | | | C | C | | |
| NC_011770 | 1133109 | C | T | 80 | 0 | 2 | 0 | NON_SYNONYMOU | MODERAT | MISSENSE | Gcg/Acg | A184T | PLES_10501 | T | T | | | | | | | | | | |
| NC_011770 | 1133146 | G | C | 78 | 0 | 4 | 0 | NON_SYNONYMOU | MODERAT | MISSENSE | gaC/gaG | D171E | PLES_10501 | | | C | C | C | C | | | | | | |
| NC_011770 | 1134322 | T | C | 80 | 0 | 12 | 0 | NON_SYNONYMOU | MODERAT | MISSENSE | tTg/tCg | L188S | PLES_10511 | C | C | C | C | C | C | C | C | C | C | C | C |
| NC_011770 | 1134759 | T | C | 85 | 0 | 2 | 0 | NON_SYNONYMOU | MODERAT | MISSENSE | Ttc/Ctc | F334L | PLES_10511 | | | | | | | C | C | | | | |
| NC_011770 | 1134762 | G | T | 86 | 0 | 2 | 0 | NON_SYNONYMOU | MODERAT | MISSENSE | Gcc/Tcc | A335S | PLES_10511 | | | | | | | T | T | | | | |
| NC_011770 | 1135819 | A | G | 85 | 0 | 1 | 0 | NON_SYNONYMOU | MODERAT | MISSENSE | Tcc/Ccc | S119P | PLES_10521 | | | | | | | G | | | | | |
| NC_011770 | 1136076 | A | G | 78 | 0 | 4 | 0 | NON_SYNONYMOU | MODERAT | MISSENSE | gTg/gCg | V33A | PLES_10521 | G | G | | | | | G | G | | | | |
| NC_011770 | 1137946 | A | G | 75 | 0 | 2 | 0 | NON_SYNONYMOU | MODERAT | MISSENSE | aAg/aGg | K531R | PLES_10531 | | | | | | | G | G | | | | |
| NC_011770 | 1139057 | A | G | 58 | 0 | 4 | 0 | NON_SYNONYMOU | MODERAT | MISSENSE | aAc/aGc | N270S | PLES_10541 | | | G | G | G | G | | | | | | |
| NC_011770 | 1139059 | G | C | 57 | 0 | 4 | 0 | NON_SYNONYMOU | MODERAT | MISSENSE | Gtg/Ctg | V271L | PLES_10541 | | | C | C | C | C | | | | | | |
| NC_011770 | 1139062 | C | T | 59 | 0 | 4 | 0 | NON_SYNONYMOU | MODERAT | MISSENSE | Ctc/Ttc | L272F | PLES_10541 | | | T | T | T | T | | | | | | |
| NC_011770 | 1139065 | G | T | 57 | 0 | 4 | 0 | NON_SYNONYMOU | MODERAT | MISSENSE | Gcc/Tcc | A273S | PLES_10541 | | | T | T | T | T | | | | | | |
| NC_011770 | 1139080 | A | G | 58 | 0 | 4 | 0 | NON_SYNONYMOU | MODERAT | MISSENSE | Agc/Ggc | S278G | PLES_10541 | | | G | G | G | G | | | | | | |
| NC_011770 | 1139082 | C | A | 62 | 0 | 4 | 0 | NON_SYNONYMOU | MODERAT | MISSENSE | agC/agA | S278R | PLES_10541 | | | A | A | A | A | | | | | | |
| NC_011770 | 1139083 | G | A | 61 | 0 | 4 | 0 | NON_SYNONYMOU | MODERAT | MISSENSE | Gtt/Att | V279I | PLES_10541 | | | A | A | A | A | | | | | | |
| NC_011770 | 1139101 | G | A | 60 | 0 | 4 | 0 | NON_SYNONYMOU | MODERAT | MISSENSE | Gac/Aac | D285N | PLES_10541 | | | A | A | A | A | | | | | | |
| NC_011770 | 1139113 | C | G | 59 | 0 | 4 | 0 | NON_SYNONYMOU | MODERAT | MISSENSE | Ctg/Gtg | L289V | PLES_10541 | | | G | G | G | G | | | | | | |
| NC_011770 | 1139120 | G | A | 56 | 0 | 4 | 0 | NON_SYNONYMOU | MODERAT | MISSENSE | cGc/cAc | R291H | PLES_10541 | | | A | A | A | A | | | | | | |
| NC_011770 | 1139152 | G | A | 83 | 0 | 2 | 0 | NON_SYNONYMOU | MODERAT | MISSENSE | Ggc/Agc | G302S | PLES_10541 | | | | | | | | | | | A | A |
| NC_011770 | 1139260 | A | G | 41 | 4 | 8 | 0 | NON_SYNONYMOU | MODERAT | MISSENSE | Acc/Gcc | T338A | PLES_10541 | G | G | . | . | . | . | G | G | G | G | G | G |
| NC_011770 | 1139369 | G | A | 63 | 0 | 4 | 0 | NON_SYNONYMOU | MODERAT | MISSENSE | aGc/aAc | S374N | PLES_10541 | | | A | A | A | A | | | | | | |
| NC_011770 | 1139593 | C | A | 81 | 0 | 4 | 0 | NON_SYNONYMOU | MODERAT | MISSENSE | Ctg/Atg | L449M | PLES_10541 | | | A | A | A | A | | | | | | |
| NC_011770 | 1139683 | C | G | 66 | 0 | 4 | 0 | NON_SYNONYMOU | MODERAT | MISSENSE | Ctg/Gtg | L479V | PLES_10541 | | | G | G | G | G | | | | | | |
| NC_011770 | 1142346 | T | G | 67 | 0 | 12 | 0 | NON_SYNONYMOU | MODERAT | MISSENSE | Tgc/Ggc | C138G | PLES_10561 | G | G | G | G | G | G | G | G | G | G | G | G |
| NC_011770 | 1142347 | G | C | 67 | 0 | 8 | 0 | NON_SYNONYMOU | MODERAT | MISSENSE | tGc/tCc | C138S | PLES_10561 | | | C | C | C | C | C | C | C | C | | |
| NC_011770 | 1142618 | G | T | 75 | 0 | 2 | 0 | NON_SYNONYMOU | MODERAT | MISSENSE | gaG/gaT | E228D | PLES_10561 | T | T | | | | | | | | | | |
| NC_011770 | 1142704 | A | C | 72 | 0 | 2 | 0 | NON_SYNONYMOU | MODERAT | MISSENSE | gAg/gCg | E257A | PLES_10561 | C | C | | | | | | | | | | |
| NC_011770 | 1143508 | G | A | 93 | 0 | 2 | 0 | NON_SYNONYMOU | MODERAT | MISSENSE | gGc/gAc | G525D | PLES_10561 | | | | | | | | | | | A | A |
| NC_011770 | 1144399 | G | A | 71 | 0 | 1 | 0 | NON_SYNONYMOU | MODERAT | MISSENSE | gGc/gAc | G822D | PLES_10561 | | | | | | | A | | | | | |
| NC_011770 | 1144452 | G | A | 74 | 0 | 4 | 0 | NON_SYNONYMOU | MODERAT | MISSENSE | Ggt/Agt | G840S | PLES_10561 | | | A | A | A | A | | | | | | |
| NC_011770 | 1145439 | G | A | 71 | 0 | 2 | 0 | NON_SYNONYMOU | MODERAT | MISSENSE | Ccg/Tcg | P543S | PLES_10571 | | | | | | | | | | | A | A |
| NC_011770 | 1145543 | T | G | 66 | 0 | 2 | 0 | NON_SYNONYMOU | MODERAT | MISSENSE | cAg/cCg | Q508P | PLES_10571 | G | G | | | | | | | | | | |
| NC_011770 | 1146888 | C | A | 64 | 0 | 4 | 0 | NON_SYNONYMOU | MODERAT | MISSENSE | Gtg/Ttg | V60L | PLES_10571 | | | A | A | A | A | | | | | | |
| NC_011770 | 1148839 | C | T | 74 | 0 | 1 | 0 | NON_SYNONYMOU | MODERAT | MISSENSE | gCc/gTc | A424V | PLES_10581 | | | | | T | | | | | | | |
| NC_011770 | 1152675 | T | G | 83 | 0 | 12 | 0 | NON_SYNONYMOU | MODERAT | MISSENSE | Tac/Gac | Y96D | PLES_10641 | G | G | G | G | G | G | G | G | G | G | G | G |
| NC_011770 | 1153472 | A | G | 84 | 0 | 8 | 0 | NON_SYNONYMOU | MODERAT | MISSENSE | Agc/Ggc | S26G | PLES_10651 | G | G | G | G | G | G | | | | | G | G |
| NC_011770 | 1154192 | A | C | 88 | 0 | 2 | 0 | NON_SYNONYMOU | MODERAT | MISSENSE | Aag/Cag | K266Q | PLES_10651 | | | | | | | C | C | | | | |
| NC_011770 | 1154274 | T | C | 79 | 0 | 12 | 0 | NON_SYNONYMOU | MODERAT | MISSENSE | gTa/gCa | V293A | PLES_10651 | C | C | C | C | C | C | C | C | C | C | C | C |
| NC_011770 | 1154767 | GGGC | G | 34 | 0 | 2 | 0 | CODON_CHANGE_ | MODERATE | | gggccg/gc | GP163A | PLES_10661 | | | | | | | G | G | | | | |
| NC_011770 | 1155313 | G | A | 62 | 0 | 6 | 0 | NON_SYNONYMOU | MODERAT | MISSENSE | cGc/cAc | R89H | PLES_10671 | | | A | A | A | A | | | A | A | | |
| NC_011770 | 1155384 | C | T | 72 | 0 | 12 | 0 | NON_SYNONYMOU | MODERAT | MISSENSE | Ctc/Ttc | L113F | PLES_10671 | T | T | T | T | T | T | T | T | T | T | T | T |
| NC_011770 | 1156270 | G | A | 80 | 0 | 12 | 0 | NON_SYNONYMOU | MODERAT | MISSENSE | gGg/gAg | G408E | PLES_10671 | A | A | A | A | A | A | A | A | A | A | A | A |
| NC_011770 | 1156411 | C | T | 73 | 0 | 1 | 0 | NON_SYNONYMOU | MODERAT | MISSENSE | cCc/cTc | P455L | PLES_10671 | T | | | | | | | | | | | |
| NC_011770 | 1156654 | C | T | 68 | 0 | 8 | 0 | NON_SYNONYMOU | MODERAT | MISSENSE | cCc/cTc | P3L | PLES_10681 | T | T | T | T | T | T | | | | | T | T |
| NC_011770 | 1156669 | C | T | 66 | 0 | 2 | 0 | NON_SYNONYMOU | MODERAT | MISSENSE | gCc/gTc | A8V | PLES_10681 | | | | | | | T | T | | | | |
| NC_011770 | 1157004 | A | G | 57 | 0 | 2 | 0 | NON_SYNONYMOU | MODERAT | MISSENSE | Agc/Ggc | S120G | PLES_10681 | | | | | | | G | G | | | | |
| NC_011770 | 1157110 | A | G | 59 | 0 | 2 | 0 | NON_SYNONYMOU | MODERAT | MISSENSE | gAc/gGc | D155G | PLES_10681 | | | | | | | G | G | | | | |
| NC_011770 | 1157491 | G | T | 66 | 0 | 2 | 0 | NON_SYNONYMOU | MODERAT | MISSENSE | cGc/cTc | R282L | PLES_10681 | | | | | | | T | T | | | | |
| NC_011770 | 1157805 | G | A | 65 | 0 | 2 | 0 | NON_SYNONYMOU | MODERAT | MISSENSE | Ggc/Agc | G387S | PLES_10681 | | | | | | | | | A | A | | |
| NC_011770 | 1158472 | A | G | 79 | 0 | 1 | 0 | NON_SYNONYMOU | MODERAT | MISSENSE | aAc/aGc | N609S | PLES_10681 | | | | | | | G | | | | | |

| | | | | | | | | | | | | | | | | | | | | | | | | | |
|---|---|---|---|---|---|---|---|---|---|---|---|---|---|---|---|---|---|---|---|---|---|---|---|---|---|
| NC_011770 | 1158511 | C | T | 77 | 0 | 2 | 0 | NON_SYNONYMOU | MODERAT | MISSENSE | cCg/cTg | P622L | PLES_10681 | T | T | | | | | | | | | | |
| NC_011770 | 1158678 | A | G | 67 | 0 | 4 | 0 | NON_SYNONYMOU | MODERAT | MISSENSE | Agc/Ggc | S678G | PLES_10681 | | | G | G | G | G | | | | | | |
| NC_011770 | 1159696 | C | T | 71 | 0 | 2 | 0 | NON_SYNONYMOU | MODERAT | MISSENSE | Ggc/Agc | G198S | PLES_10691 | | | | | | | | | | | T | T |
| NC_011770 | 1159816 | C | T | 77 | 0 | 6 | 0 | NON_SYNONYMOU | MODERAT | MISSENSE | Gcg/Acg | A158T | PLES_10691 | | | T | T | T | T | | | | | T | T |
| NC_011770 | 1160220 | T | G | 79 | 0 | 4 | 0 | NON_SYNONYMOU | MODERAT | MISSENSE | gAa/gCa | E23A | PLES_10691 | | | | | | | G | G | G | G | | |
| NC_011770 | 1160675 | A | G | 78 | 0 | 2 | 0 | NON_SYNONYMOU | MODERAT | MISSENSE | tTc/tCc | F132S | PLES_10701 | | | | | | | | | G | G | | |
| NC_011770 | 1160780 | C | T | 72 | 0 | 4 | 0 | NON_SYNONYMOU | MODERAT | MISSENSE | cGg/cAg | R97Q | PLES_10701 | | | T | T | T | T | | | | | | |
| NC_011770 | 1161455 | G | A | 70 | 0 | 4 | 0 | NON_SYNONYMOU | MODERAT | MISSENSE | aCc/aTc | T172I | PLES_10721 | | | | | | | A | A | | | A | A |
| NC_011770 | 1161887 | G | T | 65 | 0 | 2 | 0 | NON_SYNONYMOU | MODERAT | MISSENSE | aCc/aAc | T28N | PLES_10721 | | | | | | | | | | | T | T |
| NC_011770 | 1161962 | C | T | 67 | 0 | 2 | 0 | NON_SYNONYMOU | MODERAT | MISSENSE | cGg/cAg | R3Q | PLES_10721 | T | T | | | | | | | | | | |
| NC_011770 | 1162204 | C | T | 61 | 0 | 4 | 0 | NON_SYNONYMOU | MODERAT | MISSENSE | aGc/aAc | S53N | PLES_10731 | | | T | T | T | T | | | | | | |
| NC_011770 | 1162252 | C | A | 60 | 0 | 8 | 0 | NON_SYNONYMOU | MODERAT | MISSENSE | cGg/cTg | R37L | PLES_10731 | A | A | | | | | A | A | A | A | A | A |
| NC_011770 | 1162240 | T | C | 62 | 0 | 11 | 0 | NON_SYNONYMOU | MODERAT | MISSENSE | cAc/cGc | H41R | PLES_10731 | C | C | C | C | | C | C | C | C | C | C | C |
| NC_011770 | 1162054 | G | GGGGC( | 61 | 0 | 2 | 0 | CODON_INSERTIOI | MODERATE | | cca/cTCGC | P103LAP | PLES_10731 | | | GGGGCGA | | | GGGGCGA | | | | | | |
| NC_011770 | 1167558 | A | ATTC | 80 | 0 | 2 | 0 | CODON_INSERTIOI | MODERATE | | gaa/gaaGA | E317EE | PLES_10771 | | | | | | | ATTC | ATTC | | | | |
| NC_011770 | 1167594 | C | T | 80 | 0 | 2 | 0 | NON_SYNONYMOU | MODERAT | MISSENSE | Gtc/Atc | V306I | PLES_10771 | | | | | | | T | T | | | | |
| NC_011770 | 1167716 | A | G | 77 | 0 | 12 | 0 | NON_SYNONYMOU | MODERAT | MISSENSE | cTc/cCc | L265P | PLES_10771 | G | G | G | G | G | G | G | G | G | G | G | G |
| NC_011770 | 1167887 | C | T | 74 | 0 | 6 | 0 | NON_SYNONYMOU | MODERAT | MISSENSE | cGc/cAc | R208H | PLES_10771 | | | | | | | T | T | T | T | T | T |
| NC_011770 | 1168018 | G | C | 82 | 0 | 6 | 0 | NON_SYNONYMOU | MODERAT | MISSENSE | caC/caG | H164Q | PLES_10771 | | | | | | | C | C | C | C | C | C |
| NC_011770 | 1168268 | C | T | 69 | 0 | 1 | 0 | NON_SYNONYMOU | MODERAT | MISSENSE | cGc/cAc | R81H | PLES_10771 | | | | | | | T | | | | | |
| NC_011770 | 1168395 | G | A | 77 | 0 | 6 | 0 | NON_SYNONYMOU | MODERAT | MISSENSE | Cgc/Tgc | R39C | PLES_10771 | | | | | | | A | A | A | A | A | A |
| NC_011770 | 1168398 | G | C | 78 | 0 | 6 | 0 | NON_SYNONYMOU | MODERAT | MISSENSE | Cag/Gag | Q38E | PLES_10771 | | | | | | | C | C | C | C | C | C |
| NC_011770 | 1168400 | C | A | 78 | 0 | 6 | 0 | NON_SYNONYMOU | MODERAT | MISSENSE | tGg/tTg | W37L | PLES_10771 | | | | | | | A | A | A | A | A | A |
| NC_011770 | 1168113 | C | T | 87 | 0 | 1 | 0 | NON_SYNONYMOU | MODERAT | MISSENSE | Gag/Aag | E133K | PLES_10771 | T | | | | | | | | | | | |
| NC_011770 | 1168513 | A | G | 87 | 0 | 1 | 0 | NON_SYNONYMOU | MODERAT | MISSENSE | cTg/cCg | L168P | PLES_10781 | | | | | | | G | | | | | |
| NC_011770 | 1168583 | G | T | 85 | 0 | 2 | 0 | NON_SYNONYMOU | MODERAT | MISSENSE | Ctg/Atg | L145M | PLES_10781 | | | | | | | T | T | | | | |
| NC_011770 | 1168862 | A | G | 77 | 0 | 12 | 0 | NON_SYNONYMOU | MODERAT | MISSENSE | Tgc/Cgc | C52R | PLES_10781 | G | G | G | G | G | G | G | G | G | G | G | G |
| NC_011770 | 1169123 | G | C | 85 | 0 | 8 | 0 | NON_SYNONYMOU | MODERAT | MISSENSE | gCc/gGc | A292G | PLES_10791 | C | C | C | C | C | C | C | C | | | | |
| NC_011770 | 1169235 | A | G | 84 | 0 | 6 | 0 | NON_SYNONYMOU | MODERAT | MISSENSE | Tcg/Ccg | S255P | PLES_10791 | | | G | G | G | G | G | G | | | | |
| NC_011770 | 1169354 | G | A | 79 | 0 | 2 | 0 | NON_SYNONYMOU | MODERAT | MISSENSE | gCg/gTg | A215V | PLES_10791 | A | A | | | | | | | | | | |
| NC_011770 | 1169468 | G | T | 87 | 0 | 2 | 0 | NON_SYNONYMOU | MODERAT | MISSENSE | aCc/aAc | T177N | PLES_10791 | | | | | | | | | T | T | | |
| NC_011770 | 1169544 | C | T | 91 | 0 | 4 | 0 | NON_SYNONYMOU | MODERAT | MISSENSE | Ggc/Agc | G152S | PLES_10791 | | | T | T | T | T | | | | | | |
| NC_011770 | 1169247 | A | G | 89 | 0 | 1 | 0 | NON_SYNONYMOU | MODERAT | MISSENSE | Tac/Cac | Y251H | PLES_10791 | | | | | | | | G | | | | |
| NC_011770 | 1170652 | G | T | 87 | 0 | 2 | 0 | NON_SYNONYMOU | MODERAT | MISSENSE | cGc/cTc | R177L | PLES_10801 | | | | | | | T | T | | | | |
| NC_011770 | 1170736 | A | G | 92 | 0 | 6 | 0 | NON_SYNONYMOU | MODERAT | MISSENSE | aAt/aGt | N205S | PLES_10801 | G | G | | | | | | | G | G | G | G |
| NC_011770 | 1171591 | A | G | 74 | 0 | 10 | 0 | NON_SYNONYMOU | MODERAT | MISSENSE | Tat/Cat | Y144H | PLES_10811 | G | G | G | G | G | G | G | G | | | G | G |
| NC_011770 | 1171827 | T | A | 64 | 0 | 4 | 0 | NON_SYNONYMOU | MODERAT | MISSENSE | aAg/aTg | K65M | PLES_10811 | A | A | | | | | | | | | A | A |
| NC_011770 | 1171960 | G | A | 56 | 0 | 8 | 0 | NON_SYNONYMOU | MODERAT | MISSENSE | Cgc/Tgc | R21C | PLES_10811 | | | A | A | A | A | A | A | | | A | A |
| NC_011770 | 1172122 | A | G | 61 | 0 | 12 | 0 | NON_SYNONYMOU | MODERAT | MISSENSE | cTc/cCc | L316P | PLES_10821 | G | G | G | G | G | G | G | G | G | G | G | G |
| NC_011770 | 1172123 | G | A | 61 | 0 | 2 | 0 | NON_SYNONYMOU | MODERAT | MISSENSE | Ctc/Ttc | L316F | PLES_10821 | | | | | | | | | A | A | | |
| NC_011770 | 1172257 | T | A | 79 | 0 | 2 | 0 | NON_SYNONYMOU | MODERAT | MISSENSE | tAt/tTt | Y271F | PLES_10821 | | | | | | | | | A | A | | |
| NC_011770 | 1172549 | G | T | 83 | 0 | 2 | 0 | NON_SYNONYMOU | MODERAT | MISSENSE | Cgc/Agc | R174S | PLES_10821 | T | T | | | | | | | | | | |
| NC_011770 | 1172633 | C | T | 84 | 0 | 1 | 0 | NON_SYNONYMOU | MODERAT | MISSENSE | Gcc/Acc | A146T | PLES_10821 | | | | | | | T | | | | | |
| NC_011770 | 1173752 | T | TG | 73 | 0 | 1 | 0 | FRAME_SHIFT | HIGH | | cgc/Gcgc | R149A? | PLES_10831 | | | | | | | | TG | | | | |
| NC_011770 | 1173537 | G | A | 74 | 0 | 2 | 0 | NON_SYNONYMOU | MODERAT | MISSENSE | gGc/gAc | G75D | PLES_10831 | A | A | | | | | | | | | | |
| NC_011770 | 1174205 | C | T | 56 | 0 | 1 | 0 | NON_SYNONYMOU | MODERAT | MISSENSE | Cgc/Tgc | R298C | PLES_10831 | | | | | | | T | | | | | |
| NC_011770 | 1174749 | C | T | 71 | 0 | 2 | 0 | NON_SYNONYMOU | MODERAT | MISSENSE | gCg/gTg | A479V | PLES_10831 | | | | | | | T | T | | | | |
| NC_011770 | 1174805 | G | C | 78 | 0 | 2 | 0 | NON_SYNONYMOU | MODERAT | MISSENSE | aGt/aCt | S2T | PLES_10841 | | | | | | | | | C | C | | |
| NC_011770 | 1174837 | C | T | 78 | 0 | 1 | 0 | NON_SYNONYMOU | MODERAT | MISSENSE | Cgc/Tgc | R13C | PLES_10841 | | | | | | | T | | | | | |
| NC_011770 | 1175652 | T | A | 76 | 0 | 6 | 0 | NON_SYNONYMOU | MODERAT | MISSENSE | gaT/gaA | D284E | PLES_10841 | | | A | A | A | A | | | A | A | | |
| NC_011770 | 1175671 | G | A | 81 | 0 | 1 | 0 | NON_SYNONYMOU | MODERAT | MISSENSE | Gag/Aag | E291K | PLES_10841 | | | | | | | A | | | | | |
| NC_011770 | 1176683 | A | G | 80 | 0 | 12 | 0 | NON_SYNONYMOU | MODERAT | MISSENSE | cAc/cGc | H628R | PLES_10841 | G | G | G | G | G | G | G | G | G | G | G | G |
| NC_011770 | 1176713 | T | C | 85 | 0 | 12 | 0 | NON_SYNONYMOU | MODERAT | MISSENSE | gTc/gCc | V638A | PLES_10841 | C | C | C | C | C | C | C | C | C | C | C | C |
| NC_011770 | 1176838 | C | A | 92 | 0 | 2 | 0 | NON_SYNONYMOU | MODERAT | MISSENSE | Cac/Aac | H680N | PLES_10841 | A | A | | | | | | | | | | |
| NC_011770 | 1178299 | T | C | 47 | 0 | 4 | 0 | NON_SYNONYMOU | MODERAT | MISSENSE | aAg/aGg | K375R | PLES_10861 | | | C | C | C | C | | | | | | |
| NC_011770 | 1178485 | G | T | 59 | 0 | 4 | 0 | NON_SYNONYMOU | MODERAT | MISSENSE | gCg/gAg | A313E | PLES_10861 | | | T | T | T | T | | | | | | |
| NC_011770 | 1179505 | G | A | 88 | 0 | 2 | 0 | NON_SYNONYMOU | MODERAT | MISSENSE | gCc/gTc | A198V | PLES_10871 | | | | | | | | | A | A | | |
| NC_011770 | 1180482 | C | G | 97 | 0 | 4 | 0 | NON_SYNONYMOU | MODERAT | MISSENSE | gGc/gCc | G184A | PLES_10881 | | | G | G | G | G | | | | | | |
| NC_011770 | 1180626 | T | C | 98 | 0 | 6 | 0 | NON_SYNONYMOU | MODERAT | MISSENSE | cAc/cGc | H136R | PLES_10881 | | | C | C | C | C | | | C | C | | |
| NC_011770 | 1184295 | G | C | 56 | 0 | 2 | 0 | NON_SYNONYMOU | MODERAT | MISSENSE | Cgc/Ggc | R137G | PLES_10921 | C | C | | | | | | | | | | |
| NC_011770 | 1184474 | G | A | 59 | 0 | 2 | 0 | NON_SYNONYMOU | MODERAT | MISSENSE | gCg/gTg | A77V | PLES_10921 | | | | | | | | | | | A | A |
| NC_011770 | 1184568 | T | C | 58 | 0 | 6 | 0 | NON_SYNONYMOU | MODERAT | MISSENSE | Acc/Gcc | T46A | PLES_10921 | C | C | | | | | C | C | C | C | | |

| | | | | | | | | | | | | | | | | | | | | | | | | | |
|---|---|---|---|---|---|---|---|---|---|---|---|---|---|---|---|---|---|---|---|---|---|---|---|---|---|
| NC_011770 | 1184906 | C | G | 66 | 0 | 2 | 0 | NON_SYNONYMOU | MODERAT | MISSENSE | Ggc/Cgc | G187R | PLES_10931 | G | G | | | | | | | | | | |
| NC_011770 | 1184933 | C | T | 67 | 0 | 4 | 0 | NON_SYNONYMOU | MODERAT | MISSENSE | Gcc/Acc | A178T | PLES_10931 | | | T | T | T | T | | | | | | |
| NC_011770 | 1184974 | T | C | 64 | 0 | 2 | 0 | NON_SYNONYMOU | MODERAT | MISSENSE | cAc/cGc | H164R | PLES_10931 | C | C | | | | | | | | | | |
| NC_011770 | 1185268 | T | C | 70 | 0 | 12 | 0 | NON_SYNONYMOU | MODERAT | MISSENSE | aAc/aGc | N66S | PLES_10931 | C | C | C | C | C | C | C | C | C | C | C | C |
| NC_011770 | 1186223 | T | C | 50 | 0 | 1 | 0 | NON_SYNONYMOU | MODERAT | MISSENSE | Agc/Ggc | S192G | PLES_10951 | | | | | | | C | | | | | |
| NC_011770 | 1186229 | T | G | 48 | 0 | 2 | 0 | NON_SYNONYMOU | MODERAT | MISSENSE | Atg/Ctg | M190L | PLES_10951 | | | | | | | G | G | | | | |
| NC_011770 | 1186503 | C | T | 28 | 0 | 2 | 0 | NON_SYNONYMOU | MODERAT | MISSENSE | atG/atA | M98I | PLES_10951 | | | | | | | T | T | | | | |
| NC_011770 | 1186509 | T | G | 29 | 0 | 2 | 0 | NON_SYNONYMOU | MODERAT | MISSENSE | gaA/gaC | E96D | PLES_10951 | | | | | | | G | G | | | | |
| NC_011770 | 1186591 | T | C | 37 | 0 | 8 | 0 | NON_SYNONYMOU | MODERAT | MISSENSE | gAa/gGa | E69G | PLES_10951 | C | C | C | C | C | C | | | C | C | | |
| NC_011770 | 1186705 | C | A | 45 | 0 | 2 | 0 | NON_SYNONYMOU | MODERAT | MISSENSE | tGg/tTg | W31L | PLES_10951 | | | | | | | | | | | A | A |
| NC_011770 | 1187434 | T | C | 79 | 0 | 6 | 0 | NON_SYNONYMOU | MODERAT | MISSENSE | Acg/Gcg | T67A | PLES_10961 | C | C | C | C | C | C | | | | | | |
| NC_011770 | 1187520 | C | T | 84 | 0 | 2 | 0 | NON_SYNONYMOU | MODERAT | MISSENSE | cGc/cAc | R38H | PLES_10961 | | | | | | | | | T | T | | |
| NC_011770 | 1201143 | G | A | 87 | 0 | 2 | 0 | NON_SYNONYMOU | MODERAT | MISSENSE | Gtg/Atg | V23M | PLES_11071 | | | | | | | | | | | A | A |
| NC_011770 | 1201570 | C | T | 82 | 0 | 2 | 0 | NON_SYNONYMOU | MODERAT | MISSENSE | gCc/gTc | A165V | PLES_11071 | | | | | | | | | | | T | T |
| NC_011770 | 1201587 | T | G | 83 | 0 | 12 | 0 | NON_SYNONYMOU | MODERAT | MISSENSE | Tcg/Gcg | S171A | PLES_11071 | G | G | G | G | G | G | G | G | G | G | G | G |
| NC_011770 | 1201723 | C | G | 85 | 0 | 2 | 0 | NON_SYNONYMOU | MODERAT | MISSENSE | gCg/gGg | A216G | PLES_11071 | G | G | | | | | | | | | | |
| NC_011770 | 1201725 | C | T | 86 | 0 | 2 | 0 | NON_SYNONYMOU | MODERAT | MISSENSE | Cgc/Tgc | R217C | PLES_11071 | T | T | | | | | | | | | | |
| NC_011770 | 1204750 | C | A | 82 | 0 | 10 | 0 | NON_SYNONYMOU | MODERAT | MISSENSE | Gcc/Tcc | A93S | PLES_11101 | A | A | A | A | A | A | A | A | | | A | A |
| NC_011770 | 1205250 | C | T | 78 | 0 | 2 | 0 | NON_SYNONYMOU | MODERAT | MISSENSE | Ggc/Agc | G323S | PLES_11111 | | | | | | | T | T | | | | |
| NC_011770 | 1205604 | C | T | 80 | 0 | 2 | 0 | NON_SYNONYMOU | MODERAT | MISSENSE | Gcc/Acc | A205T | PLES_11111 | | | | | | | | | | | T | T |
| NC_011770 | 1205631 | C | T | 83 | 0 | 2 | 0 | NON_SYNONYMOU | MODERAT | MISSENSE | Gtc/Atc | V196I | PLES_11111 | | | | | | | | | | | T | T |
| NC_011770 | 1205660 | T | G | 79 | 0 | 4 | 0 | NON_SYNONYMOU | MODERAT | MISSENSE | gAc/gCc | D186A | PLES_11111 | | | | | | | G | G | | | G | G |
| NC_011770 | 1205666 | C | T | 82 | 0 | 2 | 0 | NON_SYNONYMOU | MODERAT | MISSENSE | cGc/cAc | R184H | PLES_11111 | | | | | | | | | | | T | T |
| NC_011770 | 1205759 | G | C | 84 | 0 | 10 | 0 | NON_SYNONYMOU | MODERAT | MISSENSE | gCc/gGc | A153G | PLES_11111 | C | C | C | C | C | C | C | C | | | C | C |
| NC_011770 | 1206700 | T | A | 78 | 0 | 6 | 0 | NON_SYNONYMOU | MODERAT | MISSENSE | Agc/Tgc | S163C | PLES_11121 | A | A | A | A | A | A | | | | | | |
| NC_011770 | 1206808 | C | T | 78 | 0 | 4 | 0 | NON_SYNONYMOU | MODERAT | MISSENSE | Gcc/Acc | A127T | PLES_11121 | | | | | | | T | T | | | T | T |
| NC_011770 | 1209176 | T | C | 67 | 0 | 4 | 0 | NON_SYNONYMOU | MODERAT | MISSENSE | gTc/gCc | V8A | PLES_11141 | | | C | C | C | C | | | | | | |
| NC_011770 | 1209265 | G | A | 74 | 0 | 2 | 0 | NON_SYNONYMOU | MODERAT | MISSENSE | Gcc/Acc | A38T | PLES_11141 | | | A | A | | | | | | | | |
| NC_011770 | 1209633 | G | T | 75 | 0 | 2 | 0 | NON_SYNONYMOU | MODERAT | MISSENSE | gaG/gaT | E160D | PLES_11141 | | | | | | | | | | | T | T |
| NC_011770 | 1210486 | A | G | 81 | 0 | 2 | 0 | NON_SYNONYMOU | MODERAT | MISSENSE | Atc/Gtc | I445V | PLES_11141 | | | | | | | | | G | G | | |
| NC_011770 | 1210697 | C | T | 96 | 0 | 2 | 0 | NON_SYNONYMOU | MODERAT | MISSENSE | gCg/gTg | A515V | PLES_11141 | | | | | | | | | T | T | | |
| NC_011770 | 1210795 | A | G | 88 | 0 | 2 | 0 | NON_SYNONYMOU | MODERAT | MISSENSE | Atc/Gtc | I548V | PLES_11141 | | | | | | | | | G | G | | |
| NC_011770 | 1210940 | G | A | 83 | 0 | 2 | 0 | NON_SYNONYMOU | MODERAT | MISSENSE | cGc/cAc | R596H | PLES_11141 | | | | | | | | | A | A | | |
| NC_011770 | 1210991 | A | G | 86 | 0 | 2 | 0 | NON_SYNONYMOU | MODERAT | MISSENSE | cAg/cGg | Q613R | PLES_11141 | | | | | | | | | G | G | | |
| NC_011770 | 1209929 | T | C | 81 | 0 | 1 | 0 | NON_SYNONYMOU | MODERAT | MISSENSE | gTg/gCg | V259A | PLES_11141 | | C | | | | | | | | | | |
| NC_011770 | 1210108 | C | T | 85 | 0 | 1 | 0 | NON_SYNONYMOU | MODERAT | MISSENSE | Ccg/Tcg | P319S | PLES_11141 | | T | | | | | | | | | | |
| NC_011770 | 1211193 | G | A | 80 | 0 | 1 | 0 | NON_SYNONYMOU | MODERAT | MISSENSE | Cgc/Tgc | R175C | PLES_11151 | | | | | | | A | | | | | |
| NC_011770 | 1211225 | C | T | 82 | 0 | 2 | 0 | NON_SYNONYMOU | MODERAT | MISSENSE | aGc/aAc | S164N | PLES_11151 | | | | | | | T | T | | | | |
| NC_011770 | 1211274 | C | T | 82 | 0 | 4 | 0 | NON_SYNONYMOU | MODERAT | MISSENSE | Gcg/Acg | A148T | PLES_11151 | | | T | T | T | T | | | | | | |
| NC_011770 | 1211396 | A | G | 76 | 0 | 2 | 0 | NON_SYNONYMOU | MODERAT | MISSENSE | aTc/aCc | I107T | PLES_11151 | G | G | | | | | | | | | | |
| NC_011770 | 1211690 | A | G | 78 | 0 | 12 | 0 | NON_SYNONYMOU | MODERAT | MISSENSE | gTc/gCc | V9A | PLES_11151 | G | G | G | G | G | G | G | G | G | G | G | G |
| NC_011770 | 1212264 | T | C | 86 | 0 | 2 | 0 | NON_SYNONYMOU | MODERAT | MISSENSE | Tcc/Ccc | S85P | PLES_11161 | C | C | | | | | | | | | | |
| NC_011770 | 1212634 | A | C | 88 | 0 | 4 | 0 | NON_SYNONYMOU | MODERAT | MISSENSE | aAg/aCg | K208T | PLES_11161 | | | C | C | C | C | | | | | | |
| NC_011770 | 1212650 | G | C | 88 | 0 | 6 | 0 | NON_SYNONYMOU | MODERAT | MISSENSE | gaG/gaC | E213D | PLES_11161 | | | C | C | C | C | | | C | C | | |
| NC_011770 | 1212775 | T | A | 82 | 0 | 2 | 0 | NON_SYNONYMOU | MODERAT | MISSENSE | cTg/cAg | L255Q | PLES_11161 | A | A | | | | | | | | | | |
| NC_011770 | 1214149 | G | C | 70 | 0 | 2 | 0 | NON_SYNONYMOU | MODERAT | MISSENSE | Gag/Cag | E72Q | PLES_11181 | | | | | | | | | C | C | | |
| NC_011770 | 1214161 | C | T | 68 | 0 | 2 | 0 | NON_SYNONYMOU | MODERAT | MISSENSE | Cgt/Tgt | R76C | PLES_11181 | | | | | | | | | T | T | | |
| NC_011770 | 1214246 | A | G | 63 | 0 | 10 | 0 | NON_SYNONYMOU | MODERAT | MISSENSE | gAc/gGc | D104G | PLES_11181 | G | G | G | G | G | G | G | G | | | G | G |
| NC_011770 | 1215634 | G | A | 87 | 0 | 8 | 0 | NON_SYNONYMOU | MODERAT | MISSENSE | cGg/cAg | R38Q | PLES_11211 | A | A | A | A | A | A | | | A | A | | |
| NC_011770 | 1215796 | G | T | 87 | 0 | 2 | 0 | NON_SYNONYMOU | MODERAT | MISSENSE | aGc/aTc | S92I | PLES_11211 | | | | | | | | | T | T | | |
| NC_011770 | 1216047 | G | A | 76 | 0 | 2 | 0 | NON_SYNONYMOU | MODERAT | MISSENSE | Gag/Aag | E176K | PLES_11211 | | | | | | | | | | | A | A |
| NC_011770 | 1216647 | G | T | 65 | 0 | 2 | 0 | NON_SYNONYMOU | MODERAT | MISSENSE | Gcg/Tcg | A106S | PLES_11221 | | | | | | | T | T | | | | |
| NC_011770 | 1217869 | C | T | 87 | 0 | 2 | 0 | NON_SYNONYMOU | MODERAT | MISSENSE | tCc/tTc | S200F | PLES_11231 | | | | | | | | | | | T | T |
| NC_011770 | 1218146 | C | T | 79 | 0 | 2 | 0 | NON_SYNONYMOU | MODERAT | MISSENSE | gCc/gTc | A12V | PLES_11241 | | | | | | | T | T | | | | |
| NC_011770 | 1218232 | A | G | 70 | 0 | 1 | 0 | NON_SYNONYMOU | MODERAT | MISSENSE | Acc/Gcc | T41A | PLES_11241 | | | | | | | | G | | | | |
| NC_011770 | 1221124 | C | T | 74 | 0 | 6 | 0 | NON_SYNONYMOU | MODERAT | MISSENSE | aGc/aAc | S132N | PLES_11261 | T | T | T | T | T | T | | | | | | |
| NC_011770 | 1221251 | C | T | 73 | 0 | 2 | 0 | NON_SYNONYMOU | MODERAT | MISSENSE | Ggc/Agc | G90S | PLES_11261 | | | | | | | T | T | | | | |
| NC_011770 | 1221756 | G | A | 87 | 0 | 1 | 0 | STOP_GAINED | HIGH | NONSENSE | Cga/Tga | R103* | PLES_11271 | | | | | | | A | | | | | |
| NC_011770 | 1221651 | G | A | 83 | 0 | 1 | 0 | NON_SYNONYMOU | MODERAT | MISSENSE | Cgc/Tgc | R138C | PLES_11271 | | | | | | | A | | | | | |
| NC_011770 | 1221809 | C | T | 81 | 0 | 10 | 0 | NON_SYNONYMOU | MODERAT | MISSENSE | cGc/cAc | R85H | PLES_11271 | T | T | T | T | T | T | T | T | | | T | T |
| NC_011770 | 1222568 | C | T | 68 | 0 | 2 | 0 | NON_SYNONYMOU | MODERAT | MISSENSE | Ccg/Tcg | P135S | PLES_11281 | | | | | | | | | | | T | T |
| NC_011770 | 1222745 | C | T | 89 | 0 | 1 | 0 | NON_SYNONYMOU | MODERAT | MISSENSE | Gag/Aag | E284K | PLES_11291 | | | | | | | T | | | | | |

| | | | | | | | | | | | | | | | | | | | | | | | | | |
|---|---|---|---|---|---|---|---|---|---|---|---|---|---|---|---|---|---|---|---|---|---|---|---|---|---|
| NC_011770 | 1222767 | C | A | 90 | 0 | 2 | 0 | NON_SYNONYMOU | MODERAT | MISSENSE | caG/caT | Q276H | PLES_11291 | A | A | | | | | | | | | | |
| NC_011770 | 1223829 | C | G | 76 | 0 | 6 | 0 | NON_SYNONYMOU | MODERAT | MISSENSE | Cag/Gag | Q38E | PLES_11301 | | | G | G | G | G | G | G | | | | |
| NC_011770 | 1224081 | G | C | 72 | 0 | 4 | 0 | NON_SYNONYMOU | MODERAT | MISSENSE | Ggc/Cgc | G122R | PLES_11301 | | | C | C | C | C | | | | | | |
| NC_011770 | 1224182 | G | A | 62 | 0 | 10 | 0 | NON_SYNONYMOU | MODERAT | MISSENSE | gaG/gaC | E155D | PLES_11301 | A | A | A | A | A | A | C | C | | | A | A |
| NC_011770 | 1224309 | C | G | 62 | 0 | 2 | 0 | NON_SYNONYMOU | MODERAT | MISSENSE | Ctc/Gtc | L198V | PLES_11301 | | | | | | | G | G | | | | |
| NC_011770 | 1224882 | G | GT | 30 | 0 | 1 | 0 | FRAME_SHIFT | HIGH | | ccc/cAcc | P10H? | PLES_11311 | | | | . | | GT | | | | | | |
| NC_011770 | 1224662 | A | AG | 71 | 0 | 4 | 0 | FRAME_SHIFT | HIGH | | cct/ccCt | P83P? | PLES_11311 | | | AG | AG | AG | AG | | | | | | |
| NC_011770 | 1224640 | A | G | 73 | 0 | 4 | 0 | NON_SYNONYMOU | MODERAT | MISSENSE | Ttc/Ctc | F91L | PLES_11311 | | | G | G | G | G | | | | | | |
| NC_011770 | 1224645 | T | C | 74 | 0 | 4 | 0 | NON_SYNONYMOU | MODERAT | MISSENSE | cAt/cGt | H89R | PLES_11311 | | | C | C | C | C | | | | | | |
| NC_011770 | 1224660 | C | T | 74 | 0 | 2 | 0 | NON_SYNONYMOU | MODERAT | MISSENSE | cGt/cAt | R84H | PLES_11311 | | | | | | | | | T | T | | |
| NC_011770 | 1224687 | A | G | 71 | 0 | 4 | 0 | NON_SYNONYMOU | MODERAT | MISSENSE | cTg/cCg | L75P | PLES_11311 | | | G | G | G | G | | | | | | |
| NC_011770 | 1224775 | T | C | 67 | 0 | 10 | 0 | NON_SYNONYMOU | MODERAT | MISSENSE | Acg/Gcg | T46A | PLES_11311 | C | C | C | C | C | C | C | C | | | C | C |
| NC_011770 | 1224784 | C | A | 71 | 0 | 2 | 0 | NON_SYNONYMOU | MODERAT | MISSENSE | Gcc/Tcc | A43S | PLES_11311 | | | | | | | | | A | A | | |
| NC_011770 | 1224793 | T | C | 69 | 0 | 4 | 0 | NON_SYNONYMOU | MODERAT | MISSENSE | Act/Gct | T40A | PLES_11311 | | | C | C | C | C | | | | | | |
| NC_011770 | 1224834 | G | A | 64 | 0 | 4 | 0 | NON_SYNONYMOU | MODERAT | MISSENSE | gCc/gTc | A26V | PLES_11311 | | | A | A | A | A | | | | | | |
| NC_011770 | 1224864 | A | C | 64 | 0 | 6 | 0 | NON_SYNONYMOU | MODERAT | MISSENSE | aTt/aGt | I16S | PLES_11311 | | | C | C | C | C | C | C | | | | |
| NC_011770 | 1224902 | G | C | 63 | 0 | 12 | 0 | NON_SYNONYMOU | MODERAT | MISSENSE | caC/caG | H3Q | PLES_11311 | C | C | C | C | C | C | C | C | C | C | C | C |
| NC_011770 | 1225100 | C | T | 66 | 0 | 4 | 0 | NON_SYNONYMOU | MODERAT | MISSENSE | aGc/aAc | S62N | PLES_11321 | | | | | | | | | T | T | T | T |
| NC_011770 | 1225263 | C | T | 62 | 0 | 4 | 0 | NON_SYNONYMOU | MODERAT | MISSENSE | Gcc/Acc | A8T | PLES_11321 | | | T | T | T | T | | | | | | |
| NC_011770 | 1227369 | C | A | 60 | 0 | 2 | 0 | NON_SYNONYMOU | MODERAT | MISSENSE | Gtc/Ttc | V189F | PLES_11341 | | | | | | | A | A | | | | |
| NC_011770 | 1227615 | C | T | 63 | 0 | 2 | 0 | NON_SYNONYMOU | MODERAT | MISSENSE | Gcc/Acc | A107T | PLES_11341 | T | T | | | | | | | | | | |
| NC_011770 | 1227693 | C | T | 59 | 0 | 4 | 0 | NON_SYNONYMOU | MODERAT | MISSENSE | Gat/Aat | D81N | PLES_11341 | | | | | | | | | T | T | T | T |
| NC_011770 | 1230440 | C | T | 71 | 0 | 2 | 0 | NON_SYNONYMOU | MODERAT | MISSENSE | Gcc/Acc | A111T | PLES_11361 | T | T | | | | | | | | | | |
| NC_011770 | 1231818 | C | T | 67 | 0 | 1 | 0 | NON_SYNONYMOU | MODERAT | MISSENSE | Ggg/Agg | G303R | PLES_11381 | | | | | | | T | | | | | |
| NC_011770 | 1232641 | G | T | 60 | 0 | 4 | 0 | NON_SYNONYMOU | MODERAT | MISSENSE | gaC/gaA | D28E | PLES_11381 | T | T | | | | | | | | | T | T |
| NC_011770 | 1238748 | C | T | 44 | 0 | 2 | 0 | NON_SYNONYMOU | MODERAT | MISSENSE | Ggc/Agc | G122S | PLES_11411 | T | T | | | | | | | | | | |
| NC_011770 | 1238928 | T | C | 52 | 0 | 2 | 0 | NON_SYNONYMOU | MODERAT | MISSENSE | Acc/Gcc | T62A | PLES_11411 | | | | | | | | | | | C | C |
| NC_011770 | 1239088 | T | G | 59 | 0 | 2 | 0 | NON_SYNONYMOU | MODERAT | MISSENSE | caA/caC | Q8H | PLES_11411 | G | G | | | | | | | | | | |
| NC_011770 | 1241661 | T | C | 91 | 0 | 2 | 0 | NON_SYNONYMOU | MODERAT | MISSENSE | gTc/gCc | V97A | PLES_11441 | | | | | | | | | C | C | | |
| NC_011770 | 1241691 | C | T | 84 | 0 | 4 | 0 | NON_SYNONYMOU | MODERAT | MISSENSE | cCc/cTc | P107L | PLES_11441 | | | T | T | T | T | | | | | | |
| NC_011770 | 1241882 | A | G | 78 | 0 | 2 | 0 | NON_SYNONYMOU | MODERAT | MISSENSE | Atc/Gtc | I171V | PLES_11441 | | | | | | | | | | | G | G |
| NC_011770 | 1241943 | A | G | 71 | 0 | 2 | 0 | NON_SYNONYMOU | MODERAT | MISSENSE | tAc/tGc | Y191C | PLES_11441 | G | G | | | | | | | | | | |
| NC_011770 | 1241987 | G | C | 69 | 0 | 2 | 0 | NON_SYNONYMOU | MODERAT | MISSENSE | Gaa/Caa | E206Q | PLES_11441 | | | | | | | | | | | C | C |
| NC_011770 | 1242404 | A | G | 77 | 0 | 2 | 0 | NON_SYNONYMOU | MODERAT | MISSENSE | gAc/gGc | D46G | PLES_11451 | G | G | | | | | | | | | | |
| NC_011770 | 1242427 | G | A | 70 | 0 | 2 | 0 | NON_SYNONYMOU | MODERAT | MISSENSE | Gcc/Acc | A54T | PLES_11451 | | | | | | | | | | | A | A |
| NC_011770 | 1242499 | A | C | 68 | 0 | 2 | 0 | NON_SYNONYMOU | MODERAT | MISSENSE | Atc/Ctc | I78L | PLES_11451 | C | C | | | | | | | | | | |
| NC_011770 | 1242576 | C | G | 63 | 0 | 6 | 0 | NON_SYNONYMOU | MODERAT | MISSENSE | agC/agG | S103R | PLES_11451 | | | | | | | G | G | G | G | G | G |
| NC_011770 | 1242758 | A | T | 66 | 0 | 2 | 0 | NON_SYNONYMOU | MODERAT | MISSENSE | cAg/cTg | Q164L | PLES_11451 | | | | | | | | | | | T | T |
| NC_011770 | 1242773 | G | A | 70 | 0 | 6 | 0 | NON_SYNONYMOU | MODERAT | MISSENSE | gGc/gAc | G169D | PLES_11451 | | | A | A | A | A | | | | | A | A |
| NC_011770 | 1242808 | A | G | 70 | 0 | 6 | 0 | NON_SYNONYMOU | MODERAT | MISSENSE | Agt/Ggt | S181G | PLES_11451 | | | G | G | G | G | | | | | G | G |
| NC_011770 | 1242895 | C | T | 72 | 0 | 2 | 0 | NON_SYNONYMOU | MODERAT | MISSENSE | Cgg/Tgg | R210W | PLES_11451 | | | | | | | T | T | | | | |
| NC_011770 | 1242951 | T | G | 68 | 0 | 8 | 0 | NON_SYNONYMOU | MODERAT | MISSENSE | atT/atG | I228M | PLES_11451 | | | G | G | G | G | G | G | | | G | G |
| NC_011770 | 1242953 | G | C | 67 | 0 | 8 | 0 | NON_SYNONYMOU | MODERAT | MISSENSE | gGg/gCg | G229A | PLES_11451 | | | C | C | C | C | C | C | | | C | C |
| NC_011770 | 1243171 | C | T | 67 | 0 | 6 | 0 | NON_SYNONYMOU | MODERAT | MISSENSE | Cca/Tca | P302S | PLES_11451 | | | T | T | T | T | T | T | | | | |
| NC_011770 | 1243824 | C | T | 76 | 0 | 1 | 0 | NON_SYNONYMOU | MODERAT | MISSENSE | Cgt/Tgt | R148C | PLES_11461 | | | | | | | T | | | | | |
| NC_011770 | 1245120 | TCC | T | 54 | 0 | 4 | 0 | FRAME_SHIFT | HIGH | | ccg/ | P210 | PLES_11471 | | | T | T | T | T | | | | | | |
| NC_011770 | 1245017 | G | A | 94 | 0 | 4 | 0 | NON_SYNONYMOU | MODERAT | MISSENSE | cGc/cAc | R175H | PLES_11471 | | | A | A | A | A | | | | | | |
| NC_011770 | 1245077 | C | A | 92 | 0 | 4 | 0 | NON_SYNONYMOU | MODERAT | MISSENSE | aCc/aAc | T195N | PLES_11471 | | | A | A | A | A | | | | | | |
| NC_011770 | 1245080 | A | G | 91 | 0 | 4 | 0 | NON_SYNONYMOU | MODERAT | MISSENSE | gAt/gGt | D196G | PLES_11471 | | | G | G | G | G | | | | | | |
| NC_011770 | 1245084 | C | G | 93 | 0 | 4 | 0 | NON_SYNONYMOU | MODERAT | MISSENSE | caC/caG | H197Q | PLES_11471 | | | G | G | G | G | | | | | | |
| NC_011770 | 1245125 | G | A | 55 | 0 | 4 | 0 | NON_SYNONYMOU | MODERAT | MISSENSE | aGg/aAg | R211K | PLES_11471 | | | A | A | A | A | | | | | | |
| NC_011770 | 1245129 | A | AGTC | 55 | 0 | 4 | 0 | CODON_INSERTIOI | MODERATE | | aag/GTCaa | K213VK | PLES_11471 | | | AGTC | AGTC | AGTC | AGTC | | | | | | |
| NC_011770 | 1245134 | G | A | 55 | 0 | 4 | 0 | NON_SYNONYMOU | MODERAT | MISSENSE | cGt/cAt | R214H | PLES_11471 | | | A | A | A | A | | | | | | |
| NC_011770 | 1245152 | TACC | T | 57 | 0 | 4 | 0 | CODON_CHANGE_ | MODERATE | | ctaccg/ctg | LP220L | PLES_11471 | | | T | T | T | T | | | | | | |
| NC_011770 | 1245173 | C | T | 57 | 0 | 4 | 0 | NON_SYNONYMOU | MODERAT | MISSENSE | gCc/gTc | A227V | PLES_11471 | | | T | T | T | T | | | | | | |
| NC_011770 | 1245728 | C | T | 81 | 0 | 2 | 0 | NON_SYNONYMOU | MODERAT | MISSENSE | Gtg/Atg | V155M | PLES_11481 | | | | | | | | | | | T | T |
| NC_011770 | 1246076 | T | C | 73 | 0 | 2 | 0 | NON_SYNONYMOU | MODERAT | MISSENSE | Acc/Gcc | T39A | PLES_11481 | | | | | | | | | C | C | | |
| NC_011770 | 1246139 | T | C | 68 | 0 | 4 | 0 | NON_SYNONYMOU | MODERAT | MISSENSE | Atc/Gtc | I18V | PLES_11481 | | | C | C | C | C | | | | | | |
| NC_011770 | 1247846 | A | AG | 33 | 6 | 4 | 0 | FRAME_SHIFT | HIGH | | cgg/Gcgg | R512A? | PLES_11491 | . | . | AG | AGG | AG | AG | . | . | | | . | . |
| NC_011770 | 1246765 | C | T | 78 | 0 | 8 | 0 | NON_SYNONYMOU | MODERAT | MISSENSE | tCc/tTc | S149F | PLES_11491 | T | T | | | | | T | T | T | T | T | T |
| NC_011770 | 1247399 | A | T | 77 | 0 | 4 | 0 | NON_SYNONYMOU | MODERAT | MISSENSE | gaA/gaT | E360D | PLES_11491 | | | | | | | T | T | | | T | T |
| NC_011770 | 1247445 | C | A | 75 | 0 | 2 | 0 | NON_SYNONYMOU | MODERAT | MISSENSE | Ccc/Acc | P376T | PLES_11491 | | | | | A | A | | | | | | |

| | | | | | | | | | | | | | | | | | | | | | | | | | |
|---|---|---|---|---|---|---|---|---|---|---|---|---|---|---|---|---|---|---|---|---|---|---|---|---|---|
| NC_011770 | 1247640 | A | G | 72 | 0 | 6 | 0 | NON_SYNONYMOU | MODERAT | MISSENSE | Atc/Gtc | I441V | PLES_11491 | G | G | | | | | G | G | | | G | G |
| NC_011770 | 1247754 | C | T | 65 | 0 | 4 | 0 | NON_SYNONYMOU | MODERAT | MISSENSE | Cgt/Tgt | R479C | PLES_11491 | | | T | T | T | T | | | | | | |
| NC_011770 | 1255707 | C | T | 73 | 0 | 1 | 0 | NON_SYNONYMOU | MODERAT | MISSENSE | gGg/gAg | G158E | PLES_11561 | | | | | | | T | | | | | |
| NC_011770 | 1256542 | G | T | 76 | 0 | 2 | 0 | NON_SYNONYMOU | MODERAT | MISSENSE | aGc/aTc | S71I | PLES_11571 | | | | | | | | | T | T | | |
| NC_011770 | 1263442 | T | C | 77 | 0 | 1 | 0 | NON_SYNONYMOU | MODERAT | MISSENSE | gTg/gCg | V36A | PLES_11661 | | | | | | | C | | | | | |
| NC_011770 | 1264240 | T | C | 69 | 0 | 12 | 0 | NON_SYNONYMOU | MODERAT | MISSENSE | aTa/aCa | I2T | PLES_11681 | C | C | C | C | C | C | C | C | C | C | C | C |
| NC_011770 | 1266745 | G | T | 19 | 0 | 4 | 0 | NON_SYNONYMOU | MODERAT | MISSENSE | Gcg/Tcg | A200S | PLES_11701 | | | T | T | T | T | | | | | | |
| NC_011770 | 1266922 | A | G | 38 | 0 | 10 | 0 | NON_SYNONYMOU | MODERAT | MISSENSE | Acc/Gcc | T259A | PLES_11701 | G | G | G | G | G | G | G | G | G | G | | |
| NC_011770 | 1273697 | C | T | 80 | 0 | 6 | 0 | NON_SYNONYMOU | MODERAT | MISSENSE | gCg/gTg | A139V | PLES_11761 | | | T | T | T | T | T | T | | | | |
| NC_011770 | 1273712 | C | T | 83 | 0 | 2 | 0 | NON_SYNONYMOU | MODERAT | MISSENSE | tCc/tTc | S144F | PLES_11761 | | | | | | | | | | | T | T |
| NC_011770 | 1273805 | G | A | 84 | 0 | 2 | 0 | NON_SYNONYMOU | MODERAT | MISSENSE | aGc/aAc | S175N | PLES_11761 | A | A | | | | | | | | | | |
| NC_011770 | 1273741 | A | G | 81 | 0 | 1 | 0 | NON_SYNONYMOU | MODERAT | MISSENSE | Atc/Gtc | I154V | PLES_11761 | G | | | | | | | | | | | |
| NC_011770 | 1276731 | G | A | 54 | 0 | 2 | 0 | NON_SYNONYMOU | MODERAT | MISSENSE | Cgc/Tgc | R81C | PLES_11791 | | | | | | | | | A | A | | |
| NC_011770 | 1280412 | A | G | 67 | 0 | 8 | 0 | NON_SYNONYMOU | MODERAT | MISSENSE | Acc/Gcc | T67A | PLES_11831 | G | G | G | G | G | G | | | G | G | | |
| NC_011770 | 1280476 | C | G | 63 | 0 | 4 | 0 | NON_SYNONYMOU | MODERAT | MISSENSE | gCc/gGc | A88G | PLES_11831 | | | G | G | G | G | | | | | | |
| NC_011770 | 1283401 | A | C | 77 | 0 | 12 | 0 | NON_SYNONYMOU | MODERAT | MISSENSE | Aag/Cag | K124Q | PLES_11851 | C | C | C | C | C | C | C | C | C | C | C | C |
| NC_011770 | 1283604 | G | T | 88 | 0 | 2 | 0 | NON_SYNONYMOU | MODERAT | MISSENSE | gaG/gaT | E191D | PLES_11851 | | | | | | | | | T | T | | |
| NC_011770 | 1283614 | C | G | 83 | 0 | 2 | 0 | NON_SYNONYMOU | MODERAT | MISSENSE | Ctc/Gtc | L195V | PLES_11851 | | | | | | | | | G | G | | |
| NC_011770 | 1284320 | G | C | 72 | 0 | 4 | 0 | NON_SYNONYMOU | MODERAT | MISSENSE | aGc/aCc | S430T | PLES_11851 | | | C | C | C | C | | | | | | |
| NC_011770 | 1284269 | C | T | 75 | 0 | 1 | 0 | NON_SYNONYMOU | MODERAT | MISSENSE | gCc/gTc | A413V | PLES_11851 | | | | | | | | | | | | T |
| NC_011770 | 1284533 | C | T | 62 | 0 | 2 | 0 | NON_SYNONYMOU | MODERAT | MISSENSE | Gcc/Acc | A110T | PLES_11861 | | | | | | | T | T | | | | |
| NC_011770 | 1284779 | G | A | 34 | 0 | 4 | 0 | NON_SYNONYMOU | MODERAT | MISSENSE | Ccg/Tcg | P28S | PLES_11861 | | | A | A | A | A | | | | | | |
| NC_011770 | 1284824 | C | T | 37 | 0 | 4 | 0 | NON_SYNONYMOU | MODERAT | MISSENSE | Gca/Aca | A13T | PLES_11861 | | | | | | | T | T | T | T | | |
| NC_011770 | 1284836 | A | C | 36 | 0 | 2 | 0 | NON_SYNONYMOU | MODERAT | MISSENSE | Tgg/Ggg | W9G | PLES_11861 | | | | | | | C | C | | | | |
| NC_011770 | 1285074 | G | A | 47 | 0 | 2 | 0 | NON_SYNONYMOU | MODERAT | MISSENSE | Ccg/Tcg | P241S | PLES_11871 | | | | | | | A | A | | | | |
| NC_011770 | 1285517 | A | G | 37 | 0 | 12 | 0 | NON_SYNONYMOU | MODERAT | MISSENSE | aTc/aCc | I93T | PLES_11871 | G | G | G | G | G | G | G | G | G | G | G | G |
| NC_011770 | 1285734 | C | T | 39 | 0 | 2 | 0 | NON_SYNONYMOU | MODERAT | MISSENSE | Gcc/Acc | A21T | PLES_11871 | T | T | | | | | | | | | | |
| NC_011770 | 1285427 | A | G | 41 | 0 | 1 | 0 | NON_SYNONYMOU | MODERAT | MISSENSE | cTg/cCg | L123P | PLES_11871 | G | | | | | | | | | | | |
| NC_011770 | 1285982 | A | G | 51 | 0 | 12 | 0 | NON_SYNONYMOU | MODERAT | MISSENSE | cAc/cGc | H6R | PLES_11881 | G | G | G | G | G | G | G | G | G | G | G | G |
| NC_011770 | 1286338 | G | A | 65 | 0 | 1 | 0 | NON_SYNONYMOU | MODERAT | MISSENSE | Gct/Act | A125T | PLES_11881 | | | | | | | A | | | | | |
| NC_011770 | 1286733 | G | GC | 81 | 0 | 2 | 0 | FRAME_SHIFT | HIGH | | atg/Catg | M84H? | PLES_11891 | | | | | GC | GC | | | | | | |
| NC_011770 | 1286555 | A | G | 76 | 0 | 6 | 0 | NON_SYNONYMOU | MODERAT | MISSENSE | aAc/aGc | N24S | PLES_11891 | | | | | | | G | G | G | G | G | G |
| NC_011770 | 1286764 | G | A | 78 | 0 | 4 | 0 | NON_SYNONYMOU | MODERAT | MISSENSE | Gtc/Atc | V94I | PLES_11891 | | | | | | | A | A | | | A | A |
| NC_011770 | 1287412 | C | T | 82 | 0 | 2 | 0 | NON_SYNONYMOU | MODERAT | MISSENSE | gCg/gTg | A132V | PLES_11901 | T | T | | | | | | | | | | |
| NC_011770 | 1287636 | G | A | 76 | 0 | 4 | 0 | NON_SYNONYMOU | MODERAT | MISSENSE | gCc/gTc | A207V | PLES_11911 | | | A | A | A | A | | | | | | |
| NC_011770 | 1287930 | T | C | 74 | 0 | 2 | 0 | NON_SYNONYMOU | MODERAT | MISSENSE | gAc/gGc | D109G | PLES_11911 | | | | | | | C | C | | | | |
| NC_011770 | 1287948 | T | C | 73 | 0 | 4 | 0 | NON_SYNONYMOU | MODERAT | MISSENSE | gAc/gGc | D103G | PLES_11911 | | | C | C | C | C | | | | | | |
| NC_011770 | 1288053 | C | A | 70 | 0 | 2 | 0 | NON_SYNONYMOU | MODERAT | MISSENSE | cGc/cTc | R68L | PLES_11911 | | | | | | | A | A | | | | |
| NC_011770 | 1288093 | C | T | 76 | 0 | 1 | 0 | NON_SYNONYMOU | MODERAT | MISSENSE | Gtg/Atg | V55M | PLES_11911 | | | | | | | T | | | | | |
| NC_011770 | 1288120 | A | G | 70 | 0 | 12 | 0 | NON_SYNONYMOU | MODERAT | MISSENSE | Tac/Cac | Y46H | PLES_11911 | G | G | G | G | G | G | G | G | G | G | G | G |
| NC_011770 | 1288249 | T | C | 78 | 0 | 12 | 0 | NON_SYNONYMOU | MODERAT | MISSENSE | Aca/Gca | T3A | PLES_11911 | C | C | C | C | C | C | C | C | C | C | C | C |
| NC_011770 | 1288252 | A | G | 79 | 0 | 2 | 0 | NON_SYNONYMOU | MODERAT | MISSENSE | Tcc/Ccc | S2P | PLES_11911 | | | | | | | G | G | | | | |
| NC_011770 | 1288391 | G | T | 79 | 0 | 4 | 0 | NON_SYNONYMOU | MODERAT | MISSENSE | Gtg/Ttg | V6L | PLES_11921 | | | T | T | T | T | | | | | | |
| NC_011770 | 1288505 | G | A | 77 | 0 | 6 | 0 | NON_SYNONYMOU | MODERAT | MISSENSE | Gcc/Acc | A44T | PLES_11921 | | | | | | | A | A | A | A | A | A |
| NC_011770 | 1288676 | G | T | 79 | 0 | 2 | 0 | NON_SYNONYMOU | MODERAT | MISSENSE | Gcg/Tcg | A101S | PLES_11921 | | | | | | | | | | | T | T |
| NC_011770 | 1289151 | A | G | 82 | 0 | 4 | 0 | NON_SYNONYMOU | MODERAT | MISSENSE | aAc/aGc | N259S | PLES_11921 | | | | | | | | | G | G | G | G |
| NC_011770 | 1290322 | C | T | 75 | 0 | 2 | 0 | NON_SYNONYMOU | MODERAT | MISSENSE | Gcc/Acc | A351T | PLES_11931 | | | T | T | | | | | | | | |
| NC_011770 | 1290610 | G | T | 85 | 0 | 6 | 0 | NON_SYNONYMOU | MODERAT | MISSENSE | Ctg/Atg | L255M | PLES_11931 | T | T | | | | | T | T | | | T | T |
| NC_011770 | 1291261 | C | A | 78 | 0 | 4 | 0 | NON_SYNONYMOU | MODERAT | MISSENSE | Gcg/Tcg | A38S | PLES_11931 | | | A | A | A | A | | | | | | |
| NC_011770 | 1291395 | T | G | 79 | 0 | 4 | 0 | NON_SYNONYMOU | MODERAT | MISSENSE | Acc/Ccc | T157P | PLES_11941 | | | G | G | G | G | | | | | | |
| NC_011770 | 1291413 | G | A | 76 | 0 | 4 | 0 | NON_SYNONYMOU | MODERAT | MISSENSE | Cgg/Tgg | R151W | PLES_11941 | | | A | A | A | A | | | | | | |
| NC_011770 | 1291497 | C | T | 78 | 0 | 4 | 0 | NON_SYNONYMOU | MODERAT | MISSENSE | Gtc/Atc | V123I | PLES_11941 | | | | | | | | | T | T | T | T |
| NC_011770 | 1291607 | G | A | 81 | 0 | 4 | 0 | NON_SYNONYMOU | MODERAT | MISSENSE | gCc/gTc | A86V | PLES_11941 | | | A | A | A | A | | | | | | |
| NC_011770 | 1292159 | T | A | 69 | 0 | 2 | 0 | NON_SYNONYMOU | MODERAT | MISSENSE | Acc/Tcc | T275S | PLES_11951 | A | A | | | | | | | | | | |
| NC_011770 | 1292213 | C | T | 70 | 0 | 6 | 0 | NON_SYNONYMOU | MODERAT | MISSENSE | Gcc/Acc | A257T | PLES_11951 | | | T | T | T | T | | | | | T | T |
| NC_011770 | 1292962 | C | T | 55 | 0 | 2 | 0 | NON_SYNONYMOU | MODERAT | MISSENSE | cGc/cAc | R7H | PLES_11951 | | | | | | | | | | | T | T |
| NC_011770 | 1292701 | C | T | 66 | 0 | 1 | 0 | NON_SYNONYMOU | MODERAT | MISSENSE | gGc/gAc | G94D | PLES_11951 | | T | | | | | | | | | | |
| NC_011770 | 1293977 | T | G | 57 | 0 | 2 | 0 | NON_SYNONYMOU | MODERAT | MISSENSE | gAc/gCc | D31A | PLES_11961 | | | | | G | G | | | | | | |
| NC_011770 | 1295826 | C | A | 56 | 0 | 2 | 0 | NON_SYNONYMOU | MODERAT | MISSENSE | Gcc/Tcc | A192S | PLES_11981 | A | A | | | | | | | | | | |
| NC_011770 | 1296047 | A | G | 69 | 0 | 8 | 0 | NON_SYNONYMOU | MODERAT | MISSENSE | gTg/gCg | V118A | PLES_11981 | G | G | G | G | G | G | | | | | G | G |
| NC_011770 | 1296083 | C | T | 70 | 0 | 6 | 0 | NON_SYNONYMOU | MODERAT | MISSENSE | gGc/gAc | G106D | PLES_11981 | | | T | T | T | T | T | T | | | | |
| NC_011770 | 1296111 | C | T | 71 | 0 | 1 | 0 | NON_SYNONYMOU | MODERAT | MISSENSE | Ggg/Agg | G97R | PLES_11981 | | | | | | | | T | | | | |

| | | | | | | | | | | | | | | | | | | | | | | | | | |
|---|---|---|---|---|---|---|---|---|---|---|---|---|---|---|---|---|---|---|---|---|---|---|---|---|---|
| NC_011770 | 1296753 | G | A | 69 | 0 | 2 | 0 | NON_SYNONYMOU | MODERAT | MISSENSE | Gtc/Atc | V83I | PLES_11991 | | | | | | | | | | | A | A |
| NC_011770 | 1296870 | T | C | 63 | 0 | 8 | 0 | NON_SYNONYMOU | MODERAT | MISSENSE | Ttc/Ctc | F122L | PLES_11991 | C | C | C | C | C | C | C | C | | | | |
| NC_011770 | 1297512 | C | T | 82 | 0 | 2 | 0 | NON_SYNONYMOU | MODERAT | MISSENSE | gGc/gAc | G302D | PLES_12001 | | | | | | | T | T | | | | |
| NC_011770 | 1297537 | G | A | 82 | 0 | 2 | 0 | NON_SYNONYMOU | MODERAT | MISSENSE | Cgc/Tgc | R294C | PLES_12001 | | | | | | | | | | | A | A |
| NC_011770 | 1297540 | C | T | 84 | 0 | 2 | 0 | NON_SYNONYMOU | MODERAT | MISSENSE | Gtc/Atc | V293I | PLES_12001 | | | | | | | | | T | T | | |
| NC_011770 | 1297674 | C | T | 99 | 0 | 2 | 0 | NON_SYNONYMOU | MODERAT | MISSENSE | cGc/cAc | R248H | PLES_12001 | | | | | | | | | | | T | T |
| NC_011770 | 1297777 | T | C | 95 | 0 | 6 | 0 | NON_SYNONYMOU | MODERAT | MISSENSE | Agc/Ggc | S214G | PLES_12001 | | | C | C | C | C | C | C | | | | |
| NC_011770 | 1298029 | G | C | 77 | 0 | 10 | 0 | NON_SYNONYMOU | MODERAT | MISSENSE | Cag/Gag | Q130E | PLES_12001 | | | C | C | C | C | C | C | C | C | C | C |
| NC_011770 | 1298035 | C | T | 80 | 0 | 6 | 0 | NON_SYNONYMOU | MODERAT | MISSENSE | Gcc/Acc | A128T | PLES_12001 | | | T | T | T | T | T | T | | | | |
| NC_011770 | 1298757 | T | G | 73 | 0 | 8 | 0 | NON_SYNONYMOU | MODERAT | MISSENSE | Atg/Ctg | M283L | PLES_12011 | G | G | G | G | G | G | G | G | | | | |
| NC_011770 | 1298846 | A | C | 63 | 0 | 2 | 0 | NON_SYNONYMOU | MODERAT | MISSENSE | gTc/gGc | V253G | PLES_12011 | C | C | | | | | | | | | | |
| NC_011770 | 1298874 | C | T | 63 | 0 | 6 | 0 | NON_SYNONYMOU | MODERAT | MISSENSE | Gcc/Acc | A244T | PLES_12011 | | | T | T | T | T | T | T | | | | |
| NC_011770 | 1298904 | C | T | 59 | 0 | 6 | 0 | NON_SYNONYMOU | MODERAT | MISSENSE | Ggc/Agc | G234S | PLES_12011 | | | T | T | T | T | T | T | | | | |
| NC_011770 | 1299006 | A | G | 54 | 0 | 12 | 0 | NON_SYNONYMOU | MODERAT | MISSENSE | Tgc/Cgc | C200R | PLES_12011 | G | G | G | G | G | G | G | G | G | G | G | G |
| NC_011770 | 1299063 | C | T | 61 | 0 | 2 | 0 | NON_SYNONYMOU | MODERAT | MISSENSE | Ggc/Agc | G181S | PLES_12011 | T | T | | | | | | | | | | |
| NC_011770 | 1299465 | T | C | 47 | 0 | 6 | 0 | NON_SYNONYMOU | MODERAT | MISSENSE | Acc/Gcc | T47A | PLES_12011 | | | C | C | C | C | C | C | | | | |
| NC_011770 | 1299599 | C | T | 55 | 0 | 6 | 0 | NON_SYNONYMOU | MODERAT | MISSENSE | aGc/aAc | S2N | PLES_12011 | | | T | T | T | T | T | T | | | | |
| NC_011770 | 1299680 | T | C | 57 | 0 | 2 | 0 | NON_SYNONYMOU | MODERAT | MISSENSE | aAc/aGc | N278S | PLES_12021 | | | | | | | C | C | | | | |
| NC_011770 | 1299777 | A | T | 61 | 0 | 12 | 0 | NON_SYNONYMOU | MODERAT | MISSENSE | Tcc/Acc | S246T | PLES_12021 | T | T | T | T | T | T | T | T | T | T | T | T |
| NC_011770 | 1300020 | C | T | 55 | 0 | 6 | 0 | NON_SYNONYMOU | MODERAT | MISSENSE | Gcc/Acc | A165T | PLES_12021 | | | T | T | T | T | T | T | | | | |
| NC_011770 | 1300281 | T | C | 54 | 0 | 6 | 0 | NON_SYNONYMOU | MODERAT | MISSENSE | Atg/Gtg | M78V | PLES_12021 | | | C | C | C | C | C | C | | | | |
| NC_011770 | 1300082 | A | G | 58 | 0 | 1 | 0 | NON_SYNONYMOU | MODERAT | MISSENSE | cTg/cCg | L144P | PLES_12021 | HP | G | | | | | | | | | | |
| NC_011770 | 1300910 | T | C | 80 | 0 | 8 | 0 | NON_SYNONYMOU | MODERAT | MISSENSE | Acg/Gcg | T239A | PLES_12031 | C | C | | | | | C | C | C | C | C | C |
| NC_011770 | 1300951 | G | A | 78 | 0 | 1 | 0 | NON_SYNONYMOU | MODERAT | MISSENSE | cCg/cTg | P225L | PLES_12031 | | | | | | | | | | | A | |
| NC_011770 | 1312085 | T | C | 42 | 0 | 4 | 0 | NON_SYNONYMOU | MODERAT | MISSENSE | Ttc/Ctc | F27L | PLES_12111 | | | | | | | | | C | C | C | C |
| NC_011770 | 1312157 | A | G | 52 | 0 | 2 | 0 | NON_SYNONYMOU | MODERAT | MISSENSE | Atc/Gtc | I51V | PLES_12111 | | | | | | | | | | | G | G |
| NC_011770 | 1312176 | G | C | 55 | 0 | 2 | 0 | NON_SYNONYMOU | MODERAT | MISSENSE | aGt/aCt | S57T | PLES_12111 | | | | | | | | | | | C | C |
| NC_011770 | 1313230 | C | G | 94 | 0 | 4 | 0 | NON_SYNONYMOU | MODERAT | MISSENSE | gaC/gaG | D408E | PLES_12111 | | | G | G | G | G | | | | | | |
| NC_011770 | 1313878 | G | A | 85 | 0 | 2 | 0 | NON_SYNONYMOU | MODERAT | MISSENSE | Gcg/Acg | A164T | PLES_12121 | | | | | | | A | A | | | | |
| NC_011770 | 1314638 | T | G | 71 | 0 | 10 | 0 | NON_SYNONYMOU | MODERAT | MISSENSE | Tcc/Gcc | S105A | PLES_12131 | G | G | G | G | G | G | | | G | G | G | G |
| NC_011770 | 1314913 | C | A | 81 | 0 | 2 | 0 | NON_SYNONYMOU | MODERAT | MISSENSE | gaC/gaA | D196E | PLES_12131 | | | | | | | | | | | A | A |
| NC_011770 | 1315893 | C | A | 73 | 0 | 4 | 0 | NON_SYNONYMOU | MODERAT | MISSENSE | agC/agA | S62R | PLES_12141 | | | A | A | A | A | | | | | | |
| NC_011770 | 1315901 | A | T | 72 | 0 | 6 | 0 | NON_SYNONYMOU | MODERAT | MISSENSE | cAg/cTg | Q65L | PLES_12141 | T | T | | | | | T | T | | | T | T |
| NC_011770 | 1316207 | G | C | 76 | 0 | 4 | 0 | NON_SYNONYMOU | MODERAT | MISSENSE | cGc/cCc | R167P | PLES_12141 | | | C | C | C | C | | | | | | |
| NC_011770 | 1317265 | A | G | 84 | 0 | 2 | 0 | NON_SYNONYMOU | MODERAT | MISSENSE | Tat/Cat | Y134H | PLES_12151 | | | | | | | | | G | G | | |
| NC_011770 | 1317483 | C | A | 74 | 0 | 2 | 0 | NON_SYNONYMOU | MODERAT | MISSENSE | cGc/cTc | R61L | PLES_12151 | | | | | | | A | A | | | | |
| NC_011770 | 1323187 | C | A | 62 | 0 | 6 | 0 | NON_SYNONYMOU | MODERAT | MISSENSE | Gcc/Tcc | A491S | PLES_12191 | | | A | A | A | A | | | | | A | A |
| NC_011770 | 1323889 | G | C | 78 | 0 | 2 | 0 | NON_SYNONYMOU | MODERAT | MISSENSE | Ccg/Gcg | P257A | PLES_12191 | | | | | | | C | C | | | | |
| NC_011770 | 1323900 | C | T | 78 | 0 | 2 | 0 | NON_SYNONYMOU | MODERAT | MISSENSE | cGc/cAc | R253H | PLES_12191 | | | | | | | | | | | T | T |
| NC_011770 | 1325136 | C | T | 55 | 0 | 2 | 0 | NON_SYNONYMOU | MODERAT | MISSENSE | cGc/cAc | R692H | PLES_12201 | | | | | | | T | T | | | | |
| NC_011770 | 1325176 | C | G | 55 | 0 | 4 | 0 | NON_SYNONYMOU | MODERAT | MISSENSE | Ggc/Cgc | G679R | PLES_12201 | | | | | | | | | G | G | G | G |
| NC_011770 | 1325341 | C | T | 50 | 0 | 2 | 0 | NON_SYNONYMOU | MODERAT | MISSENSE | Gtg/Atg | V624M | PLES_12201 | | | | | | | | | | | T | T |
| NC_011770 | 1325637 | C | T | 63 | 0 | 4 | 0 | NON_SYNONYMOU | MODERAT | MISSENSE | cGc/cAc | R525H | PLES_12201 | | | | | | | T | T | | | T | T |
| NC_011770 | 1326361 | C | T | 69 | 0 | 4 | 0 | NON_SYNONYMOU | MODERAT | MISSENSE | Gcc/Acc | A284T | PLES_12201 | | | T | T | T | T | | | | | | |
| NC_011770 | 1326394 | G | C | 69 | 0 | 2 | 0 | NON_SYNONYMOU | MODERAT | MISSENSE | Ccg/Gcg | P273A | PLES_12201 | | | | | | | | | | | C | C |
| NC_011770 | 1326422 | C | A | 72 | 0 | 4 | 0 | NON_SYNONYMOU | MODERAT | MISSENSE | gaG/gaT | E263D | PLES_12201 | | | | | | | A | A | A | A | | |
| NC_011770 | 1326471 | T | C | 68 | 0 | 6 | 0 | NON_SYNONYMOU | MODERAT | MISSENSE | gAc/gGc | D247G | PLES_12201 | | | | | | | C | C | C | C | C | C |
| NC_011770 | 1326753 | G | A | 78 | 0 | 2 | 0 | NON_SYNONYMOU | MODERAT | MISSENSE | cCg/cTg | P153L | PLES_12201 | | | | | | | A | A | | | | |
| NC_011770 | 1325847 | C | T | 74 | 0 | 1 | 0 | NON_SYNONYMOU | MODERAT | MISSENSE | tGc/tAc | C455Y | PLES_12201 | | | | | | | | | | | | T |
| NC_011770 | 1328247 | C | T | 81 | 0 | 1 | 0 | START_LOST | HIGH | MISSENSE | atG/atA | M1I | PLES_12211 | | T | | | | | | | | | | |
| NC_011770 | 1327280 | C | T | 71 | 0 | 2 | 0 | NON_SYNONYMOU | MODERAT | MISSENSE | Ggc/Agc | G324S | PLES_12211 | | | | | | | | | | | T | T |
| NC_011770 | 1327628 | C | T | 67 | 0 | 2 | 0 | NON_SYNONYMOU | MODERAT | MISSENSE | Ggt/Agt | G208S | PLES_12211 | | | | | | | T | T | | | | |
| NC_011770 | 1327741 | T | C | 67 | 0 | 2 | 0 | NON_SYNONYMOU | MODERAT | MISSENSE | aAt/aGt | N170S | PLES_12211 | | | | | | | | | | | C | C |
| NC_011770 | 1327781 | T | C | 68 | 0 | 8 | 0 | NON_SYNONYMOU | MODERAT | MISSENSE | Agc/Ggc | S157G | PLES_12211 | C | C | C | C | C | C | | | C | C | | |
| NC_011770 | 1327825 | G | C | 72 | 0 | 2 | 0 | NON_SYNONYMOU | MODERAT | MISSENSE | aCc/aGc | T142S | PLES_12211 | | | | | | | | | | | C | C |
| NC_011770 | 1327826 | T | C | 70 | 0 | 2 | 0 | NON_SYNONYMOU | MODERAT | MISSENSE | Acc/Gcc | T142A | PLES_12211 | | | | | | | C | C | | | | |
| NC_011770 | 1328210 | C | T | 83 | 0 | 6 | 0 | NON_SYNONYMOU | MODERAT | MISSENSE | Gcg/Acg | A14T | PLES_12211 | T | T | | | | | T | T | | | T | T |
| NC_011770 | 1328347 | C | T | 87 | 0 | 2 | 0 | NON_SYNONYMOU | MODERAT | MISSENSE | Gcc/Acc | A331T | PLES_12221 | | | | | | | T | T | | | | |
| NC_011770 | 1328841 | C | T | 72 | 0 | 2 | 0 | NON_SYNONYMOU | MODERAT | MISSENSE | cGg/cAg | R166Q | PLES_12221 | | | | | | | | | T | T | | |
| NC_011770 | 1328902 | C | T | 77 | 0 | 2 | 0 | NON_SYNONYMOU | MODERAT | MISSENSE | Gac/Aac | D146N | PLES_12221 | | | | | | | | | | | T | T |
| NC_011770 | 1329033 | G | T | 77 | 0 | 4 | 0 | NON_SYNONYMOU | MODERAT | MISSENSE | gCt/gAt | A102D | PLES_12221 | T | T | | | | | | | | | T | T |
| NC_011770 | 1329686 | C | T | 45 | 0 | 4 | 0 | NON_SYNONYMOU | MODERAT | MISSENSE | Gcg/Acg | A138T | PLES_12231 | T | T | | | | | T | T | | | | |

| | | | | | | | | | | | | | | | | | | | | | | | | | |
|---|---|---|---|---|---|---|---|---|---|---|---|---|---|---|---|---|---|---|---|---|---|---|---|---|---|
| NC_011770 | 1330067 | G | T | 41 | 0 | 2 | 0 | NON_SYNONYMOU | MODERAT | MISSENSE | Ctg/Atg | L11M | PLES_12231 | | | | | | | | | T | T | | |
| NC_011770 | 1331079 | C | T | 49 | 0 | 2 | 0 | NON_SYNONYMOU | MODERAT | MISSENSE | Ggc/Agc | G127S | PLES_12251 | | | | | | | T | T | | | | |
| NC_011770 | 1332280 | A | G | 80 | 0 | 8 | 0 | NON_SYNONYMOU | MODERAT | MISSENSE | Acc/Gcc | T12A | PLES_12271 | G | G | G | G | G | G | | | G | G | | |
| NC_011770 | 1332745 | C | T | 92 | 0 | 8 | 0 | NON_SYNONYMOU | MODERAT | MISSENSE | Cgc/Tgc | R167C | PLES_12271 | T | T | T | T | T | T | | | T | T | | |
| NC_011770 | 1334198 | A | T | 66 | 0 | 4 | 0 | NON_SYNONYMOU | MODERAT | MISSENSE | Acg/Tcg | T6S | PLES_12301 | | | T | T | T | T | | | | | | |
| NC_011770 | 1334324 | T | G | 62 | 0 | 4 | 0 | NON_SYNONYMOU | MODERAT | MISSENSE | Tcc/Gcc | S48A | PLES_12301 | | | G | G | G | G | | | | | | |
| NC_011770 | 1334928 | T | C | 67 | 0 | 6 | 0 | NON_SYNONYMOU | MODERAT | MISSENSE | gTc/gCc | V249A | PLES_12301 | | | C | C | C | C | C | C | | | | |
| NC_011770 | 1335713 | C | T | 81 | 0 | 4 | 0 | NON_SYNONYMOU | MODERAT | MISSENSE | Gcc/Acc | A181T | PLES_12311 | | | | | | | | | T | T | T | T |
| NC_011770 | 1335022 | A | G | 65 | 0 | 1 | 0 | NON_SYNONYMOU | MODERAT | MISSENSE | gTc/gCc | V411A | PLES_12311 | | | | | | | | G | | | | |
| NC_011770 | 1335613 | C | T | 92 | 0 | 1 | 0 | NON_SYNONYMOU | MODERAT | MISSENSE | cGc/cAc | R214H | PLES_12311 | | | | | | | | T | | | | |
| NC_011770 | 1344539 | C | T | 59 | 0 | 1 | 0 | NON_SYNONYMOU | MODERAT | MISSENSE | Gcc/Acc | A414T | PLES_12411 | | | | | | | T | | | | | |
| NC_011770 | 1351277 | T | C | 72 | 0 | 8 | 0 | NON_SYNONYMOU | MODERAT | MISSENSE | gAc/gGc | D236G | PLES_12461 | C | C | C | C | C | C | | | C | C | | |
| NC_011770 | 1351292 | G | C | 74 | 0 | 4 | 0 | NON_SYNONYMOU | MODERAT | MISSENSE | gCc/gGc | A231G | PLES_12461 | | | | | | | C | C | | | C | C |
| NC_011770 | 1351388 | C | T | 75 | 0 | 4 | 0 | NON_SYNONYMOU | MODERAT | MISSENSE | gGc/gAc | G199D | PLES_12461 | | | T | T | T | T | | | | | | |
| NC_011770 | 1351470 | G | A | 68 | 0 | 2 | 0 | NON_SYNONYMOU | MODERAT | MISSENSE | Ctt/Ttt | L172F | PLES_12461 | | | | | | | | | A | A | | |
| NC_011770 | 1351581 | T | C | 64 | 0 | 12 | 0 | NON_SYNONYMOU | MODERAT | MISSENSE | Atc/Gtc | I135V | PLES_12461 | C | C | C | C | C | C | C | C | C | C | C | C |
| NC_011770 | 1351764 | C | T | 62 | 0 | 2 | 0 | NON_SYNONYMOU | MODERAT | MISSENSE | Ggc/Agc | G74S | PLES_12461 | | | | | | | T | T | | | | |
| NC_011770 | 1351782 | C | T | 64 | 0 | 2 | 0 | NON_SYNONYMOU | MODERAT | MISSENSE | Gcc/Acc | A68T | PLES_12461 | | | | | | | | | T | T | | |
| NC_011770 | 1351983 | C | T | 59 | 0 | 2 | 0 | NON_SYNONYMOU | MODERAT | MISSENSE | Gtg/Atg | V1M | PLES_12461 | | | | | | | | | T | T | | |
| NC_011770 | 1352517 | A | AG | 78 | 0 | 1 | 0 | FRAME_SHIFT | HIGH | | ccc/cccC | P8P? | PLES_12481 | | | | | | | | | AG | | | |
| NC_011770 | 1352529 | G | A | 79 | 0 | 2 | 0 | STOP_GAINED | HIGH | NONSENSE | Cag/Tag | Q5* | PLES_12481 | | | | | | | A | A | | | | |
| NC_011770 | 1352504 | C | T | 80 | 0 | 2 | 0 | NON_SYNONYMOU | MODERAT | MISSENSE | aGc/aAc | S13N | PLES_12481 | | | | | | | | | | | T | T |
| NC_011770 | 1352526 | C | A | 67 | 0 | 4 | 0 | NON_SYNONYMOU | MODERAT | MISSENSE | Ggc/Tgc | G6C | PLES_12481 | | | | | | | A | A | A | A | | |
| NC_011770 | 1352861 | A | G | 86 | 0 | 8 | 0 | NON_SYNONYMOU | MODERAT | MISSENSE | Tcc/Ccc | S312P | PLES_12491 | | | G | G | G | G | G | G | G | G | | |
| NC_011770 | 1352938 | T | C | 93 | 0 | 6 | 0 | NON_SYNONYMOU | MODERAT | MISSENSE | cAa/cGa | Q286R | PLES_12491 | | | | | | | C | C | C | C | C | C |
| NC_011770 | 1355960 | A | G | 85 | 0 | 2 | 0 | NON_SYNONYMOU | MODERAT | MISSENSE | cAa/cGa | Q159R | PLES_12521 | | | | | | | | | | | G | G |
| NC_011770 | 1356392 | CGACCGGC | C | 90 | 0 | 1 | 0 | FRAME_SHIFT | HIGH | | gaccggcgcg | DRRG78 | PLES_12531 | | | | C | | | | | | | | |
| NC_011770 | 1357758 | C | T | 76 | 0 | 1 | 0 | STOP_GAINED | HIGH | NONSENSE | Cag/Tag | Q533* | PLES_12531 | | | | | | | T | | | | | |
| NC_011770 | 1356201 | G | A | 89 | 0 | 4 | 0 | NON_SYNONYMOU | MODERAT | MISSENSE | Ggc/Agc | G14S | PLES_12531 | | | A | A | A | A | | | | | | |
| NC_011770 | 1357431 | A | G | 80 | 0 | 1 | 0 | NON_SYNONYMOU | MODERAT | MISSENSE | Acc/Gcc | T424A | PLES_12531 | | | | | | | G | | | | | |
| NC_011770 | 1357985 | G | C | 75 | 0 | 12 | 0 | NON_SYNONYMOU | MODERAT | MISSENSE | gaG/gaC | E608D | PLES_12531 | C | C | C | C | C | C | C | C | C | C | C | C |
| NC_011770 | 1358709 | A | G | 77 | 0 | 2 | 0 | NON_SYNONYMOU | MODERAT | MISSENSE | gAg/gGg | E111G | PLES_12541 | G | G | | | | | | | | | | |
| NC_011770 | 1358814 | A | G | 85 | 0 | 2 | 0 | NON_SYNONYMOU | MODERAT | MISSENSE | aAc/aGc | N146S | PLES_12541 | | | | | | | | | | | G | G |
| NC_011770 | 1360071 | T | C | 70 | 0 | 2 | 0 | NON_SYNONYMOU | MODERAT | MISSENSE | tTg/tCg | L565S | PLES_12541 | | | | | | | | | C | C | | |
| NC_011770 | 1360458 | G | T | 73 | 0 | 4 | 0 | NON_SYNONYMOU | MODERAT | MISSENSE | aGc/aTc | S694I | PLES_12541 | | | T | T | T | T | | | | | | |
| NC_011770 | 1360617 | A | G | 76 | 0 | 1 | 0 | NON_SYNONYMOU | MODERAT | MISSENSE | cAg/cGg | Q747R | PLES_12541 | | | | | | | G | | | | | |
| NC_011770 | 1361087 | C | T | 83 | 0 | 4 | 0 | NON_SYNONYMOU | MODERAT | MISSENSE | Ccg/Tcg | P904S | PLES_12541 | | | T | T | T | T | | | | | | |
| NC_011770 | 1361337 | A | G | 66 | 0 | 2 | 0 | NON_SYNONYMOU | MODERAT | MISSENSE | cAc/cGc | H987R | PLES_12541 | | | | | | | | | G | G | | |
| NC_011770 | 1361711 | C | A | 76 | 0 | 2 | 0 | NON_SYNONYMOU | MODERAT | MISSENSE | Ctg/Atg | L1112M | PLES_12541 | | | | | | | | | | | A | A |
| NC_011770 | 1361727 | C | T | 74 | 0 | 2 | 0 | NON_SYNONYMOU | MODERAT | MISSENSE | gCg/gTg | A1117V | PLES_12541 | | | | | | | | | | | T | T |
| NC_011770 | 1361969 | G | T | 81 | 0 | 6 | 0 | NON_SYNONYMOU | MODERAT | MISSENSE | Gtc/Ttc | V1198F | PLES_12541 | | | T | T | T | T | | | | | T | T |
| NC_011770 | 1362087 | G | A | 82 | 0 | 6 | 0 | NON_SYNONYMOU | MODERAT | MISSENSE | cGc/cAc | R1237H | PLES_12541 | | | A | A | A | A | | | | | A | A |
| NC_011770 | 1362098 | G | A | 88 | 0 | 2 | 0 | NON_SYNONYMOU | MODERAT | MISSENSE | Gca/Aca | A1241T | PLES_12541 | A | A | | | | | | | | | | |
| NC_011770 | 1362405 | G | A | 90 | 0 | 2 | 0 | NON_SYNONYMOU | MODERAT | MISSENSE | cGg/cAg | R1343Q | PLES_12541 | | | | | | | | | | | A | A |
| NC_011770 | 1362557 | C | T | 80 | 0 | 2 | 0 | NON_SYNONYMOU | MODERAT | MISSENSE | Cgg/Tgg | R1394W | PLES_12541 | | | | | | | T | T | | | | |
| NC_011770 | 1362849 | T | C | 86 | 0 | 10 | 0 | NON_SYNONYMOU | MODERAT | MISSENSE | gTc/gCc | V1491A | PLES_12541 | | | C | C | C | C | C | C | C | C | C | C |
| NC_011770 | 1363197 | A | G | 76 | 0 | 2 | 0 | NON_SYNONYMOU | MODERAT | MISSENSE | cAg/cGg | Q1607R | PLES_12541 | G | G | | | | | | | | | | |
| NC_011770 | 1360370 | C | T | 72 | 0 | 1 | 0 | NON_SYNONYMOU | MODERAT | MISSENSE | Cgc/Tgc | R665C | PLES_12541 | | T | | | | | | | | | | |
| NC_011770 | 1360986 | C | T | 77 | 0 | 1 | 0 | NON_SYNONYMOU | MODERAT | MISSENSE | cCg/cTg | P870L | PLES_12541 HP | | T | | | | | | | | | | |
| NC_011770 | 1364629 | A | G | 69 | 0 | 1 | 0 | NON_SYNONYMOU | MODERAT | MISSENSE | gAc/gGc | D65G | PLES_12561 | | | | G | | | | | | | | |
| NC_011770 | 1369052 | G | A | 62 | 0 | 2 | 0 | NON_SYNONYMOU | MODERAT | MISSENSE | Ggc/Agc | G123S | PLES_12591 | | | | | | | A | A | | | | |
| NC_011770 | 1369439 | C | A | 75 | 0 | 3 | 0 | NON_SYNONYMOU | MODERAT | MISSENSE | Cgc/Agc | R252S | PLES_12591 | | | A | A | | A | | | | | | |
| NC_011770 | 1370183 | G | A | 62 | 0 | 1 | 0 | NON_SYNONYMOU | MODERAT | MISSENSE | gGg/gAg | G68E | PLES_12601 | | | | | | | A | | | | | |
| NC_011770 | 1370616 | TCGCCGGA | T | 56 | 0 | 1 | 0 | FRAME_SHIFT | HIGH | | cggtttccgg | RFPA120 | PLES_12611 | | | | | | T | | | | | | |
| NC_011770 | 1370361 | T | G | 66 | 0 | 8 | 0 | NON_SYNONYMOU | MODERAT | MISSENSE | Agc/Cgc | S209R | PLES_12611 | G | G | | | | | G | G | G | G | G | G |
| NC_011770 | 1371183 | T | TC | 44 | 0 | 1 | 0 | FRAME_SHIFT | HIGH | | atg/Catg | M55H? | PLES_12621 | | | | | | | | TC | | | | |
| NC_011770 | 1371358 | A | G | 58 | 0 | 2 | 0 | NON_SYNONYMOU | MODERAT | MISSENSE | Aca/Gca | T111A | PLES_12621 | | | | | | | | | G | G | | |
| NC_011770 | 1371359 | C | A | 59 | 0 | 2 | 0 | NON_SYNONYMOU | MODERAT | MISSENSE | aCa/aAa | T111K | PLES_12621 | | | | | | | | | A | A | | |
| NC_011770 | 1371951 | T | C | 75 | 0 | 2 | 0 | NON_SYNONYMOU | MODERAT | MISSENSE | cTc/cCc | L7P | PLES_12641 | | | | | | | | | | | C | C |
| NC_011770 | 1372169 | C | T | 73 | 0 | 2 | 0 | NON_SYNONYMOU | MODERAT | MISSENSE | Cgt/Tgt | R80C | PLES_12641 | | | | | | | | | T | T | | |
| NC_011770 | 1372197 | T | G | 70 | 0 | 6 | 0 | NON_SYNONYMOU | MODERAT | MISSENSE | gTt/gGt | V89G | PLES_12641 | | | G | G | G | G | | | | | G | G |
| NC_011770 | 1372248 | A | G | 78 | 0 | 1 | 0 | NON_SYNONYMOU | MODERAT | MISSENSE | gAc/gGc | D106G | PLES_12641 | | | | | | | G | | | | | |

| | | | | | | | | | | | | | | | | | | | | | | | | | |
|---|---|---|---|---|---|---|---|---|---|---|---|---|---|---|---|---|---|---|---|---|---|---|---|---|---|
| NC_011770 | 1372520 | T | C | 71 | 0 | 4 | 0 | NON_SYNONYMOU | MODERAT | MISSENSE | Tgc/Cgc | C197R | PLES_12641 | C | C | | | | | | | | | C | C |
| NC_011770 | 1372556 | A | G | 73 | 0 | 2 | 0 | NON_SYNONYMOU | MODERAT | MISSENSE | Acg/Gcg | T209A | PLES_12641 | | | | | | | | | G | G | | |
| NC_011770 | 1372568 | G | T | 71 | 0 | 2 | 0 | NON_SYNONYMOU | MODERAT | MISSENSE | Gtt/Ttt | V213F | PLES_12641 | | | | | | | | | | | T | T |
| NC_011770 | 1372569 | T | C | 71 | 0 | 2 | 0 | NON_SYNONYMOU | MODERAT | MISSENSE | gTt/gCt | V213A | PLES_12641 | | | | | | | | | | | C | C |
| NC_011770 | 1372788 | G | C | 69 | 0 | 4 | 0 | NON_SYNONYMOU | MODERAT | MISSENSE | aGc/aCc | S286T | PLES_12641 | C | C | | | | | | | | | C | C |
| NC_011770 | 1372808 | C | G | 71 | 0 | 4 | 0 | NON_SYNONYMOU | MODERAT | MISSENSE | Ctc/Gtc | L293V | PLES_12641 | G | G | | | | | | | | | G | G |
| NC_011770 | 1372839 | T | C | 66 | 0 | 8 | 0 | NON_SYNONYMOU | MODERAT | MISSENSE | aTg/aCg | M303T | PLES_12641 | C | C | | | | | C | C | C | C | C | C |
| NC_011770 | 1372937 | G | A | 62 | 0 | 4 | 0 | NON_SYNONYMOU | MODERAT | MISSENSE | Gcg/Acg | A336T | PLES_12641 | A | A | | | | | | | | | A | A |
| NC_011770 | 1373115 | C | T | 50 | 0 | 2 | 0 | NON_SYNONYMOU | MODERAT | MISSENSE | gCg/gTg | A395V | PLES_12641 | T | T | | | | | | | | | | |
| NC_011770 | 1373558 | A | T | 83 | 0 | 2 | 0 | NON_SYNONYMOU | MODERAT | MISSENSE | gTg/gAg | V19E | PLES_12651 | | | | | | | | | T | T | | |
| NC_011770 | 1374181 | A | G | 81 | 0 | 2 | 0 | NON_SYNONYMOU | MODERAT | MISSENSE | Aac/Gac | N120D | PLES_12661 | | | | | | | | | | | G | G |
| NC_011770 | 1374566 | T | C | 80 | 0 | 2 | 0 | NON_SYNONYMOU | MODERAT | MISSENSE | gTg/gCg | V248A | PLES_12661 | | | | | | | | | | | C | C |
| NC_011770 | 1374680 | C | T | 90 | 0 | 1 | 0 | NON_SYNONYMOU | MODERAT | MISSENSE | gCg/gTg | A286V | PLES_12661 | | | | | | | T | | | | | |
| NC_011770 | 1374943 | A | C | 81 | 0 | 2 | 0 | NON_SYNONYMOU | MODERAT | MISSENSE | Agc/Cgc | S374R | PLES_12661 | C | C | | | | | | | | | | |
| NC_011770 | 1375112 | A | G | 81 | 0 | 10 | 0 | NON_SYNONYMOU | MODERAT | MISSENSE | gAc/gGc | D430G | PLES_12661 | G | G | G | G | G | G | G | G | | | G | G |
| NC_011770 | 1375187 | A | G | 83 | 0 | 2 | 0 | NON_SYNONYMOU | MODERAT | MISSENSE | cAc/cGc | H455R | PLES_12661 | | | | | | | | | | | G | G |
| NC_011770 | 1375455 | C | G | 76 | 0 | 2 | 0 | NON_SYNONYMOU | MODERAT | MISSENSE | gaC/gaG | D544E | PLES_12661 | | | | | | | | | | | G | G |
| NC_011770 | 1375463 | A | G | 77 | 0 | 2 | 0 | NON_SYNONYMOU | MODERAT | MISSENSE | gAc/gGc | D547G | PLES_12661 | | | | | | | | | | | G | G |
| NC_011770 | 1375499 | T | C | 76 | 0 | 2 | 0 | NON_SYNONYMOU | MODERAT | MISSENSE | gTc/gCc | V559A | PLES_12661 | | | | | | | | | | | C | C |
| NC_011770 | 1375642 | G | T | 76 | 0 | 2 | 0 | NON_SYNONYMOU | MODERAT | MISSENSE | ttC/ttA | F257L | PLES_12671 | | | | | | | T | T | | | | |
| NC_011770 | 1375737 | T | C | 82 | 0 | 2 | 0 | NON_SYNONYMOU | MODERAT | MISSENSE | Atc/Gtc | I226V | PLES_12671 | | | | | | | | | | | C | C |
| NC_011770 | 1376228 | G | A | 65 | 0 | 2 | 0 | NON_SYNONYMOU | MODERAT | MISSENSE | gCc/gTc | A62V | PLES_12671 | A | A | | | | | | | | | | |
| NC_011770 | 1376990 | T | C | 76 | 0 | 2 | 0 | NON_SYNONYMOU | MODERAT | MISSENSE | cAg/cGg | Q62R | PLES_12681 | | | | | | | C | C | | | | |
| NC_011770 | 1377071 | T | A | 72 | 0 | 2 | 0 | NON_SYNONYMOU | MODERAT | MISSENSE | cAg/cTg | Q35L | PLES_12681 | | | | | | | | | A | A | | |
| NC_011770 | 1380003 | C | G | 61 | 0 | 12 | 0 | NON_SYNONYMOU | MODERAT | MISSENSE | Caa/Gaa | Q108E | PLES_12701 | G | G | G | G | G | G | G | G | G | G | G | G |
| NC_011770 | 1380343 | A | G | 72 | 0 | 2 | 0 | NON_SYNONYMOU | MODERAT | MISSENSE | aAc/aGc | N221S | PLES_12701 | | | | | | | G | G | | | | |
| NC_011770 | 1380431 | C | T | 67 | 0 | 2 | 0 | NON_SYNONYMOU | MODERAT | MISSENSE | Gca/Aca | A297T | PLES_12711 | | | | | | | T | T | | | | |
| NC_011770 | 1380575 | G | A | 66 | 0 | 2 | 0 | NON_SYNONYMOU | MODERAT | MISSENSE | Ccg/Tcg | P249S | PLES_12711 | A | A | | | | | | | | | | |
| NC_011770 | 1380641 | C | A | 62 | 0 | 2 | 0 | NON_SYNONYMOU | MODERAT | MISSENSE | Gca/Tca | A227S | PLES_12711 | | | | | | | | | A | A | | |
| NC_011770 | 1382205 | T | C | 77 | 0 | 12 | 0 | NON_SYNONYMOU | MODERAT | MISSENSE | Tgt/Cgt | C238R | PLES_12721 | C | C | C | C | C | C | C | C | C | C | C | C |
| NC_011770 | 1382326 | G | A | 78 | 0 | 4 | 0 | NON_SYNONYMOU | MODERAT | MISSENSE | cGg/cAg | R278Q | PLES_12721 | | | A | A | A | A | | | | | | |
| NC_011770 | 1382654 | C | G | 80 | 0 | 2 | 0 | NON_SYNONYMOU | MODERAT | MISSENSE | gaC/gaG | D387E | PLES_12721 | | | | | | | | | | | G | G |
| NC_011770 | 1382867 | T | A | 89 | 0 | 8 | 0 | NON_SYNONYMOU | MODERAT | MISSENSE | gaT/gaA | D458E | PLES_12721 | | | A | A | A | A | A | A | A | A | | |
| NC_011770 | 1384300 | C | T | 92 | 0 | 1 | 0 | NON_SYNONYMOU | MODERAT | MISSENSE | cCg/cTg | P342L | PLES_12731 | | | | | | | T | | | | | |
| NC_011770 | 1394024 | T | C | 76 | 0 | 12 | 0 | SYNONYMOUS_ST/ | LOW | SILENT | Ttg/Ctg | L1 | PLES_12821 | C | C | C | C | C | C | C | C | C | C | C | C |
| NC_011770 | 1394984 | GT | G | 84 | 0 | 5 | 0 | FRAME_SHIFT | HIGH | | aaa/ | K29 | PLES_12831 | | G | | | | | | | G | G | G | G |
| NC_011770 | 1395065 | TGC | T | 84 | 0 | 2 | 0 | FRAME_SHIFT | HIGH | | cgc/ | R2 | PLES_12831 | T | T | | | | | | | | | | |
| NC_011770 | 1394932 | G | A | 94 | 0 | 2 | 0 | NON_SYNONYMOU | MODERAT | MISSENSE | aCa/aTa | T47I | PLES_12831 | A | A | | | | | | | | | | |
| NC_011770 | 1394969 | T | C | 86 | 0 | 2 | 0 | NON_SYNONYMOU | MODERAT | MISSENSE | Att/Gtt | I35V | PLES_12831 | | | | | | | | | | | C | C |
| NC_011770 | 1394993 | T | C | 82 | 0 | 6 | 0 | NON_SYNONYMOU | MODERAT | MISSENSE | Ata/Gta | I27V | PLES_12831 | C | C | | | | | | | C | C | C | C |
| NC_011770 | 1395009 | G | T | 84 | 0 | 2 | 0 | NON_SYNONYMOU | MODERAT | MISSENSE | gaC/gaA | D21E | PLES_12831 | | | | | | | | | | | T | T |
| NC_011770 | 1397411 | A | C | 76 | 0 | 2 | 0 | NON_SYNONYMOU | MODERAT | MISSENSE | gAg/gCg | E193A | PLES_12851 | | | | | | | C | C | | | | |
| NC_011770 | 1397845 | G | C | 82 | 0 | 2 | 0 | NON_SYNONYMOU | MODERAT | MISSENSE | Gtg/Ctg | V68L | PLES_12861 | | | | | | | | | | | C | C |
| NC_011770 | 1397962 | A | G | 83 | 0 | 2 | 0 | NON_SYNONYMOU | MODERAT | MISSENSE | Atc/Gtc | I107V | PLES_12861 | | | | | | | G | G | | | | |
| NC_011770 | 1398057 | C | G | 85 | 0 | 2 | 0 | NON_SYNONYMOU | MODERAT | MISSENSE | gaC/gaG | D138E | PLES_12861 | | | | | | | G | G | | | | |
| NC_011770 | 1398065 | C | G | 83 | 0 | 4 | 0 | NON_SYNONYMOU | MODERAT | MISSENSE | aCc/aGc | T141S | PLES_12861 | | | | | | | G | G | | | G | G |
| NC_011770 | 1398546 | G | A | 93 | 0 | 1 | 0 | NON_SYNONYMOU | MODERAT | MISSENSE | Ggc/Agc | G115S | PLES_12871 | | | | | | | A | | | | | |
| NC_011770 | 1400108 | G | C | 75 | 0 | 2 | 0 | NON_SYNONYMOU | MODERAT | MISSENSE | Gtg/Ctg | V181L | PLES_12881 | C | C | | | | | | | | | | |
| NC_011770 | 1400136 | A | G | 76 | 0 | 2 | 0 | NON_SYNONYMOU | MODERAT | MISSENSE | gAc/gGc | D190G | PLES_12881 | | | | | | | G | G | | | | |
| NC_011770 | 1401834 | C | T | 72 | 0 | 2 | 0 | NON_SYNONYMOU | MODERAT | MISSENSE | gCg/gTg | A101V | PLES_12911 | | | | | | | | | | | T | T |
| NC_011770 | 1403269 | C | T | 68 | 0 | 1 | 0 | NON_SYNONYMOU | MODERAT | MISSENSE | Gcc/Acc | A34T | PLES_12931 | | | | | | | | T | | | | |
| NC_011770 | 1403749 | T | C | 83 | 0 | 2 | 0 | SPLICE_SITE_REGIC | LOW | SILENT | tAa/tGa | *713 | PLES_12941 | | | | | | | | | | | C | C |
| NC_011770 | 1404224 | T | G | 90 | 0 | 2 | 0 | NON_SYNONYMOU | MODERAT | MISSENSE | Acc/Ccc | T555P | PLES_12941 | | | | | G | G | | | | | | |
| NC_011770 | 1404544 | A | G | 83 | 0 | 1 | 0 | NON_SYNONYMOU | MODERAT | MISSENSE | gTa/gCa | V448A | PLES_12941 | | | | | | | G | | | | | |
| NC_011770 | 1405150 | A | G | 69 | 0 | 1 | 0 | NON_SYNONYMOU | MODERAT | MISSENSE | gTg/gCg | V246A | PLES_12941 | | | | | | | G | | | | | |
| NC_011770 | 1405316 | A | G | 90 | 0 | 1 | 0 | NON_SYNONYMOU | MODERAT | MISSENSE | Tcc/Ccc | S191P | PLES_12941 | | | | | | | G | | | | | |
| NC_011770 | 1406412 | G | A | 87 | 0 | 2 | 0 | NON_SYNONYMOU | MODERAT | MISSENSE | cGg/cAg | R106Q | PLES_12951 | | | | | | | | | | | A | A |
| NC_011770 | 1406504 | G | C | 91 | 0 | 2 | 0 | NON_SYNONYMOU | MODERAT | MISSENSE | Gcc/Ccc | A137P | PLES_12951 | | | | | | | | | | | C | C |
| NC_011770 | 1411142 | A | G | 73 | 0 | 12 | 0 | NON_SYNONYMOU | MODERAT | MISSENSE | Acc/Gcc | T157A | PLES_12991 | G | G | G | G | G | G | G | G | G | G | G | G |
| NC_011770 | 1411346 | A | G | 67 | 0 | 10 | 0 | NON_SYNONYMOU | MODERAT | MISSENSE | Acg/Gcg | T225A | PLES_12991 | G | G | G | G | G | G | | | G | G | G | G |
| NC_011770 | 1410770 | C | T | 74 | 0 | 1 | 0 | NON_SYNONYMOU | MODERAT | MISSENSE | Cgc/Tgc | R33C | PLES_12991 | | | | | | | | | | | T | |
| NC_011770 | 1413168 | T | G | 69 | 0 | 2 | 0 | NON_SYNONYMOU | MODERAT | MISSENSE | Ttg/Gtg | L146V | PLES_13021 | | | | | | | G | G | | | | |

| | | | | | | | | | | | | | | | | | | | | | | | | | |
|---|---|---|---|---|---|---|---|---|---|---|---|---|---|---|---|---|---|---|---|---|---|---|---|---|---|
| NC_011770 | 1413205 | A | G | 70 | 0 | 1 | 0 | NON_SYNONYMOU | MODERAT | MISSENSE | aAc/aGc | N158S | PLES_13021 | | | | | | | G | | | | | |
| NC_011770 | 1413315 | A | T | 74 | 0 | 4 | 0 | NON_SYNONYMOU | MODERAT | MISSENSE | Atc/Ttc | I195F | PLES_13021 | | | T | T | T | T | | | | | | |
| NC_011770 | 1413471 | C | G | 75 | 0 | 2 | 0 | NON_SYNONYMOU | MODERAT | MISSENSE | Cag/Gag | Q247E | PLES_13021 | G | G | | | | | | | | | | |
| NC_011770 | 1413484 | C | T | 69 | 0 | 2 | 0 | NON_SYNONYMOU | MODERAT | MISSENSE | gCg/gTg | A251V | PLES_13021 | T | T | | | | | | | | | | |
| NC_011770 | 1413517 | C | T | 70 | 0 | 2 | 0 | NON_SYNONYMOU | MODERAT | MISSENSE | gCg/gTg | A262V | PLES_13021 | | | | | | | | | T | T | | |
| NC_011770 | 1413552 | C | T | 72 | 0 | 2 | 0 | NON_SYNONYMOU | MODERAT | MISSENSE | Cgc/Tgc | R274C | PLES_13021 | T | T | | | | | | | | | | |
| NC_011770 | 1413589 | TTAC | T | 73 | 0 | 2 | 0 | CODON_CHANGE_ | MODERATE | | ttactg/ttg | LL10L | PLES_13031 | T | T | | | | | | | | | | |
| NC_011770 | 1413610 | G | A | 67 | 0 | 6 | 0 | NON_SYNONYMOU | MODERAT | MISSENSE | Gtc/Atc | V17I | PLES_13031 | | | A | A | A | A | | | | | A | A |
| NC_011770 | 1413634 | T | C | 67 | 0 | 12 | 0 | NON_SYNONYMOU | MODERAT | MISSENSE | Ttc/Ctc | F25L | PLES_13031 | C | C | C | C | C | C | C | C | C | C | C | C |
| NC_011770 | 1413883 | T | C | 64 | 0 | 8 | 0 | NON_SYNONYMOU | MODERAT | MISSENSE | Tcg/Ccg | S108P | PLES_13031 | C | C | C | C | C | C | | | | | C | C |
| NC_011770 | 1414362 | G | A | 68 | 0 | 10 | 0 | NON_SYNONYMOU | MODERAT | MISSENSE | gGt/gAt | G17D | PLES_13041 | A | A | A | A | A | A | | | A | A | A | A |
| NC_011770 | 1414470 | T | C | 63 | 0 | 1 | 0 | NON_SYNONYMOU | MODERAT | MISSENSE | tTg/tCg | L53S | PLES_13041 | | | | | | | C | | | | | |
| NC_011770 | 1414481 | G | C | 59 | 0 | 4 | 0 | NON_SYNONYMOU | MODERAT | MISSENSE | Gcg/Ccg | A57P | PLES_13041 | | | C | C | C | C | | | | | | |
| NC_011770 | 1414736 | C | G | 75 | 0 | 2 | 0 | NON_SYNONYMOU | MODERAT | MISSENSE | Cgg/Ggg | R142G | PLES_13041 | | | | | | | | | | | G | G |
| NC_011770 | 1414812 | C | T | 82 | 0 | 2 | 0 | NON_SYNONYMOU | MODERAT | MISSENSE | gCc/gTc | A167V | PLES_13041 | | | | | | | | | T | T | | |
| NC_011770 | 1414838 | G | A | 84 | 0 | 10 | 0 | NON_SYNONYMOU | MODERAT | MISSENSE | Gtc/Atc | V176I | PLES_13041 | A | A | A | A | A | A | A | A | A | A | | |
| NC_011770 | 1415538 | GA | G | 78 | 0 | 1 | 0 | FRAME_SHIFT | HIGH | | gag/ | E138 | PLES_13051 | | | | | | | G | | | | | |
| NC_011770 | 1415203 | T | C | 79 | 0 | 4 | 0 | NON_SYNONYMOU | MODERAT | MISSENSE | gTc/gCc | V26A | PLES_13051 | | | | | | | C | C | | | C | C |
| NC_011770 | 1415625 | C | T | 78 | 0 | 2 | 0 | NON_SYNONYMOU | MODERAT | MISSENSE | Ccg/Tcg | P167S | PLES_13051 | T | T | | | | | | | | | | |
| NC_011770 | 1415940 | C | G | 74 | 0 | 2 | 0 | NON_SYNONYMOU | MODERAT | MISSENSE | Cga/Gga | R272G | PLES_13051 | | | | | | | G | G | | | | |
| NC_011770 | 1416060 | A | G | 80 | 0 | 12 | 0 | NON_SYNONYMOU | MODERAT | MISSENSE | Atg/Gtg | M312V | PLES_13051 | G | G | G | G | G | G | G | G | G | G | G | G |
| NC_011770 | 1416422 | C | T | 80 | 0 | 2 | 0 | NON_SYNONYMOU | MODERAT | MISSENSE | Gcg/Acg | A210T | PLES_13061 | | | | | | | T | T | | | | |
| NC_011770 | 1417034 | A | G | 87 | 0 | 2 | 0 | NON_SYNONYMOU | MODERAT | MISSENSE | Tcc/Ccc | S6P | PLES_13061 | G | G | | | | | | | | | | |
| NC_011770 | 1417040 | A | G | 88 | 0 | 2 | 0 | NON_SYNONYMOU | MODERAT | MISSENSE | Tcc/Ccc | S4P | PLES_13061 | | | | | | | G | G | | | | |
| NC_011770 | 1416944 | C | T | 92 | 0 | 1 | 0 | NON_SYNONYMOU | MODERAT | MISSENSE | Gac/Aac | D36N | PLES_13061 | mexL | T | | | | | | | | | | |
| NC_011770 | 1418084 | C | G | 78 | 0 | 2 | 0 | NON_SYNONYMOU | MODERAT | MISSENSE | Ccg/Gcg | P314A | PLES_13071 | | | | | | | | | G | G | | |
| NC_011770 | 1417449 | G | A | 84 | 0 | 1 | 0 | NON_SYNONYMOU | MODERAT | MISSENSE | cGg/cAg | R102Q | PLES_13071 | | | | | | | | A | | | | |
| NC_011770 | 1418415 | G | A | 80 | 0 | 1 | 0 | NON_SYNONYMOU | MODERAT | MISSENSE | Ggc/Agc | G55S | PLES_13081 | | | | | | | A | | | | | |
| NC_011770 | 1419528 | A | G | 75 | 0 | 2 | 0 | NON_SYNONYMOU | MODERAT | MISSENSE | Agc/Ggc | S426G | PLES_13081 | | | | | | | G | G | | | | |
| NC_011770 | 1419529 | G | C | 75 | 0 | 2 | 0 | NON_SYNONYMOU | MODERAT | MISSENSE | aGc/aCc | S426T | PLES_13081 | | | | | | | C | C | | | | |
| NC_011770 | 1419750 | C | G | 74 | 0 | 4 | 0 | NON_SYNONYMOU | MODERAT | MISSENSE | Cag/Gag | Q500E | PLES_13081 | | | G | G | G | G | | | | | | |
| NC_011770 | 1419848 | T | G | 76 | 0 | 12 | 0 | NON_SYNONYMOU | MODERAT | MISSENSE | aaT/aaG | N532K | PLES_13081 | G | G | G | G | G | G | G | G | G | G | G | G |
| NC_011770 | 1420333 | A | G | 87 | 0 | 8 | 0 | NON_SYNONYMOU | MODERAT | MISSENSE | aAg/aGg | K694R | PLES_13081 | | | G | G | G | G | G | G | G | G | | |
| NC_011770 | 1420546 | A | C | 76 | 0 | 2 | 0 | NON_SYNONYMOU | MODERAT | MISSENSE | gAg/gCg | E765A | PLES_13081 | C | C | | | | | | | | | | |
| NC_011770 | 1420762 | A | C | 74 | 0 | 2 | 0 | NON_SYNONYMOU | MODERAT | MISSENSE | gAa/gCa | E837A | PLES_13081 | C | C | | | | | | | | | | |
| NC_011770 | 1421406 | C | T | 81 | 0 | 2 | 0 | NON_SYNONYMOU | MODERAT | MISSENSE | Ggc/Agc | G216S | PLES_13091 | T | T | | | | | | | | | | |
| NC_011770 | 1421450 | C | T | 86 | 0 | 2 | 0 | NON_SYNONYMOU | MODERAT | MISSENSE | aGg/aAg | R201K | PLES_13091 | T | T | | | | | | | | | | |
| NC_011770 | 1422423 | G | A | 64 | 0 | 4 | 0 | NON_SYNONYMOU | MODERAT | MISSENSE | gCc/gTc | A55V | PLES_13101 | | | A | A | A | A | | | | | | |
| NC_011770 | 1425591 | A | C | 66 | 0 | 4 | 0 | NON_SYNONYMOU | MODERAT | MISSENSE | gAg/gCg | E25A | PLES_13121 | C | C | | | | | C | C | | | | |
| NC_011770 | 1425677 | T | A | 67 | 0 | 12 | 0 | NON_SYNONYMOU | MODERAT | MISSENSE | Tgc/Agc | C54S | PLES_13121 | A | A | A | A | A | A | A | A | A | A | A | A |
| NC_011770 | 1426229 | G | A | 77 | 0 | 2 | 0 | NON_SYNONYMOU | MODERAT | MISSENSE | Gcc/Acc | A238T | PLES_13121 | | | | | | | A | A | | | | |
| NC_011770 | 1426567 | C | G | 74 | 0 | 12 | 0 | NON_SYNONYMOU | MODERAT | MISSENSE | Caa/Gaa | Q44E | PLES_13131 | G | G | G | G | G | G | G | G | G | G | G | G |
| NC_011770 | 1428187 | C | A | 76 | 0 | 4 | 0 | NON_SYNONYMOU | MODERAT | MISSENSE | gaC/gaA | D335E | PLES_13141 | | | A | A | A | A | | | | | | |
| NC_011770 | 1428293 | G | A | 80 | 0 | 10 | 0 | NON_SYNONYMOU | MODERAT | MISSENSE | Gcc/Acc | A371T | PLES_13141 | | | A | A | A | A | A | A | A | A | A | A |
| NC_011770 | 1428315 | C | T | 81 | 0 | 2 | 0 | NON_SYNONYMOU | MODERAT | MISSENSE | gCc/gTc | A378V | PLES_13141 | T | T | | | | | | | | | | |
| NC_011770 | 1428509 | C | T | 89 | 0 | 2 | 0 | NON_SYNONYMOU | MODERAT | MISSENSE | Ctc/Ttc | L443F | PLES_13141 | | | | | | | | | T | T | | |
| NC_011770 | 1428727 | G | C | 87 | 0 | 8 | 0 | NON_SYNONYMOU | MODERAT | MISSENSE | gaG/gaC | E515D | PLES_13141 | C | C | C | C | C | C | | | C | C | | |
| NC_011770 | 1428789 | A | C | 83 | 0 | 6 | 0 | NON_SYNONYMOU | MODERAT | MISSENSE | gAc/gCc | D536A | PLES_13141 | | | C | C | C | C | | | C | C | | |
| NC_011770 | 1428891 | G | A | 82 | 0 | 12 | 0 | NON_SYNONYMOU | MODERAT | MISSENSE | tGc/tAc | C570Y | PLES_13141 | A | A | A | A | A | A | A | A | A | A | A | A |
| NC_011770 | 1428948 | C | T | 82 | 0 | 4 | 0 | NON_SYNONYMOU | MODERAT | MISSENSE | gCc/gTc | A589V | PLES_13141 | | | T | T | T | T | | | | | | |
| NC_011770 | 1429004 | C | T | 79 | 0 | 4 | 0 | NON_SYNONYMOU | MODERAT | MISSENSE | Cgg/Tgg | R608W | PLES_13141 | | | T | T | T | T | | | | | | |
| NC_011770 | 1429170 | G | A | 73 | 0 | 4 | 0 | NON_SYNONYMOU | MODERAT | MISSENSE | aGc/aAc | S46N | PLES_13151 | | | A | A | A | A | | | | | | |
| NC_011770 | 1429418 | C | T | 80 | 0 | 1 | 0 | NON_SYNONYMOU | MODERAT | MISSENSE | Cat/Tat | H129Y | PLES_13151 | | | | | | | T | | | | | |
| NC_011770 | 1429431 | G | A | 77 | 0 | 6 | 0 | NON_SYNONYMOU | MODERAT | MISSENSE | gGc/gAc | G133D | PLES_13151 | | | A | A | A | A | | | A | A | | |
| NC_011770 | 1429691 | A | T | 67 | 0 | 4 | 0 | NON_SYNONYMOU | MODERAT | MISSENSE | Aca/Tca | T220S | PLES_13151 | | | | | | | T | T | T | T | | |
| NC_011770 | 1429736 | A | G | 70 | 0 | 10 | 0 | NON_SYNONYMOU | MODERAT | MISSENSE | Acg/Gcg | T235A | PLES_13151 | G | G | G | G | G | G | G | G | G | G | | |
| NC_011770 | 1430226 | C | T | 65 | 0 | 10 | 0 | NON_SYNONYMOU | MODERAT | MISSENSE | gGc/gAc | G73D | PLES_13161 | T | T | T | T | T | T | T | T | T | T | | |
| NC_011770 | 1430326 | T | C | 67 | 0 | 10 | 0 | NON_SYNONYMOU | MODERAT | MISSENSE | Acg/Gcg | T40A | PLES_13161 | C | C | C | C | C | C | C | C | C | C | | |
| NC_011770 | 1430985 | T | C | 68 | 0 | 12 | 0 | NON_SYNONYMOU | MODERAT | MISSENSE | Agc/Ggc | S221G | PLES_13171 | C | C | C | C | C | C | C | C | C | C | C | C |
| NC_011770 | 1431170 | C | T | 76 | 0 | 2 | 0 | NON_SYNONYMOU | MODERAT | MISSENSE | gGc/gAc | G159D | PLES_13171 | | | | | | | T | T | | | | |
| NC_011770 | 1431242 | C | T | 74 | 0 | 1 | 0 | NON_SYNONYMOU | MODERAT | MISSENSE | gGc/gAc | G135D | PLES_13171 | | | | | | | T | | | | | |
| NC_011770 | 1431342 | A | C | 76 | 0 | 12 | 0 | NON_SYNONYMOU | MODERAT | MISSENSE | Tgc/Ggc | C102G | PLES_13171 | C | C | C | C | C | C | C | C | C | C | C | C |

| | | | | | | | | | | | | | | | | | | | | | | | | | |
|---|---|---|---|---|---|---|---|---|---|---|---|---|---|---|---|---|---|---|---|---|---|---|---|---|---|
| NC_011770 | 1431401 | T | C | 71 | 0 | 2 | 0 | NON_SYNONYMOU | MODERAT | MISSENSE | aAc/aGc | N82S | PLES_13171 | | | | | | | | | C | C | | |
| NC_011770 | 1431519 | C | T | 78 | 0 | 4 | 0 | NON_SYNONYMOU | MODERAT | MISSENSE | Ggc/Agc | G43S | PLES_13171 | | | T | T | T | T | | | | | | |
| NC_011770 | 1431555 | C | T | 76 | 0 | 8 | 0 | NON_SYNONYMOU | MODERAT | MISSENSE | Gcc/Acc | A31T | PLES_13171 | T | T | T | T | T | T | | | T | T | | |
| NC_011770 | 1433169 | C | G | 62 | 0 | 2 | 0 | NON_SYNONYMOU | MODERAT | MISSENSE | Gtg/Ctg | V87L | PLES_13191 | | | | | | | | | | | G | G |
| NC_011770 | 1433202 | C | G | 62 | 0 | 8 | 0 | NON_SYNONYMOU | MODERAT | MISSENSE | Gtg/Ctg | V76L | PLES_13191 | G | G | G | G | G | G | | | G | G | | |
| NC_011770 | 1433421 | C | T | 60 | 0 | 4 | 0 | NON_SYNONYMOU | MODERAT | MISSENSE | Gtc/Atc | V3I | PLES_13191 | | | T | T | T | T | | | | | | |
| NC_011770 | 1433493 | C | T | 57 | 0 | 4 | 0 | NON_SYNONYMOU | MODERAT | MISSENSE | Ggc/Agc | G99S | PLES_13201 | | | T | T | T | T | | | | | | |
| NC_011770 | 1433547 | C | T | 56 | 0 | 2 | 0 | NON_SYNONYMOU | MODERAT | MISSENSE | Gag/Aag | E81K | PLES_13201 | T | T | | | | | | | | | | |
| NC_011770 | 1434677 | A | G | 17 | 10 | 2 | 0 | NON_SYNONYMOU | MODERAT | MISSENSE | gTc/gCc | V94A | PLES_13211 | . | . | . | . | . | . | . | . | . | . | G | G |
| NC_011770 | 1434698 | G | A | 18 | 10 | 2 | 0 | NON_SYNONYMOU | MODERAT | MISSENSE | aCc/aTc | T87I | PLES_13211 | . | . | . | . | . | . | . | . | . | . | A | A |
| NC_011770 | 1434743 | C | T | 19 | 10 | 2 | 0 | NON_SYNONYMOU | MODERAT | MISSENSE | aGt/aAt | S72N | PLES_13211 | . | . | . | . | . | . | . | . | . | . | T | T |
| NC_011770 | 1434774 | A | C | 16 | 10 | 2 | 0 | NON_SYNONYMOU | MODERAT | MISSENSE | Tcg/Gcg | S62A | PLES_13211 | . | . | . | . | . | . | . | . | . | . | C | C |
| NC_011770 | 1434782 | T | C | 17 | 10 | 2 | 0 | NON_SYNONYMOU | MODERAT | MISSENSE | cAc/cGc | H59R | PLES_13211 | . | . | . | . | . | . | . | . | . | . | C | C |
| NC_011770 | 1434989 | T | G | 15 | 10 | 2 | 0 | NON_SYNONYMOU | MODERAT | MISSENSE | gaA/gaC | E64D | PLES_13221 | . | . | . | . | . | . | . | . | . | . | G | G |
| NC_011770 | 1434999 | T | G | 16 | 10 | 2 | 0 | NON_SYNONYMOU | MODERAT | MISSENSE | gAa/gCa | E61A | PLES_13221 | . | . | . | . | . | . | . | . | . | . | G | G |
| NC_011770 | 1435041 | T | G | 16 | 10 | 2 | 0 | NON_SYNONYMOU | MODERAT | MISSENSE | cAg/cCg | Q47P | PLES_13221 | . | . | . | . | . | . | . | . | . | . | G | G |
| NC_011770 | 1435710 | C | CTA | 4 | 10 | 2 | 0 | FRAME_SHIFT | HIGH | | atg/atgTA | M1M? | PLES_13231 | . | . | . | . | . | . | . | . | CTA | CTA | . | . |
| NC_011770 | 1435416 | C | A | 17 | 8 | 4 | 0 | NON_SYNONYMOU | MODERAT | MISSENSE | Gcc/Tcc | A100S | PLES_13231 | . | . | . | . | . | . | . | . | A | A | A | A |
| NC_011770 | 1435419 | T | C | 17 | 8 | 4 | 0 | NON_SYNONYMOU | MODERAT | MISSENSE | Atc/Gtc | I99V | PLES_13231 | . | . | . | . | . | . | . | . | C | C | C | C |
| NC_011770 | 1435494 | T | G | 21 | 8 | 4 | 0 | NON_SYNONYMOU | MODERAT | MISSENSE | Agc/Cgc | S74R | PLES_13231 | . | . | . | . | . | . | . | . | G | G | G | G |
| NC_011770 | 1435557 | G | T | 14 | 8 | 2 | 0 | NON_SYNONYMOU | MODERAT | MISSENSE | Ctc/Atc | L53I | PLES_13231 | . | . | . | . | . | . | . | . | | | T | T |
| NC_011770 | 1435561 | C | A | 14 | 8 | 2 | 0 | NON_SYNONYMOU | MODERAT | MISSENSE | ttG/ttT | L51F | PLES_13231 | . | . | . | . | . | . | . | . | | | A | A |
| NC_011770 | 1435566 | G | C | 14 | 8 | 2 | 0 | NON_SYNONYMOU | MODERAT | MISSENSE | Cag/Gag | Q50E | PLES_13231 | . | . | . | . | . | . | . | . | | | C | C |
| NC_011770 | 1435593 | C | T | 12 | 8 | 2 | 0 | NON_SYNONYMOU | MODERAT | MISSENSE | Gcg/Acg | A41T | PLES_13231 | . | . | . | . | . | . | . | . | T | T | | |
| NC_011770 | 1435594 | T | G | 13 | 8 | 2 | 0 | NON_SYNONYMOU | MODERAT | MISSENSE | ttA/ttC | L40F | PLES_13231 | . | . | . | . | . | . | . | . | G | G | | |
| NC_011770 | 1435596 | A | T | 12 | 8 | 4 | 0 | NON_SYNONYMOU | MODERAT | MISSENSE | Tta/Ata | L40I | PLES_13231 | . | . | . | . | . | . | . | . | T | T | C | C |
| NC_011770 | 1435658 | C | T | 6 | 8 | 2 | 0 | NON_SYNONYMOU | MODERAT | MISSENSE | cGc/cAc | R19H | PLES_13231 | . | . | . | . | . | . | . | . | T | T | | |
| NC_011770 | 1435674 | T | G | 4 | 10 | 2 | 0 | NON_SYNONYMOU | MODERAT | MISSENSE | Acc/Ccc | T14P | PLES_13231 | . | . | . | . | . | . | . | . | G | G | . | . |
| NC_011770 | 1435689 | T | G | 4 | 10 | 2 | 0 | NON_SYNONYMOU | MODERAT | MISSENSE | Aat/Cat | N9H | PLES_13231 | . | . | . | . | . | . | . | . | G | G | . | . |
| NC_011770 | 1436175 | T | C | 6 | 10 | 2 | 0 | NON_SYNONYMOU | MODERAT | MISSENSE | aAg/aGg | K68R | PLES_13241 | . | . | . | . | . | . | . | . | C | C | . | . |
| NC_011770 | 1438256 | G | GCGTT | 7 | 8 | 2 | 0 | FRAME_SHIFT+STC | HIGH | | tgc/tgAAC | C7*T? | PLES_13261 | . | . | . | . | . | . | . | . | GCGTT | GCGTT | | |
| NC_011770 | 1436695 | C | A | 12 | 8 | 2 | 0 | NON_SYNONYMOU | MODERAT | MISSENSE | Gtg/Ttg | V528L | PLES_13261 | A | A | . | . | . | . | . | . | . | . | | |
| NC_011770 | 1436905 | C | T | 10 | 4 | 2 | 0 | NON_SYNONYMOU | MODERAT | MISSENSE | Gct/Act | A458T | PLES_13261 | | | . | . | | | . | . | | | T | T |
| NC_011770 | 1436913 | C | G | 4 | 10 | 2 | 0 | NON_SYNONYMOU | MODERAT | MISSENSE | aGc/aCc | S455T | PLES_13261 | . | . | . | . | . | . | . | . | G | G | . | . |
| NC_011770 | 1436914 | T | C | 4 | 10 | 2 | 0 | NON_SYNONYMOU | MODERAT | MISSENSE | Agc/Ggc | S455G | PLES_13261 | . | . | . | . | . | . | . | . | C | C | . | . |
| NC_011770 | 1436919 | C | T | 3 | 10 | 2 | 0 | NON_SYNONYMOU | MODERAT | MISSENSE | gGg/gAg | G453E | PLES_13261 | . | . | . | . | . | . | . | . | T | T | . | . |
| NC_011770 | 1436985 | T | G | 3 | 10 | 2 | 0 | NON_SYNONYMOU | MODERAT | MISSENSE | aAt/aCt | N431T | PLES_13261 | . | . | . | . | . | . | . | . | G | G | . | . |
| NC_011770 | 1436986 | T | A | 3 | 10 | 2 | 0 | NON_SYNONYMOU | MODERAT | MISSENSE | Aat/Tat | N431Y | PLES_13261 | . | . | . | . | . | . | . | . | A | A | . | . |
| NC_011770 | 1437043 | T | G | 3 | 10 | 2 | 0 | NON_SYNONYMOU | MODERAT | MISSENSE | Aac/Cac | N412H | PLES_13261 | . | . | . | . | . | . | . | . | G | G | . | . |
| NC_011770 | 1437198 | A | T | 9 | 2 | 4 | 0 | NON_SYNONYMOU | MODERAT | MISSENSE | cTg/cAg | L360Q | PLES_13261 | | | | | T | T | . | . | T | T | | |
| NC_011770 | 1437204 | G | A | 9 | 2 | 4 | 0 | NON_SYNONYMOU | MODERAT | MISSENSE | gCc/gTc | A358V | PLES_13261 | | | | | A | A | . | . | A | A | | |
| NC_011770 | 1437951 | A | G | 8 | 10 | 2 | 0 | NON_SYNONYMOU | MODERAT | MISSENSE | gTc/gCc | V109A | PLES_13261 | . | . | . | . | . | . | . | . | . | . | G | G |
| NC_011770 | 1438119 | T | C | 6 | 10 | 2 | 0 | NON_SYNONYMOU | MODERAT | MISSENSE | gAc/gGc | D53G | PLES_13261 | . | . | . | . | . | . | . | . | . | . | C | C |
| NC_011770 | 1438120 | C | T | 6 | 10 | 2 | 0 | NON_SYNONYMOU | MODERAT | MISSENSE | Gac/Aac | D53N | PLES_13261 | . | . | . | . | . | . | . | . | . | . | T | T |
| NC_011770 | 1438175 | C | A | 6 | 10 | 2 | 0 | NON_SYNONYMOU | MODERAT | MISSENSE | aaG/aaT | K34N | PLES_13261 | . | . | . | . | . | . | . | . | . | . | A | A |
| NC_011770 | 1438188 | T | C | 5 | 10 | 2 | 0 | NON_SYNONYMOU | MODERAT | MISSENSE | gAc/gGc | D30G | PLES_13261 | . | . | . | . | . | . | . | . | . | . | C | C |
| NC_011770 | 1438194 | T | A | 5 | 10 | 2 | 0 | NON_SYNONYMOU | MODERAT | MISSENSE | cAg/cTg | Q28L | PLES_13261 | . | . | . | . | . | . | . | . | . | . | A | A |
| NC_011770 | 1438196 | A | C | 5 | 10 | 2 | 0 | NON_SYNONYMOU | MODERAT | MISSENSE | gaT/gaG | D27E | PLES_13261 | . | . | . | . | . | . | . | . | . | . | C | C |
| NC_011770 | 1438201 | C | CCGA | 5 | 10 | 2 | 0 | CODON_INSERTIOI | MODERATE | | gcg/gcgTC | A25AS | PLES_13261 | . | . | . | . | . | . | . | . | . | . | CCGA | CCGA |
| NC_011770 | 1438227 | A | C | 8 | 8 | 4 | 0 | NON_SYNONYMOU | MODERAT | MISSENSE | gTc/gGc | V17G | PLES_13261 | . | . | . | . | . | . | . | . | C | C | C | C |
| NC_011770 | 1438228 | C | G | 7 | 8 | 4 | 0 | NON_SYNONYMOU | MODERAT | MISSENSE | Gtc/Ctc | V17L | PLES_13261 | . | . | . | . | . | . | . | . | G | G | G | G |
| NC_011770 | 1438260 | A | T | 6 | 8 | 2 | 0 | NON_SYNONYMOU | MODERAT | MISSENSE | aTt/aAt | I6N | PLES_13261 | . | . | . | . | . | . | . | . | . | . | T | T |
| NC_011770 | 1438261 | T | C | 6 | 8 | 2 | 0 | NON_SYNONYMOU | MODERAT | MISSENSE | Att/Gtt | I6V | PLES_13261 | . | . | . | . | . | . | . | . | . | . | C | C |
| NC_011770 | 1438262 | C | A | 6 | 8 | 2 | 0 | NON_SYNONYMOU | MODERAT | MISSENSE | aaG/aaT | K5N | PLES_13261 | . | . | . | . | . | . | . | . | . | . | A | A |
| NC_011770 | 1438267 | A | C | 7 | 8 | 2 | 0 | NON_SYNONYMOU | MODERAT | MISSENSE | Tca/Gca | S4A | PLES_13261 | . | . | . | . | . | . | . | . | C | C | | |
| NC_011770 | 1438270 | C | T | 7 | 8 | 4 | 0 | NON_SYNONYMOU | MODERAT | MISSENSE | Gaa/Aaa | E3K | PLES_13261 | . | . | . | . | . | . | . | . | T | T | T | T |
| NC_011770 | 1438280 | A | T | 8 | 8 | 2 | 0 | NON_SYNONYMOU | MODERAT | MISSENSE | cTc/cAc | L67H | PLES_13271 | . | . | . | . | . | . | . | . | | | T | T |
| NC_011770 | 1438356 | G | T | 8 | 8 | 2 | 0 | NON_SYNONYMOU | MODERAT | MISSENSE | Ctg/Atg | L42M | PLES_13271 | . | . | . | . | . | . | . | . | | | T | T |
| NC_011770 | 1438360 | A | C | 8 | 8 | 2 | 0 | NON_SYNONYMOU | MODERAT | MISSENSE | aaT/aaG | N40K | PLES_13271 | . | . | . | . | . | . | . | . | | | C | C |
| NC_011770 | 1438362 | T | C | 8 | 8 | 2 | 0 | NON_SYNONYMOU | MODERAT | MISSENSE | Aat/Gat | N40D | PLES_13271 | . | . | . | . | . | . | . | . | | | C | C |
| NC_011770 | 1438427 | G | C | 4 | 8 | 4 | 0 | NON_SYNONYMOU | MODERAT | MISSENSE | cCc/cGc | P18R | PLES_13271 | . | . | . | . | . | . | . | . | C | C | T | T |
| NC_011770 | 1438428 | G | T | 4 | 8 | 2 | 0 | NON_SYNONYMOU | MODERAT | MISSENSE | Ccc/Acc | P18T | PLES_13271 | . | . | . | . | . | . | . | . | | | T | T |

| | | | | | | | | | | | | | | | | | | | | | | | | | |
|---|---|---|---|---|---|---|---|---|---|---|---|---|---|---|---|---|---|---|---|---|---|---|---|---|---|
| NC_011770 | 1438431 | A | AGGG | 4 | 8 | 2 | 0 | CODON_INSERTIOI | MODERATE | | tgc/tgcCCC( | C16CP | PLES_13271 | . | . | . | . | . | . | . | . | | | AGGG | AGGG |
| NC_011770 | 1438433 | C | T | 3 | 9 | 1 | 0 | NON_SYNONYMOI | MODERAT | MISSENSE | tGc/tAc | C16Y | PLES_13271 | . | . | . | . | . | . | . | . | | | T | . |
| NC_011770 | 1438476 | G | A | 6 | 8 | 2 | 0 | NON_SYNONYMOI | MODERAT | MISSENSE | Cca/Tca | P2S | PLES_13271 | . | . | . | . | . | . | . | . | | | A | A |
| NC_011770 | 1438512 | G | T | 6 | 8 | 2 | 0 | STOP_GAINED | HIGH | NONSENSE | tgC/tgA | C60* | PLES_13281 | . | . | . | . | . | . | . | . | | | T | T |
| NC_011770 | 1438520 | C | A | 6 | 8 | 2 | 0 | STOP_GAINED | HIGH | NONSENSE | Gag/Tag | E58* | PLES_13281 | . | . | . | . | . | . | . | . | | | A | A |
| NC_011770 | 1438481 | A | G | 6 | 8 | 2 | 0 | NON_SYNONYMOI | MODERAT | MISSENSE | Tca/Cca | S71P | PLES_13281 | . | . | . | . | . | . | . | . | | | G | G |
| NC_011770 | 1438483 | G | T | 6 | 8 | 2 | 0 | NON_SYNONYMOI | MODERAT | MISSENSE | cCa/cAa | P70Q | PLES_13281 | . | . | . | . | . | . | . | . | | | T | T |
| NC_011770 | 1438508 | T | C | 6 | 8 | 2 | 0 | NON_SYNONYMOI | MODERAT | MISSENSE | Acg/Gcg | T62A | PLES_13281 | . | . | . | . | . | . | . | . | | | C | C |
| NC_011770 | 1438513 | C | G | 6 | 8 | 2 | 0 | NON_SYNONYMOI | MODERAT | MISSENSE | tGc/tCc | C60S | PLES_13281 | . | . | . | . | . | . | . | . | | | G | G |
| NC_011770 | 1438514 | A | T | 6 | 8 | 2 | 0 | NON_SYNONYMOI | MODERAT | MISSENSE | Tgc/Agc | C60S | PLES_13281 | . | . | . | . | . | . | . | . | | | T | T |
| NC_011770 | 1438519 | T | A | 6 | 8 | 2 | 0 | NON_SYNONYMOI | MODERAT | MISSENSE | gAg/gTg | E58V | PLES_13281 | . | . | . | . | . | . | . | . | | | A | A |
| NC_011770 | 1438525 | C | G | 5 | 8 | 2 | 0 | NON_SYNONYMOI | MODERAT | MISSENSE | aGc/aCc | S56T | PLES_13281 | . | . | . | . | . | . | . | . | | | G | G |
| NC_011770 | 1438531 | C | T | 5 | 8 | 2 | 0 | NON_SYNONYMOI | MODERAT | MISSENSE | gGc/gAc | G54D | PLES_13281 | . | . | . | . | . | . | . | . | | | T | T |
| NC_011770 | 1438580 | G | T | 7 | 8 | 2 | 0 | NON_SYNONYMOI | MODERAT | MISSENSE | Cgc/Agc | R38S | PLES_13281 | . | . | . | . | . | . | . | . | T | T | | |
| NC_011770 | 1438589 | A | C | 6 | 8 | 2 | 0 | NON_SYNONYMOI | MODERAT | MISSENSE | Tct/Gct | S35A | PLES_13281 | . | . | . | . | . | . | . | . | C | C | | |
| NC_011770 | 1439100 | C | T | 8 | 10 | 2 | 0 | NON_SYNONYMOI | MODERAT | MISSENSE | Gac/Aac | D182N | PLES_13291 | . | . | . | . | T | T | . | . | . | . | . | . |
| NC_011770 | 1439129 | C | G | 9 | 10 | 2 | 0 | NON_SYNONYMOI | MODERAT | MISSENSE | gGg/gCg | G172A | PLES_13291 | . | . | . | . | G | G | . | . | . | . | . | . |
| NC_011770 | 1439132 | G | T | 9 | 10 | 2 | 0 | NON_SYNONYMOI | MODERAT | MISSENSE | cCa/cAa | P171Q | PLES_13291 | . | . | . | . | T | T | . | . | . | . | . | . |
| NC_011770 | 1439136 | T | C | 9 | 10 | 2 | 0 | NON_SYNONYMOI | MODERAT | MISSENSE | Aac/Gac | N170D | PLES_13291 | . | . | . | . | C | C | . | . | . | . | . | . |
| NC_011770 | 1439250 | A | G | 7 | 10 | 2 | 0 | NON_SYNONYMOI | MODERAT | MISSENSE | Tac/Cac | Y132H | PLES_13291 | . | . | . | . | G | G | . | . | . | . | . | . |
| NC_011770 | 1439287 | A | C | 7 | 10 | 2 | 0 | NON_SYNONYMOI | MODERAT | MISSENSE | gaT/gaG | D119E | PLES_13291 | . | . | . | . | C | C | . | . | . | . | . | . |
| NC_011770 | 1439335 | C | A | 7 | 10 | 2 | 0 | NON_SYNONYMOI | MODERAT | MISSENSE | aaG/aaT | K103N | PLES_13291 | . | . | . | . | A | A | . | . | . | . | . | . |
| NC_011770 | 1439360 | C | T | 5 | 10 | 2 | 0 | NON_SYNONYMOI | MODERAT | MISSENSE | aGc/aAc | S95N | PLES_13291 | . | . | . | . | T | T | . | . | . | . | . | . |
| NC_011770 | 1439361 | T | C | 5 | 10 | 2 | 0 | NON_SYNONYMOI | MODERAT | MISSENSE | Agc/Ggc | S95G | PLES_13291 | . | . | . | . | C | C | . | . | . | . | . | . |
| NC_011770 | 1439367 | T | A | 5 | 10 | 2 | 0 | NON_SYNONYMOI | MODERAT | MISSENSE | Agc/Tgc | S93C | PLES_13291 | . | . | . | . | A | A | . | . | . | . | . | . |
| NC_011770 | 1439369 | G | A | 5 | 10 | 2 | 0 | NON_SYNONYMOI | MODERAT | MISSENSE | cCc/cTc | P92L | PLES_13291 | . | . | . | . | A | A | . | . | . | . | . | . |
| NC_011770 | 1439370 | G | A | 5 | 10 | 2 | 0 | NON_SYNONYMOI | MODERAT | MISSENSE | Ccc/Tcc | P92S | PLES_13291 | . | . | . | . | A | A | . | . | . | . | . | . |
| NC_011770 | 1439416 | G | C | 7 | 10 | 2 | 0 | NON_SYNONYMOI | MODERAT | MISSENSE | gaC/gaG | D76E | PLES_13291 | . | . | . | . | C | C | . | . | . | . | . | . |
| NC_011770 | 1439477 | G | T | 0 | 10 | 1 | 0 | NON_SYNONYMOI | MODERAT | MISSENSE | gCa/gAa | A56E | PLES_13291 | . | . | . | . | . | . | . | . | T | . | . | . |
| NC_011770 | 1439490 | A | G | 0 | 10 | 2 | 0 | NON_SYNONYMOI | MODERAT | MISSENSE | Ttc/Ctc | F52L | PLES_13291 | . | . | . | . | . | . | . | . | G | G | . | . |
| NC_011770 | 1439504 | A | C | 0 | 10 | 2 | 0 | NON_SYNONYMOI | MODERAT | MISSENSE | aTc/aGc | I47S | PLES_13291 | . | . | . | . | . | . | . | . | C | C | . | . |
| NC_011770 | 1439754 | T | G | 1 | 10 | 2 | 0 | NON_SYNONYMOI | MODERAT | MISSENSE | aAg/aCg | K173T | PLES_13301 | . | . | . | . | . | . | . | . | G | G | . | . |
| NC_011770 | 1439794 | A | C | 10 | 6 | 2 | 0 | NON_SYNONYMOI | MODERAT | MISSENSE | Tct/Gct | S160A | PLES_13301 | . | . | . | . | . | . | | | C | C | | |
| NC_011770 | 1439796 | A | G | 10 | 6 | 2 | 0 | NON_SYNONYMOI | MODERAT | MISSENSE | gTc/gCc | V159A | PLES_13301 | . | . | . | . | . | . | | | G | G | | |
| NC_011770 | 1439862 | A | T | 6 | 8 | 2 | 0 | NON_SYNONYMOI | MODERAT | MISSENSE | cTc/cAc | L137H | PLES_13301 | . | . | . | . | . | . | . | . | T | T | | |
| NC_011770 | 1439902 | G | T | 6 | 8 | 2 | 0 | NON_SYNONYMOI | MODERAT | MISSENSE | Ccg/Acg | P124T | PLES_13301 | . | . | . | . | . | . | . | . | T | T | | |
| NC_011770 | 1439920 | T | C | 5 | 8 | 4 | 0 | NON_SYNONYMOI | MODERAT | MISSENSE | Att/Gtt | I118V | PLES_13301 | . | . | . | . | . | . | . | . | C | C | C | C |
| NC_011770 | 1439922 | C | G | 5 | 8 | 4 | 0 | NON_SYNONYMOI | MODERAT | MISSENSE | gGc/gCc | G117A | PLES_13301 | . | . | . | . | . | . | . | . | G | G | G | G |
| NC_011770 | 1439923 | C | A | 5 | 8 | 4 | 0 | NON_SYNONYMOI | MODERAT | MISSENSE | Ggc/Tgc | G117C | PLES_13301 | . | . | . | . | . | . | . | . | A | A | A | A |
| NC_011770 | 1439970 | T | G | 6 | 8 | 4 | 0 | NON_SYNONYMOI | MODERAT | MISSENSE | gAg/gCg | E101A | PLES_13301 | . | . | . | . | . | . | . | . | G | G | G | G |
| NC_011770 | 1439998 | A | G | 6 | 8 | 4 | 0 | NON_SYNONYMOI | MODERAT | MISSENSE | Tac/Cac | Y92H | PLES_13301 | . | . | . | . | . | . | . | . | G | G | G | G |
| NC_011770 | 1440130 | A | C | 6 | 8 | 2 | 0 | NON_SYNONYMOI | MODERAT | MISSENSE | Tcg/Gcg | S48A | PLES_13301 | . | . | . | . | . | . | . | . | C | C | | |
| NC_011770 | 1440167 | C | G | 6 | 8 | 4 | 0 | NON_SYNONYMOI | MODERAT | MISSENSE | atG/atC | M35I | PLES_13301 | . | . | . | . | . | . | . | . | G | G | G | G |
| NC_011770 | 1440169 | T | A | 6 | 8 | 2 | 0 | NON_SYNONYMOI | MODERAT | MISSENSE | Atg/Ttg | M35L | PLES_13301 | . | . | . | . | . | . | . | . | A | A | | |
| NC_011770 | 1440184 | G | T | 5 | 8 | 4 | 0 | NON_SYNONYMOI | MODERAT | MISSENSE | Cgc/Agc | R30S | PLES_13301 | . | . | . | . | . | . | . | . | T | T | T | T |
| NC_011770 | 1440192 | T | C | 5 | 8 | 4 | 0 | NON_SYNONYMOI | MODERAT | MISSENSE | aAg/aGg | K27R | PLES_13301 | . | . | . | . | . | . | . | . | C | C | C | C |
| NC_011770 | 1440193 | T | G | 5 | 8 | 4 | 0 | NON_SYNONYMOI | MODERAT | MISSENSE | Aag/Cag | K27Q | PLES_13301 | . | . | . | . | . | . | . | . | G | G | G | G |
| NC_011770 | 1441438 | C | A | 6 | 8 | 2 | 0 | NON_SYNONYMOI | MODERAT | MISSENSE | Gcc/Tcc | A65S | PLES_13321 | . | . | . | . | . | . | . | . | | | A | A |
| NC_011770 | 1441439 | A | T | 7 | 8 | 2 | 0 | NON_SYNONYMOI | MODERAT | MISSENSE | gaT/gaA | D64E | PLES_13321 | . | . | . | . | . | . | . | . | T | T | | |
| NC_011770 | 1441449 | G | C | 7 | 8 | 2 | 0 | NON_SYNONYMOI | MODERAT | MISSENSE | gCc/gGc | A61G | PLES_13321 | . | . | . | . | . | . | . | . | C | C | | |
| NC_011770 | 1441453 | A | C | 6 | 8 | 2 | 0 | NON_SYNONYMOI | MODERAT | MISSENSE | Tca/Gca | S60A | PLES_13321 | . | . | . | . | . | . | . | . | C | C | | |
| NC_011770 | 1441841 | C | A | 10 | 8 | 4 | 0 | NON_SYNONYMOI | MODERAT | MISSENSE | aaG/aaT | K82N | PLES_13322 | . | . | . | . | . | . | . | . | A | A | T | T |
| NC_011770 | 1441845 | G | T | 10 | 8 | 4 | 0 | NON_SYNONYMOI | MODERAT | MISSENSE | gCc/gAc | A81D | PLES_13322 | . | . | . | . | . | . | . | . | T | T | A | A |
| NC_011770 | 1441865 | G | C | 10 | 8 | 2 | 0 | NON_SYNONYMOI | MODERAT | MISSENSE | caC/caG | H74Q | PLES_13322 | . | . | . | . | . | . | . | . | | | C | C |
| NC_011770 | 1441886 | G | C | 10 | 8 | 4 | 0 | NON_SYNONYMOI | MODERAT | MISSENSE | gaC/gaG | D67E | PLES_13322 | . | . | . | . | . | . | . | . | C | C | T | T |
| NC_011770 | 1441896 | T | C | 10 | 8 | 2 | 0 | NON_SYNONYMOI | MODERAT | MISSENSE | aAc/aGc | N64S | PLES_13322 | . | . | . | . | . | . | . | . | C | C | | |
| NC_011770 | 1441917 | G | T | 11 | 8 | 2 | 0 | NON_SYNONYMOI | MODERAT | MISSENSE | gCg/gAg | A57E | PLES_13322 | . | . | . | . | . | . | . | . | T | T | | |
| NC_011770 | 1441918 | C | T | 10 | 8 | 2 | 0 | NON_SYNONYMOI | MODERAT | MISSENSE | Gcg/Acg | A57T | PLES_13322 | . | . | . | . | . | . | . | . | T | T | | |
| NC_011770 | 1441936 | T | C | 10 | 8 | 4 | 0 | NON_SYNONYMOI | MODERAT | MISSENSE | Atc/Gtc | I51V | PLES_13322 | . | . | . | . | . | . | . | . | C | C | C | C |
| NC_011770 | 1441971 | T | C | 10 | 8 | 2 | 0 | NON_SYNONYMOI | MODERAT | MISSENSE | cAa/cGa | Q39R | PLES_13322 | . | . | . | . | . | . | . | . | | | C | C |
| NC_011770 | 1441974 | T | A | 10 | 8 | 2 | 0 | NON_SYNONYMOI | MODERAT | MISSENSE | tAc/tTc | Y38F | PLES_13322 | . | . | . | . | . | . | . | . | | | A | A |
| NC_011770 | 1441986 | A | G | 11 | 8 | 2 | 0 | NON_SYNONYMOI | MODERAT | MISSENSE | gTt/gCt | V34A | PLES_13322 | . | . | . | . | . | . | . | . | G | G | | |

| | | | | | | | | | | | | | | | | | | | | | | | | | |
|---|---|---|---|---|---|---|---|---|---|---|---|---|---|---|---|---|---|---|---|---|---|---|---|---|---|
| NC_011770 | 1442002 | T | G | 12 | 8 | 4 | 0 | NON_SYNONYMOU | MODERAT | MISSENSE | Atc/Ctc | I29L | PLES_13322 | . | . | . | . | . | . | . | . | G | G | G | G |
| NC_011770 | 1442007 | A | T | 12 | 8 | 4 | 0 | NON_SYNONYMOU | MODERAT | MISSENSE | cTg/cAg | L27Q | PLES_13322 | . | . | . | . | . | . | . | . | T | T | T | T |
| NC_011770 | 1442010 | G | A | 12 | 8 | 4 | 0 | NON_SYNONYMOU | MODERAT | MISSENSE | gCt/gTt | A26V | PLES_13322 | . | . | . | . | . | . | . | . | A | A | A | A |
| NC_011770 | 1442017 | C | T | 12 | 8 | 2 | 0 | NON_SYNONYMOU | MODERAT | MISSENSE | Gcg/Acg | A24T | PLES_13322 | . | . | . | . | . | . | . | . | T | T | | |
| NC_011770 | 1442020 | T | G | 12 | 8 | 4 | 0 | NON_SYNONYMOU | MODERAT | MISSENSE | Atg/Ctg | M23L | PLES_13322 | . | . | . | . | . | . | . | . | G | G | G | G |
| NC_011770 | 1442038 | T | G | 11 | 8 | 4 | 0 | NON_SYNONYMOU | MODERAT | MISSENSE | Aaa/Caa | K17Q | PLES_13322 | . | . | . | . | . | . | . | . | G | G | G | G |
| NC_011770 | 1442102 | CGTT | C | 11 | 8 | 2 | 0 | CODON_DELETION | MODERATE | | aac/ | N127 | PLES_13331 | . | . | . | . | . | . | . | . | C | C | | |
| NC_011770 | 1442140 | G | A | 11 | 8 | 2 | 0 | NON_SYNONYMOU | MODERAT | MISSENSE | gCc/gTc | A115V | PLES_13331 | . | . | . | . | . | . | . | . | | | A | A |
| NC_011770 | 1442361 | C | G | 10 | 8 | 4 | 0 | NON_SYNONYMOU | MODERAT | MISSENSE | agG/agC | R41S | PLES_13331 | . | . | . | . | . | . | . | . | G | G | G | G |
| NC_011770 | 1442423 | G | T | 9 | 8 | 2 | 0 | NON_SYNONYMOU | MODERAT | MISSENSE | Cgc/Agc | R21S | PLES_13331 | . | . | . | . | . | . | . | . | | | T | T |
| NC_011770 | 1442442 | C | G | 9 | 8 | 2 | 0 | NON_SYNONYMOU | MODERAT | MISSENSE | caG/caC | Q14H | PLES_13331 | . | . | . | . | . | . | . | . | | | G | G |
| NC_011770 | 1442443 | T | G | 9 | 8 | 2 | 0 | NON_SYNONYMOU | MODERAT | MISSENSE | cAg/cCg | Q14P | PLES_13331 | . | . | . | . | . | . | . | . | | | G | G |
| NC_011770 | 1442444 | G | T | 9 | 8 | 2 | 0 | NON_SYNONYMOU | MODERAT | MISSENSE | Cag/Aag | Q14K | PLES_13331 | . | . | . | . | . | . | . | . | | | T | T |
| NC_011770 | 1442478 | G | T | 10 | 8 | 2 | 0 | NON_SYNONYMOU | MODERAT | MISSENSE | caC/caA | H2Q | PLES_13331 | . | . | . | . | . | . | . | . | T | T | | |
| NC_011770 | 1442876 | A | C | 4 | 8 | 4 | 0 | NON_SYNONYMOU | MODERAT | MISSENSE | Tcg/Gcg | S262A | PLES_13341 | . | . | . | . | . | . | . | . | C | C | C | C |
| NC_011770 | 1442966 | C | A | 6 | 10 | 2 | 0 | NON_SYNONYMOU | MODERAT | MISSENSE | Gcc/Tcc | A232S | PLES_13341 | . | . | . | . | . | . | . | . | . | . | A | A |
| NC_011770 | 1442978 | T | C | 6 | 10 | 2 | 0 | NON_SYNONYMOU | MODERAT | MISSENSE | Acc/Gcc | T228A | PLES_13341 | . | . | . | . | . | . | . | . | . | . | C | C |
| NC_011770 | 1442990 | C | G | 6 | 10 | 2 | 0 | NON_SYNONYMOU | MODERAT | MISSENSE | Gta/Cta | V224L | PLES_13341 | . | . | . | . | . | . | . | . | . | . | G | G |
| NC_011770 | 1442995 | C | T | 7 | 10 | 2 | 0 | NON_SYNONYMOU | MODERAT | MISSENSE | aGa/aAa | R222K | PLES_13341 | . | . | . | . | . | . | . | . | . | . | T | T |
| NC_011770 | 1443026 | T | G | 8 | 10 | 2 | 0 | NON_SYNONYMOU | MODERAT | MISSENSE | Atc/Ctc | I212L | PLES_13341 | . | . | . | . | . | . | . | . | . | . | G | G |
| NC_011770 | 1443028 | A | G | 12 | 8 | 2 | 0 | NON_SYNONYMOU | MODERAT | MISSENSE | gTg/gCg | V211A | PLES_13341 | . | . | . | . | . | . | . | . | | | G | G |
| NC_011770 | 1443064 | G | T | 12 | 8 | 4 | 0 | NON_SYNONYMOU | MODERAT | MISSENSE | gCg/gAg | A199E | PLES_13341 | . | . | . | . | . | . | . | . | T | T | T | T |
| NC_011770 | 1443065 | C | T | 12 | 8 | 4 | 0 | NON_SYNONYMOU | MODERAT | MISSENSE | Gcg/Acg | A199T | PLES_13341 | . | . | . | . | . | . | . | . | T | T | T | T |
| NC_011770 | 1443190 | C | T | 12 | 8 | 4 | 0 | NON_SYNONYMOU | MODERAT | MISSENSE | cGc/cAc | R157H | PLES_13341 | . | . | . | . | . | . | . | . | T | T | T | T |
| NC_011770 | 1443191 | G | T | 12 | 8 | 4 | 0 | NON_SYNONYMOU | MODERAT | MISSENSE | Cgc/Agc | R157S | PLES_13341 | . | . | . | . | . | . | . | . | T | T | T | T |
| NC_011770 | 1443272 | A | C | 12 | 8 | 4 | 0 | NON_SYNONYMOU | MODERAT | MISSENSE | Tcc/Gcc | S130A | PLES_13341 | . | . | . | . | . | . | . | . | C | C | C | C |
| NC_011770 | 1443292 | G | C | 10 | 8 | 4 | 0 | NON_SYNONYMOU | MODERAT | MISSENSE | tCc/tGc | S123C | PLES_13341 | . | . | . | . | . | . | . | . | C | C | C | C |
| NC_011770 | 1443293 | A | T | 10 | 8 | 4 | 0 | NON_SYNONYMOU | MODERAT | MISSENSE | Tcc/Acc | S123T | PLES_13341 | . | . | . | . | . | . | . | . | T | T | T | T |
| NC_011770 | 1443450 | T | G | 12 | 8 | 2 | 0 | NON_SYNONYMOU | MODERAT | MISSENSE | gaA/gaC | E70D | PLES_13341 | . | . | . | . | . | . | . | . | G | G | | |
| NC_011770 | 1444627 | T | G | 3 | 10 | 2 | 0 | NON_SYNONYMOU | MODERAT | MISSENSE | aaT/aaG | N4K | PLES_13361 | . | . | . | . | . | . | . | . | G | G | . | . |
| NC_011770 | 1444714 | A | T | 2 | 10 | 2 | 0 | NON_SYNONYMOU | MODERAT | MISSENSE | gaA/gaT | E33D | PLES_13361 | . | . | . | . | . | . | . | . | T | T | . | . |
| NC_011770 | 1444746 | G | C | 3 | 10 | 2 | 0 | NON_SYNONYMOU | MODERAT | MISSENSE | aGt/aCt | S44T | PLES_13361 | . | . | . | . | . | . | . | . | C | C | . | . |
| NC_011770 | 1444754 | G | A | 3 | 10 | 2 | 0 | NON_SYNONYMOU | MODERAT | MISSENSE | Gcg/Acg | A47T | PLES_13361 | . | . | . | . | . | . | . | . | A | A | . | . |
| NC_011770 | 1445931 | C | A | 4 | 10 | 2 | 0 | NON_SYNONYMOU | MODERAT | MISSENSE | Gcg/Tcg | A171S | PLES_13381 | . | . | . | . | . | . | . | . | A | A | . | . |
| NC_011770 | 1446087 | C | T | 4 | 10 | 2 | 0 | NON_SYNONYMOU | MODERAT | MISSENSE | Gta/Ata | V119I | PLES_13381 | . | . | . | . | . | . | . | . | T | T | . | . |
| NC_011770 | 1446132 | G | T | 4 | 10 | 2 | 0 | NON_SYNONYMOU | MODERAT | MISSENSE | Ctc/Atc | L104I | PLES_13381 | . | . | . | . | . | . | . | . | T | T | . | . |
| NC_011770 | 1446153 | G | C | 5 | 10 | 2 | 0 | NON_SYNONYMOU | MODERAT | MISSENSE | Ccg/Gcg | P97A | PLES_13381 | . | . | . | . | . | . | . | . | C | C | . | . |
| NC_011770 | 1446182 | T | C | 5 | 10 | 2 | 0 | NON_SYNONYMOU | MODERAT | MISSENSE | cAc/cGc | H87R | PLES_13381 | . | . | . | . | . | . | . | . | C | C | . | . |
| NC_011770 | 1446191 | G | A | 5 | 10 | 2 | 0 | NON_SYNONYMOU | MODERAT | MISSENSE | gCc/gTc | A84V | PLES_13381 | . | . | . | . | . | . | . | . | A | A | . | . |
| NC_011770 | 1446212 | A | T | 5 | 10 | 2 | 0 | NON_SYNONYMOU | MODERAT | MISSENSE | cTg/cAg | L77Q | PLES_13381 | . | . | . | . | . | . | . | . | T | T | . | . |
| NC_011770 | 1446214 | T | G | 5 | 10 | 2 | 0 | NON_SYNONYMOU | MODERAT | MISSENSE | caA/caC | Q76H | PLES_13381 | . | . | . | . | . | . | . | . | G | G | . | . |
| NC_011770 | 1446290 | C | T | 7 | 10 | 2 | 0 | NON_SYNONYMOU | MODERAT | MISSENSE | aGg/aAg | R51K | PLES_13381 | . | . | . | . | . | . | . | . | T | T | . | . |
| NC_011770 | 1446982 | A | C | 8 | 10 | 2 | 0 | NON_SYNONYMOU | MODERAT | MISSENSE | Acg/Ccg | T11P | PLES_13391 | . | . | . | . | . | . | . | . | . | . | C | C |
| NC_011770 | 1446991 | A | G | 8 | 10 | 2 | 0 | NON_SYNONYMOU | MODERAT | MISSENSE | Acg/Gcg | T14A | PLES_13391 | . | . | . | . | . | . | . | . | . | . | G | G |
| NC_011770 | 1447301 | C | T | 11 | 10 | 2 | 0 | NON_SYNONYMOU | MODERAT | MISSENSE | gCc/gTc | A117V | PLES_13391 | . | . | . | . | . | . | . | . | . | . | T | T |
| NC_011770 | 1447306 | C | A | 11 | 10 | 2 | 0 | NON_SYNONYMOU | MODERAT | MISSENSE | Cag/Aag | Q119K | PLES_13391 | . | . | . | . | . | . | . | . | . | . | A | A |
| NC_011770 | 1447314 | T | G | 11 | 10 | 2 | 0 | NON_SYNONYMOU | MODERAT | MISSENSE | caT/caG | H121Q | PLES_13391 | . | . | . | . | . | . | . | . | . | . | G | G |
| NC_011770 | 1447322 | G | A | 12 | 10 | 2 | 0 | NON_SYNONYMOU | MODERAT | MISSENSE | cGg/cAg | R124Q | PLES_13391 | . | . | . | . | . | . | . | . | . | . | A | A |
| NC_011770 | 1447343 | C | T | 13 | 10 | 2 | 0 | NON_SYNONYMOU | MODERAT | MISSENSE | gCg/gTg | A131V | PLES_13391 | . | . | . | . | . | . | . | . | . | . | T | T |
| NC_011770 | 1447423 | A | G | 12 | 10 | 2 | 0 | NON_SYNONYMOU | MODERAT | MISSENSE | Atc/Gtc | I158V | PLES_13391 | . | . | . | . | . | . | . | . | . | . | G | G |
| NC_011770 | 1447427 | A | G | 12 | 10 | 2 | 0 | NON_SYNONYMOU | MODERAT | MISSENSE | aAc/aGc | N159S | PLES_13391 | . | . | . | . | . | . | . | . | . | . | G | G |
| NC_011770 | 1447531 | G | A | 11 | 10 | 2 | 0 | NON_SYNONYMOU | MODERAT | MISSENSE | Gtc/Atc | V194I | PLES_13391 | . | . | . | . | . | . | . | . | . | . | A | A |
| NC_011770 | 1447721 | A | G | 10 | 10 | 2 | 0 | START_LOST | HIGH | MISSENSE | Atg/Gtg | M1V | PLES_13401 | . | . | . | . | . | . | . | . | . | . | G | G |
| NC_011770 | 1447986 | T | C | 0 | 11 | 1 | 0 | NON_SYNONYMOU | MODERAT | MISSENSE | cTg/cCg | L89P | PLES_13401 | . | . | . | . | . | . | . | . | C | . | . | . |
| NC_011770 | 1448403 | G | A | 13 | 10 | 2 | 0 | NON_SYNONYMOU | MODERAT | MISSENSE | aGg/aAg | R228K | PLES_13401 | . | . | . | . | . | . | . | . | . | . | A | A |
| NC_011770 | 1448453 | G | A | 13 | 10 | 2 | 0 | NON_SYNONYMOU | MODERAT | MISSENSE | Gcc/Acc | A245T | PLES_13401 | . | . | . | . | . | . | . | . | . | . | A | A |
| NC_011770 | 1448551 | G | C | 15 | 10 | 2 | 0 | NON_SYNONYMOU | MODERAT | MISSENSE | tGt/tCt | C4S | PLES_13411 | . | . | . | . | . | . | . | . | . | . | C | C |
| NC_011770 | 1448657 | G | T | 16 | 10 | 2 | 0 | NON_SYNONYMOU | MODERAT | MISSENSE | gaG/gaT | E39D | PLES_13411 | . | . | . | . | . | . | . | . | . | . | T | T |
| NC_011770 | 1448669 | G | T | 16 | 10 | 2 | 0 | NON_SYNONYMOU | MODERAT | MISSENSE | gaG/gaT | E43D | PLES_13411 | . | . | . | . | . | . | . | . | . | . | T | T |
| NC_011770 | 1448670 | G | T | 16 | 10 | 2 | 0 | NON_SYNONYMOU | MODERAT | MISSENSE | Gcg/Tcg | A44S | PLES_13411 | . | . | . | . | . | . | . | . | . | . | T | T |
| NC_011770 | 1448829 | C | A | 16 | 10 | 2 | 0 | NON_SYNONYMOU | MODERAT | MISSENSE | Ctc/Atc | L97I | PLES_13411 | . | . | . | . | . | . | . | . | . | . | A | A |
| NC_011770 | 1448879 | G | T | 15 | 10 | 2 | 0 | NON_SYNONYMOU | MODERAT | MISSENSE | gaG/gaT | E113D | PLES_13411 | . | . | . | . | . | . | . | . | . | . | T | T |

| | | | | | | | | | | | | | | | | | | | | | | | | | |
|---|---|---|---|---|---|---|---|---|---|---|---|---|---|---|---|---|---|---|---|---|---|---|---|---|---|
| NC_011770 | 1449051 | G | T | 13 | 10 | 2 | 0 | NON_SYNONYMOU | MODERAT | MISSENSE | Gcc/Tcc | A171S | PLES_13411 | . | . | . | . | . | . | . | . | . | . | T | T |
| NC_011770 | 1449075 | G | A | 8 | 10 | 2 | 0 | NON_SYNONYMOU | MODERAT | MISSENSE | Gtg/Atg | V179M | PLES_13411 | . | . | . | . | . | . | . | . | . | . | A | A |
| NC_011770 | 1449076 | T | C | 8 | 10 | 2 | 0 | NON_SYNONYMOU | MODERAT | MISSENSE | gTg/gCg | V179A | PLES_13411 | . | . | . | . | . | . | . | . | . | . | C | C |
| NC_011770 | 1449123 | G | T | 14 | 10 | 2 | 0 | NON_SYNONYMOU | MODERAT | MISSENSE | Gcc/Tcc | A195S | PLES_13411 | . | . | . | . | . | . | . | . | . | . | T | T |
| NC_011770 | 1449259 | T | C | 17 | 10 | 2 | 0 | NON_SYNONYMOU | MODERAT | MISSENSE | gTc/gCc | V240A | PLES_13411 | . | . | . | . | . | . | . | . | . | . | C | C |
| NC_011770 | 1449599 | G | C | 15 | 10 | 2 | 0 | NON_SYNONYMOU | MODERAT | MISSENSE | cGc/cCc | R21P | PLES_13421 | . | . | . | . | . | . | . | . | . | . | C | C |
| NC_011770 | 1449691 | C | A | 14 | 10 | 2 | 0 | NON_SYNONYMOU | MODERAT | MISSENSE | Ctt/Att | L52I | PLES_13421 | . | . | . | . | . | . | . | . | . | . | A | A |
| NC_011770 | 1449715 | A | G | 14 | 10 | 2 | 0 | NON_SYNONYMOU | MODERAT | MISSENSE | Atc/Gtc | I60V | PLES_13421 | . | . | . | . | . | . | . | . | . | . | G | G |
| NC_011770 | 1449718 | G | A | 14 | 10 | 2 | 0 | NON_SYNONYMOU | MODERAT | MISSENSE | Gcg/Acg | A61T | PLES_13421 | . | . | . | . | . | . | . | . | . | . | A | A |
| NC_011770 | 1449719 | C | A | 14 | 10 | 2 | 0 | NON_SYNONYMOU | MODERAT | MISSENSE | gCg/gAg | A61E | PLES_13421 | . | . | . | . | . | . | . | . | . | . | A | A |
| NC_011770 | 1449958 | A | G | 16 | 10 | 2 | 0 | NON_SYNONYMOU | MODERAT | MISSENSE | Atc/Gtc | I141V | PLES_13421 | . | . | . | . | . | . | . | . | . | . | G | G |
| NC_011770 | 1449998 | A | C | 15 | 10 | 2 | 0 | NON_SYNONYMOU | MODERAT | MISSENSE | gAc/gCc | D154A | PLES_13421 | . | . | . | . | . | . | . | . | . | . | C | C |
| NC_011770 | 1450063 | G | T | 13 | 10 | 2 | 0 | NON_SYNONYMOU | MODERAT | MISSENSE | Gcg/Tcg | A176S | PLES_13421 | . | . | . | . | . | . | . | . | . | . | T | T |
| NC_011770 | 1450072 | A | G | 12 | 10 | 2 | 0 | NON_SYNONYMOU | MODERAT | MISSENSE | Acc/Gcc | T179A | PLES_13421 | . | . | . | . | . | . | . | . | . | . | G | G |
| NC_011770 | 1450484 | C | A | 3 | 10 | 2 | 0 | NON_SYNONYMOU | MODERAT | MISSENSE | cCt/cAt | P24H | PLES_13431 | . | . | . | . | . | . | . | . | A | A | . | . |
| NC_011770 | 1450555 | G | A | 2 | 10 | 2 | 0 | NON_SYNONYMOU | MODERAT | MISSENSE | Gcc/Acc | A48T | PLES_13431 | . | . | . | . | . | . | . | . | A | A | . | . |
| NC_011770 | 1450556 | C | T | 2 | 10 | 2 | 0 | NON_SYNONYMOU | MODERAT | MISSENSE | gCc/gTc | A48V | PLES_13431 | . | . | . | . | . | . | . | . | T | T | . | . |
| NC_011770 | 1450769 | A | C | 3 | 10 | 2 | 0 | NON_SYNONYMOU | MODERAT | MISSENSE | gAg/gCg | E119A | PLES_13431 | . | . | . | . | . | . | . | . | C | C | . | . |
| NC_011770 | 1452120 | T | A | 2 | 10 | 2 | 0 | NON_SYNONYMOU | MODERAT | MISSENSE | Tcc/Acc | S4T | PLES_13471 | . | . | . | . | A | A | . | . | . | . | . | . |
| NC_011770 | 1452168 | A | T | 0 | 11 | 1 | 0 | NON_SYNONYMOU | MODERAT | MISSENSE | Acg/Tcg | T20S | PLES_13471 | . | . | . | . | . | T | . | . | . | . | . | . |
| NC_011770 | 1452177 | C | G | 0 | 11 | 1 | 0 | NON_SYNONYMOU | MODERAT | MISSENSE | Ctg/Gtg | L23V | PLES_13471 | . | . | . | . | . | G | . | . | . | . | . | . |
| NC_011770 | 1452186 | C | A | 0 | 11 | 1 | 0 | NON_SYNONYMOU | MODERAT | MISSENSE | Ctc/Atc | L26I | PLES_13471 | . | . | . | . | . | A | . | . | . | . | . | . |
| NC_011770 | 1452219 | A | T | 0 | 10 | 2 | 0 | NON_SYNONYMOU | MODERAT | MISSENSE | Acg/Tcg | T37S | PLES_13471 | . | . | . | . | T | T | . | . | . | . | . | . |
| NC_011770 | 1452345 | C | G | 0 | 11 | 1 | 0 | NON_SYNONYMOU | MODERAT | MISSENSE | Ccg/Gcg | P79A | PLES_13471 | . | . | . | . | . | . | . | . | . | . | G | . |
| NC_011770 | 1452348 | G | A | 0 | 11 | 1 | 0 | NON_SYNONYMOU | MODERAT | MISSENSE | Gca/Aca | A80T | PLES_13471 | . | . | . | . | . | . | . | . | . | . | A | . |
| NC_011770 | 1452372 | C | A | 1 | 11 | 1 | 0 | NON_SYNONYMOU | MODERAT | MISSENSE | Cag/Aag | Q88K | PLES_13471 | . | . | . | . | . | . | . | . | . | . | A | . |
| NC_011770 | 1452444 | T | C | 3 | 7 | 1 | 0 | NON_SYNONYMOU | MODERAT | MISSENSE | Tat/Cat | Y112H | PLES_13471 | . | . | . | . | C | . | . | . | | | | |
| NC_011770 | 1452576 | T | G | 4 | 6 | 4 | 0 | NON_SYNONYMOU | MODERAT | MISSENSE | Tcg/Gcg | S156A | PLES_13471 | . | . | . | . | G | G | . | . | G | G | | |
| NC_011770 | 1452580 | G | C | 3 | 7 | 2 | 0 | NON_SYNONYMOU | MODERAT | MISSENSE | gGg/gCg | G157A | PLES_13471 | . | . | . | . | | | . | . | . | | C | C |
| NC_011770 | 1452586 | G | T | 4 | 7 | 2 | 0 | NON_SYNONYMOU | MODERAT | MISSENSE | tGg/tTg | W159L | PLES_13471 | . | . | . | . | T | T | . | . | . | | | |
| NC_011770 | 1452587 | G | C | 4 | 7 | 2 | 0 | NON_SYNONYMOU | MODERAT | MISSENSE | tgG/tgC | W159C | PLES_13471 | . | . | . | . | C | C | . | . | . | | | |
| NC_011770 | 1452591 | A | C | 3 | 7 | 3 | 0 | NON_SYNONYMOU | MODERAT | MISSENSE | Aag/Cag | K161Q | PLES_13471 | . | . | . | . | C | C | . | . | . | C | | |
| NC_011770 | 1452592 | A | G | 3 | 7 | 1 | 0 | NON_SYNONYMOU | MODERAT | MISSENSE | aAg/aGg | K161R | PLES_13471 | . | . | . | . | | | . | . | . | G | | |
| NC_011770 | 1452611 | C | T | 4 | 7 | 5 | 0 | NON_SYNONYMOU | MODERAT | MISSENSE | gaC/gaA | D167E | PLES_13471 | . | . | . | . | T | T | . | . | . | T | A | A |
| NC_011770 | 1452657 | G | A | 1 | 9 | 2 | 0 | NON_SYNONYMOU | MODERAT | MISSENSE | Ggc/Agc | G183S | PLES_13471 | . | . | . | . | . | . | . | . | . | | A | A |
| NC_011770 | 1452677 | G | C | 0 | 11 | 1 | 0 | NON_SYNONYMOU | MODERAT | MISSENSE | gaG/gaC | E189D | PLES_13471 | . | . | . | . | . | . | . | . | . | C | . | . |
| NC_011770 | 1452685 | T | C | 0 | 11 | 1 | 0 | NON_SYNONYMOU | MODERAT | MISSENSE | gTg/gCg | V192A | PLES_13471 | . | . | . | . | . | . | . | . | . | C | . | . |
| NC_011770 | 1452697 | A | G | 0 | 11 | 1 | 0 | NON_SYNONYMOU | MODERAT | MISSENSE | gAa/gGa | E196G | PLES_13471 | . | . | . | . | . | . | . | . | . | G | . | . |
| NC_011770 | 1452698 | A | T | 0 | 11 | 1 | 0 | NON_SYNONYMOU | MODERAT | MISSENSE | gaA/gaT | E196D | PLES_13471 | . | . | . | . | . | . | . | . | . | T | . | . |
| NC_011770 | 1452934 | A | G | 14 | 6 | 2 | 0 | STOP_LOST+SPLICE | HIGH | MISSENSE | Tag/Cag | *78Q | PLES_13481 | . | . | . | . | | | . | . | | | G | G |
| NC_011770 | 1453165 | T | C | 16 | 8 | 4 | 0 | START_LOST | HIGH | MISSENSE | Atg/Gtg | M1V | PLES_13481 | . | . | . | . | C | C | . | . | . | . | C | C |
| NC_011770 | 1452940 | T | C | 13 | 6 | 6 | 0 | NON_SYNONYMOU | MODERAT | MISSENSE | Agc/Ggc | S76G | PLES_13481 | . | . | . | . | C | C | . | . | C | C | C | C |
| NC_011770 | 1453023 | G | A | 13 | 6 | 6 | 0 | NON_SYNONYMOU | MODERAT | MISSENSE | gCc/gTc | A48V | PLES_13481 | . | . | . | . | A | A | . | . | A | A | A | A |
| NC_011770 | 1453035 | A | C | 15 | 6 | 2 | 0 | NON_SYNONYMOU | MODERAT | MISSENSE | cTg/cGg | L44R | PLES_13481 | . | . | . | . | | | . | . | C | C | | |
| NC_011770 | 1453061 | G | C | 15 | 6 | 2 | 0 | NON_SYNONYMOU | MODERAT | MISSENSE | atC/atG | I35M | PLES_13481 | . | . | . | . | | | . | . | C | C | | |
| NC_011770 | 1453132 | G | A | 15 | 6 | 6 | 0 | NON_SYNONYMOU | MODERAT | MISSENSE | Ccg/Tcg | P12S | PLES_13481 | . | . | . | . | A | A | . | . | A | A | A | A |
| NC_011770 | 1453147 | C | T | 14 | 8 | 4 | 0 | NON_SYNONYMOU | MODERAT | MISSENSE | Gat/Aat | D7N | PLES_13481 | . | . | . | . | T | T | . | . | . | . | T | T |
| NC_011770 | 1453149 | C | A | 14 | 8 | 4 | 0 | NON_SYNONYMOU | MODERAT | MISSENSE | cGc/cTc | R6L | PLES_13481 | . | . | . | . | A | A | . | . | . | . | A | A |
| NC_011770 | 1453483 | T | C | 24 | 8 | 2 | 0 | NON_SYNONYMOU | MODERAT | MISSENSE | Tac/Cac | Y97H | PLES_13491 | . | . | . | . | C | C | . | . | . | . | | |
| NC_011770 | 1453741 | C | A | 20 | 10 | 2 | 0 | NON_SYNONYMOU | MODERAT | MISSENSE | Cag/Aag | Q183K | PLES_13491 | . | . | . | . | A | A | . | . | . | . | . | . |
| NC_011770 | 1453792 | A | G | 21 | 8 | 2 | 0 | NON_SYNONYMOU | MODERAT | MISSENSE | Atc/Gtc | I200V | PLES_13491 | . | . | . | . | | | . | . | . | . | G | G |
| NC_011770 | 1453828 | C | A | 21 | 8 | 2 | 0 | NON_SYNONYMOU | MODERAT | MISSENSE | Cgc/Agc | R212S | PLES_13491 | . | . | . | . | A | A | . | . | . | . | | |
| NC_011770 | 1453868 | A | C | 21 | 8 | 2 | 0 | NON_SYNONYMOU | MODERAT | MISSENSE | gAc/gCc | D225A | PLES_13491 | . | . | . | . | | | . | . | . | . | C | C |
| NC_011770 | 1453895 | C | T | 21 | 8 | 4 | 0 | NON_SYNONYMOU | MODERAT | MISSENSE | gCg/gTg | A234V | PLES_13491 | . | . | . | . | T | T | . | . | . | . | T | T |
| NC_011770 | 1454029 | A | T | 21 | 8 | 2 | 0 | NON_SYNONYMOU | MODERAT | MISSENSE | cAt/cTt | H4L | PLES_13501 | . | . | . | . | | | . | . | . | . | T | T |
| NC_011770 | 1454746 | A | G | 39 | 8 | 2 | 0 | NON_SYNONYMOU | MODERAT | MISSENSE | Atc/Gtc | I58V | PLES_13511 | . | . | . | . | | | . | . | . | . | G | G |
| NC_011770 | 1455232 | A | G | 38 | 8 | 2 | 0 | NON_SYNONYMOU | MODERAT | MISSENSE | Agt/Ggt | S220G | PLES_13511 | . | . | . | . | | | . | . | . | . | G | G |
| NC_011770 | 1455233 | G | C | 38 | 8 | 2 | 0 | NON_SYNONYMOU | MODERAT | MISSENSE | aGt/aCt | S220T | PLES_13511 | . | . | . | . | | | . | . | . | . | C | C |
| NC_011770 | 1455523 | G | A | 32 | 8 | 2 | 0 | NON_SYNONYMOU | MODERAT | MISSENSE | Gac/Aac | D317N | PLES_13511 | . | . | . | . | | | . | . | . | . | A | A |
| NC_011770 | 1455718 | A | G | 27 | 8 | 2 | 0 | NON_SYNONYMOU | MODERAT | MISSENSE | Acg/Gcg | T382A | PLES_13511 | . | . | . | . | G | G | . | . | . | . | | |
| NC_011770 | 1455832 | C | A | 28 | 8 | 2 | 0 | NON_SYNONYMOU | MODERAT | MISSENSE | Ctt/Att | L420I | PLES_13511 | . | . | . | . | | | . | . | . | . | A | A |
| NC_011770 | 1456021 | G | A | 26 | 8 | 2 | 0 | NON_SYNONYMOU | MODERAT | MISSENSE | Gtc/Atc | V483I | PLES_13511 | . | . | . | . | | | . | . | . | . | A | A |

| | | | | | | | | | | | | | | | | | | | | | | | | | |
|---|---|---|---|---|---|---|---|---|---|---|---|---|---|---|---|---|---|---|---|---|---|---|---|---|---|
| NC_011770 | 1456045 | G | A | 23 | 8 | 2 | 0 | NON_SYNONYMOU | MODERAT | MISSENSE | Gtt/Att | V491I | PLES_13511 | . | . | . | . | | | . | . | . | . | A | A |
| NC_011770 | 1458447 | A | T | 19 | 10 | 2 | 0 | NON_SYNONYMOU | MODERAT | MISSENSE | Acc/Tcc | T29S | PLES_13531 | . | . | . | . | T | T | . | . | . | . | . | . |
| NC_011770 | 1458580 | G | A | 18 | 10 | 2 | 0 | NON_SYNONYMOU | MODERAT | MISSENSE | aGc/aAc | S73N | PLES_13531 | . | . | . | . | A | A | . | . | . | . | . | . |
| NC_011770 | 1459329 | A | G | 12 | 10 | 2 | 0 | NON_SYNONYMOU | MODERAT | MISSENSE | Acc/Gcc | T323A | PLES_13531 | . | . | . | . | G | G | . | . | . | . | . | . |
| NC_011770 | 1459503 | A | G | 14 | 10 | 2 | 0 | NON_SYNONYMOU | MODERAT | MISSENSE | Atc/Gtc | I381V | PLES_13531 | . | . | . | . | G | G | . | . | . | . | . | . |
| NC_011770 | 1459807 | A | T | 20 | 10 | 2 | 0 | NON_SYNONYMOU | MODERAT | MISSENSE | tAc/tTc | Y482F | PLES_13531 | . | . | . | . | T | T | . | . | . | . | . | . |
| NC_011770 | 1460082 | C | A | 19 | 10 | 2 | 0 | NON_SYNONYMOU | MODERAT | MISSENSE | Ccc/Acc | P574T | PLES_13531 | . | . | . | . | A | A | . | . | . | . | . | . |
| NC_011770 | 1460563 | C | T | 16 | 8 | 2 | 0 | NON_SYNONYMOU | MODERAT | MISSENSE | gCt/gTt | A18V | PLES_13541 | . | . | . | . | T | T | . | . | . | . | | |
| NC_011770 | 1460566 | A | G | 17 | 8 | 4 | 0 | NON_SYNONYMOU | MODERAT | MISSENSE | gAc/gGc | D19G | PLES_13541 | . | . | . | . | G | G | . | . | . | . | G | G |
| NC_011770 | 1460724 | G | A | 16 | 8 | 2 | 0 | NON_SYNONYMOU | MODERAT | MISSENSE | Gat/Aat | D72N | PLES_13541 | . | . | . | . | | | . | . | . | . | A | A |
| NC_011770 | 1460740 | C | T | 15 | 8 | 2 | 0 | NON_SYNONYMOU | MODERAT | MISSENSE | gCt/gTt | A77V | PLES_13541 | . | . | . | . | | | . | . | . | . | T | T |
| NC_011770 | 1460745 | A | G | 15 | 8 | 4 | 0 | NON_SYNONYMOU | MODERAT | MISSENSE | Acc/Gcc | T79A | PLES_13541 | . | . | . | . | G | G | . | . | . | . | G | G |
| NC_011770 | 1460784 | C | G | 15 | 8 | 2 | 0 | NON_SYNONYMOU | MODERAT | MISSENSE | Ccc/Gcc | P92A | PLES_13541 | . | . | . | . | | | . | . | . | . | G | G |
| NC_011770 | 1461034 | T | C | 6 | 10 | 2 | 0 | NON_SYNONYMOU | MODERAT | MISSENSE | Tgt/Cgt | C6R | PLES_13551 | . | . | . | . | C | C | . | . | . | . | . | . |
| NC_011770 | 1461067 | T | C | 6 | 10 | 2 | 0 | NON_SYNONYMOU | MODERAT | MISSENSE | Tat/Cat | Y17H | PLES_13551 | . | . | . | . | C | C | . | . | . | . | . | . |
| NC_011770 | 1461091 | C | G | 7 | 9 | 3 | 0 | NON_SYNONYMOU | MODERAT | MISSENSE | Cga/Gga | R25G | PLES_13551 | . | . | . | . | G | G | . | . | . | . | . | G |
| NC_011770 | 1461103 | T | C | 7 | 9 | 1 | 0 | NON_SYNONYMOU | MODERAT | MISSENSE | Tgc/Cgc | C29R | PLES_13551 | . | . | . | . | | | . | . | . | . | . | C |
| NC_011770 | 1461194 | C | G | 12 | 8 | 2 | 0 | NON_SYNONYMOU | MODERAT | MISSENSE | gCg/gGg | A59G | PLES_13551 | . | . | . | . | G | G | . | . | . | . | | |
| NC_011770 | 1461381 | A | C | 14 | 9 | 3 | 0 | NON_SYNONYMOU | MODERAT | MISSENSE | gaA/gaC | E121D | PLES_13551 | . | . | . | . | C | C | . | . | . | . | . | C |
| NC_011770 | 1461507 | T | G | 12 | 8 | 2 | 0 | NON_SYNONYMOU | MODERAT | MISSENSE | gaT/gaG | D163E | PLES_13551 | . | . | . | . | G | G | . | . | . | . | | |
| NC_011770 | 1461559 | G | A | 12 | 8 | 2 | 0 | NON_SYNONYMOU | MODERAT | MISSENSE | Gat/Aat | D181N | PLES_13551 | . | . | . | . | | | . | . | . | . | A | A |
| NC_011770 | 1461918 | G | C | 18 | 10 | 2 | 0 | NON_SYNONYMOU | MODERAT | MISSENSE | aGt/aCt | S33T | PLES_13561 | . | . | . | . | C | C | . | . | . | . | . | . |
| NC_011770 | 1462008 | T | C | 18 | 10 | 2 | 0 | NON_SYNONYMOU | MODERAT | MISSENSE | aTt/aCt | I63T | PLES_13561 | . | . | . | . | C | C | . | . | . | . | . | . |
| NC_011770 | 1462773 | C | T | 10 | 8 | 2 | 0 | NON_SYNONYMOU | MODERAT | MISSENSE | aCg/aTg | T40M | PLES_13571 | . | . | . | . | | | . | . | . | . | T | T |
| NC_011770 | 1462871 | A | G | 10 | 8 | 2 | 0 | NON_SYNONYMOU | MODERAT | MISSENSE | Aca/Gca | T73A | PLES_13571 | . | . | . | . | | | . | . | . | . | G | G |
| NC_011770 | 1462929 | G | A | 12 | 8 | 2 | 0 | NON_SYNONYMOU | MODERAT | MISSENSE | gGc/gAc | G92D | PLES_13571 | . | . | . | . | | | . | . | . | . | A | A |
| NC_011770 | 1462998 | C | T | 15 | 8 | 2 | 0 | NON_SYNONYMOU | MODERAT | MISSENSE | gCt/gTt | A115V | PLES_13571 | . | . | . | . | | | . | . | . | . | T | T |
| NC_011770 | 1463013 | G | A | 15 | 8 | 2 | 0 | NON_SYNONYMOU | MODERAT | MISSENSE | gGg/gAg | G120E | PLES_13571 | . | . | . | . | | | . | . | . | . | A | A |
| NC_011770 | 1463099 | C | A | 19 | 8 | 2 | 0 | NON_SYNONYMOU | MODERAT | MISSENSE | Ctt/Att | L149I | PLES_13571 | . | . | . | . | | | . | . | . | . | A | A |
| NC_011770 | 1463157 | A | C | 17 | 8 | 2 | 0 | NON_SYNONYMOU | MODERAT | MISSENSE | gAt/gCt | D168A | PLES_13571 | . | . | . | . | | | . | . | . | . | C | C |
| NC_011770 | 1463271 | T | A | 16 | 8 | 2 | 0 | NON_SYNONYMOU | MODERAT | MISSENSE | cTg/cAg | L206Q | PLES_13571 | . | . | . | . | | | . | . | . | . | A | A |
| NC_011770 | 1463282 | A | G | 16 | 8 | 2 | 0 | NON_SYNONYMOU | MODERAT | MISSENSE | Act/Gct | T210A | PLES_13571 | . | . | . | . | | | . | . | . | . | G | G |
| NC_011770 | 1464075 | GCTTTC | G | 9 | 8 | 2 | 0 | FRAME_SHIFT | HIGH | | ctttcc/ | LS106 | PLES_13581 | . | . | . | . | | | . | . | . | . | G | G |
| NC_011770 | 1463873 | A | G | 4 | 8 | 2 | 0 | NON_SYNONYMOU | MODERAT | MISSENSE | aAg/aGg | K38R | PLES_13581 | . | . | . | . | G | G | . | . | . | . | | |
| NC_011770 | 1464067 | A | G | 6 | 9 | 1 | 0 | NON_SYNONYMOU | MODERAT | MISSENSE | Ata/Gta | I103V | PLES_13581 | . | . | . | . | | | . | . | . | . | G | . |
| NC_011770 | 1464260 | T | A | 18 | 8 | 2 | 0 | NON_SYNONYMOU | MODERAT | MISSENSE | tTc/tAc | F167Y | PLES_13581 | . | . | . | . | A | A | . | . | . | . | | |
| NC_011770 | 1464284 | G | C | 19 | 8 | 4 | 0 | NON_SYNONYMOU | MODERAT | MISSENSE | aGc/aCc | S175T | PLES_13581 | . | . | . | . | C | C | . | . | . | . | C | C |
| NC_011770 | 1464691 | A | G | 21 | 4 | 2 | 0 | NON_SYNONYMOU | MODERAT | MISSENSE | Atc/Gtc | I311V | PLES_13581 | | | . | . | | | | | . | . | G | G |
| NC_011770 | 1464731 | C | T | 16 | 8 | 2 | 0 | NON_SYNONYMOU | MODERAT | MISSENSE | gCg/gTg | A324V | PLES_13581 | . | . | . | . | | | . | . | . | . | T | T |
| NC_011770 | 1464775 | T | G | 12 | 8 | 2 | 0 | NON_SYNONYMOU | MODERAT | MISSENSE | Tcg/Gcg | S339A | PLES_13581 | . | . | . | . | | | . | . | . | . | G | G |
| NC_011770 | 1465302 | A | T | 0 | 11 | 1 | 0 | NON_SYNONYMOU | MODERAT | MISSENSE | gaA/gaT | E514D | PLES_13581 | . | . | . | . | T | . | . | . | . | . | . | . |
| NC_011770 | 1465433 | C | G | 0 | 11 | 1 | 0 | NON_SYNONYMOU | MODERAT | MISSENSE | gCc/gGc | A558G | PLES_13581 | . | . | . | . | . | . | . | . | . | . | G | . |
| NC_011770 | 1466314 | G | A | 0 | 11 | 1 | 0 | NON_SYNONYMOU | MODERAT | MISSENSE | Gta/Ata | V15I | PLES_13591 | . | . | . | . | . | . | . | . | . | . | A | . |
| NC_011770 | 1466737 | A | G | 1 | 8 | 2 | 0 | NON_SYNONYMOU | MODERAT | MISSENSE | Atc/Gtc | I156V | PLES_13591 | . | . | . | . | | | . | . | . | . | G | G |
| NC_011770 | 1467039 | A | G | 1 | 9 | 2 | 0 | NON_SYNONYMOU | MODERAT | MISSENSE | Acg/Gcg | T79A | PLES_13601 | . | . | . | . | G | G | . | . | . | . | | . |
| NC_011770 | 1467085 | C | T | 1 | 9 | 1 | 0 | NON_SYNONYMOU | MODERAT | MISSENSE | aCc/aTc | T94I | PLES_13601 | . | . | . | . | T | . | . | . | . | . | | |
| NC_011770 | 1467087 | T | G | 1 | 9 | 3 | 0 | NON_SYNONYMOU | MODERAT | MISSENSE | Tcg/Gcg | S95A | PLES_13601 | . | . | . | . | G | . | . | . | . | . | G | G |
| NC_011770 | 1467216 | T | G | 3 | 9 | 3 | 0 | NON_SYNONYMOU | MODERAT | MISSENSE | Tcg/Gcg | S138A | PLES_13601 | . | . | . | . | G | . | . | . | . | . | G | G |
| NC_011770 | 1467423 | A | T | 0 | 11 | 1 | 0 | NON_SYNONYMOU | MODERAT | MISSENSE | Acg/Tcg | T207S | PLES_13601 | . | . | . | . | . | T | . | . | . | . | . | . |
| NC_011770 | 1467510 | C | T | 1 | 10 | 2 | 0 | NON_SYNONYMOU | MODERAT | MISSENSE | Ctc/Ttc | L236F | PLES_13601 | . | . | . | . | T | T | . | . | . | . | . | . |
| NC_011770 | 1467886 | G | A | 2 | 10 | 2 | 0 | NON_SYNONYMOU | MODERAT | MISSENSE | aGg/aAg | R361K | PLES_13601 | . | . | . | . | A | A | . | . | . | . | . | . |
| NC_011770 | 1467908 | G | C | 2 | 10 | 2 | 0 | NON_SYNONYMOU | MODERAT | MISSENSE | gaG/gaC | E368D | PLES_13601 | . | . | . | . | C | C | . | . | . | . | . | . |
| NC_011770 | 1467984 | C | T | 3 | 10 | 2 | 0 | NON_SYNONYMOU | MODERAT | MISSENSE | Ctc/Ttc | L394F | PLES_13601 | . | . | . | . | T | T | . | . | . | . | . | . |
| NC_011770 | 1468548 | T | C | 2 | 10 | 2 | 0 | NON_SYNONYMOU | MODERAT | MISSENSE | gTa/gCa | V15A | PLES_13611 | . | . | . | . | C | C | . | . | . | . | . | . |
| NC_011770 | 1468566 | C | A | 2 | 10 | 2 | 0 | NON_SYNONYMOU | MODERAT | MISSENSE | gCg/gAg | A21E | PLES_13611 | . | . | . | . | A | A | . | . | . | . | . | . |
| NC_011770 | 1468572 | G | A | 2 | 10 | 2 | 0 | NON_SYNONYMOU | MODERAT | MISSENSE | aGc/aAc | S23N | PLES_13611 | . | . | . | . | A | A | . | . | . | . | . | . |
| NC_011770 | 1468623 | G | A | 0 | 10 | 1 | 0 | NON_SYNONYMOU | MODERAT | MISSENSE | aGc/aAc | S40N | PLES_13611 | . | . | . | . | A | . | . | . | . | . | . | . |
| NC_011770 | 1468650 | A | C | 0 | 10 | 2 | 0 | NON_SYNONYMOU | MODERAT | MISSENSE | gAc/gCc | D49A | PLES_13611 | . | . | . | . | C | C | . | . | . | . | . | . |
| NC_011770 | 1468662 | A | C | 1 | 9 | 2 | 0 | NON_SYNONYMOU | MODERAT | MISSENSE | cAg/cCg | Q53P | PLES_13611 | . | . | . | . | C | C | . | . | . | . | | . |
| NC_011770 | 1468685 | A | G | 2 | 9 | 3 | 0 | NON_SYNONYMOU | MODERAT | MISSENSE | Acc/Gcc | T61A | PLES_13611 | . | . | . | . | G | G | . | . | . | . | T | . |
| NC_011770 | 1468700 | G | A | 3 | 8 | 2 | 0 | NON_SYNONYMOU | MODERAT | MISSENSE | Gtg/Atg | V66M | PLES_13611 | . | . | . | . | A | A | . | . | . | . | | |
| NC_011770 | 1468728 | G | C | 4 | 8 | 4 | 0 | NON_SYNONYMOU | MODERAT | MISSENSE | gGa/gCa | G75A | PLES_13611 | . | . | . | . | C | C | . | . | . | . | C | C |

| | | | | | | | | | | | | | | | | | | | | | | | | | |
|---|---|---|---|---|---|---|---|---|---|---|---|---|---|---|---|---|---|---|---|---|---|---|---|---|---|
| NC_011770 | 1468733 | T | G | 4 | 8 | 4 | 0 | NON_SYNONYMOU | MODERAT | MISSENSE | Tcc/Gcc | S77A | PLES_13611 | . | . | . | . | G | G | . | . | . | . | G | G |
| NC_011770 | 1468737 | A | C | 4 | 8 | 2 | 0 | NON_SYNONYMOU | MODERAT | MISSENSE | gAc/gCc | D78A | PLES_13611 | . | . | . | . | C | C | . | . | . | . | | |
| NC_011770 | 1468752 | C | T | 6 | 8 | 2 | 0 | NON_SYNONYMOU | MODERAT | MISSENSE | gCg/gTg | A83V | PLES_13611 | . | . | . | . | | | . | . | . | . | T | T |
| NC_011770 | 1468757 | A | T | 6 | 8 | 4 | 0 | NON_SYNONYMOU | MODERAT | MISSENSE | Acg/Tcg | T85S | PLES_13611 | . | . | . | . | T | T | . | . | . | . | T | T |
| NC_011770 | 1468911 | T | A | 7 | 9 | 1 | 0 | NON_SYNONYMOU | MODERAT | MISSENSE | aTt/aAt | I136N | PLES_13611 | . | . | . | . | | | . | . | . | . | A | . |
| NC_011770 | 1469729 | G | T | 15 | 8 | 2 | 0 | NON_SYNONYMOU | MODERAT | MISSENSE | Gcc/Tcc | A87S | PLES_13631 | . | . | . | . | | | . | . | . | . | T | T |
| NC_011770 | 1469947 | A | G | 18 | 8 | 2 | 0 | NON_SYNONYMOU | MODERAT | MISSENSE | Aac/Gac | N26D | PLES_13641 | . | . | . | . | | | . | . | . | . | G | G |
| NC_011770 | 1469952 | T | G | 19 | 8 | 4 | 0 | NON_SYNONYMOU | MODERAT | MISSENSE | aaT/aaG | N27K | PLES_13641 | . | . | . | . | G | G | . | . | . | . | G | G |
| NC_011770 | 1469972 | C | T | 18 | 8 | 2 | 0 | NON_SYNONYMOU | MODERAT | MISSENSE | aCc/aTc | T34I | PLES_13641 | . | . | . | . | T | T | . | . | . | . | | |
| NC_011770 | 1470017 | A | C | 16 | 8 | 4 | 0 | NON_SYNONYMOU | MODERAT | MISSENSE | aAc/aCc | N49T | PLES_13641 | . | . | . | . | C | C | . | . | . | . | C | C |
| NC_011770 | 1470074 | A | C | 14 | 8 | 4 | 0 | NON_SYNONYMOU | MODERAT | MISSENSE | cAg/cCg | Q68P | PLES_13641 | . | . | . | . | C | C | . | . | . | . | C | C |
| NC_011770 | 1470201 | C | A | 17 | 8 | 4 | 0 | NON_SYNONYMOU | MODERAT | MISSENSE | gaC/gaA | D110E | PLES_13641 | . | . | . | . | A | A | . | . | . | . | A | A |
| NC_011770 | 1470232 | T | G | 18 | 8 | 4 | 0 | NON_SYNONYMOU | MODERAT | MISSENSE | Tct/Gct | S121A | PLES_13641 | . | . | . | . | G | G | . | . | . | . | G | G |
| NC_011770 | 1470233 | C | A | 18 | 8 | 4 | 0 | NON_SYNONYMOU | MODERAT | MISSENSE | tCt/tAt | S121Y | PLES_13641 | . | . | . | . | A | A | . | . | . | . | A | A |
| NC_011770 | 1470254 | G | T | 19 | 8 | 2 | 0 | NON_SYNONYMOU | MODERAT | MISSENSE | cGc/cTc | R128L | PLES_13641 | . | . | . | . | T | T | . | . | . | . | | |
| NC_011770 | 1470283 | G | T | 26 | 8 | 4 | 0 | NON_SYNONYMOU | MODERAT | MISSENSE | Gtg/Ttg | V138L | PLES_13641 | . | . | . | . | T | T | . | . | . | . | T | T |
| NC_011770 | 1470337 | G | T | 26 | 8 | 2 | 0 | NON_SYNONYMOU | MODERAT | MISSENSE | Gcc/Tcc | A156S | PLES_13641 | . | . | . | . | | | . | . | . | . | T | T |
| NC_011770 | 1470447 | A | C | 25 | 8 | 2 | 0 | NON_SYNONYMOU | MODERAT | MISSENSE | gaA/gaC | E192D | PLES_13641 | . | . | . | . | | | . | . | . | . | C | C |
| NC_011770 | 1470528 | C | A | 28 | 8 | 4 | 0 | NON_SYNONYMOU | MODERAT | MISSENSE | gaC/gaA | D219E | PLES_13641 | . | . | . | . | A | A | . | . | . | . | A | A |
| NC_011770 | 1470789 | A | C | 38 | 8 | 2 | 0 | NON_SYNONYMOU | MODERAT | MISSENSE | gaA/gaC | E306D | PLES_13641 | . | . | . | . | | | . | . | . | . | C | C |
| NC_011770 | 1470846 | G | T | 42 | 8 | 2 | 0 | NON_SYNONYMOU | MODERAT | MISSENSE | caG/caT | Q325H | PLES_13641 | . | . | . | . | | | . | . | . | . | T | T |
| NC_011770 | 1470958 | T | A | 41 | 8 | 2 | 0 | NON_SYNONYMOU | MODERAT | MISSENSE | Tcc/Acc | S363T | PLES_13641 | . | . | . | . | A | A | . | . | . | . | | |
| NC_011770 | 1470979 | A | T | 42 | 8 | 2 | 0 | NON_SYNONYMOU | MODERAT | MISSENSE | Aca/Tca | T370S | PLES_13641 | . | . | . | . | | | . | . | . | . | T | T |
| NC_011770 | 1471169 | T | C | 40 | 8 | 2 | 0 | NON_SYNONYMOU | MODERAT | MISSENSE | gTg/gCg | V433A | PLES_13641 | . | . | . | . | C | C | . | . | . | . | | |
| NC_011770 | 1471193 | A | G | 39 | 8 | 2 | 0 | NON_SYNONYMOU | MODERAT | MISSENSE | gAc/gGc | D441G | PLES_13641 | . | . | . | . | | | . | . | . | . | G | G |
| NC_011770 | 1471346 | G | A | 37 | 8 | 2 | 0 | NON_SYNONYMOU | MODERAT | MISSENSE | cGc/cAc | R492H | PLES_13641 | . | . | . | . | | | . | . | . | . | A | A |
| NC_011770 | 1471471 | G | A | 32 | 8 | 4 | 0 | NON_SYNONYMOU | MODERAT | MISSENSE | Gtc/Atc | V534I | PLES_13641 | . | . | . | . | A | A | . | . | . | . | A | A |
| NC_011770 | 1471483 | C | A | 33 | 8 | 2 | 0 | NON_SYNONYMOU | MODERAT | MISSENSE | Ctc/Atc | L538I | PLES_13641 | . | . | . | . | | | . | . | . | . | A | A |
| NC_011770 | 1471495 | T | C | 32 | 8 | 4 | 0 | NON_SYNONYMOU | MODERAT | MISSENSE | Tcg/Ccg | S542P | PLES_13641 | . | . | . | . | C | C | . | . | . | . | C | C |
| NC_011770 | 1471587 | A | G | 21 | 8 | 4 | 0 | NON_SYNONYMOU | MODERAT | MISSENSE | Att/Gtt | I10V | PLES_13651 | . | . | . | . | G | G | . | . | . | . | G | G |
| NC_011770 | 1471589 | T | G | 20 | 8 | 4 | 0 | NON_SYNONYMOU | MODERAT | MISSENSE | atT/atG | I10M | PLES_13651 | . | . | . | . | G | G | . | . | . | . | G | G |
| NC_011770 | 1471593 | A | C | 21 | 8 | 4 | 0 | NON_SYNONYMOU | MODERAT | MISSENSE | Aaa/Caa | K12Q | PLES_13651 | . | . | . | . | C | C | . | . | . | . | C | C |
| NC_011770 | 1471594 | A | G | 20 | 8 | 4 | 0 | NON_SYNONYMOU | MODERAT | MISSENSE | aAa/aGa | K12R | PLES_13651 | . | . | . | . | G | G | . | . | . | . | G | G |
| NC_011770 | 1471595 | A | C | 20 | 8 | 4 | 0 | NON_SYNONYMOU | MODERAT | MISSENSE | aaA/aaC | K12N | PLES_13651 | . | . | . | . | C | C | . | . | . | . | C | C |
| NC_011770 | 1471668 | T | C | 24 | 8 | 2 | 0 | NON_SYNONYMOU | MODERAT | MISSENSE | Tcg/Ccg | S37P | PLES_13651 | . | . | . | . | | | . | . | . | . | C | C |
| NC_011770 | 1471714 | A | T | 21 | 8 | 2 | 0 | NON_SYNONYMOU | MODERAT | MISSENSE | gAc/gTc | D52V | PLES_13651 | . | . | . | . | | | . | . | . | . | T | T |
| NC_011770 | 1471812 | G | A | 15 | 10 | 2 | 0 | NON_SYNONYMOU | MODERAT | MISSENSE | Gac/Aac | D85N | PLES_13651 | . | . | . | . | A | A | . | . | . | . | . | . |
| NC_011770 | 1471818 | A | G | 14 | 10 | 2 | 0 | NON_SYNONYMOU | MODERAT | MISSENSE | Atc/Gtc | I87V | PLES_13651 | . | . | . | . | G | G | . | . | . | . | . | . |
| NC_011770 | 1471843 | T | C | 14 | 10 | 2 | 0 | NON_SYNONYMOU | MODERAT | MISSENSE | gTt/gCt | V95A | PLES_13651 | . | . | . | . | C | C | . | . | . | . | . | . |
| NC_011770 | 1471845 | G | A | 14 | 10 | 2 | 0 | NON_SYNONYMOU | MODERAT | MISSENSE | Ggg/Agg | G96R | PLES_13651 | . | . | . | . | A | A | . | . | . | . | . | . |
| NC_011770 | 1471936 | A | G | 14 | 10 | 2 | 0 | NON_SYNONYMOU | MODERAT | MISSENSE | gAt/gGt | D126G | PLES_13651 | . | . | . | . | G | G | . | . | . | . | . | . |
| NC_011770 | 1472116 | C | G | 2 | 10 | 2 | 0 | NON_SYNONYMOU | MODERAT | MISSENSE | aCt/aGt | T186S | PLES_13651 | . | . | . | . | G | G | . | . | . | . | . | . |
| NC_011770 | 1472702 | T | A | 9 | 8 | 2 | 0 | NON_SYNONYMOU | MODERAT | MISSENSE | Tcg/Acg | S123T | PLES_13661 | . | . | . | . | A | A | . | . | . | . | | |
| NC_011770 | 1472703 | C | G | 9 | 8 | 2 | 0 | NON_SYNONYMOU | MODERAT | MISSENSE | tCg/tGg | S123W | PLES_13661 | . | . | . | . | G | G | . | . | . | . | | |
| NC_011770 | 1472717 | C | A | 12 | 8 | 2 | 0 | NON_SYNONYMOU | MODERAT | MISSENSE | Ctc/Atc | L128I | PLES_13661 | . | . | . | . | | | . | . | . | . | A | A |
| NC_011770 | 1472784 | A | T | 12 | 8 | 2 | 0 | NON_SYNONYMOU | MODERAT | MISSENSE | cAg/cTg | Q150L | PLES_13661 | . | . | . | . | | | . | . | . | . | T | T |
| NC_011770 | 1472825 | A | G | 10 | 8 | 4 | 0 | NON_SYNONYMOU | MODERAT | MISSENSE | Agc/Ggc | S164G | PLES_13661 | . | . | . | . | G | G | . | . | . | . | G | G |
| NC_011770 | 1472833 | T | G | 9 | 8 | 2 | 0 | NON_SYNONYMOU | MODERAT | MISSENSE | gaT/gaG | D166E | PLES_13661 | . | . | . | . | G | G | . | . | . | . | | |
| NC_011770 | 1472834 | G | T | 9 | 8 | 4 | 0 | NON_SYNONYMOU | MODERAT | MISSENSE | Ggc/Tgc | G167C | PLES_13661 | . | . | . | . | T | T | . | . | . | . | T | T |
| NC_011770 | 1472835 | G | C | 10 | 8 | 4 | 0 | NON_SYNONYMOU | MODERAT | MISSENSE | gGc/gCc | G167A | PLES_13661 | . | . | . | . | C | C | . | . | . | . | C | C |
| NC_011770 | 1473201 | A | G | 5 | 10 | 2 | 0 | NON_SYNONYMOU | MODERAT | MISSENSE | Aac/Gac | N11D | PLES_13671 | . | . | . | . | G | G | . | . | . | . | . | . |
| NC_011770 | 1473202 | A | C | 5 | 10 | 2 | 0 | NON_SYNONYMOU | MODERAT | MISSENSE | aAc/aCc | N11T | PLES_13671 | . | . | . | . | C | C | . | . | . | . | . | . |
| NC_011770 | 1477263 | G | C | 52 | 0 | 2 | 0 | NON_SYNONYMOU | MODERAT | MISSENSE | aCc/aGc | T94S | PLES_13731 | | | | | | | C | C | | | | |
| NC_011770 | 1478205 | G | A | 51 | 0 | 2 | 0 | NON_SYNONYMOU | MODERAT | MISSENSE | aCt/aTt | T7I | PLES_13741 | | | | | | | | | | | A | A |
| NC_011770 | 1478501 | A | G | 38 | 0 | 6 | 0 | NON_SYNONYMOU | MODERAT | MISSENSE | Acg/Gcg | T14A | PLES_13751 | | | | | | | G | G | G | G | G | G |
| NC_011770 | 1478870 | G | A | 61 | 0 | 10 | 0 | NON_SYNONYMOU | MODERAT | MISSENSE | Gtt/Att | V137I | PLES_13751 | A | A | A | A | A | A | A | A | | | A | A |
| NC_011770 | 1479894 | G | A | 70 | 0 | 2 | 0 | NON_SYNONYMOU | MODERAT | MISSENSE | aGc/aAc | S478N | PLES_13751 | A | A | | | | | | | | | | |
| NC_011770 | 1480259 | C | T | 62 | 0 | 2 | 0 | SPLICE_SITE_REGIO | LOW | SILENT | tGa/tAa | *403 | PLES_13761 | | | | | | | | | | | T | T |
| NC_011770 | 1480289 | T | C | 57 | 0 | 2 | 0 | NON_SYNONYMOU | MODERAT | MISSENSE | gAg/gGg | E393G | PLES_13761 | C | C | | | | | | | | | | |
| NC_011770 | 1480319 | G | A | 63 | 0 | 2 | 0 | NON_SYNONYMOU | MODERAT | MISSENSE | gCa/gTa | A383V | PLES_13761 | | | | | | | | | A | A | | |
| NC_011770 | 1480490 | T | G | 69 | 0 | 2 | 0 | NON_SYNONYMOU | MODERAT | MISSENSE | cAg/cCg | Q326P | PLES_13761 | | | | | | | G | G | | | | |
| NC_011770 | 1481106 | C | T | 62 | 0 | 6 | 0 | NON_SYNONYMOU | MODERAT | MISSENSE | Gcg/Acg | A121T | PLES_13761 | | | T | T | T | T | | | | | T | T |

| | | | | | | | | | | | | | | | | | | | | | | | | | |
|---|---|---|---|---|---|---|---|---|---|---|---|---|---|---|---|---|---|---|---|---|---|---|---|---|---|
| NC_011770 | 1481457 | C | T | 81 | 0 | 1 | 0 | NON_SYNONYMOU | MODERAT | MISSENSE | Gca/Aca | A4T | PLES_13761 | | | | | | | | | | | | T |
| NC_011770 | 1491642 | T | A | 75 | 0 | 12 | 0 | NON_SYNONYMOU | MODERAT | MISSENSE | Ttg/Atg | L71M | PLES_13861 | A | A | A | A | A | A | A | A | A | A | A | A |
| NC_011770 | 1492278 | C | G | 65 | 0 | 2 | 0 | NON_SYNONYMOU | MODERAT | MISSENSE | Ctg/Gtg | L283V | PLES_13861 | | | | | | | G | G | | | | |
| NC_011770 | 1492360 | A | G | 60 | 0 | 2 | 0 | NON_SYNONYMOU | MODERAT | MISSENSE | gAa/gGa | E310G | PLES_13861 | | | | | | | | | | | G | G |
| NC_011770 | 1492368 | G | A | 59 | 0 | 2 | 0 | NON_SYNONYMOU | MODERAT | MISSENSE | Gcg/Acg | A313T | PLES_13861 | | | | | | | A | A | | | | |
| NC_011770 | 1492369 | C | T | 58 | 0 | 6 | 0 | NON_SYNONYMOU | MODERAT | MISSENSE | gCg/gTg | A313V | PLES_13861 | | | T | T | T | T | T | T | | | | |
| NC_011770 | 1494848 | CAG | C | 9 | 5 | 5 | 0 | FRAME_SHIFT | HIGH | | tcagtt/ | SV665 | PLES_13871 | C | . | . | . | . | . | C | C | | | C | C |
| NC_011770 | 1494890 | A | AT | 9 | 5 | 5 | 0 | FRAME_SHIFT | HIGH | | gac/gaTc | D679D? | PLES_13871 | AT | . | . | . | . | . | AT | AT | | | AT | AT |
| NC_011770 | 1494894 | CA | C | 9 | 5 | 5 | 0 | FRAME_SHIFT | HIGH | | aaa/ | K681 | PLES_13871 | C | . | . | . | . | . | C | C | | | C | C |
| NC_011770 | 1494867 | A | C | 6 | 10 | 2 | 0 | NON_SYNONYMOU | MODERAT | MISSENSE | aaA/aaC | K671N | PLES_13871 | . | . | . | . | . | . | . | . | C | C | . | . |
| NC_011770 | 1494880 | G | A | 9 | 5 | 5 | 0 | NON_SYNONYMOU | MODERAT | MISSENSE | Gtt/Att | V676I | PLES_13871 | A | . | . | . | . | . | A | A | | | A | A |
| NC_011770 | 1494881 | T | TTCC | 9 | 5 | 5 | 0 | CODON_INSERTIO | MODERATE | | ggc/CCTgg | G677PG | PLES_13871 | TTCC | . | . | . | . | . | TTCC | TTCC | | | TTCC | TTCC |
| NC_011770 | 1495075 | A | C | 74 | 0 | 4 | 0 | NON_SYNONYMOU | MODERAT | MISSENSE | Acc/Ccc | T741P | PLES_13871 | | | C | C | C | C | | | | | | |
| NC_011770 | 1495076 | C | A | 73 | 0 | 4 | 0 | NON_SYNONYMOU | MODERAT | MISSENSE | aCc/aAc | T741N | PLES_13871 | | | A | A | A | A | | | | | | |
| NC_011770 | 1506537 | T | C | 65 | 0 | 6 | 0 | NON_SYNONYMOU | MODERAT | MISSENSE | gTg/gCg | V74A | PLES_13971 | | | C | C | C | C | | | C | C | | |
| NC_011770 | 1506648 | A | G | 62 | 0 | 4 | 0 | NON_SYNONYMOU | MODERAT | MISSENSE | aAc/aGc | N111S | PLES_13971 | | | G | G | G | G | | | | | | |
| NC_011770 | 1506842 | A | G | 62 | 0 | 8 | 0 | NON_SYNONYMOU | MODERAT | MISSENSE | Acg/Gcg | T176A | PLES_13971 | | | G | G | G | G | G | G | | | G | G |
| NC_011770 | 1513620 | C | G | 61 | 0 | 12 | 0 | NON_SYNONYMOU | MODERAT | MISSENSE | caG/caC | Q194H | PLES_14041 | G | G | G | G | G | G | G | G | G | G | G | G |
| NC_011770 | 1513621 | T | C | 59 | 0 | 10 | 0 | NON_SYNONYMOU | MODERAT | MISSENSE | cAg/cGg | Q194R | PLES_14041 | | | C | C | C | C | C | C | C | C | C | C |
| NC_011770 | 1514459 | G | A | 77 | 0 | 2 | 0 | NON_SYNONYMOU | MODERAT | MISSENSE | Cat/Tat | H243Y | PLES_14051 | | | | | | | A | A | | | | |
| NC_011770 | 1516495 | C | T | 80 | 0 | 4 | 0 | NON_SYNONYMOU | MODERAT | MISSENSE | Ctc/Ttc | L5F | PLES_14071 | | | | | | | | | T | T | T | T |
| NC_011770 | 1516519 | A | T | 77 | 0 | 4 | 0 | NON_SYNONYMOU | MODERAT | MISSENSE | Acc/Tcc | T13S | PLES_14071 | | | | | | | | | T | T | T | T |
| NC_011770 | 1516601 | T | C | 72 | 0 | 12 | 0 | NON_SYNONYMOU | MODERAT | MISSENSE | cTc/cCc | L40P | PLES_14071 | C | C | C | C | C | C | C | C | C | C | C | C |
| NC_011770 | 1517029 | G | A | 82 | 0 | 1 | 0 | NON_SYNONYMOU | MODERAT | MISSENSE | Gat/Aat | D183N | PLES_14071 | | | | | | | A | | | | | |
| NC_011770 | 1520741 | A | G | 34 | 0 | 2 | 0 | NON_SYNONYMOU | MODERAT | MISSENSE | Acg/Gcg | T117A | PLES_14121 | | | | | | | G | G | | | | |
| NC_011770 | 1526243 | C | T | 48 | 0 | 2 | 0 | NON_SYNONYMOU | MODERAT | MISSENSE | Gcg/Acg | A224T | PLES_14161 | | | T | T | | | | | | | | |
| NC_011770 | 1526261 | G | C | 50 | 0 | 4 | 0 | NON_SYNONYMOU | MODERAT | MISSENSE | Ccc/Gcc | P218A | PLES_14161 | | | | | | | C | C | C | C | | |
| NC_011770 | 1526390 | C | T | 49 | 0 | 4 | 0 | NON_SYNONYMOU | MODERAT | MISSENSE | Ggc/Agc | G175S | PLES_14161 | T | T | | | | | | | | | T | T |
| NC_011770 | 1526825 | T | A | 37 | 0 | 2 | 0 | NON_SYNONYMOU | MODERAT | MISSENSE | Acc/Tcc | T30S | PLES_14161 | | | | | | | | | | | A | A |
| NC_011770 | 1526761 | T | C | 41 | 0 | 1 | 0 | NON_SYNONYMOU | MODERAT | MISSENSE | cAc/cGc | H51R | PLES_14161 | C | | | | | | | | | | | |
| NC_011770 | 1527283 | C | T | 70 | 0 | 2 | 0 | NON_SYNONYMOU | MODERAT | MISSENSE | cCg/cTg | P75L | PLES_14171 | | | | | | | | | | | T | T |
| NC_011770 | 1529278 | C | A | 79 | 0 | 2 | 0 | NON_SYNONYMOU | MODERAT | MISSENSE | cGg/cTg | R341L | PLES_14201 | | | | | | | | | | | A | A |
| NC_011770 | 1529411 | G | C | 75 | 0 | 2 | 0 | NON_SYNONYMOU | MODERAT | MISSENSE | Cac/Gac | H297D | PLES_14201 | | | | | | | C | C | | | | |
| NC_011770 | 1529752 | G | C | 77 | 0 | 6 | 0 | NON_SYNONYMOU | MODERAT | MISSENSE | gCc/gGc | A183G | PLES_14201 | | | C | C | C | C | | | C | C | | |
| NC_011770 | 1529753 | C | T | 75 | 0 | 1 | 0 | NON_SYNONYMOU | MODERAT | MISSENSE | Gcc/Acc | A183T | PLES_14201 | | | | T | | | | | | | | |
| NC_011770 | 1530486 | T | C | 88 | 0 | 4 | 0 | NON_SYNONYMOU | MODERAT | MISSENSE | Ata/Gta | I434V | PLES_14211 | | | C | C | C | C | | | | | | |
| NC_011770 | 1530518 | C | G | 81 | 0 | 2 | 0 | NON_SYNONYMOU | MODERAT | MISSENSE | gGt/gCt | G423A | PLES_14211 | | | | | | | G | G | | | | |
| NC_011770 | 1531539 | G | C | 68 | 0 | 2 | 0 | NON_SYNONYMOU | MODERAT | MISSENSE | Ccg/Gcg | P83A | PLES_14211 | | | | | | | | | | | C | C |
| NC_011770 | 1531122 | G | A | 75 | 0 | 1 | 0 | NON_SYNONYMOU | MODERAT | MISSENSE | Cgc/Tgc | R222C | PLES_14211 | A | | | | | | | | | | | |
| NC_011770 | 1532370 | G | C | 95 | 0 | 2 | 0 | NON_SYNONYMOU | MODERAT | MISSENSE | Ggc/Cgc | G136R | PLES_14221 | | | | | | | | | C | C | | |
| NC_011770 | 1533833 | C | G | 68 | 0 | 2 | 0 | NON_SYNONYMOU | MODERAT | MISSENSE | gaC/gaG | D623E | PLES_14221 | | | | | | | G | G | | | | |
| NC_011770 | 1533837 | A | G | 67 | 0 | 12 | 0 | NON_SYNONYMOU | MODERAT | MISSENSE | Atc/Gtc | I625V | PLES_14221 | G | G | G | G | G | G | G | G | G | G | G | G |
| NC_011770 | 1532286 | G | A | 82 | 0 | 1 | 0 | NON_SYNONYMOU | MODERAT | MISSENSE | Ggc/Agc | G108S | PLES_14221 | | | | | | | | A | | | | |
| NC_011770 | 1540722 | T | C | 77 | 0 | 1 | 0 | NON_SYNONYMOU | MODERAT | MISSENSE | Acc/Gcc | T137A | PLES_14301 | | | | | | | C | | | | | |
| NC_011770 | 1540899 | T | G | 84 | 0 | 2 | 0 | NON_SYNONYMOU | MODERAT | MISSENSE | Aat/Cat | N78H | PLES_14301 | G | G | | | | | | | | | | |
| NC_011770 | 1545046 | C | T | 79 | 0 | 1 | 0 | NON_SYNONYMOU | MODERAT | MISSENSE | Ggc/Agc | G156S | PLES_14361 | | | | | | | T | | | | | |
| NC_011770 | 1545036 | C | T | 82 | 0 | 1 | 0 | NON_SYNONYMOU | MODERAT | MISSENSE | cGg/cAg | R159Q | PLES_14361 | | | | | | | | T | | | | |
| NC_011770 | 1545819 | G | A | 81 | 0 | 1 | 0 | NON_SYNONYMOU | MODERAT | MISSENSE | Ggc/Agc | G61S | PLES_14371 | | | | | | | A | | | | | |
| NC_011770 | 1546005 | A | G | 81 | 0 | 12 | 0 | NON_SYNONYMOU | MODERAT | MISSENSE | Aac/Gac | N123D | PLES_14371 | G | G | G | G | G | G | G | G | G | G | G | G |
| NC_011770 | 1546294 | G | A | 81 | 0 | 2 | 0 | NON_SYNONYMOU | MODERAT | MISSENSE | tGc/tAc | C219Y | PLES_14371 | A | A | | | | | | | | | | |
| NC_011770 | 1547945 | C | A | 76 | 0 | 8 | 0 | NON_SYNONYMOU | MODERAT | MISSENSE | cCg/cAg | P431Q | PLES_14381 | A | A | | | | | A | A | A | A | A | A |
| NC_011770 | 1549345 | G | A | 76 | 0 | 4 | 0 | STOP_GAINED | HIGH | NONSENSE | Cag/Tag | Q451* | PLES_14391 | A | A | | | | | | | A | A | | |
| NC_011770 | 1548073 | T | C | 78 | 0 | 8 | 0 | NON_SYNONYMOU | MODERAT | MISSENSE | Atc/Gtc | I875V | PLES_14391 | C | C | | | | | C | C | C | C | C | C |
| NC_011770 | 1548154 | C | T | 80 | 0 | 4 | 0 | NON_SYNONYMOU | MODERAT | MISSENSE | Gaa/Aaa | E848K | PLES_14391 | | | T | T | T | T | | | | | | |
| NC_011770 | 1548277 | G | A | 84 | 0 | 6 | 0 | NON_SYNONYMOU | MODERAT | MISSENSE | Ccg/Tcg | P807S | PLES_14391 | A | A | | | | | A | A | A | A | | |
| NC_011770 | 1548304 | G | A | 85 | 0 | 6 | 0 | NON_SYNONYMOU | MODERAT | MISSENSE | Ctc/Ttc | L798F | PLES_14391 | | | A | A | A | A | | | | | A | A |
| NC_011770 | 1548418 | T | C | 76 | 0 | 12 | 0 | NON_SYNONYMOU | MODERAT | MISSENSE | Agt/Ggt | S760G | PLES_14391 | C | C | C | C | C | C | C | C | C | C | C | C |
| NC_011770 | 1548523 | G | A | 80 | 0 | 4 | 0 | NON_SYNONYMOU | MODERAT | MISSENSE | Cac/Tac | H725Y | PLES_14391 | | | A | A | A | A | | | | | | |
| NC_011770 | 1548556 | C | T | 80 | 0 | 6 | 0 | NON_SYNONYMOU | MODERAT | MISSENSE | Gga/Aga | G714R | PLES_14391 | T | T | | | | | T | T | T | T | | |
| NC_011770 | 1548691 | C | T | 86 | 0 | 6 | 0 | NON_SYNONYMOU | MODERAT | MISSENSE | Ggc/Agc | G669S | PLES_14391 | T | T | | | | | T | T | T | T | | |
| NC_011770 | 1548706 | A | G | 86 | 0 | 10 | 0 | NON_SYNONYMOU | MODERAT | MISSENSE | Ttc/Ctc | F664L | PLES_14391 | G | G | G | G | G | G | G | G | G | G | | |
| NC_011770 | 1548712 | C | T | 87 | 0 | 4 | 0 | NON_SYNONYMOU | MODERAT | MISSENSE | Gaa/Aaa | E662K | PLES_14391 | | | T | T | T | T | | | | | | |

| | | | | | | | | | | | | | | | | | | | | | | | | | |
|---|---|---|---|---|---|---|---|---|---|---|---|---|---|---|---|---|---|---|---|---|---|---|---|---|---|
| NC_011770 | 1548756 | G | A | 86 | 0 | 4 | 0 | NON_SYNONYMOU | MODERAT | MISSENSE | gCt/gTt | A647V | PLES_14391 | | | A | A | A | A | | | | | | |
| NC_011770 | 1548822 | A | G | 83 | 0 | 4 | 0 | NON_SYNONYMOU | MODERAT | MISSENSE | gTc/gCc | V625A | PLES_14391 | | | G | G | G | G | | | | | | |
| NC_011770 | 1548831 | G | A | 86 | 0 | 4 | 0 | NON_SYNONYMOU | MODERAT | MISSENSE | gCc/gTc | A622V | PLES_14391 | | | A | A | A | A | | | | | | |
| NC_011770 | 1548859 | C | G | 87 | 0 | 10 | 0 | NON_SYNONYMOU | MODERAT | MISSENSE | Ggg/Cgg | G613R | PLES_14391 | G | G | G | G | G | G | G | G | G | G | | |
| NC_011770 | 1548918 | T | C | 84 | 0 | 12 | 0 | NON_SYNONYMOU | MODERAT | MISSENSE | cAg/cGg | Q593R | PLES_14391 | C | C | C | C | C | C | C | C | C | C | C | C |
| NC_011770 | 1548985 | T | C | 76 | 0 | 10 | 0 | NON_SYNONYMOU | MODERAT | MISSENSE | Agc/Ggc | S571G | PLES_14391 | C | C | C | C | C | C | C | C | C | C | | |
| NC_011770 | 1549021 | T | C | 76 | 0 | 12 | 0 | NON_SYNONYMOU | MODERAT | MISSENSE | Aat/Gat | N559D | PLES_14391 | C | C | C | C | C | C | C | C | C | C | C | C |
| NC_011770 | 1549069 | C | T | 76 | 0 | 4 | 0 | NON_SYNONYMOU | MODERAT | MISSENSE | Gcc/Acc | A543T | PLES_14391 | | | T | T | T | T | | | | | | |
| NC_011770 | 1549125 | G | A | 67 | 0 | 4 | 0 | NON_SYNONYMOU | MODERAT | MISSENSE | gCc/gTc | A524V | PLES_14391 | | | A | A | A | A | | | | | | |
| NC_011770 | 1549261 | T | C | 72 | 0 | 8 | 0 | NON_SYNONYMOU | MODERAT | MISSENSE | Att/Gtt | I479V | PLES_14391 | C | C | | | | | C | C | C | C | C | C |
| NC_011770 | 1549348 | G | A | 78 | 0 | 2 | 0 | NON_SYNONYMOU | MODERAT | MISSENSE | Cgg/Tgg | R450W | PLES_14391 | | | | | | | A | A | | | | |
| NC_011770 | 1549374 | G | T | 84 | 0 | 2 | 0 | NON_SYNONYMOU | MODERAT | MISSENSE | aCt/aAt | T441N | PLES_14391 | | | | | | | T | T | | | | |
| NC_011770 | 1549381 | T | C | 81 | 0 | 2 | 0 | NON_SYNONYMOU | MODERAT | MISSENSE | Acc/Gcc | T439A | PLES_14391 | | | | | | | C | C | | | | |
| NC_011770 | 1549585 | A | G | 84 | 0 | 6 | 0 | NON_SYNONYMOU | MODERAT | MISSENSE | Tat/Cat | Y371H | PLES_14391 | | | G | G | G | G | G | G | | | | |
| NC_011770 | 1549654 | T | C | 84 | 0 | 6 | 0 | NON_SYNONYMOU | MODERAT | MISSENSE | Act/Gct | T348A | PLES_14391 | | | C | C | C | C | C | C | | | | |
| NC_011770 | 1549923 | C | T | 87 | 0 | 1 | 0 | NON_SYNONYMOU | MODERAT | MISSENSE | cGg/cAg | R258Q | PLES_14391 | | | | | | | T | | | | | |
| NC_011770 | 1549939 | T | C | 87 | 0 | 4 | 0 | NON_SYNONYMOU | MODERAT | MISSENSE | Agt/Ggt | S253G | PLES_14391 | | | C | C | C | C | | | | | | |
| NC_011770 | 1549942 | G | A | 89 | 0 | 4 | 0 | NON_SYNONYMOU | MODERAT | MISSENSE | Cgc/Tgc | R252C | PLES_14391 | | | A | A | A | A | | | | | | |
| NC_011770 | 1550080 | T | C | 84 | 0 | 4 | 0 | NON_SYNONYMOU | MODERAT | MISSENSE | Agg/Ggg | R206G | PLES_14391 | | | | | | | C | C | | | C | C |
| NC_011770 | 1550107 | G | A | 83 | 0 | 4 | 0 | NON_SYNONYMOU | MODERAT | MISSENSE | Cgc/Tgc | R197C | PLES_14391 | | | A | A | A | A | | | | | | |
| NC_011770 | 1550112 | G | A | 83 | 0 | 4 | 0 | NON_SYNONYMOU | MODERAT | MISSENSE | cCt/cTt | P195L | PLES_14391 | | | A | A | A | A | | | | | | |
| NC_011770 | 1550376 | C | T | 79 | 0 | 1 | 0 | NON_SYNONYMOU | MODERAT | MISSENSE | gGg/gAg | G107E | PLES_14391 | | | | | | | T | | | | | |
| NC_011770 | 1550403 | C | G | 81 | 0 | 2 | 0 | NON_SYNONYMOU | MODERAT | MISSENSE | gGt/gCt | G98A | PLES_14391 | | | | | | | G | G | | | | |
| NC_011770 | 1550586 | G | T | 77 | 0 | 6 | 0 | NON_SYNONYMOU | MODERAT | MISSENSE | gCg/gAg | A37E | PLES_14391 | | | T | T | T | T | T | T | | | | |
| NC_011770 | 1550641 | A | G | 78 | 0 | 8 | 0 | NON_SYNONYMOU | MODERAT | MISSENSE | Ttc/Ctc | F19L | PLES_14391 | G | G | G | G | G | G | G | G | | | | |
| NC_011770 | 1551015 | G | C | 71 | 0 | 4 | 0 | NON_SYNONYMOU | MODERAT | MISSENSE | gaC/gaG | D212E | PLES_14401 | | | C | C | C | C | | | | | | |
| NC_011770 | 1551882 | C | T | 70 | 0 | 2 | 0 | NON_SYNONYMOU | MODERAT | MISSENSE | cCg/cTg | P40L | PLES_14411 | | | | | | | | | T | T | | |
| NC_011770 | 1552059 | A | G | 76 | 0 | 8 | 0 | NON_SYNONYMOU | MODERAT | MISSENSE | cAg/cGg | Q99R | PLES_14411 | G | G | | | | | G | G | G | G | G | G |
| NC_011770 | 1552590 | A | G | 83 | 0 | 2 | 0 | NON_SYNONYMOU | MODERAT | MISSENSE | gAa/gGa | E276G | PLES_14411 | | | | | | | G | G | | | | |
| NC_011770 | 1552743 | G | A | 77 | 0 | 2 | 0 | NON_SYNONYMOU | MODERAT | MISSENSE | cGc/cAc | R327H | PLES_14411 | | | | | | | | | A | A | | |
| NC_011770 | 1553435 | C | G | 80 | 0 | 12 | 0 | NON_SYNONYMOU | MODERAT | MISSENSE | Ctg/Gtg | L558V | PLES_14411 | G | G | G | G | G | G | G | G | G | G | G | G |
| NC_011770 | 1553569 | A | C | 74 | 0 | 2 | 0 | NON_SYNONYMOU | MODERAT | MISSENSE | Atc/Ctc | I14L | PLES_14421 | | | | | | | | | C | C | | |
| NC_011770 | 1553926 | G | C | 74 | 0 | 2 | 0 | NON_SYNONYMOU | MODERAT | MISSENSE | Gac/Cac | D133H | PLES_14421 | | | | | | | | | C | C | | |
| NC_011770 | 1554015 | G | C | 75 | 0 | 4 | 0 | NON_SYNONYMOU | MODERAT | MISSENSE | gaG/gaC | E162D | PLES_14421 | | | C | C | C | C | | | | | | |
| NC_011770 | 1554133 | G | A | 75 | 0 | 6 | 0 | NON_SYNONYMOU | MODERAT | MISSENSE | Gcc/Acc | A202T | PLES_14421 | | | A | A | A | A | A | A | | | | |
| NC_011770 | 1554498 | T | G | 70 | 0 | 10 | 0 | NON_SYNONYMOU | MODERAT | MISSENSE | gaT/gaG | D323E | PLES_14421 | G | G | G | G | G | G | G | G | | | G | G |
| NC_011770 | 1554801 | C | A | 81 | 0 | 2 | 0 | NON_SYNONYMOU | MODERAT | MISSENSE | Cgc/Agc | R9S | PLES_14431 | | | | | | | | | A | A | | |
| NC_011770 | 1554975 | A | G | 84 | 0 | 6 | 0 | NON_SYNONYMOU | MODERAT | MISSENSE | Acg/Gcg | T67A | PLES_14431 | G | G | G | G | G | G | | | | | | |
| NC_011770 | 1555024 | C | T | 82 | 0 | 8 | 0 | NON_SYNONYMOU | MODERAT | MISSENSE | gCg/gTg | A83V | PLES_14431 | T | T | T | T | T | T | | | | | T | T |
| NC_011770 | 1555110 | T | C | 85 | 0 | 10 | 0 | NON_SYNONYMOU | MODERAT | MISSENSE | Tat/Cat | Y112H | PLES_14431 | C | C | C | C | C | C | | | C | C | C | C |
| NC_011770 | 1555194 | A | G | 86 | 0 | 6 | 0 | NON_SYNONYMOU | MODERAT | MISSENSE | Atc/Gtc | I140V | PLES_14431 | G | G | G | G | G | G | | | | | | |
| NC_011770 | 1555297 | G | T | 96 | 0 | 6 | 0 | NON_SYNONYMOU | MODERAT | MISSENSE | cGg/cTg | R174L | PLES_14431 | | | T | T | T | T | A | A | | | | |
| NC_011770 | 1555361 | A | T | 82 | 0 | 12 | 0 | NON_SYNONYMOU | MODERAT | MISSENSE | gaA/gaT | E195D | PLES_14431 | T | T | T | T | T | T | T | T | T | T | T | T |
| NC_011770 | 1555583 | C | T | 84 | 0 | 2 | 0 | NON_SYNONYMOU | MODERAT | MISSENSE | Ccg/Tcg | P5S | PLES_14441 | T | T | | | | | | | | | | |
| NC_011770 | 1555638 | A | T | 81 | 0 | 6 | 0 | NON_SYNONYMOU | MODERAT | MISSENSE | cAg/cTg | Q23L | PLES_14441 | T | T | | | | | G | G | G | G | | |
| NC_011770 | 1555662 | C | T | 80 | 0 | 4 | 0 | NON_SYNONYMOU | MODERAT | MISSENSE | tCg/tTg | S31L | PLES_14441 | | | T | T | T | T | | | | | | |
| NC_011770 | 1555692 | C | T | 76 | 0 | 2 | 0 | NON_SYNONYMOU | MODERAT | MISSENSE | gCg/gTg | A41V | PLES_14441 | T | T | | | | | | | | | | |
| NC_011770 | 1555766 | G | T | 75 | 0 | 2 | 0 | NON_SYNONYMOU | MODERAT | MISSENSE | Gct/Tct | A66S | PLES_14441 | | | | | | | T | T | | | | |
| NC_011770 | 1555770 | T | C | 72 | 0 | 4 | 0 | NON_SYNONYMOU | MODERAT | MISSENSE | gTc/gCc | V67A | PLES_14441 | | | | | | | C | C | C | C | | |
| NC_011770 | 1555773 | A | C | 71 | 0 | 4 | 0 | NON_SYNONYMOU | MODERAT | MISSENSE | gAg/gCg | E68A | PLES_14441 | | | | | | | C | C | C | C | | |
| NC_011770 | 1555808 | T | A | 74 | 0 | 4 | 0 | NON_SYNONYMOU | MODERAT | MISSENSE | Ttg/Atg | L80M | PLES_14441 | | | A | A | A | A | | | | | | |
| NC_011770 | 1555811 | C | T | 75 | 0 | 2 | 0 | NON_SYNONYMOU | MODERAT | MISSENSE | Cgg/Tgg | R81W | PLES_14441 | | | | | | | | | | | T | T |
| NC_011770 | 1555820 | C | G | 75 | 0 | 4 | 0 | NON_SYNONYMOU | MODERAT | MISSENSE | Cgc/Ggc | R84G | PLES_14441 | G | G | | | | | | | G | G | | |
| NC_011770 | 1555884 | G | A | 69 | 0 | 2 | 0 | NON_SYNONYMOU | MODERAT | MISSENSE | cGc/cAc | R105H | PLES_14441 | | | | | | | | | | | A | A |
| NC_011770 | 1555937 | T | C | 66 | 0 | 2 | 0 | NON_SYNONYMOU | MODERAT | MISSENSE | Tcg/Ccg | S123P | PLES_14441 | | | | | | | | | C | C | | |
| NC_011770 | 1556044 | C | G | 73 | 0 | 6 | 0 | NON_SYNONYMOU | MODERAT | MISSENSE | gaC/gaG | D158E | PLES_14441 | G | G | | | | | G | G | G | G | | |
| NC_011770 | 1556111 | T | G | 73 | 0 | 4 | 0 | NON_SYNONYMOU | MODERAT | MISSENSE | Tcg/Gcg | S181A | PLES_14441 | G | G | | | | | G | G | | | | |
| NC_011770 | 1556451 | T | C | 74 | 0 | 8 | 0 | NON_SYNONYMOU | MODERAT | MISSENSE | gTc/gCc | V294A | PLES_14441 | | | C | C | C | C | | | C | C | C | C |
| NC_011770 | 1556705 | C | A | 73 | 0 | 4 | 0 | NON_SYNONYMOU | MODERAT | MISSENSE | Cac/Aac | H379N | PLES_14441 | | | A | A | A | A | | | | | | |
| NC_011770 | 1556960 | A | C | 82 | 0 | 6 | 0 | NON_SYNONYMOU | MODERAT | MISSENSE | Atg/Ctg | M464L | PLES_14441 | C | C | | | | | C | C | | | C | C |
| NC_011770 | 1556438 | G | A | 76 | 0 | 1 | 0 | NON_SYNONYMOU | MODERAT | MISSENSE | Ggg/Agg | G290R | PLES_14441 | | | | | | | | | | | | A |
| NC_011770 | 1557240 | A | G | 72 | 0 | 12 | 0 | NON_SYNONYMOU | MODERAT | MISSENSE | Aaa/Gaa | K39E | PLES_14451 | G | G | G | G | G | G | G | G | G | G | G | G |

| | | | | | | | | | | | | | | | | | | | | | | | | | |
|---|---|---|---|---|---|---|---|---|---|---|---|---|---|---|---|---|---|---|---|---|---|---|---|---|---|
| NC_011770 | 1557414 | A | G | 72 | 0 | 2 | 0 | NON_SYNONYMOU | MODERAT | MISSENSE | Atc/Gtc | I97V | PLES_14451 | | | | | | | | | | | G | G |
| NC_011770 | 1557415 | T | C | 71 | 0 | 2 | 0 | NON_SYNONYMOU | MODERAT | MISSENSE | aTc/aCc | I97T | PLES_14451 | | | | | | | | | | | C | C |
| NC_011770 | 1557433 | T | C | 68 | 0 | 2 | 0 | NON_SYNONYMOU | MODERAT | MISSENSE | aTc/aCc | I103T | PLES_14451 | C | C | | | | | | | | | | |
| NC_011770 | 1557501 | A | G | 77 | 0 | 2 | 0 | NON_SYNONYMOU | MODERAT | MISSENSE | Atg/Gtg | M126V | PLES_14451 | G | G | | | | | | | | | | |
| NC_011770 | 1557505 | G | C | 80 | 0 | 4 | 0 | NON_SYNONYMOU | MODERAT | MISSENSE | gGc/gCc | G127A | PLES_14451 | C | C | | | | | | | C | C | | |
| NC_011770 | 1557652 | C | A | 79 | 0 | 2 | 0 | NON_SYNONYMOU | MODERAT | MISSENSE | gCc/gAc | A176D | PLES_14451 | | | | | | | | | A | A | | |
| NC_011770 | 1558060 | G | A | 75 | 0 | 6 | 0 | NON_SYNONYMOU | MODERAT | MISSENSE | gGc/gAc | G312D | PLES_14451 | | | A | A | A | A | | | A | A | | |
| NC_011770 | 1558774 | G | A | 69 | 0 | 4 | 0 | NON_SYNONYMOU | MODERAT | MISSENSE | Gtc/Atc | V131I | PLES_14461 | | | A | A | A | A | | | | | | |
| NC_011770 | 1559113 | A | G | 77 | 0 | 10 | 0 | NON_SYNONYMOU | MODERAT | MISSENSE | Atc/Gtc | I244V | PLES_14461 | | | G | G | G | G | G | G | G | G | G | G |
| NC_011770 | 1559213 | C | T | 76 | 0 | 2 | 0 | NON_SYNONYMOU | MODERAT | MISSENSE | tCg/tTg | S277L | PLES_14461 | | | | | | | | | | | T | T |
| NC_011770 | 1559405 | T | G | 88 | 0 | 2 | 0 | NON_SYNONYMOU | MODERAT | MISSENSE | gTg/gGg | V341G | PLES_14461 | | | | | | | | | | | G | G |
| NC_011770 | 1559524 | G | C | 89 | 0 | 2 | 0 | NON_SYNONYMOU | MODERAT | MISSENSE | Gcg/Ccg | A381P | PLES_14461 | | | | | | | | | | | C | C |
| NC_011770 | 1561162 | A | G | 96 | 0 | 2 | 0 | NON_SYNONYMOU | MODERAT | MISSENSE | Acc/Gcc | T102A | PLES_14481 | | | | | | | G | G | | | | |
| NC_011770 | 1567793 | A | G | 46 | 0 | 12 | 0 | NON_SYNONYMOU | MODERAT | MISSENSE | Acc/Gcc | T11A | PLES_14541 | G | G | G | G | G | G | G | G | G | G | G | G |
| NC_011770 | 1567803 | C | T | 46 | 0 | 6 | 0 | NON_SYNONYMOU | MODERAT | MISSENSE | gCg/gTg | A14V | PLES_14541 | T | T | | | | | | | A | A | A | A |
| NC_011770 | 1567882 | C | G | 55 | 0 | 2 | 0 | NON_SYNONYMOU | MODERAT | MISSENSE | gaC/gaG | D40E | PLES_14541 | | | | | | | | | G | G | | |
| NC_011770 | 1569282 | C | T | 65 | 0 | 1 | 0 | NON_SYNONYMOU | MODERAT | MISSENSE | gCc/gTc | A329V | PLES_14551 | | | | | | | | T | | | | |
| NC_011770 | 1570757 | G | A | 49 | 0 | 6 | 0 | NON_SYNONYMOU | MODERAT | MISSENSE | Ggc/Agc | G261S | PLES_14561 | | | A | A | A | A | A | A | | | | |
| NC_011770 | 1573441 | T | C | 67 | 0 | 4 | 0 | NON_SYNONYMOU | MODERAT | MISSENSE | Acc/Gcc | T125A | PLES_14571 | | | | | | | | | C | C | C | C |
| NC_011770 | 1573460 | C | A | 66 | 0 | 2 | 0 | NON_SYNONYMOU | MODERAT | MISSENSE | gaG/gaT | E118D | PLES_14571 | A | A | | | | | | | | | | |
| NC_011770 | 1573520 | G | C | 64 | 0 | 2 | 0 | NON_SYNONYMOU | MODERAT | MISSENSE | gaC/gaG | D98E | PLES_14571 | C | C | | | | | | | | | | |
| NC_011770 | 1573636 | C | T | 69 | 0 | 6 | 0 | NON_SYNONYMOU | MODERAT | MISSENSE | Gtc/Atc | V60I | PLES_14571 | | | T | T | T | T | | | | | T | T |
| NC_011770 | 1573989 | C | T | 70 | 0 | 2 | 0 | NON_SYNONYMOU | MODERAT | MISSENSE | Gcc/Acc | A152T | PLES_14581 | T | T | | | | | | | | | | |
| NC_011770 | 1574881 | G | C | 100 | 0 | 12 | 0 | NON_SYNONYMOU | MODERAT | MISSENSE | Cat/Gat | H187D | PLES_14591 | C | C | C | C | C | C | C | C | C | C | C | C |
| NC_011770 | 1576076 | G | C | 72 | 0 | 2 | 0 | NON_SYNONYMOU | MODERAT | MISSENSE | aGc/aCc | S91T | PLES_14601 | | | | | | | | | | | C | C |
| NC_011770 | 1576381 | T | G | 69 | 0 | 4 | 0 | NON_SYNONYMOU | MODERAT | MISSENSE | Tcc/Gcc | S193A | PLES_14601 | | | G | G | G | G | | | | | | |
| NC_011770 | 1576655 | C | T | 77 | 0 | 6 | 0 | NON_SYNONYMOU | MODERAT | MISSENSE | gCg/gTg | A284V | PLES_14601 | T | T | T | T | T | T | | | | | | |
| NC_011770 | 1576973 | A | G | 64 | 0 | 2 | 0 | NON_SYNONYMOU | MODERAT | MISSENSE | cAg/cGg | Q390R | PLES_14601 | | | | | | | | | | | G | G |
| NC_011770 | 1577023 | C | T | 69 | 0 | 2 | 0 | NON_SYNONYMOU | MODERAT | MISSENSE | Gtg/Atg | V49M | PLES_14611 | | | | | | | | | | | T | T |
| NC_011770 | 1581460 | C | T | 89 | 0 | 2 | 0 | NON_SYNONYMOU | MODERAT | MISSENSE | cCc/cTc | P202L | PLES_14651 | | | | | | | | | | | T | T |
| NC_011770 | 1581636 | C | T | 93 | 0 | 4 | 0 | NON_SYNONYMOU | MODERAT | MISSENSE | Cgc/Tgc | R261C | PLES_14651 | | | T | T | T | T | | | | | | |
| NC_011770 | 1581678 | G | A | 97 | 0 | 4 | 0 | NON_SYNONYMOU | MODERAT | MISSENSE | Gtc/Atc | V275I | PLES_14651 | | | A | A | A | A | | | | | | |
| NC_011770 | 1581889 | C | T | 88 | 0 | 1 | 0 | NON_SYNONYMOU | MODERAT | MISSENSE | cCg/cTg | P345L | PLES_14651 | | | | | | | | T | | | | |
| NC_011770 | 1583272 | G | C | 63 | 0 | 2 | 0 | NON_SYNONYMOU | MODERAT | MISSENSE | aCc/aGc | T181S | PLES_14661 | | | | | | | | | | | C | C |
| NC_011770 | 1583522 | T | A | 55 | 0 | 2 | 0 | NON_SYNONYMOU | MODERAT | MISSENSE | Agc/Tgc | S98C | PLES_14661 | A | A | | | | | | | | | | |
| NC_011770 | 1583941 | G | T | 45 | 0 | 2 | 0 | NON_SYNONYMOU | MODERAT | MISSENSE | gCc/gAc | A63D | PLES_14671 | | | | | | | | | T | T | | |
| NC_011770 | 1585006 | G | C | 67 | 0 | 4 | 0 | NON_SYNONYMOU | MODERAT | MISSENSE | gaG/gaC | E261D | PLES_14681 | | | C | C | C | C | | | | | | |
| NC_011770 | 1587766 | C | T | 84 | 0 | 4 | 0 | NON_SYNONYMOU | MODERAT | MISSENSE | Cgc/Tgc | R77C | PLES_14711 | | | T | T | T | T | | | | | | |
| NC_011770 | 1588252 | G | A | 85 | 0 | 1 | 0 | NON_SYNONYMOU | MODERAT | MISSENSE | Ggc/Agc | G239S | PLES_14711 | | | | | | | A | | | | | |
| NC_011770 | 1588883 | A | G | 76 | 0 | 2 | 0 | NON_SYNONYMOU | MODERAT | MISSENSE | gAc/gGc | D449G | PLES_14711 | G | G | | | | | | | | | | |
| NC_011770 | 1589467 | C | A | 62 | 0 | 2 | 0 | NON_SYNONYMOU | MODERAT | MISSENSE | Cgc/Agc | R644S | PLES_14711 | | | | | | | A | A | | | | |
| NC_011770 | 1590383 | C | T | 65 | 0 | 1 | 0 | NON_SYNONYMOU | MODERAT | MISSENSE | gCc/gTc | A949V | PLES_14711 | | | | | | | T | | | | | |
| NC_011770 | 1589081 | C | T | 74 | 0 | 1 | 0 | NON_SYNONYMOU | MODERAT | MISSENSE | gCg/gTg | A515V | PLES_14711 | | | | | | | | | | | T | |
| NC_011770 | 1593761 | T | A | 75 | 0 | 6 | 0 | NON_SYNONYMOU | MODERAT | MISSENSE | cAc/cTc | H300L | PLES_14741 | | | A | A | A | A | | | A | A | | |
| NC_011770 | 1593782 | G | T | 74 | 0 | 4 | 0 | NON_SYNONYMOU | MODERAT | MISSENSE | gCg/gAg | A293E | PLES_14741 | | | T | T | T | T | | | | | | |
| NC_011770 | 1594106 | C | T | 82 | 0 | 6 | 0 | NON_SYNONYMOU | MODERAT | MISSENSE | cGc/cAc | R185H | PLES_14741 | | | T | T | T | T | | | | | T | T |
| NC_011770 | 1594358 | G | A | 83 | 0 | 2 | 0 | NON_SYNONYMOU | MODERAT | MISSENSE | gCg/gTg | A101V | PLES_14741 | A | A | | | | | | | | | | |
| NC_011770 | 1594753 | A | G | 60 | 0 | 12 | 0 | NON_SYNONYMOU | MODERAT | MISSENSE | aTc/aCc | I106T | PLES_14751 | G | G | G | G | G | G | G | G | G | G | G | G |
| NC_011770 | 1595030 | T | C | 35 | 0 | 10 | 0 | NON_SYNONYMOU | MODERAT | MISSENSE | Atg/Gtg | M14V | PLES_14751 | | | C | C | C | C | C | C | C | C | C | C |
| NC_011770 | 1597129 | C | T | 71 | 0 | 2 | 0 | NON_SYNONYMOU | MODERAT | MISSENSE | aGc/aAc | S272N | PLES_14781 | | | | | | | | | | | T | T |
| NC_011770 | 1597947 | T | A | 57 | 0 | 2 | 0 | NON_SYNONYMOU | MODERAT | MISSENSE | cAg/cTg | Q661L | PLES_14791 | | | | | | | | | | | A | A |
| NC_011770 | 1598158 | G | T | 36 | 0 | 2 | 0 | NON_SYNONYMOU | MODERAT | MISSENSE | Ctg/Atg | L591M | PLES_14791 | | | | | | | | | T | T | | |
| NC_011770 | 1598278 | T | C | 46 | 0 | 4 | 0 | NON_SYNONYMOU | MODERAT | MISSENSE | Atc/Gtc | I551V | PLES_14791 | | | C | C | C | C | | | | | | |
| NC_011770 | 1598767 | T | C | 82 | 0 | 4 | 0 | NON_SYNONYMOU | MODERAT | MISSENSE | Atc/Gtc | I388V | PLES_14791 | | | | | | | C | C | C | C | | |
| NC_011770 | 1599690 | A | T | 88 | 0 | 2 | 0 | NON_SYNONYMOU | MODERAT | MISSENSE | tTc/tAc | F80Y | PLES_14791 | | | | | | | T | T | | | | |
| NC_011770 | 1600962 | C | G | 72 | 0 | 2 | 0 | NON_SYNONYMOU | MODERAT | MISSENSE | gaG/gaC | E376D | PLES_14811 | | | | | | | | | | | G | G |
| NC_011770 | 1601143 | G | A | 69 | 0 | 4 | 0 | NON_SYNONYMOU | MODERAT | MISSENSE | gCg/gTg | A316V | PLES_14811 | | | A | A | A | A | | | | | | |
| NC_011770 | 1601234 | T | C | 72 | 0 | 2 | 0 | NON_SYNONYMOU | MODERAT | MISSENSE | Aag/Gag | K286E | PLES_14811 | | | | | | | C | C | | | | |
| NC_011770 | 1601617 | T | C | 62 | 0 | 1 | 0 | NON_SYNONYMOU | MODERAT | MISSENSE | cAc/cGc | H158R | PLES_14811 | | | | | | | C | | | | | |
| NC_011770 | 1620386 | C | T | 82 | 0 | 4 | 0 | NON_SYNONYMOU | MODERAT | MISSENSE | Gcc/Acc | A138T | PLES_14941 | | | T | T | T | T | | | | | | |
| NC_011770 | 1620509 | C | T | 83 | 0 | 2 | 0 | NON_SYNONYMOU | MODERAT | MISSENSE | Ggc/Agc | G97S | PLES_14941 | | | | | | | | | | | T | T |
| NC_011770 | 1620340 | T | C | 84 | 0 | 1 | 0 | NON_SYNONYMOU | MODERAT | MISSENSE | cAg/cGg | Q153R | PLES_14941 | | C | | | | | | | | | | |

| | | | | | | | | | | | | | | | | | | | | | | | | | |
|---|---|---|---|---|---|---|---|---|---|---|---|---|---|---|---|---|---|---|---|---|---|---|---|---|---|
| NC_011770 | 1623314 | C | T | 53 | 0 | 2 | 0 | NON_SYNONYMOU | MODERAT | MISSENSE | gCc/gTc | A43V | PLES_14971 | T | T | | | | | | | | | | |
| NC_011770 | 1623387 | T | A | 54 | 0 | 2 | 0 | NON_SYNONYMOU | MODERAT | MISSENSE | caT/caA | H67Q | PLES_14971 | | | | | | | | | A | A | | |
| NC_011770 | 1623409 | G | T | 52 | 0 | 6 | 0 | NON_SYNONYMOU | MODERAT | MISSENSE | Gtg/Ttg | V75L | PLES_14971 | T | T | | | | | T | T | | | T | T |
| NC_011770 | 1623436 | T | G | 52 | 0 | 4 | 0 | NON_SYNONYMOU | MODERAT | MISSENSE | Tgc/Ggc | C84G | PLES_14971 | | | G | G | G | G | | | | | | |
| NC_011770 | 1623466 | C | T | 50 | 0 | 6 | 0 | NON_SYNONYMOU | MODERAT | MISSENSE | Ctc/Ttc | L94F | PLES_14971 | | | T | T | T | T | | | T | T | | |
| NC_011770 | 1624310 | T | C | 70 | 0 | 10 | 0 | NON_SYNONYMOU | MODERAT | MISSENSE | gAa/gGa | E909G | PLES_14981 | C | C | C | C | C | C | C | C | | | C | C |
| NC_011770 | 1624790 | C | T | 63 | 0 | 1 | 0 | NON_SYNONYMOU | MODERAT | MISSENSE | gGc/gAc | G749D | PLES_14981 | | | | | | | T | | | | | |
| NC_011770 | 1624898 | C | T | 62 | 0 | 4 | 0 | NON_SYNONYMOU | MODERAT | MISSENSE | gGc/gAc | G713D | PLES_14981 | | | T | T | T | T | | | | | | |
| NC_011770 | 1624908 | C | T | 62 | 0 | 10 | 0 | NON_SYNONYMOU | MODERAT | MISSENSE | Gcc/Acc | A710T | PLES_14981 | T | T | T | T | T | T | T | T | T | T | | |
| NC_011770 | 1625079 | C | T | 67 | 0 | 10 | 0 | NON_SYNONYMOU | MODERAT | MISSENSE | Gcg/Acg | A653T | PLES_14981 | T | T | T | T | T | T | T | T | T | T | | |
| NC_011770 | 1625136 | C | T | 70 | 0 | 10 | 0 | NON_SYNONYMOU | MODERAT | MISSENSE | Gcc/Acc | A634T | PLES_14981 | T | T | T | T | T | T | T | T | T | T | | |
| NC_011770 | 1625318 | T | C | 68 | 0 | 10 | 0 | NON_SYNONYMOU | MODERAT | MISSENSE | aAc/aGc | N573S | PLES_14981 | C | C | C | C | C | C | C | C | C | C | | |
| NC_011770 | 1625331 | C | A | 72 | 0 | 2 | 0 | NON_SYNONYMOU | MODERAT | MISSENSE | Gcc/Tcc | A569S | PLES_14981 | | | | | | | | | | | A | A |
| NC_011770 | 1625525 | C | T | 63 | 0 | 4 | 0 | NON_SYNONYMOU | MODERAT | MISSENSE | aGc/aAc | S504N | PLES_14981 | | | T | T | T | T | | | | | | |
| NC_011770 | 1626356 | T | A | 86 | 0 | 4 | 0 | NON_SYNONYMOU | MODERAT | MISSENSE | cAg/cTg | Q227L | PLES_14981 | | | A | A | A | A | | | | | | |
| NC_011770 | 1626389 | G | C | 88 | 0 | 4 | 0 | NON_SYNONYMOU | MODERAT | MISSENSE | aCc/aGc | T216S | PLES_14981 | | | C | C | C | C | | | | | | |
| NC_011770 | 1626532 | G | T | 82 | 0 | 2 | 0 | NON_SYNONYMOU | MODERAT | MISSENSE | aaC/aaA | N168K | PLES_14981 | T | T | | | | | | | | | | |
| NC_011770 | 1626933 | T | C | 73 | 0 | 12 | 0 | NON_SYNONYMOU | MODERAT | MISSENSE | Acc/Gcc | T35A | PLES_14981 | C | C | C | C | C | C | C | C | C | C | C | C |
| NC_011770 | 1627020 | G | A | 70 | 0 | 12 | 0 | NON_SYNONYMOU | MODERAT | MISSENSE | Ccc/Tcc | P6S | PLES_14981 | A | A | A | A | A | A | A | A | A | A | A | A |
| NC_011770 | 1627022 | C | T | 68 | 0 | 12 | 0 | NON_SYNONYMOU | MODERAT | MISSENSE | cGc/cAc | R5H | PLES_14981 | T | T | T | T | T | T | T | T | T | T | T | T |
| NC_011770 | 1627025 | G | T | 69 | 0 | 12 | 0 | NON_SYNONYMOU | MODERAT | MISSENSE | gCg/gAc | A4D | PLES_14981 | T | T | T | T | T | T | T | T | T | T | T | T |
| NC_011770 | 1626374 | G | A | 86 | 0 | 1 | 0 | NON_SYNONYMOU | MODERAT | MISSENSE | gCc/gTc | A221V | PLES_14981 | | | | | | | | | | | A | |
| NC_011770 | 1627988 | G | T | 77 | 0 | 4 | 0 | NON_SYNONYMOU | MODERAT | MISSENSE | Gcc/Tcc | A219S | PLES_14991 | | | | | | | | | T | T | T | T |
| NC_011770 | 1628066 | A | C | 77 | 0 | 2 | 0 | NON_SYNONYMOU | MODERAT | MISSENSE | Aac/Cac | N245H | PLES_14991 | | | | | | | C | C | | | | |
| NC_011770 | 1628130 | T | C | 78 | 0 | 6 | 0 | NON_SYNONYMOU | MODERAT | MISSENSE | gTc/gCc | V266A | PLES_14991 | C | C | | | | | | | C | C | C | C |
| NC_011770 | 1628154 | C | T | 75 | 0 | 2 | 0 | NON_SYNONYMOU | MODERAT | MISSENSE | gCg/gTg | A274V | PLES_14991 | | | | | | | T | T | | | | |
| NC_011770 | 1628162 | A | G | 77 | 0 | 12 | 0 | NON_SYNONYMOU | MODERAT | MISSENSE | Agc/Ggc | S277G | PLES_14991 | G | G | G | G | G | G | G | G | G | G | G | G |
| NC_011770 | 1628388 | A | G | 73 | 0 | 10 | 0 | NON_SYNONYMOU | MODERAT | MISSENSE | gAc/gGc | D352G | PLES_14991 | G | G | G | G | G | G | | | G | G | G | G |
| NC_011770 | 1628391 | G | A | 74 | 0 | 2 | 0 | NON_SYNONYMOU | MODERAT | MISSENSE | cGt/cAt | R353H | PLES_14991 | | | | | | | A | A | | | | |
| NC_011770 | 1628670 | C | T | 70 | 0 | 2 | 0 | NON_SYNONYMOU | MODERAT | MISSENSE | aCc/aTc | T446I | PLES_14991 | | | | | | | | | T | T | | |
| NC_011770 | 1629054 | A | G | 44 | 0 | 4 | 0 | NON_SYNONYMOU | MODERAT | MISSENSE | gAc/gGc | D574G | PLES_14991 | G | G | | | | | | | | | G | G |
| NC_011770 | 1629137 | C | T | 31 | 0 | 2 | 0 | NON_SYNONYMOU | MODERAT | MISSENSE | Ccg/Tcg | P602S | PLES_14991 | | | | | | | | | | | T | T |
| NC_011770 | 1629183 | G | A | 27 | 0 | 2 | 0 | NON_SYNONYMOU | MODERAT | MISSENSE | gGc/gAc | G617D | PLES_14991 | A | A | | | | | | | | | | |
| NC_011770 | 1629348 | C | T | 34 | 0 | 8 | 0 | NON_SYNONYMOU | MODERAT | MISSENSE | gCc/gTc | A672V | PLES_14991 | T | T | | | | | T | T | T | T | T | T |
| NC_011770 | 1630193 | G | T | 71 | 0 | 12 | 0 | NON_SYNONYMOU | MODERAT | MISSENSE | aGc/aTc | S77I | PLES_15011 | T | T | T | T | T | T | T | T | T | T | T | T |
| NC_011770 | 1630443 | C | G | 78 | 0 | 2 | 0 | NON_SYNONYMOU | MODERAT | MISSENSE | ttC/ttG | F160L | PLES_15011 | | | | | | | | | | | G | G |
| NC_011770 | 1630462 | A | T | 72 | 0 | 10 | 0 | NON_SYNONYMOU | MODERAT | MISSENSE | Acc/Tcc | T167S | PLES_15011 | T | T | T | T | T | T | | | T | T | T | T |
| NC_011770 | 1630606 | T | C | 67 | 0 | 1 | 0 | NON_SYNONYMOU | MODERAT | MISSENSE | Ttc/Ctc | F215L | PLES_15011 | | | | | | | C | | | | | |
| NC_011770 | 1630829 | C | T | 70 | 0 | 2 | 0 | NON_SYNONYMOU | MODERAT | MISSENSE | cCc/cTc | P289L | PLES_15011 | | | | | | | T | T | | | | |
| NC_011770 | 1631092 | C | T | 55 | 0 | 2 | 0 | NON_SYNONYMOU | MODERAT | MISSENSE | Cgg/Tgg | R377W | PLES_15011 | T | T | | | | | | | | | | |
| NC_011770 | 1635310 | G | A | 49 | 0 | 4 | 0 | NON_SYNONYMOU | MODERAT | MISSENSE | aGc/aAc | S111N | PLES_15071 | | | A | A | A | A | | | | | | |
| NC_011770 | 1635485 | C | G | 71 | 0 | 2 | 0 | NON_SYNONYMOU | MODERAT | MISSENSE | gaC/gaG | D169E | PLES_15071 | | | | | | | | | | | G | G |
| NC_011770 | 1638221 | C | T | 67 | 0 | 1 | 0 | NON_SYNONYMOU | MODERAT | MISSENSE | gCg/gTg | A30V | PLES_15101 | | | | | | | T | | | | | |
| NC_011770 | 1638833 | C | T | 85 | 0 | 4 | 0 | NON_SYNONYMOU | MODERAT | MISSENSE | gCc/gTc | A234V | PLES_15101 | | | T | T | T | T | | | | | | |
| NC_011770 | 1638907 | G | A | 84 | 0 | 4 | 0 | NON_SYNONYMOU | MODERAT | MISSENSE | Gcc/Acc | A259T | PLES_15101 | | | A | A | A | A | | | | | | |
| NC_011770 | 1639022 | C | A | 78 | 0 | 2 | 0 | NON_SYNONYMOU | MODERAT | MISSENSE | gCg/gAg | A297E | PLES_15101 | A | A | | | | | | | | | | |
| NC_011770 | 1639229 | G | T | 79 | 0 | 2 | 0 | NON_SYNONYMOU | MODERAT | MISSENSE | cGc/cTc | R366L | PLES_15101 | T | T | | | | | | | | | | |
| NC_011770 | 1640166 | A | G | 90 | 0 | 4 | 0 | NON_SYNONYMOU | MODERAT | MISSENSE | Atc/Gtc | I294V | PLES_15111 | | | G | G | G | G | | | | | | |
| NC_011770 | 1640800 | G | A | 83 | 0 | 4 | 0 | NON_SYNONYMOU | MODERAT | MISSENSE | gGt/gAt | G505D | PLES_15111 | | | A | A | A | A | | | | | | |
| NC_011770 | 1640836 | C | T | 87 | 0 | 6 | 0 | NON_SYNONYMOU | MODERAT | MISSENSE | gCc/gTc | A517V | PLES_15111 | | | T | T | T | T | T | T | | | | |
| NC_011770 | 1641092 | C | G | 83 | 0 | 10 | 0 | NON_SYNONYMOU | MODERAT | MISSENSE | gaC/gaG | D602E | PLES_15111 | G | G | G | G | G | G | | | G | G | G | G |
| NC_011770 | 1641220 | G | C | 78 | 0 | 4 | 0 | NON_SYNONYMOU | MODERAT | MISSENSE | gGc/gCc | G645A | PLES_15111 | | | C | C | C | C | | | | | | |
| NC_011770 | 1641847 | C | T | 85 | 0 | 2 | 0 | NON_SYNONYMOU | MODERAT | MISSENSE | cCc/cTc | P854L | PLES_15111 | | | | | | | | | T | T | | |
| NC_011770 | 1642047 | G | A | 82 | 0 | 4 | 0 | NON_SYNONYMOU | MODERAT | MISSENSE | Gtg/Atg | V921M | PLES_15111 | | | A | A | A | A | | | | | | |
| NC_011770 | 1642153 | C | T | 89 | 0 | 2 | 0 | NON_SYNONYMOU | MODERAT | MISSENSE | gCc/gTc | A956V | PLES_15111 | T | T | | | | | | | | | | |
| NC_011770 | 1642393 | A | G | 75 | 0 | 12 | 0 | NON_SYNONYMOU | MODERAT | MISSENSE | cAc/cGc | H1036R | PLES_15111 | G | G | G | G | G | G | G | G | G | G | G | G |
| NC_011770 | 1642425 | C | T | 78 | 0 | 2 | 0 | NON_SYNONYMOU | MODERAT | MISSENSE | Cat/Tat | H1047Y | PLES_15111 | | | | | | | T | T | | | | |
| NC_011770 | 1642580 | G | A | 76 | 0 | 4 | 0 | NON_SYNONYMOU | MODERAT | MISSENSE | Ggt/Agt | G46S | PLES_15121 | | | A | A | A | A | | | | | | |
| NC_011770 | 1642592 | A | G | 75 | 0 | 2 | 0 | NON_SYNONYMOU | MODERAT | MISSENSE | Atc/Gtc | I50V | PLES_15121 | G | G | | | | | | | | | | |
| NC_011770 | 1642623 | C | A | 75 | 0 | 6 | 0 | NON_SYNONYMOU | MODERAT | MISSENSE | gCc/gAc | A60D | PLES_15121 | | | | | | | A | A | A | A | A | A |
| NC_011770 | 1642833 | C | A | 74 | 0 | 2 | 0 | NON_SYNONYMOU | MODERAT | MISSENSE | gCg/gAg | A130E | PLES_15121 | | | | | | | A | A | | | | |
| NC_011770 | 1643279 | G | T | 71 | 0 | 6 | 0 | NON_SYNONYMOU | MODERAT | MISSENSE | Gcg/Tcg | A279S | PLES_15121 | | | T | T | T | T | T | T | | | | |

| | | | | | | | | | | | | | | | | | | | | | | | | | |
|---|---|---|---|---|---|---|---|---|---|---|---|---|---|---|---|---|---|---|---|---|---|---|---|---|---|
| NC_011770 | 1643508 | T | C | 84 | 0 | 12 | 0 | NON_SYNONYMOU | MODERAT | MISSENSE | gTg/gCg | V355A | PLES_15121 | C | C | C | C | C | C | C | C | C | C | C | C |
| NC_011770 | 1644688 | C | G | 69 | 0 | 12 | 0 | NON_SYNONYMOU | MODERAT | MISSENSE | gaC/gaG | D57E | PLES_15131 | G | G | G | G | G | G | G | G | G | G | G | G |
| NC_011770 | 1645631 | T | G | 92 | 0 | 12 | 0 | NON_SYNONYMOU | MODERAT | MISSENSE | gaT/gaG | D173E | PLES_15141 | G | G | G | G | G | G | G | G | G | G | G | G |
| NC_011770 | 1646052 | A | C | 92 | 0 | 4 | 0 | NON_SYNONYMOU | MODERAT | MISSENSE | Aag/Cag | K314Q | PLES_15141 | | | | | | | | | C | C | C | C |
| NC_011770 | 1647216 | C | T | 80 | 0 | 2 | 0 | NON_SYNONYMOU | MODERAT | MISSENSE | aCg/aTg | T6M | PLES_15161 | | | | | | | T | T | | | | |
| NC_011770 | 1647477 | T | C | 79 | 0 | 8 | 0 | NON_SYNONYMOU | MODERAT | MISSENSE | tTg/tCg | L93S | PLES_15161 | C | C | | | | | C | C | C | C | C | C |
| NC_011770 | 1647570 | G | T | 80 | 0 | 2 | 0 | NON_SYNONYMOU | MODERAT | MISSENSE | tGg/tTg | W124L | PLES_15161 | | | | | | | T | T | | | | |
| NC_011770 | 1647578 | A | G | 76 | 0 | 6 | 0 | NON_SYNONYMOU | MODERAT | MISSENSE | Acg/Gcg | T127A | PLES_15161 | | | | | | | G | G | G | G | G | G |
| NC_011770 | 1647649 | G | A | 82 | 0 | 1 | 0 | NON_SYNONYMOU | MODERAT | MISSENSE | atG/atA | M150I | PLES_15161 | | | | | | | A | | | | | |
| NC_011770 | 1648007 | G | C | 85 | 0 | 2 | 0 | NON_SYNONYMOU | MODERAT | MISSENSE | Ggc/Cgc | G270R | PLES_15161 | C | C | | | | | | | | | | |
| NC_011770 | 1648277 | G | A | 77 | 0 | 6 | 0 | NON_SYNONYMOU | MODERAT | MISSENSE | Gcc/Acc | A360T | PLES_15161 | | | A | A | A | A | A | A | | | | |
| NC_011770 | 1648536 | G | A | 88 | 0 | 4 | 0 | NON_SYNONYMOU | MODERAT | MISSENSE | gGc/gAc | G446D | PLES_15161 | | | | | | | | | A | A | A | A |
| NC_011770 | 1648598 | C | A | 84 | 0 | 8 | 0 | NON_SYNONYMOU | MODERAT | MISSENSE | Ccc/Acc | P467T | PLES_15161 | A | A | A | A | A | A | A | A | | | | |
| NC_011770 | 1648909 | G | A | 92 | 0 | 4 | 0 | NON_SYNONYMOU | MODERAT | MISSENSE | Gcc/Acc | A93T | PLES_15171 | | | A | A | A | A | | | | | | |
| NC_011770 | 1648950 | G | C | 95 | 0 | 10 | 0 | NON_SYNONYMOU | MODERAT | MISSENSE | gaG/gaC | E106D | PLES_15171 | C | C | C | C | C | C | | | C | C | C | C |
| NC_011770 | 1652807 | A | G | 103 | 0 | 4 | 0 | NON_SYNONYMOU | MODERAT | MISSENSE | gTg/gCg | V80A | PLES_15211 | G | G | | | | | G | G | | | | |
| NC_011770 | 1653220 | T | C | 84 | 0 | 6 | 0 | NON_SYNONYMOU | MODERAT | MISSENSE | Agc/Ggc | S155G | PLES_15221 | | | | | | | C | C | C | C | C | C |
| NC_011770 | 1653609 | C | T | 84 | 0 | 2 | 0 | NON_SYNONYMOU | MODERAT | MISSENSE | gGc/gAc | G25D | PLES_15221 | | | | | | | | | | | T | T |
| NC_011770 | 1653651 | C | T | 81 | 0 | 2 | 0 | NON_SYNONYMOU | MODERAT | MISSENSE | cGc/cAc | R11H | PLES_15221 | T | T | | | | | | | | | | |
| NC_011770 | 1653795 | T | C | 77 | 0 | 12 | 0 | NON_SYNONYMOU | MODERAT | MISSENSE | Atg/Gtg | M308V | PLES_15231 | C | C | C | C | C | C | C | C | C | C | C | C |
| NC_011770 | 1653819 | C | T | 76 | 0 | 2 | 0 | NON_SYNONYMOU | MODERAT | MISSENSE | Gcc/Acc | A300T | PLES_15231 | T | T | | | | | | | | | | |
| NC_011770 | 1653852 | G | C | 73 | 0 | 2 | 0 | NON_SYNONYMOU | MODERAT | MISSENSE | Ctc/Gtc | L289V | PLES_15231 | | | | | | | C | C | | | | |
| NC_011770 | 1655861 | T | C | 77 | 0 | 12 | 0 | NON_SYNONYMOU | MODERAT | MISSENSE | Aac/Gac | N395D | PLES_15241 | C | C | C | C | C | C | C | C | C | C | C | C |
| NC_011770 | 1656026 | C | T | 78 | 0 | 2 | 0 | NON_SYNONYMOU | MODERAT | MISSENSE | Gat/Aat | D340N | PLES_15241 | | | | | T | T | | | | | | |
| NC_011770 | 1656576 | T | G | 79 | 0 | 6 | 0 | NON_SYNONYMOU | MODERAT | MISSENSE | gaA/gaC | E156D | PLES_15241 | | | | | | | G | G | G | G | G | G |
| NC_011770 | 1656878 | T | C | 85 | 0 | 12 | 0 | NON_SYNONYMOU | MODERAT | MISSENSE | Ata/Gta | I56V | PLES_15241 | C | C | C | C | C | C | C | C | C | C | C | C |
| NC_011770 | 1657037 | C | T | 93 | 0 | 2 | 0 | NON_SYNONYMOU | MODERAT | MISSENSE | Gcc/Acc | A3T | PLES_15241 | T | T | | | | | | | | | | |
| NC_011770 | 1658422 | T | C | 52 | 0 | 12 | 0 | NON_SYNONYMOU | MODERAT | MISSENSE | Acg/Gcg | T116A | PLES_15271 | C | C | C | C | C | C | C | C | C | C | C | C |
| NC_011770 | 1658481 | TCCTCTA | T | 53 | 0 | 1 | 0 | CODON_CHANGE_ | MODERATE | | gtagaggaa | VEE94E | PLES_15271 | | | | | | | | | . | T | | |
| NC_011770 | 1658559 | T | C | 56 | 0 | 2 | 0 | NON_SYNONYMOU | MODERAT | MISSENSE | cAg/cGg | Q70R | PLES_15271 | | | | | | | C | C | | | | |
| NC_011770 | 1670480 | C | A | 85 | 0 | 2 | 0 | NON_SYNONYMOU | MODERAT | MISSENSE | Ctg/Atg | L100M | PLES_15381 | A | A | | | | | | | | | | |
| NC_011770 | 1670742 | C | T | 87 | 0 | 2 | 0 | NON_SYNONYMOU | MODERAT | MISSENSE | cCt/cTt | P187L | PLES_15381 | | | | | | | | | | | T | T |
| NC_011770 | 1671235 | T | C | 73 | 0 | 4 | 0 | NON_SYNONYMOU | MODERAT | MISSENSE | Tgc/Cgc | C66R | PLES_15391 | C | C | | | | | C | C | | | | |
| NC_011770 | 1673319 | A | AC | 80 | 0 | 1 | 0 | FRAME_SHIFT | HIGH | | acg/Cacg | T105H? | PLES_15411 | | | | AC | | | | | | | | |
| NC_011770 | 1673409 | A | G | 74 | 0 | 2 | 0 | NON_SYNONYMOU | MODERAT | MISSENSE | Atc/Gtc | I133V | PLES_15411 | G | G | | | | | | | | | | |
| NC_011770 | 1674439 | T | C | 90 | 0 | 12 | 0 | NON_SYNONYMOU | MODERAT | MISSENSE | gTg/gCg | V476A | PLES_15411 | C | C | C | C | C | C | C | C | C | C | C | C |
| NC_011770 | 1674517 | A | G | 90 | 0 | 2 | 0 | NON_SYNONYMOU | MODERAT | MISSENSE | cAg/cGg | Q502R | PLES_15411 | | | | | | | | | | | G | G |
| NC_011770 | 1674829 | TC | T | 79 | 0 | 4 | 0 | FRAME_SHIFT+STC | HIGH | | tga/ | *153 | PLES_15421 | | | | | | | T | T | T | T | | |
| NC_011770 | 1675552 | T | C | 78 | 0 | 4 | 0 | NON_SYNONYMOU | MODERAT | MISSENSE | gAc/gGc | D92G | PLES_15431 | C | C | | | | | C | C | | | | |
| NC_011770 | 1676171 | A | G | 81 | 0 | 12 | 0 | NON_SYNONYMOU | MODERAT | MISSENSE | gTc/gCc | V335A | PLES_15441 | G | G | G | G | G | G | G | G | G | G | G | G |
| NC_011770 | 1676276 | C | T | 90 | 0 | 2 | 0 | NON_SYNONYMOU | MODERAT | MISSENSE | cGc/cAc | R300H | PLES_15441 | | | | | | | | | | | T | T |
| NC_011770 | 1675850 | A | G | 87 | 0 | 1 | 0 | NON_SYNONYMOU | MODERAT | MISSENSE | gTc/gCc | V442A | PLES_15441 | G | | | | | | | | | | | |
| NC_011770 | 1677848 | C | T | 65 | 0 | 4 | 0 | NON_SYNONYMOU | MODERAT | MISSENSE | Gcg/Acg | A288T | PLES_15451 | T | T | | | | | T | T | | | | |
| NC_011770 | 1678019 | T | A | 56 | 0 | 4 | 0 | NON_SYNONYMOU | MODERAT | MISSENSE | Acc/Tcc | T231S | PLES_15451 | A | A | | | | | A | A | | | | |
| NC_011770 | 1678046 | C | T | 60 | 0 | 4 | 0 | NON_SYNONYMOU | MODERAT | MISSENSE | Gca/Aca | A222T | PLES_15451 | T | T | | | | | T | T | | | | |
| NC_011770 | 1678363 | T | A | 70 | 0 | 12 | 0 | NON_SYNONYMOU | MODERAT | MISSENSE | tAc/tTc | Y116F | PLES_15451 | A | A | A | A | A | A | A | A | A | A | A | A |
| NC_011770 | 1678905 | C | T | 64 | 0 | 4 | 0 | NON_SYNONYMOU | MODERAT | MISSENSE | tCt/tTt | S4F | PLES_15461 | T | T | | | | | T | T | | | | |
| NC_011770 | 1679550 | G | A | 66 | 0 | 12 | 0 | NON_SYNONYMOU | MODERAT | MISSENSE | aGc/aAc | S219N | PLES_15461 | A | A | A | A | A | A | A | A | A | A | A | A |
| NC_011770 | 1681425 | G | C | 83 | 0 | 4 | 0 | NON_SYNONYMOU | MODERAT | MISSENSE | gaC/gaG | D216E | PLES_15471 | | | | | | | | | C | C | C | C |
| NC_011770 | 1680472 | C | T | 70 | 0 | 1 | 0 | NON_SYNONYMOU | MODERAT | MISSENSE | gGt/gAt | G534D | PLES_15471 | | T | | | | | | | | | | |
| NC_011770 | 1681007 | C | T | 93 | 0 | 1 | 0 | NON_SYNONYMOU | MODERAT | MISSENSE | Gtg/Atg | V356M | PLES_15471 | | | | | | | | | | | T | |
| NC_011770 | 1682433 | C | T | 72 | 0 | 2 | 0 | NON_SYNONYMOU | MODERAT | MISSENSE | Ccg/Tcg | P49S | PLES_15481 | T | T | | | | | | | | | | |
| NC_011770 | 1683303 | C | T | 34 | 0 | 2 | 0 | NON_SYNONYMOU | MODERAT | MISSENSE | Ctc/Ttc | L339F | PLES_15481 | | | | | | | T | T | | | | |
| NC_011770 | 1683371 | G | C | 34 | 0 | 8 | 0 | NON_SYNONYMOU | MODERAT | MISSENSE | gaG/gaC | E361D | PLES_15481 | | | C | C | C | C | | | C | C | C | C |
| NC_011770 | 1683511 | T | G | 19 | 0 | 6 | 0 | NON_SYNONYMOU | MODERAT | MISSENSE | gTg/gGg | V408G | PLES_15481 | G | G | | | | | | | G | G | G | G |
| NC_011770 | 1682635 | C | T | 59 | 0 | 1 | 0 | NON_SYNONYMOU | MODERAT | MISSENSE | gCg/gTg | A116V | PLES_15481 | | | | | | | | | | | T | |
| NC_011770 | 1684397 | T | C | 22 | 10 | 2 | 0 | NON_SYNONYMOU | MODERAT | MISSENSE | Atc/Gtc | I150V | PLES_15491 | . | . | . | . | C | C | . | . | . | . | . | . |
| NC_011770 | 1684477 | C | T | 23 | 10 | 2 | 0 | NON_SYNONYMOU | MODERAT | MISSENSE | cGg/cAg | R123Q | PLES_15491 | . | . | . | . | T | T | . | . | . | . | . | . |
| NC_011770 | 1684546 | C | T | 23 | 10 | 2 | 0 | NON_SYNONYMOU | MODERAT | MISSENSE | aGt/aAt | S100N | PLES_15491 | . | . | . | . | T | T | . | . | . | . | . | . |
| NC_011770 | 1684580 | A | G | 19 | 10 | 2 | 0 | NON_SYNONYMOU | MODERAT | MISSENSE | Tct/Cct | S89P | PLES_15491 | . | . | . | . | G | G | . | . | . | . | . | . |
| NC_011770 | 1684582 | G | A | 19 | 10 | 2 | 0 | NON_SYNONYMOU | MODERAT | MISSENSE | aCg/aTg | T88M | PLES_15491 | . | . | . | . | A | A | . | . | . | . | . | . |
| NC_011770 | 1684628 | T | G | 21 | 10 | 2 | 0 | NON_SYNONYMOU | MODERAT | MISSENSE | Aag/Cag | K73Q | PLES_15491 | . | . | . | . | G | G | . | . | . | . | . | . |

| | | | | | | | | | | | | | | | | | | | | | | | | | |
|---|---|---|---|---|---|---|---|---|---|---|---|---|---|---|---|---|---|---|---|---|---|---|---|---|---|
| NC_011770 | 1684734 | T | A | 20 | 10 | 2 | 0 | NON_SYNONYMOU | MODERAT | MISSENSE | caA/caT | Q37H | PLES_15491 | . | . | . | . | A | A | . | . | . | . | . | . |
| NC_011770 | 1685598 | C | T | 18 | 10 | 2 | 0 | NON_SYNONYMOU | MODERAT | MISSENSE | gCt/gTt | A76V | PLES_15511 | . | . | . | . | T | T | . | . | . | . | . | . |
| NC_011770 | 1685809 | G | A | 13 | 10 | 2 | 0 | NON_SYNONYMOU | MODERAT | MISSENSE | Gaa/Aaa | E2K | PLES_15521 | . | . | . | . | A | A | . | . | . | . | . | . |
| NC_011770 | 1685810 | A | C | 14 | 10 | 2 | 0 | NON_SYNONYMOU | MODERAT | MISSENSE | gAa/gCa | E2A | PLES_15521 | . | . | . | . | C | C | . | . | . | . | . | . |
| NC_011770 | 1685813 | C | A | 13 | 10 | 2 | 0 | NON_SYNONYMOU | MODERAT | MISSENSE | aCc/aAc | T3N | PLES_15521 | . | . | . | . | A | A | . | . | . | . | . | . |
| NC_011770 | 1685818 | A | G | 14 | 10 | 2 | 0 | NON_SYNONYMOU | MODERAT | MISSENSE | Aac/Gac | N5D | PLES_15521 | . | . | . | . | G | G | . | . | . | . | . | . |
| NC_011770 | 1685825 | C | A | 13 | 10 | 2 | 0 | NON_SYNONYMOU | MODERAT | MISSENSE | aCg/aAg | T7K | PLES_15521 | . | . | . | . | A | A | . | . | . | . | . | . |
| NC_011770 | 1685827 | G | C | 13 | 10 | 2 | 0 | NON_SYNONYMOU | MODERAT | MISSENSE | Gcg/Ccg | A8P | PLES_15521 | . | . | . | . | C | C | . | . | . | . | . | . |
| NC_011770 | 1685836 | A | G | 14 | 10 | 2 | 0 | NON_SYNONYMOU | MODERAT | MISSENSE | Agc/Ggc | S11G | PLES_15521 | . | . | . | . | G | G | . | . | . | . | . | . |
| NC_011770 | 1685837 | G | A | 13 | 10 | 2 | 0 | NON_SYNONYMOU | MODERAT | MISSENSE | aGc/aAc | S11N | PLES_15521 | . | . | . | . | A | A | . | . | . | . | . | . |
| NC_011770 | 1685848 | C | A | 14 | 10 | 2 | 0 | NON_SYNONYMOU | MODERAT | MISSENSE | Cac/Aac | H15N | PLES_15521 | . | . | . | . | A | A | . | . | . | . | . | . |
| NC_011770 | 1685965 | A | G | 0 | 11 | 1 | 0 | NON_SYNONYMOU | MODERAT | MISSENSE | Aac/Gac | N54D | PLES_15521 | . | . | . | . | . | G | . | . | . | . | . | . |
| NC_011770 | 1685986 | G | A | 16 | 10 | 2 | 0 | NON_SYNONYMOU | MODERAT | MISSENSE | Gaa/Aaa | E61K | PLES_15521 | . | . | . | . | A | A | . | . | . | . | . | . |
| NC_011770 | 1685993 | C | T | 16 | 10 | 2 | 0 | NON_SYNONYMOU | MODERAT | MISSENSE | gCt/gTt | A63V | PLES_15521 | . | . | . | . | T | T | . | . | . | . | . | . |
| NC_011770 | 1689591 | C | A | 16 | 10 | 2 | 0 | NON_SYNONYMOU | MODERAT | MISSENSE | gaC/gaA | D2E | PLES_15561 | . | . | . | . | A | A | . | . | . | . | . | . |
| NC_011770 | 1690008 | G | T | 22 | 10 | 2 | 0 | NON_SYNONYMOU | MODERAT | MISSENSE | cGg/cTg | R7L | PLES_15571 | . | . | . | . | T | T | . | . | . | . | . | . |
| NC_011770 | 1690266 | T | G | 20 | 10 | 2 | 0 | NON_SYNONYMOU | MODERAT | MISSENSE | aTt/aGt | I93S | PLES_15571 | . | . | . | . | G | G | . | . | . | . | . | . |
| NC_011770 | 1691037 | T | A | 14 | 10 | 2 | 0 | NON_SYNONYMOU | MODERAT | MISSENSE | gaT/gaA | D76E | PLES_15591 | . | . | . | . | A | A | . | . | . | . | . | . |
| NC_011770 | 1691087 | G | A | 14 | 10 | 2 | 0 | NON_SYNONYMOU | MODERAT | MISSENSE | aGg/aAg | R93K | PLES_15591 | . | . | . | . | A | A | . | . | . | . | . | . |
| NC_011770 | 1691105 | T | C | 12 | 10 | 2 | 0 | NON_SYNONYMOU | MODERAT | MISSENSE | gTc/gCc | V99A | PLES_15591 | . | . | . | . | C | C | . | . | . | . | . | . |
| NC_011770 | 1691107 | C | A | 12 | 10 | 2 | 0 | NON_SYNONYMOU | MODERAT | MISSENSE | Ctc/Atc | L100I | PLES_15591 | . | . | . | . | A | A | . | . | . | . | . | . |
| NC_011770 | 1691129 | A | G | 12 | 10 | 2 | 0 | NON_SYNONYMOU | MODERAT | MISSENSE | cAa/cGa | Q107R | PLES_15591 | . | . | . | . | G | G | . | . | . | . | . | . |
| NC_011770 | 1691152 | G | A | 11 | 10 | 2 | 0 | NON_SYNONYMOU | MODERAT | MISSENSE | Gaa/Aaa | E115K | PLES_15591 | . | . | . | . | A | A | . | . | . | . | . | . |
| NC_011770 | 1691153 | A | C | 10 | 10 | 2 | 0 | NON_SYNONYMOU | MODERAT | MISSENSE | gAa/gCa | E115A | PLES_15591 | . | . | . | . | C | C | . | . | . | . | . | . |
| NC_011770 | 1691284 | C | G | 15 | 10 | 2 | 0 | NON_SYNONYMOU | MODERAT | MISSENSE | gaC/gaG | D32E | PLES_15601 | . | . | . | . | G | G | . | . | . | . | . | . |
| NC_011770 | 1691339 | A | C | 14 | 10 | 2 | 0 | NON_SYNONYMOU | MODERAT | MISSENSE | Aag/Cag | K51Q | PLES_15601 | . | . | . | . | C | C | . | . | . | . | . | . |
| NC_011770 | 1691726 | A | G | 19 | 10 | 2 | 0 | NON_SYNONYMOU | MODERAT | MISSENSE | Aat/Gat | N16D | PLES_15621 | . | . | . | . | G | G | . | . | . | . | . | . |
| NC_011770 | 1691783 | G | A | 17 | 10 | 2 | 0 | NON_SYNONYMOU | MODERAT | MISSENSE | Gcc/Acc | A35T | PLES_15621 | . | . | . | . | A | A | . | . | . | . | . | . |
| NC_011770 | 1692443 | C | A | 20 | 10 | 2 | 0 | NON_SYNONYMOU | MODERAT | MISSENSE | aaC/aaA | N63K | PLES_15631 | . | . | . | . | A | A | . | . | . | . | . | . |
| NC_011770 | 1692849 | C | G | 21 | 10 | 2 | 0 | NON_SYNONYMOU | MODERAT | MISSENSE | gaC/gaG | D67E | PLES_15641 | . | . | . | . | G | G | . | . | . | . | . | . |
| NC_011770 | 1693072 | T | C | 18 | 10 | 2 | 0 | NON_SYNONYMOU | MODERAT | MISSENSE | Tcg/Ccg | S142P | PLES_15641 | . | . | . | . | C | C | . | . | . | . | . | . |
| NC_011770 | 1695693 | T | TC | 6 | 10 | 2 | 0 | FRAME_SHIFT | HIGH | | tca/tcCa | S123S? | PLES_15701 | . | . | . | . | TC | TC | . | . | . | . | . | . |
| NC_011770 | 1695697 | CG | C | 6 | 10 | 2 | 0 | FRAME_SHIFT | HIGH | | gcg/ | A124 | PLES_15701 | . | . | . | . | C | C | . | . | . | . | . | . |
| NC_011770 | 1695582 | G | A | 12 | 10 | 2 | 0 | NON_SYNONYMOU | MODERAT | MISSENSE | Gtc/Atc | V86I | PLES_15701 | . | . | . | . | A | A | . | . | . | . | . | . |
| NC_011770 | 1695592 | A | ACAG | 4 | 10 | 1 | 0 | CODON_INSERTIO | MODERATE | | gcg/AGCg | A90SA | PLES_15701 | . | . | . | . | ACAG | | . | . | . | . | . | . |
| NC_011770 | 1695604 | A | C | 11 | 10 | 2 | 0 | NON_SYNONYMOU | MODERAT | MISSENSE | aAg/aCg | K93T | PLES_15701 | . | . | . | . | C | C | . | . | . | . | . | . |
| NC_011770 | 1695624 | C | A | 15 | 10 | 2 | 0 | NON_SYNONYMOU | MODERAT | MISSENSE | Ctg/Atg | L100M | PLES_15701 | . | . | . | . | A | A | . | . | . | . | . | . |
| NC_011770 | 1695657 | A | T | 13 | 10 | 2 | 0 | NON_SYNONYMOU | MODERAT | MISSENSE | Acg/Tcg | T111S | PLES_15701 | . | . | . | . | T | T | . | . | . | . | . | . |
| NC_011770 | 1695660 | G | A | 13 | 10 | 2 | 0 | NON_SYNONYMOU | MODERAT | MISSENSE | Gtc/Atc | V112I | PLES_15701 | . | . | . | . | A | A | . | . | . | . | . | . |
| NC_011770 | 1695720 | C | A | 7 | 10 | 2 | 0 | NON_SYNONYMOU | MODERAT | MISSENSE | Ctc/Atc | L132I | PLES_15701 | . | . | . | . | A | A | . | . | . | . | . | . |
| NC_011770 | 1695735 | A | G | 8 | 10 | 2 | 0 | NON_SYNONYMOU | MODERAT | MISSENSE | Agc/Ggc | S137G | PLES_15701 | . | . | . | . | G | G | . | . | . | . | . | . |
| NC_011770 | 1695736 | G | C | 8 | 10 | 2 | 0 | NON_SYNONYMOU | MODERAT | MISSENSE | aGc/aCc | S137T | PLES_15701 | . | . | . | . | C | C | . | . | . | . | . | . |
| NC_011770 | 1695751 | G | A | 8 | 10 | 2 | 0 | NON_SYNONYMOU | MODERAT | MISSENSE | cGg/cAg | R142Q | PLES_15701 | . | . | . | . | A | A | . | . | . | . | . | . |
| NC_011770 | 1695766 | C | T | 9 | 10 | 2 | 0 | NON_SYNONYMOU | MODERAT | MISSENSE | gCc/gTc | A147V | PLES_15701 | . | . | . | . | T | T | . | . | . | . | . | . |
| NC_011770 | 1698228 | G | A | 1 | 10 | 2 | 0 | NON_SYNONYMOU | MODERAT | MISSENSE | Ggt/Agt | G542S | PLES_15721 | . | . | . | . | A | A | . | . | . | . | . | . |
| NC_011770 | 1698225 | G | A | 1 | 10 | 1 | 0 | NON_SYNONYMOU | MODERAT | MISSENSE | Gta/Ata | V541I | PLES_15721 | . | . | . | . | | A | . | . | . | . | . | . |
| NC_011770 | 1699495 | T | G | 19 | 10 | 2 | 0 | NON_SYNONYMOU | MODERAT | MISSENSE | aaT/aaG | N410K | PLES_15731 | . | . | . | . | G | G | . | . | . | . | . | . |
| NC_011770 | 1699512 | G | T | 19 | 10 | 2 | 0 | NON_SYNONYMOU | MODERAT | MISSENSE | gGc/gTc | G416V | PLES_15731 | . | . | . | . | T | T | . | . | . | . | . | . |
| NC_011770 | 1699520 | A | G | 20 | 10 | 2 | 0 | NON_SYNONYMOU | MODERAT | MISSENSE | Acc/Gcc | T419A | PLES_15731 | . | . | . | . | G | G | . | . | . | . | . | . |
| NC_011770 | 1699545 | G | A | 19 | 10 | 2 | 0 | NON_SYNONYMOU | MODERAT | MISSENSE | aGc/aAc | S427N | PLES_15731 | . | . | . | . | A | A | . | . | . | . | . | . |
| NC_011770 | 1699565 | A | G | 19 | 10 | 2 | 0 | NON_SYNONYMOU | MODERAT | MISSENSE | Atc/Gtc | I434V | PLES_15731 | . | . | . | . | G | G | . | . | . | . | . | . |
| NC_011770 | 1700474 | C | T | 19 | 10 | 2 | 0 | NON_SYNONYMOU | MODERAT | MISSENSE | gCc/gTc | A208V | PLES_15741 | . | . | . | . | T | T | . | . | . | . | . | . |
| NC_011770 | 1701742 | G | C | 18 | 10 | 2 | 0 | NON_SYNONYMOU | MODERAT | MISSENSE | Gaa/Caa | E9Q | PLES_15751 | . | . | . | . | C | C | . | . | . | . | . | . |
| NC_011770 | 1701831 | T | G | 19 | 10 | 2 | 0 | NON_SYNONYMOU | MODERAT | MISSENSE | gaT/gaG | D38E | PLES_15751 | . | . | . | . | G | G | . | . | . | . | . | . |
| NC_011770 | 1702550 | G | A | 17 | 10 | 2 | 0 | NON_SYNONYMOU | MODERAT | MISSENSE | aGc/aAc | S7N | PLES_15781 | . | . | . | . | A | A | . | . | . | . | . | . |
| NC_011770 | 1702636 | G | A | 21 | 10 | 2 | 0 | NON_SYNONYMOU | MODERAT | MISSENSE | Ggt/Agt | G36S | PLES_15781 | . | . | . | . | A | A | . | . | . | . | . | . |
| NC_011770 | 1702733 | G | A | 21 | 10 | 2 | 0 | NON_SYNONYMOU | MODERAT | MISSENSE | aGc/aAc | S68N | PLES_15781 | . | . | . | . | A | A | . | . | . | . | . | . |
| NC_011770 | 1703479 | A | C | 20 | 10 | 2 | 0 | NON_SYNONYMOU | MODERAT | MISSENSE | Aac/Cac | N170H | PLES_15791 | . | . | . | . | C | C | . | . | . | . | . | . |
| NC_011770 | 1703755 | A | G | 19 | 10 | 2 | 0 | NON_SYNONYMOU | MODERAT | MISSENSE | Acc/Gcc | T262A | PLES_15791 | . | . | . | . | G | G | . | . | . | . | . | . |
| NC_011770 | 1703780 | C | A | 19 | 10 | 2 | 0 | NON_SYNONYMOU | MODERAT | MISSENSE | gCg/gAg | A270E | PLES_15791 | . | . | . | . | A | A | . | . | . | . | . | . |
| NC_011770 | 1703794 | C | A | 20 | 10 | 2 | 0 | NON_SYNONYMOU | MODERAT | MISSENSE | Ctc/Atc | L275I | PLES_15791 | . | . | . | . | A | A | . | . | . | . | . | . |
| NC_011770 | 1703912 | G | A | 17 | 10 | 2 | 0 | NON_SYNONYMOU | MODERAT | MISSENSE | aGc/aAc | S314N | PLES_15791 | . | . | . | . | A | A | . | . | . | . | . | . |

| | | | | | | | | | | | | | | | | | | | | | | | | | |
|---|---|---|---|---|---|---|---|---|---|---|---|---|---|---|---|---|---|---|---|---|---|---|---|---|---|
| NC_011770 | 1704989 | G | A | 18 | 10 | 2 | 0 | NON_SYNONYMOU | MODERAT | MISSENSE | aGc/aAc | S159N | PLES_15811 | . | . | . | . | A | A | . | . | . | . | . | . |
| NC_011770 | 1705639 | T | A | 16 | 10 | 2 | 0 | NON_SYNONYMOU | MODERAT | MISSENSE | aTg/aAg | M69K | PLES_15821 | . | . | . | . | A | A | . | . | . | . | . | . |
| NC_011770 | 1707544 | C | T | 22 | 10 | 2 | 0 | NON_SYNONYMOU | MODERAT | MISSENSE | Ccg/Tcg | P157S | PLES_15851 | . | . | . | . | T | T | . | . | . | . | . | . |
| NC_011770 | 1708124 | C | A | 22 | 10 | 2 | 0 | NON_SYNONYMOU | MODERAT | MISSENSE | gCg/gAg | A92E | PLES_15861 | . | . | . | . | A | A | . | . | . | . | . | . |
| NC_011770 | 1709165 | C | G | 27 | 8 | 2 | 0 | NON_SYNONYMOU | MODERAT | MISSENSE | Ctg/Gtg | L174V | PLES_15871 | . | . | . | . | | | . | . | G | G | . | . |
| NC_011770 | 1709202 | A | C | 28 | 8 | 2 | 0 | NON_SYNONYMOU | MODERAT | MISSENSE | aAg/aCg | K186T | PLES_15871 | . | . | . | . | | | . | . | C | C | . | . |
| NC_011770 | 1709203 | G | C | 28 | 8 | 2 | 0 | NON_SYNONYMOU | MODERAT | MISSENSE | aaG/aaC | K186N | PLES_15871 | . | . | . | . | | | . | . | C | C | . | . |
| NC_011770 | 1709725 | A | T | 37 | 8 | 2 | 0 | NON_SYNONYMOU | MODERAT | MISSENSE | gaA/gaT | E360D | PLES_15871 | . | . | . | . | | | . | . | T | T | . | . |
| NC_011770 | 1709755 | G | C | 34 | 8 | 2 | 0 | NON_SYNONYMOU | MODERAT | MISSENSE | gaG/gaC | E370D | PLES_15871 | . | . | . | . | | | . | . | C | C | . | . |
| NC_011770 | 1709756 | A | C | 34 | 8 | 2 | 0 | NON_SYNONYMOU | MODERAT | MISSENSE | Aaa/Caa | K371Q | PLES_15871 | . | . | . | . | | | . | . | C | C | . | . |
| NC_011770 | 1709757 | A | G | 34 | 8 | 2 | 0 | NON_SYNONYMOU | MODERAT | MISSENSE | aAa/aGa | K371R | PLES_15871 | . | . | . | . | | | . | . | G | G | . | . |
| NC_011770 | 1709758 | A | C | 34 | 8 | 2 | 0 | NON_SYNONYMOU | MODERAT | MISSENSE | aaA/aaC | K371N | PLES_15871 | . | . | . | . | | | . | . | C | C | . | . |
| NC_011770 | 1709795 | G | A | 25 | 8 | 2 | 0 | NON_SYNONYMOU | MODERAT | MISSENSE | Gcc/Acc | A384T | PLES_15871 | . | . | . | . | | | . | . | A | A | . | . |
| NC_011770 | 1709806 | T | G | 24 | 8 | 2 | 0 | NON_SYNONYMOU | MODERAT | MISSENSE | gaT/gaG | D387E | PLES_15871 | . | . | . | . | | | . | . | G | G | . | . |
| NC_011770 | 1709807 | C | A | 25 | 8 | 2 | 0 | NON_SYNONYMOU | MODERAT | MISSENSE | Cag/Aag | Q388K | PLES_15871 | . | . | . | . | | | . | . | A | A | . | . |
| NC_011770 | 1709808 | A | G | 25 | 8 | 2 | 0 | NON_SYNONYMOU | MODERAT | MISSENSE | cAg/cGg | Q388R | PLES_15871 | . | . | . | . | | | . | . | G | G | . | . |
| NC_011770 | 1709819 | T | A | 23 | 8 | 2 | 0 | NON_SYNONYMOU | MODERAT | MISSENSE | Tcg/Acg | S392T | PLES_15871 | . | . | . | . | | | . | . | A | A | . | . |
| NC_011770 | 1709880 | G | A | 35 | 8 | 2 | 0 | NON_SYNONYMOU | MODERAT | MISSENSE | aGc/aAc | S412N | PLES_15871 | . | . | . | . | | | . | . | A | A | . | . |
| NC_011770 | 1710080 | T | C | 19 | 9 | 1 | 0 | NON_SYNONYMOU | MODERAT | MISSENSE | Ttc/Ctc | F479L | PLES_15871 | . | . | . | . | | | . | . | C | . | . | . |
| NC_011770 | 1710089 | C | G | 19 | 8 | 2 | 0 | NON_SYNONYMOU | MODERAT | MISSENSE | Cta/Gta | L482V | PLES_15871 | . | . | . | . | | | . | . | G | G | . | . |
| NC_011770 | 1710102 | A | T | 20 | 8 | 2 | 0 | NON_SYNONYMOU | MODERAT | MISSENSE | tAt/tTt | Y486F | PLES_15871 | . | . | . | . | | | . | . | T | T | . | . |
| NC_011770 | 1710111 | G | A | 37 | 8 | 2 | 0 | NON_SYNONYMOU | MODERAT | MISSENSE | aGc/aAc | S489N | PLES_15871 | . | . | . | . | | | . | . | A | A | . | . |
| NC_011770 | 1710127 | C | G | 37 | 8 | 2 | 0 | NON_SYNONYMOU | MODERAT | MISSENSE | gaC/gaG | D494E | PLES_15871 | . | . | . | . | | | . | . | G | G | . | . |
| NC_011770 | 1710437 | T | A | 35 | 8 | 2 | 0 | NON_SYNONYMOU | MODERAT | MISSENSE | Tct/Act | S598T | PLES_15871 | . | . | . | . | | | . | . | A | A | . | . |
| NC_011770 | 1710438 | C | G | 36 | 8 | 2 | 0 | NON_SYNONYMOU | MODERAT | MISSENSE | tCt/tGt | S598C | PLES_15871 | . | . | . | . | | | . | . | G | G | . | . |
| NC_011770 | 1710459 | G | C | 41 | 8 | 2 | 0 | NON_SYNONYMOU | MODERAT | MISSENSE | aGc/aCc | S605T | PLES_15871 | . | . | . | . | | | . | . | C | C | . | . |
| NC_011770 | 1710480 | G | A | 41 | 8 | 2 | 0 | NON_SYNONYMOU | MODERAT | MISSENSE | cGc/cAc | R612H | PLES_15871 | . | . | . | . | | | . | . | A | A | . | . |
| NC_011770 | 1710497 | A | G | 38 | 8 | 2 | 0 | NON_SYNONYMOU | MODERAT | MISSENSE | Acc/Gcc | T618A | PLES_15871 | . | . | . | . | | | . | . | G | G | . | . |
| NC_011770 | 1710504 | T | A | 39 | 8 | 2 | 0 | NON_SYNONYMOU | MODERAT | MISSENSE | tTc/tAc | F620Y | PLES_15871 | . | . | . | . | | | . | . | A | A | . | . |
| NC_011770 | 1710723 | C | A | 41 | 8 | 4 | 0 | NON_SYNONYMOU | MODERAT | MISSENSE | aCc/aAc | T693N | PLES_15871 | . | . | . | . | A | A | . | . | A | A | . | . |
| NC_011770 | 1711034 | A | C | 31 | 8 | 2 | 0 | NON_SYNONYMOU | MODERAT | MISSENSE | Aag/Cag | K797Q | PLES_15871 | . | . | . | . | | | . | . | C | C | . | . |
| NC_011770 | 1711037 | G | A | 31 | 8 | 2 | 0 | NON_SYNONYMOU | MODERAT | MISSENSE | Gcg/Acg | A798T | PLES_15871 | . | . | . | . | | | . | . | A | A | . | . |
| NC_011770 | 1711038 | C | A | 31 | 8 | 2 | 0 | NON_SYNONYMOU | MODERAT | MISSENSE | gCg/gAg | A798E | PLES_15871 | . | . | . | . | | | . | . | A | A | . | . |
| NC_011770 | 1711043 | G | C | 31 | 8 | 2 | 0 | NON_SYNONYMOU | MODERAT | MISSENSE | Gaa/Caa | E800Q | PLES_15871 | . | . | . | . | | | . | . | C | C | . | . |
| NC_011770 | 1711067 | C | G | 31 | 8 | 2 | 0 | NON_SYNONYMOU | MODERAT | MISSENSE | Cag/Gag | Q808E | PLES_15871 | . | . | . | . | | | . | . | G | G | . | . |
| NC_011770 | 1711166 | G | A | 32 | 8 | 2 | 0 | NON_SYNONYMOU | MODERAT | MISSENSE | Gac/Aac | D841N | PLES_15871 | . | . | . | . | | | . | . | A | A | . | . |
| NC_011770 | 1711189 | A | C | 30 | 8 | 2 | 0 | NON_SYNONYMOU | MODERAT | MISSENSE | gaA/gaC | E848D | PLES_15871 | . | . | . | . | | | . | . | C | C | . | . |
| NC_011770 | 1711686 | G | A | 24 | 10 | 2 | 0 | NON_SYNONYMOU | MODERAT | MISSENSE | cGg/cAg | R1014Q | PLES_15871 | . | . | . | . | A | A | . | . | . | . | . | . |
| NC_011770 | 1711889 | T | G | 34 | 8 | 4 | 0 | NON_SYNONYMOU | MODERAT | MISSENSE | Tcc/Gcc | S1082A | PLES_15871 | . | . | . | . | G | G | . | . | G | G | . | . |
| NC_011770 | 1711895 | G | T | 26 | 8 | 4 | 0 | NON_SYNONYMOU | MODERAT | MISSENSE | Gcg/Tcg | A1084S | PLES_15871 | . | . | . | . | T | T | . | . | T | T | . | . |
| NC_011770 | 1711898 | G | A | 25 | 8 | 4 | 0 | NON_SYNONYMOU | MODERAT | MISSENSE | Gtc/Atc | V1085I | PLES_15871 | . | . | . | . | A | A | . | . | A | A | . | . |
| NC_011770 | 1711901 | C | T | 26 | 8 | 4 | 0 | NON_SYNONYMOU | MODERAT | MISSENSE | Cac/Tac | H1086Y | PLES_15871 | . | . | . | . | T | T | . | . | T | T | . | . |
| NC_011770 | 1711922 | C | A | 25 | 8 | 4 | 0 | NON_SYNONYMOU | MODERAT | MISSENSE | Ccc/Acc | P1093T | PLES_15871 | . | . | . | . | A | A | . | . | A | A | . | . |
| NC_011770 | 1711931 | G | A | 26 | 8 | 4 | 0 | NON_SYNONYMOU | MODERAT | MISSENSE | Gtg/Atg | V1096M | PLES_15871 | . | . | . | . | A | A | . | . | A | A | . | . |
| NC_011770 | 1711966 | C | A | 30 | 8 | 4 | 0 | NON_SYNONYMOU | MODERAT | MISSENSE | gaC/gaA | D1107E | PLES_15871 | . | . | . | . | A | A | . | . | A | A | . | . |
| NC_011770 | 1711971 | A | T | 30 | 8 | 4 | 0 | NON_SYNONYMOU | MODERAT | MISSENSE | tAc/tTc | Y1109F | PLES_15871 | . | . | . | . | T | T | . | . | T | T | . | . |
| NC_011770 | 1711973 | A | C | 31 | 8 | 4 | 0 | NON_SYNONYMOU | MODERAT | MISSENSE | Atc/Ctc | I1110L | PLES_15871 | . | . | . | . | C | C | . | . | C | C | . | . |
| NC_011770 | 1711975 | C | G | 30 | 8 | 4 | 0 | NON_SYNONYMOU | MODERAT | MISSENSE | atC/atG | I1110M | PLES_15871 | . | . | . | . | G | G | . | . | G | G | . | . |
| NC_011770 | 1716022 | T | A | 47 | 8 | 2 | 0 | STOP_GAINED | HIGH | NONSENSE | tTg/tAg | L124* | PLES_15911 | . | . | . | . | A | A | . | . | | | . | . |
| NC_011770 | 1715761 | T | C | 49 | 8 | 2 | 0 | NON_SYNONYMOU | MODERAT | MISSENSE | aTc/aCc | I37T | PLES_15911 | . | . | . | . | C | C | . | . | | | . | . |
| NC_011770 | 1715808 | G | A | 48 | 8 | 2 | 0 | NON_SYNONYMOU | MODERAT | MISSENSE | Gtc/Atc | V53I | PLES_15911 | . | . | . | . | A | A | . | . | | | . | . |
| NC_011770 | 1715832 | G | A | 45 | 8 | 2 | 0 | NON_SYNONYMOU | MODERAT | MISSENSE | Gac/Aac | D61N | PLES_15911 | . | . | . | . | A | A | . | . | | | . | . |
| NC_011770 | 1715853 | T | G | 40 | 8 | 2 | 0 | NON_SYNONYMOU | MODERAT | MISSENSE | Tcc/Gcc | S68A | PLES_15911 | . | . | . | . | G | G | . | . | | | . | . |
| NC_011770 | 1715857 | G | A | 40 | 8 | 2 | 0 | NON_SYNONYMOU | MODERAT | MISSENSE | cGg/cAg | R69Q | PLES_15911 | . | . | . | . | A | A | . | . | | | . | . |
| NC_011770 | 1715869 | G | A | 43 | 8 | 2 | 0 | NON_SYNONYMOU | MODERAT | MISSENSE | gGt/gAt | G73D | PLES_15911 | . | . | . | . | A | A | . | . | | | . | . |
| NC_011770 | 1715910 | A | G | 45 | 8 | 2 | 0 | NON_SYNONYMOU | MODERAT | MISSENSE | Acc/Gcc | T87A | PLES_15911 | . | . | . | . | G | G | . | . | | | . | . |
| NC_011770 | 1715937 | C | G | 45 | 8 | 2 | 0 | NON_SYNONYMOU | MODERAT | MISSENSE | Cag/Gag | Q96E | PLES_15911 | . | . | . | . | G | G | . | . | | | . | . |
| NC_011770 | 1715973 | T | A | 47 | 8 | 2 | 0 | NON_SYNONYMOU | MODERAT | MISSENSE | Tgc/Agc | C108S | PLES_15911 | . | . | . | . | A | A | . | . | | | . | . |
| NC_011770 | 1716021 | T | A | 47 | 8 | 2 | 0 | NON_SYNONYMOU | MODERAT | MISSENSE | Ttg/Atg | L124M | PLES_15911 | . | . | . | . | A | A | . | . | | | . | . |
| NC_011770 | 1716054 | G | A | 48 | 8 | 2 | 0 | NON_SYNONYMOU | MODERAT | MISSENSE | Gtc/Atc | V135I | PLES_15911 | . | . | . | . | A | A | . | . | | | . | . |
| NC_011770 | 1716163 | T | C | 47 | 8 | 2 | 0 | NON_SYNONYMOU | MODERAT | MISSENSE | gTc/gCc | V171A | PLES_15911 | . | . | . | . | C | C | . | . | | | . | . |
| NC_011770 | 1716222 | A | C | 43 | 8 | 2 | 0 | NON_SYNONYMOU | MODERAT | MISSENSE | Agc/Cgc | S191R | PLES_15911 | . | . | . | . | C | C | . | . | | | . | . |

| | | | | | | | | | | | | | | | | | | | | | | | | | |
|---|---|---|---|---|---|---|---|---|---|---|---|---|---|---|---|---|---|---|---|---|---|---|---|---|---|
| NC_011770 | 1716229 | A | T | 45 | 8 | 2 | 0 | NON_SYNONYMOU | MODERAT | MISSENSE | tAc/tTc | Y193F | PLES_15911 | . | . | . | . | T | T | . | . | | | . | . |
| NC_011770 | 1716238 | G | A | 44 | 8 | 2 | 0 | NON_SYNONYMOU | MODERAT | MISSENSE | cGg/cAg | R196Q | PLES_15911 | . | . | . | . | A | A | . | . | | | . | . |
| NC_011770 | 1716249 | C | G | 42 | 8 | 2 | 0 | NON_SYNONYMOU | MODERAT | MISSENSE | Ccc/Gcc | P200A | PLES_15911 | . | . | . | . | G | G | . | . | | | . | . |
| NC_011770 | 1716258 | T | C | 42 | 8 | 2 | 0 | NON_SYNONYMOU | MODERAT | MISSENSE | Tac/Cac | Y203H | PLES_15911 | . | . | . | . | C | C | . | . | | | . | . |
| NC_011770 | 1716416 | C | G | 45 | 8 | 2 | 0 | NON_SYNONYMOU | MODERAT | MISSENSE | gaC/gaG | D255E | PLES_15911 | . | . | . | . | G | G | . | . | | | . | . |
| NC_011770 | 1716486 | TATCCAAG | T | 28 | 8 | 2 | 0 | FRAME_SHIFT | HIGH | | ctatccaag/ | LSK17 | PLES_15921 | . | . | . | . | T | T | . | . | | | . | . |
| NC_011770 | 1716471 | C | T | 28 | 8 | 2 | 0 | NON_SYNONYMOU | MODERAT | MISSENSE | gCg/gTg | A12V | PLES_15921 | . | . | . | . | T | T | . | . | | | . | . |
| NC_011770 | 1716480 | T | A | 28 | 8 | 2 | 0 | NON_SYNONYMOU | MODERAT | MISSENSE | tTc/tAc | F15Y | PLES_15921 | . | . | . | . | A | A | . | . | | | . | . |
| NC_011770 | 1716482 | A | G | 29 | 8 | 2 | 0 | NON_SYNONYMOU | MODERAT | MISSENSE | Atc/Gtc | I16V | PLES_15921 | . | . | . | . | G | G | . | . | | | . | . |
| NC_011770 | 1716509 | C | A | 44 | 8 | 2 | 0 | NON_SYNONYMOU | MODERAT | MISSENSE | Ccg/Acg | P25T | PLES_15921 | . | . | . | . | A | A | . | . | | | . | . |
| NC_011770 | 1716917 | G | C | 33 | 8 | 2 | 0 | NON_SYNONYMOU | MODERAT | MISSENSE | Gag/Cag | E13Q | PLES_15931 | . | . | . | . | C | C | . | . | | | . | . |
| NC_011770 | 1716918 | A | G | 33 | 8 | 2 | 0 | NON_SYNONYMOU | MODERAT | MISSENSE | gAg/gGg | E13G | PLES_15931 | . | . | . | . | G | G | . | . | | | . | . |
| NC_011770 | 1716919 | G | C | 33 | 8 | 2 | 0 | NON_SYNONYMOU | MODERAT | MISSENSE | gaG/gaC | E13D | PLES_15931 | . | . | . | . | C | C | . | . | | | . | . |
| NC_011770 | 1716924 | T | A | 33 | 8 | 2 | 0 | NON_SYNONYMOU | MODERAT | MISSENSE | tTt/tAt | F15Y | PLES_15931 | . | . | . | . | A | A | . | . | | | . | . |
| NC_011770 | 1716956 | A | G | 34 | 8 | 2 | 0 | NON_SYNONYMOU | MODERAT | MISSENSE | Atc/Gtc | I26V | PLES_15931 | . | . | . | . | G | G | . | . | | | . | . |
| NC_011770 | 1717046 | G | A | 35 | 8 | 2 | 0 | NON_SYNONYMOU | MODERAT | MISSENSE | Gac/Aac | D56N | PLES_15931 | . | . | . | . | A | A | . | . | | | . | . |
| NC_011770 | 1717061 | A | G | 35 | 8 | 2 | 0 | NON_SYNONYMOU | MODERAT | MISSENSE | Aag/Gag | K61E | PLES_15931 | . | . | . | . | G | G | . | . | | | . | . |
| NC_011770 | 1717063 | G | C | 35 | 8 | 2 | 0 | NON_SYNONYMOU | MODERAT | MISSENSE | aaG/aaC | K61N | PLES_15931 | . | . | . | . | C | C | . | . | | | . | . |
| NC_011770 | 1717493 | T | G | 33 | 8 | 2 | 0 | NON_SYNONYMOU | MODERAT | MISSENSE | Tca/Gca | S205A | PLES_15931 | . | . | . | . | G | G | . | . | | | . | . |
| NC_011770 | 1717593 | G | A | 37 | 8 | 2 | 0 | NON_SYNONYMOU | MODERAT | MISSENSE | aGc/aAc | S238N | PLES_15931 | . | . | . | . | A | A | . | . | | | . | . |
| NC_011770 | 1717929 | G | T | 38 | 8 | 2 | 0 | NON_SYNONYMOU | MODERAT | MISSENSE | cGc/cTc | R350L | PLES_15931 | . | . | . | . | | | . | . | T | T | . | . |
| NC_011770 | 1718088 | C | T | 45 | 8 | 2 | 0 | NON_SYNONYMOU | MODERAT | MISSENSE | aCc/aTc | T403I | PLES_15931 | . | . | . | . | | | . | . | T | T | . | . |
| NC_011770 | 1718114 | G | A | 45 | 8 | 2 | 0 | NON_SYNONYMOU | MODERAT | MISSENSE | Gtc/Atc | V412I | PLES_15931 | . | . | . | . | | | . | . | A | A | . | . |
| NC_011770 | 1718388 | C | G | 42 | 8 | 2 | 0 | NON_SYNONYMOU | MODERAT | MISSENSE | aCc/aGc | T503S | PLES_15931 | . | . | . | . | | | . | . | G | G | . | . |
| NC_011770 | 1718909 | A | T | 39 | 8 | 2 | 0 | NON_SYNONYMOU | MODERAT | MISSENSE | Agc/Tgc | S677C | PLES_15931 | . | . | . | . | | | . | . | T | T | . | . |
| NC_011770 | 1718910 | G | C | 39 | 8 | 2 | 0 | NON_SYNONYMOU | MODERAT | MISSENSE | aGc/aCc | S677T | PLES_15931 | . | . | . | . | | | . | . | C | C | . | . |
| NC_011770 | 1718916 | G | A | 40 | 8 | 2 | 0 | NON_SYNONYMOU | MODERAT | MISSENSE | aGt/aAt | S679N | PLES_15931 | . | . | . | . | | | . | . | A | A | . | . |
| NC_011770 | 1718942 | A | G | 39 | 8 | 2 | 0 | NON_SYNONYMOU | MODERAT | MISSENSE | Atg/Gtg | M688V | PLES_15931 | . | . | . | . | | | . | . | G | G | . | . |
| NC_011770 | 1718944 | G | C | 39 | 8 | 2 | 0 | NON_SYNONYMOU | MODERAT | MISSENSE | atG/atC | M688I | PLES_15931 | . | . | . | . | | | . | . | C | C | . | . |
| NC_011770 | 1718966 | A | G | 48 | 8 | 2 | 0 | NON_SYNONYMOU | MODERAT | MISSENSE | Atg/Gtg | M696V | PLES_15931 | . | . | . | . | | | . | . | G | G | . | . |
| NC_011770 | 1719002 | A | C | 46 | 8 | 2 | 0 | NON_SYNONYMOU | MODERAT | MISSENSE | Aac/Cac | N708H | PLES_15931 | . | . | . | . | | | . | . | C | C | . | . |
| NC_011770 | 1719004 | C | A | 46 | 8 | 2 | 0 | NON_SYNONYMOU | MODERAT | MISSENSE | aaC/aaA | N708K | PLES_15931 | . | . | . | . | | | . | . | A | A | . | . |
| NC_011770 | 1719034 | T | A | 46 | 6 | 4 | 0 | NON_SYNONYMOU | MODERAT | MISSENSE | gaT/gaA | D718E | PLES_15931 | A | A | . | . | | | . | . | A | A | . | . |
| NC_011770 | 1719067 | G | A | 49 | 6 | 4 | 0 | NON_SYNONYMOU | MODERAT | MISSENSE | gaG/gaC | E729D | PLES_15931 | A | A | . | . | | | . | . | C | C | . | . |
| NC_011770 | 1719429 | C | CG | 12 | 9 | 1 | 0 | FRAME_SHIFT | HIGH | | tgg/Gtgg | W116V? | PLES_15941 | . | . | . | . | | | . | . | . | CG | . | . |
| NC_011770 | 1719089 | G | C | 36 | 8 | 2 | 0 | NON_SYNONYMOU | MODERAT | MISSENSE | aGt/aCt | S2T | PLES_15941 | . | . | . | . | | | . | . | C | C | . | . |
| NC_011770 | 1719094 | A | T | 36 | 8 | 2 | 0 | NON_SYNONYMOU | MODERAT | MISSENSE | Acc/Tcc | T4S | PLES_15941 | . | . | . | . | | | . | . | T | T | . | . |
| NC_011770 | 1719095 | C | A | 37 | 8 | 2 | 0 | NON_SYNONYMOU | MODERAT | MISSENSE | aCc/aAc | T4N | PLES_15941 | . | . | . | . | | | . | . | A | A | . | . |
| NC_011770 | 1719130 | G | T | 28 | 8 | 2 | 0 | NON_SYNONYMOU | MODERAT | MISSENSE | Gca/Tca | A16S | PLES_15941 | . | . | . | . | | | . | . | T | T | . | . |
| NC_011770 | 1719131 | C | T | 28 | 8 | 2 | 0 | NON_SYNONYMOU | MODERAT | MISSENSE | gCa/gTa | A16V | PLES_15941 | . | . | . | . | | | . | . | T | T | . | . |
| NC_011770 | 1719152 | G | A | 44 | 6 | 2 | 0 | NON_SYNONYMOU | MODERAT | MISSENSE | aGc/aAc | S23N | PLES_15941 | A | A | . | . | | | . | . | | | . | . |
| NC_011770 | 1719156 | A | T | 43 | 6 | 4 | 0 | NON_SYNONYMOU | MODERAT | MISSENSE | gaA/gaT | E24D | PLES_15941 | T | T | . | . | | | . | . | T | T | . | . |
| NC_011770 | 1719170 | T | G | 46 | 6 | 4 | 0 | NON_SYNONYMOU | MODERAT | MISSENSE | tTg/tGg | L29W | PLES_15941 | G | G | . | . | | | . | . | G | G | . | . |
| NC_011770 | 1719190 | C | A | 46 | 6 | 2 | 0 | NON_SYNONYMOU | MODERAT | MISSENSE | Cgc/Agc | R36S | PLES_15941 | | | . | . | | | . | . | A | A | . | . |
| NC_011770 | 1719220 | A | G | 24 | 8 | 4 | 0 | NON_SYNONYMOU | MODERAT | MISSENSE | Aca/Gca | T46A | PLES_15941 | . | . | . | . | G | G | . | . | G | G | . | . |
| NC_011770 | 1719233 | C | G | 25 | 8 | 4 | 0 | NON_SYNONYMOU | MODERAT | MISSENSE | aCc/aGc | T50S | PLES_15941 | . | . | . | . | G | G | . | . | G | G | . | . |
| NC_011770 | 1719242 | C | T | 27 | 8 | 2 | 0 | NON_SYNONYMOU | MODERAT | MISSENSE | gCc/gTc | A53V | PLES_15941 | . | . | . | . | T | T | . | . | | | . | . |
| NC_011770 | 1719271 | T | A | 25 | 8 | 2 | 0 | NON_SYNONYMOU | MODERAT | MISSENSE | Tcc/Acc | S63T | PLES_15941 | . | . | . | . | A | A | . | . | | | . | . |
| NC_011770 | 1719272 | C | G | 26 | 8 | 2 | 0 | NON_SYNONYMOU | MODERAT | MISSENSE | tCc/tGc | S63C | PLES_15941 | . | . | . | . | G | G | . | . | | | . | . |
| NC_011770 | 1719277 | T | G | 26 | 8 | 2 | 0 | NON_SYNONYMOU | MODERAT | MISSENSE | Tcc/Gcc | S65A | PLES_15941 | . | . | . | . | G | G | . | . | | | . | . |
| NC_011770 | 1719292 | C | A | 27 | 8 | 2 | 0 | NON_SYNONYMOU | MODERAT | MISSENSE | Cta/Ata | L70I | PLES_15941 | . | . | . | . | A | A | . | . | | | . | . |
| NC_011770 | 1719320 | C | G | 27 | 8 | 2 | 0 | NON_SYNONYMOU | MODERAT | MISSENSE | aCc/aGc | T79S | PLES_15941 | . | . | . | . | G | G | . | . | | | . | . |
| NC_011770 | 1719325 | G | C | 28 | 8 | 2 | 0 | NON_SYNONYMOU | MODERAT | MISSENSE | Gca/Cca | A81P | PLES_15941 | . | . | . | . | C | C | . | . | | | . | . |
| NC_011770 | 1719382 | A | G | 31 | 8 | 2 | 0 | NON_SYNONYMOU | MODERAT | MISSENSE | Acg/Gcg | T100A | PLES_15941 | . | . | . | . | G | G | . | . | | | . | . |
| NC_011770 | 1719391 | A | G | 29 | 8 | 2 | 0 | NON_SYNONYMOU | MODERAT | MISSENSE | Agc/Ggc | S103G | PLES_15941 | . | . | . | . | G | G | . | . | | | . | . |
| NC_011770 | 1719392 | G | C | 30 | 8 | 2 | 0 | NON_SYNONYMOU | MODERAT | MISSENSE | aGc/aCc | S103T | PLES_15941 | . | . | . | . | C | C | . | . | | | . | . |
| NC_011770 | 1719424 | G | A | 12 | 10 | 2 | 0 | NON_SYNONYMOU | MODERAT | MISSENSE | Gat/Aat | D114N | PLES_15941 | . | . | . | . | A | A | . | . | . | . | . | . |
| NC_011770 | 1719425 | A | G | 12 | 9 | 2 | 0 | NON_SYNONYMOU | MODERAT | MISSENSE | gAt/gGt | D114G | PLES_15941 | . | . | . | . | G | G | . | . | . | | . | . |
| NC_011770 | 1719433 | A | G | 12 | 9 | 1 | 0 | NON_SYNONYMOU | MODERAT | MISSENSE | Acg/Gcg | T117A | PLES_15941 | . | . | . | . | | | . | . | . | G | . | . |
| NC_011770 | 1719454 | T | G | 38 | 6 | 2 | 0 | NON_SYNONYMOU | MODERAT | MISSENSE | Tcg/Gcg | S124A | PLES_15941 | | | . | . | G | G | . | . | | | . | . |
| NC_011770 | 1719499 | A | G | 29 | 8 | 4 | 0 | NON_SYNONYMOU | MODERAT | MISSENSE | Acg/Gcg | T139A | PLES_15941 | G | G | . | . | . | . | . | . | G | G | . | . |
| NC_011770 | 1719527 | G | T | 28 | 8 | 4 | 0 | NON_SYNONYMOU | MODERAT | MISSENSE | gGa/gTa | G148V | PLES_15941 | T | T | . | . | . | . | . | . | T | T | . | . |

| | | | | | | | | | | | | | | | | | | | | | | | | | |
|---|---|---|---|---|---|---|---|---|---|---|---|---|---|---|---|---|---|---|---|---|---|---|---|---|---|
| NC_011770 | 1719581 | A | T | 21 | 8 | 4 | 0 | NON_SYNONYMOU | MODERAT | MISSENSE | tAc/tTc | Y166F | PLES_15941 | T | T | . | . | . | . | . | . | T | T | . | . |
| NC_011770 | 1719590 | G | T | 21 | 8 | 4 | 0 | NON_SYNONYMOU | MODERAT | MISSENSE | aGt/aTt | S169I | PLES_15941 | T | T | . | . | . | . | . | . | T | T | . | . |
| NC_011770 | 1719622 | G | T | 26 | 8 | 2 | 0 | NON_SYNONYMOU | MODERAT | MISSENSE | Gct/Tct | A180S | PLES_15941 | T | T | . | . | . | . | . | . | | | . | . |
| NC_011770 | 1719634 | A | T | 27 | 8 | 4 | 0 | NON_SYNONYMOU | MODERAT | MISSENSE | Acg/Tcg | T184S | PLES_15941 | T | T | . | . | . | . | . | . | T | T | . | . |
| NC_011770 | 1719752 | T | C | 28 | 8 | 2 | 0 | NON_SYNONYMOU | MODERAT | MISSENSE | gTg/gCg | V223A | PLES_15941 | C | C | . | . | . | . | . | . | | | . | . |
| NC_011770 | 1719811 | G | T | 27 | 8 | 4 | 0 | NON_SYNONYMOU | MODERAT | MISSENSE | Gtg/Ttg | V243L | PLES_15941 | T | T | . | . | . | . | . | . | T | T | . | . |
| NC_011770 | 1719976 | G | T | 29 | 8 | 4 | 0 | NON_SYNONYMOU | MODERAT | MISSENSE | Gca/Tca | A298S | PLES_15941 | T | T | . | . | . | . | . | . | T | T | . | . |
| NC_011770 | 1720021 | C | T | 30 | 8 | 4 | 0 | NON_SYNONYMOU | MODERAT | MISSENSE | Cct/Tct | P313S | PLES_15941 | T | T | . | . | . | . | . | . | T | T | . | . |
| NC_011770 | 1720051 | G | A | 28 | 8 | 4 | 0 | NON_SYNONYMOU | MODERAT | MISSENSE | Gcc/Acc | A323T | PLES_15941 | A | A | . | . | . | . | . | . | A | A | . | . |
| NC_011770 | 1720052 | C | G | 29 | 8 | 4 | 0 | NON_SYNONYMOU | MODERAT | MISSENSE | gCc/gGc | A323G | PLES_15941 | G | G | . | . | . | . | . | . | G | G | . | . |
| NC_011770 | 1720117 | T | C | 27 | 8 | 2 | 0 | NON_SYNONYMOU | MODERAT | MISSENSE | Ttc/Ctc | F345L | PLES_15941 | | | . | . | . | . | . | . | C | C | . | . |
| NC_011770 | 1720129 | T | G | 26 | 8 | 2 | 0 | NON_SYNONYMOU | MODERAT | MISSENSE | Tcg/Gcg | S349A | PLES_15941 | | | . | . | . | . | . | . | G | G | . | . |
| NC_011770 | 1720138 | G | T | 23 | 8 | 2 | 0 | NON_SYNONYMOU | MODERAT | MISSENSE | Gcc/Tcc | A352S | PLES_15941 | | | . | . | . | . | . | . | T | T | . | . |
| NC_011770 | 1720141 | A | G | 23 | 8 | 2 | 0 | NON_SYNONYMOU | MODERAT | MISSENSE | Agt/Ggt | S353G | PLES_15941 | | | . | . | . | . | . | . | G | G | . | . |
| NC_011770 | 1720142 | G | T | 23 | 8 | 2 | 0 | NON_SYNONYMOU | MODERAT | MISSENSE | aGt/aTt | S353I | PLES_15941 | | | . | . | . | . | . | . | T | T | . | . |
| NC_011770 | 1720467 | G | T | 21 | 8 | 2 | 0 | STOP_GAINED | HIGH | NONSENSE | Gag/Tag | E79* | PLES_15951 | T | T | . | . | . | . | . | . | | | . | . |
| NC_011770 | 1720246 | A | G | 25 | 8 | 2 | 0 | NON_SYNONYMOU | MODERAT | MISSENSE | aAa/aGa | K5R | PLES_15951 | | | . | . | . | . | . | . | G | G | . | . |
| NC_011770 | 1720284 | G | T | 26 | 8 | 2 | 0 | NON_SYNONYMOU | MODERAT | MISSENSE | Gcc/Tcc | A18S | PLES_15951 | T | T | . | . | . | . | . | . | | | . | . |
| NC_011770 | 1720351 | A | C | 26 | 8 | 2 | 0 | NON_SYNONYMOU | MODERAT | MISSENSE | gAc/gCc | D40A | PLES_15951 | C | C | . | . | . | . | . | . | | | . | . |
| NC_011770 | 1720373 | G | T | 27 | 8 | 2 | 0 | NON_SYNONYMOU | MODERAT | MISSENSE | caG/caT | Q47H | PLES_15951 | | | . | . | . | . | . | . | T | T | . | . |
| NC_011770 | 1720398 | A | G | 26 | 8 | 2 | 0 | NON_SYNONYMOU | MODERAT | MISSENSE | Acc/Gcc | T56A | PLES_15951 | | | . | . | . | . | . | . | G | G | . | . |
| NC_011770 | 1720432 | T | A | 25 | 8 | 2 | 0 | NON_SYNONYMOU | MODERAT | MISSENSE | tTt/tAt | F67Y | PLES_15951 | A | A | . | . | . | . | . | . | | | . | . |
| NC_011770 | 1720468 | A | C | 22 | 8 | 4 | 0 | NON_SYNONYMOU | MODERAT | MISSENSE | gAg/gCg | E79A | PLES_15951 | C | C | . | . | . | . | . | . | C | C | . | . |
| NC_011770 | 1720489 | A | G | 5 | 8 | 2 | 0 | NON_SYNONYMOU | MODERAT | MISSENSE | cAg/cGg | Q86R | PLES_15951 | G | G | . | . | . | . | . | . | | | . | . |
| NC_011770 | 1720687 | A | AC | 18 | 8 | 1 | 0 | FRAME_SHIFT | HIGH | | gaa/gaCa | E28D? | PLES_15961 | . | . | . | . | | AC | . | . | | | . | . |
| NC_011770 | 1720690 | CG | C | 17 | 8 | 4 | 0 | FRAME_SHIFT | HIGH | | gcg/ | A29 | PLES_15961 | . | . | . | . | C | C | . | . | GG | GG | . | . |
| NC_011770 | 1720613 | G | T | 3 | 10 | 2 | 0 | NON_SYNONYMOU | MODERAT | MISSENSE | gaG/gaT | E3D | PLES_15961 | . | . | . | . | T | T | . | . | . | . | . | . |
| NC_011770 | 1720615 | T | C | 3 | 10 | 2 | 0 | NON_SYNONYMOU | MODERAT | MISSENSE | gTg/gCg | V4A | PLES_15961 | . | . | . | . | C | C | . | . | . | . | . | . |
| NC_011770 | 1720689 | G | A | 17 | 8 | 2 | 0 | NON_SYNONYMOU | MODERAT | MISSENSE | Gcg/Acg | A29T | PLES_15961 | . | . | . | . | | | . | . | A | A | . | . |
| NC_011770 | 1720716 | G | A | 19 | 8 | 2 | 0 | NON_SYNONYMOU | MODERAT | MISSENSE | Gag/Aag | E38K | PLES_15961 | . | . | . | . | | | . | . | A | A | . | . |
| NC_011770 | 1720718 | G | T | 19 | 8 | 2 | 0 | NON_SYNONYMOU | MODERAT | MISSENSE | gaG/gaT | E38D | PLES_15961 | . | . | . | . | T | T | . | . | | | . | . |
| NC_011770 | 1720746 | G | A | 18 | 8 | 4 | 0 | NON_SYNONYMOU | MODERAT | MISSENSE | Ggc/Agc | G48S | PLES_15961 | . | . | . | . | A | A | . | . | A | A | . | . |
| NC_011770 | 1720933 | T | C | 24 | 0 | 12 | 0 | SPLICE_SITE_REGI | LOW | SILENT | tAa/tGa | *77 | PLES_15971 | C | C | C | C | C | C | C | C | C | C | C | C |
| NC_011770 | 1723735 | C | T | 72 | 0 | 2 | 0 | STOP_GAINED | HIGH | NONSENSE | tGg/tAg | W79* | PLES_15981 | | | | | T | T | | | | | | |
| NC_011770 | 1721546 | C | T | 74 | 0 | 2 | 0 | NON_SYNONYMOU | MODERAT | MISSENSE | Gcc/Acc | A809T | PLES_15981 | | | | | | | T | T | | | | |
| NC_011770 | 1721614 | G | A | 79 | 0 | 4 | 0 | NON_SYNONYMOU | MODERAT | MISSENSE | cCg/cTg | P786L | PLES_15981 | | | A | A | A | A | | | | | | |
| NC_011770 | 1722038 | G | C | 84 | 0 | 4 | 0 | NON_SYNONYMOU | MODERAT | MISSENSE | Ctg/Gtg | L645V | PLES_15981 | C | C | | | | | C | C | | | | |
| NC_011770 | 1722094 | C | T | 83 | 0 | 4 | 0 | NON_SYNONYMOU | MODERAT | MISSENSE | cGg/cAg | R626Q | PLES_15981 | | | | | | | | | T | T | T | T |
| NC_011770 | 1722836 | C | T | 95 | 0 | 1 | 0 | NON_SYNONYMOU | MODERAT | MISSENSE | Ggc/Agc | G379S | PLES_15981 | | | | | | | T | | | | | |
| NC_011770 | 1723864 | G | T | 67 | 0 | 10 | 0 | NON_SYNONYMOU | MODERAT | MISSENSE | gCa/gAa | A36E | PLES_15981 | | | T | T | T | T | T | T | T | T | T | T |
| NC_011770 | 1724111 | T | C | 38 | 0 | 8 | 0 | NON_SYNONYMOU | MODERAT | MISSENSE | gAc/gGc | D398G | PLES_15991 | | | C | C | C | C | | | C | C | C | C |
| NC_011770 | 1725257 | A | G | 68 | 0 | 1 | 0 | NON_SYNONYMOU | MODERAT | MISSENSE | cTg/cCg | L16P | PLES_15991 | | G | | | | | | | | | | |
| NC_011770 | 1725890 | C | A | 81 | 0 | 2 | 0 | NON_SYNONYMOU | MODERAT | MISSENSE | Gcg/Tcg | A390S | PLES_16001 | | | | | | | A | A | | | | |
| NC_011770 | 1726172 | C | T | 76 | 0 | 2 | 0 | NON_SYNONYMOU | MODERAT | MISSENSE | Gtc/Atc | V296I | PLES_16001 | T | T | | | | | | | | | | |
| NC_011770 | 1726708 | G | A | 61 | 0 | 1 | 0 | NON_SYNONYMOU | MODERAT | MISSENSE | cCa/cTa | P117L | PLES_16001 | | | | | | | | | | | | A |
| NC_011770 | 1727181 | C | T | 66 | 0 | 2 | 0 | NON_SYNONYMOU | MODERAT | MISSENSE | Gac/Aac | D563N | PLES_16011 | T | T | | | | | | | | | | |
| NC_011770 | 1729012 | G | A | 55 | 0 | 2 | 0 | START_LOST | HIGH | MISSENSE | atG/atA | M1I | PLES_16021 | | | | | | | | | | | A | A |
| NC_011770 | 1730195 | GGC | G | 60 | 0 | 1 | 0 | FRAME_SHIFT | HIGH | | cgc/ | R65 | PLES_16031 | G | | | | | | | | | | | |
| NC_011770 | 1734079 | A | G | 75 | 0 | 6 | 0 | NON_SYNONYMOU | MODERAT | MISSENSE | aAg/aGg | K494R | PLES_16051 | | | G | G | G | G | G | G | | | | |
| NC_011770 | 1735316 | G | A | 59 | 0 | 4 | 0 | NON_SYNONYMOU | MODERAT | MISSENSE | Gtc/Atc | V359I | PLES_16061 | | | A | A | A | A | | | | | | |
| NC_011770 | 1735643 | C | T | 57 | 0 | 2 | 0 | NON_SYNONYMOU | MODERAT | MISSENSE | gCg/gTg | A30V | PLES_16071 | T | T | | | | | | | | | | |
| NC_011770 | 1735930 | C | T | 63 | 0 | 1 | 0 | NON_SYNONYMOU | MODERAT | MISSENSE | Cgc/Tgc | R126C | PLES_16071 | | | | | | | T | | | | | |
| NC_011770 | 1736122 | G | A | 57 | 0 | 2 | 0 | NON_SYNONYMOU | MODERAT | MISSENSE | Gcc/Acc | A190T | PLES_16071 | A | A | | | | | | | | | | |
| NC_011770 | 1738222 | C | T | 65 | 0 | 4 | 0 | NON_SYNONYMOU | MODERAT | MISSENSE | cGc/cAc | R69H | PLES_16091 | | | | | | | | | T | T | T | T |
| NC_011770 | 1738236 | G | C | 65 | 0 | 6 | 0 | NON_SYNONYMOU | MODERAT | MISSENSE | caC/caG | H64Q | PLES_16091 | C | C | | | | | | | C | C | C | C |
| NC_011770 | 1739021 | A | G | 64 | 0 | 4 | 0 | NON_SYNONYMOU | MODERAT | MISSENSE | aAc/aGc | N130S | PLES_16101 | G | G | | | | | | | G | G | | |
| NC_011770 | 1739543 | T | G | 36 | 0 | 4 | 0 | NON_SYNONYMOU | MODERAT | MISSENSE | Tgc/Ggc | C8G | PLES_16111 | | | G | G | G | G | | | | | | |
| NC_011770 | 1739601 | C | T | 32 | 0 | 2 | 0 | NON_SYNONYMOU | MODERAT | MISSENSE | gCc/gTc | A27V | PLES_16111 | | | | | | | | | T | T | | |
| NC_011770 | 1740596 | A | G | 62 | 0 | 12 | 0 | NON_SYNONYMOU | MODERAT | MISSENSE | Acc/Gcc | T24A | PLES_16121 | G | G | G | G | G | G | G | G | G | G | G | G |
| NC_011770 | 1741451 | TC | T | 71 | 0 | 12 | 0 | FRAME_SHIFT | HIGH | | tcg/ | S22 | PLES_16131 | T | T | T | T | T | T | T | T | T | T | T | T |
| NC_011770 | 1741521 | A | T | 77 | 0 | 4 | 0 | NON_SYNONYMOU | MODERAT | MISSENSE | cAg/cTg | Q45L | PLES_16131 | | | T | T | T | T | | | | | | |
| NC_011770 | 1742036 | G | C | 80 | 0 | 4 | 0 | NON_SYNONYMOU | MODERAT | MISSENSE | Gtg/Ctg | V217L | PLES_16131 | | | C | C | C | C | | | | | | |

| | | | | | | | | | | | | | | | | | | | | | | | | | |
|---|---|---|---|---|---|---|---|---|---|---|---|---|---|---|---|---|---|---|---|---|---|---|---|---|---|
| NC_011770 | 1742093 | T | C | 76 | 0 | 12 | 0 | NON_SYNONYMOU | MODERAT | MISSENSE | Ttc/Ctc | F236L | PLES_16131 | C | C | C | C | C | C | C | C | C | C | C | C |
| NC_011770 | 1742607 | A | G | 58 | 0 | 1 | 0 | NON_SYNONYMOU | MODERAT | MISSENSE | tAt/tGt | Y119C | PLES_16141 | | | | | | | G | | | | | |
| NC_011770 | 1743678 | G | C | 71 | 0 | 6 | 0 | NON_SYNONYMOU | MODERAT | MISSENSE | Gag/Cag | E239Q | PLES_16151 | C | C | | | | | | | C | C | C | C |
| NC_011770 | 1743890 | C | G | 73 | 0 | 4 | 0 | NON_SYNONYMOU | MODERAT | MISSENSE | gaC/gaG | D309E | PLES_16151 | G | G | | | | | | | G | G | | |
| NC_011770 | 1745329 | G | A | 68 | 0 | 2 | 0 | NON_SYNONYMOU | MODERAT | MISSENSE | Gcg/Acg | A50T | PLES_16171 | | | | | | | | | | | A | A |
| NC_011770 | 1749685 | C | T | 77 | 0 | 1 | 0 | NON_SYNONYMOU | MODERAT | MISSENSE | Ggc/Agc | G10S | PLES_16241 | | T | | | | | | | | | | |
| NC_011770 | 1751063 | C | T | 52 | 0 | 4 | 0 | NON_SYNONYMOU | MODERAT | MISSENSE | aCc/aTc | T135I | PLES_16261 | | | T | T | T | T | | | | | | |
| NC_011770 | 1751321 | C | T | 51 | 0 | 2 | 0 | NON_SYNONYMOU | MODERAT | MISSENSE | Gcg/Acg | A280T | PLES_16271 | | | | | | | | | | | T | T |
| NC_011770 | 1751386 | C | T | 60 | 0 | 2 | 0 | NON_SYNONYMOU | MODERAT | MISSENSE | aGg/aAg | R258K | PLES_16271 | T | T | | | | | | | | | | |
| NC_011770 | 1752504 | T | C | 49 | 0 | 1 | 0 | NON_SYNONYMOU | MODERAT | MISSENSE | cTc/cCc | L84P | PLES_16281 | | | | | | | | C | | | | |
| NC_011770 | 1753299 | G | A | 70 | 0 | 4 | 0 | NON_SYNONYMOU | MODERAT | MISSENSE | Gtc/Atc | V224I | PLES_16291 | | | A | A | A | A | | | | | | |
| NC_011770 | 1753943 | G | A | 93 | 0 | 8 | 0 | NON_SYNONYMOU | MODERAT | MISSENSE | Gcc/Acc | A167T | PLES_16301 | | | A | A | A | A | | | A | A | A | A |
| NC_011770 | 1754521 | A | G | 88 | 0 | 2 | 0 | NON_SYNONYMOU | MODERAT | MISSENSE | Atc/Gtc | I101V | PLES_16311 | | | | | | | | | | | G | G |
| NC_011770 | 1755244 | C | T | 71 | 0 | 10 | 0 | NON_SYNONYMOU | MODERAT | MISSENSE | gCg/gTg | A10V | PLES_16321 | T | T | T | T | T | T | | | T | T | T | T |
| NC_011770 | 1755405 | C | T | 73 | 0 | 4 | 0 | NON_SYNONYMOU | MODERAT | MISSENSE | Ctc/Ttc | L64F | PLES_16321 | T | T | | | | | | | T | T | | |
| NC_011770 | 1755645 | C | A | 81 | 0 | 2 | 0 | NON_SYNONYMOU | MODERAT | MISSENSE | ttC/ttA | F7L | PLES_16331 | A | A | | | | | | | | | | |
| NC_011770 | 1755863 | G | A | 74 | 0 | 2 | 0 | NON_SYNONYMOU | MODERAT | MISSENSE | gGc/gAc | G80D | PLES_16331 | A | A | | | | | | | | | | |
| NC_011770 | 1756490 | A | G | 63 | 0 | 1 | 0 | NON_SYNONYMOU | MODERAT | MISSENSE | gAg/gGg | E289G | PLES_16331 | | | | | | | G | | | | | |
| NC_011770 | 1756523 | A | G | 63 | 0 | 4 | 0 | NON_SYNONYMOU | MODERAT | MISSENSE | aAg/aGg | K300R | PLES_16331 | | | G | G | G | G | | | | | | |
| NC_011770 | 1758135 | C | T | 62 | 0 | 2 | 0 | NON_SYNONYMOU | MODERAT | MISSENSE | cGc/cAc | R365H | PLES_16361 | | | | | | | | | | | T | T |
| NC_011770 | 1759906 | C | T | 72 | 0 | 1 | 0 | NON_SYNONYMOU | MODERAT | MISSENSE | Gag/Aag | E53K | PLES_16371 | | | | | | | | T | | | | |
| NC_011770 | 1761862 | G | A | 70 | 0 | 12 | 0 | NON_SYNONYMOU | MODERAT | MISSENSE | Gtc/Atc | V506I | PLES_16381 | A | A | A | A | A | A | A | A | A | A | A | A |
| NC_011770 | 1761080 | A | G | 77 | 0 | 1 | 0 | NON_SYNONYMOU | MODERAT | MISSENSE | gAc/gGc | D245G | PLES_16381 | | | | | | | | G | | | | |
| NC_011770 | 1762561 | G | A | 79 | 0 | 6 | 0 | NON_SYNONYMOU | MODERAT | MISSENSE | aGg/aAg | R134K | PLES_16391 | A | A | | | | | A | A | | | A | A |
| NC_011770 | 1763142 | G | A | 72 | 0 | 2 | 0 | NON_SYNONYMOU | MODERAT | MISSENSE | Ggc/Agc | G328S | PLES_16391 | A | A | | | | | | | | | | |
| NC_011770 | 1762426 | A | G | 68 | 0 | 1 | 0 | NON_SYNONYMOU | MODERAT | MISSENSE | aAc/aGc | N89S | PLES_16391 | | G | | | | | | | | | | |
| NC_011770 | 1763893 | A | C | 36 | 0 | 2 | 0 | NON_SYNONYMOU | MODERAT | MISSENSE | gaA/gaC | E71D | PLES_16401 | | | | | | | C | C | | | | |
| NC_011770 | 1763945 | C | G | 31 | 0 | 4 | 0 | NON_SYNONYMOU | MODERAT | MISSENSE | Ccc/Gcc | P89A | PLES_16401 | G | G | | | | | G | G | | | | |
| NC_011770 | 1763953 | C | A | 31 | 0 | 6 | 0 | NON_SYNONYMOU | MODERAT | MISSENSE | gaC/gaA | D91E | PLES_16401 | | | A | A | A | A | | | A | A | | |
| NC_011770 | 1764248 | A | G | 40 | 0 | 2 | 0 | NON_SYNONYMOU | MODERAT | MISSENSE | Acg/Gcg | T190A | PLES_16401 | | | | | | | | | G | G | | |
| NC_011770 | 1764251 | G | A | 42 | 0 | 2 | 0 | NON_SYNONYMOU | MODERAT | MISSENSE | Gcc/Acc | A191T | PLES_16401 | | | | | | | | | A | A | | |
| NC_011770 | 1764257 | G | A | 39 | 0 | 8 | 0 | NON_SYNONYMOU | MODERAT | MISSENSE | Gca/Aca | A193T | PLES_16401 | | | A | A | A | A | A | A | | | A | A |
| NC_011770 | 1764515 | A | G | 49 | 0 | 10 | 0 | NON_SYNONYMOU | MODERAT | MISSENSE | Acg/Gcg | T279A | PLES_16401 | | | G | G | G | G | G | G | G | G | G | G |
| NC_011770 | 1764818 | A | G | 63 | 0 | 2 | 0 | NON_SYNONYMOU | MODERAT | MISSENSE | Acg/Gcg | T380A | PLES_16401 | | | | | | | G | G | | | | |
| NC_011770 | 1765069 | G | A | 72 | 0 | 2 | 0 | NON_SYNONYMOU | MODERAT | MISSENSE | atG/atA | M463I | PLES_16401 | | | | | | | | | A | A | | |
| NC_011770 | 1765395 | C | T | 83 | 0 | 4 | 0 | NON_SYNONYMOU | MODERAT | MISSENSE | gCg/gTg | A572V | PLES_16401 | | | T | T | T | T | | | | | | |
| NC_011770 | 1766188 | C | T | 69 | 0 | 4 | 0 | NON_SYNONYMOU | MODERAT | MISSENSE | Ggc/Agc | G266S | PLES_16411 | T | T | | | | | | | | | T | T |
| NC_011770 | 1766536 | T | G | 66 | 0 | 6 | 0 | NON_SYNONYMOU | MODERAT | MISSENSE | Acc/Ccc | T150P | PLES_16411 | G | G | | | | | | | G | G | G | G |
| NC_011770 | 1766539 | G | T | 67 | 0 | 6 | 0 | NON_SYNONYMOU | MODERAT | MISSENSE | Ccc/Acc | P149T | PLES_16411 | T | T | | | | | | | T | T | T | T |
| NC_011770 | 1766926 | C | T | 61 | 0 | 2 | 0 | NON_SYNONYMOU | MODERAT | MISSENSE | Gag/Aag | E20K | PLES_16411 | | | | | | | T | T | | | | |
| NC_011770 | 1768525 | A | G | 79 | 0 | 1 | 0 | NON_SYNONYMOU | MODERAT | MISSENSE | Acc/Gcc | T44A | PLES_16431 | | | | | | | G | | | | | |
| NC_011770 | 1769128 | G | A | 85 | 0 | 1 | 0 | NON_SYNONYMOU | MODERAT | MISSENSE | Ggc/Agc | G245S | PLES_16431 | | | | | | | A | | | | | |
| NC_011770 | 1770060 | C | T | 81 | 0 | 4 | 0 | NON_SYNONYMOU | MODERAT | MISSENSE | gCg/gTg | A192V | PLES_16441 | | | T | T | T | T | | | | | | |
| NC_011770 | 1770105 | A | G | 82 | 0 | 12 | 0 | NON_SYNONYMOU | MODERAT | MISSENSE | cAc/cGc | H207R | PLES_16441 | G | G | G | G | G | G | G | G | G | G | G | G |
| NC_011770 | 1770422 | C | G | 75 | 0 | 8 | 0 | NON_SYNONYMOU | MODERAT | MISSENSE | Ccg/Gcg | P313A | PLES_16441 | G | G | | | | | G | G | G | G | G | G |
| NC_011770 | 1770941 | C | G | 73 | 0 | 6 | 0 | NON_SYNONYMOU | MODERAT | MISSENSE | gCc/gGc | A146G | PLES_16451 | G | G | G | G | G | G | | | | | | |
| NC_011770 | 1771504 | A | G | 66 | 0 | 6 | 0 | NON_SYNONYMOU | MODERAT | MISSENSE | Atc/Gtc | I334V | PLES_16451 | G | G | G | G | G | G | | | | | | |
| NC_011770 | 1771628 | C | T | 56 | 0 | 2 | 0 | NON_SYNONYMOU | MODERAT | MISSENSE | cGc/cAc | R130H | PLES_16461 | | | | | | | T | T | | | | |
| NC_011770 | 1772553 | C | T | 34 | 0 | 2 | 0 | NON_SYNONYMOU | MODERAT | MISSENSE | Gcc/Acc | A48T | PLES_16471 | | | | | | | T | T | | | | |
| NC_011770 | 1773604 | T | C | 63 | 0 | 2 | 0 | NON_SYNONYMOU | MODERAT | MISSENSE | Ttc/Ctc | F4L | PLES_16501 | | | | | | | C | C | | | | |
| NC_011770 | 1773766 | C | T | 65 | 0 | 2 | 0 | NON_SYNONYMOU | MODERAT | MISSENSE | Ctt/Ttt | L58F | PLES_16501 | | | | | | | T | T | | | | |
| NC_011770 | 1774028 | A | G | 54 | 0 | 10 | 0 | NON_SYNONYMOU | MODERAT | MISSENSE | Acc/Gcc | T47A | PLES_16511 | | | G | G | G | G | G | G | G | G | G | G |
| NC_011770 | 1774040 | A | C | 54 | 0 | 12 | 0 | NON_SYNONYMOU | MODERAT | MISSENSE | Aac/Cac | N51H | PLES_16511 | C | C | C | C | C | C | C | C | C | C | C | C |
| NC_011770 | 1774328 | A | G | 50 | 0 | 4 | 0 | NON_SYNONYMOU | MODERAT | MISSENSE | Atc/Gtc | I147V | PLES_16511 | | | | | | | | | G | G | G | G |
| NC_011770 | 1774314 | C | T | 51 | 0 | 1 | 0 | NON_SYNONYMOU | MODERAT | MISSENSE | gCg/gTg | A142V | PLES_16511 | | | | | | | | T | | | | |
| NC_011770 | 1774507 | G | A | 49 | 0 | 12 | 0 | NON_SYNONYMOU | MODERAT | MISSENSE | cGg/cAg | R13Q | PLES_16521 | A | A | A | A | A | A | A | A | A | A | A | A |
| NC_011770 | 1774603 | C | G | 42 | 0 | 2 | 0 | NON_SYNONYMOU | MODERAT | MISSENSE | cCg/cGg | P45R | PLES_16521 | | | | | | | G | G | | | | |
| NC_011770 | 1774690 | G | A | 38 | 0 | 8 | 0 | NON_SYNONYMOU | MODERAT | MISSENSE | aGc/aAc | S74N | PLES_16521 | | | A | A | A | A | | | A | A | A | A |
| NC_011770 | 1774711 | G | T | 39 | 0 | 2 | 0 | NON_SYNONYMOU | MODERAT | MISSENSE | gGg/gTg | G81V | PLES_16521 | T | T | | | | | | | | | | |
| NC_011770 | 1774780 | G | A | 40 | 0 | 10 | 0 | NON_SYNONYMOU | MODERAT | MISSENSE | aGc/aAc | S104N | PLES_16521 | | | A | A | A | A | A | A | A | A | A | A |
| NC_011770 | 1774801 | C | G | 43 | 0 | 10 | 0 | NON_SYNONYMOU | MODERAT | MISSENSE | aCc/aGc | T111S | PLES_16521 | | | G | G | G | G | G | G | G | G | G | G |
| NC_011770 | 1774818 | T | G | 43 | 0 | 8 | 0 | NON_SYNONYMOU | MODERAT | MISSENSE | Tct/Gct | S117A | PLES_16521 | | | G | G | G | G | | | G | G | G | G |

| | | | | | | | | | | | | | | | | | | | | | | | | | |
|---|---|---|---|---|---|---|---|---|---|---|---|---|---|---|---|---|---|---|---|---|---|---|---|---|---|
| NC_011770 | 1775060 | C | G | 39 | 0 | 8 | 0 | NON_SYNONYMOU | MODERAT | MISSENSE | gaC/gaG | D197E | PLES_16521 | | | G | G | G | G | | | G | G | G | G |
| NC_011770 | 1782567 | G | A | 61 | 0 | 2 | 0 | NON_SYNONYMOU | MODERAT | MISSENSE | cGc/cAc | R64H | PLES_16571 | | | | | A | A | | | | | | |
| NC_011770 | 1783142 | C | T | 71 | 0 | 6 | 0 | NON_SYNONYMOU | MODERAT | MISSENSE | Cat/Tat | H256Y | PLES_16571 | | | T | T | T | T | | | T | T | | |
| NC_011770 | 1783615 | C | G | 80 | 0 | 2 | 0 | NON_SYNONYMOU | MODERAT | MISSENSE | aaC/aaG | N413K | PLES_16571 | G | G | | | | | | | | | | |
| NC_011770 | 1783715 | A | G | 78 | 0 | 6 | 0 | NON_SYNONYMOU | MODERAT | MISSENSE | Acc/Gcc | T447A | PLES_16571 | | | G | G | G | G | | | G | G | | |
| NC_011770 | 1783458 | C | T | 80 | 0 | 1 | 0 | NON_SYNONYMOU | MODERAT | MISSENSE | gCc/gTc | A361V | PLES_16571 | | | | | | | | T | | | | |
| NC_011770 | 1785574 | G | A | 94 | 0 | 4 | 0 | NON_SYNONYMOU | MODERAT | MISSENSE | Ggc/Agc | G264S | PLES_16591 | | | A | A | A | A | | | | | | |
| NC_011770 | 1785943 | G | A | 89 | 0 | 12 | 0 | NON_SYNONYMOU | MODERAT | MISSENSE | Gtc/Atc | V62I | PLES_16601 | A | A | A | A | A | A | A | A | A | A | A | A |
| NC_011770 | 1785987 | C | A | 90 | 0 | 12 | 0 | NON_SYNONYMOU | MODERAT | MISSENSE | gaC/gaA | D76E | PLES_16601 | A | A | A | A | A | A | A | A | A | A | A | A |
| NC_011770 | 1787083 | A | G | 95 | 0 | 1 | 0 | NON_SYNONYMOU | MODERAT | MISSENSE | cAc/cGc | H52R | PLES_16611 | | | | | | | G | | | | | |
| NC_011770 | 1787400 | G | T | 99 | 0 | 2 | 0 | NON_SYNONYMOU | MODERAT | MISSENSE | Gtg/Ttg | V158L | PLES_16611 | | | | | | | | | | | T | T |
| NC_011770 | 1787563 | A | G | 91 | 0 | 1 | 0 | NON_SYNONYMOU | MODERAT | MISSENSE | tAc/tGc | Y212C | PLES_16611 | | | | | | | G | | | | | |
| NC_011770 | 1798522 | T | G | 50 | 0 | 2 | 0 | NON_SYNONYMOU | MODERAT | MISSENSE | caA/caC | Q102H | PLES_16701 | G | G | | | | | | | | | | |
| NC_011770 | 1799762 | A | G | 56 | 0 | 2 | 0 | NON_SYNONYMOU | MODERAT | MISSENSE | gTg/gCg | V44A | PLES_16721 | G | G | | | | | | | | | | |
| NC_011770 | 1800824 | C | T | 59 | 0 | 2 | 0 | NON_SYNONYMOU | MODERAT | MISSENSE | Ggt/Agt | G119S | PLES_16741 | | | | | | | | | T | T | | |
| NC_011770 | 1800830 | G | A | 60 | 0 | 2 | 0 | NON_SYNONYMOU | MODERAT | MISSENSE | Cgc/Tgc | R117C | PLES_16741 | | | | | | | | | A | A | | |
| NC_011770 | 1800835 | A | C | 61 | 0 | 4 | 0 | NON_SYNONYMOU | MODERAT | MISSENSE | tTg/tGg | L115W | PLES_16741 | C | C | | | | | | | C | C | | |
| NC_011770 | 1800850 | G | A | 60 | 0 | 8 | 0 | NON_SYNONYMOU | MODERAT | MISSENSE | gCc/gTc | A110V | PLES_16741 | | | A | A | A | A | A | A | | | A | A |
| NC_011770 | 1800863 | G | C | 61 | 0 | 4 | 0 | NON_SYNONYMOU | MODERAT | MISSENSE | Ctc/Gtc | L106V | PLES_16741 | C | C | | | | | | | C | C | | |
| NC_011770 | 1800899 | C | T | 60 | 0 | 2 | 0 | NON_SYNONYMOU | MODERAT | MISSENSE | Gcg/Acg | A94T | PLES_16741 | T | T | | | | | | | | | | |
| NC_011770 | 1800923 | T | G | 61 | 0 | 4 | 0 | NON_SYNONYMOU | MODERAT | MISSENSE | Atc/Ctc | I86L | PLES_16741 | | | G | G | G | G | | | | | | |
| NC_011770 | 1802935 | G | A | 67 | 0 | 2 | 0 | NON_SYNONYMOU | MODERAT | MISSENSE | Gcg/Acg | A10T | PLES_16771 | A | A | | | | | | | | | | |
| NC_011770 | 1803275 | A | G | 78 | 0 | 2 | 0 | NON_SYNONYMOU | MODERAT | MISSENSE | aAc/aGc | N123S | PLES_16771 | | | | | | | | | | | G | G |
| NC_011770 | 1803281 | C | A | 77 | 0 | 12 | 0 | NON_SYNONYMOU | MODERAT | MISSENSE | gCc/gAc | A125D | PLES_16771 | A | A | A | A | A | A | A | A | A | A | A | A |
| NC_011770 | 1803298 | C | G | 79 | 0 | 2 | 0 | NON_SYNONYMOU | MODERAT | MISSENSE | Ctc/Gtc | L131V | PLES_16771 | G | G | | | | | | | | | | |
| NC_011770 | 1803787 | G | A | 75 | 0 | 2 | 0 | NON_SYNONYMOU | MODERAT | MISSENSE | Gcg/Acg | A294T | PLES_16771 | | | | | | | | | | | A | A |
| NC_011770 | 1803797 | C | T | 74 | 0 | 2 | 0 | NON_SYNONYMOU | MODERAT | MISSENSE | gCc/gTc | A297V | PLES_16771 | | | | | | | | | | | T | T |
| NC_011770 | 1812794 | C | G | 80 | 0 | 2 | 0 | NON_SYNONYMOU | MODERAT | MISSENSE | gaG/gaC | E25D | PLES_16891 | | | | | | | | | | | G | G |
| NC_011770 | 1812937 | C | T | 84 | 0 | 2 | 0 | NON_SYNONYMOU | MODERAT | MISSENSE | Gaa/Aaa | E44K | PLES_16901 | T | T | | | | | | | | | | |
| NC_011770 | 1813636 | A | T | 81 | 0 | 2 | 0 | NON_SYNONYMOU | MODERAT | MISSENSE | caT/caA | H219Q | PLES_16921 | | | | | | | | | | | T | T |
| NC_011770 | 1813982 | G | A | 74 | 0 | 2 | 0 | NON_SYNONYMOU | MODERAT | MISSENSE | cCg/cTg | P104L | PLES_16921 | | | | | | | | | | | A | A |
| NC_011770 | 1814193 | C | T | 65 | 0 | 10 | 0 | NON_SYNONYMOU | MODERAT | MISSENSE | Gct/Act | A34T | PLES_16921 | T | T | T | T | T | T | | | T | T | T | T |
| NC_011770 | 1815651 | A | G | 12 | 10 | 2 | 0 | NON_SYNONYMOU | MODERAT | MISSENSE | Agc/Ggc | S9G | PLES_16931 | . | . | . | . | . | . | G | G | . | . | . | . |
| NC_011770 | 1815666 | C | A | 13 | 10 | 2 | 0 | NON_SYNONYMOU | MODERAT | MISSENSE | Ctt/Att | L14I | PLES_16931 | . | . | . | . | . | . | A | A | . | . | . | . |
| NC_011770 | 1815684 | A | G | 11 | 10 | 2 | 0 | NON_SYNONYMOU | MODERAT | MISSENSE | Agc/Ggc | S20G | PLES_16931 | . | . | . | . | . | . | G | G | . | . | . | . |
| NC_011770 | 1815685 | G | A | 11 | 10 | 2 | 0 | NON_SYNONYMOU | MODERAT | MISSENSE | aGc/aAc | S20N | PLES_16931 | . | . | . | . | . | . | A | A | . | . | . | . |
| NC_011770 | 1815689 | A | G | 11 | 10 | 2 | 0 | NON_SYNONYMOU | MODERAT | MISSENSE | atA/atG | I21M | PLES_16931 | . | . | . | . | . | . | G | G | . | . | . | . |
| NC_011770 | 1815690 | C | G | 11 | 10 | 2 | 0 | NON_SYNONYMOU | MODERAT | MISSENSE | Cca/Gca | P22A | PLES_16931 | . | . | . | . | . | . | G | G | . | . | . | . |
| NC_011770 | 1815691 | C | G | 11 | 10 | 2 | 0 | NON_SYNONYMOU | MODERAT | MISSENSE | cCa/cGa | P22R | PLES_16931 | . | . | . | . | . | . | G | G | . | . | . | . |
| NC_011770 | 1815696 | G | C | 11 | 10 | 2 | 0 | NON_SYNONYMOU | MODERAT | MISSENSE | Gaa/Caa | E24Q | PLES_16931 | . | . | . | . | . | . | C | C | . | . | . | . |
| NC_011770 | 1815697 | A | G | 11 | 10 | 2 | 0 | NON_SYNONYMOU | MODERAT | MISSENSE | gAa/gGa | E24G | PLES_16931 | . | . | . | . | . | . | G | G | . | . | . | . |
| NC_011770 | 1815724 | C | T | 14 | 8 | 2 | 0 | NON_SYNONYMOU | MODERAT | MISSENSE | gCg/gTg | A33V | PLES_16931 | . | . | . | . | . | . | T | T | . | . | . | . |
| NC_011770 | 1815774 | A | G | 11 | 10 | 2 | 0 | NON_SYNONYMOU | MODERAT | MISSENSE | Aaa/Gaa | K50E | PLES_16931 | . | . | . | . | . | . | G | G | . | . | . | . |
| NC_011770 | 1815780 | G | C | 9 | 10 | 2 | 0 | NON_SYNONYMOU | MODERAT | MISSENSE | Gag/Cag | E52Q | PLES_16931 | . | . | . | . | . | . | C | C | . | . | . | . |
| NC_011770 | 1815782 | G | C | 9 | 10 | 2 | 0 | NON_SYNONYMOU | MODERAT | MISSENSE | gaG/gaC | E52D | PLES_16931 | . | . | . | . | . | . | C | C | . | . | . | . |
| NC_011770 | 1815784 | A | G | 9 | 10 | 2 | 0 | NON_SYNONYMOU | MODERAT | MISSENSE | aAa/aGa | K53R | PLES_16931 | . | . | . | . | . | . | G | G | . | . | . | . |
| NC_011770 | 1815786 | A | G | 9 | 10 | 2 | 0 | NON_SYNONYMOU | MODERAT | MISSENSE | Atg/Gtg | M54V | PLES_16931 | . | . | . | . | . | . | G | G | . | . | . | . |
| NC_011770 | 1815788 | G | A | 9 | 10 | 2 | 0 | NON_SYNONYMOU | MODERAT | MISSENSE | atG/atA | M54I | PLES_16931 | . | . | . | . | . | . | A | A | . | . | . | . |
| NC_011770 | 1815811 | A | G | 12 | 10 | 2 | 0 | NON_SYNONYMOU | MODERAT | MISSENSE | aAt/aGt | N62S | PLES_16931 | . | . | . | . | . | . | G | G | . | . | . | . |
| NC_011770 | 1815850 | G | A | 12 | 10 | 2 | 0 | NON_SYNONYMOU | MODERAT | MISSENSE | gGa/gAa | G75E | PLES_16931 | . | . | . | . | . | . | A | A | . | . | . | . |
| NC_011770 | 1815856 | C | G | 12 | 10 | 2 | 0 | NON_SYNONYMOU | MODERAT | MISSENSE | aCc/aGc | T77S | PLES_16931 | . | . | . | . | . | . | G | G | . | . | . | . |
| NC_011770 | 1815946 | G | C | 12 | 10 | 2 | 0 | NON_SYNONYMOU | MODERAT | MISSENSE | gGt/gCt | G107A | PLES_16931 | . | . | . | . | . | . | C | C | . | . | . | . |
| NC_011770 | 1815988 | G | C | 14 | 8 | 4 | 0 | NON_SYNONYMOU | MODERAT | MISSENSE | gGg/gCg | G121A | PLES_16931 | C | C | . | . | . | . | C | C | . | . | . | . |
| NC_011770 | 1816107 | G | A | 20 | 6 | 2 | 0 | NON_SYNONYMOU | MODERAT | MISSENSE | Gga/Aga | G161R | PLES_16931 | | | . | . | . | . | | | . | . | A | A |
| NC_011770 | 1816126 | T | A | 19 | 6 | 4 | 0 | NON_SYNONYMOU | MODERAT | MISSENSE | tTc/tAc | F167Y | PLES_16931 | A | A | . | . | . | . | | | . | . | A | A |
| NC_011770 | 1816150 | A | G | 21 | 6 | 2 | 0 | NON_SYNONYMOU | MODERAT | MISSENSE | gAt/gGt | D175G | PLES_16931 | G | G | . | . | . | . | | | . | . | | |
| NC_011770 | 1816155 | A | G | 20 | 6 | 4 | 0 | NON_SYNONYMOU | MODERAT | MISSENSE | Agc/Ggc | S177G | PLES_16931 | G | G | . | . | . | . | | | . | . | G | G |
| NC_011770 | 1816159 | A | T | 20 | 6 | 4 | 0 | NON_SYNONYMOU | MODERAT | MISSENSE | tAc/tTc | Y178F | PLES_16931 | T | T | . | . | . | . | | | . | . | T | T |
| NC_011770 | 1816176 | G | A | 19 | 6 | 4 | 0 | NON_SYNONYMOU | MODERAT | MISSENSE | Gac/Aac | D184N | PLES_16931 | A | A | . | . | . | . | | | . | . | A | A |
| NC_011770 | 1816198 | G | C | 10 | 10 | 2 | 0 | NON_SYNONYMOU | MODERAT | MISSENSE | aGt/aCt | S191T | PLES_16931 | . | . | . | . | . | . | C | C | . | . | . | . |
| NC_011770 | 1816380 | G | A | 31 | 6 | 6 | 0 | NON_SYNONYMOU | MODERAT | MISSENSE | Gtt/Att | V252I | PLES_16931 | A | A | . | . | . | . | A | A | . | . | A | A |
| NC_011770 | 1816383 | A | G | 31 | 6 | 4 | 0 | NON_SYNONYMOU | MODERAT | MISSENSE | Agc/Ggc | S253G | PLES_16931 | G | G | . | . | . | . | | | . | . | G | G |

| | | | | | | | | | | | | | | | | | | | | | | | | | |
|---|---|---|---|---|---|---|---|---|---|---|---|---|---|---|---|---|---|---|---|---|---|---|---|---|---|
| NC_011770 | 1816393 | G | A | 31 | 6 | 6 | 0 | NON_SYNONYMOU | MODERAT | MISSENSE | cGt/cAt | R256H | PLES_16931 | A | A | . | . | . | . | A | A | . | . | A | A |
| NC_011770 | 1816460 | A | G | 34 | 6 | 2 | 0 | NON_SYNONYMOU | MODERAT | MISSENSE | atA/atG | I278M | PLES_16931 | | | . | . | . | . | G | G | . | . | | |
| NC_011770 | 1816528 | A | G | 33 | 6 | 6 | 0 | NON_SYNONYMOU | MODERAT | MISSENSE | aAc/aGc | N301S | PLES_16931 | G | G | . | . | . | . | G | G | . | . | G | G |
| NC_011770 | 1816702 | C | G | 31 | 4 | 8 | 0 | NON_SYNONYMOU | MODERAT | MISSENSE | gCc/gGc | A359G | PLES_16931 | G | G | . | . | . | . | G | G | G | G | G | G |
| NC_011770 | 1816794 | G | C | 32 | 4 | 2 | 0 | NON_SYNONYMOU | MODERAT | MISSENSE | Gaa/Caa | E390Q | PLES_16931 | | | . | . | . | . | | | C | C | | |
| NC_011770 | 1816844 | C | A | 28 | 4 | 2 | 0 | NON_SYNONYMOU | MODERAT | MISSENSE | gaC/gaA | D406E | PLES_16931 | | | . | . | . | . | | | | | A | A |
| NC_011770 | 1816846 | A | G | 28 | 4 | 2 | 0 | NON_SYNONYMOU | MODERAT | MISSENSE | cAg/cGg | Q407R | PLES_16931 | | | . | . | . | . | | | | | G | G |
| NC_011770 | 1816847 | G | C | 28 | 4 | 2 | 0 | NON_SYNONYMOU | MODERAT | MISSENSE | caG/caC | Q407H | PLES_16931 | | | . | . | . | . | | | | | C | C |
| NC_011770 | 1816906 | T | C | 28 | 4 | 8 | 0 | NON_SYNONYMOU | MODERAT | MISSENSE | gTc/gCc | V427A | PLES_16931 | C | C | . | . | . | . | C | C | C | C | C | C |
| NC_011770 | 1816947 | G | C | 27 | 4 | 2 | 0 | NON_SYNONYMOU | MODERAT | MISSENSE | Gcg/Ccg | A441P | PLES_16931 | | | . | . | . | . | | | C | C | | |
| NC_011770 | 1816962 | A | G | 26 | 4 | 2 | 0 | NON_SYNONYMOU | MODERAT | MISSENSE | Atc/Gtc | I446V | PLES_16931 | | | . | . | . | . | | | G | G | | |
| NC_011770 | 1816965 | A | G | 26 | 4 | 2 | 0 | NON_SYNONYMOU | MODERAT | MISSENSE | Aca/Gca | T447A | PLES_16931 | | | . | . | . | . | | | G | G | | |
| NC_011770 | 1816966 | C | T | 26 | 4 | 2 | 0 | NON_SYNONYMOU | MODERAT | MISSENSE | aCa/aTa | T447I | PLES_16931 | | | . | . | . | . | | | T | T | | |
| NC_011770 | 1816858 | C | G | 30 | 4 | 1 | 0 | NON_SYNONYMOU | MODERAT | MISSENSE | aCc/aGc | T411S | PLES_16931 | | | . | . | . | . | | | | | | G |
| NC_011770 | 1817847 | A | G | 36 | 4 | 2 | 0 | NON_SYNONYMOU | MODERAT | MISSENSE | cAa/cGa | Q250R | PLES_16941 | G | G | . | . | . | . | | | | | | |
| NC_011770 | 1818026 | A | G | 32 | 4 | 4 | 0 | NON_SYNONYMOU | MODERAT | MISSENSE | Acg/Gcg | T310A | PLES_16941 | G | G | . | . | . | . | | | | | G | G |
| NC_011770 | 1818065 | A | G | 26 | 4 | 2 | 0 | NON_SYNONYMOU | MODERAT | MISSENSE | Atc/Gtc | I323V | PLES_16941 | G | G | . | . | . | . | | | | | | |
| NC_011770 | 1818075 | A | G | 22 | 4 | 2 | 0 | NON_SYNONYMOU | MODERAT | MISSENSE | cAc/cGc | H326R | PLES_16941 | G | G | . | . | . | . | | | | | | |
| NC_011770 | 1818137 | A | C | 11 | 4 | 4 | 0 | NON_SYNONYMOU | MODERAT | MISSENSE | Aag/Cag | K347Q | PLES_16941 | C | C | . | . | . | . | C | C | | | | |
| NC_011770 | 1818158 | T | G | 9 | 4 | 7 | 0 | NON_SYNONYMOU | MODERAT | MISSENSE | Tcc/Gcc | S354A | PLES_16941 | G | G | . | . | . | . | G | G | G | . | G | G |
| NC_011770 | 1818304 | G | T | 26 | 4 | 2 | 0 | NON_SYNONYMOU | MODERAT | MISSENSE | gaG/gaT | E402D | PLES_16941 | | | . | . | . | . | | | T | T | | |
| NC_011770 | 1818401 | A | C | 31 | 4 | 2 | 0 | NON_SYNONYMOU | MODERAT | MISSENSE | Acc/Ccc | T435P | PLES_16941 | | | . | . | . | . | | | C | C | | |
| NC_011770 | 1818444 | A | T | 32 | 4 | 4 | 0 | NON_SYNONYMOU | MODERAT | MISSENSE | cAg/cTg | Q449L | PLES_16941 | T | T | . | . | . | . | | | T | T | | |
| NC_011770 | 1818470 | A | G | 35 | 4 | 2 | 0 | NON_SYNONYMOU | MODERAT | MISSENSE | Act/Gct | T458A | PLES_16941 | | | . | . | . | . | G | G | | | | |
| NC_011770 | 1818645 | T | G | 43 | 5 | 1 | 0 | NON_SYNONYMOU | MODERAT | MISSENSE | gTc/gGc | V516G | PLES_16941 | | | . | . | . | . | | | . | G | | |
| NC_011770 | 1818749 | T | G | 28 | 4 | 8 | 0 | NON_SYNONYMOU | MODERAT | MISSENSE | Tcg/Gcg | S551A | PLES_16941 | G | G | . | . | . | . | G | G | G | G | G | G |
| NC_011770 | 1818800 | G | A | 28 | 4 | 2 | 0 | NON_SYNONYMOU | MODERAT | MISSENSE | Gac/Aac | D568N | PLES_16941 | A | A | . | . | . | . | | | | | | |
| NC_011770 | 1820830 | C | T | 235 | 1 | 1 | 0 | STOP_GAINED | HIGH | NONSENSE | Cag/Tag | Q16* | PLES_16951 | . | T | | | | | | | | | | |
| NC_011770 | 1820885 | AG | A | 263 | 1 | 1 | 0 | FRAME_SHIFT | HIGH | | gct/ | A35 | PLES_16951 | . | A | | | | | | | | | | |
| NC_011770 | 1820836 | C | A | 238 | 1 | 1 | 0 | NON_SYNONYMOU | MODERAT | MISSENSE | Ctg/Atg | L18M | PLES_16951 | . | A | | | | | | | | | | |
| NC_011770 | 1820854 | C | T | 248 | 1 | 1 | 0 | NON_SYNONYMOU | MODERAT | MISSENSE | Cgg/Tgg | R24W | PLES_16951 | . | T | | | | | | | | | | |
| NC_011770 | 1820870 | A | C | 261 | 1 | 1 | 0 | NON_SYNONYMOU | MODERAT | MISSENSE | aAa/aCa | K29T | PLES_16951 | . | C | | | | | | | | | | |
| NC_011770 | 1820872 | G | A | 263 | 1 | 1 | 0 | NON_SYNONYMOU | MODERAT | MISSENSE | Gcg/Acg | A30T | PLES_16951 | . | A | | | | | | | | | | |
| NC_011770 | 1820900 | T | C | 291 | 0 | 2 | 0 | NON_SYNONYMOU | MODERAT | MISSENSE | tTg/tCg | L39S | PLES_16951 | C | C | | | | | | | | | | |
| NC_011770 | 1820912 | A | C | 283 | 0 | 2 | 0 | NON_SYNONYMOU | MODERAT | MISSENSE | gAa/gCa | E43A | PLES_16951 | C | C | | | | | | | | | | |
| NC_011770 | 1821333 | C | T | 353 | 2 | 2 | 0 | NON_SYNONYMOU | MODERAT | MISSENSE | Cgt/Tgt | R124C | PLES_16961 | . | . | | | | | | | T | T | | |
| NC_011770 | 1821460 | G | A | 327 | 2 | 6 | 0 | NON_SYNONYMOU | MODERAT | MISSENSE | aGg/aAg | R166K | PLES_16961 | . | . | | | A | A | A | A | A | A | | |
| NC_011770 | 1826092 | A | G | 75 | 0 | 2 | 0 | NON_SYNONYMOU | MODERAT | MISSENSE | cAc/cGc | H43R | PLES_16991 | | | | | | | | | G | G | | |
| NC_011770 | 1826159 | A | T | 77 | 0 | 2 | 0 | NON_SYNONYMOU | MODERAT | MISSENSE | gaA/gaT | E65D | PLES_16991 | T | T | | | | | | | | | | |
| NC_011770 | 1826888 | G | C | 68 | 0 | 2 | 0 | NON_SYNONYMOU | MODERAT | MISSENSE | gaG/gaC | E308D | PLES_16991 | C | C | | | | | | | | | | |
| NC_011770 | 1826899 | G | A | 67 | 0 | 12 | 0 | NON_SYNONYMOU | MODERAT | MISSENSE | aGg/aAg | R312K | PLES_16991 | A | A | A | A | A | A | A | A | A | A | A | A |
| NC_011770 | 1831832 | T | C | 82 | 0 | 2 | 0 | NON_SYNONYMOU | MODERAT | MISSENSE | Tcg/Ccg | S158P | PLES_17051 | C | C | | | | | | | | | | |
| NC_011770 | 1832614 | A | G | 95 | 0 | 2 | 0 | NON_SYNONYMOU | MODERAT | MISSENSE | gTg/gCg | V214A | PLES_17061 | | | | | | | | | | | G | G |
| NC_011770 | 1832615 | C | T | 95 | 0 | 2 | 0 | NON_SYNONYMOU | MODERAT | MISSENSE | Gtg/Atg | V214M | PLES_17061 | T | T | | | | | | | | | | |
| NC_011770 | 1835259 | G | A | 86 | 0 | 2 | 0 | NON_SYNONYMOU | MODERAT | MISSENSE | Ggc/Agc | G93S | PLES_17081 | | | | | | | | | A | A | | |
| NC_011770 | 1837298 | C | T | 84 | 0 | 2 | 0 | NON_SYNONYMOU | MODERAT | MISSENSE | gCc/gTc | A297V | PLES_17091 | | | | | | | | | | | T | T |
| NC_011770 | 1837367 | A | G | 75 | 0 | 1 | 0 | NON_SYNONYMOU | MODERAT | MISSENSE | gAg/gGg | E320G | PLES_17091 | | | | | | | G | | | | | |
| NC_011770 | 1837867 | T | C | 76 | 0 | 1 | 0 | NON_SYNONYMOU | MODERAT | MISSENSE | gTa/gCa | V38A | PLES_17101 | | | | | | | C | | | | | |
| NC_011770 | 1839104 | C | G | 79 | 0 | 2 | 0 | NON_SYNONYMOU | MODERAT | MISSENSE | Gtc/Ctc | V20L | PLES_17111 | G | G | | | | | | | | | | |
| NC_011770 | 1844786 | TGC | T | 79 | 0 | 1 | 0 | FRAME_SHIFT | HIGH | | cgcggc/ | RG443 | PLES_17181 hsbR | | | | | | | | T | | | | |
| NC_011770 | 1844987 | CG | CGG | 89 | 0 | 3 | 0 | FRAME_SHIFT | HIGH | | ggg/ | G509 | **PLES_17181 hsbR** | | **CGG** | | **C** | | | | **C** | | | | |
| NC_011770 | 1848097 | C | T | 72 | 0 | 2 | 0 | NON_SYNONYMOU | MODERAT | MISSENSE | gCg/gTg | A63V | PLES_17211 | | | | | | | | | | | T | T |
| NC_011770 | 1848283 | T | C | 66 | 0 | 1 | 0 | NON_SYNONYMOU | MODERAT | MISSENSE | gTg/gCg | V125A | PLES_17211 | | | | | | | C | | | | | |
| NC_011770 | 1848894 | G | A | 80 | 0 | 2 | 0 | NON_SYNONYMOU | MODERAT | MISSENSE | Gtc/Atc | V329I | PLES_17211 | | | | | | | A | A | | | | |
| NC_011770 | 1849347 | C | G | 73 | 0 | 12 | 0 | NON_SYNONYMOU | MODERAT | MISSENSE | Ccc/Gcc | P40A | PLES_17221 | G | G | G | G | G | G | G | G | G | G | G | G |
| NC_011770 | 1850176 | C | T | 79 | 0 | 2 | 0 | NON_SYNONYMOU | MODERAT | MISSENSE | cCg/cTg | P316L | PLES_17221 | | | | | T | T | | | | | | |
| NC_011770 | 1851325 | C | T | 77 | 0 | 4 | 0 | NON_SYNONYMOU | MODERAT | MISSENSE | cCg/cTg | P84L | PLES_17241 | | | T | T | T | T | | | | | | |
| NC_011770 | 1851477 | C | A | 88 | 0 | 2 | 0 | NON_SYNONYMOU | MODERAT | MISSENSE | Ccg/Acg | P135T | PLES_17241 | | | | | | | | | A | A | | |
| NC_011770 | 1851535 | C | T | 82 | 0 | 2 | 0 | NON_SYNONYMOU | MODERAT | MISSENSE | cCg/cTg | P154L | PLES_17241 | | | | | | | | | T | T | | |
| NC_011770 | 1851538 | T | C | 83 | 0 | 2 | 0 | NON_SYNONYMOU | MODERAT | MISSENSE | gTg/gCg | V155A | PLES_17241 | | | | | | | | | C | C | | |
| NC_011770 | 1851562 | T | C | 78 | 0 | 2 | 0 | NON_SYNONYMOU | MODERAT | MISSENSE | gTc/gCc | V163A | PLES_17241 | C | C | | | | | | | | | | |
| NC_011770 | 1851616 | A | G | 79 | 0 | 4 | 0 | NON_SYNONYMOU | MODERAT | MISSENSE | aAc/aGc | N181S | PLES_17241 | | | | | | | G | G | G | G | | |

| | | | | | | | | | | | | | | | | | | | | | | | | | |
|---|---|---|---|---|---|---|---|---|---|---|---|---|---|---|---|---|---|---|---|---|---|---|---|---|---|
| NC_011770 | 1851793 | C | A | 62 | 0 | 2 | 0 | NON_SYNONYMOU | MODERAT | MISSENSE | gCg/gAg | A240E | PLES_17241 | | | | | | | | | A | A | | |
| NC_011770 | 1852006 | A | G | 57 | 0 | 4 | 0 | NON_SYNONYMOU | MODERAT | MISSENSE | cAc/cGc | H311R | PLES_17241 | | | | | | | G | G | G | G | | |
| NC_011770 | 1852060 | C | T | 56 | 0 | 2 | 0 | NON_SYNONYMOU | MODERAT | MISSENSE | gCg/gTg | A329V | PLES_17241 | | | | | | | | | T | T | | |
| NC_011770 | 1852239 | G | A | 53 | 0 | 2 | 0 | NON_SYNONYMOU | MODERAT | MISSENSE | Gtc/Atc | V389I | PLES_17241 | A | A | | | | | | | | | | |
| NC_011770 | 1852336 | C | T | 52 | 0 | 2 | 0 | NON_SYNONYMOU | MODERAT | MISSENSE | gCt/gTt | A421V | PLES_17241 | | | | | | | | | T | T | | |
| NC_011770 | 1852654 | T | C | 57 | 0 | 12 | 0 | NON_SYNONYMOU | MODERAT | MISSENSE | cTg/cCg | L527P | PLES_17241 | C | C | C | C | C | C | C | C | C | C | C | C |
| NC_011770 | 1852719 | T | C | 54 | 0 | 10 | 0 | NON_SYNONYMOU | MODERAT | MISSENSE | Tcg/Ccg | S549P | PLES_17241 | | | C | C | C | C | C | C | C | C | C | C |
| NC_011770 | 1852746 | A | T | 55 | 0 | 4 | 0 | NON_SYNONYMOU | MODERAT | MISSENSE | Acc/Tcc | T558S | PLES_17241 | | | | | | | | | T | T | T | T |
| NC_011770 | 1852770 | G | A | 57 | 0 | 2 | 0 | NON_SYNONYMOU | MODERAT | MISSENSE | Gac/Aac | D566N | PLES_17241 | | | | | | | | | A | A | | |
| NC_011770 | 1852861 | C | T | 62 | 0 | 2 | 0 | NON_SYNONYMOU | MODERAT | MISSENSE | gCg/gTg | A596V | PLES_17241 | | | | | | | | | | | T | T |
| NC_011770 | 1852870 | C | CGGA | 60 | 0 | 3 | 0 | CODON_INSERTIO | MODERATE | | ccc/GAGcc | P602EP | PLES_17241 | | | CGGA | | CGGA | | | CGGA | | | | |
| NC_011770 | 1854560 | A | G | 84 | 0 | 1 | 0 | NON_SYNONYMOU | MODERAT | MISSENSE | aTc/aCc | I253T | PLES_17251 | | | | | | | G | | | | | |
| NC_011770 | 1855125 | C | T | 86 | 0 | 2 | 0 | NON_SYNONYMOU | MODERAT | MISSENSE | Gtc/Atc | V65I | PLES_17251 | | | | | | | | | | | T | T |
| NC_011770 | 1855275 | G | A | 82 | 0 | 12 | 0 | NON_SYNONYMOU | MODERAT | MISSENSE | Ccc/Tcc | P15S | PLES_17251 | A | A | A | A | A | A | A | A | A | A | A | A |
| NC_011770 | 1857053 | C | T | 75 | 0 | 2 | 0 | NON_SYNONYMOU | MODERAT | MISSENSE | Gtg/Atg | V297M | PLES_17281 | | | | | | | | | | | T | T |
| NC_011770 | 1857061 | G | A | 72 | 0 | 4 | 0 | NON_SYNONYMOU | MODERAT | MISSENSE | gCg/gTg | A294V | PLES_17281 | A | A | | | | | | | | | A | A |
| NC_011770 | 1857157 | T | C | 76 | 0 | 12 | 0 | NON_SYNONYMOU | MODERAT | MISSENSE | cAc/cGc | H262R | PLES_17281 | C | C | C | C | C | C | C | C | C | C | C | C |
| NC_011770 | 1857469 | G | A | 70 | 0 | 2 | 0 | NON_SYNONYMOU | MODERAT | MISSENSE | gCg/gTg | A158V | PLES_17281 | A | A | | | | | | | | | | |
| NC_011770 | 1857680 | C | T | 70 | 0 | 2 | 0 | NON_SYNONYMOU | MODERAT | MISSENSE | Gcg/Acg | A88T | PLES_17281 | | | | | | | | | | | T | T |
| NC_011770 | 1858203 | A | G | 64 | 0 | 1 | 0 | NON_SYNONYMOU | MODERAT | MISSENSE | Ttc/Ctc | F195L | PLES_17291 | | | | | | | G | | | | | |
| NC_011770 | 1861062 | C | G | 72 | 0 | 6 | 0 | NON_SYNONYMOU | MODERAT | MISSENSE | Gag/Cag | E257Q | PLES_17331 | | | G | G | G | G | | | | | G | G |
| NC_011770 | 1861218 | C | T | 74 | 0 | 2 | 0 | NON_SYNONYMOU | MODERAT | MISSENSE | Gac/Aac | D205N | PLES_17331 | | | | | | | | | T | T | | |
| NC_011770 | 1861457 | G | T | 75 | 0 | 6 | 0 | NON_SYNONYMOU | MODERAT | MISSENSE | gCa/gAa | A125E | PLES_17331 | | | T | T | T | T | T | T | | | | |
| NC_011770 | 1861712 | A | T | 65 | 0 | 1 | 0 | NON_SYNONYMOU | MODERAT | MISSENSE | gTg/gAg | V40E | PLES_17331 | | | | | | | T | | | | | |
| NC_011770 | 1861742 | C | T | 63 | 0 | 1 | 0 | NON_SYNONYMOU | MODERAT | MISSENSE | cGt/cAt | R30H | PLES_17331 | | | | | | | | | | | T | |
| NC_011770 | 1862764 | TG | T | 68 | 0 | 1 | 0 | FRAME_SHIFT | HIGH | | caa/ | Q435 | PLES_17351 | | | | | | | | T | | | | |
| NC_011770 | 1863049 | G | A | 75 | 0 | 2 | 0 | NON_SYNONYMOU | MODERAT | MISSENSE | gCg/gTg | A340V | PLES_17351 | | | | | | | | | A | A | | |
| NC_011770 | 1863502 | T | C | 71 | 0 | 2 | 0 | NON_SYNONYMOU | MODERAT | MISSENSE | cAg/cGg | Q189R | PLES_17351 | | | | | | | | | C | C | | |
| NC_011770 | 1863572 | A | C | 68 | 0 | 8 | 0 | NON_SYNONYMOU | MODERAT | MISSENSE | Tcg/Gcg | S166A | PLES_17351 | | | C | C | C | C | | | C | C | C | C |
| NC_011770 | 1863748 | C | T | 66 | 0 | 2 | 0 | NON_SYNONYMOU | MODERAT | MISSENSE | aGc/aAc | S107N | PLES_17351 | | | | | | | | | | | T | T |
| NC_011770 | 1863857 | C | G | 71 | 0 | 2 | 0 | NON_SYNONYMOU | MODERAT | MISSENSE | Gtg/Ctg | V71L | PLES_17351 | | | | | | | | | G | G | | |
| NC_011770 | 1863961 | C | T | 70 | 0 | 2 | 0 | NON_SYNONYMOU | MODERAT | MISSENSE | cGg/cAg | R36Q | PLES_17351 | | | | | | | | | T | T | | |
| NC_011770 | 1864587 | C | T | 73 | 0 | 2 | 0 | NON_SYNONYMOU | MODERAT | MISSENSE | gGc/gAc | G215D | PLES_17361 | | | | | T | T | | | | | | |
| NC_011770 | 1866792 | C | CG | 76 | 0 | 1 | 0 | FRAME_SHIFT | HIGH | | ccc/cccC | P1831P? | PLES_17371 | | | | | | | | CG | | | | |
| NC_011770 | 1865973 | C | T | 84 | 0 | 4 | 0 | NON_SYNONYMOU | MODERAT | MISSENSE | Gta/Ata | V2105I | PLES_17371 | | | T | T | T | T | | | | | | |
| NC_011770 | 1866297 | G | A | 79 | 0 | 6 | 0 | NON_SYNONYMOU | MODERAT | MISSENSE | Ctc/Ttc | L1997F | PLES_17371 | | | | | | | A | A | A | A | A | A |
| NC_011770 | 1866666 | C | T | 75 | 0 | 2 | 0 | NON_SYNONYMOU | MODERAT | MISSENSE | Gtc/Atc | V1874I | PLES_17371 | | | | | | | | | | | T | T |
| NC_011770 | 1867098 | A | C | 72 | 0 | 12 | 0 | NON_SYNONYMOU | MODERAT | MISSENSE | Tcc/Gcc | S1730A | PLES_17371 | C | C | C | C | C | C | C | C | C | C | C | C |
| NC_011770 | 1867119 | T | C | 76 | 0 | 4 | 0 | NON_SYNONYMOU | MODERAT | MISSENSE | Acc/Gcc | T1723A | PLES_17371 | C | C | | | | | | | | | C | C |
| NC_011770 | 1867482 | C | T | 77 | 0 | 2 | 0 | NON_SYNONYMOU | MODERAT | MISSENSE | Gag/Aag | E1602K | PLES_17371 | T | T | | | | | | | | | | |
| NC_011770 | 1867518 | G | A | 73 | 0 | 2 | 0 | NON_SYNONYMOU | MODERAT | MISSENSE | Cgg/Tgg | R1590W | PLES_17371 | | | | | | | | | | | A | A |
| NC_011770 | 1868096 | C | T | 78 | 0 | 10 | 0 | NON_SYNONYMOU | MODERAT | MISSENSE | cGc/cAc | R1397H | PLES_17371 | | | T | T | T | T | T | T | T | T | T | T |
| NC_011770 | 1868558 | C | T | 56 | 0 | 4 | 0 | NON_SYNONYMOU | MODERAT | MISSENSE | cGc/cAc | R1243H | PLES_17371 | | | | | | | T | T | T | T | | |
| NC_011770 | 1869231 | A | G | 60 | 0 | 12 | 0 | NON_SYNONYMOU | MODERAT | MISSENSE | Tgc/Cgc | C1019R | PLES_17371 | G | G | G | G | G | G | G | G | G | G | G | G |
| NC_011770 | 1869842 | G | A | 65 | 0 | 2 | 0 | NON_SYNONYMOU | MODERAT | MISSENSE | gCc/gTc | A815V | PLES_17371 | | | | | | | | | | | A | A |
| NC_011770 | 1870116 | T | C | 88 | 0 | 8 | 0 | NON_SYNONYMOU | MODERAT | MISSENSE | Acg/Gcg | T724A | PLES_17371 | C | C | C | C | C | C | | | | | C | C |
| NC_011770 | 1870179 | C | T | 77 | 0 | 2 | 0 | NON_SYNONYMOU | MODERAT | MISSENSE | Gcg/Acg | A703T | PLES_17371 | | | | | | | | | | | T | T |
| NC_011770 | 1870850 | G | A | 85 | 0 | 2 | 0 | NON_SYNONYMOU | MODERAT | MISSENSE | gCc/gTc | A479V | PLES_17371 | | | | | | | A | A | | | | |
| NC_011770 | 1871246 | T | C | 76 | 0 | 2 | 0 | NON_SYNONYMOU | MODERAT | MISSENSE | aAc/aGc | N347S | PLES_17371 | | | | | | | | | | | C | C |
| NC_011770 | 1871468 | A | G | 83 | 0 | 12 | 0 | NON_SYNONYMOU | MODERAT | MISSENSE | gTg/gCg | V273A | PLES_17371 | G | G | G | G | G | G | G | G | G | G | G | G |
| NC_011770 | 1872057 | C | T | 95 | 0 | 4 | 0 | NON_SYNONYMOU | MODERAT | MISSENSE | Gcg/Acg | A77T | PLES_17371 | T | T | | | | | | | | | T | T |
| NC_011770 | 1872273 | T | C | 85 | 0 | 12 | 0 | NON_SYNONYMOU | MODERAT | MISSENSE | Act/Gct | T5A | PLES_17371 | C | C | C | C | C | C | C | C | C | C | C | C |
| NC_011770 | 1865921 | A | G | 69 | 0 | 1 | 0 | NON_SYNONYMOU | MODERAT | MISSENSE | gTc/gCc | V2122A | PLES_17371 | G | | | | | | | | | | | |
| NC_011770 | 1871016 | C | T | 79 | 0 | 1 | 0 | NON_SYNONYMOU | MODERAT | MISSENSE | Gcc/Acc | A424T | PLES_17371 | | | | | | T | | | | | | |
| NC_011770 | 1873840 | C | T | 61 | 0 | 2 | 0 | NON_SYNONYMOU | MODERAT | MISSENSE | cGc/cAc | R255H | PLES_17391 | T | T | | | | | | | | | | |
| NC_011770 | 1874033 | C | T | 69 | 0 | 2 | 0 | NON_SYNONYMOU | MODERAT | MISSENSE | Gtg/Atg | V191M | PLES_17391 | | | | | | | | | T | T | | |
| NC_011770 | 1874168 | T | C | 66 | 0 | 2 | 0 | NON_SYNONYMOU | MODERAT | MISSENSE | Acc/Gcc | T146A | PLES_17391 | | | | | | | | | C | C | | |
| NC_011770 | 1875220 | C | G | 72 | 0 | 8 | 0 | NON_SYNONYMOU | MODERAT | MISSENSE | cGc/cCc | R391P | PLES_17401 | G | G | | | | | G | G | G | G | G | G |
| NC_011770 | 1875221 | G | T | 73 | 0 | 8 | 0 | NON_SYNONYMOU | MODERAT | MISSENSE | Cgc/Agc | R391S | PLES_17401 | T | T | | | | | T | T | T | T | T | T |
| NC_011770 | 1875230 | C | T | 80 | 0 | 2 | 0 | NON_SYNONYMOU | MODERAT | MISSENSE | Gcc/Acc | A388T | PLES_17401 | | | | | | | | | | | T | T |
| NC_011770 | 1875259 | G | A | 83 | 0 | 8 | 0 | NON_SYNONYMOU | MODERAT | MISSENSE | gCc/gTc | A378V | PLES_17401 | A | A | | | | | A | A | A | A | A | A |
| NC_011770 | 1875421 | C | T | 68 | 0 | 6 | 0 | NON_SYNONYMOU | MODERAT | MISSENSE | aGc/aAc | S324N | PLES_17401 | T | T | | | | | T | T | T | T | | |

| | | | | | | | | | | | | | | | | | | | | | | | | | |
|---|---|---|---|---|---|---|---|---|---|---|---|---|---|---|---|---|---|---|---|---|---|---|---|---|---|
| NC_011770 | 1875448 | G | T | 63 | 0 | 12 | 0 | NON_SYNONYMOU | MODERAT | MISSENSE | cCg/cAg | P315Q | PLES_17401 | T | T | T | T | T | T | T | T | T | T | T | T |
| NC_011770 | 1875557 | C | G | 69 | 0 | 2 | 0 | NON_SYNONYMOU | MODERAT | MISSENSE | Gtc/Ctc | V279L | PLES_17401 | | | | | | | | | | | G | G |
| NC_011770 | 1875787 | C | T | 78 | 0 | 8 | 0 | NON_SYNONYMOU | MODERAT | MISSENSE | cGc/cAc | R202H | PLES_17401 | T | T | | | | | T | T | T | T | T | T |
| NC_011770 | 1875823 | A | G | 78 | 0 | 8 | 0 | NON_SYNONYMOU | MODERAT | MISSENSE | gTc/gCc | V190A | PLES_17401 | G | G | | | | | G | G | G | G | G | G |
| NC_011770 | 1876339 | T | C | 63 | 0 | 4 | 0 | NON_SYNONYMOU | MODERAT | MISSENSE | cAa/cGa | Q18R | PLES_17401 | | | C | C | C | C | | | | | | |
| NC_011770 | 1876395 | G | C | 62 | 0 | 2 | 0 | NON_SYNONYMOU | MODERAT | MISSENSE | gCc/gGc | A290G | PLES_17411 | | | | | C | C | | | | | | |
| NC_011770 | 1878843 | A | AC | 74 | 0 | 1 | 0 | FRAME_SHIFT | HIGH | | gta/gGta | V156G? | PLES_17431 | | | | | | | | AC | | | | |
| NC_011770 | 1878997 | C | T | 69 | 0 | 2 | 0 | NON_SYNONYMOU | MODERAT | MISSENSE | Ggc/Agc | G105S | PLES_17431 | | | | | T | T | | | | | | |
| NC_011770 | 1879464 | A | G | 67 | 0 | 1 | 0 | NON_SYNONYMOU | MODERAT | MISSENSE | Agc/Ggc | S23G | PLES_17441 | | | | | | | G | | | | | |
| NC_011770 | 1879585 | C | T | 57 | 0 | 2 | 0 | NON_SYNONYMOU | MODERAT | MISSENSE | aCc/aTc | T63I | PLES_17441 | T | T | | | | | | | | | | |
| NC_011770 | 1879935 | G | A | 68 | 0 | 12 | 0 | NON_SYNONYMOU | MODERAT | MISSENSE | aCt/aTt | T43I | PLES_17451 | A | A | A | A | A | A | A | A | A | A | A | A |
| NC_011770 | 1880010 | T | C | 67 | 0 | 2 | 0 | NON_SYNONYMOU | MODERAT | MISSENSE | cAg/cGg | Q18R | PLES_17451 | | | | | | | | | C | C | | |
| NC_011770 | 1882326 | G | A | 78 | 0 | 4 | 0 | NON_SYNONYMOU | MODERAT | MISSENSE | gCc/gTc | A132V | PLES_17471 | A | A | | | | | | | | | A | A |
| NC_011770 | 1882686 | C | T | 71 | 0 | 2 | 0 | NON_SYNONYMOU | MODERAT | MISSENSE | cGc/cAc | R12H | PLES_17471 | T | T | | | | | | | | | | |
| NC_011770 | 1882871 | T | G | 62 | 0 | 2 | 0 | NON_SYNONYMOU | MODERAT | MISSENSE | gAg/gCg | E215A | PLES_17481 | G | G | | | | | | | | | | |
| NC_011770 | 1882890 | C | T | 67 | 0 | 2 | 0 | NON_SYNONYMOU | MODERAT | MISSENSE | Gct/Act | A209T | PLES_17481 | T | T | | | | | | | | | | |
| NC_011770 | 1882947 | C | T | 69 | 0 | 2 | 0 | NON_SYNONYMOU | MODERAT | MISSENSE | Gcg/Acg | A190T | PLES_17481 | | | | | | | T | T | | | | |
| NC_011770 | 1883118 | A | C | 78 | 0 | 8 | 0 | NON_SYNONYMOU | MODERAT | MISSENSE | Tcc/Gcc | S133A | PLES_17481 | | | C | C | C | C | C | C | | | C | C |
| NC_011770 | 1883138 | T | C | 74 | 0 | 8 | 0 | NON_SYNONYMOU | MODERAT | MISSENSE | aAc/aGc | N126S | PLES_17481 | C | C | C | C | C | C | C | C | | | | |
| NC_011770 | 1883282 | T | A | 86 | 0 | 2 | 0 | NON_SYNONYMOU | MODERAT | MISSENSE | gAa/gTa | E78V | PLES_17481 | | | | | | | | | | | A | A |
| NC_011770 | 1883313 | C | T | 80 | 0 | 2 | 0 | NON_SYNONYMOU | MODERAT | MISSENSE | Gcg/Acg | A68T | PLES_17481 | | | | | | | | | | | T | T |
| NC_011770 | 1883348 | T | A | 79 | 0 | 2 | 0 | NON_SYNONYMOU | MODERAT | MISSENSE | gAg/gTg | E56V | PLES_17481 | | | | | | | | | | | A | A |
| NC_011770 | 1883427 | T | C | 69 | 0 | 2 | 0 | NON_SYNONYMOU | MODERAT | MISSENSE | Acc/Gcc | T30A | PLES_17481 | | | | | | | | | | | C | C |
| NC_011770 | 1883507 | G | T | 61 | 0 | 8 | 0 | NON_SYNONYMOU | MODERAT | MISSENSE | aCa/aAa | T3K | PLES_17481 | T | T | | | | | T | T | T | T | T | T |
| NC_011770 | 1884279 | A | C | 76 | 0 | 2 | 0 | NON_SYNONYMOU | MODERAT | MISSENSE | cTg/cGg | L38R | PLES_17491 | | | | | | | | | C | C | | |
| NC_011770 | 1884522 | C | T | 76 | 0 | 2 | 0 | NON_SYNONYMOU | MODERAT | MISSENSE | Ggc/Agc | G234S | PLES_17501 | | | | | | | | | | | T | T |
| NC_011770 | 1885103 | C | G | 72 | 0 | 2 | 0 | NON_SYNONYMOU | MODERAT | MISSENSE | gGc/gCc | G40A | PLES_17501 | | | | | | | | | G | G | | |
| NC_011770 | 1885849 | G | A | 74 | 0 | 6 | 0 | NON_SYNONYMOU | MODERAT | MISSENSE | gCc/gTc | A57V | PLES_17511 | A | A | | | | | | | A | A | A | A |
| NC_011770 | 1887738 | A | G | 60 | 0 | 8 | 0 | NON_SYNONYMOU | MODERAT | MISSENSE | Ata/Gta | I240V | PLES_17531 | G | G | | | | | G | G | G | G | G | G |
| NC_011770 | 1890043 | A | G | 68 | 0 | 1 | 0 | NON_SYNONYMOU | MODERAT | MISSENSE | gTg/gCg | V75A | PLES_17541 | | | | | | | G | | | | | |
| NC_011770 | 1891147 | G | C | 81 | 0 | 8 | 0 | NON_SYNONYMOU | MODERAT | MISSENSE | ttG/ttC | L242F | PLES_17551 | C | C | C | C | C | C | C | C | | | | |
| NC_011770 | 1892816 | GGAAAT | G | 80 | 0 | 2 | 0 | FRAME_SHIFT | HIGH | | atttcc/ | IS11 | PLES_17571 | G | G | | | | | | | | | | |
| NC_011770 | 1892716 | C | T | 80 | 0 | 4 | 0 | NON_SYNONYMOU | MODERAT | MISSENSE | Gtc/Atc | V46I | PLES_17571 | | | T | T | T | T | | | | | | |
| NC_011770 | 1892740 | C | T | 83 | 0 | 6 | 0 | NON_SYNONYMOU | MODERAT | MISSENSE | Ggg/Agg | G38R | PLES_17571 | T | T | T | T | T | T | | | | | | |
| NC_011770 | 1892796 | C | T | 81 | 0 | 6 | 0 | NON_SYNONYMOU | MODERAT | MISSENSE | gGa/gAa | G19E | PLES_17571 | T | T | T | T | T | T | | | | | | |
| NC_011770 | 1892805 | C | T | 81 | 0 | 4 | 0 | NON_SYNONYMOU | MODERAT | MISSENSE | cGc/cAc | R16H | PLES_17571 | | | T | T | T | T | | | | | | |
| NC_011770 | 1896032 | C | T | 59 | 0 | 4 | 0 | NON_SYNONYMOU | MODERAT | MISSENSE | Ccc/Tcc | P21S | PLES_17591 | | | | | | | | | T | T | T | T |
| NC_011770 | 1896231 | A | C | 61 | 0 | 4 | 0 | NON_SYNONYMOU | MODERAT | MISSENSE | cAg/cCg | Q87P | PLES_17591 | | | C | C | C | C | | | | | | |
| NC_011770 | 1898496 | A | G | 60 | 0 | 12 | 0 | NON_SYNONYMOU | MODERAT | MISSENSE | Acc/Gcc | T278A | PLES_17611 | G | G | G | G | G | G | G | G | G | G | G | G |
| NC_011770 | 1898515 | A | T | 60 | 0 | 12 | 0 | NON_SYNONYMOU | MODERAT | MISSENSE | cAg/cTg | Q284L | PLES_17611 | T | T | T | T | T | T | T | T | T | T | T | T |
| NC_011770 | 1898935 | C | T | 77 | 0 | 2 | 0 | NON_SYNONYMOU | MODERAT | MISSENSE | cCg/cTg | P424L | PLES_17611 | | | T | T | | | | | | | | |
| NC_011770 | 1899755 | C | T | 82 | 0 | 1 | 0 | INTRAGENIC | MODIFIER | | | | PLES_17612 | | | | | | | | | | | | T |
| NC_011770 | 1900779 | C | T | 72 | 0 | 2 | 0 | NON_SYNONYMOU | MODERAT | MISSENSE | Gcg/Acg | A371T | PLES_17631 | | | | | | | | | T | T | | |
| NC_011770 | 1901016 | A | G | 81 | 0 | 1 | 0 | NON_SYNONYMOU | MODERAT | MISSENSE | Ttc/Ctc | F292L | PLES_17631 | | | | | | | G | | | | | |
| NC_011770 | 1902647 | C | T | 75 | 0 | 2 | 0 | NON_SYNONYMOU | MODERAT | MISSENSE | Gtg/Atg | V271M | PLES_17651 | | | | | | | T | T | | | | |
| NC_011770 | 1902697 | T | C | 74 | 0 | 4 | 0 | NON_SYNONYMOU | MODERAT | MISSENSE | aAg/aGg | K254R | PLES_17651 | | | C | C | C | C | | | | | | |
| NC_011770 | 1902748 | T | C | 69 | 0 | 4 | 0 | NON_SYNONYMOU | MODERAT | MISSENSE | gAc/gGc | D237G | PLES_17651 | | | C | C | C | C | | | | | | |
| NC_011770 | 1902764 | G | C | 73 | 0 | 8 | 0 | NON_SYNONYMOU | MODERAT | MISSENSE | Cgg/Ggg | R232G | PLES_17651 | C | C | | | | | C | C | C | C | C | C |
| NC_011770 | 1902812 | C | T | 70 | 0 | 4 | 0 | NON_SYNONYMOU | MODERAT | MISSENSE | Gtg/Atg | V216M | PLES_17651 | | | T | T | T | T | | | | | | |
| NC_011770 | 1907378 | C | CG | 55 | 0 | 1 | 0 | FRAME_SHIFT | HIGH | | atg/Gatg | M13D? | PLES_17681 | | | CG | | | | | | | | | |
| NC_011770 | 1907428 | G | GA | 53 | 0 | 6 | 0 | FRAME_SHIFT | HIGH | | ccc/Accc | P29T? | PLES_17681 | | | GA | GA | GA | GA | GA | GA | | | | |
| NC_011770 | 1907771 | C | CT | 62 | 0 | 4 | 0 | FRAME_SHIFT | HIGH | | ctc/ctTc | L143L? | PLES_17681 | | | CT | CT | CT | CT | | | | | | |
| NC_011770 | 1907843 | T | C | 62 | 0 | 4 | 0 | STOP_LOST+SPLICE | HIGH | MISSENSE | Tga/Cga | *167R | PLES_17681 | | | C | C | C | C | | | | | | |
| NC_011770 | 1907381 | A | G | 14 | 7 | 1 | 0 | NON_SYNONYMOU | MODERAT | MISSENSE | Atg/Gtg | M13V | PLES_17681 | | | . | G | . | . | | | . | . | . | . |
| NC_011770 | 1907388 | T | G | 51 | 0 | 8 | 0 | NON_SYNONYMOU | MODERAT | MISSENSE | gTg/gGg | V15G | PLES_17681 | | | G | G | G | G | | | G | G | G | G |
| NC_011770 | 1907397 | T | G | 53 | 0 | 8 | 0 | NON_SYNONYMOU | MODERAT | MISSENSE | cTg/cGg | L18R | PLES_17681 | | | G | G | G | G | | | G | G | G | G |
| NC_011770 | 1907399 | G | A | 51 | 0 | 8 | 0 | NON_SYNONYMOU | MODERAT | MISSENSE | Gcg/Acg | A19T | PLES_17681 | | | A | A | A | A | | | A | A | A | A |
| NC_011770 | 1907406 | G | T | 54 | 0 | 4 | 0 | NON_SYNONYMOU | MODERAT | MISSENSE | cGg/cTg | R21L | PLES_17681 | | | T | T | T | T | | | | | | |
| NC_011770 | 1907489 | C | T | 59 | 0 | 2 | 0 | NON_SYNONYMOU | MODERAT | MISSENSE | Ccg/Tcg | P49S | PLES_17681 | | | | | | | T | T | | | | |
| NC_011770 | 1907643 | T | G | 63 | 0 | 4 | 0 | NON_SYNONYMOU | MODERAT | MISSENSE | cTg/cGg | L100R | PLES_17681 | | | | | | | | | G | G | G | G |
| NC_011770 | 1907774 | C | T | 61 | 0 | 2 | 0 | NON_SYNONYMOU | MODERAT | MISSENSE | Ccg/Tcg | P144S | PLES_17681 | | | | | | | T | T | | | | |
| NC_011770 | 1907859 | C | T | 61 | 0 | 2 | 0 | NON_SYNONYMOU | MODERAT | MISSENSE | Cgc/Tgc | R14C | PLES_17691 | | | | | | | T | T | | | | |

| | | | | | | | | | | | | | | | | | | | | | | | | | |
|---|---|---|---|---|---|---|---|---|---|---|---|---|---|---|---|---|---|---|---|---|---|---|---|---|---|
| NC_011770 | 1907970 | A | G | 66 | 0 | 6 | 0 | NON_SYNONYMOU | MODERAT | MISSENSE | Acc/Gcc | T51A | PLES_17691 | | | | | | | G | G | G | G | G | G |
| NC_011770 | 1909693 | C | T | 73 | 0 | 2 | 0 | NON_SYNONYMOU | MODERAT | MISSENSE | cCg/cTg | P625L | PLES_17691 | T | T | | | | | | | | | | |
| NC_011770 | 1909717 | C | T | 70 | 0 | 6 | 0 | NON_SYNONYMOU | MODERAT | MISSENSE | gCc/gTc | A633V | PLES_17691 | T | T | T | T | T | T | | | | | | |
| NC_011770 | 1910695 | G | A | 76 | 0 | 2 | 0 | NON_SYNONYMOU | MODERAT | MISSENSE | gGc/gAc | G959D | PLES_17691 | A | A | | | | | | | | | | |
| NC_011770 | 1910716 | T | C | 76 | 0 | 8 | 0 | NON_SYNONYMOU | MODERAT | MISSENSE | gTg/gCg | V966A | PLES_17691 | C | C | | | | | C | C | C | C | C | C |
| NC_011770 | 1910826 | G | A | 80 | 0 | 2 | 0 | NON_SYNONYMOU | MODERAT | MISSENSE | Gac/Aac | D1003N | PLES_17691 | A | A | | | | | | | | | | |
| NC_011770 | 1910875 | C | T | 76 | 0 | 6 | 0 | NON_SYNONYMOU | MODERAT | MISSENSE | aCc/aTc | T1019I | PLES_17691 | T | T | | | | | | | T | T | T | T |
| NC_011770 | 1913778 | A | G | 87 | 0 | 12 | 0 | NON_SYNONYMOU | MODERAT | MISSENSE | Atc/Gtc | I91V | PLES_17711 | G | G | G | G | G | G | G | G | G | G | G | G |
| NC_011770 | 1914211 | C | G | 94 | 0 | 6 | 0 | NON_SYNONYMOU | MODERAT | MISSENSE | aCc/aGc | T13S | PLES_17721 | | | G | G | G | G | G | G | | | | |
| NC_011770 | 1914247 | C | A | 97 | 0 | 2 | 0 | NON_SYNONYMOU | MODERAT | MISSENSE | gCc/gAc | A25D | PLES_17721 | | | | | | | A | A | | | | |
| NC_011770 | 1914253 | A | G | 95 | 0 | 2 | 0 | NON_SYNONYMOU | MODERAT | MISSENSE | aAc/aGc | N27S | PLES_17721 | | | | | | | G | G | | | | |
| NC_011770 | 1914303 | G | A | 96 | 0 | 6 | 0 | NON_SYNONYMOU | MODERAT | MISSENSE | Gga/Aga | G44R | PLES_17721 | | | A | A | A | A | A | A | | | | |
| NC_011770 | 1914330 | A | G | 89 | 0 | 6 | 0 | NON_SYNONYMOU | MODERAT | MISSENSE | Acg/Gcg | T53A | PLES_17721 | | | G | G | G | G | G | G | | | | |
| NC_011770 | 1914333 | T | C | 90 | 0 | 6 | 0 | NON_SYNONYMOU | MODERAT | MISSENSE | Ttc/Ctc | F54L | PLES_17721 | | | C | C | C | C | C | C | | | | |
| NC_011770 | 1914336 | C | A | 90 | 0 | 6 | 0 | NON_SYNONYMOU | MODERAT | MISSENSE | Ctc/Atc | L55I | PLES_17721 | | | A | A | A | A | A | A | | | | |
| NC_011770 | 1914461 | C | A | 102 | 0 | 4 | 0 | NON_SYNONYMOU | MODERAT | MISSENSE | caC/caA | H96Q | PLES_17721 | | | A | A | A | A | | | | | | |
| NC_011770 | 1914484 | A | G | 98 | 0 | 4 | 0 | NON_SYNONYMOU | MODERAT | MISSENSE | aAc/aGc | N104S | PLES_17721 | | | G | G | G | G | | | | | | |
| NC_011770 | 1914496 | T | C | 98 | 0 | 1 | 0 | NON_SYNONYMOU | MODERAT | MISSENSE | aTc/aCc | I108T | PLES_17721 | | | | | | | C | | | | | |
| NC_011770 | 1914534 | G | A | 100 | 0 | 8 | 0 | NON_SYNONYMOU | MODERAT | MISSENSE | Gtc/Atc | V121I | PLES_17721 | A | A | A | A | A | A | A | A | | | | |
| NC_011770 | 1914926 | G | T | 64 | 0 | 4 | 0 | NON_SYNONYMOU | MODERAT | MISSENSE | gaG/gaT | E251D | PLES_17721 | | | | | | | | | T | T | T | T |
| NC_011770 | 1915087 | C | T | 69 | 0 | 2 | 0 | NON_SYNONYMOU | MODERAT | MISSENSE | gCc/gTc | A305V | PLES_17721 | | | | | | | | | | | T | T |
| NC_011770 | 1915134 | G | A | 77 | 0 | 4 | 0 | NON_SYNONYMOU | MODERAT | MISSENSE | Gtc/Atc | V321I | PLES_17721 | | | A | A | A | A | | | | | | |
| NC_011770 | 1915156 | G | C | 75 | 0 | 12 | 0 | NON_SYNONYMOU | MODERAT | MISSENSE | cGg/cCg | R328P | PLES_17721 | C | C | C | C | C | C | C | C | C | C | C | C |
| NC_011770 | 1915609 | T | G | 87 | 0 | 4 | 0 | NON_SYNONYMOU | MODERAT | MISSENSE | tTc/tGc | F479C | PLES_17721 | | | G | G | G | G | | | | | | |
| NC_011770 | 1915909 | G | A | 92 | 0 | 4 | 0 | NON_SYNONYMOU | MODERAT | MISSENSE | aGg/aAg | R579K | PLES_17721 | | | A | A | A | A | | | | | | |
| NC_011770 | 1915956 | G | A | 87 | 0 | 4 | 0 | NON_SYNONYMOU | MODERAT | MISSENSE | Gag/Aag | E595K | PLES_17721 | | | A | A | A | A | | | | | | |
| NC_011770 | 1916094 | G | A | 72 | 0 | 12 | 0 | NON_SYNONYMOU | MODERAT | MISSENSE | Gtg/Atg | V641M | PLES_17721 | A | A | A | A | A | A | A | A | A | A | A | A |
| NC_011770 | 1916130 | C | T | 66 | 0 | 8 | 0 | NON_SYNONYMOU | MODERAT | MISSENSE | Ccc/Tcc | P653S | PLES_17721 | T | T | T | T | T | T | T | T | | | | |
| NC_011770 | 1916841 | GC | G | 51 | 0 | 2 | 0 | FRAME_SHIFT | HIGH | | gcc/ | A198 | PLES_17731 | G | G | | | | | | | | | | |
| NC_011770 | 1916694 | C | T | 60 | 0 | 2 | 0 | NON_SYNONYMOU | MODERAT | MISSENSE | Cgt/Tgt | R149C | PLES_17731 | | | | | | | T | T | | | | |
| NC_011770 | 1918120 | G | A | 72 | 0 | 12 | 0 | NON_SYNONYMOU | MODERAT | MISSENSE | Gac/Aac | D6N | PLES_17741 | A | A | A | A | A | A | A | A | A | A | A | A |
| NC_011770 | 1918121 | A | G | 72 | 0 | 10 | 0 | NON_SYNONYMOU | MODERAT | MISSENSE | gAc/gGc | D6G | PLES_17741 | G | G | G | G | G | G | | | G | G | G | G |
| NC_011770 | 1918139 | A | C | 71 | 0 | 4 | 0 | NON_SYNONYMOU | MODERAT | MISSENSE | aAt/aCt | N12T | PLES_17741 | | | | | | | | | C | C | C | C |
| NC_011770 | 1918146 | T | G | 72 | 0 | 4 | 0 | NON_SYNONYMOU | MODERAT | MISSENSE | aaT/aaG | N14K | PLES_17741 | | | | | | | | | G | G | G | G |
| NC_011770 | 1918222 | G | A | 72 | 0 | 2 | 0 | NON_SYNONYMOU | MODERAT | MISSENSE | Gca/Aca | A40T | PLES_17741 | | | | | | | A | A | | | | |
| NC_011770 | 1918228 | A | G | 72 | 0 | 4 | 0 | NON_SYNONYMOU | MODERAT | MISSENSE | Atc/Gtc | I42V | PLES_17741 | | | | | | | | | G | G | G | G |
| NC_011770 | 1918249 | C | T | 71 | 0 | 4 | 0 | NON_SYNONYMOU | MODERAT | MISSENSE | Ccg/Tcg | P49S | PLES_17741 | | | | | | | | | T | T | T | T |
| NC_011770 | 1918406 | A | G | 44 | 6 | 6 | 0 | NON_SYNONYMOU | MODERAT | MISSENSE | cAt/cGt | H101R | PLES_17741 | G | G | G | G | G | G | . | . | . | . | . | . |
| NC_011770 | 1919604 | C | CAGGT | 56 | 0 | 6 | 0 | FRAME_SHIFT | HIGH | | ctc/cAGGT | L311QV? | PLES_17751 | | | | | | | CAGGT | CAGGT | CAGGT | CAGGT | CAGGT | CAGGT |
| NC_011770 | 1918704 | A | G | 40 | 6 | 6 | 0 | NON_SYNONYMOU | MODERAT | MISSENSE | Acc/Gcc | T11A | PLES_17751 | G | G | G | G | G | G | . | . | . | . | . | . |
| NC_011770 | 1918772 | G | T | 41 | 6 | 2 | 0 | NON_SYNONYMOU | MODERAT | MISSENSE | caG/caT | Q33H | PLES_17751 | T | T | | | | | . | . | . | . | . | . |
| NC_011770 | 1919268 | G | A | 56 | 0 | 6 | 0 | NON_SYNONYMOU | MODERAT | MISSENSE | Gct/Act | A199T | PLES_17751 | A | A | A | A | A | A | | | | | | |
| NC_011770 | 1919320 | A | C | 58 | 0 | 2 | 0 | NON_SYNONYMOU | MODERAT | MISSENSE | cAg/cCg | Q216P | PLES_17751 | | | C | C | | | | | | | | |
| NC_011770 | 1919364 | G | A | 65 | 0 | 6 | 0 | NON_SYNONYMOU | MODERAT | MISSENSE | Gtc/Atc | V231I | PLES_17751 | | | | | | | A | A | A | A | A | A |
| NC_011770 | 1919434 | T | C | 64 | 0 | 6 | 0 | NON_SYNONYMOU | MODERAT | MISSENSE | gTc/gCc | V254A | PLES_17751 | | | | | | | C | C | C | C | C | C |
| NC_011770 | 1919440 | A | G | 64 | 0 | 6 | 0 | NON_SYNONYMOU | MODERAT | MISSENSE | gAt/gGt | D256G | PLES_17751 | | | | | | | G | G | G | G | G | G |
| NC_011770 | 1919578 | A | G | 64 | 0 | 6 | 0 | NON_SYNONYMOU | MODERAT | MISSENSE | gAc/gGc | D302G | PLES_17751 | | | | | | | G | G | G | G | G | G |
| NC_011770 | 1919626 | T | C | 55 | 0 | 6 | 0 | NON_SYNONYMOU | MODERAT | MISSENSE | aTc/aCc | I318T | PLES_17751 | | | | | | | C | C | C | C | C | C |
| NC_011770 | 1919643 | T | G | 67 | 0 | 6 | 0 | NON_SYNONYMOU | MODERAT | MISSENSE | Tct/Gct | S324A | PLES_17751 | | | | | | | G | G | G | G | G | G |
| NC_011770 | 1919670 | G | A | 71 | 0 | 6 | 0 | NON_SYNONYMOU | MODERAT | MISSENSE | Gtt/Att | V333I | PLES_17751 | | | | | | | A | A | A | A | A | A |
| NC_011770 | 1919725 | A | G | 75 | 0 | 2 | 0 | NON_SYNONYMOU | MODERAT | MISSENSE | gAc/gGc | D351G | PLES_17751 | | | | | | | G | G | | | | |
| NC_011770 | 1919727 | A | C | 75 | 0 | 6 | 0 | NON_SYNONYMOU | MODERAT | MISSENSE | Atg/Ctg | M352L | PLES_17751 | | | | | | | C | C | C | C | C | C |
| NC_011770 | 1919769 | A | C | 73 | 0 | 6 | 0 | NON_SYNONYMOU | MODERAT | MISSENSE | Aag/Cag | K366Q | PLES_17751 | | | | | | | C | C | C | C | C | C |
| NC_011770 | 1919772 | T | G | 73 | 0 | 6 | 0 | NON_SYNONYMOU | MODERAT | MISSENSE | Tgg/Ggg | W367G | PLES_17751 | | | | | | | G | G | G | G | G | G |
| NC_011770 | 1919774 | G | T | 74 | 0 | 6 | 0 | NON_SYNONYMOU | MODERAT | MISSENSE | tgG/tgT | W367C | PLES_17751 | | | | | | | T | T | T | T | T | T |
| NC_011770 | 1919886 | T | C | 66 | 0 | 6 | 0 | NON_SYNONYMOU | MODERAT | MISSENSE | Ttt/Ctt | F405L | PLES_17751 | | | | | | | C | C | C | C | C | C |
| NC_011770 | 1919888 | T | G | 64 | 0 | 12 | 0 | NON_SYNONYMOU | MODERAT | MISSENSE | ttT/ttG | F405L | PLES_17751 | G | G | G | G | G | G | G | G | G | G | G | G |
| NC_011770 | 1919914 | A | G | 59 | 0 | 6 | 0 | NON_SYNONYMOU | MODERAT | MISSENSE | gAg/gGg | E414G | PLES_17751 | | | | | | | G | G | G | G | G | G |
| NC_011770 | 1919915 | G | T | 60 | 0 | 6 | 0 | NON_SYNONYMOU | MODERAT | MISSENSE | gaG/gaT | E414D | PLES_17751 | | | | | | | T | T | T | T | T | T |
| NC_011770 | 1919940 | G | A | 53 | 0 | 6 | 0 | NON_SYNONYMOU | MODERAT | MISSENSE | Gca/Aca | A423T | PLES_17751 | | | | | | | A | A | A | A | A | A |
| NC_011770 | 1919941 | C | A | 53 | 0 | 6 | 0 | NON_SYNONYMOU | MODERAT | MISSENSE | gCa/gAa | A423E | PLES_17751 | | | | | | | A | A | A | A | A | A |
| NC_011770 | 1919943 | C | T | 53 | 0 | 6 | 0 | NON_SYNONYMOU | MODERAT | MISSENSE | Cct/Tct | P424S | PLES_17751 | | | | | | | T | T | T | T | T | T |

| | | | | | | | | | | | | | | | | | | | | | | | | | |
|---|---|---|---|---|---|---|---|---|---|---|---|---|---|---|---|---|---|---|---|---|---|---|---|---|---|
| NC_011770 | 1919971 | A | T | 54 | 0 | 6 | 0 | NON_SYNONYMOU | MODERAT | MISSENSE | gAg/gTg | E433V | PLES_17751 | | | | | | | T | T | T | T | T | T |
| NC_011770 | 1919972 | G | C | 55 | 0 | 6 | 0 | NON_SYNONYMOU | MODERAT | MISSENSE | gaG/gaC | E433D | PLES_17751 | | | | | | | C | C | C | C | C | C |
| NC_011770 | 1919980 | T | A | 47 | 0 | 6 | 0 | NON_SYNONYMOU | MODERAT | MISSENSE | gTa/gAa | V436E | PLES_17751 | | | | | | | A | A | A | A | A | A |
| NC_011770 | 1919985 | G | A | 54 | 0 | 6 | 0 | NON_SYNONYMOU | MODERAT | MISSENSE | Gtg/Atg | V438M | PLES_17751 | | | | | | | A | A | A | A | A | A |
| NC_011770 | 1919986 | T | C | 52 | 0 | 6 | 0 | NON_SYNONYMOU | MODERAT | MISSENSE | gTg/gCg | V438A | PLES_17751 | | | | | | | C | C | C | C | C | C |
| NC_011770 | 1919991 | A | G | 52 | 0 | 6 | 0 | NON_SYNONYMOU | MODERAT | MISSENSE | Aag/Gag | K440E | PLES_17751 | | | | | | | G | G | G | G | G | G |
| NC_011770 | 1919995 | T | C | 53 | 0 | 6 | 0 | NON_SYNONYMOU | MODERAT | MISSENSE | aTt/aCt | I441T | PLES_17751 | | | | | | | C | C | C | C | C | C |
| NC_011770 | 1920216 | C | T | 72 | 0 | 2 | 0 | NON_SYNONYMOU | MODERAT | MISSENSE | Cac/Tac | H515Y | PLES_17751 | | | | | | | T | T | | | | |
| NC_011770 | 1920258 | A | G | 72 | 0 | 4 | 0 | NON_SYNONYMOU | MODERAT | MISSENSE | Aag/Gag | K529E | PLES_17751 | | | | | | | | | G | G | G | G |
| NC_011770 | 1920267 | T | C | 70 | 0 | 6 | 0 | NON_SYNONYMOU | MODERAT | MISSENSE | Tcg/Ccg | S532P | PLES_17751 | | | | | | | C | C | C | C | C | C |
| NC_011770 | 1920418 | G | A | 71 | 0 | 12 | 0 | NON_SYNONYMOU | MODERAT | MISSENSE | cGt/cAt | R582H | PLES_17751 | A | A | A | A | A | A | A | A | A | A | A | A |
| NC_011770 | 1920717 | C | T | 62 | 0 | 12 | 0 | NON_SYNONYMOU | MODERAT | MISSENSE | Cat/Tat | H682Y | PLES_17751 | T | T | T | T | T | T | T | T | T | T | T | T |
| NC_011770 | 1920996 | C | T | 60 | 0 | 2 | 0 | NON_SYNONYMOU | MODERAT | MISSENSE | Cgc/Tgc | R775C | PLES_17751 | | | | | | | T | T | | | | |
| NC_011770 | 1921249 | A | G | 64 | 0 | 8 | 0 | NON_SYNONYMOU | MODERAT | MISSENSE | cAg/cGg | Q859R | PLES_17751 | G | G | G | G | G | G | | | | | G | G |
| NC_011770 | 1921264 | C | G | 63 | 0 | 12 | 0 | NON_SYNONYMOU | MODERAT | MISSENSE | gCc/gGc | A864G | PLES_17751 | G | G | G | G | G | G | A | A | G | G | G | G |
| NC_011770 | 1921290 | C | A | 66 | 0 | 2 | 0 | NON_SYNONYMOU | MODERAT | MISSENSE | Cag/Aag | Q873K | PLES_17751 | | | | | | | | | A | A | | |
| NC_011770 | 1921369 | T | C | 64 | 0 | 2 | 0 | NON_SYNONYMOU | MODERAT | MISSENSE | Acc/Gcc | T136A | PLES_17761 | | | | | | | | | C | C | | |
| NC_011770 | 1921868 | C | G | 62 | 0 | 2 | 0 | NON_SYNONYMOU | MODERAT | MISSENSE | ttC/ttG | F5L | PLES_17771 | | | | | | | | | G | G | | |
| NC_011770 | 1923668 | T | C | 79 | 0 | 2 | 0 | NON_SYNONYMOU | MODERAT | MISSENSE | gTg/gCg | V78A | PLES_17791 | C | C | | | | | | | | | | |
| NC_011770 | 1924888 | C | A | 67 | 0 | 8 | 0 | NON_SYNONYMOU | MODERAT | MISSENSE | gCg/gAg | A133E | PLES_17801 | A | A | | | | | A | A | A | A | A | A |
| NC_011770 | 1925026 | T | C | 64 | 0 | 4 | 0 | NON_SYNONYMOU | MODERAT | MISSENSE | gTc/gCc | V179A | PLES_17801 | | | | | | | C | C | C | C | | |
| NC_011770 | 1925582 | G | A | 64 | 0 | 6 | 0 | NON_SYNONYMOU | MODERAT | MISSENSE | Gat/Aat | D34N | PLES_17811 | A | A | | | | | A | A | A | A | | |
| NC_011770 | 1926150 | G | A | 64 | 0 | 10 | 0 | NON_SYNONYMOU | MODERAT | MISSENSE | gGg/gAg | G26E | PLES_17831 | | | A | A | A | A | A | A | A | A | A | A |
| NC_011770 | 1926153 | T | G | 67 | 0 | 4 | 0 | NON_SYNONYMOU | MODERAT | MISSENSE | gTc/gGc | V27G | PLES_17831 | | | G | G | G | G | | | | | | |
| NC_011770 | 1926342 | A | G | 75 | 0 | 8 | 0 | NON_SYNONYMOU | MODERAT | MISSENSE | gAc/gGc | D90G | PLES_17831 | | | G | G | G | G | | | G | G | G | G |
| NC_011770 | 1926534 | A | C | 82 | 0 | 2 | 0 | NON_SYNONYMOU | MODERAT | MISSENSE | gAa/gCa | E154A | PLES_17831 | | | | | | | C | C | | | | |
| NC_011770 | 1927081 | C | T | 77 | 0 | 2 | 0 | NON_SYNONYMOU | MODERAT | MISSENSE | Cgc/Tgc | R53C | PLES_17841 | | | | | | | T | T | | | | |
| NC_011770 | 1927085 | A | G | 75 | 0 | 4 | 0 | NON_SYNONYMOU | MODERAT | MISSENSE | cAg/cGg | Q54R | PLES_17841 | | | | | | | | | G | G | G | G |
| NC_011770 | 1927369 | C | G | 60 | 0 | 2 | 0 | NON_SYNONYMOU | MODERAT | MISSENSE | Ctg/Gtg | L149V | PLES_17841 | G | G | | | | | | | | | | |
| NC_011770 | 1927390 | A | G | 55 | 0 | 4 | 0 | NON_SYNONYMOU | MODERAT | MISSENSE | Atg/Gtg | M156V | PLES_17841 | | | | | | | G | G | G | G | | |
| NC_011770 | 1927577 | T | C | 65 | 0 | 4 | 0 | NON_SYNONYMOU | MODERAT | MISSENSE | gTg/gCg | V218A | PLES_17841 | | | | | | | C | C | C | C | | |
| NC_011770 | 1927776 | T | A | 80 | 0 | 4 | 0 | NON_SYNONYMOU | MODERAT | MISSENSE | Tgc/Agc | C39S | PLES_17851 | | | | | | | A | A | A | A | | |
| NC_011770 | 1927846 | C | T | 84 | 0 | 4 | 0 | NON_SYNONYMOU | MODERAT | MISSENSE | cCg/cTg | P62L | PLES_17851 | | | T | T | T | T | | | | | | |
| NC_011770 | 1928239 | G | A | 75 | 0 | 6 | 0 | NON_SYNONYMOU | MODERAT | MISSENSE | cGa/cAa | R193Q | PLES_17851 | A | A | A | A | A | A | | | | | | |
| NC_011770 | 1929996 | G | A | 78 | 0 | 2 | 0 | NON_SYNONYMOU | MODERAT | MISSENSE | aCc/aTc | T84I | PLES_17871 | A | A | | | | | | | | | | |
| NC_011770 | 1930001 | T | C | 78 | 0 | 12 | 0 | NON_SYNONYMOU | MODERAT | MISSENSE | atA/atG | I82M | PLES_17871 | C | C | C | C | C | C | C | C | C | C | C | C |
| NC_011770 | 1930213 | T | C | 72 | 0 | 10 | 0 | NON_SYNONYMOU | MODERAT | MISSENSE | Atc/Gtc | I12V | PLES_17871 | C | C | C | C | C | C | C | C | C | C | | |
| NC_011770 | 1931797 | CG | C | 50 | 0 | 1 | 0 | FRAME_SHIFT | HIGH | | ggg/ | G13 | PLES_17891 | | | | | | | | C | | | | |
| NC_011770 | 1931784 | T | C | 51 | 0 | 12 | 0 | NON_SYNONYMOU | MODERAT | MISSENSE | gTc/gCc | V7A | PLES_17891 | C | C | C | C | C | C | C | C | C | C | C | C |
| NC_011770 | 1931850 | G | A | 47 | 0 | 12 | 0 | NON_SYNONYMOU | MODERAT | MISSENSE | gGc/gAc | G29D | PLES_17891 | A | A | A | A | A | A | A | A | A | A | A | A |
| NC_011770 | 1933666 | T | C | 19 | 0 | 10 | 0 | NON_SYNONYMOU | MODERAT | MISSENSE | Ttc/Ctc | F5L | PLES_17921 | C | C | C | C | C | C | C | C | C | C | | |
| NC_011770 | 1934516 | A | G | 89 | 0 | 1 | 0 | NON_SYNONYMOU | MODERAT | MISSENSE | Atg/Gtg | M73V | PLES_17941 | | | | | | | G | | | | | |
| NC_011770 | 1935038 | C | T | 70 | 0 | 4 | 0 | NON_SYNONYMOU | MODERAT | MISSENSE | gCc/gTc | A24V | PLES_17951 | | | T | T | T | T | | | | | | |
| NC_011770 | 1935403 | G | T | 75 | 0 | 2 | 0 | NON_SYNONYMOU | MODERAT | MISSENSE | Gcg/Tcg | A146S | PLES_17951 | | | | | | | | | | | T | T |
| NC_011770 | 1937380 | C | G | 80 | 0 | 12 | 0 | NON_SYNONYMOU | MODERAT | MISSENSE | Cag/Gag | Q805E | PLES_17951 | G | G | G | G | G | G | G | G | G | G | G | G |
| NC_011770 | 1937448 | G | C | 81 | 0 | 2 | 0 | NON_SYNONYMOU | MODERAT | MISSENSE | gaG/gaC | E827D | PLES_17951 | C | C | | | | | | | | | | |
| NC_011770 | 1937467 | C | T | 81 | 0 | 6 | 0 | NON_SYNONYMOU | MODERAT | MISSENSE | Cgc/Tgc | R834C | PLES_17951 | | | T | T | T | T | | | | | T | T |
| NC_011770 | 1937707 | C | T | 77 | 0 | 2 | 0 | NON_SYNONYMOU | MODERAT | MISSENSE | Cgc/Tgc | R914C | PLES_17951 | | | | | | | | | | | T | T |
| NC_011770 | 1937951 | C | T | 84 | 0 | 2 | 0 | NON_SYNONYMOU | MODERAT | MISSENSE | cCg/cTg | P995L | PLES_17951 | T | T | | | | | | | | | | |
| NC_011770 | 1937957 | C | T | 85 | 0 | 2 | 0 | NON_SYNONYMOU | MODERAT | MISSENSE | tCg/tTg | S997L | PLES_17951 | | | | | | | | | T | T | | |
| NC_011770 | 1938037 | G | A | 81 | 0 | 2 | 0 | NON_SYNONYMOU | MODERAT | MISSENSE | Gac/Aac | D1024N | PLES_17951 | A | A | | | | | | | | | | |
| NC_011770 | 1938077 | C | G | 81 | 0 | 2 | 0 | NON_SYNONYMOU | MODERAT | MISSENSE | gCa/gGa | A1037G | PLES_17951 | G | G | | | | | | | | | | |
| NC_011770 | 1938217 | G | C | 78 | 0 | 2 | 0 | NON_SYNONYMOU | MODERAT | MISSENSE | Gag/Cag | E1084Q | PLES_17951 | C | C | | | | | | | | | | |
| NC_011770 | 1939262 | T | A | 63 | 0 | 4 | 0 | NON_SYNONYMOU | MODERAT | MISSENSE | gTc/gAc | V1432D | PLES_17951 | | | | | | | A | A | A | A | | |
| NC_011770 | 1939452 | A | AC | 67 | 0 | 1 | 0 | FRAME_SHIFT | HIGH | | gtg/gGtg | V1117G? | PLES_17961 | | | | AC | | | | | | | | |
| NC_011770 | 1943363 | G | T | 95 | 0 | 2 | 0 | NON_SYNONYMOU | MODERAT | MISSENSE | tgG/tgT | W99C | PLES_17971 | | | | | | | | | | | T | T |
| NC_011770 | 1943629 | T | C | 93 | 0 | 10 | 0 | NON_SYNONYMOU | MODERAT | MISSENSE | gTg/gCg | V188A | PLES_17971 | C | C | | C | | C | C | C | C | C | C | C |
| NC_011770 | 1944142 | A | G | 71 | 0 | 10 | 0 | NON_SYNONYMOU | MODERAT | MISSENSE | gAc/gGc | D141G | PLES_17981 | G | G | G | G | G | G | G | G | G | G | | |
| NC_011770 | 1944177 | G | A | 70 | 0 | 2 | 0 | NON_SYNONYMOU | MODERAT | MISSENSE | Gtg/Atg | V153M | PLES_17981 | A | A | | | | | | | | | | |
| NC_011770 | 1944243 | C | G | 71 | 0 | 10 | 0 | NON_SYNONYMOU | MODERAT | MISSENSE | Ccg/Gcg | P175A | PLES_17981 | G | G | G | G | G | G | G | G | G | G | | |
| NC_011770 | 1944565 | T | C | 78 | 0 | 8 | 0 | NON_SYNONYMOU | MODERAT | MISSENSE | aTc/aCc | I282T | PLES_17981 | C | C | C | C | C | C | | | C | C | | |
| NC_011770 | 1945740 | A | G | 89 | 0 | 4 | 0 | NON_SYNONYMOU | MODERAT | MISSENSE | Atc/Gtc | I357V | PLES_17991 | G | G | | | | | G | G | | | | |

| | | | | | | | | | | | | | | | | | | | | | | | | | |
|---|---|---|---|---|---|---|---|---|---|---|---|---|---|---|---|---|---|---|---|---|---|---|---|---|---|
| NC_011770 | 1946394 | G | A | 79 | 0 | 4 | 0 | NON_SYNONYMOU | MODERAT | MISSENSE | Gtc/Atc | V575I | PLES_17991 | | | A | A | A | A | | | | | | |
| NC_011770 | 1946500 | T | C | 71 | 0 | 1 | 0 | NON_SYNONYMOU | MODERAT | MISSENSE | gTg/gCg | V610A | PLES_17991 | | | | | | | | C | | | | |
| NC_011770 | 1947153 | G | A | 74 | 0 | 2 | 0 | NON_SYNONYMOU | MODERAT | MISSENSE | Ggc/Agc | G48S | PLES_18001 | | | | | | | A | A | | | | |
| NC_011770 | 1947976 | T | C | 68 | 0 | 12 | 0 | NON_SYNONYMOU | MODERAT | MISSENSE | gTg/gCg | V322A | PLES_18001 | C | C | C | C | C | C | C | C | C | C | C | C |
| NC_011770 | 1948105 | A | G | 70 | 0 | 2 | 0 | NON_SYNONYMOU | MODERAT | MISSENSE | gAc/gGc | D365G | PLES_18001 | G | G | | | | | | | | | | |
| NC_011770 | 1948825 | A | G | 51 | 0 | 2 | 0 | NON_SYNONYMOU | MODERAT | MISSENSE | gAg/gGg | E605G | PLES_18001 | | | | | | | | | | | G | G |
| NC_011770 | 1948877 | A | C | 47 | 0 | 2 | 0 | NON_SYNONYMOU | MODERAT | MISSENSE | gaA/gaC | E622D | PLES_18001 | | | | | | | C | C | | | | |
| NC_011770 | 1949840 | T | A | 84 | 0 | 6 | 0 | NON_SYNONYMOU | MODERAT | MISSENSE | gaA/gaT | E307D | PLES_18031 | A | A | | | | | A | A | A | A | | |
| NC_011770 | 1949845 | T | G | 82 | 0 | 6 | 0 | NON_SYNONYMOU | MODERAT | MISSENSE | Atg/Ctg | M306L | PLES_18031 | G | G | | | | | G | G | G | G | | |
| NC_011770 | 1949929 | A | C | 79 | 0 | 2 | 0 | NON_SYNONYMOU | MODERAT | MISSENSE | Tcg/Gcg | S278A | PLES_18031 | | | | | | | C | C | | | | |
| NC_011770 | 1950493 | T | G | 78 | 0 | 12 | 0 | NON_SYNONYMOU | MODERAT | MISSENSE | Agc/Cgc | S90R | PLES_18031 | G | G | G | G | G | G | G | G | G | G | G | G |
| NC_011770 | 1953913 | T | G | 75 | 0 | 2 | 0 | NON_SYNONYMOU | MODERAT | MISSENSE | gaA/gaC | E247D | PLES_18061 | | | | | | | | | G | G | | |
| NC_011770 | 1953924 | C | T | 72 | 0 | 2 | 0 | NON_SYNONYMOU | MODERAT | MISSENSE | Gcg/Acg | A244T | PLES_18061 | | | | | | | T | T | | | | |
| NC_011770 | 1953926 | T | C | 73 | 0 | 2 | 0 | NON_SYNONYMOU | MODERAT | MISSENSE | cAa/cGa | Q243R | PLES_18061 | | | | | | | C | C | | | | |
| NC_011770 | 1953950 | C | T | 73 | 0 | 2 | 0 | NON_SYNONYMOU | MODERAT | MISSENSE | gGg/gAg | G235E | PLES_18061 | | | | | | | T | T | | | | |
| NC_011770 | 1954385 | T | C | 73 | 0 | 10 | 0 | NON_SYNONYMOU | MODERAT | MISSENSE | gAa/gGa | E90G | PLES_18061 | | | C | C | C | C | C | C | C | C | C | C |
| NC_011770 | 1954854 | C | T | 66 | 0 | 2 | 0 | NON_SYNONYMOU | MODERAT | MISSENSE | gCc/gTc | A33V | PLES_18071 | | | | | | | | | | | T | T |
| NC_011770 | 1954889 | C | G | 64 | 0 | 12 | 0 | NON_SYNONYMOU | MODERAT | MISSENSE | Ccg/Gcg | P45A | PLES_18071 | G | G | G | G | G | G | G | G | G | G | G | G |
| NC_011770 | 1954890 | C | T | 62 | 0 | 12 | 0 | NON_SYNONYMOU | MODERAT | MISSENSE | cCg/cTg | P45L | PLES_18071 | T | T | T | T | T | T | T | T | T | T | T | T |
| NC_011770 | 1955687 | G | T | 61 | 0 | 4 | 0 | NON_SYNONYMOU | MODERAT | MISSENSE | Gcg/Tcg | A88S | PLES_18081 | | | T | T | T | T | | | | | | |
| NC_011770 | 1955772 | A | T | 71 | 0 | 6 | 0 | NON_SYNONYMOU | MODERAT | MISSENSE | cAa/cTa | Q116L | PLES_18081 | | | T | T | T | T | T | T | | | | |
| NC_011770 | 1955957 | G | C | 74 | 0 | 6 | 0 | NON_SYNONYMOU | MODERAT | MISSENSE | Ctg/Gtg | L590V | PLES_18091 | | | C | C | C | C | C | C | | | | |
| NC_011770 | 1956120 | T | G | 78 | 0 | 2 | 0 | NON_SYNONYMOU | MODERAT | MISSENSE | gaA/gaC | E535D | PLES_18091 | | | | | | | G | G | | | | |
| NC_011770 | 1956359 | C | T | 82 | 0 | 4 | 0 | NON_SYNONYMOU | MODERAT | MISSENSE | Gtc/Atc | V456I | PLES_18091 | | | | | | | T | T | | | T | T |
| NC_011770 | 1956548 | C | T | 75 | 0 | 2 | 0 | NON_SYNONYMOU | MODERAT | MISSENSE | Gtc/Atc | V393I | PLES_18091 | | | | | | | | | | | T | T |
| NC_011770 | 1956628 | A | G | 72 | 0 | 2 | 0 | NON_SYNONYMOU | MODERAT | MISSENSE | gTg/gCg | V366A | PLES_18091 | | | | | | | G | G | | | | |
| NC_011770 | 1956749 | C | T | 83 | 0 | 2 | 0 | NON_SYNONYMOU | MODERAT | MISSENSE | Ggg/Agg | G326R | PLES_18091 | | | | | | | | | T | T | | |
| NC_011770 | 1956778 | A | T | 83 | 0 | 4 | 0 | NON_SYNONYMOU | MODERAT | MISSENSE | cTg/cAg | L316Q | PLES_18091 | | | T | T | T | T | | | | | | |
| NC_011770 | 1956790 | C | T | 83 | 0 | 2 | 0 | NON_SYNONYMOU | MODERAT | MISSENSE | cGc/cAc | R312H | PLES_18091 | | | | | | | | | T | T | | |
| NC_011770 | 1956818 | T | G | 83 | 0 | 2 | 0 | NON_SYNONYMOU | MODERAT | MISSENSE | Acc/Ccc | T303P | PLES_18091 | | | | | | | | | G | G | | |
| NC_011770 | 1956884 | T | C | 90 | 0 | 2 | 0 | NON_SYNONYMOU | MODERAT | MISSENSE | Agc/Ggc | S281G | PLES_18091 | | | | | | | | | C | C | | |
| NC_011770 | 1956904 | T | C | 86 | 0 | 8 | 0 | NON_SYNONYMOU | MODERAT | MISSENSE | gAc/gGc | D274G | PLES_18091 | | | C | C | C | C | C | C | C | C | | |
| NC_011770 | 1956949 | T | A | 84 | 0 | 2 | 0 | NON_SYNONYMOU | MODERAT | MISSENSE | cAg/cTg | Q259L | PLES_18091 | | | | | | | | | | | A | A |
| NC_011770 | 1957391 | C | T | 81 | 0 | 2 | 0 | NON_SYNONYMOU | MODERAT | MISSENSE | Gtc/Atc | V112I | PLES_18091 | | | | | | | | | T | T | | |
| NC_011770 | 1956503 | C | T | 81 | 0 | 1 | 0 | NON_SYNONYMOU | MODERAT | MISSENSE | Gtg/Atg | V408M | PLES_18091 | | | | | | | | | | | T | |
| NC_011770 | 1960202 | T | A | 59 | 0 | 10 | 0 | NON_SYNONYMOU | MODERAT | MISSENSE | Tgc/Agc | C72S | PLES_18111 | A | A | A | A | A | A | | | A | A | A | A |
| NC_011770 | 1960439 | G | A | 64 | 0 | 2 | 0 | NON_SYNONYMOU | MODERAT | MISSENSE | Ggc/Agc | G151S | PLES_18111 | | | | | | | A | A | | | | |
| NC_011770 | 1961233 | G | A | 78 | 0 | 2 | 0 | NON_SYNONYMOU | MODERAT | MISSENSE | Cgc/Tgc | R120C | PLES_18121 | A | A | | | | | | | | | | |
| NC_011770 | 1961550 | C | T | 90 | 0 | 4 | 0 | NON_SYNONYMOU | MODERAT | MISSENSE | cGc/cAc | R14H | PLES_18121 | T | T | | | | | | | T | T | | |
| NC_011770 | 1961832 | C | T | 95 | 0 | 2 | 0 | NON_SYNONYMOU | MODERAT | MISSENSE | Gcg/Acg | A276T | PLES_18131 | | | | | | | | | | | T | T |
| NC_011770 | 1962336 | T | C | 74 | 0 | 10 | 0 | NON_SYNONYMOU | MODERAT | MISSENSE | Acg/Gcg | T108A | PLES_18131 | C | C | C | C | C | C | C | C | | | C | C |
| NC_011770 | 1963059 | C | T | 83 | 0 | 2 | 0 | NON_SYNONYMOU | MODERAT | MISSENSE | atG/atA | M133I | PLES_18141 | T | T | | | | | | | | | | |
| NC_011770 | 1963073 | C | T | 84 | 0 | 2 | 0 | NON_SYNONYMOU | MODERAT | MISSENSE | Ggg/Agg | G129R | PLES_18141 | | | | | | | | | | | T | T |
| NC_011770 | 1963385 | C | G | 94 | 0 | 2 | 0 | NON_SYNONYMOU | MODERAT | MISSENSE | Gtc/Ctc | V25L | PLES_18141 | G | G | | | | | | | | | | |
| NC_011770 | 1964252 | A | G | 76 | 0 | 1 | 0 | NON_SYNONYMOU | MODERAT | MISSENSE | cTg/cCg | L11P | PLES_18151 | | | | | | | | G | | | | |
| NC_011770 | 1964462 | C | T | 80 | 0 | 1 | 0 | NON_SYNONYMOU | MODERAT | MISSENSE | Gag/Aag | E210K | PLES_18161 | | | | | | | T | | | | | |
| NC_011770 | 1964732 | G | T | 73 | 0 | 4 | 0 | NON_SYNONYMOU | MODERAT | MISSENSE | Ctg/Atg | L120M | PLES_18161 | | | T | T | T | T | | | | | | |
| NC_011770 | 1966169 | C | T | 76 | 0 | 4 | 0 | NON_SYNONYMOU | MODERAT | MISSENSE | Gcc/Acc | A12T | PLES_18171 | | | T | T | T | T | | | | | | |
| NC_011770 | 1967209 | A | G | 72 | 0 | 12 | 0 | NON_SYNONYMOU | MODERAT | MISSENSE | gAt/gGt | D215G | PLES_18181 | G | G | G | G | G | G | G | G | G | G | G | G |
| NC_011770 | 1967678 | C | A | 42 | 0 | 12 | 0 | NON_SYNONYMOU | MODERAT | MISSENSE | Cgc/Agc | R111S | PLES_18191 | A | A | A | A | A | A | A | A | A | A | A | A |
| NC_011770 | 1968276 | C | T | 68 | 0 | 6 | 0 | NON_SYNONYMOU | MODERAT | MISSENSE | aGg/aAg | R319K | PLES_18201 | | | | | | | T | T | T | T | T | T |
| NC_011770 | 1968849 | A | G | 68 | 0 | 1 | 0 | NON_SYNONYMOU | MODERAT | MISSENSE | gTc/gCc | V128A | PLES_18201 | | G | | | | | | | | | | |
| NC_011770 | 1972704 | A | G | 89 | 0 | 6 | 0 | NON_SYNONYMOU | MODERAT | MISSENSE | Atc/Gtc | I163V | PLES_18251 | G | G | | | | | G | G | G | G | | |
| NC_011770 | 1973302 | G | A | 78 | 0 | 2 | 0 | NON_SYNONYMOU | MODERAT | MISSENSE | Gtc/Atc | V27I | PLES_18261 | | | | | | | | | A | A | | |
| NC_011770 | 1974179 | C | T | 67 | 0 | 1 | 0 | NON_SYNONYMOU | MODERAT | MISSENSE | cCg/cTg | P319L | PLES_18261 | | | | | | | T | | | | | |
| NC_011770 | 1974358 | A | G | 68 | 0 | 12 | 0 | NON_SYNONYMOU | MODERAT | MISSENSE | Acg/Gcg | T379A | PLES_18261 | G | G | G | G | G | G | G | G | G | G | G | G |
| NC_011770 | 1974554 | T | C | 73 | 0 | 12 | 0 | NON_SYNONYMOU | MODERAT | MISSENSE | cAc/cGc | H254R | PLES_18271 | C | C | C | C | C | C | C | C | C | C | C | C |
| NC_011770 | 1974867 | C | T | 78 | 0 | 2 | 0 | NON_SYNONYMOU | MODERAT | MISSENSE | Gga/Aga | G150R | PLES_18271 | | | | | T | T | | | | | | |
| NC_011770 | 1974932 | T | C | 78 | 0 | 2 | 0 | NON_SYNONYMOU | MODERAT | MISSENSE | aAc/aGc | N128S | PLES_18271 | | | | | | | | | C | C | | |
| NC_011770 | 1975227 | C | T | 67 | 0 | 2 | 0 | NON_SYNONYMOU | MODERAT | MISSENSE | Ggc/Agc | G30S | PLES_18271 | T | T | | | | | | | | | | |
| NC_011770 | 1976498 | C | T | 85 | 0 | 2 | 0 | NON_SYNONYMOU | MODERAT | MISSENSE | Gac/Aac | D376N | PLES_18291 | | | | | | | | | | | T | T |
| NC_011770 | 1979020 | G | A | 72 | 0 | 8 | 0 | NON_SYNONYMOU | MODERAT | MISSENSE | aGg/aAg | R247K | PLES_18311 | | | A | A | A | A | | | A | A | A | A |

| | | | | | | | | | | | | | | | | | | | | | | | | | |
|---|---|---|---|---|---|---|---|---|---|---|---|---|---|---|---|---|---|---|---|---|---|---|---|---|---|
| NC_011770 | 1981466 | T | C | 64 | 0 | 12 | 0 | NON_SYNONYMOU | MODERAT | MISSENSE | Tgc/Cgc | C20R | PLES_18341 | C | C | C | C | C | C | C | C | C | C | C | C |
| NC_011770 | 1982145 | C | G | 86 | 0 | 2 | 0 | NON_SYNONYMOU | MODERAT | MISSENSE | aCg/aGg | T246R | PLES_18341 | | | | | | | | | G | G | | |
| NC_011770 | 1982622 | T | C | 72 | 0 | 10 | 0 | NON_SYNONYMOU | MODERAT | MISSENSE | gTg/gCg | V405A | PLES_18341 | C | C | C | C | C | C | C | C | C | C | | |
| NC_011770 | 1983556 | T | C | 79 | 0 | 1 | 0 | NON_SYNONYMOU | MODERAT | MISSENSE | Tac/Cac | Y118H | PLES_18351 | | C | | | | | | | | | | |
| NC_011770 | 1984428 | C | T | 60 | 0 | 2 | 0 | NON_SYNONYMOU | MODERAT | MISSENSE | gCc/gTc | A24V | PLES_18371 | | | | | | | | | | | T | T |
| NC_011770 | 1985186 | A | G | 93 | 0 | 2 | 0 | NON_SYNONYMOU | MODERAT | MISSENSE | Acc/Gcc | T277A | PLES_18371 | G | G | | | | | | | | | | |
| NC_011770 | 1986643 | T | C | 62 | 0 | 12 | 0 | NON_SYNONYMOU | MODERAT | MISSENSE | cAc/cGc | H418R | PLES_18391 | C | C | C | C | C | C | C | C | C | C | C | C |
| NC_011770 | 1987091 | T | G | 62 | 0 | 4 | 0 | NON_SYNONYMOU | MODERAT | MISSENSE | Atg/Ctg | M269L | PLES_18391 | | | G | G | G | G | | | | | | |
| NC_011770 | 1987584 | C | T | 90 | 0 | 1 | 0 | NON_SYNONYMOU | MODERAT | MISSENSE | atG/atA | M104I | PLES_18391 | | | | | | | T | | | | | |
| NC_011770 | 1988831 | C | A | 76 | 0 | 4 | 0 | NON_SYNONYMOU | MODERAT | MISSENSE | gGg/gTg | G167V | PLES_18411 | | | A | A | A | A | | | | | | |
| NC_011770 | 1989188 | C | T | 70 | 0 | 2 | 0 | NON_SYNONYMOU | MODERAT | MISSENSE | cGc/cAc | R48H | PLES_18411 | | | | | | | | | T | T | | |
| NC_011770 | 1991965 | A | C | 82 | 0 | 2 | 0 | NON_SYNONYMOU | MODERAT | MISSENSE | cTc/cGc | L204R | PLES_18451 | | | | | | | | | | | C | C |
| NC_011770 | 1992146 | C | T | 75 | 0 | 2 | 0 | NON_SYNONYMOU | MODERAT | MISSENSE | Ggc/Agc | G144S | PLES_18451 | T | T | | | | | | | | | | |
| NC_011770 | 1992418 | A | G | 80 | 0 | 12 | 0 | NON_SYNONYMOU | MODERAT | MISSENSE | gTg/gCg | V53A | PLES_18451 | G | G | G | G | G | G | G | G | G | G | G | G |
| NC_011770 | 1993563 | A | C | 74 | 0 | 6 | 0 | NON_SYNONYMOU | MODERAT | MISSENSE | Tcc/Gcc | S63A | PLES_18471 | | | | | | | C | C | C | C | C | C |
| NC_011770 | 1993651 | G | C | 80 | 0 | 6 | 0 | NON_SYNONYMOU | MODERAT | MISSENSE | gaC/gaG | D33E | PLES_18471 | | | | | | | C | C | C | C | C | C |
| NC_011770 | 1994028 | G | C | 78 | 0 | 2 | 0 | NON_SYNONYMOU | MODERAT | MISSENSE | Gat/Cat | D45H | PLES_18481 | C | C | | | | | | | | | | |
| NC_011770 | 1994761 | C | T | 63 | 0 | 4 | 0 | NON_SYNONYMOU | MODERAT | MISSENSE | Ggt/Agt | G150S | PLES_18491 | | | T | T | T | T | | | | | | |
| NC_011770 | 1994785 | G | A | 61 | 0 | 4 | 0 | NON_SYNONYMOU | MODERAT | MISSENSE | Ctc/Ttc | L142F | PLES_18491 | | | A | A | A | A | | | | | | |
| NC_011770 | 1998351 | G | T | 70 | 0 | 4 | 0 | NON_SYNONYMOU | MODERAT | MISSENSE | gaG/gaT | E178D | PLES_18521 | | | T | T | T | T | | | | | | |
| NC_011770 | 1999174 | C | T | 75 | 0 | 2 | 0 | NON_SYNONYMOU | MODERAT | MISSENSE | Gcc/Acc | A109T | PLES_18531 | | | | | | | T | T | | | | |
| NC_011770 | 1999263 | T | C | 74 | 0 | 1 | 0 | NON_SYNONYMOU | MODERAT | MISSENSE | cAc/cGc | H79R | PLES_18531 | | | | | | | C | | | | | |
| NC_011770 | 1999514 | GT | G | 78 | 0 | 1 | 0 | FRAME_SHIFT | HIGH | | acc/ | T307 | PLES_18541 | | | | G | | | | | | | | |
| NC_011770 | 2000151 | C | T | 81 | 0 | 2 | 0 | NON_SYNONYMOU | MODERAT | MISSENSE | Gac/Aac | D95N | PLES_18541 | T | T | | | | | | | | | | |
| NC_011770 | 2001162 | G | A | 81 | 0 | 2 | 0 | NON_SYNONYMOU | MODERAT | MISSENSE | gCg/gTg | A23V | PLES_18551 | A | A | | | | | | | | | | |
| NC_011770 | 2002098 | G | A | 69 | 0 | 2 | 0 | STOP_GAINED | HIGH | NONSENSE | Cag/Tag | Q92* | PLES_18561 | A | A | | | | | | | | | | |
| NC_011770 | 2001923 | A | C | 81 | 0 | 2 | 0 | NON_SYNONYMOU | MODERAT | MISSENSE | tTc/tGc | F150C | PLES_18561 | C | C | | | | | | | | | | |
| NC_011770 | 2001983 | G | A | 72 | 0 | 2 | 0 | NON_SYNONYMOU | MODERAT | MISSENSE | aCc/aTc | T130I | PLES_18561 | | | | | | | A | A | | | | |
| NC_011770 | 2002097 | T | C | 65 | 0 | 2 | 0 | NON_SYNONYMOU | MODERAT | MISSENSE | cAg/cGg | Q92R | PLES_18561 | C | C | | | | | | | | | | |
| NC_011770 | 2002223 | G | A | 70 | 0 | 6 | 0 | NON_SYNONYMOU | MODERAT | MISSENSE | gCc/gTc | A50V | PLES_18561 | | | A | A | A | A | A | A | | | | |
| NC_011770 | 2002229 | C | G | 69 | 0 | 6 | 0 | NON_SYNONYMOU | MODERAT | MISSENSE | gGc/gCc | G48A | PLES_18561 | | | G | G | G | G | G | G | | | | |
| NC_011770 | 2004057 | C | T | 74 | 0 | 2 | 0 | NON_SYNONYMOU | MODERAT | MISSENSE | aCt/aTt | T3I | PLES_18581 | | | | | | | | | T | T | | |
| NC_011770 | 2004065 | C | T | 75 | 0 | 2 | 0 | NON_SYNONYMOU | MODERAT | MISSENSE | Ccg/Tcg | P6S | PLES_18581 | | | | | | | T | T | | | | |
| NC_011770 | 2004315 | C | T | 78 | 0 | 2 | 0 | NON_SYNONYMOU | MODERAT | MISSENSE | gCc/gTc | A89V | PLES_18581 | | | | | | | | | | | T | T |
| NC_011770 | 2004564 | G | C | 70 | 0 | 2 | 0 | NON_SYNONYMOU | MODERAT | MISSENSE | Gac/Cac | D27H | PLES_18591 | | | | | | | | | C | C | | |
| NC_011770 | 2004621 | C | T | 73 | 0 | 2 | 0 | NON_SYNONYMOU | MODERAT | MISSENSE | Cgc/Tgc | R46C | PLES_18591 | T | T | | | | | | | | | | |
| NC_011770 | 2004642 | G | A | 74 | 0 | 2 | 0 | NON_SYNONYMOU | MODERAT | MISSENSE | Gaa/Aaa | E53K | PLES_18591 | | | | | | | | | | | A | A |
| NC_011770 | 2005062 | G | A | 79 | 0 | 1 | 0 | NON_SYNONYMOU | MODERAT | MISSENSE | Cgc/Tgc | R149C | PLES_18601 | | | | | | | | | | | A | |
| NC_011770 | 2005473 | G | A | 78 | 0 | 1 | 0 | NON_SYNONYMOU | MODERAT | MISSENSE | Cgc/Tgc | R12C | PLES_18601 | | | | | | | | | | | A | |
| NC_011770 | 2005889 | C | T | 75 | 0 | 1 | 0 | NON_SYNONYMOU | MODERAT | MISSENSE | gGc/gAc | G332D | PLES_18611 | | | | | | | T | | | | | |
| NC_011770 | 2006297 | C | T | 66 | 0 | 2 | 0 | NON_SYNONYMOU | MODERAT | MISSENSE | aGc/aAc | S196N | PLES_18611 | T | T | | | | | | | | | | |
| NC_011770 | 2006759 | C | T | 47 | 0 | 1 | 0 | NON_SYNONYMOU | MODERAT | MISSENSE | gGc/gAc | G42D | PLES_18611 | | | | | | | | T | | | | |
| NC_011770 | 2007282 | C | T | 52 | 0 | 2 | 0 | NON_SYNONYMOU | MODERAT | MISSENSE | Gac/Aac | D53N | PLES_18621 | T | T | | | | | | | | | | |
| NC_011770 | 2007302 | C | T | 50 | 0 | 2 | 0 | NON_SYNONYMOU | MODERAT | MISSENSE | gGc/gAc | G46D | PLES_18621 | | | | | | | T | T | | | | |
| NC_011770 | 2007877 | G | A | 66 | 0 | 2 | 0 | NON_SYNONYMOU | MODERAT | MISSENSE | gCg/gTg | A123V | PLES_18631 | | | | | | | A | A | | | | |
| NC_011770 | 2008509 | T | C | 76 | 0 | 6 | 0 | NON_SYNONYMOU | MODERAT | MISSENSE | Acg/Gcg | T37A | PLES_18641 | C | C | C | C | C | C | | | | | | |
| NC_011770 | 2008559 | A | T | 78 | 0 | 6 | 0 | NON_SYNONYMOU | MODERAT | MISSENSE | cTg/cAg | L20Q | PLES_18641 | T | T | T | T | T | T | | | | | | |
| NC_011770 | 2008602 | C | T | 80 | 0 | 6 | 0 | NON_SYNONYMOU | MODERAT | MISSENSE | Gct/Act | A6T | PLES_18641 | T | T | T | T | T | T | | | | | | |
| NC_011770 | 2008575 | C | CCAGGC | 79 | 0 | 3 | 0 | CODON_INSERTIOI | MODERATE | | ctg/ctgAG( | L14LSL | PLES_18641 (hypothetica | | CCAGGCT | | CCAGGCT | | CCAGGCT | | | | | | |
| NC_011770 | 2010059 | G | A | 84 | 0 | 2 | 0 | NON_SYNONYMOU | MODERAT | MISSENSE | Ggt/Agt | G55S | PLES_18671 | A | A | | | | | | | | | | |
| NC_011770 | 2010080 | A | G | 83 | 0 | 10 | 0 | NON_SYNONYMOU | MODERAT | MISSENSE | Atc/Gtc | I62V | PLES_18671 | G | G | G | G | G | G | G | G | G | G | | |
| NC_011770 | 2010273 | G | A | 83 | 0 | 4 | 0 | NON_SYNONYMOU | MODERAT | MISSENSE | cGg/cAg | R126Q | PLES_18671 | | | | | | | A | A | A | A | | |
| NC_011770 | 2010638 | G | T | 76 | 0 | 6 | 0 | NON_SYNONYMOU | MODERAT | MISSENSE | Gcc/Tcc | A248S | PLES_18671 | | | T | T | T | T | | | T | T | | |
| NC_011770 | 2010750 | C | T | 77 | 0 | 6 | 0 | NON_SYNONYMOU | MODERAT | MISSENSE | aCc/aTc | T285I | PLES_18671 | | | T | T | T | T | | | T | T | | |
| NC_011770 | 2011230 | A | C | 76 | 0 | 12 | 0 | NON_SYNONYMOU | MODERAT | MISSENSE | agA/agC | R150S | PLES_18681 | C | C | C | C | C | C | C | C | C | C | C | C |
| NC_011770 | 2013279 | C | T | 74 | 0 | 1 | 0 | NON_SYNONYMOU | MODERAT | MISSENSE | cCg/cTg | P267L | PLES_18701 | | | | | | | T | | | | | |
| NC_011770 | 2013453 | CCGA | C | 71 | 0 | 4 | 0 | CODON_DELETION | MODERATE | | gac/ | D328 | PLES_18701 | | | | | | | C | C | C | C | | |
| NC_011770 | 2014019 | G | A | 86 | 0 | 2 | 0 | NON_SYNONYMOU | MODERAT | MISSENSE | Gcg/Acg | A154T | PLES_18711 | | | | | | | | | | | A | A |
| NC_011770 | 2020036 | CT | C | 54 | 0 | 2 | 0 | FRAME_SHIFT | HIGH | | tac/ | Y267 | PLES_18771 | | | C | C | | | | | | | | |
| NC_011770 | 2019347 | C | T | 76 | 0 | 3 | 0 | NON_SYNONYMOU | MODERAT | MISSENSE | Cgc/Tgc | R37C | PLES_18771 | T | T | | | | | T | | | | | |
| NC_011770 | 2019359 | C | T | 77 | 0 | 4 | 0 | NON_SYNONYMOU | MODERAT | MISSENSE | Ctc/Ttc | L41F | PLES_18771 | | | T | T | T | T | | | | | | |
| NC_011770 | 2019751 | C | A | 71 | 0 | 8 | 0 | NON_SYNONYMOU | MODERAT | MISSENSE | gaC/gaA | D171E | PLES_18771 | | | A | A | A | A | A | A | | | A | A |

| | | | | | | | | | | | | | | | | | | | | | | | | | |
|---|---|---|---|---|---|---|---|---|---|---|---|---|---|---|---|---|---|---|---|---|---|---|---|---|---|
| NC_011770 | 2020133 | G | C | 60 | 0 | 2 | 0 | NON_SYNONYMOU | MODERAT | MISSENSE | Gag/Cag | E299Q | PLES_18771 | | | | | | | C | C | | | | |
| NC_011770 | 2020238 | T | C | 57 | 0 | 12 | 0 | NON_SYNONYMOU | MODERAT | MISSENSE | Tcg/Ccg | S334P | PLES_18771 | C | C | C | C | C | C | C | C | C | C | C | C |
| NC_011770 | 2020440 | T | C | 47 | 0 | 4 | 0 | NON_SYNONYMOU | MODERAT | MISSENSE | gTg/gCg | V401A | PLES_18771 | C | C | | | | | | | C | C | | |
| NC_011770 | 2022055 | G | A | 79 | 0 | 2 | 0 | NON_SYNONYMOU | MODERAT | MISSENSE | Ggc/Agc | G291S | PLES_18781 | A | A | | | | | | | | | | |
| NC_011770 | 2024547 | G | A | 81 | 0 | 2 | 0 | NON_SYNONYMOU | MODERAT | MISSENSE | gGc/gAc | G66D | PLES_18811 | | | | | | | A | A | | | | |
| NC_011770 | 2025246 | C | T | 77 | 0 | 4 | 0 | NON_SYNONYMOU | MODERAT | MISSENSE | gCc/gTc | A299V | PLES_18811 | | | T | T | T | T | | | | | | |
| NC_011770 | 2025462 | C | T | 81 | 0 | 2 | 0 | NON_SYNONYMOU | MODERAT | MISSENSE | gCg/gTg | A371V | PLES_18811 | T | T | | | | | | | | | | |
| NC_011770 | 2027963 | A | G | 73 | 0 | 8 | 0 | NON_SYNONYMOU | MODERAT | MISSENSE | aAt/aGt | N277S | PLES_18831 | | | G | G | G | G | G | G | | | G | G |
| NC_011770 | 2028251 | G | C | 77 | 0 | 12 | 0 | NON_SYNONYMOU | MODERAT | MISSENSE | Cag/Gag | Q201E | PLES_18841 | C | C | C | C | C | C | C | C | C | C | C | C |
| NC_011770 | 2031387 | A | G | 85 | 0 | 12 | 0 | NON_SYNONYMOU | MODERAT | MISSENSE | Acg/Gcg | T54A | PLES_18871 | G | G | G | G | G | G | G | G | G | G | G | G |
| NC_011770 | 2031469 | A | G | 83 | 0 | 8 | 0 | NON_SYNONYMOU | MODERAT | MISSENSE | cAg/cGg | Q81R | PLES_18871 | G | G | | | | | G | G | G | G | G | G |
| NC_011770 | 2031765 | C | A | 65 | 0 | 12 | 0 | NON_SYNONYMOU | MODERAT | MISSENSE | Cac/Aac | H180N | PLES_18871 | A | A | A | A | A | A | A | A | A | A | A | A |
| NC_011770 | 2031987 | C | T | 61 | 0 | 4 | 0 | NON_SYNONYMOU | MODERAT | MISSENSE | gGc/gAc | G144D | PLES_18881 | | | T | T | T | T | | | | | | |
| NC_011770 | 2032408 | T | C | 25 | 0 | 12 | 0 | NON_SYNONYMOU | MODERAT | MISSENSE | Atc/Gtc | I4V | PLES_18881 | C | C | C | C | C | C | C | C | C | C | C | C |
| NC_011770 | 2033874 | A | C | 74 | 0 | 12 | 0 | NON_SYNONYMOU | MODERAT | MISSENSE | gAg/gCg | E10A | PLES_18901 | C | C | C | C | C | C | C | C | C | C | C | C |
| NC_011770 | 2034023 | G | A | 85 | 0 | 4 | 0 | NON_SYNONYMOU | MODERAT | MISSENSE | Ggg/Agg | G60R | PLES_18901 | | | A | A | A | A | | | | | | |
| NC_011770 | 2034027 | C | T | 80 | 0 | 12 | 0 | NON_SYNONYMOU | MODERAT | MISSENSE | gCc/gTc | A61V | PLES_18901 | T | T | T | T | T | T | T | T | T | T | T | T |
| NC_011770 | 2034168 | C | G | 76 | 0 | 12 | 0 | NON_SYNONYMOU | MODERAT | MISSENSE | gCc/gGc | A108G | PLES_18901 | G | G | G | G | G | G | G | G | G | G | G | G |
| NC_011770 | 2035221 | C | T | 74 | 0 | 2 | 0 | START_LOST | HIGH | MISSENSE | atG/atA | M1I | PLES_18911 | | | | | | | | | | | T | T |
| NC_011770 | 2034482 | G | A | 85 | 0 | 2 | 0 | NON_SYNONYMOU | MODERAT | MISSENSE | Cgc/Tgc | R248C | PLES_18911 | | | | | | | | | | | A | A |
| NC_011770 | 2036956 | T | C | 81 | 0 | 4 | 0 | NON_SYNONYMOU | MODERAT | MISSENSE | Agc/Ggc | S229G | PLES_18931 | | | C | C | C | C | | | | | | |
| NC_011770 | 2037343 | G | A | 67 | 0 | 2 | 0 | NON_SYNONYMOU | MODERAT | MISSENSE | Cgc/Tgc | R100C | PLES_18931 | A | A | | | | | | | | | | |
| NC_011770 | 2037366 | T | C | 62 | 0 | 6 | 0 | NON_SYNONYMOU | MODERAT | MISSENSE | gAc/gGc | D92G | PLES_18931 | | | | | | | C | C | A | A | C | C |
| NC_011770 | 2037540 | T | G | 67 | 0 | 4 | 0 | NON_SYNONYMOU | MODERAT | MISSENSE | gAc/gCc | D34A | PLES_18931 | | | G | G | G | G | | | | | | |
| NC_011770 | 2037892 | G | A | 72 | 0 | 4 | 0 | NON_SYNONYMOU | MODERAT | MISSENSE | Cgt/Tgt | R165C | PLES_18941 | | | A | A | A | A | | | | | | |
| NC_011770 | 2037936 | T | C | 69 | 0 | 2 | 0 | NON_SYNONYMOU | MODERAT | MISSENSE | gAa/gGa | E150G | PLES_18941 | C | C | | | | | | | | | | |
| NC_011770 | 2038676 | C | T | 58 | 0 | 4 | 0 | NON_SYNONYMOU | MODERAT | MISSENSE | aGt/aAt | S203N | PLES_18951 | | | T | T | T | T | | | | | | |
| NC_011770 | 2038820 | A | G | 63 | 0 | 1 | 0 | NON_SYNONYMOU | MODERAT | MISSENSE | gTg/gCg | V155A | PLES_18951 | | | | | | | G | | | | | |
| NC_011770 | 2039717 | T | G | 75 | 0 | 4 | 0 | NON_SYNONYMOU | MODERAT | MISSENSE | aaA/aaC | K104N | PLES_18961 | | | G | G | G | G | | | | | | |
| NC_011770 | 2039821 | G | C | 82 | 0 | 4 | 0 | NON_SYNONYMOU | MODERAT | MISSENSE | Ctg/Gtg | L70V | PLES_18961 | | | | | | | C | C | C | C | | |
| NC_011770 | 2039906 | G | C | 82 | 0 | 4 | 0 | NON_SYNONYMOU | MODERAT | MISSENSE | gaC/gaG | D41E | PLES_18961 | | | | | | | C | C | C | C | | |
| NC_011770 | 2040939 | C | T | 55 | 0 | 2 | 0 | NON_SYNONYMOU | MODERAT | MISSENSE | Gaa/Aaa | E424K | PLES_18981 | T | T | | | | | | | | | | |
| NC_011770 | 2041511 | A | G | 69 | 0 | 8 | 0 | NON_SYNONYMOU | MODERAT | MISSENSE | aTg/aCg | M233T | PLES_18981 | G | G | | | | | G | G | G | G | G | G |
| NC_011770 | 2051404 | A | T | 80 | 0 | 6 | 0 | NON_SYNONYMOU | MODERAT | MISSENSE | gAg/gTg | E521V | PLES_19041 | | | | | | | T | T | T | T | T | T |
| NC_011770 | 2071745 | C | T | 72 | 0 | 2 | 0 | NON_SYNONYMOU | MODERAT | MISSENSE | cGc/cAc | R231H | PLES_19201 | | | | | | | | | | | T | T |
| NC_011770 | 2072001 | C | T | 72 | 0 | 2 | 0 | NON_SYNONYMOU | MODERAT | MISSENSE | Gtg/Atg | V146M | PLES_19201 | | | | | | | | | | | T | T |
| NC_011770 | 2074651 | C | T | 83 | 0 | 1 | 0 | NON_SYNONYMOU | MODERAT | MISSENSE | Ggg/Agg | G497R | PLES_19221 | | | | | | | T | | | | | |
| NC_011770 | 2074666 | T | C | 83 | 0 | 2 | 0 | NON_SYNONYMOU | MODERAT | MISSENSE | Acc/Gcc | T492A | PLES_19221 | | | | | | | | | C | C | | |
| NC_011770 | 2074738 | C | T | 89 | 0 | 2 | 0 | NON_SYNONYMOU | MODERAT | MISSENSE | Gtc/Atc | V468I | PLES_19221 | | | | | | | | | T | T | | |
| NC_011770 | 2075140 | C | T | 80 | 0 | 12 | 0 | NON_SYNONYMOU | MODERAT | MISSENSE | Ggc/Agc | G334S | PLES_19221 | T | T | T | T | T | T | T | T | T | T | T | T |
| NC_011770 | 2075284 | C | T | 66 | 0 | 2 | 0 | NON_SYNONYMOU | MODERAT | MISSENSE | Ggc/Agc | G286S | PLES_19221 | | | | | | | | | | | T | T |
| NC_011770 | 2075950 | G | T | 51 | 0 | 2 | 0 | NON_SYNONYMOU | MODERAT | MISSENSE | Cgc/Agc | R64S | PLES_19221 | | | | | | | | | T | T | | |
| NC_011770 | 2076392 | G | A | 78 | 0 | 2 | 0 | NON_SYNONYMOU | MODERAT | MISSENSE | Cct/Tct | P289S | PLES_19231 | | | | | | | | | | | A | A |
| NC_011770 | 2076571 | C | T | 69 | 0 | 2 | 0 | NON_SYNONYMOU | MODERAT | MISSENSE | aGc/aAc | S229N | PLES_19231 | | | | | | | | | | | T | T |
| NC_011770 | 2076950 | T | C | 54 | 0 | 12 | 0 | NON_SYNONYMOU | MODERAT | MISSENSE | Acg/Gcg | T103A | PLES_19231 | C | C | C | C | C | C | C | C | C | C | C | C |
| NC_011770 | 2077112 | T | C | 61 | 0 | 12 | 0 | NON_SYNONYMOU | MODERAT | MISSENSE | Aac/Gac | N49D | PLES_19231 | C | C | C | C | C | C | C | C | C | C | C | C |
| NC_011770 | 2081031 | G | A | 67 | 0 | 2 | 0 | NON_SYNONYMOU | MODERAT | MISSENSE | aGt/aAt | S138N | PLES_19261 | | | | | | | | | A | A | | |
| NC_011770 | 2081257 | A | G | 65 | 0 | 8 | 0 | NON_SYNONYMOU | MODERAT | MISSENSE | Acg/Gcg | T29A | PLES_19271 | | | G | G | G | G | | | G | G | G | G |
| NC_011770 | 2081266 | G | A | 66 | 0 | 8 | 0 | NON_SYNONYMOU | MODERAT | MISSENSE | Gtg/Atg | V32M | PLES_19271 | | | A | A | A | A | | | A | A | A | A |
| NC_011770 | 2081351 | T | C | 67 | 0 | 2 | 0 | NON_SYNONYMOU | MODERAT | MISSENSE | gTg/gCg | V60A | PLES_19271 | | | | | | | | | C | C | | |
| NC_011770 | 2081371 | C | T | 67 | 0 | 1 | 0 | NON_SYNONYMOU | MODERAT | MISSENSE | Ccg/Tcg | P67S | PLES_19271 | | | | | | | T | | | | | |
| NC_011770 | 2081836 | A | C | 59 | 0 | 4 | 0 | NON_SYNONYMOU | MODERAT | MISSENSE | Acg/Ccg | T222P | PLES_19271 | | | C | C | C | C | | | | | | |
| NC_011770 | 2081876 | C | T | 63 | 0 | 2 | 0 | NON_SYNONYMOU | MODERAT | MISSENSE | gCc/gTc | A235V | PLES_19271 | | | | | | | | | T | T | | |
| NC_011770 | 2082170 | A | T | 74 | 0 | 8 | 0 | NON_SYNONYMOU | MODERAT | MISSENSE | cAg/cTg | Q10L | PLES_19281 | | | T | T | T | T | | | T | T | T | T |
| NC_011770 | 2082505 | A | G | 63 | 0 | 12 | 0 | NON_SYNONYMOU | MODERAT | MISSENSE | Acg/Gcg | T122A | PLES_19281 | G | G | G | G | G | G | G | G | G | G | G | G |
| NC_011770 | 2083667 | A | G | 84 | 0 | 12 | 0 | NON_SYNONYMOU | MODERAT | MISSENSE | Aag/Gag | K117E | PLES_19301 | G | G | G | G | G | G | G | G | G | G | G | G |
| NC_011770 | 2083929 | T | C | 88 | 0 | 12 | 0 | NON_SYNONYMOU | MODERAT | MISSENSE | aTg/aCg | M204T | PLES_19301 | C | C | C | C | C | C | C | C | C | C | C | C |
| NC_011770 | 2084055 | G | A | 78 | 0 | 2 | 0 | NON_SYNONYMOU | MODERAT | MISSENSE | cGc/cAc | R246H | PLES_19301 | A | A | | | | | | | | | | |
| NC_011770 | 2084193 | T | A | 80 | 0 | 2 | 0 | NON_SYNONYMOU | MODERAT | MISSENSE | gTc/gAc | V292D | PLES_19301 | | | | | | | A | A | | | | |
| NC_011770 | 2084772 | T | C | 70 | 0 | 2 | 0 | NON_SYNONYMOU | MODERAT | MISSENSE | Atg/Gtg | M90V | PLES_19311 | C | C | | | | | | | | | | |
| NC_011770 | 2084792 | A | G | 72 | 0 | 2 | 0 | NON_SYNONYMOU | MODERAT | MISSENSE | gTg/gCg | V83A | PLES_19311 | G | G | | | | | | | | | | |
| NC_011770 | 2084909 | T | C | 87 | 0 | 10 | 0 | NON_SYNONYMOU | MODERAT | MISSENSE | cAc/cGc | H44R | PLES_19311 | C | C | C | C | C | C | | | C | C | C | C |

| | | | | | | | | | | | | | | | | | | | | | | | | | |
|---|---|---|---|---|---|---|---|---|---|---|---|---|---|---|---|---|---|---|---|---|---|---|---|---|---|
| NC_011770 | 2085423 | C | T | 74 | 0 | 4 | 0 | NON_SYNONYMOU | MODERAT | MISSENSE | cGc/cAc | R146H | PLES_19321 | T | T | | | | | | | T | T | | |
| NC_011770 | 2085505 | C | T | 67 | 0 | 8 | 0 | NON_SYNONYMOU | MODERAT | MISSENSE | Gcg/Acg | A119T | PLES_19321 | | | T | T | T | T | T | T | | | T | T |
| NC_011770 | 2085550 | T | C | 72 | 0 | 4 | 0 | NON_SYNONYMOU | MODERAT | MISSENSE | Atc/Gtc | I104V | PLES_19321 | C | C | | | | | | | C | C | | |
| NC_011770 | 2085591 | A | G | 78 | 0 | 4 | 0 | NON_SYNONYMOU | MODERAT | MISSENSE | gTc/gCc | V90A | PLES_19321 | G | G | | | | | | | G | G | | |
| NC_011770 | 2085616 | C | T | 77 | 0 | 4 | 0 | NON_SYNONYMOU | MODERAT | MISSENSE | Ggc/Agc | G82S | PLES_19321 | T | T | | | | | | | T | T | | |
| NC_011770 | 2085634 | T | C | 76 | 0 | 4 | 0 | NON_SYNONYMOU | MODERAT | MISSENSE | Acg/Gcg | T76A | PLES_19321 | C | C | | | | | | | C | C | | |
| NC_011770 | 2086514 | T | C | 65 | 0 | 2 | 0 | NON_SYNONYMOU | MODERAT | MISSENSE | aAa/aGa | K442R | PLES_19341 | | | | | | | | | C | C | | |
| NC_011770 | 2086521 | C | T | 64 | 0 | 4 | 0 | NON_SYNONYMOU | MODERAT | MISSENSE | Gca/Aca | A440T | PLES_19341 | T | T | | | | | | | T | T | | |
| NC_011770 | 2086542 | C | T | 64 | 0 | 2 | 0 | NON_SYNONYMOU | MODERAT | MISSENSE | Gcg/Acg | A433T | PLES_19341 | | | | | | | | | T | T | | |
| NC_011770 | 2086547 | A | G | 61 | 0 | 8 | 0 | NON_SYNONYMOU | MODERAT | MISSENSE | cTa/cCa | L431P | PLES_19341 | G | G | | | | | G | G | T | T | G | G |
| NC_011770 | 2086548 | G | T | 63 | 0 | 8 | 0 | NON_SYNONYMOU | MODERAT | MISSENSE | Cta/Ata | L431I | PLES_19341 | T | T | | | | | T | T | T | T | T | T |
| NC_011770 | 2086682 | C | T | 67 | 0 | 2 | 0 | NON_SYNONYMOU | MODERAT | MISSENSE | cGc/cAc | R386H | PLES_19341 | | | | | | | | | T | T | | |
| NC_011770 | 2086770 | T | C | 71 | 0 | 12 | 0 | NON_SYNONYMOU | MODERAT | MISSENSE | Acc/Gcc | T357A | PLES_19341 | C | C | C | C | C | C | C | C | C | C | C | C |
| NC_011770 | 2086774 | C | A | 68 | 0 | 12 | 0 | NON_SYNONYMOU | MODERAT | MISSENSE | caG/caT | Q355H | PLES_19341 | A | A | A | A | A | A | A | A | A | A | A | A |
| NC_011770 | 2087199 | T | C | 81 | 0 | 2 | 0 | NON_SYNONYMOU | MODERAT | MISSENSE | Acg/Gcg | T214A | PLES_19341 | | | | | | | | | C | C | | |
| NC_011770 | 2087211 | T | C | 80 | 0 | 4 | 0 | NON_SYNONYMOU | MODERAT | MISSENSE | Acg/Gcg | T210A | PLES_19341 | | | C | C | C | C | | | | | | |
| NC_011770 | 2087252 | G | C | 79 | 0 | 6 | 0 | NON_SYNONYMOU | MODERAT | MISSENSE | aCc/aGc | T196S | PLES_19341 | C | C | | | | | C | C | | | C | C |
| NC_011770 | 2087282 | C | T | 78 | 0 | 12 | 0 | NON_SYNONYMOU | MODERAT | MISSENSE | cGg/cAg | R186Q | PLES_19341 | T | T | T | T | T | T | T | T | T | T | T | T |
| NC_011770 | 2087569 | C | G | 75 | 0 | 2 | 0 | NON_SYNONYMOU | MODERAT | MISSENSE | agG/agC | R90S | PLES_19341 | | | | | | | | | G | G | | |
| NC_011770 | 2087570 | C | T | 76 | 0 | 8 | 0 | NON_SYNONYMOU | MODERAT | MISSENSE | aGg/aAg | R90K | PLES_19341 | | | T | T | T | T | | | T | T | T | T |
| NC_011770 | 2087651 | T | C | 77 | 0 | 2 | 0 | NON_SYNONYMOU | MODERAT | MISSENSE | cAc/cGc | H63R | PLES_19341 | | | | | | | | | | | C | C |
| NC_011770 | 2087748 | C | T | 78 | 0 | 2 | 0 | NON_SYNONYMOU | MODERAT | MISSENSE | Gcg/Acg | A31T | PLES_19341 | | | | | | | | | T | T | | |
| NC_011770 | 2087775 | C | T | 77 | 0 | 2 | 0 | NON_SYNONYMOU | MODERAT | MISSENSE | Gcg/Acg | A22T | PLES_19341 | T | T | | | | | | | | | | |
| NC_011770 | 2088011 | C | CG | 78 | 0 | 1 | 0 | FRAME_SHIFT | HIGH | | ggc/ggGc | G23G? | PLES_19351 | | | | | | | CG | | | | | |
| NC_011770 | 2088854 | AG | A | 68 | 0 | 4 | 0 | FRAME_SHIFT | HIGH | | ggt/ | G303 | PLES_19351 | | | A | A | A | A | | | | | | |
| NC_011770 | 2088154 | G | A | 71 | 0 | 8 | 0 | NON_SYNONYMOU | MODERAT | MISSENSE | Gcg/Acg | A69T | PLES_19351 | | | A | A | A | A | | | A | A | A | A |
| NC_011770 | 2088292 | G | C | 81 | 0 | 2 | 0 | NON_SYNONYMOU | MODERAT | MISSENSE | Gtg/Ctg | V115L | PLES_19351 | | | | | | | C | C | | | | |
| NC_011770 | 2088611 | A | G | 72 | 0 | 1 | 0 | NON_SYNONYMOU | MODERAT | MISSENSE | aAc/aGc | N221S | PLES_19351 | | | | | | | G | | | | | |
| NC_011770 | 2088709 | G | T | 67 | 0 | 2 | 0 | NON_SYNONYMOU | MODERAT | MISSENSE | Gcc/Tcc | A254S | PLES_19351 | | | | | | | T | T | | | | |
| NC_011770 | 2089050 | C | G | 66 | 0 | 8 | 0 | NON_SYNONYMOU | MODERAT | MISSENSE | aCc/aGc | T37S | PLES_19361 | | | G | G | G | G | | | G | G | G | G |
| NC_011770 | 2089125 | G | A | 72 | 0 | 4 | 0 | NON_SYNONYMOU | MODERAT | MISSENSE | aGc/aAc | S62N | PLES_19361 | | | A | A | A | A | | | | | | |
| NC_011770 | 2089275 | T | G | 66 | 0 | 2 | 0 | NON_SYNONYMOU | MODERAT | MISSENSE | gTc/gGc | V112G | PLES_19361 | | | | | | | G | G | | | | |
| NC_011770 | 2090049 | T | C | 85 | 0 | 2 | 0 | NON_SYNONYMOU | MODERAT | MISSENSE | Acc/Gcc | T86A | PLES_19371 | | | | | | | | | | | C | C |
| NC_011770 | 2093229 | A | C | 65 | 0 | 2 | 0 | NON_SYNONYMOU | MODERAT | MISSENSE | gAg/gCg | E140A | PLES_19401 | | | | | | | | | | | C | C |
| NC_011770 | 2095994 | A | G | 74 | 0 | 2 | 0 | NON_SYNONYMOU | MODERAT | MISSENSE | Agt/Ggt | S66G | PLES_19431 | G | G | | | | | | | | | | |
| NC_011770 | 2096231 | G | A | 76 | 0 | 2 | 0 | NON_SYNONYMOU | MODERAT | MISSENSE | Gcc/Acc | A145T | PLES_19431 | A | A | | | | | | | | | | |
| NC_011770 | 2096300 | A | G | 78 | 0 | 12 | 0 | NON_SYNONYMOU | MODERAT | MISSENSE | Atc/Gtc | I168V | PLES_19431 | G | G | G | G | G | G | G | G | G | G | G | G |
| NC_011770 | 2096660 | G | A | 89 | 0 | 4 | 0 | NON_SYNONYMOU | MODERAT | MISSENSE | Ggc/Agc | G288S | PLES_19431 | | | A | A | A | A | | | | | | |
| NC_011770 | 2101296 | C | T | 70 | 0 | 4 | 0 | NON_SYNONYMOU | MODERAT | MISSENSE | gCg/gTg | A16V | PLES_19471 | T | T | | | | | | | | | T | T |
| NC_011770 | 2103986 | A | T | 21 | 0 | 4 | 0 | NON_SYNONYMOU | MODERAT | MISSENSE | aAc/aTc | N72I | PLES_19501 | | | | | | | T | T | | | T | T |
| NC_011770 | 2104042 | C | G | 16 | 1 | 2 | 0 | NON_SYNONYMOU | MODERAT | MISSENSE | Ccg/Gcg | P91A | PLES_19501 | | . | | | | | G | G | | | | |
| NC_011770 | 2104075 | G | A | 14 | 1 | 11 | 0 | NON_SYNONYMOU | MODERAT | MISSENSE | Gcc/Acc | A102T | PLES_19501 | A | . | A | A | A | A | A | A | A | A | A | A |
| NC_011770 | 2104086 | CGCCCAG | C | 14 | 0 | 1 | 0 | CODON_CHANGE_ | MODERATE | | gcccaggcg, | AQA108A | PLES_19501 | | | | | | | | C | | | | |
| NC_011770 | 2108288 | C | T | 78 | 0 | 2 | 0 | NON_SYNONYMOU | MODERAT | MISSENSE | cCg/cTg | P97L | PLES_19541 | | | | | | | T | T | | | | |
| NC_011770 | 2120389 | T | C | 70 | 0 | 10 | 0 | NON_SYNONYMOU | MODERAT | MISSENSE | aAc/aGc | N221S | PLES_19661 | C | C | C | C | C | C | | | C | C | C | C |
| NC_011770 | 2120864 | C | A | 83 | 0 | 2 | 0 | NON_SYNONYMOU | MODERAT | MISSENSE | Gcc/Tcc | A63S | PLES_19661 | | | | | | | A | A | | | | |
| NC_011770 | 2120957 | G | C | 80 | 0 | 8 | 0 | NON_SYNONYMOU | MODERAT | MISSENSE | Cca/Gca | P32A | PLES_19661 | C | C | | | | | C | C | C | C | C | C |
| NC_011770 | 2120698 | C | T | 75 | 0 | 1 | 0 | NON_SYNONYMOU | MODERAT | MISSENSE | gGc/gAc | G118D | PLES_19661 | | | | | | | | T | | | | |
| NC_011770 | 2121222 | C | T | 79 | 0 | 6 | 0 | NON_SYNONYMOU | MODERAT | MISSENSE | cGc/cAc | R315H | PLES_19671 | T | T | | | | | T | T | | | T | T |
| NC_011770 | 2121364 | G | C | 79 | 0 | 2 | 0 | NON_SYNONYMOU | MODERAT | MISSENSE | Cgc/Ggc | R268G | PLES_19671 | | | | | | | | | C | C | | |
| NC_011770 | 2121684 | G | A | 78 | 0 | 2 | 0 | NON_SYNONYMOU | MODERAT | MISSENSE | aCc/aTc | T161I | PLES_19671 | | | | | | | | | A | A | | |
| NC_011770 | 2121865 | C | T | 72 | 0 | 4 | 0 | NON_SYNONYMOU | MODERAT | MISSENSE | Gtg/Atg | V101M | PLES_19671 | | | T | T | T | T | | | | | | |
| NC_011770 | 2125535 | C | A | 70 | 0 | 2 | 0 | NON_SYNONYMOU | MODERAT | MISSENSE | agC/agA | S260R | PLES_19691 | | | | | | | A | A | | | | |
| NC_011770 | 2125638 | C | T | 71 | 0 | 1 | 0 | NON_SYNONYMOU | MODERAT | MISSENSE | Cgg/Tgg | R295W | PLES_19691 | | | | | | | T | | | | | |
| NC_011770 | 2126176 | G | A | 74 | 0 | 2 | 0 | NON_SYNONYMOU | MODERAT | MISSENSE | cGc/cAc | R474H | PLES_19691 | | | | | | | | | | | A | A |
| NC_011770 | 2126142 | A | G | 77 | 0 | 1 | 0 | NON_SYNONYMOU | MODERAT | MISSENSE | Acc/Gcc | T463A | PLES_19691 | G | | | | | | | | | | | |
| NC_011770 | 2126363 | A | G | 80 | 0 | 8 | 0 | NON_SYNONYMOU | MODERAT | MISSENSE | cAc/cGc | H44R | PLES_19701 | G | G | | | | | G | G | G | G | G | G |
| NC_011770 | 2126450 | G | A | 78 | 0 | 2 | 0 | NON_SYNONYMOU | MODERAT | MISSENSE | gGt/gAt | G73D | PLES_19701 | | | | | | | A | A | | | | |
| NC_011770 | 2126873 | G | C | 72 | 0 | 8 | 0 | NON_SYNONYMOU | MODERAT | MISSENSE | gGc/gCc | G214A | PLES_19701 | C | C | | | | | C | C | C | C | C | C |
| NC_011770 | 2126906 | G | A | 75 | 0 | 1 | 0 | NON_SYNONYMOU | MODERAT | MISSENSE | cGt/cAt | R225H | PLES_19701 | | | | | | | A | | | | | |
| NC_011770 | 2127050 | G | A | 77 | 0 | 4 | 0 | NON_SYNONYMOU | MODERAT | MISSENSE | cGc/cAc | R273H | PLES_19701 | | | A | A | A | A | | | | | | |
| NC_011770 | 2127317 | T | G | 65 | 0 | 2 | 0 | NON_SYNONYMOU | MODERAT | MISSENSE | gAc/gCc | D261A | PLES_19711 | | | | | | | G | G | | | | |

| | | | | | | | | | | | | | | | | | | | | | | | | | |
|---|---|---|---|---|---|---|---|---|---|---|---|---|---|---|---|---|---|---|---|---|---|---|---|---|---|
| NC_011770 | 2127390 | C | T | 61 | 0 | 4 | 0 | NON_SYNONYMOU | MODERAT | MISSENSE | Gcc/Acc | A237T | PLES_19711 | T | T | | | | | T | T | | | | |
| NC_011770 | 2128064 | C | T | 36 | 0 | 4 | 0 | NON_SYNONYMOU | MODERAT | MISSENSE | cGc/cAc | R12H | PLES_19711 | | | | | | | | | T | T | T | T |
| NC_011770 | 2127969 | G | A | 40 | 0 | 1 | 0 | NON_SYNONYMOU | MODERAT | MISSENSE | Cgc/Tgc | R44C | PLES_19711 | | | | | | | | A | | | | |
| NC_011770 | 2130355 | T | C | 75 | 0 | 12 | 0 | NON_SYNONYMOU | MODERAT | MISSENSE | gTg/gCg | V90A | PLES_19741 | C | C | C | C | C | C | C | C | C | C | C | C |
| NC_011770 | 2131253 | G | C | 85 | 0 | 12 | 0 | NON_SYNONYMOU | MODERAT | MISSENSE | Gcg/Ccg | A75P | PLES_19751 | C | C | C | C | C | C | C | C | C | C | C | C |
| NC_011770 | 2131417 | T | C | 81 | 0 | 12 | 0 | NON_SYNONYMOU | MODERAT | MISSENSE | Tcg/Ccg | S31P | PLES_19761 | C | C | C | C | C | C | C | C | C | C | C | C |
| NC_011770 | 2131820 | A | G | 79 | 0 | 2 | 0 | NON_SYNONYMOU | MODERAT | MISSENSE | aAg/aGg | K165R | PLES_19761 | | | | | | | G | G | | | | |
| NC_011770 | 2138859 | T | A | 74 | 0 | 6 | 0 | NON_SYNONYMOU | MODERAT | MISSENSE | Tcc/Acc | S14T | PLES_19801 | A | A | | | | | | | A | A | A | A |
| NC_011770 | 2138872 | T | C | 75 | 0 | 8 | 0 | NON_SYNONYMOU | MODERAT | MISSENSE | tTg/tCg | L18S | PLES_19801 | C | C | | | | | C | C | C | C | C | C |
| NC_011770 | 2138923 | T | C | 80 | 0 | 12 | 0 | NON_SYNONYMOU | MODERAT | MISSENSE | tTg/tCg | L35S | PLES_19801 | C | C | C | C | C | C | C | C | C | C | C | C |
| NC_011770 | 2138935 | T | C | 75 | 0 | 12 | 0 | NON_SYNONYMOU | MODERAT | MISSENSE | gTg/gCg | V39A | PLES_19801 | C | C | C | C | C | C | C | C | C | C | C | C |
| NC_011770 | 2139255 | A | G | 84 | 0 | 8 | 0 | NON_SYNONYMOU | MODERAT | MISSENSE | Acc/Gcc | T146A | PLES_19801 | G | G | G | G | G | G | | | G | G | | |
| NC_011770 | 2139675 | G | C | 71 | 0 | 2 | 0 | NON_SYNONYMOU | MODERAT | MISSENSE | Gac/Cac | D286H | PLES_19801 | | | | | | | C | C | | | | |
| NC_011770 | 2142297 | G | C | 79 | 0 | 6 | 0 | NON_SYNONYMOU | MODERAT | MISSENSE | Gtg/Ctg | V790L | PLES_19811 | | | C | C | C | C | | | C | C | | |
| NC_011770 | 2142447 | G | T | 85 | 0 | 4 | 0 | NON_SYNONYMOU | MODERAT | MISSENSE | Ctg/Atg | L411M | PLES_19821 | T | T | | | | | | | | | T | T |
| NC_011770 | 2142520 | C | G | 90 | 0 | 4 | 0 | NON_SYNONYMOU | MODERAT | MISSENSE | gaG/gaC | E386D | PLES_19821 | G | G | | | | | | | | | G | G |
| NC_011770 | 2142687 | G | A | 89 | 0 | 2 | 0 | NON_SYNONYMOU | MODERAT | MISSENSE | Cac/Tac | H331Y | PLES_19821 | | | | | | | | | | | A | A |
| NC_011770 | 2145362 | G | T | 78 | 0 | 2 | 0 | NON_SYNONYMOU | MODERAT | MISSENSE | gCc/gAc | A42D | PLES_19841 | | | | | | | | | T | T | | |
| NC_011770 | 2148664 | G | A | 67 | 0 | 1 | 0 | STOP_GAINED | HIGH | NONSENSE | Cag/Tag | Q89* | PLES_19861 | | | | | | | | | | | A | |
| NC_011770 | 2147875 | G | A | 69 | 0 | 1 | 0 | STOP_GAINED | HIGH | NONSENSE | Cag/Tag | Q352* | PLES_19861 | | | | | | | | | | | | A |
| NC_011770 | 2147335 | C | T | 63 | 0 | 4 | 0 | NON_SYNONYMOU | MODERAT | MISSENSE | Gac/Aac | D532N | PLES_19861 | T | T | | | | | T | T | | | | |
| NC_011770 | 2147505 | T | C | 62 | 0 | 12 | 0 | NON_SYNONYMOU | MODERAT | MISSENSE | cAc/cGc | H475R | PLES_19861 | C | C | C | C | C | C | C | C | C | C | C | C |
| NC_011770 | 2147524 | C | T | 63 | 0 | 2 | 0 | NON_SYNONYMOU | MODERAT | MISSENSE | Ggc/Agc | G469S | PLES_19861 | T | T | | | | | | | | | | |
| NC_011770 | 2147781 | A | T | 66 | 0 | 4 | 0 | NON_SYNONYMOU | MODERAT | MISSENSE | tTc/tAc | F383Y | PLES_19861 | | | T | T | T | T | | | | | | |
| NC_011770 | 2147809 | C | T | 67 | 0 | 2 | 0 | NON_SYNONYMOU | MODERAT | MISSENSE | Gac/Aac | D374N | PLES_19861 | T | T | | | | | | | | | | |
| NC_011770 | 2148265 | T | C | 68 | 0 | 8 | 0 | NON_SYNONYMOU | MODERAT | MISSENSE | Acg/Gcg | T222A | PLES_19861 | C | C | | | | | C | C | C | C | C | C |
| NC_011770 | 2148315 | C | T | 69 | 0 | 6 | 0 | NON_SYNONYMOU | MODERAT | MISSENSE | gGg/gAg | G205E | PLES_19861 | T | T | | | | | | | T | T | T | T |
| NC_011770 | 2149104 | C | A | 70 | 0 | 2 | 0 | NON_SYNONYMOU | MODERAT | MISSENSE | Gcg/Tcg | A282S | PLES_19871 | | | | | | | | | | | A | A |
| NC_011770 | 2149160 | C | A | 71 | 0 | 4 | 0 | NON_SYNONYMOU | MODERAT | MISSENSE | aGc/aTc | S263I | PLES_19871 | | | A | A | A | A | | | | | | |
| NC_011770 | 2149615 | T | G | 69 | 0 | 4 | 0 | NON_SYNONYMOU | MODERAT | MISSENSE | gaA/gaC | E111D | PLES_19871 | | | | | | | | | G | G | G | G |
| NC_011770 | 2149770 | C | T | 56 | 0 | 2 | 0 | NON_SYNONYMOU | MODERAT | MISSENSE | Gtg/Atg | V60M | PLES_19871 | | | | | | | | | | | T | T |
| NC_011770 | 2149787 | G | A | 56 | 0 | 2 | 0 | NON_SYNONYMOU | MODERAT | MISSENSE | cCg/cTg | P54L | PLES_19871 | | | | | | | | | | | A | A |
| NC_011770 | 2149914 | G | A | 56 | 0 | 4 | 0 | NON_SYNONYMOU | MODERAT | MISSENSE | Ctc/Ttc | L12F | PLES_19871 | A | A | | | | | | | | | A | A |
| NC_011770 | 2150393 | CAG | C | 47 | 0 | 1 | 0 | FRAME_SHIFT | HIGH | | ctg/ | L14 | PLES_19881 | | | | C | | | | | | | | |
| NC_011770 | 2150394 | A | G | 43 | 0 | 3 | 0 | NON_SYNONYMOU | MODERAT | MISSENSE | cTg/cCg | L14P | PLES_19881 | | | G | . | G | G | | | | | | |
| NC_011770 | 2150570 | A | T | 55 | 0 | 8 | 0 | NON_SYNONYMOU | MODERAT | MISSENSE | cTg/cAg | L267Q | PLES_19891 | T | T | T | T | T | T | | | | | T | T |
| NC_011770 | 2151545 | GGC | G | 85 | 0 | 1 | 0 | FRAME_SHIFT | HIGH | | gcc/ | A272 | PLES_19901 | | | G | | | | | | | | | |
| NC_011770 | 2151531 | G | C | 86 | 0 | 2 | 0 | NON_SYNONYMOU | MODERAT | MISSENSE | gCc/gGc | A277G | PLES_19901 | | | | | | | C | C | | | | |
| NC_011770 | 2153112 | A | G | 68 | 0 | 6 | 0 | NON_SYNONYMOU | MODERAT | MISSENSE | aAg/aGg | K87R | PLES_19911 | | | G | G | G | G | | | G | G | | |
| NC_011770 | 2159300 | T | C | 91 | 0 | 2 | 0 | NON_SYNONYMOU | MODERAT | MISSENSE | aTt/aCt | I21T | PLES_19941 | | | | | | | | | C | C | | |
| NC_011770 | 2159378 | G | T | 87 | 0 | 2 | 0 | NON_SYNONYMOU | MODERAT | MISSENSE | cGt/cTt | R47L | PLES_19941 | | | | | | | T | T | | | | |
| NC_011770 | 2159426 | T | C | 92 | 0 | 12 | 0 | NON_SYNONYMOU | MODERAT | MISSENSE | cTg/cCg | L63P | PLES_19941 | C | C | C | C | C | C | C | C | C | C | C | C |
| NC_011770 | 2160410 | C | G | 92 | 0 | 8 | 0 | NON_SYNONYMOU | MODERAT | MISSENSE | tCc/tGc | S196C | PLES_19951 | | | G | G | G | G | G | G | | | G | G |
| NC_011770 | 2160791 | G | T | 72 | 0 | 2 | 0 | NON_SYNONYMOU | MODERAT | MISSENSE | cGg/cTg | R323L | PLES_19951 | | | | | | | | | T | T | | |
| NC_011770 | 2174194 | A | G | 88 | 0 | 6 | 0 | NON_SYNONYMOU | MODERAT | MISSENSE | cAg/cGg | Q104R | PLES_20041 | | | | | | | G | G | G | G | G | G |
| NC_011770 | 2174295 | G | C | 101 | 0 | 2 | 0 | NON_SYNONYMOU | MODERAT | MISSENSE | Gaa/Caa | E138Q | PLES_20041 | | | | | | | | | C | C | | |
| NC_011770 | 2174305 | C | T | 99 | 0 | 2 | 0 | NON_SYNONYMOU | MODERAT | MISSENSE | cCg/cTg | P141L | PLES_20041 | | | | | | | | | T | T | | |
| NC_011770 | 2174401 | G | A | 99 | 0 | 1 | 0 | NON_SYNONYMOU | MODERAT | MISSENSE | Gtc/Atc | V2I | PLES_20051 | | | | | | | A | | | | | |
| NC_011770 | 2174750 | A | G | 65 | 0 | 4 | 0 | NON_SYNONYMOU | MODERAT | MISSENSE | aAg/aGg | K118R | PLES_20051 | | | | | | | G | G | | | G | G |
| NC_011770 | 2174773 | G | C | 65 | 0 | 2 | 0 | NON_SYNONYMOU | MODERAT | MISSENSE | Gtg/Ctg | V126L | PLES_20051 | | | | | | | | | C | C | | |
| NC_011770 | 2175125 | C | A | 72 | 0 | 2 | 0 | NON_SYNONYMOU | MODERAT | MISSENSE | Gcc/Tcc | A591S | PLES_20061 | A | A | | | | | | | | | | |
| NC_011770 | 2175709 | G | A | 77 | 0 | 4 | 0 | NON_SYNONYMOU | MODERAT | MISSENSE | gCc/gTc | A396V | PLES_20061 | | | | | | | A | A | A | A | | |
| NC_011770 | 2176649 | G | T | 69 | 0 | 2 | 0 | NON_SYNONYMOU | MODERAT | MISSENSE | Cgt/Agt | R83S | PLES_20061 | T | T | | | | | | | | | | |
| NC_011770 | 2177320 | C | G | 83 | 0 | 2 | 0 | NON_SYNONYMOU | MODERAT | MISSENSE | Gtc/Ctc | V249L | PLES_20071 | | | | | | | | | | | G | G |
| NC_011770 | 2177682 | C | T | 80 | 0 | 4 | 0 | NON_SYNONYMOU | MODERAT | MISSENSE | cGg/cAg | R128Q | PLES_20071 | | | T | T | T | T | | | | | | |
| NC_011770 | 2177689 | G | T | 84 | 0 | 4 | 0 | NON_SYNONYMOU | MODERAT | MISSENSE | Ctg/Atg | L126M | PLES_20071 | | | T | T | T | T | | | | | | |
| NC_011770 | 2177820 | C | T | 76 | 0 | 2 | 0 | NON_SYNONYMOU | MODERAT | MISSENSE | cGc/cAc | R82H | PLES_20071 | | | | | T | T | | | | | | |
| NC_011770 | 2181056 | A | T | 81 | 0 | 6 | 0 | NON_SYNONYMOU | MODERAT | MISSENSE | aaT/aaA | N27K | PLES_20111 | T | T | T | T | T | T | | | | | | |
| NC_011770 | 2181061 | C | T | 81 | 0 | 2 | 0 | NON_SYNONYMOU | MODERAT | MISSENSE | Gca/Aca | A26T | PLES_20111 | | | | | | | | | | | T | T |
| NC_011770 | 2183613 | C | T | 78 | 0 | 1 | 0 | NON_SYNONYMOU | MODERAT | MISSENSE | Gcc/Acc | A434T | PLES_20131 | | | | | | | T | | | | | |
| NC_011770 | 2183733 | G | C | 94 | 0 | 10 | 0 | NON_SYNONYMOU | MODERAT | MISSENSE | Ccg/Gcg | P394A | PLES_20131 | | | C | C | C | C | C | C | C | C | C | C |
| NC_011770 | 2185500 | A | G | 55 | 0 | 2 | 0 | NON_SYNONYMOU | MODERAT | MISSENSE | Agc/Ggc | S94G | PLES_20141 | | | | | | | G | G | | | | |

| | | | | | | | | | | | | | | | | | | | | | | | | | |
|---|---|---|---|---|---|---|---|---|---|---|---|---|---|---|---|---|---|---|---|---|---|---|---|---|---|
| NC_011770 | 2186103 | C | T | 80 | 0 | 2 | 0 | NON_SYNONYMOU | MODERAT | MISSENSE | gCc/gTc | A38V | PLES_20151 | | | | | | | | | | | T | T |
| NC_011770 | 2186713 | C | T | 70 | 0 | 2 | 0 | NON_SYNONYMOU | MODERAT | MISSENSE | gCg/gTg | A29V | PLES_20161 | T | T | | | | | | | | | | |
| NC_011770 | 2186820 | C | A | 62 | 0 | 4 | 0 | NON_SYNONYMOU | MODERAT | MISSENSE | Cgc/Agc | R65S | PLES_20161 | A | A | | | | | A | A | | | | |
| NC_011770 | 2187267 | G | A | 91 | 0 | 2 | 0 | NON_SYNONYMOU | MODERAT | MISSENSE | Gtg/Atg | V214M | PLES_20161 | A | A | | | | | | | | | | |
| NC_011770 | 2187875 | C | A | 84 | 0 | 2 | 0 | NON_SYNONYMOU | MODERAT | MISSENSE | gaC/gaA | D416E | PLES_20161 | A | A | | | | | | | | | | |
| NC_011770 | 2188522 | C | A | 79 | 0 | 4 | 0 | NON_SYNONYMOU | MODERAT | MISSENSE | gCc/gAc | A632D | PLES_20161 | | | A | A | A | A | | | | | | |
| NC_011770 | 2188546 | G | A | 77 | 0 | 6 | 0 | NON_SYNONYMOU | MODERAT | MISSENSE | gGc/gAc | G640D | PLES_20161 | A | A | | | | | A | A | A | A | | |
| NC_011770 | 2188635 | A | G | 83 | 0 | 10 | 0 | NON_SYNONYMOU | MODERAT | MISSENSE | Agc/Ggc | S670G | PLES_20161 | G | G | G | G | G | G | G | G | G | G | | |
| NC_011770 | 2188644 | A | G | 85 | 0 | 4 | 0 | NON_SYNONYMOU | MODERAT | MISSENSE | Acc/Gcc | T673A | PLES_20161 | | | G | G | G | G | | | | | | |
| NC_011770 | 2188744 | G | T | 79 | 0 | 2 | 0 | NON_SYNONYMOU | MODERAT | MISSENSE | cGg/cTg | R706L | PLES_20161 | | | T | T | | | | | | | | |
| NC_011770 | 2187909 | G | A | 84 | 0 | 1 | 0 | NON_SYNONYMOU | MODERAT | MISSENSE | Gcc/Acc | A428T | PLES_20161 | | | | | | | | A | | | | |
| NC_011770 | 2190313 | C | T | 62 | 0 | 2 | 0 | SPLICE_SITE_REGIC | LOW | SILENT | tGa/tAa | *108 | PLES_20181 | | | | | | | | | T | T | | |
| NC_011770 | 2190325 | T | C | 60 | 0 | 4 | 0 | NON_SYNONYMOU | MODERAT | MISSENSE | cAg/cGg | Q104R | PLES_20181 | | | C | C | C | C | | | | | | |
| NC_011770 | 2190328 | T | G | 58 | 0 | 4 | 0 | NON_SYNONYMOU | MODERAT | MISSENSE | aAg/aCg | K103T | PLES_20181 | | | G | G | G | G | | | | | | |
| NC_011770 | 2190872 | G | A | 74 | 0 | 2 | 0 | NON_SYNONYMOU | MODERAT | MISSENSE | gCc/gTc | A53V | PLES_20191 | | | | | | | | | A | A | | |
| NC_011770 | 2191355 | C | CG | 74 | 0 | 1 | 0 | FRAME_SHIFT | HIGH | | cgc/cCgc | R3P? | PLES_20201 | | | | CG | | | | | | | | |
| NC_011770 | 2191157 | A | G | 71 | 0 | 1 | 0 | NON_SYNONYMOU | MODERAT | MISSENSE | gTc/gCc | V69A | PLES_20201 | | | | | | | G | | | | | |
| NC_011770 | 2192109 | T | A | 64 | 0 | 2 | 0 | NON_SYNONYMOU | MODERAT | MISSENSE | cTg/cAg | L151Q | PLES_20211 | A | A | | | | | | | | | | |
| NC_011770 | 2192607 | A | C | 45 | 0 | 2 | 0 | NON_SYNONYMOU | MODERAT | MISSENSE | gAc/gCc | D317A | PLES_20211 | C | C | | | | | | | | | | |
| NC_011770 | 2192751 | G | A | 25 | 0 | 1 | 0 | NON_SYNONYMOU | MODERAT | MISSENSE | cGc/cAc | R365H | PLES_20211 | | | | | | | A | | | | | |
| NC_011770 | 2193820 | T | C | 70 | 0 | 4 | 0 | NON_SYNONYMOU | MODERAT | MISSENSE | aAc/aGc | N224S | PLES_20221 | | | C | C | C | C | | | | | | |
| NC_011770 | 2195492 | T | G | 49 | 0 | 2 | 0 | NON_SYNONYMOU | MODERAT | MISSENSE | Atc/Ctc | I12L | PLES_20231 | | | | | | | G | G | | | | |
| NC_011770 | 2196244 | CG | CGGG | 53 | 0 | 2 | 0 | FRAME_SHIFT | HIGH | | ccc/ | P71 | PLES_20241 | CGGG | | | C | | | | | | | | |
| NC_011770 | 2195539 | G | C | 46 | 0 | 12 | 0 | NON_SYNONYMOU | MODERAT | MISSENSE | Ccc/Gcc | P307A | PLES_20241 | C | C | C | C | C | C | C | C | C | C | C | C |
| NC_011770 | 2195929 | G | A | 53 | 0 | 2 | 0 | NON_SYNONYMOU | MODERAT | MISSENSE | Cgc/Tgc | R177C | PLES_20241 | | | | | | | | | A | A | | |
| NC_011770 | 2196201 | T | C | 52 | 0 | 2 | 0 | NON_SYNONYMOU | MODERAT | MISSENSE | aAc/aGc | N86S | PLES_20241 | | | | | | | | | | | C | C |
| NC_011770 | 2196313 | A | G | 54 | 0 | 2 | 0 | NON_SYNONYMOU | MODERAT | MISSENSE | Tcg/Ccg | S49P | PLES_20241 | | | | | | | | | | | G | G |
| NC_011770 | 2196322 | A | C | 53 | 0 | 10 | 0 | NON_SYNONYMOU | MODERAT | MISSENSE | Tcc/Gcc | S46A | PLES_20241 | | | C | C | C | C | C | C | C | C | C | C |
| NC_011770 | 2196430 | T | C | 61 | 0 | 4 | 0 | NON_SYNONYMOU | MODERAT | MISSENSE | Acc/Gcc | T10A | PLES_20241 | | | | | | | C | C | | | C | C |
| NC_011770 | 2196450 | T | G | 67 | 0 | 4 | 0 | NON_SYNONYMOU | MODERAT | MISSENSE | cAg/cCg | Q3P | PLES_20241 | | | | | | | G | G | | | G | G |
| NC_011770 | 2196556 | G | A | 66 | 0 | 1 | 0 | NON_SYNONYMOU | MODERAT | MISSENSE | gCg/gTg | A172V | PLES_20251 | | | | | | | A | | | | | |
| NC_011770 | 2196660 | G | C | 66 | 0 | 4 | 0 | NON_SYNONYMOU | MODERAT | MISSENSE | gaC/gaG | D137E | PLES_20251 | | | C | C | C | C | | | | | | |
| NC_011770 | 2196722 | T | C | 66 | 0 | 4 | 0 | NON_SYNONYMOU | MODERAT | MISSENSE | Aac/Gac | N117D | PLES_20251 | | | C | C | C | C | | | | | | |
| NC_011770 | 2196731 | T | C | 65 | 0 | 4 | 0 | NON_SYNONYMOU | MODERAT | MISSENSE | Act/Gct | T114A | PLES_20251 | | | C | C | C | C | | | | | | |
| NC_011770 | 2196832 | T | A | 64 | 0 | 6 | 0 | NON_SYNONYMOU | MODERAT | MISSENSE | gAg/gTg | E80V | PLES_20251 | | | A | A | A | A | A | A | | | | |
| NC_011770 | 2196838 | C | T | 63 | 0 | 6 | 0 | NON_SYNONYMOU | MODERAT | MISSENSE | gGt/gAt | G78D | PLES_20251 | | | T | T | T | T | T | T | | | | |
| NC_011770 | 2196949 | T | G | 71 | 0 | 4 | 0 | NON_SYNONYMOU | MODERAT | MISSENSE | cAg/cCg | Q41P | PLES_20251 | | | G | G | G | G | | | | | | |
| NC_011770 | 2196953 | T | C | 72 | 0 | 4 | 0 | NON_SYNONYMOU | MODERAT | MISSENSE | Acc/Gcc | T40A | PLES_20251 | | | C | C | C | C | | | | | | |
| NC_011770 | 2196974 | A | G | 71 | 0 | 4 | 0 | NON_SYNONYMOU | MODERAT | MISSENSE | Ttc/Ctc | F33L | PLES_20251 | | | G | G | G | G | | | | | | |
| NC_011770 | 2196983 | G | T | 68 | 0 | 4 | 0 | NON_SYNONYMOU | MODERAT | MISSENSE | Ccg/Acg | P30T | PLES_20251 | | | T | T | T | T | | | | | | |
| NC_011770 | 2197051 | T | G | 68 | 0 | 2 | 0 | NON_SYNONYMOU | MODERAT | MISSENSE | gAt/gCt | D7A | PLES_20251 | | | | | | | G | G | | | | |
| NC_011770 | 2197651 | C | T | 60 | 0 | 6 | 0 | NON_SYNONYMOU | MODERAT | MISSENSE | gCg/gTg | A139V | PLES_20261 | | | T | T | T | T | T | T | | | | |
| NC_011770 | 2197978 | A | G | 77 | 0 | 4 | 0 | NON_SYNONYMOU | MODERAT | MISSENSE | cAc/cGc | H60R | PLES_20271 | | | G | G | G | G | | | | | | |
| NC_011770 | 2198073 | T | C | 77 | 0 | 4 | 0 | NON_SYNONYMOU | MODERAT | MISSENSE | Tgc/Cgc | C92R | PLES_20271 | | | C | C | C | C | | | | | | |
| NC_011770 | 2198074 | G | A | 79 | 0 | 2 | 0 | NON_SYNONYMOU | MODERAT | MISSENSE | tGc/tAc | C92Y | PLES_20271 | A | A | | | | | | | | | | |
| NC_011770 | 2203396 | C | T | 73 | 0 | 2 | 0 | NON_SYNONYMOU | MODERAT | MISSENSE | Gcc/Acc | A191T | PLES_20331 | | | | | | | T | T | | | | |
| NC_011770 | 2204584 | C | A | 69 | 0 | 8 | 0 | NON_SYNONYMOU | MODERAT | MISSENSE | Ctc/Atc | L148I | PLES_20341 | A | A | A | A | A | A | A | A | | | | |
| NC_011770 | 2204942 | A | G | 74 | 0 | 2 | 0 | NON_SYNONYMOU | MODERAT | MISSENSE | gAc/gGc | D267G | PLES_20341 | | | | | | | G | G | | | | |
| NC_011770 | 2205431 | A | G | 86 | 0 | 2 | 0 | NON_SYNONYMOU | MODERAT | MISSENSE | aAa/aGa | K430R | PLES_20341 | | | | | | | G | G | | | | |
| NC_011770 | 2205503 | G | T | 93 | 0 | 2 | 0 | NON_SYNONYMOU | MODERAT | MISSENSE | cGt/cTt | R454L | PLES_20341 | | | | | | | | | T | T | | |
| NC_011770 | 2205748 | C | T | 79 | 0 | 2 | 0 | NON_SYNONYMOU | MODERAT | MISSENSE | aCt/aTt | T2I | PLES_20351 | | | | | | | | | | | T | T |
| NC_011770 | 2206046 | G | T | 95 | 0 | 2 | 0 | NON_SYNONYMOU | MODERAT | MISSENSE | gaG/gaT | E101D | PLES_20351 | T | T | | | | | | | | | | |
| NC_011770 | 2206392 | T | C | 85 | 0 | 4 | 0 | NON_SYNONYMOU | MODERAT | MISSENSE | Ttc/Ctc | F217L | PLES_20351 | C | C | | | | | C | C | | | | |
| NC_011770 | 2206419 | C | G | 88 | 0 | 4 | 0 | NON_SYNONYMOU | MODERAT | MISSENSE | Ctg/Gtg | L226V | PLES_20351 | G | G | | | | | G | G | | | | |
| NC_011770 | 2206672 | A | C | 82 | 0 | 2 | 0 | NON_SYNONYMOU | MODERAT | MISSENSE | gAc/gCc | D310A | PLES_20351 | C | C | | | | | | | | | | |
| NC_011770 | 2206908 | C | G | 69 | 0 | 2 | 0 | NON_SYNONYMOU | MODERAT | MISSENSE | Ccg/Gcg | P389A | PLES_20351 | | | | | | | | | G | G | | |
| NC_011770 | 2208014 | G | C | 72 | 0 | 8 | 0 | NON_SYNONYMOU | MODERAT | MISSENSE | Gag/Cag | E224Q | PLES_20361 | | | C | C | C | C | | | C | C | C | C |
| NC_011770 | 2208090 | C | T | 74 | 0 | 1 | 0 | NON_SYNONYMOU | MODERAT | MISSENSE | gCc/gTc | A249V | PLES_20361 | | | | | | | T | | | | | |
| NC_011770 | 2207984 | G | A | 73 | 0 | 1 | 0 | NON_SYNONYMOU | MODERAT | MISSENSE | Gag/Aag | E214K | PLES_20361 | | | | | | | | A | | | | |
| NC_011770 | 2209118 | G | A | 73 | 0 | 2 | 0 | NON_SYNONYMOU | MODERAT | MISSENSE | cGc/cAc | R47H | PLES_20371 | | | | | | | A | A | | | | |
| NC_011770 | 2209249 | G | A | 69 | 0 | 1 | 0 | NON_SYNONYMOU | MODERAT | MISSENSE | Ggc/Agc | G91S | PLES_20371 | | | | | | | | A | | | | |
| NC_011770 | 2210014 | C | T | 70 | 0 | 1 | 0 | NON_SYNONYMOU | MODERAT | MISSENSE | Cgc/Tgc | R9C | PLES_20381 | | | | | | | T | | | | | |

| | | | | | | | | | | | | | | | | | | | | | | | | | |
|---|---|---|---|---|---|---|---|---|---|---|---|---|---|---|---|---|---|---|---|---|---|---|---|---|---|
| NC_011770 | 2211193 | T | C | 43 | 0 | 6 | 0 | NON_SYNONYMOU | MODERAT | MISSENSE | Acc/Gcc | T12A | PLES_20401 | | | C | C | C | C | C | C | | | | |
| NC_011770 | 2211198 | G | A | 42 | 0 | 12 | 0 | NON_SYNONYMOU | MODERAT | MISSENSE | cCc/cTc | P10L | PLES_20401 | A | A | A | A | A | A | A | A | A | A | A | A |
| NC_011770 | 2211168 | C | T | 43 | 0 | 1 | 0 | NON_SYNONYMOU | MODERAT | MISSENSE | gGc/gAc | G20D | PLES_20401 | | | | | | | | T | | | | |
| NC_011770 | 2211288 | A | G | 52 | 0 | 6 | 0 | NON_SYNONYMOU | MODERAT | MISSENSE | Ttc/Ctc | F642L | PLES_20411 | | | G | G | G | G | G | G | | | | |
| NC_011770 | 2212854 | C | T | 86 | 0 | 1 | 0 | NON_SYNONYMOU | MODERAT | MISSENSE | Gat/Aat | D120N | PLES_20411 | | | | | | | T | | | | | |
| NC_011770 | 2212866 | T | C | 85 | 0 | 12 | 0 | NON_SYNONYMOU | MODERAT | MISSENSE | Atc/Gtc | I116V | PLES_20411 | C | C | C | C | C | C | C | C | C | C | C | C |
| NC_011770 | 2212883 | C | T | 85 | 0 | 2 | 0 | NON_SYNONYMOU | MODERAT | MISSENSE | gGc/gAc | G110D | PLES_20411 | | | | | | | T | T | | | | |
| NC_011770 | 2212986 | T | A | 92 | 0 | 8 | 0 | NON_SYNONYMOU | MODERAT | MISSENSE | Acc/Tcc | T76S | PLES_20411 | A | A | | | | | A | A | A | A | A | A |
| NC_011770 | 2213159 | G | A | 79 | 0 | 4 | 0 | NON_SYNONYMOU | MODERAT | MISSENSE | tCc/tTc | S18F | PLES_20411 | A | A | | | | | | | A | A | | |
| NC_011770 | 2211743 | C | T | 80 | 0 | 1 | 0 | NON_SYNONYMOU | MODERAT | MISSENSE | gGc/gAc | G490D | PLES_20411 | | | | | | | | T | | | | |
| NC_011770 | 2213387 | A | T | 68 | 0 | 4 | 0 | START_LOST | HIGH | MISSENSE | aTg/aAg | M1K | PLES_20421 | | | T | T | T | T | | | | | | |
| NC_011770 | 2213277 | C | T | 70 | 0 | 2 | 0 | NON_SYNONYMOU | MODERAT | MISSENSE | Gaa/Aaa | E38K | PLES_20421 | | | | | T | T | | | | | | |
| NC_011770 | 2214437 | G | A | 69 | 0 | 4 | 0 | NON_SYNONYMOU | MODERAT | MISSENSE | cGc/cAc | R334H | PLES_20431 | | | A | A | A | A | | | | | | |
| NC_011770 | 2214539 | A | T | 67 | 0 | 4 | 0 | NON_SYNONYMOU | MODERAT | MISSENSE | cAg/cTg | Q368L | PLES_20431 | | | | | | | | | T | T | T | T |
| NC_011770 | 2214625 | A | G | 68 | 0 | 1 | 0 | NON_SYNONYMOU | MODERAT | MISSENSE | Acg/Gcg | T397A | PLES_20431 | | | | | | | G | | | | | |
| NC_011770 | 2214988 | G | A | 72 | 0 | 2 | 0 | NON_SYNONYMOU | MODERAT | MISSENSE | Gtc/Atc | V518I | PLES_20431 | | | | | | | | | A | A | | |
| NC_011770 | 2215313 | A | G | 72 | 0 | 2 | 0 | NON_SYNONYMOU | MODERAT | MISSENSE | cAg/cGg | Q626R | PLES_20431 | | | | | | | G | G | | | | |
| NC_011770 | 2213738 | A | G | 59 | 0 | 1 | 0 | NON_SYNONYMOU | MODERAT | MISSENSE | tAc/tGc | Y101C | PLES_20431 | | G | | | | | | | | | | |
| NC_011770 | 2216187 | C | A | 58 | 0 | 2 | 0 | NON_SYNONYMOU | MODERAT | MISSENSE | Ctc/Atc | L227I | PLES_20441 | | | | | | | A | A | | | | |
| NC_011770 | 2216188 | T | C | 56 | 0 | 2 | 0 | NON_SYNONYMOU | MODERAT | MISSENSE | cTc/cCc | L227P | PLES_20441 | | | | | | | | | C | C | | |
| NC_011770 | 2216955 | G | A | 81 | 0 | 12 | 0 | NON_SYNONYMOU | MODERAT | MISSENSE | aGg/aAg | R29K | PLES_20461 | A | A | A | A | A | A | A | A | A | A | A | A |
| NC_011770 | 2217473 | G | T | 97 | 0 | 2 | 0 | NON_SYNONYMOU | MODERAT | MISSENSE | Ctg/Atg | L223M | PLES_20471 | | | | | | | | | T | T | | |
| NC_011770 | 2217860 | C | T | 87 | 0 | 2 | 0 | NON_SYNONYMOU | MODERAT | MISSENSE | Ggc/Agc | G94S | PLES_20471 | | | | | | | | | T | T | | |
| NC_011770 | 2222320 | T | G | 75 | 0 | 2 | 0 | NON_SYNONYMOU | MODERAT | MISSENSE | agT/agG | S106R | PLES_20501 | G | G | | | | | | | | | | |
| NC_011770 | 2222339 | G | A | 81 | 0 | 2 | 0 | NON_SYNONYMOU | MODERAT | MISSENSE | Ggc/Agc | G113S | PLES_20501 | | | | | | | | | A | A | | |
| NC_011770 | 2225606 | A | C | 64 | 0 | 2 | 0 | NON_SYNONYMOU | MODERAT | MISSENSE | gaA/gaC | E142D | PLES_20521 | | | | | | | C | C | | | | |
| NC_011770 | 2229940 | G | T | 43 | 0 | 4 | 0 | NON_SYNONYMOU | MODERAT | MISSENSE | Gcc/Tcc | A173S | PLES_20581 | | | | | | | | | T | T | T | T |
| NC_011770 | 2230788 | CG | C | 71 | 0 | 2 | 0 | FRAME_SHIFT | HIGH | | ccc/ | P16 | PLES_20591 | C | C | | | | | | | | | | |
| NC_011770 | 2230553 | A | G | 64 | 0 | 4 | 0 | NON_SYNONYMOU | MODERAT | MISSENSE | gTg/gCg | V95A | PLES_20591 | | | G | G | G | G | | | | | | |
| NC_011770 | 2243533 | T | A | 81 | 0 | 2 | 0 | NON_SYNONYMOU | MODERAT | MISSENSE | aTg/aAg | M10K | PLES_20691 | | | | | | | A | A | | | | |
| NC_011770 | 2243922 | G | A | 87 | 0 | 2 | 0 | NON_SYNONYMOU | MODERAT | MISSENSE | Gcc/Acc | A140T | PLES_20691 | | | | | | | A | A | | | | |
| NC_011770 | 2243941 | T | C | 83 | 0 | 4 | 0 | NON_SYNONYMOU | MODERAT | MISSENSE | gTg/gCg | V146A | PLES_20691 | | | | | | | C | C | C | C | | |
| NC_011770 | 2244306 | T | G | 77 | 0 | 12 | 0 | NON_SYNONYMOU | MODERAT | MISSENSE | Tcg/Gcg | S268A | PLES_20691 | G | G | G | G | G | G | G | G | G | G | G | G |
| NC_011770 | 2244391 | T | C | 81 | 0 | 4 | 0 | NON_SYNONYMOU | MODERAT | MISSENSE | gTg/gCg | V296A | PLES_20691 | C | C | | | | | | | | | C | C |
| NC_011770 | 2246690 | T | C | 76 | 0 | 12 | 0 | NON_SYNONYMOU | MODERAT | MISSENSE | Atc/Gtc | I136V | PLES_20721 | C | C | C | C | C | C | C | C | C | C | C | C |
| NC_011770 | 2246881 | G | A | 78 | 0 | 6 | 0 | NON_SYNONYMOU | MODERAT | MISSENSE | gCc/gTc | A72V | PLES_20721 | | | A | A | A | A | A | A | | | | |
| NC_011770 | 2247055 | G | A | 78 | 0 | 2 | 0 | NON_SYNONYMOU | MODERAT | MISSENSE | gCg/gTg | A14V | PLES_20721 | | | | | A | A | | | | | | |
| NC_011770 | 2247609 | C | T | 76 | 0 | 1 | 0 | NON_SYNONYMOU | MODERAT | MISSENSE | gGc/gAc | G189D | PLES_20731 | | | | | | | T | | | | | |
| NC_011770 | 2247636 | C | T | 74 | 0 | 2 | 0 | NON_SYNONYMOU | MODERAT | MISSENSE | cGc/cAc | R180H | PLES_20731 | T | T | | | | | | | | | | |
| NC_011770 | 2247903 | T | C | 81 | 0 | 2 | 0 | NON_SYNONYMOU | MODERAT | MISSENSE | gAg/gGg | E91G | PLES_20731 | | | | | | | | | C | C | | |
| NC_011770 | 2248171 | C | T | 77 | 0 | 4 | 0 | NON_SYNONYMOU | MODERAT | MISSENSE | Gct/Act | A2T | PLES_20731 | | | T | T | T | T | | | | | | |
| NC_011770 | 2248371 | A | G | 73 | 0 | 1 | 0 | NON_SYNONYMOU | MODERAT | MISSENSE | cAc/cGc | H87R | PLES_20741 | | | | | | | | G | | | | |
| NC_011770 | 2249716 | C | T | 49 | 0 | 2 | 0 | NON_SYNONYMOU | MODERAT | MISSENSE | gCg/gTg | A121V | PLES_20751 | | | | | | | | | T | T | | |
| NC_011770 | 2250489 | G | A | 82 | 0 | 4 | 0 | NON_SYNONYMOU | MODERAT | MISSENSE | Gtc/Atc | V147I | PLES_20761 | | | | | | | A | A | | | A | A |
| NC_011770 | 2250531 | C | T | 85 | 0 | 2 | 0 | NON_SYNONYMOU | MODERAT | MISSENSE | Cct/Tct | P161S | PLES_20761 | | | | | | | | | T | T | | |
| NC_011770 | 2250850 | G | A | 83 | 0 | 4 | 0 | NON_SYNONYMOU | MODERAT | MISSENSE | gGc/gAc | G267D | PLES_20761 | | | | | | | A | A | | | A | A |
| NC_011770 | 2252239 | G | A | 59 | 0 | 1 | 0 | NON_SYNONYMOU | MODERAT | MISSENSE | Ggg/Agg | G46R | PLES_20781 | | | | | | | A | | | | | |
| NC_011770 | 2252404 | C | A | 51 | 0 | 8 | 0 | NON_SYNONYMOU | MODERAT | MISSENSE | Cgc/Agc | R101S | PLES_20781 | | | A | A | A | A | A | A | | | A | A |
| NC_011770 | 2252602 | C | G | 47 | 0 | 2 | 0 | NON_SYNONYMOU | MODERAT | MISSENSE | Cag/Gag | Q167E | PLES_20781 | | | | | | | G | G | | | | |
| NC_011770 | 2253038 | C | G | 16 | 0 | 2 | 0 | NON_SYNONYMOU | MODERAT | MISSENSE | gCa/gGa | A312G | PLES_20781 | | | | | | | G | G | | | | |
| NC_011770 | 2253650 | T | C | 38 | 0 | 2 | 0 | NON_SYNONYMOU | MODERAT | MISSENSE | gTc/gCc | V516A | PLES_20781 | | | | | | | C | C | | | | |
| NC_011770 | 2253821 | T | G | 61 | 0 | 2 | 0 | NON_SYNONYMOU | MODERAT | MISSENSE | gTt/gGt | V573G | PLES_20781 | | | | | | | G | G | | | | |
| NC_011770 | 2254115 | C | T | 69 | 0 | 2 | 0 | NON_SYNONYMOU | MODERAT | MISSENSE | gCc/gTc | A671V | PLES_20781 | T | T | | | | | | | | | | |
| NC_011770 | 2254519 | C | T | 66 | 0 | 2 | 0 | NON_SYNONYMOU | MODERAT | MISSENSE | Ccg/Tcg | P40S | PLES_20791 | | | | | | | T | T | | | | |
| NC_011770 | 2254557 | T | G | 72 | 0 | 12 | 0 | NON_SYNONYMOU | MODERAT | MISSENSE | aaT/aaG | N52K | PLES_20791 | G | G | G | G | G | G | G | G | G | G | G | G |
| NC_011770 | 2262362 | G | A | 83 | 0 | 4 | 0 | NON_SYNONYMOU | MODERAT | MISSENSE | Ggg/Agg | G28R | PLES_20871 | | | A | A | A | A | | | | | | |
| NC_011770 | 2264214 | A | G | 56 | 0 | 2 | 0 | NON_SYNONYMOU | MODERAT | MISSENSE | Acg/Gcg | T140A | PLES_20891 | G | G | | | | | | | | | | |
| NC_011770 | 2264498 | A | G | 64 | 0 | 4 | 0 | NON_SYNONYMOU | MODERAT | MISSENSE | aAg/aGg | K6R | PLES_20901 | | | G | G | G | G | | | | | | |
| NC_011770 | 2264800 | A | G | 74 | 0 | 2 | 0 | NON_SYNONYMOU | MODERAT | MISSENSE | Acc/Gcc | T107A | PLES_20901 | G | G | | | | | | | | | | |
| NC_011770 | 2265259 | T | C | 67 | 0 | 4 | 0 | NON_SYNONYMOU | MODERAT | MISSENSE | Tcc/Ccc | S260P | PLES_20901 | | | C | C | C | C | | | | | | |
| NC_011770 | 2265451 | A | G | 45 | 0 | 12 | 0 | NON_SYNONYMOU | MODERAT | MISSENSE | Ata/Gta | I324V | PLES_20901 | G | G | G | G | G | G | G | G | G | G | G | G |
| NC_011770 | 2265621 | C | A | 52 | 0 | 8 | 0 | NON_SYNONYMOU | MODERAT | MISSENSE | Gcc/Tcc | A163S | PLES_20911 | | | A | A | A | A | | | A | A | A | A |

| | | | | | | | | | | | | | | | | | | | | | | | | | |
|---|---|---|---|---|---|---|---|---|---|---|---|---|---|---|---|---|---|---|---|---|---|---|---|---|---|
| NC_011770 | 2265702 | A | G | 57 | 0 | 12 | 0 | NON_SYNONYMOU | MODERAT | MISSENSE | Tgc/Cgc | C136R | PLES_20911 | G | G | G | G | G | G | G | G | G | G | G | G |
| NC_011770 | 2265788 | G | C | 59 | 0 | 2 | 0 | NON_SYNONYMOU | MODERAT | MISSENSE | gCc/gGc | A107G | PLES_20911 | C | C | | | | | | | | | | |
| NC_011770 | 2265789 | C | T | 60 | 0 | 2 | 0 | NON_SYNONYMOU | MODERAT | MISSENSE | Gcc/Acc | A107T | PLES_20911 | T | T | | | | | | | | | | |
| NC_011770 | 2265851 | T | C | 61 | 0 | 2 | 0 | NON_SYNONYMOU | MODERAT | MISSENSE | gAa/gGa | E86G | PLES_20911 | C | C | | | | | | | | | | |
| NC_011770 | 2265879 | T | C | 62 | 0 | 2 | 0 | NON_SYNONYMOU | MODERAT | MISSENSE | Aac/Gac | N77D | PLES_20911 | C | C | | | | | | | | | | |
| NC_011770 | 2265995 | T | C | 70 | 0 | 12 | 0 | NON_SYNONYMOU | MODERAT | MISSENSE | gAg/gGg | E38G | PLES_20911 | C | C | C | C | C | C | C | C | C | C | C | C |
| NC_011770 | 2265999 | G | C | 69 | 0 | 10 | 0 | NON_SYNONYMOU | MODERAT | MISSENSE | Ccc/Gcc | P37A | PLES_20911 | | | C | C | C | C | C | C | C | C | C | C |
| NC_011770 | 2266046 | C | T | 73 | 0 | 2 | 0 | NON_SYNONYMOU | MODERAT | MISSENSE | cGc/cAc | R21H | PLES_20911 | | | | | | | T | T | | | | |
| NC_011770 | 2272528 | A | G | 84 | 0 | 12 | 0 | NON_SYNONYMOU | MODERAT | MISSENSE | Att/Gtt | I44V | PLES_21001 | G | G | G | G | G | G | G | G | G | G | G | G |
| NC_011770 | 2272579 | A | G | 85 | 0 | 2 | 0 | NON_SYNONYMOU | MODERAT | MISSENSE | Aag/Gag | K61E | PLES_21001 | G | G | | | | | | | | | | |
| NC_011770 | 2272630 | T | C | 81 | 0 | 12 | 0 | NON_SYNONYMOU | MODERAT | MISSENSE | Tcg/Ccg | S78P | PLES_21001 | C | C | C | C | C | C | C | C | C | C | C | C |
| NC_011770 | 2275704 | T | A | 82 | 0 | 12 | 0 | NON_SYNONYMOU | MODERAT | MISSENSE | gTg/gAg | V71E | PLES_21041 | A | A | A | A | A | A | A | A | A | A | A | A |
| NC_011770 | 2276479 | T | C | 84 | 0 | 2 | 0 | INTRAGENIC | MODIFIER | | | | PLES_21042 | | | | | | | | | | | C | C |
| NC_011770 | 2276514 | C | T | 89 | 0 | 2 | 0 | NON_SYNONYMOU | MODERAT | MISSENSE | cGc/cAc | R372H | PLES_21051 | T | T | | | | | | | | | | |
| NC_011770 | 2276640 | T | C | 84 | 0 | 2 | 0 | NON_SYNONYMOU | MODERAT | MISSENSE | aAt/aGt | N330S | PLES_21051 | | | | | | | | | | | C | C |
| NC_011770 | 2277244 | C | T | 91 | 0 | 2 | 0 | NON_SYNONYMOU | MODERAT | MISSENSE | Gcc/Acc | A129T | PLES_21051 | T | T | | | | | | | | | | |
| NC_011770 | 2277536 | C | G | 76 | 0 | 12 | 0 | NON_SYNONYMOU | MODERAT | MISSENSE | gaG/gaC | E31D | PLES_21051 | G | G | G | G | G | G | G | G | G | G | G | G |
| NC_011770 | 2278793 | G | C | 72 | 0 | 8 | 0 | NON_SYNONYMOU | MODERAT | MISSENSE | Gag/Cag | E98Q | PLES_21071 | | | C | C | C | C | C | C | C | C | | |
| NC_011770 | 2279555 | C | A | 87 | 0 | 2 | 0 | NON_SYNONYMOU | MODERAT | MISSENSE | gaC/gaA | D31E | PLES_21081 | A | A | | | | | | | | | | |
| NC_011770 | 2280367 | G | C | 76 | 0 | 2 | 0 | NON_SYNONYMOU | MODERAT | MISSENSE | Gtg/Ctg | V94L | PLES_21091 | C | C | | | | | | | | | | |
| NC_011770 | 2280481 | C | G | 73 | 0 | 2 | 0 | NON_SYNONYMOU | MODERAT | MISSENSE | Cgc/Ggc | R132G | PLES_21091 | G | G | | | | | | | | | | |
| NC_011770 | 2280485 | C | G | 75 | 0 | 6 | 0 | NON_SYNONYMOU | MODERAT | MISSENSE | aCc/aGc | T133S | PLES_21091 | | | | | | | G | G | G | G | G | G |
| NC_011770 | 2280637 | A | C | 74 | 0 | 4 | 0 | NON_SYNONYMOU | MODERAT | MISSENSE | Atc/Ctc | I184L | PLES_21091 | | | | | | | | | C | C | C | C |
| NC_011770 | 2285590 | A | G | 77 | 0 | 1 | 0 | NON_SYNONYMOU | MODERAT | MISSENSE | gAc/gGc | D349G | PLES_21131 | | | | | | | G | | | | | |
| NC_011770 | 2285834 | C | T | 62 | 0 | 8 | 0 | NON_SYNONYMOU | MODERAT | MISSENSE | Gcc/Acc | A310T | PLES_21141 | T | T | T | T | T | T | | | | | T | T |
| NC_011770 | 2286665 | C | G | 73 | 0 | 2 | 0 | NON_SYNONYMOU | MODERAT | MISSENSE | Gtc/Ctc | V33L | PLES_21141 | | | | | | | | | | | G | G |
| NC_011770 | 2287809 | A | AG | 71 | 0 | 1 | 0 | FRAME_SHIFT | HIGH | | ccc/cccC | P86P? | PLES_21161 | | | | | | | AG | | | | | |
| NC_011770 | 2287719 | C | T | 75 | 0 | 2 | 0 | NON_SYNONYMOU | MODERAT | MISSENSE | Gat/Aat | D117N | PLES_21161 | | | | | | | | | T | T | | |
| NC_011770 | 2287884 | C | T | 78 | 0 | 10 | 0 | NON_SYNONYMOU | MODERAT | MISSENSE | Gcg/Acg | A62T | PLES_21161 | T | T | T | T | T | T | T | T | T | T | | |
| NC_011770 | 2287945 | C | G | 74 | 0 | 2 | 0 | NON_SYNONYMOU | MODERAT | MISSENSE | gaG/gaC | E41D | PLES_21161 | G | G | | | | | | | | | | |
| NC_011770 | 2287965 | G | A | 75 | 0 | 2 | 0 | NON_SYNONYMOU | MODERAT | MISSENSE | Cac/Tac | H35Y | PLES_21161 | | | | | | | A | A | | | | |
| NC_011770 | 2288667 | C | T | 66 | 0 | 4 | 0 | NON_SYNONYMOU | MODERAT | MISSENSE | Gtc/Atc | V190I | PLES_21171 | | | T | T | T | T | | | | | | |
| NC_011770 | 2288706 | C | T | 64 | 0 | 2 | 0 | NON_SYNONYMOU | MODERAT | MISSENSE | Gcc/Acc | A177T | PLES_21171 | | | | | | | | | T | T | | |
| NC_011770 | 2288850 | C | T | 63 | 0 | 2 | 0 | NON_SYNONYMOU | MODERAT | MISSENSE | Gga/Aga | G129R | PLES_21171 | T | T | | | | | | | | | | |
| NC_011770 | 2288913 | C | T | 67 | 0 | 4 | 0 | NON_SYNONYMOU | MODERAT | MISSENSE | Ggt/Agt | G108S | PLES_21171 | | | T | T | T | T | | | | | | |
| NC_011770 | 2288951 | G | A | 71 | 0 | 1 | 0 | NON_SYNONYMOU | MODERAT | MISSENSE | gCt/gTt | A95V | PLES_21171 | | | | | | | A | | | | | |
| NC_011770 | 2288955 | G | A | 70 | 0 | 2 | 0 | NON_SYNONYMOU | MODERAT | MISSENSE | Cgc/Tgc | R94C | PLES_21171 | | | | | | | | | A | A | | |
| NC_011770 | 2289012 | G | A | 72 | 0 | 4 | 0 | NON_SYNONYMOU | MODERAT | MISSENSE | Ctc/Ttc | L75F | PLES_21171 | | | A | A | A | A | | | | | | |
| NC_011770 | 2289096 | A | G | 68 | 0 | 4 | 0 | NON_SYNONYMOU | MODERAT | MISSENSE | Tgt/Cgt | C47R | PLES_21171 | | | | | | | G | G | G | G | | |
| NC_011770 | 2289099 | C | A | 72 | 0 | 2 | 0 | NON_SYNONYMOU | MODERAT | MISSENSE | Gcc/Tcc | A46S | PLES_21171 | | | | | | | | | A | A | | |
| NC_011770 | 2289749 | A | G | 96 | 0 | 2 | 0 | NON_SYNONYMOU | MODERAT | MISSENSE | gAc/gGc | D54G | PLES_21181 | G | G | | | | | | | | | | |
| NC_011770 | 2290043 | C | T | 79 | 0 | 2 | 0 | NON_SYNONYMOU | MODERAT | MISSENSE | gCc/gTc | A152V | PLES_21181 | | | | | | | | | T | T | | |
| NC_011770 | 2290111 | C | G | 70 | 0 | 4 | 0 | NON_SYNONYMOU | MODERAT | MISSENSE | Cat/Gat | H175D | PLES_21181 | | | G | G | G | G | | | | | | |
| NC_011770 | 2290243 | G | A | 55 | 0 | 2 | 0 | NON_SYNONYMOU | MODERAT | MISSENSE | Gcc/Acc | A219T | PLES_21181 | A | A | | | | | | | | | | |
| NC_011770 | 2290474 | G | T | 58 | 0 | 2 | 0 | NON_SYNONYMOU | MODERAT | MISSENSE | Gct/Tct | A296S | PLES_21181 | | | | | | | | | T | T | | |
| NC_011770 | 2290550 | C | A | 62 | 0 | 2 | 0 | NON_SYNONYMOU | MODERAT | MISSENSE | cCg/cAg | P321Q | PLES_21181 | A | A | | | | | | | | | | |
| NC_011770 | 2290634 | T | C | 55 | 0 | 4 | 0 | NON_SYNONYMOU | MODERAT | MISSENSE | gTc/gCc | V349A | PLES_21181 | C | C | | | | | | | C | C | | |
| NC_011770 | 2294847 | C | A | 73 | 0 | 2 | 0 | NON_SYNONYMOU | MODERAT | MISSENSE | aGc/aTc | S272I | PLES_21201 | A | A | | | | | | | | | | |
| NC_011770 | 2295352 | A | G | 67 | 0 | 8 | 0 | NON_SYNONYMOU | MODERAT | MISSENSE | Ttc/Ctc | F104L | PLES_21201 | G | G | G | G | G | G | | | G | G | | |
| NC_011770 | 2295409 | C | T | 68 | 0 | 1 | 0 | NON_SYNONYMOU | MODERAT | MISSENSE | Gcc/Acc | A85T | PLES_21201 | | | | | | | T | | | | | |
| NC_011770 | 2295634 | T | C | 61 | 0 | 8 | 0 | NON_SYNONYMOU | MODERAT | MISSENSE | Acc/Gcc | T10A | PLES_21201 | C | C | C | C | C | C | | | C | C | | |
| NC_011770 | 2296017 | C | T | 49 | 0 | 8 | 0 | INTRAGENIC | MODIFIER | | | | PLES_21202 | T | T | T | T | T | T | | | T | T | | |
| NC_011770 | 2296040 | A | G | 50 | 0 | 8 | 0 | INTRAGENIC | MODIFIER | | | | PLES_21202 | G | G | G | G | G | G | | | G | G | | |
| NC_011770 | 2296041 | A | G | 50 | 0 | 8 | 0 | INTRAGENIC | MODIFIER | | | | PLES_21202 | G | G | G | G | G | G | | | G | G | | |
| NC_011770 | 2296056 | A | G | 52 | 0 | 4 | 0 | INTRAGENIC | MODIFIER | | | | PLES_21202 | G | G | | | | | | | G | G | | |
| NC_011770 | 2296310 | G | A | 36 | 0 | 2 | 0 | NON_SYNONYMOU | MODERAT | MISSENSE | Gtc/Atc | V27I | PLES_21211 | | | | | | | A | A | | | | |
| NC_011770 | 2296818 | T | C | 59 | 0 | 4 | 0 | NON_SYNONYMOU | MODERAT | MISSENSE | gTc/gCc | V196A | PLES_21211 | | | C | C | C | C | | | | | | |
| NC_011770 | 2296878 | G | A | 60 | 0 | 2 | 0 | NON_SYNONYMOU | MODERAT | MISSENSE | cGg/cAg | R216Q | PLES_21211 | | | | | | | | | A | A | | |
| NC_011770 | 2297085 | A | G | 58 | 0 | 6 | 0 | NON_SYNONYMOU | MODERAT | MISSENSE | aAc/aGc | N285S | PLES_21211 | | | G | G | G | G | | | G | G | | |
| NC_011770 | 2297093 | T | C | 58 | 0 | 10 | 0 | NON_SYNONYMOU | MODERAT | MISSENSE | Tca/Cca | S288P | PLES_21211 | C | C | C | C | C | C | C | C | C | C | | |
| NC_011770 | 2297122 | CGCG | C | 61 | 0 | 6 | 0 | CODON_CHANGE_ | MODERATE | | gcggcc/gcc | AA299A | PLES_21211 | | | C | C | C | C | | | C | C | | |
| NC_011770 | 2297136 | T | C | 62 | 0 | 6 | 0 | NON_SYNONYMOU | MODERAT | MISSENSE | gTc/gCc | V302A | PLES_21211 | | | C | C | C | C | | | C | C | | |

| | | | | | | | | | | | | | | | | | | | | | | | | | |
|---|---|---|---|---|---|---|---|---|---|---|---|---|---|---|---|---|---|---|---|---|---|---|---|---|---|
| NC_011770 | 2297153 | A | G | 58 | 0 | 6 | 0 | NON_SYNONYMOU | MODERAT | MISSENSE | Acc/Gcc | T308A | PLES_21211 | | | G | G | G | G | | | G | G | | |
| NC_011770 | 2297350 | A | G | 61 | 0 | 1 | 0 | NON_SYNONYMOU | MODERAT | MISSENSE | Atc/Gtc | I28V | PLES_21221 | | | | | | | G | | | | | |
| NC_011770 | 2297636 | A | G | 61 | 0 | 10 | 0 | NON_SYNONYMOU | MODERAT | MISSENSE | cAc/cGc | H123R | PLES_21221 | G | G | G | G | G | G | G | G | G | G | | |
| NC_011770 | 2298272 | C | T | 66 | 0 | 6 | 0 | NON_SYNONYMOU | MODERAT | MISSENSE | Ccg/Tcg | P80S | PLES_21231 | | | T | T | T | T | | | | | T | T |
| NC_011770 | 2298599 | G | C | 60 | 0 | 2 | 0 | NON_SYNONYMOU | MODERAT | MISSENSE | Gag/Cag | E189Q | PLES_21231 | | | | | | | C | C | | | | |
| NC_011770 | 2298788 | C | T | 56 | 0 | 6 | 0 | NON_SYNONYMOU | MODERAT | MISSENSE | Cgc/Tgc | R252C | PLES_21231 | | | T | T | T | T | | | | | T | T |
| NC_011770 | 2298801 | C | T | 59 | 0 | 10 | 0 | NON_SYNONYMOU | MODERAT | MISSENSE | gCg/gTg | A256V | PLES_21231 | T | T | T | T | T | T | | | T | T | T | T |
| NC_011770 | 2299169 | G | A | 60 | 0 | 4 | 0 | NON_SYNONYMOU | MODERAT | MISSENSE | Gag/Aag | E379K | PLES_21231 | | | A | A | A | A | | | | | | |
| NC_011770 | 2299333 | G | A | 66 | 0 | 4 | 0 | NON_SYNONYMOU | MODERAT | MISSENSE | gCa/gTa | A513V | PLES_21241 | | | | | | | | | A | A | C | C |
| NC_011770 | 2299355 | T | G | 64 | 0 | 2 | 0 | NON_SYNONYMOU | MODERAT | MISSENSE | Atc/Ctc | I506L | PLES_21241 | | | | | | | G | G | | | | |
| NC_011770 | 2299550 | C | T | 74 | 0 | 1 | 0 | NON_SYNONYMOU | MODERAT | MISSENSE | Ggc/Agc | G441S | PLES_21241 | | | | | | | T | | | | | |
| NC_011770 | 2302009 | A | T | 78 | 0 | 2 | 0 | NON_SYNONYMOU | MODERAT | MISSENSE | Acg/Tcg | T214S | PLES_21251 | T | T | | | | | | | | | | |
| NC_011770 | 2302753 | C | T | 67 | 0 | 2 | 0 | NON_SYNONYMOU | MODERAT | MISSENSE | Ccg/Tcg | P462S | PLES_21251 | | | | | | | T | T | | | | |
| NC_011770 | 2302780 | A | G | 62 | 0 | 12 | 0 | NON_SYNONYMOU | MODERAT | MISSENSE | Acg/Gcg | T471A | PLES_21251 | G | G | G | G | G | G | G | G | G | G | G | G |
| NC_011770 | 2303043 | C | T | 75 | 0 | 1 | 0 | NON_SYNONYMOU | MODERAT | MISSENSE | gGc/gAc | G50D | PLES_21261 | | | | | | | T | | | | | |
| NC_011770 | 2303409 | C | T | 84 | 0 | 6 | 0 | NON_SYNONYMOU | MODERAT | MISSENSE | Gtg/Atg | V154M | PLES_21271 | | | T | T | T | T | T | T | | | | |
| NC_011770 | 2303487 | C | T | 82 | 0 | 2 | 0 | NON_SYNONYMOU | MODERAT | MISSENSE | Gcg/Acg | A128T | PLES_21271 | | | | | | | | | | | T | T |
| NC_011770 | 2303580 | C | T | 89 | 0 | 4 | 0 | NON_SYNONYMOU | MODERAT | MISSENSE | Gcc/Acc | A97T | PLES_21271 | | | T | T | T | T | | | | | | |
| NC_011770 | 2303682 | C | T | 90 | 0 | 6 | 0 | NON_SYNONYMOU | MODERAT | MISSENSE | Gtc/Atc | V63I | PLES_21271 | | | T | T | T | T | | | T | T | | |
| NC_011770 | 2303838 | A | G | 83 | 0 | 1 | 0 | NON_SYNONYMOU | MODERAT | MISSENSE | Tcc/Ccc | S11P | PLES_21271 | G | | | | | | | | | | | |
| NC_011770 | 2304092 | C | CTATT | 63 | 0 | 4 | 0 | FRAME_SHIFT | HIGH | | cac/cTATT | H39LL? | PLES_21281 | CTATT | CTATT | | | | | | | CTATT | CTATT | | |
| NC_011770 | 2304116 | T | TG | 67 | 0 | 4 | 0 | FRAME_SHIFT | HIGH | | ggc/ggGc | G48G? | PLES_21281 | TG | TG | | | | | | | TG | TG | | |
| NC_011770 | 2304138 | C | T | 69 | 0 | 8 | 0 | FRAME_SHIFT | HIGH | | ccc/ccGc | P54P? | PLES_21281 | | | T | T | T | T | | | CG | CG | T | T |
| NC_011770 | 2304025 | A | G | 69 | 0 | 6 | 0 | NON_SYNONYMOU | MODERAT | MISSENSE | aaA/aaC | K16N | PLES_21281 | | | G | G | G | G | | | | | C | C |
| NC_011770 | 2304030 | A | G | 69 | 0 | 4 | 0 | NON_SYNONYMOU | MODERAT | MISSENSE | aAg/aGg | K18R | PLES_21281 | G | G | | | | | | | G | G | | |
| NC_011770 | 2304031 | G | C | 69 | 0 | 4 | 0 | NON_SYNONYMOU | MODERAT | MISSENSE | aaG/aaC | K18N | PLES_21281 | C | C | | | | | | | C | C | | |
| NC_011770 | 2304033 | T | C | 71 | 0 | 4 | 0 | NON_SYNONYMOU | MODERAT | MISSENSE | cTa/cCa | L19P | PLES_21281 | C | C | | | | | | | C | C | | |
| NC_011770 | 2304072 | T | G | 68 | 0 | 4 | 0 | NON_SYNONYMOU | MODERAT | MISSENSE | tTg/tGg | L32W | PLES_21281 | G | G | | | | | | | G | G | | |
| NC_011770 | 2304083 | A | T | 62 | 0 | 12 | 0 | NON_SYNONYMOU | MODERAT | MISSENSE | Acg/Tcg | T36S | PLES_21281 | T | T | T | T | T | T | T | T | T | T | T | T |
| NC_011770 | 2304093 | A | G | 57 | 4 | 2 | 0 | NON_SYNONYMOU | MODERAT | MISSENSE | cAc/cGc | H39R | PLES_21281 | . | . | | | | | | | . | . | G | G |
| NC_011770 | 2304132 | T | A | 68 | 0 | 4 | 0 | NON_SYNONYMOU | MODERAT | MISSENSE | cTg/cAg | L52Q | PLES_21281 | A | A | | | | | | | A | A | | |
| NC_011770 | 2304285 | T | C | 86 | 0 | 2 | 0 | NON_SYNONYMOU | MODERAT | MISSENSE | aTc/aCc | I103T | PLES_21281 | | | | | | | | | | | C | C |
| NC_011770 | 2304302 | T | C | 77 | 0 | 12 | 0 | NON_SYNONYMOU | MODERAT | MISSENSE | Tat/Cat | Y109H | PLES_21281 | C | C | C | C | C | C | C | C | C | C | C | C |
| NC_011770 | 2304582 | C | G | 72 | 0 | 6 | 0 | NON_SYNONYMOU | MODERAT | MISSENSE | gaG/gaC | E285D | PLES_21291 | | | | | | | G | G | G | G | G | G |
| NC_011770 | 2304815 | C | A | 65 | 0 | 2 | 0 | NON_SYNONYMOU | MODERAT | MISSENSE | Gcc/Tcc | A208S | PLES_21291 | | | | | | | | | | | A | A |
| NC_011770 | 2305470 | T | G | 68 | 0 | 4 | 0 | NON_SYNONYMOU | MODERAT | MISSENSE | caA/caC | Q389H | PLES_21301 | G | G | | | | | | | G | G | | |
| NC_011770 | 2306053 | A | G | 74 | 0 | 1 | 0 | NON_SYNONYMOU | MODERAT | MISSENSE | gTg/gCg | V195A | PLES_21301 | | | | | | | G | | | | | |
| NC_011770 | 2306078 | C | T | 74 | 0 | 6 | 0 | NON_SYNONYMOU | MODERAT | MISSENSE | Gtc/Atc | V187I | PLES_21301 | | | T | T | T | T | | | | | T | T |
| NC_011770 | 2306174 | T | C | 75 | 0 | 10 | 0 | NON_SYNONYMOU | MODERAT | MISSENSE | Agt/Ggt | S155G | PLES_21301 | C | C | C | C | C | C | | | C | C | C | C |
| NC_011770 | 2306125 | G | A | 79 | 0 | 1 | 0 | NON_SYNONYMOU | MODERAT | MISSENSE | gCg/gTg | A171V | PLES_21301 | | A | | | | | | | | | | |
| NC_011770 | 2307943 | A | C | 85 | 0 | 2 | 0 | NON_SYNONYMOU | MODERAT | MISSENSE | Acg/Ccg | T3P | PLES_21321 | | | | | | | | | | | C | C |
| NC_011770 | 2308405 | G | A | 82 | 0 | 2 | 0 | NON_SYNONYMOU | MODERAT | MISSENSE | Gtg/Atg | V157M | PLES_21321 | | | A | A | | | | | | | | |
| NC_011770 | 2308675 | A | C | 75 | 0 | 2 | 0 | NON_SYNONYMOU | MODERAT | MISSENSE | Tcc/Gcc | S264A | PLES_21331 | | | | | | | C | C | | | | |
| NC_011770 | 2308878 | C | T | 60 | 0 | 2 | 0 | NON_SYNONYMOU | MODERAT | MISSENSE | gGc/gAc | G196D | PLES_21331 | | | | | | | T | T | | | | |
| NC_011770 | 2309152 | C | T | 62 | 0 | 4 | 0 | NON_SYNONYMOU | MODERAT | MISSENSE | Ggc/Agc | G105S | PLES_21331 | | | T | T | T | T | | | | | | |
| NC_011770 | 2309170 | T | C | 63 | 0 | 2 | 0 | NON_SYNONYMOU | MODERAT | MISSENSE | Acc/Gcc | T99A | PLES_21331 | C | C | | | | | | | | | | |
| NC_011770 | 2309418 | T | C | 70 | 0 | 12 | 0 | NON_SYNONYMOU | MODERAT | MISSENSE | cAt/cGt | H16R | PLES_21331 | C | C | C | C | C | C | C | C | C | C | C | C |
| NC_011770 | 2309687 | C | G | 62 | 0 | 12 | 0 | NON_SYNONYMOU | MODERAT | MISSENSE | gCc/gGc | A53G | PLES_21341 | G | G | G | G | G | G | G | G | G | G | G | G |
| NC_011770 | 2309840 | A | C | 48 | 0 | 12 | 0 | NON_SYNONYMOU | MODERAT | MISSENSE | gAc/gCc | D104A | PLES_21341 | C | C | C | C | C | C | C | C | C | C | C | C |
| NC_011770 | 2310112 | C | T | 62 | 0 | 12 | 0 | NON_SYNONYMOU | MODERAT | MISSENSE | Ctc/Ttc | L195F | PLES_21341 | T | T | T | T | T | T | T | T | T | T | T | T |
| NC_011770 | 2310122 | T | A | 66 | 0 | 2 | 0 | NON_SYNONYMOU | MODERAT | MISSENSE | cTg/cAg | L198Q | PLES_21341 | | | | | | | | | | | A | A |
| NC_011770 | 2313385 | G | A | 76 | 0 | 2 | 0 | NON_SYNONYMOU | MODERAT | MISSENSE | Gag/Aag | E32K | PLES_21371 | | | | | | | | | | | A | A |
| NC_011770 | 2316949 | C | T | 73 | 0 | 2 | 0 | NON_SYNONYMOU | MODERAT | MISSENSE | Gaa/Aaa | E265K | PLES_21421 | T | T | | | | | | | | | | |
| NC_011770 | 2317131 | C | T | 73 | 0 | 2 | 0 | NON_SYNONYMOU | MODERAT | MISSENSE | cGc/cAc | R204H | PLES_21421 | T | T | | | | | | | | | | |
| NC_011770 | 2317399 | C | T | 57 | 0 | 2 | 0 | NON_SYNONYMOU | MODERAT | MISSENSE | Gca/Aca | A115T | PLES_21421 | T | T | | | | | | | | | | |
| NC_011770 | 2317948 | T | C | 32 | 0 | 2 | 0 | NON_SYNONYMOU | MODERAT | MISSENSE | gTc/gCc | V36A | PLES_21431 | | | | | | | | | | | C | C |
| NC_011770 | 2318623 | C | T | 74 | 0 | 2 | 0 | NON_SYNONYMOU | MODERAT | MISSENSE | gCc/gTc | A261V | PLES_21431 | | | | | | | | | T | T | | |
| NC_011770 | 2318758 | C | T | 74 | 0 | 2 | 0 | NON_SYNONYMOU | MODERAT | MISSENSE | gCg/gTg | A306V | PLES_21431 | | | | | | | T | T | | | | |
| NC_011770 | 2320482 | A | G | 75 | 0 | 2 | 0 | NON_SYNONYMOU | MODERAT | MISSENSE | Acc/Gcc | T523A | PLES_21441 | | | | | | | | | | | G | G |
| NC_011770 | 2319301 | G | A | 80 | 0 | 1 | 0 | NON_SYNONYMOU | MODERAT | MISSENSE | aGc/aAc | S129N | PLES_21441 | A | | | | | | | | | | | |
| NC_011770 | 2319979 | C | T | 84 | 0 | 1 | 0 | NON_SYNONYMOU | MODERAT | MISSENSE | gCg/gTg | A355V | PLES_21441 | T | | | | | | | | | | | |
| NC_011770 | 2322406 | C | T | 87 | 0 | 4 | 0 | NON_SYNONYMOU | MODERAT | MISSENSE | Cac/Tac | H184Y | PLES_21471 | T | T | | | | | | | T | T | | |

| | | | | | | | | | | | | | | | | | | | | | | | | | |
|---|---|---|---|---|---|---|---|---|---|---|---|---|---|---|---|---|---|---|---|---|---|---|---|---|---|
| NC_011770 | 2322415 | C | G | 87 | 0 | 2 | 0 | NON_SYNONYMOU | MODERAT | MISSENSE | Cag/Gag | Q187E | PLES_21471 | | | | | | | G | G | | | | |
| NC_011770 | 2324133 | G | A | 78 | 0 | 2 | 0 | NON_SYNONYMOU | MODERAT | MISSENSE | cGg/cAg | R226Q | PLES_21491 | | | | | | | | | | | A | A |
| NC_011770 | 2324400 | T | TG | 74 | 0 | 1 | 0 | FRAME_SHIFT | HIGH | | cag/cCag | Q319P? | PLES_21501 | | | | | | | | TG | | | | |
| NC_011770 | 2324514 | A | G | 72 | 0 | 12 | 0 | NON_SYNONYMOU | MODERAT | MISSENSE | gTc/gCc | V281A | PLES_21501 | G | G | G | G | G | G | G | G | G | G | G | G |
| NC_011770 | 2325115 | C | T | 90 | 0 | 2 | 0 | NON_SYNONYMOU | MODERAT | MISSENSE | Gtc/Atc | V81I | PLES_21501 | | | | | | | T | T | | | | |
| NC_011770 | 2325152 | C | A | 92 | 0 | 6 | 0 | NON_SYNONYMOU | MODERAT | MISSENSE | tgG/tgT | W68C | PLES_21501 | | | A | A | A | A | | | | | A | A |
| NC_011770 | 2325735 | T | C | 74 | 0 | 12 | 0 | NON_SYNONYMOU | MODERAT | MISSENSE | Acc/Gcc | T197A | PLES_21511 | C | C | C | C | C | C | C | C | C | C | C | C |
| NC_011770 | 2326290 | A | C | 61 | 0 | 10 | 0 | NON_SYNONYMOU | MODERAT | MISSENSE | Tgc/Ggc | C12G | PLES_21511 | C | C | C | C | C | C | C | C | C | C | | |
| NC_011770 | 2326311 | T | G | 62 | 0 | 2 | 0 | NON_SYNONYMOU | MODERAT | MISSENSE | Atc/Ctc | I5L | PLES_21511 | | | | | | | G | G | | | | |
| NC_011770 | 2326541 | T | C | 61 | 0 | 6 | 0 | NON_SYNONYMOU | MODERAT | MISSENSE | Agc/Ggc | S185G | PLES_21521 | | | C | C | C | C | C | C | | | | |
| NC_011770 | 2327398 | C | A | 78 | 0 | 2 | 0 | NON_SYNONYMOU | MODERAT | MISSENSE | gaG/gaT | E635D | PLES_21531 | | | | | | | A | A | | | | |
| NC_011770 | 2327483 | C | T | 79 | 0 | 6 | 0 | NON_SYNONYMOU | MODERAT | MISSENSE | gGg/gAg | G607E | PLES_21531 | | | T | T | T | T | T | T | | | | |
| NC_011770 | 2327484 | C | T | 79 | 0 | 6 | 0 | NON_SYNONYMOU | MODERAT | MISSENSE | Ggg/Agg | G607R | PLES_21531 | | | T | T | T | T | T | T | | | | |
| NC_011770 | 2327859 | T | G | 81 | 0 | 10 | 0 | NON_SYNONYMOU | MODERAT | MISSENSE | Atg/Ctg | M482L | PLES_21531 | | | G | G | G | G | G | G | G | G | G | G |
| NC_011770 | 2327910 | C | T | 82 | 0 | 4 | 0 | NON_SYNONYMOU | MODERAT | MISSENSE | Gcc/Acc | A465T | PLES_21531 | | | T | T | T | T | | | | | | |
| NC_011770 | 2327912 | C | A | 82 | 0 | 4 | 0 | NON_SYNONYMOU | MODERAT | MISSENSE | cGc/cTc | R464L | PLES_21531 | | | A | A | A | A | | | | | | |
| NC_011770 | 2328131 | C | G | 72 | 0 | 4 | 0 | NON_SYNONYMOU | MODERAT | MISSENSE | aGc/aCc | S391T | PLES_21531 | | | G | G | G | G | | | | | | |
| NC_011770 | 2328162 | C | A | 70 | 0 | 2 | 0 | NON_SYNONYMOU | MODERAT | MISSENSE | Gcc/Tcc | A381S | PLES_21531 | | | | | | | | | A | A | | |
| NC_011770 | 2328266 | C | T | 78 | 0 | 4 | 0 | NON_SYNONYMOU | MODERAT | MISSENSE | aGg/aAg | R346K | PLES_21531 | | | T | T | T | T | | | | | | |
| NC_011770 | 2328291 | C | T | 77 | 0 | 2 | 0 | NON_SYNONYMOU | MODERAT | MISSENSE | Gtc/Atc | V338I | PLES_21531 | | | | | | | T | T | | | | |
| NC_011770 | 2328123 | A | ATAT | 69 | 0 | 3 | 0 | CODON_INSERTIOI | MODERATE | | aac/aacAT | N393NI | PLES_21531 | | | ATAT | | ATAT | ATAT | | | | | | |
| NC_011770 | 2330162 | T | C | 78 | 0 | 2 | 0 | NON_SYNONYMOU | MODERAT | MISSENSE | gTc/gCc | V75A | PLES_21541 | | | | | | | C | C | | | | |
| NC_011770 | 2330219 | A | C | 75 | 0 | 6 | 0 | NON_SYNONYMOU | MODERAT | MISSENSE | gAg/gCg | E94A | PLES_21541 | | | | | | | C | C | C | C | C | C |
| NC_011770 | 2334439 | G | A | 59 | 0 | 2 | 0 | NON_SYNONYMOU | MODERAT | MISSENSE | Gct/Act | A159T | PLES_21581 | | | | | | | A | A | | | | |
| NC_011770 | 2334472 | C | T | 60 | 0 | 2 | 0 | NON_SYNONYMOU | MODERAT | MISSENSE | Cgc/Tgc | R170C | PLES_21581 | T | T | | | | | | | | | | |
| NC_011770 | 2334559 | C | T | 70 | 0 | 2 | 0 | NON_SYNONYMOU | MODERAT | MISSENSE | Ctc/Ttc | L199F | PLES_21581 | | | | | | | T | T | | | | |
| NC_011770 | 2334742 | A | G | 65 | 0 | 8 | 0 | NON_SYNONYMOU | MODERAT | MISSENSE | Aag/Gag | K260E | PLES_21581 | G | G | | | | | G | G | G | G | G | G |
| NC_011770 | 2334827 | A | G | 62 | 0 | 6 | 0 | NON_SYNONYMOU | MODERAT | MISSENSE | cAg/cGg | Q288R | PLES_21581 | G | G | | | | | | | G | G | G | G |
| NC_011770 | 2334832 | C | T | 63 | 0 | 6 | 0 | NON_SYNONYMOU | MODERAT | MISSENSE | Ccg/Tcg | P290S | PLES_21581 | T | T | | | | | | | T | T | T | T |
| NC_011770 | 2334844 | G | C | 65 | 0 | 6 | 0 | NON_SYNONYMOU | MODERAT | MISSENSE | Gcg/Ccg | A294P | PLES_21581 | C | C | | | | | | | C | C | C | C |
| NC_011770 | 2335403 | C | A | 69 | 0 | 4 | 0 | NON_SYNONYMOU | MODERAT | MISSENSE | cCc/cAc | P480H | PLES_21581 | | | A | A | A | A | | | | | | |
| NC_011770 | 2334202 | G | A | 68 | 0 | 1 | 0 | NON_SYNONYMOU | MODERAT | MISSENSE | Gtg/Atg | V80M | PLES_21581 | | | | | | | | | | | | A |
| NC_011770 | 2334632 | C | T | 67 | 0 | 1 | 0 | NON_SYNONYMOU | MODERAT | MISSENSE | cCc/cTc | P223L | PLES_21581 | | | | | | | | T | | | | |
| NC_011770 | 2338788 | A | G | 68 | 0 | 2 | 0 | NON_SYNONYMOU | MODERAT | MISSENSE | Aag/Gag | K84E | PLES_21621 | | | | | | | | | G | G | | |
| NC_011770 | 2338875 | G | A | 75 | 0 | 4 | 0 | NON_SYNONYMOU | MODERAT | MISSENSE | Gtg/Atg | V113M | PLES_21621 | | | A | A | A | A | | | | | | |
| NC_011770 | 2341618 | C | T | 45 | 0 | 4 | 0 | NON_SYNONYMOU | MODERAT | MISSENSE | gCg/gTg | A50V | PLES_21661 | T | T | | | | | | | | | T | T |
| NC_011770 | 2341692 | G | A | 42 | 0 | 2 | 0 | NON_SYNONYMOU | MODERAT | MISSENSE | Ggc/Agc | G75S | PLES_21661 | | | | | | | A | A | | | | |
| NC_011770 | 2341713 | G | A | 40 | 0 | 2 | 0 | NON_SYNONYMOU | MODERAT | MISSENSE | Gcc/Acc | A82T | PLES_21661 | | | | | | | A | A | | | | |
| NC_011770 | 2341716 | G | T | 40 | 0 | 2 | 0 | NON_SYNONYMOU | MODERAT | MISSENSE | Gcc/Tcc | A83S | PLES_21661 | | | | | | | T | T | | | | |
| NC_011770 | 2341798 | T | C | 47 | 0 | 2 | 0 | NON_SYNONYMOU | MODERAT | MISSENSE | tTg/tCg | L110S | PLES_21661 | | | | | | | C | C | | | | |
| NC_011770 | 2343485 | C | T | 82 | 0 | 1 | 0 | NON_SYNONYMOU | MODERAT | MISSENSE | Ctt/Ttt | L476F | PLES_21671 | | | | | | | | T | | | | |
| NC_011770 | 2344755 | A | G | 60 | 0 | 2 | 0 | NON_SYNONYMOU | MODERAT | MISSENSE | aAg/aGg | K116R | PLES_21691 | | | | | | | | | G | G | | |
| NC_011770 | 2344796 | G | A | 65 | 0 | 8 | 0 | NON_SYNONYMOU | MODERAT | MISSENSE | Ggc/Agc | G130S | PLES_21691 | A | A | A | A | A | A | | | | | A | A |
| NC_011770 | 2344880 | C | G | 66 | 0 | 8 | 0 | NON_SYNONYMOU | MODERAT | MISSENSE | Ccg/Gcg | P158A | PLES_21691 | G | G | | | | | G | G | G | G | G | G |
| NC_011770 | 2344416 | C | T | 77 | 0 | 1 | 0 | NON_SYNONYMOU | MODERAT | MISSENSE | cCa/cTa | P3L | PLES_21691 | | T | | | | | | | | | | |
| NC_011770 | 2360240 | C | A | 71 | 0 | 4 | 0 | NON_SYNONYMOU | MODERAT | MISSENSE | Cag/Aag | Q156K | PLES_21821 | | | A | A | A | A | | | | | | |
| NC_011770 | 2360242 | G | T | 73 | 0 | 4 | 0 | NON_SYNONYMOU | MODERAT | MISSENSE | caG/caT | Q156H | PLES_21821 | | | T | T | T | T | | | | | | |
| NC_011770 | 2360268 | C | T | 72 | 0 | 2 | 0 | NON_SYNONYMOU | MODERAT | MISSENSE | gCg/gTg | A165V | PLES_21821 | | | | | | | | | T | T | | |
| NC_011770 | 2361799 | C | T | 67 | 0 | 2 | 0 | NON_SYNONYMOU | MODERAT | MISSENSE | gCc/gTc | A303V | PLES_21831 | | | | | | | | | | | T | T |
| NC_011770 | 2361841 | C | T | 67 | 0 | 4 | 0 | NON_SYNONYMOU | MODERAT | MISSENSE | Gcc/Acc | A163T | PLES_21841 | | | T | T | T | T | | | | | | |
| NC_011770 | 2362041 | T | A | 78 | 0 | 2 | 0 | NON_SYNONYMOU | MODERAT | MISSENSE | cAg/cTg | Q96L | PLES_21841 | | | | | | | | | A | A | | |
| NC_011770 | 2362273 | T | G | 49 | 0 | 4 | 0 | NON_SYNONYMOU | MODERAT | MISSENSE | Acc/Ccc | T19P | PLES_21841 | | | G | G | G | G | | | | | | |
| NC_011770 | 2362557 | G | A | 65 | 0 | 4 | 0 | NON_SYNONYMOU | MODERAT | MISSENSE | Gcc/Acc | A62T | PLES_21851 | | | A | A | A | A | | | | | | |
| NC_011770 | 2362923 | T | C | 62 | 0 | 12 | 0 | NON_SYNONYMOU | MODERAT | MISSENSE | Tgg/Cgg | W184R | PLES_21851 | C | C | C | C | C | C | C | C | C | C | C | C |
| NC_011770 | 2362999 | A | G | 62 | 0 | 2 | 0 | NON_SYNONYMOU | MODERAT | MISSENSE | aAc/aGc | N209S | PLES_21851 | | | | | | | | | G | G | | |
| NC_011770 | 2363339 | G | A | 59 | 0 | 2 | 0 | NON_SYNONYMOU | MODERAT | MISSENSE | cCg/cTg | P197L | PLES_21861 | A | A | | | | | | | | | | |
| NC_011770 | 2363448 | C | T | 62 | 0 | 2 | 0 | NON_SYNONYMOU | MODERAT | MISSENSE | Gcg/Acg | A161T | PLES_21861 | | | | | | | | | T | T | | |
| NC_011770 | 2363672 | T | C | 64 | 0 | 2 | 0 | NON_SYNONYMOU | MODERAT | MISSENSE | gAt/gGt | D86G | PLES_21861 | | | | | | | C | C | | | | |
| NC_011770 | 2363694 | C | T | 64 | 0 | 10 | 0 | NON_SYNONYMOU | MODERAT | MISSENSE | Gac/Aac | D79N | PLES_21861 | T | T | T | T | T | T | T | T | T | T | | |
| NC_011770 | 2363721 | C | T | 65 | 0 | 6 | 0 | NON_SYNONYMOU | MODERAT | MISSENSE | Gcc/Acc | A70T | PLES_21861 | T | T | T | T | T | T | | | | | | |
| NC_011770 | 2363825 | T | C | 69 | 0 | 12 | 0 | NON_SYNONYMOU | MODERAT | MISSENSE | aAc/aGc | N35S | PLES_21861 | C | C | C | C | C | C | C | C | C | C | C | C |
| NC_011770 | 2363873 | T | G | 75 | 0 | 2 | 0 | NON_SYNONYMOU | MODERAT | MISSENSE | tAc/tCc | Y19S | PLES_21861 | | | | | | | | | G | G | | |

| | | | | | | | | | | | | | | | | | | | | | | | | | |
|---|---|---|---|---|---|---|---|---|---|---|---|---|---|---|---|---|---|---|---|---|---|---|---|---|---|
| NC_011770 | 2366325 | A | G | 65 | 0 | 1 | 0 | NON_SYNONYMOU | MODERAT | MISSENSE | aAc/aGc | N145S | PLES_21891 | | | | | | | G | | | | | |
| NC_011770 | 2367646 | G | T | 70 | 0 | 2 | 0 | STOP_GAINED | HIGH | NONSENSE | Gag/Tag | E280* | PLES_21901 | T | T | | | | | | | | | | |
| NC_011770 | 2366863 | T | C | 72 | 0 | 12 | 0 | NON_SYNONYMOU | MODERAT | MISSENSE | Tcg/Ccg | S19P | PLES_21901 | C | C | C | C | C | C | C | C | C | C | C | C |
| NC_011770 | 2367365 | T | A | 57 | 0 | 4 | 0 | NON_SYNONYMOU | MODERAT | MISSENSE | cTg/cAg | L186Q | PLES_21901 | A | A | | | | | | | A | A | | |
| NC_011770 | 2367379 | T | C | 61 | 0 | 2 | 0 | NON_SYNONYMOU | MODERAT | MISSENSE | Ttc/Ctc | F191L | PLES_21901 | | | | | | | | | C | C | | |
| NC_011770 | 2367386 | C | G | 60 | 0 | 2 | 0 | NON_SYNONYMOU | MODERAT | MISSENSE | gCg/gGg | A193G | PLES_21901 | | | | | | | | | | | G | G |
| NC_011770 | 2367433 | A | G | 64 | 0 | 12 | 0 | NON_SYNONYMOU | MODERAT | MISSENSE | Acg/Gcg | T209A | PLES_21901 | G | G | G | G | G | G | G | G | G | G | G | G |
| NC_011770 | 2367586 | C | A | 71 | 0 | 2 | 0 | NON_SYNONYMOU | MODERAT | MISSENSE | Ctc/Atc | L260I | PLES_21901 | | | | | | | | | | | A | A |
| NC_011770 | 2367647 | A | C | 71 | 0 | 12 | 0 | NON_SYNONYMOU | MODERAT | MISSENSE | gAg/gCg | E280A | PLES_21901 | C | C | C | C | C | C | C | C | C | C | C | C |
| NC_011770 | 2368003 | G | C | 67 | 0 | 2 | 0 | NON_SYNONYMOU | MODERAT | MISSENSE | gGg/gCg | G82A | PLES_21911 | C | C | | | | | | | | | | |
| NC_011770 | 2368263 | C | G | 69 | 0 | 2 | 0 | NON_SYNONYMOU | MODERAT | MISSENSE | Ccg/Gcg | P169A | PLES_21911 | | | G | G | | | | | | | | |
| NC_011770 | 2368435 | T | C | 74 | 0 | 10 | 0 | NON_SYNONYMOU | MODERAT | MISSENSE | cTg/cCg | L226P | PLES_21911 | C | C | C | C | C | C | C | C | | | C | C |
| NC_011770 | 2368518 | T | C | 79 | 0 | 2 | 0 | NON_SYNONYMOU | MODERAT | MISSENSE | Tcg/Ccg | S254P | PLES_21911 | | | | | | | | | | | C | C |
| NC_011770 | 2368533 | C | G | 81 | 0 | 8 | 0 | NON_SYNONYMOU | MODERAT | MISSENSE | Ctg/Gtg | L259V | PLES_21911 | G | G | A | A | A | A | A | A | | | | |
| NC_011770 | 2368596 | C | T | 83 | 0 | 6 | 0 | NON_SYNONYMOU | MODERAT | MISSENSE | Ccc/Tcc | P280S | PLES_21911 | | | T | T | T | T | T | T | | | | |
| NC_011770 | 2368708 | T | C | 87 | 0 | 6 | 0 | NON_SYNONYMOU | MODERAT | MISSENSE | gTc/gCc | V317A | PLES_21911 | C | C | C | C | C | C | | | | | | |
| NC_011770 | 2368719 | C | T | 89 | 0 | 1 | 0 | NON_SYNONYMOU | MODERAT | MISSENSE | Cca/Tca | P321S | PLES_21911 | | | | | | | T | | | | | |
| NC_011770 | 2368756 | C | T | 82 | 0 | 8 | 0 | NON_SYNONYMOU | MODERAT | MISSENSE | aCg/aTg | T333M | PLES_21911 | T | T | T | T | T | T | T | T | | | | |
| NC_011770 | 2368817 | C | G | 67 | 0 | 8 | 0 | NON_SYNONYMOU | MODERAT | MISSENSE | gaC/gaG | D353E | PLES_21911 | G | G | G | G | G | G | G | G | | | | |
| NC_011770 | 2368818 | G | C | 67 | 0 | 8 | 0 | NON_SYNONYMOU | MODERAT | MISSENSE | Gag/Cag | E354Q | PLES_21911 | C | C | C | C | C | C | C | C | | | | |
| NC_011770 | 2368882 | A | G | 74 | 0 | 4 | 0 | NON_SYNONYMOU | MODERAT | MISSENSE | cAc/cGc | H375R | PLES_21911 | G | G | | | | | G | G | | | | |
| NC_011770 | 2369479 | A | G | 81 | 0 | 10 | 0 | NON_SYNONYMOU | MODERAT | MISSENSE | gAc/gGc | D574G | PLES_21911 | G | G | G | G | G | G | | | G | G | G | G |
| NC_011770 | 2369727 | A | G | 78 | 0 | 8 | 0 | NON_SYNONYMOU | MODERAT | MISSENSE | Acg/Gcg | T657A | PLES_21911 | G | G | G | G | G | G | | | | | G | G |
| NC_011770 | 2370137 | G | T | 85 | 0 | 4 | 0 | NON_SYNONYMOU | MODERAT | MISSENSE | Gcc/Tcc | A126S | PLES_21921 | | | T | T | T | T | | | | | | |
| NC_011770 | 2370228 | G | A | 83 | 0 | 2 | 0 | NON_SYNONYMOU | MODERAT | MISSENSE | cGc/cAc | R156H | PLES_21921 | | | | | | | A | A | | | | |
| NC_011770 | 2370525 | G | A | 77 | 0 | 8 | 0 | NON_SYNONYMOU | MODERAT | MISSENSE | gGg/gAg | G255E | PLES_21921 | A | A | | | | | A | A | A | A | A | A |
| NC_011770 | 2371579 | T | C | 92 | 0 | 8 | 0 | NON_SYNONYMOU | MODERAT | MISSENSE | aAc/aGc | N473S | PLES_21941 | | | C | C | C | C | C | C | C | C | | |
| NC_011770 | 2372117 | T | C | 82 | 0 | 6 | 0 | NON_SYNONYMOU | MODERAT | MISSENSE | Acc/Gcc | T294A | PLES_21941 | | | C | C | C | C | | | C | C | | |
| NC_011770 | 2372375 | T | C | 82 | 0 | 6 | 0 | NON_SYNONYMOU | MODERAT | MISSENSE | Acc/Gcc | T208A | PLES_21941 | | | C | C | C | C | | | C | C | | |
| NC_011770 | 2372606 | T | C | 86 | 0 | 6 | 0 | NON_SYNONYMOU | MODERAT | MISSENSE | Aac/Gac | N131D | PLES_21941 | | | C | C | C | C | | | C | C | | |
| NC_011770 | 2372012 | C | T | 84 | 0 | 1 | 0 | NON_SYNONYMOU | MODERAT | MISSENSE | Gcg/Acg | A329T | PLES_21941 | | | | | | | | | | | | T |
| NC_011770 | 2373232 | G | T | 56 | 0 | 4 | 0 | NON_SYNONYMOU | MODERAT | MISSENSE | aCg/aAg | T88K | PLES_21951 | | | T | T | T | T | | | | | | |
| NC_011770 | 2373239 | G | C | 55 | 0 | 2 | 0 | NON_SYNONYMOU | MODERAT | MISSENSE | Ccg/Gcg | P86A | PLES_21951 | | | | | | | | | C | C | | |
| NC_011770 | 2373376 | T | C | 50 | 0 | 4 | 0 | NON_SYNONYMOU | MODERAT | MISSENSE | gAc/gGc | D40G | PLES_21951 | | | C | C | C | C | | | | | | |
| NC_011770 | 2373463 | A | G | 47 | 0 | 4 | 0 | NON_SYNONYMOU | MODERAT | MISSENSE | cTc/cCc | L11P | PLES_21951 | | | G | G | G | G | | | | | | |
| NC_011770 | 2373464 | G | A | 47 | 0 | 4 | 0 | NON_SYNONYMOU | MODERAT | MISSENSE | Ctc/Ttc | L11F | PLES_21951 | | | A | A | A | A | | | | | | |
| NC_011770 | 2373484 | G | A | 45 | 0 | 4 | 0 | NON_SYNONYMOU | MODERAT | MISSENSE | aCg/aTg | T4M | PLES_21951 | | | A | A | A | A | | | | | | |
| NC_011770 | 2373712 | T | C | 53 | 0 | 12 | 0 | NON_SYNONYMOU | MODERAT | MISSENSE | Aaa/Gaa | K77E | PLES_21961 | C | C | C | C | C | C | C | C | C | C | C | C |
| NC_011770 | 2374659 | C | T | 62 | 0 | 2 | 0 | NON_SYNONYMOU | MODERAT | MISSENSE | Gcg/Acg | A294T | PLES_21971 | | | | | | | | | T | T | | |
| NC_011770 | 2375151 | A | G | 67 | 0 | 1 | 0 | NON_SYNONYMOU | MODERAT | MISSENSE | Tgg/Cgg | W130R | PLES_21971 | | | | | | | G | | | | | |
| NC_011770 | 2375352 | C | T | 53 | 0 | 4 | 0 | NON_SYNONYMOU | MODERAT | MISSENSE | Gcc/Acc | A63T | PLES_21971 | | | | | | | T | T | T | T | | |
| NC_011770 | 2375490 | C | T | 58 | 0 | 1 | 0 | NON_SYNONYMOU | MODERAT | MISSENSE | Gtc/Atc | V17I | PLES_21971 | | | | | | | | T | | | | |
| NC_011770 | 2376821 | G | A | 79 | 0 | 2 | 0 | NON_SYNONYMOU | MODERAT | MISSENSE | aGg/aAg | R32K | PLES_21991 | | | | | | | | | | | A | A |
| NC_011770 | 2376862 | G | T | 76 | 0 | 2 | 0 | NON_SYNONYMOU | MODERAT | MISSENSE | Gct/Tct | A46S | PLES_21991 | | | | | | | T | T | | | | |
| NC_011770 | 2377667 | G | A | 79 | 0 | 2 | 0 | NON_SYNONYMOU | MODERAT | MISSENSE | cGg/cAg | R314Q | PLES_21991 | | | | | | | | | | | A | A |
| NC_011770 | 2378448 | A | C | 74 | 0 | 2 | 0 | NON_SYNONYMOU | MODERAT | MISSENSE | Acc/Ccc | T65P | PLES_22001 | | | | | | | C | C | | | | |
| NC_011770 | 2382021 | G | A | 75 | 0 | 10 | 0 | NON_SYNONYMOU | MODERAT | MISSENSE | gGg/gAg | G120E | PLES_22041 | A | A | A | A | A | A | A | A | A | A | | |
| NC_011770 | 2382729 | A | T | 80 | 0 | 2 | 0 | NON_SYNONYMOU | MODERAT | MISSENSE | cTg/cAg | L813Q | PLES_22061 | | | | | | | T | T | | | | |
| NC_011770 | 2383057 | C | T | 70 | 0 | 2 | 0 | NON_SYNONYMOU | MODERAT | MISSENSE | Gtc/Atc | V704I | PLES_22061 | | | | | | | | | T | T | | |
| NC_011770 | 2383074 | T | C | 68 | 0 | 12 | 0 | NON_SYNONYMOU | MODERAT | MISSENSE | cAg/cGg | Q698R | PLES_22061 | C | C | C | C | C | C | C | C | C | C | C | C |
| NC_011770 | 2383153 | C | T | 70 | 0 | 2 | 0 | NON_SYNONYMOU | MODERAT | MISSENSE | Gaa/Aaa | E672K | PLES_22061 | T | T | | | | | | | | | | |
| NC_011770 | 2383331 | A | C | 67 | 0 | 12 | 0 | NON_SYNONYMOU | MODERAT | MISSENSE | caT/caG | H612Q | PLES_22061 | C | C | C | C | C | C | C | C | C | C | C | C |
| NC_011770 | 2383738 | C | T | 73 | 0 | 2 | 0 | NON_SYNONYMOU | MODERAT | MISSENSE | Ggc/Agc | G477S | PLES_22061 | | | | | | | T | T | | | | |
| NC_011770 | 2383747 | G | T | 73 | 0 | 4 | 0 | NON_SYNONYMOU | MODERAT | MISSENSE | Ctg/Atg | L474M | PLES_22061 | | | T | T | T | T | | | | | | |
| NC_011770 | 2384013 | T | C | 72 | 0 | 12 | 0 | NON_SYNONYMOU | MODERAT | MISSENSE | aAc/aGc | N385S | PLES_22061 | C | C | C | C | C | C | C | C | C | C | C | C |
| NC_011770 | 2384170 | A | G | 83 | 0 | 2 | 0 | NON_SYNONYMOU | MODERAT | MISSENSE | Ttc/Ctc | F333L | PLES_22061 | G | G | | | | | | | | | | |
| NC_011770 | 2384481 | T | C | 84 | 0 | 2 | 0 | NON_SYNONYMOU | MODERAT | MISSENSE | aAg/aGg | K229R | PLES_22061 | | | | | | | | | C | C | | |
| NC_011770 | 2384563 | C | T | 75 | 0 | 2 | 0 | NON_SYNONYMOU | MODERAT | MISSENSE | Gtc/Atc | V202I | PLES_22061 | | | | | | | | | | | T | T |
| NC_011770 | 2384938 | C | T | 79 | 0 | 2 | 0 | NON_SYNONYMOU | MODERAT | MISSENSE | Gaa/Aaa | E77K | PLES_22061 | | | | | | | | | T | T | | |
| NC_011770 | 2384968 | C | T | 82 | 0 | 2 | 0 | NON_SYNONYMOU | MODERAT | MISSENSE | Ggt/Agt | G67S | PLES_22061 | T | T | | | | | | | | | | |
| NC_011770 | 2385135 | C | G | 67 | 0 | 4 | 0 | NON_SYNONYMOU | MODERAT | MISSENSE | gGt/gCt | G11A | PLES_22061 | | | G | G | G | G | | | | | | |
| NC_011770 | 2383515 | A | G | 65 | 0 | 3 | 0 | NON_SYNONYMOU | MODERAT | MISSENSE | gTc/gCc | V551A | PLES_22061 | G | G | | | | | | G | | | | |

| | | | | | | | | | | | | | | | | | | | | | | | | | |
|---|---|---|---|---|---|---|---|---|---|---|---|---|---|---|---|---|---|---|---|---|---|---|---|---|---|
| NC_011770 | 2385507 | C | T | 56 | 0 | 4 | 0 | NON_SYNONYMOU | MODERAT | MISSENSE | Gcc/Acc | A118T | PLES_22071 | | | T | T | T | T | | | | | | |
| NC_011770 | 2385738 | C | T | 55 | 0 | 2 | 0 | NON_SYNONYMOU | MODERAT | MISSENSE | Gtc/Atc | V41I | PLES_22071 | | | | | | | T | T | | | | |
| NC_011770 | 2388907 | C | G | 66 | 0 | 2 | 0 | NON_SYNONYMOU | MODERAT | MISSENSE | gaC/gaG | D5E | PLES_22121 | | | | | | | | | | | G | G |
| NC_011770 | 2389020 | A | G | 65 | 0 | 2 | 0 | NON_SYNONYMOU | MODERAT | MISSENSE | cAg/cGg | Q43R | PLES_22121 | | | | | | | | | G | G | | |
| NC_011770 | 2389046 | G | A | 67 | 0 | 6 | 0 | NON_SYNONYMOU | MODERAT | MISSENSE | Gcg/Acg | A52T | PLES_22121 | | | A | A | A | A | | | A | A | | |
| NC_011770 | 2389056 | C | T | 67 | 0 | 6 | 0 | NON_SYNONYMOU | MODERAT | MISSENSE | gCg/gTg | A55V | PLES_22121 | | | T | T | T | T | | | T | T | | |
| NC_011770 | 2389961 | G | C | 63 | 0 | 2 | 0 | NON_SYNONYMOU | MODERAT | MISSENSE | Gaa/Caa | E357Q | PLES_22121 | | | | | | | | | C | C | | |
| NC_011770 | 2391316 | C | T | 83 | 0 | 6 | 0 | NON_SYNONYMOU | MODERAT | MISSENSE | Ggc/Agc | G145S | PLES_22151 | | | T | T | T | T | | | T | T | | |
| NC_011770 | 2392022 | A | G | 78 | 0 | 12 | 0 | NON_SYNONYMOU | MODERAT | MISSENSE | Tgc/Cgc | C278R | PLES_22161 | G | G | G | G | G | G | G | G | G | G | G | G |
| NC_011770 | 2392619 | C | T | 79 | 0 | 8 | 0 | NON_SYNONYMOU | MODERAT | MISSENSE | Gcg/Acg | A79T | PLES_22161 | T | T | T | T | T | T | | | | | T | T |
| NC_011770 | 2392807 | A | G | 75 | 0 | 4 | 0 | NON_SYNONYMOU | MODERAT | MISSENSE | tTa/tCa | L16S | PLES_22161 | G | G | | | | | | | | | G | G |
| NC_011770 | 2392939 | C | A | 75 | 0 | 4 | 0 | NON_SYNONYMOU | MODERAT | MISSENSE | Ctg/Atg | L17M | PLES_22171 | | | A | A | A | A | | | | | | |
| NC_011770 | 2393116 | G | A | 70 | 0 | 2 | 0 | NON_SYNONYMOU | MODERAT | MISSENSE | Gtc/Atc | V76I | PLES_22171 | | | | | | | | | | | A | A |
| NC_011770 | 2393255 | G | A | 67 | 0 | 2 | 0 | NON_SYNONYMOU | MODERAT | MISSENSE | cGc/cAc | R122H | PLES_22171 | | | | | | | | | | | A | A |
| NC_011770 | 2394354 | T | C | 94 | 0 | 1 | 0 | NON_SYNONYMOU | MODERAT | MISSENSE | gAc/gGc | D48G | PLES_22181 | | | | | | | C | | | | | |
| NC_011770 | 2394000 | A | G | 97 | 0 | 1 | 0 | NON_SYNONYMOU | MODERAT | MISSENSE | gTc/gCc | V166A | PLES_22181 | | | | | | | | G | | | | |
| NC_011770 | 2394621 | G | A | 86 | 0 | 2 | 0 | NON_SYNONYMOU | MODERAT | MISSENSE | aGt/aAt | S10N | PLES_22191 | A | A | | | | | | | | | | |
| NC_011770 | 2394627 | T | G | 89 | 0 | 2 | 0 | NON_SYNONYMOU | MODERAT | MISSENSE | tTc/tGc | F12C | PLES_22191 | G | G | | | | | | | | | | |
| NC_011770 | 2394636 | C | T | 88 | 0 | 2 | 0 | NON_SYNONYMOU | MODERAT | MISSENSE | gCa/gTa | A15V | PLES_22191 | T | T | | | | | | | | | | |
| NC_011770 | 2394638 | T | C | 89 | 0 | 2 | 0 | NON_SYNONYMOU | MODERAT | MISSENSE | Tcc/Ccc | S16P | PLES_22191 | C | C | | | | | | | | | | |
| NC_011770 | 2394988 | G | T | 70 | 0 | 2 | 0 | NON_SYNONYMOU | MODERAT | MISSENSE | cGc/cTc | R27L | PLES_22201 | | | | | | | T | T | | | | |
| NC_011770 | 2395378 | C | T | 61 | 0 | 1 | 0 | NON_SYNONYMOU | MODERAT | MISSENSE | gCc/gTc | A157V | PLES_22201 | | | | | | | T | | | | | |
| NC_011770 | 2395957 | C | T | 50 | 0 | 2 | 0 | NON_SYNONYMOU | MODERAT | MISSENSE | gCg/gTg | A350V | PLES_22201 | | | | | | | T | T | | | | |
| NC_011770 | 2396316 | C | T | 60 | 0 | 1 | 0 | NON_SYNONYMOU | MODERAT | MISSENSE | Ggg/Agg | G397R | PLES_22211 | | | | | | | T | | | | | |
| NC_011770 | 2396866 | C | T | 72 | 0 | 4 | 0 | NON_SYNONYMOU | MODERAT | MISSENSE | atG/atA | M213I | PLES_22211 | | | T | T | T | T | | | | | | |
| NC_011770 | 2396874 | C | A | 73 | 0 | 4 | 0 | NON_SYNONYMOU | MODERAT | MISSENSE | Gcc/Tcc | A211S | PLES_22211 | | | A | A | A | A | | | | | | |
| NC_011770 | 2397197 | A | G | 70 | 0 | 2 | 0 | NON_SYNONYMOU | MODERAT | MISSENSE | gTa/gCa | V103A | PLES_22211 | G | G | | | | | | | | | | |
| NC_011770 | 2397890 | T | C | 77 | 0 | 1 | 0 | NON_SYNONYMOU | MODERAT | MISSENSE | Ttc/Ctc | F71L | PLES_22221 | | | | | | | C | | | | | |
| NC_011770 | 2398334 | A | G | 82 | 0 | 12 | 0 | NON_SYNONYMOU | MODERAT | MISSENSE | Aac/Gac | N219D | PLES_22221 | G | G | G | G | G | G | G | G | G | G | G | G |
| NC_011770 | 2399234 | G | A | 54 | 0 | 2 | 0 | NON_SYNONYMOU | MODERAT | MISSENSE | Ctc/Ttc | L5F | PLES_22231 | | | | | | | | | | | A | A |
| NC_011770 | 2399893 | C | T | 72 | 0 | 2 | 0 | NON_SYNONYMOU | MODERAT | MISSENSE | Ccg/Tcg | P113S | PLES_22251 | T | T | | | | | | | | | | |
| NC_011770 | 2400202 | C | T | 84 | 0 | 1 | 0 | NON_SYNONYMOU | MODERAT | MISSENSE | Ccc/Tcc | P216S | PLES_22251 | | | | | | | T | | | | | |
| NC_011770 | 2401804 | C | G | 65 | 0 | 2 | 0 | NON_SYNONYMOU | MODERAT | MISSENSE | Gag/Cag | E199Q | PLES_22261 | | | | | | | | | | | G | G |
| NC_011770 | 2402229 | A | G | 72 | 0 | 6 | 0 | NON_SYNONYMOU | MODERAT | MISSENSE | gTg/gCg | V57A | PLES_22261 | G | G | | | | | G | G | | | G | G |
| NC_011770 | 2402070 | C | T | 73 | 0 | 1 | 0 | NON_SYNONYMOU | MODERAT | MISSENSE | gGc/gAc | G110D | PLES_22261 | | | | | | | | T | | | | |
| NC_011770 | 2403130 | G | T | 94 | 0 | 2 | 0 | NON_SYNONYMOU | MODERAT | MISSENSE | Ctg/Atg | L111M | PLES_22271 | | | | | | | T | T | | | | |
| NC_011770 | 2403166 | C | T | 90 | 0 | 2 | 0 | NON_SYNONYMOU | MODERAT | MISSENSE | Gtg/Atg | V99M | PLES_22271 | | | | | | | T | T | | | | |
| NC_011770 | 2404326 | C | A | 73 | 0 | 2 | 0 | NON_SYNONYMOU | MODERAT | MISSENSE | cGg/cTg | R204L | PLES_22281 | A | A | | | | | | | | | | |
| NC_011770 | 2404614 | G | T | 58 | 0 | 4 | 0 | NON_SYNONYMOU | MODERAT | MISSENSE | aCc/aAc | T108N | PLES_22281 | T | T | | | | | | | | | T | T |
| NC_011770 | 2404625 | G | C | 60 | 0 | 4 | 0 | NON_SYNONYMOU | MODERAT | MISSENSE | gaC/gaG | D104E | PLES_22281 | | | C | C | C | C | | | | | | |
| NC_011770 | 2404873 | T | C | 56 | 0 | 8 | 0 | NON_SYNONYMOU | MODERAT | MISSENSE | Aac/Gac | N22D | PLES_22281 | C | C | | | | | C | C | C | C | C | C |
| NC_011770 | 2405166 | T | C | 70 | 0 | 6 | 0 | NON_SYNONYMOU | MODERAT | MISSENSE | Aag/Gag | K278E | PLES_22291 | | | | | | | C | C | C | C | C | C |
| NC_011770 | 2405336 | T | A | 69 | 0 | 2 | 0 | NON_SYNONYMOU | MODERAT | MISSENSE | cAg/cTg | Q221L | PLES_22291 | | | | | | | | | | | A | A |
| NC_011770 | 2405658 | G | C | 59 | 0 | 2 | 0 | NON_SYNONYMOU | MODERAT | MISSENSE | Ctg/Gtg | L114V | PLES_22291 | | | | | | | | | | | C | C |
| NC_011770 | 2405711 | C | G | 62 | 0 | 4 | 0 | NON_SYNONYMOU | MODERAT | MISSENSE | gGt/gCt | G96A | PLES_22291 | | | G | G | G | G | | | | | | |
| NC_011770 | 2405939 | G | A | 67 | 0 | 2 | 0 | NON_SYNONYMOU | MODERAT | MISSENSE | aCc/aTc | T20I | PLES_22291 | | | | | | | | | | | A | A |
| NC_011770 | 2405994 | A | T | 58 | 0 | 4 | 0 | NON_SYNONYMOU | MODERAT | MISSENSE | Tcg/Acg | S2T | PLES_22291 | | | T | T | T | T | | | | | | |
| NC_011770 | 2406095 | C | T | 54 | 0 | 2 | 0 | NON_SYNONYMOU | MODERAT | MISSENSE | Gcc/Acc | A503T | PLES_22301 | | | | | | | | | | | T | T |
| NC_011770 | 2406143 | G | A | 56 | 0 | 2 | 0 | NON_SYNONYMOU | MODERAT | MISSENSE | Cgc/Tgc | R487C | PLES_22301 | | | | | | | | | | | A | A |
| NC_011770 | 2406206 | T | A | 56 | 0 | 2 | 0 | NON_SYNONYMOU | MODERAT | MISSENSE | Atc/Ttc | I466F | PLES_22301 | | | | | | | | | | | A | A |
| NC_011770 | 2406236 | A | G | 59 | 0 | 10 | 0 | NON_SYNONYMOU | MODERAT | MISSENSE | Tcc/Ccc | S456P | PLES_22301 | G | G | G | G | G | G | G | G | | | G | G |
| NC_011770 | 2407362 | C | T | 54 | 0 | 1 | 0 | NON_SYNONYMOU | MODERAT | MISSENSE | atG/atA | M80I | PLES_22301 | | | | | | | T | | | | | |
| NC_011770 | 2407532 | C | T | 35 | 0 | 4 | 0 | NON_SYNONYMOU | MODERAT | MISSENSE | Gcg/Acg | A24T | PLES_22301 | | | T | T | T | T | | | | | | |
| NC_011770 | 2408031 | C | T | 70 | 0 | 2 | 0 | NON_SYNONYMOU | MODERAT | MISSENSE | gGc/gAc | G223D | PLES_22311 | | | | | | | T | T | | | | |
| NC_011770 | 2408530 | C | T | 87 | 0 | 1 | 0 | NON_SYNONYMOU | MODERAT | MISSENSE | Gac/Aac | D57N | PLES_22311 | | | | | | | | T | | | | |
| NC_011770 | 2408861 | G | A | 80 | 0 | 2 | 0 | NON_SYNONYMOU | MODERAT | MISSENSE | Ggc/Agc | G12S | PLES_22321 | | | | | | | | | A | A | | |
| NC_011770 | 2409017 | C | T | 69 | 0 | 2 | 0 | NON_SYNONYMOU | MODERAT | MISSENSE | Cgg/Tgg | R64W | PLES_22321 | | | | | | | | | T | T | | |
| NC_011770 | 2409038 | T | C | 69 | 0 | 4 | 0 | NON_SYNONYMOU | MODERAT | MISSENSE | Tgt/Cgt | C71R | PLES_22321 | C | C | | | | | C | C | | | | |
| NC_011770 | 2410682 | T | C | 68 | 0 | 4 | 0 | NON_SYNONYMOU | MODERAT | MISSENSE | gAc/gGc | D155G | PLES_22341 | | | | | | | C | C | C | C | | |
| NC_011770 | 2410989 | G | A | 65 | 0 | 4 | 0 | NON_SYNONYMOU | MODERAT | MISSENSE | Cgc/Tgc | R53C | PLES_22341 | | | A | A | A | A | | | | | | |
| NC_011770 | 2411004 | C | G | 64 | 0 | 6 | 0 | NON_SYNONYMOU | MODERAT | MISSENSE | Gcc/Ccc | A48P | PLES_22341 | | | G | G | G | G | G | G | | | | |
| NC_011770 | 2412456 | C | A | 56 | 0 | 4 | 0 | NON_SYNONYMOU | MODERAT | MISSENSE | Gcg/Tcg | A98S | PLES_22361 | A | A | | | | | | | A | A | | |

| | | | | | | | | | | | | | | | | | | | | | | | | | |
|---|---|---|---|---|---|---|---|---|---|---|---|---|---|---|---|---|---|---|---|---|---|---|---|---|---|
| NC_011770 | 2414796 | T | C | 80 | 0 | 2 | 0 | NON_SYNONYMOU | MODERAT | MISSENSE | gTg/gCg | V31A | PLES_22391 | | | | | | | C | C | | | | |
| NC_011770 | 2415867 | T | A | 68 | 0 | 2 | 0 | NON_SYNONYMOU | MODERAT | MISSENSE | Agc/Tgc | S731C | PLES_22411 | | | | | | | | | | | A | A |
| NC_011770 | 2415885 | C | T | 71 | 0 | 12 | 0 | NON_SYNONYMOU | MODERAT | MISSENSE | Gcc/Acc | A725T | PLES_22411 | T | T | T | T | T | T | T | T | T | T | T | T |
| NC_011770 | 2417292 | C | T | 70 | 0 | 1 | 0 | NON_SYNONYMOU | MODERAT | MISSENSE | Gcg/Acg | A256T | PLES_22411 | | | | | | | T | | | | | |
| NC_011770 | 2417593 | G | C | 93 | 0 | 2 | 0 | NON_SYNONYMOU | MODERAT | MISSENSE | caC/caG | H155Q | PLES_22411 | | | | | | | C | C | | | | |
| NC_011770 | 2417706 | G | A | 84 | 0 | 6 | 0 | NON_SYNONYMOU | MODERAT | MISSENSE | Ccc/Tcc | P118S | PLES_22411 | | | A | A | A | A | | | | | A | A |
| NC_011770 | 2417568 | C | T | 91 | 0 | 1 | 0 | NON_SYNONYMOU | MODERAT | MISSENSE | Ggc/Agc | G164S | PLES_22411 | | | | | | | | T | | | | |
| NC_011770 | 2418413 | TG | T | 75 | 0 | 1 | 0 | FRAME_SHIFT | HIGH | | gcc/ | A94 | PLES_22421 | | | | | | | T | | | | | |
| NC_011770 | 2418314 | T | C | 65 | 0 | 12 | 0 | NON_SYNONYMOU | MODERAT | MISSENSE | Tgc/Cgc | C59R | PLES_22421 | C | C | C | C | C | C | C | C | C | C | C | C |
| NC_011770 | 2418661 | G | C | 86 | 0 | 2 | 0 | NON_SYNONYMOU | MODERAT | MISSENSE | gaG/gaC | E174D | PLES_22421 | | | | | | | C | C | | | | |
| NC_011770 | 2418858 | G | A | 82 | 0 | 6 | 0 | NON_SYNONYMOU | MODERAT | MISSENSE | cGc/cAc | R240H | PLES_22421 | A | A | | | | | | | A | A | A | A |
| NC_011770 | 2418902 | A | G | 77 | 0 | 2 | 0 | NON_SYNONYMOU | MODERAT | MISSENSE | Agc/Ggc | S255G | PLES_22421 | | | | | | | G | G | | | | |
| NC_011770 | 2419106 | A | G | 71 | 0 | 2 | 0 | NON_SYNONYMOU | MODERAT | MISSENSE | gAc/gGc | D27G | PLES_22431 | | | | | | | G | G | | | | |
| NC_011770 | 2419436 | G | C | 76 | 0 | 2 | 0 | NON_SYNONYMOU | MODERAT | MISSENSE | gGa/gCa | G137A | PLES_22431 | | | | | | | C | C | | | | |
| NC_011770 | 2419920 | T | C | 69 | 0 | 6 | 0 | NON_SYNONYMOU | MODERAT | MISSENSE | Agc/Ggc | S88G | PLES_22441 | | | C | C | C | C | C | C | | | | |
| NC_011770 | 2420010 | C | T | 75 | 0 | 2 | 0 | NON_SYNONYMOU | MODERAT | MISSENSE | Gac/Aac | D58N | PLES_22441 | | | | | | | T | T | | | | |
| NC_011770 | 2420377 | T | C | 83 | 0 | 12 | 0 | NON_SYNONYMOU | MODERAT | MISSENSE | Acg/Gcg | T214A | PLES_22451 | C | C | C | C | C | C | C | C | C | C | C | C |
| NC_011770 | 2420761 | C | T | 80 | 0 | 2 | 0 | NON_SYNONYMOU | MODERAT | MISSENSE | Gcc/Acc | A86T | PLES_22451 | T | T | | | | | | | | | | |
| NC_011770 | 2422746 | G | T | 24 | 4 | 2 | 0 | NON_SYNONYMOU | MODERAT | MISSENSE | agC/agA | S137R | PLES_22481 | T | T | . | . | . | . | | | | | | |
| NC_011770 | 2422867 | G | A | 29 | 4 | 1 | 0 | NON_SYNONYMOU | MODERAT | MISSENSE | gCg/gTg | A97V | PLES_22481 | | | . | . | . | . | A | | | | | |
| NC_011770 | 2424063 | G | A | 54 | 4 | 2 | 0 | NON_SYNONYMOU | MODERAT | MISSENSE | gCa/gTa | A586V | PLES_22501 | | | . | . | . | . | A | A | | | | |
| NC_011770 | 2424117 | T | C | 51 | 4 | 4 | 0 | NON_SYNONYMOU | MODERAT | MISSENSE | gAa/gGa | E568G | PLES_22501 | | | . | . | . | . | | | C | C | C | C |
| NC_011770 | 2424694 | C | G | 42 | 4 | 8 | 0 | NON_SYNONYMOU | MODERAT | MISSENSE | Ggc/Cgc | G376R | PLES_22501 | G | G | . | . | . | . | G | G | G | G | G | G |
| NC_011770 | 2425156 | T | C | 39 | 4 | 4 | 0 | NON_SYNONYMOU | MODERAT | MISSENSE | Acg/Gcg | T222A | PLES_22501 | | | . | . | . | . | | | C | C | C | C |
| NC_011770 | 2425598 | G | T | 42 | 4 | 2 | 0 | NON_SYNONYMOU | MODERAT | MISSENSE | gaC/gaA | D74E | PLES_22501 | T | T | . | . | . | . | | | | | | |
| NC_011770 | 2427266 | G | T | 34 | 4 | 6 | 0 | NON_SYNONYMOU | MODERAT | MISSENSE | Ctg/Atg | L56M | PLES_22531 | T | T | . | . | . | . | T | T | T | T | | |
| NC_011770 | 2427579 | C | T | 31 | 4 | 8 | 0 | NON_SYNONYMOU | MODERAT | MISSENSE | cGc/cAc | R215H | PLES_22541 | T | T | . | . | . | . | T | T | T | T | T | T |
| NC_011770 | 2428488 | C | T | 47 | 0 | 4 | 0 | SPLICE_SITE_REGIC | LOW | SILENT | tGa/tAa | *137 | PLES_22551 | | | | | | | | | T | T | T | T |
| NC_011770 | 2428973 | A | G | 60 | 0 | 10 | 0 | NON_SYNONYMOU | MODERAT | MISSENSE | Tcc/Ccc | S122P | PLES_22561 | G | G | G | G | G | G | | | G | G | G | G |
| NC_011770 | 2432120 | T | C | 74 | 0 | 2 | 0 | NON_SYNONYMOU | MODERAT | MISSENSE | gTg/gCg | V19A | PLES_22581 | | | | | | | | | C | C | | |
| NC_011770 | 2432450 | G | A | 82 | 0 | 1 | 0 | NON_SYNONYMOU | MODERAT | MISSENSE | gGc/gAc | G129D | PLES_22581 | | | | | | | A | | | | | |
| NC_011770 | 2432482 | C | T | 83 | 0 | 2 | 0 | NON_SYNONYMOU | MODERAT | MISSENSE | Cgt/Tgt | R140C | PLES_22581 | | | | | | | | | | | T | T |
| NC_011770 | 2432561 | G | A | 81 | 0 | 2 | 0 | NON_SYNONYMOU | MODERAT | MISSENSE | cGc/cAc | R166H | PLES_22581 | | | | | | | A | A | | | | |
| NC_011770 | 2432581 | A | G | 84 | 0 | 12 | 0 | NON_SYNONYMOU | MODERAT | MISSENSE | Aaa/Gaa | K173E | PLES_22581 | G | G | G | G | G | G | G | G | G | G | G | G |
| NC_011770 | 2432110 | C | T | 73 | 0 | 1 | 0 | NON_SYNONYMOU | MODERAT | MISSENSE | Ccg/Tcg | P16S | PLES_22581 | | | | | | | | | | | | T |
| NC_011770 | 2433205 | C | T | 68 | 0 | 2 | 0 | NON_SYNONYMOU | MODERAT | MISSENSE | Ccg/Tcg | P119S | PLES_22591 | T | T | | | | | | | | | | |
| NC_011770 | 2434342 | A | T | 82 | 0 | 2 | 0 | NON_SYNONYMOU | MODERAT | MISSENSE | Agc/Tgc | S266C | PLES_22601 | T | T | | | | | | | | | | |
| NC_011770 | 2435208 | C | T | 77 | 0 | 2 | 0 | NON_SYNONYMOU | MODERAT | MISSENSE | Ctc/Ttc | L245F | PLES_22611 | | | | | | | T | T | | | | |
| NC_011770 | 2434557 | C | T | 84 | 0 | 1 | 0 | NON_SYNONYMOU | MODERAT | MISSENSE | Ccg/Tcg | P28S | PLES_22611 | | | | | | | | T | | | | |
| NC_011770 | 2437815 | C | T | 74 | 0 | 2 | 0 | NON_SYNONYMOU | MODERAT | MISSENSE | gCc/gTc | A5V | PLES_22651 | T | T | | | | | | | | | | |
| NC_011770 | 2438333 | G | A | 78 | 0 | 2 | 0 | NON_SYNONYMOU | MODERAT | MISSENSE | Gcc/Acc | A178T | PLES_22651 | | | | | | | | | A | A | | |
| NC_011770 | 2437985 | G | A | 76 | 0 | 1 | 0 | NON_SYNONYMOU | MODERAT | MISSENSE | Gcc/Acc | A62T | PLES_22651 | | A | | | | | | | | | | |
| NC_011770 | 2439622 | C | T | 75 | 0 | 2 | 0 | NON_SYNONYMOU | MODERAT | MISSENSE | Gag/Aag | E26K | PLES_22671 | | | | | | | | | T | T | | |
| NC_011770 | 2440644 | T | C | 56 | 0 | 8 | 0 | NON_SYNONYMOU | MODERAT | MISSENSE | cAg/cGg | Q150R | PLES_22691 | C | C | | | | | C | C | C | C | C | C |
| NC_011770 | 2440807 | C | T | 58 | 0 | 2 | 0 | NON_SYNONYMOU | MODERAT | MISSENSE | Gag/Aag | E96K | PLES_22691 | | | | | | | | | | | T | T |
| NC_011770 | 2440848 | T | C | 57 | 0 | 2 | 0 | NON_SYNONYMOU | MODERAT | MISSENSE | gAg/gGg | E82G | PLES_22691 | | | | | | | | | | | C | C |
| NC_011770 | 2440890 | T | C | 61 | 0 | 2 | 0 | NON_SYNONYMOU | MODERAT | MISSENSE | aAc/aGc | N68S | PLES_22691 | | | | | | | | | | | C | C |
| NC_011770 | 2440908 | T | C | 63 | 0 | 2 | 0 | NON_SYNONYMOU | MODERAT | MISSENSE | cAg/cGg | Q62R | PLES_22691 | | | | | | | | | | | C | C |
| NC_011770 | 2442065 | T | C | 84 | 0 | 12 | 0 | NON_SYNONYMOU | MODERAT | MISSENSE | cAc/cGc | H96R | PLES_22711 | C | C | C | C | C | C | C | C | C | C | C | C |
| NC_011770 | 2442161 | G | T | 90 | 0 | 2 | 0 | NON_SYNONYMOU | MODERAT | MISSENSE | aCc/aAc | T64N | PLES_22711 | | | | | | | | | T | T | | |
| NC_011770 | 2443046 | T | C | 78 | 0 | 1 | 0 | NON_SYNONYMOU | MODERAT | MISSENSE | cTg/cCg | L198P | PLES_22721 | | | | | | | | C | | | | |
| NC_011770 | 2443427 | C | T | 66 | 0 | 2 | 0 | NON_SYNONYMOU | MODERAT | MISSENSE | cGc/cAc | R30H | PLES_22731 | | | | | | | T | T | | | | |
| NC_011770 | 2444129 | A | G | 73 | 0 | 1 | 0 | NON_SYNONYMOU | MODERAT | MISSENSE | gAt/gGt | D123G | PLES_22741 | | | | | | | G | | | | | |
| NC_011770 | 2444652 | A | C | 84 | 0 | 4 | 0 | NON_SYNONYMOU | MODERAT | MISSENSE | gaA/gaC | E297D | PLES_22741 | | | | | | | | | C | C | C | C |
| NC_011770 | 2444711 | C | G | 86 | 0 | 4 | 0 | NON_SYNONYMOU | MODERAT | MISSENSE | aCc/aGc | T317S | PLES_22741 | | | | | | | | | G | G | G | G |
| NC_011770 | 2445276 | C | T | 85 | 0 | 1 | 0 | NON_SYNONYMOU | MODERAT | MISSENSE | Ccg/Tcg | P112S | PLES_22751 | | | | | | | T | | | | | |
| NC_011770 | 2447484 | C | T | 24 | 0 | 6 | 0 | NON_SYNONYMOU | MODERAT | MISSENSE | Gcc/Acc | A11T | PLES_22771 | T | T | | | | | | | T | T | T | T |
| NC_011770 | 2447493 | C | T | 23 | 0 | 6 | 0 | NON_SYNONYMOU | MODERAT | MISSENSE | Gcc/Acc | A8T | PLES_22771 | T | T | | | | | | | T | T | T | T |
| NC_011770 | 2447495 | T | G | 23 | 0 | 6 | 0 | NON_SYNONYMOU | MODERAT | MISSENSE | aAc/aCc | N7T | PLES_22771 | G | G | | | | | | | G | G | G | G |
| NC_011770 | 2448812 | T | C | 44 | 6 | 6 | 0 | NON_SYNONYMOU | MODERAT | MISSENSE | atA/atG | I377M | PLES_22791 | . | . | C | C | C | C | C | C | . | . | . | . |
| NC_011770 | 2450979 | C | T | 67 | 0 | 2 | 0 | NON_SYNONYMOU | MODERAT | MISSENSE | Ccc/Tcc | P4S | PLES_22801 | | | | | | | | | T | T | | |
| NC_011770 | 2450994 | G | A | 67 | 0 | 2 | 0 | NON_SYNONYMOU | MODERAT | MISSENSE | Gcc/Acc | A9T | PLES_22801 | | | | | | | | | A | A | | |

| | | | | | | | | | | | | | | | | | | | | | | | | | |
|---|---|---|---|---|---|---|---|---|---|---|---|---|---|---|---|---|---|---|---|---|---|---|---|---|---|
| NC_011770 | 2451014 | G | GCTTTC | 68 | 0 | 1 | 0 | CODON_CHANGE_ | MODERATE | | tgc/tCTCT | C17SLC | PLES_22801 | | | | | | | | | . | GCTTTCT | | |
| NC_011770 | 2451019 | G | C | 65 | 0 | 2 | 0 | NON_SYNONYMOU | MODERAT | MISSENSE | tGc/tCc | C17S | PLES_22801 | | | | | | | | | C | C | | |
| NC_011770 | 2451402 | A | G | 80 | 0 | 2 | 0 | NON_SYNONYMOU | MODERAT | MISSENSE | Agc/Ggc | S145G | PLES_22801 | | | | | | | | | G | G | | |
| NC_011770 | 2451663 | G | A | 91 | 0 | 2 | 0 | NON_SYNONYMOU | MODERAT | MISSENSE | Gtc/Atc | V232I | PLES_22801 | | | | | | | | | A | A | | |
| NC_011770 | 2451706 | C | G | 86 | 0 | 2 | 0 | NON_SYNONYMOU | MODERAT | MISSENSE | aCc/aGc | T246S | PLES_22801 | | | | | | | | | G | G | | |
| NC_011770 | 2452109 | C | T | 85 | 0 | 2 | 0 | NON_SYNONYMOU | MODERAT | MISSENSE | Cgc/Tgc | R9C | PLES_22811 | | | | | | | T | T | | | | |
| NC_011770 | 2452119 | C | T | 85 | 0 | 4 | 0 | NON_SYNONYMOU | MODERAT | MISSENSE | gCg/gTg | A12V | PLES_22811 | T | T | | | | | | | T | T | | |
| NC_011770 | 2452133 | G | T | 84 | 0 | 2 | 0 | NON_SYNONYMOU | MODERAT | MISSENSE | Gcc/Tcc | A17S | PLES_22811 | | | | | | | | | T | T | | |
| NC_011770 | 2452261 | C | G | 86 | 0 | 2 | 0 | NON_SYNONYMOU | MODERAT | MISSENSE | atC/atG | I59M | PLES_22811 | G | G | | | | | | | | | | |
| NC_011770 | 2452277 | G | T | 87 | 0 | 4 | 0 | NON_SYNONYMOU | MODERAT | MISSENSE | Gtg/Ttg | V65L | PLES_22811 | | | | | | | T | T | | | T | T |
| NC_011770 | 2452484 | G | C | 75 | 0 | 4 | 0 | NON_SYNONYMOU | MODERAT | MISSENSE | Gag/Cag | E134Q | PLES_22811 | C | C | | | | | | | C | C | | |
| NC_011770 | 2452634 | A | G | 71 | 0 | 2 | 0 | NON_SYNONYMOU | MODERAT | MISSENSE | Atg/Gtg | M184V | PLES_22811 | G | G | | | | | | | | | | |
| NC_011770 | 2452857 | A | T | 60 | 0 | 2 | 0 | NON_SYNONYMOU | MODERAT | MISSENSE | gAc/gTc | D28V | PLES_22821 | | | | | | | T | T | | | | |
| NC_011770 | 2452865 | A | G | 64 | 0 | 2 | 0 | NON_SYNONYMOU | MODERAT | MISSENSE | Agc/Ggc | S31G | PLES_22821 | G | G | | | | | | | | | | |
| NC_011770 | 2452895 | A | T | 54 | 0 | 4 | 0 | NON_SYNONYMOU | MODERAT | MISSENSE | Atc/Ttc | I41F | PLES_22821 | T | T | | | | | | | T | T | | |
| NC_011770 | 2452911 | C | T | 56 | 0 | 6 | 0 | NON_SYNONYMOU | MODERAT | MISSENSE | gCc/gTc | A46V | PLES_22821 | T | T | | | | | T | T | T | T | | |
| NC_011770 | 2452943 | A | C | 54 | 0 | 2 | 0 | NON_SYNONYMOU | MODERAT | MISSENSE | Act/Cct | T57P | PLES_22821 | | | | | | | C | C | | | | |
| NC_011770 | 2453009 | T | A | 52 | 0 | 12 | 0 | NON_SYNONYMOU | MODERAT | MISSENSE | Tcg/Acg | S79T | PLES_22821 | A | A | A | A | A | A | A | A | A | A | A | A |
| NC_011770 | 2453022 | C | T | 50 | 0 | 4 | 0 | NON_SYNONYMOU | MODERAT | MISSENSE | gCg/gTg | A83V | PLES_22821 | T | T | | | | | | | T | T | | |
| NC_011770 | 2453422 | A | T | 6 | 8 | 4 | 0 | NON_SYNONYMOU | MODERAT | MISSENSE | gAg/gTg | E10V | PLES_22831 | T | T | . | . | . | . | . | . | T | T | . | . |
| NC_011770 | 2453544 | T | C | 8 | 8 | 4 | 0 | NON_SYNONYMOU | MODERAT | MISSENSE | Ttt/Ctt | F51L | PLES_22831 | C | C | . | . | . | . | . | . | C | C | . | . |
| NC_011770 | 2453790 | T | C | 10 | 8 | 4 | 0 | NON_SYNONYMOU | MODERAT | MISSENSE | Tat/Cat | Y133H | PLES_22831 | C | C | . | . | . | . | . | . | C | C | . | . |
| NC_011770 | 2454483 | G | C | 11 | 8 | 2 | 0 | NON_SYNONYMOU | MODERAT | MISSENSE | Gtc/Ctc | V364L | PLES_22831 | | | . | . | . | . | . | . | C | C | . | . |
| NC_011770 | 2454615 | C | G | 9 | 8 | 2 | 0 | NON_SYNONYMOU | MODERAT | MISSENSE | Ctc/Gtc | L408V | PLES_22831 | | | . | . | . | . | . | . | G | G | . | . |
| NC_011770 | 2454922 | C | T | 9 | 8 | 2 | 0 | NON_SYNONYMOU | MODERAT | MISSENSE | gCg/gTg | A510V | PLES_22831 | | | . | . | . | . | . | . | T | T | . | . |
| NC_011770 | 2455849 | G | T | 68 | 0 | 2 | 0 | NON_SYNONYMOU | MODERAT | MISSENSE | Ccg/Acg | P14T | PLES_22841 | T | T | | | | | | | | | | |
| NC_011770 | 2457265 | C | T | 36 | 0 | 1 | 0 | STOP_GAINED | HIGH | NONSENSE | tgG/tgA | W46* | PLES_22851 | | T | | | | | | | | | | |
| NC_011770 | 2456441 | C | T | 68 | 0 | 4 | 0 | NON_SYNONYMOU | MODERAT | MISSENSE | cGc/cAc | R321H | PLES_22851 | | | T | T | T | T | | | | | | |
| NC_011770 | 2456801 | G | A | 63 | 0 | 2 | 0 | NON_SYNONYMOU | MODERAT | MISSENSE | aCc/aTc | T201I | PLES_22851 | | | | | | | A | A | | | | |
| NC_011770 | 2456960 | T | G | 60 | 0 | 2 | 0 | NON_SYNONYMOU | MODERAT | MISSENSE | gAc/gCc | D148A | PLES_22851 | | | G | G | | | | | | | | |
| NC_011770 | 2457384 | G | T | 29 | 0 | 2 | 0 | NON_SYNONYMOU | MODERAT | MISSENSE | Ctt/Att | L7I | PLES_22851 | | | | | | | | | T | T | | |
| NC_011770 | 2458468 | C | T | 78 | 0 | 2 | 0 | NON_SYNONYMOU | MODERAT | MISSENSE | Ctc/Ttc | L275F | PLES_22861 | | | | | | | | | T | T | | |
| NC_011770 | 2458892 | A | T | 79 | 0 | 2 | 0 | NON_SYNONYMOU | MODERAT | MISSENSE | gAg/gTg | E416V | PLES_22861 | | | | | | | | | | | T | T |
| NC_011770 | 2459072 | C | T | 77 | 0 | 1 | 0 | NON_SYNONYMOU | MODERAT | MISSENSE | gCc/gTc | A476V | PLES_22861 | | | | | | | | T | | | | |
| NC_011770 | 2460771 | A | T | 64 | 0 | 12 | 0 | NON_SYNONYMOU | MODERAT | MISSENSE | Tcg/Acg | S89T | PLES_22881 | T | T | T | T | T | T | T | T | T | T | T | T |
| NC_011770 | 2461188 | T | C | 69 | 0 | 10 | 0 | NON_SYNONYMOU | MODERAT | MISSENSE | Aac/Gac | N61D | PLES_22891 | | | C | C | C | C | C | C | C | C | C | C |
| NC_011770 | 2461194 | G | A | 69 | 0 | 2 | 0 | NON_SYNONYMOU | MODERAT | MISSENSE | Ccg/Tcg | P59S | PLES_22891 | | | | | | | | | A | A | | |
| NC_011770 | 2462259 | A | G | 79 | 0 | 1 | 0 | NON_SYNONYMOU | MODERAT | MISSENSE | cTg/cCg | L549P | PLES_22911 | | | | | | | G | | | | | |
| NC_011770 | 2462383 | C | G | 74 | 0 | 2 | 0 | NON_SYNONYMOU | MODERAT | MISSENSE | Gaa/Caa | E508Q | PLES_22911 | | | | | | | G | G | | | | |
| NC_011770 | 2462485 | C | T | 65 | 0 | 2 | 0 | NON_SYNONYMOU | MODERAT | MISSENSE | Gcc/Acc | A474T | PLES_22911 | | | | | | | | | T | T | | |
| NC_011770 | 2462575 | C | A | 68 | 0 | 2 | 0 | NON_SYNONYMOU | MODERAT | MISSENSE | Gcc/Tcc | A444S | PLES_22911 | | | | | | | | | A | A | | |
| NC_011770 | 2463007 | T | C | 70 | 0 | 12 | 0 | NON_SYNONYMOU | MODERAT | MISSENSE | Aat/Gat | N300D | PLES_22911 | C | C | C | C | C | C | C | C | C | C | C | C |
| NC_011770 | 2463091 | G | C | 70 | 0 | 4 | 0 | NON_SYNONYMOU | MODERAT | MISSENSE | Ctc/Gtc | L272V | PLES_22911 | | | | | | | C | C | | | C | C |
| NC_011770 | 2463193 | T | A | 76 | 0 | 2 | 0 | NON_SYNONYMOU | MODERAT | MISSENSE | Aac/Tac | N238Y | PLES_22911 | A | A | | | | | | | | | | |
| NC_011770 | 2463646 | C | T | 69 | 0 | 2 | 0 | NON_SYNONYMOU | MODERAT | MISSENSE | Gcc/Acc | A87T | PLES_22911 | | | | | | | T | T | | | | |
| NC_011770 | 2463808 | G | T | 73 | 0 | 2 | 0 | NON_SYNONYMOU | MODERAT | MISSENSE | Ccc/Acc | P33T | PLES_22911 | | | | | | | | | | | T | T |
| NC_011770 | 2463264 | G | A | 88 | 0 | 1 | 0 | NON_SYNONYMOU | MODERAT | MISSENSE | cCg/cTg | P214L | PLES_22911 | | | | | | | | | | | | A |
| NC_011770 | 2464443 | C | T | 15 | 0 | 2 | 0 | NON_SYNONYMOU | MODERAT | MISSENSE | Gtc/Atc | V67I | PLES_22921 | | | | | | | T | T | | | | |
| NC_011770 | 2467171 | A | G | 62 | 0 | 10 | 0 | NON_SYNONYMOU | MODERAT | MISSENSE | Acg/Gcg | T178A | PLES_22961 | G | G | G | G | G | G | | | G | G | G | G |
| NC_011770 | 2467367 | T | C | 50 | 0 | 2 | 0 | NON_SYNONYMOU | MODERAT | MISSENSE | gTg/gCg | V243A | PLES_22961 | | | | | | | C | C | | | | |
| NC_011770 | 2467517 | T | C | 60 | 0 | 6 | 0 | NON_SYNONYMOU | MODERAT | MISSENSE | gTg/gCg | V293A | PLES_22961 | | | | | | | C | C | C | C | C | C |
| NC_011770 | 2467562 | C | T | 58 | 0 | 2 | 0 | NON_SYNONYMOU | MODERAT | MISSENSE | cCc/cTc | P308L | PLES_22961 | | | | | | | | | | | T | T |
| NC_011770 | 2467382 | G | A | 56 | 0 | 1 | 0 | NON_SYNONYMOU | MODERAT | MISSENSE | gGt/gAt | G248D | PLES_22961 | | A | | | | | | | | | | |
| NC_011770 | 2467681 | A | AC | 49 | 0 | 1 | 0 | FRAME_SHIFT | HIGH | | cgg/cCgg | R13P? | PLES_22971 | AC | | | | | | | | | | | |
| NC_011770 | 2467918 | C | G | 53 | 0 | 2 | 0 | NON_SYNONYMOU | MODERAT | MISSENSE | Ctg/Gtg | L90V | PLES_22971 | G | G | | | | | | | | | | |
| NC_011770 | 2468500 | A | G | 52 | 0 | 2 | 0 | NON_SYNONYMOU | MODERAT | MISSENSE | Acc/Gcc | T284A | PLES_22971 | G | G | | | | | | | | | | |
| NC_011770 | 2467819 | G | A | 59 | 0 | 1 | 0 | NON_SYNONYMOU | MODERAT | MISSENSE | Gcc/Acc | A57T | PLES_22971 | | A | | | | | | | | | | |
| NC_011770 | 2468663 | C | T | 70 | 0 | 6 | 0 | NON_SYNONYMOU | MODERAT | MISSENSE | Gtc/Atc | V427I | PLES_22981 | | | T | T | T | T | T | T | | | | |
| NC_011770 | 2469344 | C | T | 75 | 0 | 2 | 0 | NON_SYNONYMOU | MODERAT | MISSENSE | Ggc/Agc | G200S | PLES_22981 | | | | | | | | | T | T | | |
| NC_011770 | 2469883 | G | C | 21 | 0 | 2 | 0 | NON_SYNONYMOU | MODERAT | MISSENSE | cCc/cGc | P20R | PLES_22981 | | | | | | | | | | | C | C |
| NC_011770 | 2470654 | C | T | 52 | 0 | 6 | 0 | NON_SYNONYMOU | MODERAT | MISSENSE | Ccg/Tcg | P88S | PLES_22991 | T | T | | | | | | | T | T | T | T |
| NC_011770 | 2470995 | T | C | 47 | 0 | 2 | 0 | NON_SYNONYMOU | MODERAT | MISSENSE | gTg/gCg | V54A | PLES_23001 | | | | | | | | | C | C | | |

| | | | | | | | | | | | | | | | | | | | | | | | | | |
|---|---|---|---|---|---|---|---|---|---|---|---|---|---|---|---|---|---|---|---|---|---|---|---|---|---|
| NC_011770 | 2471687 | C | G | 68 | 0 | 2 | 0 | NON_SYNONYMOU | MODERAT | MISSENSE | Ccc/Gcc | P23A | PLES_23011 | G | G | | | | | | | | | | |
| NC_011770 | 2471969 | A | G | 80 | 0 | 1 | 0 | NON_SYNONYMOU | MODERAT | MISSENSE | Agc/Ggc | S117G | PLES_23011 | | | | | | | G | | | | | |
| NC_011770 | 2472970 | T | C | 70 | 0 | 8 | 0 | NON_SYNONYMOU | MODERAT | MISSENSE | cAg/cGg | Q318R | PLES_23041 | | | C | C | C | C | C | C | | | C | C |
| NC_011770 | 2473112 | G | A | 73 | 0 | 2 | 0 | NON_SYNONYMOU | MODERAT | MISSENSE | Cgc/Tgc | R271C | PLES_23041 | | | | | | | A | A | | | | |
| NC_011770 | 2473117 | T | C | 73 | 0 | 10 | 0 | NON_SYNONYMOU | MODERAT | MISSENSE | cAg/cGg | Q269R | PLES_23041 | C | C | C | C | C | C | C | C | | | C | C |
| NC_011770 | 2473505 | T | C | 49 | 0 | 8 | 0 | NON_SYNONYMOU | MODERAT | MISSENSE | Aat/Gat | N140D | PLES_23041 | | | C | C | C | C | C | C | | | C | C |
| NC_011770 | 2473741 | A | G | 40 | 0 | 12 | 0 | NON_SYNONYMOU | MODERAT | MISSENSE | gTc/gCc | V61A | PLES_23041 | G | G | G | G | G | G | G | G | G | G | G | G |
| NC_011770 | 2474297 | T | C | 60 | 0 | 2 | 0 | NON_SYNONYMOU | MODERAT | MISSENSE | gTc/gCc | V24A | PLES_23051 | | | | | | | | | C | C | | |
| NC_011770 | 2474357 | G | A | 66 | 0 | 2 | 0 | NON_SYNONYMOU | MODERAT | MISSENSE | gGc/gAc | G44D | PLES_23051 | | | | | | | | | A | A | | |
| NC_011770 | 2475043 | T | C | 81 | 0 | 12 | 0 | NON_SYNONYMOU | MODERAT | MISSENSE | aTg/aCg | M155T | PLES_23061 | C | C | C | C | C | C | C | C | C | C | C | C |
| NC_011770 | 2475114 | C | T | 76 | 0 | 2 | 0 | NON_SYNONYMOU | MODERAT | MISSENSE | Cgc/Tgc | R179C | PLES_23061 | T | T | | | | | | | | | | |
| NC_011770 | 2475521 | A | C | 89 | 0 | 12 | 0 | NON_SYNONYMOU | MODERAT | MISSENSE | gaA/gaC | E47D | PLES_23071 | C | C | C | C | C | C | C | C | C | C | C | C |
| NC_011770 | 2475946 | C | T | 87 | 0 | 10 | 0 | NON_SYNONYMOU | MODERAT | MISSENSE | aGg/aAg | R160K | PLES_23081 | T | T | T | T | T | T | | | T | T | T | T |
| NC_011770 | 2475971 | C | T | 89 | 0 | 2 | 0 | NON_SYNONYMOU | MODERAT | MISSENSE | Gcc/Acc | A152T | PLES_23081 | | | | | | | T | T | | | | |
| NC_011770 | 2476412 | A | G | 89 | 0 | 12 | 0 | NON_SYNONYMOU | MODERAT | MISSENSE | Tct/Cct | S5P | PLES_23081 | G | G | G | G | G | G | G | G | G | G | G | G |
| NC_011770 | 2476509 | G | A | 91 | 0 | 2 | 0 | NON_SYNONYMOU | MODERAT | MISSENSE | aCc/aTc | T297I | PLES_23091 | | | | | | | A | A | | | | |
| NC_011770 | 2476510 | T | C | 90 | 0 | 2 | 0 | NON_SYNONYMOU | MODERAT | MISSENSE | Acc/Gcc | T297A | PLES_23091 | | | | | | | C | C | | | | |
| NC_011770 | 2477178 | C | T | 79 | 0 | 4 | 0 | NON_SYNONYMOU | MODERAT | MISSENSE | cGc/cAc | R74H | PLES_23091 | | | T | T | T | T | | | | | | |
| NC_011770 | 2477829 | C | T | 83 | 0 | 2 | 0 | NON_SYNONYMOU | MODERAT | MISSENSE | cCc/cTc | P115L | PLES_23101 | | | | | | | T | T | | | | |
| NC_011770 | 2478018 | G | T | 80 | 0 | 2 | 0 | NON_SYNONYMOU | MODERAT | MISSENSE | gGc/gTc | G178V | PLES_23101 | | | | | | | T | T | | | | |
| NC_011770 | 2478062 | G | A | 89 | 0 | 2 | 0 | NON_SYNONYMOU | MODERAT | MISSENSE | Gac/Aac | D193N | PLES_23101 | A | A | | | | | | | | | | |
| NC_011770 | 2478595 | C | A | 65 | 0 | 2 | 0 | NON_SYNONYMOU | MODERAT | MISSENSE | Ctc/Atc | L63I | PLES_23111 | | | | | | | A | A | | | | |
| NC_011770 | 2479568 | T | C | 67 | 0 | 12 | 0 | NON_SYNONYMOU | MODERAT | MISSENSE | Acc/Gcc | T162A | PLES_23121 | C | C | C | C | C | C | C | C | C | C | C | C |
| NC_011770 | 2479610 | T | C | 65 | 0 | 8 | 0 | NON_SYNONYMOU | MODERAT | MISSENSE | Agc/Ggc | S148G | PLES_23121 | C | C | C | C | C | C | | | C | C | | |
| NC_011770 | 2479916 | C | T | 65 | 0 | 2 | 0 | NON_SYNONYMOU | MODERAT | MISSENSE | Gcg/Acg | A46T | PLES_23121 | | | | | | | | | | | T | T |
| NC_011770 | 2480024 | T | C | 63 | 0 | 12 | 0 | NON_SYNONYMOU | MODERAT | MISSENSE | Atc/Gtc | I10V | PLES_23121 | C | C | C | C | C | C | C | C | C | C | C | C |
| NC_011770 | 2481712 | T | C | 93 | 0 | 2 | 0 | NON_SYNONYMOU | MODERAT | MISSENSE | Tac/Cac | Y110H | PLES_23161 | | | | | | | C | C | | | | |
| NC_011770 | 2481777 | C | T | 100 | 0 | 2 | 0 | NON_SYNONYMOU | MODERAT | MISSENSE | Gac/Aac | D138N | PLES_23171 | T | T | | | | | | | | | | |
| NC_011770 | 2481981 | C | T | 98 | 0 | 4 | 0 | NON_SYNONYMOU | MODERAT | MISSENSE | Gcc/Acc | A70T | PLES_23171 | | | | | | | T | T | | | T | T |
| NC_011770 | 2482371 | CGT | C | 64 | 0 | 4 | 0 | FRAME_SHIFT | HIGH | | aac/ | N402 | PLES_23181 | | | C | C | C | C | | | | | | |
| NC_011770 | 2482387 | G | GCA | 55 | 3 | 1 | 0 | FRAME_SHIFT | HIGH | | ttc/ttTGc | F397F? | PLES_23181 | | | GCA | . | . | . | | | | | | |
| NC_011770 | 2482380 | C | A | 62 | 0 | 4 | 0 | NON_SYNONYMOU | MODERAT | MISSENSE | Gtc/Ttc | V400F | PLES_23181 | | | A | A | A | A | | | | | | |
| NC_011770 | 2482385 | A | G | 54 | 3 | 1 | 0 | NON_SYNONYMOU | MODERAT | MISSENSE | cTg/cCg | L398P | PLES_23181 | | | G | . | . | . | | | | | | |
| NC_011770 | 2483325 | A | G | 50 | 0 | 12 | 0 | NON_SYNONYMOU | MODERAT | MISSENSE | Ttg/Atg | L85M | PLES_23181 | G | G | T | T | T | T | G | G | G | G | G | G |
| NC_011770 | 2483333 | G | A | 47 | 0 | 4 | 0 | NON_SYNONYMOU | MODERAT | MISSENSE | gCg/gTg | A82V | PLES_23181 | | | A | A | A | A | | | | | | |
| NC_011770 | 2483448 | G | A | 43 | 4 | 8 | 0 | NON_SYNONYMOU | MODERAT | MISSENSE | Cac/Tac | H44Y | PLES_23181 | A | A | . | . | . | . | A | A | A | A | A | A |
| NC_011770 | 2484170 | G | A | 68 | 0 | 10 | 0 | NON_SYNONYMOU | MODERAT | MISSENSE | Ggg/Agg | G77R | PLES_23191 | | | A | A | A | A | A | A | A | A | A | A |
| NC_011770 | 2485352 | C | T | 80 | 0 | 2 | 0 | START_LOST | HIGH | MISSENSE | atG/atA | M1I | PLES_23201 | | | | | | | | | | | T | T |
| NC_011770 | 2485580 | G | C | 67 | 0 | 2 | 0 | NON_SYNONYMOU | MODERAT | MISSENSE | Gtg/Ctg | V42L | PLES_23211 | | | | | | | C | C | | | | |
| NC_011770 | 2485844 | C | T | 49 | 0 | 1 | 0 | NON_SYNONYMOU | MODERAT | MISSENSE | Cgc/Tgc | R130C | PLES_23211 | | | | | | | T | | | | | |
| NC_011770 | 2486099 | A | G | 36 | 0 | 8 | 0 | NON_SYNONYMOU | MODERAT | MISSENSE | Acc/Gcc | T21A | PLES_23221 | G | G | G | G | G | G | | | | | G | G |
| NC_011770 | 2486229 | G | A | 30 | 0 | 1 | 0 | NON_SYNONYMOU | MODERAT | MISSENSE | cGc/cAc | R64H | PLES_23221 | | | | | | | | | | | | A |
| NC_011770 | 2487521 | C | T | 41 | 0 | 2 | 0 | NON_SYNONYMOU | MODERAT | MISSENSE | Gct/Act | A19T | PLES_23241 | | | | | | | | | | | T | T |
| NC_011770 | 2487929 | TGGA | T | 59 | 0 | 2 | 0 | CODON_CHANGE_ | MODERATE | | gaggcg/gc | EA60A | PLES_23251 | | | | | | | | | | | T | T |
| NC_011770 | 2488031 | C | T | 59 | 0 | 4 | 0 | NON_SYNONYMOU | MODERAT | MISSENSE | Gtg/Atg | V106M | PLES_23261 | | | T | T | T | T | | | | | | |
| NC_011770 | 2488048 | A | G | 58 | 0 | 2 | 0 | NON_SYNONYMOU | MODERAT | MISSENSE | tTc/tCc | F100S | PLES_23261 | | | | | G | G | | | | | | |
| NC_011770 | 2489376 | C | T | 53 | 0 | 8 | 0 | NON_SYNONYMOU | MODERAT | MISSENSE | gCc/gTc | A137V | PLES_23281 | | | T | T | T | T | | | T | T | T | T |
| NC_011770 | 2490221 | C | T | 76 | 0 | 1 | 0 | STOP_GAINED | HIGH | NONSENSE | tgG/tgA | W184* | PLES_23291 | T | | | | | | | | | | | |
| NC_011770 | 2491774 | C | T | 69 | 0 | 2 | 0 | NON_SYNONYMOU | MODERAT | MISSENSE | Gcg/Acg | A82T | PLES_23311 | | | | | | | | | T | T | | |
| NC_011770 | 2491576 | C | T | 70 | 0 | 1 | 0 | NON_SYNONYMOU | MODERAT | MISSENSE | Gaa/Aaa | E148K | PLES_23311 | | | | | | | | T | | | | |
| NC_011770 | 2493145 | A | G | 85 | 0 | 4 | 0 | NON_SYNONYMOU | MODERAT | MISSENSE | Atc/Gtc | I8V | PLES_23331 | | | G | G | G | G | | | | | | |
| NC_011770 | 2493194 | C | T | 78 | 0 | 2 | 0 | NON_SYNONYMOU | MODERAT | MISSENSE | gCc/gTc | A24V | PLES_23331 | | | | | | | | | T | T | | |
| NC_011770 | 2493373 | A | T | 93 | 0 | 2 | 0 | NON_SYNONYMOU | MODERAT | MISSENSE | Atc/Ttc | I84F | PLES_23331 | | | | | | | T | T | | | | |
| NC_011770 | 2493870 | A | C | 75 | 0 | 12 | 0 | NON_SYNONYMOU | MODERAT | MISSENSE | gaA/gaC | E249D | PLES_23331 | C | C | C | C | C | C | C | C | C | C | C | C |
| NC_011770 | 2494089 | C | T | 73 | 0 | 1 | 0 | STOP_GAINED | HIGH | NONSENSE | Cag/Tag | Q44* | PLES_23341 | | | | | | | T | | | | | |
| NC_011770 | 2495277 | C | T | 79 | 0 | 2 | 0 | NON_SYNONYMOU | MODERAT | MISSENSE | Ggc/Agc | G106S | PLES_23371 | | | | | | | | | | | T | T |
| NC_011770 | 2495929 | A | G | 75 | 0 | 4 | 0 | NON_SYNONYMOU | MODERAT | MISSENSE | cAc/cGc | H49R | PLES_23381 | | | | | | | G | G | | | G | G |
| NC_011770 | 2496531 | T | C | 72 | 0 | 12 | 0 | NON_SYNONYMOU | MODERAT | MISSENSE | Tgc/Cgc | C250R | PLES_23381 | C | C | C | C | C | C | C | C | C | C | C | C |
| NC_011770 | 2505794 | C | T | 44 | 2 | 4 | 0 | NON_SYNONYMOU | MODERAT | MISSENSE | Cca/Tca | P104S | PLES_23491 | | | | | | | T | T | T | T | . | . |
| NC_011770 | 2508677 | G | A | 5 | 10 | 2 | 0 | NON_SYNONYMOU | MODERAT | MISSENSE | Gcc/Acc | A648T | PLES_23501 | A | A | . | . | . | . | . | . | . | . | . | . |
| NC_011770 | 2512154 | C | A | 8 | 9 | 1 | 0 | NON_SYNONYMOU | MODERAT | MISSENSE | Ggt/Tgt | G209C | PLES_23531 | | | . | . | . | A | . | . | . | . | . | . |
| NC_011770 | 2512193 | T | C | 19 | 8 | 2 | 0 | NON_SYNONYMOU | MODERAT | MISSENSE | Agc/Ggc | S196G | PLES_23531 | | | . | . | C | C | . | . | . | . | . | . |

| | | | | | | | | | | | | | | | | | | | | | | | | | |
|---|---|---|---|---|---|---|---|---|---|---|---|---|---|---|---|---|---|---|---|---|---|---|---|---|---|
| NC_011770 | 2512490 | T | C | 17 | 8 | 2 | 0 | NON_SYNONYMOU | MODERAT | MISSENSE | Atc/Gtc | I97V | PLES_23531 | | | . | . | C | C | . | . | . | . | . | . |
| NC_011770 | 2512494 | C | A | 16 | 8 | 2 | 0 | NON_SYNONYMOU | MODERAT | MISSENSE | atG/atT | M95I | PLES_23531 | | | . | . | A | A | . | . | . | . | . | . |
| NC_011770 | 2512496 | T | C | 16 | 8 | 2 | 0 | NON_SYNONYMOU | MODERAT | MISSENSE | Atg/Gtg | M95V | PLES_23531 | | | . | . | C | C | . | . | . | . | . | . |
| NC_011770 | 2512508 | T | A | 16 | 8 | 2 | 0 | NON_SYNONYMOU | MODERAT | MISSENSE | Acc/Tcc | T91S | PLES_23531 | | | . | . | A | A | . | . | . | . | . | . |
| NC_011770 | 2512593 | G | T | 19 | 8 | 2 | 0 | NON_SYNONYMOU | MODERAT | MISSENSE | gaC/gaA | D62E | PLES_23531 | | | . | . | T | T | . | . | . | . | . | . |
| NC_011770 | 2512604 | T | C | 20 | 8 | 2 | 0 | NON_SYNONYMOU | MODERAT | MISSENSE | Agc/Ggc | S59G | PLES_23531 | | | . | . | C | C | . | . | . | . | . | . |
| NC_011770 | 2512718 | T | C | 21 | 8 | 2 | 0 | NON_SYNONYMOU | MODERAT | MISSENSE | Atc/Gtc | I21V | PLES_23531 | | | . | . | C | C | . | . | . | . | . | . |
| NC_011770 | 2512921 | G | T | 17 | 8 | 2 | 0 | NON_SYNONYMOU | MODERAT | MISSENSE | gaC/gaA | D312E | PLES_23541 | | | . | . | T | T | . | . | . | . | . | . |
| NC_011770 | 2512926 | T | G | 17 | 8 | 2 | 0 | NON_SYNONYMOU | MODERAT | MISSENSE | Aag/Cag | K311Q | PLES_23541 | | | . | . | G | G | . | . | . | . | . | . |
| NC_011770 | 2512932 | T | G | 17 | 8 | 2 | 0 | NON_SYNONYMOU | MODERAT | MISSENSE | Atg/Ctg | M309L | PLES_23541 | | | . | . | G | G | . | . | . | . | . | . |
| NC_011770 | 2512941 | T | C | 17 | 8 | 2 | 0 | NON_SYNONYMOU | MODERAT | MISSENSE | Atc/Gtc | I306V | PLES_23541 | | | . | . | C | C | . | . | . | . | . | . |
| NC_011770 | 2512947 | A | G | 17 | 8 | 2 | 0 | NON_SYNONYMOU | MODERAT | MISSENSE | Ttc/Ctc | F304L | PLES_23541 | | | . | . | G | G | . | . | . | . | . | . |
| NC_011770 | 2512952 | C | G | 17 | 8 | 2 | 0 | NON_SYNONYMOU | MODERAT | MISSENSE | aGc/aCc | S302T | PLES_23541 | | | . | . | G | G | . | . | . | . | . | . |
| NC_011770 | 2513157 | C | G | 4 | 9 | 1 | 0 | NON_SYNONYMOU | MODERAT | MISSENSE | Gag/Cag | E234Q | PLES_23541 | | | . | . | . | G | . | . | . | . | . | . |
| NC_011770 | 2513234 | A | T | 19 | 8 | 2 | 0 | NON_SYNONYMOU | MODERAT | MISSENSE | cTg/cAg | L208Q | PLES_23541 | | | . | . | T | T | . | . | . | . | . | . |
| NC_011770 | 2513412 | T | A | 23 | 8 | 2 | 0 | NON_SYNONYMOU | MODERAT | MISSENSE | Agg/Tgg | R149W | PLES_23541 | | | . | . | A | A | . | . | . | . | . | . |
| NC_011770 | 2513520 | T | C | 19 | 8 | 2 | 0 | NON_SYNONYMOU | MODERAT | MISSENSE | Aat/Gat | N113D | PLES_23541 | | | . | . | C | C | . | . | . | . | . | . |
| NC_011770 | 2513559 | G | T | 20 | 8 | 2 | 0 | NON_SYNONYMOU | MODERAT | MISSENSE | Cac/Aac | H100N | PLES_23541 | | | . | . | T | T | . | . | . | . | . | . |
| NC_011770 | 2513633 | G | T | 17 | 8 | 2 | 0 | NON_SYNONYMOU | MODERAT | MISSENSE | gCc/gAc | A75D | PLES_23541 | | | . | . | T | T | . | . | . | . | . | . |
| NC_011770 | 2513639 | G | A | 17 | 8 | 2 | 0 | NON_SYNONYMOU | MODERAT | MISSENSE | cCc/cTc | P73L | PLES_23541 | | | . | . | A | A | . | . | . | . | . | . |
| NC_011770 | 2513682 | G | T | 15 | 8 | 2 | 0 | NON_SYNONYMOU | MODERAT | MISSENSE | Ctg/Atg | L59M | PLES_23541 | | | . | . | T | T | . | . | . | . | . | . |
| NC_011770 | 2514009 | C | G | 17 | 8 | 2 | 0 | NON_SYNONYMOU | MODERAT | MISSENSE | atG/atC | M298I | PLES_23551 | | | . | . | G | G | . | . | . | . | . | . |
| NC_011770 | 2514011 | T | G | 16 | 8 | 2 | 0 | NON_SYNONYMOU | MODERAT | MISSENSE | Atg/Ctg | M298L | PLES_23551 | | | . | . | G | G | . | . | . | . | . | . |
| NC_011770 | 2514021 | A | C | 15 | 8 | 2 | 0 | NON_SYNONYMOU | MODERAT | MISSENSE | aaT/aaG | N294K | PLES_23551 | | | . | . | C | C | . | . | . | . | . | . |
| NC_011770 | 2514022 | T | G | 15 | 8 | 2 | 0 | NON_SYNONYMOU | MODERAT | MISSENSE | aAt/aCt | N294T | PLES_23551 | | | . | . | G | G | . | . | . | . | . | . |
| NC_011770 | 2514052 | A | T | 14 | 8 | 2 | 0 | NON_SYNONYMOU | MODERAT | MISSENSE | tTc/tAc | F284Y | PLES_23551 | | | . | . | T | T | . | . | . | . | . | . |
| NC_011770 | 2514209 | G | T | 6 | 9 | 1 | 0 | NON_SYNONYMOU | MODERAT | MISSENSE | Ctg/Atg | L232M | PLES_23551 | | | . | . | . | T | . | . | . | . | . | . |
| NC_011770 | 2514265 | G | T | 23 | 8 | 2 | 0 | NON_SYNONYMOU | MODERAT | MISSENSE | gCa/gAa | A213E | PLES_23551 | | | . | . | T | T | . | . | . | . | . | . |
| NC_011770 | 2514266 | C | G | 23 | 8 | 2 | 0 | NON_SYNONYMOU | MODERAT | MISSENSE | Gca/Cca | A213P | PLES_23551 | | | . | . | G | G | . | . | . | . | . | . |
| NC_011770 | 2514329 | C | G | 20 | 8 | 2 | 0 | NON_SYNONYMOU | MODERAT | MISSENSE | Gtg/Ctg | V192L | PLES_23551 | | | . | . | G | G | . | . | . | . | . | . |
| NC_011770 | 2514330 | C | A | 20 | 8 | 2 | 0 | NON_SYNONYMOU | MODERAT | MISSENSE | aaG/aaT | K191N | PLES_23551 | | | . | . | A | A | . | . | . | . | . | . |
| NC_011770 | 2514332 | T | C | 20 | 8 | 2 | 0 | NON_SYNONYMOU | MODERAT | MISSENSE | Aag/Gag | K191E | PLES_23551 | | | . | . | C | C | . | . | . | . | . | . |
| NC_011770 | 2514334 | G | C | 20 | 8 | 2 | 0 | NON_SYNONYMOU | MODERAT | MISSENSE | tCc/tGc | S190C | PLES_23551 | | | . | . | C | C | . | . | . | . | . | . |
| NC_011770 | 2514335 | A | C | 20 | 8 | 2 | 0 | NON_SYNONYMOU | MODERAT | MISSENSE | Tcc/Gcc | S190A | PLES_23551 | | | . | . | C | C | . | . | . | . | . | . |
| NC_011770 | 2514349 | T | C | 22 | 8 | 2 | 0 | NON_SYNONYMOU | MODERAT | MISSENSE | aAc/aGc | N185S | PLES_23551 | | | . | . | C | C | . | . | . | . | . | . |
| NC_011770 | 2514392 | C | T | 21 | 8 | 2 | 0 | NON_SYNONYMOU | MODERAT | MISSENSE | Gtc/Atc | V171I | PLES_23551 | | | . | . | T | T | . | . | . | . | . | . |
| NC_011770 | 2514422 | C | T | 12 | 8 | 2 | 0 | NON_SYNONYMOU | MODERAT | MISSENSE | Gta/Ata | V161I | PLES_23551 | | | . | . | T | T | . | . | . | . | . | . |
| NC_011770 | 2514627 | C | A | 8 | 8 | 2 | 0 | NON_SYNONYMOU | MODERAT | MISSENSE | gaG/gaT | E92D | PLES_23551 | | | . | . | A | A | . | . | . | . | . | . |
| NC_011770 | 2514631 | C | T | 8 | 8 | 2 | 0 | NON_SYNONYMOU | MODERAT | MISSENSE | cGt/cAt | R91H | PLES_23551 | | | . | . | T | T | . | . | . | . | . | . |
| NC_011770 | 2514632 | G | T | 8 | 8 | 2 | 0 | NON_SYNONYMOU | MODERAT | MISSENSE | Cgt/Agt | R91S | PLES_23551 | | | . | . | T | T | . | . | . | . | . | . |
| NC_011770 | 2514654 | G | C | 6 | 9 | 1 | 0 | NON_SYNONYMOU | MODERAT | MISSENSE | agC/agG | S83R | PLES_23551 | | | . | . | . | C | . | . | . | . | . | . |
| NC_011770 | 2514694 | A | T | 12 | 8 | 2 | 0 | NON_SYNONYMOU | MODERAT | MISSENSE | tTc/tAc | F70Y | PLES_23551 | | | . | . | T | T | . | . | . | . | . | . |
| NC_011770 | 2514728 | T | C | 12 | 8 | 2 | 0 | NON_SYNONYMOU | MODERAT | MISSENSE | Acc/Gcc | T59A | PLES_23551 | | | . | . | C | C | . | . | . | . | . | . |
| NC_011770 | 2514745 | A | G | 12 | 8 | 2 | 0 | NON_SYNONYMOU | MODERAT | MISSENSE | gTc/gCc | V53A | PLES_23551 | | | . | . | G | G | . | . | . | . | . | . |
| NC_011770 | 2514747 | T | G | 11 | 8 | 2 | 0 | NON_SYNONYMOU | MODERAT | MISSENSE | gaA/gaC | E52D | PLES_23551 | | | . | . | G | G | . | . | . | . | . | . |
| NC_011770 | 2516900 | T | C | 4 | 10 | 2 | 0 | NON_SYNONYMOU | MODERAT | MISSENSE | cAt/cGt | H95R | PLES_23561 | C | C | . | . | . | . | . | . | . | . | . | . |
| NC_011770 | 2521968 | TC | T | 4 | 10 | 2 | 0 | FRAME_SHIFT | HIGH | | gaa/ | E3 | PLES_23571 | T | T | . | . | . | . | . | . | . | . | . | . |
| NC_011770 | 2517414 | G | A | 5 | 10 | 2 | 0 | NON_SYNONYMOU | MODERAT | MISSENSE | aCa/aTa | T1521I | PLES_23571 | A | A | . | . | . | . | . | . | . | . | . | . |
| NC_011770 | 2517766 | G | A | 6 | 10 | 2 | 0 | NON_SYNONYMOU | MODERAT | MISSENSE | Ccc/Tcc | P1404S | PLES_23571 | A | A | . | . | . | . | . | . | . | . | . | . |
| NC_011770 | 2517990 | A | G | 6 | 10 | 2 | 0 | NON_SYNONYMOU | MODERAT | MISSENSE | gTt/gCt | V1329A | PLES_23571 | G | G | . | . | . | . | . | . | . | . | . | . |
| NC_011770 | 2518002 | C | T | 6 | 10 | 2 | 0 | NON_SYNONYMOU | MODERAT | MISSENSE | gGt/gAt | G1325D | PLES_23571 | T | T | . | . | . | . | . | . | . | . | . | . |
| NC_011770 | 2518014 | C | T | 6 | 10 | 2 | 0 | NON_SYNONYMOU | MODERAT | MISSENSE | aGt/aAt | S1321N | PLES_23571 | T | T | . | . | . | . | . | . | . | . | . | . |
| NC_011770 | 2518026 | C | A | 5 | 10 | 2 | 0 | NON_SYNONYMOU | MODERAT | MISSENSE | cGg/cTg | R1317L | PLES_23571 | A | A | . | . | . | . | . | . | . | . | . | . |
| NC_011770 | 2518038 | A | C | 5 | 10 | 2 | 0 | NON_SYNONYMOU | MODERAT | MISSENSE | cTc/cGc | L1313R | PLES_23571 | C | C | . | . | . | . | . | . | . | . | . | . |
| NC_011770 | 2518063 | T | C | 5 | 10 | 2 | 0 | NON_SYNONYMOU | MODERAT | MISSENSE | Atc/Gtc | I1305V | PLES_23571 | C | C | . | . | . | . | . | . | . | . | . | . |
| NC_011770 | 2518086 | C | T | 5 | 10 | 2 | 0 | NON_SYNONYMOU | MODERAT | MISSENSE | gGg/gAg | G1297E | PLES_23571 | T | T | . | . | . | . | . | . | . | . | . | . |
| NC_011770 | 2518167 | C | T | 5 | 10 | 2 | 0 | NON_SYNONYMOU | MODERAT | MISSENSE | gGg/gAg | G1270E | PLES_23571 | T | T | . | . | . | . | . | . | . | . | . | . |
| NC_011770 | 2518181 | A | T | 5 | 10 | 2 | 0 | NON_SYNONYMOU | MODERAT | MISSENSE | caT/caA | H1265Q | PLES_23571 | T | T | . | . | . | . | . | . | . | . | . | . |
| NC_011770 | 2518849 | T | C | 5 | 10 | 2 | 0 | NON_SYNONYMOU | MODERAT | MISSENSE | Acc/Gcc | T1043A | PLES_23571 | C | C | . | . | . | . | . | . | . | . | . | . |
| NC_011770 | 2518861 | G | C | 5 | 10 | 2 | 0 | NON_SYNONYMOU | MODERAT | MISSENSE | Caa/Gaa | Q1039E | PLES_23571 | C | C | . | . | . | . | . | . | . | . | . | . |
| NC_011770 | 2519088 | G | A | 3 | 10 | 2 | 0 | NON_SYNONYMOU | MODERAT | MISSENSE | tCg/tTg | S963L | PLES_23571 | A | A | . | . | . | . | . | . | . | . | . | . |
| NC_011770 | 2519089 | A | T | 3 | 10 | 2 | 0 | NON_SYNONYMOU | MODERAT | MISSENSE | Tcg/Acg | S963T | PLES_23571 | T | T | . | . | . | . | . | . | . | . | . | . |

| | | | | | | | | | | | | | | | | | | | | | | | | | |
|---|---|---|---|---|---|---|---|---|---|---|---|---|---|---|---|---|---|---|---|---|---|---|---|---|---|
| NC_011770 | 2519094 | T | A | 3 | 10 | 2 | 0 | NON_SYNONYMOU | MODERAT | MISSENSE | tAc/tTc | Y961F | PLES_23571 | A | A | . | . | . | . | . | . | . | . | . | . |
| NC_011770 | 2519536 | C | T | 2 | 10 | 2 | 0 | NON_SYNONYMOU | MODERAT | MISSENSE | Ggt/Agt | G814S | PLES_23571 | T | T | . | . | . | . | . | . | . | . | . | . |
| NC_011770 | 2519545 | G | T | 2 | 10 | 2 | 0 | NON_SYNONYMOU | MODERAT | MISSENSE | Cgc/Agc | R811S | PLES_23571 | T | T | . | . | . | . | . | . | . | . | . | . |
| NC_011770 | 2519747 | G | T | 3 | 10 | 2 | 0 | NON_SYNONYMOU | MODERAT | MISSENSE | aaC/aaA | N743K | PLES_23571 | T | T | . | . | . | . | . | . | . | . | . | . |
| NC_011770 | 2519749 | T | C | 3 | 10 | 2 | 0 | NON_SYNONYMOU | MODERAT | MISSENSE | Aac/Gac | N743D | PLES_23571 | C | C | . | . | . | . | . | . | . | . | . | . |
| NC_011770 | 2520007 | C | T | 5 | 10 | 2 | 0 | NON_SYNONYMOU | MODERAT | MISSENSE | Gcc/Acc | A657T | PLES_23571 | T | T | . | . | . | . | . | . | . | . | . | . |
| NC_011770 | 2520141 | T | C | 5 | 10 | 2 | 0 | NON_SYNONYMOU | MODERAT | MISSENSE | tAt/tGt | Y612C | PLES_23571 | C | C | . | . | . | . | . | . | . | . | . | . |
| NC_011770 | 2520142 | A | T | 5 | 10 | 2 | 0 | NON_SYNONYMOU | MODERAT | MISSENSE | Tat/Aat | Y612N | PLES_23571 | T | T | . | . | . | . | . | . | . | . | . | . |
| NC_011770 | 2520148 | G | A | 5 | 10 | 2 | 0 | NON_SYNONYMOU | MODERAT | MISSENSE | Cac/Tac | H610Y | PLES_23571 | A | A | . | . | . | . | . | . | . | . | . | . |
| NC_011770 | 2520193 | C | G | 5 | 10 | 2 | 0 | NON_SYNONYMOU | MODERAT | MISSENSE | Gaa/Caa | E595Q | PLES_23571 | G | G | . | . | . | . | . | . | . | . | . | . |
| NC_011770 | 2520236 | C | T | 4 | 10 | 2 | 0 | NON_SYNONYMOU | MODERAT | MISSENSE | atG/atA | M580I | PLES_23571 | T | T | . | . | . | . | . | . | . | . | . | . |
| NC_011770 | 2520238 | T | A | 4 | 10 | 2 | 0 | NON_SYNONYMOU | MODERAT | MISSENSE | Atg/Ttg | M580L | PLES_23571 | A | A | . | . | . | . | . | . | . | . | . | . |
| NC_011770 | 2520247 | G | T | 4 | 10 | 2 | 0 | NON_SYNONYMOU | MODERAT | MISSENSE | Ctc/Atc | L577I | PLES_23571 | T | T | . | . | . | . | . | . | . | . | . | . |
| NC_011770 | 2520253 | T | G | 4 | 10 | 2 | 0 | NON_SYNONYMOU | MODERAT | MISSENSE | Agt/Cgt | S575R | PLES_23571 | G | G | . | . | . | . | . | . | . | . | . | . |
| NC_011770 | 2520313 | C | T | 6 | 10 | 2 | 0 | NON_SYNONYMOU | MODERAT | MISSENSE | Gtc/Atc | V555I | PLES_23571 | T | T | . | . | . | . | . | . | . | . | . | . |
| NC_011770 | 2520330 | G | A | 5 | 10 | 2 | 0 | NON_SYNONYMOU | MODERAT | MISSENSE | gCc/gTc | A549V | PLES_23571 | A | A | . | . | . | . | . | . | . | . | . | . |
| NC_011770 | 2520331 | C | A | 5 | 10 | 2 | 0 | NON_SYNONYMOU | MODERAT | MISSENSE | Gcc/Tcc | A549S | PLES_23571 | A | A | . | . | . | . | . | . | . | . | . | . |
| NC_011770 | 2520363 | G | A | 5 | 10 | 2 | 0 | NON_SYNONYMOU | MODERAT | MISSENSE | gCt/gTt | A538V | PLES_23571 | A | A | . | . | . | . | . | . | . | . | . | . |
| NC_011770 | 2520365 | T | A | 5 | 10 | 2 | 0 | NON_SYNONYMOU | MODERAT | MISSENSE | gaA/gaT | E537D | PLES_23571 | A | A | . | . | . | . | . | . | . | . | . | . |
| NC_011770 | 2520405 | A | G | 6 | 10 | 2 | 0 | NON_SYNONYMOU | MODERAT | MISSENSE | gTt/gCt | V524A | PLES_23571 | G | G | . | . | . | . | . | . | . | . | . | . |
| NC_011770 | 2520436 | G | A | 6 | 10 | 2 | 0 | NON_SYNONYMOU | MODERAT | MISSENSE | Ccc/Tcc | P514S | PLES_23571 | A | A | . | . | . | . | . | . | . | . | . | . |
| NC_011770 | 2520459 | G | A | 5 | 10 | 2 | 0 | NON_SYNONYMOU | MODERAT | MISSENSE | gCc/gTc | A506V | PLES_23571 | A | A | . | . | . | . | . | . | . | . | . | . |
| NC_011770 | 2520538 | C | T | 5 | 10 | 2 | 0 | NON_SYNONYMOU | MODERAT | MISSENSE | Ggc/Agc | G480S | PLES_23571 | T | T | . | . | . | . | . | . | . | . | . | . |
| NC_011770 | 2520547 | A | G | 5 | 10 | 2 | 0 | NON_SYNONYMOU | MODERAT | MISSENSE | Tgc/Cgc | C477R | PLES_23571 | G | G | . | . | . | . | . | . | . | . | . | . |
| NC_011770 | 2520561 | T | C | 5 | 10 | 2 | 0 | NON_SYNONYMOU | MODERAT | MISSENSE | cAg/cGg | Q472R | PLES_23571 | C | C | . | . | . | . | . | . | . | . | . | . |
| NC_011770 | 2520568 | T | C | 6 | 10 | 2 | 0 | NON_SYNONYMOU | MODERAT | MISSENSE | Aca/Gca | T470A | PLES_23571 | C | C | . | . | . | . | . | . | . | . | . | . |
| NC_011770 | 2520570 | C | A | 5 | 10 | 2 | 0 | NON_SYNONYMOU | MODERAT | MISSENSE | gGc/gTc | G469V | PLES_23571 | A | A | . | . | . | . | . | . | . | . | . | . |
| NC_011770 | 2520689 | G | T | 6 | 10 | 2 | 0 | NON_SYNONYMOU | MODERAT | MISSENSE | caC/caA | H429Q | PLES_23571 | T | T | . | . | . | . | . | . | . | . | . | . |
| NC_011770 | 2520775 | C | T | 5 | 10 | 2 | 0 | NON_SYNONYMOU | MODERAT | MISSENSE | Ggg/Agg | G401R | PLES_23571 | T | T | . | . | . | . | . | . | . | . | . | . |
| NC_011770 | 2520862 | A | G | 4 | 10 | 2 | 0 | NON_SYNONYMOU | MODERAT | MISSENSE | Tca/Cca | S372P | PLES_23571 | G | G | . | . | . | . | . | . | . | . | . | . |
| NC_011770 | 2520865 | G | C | 4 | 10 | 2 | 0 | NON_SYNONYMOU | MODERAT | MISSENSE | Ccg/Gcg | P371A | PLES_23571 | C | C | . | . | . | . | . | . | . | . | . | . |
| NC_011770 | 2521156 | T | C | 6 | 10 | 2 | 0 | NON_SYNONYMOU | MODERAT | MISSENSE | Atc/Gtc | I274V | PLES_23571 | C | C | . | . | . | . | . | . | . | . | . | . |
| NC_011770 | 2521206 | C | T | 5 | 10 | 2 | 0 | NON_SYNONYMOU | MODERAT | MISSENSE | cGt/cAt | R257H | PLES_23571 | T | T | . | . | . | . | . | . | . | . | . | . |
| NC_011770 | 2521233 | T | C | 6 | 10 | 2 | 0 | NON_SYNONYMOU | MODERAT | MISSENSE | cAg/cGg | Q248R | PLES_23571 | C | C | . | . | . | . | . | . | . | . | . | . |
| NC_011770 | 2521330 | T | C | 5 | 10 | 2 | 0 | NON_SYNONYMOU | MODERAT | MISSENSE | Aaa/Gaa | K216E | PLES_23571 | C | C | . | . | . | . | . | . | . | . | . | . |
| NC_011770 | 2521534 | C | T | 4 | 10 | 2 | 0 | NON_SYNONYMOU | MODERAT | MISSENSE | Gtc/Atc | V148I | PLES_23571 | T | T | . | . | . | . | . | . | . | . | . | . |
| NC_011770 | 2521569 | T | C | 4 | 10 | 2 | 0 | NON_SYNONYMOU | MODERAT | MISSENSE | aAa/aGa | K136R | PLES_23571 | C | C | . | . | . | . | . | . | . | . | . | . |
| NC_011770 | 2521813 | C | T | 4 | 10 | 2 | 0 | NON_SYNONYMOU | MODERAT | MISSENSE | Gtt/Att | V55I | PLES_23571 | T | T | . | . | . | . | . | . | . | . | . | . |
| NC_011770 | 2522392 | A | C | 5 | 10 | 2 | 0 | NON_SYNONYMOU | MODERAT | MISSENSE | Acg/Ccg | T74P | PLES_23581 | C | C | . | . | . | . | . | . | . | . | . | . |
| NC_011770 | 2522396 | A | C | 5 | 10 | 2 | 0 | NON_SYNONYMOU | MODERAT | MISSENSE | aAt/aCt | N75T | PLES_23581 | C | C | . | . | . | . | . | . | . | . | . | . |
| NC_011770 | 2522426 | G | A | 5 | 10 | 2 | 0 | NON_SYNONYMOU | MODERAT | MISSENSE | gGg/gAg | G85E | PLES_23581 | A | A | . | . | . | . | . | . | . | . | . | . |
| NC_011770 | 2522491 | T | G | 5 | 10 | 2 | 0 | NON_SYNONYMOU | MODERAT | MISSENSE | Ttg/Gtg | L107V | PLES_23581 | G | G | . | . | . | . | . | . | . | . | . | . |
| NC_011770 | 2522510 | T | C | 5 | 10 | 2 | 0 | NON_SYNONYMOU | MODERAT | MISSENSE | tTt/tCt | F113S | PLES_23581 | C | C | . | . | . | . | . | . | . | . | . | . |
| NC_011770 | 2522529 | A | T | 5 | 10 | 2 | 0 | NON_SYNONYMOU | MODERAT | MISSENSE | gaA/gaT | E119D | PLES_23581 | T | T | . | . | . | . | . | . | . | . | . | . |
| NC_011770 | 2522581 | G | A | 4 | 10 | 2 | 0 | NON_SYNONYMOU | MODERAT | MISSENSE | Ggc/Agc | G137S | PLES_23581 | A | A | . | . | . | . | . | . | . | . | . | . |
| NC_011770 | 2522612 | A | G | 2 | 10 | 2 | 0 | NON_SYNONYMOU | MODERAT | MISSENSE | cAc/cGc | H147R | PLES_23581 | G | G | . | . | . | . | . | . | . | . | . | . |
| NC_011770 | 2522626 | A | G | 2 | 10 | 2 | 0 | NON_SYNONYMOU | MODERAT | MISSENSE | Att/Gtt | I152V | PLES_23581 | G | G | . | . | . | . | . | . | . | . | . | . |
| NC_011770 | 2522643 | G | C | 2 | 10 | 2 | 0 | NON_SYNONYMOU | MODERAT | MISSENSE | ttG/ttC | L157F | PLES_23581 | C | C | . | . | . | . | . | . | . | . | . | . |
| NC_011770 | 2522707 | T | C | 4 | 10 | 2 | 0 | NON_SYNONYMOU | MODERAT | MISSENSE | Ttc/Ctc | F179L | PLES_23581 | C | C | . | . | . | . | . | . | . | . | . | . |
| NC_011770 | 2522728 | G | A | 3 | 10 | 2 | 0 | NON_SYNONYMOU | MODERAT | MISSENSE | Gcc/Acc | A186T | PLES_23581 | A | A | . | . | . | . | . | . | . | . | . | . |
| NC_011770 | 2522729 | C | G | 3 | 10 | 2 | 0 | NON_SYNONYMOU | MODERAT | MISSENSE | gCc/gGc | A186G | PLES_23581 | G | G | . | . | . | . | . | . | . | . | . | . |
| NC_011770 | 2522735 | T | C | 3 | 10 | 2 | 0 | NON_SYNONYMOU | MODERAT | MISSENSE | gTc/gCc | V188A | PLES_23581 | C | C | . | . | . | . | . | . | . | . | . | . |
| NC_011770 | 2522747 | G | A | 4 | 10 | 2 | 0 | NON_SYNONYMOU | MODERAT | MISSENSE | gGa/gAa | G192E | PLES_23581 | A | A | . | . | . | . | . | . | . | . | . | . |
| NC_011770 | 2522872 | G | C | 4 | 10 | 2 | 0 | NON_SYNONYMOU | MODERAT | MISSENSE | Gag/Cag | E234Q | PLES_23581 | C | C | . | . | . | . | . | . | . | . | . | . |
| NC_011770 | 2522876 | A | G | 4 | 10 | 2 | 0 | NON_SYNONYMOU | MODERAT | MISSENSE | gAg/gGg | E235G | PLES_23581 | G | G | . | . | . | . | . | . | . | . | . | . |
| NC_011770 | 2522877 | G | C | 4 | 10 | 2 | 0 | NON_SYNONYMOU | MODERAT | MISSENSE | gaG/gaC | E235D | PLES_23581 | C | C | . | . | . | . | . | . | . | . | . | . |
| NC_011770 | 2522998 | T | C | 3 | 10 | 2 | 0 | NON_SYNONYMOU | MODERAT | MISSENSE | Tct/Cct | S276P | PLES_23581 | C | C | . | . | . | . | . | . | . | . | . | . |
| NC_011770 | 2523067 | T | A | 3 | 10 | 2 | 0 | NON_SYNONYMOU | MODERAT | MISSENSE | Tcc/Acc | S299T | PLES_23581 | A | A | . | . | . | . | . | . | . | . | . | . |
| NC_011770 | 2523068 | C | T | 3 | 10 | 2 | 0 | NON_SYNONYMOU | MODERAT | MISSENSE | tCc/tTc | S299F | PLES_23581 | T | T | . | . | . | . | . | . | . | . | . | . |
| NC_011770 | 2523557 | A | T | 5 | 10 | 2 | 0 | NON_SYNONYMOU | MODERAT | MISSENSE | aaA/aaT | K17N | PLES_23591 | T | T | . | . | . | . | . | . | . | . | . | . |
| NC_011770 | 2523571 | C | T | 5 | 10 | 2 | 0 | NON_SYNONYMOU | MODERAT | MISSENSE | aCc/aTc | T22I | PLES_23591 | T | T | . | . | . | . | . | . | . | . | . | . |
| NC_011770 | 2523575 | T | G | 5 | 10 | 2 | 0 | NON_SYNONYMOU | MODERAT | MISSENSE | gaT/gaG | D23E | PLES_23591 | G | G | . | . | . | . | . | . | . | . | . | . |

| | | | | | | | | | | | | | | | | | | | | | | | | | |
|---|---|---|---|---|---|---|---|---|---|---|---|---|---|---|---|---|---|---|---|---|---|---|---|---|---|
| NC_011770 | 2523607 | C | T | 5 | 10 | 2 | 0 | NON_SYNONYMOU | MODERAT | MISSENSE | gCc/gTc | A34V | PLES_23591 | T | T | . | . | . | . | . | . | . | . | . | . |
| NC_011770 | 2523621 | G | A | 5 | 10 | 2 | 0 | NON_SYNONYMOU | MODERAT | MISSENSE | Gtc/Atc | V39I | PLES_23591 | A | A | . | . | . | . | . | . | . | . | . | . |
| NC_011770 | 2523631 | T | A | 4 | 10 | 2 | 0 | NON_SYNONYMOU | MODERAT | MISSENSE | gTg/gAg | V42E | PLES_23591 | A | A | . | . | . | . | . | . | . | . | . | . |
| NC_011770 | 2523978 | G | A | 8 | 10 | 2 | 0 | NON_SYNONYMOU | MODERAT | MISSENSE | Gcc/Acc | A158T | PLES_23591 | A | A | . | . | . | . | . | . | . | . | . | . |
| NC_011770 | 2524140 | G | A | 7 | 10 | 2 | 0 | NON_SYNONYMOU | MODERAT | MISSENSE | Gcc/Acc | A212T | PLES_23591 | A | A | . | . | . | . | . | . | . | . | . | . |
| NC_011770 | 2524533 | T | G | 5 | 10 | 2 | 0 | NON_SYNONYMOU | MODERAT | MISSENSE | Tcc/Gcc | S343A | PLES_23591 | G | G | . | . | . | . | . | . | . | . | . | . |
| NC_011770 | 2524635 | G | A | 6 | 10 | 2 | 0 | NON_SYNONYMOU | MODERAT | MISSENSE | Gtc/Atc | V377I | PLES_23591 | A | A | . | . | . | . | . | . | . | . | . | . |
| NC_011770 | 2524663 | C | G | 6 | 10 | 2 | 0 | NON_SYNONYMOU | MODERAT | MISSENSE | gCg/gGg | A386G | PLES_23591 | G | G | . | . | . | . | . | . | . | . | . | . |
| NC_011770 | 2524827 | G | C | 6 | 10 | 2 | 0 | NON_SYNONYMOU | MODERAT | MISSENSE | Gcc/Ccc | A441P | PLES_23591 | C | C | . | . | . | . | . | . | . | . | . | . |
| NC_011770 | 2525049 | A | T | 4 | 10 | 2 | 0 | NON_SYNONYMOU | MODERAT | MISSENSE | Agc/Tgc | S515C | PLES_23591 | T | T | . | . | . | . | . | . | . | . | . | . |
| NC_011770 | 2525056 | A | G | 4 | 10 | 2 | 0 | NON_SYNONYMOU | MODERAT | MISSENSE | aAg/aGg | K517R | PLES_23591 | G | G | . | . | . | . | . | . | . | . | . | . |
| NC_011770 | 2525110 | T | C | 4 | 10 | 2 | 0 | NON_SYNONYMOU | MODERAT | MISSENSE | aTc/aCc | I535T | PLES_23591 | C | C | . | . | . | . | . | . | . | . | . | . |
| NC_011770 | 2525134 | G | A | 4 | 10 | 2 | 0 | NON_SYNONYMOU | MODERAT | MISSENSE | gGt/gAt | G543D | PLES_23591 | A | A | . | . | . | . | . | . | . | . | . | . |
| NC_011770 | 2527138 | G | T | 3 | 10 | 2 | 0 | NON_SYNONYMOU | MODERAT | MISSENSE | gaG/gaT | E503D | PLES_23601 | T | T | . | . | . | . | . | . | . | . | . | . |
| NC_011770 | 2528473 | C | A | 4 | 10 | 2 | 0 | NON_SYNONYMOU | MODERAT | MISSENSE | gCc/gAc | A9D | PLES_23611 | A | A | . | . | . | . | . | . | . | . | . | . |
| NC_011770 | 2528475 | A | G | 4 | 10 | 2 | 0 | NON_SYNONYMOU | MODERAT | MISSENSE | Acc/Gcc | T10A | PLES_23611 | G | G | . | . | . | . | . | . | . | . | . | . |
| NC_011770 | 2528483 | T | G | 4 | 10 | 2 | 0 | NON_SYNONYMOU | MODERAT | MISSENSE | atT/atG | I12M | PLES_23611 | G | G | . | . | . | . | . | . | . | . | . | . |
| NC_011770 | 2528485 | C | T | 4 | 10 | 2 | 0 | NON_SYNONYMOU | MODERAT | MISSENSE | gCc/gTc | A13V | PLES_23611 | T | T | . | . | . | . | . | . | . | . | . | . |
| NC_011770 | 2528488 | G | A | 3 | 10 | 2 | 0 | NON_SYNONYMOU | MODERAT | MISSENSE | gGc/gAc | G14D | PLES_23611 | A | A | . | . | . | . | . | . | . | . | . | . |
| NC_011770 | 2528612 | T | G | 5 | 10 | 2 | 0 | NON_SYNONYMOU | MODERAT | MISSENSE | gaT/gaG | D55E | PLES_23611 | G | G | . | . | . | . | . | . | . | . | . | . |
| NC_011770 | 2529018 | T | G | 5 | 10 | 2 | 0 | NON_SYNONYMOU | MODERAT | MISSENSE | Tca/Gca | S191A | PLES_23611 | G | G | . | . | . | . | . | . | . | . | . | . |
| NC_011770 | 2529418 | T | C | 6 | 10 | 2 | 0 | NON_SYNONYMOU | MODERAT | MISSENSE | aTa/aCa | I324T | PLES_23611 | C | C | . | . | . | . | . | . | . | . | . | . |
| NC_011770 | 2529691 | A | T | 8 | 10 | 2 | 0 | NON_SYNONYMOU | MODERAT | MISSENSE | cAg/cTg | Q415L | PLES_23611 | T | T | . | . | . | . | . | . | . | . | . | . |
| NC_011770 | 2529715 | G | A | 7 | 10 | 2 | 0 | NON_SYNONYMOU | MODERAT | MISSENSE | aGa/aAa | R423K | PLES_23611 | A | A | . | . | . | . | . | . | . | . | . | . |
| NC_011770 | 2529783 | G | A | 7 | 10 | 2 | 0 | NON_SYNONYMOU | MODERAT | MISSENSE | Gac/Aac | D446N | PLES_23611 | A | A | . | . | . | . | . | . | . | . | . | . |
| NC_011770 | 2530063 | A | G | 7 | 10 | 2 | 0 | NON_SYNONYMOU | MODERAT | MISSENSE | aAt/aGt | N539S | PLES_23611 | G | G | . | . | . | . | . | . | . | . | . | . |
| NC_011770 | 2530198 | A | C | 6 | 10 | 2 | 0 | NON_SYNONYMOU | MODERAT | MISSENSE | aAc/aCc | N584T | PLES_23611 | C | C | . | . | . | . | . | . | . | . | . | . |
| NC_011770 | 2530222 | G | A | 5 | 10 | 2 | 0 | NON_SYNONYMOU | MODERAT | MISSENSE | aGt/aAt | S592N | PLES_23611 | A | A | . | . | . | . | . | . | . | . | . | . |
| NC_011770 | 2530262 | A | G | 6 | 10 | 2 | 0 | NON_SYNONYMOU | MODERAT | MISSENSE | atA/atG | I605M | PLES_23611 | G | G | . | . | . | . | . | . | . | . | . | . |
| NC_011770 | 2530323 | C | T | 7 | 10 | 2 | 0 | NON_SYNONYMOU | MODERAT | MISSENSE | Cac/Tac | H626Y | PLES_23611 | T | T | . | . | . | . | . | . | . | . | . | . |
| NC_011770 | 2530341 | C | T | 6 | 10 | 2 | 0 | NON_SYNONYMOU | MODERAT | MISSENSE | Cgt/Tgt | R632C | PLES_23611 | T | T | . | . | . | . | . | . | . | . | . | . |
| NC_011770 | 2530451 | G | C | 7 | 10 | 2 | 0 | NON_SYNONYMOU | MODERAT | MISSENSE | gaG/gaC | E668D | PLES_23611 | C | C | . | . | . | . | . | . | . | . | . | . |
| NC_011770 | 2530482 | A | G | 6 | 10 | 2 | 0 | NON_SYNONYMOU | MODERAT | MISSENSE | Act/Gct | T679A | PLES_23611 | G | G | . | . | . | . | . | . | . | . | . | . |
| NC_011770 | 2530524 | A | G | 7 | 10 | 2 | 0 | NON_SYNONYMOU | MODERAT | MISSENSE | Act/Gct | T693A | PLES_23611 | G | G | . | . | . | . | . | . | . | . | . | . |
| NC_011770 | 2530560 | A | G | 6 | 10 | 2 | 0 | NON_SYNONYMOU | MODERAT | MISSENSE | Agt/Ggt | S705G | PLES_23611 | G | G | . | . | . | . | . | . | . | . | . | . |
| NC_011770 | 2530561 | G | A | 6 | 10 | 2 | 0 | NON_SYNONYMOU | MODERAT | MISSENSE | aGt/aAt | S705N | PLES_23611 | A | A | . | . | . | . | . | . | . | . | . | . |
| NC_011770 | 2530562 | T | A | 6 | 10 | 2 | 0 | NON_SYNONYMOU | MODERAT | MISSENSE | agT/agA | S705R | PLES_23611 | A | A | . | . | . | . | . | . | . | . | . | . |
| NC_011770 | 2530578 | A | G | 6 | 10 | 2 | 0 | NON_SYNONYMOU | MODERAT | MISSENSE | Aat/Gat | N711D | PLES_23611 | G | G | . | . | . | . | . | . | . | . | . | . |
| NC_011770 | 2530641 | G | C | 6 | 10 | 2 | 0 | NON_SYNONYMOU | MODERAT | MISSENSE | Gcc/Ccc | A732P | PLES_23611 | C | C | . | . | . | . | . | . | . | . | . | . |
| NC_011770 | 2530662 | G | C | 6 | 10 | 2 | 0 | NON_SYNONYMOU | MODERAT | MISSENSE | Gtg/Ctg | V739L | PLES_23611 | C | C | . | . | . | . | . | . | . | . | . | . |
| NC_011770 | 2530673 | T | G | 6 | 10 | 2 | 0 | NON_SYNONYMOU | MODERAT | MISSENSE | gaT/gaG | D742E | PLES_23611 | G | G | . | . | . | . | . | . | . | . | . | . |
| NC_011770 | 2530708 | A | G | 6 | 9 | 2 | 0 | NON_SYNONYMOU | MODERAT | MISSENSE | aAc/aGc | N754S | PLES_23611 | G | G | . | . | . | . | . | . | . | . | . | . |
| NC_011770 | 2530742 | T | G | 6 | 10 | 2 | 0 | NON_SYNONYMOU | MODERAT | MISSENSE | caT/caG | H765Q | PLES_23611 | G | G | . | . | . | . | . | . | . | . | . | . |
| NC_011770 | 2530812 | G | A | 5 | 10 | 2 | 0 | NON_SYNONYMOU | MODERAT | MISSENSE | Gcg/Acg | A789T | PLES_23611 | A | A | . | . | . | . | . | . | . | . | . | . |
| NC_011770 | 2530846 | G | C | 5 | 10 | 2 | 0 | NON_SYNONYMOU | MODERAT | MISSENSE | aGc/aCc | S800T | PLES_23611 | C | C | . | . | . | . | . | . | . | . | . | . |
| NC_011770 | 2530869 | A | G | 4 | 10 | 2 | 0 | NON_SYNONYMOU | MODERAT | MISSENSE | Acc/Gcc | T808A | PLES_23611 | G | G | . | . | . | . | . | . | . | . | . | . |
| NC_011770 | 2530876 | C | T | 4 | 10 | 2 | 0 | NON_SYNONYMOU | MODERAT | MISSENSE | tCt/tTt | S810F | PLES_23611 | T | T | . | . | . | . | . | . | . | . | . | . |
| NC_011770 | 2530900 | G | A | 6 | 10 | 2 | 0 | NON_SYNONYMOU | MODERAT | MISSENSE | aGg/aAg | R818K | PLES_23611 | A | A | . | . | . | . | . | . | . | . | . | . |
| NC_011770 | 2530906 | C | G | 6 | 10 | 2 | 0 | NON_SYNONYMOU | MODERAT | MISSENSE | gCt/gGt | A820G | PLES_23611 | G | G | . | . | . | . | . | . | . | . | . | . |
| NC_011770 | 2530984 | T | G | 6 | 10 | 2 | 0 | NON_SYNONYMOU | MODERAT | MISSENSE | tTg/tGg | L846W | PLES_23611 | G | G | . | . | . | . | . | . | . | . | . | . |
| NC_011770 | 2530988 | C | A | 6 | 10 | 2 | 0 | NON_SYNONYMOU | MODERAT | MISSENSE | caC/caA | H847Q | PLES_23611 | A | A | . | . | . | . | . | . | . | . | . | . |
| NC_011770 | 2531043 | T | A | 5 | 10 | 2 | 0 | NON_SYNONYMOU | MODERAT | MISSENSE | Ttc/Atc | F866I | PLES_23611 | A | A | . | . | . | . | . | . | . | . | . | . |
| NC_011770 | 2531062 | A | G | 6 | 10 | 2 | 0 | NON_SYNONYMOU | MODERAT | MISSENSE | cAt/cGt | H872R | PLES_23611 | G | G | . | . | . | . | . | . | . | . | . | . |
| NC_011770 | 2531110 | G | A | 5 | 10 | 2 | 0 | NON_SYNONYMOU | MODERAT | MISSENSE | aGg/aAg | R888K | PLES_23611 | A | A | . | . | . | . | . | . | . | . | . | . |
| NC_011770 | 2531199 | A | C | 5 | 10 | 2 | 0 | NON_SYNONYMOU | MODERAT | MISSENSE | Agt/Cgt | S918R | PLES_23611 | C | C | . | . | . | . | . | . | . | . | . | . |
| NC_011770 | 2531200 | G | A | 5 | 10 | 2 | 0 | NON_SYNONYMOU | MODERAT | MISSENSE | aGt/aAt | S918N | PLES_23611 | A | A | . | . | . | . | . | . | . | . | . | . |
| NC_011770 | 2531301 | A | G | 6 | 10 | 2 | 0 | NON_SYNONYMOU | MODERAT | MISSENSE | Aac/Gac | N952D | PLES_23611 | G | G | . | . | . | . | . | . | . | . | . | . |
| NC_011770 | 2531505 | A | G | 5 | 10 | 2 | 0 | NON_SYNONYMOU | MODERAT | MISSENSE | Acc/Gcc | T1020A | PLES_23611 | G | G | . | . | . | . | . | . | . | . | . | . |
| NC_011770 | 2531512 | A | C | 5 | 10 | 2 | 0 | NON_SYNONYMOU | MODERAT | MISSENSE | gAt/gCt | D1022A | PLES_23611 | C | C | . | . | . | . | . | . | . | . | . | . |
| NC_011770 | 2531529 | A | G | 4 | 10 | 2 | 0 | NON_SYNONYMOU | MODERAT | MISSENSE | Agc/Ggc | S1028G | PLES_23611 | G | G | . | . | . | . | . | . | . | . | . | . |
| NC_011770 | 2531530 | G | A | 4 | 10 | 2 | 0 | NON_SYNONYMOU | MODERAT | MISSENSE | aGc/aAc | S1028N | PLES_23611 | A | A | . | . | . | . | . | . | . | . | . | . |
| NC_011770 | 2531532 | A | G | 4 | 10 | 2 | 0 | NON_SYNONYMOU | MODERAT | MISSENSE | Aag/Gag | K1029E | PLES_23611 | G | G | . | . | . | . | . | . | . | . | . | . |

| | | | | | | | | | | | | | | | | | | | | | | | | | |
|---|---|---|---|---|---|---|---|---|---|---|---|---|---|---|---|---|---|---|---|---|---|---|---|---|---|
| NC_011770 | 2531537 | G | A | 4 | 10 | 2 | 0 | NON_SYNONYMOU | MODERAT | MISSENSE | atG/atA | M1030I | PLES_23611 | A | A | . | . | . | . | . | . | . | . | . | . |
| NC_011770 | 2531609 | T | A | 4 | 10 | 2 | 0 | NON_SYNONYMOU | MODERAT | MISSENSE | gaT/gaA | D1054E | PLES_23611 | A | A | . | . | . | . | . | . | . | . | . | . |
| NC_011770 | 2531913 | C | T | 7 | 10 | 2 | 0 | NON_SYNONYMOU | MODERAT | MISSENSE | Cat/Tat | H1156Y | PLES_23611 | T | T | . | . | . | . | . | . | . | . | . | . |
| NC_011770 | 2538888 | G | T | 6 | 10 | 2 | 0 | NON_SYNONYMOU | MODERAT | MISSENSE | Ccc/Acc | P355T | PLES_23651 | T | T | . | . | . | . | . | . | . | . | . | . |
| NC_011770 | 2539643 | G | T | 7 | 8 | 2 | 0 | NON_SYNONYMOU | MODERAT | MISSENSE | aCc/aAc | T103N | PLES_23651 | | | . | . | T | T | . | . | . | . | . | . |
| NC_011770 | 2542377 | G | T | 8 | 6 | 2 | 0 | NON_SYNONYMOU | MODERAT | MISSENSE | Ctg/Atg | L149M | PLES_23681 | | | T | T | . | . | . | . | . | . | | |
| NC_011770 | 2542388 | G | A | 7 | 6 | 2 | 0 | NON_SYNONYMOU | MODERAT | MISSENSE | gCa/gTa | A145V | PLES_23681 | | | A | A | . | . | . | . | . | . | | |
| NC_011770 | 2542389 | C | G | 7 | 6 | 2 | 0 | NON_SYNONYMOU | MODERAT | MISSENSE | Gca/Cca | A145P | PLES_23681 | | | | | . | . | . | . | . | . | G | G |
| NC_011770 | 2542394 | C | G | 7 | 6 | 4 | 0 | NON_SYNONYMOU | MODERAT | MISSENSE | aGc/aCc | S143T | PLES_23681 | | | G | G | . | . | . | . | . | . | G | G |
| NC_011770 | 2542410 | G | C | 8 | 7 | 1 | 0 | NON_SYNONYMOU | MODERAT | MISSENSE | Cag/Gag | Q138E | PLES_23681 | | | | | C | . | . | . | . | . | . | . |
| NC_011770 | 2542497 | A | C | 9 | 8 | 2 | 0 | NON_SYNONYMOU | MODERAT | MISSENSE | Tcc/Gcc | S109A | PLES_23681 | | | C | C | . | . | . | . | . | . | . | . |
| NC_011770 | 2542559 | G | T | 8 | 8 | 2 | 0 | NON_SYNONYMOU | MODERAT | MISSENSE | aCc/aAc | T88N | PLES_23681 | | | T | T | . | . | . | . | . | . | . | . |
| NC_011770 | 2542560 | T | C | 8 | 8 | 2 | 0 | NON_SYNONYMOU | MODERAT | MISSENSE | Acc/Gcc | T88A | PLES_23681 | | | C | C | . | . | . | . | . | . | . | . |
| NC_011770 | 2543298 | A | C | 8 | 8 | 2 | 0 | NON_SYNONYMOU | MODERAT | MISSENSE | Act/Cct | T92P | PLES_23701 | | | . | . | . | . | . | . | C | C | . | . |
| NC_011770 | 2543432 | G | C | 8 | 8 | 2 | 0 | NON_SYNONYMOU | MODERAT | MISSENSE | agG/agC | R136S | PLES_23701 | | | . | . | . | . | . | . | C | C | . | . |
| NC_011770 | 2544752 | G | A | 35 | 2 | 4 | 0 | NON_SYNONYMOU | MODERAT | MISSENSE | Cca/Tca | P104S | PLES_23711 | | | | | | | A | A | A | A | . | . |
| NC_011770 | 2545247 | G | T | 29 | 2 | 2 | 0 | NON_SYNONYMOU | MODERAT | MISSENSE | ttC/ttA | F37L | PLES_23721 | | | | | | | | | T | T | . | . |
| NC_011770 | 2545321 | G | A | 19 | 4 | 6 | 0 | NON_SYNONYMOU | MODERAT | MISSENSE | Ctc/Ttc | L13F | PLES_23721 | | | . | . | A | A | A | A | A | A | . | . |
| NC_011770 | 2548858 | C | G | 10 | 8 | 2 | 0 | NON_SYNONYMOU | MODERAT | MISSENSE | Ctc/Gtc | L34V | PLES_23751 | | | . | . | . | . | . | . | . | . | G | G |
| NC_011770 | 2548873 | T | A | 10 | 8 | 2 | 0 | NON_SYNONYMOU | MODERAT | MISSENSE | Tac/Aac | Y39N | PLES_23751 | | | . | . | . | . | . | . | . | . | A | A |
| NC_011770 | 2548874 | A | G | 10 | 8 | 2 | 0 | NON_SYNONYMOU | MODERAT | MISSENSE | tAc/tGc | Y39C | PLES_23751 | | | . | . | . | . | . | . | . | . | G | G |
| NC_011770 | 2548964 | T | C | 8 | 6 | 2 | 0 | NON_SYNONYMOU | MODERAT | MISSENSE | tTg/tCg | L69S | PLES_23751 | | | . | | . | | . | . | . | . | C | C |
| NC_011770 | 2549000 | G | A | 7 | 6 | 2 | 0 | NON_SYNONYMOU | MODERAT | MISSENSE | gGc/gAc | G81D | PLES_23751 | | | . | . | | | . | . | . | . | A | A |
| NC_011770 | 2549015 | G | A | 6 | 9 | 1 | 0 | NON_SYNONYMOU | MODERAT | MISSENSE | aGa/aAa | R86K | PLES_23751 | | | . | . | . | . | . | . | . | . | . | A |
| NC_011770 | 2549308 | T | C | 7 | 2 | 8 | 0 | INTRAGENIC | MODIFIER | | | | PLES_23761 | C | C | C | . | C | C | | | C | C | C | . |
| NC_011770 | 2549313 | C | A | 7 | 2 | 10 | 0 | INTRAGENIC | MODIFIER | | | | PLES_23761 | A | A | A | . | A | A | A | A | A | A | A | . |
| NC_011770 | 2549754 | G | A | 47 | 0 | 2 | 0 | INTRAGENIC | MODIFIER | | | | PLES_23761 | | | | | | | A | A | | | | |
| NC_011770 | 2549755 | T | C | 47 | 0 | 2 | 0 | INTRAGENIC | MODIFIER | | | | PLES_23761 | | | | | | | C | C | | | | |
| NC_011770 | 2549807 | AGGC | A | 46 | 0 | 2 | 0 | INTRAGENIC | MODIFIER | | | | PLES_23761 | A | A | | | | | | | | | | |
| NC_011770 | 2549837 | G | A | 47 | 0 | 2 | 0 | INTRAGENIC | MODIFIER | | | | PLES_23761 | | | | | | | | | A | A | | |
| NC_011770 | 2549892 | G | T | 50 | 0 | 4 | 0 | INTRAGENIC | MODIFIER | | | | PLES_23761 | | | | | | | T | T | | | T | T |
| NC_011770 | 2549997 | T | G | 55 | 0 | 4 | 0 | INTRAGENIC | MODIFIER | | | | PLES_23761 | | | | | | | G | G | G | G | | |
| NC_011770 | 2550091 | C | T | 58 | 0 | 4 | 0 | INTRAGENIC | MODIFIER | | | | PLES_23761 | | | T | T | T | T | | | | | | |
| NC_011770 | 2550134 | A | T | 61 | 0 | 6 | 0 | INTRAGENIC | MODIFIER | | | | PLES_23761 | T | T | | | | | | | T | T | T | T |
| NC_011770 | 2550135 | C | G | 63 | 0 | 6 | 0 | INTRAGENIC | MODIFIER | | | | PLES_23761 | G | G | | | | | | | G | G | G | G |
| NC_011770 | 2550143 | T | C | 58 | 0 | 2 | 0 | INTRAGENIC | MODIFIER | | | | PLES_23761 | | | | | | | | | C | C | | |
| NC_011770 | 2550148 | T | C | 59 | 0 | 4 | 0 | INTRAGENIC | MODIFIER | | | | PLES_23761 | | | C | C | C | C | | | | | | |
| NC_011770 | 2550168 | A | G | 60 | 0 | 12 | 0 | INTRAGENIC | MODIFIER | | | | PLES_23761 | G | G | G | G | G | G | G | G | G | G | G | G |
| NC_011770 | 2550175 | C | T | 58 | 0 | 6 | 0 | INTRAGENIC | MODIFIER | | | | PLES_23761 | T | T | | | | | | | T | T | T | T |
| NC_011770 | 2550184 | G | T | 57 | 0 | 10 | 0 | INTRAGENIC | MODIFIER | | | | PLES_23761 | T | T | T | T | T | T | T | T | | | T | T |
| NC_011770 | 2550198 | T | C | 58 | 0 | 4 | 0 | INTRAGENIC | MODIFIER | | | | PLES_23761 | C | C | | | | | | | | | C | C |
| NC_011770 | 2550202 | G | A | 54 | 0 | 2 | 0 | INTRAGENIC | MODIFIER | | | | PLES_23761 | | | | | | | | | A | A | | |
| NC_011770 | 2550203 | C | CG | 53 | 0 | 10 | 0 | INTRAGENIC | MODIFIER | | | | PLES_23761 | CG | CG | CG | CG | CG | CG | CG | CG | . | . | CG | CG |
| NC_011770 | 2550226 | A | G | 56 | 0 | 4 | 0 | INTRAGENIC | MODIFIER | | | | PLES_23761 | | | G | G | G | G | | | | | | |
| NC_011770 | 2550280 | C | T | 56 | 0 | 6 | 0 | INTRAGENIC | MODIFIER | | | | PLES_23761 | | | T | T | T | T | T | T | | | | |
| NC_011770 | 2550281 | T | A | 56 | 0 | 6 | 0 | INTRAGENIC | MODIFIER | | | | PLES_23761 | | | A | A | A | A | A | A | | | | |
| NC_011770 | 2550321 | C | T | 57 | 0 | 4 | 0 | INTRAGENIC | MODIFIER | | | | PLES_23761 | T | T | | | | | | | | | T | T |
| NC_011770 | 2550383 | C | T | 56 | 0 | 2 | 0 | INTRAGENIC | MODIFIER | | | | PLES_23761 | | | | | | | | | | | T | T |
| NC_011770 | 2550385 | T | A | 55 | 0 | 2 | 0 | INTRAGENIC | MODIFIER | | | | PLES_23761 | | | | | | | | | | | A | A |
| NC_011770 | 2550401 | G | A | 58 | 0 | 2 | 0 | INTRAGENIC | MODIFIER | | | | PLES_23761 | | | | | | | | | A | A | | |
| NC_011770 | 2550464 | T | C | 61 | 0 | 8 | 0 | INTRAGENIC | MODIFIER | | | | PLES_23761 | | | C | C | C | C | | | C | C | C | C |
| NC_011770 | 2550466 | G | A | 61 | 0 | 8 | 0 | INTRAGENIC | MODIFIER | | | | PLES_23761 | | | A | A | A | A | | | A | A | A | A |
| NC_011770 | 2550473 | G | A | 60 | 0 | 8 | 0 | INTRAGENIC | MODIFIER | | | | PLES_23761 | | | A | A | A | A | | | A | A | A | A |
| NC_011770 | 2550547 | C | T | 57 | 0 | 2 | 0 | INTRAGENIC | MODIFIER | | | | PLES_23761 | | | T | T | | | | | | | | |
| NC_011770 | 2550548 | G | A | 60 | 0 | 2 | 0 | INTRAGENIC | MODIFIER | | | | PLES_23761 | | | | | | | A | A | | | | |
| NC_011770 | 2550569 | A | G | 58 | 0 | 12 | 0 | INTRAGENIC | MODIFIER | | | | PLES_23761 | G | G | G | G | G | G | G | G | G | G | G | G |
| NC_011770 | 2550658 | G | T | 60 | 0 | 8 | 0 | INTRAGENIC | MODIFIER | | | | PLES_23761 | | | T | T | T | T | | | T | T | T | T |
| NC_011770 | 2550662 | G | A | 59 | 0 | 8 | 0 | INTRAGENIC | MODIFIER | | | | PLES_23761 | | | A | A | A | A | | | A | A | A | A |
| NC_011770 | 2550711 | T | G | 59 | 0 | 2 | 0 | NON_SYNONYMOU | MODERAT | MISSENSE | Acc/Ccc | T2088P | PLES_23771 | G | G | | | | | | | | | | |
| NC_011770 | 2550759 | C | T | 54 | 0 | 8 | 0 | NON_SYNONYMOU | MODERAT | MISSENSE | Gca/Aca | A2072T | PLES_23771 | | | T | T | T | T | | | T | T | T | T |
| NC_011770 | 2550813 | G | A | 57 | 0 | 2 | 0 | NON_SYNONYMOU | MODERAT | MISSENSE | Cac/Tac | H2054Y | PLES_23771 | | | | | | | A | A | | | | |
| NC_011770 | 2551473 | T | C | 80 | 0 | 8 | 0 | NON_SYNONYMOU | MODERAT | MISSENSE | Acg/Gcg | T1834A | PLES_23771 | C | C | C | C | C | C | | | C | C | | |

| | | | | | | | | | | | | | | | | | | | | | | | | | |
|---|---|---|---|---|---|---|---|---|---|---|---|---|---|---|---|---|---|---|---|---|---|---|---|---|---|
| NC_011770 | 2551573 | G | T | 82 | 0 | 12 | 0 | NON_SYNONYMOU | MODERAT | MISSENSE | caC/caA | H1800Q | PLES_23771 | T | T | T | T | T | T | T | T | T | T | T | T |
| NC_011770 | 2551902 | C | T | 75 | 0 | 2 | 0 | NON_SYNONYMOU | MODERAT | MISSENSE | Gac/Aac | D1691N | PLES_23771 | | | | | | | | | | | T | T |
| NC_011770 | 2552316 | C | T | 78 | 0 | 2 | 0 | NON_SYNONYMOU | MODERAT | MISSENSE | Gat/Aat | D1553N | PLES_23771 | | | | | | | | | | | T | T |
| NC_011770 | 2552339 | C | T | 76 | 0 | 2 | 0 | NON_SYNONYMOU | MODERAT | MISSENSE | aGg/aAg | R1545K | PLES_23771 | | | | | | | | | T | T | | |
| NC_011770 | 2552453 | T | G | 77 | 0 | 12 | 0 | NON_SYNONYMOU | MODERAT | MISSENSE | gAa/gCa | E1507A | PLES_23771 | G | G | G | G | G | G | G | G | G | G | G | G |
| NC_011770 | 2552459 | G | A | 77 | 0 | 6 | 0 | NON_SYNONYMOU | MODERAT | MISSENSE | gCc/gTc | A1505V | PLES_23771 | | | A | A | A | A | | | A | A | | |
| NC_011770 | 2552515 | C | G | 74 | 0 | 10 | 0 | NON_SYNONYMOU | MODERAT | MISSENSE | gaG/gaC | E1486D | PLES_23771 | | | G | G | G | G | G | G | G | G | G | G |
| NC_011770 | 2552534 | C | T | 73 | 0 | 2 | 0 | NON_SYNONYMOU | MODERAT | MISSENSE | gGc/gAc | G1480D | PLES_23771 | | | | | | | T | T | | | | |
| NC_011770 | 2552535 | C | T | 73 | 0 | 2 | 0 | NON_SYNONYMOU | MODERAT | MISSENSE | Ggc/Agc | G1480S | PLES_23771 | | | | | | | T | T | | | | |
| NC_011770 | 2552834 | G | A | 69 | 0 | 2 | 0 | NON_SYNONYMOU | MODERAT | MISSENSE | aCc/aTc | T1380I | PLES_23771 | A | A | | | | | | | | | | |
| NC_011770 | 2552835 | T | C | 67 | 0 | 2 | 0 | NON_SYNONYMOU | MODERAT | MISSENSE | Acc/Gcc | T1380A | PLES_23771 | C | C | | | | | | | | | | |
| NC_011770 | 2552840 | A | G | 66 | 0 | 2 | 0 | NON_SYNONYMOU | MODERAT | MISSENSE | aTc/aCc | I1378T | PLES_23771 | G | G | | | | | | | | | | |
| NC_011770 | 2553071 | C | T | 61 | 0 | 2 | 0 | NON_SYNONYMOU | MODERAT | MISSENSE | cGc/cAc | R1301H | PLES_23771 | | | | | | | | | T | T | | |
| NC_011770 | 2553123 | C | T | 62 | 0 | 2 | 0 | NON_SYNONYMOU | MODERAT | MISSENSE | Gcc/Acc | A1284T | PLES_23771 | | | | | | | | | T | T | | |
| NC_011770 | 2553194 | T | C | 62 | 0 | 12 | 0 | NON_SYNONYMOU | MODERAT | MISSENSE | cAg/cGg | Q1260R | PLES_23771 | C | C | C | C | C | C | C | C | C | C | C | C |
| NC_011770 | 2553229 | C | A | 65 | 0 | 2 | 0 | NON_SYNONYMOU | MODERAT | MISSENSE | caG/caT | Q1248H | PLES_23771 | | | | | | | A | A | | | | |
| NC_011770 | 2553279 | T | C | 67 | 0 | 10 | 0 | NON_SYNONYMOU | MODERAT | MISSENSE | Acg/Gcg | T1232A | PLES_23771 | C | C | C | C | C | C | C | C | C | C | | |
| NC_011770 | 2553339 | T | G | 72 | 0 | 2 | 0 | NON_SYNONYMOU | MODERAT | MISSENSE | Aat/Cat | N1212H | PLES_23771 | | | | | | | G | G | | | | |
| NC_011770 | 2553481 | T | G | 67 | 0 | 2 | 0 | NON_SYNONYMOU | MODERAT | MISSENSE | caA/caC | Q1164H | PLES_23771 | G | G | | | | | | | | | | |
| NC_011770 | 2553510 | C | T | 66 | 0 | 1 | 0 | NON_SYNONYMOU | MODERAT | MISSENSE | Gcc/Acc | A1155T | PLES_23771 | | | | | | | T | | | | | |
| NC_011770 | 2553542 | C | T | 66 | 0 | 2 | 0 | NON_SYNONYMOU | MODERAT | MISSENSE | cGc/cAc | R1144H | PLES_23771 | | | | | | | T | T | | | | |
| NC_011770 | 2553816 | C | T | 71 | 0 | 4 | 0 | NON_SYNONYMOU | MODERAT | MISSENSE | Gcc/Acc | A1053T | PLES_23771 | | | | | | | T | T | | | T | T |
| NC_011770 | 2553826 | T | G | 69 | 0 | 10 | 0 | NON_SYNONYMOU | MODERAT | MISSENSE | gaA/gaC | E1049D | PLES_23771 | | | G | G | G | G | G | G | G | G | G | G |
| NC_011770 | 2554367 | G | C | 71 | 0 | 6 | 0 | NON_SYNONYMOU | MODERAT | MISSENSE | gCt/gGt | A869G | PLES_23771 | | | C | C | C | C | | | C | C | | |
| NC_011770 | 2554836 | G | C | 82 | 0 | 4 | 0 | NON_SYNONYMOU | MODERAT | MISSENSE | Cac/Gac | H713D | PLES_23771 | | | | | | | C | C | C | C | | |
| NC_011770 | 2555137 | C | G | 81 | 0 | 2 | 0 | NON_SYNONYMOU | MODERAT | MISSENSE | caG/caC | Q612H | PLES_23771 | | | | | | | | | G | G | | |
| NC_011770 | 2555727 | A | G | 83 | 0 | 1 | 0 | NON_SYNONYMOU | MODERAT | MISSENSE | Tgg/Cgg | W416R | PLES_23771 | | | | | | | G | | | | | |
| NC_011770 | 2556077 | T | C | 67 | 0 | 12 | 0 | NON_SYNONYMOU | MODERAT | MISSENSE | cAc/cGc | H299R | PLES_23771 | C | C | C | C | C | C | C | C | C | C | C | C |
| NC_011770 | 2556391 | G | C | 79 | 0 | 6 | 0 | NON_SYNONYMOU | MODERAT | MISSENSE | gaC/gaG | D194E | PLES_23771 | C | C | C | C | C | C | | | | | | |
| NC_011770 | 2556395 | A | G | 75 | 0 | 12 | 0 | NON_SYNONYMOU | MODERAT | MISSENSE | gTc/gCc | V193A | PLES_23771 | G | G | G | G | G | G | G | G | G | G | G | G |
| NC_011770 | 2556443 | T | C | 77 | 0 | 2 | 0 | NON_SYNONYMOU | MODERAT | MISSENSE | gAc/gGc | D177G | PLES_23771 | C | C | | | | | | | | | | |
| NC_011770 | 2556567 | T | C | 72 | 0 | 2 | 0 | NON_SYNONYMOU | MODERAT | MISSENSE | Agc/Ggc | S136G | PLES_23771 | | | | | | | | | C | C | | |
| NC_011770 | 2553728 | T | C | 75 | 0 | 1 | 0 | NON_SYNONYMOU | MODERAT | MISSENSE | cAc/cGc | H1082R | PLES_23771 | C | | | | | | | | | | | |
| NC_011770 | 2557029 | T | G | 68 | 0 | 8 | 0 | NON_SYNONYMOU | MODERAT | MISSENSE | gaA/gaC | E199D | PLES_23781 | | | G | G | G | G | | | G | G | G | G |
| NC_011770 | 2557109 | G | A | 66 | 0 | 12 | 0 | NON_SYNONYMOU | MODERAT | MISSENSE | Ctc/Ttc | L173F | PLES_23781 | A | A | A | A | A | A | A | A | A | A | A | A |
| NC_011770 | 2557357 | T | C | 62 | 0 | 6 | 0 | NON_SYNONYMOU | MODERAT | MISSENSE | cAg/cGg | Q90R | PLES_23781 | | | C | C | C | C | | | C | C | | |
| NC_011770 | 2558538 | G | A | 67 | 0 | 2 | 0 | STOP_GAINED | HIGH | NONSENSE | Cag/Tag | Q325* | PLES_23791 | | | A | A | | | | | | | | |
| NC_011770 | 2557679 | C | T | 63 | 0 | 2 | 0 | NON_SYNONYMOU | MODERAT | MISSENSE | cGc/cAc | R611H | PLES_23791 | | | | | | | | | | | T | T |
| NC_011770 | 2558153 | A | G | 68 | 0 | 1 | 0 | NON_SYNONYMOU | MODERAT | MISSENSE | cTc/cCc | L453P | PLES_23791 | | | | | | | G | | | | | |
| NC_011770 | 2558751 | C | T | 65 | 0 | 2 | 0 | NON_SYNONYMOU | MODERAT | MISSENSE | Gcc/Acc | A254T | PLES_23791 | | | | | | | T | T | | | | |
| NC_011770 | 2558822 | T | C | 65 | 0 | 12 | 0 | NON_SYNONYMOU | MODERAT | MISSENSE | cAg/cGg | Q230R | PLES_23791 | C | C | C | C | C | C | C | C | C | C | C | C |
| NC_011770 | 2559267 | T | C | 76 | 0 | 4 | 0 | NON_SYNONYMOU | MODERAT | MISSENSE | Acc/Gcc | T82A | PLES_23791 | C | C | | | | | C | C | | | | |
| NC_011770 | 2559588 | C | T | 75 | 0 | 6 | 0 | SPLICE_SITE_REGIC | LOW | SILENT | tGa/tAa | *180 | PLES_23801 | | | T | T | T | T | | | T | T | | |
| NC_011770 | 2559681 | C | T | 72 | 0 | 2 | 0 | NON_SYNONYMOU | MODERAT | MISSENSE | gGc/gAc | G149D | PLES_23801 | | | | | | | | | | | T | T |
| NC_011770 | 2559694 | G | A | 70 | 0 | 6 | 0 | NON_SYNONYMOU | MODERAT | MISSENSE | Ccg/Tcg | P145S | PLES_23801 | | | A | A | A | A | | | A | A | | |
| NC_011770 | 2559784 | C | A | 62 | 0 | 2 | 0 | NON_SYNONYMOU | MODERAT | MISSENSE | Gcc/Tcc | A115S | PLES_23801 | A | A | | | | | | | | | | |
| NC_011770 | 2559982 | G | T | 48 | 0 | 8 | 0 | NON_SYNONYMOU | MODERAT | MISSENSE | Cgc/Agc | R49S | PLES_23801 | | | T | T | T | T | | | T | T | T | T |
| NC_011770 | 2559999 | A | C | 48 | 0 | 6 | 0 | NON_SYNONYMOU | MODERAT | MISSENSE | gTc/gGc | V43G | PLES_23801 | C | C | | | | | C | C | | | C | C |
| NC_011770 | 2560076 | CAGGAGA | C | 41 | 0 | 2 | 0 | CODON_CHANGE_ | MODERATE | | cttctcctg/c | LLL15L | PLES_23801 | C | C | | | | | | | | | | |
| NC_011770 | 2560296 | C | A | 45 | 0 | 8 | 0 | NON_SYNONYMOU | MODERAT | MISSENSE | Ggc/Tgc | G62C | PLES_23811 | | | A | A | A | A | | | A | A | A | A |
| NC_011770 | 2560298 | C | A | 44 | 0 | 8 | 0 | NON_SYNONYMOU | MODERAT | MISSENSE | cGc/cTc | R61L | PLES_23811 | | | A | A | A | A | | | A | A | A | A |
| NC_011770 | 2560305 | C | T | 51 | 0 | 10 | 0 | NON_SYNONYMOU | MODERAT | MISSENSE | Gtt/Att | V59I | PLES_23811 | | | T | T | T | T | T | T | T | T | T | T |
| NC_011770 | 2560389 | G | C | 54 | 0 | 12 | 0 | NON_SYNONYMOU | MODERAT | MISSENSE | Ccc/Gcc | P31A | PLES_23811 | C | C | C | C | C | C | C | C | C | C | C | C |
| NC_011770 | 2561375 | C | T | 74 | 0 | 4 | 0 | NON_SYNONYMOU | MODERAT | MISSENSE | aGc/aAc | S13N | PLES_23831 | | | T | T | T | T | | | | | | |
| NC_011770 | 2561664 | C | G | 77 | 0 | 4 | 0 | NON_SYNONYMOU | MODERAT | MISSENSE | gaC/gaG | D20E | PLES_23841 | G | G | | | | | | | | | G | G |
| NC_011770 | 2561687 | T | G | 77 | 0 | 2 | 0 | NON_SYNONYMOU | MODERAT | MISSENSE | gTg/gGg | V28G | PLES_23841 | G | G | | | | | | | | | | |
| NC_011770 | 2561698 | G | A | 76 | 0 | 6 | 0 | NON_SYNONYMOU | MODERAT | MISSENSE | Ggg/Agg | G32R | PLES_23841 | | | A | A | A | A | | | A | A | | |
| NC_011770 | 2561819 | G | C | 75 | 0 | 2 | 0 | NON_SYNONYMOU | MODERAT | MISSENSE | gGc/gCc | G72A | PLES_23841 | | | | | | | C | C | | | | |
| NC_011770 | 2561838 | C | A | 73 | 0 | 2 | 0 | NON_SYNONYMOU | MODERAT | MISSENSE | gaC/gaA | D78E | PLES_23841 | | | | | | | A | A | | | | |
| NC_011770 | 2562014 | T | C | 74 | 0 | 12 | 0 | NON_SYNONYMOU | MODERAT | MISSENSE | gTc/gCc | V137A | PLES_23841 | C | C | C | C | C | C | C | C | C | C | C | C |
| NC_011770 | 2562049 | A | G | 78 | 0 | 4 | 0 | NON_SYNONYMOU | MODERAT | MISSENSE | Acc/Gcc | T149A | PLES_23841 | G | G | | | | | G | G | | | | |
| NC_011770 | 2562514 | C | T | 80 | 0 | 1 | 0 | NON_SYNONYMOU | MODERAT | MISSENSE | Ggg/Agg | G210R | PLES_23851 | | | | | | | | T | | | | |

| | | | | | | | | | | | | | | | | | | | | | | | | | |
|---|---|---|---|---|---|---|---|---|---|---|---|---|---|---|---|---|---|---|---|---|---|---|---|---|---|
| NC_011770 | 2563282 | AC | A | 76 | 0 | 3 | 0 | FRAME_SHIFT | HIGH | | ggg/ | G149 | PLES_23861 | | | | | | | A | | | | CC | CC |
| NC_011770 | 2563276 | G | A | 79 | 0 | 2 | 0 | NON_SYNONYMOU | MODERAT | MISSENSE | Ccc/Tcc | P152S | PLES_23861 | | | | | | | | | | | A | A |
| NC_011770 | 2563284 | C | G | 67 | 1 | 2 | 0 | NON_SYNONYMOU | MODERAT | MISSENSE | gGg/gCg | G149A | PLES_23861 | G | G | | | | | . | | | | | |
| NC_011770 | 2563585 | C | T | 69 | 0 | 2 | 0 | NON_SYNONYMOU | MODERAT | MISSENSE | Gtc/Atc | V49I | PLES_23861 | | | | | | | | | | | T | T |
| NC_011770 | 2563603 | C | T | 71 | 0 | 2 | 0 | NON_SYNONYMOU | MODERAT | MISSENSE | Gtg/Atg | V43M | PLES_23861 | | | | | | | | | | | T | T |
| NC_011770 | 2565787 | C | T | 50 | 0 | 4 | 0 | NON_SYNONYMOU | MODERAT | MISSENSE | gCg/gTg | A290V | PLES_23881 | | | T | T | T | T | | | | | | |
| NC_011770 | 2566387 | T | C | 75 | 0 | 2 | 0 | NON_SYNONYMOU | MODERAT | MISSENSE | Atc/Gtc | I41V | PLES_23891 | | | | | C | C | | | | | | |
| NC_011770 | 2567788 | C | T | 83 | 0 | 8 | 0 | NON_SYNONYMOU | MODERAT | MISSENSE | aGg/aAg | R349K | PLES_23901 | | | T | T | T | T | | | T | T | T | T |
| NC_011770 | 2568001 | C | T | 67 | 0 | 4 | 0 | NON_SYNONYMOU | MODERAT | MISSENSE | cGc/cAc | R278H | PLES_23901 | | | T | T | T | T | | | | | | |
| NC_011770 | 2569790 | C | T | 54 | 0 | 2 | 0 | NON_SYNONYMOU | MODERAT | MISSENSE | cGc/cAc | R263H | PLES_23921 | | | | | | | | | T | T | | |
| NC_011770 | 2570082 | C | T | 44 | 0 | 4 | 0 | NON_SYNONYMOU | MODERAT | MISSENSE | Ggc/Agc | G166S | PLES_23921 | | | T | T | T | T | | | | | | |
| NC_011770 | 2570177 | G | C | 45 | 0 | 2 | 0 | NON_SYNONYMOU | MODERAT | MISSENSE | gCc/gGc | A134G | PLES_23921 | C | C | | | | | | | | | | |
| NC_011770 | 2570328 | C | T | 41 | 0 | 12 | 0 | NON_SYNONYMOU | MODERAT | MISSENSE | Gtc/Atc | V84I | PLES_23921 | T | T | T | T | T | T | T | T | T | T | T | T |
| NC_011770 | 2570388 | C | T | 40 | 0 | 8 | 0 | NON_SYNONYMOU | MODERAT | MISSENSE | Gcg/Acg | A64T | PLES_23921 | | | T | T | T | T | | | T | T | T | T |
| NC_011770 | 2571386 | C | G | 73 | 0 | 2 | 0 | NON_SYNONYMOU | MODERAT | MISSENSE | Gtc/Ctc | V235L | PLES_23931 | G | G | | | | | | | | | | |
| NC_011770 | 2571764 | C | A | 72 | 0 | 2 | 0 | NON_SYNONYMOU | MODERAT | MISSENSE | Gtg/Ttg | V109L | PLES_23931 | | | | | | | | | A | A | | |
| NC_011770 | 2572565 | C | T | 85 | 0 | 4 | 0 | NON_SYNONYMOU | MODERAT | MISSENSE | Gcg/Acg | A113T | PLES_23941 | | | T | T | T | T | | | | | | |
| NC_011770 | 2572580 | C | T | 83 | 0 | 4 | 0 | NON_SYNONYMOU | MODERAT | MISSENSE | Gtc/Atc | V108I | PLES_23941 | T | T | | | | | | | T | T | | |
| NC_011770 | 2572742 | T | C | 77 | 0 | 12 | 0 | NON_SYNONYMOU | MODERAT | MISSENSE | Acc/Gcc | T54A | PLES_23941 | C | C | C | C | C | C | C | C | C | C | C | C |
| NC_011770 | 2574602 | C | G | 82 | 0 | 2 | 0 | NON_SYNONYMOU | MODERAT | MISSENSE | gCc/gGc | A183G | PLES_23961 | | | | | | | G | G | | | | |
| NC_011770 | 2574604 | C | T | 82 | 0 | 4 | 0 | NON_SYNONYMOU | MODERAT | MISSENSE | Cgg/Tgg | R184W | PLES_23961 | | | T | T | T | T | | | | | | |
| NC_011770 | 2574623 | G | A | 82 | 0 | 2 | 0 | NON_SYNONYMOU | MODERAT | MISSENSE | cGt/cAt | R190H | PLES_23961 | | | | | | | | | | | A | A |
| NC_011770 | 2575007 | G | A | 92 | 0 | 6 | 0 | NON_SYNONYMOU | MODERAT | MISSENSE | aGc/aAc | S318N | PLES_23961 | A | A | | | | | | | A | A | A | A |
| NC_011770 | 2575621 | G | C | 89 | 0 | 2 | 0 | NON_SYNONYMOU | MODERAT | MISSENSE | Gag/Cag | E107Q | PLES_23971 | C | C | | | | | | | | | | |
| NC_011770 | 2576246 | A | G | 82 | 0 | 2 | 0 | NON_SYNONYMOU | MODERAT | MISSENSE | Agc/Ggc | S12G | PLES_23981 | G | G | | | | | | | | | | |
| NC_011770 | 2577144 | A | C | 71 | 0 | 12 | 0 | NON_SYNONYMOU | MODERAT | MISSENSE | gAc/gCc | D163A | PLES_23991 | C | C | C | C | C | C | C | C | C | C | C | C |
| NC_011770 | 2578124 | A | G | 83 | 0 | 1 | 0 | NON_SYNONYMOU | MODERAT | MISSENSE | gAg/gGg | E58G | PLES_24001 | | | | | | | G | | | | | |
| NC_011770 | 2579319 | C | T | 53 | 0 | 4 | 0 | NON_SYNONYMOU | MODERAT | MISSENSE | gGc/gAc | G60D | PLES_24011 | | | T | T | T | T | | | | | | |
| NC_011770 | 2579895 | C | T | 32 | 0 | 4 | 0 | NON_SYNONYMOU | MODERAT | MISSENSE | cGt/cAt | R30H | PLES_24021 | | | | | | | | | T | T | T | T |
| NC_011770 | 2583287 | C | G | 46 | 0 | 4 | 0 | NON_SYNONYMOU | MODERAT | MISSENSE | Gcg/Ccg | A544P | PLES_24051 | | | G | G | G | G | | | | | | |
| NC_011770 | 2583329 | T | C | 46 | 0 | 6 | 0 | NON_SYNONYMOU | MODERAT | MISSENSE | Atc/Gtc | I530V | PLES_24051 | C | C | | | | | | | C | C | C | C |
| NC_011770 | 2584084 | G | A | 43 | 0 | 2 | 0 | NON_SYNONYMOU | MODERAT | MISSENSE | aCc/aTc | T278I | PLES_24051 | A | A | | | | | | | | | | |
| NC_011770 | 2584651 | A | G | 61 | 0 | 1 | 0 | NON_SYNONYMOU | MODERAT | MISSENSE | cTg/cCg | L89P | PLES_24051 | | | | | | | | G | | | | |
| NC_011770 | 2585960 | A | C | 82 | 0 | 4 | 0 | NON_SYNONYMOU | MODERAT | MISSENSE | gAc/gCc | D146A | PLES_24061 | | | C | C | C | C | | | | | | |
| NC_011770 | 2585924 | A | G | 89 | 0 | 1 | 0 | NON_SYNONYMOU | MODERAT | MISSENSE | tAc/tGc | Y134C | PLES_24061 | | | | | | | | G | | | | |
| NC_011770 | 2586393 | G | A | 70 | 0 | 2 | 0 | NON_SYNONYMOU | MODERAT | MISSENSE | Gtc/Atc | V45I | PLES_24071 | | | | | | | A | A | | | | |
| NC_011770 | 2587399 | G | T | 75 | 0 | 4 | 0 | NON_SYNONYMOU | MODERAT | MISSENSE | Gtt/Ttt | V230F | PLES_24081 | T | T | | | | | | | | | T | T |
| NC_011770 | 2587574 | T | C | 78 | 0 | 2 | 0 | NON_SYNONYMOU | MODERAT | MISSENSE | gTc/gCc | V288A | PLES_24081 | | | | | | | | | C | C | | |
| NC_011770 | 2587801 | AGCC | A | 80 | 0 | 2 | 0 | CODON_CHANGE_ | MODERATE | | ccgccc/ccc | PP31P | PLES_24091 | A | A | | | | | | | | | | |
| NC_011770 | 2588718 | A | G | 87 | 0 | 2 | 0 | NON_SYNONYMOU | MODERAT | MISSENSE | Acc/Gcc | T334A | PLES_24091 | | | | | | | | | G | G | | |
| NC_011770 | 2590149 | C | T | 77 | 0 | 1 | 0 | NON_SYNONYMOU | MODERAT | MISSENSE | Gcg/Acg | A102T | PLES_24121 | | T | | | | | | | | | | |
| NC_011770 | 2590981 | G | A | 66 | 0 | 4 | 0 | STOP_GAINED | HIGH | NONSENSE | tgG/tgA | W87* | PLES_24131 | A | A | | | | | | | | | A | A |
| NC_011770 | 2590979 | T | C | 66 | 0 | 4 | 0 | NON_SYNONYMOU | MODERAT | MISSENSE | Tgg/Cgg | W87R | PLES_24131 | C | C | | | | | | | | | C | C |
| NC_011770 | 2591013 | C | G | 67 | 0 | 2 | 0 | NON_SYNONYMOU | MODERAT | MISSENSE | gCg/gGg | A98G | PLES_24131 | | | | | | | | | | | G | G |
| NC_011770 | 2591079 | T | C | 62 | 0 | 4 | 0 | NON_SYNONYMOU | MODERAT | MISSENSE | gTc/gCc | V120A | PLES_24131 | C | C | | | | | | | | | C | C |
| NC_011770 | 2594004 | T | C | 72 | 0 | 4 | 0 | NON_SYNONYMOU | MODERAT | MISSENSE | Atc/Gtc | I111V | PLES_24151 | | | C | C | C | C | | | | | | |
| NC_011770 | 2594115 | C | T | 64 | 0 | 4 | 0 | NON_SYNONYMOU | MODERAT | MISSENSE | Gcc/Acc | A74T | PLES_24151 | | | T | T | T | T | | | | | | |
| NC_011770 | 2593850 | C | T | 75 | 0 | 1 | 0 | NON_SYNONYMOU | MODERAT | MISSENSE | cGc/cAc | R162H | PLES_24151 | | T | | | | | | | | | | |
| NC_011770 | 2594298 | C | T | 75 | 0 | 1 | 0 | NON_SYNONYMOU | MODERAT | MISSENSE | Gcc/Acc | A13T | PLES_24151 | | | | T | | | | | | | | |
| NC_011770 | 2599010 | T | TC | 79 | 0 | 2 | 0 | FRAME_SHIFT | HIGH | | atg/Catg | M38H? | PLES_24191 | | | | | | | | | | | TC | TC |
| NC_011770 | 2600025 | CG | C | 48 | 0 | 1 | 0 | FRAME_SHIFT | HIGH | | ggc/ | G376 | PLES_24191 | | | | | | | C | | | | | |
| NC_011770 | 2598953 | G | T | 81 | 0 | 2 | 0 | NON_SYNONYMOU | MODERAT | MISSENSE | tGg/tTg | W17L | PLES_24191 | | | | | | | | | T | T | | |
| NC_011770 | 2599228 | T | C | 77 | 0 | 2 | 0 | NON_SYNONYMOU | MODERAT | MISSENSE | Tat/Cat | Y109H | PLES_24191 | | | | | | | | | C | C | | |
| NC_011770 | 2600287 | A | G | 64 | 0 | 2 | 0 | NON_SYNONYMOU | MODERAT | MISSENSE | Atc/Gtc | I462V | PLES_24191 | | | | | | | | | G | G | | |
| NC_011770 | 2600289 | C | G | 67 | 0 | 2 | 0 | NON_SYNONYMOU | MODERAT | MISSENSE | atC/atG | I462M | PLES_24191 | | | | | | | | | G | G | | |
| NC_011770 | 2600561 | C | G | 78 | 0 | 8 | 0 | NON_SYNONYMOU | MODERAT | MISSENSE | aCc/aGc | T553S | PLES_24191 | G | G | G | G | G | G | | | | | G | G |
| NC_011770 | 2600825 | A | G | 73 | 0 | 2 | 0 | NON_SYNONYMOU | MODERAT | MISSENSE | aAc/aGc | N641S | PLES_24191 | | | | | | | G | G | | | | |
| NC_011770 | 2604356 | A | AC | 77 | 0 | 1 | 0 | FRAME_SHIFT | HIGH | | ctg/cCtg | L1077P? | PLES_24201 | | | | | | | | AC | | | | |
| NC_011770 | 2601278 | G | A | 60 | 0 | 2 | 0 | NON_SYNONYMOU | MODERAT | MISSENSE | Gtc/Atc | V49I | PLES_24201 | | | | | | | | | | | A | A |
| NC_011770 | 2601573 | A | C | 61 | 0 | 2 | 0 | NON_SYNONYMOU | MODERAT | MISSENSE | gAa/gCa | E147A | PLES_24201 | | | | | | | | | C | C | | |
| NC_011770 | 2601616 | A | C | 61 | 0 | 2 | 0 | NON_SYNONYMOU | MODERAT | MISSENSE | gaA/gaC | E161D | PLES_24201 | | | | | | | | | C | C | | |
| NC_011770 | 2602130 | A | G | 70 | 0 | 2 | 0 | NON_SYNONYMOU | MODERAT | MISSENSE | Acc/Gcc | T333A | PLES_24201 | | | | | | | | | | | G | G |

| | | | | | | | | | | | | | | | | | | | | | | | | | |
|---|---|---|---|---|---|---|---|---|---|---|---|---|---|---|---|---|---|---|---|---|---|---|---|---|---|
| NC_011770 | 2602545 | C | T | 62 | 0 | 1 | 0 | NON_SYNONYMOU | MODERAT | MISSENSE | cCc/cTc | P471L | PLES_24201 | | | | | | | T | | | | | |
| NC_011770 | 2602809 | C | T | 75 | 0 | 2 | 0 | NON_SYNONYMOU | MODERAT | MISSENSE | aCc/aTc | T559I | PLES_24201 | | | | | | | T | T | | | | |
| NC_011770 | 2602872 | T | C | 77 | 0 | 4 | 0 | NON_SYNONYMOU | MODERAT | MISSENSE | gTg/gCg | V580A | PLES_24201 | | | | | | | C | C | C | C | | |
| NC_011770 | 2602903 | G | C | 78 | 0 | 2 | 0 | NON_SYNONYMOU | MODERAT | MISSENSE | ttG/ttC | L590F | PLES_24201 | C | C | | | | | | | | | | |
| NC_011770 | 2603171 | C | T | 80 | 0 | 2 | 0 | NON_SYNONYMOU | MODERAT | MISSENSE | Cgt/Tgt | R680C | PLES_24201 | | | | | | | | | | | T | T |
| NC_011770 | 2603514 | G | A | 85 | 0 | 2 | 0 | NON_SYNONYMOU | MODERAT | MISSENSE | cGc/cAc | R794H | PLES_24201 | | | | | | | A | A | | | | |
| NC_011770 | 2603640 | T | C | 84 | 0 | 12 | 0 | NON_SYNONYMOU | MODERAT | MISSENSE | cTg/cCg | L836P | PLES_24201 | C | C | C | C | C | C | C | C | C | C | C | C |
| NC_011770 | 2603831 | G | A | 74 | 0 | 10 | 0 | NON_SYNONYMOU | MODERAT | MISSENSE | Gcg/Acg | A900T | PLES_24201 | | | A | A | A | A | A | A | A | A | A | A |
| NC_011770 | 2603841 | A | G | 79 | 0 | 12 | 0 | NON_SYNONYMOU | MODERAT | MISSENSE | aAc/aGc | N903S | PLES_24201 | G | G | G | G | G | G | G | G | G | G | G | G |
| NC_011770 | 2603948 | G | A | 85 | 0 | 2 | 0 | NON_SYNONYMOU | MODERAT | MISSENSE | Gcc/Acc | A939T | PLES_24201 | | | | | | | | | A | A | | |
| NC_011770 | 2604393 | G | A | 74 | 0 | 6 | 0 | NON_SYNONYMOU | MODERAT | MISSENSE | cGt/cAt | R1087H | PLES_24201 | A | A | A | A | A | A | | | | | | |
| NC_011770 | 2605037 | A | G | 70 | 0 | 1 | 0 | NON_SYNONYMOU | MODERAT | MISSENSE | Atc/Gtc | I1302V | PLES_24201 | | | | | | | G | | | | | |
| NC_011770 | 2605948 | C | T | 64 | 0 | 2 | 0 | NON_SYNONYMOU | MODERAT | MISSENSE | Ccg/Tcg | P171S | PLES_24211 | | | | | | | | | T | T | | |
| NC_011770 | 2607503 | G | C | 52 | 0 | 2 | 0 | NON_SYNONYMOU | MODERAT | MISSENSE | gaC/gaG | D49E | PLES_24221 | C | C | | | | | | | | | | |
| NC_011770 | 2607609 | C | G | 50 | 0 | 2 | 0 | NON_SYNONYMOU | MODERAT | MISSENSE | gGc/gCc | G14A | PLES_24221 | G | G | | | | | | | | | | |
| NC_011770 | 2607836 | G | T | 27 | 0 | 2 | 0 | NON_SYNONYMOU | MODERAT | MISSENSE | gCc/gAc | A309D | PLES_24231 | T | T | | | | | | | | | | |
| NC_011770 | 2607903 | C | T | 32 | 0 | 2 | 0 | NON_SYNONYMOU | MODERAT | MISSENSE | Gcc/Acc | A287T | PLES_24231 | | | | | | | | | | | T | T |
| NC_011770 | 2608230 | C | T | 58 | 0 | 6 | 0 | NON_SYNONYMOU | MODERAT | MISSENSE | Gag/Aag | E178K | PLES_24231 | | | | | | | T | T | T | T | T | T |
| NC_011770 | 2608878 | C | T | 62 | 0 | 2 | 0 | NON_SYNONYMOU | MODERAT | MISSENSE | cGg/cAg | R294Q | PLES_24241 | | | | | | | | | | | T | T |
| NC_011770 | 2609083 | T | C | 65 | 0 | 4 | 0 | NON_SYNONYMOU | MODERAT | MISSENSE | Aac/Gac | N226D | PLES_24241 | C | C | | | | | C | C | | | | |
| NC_011770 | 2609118 | T | C | 65 | 0 | 2 | 0 | NON_SYNONYMOU | MODERAT | MISSENSE | gAc/gGc | D214G | PLES_24241 | | | C | C | | | | | | | | |
| NC_011770 | 2609182 | C | T | 65 | 0 | 2 | 0 | NON_SYNONYMOU | MODERAT | MISSENSE | Gcg/Acg | A193T | PLES_24241 | T | T | | | | | | | | | | |
| NC_011770 | 2609365 | G | A | 64 | 0 | 2 | 0 | NON_SYNONYMOU | MODERAT | MISSENSE | Cac/Tac | H132Y | PLES_24241 | | | | | | | | | A | A | | |
| NC_011770 | 2609427 | G | C | 64 | 0 | 4 | 0 | NON_SYNONYMOU | MODERAT | MISSENSE | aCa/aGa | T111R | PLES_24241 | C | C | | | | | | | | | C | C |
| NC_011770 | 2609604 | A | G | 71 | 0 | 2 | 0 | NON_SYNONYMOU | MODERAT | MISSENSE | gTa/gCa | V52A | PLES_24241 | | | | | | | G | G | | | | |
| NC_011770 | 2609653 | C | T | 76 | 0 | 2 | 0 | NON_SYNONYMOU | MODERAT | MISSENSE | Gcc/Acc | A36T | PLES_24241 | | | | | | | T | T | | | | |
| NC_011770 | 2609698 | C | A | 84 | 0 | 2 | 0 | NON_SYNONYMOU | MODERAT | MISSENSE | Gtc/Ttc | V21F | PLES_24241 | | | | | | | A | A | | | | |
| NC_011770 | 2609724 | T | C | 82 | 0 | 12 | 0 | NON_SYNONYMOU | MODERAT | MISSENSE | gAc/gGc | D12G | PLES_24241 | C | C | C | C | C | C | C | C | C | C | C | C |
| NC_011770 | 2609158 | C | T | 66 | 0 | 1 | 0 | NON_SYNONYMOU | MODERAT | MISSENSE | Gcc/Acc | A201T | PLES_24241 | | | | | | | | T | | | | |
| NC_011770 | 2609991 | C | T | 88 | 0 | 2 | 0 | NON_SYNONYMOU | MODERAT | MISSENSE | gCa/gTa | A3V | PLES_24251 | | | | | | | T | T | | | | |
| NC_011770 | 2610060 | A | G | 84 | 0 | 10 | 0 | NON_SYNONYMOU | MODERAT | MISSENSE | cAg/cGg | Q26R | PLES_24251 | G | G | G | G | G | G | G | G | G | G | | |
| NC_011770 | 2610219 | A | G | 79 | 0 | 4 | 0 | NON_SYNONYMOU | MODERAT | MISSENSE | gAg/gGg | E79G | PLES_24251 | | | G | G | G | G | | | | | | |
| NC_011770 | 2610416 | C | T | 82 | 0 | 4 | 0 | NON_SYNONYMOU | MODERAT | MISSENSE | Ccg/Tcg | P145S | PLES_24251 | T | T | | | | | | | T | T | | |
| NC_011770 | 2610498 | G | A | 77 | 0 | 10 | 0 | NON_SYNONYMOU | MODERAT | MISSENSE | aGt/aAt | S172N | PLES_24251 | A | A | A | A | A | A | A | A | A | A | | |
| NC_011770 | 2611536 | T | C | 79 | 0 | 4 | 0 | NON_SYNONYMOU | MODERAT | MISSENSE | Tcc/Ccc | S71P | PLES_24261 | | | C | C | C | C | | | | | | |
| NC_011770 | 2612147 | C | T | 65 | 0 | 4 | 0 | NON_SYNONYMOU | MODERAT | MISSENSE | cCg/cTg | P10L | PLES_24271 | | | | | | | T | T | T | T | | |
| NC_011770 | 2612248 | C | T | 65 | 0 | 2 | 0 | NON_SYNONYMOU | MODERAT | MISSENSE | Ctt/Ttt | L44F | PLES_24271 | | | | | | | T | T | | | | |
| NC_011770 | 2612602 | A | G | 87 | 0 | 8 | 0 | NON_SYNONYMOU | MODERAT | MISSENSE | Acc/Gcc | T162A | PLES_24271 | G | G | G | G | G | G | | | | | G | G |
| NC_011770 | 2613562 | C | G | 95 | 0 | 2 | 0 | NON_SYNONYMOU | MODERAT | MISSENSE | Cag/Gag | Q482E | PLES_24271 | G | G | | | | | | | | | | |
| NC_011770 | 2614545 | C | T | 73 | 0 | 4 | 0 | NON_SYNONYMOU | MODERAT | MISSENSE | gCg/gTg | A232V | PLES_24281 | | | T | T | T | T | | | | | | |
| NC_011770 | 2614697 | C | G | 60 | 0 | 2 | 0 | NON_SYNONYMOU | MODERAT | MISSENSE | Ctg/Gtg | L283V | PLES_24281 | G | G | | | | | | | | | | |
| NC_011770 | 2614914 | C | T | 65 | 0 | 2 | 0 | NON_SYNONYMOU | MODERAT | MISSENSE | gCg/gTg | A355V | PLES_24281 | | | | | | | | | | | T | T |
| NC_011770 | 2615718 | G | A | 97 | 0 | 2 | 0 | NON_SYNONYMOU | MODERAT | MISSENSE | gGt/gAt | G75D | PLES_24301 | A | A | | | | | | | | | | |
| NC_011770 | 2615987 | T | C | 87 | 0 | 4 | 0 | NON_SYNONYMOU | MODERAT | MISSENSE | tTc/tCc | F28S | PLES_24311 | | | C | C | C | C | | | | | | |
| NC_011770 | 2616058 | T | C | 81 | 0 | 12 | 0 | NON_SYNONYMOU | MODERAT | MISSENSE | Tgg/Cgg | W52R | PLES_24311 | C | C | C | C | C | C | C | C | C | C | C | C |
| NC_011770 | 2616116 | T | C | 84 | 0 | 10 | 0 | NON_SYNONYMOU | MODERAT | MISSENSE | gTc/gCc | V71A | PLES_24311 | | | C | C | C | C | C | C | C | C | C | C |
| NC_011770 | 2616466 | G | A | 94 | 0 | 2 | 0 | NON_SYNONYMOU | MODERAT | MISSENSE | aGc/aAc | S47N | PLES_24321 | A | A | | | | | | | | | | |
| NC_011770 | 2616754 | G | T | 81 | 0 | 2 | 0 | NON_SYNONYMOU | MODERAT | MISSENSE | cGc/cTc | R143L | PLES_24321 | T | T | | | | | | | | | | |
| NC_011770 | 2616775 | A | G | 82 | 0 | 2 | 0 | NON_SYNONYMOU | MODERAT | MISSENSE | cAg/cGg | Q150R | PLES_24321 | | | G | G | | | | | | | | |
| NC_011770 | 2616781 | A | T | 84 | 0 | 2 | 0 | NON_SYNONYMOU | MODERAT | MISSENSE | cAc/cTc | H152L | PLES_24321 | T | T | | | | | | | | | | |
| NC_011770 | 2616840 | G | A | 86 | 0 | 2 | 0 | NON_SYNONYMOU | MODERAT | MISSENSE | Gcg/Acg | A172T | PLES_24321 | A | A | | | | | | | | | | |
| NC_011770 | 2616925 | C | T | 87 | 0 | 2 | 0 | NON_SYNONYMOU | MODERAT | MISSENSE | Cgc/Tgc | R7C | PLES_24331 | T | T | | | | | | | | | | |
| NC_011770 | 2617067 | C | T | 91 | 0 | 10 | 0 | NON_SYNONYMOU | MODERAT | MISSENSE | gCg/gTg | A54V | PLES_24331 | | | T | T | T | T | T | T | T | T | T | T |
| NC_011770 | 2617273 | G | A | 81 | 0 | 6 | 0 | NON_SYNONYMOU | MODERAT | MISSENSE | Gtt/Att | V123I | PLES_24331 | A | A | A | A | A | A | | | | | | |
| NC_011770 | 2617978 | C | T | 82 | 0 | 4 | 0 | NON_SYNONYMOU | MODERAT | MISSENSE | Ccg/Tcg | P358S | PLES_24331 | | | T | T | T | T | | | | | | |
| NC_011770 | 2618798 | C | T | 77 | 0 | 2 | 0 | STOP_GAINED | HIGH | NONSENSE | Cag/Tag | Q257* | PLES_24341 | T | T | | | | | | | | | | |
| NC_011770 | 2618747 | A | T | 77 | 0 | 2 | 0 | NON_SYNONYMOU | MODERAT | MISSENSE | Acg/Tcg | T240S | PLES_24341 | T | T | | | | | | | | | | |
| NC_011770 | 2618877 | G | C | 74 | 0 | 10 | 0 | NON_SYNONYMOU | MODERAT | MISSENSE | gGc/gCc | G283A | PLES_24341 | C | C | C | C | C | C | C | C | C | C | | |
| NC_011770 | 2619083 | C | G | 78 | 0 | 2 | 0 | NON_SYNONYMOU | MODERAT | MISSENSE | cCa/cGa | P23R | PLES_24351 | | | | | G | G | | | | | | |
| NC_011770 | 2619259 | A | G | 79 | 0 | 2 | 0 | NON_SYNONYMOU | MODERAT | MISSENSE | Aag/Gag | K82E | PLES_24351 | | | | | | | | | | | G | G |
| NC_011770 | 2621353 | C | T | 72 | 0 | 10 | 0 | SPLICE_SITE_REGIC | LOW | SILENT | tGa/tAa | *518 | PLES_24391 | T | T | T | T | T | T | | | T | T | T | T |
| NC_011770 | 2621467 | C | T | 68 | 0 | 4 | 0 | NON_SYNONYMOU | MODERAT | MISSENSE | cGc/cAc | R480H | PLES_24391 | | | | | | | T | T | | | T | T |

| | | | | | | | | | | | | | | | | | | | | | | | | | |
|---|---|---|---|---|---|---|---|---|---|---|---|---|---|---|---|---|---|---|---|---|---|---|---|---|---|
| NC_011770 | 2621555 | T | C | 64 | 0 | 12 | 0 | NON_SYNONYMOU | MODERAT | MISSENSE | Acg/Gcg | T451A | PLES_24391 | C | C | C | C | C | C | C | C | C | C | C | C |
| NC_011770 | 2621708 | C | T | 70 | 0 | 2 | 0 | NON_SYNONYMOU | MODERAT | MISSENSE | Gcc/Acc | A400T | PLES_24391 | | | | | | | T | T | | | | |
| NC_011770 | 2622271 | G | C | 78 | 0 | 2 | 0 | NON_SYNONYMOU | MODERAT | MISSENSE | gCc/gGc | A212G | PLES_24391 | | | | | | | | | | | C | C |
| NC_011770 | 2622640 | C | T | 85 | 0 | 2 | 0 | NON_SYNONYMOU | MODERAT | MISSENSE | cGc/cAc | R89H | PLES_24391 | T | T | | | | | | | | | | |
| NC_011770 | 2624525 | G | A | 69 | 0 | 2 | 0 | NON_SYNONYMOU | MODERAT | MISSENSE | gGt/gAt | G77D | PLES_24411 | | | | | | | A | A | | | | |
| NC_011770 | 2624847 | C | T | 65 | 0 | 4 | 0 | NON_SYNONYMOU | MODERAT | MISSENSE | gCg/gTg | A103V | PLES_24421 | | | T | T | T | T | | | | | | |
| NC_011770 | 2625248 | A | G | 62 | 0 | 2 | 0 | NON_SYNONYMOU | MODERAT | MISSENSE | Acg/Gcg | T237A | PLES_24421 | | | | | | | G | G | | | | |
| NC_011770 | 2625435 | C | T | 54 | 0 | 2 | 0 | NON_SYNONYMOU | MODERAT | MISSENSE | gCg/gTg | A299V | PLES_24421 | | | | | | | T | T | | | | |
| NC_011770 | 2626757 | T | C | 71 | 0 | 12 | 0 | NON_SYNONYMOU | MODERAT | MISSENSE | Aaa/Gaa | K41E | PLES_24431 | C | C | C | C | C | C | C | C | C | C | C | C |
| NC_011770 | 2626841 | C | T | 67 | 0 | 2 | 0 | NON_SYNONYMOU | MODERAT | MISSENSE | Gac/Aac | D13N | PLES_24431 | | | | | | | T | T | | | | |
| NC_011770 | 2627212 | C | T | 68 | 0 | 2 | 0 | NON_SYNONYMOU | MODERAT | MISSENSE | Gtg/Atg | V259M | PLES_24441 | | | | | | | T | T | | | | |
| NC_011770 | 2627937 | C | T | 57 | 0 | 2 | 0 | NON_SYNONYMOU | MODERAT | MISSENSE | cGc/cAc | R17H | PLES_24441 | | | | | | | | | | | T | T |
| NC_011770 | 2630784 | A | G | 75 | 0 | 10 | 0 | NON_SYNONYMOU | MODERAT | MISSENSE | Atc/Gtc | I432V | PLES_24481 | G | G | G | G | G | G | G | G | G | G | | |
| NC_011770 | 2630793 | C | T | 74 | 0 | 2 | 0 | NON_SYNONYMOU | MODERAT | MISSENSE | Ccg/Tcg | P435S | PLES_24481 | | | | | | | | | | | T | T |
| NC_011770 | 2630934 | A | G | 73 | 0 | 6 | 0 | NON_SYNONYMOU | MODERAT | MISSENSE | Agt/Ggt | S6G | PLES_24491 | | | G | G | G | G | | | | | G | G |
| NC_011770 | 2631034 | T | C | 67 | 0 | 10 | 0 | NON_SYNONYMOU | MODERAT | MISSENSE | cTc/cCc | L39P | PLES_24491 | C | C | C | C | C | C | C | C | | | C | C |
| NC_011770 | 2631108 | G | A | 63 | 0 | 12 | 0 | NON_SYNONYMOU | MODERAT | MISSENSE | Gca/Aca | A64T | PLES_24491 | A | A | A | A | A | A | A | A | A | A | A | A |
| NC_011770 | 2635551 | A | G | 70 | 0 | 6 | 0 | NON_SYNONYMOU | MODERAT | MISSENSE | Agc/Ggc | S160G | PLES_24521 | | | G | G | G | G | | | G | G | | |
| NC_011770 | 2635596 | T | C | 73 | 0 | 1 | 0 | NON_SYNONYMOU | MODERAT | MISSENSE | Tgg/Cgg | W175R | PLES_24521 | | | | | | | C | | | | | |
| NC_011770 | 2636208 | T | A | 60 | 0 | 12 | 0 | NON_SYNONYMOU | MODERAT | MISSENSE | Tcc/Acc | S379T | PLES_24521 | A | A | A | A | A | A | A | A | A | A | A | A |
| NC_011770 | 2636496 | A | G | 66 | 0 | 1 | 0 | NON_SYNONYMOU | MODERAT | MISSENSE | Acc/Gcc | T475A | PLES_24521 | | | | | | | G | | | | | |
| NC_011770 | 2635315 | C | T | 74 | 0 | 1 | 0 | NON_SYNONYMOU | MODERAT | MISSENSE | gCc/gTc | A81V | PLES_24521 | | | | | | | | T | | | | |
| NC_011770 | 2639601 | T | C | 64 | 0 | 4 | 0 | NON_SYNONYMOU | MODERAT | MISSENSE | Atg/Gtg | M253V | PLES_24551 | C | C | | | | | | | C | C | | |
| NC_011770 | 2639673 | C | T | 64 | 0 | 4 | 0 | NON_SYNONYMOU | MODERAT | MISSENSE | Gcc/Acc | A229T | PLES_24551 | | | T | T | T | T | | | | | | |
| NC_011770 | 2640051 | C | T | 45 | 0 | 2 | 0 | NON_SYNONYMOU | MODERAT | MISSENSE | Gtc/Atc | V103I | PLES_24551 | | | | | | | T | T | | | | |
| NC_011770 | 2640340 | T | A | 30 | 0 | 8 | 0 | NON_SYNONYMOU | MODERAT | MISSENSE | gaA/gaT | E6D | PLES_24551 | A | A | A | A | A | A | | | | | A | A |
| NC_011770 | 2656021 | T | TG | 65 | 0 | 10 | 0 | FRAME_SHIFT | HIGH | | cca/ccCa | P183P? | PLES_24691 | | . | TG | TG | TG | TG | TG | TG | TA | TA | TA | TA |
| NC_011770 | 2655858 | C | T | 68 | 0 | 1 | 0 | NON_SYNONYMOU | MODERAT | MISSENSE | Ggc/Agc | G238S | PLES_24691 | | | | | | | T | | | | | |
| NC_011770 | 2655894 | G | A | 72 | 0 | 2 | 0 | NON_SYNONYMOU | MODERAT | MISSENSE | Cca/Tca | P226S | PLES_24691 | | | | | | | | | | | A | A |
| NC_011770 | 2655905 | G | A | 68 | 0 | 2 | 0 | NON_SYNONYMOU | MODERAT | MISSENSE | gCa/gTa | A222V | PLES_24691 | | | | | | | | | | | A | A |
| NC_011770 | 2655906 | C | T | 69 | 0 | 2 | 0 | NON_SYNONYMOU | MODERAT | MISSENSE | Gca/Aca | A222T | PLES_24691 | | | | | | | | | | | T | T |
| NC_011770 | 2655968 | T | C | 65 | 0 | 2 | 0 | NON_SYNONYMOU | MODERAT | MISSENSE | aAa/aGa | K201R | PLES_24691 | | | | | | | | | C | C | | |
| NC_011770 | 2655980 | G | C | 63 | 0 | 2 | 0 | NON_SYNONYMOU | MODERAT | MISSENSE | gCg/gGg | A197G | PLES_24691 | | | | | | | C | C | | | | |
| NC_011770 | 2655982 | G | T | 61 | 0 | 10 | 0 | NON_SYNONYMOU | MODERAT | MISSENSE | gaC/gaA | D196E | PLES_24691 | | | T | T | T | T | T | T | T | T | T | T |
| NC_011770 | 2656037 | A | T | 61 | 0 | 4 | 0 | NON_SYNONYMOU | MODERAT | MISSENSE | gTc/gAc | V178D | PLES_24691 | | | | | | | | | T | T | T | T |
| NC_011770 | 2656383 | C | G | 74 | 0 | 4 | 0 | NON_SYNONYMOU | MODERAT | MISSENSE | Gag/Cag | E63Q | PLES_24691 | | | G | G | G | G | | | | | | |
| NC_011770 | 2656452 | T | A | 74 | 0 | 4 | 0 | NON_SYNONYMOU | MODERAT | MISSENSE | Acg/Tcg | T40S | PLES_24691 | | | | | | | A | A | A | A | | |
| NC_011770 | 2658467 | A | G | 64 | 0 | 2 | 0 | NON_SYNONYMOU | MODERAT | MISSENSE | gTg/gCg | V67A | PLES_24701 | | | | | | | | | | | G | G |
| NC_011770 | 2657939 | C | T | 66 | 0 | 1 | 0 | NON_SYNONYMOU | MODERAT | MISSENSE | gGc/gAc | G243D | PLES_24701 | | | | | | | | T | | | | |
| NC_011770 | 2661235 | C | T | 64 | 0 | 2 | 0 | NON_SYNONYMOU | MODERAT | MISSENSE | Gac/Aac | D208N | PLES_24721 | T | T | | | | | | | | | | |
| NC_011770 | 2661532 | C | T | 68 | 0 | 10 | 0 | NON_SYNONYMOU | MODERAT | MISSENSE | Gcc/Acc | A109T | PLES_24721 | T | T | T | T | T | T | T | T | T | T | | |
| NC_011770 | 2661711 | A | C | 59 | 0 | 12 | 0 | NON_SYNONYMOU | MODERAT | MISSENSE | aTc/aGc | I49S | PLES_24721 | C | C | C | C | C | C | C | C | C | C | C | C |
| NC_011770 | 2662094 | C | T | 70 | 0 | 8 | 0 | NON_SYNONYMOU | MODERAT | MISSENSE | aGc/aAc | S127N | PLES_24731 | T | T | T | T | T | T | T | T | | | | |
| NC_011770 | 2662316 | C | T | 63 | 0 | 4 | 0 | NON_SYNONYMOU | MODERAT | MISSENSE | cGc/cAc | R53H | PLES_24731 | | | T | T | T | T | | | | | | |
| NC_011770 | 2662376 | A | G | 70 | 0 | 2 | 0 | NON_SYNONYMOU | MODERAT | MISSENSE | cTg/cCg | L33P | PLES_24731 | | | | | | | | | G | G | | |
| NC_011770 | 2662410 | A | G | 74 | 0 | 2 | 0 | NON_SYNONYMOU | MODERAT | MISSENSE | Tgc/Cgc | C22R | PLES_24731 | | | | | | | | | G | G | | |
| NC_011770 | 2662464 | C | T | 76 | 0 | 2 | 0 | NON_SYNONYMOU | MODERAT | MISSENSE | Gcc/Acc | A4T | PLES_24731 | | | | | | | | | T | T | | |
| NC_011770 | 2662860 | C | G | 78 | 0 | 2 | 0 | NON_SYNONYMOU | MODERAT | MISSENSE | Gcc/Ccc | A16P | PLES_24741 | | | | | | | G | G | | | | |
| NC_011770 | 2663180 | C | A | 79 | 0 | 2 | 0 | NON_SYNONYMOU | MODERAT | MISSENSE | gaG/gaT | E296D | PLES_24751 | | | | | | | | | | | A | A |
| NC_011770 | 2663905 | T | C | 51 | 0 | 12 | 0 | NON_SYNONYMOU | MODERAT | MISSENSE | Atg/Gtg | M55V | PLES_24751 | C | C | C | C | C | C | C | C | C | C | C | C |
| NC_011770 | 2666980 | C | T | 72 | 0 | 4 | 0 | NON_SYNONYMOU | MODERAT | MISSENSE | gGc/gAc | G36D | PLES_24781 | | | T | T | T | T | | | | | | |
| NC_011770 | 2668396 | C | G | 21 | 0 | 4 | 0 | NON_SYNONYMOU | MODERAT | MISSENSE | aGc/aCc | S110T | PLES_24801 | | | | | | | G | G | | | G | G |
| NC_011770 | 2668475 | C | T | 24 | 0 | 2 | 0 | NON_SYNONYMOU | MODERAT | MISSENSE | Gta/Ata | V84I | PLES_24801 | | | | | | | | | T | T | | |
| NC_011770 | 2676780 | T | C | 92 | 0 | 6 | 0 | NON_SYNONYMOU | MODERAT | MISSENSE | gAc/gGc | D76G | PLES_24871 | C | C | C | C | C | C | | | | | | |
| NC_011770 | 2686231 | GGC | G | 77 | 0 | 1 | 0 | FRAME_SHIFT | HIGH | | gcgcca/ | AP309 | PLES_24951 | | | | G | | | | | | | | |
| NC_011770 | 2686408 | C | A | 87 | 0 | 2 | 0 | NON_SYNONYMOU | MODERAT | MISSENSE | cGg/cTg | R251L | PLES_24951 | A | A | | | | | | | | | | |
| NC_011770 | 2686540 | T | A | 91 | 0 | 2 | 0 | NON_SYNONYMOU | MODERAT | MISSENSE | gAc/gTc | D207V | PLES_24951 | | | | | | | | | A | A | | |
| NC_011770 | 2686855 | A | T | 83 | 0 | 2 | 0 | NON_SYNONYMOU | MODERAT | MISSENSE | cTg/cAg | L102Q | PLES_24951 | | | | | | | T | T | | | | |
| NC_011770 | 2686979 | C | T | 84 | 0 | 12 | 0 | NON_SYNONYMOU | MODERAT | MISSENSE | Gtc/Atc | V61I | PLES_24951 | T | T | T | T | T | T | T | T | T | T | T | T |
| NC_011770 | 2688320 | A | G | 87 | 0 | 8 | 0 | NON_SYNONYMOU | MODERAT | MISSENSE | aTg/aCg | M5T | PLES_24971 | G | G | | | | | G | G | G | G | G | G |
| NC_011770 | 2688556 | G | A | 92 | 0 | 4 | 0 | NON_SYNONYMOU | MODERAT | MISSENSE | cCc/cTc | P27L | PLES_24981 | | | | | | | A | A | A | A | | |
| NC_011770 | 2688415 | T | C | 88 | 0 | 1 | 0 | NON_SYNONYMOU | MODERAT | MISSENSE | gAc/gGc | D74G | PLES_24981 | C | | | | | | | | | | | |

| | | | | | | | | | | | | | | | | | | | | | | | | | |
|---|---|---|---|---|---|---|---|---|---|---|---|---|---|---|---|---|---|---|---|---|---|---|---|---|---|
| NC_011770 | 2688836 | T | A | 72 | 0 | 8 | 0 | NON_SYNONYMOU | MODERAT | MISSENSE | caA/caT | Q53H | PLES_24991 | A | A | | | | | A | A | A | A | A | A |
| NC_011770 | 2689066 | T | A | 55 | 0 | 2 | 0 | NON_SYNONYMOU | MODERAT | MISSENSE | gaA/gaT | E107D | PLES_25001 | | | | | | | | | A | A | | |
| NC_011770 | 2690135 | C | T | 65 | 0 | 2 | 0 | NON_SYNONYMOU | MODERAT | MISSENSE | Gcc/Acc | A11T | PLES_25011 | | | | | | | | | T | T | | |
| NC_011770 | 2690342 | T | A | 28 | 0 | 12 | 0 | INTRAGENIC | MODIFIER | | | | PLES_25012 | A | A | A | A | A | A | A | A | A | A | A | A |
| NC_011770 | 2692220 | C | A | 9 | 8 | 2 | 0 | STOP_GAINED | HIGH | NONSENSE | Gaa/Taa | E526* | PLES_25041 | . | . | . | . | . | . | . | . | | | A | A |
| NC_011770 | 2693775 | G | GCGTT | 9 | 8 | 2 | 0 | FRAME_SHIFT+STC | HIGH | | tgc/tgAAC | C7*T? | PLES_25041 | . | . | . | . | . | . | . | . | GCGTT | GCGTT | | |
| NC_011770 | 2691986 | G | T | 8 | 8 | 2 | 0 | NON_SYNONYMOU | MODERAT | MISSENSE | Ctg/Atg | L604M | PLES_25041 | . | . | T | T | . | . | . | . | . | . | | |
| NC_011770 | 2691997 | G | A | 3 | 10 | 2 | 0 | NON_SYNONYMOU | MODERAT | MISSENSE | gCg/gTg | A600V | PLES_25041 | . | . | A | A | . | . | . | . | . | . | . | . |
| NC_011770 | 2692003 | C | G | 3 | 10 | 2 | 0 | NON_SYNONYMOU | MODERAT | MISSENSE | aGc/aCc | S598T | PLES_25041 | . | . | G | G | . | . | . | . | . | . | . | . |
| NC_011770 | 2692121 | C | G | 3 | 10 | 2 | 0 | NON_SYNONYMOU | MODERAT | MISSENSE | Gag/Cag | E559Q | PLES_25041 | . | . | G | G | . | . | . | . | . | . | . | . |
| NC_011770 | 2692135 | G | T | 3 | 10 | 2 | 0 | NON_SYNONYMOU | MODERAT | MISSENSE | gCg/gAg | A554E | PLES_25041 | . | . | T | T | . | . | . | . | . | . | . | . |
| NC_011770 | 2692148 | C | G | 8 | 8 | 2 | 0 | NON_SYNONYMOU | MODERAT | MISSENSE | Gag/Cag | E550Q | PLES_25041 | . | . | | | . | . | . | . | . | . | G | G |
| NC_011770 | 2692168 | G | T | 10 | 4 | 4 | 0 | NON_SYNONYMOU | MODERAT | MISSENSE | aCc/aAc | T543N | PLES_25041 | | | T | T | | | . | . | . | . | T | T |
| NC_011770 | 2692169 | T | C | 10 | 4 | 4 | 0 | NON_SYNONYMOU | MODERAT | MISSENSE | Acc/Gcc | T543A | PLES_25041 | | | C | C | | | . | . | . | . | C | C |
| NC_011770 | 2692172 | T | C | 11 | 4 | 2 | 0 | NON_SYNONYMOU | MODERAT | MISSENSE | Atc/Gtc | I542V | PLES_25041 | | | | | | | . | . | . | . | C | C |
| NC_011770 | 2692219 | T | G | 10 | 8 | 2 | 0 | NON_SYNONYMOU | MODERAT | MISSENSE | gAa/gCa | E526A | PLES_25041 | . | . | . | . | . | . | . | . | | | G | G |
| NC_011770 | 2692221 | C | G | 10 | 8 | 2 | 0 | NON_SYNONYMOU | MODERAT | MISSENSE | aaG/aaC | K525N | PLES_25041 | . | . | . | . | . | . | . | . | | | G | G |
| NC_011770 | 2692424 | C | T | 7 | 4 | 2 | 0 | NON_SYNONYMOU | MODERAT | MISSENSE | Gct/Act | A458T | PLES_25041 | | | . | . | | | . | . | | | T | T |
| NC_011770 | 2692432 | C | G | 3 | 10 | 2 | 0 | NON_SYNONYMOU | MODERAT | MISSENSE | aGc/aCc | S455T | PLES_25041 | . | . | . | . | . | . | . | . | G | G | . | . |
| NC_011770 | 2692433 | T | C | 3 | 10 | 2 | 0 | NON_SYNONYMOU | MODERAT | MISSENSE | Agc/Ggc | S455G | PLES_25041 | . | . | . | . | . | . | . | . | C | C | . | . |
| NC_011770 | 2692438 | C | T | 3 | 10 | 2 | 0 | NON_SYNONYMOU | MODERAT | MISSENSE | gGg/gAg | G453E | PLES_25041 | . | . | . | . | . | . | . | . | T | T | . | . |
| NC_011770 | 2692504 | T | G | 4 | 10 | 2 | 0 | NON_SYNONYMOU | MODERAT | MISSENSE | aAt/aCt | N431T | PLES_25041 | . | . | . | . | . | . | . | . | G | G | . | . |
| NC_011770 | 2692505 | T | A | 4 | 10 | 2 | 0 | NON_SYNONYMOU | MODERAT | MISSENSE | Aat/Tat | N431Y | PLES_25041 | . | . | . | . | . | . | . | . | A | A | . | . |
| NC_011770 | 2692562 | T | G | 4 | 10 | 2 | 0 | NON_SYNONYMOU | MODERAT | MISSENSE | Aac/Cac | N412H | PLES_25041 | . | . | . | . | . | . | . | . | G | G | . | . |
| NC_011770 | 2692717 | A | T | 11 | 3 | 3 | 0 | NON_SYNONYMOU | MODERAT | MISSENSE | cTg/cAg | L360Q | PLES_25041 | | | | | . | T | . | . | T | T | | |
| NC_011770 | 2692723 | G | A | 11 | 3 | 3 | 0 | NON_SYNONYMOU | MODERAT | MISSENSE | gCc/gTc | A358V | PLES_25041 | | | | | . | A | . | . | A | A | | |
| NC_011770 | 2693470 | A | G | 8 | 10 | 2 | 0 | NON_SYNONYMOU | MODERAT | MISSENSE | gTc/gCc | V109A | PLES_25041 | . | . | . | . | . | . | . | . | . | . | G | G |
| NC_011770 | 2693638 | T | C | 5 | 10 | 2 | 0 | NON_SYNONYMOU | MODERAT | MISSENSE | gAc/gGc | D53G | PLES_25041 | . | . | . | . | . | . | . | . | . | . | C | C |
| NC_011770 | 2693639 | C | T | 5 | 10 | 2 | 0 | NON_SYNONYMOU | MODERAT | MISSENSE | Gac/Aac | D53N | PLES_25041 | . | . | . | . | . | . | . | . | . | . | T | T |
| NC_011770 | 2693694 | C | A | 6 | 10 | 2 | 0 | NON_SYNONYMOU | MODERAT | MISSENSE | aaG/aaT | K34N | PLES_25041 | . | . | . | . | . | . | . | . | . | . | A | A |
| NC_011770 | 2693707 | T | C | 5 | 10 | 2 | 0 | NON_SYNONYMOU | MODERAT | MISSENSE | gAc/gGc | D30G | PLES_25041 | . | . | . | . | . | . | . | . | . | . | C | C |
| NC_011770 | 2693713 | T | A | 5 | 10 | 2 | 0 | NON_SYNONYMOU | MODERAT | MISSENSE | cAg/cTg | Q28L | PLES_25041 | . | . | . | . | . | . | . | . | . | . | A | A |
| NC_011770 | 2693715 | A | C | 5 | 10 | 2 | 0 | NON_SYNONYMOU | MODERAT | MISSENSE | gaT/gaG | D27E | PLES_25041 | . | . | . | . | . | . | . | . | . | . | C | C |
| NC_011770 | 2693720 | C | CCGA | 5 | 10 | 2 | 0 | CODON_INSERTIO | MODERATE | | gcg/gcgTC | A25AS | PLES_25041 | . | . | . | . | . | . | . | . | . | . | CCGA | CCGA |
| NC_011770 | 2693746 | A | C | 9 | 8 | 4 | 0 | NON_SYNONYMOU | MODERAT | MISSENSE | gTc/gGc | V17G | PLES_25041 | . | . | . | . | . | . | . | . | C | C | C | C |
| NC_011770 | 2693747 | C | G | 9 | 8 | 4 | 0 | NON_SYNONYMOU | MODERAT | MISSENSE | Gtc/Ctc | V17L | PLES_25041 | . | . | . | . | . | . | . | . | G | G | G | G |
| NC_011770 | 2693779 | A | T | 8 | 8 | 2 | 0 | NON_SYNONYMOU | MODERAT | MISSENSE | aTt/aAt | I6N | PLES_25041 | . | . | . | . | . | . | . | . | . | . | T | T |
| NC_011770 | 2693780 | T | C | 8 | 8 | 2 | 0 | NON_SYNONYMOU | MODERAT | MISSENSE | Att/Gtt | I6V | PLES_25041 | . | . | . | . | . | . | . | . | . | . | C | C |
| NC_011770 | 2693781 | C | A | 8 | 8 | 2 | 0 | NON_SYNONYMOU | MODERAT | MISSENSE | aaG/aaT | K5N | PLES_25041 | . | . | . | . | . | . | . | . | . | . | A | A |
| NC_011770 | 2693786 | A | C | 9 | 8 | 2 | 0 | NON_SYNONYMOU | MODERAT | MISSENSE | Tca/Gca | S4A | PLES_25041 | . | . | . | . | . | . | . | . | C | C | | |
| NC_011770 | 2693789 | C | T | 9 | 8 | 4 | 0 | NON_SYNONYMOU | MODERAT | MISSENSE | Gaa/Aaa | E3K | PLES_25041 | . | . | . | . | . | . | . | . | T | T | T | T |
| NC_011770 | 2693799 | A | T | 11 | 8 | 2 | 0 | NON_SYNONYMOU | MODERAT | MISSENSE | cTc/cAc | L67H | PLES_25051 | . | . | . | . | . | . | . | . | | | T | T |
| NC_011770 | 2693875 | G | T | 8 | 8 | 2 | 0 | NON_SYNONYMOU | MODERAT | MISSENSE | Ctg/Atg | L42M | PLES_25051 | . | . | . | . | . | . | . | . | | | T | T |
| NC_011770 | 2693879 | A | C | 8 | 8 | 2 | 0 | NON_SYNONYMOU | MODERAT | MISSENSE | aaT/aaG | N40K | PLES_25051 | . | . | . | . | . | . | . | . | | | C | C |
| NC_011770 | 2693881 | T | C | 8 | 8 | 2 | 0 | NON_SYNONYMOU | MODERAT | MISSENSE | Aat/Gat | N40D | PLES_25051 | . | . | . | . | . | . | . | . | | | C | C |
| NC_011770 | 2693946 | G | C | 5 | 8 | 4 | 0 | NON_SYNONYMOU | MODERAT | MISSENSE | cCc/cGc | P18R | PLES_25051 | . | . | . | . | . | . | . | . | C | C | T | T |
| NC_011770 | 2693947 | G | T | 5 | 8 | 2 | 0 | NON_SYNONYMOU | MODERAT | MISSENSE | Ccc/Acc | P18T | PLES_25051 | . | . | . | . | . | . | . | . | | | T | T |
| NC_011770 | 2693950 | A | AGGG | 4 | 8 | 2 | 0 | CODON_INSERTIO | MODERATE | | tgc/tgcCCC | C16CP | PLES_25051 | . | . | . | . | . | . | . | . | | | AGGG | AGGG |
| NC_011770 | 2693995 | G | A | 7 | 8 | 2 | 0 | NON_SYNONYMOU | MODERAT | MISSENSE | Cca/Tca | P2S | PLES_25051 | . | . | . | . | . | . | . | . | | | A | A |
| NC_011770 | 2694031 | G | T | 7 | 8 | 2 | 0 | STOP_GAINED | HIGH | NONSENSE | tgC/tgA | C60* | PLES_25061 | . | . | . | . | . | . | . | . | | | T | T |
| NC_011770 | 2694039 | C | A | 6 | 8 | 2 | 0 | STOP_GAINED | HIGH | NONSENSE | Gag/Tag | E58* | PLES_25061 | . | . | . | . | . | . | . | . | | | A | A |
| NC_011770 | 2694000 | A | G | 6 | 8 | 2 | 0 | NON_SYNONYMOU | MODERAT | MISSENSE | Tca/Cca | S71P | PLES_25061 | . | . | . | . | . | . | . | . | | | G | G |
| NC_011770 | 2694002 | G | T | 6 | 8 | 2 | 0 | NON_SYNONYMOU | MODERAT | MISSENSE | cCa/cAa | P70Q | PLES_25061 | . | . | . | . | . | . | . | . | | | T | T |
| NC_011770 | 2694027 | T | C | 7 | 8 | 2 | 0 | NON_SYNONYMOU | MODERAT | MISSENSE | Acg/Gcg | T62A | PLES_25061 | . | . | . | . | . | . | . | . | | | C | C |
| NC_011770 | 2694032 | C | G | 7 | 8 | 2 | 0 | NON_SYNONYMOU | MODERAT | MISSENSE | tGc/tCc | C60S | PLES_25061 | . | . | . | . | . | . | . | . | | | G | G |
| NC_011770 | 2694033 | A | T | 7 | 8 | 2 | 0 | NON_SYNONYMOU | MODERAT | MISSENSE | Tgc/Agc | C60S | PLES_25061 | . | . | . | . | . | . | . | . | | | T | T |
| NC_011770 | 2694038 | T | A | 6 | 8 | 2 | 0 | NON_SYNONYMOU | MODERAT | MISSENSE | gAg/gTg | E58V | PLES_25061 | . | . | . | . | . | . | . | . | | | A | A |
| NC_011770 | 2694044 | C | G | 6 | 8 | 2 | 0 | NON_SYNONYMOU | MODERAT | MISSENSE | aGc/aCc | S56T | PLES_25061 | . | . | . | . | . | . | . | . | | | G | G |
| NC_011770 | 2694050 | C | T | 6 | 8 | 2 | 0 | NON_SYNONYMOU | MODERAT | MISSENSE | gGc/gAc | G54D | PLES_25061 | . | . | . | . | . | . | . | . | | | T | T |
| NC_011770 | 2694099 | G | T | 8 | 8 | 2 | 0 | NON_SYNONYMOU | MODERAT | MISSENSE | Cgc/Agc | R38S | PLES_25061 | . | . | . | . | . | . | . | . | T | T | | |
| NC_011770 | 2694108 | A | C | 8 | 8 | 2 | 0 | NON_SYNONYMOU | MODERAT | MISSENSE | Tct/Gct | S35A | PLES_25061 | . | . | . | . | . | . | . | . | C | C | | |
| NC_011770 | 2694619 | C | T | 9 | 10 | 2 | 0 | NON_SYNONYMOU | MODERAT | MISSENSE | Gac/Aac | D205N | PLES_25071 | . | . | . | . | T | T | . | . | . | . | . | . |

| | | | | | | | | | | | | | | | | | | | | | | | | | |
|---|---|---|---|---|---|---|---|---|---|---|---|---|---|---|---|---|---|---|---|---|---|---|---|---|---|
| NC_011770 | 2694648 | C | G | 9 | 10 | 2 | 0 | NON_SYNONYMOU | MODERAT | MISSENSE | gGg/gCg | G195A | PLES_25071 | . | . | . | . | G | G | . | . | . | . | . | . |
| NC_011770 | 2694651 | G | T | 9 | 10 | 2 | 0 | NON_SYNONYMOU | MODERAT | MISSENSE | cCa/cAa | P194Q | PLES_25071 | . | . | . | . | T | T | . | . | . | . | . | . |
| NC_011770 | 2694655 | T | C | 10 | 10 | 2 | 0 | NON_SYNONYMOU | MODERAT | MISSENSE | Aac/Gac | N193D | PLES_25071 | . | . | . | . | C | C | . | . | . | . | . | . |
| NC_011770 | 2694769 | A | G | 8 | 10 | 2 | 0 | NON_SYNONYMOU | MODERAT | MISSENSE | Tac/Cac | Y155H | PLES_25071 | . | . | . | . | G | G | . | . | . | . | . | . |
| NC_011770 | 2694806 | A | C | 8 | 10 | 2 | 0 | NON_SYNONYMOU | MODERAT | MISSENSE | gaT/gaG | D142E | PLES_25071 | . | . | . | . | C | C | . | . | . | . | . | . |
| NC_011770 | 2694854 | C | A | 7 | 10 | 2 | 0 | NON_SYNONYMOU | MODERAT | MISSENSE | aaG/aaT | K126N | PLES_25071 | . | . | . | . | A | A | . | . | . | . | . | . |
| NC_011770 | 2694879 | C | T | 6 | 10 | 2 | 0 | NON_SYNONYMOU | MODERAT | MISSENSE | aGc/aAc | S118N | PLES_25071 | . | . | . | . | T | T | . | . | . | . | . | . |
| NC_011770 | 2694880 | T | C | 6 | 10 | 2 | 0 | NON_SYNONYMOU | MODERAT | MISSENSE | Agc/Ggc | S118G | PLES_25071 | . | . | . | . | C | C | . | . | . | . | . | . |
| NC_011770 | 2694886 | T | A | 6 | 10 | 2 | 0 | NON_SYNONYMOU | MODERAT | MISSENSE | Agc/Tgc | S116C | PLES_25071 | . | . | . | . | A | A | . | . | . | . | . | . |
| NC_011770 | 2694888 | G | A | 6 | 10 | 2 | 0 | NON_SYNONYMOU | MODERAT | MISSENSE | cCc/cTc | P115L | PLES_25071 | . | . | . | . | A | A | . | . | . | . | . | . |
| NC_011770 | 2694889 | G | A | 6 | 10 | 2 | 0 | NON_SYNONYMOU | MODERAT | MISSENSE | Ccc/Tcc | P115S | PLES_25071 | . | . | . | . | A | A | . | . | . | . | . | . |
| NC_011770 | 2694935 | G | C | 7 | 10 | 2 | 0 | NON_SYNONYMOU | MODERAT | MISSENSE | gaC/gaG | D99E | PLES_25071 | . | . | . | . | C | C | . | . | . | . | . | . |
| NC_011770 | 2694996 | G | T | 1 | 10 | 2 | 0 | NON_SYNONYMOU | MODERAT | MISSENSE | gCa/gAa | A79E | PLES_25071 | . | . | . | . | . | . | . | . | T | T | . | . |
| NC_011770 | 2695009 | A | G | 1 | 10 | 2 | 0 | NON_SYNONYMOU | MODERAT | MISSENSE | Ttc/Ctc | F75L | PLES_25071 | . | . | . | . | . | . | . | . | G | G | . | . |
| NC_011770 | 2695023 | A | C | 1 | 10 | 2 | 0 | NON_SYNONYMOU | MODERAT | MISSENSE | aTc/aGc | I70S | PLES_25071 | . | . | . | . | . | . | . | . | C | C | . | . |
| NC_011770 | 2695273 | T | G | 2 | 10 | 2 | 0 | NON_SYNONYMOU | MODERAT | MISSENSE | aAg/aCg | K173T | PLES_25081 | . | . | . | . | . | . | . | . | G | G | . | . |
| NC_011770 | 2695313 | A | C | 12 | 6 | 2 | 0 | NON_SYNONYMOU | MODERAT | MISSENSE | Tct/Gct | S160A | PLES_25081 | . | . | . | . | . | . | | | C | C | | |
| NC_011770 | 2695315 | A | G | 12 | 6 | 2 | 0 | NON_SYNONYMOU | MODERAT | MISSENSE | gTc/gCc | V159A | PLES_25081 | . | . | . | . | . | . | | | G | G | | |
| NC_011770 | 2695316 | C | A | 11 | 6 | 2 | 0 | NON_SYNONYMOU | MODERAT | MISSENSE | Gtc/Ttc | V159F | PLES_25081 | . | . | . | . | . | . | | | A | A | | |
| NC_011770 | 2695349 | T | G | 6 | 8 | 4 | 0 | NON_SYNONYMOU | MODERAT | MISSENSE | Aaa/Caa | K148Q | PLES_25081 | . | . | . | . | . | . | . | . | G | G | G | G |
| NC_011770 | 2695381 | A | T | 6 | 8 | 2 | 0 | NON_SYNONYMOU | MODERAT | MISSENSE | cTc/cAc | L137H | PLES_25081 | . | . | . | . | . | . | . | . | T | T | | |
| NC_011770 | 2695421 | G | T | 5 | 8 | 2 | 0 | NON_SYNONYMOU | MODERAT | MISSENSE | Ccg/Acg | P124T | PLES_25081 | . | . | . | . | . | . | . | . | T | T | | |
| NC_011770 | 2695439 | T | C | 4 | 8 | 4 | 0 | NON_SYNONYMOU | MODERAT | MISSENSE | Att/Gtt | I118V | PLES_25081 | . | . | . | . | . | . | . | . | C | C | C | C |
| NC_011770 | 2695441 | C | G | 4 | 8 | 4 | 0 | NON_SYNONYMOU | MODERAT | MISSENSE | gGc/gCc | G117A | PLES_25081 | . | . | . | . | . | . | . | . | G | G | G | G |
| NC_011770 | 2695442 | C | A | 4 | 8 | 4 | 0 | NON_SYNONYMOU | MODERAT | MISSENSE | Ggc/Tgc | G117C | PLES_25081 | . | . | . | . | . | . | . | . | A | A | A | A |
| NC_011770 | 2695489 | T | G | 5 | 8 | 4 | 0 | NON_SYNONYMOU | MODERAT | MISSENSE | gAg/gCg | E101A | PLES_25081 | . | . | . | . | . | . | . | . | G | G | G | G |
| NC_011770 | 2695517 | A | G | 6 | 8 | 4 | 0 | NON_SYNONYMOU | MODERAT | MISSENSE | Tac/Cac | Y92H | PLES_25081 | . | . | . | . | . | . | . | . | G | G | G | G |
| NC_011770 | 2695649 | A | C | 7 | 8 | 2 | 0 | NON_SYNONYMOU | MODERAT | MISSENSE | Tcg/Gcg | S48A | PLES_25081 | . | . | . | . | . | . | . | . | C | C | | |
| NC_011770 | 2695686 | C | G | 5 | 8 | 4 | 0 | NON_SYNONYMOU | MODERAT | MISSENSE | atG/atC | M35I | PLES_25081 | . | . | . | . | . | . | . | . | G | G | G | G |
| NC_011770 | 2695688 | T | A | 5 | 8 | 2 | 0 | NON_SYNONYMOU | MODERAT | MISSENSE | Atg/Ttg | M35L | PLES_25081 | . | . | . | . | . | . | . | . | A | A | | |
| NC_011770 | 2695703 | G | T | 5 | 8 | 4 | 0 | NON_SYNONYMOU | MODERAT | MISSENSE | Cgc/Agc | R30S | PLES_25081 | . | . | . | . | . | . | . | . | T | T | T | T |
| NC_011770 | 2695711 | T | C | 5 | 8 | 4 | 0 | NON_SYNONYMOU | MODERAT | MISSENSE | aAg/aGg | K27R | PLES_25081 | . | . | . | . | . | . | . | . | C | C | C | C |
| NC_011770 | 2695712 | T | G | 5 | 8 | 4 | 0 | NON_SYNONYMOU | MODERAT | MISSENSE | Aag/Cag | K27Q | PLES_25081 | . | . | . | . | . | . | . | . | G | G | G | G |
| NC_011770 | 2696957 | C | A | 7 | 8 | 2 | 0 | NON_SYNONYMOU | MODERAT | MISSENSE | Gcc/Tcc | A65S | PLES_25101 | . | . | . | . | . | . | . | . | | | A | A |
| NC_011770 | 2696958 | A | T | 6 | 8 | 2 | 0 | NON_SYNONYMOU | MODERAT | MISSENSE | gaT/gaA | D64E | PLES_25101 | . | . | . | . | . | . | . | . | T | T | | |
| NC_011770 | 2696968 | G | C | 7 | 8 | 2 | 0 | NON_SYNONYMOU | MODERAT | MISSENSE | gCc/gGc | A61G | PLES_25101 | . | . | . | . | . | . | . | . | C | C | | |
| NC_011770 | 2696972 | A | C | 6 | 8 | 2 | 0 | NON_SYNONYMOU | MODERAT | MISSENSE | Tca/Gca | S60A | PLES_25101 | . | . | . | . | . | . | . | . | C | C | | |
| NC_011770 | 2697360 | C | A | 10 | 8 | 4 | 0 | NON_SYNONYMOU | MODERAT | MISSENSE | aaG/aaT | K82N | PLES_25111 | . | . | . | . | . | . | . | . | A | A | T | T |
| NC_011770 | 2697364 | G | T | 9 | 8 | 4 | 0 | NON_SYNONYMOU | MODERAT | MISSENSE | gCc/gAc | A81D | PLES_25111 | . | . | . | . | . | . | . | . | T | T | A | A |
| NC_011770 | 2697384 | G | C | 10 | 8 | 2 | 0 | NON_SYNONYMOU | MODERAT | MISSENSE | caC/caG | H74Q | PLES_25111 | . | . | . | . | . | . | . | . | | | C | C |
| NC_011770 | 2697405 | G | C | 10 | 8 | 4 | 0 | NON_SYNONYMOU | MODERAT | MISSENSE | gaC/gaG | D67E | PLES_25111 | . | . | . | . | . | . | . | . | C | C | T | T |
| NC_011770 | 2697436 | G | T | 11 | 8 | 2 | 0 | NON_SYNONYMOU | MODERAT | MISSENSE | gCg/gAg | A57E | PLES_25111 | . | . | . | . | . | . | . | . | T | T | | |
| NC_011770 | 2697437 | C | T | 10 | 8 | 2 | 0 | NON_SYNONYMOU | MODERAT | MISSENSE | Gcg/Acg | A57T | PLES_25111 | . | . | . | . | . | . | . | . | T | T | | |
| NC_011770 | 2697455 | T | C | 11 | 8 | 4 | 0 | NON_SYNONYMOU | MODERAT | MISSENSE | Atc/Gtc | I51V | PLES_25111 | . | . | . | . | . | . | . | . | C | C | C | C |
| NC_011770 | 2697490 | T | C | 10 | 8 | 2 | 0 | NON_SYNONYMOU | MODERAT | MISSENSE | cAa/cGa | Q39R | PLES_25111 | . | . | . | . | . | . | . | . | | | C | C |
| NC_011770 | 2697493 | T | A | 10 | 8 | 2 | 0 | NON_SYNONYMOU | MODERAT | MISSENSE | tAc/tTc | Y38F | PLES_25111 | . | . | . | . | . | . | . | . | | | A | A |
| NC_011770 | 2697505 | A | G | 11 | 8 | 2 | 0 | NON_SYNONYMOU | MODERAT | MISSENSE | gTt/gCt | V34A | PLES_25111 | . | . | . | . | . | . | . | . | G | G | | |
| NC_011770 | 2697521 | T | G | 12 | 8 | 4 | 0 | NON_SYNONYMOU | MODERAT | MISSENSE | Atc/Ctc | I29L | PLES_25111 | . | . | . | . | . | . | . | . | G | G | G | G |
| NC_011770 | 2697526 | A | T | 12 | 8 | 4 | 0 | NON_SYNONYMOU | MODERAT | MISSENSE | cTg/cAg | L27Q | PLES_25111 | . | . | . | . | . | . | . | . | T | T | T | T |
| NC_011770 | 2697529 | G | A | 12 | 8 | 4 | 0 | NON_SYNONYMOU | MODERAT | MISSENSE | gCt/gTt | A26V | PLES_25111 | . | . | . | . | . | . | . | . | A | A | A | A |
| NC_011770 | 2697536 | C | T | 12 | 8 | 2 | 0 | NON_SYNONYMOU | MODERAT | MISSENSE | Gcg/Acg | A24T | PLES_25111 | . | . | . | . | . | . | . | . | T | T | | |
| NC_011770 | 2697539 | T | G | 12 | 8 | 4 | 0 | NON_SYNONYMOU | MODERAT | MISSENSE | Atg/Ctg | M23L | PLES_25111 | . | . | . | . | . | . | . | . | G | G | G | G |
| NC_011770 | 2697557 | T | G | 11 | 8 | 4 | 0 | NON_SYNONYMOU | MODERAT | MISSENSE | Aaa/Caa | K17Q | PLES_25111 | . | . | . | . | . | . | . | . | G | G | G | G |
| NC_011770 | 2697621 | CGTT | C | 11 | 8 | 2 | 0 | CODON_DELETION | MODERATE | | aac/ | N127 | PLES_25121 | . | . | . | . | . | . | . | . | C | C | | |
| NC_011770 | 2697659 | G | A | 12 | 8 | 2 | 0 | NON_SYNONYMOU | MODERAT | MISSENSE | gCc/gTc | A115V | PLES_25121 | . | . | . | . | . | . | . | . | | | A | A |
| NC_011770 | 2697880 | C | G | 10 | 8 | 4 | 0 | NON_SYNONYMOU | MODERAT | MISSENSE | agG/agC | R41S | PLES_25121 | . | . | . | . | . | . | . | . | G | G | G | G |
| NC_011770 | 2697942 | G | T | 9 | 8 | 2 | 0 | NON_SYNONYMOU | MODERAT | MISSENSE | Cgc/Agc | R21S | PLES_25121 | . | . | . | . | . | . | . | . | | | T | T |
| NC_011770 | 2697961 | C | G | 8 | 8 | 2 | 0 | NON_SYNONYMOU | MODERAT | MISSENSE | caG/caC | Q14H | PLES_25121 | . | . | . | . | . | . | . | . | | | G | G |
| NC_011770 | 2697962 | T | G | 8 | 8 | 2 | 0 | NON_SYNONYMOU | MODERAT | MISSENSE | cAg/cCg | Q14P | PLES_25121 | . | . | . | . | . | . | . | . | | | G | G |
| NC_011770 | 2697963 | G | T | 8 | 8 | 2 | 0 | NON_SYNONYMOU | MODERAT | MISSENSE | Cag/Aag | Q14K | PLES_25121 | . | . | . | . | . | . | . | . | | | T | T |
| NC_011770 | 2697997 | G | T | 8 | 9 | 1 | 0 | NON_SYNONYMOU | MODERAT | MISSENSE | caC/caA | H2Q | PLES_25121 | . | . | . | . | . | . | . | . | T | . | | |
| NC_011770 | 2698395 | A | C | 3 | 9 | 3 | 0 | NON_SYNONYMOU | MODERAT | MISSENSE | Tcg/Gcg | S262A | PLES_25131 | . | . | . | . | . | . | . | . | . | C | C | C |

| | | | | | | | | | | | | | | | | | | | | | | | | | |
|---|---|---|---|---|---|---|---|---|---|---|---|---|---|---|---|---|---|---|---|---|---|---|---|---|---|
| NC_011770 | 2698485 | C | A | 6 | 10 | 2 | 0 | NON_SYNONYMOU | MODERAT | MISSENSE | Gcc/Tcc | A232S | PLES_25131 | . | . | . | . | . | . | . | . | . | . | A | A |
| NC_011770 | 2698497 | T | C | 6 | 10 | 2 | 0 | NON_SYNONYMOU | MODERAT | MISSENSE | Acc/Gcc | T228A | PLES_25131 | . | . | . | . | . | . | . | . | . | . | C | C |
| NC_011770 | 2698509 | C | G | 7 | 10 | 2 | 0 | NON_SYNONYMOU | MODERAT | MISSENSE | Gta/Cta | V224L | PLES_25131 | . | . | . | . | . | . | . | . | . | . | G | G |
| NC_011770 | 2698514 | C | T | 7 | 10 | 2 | 0 | NON_SYNONYMOU | MODERAT | MISSENSE | aGa/aAa | R222K | PLES_25131 | . | . | . | . | . | . | . | . | . | . | T | T |
| NC_011770 | 2698545 | T | G | 7 | 9 | 2 | 0 | NON_SYNONYMOU | MODERAT | MISSENSE | Atc/Ctc | I212L | PLES_25131 | . | . | . | . | . | . | . | . | . | | G | G |
| NC_011770 | 2698547 | A | G | 10 | 8 | 2 | 0 | NON_SYNONYMOU | MODERAT | MISSENSE | gTg/gCg | V211A | PLES_25131 | . | . | . | . | . | . | . | . | | | G | G |
| NC_011770 | 2698583 | G | T | 11 | 8 | 4 | 0 | NON_SYNONYMOU | MODERAT | MISSENSE | gCg/gAg | A199E | PLES_25131 | . | . | . | . | . | . | . | . | T | T | T | T |
| NC_011770 | 2698584 | C | T | 11 | 8 | 4 | 0 | NON_SYNONYMOU | MODERAT | MISSENSE | Gcg/Acg | A199T | PLES_25131 | . | . | . | . | . | . | . | . | T | T | T | T |
| NC_011770 | 2698709 | C | T | 10 | 8 | 4 | 0 | NON_SYNONYMOU | MODERAT | MISSENSE | cGc/cAc | R157H | PLES_25131 | . | . | . | . | . | . | . | . | T | T | T | T |
| NC_011770 | 2698710 | G | T | 10 | 8 | 4 | 0 | NON_SYNONYMOU | MODERAT | MISSENSE | Cgc/Agc | R157S | PLES_25131 | . | . | . | . | . | . | . | . | T | T | T | T |
| NC_011770 | 2698791 | A | C | 11 | 8 | 4 | 0 | NON_SYNONYMOU | MODERAT | MISSENSE | Tcc/Gcc | S130A | PLES_25131 | . | . | . | . | . | . | . | . | C | C | C | C |
| NC_011770 | 2698811 | G | C | 8 | 8 | 4 | 0 | NON_SYNONYMOU | MODERAT | MISSENSE | tCc/tGc | S123C | PLES_25131 | . | . | . | . | . | . | . | . | C | C | C | C |
| NC_011770 | 2698812 | A | T | 8 | 8 | 4 | 0 | NON_SYNONYMOU | MODERAT | MISSENSE | Tcc/Acc | S123T | PLES_25131 | . | . | . | . | . | . | . | . | T | T | T | T |
| NC_011770 | 2698969 | T | G | 10 | 8 | 2 | 0 | NON_SYNONYMOU | MODERAT | MISSENSE | gaA/gaC | E70D | PLES_25131 | . | . | . | . | . | . | . | . | G | G | | |
| NC_011770 | 2703347 | A | T | 7 | 10 | 2 | 0 | NON_SYNONYMOU | MODERAT | MISSENSE | Acc/Tcc | T12S | PLES_25191 | T | T | . | . | . | . | . | . | . | . | . | . |
| NC_011770 | 2703456 | A | G | 8 | 10 | 2 | 0 | NON_SYNONYMOU | MODERAT | MISSENSE | gAc/gGc | D48G | PLES_25191 | G | G | . | . | . | . | . | . | . | . | . | . |
| NC_011770 | 2703485 | A | C | 7 | 10 | 2 | 0 | NON_SYNONYMOU | MODERAT | MISSENSE | Atc/Ctc | I58L | PLES_25191 | C | C | . | . | . | . | . | . | . | . | . | . |
| NC_011770 | 2703513 | C | A | 10 | 10 | 2 | 0 | NON_SYNONYMOU | MODERAT | MISSENSE | gCc/gAc | A67D | PLES_25191 | A | A | . | . | . | . | . | . | . | . | . | . |
| NC_011770 | 2703541 | A | T | 10 | 10 | 2 | 0 | NON_SYNONYMOU | MODERAT | MISSENSE | aaA/aaT | K76N | PLES_25191 | T | T | . | . | . | . | . | . | . | . | . | . |
| NC_011770 | 2703560 | G | C | 10 | 10 | 2 | 0 | NON_SYNONYMOU | MODERAT | MISSENSE | Gag/Cag | E83Q | PLES_25191 | C | C | . | . | . | . | . | . | . | . | . | . |
| NC_011770 | 2704163 | C | T | 7 | 10 | 2 | 0 | NON_SYNONYMOU | MODERAT | MISSENSE | Cac/Tac | H57Y | PLES_25201 | T | T | . | . | . | . | . | . | . | . | . | . |
| NC_011770 | 2704229 | G | T | 7 | 10 | 2 | 0 | NON_SYNONYMOU | MODERAT | MISSENSE | Gtg/Ttg | V79L | PLES_25201 | T | T | . | . | . | . | . | . | . | . | . | . |
| NC_011770 | 2705983 | T | C | 15 | 10 | 2 | 0 | NON_SYNONYMOU | MODERAT | MISSENSE | Tcg/Ccg | S242P | PLES_25221 | . | . | . | . | C | C | . | . | . | . | . | . |
| NC_011770 | 2706042 | C | A | 16 | 10 | 2 | 0 | NON_SYNONYMOU | MODERAT | MISSENSE | gaC/gaA | D261E | PLES_25221 | . | . | . | . | A | A | . | . | . | . | . | . |
| NC_011770 | 2708389 | T | C | 22 | 10 | 2 | 0 | NON_SYNONYMOU | MODERAT | MISSENSE | Tcc/Ccc | S100P | PLES_25241 | . | . | . | . | C | C | . | . | . | . | . | . |
| NC_011770 | 2708545 | G | C | 19 | 10 | 2 | 0 | NON_SYNONYMOU | MODERAT | MISSENSE | Gaa/Caa | E152Q | PLES_25241 | . | . | . | . | C | C | . | . | . | . | . | . |
| NC_011770 | 2710184 | T | A | 6 | 10 | 2 | 0 | NON_SYNONYMOU | MODERAT | MISSENSE | gaT/gaA | D60E | PLES_25271 | . | . | . | . | A | A | . | . | . | . | . | . |
| NC_011770 | 2710188 | G | A | 6 | 10 | 2 | 0 | NON_SYNONYMOU | MODERAT | MISSENSE | Ggc/Agc | G62S | PLES_25271 | . | . | . | . | A | A | . | . | . | . | . | . |
| NC_011770 | 2710189 | G | A | 6 | 10 | 2 | 0 | NON_SYNONYMOU | MODERAT | MISSENSE | gGc/gAc | G62D | PLES_25271 | . | . | . | . | A | A | . | . | . | . | . | . |
| NC_011770 | 2710191 | G | A | 6 | 10 | 2 | 0 | NON_SYNONYMOU | MODERAT | MISSENSE | Gcc/Acc | A63T | PLES_25271 | . | . | . | . | A | A | . | . | . | . | . | . |
| NC_011770 | 2710192 | C | A | 6 | 10 | 2 | 0 | NON_SYNONYMOU | MODERAT | MISSENSE | gCc/gAc | A63D | PLES_25271 | . | . | . | . | A | A | . | . | . | . | . | . |
| NC_011770 | 2710765 | T | G | 13 | 8 | 4 | 0 | NON_SYNONYMOU | MODERAT | MISSENSE | Tca/Gca | S45A | PLES_25281 | G | G | . | . | G | G | . | . | . | . | . | . |
| NC_011770 | 2710770 | G | C | 13 | 8 | 2 | 0 | NON_SYNONYMOU | MODERAT | MISSENSE | aaG/aaC | K46N | PLES_25281 | C | C | . | . | | | . | . | . | . | . | . |
| NC_011770 | 2710838 | C | A | 9 | 8 | 2 | 0 | NON_SYNONYMOU | MODERAT | MISSENSE | aCg/aAg | T69K | PLES_25281 | | | . | . | A | A | . | . | . | . | . | . |
| NC_011770 | 2710849 | C | T | 9 | 8 | 2 | 0 | NON_SYNONYMOU | MODERAT | MISSENSE | Cac/Tac | H73Y | PLES_25281 | | | . | . | T | T | . | . | . | . | . | . |
| NC_011770 | 2710850 | A | T | 9 | 8 | 2 | 0 | NON_SYNONYMOU | MODERAT | MISSENSE | cAc/cTc | H73L | PLES_25281 | | | . | . | T | T | . | . | . | . | . | . |
| NC_011770 | 2710852 | A | G | 9 | 8 | 2 | 0 | NON_SYNONYMOU | MODERAT | MISSENSE | Atc/Gtc | I74V | PLES_25281 | | | . | . | G | G | . | . | . | . | . | . |
| NC_011770 | 2710854 | C | G | 9 | 8 | 2 | 0 | NON_SYNONYMOU | MODERAT | MISSENSE | atC/atG | I74M | PLES_25281 | | | . | . | G | G | . | . | . | . | . | . |
| NC_011770 | 2711186 | G | A | 24 | 8 | 2 | 0 | NON_SYNONYMOU | MODERAT | MISSENSE | cGt/cAt | R185H | PLES_25281 | A | A | . | . | | | . | . | . | . | . | . |
| NC_011770 | 2711420 | A | T | 28 | 8 | 4 | 0 | NON_SYNONYMOU | MODERAT | MISSENSE | gaA/gaT | E50D | PLES_25291 | T | T | . | . | T | T | . | . | . | . | . | . |
| NC_011770 | 2711445 | T | C | 29 | 8 | 4 | 0 | NON_SYNONYMOU | MODERAT | MISSENSE | Tac/Cac | Y59H | PLES_25291 | C | C | . | . | C | C | . | . | . | . | . | . |
| NC_011770 | 2711476 | A | G | 31 | 8 | 4 | 0 | NON_SYNONYMOU | MODERAT | MISSENSE | aAg/aGg | K69R | PLES_25291 | G | G | . | . | G | G | . | . | . | . | . | . |
| NC_011770 | 2711529 | G | A | 29 | 8 | 2 | 0 | NON_SYNONYMOU | MODERAT | MISSENSE | Gtg/Atg | V87M | PLES_25291 | A | A | . | . | | | . | . | . | . | . | . |
| NC_011770 | 2711615 | C | A | 29 | 8 | 2 | 0 | NON_SYNONYMOU | MODERAT | MISSENSE | gaC/gaA | D115E | PLES_25291 | A | A | . | . | | | . | . | . | . | . | . |
| NC_011770 | 2711644 | G | A | 30 | 8 | 2 | 0 | NON_SYNONYMOU | MODERAT | MISSENSE | aGg/aAg | R125K | PLES_25291 | A | A | . | . | | | . | . | . | . | . | . |
| NC_011770 | 2711657 | A | C | 30 | 8 | 2 | 0 | NON_SYNONYMOU | MODERAT | MISSENSE | gaA/gaC | E129D | PLES_25291 | | | . | . | C | C | . | . | . | . | . | . |
| NC_011770 | 2711871 | G | T | 0 | 11 | 1 | 0 | NON_SYNONYMOU | MODERAT | MISSENSE | Gcc/Tcc | A201S | PLES_25291 | . | . | . | . | . | T | . | . | . | . | . | . |
| NC_011770 | 2712563 | C | T | 5 | 10 | 2 | 0 | INTRAGENIC | MODIFIER | | | | PLES_25292 | T | T | . | . | . | . | . | . | . | . | . | . |
| NC_011770 | 2712572 | A | G | 4 | 10 | 2 | 0 | INTRAGENIC | MODIFIER | | | | PLES_25292 | G | G | . | . | . | . | . | . | . | . | . | . |
| NC_011770 | 2712606 | T | C | 13 | 8 | 2 | 0 | INTRAGENIC | MODIFIER | | | | PLES_25292 | C | C | . | . | | | . | . | . | . | . | . |
| NC_011770 | 2712670 | G | A | 6 | 10 | 2 | 0 | INTRAGENIC | MODIFIER | | | | PLES_25293 | A | A | . | . | . | . | . | . | . | . | . | . |
| NC_011770 | 2712676 | C | T | 6 | 10 | 2 | 0 | INTRAGENIC | MODIFIER | | | | PLES_25293 | T | T | . | . | . | . | . | . | . | . | . | . |
| NC_011770 | 2712691 | G | A | 6 | 10 | 2 | 0 | INTRAGENIC | MODIFIER | | | | PLES_25293 | A | A | . | . | . | . | . | . | . | . | . | . |
| NC_011770 | 2713013 | T | C | 17 | 8 | 2 | 0 | NON_SYNONYMOU | MODERAT | MISSENSE | gTc/gCc | V59A | PLES_25311 | | | . | . | C | C | . | . | . | . | . | . |
| NC_011770 | 2713016 | T | C | 16 | 8 | 2 | 0 | NON_SYNONYMOU | MODERAT | MISSENSE | gTg/gCg | V60A | PLES_25311 | | | . | . | C | C | . | . | . | . | . | . |
| NC_011770 | 2713094 | G | T | 14 | 8 | 2 | 0 | NON_SYNONYMOU | MODERAT | MISSENSE | tGt/tTt | C86F | PLES_25311 | | | . | . | T | T | . | . | . | . | . | . |
| NC_011770 | 2714427 | G | A | 24 | 8 | 2 | 0 | NON_SYNONYMOU | MODERAT | MISSENSE | aGg/aAg | R4K | PLES_25361 | A | A | . | . | | | . | . | . | . | . | . |
| NC_011770 | 2714428 | G | T | 24 | 8 | 2 | 0 | NON_SYNONYMOU | MODERAT | MISSENSE | agG/agT | R4S | PLES_25361 | | | . | . | T | T | . | . | . | . | . | . |
| NC_011770 | 2714430 | C | T | 24 | 8 | 2 | 0 | NON_SYNONYMOU | MODERAT | MISSENSE | gCg/gTg | A5V | PLES_25361 | | | . | . | T | T | . | . | . | . | . | . |
| NC_011770 | 2714442 | A | G | 25 | 8 | 2 | 0 | NON_SYNONYMOU | MODERAT | MISSENSE | cAt/cGt | H9R | PLES_25361 | G | G | . | . | | | . | . | . | . | . | . |
| NC_011770 | 2714451 | G | A | 24 | 8 | 2 | 0 | NON_SYNONYMOU | MODERAT | MISSENSE | gGc/gAc | G12D | PLES_25361 | A | A | . | . | | | . | . | . | . | . | . |
| NC_011770 | 2714454 | A | G | 24 | 8 | 2 | 0 | NON_SYNONYMOU | MODERAT | MISSENSE | cAg/cGg | Q13R | PLES_25361 | G | G | . | . | | | . | . | . | . | . | . |

| | | | | | | | | | | | | | | | | | | | | | | | | | |
|---|---|---|---|---|---|---|---|---|---|---|---|---|---|---|---|---|---|---|---|---|---|---|---|---|---|
| NC_011770 | 2714463 | C | T | 20 | 9 | 1 | 0 | NON_SYNONYMOU | MODERAT | MISSENSE | cCa/cTa | P16L | PLES_25361 | T | . | . | . | | | . | . | . | . | . | . |
| NC_011770 | 2714502 | T | C | 20 | 10 | 2 | 0 | NON_SYNONYMOU | MODERAT | MISSENSE | tTa/tCa | L29S | PLES_25361 | . | . | . | . | C | C | . | . | . | . | . | . |
| NC_011770 | 2714660 | C | G | 24 | 8 | 4 | 0 | NON_SYNONYMOU | MODERAT | MISSENSE | Cca/Gca | P82A | PLES_25361 | G | G | . | . | G | G | . | . | . | . | . | . |
| NC_011770 | 2714675 | G | A | 25 | 8 | 2 | 0 | NON_SYNONYMOU | MODERAT | MISSENSE | Gtg/Atg | V87M | PLES_25361 | | | . | . | A | A | . | . | . | . | . | . |
| NC_011770 | 2714684 | A | G | 25 | 8 | 4 | 0 | NON_SYNONYMOU | MODERAT | MISSENSE | Atc/Gtc | I90V | PLES_25361 | G | G | . | . | G | G | . | . | . | . | . | . |
| NC_011770 | 2714765 | C | T | 23 | 8 | 2 | 0 | NON_SYNONYMOU | MODERAT | MISSENSE | Cac/Tac | H117Y | PLES_25361 | T | T | . | . | | | . | . | . | . | . | . |
| NC_011770 | 2714805 | C | A | 21 | 8 | 2 | 0 | NON_SYNONYMOU | MODERAT | MISSENSE | gCa/gAa | A130E | PLES_25361 | A | A | . | . | | | . | . | . | . | . | . |
| NC_011770 | 2714814 | A | G | 21 | 8 | 2 | 0 | NON_SYNONYMOU | MODERAT | MISSENSE | aAa/aGa | K133R | PLES_25361 | G | G | . | . | | | . | . | . | . | . | . |
| NC_011770 | 2715067 | T | C | 19 | 8 | 2 | 0 | NON_SYNONYMOU | MODERAT | MISSENSE | Ttc/Ctc | F13L | PLES_25371 | C | C | . | . | | | . | . | . | . | . | . |
| NC_011770 | 2715132 | G | T | 14 | 10 | 2 | 0 | NON_SYNONYMOU | MODERAT | MISSENSE | gaG/gaT | E34D | PLES_25371 | . | . | . | . | T | T | . | . | . | . | . | . |
| NC_011770 | 2715271 | C | A | 18 | 8 | 2 | 0 | NON_SYNONYMOU | MODERAT | MISSENSE | Cgt/Agt | R81S | PLES_25371 | A | A | . | . | | | . | . | . | . | . | . |
| NC_011770 | 2715274 | G | T | 18 | 8 | 2 | 0 | NON_SYNONYMOU | MODERAT | MISSENSE | Gcc/Tcc | A82S | PLES_25371 | T | T | . | . | | | . | . | . | . | . | . |
| NC_011770 | 2715283 | A | T | 17 | 8 | 2 | 0 | NON_SYNONYMOU | MODERAT | MISSENSE | Aca/Tca | T85S | PLES_25371 | T | T | . | . | | | . | . | . | . | . | . |
| NC_011770 | 2715387 | G | T | 23 | 8 | 2 | 0 | NON_SYNONYMOU | MODERAT | MISSENSE | gaG/gaT | E119D | PLES_25371 | T | T | . | . | | | . | . | . | . | . | . |
| NC_011770 | 2715404 | A | C | 22 | 8 | 2 | 0 | NON_SYNONYMOU | MODERAT | MISSENSE | gAt/gCt | D125A | PLES_25371 | C | C | . | . | | | . | . | . | . | . | . |
| NC_011770 | 2715458 | A | G | 25 | 8 | 4 | 0 | NON_SYNONYMOU | MODERAT | MISSENSE | Agt/Ggt | S16G | PLES_25381 | G | G | . | . | G | G | . | . | . | . | . | . |
| NC_011770 | 2715465 | G | C | 24 | 8 | 2 | 0 | NON_SYNONYMOU | MODERAT | MISSENSE | gGt/gCt | G18A | PLES_25381 | | | . | . | C | C | . | . | . | . | . | . |
| NC_011770 | 2715491 | C | T | 23 | 8 | 2 | 0 | NON_SYNONYMOU | MODERAT | MISSENSE | Ctt/Ttt | L27F | PLES_25381 | T | T | . | . | | | . | . | . | . | . | . |
| NC_011770 | 2715523 | C | A | 23 | 8 | 2 | 0 | NON_SYNONYMOU | MODERAT | MISSENSE | gaC/gaA | D37E | PLES_25381 | A | A | . | . | | | . | . | . | . | . | . |
| NC_011770 | 2715530 | G | A | 23 | 8 | 2 | 0 | NON_SYNONYMOU | MODERAT | MISSENSE | Ggc/Agc | G40S | PLES_25381 | | | . | . | A | A | . | . | . | . | . | . |
| NC_011770 | 2715531 | G | A | 23 | 8 | 2 | 0 | NON_SYNONYMOU | MODERAT | MISSENSE | gGc/gAc | G40D | PLES_25381 | A | A | . | . | | | . | . | . | . | . | . |
| NC_011770 | 2715551 | G | T | 23 | 8 | 2 | 0 | NON_SYNONYMOU | MODERAT | MISSENSE | Gtt/Ttt | V47F | PLES_25381 | | | . | . | T | T | . | . | . | . | . | . |
| NC_011770 | 2715626 | G | T | 21 | 8 | 2 | 0 | NON_SYNONYMOU | MODERAT | MISSENSE | Gct/Tct | A72S | PLES_25381 | T | T | . | . | | | . | . | . | . | . | . |
| NC_011770 | 2716259 | C | T | 26 | 8 | 2 | 0 | NON_SYNONYMOU | MODERAT | MISSENSE | Ccg/Tcg | P283S | PLES_25381 | | | . | . | T | T | . | . | . | . | . | . |
| NC_011770 | 2716467 | A | C | 27 | 8 | 4 | 0 | NON_SYNONYMOU | MODERAT | MISSENSE | aAg/aCg | K352T | PLES_25381 | C | C | . | . | C | C | . | . | . | . | . | . |
| NC_011770 | 2716500 | T | C | 25 | 8 | 2 | 0 | NON_SYNONYMOU | MODERAT | MISSENSE | aTg/aCg | M363T | PLES_25381 | | | . | . | C | C | . | . | . | . | . | . |
| NC_011770 | 2716656 | T | C | 26 | 8 | 4 | 0 | NON_SYNONYMOU | MODERAT | MISSENSE | aTc/aCc | I415T | PLES_25381 | C | C | . | . | C | C | . | . | . | . | . | . |
| NC_011770 | 2716740 | C | T | 25 | 8 | 4 | 0 | NON_SYNONYMOU | MODERAT | MISSENSE | gCc/gTc | A443V | PLES_25381 | T | T | . | . | T | T | . | . | . | . | . | . |
| NC_011770 | 2717309 | T | G | 28 | 8 | 4 | 0 | NON_SYNONYMOU | MODERAT | MISSENSE | Tct/Gct | S18A | PLES_25401 | G | G | . | . | G | G | . | . | . | . | . | . |
| NC_011770 | 2717327 | A | G | 25 | 8 | 4 | 0 | NON_SYNONYMOU | MODERAT | MISSENSE | Aag/Gag | K24E | PLES_25401 | G | G | . | . | G | G | . | . | . | . | . | . |
| NC_011770 | 2717328 | A | G | 26 | 8 | 4 | 0 | NON_SYNONYMOU | MODERAT | MISSENSE | aAg/aGg | K24R | PLES_25401 | G | G | . | . | G | G | . | . | . | . | . | . |
| NC_011770 | 2717825 | A | G | 25 | 8 | 4 | 0 | NON_SYNONYMOU | MODERAT | MISSENSE | Att/Gtt | I190V | PLES_25401 | G | G | . | . | G | G | . | . | . | . | . | . |
| NC_011770 | 2718044 | G | A | 28 | 8 | 4 | 0 | NON_SYNONYMOU | MODERAT | MISSENSE | Gtt/Att | V263I | PLES_25401 | A | A | . | . | A | A | . | . | . | . | . | . |
| NC_011770 | 2718497 | G | A | 21 | 8 | 4 | 0 | NON_SYNONYMOU | MODERAT | MISSENSE | Gcg/Acg | A414T | PLES_25401 | A | A | . | . | A | A | . | . | . | . | . | . |
| NC_011770 | 2718763 | T | G | 23 | 8 | 2 | 0 | NON_SYNONYMOU | MODERAT | MISSENSE | Tct/Gct | S38A | PLES_25411 | G | G | . | . | | | . | . | . | . | . | . |
| NC_011770 | 2718781 | T | A | 23 | 8 | 4 | 0 | NON_SYNONYMOU | MODERAT | MISSENSE | Tcc/Acc | S44T | PLES_25411 | A | A | . | . | A | A | . | . | . | . | . | . |
| NC_011770 | 2718868 | T | G | 21 | 8 | 4 | 0 | NON_SYNONYMOU | MODERAT | MISSENSE | Tcc/Gcc | S73A | PLES_25411 | G | G | . | . | G | G | . | . | . | . | . | . |
| NC_011770 | 2718877 | A | G | 22 | 8 | 4 | 0 | NON_SYNONYMOU | MODERAT | MISSENSE | Agc/Ggc | S76G | PLES_25411 | G | G | . | . | G | G | . | . | . | . | . | . |
| NC_011770 | 2718883 | A | G | 22 | 8 | 4 | 0 | NON_SYNONYMOU | MODERAT | MISSENSE | Atc/Gtc | I78V | PLES_25411 | G | G | . | . | G | G | . | . | . | . | . | . |
| NC_011770 | 2718893 | C | T | 21 | 8 | 2 | 0 | NON_SYNONYMOU | MODERAT | MISSENSE | aCc/aTc | T81I | PLES_25411 | | | . | . | T | T | . | . | . | . | . | . |
| NC_011770 | 2718977 | C | T | 24 | 8 | 2 | 0 | NON_SYNONYMOU | MODERAT | MISSENSE | aCg/aTg | T109M | PLES_25411 | T | T | . | . | | | . | . | . | . | . | . |
| NC_011770 | 2719575 | G | A | 30 | 8 | 2 | 0 | NON_SYNONYMOU | MODERAT | MISSENSE | Gcc/Acc | A13T | PLES_25421 | | | . | . | A | A | . | . | . | . | . | . |
| NC_011770 | 2719657 | A | G | 33 | 8 | 4 | 0 | NON_SYNONYMOU | MODERAT | MISSENSE | aAc/aGc | N40S | PLES_25421 | G | G | . | . | G | G | . | . | . | . | . | . |
| NC_011770 | 2719695 | G | A | 32 | 8 | 2 | 0 | NON_SYNONYMOU | MODERAT | MISSENSE | Gct/Act | A53T | PLES_25421 | A | A | . | . | | | . | . | . | . | . | . |
| NC_011770 | 2719696 | C | A | 32 | 8 | 2 | 0 | NON_SYNONYMOU | MODERAT | MISSENSE | gCt/gAt | A53D | PLES_25421 | A | A | . | . | | | . | . | . | . | . | . |
| NC_011770 | 2719905 | G | T | 26 | 8 | 2 | 0 | NON_SYNONYMOU | MODERAT | MISSENSE | Gcc/Tcc | A123S | PLES_25421 | T | T | . | . | | | . | . | . | . | . | . |
| NC_011770 | 2720064 | A | G | 29 | 8 | 2 | 0 | NON_SYNONYMOU | MODERAT | MISSENSE | Acc/Gcc | T176A | PLES_25421 | | | . | . | G | G | . | . | . | . | . | . |
| NC_011770 | 2720088 | G | T | 28 | 8 | 4 | 0 | NON_SYNONYMOU | MODERAT | MISSENSE | Gcc/Tcc | A184S | PLES_25421 | T | T | . | . | A | A | . | . | . | . | . | . |
| NC_011770 | 2720089 | C | T | 28 | 8 | 2 | 0 | NON_SYNONYMOU | MODERAT | MISSENSE | gCc/gTc | A184V | PLES_25421 | T | T | . | . | | | . | . | . | . | . | . |
| NC_011770 | 2720463 | A | G | 24 | 8 | 4 | 0 | NON_SYNONYMOU | MODERAT | MISSENSE | Agc/Ggc | S309G | PLES_25421 | G | G | . | . | G | G | . | . | . | . | . | . |
| NC_011770 | 2721092 | G | A | 24 | 8 | 2 | 0 | NON_SYNONYMOU | MODERAT | MISSENSE | aGg/aAg | R67K | PLES_25431 | A | A | . | . | | | . | . | . | . | . | . |
| NC_011770 | 2721095 | C | T | 24 | 8 | 2 | 0 | NON_SYNONYMOU | MODERAT | MISSENSE | gCc/gTc | A68V | PLES_25431 | T | T | . | . | | | . | . | . | . | . | . |
| NC_011770 | 2721107 | G | A | 24 | 8 | 2 | 0 | NON_SYNONYMOU | MODERAT | MISSENSE | aGa/aAa | R72K | PLES_25431 | | | . | . | A | A | . | . | . | . | . | . |
| NC_011770 | 2721113 | T | C | 23 | 8 | 4 | 0 | NON_SYNONYMOU | MODERAT | MISSENSE | gTc/gCc | V74A | PLES_25431 | C | C | . | . | C | C | . | . | . | . | . | . |
| NC_011770 | 2721131 | A | C | 22 | 8 | 2 | 0 | NON_SYNONYMOU | MODERAT | MISSENSE | aAc/aCc | N80T | PLES_25431 | C | C | . | . | | | . | . | . | . | . | . |
| NC_011770 | 2721132 | C | G | 23 | 8 | 2 | 0 | NON_SYNONYMOU | MODERAT | MISSENSE | aaC/aaG | N80K | PLES_25431 | G | G | . | . | | | . | . | . | . | . | . |
| NC_011770 | 2721374 | G | T | 17 | 10 | 2 | 0 | NON_SYNONYMOU | MODERAT | MISSENSE | cGc/cTc | R161L | PLES_25431 | . | . | . | . | T | T | . | . | . | . | . | . |
| NC_011770 | 2721416 | G | C | 21 | 8 | 4 | 0 | NON_SYNONYMOU | MODERAT | MISSENSE | aGt/aCt | S175T | PLES_25431 | C | C | . | . | C | C | . | . | . | . | . | . |
| NC_011770 | 2721417 | T | A | 21 | 8 | 2 | 0 | NON_SYNONYMOU | MODERAT | MISSENSE | agT/agA | S175R | PLES_25431 | A | A | . | . | | | . | . | . | . | . | . |
| NC_011770 | 2721448 | C | A | 22 | 8 | 2 | 0 | NON_SYNONYMOU | MODERAT | MISSENSE | Cgc/Agc | R186S | PLES_25431 | A | A | . | . | | | . | . | . | . | . | . |
| NC_011770 | 2721462 | A | C | 24 | 8 | 2 | 0 | NON_SYNONYMOU | MODERAT | MISSENSE | gaA/gaC | E190D | PLES_25431 | C | C | . | . | | | . | . | . | . | . | . |
| NC_011770 | 2721464 | T | C | 23 | 8 | 2 | 0 | NON_SYNONYMOU | MODERAT | MISSENSE | gTg/gCg | V191A | PLES_25431 | C | C | . | . | | | . | . | . | . | . | . |

| | | | | | | | | | | | | | | | | | | | | | | | | | |
|---|---|---|---|---|---|---|---|---|---|---|---|---|---|---|---|---|---|---|---|---|---|---|---|---|---|
| NC_011770 | 2721476 | A | C | 24 | 8 | 2 | 0 | NON_SYNONYMOU | MODERAT | MISSENSE | cAt/cCt | H195P | PLES_25431 | C | C | . | . | | | . | . | . | . | . | . |
| NC_011770 | 2721636 | T | C | 23 | 8 | 2 | 0 | NON_SYNONYMOU | MODERAT | MISSENSE | gTa/gCa | V56A | PLES_25441 | C | C | . | . | | | . | . | . | . | . | . |
| NC_011770 | 2721641 | A | G | 23 | 8 | 2 | 0 | NON_SYNONYMOU | MODERAT | MISSENSE | Agc/Ggc | S58G | PLES_25441 | | | . | . | G | G | . | . | . | . | . | . |
| NC_011770 | 2721656 | A | C | 23 | 8 | 2 | 0 | NON_SYNONYMOU | MODERAT | MISSENSE | Att/Ctt | I63L | PLES_25441 | C | C | . | . | | | . | . | . | . | . | . |
| NC_011770 | 2721721 | A | T | 25 | 8 | 4 | 0 | NON_SYNONYMOU | MODERAT | MISSENSE | gaA/gaT | E84D | PLES_25441 | T | T | . | . | G | G | . | . | . | . | . | . |
| NC_011770 | 2721986 | C | A | 3 | 9 | 3 | 0 | NON_SYNONYMOU | MODERAT | MISSENSE | Cgc/Agc | R66S | PLES_25451 | A | A | . | . | . | A | . | . | . | . | . | . |
| NC_011770 | 2722015 | T | G | 15 | 8 | 4 | 0 | NON_SYNONYMOU | MODERAT | MISSENSE | gaT/gaG | D75E | PLES_25451 | G | G | . | . | A | A | . | . | . | . | . | . |
| NC_011770 | 2722040 | A | G | 12 | 8 | 4 | 0 | NON_SYNONYMOU | MODERAT | MISSENSE | Att/Gtt | I84V | PLES_25451 | G | G | . | . | G | G | . | . | . | . | . | . |
| NC_011770 | 2722043 | C | A | 13 | 8 | 4 | 0 | NON_SYNONYMOU | MODERAT | MISSENSE | Ctt/Att | L85I | PLES_25451 | A | A | . | . | A | A | . | . | . | . | . | . |
| NC_011770 | 2722089 | C | T | 16 | 8 | 4 | 0 | NON_SYNONYMOU | MODERAT | MISSENSE | gCc/gTc | A100V | PLES_25451 | T | T | . | . | T | T | . | . | . | . | . | . |
| NC_011770 | 2722143 | A | G | 4 | 10 | 2 | 0 | NON_SYNONYMOU | MODERAT | MISSENSE | cAg/cGg | Q118R | PLES_25451 | . | . | . | . | G | G | . | . | . | . | . | . |
| NC_011770 | 2723373 | T | A | 0 | 11 | 1 | 0 | NON_SYNONYMOU | MODERAT | MISSENSE | tTt/tAt | F21Y | PLES_25471 | . | . | . | . | . | A | . | . | . | . | . | . |
| NC_011770 | 2723651 | A | G | 10 | 8 | 2 | 0 | NON_SYNONYMOU | MODERAT | MISSENSE | Acg/Gcg | T114A | PLES_25471 | | | . | . | G | G | . | . | . | . | . | . |
| NC_011770 | 2723805 | G | A | 14 | 10 | 2 | 0 | NON_SYNONYMOU | MODERAT | MISSENSE | cGg/cAg | R165Q | PLES_25471 | . | . | . | . | A | A | . | . | . | . | . | . |
| NC_011770 | 2723813 | C | A | 14 | 10 | 2 | 0 | NON_SYNONYMOU | MODERAT | MISSENSE | Cat/Aat | H168N | PLES_25471 | . | . | . | . | A | A | . | . | . | . | . | . |
| NC_011770 | 2723815 | T | G | 14 | 10 | 2 | 0 | NON_SYNONYMOU | MODERAT | MISSENSE | caT/caG | H168Q | PLES_25471 | . | . | . | . | G | G | . | . | . | . | . | . |
| NC_011770 | 2723843 | T | G | 17 | 10 | 2 | 0 | NON_SYNONYMOU | MODERAT | MISSENSE | Tcc/Gcc | S178A | PLES_25471 | . | . | . | . | G | G | . | . | . | . | . | . |
| NC_011770 | 2723915 | G | C | 24 | 8 | 2 | 0 | NON_SYNONYMOU | MODERAT | MISSENSE | Gtg/Ctg | V202L | PLES_25471 | C | C | . | . | | | . | . | . | . | . | . |
| NC_011770 | 2723916 | T | G | 23 | 8 | 2 | 0 | NON_SYNONYMOU | MODERAT | MISSENSE | gTg/gGg | V202G | PLES_25471 | G | G | . | . | | | . | . | . | . | . | . |
| NC_011770 | 2724012 | C | T | 22 | 8 | 4 | 0 | NON_SYNONYMOU | MODERAT | MISSENSE | gCc/gTc | A234V | PLES_25471 | T | T | . | . | T | T | . | . | . | . | . | . |
| NC_011770 | 2724051 | C | A | 24 | 8 | 2 | 0 | NON_SYNONYMOU | MODERAT | MISSENSE | aCt/aAt | T247N | PLES_25471 | A | A | . | . | | | . | . | . | . | . | . |
| NC_011770 | 2724169 | C | G | 23 | 8 | 2 | 0 | NON_SYNONYMOU | MODERAT | MISSENSE | aCg/aGg | T12R | PLES_25481 | G | G | . | . | | | . | . | . | . | . | . |
| NC_011770 | 2724195 | C | G | 24 | 8 | 2 | 0 | NON_SYNONYMOU | MODERAT | MISSENSE | Ctg/Gtg | L21V | PLES_25481 | G | G | . | . | | | . | . | . | . | . | . |
| NC_011770 | 2724232 | A | G | 25 | 8 | 2 | 0 | NON_SYNONYMOU | MODERAT | MISSENSE | cAg/cGg | Q33R | PLES_25481 | | | . | . | G | G | . | . | . | . | . | . |
| NC_011770 | 2724240 | A | C | 25 | 8 | 2 | 0 | NON_SYNONYMOU | MODERAT | MISSENSE | Att/Ctt | I36L | PLES_25481 | | | . | . | C | C | . | . | . | . | . | . |
| NC_011770 | 2724265 | C | A | 25 | 8 | 2 | 0 | NON_SYNONYMOU | MODERAT | MISSENSE | gCc/gAc | A44D | PLES_25481 | A | A | . | . | | | . | . | . | . | . | . |
| NC_011770 | 2724355 | C | A | 26 | 8 | 4 | 0 | NON_SYNONYMOU | MODERAT | MISSENSE | gCt/gAt | A74D | PLES_25481 | A | A | . | . | A | A | . | . | . | . | . | . |
| NC_011770 | 2724364 | G | A | 26 | 8 | 4 | 0 | NON_SYNONYMOU | MODERAT | MISSENSE | aGg/aAg | R77K | PLES_25481 | A | A | . | . | A | A | . | . | . | . | . | . |
| NC_011770 | 2724376 | G | T | 27 | 8 | 2 | 0 | NON_SYNONYMOU | MODERAT | MISSENSE | cGc/cTc | R81L | PLES_25481 | T | T | . | . | | | . | . | . | . | . | . |
| NC_011770 | 2724498 | G | A | 5 | 10 | 2 | 0 | NON_SYNONYMOU | MODERAT | MISSENSE | Gtc/Atc | V122I | PLES_25481 | A | A | . | . | . | . | . | . | . | . | . | . |
| NC_011770 | 2724681 | T | G | 17 | 8 | 4 | 0 | NON_SYNONYMOU | MODERAT | MISSENSE | Ttc/Gtc | F183V | PLES_25481 | G | G | . | . | G | G | . | . | . | . | . | . |
| NC_011770 | 2725198 | T | A | 15 | 8 | 2 | 0 | NON_SYNONYMOU | MODERAT | MISSENSE | tTt/tAt | F6Y | PLES_25501 | A | A | . | . | | | . | . | . | . | . | . |
| NC_011770 | 2725216 | G | A | 14 | 8 | 2 | 0 | NON_SYNONYMOU | MODERAT | MISSENSE | aGc/aAc | S12N | PLES_25501 | A | A | . | . | | | . | . | . | . | . | . |
| NC_011770 | 2725219 | T | C | 14 | 8 | 4 | 0 | NON_SYNONYMOU | MODERAT | MISSENSE | cTc/cCc | L13P | PLES_25501 | C | C | . | . | C | C | . | . | . | . | . | . |
| NC_011770 | 2725275 | G | T | 13 | 8 | 2 | 0 | NON_SYNONYMOU | MODERAT | MISSENSE | Gcc/Tcc | A32S | PLES_25501 | | | . | . | T | T | . | . | . | . | . | . |
| NC_011770 | 2725350 | C | T | 14 | 8 | 2 | 0 | NON_SYNONYMOU | MODERAT | MISSENSE | Cgc/Tgc | R57C | PLES_25501 | T | T | . | . | | | . | . | . | . | . | . |
| NC_011770 | 2725374 | A | C | 15 | 8 | 2 | 0 | NON_SYNONYMOU | MODERAT | MISSENSE | Atc/Ctc | I65L | PLES_25501 | | | . | . | C | C | . | . | . | . | . | . |
| NC_011770 | 2725426 | A | T | 14 | 8 | 2 | 0 | NON_SYNONYMOU | MODERAT | MISSENSE | aAg/aTg | K82M | PLES_25501 | T | T | . | . | | | . | . | . | . | . | . |
| NC_011770 | 2726370 | T | C | 6 | 10 | 2 | 0 | NON_SYNONYMOU | MODERAT | MISSENSE | Tac/Cac | Y27H | PLES_25521 | C | C | . | . | . | . | . | . | . | . | . | . |
| NC_011770 | 2726671 | T | A | 9 | 8 | 2 | 0 | NON_SYNONYMOU | MODERAT | MISSENSE | gTt/gAt | V127D | PLES_25521 | | | . | . | A | A | . | . | . | . | . | . |
| NC_011770 | 2726805 | T | G | 7 | 10 | 2 | 0 | NON_SYNONYMOU | MODERAT | MISSENSE | Tcc/Gcc | S172A | PLES_25521 | G | G | . | . | . | . | . | . | . | . | . | . |
| NC_011770 | 2726955 | G | A | 8 | 10 | 2 | 0 | NON_SYNONYMOU | MODERAT | MISSENSE | Gcc/Acc | A45T | PLES_25531 | A | A | . | . | . | . | . | . | . | . | . | . |
| NC_011770 | 2727043 | G | A | 8 | 10 | 2 | 0 | NON_SYNONYMOU | MODERAT | MISSENSE | aGg/aAg | R74K | PLES_25531 | A | A | . | . | . | . | . | . | . | . | . | . |
| NC_011770 | 2727175 | T | C | 8 | 10 | 2 | 0 | NON_SYNONYMOU | MODERAT | MISSENSE | gTg/gCg | V118A | PLES_25531 | C | C | . | . | . | . | . | . | . | . | . | . |
| NC_011770 | 2727368 | C | A | 5 | 10 | 2 | 0 | NON_SYNONYMOU | MODERAT | MISSENSE | Cat/Aat | H61N | PLES_25541 | A | A | . | . | . | . | . | . | . | . | . | . |
| NC_011770 | 2727423 | T | C | 4 | 10 | 2 | 0 | NON_SYNONYMOU | MODERAT | MISSENSE | gTa/gCa | V79A | PLES_25541 | C | C | . | . | . | . | . | . | . | . | . | . |
| NC_011770 | 2737969 | C | T | 2 | 10 | 2 | 0 | NON_SYNONYMOU | MODERAT | MISSENSE | gCg/gTg | A41V | PLES_25621 | T | T | . | . | . | . | . | . | . | . | . | . |
| NC_011770 | 2738080 | G | T | 2 | 10 | 2 | 0 | NON_SYNONYMOU | MODERAT | MISSENSE | cGc/cTc | R78L | PLES_25621 | T | T | . | . | . | . | . | . | . | . | . | . |
| NC_011770 | 2738083 | A | G | 2 | 10 | 2 | 0 | NON_SYNONYMOU | MODERAT | MISSENSE | gAt/gGt | D79G | PLES_25621 | G | G | . | . | . | . | . | . | . | . | . | . |
| NC_011770 | 2738206 | T | A | 2 | 10 | 2 | 0 | NON_SYNONYMOU | MODERAT | MISSENSE | tTc/tAc | F120Y | PLES_25621 | A | A | . | . | . | . | . | . | . | . | . | . |
| NC_011770 | 2738286 | G | C | 3 | 10 | 2 | 0 | NON_SYNONYMOU | MODERAT | MISSENSE | Gcg/Ccg | A147P | PLES_25621 | C | C | . | . | . | . | . | . | . | . | . | . |
| NC_011770 | 2738287 | C | A | 3 | 10 | 2 | 0 | NON_SYNONYMOU | MODERAT | MISSENSE | gCg/gAg | A147E | PLES_25621 | A | A | . | . | . | . | . | . | . | . | . | . |
| NC_011770 | 2738346 | G | T | 3 | 10 | 2 | 0 | NON_SYNONYMOU | MODERAT | MISSENSE | Gcc/Tcc | A167S | PLES_25621 | T | T | . | . | . | . | . | . | . | . | . | . |
| NC_011770 | 2738350 | G | C | 3 | 10 | 2 | 0 | NON_SYNONYMOU | MODERAT | MISSENSE | aGt/aCt | S168T | PLES_25621 | C | C | . | . | . | . | . | . | . | . | . | . |
| NC_011770 | 2738351 | T | G | 3 | 10 | 2 | 0 | NON_SYNONYMOU | MODERAT | MISSENSE | agT/agG | S168R | PLES_25621 | G | G | . | . | . | . | . | . | . | . | . | . |
| NC_011770 | 2741043 | G | T | 60 | 0 | 2 | 0 | NON_SYNONYMOU | MODERAT | MISSENSE | aCg/aAg | T427K | PLES_25671 | | | | | | | | | | | T | T |
| NC_011770 | 2741249 | T | G | 68 | 0 | 4 | 0 | NON_SYNONYMOU | MODERAT | MISSENSE | ttA/ttC | L358F | PLES_25671 | | | | | | | G | G | G | G | | |
| NC_011770 | 2741258 | G | C | 68 | 0 | 2 | 0 | NON_SYNONYMOU | MODERAT | MISSENSE | gaC/gaG | D355E | PLES_25671 | | | | | | | C | C | | | | |
| NC_011770 | 2741905 | C | T | 79 | 0 | 2 | 0 | NON_SYNONYMOU | MODERAT | MISSENSE | Gat/Aat | D140N | PLES_25671 | | | | | | | | | | | T | T |
| NC_011770 | 2742078 | A | T | 62 | 0 | 4 | 0 | NON_SYNONYMOU | MODERAT | MISSENSE | cTc/cAc | L82H | PLES_25671 | T | T | | | | | | | T | T | | |
| NC_011770 | 2742115 | T | C | 62 | 0 | 12 | 0 | NON_SYNONYMOU | MODERAT | MISSENSE | Agc/Ggc | S70G | PLES_25671 | C | C | C | C | C | C | C | C | C | C | C | C |
| NC_011770 | 2742217 | C | T | 56 | 0 | 2 | 0 | NON_SYNONYMOU | MODERAT | MISSENSE | Gcc/Acc | A36T | PLES_25671 | | | | | | | T | T | | | | |

| | | | | | | | | | | | | | | | | | | | | | | | | | |
|---|---|---|---|---|---|---|---|---|---|---|---|---|---|---|---|---|---|---|---|---|---|---|---|---|---|
| NC_011770 | 2742579 | GC | G | 35 | 0 | 1 | 0 | FRAME_SHIFT | HIGH | | ggg/ | G115 | PLES_25681 | | | | | | | | G | | | | |
| NC_011770 | 2742801 | C | T | 16 | 0 | 2 | 0 | NON_SYNONYMOU | MODERAT | MISSENSE | Gtg/Atg | V42M | PLES_25681 | | | | | | | | | T | T | | |
| NC_011770 | 2742831 | C | T | 17 | 0 | 2 | 0 | NON_SYNONYMOU | MODERAT | MISSENSE | Gcc/Acc | A32T | PLES_25681 | | | | | | | | | | | T | T |
| NC_011770 | 2743292 | G | T | 60 | 0 | 2 | 0 | NON_SYNONYMOU | MODERAT | MISSENSE | Gcc/Tcc | A86S | PLES_25691 | | | | | | | | | | | T | T |
| NC_011770 | 2743557 | A | G | 64 | 0 | 2 | 0 | NON_SYNONYMOU | MODERAT | MISSENSE | gAc/gGc | D174G | PLES_25691 | | | | | | | G | G | | | | |
| NC_011770 | 2743728 | C | T | 69 | 0 | 2 | 0 | NON_SYNONYMOU | MODERAT | MISSENSE | gCc/gTc | A231V | PLES_25691 | | | | | | | | | | | T | T |
| NC_011770 | 2743761 | C | T | 69 | 0 | 2 | 0 | NON_SYNONYMOU | MODERAT | MISSENSE | cCg/cTg | P242L | PLES_25691 | | | | | | | T | T | | | | |
| NC_011770 | 2744913 | T | C | 69 | 0 | 10 | 0 | NON_SYNONYMOU | MODERAT | MISSENSE | cTg/cCg | L214P | PLES_25701 | C | C | C | C | C | C | C | C | | | C | C |
| NC_011770 | 2745202 | G | C | 66 | 0 | 2 | 0 | NON_SYNONYMOU | MODERAT | MISSENSE | gaG/gaC | E310D | PLES_25701 | | | | | | | | | | | C | C |
| NC_011770 | 2745287 | C | T | 67 | 0 | 1 | 0 | NON_SYNONYMOU | MODERAT | MISSENSE | Ccc/Tcc | P339S | PLES_25701 | | | | | | | T | | | | | |
| NC_011770 | 2744565 | T | C | 73 | 0 | 1 | 0 | NON_SYNONYMOU | MODERAT | MISSENSE | gTa/gCa | V98A | PLES_25701 | | | | | | | | C | | | | |
| NC_011770 | 2745624 | A | G | 69 | 0 | 2 | 0 | NON_SYNONYMOU | MODERAT | MISSENSE | aAg/aGg | K92R | PLES_25711 | | | | | | | | | G | G | | |
| NC_011770 | 2745750 | C | T | 67 | 0 | 1 | 0 | NON_SYNONYMOU | MODERAT | MISSENSE | cCc/cTc | P134L | PLES_25711 | | | | | | | T | | | | | |
| NC_011770 | 2746031 | T | C | 68 | 0 | 12 | 0 | NON_SYNONYMOU | MODERAT | MISSENSE | Ttc/Ctc | F228L | PLES_25711 | C | C | C | C | C | C | C | C | C | C | C | C |
| NC_011770 | 2747039 | C | T | 73 | 0 | 2 | 0 | NON_SYNONYMOU | MODERAT | MISSENSE | Cgc/Tgc | R246C | PLES_25721 | T | T | | | | | | | | | | |
| NC_011770 | 2747587 | A | G | 72 | 0 | 2 | 0 | NON_SYNONYMOU | MODERAT | MISSENSE | gAg/gGg | E66G | PLES_25731 | G | G | | | | | | | | | | |
| NC_011770 | 2747806 | G | C | 82 | 0 | 4 | 0 | NON_SYNONYMOU | MODERAT | MISSENSE | gGt/gCt | G139A | PLES_25731 | | | | | | | | | C | C | C | C |
| NC_011770 | 2748189 | G | A | 88 | 0 | 2 | 0 | NON_SYNONYMOU | MODERAT | MISSENSE | Gtc/Atc | V267I | PLES_25731 | | | | | | | A | A | | | | |
| NC_011770 | 2748271 | G | A | 83 | 0 | 10 | 0 | NON_SYNONYMOU | MODERAT | MISSENSE | gGg/gAg | G294E | PLES_25731 | A | A | A | A | A | A | A | A | A | A | | |
| NC_011770 | 2749349 | C | T | 71 | 0 | 1 | 0 | STOP_GAINED | HIGH | NONSENSE | Cag/Tag | Q234* | PLES_25741 | | T | | | | | | | | | | |
| NC_011770 | 2749260 | C | G | 75 | 0 | 6 | 0 | NON_SYNONYMOU | MODERAT | MISSENSE | aCc/aGc | T204S | PLES_25741 | G | G | G | G | G | G | | | | | | |
| NC_011770 | 2749828 | GC | G | 54 | 0 | 1 | 0 | FRAME_SHIFT | HIGH | | ctc/ | L63 | PLES_25751 | | | | | | | G | | | | | |
| NC_011770 | 2749726 | C | A | 54 | 0 | 4 | 0 | NON_SYNONYMOU | MODERAT | MISSENSE | Cgc/Agc | R27S | PLES_25751 | | | A | A | A | A | | | | | | |
| NC_011770 | 2749788 | C | G | 55 | 0 | 2 | 0 | NON_SYNONYMOU | MODERAT | MISSENSE | gaC/gaG | D47E | PLES_25751 | | | | | | | | | G | G | | |
| NC_011770 | 2750582 | C | A | 53 | 0 | 2 | 0 | NON_SYNONYMOU | MODERAT | MISSENSE | gCg/gAg | A312E | PLES_25751 | | | | | A | A | | | | | | |
| NC_011770 | 2784571 | C | T | 80 | 0 | 2 | 0 | NON_SYNONYMOU | MODERAT | MISSENSE | Gcc/Acc | A274T | PLES_25961 | | | | | | | T | T | | | | |
| NC_011770 | 2784805 | G | A | 83 | 0 | 2 | 0 | NON_SYNONYMOU | MODERAT | MISSENSE | Cac/Tac | H196Y | PLES_25961 | | | | | | | A | A | | | | |
| NC_011770 | 2784835 | A | G | 84 | 0 | 2 | 0 | NON_SYNONYMOU | MODERAT | MISSENSE | Tcc/Ccc | S186P | PLES_25961 | G | G | | | | | | | | | | |
| NC_011770 | 2785245 | T | C | 88 | 0 | 2 | 0 | NON_SYNONYMOU | MODERAT | MISSENSE | aAg/aGg | K49R | PLES_25961 | | | | | | | C | C | | | | |
| NC_011770 | 2785246 | T | G | 89 | 0 | 2 | 0 | NON_SYNONYMOU | MODERAT | MISSENSE | Aag/Cag | K49Q | PLES_25961 | | | | | | | G | G | | | | |
| NC_011770 | 2785730 | G | T | 75 | 0 | 2 | 0 | NON_SYNONYMOU | MODERAT | MISSENSE | aGa/aTa | R63I | PLES_25971 | | | | | | | | | T | T | | |
| NC_011770 | 2785738 | G | A | 71 | 0 | 2 | 0 | NON_SYNONYMOU | MODERAT | MISSENSE | Gtc/Atc | V66I | PLES_25971 | | | | | | | | | A | A | | |
| NC_011770 | 2787362 | C | CG | 71 | 0 | 1 | 0 | FRAME_SHIFT | HIGH | | aac/Gaac | N301E? | PLES_25981 | | | | | CG | | | | | | | |
| NC_011770 | 2787143 | C | CGG | 76 | 0 | 1 | 0 | FRAME_SHIFT | HIGH | | ggc/ggGGc | G226G? | PLES_25981 | | | | | | | CGG | | | | | |
| NC_011770 | 2786478 | G | A | 74 | 0 | 4 | 0 | NON_SYNONYMOU | MODERAT | MISSENSE | Gta/Ata | V4I | PLES_25981 | | | A | A | A | A | | | | | | |
| NC_011770 | 2786581 | C | T | 68 | 0 | 12 | 0 | NON_SYNONYMOU | MODERAT | MISSENSE | gCg/gTg | A38V | PLES_25981 | T | T | T | T | T | T | T | T | T | T | T | T |
| NC_011770 | 2786738 | C | G | 67 | 0 | 4 | 0 | NON_SYNONYMOU | MODERAT | MISSENSE | gaC/gaG | D90E | PLES_25981 | | | G | G | G | G | | | | | | |
| NC_011770 | 2786823 | C | A | 67 | 0 | 4 | 0 | NON_SYNONYMOU | MODERAT | MISSENSE | Cct/Act | P119T | PLES_25981 | | | A | A | A | A | | | | | | |
| NC_011770 | 2786839 | G | T | 72 | 0 | 4 | 0 | NON_SYNONYMOU | MODERAT | MISSENSE | cGa/cTa | R124L | PLES_25981 | | | T | T | T | T | | | | | | |
| NC_011770 | 2786847 | A | G | 75 | 0 | 6 | 0 | NON_SYNONYMOU | MODERAT | MISSENSE | Acg/Gcg | T127A | PLES_25981 | G | G | | | | | G | G | | | G | G |
| NC_011770 | 2786905 | A | G | 79 | 0 | 6 | 0 | NON_SYNONYMOU | MODERAT | MISSENSE | aAg/aGg | K146R | PLES_25981 | G | G | | | | | G | G | | | G | G |
| NC_011770 | 2787297 | A | C | 73 | 0 | 2 | 0 | NON_SYNONYMOU | MODERAT | MISSENSE | Agc/Cgc | S277R | PLES_25981 | C | C | | | | | | | | | | |
| NC_011770 | 2787298 | G | A | 75 | 0 | 2 | 0 | NON_SYNONYMOU | MODERAT | MISSENSE | aGc/aAc | S277N | PLES_25981 | A | A | | | | | | | | | | |
| NC_011770 | 2787717 | C | T | 69 | 0 | 2 | 0 | NON_SYNONYMOU | MODERAT | MISSENSE | Cgt/Tgt | R417C | PLES_25981 | T | T | | | | | | | | | | |
| NC_011770 | 2788399 | G | A | 65 | 0 | 2 | 0 | NON_SYNONYMOU | MODERAT | MISSENSE | aGc/aAc | S644N | PLES_25981 | A | A | | | | | | | | | | |
| NC_011770 | 2786770 | T | A | 63 | 0 | 1 | 0 | NON_SYNONYMOU | MODERAT | MISSENSE | cTg/cAg | L101Q | PLES_25981 | | | | | A | | | | | | | |
| NC_011770 | 2789258 | C | G | 53 | 0 | 2 | 0 | NON_SYNONYMOU | MODERAT | MISSENSE | gCc/gGc | A42G | PLES_25991 | G | G | | | | | | | | | | |
| NC_011770 | 2797883 | GAA | G | 28 | 6 | 4 | 0 | FRAME_SHIFT | HIGH | | aag/ | K317 | PLES_26051 | G | G | G | G | . | . | . | . | . | . | | |
| NC_011770 | 2797887 | T | TCC | 28 | 6 | 4 | 0 | FRAME_SHIFT | HIGH | | tac/tCCac | Y318S? | PLES_26051 | TCC | TCC | TCC | TCC | . | . | . | . | . | . | | |
| NC_011770 | 2796970 | G | A | 12 | 10 | 2 | 0 | NON_SYNONYMOU | MODERAT | MISSENSE | aGa/aAa | R12K | PLES_26051 | . | . | . | . | . | . | . | . | . | . | A | A |
| NC_011770 | 2797050 | A | C | 12 | 10 | 2 | 0 | NON_SYNONYMOU | MODERAT | MISSENSE | Atc/Ctc | I39L | PLES_26051 | . | . | . | . | . | . | . | . | . | . | C | C |
| NC_011770 | 2797051 | T | A | 12 | 10 | 2 | 0 | NON_SYNONYMOU | MODERAT | MISSENSE | aTc/aAc | I39N | PLES_26051 | . | . | . | . | . | . | . | . | . | . | A | A |
| NC_011770 | 2797086 | A | C | 30 | 6 | 2 | 0 | NON_SYNONYMOU | MODERAT | MISSENSE | Atg/Ctg | M51L | PLES_26051 | | | | | . | . | . | . | . | . | C | C |
| NC_011770 | 2797088 | G | C | 30 | 6 | 4 | 0 | NON_SYNONYMOU | MODERAT | MISSENSE | atG/atC | M51I | PLES_26051 | C | C | C | C | . | . | . | . | . | . | | |
| NC_011770 | 2797090 | C | G | 30 | 6 | 2 | 0 | NON_SYNONYMOU | MODERAT | MISSENSE | tCc/tGc | S52C | PLES_26051 | | | | | . | . | . | . | . | . | G | G |
| NC_011770 | 2797131 | G | A | 9 | 10 | 2 | 0 | NON_SYNONYMOU | MODERAT | MISSENSE | Gcc/Acc | A66T | PLES_26051 | . | . | . | . | . | . | . | . | . | . | A | A |
| NC_011770 | 2797132 | C | T | 9 | 10 | 2 | 0 | NON_SYNONYMOU | MODERAT | MISSENSE | gCc/gTc | A66V | PLES_26051 | . | . | . | . | . | . | . | . | . | . | T | T |
| NC_011770 | 2797153 | T | C | 8 | 10 | 2 | 0 | NON_SYNONYMOU | MODERAT | MISSENSE | gTt/gCt | V73A | PLES_26051 | . | . | . | . | . | . | . | . | . | . | C | C |
| NC_011770 | 2797165 | A | G | 8 | 10 | 2 | 0 | NON_SYNONYMOU | MODERAT | MISSENSE | aAg/aGg | K77R | PLES_26051 | . | . | . | . | . | . | . | . | . | . | G | G |
| NC_011770 | 2797182 | A | G | 7 | 10 | 2 | 0 | NON_SYNONYMOU | MODERAT | MISSENSE | Atc/Gtc | I83V | PLES_26051 | . | . | . | . | . | . | . | . | . | . | G | G |
| NC_011770 | 2797197 | C | A | 7 | 10 | 2 | 0 | NON_SYNONYMOU | MODERAT | MISSENSE | Cgc/Agc | R88S | PLES_26051 | . | . | . | . | . | . | . | . | . | . | A | A |
| NC_011770 | 2797198 | G | A | 7 | 10 | 2 | 0 | NON_SYNONYMOU | MODERAT | MISSENSE | cGc/cAc | R88H | PLES_26051 | . | . | . | . | . | . | . | . | . | . | A | A |

| | | | | | | | | | | | | | | | | | | | | | | | | | |
|---|---|---|---|---|---|---|---|---|---|---|---|---|---|---|---|---|---|---|---|---|---|---|---|---|---|
| NC_011770 | 2797210 | C | T | 29 | 6 | 4 | 0 | NON_SYNONYMOU | MODERAT | MISSENSE | gCc/gTc | A92V | PLES_26051 | T | T | T | T | . | . | . | . | . | . | | |
| NC_011770 | 2797222 | G | C | 29 | 6 | 4 | 0 | NON_SYNONYMOU | MODERAT | MISSENSE | gGc/gCc | G96A | PLES_26051 | C | C | C | C | . | . | . | . | . | . | | |
| NC_011770 | 2797226 | A | C | 30 | 6 | 4 | 0 | NON_SYNONYMOU | MODERAT | MISSENSE | gaA/gaC | E97D | PLES_26051 | C | C | C | C | . | . | . | . | . | . | | |
| NC_011770 | 2797239 | A | G | 32 | 6 | 4 | 0 | NON_SYNONYMOU | MODERAT | MISSENSE | Aac/Gac | N102D | PLES_26051 | G | G | G | G | . | . | . | . | . | . | | |
| NC_011770 | 2797240 | A | C | 33 | 6 | 6 | 0 | NON_SYNONYMOU | MODERAT | MISSENSE | aAc/aCc | N102T | PLES_26051 | C | C | C | C | . | . | . | . | . | . | C | C |
| NC_011770 | 2797278 | C | A | 35 | 6 | 4 | 0 | NON_SYNONYMOU | MODERAT | MISSENSE | Ctc/Atc | L115I | PLES_26051 | A | A | A | A | . | . | . | . | . | . | | |
| NC_011770 | 2797290 | C | A | 32 | 6 | 4 | 0 | NON_SYNONYMOU | MODERAT | MISSENSE | Cgt/Agt | R119S | PLES_26051 | A | A | A | A | . | . | . | . | . | . | | |
| NC_011770 | 2797293 | C | A | 32 | 6 | 4 | 0 | NON_SYNONYMOU | MODERAT | MISSENSE | Cag/Aag | Q120K | PLES_26051 | A | A | A | A | . | . | . | . | . | . | | |
| NC_011770 | 2797295 | G | C | 32 | 6 | 6 | 0 | NON_SYNONYMOU | MODERAT | MISSENSE | caG/caC | Q120H | PLES_26051 | C | C | C | C | . | . | . | . | . | . | A | A |
| NC_011770 | 2797345 | C | G | 35 | 6 | 4 | 0 | NON_SYNONYMOU | MODERAT | MISSENSE | aCc/aGc | T137S | PLES_26051 | G | G | G | G | . | . | . | . | . | . | | |
| NC_011770 | 2797350 | G | A | 32 | 6 | 4 | 0 | NON_SYNONYMOU | MODERAT | MISSENSE | Ggg/Agg | G139R | PLES_26051 | A | A | A | A | . | . | . | . | . | . | | |
| NC_011770 | 2797351 | G | A | 33 | 6 | 4 | 0 | NON_SYNONYMOU | MODERAT | MISSENSE | gGg/gAg | G139E | PLES_26051 | A | A | A | A | . | . | . | . | . | . | | |
| NC_011770 | 2797353 | A | C | 33 | 6 | 4 | 0 | NON_SYNONYMOU | MODERAT | MISSENSE | Aag/Cag | K140Q | PLES_26051 | C | C | C | C | . | . | . | . | . | . | | |
| NC_011770 | 2797354 | A | G | 33 | 6 | 2 | 0 | NON_SYNONYMOU | MODERAT | MISSENSE | aAg/aGg | K140R | PLES_26051 | | | | | . | . | . | . | . | . | G | G |
| NC_011770 | 2797389 | G | C | 37 | 6 | 6 | 0 | NON_SYNONYMOU | MODERAT | MISSENSE | Gtg/Ctg | V152L | PLES_26051 | C | C | C | C | . | . | . | . | . | . | C | C |
| NC_011770 | 2797398 | G | C | 36 | 6 | 6 | 0 | NON_SYNONYMOU | MODERAT | MISSENSE | Gtg/Ctg | V155L | PLES_26051 | C | C | C | C | . | . | . | . | . | . | T | T |
| NC_011770 | 2797404 | G | A | 34 | 6 | 4 | 0 | NON_SYNONYMOU | MODERAT | MISSENSE | Gcg/Acg | A157T | PLES_26051 | A | A | A | A | . | . | . | . | . | . | | |
| NC_011770 | 2797405 | C | G | 35 | 6 | 4 | 0 | NON_SYNONYMOU | MODERAT | MISSENSE | gCg/gGg | A157G | PLES_26051 | G | G | G | G | . | . | . | . | . | . | | |
| NC_011770 | 2797413 | A | G | 36 | 6 | 4 | 0 | NON_SYNONYMOU | MODERAT | MISSENSE | Aaa/Gaa | K160E | PLES_26051 | G | G | G | G | . | . | . | . | . | . | | |
| NC_011770 | 2797455 | T | C | 29 | 6 | 4 | 0 | NON_SYNONYMOU | MODERAT | MISSENSE | Ttc/Ctc | F174L | PLES_26051 | C | C | C | C | . | . | . | . | . | . | | |
| NC_011770 | 2797457 | C | G | 30 | 6 | 4 | 0 | NON_SYNONYMOU | MODERAT | MISSENSE | ttC/ttG | F174L | PLES_26051 | G | G | G | G | . | . | . | . | . | . | | |
| NC_011770 | 2797458 | A | G | 30 | 6 | 4 | 0 | NON_SYNONYMOU | MODERAT | MISSENSE | Aac/Gac | N175D | PLES_26051 | G | G | G | G | . | . | . | . | . | . | | |
| NC_011770 | 2797459 | A | G | 30 | 6 | 4 | 0 | NON_SYNONYMOU | MODERAT | MISSENSE | aAc/aGc | N175S | PLES_26051 | G | G | G | G | . | . | . | . | . | . | | |
| NC_011770 | 2797460 | C | G | 30 | 6 | 4 | 0 | NON_SYNONYMOU | MODERAT | MISSENSE | aaC/aaG | N175K | PLES_26051 | G | G | G | G | . | . | . | . | . | . | | |
| NC_011770 | 2797464 | A | T | 30 | 6 | 6 | 0 | NON_SYNONYMOU | MODERAT | MISSENSE | Atg/Ttg | M177L | PLES_26051 | T | T | T | T | . | . | . | . | . | . | T | T |
| NC_011770 | 2797489 | C | T | 31 | 6 | 4 | 0 | NON_SYNONYMOU | MODERAT | MISSENSE | gCc/gTc | A185V | PLES_26051 | T | T | T | T | . | . | . | . | . | . | | |
| NC_011770 | 2797504 | T | C | 32 | 6 | 4 | 0 | NON_SYNONYMOU | MODERAT | MISSENSE | gTc/gCc | V190A | PLES_26051 | C | C | C | C | . | . | . | . | . | . | | |
| NC_011770 | 2797515 | G | T | 32 | 6 | 4 | 0 | NON_SYNONYMOU | MODERAT | MISSENSE | Gcg/Tcg | A194S | PLES_26051 | T | T | T | T | . | . | . | . | . | . | | |
| NC_011770 | 2797554 | G | A | 31 | 6 | 4 | 0 | NON_SYNONYMOU | MODERAT | MISSENSE | Gcc/Acc | A207T | PLES_26051 | A | A | A | A | . | . | . | . | . | . | | |
| NC_011770 | 2797555 | C | A | 31 | 6 | 4 | 0 | NON_SYNONYMOU | MODERAT | MISSENSE | gCc/gAc | A207D | PLES_26051 | A | A | A | A | . | . | . | . | . | . | | |
| NC_011770 | 2797584 | C | A | 33 | 6 | 2 | 0 | NON_SYNONYMOU | MODERAT | MISSENSE | Ctg/Atg | L217M | PLES_26051 | | | | | . | . | . | . | . | . | A | A |
| NC_011770 | 2797601 | G | C | 33 | 6 | 4 | 0 | NON_SYNONYMOU | MODERAT | MISSENSE | caG/caC | Q222H | PLES_26051 | C | C | C | C | . | . | . | . | . | . | | |
| NC_011770 | 2797612 | G | T | 34 | 6 | 6 | 0 | NON_SYNONYMOU | MODERAT | MISSENSE | cGc/cTc | R226L | PLES_26051 | T | T | T | T | . | . | . | . | . | . | T | T |
| NC_011770 | 2797647 | G | A | 35 | 6 | 4 | 0 | NON_SYNONYMOU | MODERAT | MISSENSE | Gtc/Atc | V238I | PLES_26051 | A | A | A | A | . | . | . | . | . | . | | |
| NC_011770 | 2797698 | G | A | 25 | 6 | 4 | 0 | NON_SYNONYMOU | MODERAT | MISSENSE | Gag/Aag | E255K | PLES_26051 | A | A | A | A | . | . | . | . | . | . | | |
| NC_011770 | 2797699 | A | C | 26 | 6 | 4 | 0 | NON_SYNONYMOU | MODERAT | MISSENSE | gAg/gCg | E255A | PLES_26051 | C | C | C | C | . | . | . | . | . | . | | |
| NC_011770 | 2797700 | G | C | 26 | 6 | 6 | 0 | NON_SYNONYMOU | MODERAT | MISSENSE | gaG/gaC | E255D | PLES_26051 | C | C | C | C | . | . | . | . | . | . | A | A |
| NC_011770 | 2797706 | G | C | 26 | 6 | 6 | 0 | NON_SYNONYMOU | MODERAT | MISSENSE | gaG/gaC | E257D | PLES_26051 | C | C | C | C | . | . | . | . | . | . | A | A |
| NC_011770 | 2797722 | C | A | 33 | 6 | 2 | 0 | NON_SYNONYMOU | MODERAT | MISSENSE | Cgc/Agc | R263S | PLES_26051 | | | | | . | . | . | . | . | . | A | A |
| NC_011770 | 2797749 | G | A | 34 | 6 | 6 | 0 | NON_SYNONYMOU | MODERAT | MISSENSE | Gtc/Atc | V272I | PLES_26051 | A | A | A | A | . | . | . | . | . | . | A | A |
| NC_011770 | 2797830 | C | G | 35 | 6 | 4 | 0 | NON_SYNONYMOU | MODERAT | MISSENSE | Ctg/Gtg | L299V | PLES_26051 | G | G | G | G | . | . | . | . | . | . | | |
| NC_011770 | 2797872 | C | A | 28 | 6 | 4 | 0 | NON_SYNONYMOU | MODERAT | MISSENSE | Cgt/Agt | R313S | PLES_26051 | A | A | A | A | . | . | . | . | . | . | | |
| NC_011770 | 2797896 | T | A | 19 | 7 | 3 | 0 | NON_SYNONYMOU | MODERAT | MISSENSE | Tcg/Acg | S321T | PLES_26051 | . | A | A | A | . | . | . | . | . | . | | |
| NC_011770 | 2797944 | A | G | 34 | 6 | 4 | 0 | NON_SYNONYMOU | MODERAT | MISSENSE | Aag/Gag | K337E | PLES_26051 | G | G | G | G | . | . | . | . | . | . | | |
| NC_011770 | 2797945 | A | C | 34 | 6 | 4 | 0 | NON_SYNONYMOU | MODERAT | MISSENSE | aAg/aCg | K337T | PLES_26051 | C | C | C | C | . | . | . | . | . | . | | |
| NC_011770 | 2797950 | G | T | 33 | 6 | 4 | 0 | NON_SYNONYMOU | MODERAT | MISSENSE | Gtg/Ttg | V339L | PLES_26051 | T | T | T | T | . | . | . | . | . | . | | |
| NC_011770 | 2797953 | C | A | 34 | 6 | 4 | 0 | NON_SYNONYMOU | MODERAT | MISSENSE | Cag/Aag | Q340K | PLES_26051 | A | A | A | A | . | . | . | . | . | . | | |
| NC_011770 | 2797957 | T | G | 34 | 6 | 4 | 0 | NON_SYNONYMOU | MODERAT | MISSENSE | cTc/cGc | L341R | PLES_26051 | G | G | G | G | . | . | . | . | . | . | | |
| NC_011770 | 2797972 | G | A | 33 | 6 | 4 | 0 | NON_SYNONYMOU | MODERAT | MISSENSE | gGg/gAg | G346E | PLES_26051 | A | A | A | A | . | . | . | . | . | . | | |
| NC_011770 | 2797975 | G | A | 34 | 6 | 4 | 0 | NON_SYNONYMOU | MODERAT | MISSENSE | cGc/cAc | R347H | PLES_26051 | A | A | A | A | . | . | . | . | . | . | | |
| NC_011770 | 2798091 | G | T | 39 | 6 | 2 | 0 | NON_SYNONYMOU | MODERAT | MISSENSE | Gcc/Tcc | A386S | PLES_26051 | | | | | . | . | . | . | . | . | T | T |
| NC_011770 | 2798175 | G | C | 17 | 6 | 4 | 0 | NON_SYNONYMOU | MODERAT | MISSENSE | Gaa/Caa | E414Q | PLES_26051 | C | C | C | C | . | . | . | . | . | . | | |
| NC_011770 | 2798209 | G | A | 16 | 6 | 4 | 0 | NON_SYNONYMOU | MODERAT | MISSENSE | cGt/cAt | R425H | PLES_26051 | A | A | A | A | . | . | . | . | . | . | | |
| NC_011770 | 2798216 | C | G | 15 | 9 | 1 | 0 | NON_SYNONYMOU | MODERAT | MISSENSE | gaC/gaG | D427E | PLES_26051 | . | . | G | . | . | . | . | . | . | . | | |
| NC_011770 | 2798221 | T | C | 15 | 8 | 2 | 0 | NON_SYNONYMOU | MODERAT | MISSENSE | gTg/gCg | V429A | PLES_26051 | . | . | | | . | . | . | . | . | . | C | C |
| NC_011770 | 2798248 | T | C | 11 | 10 | 2 | 0 | NON_SYNONYMOU | MODERAT | MISSENSE | gTg/gCg | V438A | PLES_26051 | . | . | . | . | . | . | . | . | . | . | C | C |
| NC_011770 | 2798254 | G | A | 1 | 10 | 2 | 0 | NON_SYNONYMOU | MODERAT | MISSENSE | gGc/gAc | G440D | PLES_26051 | . | . | . | . | . | . | . | . | . | . | A | A |
| NC_011770 | 2798259 | G | A | 1 | 10 | 2 | 0 | NON_SYNONYMOU | MODERAT | MISSENSE | Gtc/Atc | V442I | PLES_26051 | . | . | . | . | . | . | . | . | . | . | A | A |
| NC_011770 | 2798260 | T | C | 1 | 10 | 2 | 0 | NON_SYNONYMOU | MODERAT | MISSENSE | gTc/gCc | V442A | PLES_26051 | . | . | . | . | . | . | . | . | . | . | C | C |
| NC_011770 | 2798262 | T | C | 1 | 10 | 2 | 0 | NON_SYNONYMOU | MODERAT | MISSENSE | Tcc/Ccc | S443P | PLES_26051 | . | . | . | . | . | . | . | . | . | . | C | C |
| NC_011770 | 2798263 | C | T | 1 | 10 | 2 | 0 | NON_SYNONYMOU | MODERAT | MISSENSE | tCc/tTc | S443F | PLES_26051 | . | . | . | . | . | . | . | . | . | . | T | T |
| NC_011770 | 2798265 | A | G | 1 | 10 | 2 | 0 | NON_SYNONYMOU | MODERAT | MISSENSE | Acg/Gcg | T444A | PLES_26051 | . | . | . | . | . | . | . | . | . | . | G | G |

| | | | | | | | | | | | | | | | | | | | | | | | | | |
|---|---|---|---|---|---|---|---|---|---|---|---|---|---|---|---|---|---|---|---|---|---|---|---|---|---|
| NC_011770 | 2798269 | C | T | 1 | 10 | 2 | 0 | NON_SYNONYMOU | MODERAT | MISSENSE | aCt/aTt | T445I | PLES_26051 | . | . | . | . | . | . | . | . | . | . | T | T |
| NC_011770 | 2798280 | G | A | 1 | 10 | 2 | 0 | NON_SYNONYMOU | MODERAT | MISSENSE | Gtt/Att | V449I | PLES_26051 | . | . | . | . | . | . | . | . | . | . | A | A |
| NC_011770 | 2798281 | T | C | 1 | 10 | 2 | 0 | NON_SYNONYMOU | MODERAT | MISSENSE | gTt/gCt | V449A | PLES_26051 | . | . | . | . | . | . | . | . | . | . | C | C |
| NC_011770 | 2798283 | G | C | 1 | 10 | 1 | 0 | NON_SYNONYMOU | MODERAT | MISSENSE | Gtc/Ctc | V450L | PLES_26051 | . | . | . | . | . | . | . | . | . | . | C | . |
| NC_011770 | 2798284 | T | C | 1 | 10 | 2 | 0 | NON_SYNONYMOU | MODERAT | MISSENSE | gTc/gCc | V450A | PLES_26051 | . | . | . | . | . | . | . | . | . | . | C | C |
| NC_011770 | 2798332 | A | G | 9 | 10 | 2 | 0 | NON_SYNONYMOU | MODERAT | MISSENSE | aAc/aGc | N466S | PLES_26051 | . | . | . | . | . | . | . | . | . | . | G | G |
| NC_011770 | 2798347 | T | C | 10 | 10 | 2 | 0 | NON_SYNONYMOU | MODERAT | MISSENSE | aTg/aCg | M471T | PLES_26051 | . | . | . | . | . | . | . | . | . | . | C | C |
| NC_011770 | 2798382 | G | A | 11 | 9 | 3 | 0 | NON_SYNONYMOU | MODERAT | MISSENSE | Gtg/Atg | V483M | PLES_26051 | . | . | . | A | . | . | . | . | . | . | A | A |
| NC_011770 | 2798383 | T | C | 11 | 9 | 3 | 0 | NON_SYNONYMOU | MODERAT | MISSENSE | gTg/gCg | V483A | PLES_26051 | . | . | . | C | . | . | . | . | . | . | C | C |
| NC_011770 | 2798410 | C | T | 12 | 9 | 1 | 0 | NON_SYNONYMOU | MODERAT | MISSENSE | gCg/gTg | A492V | PLES_26051 | . | . | . | T | . | . | . | . | . | . | | |
| NC_011770 | 2798439 | G | A | 30 | 6 | 2 | 0 | NON_SYNONYMOU | MODERAT | MISSENSE | Gcc/Acc | A502T | PLES_26051 | | | | | . | . | . | . | . | . | A | A |
| NC_011770 | 2798458 | A | G | 28 | 6 | 6 | 0 | NON_SYNONYMOU | MODERAT | MISSENSE | aAc/aGc | N508S | PLES_26051 | G | G | G | G | . | . | . | . | . | . | G | G |
| NC_011770 | 2798461 | A | G | 28 | 6 | 6 | 0 | NON_SYNONYMOU | MODERAT | MISSENSE | aAt/aGt | N509S | PLES_26051 | G | G | G | G | . | . | . | . | . | . | G | G |
| NC_011770 | 2798649 | T | C | 19 | 6 | 4 | 0 | NON_SYNONYMOU | MODERAT | MISSENSE | Tcg/Ccg | S572P | PLES_26051 | C | C | C | C | . | . | . | . | . | . | | |
| NC_011770 | 2798677 | A | G | 8 | 9 | 1 | 0 | NON_SYNONYMOU | MODERAT | MISSENSE | cAg/cGg | Q581R | PLES_26051 | G | . | . | . | . | . | . | . | . | . | | |
| NC_011770 | 2798678 | G | C | 9 | 9 | 1 | 0 | NON_SYNONYMOU | MODERAT | MISSENSE | caG/caC | Q581H | PLES_26051 | C | . | . | . | . | . | . | . | . | . | | |
| NC_011770 | 2798749 | G | A | 7 | 10 | 2 | 0 | NON_SYNONYMOU | MODERAT | MISSENSE | cGg/cAg | R605Q | PLES_26051 | . | . | . | . | . | . | . | . | . | . | A | A |
| NC_011770 | 2798754 | C | A | 7 | 10 | 2 | 0 | NON_SYNONYMOU | MODERAT | MISSENSE | Ctc/Atc | L607I | PLES_26051 | . | . | . | . | . | . | . | . | . | . | A | A |
| NC_011770 | 2798775 | C | T | 6 | 10 | 2 | 0 | NON_SYNONYMOU | MODERAT | MISSENSE | Cat/Tat | H614Y | PLES_26051 | . | . | . | . | . | . | . | . | . | . | T | T |
| NC_011770 | 2798784 | G | A | 5 | 10 | 2 | 0 | NON_SYNONYMOU | MODERAT | MISSENSE | Gtg/Atg | V617M | PLES_26051 | . | . | . | . | . | . | . | . | . | . | A | A |
| NC_011770 | 2798796 | G | A | 5 | 10 | 2 | 0 | NON_SYNONYMOU | MODERAT | MISSENSE | Gtt/Att | V621I | PLES_26051 | . | . | . | . | . | . | . | . | . | . | A | A |
| NC_011770 | 2798805 | C | T | 4 | 10 | 2 | 0 | NON_SYNONYMOU | MODERAT | MISSENSE | Cgg/Tgg | R624W | PLES_26051 | . | . | . | . | . | . | . | . | . | . | T | T |
| NC_011770 | 2798806 | G | A | 4 | 10 | 2 | 0 | NON_SYNONYMOU | MODERAT | MISSENSE | cGg/cAg | R624Q | PLES_26051 | . | . | . | . | . | . | . | . | . | . | A | A |
| NC_011770 | 2822348 | C | A | 4 | 8 | 4 | 0 | NON_SYNONYMOU | MODERAT | MISSENSE | aCc/aAc | T787N | PLES_26261 | A | A | A | A | . | . | . | . | . | . | . | . |
| NC_011770 | 2840057 | G | C | 60 | 4 | 2 | 0 | NON_SYNONYMOU | MODERAT | MISSENSE | gCg/gGg | A290G | PLES_26421 | | | | | | | . | . | . | . | C | C |
| NC_011770 | 2840109 | C | G | 54 | 6 | 2 | 0 | NON_SYNONYMOU | MODERAT | MISSENSE | Gag/Cag | E273Q | PLES_26421 | | | | | . | . | . | . | . | . | G | G |
| NC_011770 | 2840136 | T | C | 54 | 6 | 6 | 0 | NON_SYNONYMOU | MODERAT | MISSENSE | Aac/Gac | N264D | PLES_26421 | C | C | C | C | . | . | . | . | . | . | C | C |
| NC_011770 | 2840139 | C | G | 54 | 6 | 6 | 0 | NON_SYNONYMOU | MODERAT | MISSENSE | Gtc/Ctc | V263L | PLES_26421 | G | G | G | G | . | . | . | . | . | . | G | G |
| NC_011770 | 2840157 | A | C | 57 | 6 | 6 | 0 | NON_SYNONYMOU | MODERAT | MISSENSE | Tcg/Gcg | S257A | PLES_26421 | C | C | C | C | . | . | . | . | . | . | C | C |
| NC_011770 | 2840250 | C | T | 45 | 5 | 1 | 0 | NON_SYNONYMOU | MODERAT | MISSENSE | Gtt/Att | V226I | PLES_26421 | | | | | T | . | . | . | . | . | | |
| NC_011770 | 2840294 | G | C | 36 | 6 | 2 | 0 | NON_SYNONYMOU | MODERAT | MISSENSE | gCg/gGg | A211G | PLES_26421 | C | C | | | . | . | . | . | . | . | | |
| NC_011770 | 2840303 | G | T | 36 | 6 | 2 | 0 | NON_SYNONYMOU | MODERAT | MISSENSE | tCc/tAc | S208Y | PLES_26421 | T | T | | | . | . | . | . | . | . | | |
| NC_011770 | 2840304 | A | C | 36 | 6 | 2 | 0 | NON_SYNONYMOU | MODERAT | MISSENSE | Tcc/Gcc | S208A | PLES_26421 | C | C | | | . | . | . | . | . | . | | |
| NC_011770 | 2840330 | A | G | 7 | 6 | 2 | 0 | NON_SYNONYMOU | MODERAT | MISSENSE | cTg/cCg | L199P | PLES_26421 | G | G | | | . | . | . | . | . | . | | |
| NC_011770 | 2840331 | G | C | 7 | 6 | 2 | 0 | NON_SYNONYMOU | MODERAT | MISSENSE | Ctg/Gtg | L199V | PLES_26421 | C | C | | | . | . | . | . | . | . | | |
| NC_011770 | 2840340 | T | C | 13 | 6 | 6 | 0 | NON_SYNONYMOU | MODERAT | MISSENSE | Aat/Gat | N196D | PLES_26421 | C | C | C | C | . | . | . | . | . | . | C | C |
| NC_011770 | 2840344 | C | G | 14 | 6 | 2 | 0 | NON_SYNONYMOU | MODERAT | MISSENSE | ttG/ttC | L194F | PLES_26421 | G | G | | | . | . | . | . | . | . | | |
| NC_011770 | 2840350 | C | G | 13 | 6 | 6 | 0 | NON_SYNONYMOU | MODERAT | MISSENSE | caG/caC | Q192H | PLES_26421 | G | G | G | G | . | . | . | . | . | . | G | G |
| NC_011770 | 2840351 | T | C | 14 | 6 | 6 | 0 | NON_SYNONYMOU | MODERAT | MISSENSE | cAg/cGg | Q192R | PLES_26421 | C | C | C | C | . | . | . | . | . | . | C | C |
| NC_011770 | 2840426 | C | T | 62 | 6 | 6 | 0 | NON_SYNONYMOU | MODERAT | MISSENSE | cGc/cAc | R167H | PLES_26421 | T | T | T | T | . | . | . | . | . | . | T | T |
| NC_011770 | 2840439 | G | T | 62 | 5 | 1 | 0 | NON_SYNONYMOU | MODERAT | MISSENSE | Cac/Aac | H163N | PLES_26421 | | | | | . | T | . | . | . | . | | |
| NC_011770 | 2840493 | C | T | 70 | 6 | 2 | 0 | NON_SYNONYMOU | MODERAT | MISSENSE | Gtg/Atg | V145M | PLES_26421 | T | T | | | . | . | . | . | . | . | | |
| NC_011770 | 2840550 | C | A | 72 | 6 | 6 | 0 | NON_SYNONYMOU | MODERAT | MISSENSE | Gcc/Tcc | A126S | PLES_26421 | A | A | A | A | . | . | . | . | . | . | A | A |
| NC_011770 | 2840614 | G | C | 27 | 7 | 3 | 0 | NON_SYNONYMOU | MODERAT | MISSENSE | agC/agG | S104R | PLES_26421 | C | . | C | C | . | . | . | . | . | . | | |
| NC_011770 | 2840615 | C | G | 27 | 7 | 3 | 0 | NON_SYNONYMOU | MODERAT | MISSENSE | aGc/aCc | S104T | PLES_26421 | G | . | G | G | . | . | . | . | . | . | | |
| NC_011770 | 2840616 | T | C | 26 | 7 | 3 | 0 | NON_SYNONYMOU | MODERAT | MISSENSE | Agc/Ggc | S104G | PLES_26421 | C | . | C | C | . | . | . | . | . | . | | |
| NC_011770 | 2840634 | G | A | 26 | 7 | 3 | 0 | NON_SYNONYMOU | MODERAT | MISSENSE | Ctc/Ttc | L98F | PLES_26421 | A | . | A | A | . | . | . | . | . | . | | |
| NC_011770 | 2840675 | G | C | 26 | 7 | 1 | 0 | NON_SYNONYMOU | MODERAT | MISSENSE | gCg/gGg | A84G | PLES_26421 | C | . | | | . | . | . | . | . | . | | |
| NC_011770 | 2840676 | C | T | 25 | 7 | 1 | 0 | NON_SYNONYMOU | MODERAT | MISSENSE | Gcg/Acg | A84T | PLES_26421 | T | . | | | . | . | . | . | . | . | | |
| NC_011770 | 2840693 | G | T | 27 | 7 | 3 | 0 | NON_SYNONYMOU | MODERAT | MISSENSE | gCc/gAc | A78D | PLES_26421 | T | . | T | T | . | . | . | . | . | . | | |
| NC_011770 | 2840694 | C | G | 26 | 7 | 3 | 0 | NON_SYNONYMOU | MODERAT | MISSENSE | Gcc/Ccc | A78P | PLES_26421 | G | . | G | G | . | . | . | . | . | . | | |
| NC_011770 | 2840714 | C | T | 48 | 6 | 2 | 0 | NON_SYNONYMOU | MODERAT | MISSENSE | cGc/cAc | R71H | PLES_26421 | T | T | | | . | . | . | . | . | . | | |
| NC_011770 | 2840727 | C | T | 46 | 6 | 6 | 0 | NON_SYNONYMOU | MODERAT | MISSENSE | Gcg/Acg | A67T | PLES_26421 | T | T | T | T | . | . | . | . | . | . | T | T |
| NC_011770 | 2840739 | A | T | 47 | 6 | 6 | 0 | NON_SYNONYMOU | MODERAT | MISSENSE | Tcc/Acc | S63T | PLES_26421 | T | T | T | T | . | . | . | . | . | . | T | T |
| NC_011770 | 2840750 | T | C | 45 | 6 | 6 | 0 | NON_SYNONYMOU | MODERAT | MISSENSE | aAg/aGg | K59R | PLES_26421 | C | C | C | C | . | . | . | . | . | . | C | C |
| NC_011770 | 2840751 | T | G | 45 | 6 | 4 | 0 | NON_SYNONYMOU | MODERAT | MISSENSE | Aag/Cag | K59Q | PLES_26421 | G | G | G | G | . | . | . | . | . | . | | |
| NC_011770 | 2840841 | T | G | 42 | 6 | 6 | 0 | NON_SYNONYMOU | MODERAT | MISSENSE | Atg/Ctg | M29L | PLES_26421 | G | G | G | G | . | . | . | . | . | . | G | G |
| NC_011770 | 2840880 | GCGC | G | 44 | 6 | 4 | 0 | CODON_DELETION | MODERATE | | gcg/ | A15 | PLES_26421 | G | G | G | G | . | . | . | . | . | . | | |
| NC_011770 | 2841431 | G | GC | 16 | 7 | 1 | 0 | FRAME_SHIFT | HIGH | | ttg/Cttg | L65L? | PLES_26431 | . | | | | . | . | . | . | . | . | | GC |
| NC_011770 | 2841550 | G | GAT | 33 | 6 | 4 | 0 | FRAME_SHIFT | HIGH | | cct/ATcct | P104I? | PLES_26431 | GAT | GAT | GAT | GAT | . | . | . | . | . | . | | |
| NC_011770 | 2841552 | CTG | C | 33 | 6 | 4 | 0 | FRAME_SHIFT | HIGH | | cctgcc/ | PA104 | PLES_26431 | C | C | C | C | . | . | . | . | . | . | | |
| NC_011770 | 2841249 | C | A | 41 | 6 | 4 | 0 | NON_SYNONYMOU | MODERAT | MISSENSE | aCt/aAt | T3N | PLES_26431 | A | A | A | A | . | . | . | . | . | . | | |

| | | | | | | | | | | | | | | | | | | | | | | | | | |
|---|---|---|---|---|---|---|---|---|---|---|---|---|---|---|---|---|---|---|---|---|---|---|---|---|---|
| NC_011770 | 2841273 | G | A | 39 | 6 | 4 | 0 | NON_SYNONYMOU | MODERAT | MISSENSE | cGg/cAg | R11Q | PLES_26431 | A | A | A | A | . | . | . | . | . | . | | |
| NC_011770 | 2841338 | C | A | 38 | 6 | 6 | 0 | NON_SYNONYMOU | MODERAT | MISSENSE | Ccg/Acg | P33T | PLES_26431 | A | A | A | A | . | . | . | . | . | . | A | A |
| NC_011770 | 2841368 | A | G | 40 | 6 | 6 | 0 | NON_SYNONYMOU | MODERAT | MISSENSE | Atc/Gtc | I43V | PLES_26431 | G | G | G | G | . | . | . | . | . | . | G | G |
| NC_011770 | 2841393 | C | A | 38 | 6 | 4 | 0 | NON_SYNONYMOU | MODERAT | MISSENSE | aCg/aAg | T51K | PLES_26431 | A | A | A | A | . | . | . | . | . | . | | |
| NC_011770 | 2841405 | A | G | 36 | 6 | 4 | 0 | NON_SYNONYMOU | MODERAT | MISSENSE | aAc/aGc | N55S | PLES_26431 | G | G | G | G | . | . | . | . | . | . | | |
| NC_011770 | 2841414 | G | C | 17 | 6 | 4 | 0 | NON_SYNONYMOU | MODERAT | MISSENSE | gGt/gCt | G58A | PLES_26431 | C | C | C | C | . | . | . | . | . | . | | |
| NC_011770 | 2841425 | G | C | 16 | 7 | 3 | 0 | NON_SYNONYMOU | MODERAT | MISSENSE | Gcg/Ccg | A62P | PLES_26431 | . | C | C | C | . | . | . | . | . | . | | |
| NC_011770 | 2841426 | C | A | 16 | 7 | 5 | 0 | NON_SYNONYMOU | MODERAT | MISSENSE | gCg/gAg | A62E | PLES_26431 | . | A | A | A | . | . | . | . | . | . | G | G |
| NC_011770 | 2841428 | C | G | 16 | 7 | 3 | 0 | NON_SYNONYMOU | MODERAT | MISSENSE | Cag/Gag | Q63E | PLES_26431 | . | G | G | G | . | . | . | . | . | . | | |
| NC_011770 | 2841437 | C | G | 17 | 7 | 3 | 0 | NON_SYNONYMOU | MODERAT | MISSENSE | Cag/Gag | Q66E | PLES_26431 | . | G | G | G | . | . | . | . | . | . | | |
| NC_011770 | 2841450 | C | G | 20 | 7 | 3 | 0 | NON_SYNONYMOU | MODERAT | MISSENSE | gCc/gGc | A70G | PLES_26431 | . | G | G | G | . | . | . | . | . | . | | |
| NC_011770 | 2841461 | T | G | 19 | 7 | 5 | 0 | NON_SYNONYMOU | MODERAT | MISSENSE | Tcc/Gcc | S74A | PLES_26431 | . | G | G | G | . | . | . | . | . | . | G | G |
| NC_011770 | 2841466 | C | G | 21 | 7 | 3 | 0 | NON_SYNONYMOU | MODERAT | MISSENSE | gaC/gaG | D75E | PLES_26431 | . | G | G | G | . | . | . | . | . | . | | |
| NC_011770 | 2841476 | A | G | 20 | 7 | 5 | 0 | NON_SYNONYMOU | MODERAT | MISSENSE | Aca/Gca | T79A | PLES_26431 | . | G | G | G | . | . | . | . | . | . | G | G |
| NC_011770 | 2841482 | G | A | 20 | 7 | 5 | 0 | NON_SYNONYMOU | MODERAT | MISSENSE | Gct/Act | A81T | PLES_26431 | . | A | A | A | . | . | . | . | . | . | A | A |
| NC_011770 | 2841483 | C | G | 20 | 7 | 5 | 0 | NON_SYNONYMOU | MODERAT | MISSENSE | gCt/gGt | A81G | PLES_26431 | . | G | G | G | . | . | . | . | . | . | A | A |
| NC_011770 | 2841512 | T | G | 31 | 6 | 6 | 0 | NON_SYNONYMOU | MODERAT | MISSENSE | Ttg/Gtg | L91V | PLES_26431 | G | G | G | G | . | . | . | . | . | . | G | G |
| NC_011770 | 2841513 | T | C | 32 | 6 | 4 | 0 | NON_SYNONYMOU | MODERAT | MISSENSE | tTg/tCg | L91S | PLES_26431 | C | C | C | C | . | . | . | . | . | . | | |
| NC_011770 | 2841524 | G | A | 34 | 6 | 6 | 0 | NON_SYNONYMOU | MODERAT | MISSENSE | Gcc/Acc | A95T | PLES_26431 | A | A | A | A | . | . | . | . | . | . | A | A |
| NC_011770 | 2841539 | G | C | 34 | 6 | 6 | 0 | NON_SYNONYMOU | MODERAT | MISSENSE | Gac/Cac | D100H | PLES_26431 | C | C | C | C | . | . | . | . | . | . | C | C |
| NC_011770 | 2841548 | C | G | 34 | 6 | 4 | 0 | NON_SYNONYMOU | MODERAT | MISSENSE | Ccg/Gcg | P103A | PLES_26431 | G | G | G | G | . | . | . | . | . | . | | |
| NC_011770 | 2841554 | G | C | 34 | 6 | 1 | 0 | NON_SYNONYMOU | MODERAT | MISSENSE | Gcc/Ccc | A105P | PLES_26431 | . | . | . | C | . | . | . | . | . | . | | |
| NC_011770 | 2841561 | T | A | 37 | 6 | 4 | 0 | NON_SYNONYMOU | MODERAT | MISSENSE | tTc/tAc | F107Y | PLES_26431 | A | A | A | A | . | . | . | . | . | . | | |
| NC_011770 | 2841797 | G | A | 49 | 6 | 4 | 0 | NON_SYNONYMOU | MODERAT | MISSENSE | Gcg/Acg | A186T | PLES_26431 | A | A | A | A | . | . | . | . | . | . | | |
| NC_011770 | 2841438 | AG | CG | 16 | 7 | 4 | 0 | FRAME_SHIFT | HIGH | | cag/ | Q66 | PLES_26431 (i | . | **CG** | **CG** | **CG** | . | . | . | . | . | . | . | **A** |
| NC_011770 | 2841888 | A | G | 45 | 6 | 4 | 0 | NON_SYNONYMOU | MODERAT | MISSENSE | cAc/cGc | H3R | PLES_26441 | G | G | G | G | . | . | . | . | . | . | | |
| NC_011770 | 2841900 | C | T | 47 | 6 | 4 | 0 | NON_SYNONYMOU | MODERAT | MISSENSE | tCg/tTg | S7L | PLES_26441 | T | T | T | T | . | . | . | . | . | . | | |
| NC_011770 | 2841983 | A | G | 45 | 6 | 4 | 0 | NON_SYNONYMOU | MODERAT | MISSENSE | Agc/Ggc | S35G | PLES_26441 | G | G | G | G | . | . | . | . | . | . | | |
| NC_011770 | 2842010 | G | C | 42 | 6 | 4 | 0 | NON_SYNONYMOU | MODERAT | MISSENSE | Ggg/Cgg | G44R | PLES_26441 | C | C | C | C | . | . | . | . | . | . | | |
| NC_011770 | 2842013 | G | A | 42 | 6 | 4 | 0 | NON_SYNONYMOU | MODERAT | MISSENSE | Gcc/Acc | A45T | PLES_26441 | A | A | A | A | . | . | . | . | . | . | | |
| NC_011770 | 2842092 | C | A | 40 | 6 | 2 | 0 | NON_SYNONYMOU | MODERAT | MISSENSE | gCg/gAg | A71E | PLES_26441 | | | | | . | . | . | . | . | . | A | A |
| NC_011770 | 2842100 | C | A | 16 | 10 | 2 | 0 | NON_SYNONYMOU | MODERAT | MISSENSE | Caa/Aaa | Q74K | PLES_26441 | . | . | . | . | . | . | . | . | . | . | A | A |
| NC_011770 | 2842127 | C | G | 30 | 6 | 4 | 0 | NON_SYNONYMOU | MODERAT | MISSENSE | Caa/Gaa | Q83E | PLES_26441 | G | G | G | G | . | . | . | . | . | . | | |
| NC_011770 | 2842129 | A | G | 30 | 6 | 6 | 0 | NON_SYNONYMOU | MODERAT | MISSENSE | caA/caC | Q83H | PLES_26441 | G | G | G | G | . | . | . | . | . | . | C | C |
| NC_011770 | 2842130 | G | A | 29 | 6 | 4 | 0 | NON_SYNONYMOU | MODERAT | MISSENSE | Gcc/Acc | A84T | PLES_26441 | A | A | A | A | . | . | . | . | . | . | | |
| NC_011770 | 2842131 | C | G | 29 | 6 | 4 | 0 | NON_SYNONYMOU | MODERAT | MISSENSE | gCc/gGc | A84G | PLES_26441 | G | G | G | G | . | . | . | . | . | . | | |
| NC_011770 | 2842136 | A | C | 30 | 6 | 4 | 0 | NON_SYNONYMOU | MODERAT | MISSENSE | Atg/Ctg | M86L | PLES_26441 | C | C | C | C | . | . | . | . | . | . | | |
| NC_011770 | 2842138 | G | C | 30 | 6 | 4 | 0 | NON_SYNONYMOU | MODERAT | MISSENSE | atG/atC | M86I | PLES_26441 | C | C | C | C | . | . | . | . | . | . | | |
| NC_011770 | 2842139 | G | T | 30 | 6 | 4 | 0 | NON_SYNONYMOU | MODERAT | MISSENSE | Gtg/Ttg | V87L | PLES_26441 | T | T | T | T | . | . | . | . | . | . | | |
| NC_011770 | 2842157 | A | C | 30 | 6 | 6 | 0 | NON_SYNONYMOU | MODERAT | MISSENSE | Atg/Ctg | M93L | PLES_26441 | C | C | C | C | . | . | . | . | . | . | C | C |
| NC_011770 | 2842169 | T | C | 30 | 6 | 6 | 0 | NON_SYNONYMOU | MODERAT | MISSENSE | Ttt/Ctt | F97L | PLES_26441 | C | C | C | C | . | . | . | . | . | . | C | C |
| NC_011770 | 2842170 | T | G | 30 | 6 | 4 | 0 | NON_SYNONYMOU | MODERAT | MISSENSE | tTt/tGt | F97C | PLES_26441 | G | G | G | G | . | . | . | . | . | . | | |
| NC_011770 | 2842171 | T | G | 31 | 6 | 6 | 0 | NON_SYNONYMOU | MODERAT | MISSENSE | ttT/ttG | F97L | PLES_26441 | G | G | G | G | . | . | . | . | . | . | C | C |
| NC_011770 | 2842190 | C | G | 42 | 6 | 4 | 0 | NON_SYNONYMOU | MODERAT | MISSENSE | Cag/Gag | Q104E | PLES_26441 | G | G | G | G | . | . | . | . | . | . | | |
| NC_011770 | 2842191 | A | C | 41 | 6 | 2 | 0 | NON_SYNONYMOU | MODERAT | MISSENSE | cAg/cCg | Q104P | PLES_26441 | | | | | . | . | . | . | . | . | C | C |
| NC_011770 | 2842263 | G | A | 42 | 6 | 6 | 0 | NON_SYNONYMOU | MODERAT | MISSENSE | cGg/cAg | R128Q | PLES_26441 | A | A | A | A | . | . | . | . | . | . | A | A |
| NC_011770 | 2842289 | C | G | 40 | 6 | 2 | 0 | NON_SYNONYMOU | MODERAT | MISSENSE | Ctc/Gtc | L137V | PLES_26441 | | | | | . | . | . | . | . | . | G | G |
| NC_011770 | 2842307 | G | C | 39 | 6 | 6 | 0 | NON_SYNONYMOU | MODERAT | MISSENSE | Gcg/Ccg | A143P | PLES_26441 | C | C | C | C | . | . | . | . | . | . | C | C |
| NC_011770 | 2842355 | A | G | 45 | 6 | 2 | 0 | NON_SYNONYMOU | MODERAT | MISSENSE | Aag/Gag | K159E | PLES_26441 | | | | | . | . | . | . | . | . | G | G |
| NC_011770 | 2842356 | A | C | 45 | 6 | 2 | 0 | NON_SYNONYMOU | MODERAT | MISSENSE | aAg/aCg | K159T | PLES_26441 | | | | | . | . | . | . | . | . | C | C |
| NC_011770 | 2842383 | G | C | 46 | 6 | 4 | 0 | NON_SYNONYMOU | MODERAT | MISSENSE | gGa/gCa | G168A | PLES_26441 | C | C | C | C | . | . | . | . | . | . | | |
| NC_011770 | 2842385 | G | A | 47 | 6 | 2 | 0 | NON_SYNONYMOU | MODERAT | MISSENSE | Gcg/Acg | A169T | PLES_26441 | | | | | . | . | . | . | . | . | A | A |
| NC_011770 | 2842394 | G | A | 45 | 6 | 2 | 0 | NON_SYNONYMOU | MODERAT | MISSENSE | Ggc/Agc | G172S | PLES_26441 | | | | | . | . | . | . | . | . | A | A |
| NC_011770 | 2842400 | A | G | 43 | 6 | 6 | 0 | NON_SYNONYMOU | MODERAT | MISSENSE | Atc/Gtc | I174V | PLES_26441 | G | G | G | G | . | . | . | . | . | . | G | G |
| NC_011770 | 2842405 | G | T | 47 | 6 | 4 | 0 | NON_SYNONYMOU | MODERAT | MISSENSE | gaG/gaT | E175D | PLES_26441 | T | T | T | T | . | . | . | . | . | . | | |
| NC_011770 | 2842449 | T | C | 42 | 6 | 6 | 0 | NON_SYNONYMOU | MODERAT | MISSENSE | gTt/gCt | V190A | PLES_26441 | C | C | C | C | . | . | . | . | . | . | C | C |
| NC_011770 | 2842460 | A | G | 42 | 6 | 4 | 0 | NON_SYNONYMOU | MODERAT | MISSENSE | Atc/Gtc | I194V | PLES_26441 | G | G | G | G | . | . | . | . | . | . | | |
| NC_011770 | 2842491 | G | A | 43 | 6 | 6 | 0 | NON_SYNONYMOU | MODERAT | MISSENSE | gGg/gAg | G204E | PLES_26441 | A | A | A | A | . | . | . | . | . | . | A | A |
| NC_011770 | 2842617 | C | T | 37 | 6 | 4 | 0 | NON_SYNONYMOU | MODERAT | MISSENSE | aCc/aTc | T7I | PLES_26451 | T | T | T | T | . | . | . | . | . | . | | |
| NC_011770 | 2842643 | G | A | 32 | 6 | 6 | 0 | NON_SYNONYMOU | MODERAT | MISSENSE | Gcc/Acc | A16T | PLES_26451 | A | A | A | A | . | . | . | . | . | . | A | A |
| NC_011770 | 2842665 | C | T | 31 | 6 | 2 | 0 | NON_SYNONYMOU | MODERAT | MISSENSE | gCc/gTc | A23V | PLES_26451 | | | | | . | . | . | . | . | . | T | T |
| NC_011770 | 2842676 | C | G | 30 | 6 | 6 | 0 | NON_SYNONYMOU | MODERAT | MISSENSE | Cgc/Ggc | R27G | PLES_26451 | G | G | G | G | . | . | . | . | . | . | G | G |

| | | | | | | | | | | | | | | | | | | | | | | | | | |
|---|---|---|---|---|---|---|---|---|---|---|---|---|---|---|---|---|---|---|---|---|---|---|---|---|---|
| NC_011770 | 2842689 | C | T | 29 | 6 | 2 | 0 | NON_SYNONYMOU | MODERAT | MISSENSE | gCg/gTg | A31V | PLES_26451 | | | | | . | . | . | . | . | . | T | T |
| NC_011770 | 2842715 | A | G | 9 | 8 | 4 | 0 | NON_SYNONYMOU | MODERAT | MISSENSE | Atg/Gtg | M40V | PLES_26451 | G | G | G | G | . | . | . | . | . | . | . | . |
| NC_011770 | 2842721 | C | G | 9 | 8 | 4 | 0 | NON_SYNONYMOU | MODERAT | MISSENSE | Cga/Gga | R42G | PLES_26451 | G | G | G | G | . | . | . | . | . | . | . | . |
| NC_011770 | 2842722 | G | C | 8 | 8 | 4 | 0 | NON_SYNONYMOU | MODERAT | MISSENSE | cGa/cCa | R42P | PLES_26451 | C | C | C | C | . | . | . | . | . | . | . | . |
| NC_011770 | 2842728 | G | T | 8 | 8 | 4 | 0 | NON_SYNONYMOU | MODERAT | MISSENSE | gGc/gTc | G44V | PLES_26451 | T | T | T | T | . | . | . | . | . | . | . | . |
| NC_011770 | 2842752 | C | T | 11 | 8 | 4 | 0 | NON_SYNONYMOU | MODERAT | MISSENSE | gCc/gTc | A52V | PLES_26451 | T | T | T | T | . | . | . | . | . | . | . | . |
| NC_011770 | 2842760 | G | T | 18 | 6 | 4 | 0 | NON_SYNONYMOU | MODERAT | MISSENSE | Gct/Tct | A55S | PLES_26451 | T | T | T | T | . | . | . | . | . | . | | |
| NC_011770 | 2842766 | C | G | 18 | 6 | 4 | 0 | NON_SYNONYMOU | MODERAT | MISSENSE | Ctg/Gtg | L57V | PLES_26451 | G | G | G | G | . | . | . | . | . | . | | |
| NC_011770 | 2842772 | G | T | 18 | 6 | 4 | 0 | NON_SYNONYMOU | MODERAT | MISSENSE | Gtg/Ttg | V59L | PLES_26451 | T | T | T | T | . | . | . | . | . | . | | |
| NC_011770 | 2842773 | T | C | 20 | 6 | 2 | 0 | NON_SYNONYMOU | MODERAT | MISSENSE | gTg/gCg | V59A | PLES_26451 | | | | | . | . | . | . | . | . | C | C |
| NC_011770 | 2842785 | T | C | 20 | 6 | 2 | 0 | NON_SYNONYMOU | MODERAT | MISSENSE | gTt/gCt | V63A | PLES_26451 | | | | | . | . | . | . | . | . | C | C |
| NC_011770 | 2842808 | A | T | 22 | 6 | 6 | 0 | NON_SYNONYMOU | MODERAT | MISSENSE | Atg/Ttg | M71L | PLES_26451 | T | T | T | T | . | . | . | . | . | . | T | T |
| NC_011770 | 2842821 | C | T | 22 | 6 | 6 | 0 | NON_SYNONYMOU | MODERAT | MISSENSE | gCg/gTg | A75V | PLES_26451 | T | T | T | T | . | . | . | . | . | . | T | T |
| NC_011770 | 2842824 | C | T | 22 | 6 | 2 | 0 | NON_SYNONYMOU | MODERAT | MISSENSE | gCg/gTg | A76V | PLES_26451 | | | | | . | . | . | . | . | . | T | T |
| NC_011770 | 2842830 | G | C | 24 | 6 | 4 | 0 | NON_SYNONYMOU | MODERAT | MISSENSE | gGa/gCa | G78A | PLES_26451 | C | C | C | C | . | . | . | . | . | . | | |
| NC_011770 | 2842858 | T | C | 23 | 6 | 6 | 0 | NON_SYNONYMOU | MODERAT | MISSENSE | gaT/gaA | D87E | PLES_26451 | C | C | C | C | . | . | . | . | . | . | A | A |
| NC_011770 | 2843253 | T | TGC | 13 | 6 | 6 | 0 | FRAME_SHIFT | HIGH | | tca/tcaGC | S414S? | PLES_26461 | TGC | TGC | TGC | TGC | . | . | . | . | . | . | TGC | TGC |
| NC_011770 | 2843255 | GAA | G | 13 | 6 | 6 | 0 | FRAME_SHIFT | HIGH | | gattca/ | DS413 | PLES_26461 | G | G | G | G | . | . | . | . | . | . | G | G |
| NC_011770 | 2843711 | G | C | 26 | 6 | 4 | 0 | STOP_GAINED | HIGH | NONSENSE | tCa/tGa | S262* | PLES_26461 | C | C | C | C | . | . | . | . | . | . | | |
| NC_011770 | 2843052 | G | A | 6 | 7 | 3 | 0 | NON_SYNONYMOU | MODERAT | MISSENSE | Ctt/Ttt | L482F | PLES_26461 | A | . | A | A | . | . | . | . | . | . | | |
| NC_011770 | 2843063 | A | G | 6 | 7 | 3 | 0 | NON_SYNONYMOU | MODERAT | MISSENSE | gTc/gCc | V478A | PLES_26461 | G | . | G | G | . | . | . | . | . | . | | |
| NC_011770 | 2843138 | A | G | 12 | 6 | 4 | 0 | NON_SYNONYMOU | MODERAT | MISSENSE | aTg/aCg | M453T | PLES_26461 | G | G | G | G | . | . | . | . | . | . | | |
| NC_011770 | 2843139 | T | C | 12 | 6 | 4 | 0 | NON_SYNONYMOU | MODERAT | MISSENSE | Atg/Gtg | M453V | PLES_26461 | C | C | C | C | . | . | . | . | . | . | | |
| NC_011770 | 2843291 | G | A | 21 | 6 | 4 | 0 | NON_SYNONYMOU | MODERAT | MISSENSE | aCg/aTg | T402M | PLES_26461 | A | A | A | A | . | . | . | . | . | . | | |
| NC_011770 | 2843292 | T | C | 22 | 6 | 4 | 0 | NON_SYNONYMOU | MODERAT | MISSENSE | Acg/Gcg | T402A | PLES_26461 | C | C | C | C | . | . | . | . | . | . | | |
| NC_011770 | 2843407 | G | C | 24 | 6 | 2 | 0 | NON_SYNONYMOU | MODERAT | MISSENSE | atC/atG | I363M | PLES_26461 | | | | | . | . | . | . | . | . | C | C |
| NC_011770 | 2843409 | T | C | 24 | 6 | 6 | 0 | NON_SYNONYMOU | MODERAT | MISSENSE | Atc/Gtc | I363V | PLES_26461 | C | C | C | C | . | . | . | . | . | . | C | C |
| NC_011770 | 2843421 | A | C | 22 | 6 | 2 | 0 | NON_SYNONYMOU | MODERAT | MISSENSE | Tcc/Gcc | S359A | PLES_26461 | | | | | . | . | . | . | . | . | C | C |
| NC_011770 | 2843427 | G | T | 22 | 6 | 4 | 0 | NON_SYNONYMOU | MODERAT | MISSENSE | Cgc/Agc | R357S | PLES_26461 | T | T | T | T | . | . | . | . | . | . | | |
| NC_011770 | 2843457 | T | G | 23 | 6 | 2 | 0 | NON_SYNONYMOU | MODERAT | MISSENSE | Att/Ctt | I347L | PLES_26461 | | | | | . | . | . | . | . | . | G | G |
| NC_011770 | 2843548 | G | C | 26 | 6 | 2 | 0 | NON_SYNONYMOU | MODERAT | MISSENSE | agC/agG | S316R | PLES_26461 | | | | | . | . | . | . | . | . | C | C |
| NC_011770 | 2843549 | C | G | 26 | 6 | 2 | 0 | NON_SYNONYMOU | MODERAT | MISSENSE | aGc/aCc | S316T | PLES_26461 | | | | | . | . | . | . | . | . | G | G |
| NC_011770 | 2843550 | T | A | 26 | 6 | 2 | 0 | NON_SYNONYMOU | MODERAT | MISSENSE | Agc/Tgc | S316C | PLES_26461 | | | | | . | . | . | . | . | . | A | A |
| NC_011770 | 2843589 | C | T | 31 | 6 | 2 | 0 | NON_SYNONYMOU | MODERAT | MISSENSE | Gcc/Acc | A303T | PLES_26461 | | | | | . | . | . | . | . | . | T | T |
| NC_011770 | 2843595 | T | C | 32 | 6 | 2 | 0 | NON_SYNONYMOU | MODERAT | MISSENSE | Atc/Gtc | I301V | PLES_26461 | | | | | . | . | . | . | . | . | C | C |
| NC_011770 | 2843616 | C | T | 31 | 6 | 2 | 0 | NON_SYNONYMOU | MODERAT | MISSENSE | Gcc/Acc | A294T | PLES_26461 | | | | | . | . | . | . | . | . | T | T |
| NC_011770 | 2843651 | T | C | 32 | 6 | 4 | 0 | NON_SYNONYMOU | MODERAT | MISSENSE | aAt/aGt | N282S | PLES_26461 | C | C | C | C | . | . | . | . | . | . | | |
| NC_011770 | 2843660 | G | C | 30 | 6 | 6 | 0 | NON_SYNONYMOU | MODERAT | MISSENSE | aCc/aGc | T279S | PLES_26461 | C | C | C | C | . | . | . | . | . | . | C | C |
| NC_011770 | 2843712 | A | T | 25 | 6 | 6 | 0 | NON_SYNONYMOU | MODERAT | MISSENSE | Tca/Aca | S262T | PLES_26461 | T | T | T | T | . | . | . | . | . | . | C | C |
| NC_011770 | 2843723 | C | T | 35 | 5 | 1 | 0 | NON_SYNONYMOU | MODERAT | MISSENSE | aGg/aAg | R258K | PLES_26461 | | | | | . | . | . | T | . | . | | |
| NC_011770 | 2843747 | T | C | 34 | 6 | 6 | 0 | NON_SYNONYMOU | MODERAT | MISSENSE | aAt/aGt | N250S | PLES_26461 | C | C | C | C | . | . | . | . | . | . | C | C |
| NC_011770 | 2843780 | T | C | 34 | 6 | 4 | 0 | NON_SYNONYMOU | MODERAT | MISSENSE | gAc/gGc | D239G | PLES_26461 | C | C | C | C | . | . | . | . | . | . | | |
| NC_011770 | 2843781 | C | T | 34 | 6 | 4 | 0 | NON_SYNONYMOU | MODERAT | MISSENSE | Gac/Aac | D239N | PLES_26461 | T | T | T | T | . | . | . | . | . | . | | |
| NC_011770 | 2843795 | G | T | 32 | 6 | 4 | 0 | NON_SYNONYMOU | MODERAT | MISSENSE | gCa/gAa | A234E | PLES_26461 | T | T | T | T | . | . | . | . | . | . | | |
| NC_011770 | 2843796 | C | A | 31 | 6 | 2 | 0 | NON_SYNONYMOU | MODERAT | MISSENSE | Gca/Tca | A234S | PLES_26461 | | | | | . | . | . | . | . | . | A | A |
| NC_011770 | 2843805 | C | G | 31 | 6 | 4 | 0 | NON_SYNONYMOU | MODERAT | MISSENSE | Gcg/Ccg | A231P | PLES_26461 | G | G | G | G | . | . | . | . | . | . | | |
| NC_011770 | 2843821 | G | T | 35 | 6 | 4 | 0 | NON_SYNONYMOU | MODERAT | MISSENSE | agC/agA | S225R | PLES_26461 | T | T | T | T | . | . | . | . | . | . | | |
| NC_011770 | 2843822 | C | G | 35 | 6 | 4 | 0 | NON_SYNONYMOU | MODERAT | MISSENSE | aGc/aCc | S225T | PLES_26461 | G | G | G | G | . | . | . | . | . | . | | |
| NC_011770 | 2843837 | C | T | 33 | 6 | 4 | 0 | NON_SYNONYMOU | MODERAT | MISSENSE | gGt/gAt | G220D | PLES_26461 | T | T | T | T | . | . | . | . | . | . | | |
| NC_011770 | 2843838 | C | T | 34 | 6 | 4 | 0 | NON_SYNONYMOU | MODERAT | MISSENSE | Ggt/Agt | G220S | PLES_26461 | T | T | T | T | . | . | . | . | . | . | | |
| NC_011770 | 2843856 | G | T | 35 | 6 | 2 | 0 | NON_SYNONYMOU | MODERAT | MISSENSE | Cgc/Agc | R214S | PLES_26461 | | | | | . | . | . | . | . | . | T | T |
| NC_011770 | 2843880 | G | T | 24 | 7 | 1 | 0 | NON_SYNONYMOU | MODERAT | MISSENSE | Cac/Aac | H206N | PLES_26461 | | | | | . | . | . | . | . | . | T | . |
| NC_011770 | 2843899 | A | C | 22 | 8 | 4 | 0 | NON_SYNONYMOU | MODERAT | MISSENSE | gaT/gaG | D199E | PLES_26461 | C | C | C | C | . | . | . | . | . | . | . | . |
| NC_011770 | 2843901 | C | G | 21 | 8 | 4 | 0 | NON_SYNONYMOU | MODERAT | MISSENSE | Gat/Cat | D199H | PLES_26461 | G | G | G | G | . | . | . | . | . | . | . | . |
| NC_011770 | 2843988 | C | T | 33 | 6 | 2 | 0 | NON_SYNONYMOU | MODERAT | MISSENSE | Gac/Aac | D170N | PLES_26461 | | | | | . | . | . | . | . | . | T | T |
| NC_011770 | 2844013 | C | G | 35 | 6 | 6 | 0 | NON_SYNONYMOU | MODERAT | MISSENSE | agG/agC | R161S | PLES_26461 | G | G | G | G | . | . | . | . | . | . | G | G |
| NC_011770 | 2844063 | T | C | 35 | 6 | 4 | 0 | NON_SYNONYMOU | MODERAT | MISSENSE | Agc/Ggc | S145G | PLES_26461 | C | C | C | C | . | . | . | . | . | . | | |
| NC_011770 | 2844119 | G | C | 33 | 6 | 2 | 0 | NON_SYNONYMOU | MODERAT | MISSENSE | tCg/tGg | S126W | PLES_26461 | | | | | . | . | . | . | . | . | C | C |
| NC_011770 | 2844120 | A | T | 34 | 6 | 2 | 0 | NON_SYNONYMOU | MODERAT | MISSENSE | Tcg/Acg | S126T | PLES_26461 | | | | | . | . | . | . | . | . | T | T |
| NC_011770 | 2844160 | T | A | 35 | 6 | 6 | 0 | NON_SYNONYMOU | MODERAT | MISSENSE | gaA/gaT | E112D | PLES_26461 | A | A | A | A | . | . | . | . | . | . | A | A |
| NC_011770 | 2844164 | G | C | 35 | 6 | 4 | 0 | NON_SYNONYMOU | MODERAT | MISSENSE | aCg/aGg | T111R | PLES_26461 | C | C | C | C | . | . | . | . | . | . | | |
| NC_011770 | 2844165 | T | C | 35 | 6 | 4 | 0 | NON_SYNONYMOU | MODERAT | MISSENSE | Acg/Gcg | T111A | PLES_26461 | C | C | C | C | . | . | . | . | . | . | | |

| | | | | | | | | | | | | | | | | | | | | | | | | | |
|---|---|---|---|---|---|---|---|---|---|---|---|---|---|---|---|---|---|---|---|---|---|---|---|---|---|
| NC_011770 | 2844174 | G | C | 36 | 6 | 4 | 0 | NON_SYNONYMOU | MODERAT | MISSENSE | Ccg/Gcg | P108A | PLES_26461 | C | C | C | C | . | . | . | . | . | . | | |
| NC_011770 | 2844179 | C | T | 37 | 6 | 2 | 0 | NON_SYNONYMOU | MODERAT | MISSENSE | aGc/aAc | S106N | PLES_26461 | | | | | . | . | . | . | . | . | T | T |
| NC_011770 | 2844224 | C | T | 37 | 6 | 6 | 0 | NON_SYNONYMOU | MODERAT | MISSENSE | aGc/aAc | S91N | PLES_26461 | T | T | T | T | . | . | . | . | . | . | T | T |
| NC_011770 | 2844307 | A | G | 32 | 6 | 6 | 0 | NON_SYNONYMOU | MODERAT | MISSENSE | agT/agG | S63R | PLES_26461 | G | G | G | G | . | . | . | . | . | . | C | C |
| NC_011770 | 2844308 | C | G | 33 | 6 | 2 | 0 | NON_SYNONYMOU | MODERAT | MISSENSE | aGt/aCt | S63T | PLES_26461 | | | | | . | . | . | . | . | . | G | G |
| NC_011770 | 2844309 | T | A | 33 | 6 | 2 | 0 | NON_SYNONYMOU | MODERAT | MISSENSE | Agt/Tgt | S63C | PLES_26461 | | | | | . | . | . | . | . | . | A | A |
| NC_011770 | 2844372 | C | T | 34 | 6 | 6 | 0 | NON_SYNONYMOU | MODERAT | MISSENSE | Gtc/Atc | V42I | PLES_26461 | T | T | T | T | . | . | . | . | . | . | T | T |
| NC_011770 | 2844453 | G | A | 37 | 6 | 2 | 0 | NON_SYNONYMOU | MODERAT | MISSENSE | Ctc/Ttc | L15F | PLES_26461 | | | | | . | . | . | . | . | . | A | A |
| NC_011770 | 2843722 | C | G | 35 | 5 | 3 | 0 | NON_SYNONYMOU | MODERAT | MISSENSE | agG/agC | R258S | PLES_26461 | G | | G | G | . | . | . | | . | . | | |
| NC_011770 | 2844865 | CAT | C | 16 | 7 | 1 | 0 | FRAME_SHIFT | HIGH | | atg/ | M6 | PLES_26471 | | | | | . | . | . | . | . | . | . | C |
| NC_011770 | 2844873 | C | CAT | 17 | 7 | 1 | 0 | FRAME_SHIFT | HIGH | | ccg/ccgAT | P3P? | PLES_26471 | | | | | . | . | . | . | . | . | . | CAT |
| NC_011770 | 2844561 | C | T | 32 | 6 | 4 | 0 | NON_SYNONYMOU | MODERAT | MISSENSE | Gtc/Atc | V108I | PLES_26471 | T | T | T | T | . | . | . | . | . | . | | |
| NC_011770 | 2844570 | C | T | 19 | 7 | 1 | 0 | NON_SYNONYMOU | MODERAT | MISSENSE | Gtc/Atc | V105I | PLES_26471 | | | | | . | . | . | . | . | . | T | . |
| NC_011770 | 2844819 | T | C | 8 | 10 | 2 | 0 | NON_SYNONYMOU | MODERAT | MISSENSE | Acg/Gcg | T22A | PLES_26471 | . | . | . | . | . | . | . | . | . | . | C | C |
| NC_011770 | 2844836 | C | T | 21 | 6 | 2 | 0 | NON_SYNONYMOU | MODERAT | MISSENSE | aGc/aAc | S16N | PLES_26471 | | | | | . | . | . | . | . | . | T | T |
| NC_011770 | 2844840 | A | T | 21 | 6 | 4 | 0 | NON_SYNONYMOU | MODERAT | MISSENSE | Ttc/Atc | F15I | PLES_26471 | T | T | T | T | . | . | . | . | . | . | | |
| NC_011770 | 2844841 | C | G | 21 | 6 | 4 | 0 | NON_SYNONYMOU | MODERAT | MISSENSE | ttG/ttC | L14F | PLES_26471 | G | G | G | G | . | . | . | . | . | . | | |
| NC_011770 | 2844851 | A | G | 13 | 7 | 1 | 0 | NON_SYNONYMOU | MODERAT | MISSENSE | cTg/cCg | L11P | PLES_26471 | | | | | . | . | . | . | . | . | . | G |
| NC_011770 | 2844854 | G | C | 13 | 7 | 1 | 0 | NON_SYNONYMOU | MODERAT | MISSENSE | cCg/cGg | P10R | PLES_26471 | | | | | . | . | . | . | . | . | . | C |
| NC_011770 | 2844857 | C | A | 13 | 7 | 1 | 0 | NON_SYNONYMOU | MODERAT | MISSENSE | cGg/cTg | R9L | PLES_26471 | | | | | . | . | . | . | . | . | . | A |
| NC_011770 | 2844870 | G | A | 16 | 7 | 1 | 0 | NON_SYNONYMOU | MODERAT | MISSENSE | Cgc/Tgc | R5C | PLES_26471 | | | | | . | . | . | . | . | . | . | A |
| NC_011770 | 2844872 | G | T | 15 | 8 | 4 | 0 | NON_SYNONYMOU | MODERAT | MISSENSE | gCg/gAg | A4E | PLES_26471 | T | T | T | T | . | . | . | . | . | . | . | . |
| NC_011770 | 2844875 | G | A | 17 | 7 | 1 | 0 | NON_SYNONYMOU | MODERAT | MISSENSE | cCg/cTg | P3L | PLES_26471 | | | | | . | . | . | . | . | . | . | A |
| NC_011770 | 2844876 | G | C | 29 | 6 | 4 | 0 | NON_SYNONYMOU | MODERAT | MISSENSE | Ccg/Gcg | P3A | PLES_26471 | C | C | C | C | . | . | . | . | . | . | | |
| NC_011770 | 2845323 | CTT | C | 16 | 8 | 2 | 0 | FRAME_SHIFT | HIGH | | aag/ | K318 | PLES_26481 | . | . | C | C | . | . | . | . | . | . | | |
| NC_011770 | 2844890 | T | C | 28 | 6 | 6 | 0 | NON_SYNONYMOU | MODERAT | MISSENSE | Agt/Ggt | S463G | PLES_26481 | C | C | C | C | . | . | . | . | . | . | C | C |
| NC_011770 | 2844911 | G | A | 31 | 6 | 6 | 0 | NON_SYNONYMOU | MODERAT | MISSENSE | Ctc/Ttc | L456F | PLES_26481 | A | A | A | A | . | . | . | . | . | . | A | A |
| NC_011770 | 2844974 | C | T | 33 | 6 | 6 | 0 | NON_SYNONYMOU | MODERAT | MISSENSE | Gcg/Acg | A435T | PLES_26481 | T | T | T | T | . | . | . | . | . | . | T | T |
| NC_011770 | 2845103 | G | A | 28 | 6 | 4 | 0 | NON_SYNONYMOU | MODERAT | MISSENSE | Ctt/Ttt | L392F | PLES_26481 | A | A | A | A | . | . | . | . | . | . | | |
| NC_011770 | 2845106 | C | G | 29 | 6 | 2 | 0 | NON_SYNONYMOU | MODERAT | MISSENSE | Gaa/Caa | E391Q | PLES_26481 | | | | | . | . | . | . | . | . | G | G |
| NC_011770 | 2845184 | C | A | 33 | 6 | 4 | 0 | NON_SYNONYMOU | MODERAT | MISSENSE | Gca/Tca | A365S | PLES_26481 | A | A | A | A | . | . | . | . | . | . | | |
| NC_011770 | 2845210 | T | C | 36 | 6 | 4 | 0 | NON_SYNONYMOU | MODERAT | MISSENSE | cAg/cGg | Q356R | PLES_26481 | C | C | C | C | . | . | . | . | . | . | | |
| NC_011770 | 2845214 | A | C | 35 | 6 | 4 | 0 | NON_SYNONYMOU | MODERAT | MISSENSE | Tcg/Gcg | S355A | PLES_26481 | C | C | C | C | . | . | . | . | . | . | | |
| NC_011770 | 2845223 | G | C | 38 | 6 | 4 | 0 | NON_SYNONYMOU | MODERAT | MISSENSE | Cac/Gac | H352D | PLES_26481 | C | C | C | C | . | . | . | . | . | . | | |
| NC_011770 | 2845280 | C | T | 16 | 7 | 3 | 0 | NON_SYNONYMOU | MODERAT | MISSENSE | Gcg/Acg | A333T | PLES_26481 | . | T | T | T | . | . | . | . | . | . | | |
| NC_011770 | 2845306 | G | A | 15 | 8 | 2 | 0 | NON_SYNONYMOU | MODERAT | MISSENSE | cCg/cTg | P324L | PLES_26481 | . | . | A | A | . | . | . | . | . | . | | |
| NC_011770 | 2845307 | G | A | 16 | 8 | 2 | 0 | NON_SYNONYMOU | MODERAT | MISSENSE | Ccg/Tcg | P324S | PLES_26481 | . | . | A | A | . | . | . | . | . | . | | |
| NC_011770 | 2845328 | T | A | 15 | 8 | 4 | 0 | NON_SYNONYMOU | MODERAT | MISSENSE | Acc/Tcc | T317S | PLES_26481 | . | . | A | A | . | . | . | . | . | . | A | A |
| NC_011770 | 2845335 | G | T | 17 | 8 | 2 | 0 | NON_SYNONYMOU | MODERAT | MISSENSE | gaC/gaA | D314E | PLES_26481 | . | . | T | T | . | . | . | . | . | . | | |
| NC_011770 | 2845343 | G | A | 17 | 8 | 2 | 0 | NON_SYNONYMOU | MODERAT | MISSENSE | Ctc/Ttc | L312F | PLES_26481 | . | . | A | A | . | . | . | . | . | . | | |
| NC_011770 | 2845345 | G | A | 16 | 8 | 2 | 0 | NON_SYNONYMOU | MODERAT | MISSENSE | gCg/gTg | A311V | PLES_26481 | . | . | A | A | . | . | . | . | . | . | | |
| NC_011770 | 2845349 | G | C | 17 | 8 | 2 | 0 | NON_SYNONYMOU | MODERAT | MISSENSE | Cag/Gag | Q310E | PLES_26481 | . | . | C | C | . | . | . | . | . | . | | |
| NC_011770 | 2845359 | G | T | 36 | 6 | 4 | 0 | NON_SYNONYMOU | MODERAT | MISSENSE | gaC/gaA | D306E | PLES_26481 | T | T | T | T | . | . | . | . | . | . | | |
| NC_011770 | 2845402 | G | A | 36 | 6 | 6 | 0 | NON_SYNONYMOU | MODERAT | MISSENSE | aCg/aTg | T292M | PLES_26481 | A | A | A | A | . | . | . | . | . | . | A | A |
| NC_011770 | 2845403 | T | C | 36 | 6 | 6 | 0 | NON_SYNONYMOU | MODERAT | MISSENSE | Acg/Gcg | T292A | PLES_26481 | C | C | C | C | . | . | . | . | . | . | C | C |
| NC_011770 | 2845424 | C | A | 37 | 6 | 4 | 0 | NON_SYNONYMOU | MODERAT | MISSENSE | Gcc/Tcc | A285S | PLES_26481 | A | A | A | A | . | . | . | . | . | . | | |
| NC_011770 | 2845441 | T | A | 20 | 6 | 4 | 0 | NON_SYNONYMOU | MODERAT | MISSENSE | gAg/gTg | E279V | PLES_26481 | A | A | A | A | . | . | . | . | . | . | | |
| NC_011770 | 2845445 | T | G | 20 | 6 | 3 | 0 | NON_SYNONYMOU | MODERAT | MISSENSE | Aag/Cag | K278Q | PLES_26481 | G | . | G | G | . | . | . | . | . | . | | |
| NC_011770 | 2845474 | A | G | 19 | 6 | 4 | 0 | NON_SYNONYMOU | MODERAT | MISSENSE | gTc/gCc | V268A | PLES_26481 | G | G | G | G | . | . | . | . | . | . | | |
| NC_011770 | 2845475 | C | T | 19 | 6 | 6 | 0 | NON_SYNONYMOU | MODERAT | MISSENSE | Gtc/Atc | V268I | PLES_26481 | T | T | T | T | . | . | . | . | . | . | T | T |
| NC_011770 | 2845480 | G | T | 18 | 6 | 4 | 0 | NON_SYNONYMOU | MODERAT | MISSENSE | gCt/gAt | A266D | PLES_26481 | T | T | T | T | . | . | . | . | . | . | | |
| NC_011770 | 2845481 | C | G | 19 | 6 | 4 | 0 | NON_SYNONYMOU | MODERAT | MISSENSE | Gct/Cct | A266P | PLES_26481 | G | G | G | G | . | . | . | . | . | . | | |
| NC_011770 | 2845490 | A | T | 19 | 6 | 2 | 0 | NON_SYNONYMOU | MODERAT | MISSENSE | Tcg/Acg | S263T | PLES_26481 | | | | | . | . | . | . | . | . | T | T |
| NC_011770 | 2845515 | G | C | 17 | 6 | 6 | 0 | NON_SYNONYMOU | MODERAT | MISSENSE | agC/agG | S254R | PLES_26481 | C | C | C | C | . | . | . | . | . | . | C | C |
| NC_011770 | 2845516 | C | G | 17 | 6 | 6 | 0 | NON_SYNONYMOU | MODERAT | MISSENSE | aGc/aCc | S254T | PLES_26481 | G | G | G | G | . | . | . | . | . | . | G | G |
| NC_011770 | 2845517 | T | C | 17 | 6 | 4 | 0 | NON_SYNONYMOU | MODERAT | MISSENSE | Agc/Ggc | S254G | PLES_26481 | C | C | C | C | . | . | . | . | . | . | | |
| NC_011770 | 2845526 | C | T | 18 | 6 | 3 | 0 | NON_SYNONYMOU | MODERAT | MISSENSE | Gcc/Acc | A251T | PLES_26481 | . | T | T | T | . | . | . | . | . | . | | |
| NC_011770 | 2845529 | G | A | 18 | 6 | 4 | 0 | NON_SYNONYMOU | MODERAT | MISSENSE | Ccg/Tcg | P250S | PLES_26481 | A | A | A | A | . | . | . | . | . | . | | |
| NC_011770 | 2845538 | A | C | 18 | 6 | 2 | 0 | NON_SYNONYMOU | MODERAT | MISSENSE | Tcc/Gcc | S247A | PLES_26481 | | | | | . | . | . | . | . | . | C | C |
| NC_011770 | 2845605 | C | G | 28 | 6 | 4 | 0 | NON_SYNONYMOU | MODERAT | MISSENSE | aaG/aaC | K224N | PLES_26481 | G | G | G | G | . | . | . | . | . | . | | |
| NC_011770 | 2845606 | T | C | 28 | 6 | 4 | 0 | NON_SYNONYMOU | MODERAT | MISSENSE | aAg/aGg | K224R | PLES_26481 | C | C | C | C | . | . | . | . | . | . | | |
| NC_011770 | 2845607 | T | G | 28 | 6 | 4 | 0 | NON_SYNONYMOU | MODERAT | MISSENSE | Aag/Cag | K224Q | PLES_26481 | G | G | G | G | . | . | . | . | . | . | | |

| | | | | | | | | | | | | | | | | | | | | | | | | | |
|---|---|---|---|---|---|---|---|---|---|---|---|---|---|---|---|---|---|---|---|---|---|---|---|---|---|
| NC_011770 | 2845616 | G | A | 27 | 6 | 4 | 0 | NON_SYNONYMOU | MODERAT | MISSENSE | Ccc/Tcc | P221S | PLES_26481 | A | A | A | A | . | . | . | . | . | . | | |
| NC_011770 | 2845625 | A | C | 26 | 6 | 4 | 0 | NON_SYNONYMOU | MODERAT | MISSENSE | Tcc/Gcc | S218A | PLES_26481 | C | C | C | C | . | . | . | . | . | . | | |
| NC_011770 | 2845694 | T | C | 35 | 6 | 4 | 0 | NON_SYNONYMOU | MODERAT | MISSENSE | Atc/Gtc | I195V | PLES_26481 | C | C | C | C | . | . | . | . | . | . | | |
| NC_011770 | 2845711 | G | T | 18 | 6 | 3 | 0 | NON_SYNONYMOU | MODERAT | MISSENSE | aCg/aAg | T189K | PLES_26481 | T | T | T | . | . | . | . | . | . | . | | |
| NC_011770 | 2845717 | G | C | 18 | 6 | 4 | 0 | NON_SYNONYMOU | MODERAT | MISSENSE | tCg/tGg | S187W | PLES_26481 | C | C | C | C | . | . | . | . | . | . | | |
| NC_011770 | 2845718 | A | C | 17 | 6 | 4 | 0 | NON_SYNONYMOU | MODERAT | MISSENSE | Tcg/Gcg | S187A | PLES_26481 | C | C | C | C | . | . | . | . | . | . | | |
| NC_011770 | 2845739 | G | T | 18 | 6 | 6 | 0 | NON_SYNONYMOU | MODERAT | MISSENSE | Ctg/Atg | L180M | PLES_26481 | T | T | T | T | . | . | . | . | . | . | A | A |
| NC_011770 | 2845821 | G | C | 36 | 6 | 2 | 0 | NON_SYNONYMOU | MODERAT | MISSENSE | agC/agG | S152R | PLES_26481 | | | | | . | . | . | . | . | . | C | C |
| NC_011770 | 2845822 | C | G | 36 | 6 | 6 | 0 | NON_SYNONYMOU | MODERAT | MISSENSE | aGc/aCc | S152T | PLES_26481 | G | G | G | G | . | . | . | . | . | . | G | G |
| NC_011770 | 2845823 | T | A | 35 | 6 | 6 | 0 | NON_SYNONYMOU | MODERAT | MISSENSE | Agc/Tgc | S152C | PLES_26481 | A | A | A | A | . | . | . | . | . | . | A | A |
| NC_011770 | 2845853 | C | T | 49 | 3 | 3 | 0 | NON_SYNONYMOU | MODERAT | MISSENSE | Gtg/Atg | V142M | PLES_26481 | | | | | T | T | . | T | . | . | | |
| NC_011770 | 2845865 | G | T | 49 | 3 | 3 | 0 | NON_SYNONYMOU | MODERAT | MISSENSE | Ctc/Atc | L138I | PLES_26481 | | | | | T | T | . | T | . | . | | |
| NC_011770 | 2845979 | T | C | 19 | 6 | 2 | 0 | NON_SYNONYMOU | MODERAT | MISSENSE | Att/Gtt | I100V | PLES_26481 | | | | | . | . | . | . | . | . | C | C |
| NC_011770 | 2845995 | C | G | 23 | 6 | 4 | 0 | NON_SYNONYMOU | MODERAT | MISSENSE | aaG/aaC | K94N | PLES_26481 | G | G | G | G | . | . | . | . | . | . | | |
| NC_011770 | 2845996 | T | C | 23 | 6 | 4 | 0 | NON_SYNONYMOU | MODERAT | MISSENSE | aAg/aGg | K94R | PLES_26481 | C | C | C | C | . | . | . | . | . | . | | |
| NC_011770 | 2845997 | T | G | 23 | 6 | 4 | 0 | NON_SYNONYMOU | MODERAT | MISSENSE | Aag/Cag | K94Q | PLES_26481 | G | G | G | G | . | . | . | . | . | . | | |
| NC_011770 | 2846063 | T | G | 34 | 6 | 2 | 0 | NON_SYNONYMOU | MODERAT | MISSENSE | Atc/Ctc | I72L | PLES_26481 | | | | | . | . | . | . | . | . | G | G |
| NC_011770 | 2846101 | T | C | 31 | 6 | 6 | 0 | NON_SYNONYMOU | MODERAT | MISSENSE | aAc/aGc | N59S | PLES_26481 | C | C | C | C | . | . | . | . | . | . | C | C |
| NC_011770 | 2846138 | T | G | 15 | 8 | 2 | 0 | NON_SYNONYMOU | MODERAT | MISSENSE | Atc/Ctc | I47L | PLES_26481 | . | . | G | G | . | . | . | . | . | . | | |
| NC_011770 | 2846141 | C | A | 14 | 8 | 2 | 0 | NON_SYNONYMOU | MODERAT | MISSENSE | Gtc/Ttc | V46F | PLES_26481 | . | . | A | A | . | . | . | . | . | . | | |
| NC_011770 | 2846143 | C | G | 14 | 8 | 4 | 0 | NON_SYNONYMOU | MODERAT | MISSENSE | aGc/aCc | S45T | PLES_26481 | . | . | G | G | . | . | . | . | . | . | T | T |
| NC_011770 | 2846144 | T | G | 14 | 8 | 2 | 0 | NON_SYNONYMOU | MODERAT | MISSENSE | Agc/Cgc | S45R | PLES_26481 | . | . | G | G | . | . | . | . | . | . | | |
| NC_011770 | 2846152 | G | T | 14 | 8 | 2 | 0 | NON_SYNONYMOU | MODERAT | MISSENSE | aCc/aAc | T42N | PLES_26481 | . | . | T | T | . | . | . | . | . | . | | |
| NC_011770 | 2846422 | G | T | 24 | 6 | 4 | 0 | STOP_GAINED | HIGH | NONSENSE | tCg/tAg | S271* | PLES_26491 | T | T | T | T | . | . | . | . | . | . | | |
| NC_011770 | 2846391 | G | C | 23 | 6 | 4 | 0 | NON_SYNONYMOU | MODERAT | MISSENSE | ttC/ttG | F281L | PLES_26491 | C | C | C | C | . | . | . | . | . | . | | |
| NC_011770 | 2846392 | A | G | 23 | 6 | 4 | 0 | NON_SYNONYMOU | MODERAT | MISSENSE | tTc/tCc | F281S | PLES_26491 | G | G | G | G | . | . | . | . | . | . | | |
| NC_011770 | 2846393 | A | T | 23 | 6 | 4 | 0 | NON_SYNONYMOU | MODERAT | MISSENSE | Ttc/Atc | F281I | PLES_26491 | T | T | T | T | . | . | . | . | . | . | | |
| NC_011770 | 2846398 | C | G | 23 | 6 | 2 | 0 | NON_SYNONYMOU | MODERAT | MISSENSE | aGc/aCc | S279T | PLES_26491 | | | | | . | . | . | . | . | . | G | G |
| NC_011770 | 2846401 | C | T | 22 | 6 | 4 | 0 | NON_SYNONYMOU | MODERAT | MISSENSE | cGc/cAc | R278H | PLES_26491 | T | T | T | T | . | . | . | . | . | . | | |
| NC_011770 | 2846413 | G | A | 24 | 6 | 4 | 0 | NON_SYNONYMOU | MODERAT | MISSENSE | aCg/aTg | T274M | PLES_26491 | A | A | A | A | . | . | . | . | . | . | | |
| NC_011770 | 2846414 | T | C | 24 | 6 | 4 | 0 | NON_SYNONYMOU | MODERAT | MISSENSE | Acg/Gcg | T274A | PLES_26491 | C | C | C | C | . | . | . | . | . | . | | |
| NC_011770 | 2846420 | A | T | 24 | 6 | 4 | 0 | NON_SYNONYMOU | MODERAT | MISSENSE | Tcc/Acc | S272T | PLES_26491 | T | T | T | T | . | . | . | . | . | . | | |
| NC_011770 | 2846423 | A | T | 24 | 6 | 4 | 0 | NON_SYNONYMOU | MODERAT | MISSENSE | Tcg/Acg | S271T | PLES_26491 | T | T | T | T | . | . | . | . | . | . | | |
| NC_011770 | 2846431 | A | C | 24 | 6 | 4 | 0 | NON_SYNONYMOU | MODERAT | MISSENSE | gTc/gGc | V268G | PLES_26491 | C | C | C | C | . | . | . | . | . | . | | |
| NC_011770 | 2846432 | C | G | 24 | 6 | 4 | 0 | NON_SYNONYMOU | MODERAT | MISSENSE | Gtc/Ctc | V268L | PLES_26491 | G | G | G | G | . | . | . | . | . | . | | |
| NC_011770 | 2846475 | C | T | 64 | 3 | 3 | 0 | NON_SYNONYMOU | MODERAT | MISSENSE | atG/atA | M253I | PLES_26491 | | | | | T | T | . | T | . | . | | |
| NC_011770 | 2846476 | A | T | 63 | 3 | 3 | 0 | NON_SYNONYMOU | MODERAT | MISSENSE | aTg/aAg | M253K | PLES_26491 | | | | | T | T | . | T | . | . | | |
| NC_011770 | 2846482 | G | T | 61 | 3 | 3 | 0 | NON_SYNONYMOU | MODERAT | MISSENSE | gCg/gAg | A251E | PLES_26491 | | | | | T | T | . | T | . | . | | |
| NC_011770 | 2846510 | A | C | 38 | 6 | 6 | 0 | NON_SYNONYMOU | MODERAT | MISSENSE | Tcc/Gcc | S242A | PLES_26491 | C | C | C | C | . | . | . | . | . | . | C | C |
| NC_011770 | 2846538 | G | C | 41 | 6 | 2 | 0 | NON_SYNONYMOU | MODERAT | MISSENSE | atC/atG | I232M | PLES_26491 | | | | | . | . | . | . | . | . | C | C |
| NC_011770 | 2846540 | T | A | 41 | 6 | 2 | 0 | NON_SYNONYMOU | MODERAT | MISSENSE | Atc/Ttc | I232F | PLES_26491 | | | | | . | . | . | . | . | . | A | A |
| NC_011770 | 2846594 | C | A | 31 | 5 | 7 | 0 | NON_SYNONYMOU | MODERAT | MISSENSE | Gcc/Tcc | A214S | PLES_26491 | A | A | A | A | T | . | . | . | . | . | A | A |
| NC_011770 | 2846599 | A | T | 33 | 4 | 8 | 0 | NON_SYNONYMOU | MODERAT | MISSENSE | tTc/tAc | F212Y | PLES_26491 | T | T | T | T | T | T | . | . | . | . | T | T |
| NC_011770 | 2846683 | C | A | 33 | 6 | 6 | 0 | NON_SYNONYMOU | MODERAT | MISSENSE | cGg/cTg | R184L | PLES_26491 | A | A | A | A | . | . | . | . | . | . | A | A |
| NC_011770 | 2846711 | C | T | 38 | 6 | 4 | 0 | NON_SYNONYMOU | MODERAT | MISSENSE | Gcc/Acc | A175T | PLES_26491 | T | T | T | T | . | . | . | . | . | . | | |
| NC_011770 | 2846792 | C | G | 32 | 6 | 4 | 0 | NON_SYNONYMOU | MODERAT | MISSENSE | Gag/Cag | E148Q | PLES_26491 | G | G | G | G | . | . | . | . | . | . | | |
| NC_011770 | 2846815 | A | G | 32 | 6 | 4 | 0 | NON_SYNONYMOU | MODERAT | MISSENSE | gTc/gCc | V140A | PLES_26491 | G | G | G | G | . | . | . | . | . | . | | |
| NC_011770 | 2846836 | G | C | 38 | 6 | 6 | 0 | NON_SYNONYMOU | MODERAT | MISSENSE | gCa/gGa | A133G | PLES_26491 | C | C | C | C | . | . | . | . | . | . | A | A |
| NC_011770 | 2846860 | G | C | 39 | 6 | 4 | 0 | NON_SYNONYMOU | MODERAT | MISSENSE | gCg/gGg | A125G | PLES_26491 | C | C | C | C | . | . | . | . | . | . | | |
| NC_011770 | 2846861 | C | T | 40 | 6 | 4 | 0 | NON_SYNONYMOU | MODERAT | MISSENSE | Gcg/Acg | A125T | PLES_26491 | T | T | T | T | . | . | . | . | . | . | | |
| NC_011770 | 2846881 | C | T | 42 | 6 | 6 | 0 | NON_SYNONYMOU | MODERAT | MISSENSE | aGc/aAc | S118N | PLES_26491 | T | T | T | T | . | . | . | . | . | . | T | T |
| NC_011770 | 2846939 | G | C | 34 | 6 | 2 | 0 | NON_SYNONYMOU | MODERAT | MISSENSE | Ccc/Gcc | P99A | PLES_26491 | | | | | . | . | . | . | . | . | C | C |
| NC_011770 | 2846953 | T | C | 34 | 6 | 2 | 0 | NON_SYNONYMOU | MODERAT | MISSENSE | cAg/cGg | Q94R | PLES_26491 | | | | | . | . | . | . | . | . | C | C |
| NC_011770 | 2846986 | G | C | 6 | 7 | 5 | 0 | NON_SYNONYMOU | MODERAT | MISSENSE | tCg/tGg | S83W | PLES_26491 | C | . | C | C | . | . | . | . | . | . | C | C |
| NC_011770 | 2846987 | A | T | 6 | 7 | 5 | 0 | NON_SYNONYMOU | MODERAT | MISSENSE | Tcg/Acg | S83T | PLES_26491 | T | . | T | T | . | . | . | . | . | . | T | T |
| NC_011770 | 2846992 | G | C | 6 | 7 | 5 | 0 | NON_SYNONYMOU | MODERAT | MISSENSE | tCg/tGg | S81W | PLES_26491 | C | . | C | C | . | . | . | . | . | . | C | C |
| NC_011770 | 2846993 | A | T | 6 | 7 | 5 | 0 | NON_SYNONYMOU | MODERAT | MISSENSE | Tcg/Acg | S81T | PLES_26491 | T | . | T | T | . | . | . | . | . | . | T | T |
| NC_011770 | 2847012 | G | C | 15 | 6 | 4 | 0 | NON_SYNONYMOU | MODERAT | MISSENSE | atC/atG | I74M | PLES_26491 | C | C | C | C | . | . | . | . | . | . | | |
| NC_011770 | 2847014 | T | C | 15 | 6 | 4 | 0 | NON_SYNONYMOU | MODERAT | MISSENSE | Atc/Gtc | I74V | PLES_26491 | C | C | C | C | . | . | . | . | . | . | | |
| NC_011770 | 2847031 | A | G | 16 | 6 | 4 | 0 | NON_SYNONYMOU | MODERAT | MISSENSE | aTc/aCc | I68T | PLES_26491 | G | G | G | G | . | . | . | . | . | . | | |
| NC_011770 | 2847032 | T | C | 16 | 6 | 2 | 0 | NON_SYNONYMOU | MODERAT | MISSENSE | Atc/Gtc | I68V | PLES_26491 | | | | | . | . | . | . | . | . | C | C |
| NC_011770 | 2847055 | G | C | 30 | 6 | 2 | 0 | NON_SYNONYMOU | MODERAT | MISSENSE | aCg/aGg | T60R | PLES_26491 | | | | | . | . | . | . | . | . | C | C |

| | | | | | | | | | | | | | | | | | | | | | | | | | |
|---|---|---|---|---|---|---|---|---|---|---|---|---|---|---|---|---|---|---|---|---|---|---|---|---|---|
| NC_011770 | 2847056 | T | C | 30 | 6 | 2 | 0 | NON_SYNONYMOU | MODERAT | MISSENSE | Acg/Gcg | T60A | PLES_26491 | | | | | . | . | . | . | . | . | C | C |
| NC_011770 | 2847104 | G | A | 9 | 9 | 2 | 0 | NON_SYNONYMOU | MODERAT | MISSENSE | Ctc/Ttc | L44F | PLES_26491 | . | . | | . | . | . | . | . | . | . | A | A |
| NC_011770 | 2847127 | C | G | 6 | 9 | 3 | 0 | NON_SYNONYMOU | MODERAT | MISSENSE | gGc/gCc | G36A | PLES_26491 | . | . | G | . | . | . | . | . | . | . | G | G |
| NC_011770 | 2847128 | C | A | 6 | 9 | 1 | 0 | NON_SYNONYMOU | MODERAT | MISSENSE | Ggc/Tgc | G36C | PLES_26491 | . | . | A | . | . | . | . | . | . | . | | |
| NC_011770 | 2847129 | G | C | 6 | 9 | 3 | 0 | NON_SYNONYMOU | MODERAT | MISSENSE | atC/atG | I35M | PLES_26491 | . | . | C | . | . | . | . | . | . | . | A | A |
| NC_011770 | 2847131 | T | C | 6 | 9 | 3 | 0 | NON_SYNONYMOU | MODERAT | MISSENSE | Atc/Gtc | I35V | PLES_26491 | . | . | C | . | . | . | . | . | . | . | C | C |
| NC_011770 | 2847137 | T | C | 6 | 9 | 2 | 0 | NON_SYNONYMOU | MODERAT | MISSENSE | Acc/Gcc | T33A | PLES_26491 | . | . | | . | . | . | . | . | . | . | C | C |
| NC_011770 | 2847140 | C | T | 6 | 9 | 2 | 0 | NON_SYNONYMOU | MODERAT | MISSENSE | Gcg/Acg | A32T | PLES_26491 | . | . | | . | . | . | . | . | . | . | T | T |
| NC_011770 | 2847149 | G | A | 6 | 9 | 3 | 0 | NON_SYNONYMOU | MODERAT | MISSENSE | Ctc/Ttc | L29F | PLES_26491 | . | . | A | . | . | . | . | . | . | . | A | A |
| NC_011770 | 2847173 | G | T | 35 | 6 | 4 | 0 | NON_SYNONYMOU | MODERAT | MISSENSE | Ctc/Atc | L21I | PLES_26491 | T | T | T | T | . | . | . | . | . | . | | |
| NC_011770 | 2847193 | C | G | 34 | 6 | 4 | 0 | NON_SYNONYMOU | MODERAT | MISSENSE | gGc/gCc | G14A | PLES_26491 | G | G | G | G | . | . | . | . | . | . | | |
| NC_011770 | 2847196 | G | A | 34 | 6 | 4 | 0 | NON_SYNONYMOU | MODERAT | MISSENSE | gCt/gTt | A13V | PLES_26491 | A | A | A | A | . | . | . | . | . | . | | |
| NC_011770 | 2847203 | G | T | 35 | 6 | 2 | 0 | NON_SYNONYMOU | MODERAT | MISSENSE | Ctg/Atg | L11M | PLES_26491 | | | | | . | . | . | . | . | . | T | T |
| NC_011770 | 2847212 | T | G | 35 | 6 | 4 | 0 | NON_SYNONYMOU | MODERAT | MISSENSE | Atg/Ctg | M8L | PLES_26491 | G | G | G | G | . | . | . | . | . | . | | |
| NC_011770 | 2847226 | C | CGAT | 15 | 6 | 4 | 0 | CODON_INSERTION | MODERATE | | cga/cATCg | R3HR | PLES_26491 | CGAT | CGAT | CGAT | CGAT | . | . | . | . | . | . | | |
| NC_011770 | 2847233 | C | T | 34 | 6 | 2 | 0 | NON_SYNONYMOU | MODERAT | MISSENSE | Gtg/Atg | V1M | PLES_26491 | | | | | . | . | . | . | . | . | T | T |
| NC_011770 | 2847249 | T | C | 33 | 6 | 2 | 0 | NON_SYNONYMOU | MODERAT | MISSENSE | cAc/cGc | H125R | PLES_26501 | | | | | . | . | . | . | . | . | C | C |
| NC_011770 | 2847303 | T | A | 14 | 7 | 3 | 0 | NON_SYNONYMOU | MODERAT | MISSENSE | tAt/tTt | Y107F | PLES_26501 | A | . | A | A | . | . | . | . | . | . | | |
| NC_011770 | 2847309 | C | T | 11 | 9 | 3 | 0 | NON_SYNONYMOU | MODERAT | MISSENSE | aGg/aAg | R105K | PLES_26501 | . | . | T | . | . | . | . | . | . | . | T | T |
| NC_011770 | 2847331 | G | T | 12 | 10 | 2 | 0 | NON_SYNONYMOU | MODERAT | MISSENSE | Ctc/Atc | L98I | PLES_26501 | . | . | . | . | . | . | . | . | . | . | T | T |
| NC_011770 | 2847411 | C | T | 11 | 10 | 2 | 0 | NON_SYNONYMOU | MODERAT | MISSENSE | gGc/gAc | G71D | PLES_26501 | . | . | . | . | . | . | . | . | . | . | T | T |
| NC_011770 | 2847412 | C | G | 11 | 10 | 2 | 0 | NON_SYNONYMOU | MODERAT | MISSENSE | Ggc/Cgc | G71R | PLES_26501 | . | . | . | . | . | . | . | . | . | . | G | G |
| NC_011770 | 2847432 | A | G | 13 | 10 | 2 | 0 | NON_SYNONYMOU | MODERAT | MISSENSE | aTc/aCc | I64T | PLES_26501 | . | . | . | . | . | . | . | . | . | . | G | G |
| NC_011770 | 2847439 | T | C | 13 | 10 | 2 | 0 | NON_SYNONYMOU | MODERAT | MISSENSE | Acg/Gcg | T62A | PLES_26501 | . | . | . | . | . | . | . | . | . | . | C | C |
| NC_011770 | 2847453 | T | G | 12 | 10 | 2 | 0 | NON_SYNONYMOU | MODERAT | MISSENSE | gAc/gCc | D57A | PLES_26501 | . | . | . | . | . | . | . | . | . | . | G | G |
| NC_011770 | 2847457 | T | C | 13 | 10 | 2 | 0 | NON_SYNONYMOU | MODERAT | MISSENSE | Acc/Gcc | T56A | PLES_26501 | . | . | . | . | . | . | . | . | . | . | C | C |
| NC_011770 | 2847465 | C | T | 13 | 10 | 2 | 0 | NON_SYNONYMOU | MODERAT | MISSENSE | gGa/gAa | G53E | PLES_26501 | . | . | . | . | . | . | . | . | . | . | T | T |
| NC_011770 | 2847492 | T | C | 13 | 10 | 2 | 0 | NON_SYNONYMOU | MODERAT | MISSENSE | cAc/cGc | H44R | PLES_26501 | . | . | . | . | . | . | . | . | . | . | C | C |
| NC_011770 | 2847546 | C | T | 26 | 6 | 2 | 0 | NON_SYNONYMOU | MODERAT | MISSENSE | cGc/cAc | R26H | PLES_26501 | | | | | . | . | . | . | . | . | T | T |
| NC_011770 | 2847565 | C | G | 26 | 6 | 2 | 0 | NON_SYNONYMOU | MODERAT | MISSENSE | Gtg/Ctg | V20L | PLES_26501 | | | | | . | . | . | . | . | . | G | G |
| NC_011770 | 2847585 | G | A | 25 | 6 | 2 | 0 | NON_SYNONYMOU | MODERAT | MISSENSE | gCc/gTc | A13V | PLES_26501 | | | | | . | . | . | . | . | . | A | A |
| NC_011770 | 2847607 | C | A | 21 | 6 | 6 | 0 | NON_SYNONYMOU | MODERAT | MISSENSE | Gcc/Tcc | A6S | PLES_26501 | A | A | A | A | . | . | . | . | . | . | A | A |
| NC_011770 | 2847610 | C | T | 22 | 6 | 4 | 0 | NON_SYNONYMOU | MODERAT | MISSENSE | Gca/Aca | A5T | PLES_26501 | T | T | T | T | . | . | . | . | . | . | | |
| NC_011770 | 2848072 | G | T | 0 | 11 | 1 | 0 | NON_SYNONYMOU | MODERAT | MISSENSE | Cat/Aat | H88N | PLES_26511 | . | . | T | . | . | . | . | . | . | . | . | . |
| NC_011770 | 2848090 | T | A | 28 | 6 | 6 | 0 | NON_SYNONYMOU | MODERAT | MISSENSE | Atg/Ttg | M82L | PLES_26511 | A | A | A | A | . | . | . | . | . | . | A | A |
| NC_011770 | 2848111 | C | G | 28 | 6 | 6 | 0 | NON_SYNONYMOU | MODERAT | MISSENSE | Gag/Cag | E75Q | PLES_26511 | G | G | G | G | . | . | . | . | . | . | G | G |
| NC_011770 | 2848123 | T | C | 11 | 8 | 4 | 0 | NON_SYNONYMOU | MODERAT | MISSENSE | Aac/Gac | N71D | PLES_26511 | C | C | C | C | . | . | . | . | . | . | . | . |
| NC_011770 | 2848134 | A | G | 9 | 8 | 4 | 0 | NON_SYNONYMOU | MODERAT | MISSENSE | aTc/aCc | I67T | PLES_26511 | G | G | G | G | . | . | . | . | . | . | . | . |
| NC_011770 | 2848135 | T | C | 9 | 8 | 4 | 0 | NON_SYNONYMOU | MODERAT | MISSENSE | Atc/Gtc | I67V | PLES_26511 | C | C | C | C | . | . | . | . | . | . | . | . |
| NC_011770 | 2848144 | C | T | 9 | 8 | 4 | 0 | NON_SYNONYMOU | MODERAT | MISSENSE | Gtc/Atc | V64I | PLES_26511 | T | T | T | T | . | . | . | . | . | . | . | . |
| NC_011770 | 2848161 | G | A | 9 | 8 | 4 | 0 | NON_SYNONYMOU | MODERAT | MISSENSE | gCc/gTc | A58V | PLES_26511 | A | A | A | A | . | . | . | . | . | . | . | . |
| NC_011770 | 2848171 | G | C | 9 | 8 | 4 | 0 | NON_SYNONYMOU | MODERAT | MISSENSE | Ctg/Gtg | L55V | PLES_26511 | C | C | C | C | . | . | . | . | . | . | . | . |
| NC_011770 | 2848234 | G | T | 4 | 8 | 4 | 0 | NON_SYNONYMOU | MODERAT | MISSENSE | Cag/Aag | Q34K | PLES_26511 | T | T | T | T | . | . | . | . | . | . | . | . |
| NC_011770 | 2848245 | G | A | 8 | 6 | 2 | 0 | NON_SYNONYMOU | MODERAT | MISSENSE | gCg/gTg | A30V | PLES_26511 | | | | | . | . | . | . | . | . | A | A |
| NC_011770 | 2848257 | C | T | 8 | 6 | 6 | 0 | NON_SYNONYMOU | MODERAT | MISSENSE | cGg/cAg | R26Q | PLES_26511 | T | T | T | T | . | . | . | . | . | . | T | T |
| NC_011770 | 2848306 | C | G | 12 | 8 | 4 | 0 | NON_SYNONYMOU | MODERAT | MISSENSE | Gtg/Ctg | V10L | PLES_26511 | G | G | G | G | . | . | . | . | . | . | . | . |
| NC_011770 | 2848320 | G | A | 13 | 8 | 4 | 0 | NON_SYNONYMOU | MODERAT | MISSENSE | gCc/gTc | A5V | PLES_26511 | A | A | A | A | . | . | . | . | . | . | . | . |
| NC_011770 | 2848327 | G | A | 13 | 8 | 4 | 0 | NON_SYNONYMOU | MODERAT | MISSENSE | Ctc/Ttc | L3F | PLES_26511 | A | A | A | A | . | . | . | . | . | . | . | . |
| NC_011770 | 2849034 | C | A | 20 | 6 | 4 | 0 | STOP_GAINED | HIGH | NONSENSE | Gag/Tag | E90* | PLES_26521 | A | A | A | A | . | . | . | . | . | . | | |
| NC_011770 | 2848568 | G | C | 8 | 6 | 2 | 0 | NON_SYNONYMOU | MODERAT | MISSENSE | gCc/gGc | A245G | PLES_26521 | | | | | . | . | . | . | . | . | C | C |
| NC_011770 | 2848569 | C | T | 7 | 6 | 2 | 0 | NON_SYNONYMOU | MODERAT | MISSENSE | Gcc/Acc | A245T | PLES_26521 | | | | | . | . | . | . | . | . | T | T |
| NC_011770 | 2848572 | G | T | 4 | 10 | 2 | 0 | NON_SYNONYMOU | MODERAT | MISSENSE | Ccc/Acc | P244T | PLES_26521 | . | . | . | . | . | . | . | . | . | . | T | T |
| NC_011770 | 2848575 | A | G | 4 | 10 | 2 | 0 | NON_SYNONYMOU | MODERAT | MISSENSE | Tcg/Ccg | S243P | PLES_26521 | . | . | . | . | . | . | . | . | . | . | G | G |
| NC_011770 | 2848577 | CCCGGCGC | C | 4 | 10 | 2 | 0 | CODON_CHANGE_ | MODERATE | | gaaatggcca | EMASDAP | PLES_26521 | . | . | . | . | . | . | . | . | . | . | C | C |
| NC_011770 | 2848614 | G | T | 32 | 6 | 6 | 0 | NON_SYNONYMOU | MODERAT | MISSENSE | Ctg/Atg | L230M | PLES_26521 | T | T | T | T | . | . | . | . | . | . | T | T |
| NC_011770 | 2848671 | G | T | 34 | 6 | 4 | 0 | NON_SYNONYMOU | MODERAT | MISSENSE | Cag/Aag | Q211K | PLES_26521 | T | T | T | T | . | . | . | . | . | . | | |
| NC_011770 | 2848680 | G | C | 34 | 6 | 4 | 0 | NON_SYNONYMOU | MODERAT | MISSENSE | Cag/Gag | Q208E | PLES_26521 | C | C | C | C | . | . | . | . | . | . | | |
| NC_011770 | 2848688 | A | T | 38 | 6 | 4 | 0 | NON_SYNONYMOU | MODERAT | MISSENSE | cTc/cAc | L205H | PLES_26521 | T | T | T | T | . | . | . | . | . | . | | |
| NC_011770 | 2848723 | C | G | 21 | 6 | 4 | 0 | NON_SYNONYMOU | MODERAT | MISSENSE | aaG/aaC | K193N | PLES_26521 | G | G | G | G | . | . | . | . | . | . | | |
| NC_011770 | 2848724 | T | C | 21 | 6 | 4 | 0 | NON_SYNONYMOU | MODERAT | MISSENSE | aAg/aGg | K193R | PLES_26521 | C | C | C | C | . | . | . | . | . | . | | |
| NC_011770 | 2848740 | C | T | 19 | 6 | 4 | 0 | NON_SYNONYMOU | MODERAT | MISSENSE | Gcc/Acc | A188T | PLES_26521 | T | T | T | T | . | . | . | . | . | . | | |
| NC_011770 | 2848746 | C | T | 19 | 6 | 4 | 0 | NON_SYNONYMOU | MODERAT | MISSENSE | Gcg/Acg | A186T | PLES_26521 | T | T | T | T | . | . | . | . | . | . | | |

| | | | | | | | | | | | | | | | | | | | | | | | | | |
|---|---|---|---|---|---|---|---|---|---|---|---|---|---|---|---|---|---|---|---|---|---|---|---|---|---|
| NC_011770 | 2848755 | C | G | 14 | 6 | 4 | 0 | NON_SYNONYMOU | MODERAT | MISSENSE | Gcc/Ccc | A183P | PLES_26521 | G | G | G | G | . | . | . | . | . | . | | |
| NC_011770 | 2848757 | A | G | 14 | 6 | 6 | 0 | NON_SYNONYMOU | MODERAT | MISSENSE | gTg/gCg | V182A | PLES_26521 | G | G | G | G | . | . | . | . | . | . | G | G |
| NC_011770 | 2848759 | G | C | 13 | 6 | 6 | 0 | NON_SYNONYMOU | MODERAT | MISSENSE | aaC/aaG | N181K | PLES_26521 | C | C | C | C | . | . | . | . | . | . | A | A |
| NC_011770 | 2848760 | T | C | 14 | 6 | 2 | 0 | NON_SYNONYMOU | MODERAT | MISSENSE | aAc/aGc | N181S | PLES_26521 | | | | | . | . | . | . | . | . | C | C |
| NC_011770 | 2848761 | T | C | 14 | 6 | 4 | 0 | NON_SYNONYMOU | MODERAT | MISSENSE | Aac/Gac | N181D | PLES_26521 | C | C | C | C | . | . | . | . | . | . | | |
| NC_011770 | 2848767 | G | T | 14 | 6 | 2 | 0 | NON_SYNONYMOU | MODERAT | MISSENSE | Cgc/Agc | R179S | PLES_26521 | | | | | . | . | . | . | . | . | T | T |
| NC_011770 | 2848779 | T | G | 17 | 6 | 4 | 0 | NON_SYNONYMOU | MODERAT | MISSENSE | Atg/Ctg | M175L | PLES_26521 | G | G | G | G | . | . | . | . | . | . | | |
| NC_011770 | 2848790 | A | C | 17 | 6 | 2 | 0 | NON_SYNONYMOU | MODERAT | MISSENSE | aTc/aGc | I171S | PLES_26521 | | | | | . | . | . | . | . | . | C | C |
| NC_011770 | 2848797 | A | G | 17 | 6 | 6 | 0 | NON_SYNONYMOU | MODERAT | MISSENSE | Tca/Cca | S169P | PLES_26521 | G | G | G | G | . | . | . | . | . | . | G | G |
| NC_011770 | 2848802 | G | A | 17 | 6 | 4 | 0 | NON_SYNONYMOU | MODERAT | MISSENSE | aCa/aTa | T167I | PLES_26521 | A | A | A | A | . | . | . | . | . | . | | |
| NC_011770 | 2848803 | T | G | 17 | 6 | 6 | 0 | NON_SYNONYMOU | MODERAT | MISSENSE | Aca/Cca | T167P | PLES_26521 | G | G | G | G | . | . | . | . | . | . | A | A |
| NC_011770 | 2848804 | T | G | 18 | 6 | 4 | 0 | NON_SYNONYMOU | MODERAT | MISSENSE | gaA/gaC | E166D | PLES_26521 | G | G | G | G | . | . | . | . | . | . | | |
| NC_011770 | 2848820 | C | T | 20 | 6 | 4 | 0 | NON_SYNONYMOU | MODERAT | MISSENSE | gGc/gAc | G161D | PLES_26521 | T | T | T | T | . | . | . | . | . | . | | |
| NC_011770 | 2848822 | A | C | 20 | 6 | 4 | 0 | NON_SYNONYMOU | MODERAT | MISSENSE | gaT/gaG | D160E | PLES_26521 | C | C | C | C | . | . | . | . | . | . | | |
| NC_011770 | 2848841 | T | C | 34 | 6 | 2 | 0 | NON_SYNONYMOU | MODERAT | MISSENSE | gAc/gGc | D154G | PLES_26521 | | | | | . | . | . | . | . | . | C | C |
| NC_011770 | 2848872 | C | T | 33 | 6 | 2 | 0 | NON_SYNONYMOU | MODERAT | MISSENSE | Gtc/Atc | V144I | PLES_26521 | | | | | . | . | . | . | . | . | T | T |
| NC_011770 | 2848889 | G | A | 29 | 6 | 2 | 0 | NON_SYNONYMOU | MODERAT | MISSENSE | gCg/gTg | A138V | PLES_26521 | | | | | . | . | . | . | . | . | A | A |
| NC_011770 | 2848892 | T | C | 29 | 6 | 2 | 0 | NON_SYNONYMOU | MODERAT | MISSENSE | cAt/cGt | H137R | PLES_26521 | | | C | C | . | . | . | . | . | . | | |
| NC_011770 | 2848902 | C | A | 28 | 6 | 4 | 0 | NON_SYNONYMOU | MODERAT | MISSENSE | Gcg/Tcg | A134S | PLES_26521 | A | A | A | A | . | . | . | . | . | . | | |
| NC_011770 | 2848907 | G | C | 28 | 6 | 4 | 0 | NON_SYNONYMOU | MODERAT | MISSENSE | gCc/gGc | A132G | PLES_26521 | C | C | C | C | . | . | . | . | . | . | | |
| NC_011770 | 2848919 | G | A | 27 | 6 | 2 | 0 | NON_SYNONYMOU | MODERAT | MISSENSE | gCc/gTc | A128V | PLES_26521 | | | | | . | . | . | . | . | . | A | A |
| NC_011770 | 2848926 | G | T | 30 | 6 | 4 | 0 | NON_SYNONYMOU | MODERAT | MISSENSE | Cgt/Agt | R126S | PLES_26521 | T | T | T | T | . | . | . | . | . | . | | |
| NC_011770 | 2848955 | G | T | 29 | 6 | 2 | 0 | NON_SYNONYMOU | MODERAT | MISSENSE | gCg/gAg | A116E | PLES_26521 | | | | | . | . | . | . | . | . | T | T |
| NC_011770 | 2848966 | C | G | 29 | 6 | 4 | 0 | NON_SYNONYMOU | MODERAT | MISSENSE | caG/caC | Q112H | PLES_26521 | G | G | G | G | . | . | . | . | . | . | | |
| NC_011770 | 2848967 | T | A | 28 | 6 | 4 | 0 | NON_SYNONYMOU | MODERAT | MISSENSE | cAg/cTg | Q112L | PLES_26521 | A | A | A | A | . | . | . | . | . | . | | |
| NC_011770 | 2848995 | T | C | 19 | 6 | 4 | 0 | NON_SYNONYMOU | MODERAT | MISSENSE | Acc/Gcc | T103A | PLES_26521 | C | C | C | C | . | . | . | . | . | . | | |
| NC_011770 | 2849000 | T | C | 21 | 6 | 4 | 0 | NON_SYNONYMOU | MODERAT | MISSENSE | aAc/aGc | N101S | PLES_26521 | C | C | C | C | . | . | . | . | . | . | | |
| NC_011770 | 2849001 | T | C | 21 | 6 | 4 | 0 | NON_SYNONYMOU | MODERAT | MISSENSE | Aac/Gac | N101D | PLES_26521 | C | C | C | C | . | . | . | . | . | . | | |
| NC_011770 | 2849003 | C | G | 19 | 6 | 4 | 0 | NON_SYNONYMOU | MODERAT | MISSENSE | gGc/gCc | G100A | PLES_26521 | G | G | G | G | . | . | . | . | . | . | | |
| NC_011770 | 2849007 | C | T | 21 | 5 | 2 | 0 | NON_SYNONYMOU | MODERAT | MISSENSE | Gtc/Atc | V99I | PLES_26521 | | | | | . | . | . | . | . | . | T | T |
| NC_011770 | 2849021 | T | A | 21 | 5 | 4 | 0 | NON_SYNONYMOU | MODERAT | MISSENSE | cAg/cTg | Q94L | PLES_26521 | A | A | A | A | . | . | . | . | . | . | | |
| NC_011770 | 2849022 | G | C | 22 | 5 | 4 | 0 | NON_SYNONYMOU | MODERAT | MISSENSE | Cag/Gag | Q94E | PLES_26521 | C | C | C | C | . | . | . | . | . | . | | |
| NC_011770 | 2849032 | C | G | 20 | 6 | 4 | 0 | NON_SYNONYMOU | MODERAT | MISSENSE | gaG/gaC | E90D | PLES_26521 | G | G | G | G | . | . | . | . | . | . | | |
| NC_011770 | 2849033 | T | G | 20 | 6 | 4 | 0 | NON_SYNONYMOU | MODERAT | MISSENSE | gAg/gCg | E90A | PLES_26521 | G | G | G | G | . | . | . | . | . | . | | |
| NC_011770 | 2849088 | C | T | 34 | 6 | 2 | 0 | NON_SYNONYMOU | MODERAT | MISSENSE | Gaa/Aaa | E72K | PLES_26521 | | | | | . | . | . | . | . | . | T | T |
| NC_011770 | 2849102 | T | A | 30 | 6 | 2 | 0 | NON_SYNONYMOU | MODERAT | MISSENSE | cAg/cTg | Q67L | PLES_26521 | | | | | . | . | . | . | . | . | A | A |
| NC_011770 | 2849123 | T | G | 13 | 9 | 2 | 0 | NON_SYNONYMOU | MODERAT | MISSENSE | cAg/cCg | Q60P | PLES_26521 | . | . | | . | . | . | . | . | . | . | G | G |
| NC_011770 | 2849127 | C | T | 12 | 9 | 1 | 0 | NON_SYNONYMOU | MODERAT | MISSENSE | Gcg/Acg | A59T | PLES_26521 | . | . | T | . | . | . | . | . | . | . | | |
| NC_011770 | 2849129 | G | A | 12 | 9 | 3 | 0 | NON_SYNONYMOU | MODERAT | MISSENSE | aCg/aTg | T58M | PLES_26521 | . | . | A | . | . | . | . | . | . | . | A | A |
| NC_011770 | 2849130 | T | C | 13 | 9 | 1 | 0 | NON_SYNONYMOU | MODERAT | MISSENSE | Acg/Gcg | T58A | PLES_26521 | . | . | C | . | . | . | . | . | . | . | | |
| NC_011770 | 2849142 | C | T | 8 | 10 | 2 | 0 | NON_SYNONYMOU | MODERAT | MISSENSE | Gcg/Acg | A54T | PLES_26521 | . | . | . | . | . | . | . | . | . | . | T | T |
| NC_011770 | 2849144 | C | G | 8 | 10 | 2 | 0 | NON_SYNONYMOU | MODERAT | MISSENSE | cGc/cCc | R53P | PLES_26521 | . | . | . | . | . | . | . | . | . | . | G | G |
| NC_011770 | 2849148 | A | G | 8 | 10 | 2 | 0 | NON_SYNONYMOU | MODERAT | MISSENSE | Tcg/Ccg | S52P | PLES_26521 | . | . | . | . | . | . | . | . | . | . | G | G |
| NC_011770 | 2849150 | G | T | 8 | 10 | 2 | 0 | NON_SYNONYMOU | MODERAT | MISSENSE | cCg/cAg | P51Q | PLES_26521 | . | . | . | . | . | . | . | . | . | . | T | T |
| NC_011770 | 2849154 | C | G | 8 | 10 | 2 | 0 | NON_SYNONYMOU | MODERAT | MISSENSE | Gcg/Ccg | A50P | PLES_26521 | . | . | . | . | . | . | . | . | . | . | G | G |
| NC_011770 | 2849157 | A | G | 8 | 10 | 2 | 0 | NON_SYNONYMOU | MODERAT | MISSENSE | Tca/Cca | S49P | PLES_26521 | . | . | . | . | . | . | . | . | . | . | G | G |
| NC_011770 | 2849172 | T | A | 14 | 10 | 2 | 0 | NON_SYNONYMOU | MODERAT | MISSENSE | Acg/Tcg | T44S | PLES_26521 | . | . | . | . | . | . | . | . | . | . | A | A |
| NC_011770 | 2849210 | G | A | 39 | 6 | 6 | 0 | NON_SYNONYMOU | MODERAT | MISSENSE | gCg/gTg | A31V | PLES_26521 | A | A | A | A | . | . | . | . | . | . | A | A |
| NC_011770 | 2849222 | A | G | 41 | 6 | 2 | 0 | NON_SYNONYMOU | MODERAT | MISSENSE | gTa/gCa | V27A | PLES_26521 | | | | | . | . | . | . | . | . | G | G |
| NC_011770 | 2849235 | G | T | 40 | 6 | 4 | 0 | NON_SYNONYMOU | MODERAT | MISSENSE | Cgc/Agc | R23S | PLES_26521 | T | T | T | T | . | . | . | . | . | . | | |
| NC_011770 | 2849237 | G | A | 40 | 6 | 4 | 0 | NON_SYNONYMOU | MODERAT | MISSENSE | cCt/cTt | P22L | PLES_26521 | A | A | A | A | . | . | . | . | . | . | | |
| NC_011770 | 2849244 | A | C | 41 | 6 | 2 | 0 | NON_SYNONYMOU | MODERAT | MISSENSE | Tgg/Ggg | W20G | PLES_26521 | | | | | . | . | . | . | . | . | C | C |
| NC_011770 | 2849246 | C | T | 41 | 6 | 2 | 0 | NON_SYNONYMOU | MODERAT | MISSENSE | cGc/cAc | R19H | PLES_26521 | | | | | . | . | . | . | . | . | T | T |
| NC_011770 | 2849247 | G | T | 41 | 6 | 2 | 0 | NON_SYNONYMOU | MODERAT | MISSENSE | Cgc/Agc | R19S | PLES_26521 | | | | | . | . | . | . | . | . | T | T |
| NC_011770 | 2849469 | TAC | T | 34 | 6 | 2 | 0 | FRAME_SHIFT | HIGH | | ctgtac/ | LY910 | PLES_26531 | T | | T | | . | . | . | . | . | . | | |
| NC_011770 | 2849318 | C | CGGCAC | 39 | 6 | 1 | 0 | FRAME_SHIFT+STC | HIGH | | tcg/tcCTG/ | S961S*IC? | PLES_26531 | CGGCAGATTCAG | | | | . | . | . | . | . | . | | |
| NC_011770 | 2849315 | T | G | 38 | 6 | 4 | 0 | STOP_LOST+SPLICE | HIGH | MISSENSE | taA/taC | *962Y | PLES_26531 | G | G | G | G | . | . | . | . | . | . | | |
| NC_011770 | 2849477 | C | CTT | 34 | 6 | 6 | 0 | FRAME_SHIFT | HIGH | | ccg/ccAAg | P908P? | PLES_26531 | CTT | CTT | CTT | CTT | . | . | . | . | . | . | G | G |
| NC_011770 | 2849642 | G | GGCAC | 28 | 6 | 4 | 0 | FRAME_SHIFT | HIGH | | atc/atGTG | I853MC? | PLES_26531 | GGCAC | GGCAC | GGCAC | GGCAC | . | . | . | . | . | . | | |
| NC_011770 | 2851563 | G | GTCCGC | 33 | 6 | 4 | 0 | FRAME_SHIFT | HIGH | | act/aGCGC | T213SG? | PLES_26531 | GTCCGC | GTCCGC | GTCCGC | GTCCGC | . | . | . | . | . | . | | |
| NC_011770 | 2852155 | TGGAGC | T | 3 | 7 | 1 | 0 | FRAME_SHIFT | HIGH | | ggctcc/ | GS14 | PLES_26531 | . | . | . | . | . | . | . | . | . | . | T | . |
| NC_011770 | 2849320 | A | G | 21 | 8 | 2 | 0 | NON_SYNONYMOU | MODERAT | MISSENSE | Tcg/Ccg | S961P | PLES_26531 | . | . | G | G | . | . | . | . | . | . | | |

| | | | | | | | | | | | | | | | | | | | | | | | | | |
|---|---|---|---|---|---|---|---|---|---|---|---|---|---|---|---|---|---|---|---|---|---|---|---|---|---|
| NC_011770 | 2849332 | C | T | 29 | 6 | 2 | 0 | NON_SYNONYMOU | MODERAT | MISSENSE | Gcc/Acc | A957T | PLES_26531 | T | T | | | . | . | . | . | . | . | | |
| NC_011770 | 2849355 | T | C | 40 | 6 | 4 | 0 | NON_SYNONYMOU | MODERAT | MISSENSE | cAg/cGg | Q949R | PLES_26531 | C | C | C | C | . | . | . | . | . | . | | |
| NC_011770 | 2849387 | T | A | 39 | 6 | 4 | 0 | NON_SYNONYMOU | MODERAT | MISSENSE | gaA/gaT | E938D | PLES_26531 | A | A | A | A | . | . | . | . | . | . | | |
| NC_011770 | 2849397 | G | C | 38 | 6 | 4 | 0 | NON_SYNONYMOU | MODERAT | MISSENSE | aCc/aGc | T935S | PLES_26531 | C | C | C | C | . | . | . | . | . | . | | |
| NC_011770 | 2849400 | C | A | 38 | 6 | 4 | 0 | NON_SYNONYMOU | MODERAT | MISSENSE | tGc/tTc | C934F | PLES_26531 | A | A | A | A | . | . | . | . | . | . | | |
| NC_011770 | 2849401 | A | C | 37 | 6 | 4 | 0 | NON_SYNONYMOU | MODERAT | MISSENSE | Tgc/Ggc | C934G | PLES_26531 | C | C | C | C | . | . | . | . | . | . | | |
| NC_011770 | 2849407 | A | G | 41 | 6 | 4 | 0 | NON_SYNONYMOU | MODERAT | MISSENSE | Tac/Cac | Y932H | PLES_26531 | G | G | G | G | . | . | . | . | . | . | | |
| NC_011770 | 2849408 | C | G | 42 | 6 | 2 | 0 | NON_SYNONYMOU | MODERAT | MISSENSE | caG/caC | Q931H | PLES_26531 | | | | | . | . | . | . | . | . | G | G |
| NC_011770 | 2849461 | G | A | 33 | 6 | 2 | 0 | NON_SYNONYMOU | MODERAT | MISSENSE | Ctc/Ttc | L914F | PLES_26531 | | | | | . | . | . | . | . | . | A | A |
| NC_011770 | 2849467 | G | A | 33 | 6 | 2 | 0 | NON_SYNONYMOU | MODERAT | MISSENSE | Ctc/Ttc | L912F | PLES_26531 | | | | | . | . | . | . | . | . | A | A |
| NC_011770 | 2849475 | G | C | 15 | 7 | 2 | 0 | NON_SYNONYMOU | MODERAT | MISSENSE | tCc/tGc | S909C | PLES_26531 | | . | | | . | . | . | . | . | . | C | C |
| NC_011770 | 2849476 | A | T | 32 | 6 | 2 | 0 | NON_SYNONYMOU | MODERAT | MISSENSE | Tcc/Acc | S909T | PLES_26531 | | | | | . | . | . | . | . | . | T | T |
| NC_011770 | 2849633 | G | C | 14 | 10 | 2 | 0 | NON_SYNONYMOU | MODERAT | MISSENSE | agC/agG | S856R | PLES_26531 | . | . | . | . | . | . | . | . | . | . | C | C |
| NC_011770 | 2849634 | C | G | 14 | 10 | 2 | 0 | NON_SYNONYMOU | MODERAT | MISSENSE | aGc/aCc | S856T | PLES_26531 | . | . | . | . | . | . | . | . | . | . | G | G |
| NC_011770 | 2849635 | T | A | 14 | 10 | 2 | 0 | NON_SYNONYMOU | MODERAT | MISSENSE | Agc/Tgc | S856C | PLES_26531 | . | . | . | . | . | . | . | . | . | . | A | A |
| NC_011770 | 2849644 | T | C | 28 | 6 | 4 | 0 | NON_SYNONYMOU | MODERAT | MISSENSE | Atc/Gtc | I853V | PLES_26531 | C | C | C | C | . | . | . | . | . | . | | |
| NC_011770 | 2849735 | C | G | 33 | 6 | 6 | 0 | NON_SYNONYMOU | MODERAT | MISSENSE | agG/agC | R822S | PLES_26531 | G | G | G | G | . | . | . | . | . | . | G | G |
| NC_011770 | 2849818 | T | C | 36 | 6 | 4 | 0 | NON_SYNONYMOU | MODERAT | MISSENSE | Atc/Gtc | I795V | PLES_26531 | C | C | C | C | . | . | . | . | . | . | | |
| NC_011770 | 2850085 | C | T | 31 | 6 | 2 | 0 | NON_SYNONYMOU | MODERAT | MISSENSE | Gcc/Acc | A706T | PLES_26531 | | | | | . | . | . | . | . | . | T | T |
| NC_011770 | 2850128 | G | T | 31 | 6 | 6 | 0 | NON_SYNONYMOU | MODERAT | MISSENSE | gaC/gaA | D691E | PLES_26531 | T | T | T | T | . | . | . | . | . | . | C | C |
| NC_011770 | 2850282 | G | T | 33 | 6 | 2 | 0 | NON_SYNONYMOU | MODERAT | MISSENSE | gCg/gAg | A640E | PLES_26531 | | | | | . | . | . | . | . | . | T | T |
| NC_011770 | 2850291 | G | A | 33 | 6 | 2 | 0 | NON_SYNONYMOU | MODERAT | MISSENSE | gCc/gTc | A637V | PLES_26531 | | | | | . | . | . | . | . | . | A | A |
| NC_011770 | 2850298 | C | A | 30 | 6 | 2 | 0 | NON_SYNONYMOU | MODERAT | MISSENSE | Gcc/Tcc | A635S | PLES_26531 | | | | | . | . | . | . | . | . | A | A |
| NC_011770 | 2850304 | G | C | 30 | 6 | 2 | 0 | NON_SYNONYMOU | MODERAT | MISSENSE | Cca/Gca | P633A | PLES_26531 | | | | | . | . | . | . | . | . | C | C |
| NC_011770 | 2850309 | G | A | 29 | 6 | 6 | 0 | NON_SYNONYMOU | MODERAT | MISSENSE | cCg/cTg | P631L | PLES_26531 | A | A | A | A | . | . | . | . | . | . | A | A |
| NC_011770 | 2850310 | G | A | 30 | 6 | 2 | 0 | NON_SYNONYMOU | MODERAT | MISSENSE | Ccg/Tcg | P631S | PLES_26531 | | | | | . | . | . | . | . | . | A | A |
| NC_011770 | 2850311 | G | T | 30 | 6 | 2 | 0 | NON_SYNONYMOU | MODERAT | MISSENSE | gaC/gaA | D630E | PLES_26531 | | | | | . | . | . | . | . | . | T | T |
| NC_011770 | 2850349 | C | T | 35 | 6 | 2 | 0 | NON_SYNONYMOU | MODERAT | MISSENSE | Gac/Aac | D618N | PLES_26531 | | | | | . | . | . | . | . | . | T | T |
| NC_011770 | 2850446 | C | G | 0 | 10 | 1 | 0 | NON_SYNONYMOU | MODERAT | MISSENSE | atG/atC | M585I | PLES_26531 | . | . | . | . | . | . | . | . | . | . | . | G |
| NC_011770 | 2850447 | A | C | 0 | 10 | 1 | 0 | NON_SYNONYMOU | MODERAT | MISSENSE | aTg/aGg | M585R | PLES_26531 | . | . | . | . | . | . | . | . | . | . | . | C |
| NC_011770 | 2850448 | T | G | 0 | 11 | 1 | 0 | NON_SYNONYMOU | MODERAT | MISSENSE | Atg/Ctg | M585L | PLES_26531 | . | . | . | . | . | . | . | . | . | . | . | G |
| NC_011770 | 2850451 | G | GCTT | 13 | 10 | 2 | 0 | CODON_INSERTIOI | MODERATE | | gca/gcaAA | A583AK | PLES_26531 | . | . | . | . | . | . | . | . | . | . | GCTT | GCTT |
| NC_011770 | 2850555 | C | G | 35 | 6 | 4 | 0 | NON_SYNONYMOU | MODERAT | MISSENSE | aGc/aCc | S549T | PLES_26531 | G | G | G | G | . | . | . | . | . | . | | |
| NC_011770 | 2850556 | T | A | 34 | 6 | 4 | 0 | NON_SYNONYMOU | MODERAT | MISSENSE | Agc/Tgc | S549C | PLES_26531 | A | A | A | A | . | . | . | . | . | . | | |
| NC_011770 | 2850616 | G | T | 23 | 6 | 4 | 0 | NON_SYNONYMOU | MODERAT | MISSENSE | Cgc/Agc | R529S | PLES_26531 | T | T | T | T | . | . | . | . | . | . | | |
| NC_011770 | 2850618 | C | T | 23 | 6 | 6 | 0 | NON_SYNONYMOU | MODERAT | MISSENSE | aGc/aAc | S528N | PLES_26531 | T | T | T | T | . | . | . | . | . | . | T | T |
| NC_011770 | 2850622 | G | A | 23 | 6 | 4 | 0 | NON_SYNONYMOU | MODERAT | MISSENSE | Ctc/Ttc | L527F | PLES_26531 | A | A | A | A | . | . | . | . | . | . | | |
| NC_011770 | 2850659 | C | G | 30 | 6 | 2 | 0 | NON_SYNONYMOU | MODERAT | MISSENSE | atG/atC | M514I | PLES_26531 | | | | | . | . | . | . | . | . | G | G |
| NC_011770 | 2850661 | T | A | 31 | 6 | 2 | 0 | NON_SYNONYMOU | MODERAT | MISSENSE | Atg/Ttg | M514L | PLES_26531 | | | | | . | . | . | . | . | . | A | A |
| NC_011770 | 2850666 | A | T | 31 | 6 | 2 | 0 | NON_SYNONYMOU | MODERAT | MISSENSE | aTc/aAc | I512N | PLES_26531 | | | | | . | . | . | . | . | . | T | T |
| NC_011770 | 2850690 | G | C | 28 | 6 | 2 | 0 | NON_SYNONYMOU | MODERAT | MISSENSE | gCt/gGt | A504G | PLES_26531 | | | | | . | . | . | . | . | . | C | C |
| NC_011770 | 2850691 | C | T | 28 | 6 | 2 | 0 | NON_SYNONYMOU | MODERAT | MISSENSE | Gct/Act | A504T | PLES_26531 | | | | | . | . | . | . | . | . | T | T |
| NC_011770 | 2850702 | A | G | 30 | 6 | 2 | 0 | NON_SYNONYMOU | MODERAT | MISSENSE | gTg/gCg | V500A | PLES_26531 | | | | | . | . | . | . | . | . | G | G |
| NC_011770 | 2850720 | G | A | 31 | 6 | 2 | 0 | NON_SYNONYMOU | MODERAT | MISSENSE | gCg/gTg | A494V | PLES_26531 | | | | | . | . | . | . | . | . | A | A |
| NC_011770 | 2850739 | G | T | 35 | 5 | 1 | 0 | NON_SYNONYMOU | MODERAT | MISSENSE | Ctg/Atg | L488M | PLES_26531 | | | | | T | . | . | . | . | . | | |
| NC_011770 | 2850747 | T | A | 32 | 5 | 2 | 0 | NON_SYNONYMOU | MODERAT | MISSENSE | tAc/tTc | Y485F | PLES_26531 | | | | | | . | . | . | . | . | A | A |
| NC_011770 | 2850752 | C | G | 31 | 5 | 2 | 0 | NON_SYNONYMOU | MODERAT | MISSENSE | aaG/aaC | K483N | PLES_26531 | | | | | | . | . | . | . | . | G | G |
| NC_011770 | 2850753 | T | C | 30 | 6 | 4 | 0 | NON_SYNONYMOU | MODERAT | MISSENSE | aAg/aGg | K483R | PLES_26531 | C | C | C | C | . | . | . | . | . | . | | |
| NC_011770 | 2850754 | T | G | 29 | 6 | 4 | 0 | NON_SYNONYMOU | MODERAT | MISSENSE | Aag/Cag | K483Q | PLES_26531 | G | G | G | G | . | . | . | . | . | . | | |
| NC_011770 | 2850766 | G | T | 29 | 6 | 4 | 0 | NON_SYNONYMOU | MODERAT | MISSENSE | Cag/Aag | Q479K | PLES_26531 | T | T | T | T | . | . | . | . | . | . | | |
| NC_011770 | 2850768 | C | G | 30 | 6 | 2 | 0 | NON_SYNONYMOU | MODERAT | MISSENSE | gGc/gCc | G478A | PLES_26531 | | | | | . | . | . | . | . | . | G | G |
| NC_011770 | 2850980 | C | A | 26 | 6 | 4 | 0 | NON_SYNONYMOU | MODERAT | MISSENSE | caG/caT | Q407H | PLES_26531 | A | A | A | A | . | . | . | . | . | . | | |
| NC_011770 | 2850992 | C | G | 23 | 6 | 6 | 0 | NON_SYNONYMOU | MODERAT | MISSENSE | ttG/ttC | L403F | PLES_26531 | G | G | G | G | . | . | . | . | . | . | G | G |
| NC_011770 | 2851051 | C | G | 22 | 7 | 1 | 0 | NON_SYNONYMOU | MODERAT | MISSENSE | Gat/Cat | D384H | PLES_26531 | | | | | . | . | . | . | . | . | . | G |
| NC_011770 | 2851054 | C | A | 21 | 7 | 5 | 0 | NON_SYNONYMOU | MODERAT | MISSENSE | Gcg/Tcg | A383S | PLES_26531 | A | A | A | A | . | . | . | . | . | . | . | A |
| NC_011770 | 2851063 | C | T | 21 | 7 | 1 | 0 | NON_SYNONYMOU | MODERAT | MISSENSE | Gtc/Atc | V380I | PLES_26531 | | | | | . | . | . | . | . | . | . | T |
| NC_011770 | 2851096 | G | T | 25 | 7 | 5 | 0 | NON_SYNONYMOU | MODERAT | MISSENSE | Ctg/Atg | L369M | PLES_26531 | T | T | T | T | . | . | . | . | . | . | . | T |
| NC_011770 | 2851098 | A | G | 26 | 7 | 1 | 0 | NON_SYNONYMOU | MODERAT | MISSENSE | tTg/tCg | L368S | PLES_26531 | | | | | . | . | . | . | . | . | . | G |
| NC_011770 | 2851099 | A | T | 26 | 7 | 1 | 0 | NON_SYNONYMOU | MODERAT | MISSENSE | Ttg/Atg | L368M | PLES_26531 | | | | | . | . | . | . | . | . | . | T |
| NC_011770 | 2851204 | T | C | 40 | 6 | 2 | 0 | NON_SYNONYMOU | MODERAT | MISSENSE | Atc/Gtc | I333V | PLES_26531 | | | | | . | . | . | . | . | . | C | C |
| NC_011770 | 2851240 | T | C | 36 | 6 | 4 | 0 | NON_SYNONYMOU | MODERAT | MISSENSE | Acc/Gcc | T321A | PLES_26531 | C | C | C | C | . | . | . | . | . | . | | |
| NC_011770 | 2851244 | G | T | 36 | 6 | 4 | 0 | NON_SYNONYMOU | MODERAT | MISSENSE | caC/caA | H319Q | PLES_26531 | T | T | T | T | . | . | . | . | . | . | | |

| | | | | | | | | | | | | | | | | | | | | | | | | | |
|---|---|---|---|---|---|---|---|---|---|---|---|---|---|---|---|---|---|---|---|---|---|---|---|---|---|
| NC_011770 | 2851248 | T | G | 35 | 6 | 4 | 0 | NON_SYNONYMOU | MODERAT | MISSENSE | gAg/gCg | E318A | PLES_26531 | G | G | G | G | . | . | . | . | . | . | | |
| NC_011770 | 2851402 | T | G | 18 | 7 | 4 | 0 | NON_SYNONYMOU | MODERAT | MISSENSE | Atg/Ctg | M267L | PLES_26531 | G | G | G | G | . | . | . | . | . | . | | . |
| NC_011770 | 2851405 | C | T | 18 | 7 | 5 | 0 | NON_SYNONYMOU | MODERAT | MISSENSE | Gcg/Acg | A266T | PLES_26531 | T | T | T | T | . | . | . | . | . | . | T | . |
| NC_011770 | 2851518 | T | C | 42 | 6 | 4 | 0 | NON_SYNONYMOU | MODERAT | MISSENSE | aAc/aGc | N228S | PLES_26531 | C | C | C | C | . | . | . | . | . | . | | |
| NC_011770 | 2851519 | T | C | 42 | 6 | 4 | 0 | NON_SYNONYMOU | MODERAT | MISSENSE | Aac/Gac | N228D | PLES_26531 | C | C | C | C | . | . | . | . | . | . | | |
| NC_011770 | 2851551 | C | T | 42 | 6 | 6 | 0 | NON_SYNONYMOU | MODERAT | MISSENSE | aGc/aAc | S217N | PLES_26531 | T | T | T | T | . | . | . | . | . | . | T | T |
| NC_011770 | 2851564 | T | C | 17 | 10 | 2 | 0 | NON_SYNONYMOU | MODERAT | MISSENSE | Act/Gct | T213A | PLES_26531 | . | . | . | . | . | . | . | . | . | . | C | C |
| NC_011770 | 2851577 | G | C | 37 | 6 | 4 | 0 | NON_SYNONYMOU | MODERAT | MISSENSE | atC/atG | I208M | PLES_26531 | C | C | C | C | . | . | . | . | . | . | | |
| NC_011770 | 2851579 | T | C | 36 | 6 | 6 | 0 | NON_SYNONYMOU | MODERAT | MISSENSE | Atc/Gtc | I208V | PLES_26531 | C | C | C | C | . | . | . | . | . | . | C | C |
| NC_011770 | 2851691 | C | G | 46 | 6 | 2 | 0 | NON_SYNONYMOU | MODERAT | MISSENSE | ttG/ttC | L170F | PLES_26531 | | | | | . | . | . | . | . | . | G | G |
| NC_011770 | 2851714 | A | T | 45 | 6 | 2 | 0 | NON_SYNONYMOU | MODERAT | MISSENSE | Tcg/Acg | S163T | PLES_26531 | | | | | . | . | . | . | . | . | T | T |
| NC_011770 | 2851717 | C | T | 47 | 6 | 2 | 0 | NON_SYNONYMOU | MODERAT | MISSENSE | Ggc/Agc | G162S | PLES_26531 | | | | | . | . | . | . | . | . | T | T |
| NC_011770 | 2851829 | A | G | 43 | 6 | 6 | 0 | NON_SYNONYMOU | MODERAT | MISSENSE | agT/agG | S124R | PLES_26531 | G | G | G | G | . | . | . | . | . | . | C | C |
| NC_011770 | 2851830 | C | G | 43 | 6 | 2 | 0 | NON_SYNONYMOU | MODERAT | MISSENSE | aGt/aCt | S124T | PLES_26531 | | | | | . | . | . | . | . | . | G | G |
| NC_011770 | 2851831 | T | A | 43 | 6 | 2 | 0 | NON_SYNONYMOU | MODERAT | MISSENSE | Agt/Tgt | S124C | PLES_26531 | | | | | . | . | . | . | . | . | A | A |
| NC_011770 | 2851965 | C | T | 39 | 6 | 6 | 0 | NON_SYNONYMOU | MODERAT | MISSENSE | gGg/gAg | G79E | PLES_26531 | T | T | T | T | . | . | . | . | . | . | G | G |
| NC_011770 | 2851966 | C | G | 40 | 6 | 2 | 0 | NON_SYNONYMOU | MODERAT | MISSENSE | Ggg/Cgg | G79R | PLES_26531 | | | | | . | . | . | . | . | . | G | G |
| NC_011770 | 2851998 | A | G | 41 | 6 | 2 | 0 | NON_SYNONYMOU | MODERAT | MISSENSE | gTg/gCg | V68A | PLES_26531 | | | | | . | . | . | . | . | . | G | G |
| NC_011770 | 2852007 | G | T | 39 | 6 | 2 | 0 | NON_SYNONYMOU | MODERAT | MISSENSE | cCg/cAg | P65Q | PLES_26531 | | | | | . | . | . | . | . | . | T | T |
| NC_011770 | 2852023 | G | T | 41 | 6 | 2 | 0 | NON_SYNONYMOU | MODERAT | MISSENSE | Ctg/Atg | L60M | PLES_26531 | | | | | . | . | . | . | . | . | T | T |
| NC_011770 | 2852052 | T | C | 39 | 6 | 2 | 0 | NON_SYNONYMOU | MODERAT | MISSENSE | aAg/aGg | K50R | PLES_26531 | | | | | . | . | . | . | . | . | C | C |
| NC_011770 | 2852060 | G | C | 38 | 6 | 6 | 0 | NON_SYNONYMOU | MODERAT | MISSENSE | caC/caG | H47Q | PLES_26531 | C | C | C | C | . | . | . | . | . | . | C | C |
| NC_011770 | 2852068 | C | T | 33 | 6 | 2 | 0 | NON_SYNONYMOU | MODERAT | MISSENSE | Gca/Aca | A45T | PLES_26531 | | | | | . | . | . | . | . | . | T | T |
| NC_011770 | 2852071 | A | C | 33 | 6 | 6 | 0 | NON_SYNONYMOU | MODERAT | MISSENSE | Tcg/Gcg | S44A | PLES_26531 | C | C | C | C | . | . | . | . | . | . | C | C |
| NC_011770 | 2852083 | G | C | 40 | 6 | 2 | 0 | NON_SYNONYMOU | MODERAT | MISSENSE | Ccc/Gcc | P40A | PLES_26531 | | | | | . | . | . | . | . | . | C | C |
| NC_011770 | 2852125 | T | C | 41 | 6 | 6 | 0 | NON_SYNONYMOU | MODERAT | MISSENSE | Agc/Ggc | S26G | PLES_26531 | C | C | C | C | . | . | . | . | . | . | C | C |
| NC_011770 | 2852158 | A | C | 2 | 8 | 4 | 0 | NON_SYNONYMOU | MODERAT | MISSENSE | Tcc/Gcc | S15A | PLES_26531 | C | C | C | C | . | . | . | . | . | . | . | . |
| NC_011770 | 2852161 | C | T | 2 | 8 | 4 | 0 | NON_SYNONYMOU | MODERAT | MISSENSE | Ggc/Agc | G14S | PLES_26531 | T | T | T | T | . | . | . | . | . | . | . | . |
| NC_011770 | 2852163 | G | A | 2 | 8 | 4 | 0 | NON_SYNONYMOU | MODERAT | MISSENSE | gCc/gTc | A13V | PLES_26531 | A | A | A | A | . | . | . | . | . | . | . | . |
| NC_011770 | 2852166 | C | G | 2 | 8 | 4 | 0 | NON_SYNONYMOU | MODERAT | MISSENSE | gGc/gCc | G12A | PLES_26531 | G | G | G | G | . | . | . | . | . | . | . | . |
| NC_011770 | 2852097 | C | T | 46 | 4 | 3 | 0 | NON_SYNONYMOU | MODERAT | MISSENSE | cGc/cAc | R35H | PLES_26531 | | T | T | T | . | . | . | . | | | | |
| NC_011770 | 2853503 | T | TCCTC | 29 | 6 | 2 | 0 | FRAME_SHIFT | HIGH | | gca/gcGAC | A179AR? | PLES_26541 | | | | | . | . | . | . | . | . | TCCTC | TCCTC |
| NC_011770 | 2852624 | A | G | 25 | 6 | 6 | 0 | NON_SYNONYMOU | MODERAT | MISSENSE | gaT/gaA | D472E | PLES_26541 | G | G | G | G | . | . | . | . | . | . | T | T |
| NC_011770 | 2852784 | G | T | 21 | 7 | 1 | 0 | NON_SYNONYMOU | MODERAT | MISSENSE | gCc/gAc | A419D | PLES_26541 | | | | | . | . | . | . | . | . | T | . |
| NC_011770 | 2852788 | A | C | 21 | 7 | 1 | 0 | NON_SYNONYMOU | MODERAT | MISSENSE | Tcg/Gcg | S418A | PLES_26541 | | | | | . | . | . | . | . | . | C | . |
| NC_011770 | 2852816 | C | G | 21 | 7 | 1 | 0 | NON_SYNONYMOU | MODERAT | MISSENSE | caG/caC | Q408H | PLES_26541 | | | | | . | . | . | . | . | . | G | . |
| NC_011770 | 2852817 | T | C | 21 | 7 | 1 | 0 | NON_SYNONYMOU | MODERAT | MISSENSE | cAg/cGg | Q408R | PLES_26541 | | | | | . | . | . | . | . | . | C | . |
| NC_011770 | 2852831 | C | A | 22 | 7 | 1 | 0 | NON_SYNONYMOU | MODERAT | MISSENSE | caG/caT | Q403H | PLES_26541 | | | | | . | . | . | . | . | . | A | . |
| NC_011770 | 2852833 | G | T | 21 | 7 | 1 | 0 | NON_SYNONYMOU | MODERAT | MISSENSE | Cag/Aag | Q403K | PLES_26541 | | | | | . | . | . | . | . | . | T | . |
| NC_011770 | 2852839 | T | A | 21 | 7 | 1 | 0 | NON_SYNONYMOU | MODERAT | MISSENSE | Acc/Tcc | T401S | PLES_26541 | | | | | . | . | . | . | . | . | A | . |
| NC_011770 | 2852850 | G | C | 20 | 6 | 2 | 0 | NON_SYNONYMOU | MODERAT | MISSENSE | aCc/aGc | T397S | PLES_26541 | | | | | . | . | . | . | . | . | C | C |
| NC_011770 | 2852869 | A | C | 26 | 6 | 2 | 0 | NON_SYNONYMOU | MODERAT | MISSENSE | Tcg/Gcg | S391A | PLES_26541 | | | | | . | . | . | . | . | . | C | C |
| NC_011770 | 2852886 | C | T | 26 | 6 | 2 | 0 | NON_SYNONYMOU | MODERAT | MISSENSE | cGc/cAc | R385H | PLES_26541 | | | | | . | . | . | . | . | . | T | T |
| NC_011770 | 2852900 | C | G | 28 | 6 | 2 | 0 | NON_SYNONYMOU | MODERAT | MISSENSE | gaG/gaC | E380D | PLES_26541 | | | | | . | . | . | . | . | . | G | G |
| NC_011770 | 2853007 | G | A | 29 | 6 | 2 | 0 | NON_SYNONYMOU | MODERAT | MISSENSE | Cac/Tac | H345Y | PLES_26541 | | | | | . | . | . | . | . | . | A | A |
| NC_011770 | 2853031 | C | CTCCAC | 26 | 6 | 1 | 0 | CODON_INSERTIO | MODERATE | | gac/gacTC | D336DSQS | PLES_26541 | | | | | . | . | . | . | . | . | CTCCACTT | . |
| NC_011770 | 2853062 | C | G | 30 | 6 | 2 | 0 | NON_SYNONYMOU | MODERAT | MISSENSE | gaG/gaC | E326D | PLES_26541 | | | | | . | . | . | . | . | . | G | G |
| NC_011770 | 2853100 | T | G | 31 | 6 | 2 | 0 | NON_SYNONYMOU | MODERAT | MISSENSE | Aac/Cac | N314H | PLES_26541 | | | | | . | . | . | . | . | . | G | G |
| NC_011770 | 2853229 | G | A | 29 | 6 | 2 | 0 | NON_SYNONYMOU | MODERAT | MISSENSE | Ccg/Tcg | P271S | PLES_26541 | | | | | . | . | . | . | . | . | A | A |
| NC_011770 | 2853352 | G | C | 31 | 6 | 2 | 0 | NON_SYNONYMOU | MODERAT | MISSENSE | Ccg/Gcg | P230A | PLES_26541 | | | | | . | . | . | . | . | . | C | C |
| NC_011770 | 2853366 | A | C | 29 | 6 | 4 | 0 | NON_SYNONYMOU | MODERAT | MISSENSE | gTc/gGc | V225G | PLES_26541 | C | C | C | C | . | . | . | . | . | . | | |
| NC_011770 | 2853373 | A | C | 29 | 6 | 6 | 0 | NON_SYNONYMOU | MODERAT | MISSENSE | Tct/Gct | S223A | PLES_26541 | C | C | C | C | . | . | . | . | . | . | C | C |
| NC_011770 | 2853418 | G | A | 34 | 6 | 2 | 0 | NON_SYNONYMOU | MODERAT | MISSENSE | Ctc/Ttc | L208F | PLES_26541 | | | | | . | . | . | . | . | . | A | A |
| NC_011770 | 2853463 | C | T | 31 | 6 | 2 | 0 | NON_SYNONYMOU | MODERAT | MISSENSE | Ggc/Agc | G193S | PLES_26541 | | | | | . | . | . | . | . | . | T | T |
| NC_011770 | 2853466 | T | C | 31 | 6 | 2 | 0 | NON_SYNONYMOU | MODERAT | MISSENSE | Agc/Ggc | S192G | PLES_26541 | | | | | . | . | . | . | . | . | C | C |
| NC_011770 | 2853469 | C | T | 31 | 6 | 2 | 0 | NON_SYNONYMOU | MODERAT | MISSENSE | Gcg/Acg | A191T | PLES_26541 | | | | | . | . | . | . | . | . | T | T |
| NC_011770 | 2853478 | C | T | 30 | 6 | 2 | 0 | NON_SYNONYMOU | MODERAT | MISSENSE | Ggt/Agt | G188S | PLES_26541 | | | | | . | . | . | . | . | . | T | T |
| NC_011770 | 2853481 | C | T | 31 | 6 | 2 | 0 | NON_SYNONYMOU | MODERAT | MISSENSE | Ggc/Agc | G187S | PLES_26541 | | | | | . | . | . | . | . | . | T | T |
| NC_011770 | 2853498 | G | T | 17 | 7 | 1 | 0 | NON_SYNONYMOU | MODERAT | MISSENSE | cCc/cAc | P181H | PLES_26541 | | | | | . | . | . | . | . | . | T | . |
| NC_011770 | 2853499 | G | T | 17 | 7 | 1 | 0 | NON_SYNONYMOU | MODERAT | MISSENSE | Ccc/Acc | P181T | PLES_26541 | | | | | . | . | . | . | . | . | T | . |
| NC_011770 | 2853558 | C | T | 32 | 6 | 2 | 0 | NON_SYNONYMOU | MODERAT | MISSENSE | gGc/gAc | G161D | PLES_26541 | | | | | . | . | . | . | . | . | T | T |
| NC_011770 | 2853560 | C | A | 32 | 6 | 2 | 0 | NON_SYNONYMOU | MODERAT | MISSENSE | caG/caT | Q160H | PLES_26541 | | | | | . | . | . | . | . | . | A | A |

| | | | | | | | | | | | | | | | | | | | | | | | | | |
|---|---|---|---|---|---|---|---|---|---|---|---|---|---|---|---|---|---|---|---|---|---|---|---|---|---|
| NC_011770 | 2853561 | T | C | 32 | 6 | 2 | 0 | NON_SYNONYMOU | MODERAT | MISSENSE | cAg/cGg | Q160R | PLES_26541 | | | | | . | . | . | . | . | . | C | C |
| NC_011770 | 2853703 | A | T | 34 | 6 | 2 | 0 | NON_SYNONYMOU | MODERAT | MISSENSE | Tcc/Acc | S113T | PLES_26541 | | | | | . | . | . | . | . | . | T | T |
| NC_011770 | 2853713 | T | G | 34 | 6 | 2 | 0 | NON_SYNONYMOU | MODERAT | MISSENSE | gaA/gaC | E109D | PLES_26541 | | | | | . | . | . | . | . | . | G | G |
| NC_011770 | 2853902 | C | A | 35 | 6 | 4 | 0 | NON_SYNONYMOU | MODERAT | MISSENSE | gaG/gaT | E46D | PLES_26541 | A | A | A | A | . | . | . | . | . | . | | |
| NC_011770 | 2853903 | T | G | 35 | 6 | 2 | 0 | NON_SYNONYMOU | MODERAT | MISSENSE | gAg/gCg | E46A | PLES_26541 | | | | | . | . | . | . | . | . | G | G |
| NC_011770 | 2853916 | G | T | 35 | 6 | 2 | 0 | NON_SYNONYMOU | MODERAT | MISSENSE | Cag/Aag | Q42K | PLES_26541 | | | | | . | . | . | . | . | . | T | T |
| NC_011770 | 2853924 | T | G | 31 | 6 | 2 | 0 | NON_SYNONYMOU | MODERAT | MISSENSE | aAc/aCc | N39T | PLES_26541 | | | | | . | . | . | . | . | . | G | G |
| NC_011770 | 2853925 | T | C | 31 | 6 | 2 | 0 | NON_SYNONYMOU | MODERAT | MISSENSE | Aac/Gac | N39D | PLES_26541 | | | | | . | . | . | . | . | . | C | C |
| NC_011770 | 2853934 | C | T | 30 | 6 | 2 | 0 | NON_SYNONYMOU | MODERAT | MISSENSE | Gcg/Acg | A36T | PLES_26541 | | | | | . | . | . | . | . | . | T | T |
| NC_011770 | 2853944 | C | CGAA | 31 | 6 | 2 | 0 | CODON_CHANGE_ | MODERATE | | acg/acTTC | T32TS | PLES_26541 | | | | | . | . | . | . | . | . | CGAA | CGAA |
| NC_011770 | 2853953 | TCCA | T | 30 | 6 | 2 | 0 | CODON_CHANGE_ | MODERATE | | ggtgga/gg | GG28G | PLES_26541 | | | | | . | . | . | . | . | . | T | T |
| NC_011770 | 2853975 | C | G | 34 | 6 | 2 | 0 | NON_SYNONYMOU | MODERAT | MISSENSE | gGt/gCt | G22A | PLES_26541 | | | | | . | . | . | . | . | . | G | G |
| NC_011770 | 2853992 | G | C | 33 | 6 | 2 | 0 | NON_SYNONYMOU | MODERAT | MISSENSE | atC/atG | I16M | PLES_26541 | | | | | . | . | . | . | . | . | C | C |
| NC_011770 | 2853994 | T | C | 32 | 6 | 2 | 0 | NON_SYNONYMOU | MODERAT | MISSENSE | Atc/Gtc | I16V | PLES_26541 | | | | | . | . | . | . | . | . | C | C |
| NC_011770 | 2854981 | T | C | 41 | 6 | 2 | 0 | NON_SYNONYMOU | MODERAT | MISSENSE | Agt/Ggt | S217G | PLES_26551 | | | | | . | . | . | . | . | . | C | C |
| NC_011770 | 2854984 | A | G | 42 | 6 | 4 | 0 | NON_SYNONYMOU | MODERAT | MISSENSE | Tcg/Ccg | S216P | PLES_26551 | G | G | G | G | . | . | . | . | . | . | | |
| NC_011770 | 2854996 | C | T | 41 | 6 | 6 | 0 | NON_SYNONYMOU | MODERAT | MISSENSE | Gcg/Acg | A212T | PLES_26551 | T | T | T | T | . | . | . | . | . | . | T | T |
| NC_011770 | 2855008 | C | T | 41 | 6 | 2 | 0 | NON_SYNONYMOU | MODERAT | MISSENSE | Gca/Aca | A208T | PLES_26551 | | | | | . | . | . | . | . | . | T | T |
| NC_011770 | 2855030 | T | G | 41 | 6 | 6 | 0 | NON_SYNONYMOU | MODERAT | MISSENSE | gaA/gaC | E200D | PLES_26551 | G | G | G | G | . | . | . | . | . | . | C | C |
| NC_011770 | 2855096 | G | C | 36 | 6 | 2 | 0 | NON_SYNONYMOU | MODERAT | MISSENSE | atC/atG | I178M | PLES_26551 | | | | | . | . | . | . | . | . | C | C |
| NC_011770 | 2855098 | T | C | 37 | 6 | 2 | 0 | NON_SYNONYMOU | MODERAT | MISSENSE | Atc/Gtc | I178V | PLES_26551 | | | | | . | . | . | . | . | . | C | C |
| NC_011770 | 2855193 | G | A | 40 | 6 | 2 | 0 | NON_SYNONYMOU | MODERAT | MISSENSE | aCc/aTc | T146I | PLES_26551 | | | | | . | . | . | . | . | . | A | A |
| NC_011770 | 2855194 | T | C | 40 | 6 | 2 | 0 | NON_SYNONYMOU | MODERAT | MISSENSE | Acc/Gcc | T146A | PLES_26551 | | | | | . | . | . | . | . | . | C | C |
| NC_011770 | 2855280 | G | C | 40 | 6 | 2 | 0 | NON_SYNONYMOU | MODERAT | MISSENSE | tCc/tGc | S117C | PLES_26551 | | | | | . | . | . | . | . | . | C | C |
| NC_011770 | 2855281 | A | T | 38 | 6 | 2 | 0 | NON_SYNONYMOU | MODERAT | MISSENSE | Tcc/Acc | S117T | PLES_26551 | | | | | . | . | . | . | . | . | T | T |
| NC_011770 | 2855362 | A | C | 39 | 6 | 2 | 0 | NON_SYNONYMOU | MODERAT | MISSENSE | Tcg/Gcg | S90A | PLES_26551 | | | | | . | . | . | . | . | . | C | C |
| NC_011770 | 2855461 | G | T | 41 | 6 | 2 | 0 | NON_SYNONYMOU | MODERAT | MISSENSE | Cgc/Agc | R57S | PLES_26551 | | | | | . | . | . | . | . | . | T | T |
| NC_011770 | 2855551 | T | C | 40 | 6 | 4 | 0 | NON_SYNONYMOU | MODERAT | MISSENSE | Att/Gtt | I27V | PLES_26551 | C | C | C | C | . | . | . | . | . | . | | |
| NC_011770 | 2855576 | C | A | 38 | 6 | 2 | 0 | NON_SYNONYMOU | MODERAT | MISSENSE | ttG/ttT | L18F | PLES_26551 | | | | | . | . | . | . | . | . | A | A |
| NC_011770 | 2855633 | GCCCCCCT | G | 0 | 10 | 1 | 0 | CODON_CHANGE_ | MODERATE | | gttgcaagg | VARGA105 | PLES_26561 | . | . | . | . | . | . | . | . | . | . | G | . |
| NC_011770 | 2855661 | A | T | 34 | 6 | 2 | 0 | NON_SYNONYMOU | MODERAT | MISSENSE | Tcc/Acc | S100T | PLES_26561 | | | | | . | . | . | . | . | . | T | T |
| NC_011770 | 2855851 | G | C | 32 | 6 | 6 | 0 | NON_SYNONYMOU | MODERAT | MISSENSE | atC/atG | I36M | PLES_26561 | C | C | C | C | . | . | . | . | . | . | T | T |
| NC_011770 | 2855853 | T | G | 33 | 6 | 6 | 0 | NON_SYNONYMOU | MODERAT | MISSENSE | Atc/Ctc | I36L | PLES_26561 | G | G | G | G | . | . | . | . | . | . | G | G |
| NC_011770 | 2856256 | C | G | 36 | 6 | 6 | 0 | NON_SYNONYMOU | MODERAT | MISSENSE | agG/agC | R51S | PLES_26571 | G | G | G | G | . | . | . | . | . | . | G | G |
| NC_011770 | 2856335 | G | A | 36 | 6 | 2 | 0 | NON_SYNONYMOU | MODERAT | MISSENSE | gCg/gTg | A25V | PLES_26571 | | | | | . | . | . | . | . | . | A | A |
| NC_011770 | 2856363 | T | C | 37 | 6 | 6 | 0 | NON_SYNONYMOU | MODERAT | MISSENSE | Atg/Gtg | M16V | PLES_26571 | C | C | C | C | . | . | . | . | . | . | C | C |
| NC_011770 | 2856381 | G | A | 38 | 6 | 4 | 0 | NON_SYNONYMOU | MODERAT | MISSENSE | Ctc/Ttc | L10F | PLES_26571 | A | A | A | A | . | . | . | . | . | . | | |
| NC_011770 | 2856389 | G | T | 39 | 6 | 2 | 0 | NON_SYNONYMOU | MODERAT | MISSENSE | aCt/aAt | T7N | PLES_26571 | | | | | . | . | . | . | . | . | T | T |
| NC_011770 | 2856395 | A | T | 38 | 6 | 4 | 0 | NON_SYNONYMOU | MODERAT | MISSENSE | aTc/aAc | I5N | PLES_26571 | T | T | T | T | . | . | . | . | . | . | | |
| NC_011770 | 2856403 | G | T | 40 | 6 | 2 | 0 | NON_SYNONYMOU | MODERAT | MISSENSE | caC/caA | H2Q | PLES_26571 | | | | | . | . | . | . | . | . | T | T |
| NC_011770 | 2856444 | C | G | 39 | 6 | 2 | 0 | NON_SYNONYMOU | MODERAT | MISSENSE | aGt/aCt | S72T | PLES_26581 | | | | | . | . | . | . | . | . | G | G |
| NC_011770 | 2856539 | G | C | 34 | 6 | 2 | 0 | NON_SYNONYMOU | MODERAT | MISSENSE | atC/atG | I40M | PLES_26581 | | | | | . | . | . | . | . | . | C | C |
| NC_011770 | 2856574 | A | G | 34 | 6 | 4 | 0 | NON_SYNONYMOU | MODERAT | MISSENSE | Ttc/Ctc | F29L | PLES_26581 | G | G | G | G | . | . | . | . | . | . | | |
| NC_011770 | 2856628 | C | T | 34 | 6 | 2 | 0 | NON_SYNONYMOU | MODERAT | MISSENSE | Gcc/Acc | A11T | PLES_26581 | | | T | T | . | . | . | . | . | . | | |
| NC_011770 | 2856957 | G | A | 31 | 6 | 2 | 0 | STOP_GAINED | HIGH | NONSENSE | Cag/Tag | Q28* | PLES_26591 | | | | | . | . | . | . | . | . | A | A |
| NC_011770 | 2856659 | T | G | 33 | 6 | 2 | 0 | NON_SYNONYMOU | MODERAT | MISSENSE | cAa/cCa | Q127P | PLES_26591 | | | | | . | . | . | . | . | . | G | G |
| NC_011770 | 2856668 | C | T | 31 | 6 | 2 | 0 | NON_SYNONYMOU | MODERAT | MISSENSE | gGg/gAg | G124E | PLES_26591 | | | | | . | . | . | . | . | . | T | T |
| NC_011770 | 2856681 | C | G | 28 | 6 | 2 | 0 | NON_SYNONYMOU | MODERAT | MISSENSE | Gcg/Ccg | A120P | PLES_26591 | | | | | . | . | . | . | . | . | G | G |
| NC_011770 | 2856686 | A | G | 27 | 6 | 2 | 0 | NON_SYNONYMOU | MODERAT | MISSENSE | cTg/cCg | L118P | PLES_26591 | | | | | . | . | . | . | . | . | G | G |
| NC_011770 | 2856692 | T | G | 26 | 6 | 2 | 0 | NON_SYNONYMOU | MODERAT | MISSENSE | gAg/gCg | E116A | PLES_26591 | | | | | . | . | . | . | . | . | G | G |
| NC_011770 | 2856693 | C | G | 28 | 6 | 2 | 0 | NON_SYNONYMOU | MODERAT | MISSENSE | Gag/Cag | E116Q | PLES_26591 | | | | | . | . | . | . | . | . | G | G |
| NC_011770 | 2856695 | G | A | 29 | 6 | 2 | 0 | NON_SYNONYMOU | MODERAT | MISSENSE | gCc/gTc | A115V | PLES_26591 | | | | | . | . | . | . | . | . | A | A |
| NC_011770 | 2856698 | G | A | 28 | 6 | 2 | 0 | NON_SYNONYMOU | MODERAT | MISSENSE | gCg/gTg | A114V | PLES_26591 | | | | | . | . | . | . | . | . | A | A |
| NC_011770 | 2856770 | A | G | 30 | 6 | 2 | 0 | NON_SYNONYMOU | MODERAT | MISSENSE | tTt/tCt | F90S | PLES_26591 | | | | | . | . | . | . | . | . | G | G |
| NC_011770 | 2856771 | A | C | 30 | 6 | 2 | 0 | NON_SYNONYMOU | MODERAT | MISSENSE | Ttt/Gtt | F90V | PLES_26591 | | | | | . | . | . | . | . | . | C | C |
| NC_011770 | 2856956 | T | G | 31 | 6 | 2 | 0 | NON_SYNONYMOU | MODERAT | MISSENSE | cAg/cCg | Q28P | PLES_26591 | | | | | . | . | . | . | . | . | G | G |
| NC_011770 | 2856972 | G | C | 30 | 6 | 2 | 0 | NON_SYNONYMOU | MODERAT | MISSENSE | Ctg/Gtg | L23V | PLES_26591 | | | | | . | . | . | . | . | . | C | C |
| NC_011770 | 2857029 | A | G | 32 | 6 | 2 | 0 | NON_SYNONYMOU | MODERAT | MISSENSE | Tcg/Ccg | S4P | PLES_26591 | | | | | . | . | . | . | . | . | G | G |
| NC_011770 | 2857035 | C | A | 32 | 6 | 2 | 0 | NON_SYNONYMOU | MODERAT | MISSENSE | Gtg/Ttg | V2L | PLES_26591 | | | | | . | . | . | . | . | . | A | A |
| NC_011770 | 2859041 | C | T | 27 | 8 | 4 | 0 | NON_SYNONYMOU | MODERAT | MISSENSE | gCg/gTg | A247V | PLES_26621 | T | T | T | T | . | . | . | . | . | . | . | . |
| NC_011770 | 2859198 | A | C | 27 | 8 | 2 | 0 | NON_SYNONYMOU | MODERAT | MISSENSE | caA/caC | Q299H | PLES_26621 | C | C | | | . | . | . | . | . | . | . | . |
| NC_011770 | 2860691 | G | C | 30 | 8 | 4 | 0 | NON_SYNONYMOU | MODERAT | MISSENSE | gGc/gCc | G171A | PLES_26641 | C | C | C | C | . | . | . | . | . | . | . | . |

| | | | | | | | | | | | | | | | | | | | | | | | | | |
|---|---|---|---|---|---|---|---|---|---|---|---|---|---|---|---|---|---|---|---|---|---|---|---|---|---|
| NC_011770 | 2862598 | A | G | 21 | 8 | 4 | 0 | NON_SYNONYMOU | MODERAT | MISSENSE | aAc/aGc | N234S | PLES_26661 | G | G | G | G | . | . | . | . | . | . | . | . |
| NC_011770 | 2864377 | A | C | 27 | 8 | 2 | 0 | NON_SYNONYMOU | MODERAT | MISSENSE | cAc/cCc | H827P | PLES_26661 | C | C | | | . | . | . | . | . | . | . | . |
| NC_011770 | 2865274 | G | A | 28 | 6 | 4 | 0 | NON_SYNONYMOU | MODERAT | MISSENSE | gCa/gTa | A728V | PLES_26671 | A | A | A | A | . | . | . | . | . | . | | |
| NC_011770 | 2865281 | C | CGTT | 26 | 6 | 4 | 0 | CODON_INSERTIO | MODERATE | | ggc/ggcAA | G725GN | PLES_26671 | CGTT | CGTT | CGTT | CGTT | . | . | . | . | . | . | | |
| NC_011770 | 2865283 | C | T | 10 | 8 | 2 | 0 | NON_SYNONYMOU | MODERAT | MISSENSE | gGc/gAc | G725D | PLES_26671 | . | . | | | . | . | . | . | . | . | T | T |
| NC_011770 | 2865286 | G | C | 11 | 8 | 2 | 0 | NON_SYNONYMOU | MODERAT | MISSENSE | aCc/aGc | T724S | PLES_26671 | . | . | | | . | . | . | . | . | . | C | C |
| NC_011770 | 2865287 | T | A | 11 | 7 | 5 | 0 | NON_SYNONYMOU | MODERAT | MISSENSE | Acc/Tcc | T724S | PLES_26671 | . | A | A | A | . | . | . | . | . | . | C | C |
| NC_011770 | 2865290 | T | C | 18 | 6 | 4 | 0 | NON_SYNONYMOU | MODERAT | MISSENSE | Agc/Ggc | S723G | PLES_26671 | C | C | C | C | . | . | . | . | . | . | | |
| NC_011770 | 2865293 | C | G | 30 | 6 | 2 | 0 | NON_SYNONYMOU | MODERAT | MISSENSE | Gac/Cac | D722H | PLES_26671 | | | | | . | . | . | . | . | . | G | G |
| NC_011770 | 2865299 | C | T | 29 | 6 | 2 | 0 | NON_SYNONYMOU | MODERAT | MISSENSE | Gcc/Acc | A720T | PLES_26671 | | | | | . | . | . | . | . | . | T | T |
| NC_011770 | 2865301 | C | T | 30 | 6 | 2 | 0 | NON_SYNONYMOU | MODERAT | MISSENSE | gGc/gAc | G719D | PLES_26671 | | | | | . | . | . | . | . | . | T | T |
| NC_011770 | 2865316 | T | C | 33 | 6 | 6 | 0 | NON_SYNONYMOU | MODERAT | MISSENSE | gAc/gGc | D714G | PLES_26671 | C | C | C | C | . | . | . | . | . | . | G | G |
| NC_011770 | 2865329 | G | C | 34 | 6 | 4 | 0 | NON_SYNONYMOU | MODERAT | MISSENSE | Cca/Gca | P710A | PLES_26671 | C | C | C | C | . | . | . | . | . | . | | |
| NC_011770 | 2865368 | G | C | 33 | 6 | 6 | 0 | NON_SYNONYMOU | MODERAT | MISSENSE | Cat/Gat | H697D | PLES_26671 | C | C | C | C | . | . | . | . | . | . | C | C |
| NC_011770 | 2865370 | G | T | 33 | 6 | 6 | 0 | NON_SYNONYMOU | MODERAT | MISSENSE | cCg/cAg | P696Q | PLES_26671 | T | T | T | T | . | . | . | . | . | . | T | T |
| NC_011770 | 2865380 | C | G | 31 | 6 | 6 | 0 | NON_SYNONYMOU | MODERAT | MISSENSE | Gca/Cca | A693P | PLES_26671 | G | G | G | G | . | . | . | . | . | . | G | G |
| NC_011770 | 2865458 | T | C | 37 | 6 | 4 | 0 | NON_SYNONYMOU | MODERAT | MISSENSE | Acg/Gcg | T667A | PLES_26671 | C | C | C | C | . | . | . | . | . | . | | |
| NC_011770 | 2865484 | G | A | 36 | 6 | 2 | 0 | NON_SYNONYMOU | MODERAT | MISSENSE | gCg/gTg | A658V | PLES_26671 | | | | | . | . | . | . | . | . | A | A |
| NC_011770 | 2865485 | C | G | 36 | 6 | 2 | 0 | NON_SYNONYMOU | MODERAT | MISSENSE | Gcg/Ccg | A658P | PLES_26671 | | | | | . | . | . | . | . | . | G | G |
| NC_011770 | 2865593 | C | A | 38 | 6 | 6 | 0 | NON_SYNONYMOU | MODERAT | MISSENSE | Gcc/Tcc | A622S | PLES_26671 | A | A | A | A | . | . | . | . | . | . | A | A |
| NC_011770 | 2865885 | A | C | 40 | 6 | 2 | 0 | NON_SYNONYMOU | MODERAT | MISSENSE | atT/atG | I524M | PLES_26671 | | | | | . | . | . | . | . | . | C | C |
| NC_011770 | 2865887 | T | C | 40 | 6 | 2 | 0 | NON_SYNONYMOU | MODERAT | MISSENSE | Att/Gtt | I524V | PLES_26671 | | | | | . | . | . | . | . | . | C | C |
| NC_011770 | 2865903 | C | G | 41 | 6 | 2 | 0 | NON_SYNONYMOU | MODERAT | MISSENSE | ttG/ttC | L518F | PLES_26671 | | | | | . | . | . | . | . | . | G | G |
| NC_011770 | 2865961 | C | T | 38 | 6 | 2 | 0 | NON_SYNONYMOU | MODERAT | MISSENSE | gGc/gAc | G499D | PLES_26671 | | | | | . | . | . | . | . | . | T | T |
| NC_011770 | 2866058 | C | T | 52 | 4 | 2 | 0 | NON_SYNONYMOU | MODERAT | MISSENSE | Gcg/Acg | A467T | PLES_26671 | | | | | T | T | . | . | . | . | | |
| NC_011770 | 2866074 | C | G | 52 | 4 | 2 | 0 | NON_SYNONYMOU | MODERAT | MISSENSE | ttG/ttC | L461F | PLES_26671 | | | | | G | G | . | . | . | . | | |
| NC_011770 | 2866101 | C | G | 65 | 4 | 2 | 0 | NON_SYNONYMOU | MODERAT | MISSENSE | agG/agC | R452S | PLES_26671 | . | . | . | . | G | G | . | . | . | . | | |
| NC_011770 | 2866141 | G | T | 41 | 4 | 2 | 0 | NON_SYNONYMOU | MODERAT | MISSENSE | gCc/gAc | A439D | PLES_26671 | | | | | T | T | . | . | . | . | | |
| NC_011770 | 2866142 | C | T | 40 | 4 | 2 | 0 | NON_SYNONYMOU | MODERAT | MISSENSE | Gcc/Acc | A439T | PLES_26671 | | | | | T | T | . | . | . | . | | |
| NC_011770 | 2866151 | A | C | 41 | 4 | 2 | 0 | NON_SYNONYMOU | MODERAT | MISSENSE | Tcc/Gcc | S436A | PLES_26671 | | | | | C | C | . | . | . | . | | |
| NC_011770 | 2866163 | C | A | 40 | 4 | 2 | 0 | NON_SYNONYMOU | MODERAT | MISSENSE | Gcg/Tcg | A432S | PLES_26671 | | | | | A | A | . | . | . | . | | |
| NC_011770 | 2866170 | C | A | 41 | 4 | 4 | 0 | NON_SYNONYMOU | MODERAT | MISSENSE | caG/caT | Q429H | PLES_26671 | A | A | A | A | | | . | . | . | . | | |
| NC_011770 | 2866172 | G | C | 41 | 4 | 2 | 0 | NON_SYNONYMOU | MODERAT | MISSENSE | Cag/Gag | Q429E | PLES_26671 | | | | | C | C | . | . | . | . | | |
| NC_011770 | 2866175 | C | A | 41 | 4 | 2 | 0 | NON_SYNONYMOU | MODERAT | MISSENSE | Gcc/Tcc | A428S | PLES_26671 | | | | | A | A | . | . | . | . | | |
| NC_011770 | 2866180 | T | C | 41 | 4 | 2 | 0 | NON_SYNONYMOU | MODERAT | MISSENSE | aAg/aGg | K426R | PLES_26671 | | | | | C | C | . | . | . | . | | |
| NC_011770 | 2866181 | T | G | 40 | 4 | 2 | 0 | NON_SYNONYMOU | MODERAT | MISSENSE | Aag/Cag | K426Q | PLES_26671 | | | | | G | G | . | . | . | . | | |
| NC_011770 | 2866186 | C | G | 41 | 4 | 2 | 0 | NON_SYNONYMOU | MODERAT | MISSENSE | aGc/aCc | S424T | PLES_26671 | | | | | G | G | . | . | . | . | | |
| NC_011770 | 2866353 | C | G | 37 | 6 | 2 | 0 | NON_SYNONYMOU | MODERAT | MISSENSE | agG/agC | R368S | PLES_26671 | | | | | . | . | . | . | . | . | G | G |
| NC_011770 | 2866368 | C | G | 50 | 4 | 2 | 0 | NON_SYNONYMOU | MODERAT | MISSENSE | gaG/gaC | E363D | PLES_26671 | | | | | G | G | . | . | . | . | | |
| NC_011770 | 2866381 | G | C | 49 | 4 | 2 | 0 | NON_SYNONYMOU | MODERAT | MISSENSE | gCc/gGc | A359G | PLES_26671 | | | | | C | C | . | . | . | . | | |
| NC_011770 | 2866607 | T | C | 82 | 4 | 2 | 0 | NON_SYNONYMOU | MODERAT | MISSENSE | Atc/Gtc | I284V | PLES_26671 | | | | | C | C | . | . | . | . | | |
| NC_011770 | 2866721 | G | T | 38 | 6 | 2 | 0 | NON_SYNONYMOU | MODERAT | MISSENSE | Cag/Aag | Q246K | PLES_26671 | | | | | . | . | . | . | . | . | T | T |
| NC_011770 | 2866804 | T | C | 39 | 6 | 2 | 0 | NON_SYNONYMOU | MODERAT | MISSENSE | aAt/aGt | N218S | PLES_26671 | | | | | . | . | . | . | . | . | C | C |
| NC_011770 | 2866807 | C | T | 39 | 6 | 2 | 0 | NON_SYNONYMOU | MODERAT | MISSENSE | aGg/aAg | R217K | PLES_26671 | | | | | . | . | . | . | . | . | T | T |
| NC_011770 | 2866883 | A | T | 50 | 4 | 2 | 0 | NON_SYNONYMOU | MODERAT | MISSENSE | Tcg/Acg | S192T | PLES_26671 | | | | | T | T | . | . | . | . | | |
| NC_011770 | 2866947 | C | G | 39 | 6 | 2 | 0 | NON_SYNONYMOU | MODERAT | MISSENSE | agG/agC | R170S | PLES_26671 | | | | | . | . | . | . | . | . | G | G |
| NC_011770 | 2866996 | G | A | 34 | 6 | 4 | 0 | NON_SYNONYMOU | MODERAT | MISSENSE | gCg/gTg | A154V | PLES_26671 | A | A | A | A | . | . | . | . | . | . | | |
| NC_011770 | 2867014 | T | G | 29 | 6 | 2 | 0 | NON_SYNONYMOU | MODERAT | MISSENSE | aAg/aCg | K148T | PLES_26671 | | | | | . | . | . | . | . | . | G | G |
| NC_011770 | 2867015 | T | C | 29 | 6 | 2 | 0 | NON_SYNONYMOU | MODERAT | MISSENSE | Aag/Gag | K148E | PLES_26671 | | | | | . | . | . | . | . | . | C | C |
| NC_011770 | 2867018 | G | A | 29 | 6 | 2 | 0 | NON_SYNONYMOU | MODERAT | MISSENSE | Ccg/Tcg | P147S | PLES_26671 | | | | | . | . | . | . | . | . | A | A |
| NC_011770 | 2867034 | C | G | 29 | 6 | 2 | 0 | NON_SYNONYMOU | MODERAT | MISSENSE | ttG/ttC | L141F | PLES_26671 | | | | | . | . | . | . | . | . | G | G |
| NC_011770 | 2867072 | T | C | 34 | 6 | 6 | 0 | NON_SYNONYMOU | MODERAT | MISSENSE | Acc/Gcc | T129A | PLES_26671 | C | C | C | C | . | . | . | . | . | . | C | C |
| NC_011770 | 2867177 | C | T | 35 | 6 | 2 | 0 | NON_SYNONYMOU | MODERAT | MISSENSE | Gtg/Atg | V94M | PLES_26671 | | | | | . | . | . | . | . | . | T | T |
| NC_011770 | 2867231 | G | A | 37 | 6 | 2 | 0 | NON_SYNONYMOU | MODERAT | MISSENSE | Ccg/Tcg | P76S | PLES_26671 | | | | | . | . | . | . | . | . | A | A |
| NC_011770 | 2867265 | G | C | 34 | 6 | 2 | 0 | NON_SYNONYMOU | MODERAT | MISSENSE | atC/atG | I64M | PLES_26671 | | | | | . | . | . | . | . | . | C | C |
| NC_011770 | 2867267 | T | C | 34 | 6 | 2 | 0 | NON_SYNONYMOU | MODERAT | MISSENSE | Atc/Gtc | I64V | PLES_26671 | | | | | . | . | . | . | . | . | C | C |
| NC_011770 | 2867324 | T | C | 32 | 6 | 2 | 0 | NON_SYNONYMOU | MODERAT | MISSENSE | Atc/Gtc | I45V | PLES_26671 | | | | | . | . | . | . | . | . | C | C |
| NC_011770 | 2867333 | C | A | 33 | 6 | 2 | 0 | NON_SYNONYMOU | MODERAT | MISSENSE | Gcc/Tcc | A42S | PLES_26671 | | | | | . | . | . | . | . | . | A | A |
| NC_011770 | 2867964 | T | C | 9 | 6 | 2 | 0 | START_LOST | HIGH | MISSENSE | Atg/Gtg | M1V | PLES_26681 | | | | | . | . | . | . | . | . | C | C |
| NC_011770 | 2867589 | C | T | 25 | 6 | 4 | 0 | NON_SYNONYMOU | MODERAT | MISSENSE | Ggc/Agc | G126S | PLES_26681 | T | T | T | T | . | . | . | . | . | . | | |
| NC_011770 | 2867645 | G | A | 20 | 6 | 2 | 0 | NON_SYNONYMOU | MODERAT | MISSENSE | gCc/gTc | A107V | PLES_26681 | | | | | . | . | . | . | . | . | A | A |
| NC_011770 | 2867648 | G | C | 20 | 6 | 2 | 0 | NON_SYNONYMOU | MODERAT | MISSENSE | gCc/gGc | A106G | PLES_26681 | | | | | . | . | . | . | . | . | C | C |

| | | | | | | | | | | | | | | | | | | | | | | | | | |
|---|---|---|---|---|---|---|---|---|---|---|---|---|---|---|---|---|---|---|---|---|---|---|---|---|---|
| NC_011770 | 2867666 | G | A | 19 | 6 | 4 | 0 | NON_SYNONYMOU | MODERAT | MISSENSE | gCc/gTc | A100V | PLES_26681 | A | A | A | A | . | . | . | . | . | . | | |
| NC_011770 | 2867677 | C | G | 18 | 6 | 2 | 0 | NON_SYNONYMOU | MODERAT | MISSENSE | gaG/gaC | E96D | PLES_26681 | | | | | . | . | . | . | . | . | G | G |
| NC_011770 | 2867696 | A | G | 17 | 6 | 4 | 0 | NON_SYNONYMOU | MODERAT | MISSENSE | gTg/gCg | V90A | PLES_26681 | G | G | G | G | . | . | . | . | . | . | | |
| NC_011770 | 2867759 | G | C | 15 | 6 | 4 | 0 | NON_SYNONYMOU | MODERAT | MISSENSE | gCc/gGc | A69G | PLES_26681 | C | C | C | C | . | . | . | . | . | . | | |
| NC_011770 | 2867760 | C | T | 14 | 6 | 4 | 0 | NON_SYNONYMOU | MODERAT | MISSENSE | Gcc/Acc | A69T | PLES_26681 | T | T | T | T | . | . | . | . | . | . | | |
| NC_011770 | 2867795 | C | G | 12 | 7 | 5 | 0 | NON_SYNONYMOU | MODERAT | MISSENSE | cGg/cCg | R57P | PLES_26681 | G | . | G | G | . | . | . | . | . | . | G | G |
| NC_011770 | 2867810 | T | C | 10 | 7 | 3 | 0 | NON_SYNONYMOU | MODERAT | MISSENSE | cAg/cGg | Q52R | PLES_26681 | C | . | C | C | . | . | . | . | . | . | | |
| NC_011770 | 2867826 | A | G | 8 | 9 | 3 | 0 | NON_SYNONYMOU | MODERAT | MISSENSE | Tcc/Ccc | S47P | PLES_26681 | . | . | G | . | . | . | . | . | . | . | G | G |
| NC_011770 | 2867834 | G | C | 6 | 9 | 2 | 0 | NON_SYNONYMOU | MODERAT | MISSENSE | gCg/gGg | A44G | PLES_26681 | . | . | | . | . | . | . | . | . | . | C | C |
| NC_011770 | 2867840 | G | A | 6 | 8 | 2 | 0 | NON_SYNONYMOU | MODERAT | MISSENSE | gCa/gTa | A42V | PLES_26681 | | . | | . | . | . | . | . | . | . | A | A |
| NC_011770 | 2867844 | T | C | 7 | 7 | 2 | 0 | NON_SYNONYMOU | MODERAT | MISSENSE | Aac/Gac | N41D | PLES_26681 | | | | . | . | . | . | . | . | . | C | C |
| NC_011770 | 2867853 | G | C | 11 | 7 | 2 | 0 | NON_SYNONYMOU | MODERAT | MISSENSE | Ccg/Gcg | P38A | PLES_26681 | | | | . | . | . | . | . | . | . | C | C |
| NC_011770 | 2867923 | C | A | 16 | 6 | 2 | 0 | NON_SYNONYMOU | MODERAT | MISSENSE | gaG/gaT | E14D | PLES_26681 | | | | | . | . | . | . | . | . | A | A |
| NC_011770 | 2867955 | A | C | 16 | 6 | 2 | 0 | NON_SYNONYMOU | MODERAT | MISSENSE | Tca/Gca | S4A | PLES_26681 | | | | | . | . | . | . | . | . | C | C |
| NC_011770 | 2868574 | ATG | A | 10 | 8 | 4 | 0 | FRAME_SHIFT+STA | HIGH | | atg/ | M1 | PLES_26691 | A | A | A | A | . | . | . | . | . | . | . | . |
| NC_011770 | 2868046 | G | T | 19 | 6 | 2 | 0 | NON_SYNONYMOU | MODERAT | MISSENSE | gCc/gAc | A177D | PLES_26691 | | | | | . | . | . | . | . | . | T | T |
| NC_011770 | 2868047 | C | T | 19 | 6 | 6 | 0 | NON_SYNONYMOU | MODERAT | MISSENSE | Gcc/Acc | A177T | PLES_26691 | T | T | T | T | . | . | . | . | . | . | T | T |
| NC_011770 | 2868052 | C | T | 20 | 6 | 2 | 0 | NON_SYNONYMOU | MODERAT | MISSENSE | cGa/cAa | R175Q | PLES_26691 | | | | | . | . | . | . | . | . | T | T |
| NC_011770 | 2868056 | T | C | 20 | 6 | 2 | 0 | NON_SYNONYMOU | MODERAT | MISSENSE | Acg/Gcg | T174A | PLES_26691 | | | | | . | . | . | . | . | . | C | C |
| NC_011770 | 2868112 | T | G | 28 | 5 | 1 | 0 | NON_SYNONYMOU | MODERAT | MISSENSE | gAg/gCg | E155A | PLES_26691 | | | | | . | G | . | . | . | . | | |
| NC_011770 | 2868167 | G | A | 27 | 6 | 2 | 0 | NON_SYNONYMOU | MODERAT | MISSENSE | Ccc/Tcc | P137S | PLES_26691 | | | | | . | . | . | . | . | . | A | A |
| NC_011770 | 2868180 | C | G | 28 | 6 | 2 | 0 | NON_SYNONYMOU | MODERAT | MISSENSE | aaG/aaC | K132N | PLES_26691 | | | | | . | . | . | . | . | . | G | G |
| NC_011770 | 2868200 | C | T | 27 | 6 | 2 | 0 | NON_SYNONYMOU | MODERAT | MISSENSE | Gtt/Att | V126I | PLES_26691 | | | | | . | . | . | . | . | . | T | T |
| NC_011770 | 2868239 | C | T | 29 | 6 | 2 | 0 | NON_SYNONYMOU | MODERAT | MISSENSE | Ggc/Agc | G113S | PLES_26691 | | | | | . | . | . | . | . | . | T | T |
| NC_011770 | 2868277 | C | A | 33 | 6 | 2 | 0 | NON_SYNONYMOU | MODERAT | MISSENSE | gGc/gTc | G100V | PLES_26691 | | | | | . | . | . | . | . | . | A | A |
| NC_011770 | 2868311 | C | T | 32 | 6 | 2 | 0 | NON_SYNONYMOU | MODERAT | MISSENSE | Gcc/Acc | A89T | PLES_26691 | | | | | . | . | . | . | . | . | T | T |
| NC_011770 | 2868335 | C | A | 32 | 6 | 2 | 0 | NON_SYNONYMOU | MODERAT | MISSENSE | Gct/Tct | A81S | PLES_26691 | | | | | . | . | . | . | . | . | A | A |
| NC_011770 | 2868407 | G | T | 33 | 6 | 6 | 0 | NON_SYNONYMOU | MODERAT | MISSENSE | Ctg/Atg | L57M | PLES_26691 | T | T | T | T | . | . | . | . | . | . | T | T |
| NC_011770 | 2868425 | T | C | 35 | 6 | 2 | 0 | NON_SYNONYMOU | MODERAT | MISSENSE | Atc/Gtc | I51V | PLES_26691 | | | | | . | . | . | . | . | . | C | C |
| NC_011770 | 2868478 | C | T | 35 | 6 | 2 | 0 | NON_SYNONYMOU | MODERAT | MISSENSE | cGc/cAc | R33H | PLES_26691 | | | | | . | . | . | . | . | . | T | T |
| NC_011770 | 2868512 | C | T | 32 | 6 | 2 | 0 | NON_SYNONYMOU | MODERAT | MISSENSE | Gtt/Att | V22I | PLES_26691 | | | | | . | . | . | . | . | . | T | T |
| NC_011770 | 2868523 | A | T | 12 | 6 | 6 | 0 | NON_SYNONYMOU | MODERAT | MISSENSE | cTg/cAg | L18Q | PLES_26691 | T | T | T | T | . | . | . | . | . | . | T | T |
| NC_011770 | 2868526 | G | A | 12 | 6 | 2 | 0 | NON_SYNONYMOU | MODERAT | MISSENSE | gCc/gTc | A17V | PLES_26691 | | | | | . | . | . | . | . | . | A | A |
| NC_011770 | 2868527 | C | G | 12 | 6 | 4 | 0 | NON_SYNONYMOU | MODERAT | MISSENSE | Gcc/Ccc | A17P | PLES_26691 | G | G | G | G | . | . | . | . | . | . | | |
| NC_011770 | 2868530 | C | T | 11 | 6 | 4 | 0 | NON_SYNONYMOU | MODERAT | MISSENSE | Gcc/Acc | A16T | PLES_26691 | T | T | T | T | . | . | . | . | . | . | | |
| NC_011770 | 2868532 | A | C | 12 | 6 | 6 | 0 | NON_SYNONYMOU | MODERAT | MISSENSE | gTt/gGt | V15G | PLES_26691 | C | C | C | C | . | . | . | . | . | . | C | C |
| NC_011770 | 2868533 | C | T | 12 | 6 | 6 | 0 | NON_SYNONYMOU | MODERAT | MISSENSE | Gtt/Att | V15I | PLES_26691 | T | T | T | T | . | . | . | . | . | . | A | A |
| NC_011770 | 2868538 | T | C | 14 | 6 | 2 | 0 | NON_SYNONYMOU | MODERAT | MISSENSE | tAt/tGt | Y13C | PLES_26691 | | | | | . | . | . | . | . | . | C | C |
| NC_011770 | 2868542 | G | C | 13 | 6 | 4 | 0 | NON_SYNONYMOU | MODERAT | MISSENSE | Ctg/Gtg | L12V | PLES_26691 | C | C | C | C | . | . | . | . | . | . | | |
| NC_011770 | 2868548 | C | T | 14 | 6 | 2 | 0 | NON_SYNONYMOU | MODERAT | MISSENSE | Gcc/Acc | A10T | PLES_26691 | | | | | . | . | . | . | . | . | T | T |
| NC_011770 | 2868551 | T | A | 13 | 6 | 6 | 0 | NON_SYNONYMOU | MODERAT | MISSENSE | Act/Tct | T9S | PLES_26691 | A | A | A | A | . | . | . | . | . | . | C | C |
| NC_011770 | 2868103 | G | A | 29 | 5 | 3 | 0 | NON_SYNONYMOU | MODERAT | MISSENSE | gCc/gTc | A158V | PLES_26691 | | A | A | A | . | | . | . | . | . | | |
| NC_011770 | 2868690 | G | C | 38 | 6 | 2 | 0 | NON_SYNONYMOU | MODERAT | MISSENSE | gCc/gGc | A209G | PLES_26701 | | | | | . | . | . | . | . | . | C | C |
| NC_011770 | 2868691 | C | T | 37 | 6 | 2 | 0 | NON_SYNONYMOU | MODERAT | MISSENSE | Gcc/Acc | A209T | PLES_26701 | | | | | . | . | . | . | . | . | T | T |
| NC_011770 | 2868782 | C | A | 37 | 6 | 4 | 0 | NON_SYNONYMOU | MODERAT | MISSENSE | gaG/gaT | E178D | PLES_26701 | A | A | A | A | . | . | . | . | . | . | | |
| NC_011770 | 2868854 | C | G | 31 | 6 | 4 | 0 | NON_SYNONYMOU | MODERAT | MISSENSE | atG/atC | M154I | PLES_26701 | G | G | G | G | . | . | . | . | . | . | | |
| NC_011770 | 2868855 | A | G | 31 | 6 | 4 | 0 | NON_SYNONYMOU | MODERAT | MISSENSE | aTg/aCg | M154T | PLES_26701 | G | G | G | G | . | . | . | . | . | . | | |
| NC_011770 | 2868863 | G | T | 31 | 6 | 4 | 0 | NON_SYNONYMOU | MODERAT | MISSENSE | agC/agA | S151R | PLES_26701 | T | T | T | T | . | . | . | . | . | . | | |
| NC_011770 | 2868874 | A | C | 12 | 8 | 4 | 0 | NON_SYNONYMOU | MODERAT | MISSENSE | Ttg/Gtg | L148V | PLES_26701 | C | C | C | C | . | . | . | . | . | . | . | . |
| NC_011770 | 2868878 | ACTGCTGG | A | 5 | 8 | 3 | 0 | CODON_CHANGE_ | MODERATE | | ggcccatcca | GPSSS142( | PLES_26701 | A | . | A | A | . | . | . | . | . | . | . | . |
| NC_011770 | 2868895 | C | T | 34 | 6 | 2 | 0 | NON_SYNONYMOU | MODERAT | MISSENSE | Gcc/Acc | A141T | PLES_26701 | | | | | . | . | . | . | . | . | T | T |
| NC_011770 | 2868903 | T | C | 34 | 6 | 2 | 0 | NON_SYNONYMOU | MODERAT | MISSENSE | aAg/aGg | K138R | PLES_26701 | | | | | . | . | . | . | . | . | C | C |
| NC_011770 | 2868904 | T | C | 35 | 6 | 2 | 0 | NON_SYNONYMOU | MODERAT | MISSENSE | Aag/Gag | K138E | PLES_26701 | | | | | . | . | . | . | . | . | C | C |
| NC_011770 | 2869138 | C | T | 28 | 6 | 4 | 0 | NON_SYNONYMOU | MODERAT | MISSENSE | Gca/Aca | A60T | PLES_26701 | T | T | T | T | . | . | . | . | . | . | | |
| NC_011770 | 2869154 | C | G | 24 | 6 | 2 | 0 | NON_SYNONYMOU | MODERAT | MISSENSE | ttG/ttC | L54F | PLES_26701 | | | | | . | . | . | . | . | . | G | G |
| NC_011770 | 2869163 | T | G | 24 | 6 | 2 | 0 | NON_SYNONYMOU | MODERAT | MISSENSE | gaA/gaC | E51D | PLES_26701 | | | | | . | . | . | . | . | . | G | G |
| NC_011770 | 2869165 | C | T | 24 | 6 | 2 | 0 | NON_SYNONYMOU | MODERAT | MISSENSE | Gaa/Aaa | E51K | PLES_26701 | | | | | . | . | . | . | . | . | T | T |
| NC_011770 | 2869167 | G | C | 24 | 6 | 6 | 0 | NON_SYNONYMOU | MODERAT | MISSENSE | aCc/aGc | T50S | PLES_26701 | C | C | C | C | . | . | . | . | . | . | C | C |
| NC_011770 | 2869173 | A | G | 23 | 6 | 4 | 0 | NON_SYNONYMOU | MODERAT | MISSENSE | gTg/gCg | V48A | PLES_26701 | G | G | G | G | . | . | . | . | . | . | | |
| NC_011770 | 2869234 | A | G | 21 | 8 | 4 | 0 | NON_SYNONYMOU | MODERAT | MISSENSE | Tcg/Ccg | S28P | PLES_26701 | G | G | G | G | . | . | . | . | . | . | . | . |
| NC_011770 | 2869237 | T | C | 20 | 8 | 4 | 0 | NON_SYNONYMOU | MODERAT | MISSENSE | Acc/Gcc | T27A | PLES_26701 | C | C | C | C | . | . | . | . | . | . | . | . |
| NC_011770 | 2869260 | G | A | 18 | 8 | 4 | 0 | NON_SYNONYMOU | MODERAT | MISSENSE | gCc/gTc | A19V | PLES_26701 | A | A | A | A | . | . | . | . | . | . | . | . |

| | | | | | | | | | | | | | | | | | | | | | | | | | |
|---|---|---|---|---|---|---|---|---|---|---|---|---|---|---|---|---|---|---|---|---|---|---|---|---|---|
| NC_011770 | 2869261 | C | T | 18 | 8 | 4 | 0 | NON_SYNONYMOU | MODERAT | MISSENSE | Gcc/Acc | A19T | PLES_26701 | T | T | T | T | . | . | . | . | . | . | . | . |
| NC_011770 | 2869272 | G | A | 18 | 8 | 4 | 0 | NON_SYNONYMOU | MODERAT | MISSENSE | tCg/tTg | S15L | PLES_26701 | A | A | A | A | . | . | . | . | . | . | . | . |
| NC_011770 | 2869273 | A | C | 18 | 8 | 4 | 0 | NON_SYNONYMOU | MODERAT | MISSENSE | Tcg/Gcg | S15A | PLES_26701 | C | C | C | C | . | . | . | . | . | . | . | . |
| NC_011770 | 2869282 | C | A | 18 | 8 | 4 | 0 | NON_SYNONYMOU | MODERAT | MISSENSE | Gtg/Ttg | V12L | PLES_26701 | A | A | A | A | . | . | . | . | . | . | . | . |
| NC_011770 | 2869285 | G | C | 18 | 8 | 4 | 0 | NON_SYNONYMOU | MODERAT | MISSENSE | Ctg/Gtg | L11V | PLES_26701 | C | C | C | C | . | . | . | . | . | . | . | . |
| NC_011770 | 2869288 | C | G | 18 | 8 | 4 | 0 | NON_SYNONYMOU | MODERAT | MISSENSE | Gtc/Ctc | V10L | PLES_26701 | G | G | G | G | . | . | . | . | . | . | . | . |
| NC_011770 | 2869293 | A | G | 19 | 8 | 4 | 0 | NON_SYNONYMOU | MODERAT | MISSENSE | tTc/tCc | F8S | PLES_26701 | G | G | G | G | . | . | . | . | . | . | . | . |
| NC_011770 | 2869300 | G | T | 28 | 6 | 2 | 0 | NON_SYNONYMOU | MODERAT | MISSENSE | Ctc/Atc | L6I | PLES_26701 | | | | | . | . | . | . | . | . | T | T |
| NC_011770 | 2869350 | G | A | 28 | 6 | 2 | 0 | NON_SYNONYMOU | MODERAT | MISSENSE | aCc/aTc | T208I | PLES_26711 | | | | | . | . | . | . | . | . | A | A |
| NC_011770 | 2869355 | G | T | 27 | 6 | 2 | 0 | NON_SYNONYMOU | MODERAT | MISSENSE | aaC/aaA | N206K | PLES_26711 | | | | | . | . | . | . | . | . | T | T |
| NC_011770 | 2869357 | T | C | 27 | 6 | 2 | 0 | NON_SYNONYMOU | MODERAT | MISSENSE | Aac/Gac | N206D | PLES_26711 | | | | | . | . | . | . | . | . | C | C |
| NC_011770 | 2869378 | T | C | 26 | 6 | 2 | 0 | NON_SYNONYMOU | MODERAT | MISSENSE | Acc/Gcc | T199A | PLES_26711 | | | | | . | . | . | . | . | . | C | C |
| NC_011770 | 2869380 | T | C | 27 | 6 | 2 | 0 | NON_SYNONYMOU | MODERAT | MISSENSE | aAc/aGc | N198S | PLES_26711 | | | | | . | . | . | . | . | . | C | C |
| NC_011770 | 2869385 | C | A | 26 | 6 | 2 | 0 | NON_SYNONYMOU | MODERAT | MISSENSE | gaG/gaT | E196D | PLES_26711 | | | | | . | . | . | . | . | . | A | A |
| NC_011770 | 2869446 | G | T | 28 | 6 | 2 | 0 | NON_SYNONYMOU | MODERAT | MISSENSE | gCc/gAc | A176D | PLES_26711 | | | | | . | . | . | . | . | . | T | T |
| NC_011770 | 2869452 | G | A | 29 | 6 | 4 | 0 | NON_SYNONYMOU | MODERAT | MISSENSE | aCg/aTg | T174M | PLES_26711 | A | A | A | A | . | . | . | . | . | . | | |
| NC_011770 | 2869464 | A | G | 27 | 6 | 6 | 0 | NON_SYNONYMOU | MODERAT | MISSENSE | aTc/aCc | I170T | PLES_26711 | G | G | G | G | . | . | . | . | . | . | C | C |
| NC_011770 | 2869488 | C | A | 26 | 6 | 2 | 0 | NON_SYNONYMOU | MODERAT | MISSENSE | cGc/cTc | R162L | PLES_26711 | | | | | . | . | . | . | . | . | A | A |
| NC_011770 | 2869519 | C | T | 23 | 6 | 2 | 0 | NON_SYNONYMOU | MODERAT | MISSENSE | Gtc/Atc | V152I | PLES_26711 | | | | | . | . | . | . | . | . | T | T |
| NC_011770 | 2869539 | A | G | 17 | 6 | 6 | 0 | NON_SYNONYMOU | MODERAT | MISSENSE | cTg/cCg | L145P | PLES_26711 | G | G | G | G | . | . | . | . | . | . | G | G |
| NC_011770 | 2869546 | T | C | 17 | 6 | 6 | 0 | NON_SYNONYMOU | MODERAT | MISSENSE | Acc/Gcc | T143A | PLES_26711 | C | C | C | C | . | . | . | . | . | . | G | G |
| NC_011770 | 2869549 | TGGA | T | 17 | 6 | 2 | 0 | CODON_DELETION | MODERATE | | tcc/ | S141 | PLES_26711 | | | | | . | . | . | . | . | . | T | T |
| NC_011770 | 2869561 | A | C | 16 | 6 | 6 | 0 | NON_SYNONYMOU | MODERAT | MISSENSE | Tcc/Gcc | S138A | PLES_26711 | C | C | C | C | . | . | . | . | . | . | G | G |
| NC_011770 | 2869563 | G | T | 16 | 6 | 2 | 0 | NON_SYNONYMOU | MODERAT | MISSENSE | aCg/aAg | T137K | PLES_26711 | | | | | . | . | . | . | . | . | T | T |
| NC_011770 | 2869564 | T | G | 16 | 6 | 4 | 0 | NON_SYNONYMOU | MODERAT | MISSENSE | Acg/Ccg | T137P | PLES_26711 | G | G | G | G | . | . | . | . | . | . | | |
| NC_011770 | 2869567 | G | T | 16 | 6 | 2 | 0 | NON_SYNONYMOU | MODERAT | MISSENSE | Cgc/Agc | R136S | PLES_26711 | | | | | . | . | . | . | . | . | T | T |
| NC_011770 | 2869573 | C | T | 16 | 6 | 2 | 0 | NON_SYNONYMOU | MODERAT | MISSENSE | Gca/Aca | A134T | PLES_26711 | | | | | . | . | . | . | . | . | T | T |
| NC_011770 | 2869578 | C | G | 16 | 6 | 6 | 0 | NON_SYNONYMOU | MODERAT | MISSENSE | cGg/cCg | R132P | PLES_26711 | G | G | G | G | . | . | . | . | . | . | G | G |
| NC_011770 | 2869591 | C | T | 16 | 6 | 2 | 0 | NON_SYNONYMOU | MODERAT | MISSENSE | Gca/Aca | A128T | PLES_26711 | | | | | . | . | . | . | . | . | T | T |
| NC_011770 | 2869594 | T | C | 16 | 6 | 6 | 0 | NON_SYNONYMOU | MODERAT | MISSENSE | Acc/Gcc | T127A | PLES_26711 | C | C | C | C | . | . | . | . | . | . | C | C |
| NC_011770 | 2869609 | C | T | 16 | 6 | 6 | 0 | NON_SYNONYMOU | MODERAT | MISSENSE | Gcc/Acc | A122T | PLES_26711 | T | T | T | T | . | . | . | . | . | . | T | T |
| NC_011770 | 2869618 | C | T | 16 | 6 | 6 | 0 | NON_SYNONYMOU | MODERAT | MISSENSE | Gcg/Acg | A119T | PLES_26711 | T | T | T | T | . | . | . | . | . | . | T | T |
| NC_011770 | 2869633 | C | T | 13 | 6 | 6 | 0 | NON_SYNONYMOU | MODERAT | MISSENSE | Gtc/Atc | V114I | PLES_26711 | T | T | T | T | . | . | . | . | . | . | T | T |
| NC_011770 | 2869675 | C | T | 7 | 7 | 1 | 0 | NON_SYNONYMOU | MODERAT | MISSENSE | Ggc/Agc | G100S | PLES_26711 | | | | | . | . | . | . | . | . | T | . |
| NC_011770 | 2869681 | G | A | 7 | 7 | 1 | 0 | NON_SYNONYMOU | MODERAT | MISSENSE | Ctc/Ttc | L98F | PLES_26711 | | | | | . | . | . | . | . | . | A | . |
| NC_011770 | 2869684 | A | T | 7 | 7 | 1 | 0 | NON_SYNONYMOU | MODERAT | MISSENSE | Tct/Act | S97T | PLES_26711 | | | | | . | . | . | . | . | . | T | . |
| NC_011770 | 2869699 | C | A | 13 | 7 | 1 | 0 | NON_SYNONYMOU | MODERAT | MISSENSE | Gcc/Tcc | A92S | PLES_26711 | | | | | . | . | . | . | . | . | A | . |
| NC_011770 | 2869717 | T | C | 22 | 6 | 4 | 0 | NON_SYNONYMOU | MODERAT | MISSENSE | Att/Gtt | I86V | PLES_26711 | C | C | C | C | . | . | . | . | . | . | | |
| NC_011770 | 2869743 | T | G | 15 | 6 | 6 | 0 | NON_SYNONYMOU | MODERAT | MISSENSE | gAc/gCc | D77A | PLES_26711 | G | G | G | G | . | . | . | . | . | . | G | G |
| NC_011770 | 2869838 | C | A | 22 | 6 | 6 | 0 | NON_SYNONYMOU | MODERAT | MISSENSE | gaG/gaT | E45D | PLES_26711 | A | A | A | A | . | . | . | . | . | . | T | T |
| NC_011770 | 2869843 | G | C | 21 | 6 | 2 | 0 | NON_SYNONYMOU | MODERAT | MISSENSE | Cag/Gag | Q44E | PLES_26711 | | | | | . | . | . | . | . | . | C | C |
| NC_011770 | 2869861 | G | T | 6 | 10 | 2 | 0 | NON_SYNONYMOU | MODERAT | MISSENSE | Ctc/Atc | L38I | PLES_26711 | . | . | . | . | . | . | . | . | . | . | T | T |
| NC_011770 | 2869963 | G | A | 0 | 11 | 1 | 0 | NON_SYNONYMOU | MODERAT | MISSENSE | Ccg/Tcg | P4S | PLES_26711 | . | . | A | . | . | . | . | . | . | . | . | . |
| NC_011770 | 2870429 | A | G | 21 | 6 | 6 | 0 | START_LOST | HIGH | MISSENSE | tTg/tCg | L1S | PLES_26721 | G | G | G | G | . | . | . | . | . | . | G | G |
| NC_011770 | 2869991 | A | G | 12 | 8 | 4 | 0 | NON_SYNONYMOU | MODERAT | MISSENSE | aTg/aCg | M147T | PLES_26721 | G | G | G | G | . | . | . | . | . | . | . | . |
| NC_011770 | 2869992 | T | C | 12 | 8 | 4 | 0 | NON_SYNONYMOU | MODERAT | MISSENSE | Atg/Gtg | M147V | PLES_26721 | C | C | C | C | . | . | . | . | . | . | . | . |
| NC_011770 | 2870019 | C | T | 13 | 8 | 4 | 0 | NON_SYNONYMOU | MODERAT | MISSENSE | Gcg/Acg | A138T | PLES_26721 | T | T | T | T | . | . | . | . | . | . | . | . |
| NC_011770 | 2870030 | G | A | 13 | 8 | 4 | 0 | NON_SYNONYMOU | MODERAT | MISSENSE | cCc/cTc | P134L | PLES_26721 | A | A | A | A | . | . | . | . | . | . | . | . |
| NC_011770 | 2870036 | T | C | 14 | 8 | 4 | 0 | NON_SYNONYMOU | MODERAT | MISSENSE | gAg/gGg | E132G | PLES_26721 | C | C | C | C | . | . | . | . | . | . | . | . |
| NC_011770 | 2870074 | C | A | 27 | 6 | 4 | 0 | NON_SYNONYMOU | MODERAT | MISSENSE | gaG/gaT | E119D | PLES_26721 | A | A | A | A | . | . | . | . | . | . | | |
| NC_011770 | 2870121 | C | A | 26 | 6 | 4 | 0 | NON_SYNONYMOU | MODERAT | MISSENSE | Gcc/Tcc | A104S | PLES_26721 | A | A | A | A | . | . | . | . | . | . | | |
| NC_011770 | 2870139 | G | T | 26 | 6 | 2 | 0 | NON_SYNONYMOU | MODERAT | MISSENSE | Ctc/Atc | L98I | PLES_26721 | | | | | . | . | . | . | . | . | T | T |
| NC_011770 | 2870148 | C | G | 26 | 6 | 4 | 0 | NON_SYNONYMOU | MODERAT | MISSENSE | Gat/Cat | D95H | PLES_26721 | G | G | G | G | . | . | . | . | . | . | | |
| NC_011770 | 2870180 | A | G | 23 | 6 | 6 | 0 | NON_SYNONYMOU | MODERAT | MISSENSE | gTa/gCa | V84A | PLES_26721 | G | G | G | G | . | . | . | . | . | . | G | G |
| NC_011770 | 2870229 | G | C | 23 | 6 | 4 | 0 | NON_SYNONYMOU | MODERAT | MISSENSE | Cag/Gag | Q68E | PLES_26721 | C | C | C | C | . | . | . | . | . | . | | |
| NC_011770 | 2870238 | C | T | 23 | 6 | 2 | 0 | NON_SYNONYMOU | MODERAT | MISSENSE | Ggc/Agc | G65S | PLES_26721 | | | | | . | . | . | . | . | . | T | T |
| NC_011770 | 2870268 | G | T | 24 | 6 | 4 | 0 | NON_SYNONYMOU | MODERAT | MISSENSE | Ctt/Att | L55I | PLES_26721 | T | T | T | T | . | . | . | . | . | . | | |
| NC_011770 | 2870285 | G | A | 24 | 6 | 2 | 0 | NON_SYNONYMOU | MODERAT | MISSENSE | gCc/gTc | A49V | PLES_26721 | | | | | . | . | . | . | . | . | A | A |
| NC_011770 | 2870297 | T | G | 23 | 6 | 2 | 0 | NON_SYNONYMOU | MODERAT | MISSENSE | gAg/gCg | E45A | PLES_26721 | | | | | . | . | . | . | . | . | G | G |
| NC_011770 | 2870348 | A | G | 25 | 6 | 6 | 0 | NON_SYNONYMOU | MODERAT | MISSENSE | aTg/aCg | M28T | PLES_26721 | G | G | G | G | . | . | . | . | . | . | G | G |
| NC_011770 | 2870378 | G | C | 21 | 6 | 6 | 0 | NON_SYNONYMOU | MODERAT | MISSENSE | gCc/gGc | A18G | PLES_26721 | C | C | C | C | . | . | . | . | . | . | C | C |
| NC_011770 | 2870406 | A | G | 19 | 6 | 6 | 0 | NON_SYNONYMOU | MODERAT | MISSENSE | Tcc/Ccc | S9P | PLES_26721 | G | G | G | G | . | . | . | . | . | . | G | G |

| | | | | | | | | | | | | | | | | | | | | | | | | | |
|---|---|---|---|---|---|---|---|---|---|---|---|---|---|---|---|---|---|---|---|---|---|---|---|---|---|
| NC_011770 | 2870427 | C | T | 21 | 6 | 2 | 0 | NON_SYNONYMOU | MODERAT | MISSENSE | Gcc/Acc | A2T | PLES_26721 | | | | | . | . | . | . | . | . | T | T |
| NC_011770 | 2870874 | G | A | 42 | 4 | 2 | 0 | NON_SYNONYMOU | MODERAT | MISSENSE | Ctc/Ttc | L705F | PLES_26731 | | | | | A | A | . | . | . | . | | |
| NC_011770 | 2870876 | T | A | 42 | 4 | 2 | 0 | NON_SYNONYMOU | MODERAT | MISSENSE | tAc/tTc | Y704F | PLES_26731 | | | | | A | A | . | . | . | . | | |
| NC_011770 | 2870878 | G | C | 42 | 4 | 2 | 0 | NON_SYNONYMOU | MODERAT | MISSENSE | atC/atG | I703M | PLES_26731 | | | | | C | C | . | . | . | . | | |
| NC_011770 | 2870880 | T | C | 41 | 4 | 2 | 0 | NON_SYNONYMOU | MODERAT | MISSENSE | Atc/Gtc | I703V | PLES_26731 | | | | | C | C | . | . | . | . | | |
| NC_011770 | 2870893 | C | G | 41 | 4 | 4 | 0 | NON_SYNONYMOU | MODERAT | MISSENSE | caG/caC | Q698H | PLES_26731 | | | | | G | G | . | . | . | . | T | T |
| NC_011770 | 2870937 | G | A | 42 | 4 | 2 | 0 | NON_SYNONYMOU | MODERAT | MISSENSE | Ctc/Ttc | L684F | PLES_26731 | | | | | A | A | . | . | . | . | | |
| NC_011770 | 2870940 | A | T | 43 | 4 | 2 | 0 | NON_SYNONYMOU | MODERAT | MISSENSE | Tcg/Acg | S683T | PLES_26731 | | | | | T | T | . | . | . | . | | |
| NC_011770 | 2870958 | A | T | 60 | 4 | 2 | 0 | NON_SYNONYMOU | MODERAT | MISSENSE | Tcg/Acg | S677T | PLES_26731 | | | | | T | T | . | . | . | . | | |
| NC_011770 | 2870964 | T | G | 64 | 4 | 2 | 0 | NON_SYNONYMOU | MODERAT | MISSENSE | Atg/Ctg | M675L | PLES_26731 | | | | | G | G | . | . | . | . | | |
| NC_011770 | 2870969 | A | G | 64 | 4 | 2 | 0 | NON_SYNONYMOU | MODERAT | MISSENSE | gTg/gCg | V673A | PLES_26731 | | | | | G | G | . | . | . | . | | |
| NC_011770 | 2870983 | C | G | 67 | 4 | 2 | 0 | NON_SYNONYMOU | MODERAT | MISSENSE | gaG/gaC | E668D | PLES_26731 | | | | | G | G | . | . | . | . | | |
| NC_011770 | 2871149 | G | C | 53 | 4 | 2 | 0 | NON_SYNONYMOU | MODERAT | MISSENSE | tCg/tGg | S613W | PLES_26731 | | | | | | | . | . | . | . | C | C |
| NC_011770 | 2871150 | A | T | 52 | 4 | 2 | 0 | NON_SYNONYMOU | MODERAT | MISSENSE | Tcg/Acg | S613T | PLES_26731 | | | | | | | . | . | . | . | T | T |
| NC_011770 | 2871170 | G | C | 65 | 4 | 2 | 0 | NON_SYNONYMOU | MODERAT | MISSENSE | aCc/aGc | T606S | PLES_26731 | | | | | C | C | . | . | . | . | | |
| NC_011770 | 2871204 | G | T | 64 | 4 | 2 | 0 | NON_SYNONYMOU | MODERAT | MISSENSE | Cag/Aag | Q595K | PLES_26731 | | | | | T | T | . | . | . | . | | |
| NC_011770 | 2871250 | G | T | 45 | 6 | 4 | 0 | NON_SYNONYMOU | MODERAT | MISSENSE | agC/agA | S579R | PLES_26731 | T | T | T | T | . | . | . | . | . | . | | |
| NC_011770 | 2871251 | C | G | 46 | 6 | 4 | 0 | NON_SYNONYMOU | MODERAT | MISSENSE | aGc/aCc | S579T | PLES_26731 | G | G | G | G | . | . | . | . | . | . | | |
| NC_011770 | 2871253 | G | C | 45 | 6 | 4 | 0 | NON_SYNONYMOU | MODERAT | MISSENSE | atC/atG | I578M | PLES_26731 | C | C | C | C | . | . | . | . | . | . | | |
| NC_011770 | 2871255 | T | C | 45 | 6 | 4 | 0 | NON_SYNONYMOU | MODERAT | MISSENSE | Atc/Gtc | I578V | PLES_26731 | C | C | C | C | . | . | . | . | . | . | | |
| NC_011770 | 2871379 | C | G | 94 | 4 | 2 | 0 | NON_SYNONYMOU | MODERAT | MISSENSE | ttG/ttC | L536F | PLES_26731 | | | | | G | G | . | . | . | . | | |
| NC_011770 | 2871510 | C | T | 65 | 4 | 2 | 0 | NON_SYNONYMOU | MODERAT | MISSENSE | Gtc/Atc | V493I | PLES_26731 | | | | | | | . | . | . | . | T | T |
| NC_011770 | 2871786 | G | A | 54 | 6 | 6 | 0 | NON_SYNONYMOU | MODERAT | MISSENSE | Cac/Tac | H401Y | PLES_26731 | A | A | A | A | . | . | . | . | . | . | A | A |
| NC_011770 | 2871800 | C | G | 54 | 5 | 1 | 0 | NON_SYNONYMOU | MODERAT | MISSENSE | gGg/gCg | G396A | PLES_26731 | | | | | G | . | . | . | . | . | | |
| NC_011770 | 2871814 | C | G | 65 | 4 | 2 | 0 | NON_SYNONYMOU | MODERAT | MISSENSE | ttG/ttC | L391F | PLES_26731 | | | | | G | G | . | . | . | . | | |
| NC_011770 | 2871883 | A | C | 46 | 4 | 2 | 0 | NON_SYNONYMOU | MODERAT | MISSENSE | agT/agG | S368R | PLES_26731 | | | | | C | C | . | . | . | . | | |
| NC_011770 | 2871884 | C | G | 46 | 4 | 2 | 0 | NON_SYNONYMOU | MODERAT | MISSENSE | aGt/aCt | S368T | PLES_26731 | | | | | G | G | . | . | . | . | | |
| NC_011770 | 2871885 | T | A | 45 | 4 | 2 | 0 | NON_SYNONYMOU | MODERAT | MISSENSE | Agt/Tgt | S368C | PLES_26731 | | | | | A | A | . | . | . | . | | |
| NC_011770 | 2871891 | C | T | 46 | 4 | 2 | 0 | NON_SYNONYMOU | MODERAT | MISSENSE | Ggg/Agg | G366R | PLES_26731 | | | | | T | T | . | . | . | . | | |
| NC_011770 | 2871954 | G | T | 47 | 4 | 2 | 0 | NON_SYNONYMOU | MODERAT | MISSENSE | Cgc/Agc | R345S | PLES_26731 | | | T | T | | | . | . | . | . | | |
| NC_011770 | 2871969 | A | T | 47 | 4 | 2 | 0 | NON_SYNONYMOU | MODERAT | MISSENSE | Tcc/Acc | S340T | PLES_26731 | | | | | T | T | . | . | . | . | | |
| NC_011770 | 2872003 | C | G | 48 | 4 | 2 | 0 | NON_SYNONYMOU | MODERAT | MISSENSE | atG/atC | M328I | PLES_26731 | | | | | G | G | . | . | . | . | | |
| NC_011770 | 2872077 | G | A | 42 | 6 | 4 | 0 | NON_SYNONYMOU | MODERAT | MISSENSE | Ctc/Ttc | L304F | PLES_26731 | A | A | A | A | . | . | . | . | . | . | | |
| NC_011770 | 2872163 | T | G | 35 | 6 | 2 | 0 | NON_SYNONYMOU | MODERAT | MISSENSE | gAg/gCg | E275A | PLES_26731 | | | | | . | . | . | . | . | . | G | G |
| NC_011770 | 2872272 | C | T | 37 | 6 | 6 | 0 | NON_SYNONYMOU | MODERAT | MISSENSE | Ggc/Agc | G239S | PLES_26731 | T | T | T | T | . | . | . | . | . | . | T | T |
| NC_011770 | 2872286 | C | T | 38 | 6 | 6 | 0 | NON_SYNONYMOU | MODERAT | MISSENSE | tGc/tAc | C234Y | PLES_26731 | T | T | T | T | . | . | . | . | . | . | T | T |
| NC_011770 | 2872291 | C | A | 37 | 6 | 4 | 0 | NON_SYNONYMOU | MODERAT | MISSENSE | gaG/gaT | E232D | PLES_26731 | A | A | A | A | . | . | . | . | . | . | | |
| NC_011770 | 2872299 | G | C | 38 | 6 | 2 | 0 | NON_SYNONYMOU | MODERAT | MISSENSE | Caa/Gaa | Q230E | PLES_26731 | | | | | . | . | . | . | . | . | C | C |
| NC_011770 | 2872308 | T | C | 38 | 6 | 6 | 0 | NON_SYNONYMOU | MODERAT | MISSENSE | Atc/Gtc | I227V | PLES_26731 | C | C | C | C | . | . | . | . | . | . | C | C |
| NC_011770 | 2872394 | G | T | 39 | 6 | 2 | 0 | NON_SYNONYMOU | MODERAT | MISSENSE | aCg/aAg | T198K | PLES_26731 | | | | | . | . | . | . | . | . | T | T |
| NC_011770 | 2872395 | T | G | 39 | 6 | 2 | 0 | NON_SYNONYMOU | MODERAT | MISSENSE | Acg/Ccg | T198P | PLES_26731 | | | | | . | . | . | . | . | . | G | G |
| NC_011770 | 2872512 | G | A | 34 | 6 | 2 | 0 | NON_SYNONYMOU | MODERAT | MISSENSE | Ctc/Ttc | L159F | PLES_26731 | | | | | . | . | . | . | . | . | A | A |
| NC_011770 | 2872562 | T | A | 40 | 5 | 1 | 0 | NON_SYNONYMOU | MODERAT | MISSENSE | cAg/cTg | Q142L | PLES_26731 | | | | | A | . | . | . | . | . | | |
| NC_011770 | 2872650 | T | C | 39 | 6 | 4 | 0 | NON_SYNONYMOU | MODERAT | MISSENSE | Acg/Gcg | T113A | PLES_26731 | C | C | C | C | . | . | . | . | . | . | | |
| NC_011770 | 2872763 | G | T | 40 | 6 | 2 | 0 | NON_SYNONYMOU | MODERAT | MISSENSE | cCa/cAa | P75Q | PLES_26731 | | | | | . | . | . | . | . | . | T | T |
| NC_011770 | 2872794 | C | T | 35 | 6 | 2 | 0 | NON_SYNONYMOU | MODERAT | MISSENSE | Gtg/Atg | V65M | PLES_26731 | | | | | . | . | . | . | . | . | T | T |
| NC_011770 | 2872913 | G | T | 43 | 6 | 4 | 0 | NON_SYNONYMOU | MODERAT | MISSENSE | tCc/tAc | S25Y | PLES_26731 | T | T | T | T | . | . | . | . | . | . | | |
| NC_011770 | 2872914 | A | T | 43 | 6 | 4 | 0 | NON_SYNONYMOU | MODERAT | MISSENSE | Tcc/Acc | S25T | PLES_26731 | T | T | T | T | . | . | . | . | . | . | | |
| NC_011770 | 2872950 | C | T | 46 | 6 | 4 | 0 | NON_SYNONYMOU | MODERAT | MISSENSE | Gcg/Acg | A13T | PLES_26731 | T | T | T | T | . | . | . | . | . | . | | |
| NC_011770 | 2872953 | A | C | 46 | 6 | 6 | 0 | NON_SYNONYMOU | MODERAT | MISSENSE | Tcc/Gcc | S12A | PLES_26731 | C | C | C | C | . | . | . | . | . | . | C | C |
| NC_011770 | 2872964 | G | T | 48 | 6 | 2 | 0 | NON_SYNONYMOU | MODERAT | MISSENSE | aCt/aAt | T8N | PLES_26731 | | | | | . | . | . | . | . | . | T | T |
| NC_011770 | 2871834 | A | C | 65 | 4 | 1 | 0 | NON_SYNONYMOU | MODERAT | MISSENSE | Ttg/Gtg | L385V | PLES_26731 | | | | | | C | . | . | . | . | | |
| NC_011770 | 2873251 | T | TGG | 32 | 6 | 4 | 0 | FRAME_SHIFT | HIGH | | acg/acgCC | T59T? | PLES_26741 | TGG | TGG | TGG | TGG | . | . | . | . | . | . | | |
| NC_011770 | 2873148 | A | G | 38 | 6 | 4 | 0 | NON_SYNONYMOU | MODERAT | MISSENSE | gTc/gCc | V94A | PLES_26741 | G | G | G | G | . | . | . | . | . | . | | |
| NC_011770 | 2873154 | G | A | 35 | 6 | 4 | 0 | NON_SYNONYMOU | MODERAT | MISSENSE | gCa/gTa | A92V | PLES_26741 | A | A | A | A | . | . | . | . | . | . | | |
| NC_011770 | 2873155 | C | T | 35 | 6 | 4 | 0 | NON_SYNONYMOU | MODERAT | MISSENSE | Gca/Aca | A92T | PLES_26741 | T | T | T | T | . | . | . | . | . | . | | |
| NC_011770 | 2873157 | C | T | 35 | 6 | 2 | 0 | NON_SYNONYMOU | MODERAT | MISSENSE | tGc/tAc | C91Y | PLES_26741 | | | | | . | . | . | . | . | . | T | T |
| NC_011770 | 2873158 | A | T | 35 | 6 | 6 | 0 | NON_SYNONYMOU | MODERAT | MISSENSE | Tgc/Agc | C91S | PLES_26741 | T | T | T | T | . | . | . | . | . | . | G | G |
| NC_011770 | 2873163 | T | C | 34 | 6 | 4 | 0 | NON_SYNONYMOU | MODERAT | MISSENSE | cAg/cGg | Q89R | PLES_26741 | C | C | C | C | . | . | . | . | . | . | | |
| NC_011770 | 2873164 | G | C | 35 | 6 | 6 | 0 | NON_SYNONYMOU | MODERAT | MISSENSE | Cag/Gag | Q89E | PLES_26741 | C | C | C | C | . | . | . | . | . | . | C | C |
| NC_011770 | 2873167 | A | T | 35 | 6 | 6 | 0 | NON_SYNONYMOU | MODERAT | MISSENSE | Tcg/Acg | S88T | PLES_26741 | T | T | T | T | . | . | . | . | . | . | C | C |
| NC_011770 | 2873176 | C | T | 44 | 6 | 4 | 0 | NON_SYNONYMOU | MODERAT | MISSENSE | Gtc/Atc | V85I | PLES_26741 | T | T | T | T | . | . | . | . | . | . | | |

| | | | | | | | | | | | | | | | | | | | | | | | | | |
|---|---|---|---|---|---|---|---|---|---|---|---|---|---|---|---|---|---|---|---|---|---|---|---|---|---|
| NC_011770 | 2873217 | T | G | 46 | 6 | 2 | 0 | NON_SYNONYMOU | MODERAT | MISSENSE | gAa/gCa | E71A | PLES_26741 | | | | | . | . | . | . | . | . | G | G |
| NC_011770 | 2873224 | A | G | 44 | 6 | 4 | 0 | NON_SYNONYMOU | MODERAT | MISSENSE | Tcg/Ccg | S69P | PLES_26741 | G | G | G | G | . | . | . | . | . | . | | |
| NC_011770 | 2873244 | C | G | 30 | 6 | 6 | 0 | NON_SYNONYMOU | MODERAT | MISSENSE | aGc/aCc | S62T | PLES_26741 | G | G | G | G | . | . | . | . | . | . | G | G |
| NC_011770 | 2873245 | T | A | 30 | 6 | 4 | 0 | NON_SYNONYMOU | MODERAT | MISSENSE | Agc/Tgc | S62C | PLES_26741 | A | A | A | A | . | . | . | . | . | . | | |
| NC_011770 | 2873254 | T | C | 21 | 9 | 2 | 0 | NON_SYNONYMOU | MODERAT | MISSENSE | Acg/Gcg | T59A | PLES_26741 | . | . | | . | . | . | . | . | . | . | C | C |
| NC_011770 | 2873256 | T | C | 21 | 9 | 1 | 0 | NON_SYNONYMOU | MODERAT | MISSENSE | cAg/cGg | Q58R | PLES_26741 | . | . | C | . | . | . | . | . | . | . | | |
| NC_011770 | 2873257 | G | C | 21 | 7 | 3 | 0 | NON_SYNONYMOU | MODERAT | MISSENSE | Cag/Gag | Q58E | PLES_26741 | . | C | C | C | . | . | . | . | . | . | | |
| NC_011770 | 2873268 | A | G | 39 | 6 | 4 | 0 | NON_SYNONYMOU | MODERAT | MISSENSE | aTc/aCc | I54T | PLES_26741 | G | G | G | G | . | . | . | . | . | . | | |
| NC_011770 | 2873269 | T | C | 39 | 6 | 6 | 0 | NON_SYNONYMOU | MODERAT | MISSENSE | Atc/Gtc | I54V | PLES_26741 | C | C | C | C | . | . | . | . | . | . | C | C |
| NC_011770 | 2873271 | T | G | 38 | 6 | 4 | 0 | NON_SYNONYMOU | MODERAT | MISSENSE | aAc/aCc | N53T | PLES_26741 | G | G | G | G | . | . | . | . | . | . | | |
| NC_011770 | 2873296 | T | C | 47 | 6 | 4 | 0 | NON_SYNONYMOU | MODERAT | MISSENSE | Acc/Gcc | T45A | PLES_26741 | C | C | C | C | . | . | . | . | . | . | | |
| NC_011770 | 2873335 | C | A | 45 | 6 | 2 | 0 | NON_SYNONYMOU | MODERAT | MISSENSE | Gcg/Tcg | A32S | PLES_26741 | | | | | . | . | . | . | . | . | A | A |
| NC_011770 | 2873337 | C | T | 44 | 6 | 4 | 0 | NON_SYNONYMOU | MODERAT | MISSENSE | cGc/cAc | R31H | PLES_26741 | T | T | T | T | . | . | . | . | . | . | | |
| NC_011770 | 2873338 | G | T | 44 | 6 | 4 | 0 | NON_SYNONYMOU | MODERAT | MISSENSE | Cgc/Agc | R31S | PLES_26741 | T | T | T | T | . | . | . | . | . | . | | |
| NC_011770 | 2873341 | C | A | 44 | 6 | 4 | 0 | NON_SYNONYMOU | MODERAT | MISSENSE | Gtc/Ttc | V30F | PLES_26741 | A | A | A | A | . | . | . | . | . | . | | |
| NC_011770 | 2873350 | C | T | 42 | 6 | 4 | 0 | NON_SYNONYMOU | MODERAT | MISSENSE | Gac/Aac | D27N | PLES_26741 | T | T | T | T | . | . | . | . | . | . | | |
| NC_011770 | 2873353 | C | T | 43 | 6 | 6 | 0 | NON_SYNONYMOU | MODERAT | MISSENSE | Gta/Ata | V26I | PLES_26741 | T | T | T | T | . | . | . | . | . | . | T | T |
| NC_011770 | 2873358 | C | T | 47 | 6 | 2 | 0 | NON_SYNONYMOU | MODERAT | MISSENSE | aGa/aAa | R24K | PLES_26741 | | | | | . | . | . | . | . | . | T | T |
| NC_011770 | 2873389 | C | T | 47 | 6 | 4 | 0 | NON_SYNONYMOU | MODERAT | MISSENSE | Gcg/Acg | A14T | PLES_26741 | T | T | T | T | . | . | . | . | . | . | | |
| NC_011770 | 2873394 | C | T | 47 | 6 | 2 | 0 | NON_SYNONYMOU | MODERAT | MISSENSE | cGg/cAg | R12Q | PLES_26741 | | | | | . | . | . | . | . | . | T | T |
| NC_011770 | 2873396 | C | G | 46 | 6 | 4 | 0 | NON_SYNONYMOU | MODERAT | MISSENSE | caG/caC | Q11H | PLES_26741 | G | G | G | G | . | . | . | . | . | . | | |
| NC_011770 | 2873550 | C | T | 50 | 6 | 2 | 0 | NON_SYNONYMOU | MODERAT | MISSENSE | Gtc/Atc | V97I | PLES_26751 | | | | | . | . | . | . | . | . | T | T |
| NC_011770 | 2873787 | C | T | 50 | 6 | 4 | 0 | NON_SYNONYMOU | MODERAT | MISSENSE | Gca/Aca | A18T | PLES_26751 | T | T | T | T | . | . | . | . | . | . | | |
| NC_011770 | 2873835 | T | A | 48 | 6 | 2 | 0 | NON_SYNONYMOU | MODERAT | MISSENSE | Atc/Ttc | I2F | PLES_26751 | | | | | . | . | . | . | . | . | A | A |
| NC_011770 | 2873921 | C | G | 50 | 6 | 4 | 0 | NON_SYNONYMOU | MODERAT | MISSENSE | Gcg/Ccg | A360P | PLES_26761 | G | G | G | G | . | . | . | . | . | . | | |
| NC_011770 | 2874080 | T | G | 48 | 6 | 6 | 0 | NON_SYNONYMOU | MODERAT | MISSENSE | Aag/Cag | K307Q | PLES_26761 | G | G | G | G | . | . | . | . | . | . | G | G |
| NC_011770 | 2874136 | C | T | 45 | 6 | 6 | 0 | NON_SYNONYMOU | MODERAT | MISSENSE | cGc/cAc | R288H | PLES_26761 | T | T | T | T | . | . | . | . | . | . | T | T |
| NC_011770 | 2874179 | C | T | 41 | 6 | 4 | 0 | NON_SYNONYMOU | MODERAT | MISSENSE | Gct/Act | A274T | PLES_26761 | T | T | T | T | . | . | . | . | . | . | | |
| NC_011770 | 2874196 | T | A | 43 | 6 | 6 | 0 | NON_SYNONYMOU | MODERAT | MISSENSE | cAg/cTg | Q268L | PLES_26761 | A | A | A | A | . | . | . | . | . | . | A | A |
| NC_011770 | 2874207 | C | A | 43 | 6 | 6 | 0 | NON_SYNONYMOU | MODERAT | MISSENSE | ttG/ttT | L264F | PLES_26761 | A | A | A | A | . | . | . | . | . | . | G | G |
| NC_011770 | 2874283 | G | A | 48 | 6 | 2 | 0 | NON_SYNONYMOU | MODERAT | MISSENSE | gCc/gTc | A239V | PLES_26761 | | | | | . | . | . | . | . | . | A | A |
| NC_011770 | 2874354 | T | A | 42 | 6 | 6 | 0 | NON_SYNONYMOU | MODERAT | MISSENSE | caA/caT | Q215H | PLES_26761 | A | A | A | A | . | . | . | . | . | . | A | A |
| NC_011770 | 2874355 | T | A | 42 | 6 | 6 | 0 | NON_SYNONYMOU | MODERAT | MISSENSE | cAa/cTa | Q215L | PLES_26761 | A | A | A | A | . | . | . | . | . | . | A | A |
| NC_011770 | 2874403 | A | G | 39 | 6 | 6 | 0 | NON_SYNONYMOU | MODERAT | MISSENSE | gTg/gCg | V199A | PLES_26761 | G | G | G | G | . | . | . | . | . | . | G | G |
| NC_011770 | 2874406 | T | A | 39 | 6 | 4 | 0 | NON_SYNONYMOU | MODERAT | MISSENSE | cAg/cTg | Q198L | PLES_26761 | A | A | A | A | . | . | . | . | . | . | | |
| NC_011770 | 2874557 | C | T | 46 | 6 | 2 | 0 | NON_SYNONYMOU | MODERAT | MISSENSE | Gtg/Atg | V148M | PLES_26761 | | | | | . | . | . | . | . | . | T | T |
| NC_011770 | 2874584 | T | A | 46 | 6 | 6 | 0 | NON_SYNONYMOU | MODERAT | MISSENSE | Acg/Tcg | T139S | PLES_26761 | A | A | A | A | . | . | . | . | . | . | A | A |
| NC_011770 | 2874713 | C | T | 44 | 6 | 4 | 0 | NON_SYNONYMOU | MODERAT | MISSENSE | Gtc/Atc | V96I | PLES_26761 | T | T | T | T | . | . | . | . | . | . | | |
| NC_011770 | 2874813 | C | T | 40 | 6 | 6 | 0 | NON_SYNONYMOU | MODERAT | MISSENSE | gaG/gaC | E62D | PLES_26761 | T | T | T | T | . | . | . | . | . | . | G | G |
| NC_011770 | 2874830 | G | A | 41 | 6 | 2 | 0 | NON_SYNONYMOU | MODERAT | MISSENSE | Cat/Tat | H57Y | PLES_26761 | | | | | . | . | . | . | . | . | A | A |
| NC_011770 | 2874833 | C | A | 39 | 6 | 4 | 0 | NON_SYNONYMOU | MODERAT | MISSENSE | Gcc/Tcc | A56S | PLES_26761 | A | A | A | A | . | . | . | . | . | . | | |
| NC_011770 | 2874946 | A | T | 35 | 6 | 6 | 0 | NON_SYNONYMOU | MODERAT | MISSENSE | tTc/tAc | F18Y | PLES_26761 | T | T | T | T | . | . | . | . | . | . | T | T |
| NC_011770 | 2874961 | A | G | 37 | 6 | 6 | 0 | NON_SYNONYMOU | MODERAT | MISSENSE | gTc/gCc | V13A | PLES_26761 | G | G | G | G | . | . | . | . | . | . | G | G |
| NC_011770 | 2875098 | C | T | 39 | 6 | 2 | 0 | NON_SYNONYMOU | MODERAT | MISSENSE | Gcc/Acc | A205T | PLES_26771 | | | | | . | . | . | . | . | . | T | T |
| NC_011770 | 2875154 | G | C | 35 | 6 | 4 | 0 | NON_SYNONYMOU | MODERAT | MISSENSE | aCc/aGc | T186S | PLES_26771 | C | C | C | C | . | . | . | . | . | . | | |
| NC_011770 | 2875235 | G | C | 36 | 6 | 6 | 0 | NON_SYNONYMOU | MODERAT | MISSENSE | aCc/aGc | T159S | PLES_26771 | C | C | C | C | . | . | . | . | . | . | C | C |
| NC_011770 | 2875307 | A | G | 35 | 6 | 4 | 0 | NON_SYNONYMOU | MODERAT | MISSENSE | aTg/aCg | M135T | PLES_26771 | G | G | G | G | . | . | . | . | . | . | | |
| NC_011770 | 2875373 | T | C | 30 | 6 | 4 | 0 | NON_SYNONYMOU | MODERAT | MISSENSE | aAg/aGg | K113R | PLES_26771 | C | C | C | C | . | . | . | . | . | . | | |
| NC_011770 | 2875453 | G | C | 27 | 6 | 2 | 0 | NON_SYNONYMOU | MODERAT | MISSENSE | atC/atG | I86M | PLES_26771 | | | | | . | . | . | . | . | . | C | C |
| NC_011770 | 2875521 | C | T | 28 | 6 | 4 | 0 | NON_SYNONYMOU | MODERAT | MISSENSE | Gtc/Atc | V64I | PLES_26771 | T | T | T | T | . | . | . | . | . | . | | |
| NC_011770 | 2875679 | C | T | 28 | 6 | 4 | 0 | NON_SYNONYMOU | MODERAT | MISSENSE | cGc/cAc | R11H | PLES_26771 | T | T | T | T | . | . | . | . | . | . | | |
| NC_011770 | 2875682 | T | A | 29 | 6 | 4 | 0 | NON_SYNONYMOU | MODERAT | MISSENSE | tAc/tTc | Y10F | PLES_26771 | A | A | A | A | . | . | . | . | . | . | | |
| NC_011770 | 2875701 | T | G | 25 | 6 | 4 | 0 | NON_SYNONYMOU | MODERAT | MISSENSE | Aac/Cac | N4H | PLES_26771 | G | G | G | G | . | . | . | . | . | . | | |
| NC_011770 | 2875706 | G | T | 26 | 6 | 4 | 0 | NON_SYNONYMOU | MODERAT | MISSENSE | cCc/cAc | P2H | PLES_26771 | T | T | T | T | . | . | . | . | . | . | | |
| NC_011770 | 2875509 | G | C | 35 | 3 | 1 | 0 | NON_SYNONYMOU | MODERAT | MISSENSE | Cag/Gag | Q68E | PLES_26771 | | C | | | | | . | | . | . | | |
| NC_011770 | 2875831 | G | C | 17 | 10 | 2 | 0 | NON_SYNONYMOU | MODERAT | MISSENSE | Ccg/Gcg | P73A | PLES_26781 | . | . | . | . | . | . | . | . | . | . | C | C |
| NC_011770 | 2875839 | G | A | 16 | 10 | 2 | 0 | NON_SYNONYMOU | MODERAT | MISSENSE | gCg/gTg | A70V | PLES_26781 | . | . | . | . | . | . | . | . | . | . | A | A |
| NC_011770 | 2875843 | A | C | 16 | 10 | 2 | 0 | NON_SYNONYMOU | MODERAT | MISSENSE | Tcg/Gcg | S69A | PLES_26781 | . | . | . | . | . | . | . | . | . | . | C | C |
| NC_011770 | 2875862 | T | G | 17 | 10 | 2 | 0 | NON_SYNONYMOU | MODERAT | MISSENSE | gaA/gaC | E62D | PLES_26781 | . | . | . | . | . | . | . | . | . | . | G | G |
| NC_011770 | 2875919 | G | C | 19 | 10 | 2 | 0 | NON_SYNONYMOU | MODERAT | MISSENSE | gaC/gaG | D43E | PLES_26781 | . | . | . | . | . | . | . | . | . | . | C | C |
| NC_011770 | 2876007 | G | T | 12 | 10 | 2 | 0 | NON_SYNONYMOU | MODERAT | MISSENSE | gCa/gAa | A14E | PLES_26781 | . | . | . | . | . | . | . | . | . | . | T | T |
| NC_011770 | 2876008 | C | T | 13 | 10 | 2 | 0 | NON_SYNONYMOU | MODERAT | MISSENSE | Gca/Aca | A14T | PLES_26781 | . | . | . | . | . | . | . | . | . | . | T | T |

| | | | | | | | | | | | | | | | | | | | | | | | | | |
|---|---|---|---|---|---|---|---|---|---|---|---|---|---|---|---|---|---|---|---|---|---|---|---|---|---|
| NC_011770 | 2876020 | G | C | 13 | 10 | 2 | 0 | NON_SYNONYMOU | MODERAT | MISSENSE | Ccc/Gcc | P10A | PLES_26781 | . | . | . | . | . | . | . | . | . | . | C | C |
| NC_011770 | 2876029 | T | C | 13 | 10 | 2 | 0 | NON_SYNONYMOU | MODERAT | MISSENSE | Acg/Gcg | T7A | PLES_26781 | . | . | . | . | . | . | . | . | . | . | C | C |
| NC_011770 | 2876087 | A | G | 12 | 10 | 2 | 0 | NON_SYNONYMOU | MODERAT | MISSENSE | Tcc/Ccc | S129P | PLES_26791 | . | . | . | . | . | . | . | . | . | . | G | G |
| NC_011770 | 2876093 | C | T | 13 | 10 | 2 | 0 | NON_SYNONYMOU | MODERAT | MISSENSE | Gcg/Acg | A127T | PLES_26791 | . | . | . | . | . | . | . | . | . | . | T | T |
| NC_011770 | 2876096 | A | C | 12 | 10 | 2 | 0 | NON_SYNONYMOU | MODERAT | MISSENSE | Tcc/Gcc | S126A | PLES_26791 | . | . | . | . | . | . | . | . | . | . | C | C |
| NC_011770 | 2876098 | G | A | 12 | 10 | 2 | 0 | NON_SYNONYMOU | MODERAT | MISSENSE | gCa/gTa | A125V | PLES_26791 | . | . | . | . | . | . | . | . | . | . | A | A |
| NC_011770 | 2876195 | T | A | 19 | 10 | 2 | 0 | NON_SYNONYMOU | MODERAT | MISSENSE | Acg/Tcg | T93S | PLES_26791 | . | . | . | . | . | . | . | . | . | . | A | A |
| NC_011770 | 2876228 | A | G | 18 | 10 | 2 | 0 | NON_SYNONYMOU | MODERAT | MISSENSE | Tcc/Ccc | S82P | PLES_26791 | . | . | . | . | . | . | . | . | . | . | G | G |
| NC_011770 | 2876245 | C | T | 16 | 10 | 2 | 0 | NON_SYNONYMOU | MODERAT | MISSENSE | gGc/gAc | G76D | PLES_26791 | . | . | . | . | . | . | . | . | . | . | T | T |
| NC_011770 | 2876246 | C | T | 16 | 10 | 2 | 0 | NON_SYNONYMOU | MODERAT | MISSENSE | Ggc/Agc | G76S | PLES_26791 | . | . | . | . | . | . | . | . | . | . | T | T |
| NC_011770 | 2876323 | A | G | 19 | 10 | 2 | 0 | NON_SYNONYMOU | MODERAT | MISSENSE | gTc/gCc | V50A | PLES_26791 | . | . | . | . | . | . | . | . | . | . | G | G |
| NC_011770 | 2876326 | T | C | 18 | 10 | 2 | 0 | NON_SYNONYMOU | MODERAT | MISSENSE | aAg/aGg | K49R | PLES_26791 | . | . | . | . | . | . | . | . | . | . | C | C |
| NC_011770 | 2876452 | T | G | 16 | 10 | 2 | 0 | NON_SYNONYMOU | MODERAT | MISSENSE | aAc/aCc | N7T | PLES_26791 | . | . | . | . | . | . | . | . | . | . | G | G |
| NC_011770 | 2876453 | T | G | 16 | 10 | 2 | 0 | NON_SYNONYMOU | MODERAT | MISSENSE | Aac/Cac | N7H | PLES_26791 | . | . | . | . | . | . | . | . | . | . | G | G |
| NC_011770 | 2876798 | G | C | 11 | 6 | 6 | 0 | STOP_GAINED | HIGH | NONSENSE | taC/taG | Y40* | PLES_26801 | C | C | C | C | . | . | . | . | . | . | C | C |
| NC_011770 | 2876807 | G | C | 11 | 6 | 6 | 0 | STOP_GAINED | HIGH | NONSENSE | taC/taG | Y37* | PLES_26801 | C | C | C | C | . | . | . | . | . | . | C | C |
| NC_011770 | 2876703 | T | C | 30 | 6 | 4 | 0 | NON_SYNONYMOU | MODERAT | MISSENSE | gAc/gGc | D72G | PLES_26801 | C | C | C | C | . | . | . | . | . | . | | |
| NC_011770 | 2876716 | C | T | 13 | 10 | 2 | 0 | NON_SYNONYMOU | MODERAT | MISSENSE | Gcc/Acc | A68T | PLES_26801 | . | . | . | . | . | . | . | . | . | . | T | T |
| NC_011770 | 2876725 | C | G | 18 | 6 | 4 | 0 | NON_SYNONYMOU | MODERAT | MISSENSE | Gaa/Caa | E65Q | PLES_26801 | G | G | G | G | . | . | . | . | . | . | | |
| NC_011770 | 2876732 | C | A | 7 | 6 | 2 | 0 | NON_SYNONYMOU | MODERAT | MISSENSE | caG/caT | Q62H | PLES_26801 | | | | | . | . | . | . | . | . | A | A |
| NC_011770 | 2876736 | G | A | 7 | 6 | 2 | 0 | NON_SYNONYMOU | MODERAT | MISSENSE | gCt/gTt | A61V | PLES_26801 | | | | | . | . | . | . | . | . | A | A |
| NC_011770 | 2876744 | C | G | 12 | 6 | 2 | 0 | NON_SYNONYMOU | MODERAT | MISSENSE | gaG/gaC | E58D | PLES_26801 | | | | | . | . | . | . | . | . | G | G |
| NC_011770 | 2876745 | T | C | 12 | 6 | 2 | 0 | NON_SYNONYMOU | MODERAT | MISSENSE | gAg/gGg | E58G | PLES_26801 | | | | | . | . | . | . | . | . | C | C |
| NC_011770 | 2876746 | C | T | 12 | 6 | 2 | 0 | NON_SYNONYMOU | MODERAT | MISSENSE | Gag/Aag | E58K | PLES_26801 | | | | | . | . | . | . | . | . | T | T |
| NC_011770 | 2876747 | A | T | 12 | 6 | 6 | 0 | NON_SYNONYMOU | MODERAT | MISSENSE | gaT/gaA | D57E | PLES_26801 | T | T | T | T | . | . | . | . | . | . | T | T |
| NC_011770 | 2876760 | T | C | 12 | 6 | 6 | 0 | NON_SYNONYMOU | MODERAT | MISSENSE | aAt/aGt | N53S | PLES_26801 | C | C | C | C | . | . | . | . | . | . | C | C |
| NC_011770 | 2876778 | G | T | 12 | 6 | 6 | 0 | NON_SYNONYMOU | MODERAT | MISSENSE | gCg/gAg | A47E | PLES_26801 | T | T | T | T | . | . | . | . | . | . | T | T |
| NC_011770 | 2876799 | T | C | 11 | 6 | 6 | 0 | NON_SYNONYMOU | MODERAT | MISSENSE | tAc/tGc | Y40C | PLES_26801 | C | C | C | C | . | . | . | . | . | . | C | C |
| NC_011770 | 2876808 | T | C | 11 | 6 | 6 | 0 | NON_SYNONYMOU | MODERAT | MISSENSE | tAc/tGc | Y37C | PLES_26801 | C | C | C | C | . | . | . | . | . | . | C | C |
| NC_011770 | 2876838 | G | T | 8 | 6 | 6 | 0 | NON_SYNONYMOU | MODERAT | MISSENSE | gCa/gAa | A27E | PLES_26801 | T | T | T | T | . | . | . | . | . | . | T | T |
| NC_011770 | 2876848 | C | T | 8 | 6 | 5 | 0 | NON_SYNONYMOU | MODERAT | MISSENSE | Gcg/Acg | A24T | PLES_26801 | T | T | T | T | . | . | . | . | . | . | . | T |
| NC_011770 | 2876857 | G | C | 8 | 6 | 6 | 0 | NON_SYNONYMOU | MODERAT | MISSENSE | Ctt/Gtt | L21V | PLES_26801 | C | C | C | C | . | . | . | . | . | . | C | C |
| NC_011770 | 2876859 | T | G | 8 | 6 | 6 | 0 | NON_SYNONYMOU | MODERAT | MISSENSE | gAc/gCc | D20A | PLES_26801 | G | G | G | G | . | . | . | . | . | . | G | G |
| NC_011770 | 2876872 | G | T | 29 | 6 | 2 | 0 | NON_SYNONYMOU | MODERAT | MISSENSE | Ccc/Acc | P16T | PLES_26801 | | | | | . | . | . | . | . | . | T | T |
| NC_011770 | 2876877 | T | C | 29 | 6 | 4 | 0 | NON_SYNONYMOU | MODERAT | MISSENSE | gAt/gGt | D14G | PLES_26801 | C | C | C | C | . | . | . | . | . | . | | |
| NC_011770 | 2876884 | T | A | 29 | 6 | 6 | 0 | NON_SYNONYMOU | MODERAT | MISSENSE | Acg/Tcg | T12S | PLES_26801 | A | A | A | A | . | . | . | . | . | . | C | C |
| NC_011770 | 2876904 | G | T | 33 | 6 | 4 | 0 | NON_SYNONYMOU | MODERAT | MISSENSE | gCt/gAt | A5D | PLES_26801 | T | T | T | T | . | . | . | . | . | . | | |
| NC_011770 | 2876905 | C | T | 34 | 6 | 2 | 0 | NON_SYNONYMOU | MODERAT | MISSENSE | Gct/Act | A5T | PLES_26801 | | | | | . | . | . | . | . | . | T | T |
| NC_011770 | 2877023 | T | G | 33 | 6 | 6 | 0 | NON_SYNONYMOU | MODERAT | MISSENSE | Aaa/Caa | K227Q | PLES_26811 | G | G | G | G | . | . | . | . | . | . | G | G |
| NC_011770 | 2877025 | G | A | 33 | 6 | 6 | 0 | NON_SYNONYMOU | MODERAT | MISSENSE | cCa/cTa | P226L | PLES_26811 | A | A | A | A | . | . | . | . | . | . | A | A |
| NC_011770 | 2877078 | C | A | 32 | 6 | 6 | 0 | NON_SYNONYMOU | MODERAT | MISSENSE | gaG/gaT | E208D | PLES_26811 | A | A | A | A | . | . | . | . | . | . | T | T |
| NC_011770 | 2877110 | T | C | 33 | 6 | 6 | 0 | NON_SYNONYMOU | MODERAT | MISSENSE | Acc/Gcc | T198A | PLES_26811 | C | C | C | C | . | . | . | . | . | . | C | C |
| NC_011770 | 2877144 | C | A | 32 | 6 | 4 | 0 | NON_SYNONYMOU | MODERAT | MISSENSE | gaG/gaT | E186D | PLES_26811 | A | A | A | A | . | . | . | . | . | . | | |
| NC_011770 | 2877173 | G | A | 24 | 6 | 4 | 0 | NON_SYNONYMOU | MODERAT | MISSENSE | Ctc/Ttc | L177F | PLES_26811 | A | A | A | A | . | . | . | . | . | . | | |
| NC_011770 | 2877178 | G | A | 24 | 6 | 4 | 0 | NON_SYNONYMOU | MODERAT | MISSENSE | gCc/gTc | A175V | PLES_26811 | A | A | A | A | . | . | . | . | . | . | | |
| NC_011770 | 2877186 | G | C | 24 | 6 | 6 | 0 | NON_SYNONYMOU | MODERAT | MISSENSE | gaC/gaG | D172E | PLES_26811 | C | C | C | C | . | . | . | . | . | . | A | A |
| NC_011770 | 2877215 | T | C | 28 | 6 | 6 | 0 | NON_SYNONYMOU | MODERAT | MISSENSE | Acg/Gcg | T163A | PLES_26811 | C | C | C | C | . | . | . | . | . | . | C | C |
| NC_011770 | 2877222 | CGGT | C | 27 | 6 | 2 | 0 | CODON_CHANGE_ | MODERATE | | ccaccg/ccg | PP159P | PLES_26811 | | | | | . | . | . | . | . | . | C | C |
| NC_011770 | 2877224 | G | T | 17 | 8 | 4 | 0 | NON_SYNONYMOU | MODERAT | MISSENSE | Ccg/Acg | P160T | PLES_26811 | T | T | T | T | . | . | . | . | . | . | . | . |
| NC_011770 | 2877275 | C | T | 30 | 6 | 2 | 0 | NON_SYNONYMOU | MODERAT | MISSENSE | Gcg/Acg | A143T | PLES_26811 | | | | | . | . | . | . | . | . | T | T |
| NC_011770 | 2877299 | G | A | 30 | 6 | 2 | 0 | NON_SYNONYMOU | MODERAT | MISSENSE | Ccc/Tcc | P135S | PLES_26811 | | | | | . | . | . | . | . | . | A | A |
| NC_011770 | 2877314 | T | C | 29 | 6 | 6 | 0 | NON_SYNONYMOU | MODERAT | MISSENSE | Acc/Gcc | T130A | PLES_26811 | C | C | C | C | . | . | . | . | . | . | C | C |
| NC_011770 | 2877331 | G | T | 30 | 6 | 4 | 0 | NON_SYNONYMOU | MODERAT | MISSENSE | gCa/gAa | A124E | PLES_26811 | T | T | T | T | . | . | . | . | . | . | | |
| NC_011770 | 2877332 | C | T | 30 | 6 | 2 | 0 | NON_SYNONYMOU | MODERAT | MISSENSE | Gca/Aca | A124T | PLES_26811 | | | | | . | . | . | . | . | . | T | T |
| NC_011770 | 2877344 | C | T | 30 | 6 | 6 | 0 | NON_SYNONYMOU | MODERAT | MISSENSE | Gcg/Acg | A120T | PLES_26811 | T | T | T | T | . | . | . | . | . | . | T | T |
| NC_011770 | 2877349 | T | A | 30 | 6 | 4 | 0 | NON_SYNONYMOU | MODERAT | MISSENSE | cAa/cTa | Q118L | PLES_26811 | A | A | A | A | . | . | . | . | . | . | | |
| NC_011770 | 2877351 | C | A | 30 | 6 | 2 | 0 | NON_SYNONYMOU | MODERAT | MISSENSE | gaG/gaT | E117D | PLES_26811 | | | | | . | . | . | . | . | . | A | A |
| NC_011770 | 2877372 | G | T | 30 | 6 | 6 | 0 | NON_SYNONYMOU | MODERAT | MISSENSE | gaC/gaA | D110E | PLES_26811 | T | T | T | T | . | . | . | . | . | . | T | T |
| NC_011770 | 2877380 | T | G | 30 | 6 | 2 | 0 | NON_SYNONYMOU | MODERAT | MISSENSE | Acc/Ccc | T108P | PLES_26811 | | | | | . | . | . | . | . | . | G | G |
| NC_011770 | 2877445 | C | T | 11 | 8 | 4 | 0 | NON_SYNONYMOU | MODERAT | MISSENSE | aGc/aAc | S86N | PLES_26811 | T | T | T | T | . | . | . | . | . | . | . | . |
| NC_011770 | 2877453 | G | C | 11 | 8 | 4 | 0 | NON_SYNONYMOU | MODERAT | MISSENSE | atC/atG | I83M | PLES_26811 | C | C | C | C | . | . | . | . | . | . | . | . |
| NC_011770 | 2877455 | T | C | 11 | 8 | 4 | 0 | NON_SYNONYMOU | MODERAT | MISSENSE | Atc/Gtc | I83V | PLES_26811 | C | C | C | C | . | . | . | . | . | . | . | . |

| | | | | | | | | | | | | | | | | | | | | | | | | | |
|---|---|---|---|---|---|---|---|---|---|---|---|---|---|---|---|---|---|---|---|---|---|---|---|---|---|
| NC_011770 | 2877466 | A | G | 11 | 8 | 4 | 0 | NON_SYNONYMOU | MODERAT | MISSENSE | aTg/aCg | M79T | PLES_26811 | G | G | G | G | . | . | . | . | . | . | . | . |
| NC_011770 | 2877467 | T | C | 11 | 8 | 4 | 0 | NON_SYNONYMOU | MODERAT | MISSENSE | Atg/Gtg | M79V | PLES_26811 | C | C | C | C | . | . | . | . | . | . | . | . |
| NC_011770 | 2877475 | C | G | 11 | 8 | 4 | 0 | NON_SYNONYMOU | MODERAT | MISSENSE | gGc/gCc | G76A | PLES_26811 | G | G | G | G | . | . | . | . | . | . | . | . |
| NC_011770 | 2877494 | G | C | 15 | 8 | 4 | 0 | NON_SYNONYMOU | MODERAT | MISSENSE | Ccg/Gcg | P70A | PLES_26811 | C | C | C | C | . | . | . | . | . | . | . | . |
| NC_011770 | 2877504 | C | G | 15 | 8 | 4 | 0 | NON_SYNONYMOU | MODERAT | MISSENSE | agG/agC | R66S | PLES_26811 | G | G | G | G | . | . | . | . | . | . | . | . |
| NC_011770 | 2877512 | T | C | 15 | 8 | 4 | 0 | NON_SYNONYMOU | MODERAT | MISSENSE | Atc/Gtc | I64V | PLES_26811 | C | C | C | C | . | . | . | . | . | . | . | . |
| NC_011770 | 2877516 | T | A | 15 | 8 | 4 | 0 | NON_SYNONYMOU | MODERAT | MISSENSE | gaA/gaT | E62D | PLES_26811 | A | A | A | A | . | . | . | . | . | . | . | . |
| NC_011770 | 2877557 | G | C | 15 | 8 | 4 | 0 | NON_SYNONYMOU | MODERAT | MISSENSE | Ccg/Gcg | P49A | PLES_26811 | C | C | C | C | . | . | . | . | . | . | . | . |
| NC_011770 | 2877568 | T | G | 7 | 8 | 4 | 0 | NON_SYNONYMOU | MODERAT | MISSENSE | gAg/gCg | E45A | PLES_26811 | G | G | G | G | . | . | . | . | . | . | . | . |
| NC_011770 | 2877574 | G | T | 7 | 8 | 4 | 0 | NON_SYNONYMOU | MODERAT | MISSENSE | aCt/aAt | T43N | PLES_26811 | T | T | T | T | . | . | . | . | . | . | . | . |
| NC_011770 | 2877575 | T | C | 7 | 8 | 4 | 0 | NON_SYNONYMOU | MODERAT | MISSENSE | Act/Gct | T43A | PLES_26811 | C | C | C | C | . | . | . | . | . | . | . | . |
| NC_011770 | 2877581 | G | T | 7 | 8 | 4 | 0 | NON_SYNONYMOU | MODERAT | MISSENSE | Ctc/Atc | L41I | PLES_26811 | T | T | T | T | . | . | . | . | . | . | . | . |
| NC_011770 | 2877583 | T | G | 8 | 8 | 4 | 0 | NON_SYNONYMOU | MODERAT | MISSENSE | cAc/cCc | H40P | PLES_26811 | G | G | G | G | . | . | . | . | . | . | . | . |
| NC_011770 | 2877590 | C | A | 8 | 8 | 4 | 0 | NON_SYNONYMOU | MODERAT | MISSENSE | Gcg/Tcg | A38S | PLES_26811 | A | A | A | A | . | . | . | . | . | . | . | . |
| NC_011770 | 2878185 | T | C | 5 | 10 | 2 | 0 | NON_SYNONYMOU | MODERAT | MISSENSE | Agc/Ggc | S147G | PLES_26821 | . | . | C | C | . | . | . | . | . | . | . | . |
| NC_011770 | 2878213 | G | C | 4 | 10 | 2 | 0 | NON_SYNONYMOU | MODERAT | MISSENSE | atC/atG | I137M | PLES_26821 | . | . | C | C | . | . | . | . | . | . | . | . |
| NC_011770 | 2878215 | T | C | 5 | 10 | 2 | 0 | NON_SYNONYMOU | MODERAT | MISSENSE | Atc/Gtc | I137V | PLES_26821 | . | . | C | C | . | . | . | . | . | . | . | . |
| NC_011770 | 2878223 | G | A | 5 | 10 | 2 | 0 | NON_SYNONYMOU | MODERAT | MISSENSE | aCc/aTc | T134I | PLES_26821 | . | . | A | A | . | . | . | . | . | . | . | . |
| NC_011770 | 2878224 | T | C | 4 | 10 | 2 | 0 | NON_SYNONYMOU | MODERAT | MISSENSE | Acc/Gcc | T134A | PLES_26821 | . | . | C | C | . | . | . | . | . | . | . | . |
| NC_011770 | 2878249 | G | C | 6 | 10 | 2 | 0 | NON_SYNONYMOU | MODERAT | MISSENSE | caC/caG | H125Q | PLES_26821 | . | . | C | C | . | . | . | . | . | . | . | . |
| NC_011770 | 2878307 | C | G | 2 | 10 | 2 | 0 | NON_SYNONYMOU | MODERAT | MISSENSE | aGc/aCc | S106T | PLES_26821 | . | . | . | . | . | . | . | . | . | . | G | G |
| NC_011770 | 2878308 | T | A | 2 | 10 | 2 | 0 | NON_SYNONYMOU | MODERAT | MISSENSE | Agc/Tgc | S106C | PLES_26821 | . | . | . | . | . | . | . | . | . | . | A | A |
| NC_011770 | 2878318 | G | C | 2 | 10 | 2 | 0 | NON_SYNONYMOU | MODERAT | MISSENSE | atC/atG | I102M | PLES_26821 | . | . | . | . | . | . | . | . | . | . | C | C |
| NC_011770 | 2878320 | T | C | 2 | 10 | 2 | 0 | NON_SYNONYMOU | MODERAT | MISSENSE | Atc/Gtc | I102V | PLES_26821 | . | . | . | . | . | . | . | . | . | . | C | C |
| NC_011770 | 2878336 | G | C | 2 | 10 | 2 | 0 | NON_SYNONYMOU | MODERAT | MISSENSE | gaC/gaG | D96E | PLES_26821 | . | . | . | . | . | . | . | . | . | . | C | C |
| NC_011770 | 2878337 | T | G | 2 | 10 | 2 | 0 | NON_SYNONYMOU | MODERAT | MISSENSE | gAc/gCc | D96A | PLES_26821 | . | . | . | . | . | . | . | . | . | . | G | G |
| NC_011770 | 2878807 | CT | C | 18 | 6 | 4 | 0 | FRAME_SHIFT | HIGH | | caa/ | Q320 | PLES_26831 | C | C | C | C | . | . | . | . | . | . | | |
| NC_011770 | 2879709 | C | CA | 32 | 6 | 2 | 0 | FRAME_SHIFT | HIGH | | cgg/cTgg | R20L? | PLES_26831 | | | | | . | . | . | . | . | . | CA | CA |
| NC_011770 | 2879710 | G | GA | 15 | 6 | 5 | 0 | FRAME_SHIFT | HIGH | | ggc/ggcT | G19G? | PLES_26831 | GA | GA | GA | GA | . | . | . | . | . | . | . | A |
| NC_011770 | 2879726 | G | GA | 32 | 6 | 6 | 0 | FRAME_SHIFT | HIGH | | ttc/ttTc | F14F? | PLES_26831 | GA | GA | GA | GA | . | . | . | . | . | . | GA | GA |
| NC_011770 | 2878804 | G | C | 19 | 6 | 6 | 0 | NON_SYNONYMOU | MODERAT | MISSENSE | Cag/Gag | Q322E | PLES_26831 | C | C | C | C | . | . | . | . | . | . | C | C |
| NC_011770 | 2878858 | C | T | 13 | 6 | 4 | 0 | NON_SYNONYMOU | MODERAT | MISSENSE | Gat/Aat | D304N | PLES_26831 | T | T | T | T | . | . | . | . | . | . | | |
| NC_011770 | 2878860 | T | G | 17 | 6 | 4 | 0 | NON_SYNONYMOU | MODERAT | MISSENSE | gAc/gCc | D303A | PLES_26831 | G | G | G | G | . | . | . | . | . | . | | |
| NC_011770 | 2878870 | A | G | 18 | 6 | 2 | 0 | NON_SYNONYMOU | MODERAT | MISSENSE | Tcc/Ccc | S300P | PLES_26831 | | | | | . | . | . | . | . | . | G | G |
| NC_011770 | 2878876 | A | G | 19 | 6 | 6 | 0 | NON_SYNONYMOU | MODERAT | MISSENSE | Tcc/Ccc | S298P | PLES_26831 | G | G | G | G | . | . | . | . | . | . | G | G |
| NC_011770 | 2878890 | G | T | 20 | 6 | 2 | 0 | NON_SYNONYMOU | MODERAT | MISSENSE | cCg/cAg | P293Q | PLES_26831 | | | | | . | . | . | . | . | . | T | T |
| NC_011770 | 2878894 | G | C | 20 | 6 | 6 | 0 | NON_SYNONYMOU | MODERAT | MISSENSE | Ccg/Gcg | P292A | PLES_26831 | C | C | C | C | . | . | . | . | . | . | A | A |
| NC_011770 | 2878904 | T | A | 21 | 6 | 4 | 0 | NON_SYNONYMOU | MODERAT | MISSENSE | caA/caT | Q288H | PLES_26831 | A | A | A | A | . | . | . | . | . | . | | |
| NC_011770 | 2878963 | C | T | 29 | 6 | 2 | 0 | NON_SYNONYMOU | MODERAT | MISSENSE | Gtc/Atc | V269I | PLES_26831 | | | | | . | . | . | . | . | . | T | T |
| NC_011770 | 2878986 | G | C | 28 | 6 | 2 | 0 | NON_SYNONYMOU | MODERAT | MISSENSE | gCc/gGc | A261G | PLES_26831 | | | | | . | . | . | . | . | . | C | C |
| NC_011770 | 2879001 | T | C | 31 | 6 | 2 | 0 | NON_SYNONYMOU | MODERAT | MISSENSE | cAc/cGc | H256R | PLES_26831 | | | | | . | . | . | . | . | . | C | C |
| NC_011770 | 2879049 | C | T | 33 | 6 | 2 | 0 | NON_SYNONYMOU | MODERAT | MISSENSE | cGt/cAt | R240H | PLES_26831 | | | | | . | . | . | . | . | . | T | T |
| NC_011770 | 2879050 | G | T | 33 | 6 | 2 | 0 | NON_SYNONYMOU | MODERAT | MISSENSE | Cgt/Agt | R240S | PLES_26831 | | | | | . | . | . | . | . | . | T | T |
| NC_011770 | 2879083 | C | T | 35 | 6 | 6 | 0 | NON_SYNONYMOU | MODERAT | MISSENSE | Gtc/Atc | V229I | PLES_26831 | T | T | T | T | . | . | . | . | . | . | T | T |
| NC_011770 | 2879153 | C | T | 37 | 6 | 6 | 0 | NON_SYNONYMOU | MODERAT | MISSENSE | gaG/gaC | E205D | PLES_26831 | T | T | T | T | . | . | . | . | . | . | G | G |
| NC_011770 | 2879163 | T | C | 38 | 5 | 2 | 0 | NON_SYNONYMOU | MODERAT | MISSENSE | gAt/gGt | D202G | PLES_26831 | | | | | . | . | . | . | . | . | C | C |
| NC_011770 | 2879188 | C | T | 28 | 6 | 4 | 0 | NON_SYNONYMOU | MODERAT | MISSENSE | Gtc/Atc | V194I | PLES_26831 | T | T | T | T | . | . | . | . | . | . | | |
| NC_011770 | 2879190 | G | C | 24 | 6 | 2 | 0 | NON_SYNONYMOU | MODERAT | MISSENSE | aCc/aGc | T193S | PLES_26831 | | | | | . | . | . | . | . | . | C | C |
| NC_011770 | 2879191 | T | A | 24 | 6 | 6 | 0 | NON_SYNONYMOU | MODERAT | MISSENSE | Acc/Tcc | T193S | PLES_26831 | A | A | A | A | . | . | . | . | . | . | A | A |
| NC_011770 | 2879206 | A | G | 24 | 6 | 6 | 0 | NON_SYNONYMOU | MODERAT | MISSENSE | Tcg/Ccg | S188P | PLES_26831 | G | G | G | G | . | . | . | . | . | . | G | G |
| NC_011770 | 2879214 | G | T | 24 | 6 | 6 | 0 | NON_SYNONYMOU | MODERAT | MISSENSE | gCt/gAt | A185D | PLES_26831 | T | T | T | T | . | . | . | . | . | . | T | T |
| NC_011770 | 2879235 | A | G | 33 | 6 | 6 | 0 | NON_SYNONYMOU | MODERAT | MISSENSE | gTg/gCg | V178A | PLES_26831 | G | G | G | G | . | . | . | . | . | . | G | G |
| NC_011770 | 2879236 | C | G | 33 | 6 | 6 | 0 | NON_SYNONYMOU | MODERAT | MISSENSE | Gtg/Ctg | V178L | PLES_26831 | G | G | G | G | . | . | . | . | . | . | G | G |
| NC_011770 | 2879241 | T | C | 33 | 6 | 2 | 0 | NON_SYNONYMOU | MODERAT | MISSENSE | aAt/aGt | N176S | PLES_26831 | | | | | . | . | . | . | . | . | C | C |
| NC_011770 | 2879281 | C | T | 35 | 6 | 2 | 0 | NON_SYNONYMOU | MODERAT | MISSENSE | Gag/Aag | E163K | PLES_26831 | | | | | . | . | . | . | . | . | T | T |
| NC_011770 | 2879303 | G | C | 34 | 6 | 4 | 0 | NON_SYNONYMOU | MODERAT | MISSENSE | gaC/gaG | D155E | PLES_26831 | C | C | C | C | . | . | . | . | . | . | | |
| NC_011770 | 2879308 | T | G | 33 | 6 | 2 | 0 | NON_SYNONYMOU | MODERAT | MISSENSE | Atc/Ctc | I154L | PLES_26831 | | | | | . | . | . | . | . | . | G | G |
| NC_011770 | 2879310 | A | G | 33 | 6 | 2 | 0 | NON_SYNONYMOU | MODERAT | MISSENSE | cTg/cCg | L153P | PLES_26831 | | | | | . | . | . | . | . | . | G | G |
| NC_011770 | 2879353 | G | C | 32 | 6 | 2 | 0 | NON_SYNONYMOU | MODERAT | MISSENSE | Cag/Gag | Q139E | PLES_26831 | | | | | . | . | . | . | . | . | C | C |
| NC_011770 | 2879360 | C | G | 33 | 6 | 2 | 0 | NON_SYNONYMOU | MODERAT | MISSENSE | caG/caC | Q136H | PLES_26831 | | | | | . | . | . | . | . | . | G | G |
| NC_011770 | 2879361 | T | C | 33 | 6 | 2 | 0 | NON_SYNONYMOU | MODERAT | MISSENSE | cAg/cGg | Q136R | PLES_26831 | | | | | . | . | . | . | . | . | C | C |
| NC_011770 | 2879385 | A | G | 35 | 6 | 6 | 0 | NON_SYNONYMOU | MODERAT | MISSENSE | gTg/gCg | V128A | PLES_26831 | G | G | G | G | . | . | . | . | . | . | G | G |

| | | | | | | | | | | | | | | | | | | | | | | | | | |
|---|---|---|---|---|---|---|---|---|---|---|---|---|---|---|---|---|---|---|---|---|---|---|---|---|---|
| NC_011770 | 2879437 | G | A | 36 | 6 | 4 | 0 | NON_SYNONYMOU | MODERAT | MISSENSE | Ccc/Tcc | P111S | PLES_26831 | A | A | A | A | . | . | . | . | . | . | | |
| NC_011770 | 2879468 | C | G | 36 | 6 | 2 | 0 | NON_SYNONYMOU | MODERAT | MISSENSE | aaG/aaC | K100N | PLES_26831 | | | | | . | . | . | . | . | . | G | G |
| NC_011770 | 2879548 | T | C | 37 | 6 | 2 | 0 | NON_SYNONYMOU | MODERAT | MISSENSE | Atc/Gtc | I74V | PLES_26831 | | | | | . | . | . | . | . | . | C | C |
| NC_011770 | 2879614 | T | G | 36 | 6 | 4 | 0 | NON_SYNONYMOU | MODERAT | MISSENSE | Atc/Ctc | I52L | PLES_26831 | G | G | G | G | . | . | . | . | . | . | | |
| NC_011770 | 2879688 | T | A | 32 | 6 | 6 | 0 | NON_SYNONYMOU | MODERAT | MISSENSE | cAa/cTa | Q27L | PLES_26831 | A | A | A | A | . | . | . | . | . | . | A | A |
| NC_011770 | 2879736 | G | A | 31 | 6 | 6 | 0 | NON_SYNONYMOU | MODERAT | MISSENSE | tCa/tTa | S11L | PLES_26831 | A | A | A | A | . | . | . | . | . | . | A | A |
| NC_011770 | 2879751 | T | C | 30 | 6 | 2 | 0 | NON_SYNONYMOU | MODERAT | MISSENSE | cAa/cGa | Q6R | PLES_26831 | | | | | . | . | . | . | . | . | C | C |
| NC_011770 | 2880666 | G | C | 5 | 8 | 4 | 0 | NON_SYNONYMOU | MODERAT | MISSENSE | Ctg/Gtg | L58V | PLES_26851 | C | C | C | C | . | . | . | . | . | . | . | . |
| NC_011770 | 2880692 | G | C | 5 | 8 | 4 | 0 | NON_SYNONYMOU | MODERAT | MISSENSE | gCg/gGg | A49G | PLES_26851 | C | C | C | C | . | . | . | . | . | . | . | . |
| NC_011770 | 2880716 | C | G | 4 | 8 | 4 | 0 | NON_SYNONYMOU | MODERAT | MISSENSE | cGc/cCc | R41P | PLES_26851 | G | G | G | G | . | . | . | . | . | . | . | . |
| NC_011770 | 2880717 | G | C | 4 | 8 | 4 | 0 | NON_SYNONYMOU | MODERAT | MISSENSE | Cgc/Ggc | R41G | PLES_26851 | C | C | C | C | . | . | . | . | . | . | . | . |
| NC_011770 | 2880720 | C | G | 4 | 8 | 4 | 0 | NON_SYNONYMOU | MODERAT | MISSENSE | Ggc/Cgc | G40R | PLES_26851 | G | G | G | G | . | . | . | . | . | . | . | . |
| NC_011770 | 2880727 | G | T | 4 | 8 | 4 | 0 | NON_SYNONYMOU | MODERAT | MISSENSE | caC/caA | H37Q | PLES_26851 | T | T | T | T | . | . | . | . | . | . | . | . |
| NC_011770 | 2880729 | G | C | 4 | 8 | 4 | 0 | NON_SYNONYMOU | MODERAT | MISSENSE | Cac/Gac | H37D | PLES_26851 | C | C | C | C | . | . | . | . | . | . | . | . |
| NC_011770 | 2880788 | G | T | 12 | 6 | 6 | 0 | NON_SYNONYMOU | MODERAT | MISSENSE | gCa/gAa | A17E | PLES_26851 | T | T | T | T | . | . | . | . | . | . | T | T |
| NC_011770 | 2880789 | C | T | 12 | 6 | 6 | 0 | NON_SYNONYMOU | MODERAT | MISSENSE | Gca/Aca | A17T | PLES_26851 | T | T | T | T | . | . | . | . | . | . | T | T |
| NC_011770 | 2880791 | T | A | 12 | 6 | 6 | 0 | NON_SYNONYMOU | MODERAT | MISSENSE | cAg/cTg | Q16L | PLES_26851 | A | A | A | A | . | . | . | . | . | . | A | A |
| NC_011770 | 2880792 | G | C | 12 | 6 | 6 | 0 | NON_SYNONYMOU | MODERAT | MISSENSE | Cag/Gag | Q16E | PLES_26851 | C | C | C | C | . | . | . | . | . | . | C | C |
| NC_011770 | 2880818 | C | T | 20 | 6 | 4 | 0 | NON_SYNONYMOU | MODERAT | MISSENSE | cGt/cAt | R7H | PLES_26851 | T | T | T | T | . | . | . | . | . | . | | |
| NC_011770 | 2885458 | G | T | 42 | 6 | 2 | 0 | STOP_GAINED | HIGH | NONSENSE | tCg/tAg | S127* | PLES_26911 | | | | | . | . | . | . | . | . | T | T |
| NC_011770 | 2885074 | T | A | 31 | 6 | 6 | 0 | NON_SYNONYMOU | MODERAT | MISSENSE | cAg/cTg | Q255L | PLES_26911 | A | A | A | A | . | . | . | . | . | . | A | A |
| NC_011770 | 2885172 | GTTC | G | 12 | 8 | 3 | 0 | CODON_CHANGE_ | MODERATE | | cagaac/cac | QN221H | PLES_26911 | . | . | G | . | . | . | . | . | . | . | G | G |
| NC_011770 | 2885223 | G | C | 38 | 6 | 4 | 0 | NON_SYNONYMOU | MODERAT | MISSENSE | atC/atG | I205M | PLES_26911 | C | C | C | C | . | . | . | . | . | . | | |
| NC_011770 | 2885225 | T | C | 37 | 6 | 4 | 0 | NON_SYNONYMOU | MODERAT | MISSENSE | Atc/Gtc | I205V | PLES_26911 | C | C | C | C | . | . | . | . | . | . | | |
| NC_011770 | 2885233 | C | T | 40 | 6 | 2 | 0 | NON_SYNONYMOU | MODERAT | MISSENSE | cGg/cAg | R202Q | PLES_26911 | | | | | . | . | . | . | . | . | T | T |
| NC_011770 | 2885249 | C | T | 41 | 6 | 2 | 0 | NON_SYNONYMOU | MODERAT | MISSENSE | Gcg/Acg | A197T | PLES_26911 | | | | | . | . | . | . | . | . | T | T |
| NC_011770 | 2885293 | T | C | 45 | 6 | 2 | 0 | NON_SYNONYMOU | MODERAT | MISSENSE | aAg/aGg | K182R | PLES_26911 | | | | | . | . | . | . | . | . | C | C |
| NC_011770 | 2885302 | G | T | 44 | 6 | 2 | 0 | NON_SYNONYMOU | MODERAT | MISSENSE | aCg/aAg | T179K | PLES_26911 | | | | | . | . | . | . | . | . | T | T |
| NC_011770 | 2885318 | T | C | 45 | 6 | 2 | 0 | NON_SYNONYMOU | MODERAT | MISSENSE | Acc/Gcc | T174A | PLES_26911 | | | | | . | . | . | . | . | . | C | C |
| NC_011770 | 2885360 | G | A | 47 | 6 | 2 | 0 | NON_SYNONYMOU | MODERAT | MISSENSE | Cgc/Tgc | R160C | PLES_26911 | | | | | . | . | . | . | . | . | A | A |
| NC_011770 | 2885369 | G | C | 45 | 6 | 6 | 0 | NON_SYNONYMOU | MODERAT | MISSENSE | Cgc/Ggc | R157G | PLES_26911 | C | C | C | C | . | . | . | . | . | . | C | C |
| NC_011770 | 2885399 | C | G | 46 | 6 | 4 | 0 | NON_SYNONYMOU | MODERAT | MISSENSE | Gag/Cag | E147Q | PLES_26911 | G | G | G | G | . | . | . | . | . | . | | |
| NC_011770 | 2885431 | G | T | 44 | 6 | 6 | 0 | NON_SYNONYMOU | MODERAT | MISSENSE | aCg/aAg | T136K | PLES_26911 | T | T | T | T | . | . | . | . | . | . | T | T |
| NC_011770 | 2885432 | T | C | 46 | 6 | 6 | 0 | NON_SYNONYMOU | MODERAT | MISSENSE | Acg/Gcg | T136A | PLES_26911 | C | C | C | C | . | . | . | . | . | . | C | C |
| NC_011770 | 2885459 | A | T | 42 | 6 | 2 | 0 | NON_SYNONYMOU | MODERAT | MISSENSE | Tcg/Acg | S127T | PLES_26911 | | | | | . | . | . | . | . | . | T | T |
| NC_011770 | 2885473 | C | G | 26 | 6 | 4 | 0 | NON_SYNONYMOU | MODERAT | MISSENSE | aGg/aCg | R122T | PLES_26911 | G | G | G | G | . | . | . | . | . | . | | |
| NC_011770 | 2885476 | G | T | 27 | 6 | 4 | 0 | NON_SYNONYMOU | MODERAT | MISSENSE | aCg/aAg | T121K | PLES_26911 | T | T | T | T | . | . | . | . | . | . | | |
| NC_011770 | 2885480 | T | C | 27 | 6 | 4 | 0 | NON_SYNONYMOU | MODERAT | MISSENSE | Atc/Gtc | I120V | PLES_26911 | C | C | C | C | . | . | . | . | . | . | | |
| NC_011770 | 2885491 | G | A | 28 | 6 | 4 | 0 | NON_SYNONYMOU | MODERAT | MISSENSE | aCg/aTg | T116M | PLES_26911 | A | A | A | A | . | . | . | . | . | . | | |
| NC_011770 | 2885506 | T | A | 27 | 6 | 4 | 0 | NON_SYNONYMOU | MODERAT | MISSENSE | gAg/gTg | E111V | PLES_26911 | A | A | A | A | . | . | . | . | . | . | | |
| NC_011770 | 2885507 | C | T | 28 | 6 | 6 | 0 | NON_SYNONYMOU | MODERAT | MISSENSE | Gag/Aag | E111K | PLES_26911 | T | T | T | T | . | . | . | . | . | . | T | T |
| NC_011770 | 2885515 | A | C | 26 | 6 | 4 | 0 | NON_SYNONYMOU | MODERAT | MISSENSE | aTc/aGc | I108S | PLES_26911 | C | C | C | C | . | . | . | . | . | . | | |
| NC_011770 | 2885516 | T | G | 26 | 6 | 6 | 0 | NON_SYNONYMOU | MODERAT | MISSENSE | Atc/Ctc | I108L | PLES_26911 | G | G | G | G | . | . | . | . | . | . | C | C |
| NC_011770 | 2885518 | A | T | 25 | 6 | 4 | 0 | NON_SYNONYMOU | MODERAT | MISSENSE | gTc/gAc | V107D | PLES_26911 | T | T | T | T | . | . | . | . | . | . | | |
| NC_011770 | 2885593 | G | T | 40 | 6 | 6 | 0 | NON_SYNONYMOU | MODERAT | MISSENSE | gCc/gAc | A82D | PLES_26911 | T | T | T | T | . | . | . | . | . | . | C | C |
| NC_011770 | 2885594 | C | T | 41 | 6 | 2 | 0 | NON_SYNONYMOU | MODERAT | MISSENSE | Gcc/Acc | A82T | PLES_26911 | | | | | . | . | . | . | . | . | T | T |
| NC_011770 | 2885613 | G | C | 47 | 6 | 4 | 0 | NON_SYNONYMOU | MODERAT | MISSENSE | agC/agG | S75R | PLES_26911 | C | C | C | C | . | . | . | . | . | . | | |
| NC_011770 | 2885614 | C | T | 46 | 6 | 6 | 0 | NON_SYNONYMOU | MODERAT | MISSENSE | aGc/aAc | S75N | PLES_26911 | T | T | T | T | . | . | . | . | . | . | T | T |
| NC_011770 | 2885618 | G | A | 47 | 6 | 4 | 0 | NON_SYNONYMOU | MODERAT | MISSENSE | Ccc/Tcc | P74S | PLES_26911 | A | A | A | A | . | . | . | . | . | . | | |
| NC_011770 | 2885638 | G | C | 47 | 6 | 4 | 0 | NON_SYNONYMOU | MODERAT | MISSENSE | tCc/tGc | S67C | PLES_26911 | C | C | C | C | . | . | . | . | . | . | | |
| NC_011770 | 2885639 | A | T | 46 | 6 | 4 | 0 | NON_SYNONYMOU | MODERAT | MISSENSE | Tcc/Acc | S67T | PLES_26911 | T | T | T | T | . | . | . | . | . | . | | |
| NC_011770 | 2885674 | G | A | 47 | 6 | 4 | 0 | NON_SYNONYMOU | MODERAT | MISSENSE | gCg/gTg | A55V | PLES_26911 | A | A | A | A | . | . | . | . | . | . | | |
| NC_011770 | 2885675 | C | T | 47 | 6 | 4 | 0 | NON_SYNONYMOU | MODERAT | MISSENSE | Gcg/Acg | A55T | PLES_26911 | T | T | T | T | . | . | . | . | . | . | | |
| NC_011770 | 2885678 | C | A | 49 | 6 | 2 | 0 | NON_SYNONYMOU | MODERAT | MISSENSE | Gcc/Tcc | A54S | PLES_26911 | | | | | . | . | . | . | . | . | A | A |
| NC_011770 | 2885693 | C | T | 51 | 6 | 2 | 0 | NON_SYNONYMOU | MODERAT | MISSENSE | Gac/Aac | D49N | PLES_26911 | | | | | . | . | . | . | . | . | T | T |
| NC_011770 | 2885711 | T | C | 49 | 6 | 4 | 0 | NON_SYNONYMOU | MODERAT | MISSENSE | Atc/Gtc | I43V | PLES_26911 | C | C | C | C | . | . | . | . | . | . | | |
| NC_011770 | 2885734 | A | T | 44 | 6 | 6 | 0 | NON_SYNONYMOU | MODERAT | MISSENSE | cTg/cAg | L35Q | PLES_26911 | T | T | T | T | . | . | . | . | . | . | T | T |
| NC_011770 | 2885332 | C | T | 45 | 6 | 1 | 0 | NON_SYNONYMOU | MODERAT | MISSENSE | gGt/gAt | G169D | PLES_26911 | | | | | . | . | . | . | . | . | T | |
| NC_011770 | 2886134 | G | C | 42 | 6 | 6 | 0 | NON_SYNONYMOU | MODERAT | MISSENSE | Caa/Gaa | Q91E | PLES_26921 | C | C | C | C | . | . | . | . | . | . | C | C |
| NC_011770 | 2886236 | C | T | 46 | 6 | 6 | 0 | NON_SYNONYMOU | MODERAT | MISSENSE | Gtg/Atg | V57M | PLES_26921 | T | T | T | T | . | . | . | . | . | . | T | T |
| NC_011770 | 2886262 | G | T | 49 | 6 | 4 | 0 | NON_SYNONYMOU | MODERAT | MISSENSE | gCc/gAc | A48D | PLES_26921 | T | T | T | T | . | . | . | . | . | . | | |
| NC_011770 | 2886327 | T | G | 45 | 6 | 6 | 0 | NON_SYNONYMOU | MODERAT | MISSENSE | gaA/gaC | E26D | PLES_26921 | G | G | G | G | . | . | . | . | . | . | G | G |

| | | | | | | | | | | | | | | | | | | | | | | | | | |
|---|---|---|---|---|---|---|---|---|---|---|---|---|---|---|---|---|---|---|---|---|---|---|---|---|---|
| NC_011770 | 2886335 | T | C | 43 | 6 | 2 | 0 | NON_SYNONYMOU | MODERAT | MISSENSE | Agc/Ggc | S24G | PLES_26921 | | | | | . | . | . | . | . | . | C | C |
| NC_011770 | 2886365 | G | T | 15 | 9 | 1 | 0 | NON_SYNONYMOU | MODERAT | MISSENSE | Cac/Aac | H14N | PLES_26921 | . | . | . | T | . | . | . | . | . | . | | |
| NC_011770 | 2886591 | G | T | 35 | 6 | 2 | 0 | NON_SYNONYMOU | MODERAT | MISSENSE | Ctt/Att | L217I | PLES_26931 | | | | | . | . | . | . | . | . | T | T |
| NC_011770 | 2886594 | T | C | 30 | 6 | 2 | 0 | NON_SYNONYMOU | MODERAT | MISSENSE | Atc/Gtc | I216V | PLES_26931 | | | | | . | . | . | . | . | . | C | C |
| NC_011770 | 2886595 | C | G | 30 | 6 | 6 | 0 | NON_SYNONYMOU | MODERAT | MISSENSE | caG/caC | Q215H | PLES_26931 | G | G | G | G | . | . | . | . | . | . | G | G |
| NC_011770 | 2886597 | G | T | 30 | 6 | 6 | 0 | NON_SYNONYMOU | MODERAT | MISSENSE | Cag/Aag | Q215K | PLES_26931 | T | T | T | T | . | . | . | . | . | . | T | T |
| NC_011770 | 2886604 | C | G | 31 | 6 | 6 | 0 | NON_SYNONYMOU | MODERAT | MISSENSE | gaG/gaC | E212D | PLES_26931 | G | G | G | G | . | . | . | . | . | . | G | G |
| NC_011770 | 2886623 | C | T | 32 | 6 | 4 | 0 | NON_SYNONYMOU | MODERAT | MISSENSE | aGc/aAc | S206N | PLES_26931 | T | T | T | T | . | . | . | . | . | . | | |
| NC_011770 | 2886624 | T | C | 32 | 6 | 4 | 0 | NON_SYNONYMOU | MODERAT | MISSENSE | Agc/Ggc | S206G | PLES_26931 | C | C | C | C | . | . | . | . | . | . | | |
| NC_011770 | 2886653 | G | C | 18 | 6 | 6 | 0 | NON_SYNONYMOU | MODERAT | MISSENSE | gCc/gGc | A196G | PLES_26931 | C | C | C | C | . | . | . | . | . | . | C | C |
| NC_011770 | 2886654 | C | T | 17 | 6 | 6 | 0 | NON_SYNONYMOU | MODERAT | MISSENSE | Gcc/Acc | A196T | PLES_26931 | T | T | T | T | . | . | . | . | . | . | T | T |
| NC_011770 | 2886665 | A | G | 18 | 6 | 6 | 0 | NON_SYNONYMOU | MODERAT | MISSENSE | gTc/gCc | V192A | PLES_26931 | G | G | G | G | . | . | . | . | . | . | G | G |
| NC_011770 | 2886669 | C | A | 18 | 6 | 6 | 0 | NON_SYNONYMOU | MODERAT | MISSENSE | Gac/Tac | D191Y | PLES_26931 | A | A | A | A | . | . | . | . | . | . | G | G |
| NC_011770 | 2886670 | C | T | 18 | 6 | 6 | 0 | NON_SYNONYMOU | MODERAT | MISSENSE | caG/caC | Q190H | PLES_26931 | T | T | T | T | . | . | . | . | . | . | G | G |
| NC_011770 | 2886672 | G | C | 18 | 6 | 6 | 0 | NON_SYNONYMOU | MODERAT | MISSENSE | Cag/Gag | Q190E | PLES_26931 | C | C | C | C | . | . | . | . | . | . | C | C |
| NC_011770 | 2886678 | G | T | 18 | 6 | 6 | 0 | NON_SYNONYMOU | MODERAT | MISSENSE | Cac/Aac | H188N | PLES_26931 | T | T | T | T | . | . | . | . | . | . | T | T |
| NC_011770 | 2886681 | G | T | 18 | 6 | 6 | 0 | NON_SYNONYMOU | MODERAT | MISSENSE | Cag/Aag | Q187K | PLES_26931 | T | T | T | T | . | . | . | . | . | . | T | T |
| NC_011770 | 2886711 | C | T | 33 | 6 | 2 | 0 | NON_SYNONYMOU | MODERAT | MISSENSE | Gtc/Atc | V177I | PLES_26931 | | | | | . | . | . | . | . | . | T | T |
| NC_011770 | 2886735 | T | C | 31 | 6 | 6 | 0 | NON_SYNONYMOU | MODERAT | MISSENSE | Att/Gtt | I169V | PLES_26931 | C | C | C | C | . | . | . | . | . | . | C | C |
| NC_011770 | 2886809 | C | T | 31 | 6 | 6 | 0 | NON_SYNONYMOU | MODERAT | MISSENSE | cGc/cAc | R144H | PLES_26931 | T | T | T | T | . | . | . | . | . | . | T | T |
| NC_011770 | 2886813 | T | A | 32 | 6 | 4 | 0 | NON_SYNONYMOU | MODERAT | MISSENSE | Agc/Tgc | S143C | PLES_26931 | A | A | A | A | . | . | . | . | . | . | | |
| NC_011770 | 2886831 | T | C | 33 | 6 | 4 | 0 | NON_SYNONYMOU | MODERAT | MISSENSE | Atc/Gtc | I137V | PLES_26931 | C | C | C | C | . | . | . | . | . | . | | |
| NC_011770 | 2886883 | T | G | 28 | 6 | 4 | 0 | NON_SYNONYMOU | MODERAT | MISSENSE | gaA/gaC | E119D | PLES_26931 | G | G | G | G | . | . | . | . | . | . | | |
| NC_011770 | 2886909 | T | C | 27 | 6 | 2 | 0 | NON_SYNONYMOU | MODERAT | MISSENSE | Acc/Gcc | T111A | PLES_26931 | | | | | . | . | . | . | . | . | C | C |
| NC_011770 | 2886918 | T | C | 24 | 6 | 6 | 0 | NON_SYNONYMOU | MODERAT | MISSENSE | Att/Gtt | I108V | PLES_26931 | C | C | C | C | . | . | . | . | . | . | C | C |
| NC_011770 | 2886924 | C | T | 29 | 6 | 2 | 0 | NON_SYNONYMOU | MODERAT | MISSENSE | Gtc/Atc | V106I | PLES_26931 | | | | | . | . | . | . | . | . | T | T |
| NC_011770 | 2886939 | A | G | 30 | 6 | 4 | 0 | NON_SYNONYMOU | MODERAT | MISSENSE | Ttc/Ctc | F101L | PLES_26931 | G | G | G | G | . | . | . | . | . | . | | |
| NC_011770 | 2886978 | T | C | 14 | 8 | 4 | 0 | NON_SYNONYMOU | MODERAT | MISSENSE | Atg/Gtg | M88V | PLES_26931 | C | C | C | C | . | . | . | . | . | . | . | . |
| NC_011770 | 2887128 | G | T | 14 | 7 | 5 | 0 | NON_SYNONYMOU | MODERAT | MISSENSE | Ccg/Acg | P38T | PLES_26931 | T | T | T | T | . | . | . | . | . | . | C | . |
| NC_011770 | 2887136 | T | G | 25 | 6 | 6 | 0 | NON_SYNONYMOU | MODERAT | MISSENSE | aAc/aCc | N35T | PLES_26931 | G | G | G | G | . | . | . | . | . | . | A | A |
| NC_011770 | 2887137 | T | A | 26 | 6 | 4 | 0 | NON_SYNONYMOU | MODERAT | MISSENSE | Aac/Tac | N35Y | PLES_26931 | A | A | A | A | . | . | . | . | . | . | | |
| NC_011770 | 2887162 | C | G | 29 | 6 | 2 | 0 | NON_SYNONYMOU | MODERAT | MISSENSE | gaG/gaC | E26D | PLES_26931 | | | | | . | . | . | . | . | . | G | G |
| NC_011770 | 2887163 | T | C | 29 | 6 | 4 | 0 | NON_SYNONYMOU | MODERAT | MISSENSE | gAg/gGg | E26G | PLES_26931 | C | C | C | C | . | . | . | . | . | . | | |
| NC_011770 | 2887170 | T | A | 30 | 6 | 6 | 0 | NON_SYNONYMOU | MODERAT | MISSENSE | Agc/Tgc | S24C | PLES_26931 | A | A | A | A | . | . | . | . | . | . | C | C |
| NC_011770 | 2887175 | G | C | 31 | 6 | 6 | 0 | NON_SYNONYMOU | MODERAT | MISSENSE | tCc/tGc | S22C | PLES_26931 | C | C | C | C | . | . | . | . | . | . | C | C |
| NC_011770 | 2887191 | T | C | 30 | 6 | 4 | 0 | NON_SYNONYMOU | MODERAT | MISSENSE | Atc/Gtc | I17V | PLES_26931 | C | C | C | C | . | . | . | . | . | . | | |
| NC_011770 | 2887193 | T | C | 30 | 6 | 4 | 0 | NON_SYNONYMOU | MODERAT | MISSENSE | aAa/aGa | K16R | PLES_26931 | C | C | C | C | . | . | . | . | . | . | | |
| NC_011770 | 2887215 | G | C | 30 | 6 | 4 | 0 | NON_SYNONYMOU | MODERAT | MISSENSE | Cta/Gta | L9V | PLES_26931 | C | C | C | C | . | . | . | . | . | . | | |
| NC_011770 | 2887228 | A | T | 28 | 6 | 2 | 0 | NON_SYNONYMOU | MODERAT | MISSENSE | aaT/aaA | N4K | PLES_26931 | | | | | . | . | . | . | . | . | T | T |
| NC_011770 | 2887236 | A | T | 27 | 6 | 4 | 0 | NON_SYNONYMOU | MODERAT | MISSENSE | Tcc/Acc | S2T | PLES_26931 | T | T | T | T | . | . | . | . | . | . | | |
| NC_011770 | 2887472 | C | G | 11 | 10 | 2 | 0 | NON_SYNONYMOU | MODERAT | MISSENSE | Gtg/Ctg | V180L | PLES_26941 | . | . | . | . | . | . | . | . | . | . | G | G |
| NC_011770 | 2887479 | C | G | 11 | 10 | 2 | 0 | NON_SYNONYMOU | MODERAT | MISSENSE | aaG/aaC | K177N | PLES_26941 | . | . | . | . | . | . | . | . | . | . | G | G |
| NC_011770 | 2887520 | A | T | 7 | 10 | 2 | 0 | NON_SYNONYMOU | MODERAT | MISSENSE | Tcc/Acc | S164T | PLES_26941 | . | . | . | . | . | . | . | . | . | . | T | T |
| NC_011770 | 2887521 | G | T | 7 | 10 | 2 | 0 | NON_SYNONYMOU | MODERAT | MISSENSE | agC/agA | S163R | PLES_26941 | . | . | . | . | . | . | . | . | . | . | T | T |
| NC_011770 | 2887526 | T | A | 7 | 10 | 2 | 0 | NON_SYNONYMOU | MODERAT | MISSENSE | Atc/Ttc | I162F | PLES_26941 | . | . | . | . | . | . | . | . | . | . | A | A |
| NC_011770 | 2887529 | A | C | 7 | 10 | 2 | 0 | NON_SYNONYMOU | MODERAT | MISSENSE | Ttc/Gtc | F161V | PLES_26941 | . | . | . | . | . | . | . | . | . | . | C | C |
| NC_011770 | 2887531 | C | T | 7 | 10 | 2 | 0 | NON_SYNONYMOU | MODERAT | MISSENSE | cGc/cAc | R160H | PLES_26941 | . | . | . | . | . | . | . | . | . | . | T | T |
| NC_011770 | 2887628 | G | T | 13 | 10 | 2 | 0 | NON_SYNONYMOU | MODERAT | MISSENSE | Cac/Aac | H128N | PLES_26941 | . | . | . | . | . | . | . | . | . | . | T | T |
| NC_011770 | 2887747 | A | G | 1 | 10 | 2 | 0 | NON_SYNONYMOU | MODERAT | MISSENSE | gTg/gCg | V88A | PLES_26941 | . | . | . | . | . | . | . | . | . | . | G | G |
| NC_011770 | 2887756 | A | C | 1 | 10 | 2 | 0 | NON_SYNONYMOU | MODERAT | MISSENSE | cTg/cGg | L85R | PLES_26941 | . | . | . | . | . | . | . | . | . | . | C | C |
| NC_011770 | 2887783 | C | T | 1 | 10 | 2 | 0 | NON_SYNONYMOU | MODERAT | MISSENSE | gGg/gAg | G76E | PLES_26941 | . | . | . | . | . | . | . | . | . | . | T | T |
| NC_011770 | 2887826 | C | T | 1 | 10 | 2 | 0 | NON_SYNONYMOU | MODERAT | MISSENSE | Gac/Aac | D62N | PLES_26941 | . | . | . | . | . | . | . | . | . | . | T | T |
| NC_011770 | 2887912 | C | T | 15 | 10 | 2 | 0 | NON_SYNONYMOU | MODERAT | MISSENSE | aGc/aAc | S33N | PLES_26941 | . | . | . | . | . | . | . | . | . | . | T | T |
| NC_011770 | 2887946 | G | T | 16 | 10 | 2 | 0 | NON_SYNONYMOU | MODERAT | MISSENSE | Cag/Aag | Q22K | PLES_26941 | . | . | . | . | . | . | . | . | . | . | T | T |
| NC_011770 | 2887967 | C | T | 15 | 10 | 2 | 0 | NON_SYNONYMOU | MODERAT | MISSENSE | Gac/Aac | D15N | PLES_26941 | . | . | . | . | . | . | . | . | . | . | T | T |
| NC_011770 | 2887987 | G | C | 14 | 10 | 2 | 0 | NON_SYNONYMOU | MODERAT | MISSENSE | gCc/gGc | A8G | PLES_26941 | . | . | . | . | . | . | . | . | . | . | C | C |
| NC_011770 | 2888212 | G | C | 39 | 6 | 6 | 0 | NON_SYNONYMOU | MODERAT | MISSENSE | Ctg/Gtg | L125V | PLES_26951 | C | C | C | C | . | . | . | . | . | . | C | C |
| NC_011770 | 2888233 | G | T | 39 | 6 | 6 | 0 | NON_SYNONYMOU | MODERAT | MISSENSE | Cag/Aag | Q118K | PLES_26951 | T | T | T | T | . | . | . | . | . | . | T | T |
| NC_011770 | 2888242 | G | C | 40 | 6 | 2 | 0 | NON_SYNONYMOU | MODERAT | MISSENSE | Caa/Gaa | Q115E | PLES_26951 | | | | | . | . | . | . | . | . | C | C |
| NC_011770 | 2888273 | T | G | 43 | 6 | 2 | 0 | NON_SYNONYMOU | MODERAT | MISSENSE | caA/caC | Q104H | PLES_26951 | | | | | . | . | . | . | . | . | G | G |
| NC_011770 | 2888280 | T | C | 41 | 6 | 2 | 0 | NON_SYNONYMOU | MODERAT | MISSENSE | cAc/cGc | H102R | PLES_26951 | | | | | . | . | . | . | . | . | C | C |
| NC_011770 | 2888329 | A | C | 42 | 6 | 2 | 0 | NON_SYNONYMOU | MODERAT | MISSENSE | Ttc/Gtc | F86V | PLES_26951 | | | | | . | . | . | . | . | . | C | C |

| | | | | | | | | | | | | | | | | | | | | | | | | | |
|---|---|---|---|---|---|---|---|---|---|---|---|---|---|---|---|---|---|---|---|---|---|---|---|---|---|
| NC_011770 | 2888385 | G | A | 38 | 6 | 2 | 0 | NON_SYNONYMOU | MODERAT | MISSENSE | gCc/gTc | A67V | PLES_26951 | | | | | . | . | . | . | . | . | A | A |
| NC_011770 | 2888386 | C | G | 39 | 6 | 2 | 0 | NON_SYNONYMOU | MODERAT | MISSENSE | Gcc/Ccc | A67P | PLES_26951 | | | | | . | . | . | . | . | . | G | G |
| NC_011770 | 2888418 | C | G | 41 | 6 | 2 | 0 | NON_SYNONYMOU | MODERAT | MISSENSE | aGc/aCc | S56T | PLES_26951 | | | | | . | . | . | . | . | . | G | G |
| NC_011770 | 2888464 | G | C | 38 | 6 | 2 | 0 | NON_SYNONYMOU | MODERAT | MISSENSE | Caa/Gaa | Q41E | PLES_26951 | | | | | . | . | . | . | . | . | C | C |
| NC_011770 | 2888496 | A | G | 42 | 6 | 2 | 0 | NON_SYNONYMOU | MODERAT | MISSENSE | gTc/gCc | V30A | PLES_26951 | | | | | . | . | . | . | . | . | G | G |
| NC_011770 | 2888565 | T | C | 38 | 6 | 2 | 0 | NON_SYNONYMOU | MODERAT | MISSENSE | cAg/cGg | Q7R | PLES_26951 | | | | | . | . | . | . | . | . | C | C |
| NC_011770 | 2888833 | GA | G | 43 | 6 | 2 | 0 | FRAME_SHIFT | HIGH | | ttt/ | F62 | PLES_26961 | | | | | . | . | . | . | . | . | G | G |
| NC_011770 | 2888907 | GT | G | 39 | 6 | 2 | 0 | FRAME_SHIFT | HIGH | | acc/ | T38 | PLES_26961 | | | | | . | . | . | . | . | . | G | G |
| NC_011770 | 2888935 | TG | T | 38 | 6 | 2 | 0 | FRAME_SHIFT | HIGH | | gcc/ | A28 | PLES_26961 | | | | | . | . | . | . | . | . | T | T |
| NC_011770 | 2888978 | GA | G | 37 | 6 | 2 | 0 | FRAME_SHIFT | HIGH | | ttc/ | F14 | PLES_26961 | | | | | . | . | . | . | . | . | G | G |
| NC_011770 | 2888997 | TG | T | 37 | 6 | 2 | 0 | FRAME_SHIFT | HIGH | | cat/ | H8 | PLES_26961 | | | | | . | . | . | . | . | . | T | T |
| NC_011770 | 2889016 | AC | A | 38 | 6 | 2 | 0 | FRAME_SHIFT | HIGH | | gtg/ | V1 | PLES_26961 | | | | | . | . | . | . | . | . | A | A |
| NC_011770 | 2888613 | G | A | 41 | 6 | 2 | 0 | NON_SYNONYMOU | MODERAT | MISSENSE | aCa/aTa | T136I | PLES_26961 | | | | | . | . | . | . | . | . | A | A |
| NC_011770 | 2888680 | G | C | 40 | 6 | 2 | 0 | NON_SYNONYMOU | MODERAT | MISSENSE | Cag/Gag | Q114E | PLES_26961 | | | | | . | . | . | . | . | . | C | C |
| NC_011770 | 2888683 | A | G | 40 | 6 | 2 | 0 | NON_SYNONYMOU | MODERAT | MISSENSE | Tgc/Cgc | C113R | PLES_26961 | | | | | . | . | . | . | . | . | G | G |
| NC_011770 | 2888744 | A | T | 42 | 6 | 2 | 0 | NON_SYNONYMOU | MODERAT | MISSENSE | caT/caA | H92Q | PLES_26961 | | | | | . | . | . | . | . | . | T | T |
| NC_011770 | 2888752 | T | C | 41 | 6 | 2 | 0 | NON_SYNONYMOU | MODERAT | MISSENSE | Atc/Gtc | I90V | PLES_26961 | | | | | . | . | . | . | . | . | C | C |
| NC_011770 | 2888779 | G | A | 43 | 6 | 2 | 0 | NON_SYNONYMOU | MODERAT | MISSENSE | Cat/Tat | H81Y | PLES_26961 | | | | | . | . | . | . | . | . | A | A |
| NC_011770 | 2888781 | A | G | 43 | 6 | 2 | 0 | NON_SYNONYMOU | MODERAT | MISSENSE | aTc/aCc | I80T | PLES_26961 | | | | | . | . | . | . | . | . | G | G |
| NC_011770 | 2888848 | C | T | 42 | 6 | 2 | 0 | NON_SYNONYMOU | MODERAT | MISSENSE | Gag/Aag | E58K | PLES_26961 | | | | | . | . | . | . | . | . | T | T |
| NC_011770 | 2888885 | G | T | 25 | 7 | 1 | 0 | NON_SYNONYMOU | MODERAT | MISSENSE | ttC/ttA | F45L | PLES_26961 | | | | | . | . | . | . | . | . | . | T |
| NC_011770 | 2888914 | G | T | 38 | 6 | 2 | 0 | NON_SYNONYMOU | MODERAT | MISSENSE | Cgc/Agc | R36S | PLES_26961 | | | | | . | . | . | . | . | . | T | T |
| NC_011770 | 2888916 | G | A | 38 | 6 | 2 | 0 | NON_SYNONYMOU | MODERAT | MISSENSE | aCg/aTg | T35M | PLES_26961 | | | | | . | . | . | . | . | . | A | A |
| NC_011770 | 2888925 | C | A | 39 | 6 | 2 | 0 | NON_SYNONYMOU | MODERAT | MISSENSE | aGt/aTt | S32I | PLES_26961 | | | | | . | . | . | . | . | . | A | A |
| NC_011770 | 2888946 | C | T | 35 | 6 | 2 | 0 | NON_SYNONYMOU | MODERAT | MISSENSE | cGc/cAc | R25H | PLES_26961 | | | | | . | . | . | . | . | . | T | T |
| NC_011770 | 2888952 | G | A | 34 | 6 | 2 | 0 | NON_SYNONYMOU | MODERAT | MISSENSE | gCt/gTt | A23V | PLES_26961 | | | | | . | . | . | . | . | . | A | A |
| NC_011770 | 2888958 | A | G | 32 | 6 | 2 | 0 | NON_SYNONYMOU | MODERAT | MISSENSE | tTc/tCc | F21S | PLES_26961 | | | | | . | . | . | . | . | . | G | G |
| NC_011770 | 2888961 | G | A | 33 | 6 | 2 | 0 | NON_SYNONYMOU | MODERAT | MISSENSE | tCt/tTt | S20F | PLES_26961 | | | | | . | . | . | . | . | . | A | A |
| NC_011770 | 2888962 | A | G | 32 | 6 | 2 | 0 | NON_SYNONYMOU | MODERAT | MISSENSE | Tct/Cct | S20P | PLES_26961 | | | | | . | . | . | . | . | . | G | G |
| NC_011770 | 2888964 | T | C | 31 | 6 | 2 | 0 | NON_SYNONYMOU | MODERAT | MISSENSE | gAt/gGt | D19G | PLES_26961 | | | | | . | . | . | . | . | . | C | C |
| NC_011770 | 2888983 | T | C | 24 | 7 | 1 | 0 | NON_SYNONYMOU | MODERAT | MISSENSE | Agt/Ggt | S13G | PLES_26961 | | | | | . | . | . | . | . | . | C | . |
| NC_011770 | 2889196 | G | T | 24 | 6 | 2 | 0 | NON_SYNONYMOU | MODERAT | MISSENSE | gCc/gAc | A91D | PLES_26971 | | | | | . | . | . | . | . | . | T | T |
| NC_011770 | 2889294 | G | C | 27 | 6 | 2 | 0 | NON_SYNONYMOU | MODERAT | MISSENSE | aaC/aaG | N58K | PLES_26971 | | | | | . | . | . | . | . | . | C | C |
| NC_011770 | 2889986 | C | T | 28 | 8 | 1 | 0 | NON_SYNONYMOU | MODERAT | MISSENSE | Ggc/Agc | G107S | PLES_26981 | | T | | | . | . | . | . | . | . | . | . |
| NC_011770 | 2897496 | G | T | 34 | 6 | 2 | 0 | STOP_GAINED | HIGH | NONSENSE | tCg/tAg | S227* | PLES_27011 | | | | | . | . | . | . | . | . | T | T |
| NC_011770 | 2896196 | G | C | 28 | 6 | 2 | 0 | NON_SYNONYMOU | MODERAT | MISSENSE | agC/agG | S660R | PLES_27011 | | | | | . | . | . | . | . | . | C | C |
| NC_011770 | 2896197 | C | G | 27 | 6 | 2 | 0 | NON_SYNONYMOU | MODERAT | MISSENSE | aGc/aCc | S660T | PLES_27011 | | | | | . | . | . | . | . | . | G | G |
| NC_011770 | 2896272 | C | G | 30 | 6 | 2 | 0 | NON_SYNONYMOU | MODERAT | MISSENSE | aGc/aCc | S635T | PLES_27011 | | | | | . | . | . | . | . | . | G | G |
| NC_011770 | 2896309 | C | G | 28 | 6 | 2 | 0 | NON_SYNONYMOU | MODERAT | MISSENSE | Gca/Cca | A623P | PLES_27011 | | | | | . | . | . | . | . | . | G | G |
| NC_011770 | 2896316 | T | G | 27 | 6 | 2 | 0 | NON_SYNONYMOU | MODERAT | MISSENSE | caA/caC | Q620H | PLES_27011 | | | | | . | . | . | . | . | . | G | G |
| NC_011770 | 2896342 | C | T | 28 | 6 | 2 | 0 | NON_SYNONYMOU | MODERAT | MISSENSE | Gcc/Acc | A612T | PLES_27011 | | | | | . | . | . | . | . | . | T | T |
| NC_011770 | 2896480 | C | T | 30 | 6 | 2 | 0 | NON_SYNONYMOU | MODERAT | MISSENSE | Gcg/Acg | A566T | PLES_27011 | | | | | . | . | . | . | . | . | T | T |
| NC_011770 | 2896558 | C | T | 28 | 6 | 2 | 0 | NON_SYNONYMOU | MODERAT | MISSENSE | Gct/Act | A540T | PLES_27011 | | | | | . | . | . | . | . | . | T | T |
| NC_011770 | 2896559 | G | C | 28 | 6 | 2 | 0 | NON_SYNONYMOU | MODERAT | MISSENSE | atC/atG | I539M | PLES_27011 | | | | | . | . | . | . | . | . | C | C |
| NC_011770 | 2896561 | T | G | 28 | 6 | 2 | 0 | NON_SYNONYMOU | MODERAT | MISSENSE | Atc/Ctc | I539L | PLES_27011 | | | | | . | . | . | . | . | . | G | G |
| NC_011770 | 2896570 | C | T | 28 | 6 | 2 | 0 | NON_SYNONYMOU | MODERAT | MISSENSE | Ggc/Agc | G536S | PLES_27011 | | | | | . | . | . | . | . | . | T | T |
| NC_011770 | 2896701 | T | C | 22 | 6 | 2 | 0 | NON_SYNONYMOU | MODERAT | MISSENSE | aAa/aGa | K492R | PLES_27011 | | | | | . | . | . | . | . | . | C | C |
| NC_011770 | 2896702 | T | G | 23 | 6 | 2 | 0 | NON_SYNONYMOU | MODERAT | MISSENSE | Aaa/Caa | K492Q | PLES_27011 | | | | | . | . | . | . | . | . | G | G |
| NC_011770 | 2896929 | G | T | 35 | 6 | 2 | 0 | NON_SYNONYMOU | MODERAT | MISSENSE | aCt/aAt | T416N | PLES_27011 | | | | | . | . | . | . | . | . | T | T |
| NC_011770 | 2896975 | A | C | 36 | 6 | 2 | 0 | NON_SYNONYMOU | MODERAT | MISSENSE | Tcg/Gcg | S401A | PLES_27011 | | | | | . | . | . | . | . | . | C | C |
| NC_011770 | 2897017 | C | T | 37 | 6 | 2 | 0 | NON_SYNONYMOU | MODERAT | MISSENSE | Gcc/Acc | A387T | PLES_27011 | | | | | . | . | . | . | . | . | T | T |
| NC_011770 | 2897024 | C | G | 38 | 6 | 2 | 0 | NON_SYNONYMOU | MODERAT | MISSENSE | aaG/aaC | K384N | PLES_27011 | | | | | . | . | . | . | . | . | G | G |
| NC_011770 | 2897065 | T | A | 33 | 6 | 2 | 0 | NON_SYNONYMOU | MODERAT | MISSENSE | Acg/Tcg | T371S | PLES_27011 | | | | | . | . | . | . | . | . | A | A |
| NC_011770 | 2897073 | C | G | 33 | 6 | 2 | 0 | NON_SYNONYMOU | MODERAT | MISSENSE | gGc/gCc | G368A | PLES_27011 | | | | | . | . | . | . | . | . | G | G |
| NC_011770 | 2897101 | A | T | 35 | 6 | 2 | 0 | NON_SYNONYMOU | MODERAT | MISSENSE | Tcg/Acg | S359T | PLES_27011 | | | | | . | . | . | . | . | . | T | T |
| NC_011770 | 2897107 | C | T | 38 | 6 | 2 | 0 | NON_SYNONYMOU | MODERAT | MISSENSE | Gac/Aac | D357N | PLES_27011 | | | | | . | . | . | . | . | . | T | T |
| NC_011770 | 2897115 | C | T | 40 | 6 | 2 | 0 | NON_SYNONYMOU | MODERAT | MISSENSE | gGc/gAc | G354D | PLES_27011 | | | | | . | . | . | . | . | . | T | T |
| NC_011770 | 2897125 | A | G | 38 | 6 | 2 | 0 | NON_SYNONYMOU | MODERAT | MISSENSE | Tcg/Ccg | S351P | PLES_27011 | | | | | . | . | . | . | . | . | G | G |
| NC_011770 | 2897133 | G | A | 40 | 6 | 2 | 0 | NON_SYNONYMOU | MODERAT | MISSENSE | tCg/tTg | S348L | PLES_27011 | | | | | . | . | . | . | . | . | A | A |
| NC_011770 | 2897149 | G | C | 41 | 6 | 2 | 0 | NON_SYNONYMOU | MODERAT | MISSENSE | Ctc/Gtc | L343V | PLES_27011 | | | | | . | . | . | . | . | . | C | C |
| NC_011770 | 2897261 | C | G | 36 | 6 | 2 | 0 | NON_SYNONYMOU | MODERAT | MISSENSE | gaG/gaC | E305D | PLES_27011 | | | | | . | . | . | . | . | . | G | G |
| NC_011770 | 2897295 | T | C | 37 | 6 | 2 | 0 | NON_SYNONYMOU | MODERAT | MISSENSE | aAg/aGg | K294R | PLES_27011 | | | | | . | . | . | . | . | . | C | C |

| | | | | | | | | | | | | | | | | | | | | | | | | | |
|---|---|---|---|---|---|---|---|---|---|---|---|---|---|---|---|---|---|---|---|---|---|---|---|---|---|
| NC_011770 | 2897371 | A | C | 32 | 6 | 2 | 0 | NON_SYNONYMOU | MODERAT | MISSENSE | Tcg/Gcg | S269A | PLES_27011 | | | | | . | . | . | . | . | . | C | C |
| NC_011770 | 2897378 | G | C | 33 | 6 | 2 | 0 | NON_SYNONYMOU | MODERAT | MISSENSE | caC/caG | H266Q | PLES_27011 | | | | | . | . | . | . | . | . | C | C |
| NC_011770 | 2897388 | C | T | 32 | 6 | 2 | 0 | NON_SYNONYMOU | MODERAT | MISSENSE | gGc/gAc | G263D | PLES_27011 | | | | | . | . | . | . | . | . | T | T |
| NC_011770 | 2897409 | A | G | 30 | 6 | 2 | 0 | NON_SYNONYMOU | MODERAT | MISSENSE | aTg/aCg | M256T | PLES_27011 | | | | | . | . | . | . | . | . | G | G |
| NC_011770 | 2897410 | T | C | 31 | 6 | 2 | 0 | NON_SYNONYMOU | MODERAT | MISSENSE | Atg/Gtg | M256V | PLES_27011 | | | | | . | . | . | . | . | . | C | C |
| NC_011770 | 2897413 | T | C | 31 | 6 | 2 | 0 | NON_SYNONYMOU | MODERAT | MISSENSE | Acc/Gcc | T255A | PLES_27011 | | | | | . | . | . | . | . | . | C | C |
| NC_011770 | 2897414 | C | G | 31 | 6 | 2 | 0 | NON_SYNONYMOU | MODERAT | MISSENSE | caG/caC | Q254H | PLES_27011 | | | | | . | . | . | . | . | . | G | G |
| NC_011770 | 2897425 | C | T | 32 | 6 | 2 | 0 | NON_SYNONYMOU | MODERAT | MISSENSE | Gtc/Atc | V251I | PLES_27011 | | | | | . | . | . | . | . | . | T | T |
| NC_011770 | 2897433 | C | T | 36 | 6 | 2 | 0 | NON_SYNONYMOU | MODERAT | MISSENSE | cGg/cAg | R248Q | PLES_27011 | | | | | . | . | . | . | . | . | T | T |
| NC_011770 | 2897465 | A | T | 34 | 6 | 2 | 0 | NON_SYNONYMOU | MODERAT | MISSENSE | gaT/gaA | D237E | PLES_27011 | | | | | . | . | . | . | . | . | T | T |
| NC_011770 | 2897467 | C | T | 34 | 6 | 2 | 0 | NON_SYNONYMOU | MODERAT | MISSENSE | Gat/Aat | D237N | PLES_27011 | | | | | . | . | . | . | . | . | T | T |
| NC_011770 | 2897497 | A | G | 33 | 6 | 2 | 0 | NON_SYNONYMOU | MODERAT | MISSENSE | Tcg/Ccg | S227P | PLES_27011 | | | | | . | . | . | . | . | . | G | G |
| NC_011770 | 2897508 | A | G | 34 | 6 | 2 | 0 | NON_SYNONYMOU | MODERAT | MISSENSE | tTc/tCc | F223S | PLES_27011 | | | | | . | . | . | . | . | . | G | G |
| NC_011770 | 2897520 | G | T | 35 | 6 | 2 | 0 | NON_SYNONYMOU | MODERAT | MISSENSE | gCc/gAc | A219D | PLES_27011 | | | | | . | . | . | . | . | . | T | T |
| NC_011770 | 2897550 | C | G | 34 | 6 | 2 | 0 | NON_SYNONYMOU | MODERAT | MISSENSE | cGc/cCc | R209P | PLES_27011 | | | | | . | . | . | . | . | . | G | G |
| NC_011770 | 2897551 | G | C | 34 | 6 | 2 | 0 | NON_SYNONYMOU | MODERAT | MISSENSE | Cgc/Ggc | R209G | PLES_27011 | | | | | . | . | . | . | . | . | C | C |
| NC_011770 | 2897554 | C | T | 34 | 6 | 2 | 0 | NON_SYNONYMOU | MODERAT | MISSENSE | Gcg/Acg | A208T | PLES_27011 | | | | | . | . | . | . | . | . | T | T |
| NC_011770 | 2897716 | G | A | 36 | 6 | 2 | 0 | NON_SYNONYMOU | MODERAT | MISSENSE | Cac/Tac | H154Y | PLES_27011 | | | | | . | . | . | . | . | . | A | A |
| NC_011770 | 2897725 | A | G | 36 | 6 | 2 | 0 | NON_SYNONYMOU | MODERAT | MISSENSE | Tcg/Ccg | S151P | PLES_27011 | | | | | . | . | . | . | . | . | G | G |
| NC_011770 | 2897831 | C | G | 41 | 6 | 2 | 0 | NON_SYNONYMOU | MODERAT | MISSENSE | gaG/gaC | E115D | PLES_27011 | | | | | . | . | . | . | . | . | G | G |
| NC_011770 | 2897836 | C | T | 41 | 6 | 2 | 0 | NON_SYNONYMOU | MODERAT | MISSENSE | Gtg/Atg | V114M | PLES_27011 | | | | | . | . | . | . | . | . | T | T |
| NC_011770 | 2897891 | G | C | 40 | 6 | 2 | 0 | NON_SYNONYMOU | MODERAT | MISSENSE | caC/caG | H95Q | PLES_27011 | | | | | . | . | . | . | . | . | C | C |
| NC_011770 | 2897995 | C | G | 33 | 6 | 2 | 0 | NON_SYNONYMOU | MODERAT | MISSENSE | Gtt/Ctt | V61L | PLES_27011 | | | | | . | . | . | . | . | . | G | G |
| NC_011770 | 2898012 | T | G | 30 | 6 | 2 | 0 | NON_SYNONYMOU | MODERAT | MISSENSE | cAg/cCg | Q55P | PLES_27011 | | | | | . | . | . | . | . | . | G | G |
| NC_011770 | 2898013 | G | C | 30 | 6 | 2 | 0 | NON_SYNONYMOU | MODERAT | MISSENSE | Cag/Gag | Q55E | PLES_27011 | | | | | . | . | . | . | . | . | C | C |
| NC_011770 | 2898015 | T | G | 31 | 6 | 2 | 0 | NON_SYNONYMOU | MODERAT | MISSENSE | gAg/gCg | E54A | PLES_27011 | | | | | . | . | . | . | . | . | G | G |
| NC_011770 | 2898024 | G | A | 31 | 6 | 2 | 0 | NON_SYNONYMOU | MODERAT | MISSENSE | gCc/gTc | A51V | PLES_27011 | | | | | . | . | . | . | . | . | A | A |
| NC_011770 | 2898034 | G | C | 30 | 6 | 2 | 0 | NON_SYNONYMOU | MODERAT | MISSENSE | Ccg/Gcg | P48A | PLES_27011 | | | | | . | . | . | . | . | . | C | C |
| NC_011770 | 2898476 | T | C | 28 | 6 | 2 | 0 | NON_SYNONYMOU | MODERAT | MISSENSE | Acc/Gcc | T142A | PLES_27021 | | | | | . | . | . | . | . | . | C | C |
| NC_011770 | 2898503 | T | C | 30 | 6 | 2 | 0 | NON_SYNONYMOU | MODERAT | MISSENSE | Acc/Gcc | T133A | PLES_27021 | | | | | . | . | . | . | . | . | C | C |
| NC_011770 | 2898596 | C | T | 32 | 6 | 2 | 0 | NON_SYNONYMOU | MODERAT | MISSENSE | Gtc/Atc | V102I | PLES_27021 | | | | | . | . | . | . | . | . | T | T |
| NC_011770 | 2898639 | G | C | 33 | 6 | 2 | 0 | NON_SYNONYMOU | MODERAT | MISSENSE | gaC/gaG | D87E | PLES_27021 | | | | | . | . | . | . | . | . | C | C |
| NC_011770 | 2898659 | C | T | 33 | 6 | 2 | 0 | NON_SYNONYMOU | MODERAT | MISSENSE | Gtc/Atc | V81I | PLES_27021 | | | | | . | . | . | . | . | . | T | T |
| NC_011770 | 2898690 | C | G | 31 | 6 | 2 | 0 | NON_SYNONYMOU | MODERAT | MISSENSE | caG/caC | Q70H | PLES_27021 | | | | | . | . | . | . | . | . | G | G |
| NC_011770 | 2898692 | G | C | 31 | 6 | 2 | 0 | NON_SYNONYMOU | MODERAT | MISSENSE | Cag/Gag | Q70E | PLES_27021 | | | | | . | . | . | . | . | . | C | C |
| NC_011770 | 2898695 | G | T | 31 | 6 | 2 | 0 | NON_SYNONYMOU | MODERAT | MISSENSE | Cag/Aag | Q69K | PLES_27021 | | | | | . | . | . | . | . | . | T | T |
| NC_011770 | 2898700 | C | T | 31 | 6 | 2 | 0 | NON_SYNONYMOU | MODERAT | MISSENSE | cGc/cAc | R67H | PLES_27021 | | | | | . | . | . | . | . | . | T | T |
| NC_011770 | 2898702 | G | C | 30 | 6 | 2 | 0 | NON_SYNONYMOU | MODERAT | MISSENSE | gaC/gaG | D66E | PLES_27021 | | | | | . | . | . | . | . | . | C | C |
| NC_011770 | 2898712 | T | C | 35 | 6 | 2 | 0 | NON_SYNONYMOU | MODERAT | MISSENSE | cAc/cGc | H63R | PLES_27021 | | | | | . | . | . | . | . | . | C | C |
| NC_011770 | 2898722 | G | T | 37 | 6 | 2 | 0 | NON_SYNONYMOU | MODERAT | MISSENSE | Ctc/Atc | L60I | PLES_27021 | | | | | . | . | . | . | . | . | T | T |
| NC_011770 | 2898759 | C | G | 33 | 6 | 2 | 0 | NON_SYNONYMOU | MODERAT | MISSENSE | gaG/gaC | E47D | PLES_27021 | | | | | . | . | . | . | . | . | G | G |
| NC_011770 | 2898760 | T | G | 32 | 6 | 2 | 0 | NON_SYNONYMOU | MODERAT | MISSENSE | gAg/gCg | E47A | PLES_27021 | | | | | . | . | . | . | . | . | G | G |
| NC_011770 | 2898761 | C | T | 32 | 6 | 2 | 0 | NON_SYNONYMOU | MODERAT | MISSENSE | Gag/Aag | E47K | PLES_27021 | | | | | . | . | . | . | . | . | T | T |
| NC_011770 | 2898766 | C | T | 32 | 6 | 2 | 0 | NON_SYNONYMOU | MODERAT | MISSENSE | gGc/gAc | G45D | PLES_27021 | | | | | . | . | . | . | . | . | T | T |
| NC_011770 | 2898772 | G | T | 33 | 6 | 2 | 0 | NON_SYNONYMOU | MODERAT | MISSENSE | aCc/aAc | T43N | PLES_27021 | | | | | . | . | . | . | . | . | T | T |
| NC_011770 | 2898818 | G | T | 36 | 6 | 2 | 0 | NON_SYNONYMOU | MODERAT | MISSENSE | Cgc/Agc | R28S | PLES_27021 | | | | | . | . | . | . | . | . | T | T |
| NC_011770 | 2898822 | A | T | 37 | 6 | 2 | 0 | NON_SYNONYMOU | MODERAT | MISSENSE | gaT/gaA | D26E | PLES_27021 | | | | | . | . | . | . | . | . | T | T |
| NC_011770 | 2898883 | A | G | 33 | 6 | 2 | 0 | NON_SYNONYMOU | MODERAT | MISSENSE | aTc/aCc | I6T | PLES_27021 | | | | | . | . | . | . | . | . | G | G |
| NC_011770 | 2898884 | T | A | 34 | 6 | 2 | 0 | NON_SYNONYMOU | MODERAT | MISSENSE | Atc/Ttc | I6F | PLES_27021 | | | | | . | . | . | . | . | . | A | A |
| NC_011770 | 2898888 | T | A | 35 | 6 | 2 | 0 | NON_SYNONYMOU | MODERAT | MISSENSE | caA/caT | Q4H | PLES_27021 | | | | | . | . | . | . | . | . | A | A |
| NC_011770 | 2898987 | C | T | 38 | 6 | 2 | 0 | NON_SYNONYMOU | MODERAT | MISSENSE | Gtg/Atg | V172M | PLES_27031 | | | | | . | . | . | . | . | . | T | T |
| NC_011770 | 2899005 | C | T | 37 | 6 | 2 | 0 | NON_SYNONYMOU | MODERAT | MISSENSE | Gcg/Acg | A166T | PLES_27031 | | | | | . | . | . | . | . | . | T | T |
| NC_011770 | 2899013 | C | T | 36 | 6 | 2 | 0 | NON_SYNONYMOU | MODERAT | MISSENSE | cGc/cAc | R163H | PLES_27031 | | | | | . | . | . | . | . | . | T | T |
| NC_011770 | 2899023 | T | C | 36 | 6 | 2 | 0 | NON_SYNONYMOU | MODERAT | MISSENSE | Acc/Gcc | T160A | PLES_27031 | | | | | . | . | . | . | . | . | C | C |
| NC_011770 | 2899026 | T | C | 37 | 6 | 2 | 0 | NON_SYNONYMOU | MODERAT | MISSENSE | Aac/Gac | N159D | PLES_27031 | | | | | . | . | . | . | . | . | C | C |
| NC_011770 | 2899035 | G | C | 36 | 6 | 2 | 0 | NON_SYNONYMOU | MODERAT | MISSENSE | Cag/Gag | Q156E | PLES_27031 | | | | | . | . | . | . | . | . | C | C |
| NC_011770 | 2899106 | A | G | 38 | 6 | 2 | 0 | NON_SYNONYMOU | MODERAT | MISSENSE | aTg/aCg | M132T | PLES_27031 | | | | | . | . | . | . | . | . | G | G |
| NC_011770 | 2899110 | C | T | 37 | 6 | 2 | 0 | NON_SYNONYMOU | MODERAT | MISSENSE | Gtc/Atc | V131I | PLES_27031 | | | | | . | . | . | . | . | . | T | T |
| NC_011770 | 2899127 | T | G | 39 | 6 | 2 | 0 | NON_SYNONYMOU | MODERAT | MISSENSE | gAg/gCg | E125A | PLES_27031 | | | | | . | . | . | . | . | . | G | G |
| NC_011770 | 2899139 | T | G | 40 | 6 | 2 | 0 | NON_SYNONYMOU | MODERAT | MISSENSE | cAc/cCc | H121P | PLES_27031 | | | | | . | . | . | . | . | . | G | G |
| NC_011770 | 2899140 | G | C | 39 | 6 | 2 | 0 | NON_SYNONYMOU | MODERAT | MISSENSE | Cac/Gac | H121D | PLES_27031 | | | | | . | . | . | . | . | . | C | C |
| NC_011770 | 2899163 | G | C | 38 | 6 | 2 | 0 | NON_SYNONYMOU | MODERAT | MISSENSE | cCc/cGc | P113R | PLES_27031 | | | | | . | . | . | . | . | . | C | C |

| | | | | | | | | | | | | | | | | | | | | | | | | | |
|---|---|---|---|---|---|---|---|---|---|---|---|---|---|---|---|---|---|---|---|---|---|---|---|---|---|
| NC_011770 | 2899164 | G | T | 37 | 6 | 2 | 0 | NON_SYNONYMOU | MODERAT | MISSENSE | Ccc/Acc | P113T | PLES_27031 | | | | | . | . | . | . | . | . | T | T |
| NC_011770 | 2899171 | C | G | 39 | 6 | 2 | 0 | NON_SYNONYMOU | MODERAT | MISSENSE | atG/atC | M110I | PLES_27031 | | | | | . | . | . | . | . | . | G | G |
| NC_011770 | 2899181 | T | C | 42 | 6 | 2 | 0 | NON_SYNONYMOU | MODERAT | MISSENSE | cAg/cGg | Q107R | PLES_27031 | | | | | . | . | . | . | . | . | C | C |
| NC_011770 | 2899217 | T | C | 41 | 6 | 2 | 0 | NON_SYNONYMOU | MODERAT | MISSENSE | aAg/aGg | K95R | PLES_27031 | | | | | . | . | . | . | . | . | C | C |
| NC_011770 | 2899235 | G | A | 26 | 6 | 1 | 0 | NON_SYNONYMOU | MODERAT | MISSENSE | aCc/aTc | T89I | PLES_27031 | | | | | . | . | . | . | . | . | A | . |
| NC_011770 | 2899236 | T | C | 26 | 6 | 2 | 0 | NON_SYNONYMOU | MODERAT | MISSENSE | Acc/Gcc | T89A | PLES_27031 | | | | | . | . | . | . | . | . | C | C |
| NC_011770 | 2899272 | C | T | 43 | 6 | 2 | 0 | NON_SYNONYMOU | MODERAT | MISSENSE | Gtc/Atc | V77I | PLES_27031 | | | | | . | . | . | . | . | . | T | T |
| NC_011770 | 2899307 | G | T | 40 | 6 | 2 | 0 | NON_SYNONYMOU | MODERAT | MISSENSE | cCa/cAa | P65Q | PLES_27031 | | | | | . | . | . | . | . | . | T | T |
| NC_011770 | 2899308 | G | C | 40 | 6 | 2 | 0 | NON_SYNONYMOU | MODERAT | MISSENSE | Cca/Gca | P65A | PLES_27031 | | | | | . | . | . | . | . | . | C | C |
| NC_011770 | 2899650 | C | T | 32 | 6 | 2 | 0 | NON_SYNONYMOU | MODERAT | MISSENSE | atG/atA | M213I | PLES_27041 | | | | | . | . | . | . | . | . | T | T |
| NC_011770 | 2899651 | A | G | 31 | 6 | 2 | 0 | NON_SYNONYMOU | MODERAT | MISSENSE | aTg/aCg | M213T | PLES_27041 | | | | | . | . | . | . | . | . | G | G |
| NC_011770 | 2899665 | G | C | 34 | 6 | 2 | 0 | NON_SYNONYMOU | MODERAT | MISSENSE | gaC/gaG | D208E | PLES_27041 | | | | | . | . | . | . | . | . | C | C |
| NC_011770 | 2899684 | A | T | 33 | 6 | 2 | 0 | NON_SYNONYMOU | MODERAT | MISSENSE | tTc/tAc | F202Y | PLES_27041 | | | | | . | . | . | . | . | . | T | T |
| NC_011770 | 2899699 | G | A | 30 | 6 | 2 | 0 | NON_SYNONYMOU | MODERAT | MISSENSE | gCa/gTa | A197V | PLES_27041 | | | | | . | . | . | . | . | . | A | A |
| NC_011770 | 2899770 | G | C | 32 | 6 | 2 | 0 | NON_SYNONYMOU | MODERAT | MISSENSE | agC/agG | S173R | PLES_27041 | | | | | . | . | . | . | . | . | C | C |
| NC_011770 | 2899771 | C | G | 32 | 6 | 2 | 0 | NON_SYNONYMOU | MODERAT | MISSENSE | aGc/aCc | S173T | PLES_27041 | | | | | . | . | . | . | . | . | G | G |
| NC_011770 | 2899772 | T | A | 32 | 6 | 2 | 0 | NON_SYNONYMOU | MODERAT | MISSENSE | Agc/Tgc | S173C | PLES_27041 | | | | | . | . | . | . | . | . | A | A |
| NC_011770 | 2899825 | G | C | 31 | 6 | 2 | 0 | NON_SYNONYMOU | MODERAT | MISSENSE | tCg/tGg | S155W | PLES_27041 | | | | | . | . | . | . | . | . | C | C |
| NC_011770 | 2899826 | A | T | 31 | 6 | 2 | 0 | NON_SYNONYMOU | MODERAT | MISSENSE | Tcg/Acg | S155T | PLES_27041 | | | | | . | . | . | . | . | . | T | T |
| NC_011770 | 2899916 | G | A | 38 | 6 | 2 | 0 | NON_SYNONYMOU | MODERAT | MISSENSE | Ctc/Ttc | L125F | PLES_27041 | A | A | | | . | . | . | . | . | . | | |
| NC_011770 | 2899932 | C | G | 37 | 6 | 2 | 0 | NON_SYNONYMOU | MODERAT | MISSENSE | ttG/ttC | L119F | PLES_27041 | | | | | . | . | . | . | . | . | G | G |
| NC_011770 | 2899975 | C | G | 39 | 6 | 2 | 0 | NON_SYNONYMOU | MODERAT | MISSENSE | aGc/aCc | S105T | PLES_27041 | G | G | | | . | . | . | . | . | . | | |
| NC_011770 | 2899977 | C | G | 36 | 6 | 2 | 0 | NON_SYNONYMOU | MODERAT | MISSENSE | ttG/ttC | L104F | PLES_27041 | | | | | . | . | . | . | . | . | G | G |
| NC_011770 | 2899985 | C | G | 36 | 6 | 2 | 0 | NON_SYNONYMOU | MODERAT | MISSENSE | Gcg/Ccg | A102P | PLES_27041 | | | | | . | . | . | . | . | . | G | G |
| NC_011770 | 2899994 | C | T | 35 | 6 | 2 | 0 | NON_SYNONYMOU | MODERAT | MISSENSE | Ggc/Agc | G99S | PLES_27041 | | | | | . | . | . | . | . | . | T | T |
| NC_011770 | 2900016 | C | G | 37 | 6 | 2 | 0 | NON_SYNONYMOU | MODERAT | MISSENSE | gaG/gaC | E91D | PLES_27041 | G | G | | | . | . | . | . | . | . | | |
| NC_011770 | 2900027 | A | T | 37 | 6 | 2 | 0 | NON_SYNONYMOU | MODERAT | MISSENSE | Tcg/Acg | S88T | PLES_27041 | T | T | | | . | . | . | . | . | . | | |
| NC_011770 | 2900038 | G | A | 37 | 6 | 2 | 0 | NON_SYNONYMOU | MODERAT | MISSENSE | aCc/aTc | T84I | PLES_27041 | | | | | . | . | . | . | . | . | A | A |
| NC_011770 | 2900039 | T | C | 37 | 6 | 2 | 0 | NON_SYNONYMOU | MODERAT | MISSENSE | Acc/Gcc | T84A | PLES_27041 | | | | | . | . | . | . | . | . | C | C |
| NC_011770 | 2900041 | C | T | 38 | 6 | 2 | 0 | NON_SYNONYMOU | MODERAT | MISSENSE | aGg/aAg | R83K | PLES_27041 | | | | | . | . | . | . | . | . | T | T |
| NC_011770 | 2900058 | T | G | 36 | 6 | 4 | 0 | NON_SYNONYMOU | MODERAT | MISSENSE | gaA/gaC | E77D | PLES_27041 | G | G | | | . | . | . | . | . | . | G | G |
| NC_011770 | 2900070 | C | A | 35 | 6 | 2 | 0 | NON_SYNONYMOU | MODERAT | MISSENSE | ttG/ttT | L73F | PLES_27041 | A | A | | | . | . | . | . | . | . | | |
| NC_011770 | 2900085 | G | C | 32 | 6 | 2 | 0 | NON_SYNONYMOU | MODERAT | MISSENSE | agC/agG | S68R | PLES_27041 | | | | | . | . | . | . | . | . | C | C |
| NC_011770 | 2900086 | C | G | 33 | 6 | 2 | 0 | NON_SYNONYMOU | MODERAT | MISSENSE | aGc/aCc | S68T | PLES_27041 | | | | | . | . | . | . | . | . | G | G |
| NC_011770 | 2900087 | T | A | 32 | 6 | 2 | 0 | NON_SYNONYMOU | MODERAT | MISSENSE | Agc/Tgc | S68C | PLES_27041 | | | | | . | . | . | . | . | . | A | A |
| NC_011770 | 2900116 | T | C | 30 | 6 | 2 | 0 | NON_SYNONYMOU | MODERAT | MISSENSE | aAg/aGg | K58R | PLES_27041 | | | | | . | . | . | . | . | . | C | C |
| NC_011770 | 2900117 | T | G | 30 | 6 | 2 | 0 | NON_SYNONYMOU | MODERAT | MISSENSE | Aag/Cag | K58Q | PLES_27041 | | | | | . | . | . | . | . | . | G | G |
| NC_011770 | 2900131 | A | T | 30 | 6 | 2 | 0 | NON_SYNONYMOU | MODERAT | MISSENSE | gTg/gAg | V53E | PLES_27041 | | | | | . | . | . | . | . | . | T | T |
| NC_011770 | 2900132 | C | G | 31 | 6 | 2 | 0 | NON_SYNONYMOU | MODERAT | MISSENSE | Gtg/Ctg | V53L | PLES_27041 | | | | | . | . | . | . | . | . | G | G |
| NC_011770 | 2900275 | TTGC | T | 30 | 6 | 4 | 0 | CODON_CHANGE_ | MODERATE | | agcaac/aa | SN4N | PLES_27041 | | | T | T | . | . | . | . | . | . | T | T |
| NC_011770 | 2900748 | C | G | 20 | 8 | 2 | 0 | NON_SYNONYMOU | MODERAT | MISSENSE | gaG/gaC | E403D | PLES_27051 | | | G | G | . | . | . | . | . | . | . | . |
| NC_011770 | 2900875 | C | T | 36 | 6 | 2 | 0 | NON_SYNONYMOU | MODERAT | MISSENSE | cGc/cAc | R361H | PLES_27051 | | | | | . | . | . | . | . | . | T | T |
| NC_011770 | 2900877 | C | G | 35 | 6 | 4 | 0 | NON_SYNONYMOU | MODERAT | MISSENSE | atG/atC | M360I | PLES_27051 | | | G | G | . | . | . | . | . | . | G | G |
| NC_011770 | 2900931 | C | A | 21 | 8 | 2 | 0 | NON_SYNONYMOU | MODERAT | MISSENSE | aaG/aaT | K342N | PLES_27051 | | | . | . | . | . | . | . | . | . | A | A |
| NC_011770 | 2900932 | T | C | 21 | 8 | 2 | 0 | NON_SYNONYMOU | MODERAT | MISSENSE | aAg/aGg | K342R | PLES_27051 | | | . | . | . | . | . | . | . | . | C | C |
| NC_011770 | 2900936 | C | T | 21 | 8 | 2 | 0 | NON_SYNONYMOU | MODERAT | MISSENSE | Ggc/Agc | G341S | PLES_27051 | | | . | . | . | . | . | . | . | . | T | T |
| NC_011770 | 2900952 | T | G | 24 | 6 | 2 | 0 | NON_SYNONYMOU | MODERAT | MISSENSE | gaA/gaC | E335D | PLES_27051 | | | | | . | . | . | . | . | . | G | G |
| NC_011770 | 2901002 | T | C | 17 | 8 | 2 | 0 | NON_SYNONYMOU | MODERAT | MISSENSE | Atg/Gtg | M319V | PLES_27051 | | | . | . | . | . | . | . | . | . | C | C |
| NC_011770 | 2901004 | T | C | 18 | 8 | 2 | 0 | NON_SYNONYMOU | MODERAT | MISSENSE | aAg/aGg | K318R | PLES_27051 | | | . | . | . | . | . | . | . | . | C | C |
| NC_011770 | 2901014 | C | T | 18 | 8 | 2 | 0 | NON_SYNONYMOU | MODERAT | MISSENSE | Gcc/Acc | A315T | PLES_27051 | | | . | . | . | . | . | . | . | . | T | T |
| NC_011770 | 2901038 | C | T | 21 | 8 | 2 | 0 | NON_SYNONYMOU | MODERAT | MISSENSE | Gcg/Acg | A307T | PLES_27051 | | | . | . | . | . | . | . | . | . | T | T |
| NC_011770 | 2901101 | C | T | 39 | 6 | 4 | 0 | NON_SYNONYMOU | MODERAT | MISSENSE | Gat/Aat | D286N | PLES_27051 | | | T | T | . | . | . | . | . | . | T | T |
| NC_011770 | 2901102 | A | C | 40 | 6 | 2 | 0 | NON_SYNONYMOU | MODERAT | MISSENSE | atT/atG | I285M | PLES_27051 | | | C | C | . | . | . | . | . | . | | |
| NC_011770 | 2901127 | T | G | 45 | 6 | 2 | 0 | NON_SYNONYMOU | MODERAT | MISSENSE | gAc/gCc | D277A | PLES_27051 | | | | | . | . | . | . | . | . | G | G |
| NC_011770 | 2901137 | A | G | 44 | 6 | 6 | 0 | NON_SYNONYMOU | MODERAT | MISSENSE | Tcg/Ccg | S274P | PLES_27051 | G | G | G | G | . | . | . | . | . | . | G | G |
| NC_011770 | 2901149 | C | A | 43 | 6 | 2 | 0 | NON_SYNONYMOU | MODERAT | MISSENSE | Gct/Tct | A270S | PLES_27051 | | | | | . | . | . | . | . | . | A | A |
| NC_011770 | 2901151 | C | G | 44 | 6 | 2 | 0 | NON_SYNONYMOU | MODERAT | MISSENSE | gGc/gCc | G269A | PLES_27051 | | | | | . | . | . | . | . | . | G | G |
| NC_011770 | 2901158 | G | A | 45 | 6 | 2 | 0 | NON_SYNONYMOU | MODERAT | MISSENSE | Cca/Tca | P267S | PLES_27051 | | | A | A | . | . | . | . | . | . | | |
| NC_011770 | 2901170 | G | C | 46 | 6 | 2 | 0 | NON_SYNONYMOU | MODERAT | MISSENSE | Ccg/Gcg | P263A | PLES_27051 | | | | | . | . | . | . | . | . | C | C |
| NC_011770 | 2901183 | T | A | 45 | 6 | 2 | 0 | NON_SYNONYMOU | MODERAT | MISSENSE | gaA/gaT | E258D | PLES_27051 | | | A | A | . | . | . | . | . | . | | |
| NC_011770 | 2901196 | C | T | 41 | 6 | 4 | 0 | NON_SYNONYMOU | MODERAT | MISSENSE | cGc/cAc | R254H | PLES_27051 | | | T | T | . | . | . | . | . | . | T | T |
| NC_011770 | 2901197 | G | C | 40 | 6 | 4 | 0 | NON_SYNONYMOU | MODERAT | MISSENSE | Cgc/Ggc | R254G | PLES_27051 | | | C | C | . | . | . | . | . | . | C | C |

| | | | | | | | | | | | | | | | | | | | | | | | | | |
|---|---|---|---|---|---|---|---|---|---|---|---|---|---|---|---|---|---|---|---|---|---|---|---|---|---|
| NC_011770 | 2901202 | C | G | 41 | 6 | 4 | 0 | NON_SYNONYMOU | MODERAT | MISSENSE | gGc/gCc | G252A | PLES_27051 | | | G | G | . | . | . | . | . | . | G | G |
| NC_011770 | 2901203 | C | A | 41 | 6 | 2 | 0 | NON_SYNONYMOU | MODERAT | MISSENSE | Ggc/Tgc | G252C | PLES_27051 | | | A | A | . | . | . | . | . | . | | |
| NC_011770 | 2901221 | C | G | 42 | 6 | 4 | 0 | NON_SYNONYMOU | MODERAT | MISSENSE | Gcg/Ccg | A246P | PLES_27051 | | | G | G | . | . | . | . | . | . | A | A |
| NC_011770 | 2901224 | G | C | 42 | 6 | 4 | 0 | NON_SYNONYMOU | MODERAT | MISSENSE | Ccc/Gcc | P245A | PLES_27051 | | | C | C | . | . | . | . | . | . | C | C |
| NC_011770 | 2901244 | A | G | 40 | 6 | 2 | 0 | NON_SYNONYMOU | MODERAT | MISSENSE | gTc/gCc | V238A | PLES_27051 | | | | | . | . | . | . | . | . | G | G |
| NC_011770 | 2901247 | G | C | 41 | 6 | 2 | 0 | NON_SYNONYMOU | MODERAT | MISSENSE | gCg/gGg | A237G | PLES_27051 | | | C | C | . | . | . | . | . | . | | |
| NC_011770 | 2901248 | C | T | 41 | 6 | 2 | 0 | NON_SYNONYMOU | MODERAT | MISSENSE | Gcg/Acg | A237T | PLES_27051 | | | T | T | . | . | . | . | . | . | | |
| NC_011770 | 2901254 | C | T | 41 | 6 | 2 | 0 | NON_SYNONYMOU | MODERAT | MISSENSE | Gag/Aag | E235K | PLES_27051 | | | T | T | . | . | . | . | . | . | | |
| NC_011770 | 2901272 | A | C | 27 | 8 | 2 | 0 | NON_SYNONYMOU | MODERAT | MISSENSE | Tcc/Gcc | S229A | PLES_27051 | | | . | . | . | . | . | . | . | . | C | C |
| NC_011770 | 2901289 | T | C | 24 | 8 | 2 | 0 | NON_SYNONYMOU | MODERAT | MISSENSE | gAc/gGc | D223G | PLES_27051 | | | . | . | . | . | . | . | . | . | C | C |
| NC_011770 | 2901290 | C | T | 24 | 8 | 2 | 0 | NON_SYNONYMOU | MODERAT | MISSENSE | Gac/Aac | D223N | PLES_27051 | | | . | . | . | . | . | . | . | . | T | T |
| NC_011770 | 2901354 | G | T | 25 | 8 | 2 | 0 | NON_SYNONYMOU | MODERAT | MISSENSE | gaC/gaA | D201E | PLES_27051 | | | . | . | . | . | . | . | . | . | T | T |
| NC_011770 | 2901368 | G | T | 24 | 8 | 2 | 0 | NON_SYNONYMOU | MODERAT | MISSENSE | Cag/Aag | Q197K | PLES_27051 | | | . | . | . | . | . | . | . | . | T | T |
| NC_011770 | 2901385 | G | A | 20 | 8 | 4 | 0 | NON_SYNONYMOU | MODERAT | MISSENSE | gCg/gTg | A191V | PLES_27051 | A | A | . | . | . | . | . | . | . | . | A | A |
| NC_011770 | 2901389 | C | T | 36 | 6 | 2 | 0 | NON_SYNONYMOU | MODERAT | MISSENSE | Gtg/Atg | V190M | PLES_27051 | | | | | . | . | . | . | . | . | T | T |
| NC_011770 | 2901419 | A | C | 36 | 6 | 2 | 0 | NON_SYNONYMOU | MODERAT | MISSENSE | Tcg/Gcg | S180A | PLES_27051 | | | C | C | . | . | . | . | . | . | | |
| NC_011770 | 2901433 | G | T | 38 | 6 | 2 | 0 | NON_SYNONYMOU | MODERAT | MISSENSE | gCc/gAc | A175D | PLES_27051 | | | T | T | . | . | . | . | . | . | | |
| NC_011770 | 2901446 | C | G | 38 | 6 | 2 | 0 | NON_SYNONYMOU | MODERAT | MISSENSE | Gcc/Ccc | A171P | PLES_27051 | | | G | G | . | . | . | . | . | . | | |
| NC_011770 | 2901645 | C | A | 32 | 6 | 4 | 0 | NON_SYNONYMOU | MODERAT | MISSENSE | ttG/ttT | L104F | PLES_27051 | | | A | A | . | . | . | . | . | . | A | A |
| NC_011770 | 2901674 | C | T | 33 | 6 | 2 | 0 | NON_SYNONYMOU | MODERAT | MISSENSE | Gtc/Atc | V95I | PLES_27051 | | | | | . | . | . | . | . | . | T | T |
| NC_011770 | 2901681 | G | C | 35 | 6 | 2 | 0 | NON_SYNONYMOU | MODERAT | MISSENSE | gaC/gaG | D92E | PLES_27051 | | | C | C | . | . | . | . | . | . | | |
| NC_011770 | 2901695 | A | T | 34 | 6 | 4 | 0 | NON_SYNONYMOU | MODERAT | MISSENSE | Ttg/Atg | L88M | PLES_27051 | | | T | T | . | . | . | . | . | . | G | G |
| NC_011770 | 2901766 | C | G | 35 | 6 | 2 | 0 | NON_SYNONYMOU | MODERAT | MISSENSE | aGc/aCc | S64T | PLES_27051 | | | | | . | . | . | . | . | . | G | G |
| NC_011770 | 2901836 | G | C | 6 | 8 | 2 | 0 | NON_SYNONYMOU | MODERAT | MISSENSE | Ctg/Gtg | L41V | PLES_27051 | | | . | . | . | . | . | . | . | . | C | C |
| NC_011770 | 2901839 | A | G | 6 | 8 | 2 | 0 | NON_SYNONYMOU | MODERAT | MISSENSE | Tcc/Ccc | S40P | PLES_27051 | | | . | . | . | . | . | . | . | . | G | G |
| NC_011770 | 2901851 | C | T | 6 | 8 | 2 | 0 | NON_SYNONYMOU | MODERAT | MISSENSE | Ggc/Agc | G36S | PLES_27051 | | | . | . | . | . | . | . | . | . | T | T |
| NC_011770 | 2901854 | A | G | 6 | 8 | 2 | 0 | NON_SYNONYMOU | MODERAT | MISSENSE | Tct/Cct | S35P | PLES_27051 | | | . | . | . | . | . | . | . | . | G | G |
| NC_011770 | 2901860 | A | G | 15 | 8 | 2 | 0 | NON_SYNONYMOU | MODERAT | MISSENSE | Tcc/Ccc | S33P | PLES_27051 | | | . | . | . | . | . | . | . | . | G | G |
| NC_011770 | 2901911 | T | C | 17 | 8 | 2 | 0 | NON_SYNONYMOU | MODERAT | MISSENSE | Atc/Gtc | I16V | PLES_27051 | | | . | . | . | . | . | . | . | . | C | C |
| NC_011770 | 2901941 | T | C | 19 | 7 | 1 | 0 | NON_SYNONYMOU | MODERAT | MISSENSE | Acc/Gcc | T6A | PLES_27051 | | | . | C | . | . | . | . | . | . | | |
| NC_011770 | 2901961 | T | C | 28 | 6 | 4 | 0 | STOP_LOST+SPLICE | HIGH | MISSENSE | tAg/tGg | *187W | PLES_27061 | | | C | C | . | . | . | . | . | . | C | C |
| NC_011770 | 2902021 | G | T | 32 | 6 | 2 | 0 | STOP_GAINED | HIGH | NONSENSE | tCg/tAg | S167* | PLES_27061 | | | T | T | . | . | . | . | . | . | | |
| NC_011770 | 2901960 | C | T | 28 | 6 | 4 | 0 | SPLICE_SITE_REGIO | LOW | SILENT | taG/taA | *187 | PLES_27061 | | | T | T | . | . | . | . | . | . | T | T |
| NC_011770 | 2901980 | C | T | 32 | 6 | 2 | 0 | NON_SYNONYMOU | MODERAT | MISSENSE | Gtc/Atc | V181I | PLES_27061 | | | T | T | . | . | . | . | . | . | | |
| NC_011770 | 2901985 | T | C | 32 | 6 | 2 | 0 | NON_SYNONYMOU | MODERAT | MISSENSE | aAc/aGc | N179S | PLES_27061 | | | | | . | . | . | . | . | . | C | C |
| NC_011770 | 2901988 | C | T | 32 | 6 | 4 | 0 | NON_SYNONYMOU | MODERAT | MISSENSE | aGg/aAg | R178K | PLES_27061 | | | T | T | . | . | . | . | . | . | T | T |
| NC_011770 | 2901995 | C | A | 32 | 6 | 2 | 0 | NON_SYNONYMOU | MODERAT | MISSENSE | Gcg/Tcg | A176S | PLES_27061 | | | | | . | . | . | . | . | . | A | A |
| NC_011770 | 2902022 | A | C | 31 | 6 | 2 | 0 | NON_SYNONYMOU | MODERAT | MISSENSE | Tcg/Gcg | S167A | PLES_27061 | | | C | C | . | . | . | . | . | . | | |
| NC_011770 | 2902046 | C | T | 32 | 6 | 2 | 0 | NON_SYNONYMOU | MODERAT | MISSENSE | Gcc/Acc | A159T | PLES_27061 | | | | | . | . | . | . | . | . | T | T |
| NC_011770 | 2902058 | T | C | 32 | 6 | 2 | 0 | NON_SYNONYMOU | MODERAT | MISSENSE | Att/Gtt | I155V | PLES_27061 | | | C | C | . | . | . | . | . | . | | |
| NC_011770 | 2902077 | A | C | 35 | 6 | 2 | 0 | NON_SYNONYMOU | MODERAT | MISSENSE | gaT/gaG | D148E | PLES_27061 | | | C | C | . | . | . | . | . | . | | |
| NC_011770 | 2902094 | T | C | 35 | 6 | 2 | 0 | NON_SYNONYMOU | MODERAT | MISSENSE | Agc/Ggc | S143G | PLES_27061 | | | | | . | . | . | . | . | . | C | C |
| NC_011770 | 2902110 | T | G | 33 | 6 | 2 | 0 | NON_SYNONYMOU | MODERAT | MISSENSE | caA/caC | Q137H | PLES_27061 | | | G | G | . | . | . | . | . | . | | |
| NC_011770 | 2902111 | T | C | 33 | 6 | 2 | 0 | NON_SYNONYMOU | MODERAT | MISSENSE | cAa/cGa | Q137R | PLES_27061 | | | C | C | . | . | . | . | . | . | | |
| NC_011770 | 2902135 | G | T | 30 | 6 | 4 | 0 | NON_SYNONYMOU | MODERAT | MISSENSE | aCc/aAc | T129N | PLES_27061 | | | T | T | . | . | . | . | . | . | T | T |
| NC_011770 | 2902136 | T | C | 31 | 6 | 2 | 0 | NON_SYNONYMOU | MODERAT | MISSENSE | Acc/Gcc | T129A | PLES_27061 | | | C | C | . | . | . | . | . | . | | |
| NC_011770 | 2902184 | T | C | 28 | 6 | 2 | 0 | NON_SYNONYMOU | MODERAT | MISSENSE | Atc/Gtc | I113V | PLES_27061 | | | C | C | . | . | . | . | . | . | | |
| NC_011770 | 2902234 | A | G | 29 | 6 | 2 | 0 | NON_SYNONYMOU | MODERAT | MISSENSE | gTc/gCc | V96A | PLES_27061 | | | G | G | . | . | . | . | . | . | | |
| NC_011770 | 2902308 | G | T | 33 | 6 | 4 | 0 | NON_SYNONYMOU | MODERAT | MISSENSE | aaC/aaA | N71K | PLES_27061 | | | T | T | . | . | . | . | . | . | C | C |
| NC_011770 | 2902321 | T | C | 31 | 6 | 4 | 0 | NON_SYNONYMOU | MODERAT | MISSENSE | cAg/cGg | Q67R | PLES_27061 | | | C | C | . | . | . | . | . | . | C | C |
| NC_011770 | 2902343 | T | A | 33 | 6 | 2 | 0 | NON_SYNONYMOU | MODERAT | MISSENSE | Acg/Tcg | T60S | PLES_27061 | | | A | A | . | . | . | . | . | . | | |
| NC_011770 | 2902363 | C | T | 34 | 6 | 2 | 0 | NON_SYNONYMOU | MODERAT | MISSENSE | aGc/aAc | S53N | PLES_27061 | | | | | . | . | . | . | . | . | T | T |
| NC_011770 | 2902417 | G | T | 23 | 6 | 4 | 0 | NON_SYNONYMOU | MODERAT | MISSENSE | gCc/gAc | A35D | PLES_27061 | | | T | T | . | . | . | . | . | . | T | T |
| NC_011770 | 2902435 | G | A | 26 | 6 | 2 | 0 | NON_SYNONYMOU | MODERAT | MISSENSE | gCc/gTc | A29V | PLES_27061 | | | | | . | . | . | . | . | . | A | A |
| NC_011770 | 2902436 | C | G | 26 | 6 | 2 | 0 | NON_SYNONYMOU | MODERAT | MISSENSE | Gcc/Ccc | A29P | PLES_27061 | | | | | . | . | . | . | . | . | G | G |
| NC_011770 | 2902508 | G | T | 19 | 6 | 2 | 0 | NON_SYNONYMOU | MODERAT | MISSENSE | Cat/Aat | H5N | PLES_27061 | | | T | T | . | . | . | . | . | . | | |
| NC_011770 | 2902511 | A | G | 19 | 6 | 4 | 0 | NON_SYNONYMOU | MODERAT | MISSENSE | Tcg/Ccg | S4P | PLES_27061 | | | G | G | . | . | . | . | . | . | G | G |
| NC_011770 | 2902514 | T | C | 19 | 6 | 4 | 0 | NON_SYNONYMOU | MODERAT | MISSENSE | Aca/Gca | T3A | PLES_27061 | | | C | C | . | . | . | . | . | . | C | C |
| NC_011770 | 2902553 | G | GA | 9 | 8 | 2 | 0 | FRAME_SHIFT | HIGH | | acc/acTc | T554T? | PLES_27071 | | | GA | GA | . | . | . | . | . | . | . | . |
| NC_011770 | 2902552 | T | A | 9 | 8 | 2 | 0 | NON_SYNONYMOU | MODERAT | MISSENSE | Acg/Tcg | T555S | PLES_27071 | | | A | A | . | . | . | . | . | . | . | . |
| NC_011770 | 2902555 | T | C | 9 | 8 | 2 | 0 | NON_SYNONYMOU | MODERAT | MISSENSE | Acc/Gcc | T554A | PLES_27071 | | | C | C | . | . | . | . | . | . | . | . |
| NC_011770 | 2902825 | C | A | 7 | 8 | 2 | 0 | NON_SYNONYMOU | MODERAT | MISSENSE | Gcc/Tcc | A464S | PLES_27071 | | | A | A | . | . | . | . | . | . | . | . |

| | | | | | | | | | | | | | | | | | | | | | | | | | |
|---|---|---|---|---|---|---|---|---|---|---|---|---|---|---|---|---|---|---|---|---|---|---|---|---|---|
| NC_011770 | 2902837 | T | C | 15 | 6 | 2 | 0 | NON_SYNONYMOU | MODERAT | MISSENSE | Atc/Gtc | I460V | PLES_27071 | | | C | C | . | . | . | . | . | . | | |
| NC_011770 | 2902842 | T | G | 15 | 6 | 2 | 0 | NON_SYNONYMOU | MODERAT | MISSENSE | gAg/gCg | E458A | PLES_27071 | | | G | G | . | . | . | . | . | . | | |
| NC_011770 | 2902851 | C | T | 17 | 6 | 6 | 0 | NON_SYNONYMOU | MODERAT | MISSENSE | gGc/gAc | G455D | PLES_27071 | T | T | T | T | . | . | . | . | . | . | T | T |
| NC_011770 | 2902854 | G | A | 17 | 6 | 2 | 0 | NON_SYNONYMOU | MODERAT | MISSENSE | gCa/gTa | A454V | PLES_27071 | | | A | A | . | . | . | . | . | . | | |
| NC_011770 | 2902894 | T | C | 29 | 6 | 2 | 0 | NON_SYNONYMOU | MODERAT | MISSENSE | Atc/Gtc | I441V | PLES_27071 | | | C | C | . | . | . | . | . | . | | |
| NC_011770 | 2902904 | A | G | 21 | 6 | 4 | 0 | NON_SYNONYMOU | MODERAT | MISSENSE | gaT/gaG | D437E | PLES_27071 | | | G | G | . | . | . | . | . | . | C | C |
| NC_011770 | 2902905 | T | C | 20 | 6 | 2 | 0 | NON_SYNONYMOU | MODERAT | MISSENSE | gAt/gGt | D437G | PLES_27071 | | | C | C | . | . | . | . | . | . | | |
| NC_011770 | 2902909 | G | C | 22 | 6 | 2 | 0 | NON_SYNONYMOU | MODERAT | MISSENSE | Ccc/Gcc | P436A | PLES_27071 | | | | | . | . | . | . | . | . | C | C |
| NC_011770 | 2902912 | T | C | 22 | 6 | 2 | 0 | NON_SYNONYMOU | MODERAT | MISSENSE | Acg/Gcg | T435A | PLES_27071 | | | C | C | . | . | . | . | . | . | | |
| NC_011770 | 2902918 | G | C | 22 | 6 | 2 | 0 | NON_SYNONYMOU | MODERAT | MISSENSE | Ctg/Gtg | L433V | PLES_27071 | | | | | . | . | . | . | . | . | C | C |
| NC_011770 | 2902924 | C | G | 22 | 6 | 4 | 0 | NON_SYNONYMOU | MODERAT | MISSENSE | Gcc/Ccc | A431P | PLES_27071 | | | G | G | . | . | . | . | . | . | G | G |
| NC_011770 | 2902933 | A | G | 25 | 6 | 2 | 0 | NON_SYNONYMOU | MODERAT | MISSENSE | Tac/Cac | Y428H | PLES_27071 | | | G | G | . | . | . | . | . | . | | |
| NC_011770 | 2902939 | C | T | 24 | 6 | 2 | 0 | NON_SYNONYMOU | MODERAT | MISSENSE | Gtc/Atc | V426I | PLES_27071 | | | T | T | . | . | . | . | . | . | | |
| NC_011770 | 2902992 | C | T | 31 | 6 | 2 | 0 | NON_SYNONYMOU | MODERAT | MISSENSE | aGc/aAc | S408N | PLES_27071 | | | T | T | . | . | . | . | . | . | | |
| NC_011770 | 2903007 | G | A | 33 | 6 | 2 | 0 | NON_SYNONYMOU | MODERAT | MISSENSE | cCc/cTc | P403L | PLES_27071 | | | | | . | . | . | . | . | . | A | A |
| NC_011770 | 2903065 | C | G | 30 | 6 | 2 | 0 | NON_SYNONYMOU | MODERAT | MISSENSE | Gaa/Caa | E384Q | PLES_27071 | | | G | G | . | . | . | . | . | . | | |
| NC_011770 | 2903353 | G | T | 28 | 6 | 2 | 0 | NON_SYNONYMOU | MODERAT | MISSENSE | Ctg/Atg | L288M | PLES_27071 | | | | | . | . | . | . | . | . | T | T |
| NC_011770 | 2903376 | C | T | 28 | 6 | 4 | 0 | NON_SYNONYMOU | MODERAT | MISSENSE | gGc/gAc | G280D | PLES_27071 | | | T | T | . | . | . | . | . | . | T | T |
| NC_011770 | 2903380 | T | C | 28 | 6 | 2 | 0 | NON_SYNONYMOU | MODERAT | MISSENSE | Atc/Gtc | I279V | PLES_27071 | | | | | . | . | . | . | . | . | C | C |
| NC_011770 | 2903389 | G | T | 32 | 6 | 2 | 0 | NON_SYNONYMOU | MODERAT | MISSENSE | Ctg/Atg | L276M | PLES_27071 | | | | | . | . | . | . | . | . | T | T |
| NC_011770 | 2903395 | A | C | 33 | 6 | 2 | 0 | NON_SYNONYMOU | MODERAT | MISSENSE | Tcc/Gcc | S274A | PLES_27071 | | | | | . | . | . | . | . | . | C | C |
| NC_011770 | 2903431 | G | C | 17 | 7 | 1 | 0 | NON_SYNONYMOU | MODERAT | MISSENSE | Ccg/Gcg | P262A | PLES_27071 | | | | | . | . | . | . | . | . | . | C |
| NC_011770 | 2903434 | C | A | 18 | 7 | 3 | 0 | NON_SYNONYMOU | MODERAT | MISSENSE | Gcg/Tcg | A261S | PLES_27071 | | | A | A | . | . | . | . | . | . | . | A |
| NC_011770 | 2903445 | C | G | 18 | 8 | 2 | 0 | NON_SYNONYMOU | MODERAT | MISSENSE | gGc/gCc | G257A | PLES_27071 | | | G | G | . | . | . | . | . | . | . | . |
| NC_011770 | 2903464 | C | A | 18 | 8 | 2 | 0 | NON_SYNONYMOU | MODERAT | MISSENSE | Gcg/Tcg | A251S | PLES_27071 | | | A | A | . | . | . | . | . | . | . | . |
| NC_011770 | 2903511 | T | C | 19 | 8 | 2 | 0 | NON_SYNONYMOU | MODERAT | MISSENSE | gAc/gGc | D235G | PLES_27071 | | | C | C | . | . | . | . | . | . | . | . |
| NC_011770 | 2903539 | T | C | 18 | 8 | 2 | 0 | NON_SYNONYMOU | MODERAT | MISSENSE | Aaa/Gaa | K226E | PLES_27071 | | | C | C | . | . | . | . | . | . | . | . |
| NC_011770 | 2903542 | A | T | 18 | 8 | 2 | 0 | NON_SYNONYMOU | MODERAT | MISSENSE | Tgc/Agc | C225S | PLES_27071 | | | T | T | . | . | . | . | . | . | . | . |
| NC_011770 | 2903547 | G | T | 19 | 8 | 2 | 0 | NON_SYNONYMOU | MODERAT | MISSENSE | aCg/aAg | T223K | PLES_27071 | | | T | T | . | . | . | . | . | . | . | . |
| NC_011770 | 2903586 | A | T | 29 | 6 | 2 | 0 | NON_SYNONYMOU | MODERAT | MISSENSE | cTg/cAg | L210Q | PLES_27071 | T | T | | | . | . | . | . | . | . | | |
| NC_011770 | 2903628 | C | T | 33 | 6 | 2 | 0 | NON_SYNONYMOU | MODERAT | MISSENSE | cGc/cAc | R196H | PLES_27071 | | | | | . | . | . | . | . | . | T | T |
| NC_011770 | 2903640 | G | T | 36 | 6 | 2 | 0 | NON_SYNONYMOU | MODERAT | MISSENSE | gCc/gAc | A192D | PLES_27071 | | | | | . | . | . | . | . | . | T | T |
| NC_011770 | 2903641 | C | T | 36 | 6 | 2 | 0 | NON_SYNONYMOU | MODERAT | MISSENSE | Gcc/Acc | A192T | PLES_27071 | | | | | . | . | . | . | . | . | T | T |
| NC_011770 | 2903649 | G | C | 35 | 6 | 4 | 0 | NON_SYNONYMOU | MODERAT | MISSENSE | aCc/aGc | T189S | PLES_27071 | | | C | C | . | . | . | . | . | . | C | C |
| NC_011770 | 2903661 | C | T | 34 | 6 | 4 | 0 | NON_SYNONYMOU | MODERAT | MISSENSE | cGg/cAg | R185Q | PLES_27071 | | | T | T | . | . | . | . | . | . | T | T |
| NC_011770 | 2903685 | G | C | 34 | 6 | 2 | 0 | NON_SYNONYMOU | MODERAT | MISSENSE | gCg/gGg | A177G | PLES_27071 | | | C | C | . | . | . | . | . | . | | |
| NC_011770 | 2903719 | C | T | 29 | 6 | 4 | 0 | NON_SYNONYMOU | MODERAT | MISSENSE | Gcc/Acc | A166T | PLES_27071 | | | T | T | . | . | . | . | . | . | T | T |
| NC_011770 | 2903722 | C | T | 29 | 6 | 4 | 0 | NON_SYNONYMOU | MODERAT | MISSENSE | Gcc/Acc | A165T | PLES_27071 | | | T | T | . | . | . | . | . | . | T | T |
| NC_011770 | 2903725 | C | T | 30 | 6 | 2 | 0 | NON_SYNONYMOU | MODERAT | MISSENSE | Ggc/Agc | G164S | PLES_27071 | | | T | T | . | . | . | . | . | . | | |
| NC_011770 | 2903733 | T | G | 31 | 6 | 2 | 0 | NON_SYNONYMOU | MODERAT | MISSENSE | cAg/cCg | Q161P | PLES_27071 | | | G | G | . | . | . | . | . | . | | |
| NC_011770 | 2903734 | G | C | 31 | 6 | 2 | 0 | NON_SYNONYMOU | MODERAT | MISSENSE | Cag/Gag | Q161E | PLES_27071 | | | C | C | . | . | . | . | . | . | | |
| NC_011770 | 2903788 | G | A | 41 | 6 | 2 | 0 | NON_SYNONYMOU | MODERAT | MISSENSE | Ctc/Ttc | L143F | PLES_27071 | | | A | A | . | . | . | . | . | . | | |
| NC_011770 | 2903815 | G | A | 31 | 6 | 2 | 0 | NON_SYNONYMOU | MODERAT | MISSENSE | Ctt/Ttt | L134F | PLES_27071 | | | | | . | . | . | . | . | . | A | A |
| NC_011770 | 2903829 | G | A | 30 | 6 | 2 | 0 | NON_SYNONYMOU | MODERAT | MISSENSE | gCg/gTg | A129V | PLES_27071 | | | | | . | . | . | . | . | . | A | A |
| NC_011770 | 2903830 | C | T | 30 | 6 | 2 | 0 | NON_SYNONYMOU | MODERAT | MISSENSE | Gcg/Acg | A129T | PLES_27071 | | | | | . | . | . | . | . | . | T | T |
| NC_011770 | 2903836 | T | C | 30 | 6 | 2 | 0 | NON_SYNONYMOU | MODERAT | MISSENSE | Atc/Gtc | I127V | PLES_27071 | | | | | . | . | . | . | . | . | C | C |
| NC_011770 | 2903846 | C | G | 28 | 6 | 2 | 0 | NON_SYNONYMOU | MODERAT | MISSENSE | gaG/gaC | E123D | PLES_27071 | | | | | . | . | . | . | . | . | G | G |
| NC_011770 | 2903847 | T | G | 28 | 6 | 4 | 0 | NON_SYNONYMOU | MODERAT | MISSENSE | gAg/gCg | E123A | PLES_27071 | | | G | G | . | . | . | . | . | . | C | C |
| NC_011770 | 2903848 | C | T | 28 | 6 | 2 | 0 | NON_SYNONYMOU | MODERAT | MISSENSE | Gag/Aag | E123K | PLES_27071 | | | | | . | . | . | . | . | . | T | T |
| NC_011770 | 2903875 | T | C | 36 | 6 | 2 | 0 | NON_SYNONYMOU | MODERAT | MISSENSE | Ata/Gta | I114V | PLES_27071 | | | | | . | . | . | . | . | . | C | C |
| NC_011770 | 2903891 | T | G | 35 | 6 | 4 | 0 | NON_SYNONYMOU | MODERAT | MISSENSE | gaA/gaC | E108D | PLES_27071 | | | G | G | . | . | . | . | . | . | G | G |
| NC_011770 | 2903905 | A | T | 36 | 6 | 2 | 0 | NON_SYNONYMOU | MODERAT | MISSENSE | Tcg/Acg | S104T | PLES_27071 | | | T | T | . | . | . | . | . | . | | |
| NC_011770 | 2903963 | G | C | 36 | 6 | 2 | 0 | NON_SYNONYMOU | MODERAT | MISSENSE | gaC/gaG | D84E | PLES_27071 | | | C | C | . | . | . | . | . | . | | |
| NC_011770 | 2903964 | T | G | 33 | 6 | 4 | 0 | NON_SYNONYMOU | MODERAT | MISSENSE | gAc/gCc | D84A | PLES_27071 | | | G | G | . | . | . | . | . | . | G | G |
| NC_011770 | 2903965 | C | A | 33 | 6 | 2 | 0 | NON_SYNONYMOU | MODERAT | MISSENSE | Gac/Tac | D84Y | PLES_27071 | | | | | . | . | . | . | . | . | A | A |
| NC_011770 | 2903975 | C | A | 32 | 6 | 4 | 0 | NON_SYNONYMOU | MODERAT | MISSENSE | agG/agT | R80S | PLES_27071 | | | A | A | . | . | . | . | . | . | T | T |
| NC_011770 | 2904035 | C | A | 37 | 6 | 2 | 0 | NON_SYNONYMOU | MODERAT | MISSENSE | gaG/gaT | E60D | PLES_27071 | | | A | A | . | . | . | . | . | . | | |
| NC_011770 | 2904046 | G | A | 36 | 6 | 2 | 0 | NON_SYNONYMOU | MODERAT | MISSENSE | Ccg/Tcg | P57S | PLES_27071 | | | A | A | . | . | . | . | . | . | | |
| NC_011770 | 2904056 | C | T | 30 | 6 | 4 | 0 | NON_SYNONYMOU | MODERAT | MISSENSE | atG/atA | M53I | PLES_27071 | | | T | T | . | . | . | . | . | . | T | T |
| NC_011770 | 2904057 | A | T | 30 | 6 | 4 | 0 | NON_SYNONYMOU | MODERAT | MISSENSE | aTg/aAg | M53K | PLES_27071 | | | T | T | . | . | . | . | . | . | T | T |
| NC_011770 | 2904061 | G | T | 30 | 6 | 2 | 0 | NON_SYNONYMOU | MODERAT | MISSENSE | Cgc/Agc | R52S | PLES_27071 | | | | | . | . | . | . | . | . | T | T |
| NC_011770 | 2904136 | C | T | 35 | 6 | 2 | 0 | NON_SYNONYMOU | MODERAT | MISSENSE | Gcc/Acc | A27T | PLES_27071 | | | | | . | . | . | . | . | . | T | T |

| | | | | | | | | | | | | | | | | | | | | | | | |
|---|---|---|---|---|---|---|---|---|---|---|---|---|---|---|---|---|---|---|---|---|---|---|---|
| NC_011770 | 2904139 | T | A | 34 | 6 | 2 | 0 | NON_SYNONYMOU | MODERAT | MISSENSE | Acg/Tcg | T26S | PLES_27071 | | | . | . | . | . | . | . | A | A |
| NC_011770 | 2904145 | C | A | 33 | 6 | 4 | 0 | NON_SYNONYMOU | MODERAT | MISSENSE | Gcg/Tcg | A24S | PLES_27071 | A | A | . | . | . | . | . | . | A | A |
| NC_011770 | 2904191 | G | C | 34 | 6 | 2 | 0 | NON_SYNONYMOU | MODERAT | MISSENSE | gaC/gaG | D8E | PLES_27071 | | | . | . | . | . | . | . | C | C |
| NC_011770 | 2904193 | C | G | 34 | 6 | 2 | 0 | NON_SYNONYMOU | MODERAT | MISSENSE | Gac/Cac | D8H | PLES_27071 | | | . | . | . | . | . | . | G | G |
| NC_011770 | 2904203 | C | G | 31 | 6 | 2 | 0 | NON_SYNONYMOU | MODERAT | MISSENSE | atG/atC | M4I | PLES_27071 | | | . | . | . | . | . | . | G | G |
| NC_011770 | 2904206 | G | C | 31 | 6 | 4 | 0 | NON_SYNONYMOU | MODERAT | MISSENSE | gaC/gaG | D3E | PLES_27071 | C | C | . | . | . | . | . | . | C | C |
| NC_011770 | 2904476 | T | C | 32 | 6 | 2 | 0 | NON_SYNONYMOU | MODERAT | MISSENSE | cAt/cGt | H287R | PLES_27081 | C | C | . | . | . | . | . | . | | |
| NC_011770 | 2904480 | G | A | 30 | 6 | 4 | 0 | NON_SYNONYMOU | MODERAT | MISSENSE | Ccc/Tcc | P286S | PLES_27081 | A | A | . | . | . | . | . | . | A | A |
| NC_011770 | 2904489 | G | C | 18 | 8 | 2 | 0 | NON_SYNONYMOU | MODERAT | MISSENSE | Cct/Gct | P283A | PLES_27081 | . | . | . | . | . | . | . | . | C | C |
| NC_011770 | 2904558 | T | C | 15 | 8 | 2 | 0 | NON_SYNONYMOU | MODERAT | MISSENSE | Agc/Ggc | S260G | PLES_27081 | C | C | . | . | . | . | . | . | . | . |
| NC_011770 | 2904567 | C | T | 17 | 6 | 2 | 0 | NON_SYNONYMOU | MODERAT | MISSENSE | Gcg/Acg | A257T | PLES_27081 | T | T | . | . | . | . | . | . | | |
| NC_011770 | 2904590 | G | A | 27 | 6 | 2 | 0 | NON_SYNONYMOU | MODERAT | MISSENSE | cCt/cTt | P249L | PLES_27081 | | | . | . | . | . | . | . | A | A |
| NC_011770 | 2904618 | G | A | 31 | 6 | 4 | 0 | NON_SYNONYMOU | MODERAT | MISSENSE | Cac/Tac | H240Y | PLES_27081 | A | A | . | . | . | . | . | . | A | A |
| NC_011770 | 2904639 | G | T | 36 | 6 | 4 | 0 | NON_SYNONYMOU | MODERAT | MISSENSE | Ctg/Atg | L233M | PLES_27081 | T | T | . | . | . | . | . | . | A | A |
| NC_011770 | 2904689 | G | C | 39 | 6 | 2 | 0 | NON_SYNONYMOU | MODERAT | MISSENSE | gCc/gGc | A216G | PLES_27081 | C | C | . | . | . | . | . | . | | |
| NC_011770 | 2904708 | C | T | 34 | 6 | 2 | 0 | NON_SYNONYMOU | MODERAT | MISSENSE | Gcc/Acc | A210T | PLES_27081 | T | T | . | . | . | . | . | . | | |
| NC_011770 | 2904710 | C | T | 34 | 6 | 4 | 0 | NON_SYNONYMOU | MODERAT | MISSENSE | aGc/aAc | S209N | PLES_27081 | T | T | . | . | . | . | . | . | T | T |
| NC_011770 | 2904711 | T | G | 33 | 6 | 4 | 0 | NON_SYNONYMOU | MODERAT | MISSENSE | Agc/Cgc | S209R | PLES_27081 | G | G | . | . | . | . | . | . | G | G |
| NC_011770 | 2904728 | C | T | 33 | 6 | 2 | 0 | NON_SYNONYMOU | MODERAT | MISSENSE | cGc/cAc | R203H | PLES_27081 | T | T | . | . | . | . | . | . | | |
| NC_011770 | 2904747 | G | T | 40 | 6 | 2 | 0 | NON_SYNONYMOU | MODERAT | MISSENSE | Ctg/Atg | L197M | PLES_27081 | T | T | . | . | . | . | . | . | | |
| NC_011770 | 2904914 | T | C | 41 | 6 | 2 | 0 | NON_SYNONYMOU | MODERAT | MISSENSE | gAt/gGt | D141G | PLES_27081 | C | C | . | . | . | . | . | . | | |
| NC_011770 | 2904915 | C | T | 42 | 6 | 2 | 0 | NON_SYNONYMOU | MODERAT | MISSENSE | Gat/Aat | D141N | PLES_27081 | | | . | . | . | . | . | . | T | T |
| NC_011770 | 2904998 | G | A | 32 | 6 | 2 | 0 | NON_SYNONYMOU | MODERAT | MISSENSE | gCa/gTa | A113V | PLES_27081 | | | . | . | . | . | . | . | A | A |
| NC_011770 | 2905038 | C | T | 42 | 6 | 2 | 0 | NON_SYNONYMOU | MODERAT | MISSENSE | Gcg/Acg | A100T | PLES_27081 | | | . | . | . | . | . | . | T | T |
| NC_011770 | 2905077 | C | T | 43 | 6 | 4 | 0 | NON_SYNONYMOU | MODERAT | MISSENSE | Gac/Aac | D87N | PLES_27081 | T | T | . | . | . | . | . | . | T | T |
| NC_011770 | 2905100 | G | T | 46 | 6 | 2 | 0 | NON_SYNONYMOU | MODERAT | MISSENSE | gCg/gAg | A79E | PLES_27081 | | | . | . | . | . | . | . | T | T |
| NC_011770 | 2905106 | A | T | 46 | 6 | 2 | 0 | NON_SYNONYMOU | MODERAT | MISSENSE | aTc/aAc | I77N | PLES_27081 | | | . | . | . | . | . | . | T | T |
| NC_011770 | 2905124 | T | A | 44 | 6 | 2 | 0 | NON_SYNONYMOU | MODERAT | MISSENSE | cAg/cTg | Q71L | PLES_27081 | | | . | . | . | . | . | . | A | A |
| NC_011770 | 2905127 | C | T | 45 | 6 | 2 | 0 | NON_SYNONYMOU | MODERAT | MISSENSE | gGc/gAc | G70D | PLES_27081 | T | T | . | . | . | . | . | . | | |
| NC_011770 | 2905133 | T | C | 44 | 6 | 2 | 0 | NON_SYNONYMOU | MODERAT | MISSENSE | gAc/gGc | D68G | PLES_27081 | C | C | . | . | . | . | . | . | | |
| NC_011770 | 2905180 | C | G | 41 | 6 | 4 | 0 | NON_SYNONYMOU | MODERAT | MISSENSE | caG/caC | Q52H | PLES_27081 | G | G | . | . | . | . | . | . | G | G |
| NC_011770 | 2905184 | G | C | 40 | 6 | 2 | 0 | NON_SYNONYMOU | MODERAT | MISSENSE | aCc/aGc | T51S | PLES_27081 | | | . | . | . | . | . | . | C | C |
| NC_011770 | 2905185 | T | C | 40 | 6 | 4 | 0 | NON_SYNONYMOU | MODERAT | MISSENSE | Acc/Gcc | T51A | PLES_27081 | C | C | . | . | . | . | . | . | C | C |
| NC_011770 | 2905189 | T | A | 40 | 6 | 4 | 0 | NON_SYNONYMOU | MODERAT | MISSENSE | gaA/gaT | E49D | PLES_27081 | A | A | . | . | . | . | . | . | C | C |
| NC_011770 | 2905478 | CGCTT | C | 42 | 6 | 2 | 0 | FRAME_SHIFT | HIGH | | aagcgc/ | KR37 | PLES_27091 | | | . | . | . | . | . | . | C | C |
| NC_011770 | 2905563 | TCG | GCG | 41 | 6 | 4 | 0 | FRAME_SHIFT | HIGH | | ccg/ | P9 | PLES_27091 | GCG | GCG | . | . | . | . | . | . | T | T |
| NC_011770 | 2905570 | G | C | 42 | 6 | 4 | 0 | FRAME_SHIFT | HIGH | | ctc/ctGc | L7L? | PLES_27091 | C | C | . | . | . | . | . | . | GC | GC |
| NC_011770 | 2905383 | T | C | 43 | 6 | 2 | 0 | NON_SYNONYMOU | MODERAT | MISSENSE | Acc/Gcc | T70A | PLES_27091 | | | . | . | . | . | . | . | C | C |
| NC_011770 | 2905389 | G | T | 43 | 6 | 4 | 0 | NON_SYNONYMOU | MODERAT | MISSENSE | Cac/Aac | H68N | PLES_27091 | T | T | . | . | . | . | . | . | T | T |
| NC_011770 | 2905395 | G | C | 44 | 6 | 2 | 0 | NON_SYNONYMOU | MODERAT | MISSENSE | Ctc/Gtc | L66V | PLES_27091 | | | . | . | . | . | . | . | C | C |
| NC_011770 | 2905425 | T | C | 45 | 6 | 2 | 0 | NON_SYNONYMOU | MODERAT | MISSENSE | Atc/Gtc | I56V | PLES_27091 | C | C | . | . | . | . | . | . | | |
| NC_011770 | 2905467 | G | T | 42 | 6 | 2 | 0 | NON_SYNONYMOU | MODERAT | MISSENSE | Cag/Aag | Q42K | PLES_27091 | | | . | . | . | . | . | . | T | T |
| NC_011770 | 2905475 | T | A | 42 | 6 | 2 | 0 | NON_SYNONYMOU | MODERAT | MISSENSE | cAg/cTg | Q39L | PLES_27091 | | | . | . | . | . | . | . | A | A |
| NC_011770 | 2905527 | C | T | 47 | 6 | 2 | 0 | NON_SYNONYMOU | MODERAT | MISSENSE | Gcg/Acg | A22T | PLES_27091 | | | . | . | . | . | . | . | T | T |
| NC_011770 | 2905574 | A | G | 44 | 6 | 2 | 0 | NON_SYNONYMOU | MODERAT | MISSENSE | gTa/gCa | V6A | PLES_27091 | | | . | . | . | . | . | . | G | G |
| NC_011770 | 2905580 | G | A | 41 | 6 | 2 | 0 | NON_SYNONYMOU | MODERAT | MISSENSE | aCc/aTc | T4I | PLES_27091 | A | A | . | . | . | . | . | . | | |
| NC_011770 | 2905581 | T | C | 40 | 6 | 2 | 0 | NON_SYNONYMOU | MODERAT | MISSENSE | Acc/Gcc | T4A | PLES_27091 | C | C | . | . | . | . | . | . | | |
| NC_011770 | 2906257 | C | A | 27 | 8 | 2 | 0 | STOP_GAINED | HIGH | NONSENSE | Gga/Tga | G67* | PLES_27101 | A | A | . | . | . | . | . | . | . | . |
| NC_011770 | 2906366 | CATCGGGA | C | 8 | 8 | 2 | 0 | FRAME_SHIFT | HIGH | | ctcccgatg/ | LPM28 | PLES_27101 | C | C | . | . | . | . | . | . | . | . |
| NC_011770 | 2905716 | C | G | 21 | 8 | 2 | 0 | NON_SYNONYMOU | MODERAT | MISSENSE | aGc/aCc | S247T | PLES_27101 | G | G | . | . | . | . | . | . | . | . |
| NC_011770 | 2905717 | T | C | 21 | 8 | 2 | 0 | NON_SYNONYMOU | MODERAT | MISSENSE | Agc/Ggc | S247G | PLES_27101 | C | C | . | . | . | . | . | . | . | . |
| NC_011770 | 2905732 | A | T | 22 | 8 | 2 | 0 | NON_SYNONYMOU | MODERAT | MISSENSE | Tac/Aac | Y242N | PLES_27101 | T | T | . | . | . | . | . | . | . | . |
| NC_011770 | 2905734 | C | T | 22 | 8 | 2 | 0 | NON_SYNONYMOU | MODERAT | MISSENSE | aGg/aAg | R241K | PLES_27101 | T | T | . | . | . | . | . | . | . | . |
| NC_011770 | 2905737 | T | C | 22 | 8 | 2 | 0 | NON_SYNONYMOU | MODERAT | MISSENSE | aAg/aGg | K240R | PLES_27101 | C | C | . | . | . | . | . | . | . | . |
| NC_011770 | 2905804 | C | G | 31 | 8 | 2 | 0 | NON_SYNONYMOU | MODERAT | MISSENSE | Gtc/Ctc | V218L | PLES_27101 | G | G | . | . | . | . | . | . | . | . |
| NC_011770 | 2905813 | A | T | 29 | 8 | 2 | 0 | NON_SYNONYMOU | MODERAT | MISSENSE | Tct/Act | S215T | PLES_27101 | T | T | . | . | . | . | . | . | . | . |
| NC_011770 | 2905819 | G | T | 29 | 8 | 2 | 0 | NON_SYNONYMOU | MODERAT | MISSENSE | Ctt/Att | L213I | PLES_27101 | T | T | . | . | . | . | . | . | . | . |
| NC_011770 | 2905870 | T | C | 15 | 8 | 2 | 0 | NON_SYNONYMOU | MODERAT | MISSENSE | Agc/Ggc | S196G | PLES_27101 | C | C | . | . | . | . | . | . | . | . |
| NC_011770 | 2905925 | T | A | 16 | 8 | 2 | 0 | NON_SYNONYMOU | MODERAT | MISSENSE | gaA/gaT | E177D | PLES_27101 | A | A | . | . | . | . | . | . | . | . |
| NC_011770 | 2905935 | T | G | 30 | 8 | 2 | 0 | NON_SYNONYMOU | MODERAT | MISSENSE | gAg/gCg | E174A | PLES_27101 | G | G | . | . | . | . | . | . | . | . |
| NC_011770 | 2905963 | A | C | 28 | 8 | 2 | 0 | NON_SYNONYMOU | MODERAT | MISSENSE | Tcg/Gcg | S165A | PLES_27101 | C | C | . | . | . | . | . | . | . | . |
| NC_011770 | 2906080 | T | G | 14 | 8 | 2 | 0 | NON_SYNONYMOU | MODERAT | MISSENSE | Atg/Ctg | M126L | PLES_27101 | G | G | . | . | . | . | . | . | . | . |

| | | | | | | | | | | | | | | | | | | | | | | | | | |
|---|---|---|---|---|---|---|---|---|---|---|---|---|---|---|---|---|---|---|---|---|---|---|---|---|---|
| NC_011770 | 2906088 | A | G | 15 | 8 | 2 | 0 | NON_SYNONYMOU | MODERAT | MISSENSE | tTc/tCc | F123S | PLES_27101 | | | G | G | . | . | . | . | . | . | . | . |
| NC_011770 | 2906110 | C | T | 14 | 8 | 2 | 0 | NON_SYNONYMOU | MODERAT | MISSENSE | Ggc/Agc | G116S | PLES_27101 | | | T | T | . | . | . | . | . | . | . | . |
| NC_011770 | 2906151 | G | T | 22 | 6 | 2 | 0 | NON_SYNONYMOU | MODERAT | MISSENSE | aCc/aAc | T102N | PLES_27101 | | | | | . | . | . | . | . | . | T | T |
| NC_011770 | 2906152 | T | G | 21 | 6 | 2 | 0 | NON_SYNONYMOU | MODERAT | MISSENSE | Acc/Ccc | T102P | PLES_27101 | | | | | . | . | . | . | . | . | G | G |
| NC_011770 | 2906185 | C | T | 18 | 6 | 2 | 0 | NON_SYNONYMOU | MODERAT | MISSENSE | Gtt/Att | V91I | PLES_27101 | | | T | T | . | . | . | . | . | . | | |
| NC_011770 | 2906196 | A | G | 10 | 8 | 2 | 0 | NON_SYNONYMOU | MODERAT | MISSENSE | cTg/cCg | L87P | PLES_27101 | | | G | G | . | . | . | . | . | . | . | . |
| NC_011770 | 2906202 | T | C | 10 | 8 | 2 | 0 | NON_SYNONYMOU | MODERAT | MISSENSE | aAt/aGt | N85S | PLES_27101 | | | C | C | . | . | . | . | . | . | . | . |
| NC_011770 | 2906206 | G | C | 10 | 8 | 2 | 0 | NON_SYNONYMOU | MODERAT | MISSENSE | Ctc/Gtc | L84V | PLES_27101 | | | C | C | . | . | . | . | . | . | . | . |
| NC_011770 | 2906210 | G | C | 10 | 8 | 2 | 0 | NON_SYNONYMOU | MODERAT | MISSENSE | aaC/aaG | N82K | PLES_27101 | | | C | C | . | . | . | . | . | . | . | . |
| NC_011770 | 2906212 | T | C | 10 | 8 | 2 | 0 | NON_SYNONYMOU | MODERAT | MISSENSE | Aac/Gac | N82D | PLES_27101 | | | C | C | . | . | . | . | . | . | . | . |
| NC_011770 | 2906244 | T | A | 27 | 8 | 2 | 0 | NON_SYNONYMOU | MODERAT | MISSENSE | tAc/tTc | Y71F | PLES_27101 | | | A | A | . | . | . | . | . | . | . | . |
| NC_011770 | 2906256 | C | G | 27 | 8 | 2 | 0 | NON_SYNONYMOU | MODERAT | MISSENSE | gGa/gCa | G67A | PLES_27101 | | | G | G | . | . | . | . | . | . | . | . |
| NC_011770 | 2906290 | G | A | 24 | 8 | 2 | 0 | NON_SYNONYMOU | MODERAT | MISSENSE | Cac/Tac | H56Y | PLES_27101 | | | A | A | . | . | . | . | . | . | . | . |
| NC_011770 | 2906320 | T | C | 8 | 8 | 2 | 0 | NON_SYNONYMOU | MODERAT | MISSENSE | Aac/Gac | N46D | PLES_27101 | | | C | C | . | . | . | . | . | . | . | . |
| NC_011770 | 2906326 | A | T | 7 | 8 | 2 | 0 | NON_SYNONYMOU | MODERAT | MISSENSE | Tcg/Acg | S44T | PLES_27101 | | | T | T | . | . | . | . | . | . | . | . |
| NC_011770 | 2906328 | C | T | 8 | 8 | 2 | 0 | NON_SYNONYMOU | MODERAT | MISSENSE | cGg/cAg | R43Q | PLES_27101 | | | T | T | . | . | . | . | . | . | . | . |
| NC_011770 | 2906337 | A | C | 7 | 8 | 2 | 0 | NON_SYNONYMOU | MODERAT | MISSENSE | gTc/gGc | V40G | PLES_27101 | | | C | C | . | . | . | . | . | . | . | . |
| NC_011770 | 2906338 | C | A | 8 | 8 | 2 | 0 | NON_SYNONYMOU | MODERAT | MISSENSE | Gtc/Ttc | V40F | PLES_27101 | | | A | A | . | . | . | . | . | . | . | . |
| NC_011770 | 2906356 | T | A | 8 | 8 | 2 | 0 | NON_SYNONYMOU | MODERAT | MISSENSE | Acg/Tcg | T34S | PLES_27101 | | | A | A | . | . | . | . | . | . | . | . |
| NC_011770 | 2906404 | C | A | 29 | 6 | 2 | 0 | NON_SYNONYMOU | MODERAT | MISSENSE | Gcc/Tcc | A18S | PLES_27101 | | | | | . | . | . | . | . | . | A | A |
| NC_011770 | 2908719 | A | G | 63 | 0 | 1 | 0 | NON_SYNONYMOU | MODERAT | MISSENSE | gAg/gGg | E54G | PLES_27111 | | | | | | | G | | | | | |
| NC_011770 | 2908790 | G | A | 68 | 0 | 1 | 0 | NON_SYNONYMOU | MODERAT | MISSENSE | Ggc/Agc | G78S | PLES_27111 | | | | | | | A | | | | | |
| NC_011770 | 2909231 | C | G | 76 | 0 | 2 | 0 | NON_SYNONYMOU | MODERAT | MISSENSE | Ccc/Gcc | P225A | PLES_27111 | | | | | | | | | G | G | | |
| NC_011770 | 2909318 | G | A | 75 | 0 | 12 | 0 | NON_SYNONYMOU | MODERAT | MISSENSE | Ggc/Agc | G254S | PLES_27111 | A | A | A | A | A | A | A | A | A | A | A | A |
| NC_011770 | 2909383 | G | T | 71 | 0 | 8 | 0 | NON_SYNONYMOU | MODERAT | MISSENSE | gaG/gaT | E275D | PLES_27111 | T | T | T | T | T | T | | | T | T | | |
| NC_011770 | 2909480 | C | G | 74 | 0 | 2 | 0 | NON_SYNONYMOU | MODERAT | MISSENSE | Ctg/Gtg | L308V | PLES_27111 | | | | | | | | | G | G | | |
| NC_011770 | 2909487 | A | G | 74 | 0 | 2 | 0 | NON_SYNONYMOU | MODERAT | MISSENSE | aAc/aGc | N310S | PLES_27111 | | | | | | | | | G | G | | |
| NC_011770 | 2909496 | G | A | 73 | 0 | 2 | 0 | NON_SYNONYMOU | MODERAT | MISSENSE | aGg/aAg | R313K | PLES_27111 | | | | | | | | | A | A | | |
| NC_011770 | 2909527 | C | A | 75 | 0 | 2 | 0 | NON_SYNONYMOU | MODERAT | MISSENSE | gaC/gaA | D323E | PLES_27111 | | | | | | | | | A | A | | |
| NC_011770 | 2909553 | G | A | 72 | 0 | 6 | 0 | NON_SYNONYMOU | MODERAT | MISSENSE | cGg/cAg | R332Q | PLES_27111 | | | A | A | A | A | | | | | A | A |
| NC_011770 | 2909610 | A | G | 70 | 0 | 6 | 0 | NON_SYNONYMOU | MODERAT | MISSENSE | gAg/gGg | E351G | PLES_27111 | | | G | G | G | G | | | | | G | G |
| NC_011770 | 2909693 | G | A | 68 | 0 | 6 | 0 | NON_SYNONYMOU | MODERAT | MISSENSE | Gcc/Acc | A379T | PLES_27111 | | | A | A | A | A | | | | | A | A |
| NC_011770 | 2910039 | A | G | 61 | 0 | 2 | 0 | NON_SYNONYMOU | MODERAT | MISSENSE | cAg/cGg | Q494R | PLES_27111 | | | | | | | | | | | G | G |
| NC_011770 | 2910270 | A | C | 42 | 0 | 2 | 0 | NON_SYNONYMOU | MODERAT | MISSENSE | gAc/gCc | D571A | PLES_27111 | C | C | | | | | | | | | | |
| NC_011770 | 2910272 | A | G | 42 | 0 | 10 | 0 | NON_SYNONYMOU | MODERAT | MISSENSE | Act/Gct | T572A | PLES_27111 | G | G | G | G | G | G | G | G | | | G | G |
| NC_011770 | 2910613 | A | C | 30 | 0 | 6 | 0 | NON_SYNONYMOU | MODERAT | MISSENSE | caA/caC | Q685H | PLES_27111 | C | C | | | | | C | C | | | C | C |
| NC_011770 | 2910986 | A | T | 42 | 0 | 2 | 0 | NON_SYNONYMOU | MODERAT | MISSENSE | Agc/Tgc | S810C | PLES_27111 | | | | | | | | | T | T | | |
| NC_011770 | 2911032 | A | G | 43 | 0 | 12 | 0 | NON_SYNONYMOU | MODERAT | MISSENSE | cAg/cGg | Q825R | PLES_27111 | G | G | G | G | G | G | G | G | G | G | G | G |
| NC_011770 | 2911791 | A | G | 62 | 0 | 2 | 0 | NON_SYNONYMOU | MODERAT | MISSENSE | gAa/gGa | E86G | PLES_27121 | | | | | | | | | G | G | | |
| NC_011770 | 2911992 | A | G | 59 | 0 | 8 | 0 | NON_SYNONYMOU | MODERAT | MISSENSE | gAt/gGt | D153G | PLES_27121 | | | G | G | G | G | | | G | G | G | G |
| NC_011770 | 2911994 | A | G | 58 | 0 | 8 | 0 | NON_SYNONYMOU | MODERAT | MISSENSE | Agc/Ggc | S154G | PLES_27121 | | | G | G | G | G | | | G | G | G | G |
| NC_011770 | 2912003 | G | C | 58 | 0 | 8 | 0 | NON_SYNONYMOU | MODERAT | MISSENSE | Gcg/Ccg | A157P | PLES_27121 | | | C | C | C | C | | | C | C | C | C |
| NC_011770 | 2912036 | T | C | 60 | 0 | 8 | 0 | NON_SYNONYMOU | MODERAT | MISSENSE | Tcc/Ccc | S168P | PLES_27121 | | | C | C | C | C | | | C | C | C | C |
| NC_011770 | 2911682 | C | T | 67 | 0 | 1 | 0 | NON_SYNONYMOU | MODERAT | MISSENSE | Cgg/Tgg | R50W | PLES_27121 | | T | | | | | | | | | | |
| NC_011770 | 2912985 | C | T | 83 | 0 | 4 | 0 | NON_SYNONYMOU | MODERAT | MISSENSE | gaG/gaT | E293D | PLES_27131 | T | T | | | | | | | | | A | A |
| NC_011770 | 2913073 | T | G | 87 | 0 | 2 | 0 | NON_SYNONYMOU | MODERAT | MISSENSE | gAc/gCc | D264A | PLES_27131 | | | | | | | G | G | | | | |
| NC_011770 | 2913422 | C | T | 83 | 0 | 2 | 0 | NON_SYNONYMOU | MODERAT | MISSENSE | Gcc/Acc | A148T | PLES_27131 | | | | | | | T | T | | | | |
| NC_011770 | 2913769 | C | T | 79 | 0 | 4 | 0 | NON_SYNONYMOU | MODERAT | MISSENSE | gGc/gAc | G32D | PLES_27131 | | | T | T | T | T | | | | | | |
| NC_011770 | 2914413 | C | T | 80 | 0 | 6 | 0 | NON_SYNONYMOU | MODERAT | MISSENSE | Ctc/Ttc | L43F | PLES_27141 | T | T | | | | | T | T | T | T | | |
| NC_011770 | 2914809 | C | A | 67 | 0 | 4 | 0 | NON_SYNONYMOU | MODERAT | MISSENSE | Ccg/Acg | P175T | PLES_27141 | | | A | A | A | A | | | | | | |
| NC_011770 | 2914824 | A | G | 65 | 0 | 4 | 0 | NON_SYNONYMOU | MODERAT | MISSENSE | Acc/Gcc | T180A | PLES_27141 | | | G | G | G | G | | | | | | |
| NC_011770 | 2914303 | T | C | 81 | 0 | 1 | 0 | NON_SYNONYMOU | MODERAT | MISSENSE | cTg/cCg | L6P | PLES_27141 | C | | | | | | | | | | | |
| NC_011770 | 2915226 | G | A | 75 | 0 | 4 | 0 | NON_SYNONYMOU | MODERAT | MISSENSE | Gcc/Acc | A9T | PLES_27151 | A | A | | | | | | | | | A | A |
| NC_011770 | 2915318 | C | G | 78 | 0 | 4 | 0 | NON_SYNONYMOU | MODERAT | MISSENSE | gaC/gaG | D39E | PLES_27151 | G | G | | | | | | | | | G | G |
| NC_011770 | 2915785 | C | T | 76 | 0 | 4 | 0 | NON_SYNONYMOU | MODERAT | MISSENSE | gCc/gTc | A195V | PLES_27151 | | | T | T | T | T | | | | | | |
| NC_011770 | 2915817 | A | G | 77 | 0 | 12 | 0 | NON_SYNONYMOU | MODERAT | MISSENSE | Acc/Gcc | T206A | PLES_27151 | G | G | G | G | G | G | G | G | G | G | G | G |
| NC_011770 | 2916368 | G | A | 76 | 0 | 4 | 0 | NON_SYNONYMOU | MODERAT | MISSENSE | gCc/gTc | A98V | PLES_27161 | | | | | | | | | A | A | A | A |
| NC_011770 | 2916602 | T | C | 79 | 0 | 2 | 0 | NON_SYNONYMOU | MODERAT | MISSENSE | gAa/gGa | E20G | PLES_27161 | | | | | | | | | C | C | | |
| NC_011770 | 2916965 | C | T | 67 | 0 | 2 | 0 | NON_SYNONYMOU | MODERAT | MISSENSE | Ggc/Agc | G50S | PLES_27171 | T | T | | | | | | | | | | |
| NC_011770 | 2917091 | G | A | 71 | 0 | 1 | 0 | NON_SYNONYMOU | MODERAT | MISSENSE | Cgc/Tgc | R8C | PLES_27171 | | | | | | | | | | | | A |
| NC_011770 | 2917440 | A | G | 72 | 0 | 10 | 0 | NON_SYNONYMOU | MODERAT | MISSENSE | Aag/Gag | K66E | PLES_27181 | | | G | G | G | G | G | G | G | G | G | G |
| NC_011770 | 2917585 | C | T | 70 | 0 | 2 | 0 | NON_SYNONYMOU | MODERAT | MISSENSE | gCg/gTg | A114V | PLES_27181 | | | | | | | | | | | T | T |

| | | | | | | | | | | | | | | | | | | | | | | | | | |
|---|---|---|---|---|---|---|---|---|---|---|---|---|---|---|---|---|---|---|---|---|---|---|---|---|---|
| NC_011770 | 2917589 | T | A | 70 | 0 | 2 | 0 | NON_SYNONYMOU | MODERAT | MISSENSE | ttT/ttA | F115L | PLES_27181 | | | | | | | | | A | A | | |
| NC_011770 | 2917663 | G | A | 70 | 0 | 6 | 0 | NON_SYNONYMOU | MODERAT | MISSENSE | aGc/aAc | S140N | PLES_27181 | | | A | A | A | A | A | A | | | | |
| NC_011770 | 2917681 | A | G | 75 | 0 | 6 | 0 | NON_SYNONYMOU | MODERAT | MISSENSE | gAc/gGc | D146G | PLES_27181 | | | G | G | G | G | G | G | | | | |
| NC_011770 | 2917866 | G | C | 91 | 0 | 2 | 0 | NON_SYNONYMOU | MODERAT | MISSENSE | Gac/Cac | D208H | PLES_27181 | | | | | | | | | | | C | C |
| NC_011770 | 2918223 | T | C | 71 | 0 | 2 | 0 | NON_SYNONYMOU | MODERAT | MISSENSE | aAg/aGg | K196R | PLES_27191 | | | | | | | | | C | C | | |
| NC_011770 | 2920989 | G | A | 65 | 0 | 2 | 0 | NON_SYNONYMOU | MODERAT | MISSENSE | Gtc/Atc | V192I | PLES_27211 | A | A | | | | | | | | | | |
| NC_011770 | 2921031 | G | C | 66 | 0 | 6 | 0 | NON_SYNONYMOU | MODERAT | MISSENSE | Ggt/Cgt | G206R | PLES_27211 | C | C | | | | | C | C | C | C | | |
| NC_011770 | 2921043 | A | C | 65 | 0 | 6 | 0 | NON_SYNONYMOU | MODERAT | MISSENSE | Aag/Cag | K210Q | PLES_27211 | C | C | | | | | C | C | C | C | | |
| NC_011770 | 2921064 | G | T | 63 | 0 | 6 | 0 | NON_SYNONYMOU | MODERAT | MISSENSE | Gcc/Tcc | A217S | PLES_27211 | T | T | | | | | T | T | T | T | | |
| NC_011770 | 2921121 | G | A | 67 | 0 | 2 | 0 | NON_SYNONYMOU | MODERAT | MISSENSE | Gac/Aac | D236N | PLES_27211 | A | A | | | | | | | | | | |
| NC_011770 | 2921262 | G | C | 70 | 0 | 4 | 0 | NON_SYNONYMOU | MODERAT | MISSENSE | Gac/Cac | D283H | PLES_27211 | C | C | | | | | | | C | C | | |
| NC_011770 | 2921600 | G | C | 71 | 0 | 4 | 0 | NON_SYNONYMOU | MODERAT | MISSENSE | gaG/gaC | E395D | PLES_27211 | C | C | | | | | | | | | C | C |
| NC_011770 | 2923473 | C | T | 71 | 0 | 2 | 0 | STOP_GAINED | HIGH | NONSENSE | Cga/Tga | R7* | PLES_27231 | T | T | | | | | | | | | | |
| NC_011770 | 2923687 | C | T | 69 | 0 | 2 | 0 | NON_SYNONYMOU | MODERAT | MISSENSE | gCc/gTc | A78V | PLES_27231 | T | T | | | | | | | | | | |
| NC_011770 | 2924008 | C | A | 73 | 0 | 2 | 0 | NON_SYNONYMOU | MODERAT | MISSENSE | gCg/gAg | A185E | PLES_27231 | | | A | A | | | | | | | | |
| NC_011770 | 2924472 | G | C | 61 | 0 | 2 | 0 | NON_SYNONYMOU | MODERAT | MISSENSE | Gtc/Ctc | V340L | PLES_27231 | | | | | | | | | | | C | C |
| NC_011770 | 2924638 | A | G | 40 | 0 | 1 | 0 | NON_SYNONYMOU | MODERAT | MISSENSE | gAc/gGc | D395G | PLES_27231 | | | | | | | G | | | | | |
| NC_011770 | 2924395 | A | G | 70 | 0 | 1 | 0 | NON_SYNONYMOU | MODERAT | MISSENSE | cAg/cGg | Q314R | PLES_27231 | | | | | | | | G | | | | |
| NC_011770 | 2926641 | A | G | 74 | 0 | 2 | 0 | NON_SYNONYMOU | MODERAT | MISSENSE | Aga/Gga | R20G | PLES_27261 | | | | | | | | | G | G | | |
| NC_011770 | 2926878 | T | C | 80 | 0 | 12 | 0 | NON_SYNONYMOU | MODERAT | MISSENSE | Tgc/Cgc | C99R | PLES_27261 | C | C | C | C | C | C | C | C | C | C | C | C |
| NC_011770 | 2926894 | T | C | 83 | 0 | 4 | 0 | NON_SYNONYMOU | MODERAT | MISSENSE | gTg/gCg | V104A | PLES_27261 | C | C | | | | | | | C | C | | |
| NC_011770 | 2927355 | C | T | 78 | 0 | 2 | 0 | NON_SYNONYMOU | MODERAT | MISSENSE | Ggc/Agc | G487S | PLES_27271 | T | T | | | | | | | | | | |
| NC_011770 | 2927540 | C | T | 77 | 0 | 4 | 0 | NON_SYNONYMOU | MODERAT | MISSENSE | aGg/aAg | R425K | PLES_27271 | | | T | T | T | T | | | | | | |
| NC_011770 | 2928026 | C | T | 72 | 0 | 1 | 0 | NON_SYNONYMOU | MODERAT | MISSENSE | tGc/tAc | C263Y | PLES_27271 | | | | | | | T | | | | | |
| NC_011770 | 2928239 | C | T | 72 | 0 | 2 | 0 | NON_SYNONYMOU | MODERAT | MISSENSE | gGc/gAc | G192D | PLES_27271 | | | | | | | | | | | T | T |
| NC_011770 | 2928515 | C | T | 67 | 0 | 12 | 0 | NON_SYNONYMOU | MODERAT | MISSENSE | gGc/gAc | G100D | PLES_27271 | T | T | T | T | T | T | T | T | T | T | T | T |
| NC_011770 | 2927699 | G | A | 70 | 0 | 1 | 0 | NON_SYNONYMOU | MODERAT | MISSENSE | gCg/gTg | A372V | PLES_27271 | | | | | | | | | | | | A |
| NC_011770 | 2929082 | C | T | 41 | 0 | 4 | 0 | NON_SYNONYMOU | MODERAT | MISSENSE | Ctc/Ttc | L5F | PLES_27281 | | | | | | | T | T | T | T | | |
| NC_011770 | 2929155 | T | A | 44 | 0 | 4 | 0 | NON_SYNONYMOU | MODERAT | MISSENSE | tTc/tAc | F29Y | PLES_27281 | | | | | | | A | A | A | A | | |
| NC_011770 | 2929175 | C | T | 39 | 0 | 2 | 0 | NON_SYNONYMOU | MODERAT | MISSENSE | Cct/Tct | P36S | PLES_27281 | | | | | | | | | | | T | T |
| NC_011770 | 2929226 | T | G | 42 | 0 | 12 | 0 | NON_SYNONYMOU | MODERAT | MISSENSE | Ttt/Gtt | F53V | PLES_27281 | G | G | G | G | G | G | G | G | G | G | G | G |
| NC_011770 | 2929228 | T | G | 42 | 0 | 12 | 0 | NON_SYNONYMOU | MODERAT | MISSENSE | ttT/ttG | F53L | PLES_27281 | G | G | G | G | G | G | G | G | G | G | G | G |
| NC_011770 | 2929725 | T | C | 66 | 0 | 2 | 0 | NON_SYNONYMOU | MODERAT | MISSENSE | cAt/cGt | H431R | PLES_27291 | | | | | C | C | | | | | | |
| NC_011770 | 2929740 | C | T | 70 | 0 | 6 | 0 | NON_SYNONYMOU | MODERAT | MISSENSE | aGa/aAa | R426K | PLES_27291 | T | T | T | T | T | T | | | | | | |
| NC_011770 | 2929759 | T | C | 67 | 0 | 6 | 0 | NON_SYNONYMOU | MODERAT | MISSENSE | Atc/Gtc | I420V | PLES_27291 | C | C | C | C | C | C | | | | | | |
| NC_011770 | 2929793 | G | C | 71 | 0 | 4 | 0 | NON_SYNONYMOU | MODERAT | MISSENSE | agC/agG | S408R | PLES_27291 | | | C | C | C | C | | | | | | |
| NC_011770 | 2929852 | A | C | 73 | 0 | 2 | 0 | NON_SYNONYMOU | MODERAT | MISSENSE | Tac/Gac | Y389D | PLES_27291 | C | C | | | | | | | | | | |
| NC_011770 | 2930122 | C | T | 68 | 0 | 6 | 0 | NON_SYNONYMOU | MODERAT | MISSENSE | Gtg/Atg | V299M | PLES_27291 | T | T | T | T | T | T | | | | | | |
| NC_011770 | 2930531 | C | G | 69 | 0 | 10 | 0 | NON_SYNONYMOU | MODERAT | MISSENSE | caG/caC | Q162H | PLES_27291 | G | G | G | G | G | G | G | G | G | G | | |
| NC_011770 | 2930667 | C | T | 69 | 0 | 2 | 0 | NON_SYNONYMOU | MODERAT | MISSENSE | cGc/cAc | R117H | PLES_27291 | | | | | | | T | T | | | | |
| NC_011770 | 2931575 | C | G | 66 | 0 | 6 | 0 | NON_SYNONYMOU | MODERAT | MISSENSE | gCc/gGc | A68G | PLES_27301 | G | G | G | G | G | G | | | | | | |
| NC_011770 | 2933400 | T | C | 82 | 0 | 6 | 0 | NON_SYNONYMOU | MODERAT | MISSENSE | Acg/Gcg | T107A | PLES_27311 | C | C | C | C | C | C | | | | | | |
| NC_011770 | 2934114 | C | G | 76 | 0 | 4 | 0 | NON_SYNONYMOU | MODERAT | MISSENSE | gaC/gaG | D76E | PLES_27321 | G | G | | | | | | | | | G | G |
| NC_011770 | 2934169 | G | A | 74 | 0 | 4 | 0 | NON_SYNONYMOU | MODERAT | MISSENSE | Gaa/Aaa | E95K | PLES_27321 | | | A | A | A | A | | | | | | |
| NC_011770 | 2935278 | T | G | 85 | 0 | 12 | 0 | NON_SYNONYMOU | MODERAT | MISSENSE | Tcc/Gcc | S161A | PLES_27341 | G | G | G | G | G | G | G | G | G | G | G | G |
| NC_011770 | 2936100 | C | T | 92 | 0 | 1 | 0 | NON_SYNONYMOU | MODERAT | MISSENSE | Ggc/Agc | G86S | PLES_27361 | | | | | | | T | | | | | |
| NC_011770 | 2936587 | C | G | 97 | 0 | 4 | 0 | NON_SYNONYMOU | MODERAT | MISSENSE | cCg/cGg | P4R | PLES_27371 | | | G | G | G | G | | | | | | |
| NC_011770 | 2936665 | G | A | 94 | 0 | 2 | 0 | NON_SYNONYMOU | MODERAT | MISSENSE | cGt/cAt | R30H | PLES_27371 | | | | | | | | | | | A | A |
| NC_011770 | 2937012 | G | T | 87 | 0 | 6 | 0 | NON_SYNONYMOU | MODERAT | MISSENSE | Gcg/Tcg | A146S | PLES_27371 | | | T | T | T | T | | | | | T | T |
| NC_011770 | 2937118 | G | C | 89 | 0 | 2 | 0 | NON_SYNONYMOU | MODERAT | MISSENSE | gGg/gCg | G181A | PLES_27371 | | | | | | | | | C | C | | |
| NC_011770 | 2937855 | G | A | 77 | 0 | 2 | 0 | NON_SYNONYMOU | MODERAT | MISSENSE | Gcg/Acg | A427T | PLES_27371 | A | A | | | | | | | | | | |
| NC_011770 | 2938159 | A | G | 70 | 0 | 8 | 0 | NON_SYNONYMOU | MODERAT | MISSENSE | aAg/aGg | K528R | PLES_27371 | G | G | G | G | G | G | | | G | G | | |
| NC_011770 | 2938221 | A | G | 66 | 0 | 2 | 0 | NON_SYNONYMOU | MODERAT | MISSENSE | Atc/Gtc | I549V | PLES_27371 | G | G | | | | | | | | | | |
| NC_011770 | 2938263 | C | T | 62 | 0 | 2 | 0 | NON_SYNONYMOU | MODERAT | MISSENSE | Cgc/Tgc | R563C | PLES_27371 | | | | | | | | | | | T | T |
| NC_011770 | 2938439 | C | G | 52 | 0 | 12 | 0 | NON_SYNONYMOU | MODERAT | MISSENSE | Cag/Gag | Q34E | PLES_27381 | G | G | G | G | G | G | G | G | G | G | G | G |
| NC_011770 | 2938965 | T | C | 79 | 0 | 10 | 0 | NON_SYNONYMOU | MODERAT | MISSENSE | gTc/gCc | V209A | PLES_27381 | | | C | C | C | C | C | C | C | C | C | C |
| NC_011770 | 2940128 | C | T | 79 | 0 | 2 | 0 | NON_SYNONYMOU | MODERAT | MISSENSE | Cgc/Tgc | R176C | PLES_27391 | | | | | | | | | T | T | | |
| NC_011770 | 2940266 | C | T | 74 | 0 | 2 | 0 | NON_SYNONYMOU | MODERAT | MISSENSE | Cgc/Tgc | R222C | PLES_27391 | | | | | | | | | T | T | | |
| NC_011770 | 2940402 | G | A | 75 | 0 | 2 | 0 | NON_SYNONYMOU | MODERAT | MISSENSE | aGc/aAc | S267N | PLES_27391 | A | A | | | | | | | | | | |
| NC_011770 | 2940980 | C | T | 62 | 0 | 2 | 0 | NON_SYNONYMOU | MODERAT | MISSENSE | Cac/Tac | H460Y | PLES_27391 | | | | | | | | | | | T | T |
| NC_011770 | 2941082 | C | T | 64 | 0 | 2 | 0 | NON_SYNONYMOU | MODERAT | MISSENSE | Ccg/Tcg | P494S | PLES_27391 | T | T | | | | | | | | | | |
| NC_011770 | 2941311 | T | C | 63 | 0 | 2 | 0 | NON_SYNONYMOU | MODERAT | MISSENSE | Ttc/Ctc | F8L | PLES_27401 | C | C | | | | | | | | | | |

| | | | | | | | | | | | | | | | | | | | | | | | | | |
|---|---|---|---|---|---|---|---|---|---|---|---|---|---|---|---|---|---|---|---|---|---|---|---|---|---|
| NC_011770 | 2941416 | G | A | 65 | 0 | 2 | 0 | NON_SYNONYMOU | MODERAT | MISSENSE | Gcg/Acg | A43T | PLES_27401 | | | | | | | | | | | A | A |
| NC_011770 | 2941483 | A | G | 67 | 0 | 4 | 0 | NON_SYNONYMOU | MODERAT | MISSENSE | gAc/gGc | D65G | PLES_27401 | G | G | | | | | G | G | | | | |
| NC_011770 | 2941503 | A | G | 68 | 0 | 12 | 0 | NON_SYNONYMOU | MODERAT | MISSENSE | Act/Gct | T72A | PLES_27401 | G | G | G | G | G | G | G | G | G | G | G | G |
| NC_011770 | 2942289 | C | T | 65 | 0 | 2 | 0 | NON_SYNONYMOU | MODERAT | MISSENSE | gCg/gTg | A72V | PLES_27411 | | | | | | | | | T | T | | |
| NC_011770 | 2942706 | G | A | 74 | 0 | 2 | 0 | NON_SYNONYMOU | MODERAT | MISSENSE | gGc/gAc | G211D | PLES_27411 | | | | | | | | | | | A | A |
| NC_011770 | 2944071 | C | T | 77 | 0 | 4 | 0 | NON_SYNONYMOU | MODERAT | MISSENSE | Cgc/Tgc | R260C | PLES_27421 | | | T | T | T | T | | | | | | |
| NC_011770 | 2944084 | A | T | 82 | 0 | 2 | 0 | NON_SYNONYMOU | MODERAT | MISSENSE | cAt/cTt | H264L | PLES_27421 | | | | | | | | | T | T | | |
| NC_011770 | 2945710 | C | A | 74 | 0 | 2 | 0 | NON_SYNONYMOU | MODERAT | MISSENSE | Cat/Aat | H23N | PLES_27441 | | | | | | | | | | | A | A |
| NC_011770 | 2947255 | T | C | 78 | 0 | 12 | 0 | NON_SYNONYMOU | MODERAT | MISSENSE | Acg/Gcg | T285A | PLES_27451 | C | C | C | C | C | C | C | C | C | C | C | C |
| NC_011770 | 2947617 | T | G | 81 | 0 | 2 | 0 | NON_SYNONYMOU | MODERAT | MISSENSE | cAc/cCc | H164P | PLES_27451 | | | | | | | G | G | | | | |
| NC_011770 | 2947619 | C | G | 81 | 0 | 2 | 0 | NON_SYNONYMOU | MODERAT | MISSENSE | aaG/aaC | K163N | PLES_27451 | G | G | | | | | | | | | | |
| NC_011770 | 2947639 | T | C | 82 | 0 | 12 | 0 | NON_SYNONYMOU | MODERAT | MISSENSE | Atg/Gtg | M157V | PLES_27451 | C | C | C | C | C | C | C | C | C | C | C | C |
| NC_011770 | 2948370 | C | T | 70 | 0 | 2 | 0 | NON_SYNONYMOU | MODERAT | MISSENSE | Ggc/Agc | G428S | PLES_27461 | T | T | | | | | | | | | | |
| NC_011770 | 2948459 | A | G | 74 | 0 | 1 | 0 | NON_SYNONYMOU | MODERAT | MISSENSE | gTg/gCg | V398A | PLES_27461 | | | | | | | G | | | | | |
| NC_011770 | 2949011 | G | A | 70 | 0 | 4 | 0 | NON_SYNONYMOU | MODERAT | MISSENSE | aCc/aTc | T214I | PLES_27461 | | | A | A | A | A | | | | | | |
| NC_011770 | 2949048 | G | C | 72 | 0 | 2 | 0 | NON_SYNONYMOU | MODERAT | MISSENSE | Ctg/Gtg | L202V | PLES_27461 | C | C | | | | | | | | | | |
| NC_011770 | 2949303 | C | T | 70 | 0 | 8 | 0 | NON_SYNONYMOU | MODERAT | MISSENSE | Gcc/Acc | A117T | PLES_27461 | T | T | T | T | T | T | | | | | T | T |
| NC_011770 | 2949372 | G | T | 71 | 0 | 2 | 0 | NON_SYNONYMOU | MODERAT | MISSENSE | Ctc/Atc | L94I | PLES_27461 | T | T | | | | | | | | | | |
| NC_011770 | 2949838 | T | C | 71 | 0 | 2 | 0 | NON_SYNONYMOU | MODERAT | MISSENSE | gAc/gGc | D260G | PLES_27471 | | | | | | | | | C | C | | |
| NC_011770 | 2949996 | T | G | 75 | 0 | 10 | 0 | NON_SYNONYMOU | MODERAT | MISSENSE | gaA/gaC | E207D | PLES_27471 | G | G | G | G | G | G | G | G | | | G | G |
| NC_011770 | 2949997 | T | C | 72 | 0 | 2 | 0 | NON_SYNONYMOU | MODERAT | MISSENSE | gAa/gGa | E207G | PLES_27471 | | | | | | | C | C | | | | |
| NC_011770 | 2950097 | C | T | 79 | 0 | 10 | 0 | NON_SYNONYMOU | MODERAT | MISSENSE | Ggt/Agt | G174S | PLES_27471 | T | T | T | T | T | T | | | T | T | T | T |
| NC_011770 | 2950136 | T | C | 76 | 0 | 10 | 0 | NON_SYNONYMOU | MODERAT | MISSENSE | Atc/Gtc | I161V | PLES_27471 | C | C | C | C | C | C | | | C | C | C | C |
| NC_011770 | 2950257 | G | C | 70 | 0 | 8 | 0 | NON_SYNONYMOU | MODERAT | MISSENSE | gaC/gaG | D120E | PLES_27471 | C | C | C | C | C | C | | | | | C | C |
| NC_011770 | 2950612 | T | G | 66 | 0 | 2 | 0 | NON_SYNONYMOU | MODERAT | MISSENSE | gAt/gCt | D2A | PLES_27471 | G | G | | | | | | | | | | |
| NC_011770 | 2951148 | C | CTGAG | 38 | 0 | 2 | 0 | FRAME_SHIFT+STC | HIGH | | gac/gaGTC | D140E*? | PLES_27481 | | | | | | | | | CTGAG | CTGAG | | |
| NC_011770 | 2950906 | G | T | 22 | 0 | 2 | 0 | NON_SYNONYMOU | MODERAT | MISSENSE | Gcg/Tcg | A63S | PLES_27481 | | | | | | | | | T | T | | |
| NC_011770 | 2950954 | G | C | 21 | 0 | 4 | 0 | NON_SYNONYMOU | MODERAT | MISSENSE | Gtc/Ctc | V79L | PLES_27481 | | | | | | | C | C | | | C | C |
| NC_011770 | 2951002 | G | A | 25 | 0 | 2 | 0 | NON_SYNONYMOU | MODERAT | MISSENSE | Gtc/Atc | V95I | PLES_27481 | | | | | | | | | A | A | | |
| NC_011770 | 2951128 | C | G | 32 | 0 | 2 | 0 | NON_SYNONYMOU | MODERAT | MISSENSE | Ccg/Gcg | P137A | PLES_27481 | | | | | | | | | | | G | G |
| NC_011770 | 2951131 | C | T | 33 | 0 | 2 | 0 | NON_SYNONYMOU | MODERAT | MISSENSE | Cgg/Tgg | R138W | PLES_27481 | | | | | | | | | T | T | | |
| NC_011770 | 2951141 | G | A | 36 | 0 | 6 | 0 | NON_SYNONYMOU | MODERAT | MISSENSE | aGt/aAt | S141N | PLES_27481 | | | | | | | A | A | A | A | A | A |
| NC_011770 | 2952272 | G | A | 71 | 0 | 2 | 0 | NON_SYNONYMOU | MODERAT | MISSENSE | Gcc/Acc | A19T | PLES_27501 | A | A | | | | | | | | | | |
| NC_011770 | 2952314 | T | A | 74 | 0 | 6 | 0 | NON_SYNONYMOU | MODERAT | MISSENSE | Tcc/Acc | S33T | PLES_27501 | | | A | A | A | A | A | A | | | | |
| NC_011770 | 2952646 | G | C | 77 | 0 | 12 | 0 | NON_SYNONYMOU | MODERAT | MISSENSE | caG/caC | Q143H | PLES_27501 | C | C | C | C | C | C | C | C | C | C | C | C |
| NC_011770 | 2952882 | T | C | 80 | 0 | 2 | 0 | NON_SYNONYMOU | MODERAT | MISSENSE | gTg/gCg | V222A | PLES_27501 | | | | | | | | | C | C | | |
| NC_011770 | 2954146 | G | A | 39 | 0 | 10 | 0 | NON_SYNONYMOU | MODERAT | MISSENSE | cGg/cAg | R361Q | PLES_27511 | A | A | A | A | A | A | A | A | A | A | | |
| NC_011770 | 2954197 | A | G | 41 | 0 | 1 | 0 | NON_SYNONYMOU | MODERAT | MISSENSE | tAc/tGc | Y378C | PLES_27511 | | | | | | | G | | | | | |
| NC_011770 | 2954888 | A | G | 47 | 0 | 4 | 0 | NON_SYNONYMOU | MODERAT | MISSENSE | Acc/Gcc | T30A | PLES_27521 | | | | | | | | | G | G | G | G |
| NC_011770 | 2954910 | G | A | 46 | 0 | 2 | 0 | NON_SYNONYMOU | MODERAT | MISSENSE | gGc/gAc | G37D | PLES_27521 | | | | | A | A | | | | | | |
| NC_011770 | 2954973 | A | G | 46 | 0 | 6 | 0 | NON_SYNONYMOU | MODERAT | MISSENSE | cAg/cGg | Q58R | PLES_27521 | G | G | | | | | G | G | G | G | | |
| NC_011770 | 2955042 | C | G | 43 | 0 | 4 | 0 | NON_SYNONYMOU | MODERAT | MISSENSE | gCc/gGc | A81G | PLES_27521 | | | G | G | G | G | | | | | | |
| NC_011770 | 2955054 | G | A | 43 | 0 | 4 | 0 | NON_SYNONYMOU | MODERAT | MISSENSE | gGg/gAg | G85E | PLES_27521 | A | A | | | | | A | A | | | | |
| NC_011770 | 2955111 | C | T | 41 | 0 | 2 | 0 | NON_SYNONYMOU | MODERAT | MISSENSE | cCg/cTg | P104L | PLES_27521 | | | | | | | | | T | T | | |
| NC_011770 | 2955129 | G | A | 39 | 0 | 12 | 0 | NON_SYNONYMOU | MODERAT | MISSENSE | gGc/gAc | G110D | PLES_27521 | A | A | A | A | A | A | A | A | A | A | A | A |
| NC_011770 | 2955393 | G | A | 44 | 0 | 2 | 0 | NON_SYNONYMOU | MODERAT | MISSENSE | gGc/gAc | G198D | PLES_27521 | | | | | | | | | | | A | A |
| NC_011770 | 2955437 | A | C | 42 | 0 | 10 | 0 | NON_SYNONYMOU | MODERAT | MISSENSE | Aag/Cag | K213Q | PLES_27521 | C | C | C | C | C | C | C | C | C | C | | |
| NC_011770 | 2955627 | A | G | 28 | 0 | 8 | 0 | NON_SYNONYMOU | MODERAT | MISSENSE | aAc/aGc | N276S | PLES_27521 | | | G | G | G | G | | | G | G | G | G |
| NC_011770 | 2956203 | C | A | 57 | 0 | 2 | 0 | NON_SYNONYMOU | MODERAT | MISSENSE | gCg/gAg | A468E | PLES_27521 | | | | | | | | | | | A | A |
| NC_011770 | 2956827 | G | A | 65 | 0 | 2 | 0 | NON_SYNONYMOU | MODERAT | MISSENSE | aGg/aAg | R676K | PLES_27521 | | | | | | | | | A | A | | |
| NC_011770 | 2956860 | A | G | 64 | 0 | 12 | 0 | NON_SYNONYMOU | MODERAT | MISSENSE | gAc/gGc | D687G | PLES_27521 | G | G | G | G | G | G | G | G | G | G | G | G |
| NC_011770 | 2956863 | C | G | 68 | 0 | 2 | 0 | NON_SYNONYMOU | MODERAT | MISSENSE | gCc/gGc | A688G | PLES_27521 | | | | | | | | | G | G | | |
| NC_011770 | 2957539 | C | G | 58 | 0 | 8 | 0 | NON_SYNONYMOU | MODERAT | MISSENSE | gaC/gaG | D913E | PLES_27521 | | | G | G | G | G | G | G | G | G | | |
| NC_011770 | 2957766 | A | C | 65 | 0 | 2 | 0 | NON_SYNONYMOU | MODERAT | MISSENSE | gAg/gCg | E989A | PLES_27521 | | | | | | | | | | | C | C |
| NC_011770 | 2957772 | A | G | 64 | 0 | 4 | 0 | NON_SYNONYMOU | MODERAT | MISSENSE | cAg/cGg | Q991R | PLES_27521 | | | | | | | | | G | G | G | G |
| NC_011770 | 2958671 | A | G | 70 | 0 | 2 | 0 | NON_SYNONYMOU | MODERAT | MISSENSE | cAg/cGg | Q24R | PLES_27531 | G | G | | | | | | | | | | |
| NC_011770 | 2959201 | T | G | 57 | 0 | 8 | 0 | NON_SYNONYMOU | MODERAT | MISSENSE | Tcc/Gcc | S201A | PLES_27531 | G | G | | | | | G | G | G | G | G | G |
| NC_011770 | 2959222 | G | A | 59 | 0 | 4 | 0 | NON_SYNONYMOU | MODERAT | MISSENSE | Gac/Aac | D208N | PLES_27531 | | | A | A | A | A | | | | | | |
| NC_011770 | 2959539 | G | A | 74 | 0 | 1 | 0 | NON_SYNONYMOU | MODERAT | MISSENSE | Ggc/Agc | G72S | PLES_27541 | | | | | | | A | | | | | |
| NC_011770 | 2959866 | C | G | 77 | 0 | 2 | 0 | NON_SYNONYMOU | MODERAT | MISSENSE | Cag/Gag | Q181E | PLES_27541 | | | | | | | | | G | G | | |
| NC_011770 | 2960409 | A | T | 73 | 0 | 2 | 0 | NON_SYNONYMOU | MODERAT | MISSENSE | Agc/Tgc | S362C | PLES_27541 | T | T | | | | | | | | | | |
| NC_011770 | 2960497 | C | T | 66 | 0 | 2 | 0 | NON_SYNONYMOU | MODERAT | MISSENSE | gCc/gTc | A391V | PLES_27541 | T | T | | | | | | | | | | |

| | | | | | | | | | | | | | | | | | | | | | | | | | |
|---|---|---|---|---|---|---|---|---|---|---|---|---|---|---|---|---|---|---|---|---|---|---|---|---|---|
| NC_011770 | 2960788 | C | T | 62 | 0 | 1 | 0 | NON_SYNONYMOU | MODERAT | MISSENSE | tCc/tTc | S488F | PLES_27541 | | | | | | | T | | | | | |
| NC_011770 | 2962146 | T | C | 74 | 0 | 2 | 0 | NON_SYNONYMOU | MODERAT | MISSENSE | gTc/gCc | V342A | PLES_27551 | | | | | | | | | C | C | | |
| NC_011770 | 2962158 | T | C | 77 | 0 | 2 | 0 | NON_SYNONYMOU | MODERAT | MISSENSE | gTc/gCc | V346A | PLES_27551 | C | C | | | | | | | | | | |
| NC_011770 | 2962666 | A | G | 83 | 0 | 6 | 0 | NON_SYNONYMOU | MODERAT | MISSENSE | cAc/cGc | H80R | PLES_27561 | | | G | G | G | G | | | | | G | G |
| NC_011770 | 2962761 | C | A | 85 | 0 | 6 | 0 | NON_SYNONYMOU | MODERAT | MISSENSE | Cac/Aac | H112N | PLES_27561 | A | A | A | A | A | A | | | | | | |
| NC_011770 | 2962774 | A | G | 80 | 0 | 10 | 0 | NON_SYNONYMOU | MODERAT | MISSENSE | cAc/cGc | H116R | PLES_27561 | G | G | G | G | G | G | G | G | | | G | G |
| NC_011770 | 2963136 | A | G | 92 | 0 | 2 | 0 | NON_SYNONYMOU | MODERAT | MISSENSE | aAg/aGg | K82R | PLES_27571 | | | | | | | | | | | G | G |
| NC_011770 | 2963252 | A | G | 87 | 0 | 1 | 0 | NON_SYNONYMOU | MODERAT | MISSENSE | Acc/Gcc | T121A | PLES_27571 | | | | | | | G | | | | | |
| NC_011770 | 2966181 | A | G | 65 | 0 | 1 | 0 | NON_SYNONYMOU | MODERAT | MISSENSE | Agc/Ggc | S127G | PLES_27601 | | | | | | | G | | | | | |
| NC_011770 | 2966422 | T | C | 65 | 0 | 1 | 0 | NON_SYNONYMOU | MODERAT | MISSENSE | gTc/gCc | V207A | PLES_27601 | | | | | | | C | | | | | |
| NC_011770 | 2969057 | C | A | 73 | 0 | 2 | 0 | NON_SYNONYMOU | MODERAT | MISSENSE | Gcg/Tcg | A357S | PLES_27631 | | | | | | | | | A | A | | |
| NC_011770 | 2969210 | C | T | 72 | 0 | 2 | 0 | NON_SYNONYMOU | MODERAT | MISSENSE | Gcc/Acc | A306T | PLES_27631 | | | | | | | T | T | | | | |
| NC_011770 | 2969591 | T | A | 66 | 0 | 2 | 0 | NON_SYNONYMOU | MODERAT | MISSENSE | Acc/Tcc | T179S | PLES_27631 | A | A | | | | | | | | | | |
| NC_011770 | 2969780 | G | T | 64 | 0 | 2 | 0 | NON_SYNONYMOU | MODERAT | MISSENSE | Cgc/Agc | R116S | PLES_27631 | | | | | | | T | T | | | | |
| NC_011770 | 2969938 | C | T | 61 | 0 | 4 | 0 | NON_SYNONYMOU | MODERAT | MISSENSE | cGg/cAg | R63Q | PLES_27631 | | | T | T | T | T | | | | | | |
| NC_011770 | 2970707 | AG | A | 68 | 0 | 2 | 0 | FRAME_SHIFT | HIGH | | ctg/ | L314 | PLES_27641 | A | A | | | | | | | | | | |
| NC_011770 | 2970424 | T | G | 61 | 0 | 12 | 0 | NON_SYNONYMOU | MODERAT | MISSENSE | caA/caC | Q408H | PLES_27641 | G | G | G | G | G | G | G | G | G | G | G | G |
| NC_011770 | 2971253 | C | T | 64 | 0 | 2 | 0 | NON_SYNONYMOU | MODERAT | MISSENSE | cGg/cAg | R132Q | PLES_27641 | | | | | | | | | T | T | | |
| NC_011770 | 2971978 | G | T | 67 | 0 | 2 | 0 | NON_SYNONYMOU | MODERAT | MISSENSE | gCg/gAg | A368E | PLES_27651 | T | T | | | | | | | | | | |
| NC_011770 | 2971993 | C | T | 66 | 0 | 2 | 0 | NON_SYNONYMOU | MODERAT | MISSENSE | gGc/gAc | G363D | PLES_27651 | | | | | | | | | | | T | T |
| NC_011770 | 2972059 | C | T | 69 | 0 | 8 | 0 | NON_SYNONYMOU | MODERAT | MISSENSE | aGc/aAc | S341N | PLES_27651 | | | T | T | T | T | T | T | | | T | T |
| NC_011770 | 2972812 | G | A | 62 | 0 | 4 | 0 | NON_SYNONYMOU | MODERAT | MISSENSE | gCc/gTc | A90V | PLES_27651 | A | A | | | | | | | | | A | A |
| NC_011770 | 2972968 | C | T | 64 | 0 | 4 | 0 | NON_SYNONYMOU | MODERAT | MISSENSE | aGc/aAc | S38N | PLES_27651 | T | T | | | | | | | | | T | T |
| NC_011770 | 2973827 | A | G | 81 | 0 | 2 | 0 | NON_SYNONYMOU | MODERAT | MISSENSE | aAg/aGg | K164R | PLES_27661 | | | | | | | | | | | G | G |
| NC_011770 | 2973849 | A | T | 80 | 0 | 4 | 0 | NON_SYNONYMOU | MODERAT | MISSENSE | gaA/gaT | E171D | PLES_27661 | | | T | T | T | T | | | | | | |
| NC_011770 | 2974030 | C | G | 73 | 0 | 4 | 0 | NON_SYNONYMOU | MODERAT | MISSENSE | Cag/Gag | Q232E | PLES_27661 | | | G | G | G | G | | | | | | |
| NC_011770 | 2974111 | A | C | 75 | 0 | 2 | 0 | NON_SYNONYMOU | MODERAT | MISSENSE | Aag/Cag | K259Q | PLES_27661 | | | | | | | | | | | C | C |
| NC_011770 | 2974117 | A | G | 75 | 0 | 12 | 0 | NON_SYNONYMOU | MODERAT | MISSENSE | Acg/Gcg | T261A | PLES_27661 | G | G | G | G | G | G | G | G | G | G | G | G |
| NC_011770 | 2974905 | C | T | 65 | 0 | 1 | 0 | STOP_GAINED | HIGH | NONSENSE | Cag/Tag | Q98* | PLES_27671 | | | | | | | | T | | | | |
| NC_011770 | 2979342 | T | C | 85 | 0 | 2 | 0 | NON_SYNONYMOU | MODERAT | MISSENSE | gTg/gCg | V534A | PLES_27681 | | | | | | | | | | | C | C |
| NC_011770 | 2979344 | A | C | 85 | 0 | 2 | 0 | NON_SYNONYMOU | MODERAT | MISSENSE | Atc/Ctc | I535L | PLES_27681 | C | C | | | | | | | | | | |
| NC_011770 | 2980104 | A | G | 88 | 0 | 2 | 0 | NON_SYNONYMOU | MODERAT | MISSENSE | cAc/cGc | H788R | PLES_27681 | G | G | | | | | | | | | | |
| NC_011770 | 2980784 | T | C | 77 | 0 | 12 | 0 | NON_SYNONYMOU | MODERAT | MISSENSE | Tgc/Cgc | C1015R | PLES_27681 | C | C | C | C | C | C | C | C | C | C | C | C |
| NC_011770 | 2981216 | T | G | 46 | 0 | 8 | 0 | NON_SYNONYMOU | MODERAT | MISSENSE | tTg/tGg | L123W | PLES_27691 | | | G | G | G | G | | | G | G | G | G |
| NC_011770 | 2981217 | G | C | 45 | 0 | 8 | 0 | NON_SYNONYMOU | MODERAT | MISSENSE | ttG/ttC | L123F | PLES_27691 | | | C | C | C | C | | | C | C | C | C |
| NC_011770 | 2981225 | G | A | 44 | 0 | 8 | 0 | NON_SYNONYMOU | MODERAT | MISSENSE | gGa/gAa | G126E | PLES_27691 | | | A | A | A | A | | | A | A | A | A |
| NC_011770 | 2981247 | C | CGGT | 45 | 0 | 8 | 0 | CODON_CHANGE_ | MODERATE | | ggc/ggTGG | G134GG | PLES_27691 | | | CGGT | CGGT | CGGT | CGGT | | | CGGT | CGGT | CGGT | CGGT |
| NC_011770 | 2981267 | C | A | 70 | 0 | 8 | 0 | NON_SYNONYMOU | MODERAT | MISSENSE | aCc/aAc | T140N | PLES_27691 | | | A | A | A | A | | | A | A | A | A |
| NC_011770 | 2981282 | A | G | 74 | 0 | 8 | 0 | NON_SYNONYMOU | MODERAT | MISSENSE | aAc/aGc | N145S | PLES_27691 | | | G | G | G | G | | | G | G | G | G |
| NC_011770 | 2981287 | A | G | 77 | 0 | 8 | 0 | NON_SYNONYMOU | MODERAT | MISSENSE | Agt/Ggt | S147G | PLES_27691 | | | G | G | G | G | | | G | G | G | G |
| NC_011770 | 2982480 | T | A | 72 | 0 | 2 | 0 | START_LOST | HIGH | MISSENSE | aTg/aAg | M1K | PLES_27701 | | | | | | | | | A | A | | |
| NC_011770 | 2982671 | T | G | 79 | 0 | 2 | 0 | NON_SYNONYMOU | MODERAT | MISSENSE | Tgc/Ggc | C65G | PLES_27701 | | | | | | | G | G | | | | |
| NC_011770 | 2982869 | C | T | 74 | 0 | 2 | 0 | NON_SYNONYMOU | MODERAT | MISSENSE | cGc/cAc | R424H | PLES_27711 | T | T | | | | | | | | | | |
| NC_011770 | 2983082 | C | T | 69 | 0 | 6 | 0 | NON_SYNONYMOU | MODERAT | MISSENSE | aGc/aAc | S353N | PLES_27711 | T | T | | | | | | | T | T | T | T |
| NC_011770 | 2983296 | C | T | 64 | 0 | 8 | 0 | NON_SYNONYMOU | MODERAT | MISSENSE | Ggc/Agc | G282S | PLES_27711 | T | T | T | T | T | T | T | T | | | | |
| NC_011770 | 2983418 | C | G | 63 | 0 | 2 | 0 | NON_SYNONYMOU | MODERAT | MISSENSE | gGc/gCc | G241A | PLES_27711 | G | G | | | | | | | | | | |
| NC_011770 | 2983482 | T | C | 63 | 0 | 2 | 0 | NON_SYNONYMOU | MODERAT | MISSENSE | Atc/Gtc | I220V | PLES_27711 | C | C | | | | | | | | | | |
| NC_011770 | 2999621 | C | T | 80 | 0 | 2 | 0 | NON_SYNONYMOU | MODERAT | MISSENSE | gCg/gTg | A39V | PLES_27841 | | | | | | | | | | | T | T |
| NC_011770 | 2999644 | G | C | 84 | 0 | 4 | 0 | NON_SYNONYMOU | MODERAT | MISSENSE | Gag/Cag | E47Q | PLES_27841 | | | C | C | C | C | | | | | | |
| NC_011770 | 3004292 | T | C | 58 | 0 | 10 | 0 | NON_SYNONYMOU | MODERAT | MISSENSE | Acc/Gcc | T19A | PLES_27891 | C | C | C | C | C | C | C | C | C | C | | |
| NC_011770 | 3006500 | C | T | 74 | 0 | 1 | 0 | NON_SYNONYMOU | MODERAT | MISSENSE | gCg/gTg | A65V | PLES_27911 | | | | | | | T | | | | | |
| NC_011770 | 3007006 | C | T | 52 | 0 | 2 | 0 | NON_SYNONYMOU | MODERAT | MISSENSE | atG/atA | M392I | PLES_27921 | | | | | | | | | | | T | T |
| NC_011770 | 3007218 | C | G | 66 | 0 | 4 | 0 | NON_SYNONYMOU | MODERAT | MISSENSE | Gtc/Ctc | V322L | PLES_27921 | | | G | G | G | G | | | | | | |
| NC_011770 | 3007683 | G | C | 69 | 0 | 12 | 0 | NON_SYNONYMOU | MODERAT | MISSENSE | Ctg/Gtg | L167V | PLES_27921 | C | C | C | C | C | C | C | C | C | C | C | C |
| NC_011770 | 3007854 | C | T | 63 | 0 | 4 | 0 | NON_SYNONYMOU | MODERAT | MISSENSE | Gca/Aca | A110T | PLES_27921 | | | T | T | T | T | | | | | | |
| NC_011770 | 3008513 | A | C | 69 | 0 | 2 | 0 | NON_SYNONYMOU | MODERAT | MISSENSE | gAc/gCc | D125A | PLES_27931 | | | | | | | | | C | C | | |
| NC_011770 | 3008570 | T | C | 66 | 0 | 12 | 0 | NON_SYNONYMOU | MODERAT | MISSENSE | tTc/tCc | F144S | PLES_27931 | C | C | C | C | C | C | C | C | C | C | C | C |
| NC_011770 | 3009050 | T | C | 65 | 0 | 4 | 0 | NON_SYNONYMOU | MODERAT | MISSENSE | aTg/aCg | M304T | PLES_27931 | | | C | C | C | C | | | | | | |
| NC_011770 | 3009620 | C | G | 79 | 0 | 2 | 0 | NON_SYNONYMOU | MODERAT | MISSENSE | gCg/gGg | A494G | PLES_27931 | | | | | | | | | | | G | G |
| NC_011770 | 3010075 | A | G | 77 | 0 | 4 | 0 | NON_SYNONYMOU | MODERAT | MISSENSE | gTc/gCc | V369A | PLES_27951 | G | G | | | | | | | G | G | | |
| NC_011770 | 3010917 | C | G | 64 | 0 | 12 | 0 | NON_SYNONYMOU | MODERAT | MISSENSE | caG/caC | Q88H | PLES_27951 | G | G | G | G | G | G | G | G | G | G | G | G |
| NC_011770 | 3010931 | A | G | 64 | 0 | 1 | 0 | NON_SYNONYMOU | MODERAT | MISSENSE | Tgg/Cgg | W84R | PLES_27951 | | | | | | | G | | | | | |

| | | | | | | | | | | | | | | | | | | | | | | | | | |
|---|---|---|---|---|---|---|---|---|---|---|---|---|---|---|---|---|---|---|---|---|---|---|---|---|---|
| NC_011770 | 3010950 | G | C | 69 | 0 | 10 | 0 | NON_SYNONYMOU | MODERAT | MISSENSE | ttC/ttG | F77L | PLES_27951 | C | C | C | C | C | C | | | C | C | C | C |
| NC_011770 | 3011143 | G | C | 64 | 0 | 4 | 0 | NON_SYNONYMOU | MODERAT | MISSENSE | aCc/aGc | T13S | PLES_27951 | | | | | | | C | C | | | C | C |
| NC_011770 | 3011311 | G | C | 71 | 0 | 8 | 0 | NON_SYNONYMOU | MODERAT | MISSENSE | Ccg/Gcg | P108A | PLES_27961 | | | C | C | C | C | C | C | C | C | | |
| NC_011770 | 3011374 | C | T | 71 | 0 | 2 | 0 | NON_SYNONYMOU | MODERAT | MISSENSE | Gcc/Acc | A87T | PLES_27961 | | | | | | | | | T | T | | |
| NC_011770 | 3011619 | A | G | 67 | 0 | 2 | 0 | NON_SYNONYMOU | MODERAT | MISSENSE | aTc/aCc | I5T | PLES_27961 | | | | | | | | | | | G | G |
| NC_011770 | 3012078 | C | T | 77 | 0 | 4 | 0 | NON_SYNONYMOU | MODERAT | MISSENSE | Gag/Aag | E69K | PLES_27971 | | | T | T | T | T | | | | | | |
| NC_011770 | 3012153 | G | C | 74 | 0 | 2 | 0 | NON_SYNONYMOU | MODERAT | MISSENSE | Ccc/Gcc | P44A | PLES_27971 | | | | | | | C | C | | | | |
| NC_011770 | 3012914 | A | G | 59 | 0 | 6 | 0 | NON_SYNONYMOU | MODERAT | MISSENSE | Acg/Gcg | T169A | PLES_27981 | | | | | | | G | G | G | G | G | G |
| NC_011770 | 3013151 | T | G | 54 | 0 | 10 | 0 | NON_SYNONYMOU | MODERAT | MISSENSE | Tcc/Gcc | S248A | PLES_27981 | G | G | G | G | G | G | G | G | | | G | G |
| NC_011770 | 3022063 | A | G | 54 | 0 | 12 | 0 | NON_SYNONYMOU | MODERAT | MISSENSE | atA/atG | I271M | PLES_28051 | G | G | G | G | G | G | G | G | G | G | G | G |
| NC_011770 | 3022490 | C | T | 72 | 0 | 4 | 0 | NON_SYNONYMOU | MODERAT | MISSENSE | Gtg/Atg | V72M | PLES_28061 | | | T | T | T | T | | | | | | |
| NC_011770 | 3022870 | C | T | 78 | 0 | 10 | 0 | NON_SYNONYMOU | MODERAT | MISSENSE | Gcc/Acc | A244T | PLES_28071 | T | T | T | T | T | T | T | T | | | T | T |
| NC_011770 | 3023227 | C | T | 69 | 0 | 2 | 0 | NON_SYNONYMOU | MODERAT | MISSENSE | Gcc/Acc | A125T | PLES_28071 | | | | | | | | | | | T | T |
| NC_011770 | 3023560 | C | T | 78 | 0 | 2 | 0 | NON_SYNONYMOU | MODERAT | MISSENSE | Gcc/Acc | A14T | PLES_28071 | | | | | | | T | T | | | | |
| NC_011770 | 3023685 | T | C | 82 | 0 | 6 | 0 | NON_SYNONYMOU | MODERAT | MISSENSE | Aca/Gca | T236A | PLES_28081 | | | C | C | C | C | | | | | C | C |
| NC_011770 | 3023916 | C | T | 65 | 0 | 2 | 0 | NON_SYNONYMOU | MODERAT | MISSENSE | Gcc/Acc | A159T | PLES_28081 | | | | | | | | | T | T | | |
| NC_011770 | 3024038 | C | T | 69 | 0 | 2 | 0 | NON_SYNONYMOU | MODERAT | MISSENSE | aGc/aAc | S118N | PLES_28081 | | | | | | | | | | | T | T |
| NC_011770 | 3024578 | G | A | 77 | 0 | 2 | 0 | NON_SYNONYMOU | MODERAT | MISSENSE | Gcc/Acc | A31T | PLES_28091 | | | | | A | A | | | | | | |
| NC_011770 | 3025017 | G | A | 81 | 0 | 4 | 0 | NON_SYNONYMOU | MODERAT | MISSENSE | Ccg/Tcg | P87S | PLES_28111 | | | | | | | A | A | A | A | | |
| NC_011770 | 3025131 | G | A | 77 | 0 | 8 | 0 | NON_SYNONYMOU | MODERAT | MISSENSE | Cca/Tca | P49S | PLES_28111 | | | A | A | A | A | A | A | A | A | | |
| NC_011770 | 3026477 | C | T | 68 | 0 | 2 | 0 | NON_SYNONYMOU | MODERAT | MISSENSE | Ccc/Tcc | P137S | PLES_28131 | | | | | | | | | | | T | T |
| NC_011770 | 3026543 | G | A | 70 | 0 | 2 | 0 | NON_SYNONYMOU | MODERAT | MISSENSE | Gtg/Atg | V159M | PLES_28131 | | | | | | | A | A | | | | |
| NC_011770 | 3027054 | G | A | 58 | 0 | 4 | 0 | NON_SYNONYMOU | MODERAT | MISSENSE | gGg/gAg | G329E | PLES_28131 | | | A | A | A | A | | | | | | |
| NC_011770 | 3027492 | A | C | 69 | 0 | 8 | 0 | NON_SYNONYMOU | MODERAT | MISSENSE | aAt/aCt | N103T | PLES_28141 | C | C | C | C | C | C | | | | | C | C |
| NC_011770 | 3027528 | C | A | 66 | 0 | 4 | 0 | NON_SYNONYMOU | MODERAT | MISSENSE | gCg/gAg | A115E | PLES_28141 | | | | | | | A | A | A | A | | |
| NC_011770 | 3028418 | C | T | 75 | 0 | 2 | 0 | NON_SYNONYMOU | MODERAT | MISSENSE | aCg/aTg | T195M | PLES_28151 | T | T | | | | | | | | | | |
| NC_011770 | 3028905 | C | T | 45 | 0 | 2 | 0 | NON_SYNONYMOU | MODERAT | MISSENSE | Gag/Aag | E440K | PLES_28161 | | | | | | | T | T | | | | |
| NC_011770 | 3029070 | T | C | 51 | 0 | 2 | 0 | NON_SYNONYMOU | MODERAT | MISSENSE | Acc/Gcc | T385A | PLES_28161 | | | | | | | | | C | C | | |
| NC_011770 | 3029454 | C | T | 68 | 0 | 1 | 0 | NON_SYNONYMOU | MODERAT | MISSENSE | Gcc/Acc | A257T | PLES_28161 | | | | | | | T | | | | | |
| NC_011770 | 3029726 | A | G | 82 | 0 | 1 | 0 | NON_SYNONYMOU | MODERAT | MISSENSE | cTg/cCg | L166P | PLES_28161 | | | | | | | | | | | G | |
| NC_011770 | 3031510 | C | T | 55 | 0 | 2 | 0 | NON_SYNONYMOU | MODERAT | MISSENSE | aCg/aTg | T150M | PLES_28181 | | | | | | | | | | | T | T |
| NC_011770 | 3031936 | C | T | 61 | 0 | 2 | 0 | NON_SYNONYMOU | MODERAT | MISSENSE | gCc/gTc | A292V | PLES_28181 | | | | | | | T | T | | | | |
| NC_011770 | 3032817 | A | C | 71 | 0 | 12 | 0 | NON_SYNONYMOU | MODERAT | MISSENSE | Agc/Cgc | S586R | PLES_28181 | C | C | C | C | C | C | C | C | C | C | C | C |
| NC_011770 | 3033008 | T | C | 85 | 0 | 12 | 0 | NON_SYNONYMOU | MODERAT | MISSENSE | Tgc/Cgc | C62R | PLES_28191 | C | C | C | C | C | C | C | C | C | C | C | C |
| NC_011770 | 3033047 | C | G | 92 | 0 | 2 | 0 | NON_SYNONYMOU | MODERAT | MISSENSE | Ctg/Gtg | L75V | PLES_28191 | G | G | | | | | | | | | | |
| NC_011770 | 3033258 | C | G | 86 | 0 | 2 | 0 | NON_SYNONYMOU | MODERAT | MISSENSE | gCc/gGc | A145G | PLES_28191 | G | G | | | | | | | | | | |
| NC_011770 | 3033349 | G | A | 81 | 0 | 4 | 0 | NON_SYNONYMOU | MODERAT | MISSENSE | atG/atA | M175I | PLES_28191 | | | A | A | A | A | | | | | | |
| NC_011770 | 3034787 | G | C | 70 | 1 | 1 | 0 | NON_SYNONYMOU | MODERAT | MISSENSE | gGc/gCc | G96A | PLES_28211 | . | C | | | | | | | | | | |
| NC_011770 | 3035418 | G | C | 77 | 0 | 2 | 0 | NON_SYNONYMOU | MODERAT | MISSENSE | gaG/gaC | E306D | PLES_28211 | C | C | | | | | | | | | | |
| NC_011770 | 3035359 | G | A | 76 | 0 | 1 | 0 | NON_SYNONYMOU | MODERAT | MISSENSE | Gct/Act | A287T | PLES_28211 | | | | | | | | | | | A | |
| NC_011770 | 3036244 | A | G | 60 | 0 | 2 | 0 | NON_SYNONYMOU | MODERAT | MISSENSE | aTc/aCc | I162T | PLES_28221 | | | | | | | | | | | G | G |
| NC_011770 | 3036454 | G | A | 60 | 0 | 4 | 0 | NON_SYNONYMOU | MODERAT | MISSENSE | gCc/gTc | A92V | PLES_28221 | | | A | A | A | A | | | | | | |
| NC_011770 | 3036715 | G | A | 54 | 0 | 12 | 0 | NON_SYNONYMOU | MODERAT | MISSENSE | aCc/aTc | T5I | PLES_28221 | A | A | A | A | A | A | A | A | A | A | A | A |
| NC_011770 | 3037174 | G | T | 64 | 0 | 2 | 0 | NON_SYNONYMOU | MODERAT | MISSENSE | Cgc/Agc | R81S | PLES_28231 | T | T | | | | | | | | | | |
| NC_011770 | 3037371 | A | G | 72 | 0 | 2 | 0 | NON_SYNONYMOU | MODERAT | MISSENSE | gTg/gCg | V15A | PLES_28231 | | | | | | | | | | | G | G |
| NC_011770 | 3038243 | CGCCG | C | 66 | 0 | 1 | 0 | FRAME_SHIFT | HIGH | | ccggcg/ | PA176 | PLES_28241 | | | | | | | C | | | | | |
| NC_011770 | 3037437 | A | G | 66 | 0 | 12 | 0 | NON_SYNONYMOU | MODERAT | MISSENSE | gTc/gCc | V446A | PLES_28241 | G | G | G | G | G | G | G | G | G | G | G | G |
| NC_011770 | 3037438 | C | T | 64 | 0 | 12 | 0 | NON_SYNONYMOU | MODERAT | MISSENSE | Gtc/Atc | V446I | PLES_28241 | T | T | T | T | T | T | T | T | T | T | T | T |
| NC_011770 | 3037441 | A | G | 65 | 0 | 12 | 0 | NON_SYNONYMOU | MODERAT | MISSENSE | Tcc/Ccc | S445P | PLES_28241 | G | G | G | G | G | G | G | G | G | G | G | G |
| NC_011770 | 3037647 | T | A | 63 | 0 | 2 | 0 | NON_SYNONYMOU | MODERAT | MISSENSE | gAt/gTt | D376V | PLES_28241 | A | A | | | | | | | | | | |
| NC_011770 | 3037834 | G | T | 63 | 0 | 2 | 0 | NON_SYNONYMOU | MODERAT | MISSENSE | Ctc/Atc | L314I | PLES_28241 | | | | | | | | | T | T | | |
| NC_011770 | 3038068 | C | G | 60 | 0 | 12 | 0 | NON_SYNONYMOU | MODERAT | MISSENSE | Ggc/Cgc | G236R | PLES_28241 | G | G | G | G | G | G | G | G | G | G | G | G |
| NC_011770 | 3038211 | C | T | 65 | 0 | 2 | 0 | NON_SYNONYMOU | MODERAT | MISSENSE | gGc/gAc | G188D | PLES_28241 | T | T | | | | | | | | | | |
| NC_011770 | 3038404 | A | G | 68 | 0 | 6 | 0 | NON_SYNONYMOU | MODERAT | MISSENSE | Ttc/Ctc | F124L | PLES_28241 | | | G | G | G | G | G | G | | | | |
| NC_011770 | 3038430 | G | A | 67 | 0 | 12 | 0 | NON_SYNONYMOU | MODERAT | MISSENSE | gCg/gTg | A115V | PLES_28241 | A | A | A | A | A | A | A | A | A | A | A | A |
| NC_011770 | 3038858 | A | G | 50 | 0 | 2 | 0 | NON_SYNONYMOU | MODERAT | MISSENSE | gTg/gCg | V231A | PLES_28251 | G | G | | | | | | | | | | |
| NC_011770 | 3038877 | C | T | 50 | 0 | 2 | 0 | NON_SYNONYMOU | MODERAT | MISSENSE | Ggc/Agc | G225S | PLES_28251 | | | | | | | | | T | T | | |
| NC_011770 | 3039240 | G | C | 53 | 0 | 12 | 0 | NON_SYNONYMOU | MODERAT | MISSENSE | Caa/Gaa | Q104E | PLES_28251 | C | C | C | C | C | C | C | C | C | C | C | C |
| NC_011770 | 3039545 | T | C | 54 | 0 | 6 | 0 | NON_SYNONYMOU | MODERAT | MISSENSE | aAc/aGc | N2S | PLES_28251 | | | C | C | C | C | | | | | C | C |
| NC_011770 | 3046525 | A | G | 39 | 0 | 2 | 0 | NON_SYNONYMOU | MODERAT | MISSENSE | Acg/Gcg | T180A | PLES_28311 | | | | | | | | | | | G | G |
| NC_011770 | 3047087 | T | C | 74 | 0 | 2 | 0 | NON_SYNONYMOU | MODERAT | MISSENSE | gTc/gCc | V367A | PLES_28311 | C | C | | | | | | | | | | |
| NC_011770 | 3046273 | G | A | 46 | 0 | 1 | 0 | NON_SYNONYMOU | MODERAT | MISSENSE | Gcc/Acc | A96T | PLES_28311 | | | | | | | | A | | | | |

| | | | | | | | | | | | | | | | | | | | | | | | | | |
|---|---|---|---|---|---|---|---|---|---|---|---|---|---|---|---|---|---|---|---|---|---|---|---|---|---|
| NC_011770 | 3047492 | A | G | 66 | 0 | 2 | 0 | NON_SYNONYMOU | MODERAT | MISSENSE | Agc/Ggc | S104G | PLES_28321 | | | | | | | | | | | G | G |
| NC_011770 | 3047202 | C | T | 74 | 0 | 1 | 0 | NON_SYNONYMOU | MODERAT | MISSENSE | gCa/gTa | A7V | PLES_28321 | | | | | | | | | | | T | |
| NC_011770 | 3048282 | A | G | 39 | 0 | 12 | 0 | NON_SYNONYMOU | MODERAT | MISSENSE | gAc/gGc | D88G | PLES_28331 | G | G | G | G | G | G | G | G | G | G | G | G |
| NC_011770 | 3048515 | G | A | 23 | 0 | 2 | 0 | NON_SYNONYMOU | MODERAT | MISSENSE | Gcg/Acg | A166T | PLES_28331 | | | | | | | | | A | A | | |
| NC_011770 | 3048861 | C | G | 17 | 6 | 2 | 0 | NON_SYNONYMOU | MODERAT | MISSENSE | aCc/aGc | T281S | PLES_28331 | . | . | | | | | . | . | . | . | G | G |
| NC_011770 | 3049222 | C | G | 13 | 4 | 8 | 0 | NON_SYNONYMOU | MODERAT | MISSENSE | caC/caG | H401Q | PLES_28331 | G | G | G | G | G | G | . | . | . | . | G | G |
| NC_011770 | 3049235 | C | A | 14 | 4 | 5 | 0 | NON_SYNONYMOU | MODERAT | MISSENSE | Cat/Aat | H406N | PLES_28331 | . | A | A | A | A | A | . | . | . | . | | |
| NC_011770 | 3062265 | T | TC | 1 | 10 | 2 | 0 | FRAME_SHIFT | HIGH | | ttg/tCtg | L4146S? | PLES_28341 | . | . | . | . | TC | TC | . | . | . | . | . | . |
| NC_011770 | 3049983 | A | G | 78 | 0 | 4 | 0 | NON_SYNONYMOU | MODERAT | MISSENSE | Atc/Gtc | I52V | PLES_28341 | G | G | | | | | | | | | G | G |
| NC_011770 | 3050008 | G | A | 80 | 0 | 2 | 0 | NON_SYNONYMOU | MODERAT | MISSENSE | gGc/gAc | G60D | PLES_28341 | | | | | | | | | A | A | | |
| NC_011770 | 3050634 | A | C | 36 | 0 | 4 | 0 | NON_SYNONYMOU | MODERAT | MISSENSE | Aag/Cag | K269Q | PLES_28341 | C | C | | | | | | | C | C | | |
| NC_011770 | 3050635 | A | G | 35 | 0 | 4 | 0 | NON_SYNONYMOU | MODERAT | MISSENSE | aAg/aGg | K269R | PLES_28341 | G | G | | | | | | | G | G | | |
| NC_011770 | 3050848 | C | T | 50 | 0 | 4 | 0 | NON_SYNONYMOU | MODERAT | MISSENSE | gCg/gTg | A340V | PLES_28341 | T | T | | | | | | | T | T | | |
| NC_011770 | 3051072 | G | A | 51 | 0 | 2 | 0 | NON_SYNONYMOU | MODERAT | MISSENSE | Gac/Aac | D415N | PLES_28341 | | | | | | | | | | | A | A |
| NC_011770 | 3051219 | G | A | 35 | 3 | 3 | 0 | NON_SYNONYMOU | MODERAT | MISSENSE | Gtc/Atc | V464I | PLES_28341 | | | | | | | . | . | A | . | A | A |
| NC_011770 | 3051233 | A | T | 35 | 5 | 1 | 0 | NON_SYNONYMOU | MODERAT | MISSENSE | gaA/gaT | E468D | PLES_28341 | | | | | | | . | . | . | . | T | . |
| NC_011770 | 3051237 | C | G | 36 | 5 | 1 | 0 | NON_SYNONYMOU | MODERAT | MISSENSE | Cag/Gag | Q470E | PLES_28341 | | | | | | | . | . | . | . | G | . |
| NC_011770 | 3051373 | T | C | 38 | 6 | 6 | 0 | NON_SYNONYMOU | MODERAT | MISSENSE | gTg/gCg | V515A | PLES_28341 | C | C | C | C | C | C | . | . | . | . | . | . |
| NC_011770 | 3051640 | G | C | 39 | 6 | 2 | 0 | NON_SYNONYMOU | MODERAT | MISSENSE | gGg/gCg | G604A | PLES_28341 | C | C | | | | | . | . | . | . | . | . |
| NC_011770 | 3052273 | G | A | 52 | 6 | 2 | 0 | NON_SYNONYMOU | MODERAT | MISSENSE | gGg/gAg | G815E | PLES_28341 | A | A | | | | | . | . | . | . | . | . |
| NC_011770 | 3052440 | A | G | 38 | 6 | 4 | 0 | NON_SYNONYMOU | MODERAT | MISSENSE | Acc/Gcc | T871A | PLES_28341 | | | G | G | G | G | . | . | . | . | . | . |
| NC_011770 | 3052495 | T | C | 40 | 6 | 2 | 0 | NON_SYNONYMOU | MODERAT | MISSENSE | gTa/gCa | V889A | PLES_28341 | C | C | | | | | . | . | . | . | . | . |
| NC_011770 | 3052995 | C | T | 31 | 6 | 6 | 0 | NON_SYNONYMOU | MODERAT | MISSENSE | Cgg/Tgg | R1056W | PLES_28341 | T | T | T | T | T | T | . | . | . | . | . | . |
| NC_011770 | 3053086 | T | C | 25 | 6 | 6 | 0 | NON_SYNONYMOU | MODERAT | MISSENSE | gTg/gCg | V1086A | PLES_28341 | C | C | C | C | C | C | . | . | . | . | . | . |
| NC_011770 | 3053518 | G | A | 42 | 6 | 2 | 0 | NON_SYNONYMOU | MODERAT | MISSENSE | gGg/gAg | G1230E | PLES_28341 | A | A | | | | | . | . | . | . | . | . |
| NC_011770 | 3053866 | A | T | 50 | 6 | 2 | 0 | NON_SYNONYMOU | MODERAT | MISSENSE | gAc/gTc | D1346V | PLES_28341 | T | T | | | | | . | . | . | . | . | . |
| NC_011770 | 3054279 | G | A | 50 | 6 | 2 | 0 | NON_SYNONYMOU | MODERAT | MISSENSE | Ggt/Agt | G1484S | PLES_28341 | A | A | | | | | . | . | . | . | . | . |
| NC_011770 | 3055209 | G | A | 44 | 6 | 6 | 0 | NON_SYNONYMOU | MODERAT | MISSENSE | Ggc/Agc | G1794S | PLES_28341 | A | A | A | A | A | A | . | . | . | . | . | . |
| NC_011770 | 3055467 | G | A | 42 | 6 | 2 | 0 | NON_SYNONYMOU | MODERAT | MISSENSE | Gga/Aga | G1880R | PLES_28341 | A | A | | | | | . | . | . | . | . | . |
| NC_011770 | 3055672 | C | G | 34 | 6 | 4 | 0 | NON_SYNONYMOU | MODERAT | MISSENSE | aCt/aGt | T1948S | PLES_28341 | | | G | G | G | G | . | . | . | . | . | . |
| NC_011770 | 3055740 | C | G | 33 | 6 | 2 | 0 | NON_SYNONYMOU | MODERAT | MISSENSE | Ctg/Gtg | L1971V | PLES_28341 | G | G | | | | | . | . | . | . | . | . |
| NC_011770 | 3056112 | G | A | 47 | 6 | 2 | 0 | NON_SYNONYMOU | MODERAT | MISSENSE | Gat/Aat | D2095N | PLES_28341 | A | A | | | | | . | . | . | . | . | . |
| NC_011770 | 3056155 | G | C | 47 | 6 | 4 | 0 | NON_SYNONYMOU | MODERAT | MISSENSE | gGt/gCt | G2109A | PLES_28341 | | | C | C | C | C | . | . | . | . | . | . |
| NC_011770 | 3056173 | G | A | 47 | 6 | 4 | 0 | NON_SYNONYMOU | MODERAT | MISSENSE | cGc/cAc | R2115H | PLES_28341 | | | A | A | A | A | . | . | . | . | . | . |
| NC_011770 | 3056659 | A | T | 47 | 6 | 6 | 0 | NON_SYNONYMOU | MODERAT | MISSENSE | cAg/cTg | Q2277L | PLES_28341 | T | T | T | T | T | T | . | . | . | . | . | . |
| NC_011770 | 3057318 | G | A | 46 | 6 | 6 | 0 | NON_SYNONYMOU | MODERAT | MISSENSE | Gca/Aca | A2497T | PLES_28341 | A | A | A | A | A | A | . | . | . | . | . | . |
| NC_011770 | 3057506 | G | T | 42 | 6 | 4 | 0 | NON_SYNONYMOU | MODERAT | MISSENSE | caG/caT | Q2559H | PLES_28341 | | | T | T | T | T | . | . | . | . | . | . |
| NC_011770 | 3057627 | A | G | 40 | 6 | 2 | 0 | NON_SYNONYMOU | MODERAT | MISSENSE | Acg/Gcg | T2600A | PLES_28341 | G | G | | | | | . | . | . | . | . | . |
| NC_011770 | 3057628 | C | T | 40 | 6 | 2 | 0 | NON_SYNONYMOU | MODERAT | MISSENSE | aCg/aTg | T2600M | PLES_28341 | T | T | | | | | . | . | . | . | . | . |
| NC_011770 | 3058116 | A | C | 70 | 0 | 4 | 0 | NON_SYNONYMOU | MODERAT | MISSENSE | Aag/Cag | K2763Q | PLES_28341 | | | | | | | C | C | C | C | | |
| NC_011770 | 3058189 | C | T | 69 | 2 | 5 | 0 | NON_SYNONYMOU | MODERAT | MISSENSE | gCg/gTg | A2787V | PLES_28341 | T | T | | | | | . | T | . | . | T | T |
| NC_011770 | 3058278 | G | A | 54 | 1 | 4 | 0 | NON_SYNONYMOU | MODERAT | MISSENSE | Gtc/Atc | V2817I | PLES_28341 | | | A | A | A | A | | | . | | | |
| NC_011770 | 3058299 | G | T | 51 | 1 | 4 | 0 | NON_SYNONYMOU | MODERAT | MISSENSE | Gac/Tac | D2824Y | PLES_28341 | | | T | T | T | T | | | . | | | |
| NC_011770 | 3058353 | A | G | 49 | 0 | 4 | 0 | NON_SYNONYMOU | MODERAT | MISSENSE | Acc/Gcc | T2842A | PLES_28341 | | | G | G | G | G | | | | | | |
| NC_011770 | 3058450 | C | T | 54 | 0 | 4 | 0 | NON_SYNONYMOU | MODERAT | MISSENSE | gCg/gTg | A2874V | PLES_28341 | | | T | T | T | T | | | | | | |
| NC_011770 | 3058549 | A | C | 51 | 0 | 6 | 0 | NON_SYNONYMOU | MODERAT | MISSENSE | gAc/gCc | D2907A | PLES_28341 | C | C | | | | | C | C | C | C | | |
| NC_011770 | 3058594 | G | C | 51 | 0 | 4 | 0 | NON_SYNONYMOU | MODERAT | MISSENSE | gGc/gCc | G2922A | PLES_28341 | C | C | | | | | C | C | | | | |
| NC_011770 | 3058611 | G | A | 41 | 0 | 4 | 0 | NON_SYNONYMOU | MODERAT | MISSENSE | Gcc/Acc | A2928T | PLES_28341 | | | A | A | A | A | | | | | | |
| NC_011770 | 3058729 | T | C | 41 | 3 | 9 | 0 | NON_SYNONYMOU | MODERAT | MISSENSE | aTc/aCc | I2967T | PLES_28341 | C | C | C | C | C | C | . | . | . | C | C | C |
| NC_011770 | 3058819 | T | C | 58 | 2 | 8 | 0 | NON_SYNONYMOU | MODERAT | MISSENSE | gTc/gCc | V2997A | PLES_28341 | C | C | C | C | C | C | . | . | C | C | | |
| NC_011770 | 3059100 | A | G | 54 | 0 | 4 | 0 | NON_SYNONYMOU | MODERAT | MISSENSE | Atc/Gtc | I3091V | PLES_28341 | | | | | | | | | G | G | G | G |
| NC_011770 | 3059458 | G | A | 61 | 0 | 2 | 0 | NON_SYNONYMOU | MODERAT | MISSENSE | gGc/gAc | G3210D | PLES_28341 | | | | | | | | | A | A | | |
| NC_011770 | 3059468 | A | C | 62 | 0 | 10 | 0 | NON_SYNONYMOU | MODERAT | MISSENSE | gaA/gaC | E3213D | PLES_28341 | | | C | C | C | C | C | C | C | C | C | C |
| NC_011770 | 3059508 | C | T | 59 | 0 | 4 | 0 | NON_SYNONYMOU | MODERAT | MISSENSE | Cgg/Tgg | R3227W | PLES_28341 | | | T | T | T | T | | | | | | |
| NC_011770 | 3059581 | T | C | 59 | 0 | 6 | 0 | NON_SYNONYMOU | MODERAT | MISSENSE | gTc/gCc | V3251A | PLES_28341 | | | | | | | C | C | C | C | C | C |
| NC_011770 | 3059787 | A | G | 58 | 0 | 2 | 0 | NON_SYNONYMOU | MODERAT | MISSENSE | Acc/Gcc | T3320A | PLES_28341 | | | | | | | | | G | G | | |
| NC_011770 | 3059890 | C | G | 55 | 0 | 10 | 0 | NON_SYNONYMOU | MODERAT | MISSENSE | aCc/aGc | T3354S | PLES_28341 | | | G | G | G | G | G | G | G | G | G | G |
| NC_011770 | 3059892 | A | G | 54 | 0 | 10 | 0 | NON_SYNONYMOU | MODERAT | MISSENSE | Acc/Gcc | T3355A | PLES_28341 | | | G | G | G | G | G | G | G | G | G | G |
| NC_011770 | 3059941 | A | G | 57 | 0 | 4 | 0 | NON_SYNONYMOU | MODERAT | MISSENSE | gAc/gGc | D3371G | PLES_28341 | | | | | | | G | G | | | G | G |
| NC_011770 | 3059959 | T | G | 54 | 0 | 6 | 0 | NON_SYNONYMOU | MODERAT | MISSENSE | gTc/gGc | V3377G | PLES_28341 | G | G | | | | | G | G | | | G | G |
| NC_011770 | 3060079 | G | A | 60 | 0 | 2 | 0 | NON_SYNONYMOU | MODERAT | MISSENSE | cGg/cAg | R3417Q | PLES_28341 | | | | | | | | | | | A | A |
| NC_011770 | 3060321 | G | A | 45 | 0 | 2 | 0 | NON_SYNONYMOU | MODERAT | MISSENSE | Ggc/Agc | G3498S | PLES_28341 | | | A | A | | | | | | | | |

| | | | | | | | | | | | | | | | | | | | | | | | | | |
|---|---|---|---|---|---|---|---|---|---|---|---|---|---|---|---|---|---|---|---|---|---|---|---|---|---|
| NC_011770 | 3060616 | G | A | 50 | 0 | 2 | 0 | NON_SYNONYMOU | MODERAT | MISSENSE | gGc/gAc | G3596D | PLES_28341 | | | | | | | A | A | | | | |
| NC_011770 | 3060874 | C | G | 42 | 0 | 2 | 0 | NON_SYNONYMOU | MODERAT | MISSENSE | gCc/gGc | A3682G | PLES_28341 | | | | | | | | | | | G | G |
| NC_011770 | 3060962 | G | C | 32 | 0 | 4 | 0 | NON_SYNONYMOU | MODERAT | MISSENSE | ttG/ttC | L3711F | PLES_28341 | | | | | | | C | C | | | C | C |
| NC_011770 | 3061059 | G | T | 23 | 3 | 2 | 0 | NON_SYNONYMOU | MODERAT | MISSENSE | Gcc/Tcc | A3744S | PLES_28341 | | | | | | | | . | . | . | T | T |
| NC_011770 | 3062270 | C | G | 1 | 9 | 3 | 0 | NON_SYNONYMOU | MODERAT | MISSENSE | aaC/aaG | N4147K | PLES_28341 | . | . | G | . | G | G | . | . | . | . | . | . |
| NC_011770 | 3062463 | A | G | 42 | 6 | 6 | 0 | NON_SYNONYMOU | MODERAT | MISSENSE | Aca/Gca | T4212A | PLES_28341 | G | G | G | G | G | G | . | . | . | . | . | . |
| NC_011770 | 3062473 | G | A | 41 | 6 | 6 | 0 | NON_SYNONYMOU | MODERAT | MISSENSE | gGc/gAc | G4215D | PLES_28341 | A | A | A | A | A | A | . | . | . | . | . | . |
| NC_011770 | 3062532 | A | G | 41 | 6 | 6 | 0 | NON_SYNONYMOU | MODERAT | MISSENSE | Acc/Gcc | T4235A | PLES_28341 | G | G | G | G | G | G | . | . | . | . | . | . |
| NC_011770 | 3062661 | G | A | 51 | 6 | 4 | 0 | NON_SYNONYMOU | MODERAT | MISSENSE | Ggt/Agt | G4278S | PLES_28341 | | | A | A | A | A | . | . | . | . | . | . |
| NC_011770 | 3062728 | G | C | 55 | 6 | 2 | 0 | NON_SYNONYMOU | MODERAT | MISSENSE | gGt/gCt | G4300A | PLES_28341 | C | C | | | | | . | . | . | . | . | . |
| NC_011770 | 3062769 | A | G | 55 | 6 | 6 | 0 | NON_SYNONYMOU | MODERAT | MISSENSE | Att/Gtt | I4314V | PLES_28341 | G | G | G | G | G | G | . | . | . | . | . | . |
| NC_011770 | 3062782 | C | T | 53 | 6 | 6 | 0 | NON_SYNONYMOU | MODERAT | MISSENSE | gCg/gTg | A4318V | PLES_28341 | T | T | T | T | T | T | . | . | . | . | . | . |
| NC_011770 | 3062790 | C | T | 56 | 6 | 4 | 0 | NON_SYNONYMOU | MODERAT | MISSENSE | Cac/Tac | H4321Y | PLES_28341 | | | T | T | T | T | . | . | . | . | . | . |
| NC_011770 | 3062881 | T | G | 56 | 6 | 2 | 0 | NON_SYNONYMOU | MODERAT | MISSENSE | gTc/gGc | V4351G | PLES_28341 | G | G | | | | | . | . | . | . | . | . |
| NC_011770 | 3062917 | G | A | 53 | 6 | 6 | 0 | NON_SYNONYMOU | MODERAT | MISSENSE | gGt/gAt | G4363D | PLES_28341 | A | A | A | A | A | A | . | . | . | . | . | . |
| NC_011770 | 3063006 | G | C | 55 | 6 | 6 | 0 | NON_SYNONYMOU | MODERAT | MISSENSE | Gtt/Ctt | V4393L | PLES_28341 | C | C | C | C | C | C | . | . | . | . | . | . |
| NC_011770 | 3063087 | G | A | 56 | 6 | 4 | 0 | NON_SYNONYMOU | MODERAT | MISSENSE | Gcc/Acc | A4420T | PLES_28341 | | | A | A | A | A | . | . | . | . | . | . |
| NC_011770 | 3063091 | C | T | 52 | 6 | 4 | 0 | NON_SYNONYMOU | MODERAT | MISSENSE | aCa/aTa | T4421I | PLES_28341 | | | T | T | T | T | . | . | . | . | . | . |
| NC_011770 | 3065067 | G | A | 0 | 10 | 2 | 0 | NON_SYNONYMOU | MODERAT | MISSENSE | Gtg/Atg | V5080M | PLES_28341 | . | . | A | . | A | . | . | . | . | . | . | . |
| NC_011770 | 3065586 | A | G | 37 | 6 | 6 | 0 | NON_SYNONYMOU | MODERAT | MISSENSE | Aat/Gat | N5253D | PLES_28341 | G | G | G | G | G | G | . | . | . | . | . | . |
| NC_011770 | 3065644 | T | C | 42 | 6 | 2 | 0 | NON_SYNONYMOU | MODERAT | MISSENSE | aTc/aCc | I5272T | PLES_28341 | C | C | | | | | . | . | . | . | . | . |
| NC_011770 | 3065667 | G | A | 42 | 6 | 2 | 0 | NON_SYNONYMOU | MODERAT | MISSENSE | Gtg/Atg | V5280M | PLES_28341 | A | A | | | | | . | . | . | . | . | . |
| NC_011770 | 3065815 | C | A | 51 | 6 | 6 | 0 | NON_SYNONYMOU | MODERAT | MISSENSE | cCg/cAg | P5329Q | PLES_28341 | A | A | A | A | A | A | . | . | . | . | . | . |
| NC_011770 | 3066286 | G | A | 15 | 6 | 1 | 0 | NON_SYNONYMOU | MODERAT | MISSENSE | aGg/aAg | R5486K | PLES_28341 | A | . | | | | | . | . | . | . | . | . |
| NC_011770 | 3051061 | G | C | 48 | 0 | 9 | 0 | NON_SYNONYMOU | MODERAT | MISSENSE | gGc/gCc | G411A | PLES_28341 | | | C | C | | C | C | C | C | C | C | C |
| NC_011770 | 3064576 | A | G | 40 | 6 | 1 | 0 | NON_SYNONYMOU | MODERAT | MISSENSE | gAc/gGc | D4916G | PLES_28341 | | G | | | | | . | . | . | . | . | . |
| NC_011770 | 3057873 | G | A | 33 | 0 | 1 | 0 | NON_SYNONYMOU | MODERAT | MISSENSE | Ggc/Agc | G2682S | PLES_28341 | | | | A | | | | | | | | |
| NC_011770 | 3067633 | T | G | 4 | 10 | 2 | 0 | NON_SYNONYMOU | MODERAT | MISSENSE | aTc/aGc | I38S | PLES_28351 | G | G | . | . | . | . | . | . | . | . | . | . |
| NC_011770 | 3067833 | A | T | 4 | 10 | 2 | 0 | NON_SYNONYMOU | MODERAT | MISSENSE | Atg/Ttg | M105L | PLES_28351 | T | T | . | . | . | . | . | . | . | . | . | . |
| NC_011770 | 3067864 | G | A | 5 | 10 | 2 | 0 | NON_SYNONYMOU | MODERAT | MISSENSE | gGt/gAt | G115D | PLES_28351 | A | A | . | . | . | . | . | . | . | . | . | . |
| NC_011770 | 3072723 | C | T | 9 | 6 | 6 | 0 | NON_SYNONYMOU | MODERAT | MISSENSE | gGc/gAc | G638D | PLES_28371 | . | . | . | . | . | . | T | T | T | T | T | T |
| NC_011770 | 3072934 | C | T | 38 | 4 | 2 | 0 | NON_SYNONYMOU | MODERAT | MISSENSE | Gac/Aac | D568N | PLES_28371 | T | T | . | . | . | . | | | | | | |
| NC_011770 | 3073089 | C | A | 24 | 4 | 6 | 0 | NON_SYNONYMOU | MODERAT | MISSENSE | gGc/gTc | G516V | PLES_28371 | A | A | . | . | . | . | A | A | | | A | A |
| NC_011770 | 3073264 | T | C | 31 | 4 | 2 | 0 | NON_SYNONYMOU | MODERAT | MISSENSE | Act/Gct | T458A | PLES_28371 | | | . | . | . | . | C | C | | | | |
| NC_011770 | 3073290 | T | A | 33 | 4 | 4 | 0 | NON_SYNONYMOU | MODERAT | MISSENSE | cAg/cTg | Q449L | PLES_28371 | A | A | . | . | . | . | | | A | A | | |
| NC_011770 | 3073333 | T | G | 37 | 4 | 2 | 0 | NON_SYNONYMOU | MODERAT | MISSENSE | Acc/Ccc | T435P | PLES_28371 | | | . | . | . | . | | | G | G | | |
| NC_011770 | 3073430 | C | A | 42 | 4 | 2 | 0 | NON_SYNONYMOU | MODERAT | MISSENSE | gaG/gaT | E402D | PLES_28371 | | | . | . | . | . | | | A | A | | |
| NC_011770 | 3073597 | T | G | 56 | 4 | 4 | 0 | NON_SYNONYMOU | MODERAT | MISSENSE | Aag/Cag | K347Q | PLES_28371 | G | G | . | . | . | . | G | G | | | | |
| NC_011770 | 3073659 | T | C | 44 | 4 | 2 | 0 | NON_SYNONYMOU | MODERAT | MISSENSE | cAc/cGc | H326R | PLES_28371 | C | C | . | . | . | . | | | | | | |
| NC_011770 | 3073669 | T | C | 46 | 4 | 2 | 0 | NON_SYNONYMOU | MODERAT | MISSENSE | Atc/Gtc | I323V | PLES_28371 | C | C | . | . | . | . | | | | | | |
| NC_011770 | 3073708 | T | C | 41 | 4 | 4 | 0 | NON_SYNONYMOU | MODERAT | MISSENSE | Acg/Gcg | T310A | PLES_28371 | C | C | . | . | . | . | | | | | C | C |
| NC_011770 | 3073887 | T | C | 34 | 4 | 2 | 0 | NON_SYNONYMOU | MODERAT | MISSENSE | cAa/cGa | Q250R | PLES_28371 | C | C | . | . | . | . | | | | | | |
| NC_011770 | 3074161 | A | G | 14 | 8 | 4 | 0 | NON_SYNONYMOU | MODERAT | MISSENSE | Tgc/Cgc | C159R | PLES_28371 | . | . | . | . | . | . | . | . | G | G | G | G |
| NC_011770 | 3074586 | A | C | 6 | 7 | 2 | 0 | NON_SYNONYMOU | MODERAT | MISSENSE | cTc/cGc | L17R | PLES_28371 | . | . | . | . | . | . | . | | | | C | C |
| NC_011770 | 3074587 | G | A | 6 | 7 | 2 | 0 | NON_SYNONYMOU | MODERAT | MISSENSE | Ctc/Ttc | L17F | PLES_28371 | . | . | . | . | . | . | . | . | | | A | A |
| NC_011770 | 3074828 | A | G | 21 | 8 | 4 | 0 | NON_SYNONYMOU | MODERAT | MISSENSE | gTc/gCc | V241A | PLES_28381 | . | . | . | . | . | . | G | G | . | . | G | G |
| NC_011770 | 3074960 | C | T | 23 | 8 | 2 | 0 | NON_SYNONYMOU | MODERAT | MISSENSE | cGc/cAc | R197H | PLES_28381 | . | . | . | . | . | . | | | . | . | T | T |
| NC_011770 | 3075246 | G | C | 23 | 8 | 2 | 0 | NON_SYNONYMOU | MODERAT | MISSENSE | Cgc/Ggc | R102G | PLES_28381 | . | . | . | . | . | . | | | . | . | C | C |
| NC_011770 | 3075398 | C | G | 24 | 8 | 4 | 0 | NON_SYNONYMOU | MODERAT | MISSENSE | aGc/aCc | S51T | PLES_28381 | . | . | . | . | . | . | G | G | . | . | G | G |
| NC_011770 | 3075485 | T | C | 25 | 8 | 2 | 0 | NON_SYNONYMOU | MODERAT | MISSENSE | aAg/aGg | K22R | PLES_28381 | . | . | . | . | . | . | | | . | . | C | C |
| NC_011770 | 3077302 | C | T | 29 | 8 | 2 | 0 | START_LOST | HIGH | MISSENSE | atG/atA | M1I | PLES_28391 | . | . | . | . | . | . | T | T | . | . | | |
| NC_011770 | 3076487 | A | G | 36 | 8 | 4 | 0 | NON_SYNONYMOU | MODERAT | MISSENSE | aTc/aCc | I273T | PLES_28391 | . | . | . | . | . | . | G | G | . | . | G | G |
| NC_011770 | 3076662 | G | C | 35 | 8 | 2 | 0 | NON_SYNONYMOU | MODERAT | MISSENSE | Caa/Gaa | Q215E | PLES_28391 | . | . | . | . | . | . | | | . | . | C | C |
| NC_011770 | 3077111 | T | C | 31 | 8 | 2 | 0 | NON_SYNONYMOU | MODERAT | MISSENSE | aAg/aGg | K65R | PLES_28391 | . | . | . | . | . | . | | | . | . | C | C |
| NC_011770 | 3077274 | T | C | 28 | 8 | 4 | 0 | NON_SYNONYMOU | MODERAT | MISSENSE | Acc/Gcc | T11A | PLES_28391 | . | . | . | . | . | . | C | C | . | . | C | C |
| NC_011770 | 3077925 | C | A | 17 | 6 | 4 | 0 | NON_SYNONYMOU | MODERAT | MISSENSE | gGt/gTt | G177V | PLES_28401 | | | . | . | . | . | A | A | . | . | A | A |
| NC_011770 | 3078015 | G | A | 63 | 0 | 4 | 0 | NON_SYNONYMOU | MODERAT | MISSENSE | aCg/aTg | T147M | PLES_28401 | A | A | | | | | A | A | | | | |
| NC_011770 | 3078063 | C | G | 62 | 0 | 12 | 0 | NON_SYNONYMOU | MODERAT | MISSENSE | gGg/gCg | G131A | PLES_28401 | G | G | G | G | G | G | G | G | G | G | G | G |
| NC_011770 | 3078186 | C | T | 61 | 0 | 1 | 0 | NON_SYNONYMOU | MODERAT | MISSENSE | gGc/gAc | G90D | PLES_28401 | | | | | | | T | | | | | |
| NC_011770 | 3078195 | C | T | 60 | 0 | 2 | 0 | NON_SYNONYMOU | MODERAT | MISSENSE | cGc/cAc | R87H | PLES_28401 | T | T | | | | | | | | | | |
| NC_011770 | 3078231 | T | A | 64 | 0 | 4 | 0 | NON_SYNONYMOU | MODERAT | MISSENSE | cAg/cTg | Q75L | PLES_28401 | | | A | A | A | A | | | | | | |
| NC_011770 | 3078459 | G | A | 73 | 0 | 2 | 0 | NON_SYNONYMOU | MODERAT | MISSENSE | Cca/Tca | P282S | PLES_28411 | | | | | | | | | A | A | | |

| | | | | | | | | | | | | | | | | | | | | | | | | | |
|---|---|---|---|---|---|---|---|---|---|---|---|---|---|---|---|---|---|---|---|---|---|---|---|---|---|
| NC_011770 | 3079218 | C | T | 74 | 0 | 4 | 0 | NON_SYNONYMOU | MODERAT | MISSENSE | Gcc/Acc | A29T | PLES_28411 | | | T | T | T | T | | | | | | |
| NC_011770 | 3079289 | A | G | 74 | 0 | 1 | 0 | NON_SYNONYMOU | MODERAT | MISSENSE | cTg/cCg | L5P | PLES_28411 | | G | | | | | | | | | | |
| NC_011770 | 3079767 | C | A | 34 | 0 | 4 | 0 | NON_SYNONYMOU | MODERAT | MISSENSE | Ctc/Atc | L12I | PLES_28431 | | | A | A | A | A | | | | | | |
| NC_011770 | 3079770 | A | T | 33 | 0 | 2 | 0 | NON_SYNONYMOU | MODERAT | MISSENSE | Aac/Tac | N13Y | PLES_28431 | | | | | | | | | T | T | | |
| NC_011770 | 3080494 | T | C | 54 | 0 | 12 | 0 | NON_SYNONYMOU | MODERAT | MISSENSE | aTg/aCg | M254T | PLES_28431 | C | C | C | C | C | C | C | C | C | C | C | C |
| NC_011770 | 3080629 | G | C | 60 | 0 | 8 | 0 | NON_SYNONYMOU | MODERAT | MISSENSE | aGc/aCc | S299T | PLES_28431 | C | C | | | | | C | C | C | C | C | C |
| NC_011770 | 3080660 | G | C | 60 | 0 | 2 | 0 | NON_SYNONYMOU | MODERAT | MISSENSE | ttG/ttC | L309F | PLES_28431 | C | C | | | | | | | | | | |
| NC_011770 | 3080664 | C | T | 62 | 0 | 2 | 0 | NON_SYNONYMOU | MODERAT | MISSENSE | Ctc/Ttc | L311F | PLES_28431 | | | | | | | | | T | T | | |
| NC_011770 | 3081121 | T | C | 69 | 0 | 4 | 0 | NON_SYNONYMOU | MODERAT | MISSENSE | gTg/gCg | V463A | PLES_28431 | | | | | | | C | C | C | C | | |
| NC_011770 | 3081214 | G | A | 69 | 0 | 1 | 0 | NON_SYNONYMOU | MODERAT | MISSENSE | gGc/gAc | G494D | PLES_28431 | | | | | | | | A | | | | |
| NC_011770 | 3081533 | A | G | 66 | 0 | 12 | 0 | NON_SYNONYMOU | MODERAT | MISSENSE | cAc/cGc | H6R | PLES_28441 | G | G | G | G | G | G | G | G | G | G | G | G |
| NC_011770 | 3081722 | T | C | 82 | 0 | 1 | 0 | NON_SYNONYMOU | MODERAT | MISSENSE | cTc/cCc | L69P | PLES_28441 | | | | | | | C | | | | | |
| NC_011770 | 3081799 | G | A | 82 | 0 | 2 | 0 | NON_SYNONYMOU | MODERAT | MISSENSE | Ggc/Agc | G95S | PLES_28441 | A | A | | | | | | | | | | |
| NC_011770 | 3082160 | A | G | 79 | 0 | 12 | 0 | NON_SYNONYMOU | MODERAT | MISSENSE | gAc/gGc | D215G | PLES_28441 | G | G | G | G | G | G | G | G | G | G | G | G |
| NC_011770 | 3082255 | C | G | 84 | 0 | 2 | 0 | NON_SYNONYMOU | MODERAT | MISSENSE | Cag/Gag | Q247E | PLES_28441 | | | | | | | | | | | G | G |
| NC_011770 | 3082267 | A | G | 81 | 0 | 8 | 0 | NON_SYNONYMOU | MODERAT | MISSENSE | Atg/Gtg | M251V | PLES_28441 | G | G | | | | | G | G | G | G | G | G |
| NC_011770 | 3082370 | T | C | 87 | 0 | 4 | 0 | NON_SYNONYMOU | MODERAT | MISSENSE | gTg/gCg | V285A | PLES_28441 | C | C | | | | | C | C | | | | |
| NC_011770 | 3083753 | A | G | 73 | 0 | 8 | 0 | NON_SYNONYMOU | MODERAT | MISSENSE | aAc/aGc | N413S | PLES_28451 | G | G | | | | | G | G | G | G | G | G |
| NC_011770 | 3084207 | C | T | 68 | 0 | 10 | 0 | NON_SYNONYMOU | MODERAT | MISSENSE | cGc/cAc | R532H | PLES_28461 | T | T | T | T | T | T | | | T | T | T | T |
| NC_011770 | 3084886 | C | T | 67 | 0 | 2 | 0 | NON_SYNONYMOU | MODERAT | MISSENSE | Gcc/Acc | A306T | PLES_28461 | | | | | | | T | T | | | | |
| NC_011770 | 3085492 | C | T | 61 | 0 | 1 | 0 | NON_SYNONYMOU | MODERAT | MISSENSE | Ggc/Agc | G104S | PLES_28461 | | | | | | | T | | | | | |
| NC_011770 | 3085588 | C | T | 59 | 0 | 2 | 0 | NON_SYNONYMOU | MODERAT | MISSENSE | Gtc/Atc | V72I | PLES_28461 | | | | | | | | | | | T | T |
| NC_011770 | 3084204 | C | T | 70 | 0 | 1 | 0 | NON_SYNONYMOU | MODERAT | MISSENSE | cGg/cAg | R533Q | PLES_28461 | | | | | | | | | | | T | |
| NC_011770 | 3085943 | T | C | 55 | 0 | 2 | 0 | NON_SYNONYMOU | MODERAT | MISSENSE | cTg/cCg | L7P | PLES_28471 | | | | | | | | | C | C | | |
| NC_011770 | 3086465 | C | T | 62 | 0 | 2 | 0 | NON_SYNONYMOU | MODERAT | MISSENSE | cCg/cTg | P181L | PLES_28471 | | | | | | | | | T | T | | |
| NC_011770 | 3095235 | A | G | 69 | 0 | 4 | 0 | NON_SYNONYMOU | MODERAT | MISSENSE | aTg/aCg | M35T | PLES_28531 | | | | | | | G | G | | | G | G |
| NC_011770 | 3095552 | AG | AGGGG | 65 | 0 | 3 | 0 | FRAME_SHIFT | HIGH | | ctg/ | L249 | PLES_28541 | AGGGG | . | | A | | | | A | | | | |
| NC_011770 | 3095360 | T | C | 63 | 0 | 6 | 0 | NON_SYNONYMOU | MODERAT | MISSENSE | cAc/cGc | H313R | PLES_28541 | | | | | | | C | C | C | C | C | C |
| NC_011770 | 3095435 | C | T | 55 | 0 | 2 | 0 | NON_SYNONYMOU | MODERAT | MISSENSE | cGt/cAt | R288H | PLES_28541 | | | | | T | T | | | | | | |
| NC_011770 | 3095484 | G | A | 63 | 0 | 2 | 0 | NON_SYNONYMOU | MODERAT | MISSENSE | Ctt/Ttt | L272F | PLES_28541 | | | | | | | | | | | A | A |
| NC_011770 | 3096008 | T | A | 74 | 0 | 2 | 0 | NON_SYNONYMOU | MODERAT | MISSENSE | cAg/cTg | Q97L | PLES_28541 | | | | | | | | | A | A | | |
| NC_011770 | 3096081 | A | G | 77 | 0 | 12 | 0 | NON_SYNONYMOU | MODERAT | MISSENSE | Ttc/Ctc | F73L | PLES_28541 | G | G | G | G | G | G | G | G | G | G | G | G |
| NC_011770 | 3097267 | A | G | 65 | 0 | 1 | 0 | NON_SYNONYMOU | MODERAT | MISSENSE | Agc/Ggc | S95G | PLES_28551 | | | | | | | G | | | | | |
| NC_011770 | 3097301 | A | G | 66 | 0 | 6 | 0 | NON_SYNONYMOU | MODERAT | MISSENSE | gAc/gGc | D106G | PLES_28551 | G | G | G | G | G | G | | | | | | |
| NC_011770 | 3097315 | A | G | 65 | 0 | 10 | 0 | NON_SYNONYMOU | MODERAT | MISSENSE | Acc/Gcc | T111A | PLES_28551 | G | G | G | G | G | G | G | G | | | G | G |
| NC_011770 | 3097378 | A | G | 66 | 0 | 6 | 0 | NON_SYNONYMOU | MODERAT | MISSENSE | Acc/Gcc | T132A | PLES_28551 | G | G | G | G | G | G | | | | | | |
| NC_011770 | 3097552 | T | C | 67 | 0 | 2 | 0 | NON_SYNONYMOU | MODERAT | MISSENSE | Ttc/Ctc | F190L | PLES_28551 | | | | | | | | | C | C | | |
| NC_011770 | 3097565 | G | A | 70 | 0 | 2 | 0 | NON_SYNONYMOU | MODERAT | MISSENSE | gGg/gAg | G194E | PLES_28551 | | | | | | | A | A | | | | |
| NC_011770 | 3097592 | T | C | 69 | 0 | 6 | 0 | NON_SYNONYMOU | MODERAT | MISSENSE | gTt/gCt | V203A | PLES_28551 | C | C | C | C | C | C | | | | | | |
| NC_011770 | 3097849 | C | G | 67 | 0 | 2 | 0 | NON_SYNONYMOU | MODERAT | MISSENSE | Cgt/Ggt | R289G | PLES_28551 | | | | | | | G | G | | | | |
| NC_011770 | 3098261 | G | A | 56 | 0 | 6 | 0 | NON_SYNONYMOU | MODERAT | MISSENSE | cGt/cAt | R426H | PLES_28551 | A | A | A | A | A | A | | | | | | |
| NC_011770 | 3098329 | T | G | 55 | 0 | 8 | 0 | NON_SYNONYMOU | MODERAT | MISSENSE | Tcg/Gcg | S449A | PLES_28551 | | | G | G | G | G | G | G | | | G | G |
| NC_011770 | 3098962 | G | A | 82 | 0 | 2 | 0 | NON_SYNONYMOU | MODERAT | MISSENSE | Gcg/Acg | A660T | PLES_28551 | | | | | | | A | A | | | | |
| NC_011770 | 3098576 | A | G | 72 | 0 | 1 | 0 | NON_SYNONYMOU | MODERAT | MISSENSE | tAc/tGc | Y531C | PLES_28551 | | | | | | | | G | | | | |
| NC_011770 | 3099082 | T | C | 85 | 0 | 12 | 0 | NON_SYNONYMOU | MODERAT | MISSENSE | gTg/gCg | V29A | PLES_28561 | C | C | C | C | C | C | C | C | C | C | C | C |
| NC_011770 | 3099084 | C | T | 86 | 0 | 2 | 0 | NON_SYNONYMOU | MODERAT | MISSENSE | Ccg/Tcg | P30S | PLES_28561 | | | | | | | T | T | | | | |
| NC_011770 | 3099567 | C | A | 82 | 0 | 2 | 0 | NON_SYNONYMOU | MODERAT | MISSENSE | Cat/Aat | H191N | PLES_28561 | | | | | | | | | A | A | | |
| NC_011770 | 3099795 | A | G | 77 | 0 | 4 | 0 | NON_SYNONYMOU | MODERAT | MISSENSE | Atc/Gtc | I267V | PLES_28561 | | | G | G | G | G | | | | | | |
| NC_011770 | 3099978 | C | T | 82 | 0 | 2 | 0 | NON_SYNONYMOU | MODERAT | MISSENSE | Ctc/Ttc | L328F | PLES_28561 | | | | | | | | | T | T | | |
| NC_011770 | 3100002 | G | A | 86 | 0 | 2 | 0 | NON_SYNONYMOU | MODERAT | MISSENSE | Ggg/Agg | G336R | PLES_28561 | | | | | | | | | A | A | | |
| NC_011770 | 3099064 | A | G | 68 | 0 | 7 | 0 | NON_SYNONYMOU | MODERAT | MISSENSE | gAt/gGt | D23G | PLES_28561 | G | G | | G | G | G | | | G | G | | |
| NC_011770 | 3100029 | G | A | 85 | 0 | 2 | 0 | NON_SYNONYMOU | MODERAT | MISSENSE | cGc/cAc | R6H | PLES_28571 | | | | | | | | | | | A | A |
| NC_011770 | 3100041 | G | A | 88 | 0 | 2 | 0 | NON_SYNONYMOU | MODERAT | MISSENSE | cGc/cAc | R10H | PLES_28571 | | | | | | | | | A | A | | |
| NC_011770 | 3100128 | A | G | 79 | 0 | 2 | 0 | NON_SYNONYMOU | MODERAT | MISSENSE | aAc/aGc | N39S | PLES_28571 | | | | | | | G | G | | | | |
| NC_011770 | 3100601 | G | A | 56 | 0 | 2 | 0 | NON_SYNONYMOU | MODERAT | MISSENSE | Ggc/Agc | G197S | PLES_28571 | | | | | | | | | | | A | A |
| NC_011770 | 3100838 | C | G | 40 | 0 | 2 | 0 | NON_SYNONYMOU | MODERAT | MISSENSE | Ctc/Gtc | L276V | PLES_28571 | | | | | | | | | G | G | | |
| NC_011770 | 3100931 | C | G | 32 | 0 | 12 | 0 | NON_SYNONYMOU | MODERAT | MISSENSE | Ccc/Gcc | P307A | PLES_28571 | G | G | G | G | G | G | G | G | G | G | G | G |
| NC_011770 | 3101871 | A | C | 83 | 0 | 2 | 0 | NON_SYNONYMOU | MODERAT | MISSENSE | Atc/Ctc | I68L | PLES_28591 | | | | | | | C | C | | | | |
| NC_011770 | 3102186 | G | C | 78 | 0 | 8 | 0 | NON_SYNONYMOU | MODERAT | MISSENSE | Gcg/Ccg | A173P | PLES_28591 | | | C | C | C | C | | | C | C | C | C |
| NC_011770 | 3102307 | T | C | 75 | 0 | 12 | 0 | NON_SYNONYMOU | MODERAT | MISSENSE | gTg/gCg | V213A | PLES_28591 | C | C | C | C | C | C | C | C | C | C | C | C |
| NC_011770 | 3103104 | G | A | 64 | 0 | 2 | 0 | NON_SYNONYMOU | MODERAT | MISSENSE | Ggt/Agt | G479S | PLES_28591 | | | | | | | A | A | | | | |
| NC_011770 | 3103587 | C | A | 69 | 0 | 4 | 0 | NON_SYNONYMOU | MODERAT | MISSENSE | Cgt/Agt | R640S | PLES_28591 | | | A | A | A | A | | | | | | |

| | | | | | | | | | | | | | | | | | | | | | | | | | |
|---|---|---|---|---|---|---|---|---|---|---|---|---|---|---|---|---|---|---|---|---|---|---|---|---|---|
| NC_011770 | 3102291 | G | A | 79 | 0 | 1 | 0 | NON_SYNONYMOU | MODERAT | MISSENSE | Gtg/Atg | V208M | PLES_28591 | | | | | | | | | | | | A |
| NC_011770 | 3103857 | C | T | 66 | 0 | 4 | 0 | NON_SYNONYMOU | MODERAT | MISSENSE | gCa/gTa | A69V | PLES_28601 | | | T | T | T | T | | | | | | |
| NC_011770 | 3104347 | T | C | 45 | 0 | 6 | 0 | NON_SYNONYMOU | MODERAT | MISSENSE | Acc/Gcc | T86A | PLES_28611 | C | C | | | | | | | C | C | C | C |
| NC_011770 | 3105827 | T | C | 61 | 0 | 2 | 0 | NON_SYNONYMOU | MODERAT | MISSENSE | Acg/Gcg | T48A | PLES_28631 | C | C | | | | | | | | | | |
| NC_011770 | 3105847 | C | T | 62 | 0 | 2 | 0 | NON_SYNONYMOU | MODERAT | MISSENSE | gGc/gAc | G41D | PLES_28631 | T | T | | | | | | | | | | |
| NC_011770 | 3105932 | G | C | 61 | 0 | 2 | 0 | NON_SYNONYMOU | MODERAT | MISSENSE | Cgc/Ggc | R13G | PLES_28631 | C | C | | | | | | | | | | |
| NC_011770 | 3105945 | G | C | 63 | 0 | 2 | 0 | NON_SYNONYMOU | MODERAT | MISSENSE | ttC/ttG | F8L | PLES_28631 | C | C | | | | | | | | | | |
| NC_011770 | 3106047 | T | C | 67 | 0 | 2 | 0 | NON_SYNONYMOU | MODERAT | MISSENSE | Ttc/Ctc | F6L | PLES_28641 | C | C | | | | | | | | | | |
| NC_011770 | 3106476 | C | G | 84 | 0 | 12 | 0 | NON_SYNONYMOU | MODERAT | MISSENSE | Ctc/Gtc | L149V | PLES_28641 | G | G | G | G | G | G | G | G | G | G | G | G |
| NC_011770 | 3107067 | C | T | 90 | 0 | 2 | 0 | NON_SYNONYMOU | MODERAT | MISSENSE | Cgc/Tgc | R346C | PLES_28641 | | | | | | | T | T | | | | |
| NC_011770 | 3107117 | G | C | 87 | 0 | 2 | 0 | NON_SYNONYMOU | MODERAT | MISSENSE | caG/caC | Q362H | PLES_28641 | | | | | | | C | C | | | | |
| NC_011770 | 3107408 | C | A | 78 | 0 | 2 | 0 | NON_SYNONYMOU | MODERAT | MISSENSE | gaC/gaA | D459E | PLES_28641 | | | | | | | | | | | A | A |
| NC_011770 | 3107487 | A | G | 71 | 0 | 2 | 0 | NON_SYNONYMOU | MODERAT | MISSENSE | Atc/Gtc | I486V | PLES_28641 | G | G | | | | | | | | | | |
| NC_011770 | 3107805 | G | A | 69 | 0 | 4 | 0 | NON_SYNONYMOU | MODERAT | MISSENSE | Gct/Act | A592T | PLES_28641 | | | A | A | A | A | | | | | | |
| NC_011770 | 3107808 | G | T | 68 | 0 | 6 | 0 | NON_SYNONYMOU | MODERAT | MISSENSE | Gcg/Tcg | A593S | PLES_28641 | T | T | T | T | T | T | | | | | | |
| NC_011770 | 3108000 | G | A | 80 | 0 | 2 | 0 | NON_SYNONYMOU | MODERAT | MISSENSE | Gcc/Acc | A657T | PLES_28641 | A | A | | | | | | | | | | |
| NC_011770 | 3108616 | A | C | 62 | 0 | 2 | 0 | NON_SYNONYMOU | MODERAT | MISSENSE | gAg/gCg | E72A | PLES_28651 | | | | | | | | | C | C | | |
| NC_011770 | 3109102 | A | G | 59 | 0 | 10 | 0 | NON_SYNONYMOU | MODERAT | MISSENSE | cAc/cGc | H234R | PLES_28651 | G | G | G | G | G | G | | | G | G | G | G |
| NC_011770 | 3109395 | G | T | 80 | 0 | 4 | 0 | NON_SYNONYMOU | MODERAT | MISSENSE | Gtg/Ttg | V332L | PLES_28651 | T | T | | | | | T | T | | | | |
| NC_011770 | 3109473 | T | G | 78 | 0 | 4 | 0 | NON_SYNONYMOU | MODERAT | MISSENSE | caA/caC | Q72H | PLES_28661 | G | G | | | | | G | G | | | | |
| NC_011770 | 3110836 | G | A | 73 | 0 | 2 | 0 | NON_SYNONYMOU | MODERAT | MISSENSE | gGc/gAc | G296D | PLES_28671 | | | | | | | | | A | A | | |
| NC_011770 | 3111009 | T | C | 69 | 2 | 6 | 0 | NON_SYNONYMOU | MODERAT | MISSENSE | gTc/gCc | V20A | PLES_28681 | C | C | | | | | C | C | . | . | C | C |
| NC_011770 | 3111270 | A | C | 70 | 2 | 4 | 0 | NON_SYNONYMOU | MODERAT | MISSENSE | gAg/gCg | E107A | PLES_28681 | C | C | | | | | | | . | . | C | C |
| NC_011770 | 3126862 | T | G | 48 | 4 | 8 | 0 | NON_SYNONYMOU | MODERAT | MISSENSE | caA/caC | Q199H | PLES_28721 | G | G | . | . | . | . | G | G | G | G | G | G |
| NC_011770 | 3128234 | C | T | 68 | 0 | 6 | 0 | NON_SYNONYMOU | MODERAT | MISSENSE | Ggc/Agc | G186S | PLES_28731 | T | T | T | T | T | T | | | | | | |
| NC_011770 | 3128588 | T | C | 72 | 0 | 12 | 0 | NON_SYNONYMOU | MODERAT | MISSENSE | Acc/Gcc | T68A | PLES_28731 | C | C | C | C | C | C | C | C | C | C | C | C |
| NC_011770 | 3128597 | C | T | 72 | 0 | 12 | 0 | NON_SYNONYMOU | MODERAT | MISSENSE | Ggc/Agc | G65S | PLES_28731 | T | T | T | T | T | T | T | T | T | T | T | T |
| NC_011770 | 3128731 | C | T | 48 | 4 | 8 | 0 | NON_SYNONYMOU | MODERAT | MISSENSE | gGg/gAg | G20E | PLES_28731 | T | T | T | T | T | T | T | T | . | . | . | . |
| NC_011770 | 3130357 | C | T | 63 | 0 | 4 | 0 | NON_SYNONYMOU | MODERAT | MISSENSE | Gag/Aag | E186K | PLES_28751 | | | T | T | T | T | | | | | | |
| NC_011770 | 3130684 | T | A | 53 | 0 | 2 | 0 | NON_SYNONYMOU | MODERAT | MISSENSE | Acc/Tcc | T77S | PLES_28751 | | | | | | | A | A | | | | |
| NC_011770 | 3130687 | C | T | 52 | 0 | 2 | 0 | NON_SYNONYMOU | MODERAT | MISSENSE | Ggc/Agc | G76S | PLES_28751 | | | | | | | T | T | | | | |
| NC_011770 | 3130810 | T | C | 43 | 0 | 10 | 0 | NON_SYNONYMOU | MODERAT | MISSENSE | Acc/Gcc | T35A | PLES_28751 | C | C | C | C | C | C | C | C | C | C | | |
| NC_011770 | 3131190 | C | T | 44 | 0 | 2 | 0 | NON_SYNONYMOU | MODERAT | MISSENSE | Gtc/Atc | V392I | PLES_28761 | T | T | | | | | | | | | | |
| NC_011770 | 3131490 | C | T | 60 | 0 | 4 | 0 | NON_SYNONYMOU | MODERAT | MISSENSE | Ggc/Agc | G292S | PLES_28761 | | | T | T | T | T | | | | | | |
| NC_011770 | 3131496 | C | T | 61 | 0 | 4 | 0 | NON_SYNONYMOU | MODERAT | MISSENSE | Gac/Aac | D290N | PLES_28761 | T | T | | | | | T | T | | | | |
| NC_011770 | 3131594 | T | G | 70 | 0 | 2 | 0 | NON_SYNONYMOU | MODERAT | MISSENSE | gAg/gCg | E257A | PLES_28761 | | | | | | | | | G | G | | |
| NC_011770 | 3131696 | C | T | 78 | 0 | 2 | 0 | NON_SYNONYMOU | MODERAT | MISSENSE | gGc/gAc | G223D | PLES_28761 | T | T | | | | | | | | | | |
| NC_011770 | 3131795 | C | T | 71 | 0 | 2 | 0 | NON_SYNONYMOU | MODERAT | MISSENSE | cGc/cAc | R190H | PLES_28761 | T | T | | | | | | | | | | |
| NC_011770 | 3131934 | T | C | 72 | 0 | 12 | 0 | NON_SYNONYMOU | MODERAT | MISSENSE | Acg/Gcg | T144A | PLES_28761 | C | C | C | C | C | C | C | C | C | C | C | C |
| NC_011770 | 3132057 | T | C | 74 | 0 | 2 | 0 | NON_SYNONYMOU | MODERAT | MISSENSE | Atc/Gtc | I103V | PLES_28761 | | | | | | | | | C | C | | |
| NC_011770 | 3132070 | G | T | 73 | 0 | 2 | 0 | NON_SYNONYMOU | MODERAT | MISSENSE | gaC/gaA | D98E | PLES_28761 | T | T | | | | | | | | | | |
| NC_011770 | 3132177 | G | T | 69 | 0 | 10 | 0 | NON_SYNONYMOU | MODERAT | MISSENSE | Ctc/Atc | L63I | PLES_28761 | T | T | T | T | T | T | T | T | | | T | T |
| NC_011770 | 3132191 | G | A | 72 | 0 | 2 | 0 | NON_SYNONYMOU | MODERAT | MISSENSE | aCg/aTg | T58M | PLES_28761 | A | A | | | | | | | | | | |
| NC_011770 | 3132204 | C | T | 73 | 0 | 1 | 0 | NON_SYNONYMOU | MODERAT | MISSENSE | Gtc/Atc | V54I | PLES_28761 | | | | | | | T | | | | | |
| NC_011770 | 3132212 | G | T | 70 | 0 | 4 | 0 | NON_SYNONYMOU | MODERAT | MISSENSE | cCg/cAg | P51Q | PLES_28761 | | | T | T | T | T | | | | | | |
| NC_011770 | 3132296 | A | G | 62 | 2 | 10 | 0 | NON_SYNONYMOU | MODERAT | MISSENSE | cTc/cCc | L23P | PLES_28761 | G | G | G | G | G | G | G | G | . | . | G | G |
| NC_011770 | 3132297 | G | T | 61 | 2 | 10 | 0 | NON_SYNONYMOU | MODERAT | MISSENSE | Ctc/Atc | L23I | PLES_28761 | T | T | T | T | T | T | T | T | . | . | T | T |
| NC_011770 | 3132347 | G | A | 73 | 2 | 10 | 0 | NON_SYNONYMOU | MODERAT | MISSENSE | aCg/aTg | T6M | PLES_28761 | A | A | A | A | A | A | A | A | . | . | A | A |
| NC_011770 | 3132072 | C | T | 75 | 0 | 1 | 0 | NON_SYNONYMOU | MODERAT | MISSENSE | Gac/Aac | D98N | PLES_28761 | | | | | | | | | | | | T |
| NC_011770 | 3132446 | C | T | 81 | 0 | 4 | 0 | SPLICE_SITE_REGIC | LOW | SILENT | tGa/tAa | *227 | PLES_28771 | | | T | T | T | T | | | | | | |
| NC_011770 | 3134540 | A | G | 90 | 0 | 2 | 0 | NON_SYNONYMOU | MODERAT | MISSENSE | Atg/Gtg | M120V | PLES_28791 | | | G | G | | | | | | | | |
| NC_011770 | 3135090 | T | C | 75 | 0 | 12 | 0 | NON_SYNONYMOU | MODERAT | MISSENSE | gTc/gCc | V303A | PLES_28791 | C | C | C | C | C | C | C | C | C | C | C | C |
| NC_011770 | 3135111 | C | A | 73 | 0 | 2 | 0 | NON_SYNONYMOU | MODERAT | MISSENSE | gCg/gAg | A310E | PLES_28791 | | | | | | | | | | | A | A |
| NC_011770 | 3137220 | A | G | 65 | 0 | 2 | 0 | NON_SYNONYMOU | MODERAT | MISSENSE | Acc/Gcc | T34A | PLES_28811 | G | G | | | | | | | | | | |
| NC_011770 | 3137307 | C | G | 68 | 0 | 10 | 0 | NON_SYNONYMOU | MODERAT | MISSENSE | Ctc/Gtc | L63V | PLES_28811 | G | G | G | G | G | G | G | G | | | G | G |
| NC_011770 | 3137320 | G | T | 70 | 0 | 2 | 0 | NON_SYNONYMOU | MODERAT | MISSENSE | tGc/tTc | C67F | PLES_28811 | T | T | | | | | | | | | | |
| NC_011770 | 3137341 | GCGCGGCA | G | 69 | 0 | 1 | 0 | CODON_CHANGE_ | MODERATE | | gcagtcgtcg | AVVAA76A | PLES_28811 | G | | | | | | | | | | | |
| NC_011770 | 3137588 | T | C | 76 | 0 | 2 | 0 | NON_SYNONYMOU | MODERAT | MISSENSE | gTc/gCc | V17A | PLES_28821 | C | C | | | | | | | | | | |
| NC_011770 | 3138580 | G | A | 74 | 0 | 6 | 0 | NON_SYNONYMOU | MODERAT | MISSENSE | Gac/Aac | D348N | PLES_28821 | A | A | | | | | A | A | | | A | A |
| NC_011770 | 3138850 | G | A | 68 | 0 | 4 | 0 | NON_SYNONYMOU | MODERAT | MISSENSE | Gcc/Acc | A438T | PLES_28821 | A | A | | | | | A | A | | | | |
| NC_011770 | 3140672 | A | G | 86 | 0 | 1 | 0 | NON_SYNONYMOU | MODERAT | MISSENSE | tAc/tGc | Y50C | PLES_28841 | | | | | | | G | | | | | |
| NC_011770 | 3141365 | G | A | 53 | 0 | 10 | 0 | NON_SYNONYMOU | MODERAT | MISSENSE | gGa/gAa | G208E | PLES_28851 | A | A | A | A | A | A | | | A | A | A | A |

| | | | | | | | | | | | | | | | | | | | | | | | | | |
|---|---|---|---|---|---|---|---|---|---|---|---|---|---|---|---|---|---|---|---|---|---|---|---|---|---|
| NC_011770 | 3141482 | T | C | 49 | 0 | 12 | 0 | NON_SYNONYMOU | MODERAT | MISSENSE | gTg/gCg | V247A | PLES_28851 | C | C | C | C | C | C | C | C | C | C | C | C |
| NC_011770 | 3141616 | T | G | 56 | 0 | 2 | 0 | NON_SYNONYMOU | MODERAT | MISSENSE | Aaa/Caa | K304Q | PLES_28861 | G | G | | | | | | | | | | |
| NC_011770 | 3141700 | C | T | 67 | 0 | 2 | 0 | NON_SYNONYMOU | MODERAT | MISSENSE | Gta/Ata | V276I | PLES_28861 | | | | | | | | | | | T | T |
| NC_011770 | 3141813 | T | C | 66 | 0 | 2 | 0 | NON_SYNONYMOU | MODERAT | MISSENSE | aAt/aGt | N238S | PLES_28861 | | | | | | | C | C | | | | |
| NC_011770 | 3142393 | T | G | 62 | 0 | 4 | 0 | NON_SYNONYMOU | MODERAT | MISSENSE | Acc/Ccc | T45P | PLES_28861 | | | G | G | G | G | | | | | | |
| NC_011770 | 3142422 | T | G | 61 | 0 | 10 | 0 | NON_SYNONYMOU | MODERAT | MISSENSE | cAa/cCa | Q35P | PLES_28861 | G | G | G | G | G | G | | | G | G | G | G |
| NC_011770 | 3142444 | G | A | 57 | 0 | 4 | 0 | NON_SYNONYMOU | MODERAT | MISSENSE | Ccg/Tcg | P28S | PLES_28861 | A | A | | | | | | | | | A | A |
| NC_011770 | 3143440 | G | A | 73 | 0 | 2 | 0 | NON_SYNONYMOU | MODERAT | MISSENSE | gCg/gTg | A247V | PLES_28881 | | | | | | | | | A | A | | |
| NC_011770 | 3143704 | A | G | 87 | 0 | 2 | 0 | NON_SYNONYMOU | MODERAT | MISSENSE | gTa/gCa | V159A | PLES_28881 | | | | | | | | | G | G | | |
| NC_011770 | 3143813 | C | T | 81 | 0 | 2 | 0 | NON_SYNONYMOU | MODERAT | MISSENSE | Gcg/Acg | A123T | PLES_28881 | T | T | | | | | | | | | | |
| NC_011770 | 3144129 | C | G | 83 | 0 | 6 | 0 | NON_SYNONYMOU | MODERAT | MISSENSE | gaG/gaC | E17D | PLES_28881 | | | G | G | G | G | G | G | | | | |
| NC_011770 | 3144943 | T | G | 76 | 0 | 4 | 0 | NON_SYNONYMOU | MODERAT | MISSENSE | Aag/Cag | K63Q | PLES_28891 | | | G | G | G | G | | | | | | |
| NC_011770 | 3145170 | G | T | 72 | 0 | 2 | 0 | NON_SYNONYMOU | MODERAT | MISSENSE | Cgc/Agc | R185S | PLES_28901 | | | | | | | T | T | | | | |
| NC_011770 | 3145257 | T | C | 56 | 0 | 4 | 0 | NON_SYNONYMOU | MODERAT | MISSENSE | Acc/Gcc | T156A | PLES_28901 | | | | | | | C | C | | | C | C |
| NC_011770 | 3145259 | T | C | 55 | 0 | 6 | 0 | NON_SYNONYMOU | MODERAT | MISSENSE | gAc/gGc | D155G | PLES_28901 | | | | | | | C | C | C | C | C | C |
| NC_011770 | 3145331 | C | T | 51 | 0 | 2 | 0 | NON_SYNONYMOU | MODERAT | MISSENSE | gGg/gAg | G131E | PLES_28901 | | | | | | | T | T | | | | |
| NC_011770 | 3145380 | C | T | 47 | 0 | 2 | 0 | NON_SYNONYMOU | MODERAT | MISSENSE | Ggc/Agc | G115S | PLES_28901 | | | | | | | | | | | T | T |
| NC_011770 | 3145382 | C | T | 47 | 0 | 4 | 0 | NON_SYNONYMOU | MODERAT | MISSENSE | cGc/cAc | R114H | PLES_28901 | | | | | | | T | T | | | T | T |
| NC_011770 | 3145383 | G | A | 47 | 0 | 4 | 0 | NON_SYNONYMOU | MODERAT | MISSENSE | Cgc/Tgc | R114C | PLES_28901 | | | | | | | A | A | | | A | A |
| NC_011770 | 3145548 | C | G | 69 | 0 | 2 | 0 | NON_SYNONYMOU | MODERAT | MISSENSE | Gcc/Ccc | A59P | PLES_28901 | | | | | | | G | G | | | | |
| NC_011770 | 3145994 | T | G | 44 | 0 | 10 | 0 | NON_SYNONYMOU | MODERAT | MISSENSE | Atg/Ctg | M19L | PLES_28911 | G | G | G | G | G | G | G | G | | | G | G |
| NC_011770 | 3146348 | G | T | 57 | 0 | 4 | 0 | NON_SYNONYMOU | MODERAT | MISSENSE | gaC/gaA | D79E | PLES_28921 | | | T | T | T | T | | | | | | |
| NC_011770 | 3146370 | A | C | 21 | 0 | 10 | 0 | NON_SYNONYMOU | MODERAT | MISSENSE | aTg/aGg | M72R | PLES_28921 | | | C | C | C | C | C | C | C | C | C | C |
| NC_011770 | 3146371 | T | A | 21 | 0 | 10 | 0 | NON_SYNONYMOU | MODERAT | MISSENSE | Atg/Ttg | M72L | PLES_28921 | | | A | A | A | A | A | A | A | A | A | A |
| NC_011770 | 3146382 | C | T | 21 | 0 | 10 | 0 | NON_SYNONYMOU | MODERAT | MISSENSE | cGc/cAc | R68H | PLES_28921 | | | T | T | T | T | T | T | T | T | T | T |
| NC_011770 | 3146383 | G | T | 21 | 0 | 10 | 0 | NON_SYNONYMOU | MODERAT | MISSENSE | Cgc/Agc | R68S | PLES_28921 | | | T | T | T | T | T | T | T | T | T | T |
| NC_011770 | 3146407 | T | C | 20 | 0 | 10 | 0 | NON_SYNONYMOU | MODERAT | MISSENSE | Atc/Gtc | I60V | PLES_28921 | | | C | C | C | C | C | C | C | C | C | C |
| NC_011770 | 3146415 | G | A | 21 | 0 | 10 | 0 | NON_SYNONYMOU | MODERAT | MISSENSE | gCg/gTg | A57V | PLES_28921 | | | A | A | A | A | A | A | A | A | A | A |
| NC_011770 | 3146416 | C | T | 20 | 0 | 10 | 0 | NON_SYNONYMOU | MODERAT | MISSENSE | Gcg/Acg | A57T | PLES_28921 | | | T | T | T | T | T | T | T | T | T | T |
| NC_011770 | 3146439 | G | C | 20 | 0 | 10 | 0 | NON_SYNONYMOU | MODERAT | MISSENSE | aCc/aGc | T49S | PLES_28921 | | | C | C | C | C | C | C | C | C | C | C |
| NC_011770 | 3146445 | C | G | 20 | 0 | 10 | 0 | NON_SYNONYMOU | MODERAT | MISSENSE | cGc/cCc | R47P | PLES_28921 | | | G | G | G | G | G | G | G | G | G | G |
| NC_011770 | 3146446 | G | A | 20 | 0 | 10 | 0 | NON_SYNONYMOU | MODERAT | MISSENSE | Cgc/Tgc | R47C | PLES_28921 | | | A | A | A | A | A | A | A | A | A | A |
| NC_011770 | 3146458 | C | T | 20 | 0 | 10 | 0 | NON_SYNONYMOU | MODERAT | MISSENSE | Gtc/Atc | V43I | PLES_28921 | | | T | T | T | T | T | T | T | T | T | T |
| NC_011770 | 3147087 | C | G | 45 | 4 | 4 | 0 | STOP_GAINED | HIGH | NONSENSE | taC/taG | Y97* | PLES_28931 | G | G | | | | | . | . | G | G | . | . |
| NC_011770 | 3147244 | C | T | 62 | 0 | 4 | 0 | STOP_GAINED | HIGH | NONSENSE | Cag/Tag | Q150* | PLES_28931 | | | | | | | T | T | | | T | T |
| NC_011770 | 3147328 | C | T | 74 | 0 | 4 | 0 | STOP_GAINED | HIGH | NONSENSE | Caa/Taa | Q178* | PLES_28931 | T | T | | | | | | | T | T | | |
| NC_011770 | 3146806 | G | A | 71 | 0 | 2 | 0 | NON_SYNONYMOU | MODERAT | MISSENSE | Gcg/Acg | A4T | PLES_28931 | | | | | | | | | | | A | A |
| NC_011770 | 3146833 | C | G | 70 | 0 | 4 | 0 | NON_SYNONYMOU | MODERAT | MISSENSE | Cgg/Ggg | R13G | PLES_28931 | | | | | | | G | G | | | G | G |
| NC_011770 | 3146839 | G | A | 69 | 0 | 4 | 0 | NON_SYNONYMOU | MODERAT | MISSENSE | Gcg/Acg | A15T | PLES_28931 | | | | | | | A | A | | | A | A |
| NC_011770 | 3146857 | C | G | 63 | 0 | 8 | 0 | NON_SYNONYMOU | MODERAT | MISSENSE | Ctt/Gtt | L21V | PLES_28931 | G | G | | | | | G | G | G | G | G | G |
| NC_011770 | 3146858 | T | G | 64 | 0 | 8 | 0 | NON_SYNONYMOU | MODERAT | MISSENSE | cTt/cGt | L21R | PLES_28931 | G | G | | | | | G | G | G | G | G | G |
| NC_011770 | 3146860 | G | A | 63 | 0 | 4 | 0 | NON_SYNONYMOU | MODERAT | MISSENSE | Ggc/Agc | G22S | PLES_28931 | A | A | | | | | | | A | A | | |
| NC_011770 | 3146863 | T | C | 64 | 0 | 4 | 0 | NON_SYNONYMOU | MODERAT | MISSENSE | Ttc/Ctc | F23L | PLES_28931 | C | C | | | | | | | C | C | | |
| NC_011770 | 3146870 | T | C | 66 | 0 | 4 | 0 | NON_SYNONYMOU | MODERAT | MISSENSE | cTt/cCt | L25P | PLES_28931 | C | C | | | | | | | C | C | | |
| NC_011770 | 3146875 | C | G | 66 | 0 | 6 | 0 | NON_SYNONYMOU | MODERAT | MISSENSE | Cgt/Ggt | R27G | PLES_28931 | G | G | | | | | | | G | G | G | G |
| NC_011770 | 3146876 | G | T | 72 | 0 | 4 | 0 | NON_SYNONYMOU | MODERAT | MISSENSE | cGt/cTt | R27L | PLES_28931 | T | T | | | | | | | T | T | | |
| NC_011770 | 3146878 | A | G | 72 | 0 | 4 | 0 | NON_SYNONYMOU | MODERAT | MISSENSE | Ata/Gta | I28V | PLES_28931 | G | G | | | | | | | G | G | | |
| NC_011770 | 3146881 | G | A | 72 | 0 | 4 | 0 | NON_SYNONYMOU | MODERAT | MISSENSE | Ggg/Agg | G29R | PLES_28931 | | | | | | | A | A | | | A | A |
| NC_011770 | 3146884 | G | A | 71 | 0 | 4 | 0 | NON_SYNONYMOU | MODERAT | MISSENSE | Gtt/Att | V30I | PLES_28931 | A | A | | | | | | | A | A | | |
| NC_011770 | 3146887 | G | C | 68 | 0 | 4 | 0 | NON_SYNONYMOU | MODERAT | MISSENSE | Gat/Cat | D31H | PLES_28931 | | | | | | | C | C | | | C | C |
| NC_011770 | 3146889 | T | G | 68 | 0 | 4 | 0 | NON_SYNONYMOU | MODERAT | MISSENSE | gaT/gaG | D31E | PLES_28931 | | | | | | | G | G | | | G | G |
| NC_011770 | 3146893 | C | T | 68 | 0 | 8 | 0 | NON_SYNONYMOU | MODERAT | MISSENSE | Cgc/Tgc | R33C | PLES_28931 | T | T | | | | | G | G | T | T | G | G |
| NC_011770 | 3146903 | T | A | 73 | 0 | 4 | 0 | NON_SYNONYMOU | MODERAT | MISSENSE | gTa/gAa | V36E | PLES_28931 | | | | | | | A | A | | | A | A |
| NC_011770 | 3146908 | G | C | 74 | 0 | 4 | 0 | NON_SYNONYMOU | MODERAT | MISSENSE | Gag/Cag | E38Q | PLES_28931 | C | C | | | | | | | C | C | | |
| NC_011770 | 3146911 | G | C | 72 | 0 | 8 | 0 | NON_SYNONYMOU | MODERAT | MISSENSE | Ggg/Cgg | G39R | PLES_28931 | C | C | | | | | C | C | C | C | C | C |
| NC_011770 | 3146914 | G | A | 71 | 0 | 4 | 0 | NON_SYNONYMOU | MODERAT | MISSENSE | Gta/Ata | V40I | PLES_28931 | | | | | | | A | A | | | A | A |
| NC_011770 | 3146926 | G | T | 68 | 0 | 4 | 0 | NON_SYNONYMOU | MODERAT | MISSENSE | Gcc/Tcc | A44S | PLES_28931 | | | | | | | T | T | | | T | T |
| NC_011770 | 3146929 | T | A | 68 | 0 | 8 | 0 | NON_SYNONYMOU | MODERAT | MISSENSE | Ttc/Atc | F45I | PLES_28931 | A | A | | | | | G | G | A | A | G | G |
| NC_011770 | 3146930 | T | G | 68 | 0 | 4 | 0 | NON_SYNONYMOU | MODERAT | MISSENSE | tTc/tGc | F45C | PLES_28931 | | | | | | | G | G | | | G | G |
| NC_011770 | 3146931 | C | A | 69 | 0 | 4 | 0 | NON_SYNONYMOU | MODERAT | MISSENSE | ttC/ttA | F45L | PLES_28931 | A | A | | | | | | | A | A | | |
| NC_011770 | 3146932 | C | G | 69 | 0 | 4 | 0 | NON_SYNONYMOU | MODERAT | MISSENSE | Cac/Gac | H46D | PLES_28931 | G | G | | | | | | | G | G | | |
| NC_011770 | 3146935 | G | A | 70 | 0 | 4 | 0 | NON_SYNONYMOU | MODERAT | MISSENSE | Gta/Ata | V47I | PLES_28931 | A | A | | | | | | | A | A | | |

| | | | | | | | | | | | | | | | | | | | | | | | | | |
|---|---|---|---|---|---|---|---|---|---|---|---|---|---|---|---|---|---|---|---|---|---|---|---|---|---|
| NC_011770 | 3146936 | T | A | 68 | 0 | 4 | 0 | NON_SYNONYMOU | MODERAT | MISSENSE | gTa/gAa | V47E | PLES_28931 | | | | | | | A | A | | | A | A |
| NC_011770 | 3146938 | G | C | 69 | 0 | 4 | 0 | NON_SYNONYMOU | MODERAT | MISSENSE | Gat/Cat | D48H | PLES_28931 | | | | | | | C | C | | | C | C |
| NC_011770 | 3146940 | T | G | 68 | 0 | 4 | 0 | NON_SYNONYMOU | MODERAT | MISSENSE | gaT/gaG | D48E | PLES_28931 | G | G | | | | | | | G | G | | |
| NC_011770 | 3146959 | T | G | 70 | 0 | 4 | 0 | NON_SYNONYMOU | MODERAT | MISSENSE | Tgg/Ggg | W55G | PLES_28931 | | | | | | | G | G | | | G | G |
| NC_011770 | 3146966 | G | A | 41 | 6 | 2 | 0 | NON_SYNONYMOU | MODERAT | MISSENSE | gGa/gAa | G57E | PLES_28931 | . | . | | | | | A | A | . | . | . | . |
| NC_011770 | 3147041 | G | T | 36 | 8 | 4 | 0 | NON_SYNONYMOU | MODERAT | MISSENSE | cGc/cTc | R82L | PLES_28931 | . | . | T | T | T | T | . | . | . | . | . | . |
| NC_011770 | 3147085 | T | C | 46 | 4 | 4 | 0 | NON_SYNONYMOU | MODERAT | MISSENSE | Tac/Cac | Y97H | PLES_28931 | C | C | | | | | . | . | C | C | . | . |
| NC_011770 | 3147086 | A | G | 46 | 4 | 4 | 0 | NON_SYNONYMOU | MODERAT | MISSENSE | tAc/tGc | Y97C | PLES_28931 | G | G | | | | | . | . | G | G | . | . |
| NC_011770 | 3147100 | A | G | 48 | 4 | 4 | 0 | NON_SYNONYMOU | MODERAT | MISSENSE | Aag/Gag | K102E | PLES_28931 | G | G | | | | | . | . | G | G | . | . |
| NC_011770 | 3147103 | G | A | 69 | 0 | 4 | 0 | NON_SYNONYMOU | MODERAT | MISSENSE | Ggt/Agt | G103S | PLES_28931 | A | A | | | | | | | A | A | | |
| NC_011770 | 3147112 | A | G | 70 | 0 | 4 | 0 | NON_SYNONYMOU | MODERAT | MISSENSE | Att/Gtt | I106V | PLES_28931 | | | | | | | G | G | | | G | G |
| NC_011770 | 3147121 | G | A | 70 | 0 | 4 | 0 | NON_SYNONYMOU | MODERAT | MISSENSE | Gag/Aag | E109K | PLES_28931 | | | | | | | A | A | | | A | A |
| NC_011770 | 3147124 | A | G | 70 | 0 | 8 | 0 | NON_SYNONYMOU | MODERAT | MISSENSE | Aat/Gat | N110D | PLES_28931 | G | G | | | | | G | G | G | G | G | G |
| NC_011770 | 3147127 | C | G | 69 | 0 | 4 | 0 | NON_SYNONYMOU | MODERAT | MISSENSE | Cgc/Ggc | R111G | PLES_28931 | | | | | | | G | G | | | G | G |
| NC_011770 | 3147151 | C | G | 73 | 0 | 8 | 0 | NON_SYNONYMOU | MODERAT | MISSENSE | Cgt/Ggt | R119G | PLES_28931 | G | G | | | | | G | G | G | G | G | G |
| NC_011770 | 3147160 | A | G | 72 | 0 | 4 | 0 | NON_SYNONYMOU | MODERAT | MISSENSE | Aat/Gat | N122D | PLES_28931 | G | G | | | | | | | G | G | | |
| NC_011770 | 3147163 | G | C | 71 | 0 | 8 | 0 | NON_SYNONYMOU | MODERAT | MISSENSE | Gat/Cat | D123H | PLES_28931 | C | C | | | | | C | C | C | C | C | C |
| NC_011770 | 3147165 | T | C | 71 | 0 | 8 | 0 | NON_SYNONYMOU | MODERAT | MISSENSE | gaT/gaG | D123E | PLES_28931 | C | C | | | | | G | G | C | C | G | G |
| NC_011770 | 3147190 | A | G | 71 | 0 | 8 | 0 | NON_SYNONYMOU | MODERAT | MISSENSE | Acc/Gcc | T132A | PLES_28931 | G | G | | | | | G | G | G | G | G | G |
| NC_011770 | 3147202 | A | G | 70 | 0 | 4 | 0 | NON_SYNONYMOU | MODERAT | MISSENSE | Atg/Gtg | M136V | PLES_28931 | G | G | | | | | | | G | G | | |
| NC_011770 | 3147208 | G | A | 71 | 0 | 4 | 0 | NON_SYNONYMOU | MODERAT | MISSENSE | Gtc/Atc | V138I | PLES_28931 | A | A | | | | | | | A | A | | |
| NC_011770 | 3147211 | G | C | 71 | 0 | 4 | 0 | NON_SYNONYMOU | MODERAT | MISSENSE | Gcc/Ccc | A139P | PLES_28931 | C | C | | | | | | | C | C | | |
| NC_011770 | 3147223 | G | C | 73 | 0 | 4 | 0 | NON_SYNONYMOU | MODERAT | MISSENSE | Gcg/Ccg | A143P | PLES_28931 | | | | | | | C | C | | | C | C |
| NC_011770 | 3147226 | G | C | 72 | 0 | 4 | 0 | NON_SYNONYMOU | MODERAT | MISSENSE | Ggc/Cgc | G144R | PLES_28931 | C | C | | | | | | | C | C | | |
| NC_011770 | 3147236 | C | T | 68 | 0 | 4 | 0 | NON_SYNONYMOU | MODERAT | MISSENSE | gCt/gTt | A147V | PLES_28931 | | | | | | | T | T | | | T | T |
| NC_011770 | 3147238 | G | A | 68 | 0 | 2 | 0 | NON_SYNONYMOU | MODERAT | MISSENSE | Gat/Aat | D148N | PLES_28931 | | | | | | | | | | | A | A |
| NC_011770 | 3147247 | G | T | 61 | 0 | 4 | 0 | NON_SYNONYMOU | MODERAT | MISSENSE | Gcg/Tcg | A151S | PLES_28931 | | | | | | | T | T | | | T | T |
| NC_011770 | 3147250 | T | G | 62 | 0 | 4 | 0 | NON_SYNONYMOU | MODERAT | MISSENSE | Tgg/Ggg | W152G | PLES_28931 | | | | | | | G | G | | | G | G |
| NC_011770 | 3147253 | C | T | 62 | 0 | 8 | 0 | NON_SYNONYMOU | MODERAT | MISSENSE | Ctt/Ttt | L153F | PLES_28931 | T | T | | | | | T | T | T | T | T | T |
| NC_011770 | 3147254 | T | C | 62 | 0 | 4 | 0 | NON_SYNONYMOU | MODERAT | MISSENSE | cTt/cCt | L153P | PLES_28931 | C | C | | | | | | | C | C | | |
| NC_011770 | 3147259 | G | A | 63 | 0 | 4 | 0 | NON_SYNONYMOU | MODERAT | MISSENSE | Gaa/Aaa | E155K | PLES_28931 | | | | | | | A | A | | | A | A |
| NC_011770 | 3147262 | C | A | 62 | 0 | 8 | 0 | NON_SYNONYMOU | MODERAT | MISSENSE | Ccc/Acc | P156T | PLES_28931 | A | A | | | | | G | G | A | A | G | G |
| NC_011770 | 3147265 | T | C | 62 | 0 | 8 | 0 | NON_SYNONYMOU | MODERAT | MISSENSE | Ttt/Ctt | F157L | PLES_28931 | C | C | | | | | C | C | C | C | C | C |
| NC_011770 | 3147280 | G | A | 72 | 0 | 8 | 0 | NON_SYNONYMOU | MODERAT | MISSENSE | Gta/Ata | V162I | PLES_28931 | A | A | | | | | A | A | A | A | A | A |
| NC_011770 | 3147292 | G | A | 71 | 0 | 8 | 0 | NON_SYNONYMOU | MODERAT | MISSENSE | Gcc/Acc | A166T | PLES_28931 | A | A | | | | | C | C | A | A | C | C |
| NC_011770 | 3147301 | C | G | 72 | 0 | 4 | 0 | NON_SYNONYMOU | MODERAT | MISSENSE | Cag/Gag | Q169E | PLES_28931 | | | | | | | G | G | | | G | G |
| NC_011770 | 3147307 | G | T | 71 | 0 | 4 | 0 | NON_SYNONYMOU | MODERAT | MISSENSE | Ggc/Tgc | G171C | PLES_28931 | | | | | | | T | T | | | T | T |
| NC_011770 | 3147316 | C | G | 72 | 0 | 4 | 0 | NON_SYNONYMOU | MODERAT | MISSENSE | Cgg/Ggg | R174G | PLES_28931 | | | | | | | G | G | | | G | G |
| NC_011770 | 3147319 | C | G | 72 | 0 | 4 | 0 | NON_SYNONYMOU | MODERAT | MISSENSE | Cag/Gag | Q175E | PLES_28931 | | | | | | | G | G | | | G | G |
| NC_011770 | 3147322 | G | T | 73 | 0 | 4 | 0 | NON_SYNONYMOU | MODERAT | MISSENSE | Gcc/Tcc | A176S | PLES_28931 | T | T | | | | | | | T | T | | |
| NC_011770 | 3147334 | A | G | 74 | 0 | 8 | 0 | NON_SYNONYMOU | MODERAT | MISSENSE | Aac/Gac | N180D | PLES_28931 | G | G | | | | | G | G | G | G | G | G |
| NC_011770 | 3147339 | G | T | 75 | 0 | 4 | 0 | NON_SYNONYMOU | MODERAT | MISSENSE | caG/caT | Q181H | PLES_28931 | | | | | | | T | T | | | T | T |
| NC_011770 | 3147352 | A | G | 76 | 0 | 8 | 0 | NON_SYNONYMOU | MODERAT | MISSENSE | Act/Gct | T186A | PLES_28931 | G | G | | | | | G | G | G | G | G | G |
| NC_011770 | 3147361 | A | G | 73 | 0 | 8 | 0 | NON_SYNONYMOU | MODERAT | MISSENSE | Att/Gtt | I189V | PLES_28931 | G | G | | | | | G | G | G | G | G | G |
| NC_011770 | 3147364 | G | A | 74 | 0 | 4 | 0 | NON_SYNONYMOU | MODERAT | MISSENSE | Gtt/Att | V190I | PLES_28931 | A | A | | | | | | | A | A | | |
| NC_011770 | 3147373 | A | C | 72 | 0 | 4 | 0 | NON_SYNONYMOU | MODERAT | MISSENSE | Aac/Cac | N193H | PLES_28931 | C | C | | | | | | | C | C | | |
| NC_011770 | 3147385 | G | T | 71 | 0 | 4 | 0 | NON_SYNONYMOU | MODERAT | MISSENSE | Gcc/Tcc | A197S | PLES_28931 | | | | | | | T | T | | | T | T |
| NC_011770 | 3147397 | A | G | 68 | 0 | 4 | 0 | NON_SYNONYMOU | MODERAT | MISSENSE | Acg/Gcg | T201A | PLES_28931 | G | G | | | | | | | G | G | | |
| NC_011770 | 3147514 | A | G | 35 | 4 | 4 | 0 | NON_SYNONYMOU | MODERAT | MISSENSE | Aaa/Gaa | K240E | PLES_28931 | . | . | | | | | G | G | . | . | G | G |
| NC_011770 | 3147622 | G | T | 43 | 4 | 4 | 0 | NON_SYNONYMOU | MODERAT | MISSENSE | Ggg/Tgg | G276W | PLES_28931 | T | T | | | | | . | . | T | T | . | . |
| NC_011770 | 3147646 | A | G | 53 | 2 | 6 | 0 | NON_SYNONYMOU | MODERAT | MISSENSE | Att/Gtt | I284V | PLES_28931 | G | G | | | | | G | G | G | G | . | . |
| NC_011770 | 3147652 | G | C | 63 | 0 | 8 | 0 | NON_SYNONYMOU | MODERAT | MISSENSE | Gag/Cag | E286Q | PLES_28931 | C | C | | | | | C | C | C | C | C | C |
| NC_011770 | 3147679 | T | C | 65 | 0 | 8 | 0 | NON_SYNONYMOU | MODERAT | MISSENSE | Tgc/Cgc | C295R | PLES_28931 | C | C | | | | | C | C | C | C | C | C |
| NC_011770 | 3147696 | C | G | 63 | 0 | 4 | 0 | NON_SYNONYMOU | MODERAT | MISSENSE | gaC/gaG | D300E | PLES_28931 | G | G | | | | | | | G | G | | |
| NC_011770 | 3195454 | A | AG | 44 | 0 | 8 | 0 | FRAME_SHIFT | HIGH | | aag/aagC | K347K? | PLES_29091 | AG | AG | | | | | AG | AG | AG | AG | AG | AG |
| NC_011770 | 3195459 | GA | G | 27 | 4 | 3 | 0 | FRAME_SHIFT | HIGH | | tcg/ | S346 | PLES_29091 | . | . | | | | | G | . | . | . | G | G |
| NC_011770 | 3196390 | T | TTC | 31 | 1 | 5 | 0 | FRAME_SHIFT | HIGH | | agc/agcGA | S35S? | PLES_29091 | TTC | . | | | | | TTC | TTC | | | TTC | TTC |
| NC_011770 | 3195192 | A | G | 61 | 0 | 8 | 0 | NON_SYNONYMOU | MODERAT | MISSENSE | gTc/gCc | V435A | PLES_29091 | G | G | | | | | G | G | G | G | G | G |
| NC_011770 | 3195249 | A | G | 70 | 0 | 8 | 0 | NON_SYNONYMOU | MODERAT | MISSENSE | aTc/aCc | I416T | PLES_29091 | G | G | | | | | G | G | G | G | G | G |
| NC_011770 | 3195276 | T | A | 71 | 0 | 8 | 0 | NON_SYNONYMOU | MODERAT | MISSENSE | aAc/aTc | N407I | PLES_29091 | A | A | | | | | A | A | A | A | A | A |
| NC_011770 | 3195304 | G | C | 71 | 0 | 8 | 0 | NON_SYNONYMOU | MODERAT | MISSENSE | Ctg/Gtg | L398V | PLES_29091 | C | C | | | | | C | C | C | C | C | C |
| NC_011770 | 3195330 | G | C | 73 | 0 | 8 | 0 | NON_SYNONYMOU | MODERAT | MISSENSE | gCc/gGc | A389G | PLES_29091 | C | C | | | | | C | C | C | C | C | C |

| | | | | | | | | | | | | | | | | | | | | | | | | | |
|---|---|---|---|---|---|---|---|---|---|---|---|---|---|---|---|---|---|---|---|---|---|---|---|---|---|
| NC_011770 | 3195331 | C | T | 73 | 0 | 8 | 0 | NON_SYNONYMOU | MODERAT | MISSENSE | Gcc/Acc | A389T | PLES_29091 | T | T | | | | | T | T | T | T | T | T |
| NC_011770 | 3195436 | T | C | 45 | 0 | 8 | 0 | NON_SYNONYMOU | MODERAT | MISSENSE | Acg/Gcg | T354A | PLES_29091 | C | C | | | | | C | C | C | C | C | C |
| NC_011770 | 3195475 | C | T | 64 | 0 | 8 | 0 | NON_SYNONYMOU | MODERAT | MISSENSE | Gct/Act | A341T | PLES_29091 | T | T | | | | | T | T | T | T | T | T |
| NC_011770 | 3195514 | T | C | 63 | 0 | 8 | 0 | NON_SYNONYMOU | MODERAT | MISSENSE | Agc/Ggc | S328G | PLES_29091 | C | C | | | | | C | C | C | C | C | C |
| NC_011770 | 3195539 | G | C | 48 | 0 | 8 | 0 | NON_SYNONYMOU | MODERAT | MISSENSE | atC/atG | I319M | PLES_29091 | C | C | | | | | C | C | C | C | C | C |
| NC_011770 | 3195540 | A | G | 47 | 0 | 8 | 0 | NON_SYNONYMOU | MODERAT | MISSENSE | aTc/aCc | I319T | PLES_29091 | G | G | | | | | G | G | G | G | G | G |
| NC_011770 | 3195541 | T | C | 47 | 0 | 8 | 0 | NON_SYNONYMOU | MODERAT | MISSENSE | Atc/Gtc | I319V | PLES_29091 | C | C | | | | | C | C | C | C | C | C |
| NC_011770 | 3195553 | T | C | 51 | 0 | 4 | 0 | NON_SYNONYMOU | MODERAT | MISSENSE | Aat/Gat | N315D | PLES_29091 | C | C | | | | | | | C | C | | |
| NC_011770 | 3195560 | T | G | 55 | 0 | 8 | 0 | NON_SYNONYMOU | MODERAT | MISSENSE | gaA/gaC | E312D | PLES_29091 | G | G | | | | | G | G | G | G | G | G |
| NC_011770 | 3195649 | G | C | 48 | 0 | 8 | 0 | NON_SYNONYMOU | MODERAT | MISSENSE | Cag/Gag | Q283E | PLES_29091 | C | C | | | | | C | C | C | C | C | C |
| NC_011770 | 3195651 | C | T | 48 | 0 | 8 | 0 | NON_SYNONYMOU | MODERAT | MISSENSE | cGg/cAg | R282Q | PLES_29091 | T | T | | | | | T | T | T | T | T | T |
| NC_011770 | 3195666 | T | C | 49 | 0 | 8 | 0 | NON_SYNONYMOU | MODERAT | MISSENSE | cAc/cGc | H277R | PLES_29091 | C | C | | | | | C | C | C | C | C | C |
| NC_011770 | 3195669 | C | T | 49 | 0 | 8 | 0 | NON_SYNONYMOU | MODERAT | MISSENSE | aGa/aAa | R276K | PLES_29091 | T | T | | | | | T | T | T | T | T | T |
| NC_011770 | 3195753 | C | G | 56 | 0 | 8 | 0 | NON_SYNONYMOU | MODERAT | MISSENSE | gGc/gCc | G248A | PLES_29091 | G | G | | | | | G | G | G | G | G | G |
| NC_011770 | 3195754 | C | T | 56 | 0 | 8 | 0 | NON_SYNONYMOU | MODERAT | MISSENSE | Ggc/Agc | G248S | PLES_29091 | T | T | | | | | T | T | T | T | T | T |
| NC_011770 | 3195802 | G | C | 62 | 0 | 8 | 0 | NON_SYNONYMOU | MODERAT | MISSENSE | Caa/Gaa | Q232E | PLES_29091 | C | C | | | | | C | C | C | C | C | C |
| NC_011770 | 3195841 | G | C | 56 | 0 | 8 | 0 | NON_SYNONYMOU | MODERAT | MISSENSE | Ccg/Gcg | P219A | PLES_29091 | C | C | | | | | C | C | C | C | C | C |
| NC_011770 | 3195847 | G | C | 54 | 0 | 6 | 0 | NON_SYNONYMOU | MODERAT | MISSENSE | Ccg/Gcg | P217A | PLES_29091 | C | C | | | | | C | C | | | C | C |
| NC_011770 | 3195850 | A | T | 53 | 0 | 6 | 0 | NON_SYNONYMOU | MODERAT | MISSENSE | Tcg/Acg | S216T | PLES_29091 | T | T | | | | | T | T | | | T | T |
| NC_011770 | 3195877 | C | A | 45 | 0 | 8 | 0 | NON_SYNONYMOU | MODERAT | MISSENSE | Gct/Tct | A207S | PLES_29091 | A | A | | | | | A | A | A | A | A | A |
| NC_011770 | 3195880 | C | A | 45 | 0 | 8 | 0 | NON_SYNONYMOU | MODERAT | MISSENSE | Gca/Tca | A206S | PLES_29091 | A | A | | | | | A | A | A | A | A | A |
| NC_011770 | 3195886 | A | C | 52 | 0 | 2 | 0 | NON_SYNONYMOU | MODERAT | MISSENSE | Tcc/Gcc | S204A | PLES_29091 | | | | | | | | | C | C | | |
| NC_011770 | 3195891 | C | T | 53 | 0 | 8 | 0 | NON_SYNONYMOU | MODERAT | MISSENSE | aGt/aAt | S202N | PLES_29091 | T | T | | | | | T | T | T | T | T | T |
| NC_011770 | 3195892 | T | C | 52 | 0 | 8 | 0 | NON_SYNONYMOU | MODERAT | MISSENSE | Agt/Ggt | S202G | PLES_29091 | C | C | | | | | C | C | C | C | C | C |
| NC_011770 | 3195909 | T | C | 59 | 0 | 2 | 0 | NON_SYNONYMOU | MODERAT | MISSENSE | aAc/aGc | N196S | PLES_29091 | | | | | | | | | C | C | | |
| NC_011770 | 3195945 | C | T | 55 | 0 | 8 | 0 | NON_SYNONYMOU | MODERAT | MISSENSE | cGc/cAc | R184H | PLES_29091 | T | T | | | | | T | T | T | T | T | T |
| NC_011770 | 3195946 | G | T | 55 | 0 | 8 | 0 | NON_SYNONYMOU | MODERAT | MISSENSE | Cgc/Agc | R184S | PLES_29091 | T | T | | | | | T | T | T | T | T | T |
| NC_011770 | 3196003 | C | T | 60 | 0 | 8 | 0 | NON_SYNONYMOU | MODERAT | MISSENSE | Gta/Ata | V165I | PLES_29091 | T | T | | | | | T | T | T | T | T | T |
| NC_011770 | 3196059 | G | C | 54 | 0 | 8 | 0 | NON_SYNONYMOU | MODERAT | MISSENSE | gCt/gGt | A146G | PLES_29091 | C | C | | | | | C | C | C | C | C | C |
| NC_011770 | 3196063 | T | C | 53 | 0 | 8 | 0 | NON_SYNONYMOU | MODERAT | MISSENSE | Atc/Gtc | I145V | PLES_29091 | C | C | | | | | C | C | C | C | C | C |
| NC_011770 | 3196065 | C | T | 53 | 0 | 8 | 0 | NON_SYNONYMOU | MODERAT | MISSENSE | cGg/cAg | R144Q | PLES_29091 | T | T | | | | | T | T | T | T | T | T |
| NC_011770 | 3196074 | A | T | 57 | 0 | 2 | 0 | NON_SYNONYMOU | MODERAT | MISSENSE | tTt/tAt | F141Y | PLES_29091 | T | T | | | | | | | | | | |
| NC_011770 | 3196110 | G | A | 54 | 0 | 8 | 0 | NON_SYNONYMOU | MODERAT | MISSENSE | aCc/aTc | T129I | PLES_29091 | A | A | | | | | A | A | A | A | A | A |
| NC_011770 | 3196133 | T | G | 46 | 0 | 8 | 0 | NON_SYNONYMOU | MODERAT | MISSENSE | gaA/gaC | E121D | PLES_29091 | G | G | | | | | G | G | G | G | G | G |
| NC_011770 | 3196146 | A | G | 45 | 0 | 8 | 0 | NON_SYNONYMOU | MODERAT | MISSENSE | cTg/cCg | L117P | PLES_29091 | G | G | | | | | G | G | G | G | G | G |
| NC_011770 | 3196165 | A | C | 42 | 0 | 8 | 0 | NON_SYNONYMOU | MODERAT | MISSENSE | Tcc/Gcc | S111A | PLES_29091 | C | C | | | | | C | C | C | C | C | C |
| NC_011770 | 3196176 | C | T | 42 | 0 | 8 | 0 | NON_SYNONYMOU | MODERAT | MISSENSE | aGc/aAc | S107N | PLES_29091 | T | T | | | | | T | T | T | T | T | T |
| NC_011770 | 3196243 | T | C | 27 | 3 | 5 | 0 | NON_SYNONYMOU | MODERAT | MISSENSE | Atc/Gtc | I85V | PLES_29091 | . | . | | | | | C | C | C | C | C | . |
| NC_011770 | 3196252 | G | A | 26 | 5 | 2 | 0 | NON_SYNONYMOU | MODERAT | MISSENSE | Ctt/Ttt | L82F | PLES_29091 | . | . | | | | | . | | A | A | . | . |
| NC_011770 | 3196317 | T | C | 25 | 6 | 2 | 0 | NON_SYNONYMOU | MODERAT | MISSENSE | aAg/aGg | K60R | PLES_29091 | . | . | | | | | . | . | C | C | . | . |
| NC_011770 | 3196318 | T | G | 25 | 6 | 2 | 0 | NON_SYNONYMOU | MODERAT | MISSENSE | Aag/Cag | K60Q | PLES_29091 | . | . | | | | | . | . | G | G | . | . |
| NC_011770 | 3196340 | G | C | 39 | 0 | 8 | 0 | NON_SYNONYMOU | MODERAT | MISSENSE | gaC/gaG | D52E | PLES_29091 | C | C | | | | | C | C | C | C | C | C |
| NC_011770 | 3196343 | C | A | 38 | 0 | 8 | 0 | NON_SYNONYMOU | MODERAT | MISSENSE | caG/caT | Q51H | PLES_29091 | A | A | | | | | A | A | G | G | A | A |
| NC_011770 | 3196363 | T | G | 41 | 0 | 2 | 0 | NON_SYNONYMOU | MODERAT | MISSENSE | Aag/Cag | K45Q | PLES_29091 | | | | | | | | | G | G | | |
| NC_011770 | 3196384 | T | C | 28 | 6 | 2 | 0 | NON_SYNONYMOU | MODERAT | MISSENSE | Agc/Ggc | S38G | PLES_29091 | . | . | | | | | . | . | C | C | . | . |
| NC_011770 | 3196405 | C | T | 29 | 3 | 5 | 0 | NON_SYNONYMOU | MODERAT | MISSENSE | Gct/Act | A31T | PLES_29091 | T | . | | | | | T | T | . | . | T | T |
| NC_011770 | 3196425 | G | C | 33 | 2 | 6 | 0 | NON_SYNONYMOU | MODERAT | MISSENSE | gCa/gGa | A24G | PLES_29091 | C | C | | | | | C | C | . | . | C | C |
| NC_011770 | 3196444 | A | G | 34 | 2 | 6 | 0 | NON_SYNONYMOU | MODERAT | MISSENSE | Ttc/Ctc | F18L | PLES_29091 | G | G | | | | | G | G | . | . | G | G |
| NC_011770 | 3196447 | G | A | 33 | 2 | 6 | 0 | NON_SYNONYMOU | MODERAT | MISSENSE | Ctc/Ttc | L17F | PLES_29091 | A | A | | | | | A | A | . | . | A | A |
| NC_011770 | 3196460 | T | G | 37 | 2 | 6 | 0 | NON_SYNONYMOU | MODERAT | MISSENSE | ttA/ttC | L12F | PLES_29091 | G | G | | | | | G | G | . | . | G | G |
| NC_011770 | 3196471 | C | T | 36 | 2 | 1 | 0 | NON_SYNONYMOU | MODERAT | MISSENSE | Gcc/Acc | A9T | PLES_29091 | | | | | | | T | | . | . | | |
| NC_011770 | 3196477 | T | G | 37 | 2 | 6 | 0 | NON_SYNONYMOU | MODERAT | MISSENSE | Agc/Cgc | S7R | PLES_29091 | G | G | | | | | G | G | . | . | G | G |
| NC_011770 | 3196486 | G | T | 38 | 2 | 6 | 0 | NON_SYNONYMOU | MODERAT | MISSENSE | Cca/Aca | P4T | PLES_29091 | T | T | | | | | T | T | . | . | T | T |
| NC_011770 | 3196435 | GAGT | G | 34 | 2 | 4 | 0 | CODON_DELETION | MODERATE | | act/ | T20 | PLES_29091 | G | . | | | | | | G | . | . | G | G |
| NC_011770 | 3201098 | T | G | 64 | 0 | 8 | 0 | NON_SYNONYMOU | MODERAT | MISSENSE | caA/caC | Q240H | PLES_29121 | G | G | G | G | G | G | | | G | G | | |
| NC_011770 | 3201324 | A | G | 60 | 0 | 12 | 0 | NON_SYNONYMOU | MODERAT | MISSENSE | gTa/gCa | V165A | PLES_29121 | G | G | G | G | G | G | G | G | G | G | G | G |
| NC_011770 | 3201450 | G | A | 65 | 0 | 2 | 0 | NON_SYNONYMOU | MODERAT | MISSENSE | aCc/aTc | T123I | PLES_29121 | | | | | | | | | | | A | A |
| NC_011770 | 3200643 | A | G | 60 | 0 | 1 | 0 | NON_SYNONYMOU | MODERAT | MISSENSE | tTc/tCc | F392S | PLES_29121 | | | | | | | | G | | | | |
| NC_011770 | 3202910 | G | A | 88 | 0 | 4 | 0 | NON_SYNONYMOU | MODERAT | MISSENSE | gCc/gTc | A28V | PLES_29131 | | | A | A | A | A | | | | | | |
| NC_011770 | 3203135 | C | G | 82 | 0 | 4 | 0 | NON_SYNONYMOU | MODERAT | MISSENSE | gaG/gaC | E318D | PLES_29141 | | | | | | | | | G | G | G | G |
| NC_011770 | 3203177 | G | C | 82 | 0 | 2 | 0 | NON_SYNONYMOU | MODERAT | MISSENSE | gaC/gaG | D304E | PLES_29141 | | | | | | | | | C | C | | |
| NC_011770 | 3203187 | C | T | 83 | 0 | 2 | 0 | NON_SYNONYMOU | MODERAT | MISSENSE | aGc/aAc | S301N | PLES_29141 | | | | | | | | | T | T | | |

| | | | | | | | | | | | | | | | | | | | | | | | | | |
|---|---|---|---|---|---|---|---|---|---|---|---|---|---|---|---|---|---|---|---|---|---|---|---|---|---|
| NC_011770 | 3203195 | G | C | 79 | 0 | 2 | 0 | NON_SYNONYMOU | MODERAT | MISSENSE | gaC/gaG | D298E | PLES_29141 | | | | | | | | | C | C | | |
| NC_011770 | 3203238 | C | CTGG | 73 | 0 | 2 | 0 | CODON_INSERTIO | MODERATE | | agc/aCCAg | S284TS | PLES_29141 | | | | | | | | | CTGG | CTGG | | |
| NC_011770 | 3203241 | C | T | 72 | 0 | 2 | 0 | NON_SYNONYMOU | MODERAT | MISSENSE | aGg/aAg | R283K | PLES_29141 | | | | | | | | | T | T | | |
| NC_011770 | 3203247 | C | G | 75 | 0 | 2 | 0 | NON_SYNONYMOU | MODERAT | MISSENSE | gGc/gCc | G281A | PLES_29141 | | | | | | | | | G | G | | |
| NC_011770 | 3203248 | C | T | 76 | 0 | 2 | 0 | NON_SYNONYMOU | MODERAT | MISSENSE | Ggc/Agc | G281S | PLES_29141 | | | | | | | | | T | T | | |
| NC_011770 | 3203262 | G | T | 76 | 0 | 2 | 0 | NON_SYNONYMOU | MODERAT | MISSENSE | gCt/gAt | A276D | PLES_29141 | | | | | | | | | T | T | | |
| NC_011770 | 3203263 | C | G | 75 | 0 | 2 | 0 | NON_SYNONYMOU | MODERAT | MISSENSE | Gct/Cct | A276P | PLES_29141 | | | | | | | | | G | G | | |
| NC_011770 | 3203264 | G | C | 76 | 0 | 2 | 0 | NON_SYNONYMOU | MODERAT | MISSENSE | gaC/gaG | D275E | PLES_29141 | | | | | | | | | C | C | | |
| NC_011770 | 3203296 | T | G | 83 | 0 | 2 | 0 | NON_SYNONYMOU | MODERAT | MISSENSE | Aag/Cag | K265Q | PLES_29141 | | | | | | | | | G | G | | |
| NC_011770 | 3203299 | A | C | 82 | 0 | 2 | 0 | NON_SYNONYMOU | MODERAT | MISSENSE | Tcc/Gcc | S264A | PLES_29141 | | | | | | | | | C | C | | |
| NC_011770 | 3203320 | C | T | 77 | 0 | 2 | 0 | NON_SYNONYMOU | MODERAT | MISSENSE | Gta/Ata | V257I | PLES_29141 | | | | | | | | | T | T | | |
| NC_011770 | 3203347 | T | C | 76 | 0 | 2 | 0 | NON_SYNONYMOU | MODERAT | MISSENSE | Acg/Gcg | T248A | PLES_29141 | | | | | | | | | C | C | | |
| NC_011770 | 3203543 | G | C | 79 | 0 | 2 | 0 | NON_SYNONYMOU | MODERAT | MISSENSE | aaC/aaG | N182K | PLES_29141 | | | | | | | | | C | C | | |
| NC_011770 | 3203553 | T | C | 81 | 0 | 2 | 0 | NON_SYNONYMOU | MODERAT | MISSENSE | aAg/aGg | K179R | PLES_29141 | | | | | | | | | C | C | | |
| NC_011770 | 3203573 | G | C | 79 | 0 | 2 | 0 | NON_SYNONYMOU | MODERAT | MISSENSE | agC/agG | S172R | PLES_29141 | | | | | | | | | C | C | | |
| NC_011770 | 3203574 | C | G | 77 | 0 | 2 | 0 | NON_SYNONYMOU | MODERAT | MISSENSE | aGc/aCc | S172T | PLES_29141 | | | | | | | | | G | G | | |
| NC_011770 | 3203606 | C | G | 80 | 0 | 2 | 0 | NON_SYNONYMOU | MODERAT | MISSENSE | agG/agC | R161S | PLES_29141 | | | | | | | | | G | G | | |
| NC_011770 | 3203607 | C | T | 79 | 0 | 2 | 0 | NON_SYNONYMOU | MODERAT | MISSENSE | aGg/aAg | R161K | PLES_29141 | | | | | | | | | T | T | | |
| NC_011770 | 3203616 | G | A | 81 | 0 | 2 | 0 | NON_SYNONYMOU | MODERAT | MISSENSE | tCc/tTc | S158F | PLES_29141 | | | | | | | | | A | A | | |
| NC_011770 | 3203617 | A | C | 80 | 0 | 2 | 0 | NON_SYNONYMOU | MODERAT | MISSENSE | Tcc/Gcc | S158A | PLES_29141 | | | | | | | | | C | C | | |
| NC_011770 | 3203634 | T | C | 78 | 0 | 2 | 0 | NON_SYNONYMOU | MODERAT | MISSENSE | aAg/aGg | K152R | PLES_29141 | | | | | | | | | C | C | | |
| NC_011770 | 3203654 | C | G | 77 | 0 | 2 | 0 | NON_SYNONYMOU | MODERAT | MISSENSE | caG/caC | Q145H | PLES_29141 | | | | | | | | | G | G | | |
| NC_011770 | 3203660 | A | C | 78 | 0 | 2 | 0 | NON_SYNONYMOU | MODERAT | MISSENSE | aaT/aaG | N143K | PLES_29141 | | | | | | | | | C | C | | |
| NC_011770 | 3203806 | C | T | 67 | 2 | 2 | 0 | NON_SYNONYMOU | MODERAT | MISSENSE | Gcc/Acc | A95T | PLES_29141 | | | | | | | T | T | . | . | | |
| NC_011770 | 3203851 | A | G | 65 | 2 | 8 | 0 | NON_SYNONYMOU | MODERAT | MISSENSE | Ttc/Ctc | F80L | PLES_29141 | | | G | G | G | G | G | G | . | . | G | G |
| NC_011770 | 3203865 | G | C | 65 | 2 | 4 | 0 | NON_SYNONYMOU | MODERAT | MISSENSE | gCc/gGc | A75G | PLES_29141 | | | C | C | C | C | | | . | . | | |
| NC_011770 | 3203916 | G | T | 70 | 1 | 1 | 0 | NON_SYNONYMOU | MODERAT | MISSENSE | gCg/gAg | A58E | PLES_29141 | | | | | | | | | T | . | | |
| NC_011770 | 3203917 | C | G | 69 | 1 | 1 | 0 | NON_SYNONYMOU | MODERAT | MISSENSE | Gcg/Ccg | A58P | PLES_29141 | | | | | | | | | G | . | | |
| NC_011770 | 3203934 | T | G | 74 | 0 | 2 | 0 | NON_SYNONYMOU | MODERAT | MISSENSE | gAa/gCa | E52A | PLES_29141 | | | | | | | | | G | G | | |
| NC_011770 | 3203946 | G | T | 74 | 0 | 2 | 0 | NON_SYNONYMOU | MODERAT | MISSENSE | gCc/gAc | A48D | PLES_29141 | | | | | | | | | T | T | | |
| NC_011770 | 3203964 | G | C | 74 | 0 | 2 | 0 | NON_SYNONYMOU | MODERAT | MISSENSE | gCc/gGc | A42G | PLES_29141 | | | | | | | | | C | C | | |
| NC_011770 | 3203965 | C | T | 73 | 0 | 2 | 0 | NON_SYNONYMOU | MODERAT | MISSENSE | Gcc/Acc | A42T | PLES_29141 | | | | | | | | | T | T | | |
| NC_011770 | 3203978 | C | A | 75 | 0 | 2 | 0 | NON_SYNONYMOU | MODERAT | MISSENSE | caG/caT | Q37H | PLES_29141 | | | | | | | | | A | A | | |
| NC_011770 | 3203979 | T | C | 76 | 0 | 2 | 0 | NON_SYNONYMOU | MODERAT | MISSENSE | cAg/cGg | Q37R | PLES_29141 | | | | | | | | | C | C | | |
| NC_011770 | 3203988 | C | G | 76 | 0 | 2 | 0 | NON_SYNONYMOU | MODERAT | MISSENSE | gGc/gCc | G34A | PLES_29141 | | | | | | | | | G | G | | |
| NC_011770 | 3204003 | G | C | 81 | 0 | 2 | 0 | NON_SYNONYMOU | MODERAT | MISSENSE | tCc/tGc | S29C | PLES_29141 | | | | | | | | | C | C | | |
| NC_011770 | 3204004 | A | T | 76 | 0 | 2 | 0 | NON_SYNONYMOU | MODERAT | MISSENSE | Tcc/Acc | S29T | PLES_29141 | | | | | | | | | T | T | | |
| NC_011770 | 3204006 | A | C | 75 | 0 | 2 | 0 | NON_SYNONYMOU | MODERAT | MISSENSE | cTc/cGc | L28R | PLES_29141 | | | | | | | | | C | C | | |
| NC_011770 | 3204007 | G | A | 77 | 0 | 2 | 0 | NON_SYNONYMOU | MODERAT | MISSENSE | Ctc/Ttc | L28F | PLES_29141 | | | | | | | | | A | A | | |
| NC_011770 | 3204055 | G | A | 78 | 0 | 2 | 0 | NON_SYNONYMOU | MODERAT | MISSENSE | Ctt/Ttt | L12F | PLES_29141 | | | | | | | | | A | A | | |
| NC_011770 | 3204075 | G | A | 72 | 0 | 2 | 0 | NON_SYNONYMOU | MODERAT | MISSENSE | tCt/tTt | S5F | PLES_29141 | | | | | | | | | A | A | | |
| NC_011770 | 3204076 | A | G | 71 | 0 | 2 | 0 | NON_SYNONYMOU | MODERAT | MISSENSE | Tct/Cct | S5P | PLES_29141 | | | | | | | | | G | G | | |
| NC_011770 | 3204079 | G | A | 73 | 0 | 2 | 0 | NON_SYNONYMOU | MODERAT | MISSENSE | Ccc/Tcc | P4S | PLES_29141 | | | | | | | | | | | A | A |
| NC_011770 | 3204311 | C | G | 69 | 0 | 4 | 0 | NON_SYNONYMOU | MODERAT | MISSENSE | caC/caG | H20Q | PLES_29151 | | | | | | | | | G | G | T | T |
| NC_011770 | 3204373 | A | G | 70 | 0 | 2 | 0 | NON_SYNONYMOU | MODERAT | MISSENSE | cAg/cGg | Q41R | PLES_29151 | | | | | | | | | G | G | | |
| NC_011770 | 3204387 | A | C | 70 | 0 | 2 | 0 | NON_SYNONYMOU | MODERAT | MISSENSE | Aag/Cag | K46Q | PLES_29151 | | | | | | | | | C | C | | |
| NC_011770 | 3204388 | A | G | 71 | 0 | 2 | 0 | NON_SYNONYMOU | MODERAT | MISSENSE | aAg/aGg | K46R | PLES_29151 | | | | | | | | | G | G | | |
| NC_011770 | 3204691 | A | G | 82 | 0 | 2 | 0 | NON_SYNONYMOU | MODERAT | MISSENSE | aAg/aGg | K147R | PLES_29151 | | | | | | | | | G | G | | |
| NC_011770 | 3208881 | C | CG | 62 | 0 | 1 | 0 | FRAME_SHIFT | HIGH | | gag/gGag | E34G? | PLES_29181 | | | | | | | CG | | | | | |
| NC_011770 | 3209218 | C | T | 65 | 0 | 2 | 0 | NON_SYNONYMOU | MODERAT | MISSENSE | Cgg/Tgg | R23W | PLES_29191 | T | T | | | | | | | | | | |
| NC_011770 | 3209630 | A | C | 55 | 0 | 2 | 0 | NON_SYNONYMOU | MODERAT | MISSENSE | gAg/gCg | E160A | PLES_29191 | | | | | | | | | | | C | C |
| NC_011770 | 3210035 | C | T | 65 | 0 | 8 | 0 | NON_SYNONYMOU | MODERAT | MISSENSE | gCc/gTc | A295V | PLES_29191 | | | T | T | T | T | | | T | T | T | T |
| NC_011770 | 3212094 | C | T | 72 | 0 | 2 | 0 | NON_SYNONYMOU | MODERAT | MISSENSE | Ccc/Tcc | P56S | PLES_29221 | | | | | | | | | T | T | | |
| NC_011770 | 3213810 | G | A | 79 | 0 | 4 | 0 | NON_SYNONYMOU | MODERAT | MISSENSE | Gtg/Atg | V305M | PLES_29241 | | | A | A | A | A | | | | | | |
| NC_011770 | 3213811 | T | C | 77 | 0 | 2 | 0 | NON_SYNONYMOU | MODERAT | MISSENSE | gTg/gCg | V305A | PLES_29241 | | | | | | | C | C | | | | |
| NC_011770 | 3213940 | G | A | 71 | 0 | 8 | 0 | NON_SYNONYMOU | MODERAT | MISSENSE | aGg/aAg | R348K | PLES_29241 | | | A | A | A | A | | | A | A | A | A |
| NC_011770 | 3213943 | G | A | 74 | 0 | 2 | 0 | NON_SYNONYMOU | MODERAT | MISSENSE | gGc/gAc | G349D | PLES_29241 | | | | | | | | | A | A | | |
| NC_011770 | 3213951 | A | G | 71 | 0 | 4 | 0 | NON_SYNONYMOU | MODERAT | MISSENSE | Atc/Gtc | I352V | PLES_29241 | | | G | G | G | G | | | | | | |
| NC_011770 | 3213961 | C | T | 71 | 0 | 1 | 0 | NON_SYNONYMOU | MODERAT | MISSENSE | gCc/gTc | A355V | PLES_29241 | | | | | | | T | | | | | |
| NC_011770 | 3214570 | G | A | 60 | 0 | 4 | 0 | NON_SYNONYMOU | MODERAT | MISSENSE | gGc/gAc | G558D | PLES_29241 | | | A | A | A | A | | | | | | |
| NC_011770 | 3214770 | A | G | 55 | 0 | 2 | 0 | NON_SYNONYMOU | MODERAT | MISSENSE | Atc/Gtc | I625V | PLES_29241 | | | | | | | | | | | G | G |
| NC_011770 | 3215061 | A | G | 67 | 0 | 4 | 0 | NON_SYNONYMOU | MODERAT | MISSENSE | Aag/Gag | K722E | PLES_29241 | | | G | G | G | G | | | | | | |

| | | | | | | | | | | | | | | | | | | | | | | | | | |
|---|---|---|---|---|---|---|---|---|---|---|---|---|---|---|---|---|---|---|---|---|---|---|---|---|---|
| NC_011770 | 3216177 | CG | C | 67 | 0 | 2 | 0 | FRAME_SHIFT | HIGH | | cgg/ | R243 | PLES_29251 | | | C | C | | | | | | | | |
| NC_011770 | 3215749 | G | C | 69 | 0 | 4 | 0 | NON_SYNONYMOU | MODERAT | MISSENSE | Caa/Gaa | Q386E | PLES_29251 | | | C | C | C | C | | | | | | |
| NC_011770 | 3215857 | C | T | 63 | 0 | 2 | 0 | NON_SYNONYMOU | MODERAT | MISSENSE | Gcg/Acg | A350T | PLES_29251 | | | | | | | | | T | T | | |
| NC_011770 | 3216400 | C | T | 76 | 0 | 2 | 0 | NON_SYNONYMOU | MODERAT | MISSENSE | Gtc/Atc | V169I | PLES_29251 | | | | | | | T | T | | | | |
| NC_011770 | 3216438 | C | T | 79 | 0 | 6 | 0 | NON_SYNONYMOU | MODERAT | MISSENSE | aGa/aAa | R156K | PLES_29251 | | | | | | | T | T | T | T | T | T |
| NC_011770 | 3216654 | C | T | 71 | 0 | 4 | 0 | NON_SYNONYMOU | MODERAT | MISSENSE | cGg/cAg | R84Q | PLES_29251 | | | | | | | T | T | T | T | | |
| NC_011770 | 3216667 | G | A | 77 | 0 | 12 | 0 | NON_SYNONYMOU | MODERAT | MISSENSE | Cac/Tac | H80Y | PLES_29251 | A | A | A | A | A | A | A | A | A | A | A | A |
| NC_011770 | 3216769 | T | C | 80 | 0 | 4 | 0 | NON_SYNONYMOU | MODERAT | MISSENSE | Agc/Ggc | S46G | PLES_29251 | | | | | | | C | C | C | C | | |
| NC_011770 | 3216889 | C | T | 73 | 0 | 2 | 0 | NON_SYNONYMOU | MODERAT | MISSENSE | Gcg/Acg | A6T | PLES_29251 | | | | | | | T | T | | | | |
| NC_011770 | 3217052 | A | G | 76 | 0 | 2 | 0 | NON_SYNONYMOU | MODERAT | MISSENSE | gTg/gCg | V209A | PLES_29261 | | | | | | | | | G | G | | |
| NC_011770 | 3217494 | C | G | 83 | 0 | 2 | 0 | NON_SYNONYMOU | MODERAT | MISSENSE | Gtc/Ctc | V62L | PLES_29261 | | | | | | | | | G | G | | |
| NC_011770 | 3217502 | A | G | 81 | 0 | 12 | 0 | NON_SYNONYMOU | MODERAT | MISSENSE | gTc/gCc | V59A | PLES_29261 | G | G | G | G | G | G | G | G | G | G | G | G |
| NC_011770 | 3218329 | G | C | 72 | 0 | 2 | 0 | NON_SYNONYMOU | MODERAT | MISSENSE | Gtg/Ctg | V126L | PLES_29271 | | | | | | | C | C | | | | |
| NC_011770 | 3218421 | T | C | 78 | 0 | 4 | 0 | NON_SYNONYMOU | MODERAT | MISSENSE | aAg/aGg | K163R | PLES_29281 | | | | | | | | | C | C | C | C |
| NC_011770 | 3218538 | T | C | 91 | 0 | 4 | 0 | NON_SYNONYMOU | MODERAT | MISSENSE | tAc/tGc | Y124C | PLES_29281 | | | C | C | C | C | | | | | | |
| NC_011770 | 3218608 | A | G | 85 | 0 | 6 | 0 | NON_SYNONYMOU | MODERAT | MISSENSE | Ttc/Ctc | F101L | PLES_29281 | | | G | G | G | G | G | G | | | | |
| NC_011770 | 3218881 | C | T | 83 | 0 | 1 | 0 | NON_SYNONYMOU | MODERAT | MISSENSE | Gtt/Att | V10I | PLES_29281 | | | | | | | | T | | | | |
| NC_011770 | 3220169 | AGACG | A | 16 | 8 | 2 | 0 | FRAME_SHIFT | HIGH | | ggcgtc/ | GV251 | PLES_29291 | . | . | A | . | . | . | A | . | . | . | | |
| NC_011770 | 3220526 | G | A | 36 | 0 | 10 | 0 | STOP_GAINED | HIGH | NONSENSE | Cag/Tag | Q134* | PLES_29291 | A | A | A | A | A | A | A | A | A | A | | |
| NC_011770 | 3220550 | GCT | G | 36 | 0 | 10 | 0 | FRAME_SHIFT | HIGH | | cag/ | Q125 | PLES_29291 | G | G | G | G | G | G | G | G | G | G | | |
| NC_011770 | 3220554 | G | GCC | 16 | 6 | 3 | 0 | FRAME_SHIFT | HIGH | VgrG3 | gac/gaGGc | D124E? | PLES_29291 | . | . | . | GCC | . | GCC | GCC | . | . | . | | |
| NC_011770 | 3218948 | G | T | 84 | 0 | 12 | 0 | NON_SYNONYMOU | MODERAT | MISSENSE | Ctc/Atc | L660I | PLES_29291 | T | T | T | T | T | T | T | T | T | T | T | T |
| NC_011770 | 3219608 | C | T | 78 | 0 | 2 | 0 | NON_SYNONYMOU | MODERAT | MISSENSE | Gcc/Acc | A440T | PLES_29291 | | | | | | | | | T | T | | |
| NC_011770 | 3219803 | T | G | 85 | 0 | 4 | 0 | NON_SYNONYMOU | MODERAT | MISSENSE | Atc/Ctc | I375L | PLES_29291 | | | | | | | G | G | G | G | | |
| NC_011770 | 3219841 | C | T | 80 | 0 | 4 | 0 | NON_SYNONYMOU | MODERAT | MISSENSE | gGc/gAc | G362D | PLES_29291 | | | T | T | T | T | | | | | | |
| NC_011770 | 3220084 | G | T | 84 | 0 | 10 | 0 | NON_SYNONYMOU | MODERAT | MISSENSE | aCg/aAg | T281K | PLES_29291 | T | T | T | T | T | T | T | T | T | T | | |
| NC_011770 | 3220107 | G | C | 74 | 0 | 10 | 0 | NON_SYNONYMOU | MODERAT | MISSENSE | atC/atG | I273M | PLES_29291 | C | C | C | C | C | C | C | C | C | C | | |
| NC_011770 | 3220109 | T | C | 75 | 0 | 10 | 0 | NON_SYNONYMOU | MODERAT | MISSENSE | Atc/Gtc | I273V | PLES_29291 | C | C | C | C | C | C | C | C | C | C | | |
| NC_011770 | 3220125 | G | C | 17 | 5 | 4 | 0 | NON_SYNONYMOU | MODERAT | MISSENSE | agC/agG | S267R | PLES_29291 | . | . | C | C | . | . | C | C | . | . | | |
| NC_011770 | 3220126 | C | G | 17 | 5 | 5 | 0 | NON_SYNONYMOU | MODERAT | MISSENSE | aGc/aCc | S267T | PLES_29291 | . | . | G | G | G | . | G | G | . | . | | |
| NC_011770 | 3220127 | T | C | 17 | 5 | 5 | 0 | NON_SYNONYMOU | MODERAT | MISSENSE | Agc/Ggc | S267G | PLES_29291 | . | . | C | C | C | . | C | C | . | . | | |
| NC_011770 | 3220134 | C | G | 17 | 5 | 5 | 0 | NON_SYNONYMOU | MODERAT | MISSENSE | caG/caC | Q264H | PLES_29291 | . | . | G | G | G | . | G | G | . | . | | |
| NC_011770 | 3220135 | T | C | 17 | 5 | 5 | 0 | NON_SYNONYMOU | MODERAT | MISSENSE | cAg/cGg | Q264R | PLES_29291 | . | . | C | C | C | . | C | C | . | . | | |
| NC_011770 | 3220138 | G | C | 17 | 5 | 4 | 0 | NON_SYNONYMOU | MODERAT | MISSENSE | tCc/tGc | S263C | PLES_29291 | . | . | C | C | . | . | C | C | . | . | | |
| NC_011770 | 3220139 | A | C | 17 | 5 | 6 | 0 | NON_SYNONYMOU | MODERAT | MISSENSE | Tcc/Gcc | S263A | PLES_29291 | . | . | C | C | C | . | C | . | . | . | C | C |
| NC_011770 | 3220166 | G | C | 16 | 8 | 2 | 0 | NON_SYNONYMOU | MODERAT | MISSENSE | Cag/Gag | Q254E | PLES_29291 | . | . | C | . | . | . | C | . | . | . | | |
| NC_011770 | 3220187 | A | C | 15 | 8 | 2 | 0 | NON_SYNONYMOU | MODERAT | MISSENSE | Tcc/Gcc | S247A | PLES_29291 | . | . | C | . | . | . | C | . | . | . | | |
| NC_011770 | 3220189 | A | T | 15 | 8 | 2 | 0 | NON_SYNONYMOU | MODERAT | MISSENSE | gTt/gAt | V246D | PLES_29291 | . | . | T | . | . | . | T | . | . | . | | |
| NC_011770 | 3220196 | C | G | 15 | 8 | 2 | 0 | NON_SYNONYMOU | MODERAT | MISSENSE | Gag/Cag | E244Q | PLES_29291 | . | . | G | . | . | . | G | . | . | . | | |
| NC_011770 | 3220220 | C | T | 16 | 8 | 2 | 0 | NON_SYNONYMOU | MODERAT | MISSENSE | Ggc/Agc | G236S | PLES_29291 | . | . | T | . | . | . | T | . | . | . | | |
| NC_011770 | 3220262 | A | C | 82 | 0 | 10 | 0 | NON_SYNONYMOU | MODERAT | MISSENSE | Tcc/Gcc | S222A | PLES_29291 | C | C | C | C | C | C | C | C | C | C | | |
| NC_011770 | 3220277 | T | A | 82 | 0 | 10 | 0 | NON_SYNONYMOU | MODERAT | MISSENSE | Agc/Tgc | S217C | PLES_29291 | A | A | A | A | A | A | A | A | A | A | | |
| NC_011770 | 3220295 | C | T | 71 | 0 | 10 | 0 | NON_SYNONYMOU | MODERAT | MISSENSE | Gtg/Atg | V211M | PLES_29291 | T | T | T | T | T | T | T | T | T | T | | |
| NC_011770 | 3220299 | C | G | 71 | 0 | 10 | 0 | NON_SYNONYMOU | MODERAT | MISSENSE | caG/caC | Q209H | PLES_29291 | G | G | G | G | G | G | G | G | G | G | | |
| NC_011770 | 3220321 | C | T | 57 | 0 | 10 | 0 | NON_SYNONYMOU | MODERAT | MISSENSE | gGa/gAa | G202E | PLES_29291 | T | T | T | T | T | T | T | T | T | T | | |
| NC_011770 | 3220327 | C | T | 57 | 0 | 10 | 0 | NON_SYNONYMOU | MODERAT | MISSENSE | gGc/gAc | G200D | PLES_29291 | T | T | T | T | T | T | T | T | T | T | | |
| NC_011770 | 3220337 | C | G | 57 | 0 | 10 | 0 | NON_SYNONYMOU | MODERAT | MISSENSE | Gcc/Ccc | A197P | PLES_29291 | G | G | G | G | G | G | G | G | G | G | | |
| NC_011770 | 3220343 | G | T | 56 | 0 | 10 | 0 | NON_SYNONYMOU | MODERAT | MISSENSE | Caa/Aaa | Q195K | PLES_29291 | T | T | T | T | T | T | T | T | C | C | | |
| NC_011770 | 3220408 | G | A | 66 | 0 | 10 | 0 | NON_SYNONYMOU | MODERAT | MISSENSE | gCc/gTc | A173V | PLES_29291 | A | A | A | A | A | A | A | A | A | A | | |
| NC_011770 | 3220411 | G | T | 66 | 0 | 10 | 0 | NON_SYNONYMOU | MODERAT | MISSENSE | gCc/gAc | A172D | PLES_29291 | T | T | T | T | T | T | T | T | T | T | | |
| NC_011770 | 3220417 | G | A | 66 | 0 | 10 | 0 | NON_SYNONYMOU | MODERAT | MISSENSE | aCc/aTc | T170I | PLES_29291 | A | A | A | A | A | A | A | A | A | A | | |
| NC_011770 | 3220437 | G | T | 74 | 0 | 10 | 0 | NON_SYNONYMOU | MODERAT | MISSENSE | gaC/gaA | D163E | PLES_29291 | T | T | T | T | T | T | T | T | T | T | | |
| NC_011770 | 3220469 | G | A | 69 | 0 | 10 | 0 | NON_SYNONYMOU | MODERAT | MISSENSE | Ctc/Ttc | L153F | PLES_29291 | A | A | A | A | A | A | A | A | A | A | | |
| NC_011770 | 3220474 | C | G | 69 | 0 | 10 | 0 | NON_SYNONYMOU | MODERAT | MISSENSE | aGc/aCc | S151T | PLES_29291 | G | G | G | G | G | G | G | G | G | G | | |
| NC_011770 | 3220508 | G | C | 55 | 0 | 10 | 0 | NON_SYNONYMOU | MODERAT | MISSENSE | Cag/Gag | Q140E | PLES_29291 | C | C | C | C | C | C | C | C | C | C | | |
| NC_011770 | 3220519 | C | G | 36 | 0 | 10 | 0 | NON_SYNONYMOU | MODERAT | MISSENSE | aGc/aCc | S136T | PLES_29291 | G | G | G | G | G | G | G | G | G | G | | |
| NC_011770 | 3220525 | T | G | 36 | 0 | 10 | 0 | NON_SYNONYMOU | MODERAT | MISSENSE | cAg/cCg | Q134P | PLES_29291 | G | G | G | G | G | G | G | G | G | G | | |
| NC_011770 | 3220532 | G | T | 37 | 0 | 10 | 0 | NON_SYNONYMOU | MODERAT | MISSENSE | Cag/Aag | Q132K | PLES_29291 | T | T | T | T | T | T | T | T | T | T | | |
| NC_011770 | 3220534 | T | A | 36 | 0 | 10 | 0 | NON_SYNONYMOU | MODERAT | MISSENSE | tAc/tTc | Y131F | PLES_29291 | A | A | A | A | A | A | A | A | A | A | | |
| NC_011770 | 3220540 | C | G | 36 | 0 | 10 | 0 | NON_SYNONYMOU | MODERAT | MISSENSE | aGc/aCc | S129T | PLES_29291 | G | G | G | G | G | G | G | G | G | G | | |
| NC_011770 | 3220562 | C | T | 35 | 0 | 10 | 0 | NON_SYNONYMOU | MODERAT | MISSENSE | Gtc/Atc | V122I | PLES_29291 | T | T | T | T | T | T | T | T | T | T | | |
| NC_011770 | 3220564 | T | G | 35 | 0 | 10 | 0 | NON_SYNONYMOU | MODERAT | MISSENSE | cAg/cCg | Q121P | PLES_29291 | G | G | G | G | G | G | G | G | G | G | | |

| | | | | | | | | | | | | | | | | | | | | | | | | | |
|---|---|---|---|---|---|---|---|---|---|---|---|---|---|---|---|---|---|---|---|---|---|---|---|---|---|
| NC_011770 | 3220565 | G | T | 35 | 0 | 10 | 0 | NON_SYNONYMOU | MODERAT | MISSENSE | Cag/Aag | Q121K | PLES_29291 | T | T | T | T | T | T | T | T | T | T | | |
| NC_011770 | 3223228 | GGC | G | 69 | 0 | 1 | 0 | FRAME_SHIFT | HIGH | | ccgccc/ | PP665 | PLES_29321 | | | | | | | | G | | | | |
| NC_011770 | 3223370 | A | G | 66 | 0 | 12 | 0 | NON_SYNONYMOU | MODERAT | MISSENSE | Tcc/Ccc | S619P | PLES_29321 | G | G | G | G | G | G | G | G | G | G | G | G |
| NC_011770 | 3223744 | G | A | 66 | 0 | 2 | 0 | NON_SYNONYMOU | MODERAT | MISSENSE | aCc/aTc | T494I | PLES_29321 | | | | | | | | | A | A | | |
| NC_011770 | 3223843 | C | G | 74 | 0 | 8 | 0 | NON_SYNONYMOU | MODERAT | MISSENSE | aGc/aCc | S461T | PLES_29321 | | | G | G | G | G | | | G | G | G | G |
| NC_011770 | 3224704 | T | G | 75 | 0 | 4 | 0 | NON_SYNONYMOU | MODERAT | MISSENSE | cAg/cCg | Q174P | PLES_29321 | G | G | | | | | | | G | G | | |
| NC_011770 | 3224753 | C | G | 76 | 0 | 6 | 0 | NON_SYNONYMOU | MODERAT | MISSENSE | Gcc/Ccc | A158P | PLES_29321 | G | G | | | | | G | G | G | G | | |
| NC_011770 | 3225276 | C | T | 78 | 0 | 4 | 0 | NON_SYNONYMOU | MODERAT | MISSENSE | Ggc/Agc | G323S | PLES_29331 | T | T | | | | | | | | | T | T |
| NC_011770 | 3225285 | C | T | 79 | 0 | 4 | 0 | NON_SYNONYMOU | MODERAT | MISSENSE | Gtc/Atc | V320I | PLES_29331 | | | T | T | T | T | | | | | | |
| NC_011770 | 3225324 | C | T | 83 | 0 | 4 | 0 | NON_SYNONYMOU | MODERAT | MISSENSE | Gca/Aca | A307T | PLES_29331 | | | T | T | T | T | | | | | | |
| NC_011770 | 3225456 | C | A | 75 | 0 | 4 | 0 | NON_SYNONYMOU | MODERAT | MISSENSE | Gcc/Tcc | A263S | PLES_29331 | | | A | A | A | A | | | | | | |
| NC_011770 | 3226083 | G | A | 87 | 0 | 4 | 0 | NON_SYNONYMOU | MODERAT | MISSENSE | Cgc/Tgc | R54C | PLES_29331 | | | A | A | A | A | | | | | | |
| NC_011770 | 3226543 | G | C | 78 | 0 | 4 | 0 | NON_SYNONYMOU | MODERAT | MISSENSE | gCc/gGc | A486G | PLES_29341 | | | C | C | C | C | | | | | | |
| NC_011770 | 3226915 | A | G | 84 | 0 | 12 | 0 | NON_SYNONYMOU | MODERAT | MISSENSE | cTg/cCg | L362P | PLES_29341 | G | G | G | G | G | G | G | G | G | G | G | G |
| NC_011770 | 3227321 | G | A | 62 | 0 | 2 | 0 | NON_SYNONYMOU | MODERAT | MISSENSE | Cgc/Tgc | R227C | PLES_29341 | A | A | | | | | | | | | | |
| NC_011770 | 3227369 | T | C | 73 | 0 | 2 | 0 | NON_SYNONYMOU | MODERAT | MISSENSE | Acc/Gcc | T211A | PLES_29341 | C | C | | | | | | | | | | |
| NC_011770 | 3227554 | T | G | 74 | 0 | 2 | 0 | NON_SYNONYMOU | MODERAT | MISSENSE | aAc/aCc | N149T | PLES_29341 | G | G | | | | | | | | | | |
| NC_011770 | 3227680 | C | T | 63 | 0 | 10 | 0 | NON_SYNONYMOU | MODERAT | MISSENSE | aGc/aAc | S107N | PLES_29341 | T | T | T | T | T | T | T | T | T | T | | |
| NC_011770 | 3226670 | C | T | 79 | 0 | 1 | 0 | NON_SYNONYMOU | MODERAT | MISSENSE | Gtc/Atc | V444I | PLES_29341 | | | | | | | | | | | T | |
| NC_011770 | 3228235 | G | T | 73 | 0 | 2 | 0 | NON_SYNONYMOU | MODERAT | MISSENSE | gaC/gaA | D58E | PLES_29351 | T | T | | | | | | | | | | |
| NC_011770 | 3228335 | T | C | 71 | 0 | 12 | 0 | NON_SYNONYMOU | MODERAT | MISSENSE | cAg/cGg | Q25R | PLES_29351 | C | C | C | C | C | C | C | C | C | C | C | C |
| NC_011770 | 3228498 | C | T | 87 | 0 | 1 | 0 | NON_SYNONYMOU | MODERAT | MISSENSE | Gcg/Acg | A141T | PLES_29361 | | | | | | | | T | | | | |
| NC_011770 | 3230326 | C | T | 80 | 0 | 1 | 0 | NON_SYNONYMOU | MODERAT | MISSENSE | Gtc/Atc | V51I | PLES_29371 | | | | | | | T | | | | | |
| NC_011770 | 3231041 | C | T | 80 | 0 | 2 | 0 | NON_SYNONYMOU | MODERAT | MISSENSE | Gcc/Acc | A2T | PLES_29381 | | | | | | | | | | | T | T |
| NC_011770 | 3231286 | T | C | 87 | 0 | 2 | 0 | NON_SYNONYMOU | MODERAT | MISSENSE | Ttc/Ctc | F12L | PLES_29391 | | | | | | | | | | | C | C |
| NC_011770 | 3231295 | C | G | 84 | 0 | 10 | 0 | NON_SYNONYMOU | MODERAT | MISSENSE | Ccg/Gcg | P15A | PLES_29391 | G | G | G | G | G | G | G | G | G | G | | |
| NC_011770 | 3231721 | A | G | 87 | 0 | 6 | 0 | NON_SYNONYMOU | MODERAT | MISSENSE | Acg/Gcg | T157A | PLES_29391 | | | G | G | G | G | G | G | | | | |
| NC_011770 | 3231725 | G | C | 89 | 0 | 6 | 0 | NON_SYNONYMOU | MODERAT | MISSENSE | cGt/cCt | R158P | PLES_29391 | | | C | C | C | C | C | C | | | | |
| NC_011770 | 3231892 | T | C | 84 | 0 | 6 | 0 | NON_SYNONYMOU | MODERAT | MISSENSE | aTc/aCc | I35T | PLES_29401 | | | C | C | C | C | C | C | | | | |
| NC_011770 | 3232353 | A | G | 71 | 0 | 4 | 0 | NON_SYNONYMOU | MODERAT | MISSENSE | Atc/Gtc | I189V | PLES_29401 | | | G | G | G | G | | | | | | |
| NC_011770 | 3232355 | C | G | 72 | 0 | 4 | 0 | NON_SYNONYMOU | MODERAT | MISSENSE | atC/atG | I189M | PLES_29401 | | | G | G | G | G | | | | | | |
| NC_011770 | 3232383 | G | A | 71 | 0 | 2 | 0 | NON_SYNONYMOU | MODERAT | MISSENSE | Gtg/Atg | V199M | PLES_29401 | | | | | | | | | A | A | | |
| NC_011770 | 3232482 | A | G | 72 | 0 | 4 | 0 | NON_SYNONYMOU | MODERAT | MISSENSE | Atc/Gtc | I232V | PLES_29401 | | | G | G | G | G | | | | | | |
| NC_011770 | 3232548 | G | A | 69 | 0 | 2 | 0 | NON_SYNONYMOU | MODERAT | MISSENSE | Gcg/Acg | A254T | PLES_29401 | | | | | | | | | | | A | A |
| NC_011770 | 3232753 | A | G | 64 | 0 | 2 | 0 | NON_SYNONYMOU | MODERAT | MISSENSE | gAa/gGa | E322G | PLES_29401 | | | | | | | | | | | G | G |
| NC_011770 | 3232926 | C | G | 68 | 0 | 4 | 0 | NON_SYNONYMOU | MODERAT | MISSENSE | Ccg/Gcg | P380A | PLES_29401 | | | | | | | G | G | | | G | G |
| NC_011770 | 3233005 | C | T | 65 | 0 | 2 | 0 | NON_SYNONYMOU | MODERAT | MISSENSE | gCc/gTc | A406V | PLES_29401 | | | | | | | | | | | T | T |
| NC_011770 | 3233181 | A | C | 80 | 0 | 4 | 0 | NON_SYNONYMOU | MODERAT | MISSENSE | gAg/gCg | E15A | PLES_29411 | | | | | | | C | C | | | C | C |
| NC_011770 | 3233270 | C | T | 80 | 0 | 4 | 0 | NON_SYNONYMOU | MODERAT | MISSENSE | Cgg/Tgg | R45W | PLES_29411 | | | T | T | T | T | | | | | | |
| NC_011770 | 3233452 | C | A | 80 | 0 | 2 | 0 | NON_SYNONYMOU | MODERAT | MISSENSE | gaC/gaA | D105E | PLES_29411 | | | | | | | A | A | | | | |
| NC_011770 | 3233879 | G | C | 81 | 0 | 4 | 0 | NON_SYNONYMOU | MODERAT | MISSENSE | Gcg/Ccg | A248P | PLES_29411 | | | C | C | C | C | | | | | | |
| NC_011770 | 3234112 | G | T | 75 | 0 | 10 | 0 | NON_SYNONYMOU | MODERAT | MISSENSE | Gcc/Tcc | A74S | PLES_29421 | | | T | T | T | T | T | T | T | T | T | T |
| NC_011770 | 3234383 | A | T | 70 | 0 | 8 | 0 | NON_SYNONYMOU | MODERAT | MISSENSE | tAc/tTc | Y164F | PLES_29421 | | | T | T | T | T | T | T | T | T | | |
| NC_011770 | 3234436 | A | G | 78 | 0 | 8 | 0 | NON_SYNONYMOU | MODERAT | MISSENSE | Act/Gct | T182A | PLES_29421 | | | G | G | G | G | G | G | G | G | | |
| NC_011770 | 3234442 | A | T | 79 | 0 | 8 | 0 | NON_SYNONYMOU | MODERAT | MISSENSE | Atg/Ttg | M184L | PLES_29421 | | | T | T | T | T | T | T | T | T | | |
| NC_011770 | 3234798 | A | T | 75 | 0 | 2 | 0 | NON_SYNONYMOU | MODERAT | MISSENSE | gaA/gaT | E302D | PLES_29421 | | | | | | | T | T | | | | |
| NC_011770 | 3235499 | C | G | 77 | 0 | 2 | 0 | NON_SYNONYMOU | MODERAT | MISSENSE | gCc/gGc | A536G | PLES_29421 | | | | | | | | | G | G | | |
| NC_011770 | 3235879 | A | G | 75 | 0 | 2 | 0 | NON_SYNONYMOU | MODERAT | MISSENSE | Aac/Gac | N663D | PLES_29421 | | | | | | | | | G | G | | |
| NC_011770 | 3236455 | G | C | 83 | 0 | 2 | 0 | NON_SYNONYMOU | MODERAT | MISSENSE | Gcg/Ccg | A855P | PLES_29421 | C | C | | | | | | | | | | |
| NC_011770 | 3237025 | C | A | 83 | 0 | 6 | 0 | NON_SYNONYMOU | MODERAT | MISSENSE | Caa/Aaa | Q1045K | PLES_29421 | A | A | | | | | | | A | A | A | A |
| NC_011770 | 3237052 | G | A | 81 | 0 | 2 | 0 | NON_SYNONYMOU | MODERAT | MISSENSE | Gtg/Atg | V1054M | PLES_29421 | | | | | | | | | | | A | A |
| NC_011770 | 3237686 | C | T | 67 | 0 | 2 | 0 | NON_SYNONYMOU | MODERAT | MISSENSE | gCc/gTc | A1265V | PLES_29421 | | | | | | | T | T | | | | |
| NC_011770 | 3234287 | G | A | 72 | 0 | 1 | 0 | NON_SYNONYMOU | MODERAT | MISSENSE | aGg/aAg | R132K | PLES_29421 | | | | | | | | | | | A | |
| NC_011770 | 3235750 | G | A | 75 | 0 | 1 | 0 | NON_SYNONYMOU | MODERAT | MISSENSE | Gaa/Aaa | E620K | PLES_29421 | | | | | | | | | | | A | |
| NC_011770 | 3234665 | G | A | 67 | 0 | 1 | 0 | NON_SYNONYMOU | MODERAT | MISSENSE | cGc/cAc | R258H | PLES_29421 | | | | | | | | A | | | | |
| NC_011770 | 3234718 | G | A | 74 | 0 | 1 | 0 | NON_SYNONYMOU | MODERAT | MISSENSE | Ggc/Agc | G276S | PLES_29421 | | | | | | | | A | | | | |
| NC_011770 | 3238805 | A | G | 47 | 0 | 6 | 0 | SPLICE_SITE_REGIC | LOW | SILENT | taA/taG | *367 | PLES_29431 | G | G | G | G | G | G | | | | | | |
| NC_011770 | 3237735 | T | C | 72 | 0 | 2 | 0 | NON_SYNONYMOU | MODERAT | MISSENSE | Tcc/Ccc | S11P | PLES_29431 | C | C | | | | | | | | | | |
| NC_011770 | 3237774 | C | G | 68 | 0 | 4 | 0 | NON_SYNONYMOU | MODERAT | MISSENSE | Ccg/Gcg | P24A | PLES_29431 | | | G | G | G | G | | | | | | |
| NC_011770 | 3238369 | T | A | 54 | 0 | 6 | 0 | NON_SYNONYMOU | MODERAT | MISSENSE | cTg/cAg | L222Q | PLES_29431 | A | A | | | | | A | A | | | A | A |
| NC_011770 | 3238500 | T | C | 44 | 0 | 6 | 0 | NON_SYNONYMOU | MODERAT | MISSENSE | Tcc/Ccc | S266P | PLES_29431 | C | C | C | C | C | C | | | | | | |
| NC_011770 | 3238536 | A | G | 45 | 0 | 4 | 0 | NON_SYNONYMOU | MODERAT | MISSENSE | Agc/Ggc | S278G | PLES_29431 | G | G | | | | | | | | | G | G |

| | | | | | | | | | | | | | | | | | | | | | | | | | |
|---|---|---|---|---|---|---|---|---|---|---|---|---|---|---|---|---|---|---|---|---|---|---|---|---|---|
| NC_011770 | 3238537 | G | C | 45 | 0 | 4 | 0 | NON_SYNONYMOU | MODERAT | MISSENSE | aGc/aCc | S278T | PLES_29431 | C | C | | | | | | | | | C | C |
| NC_011770 | 3238548 | G | A | 45 | 0 | 6 | 0 | NON_SYNONYMOU | MODERAT | MISSENSE | Gcg/Acg | A282T | PLES_29431 | | | A | A | A | A | | | A | A | | |
| NC_011770 | 3238566 | G | C | 50 | 0 | 2 | 0 | NON_SYNONYMOU | MODERAT | MISSENSE | Gtc/Ctc | V288L | PLES_29431 | | | | | | | | | C | C | | |
| NC_011770 | 3238600 | G | C | 51 | 0 | 12 | 0 | NON_SYNONYMOU | MODERAT | MISSENSE | gGg/gCg | G299A | PLES_29431 | C | C | C | C | C | C | C | C | C | C | C | C |
| NC_011770 | 3239735 | G | A | 66 | 0 | 6 | 0 | NON_SYNONYMOU | MODERAT | MISSENSE | aGc/aAc | S274N | PLES_29441 | | | | | | | A | A | A | A | A | A |
| NC_011770 | 3240126 | C | T | 68 | 0 | 2 | 0 | NON_SYNONYMOU | MODERAT | MISSENSE | gCt/gTt | A5V | PLES_29451 | | | | | | | | | T | T | | |
| NC_011770 | 3240146 | C | T | 66 | 0 | 2 | 0 | NON_SYNONYMOU | MODERAT | MISSENSE | Ccg/Tcg | P12S | PLES_29451 | | | | | | | T | T | | | | |
| NC_011770 | 3240321 | G | A | 55 | 0 | 2 | 0 | NON_SYNONYMOU | MODERAT | MISSENSE | cGg/cAg | R70Q | PLES_29451 | | | | | | | | | | | A | A |
| NC_011770 | 3240449 | T | C | 63 | 0 | 6 | 0 | NON_SYNONYMOU | MODERAT | MISSENSE | Tat/Cat | Y113H | PLES_29451 | | | | | | | C | C | C | C | C | C |
| NC_011770 | 3242752 | C | T | 63 | 0 | 4 | 0 | NON_SYNONYMOU | MODERAT | MISSENSE | Cgc/Tgc | R116C | PLES_29481 | | | T | T | T | T | | | | | | |
| NC_011770 | 3243278 | C | G | 78 | 0 | 4 | 0 | NON_SYNONYMOU | MODERAT | MISSENSE | gCg/gGg | A291G | PLES_29481 | | | G | G | G | G | | | | | | |
| NC_011770 | 3243517 | C | G | 72 | 0 | 2 | 0 | NON_SYNONYMOU | MODERAT | MISSENSE | Ctg/Gtg | L371V | PLES_29481 | | | | | | | | | | | G | G |
| NC_011770 | 3243529 | C | A | 64 | 0 | 12 | 0 | NON_SYNONYMOU | MODERAT | MISSENSE | Ctc/Atc | L375I | PLES_29481 | A | A | A | A | A | A | A | A | A | A | A | A |
| NC_011770 | 3243604 | G | A | 62 | 0 | 2 | 0 | NON_SYNONYMOU | MODERAT | MISSENSE | cGt/cAt | R6H | PLES_29491 | | | | | | | | | A | A | | |
| NC_011770 | 3243708 | G | A | 67 | 0 | 2 | 0 | NON_SYNONYMOU | MODERAT | MISSENSE | Gaa/Aaa | E41K | PLES_29491 | A | A | | | | | | | | | | |
| NC_011770 | 3243820 | A | C | 71 | 0 | 2 | 0 | NON_SYNONYMOU | MODERAT | MISSENSE | aAc/aCc | N78T | PLES_29491 | | | | | | | | | C | C | | |
| NC_011770 | 3243895 | A | G | 74 | 0 | 2 | 0 | NON_SYNONYMOU | MODERAT | MISSENSE | aAg/aGg | K103R | PLES_29491 | | | | | | | G | G | | | | |
| NC_011770 | 3244065 | C | A | 65 | 0 | 2 | 0 | NON_SYNONYMOU | MODERAT | MISSENSE | Ccc/Acc | P160T | PLES_29491 | A | A | | | | | | | | | | |
| NC_011770 | 3244456 | G | A | 66 | 0 | 2 | 0 | NON_SYNONYMOU | MODERAT | MISSENSE | aGc/aAc | S290N | PLES_29491 | | | | | | | | | A | A | | |
| NC_011770 | 3245353 | T | C | 75 | 0 | 12 | 0 | NON_SYNONYMOU | MODERAT | MISSENSE | gTc/gCc | V155A | PLES_29501 | C | C | C | C | C | C | C | C | C | C | C | C |
| NC_011770 | 3245578 | C | T | 72 | 0 | 2 | 0 | NON_SYNONYMOU | MODERAT | MISSENSE | gCg/gTg | A230V | PLES_29501 | | | | | | | | | | | T | T |
| NC_011770 | 3245581 | C | T | 72 | 0 | 2 | 0 | NON_SYNONYMOU | MODERAT | MISSENSE | gCa/gTa | A231V | PLES_29501 | | | | | | | | | | | T | T |
| NC_011770 | 3245586 | A | G | 73 | 0 | 2 | 0 | NON_SYNONYMOU | MODERAT | MISSENSE | Acg/Gcg | T233A | PLES_29501 | | | | | | | | | | | G | G |
| NC_011770 | 3245626 | A | G | 71 | 0 | 2 | 0 | NON_SYNONYMOU | MODERAT | MISSENSE | cAc/cGc | H246R | PLES_29501 | | | | | | | | | | | G | G |
| NC_011770 | 3245677 | C | T | 74 | 0 | 2 | 0 | NON_SYNONYMOU | MODERAT | MISSENSE | gCg/gTg | A263V | PLES_29501 | | | | | | | | | | | T | T |
| NC_011770 | 3246295 | T | G | 73 | 0 | 12 | 0 | NON_SYNONYMOU | MODERAT | MISSENSE | Agc/Cgc | S315R | PLES_29511 | G | G | G | G | G | G | G | G | G | G | G | G |
| NC_011770 | 3246579 | C | T | 70 | 0 | 2 | 0 | NON_SYNONYMOU | MODERAT | MISSENSE | aGc/aAc | S220N | PLES_29511 | T | T | | | | | | | | | | |
| NC_011770 | 3246912 | T | A | 71 | 0 | 2 | 0 | NON_SYNONYMOU | MODERAT | MISSENSE | aAg/aTg | K109M | PLES_29511 | | | A | A | | | | | | | | |
| NC_011770 | 3246921 | T | C | 68 | 0 | 2 | 0 | NON_SYNONYMOU | MODERAT | MISSENSE | gAg/gGg | E106G | PLES_29511 | | | | | | | C | C | | | | |
| NC_011770 | 3247195 | G | A | 76 | 0 | 4 | 0 | NON_SYNONYMOU | MODERAT | MISSENSE | Ctc/Ttc | L15F | PLES_29511 | A | A | | | | | | | A | A | | |
| NC_011770 | 3248305 | G | C | 69 | 0 | 2 | 0 | NON_SYNONYMOU | MODERAT | MISSENSE | gCc/gGc | A286G | PLES_29531 | C | C | | | | | | | | | | |
| NC_011770 | 3248792 | A | G | 75 | 0 | 6 | 0 | NON_SYNONYMOU | MODERAT | MISSENSE | Tgg/Cgg | W124R | PLES_29531 | G | G | | | | | G | G | G | G | | |
| NC_011770 | 3248893 | T | C | 78 | 0 | 2 | 0 | NON_SYNONYMOU | MODERAT | MISSENSE | gAc/gGc | D90G | PLES_29531 | | | | | | | | | C | C | | |
| NC_011770 | 3248989 | A | C | 80 | 0 | 2 | 0 | NON_SYNONYMOU | MODERAT | MISSENSE | tTc/tGc | F58C | PLES_29531 | | | | | | | | | C | C | | |
| NC_011770 | 3249066 | G | C | 79 | 0 | 2 | 0 | NON_SYNONYMOU | MODERAT | MISSENSE | caC/caG | H32Q | PLES_29531 | | | | | | | | | | | C | C |
| NC_011770 | 3249155 | C | T | 81 | 0 | 2 | 0 | NON_SYNONYMOU | MODERAT | MISSENSE | Gcc/Acc | A3T | PLES_29531 | | | | | | | | | T | T | | |
| NC_011770 | 3249488 | C | G | 66 | 0 | 2 | 0 | NON_SYNONYMOU | MODERAT | MISSENSE | aaG/aaC | K155N | PLES_29541 | | | | | | | | | | | G | G |
| NC_011770 | 3249675 | T | C | 58 | 0 | 2 | 0 | NON_SYNONYMOU | MODERAT | MISSENSE | cAg/cGg | Q93R | PLES_29541 | C | C | | | | | | | | | | |
| NC_011770 | 3249985 | T | C | 53 | 0 | 12 | 0 | SPLICE_SITE_REGIC | LOW | SILENT | tAa/tGa | *463 | PLES_29551 | C | C | C | C | C | C | C | C | C | C | C | C |
| NC_011770 | 3250397 | G | C | 69 | 0 | 4 | 0 | NON_SYNONYMOU | MODERAT | MISSENSE | Ctc/Gtc | L326V | PLES_29551 | | | C | C | C | C | | | | | | |
| NC_011770 | 3250940 | T | G | 75 | 0 | 2 | 0 | NON_SYNONYMOU | MODERAT | MISSENSE | Aag/Cag | K145Q | PLES_29551 | | | | | | | G | G | | | | |
| NC_011770 | 3251595 | G | A | 70 | 0 | 2 | 0 | NON_SYNONYMOU | MODERAT | MISSENSE | gCg/gTg | A338V | PLES_29561 | | | | | | | A | A | | | | |
| NC_011770 | 3251754 | G | T | 69 | 0 | 6 | 0 | NON_SYNONYMOU | MODERAT | MISSENSE | gCc/gAc | A285D | PLES_29561 | T | T | | | | | T | T | | | T | T |
| NC_011770 | 3252160 | G | A | 61 | 0 | 4 | 0 | NON_SYNONYMOU | MODERAT | MISSENSE | Cgc/Tgc | R150C | PLES_29561 | | | A | A | A | A | | | | | | |
| NC_011770 | 3253817 | G | A | 46 | 0 | 1 | 0 | STOP_GAINED | HIGH | NONSENSE | Cag/Tag | Q13* | PLES_29571 | | | | | | | | A | | | | |
| NC_011770 | 3252664 | T | C | 69 | 0 | 4 | 0 | NON_SYNONYMOU | MODERAT | MISSENSE | gAc/gGc | D397G | PLES_29571 | | | C | C | C | C | | | | | | |
| NC_011770 | 3253832 | T | C | 45 | 0 | 2 | 0 | NON_SYNONYMOU | MODERAT | MISSENSE | Aag/Gag | K8E | PLES_29571 | | | | | | | C | C | | | | |
| NC_011770 | 3255023 | C | G | 81 | 0 | 4 | 0 | NON_SYNONYMOU | MODERAT | MISSENSE | aCc/aGc | T251S | PLES_29581 | | | G | G | G | G | | | | | | |
| NC_011770 | 3259611 | A | AG | 58 | 0 | 1 | 0 | FRAME_SHIFT | HIGH | | ctg/cCtg | L343P? | PLES_29621 | | . | | | | | | AG | | | | |
| NC_011770 | 3259590 | G | T | 57 | 0 | 6 | 0 | NON_SYNONYMOU | MODERAT | MISSENSE | gCg/gAg | A350E | PLES_29621 | | | | | | | T | T | T | T | T | T |
| NC_011770 | 3260316 | C | T | 66 | 0 | 4 | 0 | NON_SYNONYMOU | MODERAT | MISSENSE | cGc/cAc | R108H | PLES_29621 | | | T | T | T | T | | | | | | |
| NC_011770 | 3260458 | C | T | 76 | 0 | 2 | 0 | NON_SYNONYMOU | MODERAT | MISSENSE | Gcc/Acc | A61T | PLES_29621 | T | T | | | | | | | | | | |
| NC_011770 | 3260344 | C | T | 68 | 0 | 1 | 0 | NON_SYNONYMOU | MODERAT | MISSENSE | Gcc/Acc | A99T | PLES_29621 | | T | | | | | | | | | | |
| NC_011770 | 3261192 | T | C | 78 | 0 | 2 | 0 | NON_SYNONYMOU | MODERAT | MISSENSE | aAg/aGg | K107R | PLES_29631 | | | | | | | C | C | | | | |
| NC_011770 | 3261241 | C | T | 71 | 0 | 1 | 0 | NON_SYNONYMOU | MODERAT | MISSENSE | Gcg/Acg | A91T | PLES_29631 | | | | | | | | | | | | T |
| NC_011770 | 3262097 | C | T | 46 | 0 | 4 | 0 | NON_SYNONYMOU | MODERAT | MISSENSE | Ggc/Agc | G120S | PLES_29641 | | | | | | | T | T | | | T | T |
| NC_011770 | 3262259 | C | T | 38 | 0 | 2 | 0 | NON_SYNONYMOU | MODERAT | MISSENSE | Gcc/Acc | A66T | PLES_29641 | | | | | | | T | T | | | | |
| NC_011770 | 3263681 | A | C | 71 | 0 | 8 | 0 | NON_SYNONYMOU | MODERAT | MISSENSE | Tcg/Gcg | S54A | PLES_29651 | C | C | C | C | C | C | | | | | C | C |
| NC_011770 | 3265062 | C | T | 37 | 0 | 4 | 0 | NON_SYNONYMOU | MODERAT | MISSENSE | cGg/cAg | R445Q | PLES_29671 | | | | | | | T | T | T | T | | |
| NC_011770 | 3265071 | A | T | 39 | 0 | 4 | 0 | NON_SYNONYMOU | MODERAT | MISSENSE | gTc/gAc | V442D | PLES_29671 | | | | | | | T | T | T | T | | |
| NC_011770 | 3266176 | T | C | 74 | 0 | 2 | 0 | NON_SYNONYMOU | MODERAT | MISSENSE | Atc/Gtc | I74V | PLES_29671 | | | | | | | | | | | C | C |
| NC_011770 | 3266298 | T | G | 75 | 0 | 2 | 0 | NON_SYNONYMOU | MODERAT | MISSENSE | gAc/gCc | D33A | PLES_29671 | G | G | | | | | | | | | | |

| | | | | | | | | | | | | | | | | | | | | | | | | | |
|---|---|---|---|---|---|---|---|---|---|---|---|---|---|---|---|---|---|---|---|---|---|---|---|---|---|
| NC_011770 | 3266638 | T | A | 88 | 0 | 2 | 0 | NON_SYNONYMOU | MODERAT | MISSENSE | gAc/gTc | D717V | PLES_29681 | | | A | A | | | | | | | | |
| NC_011770 | 3266726 | C | T | 91 | 0 | 2 | 0 | NON_SYNONYMOU | MODERAT | MISSENSE | Ggc/Agc | G688S | PLES_29681 | T | T | | | | | | | | | | |
| NC_011770 | 3266732 | A | G | 91 | 0 | 10 | 0 | NON_SYNONYMOU | MODERAT | MISSENSE | Ttc/Ctc | F686L | PLES_29681 | | | G | G | G | G | G | G | G | G | G | G |
| NC_011770 | 3266801 | C | T | 90 | 0 | 10 | 0 | NON_SYNONYMOU | MODERAT | MISSENSE | Gcc/Acc | A663T | PLES_29681 | | | T | T | T | T | T | T | T | T | T | T |
| NC_011770 | 3268201 | A | G | 73 | 0 | 2 | 0 | NON_SYNONYMOU | MODERAT | MISSENSE | gTt/gCt | V196A | PLES_29681 | G | G | | | | | | | | | | |
| NC_011770 | 3270023 | C | G | 88 | 0 | 10 | 0 | NON_SYNONYMOU | MODERAT | MISSENSE | Cgc/Ggc | R27G | PLES_29701 | G | G | G | G | G | G | G | G | G | G | | |
| NC_011770 | 3270293 | T | C | 71 | 0 | 8 | 0 | NON_SYNONYMOU | MODERAT | MISSENSE | Ttc/Ctc | F117L | PLES_29701 | | | C | C | C | C | C | C | C | C | | |
| NC_011770 | 3270410 | A | C | 73 | 0 | 2 | 0 | NON_SYNONYMOU | MODERAT | MISSENSE | Agc/Cgc | S156R | PLES_29701 | | | | | | | C | C | | | | |
| NC_011770 | 3270482 | T | G | 83 | 0 | 4 | 0 | NON_SYNONYMOU | MODERAT | MISSENSE | Tcc/Gcc | S180A | PLES_29701 | G | G | | | | | G | G | | | | |
| NC_011770 | 3270491 | A | G | 83 | 0 | 8 | 0 | NON_SYNONYMOU | MODERAT | MISSENSE | Agc/Ggc | S183G | PLES_29701 | G | G | G | G | G | G | G | G | | | | |
| NC_011770 | 3272285 | GC | G | 89 | 0 | 2 | 0 | FRAME_SHIFT | HIGH | | ggc/ | G61 | PLES_29711 | G | G | | | | | | | | | | |
| NC_011770 | 3272621 | T | G | 72 | 0 | 8 | 0 | NON_SYNONYMOU | MODERAT | MISSENSE | cTt/cGt | L17R | PLES_29721 | G | G | G | G | G | G | G | G | | | | |
| NC_011770 | 3273562 | G | GC | 62 | 0 | 1 | 0 | FRAME_SHIFT | HIGH | | cac/cCac | H128P? | PLES_29731 | | | | | | | | GC | | | | |
| NC_011770 | 3273296 | G | C | 65 | 0 | 2 | 0 | NON_SYNONYMOU | MODERAT | MISSENSE | Gtg/Ctg | V39L | PLES_29731 | | | | | | | | | | | C | C |
| NC_011770 | 3273343 | C | G | 62 | 0 | 2 | 0 | NON_SYNONYMOU | MODERAT | MISSENSE | gaC/gaG | D54E | PLES_29731 | | | | | | | | | | | G | G |
| NC_011770 | 3274088 | A | G | 58 | 0 | 12 | 0 | NON_SYNONYMOU | MODERAT | MISSENSE | Atg/Gtg | M303V | PLES_29731 | G | G | G | G | G | G | G | G | G | G | G | G |
| NC_011770 | 3274178 | G | A | 60 | 0 | 1 | 0 | NON_SYNONYMOU | MODERAT | MISSENSE | Gtc/Atc | V333I | PLES_29731 | | | | | | | | | | | | A |
| NC_011770 | 3274591 | G | C | 55 | 0 | 12 | 0 | NON_SYNONYMOU | MODERAT | MISSENSE | Ggc/Cgc | G116R | PLES_29741 | C | C | C | C | C | C | C | C | C | C | C | C |
| NC_011770 | 3275853 | A | G | 89 | 0 | 4 | 0 | NON_SYNONYMOU | MODERAT | MISSENSE | gAg/gGg | E269G | PLES_29751 | | | | | | | G | G | G | G | | |
| NC_011770 | 3276221 | C | G | 84 | 0 | 2 | 0 | NON_SYNONYMOU | MODERAT | MISSENSE | Cag/Gag | Q392E | PLES_29751 | | | | | | | | | G | G | | |
| NC_011770 | 3276539 | C | T | 75 | 0 | 2 | 0 | NON_SYNONYMOU | MODERAT | MISSENSE | gCg/gTg | A99V | PLES_29761 | | | | | | | | | | | T | T |
| NC_011770 | 3277166 | T | TC | 6 | 5 | 4 | 0 | FRAME_SHIFT | HIGH | | tta/tCta | L15S? | PLES_29771 | . | . | TC | TC | TC | TC | . | . | . | | | |
| NC_011770 | 3277167 | T | A | 2 | 9 | 3 | 0 | STOP_GAINED | HIGH | NONSENSE | tTa/tAa | L15* | PLES_29771 | . | . | . | . | . | . | . | . | . | A | A | A |
| NC_011770 | 3277169 | C | T | 1 | 9 | 3 | 0 | STOP_GAINED | HIGH | NONSENSE | Cga/Tga | R16* | PLES_29771 | . | . | . | . | . | . | . | . | . | T | T | T |
| NC_011770 | 3277174 | A | AT | 7 | 5 | 6 | 0 | FRAME_SHIFT | HIGH | | gct/Tgct | A18C? | PLES_29771 | . | . | AT | AT | AT | AT | . | . | . | AT | . | AT |
| NC_011770 | 3277215 | C | G | 14 | 2 | 6 | 0 | NON_SYNONYMOU | MODERAT | MISSENSE | gCt/gGt | A31G | PLES_29771 | G | G | . | . | . | . | . | . | G | G | G | G |
| NC_011770 | 3279405 | G | C | 87 | 0 | 2 | 0 | NON_SYNONYMOU | MODERAT | MISSENSE | gCc/gGc | A317G | PLES_29791 | | | | | | | | | C | C | | |
| NC_011770 | 3280251 | C | T | 61 | 0 | 2 | 0 | NON_SYNONYMOU | MODERAT | MISSENSE | cGc/cAc | R35H | PLES_29791 | | | | | | | | | | | T | T |
| NC_011770 | 3280282 | C | T | 61 | 0 | 6 | 0 | NON_SYNONYMOU | MODERAT | MISSENSE | Gcc/Acc | A25T | PLES_29791 | | | T | T | T | T | T | T | | | | |
| NC_011770 | 3280417 | T | C | 51 | 0 | 2 | 0 | NON_SYNONYMOU | MODERAT | MISSENSE | Atc/Gtc | I410V | PLES_29801 | C | C | | | | | | | | | | |
| NC_011770 | 3280651 | T | C | 59 | 0 | 2 | 0 | NON_SYNONYMOU | MODERAT | MISSENSE | Atc/Gtc | I332V | PLES_29801 | C | C | | | | | | | | | | |
| NC_011770 | 3280678 | C | T | 58 | 0 | 2 | 0 | NON_SYNONYMOU | MODERAT | MISSENSE | Gac/Aac | D323N | PLES_29801 | T | T | | | | | | | | | | |
| NC_011770 | 3282386 | C | T | 60 | 0 | 2 | 0 | NON_SYNONYMOU | MODERAT | MISSENSE | Gtc/Atc | V368I | PLES_29811 | T | T | | | | | | | | | | |
| NC_011770 | 3282983 | T | C | 58 | 0 | 1 | 0 | NON_SYNONYMOU | MODERAT | MISSENSE | Acc/Gcc | T169A | PLES_29811 | | | | | | | C | | | | | |
| NC_011770 | 3283012 | T | C | 57 | 0 | 10 | 0 | NON_SYNONYMOU | MODERAT | MISSENSE | cAc/cGc | H159R | PLES_29811 | C | C | C | C | C | C | C | C | C | C | | |
| NC_011770 | 3283370 | G | A | 51 | 0 | 2 | 0 | NON_SYNONYMOU | MODERAT | MISSENSE | Cat/Tat | H40Y | PLES_29811 | | | | | | | A | A | | | | |
| NC_011770 | 3284328 | A | G | 76 | 0 | 1 | 0 | NON_SYNONYMOU | MODERAT | MISSENSE | cTg/cCg | L282P | PLES_29831 | | G | | | | | | | | | | |
| NC_011770 | 3285304 | C | T | 67 | 0 | 2 | 0 | NON_SYNONYMOU | MODERAT | MISSENSE | Gcc/Acc | A163T | PLES_29841 | | | | | | | T | T | | | | |
| NC_011770 | 3287983 | A | G | 58 | 0 | 6 | 0 | NON_SYNONYMOU | MODERAT | MISSENSE | gTt/gCt | V5A | PLES_29861 | | | G | G | G | G | | | | | G | G |
| NC_011770 | 3288334 | T | C | 69 | 0 | 10 | 0 | NON_SYNONYMOU | MODERAT | MISSENSE | Atc/Gtc | I332V | PLES_29871 | C | C | C | C | C | C | C | C | | | C | C |
| NC_011770 | 3288723 | G | A | 71 | 0 | 2 | 0 | NON_SYNONYMOU | MODERAT | MISSENSE | gCa/gTa | A202V | PLES_29871 | | | | | | | | | A | A | | |
| NC_011770 | 3289441 | G | A | 81 | 0 | 2 | 0 | NON_SYNONYMOU | MODERAT | MISSENSE | aCg/aTg | T295M | PLES_29881 | | | | | | | | | | | A | A |
| NC_011770 | 3289550 | T | C | 78 | 0 | 4 | 0 | NON_SYNONYMOU | MODERAT | MISSENSE | Acc/Gcc | T259A | PLES_29881 | | | | | | | C | C | C | C | | |
| NC_011770 | 3289714 | C | T | 65 | 0 | 2 | 0 | NON_SYNONYMOU | MODERAT | MISSENSE | cGc/cAc | R204H | PLES_29881 | T | T | | | | | | | | | | |
| NC_011770 | 3290299 | A | G | 67 | 0 | 1 | 0 | NON_SYNONYMOU | MODERAT | MISSENSE | gTc/gCc | V9A | PLES_29881 | | | | | | | G | | | | | |
| NC_011770 | 3290553 | CG | C | 46 | 0 | 1 | 0 | FRAME_SHIFT | HIGH | | ggg/ | G40 | PLES_29891 | C | | | | | | | | | | | |
| NC_011770 | 3290481 | A | C | 51 | 0 | 4 | 0 | NON_SYNONYMOU | MODERAT | MISSENSE | Aat/Cat | N15H | PLES_29891 | | | C | C | C | C | | | | | | |
| NC_011770 | 3290616 | G | A | 47 | 0 | 2 | 0 | NON_SYNONYMOU | MODERAT | MISSENSE | Ggc/Agc | G60S | PLES_29891 | | | | | | | | | | | A | A |
| NC_011770 | 3290844 | A | G | 45 | 0 | 12 | 0 | NON_SYNONYMOU | MODERAT | MISSENSE | Agc/Ggc | S136G | PLES_29891 | G | G | G | G | G | G | G | G | G | G | G | G |
| NC_011770 | 3291331 | T | C | 53 | 0 | 6 | 0 | NON_SYNONYMOU | MODERAT | MISSENSE | gTg/gCg | V298A | PLES_29891 | | | C | C | C | C | | | | | C | C |
| NC_011770 | 3291508 | G | A | 63 | 0 | 4 | 0 | NON_SYNONYMOU | MODERAT | MISSENSE | cGc/cAc | R357H | PLES_29891 | | | A | A | A | A | | | | | | |
| NC_011770 | 3291523 | C | T | 62 | 0 | 12 | 0 | NON_SYNONYMOU | MODERAT | MISSENSE | gCg/gTg | A362V | PLES_29891 | T | T | T | T | T | T | T | T | T | T | T | T |
| NC_011770 | 3291571 | A | G | 69 | 0 | 2 | 0 | NON_SYNONYMOU | MODERAT | MISSENSE | aAc/aGc | N378S | PLES_29891 | | | | | | | G | G | | | | |
| NC_011770 | 3291573 | G | C | 69 | 0 | 2 | 0 | NON_SYNONYMOU | MODERAT | MISSENSE | Gac/Cac | D379H | PLES_29891 | C | C | | | | | | | | | | |
| NC_011770 | 3291959 | A | T | 59 | 0 | 2 | 0 | NON_SYNONYMOU | MODERAT | MISSENSE | Acc/Tcc | T116S | PLES_29901 | T | T | | | | | | | | | | |
| NC_011770 | 3292115 | C | A | 79 | 0 | 2 | 0 | NON_SYNONYMOU | MODERAT | MISSENSE | Ctg/Atg | L168M | PLES_29901 | | | | | | | | | | | A | A |
| NC_011770 | 3292479 | C | T | 92 | 0 | 2 | 0 | NON_SYNONYMOU | MODERAT | MISSENSE | gCc/gTc | A289V | PLES_29901 | | | | | | | | | T | T | | |
| NC_011770 | 3292710 | T | C | 88 | 0 | 1 | 0 | NON_SYNONYMOU | MODERAT | MISSENSE | gTg/gCg | V366A | PLES_29901 | | | | | | | C | | | | | |
| NC_011770 | 3292744 | G | C | 91 | 0 | 2 | 0 | NON_SYNONYMOU | MODERAT | MISSENSE | tgG/tgC | W377C | PLES_29901 | C | C | | | | | | | | | | |
| NC_011770 | 3292776 | C | T | 84 | 0 | 8 | 0 | NON_SYNONYMOU | MODERAT | MISSENSE | gCg/gTg | A388V | PLES_29901 | | | T | T | T | T | T | T | | | T | T |
| NC_011770 | 3292782 | C | T | 93 | 0 | 4 | 0 | NON_SYNONYMOU | MODERAT | MISSENSE | gCc/gTc | A390V | PLES_29901 | | | | | | | T | T | | | T | T |
| NC_011770 | 3292836 | C | T | 95 | 0 | 2 | 0 | NON_SYNONYMOU | MODERAT | MISSENSE | gCa/gTa | A408V | PLES_29901 | | | | | | | | | | | T | T |

| | | | | | | | | | | | | | | | | | | | | | | | | | |
|---|---|---|---|---|---|---|---|---|---|---|---|---|---|---|---|---|---|---|---|---|---|---|---|---|---|
| NC_011770 | 3292841 | C | T | 92 | 0 | 6 | 0 | NON_SYNONYMOU | MODERAT | MISSENSE | Ccc/Tcc | P410S | PLES_29901 | | | | | | | T | T | T | T | T | T |
| NC_011770 | 3293310 | T | C | 99 | 0 | 10 | 0 | NON_SYNONYMOU | MODERAT | MISSENSE | atA/atG | I118M | PLES_29911 | C | C | C | C | C | C | | | C | C | C | C |
| NC_011770 | 3293422 | G | A | 93 | 0 | 2 | 0 | NON_SYNONYMOU | MODERAT | MISSENSE | gCc/gTc | A81V | PLES_29911 | A | A | | | | | | | | | | |
| NC_011770 | 3293829 | C | T | 58 | 0 | 1 | 0 | NON_SYNONYMOU | MODERAT | MISSENSE | Gac/Aac | D167N | PLES_29921 | | | | | | | T | | | | | |
| NC_011770 | 3294866 | G | T | 69 | 0 | 2 | 0 | NON_SYNONYMOU | MODERAT | MISSENSE | Gtt/Ttt | V13F | PLES_29941 | | | | | | | T | T | | | | |
| NC_011770 | 3295145 | A | G | 89 | 0 | 8 | 0 | NON_SYNONYMOU | MODERAT | MISSENSE | Atc/Gtc | I106V | PLES_29941 | G | G | G | G | G | G | G | G | | | | |
| NC_011770 | 3295661 | G | C | 86 | 0 | 4 | 0 | NON_SYNONYMOU | MODERAT | MISSENSE | Gcc/Ccc | A278P | PLES_29941 | | | C | C | C | C | | | | | | |
| NC_011770 | 3295764 | C | A | 95 | 0 | 4 | 0 | NON_SYNONYMOU | MODERAT | MISSENSE | Cgc/Agc | R3S | PLES_29951 | | | A | A | A | A | | | | | | |
| NC_011770 | 3296469 | G | A | 68 | 0 | 4 | 0 | NON_SYNONYMOU | MODERAT | MISSENSE | Gcc/Acc | A238T | PLES_29951 | | | A | A | A | A | | | | | | |
| NC_011770 | 3296517 | G | T | 66 | 0 | 2 | 0 | NON_SYNONYMOU | MODERAT | MISSENSE | Gtc/Ttc | V254F | PLES_29951 | | | | | | | T | T | | | | |
| NC_011770 | 3296644 | A | G | 60 | 0 | 6 | 0 | NON_SYNONYMOU | MODERAT | MISSENSE | cAg/cGg | Q296R | PLES_29951 | G | G | | | | | G | G | G | G | | |
| NC_011770 | 3295846 | C | T | 93 | 0 | 1 | 0 | NON_SYNONYMOU | MODERAT | MISSENSE | gCg/gTg | A30V | PLES_29951 | | T | | | | | | | | | | |
| NC_011770 | 3296812 | T | C | 60 | 0 | 2 | 0 | NON_SYNONYMOU | MODERAT | MISSENSE | Tcc/Ccc | S9P | PLES_29961 | | | | | | | C | C | | | | |
| NC_011770 | 3297143 | G | A | 51 | 0 | 2 | 0 | NON_SYNONYMOU | MODERAT | MISSENSE | gGc/gAc | G119D | PLES_29961 | | | | | A | A | | | | | | |
| NC_011770 | 3297151 | A | G | 52 | 0 | 2 | 0 | NON_SYNONYMOU | MODERAT | MISSENSE | Acg/Gcg | T122A | PLES_29961 | G | G | | | | | | | | | | |
| NC_011770 | 3297529 | C | G | 68 | 0 | 2 | 0 | NON_SYNONYMOU | MODERAT | MISSENSE | Ccg/Gcg | P248A | PLES_29961 | G | G | | | | | | | | | | |
| NC_011770 | 3297593 | C | T | 70 | 0 | 2 | 0 | NON_SYNONYMOU | MODERAT | MISSENSE | gCg/gTg | A269V | PLES_29961 | T | T | | | | | | | | | | |
| NC_011770 | 3297602 | C | T | 70 | 0 | 2 | 0 | NON_SYNONYMOU | MODERAT | MISSENSE | aCc/aTc | T272I | PLES_29961 | | | | | | | T | T | | | | |
| NC_011770 | 3297689 | G | A | 70 | 0 | 2 | 0 | NON_SYNONYMOU | MODERAT | MISSENSE | Gtc/Atc | V13I | PLES_29971 | A | A | | | | | | | | | | |
| NC_011770 | 3297716 | T | C | 70 | 0 | 2 | 0 | NON_SYNONYMOU | MODERAT | MISSENSE | Tcc/Ccc | S22P | PLES_29971 | C | C | | | | | | | | | | |
| NC_011770 | 3297735 | C | A | 74 | 0 | 2 | 0 | NON_SYNONYMOU | MODERAT | MISSENSE | cCc/cAc | P28H | PLES_29971 | | | | | | | A | A | | | | |
| NC_011770 | 3297740 | A | G | 69 | 0 | 2 | 0 | NON_SYNONYMOU | MODERAT | MISSENSE | Agc/Ggc | S30G | PLES_29971 | G | G | | | | | | | | | | |
| NC_011770 | 3299233 | G | A | 74 | 0 | 4 | 0 | NON_SYNONYMOU | MODERAT | MISSENSE | aGc/aAc | S202N | PLES_29981 | | | A | A | A | A | | | | | | |
| NC_011770 | 3299412 | T | C | 67 | 0 | 2 | 0 | NON_SYNONYMOU | MODERAT | MISSENSE | gTg/gCg | V24A | PLES_29991 | | | | | | | C | C | | | | |
| NC_011770 | 3299675 | G | T | 56 | 0 | 10 | 0 | NON_SYNONYMOU | MODERAT | MISSENSE | Gcc/Tcc | A112S | PLES_29991 | T | T | T | T | T | T | T | T | T | T | | |
| NC_011770 | 3299789 | C | T | 48 | 0 | 2 | 0 | NON_SYNONYMOU | MODERAT | MISSENSE | Cgc/Tgc | R150C | PLES_29991 | | | | | | | | | | | T | T |
| NC_011770 | 3299834 | C | T | 50 | 0 | 2 | 0 | NON_SYNONYMOU | MODERAT | MISSENSE | Ccc/Tcc | P165S | PLES_29991 | | | | | | | | | | | T | T |
| NC_011770 | 3300039 | G | A | 57 | 0 | 12 | 0 | NON_SYNONYMOU | MODERAT | MISSENSE | cGg/cAg | R233Q | PLES_29991 | A | A | A | A | A | A | A | A | A | A | A | A |
| NC_011770 | 3300936 | C | T | 70 | 0 | 2 | 0 | NON_SYNONYMOU | MODERAT | MISSENSE | gCg/gTg | A532V | PLES_29991 | | | | | | | T | T | | | | |
| NC_011770 | 3301509 | A | G | 72 | 0 | 12 | 0 | NON_SYNONYMOU | MODERAT | MISSENSE | cAt/cGt | H723R | PLES_29991 | G | G | G | G | G | G | G | G | G | G | G | G |
| NC_011770 | 3301521 | C | G | 72 | 0 | 2 | 0 | NON_SYNONYMOU | MODERAT | MISSENSE | aCc/aGc | T727S | PLES_29991 | | | | | | | | | | | G | G |
| NC_011770 | 3302337 | G | A | 65 | 0 | 4 | 0 | NON_SYNONYMOU | MODERAT | MISSENSE | cGc/cAc | R999H | PLES_29991 | | | A | A | A | A | | | | | | |
| NC_011770 | 3302354 | G | A | 61 | 0 | 2 | 0 | NON_SYNONYMOU | MODERAT | MISSENSE | Gag/Aag | E1005K | PLES_29991 | | | | | | | | | | | A | A |
| NC_011770 | 3302466 | C | T | 56 | 0 | 2 | 0 | NON_SYNONYMOU | MODERAT | MISSENSE | gCc/gTc | A1042V | PLES_29991 | | | | | | | T | T | | | | |
| NC_011770 | 3302891 | G | C | 74 | 0 | 2 | 0 | NON_SYNONYMOU | MODERAT | MISSENSE | Gtg/Ctg | V1184L | PLES_29991 | C | C | | | | | | | | | | |
| NC_011770 | 3302967 | T | C | 71 | 0 | 10 | 0 | NON_SYNONYMOU | MODERAT | MISSENSE | gTc/gCc | V1209A | PLES_29991 | C | C | C | C | C | C | | | C | C | C | C |
| NC_011770 | 3303047 | C | T | 72 | 0 | 4 | 0 | NON_SYNONYMOU | MODERAT | MISSENSE | Cct/Tct | P1236S | PLES_29991 | | | T | T | T | T | | | | | | |
| NC_011770 | 3303051 | T | C | 71 | 0 | 6 | 0 | NON_SYNONYMOU | MODERAT | MISSENSE | aTt/aCt | I1237T | PLES_29991 | | | C | C | C | C | | | C | C | | |
| NC_011770 | 3301022 | G | A | 68 | 0 | 1 | 0 | NON_SYNONYMOU | MODERAT | MISSENSE | Ggc/Agc | G561S | PLES_29991 | | | | | | | | A | | | | |
| NC_011770 | 3304180 | C | T | 76 | 0 | 8 | 0 | NON_SYNONYMOU | MODERAT | MISSENSE | Ccg/Tcg | P354S | PLES_30001 | T | T | T | T | T | T | | | | | T | T |
| NC_011770 | 3304452 | G | A | 83 | 0 | 2 | 0 | NON_SYNONYMOU | MODERAT | MISSENSE | Gta/Ata | V83I | PLES_30011 | | | | | | | | | | | A | A |
| NC_011770 | 3304995 | G | A | 79 | 0 | 4 | 0 | NON_SYNONYMOU | MODERAT | MISSENSE | Ggc/Agc | G264S | PLES_30011 | | | A | A | A | A | | | | | | |
| NC_011770 | 3310257 | C | G | 74 | 0 | 2 | 0 | STOP_GAINED | HIGH | NONSENSE | taC/taG | Y1671* | PLES_30021 | | | | | | | | | | | G | G |
| NC_011770 | 3305395 | C | T | 74 | 0 | 2 | 0 | NON_SYNONYMOU | MODERAT | MISSENSE | Ccg/Tcg | P51S | PLES_30021 | | | T | T | | | | | | | | |
| NC_011770 | 3305627 | C | T | 64 | 0 | 4 | 0 | NON_SYNONYMOU | MODERAT | MISSENSE | cCg/cTg | P128L | PLES_30021 | | | T | T | T | T | | | | | | |
| NC_011770 | 3305695 | C | G | 60 | 0 | 2 | 0 | NON_SYNONYMOU | MODERAT | MISSENSE | Ccg/Gcg | P151A | PLES_30021 | | | | | | | G | G | | | | |
| NC_011770 | 3306201 | G | T | 52 | 0 | 4 | 0 | NON_SYNONYMOU | MODERAT | MISSENSE | caG/caT | Q319H | PLES_30021 | | | T | T | T | T | | | | | | |
| NC_011770 | 3306619 | G | T | 59 | 0 | 2 | 0 | NON_SYNONYMOU | MODERAT | MISSENSE | Gcc/Tcc | A459S | PLES_30021 | | | | | | | | | T | T | | |
| NC_011770 | 3306682 | T | C | 55 | 0 | 12 | 0 | NON_SYNONYMOU | MODERAT | MISSENSE | Tgc/Cgc | C480R | PLES_30021 | C | C | C | C | C | C | C | C | C | C | C | C |
| NC_011770 | 3306929 | C | A | 61 | 0 | 6 | 0 | NON_SYNONYMOU | MODERAT | MISSENSE | gCc/gAc | A562D | PLES_30021 | | | A | A | A | A | | | | | A | A |
| NC_011770 | 3306959 | G | C | 62 | 0 | 12 | 0 | NON_SYNONYMOU | MODERAT | MISSENSE | aGg/aCg | R572T | PLES_30021 | C | C | C | C | C | C | C | C | C | C | C | C |
| NC_011770 | 3307583 | C | T | 61 | 0 | 1 | 0 | NON_SYNONYMOU | MODERAT | MISSENSE | aCc/aTc | T780I | PLES_30021 | | | | | | | T | | | | | |
| NC_011770 | 3307930 | G | A | 61 | 0 | 2 | 0 | NON_SYNONYMOU | MODERAT | MISSENSE | Ggg/Agg | G896R | PLES_30021 | | | | | | | | | | | A | A |
| NC_011770 | 3307931 | G | A | 60 | 0 | 4 | 0 | NON_SYNONYMOU | MODERAT | MISSENSE | gGg/gAg | G896E | PLES_30021 | | | | | | | | | A | A | A | A |
| NC_011770 | 3308486 | G | A | 45 | 0 | 2 | 0 | NON_SYNONYMOU | MODERAT | MISSENSE | gGc/gAc | G1081D | PLES_30021 | A | A | | | | | | | | | | |
| NC_011770 | 3308791 | C | G | 54 | 0 | 2 | 0 | NON_SYNONYMOU | MODERAT | MISSENSE | Ctc/Gtc | L1183V | PLES_30021 | | | | | | | G | G | | | | |
| NC_011770 | 3308980 | A | G | 50 | 0 | 10 | 0 | NON_SYNONYMOU | MODERAT | MISSENSE | Acg/Gcg | T1246A | PLES_30021 | G | G | G | G | G | G | | | G | G | G | G |
| NC_011770 | 3309067 | A | G | 50 | 0 | 8 | 0 | NON_SYNONYMOU | MODERAT | MISSENSE | Atc/Gtc | I1275V | PLES_30021 | G | G | | | | | G | G | G | G | G | G |
| NC_011770 | 3309235 | G | C | 66 | 0 | 2 | 0 | NON_SYNONYMOU | MODERAT | MISSENSE | Gac/Cac | D1331H | PLES_30021 | C | C | | | | | | | | | | |
| NC_011770 | 3309236 | A | T | 63 | 0 | 12 | 0 | NON_SYNONYMOU | MODERAT | MISSENSE | gAc/gTc | D1331V | PLES_30021 | T | T | C | C | C | C | C | C | C | C | C | C |
| NC_011770 | 3309239 | C | G | 65 | 0 | 2 | 0 | NON_SYNONYMOU | MODERAT | MISSENSE | gCg/gGg | A1332G | PLES_30021 | G | G | | | | | | | | | | |
| NC_011770 | 3309242 | G | C | 65 | 0 | 2 | 0 | NON_SYNONYMOU | MODERAT | MISSENSE | aGc/aCc | S1333T | PLES_30021 | C | C | | | | | | | | | | |

| | | | | | | | | | | | | | | | | | | | | | | | | | |
|---|---|---|---|---|---|---|---|---|---|---|---|---|---|---|---|---|---|---|---|---|---|---|---|---|---|
| NC_011770 | 3309254 | G | A | 64 | 0 | 4 | 0 | NON_SYNONYMOU | MODERAT | MISSENSE | cGg/cAg | R1337Q | PLES_30021 | A | A | | | | | | | A | A | | |
| NC_011770 | 3309256 | G | T | 65 | 0 | 4 | 0 | NON_SYNONYMOU | MODERAT | MISSENSE | Gcg/Tcg | A1338S | PLES_30021 | T | T | | | | | | | A | A | | |
| NC_011770 | 3309257 | C | T | 66 | 0 | 2 | 0 | NON_SYNONYMOU | MODERAT | MISSENSE | gCg/gTg | A1338V | PLES_30021 | | | | | | | T | T | | | | |
| NC_011770 | 3309355 | C | T | 80 | 0 | 2 | 0 | NON_SYNONYMOU | MODERAT | MISSENSE | Ctc/Ttc | L1371F | PLES_30021 | | | | | | | | | T | T | | |
| NC_011770 | 3309433 | A | G | 72 | 0 | 8 | 0 | NON_SYNONYMOU | MODERAT | MISSENSE | Agc/Ggc | S1397G | PLES_30021 | G | G | | | | | G | G | G | G | G | G |
| NC_011770 | 3309518 | C | T | 76 | 0 | 10 | 0 | NON_SYNONYMOU | MODERAT | MISSENSE | gCg/gTg | A1425V | PLES_30021 | T | T | T | T | T | T | | | T | T | T | T |
| NC_011770 | 3309544 | A | G | 80 | 0 | 8 | 0 | NON_SYNONYMOU | MODERAT | MISSENSE | Acg/Gcg | T1434A | PLES_30021 | | | G | G | G | G | | | G | G | G | G |
| NC_011770 | 3309580 | T | C | 85 | 0 | 2 | 0 | NON_SYNONYMOU | MODERAT | MISSENSE | Tgt/Cgt | C1446R | PLES_30021 | | | | | | | C | C | | | | |
| NC_011770 | 3309593 | G | A | 82 | 0 | 2 | 0 | NON_SYNONYMOU | MODERAT | MISSENSE | gGg/gAg | G1450E | PLES_30021 | | | | | | | A | A | | | | |
| NC_011770 | 3309598 | A | G | 82 | 0 | 2 | 0 | NON_SYNONYMOU | MODERAT | MISSENSE | Aca/Gca | T1452A | PLES_30021 | | | | | | | G | G | | | | |
| NC_011770 | 3309770 | G | T | 79 | 0 | 12 | 0 | NON_SYNONYMOU | MODERAT | MISSENSE | gGg/gTg | G1509V | PLES_30021 | T | T | T | T | T | T | T | T | T | T | T | T |
| NC_011770 | 3309893 | C | G | 72 | 0 | 2 | 0 | NON_SYNONYMOU | MODERAT | MISSENSE | gCc/gGc | A1550G | PLES_30021 | | | | | | | G | G | | | | |
| NC_011770 | 3310015 | T | C | 73 | 0 | 2 | 0 | NON_SYNONYMOU | MODERAT | MISSENSE | Tcg/Ccg | S1591P | PLES_30021 | | | | | | | C | C | | | | |
| NC_011770 | 3310375 | CCGA | C | 71 | 0 | 6 | 0 | CODON_DELETION | MODERATE | | acg/ | T1712 | PLES_30021 | C | C | C | C | C | C | | | | | | |
| NC_011770 | 3310378 | A | G | 32 | 6 | 4 | 0 | NON_SYNONYMOU | MODERAT | MISSENSE | Acg/Gcg | T1712A | PLES_30021 | . | . | . | . | . | . | G | G | | | G | G |
| NC_011770 | 3310535 | A | G | 65 | 0 | 8 | 0 | NON_SYNONYMOU | MODERAT | MISSENSE | gAc/gGc | D1764G | PLES_30021 | G | G | G | G | G | G | | | | | G | G |
| NC_011770 | 3310568 | A | G | 64 | 0 | 10 | 0 | NON_SYNONYMOU | MODERAT | MISSENSE | gAg/gGg | E1775G | PLES_30021 | G | G | G | G | G | G | G | G | | | G | G |
| NC_011770 | 3310577 | C | T | 63 | 0 | 8 | 0 | NON_SYNONYMOU | MODERAT | MISSENSE | gCg/gTg | A1778V | PLES_30021 | T | T | T | T | T | T | | | | | T | T |
| NC_011770 | 3310588 | A | G | 62 | 0 | 8 | 0 | NON_SYNONYMOU | MODERAT | MISSENSE | Acg/Gcg | T1782A | PLES_30021 | G | G | G | G | G | G | | | | | G | G |
| NC_011770 | 3310817 | C | T | 52 | 0 | 2 | 0 | NON_SYNONYMOU | MODERAT | MISSENSE | gCc/gTc | A1858V | PLES_30021 | | | | | | | T | T | | | | |
| NC_011770 | 3311110 | A | G | 64 | 0 | 6 | 0 | NON_SYNONYMOU | MODERAT | MISSENSE | Atc/Gtc | I1956V | PLES_30021 | | | | | | | G | G | G | G | G | G |
| NC_011770 | 3311459 | T | C | 54 | 0 | 6 | 0 | NON_SYNONYMOU | MODERAT | MISSENSE | gTg/gCg | V2072A | PLES_30021 | | | | | | | C | C | C | C | C | C |
| NC_011770 | 3312171 | C | A | 65 | 0 | 2 | 0 | NON_SYNONYMOU | MODERAT | MISSENSE | tCc/tAc | S161Y | PLES_30031 | | | | | | | | | | | A | A |
| NC_011770 | 3311952 | G | A | 65 | 0 | 1 | 0 | NON_SYNONYMOU | MODERAT | MISSENSE | cGg/cAg | R88Q | PLES_30031 | | | | A | | | | | | | | |
| NC_011770 | 3315373 | T | G | 86 | 0 | 2 | 0 | NON_SYNONYMOU | MODERAT | MISSENSE | caT/caG | H183Q | PLES_30061 | G | G | | | | | | | | | | |
| NC_011770 | 3315794 | A | G | 81 | 0 | 12 | 0 | NON_SYNONYMOU | MODERAT | MISSENSE | Agg/Ggg | R324G | PLES_30061 | G | G | G | G | G | G | G | G | G | G | G | G |
| NC_011770 | 3316058 | A | C | 66 | 0 | 8 | 0 | NON_SYNONYMOU | MODERAT | MISSENSE | Agc/Cgc | S412R | PLES_30061 | C | C | C | C | C | C | | | | | C | C |
| NC_011770 | 3316296 | T | C | 74 | 0 | 2 | 0 | NON_SYNONYMOU | MODERAT | MISSENSE | gTt/gCt | V491A | PLES_30061 | | | | | | | C | C | | | | |
| NC_011770 | 3316389 | G | A | 67 | 0 | 1 | 0 | NON_SYNONYMOU | MODERAT | MISSENSE | cGg/cAg | R522Q | PLES_30061 | | | | | | | | | | | A | |
| NC_011770 | 3317184 | C | G | 75 | 0 | 2 | 0 | NON_SYNONYMOU | MODERAT | MISSENSE | gaC/gaG | D57E | PLES_30081 | | | | | | | G | G | | | | |
| NC_011770 | 3317332 | G | A | 75 | 0 | 4 | 0 | NON_SYNONYMOU | MODERAT | MISSENSE | Gcc/Acc | A107T | PLES_30081 | | | A | A | A | A | | | | | | |
| NC_011770 | 3317889 | A | C | 58 | 0 | 10 | 0 | NON_SYNONYMOU | MODERAT | MISSENSE | gaA/gaC | E292D | PLES_30081 | C | C | C | C | C | C | C | C | | | C | C |
| NC_011770 | 3317997 | A | C | 48 | 0 | 6 | 0 | NON_SYNONYMOU | MODERAT | MISSENSE | gaA/gaC | E328D | PLES_30081 | C | C | | | | | C | C | | | C | C |
| NC_011770 | 3318317 | G | A | 69 | 0 | 10 | 0 | NON_SYNONYMOU | MODERAT | MISSENSE | aGc/aAc | S435N | PLES_30081 | A | A | A | A | A | A | A | A | | | A | A |
| NC_011770 | 3319339 | G | A | 79 | 0 | 6 | 0 | NON_SYNONYMOU | MODERAT | MISSENSE | aGc/aAc | S34N | PLES_30101 | A | A | A | A | A | A | | | | | | |
| NC_011770 | 3320136 | C | T | 63 | 0 | 4 | 0 | NON_SYNONYMOU | MODERAT | MISSENSE | cCg/cTg | P16L | PLES_30111 | T | T | | | | | | | T | T | | |
| NC_011770 | 3320225 | G | A | 61 | 0 | 6 | 0 | NON_SYNONYMOU | MODERAT | MISSENSE | Gtt/Att | V46I | PLES_30111 | A | A | | | | | A | A | | | A | A |
| NC_011770 | 3320400 | C | T | 61 | 0 | 4 | 0 | NON_SYNONYMOU | MODERAT | MISSENSE | gCc/gTc | A104V | PLES_30111 | T | T | | | | | | | | | T | T |
| NC_011770 | 3320435 | G | A | 65 | 0 | 2 | 0 | NON_SYNONYMOU | MODERAT | MISSENSE | Gat/Aat | D116N | PLES_30111 | | | A | A | | | | | | | | |
| NC_011770 | 3320499 | C | T | 67 | 0 | 10 | 0 | NON_SYNONYMOU | MODERAT | MISSENSE | gCg/gTg | A137V | PLES_30111 | T | T | T | T | T | T | | | T | T | T | T |
| NC_011770 | 3320676 | C | T | 61 | 0 | 2 | 0 | NON_SYNONYMOU | MODERAT | MISSENSE | gCg/gTg | A196V | PLES_30111 | | | | | | | | | | | T | T |
| NC_011770 | 3321044 | G | A | 64 | 0 | 4 | 0 | NON_SYNONYMOU | MODERAT | MISSENSE | Gtg/Atg | V319M | PLES_30111 | | | A | A | A | A | | | | | | |
| NC_011770 | 3321082 | T | G | 72 | 0 | 2 | 0 | NON_SYNONYMOU | MODERAT | MISSENSE | Tcc/Gcc | S11A | PLES_30121 | | | | | | | | | | | G | G |
| NC_011770 | 3321754 | T | C | 74 | 0 | 2 | 0 | NON_SYNONYMOU | MODERAT | MISSENSE | gTg/gCg | V27A | PLES_30131 | | | | | | | | | | | C | C |
| NC_011770 | 3322981 | G | A | 71 | 0 | 8 | 0 | NON_SYNONYMOU | MODERAT | MISSENSE | gGc/gAc | G436D | PLES_30131 | A | A | | | | | A | A | A | A | A | A |
| NC_011770 | 3326225 | T | G | 46 | 0 | 4 | 0 | NON_SYNONYMOU | MODERAT | MISSENSE | Ttc/Gtc | F166V | PLES_30151 | G | G | | | | | | | | | G | G |
| NC_011770 | 3326267 | G | A | 44 | 0 | 8 | 0 | NON_SYNONYMOU | MODERAT | MISSENSE | Gcc/Acc | A180T | PLES_30151 | A | A | | | | | A | A | A | A | A | A |
| NC_011770 | 3326270 | A | T | 44 | 0 | 8 | 0 | NON_SYNONYMOU | MODERAT | MISSENSE | Acc/Tcc | T181S | PLES_30151 | T | T | | | | | T | T | T | T | T | T |
| NC_011770 | 3326447 | C | G | 63 | 0 | 2 | 0 | NON_SYNONYMOU | MODERAT | MISSENSE | Ctg/Gtg | L240V | PLES_30151 | | | | | | | | | G | G | | |
| NC_011770 | 3326676 | A | T | 58 | 0 | 4 | 0 | NON_SYNONYMOU | MODERAT | MISSENSE | cAg/cTg | Q316L | PLES_30151 | T | T | | | | | | | | | T | T |
| NC_011770 | 3326678 | G | T | 59 | 0 | 4 | 0 | NON_SYNONYMOU | MODERAT | MISSENSE | Gcg/Tcg | A317S | PLES_30151 | T | T | | | | | | | | | T | T |
| NC_011770 | 3326792 | A | C | 77 | 0 | 6 | 0 | NON_SYNONYMOU | MODERAT | MISSENSE | Agc/Cgc | S355R | PLES_30151 | C | C | | | | | C | C | | | C | C |
| NC_011770 | 3326808 | A | T | 75 | 0 | 4 | 0 | NON_SYNONYMOU | MODERAT | MISSENSE | gAg/gTg | E360V | PLES_30151 | T | T | | | | | | | | | T | T |
| NC_011770 | 3326903 | A | G | 76 | 0 | 10 | 0 | NON_SYNONYMOU | MODERAT | MISSENSE | Aag/Gag | K392E | PLES_30151 | G | G | G | G | G | G | | | G | G | G | G |
| NC_011770 | 3327491 | G | A | 81 | 0 | 10 | 0 | NON_SYNONYMOU | MODERAT | MISSENSE | Gag/Aag | E588K | PLES_30151 | A | A | A | A | A | A | A | A | A | A | | |
| NC_011770 | 3327651 | A | G | 74 | 0 | 2 | 0 | NON_SYNONYMOU | MODERAT | MISSENSE | aAg/aGg | K641R | PLES_30151 | | | | | | | G | G | | | | |
| NC_011770 | 3327716 | A | G | 76 | 0 | 12 | 0 | NON_SYNONYMOU | MODERAT | MISSENSE | Acc/Gcc | T663A | PLES_30151 | G | G | G | G | G | G | G | G | G | G | G | G |
| NC_011770 | 3328023 | A | G | 69 | 0 | 12 | 0 | NON_SYNONYMOU | MODERAT | MISSENSE | cAc/cGc | H14R | PLES_30161 | G | G | G | G | G | G | G | G | G | G | G | G |
| NC_011770 | 3328097 | C | T | 70 | 0 | 6 | 0 | NON_SYNONYMOU | MODERAT | MISSENSE | Cgc/Tgc | R39C | PLES_30161 | | | | | | | T | T | T | T | T | T |
| NC_011770 | 3328100 | G | T | 68 | 0 | 4 | 0 | NON_SYNONYMOU | MODERAT | MISSENSE | Gcc/Tcc | A40S | PLES_30161 | | | T | T | T | T | | | | | | |
| NC_011770 | 3328151 | A | G | 67 | 0 | 4 | 0 | NON_SYNONYMOU | MODERAT | MISSENSE | Acc/Gcc | T57A | PLES_30161 | | | | | | | G | G | | | G | G |
| NC_011770 | 3328380 | G | A | 65 | 0 | 4 | 0 | NON_SYNONYMOU | MODERAT | MISSENSE | cCg/cTg | P180L | PLES_30171 | | | A | A | A | A | | | | | | |

| | | | | | | | | | | | | | | | | | | | | | | | | | |
|---|---|---|---|---|---|---|---|---|---|---|---|---|---|---|---|---|---|---|---|---|---|---|---|---|---|
| NC_011770 | 3328381 | G | A | 66 | 0 | 4 | 0 | NON_SYNONYMOU | MODERAT | MISSENSE | Ccg/Tcg | P180S | PLES_30171 | A | A | | | | | | | A | A | | |
| NC_011770 | 3328396 | T | C | 67 | 0 | 4 | 0 | NON_SYNONYMOU | MODERAT | MISSENSE | Aaa/Gaa | K175E | PLES_30171 | C | C | | | | | | | C | C | | |
| NC_011770 | 3328405 | T | C | 67 | 0 | 4 | 0 | NON_SYNONYMOU | MODERAT | MISSENSE | Aac/Gac | N172D | PLES_30171 | C | C | | | | | | | C | C | | |
| NC_011770 | 3328503 | C | T | 64 | 0 | 10 | 0 | NON_SYNONYMOU | MODERAT | MISSENSE | aGc/aAc | S139N | PLES_30171 | | | T | T | T | T | T | T | T | T | T | T |
| NC_011770 | 3328513 | G | T | 62 | 0 | 10 | 0 | NON_SYNONYMOU | MODERAT | MISSENSE | Cgc/Agc | R136S | PLES_30171 | | | T | T | T | T | T | T | T | T | T | T |
| NC_011770 | 3328771 | T | G | 67 | 0 | 12 | 0 | NON_SYNONYMOU | MODERAT | MISSENSE | Aac/Cac | N50H | PLES_30171 | G | G | G | G | G | G | G | G | G | G | G | G |
| NC_011770 | 3329093 | C | T | 70 | 0 | 6 | 0 | NON_SYNONYMOU | MODERAT | MISSENSE | Ggc/Agc | G453S | PLES_30181 | T | T | | | | | T | T | T | T | | |
| NC_011770 | 3329204 | C | T | 69 | 0 | 2 | 0 | NON_SYNONYMOU | MODERAT | MISSENSE | Gac/Aac | D416N | PLES_30181 | | | | | | | | | | | T | T |
| NC_011770 | 3329456 | C | T | 67 | 0 | 4 | 0 | NON_SYNONYMOU | MODERAT | MISSENSE | Gag/Aag | E332K | PLES_30181 | | | T | T | T | T | | | | | | |
| NC_011770 | 3329624 | G | C | 72 | 0 | 12 | 0 | NON_SYNONYMOU | MODERAT | MISSENSE | Caa/Gaa | Q276E | PLES_30181 | C | C | C | C | C | C | C | C | C | C | C | C |
| NC_011770 | 3329644 | C | T | 70 | 0 | 12 | 0 | NON_SYNONYMOU | MODERAT | MISSENSE | cGa/cAa | R269Q | PLES_30181 | T | T | T | T | T | T | T | T | T | T | T | T |
| NC_011770 | 3329971 | A | G | 72 | 0 | 4 | 0 | NON_SYNONYMOU | MODERAT | MISSENSE | cTg/cCg | L160P | PLES_30181 | | | G | G | G | G | | | | | | |
| NC_011770 | 3330208 | T | C | 58 | 0 | 4 | 0 | NON_SYNONYMOU | MODERAT | MISSENSE | cAa/cGa | Q81R | PLES_30181 | | | C | C | C | C | | | | | | |
| NC_011770 | 3331503 | C | T | 66 | 0 | 2 | 0 | NON_SYNONYMOU | MODERAT | MISSENSE | aGc/aAc | S93N | PLES_30201 | T | T | | | | | | | | | | |
| NC_011770 | 3331747 | C | G | 58 | 0 | 2 | 0 | NON_SYNONYMOU | MODERAT | MISSENSE | Gcc/Ccc | A12P | PLES_30201 | G | G | | | | | | | | | | |
| NC_011770 | 3331796 | C | T | 63 | 0 | 2 | 0 | NON_SYNONYMOU | MODERAT | MISSENSE | gGc/gAc | G480D | PLES_30211 | | | | | | | | | | | T | T |
| NC_011770 | 3331907 | T | A | 70 | 0 | 2 | 0 | NON_SYNONYMOU | MODERAT | MISSENSE | tAc/tTc | Y443F | PLES_30211 | A | A | | | | | | | | | | |
| NC_011770 | 3332111 | C | A | 75 | 0 | 2 | 0 | NON_SYNONYMOU | MODERAT | MISSENSE | cGg/cTg | R375L | PLES_30211 | A | A | | | | | | | | | | |
| NC_011770 | 3332202 | A | T | 83 | 0 | 2 | 0 | NON_SYNONYMOU | MODERAT | MISSENSE | Tgg/Agg | W345R | PLES_30211 | T | T | | | | | | | | | | |
| NC_011770 | 3332994 | C | T | 88 | 0 | 4 | 0 | NON_SYNONYMOU | MODERAT | MISSENSE | Gcc/Acc | A81T | PLES_30211 | | | T | T | T | T | | | | | | |
| NC_011770 | 3333125 | C | T | 87 | 0 | 2 | 0 | NON_SYNONYMOU | MODERAT | MISSENSE | cGc/cAc | R37H | PLES_30211 | T | T | | | | | | | | | | |
| NC_011770 | 3333503 | C | T | 77 | 0 | 6 | 0 | NON_SYNONYMOU | MODERAT | MISSENSE | gGc/gAc | G56D | PLES_30221 | | | T | T | T | T | T | T | | | | |
| NC_011770 | 3334252 | G | A | 77 | 0 | 4 | 0 | NON_SYNONYMOU | MODERAT | MISSENSE | aGc/aAc | S99N | PLES_30231 | | | A | A | A | A | | | | | | |
| NC_011770 | 3334663 | G | C | 82 | 0 | 2 | 0 | NON_SYNONYMOU | MODERAT | MISSENSE | cGc/cCc | R236P | PLES_30231 | | | | | | | | | | | C | C |
| NC_011770 | 3334853 | A | C | 95 | 0 | 2 | 0 | NON_SYNONYMOU | MODERAT | MISSENSE | Tcc/Gcc | S222A | PLES_30241 | | | | | | | | | | | C | C |
| NC_011770 | 3335129 | C | G | 99 | 0 | 2 | 0 | NON_SYNONYMOU | MODERAT | MISSENSE | Gtg/Ctg | V130L | PLES_30241 | | | | | G | G | | | | | | |
| NC_011770 | 3338192 | T | G | 75 | 0 | 2 | 0 | NON_SYNONYMOU | MODERAT | MISSENSE | cAg/cCg | Q152P | PLES_30281 | | | | | | | | | G | G | | |
| NC_011770 | 3338597 | T | C | 69 | 0 | 2 | 0 | NON_SYNONYMOU | MODERAT | MISSENSE | gAc/gGc | D17G | PLES_30281 | | | | | | | | | | | C | C |
| NC_011770 | 3339039 | C | T | 72 | 0 | 1 | 0 | NON_SYNONYMOU | MODERAT | MISSENSE | cCg/cTg | P62L | PLES_30291 | | | | | | | T | | | | | |
| NC_011770 | 3339113 | T | G | 67 | 0 | 12 | 0 | NON_SYNONYMOU | MODERAT | MISSENSE | Tgc/Ggc | C87G | PLES_30291 | G | G | G | G | G | G | G | G | G | G | G | G |
| NC_011770 | 3339458 | G | A | 83 | 0 | 2 | 0 | NON_SYNONYMOU | MODERAT | MISSENSE | Gag/Aag | E202K | PLES_30291 | A | A | | | | | | | | | | |
| NC_011770 | 3339507 | G | A | 77 | 0 | 2 | 0 | NON_SYNONYMOU | MODERAT | MISSENSE | cGg/cAg | R218Q | PLES_30291 | | | | | | | | | | | A | A |
| NC_011770 | 3339785 | G | A | 81 | 0 | 2 | 0 | NON_SYNONYMOU | MODERAT | MISSENSE | Gcc/Acc | A311T | PLES_30291 | | | | | | | A | A | | | | |
| NC_011770 | 3343394 | C | T | 79 | 0 | 1 | 0 | NON_SYNONYMOU | MODERAT | MISSENSE | cGc/cAc | R138H | PLES_30341 | | | | | | | T | | | | | |
| NC_011770 | 3344079 | C | T | 91 | 0 | 4 | 0 | NON_SYNONYMOU | MODERAT | MISSENSE | Cat/Tat | H43Y | PLES_30351 | | | T | T | T | T | | | | | | |
| NC_011770 | 3345222 | C | T | 71 | 0 | 2 | 0 | NON_SYNONYMOU | MODERAT | MISSENSE | cGc/cAc | R324H | PLES_30361 | T | T | | | | | | | | | | |
| NC_011770 | 3345288 | C | T | 76 | 0 | 2 | 0 | NON_SYNONYMOU | MODERAT | MISSENSE | aGc/aAc | S302N | PLES_30361 | | | | | | | | | | | T | T |
| NC_011770 | 3345474 | C | T | 80 | 0 | 2 | 0 | NON_SYNONYMOU | MODERAT | MISSENSE | cGc/cAc | R240H | PLES_30361 | | | | | | | T | T | | | | |
| NC_011770 | 3345764 | A | T | 82 | 0 | 10 | 0 | NON_SYNONYMOU | MODERAT | MISSENSE | gaT/gaA | D143E | PLES_30361 | T | T | T | T | T | T | | | T | T | T | T |
| NC_011770 | 3345802 | C | T | 83 | 0 | 2 | 0 | NON_SYNONYMOU | MODERAT | MISSENSE | Gcc/Acc | A131T | PLES_30361 | T | T | | | | | | | | | | |
| NC_011770 | 3345934 | T | C | 77 | 0 | 12 | 0 | NON_SYNONYMOU | MODERAT | MISSENSE | Agc/Ggc | S87G | PLES_30361 | C | C | C | C | C | C | C | C | C | C | C | C |
| NC_011770 | 3345982 | G | A | 77 | 0 | 2 | 0 | NON_SYNONYMOU | MODERAT | MISSENSE | Ccc/Tcc | P71S | PLES_30361 | | | | | | | A | A | | | | |
| NC_011770 | 3346888 | G | A | 72 | 0 | 4 | 0 | NON_SYNONYMOU | MODERAT | MISSENSE | Gac/Aac | D204N | PLES_30371 | A | A | | | | | | | A | A | | |
| NC_011770 | 3347271 | A | G | 59 | 0 | 6 | 0 | NON_SYNONYMOU | MODERAT | MISSENSE | Tcg/Ccg | S433P | PLES_30381 | | | G | G | G | G | G | G | | | | |
| NC_011770 | 3347662 | G | T | 74 | 0 | 4 | 0 | NON_SYNONYMOU | MODERAT | MISSENSE | agC/agA | S302R | PLES_30381 | T | T | | | | | | | T | T | | |
| NC_011770 | 3348066 | C | T | 78 | 0 | 4 | 0 | NON_SYNONYMOU | MODERAT | MISSENSE | Gcc/Acc | A168T | PLES_30381 | T | T | | | | | | | T | T | | |
| NC_011770 | 3348069 | C | T | 80 | 0 | 2 | 0 | NON_SYNONYMOU | MODERAT | MISSENSE | Gcc/Acc | A167T | PLES_30381 | | | | | | | | | T | T | | |
| NC_011770 | 3348506 | T | G | 73 | 0 | 12 | 0 | NON_SYNONYMOU | MODERAT | MISSENSE | gAa/gCa | E21A | PLES_30381 | G | G | G | G | G | G | G | G | G | G | G | G |
| NC_011770 | 3348605 | C | T | 80 | 0 | 12 | 0 | NON_SYNONYMOU | MODERAT | MISSENSE | Gac/Aac | D584N | PLES_30391 | T | T | T | T | T | T | T | T | T | T | T | T |
| NC_011770 | 3348881 | C | T | 87 | 0 | 2 | 0 | NON_SYNONYMOU | MODERAT | MISSENSE | Gcg/Acg | A492T | PLES_30391 | | | | | | | | | | | T | T |
| NC_011770 | 3350185 | A | G | 79 | 0 | 1 | 0 | NON_SYNONYMOU | MODERAT | MISSENSE | cTg/cCg | L57P | PLES_30391 | | | | | | | G | | | | | |
| NC_011770 | 3349043 | C | A | 77 | 0 | 1 | 0 | NON_SYNONYMOU | MODERAT | MISSENSE | Gcc/Tcc | A438S | PLES_30391 | | | | | | | | A | | | | |
| NC_011770 | 3349097 | T | C | 69 | 0 | 1 | 0 | NON_SYNONYMOU | MODERAT | MISSENSE | Agc/Ggc | S420G | PLES_30391 | | | | | | | | C | | | | |
| NC_011770 | 3351246 | C | A | 60 | 0 | 2 | 0 | NON_SYNONYMOU | MODERAT | MISSENSE | Gcc/Tcc | A328S | PLES_30411 | | | | | | | A | A | | | | |
| NC_011770 | 3351296 | G | T | 55 | 0 | 12 | 0 | NON_SYNONYMOU | MODERAT | MISSENSE | gCg/gAg | A311E | PLES_30411 | T | T | T | T | T | T | T | T | T | T | T | T |
| NC_011770 | 3351419 | T | C | 56 | 0 | 6 | 0 | NON_SYNONYMOU | MODERAT | MISSENSE | gAc/gGc | D270G | PLES_30411 | | | C | C | C | C | | | | | C | C |
| NC_011770 | 3351569 | T | G | 61 | 0 | 8 | 0 | NON_SYNONYMOU | MODERAT | MISSENSE | gAg/gCg | E220A | PLES_30411 | | | G | G | G | G | G | G | | | G | G |
| NC_011770 | 3351572 | G | A | 62 | 0 | 2 | 0 | NON_SYNONYMOU | MODERAT | MISSENSE | gCc/gTc | A219V | PLES_30411 | | | | | | | A | A | | | | |
| NC_011770 | 3352464 | C | T | 58 | 0 | 12 | 0 | NON_SYNONYMOU | MODERAT | MISSENSE | Gtc/Atc | V364I | PLES_30421 | T | T | T | T | T | T | T | T | T | T | T | T |
| NC_011770 | 3354416 | C | T | 71 | 0 | 6 | 0 | NON_SYNONYMOU | MODERAT | MISSENSE | Gcg/Acg | A51T | PLES_30431 | T | T | | | | | | | T | T | T | T |
| NC_011770 | 3364728 | GC | G | 83 | 0 | 2 | 0 | FRAME_SHIFT | HIGH | | gcg/ | A88 | PLES_30521 | G | G | | | | | | | | | | |
| NC_011770 | 3363552 | T | G | 60 | 0 | 2 | 0 | NON_SYNONYMOU | MODERAT | MISSENSE | aAt/aCt | N480T | PLES_30521 | | | | | | | | | | | G | G |

| | | | | | | | | | | | | | | | | | | | | | | | | | |
|---|---|---|---|---|---|---|---|---|---|---|---|---|---|---|---|---|---|---|---|---|---|---|---|---|---|
| NC_011770 | 3364954 | C | G | 81 | 0 | 4 | 0 | NON_SYNONYMOU | MODERAT | MISSENSE | Gtc/Ctc | V13L | PLES_30521 | | | G | G | G | G | | | | | | |
| NC_011770 | 3364356 | T | C | 93 | 0 | 1 | 0 | NON_SYNONYMOU | MODERAT | MISSENSE | tAc/tGc | Y212C | PLES_30521 | | | | | | | | C | | | | |
| NC_011770 | 3365168 | C | T | 88 | 0 | 4 | 0 | NON_SYNONYMOU | MODERAT | MISSENSE | Cac/Tac | H4Y | PLES_30531 | | | T | T | T | T | | | | | | |
| NC_011770 | 3391898 | T | C | 60 | 2 | 8 | 0 | NON_SYNONYMOU | MODERAT | MISSENSE | Agc/Ggc | S231G | PLES_30751 | . | . | C | C | C | C | | | C | C | C | C |
| NC_011770 | 3392114 | C | A | 57 | 2 | 2 | 0 | NON_SYNONYMOU | MODERAT | MISSENSE | Gtg/Ttg | V159L | PLES_30751 | . | . | | | | | | | | | A | A |
| NC_011770 | 3392417 | T | C | 34 | 2 | 9 | 0 | NON_SYNONYMOU | MODERAT | MISSENSE | Atc/Gtc | I58V | PLES_30751 | . | . | | C | C | C | C | C | C | C | C | C |
| NC_011770 | 3392906 | G | GC | 16 | 6 | 6 | 0 | FRAME_SHIFT | HIGH | | acc/aGcc | T213S? | PLES_30761 | . | . | . | . | . | . | GC | GC | GC | GC | GC | GC |
| NC_011770 | 3393165 | T | C | 30 | 6 | 6 | 0 | NON_SYNONYMOU | MODERAT | MISSENSE | Acc/Gcc | T127A | PLES_30761 | . | . | . | . | . | . | C | C | C | C | C | C |
| NC_011770 | 3393423 | G | C | 30 | 6 | 6 | 0 | NON_SYNONYMOU | MODERAT | MISSENSE | Cga/Gga | R41G | PLES_30761 | . | . | . | . | . | . | C | C | C | C | C | C |
| NC_011770 | 3393865 | G | A | 27 | 6 | 1 | 0 | STOP_GAINED | HIGH | NONSENSE | Cga/Tga | R31* | PLES_30771 | . | . | . | . | . | . | | A | | | | |
| NC_011770 | 3393948 | G | T | 25 | 6 | 6 | 0 | NON_SYNONYMOU | MODERAT | MISSENSE | cCc/cAc | P3H | PLES_30771 | . | . | . | . | . | . | T | T | T | T | T | T |
| NC_011770 | 3394722 | G | A | 35 | 6 | 2 | 0 | NON_SYNONYMOU | MODERAT | MISSENSE | Gta/Ata | V132I | PLES_30781 | . | . | . | . | . | . | | | A | A | | |
| NC_011770 | 3395286 | G | C | 26 | 6 | 6 | 0 | NON_SYNONYMOU | MODERAT | MISSENSE | Gcg/Ccg | A320P | PLES_30781 | . | . | . | . | . | . | C | C | C | C | C | C |
| NC_011770 | 3395352 | G | A | 28 | 6 | 1 | 0 | NON_SYNONYMOU | MODERAT | MISSENSE | Gtc/Atc | V342I | PLES_30781 | . | . | . | . | . | . | A | | | | | |
| NC_011770 | 3396082 | G | C | 34 | 6 | 4 | 0 | NON_SYNONYMOU | MODERAT | MISSENSE | aGc/aCc | S4T | PLES_30791 | | | C | C | C | C | . | . | . | . | . | . |
| NC_011770 | 3396654 | C | T | 44 | 6 | 6 | 0 | NON_SYNONYMOU | MODERAT | MISSENSE | Gtc/Atc | V395I | PLES_30801 | T | T | T | T | T | T | . | . | . | . | . | . |
| NC_011770 | 3397181 | G | A | 37 | 6 | 2 | 0 | NON_SYNONYMOU | MODERAT | MISSENSE | aCg/aTg | T219M | PLES_30801 | A | A | | | | | . | . | . | . | . | . |
| NC_011770 | 3398209 | G | T | 37 | 6 | 2 | 0 | NON_SYNONYMOU | MODERAT | MISSENSE | Gac/Tac | D27Y | PLES_30811 | | | T | T | | | . | . | . | . | . | . |
| NC_011770 | 3399153 | C | A | 34 | 6 | 2 | 0 | NON_SYNONYMOU | MODERAT | MISSENSE | Gcg/Tcg | A298S | PLES_30821 | A | A | | | | | . | . | . | . | . | . |
| NC_011770 | 3402373 | A | G | 48 | 6 | 6 | 0 | SYNONYMOUS_ST | LOW | SILENT | Ttg/Ctg | L1 | PLES_30831 | G | G | G | G | G | G | . | . | . | . | . | . |
| NC_011770 | 3401877 | T | C | 40 | 6 | 6 | 0 | NON_SYNONYMOU | MODERAT | MISSENSE | gAc/gGc | D166G | PLES_30831 | C | C | C | C | C | C | . | . | . | . | . | . |
| NC_011770 | 3403157 | A | G | 45 | 6 | 6 | 0 | NON_SYNONYMOU | MODERAT | MISSENSE | gTa/gCa | V299A | PLES_30841 | G | G | G | G | G | G | . | . | . | . | . | . |
| NC_011770 | 3404191 | CG | C | 42 | 6 | 1 | 0 | FRAME_SHIFT | HIGH | | cgc/ | R457 | PLES_30851 | | | | C | | | . | . | . | . | . | . |
| NC_011770 | 3404935 | TC | T | 41 | 6 | 1 | 0 | FRAME_SHIFT | HIGH | | gaa/ | E209 | PLES_30851 | | | | T | | | . | . | . | . | . | . |
| NC_011770 | 3405347 | G | A | 49 | 6 | 1 | 0 | STOP_GAINED | HIGH | NONSENSE | Cag/Tag | Q72* | PLES_30851 | A | | | | | | . | . | . | . | . | . |
| NC_011770 | 3405137 | C | T | 45 | 6 | 4 | 0 | NON_SYNONYMOU | MODERAT | MISSENSE | Gtc/Atc | V142I | PLES_30851 | | | T | T | T | T | . | . | . | . | . | . |
| NC_011770 | 3406191 | G | A | 54 | 6 | 6 | 0 | NON_SYNONYMOU | MODERAT | MISSENSE | gCg/gTg | A191V | PLES_30861 | A | A | A | A | A | A | . | . | . | . | . | . |
| NC_011770 | 3406308 | G | A | 54 | 6 | 4 | 0 | NON_SYNONYMOU | MODERAT | MISSENSE | gCt/gTt | A152V | PLES_30861 | | | A | A | A | A | . | . | . | . | . | . |
| NC_011770 | 3406426 | C | T | 52 | 6 | 2 | 0 | NON_SYNONYMOU | MODERAT | MISSENSE | Gag/Aag | E113K | PLES_30861 | T | T | | | | | . | . | . | . | . | . |
| NC_011770 | 3406950 | T | C | 49 | 6 | 4 | 0 | NON_SYNONYMOU | MODERAT | MISSENSE | Acg/Gcg | T431A | PLES_30871 | | | C | C | C | C | . | . | . | . | . | . |
| NC_011770 | 3407136 | C | T | 46 | 6 | 2 | 0 | NON_SYNONYMOU | MODERAT | MISSENSE | Gtg/Atg | V369M | PLES_30871 | T | T | | | | | . | . | . | . | . | . |
| NC_011770 | 3408123 | T | C | 44 | 6 | 6 | 0 | NON_SYNONYMOU | MODERAT | MISSENSE | Aca/Gca | T40A | PLES_30871 | C | C | C | C | C | C | . | . | . | . | . | . |
| NC_011770 | 3408930 | AG | A | 46 | 6 | 1 | 0 | FRAME_SHIFT | HIGH | | ctg/ | L82 | PLES_30881 | | A | | | | | . | . | . | . | . | . |
| NC_011770 | 3408656 | T | G | 47 | 6 | 2 | 0 | NON_SYNONYMOU | MODERAT | MISSENSE | caA/caC | Q173H | PLES_30881 | G | G | | | | | . | . | . | . | . | . |
| NC_011770 | 3409845 | T | C | 51 | 6 | 2 | 0 | NON_SYNONYMOU | MODERAT | MISSENSE | Tcc/Ccc | S70P | PLES_30891 | C | C | | | | | . | . | . | . | . | . |
| NC_011770 | 3411956 | A | G | 48 | 6 | 4 | 0 | NON_SYNONYMOU | MODERAT | MISSENSE | Agc/Ggc | S405G | PLES_30901 | | | G | G | G | G | . | . | . | . | . | . |
| NC_011770 | 3412351 | T | A | 42 | 6 | 6 | 0 | NON_SYNONYMOU | MODERAT | MISSENSE | Tcg/Acg | S76T | PLES_30911 | A | A | A | A | A | A | . | . | . | . | . | . |
| NC_011770 | 3412951 | G | A | 35 | 6 | 4 | 0 | NON_SYNONYMOU | MODERAT | MISSENSE | Gcc/Acc | A276T | PLES_30911 | | | A | A | A | A | . | . | . | . | . | . |
| NC_011770 | 3415550 | CG | C | 43 | 6 | 2 | 0 | FRAME_SHIFT | HIGH | | gcg/ | A385 | PLES_30931 | | | C | C | | | . | . | . | . | . | . |
| NC_011770 | 3414903 | G | A | 45 | 6 | 6 | 0 | NON_SYNONYMOU | MODERAT | MISSENSE | Ggc/Agc | G168S | PLES_30931 | A | A | A | A | A | A | . | . | . | . | . | . |
| NC_011770 | 3415369 | T | C | 44 | 6 | 6 | 0 | NON_SYNONYMOU | MODERAT | MISSENSE | gTc/gCc | V323A | PLES_30931 | C | C | C | C | C | C | . | . | . | . | . | . |
| NC_011770 | 3415535 | G | T | 43 | 6 | 6 | 0 | NON_SYNONYMOU | MODERAT | MISSENSE | gaG/gaT | E378D | PLES_30931 | T | T | T | T | T | T | . | . | . | . | . | . |
| NC_011770 | 3415920 | C | A | 47 | 6 | 6 | 0 | NON_SYNONYMOU | MODERAT | MISSENSE | Cgc/Agc | R37S | PLES_30941 | A | A | A | A | A | A | . | . | . | . | . | . |
| NC_011770 | 3416004 | A | G | 46 | 6 | 6 | 0 | NON_SYNONYMOU | MODERAT | MISSENSE | Acc/Gcc | T65A | PLES_30941 | G | G | G | G | G | G | . | . | . | . | . | . |
| NC_011770 | 3416041 | A | T | 46 | 6 | 2 | 0 | NON_SYNONYMOU | MODERAT | MISSENSE | cAg/cTg | Q77L | PLES_30941 | T | T | | | | | . | . | . | . | . | . |
| NC_011770 | 3416619 | G | A | 38 | 6 | 2 | 0 | NON_SYNONYMOU | MODERAT | MISSENSE | Gcc/Acc | A270T | PLES_30941 | A | A | | | | | . | . | . | . | . | . |
| NC_011770 | 3417675 | G | A | 44 | 6 | 2 | 0 | NON_SYNONYMOU | MODERAT | MISSENSE | Gtc/Atc | V132I | PLES_30951 | A | A | | | | | . | . | . | . | . | . |
| NC_011770 | 3418766 | C | T | 44 | 6 | 2 | 0 | NON_SYNONYMOU | MODERAT | MISSENSE | gCg/gTg | A48V | PLES_30961 | T | T | | | | | . | . | . | . | . | . |
| NC_011770 | 3419498 | G | A | 51 | 6 | 2 | 0 | NON_SYNONYMOU | MODERAT | MISSENSE | cGc/cAc | R292H | PLES_30961 | A | A | | | | | . | . | . | . | . | . |
| NC_011770 | 3420531 | G | A | 7 | 6 | 2 | 0 | NON_SYNONYMOU | MODERAT | MISSENSE | Ccc/Tcc | P326S | PLES_30971 | A | A | | | | | . | . | . | . | . | . |
| NC_011770 | 3421457 | A | G | 43 | 6 | 2 | 0 | NON_SYNONYMOU | MODERAT | MISSENSE | tTg/tCg | L17S | PLES_30971 | G | G | | | | | . | . | . | . | . | . |
| NC_011770 | 3421497 | G | A | 42 | 6 | 1 | 0 | NON_SYNONYMOU | MODERAT | MISSENSE | Ctc/Ttc | L4F | PLES_30971 | | | | A | | | . | . | . | . | . | . |
| NC_011770 | 3421679 | C | G | 42 | 6 | 6 | 0 | NON_SYNONYMOU | MODERAT | MISSENSE | Gtg/Ctg | V50L | PLES_30981 | G | G | G | G | G | G | . | . | . | . | . | . |
| NC_011770 | 3421902 | GCCT | G | 35 | 6 | 2 | 0 | CODON_CHANGE_ | MODERATE | | gaggcc/gc | EA543A | PLES_30991 | | | G | G | | | . | . | . | . | . | . |
| NC_011770 | 3422857 | T | C | 46 | 6 | 6 | 0 | NON_SYNONYMOU | MODERAT | MISSENSE | Atc/Gtc | I226V | PLES_30991 | C | C | C | C | C | C | . | . | . | . | . | . |
| NC_011770 | 3422481 | G | A | 41 | 6 | 1 | 0 | NON_SYNONYMOU | MODERAT | MISSENSE | gCc/gTc | A351V | PLES_30991 | | A | | | | | . | . | . | . | . | . |
| NC_011770 | 3423757 | T | C | 27 | 6 | 4 | 0 | SPLICE_SITE_REGI | LOW | SILENT | taA/taG | *256 | PLES_31001 | | | C | C | C | C | . | . | . | . | . | . |
| NC_011770 | 3425248 | G | A | 39 | 6 | 2 | 0 | NON_SYNONYMOU | MODERAT | MISSENSE | aCc/aTc | T150I | PLES_31011 | A | A | | | | | . | . | . | . | . | . |
| NC_011770 | 3425402 | C | T | 37 | 6 | 4 | 0 | NON_SYNONYMOU | MODERAT | MISSENSE | Ggc/Agc | G99S | PLES_31011 | | | T | T | T | T | . | . | . | . | . | . |
| NC_011770 | 3426060 | C | G | 38 | 6 | 6 | 0 | NON_SYNONYMOU | MODERAT | MISSENSE | tGg/tCg | W15S | PLES_31021 | G | G | G | G | G | G | . | . | . | . | . | . |
| NC_011770 | 3426512 | A | G | 45 | 6 | 2 | 0 | NON_SYNONYMOU | MODERAT | MISSENSE | aTc/aCc | I201T | PLES_31031 | G | G | | | | | . | . | . | . | . | . |
| NC_011770 | 3426516 | T | C | 46 | 6 | 2 | 0 | NON_SYNONYMOU | MODERAT | MISSENSE | Agc/Ggc | S200G | PLES_31031 | C | C | | | | | . | . | . | . | . | . |

| | | | | | | | | | | | | | | | | | | | | | | | | | |
|---|---|---|---|---|---|---|---|---|---|---|---|---|---|---|---|---|---|---|---|---|---|---|---|---|---|
| NC_011770 | 3426521 | C | T | 45 | 6 | 2 | 0 | NON_SYNONYMOU | MODERAT | MISSENSE | cGc/cAc | R198H | PLES_31031 | T | T | | | | | . | . | . | . | . | . |
| NC_011770 | 3426596 | T | C | 48 | 6 | 2 | 0 | NON_SYNONYMOU | MODERAT | MISSENSE | cAc/cGc | H173R | PLES_31031 | C | C | | | | | . | . | . | . | . | . |
| NC_011770 | 3426887 | T | G | 37 | 6 | 6 | 0 | NON_SYNONYMOU | MODERAT | MISSENSE | gAg/gCg | E76A | PLES_31031 | G | G | G | G | G | G | . | . | . | . | . | . |
| NC_011770 | 3428146 | T | TGGC | 44 | 6 | 4 | 0 | CODON_CHANGE_ | MODERATE | | ggt/ggCGG | G267GG | PLES_31041 | | | TGGC | TGGC | TGGC | TGGC | . | . | . | . | . | . |
| NC_011770 | 3428759 | A | G | 43 | 6 | 6 | 0 | NON_SYNONYMOU | MODERAT | MISSENSE | Atc/Gtc | I470V | PLES_31041 | G | G | G | G | G | G | . | . | . | . | . | . |
| NC_011770 | 3428807 | C | T | 41 | 6 | 6 | 0 | NON_SYNONYMOU | MODERAT | MISSENSE | Ctt/Ttt | L486F | PLES_31041 | T | T | T | T | T | T | . | . | . | . | . | . |
| NC_011770 | 3431787 | A | AG | 41 | 6 | 1 | 0 | FRAME_SHIFT | HIGH | | ctt/cCtt | L147P? | PLES_31061 | | AG | | | | | . | . | . | . | . | . |
| NC_011770 | 3429946 | T | G | 33 | 6 | 2 | 0 | NON_SYNONYMOU | MODERAT | MISSENSE | Agc/Cgc | S761R | PLES_31061 | G | G | | | | | . | . | . | . | . | . |
| NC_011770 | 3430347 | T | C | 63 | 0 | 2 | 0 | NON_SYNONYMOU | MODERAT | MISSENSE | aAc/aGc | N627S | PLES_31061 | | | | | | | C | C | | | | |
| NC_011770 | 3431292 | C | T | 39 | 1 | 4 | 0 | NON_SYNONYMOU | MODERAT | MISSENSE | gGc/gAc | G312D | PLES_31061 | | | T | T | T | T | | | | . | | |
| NC_011770 | 3432069 | A | G | 44 | 6 | 6 | 0 | NON_SYNONYMOU | MODERAT | MISSENSE | gTg/gCg | V53A | PLES_31061 | G | G | G | G | G | G | . | . | . | . | . | . |
| NC_011770 | 3433505 | T | C | 24 | 5 | 1 | 0 | NON_SYNONYMOU | MODERAT | MISSENSE | Att/Gtt | I526V | PLES_31081 | | | | | | | C | . | . | . | . | . |
| NC_011770 | 3433801 | G | C | 56 | 0 | 2 | 0 | NON_SYNONYMOU | MODERAT | MISSENSE | gCg/gGg | A427G | PLES_31081 | | | | | | | C | C | | | | |
| NC_011770 | 3434017 | T | C | 56 | 0 | 10 | 0 | NON_SYNONYMOU | MODERAT | MISSENSE | cAc/cGc | H355R | PLES_31081 | C | C | C | C | C | C | C | C | C | C | | |
| NC_011770 | 3434627 | G | A | 61 | 0 | 4 | 0 | NON_SYNONYMOU | MODERAT | MISSENSE | Ccg/Tcg | P152S | PLES_31081 | | | A | A | A | A | | | | | | |
| NC_011770 | 3434855 | C | T | 59 | 0 | 4 | 0 | NON_SYNONYMOU | MODERAT | MISSENSE | Gcc/Acc | A76T | PLES_31081 | | | T | T | T | T | | | | | | |
| NC_011770 | 3433981 | C | T | 60 | 0 | 1 | 0 | NON_SYNONYMOU | MODERAT | MISSENSE | cGc/cAc | R367H | PLES_31081 | | | | | | | | T | | | | |
| NC_011770 | 3434602 | T | C | 62 | 0 | 1 | 0 | NON_SYNONYMOU | MODERAT | MISSENSE | aAc/aGc | N160S | PLES_31081 | | | | | | | | C | | | | |
| NC_011770 | 3436141 | C | G | 81 | 0 | 2 | 0 | NON_SYNONYMOU | MODERAT | MISSENSE | Ggg/Cgg | G22R | PLES_31091 | | | | | | | | | G | G | | |
| NC_011770 | 3437134 | C | T | 80 | 0 | 1 | 0 | NON_SYNONYMOU | MODERAT | MISSENSE | Gag/Aag | E92K | PLES_31101 | T | | | | | | | | | | | |
| NC_011770 | 3437497 | C | T | 80 | 0 | 2 | 0 | NON_SYNONYMOU | MODERAT | MISSENSE | Gcc/Acc | A422T | PLES_31111 | T | T | | | | | | | | | | |
| NC_011770 | 3437751 | A | G | 71 | 0 | 2 | 0 | NON_SYNONYMOU | MODERAT | MISSENSE | cTg/cCg | L337P | PLES_31111 | G | G | | | | | | | | | | |
| NC_011770 | 3438520 | G | A | 91 | 0 | 1 | 0 | NON_SYNONYMOU | MODERAT | MISSENSE | Cac/Tac | H81Y | PLES_31111 | | | | | | | A | | | | | |
| NC_011770 | 3439839 | G | A | 63 | 0 | 2 | 0 | STOP_GAINED | HIGH | NONSENSE | Cga/Tga | R120* | PLES_31121 | | | | | | | | | | | A | A |
| NC_011770 | 3440159 | G | T | 74 | 0 | 2 | 0 | STOP_GAINED | HIGH | NONSENSE | tCg/tAg | S13* | PLES_31121 | | | T | T | | | | | | | | |
| NC_011770 | 3439007 | T | G | 85 | 0 | 2 | 0 | NON_SYNONYMOU | MODERAT | MISSENSE | aAc/aCc | N397T | PLES_31121 | G | G | | | | | | | | | | |
| NC_011770 | 3439142 | C | T | 78 | 0 | 2 | 0 | NON_SYNONYMOU | MODERAT | MISSENSE | aGc/aAc | S352N | PLES_31121 | | | | | | | | | | | T | T |
| NC_011770 | 3439316 | C | T | 72 | 0 | 2 | 0 | NON_SYNONYMOU | MODERAT | MISSENSE | cGc/cAc | R294H | PLES_31121 | T | T | | | | | | | | | | |
| NC_011770 | 3439390 | T | A | 75 | 0 | 8 | 0 | NON_SYNONYMOU | MODERAT | MISSENSE | gaA/gaT | E269D | PLES_31121 | A | A | A | A | A | A | | | | | A | A |
| NC_011770 | 3439598 | G | A | 57 | 0 | 2 | 0 | NON_SYNONYMOU | MODERAT | MISSENSE | gCg/gTg | A200V | PLES_31121 | | | | | | | | | | | A | A |
| NC_011770 | 3439814 | T | C | 60 | 0 | 2 | 0 | NON_SYNONYMOU | MODERAT | MISSENSE | aAg/aGg | K128R | PLES_31121 | | | | | | | | | | | C | C |
| NC_011770 | 3439860 | G | A | 61 | 0 | 2 | 0 | NON_SYNONYMOU | MODERAT | MISSENSE | Cgc/Tgc | R113C | PLES_31121 | | | | | | | | | | | A | A |
| NC_011770 | 3439938 | G | T | 63 | 0 | 10 | 0 | NON_SYNONYMOU | MODERAT | MISSENSE | Ctc/Atc | L87I | PLES_31121 | T | T | T | T | T | T | T | T | T | T | | |
| NC_011770 | 3440072 | C | G | 75 | 0 | 4 | 0 | NON_SYNONYMOU | MODERAT | MISSENSE | cGc/cCc | R42P | PLES_31121 | | | G | G | G | G | | | | | | |
| NC_011770 | 3441401 | G | A | 80 | 0 | 2 | 0 | NON_SYNONYMOU | MODERAT | MISSENSE | cCc/cTc | P313L | PLES_31141 | | | | | | | A | A | | | | |
| NC_011770 | 3441742 | C | G | 84 | 0 | 2 | 0 | NON_SYNONYMOU | MODERAT | MISSENSE | gaG/gaC | E199D | PLES_31141 | | | | | | | G | G | | | | |
| NC_011770 | 3442769 | T | G | 68 | 0 | 12 | 0 | NON_SYNONYMOU | MODERAT | MISSENSE | Tcc/Gcc | S145A | PLES_31151 | G | G | G | G | G | G | G | G | G | G | G | G |
| NC_011770 | 3442919 | G | A | 65 | 0 | 6 | 0 | NON_SYNONYMOU | MODERAT | MISSENSE | Gcc/Acc | A195T | PLES_31151 | | | | | | | A | A | A | A | A | A |
| NC_011770 | 3442995 | T | C | 72 | 0 | 6 | 0 | NON_SYNONYMOU | MODERAT | MISSENSE | gTc/gCc | V220A | PLES_31151 | C | C | C | C | C | C | | | | | | |
| NC_011770 | 3443351 | G | A | 72 | 0 | 4 | 0 | NON_SYNONYMOU | MODERAT | MISSENSE | Gcc/Acc | A339T | PLES_31151 | | | A | A | A | A | | | | | | |
| NC_011770 | 3443557 | C | G | 73 | 0 | 2 | 0 | NON_SYNONYMOU | MODERAT | MISSENSE | gaC/gaG | D407E | PLES_31151 | | | | | | | | | G | G | | |
| NC_011770 | 3443716 | G | C | 69 | 0 | 8 | 0 | NON_SYNONYMOU | MODERAT | MISSENSE | caG/caC | Q460H | PLES_31151 | C | C | | | | | C | C | C | C | C | C |
| NC_011770 | 3443733 | C | T | 70 | 0 | 12 | 0 | NON_SYNONYMOU | MODERAT | MISSENSE | tCc/tTc | S466F | PLES_31151 | T | T | T | T | T | T | T | T | T | T | T | T |
| NC_011770 | 3443989 | G | A | 77 | 0 | 6 | 0 | NON_SYNONYMOU | MODERAT | MISSENSE | Gcc/Acc | A51T | PLES_31161 | | | | | | | A | A | A | A | A | A |
| NC_011770 | 3445168 | T | A | 83 | 0 | 2 | 0 | NON_SYNONYMOU | MODERAT | MISSENSE | Ttg/Atg | L69M | PLES_31171 | | | | | | | A | A | | | | |
| NC_011770 | 3445172 | C | T | 82 | 0 | 4 | 0 | NON_SYNONYMOU | MODERAT | MISSENSE | gCc/gTc | A70V | PLES_31171 | | | T | T | T | T | | | | | | |
| NC_011770 | 3445196 | G | A | 80 | 0 | 2 | 0 | NON_SYNONYMOU | MODERAT | MISSENSE | gGt/gAt | G78D | PLES_31171 | A | A | | | | | | | | | | |
| NC_011770 | 3445213 | C | T | 81 | 0 | 2 | 0 | NON_SYNONYMOU | MODERAT | MISSENSE | Cgc/Tgc | R84C | PLES_31171 | T | T | | | | | | | | | | |
| NC_011770 | 3445216 | A | G | 76 | 0 | 2 | 0 | NON_SYNONYMOU | MODERAT | MISSENSE | Acg/Gcg | T85A | PLES_31171 | | | | | | | G | G | | | | |
| NC_011770 | 3445457 | C | T | 77 | 0 | 2 | 0 | NON_SYNONYMOU | MODERAT | MISSENSE | gCc/gTc | A165V | PLES_31171 | | | | | | | | | | | T | T |
| NC_011770 | 3445539 | G | T | 84 | 0 | 4 | 0 | NON_SYNONYMOU | MODERAT | MISSENSE | Gcc/Tcc | A17S | PLES_31181 | | | T | T | T | T | | | | | | |
| NC_011770 | 3446375 | G | C | 87 | 0 | 6 | 0 | NON_SYNONYMOU | MODERAT | MISSENSE | aaG/aaC | K295N | PLES_31181 | | | | | | | C | C | C | C | C | C |
| NC_011770 | 3446907 | G | A | 77 | 0 | 8 | 0 | NON_SYNONYMOU | MODERAT | MISSENSE | Ggc/Agc | G473S | PLES_31181 | A | A | A | A | A | A | A | A | | | | |
| NC_011770 | 3450896 | C | T | 79 | 0 | 2 | 0 | NON_SYNONYMOU | MODERAT | MISSENSE | Ccg/Tcg | P137S | PLES_31231 | T | T | | | | | | | | | | |
| NC_011770 | 3450825 | A | G | 74 | 0 | 1 | 0 | NON_SYNONYMOU | MODERAT | MISSENSE | cAc/cGc | H113R | PLES_31231 | | | | | | | | G | | | | |
| NC_011770 | 3451197 | G | T | 65 | 0 | 2 | 0 | NON_SYNONYMOU | MODERAT | MISSENSE | gCg/gAg | A290E | PLES_31241 | T | T | | | | | | | | | | |
| NC_011770 | 3451263 | C | T | 67 | 0 | 2 | 0 | NON_SYNONYMOU | MODERAT | MISSENSE | gGc/gAc | G268D | PLES_31241 | | | | | | | | | | | T | T |
| NC_011770 | 3451630 | A | G | 81 | 0 | 1 | 0 | NON_SYNONYMOU | MODERAT | MISSENSE | Tat/Cat | Y146H | PLES_31241 | | | | | | | G | | | | | |
| NC_011770 | 3451689 | C | T | 78 | 0 | 2 | 0 | NON_SYNONYMOU | MODERAT | MISSENSE | cGc/cAc | R126H | PLES_31241 | | | | | | | T | T | | | | |
| NC_011770 | 3452632 | C | T | 76 | 0 | 2 | 0 | NON_SYNONYMOU | MODERAT | MISSENSE | cGc/cAc | R385H | PLES_31251 | | | | | | | | | | | T | T |
| NC_011770 | 3453166 | C | T | 61 | 0 | 2 | 0 | NON_SYNONYMOU | MODERAT | MISSENSE | gGc/gAc | G207D | PLES_31251 | | | | | | | T | T | | | | |
| NC_011770 | 3453259 | T | G | 64 | 0 | 2 | 0 | NON_SYNONYMOU | MODERAT | MISSENSE | gAc/gCc | D176A | PLES_31251 | | | | | | | | | | | G | G |

| | | | | | | | | | | | | | | | | | | | | | | | | | |
|---|---|---|---|---|---|---|---|---|---|---|---|---|---|---|---|---|---|---|---|---|---|---|---|---|---|
| NC_011770 | 3453368 | T | A | 64 | 0 | 4 | 0 | NON_SYNONYMOU | MODERAT | MISSENSE | Acg/Tcg | T140S | PLES_31251 | | | | | | | A | A | | | A | A |
| NC_011770 | 3453535 | C | T | 71 | 0 | 4 | 0 | NON_SYNONYMOU | MODERAT | MISSENSE | cGg/cAg | R84Q | PLES_31251 | | | | | | | T | T | | | T | T |
| NC_011770 | 3453689 | C | T | 69 | 0 | 8 | 0 | NON_SYNONYMOU | MODERAT | MISSENSE | Gtc/Atc | V33I | PLES_31251 | | | T | T | T | T | T | T | T | T | | |
| NC_011770 | 3453698 | C | T | 69 | 0 | 6 | 0 | NON_SYNONYMOU | MODERAT | MISSENSE | Gtc/Atc | V30I | PLES_31251 | | | T | T | T | T | | | T | T | | |
| NC_011770 | 3454292 | G | A | 74 | 0 | 2 | 0 | STOP_GAINED | HIGH | NONSENSE | Cag/Tag | Q153* | PLES_31261 | A | A | | | | | | | | | | |
| NC_011770 | 3455859 | C | T | 81 | 0 | 2 | 0 | NON_SYNONYMOU | MODERAT | MISSENSE | Gtc/Atc | V92I | PLES_31281 | | | | | | | | | T | T | | |
| NC_011770 | 3456418 | G | A | 72 | 0 | 2 | 0 | NON_SYNONYMOU | MODERAT | MISSENSE | Cgc/Tgc | R144C | PLES_31291 | | | | | | | A | A | | | | |
| NC_011770 | 3456592 | T | A | 71 | 0 | 2 | 0 | NON_SYNONYMOU | MODERAT | MISSENSE | Atg/Ttg | M86L | PLES_31291 | | | | | | | | | | | A | A |
| NC_011770 | 3456655 | C | T | 69 | 0 | 2 | 0 | NON_SYNONYMOU | MODERAT | MISSENSE | Gtc/Atc | V65I | PLES_31291 | | | | | | | | | | | T | T |
| NC_011770 | 3456741 | C | G | 67 | 0 | 6 | 0 | NON_SYNONYMOU | MODERAT | MISSENSE | aGc/aCc | S36T | PLES_31291 | G | G | | | | | G | G | | | G | G |
| NC_011770 | 3456406 | A | G | 70 | 0 | 1 | 0 | NON_SYNONYMOU | MODERAT | MISSENSE | Tcc/Ccc | S148P | PLES_31291 | | G | | | | | | | | | | |
| NC_011770 | 3460436 | C | T | 60 | 0 | 4 | 0 | NON_SYNONYMOU | MODERAT | MISSENSE | Gcc/Acc | A122T | PLES_31331 | | | T | T | T | T | | | | | | |
| NC_011770 | 3460538 | C | T | 70 | 0 | 4 | 0 | NON_SYNONYMOU | MODERAT | MISSENSE | Gcg/Acg | A88T | PLES_31331 | | | T | T | T | T | | | | | | |
| NC_011770 | 3460573 | C | T | 73 | 0 | 2 | 0 | NON_SYNONYMOU | MODERAT | MISSENSE | cGc/cAc | R76H | PLES_31331 | T | T | | | | | | | | | | |
| NC_011770 | 3460706 | T | C | 73 | 0 | 2 | 0 | NON_SYNONYMOU | MODERAT | MISSENSE | Atc/Gtc | I32V | PLES_31331 | | | | | | | C | C | | | | |
| NC_011770 | 3460757 | G | T | 63 | 0 | 12 | 0 | NON_SYNONYMOU | MODERAT | MISSENSE | Ccc/Acc | P15T | PLES_31331 | T | T | T | T | T | T | T | T | T | T | T | T |
| NC_011770 | 3463836 | G | A | 76 | 0 | 1 | 0 | NON_SYNONYMOU | MODERAT | MISSENSE | Ggc/Agc | G59S | PLES_31371 | | | | | | | A | | | | | |
| NC_011770 | 3464349 | A | T | 71 | 0 | 6 | 0 | NON_SYNONYMOU | MODERAT | MISSENSE | Acc/Tcc | T230S | PLES_31371 | T | T | | | | | | | T | T | T | T |
| NC_011770 | 3464855 | T | G | 79 | 0 | 10 | 0 | NON_SYNONYMOU | MODERAT | MISSENSE | gAc/gCc | D140A | PLES_31381 | G | G | G | G | G | G | | | G | G | G | G |
| NC_011770 | 3465168 | G | A | 70 | 0 | 10 | 0 | NON_SYNONYMOU | MODERAT | MISSENSE | Ctc/Ttc | L36F | PLES_31381 | A | A | A | A | A | A | | | A | A | A | A |
| NC_011770 | 3465440 | G | T | 54 | 0 | 4 | 0 | NON_SYNONYMOU | MODERAT | MISSENSE | Ccg/Acg | P37T | PLES_31391 | | | T | T | T | T | | | | | | |
| NC_011770 | 3465445 | G | A | 54 | 0 | 1 | 0 | NON_SYNONYMOU | MODERAT | MISSENSE | gCg/gTg | A35V | PLES_31391 | | | | | | | | | | | | A |
| NC_011770 | 3467226 | A | G | 76 | 0 | 2 | 0 | NON_SYNONYMOU | MODERAT | MISSENSE | Acc/Gcc | T153A | PLES_31411 | G | G | | | | | | | | | | |
| NC_011770 | 3468400 | C | T | 71 | 0 | 12 | 0 | NON_SYNONYMOU | MODERAT | MISSENSE | cGg/cAg | R320Q | PLES_31441 | T | T | T | T | T | T | T | T | T | T | T | T |
| NC_011770 | 3468468 | T | A | 73 | 0 | 6 | 0 | NON_SYNONYMOU | MODERAT | MISSENSE | gaA/gaT | E297D | PLES_31441 | | | A | A | A | A | A | A | | | | |
| NC_011770 | 3468556 | C | T | 73 | 0 | 6 | 0 | NON_SYNONYMOU | MODERAT | MISSENSE | cGt/cAt | R268H | PLES_31441 | | | T | T | T | T | T | T | | | | |
| NC_011770 | 3468562 | T | C | 75 | 0 | 6 | 0 | NON_SYNONYMOU | MODERAT | MISSENSE | gAc/gGc | D266G | PLES_31441 | | | C | C | C | C | C | C | | | | |
| NC_011770 | 3468568 | T | C | 74 | 0 | 12 | 0 | NON_SYNONYMOU | MODERAT | MISSENSE | gAc/gGc | D264G | PLES_31441 | C | C | C | C | C | C | C | C | C | C | C | C |
| NC_011770 | 3468725 | T | C | 69 | 0 | 2 | 0 | NON_SYNONYMOU | MODERAT | MISSENSE | Acc/Gcc | T212A | PLES_31441 | | | | | | | | | | | C | C |
| NC_011770 | 3469098 | G | T | 61 | 0 | 2 | 0 | NON_SYNONYMOU | MODERAT | MISSENSE | gaC/gaA | D87E | PLES_31441 | | | | | | | | | | | T | T |
| NC_011770 | 3469336 | C | T | 49 | 0 | 6 | 0 | NON_SYNONYMOU | MODERAT | MISSENSE | aGc/aAc | S8N | PLES_31441 | | | T | T | T | T | T | T | | | | |
| NC_011770 | 3469809 | G | C | 44 | 0 | 2 | 0 | NON_SYNONYMOU | MODERAT | MISSENSE | cGa/cCa | R89P | PLES_31451 | | | | | | | C | C | | | | |
| NC_011770 | 3470127 | A | G | 71 | 0 | 1 | 0 | NON_SYNONYMOU | MODERAT | MISSENSE | tAc/tGc | Y195C | PLES_31451 | | | | | | | G | | | | | |
| NC_011770 | 3470201 | C | T | 74 | 0 | 8 | 0 | NON_SYNONYMOU | MODERAT | MISSENSE | Ccc/Tcc | P220S | PLES_31451 | | | T | T | T | T | T | T | | | T | T |
| NC_011770 | 3470273 | C | G | 84 | 0 | 2 | 0 | NON_SYNONYMOU | MODERAT | MISSENSE | Ctc/Gtc | L244V | PLES_31451 | | | | | | | | | | | G | G |
| NC_011770 | 3470841 | G | T | 67 | 0 | 4 | 0 | NON_SYNONYMOU | MODERAT | MISSENSE | cGc/cTc | R433L | PLES_31451 | | | | | | | | | T | T | T | T |
| NC_011770 | 3470916 | A | C | 68 | 0 | 4 | 0 | NON_SYNONYMOU | MODERAT | MISSENSE | gAa/gCa | E458A | PLES_31451 | | | | | | | | | C | C | C | C |
| NC_011770 | 3471075 | G | C | 64 | 0 | 2 | 0 | NON_SYNONYMOU | MODERAT | MISSENSE | Gca/Cca | A58P | PLES_31461 | C | C | | | | | | | | | | |
| NC_011770 | 3471550 | C | T | 69 | 0 | 2 | 0 | NON_SYNONYMOU | MODERAT | MISSENSE | gCc/gTc | A216V | PLES_31461 | | | | | | | | | T | T | | |
| NC_011770 | 3471612 | G | A | 69 | 0 | 2 | 0 | NON_SYNONYMOU | MODERAT | MISSENSE | Gtc/Atc | V237I | PLES_31461 | | | | | | | | | | | A | A |
| NC_011770 | 3471846 | T | C | 74 | 0 | 6 | 0 | NON_SYNONYMOU | MODERAT | MISSENSE | Tcc/Ccc | S315P | PLES_31461 | C | C | | | | | | | C | C | C | C |
| NC_011770 | 3472798 | G | A | 74 | 0 | 1 | 0 | STOP_GAINED | HIGH | NONSENSE | tgG/tgA | W190* | PLES_31471 | | | | | | | A | | | | | |
| NC_011770 | 3472265 | A | T | 77 | 0 | 10 | 0 | NON_SYNONYMOU | MODERAT | MISSENSE | Acc/Tcc | T13S | PLES_31471 | T | T | T | T | T | T | T | T | | | T | T |
| NC_011770 | 3472498 | A | C | 75 | 0 | 6 | 0 | NON_SYNONYMOU | MODERAT | MISSENSE | ttA/ttC | L90F | PLES_31471 | | | C | C | C | C | C | C | | | | |
| NC_011770 | 3472793 | T | C | 72 | 0 | 2 | 0 | NON_SYNONYMOU | MODERAT | MISSENSE | Ttc/Ctc | F189L | PLES_31471 | | | | | | | | | C | C | | |
| NC_011770 | 3473224 | C | G | 71 | 0 | 8 | 0 | NON_SYNONYMOU | MODERAT | MISSENSE | Gcg/Ccg | A573P | PLES_31481 | | | G | G | G | G | G | G | G | G | | |
| NC_011770 | 3473277 | G | A | 72 | 0 | 2 | 0 | NON_SYNONYMOU | MODERAT | MISSENSE | gCc/gTc | A555V | PLES_31481 | | | | | | | | | | | A | A |
| NC_011770 | 3474112 | T | C | 66 | 0 | 2 | 0 | NON_SYNONYMOU | MODERAT | MISSENSE | Agt/Ggt | S277G | PLES_31481 | | | | | | | | | | | C | C |
| NC_011770 | 3474172 | T | C | 64 | 0 | 12 | 0 | NON_SYNONYMOU | MODERAT | MISSENSE | Acc/Gcc | T257A | PLES_31481 | C | C | C | C | C | C | C | C | C | C | C | C |
| NC_011770 | 3474918 | T | C | 65 | 0 | 12 | 0 | NON_SYNONYMOU | MODERAT | MISSENSE | gAg/gGg | E8G | PLES_31481 | C | C | C | C | C | C | C | C | C | C | C | C |
| NC_011770 | 3474076 | G | A | 66 | 0 | 1 | 0 | NON_SYNONYMOU | MODERAT | MISSENSE | Cgc/Tgc | R289C | PLES_31481 | | | | | | | | | | | A | |
| NC_011770 | 3475387 | A | G | 69 | 0 | 2 | 0 | NON_SYNONYMOU | MODERAT | MISSENSE | gAg/gGg | E122G | PLES_31491 | G | G | | | | | | | | | | |
| NC_011770 | 3476263 | C | T | 42 | 0 | 2 | 0 | NON_SYNONYMOU | MODERAT | MISSENSE | Ctt/Ttt | L3F | PLES_31511 | | | | | | | | | | | T | T |
| NC_011770 | 3476328 | G | T | 37 | 0 | 2 | 0 | NON_SYNONYMOU | MODERAT | MISSENSE | aaG/aaT | K24N | PLES_31511 | | | | | | | | | T | T | | |
| NC_011770 | 3476372 | T | A | 33 | 0 | 12 | 0 | NON_SYNONYMOU | MODERAT | MISSENSE | cTg/cAg | L39Q | PLES_31511 | A | A | A | A | A | A | A | A | A | A | A | A |
| NC_011770 | 3476635 | T | TTCATG | 19 | 0 | 5 | 0 | FRAME_SHIFT | HIGH | | gaa/gaCAT | E62DM? | PLES_31521 | TTCATG | . | C | C | C | C | | | | | | |
| NC_011770 | 3476972 | T | G | 56 | 0 | 12 | 0 | NON_SYNONYMOU | MODERAT | MISSENSE | caA/caC | Q85H | PLES_31531 | G | G | G | G | G | G | G | G | G | G | G | G |
| NC_011770 | 3477016 | C | G | 66 | 0 | 2 | 0 | NON_SYNONYMOU | MODERAT | MISSENSE | Gaa/Caa | E71Q | PLES_31531 | | | | | | | | | G | G | | |
| NC_011770 | 3477495 | C | T | 89 | 0 | 2 | 0 | NON_SYNONYMOU | MODERAT | MISSENSE | Gtc/Atc | V279I | PLES_31541 | | | | | | | | | | | T | T |
| NC_011770 | 3477627 | C | T | 84 | 0 | 2 | 0 | NON_SYNONYMOU | MODERAT | MISSENSE | Gcc/Acc | A235T | PLES_31541 | | | | | | | T | T | | | | |
| NC_011770 | 3477641 | G | A | 84 | 0 | 2 | 0 | NON_SYNONYMOU | MODERAT | MISSENSE | gCc/gTc | A230V | PLES_31541 | | | | | | | A | A | | | | |
| NC_011770 | 3478078 | C | G | 63 | 0 | 2 | 0 | NON_SYNONYMOU | MODERAT | MISSENSE | gaG/gaC | E84D | PLES_31541 | | | | | | | G | G | | | | |

| | | | | | | | | | | | | | | | | | | | | | | | | | |
|---|---|---|---|---|---|---|---|---|---|---|---|---|---|---|---|---|---|---|---|---|---|---|---|---|---|
| NC_011770 | 3478510 | C | G | 63 | 0 | 12 | 0 | NON_SYNONYMOU | MODERAT | MISSENSE | ttG/ttC | L98F | PLES_31551 | G | G | G | G | G | G | G | G | G | G | G | G |
| NC_011770 | 3479023 | C | A | 54 | 0 | 6 | 0 | NON_SYNONYMOU | MODERAT | MISSENSE | tGg/tTg | W22L | PLES_31561 | | | A | A | A | A | A | A | | | | |
| NC_011770 | 3479212 | A | G | 58 | 0 | 2 | 0 | NON_SYNONYMOU | MODERAT | MISSENSE | Tcg/Ccg | S119P | PLES_31571 | | | | | | | G | G | | | | |
| NC_011770 | 3480045 | T | C | 68 | 0 | 1 | 0 | NON_SYNONYMOU | MODERAT | MISSENSE | Acg/Gcg | T142A | PLES_31581 | | | | | | | C | | | | | |
| NC_011770 | 3480997 | G | A | 75 | 0 | 2 | 0 | NON_SYNONYMOU | MODERAT | MISSENSE | gCc/gTc | A189V | PLES_31591 | | | | | | | | | A | A | | |
| NC_011770 | 3481039 | C | T | 77 | 0 | 2 | 0 | NON_SYNONYMOU | MODERAT | MISSENSE | gGg/gAg | G175E | PLES_31591 | | | | | | | | | T | T | | |
| NC_011770 | 3481051 | G | C | 74 | 0 | 2 | 0 | NON_SYNONYMOU | MODERAT | MISSENSE | gCt/gGt | A171G | PLES_31591 | | | | | | | | | C | C | | |
| NC_011770 | 3484282 | C | T | 85 | 0 | 4 | 0 | NON_SYNONYMOU | MODERAT | MISSENSE | Ccg/Tcg | P40S | PLES_31621 | | | T | T | T | T | | | | | | |
| NC_011770 | 3484885 | C | A | 91 | 0 | 2 | 0 | NON_SYNONYMOU | MODERAT | MISSENSE | Ctg/Atg | L241M | PLES_31621 | | | | | | | | | A | A | | |
| NC_011770 | 3484926 | C | G | 92 | 0 | 2 | 0 | NON_SYNONYMOU | MODERAT | MISSENSE | ttC/ttG | F254L | PLES_31621 | | | | | | | | | | | G | G |
| NC_011770 | 3485347 | G | A | 66 | 0 | 2 | 0 | NON_SYNONYMOU | MODERAT | MISSENSE | Gcg/Acg | A395T | PLES_31621 | | | | | | | A | A | | | | |
| NC_011770 | 3485534 | C | T | 71 | 0 | 2 | 0 | NON_SYNONYMOU | MODERAT | MISSENSE | aCc/aTc | T457I | PLES_31621 | | | | | | | T | T | | | | |
| NC_011770 | 3485806 | G | A | 64 | 0 | 4 | 0 | NON_SYNONYMOU | MODERAT | MISSENSE | Gcg/Acg | A548T | PLES_31621 | | | A | A | A | A | | | | | | |
| NC_011770 | 3485812 | A | G | 65 | 0 | 12 | 0 | NON_SYNONYMOU | MODERAT | MISSENSE | Acc/Gcc | T550A | PLES_31621 | G | G | G | G | G | G | G | G | G | G | G | G |
| NC_011770 | 3485818 | C | T | 68 | 0 | 4 | 0 | NON_SYNONYMOU | MODERAT | MISSENSE | Cgg/Tgg | R552W | PLES_31621 | T | T | | | | | T | T | | | | |
| NC_011770 | 3484814 | G | A | 83 | 0 | 1 | 0 | NON_SYNONYMOU | MODERAT | MISSENSE | cGg/cAg | R217Q | PLES_31621 | | | | | | | | | | | | A |
| NC_011770 | 3486916 | T | A | 72 | 0 | 4 | 0 | NON_SYNONYMOU | MODERAT | MISSENSE | gaT/gaA | D336E | PLES_31631 | | | A | A | A | A | | | | | | |
| NC_011770 | 3488596 | A | G | 71 | 0 | 2 | 0 | NON_SYNONYMOU | MODERAT | MISSENSE | cAa/cGa | Q214R | PLES_31641 | | | | | | | G | G | | | | |
| NC_011770 | 3488842 | T | C | 82 | 0 | 4 | 0 | NON_SYNONYMOU | MODERAT | MISSENSE | gTc/gCc | V296A | PLES_31641 | | | C | C | C | C | | | | | | |
| NC_011770 | 3488962 | A | G | 70 | 0 | 4 | 0 | NON_SYNONYMOU | MODERAT | MISSENSE | cAt/cGt | H336R | PLES_31641 | G | G | | | | | | | | | G | G |
| NC_011770 | 3489480 | C | T | 84 | 0 | 2 | 0 | NON_SYNONYMOU | MODERAT | MISSENSE | Cgc/Tgc | R509C | PLES_31641 | | | | | | | | | T | T | | |
| NC_011770 | 3490224 | G | T | 77 | 0 | 2 | 0 | NON_SYNONYMOU | MODERAT | MISSENSE | Gac/Tac | D757Y | PLES_31641 | | | | | | | T | T | | | | |
| NC_011770 | 3490146 | G | A | 79 | 0 | 1 | 0 | NON_SYNONYMOU | MODERAT | MISSENSE | Ggg/Agg | G731R | PLES_31641 | | A | | | | | | | | | | |
| NC_011770 | 3490015 | C | T | 89 | 0 | 1 | 0 | NON_SYNONYMOU | MODERAT | MISSENSE | aCg/aTg | T687M | PLES_31641 | | | | | | | | | | | | T |
| NC_011770 | 3493647 | A | T | 73 | 0 | 2 | 0 | NON_SYNONYMOU | MODERAT | MISSENSE | aAc/aTc | N116I | PLES_31671 | T | T | | | | | | | | | | |
| NC_011770 | 3493662 | G | A | 72 | 0 | 2 | 0 | NON_SYNONYMOU | MODERAT | MISSENSE | cGc/cAc | R121H | PLES_31671 | A | A | | | | | | | | | | |
| NC_011770 | 3495030 | G | A | 86 | 0 | 2 | 0 | SPLICE_SITE_REGIC | LOW | SILENT | tGa/tAa | *416 | PLES_31681 | | | | | | | A | A | | | | |
| NC_011770 | 3494099 | G | C | 74 | 0 | 2 | 0 | NON_SYNONYMOU | MODERAT | MISSENSE | Gtc/Ctc | V106L | PLES_31681 | C | C | | | | | | | | | | |
| NC_011770 | 3494895 | G | A | 74 | 0 | 2 | 0 | NON_SYNONYMOU | MODERAT | MISSENSE | cGc/cAc | R371H | PLES_31681 | | | | | | | A | A | | | | |
| NC_011770 | 3494954 | G | A | 75 | 0 | 10 | 0 | NON_SYNONYMOU | MODERAT | MISSENSE | Ggc/Agc | G391S | PLES_31681 | A | A | A | A | A | A | | | A | A | A | A |
| NC_011770 | 3495374 | C | T | 85 | 0 | 1 | 0 | STOP_GAINED | HIGH | NONSENSE | Cag/Tag | Q114* | PLES_31691 | | | | | | | T | | | | | |
| NC_011770 | 3495304 | C | A | 85 | 0 | 2 | 0 | NON_SYNONYMOU | MODERAT | MISSENSE | gaC/gaA | D90E | PLES_31691 | | | | | | | | | A | A | | |
| NC_011770 | 3495753 | G | A | 82 | 0 | 6 | 0 | NON_SYNONYMOU | MODERAT | MISSENSE | cGc/cAc | R240H | PLES_31691 | | | A | A | A | A | | | | | A | A |
| NC_011770 | 3495764 | G | A | 82 | 0 | 6 | 0 | NON_SYNONYMOU | MODERAT | MISSENSE | Ggc/Agc | G244S | PLES_31691 | | | A | A | A | A | | | | | A | A |
| NC_011770 | 3495771 | G | A | 84 | 0 | 2 | 0 | NON_SYNONYMOU | MODERAT | MISSENSE | gGc/gAc | G246D | PLES_31691 | A | A | | | | | | | | | | |
| NC_011770 | 3495776 | A | G | 83 | 0 | 10 | 0 | NON_SYNONYMOU | MODERAT | MISSENSE | Acc/Gcc | T248A | PLES_31691 | G | G | G | G | G | G | | | G | G | G | G |
| NC_011770 | 3495809 | C | CCGGCG | 81 | 0 | 1 | 0 | CODON_INSERTIOI | MODERATE | | cgc/ACGG( | R265TAR | PLES_31691 | | | | | | | CCGGCGA | | | | | |
| NC_011770 | 3495816 | C | A | 70 | 0 | 10 | 0 | NON_SYNONYMOU | MODERAT | MISSENSE | aCg/aAg | T261K | PLES_31691 | A | A | A | A | A | A | | | A | A | A | A |
| NC_011770 | 3495917 | G | A | 80 | 0 | 2 | 0 | NON_SYNONYMOU | MODERAT | MISSENSE | Ggc/Agc | G295S | PLES_31691 | | | | | | | | | | | A | A |
| NC_011770 | 3496351 | C | T | 67 | 0 | 2 | 0 | NON_SYNONYMOU | MODERAT | MISSENSE | Ccg/Tcg | P128S | PLES_31701 | | | | | | | T | T | | | | |
| NC_011770 | 3496502 | A | G | 62 | 0 | 2 | 0 | NON_SYNONYMOU | MODERAT | MISSENSE | cAa/cGa | Q178R | PLES_31701 | | | | | | | | | | | G | G |
| NC_011770 | 3496503 | A | C | 62 | 0 | 2 | 0 | NON_SYNONYMOU | MODERAT | MISSENSE | caA/caC | Q178H | PLES_31701 | | | | | | | | | | | C | C |
| NC_011770 | 3496959 | A | G | 65 | 0 | 4 | 0 | NON_SYNONYMOU | MODERAT | MISSENSE | Acc/Gcc | T86A | PLES_31711 | G | G | | | | | G | G | | | | |
| NC_011770 | 3496965 | T | G | 67 | 0 | 2 | 0 | NON_SYNONYMOU | MODERAT | MISSENSE | Tcc/Gcc | S88A | PLES_31711 | | | | | | | | | | | G | G |
| NC_011770 | 3497068 | G | A | 68 | 0 | 2 | 0 | NON_SYNONYMOU | MODERAT | MISSENSE | cGg/cAg | R122Q | PLES_31711 | | | | | | | | | | | A | A |
| NC_011770 | 3497311 | G | A | 77 | 0 | 6 | 0 | NON_SYNONYMOU | MODERAT | MISSENSE | cGc/cAc | R203H | PLES_31711 | | | A | A | A | A | | | A | A | | |
| NC_011770 | 3497316 | T | C | 80 | 0 | 1 | 0 | NON_SYNONYMOU | MODERAT | MISSENSE | Tac/Cac | Y205H | PLES_31711 | | | | | | | C | | | | | |
| NC_011770 | 3497634 | G | A | 87 | 0 | 1 | 0 | NON_SYNONYMOU | MODERAT | MISSENSE | Gag/Aag | E311K | PLES_31711 | | | | | | | A | | | | | |
| NC_011770 | 3497653 | A | G | 84 | 0 | 12 | 0 | NON_SYNONYMOU | MODERAT | MISSENSE | cAc/cGc | H317R | PLES_31711 | G | G | G | G | G | G | G | G | G | G | G | G |
| NC_011770 | 3497658 | G | A | 82 | 0 | 4 | 0 | NON_SYNONYMOU | MODERAT | MISSENSE | Gtc/Atc | V319I | PLES_31711 | | | A | A | A | A | | | | | | |
| NC_011770 | 3498066 | G | A | 88 | 0 | 4 | 0 | NON_SYNONYMOU | MODERAT | MISSENSE | gGc/gAc | G54D | PLES_31721 | A | A | | | | | A | A | | | | |
| NC_011770 | 3498291 | A | G | 75 | 0 | 2 | 0 | NON_SYNONYMOU | MODERAT | MISSENSE | gAc/gGc | D129G | PLES_31721 | | | | | | | | | | | G | G |
| NC_011770 | 3498863 | C | T | 93 | 0 | 2 | 0 | NON_SYNONYMOU | MODERAT | MISSENSE | Cgc/Tgc | R320C | PLES_31721 | | | | | | | T | T | | | | |
| NC_011770 | 3498885 | G | A | 92 | 0 | 2 | 0 | NON_SYNONYMOU | MODERAT | MISSENSE | cGc/cAc | R327H | PLES_31721 | | | | | | | | | | | A | A |
| NC_011770 | 3501493 | C | T | 70 | 0 | 2 | 0 | NON_SYNONYMOU | MODERAT | MISSENSE | Gcg/Acg | A970T | PLES_31741 | | | | | | | | | | | T | T |
| NC_011770 | 3501534 | A | G | 77 | 0 | 1 | 0 | NON_SYNONYMOU | MODERAT | MISSENSE | gTc/gCc | V956A | PLES_31741 | | | | | | | G | | | | | |
| NC_011770 | 3501660 | T | C | 73 | 0 | 2 | 0 | NON_SYNONYMOU | MODERAT | MISSENSE | cAg/cGg | Q914R | PLES_31741 | | | | | | | | | | | C | C |
| NC_011770 | 3502545 | T | A | 50 | 0 | 2 | 0 | NON_SYNONYMOU | MODERAT | MISSENSE | aAa/aTa | K619I | PLES_31741 | | | | | | | A | A | | | | |
| NC_011770 | 3502920 | T | C | 60 | 0 | 2 | 0 | NON_SYNONYMOU | MODERAT | MISSENSE | tAc/tGc | Y494C | PLES_31741 | C | C | | | | | | | | | | |
| NC_011770 | 3504427 | C | T | 75 | 0 | 4 | 0 | NON_SYNONYMOU | MODERAT | MISSENSE | cGg/cAg | R660Q | PLES_31751 | | | T | T | T | T | | | | | | |
| NC_011770 | 3505554 | C | A | 60 | 0 | 2 | 0 | NON_SYNONYMOU | MODERAT | MISSENSE | gaG/gaT | E284D | PLES_31751 | | | | | | | A | A | | | | |
| NC_011770 | 3505736 | T | C | 59 | 0 | 2 | 0 | NON_SYNONYMOU | MODERAT | MISSENSE | Aca/Gca | T224A | PLES_31751 | C | C | | | | | | | | | | |

| | | | | | | | | | | | | | | | | | | | | | | | | | |
|---|---|---|---|---|---|---|---|---|---|---|---|---|---|---|---|---|---|---|---|---|---|---|---|---|---|
| NC_011770 | 3506123 | T | G | 66 | 0 | 10 | 0 | NON_SYNONYMOU | MODERAT | MISSENSE | Acc/Ccc | T95P | PLES_31751 | G | G | G | G | G | G | G | G | | | G | G |
| NC_011770 | 3505979 | C | T | 61 | 0 | 1 | 0 | NON_SYNONYMOU | MODERAT | MISSENSE | Gag/Aag | E143K | PLES_31751 | | | | | | | | | | | T | |
| NC_011770 | 3506599 | C | G | 63 | 0 | 2 | 0 | NON_SYNONYMOU | MODERAT | MISSENSE | Gcg/Ccg | A278P | PLES_31761 | G | G | | | | | | | | | | |
| NC_011770 | 3507220 | G | A | 87 | 0 | 1 | 0 | NON_SYNONYMOU | MODERAT | MISSENSE | Cgc/Tgc | R71C | PLES_31761 | | | | | | | A | | | | | |
| NC_011770 | 3506847 | G | A | 76 | 0 | 1 | 0 | NON_SYNONYMOU | MODERAT | MISSENSE | gCc/gTc | A195V | PLES_31761 | | | | | | | | A | | | | |
| NC_011770 | 3507596 | C | A | 75 | 0 | 2 | 0 | NON_SYNONYMOU | MODERAT | MISSENSE | Gcg/Tcg | A34S | PLES_31771 | | | | | | | | | A | A | | |
| NC_011770 | 3507978 | C | T | 84 | 0 | 6 | 0 | NON_SYNONYMOU | MODERAT | MISSENSE | Ggt/Agt | G75S | PLES_31781 | | | | | | | T | T | T | T | T | T |
| NC_011770 | 3515735 | C | G | 74 | 0 | 2 | 0 | NON_SYNONYMOU | MODERAT | MISSENSE | Gag/Cag | E22Q | PLES_31851 | | | | | | | G | G | | | | |
| NC_011770 | 3516300 | T | C | 80 | 0 | 8 | 0 | NON_SYNONYMOU | MODERAT | MISSENSE | Acc/Gcc | T15A | PLES_31861 | C | C | C | C | C | C | | | C | C | | |
| NC_011770 | 3516493 | C | T | 76 | 0 | 4 | 0 | NON_SYNONYMOU | MODERAT | MISSENSE | Ggc/Agc | G34S | PLES_31871 | T | T | | | | | T | T | | | | |
| NC_011770 | 3519481 | T | G | 79 | 0 | 2 | 0 | NON_SYNONYMOU | MODERAT | MISSENSE | Atg/Ctg | M167L | PLES_31901 | | | | | | | | | G | G | | |
| NC_011770 | 3519490 | C | T | 78 | 0 | 2 | 0 | NON_SYNONYMOU | MODERAT | MISSENSE | Gcc/Acc | A164T | PLES_31901 | | | | | | | T | T | | | | |
| NC_011770 | 3519643 | A | G | 81 | 0 | 12 | 0 | NON_SYNONYMOU | MODERAT | MISSENSE | Tct/Cct | S113P | PLES_31901 | G | G | G | G | G | G | G | G | G | G | G | G |
| NC_011770 | 3519701 | G | C | 79 | 0 | 2 | 0 | NON_SYNONYMOU | MODERAT | MISSENSE | gaC/gaG | D93E | PLES_31901 | C | C | | | | | | | | | | |
| NC_011770 | 3519782 | C | G | 67 | 0 | 8 | 0 | NON_SYNONYMOU | MODERAT | MISSENSE | gaG/gaC | E66D | PLES_31901 | | | G | G | G | G | G | G | | | G | G |
| NC_011770 | 3519931 | G | A | 68 | 0 | 8 | 0 | NON_SYNONYMOU | MODERAT | MISSENSE | Ccc/Tcc | P17S | PLES_31901 | | | A | A | A | A | A | A | | | A | A |
| NC_011770 | 3520102 | G | A | 55 | 0 | 2 | 0 | NON_SYNONYMOU | MODERAT | MISSENSE | Gcc/Acc | A8T | PLES_31911 | | | | | | | A | A | | | | |
| NC_011770 | 3521091 | G | C | 71 | 0 | 2 | 0 | NON_SYNONYMOU | MODERAT | MISSENSE | caC/caG | H444Q | PLES_31921 | | | | | | | | | | | C | C |
| NC_011770 | 3521954 | C | T | 79 | 0 | 1 | 0 | NON_SYNONYMOU | MODERAT | MISSENSE | Ggc/Agc | G157S | PLES_31921 | | | | | | | T | | | | | |
| NC_011770 | 3522229 | C | G | 77 | 0 | 2 | 0 | NON_SYNONYMOU | MODERAT | MISSENSE | aGc/aCc | S65T | PLES_31921 | | | | | | | | | | | G | G |
| NC_011770 | 3522230 | T | C | 76 | 0 | 2 | 0 | NON_SYNONYMOU | MODERAT | MISSENSE | Agc/Ggc | S65G | PLES_31921 | | | | | | | | | | | C | C |
| NC_011770 | 3523123 | T | G | 71 | 0 | 6 | 0 | NON_SYNONYMOU | MODERAT | MISSENSE | cAg/cCg | Q284P | PLES_31941 | | | G | G | G | G | | | | | G | G |
| NC_011770 | 3523336 | G | A | 82 | 0 | 2 | 0 | NON_SYNONYMOU | MODERAT | MISSENSE | gCc/gTc | A213V | PLES_31941 | A | A | | | | | | | | | | |
| NC_011770 | 3523545 | C | G | 79 | 0 | 2 | 0 | NON_SYNONYMOU | MODERAT | MISSENSE | agG/agC | R143S | PLES_31941 | | | | | G | G | | | | | | |
| NC_011770 | 3523709 | T | C | 76 | 0 | 6 | 0 | NON_SYNONYMOU | MODERAT | MISSENSE | Ata/Gta | I89V | PLES_31941 | C | C | C | C | C | C | | | | | | |
| NC_011770 | 3523967 | T | C | 64 | 0 | 4 | 0 | NON_SYNONYMOU | MODERAT | MISSENSE | Agt/Ggt | S3G | PLES_31941 | | | C | C | C | C | | | | | | |
| NC_011770 | 3531457 | C | G | 71 | 0 | 2 | 0 | NON_SYNONYMOU | MODERAT | MISSENSE | tCc/tGc | S196C | PLES_32001 | | | | | | | | | G | G | | |
| NC_011770 | 3531543 | C | A | 83 | 0 | 2 | 0 | NON_SYNONYMOU | MODERAT | MISSENSE | Ctc/Atc | L225I | PLES_32001 | A | A | | | | | | | | | | |
| NC_011770 | 3532059 | A | G | 79 | 0 | 2 | 0 | NON_SYNONYMOU | MODERAT | MISSENSE | Atc/Gtc | I397V | PLES_32001 | G | G | | | | | | | | | | |
| NC_011770 | 3533344 | T | C | 90 | 0 | 1 | 0 | NON_SYNONYMOU | MODERAT | MISSENSE | Acc/Gcc | T434A | PLES_32021 | | | | | | | C | | | | | |
| NC_011770 | 3533580 | A | G | 77 | 0 | 12 | 0 | NON_SYNONYMOU | MODERAT | MISSENSE | aTc/aCc | I355T | PLES_32021 | G | G | G | G | G | G | G | G | G | G | G | G |
| NC_011770 | 3533875 | C | T | 66 | 0 | 2 | 0 | NON_SYNONYMOU | MODERAT | MISSENSE | Gtg/Atg | V257M | PLES_32021 | | | | | | | | | | | T | T |
| NC_011770 | 3534016 | C | A | 71 | 0 | 2 | 0 | NON_SYNONYMOU | MODERAT | MISSENSE | Gcc/Tcc | A210S | PLES_32021 | | | | | | | | | | | A | A |
| NC_011770 | 3534097 | G | T | 73 | 0 | 12 | 0 | NON_SYNONYMOU | MODERAT | MISSENSE | Ctg/Atg | L183M | PLES_32021 | T | T | T | T | T | T | T | T | T | T | T | T |
| NC_011770 | 3534259 | T | C | 76 | 0 | 12 | 0 | NON_SYNONYMOU | MODERAT | MISSENSE | Aac/Gac | N129D | PLES_32021 | C | C | C | C | C | C | C | C | C | C | C | C |
| NC_011770 | 3534444 | G | T | 76 | 0 | 4 | 0 | NON_SYNONYMOU | MODERAT | MISSENSE | aCa/aAa | T67K | PLES_32021 | | | | | | | T | T | | | T | T |
| NC_011770 | 3534099 | G | A | 76 | 0 | 1 | 0 | NON_SYNONYMOU | MODERAT | MISSENSE | gCc/gTc | A182V | PLES_32021 | | | | | | | | A | | | | |
| NC_011770 | 3535247 | C | G | 72 | 0 | 2 | 0 | NON_SYNONYMOU | MODERAT | MISSENSE | caG/caC | Q352H | PLES_32031 | | | | | | | | | G | G | | |
| NC_011770 | 3535483 | G | A | 67 | 0 | 1 | 0 | NON_SYNONYMOU | MODERAT | MISSENSE | Cgc/Tgc | R274C | PLES_32031 | | | | | | | A | | | | | |
| NC_011770 | 3535927 | T | C | 67 | 0 | 2 | 0 | NON_SYNONYMOU | MODERAT | MISSENSE | Atc/Gtc | I126V | PLES_32031 | | | | | | | | | | | C | C |
| NC_011770 | 3536271 | G | A | 61 | 0 | 2 | 0 | NON_SYNONYMOU | MODERAT | MISSENSE | aCc/aTc | T11I | PLES_32031 | A | A | | | | | | | | | | |
| NC_011770 | 3537545 | G | A | 72 | 0 | 2 | 0 | NON_SYNONYMOU | MODERAT | MISSENSE | gCg/gTg | A323V | PLES_32051 | A | A | | | | | | | | | | |
| NC_011770 | 3537800 | C | T | 68 | 0 | 10 | 0 | NON_SYNONYMOU | MODERAT | MISSENSE | gGg/gAg | G238E | PLES_32051 | T | T | T | T | T | T | T | T | T | T | | |
| NC_011770 | 3538908 | G | A | 71 | 0 | 12 | 0 | NON_SYNONYMOU | MODERAT | MISSENSE | Gcc/Acc | A91T | PLES_32061 | A | A | A | A | A | A | A | A | A | A | A | A |
| NC_011770 | 3539014 | G | A | 73 | 0 | 2 | 0 | NON_SYNONYMOU | MODERAT | MISSENSE | cGg/cAg | R126Q | PLES_32061 | | | | | | | | | | | A | A |
| NC_011770 | 3539164 | C | G | 68 | 0 | 4 | 0 | NON_SYNONYMOU | MODERAT | MISSENSE | gCc/gGc | A176G | PLES_32061 | | | | | | | G | G | | | G | G |
| NC_011770 | 3539302 | A | G | 68 | 0 | 2 | 0 | NON_SYNONYMOU | MODERAT | MISSENSE | cAg/cGg | Q222R | PLES_32061 | | | | | | | | | | | G | G |
| NC_011770 | 3539403 | G | A | 74 | 0 | 4 | 0 | NON_SYNONYMOU | MODERAT | MISSENSE | Ggc/Agc | G256S | PLES_32061 | A | A | | | | | A | A | | | | |
| NC_011770 | 3541690 | G | A | 61 | 0 | 8 | 0 | NON_SYNONYMOU | MODERAT | MISSENSE | aGg/aAg | R5K | PLES_32091 | | | A | A | A | A | A | A | A | A | | |
| NC_011770 | 3541720 | C | T | 63 | 0 | 8 | 0 | NON_SYNONYMOU | MODERAT | MISSENSE | gCg/gTg | A15V | PLES_32091 | | | T | T | T | T | T | T | T | T | | |
| NC_011770 | 3541726 | T | C | 63 | 0 | 8 | 0 | NON_SYNONYMOU | MODERAT | MISSENSE | gTc/gCc | V17A | PLES_32091 | | | C | C | C | C | C | C | C | C | | |
| NC_011770 | 3541899 | G | A | 81 | 0 | 4 | 0 | NON_SYNONYMOU | MODERAT | MISSENSE | Gcg/Acg | A75T | PLES_32091 | A | A | | | | | | | | | A | A |
| NC_011770 | 3541908 | A | G | 81 | 0 | 6 | 0 | NON_SYNONYMOU | MODERAT | MISSENSE | Aac/Gac | N78D | PLES_32091 | | | G | G | G | G | | | G | G | | |
| NC_011770 | 3543591 | T | C | 78 | 0 | 2 | 0 | NON_SYNONYMOU | MODERAT | MISSENSE | Tcg/Ccg | S81P | PLES_32111 | | | | | | | C | C | | | | |
| NC_011770 | 3543594 | G | A | 77 | 0 | 2 | 0 | NON_SYNONYMOU | MODERAT | MISSENSE | Gat/Aat | D82N | PLES_32111 | | | | | | | A | A | | | | |
| NC_011770 | 3543650 | T | G | 70 | 0 | 2 | 0 | NON_SYNONYMOU | MODERAT | MISSENSE | gaT/gaG | D100E | PLES_32111 | | | | | | | G | G | | | | |
| NC_011770 | 3543690 | G | T | 73 | 0 | 2 | 0 | NON_SYNONYMOU | MODERAT | MISSENSE | Gtc/Ttc | V114F | PLES_32111 | T | T | | | | | | | | | | |
| NC_011770 | 3543736 | A | C | 68 | 0 | 2 | 0 | NON_SYNONYMOU | MODERAT | MISSENSE | cAa/cCa | Q129P | PLES_32111 | | | | | | | C | C | | | | |
| NC_011770 | 3543757 | G | A | 69 | 0 | 8 | 0 | NON_SYNONYMOU | MODERAT | MISSENSE | gGa/gAa | G136E | PLES_32111 | | | A | A | A | A | A | A | A | A | | |
| NC_011770 | 3543769 | C | T | 66 | 0 | 2 | 0 | NON_SYNONYMOU | MODERAT | MISSENSE | gCc/gTc | A140V | PLES_32111 | | | | | | | | | | | T | T |
| NC_011770 | 3543786 | A | G | 68 | 0 | 6 | 0 | NON_SYNONYMOU | MODERAT | MISSENSE | Acc/Gcc | T146A | PLES_32111 | | | G | G | G | G | | | G | G | | |
| NC_011770 | 3543826 | C | G | 65 | 0 | 6 | 0 | NON_SYNONYMOU | MODERAT | MISSENSE | cCa/cGa | P159R | PLES_32111 | | | G | G | G | G | | | G | G | | |

| | | | | | | | | | | | | | | | | | | | | | | | | | |
|---|---|---|---|---|---|---|---|---|---|---|---|---|---|---|---|---|---|---|---|---|---|---|---|---|---|
| NC_011770 | 3543850 | T | A | 64 | 0 | 6 | 0 | NON_SYNONYMOU | MODERAT | MISSENSE | cTg/cAg | L167Q | PLES_32111 | | | A | A | A | A | | | A | A | | |
| NC_011770 | 3543879 | G | A | 67 | 0 | 6 | 0 | NON_SYNONYMOU | MODERAT | MISSENSE | Gcc/Acc | A177T | PLES_32111 | | | A | A | A | A | | | A | A | | |
| NC_011770 | 3543919 | G | A | 71 | 0 | 6 | 0 | NON_SYNONYMOU | MODERAT | MISSENSE | cGc/cAc | R190H | PLES_32111 | | | A | A | A | A | | | A | A | | |
| NC_011770 | 3543942 | G | C | 71 | 0 | 4 | 0 | NON_SYNONYMOU | MODERAT | MISSENSE | Gcg/Ccg | A198P | PLES_32111 | C | C | | | | | | | | | C | C |
| NC_011770 | 3543978 | A | G | 78 | 0 | 2 | 0 | NON_SYNONYMOU | MODERAT | MISSENSE | Acc/Gcc | T210A | PLES_32111 | G | G | | | | | | | | | | |
| NC_011770 | 3544024 | T | TGCC | 35 | 4 | 2 | 0 | CODON_CHANGE_ | MODERATE | | gtc/gtGCC | V225VP | PLES_32111 | | | TGCC | . | TGCC | . | | | . | . | | |
| NC_011770 | 3544038 | G | T | 73 | 0 | 6 | 0 | NON_SYNONYMOU | MODERAT | MISSENSE | Gcg/Tcg | A230S | PLES_32111 | | | T | T | T | T | | | T | T | | |
| NC_011770 | 3544045 | G | A | 76 | 0 | 6 | 0 | NON_SYNONYMOU | MODERAT | MISSENSE | aGc/aAc | S232N | PLES_32111 | | | A | A | A | A | | | A | A | | |
| NC_011770 | 3544132 | G | C | 86 | 0 | 6 | 0 | NON_SYNONYMOU | MODERAT | MISSENSE | gGc/gCc | G261A | PLES_32111 | | | C | C | C | C | | | C | C | | |
| NC_011770 | 3544680 | A | T | 71 | 0 | 2 | 0 | NON_SYNONYMOU | MODERAT | MISSENSE | cTg/cAg | L143Q | PLES_32121 | | | | | | | | | | | T | T |
| NC_011770 | 3544791 | C | T | 74 | 0 | 6 | 0 | NON_SYNONYMOU | MODERAT | MISSENSE | aGc/aAc | S106N | PLES_32121 | | | T | T | T | T | | | T | T | | |
| NC_011770 | 3545029 | T | C | 67 | 0 | 2 | 0 | NON_SYNONYMOU | MODERAT | MISSENSE | Acc/Gcc | T27A | PLES_32121 | C | C | | | | | | | | | | |
| NC_011770 | 3545306 | C | T | 69 | 0 | 2 | 0 | NON_SYNONYMOU | MODERAT | MISSENSE | Gcg/Acg | A288T | PLES_32131 | T | T | | | | | | | | | | |
| NC_011770 | 3545968 | T | C | 78 | 0 | 1 | 0 | NON_SYNONYMOU | MODERAT | MISSENSE | tAc/tGc | Y67C | PLES_32131 | | | | | | | | C | | | | |
| NC_011770 | 3548987 | A | G | 98 | 0 | 2 | 0 | NON_SYNONYMOU | MODERAT | MISSENSE | aAc/aGc | N3S | PLES_32161 | | | | | | | G | G | | | | |
| NC_011770 | 3549451 | G | A | 88 | 0 | 2 | 0 | NON_SYNONYMOU | MODERAT | MISSENSE | Gag/Aag | E158K | PLES_32161 | A | A | | | | | | | | | | |
| NC_011770 | 3549452 | A | C | 85 | 0 | 8 | 0 | NON_SYNONYMOU | MODERAT | MISSENSE | gAg/gCg | E158A | PLES_32161 | C | C | C | C | C | C | C | C | | | | |
| NC_011770 | 3549599 | C | G | 74 | 0 | 2 | 0 | NON_SYNONYMOU | MODERAT | MISSENSE | gCg/gGg | A207G | PLES_32161 | | | | | | | | | G | G | | |
| NC_011770 | 3549601 | G | C | 74 | 0 | 2 | 0 | NON_SYNONYMOU | MODERAT | MISSENSE | Gcg/Ccg | A208P | PLES_32161 | | | | | | | C | C | | | | |
| NC_011770 | 3549937 | C | T | 67 | 0 | 2 | 0 | NON_SYNONYMOU | MODERAT | MISSENSE | gCg/gTg | A73V | PLES_32171 | | | | | | | | | T | T | | |
| NC_011770 | 3549996 | C | T | 68 | 0 | 2 | 0 | NON_SYNONYMOU | MODERAT | MISSENSE | Cac/Tac | H93Y | PLES_32171 | | | | | | | T | T | | | | |
| NC_011770 | 3549999 | A | G | 66 | 0 | 2 | 0 | NON_SYNONYMOU | MODERAT | MISSENSE | Atc/Gtc | I94V | PLES_32171 | | | | | | | G | G | | | | |
| NC_011770 | 3550042 | C | A | 62 | 0 | 4 | 0 | NON_SYNONYMOU | MODERAT | MISSENSE | gCc/gAc | A108D | PLES_32171 | | | | | | | A | A | A | A | | |
| NC_011770 | 3550102 | G | A | 62 | 0 | 2 | 0 | NON_SYNONYMOU | MODERAT | MISSENSE | cGc/cAc | R128H | PLES_32171 | | | | | | | A | A | | | | |
| NC_011770 | 3550206 | A | C | 59 | 0 | 2 | 0 | NON_SYNONYMOU | MODERAT | MISSENSE | Atc/Ctc | I163L | PLES_32171 | | | | | | | | | C | C | | |
| NC_011770 | 3550464 | G | A | 72 | 0 | 2 | 0 | NON_SYNONYMOU | MODERAT | MISSENSE | tGc/tAc | C12Y | PLES_32181 | | | | | | | | | | | A | A |
| NC_011770 | 3550544 | C | T | 72 | 0 | 2 | 0 | NON_SYNONYMOU | MODERAT | MISSENSE | Ctc/Ttc | L39F | PLES_32181 | | | | | | | T | T | | | | |
| NC_011770 | 3550644 | G | A | 66 | 0 | 4 | 0 | NON_SYNONYMOU | MODERAT | MISSENSE | cGt/cAt | R72H | PLES_32181 | | | A | A | A | A | | | | | | |
| NC_011770 | 3550737 | G | A | 54 | 0 | 2 | 0 | NON_SYNONYMOU | MODERAT | MISSENSE | gGc/gAc | G103D | PLES_32181 | | | | | | | A | A | | | | |
| NC_011770 | 3551348 | G | C | 47 | 0 | 12 | 0 | NON_SYNONYMOU | MODERAT | MISSENSE | Gtg/Ctg | V307L | PLES_32181 | C | C | C | C | C | C | C | C | C | C | C | C |
| NC_011770 | 3551811 | G | A | 57 | 0 | 2 | 0 | NON_SYNONYMOU | MODERAT | MISSENSE | Gac/Aac | D125N | PLES_32191 | | | | | | | | | | | A | A |
| NC_011770 | 3552895 | G | T | 44 | 0 | 10 | 0 | STOP_GAINED | HIGH | NONSENSE | tCg/tAg | S271* | PLES_32201 | T | T | T | T | T | T | | | T | T | T | T |
| NC_011770 | 3552044 | G | A | 49 | 0 | 4 | 0 | NON_SYNONYMOU | MODERAT | MISSENSE | Cac/Tac | H555Y | PLES_32201 | | | A | A | A | A | | | | | | |
| NC_011770 | 3552113 | C | A | 54 | 0 | 4 | 0 | NON_SYNONYMOU | MODERAT | MISSENSE | Gcc/Tcc | A532S | PLES_32201 | | | A | A | A | A | | | | | | |
| NC_011770 | 3552896 | A | C | 43 | 0 | 10 | 0 | NON_SYNONYMOU | MODERAT | MISSENSE | Tcg/Gcg | S271A | PLES_32201 | C | C | C | C | C | C | | | C | C | C | C |
| NC_011770 | 3553147 | A | G | 51 | 0 | 2 | 0 | NON_SYNONYMOU | MODERAT | MISSENSE | gTc/gCc | V187A | PLES_32201 | | | | | | | G | G | | | | |
| NC_011770 | 3553498 | A | G | 67 | 0 | 4 | 0 | NON_SYNONYMOU | MODERAT | MISSENSE | gTg/gCg | V70A | PLES_32201 | | | G | G | G | G | | | | | | |
| NC_011770 | 3553965 | T | C | 63 | 0 | 2 | 0 | NON_SYNONYMOU | MODERAT | MISSENSE | gTc/gCc | V30A | PLES_32211 | | | | | | | | | C | C | | |
| NC_011770 | 3553997 | T | G | 65 | 0 | 2 | 0 | NON_SYNONYMOU | MODERAT | MISSENSE | Tcc/Gcc | S41A | PLES_32211 | | | | | | | | | G | G | | |
| NC_011770 | 3554004 | A | C | 67 | 0 | 2 | 0 | NON_SYNONYMOU | MODERAT | MISSENSE | gAc/gCc | D43A | PLES_32211 | | | | | | | | | C | C | | |
| NC_011770 | 3554043 | T | C | 63 | 0 | 12 | 0 | NON_SYNONYMOU | MODERAT | MISSENSE | aTc/aCc | I56T | PLES_32211 | C | C | C | C | C | C | C | C | C | C | C | C |
| NC_011770 | 3554880 | G | C | 53 | 0 | 6 | 0 | NON_SYNONYMOU | MODERAT | MISSENSE | Ccc/Gcc | P187A | PLES_32221 | C | C | | | | | C | C | C | C | | |
| NC_011770 | 3555121 | T | A | 57 | 0 | 4 | 0 | NON_SYNONYMOU | MODERAT | MISSENSE | caA/caT | Q106H | PLES_32221 | A | A | | | | | A | A | | | | |
| NC_011770 | 3555260 | G | A | 57 | 0 | 10 | 0 | NON_SYNONYMOU | MODERAT | MISSENSE | gCg/gTg | A60V | PLES_32221 | | | A | A | A | A | A | A | A | A | A | A |
| NC_011770 | 3555419 | C | G | 60 | 0 | 6 | 0 | NON_SYNONYMOU | MODERAT | MISSENSE | aGt/aCt | S7T | PLES_32221 | G | G | | | | | | | G | G | G | G |
| NC_011770 | 3555427 | G | A | 63 | 0 | 4 | 0 | NON_SYNONYMOU | MODERAT | MISSENSE | tCc/tTc | S309F | PLES_32231 | | | A | A | A | A | | | | | | |
| NC_011770 | 3555685 | G | A | 59 | 0 | 2 | 0 | NON_SYNONYMOU | MODERAT | MISSENSE | tCg/tTg | S223L | PLES_32231 | | | | | | | | | A | A | | |
| NC_011770 | 3556136 | G | A | 66 | 0 | 1 | 0 | NON_SYNONYMOU | MODERAT | MISSENSE | Cgg/Tgg | R73W | PLES_32231 | | | | | | | A | | | | | |
| NC_011770 | 3557146 | C | T | 73 | 0 | 1 | 0 | STOP_GAINED | HIGH | NONSENSE | tgG/tgA | W231* | PLES_32241 | T | | | | | | | | | | | |
| NC_011770 | 3556580 | G | T | 84 | 0 | 10 | 0 | NON_SYNONYMOU | MODERAT | MISSENSE | gCg/gAg | A420E | PLES_32241 | T | T | T | T | T | T | | | T | T | T | T |
| NC_011770 | 3558943 | T | G | 91 | 0 | 6 | 0 | NON_SYNONYMOU | MODERAT | MISSENSE | aTc/aGc | I320S | PLES_32251 | G | G | | | | | G | G | G | G | | |
| NC_011770 | 3560065 | T | C | 85 | 0 | 2 | 0 | NON_SYNONYMOU | MODERAT | MISSENSE | Atc/Gtc | I303V | PLES_32271 | C | C | | | | | | | | | | |
| NC_011770 | 3560325 | G | A | 83 | 0 | 2 | 0 | NON_SYNONYMOU | MODERAT | MISSENSE | gCg/gTg | A216V | PLES_32271 | A | A | | | | | | | | | | |
| NC_011770 | 3560368 | G | A | 75 | 0 | 4 | 0 | NON_SYNONYMOU | MODERAT | MISSENSE | Cgc/Tgc | R202C | PLES_32271 | | | A | A | A | A | | | | | | |
| NC_011770 | 3561611 | CCCG | C | 75 | 0 | 2 | 0 | CODON_CHANGE_ | MODERATE | | gcgggc/gg | AG388G | PLES_32291 | | | | | | | C | C | | | | |
| NC_011770 | 3561962 | C | T | 66 | 0 | 2 | 0 | NON_SYNONYMOU | MODERAT | MISSENSE | gGg/gAg | G272E | PLES_32291 | T | T | | | | | | | | | | |
| NC_011770 | 3562061 | G | T | 60 | 0 | 4 | 0 | NON_SYNONYMOU | MODERAT | MISSENSE | gCc/gAc | A239D | PLES_32291 | | | | | | | | | T | T | T | T |
| NC_011770 | 3562208 | G | C | 62 | 0 | 2 | 0 | NON_SYNONYMOU | MODERAT | MISSENSE | cCc/cGc | P190R | PLES_32291 | | | | | | | C | C | | | | |
| NC_011770 | 3562518 | T | C | 65 | 0 | 4 | 0 | NON_SYNONYMOU | MODERAT | MISSENSE | Atc/Gtc | I87V | PLES_32291 | | | | | | | | | C | C | C | C |
| NC_011770 | 3562523 | G | A | 67 | 0 | 4 | 0 | NON_SYNONYMOU | MODERAT | MISSENSE | gCg/gTg | A85V | PLES_32291 | | | | | | | | | A | A | A | A |
| NC_011770 | 3562788 | T | A | 71 | 0 | 2 | 0 | NON_SYNONYMOU | MODERAT | MISSENSE | gAg/gTg | E419V | PLES_32301 | | | A | A | | | | | | | | |
| NC_011770 | 3562921 | A | C | 72 | 0 | 2 | 0 | NON_SYNONYMOU | MODERAT | MISSENSE | Tcc/Gcc | S375A | PLES_32301 | C | C | | | | | | | | | | |

| | | | | | | | | | | | | | | | | | | | | | | | | | |
|---|---|---|---|---|---|---|---|---|---|---|---|---|---|---|---|---|---|---|---|---|---|---|---|---|---|
| NC_011770 | 3563403 | C | A | 53 | 0 | 8 | 0 | NON_SYNONYMOU | MODERAT | MISSENSE | cGg/cTg | R214L | PLES_32301 | A | A | A | A | A | A | | | A | A | | |
| NC_011770 | 3563998 | G | GCGC | 50 | 0 | 2 | 0 | CODON_INSERTIO | MODERATE | | gcg/gcgGC | A15AA | PLES_32301 | | | | | | | GCGC | GCGC | | | | |
| NC_011770 | 3563085 | G | A | 73 | 0 | 1 | 0 | NON_SYNONYMOU | MODERAT | MISSENSE | cCg/cTg | P320L | PLES_32301 | | | | | | | | A | | | | |
| NC_011770 | 3564465 | G | T | 55 | 0 | 4 | 0 | NON_SYNONYMOU | MODERAT | MISSENSE | Cgc/Agc | R219S | PLES_32311 | | | T | T | T | T | | | | | | |
| NC_011770 | 3565224 | C | A | 82 | 0 | 2 | 0 | NON_SYNONYMOU | MODERAT | MISSENSE | Gcc/Tcc | A854S | PLES_32321 | | | | | | | | | | | A | A |
| NC_011770 | 3565460 | C | A | 82 | 0 | 2 | 0 | NON_SYNONYMOU | MODERAT | MISSENSE | aGc/aTc | S775I | PLES_32321 | A | A | | | | | | | | | | |
| NC_011770 | 3565526 | T | C | 86 | 0 | 2 | 0 | NON_SYNONYMOU | MODERAT | MISSENSE | gAc/gGc | D753G | PLES_32321 | C | C | | | | | | | | | | |
| NC_011770 | 3565556 | C | T | 83 | 0 | 2 | 0 | NON_SYNONYMOU | MODERAT | MISSENSE | cGc/cAc | R743H | PLES_32321 | | | | | | | | | T | T | | |
| NC_011770 | 3566210 | G | C | 80 | 0 | 4 | 0 | NON_SYNONYMOU | MODERAT | MISSENSE | aCc/aGc | T525S | PLES_32321 | | | C | C | C | C | | | | | | |
| NC_011770 | 3567473 | C | A | 70 | 0 | 2 | 0 | NON_SYNONYMOU | MODERAT | MISSENSE | aGc/aTc | S104I | PLES_32321 | | | | | | | A | A | | | | |
| NC_011770 | 3567494 | T | C | 69 | 0 | 4 | 0 | NON_SYNONYMOU | MODERAT | MISSENSE | cAg/cGg | Q97R | PLES_32321 | | | C | C | C | C | | | | | | |
| NC_011770 | 3567669 | G | T | 77 | 0 | 8 | 0 | NON_SYNONYMOU | MODERAT | MISSENSE | Cgc/Agc | R39S | PLES_32321 | T | T | | | | | T | T | T | T | T | T |
| NC_011770 | 3567780 | A | G | 73 | 0 | 6 | 0 | NON_SYNONYMOU | MODERAT | MISSENSE | Tcc/Ccc | S2P | PLES_32321 | | | G | G | G | G | | | | | G | G |
| NC_011770 | 3565554 | C | T | 81 | 0 | 1 | 0 | NON_SYNONYMOU | MODERAT | MISSENSE | Gga/Aga | G744R | PLES_32321 | | | | | | | | T | | | | |
| NC_011770 | 3568083 | A | G | 63 | 0 | 2 | 0 | NON_SYNONYMOU | MODERAT | MISSENSE | gTc/gCc | V214A | PLES_32331 | | | | | | | | | | | G | G |
| NC_011770 | 3569276 | C | G | 73 | 0 | 4 | 0 | NON_SYNONYMOU | MODERAT | MISSENSE | Gtc/Ctc | V30L | PLES_32341 | | | G | G | G | G | | | | | | |
| NC_011770 | 3569679 | T | G | 75 | 0 | 4 | 0 | NON_SYNONYMOU | MODERAT | MISSENSE | gAa/gCa | E195A | PLES_32351 | | | G | G | G | G | | | | | | |
| NC_011770 | 3570258 | T | C | 80 | 0 | 2 | 0 | NON_SYNONYMOU | MODERAT | MISSENSE | aAt/aGt | N2S | PLES_32351 | | | | | | | C | C | | | | |
| NC_011770 | 3570355 | T | C | 79 | 0 | 4 | 0 | NON_SYNONYMOU | MODERAT | MISSENSE | gAg/gGg | E143G | PLES_32361 | | | | | | | | | C | C | C | C |
| NC_011770 | 3570397 | C | T | 82 | 0 | 6 | 0 | NON_SYNONYMOU | MODERAT | MISSENSE | aGc/aAc | S129N | PLES_32361 | | | T | T | T | T | T | T | | | | |
| NC_011770 | 3571163 | T | C | 84 | 0 | 12 | 0 | NON_SYNONYMOU | MODERAT | MISSENSE | Ata/Gta | I494V | PLES_32371 | C | C | C | C | C | C | C | C | C | C | C | C |
| NC_011770 | 3571454 | G | A | 82 | 0 | 2 | 0 | NON_SYNONYMOU | MODERAT | MISSENSE | Cac/Tac | H397Y | PLES_32371 | | | | | | | | | A | A | | |
| NC_011770 | 3572290 | G | A | 86 | 0 | 6 | 0 | NON_SYNONYMOU | MODERAT | MISSENSE | gCc/gTc | A118V | PLES_32371 | | | A | A | A | A | | | A | A | | |
| NC_011770 | 3571931 | C | T | 81 | 0 | 1 | 0 | NON_SYNONYMOU | MODERAT | MISSENSE | Gcc/Acc | A238T | PLES_32371 | | | | | | | | | | | T | |
| NC_011770 | 3572749 | A | C | 66 | 0 | 8 | 0 | NON_SYNONYMOU | MODERAT | MISSENSE | cTt/cGt | L419R | PLES_32381 | | | C | C | C | C | | | C | C | C | C |
| NC_011770 | 3573598 | C | T | 73 | 0 | 2 | 0 | NON_SYNONYMOU | MODERAT | MISSENSE | cGg/cAg | R136Q | PLES_32381 | | | | | | | T | T | | | | |
| NC_011770 | 3576300 | G | A | 66 | 0 | 1 | 0 | NON_SYNONYMOU | MODERAT | MISSENSE | Ggc/Agc | G281S | PLES_32411 | | | | | | | A | | | | | |
| NC_011770 | 3577109 | G | C | 72 | 0 | 12 | 0 | NON_SYNONYMOU | MODERAT | MISSENSE | Gcc/Ccc | A82P | PLES_32421 | C | C | C | C | C | C | C | C | C | C | C | C |
| NC_011770 | 3577332 | G | C | 88 | 0 | 2 | 0 | NON_SYNONYMOU | MODERAT | MISSENSE | gGc/gCc | G156A | PLES_32421 | | | | | | | | | C | C | | |
| NC_011770 | 3577584 | A | G | 86 | 0 | 4 | 0 | NON_SYNONYMOU | MODERAT | MISSENSE | aAg/aGg | K240R | PLES_32421 | | | | | | | | | G | G | G | G |
| NC_011770 | 3578082 | G | T | 90 | 0 | 2 | 0 | NON_SYNONYMOU | MODERAT | MISSENSE | cGc/cTc | R406L | PLES_32421 | T | T | | | | | | | | | | |
| NC_011770 | 3578589 | C | T | 81 | 0 | 2 | 0 | NON_SYNONYMOU | MODERAT | MISSENSE | gCc/gTc | A11V | PLES_32431 | T | T | | | | | | | | | | |
| NC_011770 | 3578613 | A | G | 78 | 0 | 8 | 0 | NON_SYNONYMOU | MODERAT | MISSENSE | aAc/aGc | N19S | PLES_32431 | G | G | | | | | G | G | G | G | G | G |
| NC_011770 | 3578741 | G | A | 82 | 0 | 4 | 0 | NON_SYNONYMOU | MODERAT | MISSENSE | Gac/Aac | D62N | PLES_32431 | | | A | A | A | A | | | | | | |
| NC_011770 | 3579771 | T | C | 62 | 0 | 12 | 0 | NON_SYNONYMOU | MODERAT | MISSENSE | aTc/aCc | I405T | PLES_32431 | C | C | C | C | C | C | C | C | C | C | C | C |
| NC_011770 | 3580061 | C | G | 62 | 0 | 2 | 0 | NON_SYNONYMOU | MODERAT | MISSENSE | Ctc/Gtc | L502V | PLES_32431 | | | | | | | | | | | G | G |
| NC_011770 | 3580064 | G | A | 60 | 0 | 6 | 0 | NON_SYNONYMOU | MODERAT | MISSENSE | Gcc/Acc | A503T | PLES_32431 | | | A | A | A | A | | | | | A | A |
| NC_011770 | 3580475 | A | C | 50 | 0 | 8 | 0 | NON_SYNONYMOU | MODERAT | MISSENSE | Acc/Ccc | T2P | PLES_32441 | C | C | C | C | C | C | | | | | C | C |
| NC_011770 | 3580965 | G | A | 80 | 0 | 2 | 0 | NON_SYNONYMOU | MODERAT | MISSENSE | aGc/aAc | S165N | PLES_32441 | A | A | | | | | | | | | | |
| NC_011770 | 3581204 | A | G | 70 | 0 | 12 | 0 | NON_SYNONYMOU | MODERAT | MISSENSE | Aac/Gac | N245D | PLES_32441 | G | G | G | G | G | G | G | G | G | G | G | G |
| NC_011770 | 3581323 | G | C | 74 | 0 | 4 | 0 | NON_SYNONYMOU | MODERAT | MISSENSE | gaG/gaC | E284D | PLES_32441 | C | C | | | | | | | C | C | | |
| NC_011770 | 3581955 | A | C | 58 | 0 | 2 | 0 | NON_SYNONYMOU | MODERAT | MISSENSE | gAc/gCc | D495A | PLES_32441 | C | C | | | | | | | | | | |
| NC_011770 | 3581969 | A | C | 57 | 0 | 2 | 0 | NON_SYNONYMOU | MODERAT | MISSENSE | Acc/Ccc | T500P | PLES_32441 | C | C | | | | | | | | | | |
| NC_011770 | 3581978 | G | A | 57 | 0 | 2 | 0 | NON_SYNONYMOU | MODERAT | MISSENSE | Gac/Aac | D503N | PLES_32441 | A | A | | | | | | | | | | |
| NC_011770 | 3582021 | A | G | 56 | 0 | 8 | 0 | NON_SYNONYMOU | MODERAT | MISSENSE | gAc/gGc | D517G | PLES_32441 | G | G | G | G | G | G | G | G | | | | |
| NC_011770 | 3582036 | G | A | 58 | 0 | 2 | 0 | NON_SYNONYMOU | MODERAT | MISSENSE | cGg/cAg | R522Q | PLES_32441 | A | A | | | | | | | | | | |
| NC_011770 | 3581693 | T | C | 65 | 0 | 1 | 0 | NON_SYNONYMOU | MODERAT | MISSENSE | Tgg/Cgg | W408R | PLES_32441 | | | | | | | | C | | | | |
| NC_011770 | 3582681 | G | A | 65 | 0 | 2 | 0 | NON_SYNONYMOU | MODERAT | MISSENSE | gCt/gTt | A212V | PLES_32451 | | | | | | | A | A | | | | |
| NC_011770 | 3583084 | C | T | 69 | 0 | 2 | 0 | NON_SYNONYMOU | MODERAT | MISSENSE | Gcg/Acg | A78T | PLES_32451 | | | | | | | | | T | T | | |
| NC_011770 | 3583191 | G | A | 66 | 0 | 8 | 0 | NON_SYNONYMOU | MODERAT | MISSENSE | gCg/gTg | A42V | PLES_32451 | A | A | | | | | A | A | A | A | A | A |
| NC_011770 | 3583789 | AG | A | 50 | 0 | 3 | 0 | FRAME_SHIFT | HIGH | | ggc/ | G8 | PLES_32461 | | | | | | | | A | | | AGG | AGG |
| NC_011770 | 3583795 | G | T | 37 | 0 | 2 | 0 | NON_SYNONYMOU | MODERAT | MISSENSE | gGc/gTc | G8V | PLES_32461 | T | T | | | | | | | | | | |
| NC_011770 | 3583827 | G | T | 53 | 0 | 2 | 0 | NON_SYNONYMOU | MODERAT | MISSENSE | Gcc/Tcc | A19S | PLES_32461 | | | | | | | | | | | T | T |
| NC_011770 | 3584839 | T | G | 34 | 0 | 12 | 0 | NON_SYNONYMOU | MODERAT | MISSENSE | gaT/gaG | D208E | PLES_32471 | G | G | G | G | G | G | G | G | G | G | G | G |
| NC_011770 | 3586416 | G | C | 13 | 8 | 2 | 0 | NON_SYNONYMOU | MODERAT | MISSENSE | Ccg/Gcg | P133A | PLES_32481 | . | . | C | . | C | . | . | . | . | . | | |
| NC_011770 | 3588527 | T | G | 57 | 0 | 6 | 0 | NON_SYNONYMOU | MODERAT | MISSENSE | gAg/gCg | E843A | PLES_32501 | G | G | G | G | G | G | | | | | | |
| NC_011770 | 3588546 | G | C | 58 | 0 | 6 | 0 | NON_SYNONYMOU | MODERAT | MISSENSE | Cgc/Ggc | R837G | PLES_32501 | C | C | C | C | C | C | | | | | | |
| NC_011770 | 3588929 | C | T | 73 | 0 | 4 | 0 | NON_SYNONYMOU | MODERAT | MISSENSE | cGc/cAc | R709H | PLES_32501 | | | T | T | T | T | | | | | | |
| NC_011770 | 3589028 | C | G | 71 | 0 | 4 | 0 | NON_SYNONYMOU | MODERAT | MISSENSE | aGc/aCc | S676T | PLES_32501 | | | G | G | G | G | | | | | | |
| NC_011770 | 3589029 | T | A | 68 | 0 | 4 | 0 | NON_SYNONYMOU | MODERAT | MISSENSE | Agc/Tgc | S676C | PLES_32501 | | | A | A | A | A | | | | | | |
| NC_011770 | 3589136 | C | T | 69 | 0 | 4 | 0 | NON_SYNONYMOU | MODERAT | MISSENSE | cGc/cAc | R640H | PLES_32501 | | | T | T | T | T | | | | | | |
| NC_011770 | 3590447 | G | A | 74 | 0 | 6 | 0 | NON_SYNONYMOU | MODERAT | MISSENSE | gCg/gTg | A203V | PLES_32501 | | | A | A | A | A | | | | | A | A |

| | | | | | | | | | | | | | | | | | | | | | | | | | |
|---|---|---|---|---|---|---|---|---|---|---|---|---|---|---|---|---|---|---|---|---|---|---|---|---|---|
| NC_011770 | 3590786 | A | G | 74 | 0 | 6 | 0 | NON_SYNONYMOU | MODERAT | MISSENSE | gTg/gCg | V90A | PLES_32501 | | | G | G | G | G | | | | | G | G |
| NC_011770 | 3594238 | C | A | 69 | 0 | 12 | 0 | NON_SYNONYMOU | MODERAT | MISSENSE | gCg/gAg | A210E | PLES_32521 | A | A | A | A | A | A | A | A | A | A | A | A |
| NC_011770 | 3594820 | A | G | 74 | 0 | 10 | 0 | NON_SYNONYMOU | MODERAT | MISSENSE | gAc/gGc | D404G | PLES_32521 | | | G | G | G | G | G | G | G | G | G | G |
| NC_011770 | 3594840 | G | A | 69 | 0 | 4 | 0 | NON_SYNONYMOU | MODERAT | MISSENSE | Gcc/Acc | A411T | PLES_32521 | | | A | A | A | A | | | | | | |
| NC_011770 | 3595011 | G | A | 68 | 0 | 6 | 0 | NON_SYNONYMOU | MODERAT | MISSENSE | Gtc/Atc | V468I | PLES_32521 | | | A | A | A | A | A | A | | | | |
| NC_011770 | 3595122 | C | T | 65 | 0 | 2 | 0 | NON_SYNONYMOU | MODERAT | MISSENSE | Cac/Tac | H505Y | PLES_32521 | | | | | | | | | | | T | T |
| NC_011770 | 3595158 | A | C | 65 | 0 | 2 | 0 | NON_SYNONYMOU | MODERAT | MISSENSE | Agc/Cgc | S517R | PLES_32521 | | | | | | | | | | | C | C |
| NC_011770 | 3595458 | G | T | 81 | 0 | 2 | 0 | NON_SYNONYMOU | MODERAT | MISSENSE | Gcc/Tcc | A617S | PLES_32521 | T | T | | | | | | | | | | |
| NC_011770 | 3595599 | A | G | 77 | 0 | 12 | 0 | NON_SYNONYMOU | MODERAT | MISSENSE | Aac/Gac | N664D | PLES_32521 | G | G | G | G | G | G | G | G | G | G | G | G |
| NC_011770 | 3596558 | C | A | 77 | 0 | 2 | 0 | NON_SYNONYMOU | MODERAT | MISSENSE | Cgc/Agc | R43S | PLES_32531 | | | A | A | | | | | | | | |
| NC_011770 | 3596957 | A | G | 70 | 0 | 6 | 0 | NON_SYNONYMOU | MODERAT | MISSENSE | Acg/Gcg | T176A | PLES_32531 | G | G | | | | | | | G | G | G | G |
| NC_011770 | 3597018 | G | A | 74 | 0 | 2 | 0 | NON_SYNONYMOU | MODERAT | MISSENSE | cGc/cAc | R196H | PLES_32531 | | | | | | | | | A | A | | |
| NC_011770 | 3597278 | A | G | 81 | 0 | 12 | 0 | NON_SYNONYMOU | MODERAT | MISSENSE | Acc/Gcc | T283A | PLES_32531 | G | G | G | G | G | G | G | G | G | G | G | G |
| NC_011770 | 3597519 | T | C | 75 | 0 | 12 | 0 | NON_SYNONYMOU | MODERAT | MISSENSE | gTg/gCg | V363A | PLES_32531 | C | C | C | C | C | C | C | C | C | C | C | C |
| NC_011770 | 3597870 | C | A | 79 | 0 | 2 | 0 | NON_SYNONYMOU | MODERAT | MISSENSE | gCg/gAg | A480E | PLES_32531 | | | | | | | | | | | A | A |
| NC_011770 | 3598845 | T | C | 76 | 0 | 1 | 0 | NON_SYNONYMOU | MODERAT | MISSENSE | Tac/Cac | Y216H | PLES_32541 | | | | | | | C | | | | | |
| NC_011770 | 3598981 | C | A | 76 | 0 | 4 | 0 | NON_SYNONYMOU | MODERAT | MISSENSE | gCg/gAg | A261E | PLES_32541 | | | A | A | A | A | | | | | | |
| NC_011770 | 3599953 | G | T | 57 | 0 | 6 | 0 | NON_SYNONYMOU | MODERAT | MISSENSE | Gtc/Ttc | V197F | PLES_32551 | T | T | T | T | T | T | | | | | | |
| NC_011770 | 3600514 | T | C | 67 | 0 | 2 | 0 | NON_SYNONYMOU | MODERAT | MISSENSE | gTg/gCg | V162A | PLES_32561 | C | C | | | | | | | | | | |
| NC_011770 | 3600645 | C | T | 72 | 0 | 1 | 0 | NON_SYNONYMOU | MODERAT | MISSENSE | Cac/Tac | H206Y | PLES_32561 | | | | | | | | | | | | T |
| NC_011770 | 3604817 | T | C | 82 | 0 | 1 | 0 | NON_SYNONYMOU | MODERAT | MISSENSE | tAc/tGc | Y47C | PLES_32581 | | | | | | | C | | | | | |
| NC_011770 | 3604923 | G | A | 70 | 0 | 2 | 0 | NON_SYNONYMOU | MODERAT | MISSENSE | Cct/Tct | P12S | PLES_32581 | | | | | | | A | A | | | | |
| NC_011770 | 3605957 | G | A | 84 | 0 | 2 | 0 | NON_SYNONYMOU | MODERAT | MISSENSE | aCc/aTc | T128I | PLES_32591 | | | | | | | | | A | A | | |
| NC_011770 | 3605958 | T | C | 82 | 0 | 4 | 0 | NON_SYNONYMOU | MODERAT | MISSENSE | Acc/Gcc | T128A | PLES_32591 | | | C | C | C | C | | | | | | |
| NC_011770 | 3606063 | C | T | 73 | 0 | 4 | 0 | NON_SYNONYMOU | MODERAT | MISSENSE | Gca/Aca | A93T | PLES_32591 | | | T | T | T | T | | | | | | |
| NC_011770 | 3606536 | C | T | 70 | 0 | 12 | 0 | NON_SYNONYMOU | MODERAT | MISSENSE | atG/atA | M520I | PLES_32601 | T | T | T | T | T | T | T | T | T | T | T | T |
| NC_011770 | 3606541 | T | C | 70 | 0 | 12 | 0 | NON_SYNONYMOU | MODERAT | MISSENSE | Acg/Gcg | T519A | PLES_32601 | C | C | C | C | C | C | C | C | C | C | C | C |
| NC_011770 | 3607032 | C | T | 73 | 0 | 12 | 0 | NON_SYNONYMOU | MODERAT | MISSENSE | cGg/cAg | R355Q | PLES_32601 | T | T | T | T | T | T | T | T | T | T | T | T |
| NC_011770 | 3607485 | A | G | 65 | 0 | 2 | 0 | NON_SYNONYMOU | MODERAT | MISSENSE | aTc/aCc | I204T | PLES_32601 | G | G | | | | | | | | | | |
| NC_011770 | 3607666 | G | C | 61 | 0 | 2 | 0 | NON_SYNONYMOU | MODERAT | MISSENSE | Ctc/Gtc | L144V | PLES_32601 | | | | | | | | | | | C | C |
| NC_011770 | 3606534 | G | A | 71 | 0 | 1 | 0 | NON_SYNONYMOU | MODERAT | MISSENSE | cCg/cTg | P521L | PLES_32601 | | | | | | | | A | | | | |
| NC_011770 | 3608101 | G | T | 79 | 0 | 2 | 0 | NON_SYNONYMOU | MODERAT | MISSENSE | aCa/aAa | T338K | PLES_32611 | | | | | | | T | T | | | | |
| NC_011770 | 3609034 | C | A | 69 | 0 | 2 | 0 | NON_SYNONYMOU | MODERAT | MISSENSE | cGg/cTg | R27L | PLES_32611 | | | | | | | | | | | A | A |
| NC_011770 | 3609122 | C | T | 69 | 0 | 10 | 0 | NON_SYNONYMOU | MODERAT | MISSENSE | aGa/aAa | R356K | PLES_32621 | T | T | T | T | T | T | T | T | T | T | | |
| NC_011770 | 3609491 | G | C | 51 | 0 | 12 | 0 | NON_SYNONYMOU | MODERAT | MISSENSE | aCc/aGc | T233S | PLES_32621 | C | C | C | C | C | C | C | C | C | C | C | C |
| NC_011770 | 3609690 | T | C | 48 | 0 | 4 | 0 | NON_SYNONYMOU | MODERAT | MISSENSE | Atc/Gtc | I167V | PLES_32621 | C | C | | | | | | | | | C | C |
| NC_011770 | 3609840 | T | C | 25 | 0 | 2 | 0 | NON_SYNONYMOU | MODERAT | MISSENSE | Atc/Gtc | I117V | PLES_32621 | C | C | | | | | | | | | | |
| NC_011770 | 3610606 | G | A | 72 | 0 | 2 | 0 | STOP_GAINED | HIGH | NONSENSE | Cag/Tag | Q465* | PLES_32631 | | | | | | | | | | | A | A |
| NC_011770 | 3610694 | G | T | 71 | 0 | 2 | 0 | NON_SYNONYMOU | MODERAT | MISSENSE | gaC/gaA | D435E | PLES_32631 | T | T | | | | | | | | | | |
| NC_011770 | 3610741 | C | T | 68 | 0 | 2 | 0 | NON_SYNONYMOU | MODERAT | MISSENSE | Gtc/Atc | V420I | PLES_32631 | T | T | | | | | | | | | | |
| NC_011770 | 3611189 | C | G | 68 | 0 | 2 | 0 | NON_SYNONYMOU | MODERAT | MISSENSE | caG/caC | Q270H | PLES_32631 | G | G | | | | | | | | | | |
| NC_011770 | 3613828 | CA | C | 80 | 0 | 1 | 0 | FRAME_SHIFT | HIGH | | ggt/ | G238 | PLES_32641 | | | | | | | | C | | | | |
| NC_011770 | 3612364 | T | G | 75 | 0 | 2 | 0 | NON_SYNONYMOU | MODERAT | MISSENSE | Aag/Cag | K727Q | PLES_32641 | G | G | | | | | | | | | | |
| NC_011770 | 3613077 | G | A | 83 | 0 | 2 | 0 | NON_SYNONYMOU | MODERAT | MISSENSE | gCc/gTc | A489V | PLES_32641 | | | | | | | A | A | | | | |
| NC_011770 | 3613506 | T | C | 71 | 0 | 6 | 0 | NON_SYNONYMOU | MODERAT | MISSENSE | cAc/cGc | H346R | PLES_32641 | C | C | | | | | C | C | | | C | C |
| NC_011770 | 3614033 | G | T | 72 | 0 | 2 | 0 | NON_SYNONYMOU | MODERAT | MISSENSE | gaC/gaA | D170E | PLES_32641 | | | | | | | | | | | T | T |
| NC_011770 | 3615293 | A | G | 59 | 0 | 2 | 0 | NON_SYNONYMOU | MODERAT | MISSENSE | Tgc/Cgc | C278R | PLES_32651 | G | G | | | | | | | | | | |
| NC_011770 | 3615608 | T | C | 74 | 0 | 4 | 0 | NON_SYNONYMOU | MODERAT | MISSENSE | Agc/Ggc | S173G | PLES_32651 | | | C | C | C | C | | | | | | |
| NC_011770 | 3615799 | A | G | 79 | 0 | 2 | 0 | NON_SYNONYMOU | MODERAT | MISSENSE | gTg/gCg | V109A | PLES_32651 | G | G | | | | | | | | | | |
| NC_011770 | 3616563 | T | TG | 83 | 0 | 1 | 0 | FRAME_SHIFT | HIGH | | ggt/ggGt | G111G? | PLES_32661 | TG | | | | | | | | | | | |
| NC_011770 | 3616802 | A | G | 75 | 0 | 8 | 0 | NON_SYNONYMOU | MODERAT | MISSENSE | Acc/Gcc | T190A | PLES_32661 | G | G | | | | | G | G | G | G | G | G |
| NC_011770 | 3617003 | C | T | 94 | 0 | 2 | 0 | NON_SYNONYMOU | MODERAT | MISSENSE | Ccg/Tcg | P257S | PLES_32661 | | | | | | | | | | | T | T |
| NC_011770 | 3617339 | A | G | 74 | 0 | 6 | 0 | NON_SYNONYMOU | MODERAT | MISSENSE | Atc/Gtc | I369V | PLES_32661 | G | G | | | | | | | G | G | G | G |
| NC_011770 | 3617504 | G | T | 72 | 0 | 2 | 0 | NON_SYNONYMOU | MODERAT | MISSENSE | Gcc/Tcc | A424S | PLES_32661 | T | T | | | | | | | | | | |
| NC_011770 | 3617583 | G | T | 73 | 0 | 8 | 0 | NON_SYNONYMOU | MODERAT | MISSENSE | tGc/tTc | C450F | PLES_32661 | T | T | T | T | T | T | | | T | T | | |
| NC_011770 | 3619951 | G | A | 56 | 0 | 2 | 0 | NON_SYNONYMOU | MODERAT | MISSENSE | gCg/gTg | A291V | PLES_32701 | A | A | | | | | | | | | | |
| NC_011770 | 3620018 | T | C | 57 | 0 | 2 | 0 | NON_SYNONYMOU | MODERAT | MISSENSE | Atc/Gtc | I269V | PLES_32701 | C | C | | | | | | | | | | |
| NC_011770 | 3621232 | A | T | 62 | 0 | 2 | 0 | NON_SYNONYMOU | MODERAT | MISSENSE | Tgc/Agc | C32S | PLES_32711 | T | T | | | | | | | | | | |
| NC_011770 | 3623588 | C | T | 73 | 0 | 4 | 0 | STOP_GAINED | HIGH | NONSENSE | Cga/Tga | R459* | PLES_32721 | | | T | T | T | T | | | | | | |
| NC_011770 | 3622242 | A | G | 73 | 0 | 2 | 0 | NON_SYNONYMOU | MODERAT | MISSENSE | cAc/cGc | H10R | PLES_32721 | G | G | | | | | | | | | | |
| NC_011770 | 3622286 | A | G | 76 | 0 | 2 | 0 | NON_SYNONYMOU | MODERAT | MISSENSE | Aac/Gac | N25D | PLES_32721 | G | G | | | | | | | | | | |
| NC_011770 | 3622355 | G | A | 78 | 0 | 2 | 0 | NON_SYNONYMOU | MODERAT | MISSENSE | Gag/Aag | E48K | PLES_32721 | A | A | | | | | | | | | | |

| | | | | | | | | | | | | | | | | | | | | | | | | | |
|---|---|---|---|---|---|---|---|---|---|---|---|---|---|---|---|---|---|---|---|---|---|---|---|---|---|
| NC_011770 | 3622557 | G | A | 73 | 0 | 2 | 0 | NON_SYNONYMOU | MODERAT | MISSENSE | gGc/gAc | G115D | PLES_32721 | A | A | | | | | | | | | | |
| NC_011770 | 3622598 | C | G | 77 | 0 | 2 | 0 | NON_SYNONYMOU | MODERAT | MISSENSE | Cgc/Ggc | R129G | PLES_32721 | G | G | | | | | | | | | | |
| NC_011770 | 3623024 | T | C | 84 | 0 | 2 | 0 | NON_SYNONYMOU | MODERAT | MISSENSE | Ttc/Ctc | F271L | PLES_32721 | C | C | | | | | | | | | | |
| NC_011770 | 3623600 | G | A | 73 | 0 | 8 | 0 | NON_SYNONYMOU | MODERAT | MISSENSE | Gtc/Atc | V463I | PLES_32721 | | | A | A | A | A | | | A | A | A | A |
| NC_011770 | 3623633 | A | G | 73 | 0 | 2 | 0 | NON_SYNONYMOU | MODERAT | MISSENSE | Acg/Gcg | T474A | PLES_32721 | G | G | | | | | | | | | | |
| NC_011770 | 3624673 | A | G | 60 | 0 | 4 | 0 | NON_SYNONYMOU | MODERAT | MISSENSE | Aag/Gag | K29E | PLES_32751 | G | G | | | | | G | G | | | | |
| NC_011770 | 3624955 | A | G | 60 | 0 | 1 | 0 | NON_SYNONYMOU | MODERAT | MISSENSE | Acc/Gcc | T123A | PLES_32751 | | | | | | | G | | | | | |
| NC_011770 | 3625453 | C | T | 59 | 0 | 2 | 0 | NON_SYNONYMOU | MODERAT | MISSENSE | Cgc/Tgc | R289C | PLES_32751 | | | | | | | T | T | | | | |
| NC_011770 | 3626572 | C | G | 77 | 0 | 4 | 0 | NON_SYNONYMOU | MODERAT | MISSENSE | Cag/Gag | Q6E | PLES_32761 | | | | | | | | | G | G | G | G |
| NC_011770 | 3628436 | G | A | 81 | 0 | 2 | 0 | NON_SYNONYMOU | MODERAT | MISSENSE | Gcc/Acc | A327T | PLES_32781 | | | | | | | A | A | | | | |
| NC_011770 | 3628581 | T | C | 77 | 0 | 2 | 0 | NON_SYNONYMOU | MODERAT | MISSENSE | gTg/gCg | V375A | PLES_32781 | C | C | | | | | | | | | | |
| NC_011770 | 3628583 | T | C | 78 | 0 | 1 | 0 | NON_SYNONYMOU | MODERAT | MISSENSE | Ttc/Ctc | F376L | PLES_32781 | | | | | | | C | | | | | |
| NC_011770 | 3628599 | T | C | 80 | 0 | 2 | 0 | NON_SYNONYMOU | MODERAT | MISSENSE | aTc/aCc | I381T | PLES_32781 | | | | | | | C | C | | | | |
| NC_011770 | 3629309 | A | G | 77 | 0 | 12 | 0 | NON_SYNONYMOU | MODERAT | MISSENSE | Atc/Gtc | I618V | PLES_32781 | G | G | G | G | G | G | G | G | G | G | G | G |
| NC_011770 | 3629736 | C | G | 81 | 0 | 12 | 0 | NON_SYNONYMOU | MODERAT | MISSENSE | Ggc/Cgc | G220R | PLES_32791 | G | G | G | G | G | G | G | G | G | G | G | G |
| NC_011770 | 3631526 | C | T | 82 | 0 | 8 | 0 | NON_SYNONYMOU | MODERAT | MISSENSE | aGg/aAg | R73K | PLES_32801 | T | T | | | | | T | T | T | T | T | T |
| NC_011770 | 3632743 | T | C | 65 | 0 | 2 | 0 | NON_SYNONYMOU | MODERAT | MISSENSE | Atc/Gtc | I287V | PLES_32811 | | | | | | | C | C | | | | |
| NC_011770 | 3633211 | T | C | 74 | 0 | 2 | 0 | NON_SYNONYMOU | MODERAT | MISSENSE | Atc/Gtc | I131V | PLES_32811 | | | | | | | | | | | C | C |
| NC_011770 | 3634255 | G | A | 98 | 0 | 4 | 0 | NON_SYNONYMOU | MODERAT | MISSENSE | Cgc/Tgc | R294C | PLES_32821 | | | A | A | A | A | | | | | | |
| NC_011770 | 3635910 | G | A | 60 | 0 | 12 | 0 | NON_SYNONYMOU | MODERAT | MISSENSE | Gtc/Atc | V91I | PLES_32831 | A | A | A | A | A | A | A | A | A | A | A | A |
| NC_011770 | 3636354 | A | G | 72 | 0 | 2 | 0 | NON_SYNONYMOU | MODERAT | MISSENSE | Atc/Gtc | I239V | PLES_32831 | | | | | | | | | | | G | G |
| NC_011770 | 3636569 | G | A | 65 | 0 | 4 | 0 | NON_SYNONYMOU | MODERAT | MISSENSE | Ggc/Agc | G63S | PLES_32841 | | | | | | | | | A | A | A | A |
| NC_011770 | 3638424 | C | T | 10 | 10 | 2 | 0 | SPLICE_SITE_REGIC | LOW | SILENT | tGa/tAa | *174 | PLES_32861 | . | . | . | . | . | . | T | T | . | . | . | . |
| NC_011770 | 3638431 | C | T | 15 | 8 | 2 | 0 | NON_SYNONYMOU | MODERAT | MISSENSE | Gtt/Att | V172I | PLES_32861 | | | . | . | . | . | T | T | . | . | . | . |
| NC_011770 | 3638760 | A | G | 17 | 8 | 2 | 0 | NON_SYNONYMOU | MODERAT | MISSENSE | aTt/aCt | I62T | PLES_32861 | G | G | . | . | . | . | | | . | . | . | . |
| NC_011770 | 3639802 | C | T | 57 | 0 | 2 | 0 | NON_SYNONYMOU | MODERAT | MISSENSE | Cac/Tac | H42Y | PLES_32871 | | | | | | | | | | | T | T |
| NC_011770 | 3639811 | A | G | 57 | 0 | 6 | 0 | NON_SYNONYMOU | MODERAT | MISSENSE | Aac/Gac | N45D | PLES_32871 | | | G | G | G | G | | | | | G | G |
| NC_011770 | 3639826 | A | G | 58 | 0 | 6 | 0 | NON_SYNONYMOU | MODERAT | MISSENSE | Atc/Gtc | I50V | PLES_32871 | G | G | G | G | G | G | | | | | | |
| NC_011770 | 3640067 | C | G | 68 | 0 | 2 | 0 | NON_SYNONYMOU | MODERAT | MISSENSE | gCc/gGc | A130G | PLES_32871 | | | | | | | | | | | G | G |
| NC_011770 | 3640184 | T | C | 66 | 0 | 1 | 0 | NON_SYNONYMOU | MODERAT | MISSENSE | aTc/aCc | I169T | PLES_32871 | | | | | | | C | | | | | |
| NC_011770 | 3640196 | T | C | 65 | 0 | 2 | 0 | NON_SYNONYMOU | MODERAT | MISSENSE | tTc/tCc | F173S | PLES_32871 | C | C | | | | | | | | | | |
| NC_011770 | 3640582 | A | G | 80 | 0 | 6 | 0 | NON_SYNONYMOU | MODERAT | MISSENSE | Atc/Gtc | I302V | PLES_32871 | G | G | G | G | G | G | | | | | | |
| NC_011770 | 3640713 | G | C | 75 | 0 | 12 | 0 | NON_SYNONYMOU | MODERAT | MISSENSE | gaG/gaC | E345D | PLES_32871 | C | C | C | C | C | C | C | C | C | C | C | C |
| NC_011770 | 3640777 | G | A | 80 | 0 | 2 | 0 | NON_SYNONYMOU | MODERAT | MISSENSE | Gtc/Atc | V367I | PLES_32871 | A | A | | | | | | | | | | |
| NC_011770 | 3642276 | T | C | 65 | 0 | 6 | 0 | NON_SYNONYMOU | MODERAT | MISSENSE | Atc/Gtc | I218V | PLES_32891 | | | C | C | C | C | | | C | C | | |
| NC_011770 | 3642417 | T | C | 56 | 0 | 8 | 0 | NON_SYNONYMOU | MODERAT | MISSENSE | Acc/Gcc | T171A | PLES_32891 | C | C | C | C | C | C | | | C | C | | |
| NC_011770 | 3642659 | A | G | 75 | 0 | 12 | 0 | NON_SYNONYMOU | MODERAT | MISSENSE | gTc/gCc | V90A | PLES_32891 | G | G | G | G | G | G | G | G | G | G | G | G |
| NC_011770 | 3642771 | T | A | 74 | 0 | 2 | 0 | NON_SYNONYMOU | MODERAT | MISSENSE | Agc/Tgc | S53C | PLES_32891 | | | | | | | | | | | A | A |
| NC_011770 | 3643234 | G | C | 91 | 0 | 2 | 0 | NON_SYNONYMOU | MODERAT | MISSENSE | gCc/gGc | A418G | PLES_32901 | | | | | | | | | C | C | | |
| NC_011770 | 3643316 | T | C | 83 | 0 | 1 | 0 | NON_SYNONYMOU | MODERAT | MISSENSE | Acc/Gcc | T391A | PLES_32901 | | | | | | | C | | | | | |
| NC_011770 | 3643958 | A | G | 88 | 0 | 1 | 0 | NON_SYNONYMOU | MODERAT | MISSENSE | Tgc/Cgc | C177R | PLES_32901 | | | | | | | G | | | | | |
| NC_011770 | 3644284 | A | G | 82 | 0 | 4 | 0 | NON_SYNONYMOU | MODERAT | MISSENSE | gTc/gCc | V68A | PLES_32901 | | | G | G | G | G | | | | | | |
| NC_011770 | 3644726 | G | T | 74 | 0 | 4 | 0 | NON_SYNONYMOU | MODERAT | MISSENSE | Gcc/Tcc | A21S | PLES_32911 | | | T | T | T | T | | | | | | |
| NC_011770 | 3644739 | A | C | 79 | 0 | 4 | 0 | NON_SYNONYMOU | MODERAT | MISSENSE | aAc/aCc | N25T | PLES_32911 | | | C | C | C | C | | | | | | |
| NC_011770 | 3644898 | A | G | 69 | 0 | 4 | 0 | NON_SYNONYMOU | MODERAT | MISSENSE | aAa/aGa | K78R | PLES_32911 | | | G | G | G | G | | | | | | |
| NC_011770 | 3644995 | A | G | 62 | 0 | 4 | 0 | NON_SYNONYMOU | MODERAT | MISSENSE | Atc/Gtc | I27V | PLES_32921 | | | G | G | G | G | | | | | | |
| NC_011770 | 3645019 | G | A | 60 | 0 | 4 | 0 | NON_SYNONYMOU | MODERAT | MISSENSE | Gcc/Acc | A35T | PLES_32921 | | | A | A | A | A | | | | | | |
| NC_011770 | 3645082 | G | T | 54 | 0 | 6 | 0 | NON_SYNONYMOU | MODERAT | MISSENSE | Gca/Tca | A56S | PLES_32921 | T | T | T | T | T | T | | | | | | |
| NC_011770 | 3645252 | T | G | 29 | 4 | 4 | 0 | NON_SYNONYMOU | MODERAT | MISSENSE | gaA/gaC | E99D | PLES_32931 | G | G | . | . | . | . | | | G | G | | |
| NC_011770 | 3645478 | G | A | 49 | 0 | 4 | 0 | NON_SYNONYMOU | MODERAT | MISSENSE | aCc/aTc | T24I | PLES_32931 | | | A | A | A | A | | | | | | |
| NC_011770 | 3646190 | G | GC | 67 | 0 | 1 | 0 | FRAME_SHIFT | HIGH | | ctt/cCtt | L7P? | PLES_32951 | | GC | | | | | | | | | | |
| NC_011770 | 3646476 | T | C | 80 | 0 | 1 | 0 | NON_SYNONYMOU | MODERAT | MISSENSE | cTg/cCg | L100P | PLES_32951 | | | | | | | C | | | | | |
| NC_011770 | 3647288 | G | A | 96 | 0 | 2 | 0 | NON_SYNONYMOU | MODERAT | MISSENSE | Gcc/Acc | A189T | PLES_32961 | | | | | | | A | A | | | | |
| NC_011770 | 3647690 | G | A | 58 | 0 | 2 | 0 | NON_SYNONYMOU | MODERAT | MISSENSE | Gca/Aca | A323T | PLES_32961 | | | | | | | | | | | A | A |
| NC_011770 | 3651006 | C | T | 76 | 0 | 8 | 0 | NON_SYNONYMOU | MODERAT | MISSENSE | cGa/cAa | R375Q | PLES_33011 | T | T | | | | | T | T | T | T | T | T |
| NC_011770 | 3651019 | C | G | 72 | 0 | 8 | 0 | NON_SYNONYMOU | MODERAT | MISSENSE | Gtg/Ctg | V371L | PLES_33011 | G | G | | | | | G | G | G | G | G | G |
| NC_011770 | 3651023 | T | G | 73 | 0 | 8 | 0 | NON_SYNONYMOU | MODERAT | MISSENSE | gaA/gaC | E369D | PLES_33011 | G | G | | | | | G | G | G | G | G | G |
| NC_011770 | 3651282 | T | A | 84 | 0 | 8 | 0 | NON_SYNONYMOU | MODERAT | MISSENSE | tAc/tTc | Y283F | PLES_33011 | A | A | | | | | A | A | A | A | A | A |
| NC_011770 | 3651598 | C | T | 68 | 0 | 12 | 0 | NON_SYNONYMOU | MODERAT | MISSENSE | Gcg/Acg | A178T | PLES_33011 | T | T | T | T | T | T | T | T | T | T | T | T |
| NC_011770 | 3651832 | G | T | 69 | 0 | 4 | 0 | NON_SYNONYMOU | MODERAT | MISSENSE | Ctg/Atg | L100M | PLES_33011 | | | T | T | T | T | | | | | | |
| NC_011770 | 3651968 | G | C | 75 | 0 | 2 | 0 | NON_SYNONYMOU | MODERAT | MISSENSE | atC/atG | I54M | PLES_33011 | | | | | | | | | | | C | C |
| NC_011770 | 3652081 | C | T | 75 | 0 | 2 | 0 | NON_SYNONYMOU | MODERAT | MISSENSE | Gcc/Acc | A17T | PLES_33011 | | | | | | | | | T | T | | |

| | | | | | | | | | | | | | | | | | | | | | | | | | |
|---|---|---|---|---|---|---|---|---|---|---|---|---|---|---|---|---|---|---|---|---|---|---|---|---|---|
| NC_011770 | 3651099 | G | A | 80 | 0 | 1 | 0 | NON_SYNONYMOU | MODERAT | MISSENSE | gCg/gTg | A344V | PLES_33011 | | A | | | | | | | | | | |
| NC_011770 | 3652294 | C | A | 70 | 0 | 12 | 0 | NON_SYNONYMOU | MODERAT | MISSENSE | gaC/gaA | D4E | PLES_33021 | A | A | A | A | A | A | A | A | A | A | A | A |
| NC_011770 | 3652656 | A | G | 72 | 0 | 12 | 0 | INTRAGENIC | MODIFIER | | | | PLES_33031 | G | G | G | G | G | G | G | G | G | G | G | G |
| NC_011770 | 3652764 | G | T | 76 | 0 | 2 | 0 | INTRAGENIC | MODIFIER | | | | PLES_33031 | T | T | | | | | | | | | | |
| NC_011770 | 3652825 | CT | C | 75 | 0 | 1 | 0 | INTRAGENIC | MODIFIER | | | | PLES_33031 | | | | | | | C | | | | | |
| NC_011770 | 3652864 | A | G | 69 | 0 | 12 | 0 | INTRAGENIC | MODIFIER | | | | PLES_33031 | G | G | G | G | G | G | G | G | G | G | G | G |
| NC_011770 | 3652898 | TCCAGC | T | 65 | 0 | 2 | 0 | INTRAGENIC | MODIFIER | | | | PLES_33031 | T | T | | | | | | | | | | |
| NC_011770 | 3652969 | A | G | 65 | 0 | 12 | 0 | INTRAGENIC | MODIFIER | | | | PLES_33031 | G | G | G | G | G | G | G | G | G | G | G | G |
| NC_011770 | 3653043 | C | G | 68 | 0 | 2 | 0 | INTRAGENIC | MODIFIER | | | | PLES_33031 | | | | | G | G | | | | | | |
| NC_011770 | 3653053 | A | G | 68 | 0 | 8 | 0 | INTRAGENIC | MODIFIER | | | | PLES_33031 | | | G | G | G | G | G | G | G | G | | |
| NC_011770 | 3653018 | A | AC | 65 | 0 | 1 | 0 | INTRAGENIC | MODIFIER | | | | PLES_33031 | | | | | | | | | | AC | | |
| NC_011770 | 3654300 | G | T | 86 | 0 | 2 | 0 | NON_SYNONYMOU | MODERAT | MISSENSE | Gtg/Ttg | V331L | PLES_33041 | | | | | | | T | T | | | | |
| NC_011770 | 3654601 | T | C | 78 | 0 | 2 | 0 | NON_SYNONYMOU | MODERAT | MISSENSE | tTc/tCc | F29S | PLES_33051 | | | | | | | | | | | C | C |
| NC_011770 | 3655360 | G | A | 73 | 0 | 10 | 0 | NON_SYNONYMOU | MODERAT | MISSENSE | cGa/cAa | R282Q | PLES_33051 | A | A | A | A | A | A | | | A | A | A | A |
| NC_011770 | 3656103 | G | A | 91 | 0 | 4 | 0 | NON_SYNONYMOU | MODERAT | MISSENSE | Ggc/Agc | G530S | PLES_33051 | | | A | A | A | A | | | | | | |
| NC_011770 | 3656142 | G | A | 91 | 0 | 4 | 0 | NON_SYNONYMOU | MODERAT | MISSENSE | Gcc/Acc | A543T | PLES_33051 | | | A | A | A | A | | | | | | |
| NC_011770 | 3656230 | T | C | 86 | 0 | 2 | 0 | NON_SYNONYMOU | MODERAT | MISSENSE | aTg/aCg | M572T | PLES_33051 | | | | | | | | | C | C | | |
| NC_011770 | 3656829 | G | A | 80 | 0 | 2 | 0 | NON_SYNONYMOU | MODERAT | MISSENSE | Gcc/Acc | A772T | PLES_33051 | | | | | | | | | | | A | A |
| NC_011770 | 3657033 | C | G | 74 | 0 | 4 | 0 | NON_SYNONYMOU | MODERAT | MISSENSE | Caa/Gaa | Q840E | PLES_33051 | | | | | | | | | G | G | G | G |
| NC_011770 | 3657222 | G | A | 67 | 0 | 2 | 0 | NON_SYNONYMOU | MODERAT | MISSENSE | Gta/Ata | V903I | PLES_33051 | A | A | | | | | | | | | | |
| NC_011770 | 3657244 | T | C | 68 | 0 | 1 | 0 | NON_SYNONYMOU | MODERAT | MISSENSE | gTg/gCg | V910A | PLES_33051 | | | | | | | C | | | | | |
| NC_011770 | 3657453 | A | G | 45 | 0 | 12 | 0 | NON_SYNONYMOU | MODERAT | MISSENSE | Atc/Gtc | I980V | PLES_33051 | G | G | G | G | G | G | G | G | G | G | G | G |
| NC_011770 | 3657622 | A | C | 38 | 0 | 2 | 0 | NON_SYNONYMOU | MODERAT | MISSENSE | aAc/aCc | N1036T | PLES_33051 | | | | | | | | | | | C | C |
| NC_011770 | 3658638 | T | G | 67 | 0 | 2 | 0 | NON_SYNONYMOU | MODERAT | MISSENSE | gaA/gaC | E62D | PLES_33061 | | | | | | | G | G | | | | |
| NC_011770 | 3666405 | G | A | 52 | 0 | 2 | 0 | NON_SYNONYMOU | MODERAT | MISSENSE | gCc/gTc | A176V | PLES_33131 | | | | | | | A | A | | | | |
| NC_011770 | 3666612 | A | T | 66 | 0 | 6 | 0 | NON_SYNONYMOU | MODERAT | MISSENSE | gTg/gAg | V107E | PLES_33131 | T | T | | | | | T | T | T | T | | |
| NC_011770 | 3671237 | G | A | 74 | 0 | 1 | 0 | STOP_GAINED | HIGH | NONSENSE | tgG/tgA | W275* | PLES_33171 | | A | | | | | | | | | | |
| NC_011770 | 3670560 | G | C | 71 | 0 | 1 | 0 | NON_SYNONYMOU | MODERAT | MISSENSE | Gcc/Ccc | A50P | PLES_33171 | | | | | | | C | | | | | |
| NC_011770 | 3671070 | G | A | 62 | 0 | 8 | 0 | NON_SYNONYMOU | MODERAT | MISSENSE | Gag/Aag | E220K | PLES_33171 | | | A | A | A | A | A | A | | | A | A |
| NC_011770 | 3671106 | G | C | 63 | 0 | 6 | 0 | NON_SYNONYMOU | MODERAT | MISSENSE | Gag/Cag | E232Q | PLES_33171 | C | C | | | | | C | C | C | C | | |
| NC_011770 | 3671643 | A | C | 75 | 0 | 4 | 0 | NON_SYNONYMOU | MODERAT | MISSENSE | Aac/Cac | N411H | PLES_33171 | | | C | C | C | C | | | | | | |
| NC_011770 | 3671664 | A | T | 73 | 0 | 4 | 0 | NON_SYNONYMOU | MODERAT | MISSENSE | Att/Ttt | I418F | PLES_33171 | | | | | | | T | T | | | T | T |
| NC_011770 | 3673262 | G | T | 71 | 0 | 2 | 0 | STOP_GAINED | HIGH | NONSENSE | Gaa/Taa | E477* | PLES_33181 | | | | | | | | | T | T | | |
| NC_011770 | 3672906 | A | G | 62 | 0 | 12 | 0 | NON_SYNONYMOU | MODERAT | MISSENSE | gAc/gGc | D358G | PLES_33181 | G | G | G | G | G | G | G | G | G | G | G | G |
| NC_011770 | 3672951 | T | C | 63 | 0 | 2 | 0 | NON_SYNONYMOU | MODERAT | MISSENSE | gTc/gCc | V373A | PLES_33181 | | | | | | | C | C | | | | |
| NC_011770 | 3673002 | G | C | 68 | 0 | 10 | 0 | NON_SYNONYMOU | MODERAT | MISSENSE | cGg/cCg | R390P | PLES_33181 | | | C | C | C | C | C | C | C | C | C | C |
| NC_011770 | 3673237 | T | G | 68 | 0 | 4 | 0 | NON_SYNONYMOU | MODERAT | MISSENSE | atT/atG | I468M | PLES_33181 | | | | | | | G | G | G | G | | |
| NC_011770 | 3673248 | G | A | 70 | 0 | 2 | 0 | NON_SYNONYMOU | MODERAT | MISSENSE | aGc/aAc | S472N | PLES_33181 | | | | | | | | | A | A | | |
| NC_011770 | 3673257 | G | A | 69 | 0 | 2 | 0 | NON_SYNONYMOU | MODERAT | MISSENSE | cGt/cAt | R475H | PLES_33181 | | | | | | | | | A | A | | |
| NC_011770 | 3672707 | G | A | 70 | 0 | 1 | 0 | NON_SYNONYMOU | MODERAT | MISSENSE | Gac/Aac | D292N | PLES_33181 | | | | | | | | | | | | A |
| NC_011770 | 3673619 | GC | GCC | 72 | 0 | 2 | 0 | FRAME_SHIFT | HIGH | | ctc/ | L29 | PLES_33191 | | GCC | | | | | | | | | | |
| NC_011770 | 3674705 | GC | G | 78 | 0 | 2 | 0 | FRAME_SHIFT | HIGH | | gcc/ | A389 | PLES_33191 | | | | | | | G | G | | | | |
| NC_011770 | 3674085 | T | C | 73 | 0 | 2 | 0 | NON_SYNONYMOU | MODERAT | MISSENSE | cTc/cCc | L182P | PLES_33191 | C | C | | | | | | | | | | |
| NC_011770 | 3675992 | A | C | 58 | 0 | 2 | 0 | NON_SYNONYMOU | MODERAT | MISSENSE | gTg/gGg | V457G | PLES_33211 | | | | | | | | | | | C | C |
| NC_011770 | 3676854 | A | C | 65 | 0 | 12 | 0 | NON_SYNONYMOU | MODERAT | MISSENSE | Tcg/Gcg | S170A | PLES_33211 | C | C | C | C | C | C | C | C | C | C | C | C |
| NC_011770 | 3679695 | A | T | 67 | 0 | 12 | 0 | NON_SYNONYMOU | MODERAT | MISSENSE | gaT/gaA | D200E | PLES_33241 | T | T | T | T | T | T | T | T | T | T | T | T |
| NC_011770 | 3680945 | T | C | 78 | 0 | 10 | 0 | NON_SYNONYMOU | MODERAT | MISSENSE | Tcg/Ccg | S174P | PLES_33251 | C | C | C | C | C | C | C | C | C | C | | |
| NC_011770 | 3681012 | G | A | 81 | 0 | 1 | 0 | NON_SYNONYMOU | MODERAT | MISSENSE | cGc/cAc | R196H | PLES_33251 | | A | | | | | | | | | | |
| NC_011770 | 3682053 | G | A | 84 | 0 | 2 | 0 | NON_SYNONYMOU | MODERAT | MISSENSE | Gcc/Acc | A214T | PLES_33261 | | | | | | | | | | | A | A |
| NC_011770 | 3682057 | G | A | 82 | 0 | 6 | 0 | NON_SYNONYMOU | MODERAT | MISSENSE | gGc/gAc | G215D | PLES_33261 | | | A | A | A | A | A | A | | | | |
| NC_011770 | 3682395 | G | A | 80 | 0 | 2 | 0 | NON_SYNONYMOU | MODERAT | MISSENSE | Gcg/Acg | A328T | PLES_33261 | | | | | | | | | | | A | A |
| NC_011770 | 3682536 | G | A | 74 | 0 | 4 | 0 | NON_SYNONYMOU | MODERAT | MISSENSE | Gtg/Atg | V375M | PLES_33261 | | | | | | | A | A | A | A | | |
| NC_011770 | 3682945 | T | C | 71 | 0 | 12 | 0 | NON_SYNONYMOU | MODERAT | MISSENSE | gTc/gCc | V511A | PLES_33261 | C | C | C | C | C | C | C | C | C | C | C | C |
| NC_011770 | 3683155 | T | A | 70 | 0 | 12 | 0 | NON_SYNONYMOU | MODERAT | MISSENSE | gTa/gAa | V581E | PLES_33261 | A | A | A | A | A | A | A | A | A | A | A | A |
| NC_011770 | 3683166 | C | A | 72 | 0 | 2 | 0 | NON_SYNONYMOU | MODERAT | MISSENSE | Cag/Aag | Q585K | PLES_33261 | | | | | | | A | A | | | | |
| NC_011770 | 3684195 | G | A | 60 | 0 | 4 | 0 | NON_SYNONYMOU | MODERAT | MISSENSE | cGc/cAc | R40H | PLES_33291 | | | A | A | A | A | | | | | | |
| NC_011770 | 3684413 | T | A | 60 | 0 | 4 | 0 | NON_SYNONYMOU | MODERAT | MISSENSE | Ttc/Atc | F113I | PLES_33291 | | | A | A | A | A | | | | | | |
| NC_011770 | 3684446 | C | G | 62 | 0 | 4 | 0 | NON_SYNONYMOU | MODERAT | MISSENSE | Cag/Gag | Q124E | PLES_33291 | | | G | G | G | G | | | | | | |
| NC_011770 | 3685192 | G | C | 63 | 0 | 4 | 0 | NON_SYNONYMOU | MODERAT | MISSENSE | gGc/gCc | G143A | PLES_33301 | | | C | C | C | C | | | | | | |
| NC_011770 | 3685301 | G | C | 70 | 0 | 4 | 0 | NON_SYNONYMOU | MODERAT | MISSENSE | caG/caC | Q179H | PLES_33301 | | | C | C | C | C | | | | | | |
| NC_011770 | 3686667 | C | T | 67 | 0 | 4 | 0 | NON_SYNONYMOU | MODERAT | MISSENSE | gGc/gAc | G344D | PLES_33311 | | | T | T | T | T | | | | | | |
| NC_011770 | 3687673 | C | CCTG | 76 | 0 | 5 | 0 | CODON_INSERTIOI | MODERATE | | cag/cagCA | Q8QQ | PLES_33311 | | | | CCTG | CCTG | CCTG | | | CCTG | CCTG | | |

| | | | | | | | | | | | | | | | | | | | | | | | | | |
|---|---|---|---|---|---|---|---|---|---|---|---|---|---|---|---|---|---|---|---|---|---|---|---|---|---|
| NC_011770 | 3687880 | A | T | 74 | 0 | 12 | 0 | NON_SYNONYMOU | MODERAT | MISSENSE | cTg/cAg | L322Q | PLES_33321 | T | T | T | T | T | T | T | T | T | T | T | T |
| NC_011770 | 3688666 | T | C | 85 | 0 | 2 | 0 | NON_SYNONYMOU | MODERAT | MISSENSE | gAg/gGg | E60G | PLES_33321 | | | | | | | | | C | C | | |
| NC_011770 | 3688750 | C | G | 80 | 0 | 12 | 0 | NON_SYNONYMOU | MODERAT | MISSENSE | aGc/aCc | S32T | PLES_33321 | G | G | G | G | G | G | G | G | G | G | G | G |
| NC_011770 | 3688753 | C | T | 79 | 0 | 12 | 0 | NON_SYNONYMOU | MODERAT | MISSENSE | aGg/aAg | R31K | PLES_33321 | T | T | T | T | T | T | T | T | T | T | T | T |
| NC_011770 | 3689540 | T | C | 67 | 0 | 2 | 0 | NON_SYNONYMOU | MODERAT | MISSENSE | cAg/cGg | Q406R | PLES_33331 | C | C | | | | | | | | | | |
| NC_011770 | 3689631 | G | A | 66 | 0 | 2 | 0 | NON_SYNONYMOU | MODERAT | MISSENSE | Cgc/Tgc | R376C | PLES_33331 | | | | | | | | | | | A | A |
| NC_011770 | 3690243 | G | A | 67 | 0 | 4 | 0 | NON_SYNONYMOU | MODERAT | MISSENSE | Cgg/Tgg | R172W | PLES_33331 | A | A | | | | | A | A | | | | |
| NC_011770 | 3690614 | T | A | 72 | 0 | 4 | 0 | NON_SYNONYMOU | MODERAT | MISSENSE | tAc/tTc | Y48F | PLES_33331 | A | A | | | | | A | A | | | | |
| NC_011770 | 3695622 | C | G | 50 | 0 | 12 | 0 | NON_SYNONYMOU | MODERAT | MISSENSE | gaG/gaC | E70D | PLES_33381 | G | G | G | G | G | G | G | G | G | G | G | G |
| NC_011770 | 3698866 | G | A | 67 | 0 | 2 | 0 | NON_SYNONYMOU | MODERAT | MISSENSE | Gcc/Acc | A92T | PLES_33411 | | | | | | | | | A | A | | |
| NC_011770 | 3699164 | G | A | 77 | 0 | 1 | 0 | NON_SYNONYMOU | MODERAT | MISSENSE | gGc/gAc | G191D | PLES_33411 | | | | | | | A | | | | | |
| NC_011770 | 3699936 | C | A | 82 | 0 | 2 | 0 | START_LOST | HIGH | MISSENSE | atG/atT | M1I | PLES_33421 | | | | | | | | | | | A | A |
| NC_011770 | 3699850 | T | C | 74 | 0 | 8 | 0 | NON_SYNONYMOU | MODERAT | MISSENSE | aAc/aGc | N30S | PLES_33421 | | | C | C | C | C | C | C | | | C | C |
| NC_011770 | 3699932 | C | T | 79 | 0 | 12 | 0 | NON_SYNONYMOU | MODERAT | MISSENSE | Gtc/Atc | V3I | PLES_33421 | T | T | T | T | T | T | T | T | T | T | T | T |
| NC_011770 | 3699297 | G | GAGCGC | 78 | 0 | 8 | 0 | CODON_CHANGE_ | MODERATE | | ctc/ctGCC | L214LPL | PLES_33421 | GAGCGGC | GAGCGGC | | GAGCGGC | | GAGCGGC | GAGCGGC | GAGCGGC | GAGCGGC | GAGCGGC | | |
| NC_011770 | 3699958 | T | C | 77 | 0 | 8 | 0 | NON_SYNONYMOU | MODERAT | MISSENSE | gAc/gGc | D215G | PLES_33431 | | | C | C | C | C | C | C | | | C | C |
| NC_011770 | 3699988 | G | A | 83 | 0 | 4 | 0 | NON_SYNONYMOU | MODERAT | MISSENSE | gCg/gTg | A205V | PLES_33431 | | | A | A | A | A | | | | | | |
| NC_011770 | 3700142 | A | G | 78 | 0 | 8 | 0 | NON_SYNONYMOU | MODERAT | MISSENSE | Tgc/Cgc | C154R | PLES_33431 | | | G | G | G | G | G | G | | | G | G |
| NC_011770 | 3700441 | T | G | 74 | 0 | 2 | 0 | NON_SYNONYMOU | MODERAT | MISSENSE | gAc/gCc | D54A | PLES_33431 | | | | | | | | | G | G | | |
| NC_011770 | 3700567 | T | C | 77 | 0 | 4 | 0 | NON_SYNONYMOU | MODERAT | MISSENSE | tAc/tGc | Y12C | PLES_33431 | | | | | | | C | C | | | C | C |
| NC_011770 | 3700639 | T | C | 72 | 0 | 12 | 0 | NON_SYNONYMOU | MODERAT | MISSENSE | Atc/Gtc | I50V | PLES_33441 | C | C | C | C | C | C | C | C | C | C | C | C |
| NC_011770 | 3701808 | C | G | 84 | 0 | 2 | 0 | NON_SYNONYMOU | MODERAT | MISSENSE | gaC/gaG | D181E | PLES_33451 | | | | | | | | | G | G | | |
| NC_011770 | 3702059 | C | G | 89 | 0 | 2 | 0 | NON_SYNONYMOU | MODERAT | MISSENSE | gCc/gGc | A40G | PLES_33461 | | | | | | | | | G | G | | |
| NC_011770 | 3702079 | G | T | 87 | 0 | 2 | 0 | NON_SYNONYMOU | MODERAT | MISSENSE | Gcc/Tcc | A47S | PLES_33461 | | | | | | | | | T | T | | |
| NC_011770 | 3702094 | A | G | 88 | 0 | 12 | 0 | NON_SYNONYMOU | MODERAT | MISSENSE | Atc/Gtc | I52V | PLES_33461 | G | G | G | G | G | G | G | G | G | G | G | G |
| NC_011770 | 3702263 | C | T | 91 | 0 | 2 | 0 | NON_SYNONYMOU | MODERAT | MISSENSE | gCc/gTc | A108V | PLES_33461 | | | | | | | | | T | T | | |
| NC_011770 | 3702797 | G | A | 81 | 0 | 4 | 0 | NON_SYNONYMOU | MODERAT | MISSENSE | gGc/gAc | G286D | PLES_33461 | | | A | A | A | A | | | | | | |
| NC_011770 | 3704576 | A | AC | 69 | 0 | 1 | 0 | FRAME_SHIFT | HIGH | | agt/agGt | S292R? | PLES_33471 | | | | | | | AC | | | | | |
| NC_011770 | 3703273 | C | A | 63 | 0 | 6 | 0 | NON_SYNONYMOU | MODERAT | MISSENSE | Gcc/Tcc | A727S | PLES_33471 | | | | | | | A | A | A | A | A | A |
| NC_011770 | 3703816 | C | T | 78 | 0 | 4 | 0 | NON_SYNONYMOU | MODERAT | MISSENSE | Gcg/Acg | A546T | PLES_33471 | T | T | | | | | | | T | T | | |
| NC_011770 | 3704282 | A | T | 84 | 0 | 2 | 0 | NON_SYNONYMOU | MODERAT | MISSENSE | caT/caA | H390Q | PLES_33471 | | | | | | | | | | | T | T |
| NC_011770 | 3705595 | C | T | 91 | 0 | 2 | 0 | NON_SYNONYMOU | MODERAT | MISSENSE | cGc/cAc | R317H | PLES_33481 | T | T | | | | | | | | | | |
| NC_011770 | 3706232 | G | A | 47 | 0 | 1 | 0 | NON_SYNONYMOU | MODERAT | MISSENSE | Cgc/Tgc | R105C | PLES_33481 | | | | | | | A | | | | | |
| NC_011770 | 3707270 | T | C | 66 | 0 | 4 | 0 | NON_SYNONYMOU | MODERAT | MISSENSE | Aac/Gac | N193D | PLES_33491 | | | | | | | | | C | C | C | C |
| NC_011770 | 3710711 | T | C | 90 | 0 | 2 | 0 | NON_SYNONYMOU | MODERAT | MISSENSE | Aac/Gac | N504D | PLES_33511 | | | | | | | | | C | C | | |
| NC_011770 | 3710732 | T | C | 91 | 0 | 10 | 0 | NON_SYNONYMOU | MODERAT | MISSENSE | Agc/Ggc | S497G | PLES_33511 | | | C | C | C | C | C | C | C | C | C | C |
| NC_011770 | 3711262 | T | C | 95 | 0 | 1 | 0 | NON_SYNONYMOU | MODERAT | MISSENSE | gAc/gGc | D320G | PLES_33511 | | | | | | | C | | | | | |
| NC_011770 | 3711556 | G | T | 93 | 0 | 6 | 0 | NON_SYNONYMOU | MODERAT | MISSENSE | gCc/gAc | A222D | PLES_33511 | | | T | T | T | T | | | T | T | | |
| NC_011770 | 3711850 | A | G | 81 | 0 | 8 | 0 | NON_SYNONYMOU | MODERAT | MISSENSE | gTg/gCg | V124A | PLES_33511 | | | G | G | G | G | G | G | G | G | | |
| NC_011770 | 3712108 | C | G | 67 | 0 | 8 | 0 | NON_SYNONYMOU | MODERAT | MISSENSE | gGg/gCg | G38A | PLES_33511 | | | G | G | G | G | G | G | G | G | | |
| NC_011770 | 3710729 | A | G | 90 | 0 | 1 | 0 | NON_SYNONYMOU | MODERAT | MISSENSE | Tgc/Cgc | C498R | PLES_33511 | | | | | | | | G | | | | |
| NC_011770 | 3714812 | G | A | 57 | 0 | 2 | 0 | NON_SYNONYMOU | MODERAT | MISSENSE | gCa/gTa | A61V | PLES_33541 | | | | | | | | | | | A | A |
| NC_011770 | 3714908 | G | A | 62 | 0 | 2 | 0 | NON_SYNONYMOU | MODERAT | MISSENSE | gCc/gTc | A29V | PLES_33541 | | | | | | | | | A | A | | |
| NC_011770 | 3714974 | G | C | 62 | 0 | 12 | 0 | NON_SYNONYMOU | MODERAT | MISSENSE | cCa/cGa | P7R | PLES_33541 | C | C | C | C | C | C | C | C | C | C | C | C |
| NC_011770 | 3715241 | A | C | 58 | 0 | 2 | 0 | NON_SYNONYMOU | MODERAT | MISSENSE | gaT/gaG | D11E | PLES_33551 | | | | | | | | | C | C | | |
| NC_011770 | 3715507 | T | C | 64 | 0 | 4 | 0 | NON_SYNONYMOU | MODERAT | MISSENSE | cTg/cCg | L17P | PLES_33561 | | | C | C | C | C | | | | | | |
| NC_011770 | 3715528 | A | G | 63 | 0 | 4 | 0 | NON_SYNONYMOU | MODERAT | MISSENSE | aAc/aGc | N24S | PLES_33561 | | | G | G | G | G | | | | | | |
| NC_011770 | 3715560 | C | G | 65 | 0 | 4 | 0 | NON_SYNONYMOU | MODERAT | MISSENSE | Ccg/Gcg | P35A | PLES_33561 | | | | | | | G | G | | | G | G |
| NC_011770 | 3715600 | T | C | 66 | 0 | 4 | 0 | NON_SYNONYMOU | MODERAT | MISSENSE | gTc/gCc | V48A | PLES_33561 | | | C | C | C | C | | | | | | |
| NC_011770 | 3715611 | G | A | 65 | 0 | 4 | 0 | NON_SYNONYMOU | MODERAT | MISSENSE | Gcg/Acg | A52T | PLES_33561 | | | A | A | A | A | | | | | | |
| NC_011770 | 3715632 | A | C | 68 | 0 | 4 | 0 | NON_SYNONYMOU | MODERAT | MISSENSE | Atg/Ctg | M59L | PLES_33561 | | | C | C | C | C | | | | | | |
| NC_011770 | 3715689 | C | A | 67 | 0 | 4 | 0 | NON_SYNONYMOU | MODERAT | MISSENSE | Cct/Act | P78T | PLES_33561 | | | A | A | A | A | | | | | | |
| NC_011770 | 3715724 | C | A | 66 | 0 | 2 | 0 | NON_SYNONYMOU | MODERAT | MISSENSE | gaC/gaA | D89E | PLES_33561 | A | A | | | | | | | | | | |
| NC_011770 | 3715890 | G | A | 61 | 0 | 12 | 0 | NON_SYNONYMOU | MODERAT | MISSENSE | Gac/Aac | D145N | PLES_33561 | A | A | A | A | A | A | A | A | A | A | A | A |
| NC_011770 | 3715932 | C | G | 58 | 0 | 4 | 0 | NON_SYNONYMOU | MODERAT | MISSENSE | Cgc/Ggc | R159G | PLES_33561 | | | G | G | G | G | | | | | | |
| NC_011770 | 3716098 | A | C | 60 | 0 | 4 | 0 | NON_SYNONYMOU | MODERAT | MISSENSE | Tgc/Ggc | C85G | PLES_33571 | | | C | C | C | C | | | | | | |
| NC_011770 | 3716103 | G | A | 60 | 0 | 4 | 0 | NON_SYNONYMOU | MODERAT | MISSENSE | tCg/tTg | S83L | PLES_33571 | | | A | A | A | A | | | | | | |
| NC_011770 | 3716104 | A | C | 58 | 0 | 4 | 0 | NON_SYNONYMOU | MODERAT | MISSENSE | Tcg/Gcg | S83A | PLES_33571 | | | C | C | C | C | | | | | | |
| NC_011770 | 3716124 | G | A | 60 | 0 | 2 | 0 | NON_SYNONYMOU | MODERAT | MISSENSE | aCc/aTc | T76I | PLES_33571 | | | | | | | | | | | A | A |
| NC_011770 | 3716181 | A | G | 57 | 0 | 4 | 0 | NON_SYNONYMOU | MODERAT | MISSENSE | gTc/gCc | V57A | PLES_33571 | G | G | | | | | | | G | G | | |
| NC_011770 | 3717746 | G | A | 88 | 0 | 2 | 0 | NON_SYNONYMOU | MODERAT | MISSENSE | Ggc/Agc | G151S | PLES_33591 | | | | | A | A | | | | | | |
| NC_011770 | 3718254 | G | A | 77 | 0 | 1 | 0 | NON_SYNONYMOU | MODERAT | MISSENSE | gGc/gAc | G320D | PLES_33591 | | | | | | | A | | | | | |

| | | | | | | | | | | | | | | | | | | | | | | | | | |
|---|---|---|---|---|---|---|---|---|---|---|---|---|---|---|---|---|---|---|---|---|---|---|---|---|---|
| NC_011770 | 3719134 | T | C | 63 | 0 | 12 | 0 | NON_SYNONYMOU | MODERAT | MISSENSE | cAg/cGg | Q27R | PLES_33601 | C | C | C | C | C | C | C | C | C | C | C | C |
| NC_011770 | 3719158 | T | C | 66 | 0 | 2 | 0 | NON_SYNONYMOU | MODERAT | MISSENSE | aAc/aGc | N19S | PLES_33601 | | | | | | | C | C | | | | |
| NC_011770 | 3719493 | C | G | 63 | 0 | 2 | 0 | NON_SYNONYMOU | MODERAT | MISSENSE | Ggc/Cgc | G146R | PLES_33611 | G | G | | | | | | | | | | |
| NC_011770 | 3720214 | G | T | 80 | 0 | 2 | 0 | NON_SYNONYMOU | MODERAT | MISSENSE | Gtg/Ttg | V47L | PLES_33621 | T | T | | | | | | | | | | |
| NC_011770 | 3720940 | G | A | 68 | 0 | 1 | 0 | NON_SYNONYMOU | MODERAT | MISSENSE | Gac/Aac | D289N | PLES_33621 | | | | | | | | | . | A | | |
| NC_011770 | 3721033 | A | G | 69 | 0 | 6 | 0 | NON_SYNONYMOU | MODERAT | MISSENSE | Tgg/Cgg | W236R | PLES_33631 | | | G | G | G | G | G | G | | | | |
| NC_011770 | 3721038 | G | A | 69 | 0 | 12 | 0 | NON_SYNONYMOU | MODERAT | MISSENSE | tCg/tTg | S234L | PLES_33631 | A | A | A | A | A | A | A | A | A | A | A | A |
| NC_011770 | 3721086 | A | G | 67 | 0 | 6 | 0 | NON_SYNONYMOU | MODERAT | MISSENSE | cTg/cCg | L218P | PLES_33631 | | | G | G | G | G | G | G | | | | |
| NC_011770 | 3721155 | G | A | 66 | 0 | 12 | 0 | NON_SYNONYMOU | MODERAT | MISSENSE | gCg/gTg | A195V | PLES_33631 | A | A | A | A | A | A | A | A | A | A | A | A |
| NC_011770 | 3721260 | T | C | 62 | 0 | 12 | 0 | NON_SYNONYMOU | MODERAT | MISSENSE | cAg/cGg | Q160R | PLES_33631 | C | C | C | C | C | C | C | C | C | C | C | C |
| NC_011770 | 3721495 | G | T | 56 | 0 | 12 | 0 | NON_SYNONYMOU | MODERAT | MISSENSE | Ctc/Atc | L82I | PLES_33631 | T | T | T | T | T | T | T | T | T | T | T | T |
| NC_011770 | 3721638 | G | C | 54 | 0 | 4 | 0 | NON_SYNONYMOU | MODERAT | MISSENSE | aCc/aGc | T34S | PLES_33631 | | | C | C | C | C | | | | | | |
| NC_011770 | 3723156 | G | T | 30 | 0 | 12 | 0 | NON_SYNONYMOU | MODERAT | MISSENSE | Gcg/Tcg | A78S | PLES_33651 | T | T | T | T | T | T | T | T | T | T | T | T |
| NC_011770 | 3723220 | C | T | 27 | 0 | 12 | 0 | NON_SYNONYMOU | MODERAT | MISSENSE | gCc/gTc | A99V | PLES_33651 | T | T | T | T | T | T | T | T | T | T | T | T |
| NC_011770 | 3723280 | T | G | 34 | 0 | 12 | 0 | NON_SYNONYMOU | MODERAT | MISSENSE | gTa/gGa | V119G | PLES_33651 | G | G | G | G | G | G | G | G | G | G | G | G |
| NC_011770 | 3723682 | T | C | 59 | 0 | 12 | 0 | NON_SYNONYMOU | MODERAT | MISSENSE | gaT/gaG | D62E | PLES_33661 | C | C | C | C | C | C | G | G | C | C | C | C |
| NC_011770 | 3723768 | C | A | 60 | 0 | 12 | 0 | NON_SYNONYMOU | MODERAT | MISSENSE | gCc/gAc | A91D | PLES_33661 | A | A | A | A | A | A | A | A | A | A | A | A |
| NC_011770 | 3723812 | G | A | 68 | 0 | 2 | 0 | NON_SYNONYMOU | MODERAT | MISSENSE | Gac/Aac | D106N | PLES_33661 | | | | | A | A | | | | | | |
| NC_011770 | 3723916 | A | T | 73 | 0 | 2 | 0 | NON_SYNONYMOU | MODERAT | MISSENSE | caA/caT | Q140H | PLES_33661 | | | | | | | | | T | T | | |
| NC_011770 | 3724014 | G | A | 74 | 0 | 4 | 0 | NON_SYNONYMOU | MODERAT | MISSENSE | cGg/cAg | R173Q | PLES_33661 | | | | | | | A | A | A | A | | |
| NC_011770 | 3724633 | C | A | 59 | 0 | 4 | 0 | NON_SYNONYMOU | MODERAT | MISSENSE | Ccc/Acc | P103T | PLES_33671 | | | | | | | A | A | A | A | | |
| NC_011770 | 3724639 | G | A | 58 | 0 | 4 | 0 | NON_SYNONYMOU | MODERAT | MISSENSE | Ggc/Agc | G105S | PLES_33671 | | | | | | | A | A | A | A | | |
| NC_011770 | 3724661 | A | G | 60 | 0 | 4 | 0 | NON_SYNONYMOU | MODERAT | MISSENSE | aAc/aGc | N112S | PLES_33671 | | | | | | | G | G | G | G | | |
| NC_011770 | 3724702 | G | C | 62 | 0 | 4 | 0 | NON_SYNONYMOU | MODERAT | MISSENSE | Gtc/Ctc | V126L | PLES_33671 | | | | | | | C | C | C | C | | |
| NC_011770 | 3724615 | A | G | 58 | 0 | 1 | 0 | NON_SYNONYMOU | MODERAT | MISSENSE | Att/Gtt | I97V | PLES_33671 | | | | G | | | | | | | | |
| NC_011770 | 3724906 | C | A | 56 | 0 | 4 | 0 | NON_SYNONYMOU | MODERAT | MISSENSE | gCc/gAc | A32D | PLES_33681 | | | | | | | A | A | A | A | | |
| NC_011770 | 3724953 | G | T | 52 | 0 | 4 | 0 | NON_SYNONYMOU | MODERAT | MISSENSE | Gac/Tac | D48Y | PLES_33681 | | | T | T | T | T | | | | | | |
| NC_011770 | 3725166 | G | C | 54 | 0 | 6 | 0 | NON_SYNONYMOU | MODERAT | MISSENSE | Gtc/Ctc | V119L | PLES_33681 | C | C | | | | | C | C | C | C | | |
| NC_011770 | 3725633 | C | G | 75 | 0 | 2 | 0 | NON_SYNONYMOU | MODERAT | MISSENSE | Ctg/Gtg | L66V | PLES_33691 | | | | | | | | | | | G | G |
| NC_011770 | 3726567 | G | A | 50 | 0 | 2 | 0 | NON_SYNONYMOU | MODERAT | MISSENSE | Gcc/Acc | A16T | PLES_33701 | | | | | | | | | A | A | | |
| NC_011770 | 3727344 | G | T | 45 | 0 | 2 | 0 | NON_SYNONYMOU | MODERAT | MISSENSE | caG/caT | Q43H | PLES_33711 | T | T | | | | | | | | | | |
| NC_011770 | 3727568 | C | T | 64 | 0 | 2 | 0 | NON_SYNONYMOU | MODERAT | MISSENSE | gCg/gTg | A118V | PLES_33711 | | | | | | | T | T | | | | |
| NC_011770 | 3729070 | A | C | 72 | 0 | 2 | 0 | NON_SYNONYMOU | MODERAT | MISSENSE | Atg/Ctg | M348L | PLES_33721 | | | | | | | | | | | C | C |
| NC_011770 | 3729103 | A | C | 75 | 0 | 2 | 0 | NON_SYNONYMOU | MODERAT | MISSENSE | Aag/Cag | K359Q | PLES_33721 | | | | | | | | | | | C | C |
| NC_011770 | 3735373 | C | T | 62 | 0 | 10 | 0 | NON_SYNONYMOU | MODERAT | MISSENSE | cGg/cAg | R373Q | PLES_33781 | T | T | T | T | T | T | T | T | T | T | | |
| NC_011770 | 3735525 | A | C | 61 | 0 | 8 | 0 | NON_SYNONYMOU | MODERAT | MISSENSE | aaT/aaG | N322K | PLES_33781 | C | C | C | C | C | C | C | C | | | | |
| NC_011770 | 3735527 | T | G | 62 | 0 | 8 | 0 | NON_SYNONYMOU | MODERAT | MISSENSE | Aat/Cat | N322H | PLES_33781 | G | G | G | G | G | G | G | G | | | | |
| NC_011770 | 3735694 | T | C | 69 | 0 | 10 | 0 | NON_SYNONYMOU | MODERAT | MISSENSE | aAc/aGc | N266S | PLES_33781 | C | C | C | C | C | C | C | C | C | C | | |
| NC_011770 | 3735925 | T | C | 76 | 0 | 10 | 0 | NON_SYNONYMOU | MODERAT | MISSENSE | aAg/aGg | K189R | PLES_33781 | C | C | C | C | C | C | C | C | | | C | C |
| NC_011770 | 3736235 | G | C | 66 | 0 | 8 | 0 | NON_SYNONYMOU | MODERAT | MISSENSE | Ctg/Gtg | L86V | PLES_33781 | C | C | C | C | C | C | | | | | C | C |
| NC_011770 | 3736486 | A | G | 68 | 0 | 8 | 0 | NON_SYNONYMOU | MODERAT | MISSENSE | tTa/tCa | L2S | PLES_33781 | G | G | G | G | G | G | | | | | G | G |
| NC_011770 | 3736855 | G | A | 75 | 0 | 2 | 0 | NON_SYNONYMOU | MODERAT | MISSENSE | gCg/gTg | A420V | PLES_33791 | | | | | | | | | | | A | A |
| NC_011770 | 3737179 | G | A | 82 | 0 | 1 | 0 | NON_SYNONYMOU | MODERAT | MISSENSE | cCg/cTg | P312L | PLES_33791 | | | | | | | A | | | | | |
| NC_011770 | 3737209 | A | G | 80 | 0 | 12 | 0 | NON_SYNONYMOU | MODERAT | MISSENSE | cTg/cCg | L302P | PLES_33791 | G | G | G | G | G | G | G | G | G | G | G | G |
| NC_011770 | 3738331 | G | A | 71 | 0 | 2 | 0 | NON_SYNONYMOU | MODERAT | MISSENSE | Gtg/Atg | V10M | PLES_33801 | | | | | | | | | A | A | | |
| NC_011770 | 3738332 | T | C | 72 | 0 | 2 | 0 | NON_SYNONYMOU | MODERAT | MISSENSE | gTg/gCg | V10A | PLES_33801 | | | | | | | | | C | C | | |
| NC_011770 | 3739006 | G | A | 69 | 0 | 6 | 0 | NON_SYNONYMOU | MODERAT | MISSENSE | Ggc/Agc | G235S | PLES_33801 | A | A | | | | | A | A | | | A | A |
| NC_011770 | 3739196 | T | C | 68 | 0 | 12 | 0 | NON_SYNONYMOU | MODERAT | MISSENSE | aTc/aCc | I298T | PLES_33801 | C | C | C | C | C | C | C | C | C | C | C | C |
| NC_011770 | 3740086 | C | A | 68 | 0 | 12 | 0 | NON_SYNONYMOU | MODERAT | MISSENSE | Cat/Aat | H28N | PLES_33821 | A | A | A | A | A | A | A | A | A | A | A | A |
| NC_011770 | 3740096 | T | C | 64 | 0 | 2 | 0 | NON_SYNONYMOU | MODERAT | MISSENSE | cTg/cCg | L31P | PLES_33821 | C | C | | | | | | | | | | |
| NC_011770 | 3740143 | C | A | 71 | 0 | 2 | 0 | NON_SYNONYMOU | MODERAT | MISSENSE | Cag/Aag | Q47K | PLES_33821 | A | A | | | | | | | | | | |
| NC_011770 | 3740815 | G | C | 66 | 0 | 2 | 0 | NON_SYNONYMOU | MODERAT | MISSENSE | Gtc/Ctc | V271L | PLES_33821 | | | | | | | | | C | C | | |
| NC_011770 | 3740948 | A | G | 68 | 0 | 8 | 0 | NON_SYNONYMOU | MODERAT | MISSENSE | cAg/cGg | Q315R | PLES_33821 | G | G | G | G | G | G | | | G | G | | |
| NC_011770 | 3741089 | C | T | 70 | 0 | 2 | 0 | NON_SYNONYMOU | MODERAT | MISSENSE | gCc/gTc | A362V | PLES_33821 | | | | | | | | | | | T | T |
| NC_011770 | 3741502 | G | A | 43 | 0 | 2 | 0 | NON_SYNONYMOU | MODERAT | MISSENSE | Ggc/Agc | G500S | PLES_33821 | | | | | | | | | A | A | | |
| NC_011770 | 3741505 | G | A | 43 | 0 | 4 | 0 | NON_SYNONYMOU | MODERAT | MISSENSE | Gtc/Atc | V501I | PLES_33821 | A | A | | | | | A | A | | | | |
| NC_011770 | 3740015 | T | C | 72 | 0 | 1 | 0 | NON_SYNONYMOU | MODERAT | MISSENSE | gTg/gCg | V4A | PLES_33821 | | | | | | | | C | | | | |
| NC_011770 | 3741223 | G | A | 66 | 0 | 1 | 0 | NON_SYNONYMOU | MODERAT | MISSENSE | Gcc/Acc | A407T | PLES_33821 | | | | | | | | A | | | | |
| NC_011770 | 3742244 | G | A | 27 | 0 | 1 | 0 | NON_SYNONYMOU | MODERAT | MISSENSE | Gaa/Aaa | E56K | PLES_33831 | | | | | | | A | | | | | |
| NC_011770 | 3742418 | G | C | 30 | 0 | 6 | 0 | NON_SYNONYMOU | MODERAT | MISSENSE | Gcg/Ccg | A114P | PLES_33831 | | | C | C | C | C | | | | | C | C |
| NC_011770 | 3742763 | G | A | 15 | 0 | 2 | 0 | NON_SYNONYMOU | MODERAT | MISSENSE | Gcc/Acc | A229T | PLES_33831 | A | A | | | | | | | | | | |
| NC_011770 | 3742853 | G | C | 21 | 0 | 10 | 0 | NON_SYNONYMOU | MODERAT | MISSENSE | Gag/Cag | E259Q | PLES_33831 | C | C | C | C | C | C | C | C | | | C | C |

| | | | | | | | | | | | | | | | | | | | | | | | | | |
|---|---|---|---|---|---|---|---|---|---|---|---|---|---|---|---|---|---|---|---|---|---|---|---|---|---|
| NC_011770 | 3742928 | A | G | 22 | 0 | 2 | 0 | NON_SYNONYMOU | MODERAT | MISSENSE | Acc/Gcc | T284A | PLES_33831 | | | | | | | | | G | G | | |
| NC_011770 | 3752221 | T | C | 53 | 0 | 4 | 0 | NON_SYNONYMOU | MODERAT | MISSENSE | gAg/gGg | E109G | PLES_33891 | | | C | C | C | C | | | | | | |
| NC_011770 | 3752238 | C | G | 51 | 0 | 6 | 0 | NON_SYNONYMOU | MODERAT | MISSENSE | gaG/gaC | E103D | PLES_33891 | G | G | | | | | G | G | | | G | G |
| NC_011770 | 3752252 | T | A | 51 | 0 | 6 | 0 | NON_SYNONYMOU | MODERAT | MISSENSE | Acc/Tcc | T99S | PLES_33891 | A | A | | | | | A | A | | | A | A |
| NC_011770 | 3752270 | T | C | 53 | 0 | 2 | 0 | NON_SYNONYMOU | MODERAT | MISSENSE | Acc/Gcc | T93A | PLES_33891 | | | | | | | C | C | | | | |
| NC_011770 | 3752271 | C | A | 53 | 0 | 10 | 0 | NON_SYNONYMOU | MODERAT | MISSENSE | gaG/gaT | E92D | PLES_33891 | A | A | A | A | A | A | A | A | | | A | A |
| NC_011770 | 3752420 | C | T | 62 | 0 | 2 | 0 | NON_SYNONYMOU | MODERAT | MISSENSE | Gac/Aac | D43N | PLES_33891 | | | | | | | T | T | | | | |
| NC_011770 | 3752452 | C | T | 60 | 0 | 2 | 0 | NON_SYNONYMOU | MODERAT | MISSENSE | cGc/cAc | R32H | PLES_33891 | | | | | | | T | T | | | | |
| NC_011770 | 3754820 | G | A | 38 | 6 | 2 | 0 | NON_SYNONYMOU | MODERAT | MISSENSE | gCc/gTc | A143V | PLES_33901 | . | . | | | | | . | . | A | A | . | . |
| NC_011770 | 3753048 | GCAA | G | 29 | 6 | 5 | 0 | CODON_DELETION | MODERATE | | ttg/ | L733 | PLES_33901 | . | . | | G | G | G | . | . | G | G | . | . |
| NC_011770 | 3756241 | A | G | 32 | 6 | 6 | 0 | STOP_LOST+SPLICE | HIGH | MISSENSE | Tag/Cag | *168Q | PLES_33921 | . | . | G | G | G | G | . | . | G | G | . | . |
| NC_011770 | 3756346 | G | C | 35 | 6 | 2 | 0 | NON_SYNONYMOU | MODERAT | MISSENSE | Cgc/Ggc | R133G | PLES_33921 | . | . | | | | | . | . | C | C | . | . |
| NC_011770 | 3757088 | T | G | 44 | 0 | 2 | 0 | NON_SYNONYMOU | MODERAT | MISSENSE | Aag/Cag | K374Q | PLES_33931 | G | G | | | | | | | | | | |
| NC_011770 | 3757718 | G | C | 58 | 0 | 2 | 0 | NON_SYNONYMOU | MODERAT | MISSENSE | Ctc/Gtc | L164V | PLES_33931 | C | C | | | | | | | | | | |
| NC_011770 | 3757784 | T | C | 51 | 0 | 2 | 0 | NON_SYNONYMOU | MODERAT | MISSENSE | Aac/Gac | N142D | PLES_33931 | C | C | | | | | | | | | | |
| NC_011770 | 3757811 | G | A | 54 | 0 | 2 | 0 | NON_SYNONYMOU | MODERAT | MISSENSE | Cgc/Tgc | R133C | PLES_33931 | A | A | | | | | | | | | | |
| NC_011770 | 3757901 | T | A | 58 | 0 | 2 | 0 | NON_SYNONYMOU | MODERAT | MISSENSE | Acg/Tcg | T103S | PLES_33931 | | | | | | | | | A | A | | |
| NC_011770 | 3757993 | A | T | 60 | 0 | 2 | 0 | NON_SYNONYMOU | MODERAT | MISSENSE | tTc/tAc | F72Y | PLES_33931 | | | | | | | | | T | T | | |
| NC_011770 | 3758001 | G | T | 58 | 0 | 8 | 0 | NON_SYNONYMOU | MODERAT | MISSENSE | gaC/gaA | D69E | PLES_33931 | T | T | T | T | T | T | T | T | | | | |
| NC_011770 | 3758056 | A | G | 61 | 0 | 2 | 0 | NON_SYNONYMOU | MODERAT | MISSENSE | gTg/gCg | V51A | PLES_33931 | | | | | | | | | G | G | | |
| NC_011770 | 3758136 | A | C | 63 | 0 | 2 | 0 | NON_SYNONYMOU | MODERAT | MISSENSE | gaT/gaG | D24E | PLES_33931 | | | | | | | C | C | | | | |
| NC_011770 | 3758631 | T | A | 50 | 0 | 4 | 0 | NON_SYNONYMOU | MODERAT | MISSENSE | tTc/tAc | F18Y | PLES_33941 | A | A | | | | | | | | | A | A |
| NC_011770 | 3758637 | G | A | 48 | 0 | 2 | 0 | NON_SYNONYMOU | MODERAT | MISSENSE | cGc/cAc | R20H | PLES_33941 | | | | | | | A | A | | | | |
| NC_011770 | 3762060 | C | G | 74 | 0 | 4 | 0 | NON_SYNONYMOU | MODERAT | MISSENSE | Gtc/Ctc | V593L | PLES_33971 | | | G | G | G | G | | | | | | |
| NC_011770 | 3762279 | G | C | 72 | 0 | 10 | 0 | NON_SYNONYMOU | MODERAT | MISSENSE | Ctc/Gtc | L520V | PLES_33971 | | | C | C | C | C | C | C | C | C | C | C |
| NC_011770 | 3762678 | G | T | 72 | 0 | 2 | 0 | NON_SYNONYMOU | MODERAT | MISSENSE | Cac/Aac | H387N | PLES_33971 | | | | | | | | | | | T | T |
| NC_011770 | 3763119 | T | G | 76 | 0 | 2 | 0 | NON_SYNONYMOU | MODERAT | MISSENSE | Atc/Ctc | I240L | PLES_33971 | | | | | | | | | G | G | | |
| NC_011770 | 3763414 | C | G | 80 | 0 | 2 | 0 | NON_SYNONYMOU | MODERAT | MISSENSE | gaG/gaC | E141D | PLES_33971 | | | | | | | G | G | | | | |
| NC_011770 | 3764902 | T | C | 69 | 0 | 4 | 0 | NON_SYNONYMOU | MODERAT | MISSENSE | Atc/Gtc | I80V | PLES_33991 | | | C | C | C | C | | | | | | |
| NC_011770 | 3764952 | T | C | 71 | 0 | 2 | 0 | NON_SYNONYMOU | MODERAT | MISSENSE | aAc/aGc | N63S | PLES_33991 | | | | | | | C | C | | | | |
| NC_011770 | 3764964 | A | T | 71 | 0 | 4 | 0 | NON_SYNONYMOU | MODERAT | MISSENSE | cTg/cAg | L59Q | PLES_33991 | | | T | T | T | T | | | | | | |
| NC_011770 | 3765090 | T | A | 67 | 0 | 10 | 0 | NON_SYNONYMOU | MODERAT | MISSENSE | cAg/cTg | Q17L | PLES_33991 | A | A | A | A | A | A | A | A | A | A | | |
| NC_011770 | 3765133 | C | T | 67 | 0 | 1 | 0 | NON_SYNONYMOU | MODERAT | MISSENSE | Ggt/Agt | G3S | PLES_33991 | T | | | | | | | | | | | |
| NC_011770 | 3770932 | TG | T | 52 | 0 | 1 | 0 | FRAME_SHIFT | HIGH | | cat/ | H4 | PLES_34011 | | | | | | | | T | | | | |
| NC_011770 | 3769419 | T | C | 67 | 0 | 2 | 0 | NON_SYNONYMOU | MODERAT | MISSENSE | atA/atG | I508M | PLES_34011 | | | | | | | | | | | C | C |
| NC_011770 | 3769421 | T | C | 65 | 0 | 2 | 0 | NON_SYNONYMOU | MODERAT | MISSENSE | Ata/Gta | I508V | PLES_34011 | | | | | | | | | | | C | C |
| NC_011770 | 3769426 | C | G | 65 | 0 | 2 | 0 | NON_SYNONYMOU | MODERAT | MISSENSE | gGc/gCc | G506A | PLES_34011 | | | | | | | | | | | G | G |
| NC_011770 | 3769928 | C | T | 73 | 0 | 6 | 0 | NON_SYNONYMOU | MODERAT | MISSENSE | Gtg/Atg | V339M | PLES_34011 | | | T | T | T | T | | | | | T | T |
| NC_011770 | 3770129 | G | A | 66 | 0 | 8 | 0 | NON_SYNONYMOU | MODERAT | MISSENSE | Ccg/Tcg | P272S | PLES_34011 | A | A | A | A | A | A | | | A | A | | |
| NC_011770 | 3770431 | G | A | 64 | 0 | 2 | 0 | NON_SYNONYMOU | MODERAT | MISSENSE | aCg/aTg | T171M | PLES_34011 | A | A | | | | | | | | | | |
| NC_011770 | 3770761 | G | T | 59 | 0 | 2 | 0 | NON_SYNONYMOU | MODERAT | MISSENSE | gCc/gAc | A61D | PLES_34011 | | | | | | | T | T | | | | |
| NC_011770 | 3770762 | C | T | 57 | 0 | 2 | 0 | NON_SYNONYMOU | MODERAT | MISSENSE | Gcc/Acc | A61T | PLES_34011 | | | | | | | | | T | T | | |
| NC_011770 | 3770870 | G | A | 55 | 0 | 2 | 0 | NON_SYNONYMOU | MODERAT | MISSENSE | Ctt/Ttt | L25F | PLES_34011 | | | | | | | A | A | | | | |
| NC_011770 | 3771388 | G | A | 64 | 0 | 6 | 0 | NON_SYNONYMOU | MODERAT | MISSENSE | gaG/gaC | E93D | PLES_34021 | | | A | A | A | A | C | C | | | | |
| NC_011770 | 3771587 | G | A | 78 | 0 | 2 | 0 | NON_SYNONYMOU | MODERAT | MISSENSE | Gcc/Acc | A160T | PLES_34021 | | | | | | | | | | | A | A |
| NC_011770 | 3772312 | A | C | 77 | 0 | 2 | 0 | NON_SYNONYMOU | MODERAT | MISSENSE | gAg/gCg | E79A | PLES_34031 | C | C | | | | | | | | | | |
| NC_011770 | 3773857 | A | G | 75 | 0 | 2 | 0 | NON_SYNONYMOU | MODERAT | MISSENSE | aAg/aGg | K594R | PLES_34031 | | | | | | | | | | | G | G |
| NC_011770 | 3774079 | G | A | 77 | 0 | 2 | 0 | NON_SYNONYMOU | MODERAT | MISSENSE | aGc/aAc | S668N | PLES_34031 | | | | | | | | | | | A | A |
| NC_011770 | 3774757 | T | C | 52 | 0 | 2 | 0 | NON_SYNONYMOU | MODERAT | MISSENSE | gTg/gCg | V156A | PLES_34041 | C | C | | | | | | | | | | |
| NC_011770 | 3774838 | C | A | 54 | 0 | 10 | 0 | NON_SYNONYMOU | MODERAT | MISSENSE | gCg/gAg | A183E | PLES_34041 | A | A | A | A | A | A | | | A | A | A | A |
| NC_011770 | 3775273 | A | G | 23 | 0 | 2 | 0 | NON_SYNONYMOU | MODERAT | MISSENSE | Acc/Gcc | T31A | PLES_34051 | | | | | | | G | G | | | | |
| NC_011770 | 3775469 | G | A | 45 | 0 | 2 | 0 | NON_SYNONYMOU | MODERAT | MISSENSE | aGc/aAc | S96N | PLES_34051 | | | | | | | A | A | | | | |
| NC_011770 | 3775601 | A | G | 54 | 0 | 2 | 0 | NON_SYNONYMOU | MODERAT | MISSENSE | cAc/cGc | H140R | PLES_34051 | | | | | | | G | G | | | | |
| NC_011770 | 3775865 | A | G | 66 | 0 | 12 | 0 | NON_SYNONYMOU | MODERAT | MISSENSE | aAc/aGc | N228S | PLES_34051 | G | G | G | G | G | G | G | G | G | G | G | G |
| NC_011770 | 3776282 | G | A | 78 | 0 | 2 | 0 | NON_SYNONYMOU | MODERAT | MISSENSE | cGc/cAc | R367H | PLES_34051 | | | | | | | A | A | | | | |
| NC_011770 | 3776813 | G | C | 61 | 0 | 2 | 0 | NON_SYNONYMOU | MODERAT | MISSENSE | gGt/gCt | G73A | PLES_34061 | C | C | | | | | | | | | | |
| NC_011770 | 3777384 | C | G | 58 | 0 | 6 | 0 | NON_SYNONYMOU | MODERAT | MISSENSE | Cag/Gag | Q133E | PLES_34071 | G | G | | | | | G | G | | | G | G |
| NC_011770 | 3777658 | G | C | 74 | 0 | 2 | 0 | NON_SYNONYMOU | MODERAT | MISSENSE | gGg/gCg | G224A | PLES_34071 | | | | | | | C | C | | | | |
| NC_011770 | 3777663 | G | C | 74 | 0 | 2 | 0 | NON_SYNONYMOU | MODERAT | MISSENSE | Gtg/Ctg | V226L | PLES_34071 | C | C | | | | | | | | | | |
| NC_011770 | 3778047 | A | G | 77 | 0 | 2 | 0 | NON_SYNONYMOU | MODERAT | MISSENSE | Atc/Gtc | I354V | PLES_34071 | | | | | | | | | | | G | G |
| NC_011770 | 3778463 | G | C | 73 | 0 | 8 | 0 | NON_SYNONYMOU | MODERAT | MISSENSE | gaG/gaC | E60D | PLES_34081 | | | C | C | C | C | | | C | C | C | C |
| NC_011770 | 3778843 | C | T | 78 | 0 | 2 | 0 | NON_SYNONYMOU | MODERAT | MISSENSE | gCg/gTg | A187V | PLES_34081 | | | | | | | | | | | T | T |

| | | | | | | | | | | | | | | | | | | | | | | | | | |
|---|---|---|---|---|---|---|---|---|---|---|---|---|---|---|---|---|---|---|---|---|---|---|---|---|---|
| NC_011770 | 3779086 | C | G | 71 | 0 | 10 | 0 | NON_SYNONYMOU | MODERAT | MISSENSE | gCt/gGt | A268G | PLES_34081 | G | G | G | G | G | G | | | G | G | G | G |
| NC_011770 | 3779199 | G | A | 78 | 0 | 6 | 0 | NON_SYNONYMOU | MODERAT | MISSENSE | Gca/Aca | A306T | PLES_34081 | | | A | A | A | A | | | A | A | | |
| NC_011770 | 3779437 | A | G | 76 | 0 | 10 | 0 | NON_SYNONYMOU | MODERAT | MISSENSE | cAg/cGg | Q385R | PLES_34081 | G | G | G | G | G | G | | | G | G | G | G |
| NC_011770 | 3779532 | G | A | 72 | 0 | 8 | 0 | NON_SYNONYMOU | MODERAT | MISSENSE | Gcc/Acc | A417T | PLES_34081 | | | A | A | A | A | | | A | A | A | A |
| NC_011770 | 3779566 | T | C | 66 | 0 | 12 | 0 | NON_SYNONYMOU | MODERAT | MISSENSE | gTa/gCa | V428A | PLES_34081 | C | C | C | C | C | C | C | C | C | C | C | C |
| NC_011770 | 3779574 | T | C | 67 | 0 | 8 | 0 | NON_SYNONYMOU | MODERAT | MISSENSE | Tac/Cac | Y431H | PLES_34081 | | | C | C | C | C | | | C | C | C | C |
| NC_011770 | 3779784 | G | A | 53 | 0 | 2 | 0 | NON_SYNONYMOU | MODERAT | MISSENSE | Ggc/Agc | G501S | PLES_34081 | | | | | | | A | A | | | | |
| NC_011770 | 3780173 | A | G | 6 | 6 | 1 | 0 | NON_SYNONYMOU | MODERAT | MISSENSE | gTg/gCg | V75A | PLES_34091 | | | . | . | . | . | G | | . | . | | |
| NC_011770 | 3780215 | G | A | 7 | 6 | 2 | 0 | NON_SYNONYMOU | MODERAT | MISSENSE | gCg/gTg | A61V | PLES_34091 | | | . | . | . | . | | | . | . | A | A |
| NC_011770 | 3780239 | G | C | 32 | 0 | 2 | 0 | NON_SYNONYMOU | MODERAT | MISSENSE | aCc/aGc | T53S | PLES_34091 | | | | | | | | | | | C | C |
| NC_011770 | 3780255 | A | G | 33 | 0 | 8 | 0 | NON_SYNONYMOU | MODERAT | MISSENSE | Tgg/Cgg | W48R | PLES_34091 | | | G | G | G | G | | | G | G | G | G |
| NC_011770 | 3780419 | C | T | 21 | 6 | 2 | 0 | NON_SYNONYMOU | MODERAT | MISSENSE | aCc/aTc | T7I | PLES_34101 | T | T | . | . | . | . | | | . | . | | |
| NC_011770 | 3780518 | G | A | 25 | 6 | 6 | 0 | NON_SYNONYMOU | MODERAT | MISSENSE | aGc/aAc | S40N | PLES_34101 | A | A | . | . | . | . | A | A | . | . | A | A |
| NC_011770 | 3781472 | A | C | 56 | 0 | 6 | 0 | NON_SYNONYMOU | MODERAT | MISSENSE | aAc/aCc | N358T | PLES_34101 | | | C | C | C | C | | | C | C | | |
| NC_011770 | 3780949 | T | C | 35 | 6 | 1 | 0 | NON_SYNONYMOU | MODERAT | MISSENSE | Ttc/Ctc | F184L | PLES_34101 | | | . | . | . | . | | C | . | . | | |
| NC_011770 | 3781788 | C | T | 67 | 0 | 4 | 0 | NON_SYNONYMOU | MODERAT | MISSENSE | gCc/gTc | A19V | PLES_34111 | | | T | T | T | T | | | | | | |
| NC_011770 | 3781856 | C | T | 69 | 0 | 4 | 0 | NON_SYNONYMOU | MODERAT | MISSENSE | Cgc/Tgc | R42C | PLES_34111 | | | | | | | | | T | T | T | T |
| NC_011770 | 3782024 | G | C | 78 | 0 | 4 | 0 | NON_SYNONYMOU | MODERAT | MISSENSE | Gcg/Ccg | A98P | PLES_34111 | | | | | | | C | C | C | C | | |
| NC_011770 | 3782216 | G | C | 78 | 0 | 4 | 0 | NON_SYNONYMOU | MODERAT | MISSENSE | Gtg/Ctg | V162L | PLES_34111 | | | C | C | C | C | | | | | | |
| NC_011770 | 3783014 | G | T | 72 | 0 | 2 | 0 | NON_SYNONYMOU | MODERAT | MISSENSE | Gcc/Tcc | A3S | PLES_34131 | T | T | | | | | | | | | | |
| NC_011770 | 3783021 | A | G | 72 | 0 | 2 | 0 | NON_SYNONYMOU | MODERAT | MISSENSE | gAc/gGc | D5G | PLES_34131 | G | G | | | | | | | | | | |
| NC_011770 | 3783300 | G | T | 66 | 0 | 4 | 0 | NON_SYNONYMOU | MODERAT | MISSENSE | gGg/gTg | G98V | PLES_34131 | | | T | T | T | T | | | | | | |
| NC_011770 | 3783311 | T | C | 68 | 0 | 4 | 0 | NON_SYNONYMOU | MODERAT | MISSENSE | Tgg/Cgg | W102R | PLES_34131 | | | C | C | C | C | | | | | | |
| NC_011770 | 3783434 | G | A | 64 | 0 | 2 | 0 | NON_SYNONYMOU | MODERAT | MISSENSE | Gcc/Acc | A143T | PLES_34131 | | | | | | | A | A | | | | |
| NC_011770 | 3783749 | G | A | 54 | 0 | 2 | 0 | NON_SYNONYMOU | MODERAT | MISSENSE | Gcc/Acc | A248T | PLES_34131 | | | | | A | A | | | | | | |
| NC_011770 | 3783806 | G | T | 53 | 0 | 4 | 0 | NON_SYNONYMOU | MODERAT | MISSENSE | Gta/Tta | V267L | PLES_34131 | | | T | T | T | T | | | | | | |
| NC_011770 | 3783851 | G | A | 50 | 0 | 1 | 0 | NON_SYNONYMOU | MODERAT | MISSENSE | Gcc/Acc | A282T | PLES_34131 | | | | | | | A | | | | | |
| NC_011770 | 3784213 | ACGC | A | 36 | 0 | 2 | 0 | CODON_DELETION | MODERATE | | gcc/ | A19 | PLES_34141 | | | | | | | A | A | | | | |
| NC_011770 | 3784782 | G | A | 59 | 0 | 1 | 0 | NON_SYNONYMOU | MODERAT | MISSENSE | Gtg/Atg | V206M | PLES_34141 | | | | | | | A | | | | | |
| NC_011770 | 3785031 | G | A | 55 | 0 | 2 | 0 | NON_SYNONYMOU | MODERAT | MISSENSE | Ggc/Agc | G289S | PLES_34141 | A | A | | | | | | | | | | |
| NC_011770 | 3785457 | G | A | 60 | 0 | 2 | 0 | NON_SYNONYMOU | MODERAT | MISSENSE | Gtc/Atc | V431I | PLES_34141 | A | A | | | | | | | | | | |
| NC_011770 | 3785836 | A | C | 76 | 0 | 12 | 0 | NON_SYNONYMOU | MODERAT | MISSENSE | aAc/aCc | N557T | PLES_34141 | C | C | C | C | C | C | C | C | C | C | C | C |
| NC_011770 | 3786391 | C | T | 86 | 0 | 4 | 0 | NON_SYNONYMOU | MODERAT | MISSENSE | gCc/gTc | A742V | PLES_34141 | | | T | T | T | T | | | | | | |
| NC_011770 | 3786821 | C | T | 77 | 0 | 4 | 0 | NON_SYNONYMOU | MODERAT | MISSENSE | gCc/gTc | A67V | PLES_34151 | | | | | T | T | T | T | | | | |
| NC_011770 | 3787141 | G | A | 89 | 0 | 2 | 0 | NON_SYNONYMOU | MODERAT | MISSENSE | Gac/Aac | D174N | PLES_34151 | A | A | | | | | | | | | | |
| NC_011770 | 3787235 | T | C | 88 | 0 | 2 | 0 | NON_SYNONYMOU | MODERAT | MISSENSE | gTg/gCg | V205A | PLES_34151 | | | | | | | | | C | C | | |
| NC_011770 | 3787745 | G | A | 70 | 0 | 2 | 0 | NON_SYNONYMOU | MODERAT | MISSENSE | gGc/gAc | G375D | PLES_34151 | | | | | | | | | | | A | A |
| NC_011770 | 3787981 | G | A | 29 | 0 | 2 | 0 | NON_SYNONYMOU | MODERAT | MISSENSE | Gcc/Acc | A8T | PLES_34161 | A | A | | | | | | | | | | |
| NC_011770 | 3788108 | T | C | 35 | 0 | 12 | 0 | NON_SYNONYMOU | MODERAT | MISSENSE | gTc/gCc | V50A | PLES_34161 | C | C | C | C | C | C | C | C | C | C | C | C |
| NC_011770 | 3788110 | G | A | 35 | 0 | 2 | 0 | NON_SYNONYMOU | MODERAT | MISSENSE | Gcg/Acg | A51T | PLES_34161 | A | A | | | | | | | | | | |
| NC_011770 | 3788284 | T | C | 49 | 0 | 2 | 0 | NON_SYNONYMOU | MODERAT | MISSENSE | Ttc/Ctc | F109L | PLES_34161 | | | | | | | | | C | C | | |
| NC_011770 | 3788579 | G | A | 56 | 0 | 2 | 0 | NON_SYNONYMOU | MODERAT | MISSENSE | cGc/cAc | R207H | PLES_34161 | | | | | | | | | A | A | | |
| NC_011770 | 3788596 | A | C | 54 | 0 | 4 | 0 | NON_SYNONYMOU | MODERAT | MISSENSE | Atc/Ctc | I213L | PLES_34161 | | | C | C | C | C | | | | | | |
| NC_011770 | 3788998 | A | G | 57 | 0 | 2 | 0 | NON_SYNONYMOU | MODERAT | MISSENSE | Acc/Gcc | T347A | PLES_34161 | | | | | | | G | G | | | | |
| NC_011770 | 3789021 | C | G | 57 | 0 | 6 | 0 | NON_SYNONYMOU | MODERAT | MISSENSE | ttC/ttG | F354L | PLES_34161 | | | G | G | G | G | | | G | G | | |
| NC_011770 | 3789049 | G | A | 54 | 0 | 8 | 0 | NON_SYNONYMOU | MODERAT | MISSENSE | Gcg/Acg | A364T | PLES_34161 | A | A | A | A | A | A | A | A | | | | |
| NC_011770 | 3789071 | A | G | 57 | 0 | 12 | 0 | NON_SYNONYMOU | MODERAT | MISSENSE | aAc/aGc | N371S | PLES_34161 | G | G | G | G | G | G | G | G | G | G | G | G |
| NC_011770 | 3789100 | C | A | 63 | 0 | 2 | 0 | NON_SYNONYMOU | MODERAT | MISSENSE | Ctg/Atg | L381M | PLES_34161 | | | | | | | A | A | | | | |
| NC_011770 | 3789247 | G | A | 56 | 0 | 4 | 0 | NON_SYNONYMOU | MODERAT | MISSENSE | cGc/cAc | R27H | PLES_34171 | | | A | A | A | A | | | | | | |
| NC_011770 | 3789328 | G | C | 53 | 0 | 2 | 0 | NON_SYNONYMOU | MODERAT | MISSENSE | gGc/gCc | G54A | PLES_34171 | | | | | | | | | | | C | C |
| NC_011770 | 3789388 | T | A | 51 | 0 | 8 | 0 | NON_SYNONYMOU | MODERAT | MISSENSE | gTg/gAg | V74E | PLES_34171 | A | A | A | A | A | A | A | A | | | | |
| NC_011770 | 3789490 | G | A | 53 | 0 | 2 | 0 | NON_SYNONYMOU | MODERAT | MISSENSE | aGt/aAt | S108N | PLES_34171 | | | | | | | | | | | A | A |
| NC_011770 | 3789558 | G | A | 54 | 0 | 4 | 0 | NON_SYNONYMOU | MODERAT | MISSENSE | Gcc/Acc | A131T | PLES_34171 | A | A | | | | | A | A | | | | |
| NC_011770 | 3789652 | A | G | 58 | 0 | 12 | 0 | NON_SYNONYMOU | MODERAT | MISSENSE | gAc/gGc | D162G | PLES_34171 | G | G | G | G | G | G | G | G | G | G | G | G |
| NC_011770 | 3789711 | A | G | 62 | 0 | 8 | 0 | NON_SYNONYMOU | MODERAT | MISSENSE | Acc/Gcc | T182A | PLES_34171 | G | G | G | G | G | G | G | G | | | | |
| NC_011770 | 3790086 | G | A | 75 | 0 | 2 | 0 | NON_SYNONYMOU | MODERAT | MISSENSE | Gtc/Atc | V307I | PLES_34171 | | | | | | | | | A | A | | |
| NC_011770 | 3790191 | T | G | 65 | 0 | 12 | 0 | NON_SYNONYMOU | MODERAT | MISSENSE | Tac/Gac | Y342D | PLES_34171 | G | G | G | G | G | G | G | G | G | G | G | G |
| NC_011770 | 3790198 | C | T | 66 | 0 | 2 | 0 | NON_SYNONYMOU | MODERAT | MISSENSE | gCg/gTg | A344V | PLES_34171 | | | | | | | | | | | T | T |
| NC_011770 | 3790329 | C | T | 69 | 0 | 4 | 0 | NON_SYNONYMOU | MODERAT | MISSENSE | Ctc/Ttc | L388F | PLES_34171 | | | | | | | | | T | T | T | T |
| NC_011770 | 3790656 | G | A | 73 | 0 | 2 | 0 | NON_SYNONYMOU | MODERAT | MISSENSE | Gcg/Acg | A497T | PLES_34171 | A | A | | | | | | | | | | |
| NC_011770 | 3790711 | G | T | 69 | 0 | 2 | 0 | NON_SYNONYMOU | MODERAT | MISSENSE | cGc/cTc | R515L | PLES_34171 | | | | | | | | | T | T | | |
| NC_011770 | 3791195 | G | A | 58 | 0 | 2 | 0 | NON_SYNONYMOU | MODERAT | MISSENSE | Gcc/Acc | A116T | PLES_34181 | | | | | | | | | | | A | A |

| | | | | | | | | | | | | | | | | | | | | | | | | | |
|---|---|---|---|---|---|---|---|---|---|---|---|---|---|---|---|---|---|---|---|---|---|---|---|---|---|
| NC_011770 | 3791200 | A | C | 59 | 0 | 4 | 0 | NON_SYNONYMOU | MODERAT | MISSENSE | gaA/gaC | E117D | PLES_34181 | | | | | | | C | C | C | C | | |
| NC_011770 | 3791274 | T | G | 54 | 0 | 2 | 0 | NON_SYNONYMOU | MODERAT | MISSENSE | gTg/gGg | V142G | PLES_34181 | G | G | | | | | | | | | | |
| NC_011770 | 3791351 | T | C | 55 | 0 | 6 | 0 | NON_SYNONYMOU | MODERAT | MISSENSE | Tct/Cct | S168P | PLES_34181 | | | C | C | C | C | | | | | C | C |
| NC_011770 | 3791369 | T | C | 52 | 0 | 2 | 0 | NON_SYNONYMOU | MODERAT | MISSENSE | Tcc/Ccc | S174P | PLES_34181 | | | | | | | | | | | C | C |
| NC_011770 | 3791372 | G | A | 53 | 0 | 2 | 0 | NON_SYNONYMOU | MODERAT | MISSENSE | Gcg/Acg | A175T | PLES_34181 | | | | | | | | | | | A | A |
| NC_011770 | 3791379 | A | G | 58 | 0 | 2 | 0 | NON_SYNONYMOU | MODERAT | MISSENSE | gAc/gGc | D177G | PLES_34181 | | | | | | | | | | | G | G |
| NC_011770 | 3803054 | G | A | 69 | 0 | 2 | 0 | NON_SYNONYMOU | MODERAT | MISSENSE | Ggc/Agc | G385S | PLES_34291 | | | A | A | | | | | | | | |
| NC_011770 | 3803377 | T | C | 83 | 0 | 1 | 0 | NON_SYNONYMOU | MODERAT | MISSENSE | gTg/gCg | V85A | PLES_34301 | | | | | | | C | | | | | |
| NC_011770 | 3803740 | G | A | 81 | 0 | 6 | 0 | NON_SYNONYMOU | MODERAT | MISSENSE | cGg/cAg | R206Q | PLES_34301 | A | A | | | | | A | A | | | A | A |
| NC_011770 | 3803929 | G | A | 76 | 0 | 2 | 0 | NON_SYNONYMOU | MODERAT | MISSENSE | Gtc/Atc | V30I | PLES_34311 | A | A | | | | | | | | | | |
| NC_011770 | 3804040 | A | G | 72 | 0 | 8 | 0 | NON_SYNONYMOU | MODERAT | MISSENSE | Aag/Gag | K67E | PLES_34311 | G | G | | | | | G | G | G | G | G | G |
| NC_011770 | 3804536 | G | A | 77 | 0 | 8 | 0 | NON_SYNONYMOU | MODERAT | MISSENSE | cGc/cAc | R232H | PLES_34311 | A | A | | | | | A | A | A | A | A | A |
| NC_011770 | 3804626 | G | A | 80 | 0 | 2 | 0 | NON_SYNONYMOU | MODERAT | MISSENSE | aGc/aAc | S262N | PLES_34311 | A | A | | | | | | | | | | |
| NC_011770 | 3804919 | A | G | 76 | 0 | 8 | 0 | NON_SYNONYMOU | MODERAT | MISSENSE | Acc/Gcc | T360A | PLES_34311 | G | G | | | | | G | G | G | G | G | G |
| NC_011770 | 3804937 | A | C | 75 | 0 | 4 | 0 | NON_SYNONYMOU | MODERAT | MISSENSE | Acg/Ccg | T366P | PLES_34311 | | | | | | | | | C | C | C | C |
| NC_011770 | 3805064 | G | A | 70 | 0 | 4 | 0 | NON_SYNONYMOU | MODERAT | MISSENSE | cGt/cAt | R408H | PLES_34311 | | | | | | | | | A | A | A | A |
| NC_011770 | 3805431 | C | A | 72 | 0 | 8 | 0 | NON_SYNONYMOU | MODERAT | MISSENSE | gaC/gaA | D530E | PLES_34311 | | | A | A | A | A | A | A | | | A | A |
| NC_011770 | 3805463 | A | G | 68 | 0 | 2 | 0 | NON_SYNONYMOU | MODERAT | MISSENSE | cAg/cGg | Q541R | PLES_34311 | | | | | | | G | G | | | | |
| NC_011770 | 3805834 | C | A | 64 | 0 | 2 | 0 | NON_SYNONYMOU | MODERAT | MISSENSE | Cgc/Agc | R665S | PLES_34311 | A | A | | | | | | | | | | |
| NC_011770 | 3805876 | A | G | 60 | 0 | 2 | 0 | NON_SYNONYMOU | MODERAT | MISSENSE | Agc/Ggc | S679G | PLES_34311 | G | G | | | | | | | | | | |
| NC_011770 | 3805877 | G | A | 64 | 0 | 2 | 0 | NON_SYNONYMOU | MODERAT | MISSENSE | aGc/aAc | S679N | PLES_34311 | A | A | | | | | | | | | | |
| NC_011770 | 3805929 | C | G | 68 | 0 | 2 | 0 | NON_SYNONYMOU | MODERAT | MISSENSE | ttC/ttG | F696L | PLES_34311 | | | | | | | | | | | G | G |
| NC_011770 | 3805948 | G | A | 65 | 0 | 2 | 0 | NON_SYNONYMOU | MODERAT | MISSENSE | Gcg/Acg | A703T | PLES_34311 | A | A | | | | | | | | | | |
| NC_011770 | 3806038 | A | C | 73 | 0 | 6 | 0 | NON_SYNONYMOU | MODERAT | MISSENSE | Aac/Cac | N733H | PLES_34311 | | | C | C | C | C | | | | | C | C |
| NC_011770 | 3806117 | G | C | 73 | 0 | 2 | 0 | NON_SYNONYMOU | MODERAT | MISSENSE | cGc/cCc | R759P | PLES_34311 | C | C | | | | | | | | | | |
| NC_011770 | 3806245 | T | C | 70 | 0 | 2 | 0 | NON_SYNONYMOU | MODERAT | MISSENSE | Tct/Cct | S802P | PLES_34311 | | | | | | | | | | | C | C |
| NC_011770 | 3806514 | T | G | 65 | 0 | 6 | 0 | NON_SYNONYMOU | MODERAT | MISSENSE | Tcg/Gcg | S83A | PLES_34321 | | | G | G | G | G | | | | | G | G |
| NC_011770 | 3806527 | T | C | 72 | 0 | 1 | 0 | NON_SYNONYMOU | MODERAT | MISSENSE | cTc/cCc | L87P | PLES_34321 | | | | | | | C | | | | | |
| NC_011770 | 3806749 | A | G | 50 | 0 | 8 | 0 | NON_SYNONYMOU | MODERAT | MISSENSE | aAc/aGc | N161S | PLES_34321 | | | G | G | G | G | G | G | | | G | G |
| NC_011770 | 3806773 | T | G | 46 | 0 | 6 | 0 | NON_SYNONYMOU | MODERAT | MISSENSE | cTc/cGc | L169R | PLES_34321 | | | G | G | G | G | G | G | | | | |
| NC_011770 | 3807111 | A | G | 59 | 0 | 12 | 0 | NON_SYNONYMOU | MODERAT | MISSENSE | gAg/gGg | E37G | PLES_34331 | G | G | G | G | G | G | G | G | G | G | G | G |
| NC_011770 | 3807711 | G | A | 85 | 0 | 4 | 0 | NON_SYNONYMOU | MODERAT | MISSENSE | Gcc/Acc | A78T | PLES_34341 | | | A | A | A | A | | | | | | |
| NC_011770 | 3807901 | T | C | 83 | 0 | 2 | 0 | NON_SYNONYMOU | MODERAT | MISSENSE | aTc/aCc | I141T | PLES_34341 | | | | | | | | | C | C | | |
| NC_011770 | 3808861 | ACCTACTA( | A | 53 | 0 | 1 | 0 | FRAME_SHIFT | HIGH | | gccgtagtag | AVVG26 | PLES_34351 | | | | | | | | | . | A | | |
| NC_011770 | 3808580 | A | G | 68 | 0 | 6 | 0 | NON_SYNONYMOU | MODERAT | MISSENSE | gTa/gCa | V123A | PLES_34351 | G | G | | | | | G | G | | | G | G |
| NC_011770 | 3808923 | A | G | 46 | 0 | 2 | 0 | NON_SYNONYMOU | MODERAT | MISSENSE | Ttc/Ctc | F9L | PLES_34351 | | | | | | | | | G | G | | |
| NC_011770 | 3809545 | C | G | 39 | 2 | 2 | 0 | NON_SYNONYMOU | MODERAT | MISSENSE | gCc/gGc | A98G | PLES_34361 | | | | | | | . | . | | | G | G |
| NC_011770 | 3809710 | A | G | 49 | 2 | 10 | 0 | NON_SYNONYMOU | MODERAT | MISSENSE | gAc/gGc | D153G | PLES_34361 | G | G | G | G | G | G | . | . | G | G | G | G |
| NC_011770 | 3809949 | A | G | 48 | 2 | 10 | 0 | NON_SYNONYMOU | MODERAT | MISSENSE | Acg/Gcg | T233A | PLES_34361 | G | G | G | G | G | G | . | . | G | G | G | G |
| NC_011770 | 3812583 | A | G | 71 | 0 | 6 | 0 | NON_SYNONYMOU | MODERAT | MISSENSE | gTt/gCt | V175A | PLES_34381 | G | G | | | | | G | G | G | G | | |
| NC_011770 | 3812778 | G | A | 60 | 0 | 2 | 0 | NON_SYNONYMOU | MODERAT | MISSENSE | gCg/gTg | A110V | PLES_34381 | A | A | | | | | | | | | | |
| NC_011770 | 3812845 | G | A | 62 | 0 | 6 | 0 | NON_SYNONYMOU | MODERAT | MISSENSE | Ccg/Tcg | P88S | PLES_34381 | | | A | A | A | A | | | | | A | A |
| NC_011770 | 3813403 | T | C | 52 | 0 | 10 | 0 | NON_SYNONYMOU | MODERAT | MISSENSE | gTg/gCg | V71A | PLES_34391 | C | C | C | C | C | C | C | C | | | C | C |
| NC_011770 | 3813457 | T | C | 46 | 0 | 10 | 0 | NON_SYNONYMOU | MODERAT | MISSENSE | gTc/gCc | V89A | PLES_34391 | C | C | C | C | C | C | C | C | | | C | C |
| NC_011770 | 3813490 | G | T | 44 | 0 | 6 | 0 | NON_SYNONYMOU | MODERAT | MISSENSE | cGc/cTc | R100L | PLES_34391 | | | T | T | T | T | T | T | | | | |
| NC_011770 | 3813538 | C | A | 44 | 0 | 10 | 0 | NON_SYNONYMOU | MODERAT | MISSENSE | aCc/aAc | T116N | PLES_34391 | A | A | A | A | A | A | A | A | | | A | A |
| NC_011770 | 3813540 | A | G | 44 | 0 | 10 | 0 | NON_SYNONYMOU | MODERAT | MISSENSE | Acc/Gcc | T117A | PLES_34391 | G | G | G | G | G | G | G | G | | | G | G |
| NC_011770 | 3813755 | T | A | 52 | 0 | 2 | 0 | NON_SYNONYMOU | MODERAT | MISSENSE | aaA/aaT | K128N | PLES_34401 | | | | | | | A | A | | | | |
| NC_011770 | 3813967 | A | C | 31 | 0 | 2 | 0 | NON_SYNONYMOU | MODERAT | MISSENSE | Ttg/Gtg | L58V | PLES_34401 | | | | | | | C | C | | | | |
| NC_011770 | 3814132 | C | T | 39 | 0 | 2 | 0 | NON_SYNONYMOU | MODERAT | MISSENSE | Gca/Aca | A3T | PLES_34401 | | | | | | | T | T | | | | |
| NC_011770 | 3814192 | C | T | 46 | 0 | 4 | 0 | NON_SYNONYMOU | MODERAT | MISSENSE | Gcc/Acc | A95T | PLES_34411 | | | T | T | T | T | | | | | | |
| NC_011770 | 3814225 | G | A | 47 | 0 | 2 | 0 | NON_SYNONYMOU | MODERAT | MISSENSE | Ccg/Tcg | P84S | PLES_34411 | | | | | | | | | A | A | | |
| NC_011770 | 3814354 | G | A | 65 | 0 | 2 | 0 | NON_SYNONYMOU | MODERAT | MISSENSE | Ctc/Ttc | L41F | PLES_34411 | | | | | | | | | | | A | A |
| NC_011770 | 3814357 | T | C | 65 | 0 | 2 | 0 | NON_SYNONYMOU | MODERAT | MISSENSE | Atc/Gtc | I40V | PLES_34411 | | | | | | | | | | | C | C |
| NC_011770 | 3816101 | G | A | 70 | 0 | 1 | 0 | STOP_GAINED | HIGH | NONSENSE | tgG/tgA | W277* | PLES_34431 (PA1880) | | | | | | | | | | | | A |
| NC_011770 | 3816654 | C | A | 73 | 0 | 2 | 0 | NON_SYNONYMOU | MODERAT | MISSENSE | Ctg/Atg | L462M | PLES_34431 | | | | | | | | | A | A | | |
| NC_011770 | 3816735 | A | G | 67 | 0 | 8 | 0 | NON_SYNONYMOU | MODERAT | MISSENSE | Atc/Gtc | I489V | PLES_34431 | | | G | G | G | G | G | G | G | G | | |
| NC_011770 | 3816987 | G | A | 65 | 0 | 2 | 0 | NON_SYNONYMOU | MODERAT | MISSENSE | Gaa/Aaa | E573K | PLES_34431 | A | A | | | | | | | | | | |
| NC_011770 | 3815442 | G | A | 75 | 0 | 1 | 0 | NON_SYNONYMOU | MODERAT | MISSENSE | Gcc/Acc | A58T | PLES_34431 (PA1880) | | | | | | | | A | | | | |
| NC_011770 | 3817834 | C | A | 67 | 0 | 2 | 0 | NON_SYNONYMOU | MODERAT | MISSENSE | Cag/Aag | Q71K | PLES_34441 | | | | | | | | | | | A | A |
| NC_011770 | 3818003 | G | A | 72 | 0 | 1 | 0 | NON_SYNONYMOU | MODERAT | MISSENSE | cGc/cAc | R127H | PLES_34441 | | | | | | | A | | | | | |
| NC_011770 | 3818327 | G | A | 89 | 0 | 2 | 0 | NON_SYNONYMOU | MODERAT | MISSENSE | Gac/Aac | D21N | PLES_34451 | | | | | | | | | | | A | A |

| | | | | | | | | | | | | | | | | | | | | | | | | | |
|---|---|---|---|---|---|---|---|---|---|---|---|---|---|---|---|---|---|---|---|---|---|---|---|---|---|
| NC_011770 | 3818432 | A | G | 79 | 0 | 12 | 0 | NON_SYNONYMOU | MODERAT | MISSENSE | Agc/Ggc | S56G | PLES_34451 | G | G | G | G | G | G | G | G | G | G | G | G |
| NC_011770 | 3818585 | G | A | 83 | 0 | 4 | 0 | NON_SYNONYMOU | MODERAT | MISSENSE | Gcg/Acg | A107T | PLES_34451 | | | A | A | A | A | | | | | | |
| NC_011770 | 3818599 | C | T | 85 | 0 | 6 | 0 | NON_SYNONYMOU | MODERAT | MISSENSE | gaC/gaA | D111E | PLES_34451 | | | T | T | T | T | | | | | A | A |
| NC_011770 | 3818639 | T | A | 79 | 0 | 4 | 0 | NON_SYNONYMOU | MODERAT | MISSENSE | Tcc/Acc | S125T | PLES_34451 | | | A | A | A | A | | | | | | |
| NC_011770 | 3818819 | C | T | 84 | 0 | 2 | 0 | NON_SYNONYMOU | MODERAT | MISSENSE | Cgc/Tgc | R185C | PLES_34451 | T | T | | | | | | | | | | |
| NC_011770 | 3820321 | T | G | 90 | 0 | 4 | 0 | NON_SYNONYMOU | MODERAT | MISSENSE | gAc/gCc | D637A | PLES_34471 | | | G | G | G | G | | | | | | |
| NC_011770 | 3820559 | A | G | 79 | 0 | 12 | 0 | NON_SYNONYMOU | MODERAT | MISSENSE | Tgc/Cgc | C558R | PLES_34471 | G | G | G | G | G | G | G | G | G | G | G | G |
| NC_011770 | 3821224 | T | C | 87 | 0 | 1 | 0 | NON_SYNONYMOU | MODERAT | MISSENSE | aAc/aGc | N336S | PLES_34471 | | | | | | | C | | | | | |
| NC_011770 | 3821253 | G | C | 86 | 0 | 12 | 0 | NON_SYNONYMOU | MODERAT | MISSENSE | caC/caG | H326Q | PLES_34471 | C | C | C | C | C | C | C | C | C | C | C | C |
| NC_011770 | 3821831 | C | T | 82 | 0 | 4 | 0 | NON_SYNONYMOU | MODERAT | MISSENSE | Gtc/Atc | V134I | PLES_34471 | | | | | | | T | T | T | T | | |
| NC_011770 | 3821920 | T | C | 66 | 0 | 2 | 0 | NON_SYNONYMOU | MODERAT | MISSENSE | gAg/gGg | E104G | PLES_34471 | | | | | | | C | C | | | | |
| NC_011770 | 3821647 | T | C | 95 | 0 | 1 | 0 | NON_SYNONYMOU | MODERAT | MISSENSE | gAc/gGc | D195G | PLES_34471 | | | | | | | | C | | | | |
| NC_011770 | 3822536 | C | G | 76 | 0 | 2 | 0 | NON_SYNONYMOU | MODERAT | MISSENSE | aGc/aCc | S321T | PLES_34481 | | | | | | | G | G | | | | |
| NC_011770 | 3822543 | T | G | 77 | 0 | 2 | 0 | NON_SYNONYMOU | MODERAT | MISSENSE | Aaa/Caa | K319Q | PLES_34481 | | | | | | | G | G | | | | |
| NC_011770 | 3829744 | A | T | 38 | 0 | 3 | 0 | NON_SYNONYMOU | MODERAT | MISSENSE | tTc/tAc | F633Y | PLES_34491 | | | T | T | T | | | | | | | |
| NC_011770 | 3824080 | G | C | 72 | 0 | 2 | 0 | NON_SYNONYMOU | MODERAT | MISSENSE | tCc/tGc | S2521C | PLES_34491 | | | | | | | | | C | C | | |
| NC_011770 | 3824081 | A | T | 69 | 0 | 2 | 0 | NON_SYNONYMOU | MODERAT | MISSENSE | Tcc/Acc | S2521T | PLES_34491 | | | | | | | | | T | T | | |
| NC_011770 | 3824368 | G | A | 59 | 0 | 2 | 0 | NON_SYNONYMOU | MODERAT | MISSENSE | aCc/aTc | T2425I | PLES_34491 | | | | | A | A | | | | | | |
| NC_011770 | 3825554 | C | T | 73 | 0 | 2 | 0 | NON_SYNONYMOU | MODERAT | MISSENSE | Ggc/Agc | G2030S | PLES_34491 | | | | | | | | | T | T | | |
| NC_011770 | 3825565 | G | C | 76 | 0 | 4 | 0 | NON_SYNONYMOU | MODERAT | MISSENSE | aCc/aGc | T2026S | PLES_34491 | | | | | | | C | C | C | C | | |
| NC_011770 | 3825571 | T | C | 73 | 0 | 2 | 0 | NON_SYNONYMOU | MODERAT | MISSENSE | aAc/aGc | N2024S | PLES_34491 | | | | | | | | | C | C | | |
| NC_011770 | 3826351 | C | T | 31 | 0 | 2 | 0 | NON_SYNONYMOU | MODERAT | MISSENSE | gGc/gAc | G1764D | PLES_34491 | | | | | | | | | | | T | T |
| NC_011770 | 3826352 | C | T | 31 | 0 | 2 | 0 | NON_SYNONYMOU | MODERAT | MISSENSE | Ggc/Agc | G1764S | PLES_34491 | | | | | | | | | | | T | T |
| NC_011770 | 3826526 | A | C | 28 | 4 | 6 | 0 | NON_SYNONYMOU | MODERAT | MISSENSE | Tcc/Gcc | S1706A | PLES_34491 | C | C | . | . | . | . | | | C | C | C | C |
| NC_011770 | 3826783 | C | G | 28 | 0 | 3 | 0 | NON_SYNONYMOU | MODERAT | MISSENSE | gGc/gCc | G1620A | PLES_34491 | | | | | | | G | | | | G | G |
| NC_011770 | 3826795 | C | G | 31 | 0 | 3 | 0 | NON_SYNONYMOU | MODERAT | MISSENSE | aGc/aCc | S1616T | PLES_34491 | | | | | | | G | | | | G | G |
| NC_011770 | 3826796 | T | C | 31 | 0 | 3 | 0 | NON_SYNONYMOU | MODERAT | MISSENSE | Agc/Ggc | S1616G | PLES_34491 | | | | | | | C | | | | C | C |
| NC_011770 | 3826913 | T | C | 29 | 0 | 8 | 0 | NON_SYNONYMOU | MODERAT | MISSENSE | Aat/Gat | N1577D | PLES_34491 | | | C | C | C | C | C | C | | | C | C |
| NC_011770 | 3827030 | T | C | 46 | 0 | 4 | 0 | NON_SYNONYMOU | MODERAT | MISSENSE | Agc/Ggc | S1538G | PLES_34491 | | | | | | | C | C | | | C | C |
| NC_011770 | 3827066 | T | C | 57 | 0 | 1 | 0 | NON_SYNONYMOU | MODERAT | MISSENSE | Acc/Gcc | T1526A | PLES_34491 | | | | | | | C | | | | | |
| NC_011770 | 3827455 | A | G | 48 | 0 | 10 | 0 | NON_SYNONYMOU | MODERAT | MISSENSE | cTt/cCt | L1396P | PLES_34491 | | | G | G | G | G | G | G | G | G | G | G |
| NC_011770 | 3827456 | G | C | 48 | 0 | 8 | 0 | NON_SYNONYMOU | MODERAT | MISSENSE | Ctt/Gtt | L1396V | PLES_34491 | | | C | C | C | C | | | C | C | C | C |
| NC_011770 | 3827592 | C | G | 54 | 0 | 2 | 0 | NON_SYNONYMOU | MODERAT | MISSENSE | ttG/ttC | L1350F | PLES_34491 | | | | | | | | | | | G | G |
| NC_011770 | 3827767 | C | G | 47 | 2 | 10 | 0 | NON_SYNONYMOU | MODERAT | MISSENSE | gGc/gCc | G1292A | PLES_34491 | G | G | G | G | G | G | G | G | G | G | . | . |
| NC_011770 | 3827779 | C | G | 50 | 2 | 10 | 0 | NON_SYNONYMOU | MODERAT | MISSENSE | aGc/aCc | S1288T | PLES_34491 | G | G | G | G | G | G | G | G | G | G | . | . |
| NC_011770 | 3827954 | T | A | 39 | 2 | 2 | 0 | NON_SYNONYMOU | MODERAT | MISSENSE | Acc/Tcc | T1230S | PLES_34491 | | | | | | | A | A | | | . | . |
| NC_011770 | 3828098 | C | T | 46 | 2 | 2 | 0 | NON_SYNONYMOU | MODERAT | MISSENSE | Gcc/Acc | A1182T | PLES_34491 | | | | | | | | | T | T | . | . |
| NC_011770 | 3828176 | C | T | 43 | 2 | 2 | 0 | NON_SYNONYMOU | MODERAT | MISSENSE | Gtc/Atc | V1156I | PLES_34491 | | | | | | | | | T | T | . | . |
| NC_011770 | 3828218 | G | C | 44 | 2 | 6 | 0 | NON_SYNONYMOU | MODERAT | MISSENSE | Cag/Gag | Q1142E | PLES_34491 | C | C | C | C | C | C | | | | | . | . |
| NC_011770 | 3828298 | A | G | 42 | 0 | 2 | 0 | NON_SYNONYMOU | MODERAT | MISSENSE | gTc/gCc | V1115A | PLES_34491 | | | | | | | | | | | G | G |
| NC_011770 | 3828302 | C | G | 40 | 0 | 2 | 0 | NON_SYNONYMOU | MODERAT | MISSENSE | Gag/Cag | E1114Q | PLES_34491 | | | | | | | | | | | G | G |
| NC_011770 | 3828304 | G | C | 40 | 0 | 2 | 0 | NON_SYNONYMOU | MODERAT | MISSENSE | gCc/gGc | A1113G | PLES_34491 | | | | | | | | | | | C | C |
| NC_011770 | 3828334 | A | G | 41 | 0 | 1 | 0 | NON_SYNONYMOU | MODERAT | MISSENSE | aTc/aCc | I1103T | PLES_34491 | G | | | | | | | | | | | |
| NC_011770 | 3828443 | C | T | 38 | 0 | 2 | 0 | NON_SYNONYMOU | MODERAT | MISSENSE | Ggc/Agc | G1067S | PLES_34491 | | | | | | | T | T | | | | |
| NC_011770 | 3828740 | C | G | 48 | 0 | 2 | 0 | NON_SYNONYMOU | MODERAT | MISSENSE | Gcc/Ccc | A968P | PLES_34491 | G | G | | | | | | | | | | |
| NC_011770 | 3828826 | G | C | 52 | 0 | 2 | 0 | NON_SYNONYMOU | MODERAT | MISSENSE | aCc/aGc | T939S | PLES_34491 | | | | | | | C | C | | | | |
| NC_011770 | 3829252 | A | T | 40 | 0 | 3 | 0 | NON_SYNONYMOU | MODERAT | MISSENSE | tTc/tAc | F797Y | PLES_34491 | | | T | . | T | T | | | | | | |
| NC_011770 | 3829255 | C | G | 40 | 0 | 3 | 0 | NON_SYNONYMOU | MODERAT | MISSENSE | aGc/aCc | S796T | PLES_34491 | | | G | . | G | G | | | | | | |
| NC_011770 | 3829376 | T | C | 30 | 4 | 2 | 0 | NON_SYNONYMOU | MODERAT | MISSENSE | Atc/Gtc | I756V | PLES_34491 | | | . | . | . | . | | | | | C | C |
| NC_011770 | 3829603 | G | C | 26 | 4 | 2 | 0 | NON_SYNONYMOU | MODERAT | MISSENSE | aCc/aGc | T680S | PLES_34491 | | | . | . | . | . | | | | | C | C |
| NC_011770 | 3829669 | G | A | 26 | 4 | 2 | 0 | NON_SYNONYMOU | MODERAT | MISSENSE | tCg/tTg | S658L | PLES_34491 | | | . | . | . | . | A | A | | | | |
| NC_011770 | 3829724 | A | G | 32 | 0 | 12 | 0 | NON_SYNONYMOU | MODERAT | MISSENSE | Tcc/Ccc | S640P | PLES_34491 | G | G | C | C | C | C | G | G | G | G | G | G |
| NC_011770 | 3829725 | C | G | 33 | 0 | 4 | 0 | NON_SYNONYMOU | MODERAT | MISSENSE | ttG/ttC | L639F | PLES_34491 | | | G | G | G | G | | | | | | |
| NC_011770 | 3829727 | A | T | 29 | 0 | 3 | 0 | NON_SYNONYMOU | MODERAT | MISSENSE | Ttg/Atg | L639M | PLES_34491 | | | . | T | T | T | | | | | | |
| NC_011770 | 3829735 | C | G | 29 | 0 | 4 | 0 | NON_SYNONYMOU | MODERAT | MISSENSE | aGc/aCc | S636T | PLES_34491 | | | G | G | G | G | | | | | | |
| NC_011770 | 3829736 | T | A | 31 | 0 | 4 | 0 | NON_SYNONYMOU | MODERAT | MISSENSE | Agc/Tgc | S636C | PLES_34491 | | | A | A | A | A | | | | | | |
| NC_011770 | 3829820 | C | T | 45 | 0 | 6 | 0 | NON_SYNONYMOU | MODERAT | MISSENSE | Gct/Act | A608T | PLES_34491 | | | T | T | T | T | | | | | T | T |
| NC_011770 | 3830125 | T | C | 36 | 0 | 8 | 0 | NON_SYNONYMOU | MODERAT | MISSENSE | aAc/aGc | N506S | PLES_34491 | | | C | C | C | C | | | C | C | C | C |
| NC_011770 | 3830245 | C | T | 39 | 0 | 12 | 0 | NON_SYNONYMOU | MODERAT | MISSENSE | aGc/aAc | S466N | PLES_34491 | T | T | T | T | T | T | T | T | T | T | T | T |
| NC_011770 | 3830435 | T | C | 34 | 0 | 4 | 0 | NON_SYNONYMOU | MODERAT | MISSENSE | Atc/Gtc | I403V | PLES_34491 | | | C | C | C | C | | | | | | |
| NC_011770 | 3830488 | C | T | 37 | 0 | 4 | 0 | NON_SYNONYMOU | MODERAT | MISSENSE | aGc/aAc | S385N | PLES_34491 | | | T | T | T | T | | | | | | |
| NC_011770 | 3830494 | C | T | 37 | 0 | 4 | 0 | NON_SYNONYMOU | MODERAT | MISSENSE | gGc/gAc | G383D | PLES_34491 | | | T | T | T | T | | | | | | |

| | | | | | | | | | | | | | | | | | | | | | | | | | |
|---|---|---|---|---|---|---|---|---|---|---|---|---|---|---|---|---|---|---|---|---|---|---|---|---|---|
| NC_011770 | 3830765 | C | T | 35 | 0 | 4 | 0 | NON_SYNONYMOU | MODERAT | MISSENSE | Gtg/Atg | V293M | PLES_34491 | | | T | T | T | T | | | | | | |
| NC_011770 | 3830816 | A | C | 26 | 0 | 2 | 0 | NON_SYNONYMOU | MODERAT | MISSENSE | Tcg/Gcg | S276A | PLES_34491 | | | | | | | C | C | | | | |
| NC_011770 | 3830829 | C | G | 33 | 0 | 2 | 0 | NON_SYNONYMOU | MODERAT | MISSENSE | aaG/aaC | K271N | PLES_34491 | | | | | | | G | G | | | | |
| NC_011770 | 3830836 | A | G | 31 | 0 | 10 | 0 | NON_SYNONYMOU | MODERAT | MISSENSE | aTc/aCc | I269T | PLES_34491 | | | G | G | G | G | G | G | G | G | G | G |
| NC_011770 | 3830851 | C | T | 33 | 0 | 10 | 0 | NON_SYNONYMOU | MODERAT | MISSENSE | gGc/gAc | G264D | PLES_34491 | | | T | T | T | T | T | T | T | T | T | T |
| NC_011770 | 3830858 | T | C | 35 | 0 | 10 | 0 | NON_SYNONYMOU | MODERAT | MISSENSE | Acg/Gcg | T262A | PLES_34491 | | | C | C | C | C | C | C | C | C | C | C |
| NC_011770 | 3831536 | T | C | 50 | 0 | 6 | 0 | NON_SYNONYMOU | MODERAT | MISSENSE | Atc/Gtc | I36V | PLES_34491 | | | C | C | C | C | C | C | | | | |
| NC_011770 | 3826763 | A | T | 31 | 0 | 5 | 0 | NON_SYNONYMOU | MODERAT | MISSENSE | Tcg/Acg | S1627T | PLES_34491 | | T | | | | | T | T | | | T | T |
| NC_011770 | 3827596 | G | A | 58 | 0 | 1 | 0 | NON_SYNONYMOU | MODERAT | MISSENSE | aCc/aTc | T1349I | PLES_34491 | | | | | | | | | | | A | |
| NC_011770 | 3828299 | C | T | 40 | 0 | 1 | 0 | NON_SYNONYMOU | MODERAT | MISSENSE | Gtc/Atc | V1115I | PLES_34491 | | | | | | | | | | | T | |
| NC_011770 | 3828884 | C | G | 44 | 0 | 4 | 0 | NON_SYNONYMOU | MODERAT | MISSENSE | Gtc/Ctc | V920L | PLES_34491 | G | | | | | | G | G | | | G | |
| NC_011770 | 3829816 | G | C | 47 | 0 | 5 | 0 | NON_SYNONYMOU | MODERAT | MISSENSE | aCc/aGc | T609S | PLES_34491 | | | C | C | C | C | | | | | C | |
| NC_011770 | 3829819 | G | C | 45 | 0 | 5 | 0 | NON_SYNONYMOU | MODERAT | MISSENSE | gCt/gGt | A608G | PLES_34491 | | | C | C | C | C | | | | | C | |
| NC_011770 | 3827069 | C | T | 57 | 0 | 1 | 0 | NON_SYNONYMOU | MODERAT | MISSENSE | Gcc/Acc | A1525T | PLES_34491 | | | | | | | | | | | | T |
| NC_011770 | 3832152 | G | C | 61 | 0 | 2 | 0 | NON_SYNONYMOU | MODERAT | MISSENSE | gGt/gCt | G4A | PLES_34501 | | | | | | | C | C | | | | |
| NC_011770 | 3832179 | T | C | 58 | 0 | 2 | 0 | NON_SYNONYMOU | MODERAT | MISSENSE | cTg/cCg | L13P | PLES_34501 | | | | | | | | | | | C | C |
| NC_011770 | 3832242 | C | T | 62 | 0 | 2 | 0 | NON_SYNONYMOU | MODERAT | MISSENSE | gCt/gTt | A34V | PLES_34501 | | | | | | | T | T | | | | |
| NC_011770 | 3832272 | G | C | 61 | 0 | 2 | 0 | NON_SYNONYMOU | MODERAT | MISSENSE | cGc/cCc | R44P | PLES_34501 | | | | | | | | | | | C | C |
| NC_011770 | 3832281 | C | A | 63 | 0 | 2 | 0 | NON_SYNONYMOU | MODERAT | MISSENSE | gCg/gAg | A47E | PLES_34501 | | | | | | | | | | | A | A |
| NC_011770 | 3833138 | G | A | 82 | 0 | 1 | 0 | NON_SYNONYMOU | MODERAT | MISSENSE | Gcc/Acc | A333T | PLES_34501 | | | | | | | | A | | | | |
| NC_011770 | 3833856 | T | C | 95 | 0 | 1 | 0 | NON_SYNONYMOU | MODERAT | MISSENSE | gTg/gCg | V211A | PLES_34511 | | | | | | | C | | | | | |
| NC_011770 | 3833426 | A | G | 84 | 0 | 1 | 0 | NON_SYNONYMOU | MODERAT | MISSENSE | Act/Gct | T68A | PLES_34511 | | | | | | | | G | | | | |
| NC_011770 | 3835682 | T | G | 77 | 0 | 4 | 0 | NON_SYNONYMOU | MODERAT | MISSENSE | gAc/gCc | D138A | PLES_34531 | | | G | G | G | G | | | | | | |
| NC_011770 | 3835683 | C | T | 77 | 0 | 4 | 0 | NON_SYNONYMOU | MODERAT | MISSENSE | Gac/Aac | D138N | PLES_34531 | | | T | T | T | T | | | | | | |
| NC_011770 | 3835938 | A | G | 65 | 0 | 4 | 0 | NON_SYNONYMOU | MODERAT | MISSENSE | Tcg/Ccg | S53P | PLES_34531 | | | G | G | G | G | | | | | | |
| NC_011770 | 3836029 | C | T | 66 | 0 | 4 | 0 | NON_SYNONYMOU | MODERAT | MISSENSE | atG/atA | M22I | PLES_34531 | | | T | T | T | T | | | | | | |
| NC_011770 | 3836472 | C | T | 71 | 0 | 2 | 0 | NON_SYNONYMOU | MODERAT | MISSENSE | Gcg/Acg | A49T | PLES_34541 | | | | | | | | | | | T | T |
| NC_011770 | 3836571 | A | G | 70 | 0 | 4 | 0 | NON_SYNONYMOU | MODERAT | MISSENSE | Ttc/Ctc | F16L | PLES_34541 | | | G | G | G | G | | | | | | |
| NC_011770 | 3842296 | AGC | A | 64 | 0 | 1 | 0 | FRAME_SHIFT | HIGH | | gcgctt/ | AL16 | PLES_34581 | | | | | | | | A | | | | |
| NC_011770 | 3840790 | G | C | 85 | 0 | 2 | 0 | NON_SYNONYMOU | MODERAT | MISSENSE | gCc/gGc | A519G | PLES_34581 | | | | | | | C | C | | | | |
| NC_011770 | 3841072 | G | C | 75 | 0 | 2 | 0 | NON_SYNONYMOU | MODERAT | MISSENSE | gCc/gGc | A425G | PLES_34581 | | | | | | | | | | | C | C |
| NC_011770 | 3841949 | C | T | 55 | 0 | 2 | 0 | NON_SYNONYMOU | MODERAT | MISSENSE | Gag/Aag | E133K | PLES_34581 | T | T | | | | | | | | | | |
| NC_011770 | 3842349 | C | A | 72 | 0 | 2 | 0 | NON_SYNONYMOU | MODERAT | MISSENSE | Gtc/Ttc | V557F | PLES_34591 | A | A | | | | | | | | | | |
| NC_011770 | 3842556 | G | A | 69 | 0 | 1 | 0 | NON_SYNONYMOU | MODERAT | MISSENSE | Cgc/Tgc | R488C | PLES_34591 | | | | | | | A | | | | | |
| NC_011770 | 3842640 | G | T | 71 | 0 | 2 | 0 | NON_SYNONYMOU | MODERAT | MISSENSE | Ctg/Atg | L460M | PLES_34591 | | | | | | | | | T | T | | |
| NC_011770 | 3842882 | C | T | 72 | 0 | 6 | 0 | NON_SYNONYMOU | MODERAT | MISSENSE | gGt/gAt | G379D | PLES_34591 | T | T | T | T | T | T | | | | | | |
| NC_011770 | 3842939 | A | T | 72 | 0 | 8 | 0 | NON_SYNONYMOU | MODERAT | MISSENSE | tTc/tAc | F360Y | PLES_34591 | T | T | T | T | T | T | | | T | T | | |
| NC_011770 | 3843671 | T | C | 74 | 0 | 2 | 0 | NON_SYNONYMOU | MODERAT | MISSENSE | aAg/aGg | K116R | PLES_34591 | | | | | | | | | C | C | | |
| NC_011770 | 3843804 | T | C | 73 | 0 | 1 | 0 | NON_SYNONYMOU | MODERAT | MISSENSE | Acg/Gcg | T72A | PLES_34591 | | | | | | | C | | | | | |
| NC_011770 | 3844767 | C | T | 46 | 0 | 2 | 0 | NON_SYNONYMOU | MODERAT | MISSENSE | gCg/gTg | A206V | PLES_34601 | | | | | | | | | | | T | T |
| NC_011770 | 3847923 | C | A | 80 | 0 | 2 | 0 | NON_SYNONYMOU | MODERAT | MISSENSE | Gcc/Tcc | A129S | PLES_34641 | | | | | | | A | A | | | | |
| NC_011770 | 3847948 | C | A | 73 | 0 | 8 | 0 | NON_SYNONYMOU | MODERAT | MISSENSE | gaG/gaT | E120D | PLES_34641 | A | A | A | A | A | A | A | A | | | | |
| NC_011770 | 3848058 | C | G | 83 | 0 | 12 | 0 | NON_SYNONYMOU | MODERAT | MISSENSE | Gag/Cag | E84Q | PLES_34641 | G | G | G | G | G | G | G | G | G | G | G | G |
| NC_011770 | 3848072 | C | T | 79 | 0 | 10 | 0 | NON_SYNONYMOU | MODERAT | MISSENSE | cGc/cAc | R79H | PLES_34641 | T | T | T | T | T | T | T | T | T | T | | |
| NC_011770 | 3848075 | C | T | 80 | 0 | 10 | 0 | NON_SYNONYMOU | MODERAT | MISSENSE | gGc/gAc | G78D | PLES_34641 | T | T | T | T | T | T | T | T | T | T | | |
| NC_011770 | 3848083 | G | C | 78 | 0 | 12 | 0 | NON_SYNONYMOU | MODERAT | MISSENSE | gaC/gaG | D75E | PLES_34641 | C | C | C | C | C | C | C | C | C | C | C | C |
| NC_011770 | 3848189 | T | G | 79 | 0 | 10 | 0 | NON_SYNONYMOU | MODERAT | MISSENSE | gAg/gCg | E40A | PLES_34641 | G | G | G | G | G | G | G | G | G | G | | |
| NC_011770 | 3848745 | T | G | 83 | 0 | 2 | 0 | NON_SYNONYMOU | MODERAT | MISSENSE | cTg/cGg | L118R | PLES_34651 | | | | | | | | | | | G | G |
| NC_011770 | 3850250 | C | T | 83 | 0 | 2 | 0 | NON_SYNONYMOU | MODERAT | MISSENSE | Ctc/Ttc | L7F | PLES_34671 | T | T | | | | | | | | | | |
| NC_011770 | 3851058 | T | C | 57 | 0 | 2 | 0 | NON_SYNONYMOU | MODERAT | MISSENSE | cTg/cCg | L276P | PLES_34671 | | | | | | | C | C | | | | |
| NC_011770 | 3851106 | T | G | 60 | 0 | 10 | 0 | NON_SYNONYMOU | MODERAT | MISSENSE | gTg/gGg | V292G | PLES_34671 | G | G | G | G | G | G | G | G | | | G | G |
| NC_011770 | 3851623 | C | T | 67 | 0 | 2 | 0 | NON_SYNONYMOU | MODERAT | MISSENSE | gCc/gTc | A41V | PLES_34681 | | | | | | | | | T | T | | |
| NC_011770 | 3852041 | G | C | 91 | 0 | 2 | 0 | NON_SYNONYMOU | MODERAT | MISSENSE | atG/atC | M180I | PLES_34681 | | | | | | | | | | | C | C |
| NC_011770 | 3852645 | G | A | 80 | 0 | 2 | 0 | NON_SYNONYMOU | MODERAT | MISSENSE | Gtc/Atc | V382I | PLES_34681 | | | | | | | | | | | A | A |
| NC_011770 | 3852750 | G | A | 87 | 0 | 2 | 0 | NON_SYNONYMOU | MODERAT | MISSENSE | Gtc/Atc | V417I | PLES_34681 | | | | | | | | | | | A | A |
| NC_011770 | 3852905 | C | G | 76 | 0 | 2 | 0 | NON_SYNONYMOU | MODERAT | MISSENSE | gaC/gaG | D468E | PLES_34681 | | | | | | | | | | | G | G |
| NC_011770 | 3852927 | G | T | 73 | 0 | 2 | 0 | NON_SYNONYMOU | MODERAT | MISSENSE | Gcc/Tcc | A476S | PLES_34681 | | | | | | | T | T | | | | |
| NC_011770 | 3852971 | T | C | 68 | 0 | 2 | 0 | NON_SYNONYMOU | MODERAT | MISSENSE | Tcc/Ccc | S9P | PLES_34691 | | | | | | | | | | | C | C |
| NC_011770 | 3852981 | C | T | 66 | 0 | 2 | 0 | NON_SYNONYMOU | MODERAT | MISSENSE | gCg/gTg | A12V | PLES_34691 | | | | | | | | | | | T | T |
| NC_011770 | 3853000 | C | G | 66 | 0 | 4 | 0 | NON_SYNONYMOU | MODERAT | MISSENSE | agC/agG | S18R | PLES_34691 | G | G | | | | | | | G | G | | |
| NC_011770 | 3853537 | C | T | 70 | 0 | 2 | 0 | NON_SYNONYMOU | MODERAT | MISSENSE | gCc/gTc | A93V | PLES_34701 | T | T | | | | | | | | | | |
| NC_011770 | 3853571 | T | A | 70 | 0 | 6 | 0 | NON_SYNONYMOU | MODERAT | MISSENSE | gaT/gaA | D104E | PLES_34701 | | | A | A | A | A | | | | | A | A |

| | | | | | | | | | | | | | | | | | | | | | | | | | |
|---|---|---|---|---|---|---|---|---|---|---|---|---|---|---|---|---|---|---|---|---|---|---|---|---|---|
| NC_011770 | 3853800 | T | C | 74 | 0 | 6 | 0 | NON_SYNONYMOU | MODERAT | MISSENSE | Tcg/Ccg | S181P | PLES_34701 | | | C | C | C | C | | | | | C | C |
| NC_011770 | 3854069 | A | C | 76 | 0 | 8 | 0 | NON_SYNONYMOU | MODERAT | MISSENSE | gaA/gaC | E270D | PLES_34701 | C | C | C | C | C | C | | | | | C | C |
| NC_011770 | 3854758 | G | C | 74 | 0 | 4 | 0 | NON_SYNONYMOU | MODERAT | MISSENSE | Cta/Gta | L171V | PLES_34711 | | | C | C | C | C | | | | | | |
| NC_011770 | 3854853 | G | A | 68 | 0 | 4 | 0 | NON_SYNONYMOU | MODERAT | MISSENSE | gCg/gTg | A139V | PLES_34711 | | | | | | | A | A | A | A | | |
| NC_011770 | 3855703 | G | A | 83 | 0 | 2 | 0 | NON_SYNONYMOU | MODERAT | MISSENSE | Gcg/Acg | A51T | PLES_34721 | A | A | | | | | | | | | | |
| NC_011770 | 3855878 | C | T | 91 | 0 | 2 | 0 | SPLICE_SITE_REGIC | LOW | SILENT | taG/taA | *402 | PLES_34731 | | | | | | | | | T | T | | |
| NC_011770 | 3856264 | C | T | 89 | 0 | 2 | 0 | NON_SYNONYMOU | MODERAT | MISSENSE | Gct/Act | A274T | PLES_34731 | | | | | | | | | | | T | T |
| NC_011770 | 3856848 | T | C | 83 | 0 | 12 | 0 | NON_SYNONYMOU | MODERAT | MISSENSE | aAc/aGc | N79S | PLES_34731 | C | C | C | C | C | C | C | C | C | C | C | C |
| NC_011770 | 3856876 | A | T | 86 | 0 | 10 | 0 | NON_SYNONYMOU | MODERAT | MISSENSE | Ttg/Atg | L70M | PLES_34731 | T | T | T | T | T | T | T | T | T | T | | |
| NC_011770 | 3856900 | T | C | 83 | 0 | 10 | 0 | NON_SYNONYMOU | MODERAT | MISSENSE | Acg/Gcg | T62A | PLES_34731 | C | C | C | C | C | C | C | C | C | C | | |
| NC_011770 | 3857068 | C | T | 77 | 0 | 12 | 0 | NON_SYNONYMOU | MODERAT | MISSENSE | Ggc/Agc | G6S | PLES_34731 | T | T | T | T | T | T | T | T | T | T | T | T |
| NC_011770 | 3857241 | C | CG | 75 | 0 | 6 | 0 | FRAME_SHIFT | HIGH | | gcc/gccC | A29A? | PLES_34741 | | | CG | CG | CG | CG | | | CG | CG | | |
| NC_011770 | 3857248 | C | CG | 80 | 0 | 4 | 0 | FRAME_SHIFT | HIGH | | tcg/tcCg | S27S? | PLES_34741 | | | CG | CG | CG | CG | | | | | | |
| NC_011770 | 3857326 | C | T | 74 | 0 | 12 | 0 | START_LOST | HIGH | MISSENSE | atG/atA | M1I | PLES_34741 | T | T | T | T | T | T | T | T | T | T | T | T |
| NC_011770 | 3857327 | A | G | 73 | 0 | 12 | 0 | START_LOST | HIGH | MISSENSE | aTg/aCg | M1T | PLES_34741 | G | G | G | G | G | G | G | G | G | G | G | G |
| NC_011770 | 3857270 | A | G | 80 | 0 | 12 | 0 | NON_SYNONYMOU | MODERAT | MISSENSE | tTg/tCg | L20S | PLES_34741 | G | G | G | G | G | G | G | G | G | G | G | G |
| NC_011770 | 3857289 | T | C | 81 | 0 | 12 | 0 | NON_SYNONYMOU | MODERAT | MISSENSE | Acc/Gcc | T14A | PLES_34741 | C | C | C | C | C | C | C | C | C | C | C | C |
| NC_011770 | 3857605 | A | T | 71 | 0 | 2 | 0 | NON_SYNONYMOU | MODERAT | MISSENSE | Acc/Tcc | T92S | PLES_34751 | | | | | | | | | | | T | T |
| NC_011770 | 3857650 | C | T | 70 | 0 | 12 | 0 | NON_SYNONYMOU | MODERAT | MISSENSE | Cgc/Tgc | R107C | PLES_34751 | T | T | T | T | T | T | T | T | T | T | T | T |
| NC_011770 | 3857720 | A | G | 63 | 0 | 12 | 0 | NON_SYNONYMOU | MODERAT | MISSENSE | cAc/cGc | H130R | PLES_34751 | G | G | G | G | G | G | G | G | G | G | G | G |
| NC_011770 | 3858089 | A | G | 76 | 0 | 2 | 0 | NON_SYNONYMOU | MODERAT | MISSENSE | cAg/cGg | Q253R | PLES_34751 | | | | | | | | | | | G | G |
| NC_011770 | 3858399 | A | T | 67 | 0 | 2 | 0 | NON_SYNONYMOU | MODERAT | MISSENSE | gAc/gTc | D14V | PLES_34761 | | | | | | | T | T | | | | |
| NC_011770 | 3858470 | A | G | 75 | 0 | 2 | 0 | NON_SYNONYMOU | MODERAT | MISSENSE | Acc/Gcc | T38A | PLES_34761 | | | | | | | | | G | G | | |
| NC_011770 | 3858471 | C | T | 81 | 0 | 2 | 0 | NON_SYNONYMOU | MODERAT | MISSENSE | aCc/aTc | T38I | PLES_34761 | | | | | | | | | T | T | | |
| NC_011770 | 3858474 | G | A | 81 | 0 | 2 | 0 | NON_SYNONYMOU | MODERAT | MISSENSE | cGc/cAc | R39H | PLES_34761 | | | | | | | | | | | A | A |
| NC_011770 | 3858546 | T | C | 83 | 0 | 6 | 0 | NON_SYNONYMOU | MODERAT | MISSENSE | gTc/gCc | V63A | PLES_34761 | | | C | C | C | C | | | C | C | | |
| NC_011770 | 3858553 | G | C | 84 | 0 | 6 | 0 | NON_SYNONYMOU | MODERAT | MISSENSE | caG/caC | Q65H | PLES_34761 | | | C | C | C | C | | | C | C | | |
| NC_011770 | 3859120 | T | C | 76 | 0 | 4 | 0 | NON_SYNONYMOU | MODERAT | MISSENSE | gTg/gCg | V172A | PLES_34771 | | | | | C | C | C | C | | | | |
| NC_011770 | 3859257 | A | G | 77 | 0 | 4 | 0 | NON_SYNONYMOU | MODERAT | MISSENSE | Acc/Gcc | T218A | PLES_34771 | | | | | | | | | G | G | G | G |
| NC_011770 | 3859395 | C | G | 84 | 0 | 2 | 0 | NON_SYNONYMOU | MODERAT | MISSENSE | Ccg/Gcg | P264A | PLES_34771 | | | | | | | | | | | G | G |
| NC_011770 | 3859417 | A | G | 82 | 0 | 2 | 0 | NON_SYNONYMOU | MODERAT | MISSENSE | aAc/aGc | N271S | PLES_34771 | | | | | | | G | G | | | | |
| NC_011770 | 3859527 | G | A | 83 | 0 | 1 | 0 | NON_SYNONYMOU | MODERAT | MISSENSE | Ggc/Agc | G308S | PLES_34771 | | | | | | | A | | | | | |
| NC_011770 | 3863623 | G | C | 84 | 0 | 12 | 0 | NON_SYNONYMOU | MODERAT | MISSENSE | Gag/Cag | E145Q | PLES_34801 | C | C | C | C | C | C | C | C | C | C | C | C |
| NC_011770 | 3863662 | A | G | 86 | 0 | 4 | 0 | NON_SYNONYMOU | MODERAT | MISSENSE | Atc/Gtc | I158V | PLES_34801 | | | | | | | | | G | G | G | G |
| NC_011770 | 3868329 | C | T | 64 | 0 | 8 | 0 | NON_SYNONYMOU | MODERAT | MISSENSE | gCc/gTc | A41V | PLES_34831 | T | T | T | T | T | T | T | T | | | | |
| NC_011770 | 3868502 | G | A | 71 | 0 | 2 | 0 | NON_SYNONYMOU | MODERAT | MISSENSE | Gcc/Acc | A99T | PLES_34831 | | | | | | | A | A | | | | |
| NC_011770 | 3868626 | G | A | 66 | 0 | 12 | 0 | NON_SYNONYMOU | MODERAT | MISSENSE | gCg/gTg | A165V | PLES_34841 | A | A | A | A | A | A | A | A | A | A | A | A |
| NC_011770 | 3869632 | G | A | 64 | 0 | 2 | 0 | NON_SYNONYMOU | MODERAT | MISSENSE | Gat/Aat | D34N | PLES_34861 | | | | | | | | | A | A | | |
| NC_011770 | 3870434 | T | C | 44 | 0 | 2 | 0 | NON_SYNONYMOU | MODERAT | MISSENSE | gTg/gCg | V301A | PLES_34861 | | | | | | | | | C | C | | |
| NC_011770 | 3870536 | C | G | 42 | 0 | 10 | 0 | NON_SYNONYMOU | MODERAT | MISSENSE | gCc/gGc | A335G | PLES_34861 | | | G | G | G | G | G | G | G | G | G | G |
| NC_011770 | 3870756 | C | G | 66 | 0 | 2 | 0 | NON_SYNONYMOU | MODERAT | MISSENSE | aCc/aGc | T40S | PLES_34871 | | | | | | | G | G | | | | |
| NC_011770 | 3871106 | A | G | 80 | 0 | 2 | 0 | NON_SYNONYMOU | MODERAT | MISSENSE | tTc/tCc | F50S | PLES_34881 | G | G | | | | | | | | | | |
| NC_011770 | 3873854 | C | A | 42 | 0 | 2 | 0 | NON_SYNONYMOU | MODERAT | MISSENSE | Gcc/Tcc | A70S | PLES_34911 | | | | | | | | | A | A | | |
| NC_011770 | 3875061 | CAGATCCT( | C | 74 | 0 | 1 | 0 | FRAME_SHIFT | HIGH | | gaagggcag | EGQDL142 | PLES_34931 | C | . | | | | | | | | | | |
| NC_011770 | 3874633 | T | C | 69 | 0 | 8 | 0 | NON_SYNONYMOU | MODERAT | MISSENSE | gAc/gGc | D289G | PLES_34931 | C | C | | | | | C | C | C | C | C | C |
| NC_011770 | 3874678 | T | C | 75 | 0 | 2 | 0 | NON_SYNONYMOU | MODERAT | MISSENSE | gAg/gGg | E274G | PLES_34931 | C | C | | | | | | | | | | |
| NC_011770 | 3874693 | A | G | 77 | 0 | 2 | 0 | NON_SYNONYMOU | MODERAT | MISSENSE | aTg/aCg | M269T | PLES_34931 | G | G | | | | | | | | | | |
| NC_011770 | 3874694 | T | C | 77 | 0 | 2 | 0 | NON_SYNONYMOU | MODERAT | MISSENSE | Atg/Gtg | M269V | PLES_34931 | C | C | | | | | | | | | | |
| NC_011770 | 3874697 | G | A | 77 | 0 | 2 | 0 | NON_SYNONYMOU | MODERAT | MISSENSE | Ctc/Ttc | L268F | PLES_34931 | A | A | | | | | | | | | | |
| NC_011770 | 3874768 | T | C | 76 | 0 | 8 | 0 | NON_SYNONYMOU | MODERAT | MISSENSE | gAc/gGc | D244G | PLES_34931 | C | C | | | | | C | C | C | C | C | C |
| NC_011770 | 3874832 | A | C | 83 | 0 | 2 | 0 | NON_SYNONYMOU | MODERAT | MISSENSE | Tcc/Gcc | S223A | PLES_34931 | C | C | | | | | | | | | | |
| NC_011770 | 3874844 | C | G | 83 | 0 | 2 | 0 | NON_SYNONYMOU | MODERAT | MISSENSE | Gcg/Ccg | A219P | PLES_34931 | G | G | | | | | | | | | | |
| NC_011770 | 3874963 | T | C | 88 | 0 | 2 | 0 | NON_SYNONYMOU | MODERAT | MISSENSE | cAg/cGg | Q179R | PLES_34931 | C | C | | | | | | | | | | |
| NC_011770 | 3875059 | G | A | 78 | 0 | 1 | 0 | NON_SYNONYMOU | MODERAT | MISSENSE | gCc/gTc | A147V | PLES_34931 | | | | | | | A | | | | | |
| NC_011770 | 3875099 | T | G | 82 | 0 | 2 | 0 | NON_SYNONYMOU | MODERAT | MISSENSE | Atc/Ctc | I134L | PLES_34931 | G | G | | | | | | | | | | |
| NC_011770 | 3875438 | A | G | 52 | 0 | 2 | 0 | NON_SYNONYMOU | MODERAT | MISSENSE | Ttc/Ctc | F21L | PLES_34931 | | | | | | | | | G | G | | |
| NC_011770 | 3876419 | G | A | 79 | 0 | 1 | 0 | NON_SYNONYMOU | MODERAT | MISSENSE | gCc/gTc | A51V | PLES_34941 | | A | | | | | | | | | | |
| NC_011770 | 3877456 | C | T | 72 | 0 | 6 | 0 | NON_SYNONYMOU | MODERAT | MISSENSE | Gcc/Acc | A67T | PLES_34951 | | | T | T | T | T | | | | | T | T |
| NC_011770 | 3878191 | C | T | 71 | 0 | 1 | 0 | STOP_GAINED | HIGH | NONSENSE | Cag/Tag | Q102* | PLES_34961 | | | | | | | | T | | | | |
| NC_011770 | 3878187 | T | G | 72 | 0 | 2 | 0 | NON_SYNONYMOU | MODERAT | MISSENSE | gaT/gaG | D100E | PLES_34961 | | | | | | | | | | | G | G |
| NC_011770 | 3878386 | G | A | 70 | 0 | 2 | 0 | NON_SYNONYMOU | MODERAT | MISSENSE | Gtc/Atc | V167I | PLES_34961 | | | | | | | | | | | A | A |
| NC_011770 | 3878414 | T | A | 71 | 0 | 2 | 0 | NON_SYNONYMOU | MODERAT | MISSENSE | aTt/aAt | I176N | PLES_34961 | | | | | | | | | | | A | A |

| | | | | | | | | | | | | | | | | | | | | | | | | | |
|---|---|---|---|---|---|---|---|---|---|---|---|---|---|---|---|---|---|---|---|---|---|---|---|---|---|
| NC_011770 | 3878507 | A | G | 65 | 0 | 2 | 0 | NON_SYNONYMOU | MODERAT | MISSENSE | aAa/aGa | K207R | PLES_34961 | | | | | | | | | | | G | G |
| NC_011770 | 3879538 | A | C | 73 | 0 | 4 | 0 | NON_SYNONYMOU | MODERAT | MISSENSE | gAg/gCg | E103A | PLES_34981 | | | C | C | C | C | | | | | | |
| NC_011770 | 3879911 | C | A | 71 | 0 | 10 | 0 | NON_SYNONYMOU | MODERAT | MISSENSE | gaC/gaA | D227E | PLES_34981 | A | A | A | A | A | A | A | A | A | A | | |
| NC_011770 | 3881294 | GTTC | G | 64 | 0 | 2 | 0 | CODON_CHANGE_ | MODERATE | | ggaacg/gc | GT240A | PLES_35001 | G | G | | | | | | | | | | |
| NC_011770 | 3881625 | C | T | 73 | 0 | 2 | 0 | NON_SYNONYMOU | MODERAT | MISSENSE | Gcg/Acg | A131T | PLES_35001 | | | | | | | | | | | T | T |
| NC_011770 | 3882491 | C | A | 84 | 0 | 2 | 0 | NON_SYNONYMOU | MODERAT | MISSENSE | caC/caA | H115Q | PLES_35011 | A | A | | | | | | | | | | |
| NC_011770 | 3882577 | G | A | 89 | 0 | 1 | 0 | NON_SYNONYMOU | MODERAT | MISSENSE | gGc/gAc | G144D | PLES_35011 | | | | | | | A | | | | | |
| NC_011770 | 3882600 | C | T | 86 | 0 | 2 | 0 | NON_SYNONYMOU | MODERAT | MISSENSE | Cgc/Tgc | R152C | PLES_35011 | T | T | | | | | | | | | | |
| NC_011770 | 3884222 | G | A | 77 | 0 | 1 | 0 | STOP_GAINED | HIGH | NONSENSE | tgG/tgA | W143* | PLES_35031 | | | | | | | A | | | | | |
| NC_011770 | 3884055 | G | A | 74 | 0 | 2 | 0 | NON_SYNONYMOU | MODERAT | MISSENSE | Gcg/Acg | A88T | PLES_35031 | | | | | | | | | A | A | | |
| NC_011770 | 3884202 | C | G | 74 | 0 | 2 | 0 | NON_SYNONYMOU | MODERAT | MISSENSE | Ctc/Gtc | L137V | PLES_35031 | | | | | | | G | G | | | | |
| NC_011770 | 3885825 | A | G | 78 | 0 | 2 | 0 | NON_SYNONYMOU | MODERAT | MISSENSE | aTc/aCc | I414T | PLES_35051 | | | | | | | | | | | G | G |
| NC_011770 | 3885835 | G | A | 78 | 0 | 2 | 0 | NON_SYNONYMOU | MODERAT | MISSENSE | Cgc/Tgc | R411C | PLES_35051 | | | | | A | A | | | | | | |
| NC_011770 | 3890598 | T | A | 72 | 0 | 8 | 0 | NON_SYNONYMOU | MODERAT | MISSENSE | Agc/Tgc | S129C | PLES_35081 | | | A | A | A | A | A | A | | | A | A |
| NC_011770 | 3892358 | G | C | 74 | 0 | 12 | 0 | NON_SYNONYMOU | MODERAT | MISSENSE | Cag/Gag | Q302E | PLES_35091 | C | C | C | C | C | C | C | C | C | C | C | C |
| NC_011770 | 3891745 | G | A | 81 | 0 | 1 | 0 | NON_SYNONYMOU | MODERAT | MISSENSE | cCg/cTg | P506L | PLES_35091 | | | | A | | | | | | | | |
| NC_011770 | 3893829 | C | G | 79 | 0 | 8 | 0 | NON_SYNONYMOU | MODERAT | MISSENSE | gaC/gaG | D125E | PLES_35101 | G | G | G | G | G | G | | | | | G | G |
| NC_011770 | 3895618 | GGCGCCCT | G | 61 | 0 | 1 | 0 | FRAME_SHIFT | HIGH | | gaagagggc | EEGA87 | PLES_35131 | | | G | | | | | | | | | |
| NC_011770 | 3895227 | C | T | 57 | 0 | 2 | 0 | NON_SYNONYMOU | MODERAT | MISSENSE | Gtc/Atc | V221I | PLES_35131 | | | | | | | | | T | T | | |
| NC_011770 | 3895242 | C | T | 58 | 0 | 2 | 0 | NON_SYNONYMOU | MODERAT | MISSENSE | Gtg/Atg | V216M | PLES_35131 | | | | | T | T | | | | | | |
| NC_011770 | 3895806 | G | A | 67 | 0 | 1 | 0 | NON_SYNONYMOU | MODERAT | MISSENSE | Ccg/Tcg | P28S | PLES_35131 | | | | | | | A | | | | | |
| NC_011770 | 3896010 | C | T | 63 | 0 | 6 | 0 | NON_SYNONYMOU | MODERAT | MISSENSE | Ccc/Tcc | P8S | PLES_35141 | T | T | T | T | T | T | | | | | | |
| NC_011770 | 3896115 | G | A | 55 | 0 | 4 | 0 | NON_SYNONYMOU | MODERAT | MISSENSE | Gcc/Acc | A43T | PLES_35141 | | | | | | | A | A | A | A | | |
| NC_011770 | 3896669 | A | T | 57 | 0 | 4 | 0 | NON_SYNONYMOU | MODERAT | MISSENSE | aaA/aaT | K227N | PLES_35141 | T | T | | | | | | | | | T | T |
| NC_011770 | 3898702 | C | T | 80 | 0 | 2 | 0 | NON_SYNONYMOU | MODERAT | MISSENSE | aCc/aTc | T22I | PLES_35161 | | | | | | | | | T | T | | |
| NC_011770 | 3899172 | T | C | 70 | 0 | 10 | 0 | NON_SYNONYMOU | MODERAT | MISSENSE | Tcc/Ccc | S179P | PLES_35161 | C | C | C | C | C | C | | | C | C | C | C |
| NC_011770 | 3899590 | T | C | 78 | 0 | 2 | 0 | NON_SYNONYMOU | MODERAT | MISSENSE | aTc/aCc | I318T | PLES_35161 | C | C | | | | | | | | | | |
| NC_011770 | 3899856 | A | G | 64 | 0 | 2 | 0 | NON_SYNONYMOU | MODERAT | MISSENSE | Acc/Gcc | T407A | PLES_35161 | | | | | | | G | G | | | | |
| NC_011770 | 3900265 | G | T | 63 | 0 | 12 | 0 | NON_SYNONYMOU | MODERAT | MISSENSE | cGa/cTa | R543L | PLES_35161 | T | T | T | T | T | T | T | T | T | T | T | T |
| NC_011770 | 3902282 | A | G | 71 | 0 | 2 | 0 | NON_SYNONYMOU | MODERAT | MISSENSE | Acg/Gcg | T607A | PLES_35171 | G | G | | | | | | | | | | |
| NC_011770 | 3902603 | G | A | 68 | 0 | 2 | 0 | NON_SYNONYMOU | MODERAT | MISSENSE | Gag/Aag | E96K | PLES_35181 | | | | | A | A | | | | | | |
| NC_011770 | 3903747 | G | A | 40 | 0 | 1 | 0 | NON_SYNONYMOU | MODERAT | MISSENSE | Gcc/Acc | A115T | PLES_35191 | | | | | | | | A | | | | |
| NC_011770 | 3904823 | A | G | 67 | 0 | 12 | 0 | NON_SYNONYMOU | MODERAT | MISSENSE | aAa/aGa | K133R | PLES_35201 | G | G | G | G | G | G | G | G | G | G | G | G |
| NC_011770 | 3905164 | G | A | 62 | 0 | 2 | 0 | NON_SYNONYMOU | MODERAT | MISSENSE | Gcc/Acc | A247T | PLES_35201 | | | | | | | A | A | | | | |
| NC_011770 | 3905441 | G | A | 71 | 0 | 1 | 0 | NON_SYNONYMOU | MODERAT | MISSENSE | gGc/gAc | G339D | PLES_35201 | | | | | | | A | | | | | |
| NC_011770 | 3905462 | C | T | 69 | 0 | 2 | 0 | NON_SYNONYMOU | MODERAT | MISSENSE | aCc/aTc | T346I | PLES_35201 | T | T | | | | | | | | | | |
| NC_011770 | 3908952 | AC | A | 60 | 0 | 2 | 0 | INTRAGENIC | MODIFIER | | | | PLES_35222 | | | | | | | | | | | A | A |
| NC_011770 | 3915569 | A | G | 67 | 0 | 2 | 0 | NON_SYNONYMOU | MODERAT | MISSENSE | gAt/gGt | D47G | PLES_35281 | | | | | | | G | G | | | | |
| NC_011770 | 3915634 | A | G | 66 | 0 | 12 | 0 | NON_SYNONYMOU | MODERAT | MISSENSE | Aac/Gac | N69D | PLES_35281 | G | G | G | G | G | G | G | G | G | G | G | G |
| NC_011770 | 3915992 | C | T | 54 | 0 | 1 | 0 | NON_SYNONYMOU | MODERAT | MISSENSE | gCc/gTc | A115V | PLES_35291 | | | | | | | | | | T | | |
| NC_011770 | 3917503 | C | A | 71 | 0 | 2 | 0 | NON_SYNONYMOU | MODERAT | MISSENSE | Cgc/Agc | R383S | PLES_35301 | | | | | | | | | A | A | | |
| NC_011770 | 3917567 | A | G | 68 | 0 | 1 | 0 | NON_SYNONYMOU | MODERAT | MISSENSE | cAc/cGc | H404R | PLES_35301 | | | | | | | G | | | | | |
| NC_011770 | 3917769 | C | T | 60 | 0 | 8 | 0 | NON_SYNONYMOU | MODERAT | MISSENSE | Ccc/Tcc | P8S | PLES_35311 | | | T | T | T | T | T | T | T | T | | |
| NC_011770 | 3917890 | G | A | 71 | 0 | 4 | 0 | NON_SYNONYMOU | MODERAT | MISSENSE | cGc/cAc | R48H | PLES_35311 | | | A | A | A | A | | | | | | |
| NC_011770 | 3917993 | C | A | 75 | 0 | 2 | 0 | NON_SYNONYMOU | MODERAT | MISSENSE | caC/caA | H82Q | PLES_35311 | A | A | | | | | | | | | | |
| NC_011770 | 3918129 | T | C | 71 | 0 | 1 | 0 | NON_SYNONYMOU | MODERAT | MISSENSE | Ttc/Ctc | F128L | PLES_35311 | | | | | | | C | | | | | |
| NC_011770 | 3918259 | A | G | 69 | 0 | 8 | 0 | NON_SYNONYMOU | MODERAT | MISSENSE | aAc/aGc | N171S | PLES_35311 | G | G | G | G | G | G | | | | | G | G |
| NC_011770 | 3918330 | T | C | 66 | 0 | 2 | 0 | NON_SYNONYMOU | MODERAT | MISSENSE | Tat/Cat | Y195H | PLES_35311 | | | | | | | | | C | C | | |
| NC_011770 | 3918422 | G | C | 52 | 0 | 12 | 0 | NON_SYNONYMOU | MODERAT | MISSENSE | gaG/gaC | E225D | PLES_35311 | C | C | C | C | C | C | C | C | C | C | C | C |
| NC_011770 | 3918485 | C | A | 51 | 0 | 6 | 0 | NON_SYNONYMOU | MODERAT | MISSENSE | caC/caA | H246Q | PLES_35311 | | | A | A | A | A | | | | | A | A |
| NC_011770 | 3918513 | T | TCCGCC | 48 | 0 | 7 | 0 | CODON_INSERTIOI | MODERATE | | agc/GCCG | S258AAS | PLES_35311 | TCCGCCG | TCCGCCG | TCCGCCG | TCCGCCG | TCCGCCG | TCCGCCG | | | | | TCCGCCG | |
| NC_011770 | 3918775 | C | G | 31 | 0 | 6 | 0 | NON_SYNONYMOU | MODERAT | MISSENSE | cCa/cGa | P343R | PLES_35311 | G | G | G | G | G | G | | | | | | |
| NC_011770 | 3918795 | G | A | 29 | 0 | 8 | 0 | NON_SYNONYMOU | MODERAT | MISSENSE | Gtt/Att | V350I | PLES_35311 | A | A | A | A | A | A | | | | | A | A |
| NC_011770 | 3918849 | C | A | 19 | 0 | 8 | 0 | NON_SYNONYMOU | MODERAT | MISSENSE | Cgc/Agc | R368S | PLES_35311 | A | A | A | A | A | A | | | | | A | A |
| NC_011770 | 3918907 | G | A | 11 | 1 | 8 | 0 | NON_SYNONYMOU | MODERAT | MISSENSE | aGg/aAg | R387K | PLES_35311 | A | A | A | A | A | A | | | . | | A | A |
| NC_011770 | 3919127 | T | A | 35 | 0 | 12 | 0 | NON_SYNONYMOU | MODERAT | MISSENSE | agT/agA | S460R | PLES_35311 | A | A | A | A | A | A | A | A | A | A | A | A |
| NC_011770 | 3919320 | G | A | 51 | 0 | 2 | 0 | NON_SYNONYMOU | MODERAT | MISSENSE | Ggc/Agc | G525S | PLES_35311 | A | A | | | | | | | | | | |
| NC_011770 | 3919537 | A | G | 71 | 0 | 2 | 0 | NON_SYNONYMOU | MODERAT | MISSENSE | gAc/gGc | D597G | PLES_35311 | | | | | | | | | G | G | | |
| NC_011770 | 3925845 | G | A | 69 | 0 | 2 | 0 | NON_SYNONYMOU | MODERAT | MISSENSE | gGc/gAc | G69D | PLES_35371 | A | A | | | | | | | | | | |
| NC_011770 | 3926192 | G | A | 56 | 0 | 4 | 0 | NON_SYNONYMOU | MODERAT | MISSENSE | Ggg/Agg | G185R | PLES_35371 | | | | | | | | | A | A | A | A |
| NC_011770 | 3926535 | GC | G | 37 | 1 | 3 | 0 | FRAME_SHIFT | HIGH | | ctc/ | L16 | PLES_35381 | | G | | | | | . | | GCC | GCC | | |
| NC_011770 | 3926547 | G | A | 41 | 0 | 2 | 0 | NON_SYNONYMOU | MODERAT | MISSENSE | Gtg/Atg | V18M | PLES_35381 | A | A | | | | | | | | | | |

| | | | | | | | | | | | | | | | | | | | | | | | | | |
|---|---|---|---|---|---|---|---|---|---|---|---|---|---|---|---|---|---|---|---|---|---|---|---|---|---|
| NC_011770 | 3927889 | T | C | 65 | 0 | 4 | 0 | NON_SYNONYMOU | MODERAT | MISSENSE | gAc/gGc | D204G | PLES_35391 | | | C | C | C | C | | | | | | |
| NC_011770 | 3927898 | A | T | 62 | 0 | 4 | 0 | NON_SYNONYMOU | MODERAT | MISSENSE | cTc/cAc | L201H | PLES_35391 | | | T | T | T | T | | | | | | |
| NC_011770 | 3928237 | G | C | 74 | 0 | 2 | 0 | NON_SYNONYMOU | MODERAT | MISSENSE | cCg/cGg | P88R | PLES_35391 | C | C | | | | | | | | | | |
| NC_011770 | 3928490 | T | C | 84 | 0 | 2 | 0 | NON_SYNONYMOU | MODERAT | MISSENSE | Acc/Gcc | T4A | PLES_35391 | | | | | | | | | C | C | | |
| NC_011770 | 3928898 | A | G | 82 | 0 | 2 | 0 | NON_SYNONYMOU | MODERAT | MISSENSE | Acc/Gcc | T77A | PLES_35401 | | | G | G | | | | | | | | |
| NC_011770 | 3935352 | G | A | 65 | 0 | 2 | 0 | NON_SYNONYMOU | MODERAT | MISSENSE | cGg/cAg | R32Q | PLES_35451 | | | | | | | | | A | A | | |
| NC_011770 | 3935531 | A | C | 70 | 0 | 2 | 0 | NON_SYNONYMOU | MODERAT | MISSENSE | Aag/Cag | K92Q | PLES_35451 | | | | | | | C | C | | | | |
| NC_011770 | 3938278 | A | C | 59 | 0 | 2 | 0 | NON_SYNONYMOU | MODERAT | MISSENSE | aAg/aCg | K229T | PLES_35471 | | | | | | | | | | | C | C |
| NC_011770 | 3938284 | A | G | 60 | 0 | 2 | 0 | NON_SYNONYMOU | MODERAT | MISSENSE | gAc/gGc | D231G | PLES_35471 | | | | | | | G | G | | | | |
| NC_011770 | 3939165 | C | T | 50 | 0 | 4 | 0 | NON_SYNONYMOU | MODERAT | MISSENSE | Ctc/Ttc | L525F | PLES_35471 | | | T | T | T | T | | | | | | |
| NC_011770 | 3938595 | T | C | 41 | 0 | 1 | 0 | NON_SYNONYMOU | MODERAT | MISSENSE | Tgg/Cgg | W335R | PLES_35471 | | | | | | | | C | | | | |
| NC_011770 | 3939224 | G | A | 52 | 0 | 10 | 0 | INTRAGENIC | MODIFIER | | | | PLES_35472 | A | A | A | A | A | A | A | A | | | A | A |
| NC_011770 | 3939240 | T | G | 55 | 0 | 2 | 0 | INTRAGENIC | MODIFIER | | | | PLES_35472 | | | | | | | | | G | G | | |
| NC_011770 | 3942537 | C | A | 69 | 0 | 4 | 0 | NON_SYNONYMOU | MODERAT | MISSENSE | Ctg/Atg | L76M | PLES_35501 | | | A | A | A | A | | | | | | |
| NC_011770 | 3943710 | A | G | 64 | 0 | 8 | 0 | NON_SYNONYMOU | MODERAT | MISSENSE | Atc/Gtc | I467V | PLES_35501 | G | G | G | G | G | G | | | | | G | G |
| NC_011770 | 3943970 | A | C | 80 | 0 | 4 | 0 | NON_SYNONYMOU | MODERAT | MISSENSE | caA/caC | Q553H | PLES_35501 | | | C | C | C | C | | | | | | |
| NC_011770 | 3944208 | C | A | 77 | 0 | 2 | 0 | NON_SYNONYMOU | MODERAT | MISSENSE | Cgc/Agc | R633S | PLES_35501 | | | | | | | | | A | A | | |
| NC_011770 | 3944218 | T | G | 78 | 0 | 10 | 0 | NON_SYNONYMOU | MODERAT | MISSENSE | cTg/cGg | L636R | PLES_35501 | G | G | G | G | G | G | | | G | G | G | G |
| NC_011770 | 3943600 | T | C | 71 | 0 | 1 | 0 | NON_SYNONYMOU | MODERAT | MISSENSE | gTg/gCg | V430A | PLES_35501 | | C | | | | | | | | | | |
| NC_011770 | 3956175 | G | A | 71 | 0 | 2 | 0 | NON_SYNONYMOU | MODERAT | MISSENSE | cGc/cAc | R102H | PLES_35621 | | | | | | | | | A | A | | |
| NC_011770 | 3957684 | TG | T | 83 | 0 | 2 | 0 | FRAME_SHIFT | HIGH | | gtg/ | V96 | PLES_35631 | | | T | T | | | | | | | | |
| NC_011770 | 3958375 | C | A | 65 | 0 | 4 | 0 | NON_SYNONYMOU | MODERAT | MISSENSE | caC/caA | H325Q | PLES_35631 | A | A | | | | | | | A | A | | |
| NC_011770 | 3958520 | T | C | 69 | 0 | 12 | 0 | NON_SYNONYMOU | MODERAT | MISSENSE | gAc/gGc | D370G | PLES_35641 | C | C | C | C | C | C | C | C | C | C | C | C |
| NC_011770 | 3958714 | G | C | 74 | 0 | 8 | 0 | NON_SYNONYMOU | MODERAT | MISSENSE | caC/caG | H305Q | PLES_35641 | | | C | C | C | C | C | C | | | C | C |
| NC_011770 | 3958787 | A | C | 76 | 0 | 2 | 0 | NON_SYNONYMOU | MODERAT | MISSENSE | gTc/gGc | V281G | PLES_35641 | | | | | | | | | C | C | | |
| NC_011770 | 3958895 | T | C | 73 | 0 | 2 | 0 | NON_SYNONYMOU | MODERAT | MISSENSE | gAt/gGt | D245G | PLES_35641 | C | C | | | | | | | | | | |
| NC_011770 | 3959046 | C | T | 80 | 0 | 8 | 0 | NON_SYNONYMOU | MODERAT | MISSENSE | Gcg/Acg | A195T | PLES_35641 | | | T | T | T | T | T | T | | | T | T |
| NC_011770 | 3959310 | G | T | 83 | 0 | 2 | 0 | NON_SYNONYMOU | MODERAT | MISSENSE | Cag/Aag | Q107K | PLES_35641 | | | | | | | | | T | T | | |
| NC_011770 | 3959763 | G | T | 92 | 0 | 2 | 0 | NON_SYNONYMOU | MODERAT | MISSENSE | aCc/aAc | T493N | PLES_35651 | T | T | | | | | | | | | | |
| NC_011770 | 3960351 | T | C | 83 | 0 | 10 | 0 | NON_SYNONYMOU | MODERAT | MISSENSE | aAg/aGg | K297R | PLES_35651 | C | C | C | C | C | C | | | C | C | C | C |
| NC_011770 | 3960889 | G | A | 77 | 0 | 1 | 0 | NON_SYNONYMOU | MODERAT | MISSENSE | Ccc/Tcc | P118S | PLES_35651 | | | | | | | A | | | | | |
| NC_011770 | 3961050 | C | T | 78 | 0 | 4 | 0 | NON_SYNONYMOU | MODERAT | MISSENSE | aGg/aAg | R64K | PLES_35651 | T | T | | | | | | | T | T | | |
| NC_011770 | 3961889 | C | T | 87 | 0 | 2 | 0 | NON_SYNONYMOU | MODERAT | MISSENSE | Gta/Ata | V133I | PLES_35661 | | | | | | | | | T | T | | |
| NC_011770 | 3962185 | A | G | 88 | 0 | 1 | 0 | NON_SYNONYMOU | MODERAT | MISSENSE | cTg/cCg | L34P | PLES_35661 | | | | G | | | | | | | | |
| NC_011770 | 3964227 | C | A | 69 | 0 | 2 | 0 | NON_SYNONYMOU | MODERAT | MISSENSE | Gcc/Tcc | A778S | PLES_35691 | | | | | | | | | A | A | | |
| NC_011770 | 3968350 | G | T | 75 | 0 | 4 | 0 | NON_SYNONYMOU | MODERAT | MISSENSE | gCg/gAg | A306E | PLES_35701 | | | T | T | T | T | | | | | | |
| NC_011770 | 3968591 | C | T | 88 | 0 | 4 | 0 | NON_SYNONYMOU | MODERAT | MISSENSE | Ggc/Agc | G226S | PLES_35701 | | | | | | | | | T | T | T | T |
| NC_011770 | 3968662 | G | A | 83 | 0 | 12 | 0 | NON_SYNONYMOU | MODERAT | MISSENSE | tCc/tTc | S202F | PLES_35701 | A | A | A | A | A | A | A | A | A | A | A | A |
| NC_011770 | 3968807 | T | A | 74 | 0 | 2 | 0 | NON_SYNONYMOU | MODERAT | MISSENSE | Acc/Tcc | T154S | PLES_35701 | | | | | | | | | A | A | | |
| NC_011770 | 3972737 | T | G | 67 | 0 | 4 | 0 | NON_SYNONYMOU | MODERAT | MISSENSE | Tcc/Gcc | S95A | PLES_35741 | | | | | | | G | G | | | G | G |
| NC_011770 | 3974663 | A | G | 73 | 0 | 2 | 0 | NON_SYNONYMOU | MODERAT | MISSENSE | Tac/Cac | Y304H | PLES_35771 | | | | | | | G | G | | | | |
| NC_011770 | 3974889 | C | G | 87 | 0 | 2 | 0 | NON_SYNONYMOU | MODERAT | MISSENSE | gaG/gaC | E228D | PLES_35771 | | | | | | | | | G | G | | |
| NC_011770 | 3975406 | A | G | 76 | 0 | 2 | 0 | NON_SYNONYMOU | MODERAT | MISSENSE | tTc/tCc | F56S | PLES_35771 | | | | | | | G | G | | | | |
| NC_011770 | 3975773 | G | C | 59 | 0 | 2 | 0 | NON_SYNONYMOU | MODERAT | MISSENSE | gaC/gaG | D59E | PLES_35781 | | | | | | | C | C | | | | |
| NC_011770 | 3975939 | G | A | 58 | 0 | 2 | 0 | NON_SYNONYMOU | MODERAT | MISSENSE | cCc/cTc | P4L | PLES_35781 | | | | | | | A | A | | | | |
| NC_011770 | 3975940 | G | T | 58 | 0 | 2 | 0 | NON_SYNONYMOU | MODERAT | MISSENSE | Ccc/Acc | P4T | PLES_35781 | | | | | | | T | T | | | | |
| NC_011770 | 3975945 | A | G | 59 | 0 | 2 | 0 | NON_SYNONYMOU | MODERAT | MISSENSE | gTc/gCc | V2A | PLES_35781 | | | | | | | G | G | | | | |
| NC_011770 | 3976420 | G | A | 68 | 0 | 1 | 0 | NON_SYNONYMOU | MODERAT | MISSENSE | Gac/Aac | D52N | PLES_35791 | | | | | | | A | | | | | |
| NC_011770 | 3978213 | G | C | 80 | 0 | 2 | 0 | NON_SYNONYMOU | MODERAT | MISSENSE | Gtc/Ctc | V48L | PLES_35811 | | | | | | | | | | | C | C |
| NC_011770 | 3979172 | C | T | 48 | 0 | 4 | 0 | SPLICE_SITE_REGIO | LOW | SILENT | tGa/tAa | *169 | PLES_35831 | | | T | T | T | T | | | | | | |
| NC_011770 | 3979245 | C | T | 50 | 0 | 2 | 0 | NON_SYNONYMOU | MODERAT | MISSENSE | Ggc/Agc | G145S | PLES_35831 | | | | | | | T | T | | | | |
| NC_011770 | 3980123 | A | T | 66 | 0 | 2 | 0 | NON_SYNONYMOU | MODERAT | MISSENSE | Atc/Ttc | I76F | PLES_35841 | | | T | T | | | | | | | | |
| NC_011770 | 3980195 | C | G | 67 | 0 | 4 | 0 | NON_SYNONYMOU | MODERAT | MISSENSE | Cgc/Ggc | R100G | PLES_35841 | | | | | | | | | G | G | G | G |
| NC_011770 | 3980207 | G | A | 67 | 0 | 1 | 0 | NON_SYNONYMOU | MODERAT | MISSENSE | Gcc/Acc | A104T | PLES_35841 | | | | | | | | A | | | | |
| NC_011770 | 3980712 | G | A | 74 | 0 | 2 | 0 | NON_SYNONYMOU | MODERAT | MISSENSE | Gcc/Acc | A58T | PLES_35851 | A | A | | | | | | | | | | |
| NC_011770 | 3981155 | C | T | 63 | 0 | 2 | 0 | NON_SYNONYMOU | MODERAT | MISSENSE | Ggg/Agg | G218R | PLES_35871 | T | T | | | | | | | | | | |
| NC_011770 | 3981379 | G | A | 74 | 0 | 1 | 0 | NON_SYNONYMOU | MODERAT | MISSENSE | aCc/aTc | T143I | PLES_35871 | | | | | | | | | | | A | |
| NC_011770 | 3982590 | C | G | 73 | 0 | 2 | 0 | NON_SYNONYMOU | MODERAT | MISSENSE | gaG/gaC | E280D | PLES_35891 | G | G | | | | | | | | | | |
| NC_011770 | 3982772 | T | A | 70 | 0 | 2 | 0 | NON_SYNONYMOU | MODERAT | MISSENSE | Acg/Tcg | T220S | PLES_35891 | A | A | | | | | | | | | | |
| NC_011770 | 3982838 | C | T | 65 | 0 | 2 | 0 | NON_SYNONYMOU | MODERAT | MISSENSE | Gac/Aac | D198N | PLES_35891 | T | T | | | | | | | | | | |
| NC_011770 | 3983173 | G | A | 69 | 0 | 4 | 0 | NON_SYNONYMOU | MODERAT | MISSENSE | cCg/cTg | P86L | PLES_35891 | A | A | | | | | | | | | A | A |
| NC_011770 | 3983249 | C | T | 75 | 0 | 2 | 0 | NON_SYNONYMOU | MODERAT | MISSENSE | Gtc/Atc | V61I | PLES_35891 | | | | | | | | | T | T | | |

| | | | | | | | | | | | | | | | | | | | | | | | | | |
|---|---|---|---|---|---|---|---|---|---|---|---|---|---|---|---|---|---|---|---|---|---|---|---|---|---|
| NC_011770 | 3983537 | T | G | 72 | 0 | 12 | 0 | NON_SYNONYMOU | MODERAT | MISSENSE | gAg/gCg | E296A | PLES_35901 | G | G | G | G | G | G | G | G | G | G | G | G |
| NC_011770 | 3984709 | C | G | 49 | 0 | 4 | 0 | NON_SYNONYMOU | MODERAT | MISSENSE | Ctc/Gtc | L46V | PLES_35911 | | | G | G | G | G | | | | | | |
| NC_011770 | 3984724 | C | T | 50 | 0 | 4 | 0 | NON_SYNONYMOU | MODERAT | MISSENSE | Ctc/Ttc | L51F | PLES_35911 | | | T | T | T | T | | | | | | |
| NC_011770 | 3984779 | T | C | 50 | 0 | 12 | 0 | NON_SYNONYMOU | MODERAT | MISSENSE | gTa/gCa | V69A | PLES_35911 | C | C | C | C | C | C | C | C | C | C | C | C |
| NC_011770 | 3985207 | G | A | 60 | 0 | 8 | 0 | NON_SYNONYMOU | MODERAT | MISSENSE | Gcg/Acg | A212T | PLES_35911 | | | A | A | A | A | A | A | A | A | | |
| NC_011770 | 3985213 | A | G | 58 | 0 | 8 | 0 | NON_SYNONYMOU | MODERAT | MISSENSE | Acg/Gcg | T214A | PLES_35911 | | | G | G | G | G | G | G | G | G | | |
| NC_011770 | 3985435 | G | A | 56 | 0 | 2 | 0 | NON_SYNONYMOU | MODERAT | MISSENSE | Gcc/Acc | A288T | PLES_35911 | | | | | | | | | A | A | | |
| NC_011770 | 3986322 | C | T | 62 | 0 | 2 | 0 | NON_SYNONYMOU | MODERAT | MISSENSE | Ggc/Agc | G566S | PLES_35921 | | | T | T | | | | | | | | |
| NC_011770 | 3987054 | G | T | 71 | 0 | 2 | 0 | NON_SYNONYMOU | MODERAT | MISSENSE | Ctc/Atc | L322I | PLES_35921 | T | T | | | | | | | | | | |
| NC_011770 | 3987078 | G | A | 72 | 0 | 2 | 0 | NON_SYNONYMOU | MODERAT | MISSENSE | Cgg/Tgg | R314W | PLES_35921 | | | | | | | | | A | A | | |
| NC_011770 | 3987321 | G | T | 80 | 0 | 12 | 0 | NON_SYNONYMOU | MODERAT | MISSENSE | Ctg/Atg | L233M | PLES_35921 | T | T | T | T | T | T | T | T | T | T | T | T |
| NC_011770 | 3987404 | G | A | 81 | 0 | 4 | 0 | NON_SYNONYMOU | MODERAT | MISSENSE | gCc/gTc | A205V | PLES_35921 | | | A | A | A | A | | | | | | |
| NC_011770 | 3987562 | G | C | 75 | 0 | 2 | 0 | NON_SYNONYMOU | MODERAT | MISSENSE | atC/atG | I152M | PLES_35921 | C | C | | | | | | | | | | |
| NC_011770 | 3987564 | T | C | 72 | 0 | 2 | 0 | NON_SYNONYMOU | MODERAT | MISSENSE | Atc/Gtc | I152V | PLES_35921 | C | C | | | | | | | | | | |
| NC_011770 | 3987773 | G | A | 82 | 0 | 4 | 0 | NON_SYNONYMOU | MODERAT | MISSENSE | gCc/gTc | A82V | PLES_35921 | | | A | A | A | A | | | | | | |
| NC_011770 | 3985944 | C | T | 57 | 0 | 1 | 0 | NON_SYNONYMOU | MODERAT | MISSENSE | Gaa/Aaa | E692K | PLES_35921 | | | | | | | | | | | | T |
| NC_011770 | 3988183 | C | A | 68 | 0 | 2 | 0 | NON_SYNONYMOU | MODERAT | MISSENSE | Gcc/Tcc | A352S | PLES_35931 | | | | | | | | | | | A | A |
| NC_011770 | 3990899 | C | CG | 76 | 0 | 1 | 0 | FRAME_SHIFT | HIGH | | ggc/ggGc | G33G? | PLES_35951 | | | | | | | CG | | | | | |
| NC_011770 | 3991102 | C | T | 64 | 0 | 2 | 0 | NON_SYNONYMOU | MODERAT | MISSENSE | Ccg/Tcg | P99S | PLES_35951 | | | | | | | | | T | T | | |
| NC_011770 | 3992242 | GCT | G | 72 | 0 | 1 | 0 | FRAME_SHIFT | HIGH | | gag/ | E42 | PLES_35961 | | | | | | | | G | | | | |
| NC_011770 | 3992176 | G | A | 75 | 0 | 1 | 0 | STOP_GAINED | HIGH | NONSENSE | Cag/Tag | Q65* | PLES_35961 | | | | | | | A | | | | | |
| NC_011770 | 3992074 | T | C | 66 | 0 | 8 | 0 | NON_SYNONYMOU | MODERAT | MISSENSE | Agc/Ggc | S99G | PLES_35961 | C | C | | | | | C | C | C | C | C | C |
| NC_011770 | 3992086 | G | T | 72 | 0 | 4 | 0 | NON_SYNONYMOU | MODERAT | MISSENSE | Ctg/Atg | L95M | PLES_35961 | T | T | | | | | | | T | T | | |
| NC_011770 | 3992110 | C | T | 64 | 0 | 2 | 0 | NON_SYNONYMOU | MODERAT | MISSENSE | Gtc/Atc | V87I | PLES_35961 | T | T | | | | | | | | | | |
| NC_011770 | 3992737 | T | C | 94 | 0 | 6 | 0 | NON_SYNONYMOU | MODERAT | MISSENSE | Atc/Gtc | I172V | PLES_35971 | | | | | | | C | C | C | C | C | C |
| NC_011770 | 3994474 | C | A | 73 | 0 | 4 | 0 | NON_SYNONYMOU | MODERAT | MISSENSE | cGg/cTg | R381L | PLES_35991 | | | A | A | A | A | | | | | | |
| NC_011770 | 3995189 | C | T | 64 | 0 | 1 | 0 | NON_SYNONYMOU | MODERAT | MISSENSE | Ggg/Agg | G143R | PLES_35991 | | | | | | | | | | | | T |
| NC_011770 | 3996108 | C | T | 74 | 0 | 2 | 0 | NON_SYNONYMOU | MODERAT | MISSENSE | Gcc/Acc | A186T | PLES_36001 | | | | | | | T | T | | | | |
| NC_011770 | 3996355 | G | T | 72 | 0 | 6 | 0 | NON_SYNONYMOU | MODERAT | MISSENSE | caC/caA | H103Q | PLES_36001 | T | T | | | | | T | T | T | T | | |
| NC_011770 | 3996588 | C | T | 74 | 0 | 4 | 0 | NON_SYNONYMOU | MODERAT | MISSENSE | Ggc/Agc | G26S | PLES_36001 | | | T | T | T | T | | | | | | |
| NC_011770 | 3996605 | A | G | 75 | 0 | 12 | 0 | NON_SYNONYMOU | MODERAT | MISSENSE | gTc/gCc | V20A | PLES_36001 | G | G | G | G | G | G | G | G | G | G | G | G |
| NC_011770 | 3998448 | G | A | 71 | 0 | 1 | 0 | NON_SYNONYMOU | MODERAT | MISSENSE | Gtc/Atc | V332I | PLES_36021 | | | | | | | A | | | | | |
| NC_011770 | 3998850 | G | A | 71 | 0 | 10 | 0 | NON_SYNONYMOU | MODERAT | MISSENSE | Gtc/Atc | V466I | PLES_36021 | A | A | A | A | A | A | | | A | A | A | A |
| NC_011770 | 3998901 | C | A | 70 | 0 | 10 | 0 | NON_SYNONYMOU | MODERAT | MISSENSE | Ccc/Acc | P483T | PLES_36021 | A | A | A | A | A | A | | | A | A | A | A |
| NC_011770 | 3998908 | A | G | 70 | 0 | 10 | 0 | NON_SYNONYMOU | MODERAT | MISSENSE | gAc/gGc | D485G | PLES_36021 | G | G | G | G | G | G | | | G | G | G | G |
| NC_011770 | 3999020 | G | C | 67 | 0 | 10 | 0 | NON_SYNONYMOU | MODERAT | MISSENSE | gaG/gaC | E522D | PLES_36021 | C | C | C | C | C | C | | | C | C | C | C |
| NC_011770 | 3999498 | G | C | 72 | 0 | 2 | 0 | NON_SYNONYMOU | MODERAT | MISSENSE | Gcc/Ccc | A682P | PLES_36021 | C | C | | | | | | | | | | |
| NC_011770 | 4018901 | A | C | 61 | 0 | 6 | 0 | NON_SYNONYMOU | MODERAT | MISSENSE | cTt/cGt | L80R | PLES_36251 | C | C | | | | | C | C | | | C | C |
| NC_011770 | 4019280 | T | C | 68 | 0 | 6 | 0 | NON_SYNONYMOU | MODERAT | MISSENSE | cAg/cGg | Q77R | PLES_36261 | C | C | | | | | C | C | | | C | C |
| NC_011770 | 4019569 | G | C | 68 | 0 | 6 | 0 | NON_SYNONYMOU | MODERAT | MISSENSE | atC/atG | I103M | PLES_36271 | C | C | | | | | C | C | | | C | C |
| NC_011770 | 4021499 | G | A | 50 | 0 | 12 | 0 | NON_SYNONYMOU | MODERAT | MISSENSE | cGg/cAg | R107Q | PLES_36301 | A | A | A | A | A | A | A | A | A | A | A | A |
| NC_011770 | 4021586 | C | A | 56 | 0 | 2 | 0 | NON_SYNONYMOU | MODERAT | MISSENSE | cCg/cAg | P136Q | PLES_36301 | | | | | | | | | A | A | | |
| NC_011770 | 4021756 | A | G | 54 | 0 | 12 | 0 | NON_SYNONYMOU | MODERAT | MISSENSE | Ata/Gta | I193V | PLES_36301 | G | G | G | G | G | G | G | G | G | G | G | G |
| NC_011770 | 4022057 | T | C | 42 | 0 | 2 | 0 | NON_SYNONYMOU | MODERAT | MISSENSE | cTg/cCg | L293P | PLES_36301 | | | | | | | | | | | C | C |
| NC_011770 | 4022396 | A | G | 62 | 0 | 6 | 0 | NON_SYNONYMOU | MODERAT | MISSENSE | cAg/cGg | Q406R | PLES_36301 | G | G | | | | | G | G | | | G | G |
| NC_011770 | 4022414 | G | C | 65 | 0 | 2 | 0 | NON_SYNONYMOU | MODERAT | MISSENSE | gGg/gCg | G412A | PLES_36301 | C | C | | | | | | | | | | |
| NC_011770 | 4025874 | C | A | 73 | 0 | 4 | 0 | NON_SYNONYMOU | MODERAT | MISSENSE | cCg/cAg | P85Q | PLES_36341 | | | A | A | A | A | | | | | | |
| NC_011770 | 4027866 | C | G | 82 | 0 | 4 | 0 | NON_SYNONYMOU | MODERAT | MISSENSE | gaG/gaC | E687D | PLES_36371 | G | G | | | | | G | G | | | | |
| NC_011770 | 4027938 | T | G | 83 | 0 | 8 | 0 | NON_SYNONYMOU | MODERAT | MISSENSE | gaA/gaC | E663D | PLES_36371 | G | G | G | G | G | G | G | G | | | | |
| NC_011770 | 4028914 | T | C | 81 | 0 | 2 | 0 | NON_SYNONYMOU | MODERAT | MISSENSE | aAc/aGc | N338S | PLES_36371 | | | | | | | | | C | C | | |
| NC_011770 | 4028996 | T | C | 77 | 0 | 4 | 0 | NON_SYNONYMOU | MODERAT | MISSENSE | Acc/Gcc | T311A | PLES_36371 | | | | | | | | | C | C | C | C |
| NC_011770 | 4029007 | G | A | 76 | 0 | 2 | 0 | NON_SYNONYMOU | MODERAT | MISSENSE | cCg/cTg | P307L | PLES_36371 | A | A | | | | | | | | | | |
| NC_011770 | 4029344 | T | C | 72 | 0 | 4 | 0 | NON_SYNONYMOU | MODERAT | MISSENSE | Agc/Ggc | S195G | PLES_36371 | | | C | C | C | C | | | | | | |
| NC_011770 | 4030309 | A | T | 76 | 0 | 12 | 0 | NON_SYNONYMOU | MODERAT | MISSENSE | Tcc/Acc | S164T | PLES_36381 | T | T | T | T | T | T | T | T | T | T | T | T |
| NC_011770 | 4030419 | C | T | 80 | 0 | 2 | 0 | NON_SYNONYMOU | MODERAT | MISSENSE | cGg/cAg | R127Q | PLES_36381 | T | T | | | | | | | | | | |
| NC_011770 | 4033005 | C | T | 73 | 0 | 6 | 0 | NON_SYNONYMOU | MODERAT | MISSENSE | Gtc/Atc | V22I | PLES_36411 | | | T | T | T | T | | | T | T | | |
| NC_011770 | 4034182 | A | G | 72 | 0 | 1 | 0 | NON_SYNONYMOU | MODERAT | MISSENSE | gTc/gCc | V77A | PLES_36431 | | | | | | | | G | | | | |
| NC_011770 | 4034548 | A | G | 82 | 0 | 12 | 0 | NON_SYNONYMOU | MODERAT | MISSENSE | Tat/Cat | Y160H | PLES_36441 | G | G | G | G | G | G | G | G | G | G | G | G |
| NC_011770 | 4034650 | C | T | 85 | 0 | 2 | 0 | NON_SYNONYMOU | MODERAT | MISSENSE | Gcc/Acc | A126T | PLES_36441 | T | T | | | | | | | | | | |
| NC_011770 | 4034760 | G | T | 78 | 0 | 2 | 0 | NON_SYNONYMOU | MODERAT | MISSENSE | cCg/cAg | P89Q | PLES_36441 | | | | | | | T | T | | | | |
| NC_011770 | 4034847 | G | A | 79 | 0 | 2 | 0 | NON_SYNONYMOU | MODERAT | MISSENSE | gCc/gTc | A60V | PLES_36441 | A | A | | | | | | | | | | |
| NC_011770 | 4037774 | C | T | 44 | 0 | 10 | 0 | NON_SYNONYMOU | MODERAT | MISSENSE | gCc/gTc | A120V | PLES_36471 | | | T | T | T | T | T | T | T | T | T | T |

| | | | | | | | | | | | | | | | | | | | | | | | | | |
|---|---|---|---|---|---|---|---|---|---|---|---|---|---|---|---|---|---|---|---|---|---|---|---|---|---|
| NC_011770 | 4038040 | G | A | 49 | 0 | 4 | 0 | NON_SYNONYMOU | MODERAT | MISSENSE | Ggc/Agc | G209S | PLES_36471 | | | A | A | A | A | | | | | | |
| NC_011770 | 4038065 | A | G | 50 | 0 | 6 | 0 | NON_SYNONYMOU | MODERAT | MISSENSE | aAc/aGc | N217S | PLES_36471 | | | G | G | G | G | | | | | G | G |
| NC_011770 | 4038136 | C | A | 55 | 0 | 2 | 0 | NON_SYNONYMOU | MODERAT | MISSENSE | Cgc/Agc | R241S | PLES_36471 | | | | | | | | | A | A | | |
| NC_011770 | 4041374 | C | G | 51 | 0 | 2 | 0 | NON_SYNONYMOU | MODERAT | MISSENSE | ttC/ttG | F80L | PLES_36511 | | | | | | | G | G | | | | |
| NC_011770 | 4041609 | G | A | 71 | 0 | 2 | 0 | NON_SYNONYMOU | MODERAT | MISSENSE | cGc/cAc | R33H | PLES_36521 | | | | | | | | | A | A | | |
| NC_011770 | 4041645 | T | A | 73 | 0 | 2 | 0 | NON_SYNONYMOU | MODERAT | MISSENSE | gTg/gAg | V45E | PLES_36521 | | | | | | | | | | | A | A |
| NC_011770 | 4041746 | A | G | 81 | 0 | 4 | 0 | NON_SYNONYMOU | MODERAT | MISSENSE | Agc/Ggc | S79G | PLES_36521 | | | G | G | G | G | | | | | | |
| NC_011770 | 4042912 | C | T | 75 | 0 | 1 | 0 | NON_SYNONYMOU | MODERAT | MISSENSE | Gaa/Aaa | E103K | PLES_36541 | | | | | T | | | | | | | |
| NC_011770 | 4043507 | A | G | 77 | 0 | 4 | 0 | NON_SYNONYMOU | MODERAT | MISSENSE | gAg/gGg | E15G | PLES_36551 | | | G | G | G | G | | | | | | |
| NC_011770 | 4043707 | A | C | 89 | 0 | 2 | 0 | NON_SYNONYMOU | MODERAT | MISSENSE | Att/Ctt | I82L | PLES_36551 | | | | | | | | | | | C | C |
| NC_011770 | 4045726 | A | G | 82 | 0 | 1 | 0 | NON_SYNONYMOU | MODERAT | MISSENSE | Tac/Cac | Y1130H | PLES_36581 | | | | | | | G | | | | | |
| NC_011770 | 4046599 | T | C | 65 | 0 | 1 | 0 | NON_SYNONYMOU | MODERAT | MISSENSE | Acc/Gcc | T839A | PLES_36581 | | | | | | | C | | | | | |
| NC_011770 | 4047084 | G | A | 67 | 0 | 2 | 0 | NON_SYNONYMOU | MODERAT | MISSENSE | gCc/gTc | A677V | PLES_36581 | | | | | | | | | | | A | A |
| NC_011770 | 4047212 | C | G | 72 | 0 | 2 | 0 | NON_SYNONYMOU | MODERAT | MISSENSE | gaG/gaC | E634D | PLES_36581 | | | | | | | | | | | G | G |
| NC_011770 | 4047342 | T | C | 73 | 0 | 4 | 0 | NON_SYNONYMOU | MODERAT | MISSENSE | aAc/aGc | N591S | PLES_36581 | | | C | C | C | C | | | | | | |
| NC_011770 | 4047496 | C | T | 72 | 0 | 2 | 0 | NON_SYNONYMOU | MODERAT | MISSENSE | Gcc/Acc | A540T | PLES_36581 | | | | | | | | | | | T | T |
| NC_011770 | 4047519 | C | T | 78 | 0 | 2 | 0 | NON_SYNONYMOU | MODERAT | MISSENSE | gGc/gAc | G532D | PLES_36581 | T | T | | | | | | | | | | |
| NC_011770 | 4047626 | G | T | 73 | 0 | 8 | 0 | NON_SYNONYMOU | MODERAT | MISSENSE | gaC/gaA | D496E | PLES_36581 | | | T | T | T | T | A | A | A | A | | |
| NC_011770 | 4047658 | T | C | 76 | 0 | 8 | 0 | NON_SYNONYMOU | MODERAT | MISSENSE | Acc/Gcc | T486A | PLES_36581 | C | C | C | C | C | C | | | | | C | C |
| NC_011770 | 4047682 | G | C | 71 | 0 | 8 | 0 | NON_SYNONYMOU | MODERAT | MISSENSE | Cta/Gta | L478V | PLES_36581 | C | C | C | C | C | C | | | | | C | C |
| NC_011770 | 4048600 | G | A | 77 | 0 | 2 | 0 | NON_SYNONYMOU | MODERAT | MISSENSE | Cgc/Tgc | R172C | PLES_36581 | | | | | | | | | A | A | | |
| NC_011770 | 4048933 | G | C | 77 | 0 | 12 | 0 | NON_SYNONYMOU | MODERAT | MISSENSE | Ctc/Gtc | L61V | PLES_36581 | C | C | C | C | C | C | C | C | C | C | C | C |
| NC_011770 | 4049046 | G | A | 82 | 0 | 12 | 0 | NON_SYNONYMOU | MODERAT | MISSENSE | gCa/gTa | A23V | PLES_36581 | A | A | A | A | A | A | A | A | A | A | A | A |
| NC_011770 | 4047106 | C | T | 68 | 0 | 1 | 0 | NON_SYNONYMOU | MODERAT | MISSENSE | Ggc/Agc | G670S | PLES_36581 | | | | | | | | T | | | | |
| NC_011770 | 4051372 | A | T | 86 | 0 | 6 | 0 | NON_SYNONYMOU | MODERAT | MISSENSE | cTg/cAg | L155Q | PLES_36611 | T | T | T | T | T | T | | | | | | |
| NC_011770 | 4051384 | T | C | 86 | 0 | 2 | 0 | NON_SYNONYMOU | MODERAT | MISSENSE | cAg/cGg | Q151R | PLES_36611 | C | C | | | | | | | | | | |
| NC_011770 | 4051504 | T | C | 85 | 0 | 4 | 0 | NON_SYNONYMOU | MODERAT | MISSENSE | gAg/gGg | E111G | PLES_36611 | | | | | | | C | C | | | C | C |
| NC_011770 | 4051794 | C | T | 91 | 0 | 2 | 0 | NON_SYNONYMOU | MODERAT | MISSENSE | atG/atA | M14I | PLES_36611 | | | | | | | | | | | T | T |
| NC_011770 | 4051507 | A | G | 83 | 0 | 1 | 0 | NON_SYNONYMOU | MODERAT | MISSENSE | gTg/gCg | V110A | PLES_36611 | | | | | | | | G | | | | |
| NC_011770 | 4052022 | C | G | 85 | 0 | 2 | 0 | NON_SYNONYMOU | MODERAT | MISSENSE | cGc/cCc | R338P | PLES_36621 | | | | | | | G | G | | | | |
| NC_011770 | 4052038 | C | T | 85 | 0 | 2 | 0 | NON_SYNONYMOU | MODERAT | MISSENSE | Gcc/Acc | A333T | PLES_36621 | T | T | | | | | | | | | | |
| NC_011770 | 4052206 | T | C | 69 | 0 | 10 | 0 | NON_SYNONYMOU | MODERAT | MISSENSE | Agc/Ggc | S277G | PLES_36621 | | | C | C | C | C | C | C | C | C | C | C |
| NC_011770 | 4052350 | C | T | 76 | 0 | 2 | 0 | NON_SYNONYMOU | MODERAT | MISSENSE | Gac/Aac | D229N | PLES_36621 | T | T | | | | | | | | | | |
| NC_011770 | 4052398 | T | C | 80 | 0 | 6 | 0 | NON_SYNONYMOU | MODERAT | MISSENSE | Atg/Gtg | M213V | PLES_36621 | C | C | | | | | | | C | C | C | C |
| NC_011770 | 4052442 | C | T | 74 | 0 | 4 | 0 | NON_SYNONYMOU | MODERAT | MISSENSE | gGt/gAt | G198D | PLES_36621 | | | | | | | | | T | T | T | T |
| NC_011770 | 4052448 | C | G | 75 | 0 | 4 | 0 | NON_SYNONYMOU | MODERAT | MISSENSE | cGt/cCt | R196P | PLES_36621 | | | | | | | | | G | G | G | G |
| NC_011770 | 4052456 | T | G | 79 | 0 | 2 | 0 | NON_SYNONYMOU | MODERAT | MISSENSE | gaA/gaC | E193D | PLES_36621 | G | G | | | | | | | | | | |
| NC_011770 | 4052555 | C | A | 80 | 0 | 6 | 0 | NON_SYNONYMOU | MODERAT | MISSENSE | caG/caT | Q160H | PLES_36621 | | | A | A | A | A | | | A | A | | |
| NC_011770 | 4052704 | G | A | 82 | 0 | 4 | 0 | NON_SYNONYMOU | MODERAT | MISSENSE | Ctc/Ttc | L111F | PLES_36621 | A | A | | | | | | | | | A | A |
| NC_011770 | 4053101 | C | T | 78 | 0 | 2 | 0 | NON_SYNONYMOU | MODERAT | MISSENSE | gGc/gAc | G31D | PLES_36631 | | | | | | | T | T | | | | |
| NC_011770 | 4053135 | C | A | 79 | 0 | 2 | 0 | NON_SYNONYMOU | MODERAT | MISSENSE | Gcc/Tcc | A20S | PLES_36631 | | | | | | | | | A | A | | |
| NC_011770 | 4053307 | T | C | 79 | 0 | 2 | 0 | NON_SYNONYMOU | MODERAT | MISSENSE | Att/Gtt | I496V | PLES_36641 | | | | | | | | | | | C | C |
| NC_011770 | 4053505 | T | C | 76 | 0 | 12 | 0 | NON_SYNONYMOU | MODERAT | MISSENSE | Atg/Gtg | M430V | PLES_36641 | C | C | C | C | C | C | C | C | C | C | C | C |
| NC_011770 | 4053506 | C | A | 76 | 0 | 4 | 0 | NON_SYNONYMOU | MODERAT | MISSENSE | gaG/gaT | E429D | PLES_36641 | A | A | | | | | | | | | A | A |
| NC_011770 | 4053987 | T | C | 91 | 0 | 2 | 0 | NON_SYNONYMOU | MODERAT | MISSENSE | aAg/aGg | K269R | PLES_36641 | | | | | | | C | C | | | | |
| NC_011770 | 4054309 | A | C | 90 | 0 | 2 | 0 | NON_SYNONYMOU | MODERAT | MISSENSE | Tcg/Gcg | S162A | PLES_36641 | | | | | | | | | | | C | C |
| NC_011770 | 4054443 | C | G | 85 | 0 | 2 | 0 | NON_SYNONYMOU | MODERAT | MISSENSE | gGg/gCg | G117A | PLES_36641 | | | | | | | | | G | G | | |
| NC_011770 | 4054596 | C | T | 90 | 0 | 4 | 0 | NON_SYNONYMOU | MODERAT | MISSENSE | tGc/tAc | C66Y | PLES_36641 | T | T | | | | | T | T | | | | |
| NC_011770 | 4054708 | C | T | 85 | 0 | 6 | 0 | NON_SYNONYMOU | MODERAT | MISSENSE | Gcg/Acg | A29T | PLES_36641 | | | T | T | T | T | | | T | T | | |
| NC_011770 | 4054710 | G | A | 86 | 0 | 8 | 0 | NON_SYNONYMOU | MODERAT | MISSENSE | gCc/gTc | A28V | PLES_36641 | | | A | A | A | A | | | A | A | A | A |
| NC_011770 | 4054711 | C | T | 86 | 0 | 8 | 0 | NON_SYNONYMOU | MODERAT | MISSENSE | Gcc/Acc | A28T | PLES_36641 | | | T | T | T | T | | | T | T | T | T |
| NC_011770 | 4054987 | C | T | 82 | 0 | 8 | 0 | NON_SYNONYMOU | MODERAT | MISSENSE | gGc/gAc | G817D | PLES_36651 | T | T | T | T | T | T | | | | | T | T |
| NC_011770 | 4055051 | G | C | 82 | 0 | 6 | 0 | NON_SYNONYMOU | MODERAT | MISSENSE | Caa/Gaa | Q796E | PLES_36651 | | | C | C | C | C | | | | | C | C |
| NC_011770 | 4055156 | C | T | 76 | 0 | 2 | 0 | NON_SYNONYMOU | MODERAT | MISSENSE | Gcg/Acg | A761T | PLES_36651 | | | | | | | | | | | T | T |
| NC_011770 | 4055198 | C | A | 76 | 0 | 2 | 0 | NON_SYNONYMOU | MODERAT | MISSENSE | Gcg/Tcg | A747S | PLES_36651 | | | | | | | | | | | A | A |
| NC_011770 | 4055506 | G | C | 75 | 0 | 2 | 0 | NON_SYNONYMOU | MODERAT | MISSENSE | aCc/aGc | T644S | PLES_36651 | | | | | | | C | C | | | | |
| NC_011770 | 4055693 | C | T | 75 | 0 | 2 | 0 | NON_SYNONYMOU | MODERAT | MISSENSE | Gcg/Acg | A582T | PLES_36651 | | | | | | | T | T | | | | |
| NC_011770 | 4055915 | T | C | 79 | 0 | 4 | 0 | NON_SYNONYMOU | MODERAT | MISSENSE | Act/Gct | T508A | PLES_36651 | C | C | | | | | C | C | | | | |
| NC_011770 | 4055933 | C | T | 79 | 0 | 4 | 0 | NON_SYNONYMOU | MODERAT | MISSENSE | Ggc/Agc | G502S | PLES_36651 | T | T | | | | | T | T | | | | |
| NC_011770 | 4056055 | G | A | 76 | 0 | 4 | 0 | NON_SYNONYMOU | MODERAT | MISSENSE | gCg/gTg | A461V | PLES_36651 | | | | | | | | | A | A | A | A |
| NC_011770 | 4056089 | T | C | 81 | 0 | 8 | 0 | NON_SYNONYMOU | MODERAT | MISSENSE | Acc/Gcc | T450A | PLES_36651 | C | C | | | | | C | C | C | C | C | C |
| NC_011770 | 4056350 | C | T | 83 | 0 | 2 | 0 | NON_SYNONYMOU | MODERAT | MISSENSE | Gtt/Att | V363I | PLES_36651 | | | | | | | | | | | T | T |

| | | | | | | | | | | | | | | | | | | | | | | | | | |
|---|---|---|---|---|---|---|---|---|---|---|---|---|---|---|---|---|---|---|---|---|---|---|---|---|---|
| NC_011770 | 4056607 | A | G | 77 | 0 | 4 | 0 | NON_SYNONYMOU | MODERAT | MISSENSE | cTc/cCc | L277P | PLES_36651 | | | | | | | G | G | | | G | G |
| NC_011770 | 4058336 | G | A | 80 | 0 | 6 | 0 | NON_SYNONYMOU | MODERAT | MISSENSE | Cat/Tat | H39Y | PLES_36661 | | | A | A | A | A | | | | | A | A |
| NC_011770 | 4058342 | G | C | 79 | 0 | 8 | 0 | NON_SYNONYMOU | MODERAT | MISSENSE | Cag/Gag | Q37E | PLES_36661 | C | C | C | C | C | C | | | | | C | C |
| NC_011770 | 4058659 | C | A | 77 | 0 | 6 | 0 | NON_SYNONYMOU | MODERAT | MISSENSE | Gcg/Tcg | A446S | PLES_36671 | | | A | A | A | A | | | A | A | | |
| NC_011770 | 4058848 | C | G | 74 | 0 | 2 | 0 | NON_SYNONYMOU | MODERAT | MISSENSE | Gcg/Ccg | A383P | PLES_36671 | G | G | | | | | | | | | | |
| NC_011770 | 4058973 | A | G | 83 | 0 | 2 | 0 | NON_SYNONYMOU | MODERAT | MISSENSE | aTc/aCc | I341T | PLES_36671 | G | G | | | | | | | | | | |
| NC_011770 | 4059595 | C | G | 80 | 0 | 12 | 0 | NON_SYNONYMOU | MODERAT | MISSENSE | Gcg/Ccg | A134P | PLES_36671 | G | G | G | G | G | G | G | G | G | G | G | G |
| NC_011770 | 4059597 | C | T | 80 | 0 | 6 | 0 | NON_SYNONYMOU | MODERAT | MISSENSE | cGg/cAg | R133Q | PLES_36671 | | | T | T | T | T | | | T | T | | |
| NC_011770 | 4059673 | T | C | 78 | 0 | 2 | 0 | NON_SYNONYMOU | MODERAT | MISSENSE | Atc/Gtc | I108V | PLES_36671 | | | | | | | | | | | C | C |
| NC_011770 | 4060574 | A | G | 59 | 0 | 2 | 0 | NON_SYNONYMOU | MODERAT | MISSENSE | Tgc/Cgc | C125R | PLES_36681 | | | | | | | | | | | G | G |
| NC_011770 | 4060595 | G | A | 57 | 0 | 6 | 0 | NON_SYNONYMOU | MODERAT | MISSENSE | Cgc/Tgc | R118C | PLES_36681 | | | A | A | A | A | | | A | A | | |
| NC_011770 | 4060606 | C | T | 58 | 0 | 10 | 0 | NON_SYNONYMOU | MODERAT | MISSENSE | gGa/gAa | G114E | PLES_36681 | | | T | T | T | T | T | T | T | T | T | T |
| NC_011770 | 4060685 | C | T | 62 | 0 | 2 | 0 | NON_SYNONYMOU | MODERAT | MISSENSE | Gac/Aac | D88N | PLES_36681 | | | | | | | | | | | T | T |
| NC_011770 | 4061634 | G | C | 74 | 0 | 2 | 0 | NON_SYNONYMOU | MODERAT | MISSENSE | agC/agG | S267R | PLES_36691 | | | | | | | | | | | C | C |
| NC_011770 | 4061270 | G | A | 73 | 0 | 1 | 0 | NON_SYNONYMOU | MODERAT | MISSENSE | Ccg/Tcg | P389S | PLES_36691 | | | | | | | | A | | | | |
| NC_011770 | 4062840 | T | C | 66 | 0 | 8 | 0 | NON_SYNONYMOU | MODERAT | MISSENSE | Acg/Gcg | T44A | PLES_36701 | | | C | C | C | C | C | C | | | C | C |
| NC_011770 | 4063406 | G | A | 81 | 0 | 4 | 0 | NON_SYNONYMOU | MODERAT | MISSENSE | Cgc/Tgc | R386C | PLES_36711 | | | A | A | A | A | | | | | | |
| NC_011770 | 4064525 | C | T | 73 | 0 | 2 | 0 | NON_SYNONYMOU | MODERAT | MISSENSE | Gaa/Aaa | E13K | PLES_36711 | | | | | | | | | | | T | T |
| NC_011770 | 4066101 | C | T | 55 | 0 | 2 | 0 | NON_SYNONYMOU | MODERAT | MISSENSE | Gcc/Acc | A79T | PLES_36721 | | | | | | | | | | | T | T |
| NC_011770 | 4066148 | C | A | 61 | 0 | 4 | 0 | NON_SYNONYMOU | MODERAT | MISSENSE | tGc/tTc | C63F | PLES_36721 | | | A | A | A | A | | | | | | |
| NC_011770 | 4066248 | C | T | 70 | 0 | 4 | 0 | NON_SYNONYMOU | MODERAT | MISSENSE | Gta/Ata | V30I | PLES_36721 | | | T | T | T | T | | | | | | |
| NC_011770 | 4067040 | T | A | 67 | 0 | 4 | 0 | NON_SYNONYMOU | MODERAT | MISSENSE | gAa/gTa | E173V | PLES_36731 | | | | | | | A | A | | | C | C |
| NC_011770 | 4067554 | C | G | 68 | 0 | 2 | 0 | NON_SYNONYMOU | MODERAT | MISSENSE | Gcc/Ccc | A2P | PLES_36731 | | | | | | | | | G | G | | |
| NC_011770 | 4068171 | T | C | 68 | 0 | 4 | 0 | NON_SYNONYMOU | MODERAT | MISSENSE | Agc/Ggc | S211G | PLES_36751 | | | C | C | C | C | | | | | | |
| NC_011770 | 4068219 | C | G | 74 | 0 | 2 | 0 | NON_SYNONYMOU | MODERAT | MISSENSE | Gtg/Ctg | V195L | PLES_36751 | | | | | | | | | | | G | G |
| NC_011770 | 4068612 | T | C | 77 | 0 | 2 | 0 | NON_SYNONYMOU | MODERAT | MISSENSE | Agg/Ggg | R64G | PLES_36751 | | | | | | | | | | | C | C |
| NC_011770 | 4069210 | C | T | 78 | 0 | 2 | 0 | NON_SYNONYMOU | MODERAT | MISSENSE | gCc/gTc | A60V | PLES_36761 | | | T | T | | | | | | | | |
| NC_011770 | 4069372 | T | C | 87 | 0 | 1 | 0 | NON_SYNONYMOU | MODERAT | MISSENSE | cTg/cCg | L114P | PLES_36761 | | | | | | | C | | | | | |
| NC_011770 | 4069734 | A | G | 77 | 0 | 8 | 0 | NON_SYNONYMOU | MODERAT | MISSENSE | Act/Gct | T235A | PLES_36761 | G | G | | | | | G | G | G | G | G | G |
| NC_011770 | 4070257 | T | G | 71 | 0 | 12 | 0 | NON_SYNONYMOU | MODERAT | MISSENSE | gAg/gCg | E291A | PLES_36771 | G | G | G | G | G | G | G | G | G | G | G | G |
| NC_011770 | 4070264 | C | T | 70 | 0 | 10 | 0 | NON_SYNONYMOU | MODERAT | MISSENSE | Gcg/Acg | A289T | PLES_36771 | T | T | T | T | T | T | | | T | T | T | T |
| NC_011770 | 4070567 | C | T | 76 | 0 | 2 | 0 | NON_SYNONYMOU | MODERAT | MISSENSE | Gcc/Acc | A188T | PLES_36771 | T | T | | | | | | | | | | |
| NC_011770 | 4070711 | C | T | 75 | 0 | 2 | 0 | NON_SYNONYMOU | MODERAT | MISSENSE | Gtg/Atg | V140M | PLES_36771 | T | T | | | | | | | | | | |
| NC_011770 | 4071352 | T | A | 12 | 0 | 6 | 0 | NON_SYNONYMOU | MODERAT | MISSENSE | Agc/Tgc | S233C | PLES_36781 | | | A | A | A | A | | | | | A | A |
| NC_011770 | 4071874 | C | G | 55 | 0 | 6 | 0 | NON_SYNONYMOU | MODERAT | MISSENSE | Gtc/Ctc | V59L | PLES_36781 | | | G | G | G | G | | | | | G | G |
| NC_011770 | 4074298 | G | C | 83 | 0 | 4 | 0 | NON_SYNONYMOU | MODERAT | MISSENSE | Gcc/Ccc | A285P | PLES_36801 | C | C | | | | | | | | | C | C |
| NC_011770 | 4073542 | G | A | 43 | 0 | 1 | 0 | NON_SYNONYMOU | MODERAT | MISSENSE | Gcg/Acg | A33T | PLES_36801 | | A | | | | | | | | | | |
| NC_011770 | 4075801 | C | A | 59 | 0 | 6 | 0 | NON_SYNONYMOU | MODERAT | MISSENSE | Gcg/Tcg | A436S | PLES_36811 | | | A | A | A | A | | | A | A | | |
| NC_011770 | 4076325 | A | C | 72 | 0 | 4 | 0 | NON_SYNONYMOU | MODERAT | MISSENSE | cTc/cGc | L261R | PLES_36811 | | | C | C | C | C | | | | | | |
| NC_011770 | 4076433 | C | A | 81 | 0 | 2 | 0 | NON_SYNONYMOU | MODERAT | MISSENSE | cGg/cTg | R225L | PLES_36811 | | | | | | | | | | | A | A |
| NC_011770 | 4076595 | T | C | 81 | 0 | 4 | 0 | NON_SYNONYMOU | MODERAT | MISSENSE | cAc/cGc | H171R | PLES_36811 | | | C | C | C | C | | | | | | |
| NC_011770 | 4076781 | G | A | 78 | 0 | 4 | 0 | NON_SYNONYMOU | MODERAT | MISSENSE | gCc/gTc | A109V | PLES_36811 | | | A | A | A | A | | | | | | |
| NC_011770 | 4076900 | T | G | 80 | 0 | 8 | 0 | NON_SYNONYMOU | MODERAT | MISSENSE | gaA/gaC | E69D | PLES_36811 | | | G | G | G | G | G | G | | | G | G |
| NC_011770 | 4077081 | A | G | 73 | 0 | 2 | 0 | NON_SYNONYMOU | MODERAT | MISSENSE | cTg/cCg | L9P | PLES_36811 | | | | | | | | | | | G | G |
| NC_011770 | 4077355 | G | A | 66 | 1 | 1 | 0 | NON_SYNONYMOU | MODERAT | MISSENSE | Gcc/Acc | A17T | PLES_36821 | | | | | | | | | | | A | . |
| NC_011770 | 4077759 | G | A | 68 | 0 | 2 | 0 | NON_SYNONYMOU | MODERAT | MISSENSE | Cat/Tat | H191Y | PLES_36831 | | | | | | | A | A | | | | |
| NC_011770 | 4079200 | T | C | 61 | 0 | 12 | 0 | NON_SYNONYMOU | MODERAT | MISSENSE | Agc/Ggc | S235G | PLES_36841 | C | C | C | C | C | C | C | C | C | C | C | C |
| NC_011770 | 4079833 | G | A | 73 | 0 | 1 | 0 | NON_SYNONYMOU | MODERAT | MISSENSE | Ccg/Tcg | P24S | PLES_36841 | | | | | | | | | | | A | |
| NC_011770 | 4081159 | G | T | 68 | 0 | 2 | 0 | NON_SYNONYMOU | MODERAT | MISSENSE | gCc/gAc | A47D | PLES_36861 | | | | | | | | | T | T | | |
| NC_011770 | 4082459 | G | A | 82 | 0 | 1 | 0 | NON_SYNONYMOU | MODERAT | MISSENSE | Cgc/Tgc | R201C | PLES_36881 | | | | | | | A | | | | | |
| NC_011770 | 4082830 | T | C | 70 | 0 | 2 | 0 | NON_SYNONYMOU | MODERAT | MISSENSE | cAg/cGg | Q77R | PLES_36881 | | | | | | | | | C | C | | |
| NC_011770 | 4082992 | A | G | 68 | 0 | 12 | 0 | NON_SYNONYMOU | MODERAT | MISSENSE | gTc/gCc | V23A | PLES_36881 | G | G | G | G | G | G | G | G | G | G | G | G |
| NC_011770 | 4093069 | T | C | 82 | 0 | 12 | 0 | NON_SYNONYMOU | MODERAT | MISSENSE | Aac/Gac | N172D | PLES_36951 | C | C | C | C | C | C | C | C | C | C | C | C |
| NC_011770 | 4093150 | G | C | 83 | 0 | 12 | 0 | NON_SYNONYMOU | MODERAT | MISSENSE | Cag/Gag | Q145E | PLES_36951 | C | C | C | C | C | C | C | C | C | C | C | C |
| NC_011770 | 4093456 | G | A | 83 | 0 | 2 | 0 | NON_SYNONYMOU | MODERAT | MISSENSE | Cgc/Tgc | R43C | PLES_36951 | | | | | | | | | A | A | | |
| NC_011770 | 4093578 | A | G | 86 | 0 | 2 | 0 | NON_SYNONYMOU | MODERAT | MISSENSE | aTt/aCt | I2T | PLES_36951 | | | | | | | G | G | | | | |
| NC_011770 | 4093839 | G | A | 75 | 0 | 2 | 0 | NON_SYNONYMOU | MODERAT | MISSENSE | gCc/gTc | A222V | PLES_36961 | | | | | | | | | | | A | A |
| NC_011770 | 4094796 | G | A | 69 | 0 | 12 | 0 | NON_SYNONYMOU | MODERAT | MISSENSE | Gtc/Atc | V18I | PLES_36971 | A | A | A | A | A | A | A | A | A | A | A | A |
| NC_011770 | 4094847 | G | C | 71 | 0 | 4 | 0 | NON_SYNONYMOU | MODERAT | MISSENSE | Gaa/Caa | E35Q | PLES_36971 | | | | | | | C | C | C | C | | |
| NC_011770 | 4094894 | T | A | 67 | 0 | 2 | 0 | NON_SYNONYMOU | MODERAT | MISSENSE | gaT/gaA | D50E | PLES_36971 | | | | | | | A | A | | | | |
| NC_011770 | 4094988 | C | A | 72 | 0 | 2 | 0 | NON_SYNONYMOU | MODERAT | MISSENSE | Ctg/Atg | L82M | PLES_36971 | | | A | A | | | | | | | | |
| NC_011770 | 4095003 | A | G | 67 | 0 | 12 | 0 | NON_SYNONYMOU | MODERAT | MISSENSE | Agc/Ggc | S87G | PLES_36971 | G | G | G | G | G | G | G | G | G | G | G | G |

| | | | | | | | | | | | | | | | | | | | | | | | | | |
|---|---|---|---|---|---|---|---|---|---|---|---|---|---|---|---|---|---|---|---|---|---|---|---|---|---|
| NC_011770 | 4095337 | C | T | 75 | 0 | 12 | 0 | NON_SYNONYMOU | MODERAT | MISSENSE | aCg/aTg | T198M | PLES_36971 | T | T | T | T | T | T | T | T | T | T | T | T |
| NC_011770 | 4095488 | C | G | 72 | 0 | 12 | 0 | NON_SYNONYMOU | MODERAT | MISSENSE | caC/caG | H248Q | PLES_36971 | G | G | G | G | G | G | G | G | G | G | G | G |
| NC_011770 | 4095620 | T | C | 63 | 0 | 10 | 0 | INTRAGENIC | MODIFIER | | | | PLES_36981 | C | C | C | C | C | C | C | C | C | C | | |
| NC_011770 | 4095623 | G | A | 62 | 0 | 10 | 0 | INTRAGENIC | MODIFIER | | | | PLES_36981 | A | A | A | A | A | A | A | A | A | A | | |
| NC_011770 | 4095656 | C | G | 58 | 0 | 4 | 0 | INTRAGENIC | MODIFIER | | | | PLES_36981 | | | G | G | G | G | | | | | | |
| NC_011770 | 4095676 | G | A | 56 | 0 | 6 | 0 | INTRAGENIC | MODIFIER | | | | PLES_36981 | | | A | A | A | A | | | A | A | | |
| NC_011770 | 4095700 | G | A | 56 | 0 | 8 | 0 | INTRAGENIC | MODIFIER | | | | PLES_36981 | A | A | A | A | A | A | | | A | A | | |
| NC_011770 | 4095721 | C | T | 56 | 0 | 2 | 0 | INTRAGENIC | MODIFIER | | | | PLES_36981 | | | | | | | | | T | T | | |
| NC_011770 | 4095730 | A | C | 56 | 0 | 8 | 0 | INTRAGENIC | MODIFIER | | | | PLES_36981 | C | C | C | C | C | C | | | C | C | | |
| NC_011770 | 4095763 | G | A | 57 | 0 | 8 | 0 | INTRAGENIC | MODIFIER | | | | PLES_36981 | A | A | A | A | A | A | | | A | A | | |
| NC_011770 | 4095776 | C | G | 57 | 0 | 12 | 0 | INTRAGENIC | MODIFIER | | | | PLES_36981 | G | G | G | G | G | G | G | G | G | G | G | G |
| NC_011770 | 4095795 | A | G | 58 | 0 | 12 | 0 | INTRAGENIC | MODIFIER | | | | PLES_36981 | G | G | G | G | G | G | G | G | G | G | G | G |
| NC_011770 | 4095821 | A | G | 56 | 0 | 12 | 0 | INTRAGENIC | MODIFIER | | | | PLES_36981 | G | G | G | G | G | G | G | G | G | G | G | G |
| NC_011770 | 4095871 | C | G | 59 | 0 | 2 | 0 | INTRAGENIC | MODIFIER | | | | PLES_36981 | | | | | | | | | G | G | | |
| NC_011770 | 4095897 | T | C | 64 | 0 | 10 | 0 | INTRAGENIC | MODIFIER | | | | PLES_36981 | C | C | C | C | C | C | C | C | C | C | | |
| NC_011770 | 4095950 | C | T | 65 | 0 | 6 | 0 | INTRAGENIC | MODIFIER | | | | PLES_36981 | T | T | T | T | T | T | | | | | | |
| NC_011770 | 4096043 | C | G | 68 | 0 | 2 | 0 | INTRAGENIC | MODIFIER | | | | PLES_36981 | | | | | | | G | G | | | | |
| NC_011770 | 4096058 | A | G | 72 | 0 | 12 | 0 | INTRAGENIC | MODIFIER | | | | PLES_36981 | G | G | G | G | G | G | G | G | G | G | G | G |
| NC_011770 | 4096070 | T | C | 72 | 0 | 10 | 0 | INTRAGENIC | MODIFIER | | | | PLES_36981 | C | C | C | C | C | C | C | C | | | C | C |
| NC_011770 | 4096199 | C | T | 76 | 0 | 8 | 0 | INTRAGENIC | MODIFIER | | | | PLES_36981 | T | T | | | | | T | T | T | T | T | T |
| NC_011770 | 4096265 | G | A | 78 | 0 | 12 | 0 | INTRAGENIC | MODIFIER | | | | PLES_36981 | A | A | A | A | A | A | A | A | A | A | A | A |
| NC_011770 | 4096313 | G | A | 76 | 0 | 4 | 0 | INTRAGENIC | MODIFIER | | | | PLES_36981 | | | A | A | A | A | | | | | | |
| NC_011770 | 4096440 | G | A | 67 | 0 | 12 | 0 | INTRAGENIC | MODIFIER | | | | PLES_36981 | A | A | A | A | A | A | A | A | A | A | A | A |
| NC_011770 | 4096441 | C | T | 66 | 0 | 12 | 0 | INTRAGENIC | MODIFIER | | | | PLES_36981 | T | T | T | T | T | T | T | T | T | T | T | T |
| NC_011770 | 4096445 | A | G | 67 | 0 | 12 | 0 | INTRAGENIC | MODIFIER | | | | PLES_36981 | G | G | G | G | G | G | G | G | G | G | G | G |
| NC_011770 | 4096448 | A | C | 68 | 0 | 12 | 0 | INTRAGENIC | MODIFIER | | | | PLES_36981 | C | C | G | G | G | G | C | C | C | C | C | C |
| NC_011770 | 4096456 | A | G | 74 | 0 | 12 | 0 | INTRAGENIC | MODIFIER | | | | PLES_36981 | G | G | G | G | G | G | G | G | G | G | G | G |
| NC_011770 | 4096645 | T | C | 77 | 0 | 1 | 0 | INTRAGENIC | MODIFIER | | | | PLES_36991 | | | | | | | C | | | | | |
| NC_011770 | 4096688 | G | A | 77 | 0 | 12 | 0 | INTRAGENIC | MODIFIER | | | | PLES_36991 | A | A | A | A | A | A | A | A | A | A | A | A |
| NC_011770 | 4096744 | C | T | 76 | 0 | 12 | 0 | INTRAGENIC | MODIFIER | | | | PLES_36991 | T | T | T | T | T | T | T | T | T | T | T | T |
| NC_011770 | 4096886 | C | T | 82 | 0 | 2 | 0 | INTRAGENIC | MODIFIER | | | | PLES_36991 | | | | | | | | | T | T | | |
| NC_011770 | 4096892 | C | T | 77 | 0 | 8 | 0 | INTRAGENIC | MODIFIER | | | | PLES_36991 | | | T | T | T | T | T | T | T | T | | |
| NC_011770 | 4096930 | G | A | 71 | 0 | 4 | 0 | INTRAGENIC | MODIFIER | | | | PLES_36991 | | | A | A | A | A | | | | | | |
| NC_011770 | 4096956 | C | T | 73 | 0 | 12 | 0 | INTRAGENIC | MODIFIER | | | | PLES_36991 | T | T | T | T | T | T | T | T | T | T | T | T |
| NC_011770 | 4096962 | G | A | 73 | 0 | 12 | 0 | INTRAGENIC | MODIFIER | | | | PLES_36991 | A | A | A | A | A | A | A | A | A | A | A | A |
| NC_011770 | 4097027 | G | C | 72 | 0 | 6 | 0 | INTRAGENIC | MODIFIER | | | | PLES_36991 | | | C | C | C | C | C | C | | | | |
| NC_011770 | 4097601 | A | C | 66 | 0 | 12 | 0 | NON_SYNONYMOU | MODERAT | MISSENSE | gAc/gCc | D134A | PLES_37001 | C | C | C | C | C | C | C | C | C | C | C | C |
| NC_011770 | 4097220 | G | A | 68 | 0 | 1 | 0 | NON_SYNONYMOU | MODERAT | MISSENSE | cGc/cAc | R7H | PLES_37001 | | | | | | | | A | | | | |
| NC_011770 | 4097955 | C | T | 52 | 0 | 12 | 0 | NON_SYNONYMOU | MODERAT | MISSENSE | gCc/gTc | A31V | PLES_37011 | T | T | T | T | T | T | T | T | T | T | T | T |
| NC_011770 | 4098044 | G | GCCC | 59 | 0 | 1 | 0 | CODON_CHANGE_ | MODERATE | | ctg/cCCCtg | L63PL | PLES_37011 | | | | | | | GCCC | | | | | |
| NC_011770 | 4098548 | G | A | 77 | 0 | 12 | 0 | NON_SYNONYMOU | MODERAT | MISSENSE | Gtc/Atc | V229I | PLES_37011 | A | A | A | A | A | A | A | A | A | A | A | A |
| NC_011770 | 4099886 | G | A | 75 | 0 | 1 | 0 | NON_SYNONYMOU | MODERAT | MISSENSE | tCc/tTc | S124F | PLES_37021 | | | | | | | | | | | A | |
| NC_011770 | 4101647 | C | T | 69 | 0 | 10 | 0 | NON_SYNONYMOU | MODERAT | MISSENSE | Gcc/Acc | A85T | PLES_37041 | T | T | T | T | T | T | | | T | T | T | T |
| NC_011770 | 4101379 | G | A | 78 | 0 | 1 | 0 | NON_SYNONYMOU | MODERAT | MISSENSE | gCc/gTc | A174V | PLES_37041 | | | | | | | | A | | | | |
| NC_011770 | 4102319 | C | A | 78 | 0 | 12 | 0 | NON_SYNONYMOU | MODERAT | MISSENSE | gGc/gTc | G174V | PLES_37051 | A | A | A | A | A | A | A | A | A | A | A | A |
| NC_011770 | 4102762 | G | C | 77 | 0 | 4 | 0 | NON_SYNONYMOU | MODERAT | MISSENSE | gaC/gaG | D26E | PLES_37051 | | | C | C | C | C | | | | | | |
| NC_011770 | 4103054 | G | C | 71 | 0 | 4 | 0 | NON_SYNONYMOU | MODERAT | MISSENSE | Cca/Gca | P199A | PLES_37061 | | | C | C | C | C | | | | | | |
| NC_011770 | 4103851 | C | T | 13 | 0 | 10 | 0 | NON_SYNONYMOU | MODERAT | MISSENSE | Gca/Aca | A181T | PLES_37071 | T | T | T | T | T | T | T | T | T | T | | |
| NC_011770 | 4104298 | T | A | 56 | 0 | 12 | 0 | NON_SYNONYMOU | MODERAT | MISSENSE | Acc/Tcc | T32S | PLES_37071 | A | A | A | A | A | A | A | A | A | A | A | A |
| NC_011770 | 4104619 | T | C | 66 | 0 | 4 | 0 | NON_SYNONYMOU | MODERAT | MISSENSE | cAc/cGc | H213R | PLES_37081 | | | C | C | C | C | | | | | | |
| NC_011770 | 4104742 | T | A | 66 | 0 | 2 | 0 | NON_SYNONYMOU | MODERAT | MISSENSE | cAg/cTg | Q172L | PLES_37081 | A | A | | | | | | | | | | |
| NC_011770 | 4106021 | G | A | 87 | 0 | 2 | 0 | NON_SYNONYMOU | MODERAT | MISSENSE | gCg/gTg | A483V | PLES_37101 | | | | | | | | | | | A | A |
| NC_011770 | 4107239 | A | G | 64 | 0 | 10 | 0 | NON_SYNONYMOU | MODERAT | MISSENSE | aTt/aCt | I77T | PLES_37101 | G | G | G | G | G | G | G | G | G | G | | |
| NC_011770 | 4107822 | CG | C | 71 | 0 | 1 | 0 | FRAME_SHIFT | HIGH | | ccc/ | P79 | PLES_37111 | | | | | | | | C | | | | |
| NC_011770 | 4109725 | G | A | 59 | 0 | 1 | 0 | NON_SYNONYMOU | MODERAT | MISSENSE | Gcg/Acg | A48T | PLES_37141 | | | | | | | A | | | | | |
| NC_011770 | 4110181 | T | A | 61 | 0 | 2 | 0 | NON_SYNONYMOU | MODERAT | MISSENSE | Tcg/Acg | S200T | PLES_37141 | | | | | | | | | A | A | | |
| NC_011770 | 4110215 | A | G | 59 | 0 | 12 | 0 | NON_SYNONYMOU | MODERAT | MISSENSE | cAc/cGc | H211R | PLES_37141 | G | G | G | G | G | G | G | G | G | G | G | G |
| NC_011770 | 4110553 | G | T | 66 | 0 | 4 | 0 | NON_SYNONYMOU | MODERAT | MISSENSE | Gcg/Tcg | A324S | PLES_37141 | | | T | T | T | T | | | | | | |
| NC_011770 | 4111900 | G | A | 72 | 0 | 12 | 0 | NON_SYNONYMOU | MODERAT | MISSENSE | Gtg/Atg | V13M | PLES_37151 | A | A | A | A | A | A | A | A | A | A | A | A |
| NC_011770 | 4114032 | G | A | 66 | 0 | 1 | 0 | NON_SYNONYMOU | MODERAT | MISSENSE | Ggc/Agc | G472S | PLES_37161 | | | | | | | A | | | | | |
| NC_011770 | 4114128 | G | A | 55 | 0 | 2 | 0 | NON_SYNONYMOU | MODERAT | MISSENSE | Gcc/Acc | A504T | PLES_37161 | A | A | | | | | | | | | | |
| NC_011770 | 4114256 | C | G | 64 | 0 | 1 | 0 | NON_SYNONYMOU | MODERAT | MISSENSE | gaC/gaG | D546E | PLES_37161 | | | | | | | G | | | | | |

| | | | | | | | | | | | | | | | | | | | | | | | | | |
|---|---|---|---|---|---|---|---|---|---|---|---|---|---|---|---|---|---|---|---|---|---|---|---|---|---|
| NC_011770 | 4118817 | G | A | 59 | 0 | 2 | 0 | NON_SYNONYMOU | MODERAT | MISSENSE | cGg/cAg | R142Q | PLES_37201 | | | | | | | A | A | | | | |
| NC_011770 | 4119030 | C | A | 66 | 0 | 2 | 0 | STOP_GAINED | HIGH | NONSENSE | Gag/Tag | E152* | PLES_37211 | A | A | | | | | | | | | | |
| NC_011770 | 4119029 | T | G | 64 | 0 | 2 | 0 | NON_SYNONYMOU | MODERAT | MISSENSE | gAg/gCg | E152A | PLES_37211 | G | G | | | | | | | | | | |
| NC_011770 | 4119170 | C | T | 68 | 0 | 1 | 0 | NON_SYNONYMOU | MODERAT | MISSENSE | cGg/cAg | R105Q | PLES_37211 | | T | | | | | | | | | | |
| NC_011770 | 4119851 | G | C | 81 | 0 | 2 | 0 | NON_SYNONYMOU | MODERAT | MISSENSE | Cgg/Ggg | R246G | PLES_37221 | | | | | | | C | C | | | | |
| NC_011770 | 4120051 | T | C | 76 | 0 | 4 | 0 | NON_SYNONYMOU | MODERAT | MISSENSE | cAc/cGc | H179R | PLES_37221 | | | C | C | C | C | | | | | | |
| NC_011770 | 4120446 | C | T | 87 | 0 | 2 | 0 | NON_SYNONYMOU | MODERAT | MISSENSE | atG/atA | M47I | PLES_37221 | T | T | | | | | | | | | | |
| NC_011770 | 4120625 | T | A | 75 | 0 | 2 | 0 | NON_SYNONYMOU | MODERAT | MISSENSE | cTg/cAg | L14Q | PLES_37231 | A | A | | | | | | | | | | |
| NC_011770 | 4120996 | A | C | 71 | 0 | 2 | 0 | NON_SYNONYMOU | MODERAT | MISSENSE | Atc/Ctc | I138L | PLES_37231 | C | C | | | | | | | | | | |
| NC_011770 | 4121009 | C | T | 69 | 0 | 2 | 0 | NON_SYNONYMOU | MODERAT | MISSENSE | gCg/gTg | A142V | PLES_37231 | T | T | | | | | | | | | | |
| NC_011770 | 4121062 | G | A | 72 | 0 | 1 | 0 | NON_SYNONYMOU | MODERAT | MISSENSE | Gcc/Acc | A160T | PLES_37231 | | | | | | | A | | | | | |
| NC_011770 | 4121242 | G | C | 66 | 0 | 2 | 0 | NON_SYNONYMOU | MODERAT | MISSENSE | Gtg/Ctg | V220L | PLES_37231 | C | C | | | | | | | | | | |
| NC_011770 | 4121390 | G | A | 62 | 0 | 2 | 0 | NON_SYNONYMOU | MODERAT | MISSENSE | gGc/gAc | G269D | PLES_37231 | A | A | | | | | | | | | | |
| NC_011770 | 4121539 | A | C | 64 | 0 | 2 | 0 | NON_SYNONYMOU | MODERAT | MISSENSE | aAc/aCc | N4T | PLES_37241 | C | C | | | | | | | | | | |
| NC_011770 | 4121541 | G | A | 64 | 0 | 2 | 0 | NON_SYNONYMOU | MODERAT | MISSENSE | Gcc/Acc | A5T | PLES_37241 | A | A | | | | | | | | | | |
| NC_011770 | 4121659 | T | C | 64 | 0 | 1 | 0 | NON_SYNONYMOU | MODERAT | MISSENSE | cTg/cCg | L44P | PLES_37241 | | C | | | | | | | | | | |
| NC_011770 | 4122578 | G | A | 80 | 0 | 1 | 0 | NON_SYNONYMOU | MODERAT | MISSENSE | Gtg/Atg | V141M | PLES_37251 | A | | | | | | | | | | | |
| NC_011770 | 4123305 | T | G | 78 | 0 | 2 | 0 | NON_SYNONYMOU | MODERAT | MISSENSE | caT/caG | H220Q | PLES_37261 | G | G | | | | | | | | | | |
| NC_011770 | 4123381 | G | A | 74 | 0 | 2 | 0 | NON_SYNONYMOU | MODERAT | MISSENSE | Gaa/Aaa | E246K | PLES_37261 | A | A | | | | | | | | | | |
| NC_011770 | 4123568 | C | A | 66 | 0 | 4 | 0 | NON_SYNONYMOU | MODERAT | MISSENSE | cCg/cAg | P308Q | PLES_37261 | | | A | A | A | A | | | | | | |
| NC_011770 | 4123592 | A | G | 65 | 0 | 2 | 0 | NON_SYNONYMOU | MODERAT | MISSENSE | gAc/gGc | D316G | PLES_37261 | | | | | | | G | G | | | | |
| NC_011770 | 4123786 | C | T | 71 | 0 | 2 | 0 | NON_SYNONYMOU | MODERAT | MISSENSE | Cgc/Tgc | R381C | PLES_37261 | | | | | | | T | T | | | | |
| NC_011770 | 4123886 | A | G | 66 | 0 | 2 | 0 | NON_SYNONYMOU | MODERAT | MISSENSE | cAg/cGg | Q414R | PLES_37261 | | | | | | | G | G | | | | |
| NC_011770 | 4123888 | A | G | 65 | 0 | 8 | 0 | NON_SYNONYMOU | MODERAT | MISSENSE | Acg/Gcg | T415A | PLES_37261 | G | G | G | G | G | G | G | G | | | | |
| NC_011770 | 4124347 | G | T | 58 | 0 | 2 | 0 | NON_SYNONYMOU | MODERAT | MISSENSE | Gcc/Tcc | A568S | PLES_37261 | | | | | | | T | T | | | | |
| NC_011770 | 4124885 | A | G | 69 | 0 | 4 | 0 | NON_SYNONYMOU | MODERAT | MISSENSE | cAg/cGg | Q747R | PLES_37261 | | | G | G | G | G | | | | | | |
| NC_011770 | 4125036 | G | C | 75 | 0 | 4 | 0 | NON_SYNONYMOU | MODERAT | MISSENSE | gGg/gCg | G32A | PLES_37271 | | | C | C | C | C | | | | | | |
| NC_011770 | 4125788 | T | C | 69 | 0 | 2 | 0 | NON_SYNONYMOU | MODERAT | MISSENSE | Tcg/Ccg | S283P | PLES_37271 | | | | | | | C | C | | | | |
| NC_011770 | 4126820 | G | C | 71 | 0 | 12 | 0 | NON_SYNONYMOU | MODERAT | MISSENSE | Cgc/Ggc | R73G | PLES_37281 | C | C | C | C | C | C | C | C | C | C | C | C |
| NC_011770 | 4128286 | A | G | 68 | 0 | 12 | 0 | NON_SYNONYMOU | MODERAT | MISSENSE | gTg/gCg | V130A | PLES_37301 | G | G | G | G | G | G | G | G | G | G | G | G |
| NC_011770 | 4128365 | C | T | 67 | 0 | 2 | 0 | NON_SYNONYMOU | MODERAT | MISSENSE | Gcc/Acc | A104T | PLES_37301 | | | | | | | | | T | T | | |
| NC_011770 | 4131017 | G | A | 63 | 0 | 4 | 0 | NON_SYNONYMOU | MODERAT | MISSENSE | cCg/cTg | P327L | PLES_37321 | | | A | A | A | A | | | | | | |
| NC_011770 | 4131512 | A | G | 76 | 0 | 12 | 0 | NON_SYNONYMOU | MODERAT | MISSENSE | gTg/gCg | V162A | PLES_37321 | G | G | G | G | G | G | G | G | G | G | G | G |
| NC_011770 | 4132088 | G | C | 86 | 0 | 2 | 0 | NON_SYNONYMOU | MODERAT | MISSENSE | aCc/aGc | T131S | PLES_37331 | C | C | | | | | | | | | | |
| NC_011770 | 4132522 | G | A | 74 | 0 | 4 | 0 | NON_SYNONYMOU | MODERAT | MISSENSE | gCc/gTc | A143V | PLES_37341 | | | A | A | A | A | | | | | | |
| NC_011770 | 4132624 | A | C | 71 | 0 | 2 | 0 | NON_SYNONYMOU | MODERAT | MISSENSE | gTc/gGc | V109G | PLES_37341 | | | | | C | C | | | | | | |
| NC_011770 | 4149956 | A | G | 59 | 0 | 1 | 0 | NON_SYNONYMOU | MODERAT | MISSENSE | Tgg/Cgg | W78R | PLES_37491 | G | | | | | | | | | | | |
| NC_011770 | 4150401 | C | T | 51 | 0 | 2 | 0 | NON_SYNONYMOU | MODERAT | MISSENSE | Gtg/Atg | V216M | PLES_37501 | T | T | | | | | | | | | | |
| NC_011770 | 4150488 | T | C | 49 | 0 | 10 | 0 | NON_SYNONYMOU | MODERAT | MISSENSE | Atc/Gtc | I187V | PLES_37501 | | | C | C | C | C | C | C | C | C | C | C |
| NC_011770 | 4150502 | A | C | 55 | 0 | 2 | 0 | NON_SYNONYMOU | MODERAT | MISSENSE | gTc/gGc | V182G | PLES_37501 | | | | | | | | | C | C | | |
| NC_011770 | 4151037 | A | G | 20 | 0 | 12 | 0 | NON_SYNONYMOU | MODERAT | MISSENSE | Tac/Cac | Y4H | PLES_37501 | G | G | G | G | G | G | G | G | G | G | G | G |
| NC_011770 | 4151938 | G | A | 76 | 0 | 2 | 0 | NON_SYNONYMOU | MODERAT | MISSENSE | Cgc/Tgc | R147C | PLES_37521 | A | A | | | | | | | | | | |
| NC_011770 | 4152924 | G | A | 79 | 0 | 1 | 0 | NON_SYNONYMOU | MODERAT | MISSENSE | Gcc/Acc | A149T | PLES_37531 | | | | | | | A | | | | | |
| NC_011770 | 4155012 | G | A | 70 | 0 | 6 | 0 | NON_SYNONYMOU | MODERAT | MISSENSE | gCg/gTg | A89V | PLES_37561 | | | A | A | A | A | | | | | A | A |
| NC_011770 | 4155139 | T | C | 71 | 0 | 10 | 0 | NON_SYNONYMOU | MODERAT | MISSENSE | Ata/Gta | I47V | PLES_37561 | | | C | C | C | C | C | C | C | C | C | C |
| NC_011770 | 4155551 | C | G | 45 | 0 | 2 | 0 | NON_SYNONYMOU | MODERAT | MISSENSE | aCc/aGc | T22S | PLES_37571 | | | | | | | G | G | | | | |
| NC_011770 | 4155749 | G | A | 34 | 0 | 2 | 0 | NON_SYNONYMOU | MODERAT | MISSENSE | Ccg/Tcg | P290S | PLES_37581 | | | | | | | A | A | | | | |
| NC_011770 | 4157154 | A | G | 77 | 0 | 2 | 0 | NON_SYNONYMOU | MODERAT | MISSENSE | gAc/gGc | D54G | PLES_37591 | | | | | | | G | G | | | | |
| NC_011770 | 4158236 | A | G | 69 | 0 | 2 | 0 | NON_SYNONYMOU | MODERAT | MISSENSE | Atc/Gtc | I415V | PLES_37591 | | | | | | | | | G | G | | |
| NC_011770 | 4158291 | G | A | 55 | 0 | 6 | 0 | NON_SYNONYMOU | MODERAT | MISSENSE | gGc/gAc | G433D | PLES_37591 | | | A | A | A | A | | | A | A | | |
| NC_011770 | 4158299 | C | G | 55 | 0 | 6 | 0 | NON_SYNONYMOU | MODERAT | MISSENSE | Cgg/Ggg | R436G | PLES_37591 | | | G | G | G | G | | | G | G | | |
| NC_011770 | 4158300 | G | T | 55 | 0 | 6 | 0 | NON_SYNONYMOU | MODERAT | MISSENSE | cGg/cTg | R436L | PLES_37591 | | | T | T | T | T | | | T | T | | |
| NC_011770 | 4158302 | G | A | 55 | 0 | 6 | 0 | NON_SYNONYMOU | MODERAT | MISSENSE | Gaa/Aaa | E437K | PLES_37591 | | | A | A | A | A | | | A | A | | |
| NC_011770 | 4158309 | T | A | 54 | 0 | 6 | 0 | NON_SYNONYMOU | MODERAT | MISSENSE | cTg/cAg | L439Q | PLES_37591 | | | A | A | A | A | | | A | A | | |
| NC_011770 | 4158330 | C | G | 54 | 0 | 6 | 0 | NON_SYNONYMOU | MODERAT | MISSENSE | gCg/gGg | A446G | PLES_37591 | | | G | G | G | G | | | G | G | | |
| NC_011770 | 4158350 | T | C | 53 | 0 | 1 | 0 | NON_SYNONYMOU | MODERAT | MISSENSE | Tcc/Ccc | S453P | PLES_37591 | | | | | | | C | | | | | |
| NC_011770 | 4158474 | G | C | 55 | 0 | 6 | 0 | NON_SYNONYMOU | MODERAT | MISSENSE | Gtc/Ctc | V22L | PLES_37601 | | | C | C | C | C | | | C | C | | |
| NC_011770 | 4158537 | A | G | 50 | 0 | 2 | 0 | NON_SYNONYMOU | MODERAT | MISSENSE | Atc/Gtc | I43V | PLES_37601 | | | | | | | | | G | G | | |
| NC_011770 | 4159441 | AC | ACC | 60 | 0 | 4 | 0 | FRAME_SHIFT | HIGH | | cac/ | H225 | PLES_37611 | ACC | ACC | | | | | ACC | A | | | | |
| NC_011770 | 4159051 | A | G | 61 | 0 | 1 | 0 | NON_SYNONYMOU | MODERAT | MISSENSE | Acc/Gcc | T93A | PLES_37611 | | | | | | | G | | | | | |
| NC_011770 | 4159183 | G | A | 67 | 0 | 2 | 0 | NON_SYNONYMOU | MODERAT | MISSENSE | Gtc/Atc | V137I | PLES_37611 | | | | | | | | | | | A | A |
| NC_011770 | 4159241 | C | T | 71 | 0 | 6 | 0 | NON_SYNONYMOU | MODERAT | MISSENSE | gCc/gTc | A156V | PLES_37611 | | | T | T | T | T | | | | | T | T |

| | | | | | | | | | | | | | | | | | | | | | | | | | |
|---|---|---|---|---|---|---|---|---|---|---|---|---|---|---|---|---|---|---|---|---|---|---|---|---|---|
| NC_011770 | 4159449 | C | A | 64 | 0 | 4 | 0 | NON_SYNONYMOU | MODERAT | MISSENSE | caC/caA | H225Q | PLES_37611 | | | A | A | A | A | | | | | | |
| NC_011770 | 4159466 | G | A | 57 | 0 | 4 | 0 | NON_SYNONYMOU | MODERAT | MISSENSE | gGc/gAc | G231D | PLES_37611 | | | A | A | A | A | | | | | | |
| NC_011770 | 4159788 | A | C | 58 | 0 | 12 | 0 | NON_SYNONYMOU | MODERAT | MISSENSE | ttA/ttC | L338F | PLES_37611 | C | C | C | C | C | C | C | C | C | C | C | C |
| NC_011770 | 4158847 | G | A | 62 | 0 | 1 | 0 | NON_SYNONYMOU | MODERAT | MISSENSE | Ggg/Agg | G25R | PLES_37611 | | | | | | | | A | | | | |
| NC_011770 | 4160345 | C | A | 49 | 0 | 12 | 0 | NON_SYNONYMOU | MODERAT | MISSENSE | Ggc/Tgc | G96C | PLES_37621 | A | A | A | A | A | A | A | A | A | A | A | A |
| NC_011770 | 4160597 | C | T | 41 | 0 | 4 | 0 | NON_SYNONYMOU | MODERAT | MISSENSE | Gca/Aca | A12T | PLES_37621 | | | T | T | T | T | | | | | | |
| NC_011770 | 4161313 | C | T | 72 | 0 | 2 | 0 | NON_SYNONYMOU | MODERAT | MISSENSE | gCc/gTc | A225V | PLES_37631 | | | | | | | T | T | | | | |
| NC_011770 | 4161357 | A | G | 73 | 0 | 2 | 0 | NON_SYNONYMOU | MODERAT | MISSENSE | Atc/Gtc | I240V | PLES_37631 | | | | | | | G | G | | | | |
| NC_011770 | 4162112 | T | G | 71 | 0 | 12 | 0 | NON_SYNONYMOU | MODERAT | MISSENSE | Tcg/Gcg | S14A | PLES_37641 | G | G | G | G | G | G | G | G | G | G | G | G |
| NC_011770 | 4163042 | G | A | 77 | 0 | 2 | 0 | NON_SYNONYMOU | MODERAT | MISSENSE | Gcc/Acc | A324T | PLES_37641 | A | A | | | | | | | | | | |
| NC_011770 | 4163864 | G | A | 77 | 0 | 2 | 0 | NON_SYNONYMOU | MODERAT | MISSENSE | Cgc/Tgc | R312C | PLES_37661 | | | | | | | | | A | A | | |
| NC_011770 | 4164263 | C | G | 74 | 0 | 2 | 0 | NON_SYNONYMOU | MODERAT | MISSENSE | Gag/Cag | E179Q | PLES_37661 | | | G | G | | | | | | | | |
| NC_011770 | 4171368 | T | C | 74 | 0 | 10 | 0 | NON_SYNONYMOU | MODERAT | MISSENSE | Atc/Gtc | I115V | PLES_37701 | C | C | C | C | C | C | C | C | | | C | C |
| NC_011770 | 4179312 | G | A | 95 | 0 | 12 | 0 | NON_SYNONYMOU | MODERAT | MISSENSE | Gcc/Acc | A159T | PLES_37771 | A | A | A | A | A | A | A | A | A | A | A | A |
| NC_011770 | 4180197 | A | G | 76 | 0 | 12 | 0 | NON_SYNONYMOU | MODERAT | MISSENSE | Aac/Gac | N454D | PLES_37771 | G | G | G | G | G | G | G | G | G | G | G | G |
| NC_011770 | 4180110 | C | G | 84 | 0 | 1 | 0 | NON_SYNONYMOU | MODERAT | MISSENSE | Ctc/Gtc | L425V | PLES_37771 | | | | | | | | | | | | G |
| NC_011770 | 4180551 | G | A | 68 | 0 | 4 | 0 | NON_SYNONYMOU | MODERAT | MISSENSE | cGg/cAg | R96Q | PLES_37781 | A | A | | | | | A | A | | | | |
| NC_011770 | 4181047 | C | G | 64 | 0 | 12 | 0 | NON_SYNONYMOU | MODERAT | MISSENSE | Cgc/Ggc | R83G | PLES_37791 | G | G | G | G | G | G | G | G | G | G | G | G |
| NC_011770 | 4181282 | A | T | 72 | 0 | 2 | 0 | NON_SYNONYMOU | MODERAT | MISSENSE | tAc/tTc | Y161F | PLES_37791 | | | | | | | | | T | T | | |
| NC_011770 | 4181294 | G | A | 69 | 0 | 2 | 0 | NON_SYNONYMOU | MODERAT | MISSENSE | cGc/cAc | R165H | PLES_37791 | A | A | | | | | | | | | | |
| NC_011770 | 4181327 | G | A | 66 | 0 | 8 | 0 | NON_SYNONYMOU | MODERAT | MISSENSE | cGc/cAc | R176H | PLES_37791 | | | A | A | A | A | A | A | | | A | A |
| NC_011770 | 4181932 | G | A | 58 | 0 | 2 | 0 | NON_SYNONYMOU | MODERAT | MISSENSE | Gcc/Acc | A378T | PLES_37791 | A | A | | | | | | | | | | |
| NC_011770 | 4182574 | A | G | 66 | 0 | 2 | 0 | NON_SYNONYMOU | MODERAT | MISSENSE | Aac/Gac | N592D | PLES_37791 | G | G | | | | | | | | | | |
| NC_011770 | 4182697 | G | A | 61 | 0 | 2 | 0 | NON_SYNONYMOU | MODERAT | MISSENSE | Gcc/Acc | A633T | PLES_37791 | A | A | | | | | | | | | | |
| NC_011770 | 4182817 | A | C | 60 | 0 | 2 | 0 | NON_SYNONYMOU | MODERAT | MISSENSE | Acc/Ccc | T673P | PLES_37791 | | | | | | | | | C | C | | |
| NC_011770 | 4183404 | G | C | 83 | 0 | 2 | 0 | NON_SYNONYMOU | MODERAT | MISSENSE | Gag/Cag | E64Q | PLES_37801 | C | C | | | | | | | | | | |
| NC_011770 | 4185877 | G | C | 63 | 0 | 8 | 0 | NON_SYNONYMOU | MODERAT | MISSENSE | Cag/Gag | Q84E | PLES_37831 | | | C | C | C | C | C | C | | | C | C |
| NC_011770 | 4185984 | C | T | 61 | 0 | 6 | 0 | NON_SYNONYMOU | MODERAT | MISSENSE | cGg/cAg | R48Q | PLES_37831 | | | T | T | T | T | | | | | T | T |
| NC_011770 | 4187992 | G | A | 85 | 0 | 1 | 0 | NON_SYNONYMOU | MODERAT | MISSENSE | Ggc/Agc | G5S | PLES_37861 | | | | | | | A | | | | | |
| NC_011770 | 4188097 | T | C | 84 | 0 | 1 | 0 | NON_SYNONYMOU | MODERAT | MISSENSE | Ttc/Ctc | F40L | PLES_37861 | | | | | | | C | | | | | |
| NC_011770 | 4188544 | T | C | 68 | 0 | 4 | 0 | NON_SYNONYMOU | MODERAT | MISSENSE | Ttc/Ctc | F189L | PLES_37861 | | | C | C | C | C | | | | | | |
| NC_011770 | 4189463 | A | G | 74 | 0 | 8 | 0 | NON_SYNONYMOU | MODERAT | MISSENSE | gAc/gGc | D80G | PLES_37871 | G | G | | | | | G | G | G | G | G | G |
| NC_011770 | 4189467 | T | A | 73 | 0 | 12 | 0 | NON_SYNONYMOU | MODERAT | MISSENSE | gaT/gaA | D81E | PLES_37871 | A | A | A | A | A | A | A | A | A | A | A | A |
| NC_011770 | 4189478 | T | C | 77 | 0 | 2 | 0 | NON_SYNONYMOU | MODERAT | MISSENSE | cTg/cCg | L85P | PLES_37871 | | | | | | | | | | | C | C |
| NC_011770 | 4189567 | C | T | 82 | 0 | 4 | 0 | NON_SYNONYMOU | MODERAT | MISSENSE | Ccc/Tcc | P115S | PLES_37871 | T | T | | | | | | | | | T | T |
| NC_011770 | 4189692 | G | A | 70 | 0 | 2 | 0 | NON_SYNONYMOU | MODERAT | MISSENSE | Ggc/Agc | G36S | PLES_37881 | | | | | | | | | A | A | | |
| NC_011770 | 4190342 | C | T | 45 | 0 | 6 | 0 | NON_SYNONYMOU | MODERAT | MISSENSE | Ggc/Agc | G93S | PLES_37891 | T | T | | | | | | | T | T | T | T |
| NC_011770 | 4191782 | G | A | 79 | 0 | 2 | 0 | NON_SYNONYMOU | MODERAT | MISSENSE | Gac/Aac | D351N | PLES_37901 | | | | | | | | | A | A | | |
| NC_011770 | 4192001 | A | G | 83 | 0 | 12 | 0 | NON_SYNONYMOU | MODERAT | MISSENSE | Acc/Gcc | T424A | PLES_37901 | G | G | G | G | G | G | G | G | G | G | G | G |
| NC_011770 | 4193324 | G | A | 81 | 0 | 2 | 0 | NON_SYNONYMOU | MODERAT | MISSENSE | Ggc/Agc | G18S | PLES_37921 | A | A | | | | | | | | | | |
| NC_011770 | 4193532 | T | C | 70 | 0 | 1 | 0 | NON_SYNONYMOU | MODERAT | MISSENSE | gTc/gCc | V87A | PLES_37921 | | | | | | | | C | | | | |
| NC_011770 | 4194127 | C | A | 68 | 0 | 4 | 0 | NON_SYNONYMOU | MODERAT | MISSENSE | Gtg/Ttg | V283L | PLES_37931 | | | A | A | A | A | | | | | | |
| NC_011770 | 4194886 | A | G | 79 | 0 | 2 | 0 | NON_SYNONYMOU | MODERAT | MISSENSE | Tac/Cac | Y30H | PLES_37931 | G | G | | | | | | | | | | |
| NC_011770 | 4194654 | C | T | 76 | 0 | 1 | 0 | NON_SYNONYMOU | MODERAT | MISSENSE | cGt/cAt | R107H | PLES_37931 | | | | | | | | T | | | | |
| NC_011770 | 4203988 | A | T | 80 | 0 | 4 | 0 | NON_SYNONYMOU | MODERAT | MISSENSE | gaT/gaA | D944E | PLES_38011 | T | T | | | | | | | T | T | | |
| NC_011770 | 4204400 | T | C | 81 | 0 | 4 | 0 | NON_SYNONYMOU | MODERAT | MISSENSE | aAg/aGg | K807R | PLES_38011 | C | C | | | | | | | C | C | | |
| NC_011770 | 4204512 | T | A | 87 | 0 | 2 | 0 | NON_SYNONYMOU | MODERAT | MISSENSE | Acc/Tcc | T770S | PLES_38011 | | | | | | | A | A | | | | |
| NC_011770 | 4206054 | T | C | 69 | 0 | 2 | 0 | NON_SYNONYMOU | MODERAT | MISSENSE | Atc/Gtc | I256V | PLES_38011 | | | | | | | C | C | | | | |
| NC_011770 | 4206191 | C | T | 76 | 0 | 1 | 0 | NON_SYNONYMOU | MODERAT | MISSENSE | cGg/cAg | R210Q | PLES_38011 | | | | | | | T | | | | | |
| NC_011770 | 4206335 | T | C | 75 | 0 | 2 | 0 | NON_SYNONYMOU | MODERAT | MISSENSE | aAc/aGc | N162S | PLES_38011 | | | | | | | C | C | | | | |
| NC_011770 | 4214527 | A | G | 70 | 0 | 2 | 0 | NON_SYNONYMOU | MODERAT | MISSENSE | aAc/aGc | N236S | PLES_38071 | | | G | G | | | | | | | | |
| NC_011770 | 4215751 | C | T | 64 | 0 | 2 | 0 | NON_SYNONYMOU | MODERAT | MISSENSE | Gcc/Acc | A23T | PLES_38081 | | | | | | | | | | | T | T |
| NC_011770 | 4219195 | T | G | 86 | 0 | 2 | 0 | NON_SYNONYMOU | MODERAT | MISSENSE | Atg/Ctg | M126L | PLES_38111 | | | | | | | | | G | G | | |
| NC_011770 | 4219255 | C | T | 92 | 0 | 2 | 0 | NON_SYNONYMOU | MODERAT | MISSENSE | Gag/Aag | E106K | PLES_38111 | | | | | | | | | | | T | T |
| NC_011770 | 4223061 | G | A | 51 | 0 | 2 | 0 | NON_SYNONYMOU | MODERAT | MISSENSE | cGg/cAg | R137Q | PLES_38141 | | | | | | | | | | | A | A |
| NC_011770 | 4223129 | G | A | 56 | 0 | 4 | 0 | NON_SYNONYMOU | MODERAT | MISSENSE | Ggc/Agc | G160S | PLES_38141 | | | A | A | A | A | | | | | | |
| NC_011770 | 4224409 | C | T | 19 | 8 | 2 | 0 | NON_SYNONYMOU | MODERAT | MISSENSE | Cgt/Tgt | R167C | PLES_38151 | . | . | . | . | . | . | T | T | | | . | . |
| NC_011770 | 4225595 | C | T | 85 | 0 | 4 | 0 | NON_SYNONYMOU | MODERAT | MISSENSE | gGc/gAc | G575D | PLES_38161 | | | T | T | T | T | | | | | | |
| NC_011770 | 4225650 | A | G | 77 | 0 | 12 | 0 | NON_SYNONYMOU | MODERAT | MISSENSE | Ttc/Ctc | F557L | PLES_38161 | G | G | G | G | G | G | G | G | G | G | G | G |
| NC_011770 | 4225704 | C | T | 71 | 0 | 4 | 0 | NON_SYNONYMOU | MODERAT | MISSENSE | Gtc/Atc | V539I | PLES_38161 | | | | | | | T | T | T | T | | |
| NC_011770 | 4225829 | T | C | 79 | 0 | 1 | 0 | NON_SYNONYMOU | MODERAT | MISSENSE | tAc/tGc | Y497C | PLES_38161 | | | | | | | C | | | | | |
| NC_011770 | 4226237 | T | C | 69 | 0 | 12 | 0 | NON_SYNONYMOU | MODERAT | MISSENSE | aAg/aGg | K361R | PLES_38161 | C | C | C | C | C | C | C | C | C | C | C | C |

| | | | | | | | | | | | | | | | | | | | | | | | | | |
|---|---|---|---|---|---|---|---|---|---|---|---|---|---|---|---|---|---|---|---|---|---|---|---|---|---|
| NC_011770 | 4227554 | G | T | 63 | 0 | 2 | 0 | NON_SYNONYMOU | MODERAT | MISSENSE | Ctg/Atg | L229M | PLES_38171 | | | | | | | T | T | | | | |
| NC_011770 | 4227868 | G | C | 60 | 0 | 2 | 0 | NON_SYNONYMOU | MODERAT | MISSENSE | cCc/cGc | P124R | PLES_38171 | | | | | | | | | | | C | C |
| NC_011770 | 4227899 | G | A | 59 | 0 | 2 | 0 | NON_SYNONYMOU | MODERAT | MISSENSE | Ctt/Ttt | L114F | PLES_38171 | | | | | | | | | | | A | A |
| NC_011770 | 4227911 | C | T | 58 | 0 | 2 | 0 | NON_SYNONYMOU | MODERAT | MISSENSE | Gcg/Acg | A110T | PLES_38171 | | | | | | | T | T | | | | |
| NC_011770 | 4227956 | T | C | 57 | 0 | 2 | 0 | NON_SYNONYMOU | MODERAT | MISSENSE | Atc/Gtc | I95V | PLES_38171 | | | | | | | | | C | C | | |
| NC_011770 | 4228588 | GC | G | 70 | 0 | 1 | 0 | FRAME_SHIFT | HIGH | | acg/ | T129 | PLES_38181 | | | | | | | | G | | | | |
| NC_011770 | 4228951 | AC | A | 63 | 0 | 2 | 0 | FRAME_SHIFT | HIGH | | ggg/ | G8 | PLES_38181 | | | | | | | | | A | A | | |
| NC_011770 | 4228966 | CG | C | 70 | 0 | 2 | 0 | FRAME_SHIFT | HIGH | | ccc/ | P3 | PLES_38181 | C | C | | | | | | | | | | |
| NC_011770 | 4228354 | C | T | 64 | 0 | 2 | 0 | NON_SYNONYMOU | MODERAT | MISSENSE | Gtt/Att | V208I | PLES_38181 | | | | | | | T | T | | | | |
| NC_011770 | 4228555 | C | T | 70 | 0 | 4 | 0 | NON_SYNONYMOU | MODERAT | MISSENSE | Gcc/Acc | A141T | PLES_38181 | | | | | | | T | T | T | T | | |
| NC_011770 | 4228600 | C | T | 77 | 0 | 2 | 0 | NON_SYNONYMOU | MODERAT | MISSENSE | Gac/Aac | D126N | PLES_38181 | | | | | | | T | T | | | | |
| NC_011770 | 4228845 | T | C | 74 | 0 | 4 | 0 | NON_SYNONYMOU | MODERAT | MISSENSE | gAc/gGc | D44G | PLES_38181 | | | C | C | C | C | | | | | | |
| NC_011770 | 4228861 | T | C | 72 | 0 | 2 | 0 | NON_SYNONYMOU | MODERAT | MISSENSE | Acc/Gcc | T39A | PLES_38181 | | | | | | | | | C | C | | |
| NC_011770 | 4228879 | A | G | 69 | 0 | 2 | 0 | NON_SYNONYMOU | MODERAT | MISSENSE | Tgc/Cgc | C33R | PLES_38181 | | | | | | | | | G | G | | |
| NC_011770 | 4229269 | C | G | 51 | 0 | 12 | 0 | NON_SYNONYMOU | MODERAT | MISSENSE | Gag/Cag | E15Q | PLES_38191 | G | G | G | G | G | G | G | G | G | G | G | G |
| NC_011770 | 4231132 | A | C | 78 | 0 | 6 | 0 | NON_SYNONYMOU | MODERAT | MISSENSE | gAg/gCg | E470A | PLES_38201 | | | C | C | C | C | | | C | C | | |
| NC_011770 | 4231171 | C | A | 81 | 0 | 2 | 0 | NON_SYNONYMOU | MODERAT | MISSENSE | gGc/gTc | G288V | PLES_38211 | | | | | | | A | A | | | | |
| NC_011770 | 4232105 | C | T | 94 | 0 | 4 | 0 | NON_SYNONYMOU | MODERAT | MISSENSE | aCc/aTc | T25I | PLES_38221 | | | T | T | T | T | | | | | | |
| NC_011770 | 4232636 | A | ACGACG | 98 | 0 | 2 | 0 | CODON_CHANGE_ | MODERATE | | gag/gGCG, | E206GDE | PLES_38221 | | | | | | | | | | | ACGACGG | ACGACGG |
| NC_011770 | 4233884 | T | C | 84 | 0 | 4 | 0 | NON_SYNONYMOU | MODERAT | MISSENSE | cAc/cGc | H301R | PLES_38251 | | | C | C | C | C | | | | | | |
| NC_011770 | 4234545 | G | A | 80 | 0 | 2 | 0 | NON_SYNONYMOU | MODERAT | MISSENSE | Ccg/Tcg | P81S | PLES_38251 | A | A | | | | | | | | | | |
| NC_011770 | 4234571 | T | A | 82 | 0 | 6 | 0 | NON_SYNONYMOU | MODERAT | MISSENSE | aAg/aTg | K72M | PLES_38251 | | | A | A | A | A | A | A | | | | |
| NC_011770 | 4234590 | C | T | 83 | 0 | 2 | 0 | NON_SYNONYMOU | MODERAT | MISSENSE | Gcc/Acc | A66T | PLES_38251 | T | T | | | | | | | | | | |
| NC_011770 | 4234758 | G | T | 85 | 0 | 2 | 0 | NON_SYNONYMOU | MODERAT | MISSENSE | Cgc/Agc | R10S | PLES_38251 | | | | | | | | | | | T | T |
| NC_011770 | 4234781 | C | T | 84 | 0 | 2 | 0 | NON_SYNONYMOU | MODERAT | MISSENSE | aGc/aAc | S2N | PLES_38251 | | | | | | | T | T | | | | |
| NC_011770 | 4235186 | T | C | 73 | 0 | 1 | 0 | START_LOST | HIGH | MISSENSE | Atg/Gtg | M1V | PLES_38261 | | | | | | | | | | | | C |
| NC_011770 | 4234994 | A | C | 79 | 0 | 8 | 0 | NON_SYNONYMOU | MODERAT | MISSENSE | Tcc/Gcc | S65A | PLES_38261 | | | C | C | C | C | C | C | | | C | C |
| NC_011770 | 4235078 | C | A | 79 | 0 | 4 | 0 | NON_SYNONYMOU | MODERAT | MISSENSE | Ggc/Tgc | G37C | PLES_38261 | | | A | A | A | A | | | | | | |
| NC_011770 | 4235089 | G | A | 78 | 0 | 4 | 0 | NON_SYNONYMOU | MODERAT | MISSENSE | tCc/tTc | S33F | PLES_38261 | | | A | A | A | A | | | | | | |
| NC_011770 | 4235102 | C | T | 83 | 0 | 4 | 0 | NON_SYNONYMOU | MODERAT | MISSENSE | Ggc/Agc | G29S | PLES_38261 | | | T | T | T | T | | | | | | |
| NC_011770 | 4236546 | G | A | 58 | 0 | 2 | 0 | NON_SYNONYMOU | MODERAT | MISSENSE | Gcc/Acc | A57T | PLES_38291 | | | | | | | | | | | A | A |
| NC_011770 | 4236997 | A | G | 58 | 0 | 8 | 0 | NON_SYNONYMOU | MODERAT | MISSENSE | gAc/gGc | D207G | PLES_38291 | G | G | | | | | G | G | G | G | G | G |
| NC_011770 | 4237069 | T | G | 57 | 0 | 4 | 0 | NON_SYNONYMOU | MODERAT | MISSENSE | gTg/gGg | V231G | PLES_38291 | G | G | | | | | | | G | G | | |
| NC_011770 | 4237077 | G | A | 59 | 0 | 12 | 0 | NON_SYNONYMOU | MODERAT | MISSENSE | Ggc/Agc | G234S | PLES_38291 | A | A | A | A | A | A | A | A | A | A | A | A |
| NC_011770 | 4237222 | G | A | 66 | 0 | 2 | 0 | NON_SYNONYMOU | MODERAT | MISSENSE | aGc/aAc | S282N | PLES_38291 | A | A | | | | | | | | | | |
| NC_011770 | 4237251 | T | C | 69 | 0 | 2 | 0 | NON_SYNONYMOU | MODERAT | MISSENSE | Tcc/Ccc | S292P | PLES_38291 | C | C | | | | | | | | | | |
| NC_011770 | 4237372 | A | C | 69 | 0 | 2 | 0 | NON_SYNONYMOU | MODERAT | MISSENSE | gAc/gCc | D332A | PLES_38291 | | | | | | | | | | | C | C |
| NC_011770 | 4237396 | T | C | 73 | 0 | 1 | 0 | NON_SYNONYMOU | MODERAT | MISSENSE | gTg/gCg | V340A | PLES_38291 | | | | | | | C | | | | | |
| NC_011770 | 4237968 | G | C | 88 | 0 | 2 | 0 | NON_SYNONYMOU | MODERAT | MISSENSE | Ccc/Gcc | P285A | PLES_38301 | C | C | | | | | | | | | | |
| NC_011770 | 4237974 | C | A | 79 | 0 | 2 | 0 | NON_SYNONYMOU | MODERAT | MISSENSE | Gtc/Ttc | V283F | PLES_38301 | A | A | | | | | | | | | | |
| NC_011770 | 4238744 | T | A | 72 | 0 | 2 | 0 | NON_SYNONYMOU | MODERAT | MISSENSE | gAt/gTt | D26V | PLES_38301 | A | A | | | | | | | | | | |
| NC_011770 | 4238911 | T | C | 85 | 0 | 2 | 0 | NON_SYNONYMOU | MODERAT | MISSENSE | gAc/gGc | D226G | PLES_38311 | | | | | | | C | C | | | | |
| NC_011770 | 4238924 | C | T | 74 | 0 | 2 | 0 | NON_SYNONYMOU | MODERAT | MISSENSE | Gac/Aac | D222N | PLES_38311 | | | | | | | | | | | T | T |
| NC_011770 | 4238959 | G | T | 73 | 0 | 2 | 0 | NON_SYNONYMOU | MODERAT | MISSENSE | aCc/aAc | T210N | PLES_38311 | T | T | | | | | | | | | | |
| NC_011770 | 4238978 | C | T | 62 | 0 | 12 | 0 | NON_SYNONYMOU | MODERAT | MISSENSE | Gtc/Atc | V204I | PLES_38311 | T | T | T | T | T | T | T | T | T | T | T | T |
| NC_011770 | 4239112 | G | A | 42 | 0 | 2 | 0 | NON_SYNONYMOU | MODERAT | MISSENSE | gCc/gTc | A159V | PLES_38311 | A | A | | | | | | | | | | |
| NC_011770 | 4239121 | G | A | 41 | 0 | 6 | 0 | NON_SYNONYMOU | MODERAT | MISSENSE | gCg/gTg | A156V | PLES_38311 | | | A | A | A | A | | | A | A | | |
| NC_011770 | 4239149 | G | T | 43 | 0 | 2 | 0 | NON_SYNONYMOU | MODERAT | MISSENSE | Cgt/Agt | R147S | PLES_38311 | | | | | | | T | T | | | | |
| NC_011770 | 4239440 | C | T | 60 | 0 | 4 | 0 | NON_SYNONYMOU | MODERAT | MISSENSE | Gac/Aac | D50N | PLES_38311 | | | T | T | T | T | | | | | | |
| NC_011770 | 4239553 | C | T | 62 | 0 | 8 | 0 | NON_SYNONYMOU | MODERAT | MISSENSE | aGc/aAc | S12N | PLES_38311 | | | T | T | T | T | | | T | T | T | T |
| NC_011770 | 4239956 | T | C | 69 | 0 | 2 | 0 | NON_SYNONYMOU | MODERAT | MISSENSE | Acc/Gcc | T53A | PLES_38321 | | | | | | | | | | | C | C |
| NC_011770 | 4240011 | G | C | 69 | 0 | 2 | 0 | NON_SYNONYMOU | MODERAT | MISSENSE | gaC/gaG | D34E | PLES_38321 | | | | | | | C | C | | | | |
| NC_011770 | 4240444 | C | T | 64 | 0 | 4 | 0 | NON_SYNONYMOU | MODERAT | MISSENSE | Gcg/Acg | A187T | PLES_38331 | | | | | | | T | T | T | T | | |
| NC_011770 | 4240705 | G | A | 70 | 0 | 2 | 0 | NON_SYNONYMOU | MODERAT | MISSENSE | Cgg/Tgg | R100W | PLES_38331 | | | | | | | | | A | A | | |
| NC_011770 | 4240707 | T | C | 70 | 0 | 2 | 0 | NON_SYNONYMOU | MODERAT | MISSENSE | cAc/cGc | H99R | PLES_38331 | | | | | | | | | C | C | | |
| NC_011770 | 4240941 | A | G | 70 | 0 | 2 | 0 | NON_SYNONYMOU | MODERAT | MISSENSE | gTg/gCg | V21A | PLES_38331 | G | G | | | | | | | | | | |
| NC_011770 | 4241423 | C | T | 69 | 0 | 2 | 0 | NON_SYNONYMOU | MODERAT | MISSENSE | aCc/aTc | T119I | PLES_38341 | | | | | | | | | T | T | | |
| NC_011770 | 4242690 | GC | G | 62 | 0 | 1 | 0 | FRAME_SHIFT | HIGH | | ccc/ | P284 | PLES_38351 | | | | | | | | G | | | | |
| NC_011770 | 4242022 | C | T | 70 | 0 | 2 | 0 | NON_SYNONYMOU | MODERAT | MISSENSE | gCc/gTc | A60V | PLES_38351 | | | | | | | T | T | | | | |
| NC_011770 | 4242177 | G | A | 71 | 0 | 2 | 0 | NON_SYNONYMOU | MODERAT | MISSENSE | Gag/Aag | E112K | PLES_38351 | | | | | | | | | A | A | | |
| NC_011770 | 4242181 | C | T | 73 | 0 | 4 | 0 | NON_SYNONYMOU | MODERAT | MISSENSE | gCc/gTc | A113V | PLES_38351 | T | T | | | | | | | T | T | | |
| NC_011770 | 4242346 | A | G | 79 | 0 | 8 | 0 | NON_SYNONYMOU | MODERAT | MISSENSE | cAc/cGc | H168R | PLES_38351 | G | G | G | G | G | G | | | | | G | G |

| | | | | | | | | | | | | | | | | | | | | | | | | | |
|---|---|---|---|---|---|---|---|---|---|---|---|---|---|---|---|---|---|---|---|---|---|---|---|---|---|
| NC_011770 | 4242363 | T | G | 79 | 0 | 12 | 0 | NON_SYNONYMOU | MODERAT | MISSENSE | Tcc/Gcc | S174A | PLES_38351 | G | G | G | G | G | G | G | G | G | G | G | G |
| NC_011770 | 4242652 | G | A | 65 | 0 | 4 | 0 | NON_SYNONYMOU | MODERAT | MISSENSE | aGc/aAc | S270N | PLES_38351 | | | A | A | A | A | | | | | | |
| NC_011770 | 4243086 | G | A | 83 | 0 | 2 | 0 | NON_SYNONYMOU | MODERAT | MISSENSE | Gtc/Atc | V415I | PLES_38351 | | | | | | | | | | | A | A |
| NC_011770 | 4243271 | C | A | 80 | 0 | 2 | 0 | NON_SYNONYMOU | MODERAT | MISSENSE | ttC/ttA | F476L | PLES_38351 | | | | | | | A | A | | | | |
| NC_011770 | 4242865 | T | C | 79 | 0 | 1 | 0 | NON_SYNONYMOU | MODERAT | MISSENSE | cTg/cCg | L341P | PLES_38351 | | | | | | | | C | | | | |
| NC_011770 | 4243604 | C | A | 91 | 0 | 2 | 0 | NON_SYNONYMOU | MODERAT | MISSENSE | cGg/cTg | R463L | PLES_38361 | | | | | | | | | A | A | | |
| NC_011770 | 4243946 | C | T | 88 | 0 | 1 | 0 | NON_SYNONYMOU | MODERAT | MISSENSE | cGg/cAg | R349Q | PLES_38361 | | | | | | | T | | | | | |
| NC_011770 | 4243971 | A | G | 90 | 0 | 12 | 0 | NON_SYNONYMOU | MODERAT | MISSENSE | Tcg/Ccg | S341P | PLES_38361 | G | G | G | G | G | G | G | G | G | G | G | G |
| NC_011770 | 4244927 | C | T | 80 | 0 | 2 | 0 | NON_SYNONYMOU | MODERAT | MISSENSE | cGa/cAa | R22Q | PLES_38361 | | | | | | | | | T | T | | |
| NC_011770 | 4244942 | G | A | 80 | 0 | 2 | 0 | NON_SYNONYMOU | MODERAT | MISSENSE | cCc/cTc | P17L | PLES_38361 | | | | | | | A | A | | | | |
| NC_011770 | 4244962 | G | T | 78 | 0 | 4 | 0 | NON_SYNONYMOU | MODERAT | MISSENSE | gaC/gaA | D10E | PLES_38361 | | | T | T | T | T | | | | | | |
| NC_011770 | 4245277 | A | C | 74 | 0 | 10 | 0 | NON_SYNONYMOU | MODERAT | MISSENSE | gaT/gaG | D231E | PLES_38371 | | | C | C | C | C | C | C | C | C | C | C |
| NC_011770 | 4245792 | C | T | 74 | 0 | 2 | 0 | NON_SYNONYMOU | MODERAT | MISSENSE | Gtg/Atg | V60M | PLES_38371 | | | | | | | T | T | | | | |
| NC_011770 | 4247079 | A | C | 70 | 0 | 4 | 0 | NON_SYNONYMOU | MODERAT | MISSENSE | gaA/gaC | E35D | PLES_38391 | | | | | | | C | C | | | C | C |
| NC_011770 | 4249594 | A | G | 69 | 0 | 12 | 0 | NON_SYNONYMOU | MODERAT | MISSENSE | gAg/gGg | E34G | PLES_38421 | G | G | G | G | G | G | G | G | G | G | G | G |
| NC_011770 | 4250060 | G | A | 61 | 0 | 2 | 0 | NON_SYNONYMOU | MODERAT | MISSENSE | atG/atA | M189I | PLES_38421 | A | A | | | | | | | | | | |
| NC_011770 | 4249623 | A | T | 75 | 0 | 1 | 0 | NON_SYNONYMOU | MODERAT | MISSENSE | Atc/Ttc | I44F | PLES_38421 | | | | | | | | T | | | | |
| NC_011770 | 4255360 | G | T | 70 | 0 | 2 | 0 | NON_SYNONYMOU | MODERAT | MISSENSE | Gcc/Tcc | A56S | PLES_38461 | | | | | | | | | | | T | T |
| NC_011770 | 4255558 | G | A | 80 | 0 | 2 | 0 | NON_SYNONYMOU | MODERAT | MISSENSE | Ggc/Agc | G122S | PLES_38461 | | | | | | | A | A | | | | |
| NC_011770 | 4255943 | C | G | 102 | 0 | 2 | 0 | NON_SYNONYMOU | MODERAT | MISSENSE | aCc/aGc | T250S | PLES_38461 | | | | | | | | | G | G | | |
| NC_011770 | 4256774 | G | A | 65 | 0 | 6 | 0 | NON_SYNONYMOU | MODERAT | MISSENSE | Gcc/Acc | A7T | PLES_38471 | A | A | A | A | A | A | | | | | | |
| NC_011770 | 4257716 | G | A | 71 | 0 | 4 | 0 | NON_SYNONYMOU | MODERAT | MISSENSE | Gtc/Atc | V321I | PLES_38471 | A | A | | | | | A | A | | | | |
| NC_011770 | 4258102 | G | C | 69 | 0 | 2 | 0 | NON_SYNONYMOU | MODERAT | MISSENSE | Gtc/Ctc | V14L | PLES_38481 | | | | | | | | | | | C | C |
| NC_011770 | 4258111 | A | G | 70 | 0 | 8 | 0 | NON_SYNONYMOU | MODERAT | MISSENSE | Acc/Gcc | T17A | PLES_38481 | G | G | | | | | G | G | G | G | G | G |
| NC_011770 | 4258178 | C | T | 74 | 0 | 2 | 0 | NON_SYNONYMOU | MODERAT | MISSENSE | gCc/gTc | A39V | PLES_38481 | | | | | | | | | T | T | | |
| NC_011770 | 4258458 | G | A | 63 | 0 | 2 | 0 | NON_SYNONYMOU | MODERAT | MISSENSE | Gtt/Att | V19I | PLES_38491 | | | | | | | | | | | A | A |
| NC_011770 | 4258485 | C | T | 58 | 0 | 2 | 0 | NON_SYNONYMOU | MODERAT | MISSENSE | Ccg/Tcg | P28S | PLES_38491 | | | | | | | | | | | T | T |
| NC_011770 | 4258605 | A | G | 54 | 0 | 2 | 0 | NON_SYNONYMOU | MODERAT | MISSENSE | Att/Gtt | I68V | PLES_38491 | | | | | | | | | | | G | G |
| NC_011770 | 4258617 | T | C | 54 | 0 | 2 | 0 | NON_SYNONYMOU | MODERAT | MISSENSE | Tac/Cac | Y72H | PLES_38491 | | | | | | | | | | | C | C |
| NC_011770 | 4258674 | T | C | 48 | 0 | 4 | 0 | NON_SYNONYMOU | MODERAT | MISSENSE | Tcc/Ccc | S91P | PLES_38491 | C | C | | | | | | | | | C | C |
| NC_011770 | 4258677 | C | G | 49 | 0 | 4 | 0 | NON_SYNONYMOU | MODERAT | MISSENSE | Cac/Gac | H92D | PLES_38491 | G | G | | | | | | | | | G | G |
| NC_011770 | 4258698 | G | A | 53 | 0 | 2 | 0 | NON_SYNONYMOU | MODERAT | MISSENSE | Gcg/Acg | A99T | PLES_38491 | | | | | | | | | | | A | A |
| NC_011770 | 4258770 | A | T | 56 | 0 | 6 | 0 | NON_SYNONYMOU | MODERAT | MISSENSE | Atg/Ttg | M123L | PLES_38491 | | | T | T | T | T | | | | | G | G |
| NC_011770 | 4258806 | G | A | 59 | 0 | 6 | 0 | NON_SYNONYMOU | MODERAT | MISSENSE | Gtg/Atg | V135M | PLES_38491 | A | A | | | | | A | A | A | A | | |
| NC_011770 | 4258812 | T | G | 57 | 0 | 12 | 0 | NON_SYNONYMOU | MODERAT | MISSENSE | Tcg/Gcg | S137A | PLES_38491 | G | G | G | G | G | G | G | G | G | G | G | G |
| NC_011770 | 4260022 | G | A | 77 | 0 | 2 | 0 | NON_SYNONYMOU | MODERAT | MISSENSE | Gtc/Atc | V267I | PLES_38501 | | | | | | | | | | | A | A |
| NC_011770 | 4260846 | G | A | 73 | 0 | 4 | 0 | NON_SYNONYMOU | MODERAT | MISSENSE | gCc/gTc | A254V | PLES_38511 | | | A | A | A | A | | | | | | |
| NC_011770 | 4260886 | C | G | 73 | 0 | 2 | 0 | NON_SYNONYMOU | MODERAT | MISSENSE | Gaa/Caa | E241Q | PLES_38511 | G | G | | | | | | | | | | |
| NC_011770 | 4261422 | T | G | 73 | 0 | 12 | 0 | NON_SYNONYMOU | MODERAT | MISSENSE | aAc/aCc | N62T | PLES_38511 | G | G | G | G | G | G | G | G | G | G | G | G |
| NC_011770 | 4261477 | G | A | 67 | 0 | 2 | 0 | NON_SYNONYMOU | MODERAT | MISSENSE | Cgc/Tgc | R44C | PLES_38511 | | | | | | | | | | | A | A |
| NC_011770 | 4261486 | C | T | 68 | 0 | 2 | 0 | NON_SYNONYMOU | MODERAT | MISSENSE | Gcc/Acc | A41T | PLES_38511 | | | | | | | | | | | T | T |
| NC_011770 | 4261709 | C | A | 69 | 0 | 8 | 0 | NON_SYNONYMOU | MODERAT | MISSENSE | gGt/gTt | G112V | PLES_38521 | A | A | A | A | A | A | A | A | | | | |
| NC_011770 | 4261758 | C | A | 69 | 0 | 4 | 0 | NON_SYNONYMOU | MODERAT | MISSENSE | Ggc/Tgc | G96C | PLES_38521 | | | A | A | A | A | | | | | | |
| NC_011770 | 4261874 | G | C | 69 | 0 | 2 | 0 | NON_SYNONYMOU | MODERAT | MISSENSE | gCc/gGc | A57G | PLES_38521 | C | C | | | | | | | | | | |
| NC_011770 | 4262647 | A | G | 67 | 0 | 10 | 0 | NON_SYNONYMOU | MODERAT | MISSENSE | Acc/Gcc | T174A | PLES_38531 | G | G | G | G | G | G | G | G | | | G | G |
| NC_011770 | 4262662 | T | C | 68 | 0 | 4 | 0 | NON_SYNONYMOU | MODERAT | MISSENSE | Tac/Cac | Y179H | PLES_38531 | | | C | C | C | C | | | | | | |
| NC_011770 | 4262818 | G | A | 64 | 0 | 4 | 0 | NON_SYNONYMOU | MODERAT | MISSENSE | Gcg/Acg | A231T | PLES_38531 | | | | | | | A | A | | | A | A |
| NC_011770 | 4263549 | C | T | 47 | 0 | 4 | 0 | NON_SYNONYMOU | MODERAT | MISSENSE | Gtc/Atc | V474I | PLES_38541 | | | T | T | T | T | | | | | | |
| NC_011770 | 4266753 | C | G | 84 | 0 | 2 | 0 | NON_SYNONYMOU | MODERAT | MISSENSE | Gtg/Ctg | V87L | PLES_38561 | | | | | | | G | G | | | | |
| NC_011770 | 4266956 | G | A | 85 | 0 | 4 | 0 | NON_SYNONYMOU | MODERAT | MISSENSE | gCc/gTc | A19V | PLES_38561 | | | A | A | A | A | | | | | | |
| NC_011770 | 4267180 | A | C | 63 | 0 | 2 | 0 | INTRAGENIC | MODIFIER | | | | PLES_38562 | | | | | | | | | | | C | C |
| NC_011770 | 4267182 | G | C | 64 | 0 | 2 | 0 | INTRAGENIC | MODIFIER | | | | PLES_38562 | | | | | | | | | | | C | C |
| NC_011770 | 4267746 | G | A | 84 | 0 | 2 | 0 | NON_SYNONYMOU | MODERAT | MISSENSE | gCg/gTg | A15V | PLES_38571 | A | A | | | | | | | | | | |
| NC_011770 | 4268477 | C | A | 87 | 0 | 2 | 0 | NON_SYNONYMOU | MODERAT | MISSENSE | Ggc/Tgc | G678C | PLES_38591 | | | | | | | | | A | A | | |
| NC_011770 | 4268996 | A | C | 58 | 0 | 6 | 0 | NON_SYNONYMOU | MODERAT | MISSENSE | Ttc/Gtc | F505V | PLES_38591 | | | C | C | C | C | | | C | C | | |
| NC_011770 | 4269487 | A | C | 68 | 0 | 2 | 0 | NON_SYNONYMOU | MODERAT | MISSENSE | cTg/cGg | L341R | PLES_38591 | | | | | | | | | | | C | C |
| NC_011770 | 4269604 | G | A | 67 | 0 | 2 | 0 | NON_SYNONYMOU | MODERAT | MISSENSE | gCc/gTc | A302V | PLES_38591 | | | | | | | | | | | A | A |
| NC_011770 | 4269778 | G | A | 75 | 0 | 2 | 0 | NON_SYNONYMOU | MODERAT | MISSENSE | aCc/aTc | T244I | PLES_38591 | A | A | | | | | | | | | | |
| NC_011770 | 4270170 | ACCG | A | 70 | 0 | 1 | 0 | CODON_CHANGE_ | MODERATE | | ggcggt/ggt | GG112G | PLES_38591 | | | | A | | | | | | | | |
| NC_011770 | 4275707 | G | A | 78 | 0 | 2 | 0 | NON_SYNONYMOU | MODERAT | MISSENSE | gGg/gAg | G25E | PLES_38651 | A | A | | | | | | | | | | |
| NC_011770 | 4276225 | T | C | 62 | 0 | 10 | 0 | NON_SYNONYMOU | MODERAT | MISSENSE | cAc/cGc | H513R | PLES_38661 | C | C | C | C | C | C | | | C | C | C | C |
| NC_011770 | 4276436 | C | A | 52 | 0 | 8 | 0 | NON_SYNONYMOU | MODERAT | MISSENSE | Gcc/Tcc | A443S | PLES_38661 | A | A | A | A | A | A | | | A | A | | |

| | | | | | | | | | | | | | | | | | | | | | | | | | |
|---|---|---|---|---|---|---|---|---|---|---|---|---|---|---|---|---|---|---|---|---|---|---|---|---|---|
| NC_011770 | 4276442 | T | C | 50 | 0 | 8 | 0 | NON_SYNONYMOU | MODERAT | MISSENSE | Atc/Gtc | I441V | PLES_38661 | C | C | C | C | C | C | | | C | C | | |
| NC_011770 | 4276483 | T | C | 49 | 0 | 8 | 0 | NON_SYNONYMOU | MODERAT | MISSENSE | gAg/gGg | E427G | PLES_38661 | C | C | C | C | C | C | | | C | C | | |
| NC_011770 | 4277474 | G | C | 73 | 0 | 2 | 0 | NON_SYNONYMOU | MODERAT | MISSENSE | Ccg/Gcg | P97A | PLES_38661 | | | | | | | C | C | | | | |
| NC_011770 | 4277569 | G | A | 78 | 0 | 1 | 0 | NON_SYNONYMOU | MODERAT | MISSENSE | cCg/cTg | P65L | PLES_38661 | | | | | | | A | | | | | |
| NC_011770 | 4278112 | A | G | 86 | 0 | 2 | 0 | NON_SYNONYMOU | MODERAT | MISSENSE | Acc/Gcc | T65A | PLES_38671 | | | | | | | G | G | | | | |
| NC_011770 | 4278367 | G | A | 81 | 0 | 4 | 0 | NON_SYNONYMOU | MODERAT | MISSENSE | Gcg/Acg | A150T | PLES_38671 | | | A | A | A | A | | | | | | |
| NC_011770 | 4279333 | GA | G | 48 | 0 | 4 | 0 | FRAME_SHIFT | HIGH | | tcc/ | S323 | PLES_38691 | | | G | G | G | G | | | | | | |
| NC_011770 | 4279060 | C | A | 63 | 0 | 2 | 0 | NON_SYNONYMOU | MODERAT | MISSENSE | cGg/cTg | R414L | PLES_38691 | | | | | | | | | A | A | | |
| NC_011770 | 4279649 | T | C | 78 | 0 | 12 | 0 | NON_SYNONYMOU | MODERAT | MISSENSE | Agc/Ggc | S218G | PLES_38691 | C | C | C | C | C | C | C | C | C | C | C | C |
| NC_011770 | 4279888 | A | G | 76 | 0 | 1 | 0 | NON_SYNONYMOU | MODERAT | MISSENSE | gTg/gCg | V138A | PLES_38691 | | G | | | | | | | | | | |
| NC_011770 | 4281294 | G | A | 35 | 0 | 4 | 0 | NON_SYNONYMOU | MODERAT | MISSENSE | cGc/cAc | R296H | PLES_38701 | | | A | A | A | A | | | | | | |
| NC_011770 | 4283979 | T | A | 75 | 0 | 12 | 0 | NON_SYNONYMOU | MODERAT | MISSENSE | gTg/gAg | V155E | PLES_38731 | A | A | A | A | A | A | A | A | A | A | A | A |
| NC_011770 | 4284746 | T | C | 68 | 0 | 2 | 0 | NON_SYNONYMOU | MODERAT | MISSENSE | aAc/aGc | N55S | PLES_38741 | | | | | | | | | | | C | C |
| NC_011770 | 4285840 | C | A | 76 | 0 | 2 | 0 | NON_SYNONYMOU | MODERAT | MISSENSE | cGc/cTc | R112L | PLES_38761 | | | | | | | | | | | A | A |
| NC_011770 | 4287152 | C | A | 61 | 0 | 2 | 0 | NON_SYNONYMOU | MODERAT | MISSENSE | gCg/gAg | A98E | PLES_38781 | A | A | | | | | | | | | | |
| NC_011770 | 4287267 | C | A | 67 | 0 | 2 | 0 | NON_SYNONYMOU | MODERAT | MISSENSE | agC/agA | S136R | PLES_38781 | | | | | | | | | A | A | | |
| NC_011770 | 4287363 | G | C | 75 | 0 | 8 | 0 | NON_SYNONYMOU | MODERAT | MISSENSE | gaG/gaC | E168D | PLES_38781 | | | C | C | C | C | | | C | C | C | C |
| NC_011770 | 4287754 | T | C | 82 | 0 | 2 | 0 | NON_SYNONYMOU | MODERAT | MISSENSE | Tcg/Ccg | S299P | PLES_38781 | C | C | | | | | | | | | | |
| NC_011770 | 4287760 | C | T | 85 | 0 | 2 | 0 | NON_SYNONYMOU | MODERAT | MISSENSE | Ccg/Tcg | P301S | PLES_38781 | T | T | | | | | | | | | | |
| NC_011770 | 4287766 | C | T | 85 | 0 | 2 | 0 | NON_SYNONYMOU | MODERAT | MISSENSE | Cgc/Tgc | R303C | PLES_38781 | | | | | | | T | T | | | | |
| NC_011770 | 4289366 | G | C | 78 | 0 | 4 | 0 | NON_SYNONYMOU | MODERAT | MISSENSE | Gag/Cag | E127Q | PLES_38791 | | | C | C | C | C | | | | | | |
| NC_011770 | 4289389 | C | A | 83 | 0 | 6 | 0 | NON_SYNONYMOU | MODERAT | MISSENSE | caC/caA | H134Q | PLES_38791 | A | A | | | | | A | A | A | A | | |
| NC_011770 | 4292061 | C | T | 67 | 0 | 1 | 0 | STOP_GAINED | HIGH | NONSENSE | tGg/tAg | W10* | PLES_38801 | | | | | | | | | | | | T |
| NC_011770 | 4289535 | C | T | 92 | 0 | 2 | 0 | SPLICE_SITE_REGIC | LOW | SILENT | tGa/tAa | *852 | PLES_38801 | | | | | | | | | T | T | | |
| NC_011770 | 4290067 | C | T | 84 | 0 | 2 | 0 | NON_SYNONYMOU | MODERAT | MISSENSE | Ggc/Agc | G675S | PLES_38801 | | | | | | | | | T | T | | |
| NC_011770 | 4290375 | G | A | 84 | 0 | 1 | 0 | NON_SYNONYMOU | MODERAT | MISSENSE | aCc/aTc | T572I | PLES_38801 | | | | | | | A | | | | | |
| NC_011770 | 4290408 | G | A | 86 | 0 | 2 | 0 | NON_SYNONYMOU | MODERAT | MISSENSE | tCc/tTc | S561F | PLES_38801 | | | A | A | | | | | | | | |
| NC_011770 | 4290561 | T | C | 76 | 0 | 2 | 0 | NON_SYNONYMOU | MODERAT | MISSENSE | cAg/cGg | Q510R | PLES_38801 | | | | | | | | | C | C | | |
| NC_011770 | 4290618 | C | A | 75 | 0 | 1 | 0 | NON_SYNONYMOU | MODERAT | MISSENSE | cGc/cTc | R491L | PLES_38801 | | | | | | | A | | | | | |
| NC_011770 | 4290672 | T | C | 77 | 0 | 2 | 0 | NON_SYNONYMOU | MODERAT | MISSENSE | aAg/aGg | K473R | PLES_38801 | | | | | | | C | C | | | | |
| NC_011770 | 4290871 | T | C | 83 | 0 | 2 | 0 | NON_SYNONYMOU | MODERAT | MISSENSE | Atc/Gtc | I407V | PLES_38801 | | | | | | | C | C | | | | |
| NC_011770 | 4292336 | C | T | 74 | 0 | 2 | 0 | NON_SYNONYMOU | MODERAT | MISSENSE | cGg/cAg | R289Q | PLES_38811 | | | T | T | | | | | | | | |
| NC_011770 | 4292485 | T | G | 75 | 0 | 2 | 0 | NON_SYNONYMOU | MODERAT | MISSENSE | gaA/gaC | E239D | PLES_38811 | G | G | | | | | | | | | | |
| NC_011770 | 4292497 | G | C | 70 | 0 | 8 | 0 | NON_SYNONYMOU | MODERAT | MISSENSE | gaC/gaG | D235E | PLES_38811 | | | C | C | C | C | C | C | | | C | C |
| NC_011770 | 4292688 | C | T | 75 | 0 | 4 | 0 | NON_SYNONYMOU | MODERAT | MISSENSE | Gcc/Acc | A172T | PLES_38811 | | | T | T | T | T | | | | | | |
| NC_011770 | 4293041 | C | A | 75 | 0 | 2 | 0 | NON_SYNONYMOU | MODERAT | MISSENSE | cGc/cTc | R54L | PLES_38811 | | | | | | | | | A | A | | |
| NC_011770 | 4293379 | G | A | 71 | 0 | 2 | 0 | NON_SYNONYMOU | MODERAT | MISSENSE | gCg/gTg | A116V | PLES_38821 | | | | | | | | | A | A | | |
| NC_011770 | 4293479 | G | T | 74 | 0 | 2 | 0 | NON_SYNONYMOU | MODERAT | MISSENSE | Ctc/Atc | L83I | PLES_38821 | T | T | | | | | | | | | | |
| NC_011770 | 4294816 | G | A | 33 | 0 | 2 | 0 | NON_SYNONYMOU | MODERAT | MISSENSE | cGc/cAc | R25H | PLES_38851 | | | | | | | A | A | | | | |
| NC_011770 | 4295250 | G | C | 19 | 3 | 1 | 0 | NON_SYNONYMOU | MODERAT | MISSENSE | Gac/Cac | D170H | PLES_38851 | | | | | | | . | C | . | . | | |
| NC_011770 | 4297157 | T | C | 85 | 0 | 8 | 0 | NON_SYNONYMOU | MODERAT | MISSENSE | aAc/aGc | N14S | PLES_38871 | C | C | C | C | C | C | | | C | C | | |
| NC_011770 | 4298719 | C | G | 85 | 0 | 4 | 0 | NON_SYNONYMOU | MODERAT | MISSENSE | gaC/gaG | D87E | PLES_38891 | G | G | | | | | G | G | | | | |
| NC_011770 | 4300394 | GGCA | G | 85 | 0 | 2 | 0 | CODON_CHANGE_ | MODERATE | | gctgcc/gcc | AA17A | PLES_38901 | | | | | | | | | | | G | G |
| NC_011770 | 4300876 | A | T | 89 | 0 | 4 | 0 | NON_SYNONYMOU | MODERAT | MISSENSE | Acc/Tcc | T19S | PLES_38911 | | | T | T | T | T | | | | | | |
| NC_011770 | 4300931 | C | T | 96 | 0 | 2 | 0 | NON_SYNONYMOU | MODERAT | MISSENSE | aCc/aTc | T37I | PLES_38911 | | | | | | | | | | | T | T |
| NC_011770 | 4301825 | A | G | 63 | 0 | 12 | 0 | NON_SYNONYMOU | MODERAT | MISSENSE | cAt/cGt | H335R | PLES_38911 | G | G | G | G | G | G | G | G | G | G | G | G |
| NC_011770 | 4302106 | T | C | 39 | 0 | 12 | 0 | NON_SYNONYMOU | MODERAT | MISSENSE | Tgc/Cgc | C429R | PLES_38911 | C | C | C | C | C | C | C | C | C | C | C | C |
| NC_011770 | 4303909 | A | G | 80 | 0 | 2 | 0 | NON_SYNONYMOU | MODERAT | MISSENSE | Acg/Gcg | T249A | PLES_38931 | | | | | | | G | G | | | | |
| NC_011770 | 4304139 | A | G | 76 | 0 | 12 | 0 | NON_SYNONYMOU | MODERAT | MISSENSE | Atc/Gtc | I18V | PLES_38941 | G | G | G | G | G | G | G | G | G | G | G | G |
| NC_011770 | 4304215 | G | A | 75 | 0 | 8 | 0 | NON_SYNONYMOU | MODERAT | MISSENSE | aGc/aAc | S43N | PLES_38941 | | | A | A | A | A | A | A | A | A | | |
| NC_011770 | 4305990 | T | C | 73 | 0 | 4 | 0 | NON_SYNONYMOU | MODERAT | MISSENSE | cTc/cCc | L106P | PLES_38961 | | | C | C | C | C | | | | | | |
| NC_011770 | 4306634 | G | A | 77 | 0 | 4 | 0 | NON_SYNONYMOU | MODERAT | MISSENSE | Gtc/Atc | V135I | PLES_38971 | | | | | | | | | A | A | A | A |
| NC_011770 | 4307207 | CCCGCCGC | C | 78 | 1 | 1 | 0 | CODON_DELETION | MODERATE | | gccgcgacc/ | AAT327 | PLES_38971 | | | | | | | | | . | C | | |
| NC_011770 | 4307237 | G | A | 81 | 0 | 2 | 0 | NON_SYNONYMOU | MODERAT | MISSENSE | Gcc/Acc | A336T | PLES_38971 | | | | | | | | | A | A | | |
| NC_011770 | 4306772 | G | A | 86 | 0 | 1 | 0 | NON_SYNONYMOU | MODERAT | MISSENSE | Ggc/Agc | G181S | PLES_38971 | | | | | | | | A | | | | |
| NC_011770 | 4308875 | A | C | 65 | 0 | 2 | 0 | NON_SYNONYMOU | MODERAT | MISSENSE | aaA/aaC | K16N | PLES_38991 | | | | | | | C | C | | | | |
| NC_011770 | 4308937 | A | G | 62 | 0 | 2 | 0 | NON_SYNONYMOU | MODERAT | MISSENSE | aAc/aGc | N37S | PLES_38991 | | | | | | | G | G | | | | |
| NC_011770 | 4308976 | G | A | 65 | 0 | 2 | 0 | NON_SYNONYMOU | MODERAT | MISSENSE | aGg/aAg | R50K | PLES_38991 | | | | | | | A | A | | | | |
| NC_011770 | 4309579 | G | A | 58 | 0 | 2 | 0 | NON_SYNONYMOU | MODERAT | MISSENSE | gGc/gAc | G251D | PLES_38991 | | | | | | | A | A | | | | |
| NC_011770 | 4309645 | C | G | 57 | 0 | 6 | 0 | NON_SYNONYMOU | MODERAT | MISSENSE | aCc/aGc | T273S | PLES_38991 | G | G | | | | | G | G | G | G | | |
| NC_011770 | 4309699 | G | A | 58 | 0 | 1 | 0 | NON_SYNONYMOU | MODERAT | MISSENSE | gGc/gAc | G291D | PLES_38991 | | | | | | | A | | | | | |
| NC_011770 | 4309716 | A | G | 58 | 0 | 2 | 0 | NON_SYNONYMOU | MODERAT | MISSENSE | Atc/Gtc | I297V | PLES_38991 | G | G | | | | | | | | | | |

| | | | | | | | | | | | | | | | | | | | | | | | | | |
|---|---|---|---|---|---|---|---|---|---|---|---|---|---|---|---|---|---|---|---|---|---|---|---|---|---|
| NC_011770 | 4309854 | A | C | 54 | 0 | 2 | 0 | NON_SYNONYMOU | MODERAT | MISSENSE | Agc/Cgc | S343R | PLES_38991 | C | C | | | | | | | | | | |
| NC_011770 | 4310023 | G | A | 17 | 0 | 1 | 0 | NON_SYNONYMOU | MODERAT | MISSENSE | cGc/cAc | R399H | PLES_38991 | A | . | | | | | | | | | | |
| NC_011770 | 4312550 | C | T | 25 | 8 | 1 | 0 | NON_SYNONYMOU | MODERAT | MISSENSE | Ccc/Tcc | P400S | PLES_39001 | | T | . | . | . | . | | | . | . | . | . |
| NC_011770 | 4311412 | T | G | 98 | 0 | 2 | 0 | NON_SYNONYMOU | MODERAT | MISSENSE | agT/agG | S20R | PLES_39001 | | | | | | | G | G | | | | |
| NC_011770 | 4311479 | A | G | 93 | 0 | 10 | 0 | NON_SYNONYMOU | MODERAT | MISSENSE | Att/Gtt | I43V | PLES_39001 | | | G | G | G | G | G | G | G | G | G | G |
| NC_011770 | 4311560 | G | A | 91 | 0 | 2 | 0 | NON_SYNONYMOU | MODERAT | MISSENSE | Gag/Aag | E70K | PLES_39001 | | | | | | | A | A | | | | |
| NC_011770 | 4311563 | G | A | 90 | 0 | 2 | 0 | NON_SYNONYMOU | MODERAT | MISSENSE | Gtc/Atc | V71I | PLES_39001 | | | | | | | A | A | | | | |
| NC_011770 | 4311593 | A | C | 67 | 0 | 8 | 0 | NON_SYNONYMOU | MODERAT | MISSENSE | Atg/Ctg | M81L | PLES_39001 | | | C | C | C | C | | | C | C | C | C |
| NC_011770 | 4311597 | A | G | 66 | 0 | 8 | 0 | NON_SYNONYMOU | MODERAT | MISSENSE | gAt/gGt | D82G | PLES_39001 | | | G | G | G | G | | | G | G | G | G |
| NC_011770 | 4311598 | T | A | 65 | 0 | 8 | 0 | NON_SYNONYMOU | MODERAT | MISSENSE | gaT/gaA | D82E | PLES_39001 | | | A | A | A | A | | | A | A | A | A |
| NC_011770 | 4311605 | T | G | 65 | 0 | 8 | 0 | NON_SYNONYMOU | MODERAT | MISSENSE | Ttt/Gtt | F85V | PLES_39001 | | | G | G | G | G | | | G | G | G | G |
| NC_011770 | 4311606 | T | C | 65 | 0 | 8 | 0 | NON_SYNONYMOU | MODERAT | MISSENSE | tTt/tCt | F85S | PLES_39001 | | | C | C | C | C | | | C | C | C | C |
| NC_011770 | 4311614 | A | T | 63 | 0 | 8 | 0 | NON_SYNONYMOU | MODERAT | MISSENSE | Atc/Ttc | I88F | PLES_39001 | | | T | T | T | T | | | T | T | T | T |
| NC_011770 | 4311635 | G | T | 74 | 0 | 2 | 0 | NON_SYNONYMOU | MODERAT | MISSENSE | Gcc/Tcc | A95S | PLES_39001 | | | | | | | T | T | | | | |
| NC_011770 | 4311636 | C | A | 73 | 0 | 2 | 0 | NON_SYNONYMOU | MODERAT | MISSENSE | gCc/gAc | A95D | PLES_39001 | | | | | | | A | A | | | | |
| NC_011770 | 4311645 | G | A | 69 | 0 | 2 | 0 | NON_SYNONYMOU | MODERAT | MISSENSE | gGt/gAt | G98D | PLES_39001 | | | | | | | A | A | | | | |
| NC_011770 | 4311647 | C | G | 68 | 0 | 2 | 0 | NON_SYNONYMOU | MODERAT | MISSENSE | Cca/Gca | P99A | PLES_39001 | | | | | | | G | G | | | | |
| NC_011770 | 4311654 | C | T | 66 | 0 | 8 | 0 | NON_SYNONYMOU | MODERAT | MISSENSE | gCc/gTc | A101V | PLES_39001 | | | T | T | T | T | | | T | T | T | T |
| NC_011770 | 4311662 | G | A | 46 | 0 | 10 | 0 | NON_SYNONYMOU | MODERAT | MISSENSE | Gag/Aag | E104K | PLES_39001 | | | A | A | A | A | A | A | A | A | A | A |
| NC_011770 | 4311663 | A | G | 46 | 0 | 2 | 0 | NON_SYNONYMOU | MODERAT | MISSENSE | gAg/gGg | E104G | PLES_39001 | | | | | | | G | G | | | | |
| NC_011770 | 4311665 | C | T | 46 | 0 | 2 | 0 | NON_SYNONYMOU | MODERAT | MISSENSE | Ctc/Ttc | L105F | PLES_39001 | | | | | | | T | T | | | | |
| NC_011770 | 4311672 | A | G | 47 | 0 | 8 | 0 | NON_SYNONYMOU | MODERAT | MISSENSE | aAg/aGg | K107R | PLES_39001 | | | G | G | G | G | | | G | G | G | G |
| NC_011770 | 4311680 | C | A | 45 | 0 | 10 | 0 | NON_SYNONYMOU | MODERAT | MISSENSE | Caa/Aaa | Q110K | PLES_39001 | | | A | A | A | A | A | A | A | A | A | A |
| NC_011770 | 4311681 | A | G | 46 | 0 | 10 | 0 | NON_SYNONYMOU | MODERAT | MISSENSE | cAa/cGa | Q110R | PLES_39001 | | | G | G | G | G | G | G | G | G | G | G |
| NC_011770 | 4311731 | T | G | 77 | 0 | 8 | 0 | NON_SYNONYMOU | MODERAT | MISSENSE | Tcc/Gcc | S127A | PLES_39001 | | | G | G | G | G | | | G | G | G | G |
| NC_011770 | 4311732 | C | A | 77 | 0 | 8 | 0 | NON_SYNONYMOU | MODERAT | MISSENSE | tCc/tAc | S127Y | PLES_39001 | | | A | A | A | A | | | A | A | A | A |
| NC_011770 | 4311739 | A | C | 85 | 0 | 8 | 0 | NON_SYNONYMOU | MODERAT | MISSENSE | gaA/gaC | E129D | PLES_39001 | | | C | C | C | C | | | C | C | C | C |
| NC_011770 | 4311813 | G | A | 90 | 0 | 1 | 0 | NON_SYNONYMOU | MODERAT | MISSENSE | cGa/cAa | R154Q | PLES_39001 | | | | | | | A | | | | | |
| NC_011770 | 4311820 | T | A | 94 | 0 | 2 | 0 | NON_SYNONYMOU | MODERAT | MISSENSE | gaT/gaA | D156E | PLES_39001 | | | | | | | A | A | | | | |
| NC_011770 | 4311882 | G | C | 88 | 0 | 10 | 0 | NON_SYNONYMOU | MODERAT | MISSENSE | gGa/gCa | G177A | PLES_39001 | | | C | C | C | C | C | C | C | C | C | C |
| NC_011770 | 4311939 | G | A | 95 | 0 | 10 | 0 | NON_SYNONYMOU | MODERAT | MISSENSE | aGg/aAg | R196K | PLES_39001 | | | A | A | A | A | A | A | A | A | A | A |
| NC_011770 | 4312044 | T | G | 95 | 0 | 8 | 0 | NON_SYNONYMOU | MODERAT | MISSENSE | tTg/tGg | L231W | PLES_39001 | | | G | G | G | G | | | G | G | G | G |
| NC_011770 | 4312064 | T | C | 93 | 0 | 2 | 0 | NON_SYNONYMOU | MODERAT | MISSENSE | Tgc/Cgc | C238R | PLES_39001 | | | | | | | C | C | | | | |
| NC_011770 | 4312065 | G | A | 91 | 0 | 10 | 0 | NON_SYNONYMOU | MODERAT | MISSENSE | tGc/tAc | C238Y | PLES_39001 | | | A | A | A | A | A | A | A | A | A | A |
| NC_011770 | 4312068 | G | A | 94 | 0 | 2 | 0 | NON_SYNONYMOU | MODERAT | MISSENSE | gGt/gAt | G239D | PLES_39001 | | | | | | | | | A | A | | |
| NC_011770 | 4312085 | A | G | 95 | 0 | 2 | 0 | NON_SYNONYMOU | MODERAT | MISSENSE | Atg/Gtg | M245V | PLES_39001 | | | | | | | G | G | | | | |
| NC_011770 | 4312090 | A | T | 94 | 0 | 10 | 0 | NON_SYNONYMOU | MODERAT | MISSENSE | agA/agT | R246S | PLES_39001 | | | T | T | T | T | T | T | T | T | T | T |
| NC_011770 | 4312098 | A | G | 94 | 0 | 2 | 0 | NON_SYNONYMOU | MODERAT | MISSENSE | aAg/aGg | K249R | PLES_39001 | G | G | | | | | | | | | | |
| NC_011770 | 4312298 | A | G | 95 | 0 | 10 | 0 | NON_SYNONYMOU | MODERAT | MISSENSE | Att/Gtt | I316V | PLES_39001 | | | G | G | G | G | G | G | G | G | G | G |
| NC_011770 | 4312467 | A | G | 84 | 0 | 10 | 0 | NON_SYNONYMOU | MODERAT | MISSENSE | aAc/aGc | N372S | PLES_39001 | | | G | G | G | G | G | G | G | G | G | G |
| NC_011770 | 4312469 | A | T | 83 | 0 | 10 | 0 | NON_SYNONYMOU | MODERAT | MISSENSE | Act/Tct | T373S | PLES_39001 | | | T | T | T | T | T | T | T | T | T | T |
| NC_011770 | 4312502 | T | G | 76 | 0 | 10 | 0 | NON_SYNONYMOU | MODERAT | MISSENSE | Tca/Gca | S384A | PLES_39001 | | | G | G | G | G | G | G | G | G | G | G |
| NC_011770 | 4312535 | G | A | 25 | 8 | 2 | 0 | NON_SYNONYMOU | MODERAT | MISSENSE | Ggt/Agt | G395S | PLES_39001 | | | . | . | . | . | A | A | . | . | . | . |
| NC_011770 | 4312591 | T | G | 24 | 8 | 2 | 0 | NON_SYNONYMOU | MODERAT | MISSENSE | aaT/aaG | N413K | PLES_39001 | | | . | . | . | . | G | G | . | . | . | . |
| NC_011770 | 4312638 | C | T | 25 | 6 | 2 | 0 | NON_SYNONYMOU | MODERAT | MISSENSE | gCc/gTc | A429V | PLES_39001 | | | . | . | . | T | | | . | T | . | . |
| NC_011770 | 4312805 | A | G | 71 | 0 | 10 | 0 | NON_SYNONYMOU | MODERAT | MISSENSE | Acg/Gcg | T485A | PLES_39001 | | | G | G | G | G | G | G | G | G | G | G |
| NC_011770 | 4312942 | A | C | 77 | 0 | 2 | 0 | NON_SYNONYMOU | MODERAT | MISSENSE | aaA/aaC | K530N | PLES_39001 | | | | | C | C | | | | | | |
| NC_011770 | 4313080 | C | T | 78 | 0 | 8 | 0 | NON_SYNONYMOU | MODERAT | MISSENSE | Ctt/Ttt | L8F | PLES_39011 | | | T | T | T | T | | | T | T | T | T |
| NC_011770 | 4313095 | G | T | 77 | 0 | 10 | 0 | NON_SYNONYMOU | MODERAT | MISSENSE | Gtg/Ttg | V13L | PLES_39011 | | | T | T | T | T | T | T | T | T | T | T |
| NC_011770 | 4313149 | T | C | 78 | 0 | 8 | 0 | NON_SYNONYMOU | MODERAT | MISSENSE | Tct/Cct | S31P | PLES_39011 | | | C | C | C | C | | | C | C | C | C |
| NC_011770 | 4313222 | T | C | 73 | 0 | 2 | 0 | NON_SYNONYMOU | MODERAT | MISSENSE | gTa/gCa | V55A | PLES_39011 | C | C | | | | | | | | | | |
| NC_011770 | 4313224 | A | G | 74 | 0 | 2 | 0 | NON_SYNONYMOU | MODERAT | MISSENSE | Atc/Gtc | I56V | PLES_39011 | | | | | | | G | G | | | | |
| NC_011770 | 4313333 | G | A | 21 | 8 | 2 | 0 | NON_SYNONYMOU | MODERAT | MISSENSE | aGa/aAa | R92K | PLES_39011 | | | . | . | . | . | A | A | . | . | . | . |
| NC_011770 | 4313474 | T | C | 22 | 8 | 4 | 0 | NON_SYNONYMOU | MODERAT | MISSENSE | gTc/gCc | V139A | PLES_39011 | C | C | . | . | . | . | C | C | . | . | . | . |
| NC_011770 | 4313516 | A | G | 21 | 8 | 2 | 0 | NON_SYNONYMOU | MODERAT | MISSENSE | cAt/cGt | H153R | PLES_39011 | | | . | . | . | . | G | G | . | . | . | . |
| NC_011770 | 4313529 | A | G | 20 | 8 | 2 | 0 | NON_SYNONYMOU | MODERAT | MISSENSE | atA/atG | I157M | PLES_39011 | | | . | . | . | . | G | G | . | . | . | . |
| NC_011770 | 4313536 | G | A | 21 | 8 | 2 | 0 | NON_SYNONYMOU | MODERAT | MISSENSE | Gct/Act | A160T | PLES_39011 | | | . | . | . | . | A | A | . | . | . | . |
| NC_011770 | 4313585 | A | G | 33 | 0 | 2 | 0 | NON_SYNONYMOU | MODERAT | MISSENSE | gAg/gGg | E176G | PLES_39011 | | | | | | | G | G | | | | |
| NC_011770 | 4313607 | C | T | 36 | 0 | 10 | 0 | NON_SYNONYMOU | MODERAT | MISSENSE | caC/caA | H183Q | PLES_39011 | | | T | T | T | T | A | A | T | T | T | T |
| NC_011770 | 4313625 | T | G | 30 | 0 | 8 | 0 | NON_SYNONYMOU | MODERAT | MISSENSE | gaT/gaG | D189E | PLES_39011 | | | G | G | G | G | | | G | G | G | G |
| NC_011770 | 4313627 | G | A | 30 | 0 | 2 | 0 | NON_SYNONYMOU | MODERAT | MISSENSE | aGt/aAt | S190N | PLES_39011 | | | | | | | A | A | | | | |
| NC_011770 | 4313710 | G | T | 56 | 2 | 8 | 0 | NON_SYNONYMOU | MODERAT | MISSENSE | Gct/Tct | A218S | PLES_39011 | | | T | T | T | T | . | . | T | T | T | T |

| | | | | | | | | | | | | | | | | | | | | | | | | | |
|---|---|---|---|---|---|---|---|---|---|---|---|---|---|---|---|---|---|---|---|---|---|---|---|---|---|
| NC_011770 | 4313764 | T | G | 51 | 0 | 8 | 0 | NON_SYNONYMOU | MODERAT | MISSENSE | Tcc/Gcc | S236A | PLES_39011 | | | G | G | G | G | | | G | G | G | G |
| NC_011770 | 4313770 | T | G | 52 | 0 | 8 | 0 | NON_SYNONYMOU | MODERAT | MISSENSE | Ttt/Gtt | F238V | PLES_39011 | | | G | G | G | G | | | G | G | G | G |
| NC_011770 | 4313779 | A | G | 52 | 0 | 8 | 0 | NON_SYNONYMOU | MODERAT | MISSENSE | Atg/Gtg | M241V | PLES_39011 | | | G | G | G | G | | | G | G | G | G |
| NC_011770 | 4313781 | G | C | 52 | 0 | 8 | 0 | NON_SYNONYMOU | MODERAT | MISSENSE | atG/atC | M241I | PLES_39011 | | | C | C | C | C | | | C | C | C | C |
| NC_011770 | 4313786 | C | G | 52 | 0 | 8 | 0 | NON_SYNONYMOU | MODERAT | MISSENSE | aCa/aGa | T243R | PLES_39011 | | | G | G | G | G | | | G | G | G | G |
| NC_011770 | 4313851 | G | A | 70 | 0 | 2 | 0 | NON_SYNONYMOU | MODERAT | MISSENSE | Gat/Aat | D265N | PLES_39011 | | | | | | | | | | | A | A |
| NC_011770 | 4313867 | G | C | 64 | 0 | 10 | 0 | NON_SYNONYMOU | MODERAT | MISSENSE | gGc/gCc | G270A | PLES_39011 | | | C | C | C | C | C | C | C | C | C | C |
| NC_011770 | 4313912 | C | T | 71 | 0 | 2 | 0 | NON_SYNONYMOU | MODERAT | MISSENSE | aCc/aTc | T285I | PLES_39011 | T | T | | | | | | | | | | |
| NC_011770 | 4314022 | A | G | 79 | 0 | 2 | 0 | NON_SYNONYMOU | MODERAT | MISSENSE | Acc/Gcc | T322A | PLES_39011 | | | | | | | | | | | G | G |
| NC_011770 | 4314136 | C | G | 80 | 0 | 8 | 0 | NON_SYNONYMOU | MODERAT | MISSENSE | Cac/Gac | H360D | PLES_39011 | | | G | G | G | G | G | G | G | G | | |
| NC_011770 | 4314275 | G | C | 82 | 0 | 6 | 0 | NON_SYNONYMOU | MODERAT | MISSENSE | gGc/gCc | G21A | PLES_39021 | C | C | | | | | C | C | C | C | | |
| NC_011770 | 4314292 | G | C | 86 | 0 | 2 | 0 | NON_SYNONYMOU | MODERAT | MISSENSE | Ggc/Cgc | G27R | PLES_39021 | C | C | | | | | | | | | | |
| NC_011770 | 4315667 | T | C | 74 | 0 | 2 | 0 | NON_SYNONYMOU | MODERAT | MISSENSE | Ttc/Ctc | F171L | PLES_39041 | | | C | C | | | | | | | | |
| NC_011770 | 4316943 | C | A | 70 | 0 | 2 | 0 | START_LOST | HIGH | MISSENSE | atG/atT | M1I | PLES_39051 | | | | | | | | | A | A | | |
| NC_011770 | 4316623 | C | T | 62 | 0 | 6 | 0 | NON_SYNONYMOU | MODERAT | MISSENSE | cGg/cAg | R108Q | PLES_39051 | | | T | T | T | T | | | T | T | | |
| NC_011770 | 4316705 | G | T | 66 | 0 | 2 | 0 | NON_SYNONYMOU | MODERAT | MISSENSE | Ctg/Atg | L81M | PLES_39051 | | | | | | | T | T | | | | |
| NC_011770 | 4316818 | A | T | 69 | 0 | 2 | 0 | NON_SYNONYMOU | MODERAT | MISSENSE | cTg/cAg | L43Q | PLES_39051 | | | | | | | | | T | T | | |
| NC_011770 | 4316876 | C | T | 65 | 0 | 2 | 0 | NON_SYNONYMOU | MODERAT | MISSENSE | Gac/Aac | D24N | PLES_39051 | | | | | | | | | T | T | | |
| NC_011770 | 4319307 | A | G | 67 | 0 | 10 | 0 | STOP_LOST+SPLICE | HIGH | MISSENSE | tgA/tgG | *56W | PLES_39081 | G | G | G | G | G | G | G | G | G | G | | |
| NC_011770 | 4319150 | A | G | 68 | 0 | 10 | 0 | NON_SYNONYMOU | MODERAT | MISSENSE | cAc/cGc | H4R | PLES_39081 | G | G | G | G | G | G | G | G | G | G | | |
| NC_011770 | 4319174 | G | A | 70 | 0 | 8 | 0 | NON_SYNONYMOU | MODERAT | MISSENSE | aGc/aAc | S12N | PLES_39081 | A | A | A | A | A | A | A | A | | | | |
| NC_011770 | 4319221 | C | G | 69 | 0 | 8 | 0 | NON_SYNONYMOU | MODERAT | MISSENSE | Ctc/Gtc | L28V | PLES_39081 | G | G | G | G | G | G | G | G | | | | |
| NC_011770 | 4319230 | G | A | 69 | 0 | 10 | 0 | NON_SYNONYMOU | MODERAT | MISSENSE | Gcc/Acc | A31T | PLES_39081 | A | A | A | A | A | A | A | A | A | A | | |
| NC_011770 | 4319254 | G | A | 73 | 0 | 2 | 0 | NON_SYNONYMOU | MODERAT | MISSENSE | Ggc/Agc | G39S | PLES_39081 | A | A | | | | | | | | | | |
| NC_011770 | 4319159 | G | A | 75 | 0 | 1 | 0 | NON_SYNONYMOU | MODERAT | MISSENSE | cGg/cAg | R7Q | PLES_39081 | | | | | | | | A | | | | |
| NC_011770 | 4322449 | G | A | 63 | 0 | 2 | 0 | NON_SYNONYMOU | MODERAT | MISSENSE | Gcc/Acc | A135T | PLES_39111 | | | | | | | | | | | A | A |
| NC_011770 | 4323129 | T | C | 83 | 0 | 2 | 0 | NON_SYNONYMOU | MODERAT | MISSENSE | Ttc/Ctc | F76L | PLES_39121 | | | | | | | | | C | C | | |
| NC_011770 | 4323331 | C | G | 74 | 0 | 2 | 0 | NON_SYNONYMOU | MODERAT | MISSENSE | aCc/aGc | T143S | PLES_39121 | G | G | | | | | | | | | | |
| NC_011770 | 4324352 | G | T | 81 | 0 | 2 | 0 | NON_SYNONYMOU | MODERAT | MISSENSE | Gtg/Ttg | V88L | PLES_39131 | T | T | | | | | | | | | | |
| NC_011770 | 4324731 | A | G | 73 | 0 | 8 | 0 | NON_SYNONYMOU | MODERAT | MISSENSE | aAg/aGg | K214R | PLES_39131 | | | G | G | G | G | G | G | | | G | G |
| NC_011770 | 4325276 | A | G | 73 | 0 | 12 | 0 | NON_SYNONYMOU | MODERAT | MISSENSE | Acc/Gcc | T396A | PLES_39131 | G | G | G | G | G | G | G | G | G | G | G | G |
| NC_011770 | 4327290 | G | A | 66 | 0 | 2 | 0 | NON_SYNONYMOU | MODERAT | MISSENSE | Gcc/Acc | A45T | PLES_39151 | | | | | | | A | A | | | | |
| NC_011770 | 4327567 | T | C | 82 | 0 | 2 | 0 | NON_SYNONYMOU | MODERAT | MISSENSE | gTg/gCg | V137A | PLES_39151 | | | | | | | C | C | | | | |
| NC_011770 | 4327602 | A | G | 80 | 0 | 8 | 0 | NON_SYNONYMOU | MODERAT | MISSENSE | Acc/Gcc | T149A | PLES_39151 | | | G | G | G | G | G | G | | | G | G |
| NC_011770 | 4327659 | G | A | 84 | 0 | 2 | 0 | NON_SYNONYMOU | MODERAT | MISSENSE | Gcg/Acg | A168T | PLES_39151 | | | | | | | A | A | | | | |
| NC_011770 | 4327663 | A | T | 82 | 0 | 8 | 0 | NON_SYNONYMOU | MODERAT | MISSENSE | cAg/cTg | Q169L | PLES_39151 | | | T | T | T | T | T | T | | | T | T |
| NC_011770 | 4327833 | T | C | 76 | 0 | 2 | 0 | NON_SYNONYMOU | MODERAT | MISSENSE | Ttc/Ctc | F226L | PLES_39151 | C | C | | | | | | | | | | |
| NC_011770 | 4327891 | T | C | 73 | 0 | 2 | 0 | NON_SYNONYMOU | MODERAT | MISSENSE | gTc/gCc | V245A | PLES_39151 | C | C | | | | | | | | | | |
| NC_011770 | 4327926 | G | A | 77 | 0 | 2 | 0 | NON_SYNONYMOU | MODERAT | MISSENSE | Gcg/Acg | A257T | PLES_39151 | A | A | | | | | | | | | | |
| NC_011770 | 4328046 | G | C | 79 | 0 | 2 | 0 | NON_SYNONYMOU | MODERAT | MISSENSE | Gcg/Ccg | A297P | PLES_39151 | C | C | | | | | | | | | | |
| NC_011770 | 4328970 | A | G | 62 | 0 | 10 | 0 | NON_SYNONYMOU | MODERAT | MISSENSE | gAc/gGc | D278G | PLES_39161 | G | G | G | G | G | G | | | G | G | G | G |
| NC_011770 | 4329154 | G | T | 52 | 0 | 8 | 0 | NON_SYNONYMOU | MODERAT | MISSENSE | cGg/cTg | R60L | PLES_39171 | T | T | T | T | T | T | T | T | | | | |
| NC_011770 | 4329253 | G | A | 38 | 0 | 10 | 0 | NON_SYNONYMOU | MODERAT | MISSENSE | cGc/cAc | R93H | PLES_39171 | A | A | A | A | A | A | A | A | A | A | | |
| NC_011770 | 4329300 | G | A | 61 | 0 | 2 | 0 | NON_SYNONYMOU | MODERAT | MISSENSE | Gaa/Aaa | E109K | PLES_39171 | A | A | | | | | | | | | | |
| NC_011770 | 4329366 | A | C | 63 | 0 | 4 | 0 | NON_SYNONYMOU | MODERAT | MISSENSE | Atc/Ctc | I131L | PLES_39171 | C | C | | | | | | | | | C | C |
| NC_011770 | 4329573 | G | A | 63 | 0 | 2 | 0 | NON_SYNONYMOU | MODERAT | MISSENSE | Gcc/Acc | A200T | PLES_39171 | | | | | | | | | A | A | | |
| NC_011770 | 4330155 | G | A | 63 | 0 | 1 | 0 | NON_SYNONYMOU | MODERAT | MISSENSE | Ggc/Agc | G144S | PLES_39181 | | A | | | | | | | | | | |
| NC_011770 | 4329849 | A | G | 69 | 0 | 12 | 0 | NON_SYNONYMOU | MODERAT | MISSENSE | Atg/Gtg | M42V | PLES_39181 | G | G | G | G | G | G | G | G | G | G | G | G |
| NC_011770 | 4329933 | A | G | 65 | 0 | 2 | 0 | NON_SYNONYMOU | MODERAT | MISSENSE | Acc/Gcc | T70A | PLES_39181 | | | | | | | | | | | G | G |
| NC_011770 | 4330147 | C | T | 61 | 0 | 2 | 0 | NON_SYNONYMOU | MODERAT | MISSENSE | aCc/aTc | T141I | PLES_39181 | | | | | | | | | T | T | | |
| NC_011770 | 4330521 | G | A | 72 | 0 | 2 | 0 | NON_SYNONYMOU | MODERAT | MISSENSE | Gtc/Atc | V266I | PLES_39181 | | | | | | | | | | | A | A |
| NC_011770 | 4330621 | G | C | 68 | 0 | 2 | 0 | NON_SYNONYMOU | MODERAT | MISSENSE | gGt/gCt | G299A | PLES_39181 | C | C | | | | | | | | | | |
| NC_011770 | 4331225 | G | T | 73 | 0 | 2 | 0 | NON_SYNONYMOU | MODERAT | MISSENSE | caG/caT | Q500H | PLES_39181 | | | | | | | T | T | | | | |
| NC_011770 | 4332625 | T | G | 54 | 0 | 2 | 0 | NON_SYNONYMOU | MODERAT | MISSENSE | Agc/Cgc | S108R | PLES_39201 | | | | | | | | | | | G | G |
| NC_011770 | 4332782 | C | A | 66 | 0 | 4 | 0 | NON_SYNONYMOU | MODERAT | MISSENSE | gaG/gaT | E55D | PLES_39201 | | | | | | | | | A | A | A | A |
| NC_011770 | 4333025 | C | CT | 67 | 0 | 1 | 0 | FRAME_SHIFT | HIGH | | agc/aAgc | S406K? | PLES_39211 | | | | | | | CT | | | | | |
| NC_011770 | 4333011 | C | T | 68 | 0 | 2 | 0 | NON_SYNONYMOU | MODERAT | MISSENSE | Gcc/Acc | A411T | PLES_39211 | | | | | | | | | | | T | T |
| NC_011770 | 4333572 | C | A | 86 | 0 | 2 | 0 | NON_SYNONYMOU | MODERAT | MISSENSE | Gtg/Ttg | V224L | PLES_39211 | A | A | | | | | | | | | | |
| NC_011770 | 4333679 | T | C | 90 | 0 | 2 | 0 | NON_SYNONYMOU | MODERAT | MISSENSE | gAc/gGc | D188G | PLES_39211 | | | | | | | | | | | C | C |
| NC_011770 | 4334676 | C | A | 36 | 0 | 2 | 0 | NON_SYNONYMOU | MODERAT | MISSENSE | tCc/tAc | S30Y | PLES_39221 | | | | | | | | | | | A | A |
| NC_011770 | 4334708 | G | A | 35 | 0 | 4 | 0 | NON_SYNONYMOU | MODERAT | MISSENSE | Gcc/Acc | A41T | PLES_39221 | | | A | A | A | A | | | | | | |
| NC_011770 | 4335895 | A | G | 74 | 0 | 2 | 0 | NON_SYNONYMOU | MODERAT | MISSENSE | cAg/cGg | Q43R | PLES_39231 | | | | | | | G | G | | | | |

| | | | | | | | | | | | | | | | | | | | | | | | | | |
|---|---|---|---|---|---|---|---|---|---|---|---|---|---|---|---|---|---|---|---|---|---|---|---|---|---|
| NC_011770 | 4335930 | G | A | 74 | 0 | 2 | 0 | NON_SYNONYMOU | MODERAT | MISSENSE | Gtc/Atc | V55I | PLES_39231 | A | A | | | | | | | | | | |
| NC_011770 | 4336153 | G | A | 75 | 0 | 2 | 0 | NON_SYNONYMOU | MODERAT | MISSENSE | cGc/cAc | R129H | PLES_39231 | | | | | | | | | | | A | A |
| NC_011770 | 4336648 | A | G | 45 | 0 | 2 | 0 | NON_SYNONYMOU | MODERAT | MISSENSE | gAg/gGg | E294G | PLES_39231 | G | G | | | | | | | | | | |
| NC_011770 | 4336749 | G | A | 49 | 0 | 2 | 0 | NON_SYNONYMOU | MODERAT | MISSENSE | Gtc/Atc | V328I | PLES_39231 | | | | | | | A | A | | | | |
| NC_011770 | 4337389 | T | C | 66 | 0 | 4 | 0 | NON_SYNONYMOU | MODERAT | MISSENSE | Tat/Cat | Y149H | PLES_39241 | | | C | C | C | C | | | | | | |
| NC_011770 | 4337516 | G | A | 73 | 0 | 4 | 0 | NON_SYNONYMOU | MODERAT | MISSENSE | cGg/cAg | R191Q | PLES_39241 | | | A | A | A | A | | | | | | |
| NC_011770 | 4337791 | G | A | 73 | 0 | 2 | 0 | NON_SYNONYMOU | MODERAT | MISSENSE | Gcc/Acc | A283T | PLES_39241 | | | | | | | | | A | A | | |
| NC_011770 | 4337996 | C | T | 77 | 0 | 4 | 0 | NON_SYNONYMOU | MODERAT | MISSENSE | gCg/gTg | A351V | PLES_39241 | | | T | T | T | T | | | | | | |
| NC_011770 | 4338259 | G | A | 74 | 0 | 4 | 0 | NON_SYNONYMOU | MODERAT | MISSENSE | aGc/aAc | S19N | PLES_39251 | | | A | A | A | A | | | | | | |
| NC_011770 | 4338610 | A | G | 76 | 0 | 2 | 0 | NON_SYNONYMOU | MODERAT | MISSENSE | cAc/cGc | H136R | PLES_39251 | | | | | | | | | | | G | G |
| NC_011770 | 4338756 | C | A | 76 | 0 | 4 | 0 | NON_SYNONYMOU | MODERAT | MISSENSE | Cgc/Agc | R185S | PLES_39251 | | | A | A | A | A | | | | | | |
| NC_011770 | 4338765 | C | T | 79 | 0 | 2 | 0 | NON_SYNONYMOU | MODERAT | MISSENSE | Ccc/Tcc | P188S | PLES_39251 | | | | | | | | | T | T | | |
| NC_011770 | 4338770 | C | G | 77 | 0 | 4 | 0 | NON_SYNONYMOU | MODERAT | MISSENSE | gaC/gaG | D189E | PLES_39251 | | | G | G | G | G | | | | | | |
| NC_011770 | 4338812 | A | C | 76 | 0 | 4 | 0 | NON_SYNONYMOU | MODERAT | MISSENSE | gaA/gaC | E203D | PLES_39251 | | | C | C | C | C | | | | | | |
| NC_011770 | 4339233 | T | C | 72 | 0 | 1 | 0 | NON_SYNONYMOU | MODERAT | MISSENSE | Tgg/Cgg | W344R | PLES_39251 | | | | | | | C | | | | | |
| NC_011770 | 4339269 | C | T | 76 | 0 | 2 | 0 | NON_SYNONYMOU | MODERAT | MISSENSE | Ctc/Ttc | L356F | PLES_39251 | | | | | | | | | T | T | | |
| NC_011770 | 4339407 | T | G | 73 | 0 | 12 | 0 | NON_SYNONYMOU | MODERAT | MISSENSE | Tcc/Gcc | S402A | PLES_39251 | G | G | G | G | G | G | G | G | G | G | G | G |
| NC_011770 | 4339426 | G | A | 71 | 0 | 2 | 0 | NON_SYNONYMOU | MODERAT | MISSENSE | cGc/cAc | R408H | PLES_39251 | | | | | | | | | | | A | A |
| NC_011770 | 4339497 | T | C | 73 | 0 | 8 | 0 | NON_SYNONYMOU | MODERAT | MISSENSE | Tac/Cac | Y432H | PLES_39251 | C | C | C | C | C | C | | | | | C | C |
| NC_011770 | 4339574 | G | C | 69 | 0 | 2 | 0 | NON_SYNONYMOU | MODERAT | MISSENSE | caG/caC | Q457H | PLES_39251 | | | | | | | | | | | C | C |
| NC_011770 | 4339596 | G | A | 71 | 0 | 2 | 0 | NON_SYNONYMOU | MODERAT | MISSENSE | Gcg/Acg | A465T | PLES_39251 | | | | | | | A | A | | | | |
| NC_011770 | 4339239 | G | A | 74 | 0 | 1 | 0 | NON_SYNONYMOU | MODERAT | MISSENSE | Gga/Aga | G346R | PLES_39251 | | | | | | | | A | | | | |
| NC_011770 | 4339938 | C | T | 78 | 0 | 4 | 0 | NON_SYNONYMOU | MODERAT | MISSENSE | Gtc/Atc | V264I | PLES_39261 | | | | | | | T | T | | | T | T |
| NC_011770 | 4340007 | T | C | 77 | 0 | 4 | 0 | NON_SYNONYMOU | MODERAT | MISSENSE | Atc/Gtc | I241V | PLES_39261 | | | C | C | C | C | | | | | | |
| NC_011770 | 4340289 | C | T | 74 | 0 | 2 | 0 | NON_SYNONYMOU | MODERAT | MISSENSE | Gtc/Atc | V147I | PLES_39261 | T | T | | | | | | | | | | |
| NC_011770 | 4340466 | T | C | 73 | 0 | 1 | 0 | NON_SYNONYMOU | MODERAT | MISSENSE | Acc/Gcc | T88A | PLES_39261 | | | | | | | C | | | | | |
| NC_011770 | 4340562 | C | T | 74 | 0 | 2 | 0 | NON_SYNONYMOU | MODERAT | MISSENSE | Gcg/Acg | A56T | PLES_39261 | | | | | | | | | T | T | | |
| NC_011770 | 4341175 | C | T | 80 | 0 | 2 | 0 | NON_SYNONYMOU | MODERAT | MISSENSE | Gtg/Atg | V312M | PLES_39271 | | | | | | | | | | | T | T |
| NC_011770 | 4342724 | C | T | 87 | 0 | 1 | 0 | NON_SYNONYMOU | MODERAT | MISSENSE | tCg/tTg | S139L | PLES_39281 | | | | | | | | | | | | T |
| NC_011770 | 4343511 | C | A | 61 | 0 | 4 | 0 | NON_SYNONYMOU | MODERAT | MISSENSE | Gcg/Tcg | A9S | PLES_39291 | | | A | A | A | A | | | | | | |
| NC_011770 | 4351375 | G | A | 62 | 0 | 2 | 0 | NON_SYNONYMOU | MODERAT | MISSENSE | aGc/aAc | S139N | PLES_39391 | A | A | | | | | | | | | | |
| NC_011770 | 4351818 | G | T | 72 | 0 | 2 | 0 | NON_SYNONYMOU | MODERAT | MISSENSE | Ggc/Tgc | G287C | PLES_39391 | | | | | | | | | T | T | | |
| NC_011770 | 4352272 | C | T | 71 | 0 | 2 | 0 | NON_SYNONYMOU | MODERAT | MISSENSE | gCc/gTc | A438V | PLES_39391 | T | T | | | | | | | | | | |
| NC_011770 | 4352735 | A | G | 59 | 0 | 6 | 0 | NON_SYNONYMOU | MODERAT | MISSENSE | Aac/Gac | N101D | PLES_39401 | G | G | G | G | G | G | | | | | | |
| NC_011770 | 4352738 | C | G | 60 | 0 | 6 | 0 | NON_SYNONYMOU | MODERAT | MISSENSE | Cag/Gag | Q102E | PLES_39401 | G | G | G | G | G | G | | | | | | |
| NC_011770 | 4353662 | TGGCGGTC | T | 28 | 6 | 2 | 0 | CODON_CHANGE_ | MODERATE | | ggtcatggcg | GHGGG49 | PLES_39421 | . | . | . | . | . | . | | | T | T | | |
| NC_011770 | 4353973 | C | T | 51 | 0 | 2 | 0 | NON_SYNONYMOU | MODERAT | MISSENSE | Gtc/Atc | V240I | PLES_39431 | T | T | | | | | | | | | | |
| NC_011770 | 4354492 | G | C | 75 | 0 | 4 | 0 | NON_SYNONYMOU | MODERAT | MISSENSE | Ccg/Gcg | P67A | PLES_39431 | | | C | C | C | C | | | | | | |
| NC_011770 | 4354525 | T | C | 70 | 0 | 4 | 0 | NON_SYNONYMOU | MODERAT | MISSENSE | Acg/Gcg | T56A | PLES_39431 | | | C | C | C | C | | | | | | |
| NC_011770 | 4354570 | T | C | 77 | 0 | 4 | 0 | NON_SYNONYMOU | MODERAT | MISSENSE | Acc/Gcc | T41A | PLES_39431 | | | C | C | C | C | | | | | | |
| NC_011770 | 4354579 | C | T | 76 | 0 | 4 | 0 | NON_SYNONYMOU | MODERAT | MISSENSE | Ggc/Agc | G38S | PLES_39431 | | | T | T | T | T | | | | | | |
| NC_011770 | 4355092 | A | G | 68 | 0 | 4 | 0 | NON_SYNONYMOU | MODERAT | MISSENSE | gTg/gCg | V152A | PLES_39441 | | | G | G | G | G | | | | | | |
| NC_011770 | 4355393 | G | A | 68 | 0 | 6 | 0 | NON_SYNONYMOU | MODERAT | MISSENSE | Cat/Tat | H52Y | PLES_39441 | | | A | A | A | A | A | A | | | | |
| NC_011770 | 4355621 | G | T | 73 | 0 | 4 | 0 | NON_SYNONYMOU | MODERAT | MISSENSE | Ctg/Atg | L134M | PLES_39451 | | | T | T | T | T | | | | | | |
| NC_011770 | 4355646 | C | G | 74 | 0 | 4 | 0 | NON_SYNONYMOU | MODERAT | MISSENSE | aaG/aaC | K125N | PLES_39451 | | | G | G | G | G | | | | | | |
| NC_011770 | 4356566 | C | T | 78 | 0 | 2 | 0 | NON_SYNONYMOU | MODERAT | MISSENSE | Gat/Aat | D173N | PLES_39461 | T | T | | | | | | | | | | |
| NC_011770 | 4358147 | C | G | 86 | 0 | 2 | 0 | NON_SYNONYMOU | MODERAT | MISSENSE | Gag/Cag | E56Q | PLES_39481 | | | | | | | G | G | | | | |
| NC_011770 | 4358163 | C | A | 82 | 0 | 4 | 0 | NON_SYNONYMOU | MODERAT | MISSENSE | gaG/gaT | E50D | PLES_39481 | | | A | A | A | A | | | | | | |
| NC_011770 | 4359572 | T | C | 63 | 0 | 2 | 0 | NON_SYNONYMOU | MODERAT | MISSENSE | Acg/Gcg | T63A | PLES_39501 | | | | | | | C | C | | | | |
| NC_011770 | 4359581 | C | T | 64 | 0 | 2 | 0 | NON_SYNONYMOU | MODERAT | MISSENSE | Gcg/Acg | A60T | PLES_39501 | T | T | | | | | | | | | | |
| NC_011770 | 4362856 | T | C | 87 | 0 | 2 | 0 | NON_SYNONYMOU | MODERAT | MISSENSE | aAc/aGc | N239S | PLES_39541 | | | | | | | C | C | | | | |
| NC_011770 | 4363079 | C | T | 78 | 0 | 2 | 0 | NON_SYNONYMOU | MODERAT | MISSENSE | Ggt/Agt | G165S | PLES_39541 | | | | | | | T | T | | | | |
| NC_011770 | 4363081 | A | G | 78 | 0 | 2 | 0 | NON_SYNONYMOU | MODERAT | MISSENSE | gTt/gCt | V164A | PLES_39541 | | | | | | | G | G | | | | |
| NC_011770 | 4363100 | C | G | 79 | 0 | 2 | 0 | NON_SYNONYMOU | MODERAT | MISSENSE | Gtc/Ctc | V158L | PLES_39541 | | | | | | | G | G | | | | |
| NC_011770 | 4363139 | C | T | 85 | 0 | 2 | 0 | NON_SYNONYMOU | MODERAT | MISSENSE | Gcc/Acc | A145T | PLES_39541 | T | T | | | | | | | | | | |
| NC_011770 | 4364080 | A | C | 87 | 0 | 2 | 0 | NON_SYNONYMOU | MODERAT | MISSENSE | Tcc/Gcc | S603A | PLES_39551 | | | | | | | C | C | | | | |
| NC_011770 | 4365432 | G | A | 58 | 0 | 2 | 0 | NON_SYNONYMOU | MODERAT | MISSENSE | gCg/gTg | A152V | PLES_39551 | | | | | | | A | A | | | | |
| NC_011770 | 4365489 | T | G | 59 | 0 | 2 | 0 | NON_SYNONYMOU | MODERAT | MISSENSE | gAg/gCg | E133A | PLES_39551 | | | | | | | G | G | | | | |
| NC_011770 | 4364136 | A | G | 88 | 0 | 1 | 0 | NON_SYNONYMOU | MODERAT | MISSENSE | gTg/gCg | V584A | PLES_39551 | | | | | | | | G | | | | |
| NC_011770 | 4373322 | G | GC | 63 | 0 | 1 | 0 | FRAME_SHIFT | HIGH | | gcg/gGcg | A250G? | PLES_39621 | | | | | | | GC | | | | | |
| NC_011770 | 4373137 | C | G | 74 | 0 | 6 | 0 | NON_SYNONYMOU | MODERAT | MISSENSE | Ggc/Cgc | G312R | PLES_39621 | | | G | G | G | G | | | G | G | | |
| NC_011770 | 4373272 | C | A | 67 | 0 | 2 | 0 | NON_SYNONYMOU | MODERAT | MISSENSE | Ggg/Tgg | G267W | PLES_39621 | | | | | | | | | | | A | A |

| | | | | | | | | | | | | | | | | | | | | | | | | | |
|---|---|---|---|---|---|---|---|---|---|---|---|---|---|---|---|---|---|---|---|---|---|---|---|---|---|
| NC_011770 | 4373323 | C | T | 56 | 1 | 2 | 0 | NON_SYNONYMOU | MODERAT | MISSENSE | Gcg/Acg | A250T | PLES_39621 | | | | | | | . | | | | T | T |
| NC_011770 | 4373431 | G | C | 57 | 0 | 2 | 0 | NON_SYNONYMOU | MODERAT | MISSENSE | Ctg/Gtg | L214V | PLES_39621 | | | | | | | | | | | C | C |
| NC_011770 | 4373977 | C | G | 44 | 0 | 12 | 0 | NON_SYNONYMOU | MODERAT | MISSENSE | Gcg/Ccg | A32P | PLES_39621 | G | G | G | G | G | G | G | G | G | G | G | G |
| NC_011770 | 4374145 | G | GC | 40 | 0 | 1 | 0 | FRAME_SHIFT | HIGH | | ggc/ggGc | G405G? | PLES_39631 | | | | | | | | GC | | | | |
| NC_011770 | 4374269 | A | T | 41 | 0 | 8 | 0 | NON_SYNONYMOU | MODERAT | MISSENSE | cTg/cAg | L354Q | PLES_39631 | T | T | T | T | T | T | | | | | T | T |
| NC_011770 | 4374302 | A | G | 45 | 0 | 8 | 0 | NON_SYNONYMOU | MODERAT | MISSENSE | gTa/gCa | V353A | PLES_39631 | G | G | G | G | G | G | | | | | G | G |
| NC_011770 | 4374432 | C | G | 59 | 0 | 2 | 0 | NON_SYNONYMOU | MODERAT | MISSENSE | Gtt/Ctt | V310L | PLES_39631 | | | | | | | | | | | G | G |
| NC_011770 | 4374629 | A | C | 71 | 0 | 12 | 0 | NON_SYNONYMOU | MODERAT | MISSENSE | aTc/aGc | I244S | PLES_39631 | C | C | C | C | C | C | C | C | C | C | C | C |
| NC_011770 | 4374744 | A | C | 69 | 0 | 2 | 0 | NON_SYNONYMOU | MODERAT | MISSENSE | Tcg/Gcg | S206A | PLES_39631 | | | | | | | | | | | C | C |
| NC_011770 | 4374771 | C | A | 69 | 0 | 2 | 0 | NON_SYNONYMOU | MODERAT | MISSENSE | Gcg/Tcg | A197S | PLES_39631 | | | | | | | | | | | A | A |
| NC_011770 | 4374791 | T | G | 65 | 0 | 2 | 0 | NON_SYNONYMOU | MODERAT | MISSENSE | cAa/cCa | Q190P | PLES_39631 | | | | | | | | | | | G | G |
| NC_011770 | 4374807 | T | C | 64 | 0 | 2 | 0 | NON_SYNONYMOU | MODERAT | MISSENSE | Aaa/Gaa | K185E | PLES_39631 | | | | | | | | | | | C | C |
| NC_011770 | 4374813 | C | T | 64 | 0 | 2 | 0 | NON_SYNONYMOU | MODERAT | MISSENSE | Gcg/Acg | A183T | PLES_39631 | | | | | | | | | | | T | T |
| NC_011770 | 4375163 | G | A | 74 | 0 | 2 | 0 | NON_SYNONYMOU | MODERAT | MISSENSE | gCg/gTg | A66V | PLES_39631 | | | | | | | | | | | A | A |
| NC_011770 | 4374217 | T | TCCA | 35 | 0 | 4 | 0 | CODON_CHANGE_ | MODERATE | | gga/ggTGC | G381GG | PLES_39631 | TCCA | TCCA | | TCCA | | | | | | | TCCA | |
| NC_011770 | 4381559 | C | T | 74 | 0 | 4 | 0 | NON_SYNONYMOU | MODERAT | MISSENSE | atG/atA | M283I | PLES_39721 | | | | | | | T | T | | | T | T |
| NC_011770 | 4381717 | C | T | 65 | 0 | 2 | 0 | NON_SYNONYMOU | MODERAT | MISSENSE | Gtc/Atc | V231I | PLES_39721 | | | | | | | | | | | T | T |
| NC_011770 | 4381777 | T | G | 66 | 0 | 2 | 0 | NON_SYNONYMOU | MODERAT | MISSENSE | Acc/Ccc | T211P | PLES_39721 | | | | | | | | | | | G | G |
| NC_011770 | 4381798 | C | T | 63 | 0 | 2 | 0 | NON_SYNONYMOU | MODERAT | MISSENSE | Ggc/Agc | G204S | PLES_39721 | | | | | | | | | | | T | T |
| NC_011770 | 4381864 | T | C | 59 | 0 | 2 | 0 | NON_SYNONYMOU | MODERAT | MISSENSE | Acg/Gcg | T182A | PLES_39721 | | | | | | | | | | | C | C |
| NC_011770 | 4382032 | A | T | 60 | 0 | 6 | 0 | NON_SYNONYMOU | MODERAT | MISSENSE | Ttg/Atg | L126M | PLES_39721 | | | T | T | T | T | | | | | T | T |
| NC_011770 | 4382161 | T | C | 55 | 0 | 6 | 0 | NON_SYNONYMOU | MODERAT | MISSENSE | Agg/Ggg | R83G | PLES_39721 | | | C | C | C | C | | | | | C | C |
| NC_011770 | 4382163 | G | A | 56 | 0 | 2 | 0 | NON_SYNONYMOU | MODERAT | MISSENSE | gCc/gTc | A82V | PLES_39721 | | | | | | | A | A | | | | |
| NC_011770 | 4382187 | A | G | 54 | 0 | 8 | 0 | NON_SYNONYMOU | MODERAT | MISSENSE | gTc/gCc | V74A | PLES_39721 | | | G | G | G | G | G | G | | | G | G |
| NC_011770 | 4382192 | G | C | 55 | 0 | 6 | 0 | NON_SYNONYMOU | MODERAT | MISSENSE | gaC/gaG | D72E | PLES_39721 | | | C | C | C | C | | | | | C | C |
| NC_011770 | 4382233 | C | A | 60 | 0 | 6 | 0 | NON_SYNONYMOU | MODERAT | MISSENSE | Gcg/Tcg | A59S | PLES_39721 | A | A | A | A | A | A | | | | | | |
| NC_011770 | 4383228 | T | C | 74 | 0 | 10 | 0 | NON_SYNONYMOU | MODERAT | MISSENSE | gTc/gCc | V12A | PLES_39751 | C | C | C | C | C | C | | | C | C | C | C |
| NC_011770 | 4384403 | G | A | 84 | 0 | 1 | 0 | NON_SYNONYMOU | MODERAT | MISSENSE | cCg/cTg | P218L | PLES_39761 | | A | | | | | | | | | | |
| NC_011770 | 4384623 | G | A | 84 | 0 | 1 | 0 | NON_SYNONYMOU | MODERAT | MISSENSE | Ccg/Tcg | P145S | PLES_39761 | | | | | | | A | | | | | |
| NC_011770 | 4386884 | T | A | 94 | 0 | 4 | 0 | NON_SYNONYMOU | MODERAT | MISSENSE | Acc/Tcc | T685S | PLES_39781 | | | A | A | A | A | | | | | | |
| NC_011770 | 4387375 | C | A | 73 | 0 | 2 | 0 | NON_SYNONYMOU | MODERAT | MISSENSE | tGg/tTg | W521L | PLES_39781 | | | A | A | | | | | | | | |
| NC_011770 | 4387434 | G | T | 67 | 0 | 6 | 0 | NON_SYNONYMOU | MODERAT | MISSENSE | gaC/gaA | D501E | PLES_39781 | | | T | T | T | T | | | | | T | T |
| NC_011770 | 4387499 | G | C | 66 | 0 | 2 | 0 | NON_SYNONYMOU | MODERAT | MISSENSE | Ctc/Gtc | L480V | PLES_39781 | | | | | | | | | | | C | C |
| NC_011770 | 4388446 | A | G | 64 | 0 | 12 | 0 | NON_SYNONYMOU | MODERAT | MISSENSE | aTc/aCc | I164T | PLES_39781 | G | G | G | G | G | G | G | G | G | G | G | G |
| NC_011770 | 4388659 | G | C | 71 | 0 | 2 | 0 | NON_SYNONYMOU | MODERAT | MISSENSE | aCc/aGc | T93S | PLES_39781 | | | | | | | C | C | | | | |
| NC_011770 | 4388940 | T | C | 61 | 0 | 2 | 0 | NON_SYNONYMOU | MODERAT | MISSENSE | cAg/cGg | Q384R | PLES_39791 | | | | | | | | | | | C | C |
| NC_011770 | 4388950 | C | T | 62 | 0 | 2 | 0 | NON_SYNONYMOU | MODERAT | MISSENSE | Gcg/Acg | A381T | PLES_39791 | | | | | | | T | T | | | | |
| NC_011770 | 4389612 | C | A | 57 | 0 | 2 | 0 | NON_SYNONYMOU | MODERAT | MISSENSE | cGc/cTc | R160L | PLES_39791 | | | | | | | | | | | A | A |
| NC_011770 | 4390021 | T | C | 17 | 0 | 12 | 0 | NON_SYNONYMOU | MODERAT | MISSENSE | Agc/Ggc | S24G | PLES_39791 | C | C | C | C | C | C | C | C | C | C | C | C |
| NC_011770 | 4390280 | A | C | 10 | 2 | 1 | 0 | NON_SYNONYMOU | MODERAT | MISSENSE | Acc/Ccc | T4P | PLES_39801 | | | | | | | C | . | . | | | |
| NC_011770 | 4391223 | G | T | 86 | 0 | 2 | 0 | NON_SYNONYMOU | MODERAT | MISSENSE | cGg/cTg | R80L | PLES_39811 | | | | | | | T | T | | | | |
| NC_011770 | 4391466 | C | T | 80 | 0 | 4 | 0 | NON_SYNONYMOU | MODERAT | MISSENSE | gCg/gTg | A161V | PLES_39811 | | | | | | | T | T | | | T | T |
| NC_011770 | 4391912 | A | G | 65 | 0 | 12 | 0 | NON_SYNONYMOU | MODERAT | MISSENSE | Acc/Gcc | T310A | PLES_39811 | G | G | G | G | G | G | G | G | G | G | G | G |
| NC_011770 | 4392113 | A | C | 69 | 0 | 12 | 0 | NON_SYNONYMOU | MODERAT | MISSENSE | Atg/Ctg | M377L | PLES_39811 | C | C | C | C | C | C | C | C | C | C | C | C |
| NC_011770 | 4392509 | A | G | 80 | 0 | 6 | 0 | NON_SYNONYMOU | MODERAT | MISSENSE | Acg/Gcg | T509A | PLES_39811 | G | G | | | | | G | G | | | G | G |
| NC_011770 | 4392626 | G | A | 92 | 0 | 1 | 0 | NON_SYNONYMOU | MODERAT | MISSENSE | Gtc/Atc | V548I | PLES_39811 | | | | | | | A | | | | | |
| NC_011770 | 4396433 | G | A | 83 | 0 | 2 | 0 | NON_SYNONYMOU | MODERAT | MISSENSE | Ccg/Tcg | P487S | PLES_39851 | | | | | | | | | | | A | A |
| NC_011770 | 4397061 | G | T | 99 | 0 | 2 | 0 | NON_SYNONYMOU | MODERAT | MISSENSE | gaC/gaA | D277E | PLES_39851 | | | | | | | T | T | | | | |
| NC_011770 | 4398344 | C | T | 48 | 0 | 1 | 0 | NON_SYNONYMOU | MODERAT | MISSENSE | Gtc/Atc | V49I | PLES_39861 | | | | | | | | | | | | T |
| NC_011770 | 4400307 | T | C | 59 | 2 | 10 | 0 | NON_SYNONYMOU | MODERAT | MISSENSE | cAa/cGa | Q538R | PLES_39881 | C | C | C | C | C | C | . | . | C | C | C | C |
| NC_011770 | 4400952 | G | A | 65 | 0 | 2 | 0 | NON_SYNONYMOU | MODERAT | MISSENSE | gCg/gTg | A323V | PLES_39881 | | | | | | | | | A | A | | |
| NC_011770 | 4401057 | C | T | 68 | 0 | 2 | 0 | NON_SYNONYMOU | MODERAT | MISSENSE | aGg/aAg | R288K | PLES_39881 | | | | | | | T | T | | | | |
| NC_011770 | 4401115 | T | G | 64 | 0 | 2 | 0 | NON_SYNONYMOU | MODERAT | MISSENSE | Agc/Cgc | S269R | PLES_39881 | | | | | | | | | G | G | | |
| NC_011770 | 4401120 | T | C | 62 | 0 | 1 | 0 | NON_SYNONYMOU | MODERAT | MISSENSE | cAg/cGg | Q267R | PLES_39881 | | | | | | | C | | | | | |
| NC_011770 | 4401124 | G | A | 63 | 0 | 2 | 0 | NON_SYNONYMOU | MODERAT | MISSENSE | Cgc/Tgc | R266C | PLES_39881 | | | | | | | | | A | A | | |
| NC_011770 | 4401911 | A | C | 67 | 0 | 6 | 0 | NON_SYNONYMOU | MODERAT | MISSENSE | aaT/aaG | N3K | PLES_39881 | C | C | C | C | C | C | | | | | | |
| NC_011770 | 4406656 | A | T | 75 | 0 | 2 | 0 | NON_SYNONYMOU | MODERAT | MISSENSE | gAc/gTc | D4V | PLES_39931 | | | | | | | | | | | T | T |
| NC_011770 | 4406808 | G | A | 72 | 0 | 8 | 0 | NON_SYNONYMOU | MODERAT | MISSENSE | Gag/Aag | E55K | PLES_39931 | A | A | A | A | A | A | | | A | A | | |
| NC_011770 | 4407091 | A | G | 74 | 0 | 4 | 0 | NON_SYNONYMOU | MODERAT | MISSENSE | aAc/aGc | N3S | PLES_39941 | | | G | G | G | G | | | | | | |
| NC_011770 | 4407174 | G | A | 69 | 0 | 2 | 0 | NON_SYNONYMOU | MODERAT | MISSENSE | Gtc/Atc | V31I | PLES_39941 | A | A | | | | | | | | | | |
| NC_011770 | 4407565 | C | T | 77 | 0 | 2 | 0 | NON_SYNONYMOU | MODERAT | MISSENSE | aCc/aTc | T161I | PLES_39941 | | | | | | | | | | | T | T |
| NC_011770 | 4408269 | G | A | 90 | 0 | 1 | 0 | NON_SYNONYMOU | MODERAT | MISSENSE | Gac/Aac | D396N | PLES_39941 | | | | | | | A | | | | | |

| | | | | | | | | | | | | | | | | | | | | | | | | | |
|---|---|---|---|---|---|---|---|---|---|---|---|---|---|---|---|---|---|---|---|---|---|---|---|---|---|
| NC_011770 | 4410696 | G | A | 57 | 0 | 2 | 0 | NON_SYNONYMOU | MODERAT | MISSENSE | Gcc/Acc | A181T | PLES_39961 | | | | | | | | | | | A | A |
| NC_011770 | 4411422 | G | A | 67 | 0 | 1 | 0 | NON_SYNONYMOU | MODERAT | MISSENSE | Gag/Aag | E423K | PLES_39961 | | | | | | | | | | | | A |
| NC_011770 | 4412136 | G | T | 52 | 0 | 2 | 0 | NON_SYNONYMOU | MODERAT | MISSENSE | gaG/gaT | E122D | PLES_39971 | T | T | | | | | | | | | | |
| NC_011770 | 4412185 | G | A | 51 | 0 | 8 | 0 | NON_SYNONYMOU | MODERAT | MISSENSE | Gtc/Atc | V139I | PLES_39971 | A | A | A | A | A | A | A | A | | | | |
| NC_011770 | 4412467 | A | G | 73 | 0 | 6 | 0 | NON_SYNONYMOU | MODERAT | MISSENSE | Acc/Gcc | T233A | PLES_39971 | G | G | G | G | G | G | | | | | | |
| NC_011770 | 4412669 | A | G | 78 | 0 | 2 | 0 | NON_SYNONYMOU | MODERAT | MISSENSE | gAc/gGc | D300G | PLES_39971 | | | | | | | | | | | G | G |
| NC_011770 | 4412836 | G | A | 68 | 0 | 4 | 0 | NON_SYNONYMOU | MODERAT | MISSENSE | Gag/Aag | E356K | PLES_39971 | | | A | A | A | A | | | | | | |
| NC_011770 | 4413119 | G | A | 86 | 0 | 4 | 0 | NON_SYNONYMOU | MODERAT | MISSENSE | aGc/aAc | S450N | PLES_39971 | | | A | A | A | A | | | | | | |
| NC_011770 | 4413528 | G | A | 79 | 0 | 1 | 0 | NON_SYNONYMOU | MODERAT | MISSENSE | Gcc/Acc | A111T | PLES_39981 | | A | | | | | | | | | | |
| NC_011770 | 4413366 | C | T | 80 | 0 | 2 | 0 | NON_SYNONYMOU | MODERAT | MISSENSE | Cgc/Tgc | R57C | PLES_39981 | T | T | | | | | | | | | | |
| NC_011770 | 4413391 | C | A | 75 | 0 | 12 | 0 | NON_SYNONYMOU | MODERAT | MISSENSE | gCa/gAa | A65E | PLES_39981 | A | A | A | A | A | A | A | A | A | A | A | A |
| NC_011770 | 4413514 | A | G | 76 | 0 | 2 | 0 | NON_SYNONYMOU | MODERAT | MISSENSE | aAc/aGc | N106S | PLES_39981 | | | | | | | G | G | | | | |
| NC_011770 | 4413804 | G | T | 70 | 0 | 8 | 0 | NON_SYNONYMOU | MODERAT | MISSENSE | Gcc/Tcc | A203S | PLES_39981 | | | T | T | T | T | T | T | T | T | | |
| NC_011770 | 4413823 | T | G | 67 | 0 | 12 | 0 | NON_SYNONYMOU | MODERAT | MISSENSE | cTc/cGc | L209R | PLES_39981 | G | G | G | G | G | G | G | G | G | G | G | G |
| NC_011770 | 4414664 | G | A | 54 | 0 | 1 | 0 | NON_SYNONYMOU | MODERAT | MISSENSE | Cgc/Tgc | R162C | PLES_40001 | A | | | | | | | | | | | |
| NC_011770 | 4415315 | G | A | 65 | 0 | 4 | 0 | NON_SYNONYMOU | MODERAT | MISSENSE | Gtc/Atc | V22I | PLES_40011 | A | A | | | | | A | A | | | | |
| NC_011770 | 4415501 | C | G | 74 | 0 | 2 | 0 | NON_SYNONYMOU | MODERAT | MISSENSE | Cta/Gta | L84V | PLES_40011 | | | | | G | G | | | | | | |
| NC_011770 | 4415834 | A | G | 78 | 0 | 6 | 0 | NON_SYNONYMOU | MODERAT | MISSENSE | Agc/Ggc | S195G | PLES_40011 | | | G | G | G | G | | | G | G | | |
| NC_011770 | 4416372 | T | C | 72 | 0 | 2 | 0 | NON_SYNONYMOU | MODERAT | MISSENSE | tTc/tCc | F374S | PLES_40011 | | | | | | | | | | | C | C |
| NC_011770 | 4416635 | C | A | 85 | 0 | 2 | 0 | NON_SYNONYMOU | MODERAT | MISSENSE | Gtg/Ttg | V228L | PLES_40021 | | | | | | | | | | | A | A |
| NC_011770 | 4416659 | T | C | 81 | 0 | 4 | 0 | NON_SYNONYMOU | MODERAT | MISSENSE | Acc/Gcc | T220A | PLES_40021 | | | C | C | C | C | | | | | | |
| NC_011770 | 4417598 | C | G | 91 | 0 | 4 | 0 | NON_SYNONYMOU | MODERAT | MISSENSE | agG/agC | R337S | PLES_40031 | | | G | G | G | G | | | | | | |
| NC_011770 | 4422065 | G | GC | 66 | 0 | 1 | 0 | FRAME_SHIFT | HIGH | | gcg/gcCg | A733A? | PLES_40051 | | | | GC | | | | | | | | |
| NC_011770 | 4419956 | G | A | 71 | 0 | 2 | 0 | NON_SYNONYMOU | MODERAT | MISSENSE | Gcc/Acc | A30T | PLES_40051 | | | | | | | A | A | | | | |
| NC_011770 | 4419965 | G | A | 70 | 0 | 4 | 0 | NON_SYNONYMOU | MODERAT | MISSENSE | Gct/Act | A33T | PLES_40051 | | | A | A | A | A | | | | | | |
| NC_011770 | 4419998 | C | G | 70 | 0 | 4 | 0 | NON_SYNONYMOU | MODERAT | MISSENSE | Ccc/Gcc | P44A | PLES_40051 | | | G | G | G | G | | | | | | |
| NC_011770 | 4420941 | A | G | 74 | 0 | 12 | 0 | NON_SYNONYMOU | MODERAT | MISSENSE | aAc/aGc | N358S | PLES_40051 | G | G | G | G | G | G | G | G | G | G | G | G |
| NC_011770 | 4421018 | G | A | 73 | 0 | 2 | 0 | NON_SYNONYMOU | MODERAT | MISSENSE | Gcc/Acc | A384T | PLES_40051 | | | | | | | A | A | | | | |
| NC_011770 | 4421049 | G | A | 73 | 0 | 1 | 0 | NON_SYNONYMOU | MODERAT | MISSENSE | cGc/cAc | R394H | PLES_40051 | | | | | | | A | | | | | |
| NC_011770 | 4421435 | G | A | 54 | 0 | 2 | 0 | NON_SYNONYMOU | MODERAT | MISSENSE | Gcc/Acc | A523T | PLES_40051 | | | | | | | A | A | | | | |
| NC_011770 | 4421507 | C | A | 51 | 0 | 4 | 0 | NON_SYNONYMOU | MODERAT | MISSENSE | Ctg/Atg | L547M | PLES_40051 | | | A | A | A | A | | | | | | |
| NC_011770 | 4422257 | G | C | 71 | 0 | 4 | 0 | NON_SYNONYMOU | MODERAT | MISSENSE | Gcc/Ccc | A797P | PLES_40051 | | | C | C | C | C | | | | | | |
| NC_011770 | 4422261 | G | A | 71 | 0 | 2 | 0 | NON_SYNONYMOU | MODERAT | MISSENSE | aGc/aAc | S798N | PLES_40051 | | | | | | | A | A | | | | |
| NC_011770 | 4422468 | A | G | 70 | 0 | 8 | 0 | NON_SYNONYMOU | MODERAT | MISSENSE | gAc/gGc | D37G | PLES_40061 | G | G | | | | | G | G | G | G | G | G |
| NC_011770 | 4422548 | C | G | 75 | 0 | 6 | 0 | NON_SYNONYMOU | MODERAT | MISSENSE | Cag/Gag | Q64E | PLES_40061 | | | | | | | G | G | G | G | G | G |
| NC_011770 | 4422570 | A | G | 73 | 0 | 10 | 0 | NON_SYNONYMOU | MODERAT | MISSENSE | gAg/gGg | E71G | PLES_40061 | G | G | G | G | G | G | G | G | | | G | G |
| NC_011770 | 4422873 | A | G | 62 | 0 | 2 | 0 | NON_SYNONYMOU | MODERAT | MISSENSE | cAg/cGg | Q172R | PLES_40061 | | | | | G | G | | | | | | |
| NC_011770 | 4423083 | G | A | 53 | 0 | 2 | 0 | NON_SYNONYMOU | MODERAT | MISSENSE | cGc/cAc | R242H | PLES_40061 | A | A | | | | | | | | | | |
| NC_011770 | 4423203 | C | T | 53 | 0 | 6 | 0 | NON_SYNONYMOU | MODERAT | MISSENSE | tCg/tTg | S282L | PLES_40061 | | | T | T | T | T | T | T | | | | |
| NC_011770 | 4423462 | A | G | 43 | 0 | 12 | 0 | NON_SYNONYMOU | MODERAT | MISSENSE | Atc/Gtc | I43V | PLES_40071 | G | G | G | G | G | G | G | G | G | G | G | G |
| NC_011770 | 4423736 | C | A | 45 | 0 | 10 | 0 | NON_SYNONYMOU | MODERAT | MISSENSE | gCc/gAc | A134D | PLES_40071 | A | A | A | A | A | A | A | A | A | A | | |
| NC_011770 | 4423846 | G | A | 57 | 0 | 2 | 0 | NON_SYNONYMOU | MODERAT | MISSENSE | Gag/Aag | E171K | PLES_40071 | | | | | | | | | | | A | A |
| NC_011770 | 4423972 | C | T | 62 | 0 | 2 | 0 | NON_SYNONYMOU | MODERAT | MISSENSE | Cac/Tac | H213Y | PLES_40071 | | | | | | | | | | | T | T |
| NC_011770 | 4424021 | A | G | 66 | 0 | 4 | 0 | NON_SYNONYMOU | MODERAT | MISSENSE | gAc/gGc | D229G | PLES_40071 | | | | | | | | | G | G | G | G |
| NC_011770 | 4423397 | G | A | 44 | 0 | 1 | 0 | NON_SYNONYMOU | MODERAT | MISSENSE | cGc/cAc | R21H | PLES_40071 | | | | | | | | A | | | | |
| NC_011770 | 4425026 | C | T | 72 | 0 | 1 | 0 | STOP_GAINED | HIGH | NONSENSE | tgG/tgA | W227* | PLES_40081 | | | | | | | T | | | | | |
| NC_011770 | 4424964 | T | C | 71 | 0 | 2 | 0 | NON_SYNONYMOU | MODERAT | MISSENSE | cAg/cGg | Q248R | PLES_40081 | C | C | | | | | | | | | | |
| NC_011770 | 4425270 | G | A | 66 | 0 | 10 | 0 | NON_SYNONYMOU | MODERAT | MISSENSE | aCc/aTc | T146I | PLES_40081 | A | A | A | A | A | A | | | A | A | A | A |
| NC_011770 | 4425295 | C | G | 66 | 0 | 2 | 0 | NON_SYNONYMOU | MODERAT | MISSENSE | Gaa/Caa | E138Q | PLES_40081 | | | | | | | | | G | G | | |
| NC_011770 | 4425391 | G | C | 65 | 0 | 12 | 0 | NON_SYNONYMOU | MODERAT | MISSENSE | Cac/Gac | H106D | PLES_40081 | C | C | C | C | C | C | C | C | C | C | C | C |
| NC_011770 | 4425479 | C | G | 70 | 0 | 2 | 0 | NON_SYNONYMOU | MODERAT | MISSENSE | caG/caC | Q76H | PLES_40081 | G | G | | | | | | | | | | |
| NC_011770 | 4425526 | C | G | 67 | 0 | 12 | 0 | NON_SYNONYMOU | MODERAT | MISSENSE | Gtg/Ctg | V61L | PLES_40081 | G | G | G | G | G | G | G | G | G | G | G | G |
| NC_011770 | 4425553 | C | T | 71 | 0 | 10 | 0 | NON_SYNONYMOU | MODERAT | MISSENSE | Gcg/Acg | A52T | PLES_40081 | T | T | T | T | T | T | T | T | | | T | T |
| NC_011770 | 4425669 | G | A | 76 | 0 | 1 | 0 | NON_SYNONYMOU | MODERAT | MISSENSE | gCc/gTc | A13V | PLES_40081 | | | | | | | A | | | | | |
| NC_011770 | 4426593 | C | T | 58 | 0 | 10 | 0 | NON_SYNONYMOU | MODERAT | MISSENSE | gGc/gAc | G208D | PLES_40101 | T | T | T | T | T | T | T | T | T | T | | |
| NC_011770 | 4426718 | G | T | 68 | 0 | 2 | 0 | NON_SYNONYMOU | MODERAT | MISSENSE | gaC/gaA | D166E | PLES_40101 | T | T | | | | | | | | | | |
| NC_011770 | 4426776 | G | A | 65 | 0 | 8 | 0 | NON_SYNONYMOU | MODERAT | MISSENSE | gCg/gTg | A147V | PLES_40101 | | | A | A | A | A | A | A | A | A | | |
| NC_011770 | 4426923 | G | T | 68 | 0 | 2 | 0 | NON_SYNONYMOU | MODERAT | MISSENSE | gCg/gAg | A98E | PLES_40101 | | | | | | | | | T | T | | |
| NC_011770 | 4426939 | C | T | 66 | 0 | 4 | 0 | NON_SYNONYMOU | MODERAT | MISSENSE | Gcc/Acc | A93T | PLES_40101 | T | T | | | | | T | T | | | | |
| NC_011770 | 4427512 | T | A | 69 | 0 | 4 | 0 | NON_SYNONYMOU | MODERAT | MISSENSE | Tgc/Agc | C69S | PLES_40111 | | | A | A | A | A | | | | | | |
| NC_011770 | 4427752 | G | A | 83 | 0 | 2 | 0 | NON_SYNONYMOU | MODERAT | MISSENSE | Gac/Aac | D149N | PLES_40111 | A | A | | | | | | | | | | |
| NC_011770 | 4427507 | C | T | 72 | 0 | 1 | 0 | NON_SYNONYMOU | MODERAT | MISSENSE | aCc/aTc | T67I | PLES_40111 | | | | | | | | | | | | T |

| | | | | | | | | | | | | | | | | | | | | | | | | | |
|---|---|---|---|---|---|---|---|---|---|---|---|---|---|---|---|---|---|---|---|---|---|---|---|---|---|
| NC_011770 | 4428300 | G | A | 85 | 0 | 4 | 0 | NON_SYNONYMOU | MODERAT | MISSENSE | Ggc/Agc | G84S | PLES_40121 | | | A | A | A | A | | | | | | |
| NC_011770 | 4428439 | C | T | 70 | 0 | 8 | 0 | NON_SYNONYMOU | MODERAT | MISSENSE | gCg/gTg | A130V | PLES_40121 | T | T | T | T | T | T | | | | | T | T |
| NC_011770 | 4428441 | A | G | 70 | 0 | 8 | 0 | NON_SYNONYMOU | MODERAT | MISSENSE | Acg/Gcg | T131A | PLES_40121 | G | G | G | G | G | G | | | | | G | G |
| NC_011770 | 4428444 | C | T | 71 | 0 | 4 | 0 | NON_SYNONYMOU | MODERAT | MISSENSE | Ccg/Tcg | P132S | PLES_40121 | T | T | | | | | | | | | T | T |
| NC_011770 | 4428521 | C | T | 78 | 0 | 4 | 0 | NON_SYNONYMOU | MODERAT | MISSENSE | Gcg/Acg | A1086T | PLES_40131 | T | T | | | | | | | | | T | T |
| NC_011770 | 4428529 | G | A | 78 | 0 | 2 | 0 | NON_SYNONYMOU | MODERAT | MISSENSE | gCg/gTg | A1083V | PLES_40131 | | | | | | | | | A | A | | |
| NC_011770 | 4428649 | T | G | 89 | 0 | 2 | 0 | NON_SYNONYMOU | MODERAT | MISSENSE | aAg/aCg | K1043T | PLES_40131 | | | | | | | G | G | | | | |
| NC_011770 | 4428673 | G | A | 83 | 0 | 4 | 0 | NON_SYNONYMOU | MODERAT | MISSENSE | gCg/gTg | A1035V | PLES_40131 | A | A | | | | | | | | | A | A |
| NC_011770 | 4429177 | T | G | 75 | 0 | 4 | 0 | NON_SYNONYMOU | MODERAT | MISSENSE | gAg/gCg | E867A | PLES_40131 | | | G | G | G | G | | | | | | |
| NC_011770 | 4429183 | G | A | 76 | 0 | 2 | 0 | NON_SYNONYMOU | MODERAT | MISSENSE | gCg/gTg | A865V | PLES_40131 | | | | | | | | | A | A | | |
| NC_011770 | 4429322 | C | T | 78 | 0 | 2 | 0 | NON_SYNONYMOU | MODERAT | MISSENSE | Gag/Aag | E819K | PLES_40131 | | | | | | | T | T | | | | |
| NC_011770 | 4429327 | G | T | 78 | 0 | 2 | 0 | NON_SYNONYMOU | MODERAT | MISSENSE | gCg/gAg | A817E | PLES_40131 | | | | | | | T | T | | | | |
| NC_011770 | 4430074 | G | A | 77 | 0 | 4 | 0 | NON_SYNONYMOU | MODERAT | MISSENSE | gCa/gTa | A568V | PLES_40131 | A | A | | | | | | | | | A | A |
| NC_011770 | 4430253 | A | C | 75 | 0 | 8 | 0 | NON_SYNONYMOU | MODERAT | MISSENSE | gaT/gaG | D508E | PLES_40131 | C | C | | | | | C | C | C | C | C | C |
| NC_011770 | 4430297 | A | G | 70 | 0 | 6 | 0 | NON_SYNONYMOU | MODERAT | MISSENSE | Tct/Cct | S494P | PLES_40131 | G | G | | | | | | | G | G | G | G |
| NC_011770 | 4430516 | G | A | 76 | 0 | 2 | 0 | NON_SYNONYMOU | MODERAT | MISSENSE | Cgc/Tgc | R421C | PLES_40131 | | | | | | | | | | | A | A |
| NC_011770 | 4430911 | C | T | 68 | 0 | 2 | 0 | NON_SYNONYMOU | MODERAT | MISSENSE | cGc/cAc | R289H | PLES_40131 | | | | | | | | | T | T | | |
| NC_011770 | 4430918 | G | A | 66 | 0 | 2 | 0 | NON_SYNONYMOU | MODERAT | MISSENSE | Ccc/Tcc | P287S | PLES_40131 | A | A | | | | | | | | | | |
| NC_011770 | 4430925 | G | C | 67 | 0 | 4 | 0 | NON_SYNONYMOU | MODERAT | MISSENSE | gaC/gaG | D284E | PLES_40131 | | | | | | | | | C | C | C | C |
| NC_011770 | 4431008 | G | C | 69 | 0 | 10 | 0 | NON_SYNONYMOU | MODERAT | MISSENSE | Cag/Gag | Q257E | PLES_40131 | | | C | C | C | C | C | C | C | C | C | C |
| NC_011770 | 4431181 | T | G | 64 | 0 | 2 | 0 | NON_SYNONYMOU | MODERAT | MISSENSE | gAc/gCc | D199A | PLES_40131 | | | | | | | | | G | G | | |
| NC_011770 | 4431308 | C | A | 61 | 0 | 4 | 0 | NON_SYNONYMOU | MODERAT | MISSENSE | Gcc/Tcc | A157S | PLES_40131 | | | A | A | A | A | | | | | | |
| NC_011770 | 4431361 | C | T | 68 | 0 | 2 | 0 | NON_SYNONYMOU | MODERAT | MISSENSE | cGt/cAt | R139H | PLES_40131 | | | | | | | | | | | T | T |
| NC_011770 | 4431465 | T | G | 69 | 0 | 12 | 0 | NON_SYNONYMOU | MODERAT | MISSENSE | caA/caC | Q104H | PLES_40131 | G | G | G | G | G | G | G | G | G | G | G | G |
| NC_011770 | 4431761 | AGGG | A | 75 | 0 | 5 | 0 | CODON_DELETION | MODERATE | | ccc/ | P5 | PLES_40131 | | | | | | | A | | A | A | A | A |
| NC_011770 | 4432202 | T | G | 64 | 0 | 12 | 0 | NON_SYNONYMOU | MODERAT | MISSENSE | gaT/gaG | D74E | PLES_40141 | G | G | G | G | G | G | G | G | G | G | G | G |
| NC_011770 | 4432351 | A | G | 66 | 0 | 8 | 0 | NON_SYNONYMOU | MODERAT | MISSENSE | gAc/gGc | D124G | PLES_40141 | G | G | C | C | C | C | C | C | | | | |
| NC_011770 | 4432365 | G | A | 66 | 0 | 2 | 0 | NON_SYNONYMOU | MODERAT | MISSENSE | Gcc/Acc | A129T | PLES_40141 | A | A | | | | | | | | | | |
| NC_011770 | 4432547 | C | A | 80 | 0 | 10 | 0 | NON_SYNONYMOU | MODERAT | MISSENSE | gaC/gaA | D189E | PLES_40141 | A | A | A | A | A | A | | | A | A | A | A |
| NC_011770 | 4432647 | A | G | 67 | 0 | 2 | 0 | NON_SYNONYMOU | MODERAT | MISSENSE | Acc/Gcc | T223A | PLES_40141 | | | | | | | G | G | | | | |
| NC_011770 | 4433079 | G | C | 82 | 0 | 6 | 0 | NON_SYNONYMOU | MODERAT | MISSENSE | aaC/aaG | N60K | PLES_40151 | C | C | | | | | | | C | C | C | C |
| NC_011770 | 4433170 | A | T | 77 | 0 | 2 | 0 | NON_SYNONYMOU | MODERAT | MISSENSE | cTg/cAg | L30Q | PLES_40151 | | | | | | | | | | | T | T |
| NC_011770 | 4433548 | C | G | 72 | 0 | 12 | 0 | NON_SYNONYMOU | MODERAT | MISSENSE | Ggc/Cgc | G142R | PLES_40161 | G | G | G | G | G | G | G | G | G | G | G | G |
| NC_011770 | 4433751 | C | T | 69 | 0 | 6 | 0 | NON_SYNONYMOU | MODERAT | MISSENSE | cGg/cAg | R74Q | PLES_40161 | T | T | | | | | T | T | T | T | | |
| NC_011770 | 4433838 | C | T | 72 | 0 | 4 | 0 | NON_SYNONYMOU | MODERAT | MISSENSE | gGc/gAc | G45D | PLES_40161 | | | T | T | T | T | | | | | | |
| NC_011770 | 4433550 | G | A | 73 | 0 | 1 | 0 | NON_SYNONYMOU | MODERAT | MISSENSE | cCg/cTg | P141L | PLES_40161 | | | | | | | | A | | | | |
| NC_011770 | 4434321 | T | C | 81 | 0 | 12 | 0 | NON_SYNONYMOU | MODERAT | MISSENSE | Tcg/Ccg | S69P | PLES_40171 | C | C | C | C | C | C | C | C | C | C | C | C |
| NC_011770 | 4434742 | G | A | 88 | 0 | 4 | 0 | NON_SYNONYMOU | MODERAT | MISSENSE | cGc/cAc | R209H | PLES_40171 | | | A | A | A | A | | | | | | |
| NC_011770 | 4434752 | C | A | 86 | 0 | 4 | 0 | NON_SYNONYMOU | MODERAT | MISSENSE | gaC/gaA | D212E | PLES_40171 | | | A | A | A | A | | | | | | |
| NC_011770 | 4435641 | G | A | 83 | 0 | 2 | 0 | NON_SYNONYMOU | MODERAT | MISSENSE | Gac/Aac | D509N | PLES_40171 | | | | | | | | | A | A | | |
| NC_011770 | 4435717 | A | C | 82 | 0 | 10 | 0 | NON_SYNONYMOU | MODERAT | MISSENSE | gAg/gCg | E534A | PLES_40171 | C | C | T | T | T | T | T | T | C | C | | |
| NC_011770 | 4435719 | G | A | 83 | 0 | 2 | 0 | NON_SYNONYMOU | MODERAT | MISSENSE | Gcc/Acc | A535T | PLES_40171 | | | | | | | | | | | A | A |
| NC_011770 | 4435784 | C | G | 89 | 0 | 10 | 0 | NON_SYNONYMOU | MODERAT | MISSENSE | atG/atC | M133I | PLES_40181 | G | G | G | G | G | G | G | G | G | G | | |
| NC_011770 | 4436275 | G | A | 61 | 0 | 2 | 0 | STOP_GAINED | HIGH | NONSENSE | Caa/Taa | Q60* | PLES_40191 | | | | | | | | | | | A | A |
| NC_011770 | 4436370 | TCG | T | 64 | 0 | 2 | 0 | FRAME_SHIFT | HIGH | | cgcgag/ | RE27 | PLES_40191 | | | | | | | | | | | T | T |
| NC_011770 | 4436239 | T | C | 63 | 0 | 10 | 0 | NON_SYNONYMOU | MODERAT | MISSENSE | Acc/Gcc | T72A | PLES_40191 | | | C | C | C | C | C | C | C | C | C | C |
| NC_011770 | 4436295 | G | C | 64 | 0 | 2 | 0 | NON_SYNONYMOU | MODERAT | MISSENSE | gCc/gGc | A53G | PLES_40191 | | | | | | | | | | | C | C |
| NC_011770 | 4436311 | T | G | 64 | 0 | 2 | 0 | NON_SYNONYMOU | MODERAT | MISSENSE | Agc/Cgc | S48R | PLES_40191 | | | | | | | | | | | G | G |
| NC_011770 | 4436313 | T | G | 63 | 0 | 2 | 0 | NON_SYNONYMOU | MODERAT | MISSENSE | cAg/cCg | Q47P | PLES_40191 | | | | | | | | | | | G | G |
| NC_011770 | 4436319 | C | G | 65 | 0 | 2 | 0 | NON_SYNONYMOU | MODERAT | MISSENSE | gGc/gCc | G45A | PLES_40191 | | | | | | | | | | | G | G |
| NC_011770 | 4436322 | A | G | 64 | 0 | 2 | 0 | NON_SYNONYMOU | MODERAT | MISSENSE | gTc/gCc | V44A | PLES_40191 | | | | | | | | | | | G | G |
| NC_011770 | 4436401 | A | G | 68 | 0 | 2 | 0 | NON_SYNONYMOU | MODERAT | MISSENSE | Tgc/Cgc | C18R | PLES_40191 | | | | | | | | | | | G | G |
| NC_011770 | 4436436 | A | G | 69 | 0 | 2 | 0 | NON_SYNONYMOU | MODERAT | MISSENSE | gTa/gCa | V6A | PLES_40191 | | | | | | | | | | | G | G |
| NC_011770 | 4437972 | TGAC | T | 57 | 0 | 2 | 0 | FRAME_SHIFT+STO | HIGH | | tga/ | *285 | PLES_40201 | | | | | | | | | T | T | | |
| NC_011770 | 4437336 | G | A | 59 | 0 | 4 | 0 | NON_SYNONYMOU | MODERAT | MISSENSE | Gat/Aat | D73N | PLES_40201 | | | A | A | A | A | | | | | | |
| NC_011770 | 4437342 | G | A | 60 | 0 | 4 | 0 | NON_SYNONYMOU | MODERAT | MISSENSE | Gcc/Acc | A75T | PLES_40201 | A | A | | | | | A | A | | | | |
| NC_011770 | 4437399 | C | T | 64 | 0 | 2 | 0 | NON_SYNONYMOU | MODERAT | MISSENSE | Ctc/Ttc | L94F | PLES_40201 | | | | | | | | | | | T | T |
| NC_011770 | 4437541 | G | A | 59 | 0 | 1 | 0 | NON_SYNONYMOU | MODERAT | MISSENSE | aGc/aAc | S141N | PLES_40201 | | | | | | | | | | | | A |
| NC_011770 | 4438093 | G | C | 64 | 0 | 2 | 0 | NON_SYNONYMOU | MODERAT | MISSENSE | gCc/gGc | A249G | PLES_40211 | C | C | | | | | | | | | | |
| NC_011770 | 4438405 | T | C | 73 | 0 | 2 | 0 | NON_SYNONYMOU | MODERAT | MISSENSE | cAg/cGg | Q145R | PLES_40211 | | | | | | | | | C | C | | |
| NC_011770 | 4439195 | T | G | 84 | 0 | 2 | 0 | NON_SYNONYMOU | MODERAT | MISSENSE | caT/caG | H82Q | PLES_40221 | | | | | | | G | G | | | | |
| NC_011770 | 4440017 | C | T | 84 | 0 | 2 | 0 | STOP_GAINED | HIGH | NONSENSE | Cag/Tag | Q85* | PLES_40231 | | | | | | | | | | | T | T |

| | | | | | | | | | | | | | | | | | | | | | | | | | |
|---|---|---|---|---|---|---|---|---|---|---|---|---|---|---|---|---|---|---|---|---|---|---|---|---|---|
| NC_011770 | 4440490 | T | C | 89 | 0 | 2 | 0 | NON_SYNONYMOU | MODERAT | MISSENSE | aTt/aCt | I14T | PLES_40241 | C | C | | | | | | | | | | |
| NC_011770 | 4441153 | G | A | 91 | 0 | 2 | 0 | NON_SYNONYMOU | MODERAT | MISSENSE | gGc/gAc | G235D | PLES_40241 | A | A | | | | | | | | | | |
| NC_011770 | 4441854 | A | G | 76 | 0 | 2 | 0 | NON_SYNONYMOU | MODERAT | MISSENSE | Tcc/Ccc | S162P | PLES_40251 | | | | | G | G | | | | | | |
| NC_011770 | 4442277 | T | C | 81 | 0 | 12 | 0 | NON_SYNONYMOU | MODERAT | MISSENSE | Acc/Gcc | T21A | PLES_40251 | C | C | C | C | C | C | C | C | C | C | C | C |
| NC_011770 | 4442312 | A | G | 80 | 0 | 6 | 0 | NON_SYNONYMOU | MODERAT | MISSENSE | gTc/gCc | V9A | PLES_40251 | G | G | G | G | G | G | | | | | | |
| NC_011770 | 4443465 | GC | G | 81 | 0 | 2 | 0 | FRAME_SHIFT | HIGH | | ggc/ | G79 | PLES_40261 | | | | G | | | | G | | | | |
| NC_011770 | 4442777 | GC | G | 86 | 0 | 2 | 0 | FRAME_SHIFT | HIGH | | ggg/ | G308 | PLES_40261 | G | G | | | | | | | | | | |
| NC_011770 | 4443238 | C | G | 82 | 0 | 6 | 0 | NON_SYNONYMOU | MODERAT | MISSENSE | gGc/gCc | G155A | PLES_40261 | | | G | G | G | G | | | G | G | | |
| NC_011770 | 4443415 | G | C | 79 | 0 | 4 | 0 | NON_SYNONYMOU | MODERAT | MISSENSE | aCc/aGc | T96S | PLES_40261 | | | C | C | C | C | | | | | | |
| NC_011770 | 4443653 | C | A | 93 | 0 | 2 | 0 | NON_SYNONYMOU | MODERAT | MISSENSE | Gcc/Tcc | A17S | PLES_40261 | A | A | | | | | | | | | | |
| NC_011770 | 4442756 | G | A | 86 | 0 | 1 | 0 | NON_SYNONYMOU | MODERAT | MISSENSE | Ccc/Tcc | P316S | PLES_40261 | | | | | | | | A | | | | |
| NC_011770 | 4443797 | A | G | 88 | 0 | 6 | 0 | NON_SYNONYMOU | MODERAT | MISSENSE | gTg/gCg | V143A | PLES_40271 | | | G | G | G | G | G | G | | | | |
| NC_011770 | 4444211 | T | C | 92 | 0 | 6 | 0 | NON_SYNONYMOU | MODERAT | MISSENSE | aAg/aGg | K5R | PLES_40271 | | | | | | | C | C | C | C | C | C |
| NC_011770 | 4444217 | A | T | 91 | 0 | 6 | 0 | NON_SYNONYMOU | MODERAT | MISSENSE | gTc/gAc | V3D | PLES_40271 | | | | | | | T | T | T | T | T | T |
| NC_011770 | 4445772 | G | A | 84 | 0 | 2 | 0 | NON_SYNONYMOU | MODERAT | MISSENSE | aGg/aAg | R473K | PLES_40281 | | | | | | | | | | | A | A |
| NC_011770 | 4445774 | G | A | 85 | 0 | 2 | 0 | NON_SYNONYMOU | MODERAT | MISSENSE | Ggt/Agt | G474S | PLES_40281 | | | | | | | | | | | A | A |
| NC_011770 | 4445792 | G | A | 82 | 0 | 1 | 0 | NON_SYNONYMOU | MODERAT | MISSENSE | Gtc/Atc | V480I | PLES_40281 | | | | | | | A | | | | | |
| NC_011770 | 4445855 | G | A | 80 | 0 | 2 | 0 | NON_SYNONYMOU | MODERAT | MISSENSE | Gcc/Acc | A501T | PLES_40281 | A | A | | | | | | | | | | |
| NC_011770 | 4445892 | C | G | 79 | 0 | 2 | 0 | NON_SYNONYMOU | MODERAT | MISSENSE | aCc/aGc | T513S | PLES_40281 | | | | | | | | | G | G | | |
| NC_011770 | 4446212 | T | C | 64 | 0 | 2 | 0 | NON_SYNONYMOU | MODERAT | MISSENSE | gAc/gGc | D186G | PLES_40291 | | | | | | | | | | | C | C |
| NC_011770 | 4446231 | G | C | 62 | 0 | 2 | 0 | NON_SYNONYMOU | MODERAT | MISSENSE | Ccg/Gcg | P180A | PLES_40291 | | | | | | | | | C | C | | |
| NC_011770 | 4446233 | G | T | 63 | 0 | 2 | 0 | NON_SYNONYMOU | MODERAT | MISSENSE | aCc/aAc | T179N | PLES_40291 | | | | | | | | | | | T | T |
| NC_011770 | 4446284 | T | C | 68 | 0 | 12 | 0 | NON_SYNONYMOU | MODERAT | MISSENSE | gAc/gGc | D162G | PLES_40291 | C | C | C | C | C | C | C | C | C | C | C | C |
| NC_011770 | 4446435 | T | C | 76 | 0 | 4 | 0 | NON_SYNONYMOU | MODERAT | MISSENSE | Acg/Gcg | T112A | PLES_40291 | | | C | C | C | C | | | | | | |
| NC_011770 | 4446707 | C | G | 64 | 0 | 6 | 0 | NON_SYNONYMOU | MODERAT | MISSENSE | gGc/gCc | G21A | PLES_40291 | G | G | | | | | | | G | G | G | G |
| NC_011770 | 4447058 | G | A | 77 | 0 | 2 | 0 | NON_SYNONYMOU | MODERAT | MISSENSE | Ggc/Agc | G58S | PLES_40301 | A | A | | | | | | | | | | |
| NC_011770 | 4447407 | T | C | 66 | 0 | 4 | 0 | NON_SYNONYMOU | MODERAT | MISSENSE | gTc/gCc | V174A | PLES_40301 | C | C | | | | | | | C | C | | |
| NC_011770 | 4447409 | G | A | 66 | 0 | 4 | 0 | NON_SYNONYMOU | MODERAT | MISSENSE | Gtc/Atc | V175I | PLES_40301 | A | A | | | | | | | A | A | | |
| NC_011770 | 4447541 | A | G | 69 | 0 | 6 | 0 | NON_SYNONYMOU | MODERAT | MISSENSE | Aca/Gca | T219A | PLES_40301 | G | G | G | G | G | G | | | | | | |
| NC_011770 | 4447652 | A | G | 72 | 0 | 12 | 0 | NON_SYNONYMOU | MODERAT | MISSENSE | Acg/Gcg | T256A | PLES_40301 | G | G | G | G | G | G | G | G | G | G | G | G |
| NC_011770 | 4447892 | T | C | 64 | 0 | 12 | 0 | NON_SYNONYMOU | MODERAT | MISSENSE | Ttc/Ctc | F336L | PLES_40301 | C | C | C | C | C | C | C | C | C | C | C | C |
| NC_011770 | 4447934 | T | G | 72 | 0 | 2 | 0 | NON_SYNONYMOU | MODERAT | MISSENSE | Tcc/Gcc | S350A | PLES_40301 | | | | | | | | | G | G | | |
| NC_011770 | 4447938 | C | T | 73 | 0 | 2 | 0 | NON_SYNONYMOU | MODERAT | MISSENSE | gCc/gTc | A351V | PLES_40301 | | | | | | | | | | | T | T |
| NC_011770 | 4448128 | C | A | 77 | 0 | 2 | 0 | NON_SYNONYMOU | MODERAT | MISSENSE | ttC/ttA | F414L | PLES_40301 | A | A | | | | | | | | | | |
| NC_011770 | 4448222 | G | A | 76 | 0 | 2 | 0 | NON_SYNONYMOU | MODERAT | MISSENSE | Gcg/Acg | A446T | PLES_40301 | | | | | | | A | A | | | | |
| NC_011770 | 4449189 | C | T | 84 | 0 | 8 | 0 | NON_SYNONYMOU | MODERAT | MISSENSE | cGc/cAc | R164H | PLES_40321 | T | T | T | T | T | T | T | T | | | | |
| NC_011770 | 4449199 | G | A | 83 | 0 | 4 | 0 | NON_SYNONYMOU | MODERAT | MISSENSE | Ccg/Tcg | P161S | PLES_40321 | | | A | A | A | A | | | | | | |
| NC_011770 | 4449214 | A | G | 80 | 0 | 10 | 0 | NON_SYNONYMOU | MODERAT | MISSENSE | Tgc/Cgc | C156R | PLES_40321 | G | G | G | G | G | G | G | G | G | G | | |
| NC_011770 | 4449282 | C | A | 77 | 0 | 4 | 0 | NON_SYNONYMOU | MODERAT | MISSENSE | cGc/cTc | R133L | PLES_40321 | | | A | A | A | A | | | | | | |
| NC_011770 | 4449408 | G | C | 76 | 0 | 8 | 0 | NON_SYNONYMOU | MODERAT | MISSENSE | gCc/gGc | A91G | PLES_40321 | C | C | C | C | C | C | | | C | C | | |
| NC_011770 | 4449409 | C | T | 75 | 0 | 8 | 0 | NON_SYNONYMOU | MODERAT | MISSENSE | Gcc/Acc | A91T | PLES_40321 | T | T | T | T | T | T | | | T | T | | |
| NC_011770 | 4449432 | G | T | 74 | 0 | 8 | 0 | NON_SYNONYMOU | MODERAT | MISSENSE | gCt/gAt | A83D | PLES_40321 | T | T | T | T | T | T | | | T | T | | |
| NC_011770 | 4449451 | G | A | 78 | 0 | 2 | 0 | NON_SYNONYMOU | MODERAT | MISSENSE | Cgc/Tgc | R77C | PLES_40321 | | | A | A | | | | | | | | |
| NC_011770 | 4449480 | C | T | 81 | 0 | 2 | 0 | NON_SYNONYMOU | MODERAT | MISSENSE | gGg/gAg | G67E | PLES_40321 | | | | | | | | | | | T | T |
| NC_011770 | 4449553 | G | C | 80 | 0 | 4 | 0 | NON_SYNONYMOU | MODERAT | MISSENSE | Cgc/Ggc | R43G | PLES_40321 | | | C | C | C | C | | | | | | |
| NC_011770 | 4457478 | T | C | 81 | 0 | 4 | 0 | NON_SYNONYMOU | MODERAT | MISSENSE | Aag/Gag | K559E | PLES_40411 | | | C | C | C | C | | | | | | |
| NC_011770 | 4457576 | C | T | 79 | 0 | 4 | 0 | NON_SYNONYMOU | MODERAT | MISSENSE | cGg/cAg | R526Q | PLES_40411 | | | T | T | T | T | | | | | | |
| NC_011770 | 4457798 | A | T | 39 | 5 | 1 | 0 | NON_SYNONYMOU | MODERAT | MISSENSE | tTc/tAc | F452Y | PLES_40411 | | | . | . | . | T | | | . | . | | |
| NC_011770 | 4457825 | C | T | 85 | 0 | 6 | 0 | NON_SYNONYMOU | MODERAT | MISSENSE | gGc/gAc | G443D | PLES_40411 | | | T | T | T | T | | | T | T | | |
| NC_011770 | 4457897 | G | A | 93 | 0 | 2 | 0 | NON_SYNONYMOU | MODERAT | MISSENSE | cCc/cTc | P419L | PLES_40411 | | | | | | | | | A | A | | |
| NC_011770 | 4457900 | T | A | 89 | 0 | 4 | 0 | NON_SYNONYMOU | MODERAT | MISSENSE | tAc/tTc | Y418F | PLES_40411 | | | A | A | A | A | | | | | | |
| NC_011770 | 4457901 | A | G | 92 | 0 | 4 | 0 | NON_SYNONYMOU | MODERAT | MISSENSE | Tac/Cac | Y418H | PLES_40411 | | | G | G | G | G | | | | | | |
| NC_011770 | 4457945 | C | T | 91 | 0 | 2 | 0 | NON_SYNONYMOU | MODERAT | MISSENSE | aGc/aAc | S403N | PLES_40411 | | | | | | | | | T | T | | |
| NC_011770 | 4458068 | T | C | 86 | 0 | 6 | 0 | NON_SYNONYMOU | MODERAT | MISSENSE | cAg/cGg | Q362R | PLES_40411 | | | C | C | C | C | | | | | C | C |
| NC_011770 | 4458318 | A | G | 86 | 0 | 6 | 0 | NON_SYNONYMOU | MODERAT | MISSENSE | Ttc/Ctc | F279L | PLES_40411 | | | G | G | G | G | | | G | G | | |
| NC_011770 | 4458372 | C | T | 85 | 0 | 6 | 0 | NON_SYNONYMOU | MODERAT | MISSENSE | Gcc/Acc | A261T | PLES_40411 | | | T | T | T | T | | | T | T | | |
| NC_011770 | 4458408 | C | T | 85 | 0 | 2 | 0 | NON_SYNONYMOU | MODERAT | MISSENSE | Gac/Aac | D249N | PLES_40411 | T | T | | | | | | | | | | |
| NC_011770 | 4458525 | C | T | 75 | 0 | 6 | 0 | NON_SYNONYMOU | MODERAT | MISSENSE | Ggc/Agc | G210S | PLES_40411 | | | T | T | T | T | | | T | T | | |
| NC_011770 | 4458528 | C | A | 73 | 0 | 6 | 0 | NON_SYNONYMOU | MODERAT | MISSENSE | Gcg/Tcg | A209S | PLES_40411 | | | A | A | A | A | | | A | A | | |
| NC_011770 | 4458546 | T | C | 80 | 0 | 6 | 0 | NON_SYNONYMOU | MODERAT | MISSENSE | Acc/Gcc | T203A | PLES_40411 | | | C | C | C | C | | | C | C | | |
| NC_011770 | 4459533 | C | T | 87 | 0 | 4 | 0 | NON_SYNONYMOU | MODERAT | MISSENSE | Gcg/Acg | A671T | PLES_40421 | | | | | | | | | T | T | T | T |
| NC_011770 | 4459755 | G | A | 74 | 0 | 2 | 0 | NON_SYNONYMOU | MODERAT | MISSENSE | Cgt/Tgt | R597C | PLES_40421 | | | | | | | | | | | A | A |

| | | | | | | | | | | | | | | | | | | | | | | | | | |
|---|---|---|---|---|---|---|---|---|---|---|---|---|---|---|---|---|---|---|---|---|---|---|---|---|---|
| NC_011770 | 4460190 | C | T | 97 | 0 | 2 | 0 | NON_SYNONYMOU | MODERAT | MISSENSE | Gcg/Acg | A452T | PLES_40421 | | | | | | | | | T | T | | |
| NC_011770 | 4460282 | A | G | 101 | 0 | 12 | 0 | NON_SYNONYMOU | MODERAT | MISSENSE | gTa/gCa | V421A | PLES_40421 | G | G | G | G | G | G | G | G | G | G | G | G |
| NC_011770 | 4460519 | C | G | 94 | 0 | 2 | 0 | NON_SYNONYMOU | MODERAT | MISSENSE | gGc/gCc | G342A | PLES_40421 | | | | | | | | | G | G | | |
| NC_011770 | 4461044 | T | C | 88 | 0 | 2 | 0 | NON_SYNONYMOU | MODERAT | MISSENSE | aAt/aGt | N167S | PLES_40421 | | | | | | | | | | | C | C |
| NC_011770 | 4461254 | G | A | 99 | 0 | 2 | 0 | NON_SYNONYMOU | MODERAT | MISSENSE | cCc/cTc | P97L | PLES_40421 | | | | | | | | | | | A | A |
| NC_011770 | 4461420 | T | C | 85 | 0 | 6 | 0 | NON_SYNONYMOU | MODERAT | MISSENSE | Agc/Ggc | S42G | PLES_40421 | C | C | | | | | C | C | | | C | C |
| NC_011770 | 4462159 | G | A | 73 | 0 | 6 | 0 | NON_SYNONYMOU | MODERAT | MISSENSE | Gac/Aac | D106N | PLES_40431 | | | | | | | A | A | A | A | A | A |
| NC_011770 | 4462842 | T | G | 71 | 0 | 12 | 0 | NON_SYNONYMOU | MODERAT | MISSENSE | Tcc/Gcc | S72A | PLES_40441 | G | G | G | G | G | G | G | G | G | G | G | G |
| NC_011770 | 4462945 | G | A | 75 | 0 | 1 | 0 | NON_SYNONYMOU | MODERAT | MISSENSE | gGt/gAt | G106D | PLES_40441 | | | | | | | A | | | | | |
| NC_011770 | 4463785 | T | G | 60 | 0 | 10 | 0 | NON_SYNONYMOU | MODERAT | MISSENSE | gaT/gaG | D72E | PLES_40451 | G | G | G | G | G | G | G | G | G | G | | |
| NC_011770 | 4463810 | A | G | 59 | 0 | 10 | 0 | NON_SYNONYMOU | MODERAT | MISSENSE | Aat/Gat | N81D | PLES_40451 | G | G | G | G | G | G | G | G | G | G | | |
| NC_011770 | 4463811 | A | G | 59 | 0 | 10 | 0 | NON_SYNONYMOU | MODERAT | MISSENSE | aAt/aGt | N81S | PLES_40451 | G | G | G | G | G | G | G | G | G | G | | |
| NC_011770 | 4463843 | G | A | 69 | 0 | 2 | 0 | NON_SYNONYMOU | MODERAT | MISSENSE | Gcc/Acc | A92T | PLES_40451 | | | | | | | | | | | A | A |
| NC_011770 | 4463844 | C | G | 69 | 0 | 2 | 0 | NON_SYNONYMOU | MODERAT | MISSENSE | gCc/gGc | A92G | PLES_40451 | | | | | | | | | | | G | G |
| NC_011770 | 4463883 | T | A | 64 | 0 | 2 | 0 | NON_SYNONYMOU | MODERAT | MISSENSE | cTg/cAg | L105Q | PLES_40451 | | | | | | | | | | | A | A |
| NC_011770 | 4463889 | T | C | 65 | 0 | 12 | 0 | NON_SYNONYMOU | MODERAT | MISSENSE | gTc/gCc | V107A | PLES_40451 | C | C | C | C | C | C | C | C | C | C | C | C |
| NC_011770 | 4463903 | G | C | 62 | 0 | 2 | 0 | NON_SYNONYMOU | MODERAT | MISSENSE | Gag/Cag | E112Q | PLES_40451 | | | | | | | | | | | C | C |
| NC_011770 | 4463904 | A | G | 60 | 0 | 2 | 0 | NON_SYNONYMOU | MODERAT | MISSENSE | gAg/gGg | E112G | PLES_40451 | | | | | | | | | | | G | G |
| NC_011770 | 4463913 | A | G | 57 | 0 | 2 | 0 | NON_SYNONYMOU | MODERAT | MISSENSE | gAc/gGc | D115G | PLES_40451 | | | | | | | | | | | G | G |
| NC_011770 | 4463927 | C | T | 60 | 0 | 2 | 0 | NON_SYNONYMOU | MODERAT | MISSENSE | Cgt/Tgt | R120C | PLES_40451 | | | | | | | | | | | T | T |
| NC_011770 | 4463928 | G | A | 59 | 0 | 2 | 0 | NON_SYNONYMOU | MODERAT | MISSENSE | cGt/cAt | R120H | PLES_40451 | | | | | | | | | | | A | A |
| NC_011770 | 4463931 | A | C | 60 | 0 | 2 | 0 | NON_SYNONYMOU | MODERAT | MISSENSE | gAc/gCc | D121A | PLES_40451 | | | | | | | | | | | C | C |
| NC_011770 | 4463943 | T | C | 56 | 0 | 2 | 0 | NON_SYNONYMOU | MODERAT | MISSENSE | gTc/gCc | V125A | PLES_40451 | | | | | | | | | | | C | C |
| NC_011770 | 4463972 | A | G | 63 | 0 | 2 | 0 | NON_SYNONYMOU | MODERAT | MISSENSE | Aag/Gag | K135E | PLES_40451 | | | | | | | | | | | G | G |
| NC_011770 | 4463978 | C | T | 61 | 0 | 2 | 0 | NON_SYNONYMOU | MODERAT | MISSENSE | Ccg/Tcg | P137S | PLES_40451 | | | | | | | | | | | T | T |
| NC_011770 | 4464028 | A | C | 63 | 0 | 2 | 0 | NON_SYNONYMOU | MODERAT | MISSENSE | gaA/gaC | E153D | PLES_40451 | | | | | | | | | | | C | C |
| NC_011770 | 4464056 | C | T | 66 | 0 | 2 | 0 | NON_SYNONYMOU | MODERAT | MISSENSE | Cac/Tac | H163Y | PLES_40451 | | | | | | | | | | | T | T |
| NC_011770 | 4464329 | A | G | 68 | 0 | 12 | 0 | NON_SYNONYMOU | MODERAT | MISSENSE | Atc/Gtc | I254V | PLES_40451 | G | G | G | G | G | G | G | G | G | G | G | G |
| NC_011770 | 4464416 | A | G | 70 | 0 | 12 | 0 | NON_SYNONYMOU | MODERAT | MISSENSE | Aac/Gac | N283D | PLES_40451 | G | G | G | G | G | G | G | G | G | G | G | G |
| NC_011770 | 4464629 | C | G | 60 | 0 | 12 | 0 | NON_SYNONYMOU | MODERAT | MISSENSE | Ccc/Gcc | P354A | PLES_40451 | G | G | G | G | G | G | G | G | G | G | G | G |
| NC_011770 | 4464668 | G | A | 63 | 0 | 10 | 0 | NON_SYNONYMOU | MODERAT | MISSENSE | Ggg/Agg | G367R | PLES_40451 | A | A | A | A | A | A | A | A | A | A | | |
| NC_011770 | 4464030 | G | A | 64 | 0 | 3 | 0 | NON_SYNONYMOU | MODERAT | MISSENSE | cGa/cAa | R154Q | PLES_40451 | | | | | | | | A | | | A | A |
| NC_011770 | 4465112 | C | G | 64 | 0 | 12 | 0 | NON_SYNONYMOU | MODERAT | MISSENSE | caC/caG | H66Q | PLES_40461 | G | G | G | G | G | G | G | G | G | G | G | G |
| NC_011770 | 4465755 | G | A | 63 | 0 | 2 | 0 | NON_SYNONYMOU | MODERAT | MISSENSE | Gaa/Aaa | E281K | PLES_40461 | | | A | A | | | | | | | | |
| NC_011770 | 4465800 | G | A | 64 | 0 | 4 | 0 | NON_SYNONYMOU | MODERAT | MISSENSE | Gcc/Acc | A296T | PLES_40461 | A | A | | | | | A | A | | | | |
| NC_011770 | 4465887 | G | A | 63 | 0 | 12 | 0 | NON_SYNONYMOU | MODERAT | MISSENSE | Gcc/Acc | A325T | PLES_40461 | A | A | A | A | A | A | A | A | A | A | A | A |
| NC_011770 | 4465926 | G | C | 62 | 0 | 10 | 0 | NON_SYNONYMOU | MODERAT | MISSENSE | Gaa/Caa | E338Q | PLES_40461 | C | C | C | C | C | C | | | C | C | C | C |
| NC_011770 | 4466145 | C | T | 63 | 0 | 8 | 0 | NON_SYNONYMOU | MODERAT | MISSENSE | Cgc/Tgc | R411C | PLES_40461 | T | T | T | T | T | T | T | T | | | | |
| NC_011770 | 4466154 | A | G | 65 | 0 | 12 | 0 | NON_SYNONYMOU | MODERAT | MISSENSE | Agc/Ggc | S414G | PLES_40461 | G | G | G | G | G | G | G | G | G | G | G | G |
| NC_011770 | 4466265 | G | A | 57 | 0 | 1 | 0 | NON_SYNONYMOU | MODERAT | MISSENSE | gCg/gTg | A268V | PLES_40471 | | | | | | | A | | | | | |
| NC_011770 | 4466487 | A | G | 33 | 0 | 4 | 0 | NON_SYNONYMOU | MODERAT | MISSENSE | tTc/tCc | F194S | PLES_40471 | | | G | G | G | G | | | | | | |
| NC_011770 | 4466499 | G | A | 31 | 0 | 2 | 0 | NON_SYNONYMOU | MODERAT | MISSENSE | gCc/gTc | A190V | PLES_40471 | | | | | | | | | A | A | | |
| NC_011770 | 4466679 | C | T | 24 | 0 | 2 | 0 | NON_SYNONYMOU | MODERAT | MISSENSE | cGg/cAg | R130Q | PLES_40471 | | | | | | | T | T | | | | |
| NC_011770 | 4467512 | A | G | 64 | 0 | 12 | 0 | NON_SYNONYMOU | MODERAT | MISSENSE | Agc/Ggc | S116G | PLES_40481 | G | G | G | G | G | G | G | G | G | G | G | G |
| NC_011770 | 4467524 | G | C | 63 | 0 | 12 | 0 | NON_SYNONYMOU | MODERAT | MISSENSE | Gcg/Ccg | A120P | PLES_40481 | C | C | C | C | C | C | C | C | C | C | C | C |
| NC_011770 | 4467527 | G | A | 63 | 0 | 2 | 0 | NON_SYNONYMOU | MODERAT | MISSENSE | Gcg/Acg | A121T | PLES_40481 | | | | | | | A | A | | | | |
| NC_011770 | 4467628 | C | A | 57 | 0 | 12 | 0 | NON_SYNONYMOU | MODERAT | MISSENSE | gaC/gaA | D154E | PLES_40481 | A | A | A | A | A | A | A | A | A | A | A | A |
| NC_011770 | 4467629 | C | A | 56 | 0 | 12 | 0 | NON_SYNONYMOU | MODERAT | MISSENSE | Ccc/Acc | P155T | PLES_40481 | A | A | A | A | A | A | A | A | A | A | A | A |
| NC_011770 | 4467827 | G | A | 65 | 0 | 2 | 0 | NON_SYNONYMOU | MODERAT | MISSENSE | Gcg/Acg | A221T | PLES_40481 | A | A | | | | | | | | | | |
| NC_011770 | 4468847 | G | A | 52 | 0 | 2 | 0 | NON_SYNONYMOU | MODERAT | MISSENSE | gCg/gTg | A116V | PLES_40491 | | | | | | | A | A | | | | |
| NC_011770 | 4469166 | C | A | 52 | 0 | 2 | 0 | NON_SYNONYMOU | MODERAT | MISSENSE | Gct/Tct | A10S | PLES_40491 | | | | | | | A | A | | | | |
| NC_011770 | 4469597 | C | CG | 66 | 0 | 1 | 0 | FRAME_SHIFT | HIGH | | ctg/Gctg | L97A? | PLES_40501 | | | | | | | CG | | | | | |
| NC_011770 | 4469707 | G | A | 71 | 0 | 1 | 0 | NON_SYNONYMOU | MODERAT | MISSENSE | gGc/gAc | G132D | PLES_40501 | | A | | | | | | | | | | |
| NC_011770 | 4469706 | G | A | 70 | 0 | 1 | 0 | NON_SYNONYMOU | MODERAT | MISSENSE | Ggc/Agc | G132S | PLES_40501 | | | | | | | A | | | | | |
| NC_011770 | 4469722 | G | A | 71 | 0 | 2 | 0 | NON_SYNONYMOU | MODERAT | MISSENSE | cGc/cAc | R137H | PLES_40501 | | | | | | | A | A | | | | |
| NC_011770 | 4469901 | G | A | 76 | 0 | 2 | 0 | NON_SYNONYMOU | MODERAT | MISSENSE | Gcc/Acc | A197T | PLES_40501 | | | | | | | | | | | A | A |
| NC_011770 | 4469917 | G | A | 76 | 0 | 2 | 0 | NON_SYNONYMOU | MODERAT | MISSENSE | cGc/cAc | R202H | PLES_40501 | A | A | | | | | | | | | | |
| NC_011770 | 4470507 | G | A | 46 | 0 | 2 | 0 | NON_SYNONYMOU | MODERAT | MISSENSE | Gcc/Acc | A399T | PLES_40501 | | | | | | | | | A | A | | |
| NC_011770 | 4470950 | A | G | 21 | 0 | 12 | 0 | NON_SYNONYMOU | MODERAT | MISSENSE | Aag/Gag | K11E | PLES_40511 | G | G | G | G | G | G | G | G | G | G | G | G |
| NC_011770 | 4470969 | T | C | 26 | 0 | 1 | 0 | NON_SYNONYMOU | MODERAT | MISSENSE | cTg/cCg | L17P | PLES_40511 | | | | | | | C | | | | | |
| NC_011770 | 4471340 | G | A | 46 | 0 | 4 | 0 | NON_SYNONYMOU | MODERAT | MISSENSE | Gcc/Acc | A141T | PLES_40511 | | | | | | | A | A | | | A | A |
| NC_011770 | 4471436 | G | A | 50 | 0 | 2 | 0 | NON_SYNONYMOU | MODERAT | MISSENSE | Gcc/Acc | A173T | PLES_40511 | | | | | | | | | | | A | A |

| | | | | | | | | | | | | | | | | | | | | | | | | | |
|---|---|---|---|---|---|---|---|---|---|---|---|---|---|---|---|---|---|---|---|---|---|---|---|---|---|
| NC_011770 | 4471873 | G | A | 68 | 0 | 2 | 0 | NON_SYNONYMOU | MODERAT | MISSENSE | Gcc/Acc | A8T | PLES_40521 | | | | | | | | | | | A | A |
| NC_011770 | 4472333 | A | G | 84 | 0 | 4 | 0 | NON_SYNONYMOU | MODERAT | MISSENSE | aAg/aGg | K161R | PLES_40521 | | | | | | | G | G | | | G | G |
| NC_011770 | 4473098 | G | A | 59 | 0 | 4 | 0 | NON_SYNONYMOU | MODERAT | MISSENSE | Gcc/Acc | A123T | PLES_40531 | | | | | | | A | A | | | A | A |
| NC_011770 | 4473170 | A | G | 62 | 0 | 10 | 0 | NON_SYNONYMOU | MODERAT | MISSENSE | Acc/Gcc | T147A | PLES_40531 | G | G | G | G | G | G | G | G | | | G | G |
| NC_011770 | 4473320 | A | G | 61 | 0 | 6 | 0 | NON_SYNONYMOU | MODERAT | MISSENSE | Atc/Gtc | I197V | PLES_40531 | G | G | G | G | G | G | | | | | | |
| NC_011770 | 4473325 | C | G | 63 | 0 | 6 | 0 | NON_SYNONYMOU | MODERAT | MISSENSE | gaC/gaG | D198E | PLES_40531 | G | G | G | G | G | G | | | | | | |
| NC_011770 | 4473485 | G | A | 55 | 0 | 4 | 0 | NON_SYNONYMOU | MODERAT | MISSENSE | Gcc/Acc | A252T | PLES_40531 | | | A | A | A | A | | | | | | |
| NC_011770 | 4473507 | C | T | 60 | 0 | 2 | 0 | NON_SYNONYMOU | MODERAT | MISSENSE | gCc/gTc | A259V | PLES_40531 | | | | | | | T | T | | | | |
| NC_011770 | 4473516 | A | G | 59 | 0 | 4 | 0 | NON_SYNONYMOU | MODERAT | MISSENSE | cAg/cGg | Q262R | PLES_40531 | | | | | | | G | G | | | G | G |
| NC_011770 | 4473882 | A | G | 41 | 0 | 6 | 0 | NON_SYNONYMOU | MODERAT | MISSENSE | cAc/cGc | H384R | PLES_40531 | G | G | G | G | G | G | | | | | | |
| NC_011770 | 4474068 | G | A | 47 | 0 | 4 | 0 | NON_SYNONYMOU | MODERAT | MISSENSE | gGc/gAc | G446D | PLES_40531 | | | A | A | A | A | | | | | | |
| NC_011770 | 4474209 | C | T | 55 | 0 | 4 | 0 | NON_SYNONYMOU | MODERAT | MISSENSE | gCc/gTc | A493V | PLES_40531 | | | T | T | T | T | | | | | | |
| NC_011770 | 4474227 | G | A | 54 | 0 | 2 | 0 | NON_SYNONYMOU | MODERAT | MISSENSE | gGg/gAg | G499E | PLES_40531 | | | | | | | | | A | A | | |
| NC_011770 | 4474260 | G | T | 58 | 0 | 2 | 0 | NON_SYNONYMOU | MODERAT | MISSENSE | cGg/cTg | R510L | PLES_40531 | T | T | | | | | | | | | | |
| NC_011770 | 4474373 | G | A | 61 | 0 | 4 | 0 | NON_SYNONYMOU | MODERAT | MISSENSE | Ggc/Agc | G548S | PLES_40531 | | | A | A | A | A | | | | | | |
| NC_011770 | 4474394 | G | A | 62 | 0 | 2 | 0 | NON_SYNONYMOU | MODERAT | MISSENSE | Ggc/Agc | G555S | PLES_40531 | A | A | | | | | | | | | | |
| NC_011770 | 4474624 | C | A | 69 | 0 | 8 | 0 | NON_SYNONYMOU | MODERAT | MISSENSE | Cac/Aac | H51N | PLES_40541 | A | A | A | A | A | A | | | A | A | | |
| NC_011770 | 4474990 | G | A | 84 | 0 | 6 | 0 | NON_SYNONYMOU | MODERAT | MISSENSE | Gac/Aac | D173N | PLES_40541 | A | A | A | A | A | A | | | | | | |
| NC_011770 | 4475432 | G | C | 92 | 0 | 2 | 0 | NON_SYNONYMOU | MODERAT | MISSENSE | Gtg/Ctg | V95L | PLES_40551 | C | C | | | | | | | | | | |
| NC_011770 | 4475453 | A | G | 95 | 0 | 2 | 0 | NON_SYNONYMOU | MODERAT | MISSENSE | Acc/Gcc | T102A | PLES_40551 | G | G | | | | | | | | | | |
| NC_011770 | 4475750 | A | G | 97 | 0 | 12 | 0 | NON_SYNONYMOU | MODERAT | MISSENSE | Atc/Gtc | I201V | PLES_40551 | G | G | G | G | G | G | G | G | G | G | G | G |
| NC_011770 | 4475807 | A | G | 96 | 0 | 2 | 0 | NON_SYNONYMOU | MODERAT | MISSENSE | gAc/gGc | D6G | PLES_40561 | G | G | | | | | | | | | | |
| NC_011770 | 4476050 | A | G | 80 | 0 | 1 | 0 | NON_SYNONYMOU | MODERAT | MISSENSE | aAc/aGc | N87S | PLES_40561 | | | | | | | G | | | | | |
| NC_011770 | 4476726 | C | T | 64 | 0 | 4 | 0 | NON_SYNONYMOU | MODERAT | MISSENSE | gCc/gTc | A63V | PLES_40571 | | | T | T | T | T | | | | | | |
| NC_011770 | 4477100 | A | G | 55 | 0 | 12 | 0 | NON_SYNONYMOU | MODERAT | MISSENSE | Acc/Gcc | T188A | PLES_40571 | G | G | G | G | G | G | G | G | G | G | G | G |
| NC_011770 | 4477198 | C | G | 59 | 0 | 4 | 0 | NON_SYNONYMOU | MODERAT | MISSENSE | gaC/gaG | D220E | PLES_40571 | | | G | G | G | G | | | | | | |
| NC_011770 | 4477260 | G | A | 57 | 0 | 2 | 0 | NON_SYNONYMOU | MODERAT | MISSENSE | aGc/aAc | S241N | PLES_40571 | | | | | | | A | A | | | | |
| NC_011770 | 4477280 | C | G | 58 | 0 | 2 | 0 | NON_SYNONYMOU | MODERAT | MISSENSE | Ccc/Gcc | P248A | PLES_40571 | | | | | | | | | | | G | G |
| NC_011770 | 4477445 | C | T | 55 | 0 | 2 | 0 | NON_SYNONYMOU | MODERAT | MISSENSE | Ccg/Tcg | P303S | PLES_40571 | | | | | | | T | T | | | | |
| NC_011770 | 4478507 | C | G | 56 | 0 | 6 | 0 | NON_SYNONYMOU | MODERAT | MISSENSE | Caa/Gaa | Q262E | PLES_40581 | G | G | G | G | G | G | | | | | | |
| NC_011770 | 4478791 | C | G | 60 | 0 | 4 | 0 | NON_SYNONYMOU | MODERAT | MISSENSE | Ctc/Gtc | L20V | PLES_40591 | | | G | G | G | G | | | | | | |
| NC_011770 | 4478858 | A | C | 60 | 0 | 6 | 0 | NON_SYNONYMOU | MODERAT | MISSENSE | gAg/gCg | E42A | PLES_40591 | | | C | C | C | C | | | | | C | C |
| NC_011770 | 4478861 | C | A | 61 | 0 | 4 | 0 | NON_SYNONYMOU | MODERAT | MISSENSE | gCg/gAg | A43E | PLES_40591 | | | A | A | A | A | | | | | | |
| NC_011770 | 4478926 | A | G | 67 | 0 | 4 | 0 | NON_SYNONYMOU | MODERAT | MISSENSE | Acg/Gcg | T65A | PLES_40591 | | | G | G | G | G | | | | | | |
| NC_011770 | 4479001 | A | C | 63 | 0 | 12 | 0 | NON_SYNONYMOU | MODERAT | MISSENSE | Agt/Cgt | S90R | PLES_40591 | C | C | C | C | C | C | C | C | C | C | C | C |
| NC_011770 | 4479026 | C | T | 62 | 0 | 2 | 0 | NON_SYNONYMOU | MODERAT | MISSENSE | gCa/gTa | A98V | PLES_40591 | T | T | | | | | | | | | | |
| NC_011770 | 4479034 | C | G | 63 | 0 | 4 | 0 | NON_SYNONYMOU | MODERAT | MISSENSE | Cag/Gag | Q101E | PLES_40591 | | | G | G | G | G | | | | | | |
| NC_011770 | 4479047 | T | C | 60 | 0 | 10 | 0 | NON_SYNONYMOU | MODERAT | MISSENSE | aTa/aCa | I105T | PLES_40591 | C | C | C | C | C | C | C | C | C | C | | |
| NC_011770 | 4479122 | G | C | 63 | 0 | 4 | 0 | NON_SYNONYMOU | MODERAT | MISSENSE | cGt/cCt | R130P | PLES_40591 | | | C | C | C | C | | | | | | |
| NC_011770 | 4479365 | C | A | 60 | 0 | 2 | 0 | NON_SYNONYMOU | MODERAT | MISSENSE | gCc/gAc | A211D | PLES_40591 | | | | | | | | | | | A | A |
| NC_011770 | 4479392 | G | A | 63 | 0 | 2 | 0 | NON_SYNONYMOU | MODERAT | MISSENSE | aGc/aAc | S220N | PLES_40591 | A | A | | | | | | | | | | |
| NC_011770 | 4479697 | G | T | 80 | 0 | 2 | 0 | NON_SYNONYMOU | MODERAT | MISSENSE | Ggc/Tgc | G322C | PLES_40591 | | | | | | | | | T | T | | |
| NC_011770 | 4479793 | G | A | 77 | 0 | 4 | 0 | NON_SYNONYMOU | MODERAT | MISSENSE | Ggc/Agc | G354S | PLES_40591 | | | | | | | A | A | A | A | | |
| NC_011770 | 4479988 | T | A | 72 | 0 | 2 | 0 | NON_SYNONYMOU | MODERAT | MISSENSE | Tcg/Acg | S419T | PLES_40591 | A | A | | | | | | | | | | |
| NC_011770 | 4480046 | T | A | 70 | 0 | 2 | 0 | NON_SYNONYMOU | MODERAT | MISSENSE | cTg/cAg | L438Q | PLES_40591 | A | A | | | | | | | | | | |
| NC_011770 | 4479277 | G | A | 67 | 0 | 1 | 0 | NON_SYNONYMOU | MODERAT | MISSENSE | Gtg/Atg | V182M | PLES_40591 | | | | | | | | A | | | | |
| NC_011770 | 4480588 | C | T | 88 | 0 | 2 | 0 | NON_SYNONYMOU | MODERAT | MISSENSE | Gcc/Acc | A279T | PLES_40601 | T | T | | | | | | | | | | |
| NC_011770 | 4481170 | C | T | 75 | 0 | 2 | 0 | NON_SYNONYMOU | MODERAT | MISSENSE | Ggg/Agg | G85R | PLES_40601 | | | | | T | T | | | | | | |
| NC_011770 | 4482134 | C | A | 74 | 0 | 12 | 0 | NON_SYNONYMOU | MODERAT | MISSENSE | Ccc/Acc | P162T | PLES_40611 | A | A | A | A | A | A | A | A | A | A | A | A |
| NC_011770 | 4482174 | A | G | 74 | 0 | 12 | 0 | NON_SYNONYMOU | MODERAT | MISSENSE | cAc/cGc | H175R | PLES_40611 | G | G | G | G | G | G | G | G | G | G | G | G |
| NC_011770 | 4482672 | G | A | 63 | 0 | 2 | 0 | NON_SYNONYMOU | MODERAT | MISSENSE | gGa/gAa | G341E | PLES_40611 | | | | | | | | | | | A | A |
| NC_011770 | 4482722 | G | A | 58 | 0 | 2 | 0 | NON_SYNONYMOU | MODERAT | MISSENSE | Gcg/Acg | A358T | PLES_40611 | A | A | | | | | | | | | | |
| NC_011770 | 4491149 | C | T | 59 | 0 | 12 | 0 | NON_SYNONYMOU | MODERAT | MISSENSE | aGg/aAg | R146K | PLES_40671 | T | T | T | T | T | T | T | T | T | T | T | T |
| NC_011770 | 4491393 | C | T | 71 | 0 | 2 | 0 | NON_SYNONYMOU | MODERAT | MISSENSE | Gcg/Acg | A65T | PLES_40671 | T | T | | | | | | | | | | |
| NC_011770 | 4492603 | A | T | 30 | 0 | 2 | 0 | NON_SYNONYMOU | MODERAT | MISSENSE | Act/Tct | T2S | PLES_40681 | | | | | | | | | T | T | | |
| NC_011770 | 4492715 | G | A | 31 | 0 | 2 | 0 | NON_SYNONYMOU | MODERAT | MISSENSE | cGg/cAg | R39Q | PLES_40681 | | | | | | | | | | | A | A |
| NC_011770 | 4492925 | A | C | 36 | 0 | 8 | 0 | NON_SYNONYMOU | MODERAT | MISSENSE | gAt/gCt | D109A | PLES_40681 | C | C | C | C | C | C | C | C | | | | |
| NC_011770 | 4493287 | A | G | 62 | 0 | 2 | 0 | NON_SYNONYMOU | MODERAT | MISSENSE | Agt/Ggt | S38G | PLES_40691 | | | | | | | | | | | G | G |
| NC_011770 | 4493908 | G | T | 75 | 0 | 10 | 0 | NON_SYNONYMOU | MODERAT | MISSENSE | Gcg/Tcg | A245S | PLES_40691 | T | T | T | T | T | T | T | T | T | T | | |
| NC_011770 | 4494110 | T | TCGG | 70 | 0 | 12 | 0 | CODON_CHANGE_ | MODERATE | | gaa/gGCG | E313GE | PLES_40691 | TCGG | TCGG | TCGG | TCGG | TCGG | TCGG | TCGG | TCGG | TCGG | TCGG | TCGG | TCGG |
| NC_011770 | 4494118 | C | T | 75 | 0 | 2 | 0 | NON_SYNONYMOU | MODERAT | MISSENSE | Ccg/Tcg | P315S | PLES_40691 | | | | | | | | | | | T | T |
| NC_011770 | 4494356 | A | C | 79 | 0 | 6 | 0 | NON_SYNONYMOU | MODERAT | MISSENSE | gAc/gCc | D394A | PLES_40691 | | | C | C | C | C | | | | | C | C |

| | | | | | | | | | | | | | | | | | | | | | | | | | |
|---|---|---|---|---|---|---|---|---|---|---|---|---|---|---|---|---|---|---|---|---|---|---|---|---|---|
| NC_011770 | 4494936 | G | T | 84 | 0 | 2 | 0 | NON_SYNONYMOU | MODERAT | MISSENSE | gaG/gaT | E587D | PLES_40691 | T | T | | | | | | | | | | |
| NC_011770 | 4495388 | T | C | 71 | 0 | 2 | 0 | NON_SYNONYMOU | MODERAT | MISSENSE | tTg/tCg | L738S | PLES_40691 | | | | | | | | | | | C | C |
| NC_011770 | 4496074 | G | A | 84 | 0 | 2 | 0 | NON_SYNONYMOU | MODERAT | MISSENSE | Cgg/Tgg | R491W | PLES_40701 | | | | | | | A | A | | | | |
| NC_011770 | 4496329 | T | C | 78 | 0 | 12 | 0 | NON_SYNONYMOU | MODERAT | MISSENSE | Aag/Gag | K406E | PLES_40701 | C | C | C | C | C | C | C | C | C | C | C | C |
| NC_011770 | 4496925 | T | G | 70 | 0 | 4 | 0 | NON_SYNONYMOU | MODERAT | MISSENSE | aAg/aCg | K207T | PLES_40701 | | | G | G | G | G | | | | | | |
| NC_011770 | 4497385 | C | A | 71 | 0 | 6 | 0 | NON_SYNONYMOU | MODERAT | MISSENSE | Gcc/Tcc | A54S | PLES_40701 | A | A | | | | | | | A | A | A | A |
| NC_011770 | 4497522 | A | G | 66 | 0 | 2 | 0 | NON_SYNONYMOU | MODERAT | MISSENSE | tTc/tCc | F8S | PLES_40701 | | | | | | | G | G | | | | |
| NC_011770 | 4497525 | A | G | 66 | 0 | 2 | 0 | NON_SYNONYMOU | MODERAT | MISSENSE | gTt/gCt | V7A | PLES_40701 | G | G | | | | | | | | | | |
| NC_011770 | 4497955 | C | T | 73 | 0 | 2 | 0 | NON_SYNONYMOU | MODERAT | MISSENSE | Cgc/Tgc | R25C | PLES_40711 | | | | | | | | | T | T | | |
| NC_011770 | 4498195 | G | A | 79 | 0 | 2 | 0 | NON_SYNONYMOU | MODERAT | MISSENSE | Ggc/Agc | G105S | PLES_40711 | A | A | | | | | | | | | | |
| NC_011770 | 4498265 | A | G | 83 | 0 | 4 | 0 | NON_SYNONYMOU | MODERAT | MISSENSE | cAc/cGc | H128R | PLES_40711 | G | G | | | | | | | | | G | G |
| NC_011770 | 4498609 | C | T | 69 | 0 | 2 | 0 | NON_SYNONYMOU | MODERAT | MISSENSE | Cgg/Tgg | R243W | PLES_40711 | | | | | | | | | T | T | | |
| NC_011770 | 4499319 | A | T | 67 | 0 | 2 | 0 | NON_SYNONYMOU | MODERAT | MISSENSE | gaA/gaT | E479D | PLES_40711 | | | | | | | | | T | T | | |
| NC_011770 | 4499445 | G | A | 67 | 0 | 2 | 0 | NON_SYNONYMOU | MODERAT | MISSENSE | Gcc/Acc | A40T | PLES_40721 | | | | | | | A | A | | | | |
| NC_011770 | 4499550 | G | A | 68 | 0 | 4 | 0 | NON_SYNONYMOU | MODERAT | MISSENSE | Gtc/Atc | V75I | PLES_40721 | | | A | A | A | A | | | | | | |
| NC_011770 | 4499571 | G | A | 66 | 0 | 2 | 0 | NON_SYNONYMOU | MODERAT | MISSENSE | Gac/Aac | D82N | PLES_40721 | | | | | | | A | A | | | | |
| NC_011770 | 4499734 | C | T | 57 | 0 | 2 | 0 | NON_SYNONYMOU | MODERAT | MISSENSE | gCg/gTg | A136V | PLES_40721 | | | | | | | | | T | T | | |
| NC_011770 | 4499776 | G | A | 56 | 0 | 4 | 0 | NON_SYNONYMOU | MODERAT | MISSENSE | cGa/cAa | R150Q | PLES_40721 | | | A | A | A | A | | | | | | |
| NC_011770 | 4499875 | C | T | 60 | 0 | 4 | 0 | NON_SYNONYMOU | MODERAT | MISSENSE | gCg/gTg | A183V | PLES_40721 | T | T | | | | | | | | | T | T |
| NC_011770 | 4499911 | A | G | 64 | 0 | 6 | 0 | NON_SYNONYMOU | MODERAT | MISSENSE | aAc/aGc | N195S | PLES_40721 | G | G | | | | | | | G | G | G | G |
| NC_011770 | 4500025 | G | A | 73 | 0 | 2 | 0 | NON_SYNONYMOU | MODERAT | MISSENSE | cGc/cAc | R233H | PLES_40721 | | | | | | | A | A | | | | |
| NC_011770 | 4500397 | G | A | 67 | 0 | 1 | 0 | NON_SYNONYMOU | MODERAT | MISSENSE | cGg/cAg | R357Q | PLES_40721 | | | | | | | | A | | | | |
| NC_011770 | 4500871 | A | G | 66 | 0 | 2 | 0 | NON_SYNONYMOU | MODERAT | MISSENSE | Ata/Gta | I135V | PLES_40731 | | | | | | | | | | | G | G |
| NC_011770 | 4503048 | A | G | 51 | 0 | 12 | 0 | NON_SYNONYMOU | MODERAT | MISSENSE | cAg/cGg | Q31R | PLES_40751 | G | G | G | G | G | G | G | G | G | G | G | G |
| NC_011770 | 4503284 | C | G | 60 | 0 | 6 | 0 | NON_SYNONYMOU | MODERAT | MISSENSE | Ccg/Gcg | P110A | PLES_40751 | | | | | | | G | G | G | G | G | G |
| NC_011770 | 4503383 | G | T | 69 | 0 | 4 | 0 | NON_SYNONYMOU | MODERAT | MISSENSE | Gcg/Tcg | A143S | PLES_40751 | | | | | | | T | T | | | T | T |
| NC_011770 | 4503561 | T | A | 82 | 0 | 12 | 0 | NON_SYNONYMOU | MODERAT | MISSENSE | gAg/gTg | E60V | PLES_40761 | A | A | A | A | A | A | A | A | A | A | A | A |
| NC_011770 | 4503574 | C | A | 85 | 0 | 2 | 0 | NON_SYNONYMOU | MODERAT | MISSENSE | Gtg/Ttg | V56L | PLES_40761 | | | | | | | | | A | A | | |
| NC_011770 | 4503669 | C | CCGT | 79 | 0 | 2 | 0 | CODON_INSERTIO | MODERATE | | ggg/gACG | G24DG | PLES_40761 | | | | | | | | | CCGT | CCGT | | |
| NC_011770 | 4504909 | C | A | 57 | 0 | 8 | 0 | NON_SYNONYMOU | MODERAT | MISSENSE | gaG/gaT | E289D | PLES_40771 | A | A | A | A | A | A | | | A | A | | |
| NC_011770 | 4504932 | A | G | 58 | 0 | 8 | 0 | NON_SYNONYMOU | MODERAT | MISSENSE | Tgc/Cgc | C282R | PLES_40771 | G | G | G | G | G | G | | | G | G | | |
| NC_011770 | 4505171 | G | A | 69 | 0 | 2 | 0 | NON_SYNONYMOU | MODERAT | MISSENSE | gCg/gTg | A202V | PLES_40771 | A | A | | | | | | | | | | |
| NC_011770 | 4505364 | A | G | 72 | 0 | 8 | 0 | NON_SYNONYMOU | MODERAT | MISSENSE | Tcg/Ccg | S138P | PLES_40771 | G | G | G | G | G | G | | | | | G | G |
| NC_011770 | 4505468 | A | T | 74 | 0 | 2 | 0 | NON_SYNONYMOU | MODERAT | MISSENSE | cTg/cAg | L103Q | PLES_40771 | | | | | | | | | T | T | | |
| NC_011770 | 4505499 | C | T | 78 | 0 | 2 | 0 | NON_SYNONYMOU | MODERAT | MISSENSE | Gcc/Acc | A93T | PLES_40771 | | | | | | | | | | | T | T |
| NC_011770 | 4505702 | T | C | 82 | 0 | 4 | 0 | NON_SYNONYMOU | MODERAT | MISSENSE | gAc/gGc | D25G | PLES_40771 | C | C | | | | | C | C | | | | |
| NC_011770 | 4506632 | G | A | 78 | 0 | 2 | 0 | NON_SYNONYMOU | MODERAT | MISSENSE | Ctc/Ttc | L18F | PLES_40781 | | | | | | | | | A | A | | |
| NC_011770 | 4507032 | G | A | 76 | 0 | 2 | 0 | NON_SYNONYMOU | MODERAT | MISSENSE | aGc/aAc | S2N | PLES_40801 | | | | | | | | | | | A | A |
| NC_011770 | 4507311 | A | C | 70 | 0 | 2 | 0 | NON_SYNONYMOU | MODERAT | MISSENSE | gAg/gCg | E95A | PLES_40801 | | | | | | | | | C | C | | |
| NC_011770 | 4507784 | A | C | 52 | 0 | 2 | 0 | NON_SYNONYMOU | MODERAT | MISSENSE | Agc/Cgc | S253R | PLES_40801 | C | C | | | | | | | | | | |
| NC_011770 | 4508134 | C | T | 46 | 0 | 8 | 0 | NON_SYNONYMOU | MODERAT | MISSENSE | aCc/aTc | T64I | PLES_40811 | T | T | | | | | T | T | T | T | T | T |
| NC_011770 | 4508173 | C | T | 49 | 0 | 6 | 0 | NON_SYNONYMOU | MODERAT | MISSENSE | cCc/cTc | P77L | PLES_40811 | T | T | | | | | | | T | T | T | T |
| NC_011770 | 4508393 | G | A | 68 | 0 | 4 | 0 | NON_SYNONYMOU | MODERAT | MISSENSE | gCg/gTg | A276V | PLES_40821 | A | A | | | | | | | | | A | A |
| NC_011770 | 4508428 | C | G | 70 | 0 | 4 | 0 | NON_SYNONYMOU | MODERAT | MISSENSE | gaG/gaC | E264D | PLES_40821 | G | G | | | | | | | | | G | G |
| NC_011770 | 4508799 | C | T | 77 | 0 | 2 | 0 | NON_SYNONYMOU | MODERAT | MISSENSE | Gcc/Acc | A141T | PLES_40821 | | | | | | | | | T | T | | |
| NC_011770 | 4508817 | T | C | 74 | 0 | 12 | 0 | NON_SYNONYMOU | MODERAT | MISSENSE | Acg/Gcg | T135A | PLES_40821 | C | C | C | C | C | C | C | C | C | C | C | C |
| NC_011770 | 4509039 | G | C | 76 | 0 | 2 | 0 | NON_SYNONYMOU | MODERAT | MISSENSE | Ccg/Gcg | P61A | PLES_40821 | C | C | | | | | | | | | | |
| NC_011770 | 4509183 | C | T | 77 | 0 | 2 | 0 | NON_SYNONYMOU | MODERAT | MISSENSE | Gcc/Acc | A13T | PLES_40821 | T | T | | | | | | | | | | |
| NC_011770 | 4509189 | T | C | 74 | 0 | 2 | 0 | NON_SYNONYMOU | MODERAT | MISSENSE | Act/Gct | T11A | PLES_40821 | | | | | | | | | C | C | | |
| NC_011770 | 4509554 | C | T | 84 | 0 | 2 | 0 | NON_SYNONYMOU | MODERAT | MISSENSE | Gtc/Atc | V105I | PLES_40831 | T | T | | | | | | | | | | |
| NC_011770 | 4509674 | C | G | 82 | 0 | 6 | 0 | NON_SYNONYMOU | MODERAT | MISSENSE | Ggc/Cgc | G65R | PLES_40831 | | | G | G | G | G | | | G | G | | |
| NC_011770 | 4509859 | A | G | 79 | 0 | 10 | 0 | NON_SYNONYMOU | MODERAT | MISSENSE | aTg/aCg | M3T | PLES_40831 | | | G | G | G | G | G | G | G | G | G | G |
| NC_011770 | 4510036 | G | A | 78 | 0 | 2 | 0 | NON_SYNONYMOU | MODERAT | MISSENSE | cGc/cAc | R22H | PLES_40841 | A | A | | | | | | | | | | |
| NC_011770 | 4510492 | C | G | 75 | 0 | 10 | 0 | NON_SYNONYMOU | MODERAT | MISSENSE | aCc/aGc | T174S | PLES_40841 | G | G | G | G | G | G | | | G | G | G | G |
| NC_011770 | 4510576 | A | G | 64 | 2 | 2 | 0 | NON_SYNONYMOU | MODERAT | MISSENSE | cAg/cGg | Q202R | PLES_40841 | . | . | | | | | | | | | G | G |
| NC_011770 | 4510722 | T | A | 70 | 0 | 10 | 0 | NON_SYNONYMOU | MODERAT | MISSENSE | Act/Tct | T222S | PLES_40851 | A | A | C | C | C | C | C | C | | | C | C |
| NC_011770 | 4510776 | G | A | 72 | 2 | 2 | 0 | NON_SYNONYMOU | MODERAT | MISSENSE | Cgt/Tgt | R204C | PLES_40851 | . | . | | | | | | | | | A | A |
| NC_011770 | 4511754 | G | C | 71 | 0 | 2 | 0 | NON_SYNONYMOU | MODERAT | MISSENSE | caG/caC | Q95H | PLES_40861 | . | C | | | | | | | | | | |
| NC_011770 | 4511878 | A | C | 64 | 0 | 2 | 0 | NON_SYNONYMOU | MODERAT | MISSENSE | Acg/Ccg | T137P | PLES_40861 | C | C | | | | | | | | | | |
| NC_011770 | 4512271 | T | C | 61 | 2 | 6 | 0 | NON_SYNONYMOU | MODERAT | MISSENSE | Tcc/Ccc | S268P | PLES_40861 | . | . | C | C | C | C | | | | | C | C |
| NC_011770 | 4512286 | G | A | 64 | 2 | 1 | 0 | NON_SYNONYMOU | MODERAT | MISSENSE | Gcc/Acc | A273T | PLES_40861 | . | . | | | | | A | | | | | |
| NC_011770 | 4512562 | C | A | 54 | 0 | 8 | 0 | NON_SYNONYMOU | MODERAT | MISSENSE | aCa/aAa | T35K | PLES_40871 | A | A | | | | | A | A | A | A | A | A |

| | | | | | | | | | | | | | | | | | | | | | | | | | |
|---|---|---|---|---|---|---|---|---|---|---|---|---|---|---|---|---|---|---|---|---|---|---|---|---|---|
| NC_011770 | 4512600 | G | A | 59 | 0 | 4 | 0 | NON_SYNONYMOU | MODERAT | MISSENSE | Gac/Aac | D48N | PLES_40871 | | | A | A | A | A | | | | | | |
| NC_011770 | 4512699 | C | A | 69 | 0 | 2 | 0 | NON_SYNONYMOU | MODERAT | MISSENSE | Ccc/Acc | P81T | PLES_40871 | A | A | | | | | | | | | | |
| NC_011770 | 4513027 | C | T | 73 | 0 | 1 | 0 | NON_SYNONYMOU | MODERAT | MISSENSE | gCg/gTg | A190V | PLES_40871 | | | | | | | T | | | | | |
| NC_011770 | 4514316 | A | G | 59 | 0 | 8 | 0 | NON_SYNONYMOU | MODERAT | MISSENSE | cAa/cGa | Q6R | PLES_40881 | G | G | | | | | G | G | G | G | G | G |
| NC_011770 | 4514361 | T | C | 61 | 0 | 4 | 0 | NON_SYNONYMOU | MODERAT | MISSENSE | cTg/cCg | L21P | PLES_40881 | | | C | C | C | C | | | | | | |
| NC_011770 | 4514471 | C | G | 80 | 0 | 4 | 0 | NON_SYNONYMOU | MODERAT | MISSENSE | Cat/Gat | H58D | PLES_40881 | | | | | | | G | G | G | G | | |
| NC_011770 | 4514567 | G | A | 72 | 0 | 8 | 0 | NON_SYNONYMOU | MODERAT | MISSENSE | Gtc/Atc | V90I | PLES_40881 | A | A | | | | | A | A | A | A | A | A |
| NC_011770 | 4514568 | T | C | 71 | 0 | 8 | 0 | NON_SYNONYMOU | MODERAT | MISSENSE | gTc/gCc | V90A | PLES_40881 | C | C | | | | | C | C | C | C | C | C |
| NC_011770 | 4514700 | C | G | 85 | 0 | 2 | 0 | NON_SYNONYMOU | MODERAT | MISSENSE | aCg/aGg | T134R | PLES_40881 | | | | | | | | | | | G | G |
| NC_011770 | 4515033 | A | G | 90 | 0 | 4 | 0 | NON_SYNONYMOU | MODERAT | MISSENSE | cAg/cGg | Q245R | PLES_40881 | G | G | | | | | | | G | G | | |
| NC_011770 | 4515042 | G | A | 91 | 0 | 4 | 0 | NON_SYNONYMOU | MODERAT | MISSENSE | gGc/gAc | G248D | PLES_40881 | A | A | | | | | | | A | A | | |
| NC_011770 | 4515266 | G | A | 94 | 0 | 4 | 0 | NON_SYNONYMOU | MODERAT | MISSENSE | Gac/Aac | D323N | PLES_40881 | A | A | | | | | | | A | A | | |
| NC_011770 | 4515607 | C | G | 83 | 0 | 4 | 0 | NON_SYNONYMOU | MODERAT | MISSENSE | ttC/ttG | F436L | PLES_40881 | | | | | | | G | G | G | G | | |
| NC_011770 | 4515612 | G | A | 84 | 0 | 2 | 0 | NON_SYNONYMOU | MODERAT | MISSENSE | cGc/cAc | R438H | PLES_40881 | A | A | | | | | | | | | | |
| NC_011770 | 4515662 | A | G | 85 | 0 | 6 | 0 | NON_SYNONYMOU | MODERAT | MISSENSE | Atc/Gtc | I455V | PLES_40881 | | | | | | | G | G | G | G | G | G |
| NC_011770 | 4516149 | C | A | 81 | 0 | 2 | 0 | NON_SYNONYMOU | MODERAT | MISSENSE | gCg/gAg | A617E | PLES_40881 | A | A | | | | | | | | | | |
| NC_011770 | 4516283 | C | T | 79 | 0 | 2 | 0 | NON_SYNONYMOU | MODERAT | MISSENSE | gCc/gTc | A44V | PLES_40891 | T | T | | | | | | | | | | |
| NC_011770 | 4516318 | T | C | 77 | 0 | 2 | 0 | NON_SYNONYMOU | MODERAT | MISSENSE | Tcc/Ccc | S56P | PLES_40891 | | | | | | | | | C | C | | |
| NC_011770 | 4516508 | G | A | 74 | 0 | 12 | 0 | NON_SYNONYMOU | MODERAT | MISSENSE | cGg/cAg | R119Q | PLES_40891 | A | A | A | A | A | A | A | A | A | A | A | A |
| NC_011770 | 4516558 | G | A | 68 | 0 | 4 | 0 | NON_SYNONYMOU | MODERAT | MISSENSE | Gcg/Acg | A136T | PLES_40891 | | | | | | | | | A | A | A | A |
| NC_011770 | 4516733 | A | C | 54 | 0 | 2 | 0 | NON_SYNONYMOU | MODERAT | MISSENSE | gAc/gCc | D194A | PLES_40891 | | | | | | | | | | | C | C |
| NC_011770 | 4516837 | A | T | 61 | 0 | 2 | 0 | NON_SYNONYMOU | MODERAT | MISSENSE | Agc/Tgc | S229C | PLES_40891 | | | | | | | | | | | T | T |
| NC_011770 | 4516889 | G | T | 55 | 0 | 4 | 0 | NON_SYNONYMOU | MODERAT | MISSENSE | cGc/cTc | R246L | PLES_40891 | | | | | | | T | T | T | T | | |
| NC_011770 | 4516939 | G | C | 56 | 0 | 4 | 0 | NON_SYNONYMOU | MODERAT | MISSENSE | Gaa/Caa | E263Q | PLES_40891 | | | C | C | C | C | | | | | | |
| NC_011770 | 4517494 | C | T | 63 | 0 | 12 | 0 | NON_SYNONYMOU | MODERAT | MISSENSE | cCg/cTg | P29L | PLES_40901 | T | T | T | T | T | T | T | T | T | T | T | T |
| NC_011770 | 4518015 | A | G | 67 | 0 | 8 | 0 | NON_SYNONYMOU | MODERAT | MISSENSE | Atc/Gtc | I203V | PLES_40901 | | | G | G | G | G | G | G | | | G | G |
| NC_011770 | 4518557 | C | A | 76 | 0 | 2 | 0 | NON_SYNONYMOU | MODERAT | MISSENSE | Ccc/Acc | P180T | PLES_40911 | | | | | A | A | | | | | | |
| NC_011770 | 4518599 | G | C | 74 | 0 | 12 | 0 | NON_SYNONYMOU | MODERAT | MISSENSE | Gag/Cag | E194Q | PLES_40911 | C | C | C | C | C | C | C | C | C | C | C | C |
| NC_011770 | 4518634 | G | C | 71 | 0 | 6 | 0 | NON_SYNONYMOU | MODERAT | MISSENSE | gaG/gaC | E205D | PLES_40911 | C | C | | | | | C | C | | | C | C |
| NC_011770 | 4519214 | G | A | 67 | 0 | 4 | 0 | NON_SYNONYMOU | MODERAT | MISSENSE | aGc/aAc | S107N | PLES_40921 | | | A | A | A | A | | | | | | |
| NC_011770 | 4520173 | A | G | 68 | 0 | 1 | 0 | NON_SYNONYMOU | MODERAT | MISSENSE | Agc/Ggc | S427G | PLES_40921 | | | | | | | G | | | | | |
| NC_011770 | 4520917 | G | A | 77 | 0 | 2 | 0 | NON_SYNONYMOU | MODERAT | MISSENSE | cGc/cAc | R220H | PLES_40931 | | | | | | | | | A | A | | |
| NC_011770 | 4521341 | C | A | 83 | 0 | 4 | 0 | NON_SYNONYMOU | MODERAT | MISSENSE | gaC/gaA | D45E | PLES_40941 | A | A | | | | | A | A | | | | |
| NC_011770 | 4521366 | G | A | 83 | 0 | 6 | 0 | NON_SYNONYMOU | MODERAT | MISSENSE | Gag/Aag | E54K | PLES_40941 | | | A | A | A | A | | | A | A | | |
| NC_011770 | 4521409 | A | G | 76 | 0 | 12 | 0 | NON_SYNONYMOU | MODERAT | MISSENSE | cAc/cGc | H68R | PLES_40941 | G | G | G | G | G | G | G | G | G | G | G | G |
| NC_011770 | 4521700 | G | A | 80 | 0 | 2 | 0 | NON_SYNONYMOU | MODERAT | MISSENSE | tGc/tAc | C165Y | PLES_40941 | | | | | | | | | A | A | | |
| NC_011770 | 4522410 | G | T | 73 | 0 | 12 | 0 | NON_SYNONYMOU | MODERAT | MISSENSE | Gcg/Tcg | A402S | PLES_40941 | T | T | T | T | T | T | T | T | T | T | T | T |
| NC_011770 | 4522515 | T | C | 72 | 0 | 2 | 0 | NON_SYNONYMOU | MODERAT | MISSENSE | gTc/gCc | V8A | PLES_40951 | C | C | | | | | | | | | | |
| NC_011770 | 4522718 | G | A | 63 | 0 | 4 | 0 | NON_SYNONYMOU | MODERAT | MISSENSE | Ggc/Agc | G76S | PLES_40951 | | | A | A | A | A | | | | | | |
| NC_011770 | 4523514 | G | C | 60 | 0 | 2 | 0 | NON_SYNONYMOU | MODERAT | MISSENSE | tGg/tCg | W341S | PLES_40951 | C | C | | | | | | | | | | |
| NC_011770 | 4523519 | C | T | 58 | 0 | 2 | 0 | NON_SYNONYMOU | MODERAT | MISSENSE | Cgg/Tgg | R343W | PLES_40951 | T | T | | | | | | | | | | |
| NC_011770 | 4523604 | C | T | 63 | 0 | 2 | 0 | NON_SYNONYMOU | MODERAT | MISSENSE | gCg/gTg | A371V | PLES_40951 | | | | | | | | | T | T | | |
| NC_011770 | 4525065 | G | A | 30 | 0 | 4 | 0 | NON_SYNONYMOU | MODERAT | MISSENSE | Gtg/Atg | V8M | PLES_40971 | | | A | A | A | A | | | | | | |
| NC_011770 | 4525369 | G | A | 64 | 0 | 4 | 0 | NON_SYNONYMOU | MODERAT | MISSENSE | gGa/gAa | G109E | PLES_40971 | | | A | A | A | A | | | | | | |
| NC_011770 | 4525639 | A | G | 84 | 0 | 12 | 0 | NON_SYNONYMOU | MODERAT | MISSENSE | cAg/cGg | Q199R | PLES_40971 | G | G | G | G | G | G | G | G | G | G | G | G |
| NC_011770 | 4526714 | T | C | 71 | 0 | 4 | 0 | NON_SYNONYMOU | MODERAT | MISSENSE | Tcg/Ccg | S149P | PLES_40981 | | | C | C | C | C | | | | | | |
| NC_011770 | 4526721 | G | C | 71 | 0 | 4 | 0 | NON_SYNONYMOU | MODERAT | MISSENSE | cGg/cCg | R151P | PLES_40981 | | | C | C | C | C | | | | | | |
| NC_011770 | 4526726 | G | A | 71 | 0 | 2 | 0 | NON_SYNONYMOU | MODERAT | MISSENSE | Gtt/Att | V153I | PLES_40981 | | | | | | | | | | | A | A |
| NC_011770 | 4526727 | T | C | 71 | 0 | 2 | 0 | NON_SYNONYMOU | MODERAT | MISSENSE | gTt/gCt | V153A | PLES_40981 | | | | | | | | | | | C | C |
| NC_011770 | 4526786 | C | T | 63 | 0 | 2 | 0 | NON_SYNONYMOU | MODERAT | MISSENSE | Ccg/Tcg | P173S | PLES_40981 | | | | | | | | | | | T | T |
| NC_011770 | 4526798 | A | T | 65 | 0 | 2 | 0 | NON_SYNONYMOU | MODERAT | MISSENSE | Acc/Tcc | T177S | PLES_40981 | | | | | | | | | | | T | T |
| NC_011770 | 4526808 | A | G | 61 | 0 | 2 | 0 | NON_SYNONYMOU | MODERAT | MISSENSE | cAc/cGc | H180R | PLES_40981 | | | | | | | | | | | G | G |
| NC_011770 | 4527938 | T | C | 65 | 0 | 2 | 0 | NON_SYNONYMOU | MODERAT | MISSENSE | cTg/cCg | L16P | PLES_41001 | | | C | C | | | | | | | | |
| NC_011770 | 4528459 | T | A | 69 | 0 | 4 | 0 | NON_SYNONYMOU | MODERAT | MISSENSE | Tgg/Agg | W190R | PLES_41001 | | | | | | | | | A | A | A | A |
| NC_011770 | 4528756 | C | T | 80 | 0 | 1 | 0 | NON_SYNONYMOU | MODERAT | MISSENSE | Cgc/Tgc | R289C | PLES_41001 | | | | | | | T | | | | | |
| NC_011770 | 4529278 | C | T | 84 | 0 | 2 | 0 | NON_SYNONYMOU | MODERAT | MISSENSE | Gcg/Acg | A356T | PLES_41011 | | | | | T | T | | | | | | |
| NC_011770 | 4529464 | C | A | 97 | 0 | 2 | 0 | NON_SYNONYMOU | MODERAT | MISSENSE | Gtc/Ttc | V294F | PLES_41011 | | | | | | | | | | | A | A |
| NC_011770 | 4529550 | G | A | 93 | 0 | 2 | 0 | NON_SYNONYMOU | MODERAT | MISSENSE | gCc/gTc | A265V | PLES_41011 | A | A | | | | | | | | | | |
| NC_011770 | 4530112 | C | T | 80 | 0 | 2 | 0 | NON_SYNONYMOU | MODERAT | MISSENSE | Gac/Aac | D78N | PLES_41011 | | | | | T | T | | | | | | |
| NC_011770 | 4532400 | C | G | 87 | 0 | 12 | 0 | NON_SYNONYMOU | MODERAT | MISSENSE | gGc/gCc | G143A | PLES_41031 | G | G | G | G | G | G | G | G | G | G | G | G |
| NC_011770 | 4532632 | C | G | 86 | 0 | 2 | 0 | NON_SYNONYMOU | MODERAT | MISSENSE | Gtg/Ctg | V66L | PLES_41031 | | | | | | | G | G | | | | |
| NC_011770 | 4533325 | T | C | 73 | 0 | 2 | 0 | NON_SYNONYMOU | MODERAT | MISSENSE | Acg/Gcg | T204A | PLES_41041 | C | C | | | | | | | | | | |

| | | | | | | | | | | | | | | | | | | | | | | | | | |
|---|---|---|---|---|---|---|---|---|---|---|---|---|---|---|---|---|---|---|---|---|---|---|---|---|---|
| NC_011770 | 4533349 | G | C | 76 | 0 | 2 | 0 | NON_SYNONYMOU | MODERAT | MISSENSE | Cac/Gac | H196D | PLES_41041 | C | C | | | | | | | | | | |
| NC_011770 | 4533408 | G | T | 72 | 0 | 6 | 0 | NON_SYNONYMOU | MODERAT | MISSENSE | gCc/gAc | A176D | PLES_41041 | T | T | T | T | T | T | | | | | | |
| NC_011770 | 4534844 | G | C | 72 | 0 | 6 | 0 | NON_SYNONYMOU | MODERAT | MISSENSE | gCc/gGc | A37G | PLES_41061 | C | C | C | C | C | C | | | | | | |
| NC_011770 | 4534920 | T | C | 72 | 0 | 2 | 0 | NON_SYNONYMOU | MODERAT | MISSENSE | Acc/Gcc | T12A | PLES_41061 | C | C | | | | | | | | | | |
| NC_011770 | 4536927 | G | A | 56 | 0 | 2 | 0 | SPLICE_SITE_REGIO | LOW | SILENT | tGa/tAa | *305 | PLES_41081 | A | A | | | | | | | | | | |
| NC_011770 | 4536087 | A | C | 80 | 0 | 2 | 0 | NON_SYNONYMOU | MODERAT | MISSENSE | gAa/gCa | E25A | PLES_41081 | | | | | | | | | | | C | C |
| NC_011770 | 4537319 | G | A | 77 | 0 | 4 | 0 | NON_SYNONYMOU | MODERAT | MISSENSE | Ccc/Tcc | P94S | PLES_41091 | | | | | | | | | A | A | A | A |
| NC_011770 | 4537629 | C | T | 73 | 0 | 4 | 0 | NON_SYNONYMOU | MODERAT | MISSENSE | Gtg/Atg | V176M | PLES_41101 | | | T | T | T | T | | | | | | |
| NC_011770 | 4539001 | G | C | 86 | 0 | 4 | 0 | NON_SYNONYMOU | MODERAT | MISSENSE | gaC/gaG | D230E | PLES_41121 | | | C | C | C | C | | | | | | |
| NC_011770 | 4539047 | C | T | 89 | 0 | 4 | 0 | NON_SYNONYMOU | MODERAT | MISSENSE | cGg/cAg | R215Q | PLES_41121 | | | T | T | T | T | | | | | | |
| NC_011770 | 4539280 | C | A | 82 | 0 | 4 | 0 | NON_SYNONYMOU | MODERAT | MISSENSE | caG/caT | Q137H | PLES_41121 | | | | | | | | | A | A | A | A |
| NC_011770 | 4539644 | C | T | 88 | 0 | 4 | 0 | NON_SYNONYMOU | MODERAT | MISSENSE | cGc/cAc | R16H | PLES_41121 | | | | | | | | | T | T | T | T |
| NC_011770 | 4540184 | T | C | 88 | 0 | 1 | 0 | NON_SYNONYMOU | MODERAT | MISSENSE | gAc/gGc | D307G | PLES_41131 | | | | | | | C | | | | | |
| NC_011770 | 4541846 | C | T | 52 | 0 | 2 | 0 | NON_SYNONYMOU | MODERAT | MISSENSE | gCc/gTc | A178V | PLES_41141 | T | T | | | | | | | | | | |
| NC_011770 | 4541872 | T | C | 53 | 0 | 4 | 0 | NON_SYNONYMOU | MODERAT | MISSENSE | Ttc/Ctc | F187L | PLES_41141 | | | | | | | | | C | C | C | C |
| NC_011770 | 4543601 | G | A | 70 | 0 | 4 | 0 | SPLICE_SITE_REGIO | LOW | SILENT | taG/taA | *473 | PLES_41151 | | | A | A | A | A | | | | | | |
| NC_011770 | 4543267 | T | C | 82 | 0 | 1 | 0 | NON_SYNONYMOU | MODERAT | MISSENSE | aTc/aCc | I362T | PLES_41151 | | | | | | | C | | | | | |
| NC_011770 | 4543594 | G | C | 68 | 0 | 8 | 0 | NON_SYNONYMOU | MODERAT | MISSENSE | aGc/aCc | S471T | PLES_41151 | C | C | C | C | C | C | C | C | | | | |
| NC_011770 | 4543075 | T | C | 70 | 0 | 1 | 0 | NON_SYNONYMOU | MODERAT | MISSENSE | cTg/cCg | L298P | PLES_41151 | | | | | | | | C | | | | |
| NC_011770 | 4543685 | C | G | 68 | 0 | 4 | 0 | NON_SYNONYMOU | MODERAT | MISSENSE | gGa/gCa | G197A | PLES_41161 | G | G | | | | | G | G | | | | |
| NC_011770 | 4543847 | G | C | 79 | 0 | 10 | 0 | NON_SYNONYMOU | MODERAT | MISSENSE | tCc/tGc | S143C | PLES_41161 | C | C | C | C | C | C | C | C | | | C | C |
| NC_011770 | 4543848 | A | T | 77 | 0 | 10 | 0 | NON_SYNONYMOU | MODERAT | MISSENSE | Tcc/Acc | S143T | PLES_41161 | T | T | T | T | T | T | T | T | | | T | T |
| NC_011770 | 4546230 | G | A | 17 | 8 | 2 | 0 | NON_SYNONYMOU | MODERAT | MISSENSE | gCc/gTc | A13V | PLES_41181 | . | . | . | . | . | . | | | . | . | A | A |
| NC_011770 | 4546247 | C | G | 17 | 8 | 2 | 0 | NON_SYNONYMOU | MODERAT | MISSENSE | gaG/gaC | E7D | PLES_41181 | . | . | . | . | . | . | | | . | . | G | G |
| NC_011770 | 4546248 | T | G | 17 | 8 | 2 | 0 | NON_SYNONYMOU | MODERAT | MISSENSE | gAg/gCg | E7A | PLES_41181 | . | . | . | . | . | . | | | . | . | G | G |
| NC_011770 | 4546249 | C | G | 17 | 8 | 2 | 0 | NON_SYNONYMOU | MODERAT | MISSENSE | Gag/Cag | E7Q | PLES_41181 | . | . | . | . | . | . | | | . | . | G | G |
| NC_011770 | 4546264 | C | A | 17 | 8 | 2 | 0 | NON_SYNONYMOU | MODERAT | MISSENSE | Gcg/Tcg | A2S | PLES_41181 | . | . | . | . | . | . | | | . | . | A | A |
| NC_011770 | 4547040 | C | CA | 50 | 4 | 3 | 0 | FRAME_SHIFT | HIGH | | cgc/cgcT | R173R? | PLES_41191 | . | . | | CA | . | . | | | CA | CA | | |
| NC_011770 | 4547042 | C | CCG | 50 | 4 | 3 | 0 | FRAME_SHIFT | HIGH | | cgc/cCGgc | R173P? | PLES_41191 | . | . | . | CCG | . | . | | | CCG | CCG | | |
| NC_011770 | 4546268 | C | T | 17 | 8 | 2 | 0 | SPLICE_SITE_REGIO | LOW | SILENT | tGa/tAa | *431 | PLES_41191 | . | . | . | . | . | . | | | . | . | T | T |
| NC_011770 | 4546272 | T | C | 16 | 8 | 2 | 0 | NON_SYNONYMOU | MODERAT | MISSENSE | Atc/Gtc | I430V | PLES_41191 | . | . | . | . | . | . | | | . | . | C | C |
| NC_011770 | 4546425 | T | C | 53 | 4 | 8 | 0 | NON_SYNONYMOU | MODERAT | MISSENSE | Atc/Gtc | I379V | PLES_41191 | . | . | C | C | . | . | C | C | C | C | C | C |
| NC_011770 | 4546447 | G | C | 56 | 4 | 8 | 0 | NON_SYNONYMOU | MODERAT | MISSENSE | atC/atG | I371M | PLES_41191 | . | . | C | C | . | . | C | C | C | C | C | C |
| NC_011770 | 4546449 | T | C | 57 | 4 | 8 | 0 | NON_SYNONYMOU | MODERAT | MISSENSE | Atc/Gtc | I371V | PLES_41191 | . | . | C | C | . | . | C | C | C | C | C | C |
| NC_011770 | 4546524 | T | C | 66 | 0 | 1 | 0 | NON_SYNONYMOU | MODERAT | MISSENSE | Acc/Gcc | T346A | PLES_41191 | | | | | | | C | | | | | |
| NC_011770 | 4546590 | G | C | 36 | 5 | 3 | 0 | NON_SYNONYMOU | MODERAT | MISSENSE | Ccg/Gcg | P324A | PLES_41191 | . | . | C | C | . | . | | | . | C | | |
| NC_011770 | 4546599 | C | T | 37 | 5 | 3 | 0 | NON_SYNONYMOU | MODERAT | MISSENSE | Gtc/Atc | V321I | PLES_41191 | . | . | T | T | . | . | | | . | T | | |
| NC_011770 | 4546601 | G | C | 38 | 5 | 3 | 0 | NON_SYNONYMOU | MODERAT | MISSENSE | gCg/gGg | A320G | PLES_41191 | . | . | C | C | . | . | | | . | C | | |
| NC_011770 | 4546604 | G | T | 37 | 5 | 3 | 0 | NON_SYNONYMOU | MODERAT | MISSENSE | aCg/aAg | T319K | PLES_41191 | . | . | T | T | . | . | | | . | T | | |
| NC_011770 | 4546605 | T | C | 36 | 5 | 3 | 0 | NON_SYNONYMOU | MODERAT | MISSENSE | Acg/Gcg | T319A | PLES_41191 | . | . | C | C | . | . | | | . | C | | |
| NC_011770 | 4546610 | C | G | 36 | 5 | 3 | 0 | NON_SYNONYMOU | MODERAT | MISSENSE | aGc/aCc | S317T | PLES_41191 | . | . | G | G | . | . | | | . | G | | |
| NC_011770 | 4546611 | T | A | 36 | 5 | 3 | 0 | NON_SYNONYMOU | MODERAT | MISSENSE | Agc/Tgc | S317C | PLES_41191 | . | . | A | A | . | . | | | . | A | | |
| NC_011770 | 4546617 | G | C | 38 | 5 | 3 | 0 | NON_SYNONYMOU | MODERAT | MISSENSE | Ctg/Gtg | L315V | PLES_41191 | . | . | C | C | . | . | | | . | C | | |
| NC_011770 | 4546618 | C | A | 37 | 5 | 3 | 0 | NON_SYNONYMOU | MODERAT | MISSENSE | aaG/aaT | K314N | PLES_41191 | . | . | A | A | . | . | | | . | A | | |
| NC_011770 | 4546619 | T | C | 37 | 5 | 3 | 0 | NON_SYNONYMOU | MODERAT | MISSENSE | aAg/aGg | K314R | PLES_41191 | . | . | C | C | . | . | | | . | C | | |
| NC_011770 | 4546620 | T | G | 37 | 5 | 3 | 0 | NON_SYNONYMOU | MODERAT | MISSENSE | Aag/Cag | K314Q | PLES_41191 | . | . | G | G | . | . | | | . | G | | |
| NC_011770 | 4546624 | G | T | 37 | 5 | 3 | 0 | NON_SYNONYMOU | MODERAT | MISSENSE | agC/agA | S312R | PLES_41191 | . | . | T | T | . | . | | | . | T | | |
| NC_011770 | 4546625 | C | G | 36 | 5 | 3 | 0 | NON_SYNONYMOU | MODERAT | MISSENSE | aGc/aCc | S312T | PLES_41191 | . | . | G | G | . | . | | | . | G | | |
| NC_011770 | 4546626 | T | C | 36 | 5 | 3 | 0 | NON_SYNONYMOU | MODERAT | MISSENSE | Agc/Ggc | S312G | PLES_41191 | . | . | C | C | . | . | | | . | C | | |
| NC_011770 | 4546830 | T | A | 64 | 4 | 1 | 0 | NON_SYNONYMOU | MODERAT | MISSENSE | Acc/Tcc | T244S | PLES_41191 | . | . | | | . | . | A | | | | | |
| NC_011770 | 4547051 | T | G | 51 | 4 | 2 | 0 | NON_SYNONYMOU | MODERAT | MISSENSE | aAa/aCa | K170T | PLES_41191 | . | . | G | G | . | . | | | . | . | | |
| NC_011770 | 4547153 | T | C | 50 | 4 | 6 | 0 | NON_SYNONYMOU | MODERAT | MISSENSE | cAt/cGt | H136R | PLES_41191 | . | . | C | C | . | . | | | C | C | C | C |
| NC_011770 | 4547172 | G | T | 51 | 4 | 4 | 0 | NON_SYNONYMOU | MODERAT | MISSENSE | Ctc/Atc | L130I | PLES_41191 | . | . | T | T | . | . | | | T | T | | |
| NC_011770 | 4547211 | G | T | 53 | 4 | 4 | 0 | NON_SYNONYMOU | MODERAT | MISSENSE | Ctt/Att | L117I | PLES_41191 | . | . | T | T | . | . | | | T | T | | |
| NC_011770 | 4547259 | A | T | 52 | 4 | 8 | 0 | NON_SYNONYMOU | MODERAT | MISSENSE | Ttg/Atg | L101M | PLES_41191 | . | . | T | T | . | . | T | T | T | T | T | T |
| NC_011770 | 4547383 | C | G | 47 | 4 | 2 | 0 | NON_SYNONYMOU | MODERAT | MISSENSE | ttG/ttC | L59F | PLES_41191 | . | . | | | . | . | | | G | G | | |
| NC_011770 | 4547048 | GGTT | G | 51 | 4 | 3 | 0 | CODON_CHANGE_ | MODERATE | | aaaccg/acg | KP170T | PLES_41191 | . | . | | G | . | . | | | G | G | | |
| NC_011770 | 4546751 | A | T | 60 | 4 | 5 | 0 | NON_SYNONYMOU | MODERAT | MISSENSE | tTc/tAc | F270Y | PLES_41191 | . | . | T | T | . | . | | T | T | T | | |
| NC_011770 | 4546764 | G | T | 59 | 4 | 5 | 0 | NON_SYNONYMOU | MODERAT | MISSENSE | Ctc/Atc | L266I | PLES_41191 | . | . | T | T | . | . | | T | T | T | | |
| NC_011770 | 4547894 | G | A | 18 | 10 | 1 | 0 | NON_SYNONYMOU | MODERAT | MISSENSE | Ccc/Tcc | P396S | PLES_41201 | . | . | . | . | . | . | A | | . | . | . | . |
| NC_011770 | 4547915 | T | C | 18 | 10 | 2 | 0 | NON_SYNONYMOU | MODERAT | MISSENSE | Acc/Gcc | T389A | PLES_41201 | . | . | . | . | . | . | C | C | . | . | . | . |
| NC_011770 | 4547929 | A | G | 18 | 10 | 2 | 0 | NON_SYNONYMOU | MODERAT | MISSENSE | gTg/gCg | V384A | PLES_41201 | . | . | . | . | . | . | G | G | . | . | . | . |

| | | | | | | | | | | | | | | | | | | | | | | | | | |
|---|---|---|---|---|---|---|---|---|---|---|---|---|---|---|---|---|---|---|---|---|---|---|---|---|---|
| NC_011770 | 4548196 | C | G | 24 | 10 | 2 | 0 | NON_SYNONYMOU | MODERAT | MISSENSE | cGa/cCa | R295P | PLES_41201 | . | . | . | . | . | . | G | G | . | . | . | . |
| NC_011770 | 4548208 | A | G | 26 | 10 | 1 | 0 | NON_SYNONYMOU | MODERAT | MISSENSE | gTa/gCa | V291A | PLES_41201 | . | . | . | . | . | . | G | | . | . | . | . |
| NC_011770 | 4548229 | C | T | 25 | 10 | 1 | 0 | NON_SYNONYMOU | MODERAT | MISSENSE | gGt/gAt | G284D | PLES_41201 | . | . | . | . | . | . | T | | . | . | . | . |
| NC_011770 | 4548244 | G | A | 25 | 10 | 1 | 0 | NON_SYNONYMOU | MODERAT | MISSENSE | gCt/gTt | A279V | PLES_41201 | . | . | . | . | . | . | A | | . | . | . | . |
| NC_011770 | 4548302 | G | A | 21 | 10 | 1 | 0 | NON_SYNONYMOU | MODERAT | MISSENSE | Cct/Tct | P260S | PLES_41201 | . | . | . | . | . | . | A | | . | . | . | . |
| NC_011770 | 4548568 | C | G | 22 | 9 | 1 | 0 | NON_SYNONYMOU | MODERAT | MISSENSE | cGg/cCg | R171P | PLES_41201 | . | . | . | . | . | . | | | . | . | . | G |
| NC_011770 | 4548713 | C | T | 33 | 4 | 2 | 0 | NON_SYNONYMOU | MODERAT | MISSENSE | Ggc/Agc | G123S | PLES_41201 | . | . | | | . | . | | | | | T | T |
| NC_011770 | 4548724 | C | T | 26 | 6 | 4 | 0 | NON_SYNONYMOU | MODERAT | MISSENSE | gGg/gAg | G119E | PLES_41201 | . | . | T | T | . | . | | | . | . | T | T |
| NC_011770 | 4548732 | C | A | 28 | 6 | 4 | 0 | NON_SYNONYMOU | MODERAT | MISSENSE | aaG/aaT | K116N | PLES_41201 | . | . | A | A | . | . | | | . | . | G | G |
| NC_011770 | 4548734 | T | G | 28 | 6 | 4 | 0 | NON_SYNONYMOU | MODERAT | MISSENSE | Aag/Cag | K116Q | PLES_41201 | . | . | G | G | . | . | | | . | . | G | G |
| NC_011770 | 4548751 | T | C | 29 | 6 | 4 | 0 | NON_SYNONYMOU | MODERAT | MISSENSE | gAc/gGc | D110G | PLES_41201 | . | . | C | C | . | . | | | . | . | C | C |
| NC_011770 | 4548757 | A | T | 30 | 6 | 4 | 0 | NON_SYNONYMOU | MODERAT | MISSENSE | tTc/tAc | F108Y | PLES_41201 | . | . | T | T | . | . | | | . | . | T | T |
| NC_011770 | 4548770 | T | C | 33 | 6 | 4 | 0 | NON_SYNONYMOU | MODERAT | MISSENSE | Aag/Gag | K104E | PLES_41201 | . | . | C | C | . | . | | | . | . | C | C |
| NC_011770 | 4548778 | G | T | 30 | 6 | 4 | 0 | NON_SYNONYMOU | MODERAT | MISSENSE | gCc/gAc | A101D | PLES_41201 | . | . | T | T | . | . | | | . | . | T | T |
| NC_011770 | 4548779 | C | T | 31 | 6 | 4 | 0 | NON_SYNONYMOU | MODERAT | MISSENSE | Gcc/Acc | A101T | PLES_41201 | . | . | T | T | . | . | | | . | . | T | T |
| NC_011770 | 4548796 | A | G | 32 | 6 | 4 | 0 | NON_SYNONYMOU | MODERAT | MISSENSE | cTg/cCg | L95P | PLES_41201 | . | . | G | G | . | . | | | . | . | G | G |
| NC_011770 | 4548801 | G | C | 32 | 6 | 4 | 0 | NON_SYNONYMOU | MODERAT | MISSENSE | atC/atG | I93M | PLES_41201 | . | . | C | C | . | . | | | . | . | C | C |
| NC_011770 | 4548803 | T | G | 32 | 6 | 4 | 0 | NON_SYNONYMOU | MODERAT | MISSENSE | Atc/Ctc | I93L | PLES_41201 | . | . | G | G | . | . | | | . | . | G | G |
| NC_011770 | 4548825 | G | C | 28 | 8 | 2 | 0 | NON_SYNONYMOU | MODERAT | MISSENSE | atC/atG | I85M | PLES_41201 | . | . | C | C | . | . | | | . | . | . | . |
| NC_011770 | 4548827 | T | A | 28 | 8 | 2 | 0 | NON_SYNONYMOU | MODERAT | MISSENSE | Atc/Ttc | I85F | PLES_41201 | . | . | A | A | . | . | | | . | . | . | . |
| NC_011770 | 4548850 | A | T | 29 | 8 | 2 | 0 | NON_SYNONYMOU | MODERAT | MISSENSE | aTg/aAg | M77K | PLES_41201 | . | . | T | T | . | . | | | . | . | . | . |
| NC_011770 | 4548851 | T | G | 29 | 8 | 2 | 0 | NON_SYNONYMOU | MODERAT | MISSENSE | Atg/Ctg | M77L | PLES_41201 | . | . | G | G | . | . | | | . | . | . | . |
| NC_011770 | 4547879 | C | T | 18 | 10 | 1 | 0 | NON_SYNONYMOU | MODERAT | MISSENSE | Ggc/Agc | G401S | PLES_41201 | . | . | . | . | . | . | | T | . | . | . | . |
| NC_011770 | 4548197 | G | C | 25 | 10 | 1 | 0 | NON_SYNONYMOU | MODERAT | MISSENSE | Cga/Gga | R295G | PLES_41201 | . | . | . | . | . | . | | C | . | . | . | . |
| NC_011770 | 4549897 | A | ACCG | 0 | 10 | 1 | 0 | CODON_CHANGE_ | MODERATE | | ggt/ggCG | G223GG | PLES_41221 | . | . | . | . | . | . | ACCG | . | . | . | . | . |
| NC_011770 | 4549953 | C | T | 1 | 10 | 1 | 0 | NON_SYNONYMOU | MODERAT | MISSENSE | Ggc/Agc | G205S | PLES_41221 | . | . | . | . | . | . | T | | . | . | . | . |
| NC_011770 | 4550429 | A | G | 0 | 10 | 2 | 0 | NON_SYNONYMOU | MODERAT | MISSENSE | gTt/gCt | V46A | PLES_41221 | . | . | . | . | . | . | G | G | . | . | . | . |
| NC_011770 | 4550457 | A | T | 14 | 10 | 2 | 0 | NON_SYNONYMOU | MODERAT | MISSENSE | Ttt/Att | F37I | PLES_41221 | . | . | . | . | . | . | T | T | . | . | . | . |
| NC_011770 | 4550487 | T | A | 15 | 10 | 2 | 0 | NON_SYNONYMOU | MODERAT | MISSENSE | Acg/Tcg | T27S | PLES_41221 | . | . | . | . | . | . | A | A | . | . | . | . |
| NC_011770 | 4550496 | A | G | 15 | 10 | 2 | 0 | NON_SYNONYMOU | MODERAT | MISSENSE | Tca/Cca | S24P | PLES_41221 | . | . | . | . | . | . | G | G | . | . | . | . |
| NC_011770 | 4550516 | C | T | 17 | 10 | 2 | 0 | NON_SYNONYMOU | MODERAT | MISSENSE | gGt/gAt | G17D | PLES_41221 | . | . | . | . | . | . | T | T | . | . | . | . |
| NC_011770 | 4550522 | T | A | 17 | 10 | 2 | 0 | NON_SYNONYMOU | MODERAT | MISSENSE | tAc/tTc | Y15F | PLES_41221 | . | . | . | . | . | . | A | A | . | . | . | . |
| NC_011770 | 4550549 | G | A | 17 | 10 | 2 | 0 | NON_SYNONYMOU | MODERAT | MISSENSE | gCg/gTg | A6V | PLES_41221 | . | . | . | . | . | . | A | A | . | . | . | . |
| NC_011770 | 4550781 | A | G | 19 | 10 | 1 | 0 | NON_SYNONYMOU | MODERAT | MISSENSE | gTg/gCg | V48A | PLES_41231 | . | . | . | . | . | . | G | | . | . | . | . |
| NC_011770 | 4550787 | C | G | 14 | 10 | 2 | 0 | NON_SYNONYMOU | MODERAT | MISSENSE | gGg/gCg | G46A | PLES_41231 | . | . | . | . | . | . | G | G | . | . | . | . |
| NC_011770 | 4550788 | C | A | 14 | 10 | 2 | 0 | NON_SYNONYMOU | MODERAT | MISSENSE | Ggg/Tgg | G46W | PLES_41231 | . | . | . | . | . | . | A | A | . | . | . | . |
| NC_011770 | 4550800 | T | C | 15 | 10 | 2 | 0 | NON_SYNONYMOU | MODERAT | MISSENSE | Agc/Ggc | S42G | PLES_41231 | . | . | . | . | . | . | C | C | . | . | . | . |
| NC_011770 | 4550809 | C | T | 14 | 10 | 2 | 0 | NON_SYNONYMOU | MODERAT | MISSENSE | Gcg/Acg | A39T | PLES_41231 | . | . | . | . | . | . | T | T | . | . | . | . |
| NC_011770 | 4550817 | C | T | 14 | 10 | 2 | 0 | NON_SYNONYMOU | MODERAT | MISSENSE | gGg/gAg | G36E | PLES_41231 | . | . | . | . | . | . | T | T | . | . | . | . |
| NC_011770 | 4550836 | G | T | 20 | 10 | 2 | 0 | NON_SYNONYMOU | MODERAT | MISSENSE | Ctc/Atc | L30I | PLES_41231 | . | . | . | . | . | . | T | T | . | . | . | . |
| NC_011770 | 4550843 | T | G | 21 | 10 | 2 | 0 | NON_SYNONYMOU | MODERAT | MISSENSE | gaA/gaC | E27D | PLES_41231 | . | . | . | . | . | . | G | G | . | . | . | . |
| NC_011770 | 4550875 | C | T | 20 | 10 | 1 | 0 | NON_SYNONYMOU | MODERAT | MISSENSE | Ggc/Agc | G17S | PLES_41231 | . | . | . | . | . | . | T | | . | . | . | . |
| NC_011770 | 4550877 | G | A | 20 | 10 | 2 | 0 | NON_SYNONYMOU | MODERAT | MISSENSE | gCc/gTc | A16V | PLES_41231 | . | . | . | . | . | . | A | A | . | . | . | . |
| NC_011770 | 4551208 | C | T | 21 | 4 | 4 | 0 | NON_SYNONYMOU | MODERAT | MISSENSE | Gct/Act | A29T | PLES_41242 | . | . | | | . | . | T | T | | | T | T |
| NC_011770 | 4551223 | G | C | 19 | 4 | 2 | 0 | NON_SYNONYMOU | MODERAT | MISSENSE | Cta/Gta | L24V | PLES_41242 | . | . | | | . | . | C | C | | | | |
| NC_011770 | 4551374 | G | T | 8 | 4 | 2 | 0 | NON_SYNONYMOU | MODERAT | MISSENSE | Ccg/Acg | P124T | PLES_41251 | . | . | | | . | . | | | | | T | T |
| NC_011770 | 4551383 | C | A | 6 | 5 | 2 | 0 | NON_SYNONYMOU | MODERAT | MISSENSE | Gcc/Tcc | A121S | PLES_41251 | . | . | | | . | . | A | A | | | | . |
| NC_011770 | 4551388 | G | A | 5 | 5 | 2 | 0 | NON_SYNONYMOU | MODERAT | MISSENSE | gCg/gTg | A119V | PLES_41251 | . | . | | . | . | . | | | A | A | | |
| NC_011770 | 4551425 | T | C | 6 | 4 | 2 | 0 | NON_SYNONYMOU | MODERAT | MISSENSE | Atg/Gtg | M107V | PLES_41251 | . | . | C | C | . | . | | | | | | |
| NC_011770 | 4551478 | C | T | 15 | 4 | 2 | 0 | NON_SYNONYMOU | MODERAT | MISSENSE | cGg/cAg | R89Q | PLES_41251 | . | . | | | . | . | | | | | T | T |
| NC_011770 | 4551494 | G | T | 18 | 4 | 2 | 0 | NON_SYNONYMOU | MODERAT | MISSENSE | Caa/Aaa | Q84K | PLES_41251 | . | . | | | . | . | | | | | T | T |
| NC_011770 | 4552103 | G | A | 52 | 4 | 2 | 0 | NON_SYNONYMOU | MODERAT | MISSENSE | cCg/cTg | P59L | PLES_41261 | . | . | | | . | . | | | | | A | A |
| NC_011770 | 4552122 | A | G | 47 | 4 | 1 | 0 | NON_SYNONYMOU | MODERAT | MISSENSE | Tcc/Ccc | S53P | PLES_41261 | . | . | | | . | . | G | | | | | |
| NC_011770 | 4552125 | T | C | 50 | 4 | 8 | 0 | NON_SYNONYMOU | MODERAT | MISSENSE | Acc/Gcc | T52A | PLES_41261 | . | . | C | C | . | . | C | C | C | C | C | C |
| NC_011770 | 4552172 | A | G | 55 | 4 | 2 | 0 | NON_SYNONYMOU | MODERAT | MISSENSE | cTt/cCt | L36P | PLES_41261 | . | . | | | . | . | | | | | G | G |
| NC_011770 | 4552232 | T | C | 54 | 4 | 2 | 0 | NON_SYNONYMOU | MODERAT | MISSENSE | cAg/cGg | Q16R | PLES_41261 | . | . | | | . | . | | | C | C | | |
| NC_011770 | 4552245 | G | A | 57 | 4 | 2 | 0 | NON_SYNONYMOU | MODERAT | MISSENSE | Ctc/Ttc | L12F | PLES_41261 | . | . | A | A | . | . | | | | | | |
| NC_011770 | 4552265 | G | A | 54 | 4 | 2 | 0 | NON_SYNONYMOU | MODERAT | MISSENSE | aCc/aTc | T5I | PLES_41261 | . | . | | | . | . | | | | | A | A |
| NC_011770 | 4552267 | C | G | 54 | 4 | 4 | 0 | NON_SYNONYMOU | MODERAT | MISSENSE | aaG/aaC | K4N | PLES_41261 | . | . | | | . | . | | | G | G | G | G |
| NC_011770 | 4552269 | T | C | 54 | 4 | 2 | 0 | NON_SYNONYMOU | MODERAT | MISSENSE | Aag/Gag | K4E | PLES_41261 | . | . | | | . | . | | | | | C | C |
| NC_011770 | 4552391 | G | A | 48 | 4 | 2 | 0 | NON_SYNONYMOU | MODERAT | MISSENSE | gCg/gTg | A61V | PLES_41271 | . | . | | | . | . | | | A | A | | |
| NC_011770 | 4552416 | G | A | 49 | 4 | 2 | 0 | NON_SYNONYMOU | MODERAT | MISSENSE | Ctc/Ttc | L53F | PLES_41271 | . | . | | | . | . | | | A | A | | |

| | | | | | | | | | | | | | | | | | | | | | | | | | |
|---|---|---|---|---|---|---|---|---|---|---|---|---|---|---|---|---|---|---|---|---|---|---|---|---|---|
| NC_011770 | 4552451 | G | C | 44 | 4 | 1 | 0 | NON_SYNONYMOU | MODERAT | MISSENSE | cCc/cGc | P41R | PLES_41271 | . | . | | | . | . | C | | | | | |
| NC_011770 | 4552472 | G | A | 44 | 4 | 2 | 0 | NON_SYNONYMOU | MODERAT | MISSENSE | tCg/tTg | S34L | PLES_41271 | . | . | | | . | . | | | A | A | | |
| NC_011770 | 4552481 | T | C | 44 | 4 | 1 | 0 | NON_SYNONYMOU | MODERAT | MISSENSE | cAc/cGc | H31R | PLES_41271 | . | . | | | . | . | C | | | | | |
| NC_011770 | 4552493 | A | G | 41 | 4 | 6 | 0 | NON_SYNONYMOU | MODERAT | MISSENSE | gTc/gCc | V27A | PLES_41271 | . | . | | | . | . | G | G | G | G | G | G |
| NC_011770 | 4552523 | G | C | 36 | 4 | 3 | 0 | NON_SYNONYMOU | MODERAT | MISSENSE | gCc/gGc | A17G | PLES_41271 | . | . | | | . | . | C | | C | C | | |
| NC_011770 | 4552536 | C | T | 33 | 4 | 4 | 0 | NON_SYNONYMOU | MODERAT | MISSENSE | Gcc/Acc | A13T | PLES_41271 | . | . | T | T | . | . | | | | | T | T |
| NC_011770 | 4552542 | T | C | 32 | 4 | 2 | 0 | NON_SYNONYMOU | MODERAT | MISSENSE | Acc/Gcc | T11A | PLES_41271 | . | . | | | . | . | | | | | C | C |
| NC_011770 | 4556266 | G | A | 77 | 0 | 4 | 0 | NON_SYNONYMOU | MODERAT | MISSENSE | Gac/Aac | D154N | PLES_41301 | | | | | | | | | A | A | A | A |
| NC_011770 | 4557125 | G | A | 70 | 0 | 1 | 0 | STOP_GAINED | HIGH | NONSENSE | tgG/tgA | W189* | PLES_41311 | | | | | | | A | | | | | |
| NC_011770 | 4557919 | G | A | 50 | 0 | 2 | 0 | NON_SYNONYMOU | MODERAT | MISSENSE | gCg/gTg | A363V | PLES_41331 | | | | | | | A | A | | | | |
| NC_011770 | 4558199 | C | A | 51 | 0 | 2 | 0 | NON_SYNONYMOU | MODERAT | MISSENSE | Gcc/Tcc | A270S | PLES_41331 | A | A | | | | | | | | | | |
| NC_011770 | 4558853 | T | C | 72 | 0 | 12 | 0 | NON_SYNONYMOU | MODERAT | MISSENSE | Atc/Gtc | I52V | PLES_41331 | C | C | C | C | C | C | C | C | C | C | C | C |
| NC_011770 | 4559301 | A | T | 85 | 0 | 4 | 0 | NON_SYNONYMOU | MODERAT | MISSENSE | Tcc/Acc | S312T | PLES_41341 | | | | | | | | | T | T | T | T |
| NC_011770 | 4560437 | A | C | 66 | 0 | 10 | 0 | NON_SYNONYMOU | MODERAT | MISSENSE | Tcc/Gcc | S307A | PLES_41351 | | | C | C | C | C | C | C | C | C | C | C |
| NC_011770 | 4560628 | A | C | 69 | 0 | 2 | 0 | NON_SYNONYMOU | MODERAT | MISSENSE | aTg/aGg | M243R | PLES_41351 | | | | | | | C | C | | | | |
| NC_011770 | 4560650 | G | C | 74 | 0 | 2 | 0 | NON_SYNONYMOU | MODERAT | MISSENSE | Cgg/Ggg | R236G | PLES_41351 | C | C | | | | | | | | | | |
| NC_011770 | 4560974 | C | A | 74 | 0 | 2 | 0 | NON_SYNONYMOU | MODERAT | MISSENSE | Ggc/Tgc | G128C | PLES_41351 | | | A | A | | | | | | | | |
| NC_011770 | 4561295 | G | A | 74 | 0 | 1 | 0 | NON_SYNONYMOU | MODERAT | MISSENSE | Cgc/Tgc | R21C | PLES_41351 | | | | | | | A | | | | | |
| NC_011770 | 4561537 | A | G | 76 | 0 | 12 | 0 | NON_SYNONYMOU | MODERAT | MISSENSE | Tcc/Ccc | S149P | PLES_41361 | G | G | G | G | G | G | G | G | G | G | G | G |
| NC_011770 | 4561608 | G | A | 71 | 0 | 4 | 0 | NON_SYNONYMOU | MODERAT | MISSENSE | cCg/cTg | P125L | PLES_41361 | | | A | A | A | A | | | | | | |
| NC_011770 | 4562606 | G | A | 70 | 0 | 2 | 0 | NON_SYNONYMOU | MODERAT | MISSENSE | Gtc/Atc | V142I | PLES_41371 | | | | | | | A | A | | | | |
| NC_011770 | 4562928 | C | T | 63 | 0 | 2 | 0 | NON_SYNONYMOU | MODERAT | MISSENSE | gCc/gTc | A249V | PLES_41371 | | | | | | | T | T | | | | |
| NC_011770 | 4565143 | C | A | 73 | 0 | 4 | 0 | NON_SYNONYMOU | MODERAT | MISSENSE | gaC/gaA | D117E | PLES_41391 | | | | | | | | | A | A | A | A |
| NC_011770 | 4565309 | A | G | 65 | 0 | 12 | 0 | NON_SYNONYMOU | MODERAT | MISSENSE | Acc/Gcc | T173A | PLES_41391 | G | G | G | G | G | G | G | G | G | G | G | G |
| NC_011770 | 4565526 | G | A | 69 | 0 | 8 | 0 | NON_SYNONYMOU | MODERAT | MISSENSE | cGc/cAc | R245H | PLES_41391 | | | A | A | A | A | | | A | A | A | A |
| NC_011770 | 4565905 | A | G | 65 | 0 | 12 | 0 | NON_SYNONYMOU | MODERAT | MISSENSE | Tgc/Cgc | C1086R | PLES_41401 | G | G | G | G | G | G | G | G | G | G | G | G |
| NC_011770 | 4568113 | A | T | 62 | 0 | 2 | 0 | NON_SYNONYMOU | MODERAT | MISSENSE | Tat/Aat | Y350N | PLES_41401 | | | | | | | | | | | T | T |
| NC_011770 | 4568401 | C | G | 43 | 0 | 2 | 0 | NON_SYNONYMOU | MODERAT | MISSENSE | Gtc/Ctc | V254L | PLES_41401 | | | | | | | G | G | | | | |
| NC_011770 | 4568721 | A | G | 23 | 0 | 4 | 0 | NON_SYNONYMOU | MODERAT | MISSENSE | aTg/aCg | M147T | PLES_41401 | | | G | G | G | G | | | | | | |
| NC_011770 | 4567053 | A | G | 85 | 0 | 1 | 0 | NON_SYNONYMOU | MODERAT | MISSENSE | cTg/cCg | L703P | PLES_41401 | | | | | | | | G | | | | |
| NC_011770 | 4576858 | C | CG | 60 | 0 | 1 | 0 | FRAME_SHIFT | HIGH | | ccg/ccCg | P380P? | PLES_41501 | | | | | | | CG | | | | | |
| NC_011770 | 4577154 | C | T | 49 | 0 | 10 | 0 | NON_SYNONYMOU | MODERAT | MISSENSE | Gcc/Acc | A282T | PLES_41501 | T | T | T | T | T | T | T | T | | | T | T |
| NC_011770 | 4577156 | G | A | 51 | 0 | 1 | 0 | NON_SYNONYMOU | MODERAT | MISSENSE | cCg/cTg | P281L | PLES_41501 | | | | | | | A | | | | | |
| NC_011770 | 4577282 | T | C | 61 | 0 | 2 | 0 | NON_SYNONYMOU | MODERAT | MISSENSE | gAc/gGc | D239G | PLES_41501 | C | C | | | | | | | | | | |
| NC_011770 | 4577684 | A | G | 57 | 0 | 12 | 0 | NON_SYNONYMOU | MODERAT | MISSENSE | gTg/gCg | V105A | PLES_41501 | G | G | G | G | G | G | G | G | G | G | G | G |
| NC_011770 | 4578728 | T | C | 71 | 0 | 1 | 0 | NON_SYNONYMOU | MODERAT | MISSENSE | cTg/cCg | L92P | PLES_41511 | | | | | | | C | | | | | |
| NC_011770 | 4578959 | T | C | 68 | 0 | 1 | 0 | NON_SYNONYMOU | MODERAT | MISSENSE | aTc/aCc | I169T | PLES_41511 | | | | | | | C | | | | | |
| NC_011770 | 4579082 | C | T | 64 | 0 | 2 | 0 | NON_SYNONYMOU | MODERAT | MISSENSE | gCc/gTc | A210V | PLES_41511 | | | | | | | | | T | T | | |
| NC_011770 | 4579537 | T | C | 69 | 0 | 2 | 0 | NON_SYNONYMOU | MODERAT | MISSENSE | Agc/Ggc | S585G | PLES_41521 | | | | | | | | | | | C | C |
| NC_011770 | 4579552 | C | G | 70 | 0 | 2 | 0 | NON_SYNONYMOU | MODERAT | MISSENSE | Gca/Cca | A580P | PLES_41521 | | | | | | | G | G | | | | |
| NC_011770 | 4580257 | G | A | 69 | 0 | 4 | 0 | NON_SYNONYMOU | MODERAT | MISSENSE | Cgt/Tgt | R345C | PLES_41521 | | | A | A | A | A | | | | | | |
| NC_011770 | 4580683 | C | T | 59 | 1 | 2 | 0 | NON_SYNONYMOU | MODERAT | MISSENSE | Ggt/Agt | G203S | PLES_41521 | T | T | | | | | | . | | | | |
| NC_011770 | 4580749 | A | T | 69 | 0 | 2 | 0 | NON_SYNONYMOU | MODERAT | MISSENSE | Ttg/Atg | L181M | PLES_41521 | T | T | | | | | | | | | | |
| NC_011770 | 4580881 | C | T | 74 | 0 | 2 | 0 | NON_SYNONYMOU | MODERAT | MISSENSE | Gcc/Acc | A137T | PLES_41521 | T | T | | | | | | | | | | |
| NC_011770 | 4581039 | G | C | 70 | 0 | 2 | 0 | NON_SYNONYMOU | MODERAT | MISSENSE | gCc/gGc | A84G | PLES_41521 | C | C | | | | | | | | | | |
| NC_011770 | 4580486 | C | T | 60 | 0 | 1 | 0 | NON_SYNONYMOU | MODERAT | MISSENSE | atG/atA | M268I | PLES_41521 | | | | | | | | | | | | T |
| NC_011770 | 4582219 | TCGCC | T | 55 | 0 | 2 | 0 | FRAME_SHIFT | HIGH | | cgggcg/ | RA212 | PLES_41541 | | | T | T | | | | | | | | |
| NC_011770 | 4582647 | G | A | 78 | 0 | 1 | 0 | NON_SYNONYMOU | MODERAT | MISSENSE | cCc/cTc | P71L | PLES_41541 | | A | | | | | | | | | | |
| NC_011770 | 4582287 | C | T | 63 | 0 | 2 | 0 | NON_SYNONYMOU | MODERAT | MISSENSE | gGc/gAc | G191D | PLES_41541 | | | | | | | | | | | T | T |
| NC_011770 | 4582390 | A | G | 78 | 0 | 2 | 0 | NON_SYNONYMOU | MODERAT | MISSENSE | Tcg/Ccg | S157P | PLES_41541 | G | G | | | | | | | | | | |
| NC_011770 | 4582548 | T | C | 76 | 0 | 1 | 0 | NON_SYNONYMOU | MODERAT | MISSENSE | cAc/cGc | H104R | PLES_41541 | | | | | | | | C | | | | |
| NC_011770 | 4583043 | C | T | 83 | 0 | 6 | 0 | NON_SYNONYMOU | MODERAT | MISSENSE | Ctc/Ttc | L4F | PLES_41551 | T | T | | | | | | | T | T | T | T |
| NC_011770 | 4583058 | T | C | 83 | 0 | 8 | 0 | NON_SYNONYMOU | MODERAT | MISSENSE | Ttc/Ctc | F9L | PLES_41551 | C | C | | | | | C | C | C | C | C | C |
| NC_011770 | 4583439 | G | A | 60 | 0 | 8 | 0 | NON_SYNONYMOU | MODERAT | MISSENSE | Gtc/Atc | V136I | PLES_41551 | A | A | | | | | A | A | A | A | A | A |
| NC_011770 | 4583683 | G | A | 68 | 0 | 12 | 0 | NON_SYNONYMOU | MODERAT | MISSENSE | gGc/gAc | G217D | PLES_41551 | A | A | A | A | A | A | A | A | A | A | A | A |
| NC_011770 | 4583767 | G | A | 68 | 0 | 12 | 0 | NON_SYNONYMOU | MODERAT | MISSENSE | cGg/cAg | R245Q | PLES_41551 | A | A | A | A | A | A | A | A | A | A | A | A |
| NC_011770 | 4584790 | C | T | 71 | 0 | 2 | 0 | NON_SYNONYMOU | MODERAT | MISSENSE | Ggg/Agg | G268R | PLES_41571 | | | | | | | T | T | | | | |
| NC_011770 | 4584824 | T | G | 70 | 0 | 6 | 0 | NON_SYNONYMOU | MODERAT | MISSENSE | gaA/gaC | E256D | PLES_41571 | G | G | | | | | | | G | G | G | G |
| NC_011770 | 4585317 | T | C | 68 | 0 | 1 | 0 | NON_SYNONYMOU | MODERAT | MISSENSE | aAc/aGc | N92S | PLES_41571 | | | | | | | C | | | | | |
| NC_011770 | 4585929 | C | T | 77 | 0 | 6 | 0 | NON_SYNONYMOU | MODERAT | MISSENSE | Ctc/Ttc | L43F | PLES_41581 | | | T | T | T | T | T | T | | | | |
| NC_011770 | 4586001 | A | G | 74 | 0 | 4 | 0 | NON_SYNONYMOU | MODERAT | MISSENSE | Acc/Gcc | T67A | PLES_41581 | | | G | G | G | G | | | | | | |
| NC_011770 | 4586011 | A | G | 85 | 0 | 2 | 0 | NON_SYNONYMOU | MODERAT | MISSENSE | gAa/gGa | E70G | PLES_41581 | | | | | | | G | G | | | | |

| | | | | | | | | | | | | | | | | | | | | | | | | | |
|---|---|---|---|---|---|---|---|---|---|---|---|---|---|---|---|---|---|---|---|---|---|---|---|---|---|
| NC_011770 | 4586337 | T | C | 77 | 0 | 8 | 0 | NON_SYNONYMOU | MODERAT | MISSENSE | Tac/Cac | Y179H | PLES_41581 | | | C | C | C | C | C | C | | | C | C |
| NC_011770 | 4586515 | T | A | 82 | 0 | 2 | 0 | NON_SYNONYMOU | MODERAT | MISSENSE | aTg/aAg | M238K | PLES_41581 | | | | | | | A | A | | | | |
| NC_011770 | 4586943 | G | A | 69 | 0 | 2 | 0 | NON_SYNONYMOU | MODERAT | MISSENSE | Gtc/Atc | V381I | PLES_41581 | | | | | | | | | A | A | | |
| NC_011770 | 4587858 | C | T | 66 | 0 | 2 | 0 | NON_SYNONYMOU | MODERAT | MISSENSE | Ccg/Tcg | P686S | PLES_41581 | | | | | | | | | | | T | T |
| NC_011770 | 4587991 | T | C | 79 | 0 | 2 | 0 | NON_SYNONYMOU | MODERAT | MISSENSE | gTc/gCc | V730A | PLES_41581 | | | | | | | C | C | | | | |
| NC_011770 | 4588399 | G | A | 50 | 0 | 2 | 0 | NON_SYNONYMOU | MODERAT | MISSENSE | cGg/cAg | R866Q | PLES_41581 | | | | | | | A | A | | | | |
| NC_011770 | 4592448 | G | A | 93 | 0 | 2 | 0 | NON_SYNONYMOU | MODERAT | MISSENSE | cCc/cTc | P143L | PLES_41631 | | | | | | | A | A | | | | |
| NC_011770 | 4593536 | G | T | 83 | 0 | 2 | 0 | NON_SYNONYMOU | MODERAT | MISSENSE | cCg/cAg | P131Q | PLES_41641 | | | | | | | T | T | | | | |
| NC_011770 | 4593885 | G | C | 91 | 0 | 2 | 0 | NON_SYNONYMOU | MODERAT | MISSENSE | Cga/Gga | R15G | PLES_41641 | | | | | | | C | C | | | | |
| NC_011770 | 4599737 | G | A | 35 | 6 | 4 | 0 | STOP_GAINED | HIGH | NONSENSE | tgG/tgA | W107* | PLES_41671 | | | A | A | A | A | . | . | . | . | . | . |
| NC_011770 | 4599702 | T | C | 35 | 6 | 4 | 0 | NON_SYNONYMOU | MODERAT | MISSENSE | Ttc/Ctc | F96L | PLES_41671 | | | C | C | C | C | . | . | . | . | . | . |
| NC_011770 | 4600663 | G | A | 34 | 2 | 4 | 0 | NON_SYNONYMOU | MODERAT | MISSENSE | Gcg/Acg | A32T | PLES_41681 | | | A | A | A | A | | | . | . | | |
| NC_011770 | 4600667 | C | A | 35 | 2 | 8 | 0 | NON_SYNONYMOU | MODERAT | MISSENSE | gCa/gAa | A33E | PLES_41681 | | | A | A | A | A | A | A | . | . | A | A |
| NC_011770 | 4600687 | A | G | 35 | 2 | 8 | 0 | NON_SYNONYMOU | MODERAT | MISSENSE | Acc/Gcc | T40A | PLES_41681 | | | G | G | G | G | G | G | . | . | G | G |
| NC_011770 | 4600700 | T | C | 36 | 2 | 6 | 0 | NON_SYNONYMOU | MODERAT | MISSENSE | gTg/gCg | V44A | PLES_41681 | | | C | C | C | C | | | . | . | C | C |
| NC_011770 | 4600720 | G | A | 34 | 2 | 6 | 0 | NON_SYNONYMOU | MODERAT | MISSENSE | Gcc/Acc | A51T | PLES_41681 | | | A | A | A | A | | | . | . | A | A |
| NC_011770 | 4600798 | C | T | 34 | 2 | 4 | 0 | NON_SYNONYMOU | MODERAT | MISSENSE | Ctc/Ttc | L77F | PLES_41681 | | | T | T | T | T | | | . | . | | |
| NC_011770 | 4600823 | T | C | 32 | 2 | 2 | 0 | NON_SYNONYMOU | MODERAT | MISSENSE | gTg/gCg | V85A | PLES_41681 | | | | | | | | | . | . | C | C |
| NC_011770 | 4600895 | G | A | 31 | 2 | 4 | 0 | NON_SYNONYMOU | MODERAT | MISSENSE | cGc/cAc | R109H | PLES_41681 | | | A | A | A | A | | | . | . | | |
| NC_011770 | 4600988 | C | G | 30 | 2 | 8 | 0 | NON_SYNONYMOU | MODERAT | MISSENSE | tCc/tGc | S140C | PLES_41681 | G | G | G | G | G | G | | | . | . | G | G |
| NC_011770 | 4601001 | A | C | 31 | 2 | 2 | 0 | NON_SYNONYMOU | MODERAT | MISSENSE | caA/caC | Q144H | PLES_41681 | | | | | | | C | C | . | . | | |
| NC_011770 | 4601010 | C | A | 30 | 2 | 2 | 0 | NON_SYNONYMOU | MODERAT | MISSENSE | gaC/gaA | D147E | PLES_41681 | | | | | | | A | A | . | . | | |
| NC_011770 | 4604285 | G | A | 37 | 2 | 2 | 0 | NON_SYNONYMOU | MODERAT | MISSENSE | Gcg/Acg | A20T | PLES_41701 | | | | | . | . | A | A | | | | |
| NC_011770 | 4607040 | A | G | 68 | 0 | 4 | 0 | NON_SYNONYMOU | MODERAT | MISSENSE | Acc/Gcc | T7A | PLES_41721 | | | G | G | G | G | | | | | | |
| NC_011770 | 4607385 | G | A | 82 | 0 | 12 | 0 | NON_SYNONYMOU | MODERAT | MISSENSE | Gcc/Acc | A122T | PLES_41721 | A | A | A | A | A | A | A | A | A | A | A | A |
| NC_011770 | 4607650 | A | G | 72 | 0 | 4 | 0 | NON_SYNONYMOU | MODERAT | MISSENSE | gAt/gGt | D210G | PLES_41721 | | | G | G | G | G | | | | | | |
| NC_011770 | 4607661 | T | G | 77 | 0 | 4 | 0 | NON_SYNONYMOU | MODERAT | MISSENSE | Tcc/Gcc | S214A | PLES_41721 | | | G | G | G | G | | | | | | |
| NC_011770 | 4608744 | T | G | 76 | 0 | 2 | 0 | NON_SYNONYMOU | MODERAT | MISSENSE | cTc/cGc | L44R | PLES_41731 | | | | | | | | | G | G | | |
| NC_011770 | 4609026 | T | C | 70 | 0 | 6 | 0 | NON_SYNONYMOU | MODERAT | MISSENSE | cTg/cCg | L138P | PLES_41731 | | | | | | | C | C | C | C | C | C |
| NC_011770 | 4609197 | C | T | 75 | 0 | 1 | 0 | NON_SYNONYMOU | MODERAT | MISSENSE | cCg/cTg | P195L | PLES_41731 | | | | | | | T | | | | | |
| NC_011770 | 4609529 | A | G | 74 | 0 | 4 | 0 | NON_SYNONYMOU | MODERAT | MISSENSE | Ata/Gta | I306V | PLES_41731 | | | G | G | G | G | | | | | | |
| NC_011770 | 4609535 | G | T | 75 | 0 | 4 | 0 | NON_SYNONYMOU | MODERAT | MISSENSE | Gcc/Tcc | A308S | PLES_41731 | | | T | T | T | T | | | | | | |
| NC_011770 | 4609592 | A | C | 80 | 0 | 12 | 0 | NON_SYNONYMOU | MODERAT | MISSENSE | Atg/Ctg | M327L | PLES_41731 | C | C | C | C | C | C | C | C | C | C | C | C |
| NC_011770 | 4609807 | C | G | 83 | 0 | 4 | 0 | NON_SYNONYMOU | MODERAT | MISSENSE | gGg/gCg | G299A | PLES_41741 | | | G | G | G | G | | | | | | |
| NC_011770 | 4610437 | G | A | 81 | 0 | 2 | 0 | NON_SYNONYMOU | MODERAT | MISSENSE | gCg/gTg | A89V | PLES_41741 | | | | | | | | | | | A | A |
| NC_011770 | 4611427 | T | TG | 71 | 0 | 1 | 0 | FRAME_SHIFT | HIGH | | gtg/gGtg | V80G? | PLES_41751 | TG | | | | | | | | | | | |
| NC_011770 | 4611506 | C | T | 73 | 0 | 8 | 0 | NON_SYNONYMOU | MODERAT | MISSENSE | gCg/gTg | A104V | PLES_41751 | | | T | T | T | T | | | T | T | T | T |
| NC_011770 | 4611859 | G | A | 61 | 0 | 4 | 0 | NON_SYNONYMOU | MODERAT | MISSENSE | Gcg/Acg | A222T | PLES_41751 | | | | | | | | | A | A | A | A |
| NC_011770 | 4611928 | A | C | 58 | 0 | 10 | 0 | NON_SYNONYMOU | MODERAT | MISSENSE | Atc/Ctc | I245L | PLES_41751 | | | C | C | C | C | C | C | C | C | C | C |
| NC_011770 | 4612085 | T | A | 77 | 0 | 2 | 0 | NON_SYNONYMOU | MODERAT | MISSENSE | cTg/cAg | L297Q | PLES_41751 | | | A | A | | | | | | | | |
| NC_011770 | 4612153 | G | C | 78 | 0 | 2 | 0 | NON_SYNONYMOU | MODERAT | MISSENSE | Gtg/Ctg | V320L | PLES_41751 | C | C | | | | | | | | | | |
| NC_011770 | 4612468 | T | C | 73 | 0 | 2 | 0 | NON_SYNONYMOU | MODERAT | MISSENSE | Tgc/Cgc | C425R | PLES_41751 | C | C | | | | | | | | | | |
| NC_011770 | 4612880 | G | C | 77 | 0 | 12 | 0 | NON_SYNONYMOU | MODERAT | MISSENSE | Ggc/Cgc | G95R | PLES_41761 | C | C | C | C | C | C | C | C | C | C | C | C |
| NC_011770 | 4614372 | T | C | 58 | 0 | 8 | 0 | NON_SYNONYMOU | MODERAT | MISSENSE | aTg/aCg | M53T | PLES_41781 | C | C | C | C | C | C | | | | | C | C |
| NC_011770 | 4615103 | T | G | 57 | 0 | 10 | 0 | NON_SYNONYMOU | MODERAT | MISSENSE | Tca/Gca | S297A | PLES_41781 | G | G | G | G | G | G | G | G | | | G | G |
| NC_011770 | 4615648 | C | T | 67 | 0 | 2 | 0 | NON_SYNONYMOU | MODERAT | MISSENSE | aCc/aTc | T132I | PLES_41791 | | | | | | | | | | | T | T |
| NC_011770 | 4616277 | G | A | 68 | 0 | 2 | 0 | NON_SYNONYMOU | MODERAT | MISSENSE | Ccc/Tcc | P254S | PLES_41801 | | | | | | | | | | | A | A |
| NC_011770 | 4616370 | T | C | 67 | 0 | 2 | 0 | NON_SYNONYMOU | MODERAT | MISSENSE | Atc/Gtc | I223V | PLES_41801 | | | | | | | | | | | C | C |
| NC_011770 | 4616520 | C | T | 63 | 0 | 8 | 0 | NON_SYNONYMOU | MODERAT | MISSENSE | Gcc/Acc | A173T | PLES_41801 | T | T | T | T | T | T | T | T | | | | |
| NC_011770 | 4616727 | G | C | 76 | 0 | 8 | 0 | NON_SYNONYMOU | MODERAT | MISSENSE | Cag/Gag | Q104E | PLES_41801 | C | C | C | C | C | C | C | C | | | | |
| NC_011770 | 4616775 | T | C | 73 | 0 | 2 | 0 | NON_SYNONYMOU | MODERAT | MISSENSE | Atc/Gtc | I88V | PLES_41801 | C | C | | | | | | | | | | |
| NC_011770 | 4616946 | C | T | 65 | 0 | 4 | 0 | NON_SYNONYMOU | MODERAT | MISSENSE | Gtc/Atc | V31I | PLES_41801 | T | T | | | | | | | | | T | T |
| NC_011770 | 4617161 | C | T | 68 | 0 | 4 | 0 | NON_SYNONYMOU | MODERAT | MISSENSE | cGg/cAg | R281Q | PLES_41811 | T | T | | | | | | | | | T | T |
| NC_011770 | 4618346 | G | A | 53 | 0 | 1 | 0 | NON_SYNONYMOU | MODERAT | MISSENSE | Ggc/Agc | G67S | PLES_41821 | | | | | | | A | | | | | |
| NC_011770 | 4618610 | A | G | 60 | 0 | 12 | 0 | NON_SYNONYMOU | MODERAT | MISSENSE | Acc/Gcc | T155A | PLES_41821 | G | G | G | G | G | G | G | G | G | G | G | G |
| NC_011770 | 4618709 | G | C | 67 | 0 | 12 | 0 | NON_SYNONYMOU | MODERAT | MISSENSE | Gtg/Ctg | V188L | PLES_41821 | C | C | C | C | C | C | C | C | C | C | C | C |
| NC_011770 | 4618718 | G | A | 69 | 0 | 12 | 0 | NON_SYNONYMOU | MODERAT | MISSENSE | Gac/Aac | D191N | PLES_41821 | A | A | A | A | A | A | A | A | A | A | A | A |
| NC_011770 | 4619529 | A | C | 71 | 0 | 2 | 0 | NON_SYNONYMOU | MODERAT | MISSENSE | gTc/gGc | V115G | PLES_41831 | | | | | | | | | | | C | C |
| NC_011770 | 4619656 | G | C | 67 | 0 | 6 | 0 | NON_SYNONYMOU | MODERAT | MISSENSE | Cag/Gag | Q73E | PLES_41831 | C | C | C | C | C | C | | | | | | |
| NC_011770 | 4619724 | C | T | 64 | 0 | 12 | 0 | NON_SYNONYMOU | MODERAT | MISSENSE | gGc/gAc | G50D | PLES_41831 | T | T | T | T | T | T | T | T | T | T | T | T |
| NC_011770 | 4619779 | C | A | 65 | 0 | 4 | 0 | NON_SYNONYMOU | MODERAT | MISSENSE | Gcc/Tcc | A32S | PLES_41831 | | | | | | | A | A | A | A | | |
| NC_011770 | 4619817 | G | A | 66 | 0 | 2 | 0 | NON_SYNONYMOU | MODERAT | MISSENSE | gCt/gTt | A19V | PLES_41831 | A | A | | | | | | | | | | |

| | | | | | | | | | | | | | | | | | | | | | | | | | |
|---|---|---|---|---|---|---|---|---|---|---|---|---|---|---|---|---|---|---|---|---|---|---|---|---|---|
| NC_011770 | 4620087 | A | C | 61 | 0 | 4 | 0 | NON_SYNONYMOU | MODERAT | MISSENSE | cAc/cCc | H15P | PLES_41841 | | | C | C | C | C | | | | | | |
| NC_011770 | 4620344 | G | A | 71 | 0 | 2 | 0 | NON_SYNONYMOU | MODERAT | MISSENSE | Ggc/Agc | G101S | PLES_41841 | A | A | | | | | | | | | | |
| NC_011770 | 4620441 | A | G | 63 | 0 | 12 | 0 | NON_SYNONYMOU | MODERAT | MISSENSE | cAg/cGg | Q133R | PLES_41841 | G | G | G | G | G | G | G | G | G | G | G | G |
| NC_011770 | 4620643 | G | C | 56 | 0 | 12 | 0 | NON_SYNONYMOU | MODERAT | MISSENSE | gaG/gaC | E200D | PLES_41841 | C | C | C | C | C | C | C | C | C | C | C | C |
| NC_011770 | 4620672 | A | G | 58 | 0 | 12 | 0 | NON_SYNONYMOU | MODERAT | MISSENSE | gAc/gGc | D210G | PLES_41841 | G | G | G | G | G | G | G | G | G | G | G | G |
| NC_011770 | 4621361 | A | G | 69 | 0 | 10 | 0 | NON_SYNONYMOU | MODERAT | MISSENSE | Aac/Gac | N63D | PLES_41851 | G | G | G | G | G | G | G | G | | | G | G |
| NC_011770 | 4621683 | G | A | 83 | 0 | 2 | 0 | NON_SYNONYMOU | MODERAT | MISSENSE | cGa/cAa | R10Q | PLES_41861 | A | A | | | | | | | | | | |
| NC_011770 | 4621739 | A | T | 88 | 0 | 2 | 0 | NON_SYNONYMOU | MODERAT | MISSENSE | Atc/Ttc | I29F | PLES_41861 | | | | | | | | | T | T | | |
| NC_011770 | 4621913 | T | A | 82 | 0 | 2 | 0 | NON_SYNONYMOU | MODERAT | MISSENSE | Tgc/Agc | C87S | PLES_41861 | | | | | | | | | | | A | A |
| NC_011770 | 4622172 | C | T | 85 | 0 | 2 | 0 | NON_SYNONYMOU | MODERAT | MISSENSE | cGt/cAt | R178H | PLES_41871 | | | | | | | | | T | T | | |
| NC_011770 | 4623179 | A | G | 60 | 0 | 8 | 0 | NON_SYNONYMOU | MODERAT | MISSENSE | Atc/Gtc | I55V | PLES_41881 | G | G | G | G | G | G | | | G | G | | |
| NC_011770 | 4623351 | G | A | 59 | 0 | 2 | 0 | NON_SYNONYMOU | MODERAT | MISSENSE | aGc/aAc | S112N | PLES_41881 | | | | | | | A | A | | | | |
| NC_011770 | 4624118 | A | G | 66 | 0 | 10 | 0 | NON_SYNONYMOU | MODERAT | MISSENSE | Atc/Gtc | I368V | PLES_41881 | G | G | G | G | G | G | | | G | G | G | G |
| NC_011770 | 4624175 | C | G | 71 | 0 | 4 | 0 | NON_SYNONYMOU | MODERAT | MISSENSE | Ctg/Gtg | L387V | PLES_41881 | G | G | | | | | | | | | G | G |
| NC_011770 | 4625327 | C | A | 77 | 0 | 4 | 0 | STOP_GAINED | HIGH | NONSENSE | Gag/Tag | E110* | PLES_41901 | | | A | A | A | A | | | | | | |
| NC_011770 | 4625491 | T | C | 74 | 0 | 4 | 0 | NON_SYNONYMOU | MODERAT | MISSENSE | tAc/tGc | Y55C | PLES_41901 | | | C | C | C | C | | | | | | |
| NC_011770 | 4625720 | C | T | 71 | 0 | 2 | 0 | NON_SYNONYMOU | MODERAT | MISSENSE | atG/atA | M294I | PLES_41911 | | | | | | | | | T | T | | |
| NC_011770 | 4625881 | C | T | 66 | 0 | 6 | 0 | NON_SYNONYMOU | MODERAT | MISSENSE | Gtc/Atc | V241I | PLES_41911 | | | T | T | T | T | | | T | T | | |
| NC_011770 | 4626028 | G | A | 83 | 0 | 2 | 0 | NON_SYNONYMOU | MODERAT | MISSENSE | Cgg/Tgg | R192W | PLES_41911 | A | A | | | | | | | | | | |
| NC_011770 | 4628110 | A | G | 69 | 0 | 4 | 0 | NON_SYNONYMOU | MODERAT | MISSENSE | Aag/Gag | K98E | PLES_41931 | | | G | G | G | G | | | | | | |
| NC_011770 | 4628201 | T | C | 69 | 0 | 1 | 0 | NON_SYNONYMOU | MODERAT | MISSENSE | cTg/cCg | L128P | PLES_41931 | | | | C | | | | | | | | |
| NC_011770 | 4628451 | C | T | 80 | 0 | 6 | 0 | NON_SYNONYMOU | MODERAT | MISSENSE | Gtg/Atg | V239M | PLES_41941 | | | | | | | T | T | T | T | T | T |
| NC_011770 | 4628505 | G | T | 77 | 0 | 4 | 0 | NON_SYNONYMOU | MODERAT | MISSENSE | Ccg/Acg | P221T | PLES_41941 | | | | | | | | | T | T | T | T |
| NC_011770 | 4628783 | G | A | 71 | 0 | 4 | 0 | NON_SYNONYMOU | MODERAT | MISSENSE | gCg/gTg | A128V | PLES_41941 | | | A | A | A | A | | | | | | |
| NC_011770 | 4628789 | G | A | 69 | 0 | 4 | 0 | NON_SYNONYMOU | MODERAT | MISSENSE | aCc/aTc | T126I | PLES_41941 | | | A | A | A | A | | | | | | |
| NC_011770 | 4628891 | C | T | 66 | 0 | 8 | 0 | NON_SYNONYMOU | MODERAT | MISSENSE | gGc/gAc | G92D | PLES_41941 | T | T | | | | | T | T | T | T | T | T |
| NC_011770 | 4629131 | C | T | 74 | 0 | 2 | 0 | NON_SYNONYMOU | MODERAT | MISSENSE | cGc/cAc | R12H | PLES_41941 | | | | | | | T | T | | | | |
| NC_011770 | 4631718 | G | A | 39 | 0 | 4 | 0 | NON_SYNONYMOU | MODERAT | MISSENSE | Cgc/Tgc | R116C | PLES_41971 | | | A | A | A | A | | | | | | |
| NC_011770 | 4631798 | G | A | 43 | 0 | 4 | 0 | NON_SYNONYMOU | MODERAT | MISSENSE | cCg/cTg | P89L | PLES_41971 | | | | | | | A | A | | | A | A |
| NC_011770 | 4632513 | G | A | 82 | 0 | 2 | 0 | NON_SYNONYMOU | MODERAT | MISSENSE | Ggc/Agc | G81S | PLES_41981 | | | | | | | | | | | A | A |
| NC_011770 | 4632570 | A | G | 73 | 0 | 12 | 0 | NON_SYNONYMOU | MODERAT | MISSENSE | Acg/Gcg | T100A | PLES_41981 | G | G | G | G | G | G | G | G | G | G | G | G |
| NC_011770 | 4633668 | A | G | 86 | 0 | 2 | 0 | NON_SYNONYMOU | MODERAT | MISSENSE | aAc/aGc | N275S | PLES_41991 | G | G | | | | | | | | | | |
| NC_011770 | 4634377 | G | A | 57 | 0 | 2 | 0 | NON_SYNONYMOU | MODERAT | MISSENSE | cGc/cAc | R85H | PLES_42001 | A | A | | | | | | | | | | |
| NC_011770 | 4634887 | G | A | 62 | 0 | 2 | 0 | NON_SYNONYMOU | MODERAT | MISSENSE | Gac/Aac | D38N | PLES_42011 | | | | | | | | | | | A | A |
| NC_011770 | 4634903 | C | T | 62 | 0 | 2 | 0 | NON_SYNONYMOU | MODERAT | MISSENSE | gCc/gTc | A43V | PLES_42011 | | | | | | | T | T | | | | |
| NC_011770 | 4635022 | G | A | 62 | 0 | 2 | 0 | NON_SYNONYMOU | MODERAT | MISSENSE | Gag/Aag | E83K | PLES_42011 | | | | | | | | | | | A | A |
| NC_011770 | 4635202 | G | A | 68 | 0 | 8 | 0 | NON_SYNONYMOU | MODERAT | MISSENSE | Gtg/Atg | V143M | PLES_42011 | | | A | A | A | A | A | A | A | A | | |
| NC_011770 | 4635868 | C | G | 74 | 0 | 2 | 0 | NON_SYNONYMOU | MODERAT | MISSENSE | gaC/gaG | D101E | PLES_42021 | | | | | | | G | G | | | | |
| NC_011770 | 4635872 | G | A | 73 | 0 | 2 | 0 | NON_SYNONYMOU | MODERAT | MISSENSE | Ggc/Agc | G103S | PLES_42021 | | | | | | | | | | | A | A |
| NC_011770 | 4636033 | C | T | 79 | 0 | 2 | 0 | NON_SYNONYMOU | MODERAT | MISSENSE | Ggg/Agg | G253R | PLES_42031 | T | T | | | | | | | | | | |
| NC_011770 | 4636225 | G | A | 87 | 0 | 2 | 0 | NON_SYNONYMOU | MODERAT | MISSENSE | Cgc/Tgc | R189C | PLES_42031 | | | | | | | A | A | | | | |
| NC_011770 | 4638088 | A | G | 70 | 0 | 1 | 0 | NON_SYNONYMOU | MODERAT | MISSENSE | Acc/Gcc | T349A | PLES_42041 | | | | | G | | | | | | | |
| NC_011770 | 4637110 | G | T | 82 | 0 | 2 | 0 | NON_SYNONYMOU | MODERAT | MISSENSE | Gtc/Ttc | V23F | PLES_42041 | | | | | | | | | | | T | T |
| NC_011770 | 4637206 | C | T | 83 | 0 | 2 | 0 | NON_SYNONYMOU | MODERAT | MISSENSE | Cat/Tat | H55Y | PLES_42041 | | | | | | | T | T | | | | |
| NC_011770 | 4637621 | A | G | 69 | 0 | 2 | 0 | NON_SYNONYMOU | MODERAT | MISSENSE | gAc/gGc | D193G | PLES_42041 | | | | | | | G | G | | | | |
| NC_011770 | 4638409 | G | A | 82 | 0 | 2 | 0 | NON_SYNONYMOU | MODERAT | MISSENSE | Gtc/Atc | V456I | PLES_42041 | A | A | | | | | | | | | | |
| NC_011770 | 4638472 | A | G | 74 | 0 | 12 | 0 | NON_SYNONYMOU | MODERAT | MISSENSE | Acg/Gcg | T477A | PLES_42041 | G | G | G | G | G | G | G | G | G | G | G | G |
| NC_011770 | 4638517 | G | A | 75 | 0 | 1 | 0 | NON_SYNONYMOU | MODERAT | MISSENSE | Gcc/Acc | A492T | PLES_42041 | | | | | | | A | | | | | |
| NC_011770 | 4639190 | A | C | 67 | 0 | 2 | 0 | NON_SYNONYMOU | MODERAT | MISSENSE | aAc/aCc | N716T | PLES_42041 | | | | | | | | | | | C | C |
| NC_011770 | 4639616 | A | G | 61 | 0 | 2 | 0 | NON_SYNONYMOU | MODERAT | MISSENSE | Atg/Gtg | M49V | PLES_42051 | G | G | | | | | | | | | | |
| NC_011770 | 4639632 | A | G | 61 | 0 | 6 | 0 | NON_SYNONYMOU | MODERAT | MISSENSE | cAc/cGc | H54R | PLES_42051 | G | G | G | G | G | G | | | | | | |
| NC_011770 | 4639703 | C | G | 61 | 0 | 8 | 0 | NON_SYNONYMOU | MODERAT | MISSENSE | Cag/Gag | Q78E | PLES_42051 | G | G | G | G | G | G | | | | | G | G |
| NC_011770 | 4639722 | G | A | 63 | 0 | 2 | 0 | NON_SYNONYMOU | MODERAT | MISSENSE | gGc/gAc | G84D | PLES_42051 | A | A | | | | | | | | | | |
| NC_011770 | 4640271 | T | A | 44 | 0 | 2 | 0 | NON_SYNONYMOU | MODERAT | MISSENSE | Agc/Tgc | S436C | PLES_42061 | | | | | | | A | A | | | | |
| NC_011770 | 4640589 | G | A | 70 | 0 | 8 | 0 | NON_SYNONYMOU | MODERAT | MISSENSE | Ccg/Tcg | P330S | PLES_42061 | A | A | A | A | A | A | | | A | A | | |
| NC_011770 | 4640679 | T | C | 67 | 0 | 1 | 0 | NON_SYNONYMOU | MODERAT | MISSENSE | Acc/Gcc | T300A | PLES_42061 | | | | | | | C | | | | | |
| NC_011770 | 4642857 | T | G | 72 | 0 | 2 | 0 | NON_SYNONYMOU | MODERAT | MISSENSE | gaA/gaC | E286D | PLES_42091 | | | | | | | | | | | G | G |
| NC_011770 | 4643002 | C | T | 82 | 0 | 2 | 0 | NON_SYNONYMOU | MODERAT | MISSENSE | cGg/cAg | R238Q | PLES_42091 | | | | | | | | | | | T | T |
| NC_011770 | 4643050 | C | T | 80 | 0 | 12 | 0 | NON_SYNONYMOU | MODERAT | MISSENSE | tGt/tAt | C222Y | PLES_42091 | T | T | T | T | T | T | T | T | T | T | T | T |
| NC_011770 | 4643078 | C | T | 79 | 0 | 6 | 0 | NON_SYNONYMOU | MODERAT | MISSENSE | Gcc/Acc | A213T | PLES_42091 | T | T | | | | | | | T | T | T | T |
| NC_011770 | 4643135 | T | C | 84 | 0 | 4 | 0 | NON_SYNONYMOU | MODERAT | MISSENSE | Atc/Gtc | I194V | PLES_42091 | C | C | | | | | | | C | C | | |
| NC_011770 | 4643213 | T | C | 79 | 0 | 2 | 0 | NON_SYNONYMOU | MODERAT | MISSENSE | Aag/Gag | K168E | PLES_42091 | | | | | | | | | C | C | | |

| | | | | | | | | | | | | | | | | | | | | | | | | | |
|---|---|---|---|---|---|---|---|---|---|---|---|---|---|---|---|---|---|---|---|---|---|---|---|---|---|
| NC_011770 | 4643498 | T | C | 73 | 0 | 2 | 0 | NON_SYNONYMOU | MODERAT | MISSENSE | Agt/Ggt | S73G | PLES_42091 | | | | | | | C | C | | | | |
| NC_011770 | 4643535 | T | C | 78 | 0 | 2 | 0 | NON_SYNONYMOU | MODERAT | MISSENSE | atA/atG | I60M | PLES_42091 | | | | | | | | | | | C | C |
| NC_011770 | 4644055 | C | G | 81 | 0 | 4 | 0 | NON_SYNONYMOU | MODERAT | MISSENSE | Gag/Cag | E126Q | PLES_42101 | | | G | G | G | G | | | | | | |
| NC_011770 | 4644358 | G | T | 69 | 0 | 6 | 0 | NON_SYNONYMOU | MODERAT | MISSENSE | Cat/Aat | H25N | PLES_42101 | T | T | | | | | T | T | | | T | T |
| NC_011770 | 4644360 | A | G | 69 | 0 | 6 | 0 | NON_SYNONYMOU | MODERAT | MISSENSE | gTc/gCc | V24A | PLES_42101 | G | G | | | | | G | G | | | G | G |
| NC_011770 | 4644735 | AC | A | 70 | 0 | 2 | 0 | FRAME_SHIFT | HIGH | | ctg/ | L66 | PLES_42111 | | | | | | | | | | | A | A |
| NC_011770 | 4644569 | A | C | 64 | 0 | 2 | 0 | NON_SYNONYMOU | MODERAT | MISSENSE | aaA/aaC | K9N | PLES_42111 | | | | | | | | | C | C | | |
| NC_011770 | 4644779 | C | G | 72 | 0 | 2 | 0 | NON_SYNONYMOU | MODERAT | MISSENSE | ttC/ttG | F79L | PLES_42111 | | | | | | | | | G | G | | |
| NC_011770 | 4644633 | G | A | 67 | 0 | 1 | 0 | NON_SYNONYMOU | MODERAT | MISSENSE | Gtc/Atc | V31I | PLES_42111 | | | | A | | | | | | | | |
| NC_011770 | 4645988 | C | G | 82 | 0 | 4 | 0 | NON_SYNONYMOU | MODERAT | MISSENSE | tGc/tCc | C17S | PLES_42121 | | | G | G | G | G | | | | | | |
| NC_011770 | 4647283 | A | G | 77 | 0 | 12 | 0 | SPLICE_SITE_REGIC | LOW | SILENT | tAa/tGa | *386 | PLES_42131 | G | G | G | G | G | G | G | G | G | G | G | G |
| NC_011770 | 4646958 | C | A | 86 | 0 | 4 | 0 | NON_SYNONYMOU | MODERAT | MISSENSE | Ctg/Atg | L278M | PLES_42131 | | | A | A | A | A | | | | | | |
| NC_011770 | 4647268 | G | A | 79 | 0 | 2 | 0 | NON_SYNONYMOU | MODERAT | MISSENSE | aGg/aAg | R381K | PLES_42131 | | | | | | | | | A | A | | |
| NC_011770 | 4647515 | G | A | 72 | 0 | 1 | 0 | NON_SYNONYMOU | MODERAT | MISSENSE | Gtc/Atc | V39I | PLES_42141 | | | | | | | | A | | | | |
| NC_011770 | 4649430 | A | G | 75 | 0 | 12 | 0 | NON_SYNONYMOU | MODERAT | MISSENSE | aAc/aGc | N212S | PLES_42151 | G | G | G | G | G | G | G | G | G | G | G | G |
| NC_011770 | 4670601 | G | A | 46 | 6 | 4 | 0 | NON_SYNONYMOU | MODERAT | MISSENSE | Cgt/Tgt | R123C | PLES_42321 | . | . | A | A | A | A | . | . | | | . | . |
| NC_011770 | 4670768 | G | C | 49 | 6 | 6 | 0 | NON_SYNONYMOU | MODERAT | MISSENSE | aCc/aGc | T67S | PLES_42321 | . | . | C | C | C | C | . | . | C | C | . | . |
| NC_011770 | 4671015 | G | A | 44 | 6 | 6 | 0 | NON_SYNONYMOU | MODERAT | MISSENSE | gCg/gTg | A240V | PLES_42331 | . | . | A | A | A | A | . | . | A | A | . | . |
| NC_011770 | 4684315 | G | C | 84 | 0 | 1 | 0 | NON_SYNONYMOU | MODERAT | MISSENSE | Gcc/Ccc | A76P | PLES_42471 | | | | | | | C | | | | | |
| NC_011770 | 4689586 | C | T | 73 | 0 | 2 | 0 | NON_SYNONYMOU | MODERAT | MISSENSE | Gcg/Acg | A727T | PLES_42531 | T | T | | | | | | | | | | |
| NC_011770 | 4689596 | G | T | 73 | 0 | 4 | 0 | NON_SYNONYMOU | MODERAT | MISSENSE | gaC/gaA | D723E | PLES_42531 | | | T | T | T | T | | | | | | |
| NC_011770 | 4689660 | C | T | 74 | 0 | 4 | 0 | NON_SYNONYMOU | MODERAT | MISSENSE | aGc/aAc | S702N | PLES_42531 | | | T | T | T | T | | | | | | |
| NC_011770 | 4689667 | C | T | 75 | 0 | 4 | 0 | NON_SYNONYMOU | MODERAT | MISSENSE | Gcc/Acc | A700T | PLES_42531 | | | T | T | T | T | | | | | | |
| NC_011770 | 4690194 | A | T | 74 | 0 | 4 | 0 | NON_SYNONYMOU | MODERAT | MISSENSE | cTg/cAg | L524Q | PLES_42531 | | | T | T | T | T | | | | | | |
| NC_011770 | 4691808 | C | T | 93 | 0 | 4 | 0 | NON_SYNONYMOU | MODERAT | MISSENSE | gGg/gAg | G635E | PLES_42541 | T | T | | | | | | | | | T | T |
| NC_011770 | 4692699 | T | C | 55 | 0 | 1 | 0 | NON_SYNONYMOU | MODERAT | MISSENSE | aAc/aGc | N338S | PLES_42541 | | | | | | | C | | | | | |
| NC_011770 | 4693009 | A | G | 73 | 0 | 2 | 0 | NON_SYNONYMOU | MODERAT | MISSENSE | Tcg/Ccg | S235P | PLES_42541 | | | | | | | | | | | G | G |
| NC_011770 | 4693456 | G | A | 76 | 0 | 1 | 0 | NON_SYNONYMOU | MODERAT | MISSENSE | Ccg/Tcg | P86S | PLES_42541 | | | | | | | | | | | | A |
| NC_011770 | 4694277 | C | T | 92 | 0 | 2 | 0 | NON_SYNONYMOU | MODERAT | MISSENSE | Gac/Aac | D150N | PLES_42551 | | | | | | | T | T | | | | |
| NC_011770 | 4694890 | T | G | 77 | 0 | 6 | 0 | NON_SYNONYMOU | MODERAT | MISSENSE | Tgc/Ggc | C40G | PLES_42561 | | | | | | | G | G | G | G | G | G |
| NC_011770 | 4694908 | A | G | 79 | 0 | 12 | 0 | NON_SYNONYMOU | MODERAT | MISSENSE | Atc/Gtc | I46V | PLES_42561 | G | G | G | G | G | G | G | G | G | G | G | G |
| NC_011770 | 4694979 | A | C | 84 | 0 | 12 | 0 | NON_SYNONYMOU | MODERAT | MISSENSE | ttA/ttC | L69F | PLES_42561 | C | C | C | C | C | C | C | C | C | C | C | C |
| NC_011770 | 4695007 | A | G | 79 | 0 | 12 | 0 | NON_SYNONYMOU | MODERAT | MISSENSE | Atc/Gtc | I79V | PLES_42561 | G | G | G | G | G | G | G | G | G | G | G | G |
| NC_011770 | 4695448 | A | G | 83 | 0 | 8 | 0 | NON_SYNONYMOU | MODERAT | MISSENSE | Ttc/Ctc | F156L | PLES_42571 | G | G | G | G | G | G | | | | | G | G |
| NC_011770 | 4695456 | G | A | 85 | 0 | 2 | 0 | NON_SYNONYMOU | MODERAT | MISSENSE | gCc/gTc | A153V | PLES_42571 | A | A | | | | | | | | | | |
| NC_011770 | 4695555 | C | T | 80 | 0 | 1 | 0 | NON_SYNONYMOU | MODERAT | MISSENSE | gGc/gAc | G120D | PLES_42571 | | | | | | | T | | | | | |
| NC_011770 | 4695747 | T | C | 91 | 0 | 2 | 0 | NON_SYNONYMOU | MODERAT | MISSENSE | aAg/aGg | K56R | PLES_42571 | C | C | | | | | | | | | | |
| NC_011770 | 4696656 | A | G | 75 | 0 | 12 | 0 | NON_SYNONYMOU | MODERAT | MISSENSE | Tgc/Cgc | C75R | PLES_42591 | G | G | G | G | G | G | G | G | G | G | G | G |
| NC_011770 | 4697234 | C | T | 76 | 0 | 2 | 0 | NON_SYNONYMOU | MODERAT | MISSENSE | Gac/Aac | D55N | PLES_42601 | T | T | | | | | | | | | | |
| NC_011770 | 4697293 | A | C | 72 | 0 | 2 | 0 | NON_SYNONYMOU | MODERAT | MISSENSE | gTg/gGg | V35G | PLES_42601 | | | | | | | C | C | | | | |
| NC_011770 | 4697309 | C | A | 75 | 0 | 12 | 0 | NON_SYNONYMOU | MODERAT | MISSENSE | Gtg/Ttg | V30L | PLES_42601 | A | A | A | A | A | A | A | A | A | A | A | A |
| NC_011770 | 4698117 | G | A | 71 | 0 | 1 | 0 | NON_SYNONYMOU | MODERAT | MISSENSE | Ccc/Tcc | P130S | PLES_42611 | | | | | | | A | | | | | |
| NC_011770 | 4698490 | G | T | 61 | 0 | 2 | 0 | NON_SYNONYMOU | MODERAT | MISSENSE | caC/caA | H5Q | PLES_42611 | | | | | | | | | T | T | | |
| NC_011770 | 4698589 | G | T | 56 | 0 | 2 | 0 | NON_SYNONYMOU | MODERAT | MISSENSE | Ctg/Atg | L278M | PLES_42621 | | | | | | | | | T | T | | |
| NC_011770 | 4698769 | C | G | 52 | 0 | 2 | 0 | NON_SYNONYMOU | MODERAT | MISSENSE | Gtg/Ctg | V218L | PLES_42621 | | | | | | | G | G | | | | |
| NC_011770 | 4698934 | G | T | 61 | 0 | 2 | 0 | NON_SYNONYMOU | MODERAT | MISSENSE | Ctc/Atc | L163I | PLES_42621 | | | | | | | | | T | T | | |
| NC_011770 | 4699231 | G | A | 67 | 0 | 2 | 0 | NON_SYNONYMOU | MODERAT | MISSENSE | Ccg/Tcg | P64S | PLES_42621 | | | | | | | | | A | A | | |
| NC_011770 | 4699315 | G | A | 70 | 0 | 2 | 0 | NON_SYNONYMOU | MODERAT | MISSENSE | Ctc/Ttc | L36F | PLES_42621 | | | | | | | | | | | A | A |
| NC_011770 | 4700252 | G | T | 88 | 0 | 2 | 0 | NON_SYNONYMOU | MODERAT | MISSENSE | gaC/gaA | D135E | PLES_42651 | | | | | | | T | T | | | | |
| NC_011770 | 4700436 | A | G | 91 | 0 | 6 | 0 | NON_SYNONYMOU | MODERAT | MISSENSE | gTg/gCg | V74A | PLES_42651 | | | G | G | G | G | | | G | G | | |
| NC_011770 | 4701209 | G | A | 93 | 0 | 2 | 0 | NON_SYNONYMOU | MODERAT | MISSENSE | gCt/gTt | A315V | PLES_42661 | | | | | | | A | A | | | | |
| NC_011770 | 4701255 | T | C | 98 | 0 | 2 | 0 | NON_SYNONYMOU | MODERAT | MISSENSE | Agc/Ggc | S300G | PLES_42661 | | | | | | | | | C | C | | |
| NC_011770 | 4701783 | C | A | 102 | 0 | 2 | 0 | NON_SYNONYMOU | MODERAT | MISSENSE | Ggc/Tgc | G124C | PLES_42661 | | | | | | | | | | | A | A |
| NC_011770 | 4703451 | C | A | 74 | 0 | 12 | 0 | NON_SYNONYMOU | MODERAT | MISSENSE | tGc/tTc | C613F | PLES_42681 | A | A | A | A | A | A | A | A | A | A | A | A |
| NC_011770 | 4704762 | G | A | 69 | 0 | 1 | 0 | NON_SYNONYMOU | MODERAT | MISSENSE | gCc/gTc | A176V | PLES_42681 | | | | | | | | A | | | | |
| NC_011770 | 4705501 | GA | G | 65 | 0 | 3 | 0 | FRAME_SHIFT | HIGH | | act/ | T63 | PLES_42691 | G | G | | G | | | | | | | | |
| NC_011770 | 4705407 | G | A | 63 | 0 | 2 | 0 | NON_SYNONYMOU | MODERAT | MISSENSE | cGc/cAc | R29H | PLES_42691 | | | | | | | | | | | A | A |
| NC_011770 | 4705484 | C | G | 64 | 0 | 2 | 0 | NON_SYNONYMOU | MODERAT | MISSENSE | Ccg/Gcg | P55A | PLES_42691 | G | G | | | | | | | | | | |
| NC_011770 | 4707200 | T | C | 58 | 0 | 12 | 0 | NON_SYNONYMOU | MODERAT | MISSENSE | Agc/Ggc | S146G | PLES_42711 | C | C | C | C | C | C | C | C | C | C | C | C |
| NC_011770 | 4707493 | G | A | 71 | 0 | 1 | 0 | NON_SYNONYMOU | MODERAT | MISSENSE | aCg/aTg | T48M | PLES_42711 | | | | | | | | A | | | | |
| NC_011770 | 4709504 | G | A | 65 | 0 | 4 | 0 | NON_SYNONYMOU | MODERAT | MISSENSE | aCc/aTc | T268I | PLES_42731 | | | | | | | A | A | | | A | A |
| NC_011770 | 4711836 | C | T | 82 | 0 | 1 | 0 | NON_SYNONYMOU | MODERAT | MISSENSE | Gag/Aag | E36K | PLES_42751 | | | | | | | | | | | T | |

| | | | | | | | | | | | | | | | | | | | | | | | | | |
|---|---|---|---|---|---|---|---|---|---|---|---|---|---|---|---|---|---|---|---|---|---|---|---|---|---|
| NC_011770 | 4712305 | A | G | 66 | 0 | 4 | 0 | NON_SYNONYMOU | MODERAT | MISSENSE | Tcc/Ccc | S341P | PLES_42761 | G | G | | | | | | | G | G | | |
| NC_011770 | 4712454 | C | T | 75 | 0 | 2 | 0 | NON_SYNONYMOU | MODERAT | MISSENSE | cGg/cAg | R291Q | PLES_42761 | T | T | | | | | | | | | | |
| NC_011770 | 4712710 | C | T | 74 | 0 | 2 | 0 | NON_SYNONYMOU | MODERAT | MISSENSE | Gtc/Atc | V206I | PLES_42761 | T | T | | | | | | | | | | |
| NC_011770 | 4712734 | C | T | 71 | 0 | 2 | 0 | NON_SYNONYMOU | MODERAT | MISSENSE | Gca/Aca | A198T | PLES_42761 | T | T | | | | | | | | | | |
| NC_011770 | 4712905 | C | T | 78 | 0 | 2 | 0 | NON_SYNONYMOU | MODERAT | MISSENSE | Gca/Aca | A141T | PLES_42761 | | | | | | | | | T | T | | |
| NC_011770 | 4712911 | C | G | 75 | 0 | 12 | 0 | NON_SYNONYMOU | MODERAT | MISSENSE | Gcc/Ccc | A139P | PLES_42761 | G | G | G | G | G | G | G | G | G | G | G | G |
| NC_011770 | 4712970 | C | T | 81 | 0 | 4 | 0 | NON_SYNONYMOU | MODERAT | MISSENSE | aGc/aAc | S119N | PLES_42761 | | | | | | | T | T | | | T | T |
| NC_011770 | 4714096 | G | A | 78 | 0 | 4 | 0 | NON_SYNONYMOU | MODERAT | MISSENSE | gCc/gTc | A527V | PLES_42771 | | | A | A | A | A | | | | | | |
| NC_011770 | 4714102 | G | A | 78 | 0 | 2 | 0 | NON_SYNONYMOU | MODERAT | MISSENSE | gCa/gTa | A525V | PLES_42771 | | | | | | | | | A | A | | |
| NC_011770 | 4714174 | G | C | 79 | 0 | 2 | 0 | NON_SYNONYMOU | MODERAT | MISSENSE | aCg/aGg | T501R | PLES_42771 | | | | | | | | | C | C | | |
| NC_011770 | 4714207 | T | A | 76 | 0 | 2 | 0 | NON_SYNONYMOU | MODERAT | MISSENSE | cAg/cTg | Q490L | PLES_42771 | | | | | | | A | A | | | | |
| NC_011770 | 4714433 | T | C | 78 | 0 | 6 | 0 | NON_SYNONYMOU | MODERAT | MISSENSE | Acg/Gcg | T415A | PLES_42771 | | | | | | | C | C | C | C | C | C |
| NC_011770 | 4715045 | T | C | 86 | 0 | 6 | 0 | NON_SYNONYMOU | MODERAT | MISSENSE | Aca/Gca | T211A | PLES_42771 | | | C | C | C | C | | | C | C | | |
| NC_011770 | 4715090 | G | A | 80 | 0 | 2 | 0 | NON_SYNONYMOU | MODERAT | MISSENSE | Ccg/Tcg | P196S | PLES_42771 | | | | | | | | | A | A | | |
| NC_011770 | 4715407 | A | T | 84 | 0 | 6 | 0 | NON_SYNONYMOU | MODERAT | MISSENSE | cTg/cAg | L90Q | PLES_42771 | | | T | T | T | T | | | T | T | | |
| NC_011770 | 4715497 | G | A | 83 | 0 | 8 | 0 | NON_SYNONYMOU | MODERAT | MISSENSE | gCg/gTg | A60V | PLES_42771 | A | A | A | A | A | A | | | A | A | | |
| NC_011770 | 4715575 | A | G | 90 | 0 | 10 | 0 | NON_SYNONYMOU | MODERAT | MISSENSE | aTc/aCc | I34T | PLES_42771 | G | G | G | G | G | G | | | G | G | G | G |
| NC_011770 | 4715581 | C | T | 89 | 0 | 8 | 0 | NON_SYNONYMOU | MODERAT | MISSENSE | aGc/aAc | S32N | PLES_42771 | T | T | T | T | T | T | | | T | T | | |
| NC_011770 | 4715597 | T | C | 87 | 0 | 8 | 0 | NON_SYNONYMOU | MODERAT | MISSENSE | Acg/Gcg | T27A | PLES_42771 | C | C | C | C | C | C | | | C | C | | |
| NC_011770 | 4718643 | G | T | 46 | 0 | 2 | 0 | NON_SYNONYMOU | MODERAT | MISSENSE | gCc/gAc | A233D | PLES_42801 | | | | | | | | | T | T | | |
| NC_011770 | 4719262 | G | T | 68 | 0 | 2 | 0 | NON_SYNONYMOU | MODERAT | MISSENSE | Ccg/Acg | P27T | PLES_42801 | | | | | | | | | T | T | | |
| NC_011770 | 4719282 | A | G | 68 | 0 | 12 | 0 | NON_SYNONYMOU | MODERAT | MISSENSE | gTg/gCg | V20A | PLES_42801 | G | G | G | G | G | G | G | G | G | G | G | G |
| NC_011770 | 4719470 | C | G | 72 | 0 | 2 | 0 | NON_SYNONYMOU | MODERAT | MISSENSE | Gga/Cga | G55R | PLES_42811 | | | | | | | | | G | G | | |
| NC_011770 | 4720832 | G | A | 82 | 0 | 2 | 0 | NON_SYNONYMOU | MODERAT | MISSENSE | Gac/Aac | D108N | PLES_42831 | | | | | | | A | A | | | | |
| NC_011770 | 4720928 | T | C | 82 | 0 | 2 | 0 | NON_SYNONYMOU | MODERAT | MISSENSE | Tac/Cac | Y140H | PLES_42831 | | | | | | | C | C | | | | |
| NC_011770 | 4721325 | C | G | 88 | 0 | 6 | 0 | NON_SYNONYMOU | MODERAT | MISSENSE | gaC/gaG | D105E | PLES_42841 | | | G | G | G | G | | | G | G | | |
| NC_011770 | 4721728 | A | G | 97 | 0 | 6 | 0 | NON_SYNONYMOU | MODERAT | MISSENSE | cAg/cGg | Q82R | PLES_42851 | | | G | G | G | G | | | G | G | | |
| NC_011770 | 4722677 | A | G | 57 | 0 | 2 | 0 | NON_SYNONYMOU | MODERAT | MISSENSE | Tcc/Ccc | S170P | PLES_42871 | | | | | | | G | G | | | | |
| NC_011770 | 4722866 | C | A | 66 | 0 | 2 | 0 | NON_SYNONYMOU | MODERAT | MISSENSE | Gcc/Tcc | A107S | PLES_42871 | | | | | | | | | A | A | | |
| NC_011770 | 4722877 | G | T | 70 | 0 | 2 | 0 | NON_SYNONYMOU | MODERAT | MISSENSE | aCc/aAc | T103N | PLES_42871 | | | | | | | | | T | T | | |
| NC_011770 | 4722878 | T | C | 70 | 0 | 2 | 0 | NON_SYNONYMOU | MODERAT | MISSENSE | Acc/Gcc | T103A | PLES_42871 | | | | | | | | | C | C | | |
| NC_011770 | 4722881 | C | G | 71 | 0 | 2 | 0 | NON_SYNONYMOU | MODERAT | MISSENSE | Ggt/Cgt | G102R | PLES_42871 | | | | | | | | | G | G | | |
| NC_011770 | 4722884 | T | C | 70 | 0 | 2 | 0 | NON_SYNONYMOU | MODERAT | MISSENSE | Act/Gct | T101A | PLES_42871 | | | | | | | | | C | C | | |
| NC_011770 | 4722889 | G | A | 68 | 0 | 2 | 0 | NON_SYNONYMOU | MODERAT | MISSENSE | gCc/gTc | A99V | PLES_42871 | | | | | | | | | A | A | | |
| NC_011770 | 4722896 | T | C | 65 | 0 | 2 | 0 | NON_SYNONYMOU | MODERAT | MISSENSE | Acc/Gcc | T97A | PLES_42871 | | | | | | | | | C | C | | |
| NC_011770 | 4722914 | G | C | 70 | 0 | 2 | 0 | NON_SYNONYMOU | MODERAT | MISSENSE | Ccc/Gcc | P91A | PLES_42871 | | | | | | | | | C | C | | |
| NC_011770 | 4722917 | T | C | 65 | 0 | 2 | 0 | NON_SYNONYMOU | MODERAT | MISSENSE | Act/Gct | T90A | PLES_42871 | | | | | | | | | C | C | | |
| NC_011770 | 4722919 | G | T | 68 | 0 | 2 | 0 | NON_SYNONYMOU | MODERAT | MISSENSE | aCg/aAg | T89K | PLES_42871 | | | | | | | | | T | T | | |
| NC_011770 | 4722920 | T | G | 65 | 0 | 2 | 0 | NON_SYNONYMOU | MODERAT | MISSENSE | Acg/Ccg | T89P | PLES_42871 | | | | | | | | | G | G | | |
| NC_011770 | 4722956 | C | T | 72 | 0 | 4 | 0 | NON_SYNONYMOU | MODERAT | MISSENSE | Gtc/Atc | V77I | PLES_42871 | | | | | | | T | T | T | T | | |
| NC_011770 | 4723562 | C | A | 80 | 0 | 2 | 0 | NON_SYNONYMOU | MODERAT | MISSENSE | Cat/Aat | H22N | PLES_42881 | | | | | | | | | A | A | | |
| NC_011770 | 4724639 | TGC | T | 34 | 0 | 1 | 0 | FRAME_SHIFT | HIGH | | cgc/ | R94 | PLES_42901 | | | | T | | | | | | | | |
| NC_011770 | 4725004 | A | C | 46 | 0 | 2 | 0 | NON_SYNONYMOU | MODERAT | MISSENSE | Aag/Cag | K213Q | PLES_42901 | | | | | | | | | | | C | C |
| NC_011770 | 4728888 | T | A | 81 | 0 | 2 | 0 | NON_SYNONYMOU | MODERAT | MISSENSE | Acc/Tcc | T102S | PLES_42921 | | | | | | | | | A | A | | |
| NC_011770 | 4729013 | T | A | 79 | 0 | 6 | 0 | NON_SYNONYMOU | MODERAT | MISSENSE | cAg/cTg | Q60L | PLES_42921 | | | A | A | A | A | | | A | A | | |
| NC_011770 | 4728729 | G | A | 76 | 0 | 1 | 0 | NON_SYNONYMOU | MODERAT | MISSENSE | Cgg/Tgg | R155W | PLES_42921 | | | | | | | | A | | | | |
| NC_011770 | 4730000 | C | T | 80 | 0 | 2 | 0 | NON_SYNONYMOU | MODERAT | MISSENSE | Gag/Aag | E239K | PLES_42931 | T | T | | | | | | | | | | |
| NC_011770 | 4730206 | G | C | 82 | 0 | 8 | 0 | NON_SYNONYMOU | MODERAT | MISSENSE | aCt/aGt | T170S | PLES_42931 | | | C | C | C | C | C | C | C | C | | |
| NC_011770 | 4730289 | G | C | 93 | 0 | 2 | 0 | NON_SYNONYMOU | MODERAT | MISSENSE | atC/atG | I142M | PLES_42931 | C | C | | | | | | | | | | |
| NC_011770 | 4730365 | G | A | 90 | 0 | 4 | 0 | NON_SYNONYMOU | MODERAT | MISSENSE | gCg/gTg | A117V | PLES_42931 | A | A | | | | | | | | | A | A |
| NC_011770 | 4730675 | C | T | 95 | 0 | 8 | 0 | NON_SYNONYMOU | MODERAT | MISSENSE | Gtc/Atc | V14I | PLES_42931 | T | T | | | | | T | T | T | T | T | T |
| NC_011770 | 4731127 | G | T | 96 | 0 | 2 | 0 | NON_SYNONYMOU | MODERAT | MISSENSE | Ctc/Atc | L31I | PLES_42941 | | | | | | | T | T | | | | |
| NC_011770 | 4731511 | G | C | 85 | 0 | 6 | 0 | NON_SYNONYMOU | MODERAT | MISSENSE | Caa/Gaa | Q356E | PLES_42951 | C | C | | | | | | | C | C | C | C |
| NC_011770 | 4731615 | T | C | 83 | 0 | 2 | 0 | NON_SYNONYMOU | MODERAT | MISSENSE | cAg/cGg | Q321R | PLES_42951 | C | C | | | | | | | | | | |
| NC_011770 | 4731909 | G | C | 75 | 0 | 4 | 0 | NON_SYNONYMOU | MODERAT | MISSENSE | gCc/gGc | A223G | PLES_42951 | | | | | | | C | C | | | C | C |
| NC_011770 | 4731910 | C | T | 76 | 0 | 4 | 0 | NON_SYNONYMOU | MODERAT | MISSENSE | Gcc/Acc | A223T | PLES_42951 | | | | | | | T | T | | | T | T |
| NC_011770 | 4732399 | G | A | 69 | 0 | 12 | 0 | NON_SYNONYMOU | MODERAT | MISSENSE | Cac/Tac | H60Y | PLES_42951 | A | A | A | A | A | A | A | A | A | A | A | A |
| NC_011770 | 4733078 | GC | G | 82 | 0 | 1 | 0 | FRAME_SHIFT | HIGH | | ggg/ | G381 | PLES_42961 (amaB) | | | | | | | | G | | | | |
| NC_011770 | 4733005 | T | G | 77 | 0 | 2 | 0 | NON_SYNONYMOU | MODERAT | MISSENSE | gAg/gCg | E406A | PLES_42961 (a | G | G | | | | | | | | | | |
| NC_011770 | 4733992 | G | A | 73 | 0 | 4 | 0 | NON_SYNONYMOU | MODERAT | MISSENSE | gCc/gTc | A77V | PLES_42961 (amaB) | | | A | A | A | A | | | | | | |
| NC_011770 | 4734089 | G | A | 76 | 0 | 2 | 0 | NON_SYNONYMOU | MODERAT | MISSENSE | Ccc/Tcc | P45S | PLES_42961 (a | A | A | | | | | | | | | | |
| NC_011770 | 4734181 | A | G | 70 | 0 | 12 | 0 | NON_SYNONYMOU | MODERAT | MISSENSE | tTg/tCg | L14S | PLES_42961 (a | G | G | G | G | G | G | G | G | G | G | G | G |

| | | | | | | | | | | | | | | | | | | | | | | | | | |
|---|---|---|---|---|---|---|---|---|---|---|---|---|---|---|---|---|---|---|---|---|---|---|---|---|---|
| NC_011770 | 4734157 | G | A | 72 | 0 | 1 | 0 | NON_SYNONYMOU | MODERAT | MISSENSE | aCg/aTg | T22M | PLES_42961 (amaB) | | | | | | | | | | | | A |
| NC_011770 | 4734302 | G | A | 74 | 0 | 2 | 0 | START_LOST | HIGH | MISSENSE | atG/atA | M1I | PLES_42971 | | | A | A | | | | | | | | |
| NC_011770 | 4734775 | G | A | 66 | 0 | 6 | 0 | SPLICE_SITE_REGIC | LOW | SILENT | tGa/tAa | *159 | PLES_42971 | A | A | A | A | A | A | | | | | | |
| NC_011770 | 4734303 | G | A | 75 | 0 | 2 | 0 | NON_SYNONYMOU | MODERAT | MISSENSE | Gac/Aac | D2N | PLES_42971 | | | | | | | A | A | | | | |
| NC_011770 | 4734312 | A | C | 75 | 0 | 6 | 0 | NON_SYNONYMOU | MODERAT | MISSENSE | Aag/Cag | K5Q | PLES_42971 | C | C | C | C | C | C | | | | | | |
| NC_011770 | 4734357 | G | A | 73 | 0 | 6 | 0 | NON_SYNONYMOU | MODERAT | MISSENSE | Gtc/Atc | V20I | PLES_42971 | A | A | A | A | A | A | | | | | | |
| NC_011770 | 4734702 | A | G | 68 | 0 | 2 | 0 | NON_SYNONYMOU | MODERAT | MISSENSE | Atc/Gtc | I135V | PLES_42971 | G | G | | | | | | | | | | |
| NC_011770 | 4734745 | C | T | 67 | 0 | 6 | 0 | NON_SYNONYMOU | MODERAT | MISSENSE | cCc/cTc | P149L | PLES_42971 | T | T | T | T | T | T | | | | | | |
| NC_011770 | 4734837 | C | T | 67 | 0 | 6 | 0 | NON_SYNONYMOU | MODERAT | MISSENSE | aGc/aAc | S33N | PLES_42981 | T | T | T | T | T | T | | | | | | |
| NC_011770 | 4734924 | G | C | 69 | 0 | 6 | 0 | NON_SYNONYMOU | MODERAT | MISSENSE | cCc/cGc | P4R | PLES_42981 | | | | | | | C | C | C | C | C | C |
| NC_011770 | 4735665 | C | T | 73 | 0 | 12 | 0 | NON_SYNONYMOU | MODERAT | MISSENSE | Gcc/Acc | A247T | PLES_42991 | T | T | T | T | T | T | T | T | T | T | T | T |
| NC_011770 | 4735859 | G | T | 73 | 0 | 12 | 0 | NON_SYNONYMOU | MODERAT | MISSENSE | aCc/aAc | T182N | PLES_42991 | T | T | T | T | T | T | T | T | T | T | T | T |
| NC_011770 | 4736175 | C | T | 75 | 0 | 4 | 0 | NON_SYNONYMOU | MODERAT | MISSENSE | Gtc/Atc | V77I | PLES_42991 | | | | | | | T | T | | | T | T |
| NC_011770 | 4736517 | G | A | 75 | 0 | 1 | 0 | NON_SYNONYMOU | MODERAT | MISSENSE | Cgt/Tgt | R318C | PLES_43001 | | A | | | | | | | | | | |
| NC_011770 | 4736625 | C | T | 83 | 0 | 10 | 0 | NON_SYNONYMOU | MODERAT | MISSENSE | Gcc/Acc | A282T | PLES_43001 | T | T | T | T | T | T | T | T | | | T | T |
| NC_011770 | 4736636 | T | C | 85 | 0 | 4 | 0 | NON_SYNONYMOU | MODERAT | MISSENSE | cAg/cGg | Q278R | PLES_43001 | | | C | C | C | C | | | | | | |
| NC_011770 | 4737176 | G | C | 60 | 0 | 2 | 0 | NON_SYNONYMOU | MODERAT | MISSENSE | cCc/cGc | P98R | PLES_43001 | | | | | | | | | C | C | | |
| NC_011770 | 4737183 | T | C | 60 | 0 | 12 | 0 | NON_SYNONYMOU | MODERAT | MISSENSE | Aga/Gga | R96G | PLES_43001 | C | C | C | C | C | C | C | C | C | C | C | C |
| NC_011770 | 4737186 | T | C | 60 | 0 | 2 | 0 | NON_SYNONYMOU | MODERAT | MISSENSE | Acg/Gcg | T95A | PLES_43001 | C | C | | | | | | | | | | |
| NC_011770 | 4737576 | G | C | 63 | 0 | 2 | 0 | NON_SYNONYMOU | MODERAT | MISSENSE | Cag/Gag | Q276E | PLES_43011 | C | C | | | | | | | | | | |
| NC_011770 | 4737776 | G | T | 54 | 0 | 4 | 0 | NON_SYNONYMOU | MODERAT | MISSENSE | gCa/gAa | A209E | PLES_43011 | T | T | | | | | | | | | T | T |
| NC_011770 | 4738767 | G | A | 85 | 0 | 12 | 0 | NON_SYNONYMOU | MODERAT | MISSENSE | gCg/gTg | A274V | PLES_43021 | A | A | A | A | A | A | A | A | A | A | A | A |
| NC_011770 | 4738777 | T | G | 86 | 0 | 12 | 0 | NON_SYNONYMOU | MODERAT | MISSENSE | Act/Cct | T271P | PLES_43021 | G | G | G | G | G | G | G | G | G | G | G | G |
| NC_011770 | 4738911 | T | C | 87 | 0 | 12 | 0 | NON_SYNONYMOU | MODERAT | MISSENSE | gAa/gGa | E226G | PLES_43021 | C | C | C | C | C | C | C | C | C | C | C | C |
| NC_011770 | 4739214 | T | C | 85 | 0 | 1 | 0 | NON_SYNONYMOU | MODERAT | MISSENSE | tAc/tGc | Y125C | PLES_43021 | | | | | | | C | | | | | |
| NC_011770 | 4739583 | G | C | 61 | 0 | 8 | 0 | NON_SYNONYMOU | MODERAT | MISSENSE | aCc/aGc | T2S | PLES_43021 | C | C | | | | | C | C | C | C | C | C |
| NC_011770 | 4740351 | C | T | 73 | 0 | 2 | 0 | SPLICE_SITE_REGIC | LOW | SILENT | tGa/tAa | *371 | PLES_43041 | | | | | | | T | T | | | | |
| NC_011770 | 4740678 | C | A | 80 | 0 | 2 | 0 | NON_SYNONYMOU | MODERAT | MISSENSE | aGt/aTt | S262I | PLES_43041 | | | | | | | A | A | | | | |
| NC_011770 | 4740805 | C | T | 92 | 0 | 6 | 0 | NON_SYNONYMOU | MODERAT | MISSENSE | Gcg/Acg | A220T | PLES_43041 | | | | | | | T | T | T | T | T | T |
| NC_011770 | 4741009 | C | G | 90 | 0 | 6 | 0 | NON_SYNONYMOU | MODERAT | MISSENSE | Gaa/Caa | E152Q | PLES_43041 | | | | | | | G | G | G | G | G | G |
| NC_011770 | 4741086 | G | A | 86 | 0 | 2 | 0 | NON_SYNONYMOU | MODERAT | MISSENSE | gCg/gTg | A126V | PLES_43041 | A | A | | | | | | | | | | |
| NC_011770 | 4741317 | C | G | 72 | 0 | 12 | 0 | NON_SYNONYMOU | MODERAT | MISSENSE | aGc/aCc | S49T | PLES_43041 | G | G | G | G | G | G | G | G | G | G | G | G |
| NC_011770 | 4741664 | AC | A | 85 | 0 | 2 | 0 | FRAME_SHIFT | HIGH | | gtg/ | V100 | PLES_43051 | | | | | | | A | A | | | | |
| NC_011770 | 4741613 | T | C | 86 | 0 | 2 | 0 | NON_SYNONYMOU | MODERAT | MISSENSE | gAc/gGc | D117G | PLES_43051 | | | | | | | | | | | C | C |
| NC_011770 | 4741869 | C | T | 80 | 0 | 2 | 0 | NON_SYNONYMOU | MODERAT | MISSENSE | Gtg/Atg | V32M | PLES_43051 | T | T | | | | | | | | | | |
| NC_011770 | 4743589 | T | C | 80 | 0 | 8 | 0 | NON_SYNONYMOU | MODERAT | MISSENSE | gAg/gGg | E234G | PLES_43071 | C | C | | | | | C | C | C | C | C | C |
| NC_011770 | 4743679 | G | C | 75 | 0 | 4 | 0 | NON_SYNONYMOU | MODERAT | MISSENSE | gCc/gGc | A204G | PLES_43071 | | | C | C | C | C | | | | | | |
| NC_011770 | 4743719 | C | T | 80 | 0 | 1 | 0 | NON_SYNONYMOU | MODERAT | MISSENSE | Gac/Aac | D191N | PLES_43071 | | | | | | | T | | | | | |
| NC_011770 | 4743763 | T | G | 79 | 0 | 12 | 0 | NON_SYNONYMOU | MODERAT | MISSENSE | cAg/cCg | Q176P | PLES_43071 | G | G | G | G | G | G | G | G | G | G | G | G |
| NC_011770 | 4743811 | A | G | 78 | 0 | 12 | 0 | NON_SYNONYMOU | MODERAT | MISSENSE | aTg/aCg | M160T | PLES_43071 | G | G | G | G | G | G | G | G | G | G | G | G |
| NC_011770 | 4744156 | A | G | 86 | 0 | 8 | 0 | NON_SYNONYMOU | MODERAT | MISSENSE | gTc/gCc | V45A | PLES_43071 | G | G | | | | | G | G | G | G | G | G |
| NC_011770 | 4744198 | G | C | 80 | 0 | 8 | 0 | NON_SYNONYMOU | MODERAT | MISSENSE | aCc/aGc | T31S | PLES_43071 | C | C | | | | | C | C | C | C | C | C |
| NC_011770 | 4746636 | G | C | 86 | 0 | 4 | 0 | NON_SYNONYMOU | MODERAT | MISSENSE | gCc/gGc | A349G | PLES_43091 | | | | | | | C | C | C | C | | |
| NC_011770 | 4746906 | G | C | 84 | 0 | 4 | 0 | NON_SYNONYMOU | MODERAT | MISSENSE | gCt/gGt | A259G | PLES_43091 | C | C | | | | | C | C | | | | |
| NC_011770 | 4747044 | GCGC | G | 80 | 0 | 2 | 0 | CODON_CHANGE_ | MODERATE | | ggcgcc/gcc | GA212A | PLES_43091 | G | G | | | | | | | | | | |
| NC_011770 | 4747152 | T | C | 84 | 0 | 2 | 0 | NON_SYNONYMOU | MODERAT | MISSENSE | gAg/gGg | E177G | PLES_43091 | C | C | | | | | | | | | | |
| NC_011770 | 4747572 | T | C | 78 | 0 | 4 | 0 | NON_SYNONYMOU | MODERAT | MISSENSE | aAg/aGg | K37R | PLES_43091 | C | C | | | | | | | C | C | | |
| NC_011770 | 4747663 | C | G | 77 | 0 | 2 | 0 | NON_SYNONYMOU | MODERAT | MISSENSE | Ggc/Cgc | G7R | PLES_43091 | | | | | | | | | | | G | G |
| NC_011770 | 4748424 | A | T | 83 | 0 | 2 | 0 | NON_SYNONYMOU | MODERAT | MISSENSE | gAt/gTt | D68V | PLES_43101 | | | | | | | | | | | T | T |
| NC_011770 | 4748513 | G | C | 86 | 0 | 2 | 0 | NON_SYNONYMOU | MODERAT | MISSENSE | Gag/Cag | E98Q | PLES_43101 | | | | | C | C | | | | | | |
| NC_011770 | 4748843 | G | A | 79 | 0 | 2 | 0 | NON_SYNONYMOU | MODERAT | MISSENSE | Gtc/Atc | V208I | PLES_43101 | | | | | | | | | | | A | A |
| NC_011770 | 4749012 | G | A | 80 | 0 | 2 | 0 | NON_SYNONYMOU | MODERAT | MISSENSE | gGg/gAg | G264E | PLES_43101 | A | A | | | | | | | | | | |
| NC_011770 | 4749714 | TC | T | 79 | 0 | 1 | 0 | FRAME_SHIFT | HIGH | | gag/ | E114 | PLES_43111 | | | | | | | T | | | | | |
| NC_011770 | 4749969 | TC | T | 66 | 0 | 4 | 0 | FRAME_SHIFT | HIGH | | gag/ | E29 | PLES_43111 | | | T | T | T | T | | | | | | |
| NC_011770 | 4749063 | T | C | 77 | 0 | 2 | 0 | SPLICE_SITE_REGIC | LOW | SILENT | tAa/tGa | *331 | PLES_43111 | C | C | | | | | | | | | | |
| NC_011770 | 4749072 | G | T | 73 | 0 | 2 | 0 | NON_SYNONYMOU | MODERAT | MISSENSE | gCg/gAg | A328E | PLES_43111 | T | T | | | | | | | | | | |
| NC_011770 | 4749078 | T | C | 73 | 0 | 2 | 0 | NON_SYNONYMOU | MODERAT | MISSENSE | gAc/gGc | D326G | PLES_43111 | C | C | | | | | | | | | | |
| NC_011770 | 4749318 | T | C | 77 | 0 | 1 | 0 | NON_SYNONYMOU | MODERAT | MISSENSE | tAc/tGc | Y246C | PLES_43111 | | | | | | | C | | | | | |
| NC_011770 | 4749470 | G | T | 77 | 0 | 2 | 0 | NON_SYNONYMOU | MODERAT | MISSENSE | gaC/gaA | D195E | PLES_43111 | T | T | | | | | | | | | | |
| NC_011770 | 4749789 | C | T | 82 | 0 | 2 | 0 | NON_SYNONYMOU | MODERAT | MISSENSE | gGt/gAt | G89D | PLES_43111 | T | T | | | | | | | | | | |
| NC_011770 | 4749795 | C | A | 79 | 0 | 6 | 0 | NON_SYNONYMOU | MODERAT | MISSENSE | cGt/cTt | R87L | PLES_43111 | A | A | | | | | | | A | A | A | A |
| NC_011770 | 4749979 | C | T | 68 | 0 | 4 | 0 | NON_SYNONYMOU | MODERAT | MISSENSE | Ggg/Agg | G26R | PLES_43111 | | | T | T | T | T | | | | | | |

| | | | | | | | | | | | | | | | | | | | | | | | | | |
|---|---|---|---|---|---|---|---|---|---|---|---|---|---|---|---|---|---|---|---|---|---|---|---|---|---|
| NC_011770 | 4749981 | G | T | 67 | 0 | 2 | 0 | NON_SYNONYMOU | MODERAT | MISSENSE | gCc/gAc | A25D | PLES_43111 | | | | | | | T | T | | | | |
| NC_011770 | 4750021 | T | C | 71 | 0 | 2 | 0 | NON_SYNONYMOU | MODERAT | MISSENSE | Agc/Ggc | S12G | PLES_43111 | | | | | | | C | C | | | | |
| NC_011770 | 4751382 | T | C | 85 | 0 | 8 | 0 | NON_SYNONYMOU | MODERAT | MISSENSE | gAc/gGc | D235G | PLES_43131 | C | C | | | | | C | C | C | C | C | C |
| NC_011770 | 4751458 | C | T | 94 | 0 | 1 | 0 | NON_SYNONYMOU | MODERAT | MISSENSE | Gcc/Acc | A210T | PLES_43131 | | | | | | | T | | | | | |
| NC_011770 | 4751479 | C | G | 89 | 0 | 12 | 0 | NON_SYNONYMOU | MODERAT | MISSENSE | Ggc/Cgc | G203R | PLES_43131 | G | G | G | G | G | G | G | G | G | G | G | G |
| NC_011770 | 4751665 | T | C | 90 | 0 | 2 | 0 | NON_SYNONYMOU | MODERAT | MISSENSE | Atg/Gtg | M141V | PLES_43131 | | | | | | | C | C | | | | |
| NC_011770 | 4755896 | G | A | 80 | 0 | 1 | 0 | NON_SYNONYMOU | MODERAT | MISSENSE | Ccg/Tcg | P192S | PLES_43181 | | | | | | | | | | | | A |
| NC_011770 | 4758057 | G | A | 71 | 0 | 2 | 0 | NON_SYNONYMOU | MODERAT | MISSENSE | Ggc/Agc | G381S | PLES_43201 | | | | | | | | | A | A | | |
| NC_011770 | 4758124 | T | C | 69 | 0 | 1 | 0 | NON_SYNONYMOU | MODERAT | MISSENSE | gTc/gCc | V403A | PLES_43201 | | | | | | | C | | | | | |
| NC_011770 | 4774509 | T | C | 70 | 0 | 2 | 0 | NON_SYNONYMOU | MODERAT | MISSENSE | aAg/aGg | K124R | PLES_43341 | | | | | | | C | C | | | | |
| NC_011770 | 4774551 | T | C | 73 | 0 | 2 | 0 | NON_SYNONYMOU | MODERAT | MISSENSE | aAa/aGa | K110R | PLES_43341 | | | | | | | | | | | C | C |
| NC_011770 | 4774569 | G | A | 71 | 0 | 2 | 0 | NON_SYNONYMOU | MODERAT | MISSENSE | gCc/gTc | A104V | PLES_43341 | | | | | | | | | | | A | A |
| NC_011770 | 4774689 | G | A | 78 | 0 | 2 | 0 | NON_SYNONYMOU | MODERAT | MISSENSE | gCg/gTg | A64V | PLES_43341 | | | | | | | A | A | | | | |
| NC_011770 | 4774818 | G | T | 85 | 0 | 2 | 0 | NON_SYNONYMOU | MODERAT | MISSENSE | gCc/gAc | A21D | PLES_43341 | T | T | | | | | | | | | | |
| NC_011770 | 4775885 | TC | T | 57 | 0 | 2 | 0 | FRAME_SHIFT | HIGH | | cac/ | H143 | PLES_43351 | | | | | | | T | T | | | | |
| NC_011770 | 4775757 | A | G | 62 | 0 | 2 | 0 | NON_SYNONYMOU | MODERAT | MISSENSE | Acc/Gcc | T99A | PLES_43351 | G | G | | | | | | | | | | |
| NC_011770 | 4776508 | C | T | 89 | 0 | 2 | 0 | NON_SYNONYMOU | MODERAT | MISSENSE | gCt/gTt | A8V | PLES_43361 | | | | | | | | | | | T | T |
| NC_011770 | 4776703 | G | A | 79 | 0 | 4 | 0 | NON_SYNONYMOU | MODERAT | MISSENSE | cGg/cAg | R73Q | PLES_43361 | | | A | A | A | A | | | | | | |
| NC_011770 | 4776613 | A | G | 92 | 0 | 1 | 0 | NON_SYNONYMOU | MODERAT | MISSENSE | aAc/aGc | N43S | PLES_43361 | | | | | | | | G | | | | |
| NC_011770 | 4778245 | G | A | 67 | 0 | 2 | 0 | NON_SYNONYMOU | MODERAT | MISSENSE | gCg/gTg | A175V | PLES_43381 | | | | | | | | | A | A | | |
| NC_011770 | 4779245 | C | T | 83 | 0 | 4 | 0 | NON_SYNONYMOU | MODERAT | MISSENSE | Gcc/Acc | A112T | PLES_43391 | | | T | T | T | T | | | | | | |
| NC_011770 | 4779260 | A | G | 87 | 0 | 4 | 0 | NON_SYNONYMOU | MODERAT | MISSENSE | Ttc/Ctc | F107L | PLES_43391 | | | G | G | G | G | | | | | | |
| NC_011770 | 4779491 | A | G | 72 | 0 | 4 | 0 | NON_SYNONYMOU | MODERAT | MISSENSE | Tcc/Ccc | S30P | PLES_43391 | | | G | G | G | G | | | | | | |
| NC_011770 | 4779535 | A | G | 72 | 0 | 12 | 0 | NON_SYNONYMOU | MODERAT | MISSENSE | cTt/cCt | L15P | PLES_43391 | G | G | G | G | G | G | G | G | G | G | G | G |
| NC_011770 | 4779544 | T | A | 74 | 0 | 6 | 0 | NON_SYNONYMOU | MODERAT | MISSENSE | cAc/cTc | H12L | PLES_43391 | | | A | A | A | A | | | A | A | | |
| NC_011770 | 4780118 | A | G | 78 | 0 | 4 | 0 | NON_SYNONYMOU | MODERAT | MISSENSE | Tcc/Ccc | S120P | PLES_43401 | | | G | G | G | G | | | | | | |
| NC_011770 | 4780148 | C | A | 73 | 0 | 12 | 0 | NON_SYNONYMOU | MODERAT | MISSENSE | Gcc/Tcc | A110S | PLES_43401 | A | A | A | A | A | A | A | A | A | A | A | A |
| NC_011770 | 4784989 | C | T | 78 | 0 | 2 | 0 | START_LOST | HIGH | MISSENSE | atG/atA | M1I | PLES_43461 | | | | | | | | | | | T | T |
| NC_011770 | 4784988 | G | T | 77 | 0 | 1 | 0 | NON_SYNONYMOU | MODERAT | MISSENSE | Cgc/Agc | R2S | PLES_43461 | | | T | | | | | | | | | |
| NC_011770 | 4784681 | A | G | 81 | 0 | 12 | 0 | NON_SYNONYMOU | MODERAT | MISSENSE | tTg/tCg | L104S | PLES_43461 | G | G | G | G | G | G | G | G | G | G | G | G |
| NC_011770 | 4784972 | G | C | 74 | 0 | 12 | 0 | NON_SYNONYMOU | MODERAT | MISSENSE | gCa/gGa | A7G | PLES_43461 | C | C | C | C | C | C | C | C | C | C | C | C |
| NC_011770 | 4791598 | A | G | 31 | 0 | 1 | 0 | NON_SYNONYMOU | MODERAT | MISSENSE | cAg/cGg | Q40R | PLES_43531 | | G | | | | | | | | | | |
| NC_011770 | 4791919 | T | C | 49 | 0 | 4 | 0 | NON_SYNONYMOU | MODERAT | MISSENSE | gAc/gGc | D4G | PLES_43541 | | | C | C | C | C | | | | | | |
| NC_011770 | 4791950 | G | A | 52 | 0 | 4 | 0 | NON_SYNONYMOU | MODERAT | MISSENSE | Cgt/Tgt | R204C | PLES_43551 | | | | | | | A | A | A | A | | |
| NC_011770 | 4792018 | C | T | 59 | 0 | 4 | 0 | NON_SYNONYMOU | MODERAT | MISSENSE | cGg/cAg | R181Q | PLES_43551 | | | T | T | T | T | | | | | | |
| NC_011770 | 4792198 | C | T | 69 | 0 | 4 | 0 | NON_SYNONYMOU | MODERAT | MISSENSE | cGg/cAg | R121Q | PLES_43551 | | | T | T | T | T | | | | | | |
| NC_011770 | 4792412 | G | A | 61 | 0 | 4 | 0 | NON_SYNONYMOU | MODERAT | MISSENSE | Ccg/Tcg | P50S | PLES_43551 | | | A | A | A | A | | | | | | |
| NC_011770 | 4792426 | G | A | 66 | 0 | 4 | 0 | NON_SYNONYMOU | MODERAT | MISSENSE | aCg/aTg | T45M | PLES_43551 | | | A | A | A | A | | | | | | |
| NC_011770 | 4792471 | A | G | 64 | 0 | 4 | 0 | NON_SYNONYMOU | MODERAT | MISSENSE | gTg/gCg | V30A | PLES_43551 | | | G | G | G | G | | | | | | |
| NC_011770 | 4792480 | C | A | 65 | 0 | 4 | 0 | NON_SYNONYMOU | MODERAT | MISSENSE | gGc/gTc | G27V | PLES_43551 | | | A | A | A | A | | | | | | |
| NC_011770 | 4792531 | C | T | 69 | 0 | 4 | 0 | NON_SYNONYMOU | MODERAT | MISSENSE | gGc/gAc | G10D | PLES_43551 | | | T | T | T | T | | | | | | |
| NC_011770 | 4792550 | C | T | 75 | 0 | 4 | 0 | NON_SYNONYMOU | MODERAT | MISSENSE | Gtc/Atc | V4I | PLES_43551 | | | T | T | T | T | | | | | | |
| NC_011770 | 4792457 | G | A | 65 | 0 | 1 | 0 | NON_SYNONYMOU | MODERAT | MISSENSE | Ccg/Tcg | P35S | PLES_43551 | | | | | | | | A | | | | |
| NC_011770 | 4794521 | C | T | 75 | 0 | 2 | 0 | NON_SYNONYMOU | MODERAT | MISSENSE | aGt/aAt | S118N | PLES_43571 | | | | | | | | | T | T | | |
| NC_011770 | 4797492 | T | C | 65 | 0 | 1 | 0 | NON_SYNONYMOU | MODERAT | MISSENSE | gTc/gCc | V263A | PLES_43591 | C | | | | | | | | | | | |
| NC_011770 | 4797041 | G | A | 63 | 0 | 2 | 0 | NON_SYNONYMOU | MODERAT | MISSENSE | Gtc/Atc | V113I | PLES_43591 | | | | | | | | | A | A | | |
| NC_011770 | 4797087 | G | C | 62 | 0 | 2 | 0 | NON_SYNONYMOU | MODERAT | MISSENSE | gGc/gCc | G128A | PLES_43591 | | | | | | | C | C | | | | |
| NC_011770 | 4798086 | T | G | 54 | 0 | 2 | 0 | NON_SYNONYMOU | MODERAT | MISSENSE | aAg/aCg | K111T | PLES_43611 | G | G | | | | | | | | | | |
| NC_011770 | 4798212 | C | T | 54 | 1 | 10 | 0 | NON_SYNONYMOU | MODERAT | MISSENSE | gGa/gAa | G69E | PLES_43611 | T | T | T | T | T | T | T | T | . | | T | T |
| NC_011770 | 4798402 | A | G | 70 | 0 | 12 | 0 | NON_SYNONYMOU | MODERAT | MISSENSE | Tgc/Cgc | C6R | PLES_43611 | G | G | G | G | G | G | G | G | G | G | G | G |
| NC_011770 | 4798401 | C | T | 70 | 0 | 1 | 0 | NON_SYNONYMOU | MODERAT | MISSENSE | tGc/tAc | C6Y | PLES_43611 | | | | | | | | T | | | | |
| NC_011770 | 4798786 | G | A | 73 | 0 | 2 | 0 | NON_SYNONYMOU | MODERAT | MISSENSE | Gtc/Atc | V9I | PLES_43621 | A | A | | | | | | | | | | |
| NC_011770 | 4799541 | G | A | 80 | 0 | 2 | 0 | NON_SYNONYMOU | MODERAT | MISSENSE | Cac/Tac | H399Y | PLES_43641 | | | | | | | A | A | | | | |
| NC_011770 | 4799630 | C | T | 75 | 0 | 10 | 0 | NON_SYNONYMOU | MODERAT | MISSENSE | aGt/aAt | S369N | PLES_43641 | | | T | T | T | T | T | T | T | T | T | T |
| NC_011770 | 4799864 | T | A | 93 | 0 | 2 | 0 | NON_SYNONYMOU | MODERAT | MISSENSE | cAc/cTc | H291L | PLES_43641 | A | A | | | | | | | | | | |
| NC_011770 | 4801838 | G | A | 78 | 0 | 2 | 0 | NON_SYNONYMOU | MODERAT | MISSENSE | Gta/Ata | V11I | PLES_43671 | | | | | | | A | A | | | | |
| NC_011770 | 4801919 | A | C | 74 | 0 | 2 | 0 | NON_SYNONYMOU | MODERAT | MISSENSE | Att/Ctt | I38L | PLES_43671 | C | C | | | | | | | | | | |
| NC_011770 | 4802210 | G | C | 87 | 0 | 4 | 0 | NON_SYNONYMOU | MODERAT | MISSENSE | Gtc/Ctc | V135L | PLES_43671 | C | C | | | | | | | C | C | | |
| NC_011770 | 4803412 | T | C | 74 | 0 | 4 | 0 | NON_SYNONYMOU | MODERAT | MISSENSE | Atc/Gtc | I314V | PLES_43691 | | | C | C | C | C | | | | | | |
| NC_011770 | 4803597 | T | C | 75 | 0 | 4 | 0 | NON_SYNONYMOU | MODERAT | MISSENSE | gAa/gGa | E252G | PLES_43691 | | | C | C | C | C | | | | | | |
| NC_011770 | 4803747 | G | A | 74 | 0 | 2 | 0 | NON_SYNONYMOU | MODERAT | MISSENSE | gCg/gTg | A202V | PLES_43691 | | | | | | | A | A | | | | |
| NC_011770 | 4803802 | C | T | 69 | 0 | 4 | 0 | NON_SYNONYMOU | MODERAT | MISSENSE | Gca/Aca | A184T | PLES_43691 | | | T | T | T | T | | | | | | |

| | | | | | | | | | | | | | | | | | | | | | | | | | |
|---|---|---|---|---|---|---|---|---|---|---|---|---|---|---|---|---|---|---|---|---|---|---|---|---|---|
| NC_011770 | 4803829 | T | A | 66 | 0 | 4 | 0 | NON_SYNONYMOU | MODERAT | MISSENSE | Acc/Tcc | T175S | PLES_43691 | | | A | A | A | A | | | | | | |
| NC_011770 | 4803925 | T | C | 67 | 0 | 12 | 0 | NON_SYNONYMOU | MODERAT | MISSENSE | Aaa/Gaa | K143E | PLES_43691 | C | C | C | C | C | C | C | C | C | C | C | C |
| NC_011770 | 4804236 | C | T | 64 | 0 | 2 | 0 | NON_SYNONYMOU | MODERAT | MISSENSE | gGg/gAg | G39E | PLES_43691 | | | | | | | T | T | | | | |
| NC_011770 | 4807100 | G | A | 67 | 0 | 2 | 0 | NON_SYNONYMOU | MODERAT | MISSENSE | gCc/gTc | A173V | PLES_43731 | | | | | | | | | A | A | | |
| NC_011770 | 4807103 | T | G | 64 | 0 | 2 | 0 | NON_SYNONYMOU | MODERAT | MISSENSE | gAt/gCt | D172A | PLES_43731 | | | | | | | | | G | G | | |
| NC_011770 | 4807184 | A | T | 66 | 0 | 2 | 0 | NON_SYNONYMOU | MODERAT | MISSENSE | tTc/tAc | F145Y | PLES_43731 | T | T | | | | | | | | | | |
| NC_011770 | 4807320 | C | A | 72 | 0 | 4 | 0 | NON_SYNONYMOU | MODERAT | MISSENSE | Gcg/Tcg | A100S | PLES_43731 | | | A | A | A | A | | | | | | |
| NC_011770 | 4807881 | C | T | 84 | 0 | 4 | 0 | NON_SYNONYMOU | MODERAT | MISSENSE | gCc/gTc | A50V | PLES_43741 | | | T | T | T | T | | | | | | |
| NC_011770 | 4807890 | A | G | 85 | 0 | 4 | 0 | NON_SYNONYMOU | MODERAT | MISSENSE | aAg/aGg | K53R | PLES_43741 | | | | | | | | | G | G | G | G |
| NC_011770 | 4808056 | G | A | 92 | 0 | 4 | 0 | NON_SYNONYMOU | MODERAT | MISSENSE | Gag/Aag | E27K | PLES_43751 | | | A | A | A | A | | | | | | |
| NC_011770 | 4808176 | G | A | 93 | 0 | 2 | 0 | NON_SYNONYMOU | MODERAT | MISSENSE | Gtt/Att | V67I | PLES_43751 | | | | | | | | | A | A | | |
| NC_011770 | 4808251 | A | G | 88 | 0 | 2 | 0 | NON_SYNONYMOU | MODERAT | MISSENSE | cAc/cGc | H5R | PLES_43761 | | | | | | | | | | | G | G |
| NC_011770 | 4808275 | A | G | 91 | 0 | 2 | 0 | NON_SYNONYMOU | MODERAT | MISSENSE | gAc/gGc | D13G | PLES_43761 | | | | | | | | | | | G | G |
| NC_011770 | 4808544 | C | T | 90 | 0 | 10 | 0 | NON_SYNONYMOU | MODERAT | MISSENSE | Cct/Tct | P103S | PLES_43761 | | | T | T | T | T | T | T | T | T | T | T |
| NC_011770 | 4808652 | GA | G | 91 | 0 | 6 | 0 | FRAME_SHIFT | HIGH | | aag/ | K9 | PLES_43771 | | | G | G | G | G | | | G | G | | |
| NC_011770 | 4808775 | G | A | 82 | 0 | 2 | 0 | NON_SYNONYMOU | MODERAT | MISSENSE | Gag/Aag | E49K | PLES_43771 | | | | | | | | | A | A | | |
| NC_011770 | 4808790 | G | A | 81 | 0 | 2 | 0 | NON_SYNONYMOU | MODERAT | MISSENSE | Gcc/Acc | A54T | PLES_43771 | | | | | A | A | | | | | | |
| NC_011770 | 4808838 | A | G | 78 | 0 | 2 | 0 | NON_SYNONYMOU | MODERAT | MISSENSE | Aaa/Gaa | K70E | PLES_43771 | | | | | | | | | G | G | | |
| NC_011770 | 4809483 | G | A | 71 | 0 | 1 | 0 | STOP_GAINED | HIGH | NONSENSE | Cag/Tag | Q246* | PLES_43781 | | | | | | | | | | | A | |
| NC_011770 | 4809006 | A | G | 77 | 0 | 4 | 0 | NON_SYNONYMOU | MODERAT | MISSENSE | Ttc/Ctc | F405L | PLES_43781 | | | G | G | G | G | | | | | | |
| NC_011770 | 4809008 | A | G | 76 | 0 | 4 | 0 | NON_SYNONYMOU | MODERAT | MISSENSE | gTg/gCg | V404A | PLES_43781 | | | G | G | G | G | | | | | | |
| NC_011770 | 4809065 | T | C | 75 | 0 | 4 | 0 | NON_SYNONYMOU | MODERAT | MISSENSE | aAa/aGa | K385R | PLES_43781 | | | C | C | C | C | | | | | | |
| NC_011770 | 4809078 | A | C | 72 | 0 | 4 | 0 | NON_SYNONYMOU | MODERAT | MISSENSE | Tcg/Gcg | S381A | PLES_43781 | | | C | C | C | C | | | | | | |
| NC_011770 | 4809093 | T | C | 71 | 0 | 4 | 0 | NON_SYNONYMOU | MODERAT | MISSENSE | Atc/Gtc | I376V | PLES_43781 | | | C | C | C | C | | | | | | |
| NC_011770 | 4809115 | C | G | 61 | 0 | 4 | 0 | NON_SYNONYMOU | MODERAT | MISSENSE | gaG/gaC | E368D | PLES_43781 | | | G | G | G | G | | | | | | |
| NC_011770 | 4809125 | T | C | 62 | 0 | 4 | 0 | NON_SYNONYMOU | MODERAT | MISSENSE | aAg/aGg | K365R | PLES_43781 | | | C | C | C | C | | | | | | |
| NC_011770 | 4809135 | T | G | 63 | 0 | 4 | 0 | NON_SYNONYMOU | MODERAT | MISSENSE | Aag/Cag | K362Q | PLES_43781 | | | G | G | G | G | | | | | | |
| NC_011770 | 4809257 | T | G | 77 | 0 | 4 | 0 | NON_SYNONYMOU | MODERAT | MISSENSE | aAg/aCg | K321T | PLES_43781 | | | G | G | G | G | | | | | | |
| NC_011770 | 4809530 | T | C | 75 | 0 | 4 | 0 | NON_SYNONYMOU | MODERAT | MISSENSE | aAg/aGg | K230R | PLES_43781 | | | C | C | C | C | | | | | | |
| NC_011770 | 4809602 | G | C | 75 | 0 | 4 | 0 | NON_SYNONYMOU | MODERAT | MISSENSE | aCc/aGc | T206S | PLES_43781 | | | C | C | C | C | | | | | | |
| NC_011770 | 4809803 | T | G | 68 | 0 | 4 | 0 | NON_SYNONYMOU | MODERAT | MISSENSE | gAc/gCc | D139A | PLES_43781 | | | G | G | G | G | | | | | | |
| NC_011770 | 4809806 | T | G | 71 | 0 | 4 | 0 | NON_SYNONYMOU | MODERAT | MISSENSE | aAg/aCg | K138T | PLES_43781 | | | G | G | G | G | | | | | | |
| NC_011770 | 4809819 | C | T | 71 | 0 | 6 | 0 | NON_SYNONYMOU | MODERAT | MISSENSE | Gcg/Acg | A134T | PLES_43781 | | | T | T | T | T | T | T | | | | |
| NC_011770 | 4809885 | C | T | 75 | 0 | 4 | 0 | NON_SYNONYMOU | MODERAT | MISSENSE | Gcc/Acc | A112T | PLES_43781 | | | T | T | T | T | | | | | | |
| NC_011770 | 4809967 | A | C | 72 | 0 | 4 | 0 | NON_SYNONYMOU | MODERAT | MISSENSE | gaT/gaG | D84E | PLES_43781 | | | C | C | C | C | | | | | | |
| NC_011770 | 4809990 | G | T | 65 | 0 | 4 | 0 | NON_SYNONYMOU | MODERAT | MISSENSE | Ctg/Atg | L77M | PLES_43781 | | | T | T | T | T | | | | | | |
| NC_011770 | 4810003 | G | T | 61 | 0 | 6 | 0 | NON_SYNONYMOU | MODERAT | MISSENSE | aaC/aaA | N72K | PLES_43781 | | | T | T | T | T | C | C | | | | |
| NC_011770 | 4810005 | T | C | 62 | 0 | 4 | 0 | NON_SYNONYMOU | MODERAT | MISSENSE | Aac/Gac | N72D | PLES_43781 | | | C | C | C | C | | | | | | |
| NC_011770 | 4810008 | C | T | 63 | 0 | 4 | 0 | NON_SYNONYMOU | MODERAT | MISSENSE | Gtc/Atc | V71I | PLES_43781 | | | T | T | T | T | | | | | | |
| NC_011770 | 4810076 | G | A | 68 | 0 | 4 | 0 | NON_SYNONYMOU | MODERAT | MISSENSE | gCc/gTc | A48V | PLES_43781 | | | A | A | A | A | | | | | | |
| NC_011770 | 4810173 | G | T | 66 | 0 | 4 | 0 | NON_SYNONYMOU | MODERAT | MISSENSE | Ctc/Atc | L16I | PLES_43781 | | | T | T | T | T | | | | | | |
| NC_011770 | 4810202 | A | C | 70 | 0 | 4 | 0 | NON_SYNONYMOU | MODERAT | MISSENSE | cTt/cGt | L6R | PLES_43781 | | | C | C | C | C | | | | | | |
| NC_011770 | 4810477 | C | T | 80 | 0 | 2 | 0 | NON_SYNONYMOU | MODERAT | MISSENSE | Gta/Ata | V157I | PLES_43791 | | | | | | | T | T | | | | |
| NC_011770 | 4820655 | C | T | 79 | 0 | 8 | 0 | NON_SYNONYMOU | MODERAT | MISSENSE | gGc/gAc | G360D | PLES_43861 | | | T | T | T | T | T | T | | | T | T |
| NC_011770 | 4820751 | G | T | 84 | 0 | 8 | 0 | NON_SYNONYMOU | MODERAT | MISSENSE | gCt/gAt | A328D | PLES_43861 | T | T | T | T | T | T | | | | | T | T |
| NC_011770 | 4821192 | T | A | 87 | 0 | 12 | 0 | NON_SYNONYMOU | MODERAT | MISSENSE | gAg/gTg | E181V | PLES_43861 | A | A | A | A | A | A | A | A | A | A | A | A |
| NC_011770 | 4821528 | C | T | 86 | 0 | 2 | 0 | NON_SYNONYMOU | MODERAT | MISSENSE | cGt/cAt | R69H | PLES_43861 | | | | | | | | | T | T | | |
| NC_011770 | 4821541 | C | T | 88 | 0 | 2 | 0 | NON_SYNONYMOU | MODERAT | MISSENSE | Gag/Aag | E65K | PLES_43861 | | | | | | | | | T | T | | |
| NC_011770 | 4821684 | A | G | 91 | 0 | 12 | 0 | NON_SYNONYMOU | MODERAT | MISSENSE | gTg/gCg | V17A | PLES_43861 | G | G | G | G | G | G | G | G | G | G | G | G |
| NC_011770 | 4822049 | C | T | 61 | 0 | 2 | 0 | NON_SYNONYMOU | MODERAT | MISSENSE | gGc/gAc | G135D | PLES_43871 pirR | | | | | | | | | T | T | | |
| NC_011770 | 4822298 | C | G | 57 | 0 | 4 | 0 | NON_SYNONYMOU | MODERAT | MISSENSE | gGc/gCc | G52A | PLES_43871 pirR | | | | | | | | | G | G | G | G |
| NC_011770 | 4822057 | GCC | GCCC | 55 | 1 | 3 | 0 | FRAME_SHIFT | HIGH | | ggc/ | G132 | PLES_43871 **pirR** | | GCCC | | G | | | . | G | | | | |
| NC_011770 | 4826868 | G | A | 79 | 0 | 6 | 0 | SPLICE_SITE_REGI | LOW | SILENT | taG/taA | *142 | PLES_43901 | A | A | A | A | A | A | | | | | | |
| NC_011770 | 4826704 | A | G | 83 | 0 | 2 | 0 | NON_SYNONYMOU | MODERAT | MISSENSE | Acc/Gcc | T88A | PLES_43901 | | | | | | | G | G | | | | |
| NC_011770 | 4826827 | C | A | 83 | 0 | 6 | 0 | NON_SYNONYMOU | MODERAT | MISSENSE | Cgc/Agc | R129S | PLES_43901 | | | A | A | A | A | | | A | A | | |
| NC_011770 | 4826840 | G | A | 83 | 0 | 2 | 0 | NON_SYNONYMOU | MODERAT | MISSENSE | gGc/gAc | G133D | PLES_43901 | A | A | | | | | | | | | | |
| NC_011770 | 4826966 | G | A | 79 | 0 | 2 | 0 | NON_SYNONYMOU | MODERAT | MISSENSE | Gcc/Acc | A11T | PLES_43911 | | | | | | | | | A | A | | |
| NC_011770 | 4826975 | A | G | 81 | 0 | 6 | 0 | NON_SYNONYMOU | MODERAT | MISSENSE | Atc/Gtc | I14V | PLES_43911 | | | G | G | G | G | | | G | G | | |
| NC_011770 | 4827160 | C | A | 79 | 0 | 4 | 0 | NON_SYNONYMOU | MODERAT | MISSENSE | gaC/gaA | D75E | PLES_43911 | | | A | A | A | A | | | | | | |
| NC_011770 | 4828889 | TG | T | 65 | 2 | 1 | 0 | FRAME_SHIFT | HIGH | | gaa/ | E480 | PLES_43921 | | | | T | | | | | | | . | . |
| NC_011770 | 4827670 | T | C | 54 | 0 | 2 | 0 | NON_SYNONYMOU | MODERAT | MISSENSE | cTt/cCt | L72P | PLES_43921 | | | | | | | | | C | C | | |
| NC_011770 | 4828015 | T | C | 74 | 0 | 8 | 0 | NON_SYNONYMOU | MODERAT | MISSENSE | gTc/gCc | V187A | PLES_43921 | C | C | C | C | C | C | | | | | C | C |

| | | | | | | | | | | | | | | | | | | | | | | | | | |
|---|---|---|---|---|---|---|---|---|---|---|---|---|---|---|---|---|---|---|---|---|---|---|---|---|---|
| NC_011770 | 4828212 | A | C | 78 | 0 | 6 | 0 | NON_SYNONYMOU | MODERAT | MISSENSE | Aac/Cac | N253H | PLES_43921 | | | | | | | C | C | C | C | C | C |
| NC_011770 | 4829004 | G | A | 66 | 2 | 8 | 0 | NON_SYNONYMOU | MODERAT | MISSENSE | Ggc/Agc | G517S | PLES_43921 | A | A | A | A | A | A | | | A | A | . | . |
| NC_011770 | 4829059 | G | A | 67 | 2 | 2 | 0 | NON_SYNONYMOU | MODERAT | MISSENSE | cGc/cAc | R535H | PLES_43921 | | | | | | | A | A | | | . | . |
| NC_011770 | 4829359 | A | G | 72 | 2 | 8 | 0 | NON_SYNONYMOU | MODERAT | MISSENSE | aAc/aGc | N635S | PLES_43921 | G | G | G | G | G | G | | | G | G | . | . |
| NC_011770 | 4829377 | C | G | 74 | 2 | 6 | 0 | NON_SYNONYMOU | MODERAT | MISSENSE | aCg/aGg | T641R | PLES_43921 | G | G | | | | | G | G | G | G | . | . |
| NC_011770 | 4830984 | T | G | 69 | 0 | 2 | 0 | NON_SYNONYMOU | MODERAT | MISSENSE | tTc/tGc | F48C | PLES_43941 | | | | | | | | | | | G | G |
| NC_011770 | 4831037 | G | A | 66 | 0 | 1 | 0 | NON_SYNONYMOU | MODERAT | MISSENSE | Gtg/Atg | V66M | PLES_43941 | | | | | | | A | | | | | |
| NC_011770 | 4832488 | T | C | 66 | 0 | 6 | 0 | NON_SYNONYMOU | MODERAT | MISSENSE | cTg/cCg | L237P | PLES_43961 | | | | | | | C | C | C | C | C | C |
| NC_011770 | 4833969 | G | A | 76 | 0 | 2 | 0 | NON_SYNONYMOU | MODERAT | MISSENSE | Gcg/Acg | A731T | PLES_43961 | A | A | | | | | | | | | | |
| NC_011770 | 4835140 | T | C | 79 | 0 | 1 | 0 | NON_SYNONYMOU | MODERAT | MISSENSE | Tat/Cat | Y239H | PLES_43971 | | C | | | | | | | | | | |
| NC_011770 | 4834754 | G | A | 70 | 0 | 2 | 0 | NON_SYNONYMOU | MODERAT | MISSENSE | cGc/cAc | R110H | PLES_43971 | | | | | | | | | | | A | A |
| NC_011770 | 4835113 | C | A | 77 | 0 | 4 | 0 | NON_SYNONYMOU | MODERAT | MISSENSE | Caa/Aaa | Q230K | PLES_43971 | | | | | | | A | A | | | A | A |
| NC_011770 | 4835916 | C | T | 50 | 0 | 8 | 0 | NON_SYNONYMOU | MODERAT | MISSENSE | gCc/gTc | A16V | PLES_43981 | T | T | T | T | T | T | | | T | T | | |
| NC_011770 | 4836087 | C | G | 38 | 0 | 2 | 0 | NON_SYNONYMOU | MODERAT | MISSENSE | aCc/aGc | T73S | PLES_43981 | | | | | | | G | G | | | | |
| NC_011770 | 4836236 | G | A | 54 | 0 | 2 | 0 | NON_SYNONYMOU | MODERAT | MISSENSE | Ggc/Agc | G123S | PLES_43981 | | | | | | | A | A | | | | |
| NC_011770 | 4836392 | G | C | 57 | 0 | 2 | 0 | NON_SYNONYMOU | MODERAT | MISSENSE | Gta/Cta | V175L | PLES_43981 | | | | | | | | | C | C | | |
| NC_011770 | 4840086 | C | G | 65 | 0 | 2 | 0 | NON_SYNONYMOU | MODERAT | MISSENSE | cCg/cGg | P27R | PLES_44011 | G | G | | | | | | | | | | |
| NC_011770 | 4840585 | A | G | 30 | 0 | 4 | 0 | NON_SYNONYMOU | MODERAT | MISSENSE | aAc/aGc | N42S | PLES_44021 | | | | | | | G | G | | | G | G |
| NC_011770 | 4843150 | G | GA | 94 | 0 | 8 | 0 | FRAME_SHIFT | HIGH | | aga/aAga | R159K? | PLES_44041 | | | GA | GA | GA | GA | GA | GA | GA | GA | | |
| NC_011770 | 4842721 | C | T | 81 | 0 | 6 | 0 | NON_SYNONYMOU | MODERAT | MISSENSE | aCc/aTc | T15I | PLES_44041 | | | T | T | T | T | T | T | | | | |
| NC_011770 | 4842862 | T | G | 79 | 0 | 8 | 0 | NON_SYNONYMOU | MODERAT | MISSENSE | aTg/aGg | M62R | PLES_44041 | | | G | G | G | G | G | G | | | G | G |
| NC_011770 | 4843572 | C | A | 71 | 0 | 4 | 0 | NON_SYNONYMOU | MODERAT | MISSENSE | aGc/aTc | S40I | PLES_44051 | | | A | A | A | A | | | | | | |
| NC_011770 | 4843668 | C | T | 70 | 0 | 6 | 0 | NON_SYNONYMOU | MODERAT | MISSENSE | cGc/cAc | R8H | PLES_44051 | | | T | T | T | T | | | T | T | | |
| NC_011770 | 4843674 | C | T | 72 | 0 | 2 | 0 | NON_SYNONYMOU | MODERAT | MISSENSE | aGt/aAt | S6N | PLES_44051 | | | | | | | | | T | T | | |
| NC_011770 | 4843762 | G | GCC | 62 | 0 | 2 | 0 | FRAME_SHIFT | HIGH | | gcg/gGGcg | A162G? | PLES_44061 | GCC | | | | | | | | | | | GC |
| NC_011770 | 4843763 | C | T | 52 | 2 | 8 | 0 | NON_SYNONYMOU | MODERAT | MISSENSE | Gcg/Acg | A162T | PLES_44061 | . | | T | T | T | T | T | T | T | T | | . |
| NC_011770 | 4844104 | T | A | 65 | 0 | 2 | 0 | NON_SYNONYMOU | MODERAT | MISSENSE | gAg/gTg | E48V | PLES_44061 | | | | | | | A | A | | | | |
| NC_011770 | 4844532 | T | C | 60 | 0 | 12 | 0 | NON_SYNONYMOU | MODERAT | MISSENSE | Acg/Gcg | T28A | PLES_44071 | C | C | C | C | C | C | C | C | C | C | C | C |
| NC_011770 | 4844655 | C | T | 61 | 0 | 2 | 0 | NON_SYNONYMOU | MODERAT | MISSENSE | aGc/aAc | S109N | PLES_44081 | | | | | | | T | T | | | | |
| NC_011770 | 4845638 | T | C | 79 | 0 | 1 | 0 | NON_SYNONYMOU | MODERAT | MISSENSE | tAc/tGc | Y13C | PLES_44091 | | | | | | | | C | | | | |
| NC_011770 | 4860830 | T | C | 72 | 0 | 6 | 0 | NON_SYNONYMOU | MODERAT | MISSENSE | gTc/gCc | V62A | PLES_44221 | | | | | | | C | C | C | C | C | C |
| NC_011770 | 4860889 | G | A | 73 | 0 | 6 | 0 | NON_SYNONYMOU | MODERAT | MISSENSE | Gcg/Acg | A82T | PLES_44221 | | | | | | | A | A | A | A | A | A |
| NC_011770 | 4863166 | T | A | 87 | 0 | 2 | 0 | NON_SYNONYMOU | MODERAT | MISSENSE | Acc/Tcc | T248S | PLES_44251 | A | A | | | | | | | | | | |
| NC_011770 | 4863352 | T | C | 83 | 0 | 12 | 0 | NON_SYNONYMOU | MODERAT | MISSENSE | Acc/Gcc | T186A | PLES_44251 | C | C | C | C | C | C | C | C | C | C | C | C |
| NC_011770 | 4863451 | C | G | 79 | 0 | 10 | 0 | NON_SYNONYMOU | MODERAT | MISSENSE | Gtg/Ctg | V153L | PLES_44251 | | | G | G | G | G | G | G | G | G | G | G |
| NC_011770 | 4863514 | C | T | 75 | 0 | 6 | 0 | NON_SYNONYMOU | MODERAT | MISSENSE | Gcc/Acc | A132T | PLES_44251 | T | T | | | | | T | T | T | T | | |
| NC_011770 | 4863729 | G | A | 73 | 0 | 2 | 0 | NON_SYNONYMOU | MODERAT | MISSENSE | gCg/gTg | A60V | PLES_44251 | | | | | | | | | | | A | A |
| NC_011770 | 4866432 | A | G | 54 | 0 | 4 | 0 | INTRAGENIC | MODIFIER | | | | PLES_44282 | | | G | G | G | G | | | | | | |
| NC_011770 | 4870977 | T | C | 92 | 0 | 2 | 0 | NON_SYNONYMOU | MODERAT | MISSENSE | cAg/cGg | Q296R | PLES_44321 | C | C | | | | | | | | | | |
| NC_011770 | 4871127 | C | T | 96 | 0 | 12 | 0 | NON_SYNONYMOU | MODERAT | MISSENSE | aGt/aAt | S246N | PLES_44321 | T | T | T | T | T | T | T | T | T | T | T | T |
| NC_011770 | 4871808 | C | G | 69 | 0 | 2 | 0 | NON_SYNONYMOU | MODERAT | MISSENSE | tGc/tCc | C19S | PLES_44321 | G | G | | | | | | | | | | |
| NC_011770 | 4872288 | T | A | 62 | 0 | 6 | 0 | NON_SYNONYMOU | MODERAT | MISSENSE | Acg/Tcg | T173S | PLES_44331 | A | A | | | | | | | A | A | A | A |
| NC_011770 | 4872828 | A | G | 70 | 0 | 8 | 0 | NON_SYNONYMOU | MODERAT | MISSENSE | aTc/aCc | I397T | PLES_44341 | G | G | G | G | G | G | G | G | | | | |
| NC_011770 | 4873059 | T | C | 68 | 0 | 4 | 0 | NON_SYNONYMOU | MODERAT | MISSENSE | aAc/aGc | N320S | PLES_44341 | | | C | C | C | C | | | | | | |
| NC_011770 | 4874001 | T | C | 75 | 0 | 8 | 0 | NON_SYNONYMOU | MODERAT | MISSENSE | cAg/cGg | Q6R | PLES_44341 | C | C | C | C | C | C | C | C | | | | |
| NC_011770 | 4874087 | T | C | 76 | 0 | 8 | 0 | NON_SYNONYMOU | MODERAT | MISSENSE | Acg/Gcg | T447A | PLES_44351 | C | C | C | C | C | C | C | C | | | | |
| NC_011770 | 4874146 | C | T | 73 | 0 | 8 | 0 | NON_SYNONYMOU | MODERAT | MISSENSE | gGc/gAc | G427D | PLES_44351 | T | T | T | T | T | T | T | T | | | | |
| NC_011770 | 4874267 | C | G | 75 | 0 | 2 | 0 | NON_SYNONYMOU | MODERAT | MISSENSE | Gaa/Caa | E387Q | PLES_44351 | G | G | | | | | | | | | | |
| NC_011770 | 4874687 | G | A | 68 | 0 | 2 | 0 | NON_SYNONYMOU | MODERAT | MISSENSE | Ccc/Tcc | P247S | PLES_44351 | | | | | | | | | | | A | A |
| NC_011770 | 4875032 | C | T | 75 | 0 | 2 | 0 | NON_SYNONYMOU | MODERAT | MISSENSE | Ggc/Agc | G132S | PLES_44351 | T | T | | | | | | | | | | |
| NC_011770 | 4875349 | A | G | 81 | 0 | 12 | 0 | NON_SYNONYMOU | MODERAT | MISSENSE | gTg/gCg | V26A | PLES_44351 | G | G | G | G | G | G | G | G | G | G | G | G |
| NC_011770 | 4875386 | C | T | 77 | 0 | 2 | 0 | NON_SYNONYMOU | MODERAT | MISSENSE | Ggc/Agc | G14S | PLES_44351 | | | | | | | | | | | T | T |
| NC_011770 | 4875643 | A | C | 85 | 0 | 2 | 0 | NON_SYNONYMOU | MODERAT | MISSENSE | Tcc/Gcc | S68A | PLES_44361 | C | C | | | | | | | | | | |
| NC_011770 | 4875715 | C | T | 86 | 0 | 2 | 0 | NON_SYNONYMOU | MODERAT | MISSENSE | Ggt/Agt | G44S | PLES_44361 | | | | | | | | | T | T | | |
| NC_011770 | 4875831 | C | T | 85 | 0 | 2 | 0 | NON_SYNONYMOU | MODERAT | MISSENSE | cGt/cAt | R5H | PLES_44361 | | | | | | | | | T | T | | |
| NC_011770 | 4876113 | G | A | 80 | 0 | 2 | 0 | NON_SYNONYMOU | MODERAT | MISSENSE | Cgc/Tgc | R303C | PLES_44371 | | | | | | | A | A | | | | |
| NC_011770 | 4877633 | C | A | 64 | 0 | 2 | 0 | NON_SYNONYMOU | MODERAT | MISSENSE | cGc/cTc | R87L | PLES_44381 | A | A | | | | | | | | | | |
| NC_011770 | 4877880 | C | T | 85 | 0 | 2 | 0 | NON_SYNONYMOU | MODERAT | MISSENSE | Gct/Act | A5T | PLES_44381 | T | T | | | | | | | | | | |
| NC_011770 | 4878718 | T | C | 77 | 0 | 1 | 0 | NON_SYNONYMOU | MODERAT | MISSENSE | gTc/gCc | V237A | PLES_44391 | C | | | | | | | | | | | |
| NC_011770 | 4878472 | G | C | 84 | 0 | 2 | 0 | NON_SYNONYMOU | MODERAT | MISSENSE | aGt/aCt | S155T | PLES_44391 | C | C | | | | | | | | | | |
| NC_011770 | 4879072 | C | T | 65 | 0 | 2 | 0 | NON_SYNONYMOU | MODERAT | MISSENSE | cGc/cAc | R304H | PLES_44401 | | | | | | | | | T | T | | |
| NC_011770 | 4879693 | T | C | 70 | 0 | 1 | 0 | NON_SYNONYMOU | MODERAT | MISSENSE | cAc/cGc | H97R | PLES_44401 | | | | | | | C | | | | | |

| | | | | | | | | | | | | | | | | | | | | | | | | | |
|---|---|---|---|---|---|---|---|---|---|---|---|---|---|---|---|---|---|---|---|---|---|---|---|---|---|
| NC_011770 | 4879735 | G | A | 68 | 0 | 1 | 0 | NON_SYNONYMOU | MODERAT | MISSENSE | gCg/gTg | A83V | PLES_44401 | | | | | | | A | | | | | |
| NC_011770 | 4881055 | A | G | 63 | 0 | 12 | 0 | NON_SYNONYMOU | MODERAT | MISSENSE | cAc/cGc | H285R | PLES_44411 | G | G | G | G | G | G | G | G | G | G | G | G |
| NC_011770 | 4881238 | C | T | 60 | 0 | 10 | 0 | NON_SYNONYMOU | MODERAT | MISSENSE | gCg/gTg | A346V | PLES_44411 | T | T | T | T | T | T | | | T | T | T | T |
| NC_011770 | 4881604 | C | T | 64 | 0 | 4 | 0 | NON_SYNONYMOU | MODERAT | MISSENSE | aCc/aTc | T468I | PLES_44411 | | | T | T | T | T | | | | | | |
| NC_011770 | 4892675 | G | A | 45 | 0 | 10 | 0 | NON_SYNONYMOU | MODERAT | MISSENSE | Gca/Aca | A22T | PLES_44521 | A | A | A | A | A | A | A | A | A | A | | |
| NC_011770 | 4893248 | G | C | 58 | 0 | 4 | 0 | NON_SYNONYMOU | MODERAT | MISSENSE | Gac/Cac | D125H | PLES_44531 | | | | | | | | | C | C | C | C |
| NC_011770 | 4893515 | G | T | 70 | 0 | 2 | 0 | NON_SYNONYMOU | MODERAT | MISSENSE | Gcc/Tcc | A214S | PLES_44531 | | | | | | | T | T | | | | |
| NC_011770 | 4893790 | G | C | 75 | 0 | 2 | 0 | NON_SYNONYMOU | MODERAT | MISSENSE | Gcg/Ccg | A15P | PLES_44541 | | | | | | | C | C | | | | |
| NC_011770 | 4893791 | C | T | 75 | 0 | 4 | 0 | NON_SYNONYMOU | MODERAT | MISSENSE | gCg/gTg | A15V | PLES_44541 | | | | | | | T | T | | | T | T |
| NC_011770 | 4893832 | G | A | 69 | 0 | 2 | 0 | NON_SYNONYMOU | MODERAT | MISSENSE | Gcc/Acc | A29T | PLES_44541 | | | | | | | A | A | | | | |
| NC_011770 | 4893967 | C | A | 63 | 0 | 4 | 0 | NON_SYNONYMOU | MODERAT | MISSENSE | gaC/gaA | D7E | PLES_44551 | | | A | A | A | A | | | | | | |
| NC_011770 | 4893969 | A | G | 63 | 0 | 12 | 0 | NON_SYNONYMOU | MODERAT | MISSENSE | cAt/cGt | H8R | PLES_44551 | G | G | G | G | G | G | G | G | G | G | G | G |
| NC_011770 | 4893992 | G | A | 63 | 0 | 2 | 0 | NON_SYNONYMOU | MODERAT | MISSENSE | Gag/Aag | E16K | PLES_44551 | | | | | | | | | | | A | A |
| NC_011770 | 4894392 | G | A | 48 | 0 | 10 | 0 | NON_SYNONYMOU | MODERAT | MISSENSE | aGc/aAc | S149N | PLES_44551 | A | A | A | A | A | A | A | A | A | A | | |
| NC_011770 | 4894490 | G | A | 47 | 0 | 10 | 0 | NON_SYNONYMOU | MODERAT | MISSENSE | Gcg/Acg | A182T | PLES_44551 | A | A | A | A | A | A | A | A | A | A | | |
| NC_011770 | 4894553 | A | G | 46 | 0 | 2 | 0 | NON_SYNONYMOU | MODERAT | MISSENSE | Atc/Gtc | I203V | PLES_44551 | | | | | | | | | | | G | G |
| NC_011770 | 4894751 | A | G | 41 | 0 | 12 | 0 | NON_SYNONYMOU | MODERAT | MISSENSE | Acc/Gcc | T269A | PLES_44551 | G | G | G | G | G | G | G | G | G | G | G | G |
| NC_011770 | 4894820 | C | G | 42 | 0 | 2 | 0 | NON_SYNONYMOU | MODERAT | MISSENSE | Ctg/Gtg | L292V | PLES_44551 | | | | | G | G | | | | | | |
| NC_011770 | 4895298 | G | C | 68 | 0 | 10 | 0 | NON_SYNONYMOU | MODERAT | MISSENSE | Ccg/Gcg | P148A | PLES_44561 | C | C | C | C | C | C | C | C | C | C | | |
| NC_011770 | 4897473 | GC | G | 80 | 0 | 3 | 0 | FRAME_SHIFT | HIGH | | ggc/ | G266 | PLES_44571 | | | | G | | | G | G | | | | |
| NC_011770 | 4895861 | C | G | 81 | 0 | 2 | 0 | NON_SYNONYMOU | MODERAT | MISSENSE | Gtg/Ctg | V804L | PLES_44571 | | | | | | | | | | | G | G |
| NC_011770 | 4896833 | T | C | 84 | 0 | 2 | 0 | NON_SYNONYMOU | MODERAT | MISSENSE | Atc/Gtc | I480V | PLES_44571 | | | | | | | | | | | C | C |
| NC_011770 | 4897159 | G | A | 89 | 0 | 2 | 0 | NON_SYNONYMOU | MODERAT | MISSENSE | tCc/tTc | S371F | PLES_44571 | A | A | | | | | | | | | | |
| NC_011770 | 4897228 | G | C | 84 | 0 | 8 | 0 | NON_SYNONYMOU | MODERAT | MISSENSE | gCc/gGc | A348G | PLES_44571 | C | C | | | | | C | C | C | C | C | C |
| NC_011770 | 4897784 | C | T | 67 | 0 | 2 | 0 | NON_SYNONYMOU | MODERAT | MISSENSE | Gcc/Acc | A163T | PLES_44571 | | | | | | | | | | | T | T |
| NC_011770 | 4900081 | GC | G | 69 | 0 | 1 | 0 | FRAME_SHIFT | HIGH | | gcg/ | A92 | PLES_44581 | | | | | | | G | | | | | |
| NC_011770 | 4899325 | T | C | 70 | 0 | 12 | 0 | NON_SYNONYMOU | MODERAT | MISSENSE | cAc/cGc | H344R | PLES_44581 | C | C | C | C | C | C | C | C | C | C | C | C |
| NC_011770 | 4900735 | T | A | 79 | 0 | 2 | 0 | NON_SYNONYMOU | MODERAT | MISSENSE | gaA/gaT | E72D | PLES_44591 | A | A | | | | | | | | | | |
| NC_011770 | 4900892 | G | A | 70 | 0 | 6 | 0 | NON_SYNONYMOU | MODERAT | MISSENSE | gCc/gTc | A20V | PLES_44591 | | | | | | | A | A | A | A | A | A |
| NC_011770 | 4901874 | C | T | 88 | 0 | 2 | 0 | NON_SYNONYMOU | MODERAT | MISSENSE | Gtg/Atg | V19M | PLES_44601 | | | | | | | | | | | T | T |
| NC_011770 | 4903191 | G | A | 62 | 0 | 2 | 0 | NON_SYNONYMOU | MODERAT | MISSENSE | Ggc/Agc | G22S | PLES_44631 | | | | | | | | | | | A | A |
| NC_011770 | 4903195 | A | G | 63 | 0 | 2 | 0 | NON_SYNONYMOU | MODERAT | MISSENSE | cAc/cGc | H23R | PLES_44631 | | | | | | | | | G | G | | |
| NC_011770 | 4903369 | A | T | 70 | 0 | 2 | 0 | NON_SYNONYMOU | MODERAT | MISSENSE | tAt/tTt | Y81F | PLES_44631 | | | | | | | | | T | T | | |
| NC_011770 | 4906000 | T | C | 59 | 0 | 6 | 0 | NON_SYNONYMOU | MODERAT | MISSENSE | gTc/gCc | V113A | PLES_44651 | C | C | | | | | C | C | | | C | C |
| NC_011770 | 4907930 | G | T | 69 | 0 | 2 | 0 | NON_SYNONYMOU | MODERAT | MISSENSE | Gcc/Tcc | A7S | PLES_44671 | | | | | | | | | | | T | T |
| NC_011770 | 4908014 | G | T | 81 | 0 | 4 | 0 | NON_SYNONYMOU | MODERAT | MISSENSE | Ggt/Tgt | G35C | PLES_44671 | | | T | T | T | T | | | | | | |
| NC_011770 | 4908111 | G | A | 82 | 0 | 2 | 0 | NON_SYNONYMOU | MODERAT | MISSENSE | gGc/gAc | G67D | PLES_44671 | | | | | | | | | | | A | A |
| NC_011770 | 4908126 | C | G | 79 | 0 | 2 | 0 | NON_SYNONYMOU | MODERAT | MISSENSE | gCc/gGc | A72G | PLES_44671 | | | | | | | | | | | G | G |
| NC_011770 | 4908327 | G | A | 81 | 0 | 2 | 0 | NON_SYNONYMOU | MODERAT | MISSENSE | cGc/cAc | R139H | PLES_44671 | | | | | | | | | | | A | A |
| NC_011770 | 4908650 | G | A | 81 | 0 | 2 | 0 | NON_SYNONYMOU | MODERAT | MISSENSE | Gcc/Acc | A247T | PLES_44671 | | | | | | | A | A | | | | |
| NC_011770 | 4908851 | G | A | 82 | 0 | 2 | 0 | NON_SYNONYMOU | MODERAT | MISSENSE | Gag/Aag | E314K | PLES_44671 | | | | | | | | | A | A | | |
| NC_011770 | 4911267 | G | C | 86 | 0 | 8 | 0 | NON_SYNONYMOU | MODERAT | MISSENSE | Gcg/Ccg | A32P | PLES_44711 | | | C | C | C | C | C | C | C | C | | |
| NC_011770 | 4911686 | A | C | 64 | 0 | 2 | 0 | NON_SYNONYMOU | MODERAT | MISSENSE | gaA/gaC | E171D | PLES_44711 | | | | | | | | | | | C | C |
| NC_011770 | 4912110 | C | A | 66 | 0 | 6 | 0 | NON_SYNONYMOU | MODERAT | MISSENSE | Ctg/Atg | L313M | PLES_44711 | | | A | A | A | A | A | A | | | | |
| NC_011770 | 4912434 | G | A | 82 | 0 | 1 | 0 | NON_SYNONYMOU | MODERAT | MISSENSE | Gac/Aac | D421N | PLES_44711 | | | | | | | A | | | | | |
| NC_011770 | 4912705 | T | C | 97 | 0 | 1 | 0 | NON_SYNONYMOU | MODERAT | MISSENSE | aTg/aCg | M511T | PLES_44711 | | | | | | | C | | | | | |
| NC_011770 | 4912735 | T | C | 88 | 0 | 1 | 0 | NON_SYNONYMOU | MODERAT | MISSENSE | aTc/aCc | I521T | PLES_44711 | | | | | | | | C | | | | |
| NC_011770 | 4914874 | G | T | 66 | 0 | 4 | 0 | NON_SYNONYMOU | MODERAT | MISSENSE | aaG/aaT | K97N | PLES_44731 | | | | | | | T | T | T | T | | |
| NC_011770 | 4915187 | A | G | 79 | 0 | 4 | 0 | NON_SYNONYMOU | MODERAT | MISSENSE | Atc/Gtc | I202V | PLES_44731 | | | G | G | G | G | | | | | | |
| NC_011770 | 4915193 | C | T | 78 | 0 | 4 | 0 | NON_SYNONYMOU | MODERAT | MISSENSE | Ccg/Tcg | P204S | PLES_44731 | | | T | T | T | T | | | | | | |
| NC_011770 | 4915356 | A | G | 72 | 0 | 4 | 0 | NON_SYNONYMOU | MODERAT | MISSENSE | aAg/aGg | K258R | PLES_44731 | | | G | G | G | G | | | | | | |
| NC_011770 | 4915358 | A | G | 71 | 0 | 4 | 0 | NON_SYNONYMOU | MODERAT | MISSENSE | Agc/Ggc | S259G | PLES_44731 | | | G | G | G | G | | | | | | |
| NC_011770 | 4915359 | G | C | 74 | 0 | 4 | 0 | NON_SYNONYMOU | MODERAT | MISSENSE | aGc/aCc | S259T | PLES_44731 | | | C | C | C | C | | | | | | |
| NC_011770 | 4915367 | G | C | 75 | 0 | 2 | 0 | NON_SYNONYMOU | MODERAT | MISSENSE | Gtc/Ctc | V262L | PLES_44731 | | | | | | | | | C | C | | |
| NC_011770 | 4915433 | G | A | 77 | 0 | 4 | 0 | NON_SYNONYMOU | MODERAT | MISSENSE | Ggt/Agt | G284S | PLES_44731 | | | A | A | A | A | | | | | | |
| NC_011770 | 4915949 | G | A | 76 | 0 | 1 | 0 | NON_SYNONYMOU | MODERAT | MISSENSE | Ggt/Agt | G456S | PLES_44731 | | | | A | | | | | | | | |
| NC_011770 | 4916457 | A | G | 75 | 0 | 2 | 0 | NON_SYNONYMOU | MODERAT | MISSENSE | aAc/aGc | N625S | PLES_44731 | | | | | | | G | G | | | | |
| NC_011770 | 4920255 | G | A | 78 | 0 | 2 | 0 | NON_SYNONYMOU | MODERAT | MISSENSE | Ggc/Agc | G133S | PLES_44761 | | | | | | | A | A | | | | |
| NC_011770 | 4920610 | G | A | 78 | 0 | 2 | 0 | NON_SYNONYMOU | MODERAT | MISSENSE | aGg/aAg | R251K | PLES_44761 | | | | | | | | | | | A | A |
| NC_011770 | 4920864 | G | A | 75 | 0 | 2 | 0 | NON_SYNONYMOU | MODERAT | MISSENSE | Gtc/Atc | V336I | PLES_44761 | | | | | | | A | A | | | | |
| NC_011770 | 4921100 | G | A | 58 | 0 | 2 | 0 | NON_SYNONYMOU | MODERAT | MISSENSE | Ccg/Tcg | P315S | PLES_44771 | A | A | . | . | | | | | | | | |
| NC_011770 | 4921103 | T | C | 68 | 0 | 2 | 0 | NON_SYNONYMOU | MODERAT | MISSENSE | Acg/Gcg | T314A | PLES_44771 | | | . | . | | | | | | | C | C |

| | | | | | | | | | | | | | | | | | | | | | | | | | |
|---|---|---|---|---|---|---|---|---|---|---|---|---|---|---|---|---|---|---|---|---|---|---|---|---|---|
| NC_011770 | 4921190 | G | A | 69 | 0 | 2 | 0 | NON_SYNONYMOU | MODERAT | MISSENSE | Cgc/Tgc | R285C | PLES_44771 | | | | | | | | | | | A | A |
| NC_011770 | 4921366 | A | G | 66 | 0 | 12 | 0 | NON_SYNONYMOU | MODERAT | MISSENSE | cTg/cCg | L226P | PLES_44771 | G | G | G | G | G | G | G | G | G | G | G | G |
| NC_011770 | 4921588 | A | G | 74 | 0 | 12 | 0 | NON_SYNONYMOU | MODERAT | MISSENSE | gTg/gCg | V152A | PLES_44771 | G | G | G | G | G | G | G | G | G | G | G | G |
| NC_011770 | 4921880 | G | C | 66 | 0 | 6 | 0 | NON_SYNONYMOU | MODERAT | MISSENSE | Ccc/Gcc | P55A | PLES_44771 | | | C | C | C | C | C | C | | | | |
| NC_011770 | 4922483 | T | A | 58 | 0 | 2 | 0 | NON_SYNONYMOU | MODERAT | MISSENSE | aAg/aTg | K288M | PLES_44781 | A | A | | | | | | | | | | |
| NC_011770 | 4922902 | G | C | 50 | 0 | 6 | 0 | NON_SYNONYMOU | MODERAT | MISSENSE | gaC/gaG | D148E | PLES_44781 | | | | | | | C | C | C | C | C | C |
| NC_011770 | 4923070 | G | C | 42 | 0 | 2 | 0 | NON_SYNONYMOU | MODERAT | MISSENSE | gaC/gaG | D92E | PLES_44781 | | | | | | | | | C | C | | |
| NC_011770 | 4923573 | C | T | 63 | 0 | 2 | 0 | NON_SYNONYMOU | MODERAT | MISSENSE | cGa/cAa | R135Q | PLES_44791 | | | | | | | T | T | | | | |
| NC_011770 | 4925558 | A | G | 79 | 0 | 12 | 0 | INTRAGENIC | MODIFIER | | | | PLES_44812 | G | G | G | G | G | G | G | G | G | G | G | G |
| NC_011770 | 4925568 | G | A | 81 | 0 | 2 | 0 | INTRAGENIC | MODIFIER | | | | PLES_44812 | | | | | | | A | A | | | | |
| NC_011770 | 4929522 | G | A | 76 | 0 | 1 | 0 | NON_SYNONYMOU | MODERAT | MISSENSE | cGc/cAc | R143H | PLES_44841 | | | | | | | A | | | | | |
| NC_011770 | 4930242 | A | T | 62 | 0 | 12 | 0 | NON_SYNONYMOU | MODERAT | MISSENSE | cTg/cAg | L184Q | PLES_44851 | T | T | T | T | T | T | T | T | T | T | T | T |
| NC_011770 | 4931413 | G | T | 50 | 0 | 4 | 0 | NON_SYNONYMOU | MODERAT | MISSENSE | Ctg/Atg | L44M | PLES_44861 | T | T | | | | | T | T | | | | |
| NC_011770 | 4931602 | C | A | 19 | 8 | 4 | 0 | NON_SYNONYMOU | MODERAT | MISSENSE | Ctt/Att | L4I | PLES_44871 | . | . | A | A | A | A | . | . | . | . | . | . |
| NC_011770 | 4931681 | C | T | 21 | 8 | 4 | 0 | NON_SYNONYMOU | MODERAT | MISSENSE | aCc/aTc | T30I | PLES_44871 | . | . | T | T | T | T | . | . | . | . | . | . |
| NC_011770 | 4931716 | G | A | 20 | 8 | 4 | 0 | NON_SYNONYMOU | MODERAT | MISSENSE | Gag/Aag | E42K | PLES_44871 | . | . | A | A | A | A | . | . | . | . | . | . |
| NC_011770 | 4931823 | A | C | 18 | 8 | 4 | 0 | NON_SYNONYMOU | MODERAT | MISSENSE | agA/agC | R77S | PLES_44871 | . | . | C | C | C | C | . | . | . | . | . | . |
| NC_011770 | 4948917 | G | A | 32 | 2 | 4 | 0 | NON_SYNONYMOU | MODERAT | MISSENSE | Cca/Tca | P104S | PLES_44981 | | | | | | | A | A | A | A | . | . |
| NC_011770 | 4949412 | G | T | 37 | 2 | 2 | 0 | NON_SYNONYMOU | MODERAT | MISSENSE | ttC/ttA | F37L | PLES_44991 | | | | | | | | | T | T | . | . |
| NC_011770 | 4962807 | C | T | 90 | 0 | 12 | 0 | NON_SYNONYMOU | MODERAT | MISSENSE | gCc/gTc | A193V | PLES_45131 | T | T | T | T | T | T | T | T | T | T | T | T |
| NC_011770 | 4964280 | G | T | 81 | 0 | 6 | 0 | NON_SYNONYMOU | MODERAT | MISSENSE | Cgc/Agc | R218S | PLES_45151 | | | T | T | T | T | T | T | | | | |
| NC_011770 | 4964325 | C | T | 85 | 0 | 1 | 0 | NON_SYNONYMOU | MODERAT | MISSENSE | Gtc/Atc | V203I | PLES_45151 | | | | | | | T | | | | | |
| NC_011770 | 4964769 | T | C | 70 | 0 | 6 | 0 | NON_SYNONYMOU | MODERAT | MISSENSE | Atc/Gtc | I55V | PLES_45151 | | | C | C | C | C | | | C | C | | |
| NC_011770 | 4965521 | A | G | 69 | 0 | 2 | 0 | NON_SYNONYMOU | MODERAT | MISSENSE | cAg/cGg | Q168R | PLES_45161 | | | | | | | | | | | G | G |
| NC_011770 | 4965685 | G | A | 69 | 0 | 2 | 0 | NON_SYNONYMOU | MODERAT | MISSENSE | Gcc/Acc | A223T | PLES_45161 | | | | | | | A | A | | | | |
| NC_011770 | 4965710 | C | G | 68 | 0 | 4 | 0 | NON_SYNONYMOU | MODERAT | MISSENSE | gCt/gGt | A231G | PLES_45161 | | | | | | | G | G | | | G | G |
| NC_011770 | 4965836 | C | A | 67 | 0 | 4 | 0 | NON_SYNONYMOU | MODERAT | MISSENSE | gCc/gAc | A273D | PLES_45161 | A | A | | | | | | | A | A | | |
| NC_011770 | 4965841 | G | C | 69 | 0 | 4 | 0 | NON_SYNONYMOU | MODERAT | MISSENSE | Gcc/Ccc | A275P | PLES_45161 | C | C | | | | | | | C | C | | |
| NC_011770 | 4967155 | A | G | 70 | 2 | 6 | 0 | NON_SYNONYMOU | MODERAT | MISSENSE | atA/atG | I124M | PLES_45171 | | | G | G | G | G | G | G | . | . | | |
| NC_011770 | 4967175 | G | A | 73 | 2 | 2 | 0 | NON_SYNONYMOU | MODERAT | MISSENSE | aGc/aAc | S131N | PLES_45171 | | | | | | | | | . | . | A | A |
| NC_011770 | 4967871 | G | A | 65 | 2 | 4 | 0 | STOP_GAINED | HIGH | NONSENSE | tgG/tgA | W69* | PLES_45181 | | | A | A | A | A | | | . | . | | |
| NC_011770 | 4967857 | G | A | 65 | 2 | 10 | 0 | NON_SYNONYMOU | MODERAT | MISSENSE | Gtt/Att | V65I | PLES_45181 | A | A | A | A | A | A | A | A | . | . | A | A |
| NC_011770 | 4967981 | C | T | 57 | 2 | 6 | 0 | NON_SYNONYMOU | MODERAT | MISSENSE | gCg/gTg | A106V | PLES_45181 | | | T | T | T | T | T | T | . | . | | |
| NC_011770 | 4968306 | G | A | 34 | 2 | 2 | 0 | NON_SYNONYMOU | MODERAT | MISSENSE | Gcc/Acc | A45T | PLES_45191 | | | | | | | | | . | . | A | A |
| NC_011770 | 4968484 | T | C | 5 | 3 | 6 | 0 | NON_SYNONYMOU | MODERAT | MISSENSE | cTt/cCt | L104P | PLES_45191 | | | C | C | C | C | C | C | . | . | | . |
| NC_011770 | 4968495 | A | G | 7 | 4 | 6 | 0 | NON_SYNONYMOU | MODERAT | MISSENSE | Agc/Ggc | S108G | PLES_45191 | | | G | G | G | G | G | G | . | . | . | . |
| NC_011770 | 4969127 | A | G | 59 | 4 | 1 | 0 | NON_SYNONYMOU | MODERAT | MISSENSE | Acc/Gcc | T62A | PLES_45201 | | | | | | | G | | . | . | . | . |
| NC_011770 | 4969175 | C | T | 56 | 4 | 4 | 0 | NON_SYNONYMOU | MODERAT | MISSENSE | Cca/Tca | P78S | PLES_45201 | | | T | T | T | T | | | . | . | . | . |
| NC_011770 | 4969206 | C | T | 57 | 4 | 4 | 0 | NON_SYNONYMOU | MODERAT | MISSENSE | gCa/gTa | A88V | PLES_45201 | | | T | T | T | T | | | . | . | . | . |
| NC_011770 | 4969305 | C | T | 55 | 4 | 6 | 0 | NON_SYNONYMOU | MODERAT | MISSENSE | gCa/gTa | A121V | PLES_45201 | | | T | T | T | T | T | T | . | . | . | . |
| NC_011770 | 4969317 | T | C | 51 | 4 | 6 | 0 | NON_SYNONYMOU | MODERAT | MISSENSE | gTc/gCc | V125A | PLES_45201 | | | C | C | C | C | C | C | . | . | . | . |
| NC_011770 | 4969550 | T | G | 51 | 4 | 6 | 0 | NON_SYNONYMOU | MODERAT | MISSENSE | gaT/gaG | D48E | PLES_45211 | | | G | G | G | G | G | G | . | . | . | . |
| NC_011770 | 4969629 | T | C | 53 | 4 | 6 | 0 | NON_SYNONYMOU | MODERAT | MISSENSE | Tgg/Cgg | W75R | PLES_45211 | | | C | C | C | C | C | C | . | . | . | . |
| NC_011770 | 4969876 | G | A | 56 | 4 | 6 | 0 | NON_SYNONYMOU | MODERAT | MISSENSE | cGg/cAg | R157Q | PLES_45211 | | | A | A | A | A | A | A | . | . | . | . |
| NC_011770 | 4970119 | T | A | 62 | 4 | 6 | 0 | NON_SYNONYMOU | MODERAT | MISSENSE | aTc/aAc | I238N | PLES_45211 | | | A | A | A | A | A | A | . | . | . | . |
| NC_011770 | 4970319 | G | A | 58 | 4 | 4 | 0 | NON_SYNONYMOU | MODERAT | MISSENSE | Gtc/Atc | V305I | PLES_45211 | | | A | A | A | A | | | . | . | . | . |
| NC_011770 | 4970418 | C | A | 52 | 4 | 4 | 0 | NON_SYNONYMOU | MODERAT | MISSENSE | Ctc/Atc | L338I | PLES_45211 | | | A | A | A | A | | | . | . | . | . |
| NC_011770 | 4970434 | C | A | 55 | 4 | 6 | 0 | NON_SYNONYMOU | MODERAT | MISSENSE | gCc/gAc | A343D | PLES_45211 | | | A | A | A | A | A | A | . | . | . | . |
| NC_011770 | 4970446 | A | G | 59 | 4 | 4 | 0 | NON_SYNONYMOU | MODERAT | MISSENSE | aAc/aGc | N347S | PLES_45211 | | | G | G | G | G | | | . | . | . | . |
| NC_011770 | 4970448 | A | G | 59 | 4 | 4 | 0 | NON_SYNONYMOU | MODERAT | MISSENSE | Act/Gct | T348A | PLES_45211 | | | G | G | G | G | | | . | . | . | . |
| NC_011770 | 4970475 | T | A | 56 | 4 | 4 | 0 | NON_SYNONYMOU | MODERAT | MISSENSE | Tca/Aca | S357T | PLES_45211 | | | A | A | A | A | | | . | . | . | . |
| NC_011770 | 4970491 | C | T | 56 | 4 | 4 | 0 | NON_SYNONYMOU | MODERAT | MISSENSE | cCg/cTg | P362L | PLES_45211 | | | T | T | T | T | | | . | . | . | . |
| NC_011770 | 4971039 | C | G | 67 | 4 | 2 | 0 | NON_SYNONYMOU | MODERAT | MISSENSE | caG/caC | Q52H | PLES_45221 | | | | | | | G | G | . | . | . | . |
| NC_011770 | 4971190 | A | G | 71 | 4 | 2 | 0 | NON_SYNONYMOU | MODERAT | MISSENSE | gTt/gCt | V2A | PLES_45221 | | | | | | | G | G | . | . | . | . |
| NC_011770 | 4972563 | A | AAGCGC | 47 | 0 | 3 | 0 | FRAME_SHIFT | HIGH | | gcg/gcGC | A80ARAS? | PLES_45241 | | AAGCGCGCCTCC | | AAGCGCAGCCTCC | | | | | AAGCGCAGCCTCC | | | |
| NC_011770 | 4972296 | C | T | 31 | 0 | 2 | 0 | NON_SYNONYMOU | MODERAT | MISSENSE | cCc/cTc | P2L | PLES_45241 | | | | | | | T | T | | | | |
| NC_011770 | 4972409 | A | C | 35 | 0 | 4 | 0 | NON_SYNONYMOU | MODERAT | MISSENSE | Acc/Ccc | T40P | PLES_45241 | | | C | C | C | C | | | | | | |
| NC_011770 | 4972412 | A | G | 34 | 0 | 10 | 0 | NON_SYNONYMOU | MODERAT | MISSENSE | Aca/Gca | T41A | PLES_45241 | | | G | G | G | G | G | G | G | G | G | G |
| NC_011770 | 4972413 | C | A | 34 | 0 | 10 | 0 | NON_SYNONYMOU | MODERAT | MISSENSE | aCa/aAa | T41K | PLES_45241 | | | A | A | A | A | A | A | A | A | A | A |
| NC_011770 | 4972436 | C | T | 38 | 0 | 2 | 0 | NON_SYNONYMOU | MODERAT | MISSENSE | Ccg/Tcg | P49S | PLES_45241 | T | T | | | | | | | | | | |
| NC_011770 | 4972442 | G | A | 37 | 0 | 6 | 0 | NON_SYNONYMOU | MODERAT | MISSENSE | Gag/Aag | E51K | PLES_45241 | | | A | A | A | A | | | | | A | A |
| NC_011770 | 4972504 | C | A | 44 | 0 | 4 | 0 | NON_SYNONYMOU | MODERAT | MISSENSE | agC/agA | S71R | PLES_45241 | | | | | | | A | A | A | A | | |

| | | | | | | | | | | | | | | | | | | | | | | | | | |
|---|---|---|---|---|---|---|---|---|---|---|---|---|---|---|---|---|---|---|---|---|---|---|---|---|---|
| NC_011770 | 4972476 | T | C | 43 | 0 | 1 | 0 | NON_SYNONYMOU | MODERAT | MISSENSE | gTg/gCg | V62A | PLES_45241 | | | | | | | | C | | | | |
| NC_011770 | 4972606 | T | G | 56 | 0 | 4 | 0 | NON_SYNONYMOU | MODERAT | MISSENSE | Atg/Ctg | M135L | PLES_45251 | | | G | G | G | G | | | | | | |
| NC_011770 | 4972732 | G | T | 62 | 0 | 12 | 0 | NON_SYNONYMOU | MODERAT | MISSENSE | Cag/Aag | Q93K | PLES_45251 | T | T | T | T | T | T | T | T | T | T | T | T |
| NC_011770 | 4972887 | T | C | 66 | 0 | 1 | 0 | NON_SYNONYMOU | MODERAT | MISSENSE | gAc/gGc | D41G | PLES_45251 | | | | | | | C | | | | | |
| NC_011770 | 4973284 | T | C | 69 | 0 | 4 | 0 | NON_SYNONYMOU | MODERAT | MISSENSE | gTc/gCc | V59A | PLES_45261 | | | C | C | C | C | | | | | | |
| NC_011770 | 4973463 | A | G | 76 | 0 | 12 | 0 | NON_SYNONYMOU | MODERAT | MISSENSE | Atg/Gtg | M119V | PLES_45261 | G | G | G | G | G | G | G | G | G | G | G | G |
| NC_011770 | 4973502 | G | A | 71 | 0 | 4 | 0 | NON_SYNONYMOU | MODERAT | MISSENSE | Gcc/Acc | A132T | PLES_45261 | | | A | A | A | A | | | | | | |
| NC_011770 | 4973619 | G | A | 74 | 0 | 2 | 0 | NON_SYNONYMOU | MODERAT | MISSENSE | Gcg/Acg | A171T | PLES_45261 | A | A | | | | | | | | | | |
| NC_011770 | 4973673 | G | A | 76 | 0 | 2 | 0 | NON_SYNONYMOU | MODERAT | MISSENSE | Gtg/Atg | V189M | PLES_45261 | | | | | | | | | | | A | A |
| NC_011770 | 4974085 | G | A | 82 | 0 | 4 | 0 | NON_SYNONYMOU | MODERAT | MISSENSE | Cgc/Tgc | R279C | PLES_45271 | | | A | A | A | A | | | | | | |
| NC_011770 | 4974687 | T | A | 94 | 0 | 2 | 0 | NON_SYNONYMOU | MODERAT | MISSENSE | aAc/aTc | N78I | PLES_45271 | | | | | A | A | | | | | | |
| NC_011770 | 4975293 | C | G | 49 | 0 | 2 | 0 | NON_SYNONYMOU | MODERAT | MISSENSE | Cag/Gag | Q52E | PLES_45281 | | | G | G | | | | | | | | |
| NC_011770 | 4975449 | G | A | 41 | 0 | 2 | 0 | NON_SYNONYMOU | MODERAT | MISSENSE | Gcc/Acc | A104T | PLES_45281 | | | | | | | | | | | A | A |
| NC_011770 | 4975153 | C | T | 54 | 0 | 1 | 0 | NON_SYNONYMOU | MODERAT | MISSENSE | cCc/cTc | P5L | PLES_45281 | | | | | | | | T | | | | |
| NC_011770 | 4975883 | G | A | 65 | 0 | 1 | 0 | NON_SYNONYMOU | MODERAT | MISSENSE | Gcc/Acc | A105T | PLES_45291 | | | | | | | | | | | | A |
| NC_011770 | 4975711 | C | G | 59 | 0 | 2 | 0 | NON_SYNONYMOU | MODERAT | MISSENSE | gaC/gaG | D47E | PLES_45291 | G | G | | | | | | | | | | |
| NC_011770 | 4975820 | G | T | 65 | 0 | 6 | 0 | NON_SYNONYMOU | MODERAT | MISSENSE | Ggc/Tgc | G84C | PLES_45291 | | | T | T | T | T | T | T | | | | |
| NC_011770 | 4976163 | A | G | 69 | 0 | 2 | 0 | NON_SYNONYMOU | MODERAT | MISSENSE | gAc/gGc | D198G | PLES_45291 | | | | | | | | | G | G | | |
| NC_011770 | 4976173 | C | G | 73 | 0 | 2 | 0 | NON_SYNONYMOU | MODERAT | MISSENSE | ttC/ttG | F201L | PLES_45291 | | | | | | | | | G | G | | |
| NC_011770 | 4976484 | A | G | 81 | 0 | 12 | 0 | NON_SYNONYMOU | MODERAT | MISSENSE | cAg/cGg | Q305R | PLES_45291 | G | G | G | G | G | G | G | G | G | G | G | G |
| NC_011770 | 4976579 | G | T | 84 | 0 | 6 | 0 | NON_SYNONYMOU | MODERAT | MISSENSE | Gcc/Tcc | A337S | PLES_45291 | | | T | T | T | T | T | T | | | | |
| NC_011770 | 4976756 | A | G | 81 | 0 | 6 | 0 | NON_SYNONYMOU | MODERAT | MISSENSE | Agc/Ggc | S396G | PLES_45291 | | | G | G | G | G | G | G | | | | |
| NC_011770 | 4977654 | A | G | 86 | 0 | 4 | 0 | NON_SYNONYMOU | MODERAT | MISSENSE | aAg/aGg | K183R | PLES_45301 | | | G | G | G | G | | | | | | |
| NC_011770 | 4977977 | G | A | 77 | 0 | 4 | 0 | NON_SYNONYMOU | MODERAT | MISSENSE | Gaa/Aaa | E291K | PLES_45301 | | | | | | | A | A | A | A | | |
| NC_011770 | 4978100 | G | C | 78 | 0 | 2 | 0 | NON_SYNONYMOU | MODERAT | MISSENSE | Gtg/Ctg | V332L | PLES_45301 | | | | | | | | | | | C | C |
| NC_011770 | 4978200 | A | G | 74 | 0 | 4 | 0 | NON_SYNONYMOU | MODERAT | MISSENSE | cAg/cGg | Q365R | PLES_45301 | | | G | G | G | G | | | | | | |
| NC_011770 | 4978915 | T | C | 65 | 0 | 2 | 0 | NON_SYNONYMOU | MODERAT | MISSENSE | Ttc/Ctc | F173L | PLES_45311 | | | | | | | C | C | | | | |
| NC_011770 | 4980292 | T | A | 80 | 0 | 4 | 0 | NON_SYNONYMOU | MODERAT | MISSENSE | Tcc/Acc | S206T | PLES_45321 | | | | | | | | | A | A | A | A |
| NC_011770 | 4980344 | C | T | 70 | 0 | 6 | 0 | NON_SYNONYMOU | MODERAT | MISSENSE | cCg/cTg | P223L | PLES_45321 | | | T | T | T | T | T | T | | | | |
| NC_011770 | 4980504 | CG | C | 68 | 0 | 1 | 0 | FRAME_SHIFT | HIGH | | gcc/ | A8 | PLES_45331 | | | | | | | C | | | | | |
| NC_011770 | 4980932 | G | A | 81 | 0 | 2 | 0 | NON_SYNONYMOU | MODERAT | MISSENSE | Gtg/Atg | V150M | PLES_45331 | | | | | | | | | A | A | | |
| NC_011770 | 4981601 | G | A | 66 | 0 | 2 | 0 | NON_SYNONYMOU | MODERAT | MISSENSE | Gcg/Acg | A373T | PLES_45331 | | | | | | | A | A | | | | |
| NC_011770 | 4981715 | G | A | 68 | 0 | 2 | 0 | NON_SYNONYMOU | MODERAT | MISSENSE | Gcc/Acc | A411T | PLES_45331 | | | | | | | A | A | | | | |
| NC_011770 | 4981781 | T | G | 65 | 0 | 12 | 0 | NON_SYNONYMOU | MODERAT | MISSENSE | Tcc/Gcc | S433A | PLES_45331 | G | G | G | G | G | G | G | G | G | G | G | G |
| NC_011770 | 4980506 | GC | G | 56 | 1 | 2 | 0 | FRAME_SHIFT | HIGH | | cag/ | Q10 | PLES_45331 MntH | | | | G | | | . | GCCC | | | | |
| NC_011770 | 4982217 | T | A | 43 | 0 | 12 | 0 | NON_SYNONYMOU | MODERAT | MISSENSE | tAc/tTc | Y37F | PLES_45341 | A | A | A | A | A | A | A | A | A | A | A | A |
| NC_011770 | 4984012 | G | C | 40 | 0 | 4 | 0 | NON_SYNONYMOU | MODERAT | MISSENSE | gCc/gGc | A58G | PLES_45361 | | | | | | | C | C | | | C | C |
| NC_011770 | 4984157 | C | G | 44 | 0 | 4 | 0 | NON_SYNONYMOU | MODERAT | MISSENSE | Gtg/Ctg | V10L | PLES_45361 | G | G | | | | | G | G | | | | |
| NC_011770 | 4984160 | G | T | 44 | 0 | 2 | 0 | NON_SYNONYMOU | MODERAT | MISSENSE | Ctc/Atc | L9I | PLES_45361 | | | | | | | T | T | | | | |
| NC_011770 | 4984932 | A | C | 80 | 0 | 4 | 0 | NON_SYNONYMOU | MODERAT | MISSENSE | gAa/gCa | E63A | PLES_45371 | | | C | C | C | C | | | | | | |
| NC_011770 | 4985285 | T | A | 74 | 0 | 2 | 0 | NON_SYNONYMOU | MODERAT | MISSENSE | cAg/cTg | Q240L | PLES_45381 | A | A | | | | | | | | | | |
| NC_011770 | 4985544 | C | T | 75 | 0 | 2 | 0 | NON_SYNONYMOU | MODERAT | MISSENSE | Gcc/Acc | A154T | PLES_45381 | | | | | | | T | T | | | | |
| NC_011770 | 4985745 | T | C | 71 | 0 | 2 | 0 | NON_SYNONYMOU | MODERAT | MISSENSE | Aag/Gag | K87E | PLES_45381 | C | C | | | | | | | | | | |
| NC_011770 | 4985789 | T | C | 72 | 0 | 2 | 0 | NON_SYNONYMOU | MODERAT | MISSENSE | gAg/gGg | E72G | PLES_45381 | | | | | | | C | C | | | | |
| NC_011770 | 4985790 | C | T | 73 | 0 | 2 | 0 | NON_SYNONYMOU | MODERAT | MISSENSE | Gag/Aag | E72K | PLES_45381 | | | | | | | | | | | T | T |
| NC_011770 | 4986270 | C | T | 64 | 0 | 2 | 0 | NON_SYNONYMOU | MODERAT | MISSENSE | aGc/aAc | S69N | PLES_45391 | T | T | | | | | | | | | | |
| NC_011770 | 4986366 | T | C | 59 | 0 | 2 | 0 | NON_SYNONYMOU | MODERAT | MISSENSE | aAg/aGg | K37R | PLES_45391 | | | | | | | | | | | C | C |
| NC_011770 | 4986785 | G | A | 64 | 0 | 12 | 0 | NON_SYNONYMOU | MODERAT | MISSENSE | Ccc/Tcc | P34S | PLES_45401 | A | A | A | A | A | A | A | A | A | A | A | A |
| NC_011770 | 4986787 | G | A | 66 | 0 | 2 | 0 | NON_SYNONYMOU | MODERAT | MISSENSE | gCt/gTt | A33V | PLES_45401 | | | | | | | A | A | | | | |
| NC_011770 | 4986898 | G | C | 69 | 0 | 2 | 0 | NON_SYNONYMOU | MODERAT | MISSENSE | Cag/Gag | Q503E | PLES_45411 | C | C | | | | | | | | | | |
| NC_011770 | 4986918 | A | G | 62 | 0 | 8 | 0 | NON_SYNONYMOU | MODERAT | MISSENSE | gTc/gCc | V496A | PLES_45411 | G | G | G | G | G | G | | | | | G | G |
| NC_011770 | 4987126 | T | C | 61 | 0 | 4 | 0 | NON_SYNONYMOU | MODERAT | MISSENSE | Acc/Gcc | T427A | PLES_45411 | | | C | C | C | C | | | | | | |
| NC_011770 | 4987317 | C | A | 75 | 0 | 2 | 0 | NON_SYNONYMOU | MODERAT | MISSENSE | cGt/cTt | R363L | PLES_45411 | | | | | | | | | | | A | A |
| NC_011770 | 4987329 | A | G | 73 | 0 | 2 | 0 | NON_SYNONYMOU | MODERAT | MISSENSE | gTc/gCc | V359A | PLES_45411 | | | | | | | | | | | G | G |
| NC_011770 | 4987641 | C | G | 83 | 0 | 2 | 0 | NON_SYNONYMOU | MODERAT | MISSENSE | gGc/gCc | G255A | PLES_45411 | | | | | | | | | | | G | G |
| NC_011770 | 4987674 | T | C | 79 | 0 | 2 | 0 | NON_SYNONYMOU | MODERAT | MISSENSE | gAc/gGc | D244G | PLES_45411 | C | C | | | | | | | | | | |
| NC_011770 | 4988002 | C | T | 75 | 0 | 12 | 0 | NON_SYNONYMOU | MODERAT | MISSENSE | Gtc/Atc | V135I | PLES_45411 | T | T | T | T | T | T | T | T | T | T | T | T |
| NC_011770 | 4988109 | A | G | 56 | 0 | 2 | 0 | NON_SYNONYMOU | MODERAT | MISSENSE | gTc/gCc | V99A | PLES_45411 | | | | | | | | | | | G | G |
| NC_011770 | 4988166 | T | G | 59 | 0 | 2 | 0 | NON_SYNONYMOU | MODERAT | MISSENSE | gAg/gCg | E80A | PLES_45411 | | | | | | | | | | | G | G |
| NC_011770 | 4988182 | T | C | 60 | 0 | 2 | 0 | NON_SYNONYMOU | MODERAT | MISSENSE | Atg/Gtg | M75V | PLES_45411 | C | C | | | | | | | | | | |
| NC_011770 | 4988507 | G | A | 66 | 0 | 2 | 0 | NON_SYNONYMOU | MODERAT | MISSENSE | gCc/gTc | A143V | PLES_45421 | | | | | | | A | A | | | | |
| NC_011770 | 4988712 | G | C | 71 | 0 | 2 | 0 | NON_SYNONYMOU | MODERAT | MISSENSE | Ccg/Gcg | P75A | PLES_45421 | | | | | | | C | C | | | | |

| | | | | | | | | | | | | | | | | | | | | | | | | | |
|---|---|---|---|---|---|---|---|---|---|---|---|---|---|---|---|---|---|---|---|---|---|---|---|---|---|
| NC_011770 | 4988721 | A | T | 70 | 0 | 2 | 0 | NON_SYNONYMOU | MODERAT | MISSENSE | Tgt/Agt | C72S | PLES_45421 | | | | | | | | | T | T | | |
| NC_011770 | 4988787 | G | A | 68 | 0 | 2 | 0 | NON_SYNONYMOU | MODERAT | MISSENSE | Cgc/Tgc | R50C | PLES_45421 | | | | | | | | | A | A | | |
| NC_011770 | 4988796 | T | C | 69 | 0 | 2 | 0 | NON_SYNONYMOU | MODERAT | MISSENSE | Atc/Gtc | I47V | PLES_45421 | | | | | | | | | C | C | | |
| NC_011770 | 4988867 | T | G | 65 | 0 | 2 | 0 | NON_SYNONYMOU | MODERAT | MISSENSE | gAc/gCc | D23A | PLES_45421 | | | | | | | | | | | G | G |
| NC_011770 | 4988874 | A | T | 66 | 0 | 2 | 0 | NON_SYNONYMOU | MODERAT | MISSENSE | Tgg/Agg | W21R | PLES_45421 | | | | | | | | | | | T | T |
| NC_011770 | 4988897 | T | C | 63 | 0 | 4 | 0 | NON_SYNONYMOU | MODERAT | MISSENSE | cAc/cGc | H13R | PLES_45421 | | | | | | | | | C | C | A | A |
| NC_011770 | 4988906 | T | C | 68 | 0 | 2 | 0 | NON_SYNONYMOU | MODERAT | MISSENSE | tAc/tGc | Y10C | PLES_45421 | | | | | | | | | C | C | | |
| NC_011770 | 4988913 | T | C | 64 | 0 | 4 | 0 | NON_SYNONYMOU | MODERAT | MISSENSE | Acg/Gcg | T8A | PLES_45421 | | | | | | | | | C | C | C | C |
| NC_011770 | 4988916 | C | T | 66 | 0 | 2 | 0 | NON_SYNONYMOU | MODERAT | MISSENSE | Gcg/Acg | A7T | PLES_45421 | | | | | | | | | | | T | T |
| NC_011770 | 4988917 | A | C | 65 | 0 | 2 | 0 | NON_SYNONYMOU | MODERAT | MISSENSE | aaT/aaG | N6K | PLES_45421 | | | | | | | | | | | C | C |
| NC_011770 | 4988918 | T | C | 66 | 0 | 2 | 0 | NON_SYNONYMOU | MODERAT | MISSENSE | aAt/aGt | N6S | PLES_45421 | | | | | | | | | | | C | C |
| NC_011770 | 4988919 | T | C | 66 | 0 | 2 | 0 | NON_SYNONYMOU | MODERAT | MISSENSE | Aat/Gat | N6D | PLES_45421 | | | | | | | | | | | C | C |
| NC_011770 | 4988922 | T | C | 65 | 0 | 2 | 0 | NON_SYNONYMOU | MODERAT | MISSENSE | Acg/Gcg | T5A | PLES_45421 | | | | | | | | | | | C | C |
| NC_011770 | 4989728 | G | A | 66 | 0 | 2 | 0 | NON_SYNONYMOU | MODERAT | MISSENSE | Gcg/Acg | A126T | PLES_45431 | A | A | | | | | | | | | | |
| NC_011770 | 4989887 | G | A | 72 | 0 | 2 | 0 | NON_SYNONYMOU | MODERAT | MISSENSE | Gct/Act | A179T | PLES_45431 | | | | | | | | | A | A | | |
| NC_011770 | 4990184 | C | A | 75 | 0 | 2 | 0 | NON_SYNONYMOU | MODERAT | MISSENSE | Cgc/Agc | R278S | PLES_45431 | | | | | | | A | A | | | | |
| NC_011770 | 4990789 | G | C | 85 | 0 | 2 | 0 | NON_SYNONYMOU | MODERAT | MISSENSE | caG/caC | Q479H | PLES_45431 | | | | | | | C | C | | | | |
| NC_011770 | 4990799 | G | A | 82 | 0 | 2 | 0 | NON_SYNONYMOU | MODERAT | MISSENSE | Gcg/Acg | A483T | PLES_45431 | | | | | | | | | A | A | | |
| NC_011770 | 4992136 | A | G | 62 | 0 | 4 | 0 | NON_SYNONYMOU | MODERAT | MISSENSE | aAc/aGc | N4S | PLES_45451 | | | G | G | G | G | | | | | | |
| NC_011770 | 4992375 | A | G | 60 | 0 | 4 | 0 | NON_SYNONYMOU | MODERAT | MISSENSE | Agc/Ggc | S84G | PLES_45451 | | | G | G | G | G | | | | | | |
| NC_011770 | 4995263 | C | T | 78 | 0 | 2 | 0 | NON_SYNONYMOU | MODERAT | MISSENSE | aCc/aTc | T53I | PLES_45481 | | | | | | | T | T | | | | |
| NC_011770 | 4995851 | G | A | 78 | 0 | 2 | 0 | NON_SYNONYMOU | MODERAT | MISSENSE | gGc/gAc | G249D | PLES_45481 | A | A | | | | | | | | | | |
| NC_011770 | 4996820 | T | A | 76 | 0 | 2 | 0 | NON_SYNONYMOU | MODERAT | MISSENSE | gTg/gAg | V572E | PLES_45481 | | | | | | | | | A | A | | |
| NC_011770 | 4996901 | A | G | 70 | 0 | 2 | 0 | NON_SYNONYMOU | MODERAT | MISSENSE | aAg/aGg | K599R | PLES_45481 | G | G | | | | | | | | | | |
| NC_011770 | 4997584 | C | T | 27 | 0 | 4 | 0 | NON_SYNONYMOU | MODERAT | MISSENSE | Cgc/Tgc | R827C | PLES_45481 | | | T | T | T | T | | | | | | |
| NC_011770 | 4998215 | G | A | 46 | 0 | 2 | 0 | NON_SYNONYMOU | MODERAT | MISSENSE | Gac/Aac | D129N | PLES_45491 | | | | | | | A | A | | | | |
| NC_011770 | 4998893 | A | T | 66 | 0 | 2 | 0 | NON_SYNONYMOU | MODERAT | MISSENSE | Acc/Tcc | T355S | PLES_45491 | | | | | | | | | T | T | | |
| NC_011770 | 4999009 | C | G | 70 | 0 | 4 | 0 | NON_SYNONYMOU | MODERAT | MISSENSE | gaC/gaG | D393E | PLES_45491 | | | G | G | G | G | | | | | | |
| NC_011770 | 5000770 | G | T | 83 | 0 | 4 | 0 | NON_SYNONYMOU | MODERAT | MISSENSE | Ctg/Atg | L212M | PLES_45511 | | | T | T | T | T | | | | | | |
| NC_011770 | 5000829 | C | A | 82 | 0 | 2 | 0 | NON_SYNONYMOU | MODERAT | MISSENSE | cGt/cTt | R192L | PLES_45511 | | | | | | | | | | | A | A |
| NC_011770 | 5001546 | A | G | 80 | 0 | 1 | 0 | NON_SYNONYMOU | MODERAT | MISSENSE | Acc/Gcc | T13A | PLES_45521 | | | | | | | G | | | | | |
| NC_011770 | 5001675 | G | C | 68 | 0 | 1 | 0 | NON_SYNONYMOU | MODERAT | MISSENSE | Gcc/Ccc | A56P | PLES_45521 | | | | | | | C | | | | | |
| NC_011770 | 5001786 | G | A | 68 | 0 | 10 | 0 | NON_SYNONYMOU | MODERAT | MISSENSE | Gtc/Atc | V93I | PLES_45521 | A | A | A | A | A | A | A | A | A | A | | |
| NC_011770 | 5003425 | C | T | 97 | 0 | 2 | 0 | NON_SYNONYMOU | MODERAT | MISSENSE | Gcg/Acg | A148T | PLES_45531 | | | | | | | T | T | | | | |
| NC_011770 | 5004461 | C | T | 71 | 0 | 2 | 0 | NON_SYNONYMOU | MODERAT | MISSENSE | Gcg/Acg | A14T | PLES_45541 | | | | | | | | | T | T | | |
| NC_011770 | 5004485 | A | G | 73 | 0 | 12 | 0 | NON_SYNONYMOU | MODERAT | MISSENSE | Tcc/Ccc | S6P | PLES_45541 | G | G | G | G | G | G | G | G | G | G | G | G |
| NC_011770 | 5005214 | T | C | 72 | 0 | 4 | 0 | NON_SYNONYMOU | MODERAT | MISSENSE | gTg/gCg | V185A | PLES_45551 | | | C | C | C | C | | | | | | |
| NC_011770 | 5005408 | G | A | 75 | 0 | 2 | 0 | NON_SYNONYMOU | MODERAT | MISSENSE | Gaa/Aaa | E250K | PLES_45551 | | | A | A | | | | | | | | |
| NC_011770 | 5005850 | C | A | 56 | 0 | 12 | 0 | NON_SYNONYMOU | MODERAT | MISSENSE | gCc/gAc | A397D | PLES_45551 | A | A | A | A | A | A | A | A | A | A | A | A |
| NC_011770 | 5007312 | A | C | 70 | 0 | 12 | 0 | NON_SYNONYMOU | MODERAT | MISSENSE | ttA/ttC | L884F | PLES_45551 | C | C | C | C | C | C | C | C | C | C | C | C |
| NC_011770 | 5007331 | G | C | 73 | 0 | 2 | 0 | NON_SYNONYMOU | MODERAT | MISSENSE | Gag/Cag | E891Q | PLES_45551 | C | C | | | | | | | | | | |
| NC_011770 | 5008321 | G | A | 73 | 0 | 6 | 0 | NON_SYNONYMOU | MODERAT | MISSENSE | Cgg/Tgg | R212W | PLES_45561 | | | | | | | A | A | A | A | A | A |
| NC_011770 | 5008732 | G | A | 75 | 0 | 2 | 0 | NON_SYNONYMOU | MODERAT | MISSENSE | Cgg/Tgg | R75W | PLES_45561 | A | A | | | | | | | | | | |
| NC_011770 | 5008798 | T | C | 71 | 0 | 2 | 0 | NON_SYNONYMOU | MODERAT | MISSENSE | Atc/Gtc | I53V | PLES_45561 | | | | | | | C | C | | | | |
| NC_011770 | 5009368 | G | T | 69 | 0 | 2 | 0 | NON_SYNONYMOU | MODERAT | MISSENSE | gCc/gAc | A199D | PLES_45581 | T | T | | | | | | | | | | |
| NC_011770 | 5009708 | A | G | 79 | 0 | 2 | 0 | NON_SYNONYMOU | MODERAT | MISSENSE | Ttc/Ctc | F86L | PLES_45581 | | | | | | | G | G | | | | |
| NC_011770 | 5009794 | G | A | 78 | 0 | 2 | 0 | NON_SYNONYMOU | MODERAT | MISSENSE | gCc/gTc | A57V | PLES_45581 | | | | | | | | | A | A | | |
| NC_011770 | 5009922 | C | A | 70 | 0 | 2 | 0 | NON_SYNONYMOU | MODERAT | MISSENSE | gaG/gaT | E14D | PLES_45581 | | | | | | | | | A | A | | |
| NC_011770 | 5010271 | G | A | 80 | 0 | 1 | 0 | NON_SYNONYMOU | MODERAT | MISSENSE | cGg/cAg | R61Q | PLES_45591 | | | | | | | A | | | | | |
| NC_011770 | 5010373 | G | A | 81 | 0 | 2 | 0 | NON_SYNONYMOU | MODERAT | MISSENSE | aGc/aAc | S95N | PLES_45591 | A | A | | | | | | | | | | |
| NC_011770 | 5010376 | T | G | 81 | 0 | 2 | 0 | NON_SYNONYMOU | MODERAT | MISSENSE | gTa/gGa | V96G | PLES_45591 | | | | | | | | | G | G | | |
| NC_011770 | 5010555 | C | T | 89 | 0 | 2 | 0 | NON_SYNONYMOU | MODERAT | MISSENSE | Cat/Tat | H156Y | PLES_45591 | | | | | | | | | T | T | | |
| NC_011770 | 5010634 | C | G | 89 | 0 | 2 | 0 | NON_SYNONYMOU | MODERAT | MISSENSE | gCa/gGa | A182G | PLES_45591 | | | | | | | | | G | G | | |
| NC_011770 | 5010862 | A | G | 84 | 0 | 6 | 0 | NON_SYNONYMOU | MODERAT | MISSENSE | cAa/cGa | Q258R | PLES_45591 | | | G | G | G | G | | | | | G | G |
| NC_011770 | 5017497 | G | A | 78 | 0 | 2 | 0 | NON_SYNONYMOU | MODERAT | MISSENSE | Gac/Aac | D286N | PLES_45621 | | | | | | | A | A | | | | |
| NC_011770 | 5018037 | A | C | 67 | 0 | 2 | 0 | NON_SYNONYMOU | MODERAT | MISSENSE | Acc/Ccc | T466P | PLES_45621 | C | C | | | | | | | | | | |
| NC_011770 | 5018226 | A | G | 61 | 0 | 8 | 0 | NON_SYNONYMOU | MODERAT | MISSENSE | Agc/Ggc | S529G | PLES_45621 | G | G | | | | | G | G | G | G | G | G |
| NC_011770 | 5019827 | G | A | 56 | 0 | 2 | 0 | NON_SYNONYMOU | MODERAT | MISSENSE | gGc/gAc | G20D | PLES_45641 | A | A | | | | | | | | | | |
| NC_011770 | 5020855 | G | A | 84 | 0 | 1 | 0 | NON_SYNONYMOU | MODERAT | MISSENSE | Gag/Aag | E363K | PLES_45641 | | | | | | | A | | | | | |
| NC_011770 | 5021629 | G | A | 73 | 0 | 1 | 0 | NON_SYNONYMOU | MODERAT | MISSENSE | Ggc/Agc | G621S | PLES_45641 | | | | | | | A | | | | | |
| NC_011770 | 5022101 | G | A | 60 | 0 | 4 | 0 | NON_SYNONYMOU | MODERAT | MISSENSE | aGg/aAg | R778K | PLES_45641 | | | A | A | A | A | | | | | | |
| NC_011770 | 5023269 | T | C | 66 | 0 | 1 | 0 | NON_SYNONYMOU | MODERAT | MISSENSE | cTg/cCg | L163P | PLES_45661 | | | | C | | | | | | | | |

| | | | | | | | | | | | | | | | | | | | | | | | | | |
|---|---|---|---|---|---|---|---|---|---|---|---|---|---|---|---|---|---|---|---|---|---|---|---|---|---|
| NC_011770 | 5022809 | G | A | 60 | 0 | 4 | 0 | NON_SYNONYMOU | MODERAT | MISSENSE | Gcc/Acc | A10T | PLES_45661 | | | A | A | A | A | | | | | | |
| NC_011770 | 5022834 | T | C | 61 | 0 | 6 | 0 | NON_SYNONYMOU | MODERAT | MISSENSE | cTc/cCc | L18P | PLES_45661 | C | C | C | C | C | C | | | | | | |
| NC_011770 | 5022836 | T | A | 60 | 0 | 4 | 0 | NON_SYNONYMOU | MODERAT | MISSENSE | Tcg/Acg | S19T | PLES_45661 | | | A | A | A | A | | | | | | |
| NC_011770 | 5023080 | G | A | 48 | 0 | 4 | 0 | NON_SYNONYMOU | MODERAT | MISSENSE | gGg/gAg | G100E | PLES_45661 | | | A | A | A | A | | | | | | |
| NC_011770 | 5023116 | A | G | 60 | 0 | 4 | 0 | NON_SYNONYMOU | MODERAT | MISSENSE | cAc/cGc | H112R | PLES_45661 | | | G | G | G | G | | | | | | |
| NC_011770 | 5023364 | A | G | 64 | 0 | 4 | 0 | NON_SYNONYMOU | MODERAT | MISSENSE | Atc/Gtc | I195V | PLES_45661 | | | G | G | G | G | | | | | | |
| NC_011770 | 5024122 | G | A | 74 | 0 | 1 | 0 | NON_SYNONYMOU | MODERAT | MISSENSE | Gtg/Atg | V153M | PLES_45671 | | | | A | | | | | | | | |
| NC_011770 | 5025903 | G | A | 77 | 0 | 1 | 0 | NON_SYNONYMOU | MODERAT | MISSENSE | aGc/aAc | S275N | PLES_45691 | | | | | | | | | | | | A |
| NC_011770 | 5025476 | G | A | 82 | 0 | 1 | 0 | NON_SYNONYMOU | MODERAT | MISSENSE | Ggc/Agc | G133S | PLES_45691 | | | | | | | A | | | | | |
| NC_011770 | 5029505 | T | A | 73 | 0 | 2 | 0 | NON_SYNONYMOU | MODERAT | MISSENSE | Atg/Ttg | M37L | PLES_45741 | A | A | | | | | | | | | | |
| NC_011770 | 5029454 | C | T | 68 | 0 | 1 | 0 | NON_SYNONYMOU | MODERAT | MISSENSE | Gcg/Acg | A54T | PLES_45741 | | | | | | | | T | | | | |
| NC_011770 | 5039905 | A | G | 69 | 0 | 12 | 0 | NON_SYNONYMOU | MODERAT | MISSENSE | Acc/Gcc | T96A | PLES_45841 | G | G | G | G | G | G | G | G | G | G | G | G |
| NC_011770 | 5039965 | A | G | 70 | 0 | 12 | 0 | NON_SYNONYMOU | MODERAT | MISSENSE | Acc/Gcc | T116A | PLES_45841 | G | G | G | G | G | G | G | G | G | G | G | G |
| NC_011770 | 5040100 | C | T | 72 | 0 | 2 | 0 | NON_SYNONYMOU | MODERAT | MISSENSE | Cgc/Tgc | R161C | PLES_45841 | | | | | | | | | T | T | | |
| NC_011770 | 5041295 | G | A | 67 | 0 | 10 | 0 | NON_SYNONYMOU | MODERAT | MISSENSE | Gcc/Acc | A211T | PLES_45851 | A | A | A | A | A | A | | | A | A | A | A |
| NC_011770 | 5041516 | A | G | 70 | 0 | 10 | 0 | NON_SYNONYMOU | MODERAT | MISSENSE | gTg/gCg | V459A | PLES_45861 | G | G | G | G | G | G | | | G | G | G | G |
| NC_011770 | 5042151 | C | G | 75 | 0 | 8 | 0 | NON_SYNONYMOU | MODERAT | MISSENSE | caG/caC | Q247H | PLES_45861 | G | G | G | G | G | G | G | G | | | | |
| NC_011770 | 5042567 | T | C | 88 | 0 | 2 | 0 | NON_SYNONYMOU | MODERAT | MISSENSE | Acc/Gcc | T109A | PLES_45861 | | | | | | | | | C | C | | |
| NC_011770 | 5042590 | T | C | 79 | 0 | 2 | 0 | NON_SYNONYMOU | MODERAT | MISSENSE | aAt/aGt | N101S | PLES_45861 | | | | | | | | | C | C | | |
| NC_011770 | 5043098 | C | T | 85 | 0 | 1 | 0 | NON_SYNONYMOU | MODERAT | MISSENSE | cGg/cAg | R153Q | PLES_45871 | | | | | | | T | | | | | |
| NC_011770 | 5045490 | A | G | 66 | 0 | 4 | 0 | NON_SYNONYMOU | MODERAT | MISSENSE | Aac/Gac | N138D | PLES_45891 | | | | | | | G | G | | | G | G |
| NC_011770 | 5045491 | A | G | 66 | 0 | 4 | 0 | NON_SYNONYMOU | MODERAT | MISSENSE | aAc/aGc | N138S | PLES_45891 | | | | | | | G | G | | | G | G |
| NC_011770 | 5046432 | G | A | 76 | 0 | 2 | 0 | NON_SYNONYMOU | MODERAT | MISSENSE | Gtc/Atc | V108I | PLES_45901 | A | A | | | | | | | | | | |
| NC_011770 | 5048027 | T | C | 81 | 0 | 4 | 0 | NON_SYNONYMOU | MODERAT | MISSENSE | gTg/gCg | V479A | PLES_45911 | | | C | C | C | C | | | | | | |
| NC_011770 | 5048075 | T | C | 76 | 0 | 8 | 0 | NON_SYNONYMOU | MODERAT | MISSENSE | gTg/gCg | V495A | PLES_45911 | C | C | C | C | C | C | | | C | C | | |
| NC_011770 | 5049069 | T | C | 82 | 0 | 2 | 0 | NON_SYNONYMOU | MODERAT | MISSENSE | gTa/gCa | V323A | PLES_45921 | | | | | | | C | C | | | | |
| NC_011770 | 5050090 | C | G | 82 | 0 | 4 | 0 | NON_SYNONYMOU | MODERAT | MISSENSE | Gtg/Ctg | V265L | PLES_45941 | | | G | G | G | G | | | | | | |
| NC_011770 | 5050762 | T | C | 95 | 0 | 6 | 0 | NON_SYNONYMOU | MODERAT | MISSENSE | Atc/Gtc | I41V | PLES_45941 | | | | | | | C | C | C | C | C | C |
| NC_011770 | 5051605 | G | A | 77 | 0 | 4 | 0 | INTRAGENIC | MODIFIER | | | | PLES_45951 | | | A | A | A | A | | | | | | |
| NC_011770 | 5051773 | A | C | 68 | 0 | 6 | 0 | INTRAGENIC | MODIFIER | | | | PLES_45951 | | | | | | | C | C | C | C | C | C |
| NC_011770 | 5052871 | A | G | 65 | 0 | 12 | 0 | NON_SYNONYMOU | MODERAT | MISSENSE | Acc/Gcc | T307A | PLES_45961 | G | G | G | G | G | G | G | G | G | G | G | G |
| NC_011770 | 5053235 | C | G | 81 | 0 | 2 | 0 | NON_SYNONYMOU | MODERAT | MISSENSE | gCg/gGg | A428G | PLES_45961 | G | G | | | | | | | | | | |
| NC_011770 | 5053396 | T | C | 74 | 0 | 4 | 0 | NON_SYNONYMOU | MODERAT | MISSENSE | Ttc/Ctc | F482L | PLES_45961 | | | C | C | C | C | | | | | | |
| NC_011770 | 5053514 | G | T | 79 | 0 | 2 | 0 | NON_SYNONYMOU | MODERAT | MISSENSE | gaG/gaT | E14D | PLES_45971 | | | | | | | | | T | T | | |
| NC_011770 | 5053791 | G | A | 85 | 0 | 2 | 0 | NON_SYNONYMOU | MODERAT | MISSENSE | Gag/Aag | E107K | PLES_45971 | A | A | | | | | | | | | | |
| NC_011770 | 5054587 | T | C | 84 | 0 | 4 | 0 | NON_SYNONYMOU | MODERAT | MISSENSE | gTc/gCc | V372A | PLES_45971 | | | C | C | C | C | | | | | | |
| NC_011770 | 5055083 | G | A | 75 | 0 | 2 | 0 | NON_SYNONYMOU | MODERAT | MISSENSE | Gcc/Acc | A128T | PLES_45981 | | | | | | | A | A | | | | |
| NC_011770 | 5056401 | A | G | 80 | 0 | 4 | 0 | NON_SYNONYMOU | MODERAT | MISSENSE | Aag/Gag | K275E | PLES_45991 | G | G | | | | | | | G | G | | |
| NC_011770 | 5055844 | G | A | 70 | 0 | 1 | 0 | NON_SYNONYMOU | MODERAT | MISSENSE | cGa/cAa | R89Q | PLES_45991 | | | | | | | | A | | | | |
| NC_011770 | 5056944 | G | T | 85 | 0 | 4 | 0 | NON_SYNONYMOU | MODERAT | MISSENSE | Gcc/Tcc | A51S | PLES_46001 | T | T | | | | | | | T | T | | |
| NC_011770 | 5057050 | C | T | 79 | 0 | 4 | 0 | NON_SYNONYMOU | MODERAT | MISSENSE | gCc/gTc | A86V | PLES_46001 | | | T | T | T | T | | | | | | |
| NC_011770 | 5057387 | G | C | 96 | 0 | 12 | 0 | NON_SYNONYMOU | MODERAT | MISSENSE | gaG/gaC | E198D | PLES_46001 | C | C | C | C | C | C | C | C | C | C | C | C |
| NC_011770 | 5058209 | G | A | 82 | 0 | 1 | 0 | NON_SYNONYMOU | MODERAT | MISSENSE | Ggc/Agc | G43S | PLES_46011 | | | | | | | | A | | | | |
| NC_011770 | 5061544 | T | C | 73 | 0 | 12 | 0 | NON_SYNONYMOU | MODERAT | MISSENSE | Tgc/Cgc | C95R | PLES_46041 | C | C | C | C | C | C | C | C | C | C | C | C |
| NC_011770 | 5061632 | T | C | 81 | 0 | 2 | 0 | NON_SYNONYMOU | MODERAT | MISSENSE | gTa/gCa | V124A | PLES_46041 | | | | | | | C | C | | | | |
| NC_011770 | 5061757 | T | C | 93 | 0 | 2 | 0 | NON_SYNONYMOU | MODERAT | MISSENSE | Tat/Cat | Y166H | PLES_46041 | | | | | | | C | C | | | | |
| NC_011770 | 5061799 | C | G | 92 | 0 | 2 | 0 | NON_SYNONYMOU | MODERAT | MISSENSE | Cgt/Ggt | R180G | PLES_46041 | | | | | | | G | G | | | | |
| NC_011770 | 5061806 | T | A | 92 | 0 | 2 | 0 | NON_SYNONYMOU | MODERAT | MISSENSE | gTg/gAg | V182E | PLES_46041 | | | | | | | A | A | | | | |
| NC_011770 | 5061821 | A | G | 87 | 0 | 2 | 0 | NON_SYNONYMOU | MODERAT | MISSENSE | gAg/gGg | E187G | PLES_46041 | | | | | | | G | G | | | | |
| NC_011770 | 5061863 | A | C | 85 | 0 | 2 | 0 | NON_SYNONYMOU | MODERAT | MISSENSE | gAg/gCg | E201A | PLES_46041 | | | | | | | C | C | | | | |
| NC_011770 | 5061866 | C | A | 86 | 0 | 2 | 0 | NON_SYNONYMOU | MODERAT | MISSENSE | gCa/gAa | A202E | PLES_46041 | | | | | | | A | A | | | | |
| NC_011770 | 5061903 | A | C | 81 | 0 | 2 | 0 | NON_SYNONYMOU | MODERAT | MISSENSE | gaA/gaC | E214D | PLES_46041 | | | | | | | C | C | | | | |
| NC_011770 | 5062666 | G | A | 84 | 0 | 1 | 0 | NON_SYNONYMOU | MODERAT | MISSENSE | gCc/gTc | A24V | PLES_46061 | | A | | | | | | | | | | |
| NC_011770 | 5063086 | C | A | 82 | 0 | 4 | 0 | NON_SYNONYMOU | MODERAT | MISSENSE | Gcg/Tcg | A75S | PLES_46071 | | | A | A | A | A | | | | | | |
| NC_011770 | 5064100 | G | A | 78 | 0 | 1 | 0 | STOP_GAINED | HIGH | NONSENSE | tgG/tgA | W95* | PLES_46091 | | | | | | | | A | | | | |
| NC_011770 | 5064830 | ATCCTGCT( | A | 83 | 0 | 6 | 0 | CODON_CHANGE_ | MODERATE | | ctggtcctgc | LVLR344R | PLES_46091 | | A | A | | A | | A | | A | | | A |
| NC_011770 | 5064839 | G | A | 14 | 1 | 1 | 0 | NON_SYNONYMOU | MODERAT | MISSENSE | Gtc/Atc | V342I | PLES_46091 | | A | | | | | | | . | | | |
| NC_011770 | 5064505 | G | T | 75 | 0 | 2 | 0 | NON_SYNONYMOU | MODERAT | MISSENSE | gaG/gaT | E230D | PLES_46091 | T | T | | | | | | | | | | |
| NC_011770 | 5065529 | A | G | 80 | 0 | 12 | 0 | NON_SYNONYMOU | MODERAT | MISSENSE | gAa/gGa | E211G | PLES_46101 | G | G | G | G | G | G | G | G | G | G | G | G |
| NC_011770 | 5065625 | G | A | 72 | 0 | 12 | 0 | NON_SYNONYMOU | MODERAT | MISSENSE | gGa/gAa | G243E | PLES_46101 | A | A | A | A | A | A | A | A | A | A | A | A |
| NC_011770 | 5065860 | G | T | 69 | 0 | 12 | 0 | NON_SYNONYMOU | MODERAT | MISSENSE | Gat/Tat | D43Y | PLES_46111 | T | T | T | T | T | T | T | T | T | T | T | T |
| NC_011770 | 5065875 | G | A | 67 | 0 | 12 | 0 | NON_SYNONYMOU | MODERAT | MISSENSE | Gcc/Acc | A48T | PLES_46111 | A | A | A | A | A | A | A | A | A | A | A | A |

| | | | | | | | | | | | | | | | | | | | | | | | | | |
|---|---|---|---|---|---|---|---|---|---|---|---|---|---|---|---|---|---|---|---|---|---|---|---|---|---|
| NC_011770 | 5065908 | A | G | 67 | 0 | 12 | 0 | NON_SYNONYMOU | MODERAT | MISSENSE | Aat/Gat | N59D | PLES_46111 | G | G | G | G | G | G | G | G | G | G | G | G |
| NC_011770 | 5065918 | C | T | 66 | 0 | 12 | 0 | NON_SYNONYMOU | MODERAT | MISSENSE | gCg/gTg | A62V | PLES_46111 | T | T | T | T | T | T | T | T | T | T | T | T |
| NC_011770 | 5066082 | G | A | 69 | 0 | 8 | 0 | NON_SYNONYMOU | MODERAT | MISSENSE | cGc/cAc | R26H | PLES_46121 | A | A | A | A | A | A | | | | | A | A |
| NC_011770 | 5066160 | A | G | 64 | 0 | 12 | 0 | NON_SYNONYMOU | MODERAT | MISSENSE | cAg/cGg | Q52R | PLES_46121 | G | G | G | G | G | G | G | G | G | G | G | G |
| NC_011770 | 5066285 | G | A | 65 | 0 | 2 | 0 | NON_SYNONYMOU | MODERAT | MISSENSE | Gac/Aac | D94N | PLES_46121 | | | | | | | | | | | A | A |
| NC_011770 | 5066586 | A | T | 62 | 0 | 1 | 0 | NON_SYNONYMOU | MODERAT | MISSENSE | gAc/gTc | D194V | PLES_46121 | | | | | | | T | | | | | |
| NC_011770 | 5067612 | C | T | 55 | 0 | 4 | 0 | NON_SYNONYMOU | MODERAT | MISSENSE | cCg/cTg | P260L | PLES_46131 | | | T | T | T | T | | | | | | |
| NC_011770 | 5067616 | A | C | 55 | 0 | 4 | 0 | NON_SYNONYMOU | MODERAT | MISSENSE | gaA/gaC | E261D | PLES_46131 | | | C | C | C | C | | | | | | |
| NC_011770 | 5067623 | A | G | 56 | 0 | 4 | 0 | NON_SYNONYMOU | MODERAT | MISSENSE | Agc/Ggc | S264G | PLES_46131 | | | G | G | G | G | | | | | | |
| NC_011770 | 5067746 | G | A | 51 | 0 | 4 | 0 | NON_SYNONYMOU | MODERAT | MISSENSE | Ggt/Agt | G305S | PLES_46131 | | | A | A | A | A | | | | | | |
| NC_011770 | 5068230 | G | A | 65 | 0 | 4 | 0 | NON_SYNONYMOU | MODERAT | MISSENSE | Gcc/Acc | A92T | PLES_46141 | A | A | | | | | | | | | A | A |
| NC_011770 | 5068323 | C | G | 67 | 0 | 2 | 0 | NON_SYNONYMOU | MODERAT | MISSENSE | Cag/Gag | Q123E | PLES_46141 | | | | | | | G | G | | | | |
| NC_011770 | 5069218 | T | G | 73 | 0 | 8 | 0 | NON_SYNONYMOU | MODERAT | MISSENSE | Tcg/Gcg | S25A | PLES_46151 | G | G | | | | | G | G | G | G | G | G |
| NC_011770 | 5069433 | C | G | 68 | 0 | 8 | 0 | NON_SYNONYMOU | MODERAT | MISSENSE | gaC/gaG | D96E | PLES_46151 | G | G | | | | | G | G | G | G | G | G |
| NC_011770 | 5069605 | G | A | 59 | 0 | 6 | 0 | NON_SYNONYMOU | MODERAT | MISSENSE | Gtc/Atc | V154I | PLES_46151 | A | A | | | | | | | A | A | A | A |
| NC_011770 | 5070793 | GC | G | 62 | 0 | 1 | 0 | FRAME_SHIFT | HIGH | | gcg/ | A25 | PLES_46161 | | . | | | | | G | | | | | |
| NC_011770 | 5070435 | A | C | 52 | 0 | 2 | 0 | NON_SYNONYMOU | MODERAT | MISSENSE | gaT/gaG | D144E | PLES_46161 | | | | | | | C | C | | | | |
| NC_011770 | 5070469 | C | T | 50 | 0 | 12 | 0 | NON_SYNONYMOU | MODERAT | MISSENSE | cGc/cAc | R133H | PLES_46161 | T | T | T | T | T | T | T | T | T | T | T | T |
| NC_011770 | 5070538 | C | T | 53 | 0 | 2 | 0 | NON_SYNONYMOU | MODERAT | MISSENSE | gGg/gAg | G110E | PLES_46161 | T | T | | | | | | | | | | |
| NC_011770 | 5070604 | G | A | 58 | 0 | 2 | 0 | NON_SYNONYMOU | MODERAT | MISSENSE | tCg/tTg | S88L | PLES_46161 | | | | | | | A | A | | | | |
| NC_011770 | 5070700 | A | G | 58 | 0 | 12 | 0 | NON_SYNONYMOU | MODERAT | MISSENSE | gTc/gCc | V56A | PLES_46161 | G | G | G | G | G | G | G | G | G | G | G | G |
| NC_011770 | 5070730 | T | C | 61 | 0 | 6 | 0 | NON_SYNONYMOU | MODERAT | MISSENSE | gAg/gGg | E46G | PLES_46161 | C | C | | | | | | | C | C | C | C |
| NC_011770 | 5070750 | G | C | 60 | 0 | 4 | 0 | NON_SYNONYMOU | MODERAT | MISSENSE | aaC/aaG | N39K | PLES_46161 | | | | | | | | | C | C | C | C |
| NC_011770 | 5070827 | A | C | 67 | 0 | 8 | 0 | NON_SYNONYMOU | MODERAT | MISSENSE | Tgt/Ggt | C14G | PLES_46161 | C | C | | | | | C | C | C | C | C | C |
| NC_011770 | 5071367 | A | C | 94 | 0 | 4 | 0 | NON_SYNONYMOU | MODERAT | MISSENSE | Ttc/Gtc | F9V | PLES_46171 | | | | | | | | | C | C | C | C |
| NC_011770 | 5071372 | G | A | 97 | 0 | 2 | 0 | NON_SYNONYMOU | MODERAT | MISSENSE | aCg/aTg | T7M | PLES_46171 | A | A | | | | | | | | | | |
| NC_011770 | 5071827 | G | A | 98 | 0 | 2 | 0 | NON_SYNONYMOU | MODERAT | MISSENSE | Gtc/Atc | V33I | PLES_46181 | | | | | | | A | A | | | | |
| NC_011770 | 5071908 | G | A | 98 | 0 | 2 | 0 | NON_SYNONYMOU | MODERAT | MISSENSE | Gtc/Atc | V60I | PLES_46181 | | | | | | | A | A | | | | |
| NC_011770 | 5072159 | G | A | 92 | 0 | 6 | 0 | NON_SYNONYMOU | MODERAT | MISSENSE | Gtc/Atc | V38I | PLES_46191 | A | A | | | | | | | A | A | A | A |
| NC_011770 | 5072192 | T | C | 85 | 0 | 4 | 0 | NON_SYNONYMOU | MODERAT | MISSENSE | Ttt/Ctt | F49L | PLES_46191 | | | C | C | C | C | | | | | | |
| NC_011770 | 5072274 | A | G | 84 | 0 | 4 | 0 | NON_SYNONYMOU | MODERAT | MISSENSE | gAc/gGc | D76G | PLES_46191 | G | G | | | | | G | G | | | | |
| NC_011770 | 5072282 | G | C | 89 | 0 | 2 | 0 | NON_SYNONYMOU | MODERAT | MISSENSE | Ggc/Cgc | G79R | PLES_46191 | | | | | | | C | C | | | | |
| NC_011770 | 5072310 | A | G | 81 | 0 | 8 | 0 | NON_SYNONYMOU | MODERAT | MISSENSE | gAg/gGg | E88G | PLES_46191 | G | G | G | G | G | G | G | G | | | | |
| NC_011770 | 5072357 | A | G | 82 | 0 | 4 | 0 | NON_SYNONYMOU | MODERAT | MISSENSE | Acc/Gcc | T104A | PLES_46191 | | | | | | | | | G | G | G | G |
| NC_011770 | 5072369 | C | A | 81 | 0 | 4 | 0 | NON_SYNONYMOU | MODERAT | MISSENSE | Ctg/Atg | L108M | PLES_46191 | | | | | | | | | A | A | A | A |
| NC_011770 | 5072421 | G | A | 81 | 0 | 4 | 0 | NON_SYNONYMOU | MODERAT | MISSENSE | cGc/cAc | R125H | PLES_46191 | | | | | | | | | A | A | A | A |
| NC_011770 | 5072431 | T | A | 83 | 0 | 2 | 0 | NON_SYNONYMOU | MODERAT | MISSENSE | ttT/ttA | F128L | PLES_46191 | | | | | | | A | A | | | | |
| NC_011770 | 5072489 | T | A | 93 | 0 | 2 | 0 | NON_SYNONYMOU | MODERAT | MISSENSE | Tcc/Acc | S148T | PLES_46191 | A | A | | | | | | | | | | |
| NC_011770 | 5072511 | G | A | 90 | 0 | 4 | 0 | NON_SYNONYMOU | MODERAT | MISSENSE | cGa/cAa | R155Q | PLES_46191 | | | | | | | | | A | A | A | A |
| NC_011770 | 5072550 | A | G | 89 | 0 | 4 | 0 | NON_SYNONYMOU | MODERAT | MISSENSE | gAc/gGc | D168G | PLES_46191 | | | | | | | | | G | G | G | G |
| NC_011770 | 5073049 | T | C | 84 | 0 | 6 | 0 | NON_SYNONYMOU | MODERAT | MISSENSE | Atc/Gtc | I96V | PLES_46211 | C | C | | | | | | | C | C | C | C |
| NC_011770 | 5073582 | A | G | 89 | 0 | 12 | 0 | NON_SYNONYMOU | MODERAT | MISSENSE | Ata/Gta | I45V | PLES_46221 | G | G | G | G | G | G | G | G | G | G | G | G |
| NC_011770 | 5073622 | C | T | 85 | 0 | 2 | 0 | NON_SYNONYMOU | MODERAT | MISSENSE | aCc/aTc | T58I | PLES_46221 | T | T | | | | | | | | | | |
| NC_011770 | 5073655 | G | C | 77 | 0 | 8 | 0 | NON_SYNONYMOU | MODERAT | MISSENSE | cGg/cCg | R69P | PLES_46221 | C | C | | | | | C | C | C | C | C | C |
| NC_011770 | 5073675 | A | G | 79 | 0 | 12 | 0 | NON_SYNONYMOU | MODERAT | MISSENSE | Aac/Gac | N76D | PLES_46221 | G | G | G | G | G | G | G | G | G | G | G | G |
| NC_011770 | 5073879 | A | G | 73 | 0 | 4 | 0 | NON_SYNONYMOU | MODERAT | MISSENSE | Acc/Gcc | T144A | PLES_46221 | | | | | | | | | G | G | G | G |
| NC_011770 | 5073957 | G | A | 76 | 0 | 4 | 0 | NON_SYNONYMOU | MODERAT | MISSENSE | Gac/Aac | D170N | PLES_46221 | | | | | | | | | A | A | A | A |
| NC_011770 | 5074063 | G | C | 75 | 0 | 2 | 0 | NON_SYNONYMOU | MODERAT | MISSENSE | gGt/gCt | G205A | PLES_46221 | | | | | | | C | C | | | | |
| NC_011770 | 5074179 | G | A | 65 | 0 | 8 | 0 | NON_SYNONYMOU | MODERAT | MISSENSE | Gcg/Acg | A244T | PLES_46221 | A | A | | | | | A | A | A | A | A | A |
| NC_011770 | 5074273 | A | G | 64 | 0 | 2 | 0 | NON_SYNONYMOU | MODERAT | MISSENSE | cAg/cGg | Q275R | PLES_46221 | G | G | | | | | | | | | | |
| NC_011770 | 5076577 | C | T | 86 | 0 | 1 | 0 | STOP_GAINED | HIGH | NONSENSE | Cag/Tag | Q81* | PLES_46251 | | | T | | | | | | | | | |
| NC_011770 | 5077093 | G | T | 66 | 0 | 1 | 0 | STOP_GAINED | HIGH | NONSENSE | Gag/Tag | E253* | PLES_46251 | | | | | | | T | | | | | |
| NC_011770 | 5076371 | A | G | 87 | 0 | 8 | 0 | NON_SYNONYMOU | MODERAT | MISSENSE | gAc/gGc | D12G | PLES_46251 | G | G | | | | | G | G | G | G | G | G |
| NC_011770 | 5076611 | G | A | 81 | 0 | 8 | 0 | NON_SYNONYMOU | MODERAT | MISSENSE | cGa/cAa | R92Q | PLES_46251 | A | A | | | | | A | A | A | A | A | A |
| NC_011770 | 5076922 | A | C | 79 | 0 | 1 | 0 | NON_SYNONYMOU | MODERAT | MISSENSE | Acc/Ccc | T196P | PLES_46251 | | | | | | | | C | | | | |
| NC_011770 | 5077517 | A | G | 61 | 0 | 10 | 0 | NON_SYNONYMOU | MODERAT | MISSENSE | Att/Gtt | I52V | PLES_46261 | G | G | G | G | G | G | | | G | G | G | G |
| NC_011770 | 5077525 | A | T | 59 | 0 | 4 | 0 | NON_SYNONYMOU | MODERAT | MISSENSE | gaA/gaT | E54D | PLES_46261 | | | T | T | T | T | | | | | | |
| NC_011770 | 5077775 | T | A | 73 | 0 | 2 | 0 | NON_SYNONYMOU | MODERAT | MISSENSE | Ttc/Atc | F138I | PLES_46261 | | | | | | | | | A | A | | |
| NC_011770 | 5078088 | C | T | 67 | 0 | 6 | 0 | NON_SYNONYMOU | MODERAT | MISSENSE | gCg/gTg | A242V | PLES_46261 | | | T | T | T | T | | | | | T | T |
| NC_011770 | 5078162 | A | C | 72 | 0 | 12 | 0 | NON_SYNONYMOU | MODERAT | MISSENSE | Atg/Ctg | M267L | PLES_46261 | C | C | C | C | C | C | C | C | C | C | C | C |
| NC_011770 | 5078184 | G | T | 69 | 0 | 4 | 0 | NON_SYNONYMOU | MODERAT | MISSENSE | cGc/cTc | R274L | PLES_46261 | | | T | T | T | T | | | | | | |
| NC_011770 | 5078654 | C | G | 54 | 0 | 4 | 0 | NON_SYNONYMOU | MODERAT | MISSENSE | Ctg/Gtg | L431V | PLES_46261 | | | G | G | G | G | | | | | | |

| | | | | | | | | | | | | | | | | | | | | | | | | | |
|---|---|---|---|---|---|---|---|---|---|---|---|---|---|---|---|---|---|---|---|---|---|---|---|---|---|
| NC_011770 | 5078601 | C | T | 60 | 0 | 1 | 0 | NON_SYNONYMOU | MODERAT | MISSENSE | gCg/gTg | A413V | PLES_46261 | | | | | | | | T | | | | |
| NC_011770 | 5078985 | G | A | 61 | 0 | 2 | 0 | NON_SYNONYMOU | MODERAT | MISSENSE | Gcc/Acc | A53T | PLES_46271 | | | | | | | A | A | | | | |
| NC_011770 | 5079033 | A | G | 63 | 0 | 8 | 0 | NON_SYNONYMOU | MODERAT | MISSENSE | Atc/Gtc | I69V | PLES_46271 | G | G | | | | | G | G | G | G | G | G |
| NC_011770 | 5079418 | C | T | 77 | 0 | 2 | 0 | NON_SYNONYMOU | MODERAT | MISSENSE | gCg/gTg | A197V | PLES_46271 | | | | | | | T | T | | | | |
| NC_011770 | 5079480 | G | T | 77 | 0 | 6 | 0 | NON_SYNONYMOU | MODERAT | MISSENSE | Gcc/Tcc | A218S | PLES_46271 | | | T | T | T | T | T | T | | | | |
| NC_011770 | 5079511 | G | A | 78 | 0 | 6 | 0 | NON_SYNONYMOU | MODERAT | MISSENSE | aGc/aAc | S228N | PLES_46271 | | | A | A | A | A | A | A | | | | |
| NC_011770 | 5080069 | G | T | 86 | 0 | 6 | 0 | NON_SYNONYMOU | MODERAT | MISSENSE | gGt/gTt | G414V | PLES_46271 | | | T | T | T | T | T | T | | | | |
| NC_011770 | 5080117 | G | A | 88 | 0 | 2 | 0 | NON_SYNONYMOU | MODERAT | MISSENSE | gGa/gAa | G430E | PLES_46271 | | | | | | | A | A | | | | |
| NC_011770 | 5080350 | T | C | 80 | 0 | 12 | 0 | NON_SYNONYMOU | MODERAT | MISSENSE | Ttc/Ctc | F43L | PLES_46281 | C | C | C | C | C | C | C | C | C | C | C | C |
| NC_011770 | 5080431 | A | G | 94 | 0 | 12 | 0 | NON_SYNONYMOU | MODERAT | MISSENSE | Atc/Gtc | I70V | PLES_46281 | G | G | G | G | G | G | G | G | G | G | G | G |
| NC_011770 | 5080471 | G | A | 100 | 0 | 1 | 0 | NON_SYNONYMOU | MODERAT | MISSENSE | gGc/gAc | G83D | PLES_46281 | | | | | | | A | | | | | |
| NC_011770 | 5080506 | G | A | 97 | 0 | 4 | 0 | NON_SYNONYMOU | MODERAT | MISSENSE | Ggc/Agc | G95S | PLES_46281 | | | A | A | A | A | | | | | | |
| NC_011770 | 5080543 | G | T | 95 | 0 | 4 | 0 | NON_SYNONYMOU | MODERAT | MISSENSE | tGg/tTg | W107L | PLES_46281 | | | T | T | T | T | | | | | | |
| NC_011770 | 5081920 | T | C | 81 | 0 | 12 | 0 | NON_SYNONYMOU | MODERAT | MISSENSE | Atg/Gtg | M306V | PLES_46291 | C | C | C | C | C | C | C | C | C | C | C | C |
| NC_011770 | 5082158 | G | C | 84 | 0 | 4 | 0 | NON_SYNONYMOU | MODERAT | MISSENSE | caC/caG | H226Q | PLES_46291 | | | | | | | | | C | C | C | C |
| NC_011770 | 5082304 | G | C | 82 | 0 | 2 | 0 | NON_SYNONYMOU | MODERAT | MISSENSE | Ccg/Gcg | P178A | PLES_46291 | C | C | | | | | | | | | | |
| NC_011770 | 5082460 | C | T | 87 | 0 | 4 | 0 | NON_SYNONYMOU | MODERAT | MISSENSE | Gtc/Atc | V126I | PLES_46291 | | | T | T | T | T | | | | | | |
| NC_011770 | 5083053 | A | G | 66 | 0 | 12 | 0 | NON_SYNONYMOU | MODERAT | MISSENSE | gTg/gCg | V73A | PLES_46301 | G | G | G | G | G | G | G | G | G | G | G | G |
| NC_011770 | 5083093 | C | T | 73 | 0 | 2 | 0 | NON_SYNONYMOU | MODERAT | MISSENSE | Gcg/Acg | A60T | PLES_46301 | | | | | | | | | | | T | T |
| NC_011770 | 5083134 | C | T | 80 | 0 | 2 | 0 | NON_SYNONYMOU | MODERAT | MISSENSE | gGc/gAc | G46D | PLES_46301 | | | | | | | T | T | | | | |
| NC_011770 | 5083259 | G | C | 84 | 0 | 6 | 0 | NON_SYNONYMOU | MODERAT | MISSENSE | atC/atG | I4M | PLES_46301 | | | C | C | C | C | | | | | C | C |
| NC_011770 | 5083260 | A | G | 82 | 0 | 6 | 0 | NON_SYNONYMOU | MODERAT | MISSENSE | aTc/aCc | I4T | PLES_46301 | | | G | G | G | G | | | | | G | G |
| NC_011770 | 5083266 | T | C | 87 | 0 | 6 | 0 | NON_SYNONYMOU | MODERAT | MISSENSE | gAc/gGc | D2G | PLES_46301 | | | C | C | C | C | | | | | C | C |
| NC_011770 | 5083487 | G | A | 85 | 0 | 1 | 0 | NON_SYNONYMOU | MODERAT | MISSENSE | cCg/cTg | P243L | PLES_46311 | | | | | | | A | | | | | |
| NC_011770 | 5083550 | A | G | 71 | 0 | 12 | 0 | NON_SYNONYMOU | MODERAT | MISSENSE | gTg/gCg | V222A | PLES_46311 | G | G | G | G | G | G | G | G | G | G | G | G |
| NC_011770 | 5084190 | T | C | 66 | 0 | 8 | 0 | NON_SYNONYMOU | MODERAT | MISSENSE | Atc/Gtc | I9V | PLES_46311 | | | C | C | C | C | C | C | | | C | C |
| NC_011770 | 5084418 | T | C | 73 | 0 | 2 | 0 | NON_SYNONYMOU | MODERAT | MISSENSE | cAg/cGg | Q94R | PLES_46321 | | | | | | | | | | | C | C |
| NC_011770 | 5084636 | G | C | 78 | 0 | 4 | 0 | NON_SYNONYMOU | MODERAT | MISSENSE | gaC/gaG | D21E | PLES_46321 | | | C | C | C | C | | | | | | |
| NC_011770 | 5084690 | G | C | 86 | 0 | 2 | 0 | NON_SYNONYMOU | MODERAT | MISSENSE | gaC/gaG | D3E | PLES_46321 | | | | | | | C | C | | | | |
| NC_011770 | 5084881 | C | T | 81 | 0 | 4 | 0 | NON_SYNONYMOU | MODERAT | MISSENSE | gGc/gAc | G170D | PLES_46331 | | | T | T | T | T | | | | | | |
| NC_011770 | 5085100 | T | C | 78 | 0 | 1 | 0 | NON_SYNONYMOU | MODERAT | MISSENSE | cAg/cGg | Q97R | PLES_46331 | | | | | | | C | | | | | |
| NC_011770 | 5085137 | T | C | 76 | 0 | 8 | 0 | NON_SYNONYMOU | MODERAT | MISSENSE | Acc/Gcc | T85A | PLES_46331 | | | C | C | C | C | C | C | | | C | C |
| NC_011770 | 5085212 | C | T | 75 | 0 | 8 | 0 | NON_SYNONYMOU | MODERAT | MISSENSE | Gcc/Acc | A60T | PLES_46331 | | | T | T | T | T | T | T | | | T | T |
| NC_011770 | 5085238 | A | G | 82 | 0 | 8 | 0 | NON_SYNONYMOU | MODERAT | MISSENSE | gTg/gCg | V51A | PLES_46331 | | | G | G | G | G | G | G | | | G | G |
| NC_011770 | 5085272 | T | C | 85 | 0 | 4 | 0 | NON_SYNONYMOU | MODERAT | MISSENSE | Acc/Gcc | T40A | PLES_46331 | | | | | | | C | C | | | C | C |
| NC_011770 | 5086938 | G | A | 71 | 0 | 10 | 0 | STOP_GAINED | HIGH | NONSENSE | Cag/Tag | Q65* | PLES_46341 | A | A | A | A | A | A | A | A | A | A | | |
| NC_011770 | 5086167 | G | T | 93 | 0 | 6 | 0 | NON_SYNONYMOU | MODERAT | MISSENSE | Ctc/Atc | L322I | PLES_46341 | | | T | T | T | T | T | T | | | | |
| NC_011770 | 5086263 | C | A | 89 | 0 | 6 | 0 | NON_SYNONYMOU | MODERAT | MISSENSE | Gcg/Tcg | A290S | PLES_46341 | | | A | A | A | A | A | A | | | | |
| NC_011770 | 5086614 | C | A | 74 | 0 | 10 | 0 | NON_SYNONYMOU | MODERAT | MISSENSE | Gcg/Tcg | A173S | PLES_46341 | A | A | A | A | A | A | A | A | A | A | | |
| NC_011770 | 5086937 | T | A | 70 | 0 | 10 | 0 | NON_SYNONYMOU | MODERAT | MISSENSE | cAg/cTg | Q65L | PLES_46341 | A | A | A | A | A | A | A | A | A | A | | |
| NC_011770 | 5087521 | C | A | 46 | 0 | 2 | 0 | NON_SYNONYMOU | MODERAT | MISSENSE | Gcc/Tcc | A142S | PLES_46351 | | | | | | | | | A | A | | |
| NC_011770 | 5087555 | G | T | 49 | 0 | 2 | 0 | NON_SYNONYMOU | MODERAT | MISSENSE | gaC/gaA | D130E | PLES_46351 | | | | | | | | | | | T | T |
| NC_011770 | 5091348 | G | A | 85 | 0 | 1 | 0 | STOP_GAINED | HIGH | NONSENSE | Cag/Tag | Q181* | PLES_46381 | | | | | | | A | | | | | |
| NC_011770 | 5090490 | C | T | 70 | 0 | 2 | 0 | NON_SYNONYMOU | MODERAT | MISSENSE | Ggc/Agc | G467S | PLES_46381 | | | | | | | | | T | T | | |
| NC_011770 | 5091762 | T | G | 88 | 0 | 4 | 0 | NON_SYNONYMOU | MODERAT | MISSENSE | Acc/Ccc | T43P | PLES_46381 | G | G | | | | | G | G | | | | |
| NC_011770 | 5092222 | T | C | 70 | 0 | 4 | 0 | NON_SYNONYMOU | MODERAT | MISSENSE | Agc/Ggc | S145G | PLES_46391 | C | C | | | | | C | C | | | | |
| NC_011770 | 5092239 | T | C | 72 | 0 | 4 | 0 | NON_SYNONYMOU | MODERAT | MISSENSE | aAt/aGt | N139S | PLES_46391 | C | C | | | | | C | C | | | | |
| NC_011770 | 5092431 | A | C | 75 | 0 | 6 | 0 | NON_SYNONYMOU | MODERAT | MISSENSE | gTc/gGc | V75G | PLES_46391 | C | C | C | C | C | C | | | | | | |
| NC_011770 | 5100886 | CA | C | 32 | 4 | 1 | 0 | FRAME_SHIFT | HIGH | | ggt/ | G1567 | PLES_46401 | | | . | . | . | . | | C | | | | |
| NC_011770 | 5093668 | TGC | T | 55 | 1 | 3 | 0 | FRAME_SHIFT | HIGH | | agc/ | S3973 | PLES_46401 | | | T | . | T | T | | | | | | |
| NC_011770 | 5094249 | GC | G | 67 | 0 | 1 | 0 | FRAME_SHIFT | HIGH | | gcc/ | A3780 | PLES_46401 | | | | | | | G | | | | | |
| NC_011770 | 5097937 | G | A | 36 | 4 | 1 | 0 | STOP_GAINED | HIGH | NONSENSE | Cag/Tag | Q2551* | PLES_46401 | | | . | . | . | . | A | | | | | |
| NC_011770 | 5103635 | G | GC | 44 | 0 | 4 | 0 | FRAME_SHIFT | HIGH | | cac/caGc | H651Q? | PLES_46401 | | | GC | GC | GC | GC | | | | | | |
| NC_011770 | 5103638 | TG | T | 44 | 0 | 4 | 0 | FRAME_SHIFT | HIGH | | gca/ | A650 | PLES_46401 | | | T | T | T | T | | | | | | |
| NC_011770 | 5103740 | G | C | 57 | 0 | 4 | 0 | STOP_GAINED | HIGH | NONSENSE | taC/taG | Y616* | PLES_46401 | | | C | C | C | C | | | | | | |
| NC_011770 | 5105412 | GCAGCGC | G | 44 | 0 | 1 | 0 | CODON_CHANGE_ | MODERATE | | ggcgctgcc/ | GAA57A | PLES_46401 | | | G | | | | | | | | | |
| NC_011770 | 5093620 | G | T | 77 | 0 | 4 | 0 | NON_SYNONYMOU | MODERAT | MISSENSE | Cag/Aag | Q3990K | PLES_46401 | | | T | T | T | T | | | | | | |
| NC_011770 | 5093622 | G | C | 76 | 0 | 4 | 0 | NON_SYNONYMOU | MODERAT | MISSENSE | tCc/tGc | S3989C | PLES_46401 | | | C | C | C | C | | | | | | |
| NC_011770 | 5093623 | A | T | 76 | 0 | 4 | 0 | NON_SYNONYMOU | MODERAT | MISSENSE | Tcc/Acc | S3989T | PLES_46401 | | | T | T | T | T | | | | | | |
| NC_011770 | 5093690 | G | C | 55 | 0 | 4 | 0 | NON_SYNONYMOU | MODERAT | MISSENSE | atC/atG | I3966M | PLES_46401 | | | C | C | C | C | | | | | | |
| NC_011770 | 5093692 | T | C | 54 | 0 | 4 | 0 | NON_SYNONYMOU | MODERAT | MISSENSE | Atc/Gtc | I3966V | PLES_46401 | | | C | C | C | C | | | | | | |
| NC_011770 | 5093740 | T | C | 56 | 4 | 4 | 0 | NON_SYNONYMOU | MODERAT | MISSENSE | Aat/Gat | N3950D | PLES_46401 | | | . | . | . | . | | | C | C | C | C |

| | | | | | | | | | | | | | | | | | | | | | | | | | |
|---|---|---|---|---|---|---|---|---|---|---|---|---|---|---|---|---|---|---|---|---|---|---|---|---|---|
| NC_011770 | 5093808 | C | T | 82 | 0 | 4 | 0 | NON_SYNONYMOU | MODERAT | MISSENSE | cGc/cAc | R3927H | PLES_46401 | | | T | T | T | T | | | | | | |
| NC_011770 | 5093820 | C | G | 81 | 0 | 4 | 0 | NON_SYNONYMOU | MODERAT | MISSENSE | aGc/aCc | S3923T | PLES_46401 | | | G | G | G | G | | | | | | |
| NC_011770 | 5093848 | C | T | 85 | 0 | 4 | 0 | NON_SYNONYMOU | MODERAT | MISSENSE | Gac/Aac | D3914N | PLES_46401 | | | T | T | T | T | | | | | | |
| NC_011770 | 5094097 | G | C | 70 | 0 | 12 | 0 | NON_SYNONYMOU | MODERAT | MISSENSE | Ccg/Gcg | P3831A | PLES_46401 | C | C | C | C | C | C | C | C | C | C | C | C |
| NC_011770 | 5096650 | T | C | 44 | 4 | 8 | 0 | NON_SYNONYMOU | MODERAT | MISSENSE | Agt/Ggt | S2980G | PLES_46401 | C | C | . | . | . | . | C | C | C | C | C | C |
| NC_011770 | 5096901 | A | G | 47 | 4 | 6 | 0 | NON_SYNONYMOU | MODERAT | MISSENSE | aTc/aCc | I2896T | PLES_46401 | | | . | . | . | . | G | G | G | G | G | G |
| NC_011770 | 5097000 | A | G | 42 | 4 | 6 | 0 | NON_SYNONYMOU | MODERAT | MISSENSE | gTc/gCc | V2863A | PLES_46401 | | | . | . | . | . | G | G | G | G | G | G |
| NC_011770 | 5097120 | G | A | 44 | 0 | 4 | 0 | NON_SYNONYMOU | MODERAT | MISSENSE | aCc/aTc | T2823I | PLES_46401 | | | A | A | A | A | | | | | | |
| NC_011770 | 5097123 | C | G | 44 | 0 | 4 | 0 | NON_SYNONYMOU | MODERAT | MISSENSE | gGc/gCc | G2822A | PLES_46401 | | | G | G | G | G | | | | | | |
| NC_011770 | 5097137 | G | T | 43 | 0 | 4 | 0 | NON_SYNONYMOU | MODERAT | MISSENSE | gaC/gaA | D2817E | PLES_46401 | | | T | T | T | T | | | | | | |
| NC_011770 | 5097141 | T | A | 43 | 0 | 4 | 0 | NON_SYNONYMOU | MODERAT | MISSENSE | tAc/tTc | Y2816F | PLES_46401 | | | A | A | A | A | | | | | | |
| NC_011770 | 5097294 | T | G | 47 | 0 | 4 | 0 | NON_SYNONYMOU | MODERAT | MISSENSE | aAc/aCc | N2765T | PLES_46401 | | | G | G | G | G | | | | | | |
| NC_011770 | 5097298 | T | C | 48 | 0 | 4 | 0 | NON_SYNONYMOU | MODERAT | MISSENSE | Atc/Gtc | I2764V | PLES_46401 | | | C | C | C | C | | | | | | |
| NC_011770 | 5097310 | G | T | 48 | 0 | 4 | 0 | NON_SYNONYMOU | MODERAT | MISSENSE | Ctc/Atc | L2760I | PLES_46401 | | | T | T | T | T | | | | | | |
| NC_011770 | 5097313 | T | A | 45 | 0 | 2 | 0 | NON_SYNONYMOU | MODERAT | MISSENSE | Aac/Tac | N2759Y | PLES_46401 | | | | | | | A | A | | | | |
| NC_011770 | 5097409 | A | G | 51 | 0 | 4 | 0 | NON_SYNONYMOU | MODERAT | MISSENSE | Tac/Cac | Y2727H | PLES_46401 | | | G | G | G | G | | | | | | |
| NC_011770 | 5097475 | C | G | 64 | 0 | 4 | 0 | NON_SYNONYMOU | MODERAT | MISSENSE | Gtg/Ctg | V2705L | PLES_46401 | | | G | G | G | G | | | | | | |
| NC_011770 | 5097497 | A | T | 63 | 0 | 4 | 0 | NON_SYNONYMOU | MODERAT | MISSENSE | gaT/gaA | D2697E | PLES_46401 | | | T | T | T | T | | | | | | |
| NC_011770 | 5097516 | G | T | 65 | 0 | 4 | 0 | NON_SYNONYMOU | MODERAT | MISSENSE | aCc/aAc | T2691N | PLES_46401 | | | T | T | T | T | | | | | | |
| NC_011770 | 5097854 | G | C | 42 | 0 | 6 | 0 | NON_SYNONYMOU | MODERAT | MISSENSE | atC/atG | I2578M | PLES_46401 | | | | | | | C | C | C | C | C | C |
| NC_011770 | 5097868 | T | C | 35 | 4 | 6 | 0 | NON_SYNONYMOU | MODERAT | MISSENSE | Agc/Ggc | S2574G | PLES_46401 | | | . | . | . | . | C | C | C | C | C | C |
| NC_011770 | 5098028 | C | A | 46 | 0 | 4 | 0 | NON_SYNONYMOU | MODERAT | MISSENSE | gaG/gaT | E2520D | PLES_46401 | | | A | A | A | A | | | | | | |
| NC_011770 | 5098029 | T | C | 45 | 0 | 4 | 0 | NON_SYNONYMOU | MODERAT | MISSENSE | gAg/gGg | E2520G | PLES_46401 | | | C | C | C | C | | | | | | |
| NC_011770 | 5098030 | C | T | 48 | 0 | 4 | 0 | NON_SYNONYMOU | MODERAT | MISSENSE | Gag/Aag | E2520K | PLES_46401 | | | T | T | T | T | | | | | | |
| NC_011770 | 5098041 | G | T | 45 | 0 | 4 | 0 | NON_SYNONYMOU | MODERAT | MISSENSE | gCg/gAg | A2516E | PLES_46401 | | | T | T | T | T | | | | | | |
| NC_011770 | 5098207 | T | C | 44 | 4 | 4 | 0 | NON_SYNONYMOU | MODERAT | MISSENSE | Acc/Gcc | T2461A | PLES_46401 | | | . | . | . | . | | | C | C | C | C |
| NC_011770 | 5098348 | C | T | 40 | 4 | 4 | 0 | NON_SYNONYMOU | MODERAT | MISSENSE | Gtg/Atg | V2414M | PLES_46401 | | | . | . | . | . | | | T | T | T | T |
| NC_011770 | 5098797 | C | T | 47 | 4 | 4 | 0 | NON_SYNONYMOU | MODERAT | MISSENSE | cGc/cAc | R2264H | PLES_46401 | | | . | . | . | . | | | T | T | T | T |
| NC_011770 | 5098917 | C | T | 62 | 0 | 2 | 0 | NON_SYNONYMOU | MODERAT | MISSENSE | cGc/cAc | R2224H | PLES_46401 | T | T | | | | | | | | | | |
| NC_011770 | 5099023 | C | T | 46 | 4 | 4 | 0 | NON_SYNONYMOU | MODERAT | MISSENSE | Gcc/Acc | A2189T | PLES_46401 | | | . | . | . | . | | | T | T | T | T |
| NC_011770 | 5099198 | T | C | 44 | 4 | 4 | 0 | NON_SYNONYMOU | MODERAT | MISSENSE | atA/atG | I2130M | PLES_46401 | | | . | . | . | . | | | C | C | C | C |
| NC_011770 | 5100225 | A | G | 44 | 4 | 8 | 0 | NON_SYNONYMOU | MODERAT | MISSENSE | tTc/tCc | F1788S | PLES_46401 | G | G | . | . | . | . | G | G | G | G | G | G |
| NC_011770 | 5100676 | T | C | 38 | 4 | 2 | 0 | NON_SYNONYMOU | MODERAT | MISSENSE | Aca/Gca | T1638A | PLES_46401 | | | . | . | . | . | | | C | C | | |
| NC_011770 | 5101012 | C | T | 32 | 4 | 2 | 0 | NON_SYNONYMOU | MODERAT | MISSENSE | Ggg/Agg | G1526R | PLES_46401 | | | . | . | . | . | | | T | T | | |
| NC_011770 | 5101498 | C | T | 42 | 4 | 8 | 0 | NON_SYNONYMOU | MODERAT | MISSENSE | Gac/Aac | D1364N | PLES_46401 | T | T | . | . | . | . | T | T | T | T | T | T |
| NC_011770 | 5101846 | C | T | 48 | 4 | 2 | 0 | NON_SYNONYMOU | MODERAT | MISSENSE | Gcc/Acc | A1248T | PLES_46401 | | | . | . | . | . | | | T | T | | |
| NC_011770 | 5102038 | A | C | 46 | 4 | 2 | 0 | NON_SYNONYMOU | MODERAT | MISSENSE | Ttg/Gtg | L1184V | PLES_46401 | | | . | . | . | . | | | C | C | | |
| NC_011770 | 5102202 | G | A | 48 | 4 | 6 | 0 | NON_SYNONYMOU | MODERAT | MISSENSE | aCt/aTt | T1129I | PLES_46401 | A | A | . | . | . | . | A | A | | | A | A |
| NC_011770 | 5102374 | C | T | 47 | 4 | 2 | 0 | NON_SYNONYMOU | MODERAT | MISSENSE | Gtc/Atc | V1072I | PLES_46401 | | | . | . | . | . | | | T | T | | |
| NC_011770 | 5102415 | A | G | 45 | 4 | 2 | 0 | NON_SYNONYMOU | MODERAT | MISSENSE | aTa/aCa | I1058T | PLES_46401 | | | . | . | . | . | | | G | G | | |
| NC_011770 | 5102616 | T | C | 43 | 4 | 6 | 0 | NON_SYNONYMOU | MODERAT | MISSENSE | gAc/gGc | D991G | PLES_46401 | C | C | . | . | . | . | C | C | | | C | C |
| NC_011770 | 5102851 | C | T | 52 | 0 | 4 | 0 | NON_SYNONYMOU | MODERAT | MISSENSE | Ggc/Agc | G913S | PLES_46401 | | | T | T | T | T | | | | | | |
| NC_011770 | 5102979 | G | T | 44 | 4 | 4 | 0 | NON_SYNONYMOU | MODERAT | MISSENSE | gCc/gAc | A870D | PLES_46401 | | | . | . | . | . | T | T | T | T | | |
| NC_011770 | 5102992 | C | A | 44 | 4 | 2 | 0 | NON_SYNONYMOU | MODERAT | MISSENSE | Gcc/Tcc | A866S | PLES_46401 | | | . | . | . | . | A | A | | | | |
| NC_011770 | 5103145 | T | C | 40 | 4 | 4 | 0 | NON_SYNONYMOU | MODERAT | MISSENSE | Agc/Ggc | S815G | PLES_46401 | | | . | . | . | . | C | C | C | C | | |
| NC_011770 | 5103282 | A | G | 42 | 4 | 4 | 0 | NON_SYNONYMOU | MODERAT | MISSENSE | gTg/gCg | V769A | PLES_46401 | | | . | . | . | . | G | G | G | G | | |
| NC_011770 | 5103292 | T | C | 43 | 4 | 8 | 0 | NON_SYNONYMOU | MODERAT | MISSENSE | Acc/Gcc | T766A | PLES_46401 | C | C | . | . | . | . | C | C | C | C | C | C |
| NC_011770 | 5103415 | C | T | 43 | 4 | 2 | 0 | NON_SYNONYMOU | MODERAT | MISSENSE | Gcc/Acc | A725T | PLES_46401 | T | T | . | . | . | . | | | | | | |
| NC_011770 | 5103558 | C | A | 38 | 4 | 2 | 0 | NON_SYNONYMOU | MODERAT | MISSENSE | aGc/aTc | S677I | PLES_46401 | | | . | . | . | . | A | A | | | | |
| NC_011770 | 5103620 | G | C | 52 | 0 | 4 | 0 | NON_SYNONYMOU | MODERAT | MISSENSE | gaC/gaG | D656E | PLES_46401 | | | C | C | C | C | | | | | | |
| NC_011770 | 5103621 | T | C | 51 | 0 | 4 | 0 | NON_SYNONYMOU | MODERAT | MISSENSE | gAc/gGc | D656G | PLES_46401 | | | C | C | C | C | | | | | | |
| NC_011770 | 5103630 | T | G | 41 | 0 | 4 | 0 | NON_SYNONYMOU | MODERAT | MISSENSE | gAg/gCg | E653A | PLES_46401 | | | G | G | G | G | | | | | | |
| NC_011770 | 5103631 | C | G | 41 | 0 | 4 | 0 | NON_SYNONYMOU | MODERAT | MISSENSE | Gag/Cag | E653Q | PLES_46401 | | | G | G | G | G | | | | | | |
| NC_011770 | 5103662 | G | C | 52 | 0 | 4 | 0 | NON_SYNONYMOU | MODERAT | MISSENSE | gaC/gaG | D642E | PLES_46401 | | | C | C | C | C | | | | | | |
| NC_011770 | 5103687 | G | C | 44 | 0 | 4 | 0 | NON_SYNONYMOU | MODERAT | MISSENSE | tCg/tGg | S634W | PLES_46401 | | | C | C | C | C | | | | | | |
| NC_011770 | 5103688 | A | G | 43 | 0 | 4 | 0 | NON_SYNONYMOU | MODERAT | MISSENSE | Tcg/Ccg | S634P | PLES_46401 | | | G | G | G | G | | | | | | |
| NC_011770 | 5103690 | C | G | 45 | 0 | 4 | 0 | NON_SYNONYMOU | MODERAT | MISSENSE | gGc/gCc | G633A | PLES_46401 | | | G | G | G | G | | | | | | |
| NC_011770 | 5103691 | C | A | 44 | 0 | 4 | 0 | NON_SYNONYMOU | MODERAT | MISSENSE | Ggc/Tgc | G633C | PLES_46401 | | | A | A | A | A | | | | | | |
| NC_011770 | 5103694 | G | C | 45 | 0 | 4 | 0 | NON_SYNONYMOU | MODERAT | MISSENSE | Ctc/Gtc | L632V | PLES_46401 | | | C | C | C | C | | | | | | |
| NC_011770 | 5103714 | G | C | 53 | 0 | 4 | 0 | NON_SYNONYMOU | MODERAT | MISSENSE | tCg/tGg | S625W | PLES_46401 | | | C | C | C | C | | | | | | |
| NC_011770 | 5103715 | A | T | 53 | 0 | 4 | 0 | NON_SYNONYMOU | MODERAT | MISSENSE | Tcg/Acg | S625T | PLES_46401 | | | T | T | T | T | | | | | | |
| NC_011770 | 5103741 | T | A | 58 | 0 | 4 | 0 | NON_SYNONYMOU | MODERAT | MISSENSE | tAc/tTc | Y616F | PLES_46401 | | | A | A | A | A | | | | | | |

| | | | | | | | | | | | | | | | | | | | | | | | | | |
|---|---|---|---|---|---|---|---|---|---|---|---|---|---|---|---|---|---|---|---|---|---|---|---|---|---|
| NC_011770 | 5103742 | A | G | 57 | 0 | 4 | 0 | NON_SYNONYMOU | MODERAT | MISSENSE | Tac/Cac | Y616H | PLES_46401 | | | G | G | G | G | | | | | | |
| NC_011770 | 5103864 | C | T | 49 | 0 | 4 | 0 | NON_SYNONYMOU | MODERAT | MISSENSE | gGc/gAc | G575D | PLES_46401 | | | T | T | T | T | | | | | | |
| NC_011770 | 5103865 | C | T | 48 | 0 | 4 | 0 | NON_SYNONYMOU | MODERAT | MISSENSE | Ggc/Agc | G575S | PLES_46401 | | | T | T | T | T | | | | | | |
| NC_011770 | 5103866 | C | G | 49 | 0 | 4 | 0 | NON_SYNONYMOU | MODERAT | MISSENSE | caG/caC | Q574H | PLES_46401 | | | G | G | G | G | | | | | | |
| NC_011770 | 5103867 | T | C | 48 | 0 | 4 | 0 | NON_SYNONYMOU | MODERAT | MISSENSE | cAg/cGg | Q574R | PLES_46401 | | | C | C | C | C | | | | | | |
| NC_011770 | 5103889 | C | G | 46 | 0 | 4 | 0 | NON_SYNONYMOU | MODERAT | MISSENSE | Gtc/Ctc | V567L | PLES_46401 | | | G | G | G | G | | | | | | |
| NC_011770 | 5103924 | C | G | 44 | 0 | 4 | 0 | NON_SYNONYMOU | MODERAT | MISSENSE | aGc/aCc | S555T | PLES_46401 | | | G | G | G | G | | | | | | |
| NC_011770 | 5103954 | C | A | 38 | 1 | 3 | 0 | NON_SYNONYMOU | MODERAT | MISSENSE | cGg/cTg | R545L | PLES_46401 | | | A | . | A | A | | | | | | |
| NC_011770 | 5103962 | G | T | 37 | 1 | 3 | 0 | NON_SYNONYMOU | MODERAT | MISSENSE | gaC/gaA | D542E | PLES_46401 | | | T | . | T | T | | | | | | |
| NC_011770 | 5103969 | A | G | 34 | 1 | 3 | 0 | NON_SYNONYMOU | MODERAT | MISSENSE | gTg/gCg | V540A | PLES_46401 | | | G | . | G | G | | | | | | |
| NC_011770 | 5103986 | C | G | 36 | 1 | 3 | 0 | NON_SYNONYMOU | MODERAT | MISSENSE | caG/caC | Q534H | PLES_46401 | | | G | . | G | G | | | | | | |
| NC_011770 | 5103988 | G | T | 36 | 1 | 3 | 0 | NON_SYNONYMOU | MODERAT | MISSENSE | Cag/Aag | Q534K | PLES_46401 | | | T | . | T | T | | | | | | |
| NC_011770 | 5103993 | T | G | 34 | 1 | 3 | 0 | NON_SYNONYMOU | MODERAT | MISSENSE | aAc/aCc | N532T | PLES_46401 | | | G | . | G | G | | | | | | |
| NC_011770 | 5103994 | T | C | 34 | 1 | 3 | 0 | NON_SYNONYMOU | MODERAT | MISSENSE | Aac/Gac | N532D | PLES_46401 | | | C | . | C | C | | | | | | |
| NC_011770 | 5104007 | C | G | 51 | 0 | 4 | 0 | NON_SYNONYMOU | MODERAT | MISSENSE | gaG/gaC | E527D | PLES_46401 | | | G | G | G | G | | | | | | |
| NC_011770 | 5104021 | A | G | 52 | 0 | 4 | 0 | NON_SYNONYMOU | MODERAT | MISSENSE | Tcg/Ccg | S523P | PLES_46401 | | | G | G | G | G | | | | | | |
| NC_011770 | 5104044 | C | T | 48 | 0 | 4 | 0 | NON_SYNONYMOU | MODERAT | MISSENSE | cGc/cAc | R515H | PLES_46401 | | | T | T | T | T | | | | | | |
| NC_011770 | 5104054 | C | T | 49 | 0 | 4 | 0 | NON_SYNONYMOU | MODERAT | MISSENSE | Gtc/Atc | V512I | PLES_46401 | | | T | T | T | T | | | | | | |
| NC_011770 | 5104057 | T | G | 48 | 0 | 4 | 0 | NON_SYNONYMOU | MODERAT | MISSENSE | Aag/Cag | K511Q | PLES_46401 | | | G | G | G | G | | | | | | |
| NC_011770 | 5104062 | C | T | 47 | 0 | 10 | 0 | NON_SYNONYMOU | MODERAT | MISSENSE | aGc/aAc | S509N | PLES_46401 | T | T | T | T | T | T | T | T | | | T | T |
| NC_011770 | 5104068 | G | C | 47 | 0 | 4 | 0 | NON_SYNONYMOU | MODERAT | MISSENSE | aCc/aGc | T507S | PLES_46401 | | | C | C | C | C | | | | | | |
| NC_011770 | 5104084 | G | T | 28 | 2 | 2 | 0 | NON_SYNONYMOU | MODERAT | MISSENSE | Cag/Aag | Q502K | PLES_46401 | | | T | . | T | . | | | | | | |
| NC_011770 | 5104239 | T | A | 22 | 4 | 2 | 0 | NON_SYNONYMOU | MODERAT | MISSENSE | gAc/gTc | D450V | PLES_46401 | | | . | . | . | . | | | A | A | | |
| NC_011770 | 5104581 | G | C | 48 | 0 | 4 | 0 | NON_SYNONYMOU | MODERAT | MISSENSE | aCc/aGc | T336S | PLES_46401 | | | C | C | C | C | | | | | | |
| NC_011770 | 5104582 | T | G | 46 | 0 | 4 | 0 | NON_SYNONYMOU | MODERAT | MISSENSE | Acc/Ccc | T336P | PLES_46401 | | | G | G | G | G | | | | | | |
| NC_011770 | 5104602 | T | C | 32 | 4 | 2 | 0 | NON_SYNONYMOU | MODERAT | MISSENSE | cAg/cGg | Q329R | PLES_46401 | | | . | . | . | . | | | | | C | C |
| NC_011770 | 5104623 | C | T | 30 | 4 | 2 | 0 | NON_SYNONYMOU | MODERAT | MISSENSE | aGc/aAc | S322N | PLES_46401 | T | T | . | . | . | . | | | | | | |
| NC_011770 | 5104679 | G | C | 52 | 0 | 4 | 0 | NON_SYNONYMOU | MODERAT | MISSENSE | gaC/gaG | D303E | PLES_46401 | | | C | C | C | C | | | | | | |
| NC_011770 | 5104891 | C | T | 56 | 0 | 4 | 0 | NON_SYNONYMOU | MODERAT | MISSENSE | Gac/Aac | D233N | PLES_46401 | | | T | T | T | T | | | | | | |
| NC_011770 | 5104908 | G | C | 59 | 0 | 4 | 0 | NON_SYNONYMOU | MODERAT | MISSENSE | tCc/tGc | S227C | PLES_46401 | | | C | C | C | C | | | | | | |
| NC_011770 | 5104909 | A | T | 59 | 0 | 4 | 0 | NON_SYNONYMOU | MODERAT | MISSENSE | Tcc/Acc | S227T | PLES_46401 | | | T | T | T | T | | | | | | |
| NC_011770 | 5104932 | A | G | 66 | 0 | 4 | 0 | NON_SYNONYMOU | MODERAT | MISSENSE | gTc/gCc | V219A | PLES_46401 | | | G | G | G | G | | | | | | |
| NC_011770 | 5104936 | G | C | 69 | 0 | 4 | 0 | NON_SYNONYMOU | MODERAT | MISSENSE | Ctg/Gtg | L218V | PLES_46401 | | | C | C | C | C | | | | | | |
| NC_011770 | 5105014 | G | C | 65 | 0 | 4 | 0 | NON_SYNONYMOU | MODERAT | MISSENSE | Ctg/Gtg | L192V | PLES_46401 | | | C | C | C | C | | | | | | |
| NC_011770 | 5105130 | G | C | 40 | 4 | 4 | 0 | NON_SYNONYMOU | MODERAT | MISSENSE | gCc/gGc | A153G | PLES_46401 | | | . | . | . | . | C | C | C | C | | |
| NC_011770 | 5105134 | C | T | 41 | 4 | 2 | 0 | NON_SYNONYMOU | MODERAT | MISSENSE | Gcc/Acc | A152T | PLES_46401 | T | T | . | . | . | . | | | | | | |
| NC_011770 | 5105230 | C | T | 51 | 0 | 4 | 0 | NON_SYNONYMOU | MODERAT | MISSENSE | Gtt/Att | V120I | PLES_46401 | | | T | T | T | T | | | | | | |
| NC_011770 | 5105235 | T | G | 42 | 3 | 1 | 0 | NON_SYNONYMOU | MODERAT | MISSENSE | cAg/cCg | Q118P | PLES_46401 | | | . | . | . | G | | | | | | |
| NC_011770 | 5105319 | G | T | 61 | 0 | 4 | 0 | NON_SYNONYMOU | MODERAT | MISSENSE | gCc/gAc | A90D | PLES_46401 | | | T | T | T | T | | | | | | |
| NC_011770 | 5105320 | C | G | 62 | 0 | 4 | 0 | NON_SYNONYMOU | MODERAT | MISSENSE | Gcc/Ccc | A90P | PLES_46401 | | | G | G | G | G | | | | | | |
| NC_011770 | 5105323 | T | G | 61 | 0 | 4 | 0 | NON_SYNONYMOU | MODERAT | MISSENSE | Atg/Ctg | M89L | PLES_46401 | | | G | G | G | G | | | | | | |
| NC_011770 | 5105349 | C | T | 62 | 0 | 4 | 0 | NON_SYNONYMOU | MODERAT | MISSENSE | cGg/cAg | R80Q | PLES_46401 | | | T | T | T | T | | | | | | |
| NC_011770 | 5105377 | C | T | 43 | 0 | 4 | 0 | NON_SYNONYMOU | MODERAT | MISSENSE | Gcg/Acg | A71T | PLES_46401 | | | T | T | T | T | | | | | | |
| NC_011770 | 5105378 | C | CATG | 45 | 0 | 4 | 0 | CODON_CHANGE_ | MODERATE | | ggg/ggCAT | G70GM | PLES_46401 | | | CATG | CATG | CATG | CATG | | | | | | |
| NC_011770 | 5105404 | G | C | 55 | 0 | 4 | 0 | NON_SYNONYMOU | MODERAT | MISSENSE | Ccg/Gcg | P62A | PLES_46401 | | | C | C | C | C | | | | | | |
| NC_011770 | 5105427 | G | T | 38 | 1 | 3 | 0 | NON_SYNONYMOU | MODERAT | MISSENSE | gCg/gAg | A54E | PLES_46401 | | | T | . | T | T | | | | | | |
| NC_011770 | 5105428 | C | G | 38 | 1 | 3 | 0 | NON_SYNONYMOU | MODERAT | MISSENSE | Gcg/Ccg | A54P | PLES_46401 | | | G | . | G | G | | | | | | |
| NC_011770 | 5105541 | T | G | 40 | 4 | 2 | 0 | NON_SYNONYMOU | MODERAT | MISSENSE | tAc/tCc | Y16S | PLES_46401 | G | G | . | . | . | . | | | | | | |
| NC_011770 | 5106661 | GGTA | G | 38 | 6 | 2 | 0 | CODON_CHANGE_ | MODERATE | | actacc/acc | TT73T | PLES_46411 | . | . | . | . | . | . | G | G | | | | |
| NC_011770 | 5107155 | G | C | 41 | 4 | 2 | 0 | NON_SYNONYMOU | MODERAT | MISSENSE | aCc/aGc | T307S | PLES_46421 | C | C | . | . | . | . | | | | | | |
| NC_011770 | 5107227 | C | G | 42 | 4 | 2 | 0 | NON_SYNONYMOU | MODERAT | MISSENSE | aGc/aCc | S283T | PLES_46421 | G | G | . | . | . | . | | | | | | |
| NC_011770 | 5107237 | T | C | 43 | 4 | 4 | 0 | NON_SYNONYMOU | MODERAT | MISSENSE | Acc/Gcc | T280A | PLES_46421 | C | C | . | . | . | . | | | | | C | C |
| NC_011770 | 5107281 | T | C | 42 | 4 | 2 | 0 | NON_SYNONYMOU | MODERAT | MISSENSE | aAc/aGc | N265S | PLES_46421 | C | C | . | . | . | . | | | | | | |
| NC_011770 | 5107480 | T | C | 45 | 4 | 2 | 0 | NON_SYNONYMOU | MODERAT | MISSENSE | Acg/Gcg | T199A | PLES_46421 | C | C | . | . | . | . | | | | | | |
| NC_011770 | 5107543 | T | C | 44 | 4 | 2 | 0 | NON_SYNONYMOU | MODERAT | MISSENSE | Atc/Gtc | I178V | PLES_46421 | C | C | . | . | . | . | | | | | | |
| NC_011770 | 5107779 | C | T | 45 | 4 | 2 | 0 | NON_SYNONYMOU | MODERAT | MISSENSE | gGc/gAc | G99D | PLES_46421 | | | . | . | . | . | T | T | | | | |
| NC_011770 | 5107780 | C | T | 46 | 4 | 2 | 0 | NON_SYNONYMOU | MODERAT | MISSENSE | Ggc/Agc | G99S | PLES_46421 | | | . | . | . | . | T | T | | | | |
| NC_011770 | 5107788 | T | C | 46 | 4 | 2 | 0 | NON_SYNONYMOU | MODERAT | MISSENSE | aAc/aGc | N96S | PLES_46421 | C | C | . | . | . | . | | | | | | |
| NC_011770 | 5107791 | C | G | 45 | 4 | 4 | 0 | NON_SYNONYMOU | MODERAT | MISSENSE | aGc/aCc | S95T | PLES_46421 | G | G | . | . | . | . | G | G | | | | |
| NC_011770 | 5107800 | G | T | 46 | 4 | 2 | 0 | NON_SYNONYMOU | MODERAT | MISSENSE | aCc/aAc | T92N | PLES_46421 | T | T | . | . | . | . | | | | | | |
| NC_011770 | 5107803 | G | C | 44 | 4 | 2 | 0 | NON_SYNONYMOU | MODERAT | MISSENSE | aCc/aGc | T91S | PLES_46421 | C | C | . | . | . | . | | | | | | |
| NC_011770 | 5107804 | T | A | 40 | 4 | 2 | 0 | NON_SYNONYMOU | MODERAT | MISSENSE | Acc/Tcc | T91S | PLES_46421 | | | . | . | . | . | A | A | | | | |

| | | | | | | | | | | | | | | | | | | | | | | | | | |
|---|---|---|---|---|---|---|---|---|---|---|---|---|---|---|---|---|---|---|---|---|---|---|---|---|---|
| NC_011770 | 5107812 | C | G | 39 | 4 | 2 | 0 | NON_SYNONYMOU | MODERAT | MISSENSE | aGc/aCc | S88T | PLES_46421 | | | . | . | . | . | G | G | | | | |
| NC_011770 | 5107813 | T | A | 38 | 4 | 2 | 0 | NON_SYNONYMOU | MODERAT | MISSENSE | Agc/Tgc | S88C | PLES_46421 | | | . | . | . | . | A | A | | | | |
| NC_011770 | 5107816 | C | T | 39 | 4 | 4 | 0 | NON_SYNONYMOU | MODERAT | MISSENSE | Ggt/Agt | G87S | PLES_46421 | T | T | . | . | . | . | T | T | | | | |
| NC_011770 | 5108001 | G | T | 64 | 0 | 4 | 0 | NON_SYNONYMOU | MODERAT | MISSENSE | aCc/aAc | T25N | PLES_46421 | | | T | T | T | T | | | | | | |
| NC_011770 | 5108004 | A | ATTT | 64 | 0 | 4 | 0 | CODON_INSERTIOI | MODERATE | | gtc/gAAAt | V24EI | PLES_46421 | | | ATTT | ATTT | ATTT | ATTT | | | | | | |
| NC_011770 | 5109106 | T | C | 79 | 0 | 6 | 0 | NON_SYNONYMOU | MODERAT | MISSENSE | Agc/Ggc | S233G | PLES_46431 | | | C | C | C | C | | | C | C | | |
| NC_011770 | 5109655 | T | A | 56 | 0 | 2 | 0 | NON_SYNONYMOU | MODERAT | MISSENSE | Acc/Tcc | T50S | PLES_46431 | A | A | | | | | | | | | | |
| NC_011770 | 5110713 | G | T | 83 | 0 | 2 | 0 | NON_SYNONYMOU | MODERAT | MISSENSE | Ctg/Atg | L167M | PLES_46441 | | | | | | | T | T | | | | |
| NC_011770 | 5111078 | G | A | 81 | 0 | 6 | 0 | NON_SYNONYMOU | MODERAT | MISSENSE | cCg/cTg | P45L | PLES_46441 | | | A | A | A | A | | | A | A | | |
| NC_011770 | 5111300 | C | T | 75 | 0 | 2 | 0 | NON_SYNONYMOU | MODERAT | MISSENSE | Gcg/Acg | A774T | PLES_46451 | T | T | | | | | | | | | | |
| NC_011770 | 5111333 | C | T | 80 | 0 | 2 | 0 | NON_SYNONYMOU | MODERAT | MISSENSE | Gtc/Atc | V763I | PLES_46451 | | | | | | | T | T | | | | |
| NC_011770 | 5111350 | G | A | 80 | 0 | 2 | 0 | NON_SYNONYMOU | MODERAT | MISSENSE | cCg/cTg | P757L | PLES_46451 | | | | | A | A | | | | | | |
| NC_011770 | 5112354 | C | T | 63 | 0 | 2 | 0 | NON_SYNONYMOU | MODERAT | MISSENSE | atG/atA | M422I | PLES_46451 | | | | | | | | | T | T | | |
| NC_011770 | 5112430 | C | G | 60 | 0 | 4 | 0 | NON_SYNONYMOU | MODERAT | MISSENSE | aGc/aCc | S397T | PLES_46451 | | | G | G | G | G | | | | | | |
| NC_011770 | 5112467 | C | T | 65 | 0 | 6 | 0 | NON_SYNONYMOU | MODERAT | MISSENSE | Ggc/Agc | G385S | PLES_46451 | | | T | T | T | T | | | T | T | | |
| NC_011770 | 5113343 | T | C | 76 | 0 | 4 | 0 | NON_SYNONYMOU | MODERAT | MISSENSE | Aca/Gca | T93A | PLES_46451 | | | | | | | C | C | C | C | | |
| NC_011770 | 5113363 | C | G | 76 | 0 | 8 | 0 | NON_SYNONYMOU | MODERAT | MISSENSE | gGc/gCc | G86A | PLES_46451 | | | G | G | G | G | G | G | G | G | | |
| NC_011770 | 5113683 | A | G | 74 | 0 | 6 | 0 | NON_SYNONYMOU | MODERAT | MISSENSE | cTg/cCg | L176P | PLES_46461 | G | G | | | | | G | G | G | G | | |
| NC_011770 | 5114041 | T | C | 71 | 0 | 6 | 0 | NON_SYNONYMOU | MODERAT | MISSENSE | Atc/Gtc | I57V | PLES_46461 | | | | | | | C | C | C | C | C | C |
| NC_011770 | 5114176 | T | C | 66 | 0 | 2 | 0 | NON_SYNONYMOU | MODERAT | MISSENSE | Aat/Gat | N12D | PLES_46461 | | | | | | | | | C | C | | |
| NC_011770 | 5115991 | G | A | 84 | 0 | 1 | 0 | NON_SYNONYMOU | MODERAT | MISSENSE | Cgc/Tgc | R127C | PLES_46471 | | | | | | | | | | | | A |
| NC_011770 | 5115421 | T | C | 80 | 0 | 4 | 0 | NON_SYNONYMOU | MODERAT | MISSENSE | Atg/Gtg | M317V | PLES_46471 | C | C | | | | | | | | | C | C |
| NC_011770 | 5115496 | G | C | 84 | 0 | 2 | 0 | NON_SYNONYMOU | MODERAT | MISSENSE | Cgg/Ggg | R292G | PLES_46471 | | | | | | | | | C | C | | |
| NC_011770 | 5115759 | G | A | 74 | 0 | 2 | 0 | NON_SYNONYMOU | MODERAT | MISSENSE | aCc/aTc | T204I | PLES_46471 | | | | | | | A | A | | | | |
| NC_011770 | 5115927 | G | T | 83 | 0 | 4 | 0 | NON_SYNONYMOU | MODERAT | MISSENSE | gCc/gAc | A148D | PLES_46471 | T | T | | | | | | | | | T | T |
| NC_011770 | 5115955 | C | T | 84 | 0 | 2 | 0 | NON_SYNONYMOU | MODERAT | MISSENSE | Gcg/Acg | A139T | PLES_46471 | | | | | | | | | | | T | T |
| NC_011770 | 5117490 | G | C | 65 | 0 | 2 | 0 | NON_SYNONYMOU | MODERAT | MISSENSE | Ccc/Gcc | P127A | PLES_46501 | | | | | | | C | C | | | | |
| NC_011770 | 5117676 | C | T | 60 | 0 | 4 | 0 | NON_SYNONYMOU | MODERAT | MISSENSE | Ggt/Agt | G65S | PLES_46501 | T | T | | | | | | | T | T | | |
| NC_011770 | 5117685 | C | T | 57 | 0 | 4 | 0 | NON_SYNONYMOU | MODERAT | MISSENSE | Gcg/Acg | A62T | PLES_46501 | T | T | | | | | | | T | T | | |
| NC_011770 | 5117740 | C | A | 56 | 0 | 6 | 0 | NON_SYNONYMOU | MODERAT | MISSENSE | gaG/gaT | E43D | PLES_46501 | | | A | A | A | A | A | A | | | | |
| NC_011770 | 5118491 | C | T | 94 | 0 | 2 | 0 | NON_SYNONYMOU | MODERAT | MISSENSE | gCc/gTc | A15V | PLES_46521 | | | | | | | T | T | | | | |
| NC_011770 | 5118688 | A | T | 96 | 0 | 2 | 0 | NON_SYNONYMOU | MODERAT | MISSENSE | Acc/Tcc | T81S | PLES_46521 | T | T | | | | | | | | | | |
| NC_011770 | 5118724 | G | A | 100 | 0 | 2 | 0 | NON_SYNONYMOU | MODERAT | MISSENSE | Ggc/Agc | G93S | PLES_46521 | | | | | | | | | | | A | A |
| NC_011770 | 5119033 | G | T | 92 | 0 | 2 | 0 | NON_SYNONYMOU | MODERAT | MISSENSE | Gca/Tca | A196S | PLES_46521 | | | | | | | | | | | T | T |
| NC_011770 | 5119761 | A | G | 69 | 0 | 2 | 0 | NON_SYNONYMOU | MODERAT | MISSENSE | Ttc/Ctc | F100L | PLES_46531 | | | | | | | | | | | G | G |
| NC_011770 | 5121649 | A | G | 78 | 0 | 4 | 0 | NON_SYNONYMOU | MODERAT | MISSENSE | aAc/aGc | N33S | PLES_46561 | | | G | G | G | G | | | | | | |
| NC_011770 | 5121705 | G | A | 72 | 0 | 6 | 0 | NON_SYNONYMOU | MODERAT | MISSENSE | Gcc/Acc | A52T | PLES_46561 | A | A | | | | | A | A | A | A | | |
| NC_011770 | 5121729 | G | A | 71 | 0 | 4 | 0 | NON_SYNONYMOU | MODERAT | MISSENSE | Gcc/Acc | A60T | PLES_46561 | | | | | | | A | A | A | A | | |
| NC_011770 | 5122689 | T | G | 80 | 0 | 4 | 0 | START_LOST | HIGH | MISSENSE | aTg/aGg | M1R | PLES_46581 | | | G | G | G | G | | | | | | |
| NC_011770 | 5123364 | GC | G | 62 | 1 | 1 | 0 | FRAME_SHIFT | HIGH | | cca/ | P228 | PLES_46581 | | | | | | | . | G | | | | |
| NC_011770 | 5122802 | A | T | 87 | 0 | 2 | 0 | NON_SYNONYMOU | MODERAT | MISSENSE | Acc/Tcc | T39S | PLES_46581 | | | | | | | | | T | T | | |
| NC_011770 | 5122892 | A | G | 85 | 0 | 12 | 0 | NON_SYNONYMOU | MODERAT | MISSENSE | Ata/Gta | I69V | PLES_46581 | G | G | G | G | G | G | G | G | G | G | G | G |
| NC_011770 | 5122894 | A | G | 87 | 0 | 4 | 0 | NON_SYNONYMOU | MODERAT | MISSENSE | atA/atG | I69M | PLES_46581 | | | G | G | G | G | | | | | | |
| NC_011770 | 5122911 | A | T | 89 | 0 | 12 | 0 | NON_SYNONYMOU | MODERAT | MISSENSE | cAg/cTg | Q75L | PLES_46581 | T | T | T | T | T | T | T | T | T | T | T | T |
| NC_011770 | 5122926 | T | C | 87 | 0 | 12 | 0 | NON_SYNONYMOU | MODERAT | MISSENSE | gTg/gCg | V80A | PLES_46581 | C | C | C | C | C | C | C | C | C | C | C | C |
| NC_011770 | 5122929 | A | G | 86 | 0 | 12 | 0 | NON_SYNONYMOU | MODERAT | MISSENSE | cAc/cGc | H81R | PLES_46581 | G | G | G | G | G | G | G | G | G | G | G | G |
| NC_011770 | 5123058 | G | A | 79 | 0 | 4 | 0 | NON_SYNONYMOU | MODERAT | MISSENSE | cGt/cAt | R124H | PLES_46581 | | | A | A | A | A | | | | | | |
| NC_011770 | 5123418 | C | T | 69 | 0 | 4 | 0 | NON_SYNONYMOU | MODERAT | MISSENSE | gCc/gTc | A244V | PLES_46581 | | | T | T | T | T | | | | | | |
| NC_011770 | 5123512 | A | G | 77 | 0 | 8 | 0 | NON_SYNONYMOU | MODERAT | MISSENSE | aAc/aGc | N21S | PLES_46591 | G | G | | | | | G | G | G | G | G | G |
| NC_011770 | 5123605 | A | G | 84 | 0 | 6 | 0 | NON_SYNONYMOU | MODERAT | MISSENSE | aAg/aGg | K52R | PLES_46591 | | | G | G | G | G | G | G | | | | |
| NC_011770 | 5124027 | C | T | 80 | 0 | 4 | 0 | NON_SYNONYMOU | MODERAT | MISSENSE | Cat/Tat | H193Y | PLES_46591 | | | T | T | T | T | | | | | | |
| NC_011770 | 5124685 | C | T | 53 | 0 | 2 | 0 | NON_SYNONYMOU | MODERAT | MISSENSE | cCc/cTc | P412L | PLES_46591 | | | | | | | T | T | | | | |
| NC_011770 | 5124714 | G | A | 52 | 0 | 2 | 0 | NON_SYNONYMOU | MODERAT | MISSENSE | Gcc/Acc | A422T | PLES_46591 | | | | | | | A | A | | | | |
| NC_011770 | 5124825 | G | A | 45 | 0 | 2 | 0 | NON_SYNONYMOU | MODERAT | MISSENSE | Ggc/Agc | G459S | PLES_46591 | | | | | | | | | | | A | A |
| NC_011770 | 5128322 | G | A | 78 | 0 | 1 | 0 | INTRAGENIC | MODIFIER | | | | PLES_46602 | | | | | | | | A | | | | |
| NC_011770 | 5129397 | T | C | 112 | 0 | 1 | 0 | INTRAGENIC | MODIFIER | | | | PLES_46603 | | | | | | | C | | | | | |
| NC_011770 | 5128844 | G | A | 95 | 0 | 1 | 0 | INTRAGENIC | MODIFIER | | | | PLES_46603 | | | | | | | | A | | | | |
| NC_011770 | 5129250 | C | T | 101 | 0 | 1 | 0 | INTRAGENIC | MODIFIER | | | | PLES_46603 | | | | | | | | T | | | | |
| NC_011770 | 5136717 | G | A | 77 | 0 | 1 | 0 | NON_SYNONYMOU | MODERAT | MISSENSE | Gtg/Atg | V454M | PLES_46621 | | | | A | | | | | | | | |
| NC_011770 | 5135798 | G | C | 65 | 0 | 2 | 0 | NON_SYNONYMOU | MODERAT | MISSENSE | gaG/gaC | E147D | PLES_46621 | | | | | | | | | | | C | C |
| NC_011770 | 5136097 | C | T | 64 | 0 | 2 | 0 | NON_SYNONYMOU | MODERAT | MISSENSE | gCg/gTg | A247V | PLES_46621 | | | | | | | T | T | | | | |
| NC_011770 | 5136102 | C | A | 66 | 0 | 4 | 0 | NON_SYNONYMOU | MODERAT | MISSENSE | Ctc/Atc | L249I | PLES_46621 | | | | | | | | | A | A | A | A |

| | | | | | | | | | | | | | | | | | | | | | | | | | |
|---|---|---|---|---|---|---|---|---|---|---|---|---|---|---|---|---|---|---|---|---|---|---|---|---|---|
| NC_011770 | 5136124 | G | A | 66 | 0 | 12 | 0 | NON_SYNONYMOU | MODERAT | MISSENSE | cGa/cAa | R256Q | PLES_46621 | A | A | A | A | A | A | A | A | A | A | A | A |
| NC_011770 | 5136279 | G | A | 66 | 0 | 4 | 0 | NON_SYNONYMOU | MODERAT | MISSENSE | Gtt/Att | V308I | PLES_46621 | | | A | A | A | A | | | | | | |
| NC_011770 | 5136499 | C | T | 70 | 0 | 2 | 0 | NON_SYNONYMOU | MODERAT | MISSENSE | gCc/gTc | A381V | PLES_46621 | | | | | | | T | T | | | | |
| NC_011770 | 5136600 | G | C | 78 | 0 | 2 | 0 | NON_SYNONYMOU | MODERAT | MISSENSE | Gag/Cag | E415Q | PLES_46621 | C | C | | | | | | | | | | |
| NC_011770 | 5136660 | G | A | 77 | 0 | 4 | 0 | NON_SYNONYMOU | MODERAT | MISSENSE | Gct/Act | A435T | PLES_46621 | | | A | A | A | A | | | | | | |
| NC_011770 | 5136688 | C | T | 77 | 0 | 6 | 0 | NON_SYNONYMOU | MODERAT | MISSENSE | gCg/gTg | A444V | PLES_46621 | | | T | T | T | T | | | T | T | | |
| NC_011770 | 5136711 | G | A | 77 | 0 | 2 | 0 | NON_SYNONYMOU | MODERAT | MISSENSE | Gcc/Acc | A452T | PLES_46621 | | | | | | | | | A | A | | |
| NC_011770 | 5136868 | A | T | 77 | 0 | 6 | 0 | NON_SYNONYMOU | MODERAT | MISSENSE | gAc/gTc | D504V | PLES_46621 | | | T | T | T | T | | | T | T | | |
| NC_011770 | 5136934 | T | C | 71 | 0 | 2 | 0 | NON_SYNONYMOU | MODERAT | MISSENSE | gTc/gCc | V526A | PLES_46621 | C | C | | | | | | | | | | |
| NC_011770 | 5136948 | G | A | 70 | 0 | 2 | 0 | NON_SYNONYMOU | MODERAT | MISSENSE | Gcc/Acc | A531T | PLES_46621 | A | A | | | | | | | | | | |
| NC_011770 | 5137291 | T | A | 49 | 0 | 4 | 0 | NON_SYNONYMOU | MODERAT | MISSENSE | cTg/cAg | L645Q | PLES_46621 | | | A | A | A | A | | | | | | |
| NC_011770 | 5137294 | C | T | 48 | 0 | 12 | 0 | NON_SYNONYMOU | MODERAT | MISSENSE | gCc/gTc | A646V | PLES_46621 | T | T | T | T | T | T | T | T | T | T | T | T |
| NC_011770 | 5137573 | C | T | 45 | 0 | 2 | 0 | NON_SYNONYMOU | MODERAT | MISSENSE | gCc/gTc | A739V | PLES_46621 | T | T | | | | | | | | | | |
| NC_011770 | 5137788 | A | G | 54 | 0 | 2 | 0 | NON_SYNONYMOU | MODERAT | MISSENSE | Atc/Gtc | I811V | PLES_46621 | | | | | | | G | G | | | | |
| NC_011770 | 5137830 | T | A | 58 | 0 | 6 | 0 | NON_SYNONYMOU | MODERAT | MISSENSE | Tcg/Acg | S825T | PLES_46621 | | | A | A | A | A | A | A | | | | |
| NC_011770 | 5137858 | A | T | 58 | 0 | 2 | 0 | NON_SYNONYMOU | MODERAT | MISSENSE | gAg/gTg | E834V | PLES_46621 | | | | | | | | | T | T | | |
| NC_011770 | 5138178 | C | A | 70 | 0 | 2 | 0 | NON_SYNONYMOU | MODERAT | MISSENSE | Cgc/Agc | R941S | PLES_46621 | | | | | | | | | A | A | | |
| NC_011770 | 5138341 | C | T | 65 | 0 | 2 | 0 | NON_SYNONYMOU | MODERAT | MISSENSE | gCg/gTg | A995V | PLES_46621 | | | | | | | | | T | T | | |
| NC_011770 | 5138405 | T | G | 73 | 0 | 4 | 0 | NON_SYNONYMOU | MODERAT | MISSENSE | gaT/gaG | D1016E | PLES_46621 | G | G | | | | | G | G | | | | |
| NC_011770 | 5138520 | A | T | 73 | 0 | 2 | 0 | NON_SYNONYMOU | MODERAT | MISSENSE | Atc/Ttc | I1055F | PLES_46621 | | | | | | | | | T | T | | |
| NC_011770 | 5138725 | C | G | 80 | 0 | 2 | 0 | NON_SYNONYMOU | MODERAT | MISSENSE | aCc/aGc | T1123S | PLES_46621 | | | | | | | | | | | G | G |
| NC_011770 | 5148484 | T | C | 77 | 0 | 12 | 0 | NON_SYNONYMOU | MODERAT | MISSENSE | Atc/Gtc | I223V | PLES_46661 | C | C | C | C | C | C | C | C | C | C | C | C |
| NC_011770 | 5148582 | G | C | 80 | 0 | 4 | 0 | NON_SYNONYMOU | MODERAT | MISSENSE | gCc/gGc | A190G | PLES_46661 | | | | | | | C | C | C | C | | |
| NC_011770 | 5149457 | C | A | 71 | 0 | 2 | 0 | NON_SYNONYMOU | MODERAT | MISSENSE | Gcc/Tcc | A233S | PLES_46671 | | | | | | | A | A | | | | |
| NC_011770 | 5149973 | C | T | 84 | 0 | 2 | 0 | NON_SYNONYMOU | MODERAT | MISSENSE | Gcc/Acc | A61T | PLES_46671 | T | T | | | | | | | | | | |
| NC_011770 | 5150078 | C | T | 90 | 0 | 10 | 0 | NON_SYNONYMOU | MODERAT | MISSENSE | Gtc/Atc | V26I | PLES_46671 | T | T | T | T | T | T | | | T | T | T | T |
| NC_011770 | 5149286 | G | A | 77 | 0 | 1 | 0 | NON_SYNONYMOU | MODERAT | MISSENSE | Cgt/Tgt | R290C | PLES_46671 | | | | | | | | A | | | | |
| NC_011770 | 5150263 | A | T | 90 | 0 | 2 | 0 | NON_SYNONYMOU | MODERAT | MISSENSE | Acc/Tcc | T4S | PLES_46681 | | | | | | | T | T | | | | |
| NC_011770 | 5151644 | G | T | 80 | 0 | 4 | 0 | NON_SYNONYMOU | MODERAT | MISSENSE | cGc/cTc | R178L | PLES_46691 | T | T | | | | | T | T | | | | |
| NC_011770 | 5151650 | T | A | 79 | 0 | 2 | 0 | NON_SYNONYMOU | MODERAT | MISSENSE | cTg/cAg | L180Q | PLES_46691 | | | | | | | | | A | A | | |
| NC_011770 | 5151786 | G | C | 76 | 0 | 2 | 0 | NON_SYNONYMOU | MODERAT | MISSENSE | caG/caC | Q225H | PLES_46691 | C | C | | | | | | | | | | |
| NC_011770 | 5152648 | G | A | 63 | 0 | 1 | 0 | NON_SYNONYMOU | MODERAT | MISSENSE | gCc/gTc | A428V | PLES_46701 | A | | | | | | | | | | | |
| NC_011770 | 5152349 | C | T | 63 | 0 | 10 | 0 | NON_SYNONYMOU | MODERAT | MISSENSE | Gcg/Acg | A528T | PLES_46701 | | | T | T | T | T | T | T | T | T | T | T |
| NC_011770 | 5152427 | T | C | 64 | 0 | 2 | 0 | NON_SYNONYMOU | MODERAT | MISSENSE | Atc/Gtc | I502V | PLES_46701 | C | C | | | | | | | | | | |
| NC_011770 | 5153806 | A | G | 45 | 0 | 12 | 0 | NON_SYNONYMOU | MODERAT | MISSENSE | gTg/gCg | V42A | PLES_46701 | G | G | G | G | G | G | G | G | G | G | G | G |
| NC_011770 | 5153983 | GCGA | G | 48 | 0 | 6 | 0 | CODON_DELETION | MODERATE | | acg/ | T20 | PLES_46711 | | | G | G | G | G | | | | | G | G |
| NC_011770 | 5154019 | G | T | 52 | 0 | 2 | 0 | NON_SYNONYMOU | MODERAT | MISSENSE | Gtt/Ttt | V31F | PLES_46711 | | | | | | | | | T | T | | |
| NC_011770 | 5154158 | G | A | 51 | 0 | 2 | 0 | NON_SYNONYMOU | MODERAT | MISSENSE | cGc/cAc | R77H | PLES_46711 | | | | | | | A | A | | | | |
| NC_011770 | 5154504 | G | T | 57 | 0 | 2 | 0 | NON_SYNONYMOU | MODERAT | MISSENSE | gaG/gaT | E192D | PLES_46711 | | | | | | | T | T | | | | |
| NC_011770 | 5155260 | A | C | 65 | 0 | 2 | 0 | NON_SYNONYMOU | MODERAT | MISSENSE | Atg/Ctg | M168L | PLES_46721 | C | C | | | | | | | | | | |
| NC_011770 | 5155558 | G | A | 83 | 0 | 2 | 0 | NON_SYNONYMOU | MODERAT | MISSENSE | aGt/aAt | S267N | PLES_46721 | A | A | | | | | | | | | | |
| NC_011770 | 5155677 | A | C | 84 | 0 | 2 | 0 | NON_SYNONYMOU | MODERAT | MISSENSE | Aac/Cac | N307H | PLES_46721 | C | C | | | | | | | | | | |
| NC_011770 | 5155699 | G | A | 82 | 0 | 2 | 0 | NON_SYNONYMOU | MODERAT | MISSENSE | cGc/cAc | R314H | PLES_46721 | A | A | | | | | | | | | | |
| NC_011770 | 5159375 | T | C | 68 | 0 | 2 | 0 | NON_SYNONYMOU | MODERAT | MISSENSE | Atc/Gtc | I71V | PLES_46741 | | | | | | | | | C | C | | |
| NC_011770 | 5159527 | G | A | 71 | 0 | 2 | 0 | NON_SYNONYMOU | MODERAT | MISSENSE | tCc/tTc | S20F | PLES_46741 | | | | | | | | | A | A | | |
| NC_011770 | 5159581 | G | T | 72 | 0 | 2 | 0 | NON_SYNONYMOU | MODERAT | MISSENSE | aCg/aAg | T2K | PLES_46741 | | | | | | | | | T | T | | |
| NC_011770 | 5163042 | G | T | 65 | 0 | 6 | 0 | NON_SYNONYMOU | MODERAT | MISSENSE | Ccg/Acg | P16T | PLES_46781 | T | T | | | | | | | T | T | T | T |
| NC_011770 | 5170419 | TG | T | 78 | 0 | 10 | 0 | FRAME_SHIFT | HIGH | | ggt/ | G38 | PLES_46861 | | | T | T | T | T | T | T | T | T | T | T |
| NC_011770 | 5170435 | C | A | 74 | 0 | 10 | 0 | NON_SYNONYMOU | MODERAT | MISSENSE | gCg/gAg | A42E | PLES_46861 | | | A | A | A | A | A | A | A | A | A | A |
| NC_011770 | 5170474 | A | C | 69 | 0 | 12 | 0 | NON_SYNONYMOU | MODERAT | MISSENSE | aAc/aCc | N55T | PLES_46861 | C | C | C | C | C | C | C | C | C | C | C | C |
| NC_011770 | 5170476 | T | A | 68 | 0 | 12 | 0 | NON_SYNONYMOU | MODERAT | MISSENSE | Ttg/Atg | L56M | PLES_46861 | A | A | A | A | A | A | A | A | A | A | A | A |
| NC_011770 | 5173344 | CG | C | 61 | 0 | 1 | 0 | FRAME_SHIFT | HIGH | | gta/ | V17 | PLES_46891 | | | | | | | C | | | | | |
| NC_011770 | 5174279 | A | G | 82 | 0 | 1 | 0 | NON_SYNONYMOU | MODERAT | MISSENSE | cAc/cGc | H326R | PLES_46891 | | | | | | | | G | | | | |
| NC_011770 | 5174507 | G | C | 89 | 0 | 2 | 0 | NON_SYNONYMOU | MODERAT | MISSENSE | Gag/Cag | E33Q | PLES_46901 | C | C | | | | | | | | | | |
| NC_011770 | 5174744 | A | G | 80 | 0 | 2 | 0 | NON_SYNONYMOU | MODERAT | MISSENSE | Acc/Gcc | T112A | PLES_46901 | | | | | | | | | | | G | G |
| NC_011770 | 5174838 | G | A | 82 | 0 | 2 | 0 | NON_SYNONYMOU | MODERAT | MISSENSE | cGc/cAc | R143H | PLES_46901 | | | | | | | | | | | A | A |
| NC_011770 | 5175008 | G | A | 78 | 0 | 1 | 0 | NON_SYNONYMOU | MODERAT | MISSENSE | Gca/Aca | A200T | PLES_46901 | | | | | | | A | | | | | |
| NC_011770 | 5175080 | C | A | 86 | 0 | 4 | 0 | NON_SYNONYMOU | MODERAT | MISSENSE | Cgt/Agt | R224S | PLES_46901 | | | | | | | | | A | A | A | A |
| NC_011770 | 5175081 | G | A | 85 | 0 | 4 | 0 | NON_SYNONYMOU | MODERAT | MISSENSE | cGt/cAt | R224H | PLES_46901 | | | A | A | A | A | | | | | | |
| NC_011770 | 5175114 | A | G | 80 | 0 | 2 | 0 | NON_SYNONYMOU | MODERAT | MISSENSE | cAg/cGg | Q235R | PLES_46901 | | | | | | | | | | | G | G |
| NC_011770 | 5175162 | G | A | 74 | 0 | 2 | 0 | NON_SYNONYMOU | MODERAT | MISSENSE | cGc/cAc | R251H | PLES_46901 | | | | | | | A | A | | | | |
| NC_011770 | 5180459 | G | GC | 42 | 0 | 2 | 0 | FRAME_SHIFT | HIGH | | ccg/ccCg | P16P? | PLES_46961 | | | | | | | GC | GC | | | | |

| | | | | | | | | | | | | | | | | | | | | | | | | | |
|---|---|---|---|---|---|---|---|---|---|---|---|---|---|---|---|---|---|---|---|---|---|---|---|---|---|
| NC_011770 | 5180503 | G | A | 43 | 0 | 4 | 0 | NON_SYNONYMOU | MODERAT | MISSENSE | Gcc/Acc | A29T | PLES_46961 | | | A | A | A | A | | | | | | |
| NC_011770 | 5180716 | A | G | 72 | 0 | 6 | 0 | NON_SYNONYMOU | MODERAT | MISSENSE | Atc/Gtc | I100V | PLES_46961 | | | G | G | G | G | | | G | G | | |
| NC_011770 | 5180795 | G | A | 76 | 0 | 2 | 0 | NON_SYNONYMOU | MODERAT | MISSENSE | cGc/cAc | R126H | PLES_46961 | A | A | | | | | | | | | | |
| NC_011770 | 5181145 | C | T | 82 | 0 | 4 | 0 | NON_SYNONYMOU | MODERAT | MISSENSE | Ccg/Tcg | P243S | PLES_46961 | | | T | T | T | T | | | | | | |
| NC_011770 | 5181256 | A | T | 80 | 0 | 4 | 0 | NON_SYNONYMOU | MODERAT | MISSENSE | cAa/cTa | Q15L | PLES_46971 | | | | | | | T | T | | | T | T |
| NC_011770 | 5182576 | C | T | 61 | 0 | 2 | 0 | NON_SYNONYMOU | MODERAT | MISSENSE | Ccc/Tcc | P133S | PLES_46981 | | | | | | | | | T | T | | |
| NC_011770 | 5182960 | G | A | 57 | 0 | 8 | 0 | NON_SYNONYMOU | MODERAT | MISSENSE | Gcc/Acc | A261T | PLES_46981 | | | A | A | A | A | A | A | | | A | A |
| NC_011770 | 5183017 | A | G | 57 | 0 | 2 | 0 | NON_SYNONYMOU | MODERAT | MISSENSE | Acc/Gcc | T280A | PLES_46981 | | | | | | | | | G | G | | |
| NC_011770 | 5183123 | C | T | 59 | 0 | 2 | 0 | NON_SYNONYMOU | MODERAT | MISSENSE | gCg/gTg | A315V | PLES_46981 | | | | | | | | | T | T | | |
| NC_011770 | 5183662 | G | A | 74 | 0 | 1 | 0 | NON_SYNONYMOU | MODERAT | MISSENSE | Gcc/Acc | A495T | PLES_46981 | | | | | | | A | | | | | |
| NC_011770 | 5183947 | A | G | 79 | 0 | 2 | 0 | NON_SYNONYMOU | MODERAT | MISSENSE | gAc/gGc | D67G | PLES_46991 | | | | | | | G | G | | | | |
| NC_011770 | 5184269 | C | A | 70 | 0 | 4 | 0 | NON_SYNONYMOU | MODERAT | MISSENSE | gaC/gaA | D174E | PLES_46991 | A | A | | | | | | | A | A | | |
| NC_011770 | 5184270 | A | G | 69 | 0 | 5 | 0 | NON_SYNONYMOU | MODERAT | MISSENSE | Acc/Gcc | T175A | PLES_46991 | G | G | | | | | G | | G | G | | |
| NC_011770 | 5185107 | C | G | 64 | 0 | 4 | 0 | NON_SYNONYMOU | MODERAT | MISSENSE | aCc/aGc | T31S | PLES_47001 | | | | | | | | | G | G | G | G |
| NC_011770 | 5185370 | C | A | 77 | 0 | 4 | 0 | NON_SYNONYMOU | MODERAT | MISSENSE | Cac/Aac | H119N | PLES_47001 | | | A | A | A | A | | | | | | |
| NC_011770 | 5186923 | G | A | 74 | 0 | 2 | 0 | NON_SYNONYMOU | MODERAT | MISSENSE | Gcc/Acc | A302T | PLES_47011 | | | | | | | | | | | A | A |
| NC_011770 | 5187163 | A | C | 73 | 0 | 12 | 0 | NON_SYNONYMOU | MODERAT | MISSENSE | Aac/Cac | N382H | PLES_47011 | C | C | C | C | C | C | C | C | C | C | C | C |
| NC_011770 | 5187824 | C | G | 64 | 0 | 12 | 0 | NON_SYNONYMOU | MODERAT | MISSENSE | gaG/gaC | E125D | PLES_47031 | G | G | G | G | G | G | G | G | G | G | G | G |
| NC_011770 | 5188117 | C | T | 76 | 0 | 12 | 0 | NON_SYNONYMOU | MODERAT | MISSENSE | Gag/Aag | E28K | PLES_47031 | T | T | T | T | T | T | T | T | T | T | T | T |
| NC_011770 | 5188351 | G | A | 82 | 0 | 1 | 0 | NON_SYNONYMOU | MODERAT | MISSENSE | Cgg/Tgg | R112W | PLES_47041 | | | | | | | A | | | | | |
| NC_011770 | 5189145 | G | A | 89 | 0 | 2 | 0 | NON_SYNONYMOU | MODERAT | MISSENSE | cGc/cAc | R123H | PLES_47051 | A | A | | | | | | | | | | |
| NC_011770 | 5189795 | C | T | 77 | 0 | 1 | 0 | NON_SYNONYMOU | MODERAT | MISSENSE | Ccg/Tcg | P43S | PLES_47061 | | | | | | | | | | | T | |
| NC_011770 | 5189729 | A | T | 78 | 0 | 2 | 0 | NON_SYNONYMOU | MODERAT | MISSENSE | Acc/Tcc | T21S | PLES_47061 | | | | | | | | | | | T | T |
| NC_011770 | 5189754 | G | A | 73 | 0 | 2 | 0 | NON_SYNONYMOU | MODERAT | MISSENSE | cGc/cAc | R29H | PLES_47061 | | | | | | | A | A | | | | |
| NC_011770 | 5190309 | G | T | 69 | 0 | 2 | 0 | NON_SYNONYMOU | MODERAT | MISSENSE | cGt/cTt | R214L | PLES_47061 | T | T | | | | | | | | | | |
| NC_011770 | 5190366 | C | G | 76 | 0 | 2 | 0 | NON_SYNONYMOU | MODERAT | MISSENSE | gCc/gGc | A233G | PLES_47061 | G | G | | | | | | | | | | |
| NC_011770 | 5190443 | A | G | 81 | 0 | 12 | 0 | NON_SYNONYMOU | MODERAT | MISSENSE | Acc/Gcc | T259A | PLES_47061 | G | G | G | G | G | G | G | G | G | G | G | G |
| NC_011770 | 5192383 | A | C | 85 | 0 | 2 | 0 | NON_SYNONYMOU | MODERAT | MISSENSE | cTg/cGg | L196R | PLES_47081 | | | | | | | | | | | C | C |
| NC_011770 | 5192426 | C | A | 86 | 0 | 2 | 0 | NON_SYNONYMOU | MODERAT | MISSENSE | Gcc/Tcc | A182S | PLES_47081 | | | | | | | | | | | A | A |
| NC_011770 | 5192515 | C | T | 94 | 0 | 2 | 0 | NON_SYNONYMOU | MODERAT | MISSENSE | aGg/aAg | R152K | PLES_47081 | T | T | | | | | | | | | | |
| NC_011770 | 5192738 | C | T | 91 | 0 | 2 | 0 | NON_SYNONYMOU | MODERAT | MISSENSE | Gac/Aac | D78N | PLES_47081 | T | T | | | | | | | | | | |
| NC_011770 | 5192939 | C | T | 96 | 0 | 2 | 0 | NON_SYNONYMOU | MODERAT | MISSENSE | Gcg/Acg | A11T | PLES_47081 | T | T | | | | | | | | | | |
| NC_011770 | 5193087 | C | A | 89 | 0 | 4 | 0 | NON_SYNONYMOU | MODERAT | MISSENSE | Gcc/Tcc | A292S | PLES_47091 | | | A | A | A | A | | | | | | |
| NC_011770 | 5193222 | C | G | 81 | 0 | 2 | 0 | NON_SYNONYMOU | MODERAT | MISSENSE | Gtg/Ctg | V247L | PLES_47091 | G | G | | | | | | | | | | |
| NC_011770 | 5193776 | G | A | 79 | 0 | 2 | 0 | NON_SYNONYMOU | MODERAT | MISSENSE | gCa/gTa | A62V | PLES_47091 | A | A | | | | | | | | | | |
| NC_011770 | 5194399 | G | T | 83 | 0 | 10 | 0 | NON_SYNONYMOU | MODERAT | MISSENSE | Ctg/Atg | L340M | PLES_47101 | | | T | T | T | T | T | T | T | T | T | T |
| NC_011770 | 5196867 | G | A | 56 | 0 | 5 | 0 | NON_SYNONYMOU | MODERAT | MISSENSE | Gtc/Atc | V504I | PLES_47111 | A | A | | | | | | | A | A | | A |
| NC_011770 | 5197183 | G | C | 72 | 0 | 4 | 0 | NON_SYNONYMOU | MODERAT | MISSENSE | gCc/gGc | A351G | PLES_47121 | C | C | | | | | | | C | C | | |
| NC_011770 | 5198511 | A | C | 81 | 0 | 2 | 0 | NON_SYNONYMOU | MODERAT | MISSENSE | aaT/aaG | N56K | PLES_47131 | | | | | | | | | C | C | | |
| NC_011770 | 5198724 | A | G | 80 | 0 | 10 | 0 | NON_SYNONYMOU | MODERAT | MISSENSE | gTg/gCg | V187A | PLES_47141 | G | G | G | G | G | G | G | G | | | G | G |
| NC_011770 | 5198961 | T | G | 78 | 0 | 2 | 0 | NON_SYNONYMOU | MODERAT | MISSENSE | gAc/gCc | D108A | PLES_47141 | | | | | | | | | G | G | | |
| NC_011770 | 5199057 | T | A | 85 | 0 | 2 | 0 | NON_SYNONYMOU | MODERAT | MISSENSE | cAg/cTg | Q76L | PLES_47141 | | | | | | | | | A | A | | |
| NC_011770 | 5199527 | T | C | 77 | 0 | 12 | 0 | NON_SYNONYMOU | MODERAT | MISSENSE | Acc/Gcc | T158A | PLES_47151 | C | C | C | C | C | C | C | C | C | C | C | C |
| NC_011770 | 5199937 | G | A | 76 | 0 | 2 | 0 | NON_SYNONYMOU | MODERAT | MISSENSE | cCc/cTc | P21L | PLES_47151 | | | | | | | | | A | A | | |
| NC_011770 | 5201783 | C | A | 67 | 0 | 2 | 0 | NON_SYNONYMOU | MODERAT | MISSENSE | Gcc/Tcc | A257S | PLES_47171 | A | A | | | | | | | | | | |
| NC_011770 | 5202019 | T | C | 55 | 0 | 1 | 0 | NON_SYNONYMOU | MODERAT | MISSENSE | tAc/tGc | Y178C | PLES_47171 | | | | | | | | C | | | | |
| NC_011770 | 5203027 | C | G | 74 | 0 | 1 | 0 | NON_SYNONYMOU | MODERAT | MISSENSE | Gcg/Ccg | A58P | PLES_47181 | | | | | | | G | | | | | |
| NC_011770 | 5203153 | T | C | 80 | 0 | 12 | 0 | NON_SYNONYMOU | MODERAT | MISSENSE | Atc/Gtc | I16V | PLES_47181 | C | C | C | C | C | C | C | C | C | C | C | C |
| NC_011770 | 5203481 | G | A | 80 | 0 | 1 | 0 | STOP_GAINED | HIGH | NONSENSE | Cag/Tag | Q185* | PLES_47191 | | | | | | | A | | | | | |
| NC_011770 | 5203343 | C | T | 83 | 0 | 2 | 0 | NON_SYNONYMOU | MODERAT | MISSENSE | Gat/Aat | D231N | PLES_47191 | | | | | T | T | | | | | | |
| NC_011770 | 5203400 | T | C | 74 | 0 | 12 | 0 | NON_SYNONYMOU | MODERAT | MISSENSE | Ata/Gta | I212V | PLES_47191 | C | C | C | C | C | C | C | C | C | C | C | C |
| NC_011770 | 5204522 | T | C | 76 | 0 | 4 | 0 | NON_SYNONYMOU | MODERAT | MISSENSE | Acc/Gcc | T366A | PLES_47201 | | | C | C | C | C | | | | | | |
| NC_011770 | 5204579 | C | G | 75 | 0 | 2 | 0 | NON_SYNONYMOU | MODERAT | MISSENSE | Gtc/Ctc | V347L | PLES_47201 | G | G | | | | | | | | | | |
| NC_011770 | 5204697 | G | C | 72 | 0 | 2 | 0 | NON_SYNONYMOU | MODERAT | MISSENSE | gaC/gaG | D307E | PLES_47201 | | | | | | | C | C | | | | |
| NC_011770 | 5204809 | C | T | 74 | 0 | 2 | 0 | NON_SYNONYMOU | MODERAT | MISSENSE | cGc/cAc | R270H | PLES_47201 | T | T | | | | | | | | | | |
| NC_011770 | 5204869 | C | G | 69 | 0 | 2 | 0 | NON_SYNONYMOU | MODERAT | MISSENSE | tGt/tCt | C250S | PLES_47201 | | | | | | | | | | | G | G |
| NC_011770 | 5205251 | C | T | 65 | 0 | 4 | 0 | NON_SYNONYMOU | MODERAT | MISSENSE | Gtg/Atg | V123M | PLES_47201 | | | T | T | T | T | | | | | | |
| NC_011770 | 5205292 | G | A | 68 | 0 | 2 | 0 | NON_SYNONYMOU | MODERAT | MISSENSE | gCc/gTc | A109V | PLES_47201 | A | A | | | | | | | | | | |
| NC_011770 | 5205773 | G | A | 64 | 0 | 2 | 0 | NON_SYNONYMOU | MODERAT | MISSENSE | Ctc/Ttc | L389F | PLES_47211 | | | | | | | | | | | A | A |
| NC_011770 | 5206397 | A | C | 80 | 0 | 6 | 0 | NON_SYNONYMOU | MODERAT | MISSENSE | Tcc/Gcc | S181A | PLES_47211 | C | C | C | C | C | C | | | | | | |
| NC_011770 | 5207729 | C | T | 86 | 0 | 1 | 0 | NON_SYNONYMOU | MODERAT | MISSENSE | cGc/cAc | R156H | PLES_47221 | | | | | | | T | | | | | |
| NC_011770 | 5208149 | T | C | 71 | 0 | 2 | 0 | NON_SYNONYMOU | MODERAT | MISSENSE | gAc/gGc | D16G | PLES_47221 | | | | | | | | | C | C | | |

| | | | | | | | | | | | | | | | | | | | | | | | | | |
|---|---|---|---|---|---|---|---|---|---|---|---|---|---|---|---|---|---|---|---|---|---|---|---|---|---|
| NC_011770 | 5208619 | G | A | 79 | 0 | 1 | 0 | NON_SYNONYMOU | MODERAT | MISSENSE | tCg/tTg | S111L | PLES_47231 | | A | | | | | | | | | | |
| NC_011770 | 5209298 | C | G | 79 | 0 | 2 | 0 | NON_SYNONYMOU | MODERAT | MISSENSE | Gct/Cct | A17P | PLES_47241 | | | | | | | | | | | G | G |
| NC_011770 | 5209902 | A | C | 68 | 0 | 12 | 0 | NON_SYNONYMOU | MODERAT | MISSENSE | cTg/cGg | L199R | PLES_47251 | C | C | C | C | C | C | C | C | C | C | C | C |
| NC_011770 | 5210482 | G | A | 54 | 0 | 1 | 0 | NON_SYNONYMOU | MODERAT | MISSENSE | Ctc/Ttc | L6F | PLES_47251 | | | | | | | A | | | | | |
| NC_011770 | 5210667 | G | C | 65 | 0 | 2 | 0 | NON_SYNONYMOU | MODERAT | MISSENSE | gaC/gaG | D215E | PLES_47261 | | | | | | | | | | | C | C |
| NC_011770 | 5210934 | C | A | 66 | 0 | 2 | 0 | NON_SYNONYMOU | MODERAT | MISSENSE | atG/atT | M126I | PLES_47261 | A | A | | | | | | | | | | |
| NC_011770 | 5211871 | T | C | 74 | 0 | 12 | 0 | NON_SYNONYMOU | MODERAT | MISSENSE | cAc/cGc | H141R | PLES_47271 | C | C | C | C | C | C | C | C | C | C | C | C |
| NC_011770 | 5212729 | A | G | 87 | 0 | 2 | 0 | NON_SYNONYMOU | MODERAT | MISSENSE | Acc/Gcc | T79A | PLES_47281 | G | G | | | | | | | | | | |
| NC_011770 | 5213809 | T | C | 86 | 0 | 1 | 0 | NON_SYNONYMOU | MODERAT | MISSENSE | gTg/gCg | V187A | PLES_47291 | | | | | | | C | | | | | |
| NC_011770 | 5213821 | A | T | 88 | 0 | 2 | 0 | NON_SYNONYMOU | MODERAT | MISSENSE | cAg/cTg | Q191L | PLES_47291 | | | | | | | | | T | T | | |
| NC_011770 | 5213952 | C | T | 85 | 0 | 2 | 0 | NON_SYNONYMOU | MODERAT | MISSENSE | Cgc/Tgc | R235C | PLES_47291 | | | | | | | | | | | T | T |
| NC_011770 | 5214000 | A | C | 80 | 0 | 8 | 0 | NON_SYNONYMOU | MODERAT | MISSENSE | Act/Cct | T251P | PLES_47291 | C | C | | | | | C | C | C | C | C | C |
| NC_011770 | 5214016 | G | A | 77 | 0 | 4 | 0 | NON_SYNONYMOU | MODERAT | MISSENSE | cGc/cAc | R256H | PLES_47291 | | | A | A | A | A | | | | | | |
| NC_011770 | 5215193 | G | A | 72 | 0 | 2 | 0 | NON_SYNONYMOU | MODERAT | MISSENSE | gCg/gTg | A138V | PLES_47311 | | | | | | | | | | | A | A |
| NC_011770 | 5215499 | C | T | 74 | 0 | 8 | 0 | NON_SYNONYMOU | MODERAT | MISSENSE | gGc/gAc | G36D | PLES_47311 | T | T | | | | | T | T | T | T | T | T |
| NC_011770 | 5216317 | C | T | 71 | 0 | 2 | 0 | NON_SYNONYMOU | MODERAT | MISSENSE | gCc/gTc | A70V | PLES_47331 | | | | | | | T | T | | | | |
| NC_011770 | 5218442 | G | A | 64 | 0 | 6 | 0 | NON_SYNONYMOU | MODERAT | MISSENSE | Cat/Tat | H76Y | PLES_47351 | A | A | | | | | A | A | | | A | A |
| NC_011770 | 5218649 | G | T | 73 | 0 | 12 | 0 | NON_SYNONYMOU | MODERAT | MISSENSE | Ctc/Atc | L7I | PLES_47351 | T | T | T | T | T | T | T | T | T | T | T | T |
| NC_011770 | 5221602 | T | C | 62 | 0 | 2 | 0 | NON_SYNONYMOU | MODERAT | MISSENSE | Acg/Gcg | T222A | PLES_47381 | | | | | | | C | C | | | | |
| NC_011770 | 5222177 | C | A | 62 | 0 | 2 | 0 | NON_SYNONYMOU | MODERAT | MISSENSE | cGg/cTg | R30L | PLES_47381 | | | | | | | | | | | A | A |
| NC_011770 | 5222650 | G | A | 74 | 0 | 12 | 0 | NON_SYNONYMOU | MODERAT | MISSENSE | gGt/gAt | G32D | PLES_47391 | A | A | A | A | A | A | A | A | A | A | A | A |
| NC_011770 | 5222716 | A | T | 72 | 0 | 2 | 0 | NON_SYNONYMOU | MODERAT | MISSENSE | cAg/cTg | Q54L | PLES_47391 | | | | | | | | | | | T | T |
| NC_011770 | 5223725 | C | T | 82 | 0 | 2 | 0 | NON_SYNONYMOU | MODERAT | MISSENSE | Gtc/Atc | V29I | PLES_47401 | | | | | | | | | | | T | T |
| NC_011770 | 5224317 | G | C | 74 | 0 | 4 | 0 | NON_SYNONYMOU | MODERAT | MISSENSE | caG/caC | Q108H | PLES_47411 | C | C | | | | | | | C | C | | |
| NC_011770 | 5226141 | C | A | 78 | 0 | 4 | 0 | NON_SYNONYMOU | MODERAT | MISSENSE | gCt/gAt | A30D | PLES_47431 | A | A | | | | | | | | | A | A |
| NC_011770 | 5226261 | C | T | 78 | 0 | 2 | 0 | NON_SYNONYMOU | MODERAT | MISSENSE | gCc/gTc | A70V | PLES_47431 | T | T | | | | | | | | | | |
| NC_011770 | 5226960 | G | GC | 69 | 0 | 1 | 0 | FRAME_SHIFT | HIGH | | tgg/Ctgg | W171L? | PLES_47441 | | | | | | | GC | | | | | |
| NC_011770 | 5226567 | T | A | 68 | 0 | 2 | 0 | NON_SYNONYMOU | MODERAT | MISSENSE | Tcg/Acg | S38T | PLES_47441 | | | | | | | | | | | A | A |
| NC_011770 | 5228552 | GCCGC | G | 66 | 0 | 1 | 0 | FRAME_SHIFT | HIGH | | ccgccc/ | PP131 | PLES_47461 (bifA) | | | | | | G | | | | | | |
| NC_011770 | 5229128 | G | T | 87 | 0 | 2 | 0 | NON_SYNONYMOU | MODERAT | MISSENSE | cGg/cTg | R321L | PLES_47461 (bifA) | | | | | | | T | T | | | | |
| NC_011770 | 5229479 | T | G | 79 | 0 | 2 | 0 | NON_SYNONYMOU | MODERAT | MISSENSE | cTg/cGg | L438R | PLES_47461 (bifA) | | | | | | | | | G | G | | |
| NC_011770 | 5229269 | T | C | 87 | 0 | 1 | 0 | NON_SYNONYMOU | MODERAT | MISSENSE | gTg/gCg | V368A | PLES_47461 (bifA) | | | | | | | | C | | | | |
| NC_011770 | 5230818 | C | T | 86 | 0 | 6 | 0 | NON_SYNONYMOU | MODERAT | MISSENSE | Ggc/Agc | G106S | PLES_47471 | T | T | T | T | T | T | | | | | | |
| NC_011770 | 5231061 | C | T | 79 | 0 | 2 | 0 | NON_SYNONYMOU | MODERAT | MISSENSE | Gtg/Atg | V25M | PLES_47471 | | | | | | | | | | | T | T |
| NC_011770 | 5231410 | C | G | 71 | 0 | 2 | 0 | NON_SYNONYMOU | MODERAT | MISSENSE | Gcc/Ccc | A100P | PLES_47481 | | | G | G | | | | | | | | |
| NC_011770 | 5231419 | C | A | 71 | 0 | 2 | 0 | NON_SYNONYMOU | MODERAT | MISSENSE | Ggc/Tgc | G97C | PLES_47481 | | | | | | | | | | | A | A |
| NC_011770 | 5231443 | T | C | 70 | 0 | 2 | 0 | NON_SYNONYMOU | MODERAT | MISSENSE | Aat/Gat | N89D | PLES_47481 | | | | | | | | | | | C | C |
| NC_011770 | 5233911 | G | A | 71 | 0 | 4 | 0 | NON_SYNONYMOU | MODERAT | MISSENSE | Gtg/Atg | V170M | PLES_47501 | | | A | A | A | A | | | | | | |
| NC_011770 | 5233926 | C | A | 70 | 0 | 4 | 0 | NON_SYNONYMOU | MODERAT | MISSENSE | Ctg/Atg | L175M | PLES_47501 | | | A | A | A | A | | | | | | |
| NC_011770 | 5234339 | G | T | 73 | 0 | 4 | 0 | NON_SYNONYMOU | MODERAT | MISSENSE | gaG/gaT | E312D | PLES_47501 | | | T | T | T | T | | | | | | |
| NC_011770 | 5234542 | G | T | 71 | 0 | 4 | 0 | NON_SYNONYMOU | MODERAT | MISSENSE | cGt/cTt | R380L | PLES_47501 | | | T | T | T | T | | | | | | |
| NC_011770 | 5235682 | A | C | 77 | 0 | 6 | 0 | NON_SYNONYMOU | MODERAT | MISSENSE | gAc/gCc | D283A | PLES_47511 | C | C | C | C | C | C | | | | | | |
| NC_011770 | 5235716 | C | A | 79 | 0 | 2 | 0 | NON_SYNONYMOU | MODERAT | MISSENSE | gaC/gaA | D294E | PLES_47511 | A | A | | | | | | | | | | |
| NC_011770 | 5235681 | G | A | 77 | 0 | 1 | 0 | NON_SYNONYMOU | MODERAT | MISSENSE | Gac/Aac | D283N | PLES_47511 | | | | | | | | A | | | | |
| NC_011770 | 5235989 | T | A | 74 | 0 | 6 | 0 | NON_SYNONYMOU | MODERAT | MISSENSE | cTg/cAg | L27Q | PLES_47521 | A | A | A | A | A | A | | | | | | |
| NC_011770 | 5236198 | C | T | 71 | 0 | 2 | 0 | NON_SYNONYMOU | MODERAT | MISSENSE | Cgc/Tgc | R97C | PLES_47521 | T | T | | | | | | | | | | |
| NC_011770 | 5236303 | G | A | 72 | 0 | 2 | 0 | NON_SYNONYMOU | MODERAT | MISSENSE | Gat/Aat | D132N | PLES_47521 | | | | | | | | | A | A | | |
| NC_011770 | 5236840 | G | A | 75 | 0 | 2 | 0 | NON_SYNONYMOU | MODERAT | MISSENSE | Gcg/Acg | A311T | PLES_47521 | A | A | | | | | | | | | | |
| NC_011770 | 5236315 | G | A | 70 | 0 | 1 | 0 | NON_SYNONYMOU | MODERAT | MISSENSE | Ggc/Agc | G136S | PLES_47521 | | | | | | | | A | | | | |
| NC_011770 | 5238215 | G | A | 77 | 0 | 1 | 0 | NON_SYNONYMOU | MODERAT | MISSENSE | Gag/Aag | E348K | PLES_47531 | | | | | | | | | | | | A |
| NC_011770 | 5237859 | G | C | 65 | 0 | 4 | 0 | NON_SYNONYMOU | MODERAT | MISSENSE | gGc/gCc | G229A | PLES_47531 | | | | | | | C | C | | | C | C |
| NC_011770 | 5238135 | G | A | 77 | 0 | 6 | 0 | NON_SYNONYMOU | MODERAT | MISSENSE | cGg/cAg | R321Q | PLES_47531 | | | A | A | A | A | | | A | A | | |
| NC_011770 | 5237805 | T | TC | 60 | 0 | 1 | 0 | FRAME_SHIFT | HIGH | | ctg/cCtg | L213P? | PLES_47531 (mexV) | | | | | | | TC | | | | | |
| NC_011770 | 5239886 | A | G | 90 | 0 | 4 | 0 | NON_SYNONYMOU | MODERAT | MISSENSE | cAa/cGa | Q511R | PLES_47541 | | | | | | | | | G | G | G | G |
| NC_011770 | 5240078 | G | A | 90 | 0 | 2 | 0 | NON_SYNONYMOU | MODERAT | MISSENSE | cGc/cAc | R575H | PLES_47541 | | | | | | | A | A | | | | |
| NC_011770 | 5240164 | G | A | 88 | 0 | 2 | 0 | NON_SYNONYMOU | MODERAT | MISSENSE | Gcc/Acc | A604T | PLES_47541 | | | | | | | | | A | A | | |
| NC_011770 | 5241403 | G | T | 72 | 0 | 2 | 0 | NON_SYNONYMOU | MODERAT | MISSENSE | Gca/Tca | A1017S | PLES_47541 | | | | | | | | | T | T | | |
| NC_011770 | 5238432 | G | T | 87 | 0 | 2 | 0 | NON_SYNONYMOU | MODERAT | MISSENSE | atG/atT | M26I | PLES_47541 (mexW) | | | | | | | | | T | T | | |
| NC_011770 | 5244222 | G | A | 76 | 0 | 4 | 0 | NON_SYNONYMOU | MODERAT | MISSENSE | gCc/gTc | A93V | PLES_47581 | | | A | A | A | A | | | | | | |
| NC_011770 | 5247494 | A | T | 86 | 0 | 2 | 0 | NON_SYNONYMOU | MODERAT | MISSENSE | gAc/gTc | D249V | PLES_47611 | | | | | | | | | T | T | | |
| NC_011770 | 5246992 | G | A | 92 | 0 | 1 | 0 | NON_SYNONYMOU | MODERAT | MISSENSE | Gtc/Atc | V82I | PLES_47611 | | | | | | | | A | | | | |
| NC_011770 | 5248881 | G | A | 65 | 0 | 8 | 0 | NON_SYNONYMOU | MODERAT | MISSENSE | Gac/Aac | D146N | PLES_47631 | A | A | A | A | A | A | | | | | A | A |

| | | | | | | | | | | | | | | | | | | | | | | | | | |
|---|---|---|---|---|---|---|---|---|---|---|---|---|---|---|---|---|---|---|---|---|---|---|---|---|---|
| NC_011770 | 5249071 | G | A | 72 | 0 | 8 | 0 | NON_SYNONYMOU | MODERAT | MISSENSE | cGc/cAc | R209H | PLES_47631 | A | A | A | A | A | A | | | | | A | A |
| NC_011770 | 5248554 | G | A | 59 | 0 | 1 | 0 | NON_SYNONYMOU | MODERAT | MISSENSE | Gcc/Acc | A37T | PLES_47631 | | | | | | | | A | | | | |
| NC_011770 | 5248582 | G | A | 58 | 0 | 1 | 0 | NON_SYNONYMOU | MODERAT | MISSENSE | gGc/gAc | G46D | PLES_47631 | | | | | | | | A | | | | |
| NC_011770 | 5254737 | GC | G | 74 | 0 | 2 | 0 | FRAME_SHIFT | HIGH | | ggc/ | G318 | PLES_47701 | | | | G | | | GCCC | | | | | |
| NC_011770 | 5254891 | T | C | 67 | 0 | 12 | 0 | NON_SYNONYMOU | MODERAT | MISSENSE | cAg/cGg | Q267R | PLES_47701 | C | C | C | C | C | C | C | C | C | C | C | C |
| NC_011770 | 5255525 | C | T | 71 | 0 | 4 | 0 | NON_SYNONYMOU | MODERAT | MISSENSE | Gat/Aat | D56N | PLES_47701 | | | T | T | T | T | | | | | | |
| NC_011770 | 5255668 | A | G | 74 | 0 | 12 | 0 | NON_SYNONYMOU | MODERAT | MISSENSE | tTa/tCa | L8S | PLES_47701 | G | G | G | G | G | G | G | G | G | G | G | G |
| NC_011770 | 5255936 | G | C | 80 | 0 | 2 | 0 | NON_SYNONYMOU | MODERAT | MISSENSE | gGc/gCc | G83A | PLES_47711 | | | | | | | | | | | C | C |
| NC_011770 | 5255967 | C | A | 79 | 0 | 2 | 0 | NON_SYNONYMOU | MODERAT | MISSENSE | gaC/gaA | D93E | PLES_47711 | A | A | | | | | | | | | | |
| NC_011770 | 5257537 | G | A | 82 | 0 | 2 | 0 | NON_SYNONYMOU | MODERAT | MISSENSE | Gac/Aac | D475N | PLES_47721 | | | | | | | | | A | A | | |
| NC_011770 | 5257775 | A | G | 83 | 0 | 1 | 0 | NON_SYNONYMOU | MODERAT | MISSENSE | cAc/cGc | H554R | PLES_47721 | | | | | | | G | | | | | |
| NC_011770 | 5258758 | T | A | 56 | 0 | 2 | 0 | NON_SYNONYMOU | MODERAT | MISSENSE | Acc/Tcc | T12S | PLES_47731 | | | | | | | | | | | A | A |
| NC_011770 | 5259820 | C | A | 77 | 0 | 2 | 0 | NON_SYNONYMOU | MODERAT | MISSENSE | Cag/Aag | Q72K | PLES_47751 | A | A | | | | | | | | | | |
| NC_011770 | 5263749 | G | A | 58 | 0 | 1 | 0 | NON_SYNONYMOU | MODERAT | MISSENSE | Gct/Act | A682T | PLES_47771 | | | | | | | A | | | | | |
| NC_011770 | 5263000 | T | C | 73 | 0 | 1 | 0 | NON_SYNONYMOU | MODERAT | MISSENSE | cTg/cCg | L432P | PLES_47771 | | | | | | | | C | | | | |
| NC_011770 | 5263906 | G | A | 60 | 0 | 4 | 0 | NON_SYNONYMOU | MODERAT | MISSENSE | Gag/Aag | E34K | PLES_47781 | | | A | A | A | A | | | | | | |
| NC_011770 | 5264180 | T | C | 60 | 0 | 2 | 0 | NON_SYNONYMOU | MODERAT | MISSENSE | cTg/cCg | L125P | PLES_47781 | C | C | | | | | | | | | | |
| NC_011770 | 5264278 | T | C | 57 | 0 | 2 | 0 | NON_SYNONYMOU | MODERAT | MISSENSE | Tgc/Cgc | C158R | PLES_47781 | | | | | | | | | | | C | C |
| NC_011770 | 5264542 | G | C | 60 | 0 | 2 | 0 | NON_SYNONYMOU | MODERAT | MISSENSE | Ctc/Gtc | L288V | PLES_47791 | | | | | | | | | | | C | C |
| NC_011770 | 5264689 | C | T | 70 | 0 | 2 | 0 | NON_SYNONYMOU | MODERAT | MISSENSE | Gcc/Acc | A239T | PLES_47791 | T | T | | | | | | | | | | |
| NC_011770 | 5264696 | C | A | 70 | 0 | 2 | 0 | NON_SYNONYMOU | MODERAT | MISSENSE | gaG/gaT | E236D | PLES_47791 | | | | | | | | | A | A | | |
| NC_011770 | 5265498 | C | T | 76 | 0 | 2 | 0 | NON_SYNONYMOU | MODERAT | MISSENSE | Gcc/Acc | A199T | PLES_47801 | | | | | | | T | T | | | | |
| NC_011770 | 5265704 | G | A | 72 | 0 | 2 | 0 | NON_SYNONYMOU | MODERAT | MISSENSE | aCa/aTa | T130I | PLES_47801 | | | A | A | | | | | | | | |
| NC_011770 | 5265728 | G | A | 73 | 0 | 4 | 0 | NON_SYNONYMOU | MODERAT | MISSENSE | cCa/cTa | P122L | PLES_47801 | | | A | A | A | A | | | | | | |
| NC_011770 | 5265855 | G | A | 73 | 0 | 2 | 0 | NON_SYNONYMOU | MODERAT | MISSENSE | Ccc/Tcc | P80S | PLES_47801 | | | | | | | A | A | | | | |
| NC_011770 | 5271264 | G | T | 60 | 0 | 12 | 0 | NON_SYNONYMOU | MODERAT | MISSENSE | Ctg/Atg | L41M | PLES_47831 | T | T | T | T | T | T | T | T | T | T | T | T |
| NC_011770 | 5271808 | C | A | 51 | 0 | 2 | 0 | NON_SYNONYMOU | MODERAT | MISSENSE | aCc/aAc | T93N | PLES_47841 | | | | | | | | | A | A | | |
| NC_011770 | 5290610 | C | G | 65 | 0 | 2 | 0 | NON_SYNONYMOU | MODERAT | MISSENSE | caG/caC | Q228H | PLES_48011 | | | | | | | | | G | G | | |
| NC_011770 | 5291578 | A | C | 67 | 0 | 2 | 0 | NON_SYNONYMOU | MODERAT | MISSENSE | gaA/gaC | E53D | PLES_48021 | | | | | | | | | C | C | | |
| NC_011770 | 5291625 | A | G | 72 | 0 | 2 | 0 | NON_SYNONYMOU | MODERAT | MISSENSE | aAg/aGg | K69R | PLES_48021 | | | | | | | | | G | G | | |
| NC_011770 | 5292146 | T | A | 76 | 0 | 12 | 0 | NON_SYNONYMOU | MODERAT | MISSENSE | Tcg/Acg | S243T | PLES_48021 | A | A | A | A | A | A | A | A | A | A | A | A |
| NC_011770 | 5292462 | A | C | 69 | 0 | 2 | 0 | NON_SYNONYMOU | MODERAT | MISSENSE | cAg/cCg | Q348P | PLES_48021 | | | | | | | C | C | | | | |
| NC_011770 | 5296891 | G | A | 93 | 0 | 1 | 0 | NON_SYNONYMOU | MODERAT | MISSENSE | Ccg/Tcg | P363S | PLES_48091 | A | | | | | | | | | | | |
| NC_011770 | 5300076 | C | T | 67 | 0 | 2 | 0 | NON_SYNONYMOU | MODERAT | MISSENSE | gCc/gTc | A59V | PLES_48131 | | | | | | | T | T | | | | |
| NC_011770 | 5300129 | A | G | 69 | 0 | 2 | 0 | NON_SYNONYMOU | MODERAT | MISSENSE | Agc/Ggc | S77G | PLES_48131 | | | | | | | | | | | G | G |
| NC_011770 | 5300339 | C | G | 67 | 0 | 2 | 0 | NON_SYNONYMOU | MODERAT | MISSENSE | Cag/Gag | Q147E | PLES_48131 | | | | | | | | | | | G | G |
| NC_011770 | 5300400 | T | C | 63 | 0 | 2 | 0 | NON_SYNONYMOU | MODERAT | MISSENSE | gTg/gCg | V167A | PLES_48131 | | | | | | | | | | | C | C |
| NC_011770 | 5300595 | A | G | 62 | 0 | 1 | 0 | NON_SYNONYMOU | MODERAT | MISSENSE | tAc/tGc | Y232C | PLES_48131 | | | | | | | G | | | | | |
| NC_011770 | 5300615 | A | G | 61 | 0 | 2 | 0 | NON_SYNONYMOU | MODERAT | MISSENSE | Atg/Gtg | M239V | PLES_48131 | | | | | | | | | | | G | G |
| NC_011770 | 5300753 | A | C | 54 | 0 | 2 | 0 | NON_SYNONYMOU | MODERAT | MISSENSE | Atg/Ctg | M285L | PLES_48131 | | | | | | | | | | | C | C |
| NC_011770 | 5300795 | C | T | 48 | 0 | 2 | 0 | NON_SYNONYMOU | MODERAT | MISSENSE | Ccg/Tcg | P299S | PLES_48131 | | | | | | | T | T | | | | |
| NC_011770 | 5300846 | A | G | 48 | 0 | 2 | 0 | NON_SYNONYMOU | MODERAT | MISSENSE | Acc/Gcc | T316A | PLES_48131 | | | | | | | | | | | G | G |
| NC_011770 | 5300861 | G | A | 49 | 0 | 4 | 0 | NON_SYNONYMOU | MODERAT | MISSENSE | Gtc/Atc | V321I | PLES_48131 | A | A | | | | | A | A | | | | |
| NC_011770 | 5300885 | T | G | 49 | 0 | 6 | 0 | NON_SYNONYMOU | MODERAT | MISSENSE | Ttg/Gtg | L329V | PLES_48131 | G | G | | | | | G | G | | | G | G |
| NC_011770 | 5300762 | G | A | 50 | 0 | 1 | 0 | NON_SYNONYMOU | MODERAT | MISSENSE | Gcg/Acg | A288T | PLES_48131 | | | | | | | | A | | | | |
| NC_011770 | 5301396 | G | A | 79 | 0 | 1 | 0 | NON_SYNONYMOU | MODERAT | MISSENSE | gCc/gTc | A258V | PLES_48141 | | A | | | | | | | | | | |
| NC_011770 | 5302068 | C | T | 80 | 0 | 12 | 0 | NON_SYNONYMOU | MODERAT | MISSENSE | cGc/cAc | R34H | PLES_48141 | T | T | T | T | T | T | T | T | T | T | T | T |
| NC_011770 | 5301867 | T | C | 86 | 0 | 1 | 0 | NON_SYNONYMOU | MODERAT | MISSENSE | tAc/tGc | Y101C | PLES_48141 | | | | | | | | C | | | | |
| NC_011770 | 5303747 | A | G | 62 | 0 | 12 | 0 | NON_SYNONYMOU | MODERAT | MISSENSE | Acg/Gcg | T90A | PLES_48161 | G | G | G | G | G | G | G | G | G | G | G | G |
| NC_011770 | 5304245 | G | A | 73 | 0 | 2 | 0 | NON_SYNONYMOU | MODERAT | MISSENSE | Gcc/Acc | A256T | PLES_48161 | A | A | | | | | | | | | | |
| NC_011770 | 5304308 | G | A | 72 | 0 | 6 | 0 | NON_SYNONYMOU | MODERAT | MISSENSE | Gac/Aac | D277N | PLES_48161 | | | A | A | A | A | | | A | A | | |
| NC_011770 | 5307364 | G | C | 80 | 0 | 2 | 0 | NON_SYNONYMOU | MODERAT | MISSENSE | Cag/Gag | Q126E | PLES_48191 | | | | | | | | | | | C | C |
| NC_011770 | 5307403 | A | C | 82 | 0 | 4 | 0 | NON_SYNONYMOU | MODERAT | MISSENSE | Tcc/Gcc | S113A | PLES_48191 | | | C | C | C | C | | | | | | |
| NC_011770 | 5307555 | G | A | 85 | 0 | 2 | 0 | NON_SYNONYMOU | MODERAT | MISSENSE | gCg/gTg | A62V | PLES_48191 | | | | | | | A | A | | | | |
| NC_011770 | 5313168 | T | C | 81 | 0 | 2 | 0 | NON_SYNONYMOU | MODERAT | MISSENSE | Acc/Gcc | T87A | PLES_48241 | C | C | | | | | | | | | | |
| NC_011770 | 5319616 | T | G | 76 | 0 | 12 | 0 | NON_SYNONYMOU | MODERAT | MISSENSE | gaA/gaC | E83D | PLES_48311 | G | G | G | G | G | G | G | G | G | G | G | G |
| NC_011770 | 5319859 | G | C | 78 | 0 | 4 | 0 | NON_SYNONYMOU | MODERAT | MISSENSE | agC/agG | S2R | PLES_48311 | | | C | C | C | C | | | | | | |
| NC_011770 | 5320390 | T | C | 84 | 0 | 1 | 0 | NON_SYNONYMOU | MODERAT | MISSENSE | gAc/gGc | D40G | PLES_48321 | | | | | | C | | | | | | |
| NC_011770 | 5320711 | T | C | 75 | 0 | 12 | 0 | NON_SYNONYMOU | MODERAT | MISSENSE | Atc/Gtc | I95V | PLES_48331 | C | C | C | C | C | C | C | C | C | C | C | C |
| NC_011770 | 5321194 | T | C | 75 | 0 | 4 | 0 | NON_SYNONYMOU | MODERAT | MISSENSE | Acc/Gcc | T200A | PLES_48341 | | | C | C | C | C | | | | | | |
| NC_011770 | 5323191 | G | A | 77 | 0 | 8 | 0 | NON_SYNONYMOU | MODERAT | MISSENSE | Gtc/Atc | V60I | PLES_48361 | A | A | | | | | A | A | A | A | A | A |
| NC_011770 | 5324036 | A | C | 81 | 0 | 2 | 0 | NON_SYNONYMOU | MODERAT | MISSENSE | Agc/Cgc | S15R | PLES_48371 | | | | | | | C | C | | | | |

| | | | | | | | | | | | | | | | | | | | | | | | | | |
|---|---|---|---|---|---|---|---|---|---|---|---|---|---|---|---|---|---|---|---|---|---|---|---|---|---|
| NC_011770 | 5324519 | G | A | 90 | 0 | 4 | 0 | NON_SYNONYMOU | MODERAT | MISSENSE | Ggc/Agc | G176S | PLES_48371 | A | A | | | | | | | | | A | A |
| NC_011770 | 5325052 | T | C | 88 | 0 | 1 | 0 | NON_SYNONYMOU | MODERAT | MISSENSE | Tat/Cat | Y171H | PLES_48381 | | | | | | | C | | | | | |
| NC_011770 | 5325203 | G | A | 90 | 0 | 1 | 0 | NON_SYNONYMOU | MODERAT | MISSENSE | Gtc/Atc | V35I | PLES_48391 | | | | | | | A | | | | | |
| NC_011770 | 5329217 | G | A | 67 | 0 | 2 | 0 | NON_SYNONYMOU | MODERAT | MISSENSE | Gac/Aac | D94N | PLES_48441 | | | | | | | | | A | A | | |
| NC_011770 | 5329292 | G | A | 67 | 0 | 1 | 0 | NON_SYNONYMOU | MODERAT | MISSENSE | Gag/Aag | E119K | PLES_48441 | | | | | | | A | | | | | |
| NC_011770 | 5329900 | G | A | 58 | 0 | 4 | 0 | NON_SYNONYMOU | MODERAT | MISSENSE | Gtc/Atc | V25I | PLES_48451 | | | A | A | A | A | | | | | | |
| NC_011770 | 5330082 | G | T | 61 | 0 | 4 | 0 | NON_SYNONYMOU | MODERAT | MISSENSE | agG/agT | R85S | PLES_48451 | | | | | | | T | T | T | T | | |
| NC_011770 | 5330092 | G | T | 58 | 0 | 2 | 0 | NON_SYNONYMOU | MODERAT | MISSENSE | Ggc/Tgc | G89C | PLES_48451 | | | | | | | | | T | T | | |
| NC_011770 | 5330330 | C | G | 65 | 0 | 10 | 0 | NON_SYNONYMOU | MODERAT | MISSENSE | gGg/gCg | G267A | PLES_48461 | G | G | G | G | G | G | G | G | G | G | | |
| NC_011770 | 5330855 | G | C | 71 | 0 | 2 | 0 | NON_SYNONYMOU | MODERAT | MISSENSE | gCg/gGg | A92G | PLES_48461 | | | | | | | C | C | | | | |
| NC_011770 | 5331070 | T | G | 82 | 0 | 2 | 0 | NON_SYNONYMOU | MODERAT | MISSENSE | gaA/gaC | E20D | PLES_48461 | | | | | | | G | G | | | | |
| NC_011770 | 5331789 | AG | AGG | 73 | 0 | 3 | 0 | FRAME_SHIFT | HIGH | | ccc/ | P144 | PLES_48481 | | AGG | | | | | AGG | A | | | | |
| NC_011770 | 5333667 | G | A | 88 | 0 | 2 | 0 | NON_SYNONYMOU | MODERAT | MISSENSE | gCg/gTg | A116V | PLES_48501 | | | | | | | A | A | | | | |
| NC_011770 | 5333701 | G | T | 86 | 0 | 4 | 0 | NON_SYNONYMOU | MODERAT | MISSENSE | Cag/Aag | Q105K | PLES_48501 | | | T | T | T | T | | | | | | |
| NC_011770 | 5335945 | G | A | 84 | 0 | 2 | 0 | NON_SYNONYMOU | MODERAT | MISSENSE | Ggc/Agc | G117S | PLES_48521 | | | | | | | | | A | A | | |
| NC_011770 | 5336474 | T | C | 85 | 0 | 2 | 0 | NON_SYNONYMOU | MODERAT | MISSENSE | cAg/cGg | Q374R | PLES_48531 | C | C | | | | | | | | | | |
| NC_011770 | 5337245 | A | G | 75 | 0 | 2 | 0 | NON_SYNONYMOU | MODERAT | MISSENSE | gTg/gCg | V117A | PLES_48531 | | | | | | | G | G | | | | |
| NC_011770 | 5337700 | A | G | 63 | 0 | 4 | 0 | NON_SYNONYMOU | MODERAT | MISSENSE | gTg/gCg | V249A | PLES_48541 | G | G | | | | | G | G | | | | |
| NC_011770 | 5338117 | A | C | 66 | 0 | 2 | 0 | NON_SYNONYMOU | MODERAT | MISSENSE | gTc/gGc | V110G | PLES_48541 | | | | | | | C | C | | | | |
| NC_011770 | 5338791 | C | T | 67 | 0 | 4 | 0 | NON_SYNONYMOU | MODERAT | MISSENSE | Gtc/Atc | V1179I | PLES_48551 | | | | | | | T | T | | | T | T |
| NC_011770 | 5338817 | A | C | 60 | 0 | 2 | 0 | NON_SYNONYMOU | MODERAT | MISSENSE | gTt/gGt | V1170G | PLES_48551 | | | | | | | C | C | | | | |
| NC_011770 | 5339496 | G | A | 75 | 0 | 4 | 0 | NON_SYNONYMOU | MODERAT | MISSENSE | Cgg/Tgg | R944W | PLES_48551 | | | A | A | A | A | | | | | | |
| NC_011770 | 5339500 | C | A | 77 | 0 | 2 | 0 | NON_SYNONYMOU | MODERAT | MISSENSE | caG/caT | Q942H | PLES_48551 | | | | | | | | | | | A | A |
| NC_011770 | 5340113 | C | T | 71 | 0 | 4 | 0 | NON_SYNONYMOU | MODERAT | MISSENSE | aGc/aAc | S738N | PLES_48551 | | | | | | | T | T | T | T | | |
| NC_011770 | 5340566 | G | A | 60 | 0 | 4 | 0 | NON_SYNONYMOU | MODERAT | MISSENSE | gCc/gTc | A587V | PLES_48551 | | | | | | | A | A | A | A | | |
| NC_011770 | 5340828 | C | A | 62 | 0 | 4 | 0 | NON_SYNONYMOU | MODERAT | MISSENSE | Gct/Tct | A500S | PLES_48551 | | | A | A | A | A | | | | | | |
| NC_011770 | 5340873 | G | T | 61 | 0 | 6 | 0 | NON_SYNONYMOU | MODERAT | MISSENSE | Cgc/Agc | R485S | PLES_48551 | | | T | T | T | T | T | T | | | | |
| NC_011770 | 5341577 | A | G | 73 | 0 | 2 | 0 | NON_SYNONYMOU | MODERAT | MISSENSE | gTc/gCc | V250A | PLES_48551 | | | | | | | G | G | | | | |
| NC_011770 | 5341583 | G | T | 78 | 0 | 4 | 0 | NON_SYNONYMOU | MODERAT | MISSENSE | aCg/aAg | T248K | PLES_48551 | T | T | | | | | T | T | | | | |
| NC_011770 | 5344043 | C | T | 67 | 0 | 8 | 0 | NON_SYNONYMOU | MODERAT | MISSENSE | Gtc/Atc | V132I | PLES_48571 | T | T | T | T | T | T | T | T | | | | |
| NC_011770 | 5344164 | G | C | 76 | 0 | 10 | 0 | NON_SYNONYMOU | MODERAT | MISSENSE | gaC/gaG | D91E | PLES_48571 | C | C | C | C | C | C | C | C | C | C | | |
| NC_011770 | 5344234 | C | T | 84 | 0 | 6 | 0 | NON_SYNONYMOU | MODERAT | MISSENSE | tGt/tAt | C68Y | PLES_48571 | | | T | T | T | T | | | T | T | | |
| NC_011770 | 5344286 | A | C | 84 | 0 | 12 | 0 | NON_SYNONYMOU | MODERAT | MISSENSE | Tgc/Ggc | C51G | PLES_48571 | C | C | C | C | C | C | C | C | C | C | C | C |
| NC_011770 | 5344412 | C | T | 90 | 0 | 2 | 0 | NON_SYNONYMOU | MODERAT | MISSENSE | Gcc/Acc | A9T | PLES_48571 | | | | | | | | | | | T | T |
| NC_011770 | 5350778 | T | C | 30 | 0 | 2 | 0 | NON_SYNONYMOU | MODERAT | MISSENSE | gTc/gCc | V14A | PLES_48651 | | | | | | | | | C | C | | |
| NC_011770 | 5352085 | A | C | 72 | 0 | 2 | 0 | NON_SYNONYMOU | MODERAT | MISSENSE | gTc/gGc | V122G | PLES_48671 | C | C | | | | | | | | | | |
| NC_011770 | 5352225 | G | C | 75 | 0 | 2 | 0 | NON_SYNONYMOU | MODERAT | MISSENSE | gaC/gaG | D75E | PLES_48671 | | | | | | | | | | | C | C |
| NC_011770 | 5352274 | T | C | 69 | 0 | 1 | 0 | NON_SYNONYMOU | MODERAT | MISSENSE | cAg/cGg | Q59R | PLES_48671 | | | | | | | C | | | | | |
| NC_011770 | 5352413 | G | C | 75 | 0 | 4 | 0 | NON_SYNONYMOU | MODERAT | MISSENSE | Ctt/Gtt | L13V | PLES_48671 | | | C | C | C | C | | | | | | |
| NC_011770 | 5352489 | T | C | 66 | 0 | 10 | 0 | NON_SYNONYMOU | MODERAT | MISSENSE | Atc/Gtc | I539V | PLES_48681 | C | C | C | C | C | C | | | C | C | C | C |
| NC_011770 | 5352656 | G | T | 75 | 0 | 6 | 0 | NON_SYNONYMOU | MODERAT | MISSENSE | cCg/cAg | P483Q | PLES_48681 | | | T | T | T | T | | | | | T | T |
| NC_011770 | 5352744 | G | A | 73 | 0 | 6 | 0 | NON_SYNONYMOU | MODERAT | MISSENSE | Ccg/Tcg | P454S | PLES_48681 | | | A | A | A | A | | | | | A | A |
| NC_011770 | 5352944 | C | T | 69 | 0 | 2 | 0 | NON_SYNONYMOU | MODERAT | MISSENSE | cGg/cAg | R387Q | PLES_48681 | | | | | | | | | | | T | T |
| NC_011770 | 5352957 | A | C | 68 | 0 | 12 | 0 | NON_SYNONYMOU | MODERAT | MISSENSE | Tcg/Gcg | S383A | PLES_48681 | C | C | C | C | C | C | C | C | C | C | C | C |
| NC_011770 | 5353407 | T | C | 72 | 0 | 6 | 0 | NON_SYNONYMOU | MODERAT | MISSENSE | Atg/Gtg | M233V | PLES_48681 | | | C | C | C | C | | | | | C | C |
| NC_011770 | 5353686 | A | G | 71 | 0 | 12 | 0 | NON_SYNONYMOU | MODERAT | MISSENSE | Ttc/Ctc | F140L | PLES_48681 | G | G | G | G | G | G | G | G | G | G | G | G |
| NC_011770 | 5353803 | C | T | 69 | 0 | 4 | 0 | NON_SYNONYMOU | MODERAT | MISSENSE | Ggt/Agt | G101S | PLES_48681 | | | T | T | T | T | | | | | | |
| NC_011770 | 5354597 | G | C | 69 | 0 | 2 | 0 | NON_SYNONYMOU | MODERAT | MISSENSE | Ctc/Gtc | L1352V | PLES_48691 | | | | | | | C | C | | | | |
| NC_011770 | 5354833 | A | G | 72 | 0 | 10 | 0 | NON_SYNONYMOU | MODERAT | MISSENSE | gTc/gCc | V1273A | PLES_48691 | G | G | G | G | G | G | | | G | G | G | G |
| NC_011770 | 5354857 | C | T | 73 | 0 | 2 | 0 | NON_SYNONYMOU | MODERAT | MISSENSE | cGc/cAc | R1265H | PLES_48691 | | | | | | | | | T | T | | |
| NC_011770 | 5355625 | A | G | 76 | 0 | 4 | 0 | NON_SYNONYMOU | MODERAT | MISSENSE | gTg/gCg | V1009A | PLES_48691 | | | G | G | G | G | | | | | | |
| NC_011770 | 5355626 | C | T | 78 | 0 | 2 | 0 | NON_SYNONYMOU | MODERAT | MISSENSE | Gtg/Atg | V1009M | PLES_48691 | | | | | | | | | | | T | T |
| NC_011770 | 5357290 | T | C | 88 | 0 | 4 | 0 | NON_SYNONYMOU | MODERAT | MISSENSE | aAg/aGg | K454R | PLES_48691 | | | C | C | C | C | | | | | | |
| NC_011770 | 5357336 | C | A | 82 | 0 | 4 | 0 | NON_SYNONYMOU | MODERAT | MISSENSE | Gcc/Tcc | A439S | PLES_48691 | | | A | A | A | A | | | | | | |
| NC_011770 | 5358626 | G | A | 85 | 0 | 4 | 0 | NON_SYNONYMOU | MODERAT | MISSENSE | Cgt/Tgt | R9C | PLES_48691 | | | A | A | A | A | | | | | | |
| NC_011770 | 5359250 | C | A | 62 | 0 | 6 | 0 | NON_SYNONYMOU | MODERAT | MISSENSE | gaG/gaT | E22D | PLES_48701 | | | A | A | A | A | | | | | A | A |
| NC_011770 | 5359297 | A | T | 63 | 0 | 2 | 0 | NON_SYNONYMOU | MODERAT | MISSENSE | Ttc/Atc | F7I | PLES_48701 | | | | | | | | | T | T | | |
| NC_011770 | 5359808 | G | A | 62 | 0 | 12 | 0 | NON_SYNONYMOU | MODERAT | MISSENSE | gCc/gTc | A425V | PLES_48711 | A | A | A | A | A | A | A | A | A | A | A | A |
| NC_011770 | 5360306 | G | A | 60 | 0 | 4 | 0 | NON_SYNONYMOU | MODERAT | MISSENSE | gCa/gTa | A259V | PLES_48711 | A | A | | | | | | | | | A | A |
| NC_011770 | 5360484 | C | T | 54 | 0 | 2 | 0 | NON_SYNONYMOU | MODERAT | MISSENSE | Gtc/Atc | V200I | PLES_48711 | T | T | | | | | | | | | | |
| NC_011770 | 5360867 | T | G | 71 | 0 | 8 | 0 | NON_SYNONYMOU | MODERAT | MISSENSE | gAc/gCc | D72A | PLES_48711 | G | G | G | G | G | G | | | | | G | G |
| NC_011770 | 5361244 | T | C | 87 | 0 | 2 | 0 | NON_SYNONYMOU | MODERAT | MISSENSE | gAc/gGc | D229G | PLES_48721 | | | | | C | C | | | | | | |

| | | | | | | | | | | | | | | | | | | | | | | | | | |
|---|---|---|---|---|---|---|---|---|---|---|---|---|---|---|---|---|---|---|---|---|---|---|---|---|---|
| NC_011770 | 5361428 | C | A | 87 | 0 | 6 | 0 | NON_SYNONYMOU | MODERAT | MISSENSE | Gcc/Tcc | A168S | PLES_48721 | | | A | A | A | A | | | | | A | A |
| NC_011770 | 5365200 | G | T | 84 | 0 | 2 | 0 | NON_SYNONYMOU | MODERAT | MISSENSE | Gtc/Ttc | V6F | PLES_48761 | T | T | | | | | | | | | | |
| NC_011770 | 5366581 | C | T | 81 | 0 | 2 | 0 | NON_SYNONYMOU | MODERAT | MISSENSE | gCc/gTc | A466V | PLES_48761 | | | | | | | | | | | T | T |
| NC_011770 | 5366937 | T | C | 65 | 0 | 6 | 0 | NON_SYNONYMOU | MODERAT | MISSENSE | Ttc/Ctc | F16L | PLES_48771 | | | C | C | C | C | | | | | C | C |
| NC_011770 | 5367214 | C | G | 59 | 0 | 2 | 0 | NON_SYNONYMOU | MODERAT | MISSENSE | aCc/aGc | T108S | PLES_48771 | G | G | | | | | | | | | | |
| NC_011770 | 5367225 | G | A | 58 | 0 | 2 | 0 | NON_SYNONYMOU | MODERAT | MISSENSE | Gac/Aac | D112N | PLES_48771 | A | A | | | | | | | | | | |
| NC_011770 | 5367568 | T | C | 63 | 0 | 12 | 0 | NON_SYNONYMOU | MODERAT | MISSENSE | gTc/gCc | V226A | PLES_48771 | C | C | C | C | C | C | C | C | C | C | C | C |
| NC_011770 | 5367830 | A | G | 65 | 0 | 12 | 0 | NON_SYNONYMOU | MODERAT | MISSENSE | atA/atG | I313M | PLES_48771 | G | G | G | G | G | G | G | G | G | G | G | G |
| NC_011770 | 5367843 | T | C | 60 | 0 | 6 | 0 | NON_SYNONYMOU | MODERAT | MISSENSE | Tcg/Ccg | S318P | PLES_48771 | | | C | C | C | C | C | C | | | | |
| NC_011770 | 5367883 | C | T | 57 | 0 | 2 | 0 | NON_SYNONYMOU | MODERAT | MISSENSE | cCg/cTg | P331L | PLES_48771 | | | | | | | | | T | T | | |
| NC_011770 | 5368026 | G | A | 60 | 0 | 1 | 0 | NON_SYNONYMOU | MODERAT | MISSENSE | Ggc/Agc | G379S | PLES_48771 | | | | | | | A | | | | | |
| NC_011770 | 5368083 | G | T | 62 | 0 | 4 | 0 | NON_SYNONYMOU | MODERAT | MISSENSE | Gcc/Tcc | A398S | PLES_48771 | | | T | T | T | T | | | | | | |
| NC_011770 | 5369297 | G | A | 98 | 0 | 1 | 0 | NON_SYNONYMOU | MODERAT | MISSENSE | gCc/gTc | A160V | PLES_48781 | | | | | | | | | | | | A |
| NC_011770 | 5368874 | T | C | 82 | 0 | 2 | 0 | NON_SYNONYMOU | MODERAT | MISSENSE | cAg/cGg | Q301R | PLES_48781 | | | | | | | C | C | | | | |
| NC_011770 | 5369024 | T | A | 85 | 0 | 12 | 0 | NON_SYNONYMOU | MODERAT | MISSENSE | gAg/gTg | E251V | PLES_48781 | A | A | A | A | A | A | A | A | A | A | A | A |
| NC_011770 | 5369039 | T | C | 89 | 0 | 2 | 0 | NON_SYNONYMOU | MODERAT | MISSENSE | aAg/aGg | K246R | PLES_48781 | | | | | | | C | C | | | | |
| NC_011770 | 5369402 | A | G | 97 | 0 | 10 | 0 | NON_SYNONYMOU | MODERAT | MISSENSE | gTg/gCg | V125A | PLES_48781 | G | G | G | G | G | G | | | G | G | G | G |
| NC_011770 | 5370205 | C | T | 68 | 0 | 1 | 0 | STOP_GAINED | HIGH | NONSENSE | Cag/Tag | Q88* | PLES_48791 | | | | | | | | | | | T | |
| NC_011770 | 5370175 | G | A | 71 | 0 | 2 | 0 | NON_SYNONYMOU | MODERAT | MISSENSE | Ggc/Agc | G78S | PLES_48791 | | | | | | | | | | | A | A |
| NC_011770 | 5370379 | A | C | 56 | 0 | 12 | 0 | NON_SYNONYMOU | MODERAT | MISSENSE | Atc/Ctc | I146L | PLES_48791 | C | C | C | C | C | C | C | C | C | C | C | C |
| NC_011770 | 5371046 | C | A | 64 | 0 | 2 | 0 | NON_SYNONYMOU | MODERAT | MISSENSE | Ctg/Atg | L92M | PLES_48801 | | | | | | | | | A | A | | |
| NC_011770 | 5371868 | G | A | 79 | 0 | 2 | 0 | NON_SYNONYMOU | MODERAT | MISSENSE | Gtc/Atc | V366I | PLES_48801 | | | | | | | A | A | | | | |
| NC_011770 | 5371871 | A | G | 80 | 0 | 12 | 0 | NON_SYNONYMOU | MODERAT | MISSENSE | Acc/Gcc | T367A | PLES_48801 | G | G | G | G | G | G | G | G | G | G | G | G |
| NC_011770 | 5374163 | G | A | 63 | 0 | 1 | 0 | NON_SYNONYMOU | MODERAT | MISSENSE | Gcc/Acc | A21T | PLES_48821 | | | | | | | A | | | | | |
| NC_011770 | 5374562 | G | A | 82 | 0 | 2 | 0 | NON_SYNONYMOU | MODERAT | MISSENSE | Gac/Aac | D154N | PLES_48821 | | | | | | | A | A | | | | |
| NC_011770 | 5376675 | C | T | 89 | 0 | 1 | 0 | STOP_GAINED | HIGH | NONSENSE | Cag/Tag | Q304* | PLES_48831 | | | | | | | | T | | | | |
| NC_011770 | 5379900 | T | C | 72 | 0 | 2 | 0 | NON_SYNONYMOU | MODERAT | MISSENSE | Acg/Gcg | T155A | PLES_48871 | | | | | | | C | C | | | | |
| NC_011770 | 5380041 | A | G | 70 | 0 | 6 | 0 | NON_SYNONYMOU | MODERAT | MISSENSE | Tac/Cac | Y108H | PLES_48871 | | | G | G | G | G | | | G | G | | |
| NC_011770 | 5380047 | C | T | 70 | 0 | 2 | 0 | NON_SYNONYMOU | MODERAT | MISSENSE | Gtc/Atc | V106I | PLES_48871 | | | | | | | | | | | T | T |
| NC_011770 | 5381068 | CG | C | 81 | 0 | 2 | 0 | FRAME_SHIFT | HIGH | | ccc/ | P281 | PLES_48891 | | | | | | | | | | | C | C |
| NC_011770 | 5381242 | T | C | 76 | 0 | 8 | 0 | NON_SYNONYMOU | MODERAT | MISSENSE | Aag/Gag | K224E | PLES_48891 | | | C | C | C | C | C | C | C | C | | |
| NC_011770 | 5381263 | T | C | 77 | 0 | 2 | 0 | NON_SYNONYMOU | MODERAT | MISSENSE | Atg/Gtg | M217V | PLES_48891 | C | C | | | | | | | | | | |
| NC_011770 | 5381350 | A | G | 71 | 0 | 8 | 0 | NON_SYNONYMOU | MODERAT | MISSENSE | Ttc/Ctc | F188L | PLES_48891 | G | G | G | G | G | G | | | G | G | | |
| NC_011770 | 5381508 | G | A | 62 | 0 | 12 | 0 | NON_SYNONYMOU | MODERAT | MISSENSE | gCc/gTc | A135V | PLES_48891 | A | A | A | A | A | A | A | A | A | A | A | A |
| NC_011770 | 5382504 | G | GGC | 79 | 0 | 1 | 0 | FRAME_SHIFT | HIGH | | cgc/cgcGC | R27R? | PLES_48901 | | | | | | | GGC | | | | | |
| NC_011770 | 5381936 | G | T | 65 | 0 | 2 | 0 | NON_SYNONYMOU | MODERAT | MISSENSE | gCt/gAt | A217D | PLES_48901 | | | | | | | | | | | T | T |
| NC_011770 | 5382390 | T | C | 73 | 0 | 12 | 0 | NON_SYNONYMOU | MODERAT | MISSENSE | Ata/Gta | I66V | PLES_48901 | C | C | C | C | C | C | C | C | C | C | C | C |
| NC_011770 | 5382413 | T | G | 77 | 0 | 2 | 0 | NON_SYNONYMOU | MODERAT | MISSENSE | gAc/gCc | D58A | PLES_48901 | G | G | | | | | | | | | | |
| NC_011770 | 5382476 | T | C | 72 | 0 | 4 | 0 | NON_SYNONYMOU | MODERAT | MISSENSE | gAg/gGg | E37G | PLES_48901 | | | C | C | C | C | | | | | | |
| NC_011770 | 5382740 | C | T | 78 | 0 | 12 | 0 | NON_SYNONYMOU | MODERAT | MISSENSE | Gtc/Atc | V200I | PLES_48911 | T | T | T | T | T | T | T | T | T | T | T | T |
| NC_011770 | 5382839 | G | A | 86 | 0 | 2 | 0 | NON_SYNONYMOU | MODERAT | MISSENSE | Ccg/Tcg | P167S | PLES_48911 | | | | | | | | | A | A | | |
| NC_011770 | 5383186 | C | A | 101 | 0 | 4 | 0 | NON_SYNONYMOU | MODERAT | MISSENSE | cGg/cTg | R51L | PLES_48911 | | | A | A | A | A | | | | | | |
| NC_011770 | 5386563 | C | T | 83 | 0 | 1 | 0 | NON_SYNONYMOU | MODERAT | MISSENSE | gGg/gAg | G179E | PLES_48931 | T | | | | | | | | | | | |
| NC_011770 | 5384923 | G | C | 87 | 0 | 2 | 0 | NON_SYNONYMOU | MODERAT | MISSENSE | Ctc/Gtc | L726V | PLES_48931 | | | | | | | | | | | C | C |
| NC_011770 | 5385058 | G | T | 86 | 0 | 2 | 0 | NON_SYNONYMOU | MODERAT | MISSENSE | Ctg/Atg | L681M | PLES_48931 | T | T | | | | | | | | | | |
| NC_011770 | 5385207 | T | A | 94 | 0 | 2 | 0 | NON_SYNONYMOU | MODERAT | MISSENSE | gAc/gTc | D631V | PLES_48931 | | | | | | | A | A | | | | |
| NC_011770 | 5385318 | G | A | 87 | 0 | 4 | 0 | NON_SYNONYMOU | MODERAT | MISSENSE | gCg/gTg | A594V | PLES_48931 | | | | | | | A | A | A | A | | |
| NC_011770 | 5385379 | C | T | 84 | 0 | 2 | 0 | NON_SYNONYMOU | MODERAT | MISSENSE | Gac/Aac | D574N | PLES_48931 | | | | | | | T | T | | | | |
| NC_011770 | 5385382 | T | C | 80 | 0 | 2 | 0 | NON_SYNONYMOU | MODERAT | MISSENSE | Acc/Gcc | T573A | PLES_48931 | | | | | | | C | C | | | | |
| NC_011770 | 5385967 | T | C | 77 | 0 | 12 | 0 | NON_SYNONYMOU | MODERAT | MISSENSE | Acc/Gcc | T378A | PLES_48931 | C | C | C | C | C | C | C | C | C | C | C | C |
| NC_011770 | 5386072 | T | G | 87 | 0 | 2 | 0 | NON_SYNONYMOU | MODERAT | MISSENSE | Atc/Ctc | I343L | PLES_48931 | | | | | | | | | G | G | | |
| NC_011770 | 5386167 | T | C | 81 | 0 | 12 | 0 | NON_SYNONYMOU | MODERAT | MISSENSE | aAc/aGc | N311S | PLES_48931 | C | C | C | C | C | C | C | C | C | C | C | C |
| NC_011770 | 5386671 | A | G | 92 | 0 | 8 | 0 | NON_SYNONYMOU | MODERAT | MISSENSE | gTc/gCc | V143A | PLES_48931 | G | G | | | | | G | G | G | G | G | G |
| NC_011770 | 5386932 | G | A | 84 | 0 | 1 | 0 | NON_SYNONYMOU | MODERAT | MISSENSE | cCg/cTg | P56L | PLES_48931 | | | | | | | A | | | | | |
| NC_011770 | 5386948 | A | G | 84 | 0 | 8 | 0 | NON_SYNONYMOU | MODERAT | MISSENSE | Tcc/Ccc | S51P | PLES_48931 | G | G | | | | | G | G | G | G | G | G |
| NC_011770 | 5387451 | G | C | 69 | 0 | 6 | 0 | NON_SYNONYMOU | MODERAT | MISSENSE | Ctg/Gtg | L722V | PLES_48941 | C | C | | | | | | | C | C | C | C |
| NC_011770 | 5387454 | T | G | 69 | 0 | 2 | 0 | NON_SYNONYMOU | MODERAT | MISSENSE | Aaa/Caa | K721Q | PLES_48941 | | | | | | | G | G | | | | |
| NC_011770 | 5387708 | C | G | 67 | 0 | 6 | 0 | NON_SYNONYMOU | MODERAT | MISSENSE | aGc/aCc | S636T | PLES_48941 | G | G | | | | | | | G | G | G | G |
| NC_011770 | 5387717 | G | C | 63 | 0 | 6 | 0 | NON_SYNONYMOU | MODERAT | MISSENSE | gCg/gGg | A633G | PLES_48941 | C | C | | | | | | | C | C | C | C |
| NC_011770 | 5387885 | A | G | 57 | 6 | 6 | 0 | NON_SYNONYMOU | MODERAT | MISSENSE | gTt/gCt | V577A | PLES_48941 | . | . | G | G | G | G | G | G | . | . | . | . |
| NC_011770 | 5387971 | A | C | 58 | 6 | 2 | 0 | NON_SYNONYMOU | MODERAT | MISSENSE | gaT/gaG | D548E | PLES_48941 | . | . | | | | | C | C | . | . | . | . |
| NC_011770 | 5389019 | T | C | 54 | 6 | 1 | 0 | NON_SYNONYMOU | MODERAT | MISSENSE | tAt/tGt | Y199C | PLES_48941 | . | . | | | | | C | | . | . | . | . |

| | | | | | | | | | | | | | | | | | | | | | | | | | |
|---|---|---|---|---|---|---|---|---|---|---|---|---|---|---|---|---|---|---|---|---|---|---|---|---|---|
| NC_011770 | 5389245 | T | C | 60 | 1 | 5 | 0 | NON_SYNONYMOU | MODERAT | MISSENSE | Acc/Gcc | T124A | PLES_48941 | C | C | | | | | | | . | C | C | C |
| NC_011770 | 5389329 | G | T | 54 | 6 | 6 | 0 | NON_SYNONYMOU | MODERAT | MISSENSE | Cag/Aag | Q96K | PLES_48941 | . | . | T | T | T | T | T | T | . | . | . | . |
| NC_011770 | 5390141 | A | G | 67 | 0 | 6 | 0 | NON_SYNONYMOU | MODERAT | MISSENSE | Atc/Gtc | I104V | PLES_48951 | G | G | | | | | G | G | G | G | | |
| NC_011770 | 5390247 | G | C | 80 | 0 | 2 | 0 | NON_SYNONYMOU | MODERAT | MISSENSE | gGc/gCc | G139A | PLES_48951 | C | C | | | | | | | | | | |
| NC_011770 | 5390581 | G | A | 86 | 0 | 8 | 0 | NON_SYNONYMOU | MODERAT | MISSENSE | Gcc/Acc | A23T | PLES_48961 | A | A | | | | | A | A | A | A | A | A |
| NC_011770 | 5390696 | G | C | 94 | 0 | 8 | 0 | NON_SYNONYMOU | MODERAT | MISSENSE | aGc/aCc | S61T | PLES_48961 | C | C | | | | | C | C | C | C | C | C |
| NC_011770 | 5392281 | A | G | 97 | 0 | 2 | 0 | NON_SYNONYMOU | MODERAT | MISSENSE | Acg/Gcg | T277A | PLES_48971 | | | | | | | | | | | G | G |
| NC_011770 | 5393244 | A | G | 78 | 0 | 2 | 0 | NON_SYNONYMOU | MODERAT | MISSENSE | Aac/Gac | N598D | PLES_48971 | | | | | | | | | G | G | | |
| NC_011770 | 5393562 | C | T | 84 | 0 | 4 | 0 | NON_SYNONYMOU | MODERAT | MISSENSE | Gat/Aat | D59N | PLES_48981 | | | T | T | T | T | | | | | | |
| NC_011770 | 5393721 | C | T | 76 | 0 | 2 | 0 | NON_SYNONYMOU | MODERAT | MISSENSE | Gtt/Att | V6I | PLES_48981 | | | | | | | T | T | | | | |
| NC_011770 | 5395175 | T | C | 73 | 0 | 1 | 0 | NON_SYNONYMOU | MODERAT | MISSENSE | Tgt/Cgt | C30R | PLES_49001 | | | | | | | C | | | | | |
| NC_011770 | 5395705 | T | C | 74 | 0 | 1 | 0 | NON_SYNONYMOU | MODERAT | MISSENSE | cAg/cGg | Q595R | PLES_49011 | C | | | | | | | | | | | |
| NC_011770 | 5395756 | C | G | 81 | 0 | 2 | 0 | NON_SYNONYMOU | MODERAT | MISSENSE | gGc/gCc | G578A | PLES_49011 | | | | | | | | | G | G | | |
| NC_011770 | 5396143 | C | T | 86 | 0 | 2 | 0 | NON_SYNONYMOU | MODERAT | MISSENSE | cGc/cAc | R449H | PLES_49011 | | | | | | | T | T | | | | |
| NC_011770 | 5396746 | G | A | 78 | 0 | 8 | 0 | NON_SYNONYMOU | MODERAT | MISSENSE | gCg/gTg | A248V | PLES_49011 | A | A | A | A | A | A | | | A | A | | |
| NC_011770 | 5396747 | C | G | 80 | 0 | 8 | 0 | NON_SYNONYMOU | MODERAT | MISSENSE | Gcg/Ccg | A248P | PLES_49011 | G | G | G | G | G | G | | | G | G | | |
| NC_011770 | 5396749 | G | T | 80 | 0 | 8 | 0 | NON_SYNONYMOU | MODERAT | MISSENSE | cCg/cAg | P247Q | PLES_49011 | T | T | T | T | T | T | | | T | T | | |
| NC_011770 | 5396939 | G | T | 80 | 0 | 2 | 0 | NON_SYNONYMOU | MODERAT | MISSENSE | Ctg/Atg | L184M | PLES_49011 | | | | | | | | | | | T | T |
| NC_011770 | 5398128 | T | G | 81 | 0 | 6 | 0 | NON_SYNONYMOU | MODERAT | MISSENSE | gaA/gaC | E129D | PLES_49021 | | | G | G | G | G | | | | | G | G |
| NC_011770 | 5398418 | T | C | 83 | 0 | 4 | 0 | NON_SYNONYMOU | MODERAT | MISSENSE | Agc/Ggc | S33G | PLES_49021 | | | C | C | C | C | | | | | | |
| NC_011770 | 5399605 | C | T | 64 | 0 | 2 | 0 | NON_SYNONYMOU | MODERAT | MISSENSE | Gcg/Acg | A627T | PLES_49041 | | | | | | | | | | | T | T |
| NC_011770 | 5399619 | A | G | 62 | 0 | 12 | 0 | NON_SYNONYMOU | MODERAT | MISSENSE | gTg/gCg | V622A | PLES_49041 | G | G | G | G | G | G | G | G | G | G | G | G |
| NC_011770 | 5399622 | A | G | 62 | 0 | 12 | 0 | NON_SYNONYMOU | MODERAT | MISSENSE | gTc/gCc | V621A | PLES_49041 | G | G | G | G | G | G | G | G | G | G | G | G |
| NC_011770 | 5399634 | G | C | 64 | 0 | 10 | 0 | NON_SYNONYMOU | MODERAT | MISSENSE | aCc/aGc | T617S | PLES_49041 | C | C | C | C | C | C | C | C | C | C | | |
| NC_011770 | 5400724 | T | C | 78 | 0 | 12 | 0 | NON_SYNONYMOU | MODERAT | MISSENSE | Aca/Gca | T254A | PLES_49041 | C | C | C | C | C | C | C | C | C | C | C | C |
| NC_011770 | 5401042 | A | T | 93 | 0 | 10 | 0 | NON_SYNONYMOU | MODERAT | MISSENSE | Tcc/Acc | S148T | PLES_49041 | T | T | T | T | T | T | T | T | T | T | | |
| NC_011770 | 5401120 | A | T | 88 | 0 | 12 | 0 | NON_SYNONYMOU | MODERAT | MISSENSE | Tgc/Agc | C122S | PLES_49041 | T | T | T | T | T | T | T | T | T | T | T | T |
| NC_011770 | 5401219 | C | T | 91 | 0 | 2 | 0 | NON_SYNONYMOU | MODERAT | MISSENSE | Gtc/Atc | V89I | PLES_49041 | T | T | | | | | | | | | | |
| NC_011770 | 5403199 | CG | C | 50 | 4 | 1 | 0 | FRAME_SHIFT | HIGH | | ggc/ | G328 | PLES_49061 | | | | C | | | | | . | . | . | . |
| NC_011770 | 5403165 | G | A | 51 | 4 | 8 | 0 | NON_SYNONYMOU | MODERAT | MISSENSE | aCc/aTc | T340I | PLES_49061 | A | A | A | A | A | A | A | A | . | . | . | . |
| NC_011770 | 5403598 | A | G | 48 | 4 | 2 | 0 | NON_SYNONYMOU | MODERAT | MISSENSE | Tgg/Cgg | W196R | PLES_49061 | | | | | | | G | G | . | . | . | . |
| NC_011770 | 5403615 | C | A | 44 | 4 | 2 | 0 | NON_SYNONYMOU | MODERAT | MISSENSE | gGg/gTg | G190V | PLES_49061 | | | | | | | A | A | . | . | . | . |
| NC_011770 | 5403647 | C | T | 43 | 4 | 2 | 0 | NON_SYNONYMOU | MODERAT | MISSENSE | atG/atA | M179I | PLES_49061 | T | T | | | | | | | . | . | . | . |
| NC_011770 | 5403718 | T | C | 40 | 4 | 2 | 0 | NON_SYNONYMOU | MODERAT | MISSENSE | Agg/Ggg | R156G | PLES_49061 | C | C | | | | | | | . | . | . | . |
| NC_011770 | 5403754 | C | A | 41 | 4 | 2 | 0 | NON_SYNONYMOU | MODERAT | MISSENSE | Gcg/Tcg | A144S | PLES_49061 | A | A | | | | | | | . | . | . | . |
| NC_011770 | 5403805 | C | A | 39 | 4 | 8 | 0 | NON_SYNONYMOU | MODERAT | MISSENSE | Gtg/Ttg | V127L | PLES_49061 | A | A | G | G | G | G | G | G | . | . | . | . |
| NC_011770 | 5403814 | G | T | 35 | 4 | 6 | 0 | NON_SYNONYMOU | MODERAT | MISSENSE | Ctc/Atc | L124I | PLES_49061 | | | T | T | T | T | T | T | . | . | . | . |
| NC_011770 | 5403819 | A | G | 35 | 4 | 8 | 0 | NON_SYNONYMOU | MODERAT | MISSENSE | aTc/aCc | I122T | PLES_49061 | G | G | G | G | G | G | G | G | . | . | . | . |
| NC_011770 | 5403820 | T | C | 35 | 4 | 6 | 0 | NON_SYNONYMOU | MODERAT | MISSENSE | Atc/Gtc | I122V | PLES_49061 | | | C | C | C | C | C | C | . | . | . | . |
| NC_011770 | 5403843 | G | A | 36 | 4 | 6 | 0 | NON_SYNONYMOU | MODERAT | MISSENSE | gCc/gTc | A114V | PLES_49061 | | | A | A | A | A | A | A | . | . | . | . |
| NC_011770 | 5403877 | C | A | 35 | 4 | 4 | 0 | NON_SYNONYMOU | MODERAT | MISSENSE | Gcc/Tcc | A103S | PLES_49061 | | | A | A | A | A | | | . | . | . | . |
| NC_011770 | 5403898 | A | G | 26 | 4 | 6 | 0 | NON_SYNONYMOU | MODERAT | MISSENSE | Ttc/Ctc | F96L | PLES_49061 | | | G | G | G | G | G | G | . | . | . | . |
| NC_011770 | 5403928 | C | A | 4 | 10 | 2 | 0 | NON_SYNONYMOU | MODERAT | MISSENSE | Gca/Tca | A86S | PLES_49061 | A | A | . | . | . | . | . | . | . | . | . | . |
| NC_011770 | 5403955 | C | A | 1 | 10 | 2 | 0 | NON_SYNONYMOU | MODERAT | MISSENSE | Gtt/Ttt | V77F | PLES_49061 | A | A | . | . | . | . | . | . | . | . | . | . |
| NC_011770 | 5403964 | T | C | 1 | 10 | 2 | 0 | NON_SYNONYMOU | MODERAT | MISSENSE | Aca/Gca | T74A | PLES_49061 | C | C | . | . | . | . | . | . | . | . | . | . |
| NC_011770 | 5403966 | A | G | 1 | 10 | 2 | 0 | NON_SYNONYMOU | MODERAT | MISSENSE | gTa/gCa | V73A | PLES_49061 | G | G | . | . | . | . | . | . | . | . | . | . |
| NC_011770 | 5403968 | C | A | 1 | 10 | 2 | 0 | NON_SYNONYMOU | MODERAT | MISSENSE | caG/caT | Q72H | PLES_49061 | A | A | . | . | . | . | . | . | . | . | . | . |
| NC_011770 | 5403972 | A | G | 1 | 10 | 2 | 0 | NON_SYNONYMOU | MODERAT | MISSENSE | gTt/gCt | V71A | PLES_49061 | G | G | . | . | . | . | . | . | . | . | . | . |
| NC_011770 | 5403975 | A | T | 1 | 10 | 2 | 0 | NON_SYNONYMOU | MODERAT | MISSENSE | aTg/aAg | M70K | PLES_49061 | T | T | . | . | . | . | . | . | . | . | . | . |
| NC_011770 | 5403976 | T | C | 1 | 10 | 2 | 0 | NON_SYNONYMOU | MODERAT | MISSENSE | Atg/Gtg | M70V | PLES_49061 | C | C | . | . | . | . | . | . | . | . | . | . |
| NC_011770 | 5403982 | T | C | 1 | 10 | 2 | 0 | NON_SYNONYMOU | MODERAT | MISSENSE | Aga/Gga | R68G | PLES_49061 | C | C | . | . | . | . | . | . | . | . | . | . |
| NC_011770 | 5403985 | C | A | 1 | 10 | 2 | 0 | NON_SYNONYMOU | MODERAT | MISSENSE | Gcc/Tcc | A67S | PLES_49061 | A | A | . | . | . | . | . | . | . | . | . | . |
| NC_011770 | 5403991 | A | T | 1 | 10 | 2 | 0 | NON_SYNONYMOU | MODERAT | MISSENSE | Tca/Aca | S65T | PLES_49061 | T | T | . | . | . | . | . | . | . | . | . | . |
| NC_011770 | 5403996 | C | G | 1 | 10 | 2 | 0 | NON_SYNONYMOU | MODERAT | MISSENSE | cGt/cCt | R63P | PLES_49061 | G | G | . | . | . | . | . | . | . | . | . | . |
| NC_011770 | 5403999 | G | A | 1 | 10 | 2 | 0 | NON_SYNONYMOU | MODERAT | MISSENSE | tCt/tTt | S62F | PLES_49061 | A | A | . | . | . | . | . | . | . | . | . | . |
| NC_011770 | 5404039 | C | G | 3 | 10 | 2 | 0 | NON_SYNONYMOU | MODERAT | MISSENSE | Gtg/Ctg | V49L | PLES_49061 | G | G | . | . | . | . | . | . | . | . | . | . |
| NC_011770 | 5404075 | T | C | 4 | 10 | 2 | 0 | NON_SYNONYMOU | MODERAT | MISSENSE | Aag/Gag | K37E | PLES_49061 | C | C | . | . | . | . | . | . | . | . | . | . |
| NC_011770 | 5404081 | C | A | 4 | 10 | 2 | 0 | NON_SYNONYMOU | MODERAT | MISSENSE | Gtt/Ttt | V35F | PLES_49061 | A | A | . | . | . | . | . | . | . | . | . | . |
| NC_011770 | 5404110 | A | G | 5 | 10 | 2 | 0 | NON_SYNONYMOU | MODERAT | MISSENSE | cTt/cCt | L25P | PLES_49061 | G | G | . | . | . | . | . | . | . | . | . | . |
| NC_011770 | 5404126 | A | T | 3 | 10 | 2 | 0 | NON_SYNONYMOU | MODERAT | MISSENSE | Ttg/Atg | L20M | PLES_49061 | T | T | . | . | . | . | . | . | . | . | . | . |
| NC_011770 | 5404135 | T | C | 3 | 10 | 2 | 0 | NON_SYNONYMOU | MODERAT | MISSENSE | Att/Gtt | I17V | PLES_49061 | C | C | . | . | . | . | . | . | . | . | . | . |
| NC_011770 | 5404137 | G | A | 3 | 10 | 2 | 0 | NON_SYNONYMOU | MODERAT | MISSENSE | aCt/aTt | T16I | PLES_49061 | A | A | . | . | . | . | . | . | . | . | . | . |

| | | | | | | | | | | | | | | | | | | | | | | | | | |
|---|---|---|---|---|---|---|---|---|---|---|---|---|---|---|---|---|---|---|---|---|---|---|---|---|---|
| NC_011770 | 5404144 | A | G | 3 | 10 | 2 | 0 | NON_SYNONYMOU | MODERAT | MISSENSE | Tcg/Ccg | S14P | PLES_49061 | G | G | . | . | . | . | . | . | . | . | . | . |
| NC_011770 | 5404149 | T | C | 3 | 10 | 2 | 0 | NON_SYNONYMOU | MODERAT | MISSENSE | aAg/aGg | K12R | PLES_49061 | C | C | . | . | . | . | . | . | . | . | . | . |
| NC_011770 | 5404152 | C | T | 3 | 10 | 2 | 0 | NON_SYNONYMOU | MODERAT | MISSENSE | gGt/gAt | G11D | PLES_49061 | T | T | . | . | . | . | . | . | . | . | . | . |
| NC_011770 | 5404162 | C | T | 5 | 10 | 2 | 0 | NON_SYNONYMOU | MODERAT | MISSENSE | Gaa/Aaa | E8K | PLES_49061 | T | T | . | . | . | . | . | . | . | . | . | . |
| NC_011770 | 5404175 | A | C | 6 | 10 | 2 | 0 | NON_SYNONYMOU | MODERAT | MISSENSE | atT/atG | I3M | PLES_49061 | C | C | . | . | . | . | . | . | . | . | . | . |
| NC_011770 | 5404818 | G | A | 46 | 4 | 4 | 0 | STOP_GAINED | HIGH | NONSENSE | Caa/Taa | Q59* | PLES_49081 | | | A | A | A | A | | | . | . | . | . |
| NC_011770 | 5404822 | G | A | 42 | 4 | 6 | 0 | FRAME_SHIFT | HIGH | | tcc/tcGAC | S57ST? | PLES_49081 | A | A | GTGTC | GTGTC | GTGTC | GTGTC | | | . | . | . | . |
| NC_011770 | 5404916 | G | T | 68 | 4 | 8 | 0 | STOP_GAINED | HIGH | NONSENSE | tCa/tAa | S26* | PLES_49081 | T | T | T | T | T | T | T | T | . | . | . | . |
| NC_011770 | 5404796 | A | G | 60 | 4 | 4 | 0 | NON_SYNONYMOU | MODERAT | MISSENSE | gTt/gCt | V66A | PLES_49081 | | | G | G | G | G | | | . | . | . | . |
| NC_011770 | 5404817 | T | C | 46 | 4 | 6 | 0 | NON_SYNONYMOU | MODERAT | MISSENSE | cAa/cGa | Q59R | PLES_49081 | C | C | C | C | C | C | | | . | . | . | . |
| NC_011770 | 5404833 | G | C | 51 | 4 | 4 | 0 | NON_SYNONYMOU | MODERAT | MISSENSE | Ctt/Gtt | L54V | PLES_49081 | | | C | C | C | C | | | . | . | . | . |
| NC_011770 | 5404902 | C | T | 61 | 4 | 8 | 0 | NON_SYNONYMOU | MODERAT | MISSENSE | Ggg/Agg | G31R | PLES_49081 | T | T | T | T | T | T | T | T | . | . | . | . |
| NC_011770 | 5404904 | A | T | 60 | 4 | 8 | 0 | NON_SYNONYMOU | MODERAT | MISSENSE | cTg/cAg | L30Q | PLES_49081 | T | T | T | T | T | T | T | T | . | . | . | . |
| NC_011770 | 5409357 | G | A | 82 | 0 | 2 | 0 | NON_SYNONYMOU | MODERAT | MISSENSE | aGc/aAc | S2N | PLES_49131 | | | | | | | | | A | A | | |
| NC_011770 | 5409530 | G | A | 68 | 0 | 2 | 0 | NON_SYNONYMOU | MODERAT | MISSENSE | Gac/Aac | D60N | PLES_49131 | | | | | | | A | A | | | | |
| NC_011770 | 5410203 | T | C | 75 | 0 | 2 | 0 | NON_SYNONYMOU | MODERAT | MISSENSE | aAg/aGg | K131R | PLES_49151 | | | | | | | | | | | C | C |
| NC_011770 | 5410531 | C | G | 74 | 0 | 4 | 0 | NON_SYNONYMOU | MODERAT | MISSENSE | Gtg/Ctg | V22L | PLES_49151 | | | | | | | | | G | G | G | G |
| NC_011770 | 5410835 | T | C | 69 | 0 | 2 | 0 | NON_SYNONYMOU | MODERAT | MISSENSE | cAc/cGc | H76R | PLES_49161 | | | | | | | | | C | C | | |
| NC_011770 | 5410898 | T | C | 70 | 0 | 4 | 0 | NON_SYNONYMOU | MODERAT | MISSENSE | gAc/gGc | D55G | PLES_49161 | C | C | | | | | C | C | | | | |
| NC_011770 | 5411033 | G | C | 64 | 0 | 4 | 0 | NON_SYNONYMOU | MODERAT | MISSENSE | gCc/gGc | A10G | PLES_49161 | | | | | | | | | C | C | C | C |
| NC_011770 | 5411219 | G | T | 47 | 0 | 2 | 0 | NON_SYNONYMOU | MODERAT | MISSENSE | Ctc/Atc | L89I | PLES_49171 | | | | | | | | | T | T | | |
| NC_011770 | 5411224 | C | T | 46 | 0 | 2 | 0 | NON_SYNONYMOU | MODERAT | MISSENSE | cGg/cAg | R87Q | PLES_49171 | | | | | | | T | T | | | | |
| NC_011770 | 5411712 | G | A | 68 | 0 | 6 | 0 | NON_SYNONYMOU | MODERAT | MISSENSE | Gtc/Atc | V33I | PLES_49181 | | | A | A | A | A | | | | | A | A |
| NC_011770 | 5411796 | C | A | 78 | 0 | 2 | 0 | NON_SYNONYMOU | MODERAT | MISSENSE | Ctg/Atg | L61M | PLES_49181 | A | A | | | | | | | | | | |
| NC_011770 | 5411891 | G | T | 75 | 0 | 6 | 0 | NON_SYNONYMOU | MODERAT | MISSENSE | caG/caT | Q92H | PLES_49181 | | | T | T | T | T | | | | | T | T |
| NC_011770 | 5412621 | G | A | 78 | 0 | 6 | 0 | NON_SYNONYMOU | MODERAT | MISSENSE | cGg/cAg | R127Q | PLES_49191 | A | A | | | | | | | A | A | A | A |
| NC_011770 | 5414413 | C | T | 78 | 0 | 2 | 0 | NON_SYNONYMOU | MODERAT | MISSENSE | cGt/cAt | R326H | PLES_49221 | T | T | | | | | | | | | | |
| NC_011770 | 5414480 | G | A | 80 | 0 | 4 | 0 | NON_SYNONYMOU | MODERAT | MISSENSE | Cgc/Tgc | R304C | PLES_49221 | | | | | | | A | A | A | A | | |
| NC_011770 | 5414499 | C | G | 83 | 0 | 4 | 0 | NON_SYNONYMOU | MODERAT | MISSENSE | gaG/gaC | E297D | PLES_49221 | | | | | | | G | G | G | G | | |
| NC_011770 | 5414515 | G | A | 85 | 0 | 4 | 0 | NON_SYNONYMOU | MODERAT | MISSENSE | tCt/tTt | S292F | PLES_49221 | | | A | A | A | A | | | | | | |
| NC_011770 | 5414659 | G | C | 79 | 0 | 4 | 0 | NON_SYNONYMOU | MODERAT | MISSENSE | cCc/cGc | P244R | PLES_49221 | | | | | | | C | C | C | C | | |
| NC_011770 | 5414764 | G | A | 83 | 0 | 2 | 0 | NON_SYNONYMOU | MODERAT | MISSENSE | aCg/aTg | T209M | PLES_49221 | | | | | | | A | A | | | | |
| NC_011770 | 5414915 | G | C | 84 | 0 | 4 | 0 | NON_SYNONYMOU | MODERAT | MISSENSE | Cgc/Ggc | R159G | PLES_49221 | C | C | | | | | | | | | C | C |
| NC_011770 | 5415076 | C | T | 84 | 0 | 2 | 0 | NON_SYNONYMOU | MODERAT | MISSENSE | cGg/cAg | R105Q | PLES_49221 | | | | | | | T | T | | | | |
| NC_011770 | 5415278 | C | T | 75 | 0 | 2 | 0 | NON_SYNONYMOU | MODERAT | MISSENSE | Gca/Aca | A38T | PLES_49221 | | | | | | | | | | | T | T |
| NC_011770 | 5415299 | G | A | 70 | 0 | 2 | 0 | NON_SYNONYMOU | MODERAT | MISSENSE | Cgg/Tgg | R31W | PLES_49221 | | | | | | | A | A | | | | |
| NC_011770 | 5415337 | T | G | 72 | 0 | 12 | 0 | NON_SYNONYMOU | MODERAT | MISSENSE | aAa/aCa | K18T | PLES_49221 | G | G | G | G | G | G | G | G | G | G | G | G |
| NC_011770 | 5415368 | G | A | 70 | 0 | 10 | 0 | NON_SYNONYMOU | MODERAT | MISSENSE | Cgg/Tgg | R8W | PLES_49221 | A | A | A | A | A | A | | | A | A | A | A |
| NC_011770 | 5417510 | C | T | 60 | 0 | 2 | 0 | STOP_GAINED | HIGH | NONSENSE | Cag/Tag | Q525* | PLES_49231 | | | | | | | T | T | | | | |
| NC_011770 | 5417031 | G | A | 75 | 0 | 1 | 0 | NON_SYNONYMOU | MODERAT | MISSENSE | gGc/gAc | G365D | PLES_49231 | A | | | | | | | | | | | |
| NC_011770 | 5415962 | G | T | 76 | 0 | 12 | 0 | NON_SYNONYMOU | MODERAT | MISSENSE | Ggc/Tgc | G9C | PLES_49231 | T | T | T | T | T | T | T | T | T | T | T | T |
| NC_011770 | 5415972 | G | A | 74 | 0 | 10 | 0 | NON_SYNONYMOU | MODERAT | MISSENSE | aGt/aAt | S12N | PLES_49231 | A | A | A | A | A | A | | | A | A | A | A |
| NC_011770 | 5415983 | G | A | 71 | 0 | 2 | 0 | NON_SYNONYMOU | MODERAT | MISSENSE | Gat/Aat | D16N | PLES_49231 | | | | | A | A | | | | | | |
| NC_011770 | 5416254 | G | A | 74 | 0 | 10 | 0 | NON_SYNONYMOU | MODERAT | MISSENSE | cGg/cAg | R106Q | PLES_49231 | A | A | A | A | A | A | | | A | A | A | A |
| NC_011770 | 5416269 | T | C | 79 | 0 | 2 | 0 | NON_SYNONYMOU | MODERAT | MISSENSE | gTg/gCg | V111A | PLES_49231 | | | | | | | | | C | C | | |
| NC_011770 | 5417076 | G | C | 75 | 0 | 4 | 0 | NON_SYNONYMOU | MODERAT | MISSENSE | gGc/gCc | G380A | PLES_49231 | C | C | | | | | C | C | | | | |
| NC_011770 | 5417369 | G | A | 58 | 0 | 2 | 0 | NON_SYNONYMOU | MODERAT | MISSENSE | Gtc/Atc | V478I | PLES_49231 | | | | | | | A | A | | | | |
| NC_011770 | 5417412 | C | A | 58 | 0 | 2 | 0 | NON_SYNONYMOU | MODERAT | MISSENSE | aCg/aAg | T492K | PLES_49231 | | | | | | | A | A | | | | |
| NC_011770 | 5417448 | G | A | 57 | 0 | 2 | 0 | NON_SYNONYMOU | MODERAT | MISSENSE | aGc/aAc | S504N | PLES_49231 | A | A | | | | | | | | | | |
| NC_011770 | 5420871 | G | A | 59 | 0 | 1 | 0 | NON_SYNONYMOU | MODERAT | MISSENSE | gGt/gAt | G1083D | PLES_49241 | | | | | | | | A | | | | |
| NC_011770 | 5417658 | C | T | 69 | 0 | 2 | 0 | NON_SYNONYMOU | MODERAT | MISSENSE | gCc/gTc | A12V | PLES_49241 | | | | | | | T | T | | | | |
| NC_011770 | 5417690 | A | G | 71 | 0 | 4 | 0 | NON_SYNONYMOU | MODERAT | MISSENSE | Acc/Gcc | T23A | PLES_49241 | | | G | G | G | G | | | | | | |
| NC_011770 | 5418332 | G | A | 88 | 0 | 2 | 0 | NON_SYNONYMOU | MODERAT | MISSENSE | Gtg/Atg | V237M | PLES_49241 | A | A | | | | | | | | | | |
| NC_011770 | 5418515 | G | A | 97 | 0 | 2 | 0 | NON_SYNONYMOU | MODERAT | MISSENSE | Gaa/Aaa | E298K | PLES_49241 | A | A | | | | | | | | | | |
| NC_011770 | 5418530 | C | T | 95 | 0 | 2 | 0 | NON_SYNONYMOU | MODERAT | MISSENSE | Cgg/Tgg | R303W | PLES_49241 | | | | | | | | | | | T | T |
| NC_011770 | 5418639 | A | G | 100 | 0 | 2 | 0 | NON_SYNONYMOU | MODERAT | MISSENSE | cAt/cGt | H339R | PLES_49241 | G | G | | | | | | | | | | |
| NC_011770 | 5418815 | G | A | 94 | 0 | 2 | 0 | NON_SYNONYMOU | MODERAT | MISSENSE | Gcc/Acc | A398T | PLES_49241 | | | | | A | A | | | | | | |
| NC_011770 | 5418894 | G | A | 91 | 0 | 2 | 0 | NON_SYNONYMOU | MODERAT | MISSENSE | aGt/aAt | S424N | PLES_49241 | A | A | | | | | | | | . | | |
| NC_011770 | 5418929 | G | A | 81 | 1 | 11 | 0 | NON_SYNONYMOU | MODERAT | MISSENSE | Gat/Aat | D436N | PLES_49241 | A | A | A | A | A | A | A | A | A | . | A | A |
| NC_011770 | 5418938 | G | T | 75 | 1 | 3 | 0 | NON_SYNONYMOU | MODERAT | MISSENSE | Gcc/Tcc | A439S | PLES_49241 | T | T | | | | | | | T | . | | |
| NC_011770 | 5418975 | A | G | 83 | 2 | 2 | 0 | NON_SYNONYMOU | MODERAT | MISSENSE | aAc/aGc | N451S | PLES_49241 | G | G | | | | | | | . | . | | |
| NC_011770 | 5418977 | A | G | 75 | 2 | 2 | 0 | NON_SYNONYMOU | MODERAT | MISSENSE | Agc/Ggc | S452G | PLES_49241 | G | G | | | | | | | . | . | | |

| | | | | | | | | | | | | | | | | | | | | | | | | | |
|---|---|---|---|---|---|---|---|---|---|---|---|---|---|---|---|---|---|---|---|---|---|---|---|---|---|
| NC_011770 | 5418984 | A | G | 87 | 2 | 1 | 0 | NON_SYNONYMOU | MODERAT | MISSENSE | aAc/aGc | N454S | PLES_49241 | . | G | | | | | | | . | . | | |
| NC_011770 | 5419013 | G | A | 85 | 1 | 1 | 0 | NON_SYNONYMOU | MODERAT | MISSENSE | Gtc/Atc | V464I | PLES_49241 | . | A | | | | | | | | | | |
| NC_011770 | 5419029 | A | G | 83 | 0 | 2 | 0 | NON_SYNONYMOU | MODERAT | MISSENSE | aAc/aGc | N469S | PLES_49241 | G | G | | | | | | | | | | |
| NC_011770 | 5419031 | G | A | 87 | 0 | 2 | 0 | NON_SYNONYMOU | MODERAT | MISSENSE | Ggt/Agt | G470S | PLES_49241 | | | | | | | | | A | A | | |
| NC_011770 | 5419034 | G | C | 88 | 0 | 2 | 0 | NON_SYNONYMOU | MODERAT | MISSENSE | Gcc/Ccc | A471P | PLES_49241 | | | | | | | | | C | C | | |
| NC_011770 | 5419056 | G | A | 89 | 0 | 2 | 0 | NON_SYNONYMOU | MODERAT | MISSENSE | aGt/aAt | S478N | PLES_49241 | A | A | | | | | | | | | | |
| NC_011770 | 5419065 | C | G | 87 | 0 | 2 | 0 | NON_SYNONYMOU | MODERAT | MISSENSE | aCc/aGc | T481S | PLES_49241 | G | G | | | | | | | | | | |
| NC_011770 | 5419340 | A | G | 81 | 0 | 10 | 0 | NON_SYNONYMOU | MODERAT | MISSENSE | Atg/Gtg | M573V | PLES_49241 | G | G | G | G | G | G | G | G | G | G | | |
| NC_011770 | 5419342 | G | C | 82 | 0 | 4 | 0 | NON_SYNONYMOU | MODERAT | MISSENSE | atG/atC | M573I | PLES_49241 | | | | | | | C | C | C | C | | |
| NC_011770 | 5419422 | A | G | 75 | 0 | 6 | 0 | NON_SYNONYMOU | MODERAT | MISSENSE | aAc/aGc | N600S | PLES_49241 | G | G | | | | | G | G | G | G | | |
| NC_011770 | 5419700 | A | G | 77 | 0 | 8 | 0 | NON_SYNONYMOU | MODERAT | MISSENSE | Aac/Gac | N693D | PLES_49241 | | | G | G | G | G | G | G | G | G | | |
| NC_011770 | 5419760 | G | T | 85 | 0 | 2 | 0 | NON_SYNONYMOU | MODERAT | MISSENSE | Gcc/Tcc | A713S | PLES_49241 | | | | | | | T | T | | | | |
| NC_011770 | 5419775 | G | A | 79 | 0 | 8 | 0 | NON_SYNONYMOU | MODERAT | MISSENSE | Gcc/Acc | A718T | PLES_49241 | A | A | A | A | A | A | A | A | | | | |
| NC_011770 | 5419781 | A | G | 82 | 0 | 2 | 0 | NON_SYNONYMOU | MODERAT | MISSENSE | Atc/Gtc | I720V | PLES_49241 | | | | | | | G | G | | | | |
| NC_011770 | 5419874 | G | A | 94 | 0 | 6 | 0 | NON_SYNONYMOU | MODERAT | MISSENSE | Gac/Aac | D751N | PLES_49241 | A | A | A | A | A | A | | | | | | |
| NC_011770 | 5419878 | A | T | 90 | 0 | 12 | 0 | NON_SYNONYMOU | MODERAT | MISSENSE | cAg/cTg | Q752L | PLES_49241 | T | T | T | T | T | T | T | T | T | T | T | T |
| NC_011770 | 5419893 | T | C | 91 | 0 | 6 | 0 | NON_SYNONYMOU | MODERAT | MISSENSE | gTc/gCc | V757A | PLES_49241 | C | C | C | C | C | C | | | | | | |
| NC_011770 | 5419919 | A | C | 85 | 0 | 10 | 0 | NON_SYNONYMOU | MODERAT | MISSENSE | Aac/Cac | N766H | PLES_49241 | C | C | C | C | C | C | C | C | C | C | | |
| NC_011770 | 5419965 | C | G | 91 | 0 | 2 | 0 | NON_SYNONYMOU | MODERAT | MISSENSE | aCc/aGc | T781S | PLES_49241 | | | | | | | | | | | G | G |
| NC_011770 | 5420097 | A | G | 92 | 0 | 4 | 0 | NON_SYNONYMOU | MODERAT | MISSENSE | cAg/cGg | Q825R | PLES_49241 | | | | | | | G | G | | | G | G |
| NC_011770 | 5420198 | C | A | 94 | 0 | 2 | 0 | NON_SYNONYMOU | MODERAT | MISSENSE | Cag/Aag | Q859K | PLES_49241 | | | | | | | A | A | | | | |
| NC_011770 | 5420287 | C | A | 92 | 0 | 2 | 0 | NON_SYNONYMOU | MODERAT | MISSENSE | caC/caA | H888Q | PLES_49241 | | | | | | | | | A | A | | |
| NC_011770 | 5420543 | A | G | 83 | 0 | 12 | 0 | NON_SYNONYMOU | MODERAT | MISSENSE | Acg/Gcg | T974A | PLES_49241 | G | G | G | G | G | G | G | G | G | G | G | G |
| NC_011770 | 5420559 | C | T | 84 | 0 | 2 | 0 | NON_SYNONYMOU | MODERAT | MISSENSE | tCc/tTc | S979F | PLES_49241 | | | | | | | T | T | | | | |
| NC_011770 | 5420586 | G | A | 81 | 0 | 8 | 0 | NON_SYNONYMOU | MODERAT | MISSENSE | cGc/cAc | R988H | PLES_49241 | A | A | A | A | A | A | | | A | A | | |
| NC_011770 | 5421266 | G | A | 54 | 0 | 2 | 0 | NON_SYNONYMOU | MODERAT | MISSENSE | Gtc/Atc | V1215I | PLES_49241 | | | | | | | | | | | A | A |
| NC_011770 | 5421521 | G | A | 57 | 0 | 6 | 0 | NON_SYNONYMOU | MODERAT | MISSENSE | Gcc/Acc | A1300T | PLES_49241 | | | A | A | A | A | | | A | A | | |
| NC_011770 | 5421726 | C | G | 55 | 0 | 12 | 0 | NON_SYNONYMOU | MODERAT | MISSENSE | gCc/gGc | A1368G | PLES_49241 | G | G | G | G | G | G | G | G | G | G | G | G |
| NC_011770 | 5421788 | A | G | 53 | 0 | 2 | 0 | NON_SYNONYMOU | MODERAT | MISSENSE | Acc/Gcc | T1389A | PLES_49241 | G | G | | | | | | | | | | |
| NC_011770 | 5425325 | G | T | 80 | 0 | 2 | 0 | NON_SYNONYMOU | MODERAT | MISSENSE | gCg/gAg | A220E | PLES_49261 | T | T | | | | | | | | | | |
| NC_011770 | 5425345 | G | C | 83 | 0 | 2 | 0 | NON_SYNONYMOU | MODERAT | MISSENSE | ttC/ttG | F213L | PLES_49261 | C | C | | | | | | | | | | |
| NC_011770 | 5425517 | T | A | 68 | 0 | 2 | 0 | NON_SYNONYMOU | MODERAT | MISSENSE | cAg/cTg | Q156L | PLES_49261 | | | | | | | | | A | A | | |
| NC_011770 | 5425739 | G | A | 71 | 0 | 12 | 0 | NON_SYNONYMOU | MODERAT | MISSENSE | cCg/cTg | P82L | PLES_49261 | A | A | A | A | A | A | A | A | A | A | A | A |
| NC_011770 | 5425758 | C | T | 72 | 0 | 2 | 0 | NON_SYNONYMOU | MODERAT | MISSENSE | Gcc/Acc | A76T | PLES_49261 | | | | | | | T | T | | | | |
| NC_011770 | 5425786 | T | G | 70 | 0 | 10 | 0 | NON_SYNONYMOU | MODERAT | MISSENSE | gaA/gaC | E66D | PLES_49261 | | | G | G | G | G | G | G | G | G | G | G |
| NC_011770 | 5431523 | G | A | 77 | 0 | 8 | 0 | NON_SYNONYMOU | MODERAT | MISSENSE | gCc/gTc | A356V | PLES_49311 | | | A | A | A | A | | | A | A | A | A |
| NC_011770 | 5431535 | A | T | 79 | 0 | 2 | 0 | NON_SYNONYMOU | MODERAT | MISSENSE | aTc/aAc | I352N | PLES_49311 | | | | | | | | | | | T | T |
| NC_011770 | 5431539 | C | G | 79 | 0 | 12 | 0 | NON_SYNONYMOU | MODERAT | MISSENSE | Gcg/Ccg | A351P | PLES_49311 | G | G | G | G | G | G | G | G | G | G | G | G |
| NC_011770 | 5431766 | A | T | 94 | 0 | 6 | 0 | NON_SYNONYMOU | MODERAT | MISSENSE | cTg/cAg | L275Q | PLES_49311 | | | T | T | T | T | T | T | | | | |
| NC_011770 | 5431774 | G | T | 93 | 0 | 2 | 0 | NON_SYNONYMOU | MODERAT | MISSENSE | gaC/gaA | D272E | PLES_49311 | T | T | | | | | | | | | | |
| NC_011770 | 5432040 | C | G | 88 | 0 | 2 | 0 | NON_SYNONYMOU | MODERAT | MISSENSE | Gtc/Ctc | V184L | PLES_49311 | | | | | | | | | G | G | | |
| NC_011770 | 5432097 | G | T | 95 | 0 | 4 | 0 | NON_SYNONYMOU | MODERAT | MISSENSE | Ctc/Atc | L165I | PLES_49311 | | | | | | | T | T | T | T | | |
| NC_011770 | 5432239 | T | G | 80 | 0 | 6 | 0 | NON_SYNONYMOU | MODERAT | MISSENSE | gaA/gaC | E117D | PLES_49311 | | | G | G | G | G | | | | | G | G |
| NC_011770 | 5432243 | T | C | 84 | 0 | 6 | 0 | NON_SYNONYMOU | MODERAT | MISSENSE | aAg/aGg | K116R | PLES_49311 | | | C | C | C | C | | | | | C | C |
| NC_011770 | 5432260 | C | G | 82 | 0 | 6 | 0 | NON_SYNONYMOU | MODERAT | MISSENSE | aaG/aaC | K110N | PLES_49311 | G | G | G | G | G | G | | | | | | |
| NC_011770 | 5432285 | G | A | 79 | 0 | 4 | 0 | NON_SYNONYMOU | MODERAT | MISSENSE | aCc/aTc | T102I | PLES_49311 | A | A | | | | | | | A | A | | |
| NC_011770 | 5441438 | T | A | 75 | 0 | 2 | 0 | NON_SYNONYMOU | MODERAT | MISSENSE | cAg/cTg | Q51L | PLES_49411 | A | A | | | | | | | | | | |
| NC_011770 | 5448347 | G | A | 47 | 0 | 1 | 0 | NON_SYNONYMOU | MODERAT | MISSENSE | Ccg/Tcg | P86S | PLES_49471 | | | | | | | | A | | | | |
| NC_011770 | 5448322 | G | T | 49 | 0 | 12 | 0 | NON_SYNONYMOU | MODERAT | MISSENSE | aCg/aAg | T94K | PLES_49471 | T | T | T | T | T | T | T | T | T | T | T | T |
| NC_011770 | 5448328 | T | C | 47 | 0 | 2 | 0 | NON_SYNONYMOU | MODERAT | MISSENSE | gAa/gGa | E92G | PLES_49471 | | | | | | | | | C | C | | |
| NC_011770 | 5453317 | G | A | 77 | 0 | 1 | 0 | NON_SYNONYMOU | MODERAT | MISSENSE | Gcc/Acc | A72T | PLES_49531 | | | | | | | A | | | | | |
| NC_011770 | 5453324 | G | A | 78 | 0 | 2 | 0 | NON_SYNONYMOU | MODERAT | MISSENSE | cGc/cAc | R74H | PLES_49531 | | | | | A | A | | | | | | |
| NC_011770 | 5454179 | T | C | 97 | 0 | 10 | 0 | NON_SYNONYMOU | MODERAT | MISSENSE | aTc/aCc | I170T | PLES_49541 | C | C | C | C | C | C | | | C | C | C | C |
| NC_011770 | 5454203 | C | T | 97 | 0 | 10 | 0 | NON_SYNONYMOU | MODERAT | MISSENSE | gCg/gTg | A178V | PLES_49541 | T | T | T | T | T | T | | | T | T | T | T |
| NC_011770 | 5454619 | G | A | 81 | 0 | 2 | 0 | NON_SYNONYMOU | MODERAT | MISSENSE | Gcg/Acg | A317T | PLES_49541 | A | A | | | | | | | | | | |
| NC_011770 | 5454911 | A | G | 84 | 0 | 12 | 0 | NON_SYNONYMOU | MODERAT | MISSENSE | cAg/cGg | Q414R | PLES_49541 | G | G | G | G | G | G | G | G | G | G | G | G |
| NC_011770 | 5455063 | A | C | 73 | 0 | 2 | 0 | NON_SYNONYMOU | MODERAT | MISSENSE | Aac/Cac | N465H | PLES_49541 | | | | | | | | | | | C | C |
| NC_011770 | 5455163 | G | A | 78 | 0 | 12 | 0 | NON_SYNONYMOU | MODERAT | MISSENSE | aGg/aAg | R498K | PLES_49541 | A | A | A | A | A | A | A | A | A | A | A | A |
| NC_011770 | 5455337 | T | C | 75 | 0 | 12 | 0 | NON_SYNONYMOU | MODERAT | MISSENSE | cTg/cCg | L556P | PLES_49541 | C | C | C | C | C | C | C | C | C | C | C | C |
| NC_011770 | 5455354 | T | C | 74 | 0 | 12 | 0 | NON_SYNONYMOU | MODERAT | MISSENSE | Tcg/Ccg | S562P | PLES_49541 | C | C | C | C | C | C | C | C | C | C | C | C |
| NC_011770 | 5456583 | T | C | 90 | 0 | 10 | 0 | NON_SYNONYMOU | MODERAT | MISSENSE | Aac/Gac | N65D | PLES_49561 | C | C | C | C | C | C | | | C | C | C | C |
| NC_011770 | 5458220 | A | G | 59 | 0 | 2 | 0 | NON_SYNONYMOU | MODERAT | MISSENSE | gTg/gCg | V807A | PLES_49591 | G | G | | | | | | | | | | |

| | | | | | | | | | | | | | | | | | | | | | | | | | |
|---|---|---|---|---|---|---|---|---|---|---|---|---|---|---|---|---|---|---|---|---|---|---|---|---|---|
| NC_011770 | 5458226 | G | A | 59 | 0 | 2 | 0 | NON_SYNONYMOU | MODERAT | MISSENSE | aCc/aTc | T805I | PLES_49591 | | | | | | | | | | | A | A |
| NC_011770 | 5461040 | C | T | 77 | 0 | 2 | 0 | NON_SYNONYMOU | MODERAT | MISSENSE | Gcc/Acc | A9T | PLES_49601 | T | T | | | | | | | | | | |
| NC_011770 | 5461698 | A | T | 73 | 0 | 10 | 0 | NON_SYNONYMOU | MODERAT | MISSENSE | Acc/Tcc | T134S | PLES_49611 | T | T | T | T | T | T | T | T | | | T | T |
| NC_011770 | 5462095 | T | TC | 82 | 0 | 2 | 0 | FRAME_SHIFT | HIGH | | tcg/tcCg | S84S? | PLES_49621 | | | | | | | | | | | TC | TC |
| NC_011770 | 5463148 | G | A | 78 | 0 | 1 | 0 | NON_SYNONYMOU | MODERAT | MISSENSE | Gcc/Acc | A435T | PLES_49621 | | | | | | | | A | | | | |
| NC_011770 | 5462815 | G | T | 75 | 0 | 2 | 0 | NON_SYNONYMOU | MODERAT | MISSENSE | Gcc/Tcc | A324S | PLES_49621 | | | | | | | | | | | T | T |
| NC_011770 | 5463097 | A | G | 68 | 0 | 12 | 0 | NON_SYNONYMOU | MODERAT | MISSENSE | Acc/Gcc | T418A | PLES_49621 | G | G | G | G | G | G | G | G | G | G | G | G |
| NC_011770 | 5463625 | G | A | 91 | 0 | 6 | 0 | NON_SYNONYMOU | MODERAT | MISSENSE | Ggc/Agc | G594S | PLES_49621 | | | | | | | A | A | A | A | A | A |
| NC_011770 | 5464082 | G | C | 85 | 0 | 10 | 0 | NON_SYNONYMOU | MODERAT | MISSENSE | Ccc/Gcc | P57A | PLES_49631 | C | C | C | C | C | C | | | C | C | C | C |
| NC_011770 | 5464157 | C | T | 84 | 0 | 12 | 0 | NON_SYNONYMOU | MODERAT | MISSENSE | Gcg/Acg | A32T | PLES_49631 | T | T | T | T | T | T | T | T | T | T | T | T |
| NC_011770 | 5466711 | C | T | 73 | 0 | 2 | 0 | NON_SYNONYMOU | MODERAT | MISSENSE | Ccg/Tcg | P51S | PLES_49651 | | | | | | | T | T | | | | |
| NC_011770 | 5466976 | A | G | 80 | 0 | 2 | 0 | NON_SYNONYMOU | MODERAT | MISSENSE | gAc/gGc | D139G | PLES_49651 | | | | | | | | | G | G | | |
| NC_011770 | 5467228 | G | A | 82 | 0 | 2 | 0 | NON_SYNONYMOU | MODERAT | MISSENSE | aGc/aAc | S223N | PLES_49651 | | | | | | | | | A | A | | |
| NC_011770 | 5467402 | G | A | 83 | 0 | 2 | 0 | NON_SYNONYMOU | MODERAT | MISSENSE | aGc/aAc | S281N | PLES_49651 | A | A | | | | | | | | | | |
| NC_011770 | 5467958 | T | A | 71 | 0 | 2 | 0 | NON_SYNONYMOU | MODERAT | MISSENSE | cTc/cAc | L66H | PLES_49661 | | | | | | | A | A | | | | |
| NC_011770 | 5468060 | G | C | 68 | 0 | 2 | 0 | NON_SYNONYMOU | MODERAT | MISSENSE | gGc/gCc | G100A | PLES_49661 | | | | | | | | | C | C | | |
| NC_011770 | 5468845 | A | T | 78 | 0 | 12 | 0 | NON_SYNONYMOU | MODERAT | MISSENSE | Agc/Tgc | S362C | PLES_49661 | T | T | T | T | T | T | T | T | T | T | T | T |
| NC_011770 | 5469061 | G | A | 76 | 0 | 4 | 0 | NON_SYNONYMOU | MODERAT | MISSENSE | Gta/Ata | V17I | PLES_49671 | | | A | A | A | A | | | | | | |
| NC_011770 | 5469475 | G | C | 78 | 0 | 2 | 0 | NON_SYNONYMOU | MODERAT | MISSENSE | Ggc/Cgc | G155R | PLES_49671 | | | | | | | | | | | C | C |
| NC_011770 | 5469583 | G | A | 91 | 0 | 2 | 0 | NON_SYNONYMOU | MODERAT | MISSENSE | Gtg/Atg | V191M | PLES_49671 | | | | | | | | | A | A | | |
| NC_011770 | 5469736 | A | G | 83 | 0 | 2 | 0 | NON_SYNONYMOU | MODERAT | MISSENSE | Acc/Gcc | T242A | PLES_49671 | | | | | | | | | G | G | | |
| NC_011770 | 5470988 | A | C | 70 | 0 | 2 | 0 | NON_SYNONYMOU | MODERAT | MISSENSE | Atc/Ctc | I14L | PLES_49691 | | | | | | | | | C | C | | |
| NC_011770 | 5471004 | C | G | 71 | 0 | 4 | 0 | NON_SYNONYMOU | MODERAT | MISSENSE | aCc/aGc | T19S | PLES_49691 | | | G | G | G | G | | | | | | |
| NC_011770 | 5471025 | G | A | 69 | 0 | 2 | 0 | NON_SYNONYMOU | MODERAT | MISSENSE | aGc/aAc | S26N | PLES_49691 | | | | | | | | | A | A | | |
| NC_011770 | 5471036 | T | C | 76 | 0 | 2 | 0 | NON_SYNONYMOU | MODERAT | MISSENSE | Tac/Cac | Y30H | PLES_49691 | | | | | | | | | C | C | | |
| NC_011770 | 5471048 | A | C | 72 | 0 | 2 | 0 | NON_SYNONYMOU | MODERAT | MISSENSE | Aag/Cag | K34Q | PLES_49691 | | | | | | | | | C | C | | |
| NC_011770 | 5471160 | A | G | 70 | 0 | 12 | 0 | NON_SYNONYMOU | MODERAT | MISSENSE | cAc/cGc | H71R | PLES_49691 | G | G | G | G | G | G | G | G | G | G | G | G |
| NC_011770 | 5474017 | G | A | 58 | 0 | 2 | 0 | NON_SYNONYMOU | MODERAT | MISSENSE | Cgg/Tgg | R22W | PLES_49721 | | | | | | | | | | | A | A |
| NC_011770 | 5474056 | G | C | 55 | 0 | 2 | 0 | NON_SYNONYMOU | MODERAT | MISSENSE | Cct/Gct | P9A | PLES_49721 | | | | | | | | | | | C | C |
| NC_011770 | 5474065 | C | G | 55 | 0 | 4 | 0 | NON_SYNONYMOU | MODERAT | MISSENSE | Ggg/Cgg | G6R | PLES_49721 | | | G | G | G | G | | | | | | |
| NC_011770 | 5474070 | G | T | 53 | 0 | 12 | 0 | NON_SYNONYMOU | MODERAT | MISSENSE | cCg/cAg | P4Q | PLES_49721 | T | T | T | T | T | T | T | T | T | T | T | T |
| NC_011770 | 5474467 | T | G | 64 | 0 | 12 | 0 | NON_SYNONYMOU | MODERAT | MISSENSE | gaA/gaC | E369D | PLES_49731 | G | G | G | G | G | G | G | G | G | G | G | G |
| NC_011770 | 5477663 | G | A | 86 | 0 | 1 | 0 | NON_SYNONYMOU | MODERAT | MISSENSE | gCc/gTc | A178V | PLES_49751 | | | | | | | | A | | | | |
| NC_011770 | 5477030 | C | T | 74 | 0 | 4 | 0 | NON_SYNONYMOU | MODERAT | MISSENSE | gGc/gAc | G389D | PLES_49751 | | | | | | | | | T | T | T | T |
| NC_011770 | 5477046 | A | C | 73 | 0 | 4 | 0 | NON_SYNONYMOU | MODERAT | MISSENSE | Ttt/Gtt | F384V | PLES_49751 | | | C | C | C | C | | | | | | |
| NC_011770 | 5477174 | G | A | 81 | 0 | 4 | 0 | NON_SYNONYMOU | MODERAT | MISSENSE | cCc/cTc | P341L | PLES_49751 | | | A | A | A | A | | | | | | |
| NC_011770 | 5477531 | T | C | 78 | 0 | 2 | 0 | NON_SYNONYMOU | MODERAT | MISSENSE | cAg/cGg | Q222R | PLES_49751 | C | C | | | | | | | | | | |
| NC_011770 | 5478006 | G | A | 85 | 0 | 1 | 0 | NON_SYNONYMOU | MODERAT | MISSENSE | Ccg/Tcg | P64S | PLES_49751 | | | | | | | A | | | | | |
| NC_011770 | 5478615 | T | TCGCCA | 79 | 0 | 2 | 0 | CODON_INSERTIO | MODERATE | | gac/gGTG | D351GGD | PLES_49761 | | | | | | | TCGCCAC | TCGCCAC | | | | |
| NC_011770 | 5479606 | A | C | 84 | 0 | 8 | 0 | NON_SYNONYMOU | MODERAT | MISSENSE | Ttc/Gtc | F21V | PLES_49761 | C | C | | | | | C | C | C | C | C | C |
| NC_011770 | 5479630 | C | T | 83 | 0 | 4 | 0 | NON_SYNONYMOU | MODERAT | MISSENSE | Gcc/Acc | A13T | PLES_49761 | T | T | | | | | T | T | | | | |
| NC_011770 | 5480306 | T | C | 74 | 0 | 2 | 0 | NON_SYNONYMOU | MODERAT | MISSENSE | Atc/Gtc | I183V | PLES_49771 | | | | | | | C | C | | | | |
| NC_011770 | 5480368 | G | A | 77 | 0 | 2 | 0 | NON_SYNONYMOU | MODERAT | MISSENSE | gCg/gTg | A162V | PLES_49771 | | | | | | | | | A | A | | |
| NC_011770 | 5481490 | GC | G | 42 | 1 | 1 | 0 | FRAME_SHIFT | HIGH | | ggg/ | G15 | PLES_49781 | | | | G | | | . | | | | | |
| NC_011770 | 5480929 | C | T | 69 | 0 | 12 | 0 | NON_SYNONYMOU | MODERAT | MISSENSE | Ggc/Agc | G203S | PLES_49781 | T | T | T | T | T | T | T | T | T | T | T | T |
| NC_011770 | 5481177 | T | C | 66 | 0 | 2 | 0 | NON_SYNONYMOU | MODERAT | MISSENSE | gAg/gGg | E120G | PLES_49781 | | | | | | | | | | | C | C |
| NC_011770 | 5483842 | C | T | 65 | 0 | 4 | 0 | NON_SYNONYMOU | MODERAT | MISSENSE | gCt/gTt | A9V | PLES_49801 | | | T | T | T | T | | | | | | |
| NC_011770 | 5483941 | C | T | 64 | 0 | 4 | 0 | NON_SYNONYMOU | MODERAT | MISSENSE | cCg/cTg | P42L | PLES_49801 | T | T | | | | | | | | | T | T |
| NC_011770 | 5497280 | C | T | 83 | 0 | 4 | 0 | NON_SYNONYMOU | MODERAT | MISSENSE | Gtc/Atc | V102I | PLES_49881 | T | T | | | | | | | T | T | | |
| NC_011770 | 5498099 | T | C | 75 | 0 | 2 | 0 | NON_SYNONYMOU | MODERAT | MISSENSE | gAc/gGc | D171G | PLES_49891 | C | C | | | | | | | | | | |
| NC_011770 | 5498401 | A | G | 89 | 0 | 12 | 0 | NON_SYNONYMOU | MODERAT | MISSENSE | aaT/aaG | N70K | PLES_49891 | G | G | C | C | C | C | G | G | G | G | G | G |
| NC_011770 | 5504124 | T | C | 61 | 0 | 6 | 0 | NON_SYNONYMOU | MODERAT | MISSENSE | Tcg/Ccg | S79P | PLES_49951 | | | C | C | C | C | | | C | C | | |
| NC_011770 | 5504143 | A | G | 60 | 0 | 2 | 0 | NON_SYNONYMOU | MODERAT | MISSENSE | cAg/cGg | Q85R | PLES_49951 | G | G | | | | | | | | | | |
| NC_011770 | 5504488 | T | G | 68 | 0 | 2 | 0 | NON_SYNONYMOU | MODERAT | MISSENSE | gaA/gaC | E51D | PLES_49961 | | | | | | | | | | | G | G |
| NC_011770 | 5504840 | C | T | 83 | 0 | 4 | 0 | NON_SYNONYMOU | MODERAT | MISSENSE | Gag/Aag | E143K | PLES_49971 | | | T | T | T | T | | | | | | |
| NC_011770 | 5504897 | T | C | 84 | 0 | 2 | 0 | NON_SYNONYMOU | MODERAT | MISSENSE | Agc/Ggc | S124G | PLES_49971 | | | | | | | | | C | C | | |
| NC_011770 | 5505061 | C | G | 84 | 0 | 4 | 0 | NON_SYNONYMOU | MODERAT | MISSENSE | gGg/gCg | G69A | PLES_49971 | | | G | G | G | G | | | | | | |
| NC_011770 | 5508061 | G | A | 74 | 0 | 2 | 0 | NON_SYNONYMOU | MODERAT | MISSENSE | gCc/gTc | A208V | PLES_50001 | | | | | | | | | A | A | | |
| NC_011770 | 5508212 | G | A | 80 | 0 | 4 | 0 | NON_SYNONYMOU | MODERAT | MISSENSE | Cgc/Tgc | R158C | PLES_50001 | | | A | A | A | A | | | | | | |
| NC_011770 | 5509731 | C | T | 74 | 0 | 1 | 0 | NON_SYNONYMOU | MODERAT | MISSENSE | Gcc/Acc | A20T | PLES_50011 | | | | | | | | | | | T | |
| NC_011770 | 5508902 | G | A | 85 | 0 | 4 | 0 | NON_SYNONYMOU | MODERAT | MISSENSE | gCg/gTg | A296V | PLES_50011 | | | A | A | A | A | | | | | | |
| NC_011770 | 5509552 | A | C | 77 | 0 | 12 | 0 | NON_SYNONYMOU | MODERAT | MISSENSE | gaT/gaG | D79E | PLES_50011 | C | C | C | C | C | C | C | C | C | C | C | C |

| | | | | | | | | | | | | | | | | | | | | | | | | | |
|---|---|---|---|---|---|---|---|---|---|---|---|---|---|---|---|---|---|---|---|---|---|---|---|---|---|
| NC_011770 | 5509761 | G | A | 71 | 0 | 2 | 0 | NON_SYNONYMOU | MODERAT | MISSENSE | Ctc/Ttc | L10F | PLES_50011 | | | | | | | A | A | | | | |
| NC_011770 | 5510431 | T | G | 56 | 0 | 12 | 0 | NON_SYNONYMOU | MODERAT | MISSENSE | Ttg/Gtg | L106V | PLES_50021 | G | G | G | G | G | G | G | G | G | G | G | G |
| NC_011770 | 5510867 | T | A | 68 | 0 | 2 | 0 | NON_SYNONYMOU | MODERAT | MISSENSE | cTg/cAg | L251Q | PLES_50021 | A | A | | | | | | | | | | |
| NC_011770 | 5510890 | G | A | 70 | 0 | 10 | 0 | NON_SYNONYMOU | MODERAT | MISSENSE | Gcc/Acc | A259T | PLES_50021 | A | A | A | A | A | A | A | A | A | A | | |
| NC_011770 | 5510977 | G | A | 71 | 0 | 2 | 0 | NON_SYNONYMOU | MODERAT | MISSENSE | Gcc/Acc | A288T | PLES_50021 | A | A | | | | | | | | | | |
| NC_011770 | 5511004 | G | A | 65 | 0 | 6 | 0 | NON_SYNONYMOU | MODERAT | MISSENSE | Gcg/Acg | A297T | PLES_50021 | | | A | A | A | A | A | A | | | | |
| NC_011770 | 5512316 | A | G | 80 | 0 | 6 | 0 | NON_SYNONYMOU | MODERAT | MISSENSE | Tgc/Cgc | C158R | PLES_50031 | | | G | G | G | G | G | G | | | | |
| NC_011770 | 5512454 | C | T | 70 | 0 | 6 | 0 | NON_SYNONYMOU | MODERAT | MISSENSE | Gtc/Atc | V112I | PLES_50031 | | | T | T | T | T | T | T | | | | |
| NC_011770 | 5512537 | C | G | 73 | 0 | 10 | 0 | NON_SYNONYMOU | MODERAT | MISSENSE | gGc/gCc | G84A | PLES_50031 | G | G | G | G | G | G | G | G | | | G | G |
| NC_011770 | 5513306 | C | T | 97 | 0 | 1 | 0 | NON_SYNONYMOU | MODERAT | MISSENSE | Gcc/Acc | A245T | PLES_50041 | | T | | | | | | | | | | |
| NC_011770 | 5513337 | G | C | 98 | 0 | 2 | 0 | NON_SYNONYMOU | MODERAT | MISSENSE | gaC/gaG | D234E | PLES_50041 | C | C | | | | | | | | | | |
| NC_011770 | 5513690 | G | A | 87 | 0 | 2 | 0 | NON_SYNONYMOU | MODERAT | MISSENSE | Cgc/Tgc | R117C | PLES_50041 | A | A | | | | | | | | | | |
| NC_011770 | 5513951 | C | T | 82 | 0 | 6 | 0 | NON_SYNONYMOU | MODERAT | MISSENSE | Gtc/Atc | V30I | PLES_50041 | | | T | T | T | T | T | T | | | | |
| NC_011770 | 5514051 | T | C | 84 | 0 | 4 | 0 | NON_SYNONYMOU | MODERAT | MISSENSE | aAg/aGg | K177R | PLES_50051 | C | C | | | | | | | | | C | C |
| NC_011770 | 5514489 | T | C | 77 | 0 | 2 | 0 | NON_SYNONYMOU | MODERAT | MISSENSE | cAc/cGc | H31R | PLES_50051 | | | | | | | C | C | | | | |
| NC_011770 | 5514718 | G | A | 72 | 0 | 4 | 0 | NON_SYNONYMOU | MODERAT | MISSENSE | cCg/cTg | P896L | PLES_50061 | | | A | A | A | A | | | | | | |
| NC_011770 | 5514751 | T | C | 72 | 0 | 8 | 0 | NON_SYNONYMOU | MODERAT | MISSENSE | aAc/aGc | N885S | PLES_50061 | | | C | C | C | C | C | C | C | C | | |
| NC_011770 | 5514887 | G | T | 81 | 0 | 4 | 0 | NON_SYNONYMOU | MODERAT | MISSENSE | Ctg/Atg | L840M | PLES_50061 | T | T | | | | | | | | | T | T |
| NC_011770 | 5515277 | C | T | 75 | 0 | 8 | 0 | NON_SYNONYMOU | MODERAT | MISSENSE | Gcc/Acc | A710T | PLES_50061 | | | T | T | T | T | T | T | T | T | | |
| NC_011770 | 5516429 | A | G | 89 | 0 | 6 | 0 | NON_SYNONYMOU | MODERAT | MISSENSE | Ttc/Ctc | F326L | PLES_50061 | | | G | G | G | G | | | G | G | | |
| NC_011770 | 5516708 | T | G | 83 | 0 | 8 | 0 | NON_SYNONYMOU | MODERAT | MISSENSE | Agc/Cgc | S233R | PLES_50061 | | | G | G | G | G | G | G | G | G | | |
| NC_011770 | 5518426 | G | A | 81 | 0 | 1 | 0 | NON_SYNONYMOU | MODERAT | MISSENSE | Gcc/Acc | A238T | PLES_50071 | | | | | | | | A | | | | |
| NC_011770 | 5520865 | A | G | 80 | 0 | 1 | 0 | NON_SYNONYMOU | MODERAT | MISSENSE | cTg/cCg | L207P | PLES_50091 | | G | | | | | | | | | | |
| NC_011770 | 5519954 | C | T | 83 | 0 | 4 | 0 | NON_SYNONYMOU | MODERAT | MISSENSE | Gtg/Atg | V511M | PLES_50091 | | | T | T | T | T | | | | | | |
| NC_011770 | 5520809 | T | C | 83 | 0 | 1 | 0 | NON_SYNONYMOU | MODERAT | MISSENSE | Agc/Ggc | S226G | PLES_50091 | | | | | | | C | | | | | |
| NC_011770 | 5525643 | C | T | 65 | 0 | 1 | 0 | NON_SYNONYMOU | MODERAT | MISSENSE | cGc/cAc | R791H | PLES_50101 | | | | | | | | | | | T | |
| NC_011770 | 5522158 | C | T | 80 | 0 | 1 | 0 | NON_SYNONYMOU | MODERAT | MISSENSE | Gac/Aac | D1953N | PLES_50101 | | | | | | | | T | | | | |
| NC_011770 | 5522172 | G | C | 84 | 0 | 1 | 0 | NON_SYNONYMOU | MODERAT | MISSENSE | gCc/gGc | A1948G | PLES_50101 | | | | | | | | C | | | | |
| NC_011770 | 5522173 | C | T | 83 | 0 | 1 | 0 | NON_SYNONYMOU | MODERAT | MISSENSE | Gcc/Acc | A1948T | PLES_50101 | | | | | | | | T | | | | |
| NC_011770 | 5522193 | G | T | 86 | 0 | 1 | 0 | NON_SYNONYMOU | MODERAT | MISSENSE | aCc/aAc | T1941N | PLES_50101 | | | | | | | | T | | | | |
| NC_011770 | 5522199 | A | G | 87 | 0 | 1 | 0 | NON_SYNONYMOU | MODERAT | MISSENSE | gTg/gCg | V1939A | PLES_50101 | | | | | | | | G | | | | |
| NC_011770 | 5522205 | C | T | 91 | 0 | 1 | 0 | NON_SYNONYMOU | MODERAT | MISSENSE | gGc/gAc | G1937D | PLES_50101 | | | | | | | | T | | | | |
| NC_011770 | 5522247 | T | G | 95 | 0 | 1 | 0 | NON_SYNONYMOU | MODERAT | MISSENSE | gAc/gCc | D1923A | PLES_50101 | | | | | | | | G | | | | |
| NC_011770 | 5522355 | C | G | 91 | 0 | 1 | 0 | NON_SYNONYMOU | MODERAT | MISSENSE | gGc/gCc | G1887A | PLES_50101 | | | | | | | | G | . | | | |
| NC_011770 | 5523009 | G | C | 69 | 0 | 7 | 0 | NON_SYNONYMOU | MODERAT | MISSENSE | aCc/aGc | T1669S | PLES_50101 | C | C | | | C | C | | C | | | C | C |
| NC_011770 | 5522160 | T | C | 83 | 0 | 2 | 0 | NON_SYNONYMOU | MODERAT | MISSENSE | aAt/aGt | N1952S | PLES_50101 | | | | | | | | | C | C | | |
| NC_011770 | 5522749 | T | C | 47 | 0 | 12 | 0 | NON_SYNONYMOU | MODERAT | MISSENSE | Agt/Ggt | S1756G | PLES_50101 | C | C | C | C | C | C | C | C | C | C | C | C |
| NC_011770 | 5523138 | G | C | 23 | 0 | 4 | 0 | NON_SYNONYMOU | MODERAT | MISSENSE | gCt/gGt | A1626G | PLES_50101 | C | C | | | | | | | | | C | C |
| NC_011770 | 5523139 | C | T | 23 | 0 | 4 | 0 | NON_SYNONYMOU | MODERAT | MISSENSE | Gct/Act | A1626T | PLES_50101 | T | T | | | | | | | | | T | T |
| NC_011770 | 5523492 | G | C | 80 | 0 | 10 | 0 | NON_SYNONYMOU | MODERAT | MISSENSE | aCc/aGc | T1508S | PLES_50101 | C | C | C | C | C | C | C | C | | | C | C |
| NC_011770 | 5523621 | G | C | 32 | 0 | 4 | 0 | NON_SYNONYMOU | MODERAT | MISSENSE | gCt/gGt | A1465G | PLES_50101 | C | C | | | | | | | | | C | C |
| NC_011770 | 5523622 | C | T | 31 | 0 | 4 | 0 | NON_SYNONYMOU | MODERAT | MISSENSE | Gct/Act | A1465T | PLES_50101 | T | T | | | | | | | | | T | T |
| NC_011770 | 5524615 | G | C | 94 | 1 | 11 | 0 | NON_SYNONYMOU | MODERAT | MISSENSE | Ctc/Gtc | L1134V | PLES_50101 | C | C | C | C | C | C | C | . | C | C | C | C |
| NC_011770 | 5524728 | G | C | 61 | 0 | 6 | 0 | NON_SYNONYMOU | MODERAT | MISSENSE | tCc/tGc | S1096C | PLES_50101 | | | C | C | C | C | C | C | | | | |
| NC_011770 | 5524729 | A | T | 61 | 0 | 6 | 0 | NON_SYNONYMOU | MODERAT | MISSENSE | Tcc/Acc | S1096T | PLES_50101 | | | T | T | T | T | T | T | | | | |
| NC_011770 | 5524746 | G | C | 62 | 0 | 6 | 0 | NON_SYNONYMOU | MODERAT | MISSENSE | gCa/gGa | A1090G | PLES_50101 | | | C | C | C | C | C | C | | | | |
| NC_011770 | 5524747 | C | T | 63 | 0 | 6 | 0 | NON_SYNONYMOU | MODERAT | MISSENSE | Gca/Aca | A1090T | PLES_50101 | | | T | T | T | T | T | T | | | | |
| NC_011770 | 5524877 | G | T | 91 | 0 | 2 | 0 | NON_SYNONYMOU | MODERAT | MISSENSE | aaC/aaA | N1046K | PLES_50101 | | | | | | | T | T | | | | |
| NC_011770 | 5524917 | C | T | 94 | 0 | 4 | 0 | NON_SYNONYMOU | MODERAT | MISSENSE | aGc/aAc | S1033N | PLES_50101 | | | T | T | T | T | | | | | | |
| NC_011770 | 5525112 | C | T | 64 | 0 | 4 | 0 | NON_SYNONYMOU | MODERAT | MISSENSE | gGg/gAg | G968E | PLES_50101 | | | T | T | T | T | | | | | | |
| NC_011770 | 5526378 | T | A | 59 | 0 | 2 | 0 | NON_SYNONYMOU | MODERAT | MISSENSE | cAg/cTg | Q546L | PLES_50101 | | | | | | | | | A | A | | |
| NC_011770 | 5526402 | C | T | 57 | 0 | 4 | 0 | NON_SYNONYMOU | MODERAT | MISSENSE | aGc/aAc | S538N | PLES_50101 | T | T | | | | | | | | | T | T |
| NC_011770 | 5527089 | C | T | 71 | 0 | 4 | 0 | NON_SYNONYMOU | MODERAT | MISSENSE | aGc/aAc | S309N | PLES_50101 | T | T | | | | | | | | | T | T |
| NC_011770 | 5527444 | C | G | 65 | 0 | 2 | 0 | NON_SYNONYMOU | MODERAT | MISSENSE | Gtg/Ctg | V191L | PLES_50101 | | | | | | | | | G | G | | |
| NC_011770 | 5529579 | G | A | 60 | 0 | 12 | 0 | NON_SYNONYMOU | MODERAT | MISSENSE | gGc/gAc | G55D | PLES_50121 | A | A | A | A | A | A | A | A | A | A | A | A |
| NC_011770 | 5529887 | G | A | 66 | 0 | 2 | 0 | NON_SYNONYMOU | MODERAT | MISSENSE | Gcc/Acc | A158T | PLES_50121 | A | A | | | | | | | | | | |
| NC_011770 | 5530038 | C | T | 63 | 0 | 8 | 0 | NON_SYNONYMOU | MODERAT | MISSENSE | gCt/gTt | A208V | PLES_50121 | T | T | | | | | T | T | T | T | T | T |
| NC_011770 | 5530071 | C | G | 64 | 0 | 8 | 0 | NON_SYNONYMOU | MODERAT | MISSENSE | aCt/aGt | T219S | PLES_50121 | G | G | | | | | G | G | G | G | G | G |
| NC_011770 | 5530400 | G | A | 61 | 0 | 2 | 0 | NON_SYNONYMOU | MODERAT | MISSENSE | Gcc/Acc | A329T | PLES_50121 | | | | | | | | | A | A | | |
| NC_011770 | 5532773 | T | C | 84 | 0 | 2 | 0 | NON_SYNONYMOU | MODERAT | MISSENSE | gTg/gCg | V150A | PLES_50141 | | | | | | | | | | | C | C |
| NC_011770 | 5533047 | G | A | 72 | 0 | 1 | 0 | NON_SYNONYMOU | MODERAT | MISSENSE | Gcc/Acc | A10T | PLES_50151 | | | | | | | A | | | | | |
| NC_011770 | 5533146 | C | A | 76 | 0 | 2 | 0 | NON_SYNONYMOU | MODERAT | MISSENSE | Cgt/Agt | R43S | PLES_50151 | | | | | | | | | | | A | A |

| | | | | | | | | | | | | | | | | | | | | | | | | | |
|---|---|---|---|---|---|---|---|---|---|---|---|---|---|---|---|---|---|---|---|---|---|---|---|---|---|
| NC_011770 | 5533452 | A | G | 79 | 0 | 12 | 0 | NON_SYNONYMOU | MODERAT | MISSENSE | Acg/Gcg | T145A | PLES_50151 | G | G | G | G | G | G | G | G | G | G | G | G |
| NC_011770 | 5534290 | A | T | 92 | 0 | 2 | 0 | NON_SYNONYMOU | MODERAT | MISSENSE | cTg/cAg | L70Q | PLES_50161 | T | T | | | | | | | | | | |
| NC_011770 | 5536133 | G | A | 69 | 0 | 4 | 0 | NON_SYNONYMOU | MODERAT | MISSENSE | Gcc/Acc | A170T | PLES_50181 | | | A | A | A | A | | | | | | |
| NC_011770 | 5536385 | G | A | 65 | 0 | 4 | 0 | NON_SYNONYMOU | MODERAT | MISSENSE | Gat/Aat | D254N | PLES_50181 | | | A | A | A | A | | | | | | |
| NC_011770 | 5537892 | C | G | 74 | 0 | 4 | 0 | NON_SYNONYMOU | MODERAT | MISSENSE | Gac/Cac | D164H | PLES_50191 | G | G | | | | | | | | | G | G |
| NC_011770 | 5538234 | T | G | 75 | 0 | 6 | 0 | NON_SYNONYMOU | MODERAT | MISSENSE | Atg/Ctg | M50L | PLES_50191 | | | G | G | G | G | | | G | G | | |
| NC_011770 | 5538850 | C | T | 98 | 0 | 6 | 0 | NON_SYNONYMOU | MODERAT | MISSENSE | gCc/gTc | A15V | PLES_50201 | | | T | T | T | T | | | T | T | | |
| NC_011770 | 5538867 | G | A | 96 | 0 | 2 | 0 | NON_SYNONYMOU | MODERAT | MISSENSE | Ggc/Agc | G21S | PLES_50201 | A | A | | | | | | | | | | |
| NC_011770 | 5539633 | G | A | 77 | 0 | 4 | 0 | NON_SYNONYMOU | MODERAT | MISSENSE | gCg/gTg | A52V | PLES_50211 | | | A | A | A | A | | | | | | |
| NC_011770 | 5541058 | G | A | 55 | 0 | 2 | 0 | NON_SYNONYMOU | MODERAT | MISSENSE | gGc/gAc | G331D | PLES_50231 | | | | | | | A | A | | | | |
| NC_011770 | 5541474 | C | G | 39 | 0 | 8 | 0 | INTRAGENIC | MODIFIER | | | | PLES_50241 | G | G | | | | | G | G | G | G | G | G |
| NC_011770 | 5541478 | G | T | 37 | 0 | 8 | 0 | INTRAGENIC | MODIFIER | | | | PLES_50241 | T | T | | | | | T | T | T | T | T | T |
| NC_011770 | 5541582 | G | C | 48 | 0 | 8 | 0 | INTRAGENIC | MODIFIER | | | | PLES_50241 | C | C | | | | | C | C | C | C | C | C |
| NC_011770 | 5541714 | A | G | 60 | 0 | 8 | 0 | INTRAGENIC | MODIFIER | | | | PLES_50241 | G | G | | | | | G | G | G | G | G | G |
| NC_011770 | 5541764 | G | A | 61 | 0 | 4 | 0 | INTRAGENIC | MODIFIER | | | | PLES_50241 | A | A | | | | | | | | | A | A |
| NC_011770 | 5541766 | A | G | 60 | 0 | 4 | 0 | INTRAGENIC | MODIFIER | | | | PLES_50241 | G | G | | | | | | | | | G | G |
| NC_011770 | 5541767 | C | A | 60 | 0 | 4 | 0 | INTRAGENIC | MODIFIER | | | | PLES_50241 | A | A | | | | | | | | | A | A |
| NC_011770 | 5541834 | A | C | 64 | 0 | 4 | 0 | INTRAGENIC | MODIFIER | | | | PLES_50241 | C | C | | | | | | | | | C | C |
| NC_011770 | 5541842 | A | G | 64 | 0 | 4 | 0 | INTRAGENIC | MODIFIER | | | | PLES_50241 | G | G | | | | | | | | | G | G |
| NC_011770 | 5541861 | G | T | 64 | 0 | 2 | 0 | INTRAGENIC | MODIFIER | | | | PLES_50241 | | | | | | | | | T | T | | |
| NC_011770 | 5545189 | T | TCG | 60 | 0 | 12 | 0 | FRAME_SHIFT+STC | HIGH | | tga/tgCGa | *133C? | PLES_50271 | TCG | TCG | TCG | TCG | TCG | TCG | TCG | TCG | TCG | TCG | TCG | TCG |
| NC_011770 | 5546051 | AG | A | 69 | 0 | 2 | 0 | FRAME_SHIFT | HIGH | | ccc/ | P32 | PLES_50281 | | | | A | | | A | | | | | |
| NC_011770 | 5546244 | C | T | 79 | 0 | 2 | 0 | NON_SYNONYMOU | MODERAT | MISSENSE | Gta/Ata | V158I | PLES_50291 | | | | | | | T | T | | | | |
| NC_011770 | 5546348 | A | G | 85 | 0 | 12 | 0 | NON_SYNONYMOU | MODERAT | MISSENSE | gTg/gCg | V123A | PLES_50291 | G | G | G | G | G | G | G | G | G | G | G | G |
| NC_011770 | 5546873 | G | T | 82 | 0 | 2 | 0 | NON_SYNONYMOU | MODERAT | MISSENSE | Ccg/Acg | P127T | PLES_50301 | | | | | | | T | T | | | | |
| NC_011770 | 5547776 | A | C | 62 | 0 | 2 | 0 | NON_SYNONYMOU | MODERAT | MISSENSE | aTg/aGg | M12R | PLES_50311 | | | | | | | C | C | | | | |
| NC_011770 | 5550249 | T | C | 54 | 0 | 2 | 0 | NON_SYNONYMOU | MODERAT | MISSENSE | aTt/aCt | I5T | PLES_50341 | | | | | | | | | C | C | | |
| NC_011770 | 5551068 | A | G | 79 | 0 | 8 | 0 | NON_SYNONYMOU | MODERAT | MISSENSE | aAg/aGg | K85R | PLES_50351 | | | G | G | G | G | G | G | G | G | | |
| NC_011770 | 5551378 | C | T | 86 | 0 | 8 | 0 | NON_SYNONYMOU | MODERAT | MISSENSE | Ccg/Tcg | P11S | PLES_50361 | | | T | T | T | T | T | T | T | T | | |
| NC_011770 | 5551403 | G | A | 82 | 0 | 8 | 0 | NON_SYNONYMOU | MODERAT | MISSENSE | aGc/aAc | S19N | PLES_50361 | | | A | A | A | A | A | A | A | A | | |
| NC_011770 | 5551498 | G | A | 76 | 0 | 8 | 0 | NON_SYNONYMOU | MODERAT | MISSENSE | Gac/Aac | D51N | PLES_50361 | | | A | A | A | A | A | A | A | A | | |
| NC_011770 | 5551939 | A | G | 86 | 0 | 8 | 0 | NON_SYNONYMOU | MODERAT | MISSENSE | Agc/Ggc | S11G | PLES_50371 | | | G | G | G | G | G | G | G | G | | |
| NC_011770 | 5551969 | C | T | 87 | 0 | 6 | 0 | NON_SYNONYMOU | MODERAT | MISSENSE | Ctc/Ttc | L21F | PLES_50371 | | | T | T | T | T | T | T | | | | |
| NC_011770 | 5552162 | T | C | 76 | 0 | 2 | 0 | NON_SYNONYMOU | MODERAT | MISSENSE | aTt/aCt | I85T | PLES_50371 | | | | | | | C | C | | | | |
| NC_011770 | 5552290 | G | A | 81 | 0 | 6 | 0 | NON_SYNONYMOU | MODERAT | MISSENSE | Ggc/Agc | G128S | PLES_50371 | | | A | A | A | A | A | A | | | | |
| NC_011770 | 5552291 | G | A | 83 | 0 | 6 | 0 | NON_SYNONYMOU | MODERAT | MISSENSE | gGc/gAc | G128D | PLES_50371 | A | A | | | | | | | A | A | A | A |
| NC_011770 | 5552331 | G | C | 94 | 0 | 2 | 0 | NON_SYNONYMOU | MODERAT | MISSENSE | caG/caC | Q141H | PLES_50371 | | | | | | | | | C | C | | |
| NC_011770 | 5552546 | G | T | 89 | 0 | 6 | 0 | NON_SYNONYMOU | MODERAT | MISSENSE | aGc/aTc | S213I | PLES_50371 | T | T | | | | | A | A | | | T | T |
| NC_011770 | 5552588 | G | A | 84 | 0 | 2 | 0 | NON_SYNONYMOU | MODERAT | MISSENSE | gGc/gAc | G227D | PLES_50371 | | | | | A | A | | | | | | |
| NC_011770 | 5552591 | G | A | 86 | 0 | 2 | 0 | NON_SYNONYMOU | MODERAT | MISSENSE | cGg/cAg | R228Q | PLES_50371 | | | | | | | A | A | | | | |
| NC_011770 | 5552632 | G | T | 84 | 0 | 6 | 0 | NON_SYNONYMOU | MODERAT | MISSENSE | Gcc/Tcc | A242S | PLES_50371 | T | T | | | | | | | T | T | T | T |
| NC_011770 | 5552745 | G | A | 81 | 0 | 6 | 0 | NON_SYNONYMOU | MODERAT | MISSENSE | cGc/cAc | R11H | PLES_50381 | | | A | A | A | A | A | A | | | | |
| NC_011770 | 5552846 | G | A | 77 | 0 | 6 | 0 | NON_SYNONYMOU | MODERAT | MISSENSE | Ggc/Agc | G45S | PLES_50381 | | | A | A | A | A | A | A | | | | |
| NC_011770 | 5552853 | C | T | 78 | 0 | 12 | 0 | NON_SYNONYMOU | MODERAT | MISSENSE | gCc/gTc | A47V | PLES_50381 | T | T | T | T | T | T | T | T | T | T | T | T |
| NC_011770 | 5552919 | C | G | 84 | 0 | 4 | 0 | NON_SYNONYMOU | MODERAT | MISSENSE | gCc/gGc | A69G | PLES_50381 | | | G | G | G | G | | | | | | |
| NC_011770 | 5553668 | T | C | 94 | 0 | 2 | 0 | NON_SYNONYMOU | MODERAT | MISSENSE | Ttc/Ctc | F319L | PLES_50381 | | | | | | | C | C | | | | |
| NC_011770 | 5553828 | C | G | 98 | 0 | 4 | 0 | NON_SYNONYMOU | MODERAT | MISSENSE | aCc/aGc | T372S | PLES_50381 | G | G | | | | | | | | | G | G |
| NC_011770 | 5554124 | A | G | 94 | 0 | 4 | 0 | NON_SYNONYMOU | MODERAT | MISSENSE | Acc/Gcc | T471A | PLES_50381 | G | G | | | | | | | | | G | G |
| NC_011770 | 5554403 | C | T | 79 | 0 | 12 | 0 | NON_SYNONYMOU | MODERAT | MISSENSE | Cat/Tat | H564Y | PLES_50381 | T | T | T | T | T | T | T | T | T | T | T | T |
| NC_011770 | 5554836 | A | G | 76 | 0 | 12 | 0 | NON_SYNONYMOU | MODERAT | MISSENSE | aAc/aGc | N708S | PLES_50381 | G | G | G | G | G | G | G | G | G | G | G | G |
| NC_011770 | 5555594 | G | GCATC | 1 | 8 | 2 | 0 | FRAME_SHIFT | HIGH | | ggt/ggCAT | G171GI? | PLES_50391 ( | . | . | . | . | . | . | . | GCATC | . | . | . | GCATC |
| NC_011770 | 5555599 | CTGGG | C | 53 | 0 | 12 | 0 | FRAME_SHIFT | HIGH | | ctggga/ | LG173 | PLES_50391 ( | C | C | C | C | C | C | C | C | C | C | C | C |
| NC_011770 | 5555605 | C | G | 1 | 10 | 2 | 0 | NON_SYNONYMOU | MODERAT | MISSENSE | Ctc/Gtc | L175V | PLES_50391 | . | . | . | . | . | . | . | . | . | . | G | G |
| NC_011770 | 5556960 | C | T | 66 | 0 | 2 | 0 | NON_SYNONYMOU | MODERAT | MISSENSE | gGc/gAc | G149D | PLES_50401 | T | T | | | | | | | | | | |
| NC_011770 | 5559176 | C | A | 72 | 0 | 2 | 0 | NON_SYNONYMOU | MODERAT | MISSENSE | cGc/cTc | R149L | PLES_50421 | | | | | | | | | A | A | | |
| NC_011770 | 5559532 | G | C | 82 | 0 | 2 | 0 | NON_SYNONYMOU | MODERAT | MISSENSE | ttC/ttG | F30L | PLES_50421 | | | C | C | | | | | | | | |
| NC_011770 | 5560372 | C | T | 80 | 0 | 1 | 0 | NON_SYNONYMOU | MODERAT | MISSENSE | aCc/aTc | T113I | PLES_50431 | | | | | | | | | | | | T |
| NC_011770 | 5560461 | T | G | 86 | 0 | 12 | 0 | NON_SYNONYMOU | MODERAT | MISSENSE | Tcc/Gcc | S143A | PLES_50431 | G | G | G | G | G | G | G | G | G | G | G | G |
| NC_011770 | 5560654 | A | G | 94 | 0 | 4 | 0 | NON_SYNONYMOU | MODERAT | MISSENSE | gAc/gGc | D207G | PLES_50431 | | | G | G | G | G | | | | | | |
| NC_011770 | 5560707 | A | G | 91 | 0 | 4 | 0 | NON_SYNONYMOU | MODERAT | MISSENSE | Acc/Gcc | T225A | PLES_50431 | | | G | G | G | G | | | | | | |
| NC_011770 | 5561876 | C | T | 86 | 0 | 2 | 0 | NON_SYNONYMOU | MODERAT | MISSENSE | Cgc/Tgc | R236C | PLES_50441 | | | | | | | | | | | T | T |
| NC_011770 | 5562599 | T | C | 77 | 0 | 1 | 0 | NON_SYNONYMOU | MODERAT | MISSENSE | Ttc/Ctc | F155L | PLES_50451 | | C | | | | | | | | | | |

| | | | | | | | | | | | | | | | | | | | | | | | | | |
|---|---|---|---|---|---|---|---|---|---|---|---|---|---|---|---|---|---|---|---|---|---|---|---|---|---|
| NC_011770 | 5562372 | G | A | 76 | 0 | 2 | 0 | NON_SYNONYMOU | MODERAT | MISSENSE | cGt/cAt | R79H | PLES_50451 | | | | | | | A | A | | | | |
| NC_011770 | 5562668 | G | A | 75 | 0 | 4 | 0 | NON_SYNONYMOU | MODERAT | MISSENSE | Gcg/Acg | A178T | PLES_50451 | | | A | A | A | A | | | | | | |
| NC_011770 | 5562740 | G | A | 72 | 0 | 4 | 0 | NON_SYNONYMOU | MODERAT | MISSENSE | Ggc/Agc | G202S | PLES_50451 | | | A | A | A | A | | | | | | |
| NC_011770 | 5562830 | G | C | 70 | 0 | 4 | 0 | NON_SYNONYMOU | MODERAT | MISSENSE | Gtg/Ctg | V232L | PLES_50451 | C | C | | | | | | | | | C | C |
| NC_011770 | 5562903 | C | T | 74 | 0 | 12 | 0 | NON_SYNONYMOU | MODERAT | MISSENSE | gCg/gTg | A256V | PLES_50451 | T | T | T | T | T | T | T | T | T | T | T | T |
| NC_011770 | 5562912 | T | C | 75 | 0 | 1 | 0 | NON_SYNONYMOU | MODERAT | MISSENSE | gTc/gCc | V259A | PLES_50451 | | | | | | | C | | | | | |
| NC_011770 | 5563017 | G | A | 77 | 0 | 2 | 0 | NON_SYNONYMOU | MODERAT | MISSENSE | cGa/cAa | R294Q | PLES_50451 | | | | | | | | | A | A | | |
| NC_011770 | 5563031 | C | T | 79 | 0 | 2 | 0 | NON_SYNONYMOU | MODERAT | MISSENSE | Cca/Tca | P299S | PLES_50451 | T | T | | | | | | | | | | |
| NC_011770 | 5570323 | G | C | 80 | 0 | 2 | 0 | NON_SYNONYMOU | MODERAT | MISSENSE | gGc/gCc | G61A | PLES_50531 | | | | | | | | | C | C | | |
| NC_011770 | 5570524 | G | A | 86 | 0 | 2 | 0 | NON_SYNONYMOU | MODERAT | MISSENSE | cGc/cAc | R128H | PLES_50531 | | | | | | | | | A | A | | |
| NC_011770 | 5570931 | G | A | 82 | 0 | 2 | 0 | NON_SYNONYMOU | MODERAT | MISSENSE | Gaa/Aaa | E264K | PLES_50531 | | | | | | | | | | | A | A |
| NC_011770 | 5571223 | C | G | 93 | 0 | 4 | 0 | NON_SYNONYMOU | MODERAT | MISSENSE | gCc/gGc | A361G | PLES_50531 | | | | | | | G | G | G | G | | |
| NC_011770 | 5571508 | C | T | 87 | 0 | 2 | 0 | NON_SYNONYMOU | MODERAT | MISSENSE | gCg/gTg | A456V | PLES_50531 | | | | | | | T | T | | | | |
| NC_011770 | 5571553 | A | G | 88 | 0 | 4 | 0 | NON_SYNONYMOU | MODERAT | MISSENSE | gAc/gGc | D471G | PLES_50531 | | | G | G | G | G | | | | | | |
| NC_011770 | 5571612 | G | A | 88 | 0 | 2 | 0 | NON_SYNONYMOU | MODERAT | MISSENSE | Ggc/Agc | G491S | PLES_50531 | A | A | | | | | | | | | | |
| NC_011770 | 5571613 | G | C | 86 | 0 | 2 | 0 | NON_SYNONYMOU | MODERAT | MISSENSE | gGc/gCc | G491A | PLES_50531 | C | C | | | | | | | | | | |
| NC_011770 | 5571814 | G | A | 93 | 0 | 2 | 0 | NON_SYNONYMOU | MODERAT | MISSENSE | gGc/gAc | G558D | PLES_50531 | A | A | | | | | | | | | | |
| NC_011770 | 5574977 | G | T | 79 | 0 | 2 | 0 | NON_SYNONYMOU | MODERAT | MISSENSE | Gcc/Tcc | A123S | PLES_50571 | T | T | | | | | | | | | | |
| NC_011770 | 5574980 | G | A | 81 | 0 | 2 | 0 | NON_SYNONYMOU | MODERAT | MISSENSE | Ggt/Agt | G124S | PLES_50571 | | | | | | | A | A | | | | |
| NC_011770 | 5576493 | G | C | 74 | 0 | 2 | 0 | NON_SYNONYMOU | MODERAT | MISSENSE | aGc/aCc | S201T | PLES_50591 | | | | | C | C | | | | | | |
| NC_011770 | 5577404 | C | G | 52 | 0 | 12 | 0 | NON_SYNONYMOU | MODERAT | MISSENSE | caG/caC | Q100H | PLES_50601 | G | G | G | G | G | G | G | G | G | G | G | G |
| NC_011770 | 5577635 | C | A | 80 | 0 | 4 | 0 | NON_SYNONYMOU | MODERAT | MISSENSE | ttG/ttT | L23F | PLES_50601 | A | A | | | | | | | A | A | | |
| NC_011770 | 5579907 | A | G | 88 | 0 | 1 | 0 | NON_SYNONYMOU | MODERAT | MISSENSE | cAc/cGc | H529R | PLES_50611 | | | | | | | | G | | | | |
| NC_011770 | 5578341 | T | C | 77 | 0 | 2 | 0 | NON_SYNONYMOU | MODERAT | MISSENSE | tTg/tCg | L7S | PLES_50611 | | | | | | | | | | | C | C |
| NC_011770 | 5578962 | G | A | 85 | 0 | 2 | 0 | NON_SYNONYMOU | MODERAT | MISSENSE | aGt/aAt | S214N | PLES_50611 | | | | | | | | | | | A | A |
| NC_011770 | 5579245 | A | C | 90 | 0 | 2 | 0 | NON_SYNONYMOU | MODERAT | MISSENSE | gaA/gaC | E308D | PLES_50611 | | | | | | | | | | | C | C |
| NC_011770 | 5579334 | C | G | 83 | 0 | 4 | 0 | NON_SYNONYMOU | MODERAT | MISSENSE | aCc/aGc | T338S | PLES_50611 | | | | | | | G | G | | | G | G |
| NC_011770 | 5580155 | G | C | 87 | 0 | 2 | 0 | NON_SYNONYMOU | MODERAT | MISSENSE | Gtc/Ctc | V612L | PLES_50611 | | | | | | | | | | | C | C |
| NC_011770 | 5581561 | G | C | 84 | 0 | 2 | 0 | NON_SYNONYMOU | MODERAT | MISSENSE | cCc/cGc | P336R | PLES_50631 | | | | | | | C | C | | | | |
| NC_011770 | 5581604 | C | T | 80 | 0 | 2 | 0 | NON_SYNONYMOU | MODERAT | MISSENSE | Gcc/Acc | A322T | PLES_50631 | | | | | | | T | T | | | | |
| NC_011770 | 5581781 | G | A | 79 | 0 | 2 | 0 | NON_SYNONYMOU | MODERAT | MISSENSE | Cgc/Tgc | R263C | PLES_50631 | | | | | | | A | A | | | | |
| NC_011770 | 5582296 | G | C | 81 | 0 | 2 | 0 | NON_SYNONYMOU | MODERAT | MISSENSE | aCc/aGc | T91S | PLES_50631 | | | | | | | | | | | C | C |
| NC_011770 | 5582411 | T | C | 82 | 0 | 10 | 0 | NON_SYNONYMOU | MODERAT | MISSENSE | Acc/Gcc | T53A | PLES_50631 | | | C | C | C | C | C | C | C | C | C | C |
| NC_011770 | 5582462 | G | A | 79 | 0 | 2 | 0 | NON_SYNONYMOU | MODERAT | MISSENSE | Cgc/Tgc | R36C | PLES_50631 | | | | | | | | | | | A | A |
| NC_011770 | 5583718 | A | G | 65 | 0 | 10 | 0 | NON_SYNONYMOU | MODERAT | MISSENSE | Tcg/Ccg | S67P | PLES_50651 | | | G | G | G | G | G | G | G | G | G | G |
| NC_011770 | 5583726 | G | A | 65 | 0 | 1 | 0 | NON_SYNONYMOU | MODERAT | MISSENSE | cCg/cTg | P64L | PLES_50651 | | | | | | | A | | | | | |
| NC_011770 | 5583727 | G | A | 64 | 0 | 2 | 0 | NON_SYNONYMOU | MODERAT | MISSENSE | Ccg/Tcg | P64S | PLES_50651 | | | | | | | | | | | A | A |
| NC_011770 | 5583750 | G | A | 62 | 0 | 6 | 0 | NON_SYNONYMOU | MODERAT | MISSENSE | cCg/cTg | P56L | PLES_50651 | | | A | A | A | A | A | A | | | | |
| NC_011770 | 5583816 | A | G | 62 | 0 | 6 | 0 | NON_SYNONYMOU | MODERAT | MISSENSE | gTg/gCg | V34A | PLES_50651 | | | G | G | G | G | G | G | | | | |
| NC_011770 | 5584180 | C | T | 85 | 0 | 1 | 0 | NON_SYNONYMOU | MODERAT | MISSENSE | cGt/cAt | R178H | PLES_50661 | | | | | | | T | | | | | |
| NC_011770 | 5584313 | G | T | 84 | 0 | 2 | 0 | NON_SYNONYMOU | MODERAT | MISSENSE | Cgc/Agc | R134S | PLES_50661 | T | T | | | | | | | | | | |
| NC_011770 | 5584357 | G | A | 82 | 0 | 2 | 0 | NON_SYNONYMOU | MODERAT | MISSENSE | cCg/cTg | P119L | PLES_50661 | | | | | A | A | | | | | | |
| NC_011770 | 5584406 | C | T | 80 | 0 | 2 | 0 | NON_SYNONYMOU | MODERAT | MISSENSE | Gtg/Atg | V103M | PLES_50661 | T | T | | | | | | | | | | |
| NC_011770 | 5585747 | G | T | 82 | 0 | 2 | 0 | NON_SYNONYMOU | MODERAT | MISSENSE | Ctg/Atg | L6M | PLES_50671 | | | | | | | | | T | T | | |
| NC_011770 | 5586498 | T | C | 71 | 0 | 6 | 0 | NON_SYNONYMOU | MODERAT | MISSENSE | Acc/Gcc | T105A | PLES_50681 | | | C | C | C | C | C | C | | | | |
| NC_011770 | 5587095 | T | C | 70 | 0 | 1 | 0 | NON_SYNONYMOU | MODERAT | MISSENSE | Agc/Ggc | S175G | PLES_50691 | | | | | | | | C | | | | |
| NC_011770 | 5587266 | G | A | 78 | 0 | 2 | 0 | NON_SYNONYMOU | MODERAT | MISSENSE | Cgc/Tgc | R118C | PLES_50691 | | | | | | | | | | | A | A |
| NC_011770 | 5588764 | T | C | 84 | 0 | 1 | 0 | NON_SYNONYMOU | MODERAT | MISSENSE | cTg/cCg | L288P | PLES_50701 | | | | | | | C | | | | | |
| NC_011770 | 5588898 | A | G | 74 | 0 | 6 | 0 | NON_SYNONYMOU | MODERAT | MISSENSE | Acg/Gcg | T333A | PLES_50701 | | | G | G | G | G | G | G | | | | |
| NC_011770 | 5588899 | C | A | 75 | 0 | 2 | 0 | NON_SYNONYMOU | MODERAT | MISSENSE | aCg/aAg | T333K | PLES_50701 | | | | | | | A | A | | | | |
| NC_011770 | 5588959 | C | G | 79 | 0 | 2 | 0 | NON_SYNONYMOU | MODERAT | MISSENSE | cCg/cGg | P353R | PLES_50701 | | | | | G | G | | | | | | |
| NC_011770 | 5589309 | C | G | 79 | 0 | 2 | 0 | NON_SYNONYMOU | MODERAT | MISSENSE | gaC/gaG | D40E | PLES_50711 | | | | | | | | | | | G | G |
| NC_011770 | 5591073 | G | A | 94 | 0 | 1 | 0 | NON_SYNONYMOU | MODERAT | MISSENSE | Gac/Aac | D398N | PLES_50721 | | | | | | | A | | | | | |
| NC_011770 | 5591196 | G | A | 93 | 0 | 4 | 0 | NON_SYNONYMOU | MODERAT | MISSENSE | Gac/Aac | D439N | PLES_50721 | | | A | A | A | A | | | | | | |
| NC_011770 | 5592691 | C | T | 63 | 0 | 2 | 0 | NON_SYNONYMOU | MODERAT | MISSENSE | gCa/gTa | A937V | PLES_50721 | | | | | | | T | T | | | | |
| NC_011770 | 5594585 | T | C | 46 | 0 | 2 | 0 | NON_SYNONYMOU | MODERAT | MISSENSE | aAc/aGc | N630S | PLES_50741 | | | | | | | C | C | | | | |
| NC_011770 | 5597375 | A | C | 91 | 0 | 2 | 0 | NON_SYNONYMOU | MODERAT | MISSENSE | Ttg/Gtg | L119V | PLES_50751 | C | C | | | | | | | | | | |
| NC_011770 | 5597957 | G | A | 89 | 0 | 1 | 0 | INTRAGENIC | MODIFIER | | | | PLES_50752 | | | | | | | | A | | | | |
| NC_011770 | 5599032 | T | C | 10 | 0 | 1 | 0 | INTRAGENIC | MODIFIER | | | | PLES_50753 | | | | | | | C | | | | | |
| NC_011770 | 5598479 | G | A | 84 | 0 | 1 | 0 | INTRAGENIC | MODIFIER | | | | PLES_50753 | | | | | | | | A | | | | |
| NC_011770 | 5598885 | C | T | 33 | 0 | 1 | 0 | INTRAGENIC | MODIFIER | | | | PLES_50753 | | | | | | | | T | | | | |
| NC_011770 | 5603639 | G | T | 80 | 0 | 8 | 0 | NON_SYNONYMOU | MODERAT | MISSENSE | cCc/cAc | P199H | PLES_50761 | T | T | | | | | T | T | T | T | T | T |

| | | | | | | | | | | | | | | | | | | | | | | | | | |
|---|---|---|---|---|---|---|---|---|---|---|---|---|---|---|---|---|---|---|---|---|---|---|---|---|---|
| NC_011770 | 5603708 | A | G | 83 | 0 | 2 | 0 | NON_SYNONYMOU | MODERAT | MISSENSE | gTt/gCt | V176A | PLES_50761 | G | G | | | | | | | | | | |
| NC_011770 | 5603783 | A | T | 84 | 0 | 4 | 0 | NON_SYNONYMOU | MODERAT | MISSENSE | aTc/aAc | I151N | PLES_50761 | T | T | | | | | | | G | G | | |
| NC_011770 | 5603877 | T | C | 83 | 0 | 8 | 0 | NON_SYNONYMOU | MODERAT | MISSENSE | Atc/Gtc | I120V | PLES_50761 | C | C | | | | | C | C | C | C | C | C |
| NC_011770 | 5604709 | G | A | 71 | 0 | 2 | 0 | NON_SYNONYMOU | MODERAT | MISSENSE | gCg/gTg | A180V | PLES_50771 | | | | | | | A | A | | | | |
| NC_011770 | 5604907 | T | C | 76 | 0 | 2 | 0 | NON_SYNONYMOU | MODERAT | MISSENSE | aAg/aGg | K114R | PLES_50771 | | | | | | | | | | | C | C |
| NC_011770 | 5605007 | C | T | 76 | 0 | 2 | 0 | NON_SYNONYMOU | MODERAT | MISSENSE | Ggg/Agg | G81R | PLES_50771 | T | T | | | | | | | | | | |
| NC_011770 | 5605016 | T | C | 74 | 0 | 2 | 0 | NON_SYNONYMOU | MODERAT | MISSENSE | Aat/Gat | N78D | PLES_50771 | | | | | | | | | | | C | C |
| NC_011770 | 5605076 | A | G | 69 | 0 | 2 | 0 | NON_SYNONYMOU | MODERAT | MISSENSE | Tcg/Ccg | S58P | PLES_50771 | | | | | | | | | | | G | G |
| NC_011770 | 5605151 | A | G | 74 | 0 | 2 | 0 | NON_SYNONYMOU | MODERAT | MISSENSE | Ttc/Ctc | F33L | PLES_50771 | | | | | | | | | G | G | | |
| NC_011770 | 5610288 | GC | G | 2 | 5 | 1 | 0 | FRAME_SHIFT | HIGH | | ccc/ | P63 | PLES_50821 | . | . | | | | | | G | . | . | | . |
| NC_011770 | 5610759 | C | T | 32 | 0 | 1 | 0 | NON_SYNONYMOU | MODERAT | MISSENSE | atG/atA | M79I | PLES_50831 | | | | | | | | | | T | | |
| NC_011770 | 5610911 | G | T | 32 | 0 | 2 | 0 | NON_SYNONYMOU | MODERAT | MISSENSE | Cgc/Agc | R29S | PLES_50831 | | | | | | | | | T | T | | |
| NC_011770 | 5611018 | G | C | 45 | 0 | 2 | 0 | NON_SYNONYMOU | MODERAT | MISSENSE | Cag/Gag | Q253E | PLES_50841 | C | C | | | | | | | | | | |
| NC_011770 | 5611099 | C | T | 51 | 0 | 4 | 0 | NON_SYNONYMOU | MODERAT | MISSENSE | Gcc/Acc | A226T | PLES_50841 | | | T | T | T | T | | | | | | |
| NC_011770 | 5611413 | C | T | 63 | 0 | 4 | 0 | NON_SYNONYMOU | MODERAT | MISSENSE | gGc/gAc | G121D | PLES_50841 | | | T | T | T | T | | | | | | |
| NC_011770 | 5611468 | A | T | 66 | 0 | 12 | 0 | NON_SYNONYMOU | MODERAT | MISSENSE | Tcc/Acc | S103T | PLES_50841 | T | T | T | T | T | T | T | T | T | T | T | T |
| NC_011770 | 5614785 | G | A | 61 | 0 | 1 | 0 | NON_SYNONYMOU | MODERAT | MISSENSE | gGc/gAc | G153D | PLES_50861 | | | | | | | | A | | | | |
| NC_011770 | 5615432 | A | G | 72 | 0 | 4 | 0 | NON_SYNONYMOU | MODERAT | MISSENSE | Atc/Gtc | I369V | PLES_50861 | | | | | | | G | G | | | G | G |
| NC_011770 | 5617368 | G | A | 76 | 0 | 2 | 0 | NON_SYNONYMOU | MODERAT | MISSENSE | cGc/cAc | R146H | PLES_50891 | | | | | | | | | | | A | A |
| NC_011770 | 5618225 | C | G | 89 | 0 | 4 | 0 | NON_SYNONYMOU | MODERAT | MISSENSE | Gac/Cac | D286H | PLES_50901 | | | G | G | G | G | | | | | | |
| NC_011770 | 5618420 | A | G | 83 | 0 | 2 | 0 | NON_SYNONYMOU | MODERAT | MISSENSE | Ttt/Ctt | F221L | PLES_50901 | G | G | | | | | | | | | | |
| NC_011770 | 5618645 | C | T | 83 | 0 | 4 | 0 | NON_SYNONYMOU | MODERAT | MISSENSE | Gtt/Att | V146I | PLES_50901 | | | | | | | | | T | T | T | T |
| NC_011770 | 5618708 | G | C | 88 | 0 | 4 | 0 | NON_SYNONYMOU | MODERAT | MISSENSE | Cag/Gag | Q125E | PLES_50901 | C | C | | | | | C | C | | | | |
| NC_011770 | 5618734 | G | T | 88 | 0 | 4 | 0 | NON_SYNONYMOU | MODERAT | MISSENSE | aCc/aAc | T116N | PLES_50901 | T | T | | | | | T | T | | | | |
| NC_011770 | 5618891 | T | A | 79 | 0 | 4 | 0 | NON_SYNONYMOU | MODERAT | MISSENSE | Atc/Ttc | I64F | PLES_50901 | | | A | A | A | A | | | | | | |
| NC_011770 | 5618999 | C | G | 74 | 0 | 6 | 0 | NON_SYNONYMOU | MODERAT | MISSENSE | Gcg/Ccg | A28P | PLES_50901 | | | G | G | G | G | G | G | | | | |
| NC_011770 | 5622143 | C | A | 66 | 0 | 10 | 0 | NON_SYNONYMOU | MODERAT | MISSENSE | aaG/aaT | K239N | PLES_50941 | | | A | A | A | A | A | A | A | A | A | A |
| NC_011770 | 5622301 | C | A | 65 | 0 | 8 | 0 | NON_SYNONYMOU | MODERAT | MISSENSE | Gcg/Tcg | A187S | PLES_50941 | | | A | A | A | A | A | A | | | T | T |
| NC_011770 | 5622324 | T | C | 68 | 0 | 6 | 0 | NON_SYNONYMOU | MODERAT | MISSENSE | cAc/cGc | H179R | PLES_50941 | | | C | C | C | C | C | C | | | | |
| NC_011770 | 5622406 | C | T | 72 | 0 | 10 | 0 | NON_SYNONYMOU | MODERAT | MISSENSE | Gcc/Acc | A152T | PLES_50941 | T | T | T | T | T | T | T | T | | | T | T |
| NC_011770 | 5622504 | T | C | 60 | 0 | 6 | 0 | NON_SYNONYMOU | MODERAT | MISSENSE | aAc/aGc | N119S | PLES_50941 | | | C | C | C | C | C | C | | | | |
| NC_011770 | 5622687 | A | G | 50 | 0 | 12 | 0 | NON_SYNONYMOU | MODERAT | MISSENSE | cTg/cCg | L58P | PLES_50941 | G | G | T | T | T | T | T | T | G | G | G | G |
| NC_011770 | 5626091 | C | G | 79 | 0 | 2 | 0 | NON_SYNONYMOU | MODERAT | MISSENSE | caC/caG | H76Q | PLES_50971 | G | G | | | | | | | | | | |
| NC_011770 | 5626104 | A | G | 75 | 0 | 10 | 0 | NON_SYNONYMOU | MODERAT | MISSENSE | Aac/Gac | N81D | PLES_50971 | G | G | G | G | G | G | G | G | G | G | | |
| NC_011770 | 5626312 | A | T | 74 | 0 | 2 | 0 | NON_SYNONYMOU | MODERAT | MISSENSE | cAa/cTa | Q150L | PLES_50971 | T | T | | | | | | | | | | |
| NC_011770 | 5626380 | T | C | 75 | 0 | 1 | 0 | STOP_LOST+SPLICE | HIGH | MISSENSE | tgA/tgG | *150W | PLES_50981 | | | | | | | C | | | | | |
| NC_011770 | 5626415 | T | C | 73 | 0 | 10 | 0 | NON_SYNONYMOU | MODERAT | MISSENSE | Acc/Gcc | T139A | PLES_50981 | C | C | C | C | C | C | | | C | C | C | C |
| NC_011770 | 5626430 | T | C | 69 | 0 | 10 | 0 | NON_SYNONYMOU | MODERAT | MISSENSE | Acg/Gcg | T134A | PLES_50981 | C | C | C | C | C | C | | | C | C | C | C |
| NC_011770 | 5627232 | A | G | 74 | 0 | 2 | 0 | NON_SYNONYMOU | MODERAT | MISSENSE | Aag/Gag | K52E | PLES_50991 | | | | | | | G | G | | | | |
| NC_011770 | 5627256 | G | A | 72 | 0 | 2 | 0 | NON_SYNONYMOU | MODERAT | MISSENSE | Gac/Aac | D60N | PLES_50991 | A | A | | | | | | | | | | |
| NC_011770 | 5627347 | A | T | 69 | 0 | 2 | 0 | NON_SYNONYMOU | MODERAT | MISSENSE | cAc/cTc | H90L | PLES_50991 | | | | | T | T | | | | | | |
| NC_011770 | 5627441 | A | T | 69 | 0 | 4 | 0 | NON_SYNONYMOU | MODERAT | MISSENSE | gaA/gaT | E121D | PLES_50991 | | | T | T | T | T | | | | | | |
| NC_011770 | 5627509 | C | T | 71 | 0 | 2 | 0 | NON_SYNONYMOU | MODERAT | MISSENSE | gCg/gTg | A144V | PLES_50991 | | | | | | | | | | | T | T |
| NC_011770 | 5627593 | C | G | 61 | 0 | 12 | 0 | NON_SYNONYMOU | MODERAT | MISSENSE | gGg/gCg | G410A | PLES_51001 | G | G | G | G | G | G | G | G | G | G | G | G |
| NC_011770 | 5627611 | T | C | 64 | 0 | 12 | 0 | NON_SYNONYMOU | MODERAT | MISSENSE | aAg/aGg | K404R | PLES_51001 | C | C | C | C | C | C | C | C | C | C | C | C |
| NC_011770 | 5628368 | C | T | 87 | 0 | 2 | 0 | NON_SYNONYMOU | MODERAT | MISSENSE | Gtg/Atg | V152M | PLES_51001 | | | | | | | | | T | T | | |
| NC_011770 | 5629075 | G | A | 63 | 0 | 6 | 0 | NON_SYNONYMOU | MODERAT | MISSENSE | Gcc/Acc | A29T | PLES_51011 | | | A | A | A | A | | | | | A | A |
| NC_011770 | 5629098 | G | C | 67 | 0 | 2 | 0 | NON_SYNONYMOU | MODERAT | MISSENSE | caG/caC | Q36H | PLES_51011 | C | C | | | | | | | | | | |
| NC_011770 | 5629168 | C | G | 67 | 0 | 4 | 0 | NON_SYNONYMOU | MODERAT | MISSENSE | Cgc/Ggc | R60G | PLES_51011 | | | G | G | G | G | | | | | | |
| NC_011770 | 5629266 | A | C | 62 | 0 | 12 | 0 | NON_SYNONYMOU | MODERAT | MISSENSE | aaA/aaC | K92N | PLES_51011 | C | C | C | C | C | C | C | C | C | C | C | C |
| NC_011770 | 5629619 | G | A | 57 | 0 | 8 | 0 | NON_SYNONYMOU | MODERAT | MISSENSE | aGc/aAc | S210N | PLES_51011 | | | A | A | A | A | A | A | A | A | | |
| NC_011770 | 5629743 | G | C | 57 | 0 | 8 | 0 | NON_SYNONYMOU | MODERAT | MISSENSE | gaG/gaC | E251D | PLES_51011 | | | C | C | C | C | C | C | C | C | | |
| NC_011770 | 5629783 | A | G | 58 | 0 | 4 | 0 | NON_SYNONYMOU | MODERAT | MISSENSE | Agc/Ggc | S265G | PLES_51011 | | | G | G | G | G | | | | | | |
| NC_011770 | 5630042 | A | G | 56 | 0 | 12 | 0 | NON_SYNONYMOU | MODERAT | MISSENSE | Atg/Gtg | M44V | PLES_51021 | G | G | G | G | G | G | G | G | G | G | G | G |
| NC_011770 | 5630889 | A | G | 67 | 0 | 4 | 0 | NON_SYNONYMOU | MODERAT | MISSENSE | gTc/gCc | V139A | PLES_51031 | G | G | | | | | | | | | G | G |
| NC_011770 | 5631021 | T | C | 69 | 0 | 12 | 0 | NON_SYNONYMOU | MODERAT | MISSENSE | cAc/cGc | H95R | PLES_51031 | C | C | C | C | C | C | C | C | C | C | C | C |
| NC_011770 | 5631943 | GC | G | 82 | 0 | 1 | 0 | FRAME_SHIFT | HIGH | | ccc/ | P77 | PLES_51041 | | | | | | | | G | | | | |
| NC_011770 | 5631773 | T | C | 74 | 0 | 2 | 0 | NON_SYNONYMOU | MODERAT | MISSENSE | aTc/aCc | I19T | PLES_51041 | | | | | | | | | | | C | C |
| NC_011770 | 5632964 | C | T | 74 | 0 | 12 | 0 | NON_SYNONYMOU | MODERAT | MISSENSE | gCg/gTg | A416V | PLES_51041 | T | T | T | T | T | T | T | T | T | T | T | T |
| NC_011770 | 5634883 | G | A | 79 | 0 | 1 | 0 | NON_SYNONYMOU | MODERAT | MISSENSE | gCg/gTg | A377V | PLES_51071 | | A | | | | | | | | | | |
| NC_011770 | 5634865 | C | T | 77 | 0 | 4 | 0 | NON_SYNONYMOU | MODERAT | MISSENSE | cGc/cAc | R383H | PLES_51071 | | | T | T | T | T | | | | | | |
| NC_011770 | 5635265 | C | T | 74 | 0 | 2 | 0 | NON_SYNONYMOU | MODERAT | MISSENSE | Gca/Aca | A250T | PLES_51071 | | | | | | | | | T | T | | |

| | | | | | | | | | | | | | | | | | | | | | | | | | |
|---|---|---|---|---|---|---|---|---|---|---|---|---|---|---|---|---|---|---|---|---|---|---|---|---|---|
| NC_011770 | 5635871 | C | T | 77 | 0 | 2 | 0 | NON_SYNONYMOU | MODERAT | MISSENSE | Gag/Aag | E48K | PLES_51071 | | | | | | | | | | | T | T |
| NC_011770 | 5636966 | C | A | 57 | 0 | 6 | 0 | NON_SYNONYMOU | MODERAT | MISSENSE | gCc/gAc | A83D | PLES_51091 | | | A | A | A | A | | | A | A | | |
| NC_011770 | 5637086 | A | G | 52 | 0 | 1 | 0 | NON_SYNONYMOU | MODERAT | MISSENSE | cAc/cGc | H123R | PLES_51091 | | | | | | | G | | | | | |
| NC_011770 | 5637167 | G | A | 53 | 0 | 2 | 0 | NON_SYNONYMOU | MODERAT | MISSENSE | cGc/cAc | R150H | PLES_51091 | | | | | | | | | A | A | | |
| NC_011770 | 5637512 | G | A | 29 | 0 | 4 | 0 | NON_SYNONYMOU | MODERAT | MISSENSE | aGc/aAc | S265N | PLES_51091 | | | A | A | A | A | | | | | | |
| NC_011770 | 5642570 | T | C | 50 | 0 | 10 | 0 | INTRAGENIC | MODIFIER | | | | PLES_51113 | | | C | C | C | C | C | C | C | C | C | C |
| NC_011770 | 5642613 | A | G | 50 | 0 | 4 | 0 | INTRAGENIC | MODIFIER | | | | PLES_51113 | | | | | | | G | G | | | G | G |
| NC_011770 | 5651615 | G | A | 65 | 0 | 12 | 0 | NON_SYNONYMOU | MODERAT | MISSENSE | aGt/aAt | S77N | PLES_51191 | A | A | A | A | A | A | A | A | A | A | A | A |
| NC_011770 | 5651777 | G | T | 69 | 0 | 2 | 0 | NON_SYNONYMOU | MODERAT | MISSENSE | aGc/aTc | S131I | PLES_51191 | T | T | | | | | | | | | | |
| NC_011770 | 5653742 | A | T | 49 | 0 | 10 | 0 | STOP_GAINED | HIGH | NONSENSE | Aag/Tag | K451* | PLES_51201 | T | T | T | T | T | T | T | T | | | T | T |
| NC_011770 | 5653624 | AGGC | A | 58 | 0 | 4 | 0 | CODON_CHANGE_ | MODERATE | | ggcgct/gct | GA413A | PLES_51201 | | A | | | A | | A | | | | A | |
| NC_011770 | 5654288 | G | A | 78 | 0 | 1 | 0 | NON_SYNONYMOU | MODERAT | MISSENSE | Gcc/Acc | A633T | PLES_51201 | | | | | | | | A | | | | |
| NC_011770 | 5655222 | T | C | 88 | 0 | 1 | 0 | NON_SYNONYMOU | MODERAT | MISSENSE | gTg/gCg | V944A | PLES_51201 | | | | | | | | C | | | | |
| NC_011770 | 5652601 | C | G | 79 | 0 | 12 | 0 | NON_SYNONYMOU | MODERAT | MISSENSE | gaC/gaG | D70E | PLES_51201 | G | G | G | G | G | G | G | G | G | G | G | G |
| NC_011770 | 5652779 | T | G | 73 | 0 | 2 | 0 | NON_SYNONYMOU | MODERAT | MISSENSE | Tcg/Gcg | S130A | PLES_51201 | | | | | | | G | G | | | | |
| NC_011770 | 5653019 | A | G | 76 | 0 | 12 | 0 | NON_SYNONYMOU | MODERAT | MISSENSE | Atg/Gtg | M210V | PLES_51201 | G | G | G | G | G | G | G | G | G | G | G | G |
| NC_011770 | 5653460 | G | A | 60 | 0 | 10 | 0 | NON_SYNONYMOU | MODERAT | MISSENSE | Gcc/Acc | A357T | PLES_51201 | A | A | A | A | A | A | A | A | | | A | A |
| NC_011770 | 5653472 | C | A | 62 | 0 | 10 | 0 | NON_SYNONYMOU | MODERAT | MISSENSE | Cag/Aag | Q361K | PLES_51201 | A | A | A | A | A | A | A | A | | | A | A |
| NC_011770 | 5653481 | C | A | 62 | 0 | 10 | 0 | NON_SYNONYMOU | MODERAT | MISSENSE | Cgc/Agc | R364S | PLES_51201 | A | A | A | A | A | A | A | A | | | A | A |
| NC_011770 | 5653496 | G | A | 66 | 0 | 2 | 0 | NON_SYNONYMOU | MODERAT | MISSENSE | Gac/Aac | D369N | PLES_51201 | | | | | | | | | | | A | A |
| NC_011770 | 5653534 | A | C | 62 | 0 | 2 | 0 | NON_SYNONYMOU | MODERAT | MISSENSE | gaA/gaC | E381D | PLES_51201 | C | C | | | | | | | | | | |
| NC_011770 | 5653538 | G | A | 64 | 0 | 8 | 0 | NON_SYNONYMOU | MODERAT | MISSENSE | Ggc/Agc | G383S | PLES_51201 | | | A | A | A | A | A | A | | | A | A |
| NC_011770 | 5653563 | G | A | 60 | 0 | 10 | 0 | NON_SYNONYMOU | MODERAT | MISSENSE | gGc/gAc | G391D | PLES_51201 | A | A | A | A | A | A | A | A | | | A | A |
| NC_011770 | 5653565 | A | G | 60 | 0 | 10 | 0 | NON_SYNONYMOU | MODERAT | MISSENSE | Acc/Gcc | T392A | PLES_51201 | G | G | G | G | G | G | G | G | | | G | G |
| NC_011770 | 5653583 | A | G | 60 | 0 | 10 | 0 | NON_SYNONYMOU | MODERAT | MISSENSE | Aaa/Gaa | K398E | PLES_51201 | G | G | G | G | G | G | G | G | | | G | G |
| NC_011770 | 5653637 | GATA | G | 53 | 0 | 10 | 0 | CODON_CHANGE_ | MODERATE | | gataaa/gaa | DK416E | PLES_51201 | G | G | G | G | G | G | G | G | | | G | G |
| NC_011770 | 5653659 | A | G | 61 | 0 | 4 | 0 | NON_SYNONYMOU | MODERAT | MISSENSE | gAg/gGg | E423G | PLES_51201 | | | G | G | G | G | | | | | | |
| NC_011770 | 5653667 | G | A | 60 | 0 | 6 | 0 | NON_SYNONYMOU | MODERAT | MISSENSE | Gag/Aag | E426K | PLES_51201 | A | A | | | | | A | A | | | A | A |
| NC_011770 | 5653706 | G | A | 60 | 0 | 4 | 0 | NON_SYNONYMOU | MODERAT | MISSENSE | Gcc/Acc | A439T | PLES_51201 | | | | | | | A | A | | | A | A |
| NC_011770 | 5653737 | A | G | 52 | 0 | 10 | 0 | NON_SYNONYMOU | MODERAT | MISSENSE | aAc/aGc | N449S | PLES_51201 | G | G | G | G | G | G | G | G | | | G | G |
| NC_011770 | 5653743 | A | C | 51 | 0 | 10 | 0 | NON_SYNONYMOU | MODERAT | MISSENSE | aAg/aCg | K451T | PLES_51201 | C | C | C | C | C | C | C | C | | | C | C |
| NC_011770 | 5653744 | G | T | 49 | 0 | 10 | 0 | NON_SYNONYMOU | MODERAT | MISSENSE | aaG/aaT | K451N | PLES_51201 | T | T | T | T | T | T | T | T | | | T | T |
| NC_011770 | 5653761 | G | A | 57 | 0 | 8 | 0 | NON_SYNONYMOU | MODERAT | MISSENSE | gGc/gAc | G457D | PLES_51201 | | | A | A | A | A | A | A | | | A | A |
| NC_011770 | 5653772 | G | A | 57 | 0 | 12 | 0 | NON_SYNONYMOU | MODERAT | MISSENSE | Gcg/Acg | A461T | PLES_51201 | A | A | A | A | A | A | A | A | A | A | A | A |
| NC_011770 | 5653799 | G | T | 57 | 0 | 4 | 0 | NON_SYNONYMOU | MODERAT | MISSENSE | Gcc/Tcc | A470S | PLES_51201 | | | T | T | T | T | | | | | | |
| NC_011770 | 5653814 | G | A | 61 | 0 | 2 | 0 | NON_SYNONYMOU | MODERAT | MISSENSE | Gcc/Acc | A475T | PLES_51201 | | | | | | | | | A | A | | |
| NC_011770 | 5653985 | G | A | 63 | 0 | 2 | 0 | NON_SYNONYMOU | MODERAT | MISSENSE | Ggc/Agc | G532S | PLES_51201 | A | A | | | | | | | | | | |
| NC_011770 | 5654012 | G | A | 67 | 0 | 2 | 0 | NON_SYNONYMOU | MODERAT | MISSENSE | Gcc/Acc | A541T | PLES_51201 | | | | | | | A | A | | | | |
| NC_011770 | 5654312 | T | C | 76 | 0 | 2 | 0 | NON_SYNONYMOU | MODERAT | MISSENSE | Tcg/Ccg | S641P | PLES_51201 | C | C | | | | | | | | | | |
| NC_011770 | 5654409 | C | G | 76 | 0 | 2 | 0 | NON_SYNONYMOU | MODERAT | MISSENSE | gCt/gGt | A673G | PLES_51201 | | | | | | | | | G | G | | |
| NC_011770 | 5654426 | A | G | 74 | 0 | 2 | 0 | NON_SYNONYMOU | MODERAT | MISSENSE | Acc/Gcc | T679A | PLES_51201 | | | | | | | | | G | G | | |
| NC_011770 | 5654475 | A | T | 74 | 0 | 4 | 0 | NON_SYNONYMOU | MODERAT | MISSENSE | cAc/cTc | H695L | PLES_51201 | | | T | T | T | T | | | | | | |
| NC_011770 | 5655808 | A | C | 76 | 0 | 2 | 0 | NON_SYNONYMOU | MODERAT | MISSENSE | gAt/gCt | D47A | PLES_51211 | C | C | | | | | | | | | | |
| NC_011770 | 5672806 | T | C | 74 | 0 | 8 | 0 | NON_SYNONYMOU | MODERAT | MISSENSE | tAc/tGc | Y134C | PLES_51391 | C | C | | | | | C | C | C | C | C | C |
| NC_011770 | 5672807 | A | C | 74 | 0 | 8 | 0 | NON_SYNONYMOU | MODERAT | MISSENSE | Tac/Gac | Y134D | PLES_51391 | C | C | | | | | C | C | C | C | C | C |
| NC_011770 | 5673028 | A | G | 83 | 0 | 2 | 0 | NON_SYNONYMOU | MODERAT | MISSENSE | gTg/gCg | V60A | PLES_51391 | | | | | | | | | G | G | | |
| NC_011770 | 5678855 | G | A | 73 | 0 | 1 | 0 | INTRAGENIC | MODIFIER | | | | PLES_51432 | | | | | | | | | | | A | |
| NC_011770 | 5686702 | C | T | 50 | 0 | 2 | 0 | NON_SYNONYMOU | MODERAT | MISSENSE | Gcc/Acc | A69T | PLES_51511 | T | T | | | | | | | | | | |
| NC_011770 | 5686711 | C | T | 49 | 0 | 6 | 0 | NON_SYNONYMOU | MODERAT | MISSENSE | Gtg/Atg | V66M | PLES_51511 | | | T | T | T | T | T | T | | | | |
| NC_011770 | 5687216 | C | T | 75 | 0 | 1 | 0 | NON_SYNONYMOU | MODERAT | MISSENSE | Gtc/Atc | V40I | PLES_51521 | | | | | | | | T | | | | |
| NC_011770 | 5687113 | G | A | 75 | 0 | 4 | 0 | NON_SYNONYMOU | MODERAT | MISSENSE | tCc/tTc | S74F | PLES_51521 | | | A | A | A | A | | | | | | |
| NC_011770 | 5688675 | G | GC | 80 | 0 | 1 | 0 | FRAME_SHIFT | HIGH | | ggc/ggGc | G70G? | PLES_51541 | | | | GC | | | | | | | | |
| NC_011770 | 5688114 | G | T | 82 | 0 | 4 | 0 | NON_SYNONYMOU | MODERAT | MISSENSE | gaC/gaA | D257E | PLES_51541 | T | T | | | | | | | | | T | T |
| NC_011770 | 5692437 | C | T | 64 | 0 | 4 | 0 | NON_SYNONYMOU | MODERAT | MISSENSE | gCc/gTc | A59V | PLES_51571 | | | T | T | T | T | | | | | | |
| NC_011770 | 5693259 | A | G | 73 | 0 | 12 | 0 | NON_SYNONYMOU | MODERAT | MISSENSE | cAc/cGc | H333R | PLES_51571 | G | G | G | G | G | G | G | G | G | G | G | G |
| NC_011770 | 5694014 | G | A | 60 | 0 | 4 | 0 | NON_SYNONYMOU | MODERAT | MISSENSE | Gac/Aac | D585N | PLES_51571 | | | A | A | A | A | | | | | | |
| NC_011770 | 5694138 | G | A | 57 | 0 | 10 | 0 | NON_SYNONYMOU | MODERAT | MISSENSE | gGt/gAt | G626D | PLES_51571 | A | A | A | A | A | A | | | A | A | A | A |
| NC_011770 | 5694222 | G | A | 49 | 0 | 1 | 0 | NON_SYNONYMOU | MODERAT | MISSENSE | cGc/cAc | R654H | PLES_51571 | | | | | | | A | | | | | |
| NC_011770 | 5694250 | C | G | 51 | 0 | 12 | 0 | NON_SYNONYMOU | MODERAT | MISSENSE | agC/agG | S663R | PLES_51571 | G | G | G | G | G | G | G | G | G | G | G | G |
| NC_011770 | 5694338 | C | A | 48 | 0 | 10 | 0 | NON_SYNONYMOU | MODERAT | MISSENSE | Ccc/Acc | P693T | PLES_51571 | A | A | A | A | A | A | | | A | A | A | A |
| NC_011770 | 5694408 | C | T | 60 | 0 | 2 | 0 | NON_SYNONYMOU | MODERAT | MISSENSE | cCg/cTg | P716L | PLES_51571 | | | | | T | T | | | | | | |
| NC_011770 | 5694443 | G | A | 67 | 0 | 1 | 0 | NON_SYNONYMOU | MODERAT | MISSENSE | Gac/Aac | D728N | PLES_51571 | | | | | | | A | | | | | |

| | | | | | | | | | | | | | | | | | | | | | | | | | |
|---|---|---|---|---|---|---|---|---|---|---|---|---|---|---|---|---|---|---|---|---|---|---|---|---|---|
| NC_011770 | 5694821 | G | A | 76 | 0 | 2 | 0 | NON_SYNONYMOU | MODERAT | MISSENSE | Gtc/Atc | V854I | PLES_51571 | | | | | | | | | | | A | A |
| NC_011770 | 5694839 | G | A | 77 | 0 | 12 | 0 | NON_SYNONYMOU | MODERAT | MISSENSE | Gag/Aag | E860K | PLES_51571 | A | A | A | A | A | A | A | A | A | A | A | A |
| NC_011770 | 5694888 | A | G | 75 | 0 | 2 | 0 | NON_SYNONYMOU | MODERAT | MISSENSE | aAg/aGg | K876R | PLES_51571 | G | G | | | | | | | | | | |
| NC_011770 | 5694930 | A | G | 75 | 0 | 10 | 0 | NON_SYNONYMOU | MODERAT | MISSENSE | aAc/aGc | N890S | PLES_51571 | G | G | G | G | G | G | | | G | G | G | G |
| NC_011770 | 5695058 | G | A | 71 | 0 | 2 | 0 | NON_SYNONYMOU | MODERAT | MISSENSE | Gcg/Acg | A933T | PLES_51571 | | | | | | | | | A | A | | |
| NC_011770 | 5695067 | A | G | 68 | 0 | 10 | 0 | NON_SYNONYMOU | MODERAT | MISSENSE | Acg/Gcg | T936A | PLES_51571 | G | G | G | G | G | G | G | G | G | G | | |
| NC_011770 | 5695806 | C | T | 98 | 0 | 1 | 0 | NON_SYNONYMOU | MODERAT | MISSENSE | gCc/gTc | A100V | PLES_51581 | | T | | | | | | | | | | |
| NC_011770 | 5695532 | G | A | 86 | 0 | 2 | 0 | NON_SYNONYMOU | MODERAT | MISSENSE | Gat/Aat | D9N | PLES_51581 | | | | | | | | | | | A | A |
| NC_011770 | 5697478 | G | A | 80 | 0 | 2 | 0 | NON_SYNONYMOU | MODERAT | MISSENSE | cGc/cAc | R120H | PLES_51601 | A | A | | | | | | | | | | |
| NC_011770 | 5701261 | C | A | 92 | 0 | 12 | 0 | NON_SYNONYMOU | MODERAT | MISSENSE | Gcg/Tcg | A248S | PLES_51641 | A | A | A | A | A | A | A | A | A | A | A | A |
| NC_011770 | 5701347 | G | A | 89 | 0 | 2 | 0 | NON_SYNONYMOU | MODERAT | MISSENSE | aCc/aTc | T219I | PLES_51641 | | | | | | | A | A | | | | |
| NC_011770 | 5701421 | G | T | 81 | 0 | 10 | 0 | NON_SYNONYMOU | MODERAT | MISSENSE | caC/caA | H194Q | PLES_51641 | | | T | T | T | T | T | T | T | T | T | T |
| NC_011770 | 5701438 | C | G | 81 | 0 | 12 | 0 | NON_SYNONYMOU | MODERAT | MISSENSE | Gta/Cta | V189L | PLES_51641 | G | G | G | G | G | G | G | G | G | G | G | G |
| NC_011770 | 5701656 | C | A | 77 | 0 | 2 | 0 | NON_SYNONYMOU | MODERAT | MISSENSE | gGg/gTg | G116V | PLES_51641 | | | | | | | | | A | A | | |
| NC_011770 | 5701666 | A | T | 73 | 0 | 4 | 0 | NON_SYNONYMOU | MODERAT | MISSENSE | Tcg/Acg | S113T | PLES_51641 | | | | | | | | | T | T | T | T |
| NC_011770 | 5702931 | C | T | 87 | 0 | 4 | 0 | NON_SYNONYMOU | MODERAT | MISSENSE | Gtc/Atc | V19I | PLES_51651 | | | T | T | T | T | | | | | | |
| NC_011770 | 5703204 | G | A | 76 | 0 | 2 | 0 | NON_SYNONYMOU | MODERAT | MISSENSE | Cgc/Tgc | R363C | PLES_51661 | | | | | | | A | A | | | | |
| NC_011770 | 5704254 | G | C | 61 | 0 | 12 | 0 | NON_SYNONYMOU | MODERAT | MISSENSE | Ctg/Gtg | L13V | PLES_51661 | C | C | C | C | C | C | C | C | C | C | C | C |
| NC_011770 | 5704602 | A | AC | 64 | 0 | 1 | 0 | FRAME_SHIFT | HIGH | | ggt/ggGt | G14G? | PLES_51671 | | | | | | | AC | | | | | |
| NC_011770 | 5704634 | G | T | 65 | 0 | 2 | 0 | NON_SYNONYMOU | MODERAT | MISSENSE | Cgc/Agc | R4S | PLES_51671 | | | | | | | | | T | T | | |
| NC_011770 | 5704744 | G | A | 64 | 0 | 10 | 0 | NON_SYNONYMOU | MODERAT | MISSENSE | gCg/gTg | A275V | PLES_51681 | A | A | A | A | A | A | | | A | A | A | A |
| NC_011770 | 5704745 | C | G | 65 | 0 | 10 | 0 | NON_SYNONYMOU | MODERAT | MISSENSE | Gcg/Ccg | A275P | PLES_51681 | G | G | G | G | G | G | | | G | G | G | G |
| NC_011770 | 5704768 | CCGA | C | 65 | 0 | 1 | 0 | CODON_CHANGE_ | MODERATE | | atcggc/agc | IG266S | PLES_51681 | | | | | | | C | | | | | |
| NC_011770 | 5704781 | C | T | 64 | 0 | 2 | 0 | NON_SYNONYMOU | MODERAT | MISSENSE | Ggc/Agc | G263S | PLES_51681 | | | | | | | T | T | | | | |
| NC_011770 | 5705015 | C | G | 58 | 0 | 6 | 0 | NON_SYNONYMOU | MODERAT | MISSENSE | Gaa/Caa | E185Q | PLES_51681 | G | G | G | G | G | G | | | | | | |
| NC_011770 | 5705347 | A | C | 45 | 0 | 2 | 0 | NON_SYNONYMOU | MODERAT | MISSENSE | gTc/gGc | V74G | PLES_51681 | C | C | | | | | | | | | | |
| NC_011770 | 5705348 | C | T | 48 | 0 | 2 | 0 | NON_SYNONYMOU | MODERAT | MISSENSE | Gtc/Atc | V74I | PLES_51681 | | | | | | | T | T | | | | |
| NC_011770 | 5705552 | G | T | 52 | 0 | 2 | 0 | NON_SYNONYMOU | MODERAT | MISSENSE | Ctt/Att | L6I | PLES_51681 | | | | | | | T | T | | | | |
| NC_011770 | 5705927 | T | C | 54 | 0 | 2 | 0 | NON_SYNONYMOU | MODERAT | MISSENSE | cTg/cCg | L69P | PLES_51691 | C | C | | | | | | | | | | |
| NC_011770 | 5706531 | G | A | 59 | 0 | 2 | 0 | NON_SYNONYMOU | MODERAT | MISSENSE | aCc/aTc | T420I | PLES_51701 | | | | | | | A | A | | | | |
| NC_011770 | 5706673 | C | G | 73 | 0 | 2 | 0 | NON_SYNONYMOU | MODERAT | MISSENSE | Gtc/Ctc | V373L | PLES_51701 | | | | | | | G | G | | | | |
| NC_011770 | 5708479 | C | T | 90 | 0 | 2 | 0 | INTRAGENIC | MODIFIER | | | | PLES_51711 | | | | | | | | | | | T | T |
| NC_011770 | 5708695 | A | G | 72 | 0 | 8 | 0 | INTRAGENIC | MODIFIER | | | | PLES_51711 | G | G | G | G | G | G | | | | | G | G |
| NC_011770 | 5708728 | G | A | 74 | 0 | 2 | 0 | INTRAGENIC | MODIFIER | | | | PLES_51711 | A | A | | | | | | | | | | |
| NC_011770 | 5708805 | A | T | 78 | 0 | 2 | 0 | INTRAGENIC | MODIFIER | | | | PLES_51711 | T | T | | | | | | | | | | |
| NC_011770 | 5708842 | C | T | 76 | 0 | 8 | 0 | INTRAGENIC | MODIFIER | | | | PLES_51711 | T | T | T | T | T | T | | | | | T | T |
| NC_011770 | 5708858 | G | A | 75 | 0 | 4 | 0 | INTRAGENIC | MODIFIER | | | | PLES_51711 | | | A | A | A | A | | | | | | |
| NC_011770 | 5708886 | T | C | 75 | 0 | 8 | 0 | INTRAGENIC | MODIFIER | | | | PLES_51711 | C | C | C | C | C | C | | | | | C | C |
| NC_011770 | 5708911 | C | T | 79 | 0 | 2 | 0 | INTRAGENIC | MODIFIER | | | | PLES_51711 | | | | | | | | | T | T | | |
| NC_011770 | 5708923 | C | A | 77 | 0 | 2 | 0 | INTRAGENIC | MODIFIER | | | | PLES_51711 | A | A | | | | | | | | | | |
| NC_011770 | 5708932 | T | C | 73 | 0 | 4 | 0 | INTRAGENIC | MODIFIER | | | | PLES_51711 | | | | | | | C | C | C | C | | |
| NC_011770 | 5709010 | G | C | 76 | 0 | 4 | 0 | INTRAGENIC | MODIFIER | | | | PLES_51711 | | | C | C | C | C | | | | | | |
| NC_011770 | 5709130 | C | T | 66 | 0 | 4 | 0 | INTRAGENIC | MODIFIER | | | | PLES_51711 | | | T | T | T | T | | | | | | |
| NC_011770 | 5709208 | C | G | 58 | 0 | 6 | 0 | INTRAGENIC | MODIFIER | | | | PLES_51711 | | | G | G | G | G | | | | | G | G |
| NC_011770 | 5709463 | A | G | 40 | 0 | 1 | 0 | STOP_LOST+SPLICE | HIGH | MISSENSE | Tag/Cag | *334Q | PLES_51721 | | | | | | | | G | | | | |
| NC_011770 | 5709703 | C | T | 50 | 0 | 2 | 0 | NON_SYNONYMOU | MODERAT | MISSENSE | Ggc/Agc | G254S | PLES_51721 | | | | | | | | | | | T | T |
| NC_011770 | 5709945 | C | A | 66 | 0 | 2 | 0 | NON_SYNONYMOU | MODERAT | MISSENSE | cGt/cTt | R173L | PLES_51721 | A | A | | | | | | | | | | |
| NC_011770 | 5710312 | G | T | 56 | 0 | 4 | 0 | NON_SYNONYMOU | MODERAT | MISSENSE | Ctg/Atg | L51M | PLES_51721 | T | T | | | | | | | | | T | T |
| NC_011770 | 5712208 | G | A | 84 | 0 | 2 | 0 | NON_SYNONYMOU | MODERAT | MISSENSE | cGc/cAc | R82H | PLES_51751 | A | A | | | | | | | | | | |
| NC_011770 | 5712459 | C | G | 89 | 0 | 10 | 0 | NON_SYNONYMOU | MODERAT | MISSENSE | Caa/Gaa | Q166E | PLES_51751 | G | G | G | G | G | G | G | G | | | G | G |
| NC_011770 | 5712893 | A | G | 73 | 0 | 2 | 0 | NON_SYNONYMOU | MODERAT | MISSENSE | Acc/Gcc | T39A | PLES_51761 | | | | | | | | | | | G | G |
| NC_011770 | 5713005 | G | A | 83 | 0 | 1 | 0 | NON_SYNONYMOU | MODERAT | MISSENSE | gGc/gAc | G76D | PLES_51761 | | | | | | | A | | | | | |
| NC_011770 | 5713119 | A | G | 84 | 0 | 2 | 0 | NON_SYNONYMOU | MODERAT | MISSENSE | aAc/aGc | N114S | PLES_51761 | | | | | | | G | G | | | | |
| NC_011770 | 5715310 | G | A | 67 | 0 | 1 | 0 | NON_SYNONYMOU | MODERAT | MISSENSE | Gtc/Atc | V99I | PLES_51791 | | | | | | | A | | | | | |
| NC_011770 | 5715653 | G | C | 61 | 0 | 2 | 0 | NON_SYNONYMOU | MODERAT | MISSENSE | Gcc/Ccc | A56P | PLES_51801 | | | | | | | C | C | | | | |
| NC_011770 | 5715662 | G | A | 63 | 0 | 4 | 0 | NON_SYNONYMOU | MODERAT | MISSENSE | Gac/Aac | D59N | PLES_51801 | | | A | A | A | A | | | | | | |
| NC_011770 | 5715759 | G | A | 74 | 0 | 1 | 0 | NON_SYNONYMOU | MODERAT | MISSENSE | gGg/gAg | G91E | PLES_51801 | | | | | | | A | | | | | |
| NC_011770 | 5715881 | G | A | 69 | 0 | 2 | 0 | NON_SYNONYMOU | MODERAT | MISSENSE | gGc/gAc | G7D | PLES_51811 | A | A | | | | | | | | | | |
| NC_011770 | 5715917 | A | C | 66 | 0 | 2 | 0 | NON_SYNONYMOU | MODERAT | MISSENSE | gAc/gCc | D19A | PLES_51811 | | | | | | | | | C | C | | |
| NC_011770 | 5716324 | T | G | 43 | 0 | 4 | 0 | NON_SYNONYMOU | MODERAT | MISSENSE | Ttc/Gtc | F155V | PLES_51811 | | | | | | | G | G | G | G | | |
| NC_011770 | 5716776 | G | A | 35 | 0 | 2 | 0 | NON_SYNONYMOU | MODERAT | MISSENSE | Ggc/Agc | G111S | PLES_51821 | A | A | | | | | | | | | | |
| NC_011770 | 5716927 | G | A | 35 | 0 | 2 | 0 | NON_SYNONYMOU | MODERAT | MISSENSE | cGc/cAc | R161H | PLES_51821 | | | | | | | A | A | | | | |

| | | | | | | | | | | | | | | | | | | | | | | | | | |
|---|---|---|---|---|---|---|---|---|---|---|---|---|---|---|---|---|---|---|---|---|---|---|---|---|---|
| NC_011770 | 5717086 | T | A | 22 | 0 | 2 | 0 | NON_SYNONYMOU | MODERAT | MISSENSE | gTc/gAc | V214D | PLES_51821 | A | A | | | | | | | | | | |
| NC_011770 | 5717262 | C | A | 8 | 2 | 2 | 0 | STOP_GAINED | HIGH | NONSENSE | Gag/Tag | E172* | PLES_51831 | | . | | | | | A | A | . | | | |
| NC_011770 | 5717668 | AC | A | 63 | 0 | 2 | 0 | FRAME_SHIFT | HIGH | | ggt/ | G36 | PLES_51831 | A | A | | | | | | | | | | |
| NC_011770 | 5717699 | G | A | 63 | 0 | 1 | 0 | NON_SYNONYMOU | MODERAT | MISSENSE | aCc/aTc | T26I | PLES_51831 | A | | | | | | | | | | | |
| NC_011770 | 5717216 | G | A | 10 | 0 | 4 | 0 | NON_SYNONYMOU | MODERAT | MISSENSE | gCg/gTg | A187V | PLES_51831 | | | | | | | A | A | | | A | A |
| NC_011770 | 5717570 | G | C | 48 | 0 | 4 | 0 | NON_SYNONYMOU | MODERAT | MISSENSE | gCg/gGg | A69G | PLES_51831 | | | C | C | C | C | | | | | | |
| NC_011770 | 5717582 | C | T | 50 | 0 | 4 | 0 | NON_SYNONYMOU | MODERAT | MISSENSE | cGg/cAg | R65Q | PLES_51831 | | | T | T | T | T | | | | | | |
| NC_011770 | 5717635 | C | A | 57 | 0 | 2 | 0 | NON_SYNONYMOU | MODERAT | MISSENSE | tgG/tgT | W47C | PLES_51831 | | | | | | | | | A | A | | |
| NC_011770 | 5717703 | A | T | 60 | 0 | 6 | 0 | NON_SYNONYMOU | MODERAT | MISSENSE | Tcg/Acg | S25T | PLES_51831 | | | T | T | T | T | | | T | T | | |
| NC_011770 | 5717727 | C | A | 60 | 0 | 2 | 0 | NON_SYNONYMOU | MODERAT | MISSENSE | Gtc/Ttc | V17F | PLES_51831 | | | | | | | | | A | A | | |
| NC_011770 | 5717738 | C | T | 64 | 0 | 2 | 0 | NON_SYNONYMOU | MODERAT | MISSENSE | cGt/cAt | R13H | PLES_51831 | T | T | | | | | | | | | | |
| NC_011770 | 5717756 | T | C | 61 | 0 | 8 | 0 | NON_SYNONYMOU | MODERAT | MISSENSE | gAg/gGg | E7G | PLES_51831 | | | C | C | C | C | C | C | | | C | C |
| NC_011770 | 5717771 | G | A | 66 | 0 | 2 | 0 | NON_SYNONYMOU | MODERAT | MISSENSE | aCc/aTc | T2I | PLES_51831 | | | | | | | | | A | A | | |
| NC_011770 | 5718076 | A | C | 62 | 0 | 2 | 0 | NON_SYNONYMOU | MODERAT | MISSENSE | tTc/tGc | F166C | PLES_51841 | | | | | | | | | | | C | C |
| NC_011770 | 5718217 | G | A | 63 | 0 | 6 | 0 | NON_SYNONYMOU | MODERAT | MISSENSE | gCc/gTc | A119V | PLES_51841 | A | A | | | | | A | A | | | A | A |
| NC_011770 | 5718440 | C | T | 61 | 0 | 4 | 0 | NON_SYNONYMOU | MODERAT | MISSENSE | Gag/Aag | E45K | PLES_51841 | | | T | T | T | T | | | | | | |
| NC_011770 | 5718545 | T | G | 62 | 0 | 4 | 0 | NON_SYNONYMOU | MODERAT | MISSENSE | Agc/Cgc | S10R | PLES_51841 | | | G | G | G | G | | | | | | |
| NC_011770 | 5718978 | T | C | 68 | 0 | 10 | 0 | NON_SYNONYMOU | MODERAT | MISSENSE | Agc/Ggc | S87G | PLES_51851 | C | C | C | C | C | C | C | C | | | C | C |
| NC_011770 | 5719650 | C | G | 51 | 0 | 6 | 0 | NON_SYNONYMOU | MODERAT | MISSENSE | aCc/aGc | T51S | PLES_51861 | G | G | G | G | G | G | | | | | | |
| NC_011770 | 5719901 | G | A | 46 | 0 | 2 | 0 | NON_SYNONYMOU | MODERAT | MISSENSE | Ggc/Agc | G135S | PLES_51861 | A | A | | | | | | | | | | |
| NC_011770 | 5722348 | G | GC | 86 | 0 | 1 | 0 | FRAME_SHIFT | HIGH | | gcc/gGcc | A84G? | PLES_51881 | GC | | | | | | | | | | | |
| NC_011770 | 5721947 | C | T | 91 | 0 | 1 | 0 | NON_SYNONYMOU | MODERAT | MISSENSE | Gcg/Acg | A218T | PLES_51881 | | | | | | | | | | | T | |
| NC_011770 | 5721400 | C | G | 82 | 0 | 2 | 0 | NON_SYNONYMOU | MODERAT | MISSENSE | aGt/aCt | S400T | PLES_51881 | | | | | | | G | G | | | | |
| NC_011770 | 5722130 | G | T | 92 | 0 | 4 | 0 | NON_SYNONYMOU | MODERAT | MISSENSE | Ctc/Atc | L157I | PLES_51881 | | | T | T | T | T | | | | | | |
| NC_011770 | 5724022 | AG | AGG | 89 | 0 | 2 | 0 | FRAME_SHIFT | HIGH | | ctc/ | L11 | PLES_51891 | AGG | | | A | | | | | | | | |
| NC_011770 | 5723108 | C | T | 86 | 0 | 2 | 0 | NON_SYNONYMOU | MODERAT | MISSENSE | Gag/Aag | E316K | PLES_51891 | | | | | | | | | T | T | | |
| NC_011770 | 5723292 | C | A | 83 | 0 | 4 | 0 | NON_SYNONYMOU | MODERAT | MISSENSE | gaG/gaT | E254D | PLES_51891 | | | | | | | A | A | | | A | A |
| NC_011770 | 5723415 | C | G | 81 | 0 | 4 | 0 | NON_SYNONYMOU | MODERAT | MISSENSE | gaG/gaC | E213D | PLES_51891 | | | G | G | G | G | | | | | | |
| NC_011770 | 5723416 | T | G | 82 | 0 | 8 | 0 | NON_SYNONYMOU | MODERAT | MISSENSE | gAg/gCg | E213A | PLES_51891 | G | G | | | | | G | G | G | G | G | G |
| NC_011770 | 5723450 | C | G | 86 | 0 | 4 | 0 | NON_SYNONYMOU | MODERAT | MISSENSE | Gtg/Ctg | V202L | PLES_51891 | | | G | G | G | G | | | | | | |
| NC_011770 | 5724041 | G | T | 96 | 0 | 2 | 0 | NON_SYNONYMOU | MODERAT | MISSENSE | Ctg/Atg | L5M | PLES_51891 | T | T | | | | | | | | | | |
| NC_011770 | 5724678 | C | T | 78 | 0 | 6 | 0 | NON_SYNONYMOU | MODERAT | MISSENSE | gCc/gTc | A115V | PLES_51901 | | | T | T | T | T | | | T | T | | |
| NC_011770 | 5724810 | A | G | 75 | 0 | 4 | 0 | NON_SYNONYMOU | MODERAT | MISSENSE | gAc/gGc | D159G | PLES_51901 | | | G | G | G | G | | | | | | |
| NC_011770 | 5734123 | T | TG | 87 | 0 | 1 | 0 | FRAME_SHIFT | HIGH | | ccc/cccC | P28P? | PLES_51971 | | | | | | | TG | | | | | |
| NC_011770 | 5738679 | C | T | 89 | 0 | 10 | 0 | NON_SYNONYMOU | MODERAT | MISSENSE | aCt/aTt | T6I | PLES_52011 | T | T | T | T | T | T | | | T | T | T | T |
| NC_011770 | 5738787 | C | A | 79 | 0 | 2 | 0 | NON_SYNONYMOU | MODERAT | MISSENSE | gCg/gAg | A42E | PLES_52011 | | | | | | | A | A | | | | |
| NC_011770 | 5738988 | A | G | 75 | 0 | 10 | 0 | NON_SYNONYMOU | MODERAT | MISSENSE | gAc/gGc | D109G | PLES_52011 | G | G | G | G | G | G | | | G | G | G | G |
| NC_011770 | 5739345 | A | G | 61 | 0 | 10 | 0 | NON_SYNONYMOU | MODERAT | MISSENSE | cAg/cGg | Q228R | PLES_52011 | G | G | G | G | G | G | | | G | G | G | G |
| NC_011770 | 5739390 | C | T | 63 | 0 | 2 | 0 | NON_SYNONYMOU | MODERAT | MISSENSE | gCc/gTc | A243V | PLES_52011 | T | T | | | | | | | | | | |
| NC_011770 | 5739563 | G | A | 56 | 0 | 2 | 0 | NON_SYNONYMOU | MODERAT | MISSENSE | Ggc/Agc | G301S | PLES_52011 | | | | | | | | | A | A | | |
| NC_011770 | 5739891 | G | A | 32 | 0 | 2 | 0 | NON_SYNONYMOU | MODERAT | MISSENSE | cGg/cAg | R410Q | PLES_52011 | | | | | | | A | A | | | | |
| NC_011770 | 5739889 | A | T | 31 | 0 | 6 | 0 | NON_SYNONYMOU | MODERAT | MISSENSE | Acg/Tcg | T6S | PLES_52021 | | | T | T | T | T | T | T | | | | |
| NC_011770 | 5740094 | A | G | 22 | 0 | 12 | 0 | NON_SYNONYMOU | MODERAT | MISSENSE | cAc/cGc | H74R | PLES_52021 | G | G | G | G | G | G | G | G | G | G | G | G |
| NC_011770 | 5740117 | C | G | 27 | 0 | 2 | 0 | NON_SYNONYMOU | MODERAT | MISSENSE | Cga/Gga | R82G | PLES_52021 | | | | | | | | | | | G | G |
| NC_011770 | 5740283 | C | G | 47 | 0 | 2 | 0 | NON_SYNONYMOU | MODERAT | MISSENSE | aCc/aGc | T137S | PLES_52021 | G | G | | | | | | | | | | |
| NC_011770 | 5741643 | C | T | 71 | 0 | 1 | 0 | STOP_GAINED | HIGH | NONSENSE | tgG/tgA | W55* | PLES_52031 | | | | | | | | | | | | T |
| NC_011770 | 5741012 | G | C | 59 | 0 | 2 | 0 | NON_SYNONYMOU | MODERAT | MISSENSE | Ccc/Gcc | P266A | PLES_52031 | C | C | | | | | | | | | | |
| NC_011770 | 5741222 | T | C | 63 | 0 | 12 | 0 | NON_SYNONYMOU | MODERAT | MISSENSE | Acc/Gcc | T196A | PLES_52031 | C | C | C | C | C | C | C | C | C | C | C | C |
| NC_011770 | 5741369 | C | T | 66 | 0 | 4 | 0 | NON_SYNONYMOU | MODERAT | MISSENSE | Gcc/Acc | A147T | PLES_52031 | | | T | T | T | T | | | | | | |
| NC_011770 | 5741576 | G | T | 65 | 0 | 4 | 0 | NON_SYNONYMOU | MODERAT | MISSENSE | Ctt/Att | L78I | PLES_52031 | | | T | T | T | T | | | | | | |
| NC_011770 | 5741774 | A | G | 75 | 0 | 2 | 0 | NON_SYNONYMOU | MODERAT | MISSENSE | Ttc/Ctc | F12L | PLES_52031 | | | | | | | G | G | | | | |
| NC_011770 | 5742592 | C | T | 73 | 0 | 6 | 0 | NON_SYNONYMOU | MODERAT | MISSENSE | Gcg/Acg | A63T | PLES_52041 | | | T | T | T | T | T | T | | | | |
| NC_011770 | 5742981 | G | A | 73 | 0 | 6 | 0 | NON_SYNONYMOU | MODERAT | MISSENSE | Cgg/Tgg | R53W | PLES_52051 | A | A | | | | | A | A | A | A | | |
| NC_011770 | 5744145 | T | C | 72 | 0 | 1 | 0 | NON_SYNONYMOU | MODERAT | MISSENSE | Ttc/Ctc | F258L | PLES_52061 | | | | | | | | C | | | | |
| NC_011770 | 5744142 | G | A | 73 | 0 | 2 | 0 | NON_SYNONYMOU | MODERAT | MISSENSE | Gtg/Atg | V257M | PLES_52061 | | | | | | | | | | | A | A |
| NC_011770 | 5744556 | G | C | 53 | 0 | 2 | 0 | NON_SYNONYMOU | MODERAT | MISSENSE | Gtc/Ctc | V395L | PLES_52061 | | | | | | | | | C | C | | |
| NC_011770 | 5744595 | A | C | 52 | 0 | 12 | 0 | NON_SYNONYMOU | MODERAT | MISSENSE | Aag/Cag | K408Q | PLES_52061 | C | C | C | C | C | C | C | C | C | C | C | C |
| NC_011770 | 5745802 | G | C | 78 | 0 | 8 | 0 | NON_SYNONYMOU | MODERAT | MISSENSE | gaC/gaG | D311E | PLES_52071 | C | C | C | C | C | C | | | | | C | C |
| NC_011770 | 5745837 | C | T | 80 | 0 | 2 | 0 | NON_SYNONYMOU | MODERAT | MISSENSE | Gcg/Acg | A300T | PLES_52071 | T | T | | | | | | | | | | |
| NC_011770 | 5745918 | C | T | 78 | 0 | 4 | 0 | NON_SYNONYMOU | MODERAT | MISSENSE | Gcg/Acg | A273T | PLES_52071 | | | T | T | T | T | | | | | | |
| NC_011770 | 5746037 | C | T | 77 | 0 | 2 | 0 | NON_SYNONYMOU | MODERAT | MISSENSE | aGc/aAc | S233N | PLES_52071 | | | | | | | T | T | | | | |
| NC_011770 | 5746631 | A | G | 88 | 0 | 2 | 0 | NON_SYNONYMOU | MODERAT | MISSENSE | gTg/gCg | V35A | PLES_52071 | | | | | | | G | G | | | | |

| | | | | | | | | | | | | | | | | | | | | | | | | | |
|---|---|---|---|---|---|---|---|---|---|---|---|---|---|---|---|---|---|---|---|---|---|---|---|---|---|
| NC_011770 | 5746658 | C | T | 89 | 0 | 2 | 0 | NON_SYNONYMOU | MODERAT | MISSENSE | aGc/aAc | S26N | PLES_52071 | | | | | | | T | T | | | | |
| NC_011770 | 5746818 | C | T | 97 | 0 | 4 | 0 | NON_SYNONYMOU | MODERAT | MISSENSE | Gcg/Acg | A57T | PLES_52081 | | | T | T | T | T | | | | | | |
| NC_011770 | 5746890 | C | T | 92 | 0 | 2 | 0 | NON_SYNONYMOU | MODERAT | MISSENSE | Gtc/Atc | V33I | PLES_52081 | | | | | | | T | T | | | | |
| NC_011770 | 5747102 | G | C | 81 | 0 | 12 | 0 | NON_SYNONYMOU | MODERAT | MISSENSE | Ctc/Gtc | L238V | PLES_52091 | C | C | C | C | C | C | C | C | C | C | C | C |
| NC_011770 | 5747234 | C | T | 82 | 0 | 4 | 0 | NON_SYNONYMOU | MODERAT | MISSENSE | Gtg/Atg | V194M | PLES_52091 | T | T | | | | | | | | | T | T |
| NC_011770 | 5747248 | A | C | 80 | 0 | 12 | 0 | NON_SYNONYMOU | MODERAT | MISSENSE | cTc/cGc | L189R | PLES_52091 | C | C | C | C | C | C | C | C | C | C | C | C |
| NC_011770 | 5747317 | A | G | 87 | 0 | 2 | 0 | NON_SYNONYMOU | MODERAT | MISSENSE | tTc/tCc | F166S | PLES_52091 | | | | | | | | | G | G | | |
| NC_011770 | 5747626 | G | A | 72 | 0 | 2 | 0 | NON_SYNONYMOU | MODERAT | MISSENSE | cCc/cTc | P63L | PLES_52091 | | | | | | | | | | | A | A |
| NC_011770 | 5751691 | G | C | 77 | 0 | 6 | 0 | NON_SYNONYMOU | MODERAT | MISSENSE | gaC/gaG | D182E | PLES_52121 | | | C | C | C | C | | | | | C | C |
| NC_011770 | 5751776 | T | C | 72 | 0 | 12 | 0 | NON_SYNONYMOU | MODERAT | MISSENSE | tAc/tGc | Y154C | PLES_52121 | C | C | C | C | C | C | C | C | C | C | C | C |
| NC_011770 | 5751782 | A | T | 77 | 0 | 2 | 0 | NON_SYNONYMOU | MODERAT | MISSENSE | gTg/gAg | V152E | PLES_52121 | T | T | | | | | | | | | | |
| NC_011770 | 5751925 | C | G | 62 | 0 | 2 | 0 | NON_SYNONYMOU | MODERAT | MISSENSE | gaG/gaC | E104D | PLES_52121 | | | | | | | | | G | G | | |
| NC_011770 | 5752154 | C | T | 65 | 0 | 2 | 0 | NON_SYNONYMOU | MODERAT | MISSENSE | aGt/aAt | S28N | PLES_52121 | | | | | | | | | T | T | | |
| NC_011770 | 5752532 | T | G | 65 | 0 | 12 | 0 | NON_SYNONYMOU | MODERAT | MISSENSE | Tgg/Ggg | W58G | PLES_52131 | G | G | G | G | G | G | G | G | G | G | G | G |
| NC_011770 | 5752565 | G | C | 65 | 0 | 8 | 0 | NON_SYNONYMOU | MODERAT | MISSENSE | Gtc/Ctc | V69L | PLES_52131 | C | C | C | C | C | C | C | C | | | | |
| NC_011770 | 5752661 | G | A | 73 | 0 | 2 | 0 | NON_SYNONYMOU | MODERAT | MISSENSE | Ggc/Agc | G101S | PLES_52131 | A | A | | | | | | | | | | |
| NC_011770 | 5752680 | A | C | 74 | 0 | 2 | 0 | NON_SYNONYMOU | MODERAT | MISSENSE | aAg/aCg | K107T | PLES_52131 | C | C | | | | | | | | | | |
| NC_011770 | 5752730 | C | A | 74 | 0 | 8 | 0 | NON_SYNONYMOU | MODERAT | MISSENSE | Ctg/Atg | L124M | PLES_52131 | | | A | A | A | A | A | A | A | A | | |
| NC_011770 | 5755125 | C | G | 87 | 0 | 12 | 0 | NON_SYNONYMOU | MODERAT | MISSENSE | gCc/gGc | A41G | PLES_52161 | G | G | G | G | G | G | G | G | G | G | G | G |
| NC_011770 | 5755409 | G | T | 88 | 0 | 2 | 0 | NON_SYNONYMOU | MODERAT | MISSENSE | Gcc/Tcc | A136S | PLES_52161 | | | | | | | | | | | T | T |
| NC_011770 | 5755743 | C | T | 75 | 0 | 1 | 0 | NON_SYNONYMOU | MODERAT | MISSENSE | atG/atA | M208I | PLES_52171 | | | | | | | T | | | | | |
| NC_011770 | 5755772 | T | C | 72 | 0 | 8 | 0 | NON_SYNONYMOU | MODERAT | MISSENSE | Acc/Gcc | T199A | PLES_52171 | C | C | C | C | C | C | C | C | | | | |
| NC_011770 | 5755780 | C | T | 73 | 0 | 2 | 0 | NON_SYNONYMOU | MODERAT | MISSENSE | cGc/cAc | R196H | PLES_52171 | T | T | | | | | | | | | | |
| NC_011770 | 5755805 | T | C | 74 | 0 | 12 | 0 | NON_SYNONYMOU | MODERAT | MISSENSE | Atc/Gtc | I188V | PLES_52171 | C | C | C | C | C | C | C | C | C | C | C | C |
| NC_011770 | 5756306 | T | C | 78 | 0 | 12 | 0 | NON_SYNONYMOU | MODERAT | MISSENSE | Acg/Gcg | T21A | PLES_52171 | C | C | C | C | C | C | C | C | C | C | C | C |
| NC_011770 | 5757051 | A | G | 81 | 0 | 12 | 0 | NON_SYNONYMOU | MODERAT | MISSENSE | aAc/aGc | N152S | PLES_52181 | G | G | G | G | G | G | G | G | G | G | G | G |
| NC_011770 | 5757489 | C | T | 84 | 0 | 4 | 0 | NON_SYNONYMOU | MODERAT | MISSENSE | Ggg/Agg | G209R | PLES_52191 | | | T | T | T | T | | | | | | |
| NC_011770 | 5758135 | T | G | 87 | 0 | 6 | 0 | NON_SYNONYMOU | MODERAT | MISSENSE | gAg/gCg | E421A | PLES_52201 | G | G | | | | | | | G | G | G | G |
| NC_011770 | 5758634 | C | T | 86 | 0 | 2 | 0 | NON_SYNONYMOU | MODERAT | MISSENSE | Gcc/Acc | A255T | PLES_52201 | | | | | | | | | T | T | | |
| NC_011770 | 5759246 | C | T | 77 | 0 | 1 | 0 | NON_SYNONYMOU | MODERAT | MISSENSE | Gcc/Acc | A51T | PLES_52201 | | | | | | | T | | | | | |
| NC_011770 | 5759404 | C | A | 75 | 0 | 2 | 0 | NON_SYNONYMOU | MODERAT | MISSENSE | Gcc/Tcc | A261S | PLES_52211 | | | | | | | | | A | A | | |
| NC_011770 | 5759468 | C | A | 65 | 0 | 2 | 0 | NON_SYNONYMOU | MODERAT | MISSENSE | gaG/gaT | E239D | PLES_52211 | | | | | | | | | A | A | | |
| NC_011770 | 5759712 | T | C | 64 | 0 | 12 | 0 | NON_SYNONYMOU | MODERAT | MISSENSE | tAt/tGt | Y158C | PLES_52211 | C | C | C | C | C | C | C | C | C | C | C | C |
| NC_011770 | 5759859 | C | T | 72 | 0 | 2 | 0 | NON_SYNONYMOU | MODERAT | MISSENSE | cGc/cAc | R109H | PLES_52211 | T | T | | | | | | | | | | |
| NC_011770 | 5760096 | T | C | 75 | 0 | 12 | 0 | NON_SYNONYMOU | MODERAT | MISSENSE | gAc/gGc | D30G | PLES_52211 | C | C | C | C | C | C | C | C | C | C | C | C |
| NC_011770 | 5760359 | T | C | 81 | 0 | 2 | 0 | NON_SYNONYMOU | MODERAT | MISSENSE | cAc/cGc | H661R | PLES_52221 | | | | | | | | | C | C | | |
| NC_011770 | 5760444 | G | C | 80 | 0 | 10 | 0 | NON_SYNONYMOU | MODERAT | MISSENSE | Ctc/Gtc | L633V | PLES_52221 | | | C | C | C | C | C | C | C | C | C | C |
| NC_011770 | 5760747 | A | T | 86 | 0 | 10 | 0 | NON_SYNONYMOU | MODERAT | MISSENSE | Ttg/Atg | L532M | PLES_52221 | | | T | T | T | T | T | T | T | T | T | T |
| NC_011770 | 5761343 | C | G | 82 | 0 | 2 | 0 | NON_SYNONYMOU | MODERAT | MISSENSE | aGc/aCc | S333T | PLES_52221 | G | G | | | | | | | | | | |
| NC_011770 | 5762181 | T | A | 70 | 0 | 6 | 0 | NON_SYNONYMOU | MODERAT | MISSENSE | Acg/Tcg | T54S | PLES_52221 | A | A | A | A | A | A | | | | | | |
| NC_011770 | 5762213 | T | G | 72 | 0 | 4 | 0 | NON_SYNONYMOU | MODERAT | MISSENSE | gAg/gCg | E43A | PLES_52221 | | | G | G | G | G | | | | | | |
| NC_011770 | 5762494 | A | G | 90 | 0 | 2 | 0 | NON_SYNONYMOU | MODERAT | MISSENSE | Atc/Gtc | I5V | PLES_52231 | | | | | | | | | G | G | | |
| NC_011770 | 5762726 | A | G | 85 | 0 | 6 | 0 | NON_SYNONYMOU | MODERAT | MISSENSE | cAc/cGc | H82R | PLES_52231 | G | G | G | G | G | G | | | | | | |
| NC_011770 | 5763259 | G | A | 72 | 0 | 2 | 0 | NON_SYNONYMOU | MODERAT | MISSENSE | Gcg/Acg | A260T | PLES_52231 | | | | | | | | | A | A | | |
| NC_011770 | 5763377 | G | A | 70 | 0 | 4 | 0 | NON_SYNONYMOU | MODERAT | MISSENSE | aGc/aAc | S299N | PLES_52231 | | | A | A | A | A | | | | | | |
| NC_011770 | 5763379 | T | G | 67 | 0 | 12 | 0 | NON_SYNONYMOU | MODERAT | MISSENSE | Tgt/Ggt | C300G | PLES_52231 | G | G | G | G | G | G | G | G | G | G | G | G |
| NC_011770 | 5763559 | A | G | 66 | 0 | 10 | 0 | NON_SYNONYMOU | MODERAT | MISSENSE | Acg/Gcg | T360A | PLES_52231 | G | G | G | G | G | G | G | G | G | G | | |
| NC_011770 | 5769363 | C | T | 92 | 0 | 1 | 0 | STOP_GAINED | HIGH | NONSENSE | Cag/Tag | Q261* | PLES_52281 | | | | | | | T | | | | | |
| NC_011770 | 5770158 | G | A | 91 | 0 | 1 | 0 | NON_SYNONYMOU | MODERAT | MISSENSE | Gcc/Acc | A526T | PLES_52281 | | | | | | | | A | | | | |
| NC_011770 | 5770200 | A | G | 89 | 0 | 10 | 0 | NON_SYNONYMOU | MODERAT | MISSENSE | Aac/Gac | N540D | PLES_52281 | | | G | G | G | G | G | G | G | G | G | G |
| NC_011770 | 5770892 | G | A | 83 | 0 | 1 | 0 | NON_SYNONYMOU | MODERAT | MISSENSE | Gcc/Acc | A202T | PLES_52291 | | | | | | | | A | | | | |
| NC_011770 | 5770743 | T | C | 75 | 0 | 8 | 0 | NON_SYNONYMOU | MODERAT | MISSENSE | gTc/gCc | V152A | PLES_52291 | | | C | C | C | C | | | C | C | C | C |
| NC_011770 | 5770799 | C | T | 80 | 0 | 1 | 0 | NON_SYNONYMOU | MODERAT | MISSENSE | Cgc/Tgc | R171C | PLES_52291 | | | | | | | T | | | | | |
| NC_011770 | 5770974 | C | T | 80 | 0 | 2 | 0 | NON_SYNONYMOU | MODERAT | MISSENSE | gCc/gTc | A229V | PLES_52291 | | | | | | | | | | | T | T |
| NC_011770 | 5770976 | G | A | 81 | 0 | 2 | 0 | NON_SYNONYMOU | MODERAT | MISSENSE | Gag/Aag | E230K | PLES_52291 | | | | | | | A | A | | | | |
| NC_011770 | 5777129 | C | T | 81 | 0 | 2 | 0 | NON_SYNONYMOU | MODERAT | MISSENSE | gCc/gTc | A125V | PLES_52341 | | | | | | | | | T | T | | |
| NC_011770 | 5779556 | C | T | 83 | 0 | 1 | 0 | NON_SYNONYMOU | MODERAT | MISSENSE | Cat/Tat | H292Y | PLES_52361 | | | | | | | | | | | | T |
| NC_011770 | 5778744 | T | C | 68 | 0 | 12 | 0 | NON_SYNONYMOU | MODERAT | MISSENSE | gTg/gCg | V21A | PLES_52361 | C | C | C | C | C | C | C | C | C | C | C | C |
| NC_011770 | 5778870 | C | T | 67 | 0 | 6 | 0 | NON_SYNONYMOU | MODERAT | MISSENSE | gCc/gTc | A63V | PLES_52361 | T | T | | | | | T | T | | | T | T |
| NC_011770 | 5779124 | G | A | 67 | 0 | 2 | 0 | NON_SYNONYMOU | MODERAT | MISSENSE | Gcc/Acc | A148T | PLES_52361 | | | | | | | | | A | A | | |
| NC_011770 | 5779272 | G | T | 64 | 0 | 4 | 0 | NON_SYNONYMOU | MODERAT | MISSENSE | aGc/aTc | S197I | PLES_52361 | | | T | T | T | T | | | | | | |
| NC_011770 | 5779277 | G | C | 54 | 0 | 2 | 0 | NON_SYNONYMOU | MODERAT | MISSENSE | Gat/Cat | D199H | PLES_52361 | C | C | | | | | | | | | | |

| | | | | | | | | | | | | | | | | | | | | | | | | | |
|---|---|---|---|---|---|---|---|---|---|---|---|---|---|---|---|---|---|---|---|---|---|---|---|---|---|
| NC_011770 | 5779389 | G | A | 70 | 0 | 2 | 0 | NON_SYNONYMOU | MODERAT | MISSENSE | aGc/aAc | S236N | PLES_52361 | | | | | | | | | | | A | A |
| NC_011770 | 5779547 | G | A | 84 | 0 | 10 | 0 | NON_SYNONYMOU | MODERAT | MISSENSE | Gtc/Atc | V289I | PLES_52361 | A | A | A | A | A | A | A | A | A | A | | |
| NC_011770 | 5780204 | G | A | 88 | 0 | 4 | 0 | NON_SYNONYMOU | MODERAT | MISSENSE | aGa/aAa | R14K | PLES_52371 | A | A | | | | | | | | | A | A |
| NC_011770 | 5780851 | G | A | 88 | 0 | 1 | 0 | NON_SYNONYMOU | MODERAT | MISSENSE | Gcc/Acc | A230T | PLES_52371 | | | | | | | A | | | | | |
| NC_011770 | 5780910 | C | G | 84 | 0 | 2 | 0 | NON_SYNONYMOU | MODERAT | MISSENSE | caC/caG | H249Q | PLES_52371 | | | | | | | | | | | G | G |
| NC_011770 | 5788185 | A | G | 92 | 0 | 1 | 0 | NON_SYNONYMOU | MODERAT | MISSENSE | Acc/Gcc | T150A | PLES_52421 | | | | | | | G | | | | | |
| NC_011770 | 5788302 | G | A | 93 | 0 | 1 | 0 | NON_SYNONYMOU | MODERAT | MISSENSE | Ggc/Agc | G189S | PLES_52421 | | | | | | | A | | | | | |
| NC_011770 | 5788456 | C | A | 78 | 0 | 4 | 0 | NON_SYNONYMOU | MODERAT | MISSENSE | gaG/gaT | E81D | PLES_52431 | A | A | | | | | | | A | A | | |
| NC_011770 | 5788466 | G | A | 80 | 0 | 2 | 0 | NON_SYNONYMOU | MODERAT | MISSENSE | cCg/cTg | P78L | PLES_52431 | | | | | | | | | A | A | | |
| NC_011770 | 5788533 | T | C | 79 | 0 | 4 | 0 | NON_SYNONYMOU | MODERAT | MISSENSE | Atg/Gtg | M56V | PLES_52431 | C | C | | | | | | | C | C | | |
| NC_011770 | 5788535 | G | A | 82 | 0 | 4 | 0 | NON_SYNONYMOU | MODERAT | MISSENSE | gCt/gTt | A55V | PLES_52431 | A | A | | | | | | | A | A | | |
| NC_011770 | 5788554 | C | A | 80 | 0 | 2 | 0 | NON_SYNONYMOU | MODERAT | MISSENSE | Gca/Tca | A49S | PLES_52431 | | | | | | | | | A | A | | |
| NC_011770 | 5789337 | C | CT | 97 | 0 | 1 | 0 | FRAME_SHIFT | HIGH | | gtg/Tgtg | V195C? | PLES_52441 | | | CT | | | | | | | | | |
| NC_011770 | 5789696 | G | C | 70 | 0 | 4 | 0 | NON_SYNONYMOU | MODERAT | MISSENSE | gGc/gCc | G314A | PLES_52441 | | | C | C | C | C | | | | | | |
| NC_011770 | 5789926 | G | A | 61 | 0 | 1 | 0 | NON_SYNONYMOU | MODERAT | MISSENSE | Gtc/Atc | V391I | PLES_52441 | | | | | | | A | | | | | |
| NC_011770 | 5791788 | G | A | 81 | 0 | 2 | 0 | NON_SYNONYMOU | MODERAT | MISSENSE | cGg/cAg | R12Q | PLES_52461 | | | | | | | | | | | A | A |
| NC_011770 | 5791809 | C | T | 84 | 0 | 2 | 0 | NON_SYNONYMOU | MODERAT | MISSENSE | gCt/gTt | A19V | PLES_52461 | | | | | | | | | | | T | T |
| NC_011770 | 5791845 | T | C | 85 | 0 | 8 | 0 | NON_SYNONYMOU | MODERAT | MISSENSE | gTc/gCc | V31A | PLES_52461 | C | C | | | | | C | C | C | C | C | C |
| NC_011770 | 5792608 | G | A | 81 | 0 | 2 | 0 | NON_SYNONYMOU | MODERAT | MISSENSE | atG/atA | M285I | PLES_52461 | | | | | | | | | | | A | A |
| NC_011770 | 5793849 | T | A | 74 | 0 | 12 | 0 | NON_SYNONYMOU | MODERAT | MISSENSE | cTg/cAg | L9Q | PLES_52481 | A | A | A | A | A | A | A | A | A | A | A | A |
| NC_011770 | 5794052 | G | A | 82 | 0 | 4 | 0 | NON_SYNONYMOU | MODERAT | MISSENSE | Ggg/Agg | G77R | PLES_52481 | | | A | A | A | A | | | | | | |
| NC_011770 | 5794343 | A | G | 92 | 0 | 2 | 0 | NON_SYNONYMOU | MODERAT | MISSENSE | Acg/Gcg | T174A | PLES_52481 | G | G | | | | | | | | | | |
| NC_011770 | 5794970 | C | A | 40 | 0 | 10 | 0 | NON_SYNONYMOU | MODERAT | MISSENSE | gCg/gAg | A122E | PLES_52491 | A | A | A | A | A | A | | | A | A | A | A |
| NC_011770 | 5796378 | T | C | 67 | 0 | 8 | 0 | NON_SYNONYMOU | MODERAT | MISSENSE | Tac/Cac | Y53H | PLES_52521 | C | C | C | C | C | C | C | C | | | | |
| NC_011770 | 5796465 | G | A | 65 | 0 | 2 | 0 | NON_SYNONYMOU | MODERAT | MISSENSE | Ggc/Agc | G82S | PLES_52521 | | | | | | | | | | | A | A |
| NC_011770 | 5796535 | A | T | 69 | 0 | 2 | 0 | NON_SYNONYMOU | MODERAT | MISSENSE | cAg/cTg | Q105L | PLES_52521 | | | | | | | | | T | T | | |
| NC_011770 | 5800177 | G | A | 84 | 0 | 1 | 0 | NON_SYNONYMOU | MODERAT | MISSENSE | Ggc/Agc | G362S | PLES_52551 | | | | | | | | | | | | A |
| NC_011770 | 5799499 | C | A | 80 | 0 | 4 | 0 | NON_SYNONYMOU | MODERAT | MISSENSE | Ctg/Atg | L136M | PLES_52551 | | | A | A | A | A | | | | | | |
| NC_011770 | 5799668 | A | G | 72 | 0 | 12 | 0 | NON_SYNONYMOU | MODERAT | MISSENSE | cAg/cGg | Q192R | PLES_52551 | G | G | G | G | G | G | G | G | G | G | G | G |
| NC_011770 | 5799812 | C | A | 76 | 0 | 2 | 0 | NON_SYNONYMOU | MODERAT | MISSENSE | cCg/cAg | P240Q | PLES_52551 | | | | | | | | | A | A | | |
| NC_011770 | 5799911 | G | C | 82 | 0 | 4 | 0 | NON_SYNONYMOU | MODERAT | MISSENSE | aGc/aCc | S273T | PLES_52551 | | | C | C | C | C | | | | | | |
| NC_011770 | 5800093 | A | G | 77 | 0 | 2 | 0 | NON_SYNONYMOU | MODERAT | MISSENSE | Agc/Ggc | S334G | PLES_52551 | G | G | | | | | | | | | | |
| NC_011770 | 5800715 | G | A | 102 | 0 | 4 | 0 | START_LOST | HIGH | MISSENSE | atG/atA | M1I | PLES_52571 | | | A | A | A | A | | | | | | |
| NC_011770 | 5802281 | AT | A | 95 | 0 | 12 | 0 | FRAME_SHIFT+STA | HIGH | | atg/ | M1 | PLES_52581 | A | A | A | A | A | A | A | A | A | A | A | A |
| NC_011770 | 5802060 | T | C | 91 | 0 | 2 | 0 | NON_SYNONYMOU | MODERAT | MISSENSE | Acc/Gcc | T75A | PLES_52581 | C | C | | | | | | | | | | |
| NC_011770 | 5802566 | C | T | 74 | 0 | 2 | 0 | NON_SYNONYMOU | MODERAT | MISSENSE | Ccg/Tcg | P26S | PLES_52591 | | | | | | | | | T | T | | |
| NC_011770 | 5802598 | C | G | 73 | 0 | 2 | 0 | NON_SYNONYMOU | MODERAT | MISSENSE | gaC/gaG | D36E | PLES_52591 | | | | | | | | | | | G | G |
| NC_011770 | 5803013 | G | T | 84 | 0 | 12 | 0 | NON_SYNONYMOU | MODERAT | MISSENSE | Ggc/Tgc | G175C | PLES_52591 | T | T | T | T | T | T | T | T | T | T | T | T |
| NC_011770 | 5804313 | G | A | 93 | 0 | 1 | 0 | NON_SYNONYMOU | MODERAT | MISSENSE | gCc/gTc | A12V | PLES_52601 | | | | | | | | A | | | | |
| NC_011770 | 5803992 | G | C | 79 | 0 | 4 | 0 | NON_SYNONYMOU | MODERAT | MISSENSE | aCc/aGc | T119S | PLES_52601 | | | | | | | C | C | | | C | C |
| NC_011770 | 5806570 | G | A | 78 | 0 | 1 | 0 | NON_SYNONYMOU | MODERAT | MISSENSE | Gag/Aag | E167K | PLES_52641 | | | | | | | A | | | | | |
| NC_011770 | 5806917 | A | T | 76 | 0 | 4 | 0 | STOP_GAINED | HIGH | NONSENSE | tTg/tAg | L680* | PLES_52651 | | | | | | | T | T | | | T | T |
| NC_011770 | 5806905 | A | G | 77 | 0 | 2 | 0 | NON_SYNONYMOU | MODERAT | MISSENSE | gTg/gCg | V684A | PLES_52651 | | | | | | | | | | | G | G |
| NC_011770 | 5807008 | T | C | 82 | 0 | 2 | 0 | NON_SYNONYMOU | MODERAT | MISSENSE | Acg/Gcg | T650A | PLES_52651 | | | | | | | C | C | | | | |
| NC_011770 | 5807514 | T | C | 78 | 0 | 2 | 0 | NON_SYNONYMOU | MODERAT | MISSENSE | aAg/aGg | K481R | PLES_52651 | C | C | | | | | | | | | | |
| NC_011770 | 5808557 | C | G | 73 | 0 | 4 | 0 | NON_SYNONYMOU | MODERAT | MISSENSE | gaG/gaC | E133D | PLES_52651 | | | G | G | G | G | | | | | | |
| NC_011770 | 5808600 | G | A | 75 | 0 | 2 | 0 | NON_SYNONYMOU | MODERAT | MISSENSE | aCc/aTc | T119I | PLES_52651 | | | | | | | A | A | | | | |
| NC_011770 | 5808685 | C | T | 77 | 0 | 10 | 0 | NON_SYNONYMOU | MODERAT | MISSENSE | Gcg/Acg | A91T | PLES_52651 | T | T | T | T | T | T | T | T | T | T | | |
| NC_011770 | 5810082 | GCCTCGC | G | 61 | 0 | 7 | 0 | CODON_DELETION | MODERATE | | ctcgcc/ | LA13 | PLES_52671 | | | G | G | G | G | | G | G | G | | |
| NC_011770 | 5810354 | A | G | 51 | 0 | 1 | 0 | NON_SYNONYMOU | MODERAT | MISSENSE | Acc/Gcc | T99A | PLES_52671 | | | | | | | | G | | | | |
| NC_011770 | 5810100 | C | T | 20 | 1 | 2 | 0 | NON_SYNONYMOU | MODERAT | MISSENSE | gCc/gTc | A14V | PLES_52671 | | | | | T | T | | | | . | | |
| NC_011770 | 5810165 | C | A | 58 | 0 | 2 | 0 | NON_SYNONYMOU | MODERAT | MISSENSE | Cag/Aag | Q36K | PLES_52671 | | | | | | | | | | | A | A |
| NC_011770 | 5810195 | G | A | 41 | 0 | 4 | 0 | NON_SYNONYMOU | MODERAT | MISSENSE | Gcc/Acc | A46T | PLES_52671 | | | | | | | A | A | A | A | | |
| NC_011770 | 5810219 | A | G | 37 | 0 | 4 | 0 | NON_SYNONYMOU | MODERAT | MISSENSE | Atc/Gtc | I54V | PLES_52671 | | | G | G | G | G | | | | | | |
| NC_011770 | 5810691 | C | T | 76 | 0 | 2 | 0 | NON_SYNONYMOU | MODERAT | MISSENSE | Gag/Aag | E200K | PLES_52681 | T | T | | | | | | | | | | |
| NC_011770 | 5811252 | C | T | 79 | 0 | 6 | 0 | NON_SYNONYMOU | MODERAT | MISSENSE | Gtc/Atc | V13I | PLES_52681 | | | T | T | T | T | | | T | T | | |
| NC_011770 | 5811276 | C | T | 83 | 0 | 2 | 0 | NON_SYNONYMOU | MODERAT | MISSENSE | Gag/Aag | E5K | PLES_52681 | | | | | | | | | T | T | | |
| NC_011770 | 5811292 | T | C | 81 | 0 | 2 | 0 | NON_SYNONYMOU | MODERAT | MISSENSE | gAa/gGa | E203G | PLES_52691 | | | | | | | | | C | C | | |
| NC_011770 | 5811319 | C | A | 84 | 0 | 2 | 0 | NON_SYNONYMOU | MODERAT | MISSENSE | cGc/cTc | R194L | PLES_52691 | | | | | A | A | | | | | | |
| NC_011770 | 5811338 | G | C | 85 | 0 | 2 | 0 | NON_SYNONYMOU | MODERAT | MISSENSE | Ctg/Gtg | L188V | PLES_52691 | | | | | | | | | C | C | | |
| NC_011770 | 5811451 | G | A | 80 | 0 | 4 | 0 | NON_SYNONYMOU | MODERAT | MISSENSE | gCg/gTg | A150V | PLES_52691 | | | A | A | A | A | | | | | | |
| NC_011770 | 5811520 | G | C | 81 | 0 | 4 | 0 | NON_SYNONYMOU | MODERAT | MISSENSE | gCg/gGg | A127G | PLES_52691 | | | C | C | C | C | | | | | | |

| | | | | | | | | | | | | | | | | | | | | | | | | | |
|---|---|---|---|---|---|---|---|---|---|---|---|---|---|---|---|---|---|---|---|---|---|---|---|---|---|
| NC_011770 | 5811599 | G | A | 80 | 0 | 2 | 0 | NON_SYNONYMOU | MODERAT | MISSENSE | Ctc/Ttc | L101F | PLES_52691 | | | | | | | | | | | A | A |
| NC_011770 | 5811649 | C | T | 81 | 0 | 2 | 0 | NON_SYNONYMOU | MODERAT | MISSENSE | cGt/cAt | R84H | PLES_52691 | | | | | | | | | T | T | | |
| NC_011770 | 5812093 | T | C | 71 | 0 | 2 | 0 | NON_SYNONYMOU | MODERAT | MISSENSE | Atc/Gtc | I162V | PLES_52701 | | | | | | | | | | | C | C |
| NC_011770 | 5812546 | G | A | 71 | 0 | 6 | 0 | NON_SYNONYMOU | MODERAT | MISSENSE | Ctc/Ttc | L11F | PLES_52701 | A | A | | | | | | | A | A | A | A |
| NC_011770 | 5813755 | T | C | 63 | 0 | 1 | 0 | NON_SYNONYMOU | MODERAT | MISSENSE | cTg/cCg | L125P | PLES_52721 | | | | | | | C | | | | | |
| NC_011770 | 5813769 | A | G | 64 | 0 | 4 | 0 | NON_SYNONYMOU | MODERAT | MISSENSE | Atc/Gtc | I130V | PLES_52721 | | | G | G | G | G | | | | | | |
| NC_011770 | 5814096 | G | A | 81 | 0 | 2 | 0 | NON_SYNONYMOU | MODERAT | MISSENSE | Gac/Aac | D239N | PLES_52721 | | | | | | | A | A | | | | |
| NC_011770 | 5814993 | T | C | 71 | 0 | 1 | 0 | NON_SYNONYMOU | MODERAT | MISSENSE | tAc/tGc | Y363C | PLES_52731 | | | | | | | | C | | | | |
| NC_011770 | 5815539 | A | G | 61 | 0 | 1 | 0 | NON_SYNONYMOU | MODERAT | MISSENSE | gTg/gCg | V181A | PLES_52731 | | | | | | | | G | | | | |
| NC_011770 | 5817462 | G | A | 86 | 0 | 2 | 0 | NON_SYNONYMOU | MODERAT | MISSENSE | Cct/Tct | P350S | PLES_52751 | | | | | | | | | A | A | | |
| NC_011770 | 5817666 | G | C | 73 | 0 | 2 | 0 | NON_SYNONYMOU | MODERAT | MISSENSE | Cgc/Ggc | R282G | PLES_52751 | | | | | | | | | | | C | C |
| NC_011770 | 5817807 | C | T | 70 | 0 | 2 | 0 | NON_SYNONYMOU | MODERAT | MISSENSE | Gcc/Acc | A235T | PLES_52751 | | | | | | | | | T | T | | |
| NC_011770 | 5817962 | A | G | 73 | 0 | 2 | 0 | NON_SYNONYMOU | MODERAT | MISSENSE | gTc/gCc | V183A | PLES_52751 | | | | | | | G | G | | | | |
| NC_011770 | 5818289 | G | A | 63 | 0 | 4 | 0 | NON_SYNONYMOU | MODERAT | MISSENSE | gCg/gTg | A74V | PLES_52751 | | | A | A | A | A | | | | | | |
| NC_011770 | 5818427 | G | A | 65 | 0 | 4 | 0 | NON_SYNONYMOU | MODERAT | MISSENSE | gCc/gTc | A28V | PLES_52751 | | | A | A | A | A | | | | | | |
| NC_011770 | 5822061 | C | G | 69 | 0 | 10 | 0 | NON_SYNONYMOU | MODERAT | MISSENSE | Gcg/Ccg | A302P | PLES_52811 | G | G | G | G | G | G | G | G | G | G | | |
| NC_011770 | 5822199 | G | C | 80 | 0 | 2 | 0 | NON_SYNONYMOU | MODERAT | MISSENSE | Ccc/Gcc | P256A | PLES_52811 | | | | | | | | | C | C | | |
| NC_011770 | 5822295 | C | T | 72 | 0 | 2 | 0 | NON_SYNONYMOU | MODERAT | MISSENSE | Gcg/Acg | A224T | PLES_52811 | | | | | | | T | T | | | | |
| NC_011770 | 5822394 | A | G | 70 | 0 | 1 | 0 | NON_SYNONYMOU | MODERAT | MISSENSE | Ttc/Ctc | F191L | PLES_52811 | | | | | | | G | | | | | |
| NC_011770 | 5822517 | G | C | 62 | 0 | 12 | 0 | NON_SYNONYMOU | MODERAT | MISSENSE | Ctc/Gtc | L150V | PLES_52811 | C | C | C | C | C | C | C | C | C | C | C | C |
| NC_011770 | 5822608 | C | A | 51 | 0 | 4 | 0 | NON_SYNONYMOU | MODERAT | MISSENSE | tgG/tgT | W119C | PLES_52811 | | | | | | | | | A | A | A | A |
| NC_011770 | 5823396 | C | T | 59 | 0 | 4 | 0 | NON_SYNONYMOU | MODERAT | MISSENSE | cGa/cAa | R33Q | PLES_52821 | | | T | T | T | T | | | | | | |
| NC_011770 | 5826353 | G | A | 82 | 0 | 1 | 0 | NON_SYNONYMOU | MODERAT | MISSENSE | gGc/gAc | G894D | PLES_52831 | A | | | | | | | | | | | |
| NC_011770 | 5823826 | G | A | 66 | 0 | 2 | 0 | NON_SYNONYMOU | MODERAT | MISSENSE | Ggc/Agc | G52S | PLES_52831 | | | | | | | | | A | A | | |
| NC_011770 | 5823953 | A | G | 73 | 0 | 1 | 0 | NON_SYNONYMOU | MODERAT | MISSENSE | cAg/cGg | Q94R | PLES_52831 | | | | | | | G | | | | | |
| NC_011770 | 5823959 | G | A | 74 | 0 | 2 | 0 | NON_SYNONYMOU | MODERAT | MISSENSE | gGc/gAc | G96D | PLES_52831 | | | | | | | | | A | A | | |
| NC_011770 | 5824132 | G | A | 75 | 0 | 2 | 0 | NON_SYNONYMOU | MODERAT | MISSENSE | Gcc/Acc | A154T | PLES_52831 | A | A | | | | | | | | | | |
| NC_011770 | 5824576 | G | A | 81 | 0 | 2 | 0 | NON_SYNONYMOU | MODERAT | MISSENSE | Gtg/Atg | V302M | PLES_52831 | | | | | | | A | A | | | | |
| NC_011770 | 5824591 | G | A | 81 | 0 | 2 | 0 | NON_SYNONYMOU | MODERAT | MISSENSE | Gac/Aac | D307N | PLES_52831 | | | | | | | A | A | | | | |
| NC_011770 | 5824610 | T | C | 81 | 0 | 2 | 0 | NON_SYNONYMOU | MODERAT | MISSENSE | gTc/gCc | V313A | PLES_52831 | | | | | | | C | C | | | | |
| NC_011770 | 5824807 | G | A | 90 | 0 | 2 | 0 | NON_SYNONYMOU | MODERAT | MISSENSE | Gac/Aac | D379N | PLES_52831 | | | | | | | A | A | | | | |
| NC_011770 | 5824988 | C | T | 95 | 0 | 2 | 0 | NON_SYNONYMOU | MODERAT | MISSENSE | aCc/aTc | T439I | PLES_52831 | T | T | | | | | | | | | | |
| NC_011770 | 5825148 | C | G | 87 | 0 | 10 | 0 | NON_SYNONYMOU | MODERAT | MISSENSE | gaC/gaG | D492E | PLES_52831 | G | G | G | G | G | G | | | G | G | G | G |
| NC_011770 | 5828339 | G | A | 81 | 0 | 2 | 0 | NON_SYNONYMOU | MODERAT | MISSENSE | Cgc/Tgc | R427C | PLES_52851 | | | | | | | A | A | | | | |
| NC_011770 | 5829668 | A | G | 63 | 0 | 2 | 0 | NON_SYNONYMOU | MODERAT | MISSENSE | gTc/gCc | V438A | PLES_52861 | | | | | | | | | G | G | | |
| NC_011770 | 5833372 | G | A | 63 | 0 | 2 | 0 | NON_SYNONYMOU | MODERAT | MISSENSE | Gtg/Atg | V207M | PLES_52881 | | | | | | | | | | | A | A |
| NC_011770 | 5833738 | G | A | 53 | 0 | 4 | 0 | NON_SYNONYMOU | MODERAT | MISSENSE | gCc/gTc | A421V | PLES_52891 | A | A | | | | | | | | | A | A |
| NC_011770 | 5834153 | C | A | 70 | 0 | 2 | 0 | NON_SYNONYMOU | MODERAT | MISSENSE | Gcc/Tcc | A283S | PLES_52891 | | | | | | | | | A | A | | |
| NC_011770 | 5834419 | C | T | 77 | 0 | 2 | 0 | NON_SYNONYMOU | MODERAT | MISSENSE | tGc/tAc | C194Y | PLES_52891 | | | | | | | | | | | T | T |
| NC_011770 | 5834680 | G | A | 78 | 0 | 4 | 0 | NON_SYNONYMOU | MODERAT | MISSENSE | gCg/gTg | A107V | PLES_52891 | | | A | A | A | A | | | | | | |
| NC_011770 | 5834777 | T | C | 75 | 0 | 2 | 0 | NON_SYNONYMOU | MODERAT | MISSENSE | Atc/Gtc | I75V | PLES_52891 | | | | | | | | | | | C | C |
| NC_011770 | 5834951 | A | C | 75 | 0 | 4 | 0 | NON_SYNONYMOU | MODERAT | MISSENSE | Ttc/Gtc | F17V | PLES_52891 | | | | | | | C | C | C | C | | |
| NC_011770 | 5837513 | G | GC | 82 | 0 | 1 | 0 | FRAME_SHIFT | HIGH | | gcg/gGcg | A200G? | PLES_52921 | | | | | | | GC | | | | | |
| NC_011770 | 5837811 | G | A | 85 | 0 | 1 | 0 | NON_SYNONYMOU | MODERAT | MISSENSE | Cgg/Tgg | R101W | PLES_52921 | | | | | | | | A | | | | |
| NC_011770 | 5837444 | G | T | 82 | 0 | 2 | 0 | NON_SYNONYMOU | MODERAT | MISSENSE | gCg/gAg | A223E | PLES_52921 | | | | | | | | | | | T | T |
| NC_011770 | 5838867 | C | T | 74 | 0 | 2 | 0 | NON_SYNONYMOU | MODERAT | MISSENSE | Gag/Aag | E24K | PLES_52931 | | | | | | | T | T | | | | |
| NC_011770 | 5839399 | C | T | 80 | 0 | 4 | 0 | NON_SYNONYMOU | MODERAT | MISSENSE | gGg/gAg | G276E | PLES_52941 | | | T | T | T | T | | | | | | |
| NC_011770 | 5839403 | T | C | 78 | 0 | 2 | 0 | NON_SYNONYMOU | MODERAT | MISSENSE | Agc/Ggc | S275G | PLES_52941 | C | C | | | | | | | | | | |
| NC_011770 | 5839558 | G | A | 78 | 0 | 2 | 0 | NON_SYNONYMOU | MODERAT | MISSENSE | tCc/tTc | S223F | PLES_52941 | | | | | | | | | | | A | A |
| NC_011770 | 5840664 | T | C | 77 | 0 | 1 | 0 | NON_SYNONYMOU | MODERAT | MISSENSE | Atg/Gtg | M117V | PLES_52951 | | C | | | | | | | | | | |
| NC_011770 | 5842306 | G | A | 89 | 0 | 4 | 0 | NON_SYNONYMOU | MODERAT | MISSENSE | Cac/Tac | H284Y | PLES_52971 | | | A | A | A | A | | | | | | |
| NC_011770 | 5842597 | T | A | 82 | 0 | 2 | 0 | NON_SYNONYMOU | MODERAT | MISSENSE | Atc/Ttc | I187F | PLES_52971 | | | | | | | | | | | A | A |
| NC_011770 | 5842846 | T | C | 80 | 0 | 8 | 0 | NON_SYNONYMOU | MODERAT | MISSENSE | Atg/Gtg | M104V | PLES_52971 | C | C | | | | | C | C | C | C | C | C |
| NC_011770 | 5843116 | G | C | 87 | 0 | 2 | 0 | NON_SYNONYMOU | MODERAT | MISSENSE | Ctc/Gtc | L14V | PLES_52971 | C | C | | | | | | | | | | |
| NC_011770 | 5845437 | C | T | 78 | 0 | 1 | 0 | START_LOST | HIGH | MISSENSE | atG/atA | M1I | PLES_52991 | | | | | | | T | | | | | |
| NC_011770 | 5844577 | T | C | 93 | 0 | 1 | 0 | NON_SYNONYMOU | MODERAT | MISSENSE | aAg/aGg | K288R | PLES_52991 | | | | | | | C | | | | | |
| NC_011770 | 5844697 | T | C | 91 | 0 | 10 | 0 | NON_SYNONYMOU | MODERAT | MISSENSE | aAg/aGg | K248R | PLES_52991 | C | C | C | C | C | C | | | C | C | C | C |
| NC_011770 | 5846627 | A | G | 72 | 0 | 8 | 0 | NON_SYNONYMOU | MODERAT | MISSENSE | gAg/gGg | E190G | PLES_53001 | G | G | G | G | G | G | | | | | G | G |
| NC_011770 | 5846966 | G | C | 79 | 0 | 2 | 0 | NON_SYNONYMOU | MODERAT | MISSENSE | gGc/gCc | G303A | PLES_53001 | C | C | | | | | | | | | | |
| NC_011770 | 5846971 | C | G | 83 | 0 | 2 | 0 | NON_SYNONYMOU | MODERAT | MISSENSE | Cgc/Ggc | R305G | PLES_53001 | G | G | | | | | | | | | | |
| NC_011770 | 5848419 | G | C | 66 | 0 | 2 | 0 | NON_SYNONYMOU | MODERAT | MISSENSE | Gtc/Ctc | V456L | PLES_53011 | | | | | | | | | C | C | | |
| NC_011770 | 5848791 | C | T | 79 | 0 | 2 | 0 | NON_SYNONYMOU | MODERAT | MISSENSE | Gtc/Atc | V211I | PLES_53021 | | | | | | | | | T | T | | |

| | | | | | | | | | | | | | | | | | | | | | | | | | |
|---|---|---|---|---|---|---|---|---|---|---|---|---|---|---|---|---|---|---|---|---|---|---|---|---|---|
| NC_011770 | 5849106 | C | T | 82 | 0 | 2 | 0 | NON_SYNONYMOU | MODERAT | MISSENSE | Gtg/Atg | V106M | PLES_53021 | T | T | | | | | | | | | | |
| NC_011770 | 5849669 | G | C | 80 | 0 | 2 | 0 | NON_SYNONYMOU | MODERAT | MISSENSE | Ccg/Gcg | P124A | PLES_53031 | | | | | | | | | | | C | C |
| NC_011770 | 5849732 | G | A | 77 | 0 | 2 | 0 | NON_SYNONYMOU | MODERAT | MISSENSE | Cgt/Tgt | R103C | PLES_53031 | | | | | | | | | A | A | | |
| NC_011770 | 5850019 | G | T | 75 | 0 | 2 | 0 | NON_SYNONYMOU | MODERAT | MISSENSE | tCc/tAc | S7Y | PLES_53031 | | | | | | | | | T | T | | |
| NC_011770 | 5853041 | G | T | 68 | 0 | 4 | 0 | NON_SYNONYMOU | MODERAT | MISSENSE | Gcc/Tcc | A15S | PLES_53071 | | | T | T | T | T | | | | | | |
| NC_011770 | 5853606 | C | T | 66 | 0 | 6 | 0 | NON_SYNONYMOU | MODERAT | MISSENSE | gCc/gTc | A203V | PLES_53071 | | | T | T | T | T | | | T | T | | |
| NC_011770 | 5853693 | G | A | 71 | 0 | 2 | 0 | NON_SYNONYMOU | MODERAT | MISSENSE | cGc/cAc | R232H | PLES_53071 | A | A | | | | | | | | | | |
| NC_011770 | 5853824 | G | T | 67 | 0 | 2 | 0 | NON_SYNONYMOU | MODERAT | MISSENSE | Gcc/Tcc | A276S | PLES_53071 | | | | | | | T | T | | | | |
| NC_011770 | 5854821 | C | A | 84 | 0 | 12 | 0 | NON_SYNONYMOU | MODERAT | MISSENSE | Ccc/Acc | P37T | PLES_53091 | A | A | A | A | A | A | A | A | A | A | A | A |
| NC_011770 | 5855022 | A | G | 83 | 0 | 6 | 0 | NON_SYNONYMOU | MODERAT | MISSENSE | Atc/Gtc | I104V | PLES_53091 | | | G | G | G | G | | | G | G | | |
| NC_011770 | 5855158 | A | G | 74 | 0 | 2 | 0 | NON_SYNONYMOU | MODERAT | MISSENSE | gAc/gGc | D149G | PLES_53091 | G | G | | | | | | | | | | |
| NC_011770 | 5855227 | A | C | 69 | 0 | 6 | 0 | NON_SYNONYMOU | MODERAT | MISSENSE | gAg/gCg | E172A | PLES_53091 | C | C | | | | | C | C | | | C | C |
| NC_011770 | 5855432 | C | T | 86 | 0 | 4 | 0 | NON_SYNONYMOU | MODERAT | MISSENSE | Ggc/Agc | G193S | PLES_53101 | | | T | T | T | T | | | | | | |
| NC_011770 | 5855725 | G | C | 79 | 0 | 6 | 0 | NON_SYNONYMOU | MODERAT | MISSENSE | aCc/aGc | T95S | PLES_53101 | C | C | C | C | C | C | | | | | | |
| NC_011770 | 5855728 | T | C | 76 | 0 | 6 | 0 | NON_SYNONYMOU | MODERAT | MISSENSE | aAc/aGc | N94S | PLES_53101 | C | C | C | C | C | C | | | | | | |
| NC_011770 | 5855915 | T | C | 75 | 0 | 12 | 0 | NON_SYNONYMOU | MODERAT | MISSENSE | Atg/Gtg | M32V | PLES_53101 | C | C | C | C | C | C | C | C | C | C | C | C |
| NC_011770 | 5856427 | T | C | 89 | 0 | 2 | 0 | NON_SYNONYMOU | MODERAT | MISSENSE | aAg/aGg | K207R | PLES_53111 | | | | | | | C | C | | | | |
| NC_011770 | 5857307 | G | C | 60 | 0 | 2 | 0 | NON_SYNONYMOU | MODERAT | MISSENSE | aCt/aGt | T309S | PLES_53121 | C | C | | | | | | | | | | |
| NC_011770 | 5857313 | C | T | 61 | 0 | 8 | 0 | NON_SYNONYMOU | MODERAT | MISSENSE | cGg/cAg | R307Q | PLES_53121 | T | T | | | | | T | T | T | T | T | T |
| NC_011770 | 5857694 | G | C | 72 | 0 | 4 | 0 | NON_SYNONYMOU | MODERAT | MISSENSE | gCg/gGg | A180G | PLES_53121 | | | C | C | C | C | | | | | | |
| NC_011770 | 5857892 | G | A | 68 | 0 | 4 | 0 | NON_SYNONYMOU | MODERAT | MISSENSE | gCc/gTc | A114V | PLES_53121 | | | A | A | A | A | | | | | | |
| NC_011770 | 5858150 | G | A | 83 | 0 | 2 | 0 | NON_SYNONYMOU | MODERAT | MISSENSE | aCg/aTg | T28M | PLES_53121 | | | | | | | | | A | A | | |
| NC_011770 | 5858662 | CCGCCGT | C | 89 | 0 | 1 | 0 | CODON_CHANGE_ | MODERATE | | gacggcggt, | DGG688G | PLES_53131 | | | | | | | | | | C | | |
| NC_011770 | 5859016 | G | A | 76 | 0 | 2 | 0 | NON_SYNONYMOU | MODERAT | MISSENSE | cCg/cTg | P572L | PLES_53131 | | | | | | | A | A | | | | |
| NC_011770 | 5859109 | G | A | 76 | 0 | 4 | 0 | NON_SYNONYMOU | MODERAT | MISSENSE | tCg/tTg | S541L | PLES_53131 | | | A | A | A | A | | | | | | |
| NC_011770 | 5859128 | G | T | 78 | 0 | 2 | 0 | NON_SYNONYMOU | MODERAT | MISSENSE | Ctg/Atg | L535M | PLES_53131 | | | | | | | | | | | T | T |
| NC_011770 | 5859333 | G | T | 77 | 0 | 2 | 0 | NON_SYNONYMOU | MODERAT | MISSENSE | gaC/gaA | D466E | PLES_53131 | | | | | | | | | | | T | T |
| NC_011770 | 5859557 | T | C | 61 | 0 | 2 | 0 | NON_SYNONYMOU | MODERAT | MISSENSE | Agt/Ggt | S392G | PLES_53131 | C | C | | | | | | | | | | |
| NC_011770 | 5859941 | T | C | 81 | 0 | 2 | 0 | NON_SYNONYMOU | MODERAT | MISSENSE | Acc/Gcc | T264A | PLES_53131 | | | | | | | | | | | C | C |
| NC_011770 | 5860392 | G | C | 74 | 0 | 2 | 0 | NON_SYNONYMOU | MODERAT | MISSENSE | gaC/gaG | D113E | PLES_53131 | C | C | | | | | | | | | | |
| NC_011770 | 5862259 | A | G | 100 | 0 | 1 | 0 | NON_SYNONYMOU | MODERAT | MISSENSE | gTg/gCg | V326A | PLES_53141 | | | | | | | | G | | | | |
| NC_011770 | 5861018 | A | G | 74 | 0 | 2 | 0 | NON_SYNONYMOU | MODERAT | MISSENSE | Tcg/Ccg | S740P | PLES_53141 | | | | | | | | | | | G | G |
| NC_011770 | 5861045 | T | C | 76 | 0 | 2 | 0 | NON_SYNONYMOU | MODERAT | MISSENSE | Acg/Gcg | T731A | PLES_53141 | | | | | | | | | | | C | C |
| NC_011770 | 5861078 | T | G | 80 | 0 | 2 | 0 | NON_SYNONYMOU | MODERAT | MISSENSE | Aag/Cag | K720Q | PLES_53141 | | | | | | | | | | | G | G |
| NC_011770 | 5861404 | T | C | 90 | 0 | 2 | 0 | NON_SYNONYMOU | MODERAT | MISSENSE | cAc/cGc | H611R | PLES_53141 | C | C | | | | | | | | | | |
| NC_011770 | 5862430 | T | C | 93 | 0 | 2 | 0 | NON_SYNONYMOU | MODERAT | MISSENSE | gAc/gGc | D269G | PLES_53141 | | | | | | | | | C | C | | |
| NC_011770 | 5862439 | G | A | 92 | 0 | 2 | 0 | NON_SYNONYMOU | MODERAT | MISSENSE | gCg/gTg | A266V | PLES_53141 | | | | | | | | | | | A | A |
| NC_011770 | 5862614 | C | G | 95 | 0 | 6 | 0 | NON_SYNONYMOU | MODERAT | MISSENSE | Gtg/Ctg | V208L | PLES_53141 | G | G | | | | | | | G | G | G | G |
| NC_011770 | 5862616 | A | G | 92 | 0 | 6 | 0 | NON_SYNONYMOU | MODERAT | MISSENSE | cTg/cCg | L207P | PLES_53141 | G | G | | | | | | | G | G | G | G |
| NC_011770 | 5863529 | G | A | 80 | 0 | 2 | 0 | NON_SYNONYMOU | MODERAT | MISSENSE | Cgg/Tgg | R518W | PLES_53151 | A | A | | | | | | | | | | |
| NC_011770 | 5863535 | T | C | 79 | 0 | 4 | 0 | NON_SYNONYMOU | MODERAT | MISSENSE | Agg/Ggg | R516G | PLES_53151 | C | C | | | | | | | | | C | C |
| NC_011770 | 5863540 | C | T | 79 | 0 | 2 | 0 | NON_SYNONYMOU | MODERAT | MISSENSE | gGc/gAc | G514D | PLES_53151 | | | | | | | | | | | T | T |
| NC_011770 | 5863676 | C | T | 69 | 0 | 4 | 0 | NON_SYNONYMOU | MODERAT | MISSENSE | Gct/Act | A469T | PLES_53151 | T | T | | | | | | | | | T | T |
| NC_011770 | 5863684 | C | T | 66 | 0 | 4 | 0 | NON_SYNONYMOU | MODERAT | MISSENSE | gGc/gAc | G466D | PLES_53151 | T | T | | | | | | | | | T | T |
| NC_011770 | 5863712 | A | G | 63 | 0 | 2 | 0 | NON_SYNONYMOU | MODERAT | MISSENSE | Tct/Cct | S457P | PLES_53151 | | | | | | | | | | | G | G |
| NC_011770 | 5863766 | C | T | 68 | 0 | 2 | 0 | NON_SYNONYMOU | MODERAT | MISSENSE | Gtg/Atg | V439M | PLES_53151 | | | | | | | | | | | T | T |
| NC_011770 | 5863781 | T | C | 71 | 0 | 10 | 0 | NON_SYNONYMOU | MODERAT | MISSENSE | Ata/Gta | I434V | PLES_53151 | C | C | C | C | C | C | C | C | | | C | C |
| NC_011770 | 5863808 | T | C | 77 | 0 | 4 | 0 | NON_SYNONYMOU | MODERAT | MISSENSE | Acg/Gcg | T425A | PLES_53151 | C | C | | | | | | | | | C | C |
| NC_011770 | 5863862 | T | C | 83 | 0 | 2 | 0 | NON_SYNONYMOU | MODERAT | MISSENSE | Ata/Gta | I407V | PLES_53151 | | | | | | | | | C | C | | |
| NC_011770 | 5863879 | G | A | 83 | 0 | 2 | 0 | NON_SYNONYMOU | MODERAT | MISSENSE | cCg/cTg | P401L | PLES_53151 | | | | | | | | | | | A | A |
| NC_011770 | 5863881 | A | C | 83 | 0 | 2 | 0 | NON_SYNONYMOU | MODERAT | MISSENSE | gaT/gaG | D400E | PLES_53151 | | | | | | | | | | | C | C |
| NC_011770 | 5864033 | C | T | 78 | 0 | 2 | 0 | NON_SYNONYMOU | MODERAT | MISSENSE | Gag/Aag | E350K | PLES_53151 | | | | | | | | | | | T | T |
| NC_011770 | 5864055 | T | A | 74 | 0 | 2 | 0 | NON_SYNONYMOU | MODERAT | MISSENSE | gaA/gaT | E342D | PLES_53151 | | | | | | | | | | | A | A |
| NC_011770 | 5864060 | C | G | 73 | 0 | 2 | 0 | NON_SYNONYMOU | MODERAT | MISSENSE | Gag/Cag | E341Q | PLES_53151 | | | | | | | | | | | G | G |
| NC_011770 | 5864282 | C | G | 62 | 0 | 2 | 0 | NON_SYNONYMOU | MODERAT | MISSENSE | Ggc/Cgc | G267R | PLES_53151 | | | | | | | | | | | G | G |
| NC_011770 | 5864468 | C | T | 43 | 0 | 2 | 0 | NON_SYNONYMOU | MODERAT | MISSENSE | Gcc/Acc | A205T | PLES_53151 | | | | | | | | | | | T | T |
| NC_011770 | 5864516 | C | T | 46 | 0 | 2 | 0 | NON_SYNONYMOU | MODERAT | MISSENSE | Ggt/Agt | G189S | PLES_53151 | T | T | | | | | | | | | | |
| NC_011770 | 5864618 | T | C | 64 | 0 | 4 | 0 | NON_SYNONYMOU | MODERAT | MISSENSE | Atc/Gtc | I155V | PLES_53151 | C | C | | | | | | | | | C | C |
| NC_011770 | 5864669 | C | T | 70 | 0 | 2 | 0 | NON_SYNONYMOU | MODERAT | MISSENSE | Gtt/Att | V138I | PLES_53151 | T | T | | | | | | | | | | |
| NC_011770 | 5864807 | G | A | 78 | 0 | 2 | 0 | NON_SYNONYMOU | MODERAT | MISSENSE | Cgg/Tgg | R92W | PLES_53151 | A | A | | | | | | | | | | |
| NC_011770 | 5864855 | C | T | 74 | 0 | 2 | 0 | NON_SYNONYMOU | MODERAT | MISSENSE | Gct/Act | A76T | PLES_53151 | T | T | | | | | | | | | | |
| NC_011770 | 5864909 | C | T | 72 | 0 | 2 | 0 | NON_SYNONYMOU | MODERAT | MISSENSE | Gtc/Atc | V58I | PLES_53151 | T | T | | | | | | | | | | |

| | | | | | | | | | | | | | | | | | | | | | | | | | |
|---|---|---|---|---|---|---|---|---|---|---|---|---|---|---|---|---|---|---|---|---|---|---|---|---|---|
| NC_011770 | 5864936 | T | C | 71 | 0 | 4 | 0 | NON_SYNONYMOU | MODERAT | MISSENSE | Atc/Gtc | I49V | PLES_53151 | C | C | | | | | | | | | C | C |
| NC_011770 | 5868538 | A | T | 77 | 0 | 2 | 0 | NON_SYNONYMOU | MODERAT | MISSENSE | Ttt/Att | F274I | PLES_53191 | T | T | | | | | | | | | | |
| NC_011770 | 5869243 | C | A | 104 | 0 | 2 | 0 | NON_SYNONYMOU | MODERAT | MISSENSE | Gtg/Ttg | V39L | PLES_53191 | | | | | | | | | A | A | | |
| NC_011770 | 5870461 | G | A | 55 | 0 | 1 | 0 | NON_SYNONYMOU | MODERAT | MISSENSE | Ccg/Tcg | P192S | PLES_53221 | A | | | | | | | | | | | |
| NC_011770 | 5870656 | T | C | 67 | 0 | 2 | 0 | NON_SYNONYMOU | MODERAT | MISSENSE | Atc/Gtc | I127V | PLES_53221 | | | | | | | | | | | C | C |
| NC_011770 | 5870758 | T | C | 65 | 0 | 2 | 0 | NON_SYNONYMOU | MODERAT | MISSENSE | Agc/Ggc | S93G | PLES_53221 | | | | | | | | | C | C | | |
| NC_011770 | 5877057 | TG | T | 61 | 0 | 2 | 0 | FRAME_SHIFT | HIGH | | ggc/ | G33 | PLES_53261 | T | T | | | | | | | | | | |
| NC_011770 | 5877009 | T | C | 68 | 0 | 4 | 0 | NON_SYNONYMOU | MODERAT | MISSENSE | Atg/Gtg | M50V | PLES_53261 | | | | | | | C | C | | | C | C |
| NC_011770 | 5880071 | C | A | 77 | 0 | 6 | 0 | NON_SYNONYMOU | MODERAT | MISSENSE | Gcg/Tcg | A217S | PLES_53291 | A | A | | | | | A | A | A | A | | |
| NC_011770 | 5886058 | T | C | 65 | 0 | 4 | 0 | NON_SYNONYMOU | MODERAT | MISSENSE | cAc/cGc | H453R | PLES_53351 | | | | | | | C | C | C | C | | |
| NC_011770 | 5886065 | C | T | 66 | 0 | 2 | 0 | NON_SYNONYMOU | MODERAT | MISSENSE | Gcc/Acc | A451T | PLES_53351 | | | | | | | | | | | T | T |
| NC_011770 | 5886104 | T | G | 70 | 0 | 4 | 0 | NON_SYNONYMOU | MODERAT | MISSENSE | Atg/Ctg | M438L | PLES_53351 | | | | | | | G | G | G | G | | |
| NC_011770 | 5886215 | A | G | 63 | 0 | 4 | 0 | NON_SYNONYMOU | MODERAT | MISSENSE | Tgt/Cgt | C401R | PLES_53351 | | | | | | | G | G | G | G | | |
| NC_011770 | 5886260 | C | T | 66 | 0 | 2 | 0 | NON_SYNONYMOU | MODERAT | MISSENSE | Gcc/Acc | A386T | PLES_53351 | | | | | | | | | T | T | | |
| NC_011770 | 5886322 | G | A | 67 | 0 | 1 | 0 | NON_SYNONYMOU | MODERAT | MISSENSE | cCg/cTg | P365L | PLES_53351 | | | | | | | A | | | | | |
| NC_011770 | 5886536 | T | C | 64 | 0 | 2 | 0 | NON_SYNONYMOU | MODERAT | MISSENSE | Agc/Ggc | S294G | PLES_53351 | C | C | | | | | | | | | | |
| NC_011770 | 5886554 | G | A | 66 | 0 | 4 | 0 | NON_SYNONYMOU | MODERAT | MISSENSE | Cat/Tat | H288Y | PLES_53351 | | | | | | | A | A | A | A | | |
| NC_011770 | 5886571 | G | A | 64 | 0 | 4 | 0 | NON_SYNONYMOU | MODERAT | MISSENSE | gCg/gTg | A282V | PLES_53351 | | | | | | | A | A | A | A | | |
| NC_011770 | 5886595 | C | T | 60 | 0 | 4 | 0 | NON_SYNONYMOU | MODERAT | MISSENSE | cGc/cAc | R274H | PLES_53351 | | | | | | | | | T | T | T | T |
| NC_011770 | 5887093 | A | G | 53 | 0 | 10 | 0 | NON_SYNONYMOU | MODERAT | MISSENSE | gTg/gCg | V108A | PLES_53351 | G | G | G | G | G | G | G | G | G | G | | |
| NC_011770 | 5887542 | G | A | 31 | 0 | 2 | 0 | NON_SYNONYMOU | MODERAT | MISSENSE | Gcc/Acc | A16T | PLES_53361 | | | | | | | A | A | | | | |
| NC_011770 | 5888507 | C | G | 70 | 0 | 12 | 0 | NON_SYNONYMOU | MODERAT | MISSENSE | aaC/aaG | N337K | PLES_53361 | G | G | G | G | G | G | G | G | G | G | G | G |
| NC_011770 | 5889371 | C | A | 73 | 0 | 12 | 0 | NON_SYNONYMOU | MODERAT | MISSENSE | gaC/gaA | D34E | PLES_53381 | A | A | A | A | A | A | A | A | A | A | A | A |
| NC_011770 | 5889977 | C | G | 72 | 0 | 2 | 0 | NON_SYNONYMOU | MODERAT | MISSENSE | gaC/gaG | D236E | PLES_53381 | | | | | | | | | | | G | G |
| NC_011770 | 5893108 | G | A | 79 | 0 | 1 | 0 | NON_SYNONYMOU | MODERAT | MISSENSE | gGc/gAc | G254D | PLES_53411 | | | | | | | | A | | | | |
| NC_011770 | 5892595 | C | G | 70 | 0 | 2 | 0 | NON_SYNONYMOU | MODERAT | MISSENSE | gCg/gGg | A83G | PLES_53411 | | | | | | | G | G | | | | |
| NC_011770 | 5893455 | G | A | 68 | 0 | 2 | 0 | NON_SYNONYMOU | MODERAT | MISSENSE | Gcc/Acc | A370T | PLES_53411 | A | A | | | | | | | | | | |
| NC_011770 | 5893540 | C | A | 70 | 0 | 2 | 0 | NON_SYNONYMOU | MODERAT | MISSENSE | cCg/cAg | P398Q | PLES_53411 | A | A | | | | | | | | | | |
| NC_011770 | 5893849 | A | G | 68 | 0 | 10 | 0 | NON_SYNONYMOU | MODERAT | MISSENSE | cAc/cGc | H501R | PLES_53411 | | | G | G | G | G | G | G | G | G | G | G |
| NC_011770 | 5896892 | G | A | 78 | 0 | 1 | 0 | STOP_GAINED | HIGH | NONSENSE | tGg/tAg | W198* | PLES_53441 | | | | | | | A | | | | | |
| NC_011770 | 5896304 | T | C | 71 | 0 | 2 | 0 | NON_SYNONYMOU | MODERAT | MISSENSE | gTc/gCc | V2A | PLES_53441 | | | | | | | | | C | C | | |
| NC_011770 | 5896345 | C | T | 69 | 0 | 12 | 0 | NON_SYNONYMOU | MODERAT | MISSENSE | Ctc/Ttc | L16F | PLES_53441 | T | T | T | T | T | T | T | T | T | T | T | T |
| NC_011770 | 5896517 | C | A | 75 | 0 | 12 | 0 | NON_SYNONYMOU | MODERAT | MISSENSE | cCc/cAc | P73H | PLES_53441 | A | A | A | A | A | A | A | A | A | A | A | A |
| NC_011770 | 5896519 | C | A | 73 | 0 | 12 | 0 | NON_SYNONYMOU | MODERAT | MISSENSE | Ccc/Acc | P74T | PLES_53441 | A | A | A | A | A | A | A | A | A | A | A | A |
| NC_011770 | 5896751 | A | G | 83 | 0 | 12 | 0 | NON_SYNONYMOU | MODERAT | MISSENSE | aAt/aGt | N151S | PLES_53441 | G | G | G | G | G | G | G | G | G | G | G | G |
| NC_011770 | 5900185 | A | T | 71 | 0 | 12 | 0 | NON_SYNONYMOU | MODERAT | MISSENSE | gaT/gaA | D365E | PLES_53461 | T | T | T | T | T | T | T | T | T | T | T | T |
| NC_011770 | 5902361 | CCCGCGGC | C | 79 | 0 | 2 | 0 | CODON_CHANGE_ | MODERATE | | gcggcgcccg | AAPAA314 | PLES_53471 | | | | | | | | | | | C | C |
| NC_011770 | 5902884 | G | A | 46 | 0 | 1 | 0 | NON_SYNONYMOU | MODERAT | MISSENSE | tGc/tAc | C487Y | PLES_53471 | | | | | | | A | | | | | |
| NC_011770 | 5903036 | G | A | 46 | 0 | 8 | 0 | NON_SYNONYMOU | MODERAT | MISSENSE | cGc/cAc | R26H | PLES_53481 | A | A | A | A | A | A | | | | | A | A |
| NC_011770 | 5903134 | G | A | 54 | 0 | 1 | 0 | NON_SYNONYMOU | MODERAT | MISSENSE | Ggc/Agc | G59S | PLES_53481 | | | | | | | A | | | | | |
| NC_011770 | 5904152 | A | G | 67 | 0 | 10 | 0 | NON_SYNONYMOU | MODERAT | MISSENSE | gTg/gCg | V38A | PLES_53491 | G | G | G | G | G | G | G | G | | | G | G |
| NC_011770 | 5907296 | T | C | 80 | 0 | 6 | 0 | NON_SYNONYMOU | MODERAT | MISSENSE | Atc/Gtc | I20V | PLES_53511 | | | C | C | C | C | | | | | C | C |
| NC_011770 | 5907343 | C | T | 80 | 0 | 6 | 0 | NON_SYNONYMOU | MODERAT | MISSENSE | cGg/cAg | R4Q | PLES_53511 | | | T | T | T | T | | | | | T | T |
| NC_011770 | 5907559 | C | T | 92 | 0 | 1 | 0 | NON_SYNONYMOU | MODERAT | MISSENSE | Gtc/Atc | V269I | PLES_53521 | | | | | | | | | | | | T |
| NC_011770 | 5907886 | G | A | 79 | 0 | 4 | 0 | NON_SYNONYMOU | MODERAT | MISSENSE | Ccg/Tcg | P160S | PLES_53521 | A | A | | | | | A | A | | | | |
| NC_011770 | 5907922 | T | C | 80 | 0 | 4 | 0 | NON_SYNONYMOU | MODERAT | MISSENSE | Acg/Gcg | T148A | PLES_53521 | C | C | | | | | C | C | | | | |
| NC_011770 | 5910418 | A | G | 65 | 0 | 4 | 0 | NON_SYNONYMOU | MODERAT | MISSENSE | gTc/gCc | V156A | PLES_53541 | | | | | | | G | G | | | G | G |
| NC_011770 | 5910484 | T | C | 66 | 0 | 2 | 0 | NON_SYNONYMOU | MODERAT | MISSENSE | cAc/cGc | H134R | PLES_53541 | | | | | | | C | C | | | | |
| NC_011770 | 5911584 | T | G | 67 | 0 | 2 | 0 | NON_SYNONYMOU | MODERAT | MISSENSE | gAc/gCc | D84A | PLES_53551 | | | | | | | | | | | G | G |
| NC_011770 | 5911999 | C | T | 73 | 0 | 2 | 0 | NON_SYNONYMOU | MODERAT | MISSENSE | Gcc/Acc | A150T | PLES_53561 | | | | | | | T | T | | | | |
| NC_011770 | 5916659 | C | G | 34 | 0 | 8 | 0 | NON_SYNONYMOU | MODERAT | MISSENSE | gCc/gGc | A75G | PLES_53621 | G | G | G | G | G | G | | | | | G | G |
| NC_011770 | 5916664 | G | A | 35 | 0 | 8 | 0 | NON_SYNONYMOU | MODERAT | MISSENSE | Gtc/Atc | V77I | PLES_53621 | A | A | A | A | A | A | | | | | A | A |
| NC_011770 | 5916673 | A | T | 34 | 0 | 8 | 0 | NON_SYNONYMOU | MODERAT | MISSENSE | Acc/Tcc | T80S | PLES_53621 | T | T | T | T | T | T | | | | | T | T |
| NC_011770 | 5916685 | T | C | 35 | 0 | 8 | 0 | NON_SYNONYMOU | MODERAT | MISSENSE | Ttc/Ctc | F84L | PLES_53621 | C | C | C | C | C | C | | | | | C | C |
| NC_011770 | 5916695 | G | C | 35 | 0 | 8 | 0 | NON_SYNONYMOU | MODERAT | MISSENSE | cGt/cCt | R87P | PLES_53621 | C | C | C | C | C | C | | | | | C | C |
| NC_011770 | 5916697 | C | G | 34 | 0 | 8 | 0 | NON_SYNONYMOU | MODERAT | MISSENSE | Ccg/Gcg | P88A | PLES_53621 | G | G | G | G | G | G | | | | | G | G |
| NC_011770 | 5916716 | A | G | 36 | 0 | 8 | 0 | NON_SYNONYMOU | MODERAT | MISSENSE | aAc/aGc | N94S | PLES_53621 | G | G | G | G | G | G | | | | | G | G |
| NC_011770 | 5916723 | G | T | 38 | 0 | 8 | 0 | NON_SYNONYMOU | MODERAT | MISSENSE | caG/caT | Q96H | PLES_53621 | T | T | T | T | T | T | | | | | T | T |
| NC_011770 | 5916737 | C | G | 59 | 0 | 8 | 0 | NON_SYNONYMOU | MODERAT | MISSENSE | aCc/aGc | T101S | PLES_53621 | G | G | G | G | G | G | | | | | G | G |
| NC_011770 | 5916812 | G | A | 65 | 0 | 8 | 0 | NON_SYNONYMOU | MODERAT | MISSENSE | cGg/cAg | R126Q | PLES_53621 | A | A | A | A | A | A | | | | | A | A |
| NC_011770 | 5917363 | T | G | 80 | 0 | 8 | 0 | NON_SYNONYMOU | MODERAT | MISSENSE | Tat/Gat | Y310D | PLES_53621 | G | G | G | G | G | G | | | | | G | G |
| NC_011770 | 5917371 | C | G | 72 | 0 | 8 | 0 | NON_SYNONYMOU | MODERAT | MISSENSE | gaC/gaG | D312E | PLES_53621 | G | G | G | G | G | G | | | | | G | G |

| | | | | | | | | | | | | | | | | | | | | | | | | | |
|---|---|---|---|---|---|---|---|---|---|---|---|---|---|---|---|---|---|---|---|---|---|---|---|---|---|
| NC_011770 | 5917379 | G | C | 74 | 0 | 8 | 0 | NON_SYNONYMOU | MODERAT | MISSENSE | aGc/aCc | S315T | PLES_53621 | C | C | C | C | C | C | | | | | C | C |
| NC_011770 | 5917380 | C | G | 74 | 0 | 8 | 0 | NON_SYNONYMOU | MODERAT | MISSENSE | agC/agG | S315R | PLES_53621 | G | G | G | G | G | G | | | | | G | G |
| NC_011770 | 5917846 | G | T | 63 | 0 | 6 | 0 | NON_SYNONYMOU | MODERAT | MISSENSE | Gcc/Tcc | A471S | PLES_53621 | T | T | T | T | T | T | | | | | | |
| NC_011770 | 5918444 | T | C | 76 | 0 | 4 | 0 | NON_SYNONYMOU | MODERAT | MISSENSE | Agc/Ggc | S71G | PLES_53631 | | | C | C | C | C | | | | | | |
| NC_011770 | 5918485 | G | A | 83 | 0 | 4 | 0 | NON_SYNONYMOU | MODERAT | MISSENSE | gCc/gTc | A57V | PLES_53631 | | | A | A | A | A | | | | | | |
| NC_011770 | 5918605 | C | T | 87 | 0 | 2 | 0 | NON_SYNONYMOU | MODERAT | MISSENSE | cGc/cAc | R17H | PLES_53631 | T | T | | | | | | | | | | |
| NC_011770 | 5921680 | C | T | 78 | 0 | 4 | 0 | NON_SYNONYMOU | MODERAT | MISSENSE | Gcc/Acc | A681T | PLES_53661 | | | T | T | T | T | | | | | | |
| NC_011770 | 5921803 | A | G | 76 | 0 | 2 | 0 | NON_SYNONYMOU | MODERAT | MISSENSE | Ttc/Ctc | F640L | PLES_53661 | | | | | | | G | G | | | | |
| NC_011770 | 5922741 | C | T | 74 | 0 | 4 | 0 | NON_SYNONYMOU | MODERAT | MISSENSE | gGa/gAa | G327E | PLES_53661 | | | | | | | | | T | T | T | T |
| NC_011770 | 5923000 | T | C | 76 | 0 | 6 | 0 | NON_SYNONYMOU | MODERAT | MISSENSE | Aag/Gag | K241E | PLES_53661 | C | C | | | | | | | C | C | C | C |
| NC_011770 | 5923489 | G | A | 67 | 0 | 2 | 0 | NON_SYNONYMOU | MODERAT | MISSENSE | Ctc/Ttc | L78F | PLES_53661 | A | A | | | | | | | | | | |
| NC_011770 | 5923962 | G | A | 79 | 0 | 2 | 0 | NON_SYNONYMOU | MODERAT | MISSENSE | aCc/aTc | T329I | PLES_53671 | A | A | | | | | | | | | | |
| NC_011770 | 5923972 | C | T | 81 | 0 | 2 | 0 | NON_SYNONYMOU | MODERAT | MISSENSE | Gcc/Acc | A326T | PLES_53671 | | | | | | | T | T | | | | |
| NC_011770 | 5924142 | G | A | 77 | 0 | 2 | 0 | NON_SYNONYMOU | MODERAT | MISSENSE | gCc/gTc | A269V | PLES_53671 | | | | | A | A | | | | | | |
| NC_011770 | 5924151 | C | T | 80 | 0 | 2 | 0 | NON_SYNONYMOU | MODERAT | MISSENSE | cGc/cAc | R266H | PLES_53671 | T | T | | | | | | | | | | |
| NC_011770 | 5925129 | A | C | 74 | 0 | 10 | 0 | NON_SYNONYMOU | MODERAT | MISSENSE | Tcc/Gcc | S211A | PLES_53681 | C | C | C | C | C | C | | | C | C | C | C |
| NC_011770 | 5925173 | A | C | 72 | 0 | 10 | 0 | NON_SYNONYMOU | MODERAT | MISSENSE | gTc/gGc | V196G | PLES_53681 | C | C | C | C | C | C | | | C | C | C | C |
| NC_011770 | 5925180 | T | C | 73 | 0 | 2 | 0 | NON_SYNONYMOU | MODERAT | MISSENSE | Acg/Gcg | T194A | PLES_53681 | C | C | | | | | | | | | | |
| NC_011770 | 5925516 | T | G | 73 | 0 | 2 | 0 | NON_SYNONYMOU | MODERAT | MISSENSE | Agc/Cgc | S82R | PLES_53681 | G | G | | | | | | | | | | |
| NC_011770 | 5926041 | G | A | 67 | 0 | 1 | 0 | NON_SYNONYMOU | MODERAT | MISSENSE | Ccc/Tcc | P464S | PLES_53691 | | | | | | | | A | | | | |
| NC_011770 | 5926196 | G | A | 72 | 0 | 1 | 0 | NON_SYNONYMOU | MODERAT | MISSENSE | cCg/cTg | P412L | PLES_53691 | | | | | | | | A | | | | |
| NC_011770 | 5926083 | A | G | 65 | 0 | 2 | 0 | NON_SYNONYMOU | MODERAT | MISSENSE | Tat/Cat | Y450H | PLES_53691 | G | G | | | | | | | | | | |
| NC_011770 | 5926232 | T | C | 73 | 0 | 2 | 0 | NON_SYNONYMOU | MODERAT | MISSENSE | gAa/gGa | E400G | PLES_53691 | | | | | | | | | | | C | C |
| NC_011770 | 5926260 | G | A | 79 | 0 | 10 | 0 | NON_SYNONYMOU | MODERAT | MISSENSE | Cac/Tac | H391Y | PLES_53691 | A | A | A | A | A | A | A | A | A | A | | |
| NC_011770 | 5926716 | G | A | 81 | 0 | 2 | 0 | NON_SYNONYMOU | MODERAT | MISSENSE | Cgg/Tgg | R239W | PLES_53691 | | | | | A | A | | | | | | |
| NC_011770 | 5929507 | G | GC | 68 | 0 | 1 | 0 | FRAME_SHIFT | HIGH | | ggc/ggGc | G368G? | PLES_53701 | | | | | | | GC | | | | | |
| NC_011770 | 5930187 | G | A | 72 | 0 | 2 | 0 | STOP_GAINED | HIGH | NONSENSE | Caa/Taa | Q142* | PLES_53701 | | | | | | | | | A | A | | |
| NC_011770 | 5928162 | G | A | 74 | 0 | 1 | 0 | NON_SYNONYMOU | MODERAT | MISSENSE | Cgg/Tgg | R817W | PLES_53701 | | | | | | | | | | | | A |
| NC_011770 | 5927675 | G | T | 73 | 0 | 12 | 0 | NON_SYNONYMOU | MODERAT | MISSENSE | gCc/gAc | A979D | PLES_53701 | T | T | T | T | T | T | T | T | T | T | T | T |
| NC_011770 | 5927977 | G | C | 77 | 0 | 2 | 0 | NON_SYNONYMOU | MODERAT | MISSENSE | gaC/gaG | D878E | PLES_53701 | | | | | | | | | | | C | C |
| NC_011770 | 5928138 | C | A | 74 | 0 | 2 | 0 | NON_SYNONYMOU | MODERAT | MISSENSE | Gcc/Tcc | A825S | PLES_53701 | | | | | | | | | | | A | A |
| NC_011770 | 5928659 | A | G | 67 | 0 | 2 | 0 | NON_SYNONYMOU | MODERAT | MISSENSE | gTg/gCg | V651A | PLES_53701 | | | | | | | | | | | G | G |
| NC_011770 | 5929013 | C | T | 72 | 0 | 6 | 0 | NON_SYNONYMOU | MODERAT | MISSENSE | cGc/cAc | R533H | PLES_53701 | | | T | T | T | T | | | T | T | | |
| NC_011770 | 5929890 | A | G | 72 | 0 | 4 | 0 | NON_SYNONYMOU | MODERAT | MISSENSE | Tgc/Cgc | C241R | PLES_53701 | | | | | | | G | G | | | G | G |
| NC_011770 | 5929895 | G | A | 74 | 0 | 2 | 0 | NON_SYNONYMOU | MODERAT | MISSENSE | gCg/gTg | A239V | PLES_53701 | A | A | | | | | | | | | | |
| NC_011770 | 5930106 | G | C | 63 | 0 | 6 | 0 | NON_SYNONYMOU | MODERAT | MISSENSE | Ccg/Gcg | P169A | PLES_53701 | | | | | | | C | C | C | C | C | C |
| NC_011770 | 5930319 | G | C | 75 | 0 | 4 | 0 | NON_SYNONYMOU | MODERAT | MISSENSE | Cgt/Ggt | R98G | PLES_53701 | | | | | | | C | C | C | C | | |
| NC_011770 | 5930342 | C | T | 77 | 0 | 2 | 0 | NON_SYNONYMOU | MODERAT | MISSENSE | cGc/cAc | R90H | PLES_53701 | T | T | | | | | | | | | | |
| NC_011770 | 5932320 | C | T | 80 | 0 | 4 | 0 | NON_SYNONYMOU | MODERAT | MISSENSE | Gcg/Acg | A331T | PLES_53731 | | | T | T | T | T | | | | | | |
| NC_011770 | 5932458 | T | C | 89 | 0 | 10 | 0 | NON_SYNONYMOU | MODERAT | MISSENSE | Acc/Gcc | T285A | PLES_53731 | C | C | C | C | C | C | C | C | | | C | C |
| NC_011770 | 5932514 | G | T | 91 | 0 | 2 | 0 | NON_SYNONYMOU | MODERAT | MISSENSE | cCg/cAg | P266Q | PLES_53731 | | | | | | | T | T | | | | |
| NC_011770 | 5932854 | C | G | 78 | 0 | 2 | 0 | NON_SYNONYMOU | MODERAT | MISSENSE | Gac/Cac | D153H | PLES_53731 | G | G | | | | | | | | | | |
| NC_011770 | 5932996 | G | C | 80 | 0 | 8 | 0 | NON_SYNONYMOU | MODERAT | MISSENSE | gaC/gaG | D105E | PLES_53731 | C | C | A | A | A | A | | | | | A | A |
| NC_011770 | 5935371 | GC | G | 46 | 0 | 2 | 0 | FRAME_SHIFT | HIGH | | cgc/ | R632 | PLES_53741 | G | | | | | | GCC | | | | | |
| NC_011770 | 5935105 | G | A | 78 | 0 | 1 | 0 | NON_SYNONYMOU | MODERAT | MISSENSE | gGc/gAc | G541D | PLES_53741 | | | | | | | | A | | | | |
| NC_011770 | 5933958 | G | C | 63 | 0 | 8 | 0 | NON_SYNONYMOU | MODERAT | MISSENSE | Gag/Cag | E159Q | PLES_53741 | C | C | C | C | C | C | | | C | C | | |
| NC_011770 | 5934043 | A | C | 64 | 0 | 8 | 0 | NON_SYNONYMOU | MODERAT | MISSENSE | aAg/aCg | K187T | PLES_53741 | C | C | C | C | C | C | | | C | C | | |
| NC_011770 | 5934296 | G | A | 69 | 0 | 1 | 0 | NON_SYNONYMOU | MODERAT | MISSENSE | atG/atA | M271I | PLES_53741 | | | | | | | A | | | | | |
| NC_011770 | 5934330 | G | A | 73 | 0 | 2 | 0 | NON_SYNONYMOU | MODERAT | MISSENSE | Ggt/Agt | G283S | PLES_53741 | | | | | | | A | A | | | | |
| NC_011770 | 5934624 | G | C | 83 | 0 | 2 | 0 | NON_SYNONYMOU | MODERAT | MISSENSE | Gcc/Ccc | A381P | PLES_53741 | | | | | | | C | C | | | | |
| NC_011770 | 5937570 | C | A | 79 | 0 | 2 | 0 | NON_SYNONYMOU | MODERAT | MISSENSE | Gcg/Tcg | A262S | PLES_53771 | A | A | | | | | | | | | | |
| NC_011770 | 5938459 | G | T | 76 | 0 | 2 | 0 | NON_SYNONYMOU | MODERAT | MISSENSE | Gcc/Tcc | A8S | PLES_53781 | | | | | | | T | T | | | | |
| NC_011770 | 5939981 | C | T | 79 | 0 | 1 | 0 | NON_SYNONYMOU | MODERAT | MISSENSE | cCg/cTg | P384L | PLES_53791 | | | | | | | | | | | T | |
| NC_011770 | 5939225 | G | A | 76 | 0 | 8 | 0 | NON_SYNONYMOU | MODERAT | MISSENSE | gGt/gAt | G132D | PLES_53791 | A | A | | | | | A | A | A | A | A | A |
| NC_011770 | 5939919 | C | A | 78 | 0 | 2 | 0 | NON_SYNONYMOU | MODERAT | MISSENSE | gaC/gaA | D363E | PLES_53791 | | | | | | | A | A | | | | |
| NC_011770 | 5939954 | G | A | 75 | 0 | 12 | 0 | NON_SYNONYMOU | MODERAT | MISSENSE | cGa/cAa | R375Q | PLES_53791 | A | A | A | A | A | A | A | A | A | A | A | A |
| NC_011770 | 5940175 | A | G | 77 | 0 | 12 | 0 | NON_SYNONYMOU | MODERAT | MISSENSE | cAc/cGc | H58R | PLES_53801 | G | G | G | G | G | G | G | G | G | G | G | G |
| NC_011770 | 5940383 | G | C | 77 | 0 | 4 | 0 | NON_SYNONYMOU | MODERAT | MISSENSE | gaG/gaC | E127D | PLES_53801 | | | | | | | C | C | C | C | | |
| NC_011770 | 5940696 | A | G | 62 | 0 | 10 | 0 | NON_SYNONYMOU | MODERAT | MISSENSE | Acc/Gcc | T232A | PLES_53801 | G | G | G | G | G | G | | | G | G | G | G |
| NC_011770 | 5941123 | G | A | 44 | 0 | 4 | 0 | NON_SYNONYMOU | MODERAT | MISSENSE | cGc/cAc | R47H | PLES_53811 | A | A | | | | | | | | | A | A |
| NC_011770 | 5943566 | C | A | 99 | 0 | 8 | 0 | NON_SYNONYMOU | MODERAT | MISSENSE | Gcc/Tcc | A304S | PLES_53831 | A | A | A | A | A | A | | | A | A | | |
| NC_011770 | 5943631 | T | C | 96 | 0 | 2 | 0 | NON_SYNONYMOU | MODERAT | MISSENSE | aAg/aGg | K282R | PLES_53831 | | | | | | | C | C | | | | |

| | | | | | | | | | | | | | | | | | | | | | | | | | |
|---|---|---|---|---|---|---|---|---|---|---|---|---|---|---|---|---|---|---|---|---|---|---|---|---|---|
| NC_011770 | 5943962 | G | A | 94 | 0 | 2 | 0 | NON_SYNONYMOU | MODERAT | MISSENSE | Ccc/Tcc | P172S | PLES_53831 | A | A | | | | | | | | | | |
| NC_011770 | 5944025 | T | A | 90 | 0 | 2 | 0 | NON_SYNONYMOU | MODERAT | MISSENSE | Acg/Tcg | T151S | PLES_53831 | | | | | | | A | A | | | | |
| NC_011770 | 5944318 | G | C | 77 | 0 | 4 | 0 | NON_SYNONYMOU | MODERAT | MISSENSE | gCg/gGg | A53G | PLES_53831 | | | C | C | C | C | | | | | | |
| NC_011770 | 5950660 | G | A | 92 | 0 | 2 | 0 | NON_SYNONYMOU | MODERAT | MISSENSE | Gca/Aca | A112T | PLES_53871 | | | | | | | | | | | A | A |
| NC_011770 | 5950694 | C | T | 92 | 0 | 2 | 0 | NON_SYNONYMOU | MODERAT | MISSENSE | gCg/gTg | A123V | PLES_53871 | | | | | | | | | | | T | T |
| NC_011770 | 5952861 | C | T | 76 | 0 | 4 | 0 | NON_SYNONYMOU | MODERAT | MISSENSE | aGa/aAa | R78K | PLES_53891 | | | | | | | T | T | | | T | T |
| NC_011770 | 5953374 | GGCCA | G | 78 | 0 | 1 | 0 | FRAME_SHIFT | HIGH | | ctggcc/ | LA278 | PLES_53901 | | | | | | G | | | | | | |
| NC_011770 | 5954303 | A | T | 79 | 0 | 2 | 0 | NON_SYNONYMOU | MODERAT | MISSENSE | Tcg/Acg | S467T | PLES_53911 | | | | | | | | | | | T | T |
| NC_011770 | 5954336 | C | A | 79 | 0 | 2 | 0 | NON_SYNONYMOU | MODERAT | MISSENSE | Gcg/Tcg | A456S | PLES_53911 | | | | | | | | | A | A | | |
| NC_011770 | 5954762 | C | A | 68 | 0 | 2 | 0 | NON_SYNONYMOU | MODERAT | MISSENSE | Gcc/Tcc | A314S | PLES_53911 | | | | | | | A | A | | | | |
| NC_011770 | 5954839 | A | G | 65 | 0 | 4 | 0 | NON_SYNONYMOU | MODERAT | MISSENSE | aTc/aCc | I288T | PLES_53911 | | | G | G | G | G | | | | | | |
| NC_011770 | 5954960 | T | C | 55 | 0 | 4 | 0 | NON_SYNONYMOU | MODERAT | MISSENSE | Acg/Gcg | T248A | PLES_53911 | | | | | | | | | C | C | C | C |
| NC_011770 | 5955165 | A | C | 62 | 0 | 6 | 0 | NON_SYNONYMOU | MODERAT | MISSENSE | gaT/gaG | D179E | PLES_53911 | C | C | | | | | | | G | G | G | G |
| NC_011770 | 5955329 | C | T | 65 | 0 | 4 | 0 | NON_SYNONYMOU | MODERAT | MISSENSE | Gcc/Acc | A125T | PLES_53911 | | | | | | | | | T | T | T | T |
| NC_011770 | 5955572 | C | T | 72 | 0 | 2 | 0 | NON_SYNONYMOU | MODERAT | MISSENSE | Gtg/Atg | V44M | PLES_53911 | | | | | | | | | | | T | T |
| NC_011770 | 5956217 | T | C | 77 | 0 | 2 | 0 | NON_SYNONYMOU | MODERAT | MISSENSE | Acc/Gcc | T129A | PLES_53921 | | | | | | | | | | | C | C |
| NC_011770 | 5956255 | G | A | 72 | 0 | 2 | 0 | NON_SYNONYMOU | MODERAT | MISSENSE | gCg/gTg | A116V | PLES_53921 | | | | | | | | | | | A | A |
| NC_011770 | 5956273 | G | C | 72 | 0 | 2 | 0 | NON_SYNONYMOU | MODERAT | MISSENSE | gCg/gGg | A110G | PLES_53921 | | | | | | | | | | | C | C |
| NC_011770 | 5956300 | T | C | 71 | 0 | 4 | 0 | NON_SYNONYMOU | MODERAT | MISSENSE | gAc/gGc | D101G | PLES_53921 | | | | | | | C | C | | | C | C |
| NC_011770 | 5956546 | G | A | 51 | 0 | 6 | 0 | NON_SYNONYMOU | MODERAT | MISSENSE | cCg/cTg | P19L | PLES_53921 | | | A | A | A | A | | | A | A | | |
| NC_011770 | 5956550 | C | A | 51 | 0 | 2 | 0 | NON_SYNONYMOU | MODERAT | MISSENSE | Gcg/Tcg | A18S | PLES_53921 | | | | | | | | | A | A | | |
| NC_011770 | 5956768 | T | C | 68 | 0 | 6 | 0 | NON_SYNONYMOU | MODERAT | MISSENSE | Acg/Gcg | T322A | PLES_53931 | C | C | C | C | C | C | | | | | | |
| NC_011770 | 5957395 | C | T | 85 | 0 | 4 | 0 | NON_SYNONYMOU | MODERAT | MISSENSE | Gtt/Att | V113I | PLES_53931 | | | T | T | T | T | | | | | | |
| NC_011770 | 5958536 | C | G | 81 | 0 | 2 | 0 | NON_SYNONYMOU | MODERAT | MISSENSE | Gtc/Ctc | V313L | PLES_53941 | | | | | | | | | | | G | G |
| NC_011770 | 5960097 | C | T | 83 | 0 | 1 | 0 | NON_SYNONYMOU | MODERAT | MISSENSE | Gtc/Atc | V320I | PLES_53951 | | | | | | | | T | | | | |
| NC_011770 | 5959635 | C | T | 91 | 0 | 2 | 0 | NON_SYNONYMOU | MODERAT | MISSENSE | Ggc/Agc | G474S | PLES_53951 | | | | | | | | | | | T | T |
| NC_011770 | 5959692 | A | C | 86 | 0 | 12 | 0 | NON_SYNONYMOU | MODERAT | MISSENSE | Tct/Gct | S455A | PLES_53951 | C | C | C | C | C | C | C | C | C | C | C | C |
| NC_011770 | 5960228 | G | A | 79 | 0 | 2 | 0 | NON_SYNONYMOU | MODERAT | MISSENSE | cCc/cTc | P276L | PLES_53951 | A | A | | | | | | | | | | |
| NC_011770 | 5960561 | T | C | 77 | 0 | 10 | 0 | NON_SYNONYMOU | MODERAT | MISSENSE | cAc/cGc | H165R | PLES_53951 | C | C | C | C | C | C | | | C | C | C | C |
| NC_011770 | 5960600 | T | G | 77 | 0 | 2 | 0 | NON_SYNONYMOU | MODERAT | MISSENSE | gAc/gCc | D152A | PLES_53951 | | | | | | | | | | | G | G |
| NC_011770 | 5960601 | C | T | 77 | 0 | 2 | 0 | NON_SYNONYMOU | MODERAT | MISSENSE | Gac/Aac | D152N | PLES_53951 | | | | | | | | | | | T | T |
| NC_011770 | 5960690 | G | A | 83 | 0 | 2 | 0 | NON_SYNONYMOU | MODERAT | MISSENSE | gCc/gTc | A122V | PLES_53951 | | | | | | | A | A | | | | |
| NC_011770 | 5960769 | C | G | 86 | 0 | 8 | 0 | NON_SYNONYMOU | MODERAT | MISSENSE | Gag/Cag | E96Q | PLES_53951 | G | G | G | G | G | G | | | | | G | G |
| NC_011770 | 5961308 | G | C | 83 | 0 | 12 | 0 | NON_SYNONYMOU | MODERAT | MISSENSE | Cta/Gta | L168V | PLES_53961 | C | C | C | C | C | C | C | C | C | C | C | C |
| NC_011770 | 5961880 | G | A | 88 | 0 | 2 | 0 | NON_SYNONYMOU | MODERAT | MISSENSE | Cgt/Tgt | R221C | PLES_53971 | | | | | | | | | | | A | A |
| NC_011770 | 5962038 | T | G | 84 | 0 | 4 | 0 | NON_SYNONYMOU | MODERAT | MISSENSE | gAc/gCc | D168A | PLES_53971 | | | | | | | G | G | | | C | C |
| NC_011770 | 5975361 | C | T | 78 | 0 | 4 | 0 | NON_SYNONYMOU | MODERAT | MISSENSE | cCg/cTg | P17L | PLES_54061 | T | T | | | | | | | | | T | T |
| NC_011770 | 5975435 | G | A | 81 | 0 | 2 | 0 | NON_SYNONYMOU | MODERAT | MISSENSE | Gcg/Acg | A42T | PLES_54061 | | | | | | | | | | | A | A |
| NC_011770 | 5975472 | G | A | 80 | 0 | 4 | 0 | NON_SYNONYMOU | MODERAT | MISSENSE | cGg/cAg | R54Q | PLES_54061 | | | A | A | A | A | | | | | | |
| NC_011770 | 5975513 | A | G | 77 | 0 | 4 | 0 | NON_SYNONYMOU | MODERAT | MISSENSE | Acc/Gcc | T68A | PLES_54061 | G | G | | | | | | | | | G | G |
| NC_011770 | 5978241 | G | A | 81 | 0 | 1 | 0 | NON_SYNONYMOU | MODERAT | MISSENSE | Gaa/Aaa | E853K | PLES_54071 | | | | | | | | | | | A | |
| NC_011770 | 5975746 | T | C | 85 | 0 | 1 | 0 | NON_SYNONYMOU | MODERAT | MISSENSE | gTg/gCg | V21A | PLES_54071 | | | | | | | C | | | | | |
| NC_011770 | 5976078 | G | C | 98 | 0 | 2 | 0 | NON_SYNONYMOU | MODERAT | MISSENSE | Gcg/Ccg | A132P | PLES_54071 | | | | | | | | | | | C | C |
| NC_011770 | 5976526 | A | G | 97 | 0 | 2 | 0 | NON_SYNONYMOU | MODERAT | MISSENSE | aAg/aGg | K281R | PLES_54071 | | | | | | | | | G | G | | |
| NC_011770 | 5976967 | A | G | 85 | 0 | 1 | 0 | NON_SYNONYMOU | MODERAT | MISSENSE | cAc/cGc | H428R | PLES_54071 | | | | | | | G | | | | | |
| NC_011770 | 5979160 | C | T | 75 | 0 | 2 | 0 | NON_SYNONYMOU | MODERAT | MISSENSE | gGc/gAc | G269D | PLES_54091 | | | | | | | | | | | T | T |
| NC_011770 | 5979203 | G | T | 82 | 0 | 2 | 0 | NON_SYNONYMOU | MODERAT | MISSENSE | Ctg/Atg | L255M | PLES_54091 | | | | | | | T | T | | | | |
| NC_011770 | 5980708 | C | T | 70 | 0 | 1 | 0 | NON_SYNONYMOU | MODERAT | MISSENSE | Cgc/Tgc | R148C | PLES_54101 | | | | | | | | | | | | T |
| NC_011770 | 5980344 | G | T | 68 | 0 | 2 | 0 | NON_SYNONYMOU | MODERAT | MISSENSE | caG/caT | Q26H | PLES_54101 | | | | | | | | | T | T | | |
| NC_011770 | 5980531 | G | A | 69 | 0 | 1 | 0 | NON_SYNONYMOU | MODERAT | MISSENSE | Ggc/Agc | G89S | PLES_54101 | | | | | | | A | | | | | |
| NC_011770 | 5981079 | C | A | 75 | 0 | 4 | 0 | NON_SYNONYMOU | MODERAT | MISSENSE | gaC/gaA | D271E | PLES_54101 | A | A | | | | | | | | | A | A |
| NC_011770 | 5981433 | C | A | 71 | 0 | 4 | 0 | NON_SYNONYMOU | MODERAT | MISSENSE | gaC/gaA | D389E | PLES_54101 | | | A | A | A | A | | | | | | |
| NC_011770 | 5981723 | T | C | 61 | 0 | 6 | 0 | NON_SYNONYMOU | MODERAT | MISSENSE | gTc/gCc | V486A | PLES_54101 | | | C | C | C | C | | | C | C | | |
| NC_011770 | 5983232 | A | AC | 86 | 0 | 1 | 0 | FRAME_SHIFT | HIGH | | cac/cCac | H251P? | PLES_54111 | | AC | | | | | | | | | | |
| NC_011770 | 5983995 | TG | T | 65 | 0 | 2 | 0 | FRAME_SHIFT | HIGH | | gag/ | E504 | PLES_54111 | | | T | T | | | | | | | | |
| NC_011770 | 5984224 | T | C | 66 | 0 | 4 | 0 | NON_SYNONYMOU | MODERAT | MISSENSE | aTt/aCt | I10T | PLES_54121 | | | C | C | C | C | | | | | | |
| NC_011770 | 5985031 | A | G | 75 | 0 | 2 | 0 | NON_SYNONYMOU | MODERAT | MISSENSE | aAc/aGc | N279S | PLES_54121 | | | | | | | | | G | G | | |
| NC_011770 | 5987671 | G | A | 55 | 0 | 12 | 0 | NON_SYNONYMOU | MODERAT | MISSENSE | gGc/gAc | G18D | PLES_54131 | A | A | A | A | A | A | A | A | A | A | A | A |
| NC_011770 | 5987929 | C | T | 67 | 0 | 4 | 0 | NON_SYNONYMOU | MODERAT | MISSENSE | gCc/gTc | A104V | PLES_54131 | | | | | | | T | T | | | T | T |
| NC_011770 | 5988267 | G | A | 86 | 0 | 2 | 0 | NON_SYNONYMOU | MODERAT | MISSENSE | Gcc/Acc | A217T | PLES_54131 | | | | | | | A | A | | | | |
| NC_011770 | 5988763 | T | C | 84 | 0 | 2 | 0 | NON_SYNONYMOU | MODERAT | MISSENSE | cTg/cCg | L382P | PLES_54131 | | | | | | | | | | | C | C |
| NC_011770 | 5989834 | A | C | 72 | 0 | 2 | 0 | NON_SYNONYMOU | MODERAT | MISSENSE | aAg/aCg | K117T | PLES_54141 | | | | | | | C | C | | | | |

| | | | | | | | | | | | | | | | | | | | | | | | | | |
|---|---|---|---|---|---|---|---|---|---|---|---|---|---|---|---|---|---|---|---|---|---|---|---|---|---|
| NC_011770 | 5991873 | A | C | 78 | 0 | 2 | 0 | NON_SYNONYMOU | MODERAT | MISSENSE | Att/Ctt | I61L | PLES_54161 | C | C | | | | | | | | | | |
| NC_011770 | 5992107 | A | G | 79 | 0 | 2 | 0 | NON_SYNONYMOU | MODERAT | MISSENSE | Acc/Gcc | T139A | PLES_54161 | G | G | | | | | | | | | | |
| NC_011770 | 5992375 | G | A | 78 | 0 | 1 | 0 | NON_SYNONYMOU | MODERAT | MISSENSE | Gac/Aac | D40N | PLES_54171 | | A | | | | | | | | | | |
| NC_011770 | 5992516 | G | A | 85 | 0 | 4 | 0 | NON_SYNONYMOU | MODERAT | MISSENSE | Gcg/Acg | A87T | PLES_54171 | | | A | A | A | A | | | | | | |
| NC_011770 | 5992780 | G | A | 83 | 0 | 6 | 0 | NON_SYNONYMOU | MODERAT | MISSENSE | Ggc/Agc | G175S | PLES_54171 | | | A | A | A | A | A | A | | | | |
| NC_011770 | 5993562 | G | A | 86 | 0 | 1 | 0 | STOP_GAINED | HIGH | NONSENSE | Cag/Tag | Q101* | PLES_54181 | | | | | | | | | | | A | |
| NC_011770 | 5994788 | C | G | 79 | 0 | 1 | 0 | NON_SYNONYMOU | MODERAT | MISSENSE | cGc/cCc | R24P | PLES_54191 | | | | | | G | | | | | | |
| NC_011770 | 5994858 | C | A | 82 | 0 | 8 | 0 | NON_SYNONYMOU | MODERAT | MISSENSE | Gtg/Ttg | V1L | PLES_54191 | | | A | A | A | A | | | A | A | A | A |
| NC_011770 | 5996081 | G | A | 61 | 0 | 1 | 0 | NON_SYNONYMOU | MODERAT | MISSENSE | Ggc/Agc | G350S | PLES_54201 | | | | | | | | A | | | | |
| NC_011770 | 5996816 | GT | G | 85 | 0 | 2 | 0 | FRAME_SHIFT | HIGH | | aac/ | N164 | PLES_54211 | G | G | | | | | | | | | | |
| NC_011770 | 5997192 | G | GC | 72 | 0 | 1 | 0 | FRAME_SHIFT | HIGH | | gcg/gGcg | A39G? | PLES_54211 | | | | | | | GC | | | | | |
| NC_011770 | 5996418 | A | C | 65 | 0 | 4 | 0 | NON_SYNONYMOU | MODERAT | MISSENSE | cTa/cGa | L297R | PLES_54211 | C | C | | | | | C | C | | | | |
| NC_011770 | 5996611 | C | T | 71 | 0 | 2 | 0 | NON_SYNONYMOU | MODERAT | MISSENSE | Gtg/Atg | V233M | PLES_54211 | | | | | | | T | T | | | | |
| NC_011770 | 5996644 | A | C | 67 | 0 | 2 | 0 | NON_SYNONYMOU | MODERAT | MISSENSE | Tcc/Gcc | S222A | PLES_54211 | | | | | | | C | C | | | | |
| NC_011770 | 5996903 | C | G | 84 | 0 | 2 | 0 | NON_SYNONYMOU | MODERAT | MISSENSE | gaG/gaC | E135D | PLES_54211 | G | G | | | | | | | | | | |
| NC_011770 | 5997022 | G | A | 77 | 0 | 2 | 0 | NON_SYNONYMOU | MODERAT | MISSENSE | Ctc/Ttc | L96F | PLES_54211 | | | | | | | | | A | A | | |
| NC_011770 | 5997033 | T | G | 73 | 0 | 4 | 0 | NON_SYNONYMOU | MODERAT | MISSENSE | gAa/gCa | E92A | PLES_54211 | G | G | | | | | | | | | G | G |
| NC_011770 | 5997070 | C | T | 75 | 0 | 12 | 0 | NON_SYNONYMOU | MODERAT | MISSENSE | Gtc/Atc | V80I | PLES_54211 | T | T | T | T | T | T | T | T | T | T | T | T |
| NC_011770 | 5997075 | A | G | 77 | 0 | 12 | 0 | NON_SYNONYMOU | MODERAT | MISSENSE | gTc/gCc | V78A | PLES_54211 | G | G | G | G | G | G | G | G | G | G | G | G |
| NC_011770 | 5997089 | G | C | 79 | 0 | 2 | 0 | NON_SYNONYMOU | MODERAT | MISSENSE | gaC/gaG | D73E | PLES_54211 | C | C | | | | | | | | | | |
| NC_011770 | 5997605 | G | A | 71 | 0 | 2 | 0 | NON_SYNONYMOU | MODERAT | MISSENSE | Gcg/Acg | A63T | PLES_54221 | | | | | | | | | | | A | A |
| NC_011770 | 5997812 | C | A | 74 | 0 | 8 | 0 | NON_SYNONYMOU | MODERAT | MISSENSE | Ctg/Atg | L132M | PLES_54221 | | | A | A | A | A | A | A | A | A | | |
| NC_011770 | 5998193 | G | A | 70 | 0 | 2 | 0 | NON_SYNONYMOU | MODERAT | MISSENSE | Gcg/Acg | A259T | PLES_54221 | A | A | | | | | | | | | | |
| NC_011770 | 5998419 | C | A | 73 | 0 | 2 | 0 | NON_SYNONYMOU | MODERAT | MISSENSE | gCc/gAc | A334D | PLES_54221 | A | A | | | | | | | | | | |
| NC_011770 | 5998439 | C | G | 74 | 0 | 2 | 0 | NON_SYNONYMOU | MODERAT | MISSENSE | Cca/Gca | P341A | PLES_54221 | G | G | | | | | | | | | | |
| NC_011770 | 5998484 | T | C | 75 | 0 | 2 | 0 | NON_SYNONYMOU | MODERAT | MISSENSE | Ttc/Ctc | F356L | PLES_54221 | C | C | | | | | | | | | | |
| NC_011770 | 5999133 | G | A | 69 | 0 | 10 | 0 | NON_SYNONYMOU | MODERAT | MISSENSE | Ggt/Agt | G190S | PLES_54231 | A | A | A | A | A | A | | | A | A | A | A |
| NC_011770 | 5999152 | C | G | 72 | 0 | 2 | 0 | NON_SYNONYMOU | MODERAT | MISSENSE | aCc/aGc | T196S | PLES_54231 | | | | | | | | | G | G | | |
| NC_011770 | 5999260 | G | C | 67 | 0 | 2 | 0 | NON_SYNONYMOU | MODERAT | MISSENSE | aGt/aCt | S232T | PLES_54231 | C | C | | | | | | | | | | |
| NC_011770 | 5999278 | A | G | 66 | 0 | 6 | 0 | NON_SYNONYMOU | MODERAT | MISSENSE | aAc/aGc | N238S | PLES_54231 | G | G | | | G | G | | | G | G | | |
| NC_011770 | 6007433 | A | AGCGG( | 55 | 0 | 1 | 0 | CODON_CHANGE_ | MODERATE | | gct/gcCCC( | A422APA | PLES_54271 | | | | | | | | | . | AGCGGGG | | |
| NC_011770 | 6007483 | C | T | 56 | 0 | 2 | 0 | NON_SYNONYMOU | MODERAT | MISSENSE | Gca/Aca | A406T | PLES_54271 | | | | | | | | | T | T | | |
| NC_011770 | 6007489 | T | C | 51 | 0 | 2 | 0 | NON_SYNONYMOU | MODERAT | MISSENSE | Acc/Gcc | T404A | PLES_54271 | | | | | | | | | C | C | | |
| NC_011770 | 6018188 | T | C | 55 | 0 | 4 | 0 | NON_SYNONYMOU | MODERAT | MISSENSE | aAc/aGc | N410S | PLES_54361 | C | C | | | | | | | C | C | | |
| NC_011770 | 6021016 | G | A | 71 | 0 | 6 | 0 | NON_SYNONYMOU | MODERAT | MISSENSE | gCc/gTc | A236V | PLES_54381 | | | A | A | A | A | | | A | A | | |
| NC_011770 | 6021679 | G | A | 76 | 0 | 2 | 0 | NON_SYNONYMOU | MODERAT | MISSENSE | gCg/gTg | A15V | PLES_54381 | | | | | | | | | A | A | | |
| NC_011770 | 6034899 | A | G | 70 | 0 | 1 | 0 | NON_SYNONYMOU | MODERAT | MISSENSE | cAc/cGc | H42R | PLES_54491 | | | | | | | G | | | | | |
| NC_011770 | 6038236 | G | A | 78 | 0 | 2 | 0 | NON_SYNONYMOU | MODERAT | MISSENSE | Gcc/Acc | A36T | PLES_54541 | | | | | | | | | | | A | A |
| NC_011770 | 6038278 | G | A | 79 | 0 | 4 | 0 | NON_SYNONYMOU | MODERAT | MISSENSE | Gtc/Atc | V50I | PLES_54541 | | | | | | | A | A | A | A | | |
| NC_011770 | 6038318 | C | G | 79 | 0 | 2 | 0 | NON_SYNONYMOU | MODERAT | MISSENSE | gCg/gGg | A63G | PLES_54541 | G | G | | | | | | | | | | |
| NC_011770 | 6043415 | T | C | 72 | 0 | 2 | 0 | NON_SYNONYMOU | MODERAT | MISSENSE | Ttc/Ctc | F235L | PLES_54611 | | | | | | | C | C | | | | |
| NC_011770 | 6043935 | G | A | 72 | 0 | 6 | 0 | NON_SYNONYMOU | MODERAT | MISSENSE | Gac/Aac | D102N | PLES_54621 | A | A | A | A | A | A | | | | | | |
| NC_011770 | 6043959 | A | G | 72 | 0 | 10 | 0 | NON_SYNONYMOU | MODERAT | MISSENSE | Acc/Gcc | T110A | PLES_54621 | G | G | G | G | G | G | | | G | G | G | G |
| NC_011770 | 6044080 | G | A | 76 | 0 | 4 | 0 | NON_SYNONYMOU | MODERAT | MISSENSE | cGc/cAc | R150H | PLES_54621 | | | A | A | A | A | | | | | | |
| NC_011770 | 6045763 | T | C | 82 | 0 | 4 | 0 | NON_SYNONYMOU | MODERAT | MISSENSE | Tac/Cac | Y27H | PLES_54631 | | | C | C | C | C | | | | | | |
| NC_011770 | 6047516 | T | A | 81 | 0 | 2 | 0 | NON_SYNONYMOU | MODERAT | MISSENSE | Acc/Tcc | T110S | PLES_54651 | | | | | | | | | | | A | A |
| NC_011770 | 6054598 | G | A | 81 | 0 | 4 | 0 | NON_SYNONYMOU | MODERAT | MISSENSE | Cat/Tat | H123Y | PLES_54701 | | | A | A | A | A | | | | | | |
| NC_011770 | 6054685 | G | A | 80 | 0 | 2 | 0 | NON_SYNONYMOU | MODERAT | MISSENSE | Cgc/Tgc | R94C | PLES_54701 | A | A | | | | | | | | | | |
| NC_011770 | 6054948 | G | A | 78 | 0 | 2 | 0 | NON_SYNONYMOU | MODERAT | MISSENSE | cCg/cTg | P6L | PLES_54701 | A | A | | | | | | | | | | |
| NC_011770 | 6056072 | T | C | 91 | 0 | 1 | 0 | NON_SYNONYMOU | MODERAT | MISSENSE | aAc/aGc | N152S | PLES_54721 | | C | | | | | | | | | | |
| NC_011770 | 6056487 | A | C | 88 | 0 | 2 | 0 | NON_SYNONYMOU | MODERAT | MISSENSE | Tcg/Gcg | S14A | PLES_54721 | C | C | | | | | | | | | | |
| NC_011770 | 6056519 | G | A | 83 | 0 | 10 | 0 | NON_SYNONYMOU | MODERAT | MISSENSE | aCc/aTc | T3I | PLES_54721 | A | A | A | A | A | A | A | A | | | A | A |
| NC_011770 | 6056909 | G | C | 82 | 0 | 4 | 0 | NON_SYNONYMOU | MODERAT | MISSENSE | aCc/aGc | T4S | PLES_54731 | | | C | C | C | C | | | | | | |
| NC_011770 | 6057363 | G | A | 75 | 0 | 1 | 0 | NON_SYNONYMOU | MODERAT | MISSENSE | cCg/cTg | P278L | PLES_54741 | | | | A | | | | | | | | |
| NC_011770 | 6057657 | T | G | 79 | 0 | 12 | 0 | NON_SYNONYMOU | MODERAT | MISSENSE | gAc/gCc | D180A | PLES_54741 | G | G | G | G | G | G | G | G | G | G | G | G |
| NC_011770 | 6057678 | T | G | 78 | 0 | 12 | 0 | NON_SYNONYMOU | MODERAT | MISSENSE | gAg/gCg | E173A | PLES_54741 | G | G | G | G | G | G | G | G | G | G | G | G |
| NC_011770 | 6057681 | C | A | 76 | 0 | 12 | 0 | NON_SYNONYMOU | MODERAT | MISSENSE | cGc/cTc | R172L | PLES_54741 | A | A | A | A | A | A | A | A | A | A | A | A |
| NC_011770 | 6057697 | A | T | 79 | 0 | 12 | 0 | NON_SYNONYMOU | MODERAT | MISSENSE | Tcc/Acc | S167T | PLES_54741 | T | T | T | T | T | T | T | T | T | T | T | T |
| NC_011770 | 6057706 | A | G | 81 | 0 | 2 | 0 | NON_SYNONYMOU | MODERAT | MISSENSE | Ttc/Ctc | F164L | PLES_54741 | G | G | | | | | | | | | | |
| NC_011770 | 6057802 | C | T | 80 | 0 | 2 | 0 | NON_SYNONYMOU | MODERAT | MISSENSE | Ggc/Agc | G132S | PLES_54741 | T | T | | | | | | | | | | |
| NC_011770 | 6057828 | G | T | 79 | 0 | 6 | 0 | NON_SYNONYMOU | MODERAT | MISSENSE | gCc/gAc | A123D | PLES_54741 | | | T | T | T | T | | | | | T | T |
| NC_011770 | 6057886 | A | G | 79 | 0 | 12 | 0 | NON_SYNONYMOU | MODERAT | MISSENSE | Tgc/Cgc | C104R | PLES_54741 | G | G | G | G | G | G | G | G | G | G | G | G |

| | | | | | | | | | | | | | | | | | | | | | | | | | |
|---|---|---|---|---|---|---|---|---|---|---|---|---|---|---|---|---|---|---|---|---|---|---|---|---|---|
| NC_011770 | 6058299 | A | C | 87 | 0 | 4 | 0 | NON_SYNONYMOU | MODERAT | MISSENSE | Atc/Ctc | I14L | PLES_54751 | | | | | | | C | C | C | C | | |
| NC_011770 | 6059136 | A | C | 58 | 0 | 2 | 0 | NON_SYNONYMOU | MODERAT | MISSENSE | Acc/Ccc | T293P | PLES_54751 | | | | | | | | | C | C | | |
| NC_011770 | 6059175 | A | G | 63 | 0 | 12 | 0 | NON_SYNONYMOU | MODERAT | MISSENSE | Acc/Gcc | T306A | PLES_54751 | G | G | G | G | G | G | G | G | G | G | G | G |
| NC_011770 | 6059654 | CT | C | 81 | 0 | 2 | 0 | FRAME_SHIFT | HIGH | | aag/ | K89 | PLES_54761 | | | | | | | | | | | C | C |
| NC_011770 | 6059302 | G | A | 57 | 0 | 1 | 0 | NON_SYNONYMOU | MODERAT | MISSENSE | Cca/Tca | P207S | PLES_54761 | | | | | | | A | | | | | |
| NC_011770 | 6059304 | T | C | 56 | 0 | 8 | 0 | NON_SYNONYMOU | MODERAT | MISSENSE | cAc/cGc | H206R | PLES_54761 | | | C | C | C | C | C | C | C | C | | |
| NC_011770 | 6059310 | G | A | 55 | 0 | 8 | 0 | NON_SYNONYMOU | MODERAT | MISSENSE | gCg/gTg | A204V | PLES_54761 | | | A | A | A | A | A | A | A | A | | |
| NC_011770 | 6059347 | A | G | 61 | 0 | 10 | 0 | NON_SYNONYMOU | MODERAT | MISSENSE | Tat/Cat | Y192H | PLES_54761 | G | G | G | G | G | G | G | G | G | G | | |
| NC_011770 | 6059353 | C | T | 62 | 0 | 2 | 0 | NON_SYNONYMOU | MODERAT | MISSENSE | Gag/Aag | E190K | PLES_54761 | | | | | | | T | T | | | | |
| NC_011770 | 6059362 | T | G | 59 | 0 | 2 | 0 | NON_SYNONYMOU | MODERAT | MISSENSE | Aat/Cat | N187H | PLES_54761 | | | | | | | | | G | G | | |
| NC_011770 | 6059421 | C | T | 41 | 2 | 8 | 0 | NON_SYNONYMOU | MODERAT | MISSENSE | cGa/cAa | R167Q | PLES_54761 | T | T | T | T | T | T | . | . | T | T | | |
| NC_011770 | 6059529 | A | G | 73 | 0 | 10 | 0 | NON_SYNONYMOU | MODERAT | MISSENSE | gTg/gCg | V131A | PLES_54761 | G | G | G | G | G | G | G | G | G | G | | |
| NC_011770 | 6059541 | A | C | 78 | 0 | 2 | 0 | NON_SYNONYMOU | MODERAT | MISSENSE | gTt/gGt | V127G | PLES_54761 | | | | | | | C | C | | | | |
| NC_011770 | 6059598 | C | T | 83 | 0 | 2 | 0 | NON_SYNONYMOU | MODERAT | MISSENSE | aGt/aAt | S108N | PLES_54761 | T | T | | | | | | | | | | |
| NC_011770 | 6059819 | A | C | 127 | 0 | 2 | 0 | NON_SYNONYMOU | MODERAT | MISSENSE | gaT/gaG | D34E | PLES_54761 | C | C | | | | | | | | | | |
| NC_011770 | 6060168 | A | G | 31 | 6 | 2 | 0 | NON_SYNONYMOU | MODERAT | MISSENSE | Tcg/Ccg | S271P | PLES_54771 | . | . | . | . | . | . | | | G | G | | |
| NC_011770 | 6060267 | A | C | 58 | 2 | 6 | 0 | NON_SYNONYMOU | MODERAT | MISSENSE | Tcc/Gcc | S238A | PLES_54771 | . | . | C | C | C | C | | | C | C | | |
| NC_011770 | 6060273 | G | T | 59 | 2 | 2 | 0 | NON_SYNONYMOU | MODERAT | MISSENSE | Caa/Aaa | Q236K | PLES_54771 | . | . | | | | | | | T | T | | |
| NC_011770 | 6060299 | T | C | 65 | 2 | 2 | 0 | NON_SYNONYMOU | MODERAT | MISSENSE | cAc/cGc | H227R | PLES_54771 | . | . | | | | | | | C | C | | |
| NC_011770 | 6060305 | C | T | 65 | 2 | 8 | 0 | NON_SYNONYMOU | MODERAT | MISSENSE | aGg/aAg | R225K | PLES_54771 | . | . | T | T | T | T | | | T | T | T | T |
| NC_011770 | 6060348 | T | C | 47 | 6 | 2 | 0 | NON_SYNONYMOU | MODERAT | MISSENSE | Act/Gct | T211A | PLES_54771 | . | . | . | . | . | . | | | C | C | | |
| NC_011770 | 6060387 | T | C | 39 | 8 | 2 | 0 | NON_SYNONYMOU | MODERAT | MISSENSE | Aat/Gat | N198D | PLES_54771 | . | . | . | . | . | . | | | . | . | C | C |
| NC_011770 | 6060656 | C | T | 30 | 8 | 2 | 0 | NON_SYNONYMOU | MODERAT | MISSENSE | aGt/aAt | S108N | PLES_54771 | . | . | . | . | . | . | | | . | . | T | T |
| NC_011770 | 6060687 | C | T | 32 | 8 | 2 | 0 | NON_SYNONYMOU | MODERAT | MISSENSE | Ggt/Agt | G98S | PLES_54771 | . | . | . | . | . | . | T | T | . | . | | |
| NC_011770 | 6061897 | GA | G | 16 | 10 | 1 | 0 | FRAME_SHIFT | HIGH | | tct/ | S5 | PLES_54781 | . | . | . | . | . | . | G | | . | . | . | . |
| NC_011770 | 6061136 | T | G | 47 | 2 | 10 | 0 | NON_SYNONYMOU | MODERAT | MISSENSE | Acc/Ccc | T259P | PLES_54781 | . | . | G | G | G | G | G | G | G | G | G | G |
| NC_011770 | 6061226 | T | C | 51 | 2 | 4 | 0 | NON_SYNONYMOU | MODERAT | MISSENSE | Agg/Ggg | R229G | PLES_54781 | . | . | C | C | C | C | | | | | | |
| NC_011770 | 6061296 | T | G | 51 | 2 | 10 | 0 | NON_SYNONYMOU | MODERAT | MISSENSE | agA/agC | R205S | PLES_54781 | . | . | G | G | G | G | G | G | G | G | G | G |
| NC_011770 | 6061298 | T | C | 51 | 2 | 2 | 0 | NON_SYNONYMOU | MODERAT | MISSENSE | Aga/Gga | R205G | PLES_54781 | . | . | | | | | | | C | C | | |
| NC_011770 | 6061319 | G | T | 48 | 2 | 4 | 0 | NON_SYNONYMOU | MODERAT | MISSENSE | Cat/Aat | H198N | PLES_54781 | . | . | | | | | | | T | T | T | T |
| NC_011770 | 6061391 | T | C | 47 | 2 | 4 | 0 | NON_SYNONYMOU | MODERAT | MISSENSE | Acc/Gcc | T174A | PLES_54781 | . | . | | | | | | | C | C | C | C |
| NC_011770 | 6061396 | G | A | 44 | 2 | 4 | 0 | NON_SYNONYMOU | MODERAT | MISSENSE | gCg/gTg | A172V | PLES_54781 | . | . | A | A | A | A | | | | | | |
| NC_011770 | 6061400 | G | C | 41 | 2 | 4 | 0 | NON_SYNONYMOU | MODERAT | MISSENSE | Ctg/Gtg | L171V | PLES_54781 | . | . | C | C | C | C | | | | | | |
| NC_011770 | 6061465 | T | C | 39 | 2 | 2 | 0 | NON_SYNONYMOU | MODERAT | MISSENSE | aAc/aGc | N149S | PLES_54781 | . | . | | | | | | | | | C | C |
| NC_011770 | 6061522 | A | C | 39 | 2 | 4 | 0 | NON_SYNONYMOU | MODERAT | MISSENSE | gTt/gGt | V130G | PLES_54781 | . | . | C | C | C | C | | | | | | |
| NC_011770 | 6061947 | T | C | 22 | 8 | 2 | 0 | NON_SYNONYMOU | MODERAT | MISSENSE | atA/atG | I735M | PLES_54791 | . | . | . | . | . | . | | | . | . | C | C |
| NC_011770 | 6061949 | T | C | 23 | 8 | 2 | 0 | NON_SYNONYMOU | MODERAT | MISSENSE | Ata/Gta | I735V | PLES_54791 | . | . | . | . | . | . | | | . | . | C | C |
| NC_011770 | 6062135 | T | C | 63 | 0 | 2 | 0 | NON_SYNONYMOU | MODERAT | MISSENSE | Att/Gtt | I673V | PLES_54791 | | | | | | | | | | | C | C |
| NC_011770 | 6062153 | C | T | 64 | 0 | 2 | 0 | NON_SYNONYMOU | MODERAT | MISSENSE | Ggc/Agc | G667S | PLES_54791 | T | T | | | | | | | | | | |
| NC_011770 | 6062260 | G | A | 65 | 0 | 2 | 0 | NON_SYNONYMOU | MODERAT | MISSENSE | cCg/cTg | P631L | PLES_54791 | | | | | | | | | A | A | | |
| NC_011770 | 6062377 | G | A | 67 | 0 | 8 | 0 | NON_SYNONYMOU | MODERAT | MISSENSE | gCc/gTc | A592V | PLES_54791 | | | A | A | A | A | A | A | | | A | A |
| NC_011770 | 6062660 | C | T | 82 | 0 | 2 | 0 | NON_SYNONYMOU | MODERAT | MISSENSE | Ggc/Agc | G498S | PLES_54791 | | | | | | | T | T | | | | |
| NC_011770 | 6062950 | A | C | 76 | 0 | 4 | 0 | NON_SYNONYMOU | MODERAT | MISSENSE | cTg/cGg | L401R | PLES_54791 | | | C | C | C | C | | | | | | |
| NC_011770 | 6063963 | G | C | 61 | 0 | 2 | 0 | NON_SYNONYMOU | MODERAT | MISSENSE | caC/caG | H63Q | PLES_54791 | | | | | | | | | C | C | | |
| NC_011770 | 6064160 | C | G | 60 | 0 | 2 | 0 | NON_SYNONYMOU | MODERAT | MISSENSE | gaG/gaC | E787D | PLES_54801 | G | G | | | | | | | | | | |
| NC_011770 | 6064287 | A | G | 60 | 0 | 12 | 0 | NON_SYNONYMOU | MODERAT | MISSENSE | gTc/gCc | V745A | PLES_54801 | G | G | G | G | G | G | G | G | G | G | G | G |
| NC_011770 | 6064305 | C | T | 60 | 0 | 8 | 0 | NON_SYNONYMOU | MODERAT | MISSENSE | cGg/cAg | R739Q | PLES_54801 | T | T | | | | | T | T | T | T | T | T |
| NC_011770 | 6064311 | A | C | 59 | 0 | 2 | 0 | NON_SYNONYMOU | MODERAT | MISSENSE | gTc/gGc | V737G | PLES_54801 | C | C | | | | | | | | | | |
| NC_011770 | 6064315 | T | C | 64 | 0 | 2 | 0 | NON_SYNONYMOU | MODERAT | MISSENSE | Atg/Gtg | M736V | PLES_54801 | C | C | | | | | | | | | | |
| NC_011770 | 6064426 | G | A | 62 | 0 | 12 | 0 | NON_SYNONYMOU | MODERAT | MISSENSE | Ctc/Ttc | L699F | PLES_54801 | A | A | A | A | A | A | A | A | A | A | A | A |
| NC_011770 | 6064471 | C | G | 64 | 0 | 6 | 0 | NON_SYNONYMOU | MODERAT | MISSENSE | Gca/Cca | A684P | PLES_54801 | G | G | | | | | G | G | | | G | G |
| NC_011770 | 6064525 | T | C | 66 | 0 | 12 | 0 | NON_SYNONYMOU | MODERAT | MISSENSE | Agc/Ggc | S666G | PLES_54801 | C | C | C | C | C | C | C | C | C | C | C | C |
| NC_011770 | 6064528 | T | C | 64 | 0 | 4 | 0 | NON_SYNONYMOU | MODERAT | MISSENSE | Atc/Gtc | I665V | PLES_54801 | C | C | | | | | | | | | C | C |
| NC_011770 | 6064680 | C | A | 67 | 0 | 2 | 0 | NON_SYNONYMOU | MODERAT | MISSENSE | gGc/gTc | G614V | PLES_54801 | | | | | | | | | A | A | | |
| NC_011770 | 6064840 | A | G | 68 | 0 | 12 | 0 | NON_SYNONYMOU | MODERAT | MISSENSE | Tac/Cac | Y561H | PLES_54801 | G | G | G | G | G | G | G | G | G | G | G | G |
| NC_011770 | 6064845 | G | A | 67 | 0 | 2 | 0 | NON_SYNONYMOU | MODERAT | MISSENSE | gCc/gTc | A559V | PLES_54801 | | | | | | | A | A | | | | |
| NC_011770 | 6065125 | C | G | 45 | 0 | 4 | 0 | NON_SYNONYMOU | MODERAT | MISSENSE | Gtg/Ctg | V466L | PLES_54801 | | | G | G | G | G | | | | | | |
| NC_011770 | 6065401 | C | G | 63 | 0 | 2 | 0 | NON_SYNONYMOU | MODERAT | MISSENSE | Gtg/Ctg | V374L | PLES_54801 | G | G | | | | | | | | | | |
| NC_011770 | 6065403 | C | T | 62 | 0 | 4 | 0 | NON_SYNONYMOU | MODERAT | MISSENSE | cGc/cAc | R373H | PLES_54801 | | | T | T | T | T | | | | | | |
| NC_011770 | 6065409 | C | T | 63 | 0 | 6 | 0 | NON_SYNONYMOU | MODERAT | MISSENSE | aGg/aAg | R371K | PLES_54801 | | | T | T | T | T | | | | | T | T |
| NC_011770 | 6065463 | C | T | 66 | 0 | 6 | 0 | NON_SYNONYMOU | MODERAT | MISSENSE | cGt/cAt | R353H | PLES_54801 | | | T | T | T | T | | | T | T | | |
| NC_011770 | 6065493 | C | T | 68 | 0 | 6 | 0 | NON_SYNONYMOU | MODERAT | MISSENSE | cGg/cAg | R343Q | PLES_54801 | | | T | T | T | T | | | T | T | | |

| | | | | | | | | | | | | | | | | | | | | | | | | | |
|---|---|---|---|---|---|---|---|---|---|---|---|---|---|---|---|---|---|---|---|---|---|---|---|---|---|
| NC_011770 | 6065593 | C | T | 74 | 0 | 2 | 0 | NON_SYNONYMOU | MODERAT | MISSENSE | Ggc/Agc | G310S | PLES_54801 | | | | | | | | | T | T | | |
| NC_011770 | 6065637 | A | G | 79 | 0 | 12 | 0 | NON_SYNONYMOU | MODERAT | MISSENSE | gTg/gCg | V295A | PLES_54801 | G | G | G | G | G | G | G | G | G | G | G | G |
| NC_011770 | 6065778 | A | C | 79 | 0 | 10 | 0 | NON_SYNONYMOU | MODERAT | MISSENSE | gTg/gGg | V248G | PLES_54801 | C | C | C | C | C | C | C | C | | | C | C |
| NC_011770 | 6065781 | A | G | 78 | 0 | 10 | 0 | NON_SYNONYMOU | MODERAT | MISSENSE | cTc/cCc | L247P | PLES_54801 | G | G | G | G | G | G | G | G | | | G | G |
| NC_011770 | 6065797 | T | C | 81 | 0 | 10 | 0 | NON_SYNONYMOU | MODERAT | MISSENSE | Acc/Gcc | T242A | PLES_54801 | C | C | C | C | C | C | C | C | | | C | C |
| NC_011770 | 6066115 | C | G | 91 | 0 | 12 | 0 | NON_SYNONYMOU | MODERAT | MISSENSE | Gtg/Ctg | V136L | PLES_54801 | G | G | G | G | G | G | G | G | G | G | G | G |
| NC_011770 | 6066319 | G | C | 74 | 0 | 4 | 0 | NON_SYNONYMOU | MODERAT | MISSENSE | Caa/Gaa | Q68E | PLES_54801 | | | C | C | C | C | | | | | | |
| NC_011770 | 6066460 | A | G | 78 | 0 | 2 | 0 | NON_SYNONYMOU | MODERAT | MISSENSE | Ttc/Ctc | F21L | PLES_54801 | | | | | | | G | G | | | | |
| NC_011770 | 6069618 | C | A | 71 | 0 | 4 | 0 | NON_SYNONYMOU | MODERAT | MISSENSE | Gcg/Tcg | A235S | PLES_54831 | | | | | | | A | A | | | A | A |
| NC_011770 | 6070710 | G | T | 86 | 0 | 2 | 0 | NON_SYNONYMOU | MODERAT | MISSENSE | caC/caA | H146Q | PLES_54841 | | | | | | | | | T | T | | |
| NC_011770 | 6070988 | T | A | 89 | 0 | 4 | 0 | NON_SYNONYMOU | MODERAT | MISSENSE | Acc/Tcc | T54S | PLES_54841 | | | | | | | A | A | A | A | | |
| NC_011770 | 6071041 | C | G | 91 | 0 | 6 | 0 | NON_SYNONYMOU | MODERAT | MISSENSE | gGc/gCc | G36A | PLES_54841 | | | | | | | G | G | G | G | G | G |
| NC_011770 | 6073161 | A | C | 43 | 0 | 2 | 0 | NON_SYNONYMOU | MODERAT | MISSENSE | gTc/gGc | V450G | PLES_54871 | | | | | | | C | C | | | | |
| NC_011770 | 6073189 | A | C | 42 | 0 | 2 | 0 | NON_SYNONYMOU | MODERAT | MISSENSE | Ttg/Gtg | L441V | PLES_54871 | | | | | | | | | C | C | | |
| NC_011770 | 6074362 | C | T | 66 | 0 | 10 | 0 | NON_SYNONYMOU | MODERAT | MISSENSE | Gtc/Atc | V50I | PLES_54871 | T | T | T | T | T | T | T | T | T | T | | |
| NC_011770 | 6077528 | GC | G | 84 | 0 | 1 | 0 | FRAME_SHIFT | HIGH | | gcc/ | A38 | PLES_54891 | | | | G | | | | | | | | |
| NC_011770 | 6076217 | C | T | 67 | 0 | 4 | 0 | NON_SYNONYMOU | MODERAT | MISSENSE | cGt/cAt | R475H | PLES_54891 | T | T | | | | | T | T | | | | |
| NC_011770 | 6076236 | C | T | 69 | 0 | 2 | 0 | NON_SYNONYMOU | MODERAT | MISSENSE | Gat/Aat | D469N | PLES_54891 | | | | | | | | | | | T | T |
| NC_011770 | 6076467 | C | A | 85 | 0 | 4 | 0 | NON_SYNONYMOU | MODERAT | MISSENSE | Gac/Tac | D392Y | PLES_54891 | | | A | A | A | A | | | | | | |
| NC_011770 | 6076482 | G | C | 83 | 0 | 12 | 0 | NON_SYNONYMOU | MODERAT | MISSENSE | Ccg/Gcg | P387A | PLES_54891 | C | C | C | C | C | C | C | C | C | C | C | C |
| NC_011770 | 6076673 | G | T | 88 | 0 | 2 | 0 | NON_SYNONYMOU | MODERAT | MISSENSE | aCc/aAc | T323N | PLES_54891 | | | | | | | | | T | T | | |
| NC_011770 | 6077226 | T | C | 84 | 0 | 10 | 0 | NON_SYNONYMOU | MODERAT | MISSENSE | Acc/Gcc | T139A | PLES_54891 | | | C | C | C | C | C | C | C | C | C | C |
| NC_011770 | 6080037 | T | A | 80 | 0 | 4 | 0 | NON_SYNONYMOU | MODERAT | MISSENSE | Acc/Tcc | T172S | PLES_54911 | | | A | A | A | A | | | | | | |
| NC_011770 | 6080114 | T | C | 88 | 0 | 2 | 0 | NON_SYNONYMOU | MODERAT | MISSENSE | cAc/cGc | H146R | PLES_54911 | | | | | | | C | C | | | | |
| NC_011770 | 6080195 | T | C | 85 | 0 | 2 | 0 | NON_SYNONYMOU | MODERAT | MISSENSE | cAc/cGc | H119R | PLES_54911 | | | | | | | | | | | C | C |
| NC_011770 | 6080421 | T | C | 84 | 0 | 2 | 0 | NON_SYNONYMOU | MODERAT | MISSENSE | Acc/Gcc | T44A | PLES_54911 | | | | | | | | | | | C | C |
| NC_011770 | 6080430 | A | G | 82 | 0 | 2 | 0 | NON_SYNONYMOU | MODERAT | MISSENSE | Tcg/Ccg | S41P | PLES_54911 | | | | | | | | | | | G | G |
| NC_011770 | 6080508 | C | T | 88 | 0 | 2 | 0 | NON_SYNONYMOU | MODERAT | MISSENSE | Gag/Aag | E15K | PLES_54911 | | | | | | | | | T | T | | |
| NC_011770 | 6080537 | G | A | 85 | 0 | 2 | 0 | NON_SYNONYMOU | MODERAT | MISSENSE | cCg/cTg | P5L | PLES_54911 | A | A | | | | | | | | | | |
| NC_011770 | 6080687 | C | T | 85 | 0 | 2 | 0 | NON_SYNONYMOU | MODERAT | MISSENSE | aGc/aAc | S274N | PLES_54921 | | | | | | | | | | | T | T |
| NC_011770 | 6080788 | G | C | 70 | 0 | 8 | 0 | NON_SYNONYMOU | MODERAT | MISSENSE | gaC/gaG | D240E | PLES_54921 | C | C | C | C | C | C | | | | | C | C |
| NC_011770 | 6080894 | A | C | 66 | 0 | 2 | 0 | NON_SYNONYMOU | MODERAT | MISSENSE | cTg/cGg | L205R | PLES_54921 | | | | | | | | | | | C | C |
| NC_011770 | 6080993 | C | T | 66 | 0 | 2 | 0 | NON_SYNONYMOU | MODERAT | MISSENSE | cGg/cAg | R172Q | PLES_54921 | | | | | | | | | | | T | T |
| NC_011770 | 6081182 | A | C | 78 | 0 | 2 | 0 | NON_SYNONYMOU | MODERAT | MISSENSE | cTc/cGc | L109R | PLES_54921 | | | | | | | | | | | C | C |
| NC_011770 | 6081255 | T | C | 79 | 0 | 4 | 0 | NON_SYNONYMOU | MODERAT | MISSENSE | Atg/Gtg | M85V | PLES_54921 | C | C | | | | | | | | | C | C |
| NC_011770 | 6081365 | C | T | 82 | 0 | 2 | 0 | NON_SYNONYMOU | MODERAT | MISSENSE | cGc/cAc | R48H | PLES_54921 | | | | | | | | | | | T | T |
| NC_011770 | 6081368 | T | C | 81 | 0 | 2 | 0 | NON_SYNONYMOU | MODERAT | MISSENSE | gAg/gGg | E47G | PLES_54921 | | | | | | | | | | | C | C |
| NC_011770 | 6081369 | C | G | 83 | 0 | 2 | 0 | NON_SYNONYMOU | MODERAT | MISSENSE | Gag/Cag | E47Q | PLES_54921 | | | | | | | | | | | G | G |
| NC_011770 | 6081414 | T | C | 79 | 0 | 4 | 0 | NON_SYNONYMOU | MODERAT | MISSENSE | Atc/Gtc | I32V | PLES_54921 | C | C | | | | | | | | | C | C |
| NC_011770 | 6081473 | A | T | 79 | 0 | 2 | 0 | NON_SYNONYMOU | MODERAT | MISSENSE | cTg/cAg | L12Q | PLES_54921 | | | | | | | | | | | T | T |
| NC_011770 | 6082119 | T | C | 69 | 0 | 10 | 0 | NON_SYNONYMOU | MODERAT | MISSENSE | Atc/Gtc | I143V | PLES_54931 | C | C | C | C | C | C | | | C | C | C | C |
| NC_011770 | 6082749 | G | A | 82 | 0 | 2 | 0 | NON_SYNONYMOU | MODERAT | MISSENSE | Cat/Tat | H177Y | PLES_54941 | | | | | | | | | | | A | A |
| NC_011770 | 6082767 | C | T | 82 | 0 | 2 | 0 | NON_SYNONYMOU | MODERAT | MISSENSE | Ggg/Agg | G171R | PLES_54941 | | | T | T | | | | | | | | |
| NC_011770 | 6082790 | G | T | 81 | 0 | 2 | 0 | NON_SYNONYMOU | MODERAT | MISSENSE | gCc/gAc | A163D | PLES_54941 | | | | | | | | | | | T | T |
| NC_011770 | 6082838 | T | C | 78 | 0 | 2 | 0 | NON_SYNONYMOU | MODERAT | MISSENSE | gAc/gGc | D147G | PLES_54941 | | | | | | | | | | | C | C |
| NC_011770 | 6082931 | T | C | 79 | 0 | 10 | 0 | NON_SYNONYMOU | MODERAT | MISSENSE | cAg/cGg | Q116R | PLES_54941 | C | C | C | C | C | C | | | C | C | C | C |
| NC_011770 | 6083268 | G | A | 82 | 0 | 4 | 0 | NON_SYNONYMOU | MODERAT | MISSENSE | Ctc/Ttc | L4F | PLES_54941 | | | | | | | A | A | | | A | A |
| NC_011770 | 6084661 | G | A | 85 | 0 | 1 | 0 | NON_SYNONYMOU | MODERAT | MISSENSE | Ggc/Agc | G179S | PLES_54961 | | A | | | | | | | | | | |
| NC_011770 | 6084218 | A | G | 85 | 0 | 10 | 0 | NON_SYNONYMOU | MODERAT | MISSENSE | gAg/gGg | E31G | PLES_54961 | G | G | G | G | G | G | G | G | G | G | | |
| NC_011770 | 6084430 | T | C | 73 | 0 | 2 | 0 | NON_SYNONYMOU | MODERAT | MISSENSE | Tgc/Cgc | C102R | PLES_54961 | | | | | | | | | | | C | C |
| NC_011770 | 6084568 | A | G | 83 | 0 | 8 | 0 | NON_SYNONYMOU | MODERAT | MISSENSE | Aca/Gca | T148A | PLES_54961 | G | G | G | G | G | G | | | | | G | G |
| NC_011770 | 6084857 | T | A | 89 | 0 | 10 | 0 | NON_SYNONYMOU | MODERAT | MISSENSE | cTg/cAg | L244Q | PLES_54961 | A | A | A | A | A | A | | | A | A | A | A |
| NC_011770 | 6087111 | T | C | 78 | 0 | 1 | 0 | NON_SYNONYMOU | MODERAT | MISSENSE | Acc/Gcc | T7A | PLES_54991 | | | | | | | | C | | | | |
| NC_011770 | 6091144 | A | G | 69 | 0 | 4 | 0 | NON_SYNONYMOU | MODERAT | MISSENSE | gTc/gCc | V424A | PLES_55031 | | | G | G | G | G | | | | | | |
| NC_011770 | 6091580 | C | G | 74 | 0 | 2 | 0 | NON_SYNONYMOU | MODERAT | MISSENSE | Gag/Cag | E279Q | PLES_55031 | | | | | | | | | G | G | | |
| NC_011770 | 6091728 | T | A | 67 | 0 | 6 | 0 | NON_SYNONYMOU | MODERAT | MISSENSE | caA/caT | Q229H | PLES_55031 | | | A | A | A | A | | | | | A | A |
| NC_011770 | 6092177 | C | T | 65 | 0 | 2 | 0 | NON_SYNONYMOU | MODERAT | MISSENSE | Gcc/Acc | A80T | PLES_55031 | | | | | | | T | T | | | | |
| NC_011770 | 6092256 | T | C | 67 | 0 | 12 | 0 | NON_SYNONYMOU | MODERAT | MISSENSE | atA/atG | I53M | PLES_55031 | C | C | C | C | C | C | C | C | C | C | C | C |
| NC_011770 | 6095561 | ACGGCG | A | 50 | 0 | 2 | 0 | FRAME_SHIFT | HIGH | | cgccgt/ | RR151 | PLES_55041 | | | A | A | | | | | | | | |
| NC_011770 | 6095626 | CG | C | 47 | 0 | 1 | 0 | FRAME_SHIFT | HIGH | | ccc/ | P130 | PLES_55041 | | | | | | | C | | | | | |
| NC_011770 | 6094012 | C | T | 78 | 0 | 1 | 0 | NON_SYNONYMOU | MODERAT | MISSENSE | Gcc/Acc | A669T | PLES_55041 | | | | T | | | | | | | | |
| NC_011770 | 6092698 | C | T | 65 | 0 | 6 | 0 | NON_SYNONYMOU | MODERAT | MISSENSE | Gcc/Acc | A1107T | PLES_55041 | | | T | T | T | T | T | T | | | | |

| | | | | | | | | | | | | | | | | | | | | | | | | | |
|---|---|---|---|---|---|---|---|---|---|---|---|---|---|---|---|---|---|---|---|---|---|---|---|---|---|
| NC_011770 | 6092950 | G | A | 69 | 0 | 2 | 0 | NON_SYNONYMOU | MODERAT | MISSENSE | Ctc/Ttc | L1023F | PLES_55041 | | | | | | | | | | | A | A |
| NC_011770 | 6093057 | C | A | 74 | 0 | 2 | 0 | NON_SYNONYMOU | MODERAT | MISSENSE | gGg/gTg | G987V | PLES_55041 | | | | | | | | | | | A | A |
| NC_011770 | 6093852 | C | T | 79 | 0 | 6 | 0 | NON_SYNONYMOU | MODERAT | MISSENSE | aGc/aAc | S722N | PLES_55041 | | | T | T | T | T | | | | | T | T |
| NC_011770 | 6094201 | T | C | 86 | 0 | 12 | 0 | NON_SYNONYMOU | MODERAT | MISSENSE | Agc/Ggc | S606G | PLES_55041 | C | C | C | C | C | C | C | C | C | C | C | C |
| NC_011770 | 6094603 | C | T | 94 | 0 | 2 | 0 | NON_SYNONYMOU | MODERAT | MISSENSE | Gtg/Atg | V472M | PLES_55041 | | | | | | | | | T | T | | |
| NC_011770 | 6095559 | G | A | 50 | 0 | 4 | 0 | NON_SYNONYMOU | MODERAT | MISSENSE | gCc/gTc | A153V | PLES_55041 | | | A | A | A | A | | | | | | |
| NC_011770 | 6096400 | G | A | 64 | 0 | 6 | 0 | NON_SYNONYMOU | MODERAT | MISSENSE | Cgc/Tgc | R114C | PLES_55051 | A | A | | | | | A | A | A | A | | |
| NC_011770 | 6096562 | C | T | 72 | 0 | 6 | 0 | NON_SYNONYMOU | MODERAT | MISSENSE | Gcc/Acc | A60T | PLES_55051 | | | T | T | T | T | | | T | T | | |
| NC_011770 | 6102745 | C | T | 104 | 0 | 4 | 0 | NON_SYNONYMOU | MODERAT | MISSENSE | gCc/gTc | A11V | PLES_55101 | | | | | | | | | T | T | T | T |
| NC_011770 | 6102747 | A | G | 102 | 0 | 4 | 0 | NON_SYNONYMOU | MODERAT | MISSENSE | Atc/Gtc | I12V | PLES_55101 | | | | | | | | | G | G | G | G |
| NC_011770 | 6102772 | G | C | 102 | 0 | 2 | 0 | NON_SYNONYMOU | MODERAT | MISSENSE | gGc/gCc | G20A | PLES_55101 | | | | | | | | | C | C | | |
| NC_011770 | 6102822 | G | A | 106 | 0 | 2 | 0 | NON_SYNONYMOU | MODERAT | MISSENSE | Gca/Aca | A37T | PLES_55101 | | | | | | | | | | | A | A |
| NC_011770 | 6103143 | T | G | 105 | 0 | 10 | 0 | NON_SYNONYMOU | MODERAT | MISSENSE | cAg/cCg | Q727P | PLES_55111 | G | G | G | G | G | G | G | G | G | G | | |
| NC_011770 | 6103926 | A | G | 89 | 0 | 1 | 0 | NON_SYNONYMOU | MODERAT | MISSENSE | cTg/cCg | L466P | PLES_55111 | | | | | | | G | | | | | |
| NC_011770 | 6105805 | G | T | 77 | 0 | 2 | 0 | NON_SYNONYMOU | MODERAT | MISSENSE | Gcc/Tcc | A66S | PLES_55121 | | | | | | | | | T | T | | |
| NC_011770 | 6105811 | G | A | 75 | 0 | 2 | 0 | NON_SYNONYMOU | MODERAT | MISSENSE | Gcc/Acc | A68T | PLES_55121 | | | | | | | | | A | A | | |
| NC_011770 | 6105988 | G | A | 83 | 0 | 8 | 0 | NON_SYNONYMOU | MODERAT | MISSENSE | Ggc/Agc | G127S | PLES_55121 | A | A | A | A | A | A | | | | | A | A |
| NC_011770 | 6106495 | T | C | 81 | 0 | 2 | 0 | NON_SYNONYMOU | MODERAT | MISSENSE | gTg/gCg | V123A | PLES_55131 | | | | | | | C | C | | | | |
| NC_011770 | 6106680 | A | G | 85 | 0 | 2 | 0 | NON_SYNONYMOU | MODERAT | MISSENSE | Ata/Gta | I185V | PLES_55131 | | | | | | | G | G | | | | |
| NC_011770 | 6106683 | G | C | 87 | 0 | 2 | 0 | NON_SYNONYMOU | MODERAT | MISSENSE | Gag/Cag | E186Q | PLES_55131 | | | | | | | C | C | | | | |
| NC_011770 | 6110322 | C | A | 72 | 0 | 2 | 0 | NON_SYNONYMOU | MODERAT | MISSENSE | caG/caT | Q98H | PLES_55161 | | | | | | | | | A | A | | |
| NC_011770 | 6114659 | T | C | 72 | 0 | 12 | 0 | NON_SYNONYMOU | MODERAT | MISSENSE | gTa/gCa | V46A | PLES_55221 | C | C | C | C | C | C | C | C | C | C | C | C |
| NC_011770 | 6114793 | A | G | 72 | 0 | 10 | 0 | NON_SYNONYMOU | MODERAT | MISSENSE | Atc/Gtc | I91V | PLES_55221 | | | G | G | G | G | G | G | G | G | G | G |
| NC_011770 | 6114937 | T | C | 75 | 0 | 4 | 0 | NON_SYNONYMOU | MODERAT | MISSENSE | Tac/Cac | Y139H | PLES_55221 | | | | | | | C | C | | | C | C |
| NC_011770 | 6115042 | A | G | 70 | 0 | 6 | 0 | NON_SYNONYMOU | MODERAT | MISSENSE | Atc/Gtc | I174V | PLES_55221 | | | | | | | G | G | G | G | G | G |
| NC_011770 | 6115332 | A | C | 63 | 0 | 6 | 0 | NON_SYNONYMOU | MODERAT | MISSENSE | agA/agC | R270S | PLES_55221 | C | C | C | C | C | C | | | | | | |
| NC_011770 | 6115763 | GA | G | 47 | 0 | 1 | 0 | FRAME_SHIFT | HIGH | | aaa/ | K94 | PLES_55231 | | | | | | | | G | | | | |
| NC_011770 | 6115713 | C | G | 48 | 0 | 2 | 0 | NON_SYNONYMOU | MODERAT | MISSENSE | Cag/Gag | Q75E | PLES_55231 | G | G | | | | | | | | | | |
| NC_011770 | 6115992 | A | G | 74 | 0 | 12 | 0 | NON_SYNONYMOU | MODERAT | MISSENSE | Aac/Gac | N168D | PLES_55231 | G | G | G | G | G | G | G | G | G | G | G | G |
| NC_011770 | 6116214 | G | A | 68 | 0 | 6 | 0 | NON_SYNONYMOU | MODERAT | MISSENSE | Gaa/Aaa | E242K | PLES_55231 | A | A | A | A | A | A | | | | | | |
| NC_011770 | 6116272 | G | A | 61 | 0 | 2 | 0 | NON_SYNONYMOU | MODERAT | MISSENSE | cGt/cAt | R261H | PLES_55231 | | | | | | | | | | | A | A |
| NC_011770 | 6116292 | G | T | 63 | 0 | 2 | 0 | NON_SYNONYMOU | MODERAT | MISSENSE | Gcc/Tcc | A268S | PLES_55231 | | | | | | | T | T | | | | |
| NC_011770 | 6117304 | G | C | 66 | 0 | 10 | 0 | NON_SYNONYMOU | MODERAT | MISSENSE | Gcg/Ccg | A167P | PLES_55241 | | | C | C | C | C | C | C | C | C | C | C |
| NC_011770 | 6118189 | T | C | 51 | 1 | 1 | 0 | NON_SYNONYMOU | MODERAT | MISSENSE | gTc/gCc | V25A | PLES_55251 | | | | | | | | | . | C | | |
| NC_011770 | 6118700 | G | T | 43 | 0 | 2 | 0 | NON_SYNONYMOU | MODERAT | MISSENSE | caG/caT | Q195H | PLES_55251 | | | | | | | T | T | | | | |
| NC_011770 | 6119496 | T | C | 72 | 0 | 1 | 0 | NON_SYNONYMOU | MODERAT | MISSENSE | gTg/gCg | V180A | PLES_55261 | | | | | | | C | | | | | |
| NC_011770 | 6119588 | A | C | 70 | 0 | 12 | 0 | NON_SYNONYMOU | MODERAT | MISSENSE | Agc/Cgc | S211R | PLES_55261 | C | C | C | C | C | C | C | C | C | C | C | C |
| NC_011770 | 6119655 | A | G | 75 | 0 | 4 | 0 | NON_SYNONYMOU | MODERAT | MISSENSE | cAg/cGg | Q233R | PLES_55261 | G | G | | | | | | | G | G | | |
| NC_011770 | 6119784 | G | A | 87 | 0 | 4 | 0 | NON_SYNONYMOU | MODERAT | MISSENSE | aGc/aAc | S276N | PLES_55261 | A | A | | | | | | | A | A | | |
| NC_011770 | 6120434 | G | A | 98 | 0 | 4 | 0 | NON_SYNONYMOU | MODERAT | MISSENSE | Gac/Aac | D493N | PLES_55261 | A | A | | | | | | | | | A | A |
| NC_011770 | 6121953 | T | C | 92 | 0 | 2 | 0 | NON_SYNONYMOU | MODERAT | MISSENSE | Atc/Gtc | I61V | PLES_55281 | | | | | | | | | | | C | C |
| NC_011770 | 6124949 | C | A | 72 | 0 | 10 | 0 | NON_SYNONYMOU | MODERAT | MISSENSE | aaG/aaT | K7N | PLES_55321 | A | A | A | A | A | A | A | A | A | A | | |
| NC_011770 | 6126435 | C | T | 79 | 0 | 2 | 0 | NON_SYNONYMOU | MODERAT | MISSENSE | gCc/gTc | A24V | PLES_55351 | | | | | | | | | | | T | T |
| NC_011770 | 6126515 | C | A | 63 | 0 | 12 | 0 | NON_SYNONYMOU | MODERAT | MISSENSE | Ctc/Atc | L51I | PLES_55351 | A | A | A | A | A | A | T | T | A | A | A | A |
| NC_011770 | 6126570 | G | A | 72 | 0 | 10 | 0 | NON_SYNONYMOU | MODERAT | MISSENSE | gGc/gAc | G69D | PLES_55351 | A | A | A | A | A | A | | | A | A | A | A |
| NC_011770 | 6126716 | G | T | 84 | 0 | 2 | 0 | NON_SYNONYMOU | MODERAT | MISSENSE | Gcc/Tcc | A118S | PLES_55351 | | | | | | | T | T | | | | |
| NC_011770 | 6127279 | G | A | 77 | 0 | 2 | 0 | NON_SYNONYMOU | MODERAT | MISSENSE | Ggc/Agc | G83S | PLES_55361 | | | | | | | A | A | | | | |
| NC_011770 | 6127334 | G | A | 76 | 0 | 8 | 0 | NON_SYNONYMOU | MODERAT | MISSENSE | aGg/aAg | R101K | PLES_55361 | | | A | A | A | A | A | A | | | A | A |
| NC_011770 | 6127399 | A | G | 78 | 0 | 2 | 0 | NON_SYNONYMOU | MODERAT | MISSENSE | Acc/Gcc | T123A | PLES_55361 | | | | | | | | | | | G | G |
| NC_011770 | 6127403 | C | A | 77 | 0 | 2 | 0 | NON_SYNONYMOU | MODERAT | MISSENSE | aCg/aAg | T124K | PLES_55361 | | | | | | | | | | | A | A |
| NC_011770 | 6127411 | G | T | 72 | 0 | 2 | 0 | NON_SYNONYMOU | MODERAT | MISSENSE | Gcg/Tcg | A127S | PLES_55361 | | | | | | | T | T | | | | |
| NC_011770 | 6127529 | T | C | 67 | 0 | 2 | 0 | NON_SYNONYMOU | MODERAT | MISSENSE | cTg/cCg | L166P | PLES_55361 | | | | | | | | | | | C | C |
| NC_011770 | 6127633 | G | A | 73 | 0 | 2 | 0 | NON_SYNONYMOU | MODERAT | MISSENSE | Gtc/Atc | V201I | PLES_55361 | | | | | | | | | | | A | A |
| NC_011770 | 6127688 | A | C | 69 | 0 | 2 | 0 | NON_SYNONYMOU | MODERAT | MISSENSE | gAt/gCt | D219A | PLES_55361 | | | | | | | C | C | | | | |
| NC_011770 | 6127790 | A | G | 65 | 0 | 8 | 0 | NON_SYNONYMOU | MODERAT | MISSENSE | cAc/cGc | H253R | PLES_55361 | G | G | | | | | G | G | G | G | G | G |
| NC_011770 | 6127795 | G | A | 67 | 0 | 2 | 0 | NON_SYNONYMOU | MODERAT | MISSENSE | Gcc/Acc | A255T | PLES_55361 | | | | | | | | | | | A | A |
| NC_011770 | 6127900 | C | G | 60 | 0 | 2 | 0 | NON_SYNONYMOU | MODERAT | MISSENSE | Cgc/Ggc | R290G | PLES_55361 | | | | | | | | | | | G | G |
| NC_011770 | 6127918 | G | A | 60 | 0 | 2 | 0 | NON_SYNONYMOU | MODERAT | MISSENSE | Gcc/Acc | A296T | PLES_55361 | | | | | | | | | | | A | A |
| NC_011770 | 6127919 | C | T | 62 | 0 | 2 | 0 | NON_SYNONYMOU | MODERAT | MISSENSE | gCc/gTc | A296V | PLES_55361 | T | T | | | | | | | | | | |
| NC_011770 | 6127949 | C | A | 57 | 0 | 2 | 0 | NON_SYNONYMOU | MODERAT | MISSENSE | cCt/cAt | P306H | PLES_55361 | | | | | | | A | A | | | | |
| NC_011770 | 6128742 | CA | C | 24 | 0 | 1 | 0 | FRAME_SHIFT | HIGH | | acc/ | T171 | PLES_55371 | | | | | | | | C | | | | |
| NC_011770 | 6128272 | G | C | 50 | 0 | 1 | 0 | NON_SYNONYMOU | MODERAT | MISSENSE | Gga/Cga | G14R | PLES_55371 | | | | C | | | | | | | | |

| | | | | | | | | | | | | | | | | | | | | | | | | | |
|---|---|---|---|---|---|---|---|---|---|---|---|---|---|---|---|---|---|---|---|---|---|---|---|---|---|
| NC_011770 | 6128461 | G | A | 33 | 0 | 2 | 0 | NON_SYNONYMOU | MODERAT | MISSENSE | Gcc/Acc | A77T | PLES_55371 | | | | | | | | | | | A | A |
| NC_011770 | 6128653 | G | T | 14 | 0 | 2 | 0 | NON_SYNONYMOU | MODERAT | MISSENSE | Gcc/Tcc | A141S | PLES_55371 | | | | | | | | | | | T | T |
| NC_011770 | 6128696 | G | C | 18 | 0 | 11 | 0 | NON_SYNONYMOU | MODERAT | MISSENSE | gGc/gCc | G155A | PLES_55371 | . | C | C | C | C | C | C | C | C | C | C | C |
| NC_011770 | 6128723 | C | A | 22 | 0 | 2 | 0 | NON_SYNONYMOU | MODERAT | MISSENSE | cCg/cAg | P164Q | PLES_55371 | | | | | | | A | A | | | | |
| NC_011770 | 6129167 | A | G | 72 | 0 | 2 | 0 | NON_SYNONYMOU | MODERAT | MISSENSE | gAc/gGc | D312G | PLES_55371 | | | | | | | | | | | G | G |
| NC_011770 | 6129175 | A | G | 70 | 0 | 4 | 0 | NON_SYNONYMOU | MODERAT | MISSENSE | Acc/Gcc | T315A | PLES_55371 | | | | | | | G | G | | | G | G |
| NC_011770 | 6129530 | C | A | 79 | 0 | 2 | 0 | NON_SYNONYMOU | MODERAT | MISSENSE | gCc/gAc | A433D | PLES_55371 | | | | | | | | | A | A | | |
| NC_011770 | 6129603 | C | G | 87 | 0 | 2 | 0 | NON_SYNONYMOU | MODERAT | MISSENSE | atC/atG | I457M | PLES_55371 | | | | | | | G | G | | | | |
| NC_011770 | 6129943 | C | T | 90 | 0 | 2 | 0 | NON_SYNONYMOU | MODERAT | MISSENSE | Cat/Tat | H571Y | PLES_55371 | | | | | | | | | | | T | T |
| NC_011770 | 6132484 | A | G | 73 | 0 | 10 | 0 | NON_SYNONYMOU | MODERAT | MISSENSE | Ata/Gta | I198V | PLES_55401 | G | G | G | G | G | G | G | G | G | G | | |
| NC_011770 | 6132496 | G | A | 76 | 0 | 2 | 0 | NON_SYNONYMOU | MODERAT | MISSENSE | Gtg/Atg | V202M | PLES_55401 | | | | | | | | | | | A | A |
| NC_011770 | 6132533 | T | A | 77 | 0 | 4 | 0 | NON_SYNONYMOU | MODERAT | MISSENSE | cTg/cAg | L214Q | PLES_55401 | | | A | A | A | A | | | | | | |
| NC_011770 | 6132824 | C | T | 66 | 0 | 2 | 0 | NON_SYNONYMOU | MODERAT | MISSENSE | gCc/gTc | A311V | PLES_55401 | | | | | | | T | T | | | | |
| NC_011770 | 6133631 | G | A | 84 | 0 | 1 | 0 | NON_SYNONYMOU | MODERAT | MISSENSE | Cgg/Tgg | R110W | PLES_55411 | | | | | | | | | | | A | |
| NC_011770 | 6133250 | T | C | 66 | 0 | 2 | 0 | NON_SYNONYMOU | MODERAT | MISSENSE | Atc/Gtc | I237V | PLES_55411 | | | | | | | C | C | | | | |
| NC_011770 | 6133818 | C | G | 84 | 0 | 12 | 0 | NON_SYNONYMOU | MODERAT | MISSENSE | gaG/gaC | E47D | PLES_55411 | G | G | G | G | G | G | G | G | G | G | G | G |
| NC_011770 | 6134304 | C | T | 93 | 0 | 2 | 0 | NON_SYNONYMOU | MODERAT | MISSENSE | cCa/cTa | P63L | PLES_55421 | | | | | | | | | | | T | T |
| NC_011770 | 6134759 | A | G | 72 | 0 | 2 | 0 | NON_SYNONYMOU | MODERAT | MISSENSE | Aag/Gag | K215E | PLES_55421 | G | G | | | | | | | | | | |
| NC_011770 | 6135146 | G | A | 77 | 0 | 2 | 0 | NON_SYNONYMOU | MODERAT | MISSENSE | cGc/cAc | R31H | PLES_55431 | | | | | | | | | | | A | A |
| NC_011770 | 6137284 | G | A | 93 | 0 | 2 | 0 | NON_SYNONYMOU | MODERAT | MISSENSE | Gtc/Atc | V210I | PLES_55451 | A | A | | | | | | | | | | |
| NC_011770 | 6137511 | T | C | 85 | 0 | 1 | 0 | NON_SYNONYMOU | MODERAT | MISSENSE | Tac/Cac | Y55H | PLES_55461 | | | | | | | | C | | | | |
| NC_011770 | 6138560 | G | A | 71 | 0 | 4 | 0 | NON_SYNONYMOU | MODERAT | MISSENSE | Gcg/Acg | A56T | PLES_55471 | | | | | | | | | A | A | A | A |
| NC_011770 | 6138848 | G | A | 78 | 0 | 2 | 0 | NON_SYNONYMOU | MODERAT | MISSENSE | Gca/Aca | A152T | PLES_55471 | | | | | | | A | A | | | | |
| NC_011770 | 6139962 | G | A | 85 | 0 | 2 | 0 | NON_SYNONYMOU | MODERAT | MISSENSE | Gcg/Acg | A77T | PLES_55481 | | | | | | | A | A | | | | |
| NC_011770 | 6141351 | C | T | 90 | 0 | 1 | 0 | STOP_GAINED | HIGH | NONSENSE | Cag/Tag | Q382* | PLES_55491 | | | | | | | | | | | T | |
| NC_011770 | 6140221 | T | C | 87 | 0 | 2 | 0 | NON_SYNONYMOU | MODERAT | MISSENSE | cTt/cCt | L5P | PLES_55491 | | | | | | | C | C | | | | |
| NC_011770 | 6140265 | G | C | 86 | 0 | 8 | 0 | NON_SYNONYMOU | MODERAT | MISSENSE | Gtg/Ctg | V20L | PLES_55491 | C | C | C | C | C | C | C | C | | | | |
| NC_011770 | 6140314 | G | A | 83 | 0 | 4 | 0 | NON_SYNONYMOU | MODERAT | MISSENSE | cGa/cAa | R36Q | PLES_55491 | | | A | A | A | A | | | | | | |
| NC_011770 | 6140317 | T | C | 83 | 0 | 2 | 0 | NON_SYNONYMOU | MODERAT | MISSENSE | cTg/cCg | L37P | PLES_55491 | | | | | | | C | C | | | | |
| NC_011770 | 6140357 | C | A | 88 | 0 | 2 | 0 | NON_SYNONYMOU | MODERAT | MISSENSE | gaC/gaA | D50E | PLES_55491 | | | | | | | A | A | | | | |
| NC_011770 | 6140379 | A | G | 85 | 0 | 12 | 0 | NON_SYNONYMOU | MODERAT | MISSENSE | Acc/Gcc | T58A | PLES_55491 | G | G | G | G | G | G | G | G | G | G | G | G |
| NC_011770 | 6141042 | A | G | 77 | 0 | 2 | 0 | NON_SYNONYMOU | MODERAT | MISSENSE | Aac/Gac | N279D | PLES_55491 | | | | | | | G | G | | | | |
| NC_011770 | 6141396 | G | A | 88 | 0 | 4 | 0 | NON_SYNONYMOU | MODERAT | MISSENSE | Gac/Aac | D397N | PLES_55491 | | | | | | | | | A | A | A | A |
| NC_011770 | 6142407 | C | A | 93 | 0 | 10 | 0 | NON_SYNONYMOU | MODERAT | MISSENSE | aCt/aAt | T236N | PLES_55501 | | | A | A | A | A | A | A | A | A | A | A |
| NC_011770 | 6142697 | G | A | 90 | 0 | 1 | 0 | NON_SYNONYMOU | MODERAT | MISSENSE | Ggc/Agc | G333S | PLES_55501 | | | | | | | A | | | | | |
| NC_011770 | 6143725 | T | C | 75 | 0 | 1 | 0 | NON_SYNONYMOU | MODERAT | MISSENSE | cTg/cCg | L277P | PLES_55511 | | | | | | | | C | | | | |
| NC_011770 | 6143926 | T | C | 74 | 0 | 1 | 0 | NON_SYNONYMOU | MODERAT | MISSENSE | aTg/aCg | M344T | PLES_55511 | | | | | | | C | | | | | |
| NC_011770 | 6149027 | A | T | 71 | 0 | 2 | 0 | NON_SYNONYMOU | MODERAT | MISSENSE | cAa/cTa | Q258L | PLES_55571 | | | | | | | | | | | T | T |
| NC_011770 | 6149644 | G | C | 78 | 0 | 2 | 0 | NON_SYNONYMOU | MODERAT | MISSENSE | Gtg/Ctg | V464L | PLES_55571 | | | | | | | | | C | C | | |
| NC_011770 | 6149840 | C | T | 66 | 0 | 4 | 0 | NON_SYNONYMOU | MODERAT | MISSENSE | aCc/aTc | T529I | PLES_55571 | | | | | | | | | T | T | T | T |
| NC_011770 | 6152488 | C | T | 114 | 0 | 1 | 0 | NON_SYNONYMOU | MODERAT | MISSENSE | gCc/gTc | A257V | PLES_55591 | | | | | | | T | | | | | |
| NC_011770 | 6154577 | T | C | 68 | 0 | 2 | 0 | NON_SYNONYMOU | MODERAT | MISSENSE | Ttc/Ctc | F406L | PLES_55611 | C | C | | | | | | | | | | |
| NC_011770 | 6160670 | A | C | 65 | 0 | 12 | 0 | NON_SYNONYMOU | MODERAT | MISSENSE | tAt/tCt | Y2S | PLES_55661 | C | C | C | C | C | C | C | C | C | C | C | C |
| NC_011770 | 6160729 | C | T | 68 | 0 | 1 | 0 | NON_SYNONYMOU | MODERAT | MISSENSE | Cac/Tac | H22Y | PLES_55661 | | | | | | | T | | | | | |
| NC_011770 | 6161062 | A | T | 52 | 0 | 12 | 0 | NON_SYNONYMOU | MODERAT | MISSENSE | Acg/Tcg | T133S | PLES_55661 | T | T | T | T | T | T | T | T | T | T | T | T |
| NC_011770 | 6161074 | G | A | 52 | 0 | 2 | 0 | NON_SYNONYMOU | MODERAT | MISSENSE | Gcg/Acg | A137T | PLES_55661 | | | | | | | A | A | | | | |
| NC_011770 | 6161537 | G | C | 52 | 0 | 4 | 0 | NON_SYNONYMOU | MODERAT | MISSENSE | gGt/gCt | G291A | PLES_55661 | | | C | C | C | C | | | | | | |
| NC_011770 | 6162343 | C | G | 66 | 0 | 4 | 0 | NON_SYNONYMOU | MODERAT | MISSENSE | Caa/Gaa | Q560E | PLES_55661 | | | G | G | G | G | | | | | | |
| NC_011770 | 6162624 | A | AGC | 43 | 0 | 10 | 0 | FRAME_SHIFT | HIGH | | gct/gcGCt | A52A? | PLES_55671 | AGC | AGC | | AGC | AGC | AGC | AGC | AGC | . | AGC | AGC | AGC |
| NC_011770 | 6162766 | T | C | 15 | 10 | 2 | 0 | NON_SYNONYMOU | MODERAT | MISSENSE | gAg/gGg | E5G | PLES_55671 | . | . | . | . | . | . | . | . | . | . | C | C |
| NC_011770 | 6162772 | A | T | 14 | 10 | 2 | 0 | NON_SYNONYMOU | MODERAT | MISSENSE | cTc/cAc | L3H | PLES_55671 | . | . | . | . | . | . | . | . | . | . | T | T |
| NC_011770 | 6162773 | G | T | 14 | 10 | 2 | 0 | NON_SYNONYMOU | MODERAT | MISSENSE | Ctc/Atc | L3I | PLES_55671 | . | . | . | . | . | . | . | . | . | . | T | T |
| NC_011770 | 6165660 | A | G | 76 | 0 | 4 | 0 | NON_SYNONYMOU | MODERAT | MISSENSE | Acc/Gcc | T47A | PLES_55701 | | | G | G | G | G | | | | | | |
| NC_011770 | 6167401 | A | T | 87 | 0 | 4 | 0 | NON_SYNONYMOU | MODERAT | MISSENSE | gaA/gaT | E184D | PLES_55721 | | | T | T | T | T | | | | | | |
| NC_011770 | 6167592 | G | A | 83 | 0 | 4 | 0 | NON_SYNONYMOU | MODERAT | MISSENSE | gGc/gAc | G248D | PLES_55721 | | | A | A | A | A | | | | | | |
| NC_011770 | 6167606 | A | G | 84 | 0 | 4 | 0 | NON_SYNONYMOU | MODERAT | MISSENSE | Atc/Gtc | I253V | PLES_55721 | | | G | G | G | G | | | | | | |
| NC_011770 | 6167709 | A | C | 88 | 0 | 4 | 0 | NON_SYNONYMOU | MODERAT | MISSENSE | aAc/aCc | N287T | PLES_55721 | | | C | C | C | C | | | | | | |
| NC_011770 | 6167724 | A | C | 78 | 0 | 4 | 0 | NON_SYNONYMOU | MODERAT | MISSENSE | cAg/cCg | Q292P | PLES_55721 | | | C | C | C | C | | | | | | |
| NC_011770 | 6167726 | T | G | 80 | 0 | 4 | 0 | NON_SYNONYMOU | MODERAT | MISSENSE | Tct/Gct | S293A | PLES_55721 | | | G | G | G | G | | | | | | |
| NC_011770 | 6167892 | A | T | 84 | 0 | 4 | 0 | NON_SYNONYMOU | MODERAT | MISSENSE | gAt/gTt | D22V | PLES_55731 | | | T | T | T | T | | | | | | |
| NC_011770 | 6167927 | G | A | 85 | 0 | 4 | 0 | NON_SYNONYMOU | MODERAT | MISSENSE | Gtc/Atc | V34I | PLES_55731 | | | A | A | A | A | | | | | | |
| NC_011770 | 6167954 | T | A | 83 | 0 | 2 | 0 | NON_SYNONYMOU | MODERAT | MISSENSE | Tgt/Agt | C43S | PLES_55731 | A | A | | | | | | | | | | |

| | | | | | | | | | | | | | | | | | | | | | | | | | |
|---|---|---|---|---|---|---|---|---|---|---|---|---|---|---|---|---|---|---|---|---|---|---|---|---|---|
| NC_011770 | 6167960 | G | C | 83 | 0 | 4 | 0 | NON_SYNONYMOU | MODERAT | MISSENSE | Gtg/Ctg | V45L | PLES_55731 | | | C | C | C | C | | | | | | |
| NC_011770 | 6168062 | G | C | 73 | 0 | 4 | 0 | NON_SYNONYMOU | MODERAT | MISSENSE | Gag/Cag | E79Q | PLES_55731 | | | C | C | C | C | | | | | | |
| NC_011770 | 6168065 | T | G | 71 | 0 | 4 | 0 | NON_SYNONYMOU | MODERAT | MISSENSE | Tcc/Gcc | S80A | PLES_55731 | | | G | G | G | G | | | | | | |
| NC_011770 | 6168076 | C | A | 72 | 0 | 6 | 0 | NON_SYNONYMOU | MODERAT | MISSENSE | gaC/gaA | D83E | PLES_55731 | | | A | A | A | A | | | A | A | | |
| NC_011770 | 6168276 | C | T | 73 | 0 | 4 | 0 | NON_SYNONYMOU | MODERAT | MISSENSE | gCc/gTc | A150V | PLES_55731 | | | T | T | T | T | | | | | | |
| NC_011770 | 6168330 | A | G | 75 | 0 | 2 | 0 | NON_SYNONYMOU | MODERAT | MISSENSE | cAt/cGt | H168R | PLES_55731 | | | | | | | | | | | G | G |
| NC_011770 | 6168353 | G | C | 72 | 0 | 12 | 0 | NON_SYNONYMOU | MODERAT | MISSENSE | Gtg/Ctg | V176L | PLES_55731 | C | C | C | C | C | C | C | C | C | C | C | C |
| NC_011770 | 6168366 | G | A | 72 | 0 | 4 | 0 | NON_SYNONYMOU | MODERAT | MISSENSE | gGg/gAg | G180E | PLES_55731 | | | A | A | A | A | | | | | | |
| NC_011770 | 6168443 | C | G | 66 | 0 | 4 | 0 | NON_SYNONYMOU | MODERAT | MISSENSE | Cgc/Ggc | R206G | PLES_55731 | | | G | G | G | G | | | | | | |
| NC_011770 | 6168465 | G | A | 63 | 0 | 4 | 0 | NON_SYNONYMOU | MODERAT | MISSENSE | cGc/cAc | R213H | PLES_55731 | | | A | A | A | A | | | | | | |
| NC_011770 | 6168624 | A | G | 63 | 0 | 4 | 0 | NON_SYNONYMOU | MODERAT | MISSENSE | aAc/aGc | N266S | PLES_55731 | | | G | G | G | G | | | | | | |
| NC_011770 | 6168657 | C | T | 64 | 0 | 4 | 0 | NON_SYNONYMOU | MODERAT | MISSENSE | gCa/gTa | A277V | PLES_55731 | | | T | T | T | T | | | | | | |
| NC_011770 | 6168939 | G | A | 70 | 0 | 4 | 0 | NON_SYNONYMOU | MODERAT | MISSENSE | Gcg/Acg | A89T | PLES_55741 | | | A | A | A | A | | | | | | |
| NC_011770 | 6169589 | A | C | 79 | 0 | 4 | 0 | NON_SYNONYMOU | MODERAT | MISSENSE | gaA/gaC | E305D | PLES_55741 | C | C | | | | | | | C | C | | |
| NC_011770 | 6169605 | A | G | 81 | 0 | 4 | 0 | NON_SYNONYMOU | MODERAT | MISSENSE | Agc/Ggc | S311G | PLES_55741 | G | G | | | | | | | G | G | | |
| NC_011770 | 6169719 | A | G | 79 | 0 | 2 | 0 | NON_SYNONYMOU | MODERAT | MISSENSE | Agc/Ggc | S349G | PLES_55741 | | | | | | | | | | | G | G |
| NC_011770 | 6169960 | T | C | 77 | 0 | 2 | 0 | NON_SYNONYMOU | MODERAT | MISSENSE | cTg/cCg | L429P | PLES_55741 | C | C | | | | | | | | | | |
| NC_011770 | 6170368 | A | G | 76 | 0 | 2 | 0 | NON_SYNONYMOU | MODERAT | MISSENSE | aAg/aGg | K565R | PLES_55741 | G | G | | | | | | | | | | |
| NC_011770 | 6170718 | G | A | 72 | 0 | 2 | 0 | NON_SYNONYMOU | MODERAT | MISSENSE | Gaa/Aaa | E682K | PLES_55741 | | | | | | | | | | | A | A |
| NC_011770 | 6171044 | GA | G | 56 | 0 | 4 | 0 | INTRAGENIC | MODIFIER | | | | PLES_55742 | | | G | G | G | G | | | | | | |
| NC_011770 | 6171143 | C | G | 65 | 0 | 6 | 0 | INTRAGENIC | MODIFIER | | | | PLES_55742 | G | G | | | | | G | G | | | G | G |
| NC_011770 | 6171152 | C | T | 65 | 0 | 4 | 0 | INTRAGENIC | MODIFIER | | | | PLES_55742 | | | | | | | T | T | | | T | T |
| NC_011770 | 6171153 | G | GT | 65 | 0 | 2 | 0 | INTRAGENIC | MODIFIER | | | | PLES_55742 | GT | GT | | | | | | | | | | |
| NC_011770 | 6171159 | C | G | 69 | 0 | 2 | 0 | INTRAGENIC | MODIFIER | | | | PLES_55742 | G | G | | | | | | | | | | |
| NC_011770 | 6171199 | T | C | 73 | 0 | 4 | 0 | INTRAGENIC | MODIFIER | | | | PLES_55742 | | | C | C | C | C | | | | | | |
| NC_011770 | 6171247 | T | C | 74 | 0 | 4 | 0 | INTRAGENIC | MODIFIER | | | | PLES_55742 | | | | | | | C | C | | | C | C |
| NC_011770 | 6171290 | CAG | C | 76 | 0 | 4 | 0 | INTRAGENIC | MODIFIER | | | | PLES_55742 | | | C | C | C | C | | | | | | |
| NC_011770 | 6172161 | G | C | 73 | 0 | 4 | 0 | NON_SYNONYMOU | MODERAT | MISSENSE | gGc/gCc | G81A | PLES_55761 | | | C | C | C | C | | | | | | |
| NC_011770 | 6172901 | T | G | 88 | 0 | 12 | 0 | NON_SYNONYMOU | MODERAT | MISSENSE | caT/caG | H22Q | PLES_55781 | G | G | G | G | G | G | G | G | G | G | G | G |
| NC_011770 | 6173100 | T | C | 82 | 0 | 4 | 0 | NON_SYNONYMOU | MODERAT | MISSENSE | Ttc/Ctc | F89L | PLES_55781 | | | C | C | C | C | | | | | | |
| NC_011770 | 6173265 | G | C | 78 | 0 | 2 | 0 | NON_SYNONYMOU | MODERAT | MISSENSE | Gtc/Ctc | V144L | PLES_55781 | | | | | | | | | | | C | C |
| NC_011770 | 6173317 | A | T | 85 | 0 | 2 | 0 | NON_SYNONYMOU | MODERAT | MISSENSE | gAc/gTc | D161V | PLES_55781 | | | | | | | T | T | | | | |
| NC_011770 | 6173386 | C | A | 78 | 0 | 2 | 0 | NON_SYNONYMOU | MODERAT | MISSENSE | gCg/gAg | A184E | PLES_55781 | | | | | | | A | A | | | | |
| NC_011770 | 6173541 | C | T | 86 | 0 | 12 | 0 | NON_SYNONYMOU | MODERAT | MISSENSE | aGc/aAc | S93N | PLES_55791 | T | T | T | T | T | T | T | T | T | T | T | T |
| NC_011770 | 6173644 | C | T | 84 | 0 | 1 | 0 | NON_SYNONYMOU | MODERAT | MISSENSE | Ggc/Agc | G59S | PLES_55791 | | | | | | | T | | | | | |
| NC_011770 | 6173763 | G | C | 83 | 0 | 2 | 0 | NON_SYNONYMOU | MODERAT | MISSENSE | aCc/aGc | T19S | PLES_55791 | | | | | | | C | C | | | | |
| NC_011770 | 6173805 | G | A | 81 | 0 | 4 | 0 | NON_SYNONYMOU | MODERAT | MISSENSE | cCg/cTg | P5L | PLES_55791 | A | A | | | | | | | | | A | A |
| NC_011770 | 6174034 | A | G | 75 | 0 | 2 | 0 | NON_SYNONYMOU | MODERAT | MISSENSE | gTg/gCg | V324A | PLES_55801 | | | | | | | | | | | G | G |
| NC_011770 | 6174502 | T | C | 67 | 0 | 1 | 0 | NON_SYNONYMOU | MODERAT | MISSENSE | cAc/cGc | H168R | PLES_55801 | | | | | | | C | | | | | |
| NC_011770 | 6174503 | G | A | 67 | 0 | 2 | 0 | NON_SYNONYMOU | MODERAT | MISSENSE | Cac/Tac | H168Y | PLES_55801 | | | | | | | | | | | A | A |
| NC_011770 | 6174521 | C | T | 66 | 0 | 12 | 0 | NON_SYNONYMOU | MODERAT | MISSENSE | Gtg/Atg | V162M | PLES_55801 | T | T | T | T | T | T | T | T | T | T | T | T |
| NC_011770 | 6174748 | C | G | 64 | 0 | 2 | 0 | NON_SYNONYMOU | MODERAT | MISSENSE | gGc/gCc | G86A | PLES_55801 | | | | | | | | | | | G | G |
| NC_011770 | 6174782 | A | C | 61 | 0 | 2 | 0 | NON_SYNONYMOU | MODERAT | MISSENSE | Tcg/Gcg | S75A | PLES_55801 | C | C | | | | | | | | | | |
| NC_011770 | 6174860 | C | T | 74 | 0 | 4 | 0 | NON_SYNONYMOU | MODERAT | MISSENSE | Gat/Aat | D49N | PLES_55801 | | | T | T | T | T | | | | | | |
| NC_011770 | 6174958 | G | A | 73 | 0 | 2 | 0 | NON_SYNONYMOU | MODERAT | MISSENSE | gCc/gTc | A16V | PLES_55801 | | | | | | | | | | | A | A |
| NC_011770 | 6175996 | C | T | 95 | 0 | 1 | 0 | NON_SYNONYMOU | MODERAT | MISSENSE | Ggc/Agc | G276S | PLES_55811 | | | | | | | | | | | | T |
| NC_011770 | 6175368 | G | A | 91 | 0 | 2 | 0 | NON_SYNONYMOU | MODERAT | MISSENSE | gCt/gTt | A485V | PLES_55811 | | | | | | | | | | | A | A |
| NC_011770 | 6175383 | G | A | 89 | 0 | 4 | 0 | NON_SYNONYMOU | MODERAT | MISSENSE | gCg/gTg | A480V | PLES_55811 | | | | | | | A | A | A | A | | |
| NC_011770 | 6175413 | A | G | 95 | 0 | 2 | 0 | NON_SYNONYMOU | MODERAT | MISSENSE | gTg/gCg | V470A | PLES_55811 | | | | | | | | | | | G | G |
| NC_011770 | 6175738 | A | C | 88 | 0 | 10 | 0 | NON_SYNONYMOU | MODERAT | MISSENSE | Tcc/Gcc | S362A | PLES_55811 | C | C | C | C | C | C | C | C | C | C | | |
| NC_011770 | 6176830 | A | G | 96 | 0 | 6 | 0 | NON_SYNONYMOU | MODERAT | MISSENSE | gTg/gCg | V410A | PLES_55821 | | | | | | | G | G | G | G | G | G |
| NC_011770 | 6176950 | G | A | 92 | 0 | 4 | 0 | NON_SYNONYMOU | MODERAT | MISSENSE | gCg/gTg | A370V | PLES_55821 | | | A | A | A | A | | | | | | |
| NC_011770 | 6177260 | T | G | 92 | 0 | 6 | 0 | NON_SYNONYMOU | MODERAT | MISSENSE | Atg/Ctg | M267L | PLES_55821 | | | G | G | G | G | | | | | G | G |
| NC_011770 | 6177647 | C | G | 89 | 0 | 2 | 0 | NON_SYNONYMOU | MODERAT | MISSENSE | Ggt/Cgt | G138R | PLES_55821 | | | | | | | | | | | G | G |
| NC_011770 | 6177890 | T | G | 83 | 0 | 2 | 0 | NON_SYNONYMOU | MODERAT | MISSENSE | Aac/Cac | N57H | PLES_55821 | | | | | | | G | G | | | | |
| NC_011770 | 6178240 | C | A | 92 | 0 | 2 | 0 | NON_SYNONYMOU | MODERAT | MISSENSE | Ctt/Att | L14I | PLES_55831 | | | | | | | | | A | A | | |
| NC_011770 | 6180087 | A | C | 72 | 0 | 4 | 0 | NON_SYNONYMOU | MODERAT | MISSENSE | Att/Ctt | I37L | PLES_55851 | | | C | C | C | C | | | | | | |
| NC_011770 | 6180139 | G | A | 69 | 0 | 2 | 0 | NON_SYNONYMOU | MODERAT | MISSENSE | gGc/gAc | G54D | PLES_55851 | | | | | | | A | A | | | | |
| NC_011770 | 6180250 | G | A | 63 | 0 | 4 | 0 | NON_SYNONYMOU | MODERAT | MISSENSE | gGt/gAt | G91D | PLES_55851 | | | A | A | A | A | | | | | | |
| NC_011770 | 6180271 | A | G | 63 | 0 | 6 | 0 | NON_SYNONYMOU | MODERAT | MISSENSE | cAc/cGc | H98R | PLES_55851 | | | G | G | G | G | | | G | G | | |
| NC_011770 | 6180292 | A | G | 62 | 0 | 8 | 0 | NON_SYNONYMOU | MODERAT | MISSENSE | gAt/gGt | D105G | PLES_55851 | | | G | G | G | G | | | G | G | G | G |
| NC_011770 | 6183297 | T | C | 84 | 0 | 1 | 0 | NON_SYNONYMOU | MODERAT | MISSENSE | cTc/cCc | L81P | PLES_55881 | | | | | | | | C | | | | |

| | | | | | | | | | | | | | | | | | | | | | | | | | |
|---|---|---|---|---|---|---|---|---|---|---|---|---|---|---|---|---|---|---|---|---|---|---|---|---|---|
| NC_011770 | 6183801 | G | A | 81 | 0 | 1 | 0 | NON_SYNONYMOU | MODERAT | MISSENSE | cGc/cAc | R249H | PLES_55881 | | | | | | | | A | | | | |
| NC_011770 | 6183108 | C | T | 90 | 0 | 6 | 0 | NON_SYNONYMOU | MODERAT | MISSENSE | aCc/aTc | T18I | PLES_55881 | T | T | T | T | T | T | | | | | | |
| NC_011770 | 6184015 | G | T | 83 | 0 | 2 | 0 | NON_SYNONYMOU | MODERAT | MISSENSE | gCa/gAa | A93E | PLES_55891 | | | | | | | | | T | T | | |
| NC_011770 | 6184057 | G | T | 85 | 0 | 2 | 0 | NON_SYNONYMOU | MODERAT | MISSENSE | cCg/cAg | P79Q | PLES_55891 | | | | | | | | | T | T | | |
| NC_011770 | 6184285 | T | G | 80 | 0 | 2 | 0 | NON_SYNONYMOU | MODERAT | MISSENSE | gAg/gCg | E3A | PLES_55891 | | | | | | | | | G | G | | |
| NC_011770 | 6184905 | G | A | 69 | 0 | 2 | 0 | NON_SYNONYMOU | MODERAT | MISSENSE | aGc/aAc | S135N | PLES_55901 | | | | | | | | | A | A | | |
| NC_011770 | 6186170 | T | C | 74 | 0 | 2 | 0 | NON_SYNONYMOU | MODERAT | MISSENSE | Tct/Cct | S3P | PLES_55921 | | | | | | | | | C | C | | |
| NC_011770 | 6186227 | G | A | 73 | 0 | 2 | 0 | NON_SYNONYMOU | MODERAT | MISSENSE | Gcg/Acg | A22T | PLES_55921 | | | A | A | | | | | | | | |
| NC_011770 | 6186702 | A | C | 68 | 0 | 4 | 0 | NON_SYNONYMOU | MODERAT | MISSENSE | cAc/cCc | H180P | PLES_55921 | C | C | | | | | | | C | C | | |
| NC_011770 | 6186723 | T | C | 64 | 0 | 12 | 0 | NON_SYNONYMOU | MODERAT | MISSENSE | gTg/gCg | V187A | PLES_55921 | C | C | C | C | C | C | C | C | C | C | C | C |
| NC_011770 | 6187035 | C | T | 52 | 0 | 2 | 0 | NON_SYNONYMOU | MODERAT | MISSENSE | gCc/gTc | A291V | PLES_55921 | T | T | | | | | | | | | | |
| NC_011770 | 6189744 | G | A | 74 | 0 | 2 | 0 | NON_SYNONYMOU | MODERAT | MISSENSE | Gcc/Acc | A79T | PLES_55951 | | | | | | | | | A | A | | |
| NC_011770 | 6189918 | G | A | 81 | 0 | 10 | 0 | NON_SYNONYMOU | MODERAT | MISSENSE | Gac/Aac | D137N | PLES_55951 | A | A | A | A | A | A | A | A | A | A | | |
| NC_011770 | 6190281 | G | A | 79 | 0 | 2 | 0 | NON_SYNONYMOU | MODERAT | MISSENSE | Gtc/Atc | V258I | PLES_55951 | A | A | | | | | | | | | | |
| NC_011770 | 6190293 | C | G | 77 | 0 | 12 | 0 | NON_SYNONYMOU | MODERAT | MISSENSE | Cag/Gag | Q262E | PLES_55951 | G | G | G | G | G | G | G | G | G | G | G | G |
| NC_011770 | 6190318 | G | A | 77 | 0 | 2 | 0 | NON_SYNONYMOU | MODERAT | MISSENSE | gGc/gAc | G270D | PLES_55951 | | | | | | | | | | | A | A |
| NC_011770 | 6191733 | G | A | 68 | 0 | 8 | 0 | NON_SYNONYMOU | MODERAT | MISSENSE | Gcc/Acc | A742T | PLES_55951 | A | A | A | A | A | A | | | A | A | | |
| NC_011770 | 6191739 | A | G | 67 | 0 | 2 | 0 | NON_SYNONYMOU | MODERAT | MISSENSE | Agc/Ggc | S744G | PLES_55951 | | | | | | | | | | | G | G |
| NC_011770 | 6191876 | G | T | 70 | 0 | 4 | 0 | NON_SYNONYMOU | MODERAT | MISSENSE | Gcc/Tcc | A6S | PLES_55961 | | | T | T | T | T | | | | | | |
| NC_011770 | 6195745 | A | G | 66 | 0 | 8 | 0 | NON_SYNONYMOU | MODERAT | MISSENSE | Atc/Gtc | I10V | PLES_55991 | | | G | G | G | G | G | G | | | G | G |
| NC_011770 | 6195862 | C | T | 75 | 0 | 2 | 0 | NON_SYNONYMOU | MODERAT | MISSENSE | Ctc/Ttc | L49F | PLES_55991 | | | | | | | | | | | T | T |
| NC_011770 | 6196310 | T | C | 66 | 0 | 1 | 0 | NON_SYNONYMOU | MODERAT | MISSENSE | gTg/gCg | V198A | PLES_55991 | | | | | | | C | | | | | |
| NC_011770 | 6198757 | G | A | 70 | 0 | 1 | 0 | NON_SYNONYMOU | MODERAT | MISSENSE | gCg/gTg | A53V | PLES_56011 | | A | | | | | | | | | | |
| NC_011770 | 6197696 | A | C | 74 | 0 | 12 | 0 | NON_SYNONYMOU | MODERAT | MISSENSE | Ttg/Gtg | L407V | PLES_56011 | C | C | C | C | C | C | C | C | C | C | C | C |
| NC_011770 | 6198016 | G | C | 79 | 0 | 12 | 0 | NON_SYNONYMOU | MODERAT | MISSENSE | aCc/aGc | T300S | PLES_56011 | C | C | C | C | C | C | C | C | C | C | C | C |
| NC_011770 | 6198024 | T | A | 80 | 0 | 12 | 0 | NON_SYNONYMOU | MODERAT | MISSENSE | gaA/gaT | E297D | PLES_56011 | A | A | A | A | A | A | A | A | A | A | A | A |
| NC_011770 | 6198200 | G | A | 93 | 0 | 2 | 0 | NON_SYNONYMOU | MODERAT | MISSENSE | Ccc/Tcc | P239S | PLES_56011 | | | | | | | A | A | | | | |
| NC_011770 | 6198278 | T | G | 92 | 0 | 12 | 0 | NON_SYNONYMOU | MODERAT | MISSENSE | Atc/Ctc | I213L | PLES_56011 | G | G | G | G | G | G | G | G | G | G | G | G |
| NC_011770 | 6198660 | G | C | 73 | 0 | 2 | 0 | NON_SYNONYMOU | MODERAT | MISSENSE | gaC/gaG | D85E | PLES_56011 | | | | | | | C | C | | | | |
| NC_011770 | 6198682 | T | G | 66 | 0 | 2 | 0 | NON_SYNONYMOU | MODERAT | MISSENSE | gAt/gCt | D78A | PLES_56011 | | | | | | | | | G | G | | |
| NC_011770 | 6198902 | A | C | 63 | 0 | 12 | 0 | NON_SYNONYMOU | MODERAT | MISSENSE | Tcc/Gcc | S5A | PLES_56011 | C | C | C | C | C | C | C | C | C | C | C | C |
| NC_011770 | 6200670 | A | G | 81 | 0 | 2 | 0 | NON_SYNONYMOU | MODERAT | MISSENSE | Acg/Gcg | T307A | PLES_56031 | | | | | | | | | G | G | | |
| NC_011770 | 6200671 | C | T | 82 | 0 | 2 | 0 | NON_SYNONYMOU | MODERAT | MISSENSE | aCg/aTg | T307M | PLES_56031 | | | | | | | | | T | T | | |
| NC_011770 | 6201280 | G | A | 81 | 0 | 2 | 0 | NON_SYNONYMOU | MODERAT | MISSENSE | Gcg/Acg | A43T | PLES_56041 | | | | | A | A | | | | | | |
| NC_011770 | 6202160 | C | G | 61 | 0 | 12 | 0 | NON_SYNONYMOU | MODERAT | MISSENSE | aCc/aGc | T336S | PLES_56041 | G | G | G | G | G | G | G | G | G | G | G | G |
| NC_011770 | 6203548 | G | A | 47 | 0 | 1 | 0 | NON_SYNONYMOU | MODERAT | MISSENSE | gGc/gAc | G130D | PLES_56051 | | A | | | | | | | | | | |
| NC_011770 | 6203215 | T | C | 51 | 0 | 2 | 0 | NON_SYNONYMOU | MODERAT | MISSENSE | gTc/gCc | V19A | PLES_56051 | | | | | | | | | | | C | C |
| NC_011770 | 6203604 | G | A | 50 | 0 | 2 | 0 | NON_SYNONYMOU | MODERAT | MISSENSE | Gcg/Acg | A149T | PLES_56051 | | | | | | | | | | | A | A |
| NC_011770 | 6204071 | T | C | 64 | 0 | 1 | 0 | STOP_LOST+SPLICE | HIGH | MISSENSE | Tag/Cag | *110Q | PLES_56061 | | | | C | | | | | | | | |
| NC_011770 | 6203768 | G | A | 53 | 0 | 2 | 0 | NON_SYNONYMOU | MODERAT | MISSENSE | Gcc/Acc | A9T | PLES_56061 | | | | | | | | | A | A | | |
| NC_011770 | 6208898 | T | A | 82 | 0 | 2 | 0 | NON_SYNONYMOU | MODERAT | MISSENSE | Acc/Tcc | T517S | PLES_56101 | | | | | | | A | A | | | | |
| NC_011770 | 6209081 | G | C | 71 | 0 | 4 | 0 | NON_SYNONYMOU | MODERAT | MISSENSE | Cgg/Ggg | R456G | PLES_56101 | | | C | C | C | C | | | | | | |
| NC_011770 | 6209278 | A | G | 70 | 0 | 1 | 0 | NON_SYNONYMOU | MODERAT | MISSENSE | gTg/gCg | V390A | PLES_56101 | | | | | | | G | | | | | |
| NC_011770 | 6210389 | C | A | 51 | 0 | 2 | 0 | NON_SYNONYMOU | MODERAT | MISSENSE | Gtg/Ttg | V20L | PLES_56101 | A | A | | | | | | | | | | |
| NC_011770 | 6210401 | C | A | 53 | 0 | 2 | 0 | NON_SYNONYMOU | MODERAT | MISSENSE | Gtt/Ttt | V16F | PLES_56101 | | | | | | | | | A | A | | |
| NC_011770 | 6210729 | T | C | 56 | 0 | 12 | 0 | NON_SYNONYMOU | MODERAT | MISSENSE | Aag/Gag | K262E | PLES_56111 | C | C | C | C | C | C | C | C | C | C | C | C |
| NC_011770 | 6211729 | C | T | 81 | 0 | 2 | 0 | NON_SYNONYMOU | MODERAT | MISSENSE | Gcc/Acc | A256T | PLES_56121 | | | | | | | | | | | T | T |
| NC_011770 | 6213158 | G | T | 94 | 0 | 2 | 0 | NON_SYNONYMOU | MODERAT | MISSENSE | Gcg/Tcg | A190S | PLES_56131 | | | | | | | | | | | T | T |
| NC_011770 | 6213163 | CGCACCG | C | 79 | 0 | 2 | 0 | CODON_CHANGE_ | MODERATE | | gcaccgggg, | APG192G | PLES_56131 | | | | | | | C | C | | | | |
| NC_011770 | 6213389 | C | T | 87 | 0 | 6 | 0 | NON_SYNONYMOU | MODERAT | MISSENSE | Cgc/Tgc | R267C | PLES_56131 | | | T | T | T | T | T | T | | | | |
| NC_011770 | 6213720 | A | G | 78 | 0 | 12 | 0 | NON_SYNONYMOU | MODERAT | MISSENSE | cAc/cGc | H377R | PLES_56131 | G | G | G | G | G | G | G | G | G | G | G | G |
| NC_011770 | 6213923 | C | T | 77 | 0 | 1 | 0 | NON_SYNONYMOU | MODERAT | MISSENSE | gCc/gTc | A34V | PLES_56141 | | T | | | | | | | | | | |
| NC_011770 | 6214354 | G | A | 80 | 0 | 1 | 0 | NON_SYNONYMOU | MODERAT | MISSENSE | Gta/Ata | V178I | PLES_56141 | | | | | | | A | | | | | |
| NC_011770 | 6214585 | C | T | 73 | 0 | 12 | 0 | NON_SYNONYMOU | MODERAT | MISSENSE | Ctc/Ttc | L255F | PLES_56141 | T | T | T | T | T | T | T | T | T | T | T | T |
| NC_011770 | 6214701 | G | A | 70 | 0 | 1 | 0 | NON_SYNONYMOU | MODERAT | MISSENSE | Ccg/Tcg | P400S | PLES_56151 | | | | | | | | A | | | | |
| NC_011770 | 6214697 | G | A | 68 | 0 | 8 | 0 | NON_SYNONYMOU | MODERAT | MISSENSE | tCg/tTg | S401L | PLES_56151 | | | A | A | A | A | | | A | A | A | A |
| NC_011770 | 6214698 | A | T | 68 | 0 | 8 | 0 | NON_SYNONYMOU | MODERAT | MISSENSE | Tcg/Acg | S401T | PLES_56151 | | | T | T | T | T | | | T | T | T | T |
| NC_011770 | 6214722 | G | A | 70 | 0 | 4 | 0 | NON_SYNONYMOU | MODERAT | MISSENSE | Ctt/Ttt | L393F | PLES_56151 | | | A | A | A | A | | | | | | |
| NC_011770 | 6219087 | G | A | 101 | 0 | 1 | 0 | STOP_GAINED | HIGH | NONSENSE | Cag/Tag | Q140* | PLES_56191 | | | | | | | A | | | | | |
| NC_011770 | 6219619 | G | A | 86 | 0 | 4 | 0 | NON_SYNONYMOU | MODERAT | MISSENSE | cGg/cAg | R10Q | PLES_56201 | | | A | A | A | A | | | | | | |
| NC_011770 | 6219672 | C | T | 82 | 0 | 4 | 0 | NON_SYNONYMOU | MODERAT | MISSENSE | Cca/Tca | P28S | PLES_56201 | | | T | T | T | T | | | | | | |
| NC_011770 | 6220829 | T | C | 35 | 0 | 1 | 0 | NON_SYNONYMOU | MODERAT | MISSENSE | gTg/gCg | V132A | PLES_56221 | | | | | | | C | | | | | |

| | | | | | | | | | | | | | | | | | | | | | | | | | |
|---|---|---|---|---|---|---|---|---|---|---|---|---|---|---|---|---|---|---|---|---|---|---|---|---|---|
| NC_011770 | 6220967 | A | G | 47 | 0 | 12 | 0 | NON_SYNONYMOU | MODERAT | MISSENSE | cAa/cGa | Q178R | PLES_56221 | G | G | G | G | G | G | G | G | G | G | G | G |
| NC_011770 | 6221474 | A | G | 43 | 0 | 12 | 0 | NON_SYNONYMOU | MODERAT | MISSENSE | cAc/cGc | H21R | PLES_56231 | G | G | G | G | G | G | G | G | G | G | G | G |
| NC_011770 | 6221542 | T | C | 39 | 0 | 4 | 0 | NON_SYNONYMOU | MODERAT | MISSENSE | Tcc/Ccc | S44P | PLES_56231 | | | | | | | | | C | C | C | C |
| NC_011770 | 6221633 | T | C | 29 | 0 | 12 | 0 | NON_SYNONYMOU | MODERAT | MISSENSE | cTc/cCc | L74P | PLES_56231 | C | C | C | C | C | C | C | C | C | C | C | C |
| NC_011770 | 6223661 | G | A | 79 | 0 | 2 | 0 | NON_SYNONYMOU | MODERAT | MISSENSE | gCc/gTc | A700V | PLES_56251 | | | | | | | A | A | | | | |
| NC_011770 | 6224153 | T | C | 82 | 0 | 2 | 0 | NON_SYNONYMOU | MODERAT | MISSENSE | gAg/gGg | E536G | PLES_56251 | | | | | | | | | C | C | | |
| NC_011770 | 6224550 | T | C | 74 | 0 | 2 | 0 | NON_SYNONYMOU | MODERAT | MISSENSE | Aac/Gac | N404D | PLES_56251 | | | | | | | | | C | C | | |
| NC_011770 | 6224667 | G | A | 76 | 0 | 2 | 0 | NON_SYNONYMOU | MODERAT | MISSENSE | Cgg/Tgg | R365W | PLES_56251 | A | A | | | | | | | | | | |
| NC_011770 | 6224751 | G | C | 78 | 0 | 2 | 0 | NON_SYNONYMOU | MODERAT | MISSENSE | Ccg/Gcg | P337A | PLES_56251 | | | | | | | | | C | C | | |
| NC_011770 | 6225177 | C | T | 84 | 0 | 2 | 0 | NON_SYNONYMOU | MODERAT | MISSENSE | Ggc/Agc | G195S | PLES_56251 | | | | | | | | | T | T | | |
| NC_011770 | 6225669 | T | C | 72 | 0 | 4 | 0 | NON_SYNONYMOU | MODERAT | MISSENSE | Atc/Gtc | I31V | PLES_56251 | | | C | C | C | C | | | | | | |
| NC_011770 | 6226790 | C | T | 77 | 0 | 10 | 0 | NON_SYNONYMOU | MODERAT | MISSENSE | Gtc/Atc | V14I | PLES_56261 | T | T | T | T | T | T | T | T | T | T | | |
| NC_011770 | 6227082 | G | A | 88 | 0 | 2 | 0 | NON_SYNONYMOU | MODERAT | MISSENSE | cCg/cTg | P125L | PLES_56271 | | | | | | | | | A | A | | |
| NC_011770 | 6227697 | A | G | 65 | 0 | 2 | 0 | NON_SYNONYMOU | MODERAT | MISSENSE | Atc/Gtc | I30V | PLES_56281 | | | | | | | | | G | G | | |
| NC_011770 | 6228553 | G | A | 66 | 0 | 6 | 0 | NON_SYNONYMOU | MODERAT | MISSENSE | aGg/aAg | R315K | PLES_56281 | | | A | A | A | A | | | A | A | | |
| NC_011770 | 6232264 | C | T | 82 | 0 | 2 | 0 | NON_SYNONYMOU | MODERAT | MISSENSE | Gtc/Atc | V131I | PLES_56311 | T | T | | | | | | | | | | |
| NC_011770 | 6232738 | G | T | 94 | 0 | 4 | 0 | NON_SYNONYMOU | MODERAT | MISSENSE | Cag/Aag | Q662K | PLES_56321 | | | T | T | T | T | | | | | | |
| NC_011770 | 6232906 | T | G | 82 | 0 | 12 | 0 | NON_SYNONYMOU | MODERAT | MISSENSE | Atc/Ctc | I606L | PLES_56321 | G | G | G | G | G | G | G | G | G | G | G | G |
| NC_011770 | 6232975 | T | C | 77 | 0 | 12 | 0 | NON_SYNONYMOU | MODERAT | MISSENSE | Act/Gct | T583A | PLES_56321 | C | C | C | C | C | C | C | C | C | C | C | C |
| NC_011770 | 6233227 | C | T | 85 | 0 | 12 | 0 | NON_SYNONYMOU | MODERAT | MISSENSE | Gtc/Atc | V499I | PLES_56321 | T | T | T | T | T | T | T | T | T | T | T | T |
| NC_011770 | 6233316 | T | G | 85 | 0 | 8 | 0 | NON_SYNONYMOU | MODERAT | MISSENSE | aAt/aCt | N469T | PLES_56321 | | | G | G | G | G | | | G | G | G | G |
| NC_011770 | 6233364 | G | A | 81 | 0 | 2 | 0 | NON_SYNONYMOU | MODERAT | MISSENSE | gCt/gTt | A453V | PLES_56321 | | | | | | | A | A | | | | |
| NC_011770 | 6233380 | C | G | 80 | 0 | 12 | 0 | NON_SYNONYMOU | MODERAT | MISSENSE | Gtg/Ctg | V448L | PLES_56321 | G | G | G | G | G | G | G | G | G | G | G | G |
| NC_011770 | 6233392 | C | T | 85 | 0 | 12 | 0 | NON_SYNONYMOU | MODERAT | MISSENSE | Gcc/Acc | A444T | PLES_56321 | T | T | T | T | T | T | T | T | T | T | T | T |
| NC_011770 | 6233632 | C | T | 98 | 0 | 12 | 0 | NON_SYNONYMOU | MODERAT | MISSENSE | Gtc/Atc | V364I | PLES_56321 | T | T | T | T | T | T | T | T | T | T | T | T |
| NC_011770 | 6233904 | G | A | 85 | 0 | 8 | 0 | NON_SYNONYMOU | MODERAT | MISSENSE | gCt/gTt | A273V | PLES_56321 | | | A | A | A | A | | | A | A | A | A |
| NC_011770 | 6233998 | C | T | 78 | 0 | 2 | 0 | NON_SYNONYMOU | MODERAT | MISSENSE | Gac/Aac | D242N | PLES_56321 | T | T | | | | | | | | | | |
| NC_011770 | 6234081 | C | G | 77 | 0 | 4 | 0 | NON_SYNONYMOU | MODERAT | MISSENSE | gGt/gCt | G214A | PLES_56321 | G | G | | | | | G | G | | | | |
| NC_011770 | 6234102 | G | A | 77 | 0 | 8 | 0 | NON_SYNONYMOU | MODERAT | MISSENSE | gCg/gTg | A207V | PLES_56321 | | | A | A | A | A | | | A | A | A | A |
| NC_011770 | 6234168 | T | C | 78 | 0 | 12 | 0 | NON_SYNONYMOU | MODERAT | MISSENSE | aAc/aGc | N185S | PLES_56321 | C | C | C | C | C | C | C | C | C | C | C | C |
| NC_011770 | 6234208 | C | G | 83 | 0 | 4 | 0 | NON_SYNONYMOU | MODERAT | MISSENSE | Gtc/Ctc | V172L | PLES_56321 | | | G | G | G | G | | | | | | |
| NC_011770 | 6234253 | C | T | 81 | 0 | 4 | 0 | NON_SYNONYMOU | MODERAT | MISSENSE | Gtc/Atc | V157I | PLES_56321 | T | T | | | | | T | T | | | | |
| NC_011770 | 6234409 | C | A | 46 | 0 | 6 | 0 | NON_SYNONYMOU | MODERAT | MISSENSE | Gcg/Tcg | A105S | PLES_56321 | A | A | | | | | A | A | | | A | A |
| NC_011770 | 6234421 | G | C | 47 | 0 | 6 | 0 | NON_SYNONYMOU | MODERAT | MISSENSE | Caa/Gaa | Q101E | PLES_56321 | C | C | | | | | C | C | | | C | C |
| NC_011770 | 6234438 | T | G | 47 | 0 | 6 | 0 | NON_SYNONYMOU | MODERAT | MISSENSE | gAg/gCg | E95A | PLES_56321 | G | G | | | | | G | G | | | G | G |
| NC_011770 | 6234456 | G | A | 48 | 0 | 6 | 0 | NON_SYNONYMOU | MODERAT | MISSENSE | gCc/gTc | A89V | PLES_56321 | A | A | | | | | A | A | | | A | A |
| NC_011770 | 6234457 | C | T | 49 | 0 | 2 | 0 | NON_SYNONYMOU | MODERAT | MISSENSE | Gcc/Acc | A89T | PLES_56321 | | | | | | | | | | | T | T |
| NC_011770 | 6234459 | G | A | 49 | 0 | 6 | 0 | NON_SYNONYMOU | MODERAT | MISSENSE | gCg/gTg | A88V | PLES_56321 | A | A | | | | | A | A | | | A | A |
| NC_011770 | 6234468 | A | G | 48 | 0 | 6 | 0 | NON_SYNONYMOU | MODERAT | MISSENSE | gTc/gCc | V85A | PLES_56321 | G | G | | | | | G | G | | | G | G |
| NC_011770 | 6234552 | C | T | 62 | 0 | 6 | 0 | NON_SYNONYMOU | MODERAT | MISSENSE | cGg/cAg | R57Q | PLES_56321 | T | T | | | | | T | T | | | T | T |
| NC_011770 | 6234554 | A | C | 61 | 0 | 6 | 0 | NON_SYNONYMOU | MODERAT | MISSENSE | gaT/gaG | D56E | PLES_56321 | C | C | | | | | C | C | | | C | C |
| NC_011770 | 6234631 | A | T | 54 | 0 | 6 | 0 | NON_SYNONYMOU | MODERAT | MISSENSE | Ttc/Atc | F31I | PLES_56321 | T | T | | | | | T | T | | | T | T |
| NC_011770 | 6234644 | C | A | 54 | 0 | 6 | 0 | NON_SYNONYMOU | MODERAT | MISSENSE | ttG/ttT | L26F | PLES_56321 | A | A | | | | | A | A | | | A | A |
| NC_011770 | 6234709 | G | C | 43 | 0 | 6 | 0 | NON_SYNONYMOU | MODERAT | MISSENSE | Cct/Gct | P5A | PLES_56321 | C | C | | | | | C | C | | | C | C |
| NC_011770 | 6242905 | G | A | 94 | 0 | 2 | 0 | NON_SYNONYMOU | MODERAT | MISSENSE | Gtg/Atg | V118M | PLES_56391 | | | | | | | | | | | A | A |
| NC_011770 | 6243321 | C | A | 81 | 0 | 12 | 0 | NON_SYNONYMOU | MODERAT | MISSENSE | Cgc/Agc | R2S | PLES_56401 | A | A | A | A | A | A | A | A | A | A | A | A |
| NC_011770 | 6243336 | C | T | 82 | 0 | 10 | 0 | NON_SYNONYMOU | MODERAT | MISSENSE | Ccc/Tcc | P7S | PLES_56401 | T | T | T | T | T | T | | | T | T | T | T |
| NC_011770 | 6243421 | T | C | 76 | 0 | 10 | 0 | NON_SYNONYMOU | MODERAT | MISSENSE | gTg/gCg | V35A | PLES_56401 | C | C | C | C | C | C | C | C | | | C | C |
| NC_011770 | 6243714 | A | G | 82 | 0 | 12 | 0 | NON_SYNONYMOU | MODERAT | MISSENSE | Aca/Gca | T133A | PLES_56401 | G | G | G | G | G | G | G | G | G | G | G | G |
| NC_011770 | 6243869 | C | A | 82 | 0 | 2 | 0 | NON_SYNONYMOU | MODERAT | MISSENSE | gaG/gaT | E130D | PLES_56411 | | | | | | | | | A | A | | |
| NC_011770 | 6243873 | C | G | 84 | 0 | 2 | 0 | NON_SYNONYMOU | MODERAT | MISSENSE | gGt/gCt | G129A | PLES_56411 | | | | | | | G | G | | | | |
| NC_011770 | 6243919 | C | T | 77 | 0 | 4 | 0 | NON_SYNONYMOU | MODERAT | MISSENSE | Gac/Aac | D114N | PLES_56411 | | | T | T | T | T | | | | | | |
| NC_011770 | 6243963 | G | A | 70 | 0 | 4 | 0 | NON_SYNONYMOU | MODERAT | MISSENSE | gCg/gTg | A99V | PLES_56411 | | | A | A | A | A | | | | | | |
| NC_011770 | 6244122 | T | C | 65 | 0 | 12 | 0 | NON_SYNONYMOU | MODERAT | MISSENSE | cAt/cGt | H46R | PLES_56411 | C | C | C | C | C | C | C | C | C | C | C | C |
| NC_011770 | 6244221 | A | G | 65 | 0 | 12 | 0 | NON_SYNONYMOU | MODERAT | MISSENSE | gTg/gCg | V13A | PLES_56411 | G | G | G | G | G | G | G | G | G | G | G | G |
| NC_011770 | 6244402 | G | T | 63 | 0 | 2 | 0 | NON_SYNONYMOU | MODERAT | MISSENSE | Ctg/Atg | L609M | PLES_56421 | | | | | | | | | T | T | | |
| NC_011770 | 6245202 | G | A | 68 | 0 | 2 | 0 | NON_SYNONYMOU | MODERAT | MISSENSE | aCc/aTc | T342I | PLES_56421 | | | | | | | | | A | A | | |
| NC_011770 | 6245203 | T | C | 66 | 0 | 4 | 0 | NON_SYNONYMOU | MODERAT | MISSENSE | Acc/Gcc | T342A | PLES_56421 | | | C | C | C | C | | | | | | |
| NC_011770 | 6245494 | G | A | 69 | 0 | 10 | 0 | NON_SYNONYMOU | MODERAT | MISSENSE | Cac/Tac | H245Y | PLES_56421 | A | A | A | A | A | A | | | A | A | A | A |
| NC_011770 | 6245572 | C | G | 68 | 0 | 6 | 0 | NON_SYNONYMOU | MODERAT | MISSENSE | Ggc/Cgc | G219R | PLES_56421 | G | G | | | | | | | G | G | G | G |
| NC_011770 | 6245875 | C | T | 68 | 0 | 12 | 0 | NON_SYNONYMOU | MODERAT | MISSENSE | Gtc/Atc | V118I | PLES_56421 | T | T | T | T | T | T | T | T | T | T | T | T |
| NC_011770 | 6246163 | G | T | 78 | 0 | 2 | 0 | NON_SYNONYMOU | MODERAT | MISSENSE | Cta/Ata | L22I | PLES_56421 | | | | | | | T | T | | | | |

| | | | | | | | | | | | | | | | | | | | | | | | | | |
|---|---|---|---|---|---|---|---|---|---|---|---|---|---|---|---|---|---|---|---|---|---|---|---|---|---|
| NC_011770 | 6246493 | G | C | 87 | 0 | 4 | 0 | NON_SYNONYMOU | MODERAT | MISSENSE | atC/atG | I156M | PLES_56431 | C | C | | | | | | | | | C | C |
| NC_011770 | 6247246 | G | T | 77 | 0 | 2 | 0 | NON_SYNONYMOU | MODERAT | MISSENSE | Ctg/Atg | L235M | PLES_56441 | | | | | | | T | T | | | | |
| NC_011770 | 6247293 | G | A | 85 | 0 | 2 | 0 | NON_SYNONYMOU | MODERAT | MISSENSE | gCc/gTc | A219V | PLES_56441 | | | | | | | | | A | A | | |
| NC_011770 | 6247584 | A | G | 82 | 0 | 2 | 0 | NON_SYNONYMOU | MODERAT | MISSENSE | gTc/gCc | V122A | PLES_56441 | | | | | | | G | G | | | | |
| NC_011770 | 6248357 | GC | G | 79 | 0 | 1 | 0 | FRAME_SHIFT | HIGH | | ggc/ | G59 | PLES_56451 | | | | | | | | G | | | | |
| NC_011770 | 6248038 | C | T | 77 | 0 | 2 | 0 | NON_SYNONYMOU | MODERAT | MISSENSE | Gtt/Att | V166I | PLES_56451 | | | | | | | | | T | T | | |
| NC_011770 | 6248479 | T | G | 85 | 0 | 10 | 0 | NON_SYNONYMOU | MODERAT | MISSENSE | Atg/Ctg | M19L | PLES_56451 | G | G | G | G | G | G | G | G | | | G | G |
| NC_011770 | 6249776 | A | G | 83 | 0 | 12 | 0 | NON_SYNONYMOU | MODERAT | MISSENSE | aTc/aCc | I225T | PLES_56461 | G | G | G | G | G | G | G | G | G | G | G | G |
| NC_011770 | 6250151 | A | G | 69 | 0 | 4 | 0 | NON_SYNONYMOU | MODERAT | MISSENSE | gTc/gCc | V100A | PLES_56461 | | | G | G | G | G | | | | | | |
| NC_011770 | 6252345 | T | TC | 56 | 0 | 1 | 0 | FRAME_SHIFT | HIGH | | gaa/Cgaa | E67R? | PLES_56481 | | | | | | | | TC | | | | |
| NC_011770 | 6252776 | T | G | 82 | 0 | 2 | 0 | NON_SYNONYMOU | MODERAT | MISSENSE | Ttc/Gtc | F208V | PLES_56481 | | | | | | | | | | | G | G |
| NC_011770 | 6254815 | CA | C | 96 | 0 | 2 | 0 | FRAME_SHIFT | HIGH | | gtg/ | V234 | PLES_56511 | C | C | | | | | | | | | | |
| NC_011770 | 6255593 | A | C | 97 | 0 | 6 | 0 | NON_SYNONYMOU | MODERAT | MISSENSE | Tcc/Gcc | S351A | PLES_56521 | C | C | | | | | C | C | | | C | C |
| NC_011770 | 6255641 | C | T | 94 | 0 | 2 | 0 | NON_SYNONYMOU | MODERAT | MISSENSE | Gtc/Atc | V335I | PLES_56521 | | | | | | | T | T | | | | |
| NC_011770 | 6256547 | T | C | 73 | 0 | 2 | 0 | NON_SYNONYMOU | MODERAT | MISSENSE | Acc/Gcc | T33A | PLES_56521 | | | | | | | C | C | | | | |
| NC_011770 | 6262201 | T | C | 44 | 6 | 4 | 0 | NON_SYNONYMOU | MODERAT | MISSENSE | gAg/gGg | E278G | PLES_56581 | . | . | . | . | . | . | | | C | C | C | C |
| NC_011770 | 6262203 | A | T | 45 | 6 | 2 | 0 | NON_SYNONYMOU | MODERAT | MISSENSE | caT/caA | H277Q | PLES_56581 | . | . | . | . | . | . | | | T | T | | |
| NC_011770 | 6262262 | G | T | 43 | 6 | 4 | 0 | NON_SYNONYMOU | MODERAT | MISSENSE | Cag/Aag | Q258K | PLES_56581 | . | . | . | . | . | . | | | T | T | T | T |
| NC_011770 | 6262414 | A | G | 44 | 6 | 4 | 0 | NON_SYNONYMOU | MODERAT | MISSENSE | gTc/gCc | V207A | PLES_56581 | . | . | . | . | . | . | | | G | G | G | G |
| NC_011770 | 6262629 | T | G | 42 | 6 | 4 | 0 | NON_SYNONYMOU | MODERAT | MISSENSE | caA/caC | Q135H | PLES_56581 | . | . | . | . | . | . | | | G | G | A | A |
| NC_011770 | 6262666 | C | T | 41 | 6 | 4 | 0 | NON_SYNONYMOU | MODERAT | MISSENSE | aGc/aAc | S123N | PLES_56581 | . | . | . | . | . | . | | | T | T | T | T |
| NC_011770 | 6262697 | C | A | 41 | 6 | 4 | 0 | NON_SYNONYMOU | MODERAT | MISSENSE | Gct/Tct | A113S | PLES_56581 | . | . | . | . | . | . | | | A | A | A | A |
| NC_011770 | 6262791 | C | A | 38 | 6 | 4 | 0 | NON_SYNONYMOU | MODERAT | MISSENSE | atG/atT | M81I | PLES_56581 | . | . | . | . | . | . | | | A | A | A | A |
| NC_011770 | 6262832 | C | T | 38 | 6 | 2 | 0 | NON_SYNONYMOU | MODERAT | MISSENSE | Gcc/Acc | A68T | PLES_56581 | . | . | . | . | . | . | | | T | T | | |
| NC_011770 | 6262896 | T | C | 37 | 6 | 4 | 0 | NON_SYNONYMOU | MODERAT | MISSENSE | atA/atG | I46M | PLES_56581 | . | . | . | . | . | . | | | C | C | C | C |
| NC_011770 | 6262940 | T | C | 34 | 6 | 4 | 0 | NON_SYNONYMOU | MODERAT | MISSENSE | Att/Gtt | I32V | PLES_56581 | . | . | . | . | . | . | | | C | C | C | C |
| NC_011770 | 6265554 | T | C | 44 | 6 | 1 | 0 | NON_SYNONYMOU | MODERAT | MISSENSE | aAc/aGc | N327S | PLES_56591 | . | . | . | . | . | . | | C | | | | |
| NC_011770 | 6263377 | G | C | 40 | 6 | 2 | 0 | NON_SYNONYMOU | MODERAT | MISSENSE | Ccg/Gcg | P1053A | PLES_56591 | . | . | . | . | . | . | | | C | C | | |
| NC_011770 | 6263404 | G | C | 41 | 6 | 4 | 0 | NON_SYNONYMOU | MODERAT | MISSENSE | Cag/Gag | Q1044E | PLES_56591 | . | . | . | . | . | . | | | C | C | C | C |
| NC_011770 | 6263467 | T | C | 42 | 6 | 4 | 0 | NON_SYNONYMOU | MODERAT | MISSENSE | Atg/Gtg | M1023V | PLES_56591 | . | . | . | . | . | . | | | C | C | C | C |
| NC_011770 | 6263668 | T | G | 42 | 6 | 4 | 0 | NON_SYNONYMOU | MODERAT | MISSENSE | Aag/Cag | K956Q | PLES_56591 | . | . | . | . | . | . | | | G | G | G | G |
| NC_011770 | 6263772 | C | G | 34 | 6 | 2 | 0 | NON_SYNONYMOU | MODERAT | MISSENSE | cGc/cCc | R921P | PLES_56591 | . | . | . | . | . | . | | | | | G | G |
| NC_011770 | 6263927 | A | T | 40 | 6 | 2 | 0 | NON_SYNONYMOU | MODERAT | MISSENSE | caT/caA | H869Q | PLES_56591 | . | . | . | . | . | . | | | | | T | T |
| NC_011770 | 6263931 | G | A | 39 | 6 | 4 | 0 | NON_SYNONYMOU | MODERAT | MISSENSE | gCc/gTc | A868V | PLES_56591 | . | . | . | . | . | . | | | A | A | A | A |
| NC_011770 | 6264469 | C | T | 41 | 6 | 4 | 0 | NON_SYNONYMOU | MODERAT | MISSENSE | Gta/Ata | V689I | PLES_56591 | . | . | . | . | . | . | | | T | T | T | T |
| NC_011770 | 6264481 | A | G | 39 | 6 | 4 | 0 | NON_SYNONYMOU | MODERAT | MISSENSE | Tac/Cac | Y685H | PLES_56591 | . | . | . | . | . | . | | | G | G | G | G |
| NC_011770 | 6264675 | T | C | 45 | 6 | 4 | 0 | NON_SYNONYMOU | MODERAT | MISSENSE | gAc/gGc | D620G | PLES_56591 | . | . | . | . | . | . | | | C | C | C | C |
| NC_011770 | 6264835 | T | C | 45 | 6 | 4 | 0 | NON_SYNONYMOU | MODERAT | MISSENSE | Agt/Ggt | S567G | PLES_56591 | . | . | . | . | . | . | | | C | C | C | C |
| NC_011770 | 6265053 | G | A | 47 | 6 | 2 | 0 | NON_SYNONYMOU | MODERAT | MISSENSE | gCc/gTc | A494V | PLES_56591 | . | . | . | . | . | . | | | A | A | | |
| NC_011770 | 6265323 | T | A | 48 | 6 | 2 | 0 | NON_SYNONYMOU | MODERAT | MISSENSE | tAc/tTc | Y404F | PLES_56591 | . | . | . | . | . | . | | | | | A | A |
| NC_011770 | 6265418 | C | A | 50 | 6 | 4 | 0 | NON_SYNONYMOU | MODERAT | MISSENSE | caG/caT | Q372H | PLES_56591 | . | . | . | . | . | . | | | A | A | A | A |
| NC_011770 | 6265469 | G | T | 49 | 6 | 4 | 0 | NON_SYNONYMOU | MODERAT | MISSENSE | gaC/gaA | D355E | PLES_56591 | . | . | . | . | . | . | | | T | T | T | T |
| NC_011770 | 6265827 | T | G | 43 | 6 | 4 | 0 | NON_SYNONYMOU | MODERAT | MISSENSE | gAg/gCg | E236A | PLES_56591 | . | . | . | . | . | . | | | G | G | G | G |
| NC_011770 | 6265922 | T | A | 43 | 6 | 4 | 0 | NON_SYNONYMOU | MODERAT | MISSENSE | caA/caT | Q204H | PLES_56591 | . | . | . | . | . | . | | | A | A | A | A |
| NC_011770 | 6266601 | T | C | 41 | 6 | 2 | 0 | NON_SYNONYMOU | MODERAT | MISSENSE | Atc/Gtc | I669V | PLES_56601 | . | . | . | . | . | . | | | C | C | | |
| NC_011770 | 6266618 | T | C | 39 | 6 | 2 | 0 | NON_SYNONYMOU | MODERAT | MISSENSE | aAg/aGg | K663R | PLES_56601 | . | . | . | . | . | . | C | C | | | | |
| NC_011770 | 6266628 | G | A | 40 | 6 | 2 | 0 | NON_SYNONYMOU | MODERAT | MISSENSE | Ccc/Tcc | P660S | PLES_56601 | . | . | . | . | . | . | | | A | A | | |
| NC_011770 | 6266663 | G | A | 38 | 6 | 2 | 0 | NON_SYNONYMOU | MODERAT | MISSENSE | gCg/gTg | A648V | PLES_56601 | . | . | . | . | . | . | | | A | A | | |
| NC_011770 | 6266723 | T | C | 53 | 0 | 2 | 0 | NON_SYNONYMOU | MODERAT | MISSENSE | aAc/aGc | N628S | PLES_56601 | | | | | | | | | C | C | | |
| NC_011770 | 6266759 | G | A | 55 | 0 | 2 | 0 | NON_SYNONYMOU | MODERAT | MISSENSE | gCc/gTc | A616V | PLES_56601 | | | | | | | A | A | | | | |
| NC_011770 | 6267011 | T | C | 77 | 0 | 4 | 0 | NON_SYNONYMOU | MODERAT | MISSENSE | tAc/tGc | Y532C | PLES_56601 | | | C | C | C | C | | | | | | |
| NC_011770 | 6267024 | G | C | 77 | 0 | 6 | 0 | NON_SYNONYMOU | MODERAT | MISSENSE | Cag/Gag | Q528E | PLES_56601 | | | | | | | C | C | C | C | C | C |
| NC_011770 | 6267206 | G | C | 74 | 0 | 8 | 0 | NON_SYNONYMOU | MODERAT | MISSENSE | gCg/gGg | A467G | PLES_56601 | C | C | C | C | C | C | C | C | | | | |
| NC_011770 | 6267341 | T | C | 79 | 0 | 12 | 0 | NON_SYNONYMOU | MODERAT | MISSENSE | aAg/aGg | K422R | PLES_56601 | C | C | C | C | C | C | C | C | C | C | C | C |
| NC_011770 | 6267521 | G | A | 78 | 0 | 6 | 0 | NON_SYNONYMOU | MODERAT | MISSENSE | cCg/cTg | P362L | PLES_56601 | | | | | | | A | A | A | A | A | A |
| NC_011770 | 6267606 | T | C | 81 | 0 | 8 | 0 | NON_SYNONYMOU | MODERAT | MISSENSE | Acc/Gcc | T334A | PLES_56601 | C | C | C | C | C | C | C | C | | | | |
| NC_011770 | 6267732 | G | T | 76 | 0 | 2 | 0 | NON_SYNONYMOU | MODERAT | MISSENSE | Ctc/Atc | L292I | PLES_56601 | T | T | | | | | | | | | | |
| NC_011770 | 6267741 | A | G | 75 | 0 | 12 | 0 | NON_SYNONYMOU | MODERAT | MISSENSE | Tcg/Ccg | S289P | PLES_56601 | G | G | G | G | G | G | G | G | G | G | G | G |
| NC_011770 | 6268299 | G | A | 82 | 0 | 4 | 0 | NON_SYNONYMOU | MODERAT | MISSENSE | Cgc/Tgc | R103C | PLES_56601 | | | | | | | | | A | A | A | A |
| NC_011770 | 6271004 | G | T | 62 | 0 | 4 | 0 | NON_SYNONYMOU | MODERAT | MISSENSE | Gcc/Tcc | A5S | PLES_56641 | | | | | | | | | T | T | T | T |
| NC_011770 | 6275794 | A | G | 16 | 0 | 1 | 0 | NON_SYNONYMOU | MODERAT | MISSENSE | Acc/Gcc | T202A | PLES_56671 | | | | G | | | | | | | | |
| NC_011770 | 6275582 | A | G | 48 | 0 | 4 | 0 | NON_SYNONYMOU | MODERAT | MISSENSE | cAg/cGg | Q131R | PLES_56671 | | | G | G | G | G | | | | | | |

| | | | | | | | | | | | | | | | | | | | | | | | | | |
|---|---|---|---|---|---|---|---|---|---|---|---|---|---|---|---|---|---|---|---|---|---|---|---|---|---|
| NC_011770 | 6275627 | T | A | 45 | 0 | 4 | 0 | NON_SYNONYMOU | MODERAT | MISSENSE | cTg/cAg | L146Q | PLES_56671 | | | A | A | A | A | | | | | | |
| NC_011770 | 6275636 | C | T | 42 | 0 | 4 | 0 | NON_SYNONYMOU | MODERAT | MISSENSE | aCg/aTg | T149M | PLES_56671 | | | T | T | T | T | | | | | | |
| NC_011770 | 6275698 | G | A | 37 | 0 | 4 | 0 | NON_SYNONYMOU | MODERAT | MISSENSE | Gcc/Acc | A170T | PLES_56671 | | | A | A | A | A | | | | | | |
| NC_011770 | 6275784 | T | A | 16 | 0 | 4 | 0 | NON_SYNONYMOU | MODERAT | MISSENSE | gaT/gaA | D198E | PLES_56671 | | | A | A | A | A | | | | | | |
| NC_011770 | 6275797 | G | T | 14 | 0 | 12 | 0 | NON_SYNONYMOU | MODERAT | MISSENSE | Gtc/Ttc | V203F | PLES_56671 | T | T | T | T | T | T | T | T | T | T | T | T |
| NC_011770 | 6276523 | C | T | 56 | 0 | 2 | 0 | NON_SYNONYMOU | MODERAT | MISSENSE | Gct/Act | A46T | PLES_56691 | | | | | | | | | | | T | T |
| NC_011770 | 6281693 | A | G | 77 | 0 | 4 | 0 | NON_SYNONYMOU | MODERAT | MISSENSE | Ata/Gta | I178V | PLES_56761 | G | G | | | | | G | G | | | | |
| NC_011770 | 6281772 | A | G | 74 | 0 | 8 | 0 | NON_SYNONYMOU | MODERAT | MISSENSE | aAc/aGc | N204S | PLES_56761 | | | G | G | G | G | | | G | G | G | G |
| NC_011770 | 6281915 | G | T | 72 | 0 | 10 | 0 | NON_SYNONYMOU | MODERAT | MISSENSE | ttC/ttA | F382L | PLES_56771 | | | T | T | T | T | T | T | T | T | T | T |
| NC_011770 | 6283509 | A | G | 66 | 0 | 6 | 0 | NON_SYNONYMOU | MODERAT | MISSENSE | aAc/aGc | N96S | PLES_56781 | | | | | | | G | G | G | G | G | G |
| NC_011770 | 6284873 | GC | G | 62 | 0 | 2 | 0 | FRAME_SHIFT | HIGH | | ggc/ | G222 | PLES_56791 | | | | | | | G | GCC | | | | |
| NC_011770 | 6285187 | C | T | 68 | 0 | 10 | 0 | NON_SYNONYMOU | MODERAT | MISSENSE | Gtc/Atc | V118I | PLES_56791 | | | T | T | T | T | T | T | T | T | T | T |
| NC_011770 | 6285298 | T | C | 67 | 0 | 4 | 0 | NON_SYNONYMOU | MODERAT | MISSENSE | Atc/Gtc | I81V | PLES_56791 | | | C | C | C | C | | | | | | |
| NC_011770 | 6285396 | G | A | 69 | 0 | 10 | 0 | NON_SYNONYMOU | MODERAT | MISSENSE | aCg/aTg | T48M | PLES_56791 | | | A | A | A | A | A | A | A | A | A | A |
| NC_011770 | 6285402 | T | C | 70 | 0 | 10 | 0 | NON_SYNONYMOU | MODERAT | MISSENSE | aAc/aGc | N46S | PLES_56791 | | | C | C | C | C | C | C | C | C | C | C |
| NC_011770 | 6285496 | C | A | 80 | 0 | 2 | 0 | NON_SYNONYMOU | MODERAT | MISSENSE | Gtc/Ttc | V15F | PLES_56791 | A | A | | | | | | | | | | |
| NC_011770 | 6285505 | G | T | 77 | 0 | 2 | 0 | NON_SYNONYMOU | MODERAT | MISSENSE | Ctt/Att | L12I | PLES_56791 | T | T | | | | | | | | | | |
| NC_011770 | 6286147 | C | CG | 59 | 0 | 1 | 0 | FRAME_SHIFT | HIGH | | ccc/cccC | P41P? | PLES_56801 | | | | | | | | CG | | | | |
| NC_011770 | 6286659 | G | A | 60 | 0 | 4 | 0 | NON_SYNONYMOU | MODERAT | MISSENSE | Cac/Tac | H39Y | PLES_56811 | | | A | A | A | A | | | | | | |
| NC_011770 | 6290001 | C | T | 72 | 0 | 6 | 0 | NON_SYNONYMOU | MODERAT | MISSENSE | cCg/cTg | P174L | PLES_56851 | T | T | T | T | T | T | | | | | | |
| NC_011770 | 6290469 | G | A | 70 | 0 | 2 | 0 | NON_SYNONYMOU | MODERAT | MISSENSE | gGc/gAc | G330D | PLES_56851 | A | A | | | | | | | | | | |
| NC_011770 | 6290696 | A | G | 66 | 0 | 10 | 0 | NON_SYNONYMOU | MODERAT | MISSENSE | Atc/Gtc | I406V | PLES_56851 | | | G | G | G | G | G | G | G | G | G | G |
| NC_011770 | 6290961 | C | T | 73 | 0 | 12 | 0 | NON_SYNONYMOU | MODERAT | MISSENSE | aCg/aTg | T494M | PLES_56851 | T | T | T | T | T | T | T | T | T | T | T | T |
| NC_011770 | 6293079 | C | T | 85 | 0 | 2 | 0 | NON_SYNONYMOU | MODERAT | MISSENSE | Cgc/Tgc | R661C | PLES_56861 | | | | | | | | | T | T | | |
| NC_011770 | 6295241 | T | C | 72 | 0 | 2 | 0 | NON_SYNONYMOU | MODERAT | MISSENSE | Acc/Gcc | T2A | PLES_56881 | C | C | | | | | | | | | | |
| NC_011770 | 6295693 | A | C | 84 | 0 | 12 | 0 | NON_SYNONYMOU | MODERAT | MISSENSE | Atg/Ctg | M101L | PLES_56891 | C | C | C | C | C | C | C | C | C | C | C | C |
| NC_011770 | 6296170 | A | G | 90 | 0 | 2 | 0 | NON_SYNONYMOU | MODERAT | MISSENSE | Acc/Gcc | T260A | PLES_56891 | | | | | | | G | G | | | | |
| NC_011770 | 6296761 | G | A | 55 | 0 | 4 | 0 | NON_SYNONYMOU | MODERAT | MISSENSE | Gcg/Acg | A457T | PLES_56891 | | | | | | | A | A | A | A | | |
| NC_011770 | 6296813 | G | T | 58 | 0 | 2 | 0 | NON_SYNONYMOU | MODERAT | MISSENSE | cGa/cTa | R474L | PLES_56891 | T | T | | | | | | | | | | |
| NC_011770 | 6296842 | A | G | 60 | 0 | 4 | 0 | NON_SYNONYMOU | MODERAT | MISSENSE | Acg/Gcg | T484A | PLES_56891 | | | | | | | G | G | G | G | | |
| NC_011770 | 6302790 | T | C | 87 | 0 | 2 | 0 | NON_SYNONYMOU | MODERAT | MISSENSE | aTg/aCg | M44T | PLES_56931 | | | | | | | | | C | C | | |
| NC_011770 | 6304064 | G | T | 72 | 0 | 2 | 0 | NON_SYNONYMOU | MODERAT | MISSENSE | Cgc/Agc | R346S | PLES_56941 | | | | | | | | | | | T | T |
| NC_011770 | 6304129 | A | T | 62 | 0 | 2 | 0 | NON_SYNONYMOU | MODERAT | MISSENSE | gTg/gAg | V324E | PLES_56941 | T | T | | | | | | | | | | |
| NC_011770 | 6304624 | C | T | 84 | 0 | 2 | 0 | NON_SYNONYMOU | MODERAT | MISSENSE | gGc/gAc | G159D | PLES_56941 | | | | | | | | | T | T | | |
| NC_011770 | 6304883 | G | A | 78 | 0 | 1 | 0 | NON_SYNONYMOU | MODERAT | MISSENSE | Cgc/Tgc | R73C | PLES_56941 | | | | | | | A | | | | | |
| NC_011770 | 6304942 | G | T | 78 | 0 | 2 | 0 | NON_SYNONYMOU | MODERAT | MISSENSE | cCt/cAt | P53H | PLES_56941 | | | | | | | | | | | T | T |
| NC_011770 | 6310664 | C | T | 80 | 0 | 2 | 0 | NON_SYNONYMOU | MODERAT | MISSENSE | Gcg/Acg | A754T | PLES_57021 | | | | | | | | | | | T | T |
| NC_011770 | 6310765 | G | A | 73 | 0 | 2 | 0 | NON_SYNONYMOU | MODERAT | MISSENSE | gCc/gTc | A720V | PLES_57021 | | | | | | | A | A | | | | |
| NC_011770 | 6310766 | C | T | 73 | 0 | 2 | 0 | NON_SYNONYMOU | MODERAT | MISSENSE | Gcc/Acc | A720T | PLES_57021 | T | T | | | | | | | | | | |
| NC_011770 | 6311227 | C | T | 74 | 0 | 2 | 0 | NON_SYNONYMOU | MODERAT | MISSENSE | gGg/gAg | G566E | PLES_57021 | T | T | | | | | | | | | | |
| NC_011770 | 6311290 | C | T | 72 | 0 | 2 | 0 | NON_SYNONYMOU | MODERAT | MISSENSE | aGg/aAg | R545K | PLES_57021 | T | T | | | | | | | | | | |
| NC_011770 | 6311431 | A | G | 66 | 0 | 12 | 0 | NON_SYNONYMOU | MODERAT | MISSENSE | cTg/cCg | L498P | PLES_57021 | G | G | G | G | G | G | G | G | G | G | G | G |
| NC_011770 | 6311525 | C | T | 62 | 0 | 2 | 0 | NON_SYNONYMOU | MODERAT | MISSENSE | Gcc/Acc | A467T | PLES_57021 | T | T | | | | | | | | | | |
| NC_011770 | 6311576 | C | T | 61 | 0 | 2 | 0 | NON_SYNONYMOU | MODERAT | MISSENSE | Gcc/Acc | A450T | PLES_57021 | T | T | | | | | | | | | | |
| NC_011770 | 6311689 | A | T | 60 | 0 | 2 | 0 | NON_SYNONYMOU | MODERAT | MISSENSE | cTg/cAg | L412Q | PLES_57021 | T | T | | | | | | | | | | |
| NC_011770 | 6311791 | G | A | 70 | 0 | 2 | 0 | NON_SYNONYMOU | MODERAT | MISSENSE | aCg/aTg | T378M | PLES_57021 | A | A | | | | | | | | | | |
| NC_011770 | 6311866 | C | T | 69 | 0 | 2 | 0 | NON_SYNONYMOU | MODERAT | MISSENSE | cGc/cAc | R353H | PLES_57021 | | | | | | | | | | | T | T |
| NC_011770 | 6312151 | T | C | 68 | 0 | 12 | 0 | NON_SYNONYMOU | MODERAT | MISSENSE | cAt/cGt | H258R | PLES_57021 | C | C | C | C | C | C | C | C | C | C | C | C |
| NC_011770 | 6312338 | T | C | 66 | 0 | 6 | 0 | NON_SYNONYMOU | MODERAT | MISSENSE | Acg/Gcg | T196A | PLES_57021 | C | C | | | | | C | C | C | C | | |
| NC_011770 | 6314392 | C | A | 76 | 0 | 2 | 0 | NON_SYNONYMOU | MODERAT | MISSENSE | Ccg/Acg | P217T | PLES_57041 | | | | | | | | | | | A | A |
| NC_011770 | 6314399 | C | T | 77 | 0 | 2 | 0 | NON_SYNONYMOU | MODERAT | MISSENSE | aCg/aTg | T219M | PLES_57041 | | | | | | | | | | | T | T |
| NC_011770 | 6314440 | A | G | 72 | 0 | 2 | 0 | NON_SYNONYMOU | MODERAT | MISSENSE | Acc/Gcc | T233A | PLES_57041 | G | G | | | | | | | | | | |
| NC_011770 | 6314569 | A | C | 71 | 0 | 2 | 0 | NON_SYNONYMOU | MODERAT | MISSENSE | Aaa/Caa | K276Q | PLES_57041 | C | C | | | | | | | | | | |
| NC_011770 | 6315181 | A | G | 72 | 0 | 2 | 0 | NON_SYNONYMOU | MODERAT | MISSENSE | atA/atG | I11M | PLES_57051 | | | | | | | G | G | | | | |
| NC_011770 | 6315371 | G | T | 87 | 0 | 2 | 0 | NON_SYNONYMOU | MODERAT | MISSENSE | Gtc/Ttc | V75F | PLES_57051 | T | T | | | | | | | | | | |
| NC_011770 | 6315716 | G | A | 86 | 0 | 2 | 0 | NON_SYNONYMOU | MODERAT | MISSENSE | Gtc/Atc | V190I | PLES_57051 | | | | | | | A | A | | | | |
| NC_011770 | 6316059 | C | T | 91 | 0 | 2 | 0 | NON_SYNONYMOU | MODERAT | MISSENSE | gCg/gTg | A304V | PLES_57051 | | | | | | | T | T | | | | |
| NC_011770 | 6316649 | C | A | 90 | 0 | 2 | 0 | NON_SYNONYMOU | MODERAT | MISSENSE | Ccc/Acc | P501T | PLES_57051 | | | | | | | A | A | | | | |
| NC_011770 | 6316764 | G | A | 79 | 0 | 2 | 0 | NON_SYNONYMOU | MODERAT | MISSENSE | aCc/aTc | T381I | PLES_57061 | | | | | | | | | | | A | A |
| NC_011770 | 6316772 | C | G | 81 | 0 | 6 | 0 | NON_SYNONYMOU | MODERAT | MISSENSE | gaG/gaC | E378D | PLES_57061 | | | | | | | G | G | G | G | G | G |
| NC_011770 | 6317050 | G | A | 82 | 0 | 2 | 0 | NON_SYNONYMOU | MODERAT | MISSENSE | Ccg/Tcg | P286S | PLES_57061 | | | | | | | | | A | A | | |
| NC_011770 | 6317377 | T | C | 79 | 0 | 2 | 0 | NON_SYNONYMOU | MODERAT | MISSENSE | Agc/Ggc | S177G | PLES_57061 | | | | | | | C | C | | | | |

| | | | | | | | | | | | | | | | | | | | | | | | | | |
|---|---|---|---|---|---|---|---|---|---|---|---|---|---|---|---|---|---|---|---|---|---|---|---|---|---|
| NC_011770 | 6318723 | T | C | 62 | 0 | 2 | 0 | NON_SYNONYMOU | MODERAT | MISSENSE | Atc/Gtc | I275V | PLES_57071 | | | | | | | | | | | C | C |
| NC_011770 | 6320003 | A | G | 63 | 0 | 1 | 0 | NON_SYNONYMOU | MODERAT | MISSENSE | Acc/Gcc | T77A | PLES_57081 | | | | | | | G | | | | | |
| NC_011770 | 6320087 | G | A | 63 | 0 | 2 | 0 | NON_SYNONYMOU | MODERAT | MISSENSE | Ggc/Agc | G105S | PLES_57081 | | | | | | | | | A | A | | |
| NC_011770 | 6320300 | G | A | 72 | 0 | 4 | 0 | NON_SYNONYMOU | MODERAT | MISSENSE | Gtc/Atc | V176I | PLES_57081 | A | A | | | | | | | | | A | A |
| NC_011770 | 6320340 | T | C | 79 | 0 | 8 | 0 | NON_SYNONYMOU | MODERAT | MISSENSE | gTc/gCc | V189A | PLES_57081 | C | C | | | | | C | C | C | C | C | C |
| NC_011770 | 6320827 | A | C | 78 | 0 | 2 | 0 | NON_SYNONYMOU | MODERAT | MISSENSE | gaA/gaC | E351D | PLES_57081 | | | | | | | | | | | C | C |
| NC_011770 | 6320897 | G | A | 78 | 0 | 6 | 0 | NON_SYNONYMOU | MODERAT | MISSENSE | Gta/Ata | V375I | PLES_57081 | | | A | A | A | A | A | A | | | | |
| NC_011770 | 6321089 | G | T | 78 | 0 | 2 | 0 | NON_SYNONYMOU | MODERAT | MISSENSE | Gcc/Tcc | A439S | PLES_57081 | T | T | | | | | | | | | | |
| NC_011770 | 6322754 | G | A | 59 | 0 | 2 | 0 | NON_SYNONYMOU | MODERAT | MISSENSE | Gac/Aac | D109N | PLES_57121 | | | | | | | | | A | A | | |
| NC_011770 | 6322776 | A | G | 60 | 0 | 2 | 0 | NON_SYNONYMOU | MODERAT | MISSENSE | cAg/cGg | Q116R | PLES_57121 | | | | | | | | | G | G | | |
| NC_011770 | 6323741 | T | C | 44 | 0 | 2 | 0 | NON_SYNONYMOU | MODERAT | MISSENSE | Ttc/Ctc | F438L | PLES_57121 | C | C | | | | | | | | | | |
| NC_011770 | 6324535 | G | A | 71 | 0 | 1 | 0 | NON_SYNONYMOU | MODERAT | MISSENSE | gGg/gAg | G167E | PLES_57131 | A | | | | | | | | | | | |
| NC_011770 | 6331370 | G | C | 103 | 0 | 2 | 0 | NON_SYNONYMOU | MODERAT | MISSENSE | aCc/aGc | T130S | PLES_57191 | | | | | | | | | C | C | | |
| NC_011770 | 6332371 | G | A | 57 | 0 | 2 | 0 | NON_SYNONYMOU | MODERAT | MISSENSE | aGc/aAc | S130N | PLES_57201 | | | | | | | | | | | A | A |
| NC_011770 | 6332703 | A | G | 71 | 0 | 2 | 0 | NON_SYNONYMOU | MODERAT | MISSENSE | Aag/Gag | K241E | PLES_57201 | | | | | | | | | G | G | | |
| NC_011770 | 6333509 | G | A | 63 | 0 | 2 | 0 | NON_SYNONYMOU | MODERAT | MISSENSE | gCg/gTg | A235V | PLES_57211 | | | | | | | A | A | | | | |
| NC_011770 | 6333597 | C | T | 70 | 0 | 2 | 0 | NON_SYNONYMOU | MODERAT | MISSENSE | Ggc/Agc | G206S | PLES_57211 | T | T | | | | | | | | | | |
| NC_011770 | 6334191 | A | C | 79 | 0 | 10 | 0 | NON_SYNONYMOU | MODERAT | MISSENSE | Tgg/Ggg | W8G | PLES_57211 | C | C | C | C | C | C | | | C | C | C | C |
| NC_011770 | 6334884 | G | A | 68 | 0 | 4 | 0 | NON_SYNONYMOU | MODERAT | MISSENSE | cCg/cTg | P247L | PLES_57221 | | | A | A | A | A | | | | | | |
| NC_011770 | 6335049 | A | G | 77 | 0 | 12 | 0 | NON_SYNONYMOU | MODERAT | MISSENSE | gTg/gCg | V192A | PLES_57221 | G | G | G | G | G | G | G | G | G | G | G | G |
| NC_011770 | 6335300 | C | A | 82 | 0 | 4 | 0 | NON_SYNONYMOU | MODERAT | MISSENSE | gaG/gaT | E108D | PLES_57221 | | | | | | | | | A | A | A | A |
| NC_011770 | 6335397 | G | A | 78 | 0 | 2 | 0 | NON_SYNONYMOU | MODERAT | MISSENSE | gCc/gTc | A76V | PLES_57221 | | | | | | | | | | | A | A |
| NC_011770 | 6335895 | T | C | 87 | 0 | 2 | 0 | NON_SYNONYMOU | MODERAT | MISSENSE | Aat/Gat | N66D | PLES_57231 | | | | | | | C | C | | | | |
| NC_011770 | 6335944 | G | C | 86 | 0 | 2 | 0 | NON_SYNONYMOU | MODERAT | MISSENSE | gaC/gaG | D49E | PLES_57231 | | | | | | | C | C | | | | |
| NC_011770 | 6335995 | T | G | 84 | 0 | 2 | 0 | NON_SYNONYMOU | MODERAT | MISSENSE | gaA/gaC | E32D | PLES_57231 | G | G | | | | | | | | | | |
| NC_011770 | 6336508 | G | A | 96 | 0 | 1 | 0 | STOP_GAINED | HIGH | NONSENSE | tGg/tAg | W82* | PLES_57241 | | | | | | | A | | | | | |
| NC_011770 | 6336651 | C | A | 92 | 0 | 2 | 0 | NON_SYNONYMOU | MODERAT | MISSENSE | Ctg/Atg | L130M | PLES_57241 | | | | | | | | | A | A | | |
| NC_011770 | 6336977 | C | T | 86 | 0 | 4 | 0 | NON_SYNONYMOU | MODERAT | MISSENSE | Gcg/Acg | A117T | PLES_57251 | | | T | T | T | T | | | | | | |
| NC_011770 | 6337268 | G | T | 89 | 0 | 2 | 0 | NON_SYNONYMOU | MODERAT | MISSENSE | Ctt/Att | L20I | PLES_57251 | | | | | | | | | | | T | T |
| NC_011770 | 6340285 | G | T | 98 | 0 | 2 | 0 | NON_SYNONYMOU | MODERAT | MISSENSE | aGc/aTc | S21I | PLES_57301 | T | T | | | | | | | | | | |
| NC_011770 | 6340524 | C | A | 87 | 0 | 4 | 0 | NON_SYNONYMOU | MODERAT | MISSENSE | Ccg/Acg | P101T | PLES_57301 | | | A | A | A | A | | | | | | |
| NC_011770 | 6345609 | T | C | 81 | 0 | 12 | 0 | NON_SYNONYMOU | MODERAT | MISSENSE | gAc/gGc | D177G | PLES_57361 | C | C | C | C | C | C | C | C | C | C | C | C |
| NC_011770 | 6346038 | C | T | 81 | 0 | 2 | 0 | NON_SYNONYMOU | MODERAT | MISSENSE | gGc/gAc | G34D | PLES_57361 | | | | | | | T | T | | | | |
| NC_011770 | 6346643 | C | T | 65 | 0 | 4 | 0 | STOP_GAINED | HIGH | NONSENSE | tgG/tgA | W121* | PLES_57371 | | | | | | | T | T | | | T | T |
| NC_011770 | 6346516 | T | C | 66 | 0 | 2 | 0 | NON_SYNONYMOU | MODERAT | MISSENSE | Acc/Gcc | T164A | PLES_57371 | C | C | | | | | | | | | | |
| NC_011770 | 6346645 | A | G | 66 | 0 | 4 | 0 | NON_SYNONYMOU | MODERAT | MISSENSE | Tgg/Cgg | W121R | PLES_57371 | | | | | | | G | G | | | G | G |
| NC_011770 | 6346647 | G | T | 67 | 0 | 2 | 0 | NON_SYNONYMOU | MODERAT | MISSENSE | gCg/gAg | A120E | PLES_57371 | T | T | | | | | | | | | | |
| NC_011770 | 6346656 | G | A | 65 | 0 | 4 | 0 | NON_SYNONYMOU | MODERAT | MISSENSE | gCt/gTt | A117V | PLES_57371 | | | | | | | A | A | | | A | A |
| NC_011770 | 6346702 | G | A | 63 | 0 | 2 | 0 | NON_SYNONYMOU | MODERAT | MISSENSE | Cgc/Tgc | R102C | PLES_57371 | | | | | | | | | | | A | A |
| NC_011770 | 6347182 | C | T | 77 | 0 | 2 | 0 | NON_SYNONYMOU | MODERAT | MISSENSE | atG/atA | M225I | PLES_57381 | | | | | | | | | | | T | T |
| NC_011770 | 6347564 | C | T | 87 | 0 | 4 | 0 | NON_SYNONYMOU | MODERAT | MISSENSE | cGc/cAc | R98H | PLES_57381 | | | | | | | | | T | T | T | T |
| NC_011770 | 6347750 | T | C | 90 | 0 | 4 | 0 | NON_SYNONYMOU | MODERAT | MISSENSE | gAc/gGc | D36G | PLES_57381 | | | | | | | | | C | C | C | C |
| NC_011770 | 6352723 | G | A | 73 | 0 | 2 | 0 | NON_SYNONYMOU | MODERAT | MISSENSE | gCg/gTg | A80V | PLES_57421 | | | | | | | | | A | A | | |
| NC_011770 | 6352766 | C | A | 76 | 0 | 2 | 0 | NON_SYNONYMOU | MODERAT | MISSENSE | Gca/Tca | A66S | PLES_57421 | | | | | | | | | A | A | | |
| NC_011770 | 6352844 | G | A | 80 | 0 | 2 | 0 | NON_SYNONYMOU | MODERAT | MISSENSE | Ctc/Ttc | L40F | PLES_57421 | A | A | | | | | | | | | | |
| NC_011770 | 6352912 | G | A | 76 | 0 | 2 | 0 | NON_SYNONYMOU | MODERAT | MISSENSE | gCc/gTc | A17V | PLES_57421 | | | | | | | A | A | | | | |
| NC_011770 | 6354289 | C | T | 83 | 0 | 8 | 0 | NON_SYNONYMOU | MODERAT | MISSENSE | cGc/cAc | R146H | PLES_57441 | T | T | | | | | T | T | T | T | T | T |
| NC_011770 | 6354644 | C | T | 71 | 0 | 12 | 0 | NON_SYNONYMOU | MODERAT | MISSENSE | Ggc/Agc | G28S | PLES_57441 | T | T | T | T | T | T | T | T | T | T | T | T |
| NC_011770 | 6355460 | C | T | 93 | 0 | 4 | 0 | NON_SYNONYMOU | MODERAT | MISSENSE | cGg/cAg | R124Q | PLES_57471 | | | T | T | T | T | | | | | | |
| NC_011770 | 6355523 | A | T | 81 | 0 | 2 | 0 | NON_SYNONYMOU | MODERAT | MISSENSE | cTg/cAg | L103Q | PLES_57471 | | | | | | | | | | | T | T |
| NC_011770 | 6362233 | T | C | 90 | 0 | 2 | 0 | NON_SYNONYMOU | MODERAT | MISSENSE | aAg/aGg | K130R | PLES_57541 | | | | | | | C | C | | | | |
| NC_011770 | 6365096 | G | A | 99 | 0 | 4 | 0 | NON_SYNONYMOU | MODERAT | MISSENSE | Gcc/Acc | A59T | PLES_57571 | | | A | A | A | A | | | | | | |
| NC_011770 | 6366809 | T | C | 79 | 0 | 2 | 0 | NON_SYNONYMOU | MODERAT | MISSENSE | Acc/Gcc | T130A | PLES_57581 | | | | | | | | | C | C | | |
| NC_011770 | 6367448 | G | C | 72 | 0 | 2 | 0 | NON_SYNONYMOU | MODERAT | MISSENSE | gCg/gGg | A256G | PLES_57591 | | | | | | | C | C | | | | |
| NC_011770 | 6375563 | G | A | 77 | 0 | 2 | 0 | INTRAGENIC | MODIFIER | | | | PLES_57642 | | | | | | | A | A | | | | |
| NC_011770 | 6376085 | G | A | 111 | 0 | 1 | 0 | INTRAGENIC | MODIFIER | | | | PLES_57643 | | | | | | | | A | | | | |
| NC_011770 | 6376491 | C | T | 94 | 0 | 1 | 0 | INTRAGENIC | MODIFIER | | | | PLES_57643 | | | | | | | | T | | | | |
| NC_011770 | 6376638 | T | C | 100 | 0 | 1 | 0 | INTRAGENIC | MODIFIER | | | | PLES_57643 | | | | | | | C | | | | | |
| NC_011770 | 6381338 | T | C | 69 | 0 | 2 | 0 | NON_SYNONYMOU | MODERAT | MISSENSE | Act/Gct | T412A | PLES_57651 | | | | | | | | | | | C | C |
| NC_011770 | 6381364 | G | A | 69 | 0 | 2 | 0 | NON_SYNONYMOU | MODERAT | MISSENSE | cCg/cTg | P403L | PLES_57651 | A | A | | | | | | | | | | |
| NC_011770 | 6381391 | G | A | 66 | 0 | 2 | 0 | NON_SYNONYMOU | MODERAT | MISSENSE | gCg/gTg | A394V | PLES_57651 | | | | | | | | | | | A | A |
| NC_011770 | 6381416 | T | C | 65 | 0 | 2 | 0 | NON_SYNONYMOU | MODERAT | MISSENSE | Atc/Gtc | I386V | PLES_57651 | | | | | | | C | C | | | | |

| | | | | | | | | | | | | | | | | | | | | | | | | | |
|---|---|---|---|---|---|---|---|---|---|---|---|---|---|---|---|---|---|---|---|---|---|---|---|---|---|
| NC_011770 | 6381640 | T | A | 62 | 0 | 4 | 0 | NON_SYNONYMOU | MODERAT | MISSENSE | tAc/tTc | Y311F | PLES_57651 | | | | | | | A | A | | | A | A |
| NC_011770 | 6381845 | C | T | 66 | 0 | 2 | 0 | NON_SYNONYMOU | MODERAT | MISSENSE | Gtc/Atc | V243I | PLES_57651 | T | T | | | | | | | | | | |
| NC_011770 | 6392679 | A | G | 96 | 0 | 2 | 0 | NON_SYNONYMOU | MODERAT | MISSENSE | gTc/gCc | V35A | PLES_57741 | | | | | | | G | G | | | | |
| NC_011770 | 6395817 | C | A | 63 | 0 | 2 | 0 | NON_SYNONYMOU | MODERAT | MISSENSE | cGc/cTc | R110L | PLES_57771 | | | | | | | | | A | A | | |
| NC_011770 | 6396057 | G | A | 62 | 0 | 2 | 0 | NON_SYNONYMOU | MODERAT | MISSENSE | tCa/tTa | S30L | PLES_57771 | A | A | | | | | | | | | | |
| NC_011770 | 6396130 | C | T | 61 | 0 | 4 | 0 | NON_SYNONYMOU | MODERAT | MISSENSE | Gcg/Acg | A6T | PLES_57771 | | | T | T | T | T | | | | | | |
| NC_011770 | 6396295 | C | T | 65 | 0 | 4 | 0 | NON_SYNONYMOU | MODERAT | MISSENSE | gGc/gAc | G295D | PLES_57781 | | | T | T | T | T | | | | | | |
| NC_011770 | 6396299 | G | C | 66 | 0 | 2 | 0 | NON_SYNONYMOU | MODERAT | MISSENSE | Ctc/Gtc | L294V | PLES_57781 | C | C | | | | | | | | | | |
| NC_011770 | 6396646 | G | A | 69 | 0 | 2 | 0 | NON_SYNONYMOU | MODERAT | MISSENSE | tCg/tTg | S178L | PLES_57781 | A | A | | | | | | | | | | |
| NC_011770 | 6397380 | T | C | 78 | 0 | 4 | 0 | NON_SYNONYMOU | MODERAT | MISSENSE | gTg/gCg | V32A | PLES_57791 | | | C | C | C | C | | | | | | |
| NC_011770 | 6397589 | C | T | 82 | 0 | 2 | 0 | NON_SYNONYMOU | MODERAT | MISSENSE | Cac/Tac | H102Y | PLES_57791 | | | | | | | | | T | T | | |
| NC_011770 | 6397608 | C | T | 84 | 0 | 2 | 0 | NON_SYNONYMOU | MODERAT | MISSENSE | gCc/gTc | A108V | PLES_57791 | T | T | | | | | | | | | | |
| NC_011770 | 6397830 | C | T | 91 | 0 | 2 | 0 | NON_SYNONYMOU | MODERAT | MISSENSE | gCg/gTg | A182V | PLES_57791 | | | | | | | | | T | T | | |
| NC_011770 | 6397872 | A | C | 87 | 0 | 2 | 0 | NON_SYNONYMOU | MODERAT | MISSENSE | gAc/gCc | D196A | PLES_57791 | | | | | | | | | | | C | C |
| NC_011770 | 6398037 | A | T | 89 | 0 | 12 | 0 | NON_SYNONYMOU | MODERAT | MISSENSE | cAg/cTg | Q251L | PLES_57791 | T | T | T | T | T | T | T | T | T | T | T | T |
| NC_011770 | 6398317 | C | A | 80 | 0 | 6 | 0 | STOP_GAINED | HIGH | NONSENSE | Gaa/Taa | E322* | PLES_57801 | A | A | | | | | A | A | A | A | | |
| NC_011770 | 6398442 | C | T | 73 | 0 | 4 | 0 | NON_SYNONYMOU | MODERAT | MISSENSE | aGc/aAc | S280N | PLES_57801 | | | T | T | T | T | | | | | | |
| NC_011770 | 6398673 | G | A | 82 | 0 | 4 | 0 | NON_SYNONYMOU | MODERAT | MISSENSE | gCc/gTc | A203V | PLES_57801 | | | A | A | A | A | | | | | | |
| NC_011770 | 6398755 | C | T | 83 | 0 | 4 | 0 | NON_SYNONYMOU | MODERAT | MISSENSE | Ggc/Agc | G176S | PLES_57801 | T | T | | | | | T | T | | | | |
| NC_011770 | 6398961 | G | A | 82 | 0 | 4 | 0 | NON_SYNONYMOU | MODERAT | MISSENSE | gCg/gTg | A107V | PLES_57801 | | | A | A | A | A | | | | | | |
| NC_011770 | 6399294 | C | G | 93 | 0 | 4 | 0 | NON_SYNONYMOU | MODERAT | MISSENSE | Gtg/Ctg | V158L | PLES_57811 | | | G | G | G | G | | | | | | |
| NC_011770 | 6399369 | T | C | 94 | 0 | 12 | 0 | NON_SYNONYMOU | MODERAT | MISSENSE | Atc/Gtc | I133V | PLES_57811 | C | C | C | C | C | C | C | C | C | C | C | C |
| NC_011770 | 6399389 | T | G | 95 | 0 | 8 | 0 | NON_SYNONYMOU | MODERAT | MISSENSE | gAa/gCa | E126A | PLES_57811 | G | G | G | G | G | G | G | G | | | | |
| NC_011770 | 6399398 | A | G | 98 | 0 | 2 | 0 | NON_SYNONYMOU | MODERAT | MISSENSE | gTa/gCa | V123A | PLES_57811 | G | G | | | | | | | | | | |
| NC_011770 | 6399675 | C | G | 101 | 0 | 2 | 0 | NON_SYNONYMOU | MODERAT | MISSENSE | Gtg/Ctg | V31L | PLES_57811 | | | | | | | G | G | | | | |
| NC_011770 | 6399755 | A | G | 92 | 0 | 8 | 0 | NON_SYNONYMOU | MODERAT | MISSENSE | gTc/gCc | V4A | PLES_57811 | G | G | G | G | G | G | G | G | | | | |
| NC_011770 | 6399988 | G | T | 92 | 0 | 4 | 0 | NON_SYNONYMOU | MODERAT | MISSENSE | cCg/cAg | P265Q | PLES_57821 | | | | | | | | | T | T | T | T |
| NC_011770 | 6400297 | A | G | 79 | 0 | 6 | 0 | NON_SYNONYMOU | MODERAT | MISSENSE | gTc/gCc | V162A | PLES_57821 | G | G | G | G | G | G | | | | | | |
| NC_011770 | 6401061 | G | A | 65 | 0 | 2 | 0 | NON_SYNONYMOU | MODERAT | MISSENSE | aCc/aTc | T219I | PLES_57831 | A | A | | | | | | | | | | |
| NC_011770 | 6401512 | C | T | 76 | 0 | 2 | 0 | NON_SYNONYMOU | MODERAT | MISSENSE | Gtc/Atc | V69I | PLES_57831 | T | T | | | | | | | | | | |
| NC_011770 | 6402402 | G | A | 85 | 0 | 2 | 0 | NON_SYNONYMOU | MODERAT | MISSENSE | aCc/aTc | T109I | PLES_57841 | | | | | | | | | A | A | | |
| NC_011770 | 6402971 | A | G | 91 | 0 | 12 | 0 | NON_SYNONYMOU | MODERAT | MISSENSE | Agc/Ggc | S12G | PLES_57851 | G | G | G | G | G | G | G | G | G | G | G | G |
| NC_011770 | 6403641 | G | A | 99 | 0 | 2 | 0 | NON_SYNONYMOU | MODERAT | MISSENSE | cGc/cAc | R235H | PLES_57851 | | | | | | | A | A | | | | |
| NC_011770 | 6403841 | G | A | 94 | 0 | 2 | 0 | NON_SYNONYMOU | MODERAT | MISSENSE | Gcc/Acc | A302T | PLES_57851 | | | | | | | A | A | | | | |
| NC_011770 | 6404380 | G | A | 78 | 0 | 4 | 0 | NON_SYNONYMOU | MODERAT | MISSENSE | gCg/gTg | A218V | PLES_57861 | A | A | | | | | A | A | | | | |
| NC_011770 | 6405181 | A | G | 79 | 0 | 1 | 0 | NON_SYNONYMOU | MODERAT | MISSENSE | Tgg/Cgg | W195R | PLES_57871 | | | | | | | G | | | | | |
| NC_011770 | 6407162 | T | G | 55 | 0 | 8 | 0 | NON_SYNONYMOU | MODERAT | MISSENSE | gaA/gaC | E130D | PLES_57891 | G | G | G | G | G | G | G | G | | | | |
| NC_011770 | 6407188 | C | T | 55 | 0 | 4 | 0 | NON_SYNONYMOU | MODERAT | MISSENSE | Gtc/Atc | V122I | PLES_57891 | | | T | T | T | T | | | | | | |
| NC_011770 | 6407197 | T | C | 58 | 0 | 12 | 0 | NON_SYNONYMOU | MODERAT | MISSENSE | Atg/Gtg | M119V | PLES_57891 | C | C | C | C | C | C | C | C | C | C | C | C |
| NC_011770 | 6409565 | C | T | 51 | 0 | 2 | 0 | NON_SYNONYMOU | MODERAT | MISSENSE | Ccg/Tcg | P32S | PLES_57911 | T | T | | | | | | | | | | |
| NC_011770 | 6409910 | A | G | 69 | 0 | 6 | 0 | NON_SYNONYMOU | MODERAT | MISSENSE | Aac/Gac | N147D | PLES_57911 | G | G | G | G | G | G | | | | | | |
| NC_011770 | 6410854 | G | A | 79 | 0 | 1 | 0 | NON_SYNONYMOU | MODERAT | MISSENSE | Gcc/Acc | A231T | PLES_57921 | | | | | | | | A | | | | |
| NC_011770 | 6411547 | G | T | 96 | 0 | 2 | 0 | NON_SYNONYMOU | MODERAT | MISSENSE | Ggc/Tgc | G94C | PLES_57931 | | | T | T | | | | | | | | |
| NC_011770 | 6411717 | C | A | 105 | 0 | 4 | 0 | NON_SYNONYMOU | MODERAT | MISSENSE | agC/agA | S150R | PLES_57931 | | | A | A | A | A | | | | | | |
| NC_011770 | 6411722 | G | A | 104 | 0 | 2 | 0 | NON_SYNONYMOU | MODERAT | MISSENSE | cGc/cAc | R152H | PLES_57931 | A | A | | | | | | | | | | |
| NC_011770 | 6414104 | A | T | 89 | 0 | 8 | 0 | NON_SYNONYMOU | MODERAT | MISSENSE | Atg/Ttg | M43L | PLES_57951 | | | T | T | T | T | | | T | T | T | T |
| NC_011770 | 6414641 | C | G | 76 | 0 | 2 | 0 | NON_SYNONYMOU | MODERAT | MISSENSE | Ccg/Gcg | P222A | PLES_57951 | | | | | | | G | G | | | | |
| NC_011770 | 6414819 | A | C | 77 | 0 | 4 | 0 | NON_SYNONYMOU | MODERAT | MISSENSE | aAc/aCc | N281T | PLES_57951 | | | | | | | | | C | C | C | C |
| NC_011770 | 6415907 | T | G | 82 | 0 | 4 | 0 | NON_SYNONYMOU | MODERAT | MISSENSE | Ttg/Gtg | L644V | PLES_57951 | | | G | G | G | G | | | | | | |
| NC_011770 | 6416511 | T | G | 71 | 0 | 4 | 0 | NON_SYNONYMOU | MODERAT | MISSENSE | Tcc/Gcc | S70A | PLES_57961 | | | G | G | G | G | | | | | | |
| NC_011770 | 6416535 | G | A | 71 | 0 | 2 | 0 | NON_SYNONYMOU | MODERAT | MISSENSE | Gcg/Acg | A78T | PLES_57961 | A | A | | | | | | | | | | |
| NC_011770 | 6418554 | A | C | 90 | 0 | 2 | 0 | NON_SYNONYMOU | MODERAT | MISSENSE | Acc/Ccc | T111P | PLES_57981 | | | | | | | C | C | | | | |
| NC_011770 | 6419114 | T | C | 94 | 0 | 1 | 0 | NON_SYNONYMOU | MODERAT | MISSENSE | Tac/Cac | Y26H | PLES_58001 | | | | | | | C | | | | | |
| NC_011770 | 6419369 | A | G | 87 | 0 | 2 | 0 | NON_SYNONYMOU | MODERAT | MISSENSE | Agt/Ggt | S111G | PLES_58001 | | | | | | | | | G | G | | |
| NC_011770 | 6419436 | T | C | 80 | 0 | 12 | 0 | NON_SYNONYMOU | MODERAT | MISSENSE | cTt/cCt | L133P | PLES_58001 | C | C | C | C | C | C | C | C | C | C | C | C |
| NC_011770 | 6419536 | T | G | 73 | 0 | 2 | 0 | NON_SYNONYMOU | MODERAT | MISSENSE | aTc/aGc | I27S | PLES_58011 | | | | | | | | | G | G | | |
| NC_011770 | 6419547 | G | T | 75 | 0 | 2 | 0 | NON_SYNONYMOU | MODERAT | MISSENSE | Gtc/Ttc | V31F | PLES_58011 | | | | | | | | | T | T | | |
| NC_011770 | 6419572 | G | A | 80 | 0 | 2 | 0 | NON_SYNONYMOU | MODERAT | MISSENSE | aGc/aAc | S39N | PLES_58011 | | | | | | | | | A | A | | |
| NC_011770 | 6419792 | A | G | 87 | 0 | 12 | 0 | NON_SYNONYMOU | MODERAT | MISSENSE | Aag/Gag | K36E | PLES_58021 | G | G | G | G | G | G | G | G | G | G | G | G |
| NC_011770 | 6419984 | G | C | 89 | 0 | 2 | 0 | NON_SYNONYMOU | MODERAT | MISSENSE | Gag/Cag | E100Q | PLES_58021 | C | C | | | | | | | | | | |
| NC_011770 | 6420079 | G | A | 85 | 0 | 2 | 0 | NON_SYNONYMOU | MODERAT | MISSENSE | Gcc/Acc | A22T | PLES_58031 | | | | | | | | | A | A | | |
| NC_011770 | 6439133 | A | G | 73 | 0 | 2 | 0 | NON_SYNONYMOU | MODERAT | MISSENSE | aAc/aGc | N15S | PLES_58171 | | | | | | | | | | | G | G |

| | | | | | | | | | | | | | | | | | | | | | | | | | |
|---|---|---|---|---|---|---|---|---|---|---|---|---|---|---|---|---|---|---|---|---|---|---|---|---|---|
| NC_011770 | 6440519 | C | T | 65 | 0 | 4 | 0 | NON_SYNONYMOU | MODERAT | MISSENSE | Gtc/Atc | V48I | PLES_58181 | | | | | | | | | T | T | T | T |
| NC_011770 | 6440602 | A | G | 65 | 0 | 8 | 0 | NON_SYNONYMOU | MODERAT | MISSENSE | gTc/gCc | V20A | PLES_58181 | | | G | G | G | G | | | G | G | G | G |
| NC_011770 | 6444457 | T | A | 81 | 0 | 6 | 0 | NON_SYNONYMOU | MODERAT | MISSENSE | cAg/cTg | Q184L | PLES_58231 | | | A | A | A | A | | | | | A | A |
| NC_011770 | 6448136 | A | AG | 49 | 0 | 1 | 0 | FRAME_SHIFT | HIGH | | ctg/cCtg | L17P? | PLES_58251 | | | | | | | AG | | | | | |
| NC_011770 | 6447860 | G | A | 47 | 0 | 1 | 0 | NON_SYNONYMOU | MODERAT | MISSENSE | cCg/cTg | P109L | PLES_58251 | | | | | | | A | | | | | |
| NC_011770 | 6447936 | C | G | 45 | 0 | 10 | 0 | NON_SYNONYMOU | MODERAT | MISSENSE | Gtg/Ctg | V84L | PLES_58251 | G | G | G | G | G | G | G | G | | | G | G |
| NC_011770 | 6448059 | T | C | 40 | 0 | 4 | 0 | NON_SYNONYMOU | MODERAT | MISSENSE | Atc/Gtc | I43V | PLES_58251 | | | C | C | C | C | | | | | | |
| NC_011770 | 6448304 | A | G | 60 | 0 | 4 | 0 | NON_SYNONYMOU | MODERAT | MISSENSE | gTg/gCg | V465A | PLES_58261 | G | G | | | | | | | | | G | G |
| NC_011770 | 6448783 | GGCCAGCG | G | 74 | 0 | 2 | 0 | CODON_CHANGE_ | MODERATE | | gcgccgctgg | APLA302A | PLES_58261 | G | G | | | | | | | | | | |
| NC_011770 | 6448975 | G | C | 85 | 0 | 2 | 0 | NON_SYNONYMOU | MODERAT | MISSENSE | gaC/gaG | D241E | PLES_58261 | | | | | | | | | C | C | | |
| NC_011770 | 6449001 | T | G | 84 | 0 | 12 | 0 | NON_SYNONYMOU | MODERAT | MISSENSE | Atc/Ctc | I233L | PLES_58261 | G | G | G | G | G | G | G | G | G | G | G | G |
| NC_011770 | 6449853 | T | C | 69 | 0 | 2 | 0 | NON_SYNONYMOU | MODERAT | MISSENSE | gTc/gCc | V10A | PLES_58271 | | | | | | | | | C | C | | |
| NC_011770 | 6450110 | G | A | 65 | 0 | 2 | 0 | NON_SYNONYMOU | MODERAT | MISSENSE | Gaa/Aaa | E96K | PLES_58271 | | | | | | | | | A | A | | |
| NC_011770 | 6450628 | G | A | 71 | 0 | 2 | 0 | NON_SYNONYMOU | MODERAT | MISSENSE | Gcg/Acg | A116T | PLES_58281 | | | | | | | A | A | | | | |
| NC_011770 | 6450679 | G | A | 74 | 0 | 4 | 0 | NON_SYNONYMOU | MODERAT | MISSENSE | Gaa/Aaa | E133K | PLES_58281 | | | A | A | A | A | | | | | | |
| NC_011770 | 6453462 | A | AG | 94 | 0 | 1 | 0 | FRAME_SHIFT | HIGH | | ccc/cccC | P259P? | PLES_58301 | | | | | | | | AG | | | | |
| NC_011770 | 6453741 | T | C | 104 | 0 | 1 | 0 | NON_SYNONYMOU | MODERAT | MISSENSE | Aag/Gag | K167E | PLES_58301 | | | | | | | C | | | | | |
| NC_011770 | 6457145 | G | GGCGCA | 84 | 0 | 1 | 0 | CODON_INSERTIOI | MODERATE | | cgc/cgcAG | R274RSLR | PLES_58331 | GGCGCAGGCT | | | | | | | | | | | |
| NC_011770 | 6457457 | A | C | 77 | 0 | 12 | 0 | NON_SYNONYMOU | MODERAT | MISSENSE | Ttg/Gtg | L171V | PLES_58331 | C | C | C | C | C | C | C | C | C | C | C | C |
| NC_011770 | 6459278 | C | T | 66 | 0 | 2 | 0 | NON_SYNONYMOU | MODERAT | MISSENSE | gCc/gTc | A409V | PLES_58341 | | | | | | | | | | | T | T |
| NC_011770 | 6460884 | T | C | 90 | 0 | 2 | 0 | NON_SYNONYMOU | MODERAT | MISSENSE | Tgc/Cgc | C429R | PLES_58351 | | | | | | | | | C | C | | |
| NC_011770 | 6460959 | T | A | 87 | 0 | 2 | 0 | NON_SYNONYMOU | MODERAT | MISSENSE | Tgt/Agt | C454S | PLES_58351 | A | A | | | | | | | | | | |
| NC_011770 | 6462582 | G | A | 78 | 0 | 1 | 0 | STOP_GAINED | HIGH | NONSENSE | Cag/Tag | Q444* | PLES_58361 | | | | | | | | A | | | | |
| NC_011770 | 6461937 | T | C | 101 | 0 | 12 | 0 | NON_SYNONYMOU | MODERAT | MISSENSE | Atg/Gtg | M659V | PLES_58361 | C | C | C | C | C | C | C | C | C | C | C | C |
| NC_011770 | 6461939 | T | G | 101 | 0 | 4 | 0 | NON_SYNONYMOU | MODERAT | MISSENSE | gAg/gCg | E658A | PLES_58361 | | | | | | | | | G | G | G | G |
| NC_011770 | 6462137 | T | A | 98 | 0 | 2 | 0 | NON_SYNONYMOU | MODERAT | MISSENSE | aAg/aTg | K592M | PLES_58361 | | | | | | | | | | | A | A |
| NC_011770 | 6462554 | C | T | 78 | 0 | 12 | 0 | NON_SYNONYMOU | MODERAT | MISSENSE | cGg/cAg | R453Q | PLES_58361 | T | T | T | T | T | T | T | T | T | T | T | T |
| NC_011770 | 6462767 | G | A | 84 | 0 | 2 | 0 | NON_SYNONYMOU | MODERAT | MISSENSE | cCg/cTg | P382L | PLES_58361 | | | | | | | | | | | A | A |
| NC_011770 | 6463203 | T | C | 92 | 0 | 12 | 0 | NON_SYNONYMOU | MODERAT | MISSENSE | Acg/Gcg | T237A | PLES_58361 | C | C | C | C | C | C | C | C | C | C | C | C |
| NC_011770 | 6463250 | G | A | 91 | 0 | 2 | 0 | NON_SYNONYMOU | MODERAT | MISSENSE | gCc/gTc | A221V | PLES_58361 | | | | | | | A | A | | | | |
| NC_011770 | 6463357 | C | G | 81 | 0 | 10 | 0 | NON_SYNONYMOU | MODERAT | MISSENSE | gaG/gaC | E185D | PLES_58361 | G | G | G | G | G | G | G | G | G | G | | |
| NC_011770 | 6463517 | T | G | 63 | 0 | 2 | 0 | NON_SYNONYMOU | MODERAT | MISSENSE | gAa/gCa | E132A | PLES_58361 | | | | | | | | | | | G | G |
| NC_011770 | 6463599 | G | A | 61 | 0 | 4 | 0 | NON_SYNONYMOU | MODERAT | MISSENSE | Ccg/Tcg | P105S | PLES_58361 | | | A | A | A | A | | | | | | |
| NC_011770 | 6463601 | G | A | 64 | 0 | 2 | 0 | NON_SYNONYMOU | MODERAT | MISSENSE | cCg/cTg | P104L | PLES_58361 | A | A | | | | | | | | | | |
| NC_011770 | 6463724 | T | G | 67 | 0 | 2 | 0 | NON_SYNONYMOU | MODERAT | MISSENSE | gAc/gCc | D63A | PLES_58361 | | | | | | | | | | | G | G |
| NC_011770 | 6464062 | C | T | 80 | 0 | 2 | 0 | NON_SYNONYMOU | MODERAT | MISSENSE | Gtg/Atg | V922M | PLES_58371 | | | | | | | T | T | | | | |
| NC_011770 | 6464313 | G | T | 78 | 0 | 2 | 0 | NON_SYNONYMOU | MODERAT | MISSENSE | gCg/gAg | A838E | PLES_58371 | | | | | | | | | | | T | T |
| NC_011770 | 6464641 | C | G | 86 | 0 | 8 | 0 | NON_SYNONYMOU | MODERAT | MISSENSE | Gcg/Ccg | A729P | PLES_58371 | | | G | G | G | G | G | G | G | G | | |
| NC_011770 | 6464644 | A | G | 85 | 0 | 8 | 0 | NON_SYNONYMOU | MODERAT | MISSENSE | Tgc/Cgc | C728R | PLES_58371 | | | G | G | G | G | G | G | G | G | | |
| NC_011770 | 6464646 | C | G | 85 | 0 | 8 | 0 | NON_SYNONYMOU | MODERAT | MISSENSE | gGc/gCc | G727A | PLES_58371 | | | G | G | G | G | G | G | G | G | | |
| NC_011770 | 6464950 | A | C | 91 | 0 | 12 | 0 | NON_SYNONYMOU | MODERAT | MISSENSE | Tcc/Gcc | S626A | PLES_58371 | C | C | C | C | C | C | C | C | C | C | C | C |
| NC_011770 | 6464971 | T | C | 93 | 0 | 2 | 0 | NON_SYNONYMOU | MODERAT | MISSENSE | Acc/Gcc | T619A | PLES_58371 | | | | | | | C | C | | | | |
| NC_011770 | 6465885 | T | C | 81 | 0 | 4 | 0 | NON_SYNONYMOU | MODERAT | MISSENSE | gAc/gGc | D314G | PLES_58371 | | | C | C | C | C | | | | | | |
| NC_011770 | 6465891 | GCGT | G | 80 | 0 | 2 | 0 | CODON_CHANGE_ | MODERATE | | gacgcc/gcc | DA311A | PLES_58371 | | | | | | | G | G | | | | |
| NC_011770 | 6465946 | C | T | 83 | 0 | 2 | 0 | NON_SYNONYMOU | MODERAT | MISSENSE | Gac/Aac | D294N | PLES_58371 | | | | | | | T | T | | | | |
| NC_011770 | 6466647 | C | T | 82 | 0 | 2 | 0 | NON_SYNONYMOU | MODERAT | MISSENSE | aGc/aAc | S60N | PLES_58371 | | | T | T | | | | | | | | |
| NC_011770 | 6469581 | G | A | 92 | 0 | 1 | 0 | NON_SYNONYMOU | MODERAT | MISSENSE | cGc/cAc | R119H | PLES_58391 | A | | | | | | | | | | | |
| NC_011770 | 6469403 | A | C | 90 | 0 | 12 | 0 | NON_SYNONYMOU | MODERAT | MISSENSE | Acg/Ccg | T60P | PLES_58391 | C | C | C | C | C | C | C | C | C | C | C | C |
| NC_011770 | 6469416 | T | C | 89 | 0 | 8 | 0 | NON_SYNONYMOU | MODERAT | MISSENSE | cTg/cCg | L64P | PLES_58391 | C | C | C | C | C | C | C | C | | | | |
| NC_011770 | 6470370 | C | T | 81 | 0 | 2 | 0 | NON_SYNONYMOU | MODERAT | MISSENSE | Ccg/Tcg | P207S | PLES_58401 | | | | | | | T | T | | | | |
| NC_011770 | 6471088 | T | A | 81 | 0 | 2 | 0 | NON_SYNONYMOU | MODERAT | MISSENSE | gTg/gAg | V446E | PLES_58401 | A | A | | | | | | | | | | |
| NC_011770 | 6481969 | ACCAGAGC | GCCAGA | 45 | 1 | 5 | 0 | CODON_CHANGE_ | MODERATE | | ccagagccag | PEPEP222I | PLES_58501 | . | GCCAGAGCCAGAG | | A | | A | | | . | A | | A |
| NC_011770 | 6481410 | G | A | 80 | 0 | 2 | 0 | NON_SYNONYMOU | MODERAT | MISSENSE | cGg/cAg | R33Q | PLES_58501 | | | | | | | | | | | A | A |
| NC_011770 | 6481592 | T | A | 56 | 0 | 6 | 0 | NON_SYNONYMOU | MODERAT | MISSENSE | Tcg/Acg | S94T | PLES_58501 | A | A | A | A | A | A | | | | | | |
| NC_011770 | 6481619 | G | A | 53 | 0 | 4 | 0 | NON_SYNONYMOU | MODERAT | MISSENSE | Gat/Aat | D103N | PLES_58501 | | | | | | | A | A | A | A | | |
| NC_011770 | 6481718 | A | G | 43 | 0 | 12 | 0 | NON_SYNONYMOU | MODERAT | MISSENSE | Aat/Gat | N136D | PLES_58501 | G | G | G | G | G | G | G | G | G | G | G | G |
| NC_011770 | 6481887 | G | T | 55 | 0 | 4 | 0 | NON_SYNONYMOU | MODERAT | MISSENSE | aGg/aTg | R192M | PLES_58501 | | | T | T | T | T | | | | | | |
| NC_011770 | 6481949 | G | A | 50 | 0 | 2 | 0 | NON_SYNONYMOU | MODERAT | MISSENSE | Gaa/Aaa | E213K | PLES_58501 | A | A | | | | | | | | | | |
| NC_011770 | 6482185 | A | G | 50 | 0 | 2 | 0 | NON_SYNONYMOU | MODERAT | MISSENSE | atA/atG | I291M | PLES_58501 | | | | | | | | | | | G | G |
| NC_011770 | 6482226 | A | C | 44 | 0 | 2 | 0 | NON_SYNONYMOU | MODERAT | MISSENSE | gAc/gCc | D305A | PLES_58501 | | | | | | | | | | | C | C |
| NC_011770 | 6482314 | C | G | 59 | 0 | 4 | 0 | NON_SYNONYMOU | MODERAT | MISSENSE | caC/caG | H334Q | PLES_58501 | | | | | | | G | G | G | G | | |
| NC_011770 | 6482828 | G | C | 79 | 0 | 2 | 0 | NON_SYNONYMOU | MODERAT | MISSENSE | Ggc/Cgc | G506R | PLES_58501 | | | | | | | | | C | C | | |

| | | | | | | | | | | | | | | | | | | | | | | | | | |
|---|---|---|---|---|---|---|---|---|---|---|---|---|---|---|---|---|---|---|---|---|---|---|---|---|---|
| NC_011770 | 6483449 | A | G | 82 | 0 | 12 | 0 | NON_SYNONYMOU | MODERAT | MISSENSE | cAc/cGc | H170R | PLES_58511 | G | G | G | G | G | G | G | G | G | G | G | G |
| NC_011770 | 6483480 | G | C | 84 | 0 | 2 | 0 | NON_SYNONYMOU | MODERAT | MISSENSE | caG/caC | Q180H | PLES_58511 | | | | | | | | | | | C | C |
| NC_011770 | 6483494 | A | T | 81 | 0 | 8 | 0 | NON_SYNONYMOU | MODERAT | MISSENSE | gAc/gTc | D185V | PLES_58511 | T | T | T | T | T | T | T | T | | | | |
| NC_011770 | 6483542 | A | G | 82 | 0 | 12 | 0 | NON_SYNONYMOU | MODERAT | MISSENSE | cAc/cGc | H201R | PLES_58511 | G | G | G | G | G | G | G | G | G | G | G | G |
| NC_011770 | 6483698 | A | G | 86 | 0 | 1 | 0 | NON_SYNONYMOU | MODERAT | MISSENSE | cAc/cGc | H253R | PLES_58511 | | | | | | | G | | | | | |
| NC_011770 | 6484144 | T | C | 89 | 0 | 12 | 0 | NON_SYNONYMOU | MODERAT | MISSENSE | Tgc/Cgc | C402R | PLES_58511 | C | C | C | C | C | C | C | C | C | C | C | C |
| NC_011770 | 6484205 | G | A | 88 | 0 | 12 | 0 | NON_SYNONYMOU | MODERAT | MISSENSE | cGa/cAa | R422Q | PLES_58511 | A | A | A | A | A | A | A | A | A | A | A | A |
| NC_011770 | 6484216 | T | G | 87 | 0 | 10 | 0 | NON_SYNONYMOU | MODERAT | MISSENSE | Tcc/Gcc | S426A | PLES_58511 | G | G | G | G | G | G | G | G | | | G | G |
| NC_011770 | 6484651 | C | A | 84 | 0 | 8 | 0 | NON_SYNONYMOU | MODERAT | MISSENSE | gCc/gAc | A138D | PLES_58521 | A | A | A | A | A | A | | | | | A | A |
| NC_011770 | 6484908 | G | A | 83 | 0 | 2 | 0 | NON_SYNONYMOU | MODERAT | MISSENSE | Gag/Aag | E224K | PLES_58521 | | | | | | | | | A | A | | |
| NC_011770 | 6485083 | C | G | 90 | 0 | 2 | 0 | NON_SYNONYMOU | MODERAT | MISSENSE | cCc/cGc | P282R | PLES_58521 | G | G | | | | | | | | | | |
| NC_011770 | 6485201 | A | G | 94 | 0 | 8 | 0 | NON_SYNONYMOU | MODERAT | MISSENSE | Aca/Gca | T35A | PLES_58531 | G | G | | | | | G | G | G | G | G | G |
| NC_011770 | 6485817 | C | T | 64 | 0 | 6 | 0 | NON_SYNONYMOU | MODERAT | MISSENSE | gCc/gTc | A240V | PLES_58531 | | | T | T | T | T | | | | | T | T |
| NC_011770 | 6485820 | C | T | 64 | 0 | 2 | 0 | NON_SYNONYMOU | MODERAT | MISSENSE | gCc/gTc | A241V | PLES_58531 | | | | | | | | | | | T | T |
| NC_011770 | 6485870 | G | A | 58 | 0 | 6 | 0 | NON_SYNONYMOU | MODERAT | MISSENSE | Gcc/Acc | A258T | PLES_58531 | A | A | | | | | A | A | A | A | | |
| NC_011770 | 6486096 | G | C | 42 | 0 | 2 | 0 | NON_SYNONYMOU | MODERAT | MISSENSE | gGc/gCc | G333A | PLES_58531 | | | | | | | | | | | C | C |
| NC_011770 | 6486099 | C | T | 42 | 0 | 2 | 0 | NON_SYNONYMOU | MODERAT | MISSENSE | aCc/aTc | T334I | PLES_58531 | | | | | | | | | | | T | T |
| NC_011770 | 6486277 | G | A | 14 | 3 | 4 | 0 | NON_SYNONYMOU | MODERAT | MISSENSE | Gcc/Acc | A15T | PLES_58541 | . | . | A | A | A | A | | | | | . | |
| NC_011770 | 6486278 | C | T | 14 | 2 | 4 | 0 | NON_SYNONYMOU | MODERAT | MISSENSE | gCc/gTc | A15V | PLES_58541 | . | . | | | | | T | T | T | T | | |
| NC_011770 | 6486280 | A | G | 14 | 2 | 4 | 0 | NON_SYNONYMOU | MODERAT | MISSENSE | Acc/Gcc | T16A | PLES_58541 | . | . | | | | | G | G | G | G | | |
| NC_011770 | 6486289 | G | A | 13 | 4 | 4 | 0 | NON_SYNONYMOU | MODERAT | MISSENSE | Gtc/Atc | V19I | PLES_58541 | . | . | . | | | | A | A | A | A | . | |
| NC_011770 | 6486304 | G | A | 14 | 1 | 3 | 0 | NON_SYNONYMOU | MODERAT | MISSENSE | Gcc/Acc | A24T | PLES_58541 | | . | . | A | A | A | | | | | | |
| NC_011770 | 6486436 | T | C | 31 | 0 | 1 | 0 | NON_SYNONYMOU | MODERAT | MISSENSE | Tgc/Cgc | C68R | PLES_58541 | | | | | | | C | | | | | |
| NC_011770 | 6486698 | G | A | 37 | 0 | 4 | 0 | NON_SYNONYMOU | MODERAT | MISSENSE | gGc/gAc | G155D | PLES_58541 | | | A | A | A | A | | | | | | |
| NC_011770 | 6486970 | G | A | 42 | 0 | 4 | 0 | NON_SYNONYMOU | MODERAT | MISSENSE | Gcg/Acg | A246T | PLES_58541 | | | A | A | A | A | | | | | | |
| NC_011770 | 6488005 | C | T | 78 | 0 | 2 | 0 | NON_SYNONYMOU | MODERAT | MISSENSE | Ctc/Ttc | L8F | PLES_58571 | | | | | | | | | | | T | T |
| NC_011770 | 6488358 | C | T | 77 | 0 | 4 | 0 | NON_SYNONYMOU | MODERAT | MISSENSE | gCg/gTg | A18V | PLES_58581 | | | | | | | T | T | T | T | | |
| NC_011770 | 6488729 | G | A | 82 | 0 | 12 | 0 | NON_SYNONYMOU | MODERAT | MISSENSE | gGg/gAg | G31E | PLES_58591 | A | A | A | A | A | A | A | A | A | A | A | A |
| NC_011770 | 6488928 | C | G | 79 | 0 | 2 | 0 | NON_SYNONYMOU | MODERAT | MISSENSE | gaC/gaG | D97E | PLES_58591 | | | | | | | | | | | G | G |
| NC_011770 | 6490711 | T | G | 75 | 0 | 2 | 0 | NON_SYNONYMOU | MODERAT | MISSENSE | gAc/gCc | D138A | PLES_58601 | | | | | | | | | | | G | G |
| NC_011770 | 6490949 | G | C | 66 | 0 | 2 | 0 | NON_SYNONYMOU | MODERAT | MISSENSE | Ccg/Gcg | P59A | PLES_58601 | | | | | | | | | | | C | C |
| NC_011770 | 6491210 | CG | C | 67 | 0 | 1 | 0 | FRAME_SHIFT | HIGH | | gtg/ | V9 | PLES_58611 | | | | | | | | C | | | | |
| NC_011770 | 6492103 | G | A | 68 | 0 | 1 | 0 | NON_SYNONYMOU | MODERAT | MISSENSE | Ggc/Agc | G305S | PLES_58611 | | | | | | | | | | | A | |
| NC_011770 | 6492261 | G | A | 62 | 0 | 1 | 0 | NON_SYNONYMOU | MODERAT | MISSENSE | Gtc/Atc | V44I | PLES_58621 | | | | A | | | | | | | | |
| NC_011770 | 6496052 | C | T | 87 | 0 | 2 | 0 | NON_SYNONYMOU | MODERAT | MISSENSE | Ggc/Agc | G190S | PLES_58661 | | | | | | | | | | | T | T |
| NC_011770 | 6496502 | A | G | 76 | 0 | 4 | 0 | NON_SYNONYMOU | MODERAT | MISSENSE | Tgc/Cgc | C40R | PLES_58661 | | | | | | | G | G | G | G | | |
| NC_011770 | 6497888 | A | C | 77 | 0 | 4 | 0 | NON_SYNONYMOU | MODERAT | MISSENSE | Tcg/Gcg | S76A | PLES_58681 | | | | | | | C | C | C | C | | |
| NC_011770 | 6497918 | G | C | 76 | 0 | 8 | 0 | NON_SYNONYMOU | MODERAT | MISSENSE | Cga/Gga | R66G | PLES_58681 | | | C | C | C | C | C | C | C | C | | |
| NC_011770 | 6500168 | CTCGAGCA | C | 93 | 0 | 1 | 0 | CODON_DELETION | MODERATE | | gagcaccttt | EHLF72 | PLES_58701 | | | | | C | | | | | | | |
| NC_011770 | 6500754 | T | C | 94 | 0 | 4 | 0 | NON_SYNONYMOU | MODERAT | MISSENSE | aTg/aCg | M266T | PLES_58701 | | | C | C | C | C | | | | | | |
| NC_011770 | 6500790 | C | A | 90 | 0 | 10 | 0 | NON_SYNONYMOU | MODERAT | MISSENSE | gCc/gAc | A278D | PLES_58701 | | | A | A | A | A | A | A | A | A | A | A |
| NC_011770 | 6500805 | T | C | 102 | 0 | 4 | 0 | NON_SYNONYMOU | MODERAT | MISSENSE | gTc/gCc | V283A | PLES_58701 | | | | | | | C | C | C | C | | |
| NC_011770 | 6501308 | G | A | 91 | 0 | 2 | 0 | NON_SYNONYMOU | MODERAT | MISSENSE | Ggc/Agc | G451S | PLES_58701 | | | | | | | | | A | A | | |
| NC_011770 | 6503724 | G | A | 63 | 0 | 4 | 0 | NON_SYNONYMOU | MODERAT | MISSENSE | Gtc/Atc | V8I | PLES_58731 | | | A | A | A | A | | | | | | |
| NC_011770 | 6503763 | G | A | 65 | 0 | 4 | 0 | NON_SYNONYMOU | MODERAT | MISSENSE | Ggc/Agc | G21S | PLES_58731 | | | | | | | A | A | A | A | | |
| NC_011770 | 6504787 | C | G | 61 | 0 | 12 | 0 | NON_SYNONYMOU | MODERAT | MISSENSE | gCc/gGc | G402A | PLES_58741 | G | G | G | G | G | G | G | G | G | G | G | G |
| NC_011770 | 6504968 | T | C | 54 | 0 | 1 | 0 | NON_SYNONYMOU | MODERAT | MISSENSE | Acc/Gcc | T342A | PLES_58741 | putative spore mauration protein | | | C | | | | | | | | |
| NC_011770 | 6507809 | TG | T | 82 | 0 | 1 | 0 | FRAME_SHIFT | HIGH | | gag/ | E54 | PLES_58761 | | T | | | | | | | | | | |
| NC_011770 | 6507671 | A | T | 81 | 0 | 6 | 0 | NON_SYNONYMOU | MODERAT | MISSENSE | Acg/Tcg | T7S | PLES_58761 | | | T | T | T | T | | | | | T | T |
| NC_011770 | 6507699 | G | T | 80 | 0 | 2 | 0 | NON_SYNONYMOU | MODERAT | MISSENSE | cGc/cTc | R16L | PLES_58761 | | | | | | | T | T | | | | |
| NC_011770 | 6507738 | C | G | 83 | 0 | 8 | 0 | NON_SYNONYMOU | MODERAT | MISSENSE | aCg/aGg | T29R | PLES_58761 | G | G | G | G | G | G | | | | | G | G |
| NC_011770 | 6507750 | C | T | 82 | 0 | 2 | 0 | NON_SYNONYMOU | MODERAT | MISSENSE | aCg/aTg | T33M | PLES_58761 | T | T | | | | | | | | | | |
| NC_011770 | 6507759 | C | T | 82 | 0 | 2 | 0 | NON_SYNONYMOU | MODERAT | MISSENSE | cCg/cTg | P36L | PLES_58761 | | | | | | | | | | | T | T |
| NC_011770 | 6508329 | A | C | 71 | 0 | 10 | 0 | NON_SYNONYMOU | MODERAT | MISSENSE | caT/caG | H94Q | PLES_58771 | C | C | C | C | C | C | | | C | C | C | C |
| NC_011770 | 6508481 | T | C | 77 | 0 | 2 | 0 | NON_SYNONYMOU | MODERAT | MISSENSE | Aag/Gag | K44E | PLES_58771 | | | | | | | | | C | C | | |
| NC_011770 | 6508632 | T | C | 76 | 0 | 10 | 0 | SPLICE_SITE_REGIC | LOW | SILENT | taA/taG | *54 | PLES_58781 | C | C | C | C | C | C | | | C | C | C | C |
| NC_011770 | 6510947 | G | A | 76 | 0 | 2 | 0 | NON_SYNONYMOU | MODERAT | MISSENSE | Gac/Aac | D112N | PLES_58801 | A | A | | | | | | | | | | |
| NC_011770 | 6511090 | A | C | 74 | 0 | 4 | 0 | NON_SYNONYMOU | MODERAT | MISSENSE | gaA/gaC | E159D | PLES_58801 | | | C | C | C | C | | | | | | |
| NC_011770 | 6515071 | G | A | 75 | 0 | 2 | 0 | NON_SYNONYMOU | MODERAT | MISSENSE | gCc/gTc | A299V | PLES_58831 | | | | | | | | | | | A | A |
| NC_011770 | 6515104 | G | A | 72 | 0 | 2 | 0 | NON_SYNONYMOU | MODERAT | MISSENSE | cCg/cTg | P288L | PLES_58831 | | | | | | | | | | | A | A |
| NC_011770 | 6515108 | T | C | 71 | 0 | 2 | 0 | NON_SYNONYMOU | MODERAT | MISSENSE | Atg/Gtg | M287V | PLES_58831 | | | | | | | | | | | C | C |
| NC_011770 | 6515113 | G | C | 70 | 0 | 2 | 0 | NON_SYNONYMOU | MODERAT | MISSENSE | cCc/cGc | P285R | PLES_58831 | | | | | | | | | | | C | C |

| | | | | | | | | | | | | | | | | | | | | | | | | | |
|---|---|---|---|---|---|---|---|---|---|---|---|---|---|---|---|---|---|---|---|---|---|---|---|---|---|
| NC_011770 | 6515117 | A | G | 67 | 0 | 2 | 0 | NON_SYNONYMOU | MODERAT | MISSENSE | Tcg/Ccg | S284P | PLES_58831 | | | | | | | | | | | G | G |
| NC_011770 | 6515120 | G | A | 70 | 0 | 2 | 0 | NON_SYNONYMOU | MODERAT | MISSENSE | Ccg/Tcg | P283S | PLES_58831 | | | | | | | | | | | A | A |
| NC_011770 | 6515133 | T | G | 72 | 0 | 2 | 0 | NON_SYNONYMOU | MODERAT | MISSENSE | gaA/gaC | E278D | PLES_58831 | | | | | | | | | | | G | G |
| NC_011770 | 6515284 | G | C | 80 | 0 | 2 | 0 | NON_SYNONYMOU | MODERAT | MISSENSE | aCg/aGg | T228R | PLES_58831 | | | | | | | | | | | C | C |
| NC_011770 | 6515288 | G | A | 78 | 0 | 4 | 0 | NON_SYNONYMOU | MODERAT | MISSENSE | Cgc/Tgc | R227C | PLES_58831 | A | A | | | | | | | | | T | T |
| NC_011770 | 6515308 | G | T | 78 | 0 | 2 | 0 | NON_SYNONYMOU | MODERAT | MISSENSE | cCg/cAg | P220Q | PLES_58831 | | | | | | | | | | | T | T |
| NC_011770 | 6515354 | C | T | 75 | 0 | 2 | 0 | NON_SYNONYMOU | MODERAT | MISSENSE | Gcg/Acg | A205T | PLES_58831 | | | | | | | | | | | T | T |
| NC_011770 | 6515426 | T | C | 75 | 0 | 2 | 0 | NON_SYNONYMOU | MODERAT | MISSENSE | Acc/Gcc | T181A | PLES_58831 | | | | | | | | | | | C | C |
| NC_011770 | 6527674 | C | T | 96 | 0 | 2 | 0 | NON_SYNONYMOU | MODERAT | MISSENSE | Gaa/Aaa | E190K | PLES_58941 | | | | | | | T | T | | | | |
| NC_011770 | 6527731 | A | G | 102 | 0 | 12 | 0 | NON_SYNONYMOU | MODERAT | MISSENSE | Tgg/Cgg | W171R | PLES_58941 | G | G | G | G | G | G | G | G | G | G | G | G |
| NC_011770 | 6527947 | C | T | 98 | 0 | 6 | 0 | NON_SYNONYMOU | MODERAT | MISSENSE | Gcc/Acc | A99T | PLES_58941 | | | T | T | T | T | T | T | | | | |
| NC_011770 | 6528157 | G | T | 107 | 0 | 2 | 0 | NON_SYNONYMOU | MODERAT | MISSENSE | Ctg/Atg | L29M | PLES_58941 | | | | | | | | | T | T | | |
| NC_011770 | 6527433 | A | ACCGGC | 89 | 0 | 1 | 0 | FRAME_SHIFT | HIGH | | gtg/gGCCC | V270GR? | PLES_58941 (putative adhesin) | | | | | | | ACCGGC | | | | | |
| NC_011770 | 6530584 | C | G | 89 | 0 | 4 | 0 | NON_SYNONYMOU | MODERAT | MISSENSE | Caa/Gaa | Q47E | PLES_58981 | | | G | G | G | G | | | | | | |
| NC_011770 | 6530708 | G | A | 74 | 0 | 2 | 0 | NON_SYNONYMOU | MODERAT | MISSENSE | cGc/cAc | R88H | PLES_58981 | | | | | | | | | A | A | | |
| NC_011770 | 6530895 | G | A | 81 | 0 | 2 | 0 | NON_SYNONYMOU | MODERAT | MISSENSE | atG/atA | M150I | PLES_58981 | | | | | | | | | | | A | A |
| NC_011770 | 6530921 | G | A | 76 | 0 | 4 | 0 | NON_SYNONYMOU | MODERAT | MISSENSE | gGc/gAc | G159D | PLES_58981 | | | A | A | A | A | | | | | | |
| NC_011770 | 6531052 | G | A | 85 | 0 | 2 | 0 | NON_SYNONYMOU | MODERAT | MISSENSE | Gcg/Acg | A203T | PLES_58981 | | | | | | | | | | | A | A |
| NC_011770 | 6531061 | G | A | 82 | 0 | 2 | 0 | NON_SYNONYMOU | MODERAT | MISSENSE | Gtc/Atc | V206I | PLES_58981 | | | | | | | | | | | A | A |
| NC_011770 | 6533479 | A | T | 83 | 0 | 2 | 0 | NON_SYNONYMOU | MODERAT | MISSENSE | gAc/gTc | D93V | PLES_59011 | | | | | | | | | | | T | T |
| NC_011770 | 6533931 | A | G | 61 | 0 | 12 | 0 | NON_SYNONYMOU | MODERAT | MISSENSE | Atc/Gtc | I244V | PLES_59011 | G | G | G | G | G | G | G | G | G | G | G | G |
| NC_011770 | 6535596 | C | T | 67 | 0 | 2 | 0 | NON_SYNONYMOU | MODERAT | MISSENSE | cCg/cTg | P183L | PLES_59031 | | | | | | | | | | | T | T |
| NC_011770 | 6536803 | G | A | 81 | 0 | 10 | 0 | NON_SYNONYMOU | MODERAT | MISSENSE | Gtc/Atc | V369I | PLES_59041 | A | A | A | A | A | A | A | A | | | A | A |
| NC_011770 | 6537332 | G | A | 84 | 0 | 2 | 0 | NON_SYNONYMOU | MODERAT | MISSENSE | Gtc/Atc | V79I | PLES_59051 | | | | | | | | | | | A | A |
| NC_011770 | 6537461 | A | C | 86 | 0 | 2 | 0 | NON_SYNONYMOU | MODERAT | MISSENSE | Acc/Ccc | T122P | PLES_59051 | C | C | | | | | | | | | | |
| NC_011770 | 6537560 | A | G | 88 | 0 | 2 | 0 | NON_SYNONYMOU | MODERAT | MISSENSE | Agg/Ggg | R155G | PLES_59051 | | | | | | | | | | | G | G |
| NC_011770 | 6537584 | C | T | 87 | 0 | 4 | 0 | NON_SYNONYMOU | MODERAT | MISSENSE | Cgg/Tgg | R163W | PLES_59051 | | | T | T | T | T | | | | | | |
| NC_011770 | 6537731 | T | A | 85 | 0 | 2 | 0 | NON_SYNONYMOU | MODERAT | MISSENSE | Tac/Aac | Y212N | PLES_59051 | | | | | | | | | | | A | A |
| NC_011770 | 6537732 | A | G | 87 | 0 | 2 | 0 | NON_SYNONYMOU | MODERAT | MISSENSE | tAc/tGc | Y212C | PLES_59051 | | | | | | | | | | | G | G |
| NC_011770 | 6538711 | T | G | 98 | 0 | 2 | 0 | NON_SYNONYMOU | MODERAT | MISSENSE | Tgg/Ggg | W282G | PLES_59061 | | | | | | | G | G | | | | |
| NC_011770 | 6540195 | G | A | 77 | 0 | 4 | 0 | NON_SYNONYMOU | MODERAT | MISSENSE | Cgc/Tgc | R129C | PLES_59071 | | | A | A | A | A | | | | | | |
| NC_011770 | 6541574 | T | G | 67 | 0 | 2 | 0 | NON_SYNONYMOU | MODERAT | MISSENSE | Agc/Cgc | S257R | PLES_59081 | | | | | | | | | G | G | | |
| NC_011770 | 6542443 | T | C | 84 | 0 | 12 | 0 | NON_SYNONYMOU | MODERAT | MISSENSE | tTc/tCc | F7S | PLES_59091 | C | C | C | C | C | C | C | C | C | C | C | C |
| NC_011770 | 6542571 | G | A | 90 | 0 | 2 | 0 | NON_SYNONYMOU | MODERAT | MISSENSE | Gcg/Acg | A50T | PLES_59091 | A | A | | | | | | | | | | |
| NC_011770 | 6542745 | G | A | 80 | 0 | 2 | 0 | NON_SYNONYMOU | MODERAT | MISSENSE | Gtc/Atc | V108I | PLES_59091 | | | | | | | | | | | A | A |
| NC_011770 | 6542995 | C | T | 86 | 0 | 2 | 0 | NON_SYNONYMOU | MODERAT | MISSENSE | gCc/gTc | A191V | PLES_59091 | | | | | | | | | | | T | T |
| NC_011770 | 6543109 | G | A | 80 | 0 | 4 | 0 | NON_SYNONYMOU | MODERAT | MISSENSE | gGc/gAc | G229D | PLES_59091 | | | A | A | A | A | | | | | | |
| NC_011770 | 6543118 | C | T | 83 | 0 | 4 | 0 | NON_SYNONYMOU | MODERAT | MISSENSE | aCc/aTc | T232I | PLES_59091 | T | T | | | | | | | | | T | T |
| NC_011770 | 6543217 | T | C | 86 | 0 | 8 | 0 | NON_SYNONYMOU | MODERAT | MISSENSE | gTg/gCg | V265A | PLES_59091 | | | C | C | C | C | C | C | | | C | C |
| NC_011770 | 6543259 | A | C | 85 | 0 | 4 | 0 | NON_SYNONYMOU | MODERAT | MISSENSE | gAt/gCt | D279A | PLES_59091 | | | | | | | C | C | | | C | C |
| NC_011770 | 6543868 | A | G | 90 | 0 | 1 | 0 | NON_SYNONYMOU | MODERAT | MISSENSE | Acc/Gcc | T166A | PLES_59101 | G | | | | | | | | | | | |
| NC_011770 | 6543418 | A | G | 89 | 0 | 8 | 0 | NON_SYNONYMOU | MODERAT | MISSENSE | Acc/Gcc | T16A | PLES_59101 | | | G | G | G | G | G | G | | | G | G |
| NC_011770 | 6543620 | G | A | 83 | 0 | 4 | 0 | NON_SYNONYMOU | MODERAT | MISSENSE | aGa/aAa | R83K | PLES_59101 | | | A | A | A | A | | | | | | |
| NC_011770 | 6543699 | C | G | 84 | 0 | 4 | 0 | NON_SYNONYMOU | MODERAT | MISSENSE | gaC/gaG | D109E | PLES_59101 | G | G | | | | | | | G | G | | |
| NC_011770 | 6543872 | G | A | 91 | 0 | 4 | 0 | NON_SYNONYMOU | MODERAT | MISSENSE | cGc/cAc | R167H | PLES_59101 | A | A | | | | | | | A | A | | |
| NC_011770 | 6544028 | T | C | 81 | 0 | 1 | 0 | NON_SYNONYMOU | MODERAT | MISSENSE | gTg/gCg | V219A | PLES_59101 | | | | | | | C | | | | | |
| NC_011770 | 6544403 | T | C | 79 | 0 | 2 | 0 | NON_SYNONYMOU | MODERAT | MISSENSE | cAc/cGc | H91R | PLES_59111 | | | | | | | C | C | | | | |
| NC_011770 | 6544478 | C | T | 82 | 0 | 12 | 0 | NON_SYNONYMOU | MODERAT | MISSENSE | cGg/cAg | R66Q | PLES_59111 | T | T | T | T | T | T | T | T | T | T | T | T |
| NC_011770 | 6546827 | T | TC | 43 | 4 | 2 | 0 | FRAME_SHIFT+SPL | HIGH | | tag/tGag | *195*? | PLES_59131 | . | . | | | | | . | TC | | | TC | . |
| NC_011770 | 6546852 | A | C | 77 | 0 | 6 | 0 | NON_SYNONYMOU | MODERAT | MISSENSE | Ttg/Gtg | L187V | PLES_59131 | | | C | C | C | C | | | C | C | | |
| NC_011770 | 6547174 | C | A | 87 | 0 | 2 | 0 | NON_SYNONYMOU | MODERAT | MISSENSE | gaG/gaT | E79D | PLES_59131 | | | | | | | | | A | A | | |
| NC_011770 | 6547242 | T | C | 86 | 0 | 4 | 0 | NON_SYNONYMOU | MODERAT | MISSENSE | Acg/Gcg | T57A | PLES_59131 | | | | | | | C | C | | | C | C |
| NC_011770 | 6547795 | C | T | 66 | 0 | 4 | 0 | NON_SYNONYMOU | MODERAT | MISSENSE | Gcc/Acc | A441T | PLES_59141 | T | T | | | | | | | T | T | | |
| NC_011770 | 6548570 | G | T | 76 | 0 | 4 | 0 | NON_SYNONYMOU | MODERAT | MISSENSE | gaC/gaA | D182E | PLES_59141 | | | T | T | T | T | | | | | | |
| NC_011770 | 6550003 | C | CG | 78 | 0 | 2 | 0 | FRAME_SHIFT | HIGH | | ccc/cccC | P19P? | PLES_59151 | | | | | | | | | | | CG | CG |
| NC_011770 | 6549986 | T | A | 78 | 0 | 2 | 0 | NON_SYNONYMOU | MODERAT | MISSENSE | agA/agT | R25S | PLES_59151 | A | A | | | | | | | | | | |
| NC_011770 | 6550298 | C | G | 80 | 0 | 4 | 0 | NON_SYNONYMOU | MODERAT | MISSENSE | gaC/gaG | D54E | PLES_59161 | | | G | G | G | G | | | | | | |
| NC_011770 | 6550470 | T | C | 69 | 2 | 4 | 0 | NON_SYNONYMOU | MODERAT | MISSENSE | Tac/Cac | Y112H | PLES_59161 | | | C | C | C | C | | | | | . | . |
| NC_011770 | 6550744 | G | A | 82 | 0 | 2 | 0 | NON_SYNONYMOU | MODERAT | MISSENSE | aGc/aAc | S203N | PLES_59161 | A | A | | | | | | | | | | |
| NC_011770 | 6550800 | G | A | 75 | 0 | 10 | 0 | NON_SYNONYMOU | MODERAT | MISSENSE | Gac/Aac | D222N | PLES_59161 | A | A | A | A | A | A | A | A | A | A | | |
| NC_011770 | 6552271 | G | A | 76 | 0 | 1 | 0 | NON_SYNONYMOU | MODERAT | MISSENSE | gCc/gTc | A293V | PLES_59181 | | | | | | | | A | | | | |
| NC_011770 | 6552113 | C | G | 77 | 0 | 2 | 0 | NON_SYNONYMOU | MODERAT | MISSENSE | Gcg/Ccg | A346P | PLES_59181 | | | | | | | | | | | G | G |

| | | | | | | | | | | | | | | | | | | | | | | | | | |
|---|---|---|---|---|---|---|---|---|---|---|---|---|---|---|---|---|---|---|---|---|---|---|---|---|---|
| NC_011770 | 6552161 | A | G | 76 | 0 | 1 | 0 | NON_SYNONYMOU | MODERAT | MISSENSE | Tgg/Cgg | W330R | PLES_59181 | | | | | | | G | | | | | |
| NC_011770 | 6552364 | C | T | 73 | 0 | 2 | 0 | NON_SYNONYMOU | MODERAT | MISSENSE | aGc/aAc | S262N | PLES_59181 | | | | | | | | | | | T | T |
| NC_011770 | 6552479 | T | C | 75 | 0 | 2 | 0 | NON_SYNONYMOU | MODERAT | MISSENSE | Acg/Gcg | T224A | PLES_59181 | | | | | | | | | | | C | C |
| NC_011770 | 6553208 | A | G | 100 | 0 | 12 | 0 | NON_SYNONYMOU | MODERAT | MISSENSE | tTc/tCc | F432S | PLES_59191 | G | G | G | G | G | G | G | G | G | G | G | G |
| NC_011770 | 6553368 | G | A | 93 | 0 | 2 | 0 | NON_SYNONYMOU | MODERAT | MISSENSE | Ccc/Tcc | P379S | PLES_59191 | | | | | | | | | | | A | A |
| NC_011770 | 6553431 | G | A | 102 | 0 | 2 | 0 | NON_SYNONYMOU | MODERAT | MISSENSE | Cac/Tac | H358Y | PLES_59191 | | | | | | | A | A | | | | |
| NC_011770 | 6553479 | G | A | 104 | 0 | 2 | 0 | NON_SYNONYMOU | MODERAT | MISSENSE | Ccc/Tcc | P342S | PLES_59191 | | | | | | | A | A | | | | |
| NC_011770 | 6553500 | T | C | 98 | 0 | 2 | 0 | NON_SYNONYMOU | MODERAT | MISSENSE | Acc/Gcc | T335A | PLES_59191 | | | | | | | | | | | C | C |
| NC_011770 | 6553902 | T | C | 95 | 0 | 4 | 0 | NON_SYNONYMOU | MODERAT | MISSENSE | Agc/Ggc | S201G | PLES_59191 | C | C | | | | | | | C | C | | |
| NC_011770 | 6554265 | G | A | 91 | 0 | 4 | 0 | NON_SYNONYMOU | MODERAT | MISSENSE | Ccg/Tcg | P80S | PLES_59191 | | | A | A | A | A | | | | | | |
| NC_011770 | 6554363 | G | T | 77 | 0 | 4 | 0 | NON_SYNONYMOU | MODERAT | MISSENSE | gCg/gAg | A47E | PLES_59191 | | | | | | | T | T | | | T | T |
| NC_011770 | 6554753 | G | T | 74 | 0 | 2 | 0 | NON_SYNONYMOU | MODERAT | MISSENSE | Gcc/Tcc | A32S | PLES_59201 | | | | | | | | | | | T | T |
| NC_011770 | 6555264 | C | G | 72 | 0 | 4 | 0 | NON_SYNONYMOU | MODERAT | MISSENSE | gCc/gGc | A202G | PLES_59201 | G | G | | | | | | | G | G | | |
| NC_011770 | 6555859 | G | A | 76 | 0 | 4 | 0 | NON_SYNONYMOU | MODERAT | MISSENSE | cGa/cAa | R118Q | PLES_59211 | A | A | | | | | A | A | | | | |
| NC_011770 | 6556075 | C | T | 76 | 0 | 2 | 0 | NON_SYNONYMOU | MODERAT | MISSENSE | gCc/gTc | A190V | PLES_59211 | T | T | | | | | | | | | | |
| NC_011770 | 6557410 | T | A | 78 | 0 | 12 | 0 | NON_SYNONYMOU | MODERAT | MISSENSE | gAc/gTc | D226V | PLES_59241 | A | A | A | A | A | A | A | A | A | A | A | A |
| NC_011770 | 6557547 | C | T | 81 | 0 | 2 | 0 | NON_SYNONYMOU | MODERAT | MISSENSE | atG/atA | M180I | PLES_59241 | T | T | | | | | | | | | | |
| NC_011770 | 6560182 | G | A | 76 | 0 | 1 | 0 | NON_SYNONYMOU | MODERAT | MISSENSE | Gac/Aac | D579N | PLES_59251 | | | A | | | | | | | | | |
| NC_011770 | 6558859 | A | G | 75 | 0 | 12 | 0 | NON_SYNONYMOU | MODERAT | MISSENSE | Agc/Ggc | S138G | PLES_59251 | G | G | G | G | G | G | G | G | G | G | G | G |
| NC_011770 | 6559645 | G | A | 73 | 0 | 1 | 0 | NON_SYNONYMOU | MODERAT | MISSENSE | Ggc/Agc | G400S | PLES_59251 | | | | | | | A | | | | | |
| NC_011770 | 6561424 | C | T | 72 | 0 | 2 | 0 | NON_SYNONYMOU | MODERAT | MISSENSE | Ctc/Ttc | L302F | PLES_59261 | | | | | | | T | T | | | | |
| NC_011770 | 6563569 | T | C | 64 | 0 | 2 | 0 | NON_SYNONYMOU | MODERAT | MISSENSE | aAc/aGc | N256S | PLES_59281 | | | | | | | C | C | | | | |
| NC_011770 | 6563634 | G | C | 64 | 0 | 2 | 0 | NON_SYNONYMOU | MODERAT | MISSENSE | ttC/ttG | F234L | PLES_59281 | | | | | | | C | C | | | | |
| NC_011770 | 6563709 | C | G | 64 | 0 | 2 | 0 | NON_SYNONYMOU | MODERAT | MISSENSE | gaG/gaC | E209D | PLES_59281 | | | | | | | G | G | | | | |
| NC_011770 | 6563858 | T | C | 65 | 0 | 8 | 0 | NON_SYNONYMOU | MODERAT | MISSENSE | Acg/Gcg | T160A | PLES_59281 | | | C | C | C | C | C | C | | | C | C |
| NC_011770 | 6564025 | A | T | 71 | 0 | 2 | 0 | NON_SYNONYMOU | MODERAT | MISSENSE | cTg/cAg | L104Q | PLES_59281 | T | T | | | | | | | | | | |
| NC_011770 | 6564574 | G | A | 87 | 0 | 2 | 0 | NON_SYNONYMOU | MODERAT | MISSENSE | cGc/cAc | R46H | PLES_59291 | A | A | | | | | | | | | | |
| NC_011770 | 6565121 | G | T | 87 | 0 | 12 | 0 | NON_SYNONYMOU | MODERAT | MISSENSE | gaC/gaA | D160E | PLES_59301 | T | T | T | T | T | T | T | T | T | T | T | T |
| NC_011770 | 6565308 | G | C | 82 | 0 | 4 | 0 | NON_SYNONYMOU | MODERAT | MISSENSE | gCc/gGc | A98G | PLES_59301 | | | C | C | C | C | | | | | | |
| NC_011770 | 6566727 | G | C | 73 | 0 | 8 | 0 | NON_SYNONYMOU | MODERAT | MISSENSE | Ctg/Gtg | L25V | PLES_59311 | C | C | | | | | C | C | C | C | C | C |
| NC_011770 | 6566914 | G | T | 64 | 0 | 2 | 0 | NON_SYNONYMOU | MODERAT | MISSENSE | cCg/cAg | P102Q | PLES_59321 | | | | | | | | | T | T | | |
| NC_011770 | 6566921 | C | T | 64 | 0 | 2 | 0 | NON_SYNONYMOU | MODERAT | MISSENSE | Ggc/Agc | G100S | PLES_59321 | T | T | | | | | | | | | | |
| NC_011770 | 6567167 | C | T | 54 | 0 | 4 | 0 | NON_SYNONYMOU | MODERAT | MISSENSE | Gcc/Acc | A18T | PLES_59321 | | | T | T | T | T | | | | | | |
| NC_011770 | 6567347 | G | T | 77 | 0 | 2 | 0 | NON_SYNONYMOU | MODERAT | MISSENSE | cGc/cTc | R6L | PLES_59331 | | | | | | | T | T | | | | |
| NC_011770 | 6567530 | C | G | 81 | 0 | 2 | 0 | NON_SYNONYMOU | MODERAT | MISSENSE | gCg/gGg | A67G | PLES_59331 | G | G | | | | | | | | | | |
| NC_011770 | 6567647 | G | A | 85 | 0 | 2 | 0 | NON_SYNONYMOU | MODERAT | MISSENSE | aGc/aAc | S106N | PLES_59331 | | | | | | | | | A | A | | |
| NC_011770 | 6567649 | A | G | 83 | 0 | 8 | 0 | NON_SYNONYMOU | MODERAT | MISSENSE | Atc/Gtc | I107V | PLES_59331 | G | G | | | | | G | G | G | G | G | G |
| NC_011770 | 6567736 | G | A | 74 | 0 | 2 | 0 | NON_SYNONYMOU | MODERAT | MISSENSE | Gct/Act | A136T | PLES_59331 | | | | | | | | | A | A | | |
| NC_011770 | 6569306 | G | A | 85 | 0 | 1 | 0 | NON_SYNONYMOU | MODERAT | MISSENSE | Gcc/Acc | A97T | PLES_59351 | | | | | | | | A | | | | |
| NC_011770 | 6569249 | T | C | 87 | 0 | 2 | 0 | NON_SYNONYMOU | MODERAT | MISSENSE | Tgc/Cgc | C78R | PLES_59351 | C | C | | | | | | | | | | |
| NC_011770 | 6569280 | A | C | 88 | 0 | 8 | 0 | NON_SYNONYMOU | MODERAT | MISSENSE | gAc/gCc | D88A | PLES_59351 | | | C | C | C | C | | | C | C | C | C |
| NC_011770 | 6569645 | A | G | 96 | 0 | 2 | 0 | NON_SYNONYMOU | MODERAT | MISSENSE | Atc/Gtc | I210V | PLES_59351 | | | | | | | | | | | G | G |
| NC_011770 | 6569837 | A | G | 90 | 0 | 2 | 0 | NON_SYNONYMOU | MODERAT | MISSENSE | Ata/Gta | I274V | PLES_59351 | G | G | | | | | | | | | | |
| NC_011770 | 6572197 | C | T | 77 | 0 | 2 | 0 | NON_SYNONYMOU | MODERAT | MISSENSE | Gac/Aac | D302N | PLES_59381 | T | T | | | | | | | | | | |
| NC_011770 | 6572427 | C | T | 67 | 0 | 2 | 0 | NON_SYNONYMOU | MODERAT | MISSENSE | gGt/gAt | G225D | PLES_59381 | | | | | | | | | T | T | | |
| NC_011770 | 6572430 | C | G | 64 | 0 | 8 | 0 | NON_SYNONYMOU | MODERAT | MISSENSE | aGc/aCc | S224T | PLES_59381 | | | G | G | G | G | | | G | G | G | G |
| NC_011770 | 6572431 | T | C | 62 | 0 | 6 | 0 | NON_SYNONYMOU | MODERAT | MISSENSE | Agc/Ggc | S224G | PLES_59381 | | | C | C | C | C | | | C | C | | |
| NC_011770 | 6572871 | A | G | 53 | 0 | 2 | 0 | NON_SYNONYMOU | MODERAT | MISSENSE | gTg/gCg | V77A | PLES_59381 | | | | | | | | | | | G | G |
| NC_011770 | 6572898 | A | G | 52 | 0 | 12 | 0 | NON_SYNONYMOU | MODERAT | MISSENSE | aTc/aCc | I68T | PLES_59381 | G | G | G | G | G | G | G | G | G | G | G | G |
| NC_011770 | 6573036 | A | G | 56 | 0 | 12 | 0 | NON_SYNONYMOU | MODERAT | MISSENSE | tTg/tCg | L22S | PLES_59381 | G | G | G | G | G | G | G | G | G | G | G | G |
| NC_011770 | 6573052 | C | T | 57 | 0 | 4 | 0 | NON_SYNONYMOU | MODERAT | MISSENSE | Gtc/Atc | V17I | PLES_59381 | | | T | T | T | T | | | | | | |
| NC_011770 | 6573200 | T | C | 50 | 0 | 2 | 0 | NON_SYNONYMOU | MODERAT | MISSENSE | gAc/gGc | D129G | PLES_59391 | | | | | | | | | | | C | C |
| NC_011770 | 6573233 | G | T | 45 | 0 | 1 | 0 | NON_SYNONYMOU | MODERAT | MISSENSE | cCg/cAg | P118Q | PLES_59391 | | | | | | | T | | | | | |
| NC_011770 | 6573242 | G | C | 46 | 0 | 2 | 0 | NON_SYNONYMOU | MODERAT | MISSENSE | cCt/cGt | P115R | PLES_59391 | | | | | | | C | C | | | | |
| NC_011770 | 6573354 | C | A | 38 | 0 | 2 | 0 | NON_SYNONYMOU | MODERAT | MISSENSE | Gtg/Ttg | V78L | PLES_59391 | | | | | | | | | | | A | A |
| NC_011770 | 6573362 | T | C | 39 | 0 | 2 | 0 | NON_SYNONYMOU | MODERAT | MISSENSE | cAc/cGc | H75R | PLES_59391 | | | | | | | | | | | C | C |
| NC_011770 | 6573644 | G | A | 62 | 0 | 2 | 0 | NON_SYNONYMOU | MODERAT | MISSENSE | Ctc/Ttc | L655F | PLES_59401 | | | | | | | | | | | A | A |
| NC_011770 | 6573776 | C | T | 65 | 0 | 12 | 0 | NON_SYNONYMOU | MODERAT | MISSENSE | Gtc/Atc | V611I | PLES_59401 | T | T | T | T | T | T | T | T | T | T | T | T |
| NC_011770 | 6574620 | A | C | 83 | 0 | 2 | 0 | NON_SYNONYMOU | MODERAT | MISSENSE | gaT/gaG | D329E | PLES_59401 | | | | | | | | | | | C | C |
| NC_011770 | 6575186 | A | C | 88 | 0 | 2 | 0 | NON_SYNONYMOU | MODERAT | MISSENSE | Ttc/Gtc | F141V | PLES_59401 | | | | | | | C | C | | | | |
| NC_011770 | 6575332 | C | G | 77 | 0 | 2 | 0 | NON_SYNONYMOU | MODERAT | MISSENSE | gGc/gCc | G92A | PLES_59401 | | | | | | | | | | | G | G |
| NC_011770 | 6575411 | C | G | 82 | 0 | 2 | 0 | NON_SYNONYMOU | MODERAT | MISSENSE | Ggt/Cgt | G66R | PLES_59401 | | | | | | | | | | | G | G |

| | | | | | | | | | | | | | | | | | | | | | | | | | |
|---|---|---|---|---|---|---|---|---|---|---|---|---|---|---|---|---|---|---|---|---|---|---|---|---|---|
| NC_011770 | 6575775 | T | C | 85 | 0 | 1 | 0 | NON_SYNONYMOU | MODERAT | MISSENSE | gAc/gGc | D289G | PLES_59411 | | | | | | | | C | | | | |
| NC_011770 | 6576579 | G | A | 80 | 0 | 6 | 0 | NON_SYNONYMOU | MODERAT | MISSENSE | gCg/gTg | A21V | PLES_59411 | | | A | A | A | A | | | A | A | | |
| NC_011770 | 6577230 | C | A | 87 | 0 | 8 | 0 | NON_SYNONYMOU | MODERAT | MISSENSE | gaG/gaT | E253D | PLES_59421 | A | A | | | | | A | A | A | A | A | A |
| NC_011770 | 6577352 | A | G | 81 | 0 | 12 | 0 | NON_SYNONYMOU | MODERAT | MISSENSE | Ttc/Ctc | F213L | PLES_59421 | G | G | G | G | G | G | G | G | G | G | G | G |
| NC_011770 | 6577358 | T | C | 77 | 0 | 12 | 0 | NON_SYNONYMOU | MODERAT | MISSENSE | Atc/Gtc | I211V | PLES_59421 | C | C | C | C | C | C | C | C | C | C | C | C |
| NC_011770 | 6578367 | C | G | 94 | 0 | 4 | 0 | NON_SYNONYMOU | MODERAT | MISSENSE | Ctg/Gtg | L48V | PLES_59431 | | | G | G | G | G | | | | | | |
| NC_011770 | 6578413 | A | G | 86 | 0 | 2 | 0 | NON_SYNONYMOU | MODERAT | MISSENSE | gAc/gGc | D63G | PLES_59431 | | | | | | | | | | | G | G |
| NC_011770 | 6578469 | G | C | 87 | 0 | 8 | 0 | NON_SYNONYMOU | MODERAT | MISSENSE | Gat/Cat | D82H | PLES_59431 | C | C | C | C | C | C | | | C | C | | |
| NC_011770 | 6580265 | G | GC | 94 | 0 | 1 | 0 | FRAME_SHIFT | HIGH | | ctg/cCtg | L355P? | PLES_59441 | | | | | | | GC | | | | | |
| NC_011770 | 6579738 | G | A | 91 | 0 | 1 | 0 | NON_SYNONYMOU | MODERAT | MISSENSE | Ggt/Agt | G178S | PLES_59441 | | | | | | | | | | | A | |
| NC_011770 | 6579468 | G | A | 77 | 0 | 2 | 0 | NON_SYNONYMOU | MODERAT | MISSENSE | Gtc/Atc | V88I | PLES_59441 | | | | | | | | | | | A | A |
| NC_011770 | 6579714 | G | A | 83 | 0 | 2 | 0 | NON_SYNONYMOU | MODERAT | MISSENSE | Gcg/Acg | A170T | PLES_59441 | | | | | | | | | | | A | A |
| NC_011770 | 6579726 | A | T | 91 | 0 | 2 | 0 | NON_SYNONYMOU | MODERAT | MISSENSE | Atg/Ttg | M174L | PLES_59441 | | | | | | | | | T | T | | |
| NC_011770 | 6579954 | G | A | 99 | 0 | 6 | 0 | NON_SYNONYMOU | MODERAT | MISSENSE | Gtc/Atc | V250I | PLES_59441 | | | A | A | A | A | | | | | A | A |
| NC_011770 | 6580015 | G | A | 97 | 0 | 4 | 0 | NON_SYNONYMOU | MODERAT | MISSENSE | cGa/cAa | R270Q | PLES_59441 | | | A | A | A | A | | | | | | |
| NC_011770 | 6580294 | A | G | 92 | 0 | 4 | 0 | NON_SYNONYMOU | MODERAT | MISSENSE | aAg/aGg | K363R | PLES_59441 | | | G | G | G | G | | | | | | |
| NC_011770 | 6580317 | G | A | 97 | 0 | 2 | 0 | NON_SYNONYMOU | MODERAT | MISSENSE | Gta/Ata | V371I | PLES_59441 | A | A | | | | | | | | | | |
| NC_011770 | 6580350 | C | T | 96 | 0 | 2 | 0 | NON_SYNONYMOU | MODERAT | MISSENSE | Ctc/Ttc | L382F | PLES_59441 | | | | | | | | | | | T | T |
| NC_011770 | 6583488 | CG | C | 80 | 0 | 2 | 0 | FRAME_SHIFT | HIGH | | ccc/ | P63 | PLES_59471 | | | | | | | C | C | | | | |
| NC_011770 | 6583191 | T | C | 77 | 0 | 10 | 0 | NON_SYNONYMOU | MODERAT | MISSENSE | Aat/Gat | N163D | PLES_59471 | C | C | C | C | C | C | C | C | C | C | | |
| NC_011770 | 6583232 | G | T | 71 | 0 | 10 | 0 | NON_SYNONYMOU | MODERAT | MISSENSE | gCc/gAc | A149D | PLES_59471 | T | T | T | T | T | T | T | T | T | T | | |
| NC_011770 | 6583323 | C | T | 77 | 0 | 10 | 0 | NON_SYNONYMOU | MODERAT | MISSENSE | Gcg/Acg | A119T | PLES_59471 | T | T | T | T | T | T | T | T | T | T | | |
| NC_011770 | 6583365 | C | T | 75 | 0 | 6 | 0 | NON_SYNONYMOU | MODERAT | MISSENSE | Gcc/Acc | A105T | PLES_59471 | | | T | T | T | T | | | | | T | T |
| NC_011770 | 6583632 | C | A | 79 | 0 | 12 | 0 | NON_SYNONYMOU | MODERAT | MISSENSE | Ggg/Tgg | G16W | PLES_59471 | A | A | A | A | A | A | A | A | A | A | A | A |
| NC_011770 | 6583659 | A | G | 80 | 0 | 8 | 0 | NON_SYNONYMOU | MODERAT | MISSENSE | Tgg/Cgg | W7R | PLES_59471 | G | G | G | G | G | G | | | G | G | | |
| NC_011770 | 6597213 | A | C | 83 | 0 | 10 | 0 | INTRAGENIC | MODIFIER | | | | PLES_59621 | C | C | C | C | C | C | C | C | | | C | C |
| NC_011770 | 6597234 | T | C | 84 | 0 | 10 | 0 | INTRAGENIC | MODIFIER | | | | PLES_59621 | C | C | C | C | C | C | C | C | | | C | C |
| NC_011770 | 6600460 | T | C | 104 | 0 | 2 | 0 | NON_SYNONYMOU | MODERAT | MISSENSE | cAc/cGc | H153R | PLES_59641 | C | C | | | | | | | | | | |
| NC_011770 | 1423193 | T | C | 90 | 0 | 6 | 0 | NON_SYNONYMOU | MODERAT | MISSENSE | gAc/gGc | D681G | plsB | C | C | C | C | C | C | | | | | | |
| NC_011770 | 2267088 | A | G | 83 | 0 | 12 | 0 | NON_SYNONYMOU | MODERAT | MISSENSE | Aaa/Gaa | K36E | plsX | G | G | G | G | G | G | G | G | G | G | G | G |
| NC_011770 | 2758105 | C | T | 17 | 10 | 2 | 0 | INTRAGENIC | MODIFIER | | | | pltB | . | . | . | . | . | . | . | . | . | . | T | T |
| NC_011770 | 2758129 | C | T | 16 | 10 | 2 | 0 | INTRAGENIC | MODIFIER | | | | pltB | . | . | . | . | . | . | . | . | . | . | T | T |
| NC_011770 | 2758190 | T | G | 15 | 10 | 2 | 0 | INTRAGENIC | MODIFIER | | | | pltB | . | . | . | . | . | . | . | . | . | . | G | G |
| NC_011770 | 2758723 | G | A | 11 | 10 | 2 | 0 | INTRAGENIC | MODIFIER | | | | pltB | . | . | . | . | . | . | . | . | . | . | A | A |
| NC_011770 | 2758855 | T | C | 14 | 10 | 2 | 0 | INTRAGENIC | MODIFIER | | | | pltB | . | . | . | . | . | . | . | . | . | . | C | C |
| NC_011770 | 2759223 | G | A | 14 | 10 | 2 | 0 | INTRAGENIC | MODIFIER | | | | pltB | . | . | . | . | . | . | . | . | . | . | A | A |
| NC_011770 | 2760129 | G | A | 12 | 10 | 2 | 0 | INTRAGENIC | MODIFIER | | | | pltB | . | . | . | . | . | . | . | . | . | . | A | A |
| NC_011770 | 2760361 | T | C | 13 | 10 | 2 | 0 | INTRAGENIC | MODIFIER | | | | pltB | . | . | . | . | . | . | . | . | . | . | C | C |
| NC_011770 | 2760580 | C | T | 15 | 10 | 2 | 0 | INTRAGENIC | MODIFIER | | | | pltB | . | . | . | . | . | . | . | . | . | . | T | T |
| NC_011770 | 2760907 | G | A | 11 | 10 | 2 | 0 | INTRAGENIC | MODIFIER | | | | pltB | . | . | . | . | . | . | . | . | . | . | A | A |
| NC_011770 | 2761064 | G | A | 12 | 10 | 2 | 0 | INTRAGENIC | MODIFIER | | | | pltB | . | . | . | . | . | . | . | . | . | . | A | A |
| NC_011770 | 2761065 | G | A | 12 | 10 | 2 | 0 | INTRAGENIC | MODIFIER | | | | pltB | . | . | . | . | . | . | . | . | . | . | A | A |
| NC_011770 | 2762314 | T | C | 14 | 10 | 2 | 0 | INTRAGENIC | MODIFIER | | | | pltB | . | . | . | . | . | . | . | . | . | . | C | C |
| NC_011770 | 2762960 | G | A | 10 | 10 | 2 | 0 | INTRAGENIC | MODIFIER | | | | pltB | . | . | . | . | . | . | . | . | . | . | A | A |
| NC_011770 | 2763767 | T | TGGTGG | 7 | 10 | 1 | 0 | INTRAGENIC | MODIFIER | | | | pltB | . | . | . | . | . | . | . | . | . | . | TGGTGG | |
| NC_011770 | 2767187 | T | G | 17 | 10 | 2 | 0 | NON_SYNONYMOU | MODERAT | MISSENSE | aTg/aGg | M681R | pltC | . | . | . | . | . | . | . | . | . | . | G | G |
| NC_011770 | 2768779 | C | G | 16 | 10 | 2 | 0 | NON_SYNONYMOU | MODERAT | MISSENSE | Ccc/Gcc | P1212A | pltC | . | . | . | . | . | . | . | . | . | . | G | G |
| NC_011770 | 2768872 | A | T | 15 | 10 | 2 | 0 | NON_SYNONYMOU | MODERAT | MISSENSE | Acc/Tcc | T1243S | pltC | . | . | . | . | . | . | . | . | . | . | T | T |
| NC_011770 | 2769353 | A | G | 8 | 10 | 2 | 0 | NON_SYNONYMOU | MODERAT | MISSENSE | cAg/cGg | Q1403R | pltC | . | . | . | . | . | . | . | . | . | . | G | G |
| NC_011770 | 2770536 | C | T | 19 | 10 | 2 | 0 | NON_SYNONYMOU | MODERAT | MISSENSE | gCc/gTc | A39V | pltD | . | . | . | . | . | . | . | . | . | . | T | T |
| NC_011770 | 2770586 | A | G | 19 | 10 | 2 | 0 | NON_SYNONYMOU | MODERAT | MISSENSE | Att/Gtt | I56V | pltD | . | . | . | . | . | . | . | . | . | . | G | G |
| NC_011770 | 2772500 | C | G | 11 | 10 | 2 | 0 | NON_SYNONYMOU | MODERAT | MISSENSE | atC/atG | I147M | pltE | . | . | . | . | . | . | . | . | . | . | G | G |
| NC_011770 | 2774519 | T | C | 13 | 10 | 2 | 0 | NON_SYNONYMOU | MODERAT | MISSENSE | Tca/Cca | S444P | pltF | . | . | . | . | . | . | . | . | . | . | C | C |
| NC_011770 | 2779054 | A | C | 12 | 10 | 2 | 0 | NON_SYNONYMOU | MODERAT | MISSENSE | gAc/gCc | D5A | pltJ | . | . | . | . | . | . | . | . | . | . | C | C |
| NC_011770 | 2780866 | T | C | 19 | 10 | 2 | 0 | NON_SYNONYMOU | MODERAT | MISSENSE | Tcg/Ccg | S230P | PltK | . | . | . | . | . | . | . | . | . | . | C | C |
| NC_011770 | 2781026 | C | G | 16 | 10 | 2 | 0 | NON_SYNONYMOU | MODERAT | MISSENSE | gCc/gGc | A283G | PltK | . | . | . | . | . | . | . | . | . | . | G | G |
| NC_011770 | 2781487 | A | G | 21 | 10 | 2 | 0 | NON_SYNONYMOU | MODERAT | MISSENSE | cAa/cGa | Q64R | pltN | . | . | . | . | . | . | . | . | . | . | G | G |
| NC_011770 | 2781741 | C | T | 18 | 10 | 2 | 0 | NON_SYNONYMOU | MODERAT | MISSENSE | Ccg/Tcg | P149S | pltN | . | . | . | . | . | . | . | . | . | . | T | T |
| NC_011770 | 5698302 | T | G | 65 | 0 | 12 | 0 | NON_SYNONYMOU | MODERAT | MISSENSE | Tat/Gat | Y104D | pmrA | G | G | G | G | G | G | G | G | G | G | G | G |
| NC_011770 | 5699795 | G | A | 81 | 0 | 1 | 0 | NON_SYNONYMOU | MODERAT | MISSENSE | Ggc/Agc | G372S | pmrB | | | | | | | | A | | | | |
| NC_011770 | 5698685 | T | C | 60 | 0 | 4 | 0 | NON_SYNONYMOU | MODERAT | MISSENSE | Tcc/Ccc | S2P | pmrB | | | C | C | C | C | | | | | | |
| NC_011770 | 5698691 | G | A | 57 | 0 | 4 | 0 | NON_SYNONYMOU | MODERAT | MISSENSE | Gcc/Acc | A4T | pmrB | | | A | A | A | A | | | | | | |

| | | | | | | | | | | | | | | | | | | | | | | | | | |
|---|---|---|---|---|---|---|---|---|---|---|---|---|---|---|---|---|---|---|---|---|---|---|---|---|---|
| NC_011770 | 5698698 | T | C | 57 | 0 | 4 | 0 | NON_SYNONYMOU | MODERAT | MISSENSE | gTc/gCc | V6A | pmrB | | | C | C | C | C | | | | | | |
| NC_011770 | 5698724 | G | A | 59 | 0 | 4 | 0 | NON_SYNONYMOU | MODERAT | MISSENSE | Gtc/Atc | V15I | pmrB | | | A | A | A | A | | | | | | |
| NC_011770 | 5698880 | G | A | 61 | 0 | 2 | 0 | NON_SYNONYMOU | MODERAT | MISSENSE | Gcc/Acc | A67T | pmrB | | | | | | | | | | | A | A |
| NC_011770 | 5698883 | G | A | 61 | 0 | 4 | 0 | NON_SYNONYMOU | MODERAT | MISSENSE | Ggc/Agc | G68S | pmrB | | | A | A | A | A | | | | | | |
| NC_011770 | 5699183 | T | C | 68 | 0 | 1 | 0 | NON_SYNONYMOU | MODERAT | MISSENSE | Ttc/Ctc | F168L | pmrB | | | | | | | C | | | | | |
| NC_011770 | 4991398 | G | A | 75 | 0 | 1 | 0 | NON_SYNONYMOU | MODERAT | MISSENSE | cCg/cTg | P204L | pmtA | | | | | | | A | | | | | |
| NC_011770 | 4991779 | G | A | 73 | 0 | 1 | 0 | NON_SYNONYMOU | MODERAT | MISSENSE | cCg/cTg | P77L | pmtA | | | | | | | A | | | | | |
| NC_011770 | 5850214 | C | G | 75 | 0 | 2 | 0 | NON_SYNONYMOU | MODERAT | MISSENSE | Ccg/Gcg | P23A | pncA | G | G | | | | | | | | | | |
| NC_011770 | 5850386 | T | A | 76 | 0 | 4 | 0 | NON_SYNONYMOU | MODERAT | MISSENSE | cTg/cAg | L80Q | pncA | | | A | A | A | A | | | | | | |
| NC_011770 | 5850398 | G | A | 81 | 0 | 2 | 0 | NON_SYNONYMOU | MODERAT | MISSENSE | aGc/aAc | S84N | pncA | | | | | | | | | A | A | | |
| NC_011770 | 5851861 | C | A | 75 | 0 | 2 | 0 | NON_SYNONYMOU | MODERAT | MISSENSE | cCg/cAg | P351Q | pncB1 | | | | | | | | | A | A | | |
| NC_011770 | 223054 | A | G | 79 | 0 | 4 | 0 | NON_SYNONYMOU | MODERAT | MISSENSE | aAc/aGc | N42S | pntAA | | | G | G | G | G | | | | | | |
| NC_011770 | 224019 | G | A | 96 | 0 | 2 | 0 | NON_SYNONYMOU | MODERAT | MISSENSE | Gac/Aac | D364N | pntAA | | | | | | | | | | | A | A |
| NC_011770 | 224778 | C | T | 85 | 0 | 1 | 0 | NON_SYNONYMOU | MODERAT | MISSENSE | cCg/cTg | P116L | pntB | | | | | | | T | | | | | |
| NC_011770 | 224789 | A | G | 86 | 0 | 12 | 0 | NON_SYNONYMOU | MODERAT | MISSENSE | Agg/Ggg | R120G | pntB | G | G | G | G | G | G | G | G | G | G | G | G |
| NC_011770 | 272946 | A | C | 77 | 0 | 2 | 0 | NON_SYNONYMOU | MODERAT | MISSENSE | Tcc/Gcc | S249A | pobA | | | | | | | | | C | C | | |
| NC_011770 | 273009 | A | T | 78 | 0 | 2 | 0 | NON_SYNONYMOU | MODERAT | MISSENSE | Tcg/Acg | S228T | pobA | | | | | | | | | T | T | | |
| NC_011770 | 6519813 | C | G | 74 | 0 | 4 | 0 | NON_SYNONYMOU | MODERAT | MISSENSE | gaG/gaC | E876D | polA | | | G | G | G | G | | | | | | |
| NC_011770 | 6519872 | G | T | 82 | 0 | 4 | 0 | NON_SYNONYMOU | MODERAT | MISSENSE | Ctc/Atc | L857I | polA | | | T | T | T | T | | | | | | |
| NC_011770 | 6520297 | C | T | 101 | 0 | 12 | 0 | NON_SYNONYMOU | MODERAT | MISSENSE | gGc/gAc | G715D | polA | T | T | T | T | T | T | T | T | T | T | T | T |
| NC_011770 | 6520344 | C | T | 107 | 0 | 1 | 0 | NON_SYNONYMOU | MODERAT | MISSENSE | atG/atA | M699I | polA | | | | | | | T | | | | | |
| NC_011770 | 6521254 | T | C | 100 | 0 | 4 | 0 | NON_SYNONYMOU | MODERAT | MISSENSE | aAg/aGg | K396R | polA | | | C | C | C | C | | | | | | |
| NC_011770 | 6521393 | C | A | 102 | 0 | 4 | 0 | NON_SYNONYMOU | MODERAT | MISSENSE | Gtg/Ttg | V350L | polA | | | A | A | A | A | | | | | | |
| NC_011770 | 6521644 | T | C | 82 | 0 | 12 | 0 | NON_SYNONYMOU | MODERAT | MISSENSE | gAc/gGc | D266G | polA | C | C | C | C | C | C | C | C | C | C | C | C |
| NC_011770 | 6521813 | C | T | 89 | 0 | 4 | 0 | NON_SYNONYMOU | MODERAT | MISSENSE | Gag/Aag | E210K | polA | | | T | T | T | T | | | | | | |
| NC_011770 | 6522064 | T | C | 71 | 0 | 4 | 0 | NON_SYNONYMOU | MODERAT | MISSENSE | gAc/gGc | D126G | polA | | | C | C | C | C | | | | | | |
| NC_011770 | 3811742 | G | T | 79 | 0 | 6 | 0 | NON_SYNONYMOU | MODERAT | MISSENSE | Ctg/Atg | L255M | polB | T | T | | | | | T | T | T | T | | |
| NC_011770 | 3811978 | C | T | 82 | 0 | 2 | 0 | NON_SYNONYMOU | MODERAT | MISSENSE | gGc/gAc | G176D | polB | T | T | | | | | | | | | | |
| NC_011770 | 3812297 | C | T | 68 | 0 | 4 | 0 | NON_SYNONYMOU | MODERAT | MISSENSE | Gcg/Acg | A70T | polB | | | T | T | T | T | | | | | | |
| NC_011770 | 6016340 | A | C | 86 | 0 | 2 | 0 | NON_SYNONYMOU | MODERAT | MISSENSE | Acc/Ccc | T256P | ponA | | | | | | | | | | | C | C |
| NC_011770 | 6016736 | C | T | 86 | 0 | 2 | 0 | NON_SYNONYMOU | MODERAT | MISSENSE | Ccg/Tcg | P388S | ponA | | | | | | | T | T | | | | |
| NC_011770 | 6017360 | G | C | 85 | 0 | 2 | 0 | NON_SYNONYMOU | MODERAT | MISSENSE | Gga/Cga | G596R | ponA | | | C | C | | | | | | | | |
| NC_011770 | 6017417 | G | GCGC | 83 | 0 | 8 | 0 | CODON_INSERTIO | MODERATE | | gat/CCGga | D616PD | ponA | GCGC | GCGC | | | | | GCGC | GCGC | GCGC | GCGC | GCGC | GCGC |
| NC_011770 | 4013563 | T | C | 64 | 0 | 2 | 0 | NON_SYNONYMOU | MODERAT | MISSENSE | aAc/aGc | N329S | popB | | | | | | | C | C | | | | |
| NC_011770 | 4013195 | A | G | 66 | 0 | 2 | 0 | NON_SYNONYMOU | MODERAT | MISSENSE | gTg/gCg | V57A | popD | | | | | | | G | G | | | | |
| NC_011770 | 4013241 | G | A | 65 | 0 | 2 | 0 | NON_SYNONYMOU | MODERAT | MISSENSE | Ccg/Tcg | P42S | popD | | | | | | | A | A | | | | |
| NC_011770 | 4013321 | A | G | 67 | 0 | 2 | 0 | NON_SYNONYMOU | MODERAT | MISSENSE | aTc/aCc | I15T | popD | | | | | | | G | G | | | | |
| NC_011770 | 4020589 | G | A | 60 | 0 | 2 | 0 | NON_SYNONYMOU | MODERAT | MISSENSE | tCg/tTg | S134L | popN | | | | | | | A | A | | | | |
| NC_011770 | 4020724 | G | A | 53 | 0 | 2 | 0 | NON_SYNONYMOU | MODERAT | MISSENSE | cCg/cTg | P89L | popN | | | | | | | A | A | | | | |
| NC_011770 | 4020899 | C | T | 59 | 0 | 2 | 0 | NON_SYNONYMOU | MODERAT | MISSENSE | Ggc/Agc | G31S | popN | | | | | T | T | | | | | | |
| NC_011770 | 4020916 | G | A | 57 | 0 | 2 | 0 | NON_SYNONYMOU | MODERAT | MISSENSE | gCt/gTt | A25V | popN | | | | | | | A | A | | | | |
| NC_011770 | 4020932 | C | G | 56 | 1 | 3 | 0 | NON_SYNONYMOU | MODERAT | MISSENSE | Gcc/Ccc | A20P | popN | | | G | . | G | G | | | | | | |
| NC_011770 | 4020595 | G | A | 62 | 0 | 1 | 0 | NON_SYNONYMOU | MODERAT | MISSENSE | gCg/gTg | A132V | popN | | | | | | | | A | | | | |
| NC_011770 | 1539213 | AG | A | 69 | 0 | 1 | 0 | FRAME_SHIFT | HIGH | | ccc/ | P4 | potA | | | | | | | | A | | | | |
| NC_011770 | 1538899 | C | A | 75 | 0 | 4 | 0 | NON_SYNONYMOU | MODERAT | MISSENSE | aaG/aaT | K109N | potA | | | A | A | A | A | | | | | | |
| NC_011770 | 1539038 | C | T | 70 | 0 | 2 | 0 | NON_SYNONYMOU | MODERAT | MISSENSE | cGc/cAc | R63H | potA | | | | | | | | | | | T | T |
| NC_011770 | 1537846 | T | A | 88 | 0 | 2 | 0 | NON_SYNONYMOU | MODERAT | MISSENSE | tAc/tTc | Y95F | potB | | | | | | | | | | | A | A |
| NC_011770 | 1537855 | A | G | 86 | 0 | 1 | 0 | NON_SYNONYMOU | MODERAT | MISSENSE | cTg/cCg | L92P | potB | | G | | | | | | | | | | |
| NC_011770 | 1537820 | C | T | 84 | 0 | 1 | 0 | NON_SYNONYMOU | MODERAT | MISSENSE | Gcc/Acc | A104T | potB | | | | | | | | T | | | | |
| NC_011770 | 1536732 | C | T | 79 | 0 | 2 | 0 | NON_SYNONYMOU | MODERAT | MISSENSE | cGc/cAc | R171H | potC | | | | | | | | | T | T | | |
| NC_011770 | 1536003 | T | A | 76 | 0 | 2 | 0 | NON_SYNONYMOU | MODERAT | MISSENSE | Acc/Tcc | T128S | potD | | | | | | | | | | | A | A |
| NC_011770 | 1536080 | T | C | 79 | 0 | 10 | 0 | NON_SYNONYMOU | MODERAT | MISSENSE | aAc/aGc | N102S | potD | | | C | C | C | C | C | C | C | C | C | C |
| NC_011770 | 1536147 | C | T | 76 | 0 | 2 | 0 | NON_SYNONYMOU | MODERAT | MISSENSE | Gcg/Acg | A80T | potD | | | | | | | | | T | T | | |
| NC_011770 | 1536273 | G | C | 70 | 0 | 2 | 0 | NON_SYNONYMOU | MODERAT | MISSENSE | Ctg/Gtg | L38V | potD | C | C | | | | | | | | | | |
| NC_011770 | 6301042 | C | T | 89 | 0 | 1 | 0 | STOP_GAINED | HIGH | NONSENSE | Cag/Tag | Q89* | poxB | | | | | | | | T | | | | |
| NC_011770 | 6300992 | G | A | 90 | 0 | 1 | 0 | NON_SYNONYMOU | MODERAT | MISSENSE | gGc/gAc | G72D | poxB | | | | | | | A | | | | | |
| NC_011770 | 6301115 | A | G | 85 | 0 | 1 | 0 | NON_SYNONYMOU | MODERAT | MISSENSE | cAg/cGg | Q113R | poxB | | | | | | | G | | | | | |
| NC_011770 | 6301354 | G | A | 82 | 0 | 2 | 0 | NON_SYNONYMOU | MODERAT | MISSENSE | Gtc/Atc | V193I | poxB | A | A | | | | | | | | | | |
| NC_011770 | 6301423 | G | A | 82 | 0 | 2 | 0 | NON_SYNONYMOU | MODERAT | MISSENSE | Gac/Aac | D216N | poxB | A | A | | | | | | | | | | |
| NC_011770 | 6301924 | G | C | 81 | 0 | 4 | 0 | NON_SYNONYMOU | MODERAT | MISSENSE | Gcg/Ccg | A383P | poxB | | | C | C | C | C | | | | | | |
| NC_011770 | 6302449 | C | T | 72 | 0 | 2 | 0 | NON_SYNONYMOU | MODERAT | MISSENSE | Cgc/Tgc | R558C | poxB | | | | | | | T | T | | | | |
| NC_011770 | 1407877 | TG | T | 80 | 0 | 1 | 0 | FRAME_SHIFT | HIGH | | ggc/ | G253 | ppc | | | | | | | | T | | | | |

| | | | | | | | | | | | | | | | | | | | | | | | | | |
|---|---|---|---|---|---|---|---|---|---|---|---|---|---|---|---|---|---|---|---|---|---|---|---|---|---|
| NC_011770 | 1407852 | G | A | 77 | 0 | 4 | 0 | NON_SYNONYMOU | MODERAT | MISSENSE | Gcg/Acg | A243T | ppc | | | A | A | A | A | | | | | | |
| NC_011770 | 1408048 | A | G | 75 | 0 | 10 | 0 | NON_SYNONYMOU | MODERAT | MISSENSE | aAc/aGc | N308S | ppc | G | G | G | G | G | G | G | G | | | G | G |
| NC_011770 | 1409130 | A | G | 59 | 0 | 2 | 0 | NON_SYNONYMOU | MODERAT | MISSENSE | Agg/Ggg | R669G | ppc | | | | | | | G | G | | | | |
| NC_011770 | 1409489 | G | T | 57 | 0 | 2 | 0 | NON_SYNONYMOU | MODERAT | MISSENSE | gaG/gaT | E788D | ppc | | | | | | | | | | | T | T |
| NC_011770 | 1409662 | C | T | 36 | 0 | 2 | 0 | NON_SYNONYMOU | MODERAT | MISSENSE | gCc/gTc | A846V | ppc | | | | | | | | | | | T | T |
| NC_011770 | 1409680 | G | A | 32 | 0 | 10 | 0 | NON_SYNONYMOU | MODERAT | MISSENSE | cGg/cAg | R852Q | ppc | A | A | A | A | A | A | A | A | | | A | A |
| NC_011770 | 1409227 | A | G | 58 | 0 | 1 | 0 | NON_SYNONYMOU | MODERAT | MISSENSE | aAg/aGg | K701R | ppc | | | | | | | | G | | | | |
| NC_011770 | 1988428 | C | A | 78 | 0 | 2 | 0 | NON_SYNONYMOU | MODERAT | MISSENSE | Gcc/Tcc | A32S | ppiA | | | | | | | | | A | A | | |
| NC_011770 | 3925208 | C | T | 61 | 0 | 2 | 0 | NON_SYNONYMOU | MODERAT | MISSENSE | gCg/gTg | A24V | ppiB | | | | | | | T | T | | | | |
| NC_011770 | 819897 | T | A | 66 | 0 | 4 | 0 | NON_SYNONYMOU | MODERAT | MISSENSE | Tcc/Acc | S39T | ppiC2 | | | | | | | | | A | A | A | A |
| NC_011770 | 3907145 | A | G | 55 | 0 | 12 | 0 | NON_SYNONYMOU | MODERAT | MISSENSE | Ttc/Ctc | F559L | ppiD | G | G | G | G | G | G | G | G | G | G | G | G |
| NC_011770 | 3907218 | G | C | 59 | 0 | 12 | 0 | NON_SYNONYMOU | MODERAT | MISSENSE | caC/caG | H534Q | ppiD | C | C | C | C | C | C | C | C | C | C | C | C |
| NC_011770 | 3907487 | C | G | 62 | 0 | 12 | 0 | NON_SYNONYMOU | MODERAT | MISSENSE | Gcc/Ccc | A445P | ppiD | G | G | G | G | G | G | G | G | G | G | G | G |
| NC_011770 | 3908199 | C | G | 77 | 0 | 8 | 0 | NON_SYNONYMOU | MODERAT | MISSENSE | gaG/gaC | E207D | ppiD | G | G | | | | | G | G | G | G | G | G |
| NC_011770 | 3908559 | T | G | 72 | 0 | 2 | 0 | NON_SYNONYMOU | MODERAT | MISSENSE | gaA/gaC | E87D | ppiD | | | | | | | | | | | G | G |
| NC_011770 | 6239313 | C | T | 69 | 0 | 1 | 0 | NON_SYNONYMOU | MODERAT | MISSENSE | cGc/cAc | R404H | ppk | | | | T | | | | | | | | |
| NC_011770 | 88060 | T | C | 18 | 0 | 10 | 0 | NON_SYNONYMOU | MODERAT | MISSENSE | Agc/Ggc | S376G | ppkA | C | C | C | C | C | C | C | C | | | C | C |
| NC_011770 | 88107 | G | A | 14 | 0 | 2 | 0 | NON_SYNONYMOU | MODERAT | MISSENSE | gCg/gTg | A360V | ppkA | A | A | | | | | | | | | | |
| NC_011770 | 88863 | A | G | 74 | 0 | 2 | 0 | NON_SYNONYMOU | MODERAT | MISSENSE | gTc/gCc | V108A | ppkA | G | G | | | | | | | | | | |
| NC_011770 | 88872 | A | T | 72 | 0 | 2 | 0 | NON_SYNONYMOU | MODERAT | MISSENSE | cTg/cAg | L105Q | ppkA | T | T | | | | | | | | | | |
| NC_011770 | 5156305 | C | T | 56 | 0 | 6 | 0 | NON_SYNONYMOU | MODERAT | MISSENSE | gGc/gAc | G922D | pprA | | | T | T | T | T | | | | | T | T |
| NC_011770 | 5156317 | GGCA | G | 55 | 0 | 8 | 0 | CODON_CHANGE_ | MODERATE | | ctgccc/ccc | LP917P | pprA | | | G | G | G | G | G | G | | | G | G |
| NC_011770 | 5156449 | A | G | 58 | 0 | 8 | 0 | NON_SYNONYMOU | MODERAT | MISSENSE | gTg/gCg | V874A | pprA | | | G | G | G | G | G | G | | | G | G |
| NC_011770 | 5156573 | T | C | 64 | 0 | 8 | 0 | NON_SYNONYMOU | MODERAT | MISSENSE | Att/Gtt | I833V | pprA | | | C | C | C | C | C | C | | | C | C |
| NC_011770 | 5156593 | A | G | 63 | 0 | 8 | 0 | NON_SYNONYMOU | MODERAT | MISSENSE | cTt/cCt | L826P | pprA | | | G | G | G | G | G | G | | | G | G |
| NC_011770 | 5156765 | T | C | 62 | 0 | 8 | 0 | NON_SYNONYMOU | MODERAT | MISSENSE | Acc/Gcc | T769A | pprA | | | C | C | C | C | C | C | | | C | C |
| NC_011770 | 5156846 | T | G | 62 | 0 | 8 | 0 | NON_SYNONYMOU | MODERAT | MISSENSE | Atg/Ctg | M742L | pprA | | | G | G | G | G | G | G | | | G | G |
| NC_011770 | 5157161 | G | A | 62 | 0 | 6 | 0 | NON_SYNONYMOU | MODERAT | MISSENSE | Cgc/Tgc | R637C | pprA | | | A | A | A | A | | | | | A | A |
| NC_011770 | 5157787 | G | A | 66 | 0 | 2 | 0 | NON_SYNONYMOU | MODERAT | MISSENSE | gCa/gTa | A428V | pprA | | | | | | | A | A | | | | |
| NC_011770 | 5157980 | G | A | 67 | 0 | 6 | 0 | NON_SYNONYMOU | MODERAT | MISSENSE | Ctc/Ttc | L364F | pprA | | | A | A | A | A | | | | | A | A |
| NC_011770 | 5158167 | G | C | 68 | 0 | 2 | 0 | NON_SYNONYMOU | MODERAT | MISSENSE | caC/caG | H301Q | pprA | | | | | | | C | C | | | | |
| NC_011770 | 5158316 | C | T | 65 | 0 | 6 | 0 | NON_SYNONYMOU | MODERAT | MISSENSE | Gcg/Acg | A252T | pprA | T | T | | | | | | | T | T | T | T |
| NC_011770 | 5158663 | T | C | 69 | 0 | 12 | 0 | NON_SYNONYMOU | MODERAT | MISSENSE | aAc/aGc | N136S | pprA | C | C | C | C | C | C | C | C | C | C | C | C |
| NC_011770 | 5158774 | T | A | 66 | 0 | 2 | 0 | NON_SYNONYMOU | MODERAT | MISSENSE | gAt/gTt | D99V | pprA | | | | | | | | | A | A | | |
| NC_011770 | 5159000 | C | A | 78 | 0 | 8 | 0 | NON_SYNONYMOU | MODERAT | MISSENSE | Gcc/Tcc | A24S | pprA | A | A | A | A | A | A | A | A | | | | |
| NC_011770 | 5160822 | T | C | 75 | 0 | 2 | 0 | NON_SYNONYMOU | MODERAT | MISSENSE | Tcg/Ccg | S191P | pprB | | | | | | | | | C | C | | |
| NC_011770 | 3952752 | A | C | 73 | 0 | 12 | 0 | NON_SYNONYMOU | MODERAT | MISSENSE | Ttc/Gtc | F432V | ppsA | C | C | C | C | C | C | C | C | C | C | C | C |
| NC_011770 | 6237887 | G | A | 94 | 0 | 2 | 0 | NON_SYNONYMOU | MODERAT | MISSENSE | gGc/gAc | G361D | ppx | | | | | | | | | A | A | | |
| NC_011770 | 6238097 | A | C | 90 | 0 | 4 | 0 | NON_SYNONYMOU | MODERAT | MISSENSE | gAg/gCg | E431A | ppx | | | | | | | C | C | C | C | | |
| NC_011770 | 3693396 | T | G | 75 | 0 | 2 | 0 | NON_SYNONYMOU | MODERAT | MISSENSE | gAg/gCg | E145A | pqqB | G | G | | | | | | | | | | |
| NC_011770 | 3693689 | T | G | 71 | 0 | 12 | 0 | NON_SYNONYMOU | MODERAT | MISSENSE | gaA/gaC | E47D | pqqB | G | G | G | G | G | G | G | G | G | G | G | G |
| NC_011770 | 3692794 | T | C | 97 | 0 | 4 | 0 | NON_SYNONYMOU | MODERAT | MISSENSE | Agt/Ggt | S38G | pqqC | | | C | C | C | C | | | | | | |
| NC_011770 | 3690906 | G | A | 65 | 0 | 4 | 0 | NON_SYNONYMOU | MODERAT | MISSENSE | gCc/gTc | A334V | pqqE | | | A | A | A | A | | | | | | |
| NC_011770 | 3708178 | G | A | 57 | 0 | 2 | 0 | NON_SYNONYMOU | MODERAT | MISSENSE | Ctc/Ttc | L755F | pqqF | | | | | | | | | | | A | A |
| NC_011770 | 3708322 | C | T | 57 | 0 | 2 | 0 | NON_SYNONYMOU | MODERAT | MISSENSE | Gaa/Aaa | E707K | pqqF | | | | | | | T | T | | | | |
| NC_011770 | 3708342 | C | T | 55 | 0 | 6 | 0 | NON_SYNONYMOU | MODERAT | MISSENSE | cGa/cAa | R700Q | pqqF | | | T | T | T | T | | | | | T | T |
| NC_011770 | 3708369 | G | A | 62 | 0 | 2 | 0 | NON_SYNONYMOU | MODERAT | MISSENSE | gCc/gTc | A691V | pqqF | | | | | | | | | | | A | A |
| NC_011770 | 3708585 | T | G | 59 | 0 | 2 | 0 | NON_SYNONYMOU | MODERAT | MISSENSE | cAg/cCg | Q619P | pqqF | | | | | | | | | | | G | G |
| NC_011770 | 3708589 | T | A | 58 | 0 | 2 | 0 | NON_SYNONYMOU | MODERAT | MISSENSE | Acg/Tcg | T618S | pqqF | | | | | | | | | | | A | A |
| NC_011770 | 3708631 | C | T | 62 | 0 | 6 | 0 | NON_SYNONYMOU | MODERAT | MISSENSE | Gtg/Atg | V604M | pqqF | T | T | T | T | T | T | | | | | | |
| NC_011770 | 3708675 | T | G | 57 | 0 | 2 | 0 | NON_SYNONYMOU | MODERAT | MISSENSE | gAg/gCg | E589A | pqqF | | | | | | | | | | | G | G |
| NC_011770 | 3708705 | A | G | 57 | 0 | 2 | 0 | NON_SYNONYMOU | MODERAT | MISSENSE | aTg/aCg | M579T | pqqF | | | | | | | | | G | G | | |
| NC_011770 | 3708760 | G | C | 57 | 0 | 2 | 0 | NON_SYNONYMOU | MODERAT | MISSENSE | Cgg/Ggg | R561G | pqqF | | | | | | | C | C | | | | |
| NC_011770 | 3708796 | A | G | 62 | 0 | 12 | 0 | NON_SYNONYMOU | MODERAT | MISSENSE | Ttc/Ctc | F549L | pqqF | G | G | G | G | G | G | G | G | G | G | G | G |
| NC_011770 | 3708799 | C | G | 63 | 0 | 12 | 0 | NON_SYNONYMOU | MODERAT | MISSENSE | Gta/Cta | V548L | pqqF | G | G | G | G | G | G | G | G | G | G | G | G |
| NC_011770 | 3708855 | C | T | 60 | 0 | 2 | 0 | NON_SYNONYMOU | MODERAT | MISSENSE | cGg/cAg | R529Q | pqqF | | | | | | | | | T | T | | |
| NC_011770 | 3708882 | C | T | 61 | 0 | 4 | 0 | NON_SYNONYMOU | MODERAT | MISSENSE | aGc/aAc | S520N | pqqF | T | T | | | | | T | T | | | | |
| NC_011770 | 3708898 | G | A | 62 | 0 | 4 | 0 | NON_SYNONYMOU | MODERAT | MISSENSE | Ccc/Tcc | P515S | pqqF | | | A | A | A | A | | | | | | |
| NC_011770 | 3708970 | T | A | 58 | 0 | 4 | 0 | NON_SYNONYMOU | MODERAT | MISSENSE | Agc/Tgc | S491C | pqqF | | | A | A | A | A | | | | | | |
| NC_011770 | 3709003 | A | C | 59 | 0 | 12 | 0 | NON_SYNONYMOU | MODERAT | MISSENSE | Tgc/Ggc | C480G | pqqF | C | C | C | C | C | C | C | C | C | C | C | C |
| NC_011770 | 3709080 | A | G | 60 | 0 | 12 | 0 | NON_SYNONYMOU | MODERAT | MISSENSE | gTa/gCa | V454A | pqqF | G | G | G | G | G | G | G | G | G | G | G | G |
| NC_011770 | 3709143 | G | A | 60 | 0 | 12 | 0 | NON_SYNONYMOU | MODERAT | MISSENSE | tCg/tTg | S433L | pqqF | A | A | A | A | A | A | A | A | A | A | A | A |

| | | | | | | | | | | | | | | | | | | | | | | | | | |
|---|---|---|---|---|---|---|---|---|---|---|---|---|---|---|---|---|---|---|---|---|---|---|---|---|---|
| NC_011770 | 3709180 | T | C | 58 | 0 | 10 | 0 | NON_SYNONYMOU | MODERAT | MISSENSE | Acc/Gcc | T421A | pqqF | C | C | C | C | C | C | C | C | C | C | | |
| NC_011770 | 3709293 | C | T | 53 | 0 | 2 | 0 | NON_SYNONYMOU | MODERAT | MISSENSE | cGc/cAc | R383H | pqqF | | | | | | | | | | | T | T |
| NC_011770 | 3709392 | C | T | 49 | 0 | 2 | 0 | NON_SYNONYMOU | MODERAT | MISSENSE | cGg/cAg | R350Q | pqqF | | | | | | | | | | | T | T |
| NC_011770 | 3709431 | G | T | 50 | 0 | 4 | 0 | NON_SYNONYMOU | MODERAT | MISSENSE | gCc/gAc | A337D | pqqF | T | T | | | | | | | T | T | | |
| NC_011770 | 3709471 | C | T | 50 | 0 | 4 | 0 | NON_SYNONYMOU | MODERAT | MISSENSE | Gcg/Acg | A324T | pqqF | | | T | T | T | T | | | | | | |
| NC_011770 | 3709548 | G | A | 47 | 0 | 4 | 0 | NON_SYNONYMOU | MODERAT | MISSENSE | gCg/gTg | A298V | pqqF | | | | | | | A | A | | | A | A |
| NC_011770 | 3709629 | T | A | 50 | 0 | 4 | 0 | NON_SYNONYMOU | MODERAT | MISSENSE | cAg/cTg | Q271L | pqqF | | | | | | | A | A | | | A | A |
| NC_011770 | 3709650 | T | C | 48 | 0 | 4 | 0 | NON_SYNONYMOU | MODERAT | MISSENSE | gAg/gGg | E264G | pqqF | | | | | | | C | C | | | C | C |
| NC_011770 | 3709755 | C | T | 49 | 0 | 2 | 0 | NON_SYNONYMOU | MODERAT | MISSENSE | aGt/aAt | S229N | pqqF | | | | | | | | | T | T | | |
| NC_011770 | 3710088 | C | T | 68 | 0 | 2 | 0 | NON_SYNONYMOU | MODERAT | MISSENSE | cGc/cAc | R118H | pqqF | | | | | | | | | T | T | | |
| NC_011770 | 3710371 | G | C | 73 | 0 | 2 | 0 | NON_SYNONYMOU | MODERAT | MISSENSE | Ctg/Gtg | L24V | pqqF | | | | | | | | | C | C | | |
| NC_011770 | 4767839 | A | G | 82 | 0 | 12 | 0 | NON_SYNONYMOU | MODERAT | MISSENSE | Tgt/Cgt | C354R | pqsA | G | G | G | G | G | G | G | G | G | G | G | G |
| NC_011770 | 4766753 | G | A | 86 | 0 | 4 | 0 | NON_SYNONYMOU | MODERAT | MISSENSE | cCt/cTt | P200L | pqsB | | | | | | | A | A | | | A | A |
| NC_011770 | 4766885 | C | T | 83 | 0 | 2 | 0 | NON_SYNONYMOU | MODERAT | MISSENSE | gGc/gAc | G156D | pqsB | | | | | | | | | T | T | | |
| NC_011770 | 4767000 | C | T | 89 | 0 | 2 | 0 | NON_SYNONYMOU | MODERAT | MISSENSE | Ggc/Agc | G118S | pqsB | | | | | | | T | T | | | | |
| NC_011770 | 4767203 | G | A | 92 | 0 | 4 | 0 | NON_SYNONYMOU | MODERAT | MISSENSE | gCg/gTg | A50V | pqsB | A | A | | | | | | | A | A | | |
| NC_011770 | 4766414 | T | C | 84 | 0 | 2 | 0 | NON_SYNONYMOU | MODERAT | MISSENSE | Agc/Ggc | S32G | pqsC | | | | | | | | | C | C | | |
| NC_011770 | 4765059 | C | G | 87 | 0 | 2 | 0 | NON_SYNONYMOU | MODERAT | MISSENSE | ttG/ttC | L120F | pqsD | G | G | | | | | | | | | | |
| NC_011770 | 4764356 | T | C | 64 | 0 | 1 | 0 | NON_SYNONYMOU | MODERAT | MISSENSE | gAc/gGc | D19G | pqsE | C | | | | | | | | | | | |
| NC_011770 | 2792026 | A | AG | 74 | 0 | 1 | 0 | FRAME_SHIFT | HIGH | | gcc/gGcc | A8G? | pqsH | | | | | | | AG | | | | | |
| NC_011770 | 2792281 | A | G | 78 | 0 | 4 | 0 | NON_SYNONYMOU | MODERAT | MISSENSE | aAt/aGt | N91S | pqsH | G | G | | | | | | | G | G | | |
| NC_011770 | 2793049 | C | T | 85 | 0 | 2 | 0 | NON_SYNONYMOU | MODERAT | MISSENSE | cCc/cTc | P347L | pqsH | | | | | | | T | T | | | | |
| NC_011770 | 805213 | G | C | 86 | 0 | 2 | 0 | NON_SYNONYMOU | MODERAT | MISSENSE | Cat/Gat | H75D | pqsL | | | | | | | | | | | C | C |
| NC_011770 | 805347 | C | T | 83 | 0 | 4 | 0 | NON_SYNONYMOU | MODERAT | MISSENSE | cGc/cAc | R30H | pqsL | | | T | T | T | T | | | | | | |
| NC_011770 | 5475865 | T | G | 53 | 0 | 4 | 0 | NON_SYNONYMOU | MODERAT | MISSENSE | Aag/Cag | K153Q | pra | | | G | G | G | G | | | | | | |
| NC_011770 | 5476137 | T | G | 67 | 0 | 4 | 0 | NON_SYNONYMOU | MODERAT | MISSENSE | aAg/aCg | K62T | pra | | | G | G | G | G | | | | | | |
| NC_011770 | 1959382 | C | T | 70 | 0 | 4 | 0 | NON_SYNONYMOU | MODERAT | MISSENSE | Gcc/Acc | A160T | prc | | | T | T | T | T | | | | | | |
| NC_011770 | 1958238 | T | C | 76 | 0 | 1 | 0 | NON_SYNONYMOU | MODERAT | MISSENSE | gAc/gGc | D541G | prc | C | | | | | | | | | | | |
| NC_011770 | 5568271 | C | T | 70 | 0 | 10 | 0 | NON_SYNONYMOU | MODERAT | MISSENSE | Gtc/Atc | V148I | prfA | T | T | T | T | T | T | T | T | | | T | T |
| NC_011770 | 6494939 | T | C | 70 | 0 | 8 | 0 | NON_SYNONYMOU | MODERAT | MISSENSE | cAg/cGg | Q182R | prfH | C | C | | | | | C | C | C | C | C | C |
| NC_011770 | 6494952 | C | T | 69 | 0 | 2 | 0 | NON_SYNONYMOU | MODERAT | MISSENSE | Gcg/Acg | A178T | prfH | | | | | | | | | | | T | T |
| NC_011770 | 6495104 | G | C | 80 | 0 | 2 | 0 | NON_SYNONYMOU | MODERAT | MISSENSE | aCc/aGc | T127S | prfH | C | C | | | | | | | | | | |
| NC_011770 | 6495249 | T | G | 80 | 0 | 12 | 0 | NON_SYNONYMOU | MODERAT | MISSENSE | Acc/Ccc | T79P | prfH | G | G | G | G | G | G | G | G | G | G | G | G |
| NC_011770 | 6495278 | T | C | 80 | 0 | 12 | 0 | NON_SYNONYMOU | MODERAT | MISSENSE | cAc/cGc | H69R | prfH | C | C | C | C | C | C | C | C | C | C | C | C |
| NC_011770 | 6495302 | A | G | 79 | 0 | 12 | 0 | NON_SYNONYMOU | MODERAT | MISSENSE | aTt/aCt | I61T | prfH | G | G | G | G | G | G | G | G | G | G | G | G |
| NC_011770 | 6495354 | T | C | 78 | 0 | 12 | 0 | NON_SYNONYMOU | MODERAT | MISSENSE | Acc/Gcc | T44A | prfH | C | C | C | C | C | C | C | C | C | C | C | C |
| NC_011770 | 6495366 | G | T | 76 | 0 | 12 | 0 | NON_SYNONYMOU | MODERAT | MISSENSE | Ctc/Atc | L40I | prfH | T | T | T | T | T | T | T | T | T | T | T | T |
| NC_011770 | 6022328 | G | C | 82 | 0 | 2 | 0 | NON_SYNONYMOU | MODERAT | MISSENSE | gaG/gaC | E52D | priA | | | | | | | | | | | C | C |
| NC_011770 | 6022662 | C | G | 87 | 0 | 2 | 0 | NON_SYNONYMOU | MODERAT | MISSENSE | Cag/Gag | Q164E | priA | G | G | | | | | | | | | | |
| NC_011770 | 6022731 | A | T | 76 | 0 | 2 | 0 | NON_SYNONYMOU | MODERAT | MISSENSE | Agc/Tgc | S187C | priA | | | | | | | | | T | T | | |
| NC_011770 | 6022906 | C | G | 82 | 0 | 2 | 0 | NON_SYNONYMOU | MODERAT | MISSENSE | aCc/aGc | T245S | priA | | | | | | | G | G | | | | |
| NC_011770 | 6023643 | G | C | 92 | 0 | 2 | 0 | NON_SYNONYMOU | MODERAT | MISSENSE | Gta/Cta | V491L | priA | C | C | | | | | | | | | | |
| NC_011770 | 79426 | T | C | 84 | 0 | 1 | 0 | NON_SYNONYMOU | MODERAT | MISSENSE | Tcc/Ccc | S544P | prlC | | | | | | | C | | | | | |
| NC_011770 | 5777710 | G | A | 76 | 0 | 1 | 0 | STOP_GAINED | HIGH | NONSENSE | tGg/tAg | W3* | prmA | | | | | | | | A | | | | |
| NC_011770 | 5778490 | A | C | 68 | 0 | 10 | 0 | NON_SYNONYMOU | MODERAT | MISSENSE | gAg/gCg | E263A | prmA | C | C | C | C | C | C | C | C | C | C | | |
| NC_011770 | 5778502 | C | A | 67 | 0 | 2 | 0 | NON_SYNONYMOU | MODERAT | MISSENSE | gCg/gAg | A267E | prmA | | | | | | | A | A | | | | |
| NC_011770 | 1049705 | T | A | 79 | 0 | 12 | 0 | NON_SYNONYMOU | MODERAT | MISSENSE | cTc/cAc | L51H | proA | A | A | A | A | A | A | A | A | A | A | A | A |
| NC_011770 | 1050199 | A | G | 90 | 0 | 2 | 0 | NON_SYNONYMOU | MODERAT | MISSENSE | Atc/Gtc | I216V | proA | | | | | | | G | G | | | | |
| NC_011770 | 1050367 | A | G | 81 | 0 | 2 | 0 | NON_SYNONYMOU | MODERAT | MISSENSE | Atc/Gtc | I272V | proA | | | | | | | | | | | G | G |
| NC_011770 | 1050377 | G | A | 81 | 0 | 4 | 0 | NON_SYNONYMOU | MODERAT | MISSENSE | cGg/cAg | R275Q | proA | | | A | A | A | A | | | | | | |
| NC_011770 | 1050380 | T | C | 82 | 0 | 4 | 0 | NON_SYNONYMOU | MODERAT | MISSENSE | gTg/gCg | V276A | proA | | | | | | | | | C | C | C | C |
| NC_011770 | 1050815 | C | T | 79 | 0 | 4 | 0 | NON_SYNONYMOU | MODERAT | MISSENSE | aCc/aTc | T421I | proA | | | T | T | T | T | | | | | | |
| NC_011770 | 5448979 | C | T | 59 | 0 | 1 | 0 | NON_SYNONYMOU | MODERAT | MISSENSE | Ggc/Agc | G252S | proB | | | | | | | T | | | | | |
| NC_011770 | 428765 | G | A | 89 | 0 | 4 | 0 | NON_SYNONYMOU | MODERAT | MISSENSE | Ctc/Ttc | L210F | proC | | | | | | | | | A | A | A | A |
| NC_011770 | 4795494 | C | A | 59 | 0 | 1 | 0 | NON_SYNONYMOU | MODERAT | MISSENSE | Cag/Aag | Q172K | proS | | | | | | | A | | | | | |
| NC_011770 | 4795837 | C | T | 74 | 0 | 2 | 0 | NON_SYNONYMOU | MODERAT | MISSENSE | gCc/gTc | A286V | proS | | | | | | | | | | | T | T |
| NC_011770 | 4796257 | G | A | 73 | 0 | 4 | 0 | NON_SYNONYMOU | MODERAT | MISSENSE | aGc/aAc | S426N | proS | | | A | A | A | A | | | | | | |
| NC_011770 | 4796596 | G | A | 61 | 0 | 4 | 0 | NON_SYNONYMOU | MODERAT | MISSENSE | aGc/aAc | S539N | proS | | | A | A | A | A | | | | | | |
| NC_011770 | 4992955 | G | T | 79 | 0 | 2 | 0 | NON_SYNONYMOU | MODERAT | MISSENSE | Gcc/Tcc | A33S | prpB | | | | | | | | | T | T | | |
| NC_011770 | 4993310 | G | A | 85 | 0 | 2 | 0 | NON_SYNONYMOU | MODERAT | MISSENSE | aGc/aAc | S151N | prpB | A | A | | | | | | | | | | |
| NC_011770 | 4993630 | C | T | 88 | 0 | 2 | 0 | NON_SYNONYMOU | MODERAT | MISSENSE | Cgc/Tgc | R258C | prpB | | | | | | | | | T | T | | |
| NC_011770 | 4999246 | A | G | 80 | 0 | 4 | 0 | NON_SYNONYMOU | MODERAT | MISSENSE | Acg/Gcg | T32A | prpD | | | G | G | G | G | | | | | | |

| | | | | | | | | | | | | | | | | | | | | | | | | | |
|---|---|---|---|---|---|---|---|---|---|---|---|---|---|---|---|---|---|---|---|---|---|---|---|---|---|
| NC_011770 | 4999967 | C | T | 79 | 0 | 2 | 0 | NON_SYNONYMOU | MODERAT | MISSENSE | aCc/aTc | T272I | prpD | | | | | | | | | T | T | | |
| NC_011770 | 4999975 | G | A | 80 | 0 | 2 | 0 | NON_SYNONYMOU | MODERAT | MISSENSE | Gag/Aag | E275K | prpD | | | | | | | | | A | A | | |
| NC_011770 | 4999996 | T | C | 83 | 0 | 2 | 0 | NON_SYNONYMOU | MODERAT | MISSENSE | Ttc/Ctc | F282L | prpD | | | | | | | | | | | C | C |
| NC_011770 | 5000116 | A | G | 81 | 0 | 12 | 0 | NON_SYNONYMOU | MODERAT | MISSENSE | Aac/Gac | N322D | prpD | G | G | G | G | G | G | G | G | G | G | G | G |
| NC_011770 | 5618097 | C | A | 73 | 0 | 1 | 0 | INTRAGENIC | MODIFIER | | | | PrrF2 | | | A | | | | | | | | | |
| NC_011770 | 5618046 | G | A | 60 | 0 | 1 | 0 | INTRAGENIC | MODIFIER | | | | PrrF2 | | | | | | | | | | | | A |
| NC_011770 | 5618027 | A | T | 59 | 1 | 2 | 0 | INTRAGENIC | MODIFIER | | | | PrrF2 | T | T | | | | | . | | | | | |
| NC_011770 | 5618042 | G | A | 59 | 0 | 1 | 0 | INTRAGENIC | MODIFIER | | | | PrrF2 | | | | | | | A | | | | | |
| NC_011770 | 5618043 | C | T | 59 | 0 | 1 | 0 | INTRAGENIC | MODIFIER | | | | PrrF2 | | | | | | | T | | | | | |
| NC_011770 | 5618044 | C | T | 59 | 0 | 2 | 0 | INTRAGENIC | MODIFIER | | | | PrrF2 | | | | | | | | | | | T | T |
| NC_011770 | 5618048 | A | C | 61 | 0 | 1 | 0 | INTRAGENIC | MODIFIER | | | | PrrF2 | | | | | | | C | | | | | |
| NC_011770 | 5618062 | A | AC | 63 | 0 | 1 | 0 | INTRAGENIC | MODIFIER | | | | PrrF2 | | | | | | | AC | | | | | |
| NC_011770 | 5618110 | G | GT | 73 | 0 | 1 | 0 | INTRAGENIC | MODIFIER | | | | PrrF2 | | | | | | | GT | | | | | |
| NC_011770 | 5618114 | C | T | 69 | 0 | 1 | 0 | INTRAGENIC | MODIFIER | | | | PrrF2 | | | | | | | T | | | | | |
| NC_011770 | 5574328 | A | G | 87 | 0 | 1 | 0 | NON_SYNONYMOU | MODERAT | MISSENSE | aAc/aGc | N259S | prs | | | | | | | | G | | | | |
| NC_011770 | 664233 | T | C | 82 | 0 | 2 | 0 | START_LOST | HIGH | MISSENSE | Atg/Gtg | M1V | prtN | | | | | | | | | C | C | | |
| NC_011770 | 663963 | T | C | 65 | 0 | 6 | 0 | NON_SYNONYMOU | MODERAT | MISSENSE | Agc/Ggc | S91G | prtN | | | C | C | C | C | | | | | C | C |
| NC_011770 | 664211 | C | G | 81 | 0 | 2 | 0 | NON_SYNONYMOU | MODERAT | MISSENSE | aGc/aCc | S8T | prtN | | | | | | | | | G | G | | |
| NC_011770 | 664224 | A | T | 80 | 0 | 2 | 0 | NON_SYNONYMOU | MODERAT | MISSENSE | Tcc/Acc | S4T | prtN | | | | | | | | | T | T | | |
| NC_011770 | 664352 | C | G | 83 | 0 | 2 | 0 | NON_SYNONYMOU | MODERAT | MISSENSE | gGc/gCc | G251A | prtR | | | | | | | | | G | G | | |
| NC_011770 | 4008738 | G | A | 60 | 0 | 2 | 0 | NON_SYNONYMOU | MODERAT | MISSENSE | gCt/gTt | A135V | pscB | | | | | | | | | | | A | A |
| NC_011770 | 4008945 | A | G | 65 | 0 | 12 | 0 | NON_SYNONYMOU | MODERAT | MISSENSE | gTg/gCg | V66A | pscB | G | G | G | G | G | G | G | G | G | G | G | G |
| NC_011770 | 4008960 | T | A | 69 | 0 | 2 | 0 | NON_SYNONYMOU | MODERAT | MISSENSE | cAg/cTg | Q61L | pscB | | | | | | | | | A | A | | |
| NC_011770 | 4006958 | T | C | 86 | 0 | 2 | 0 | NON_SYNONYMOU | MODERAT | MISSENSE | Aag/Gag | K588E | pscC | | | C | C | | | | | | | | |
| NC_011770 | 4007210 | C | T | 84 | 0 | 12 | 0 | NON_SYNONYMOU | MODERAT | MISSENSE | Gtc/Atc | V504I | pscC | T | T | T | T | T | T | T | T | T | T | T | T |
| NC_011770 | 4007749 | T | C | 66 | 0 | 4 | 0 | NON_SYNONYMOU | MODERAT | MISSENSE | aAc/aGc | N324S | pscC | C | C | | | | | | | C | C | | |
| NC_011770 | 4008332 | G | C | 68 | 0 | 8 | 0 | NON_SYNONYMOU | MODERAT | MISSENSE | Ccg/Gcg | P130A | pscC | C | C | C | C | C | C | C | C | | | | |
| NC_011770 | 4006434 | A | T | 81 | 0 | 2 | 0 | NON_SYNONYMOU | MODERAT | MISSENSE | cTg/cAg | L161Q | pscD | | | | | | | | | T | T | | |
| NC_011770 | 4006507 | C | T | 77 | 0 | 2 | 0 | NON_SYNONYMOU | MODERAT | MISSENSE | Gcg/Acg | A137T | pscD | | | | | | | T | T | | | | |
| NC_011770 | 4006636 | T | C | 77 | 0 | 12 | 0 | NON_SYNONYMOU | MODERAT | MISSENSE | Aca/Gca | T94A | pscD | C | C | C | C | C | C | C | C | C | C | C | C |
| NC_011770 | 4006842 | G | A | 81 | 0 | 2 | 0 | NON_SYNONYMOU | MODERAT | MISSENSE | gCg/gTg | A25V | pscD | A | A | | | | | | | | | | |
| NC_011770 | 4005339 | G | A | 85 | 0 | 1 | 0 | NON_SYNONYMOU | MODERAT | MISSENSE | gCc/gTc | A37V | pscF | | | | | | | A | | | | | |
| NC_011770 | 4005350 | G | T | 80 | 0 | 6 | 0 | NON_SYNONYMOU | MODERAT | MISSENSE | gaC/gaA | D33E | pscF | | | | | | | T | T | T | T | T | T |
| NC_011770 | 4005167 | C | G | 77 | 0 | 2 | 0 | NON_SYNONYMOU | MODERAT | MISSENSE | Gaa/Caa | E8Q | pscG | G | G | | | | | | | | | | |
| NC_011770 | 4004679 | T | C | 70 | 0 | 10 | 0 | NON_SYNONYMOU | MODERAT | MISSENSE | Agc/Ggc | S56G | pscH | | | C | C | C | C | C | C | C | C | C | C |
| NC_011770 | 4004723 | G | T | 70 | 0 | 2 | 0 | NON_SYNONYMOU | MODERAT | MISSENSE | cCg/cAg | P41Q | pscH | | | | | | | | | T | T | | |
| NC_011770 | 4004221 | C | T | 78 | 0 | 4 | 0 | NON_SYNONYMOU | MODERAT | MISSENSE | Gtg/Atg | V65M | pscI | | | | | | | T | T | T | T | | |
| NC_011770 | 4004059 | T | C | 75 | 0 | 10 | 0 | NON_SYNONYMOU | MODERAT | MISSENSE | gAt/gGt | D7G | pscJ | C | C | C | C | C | C | C | C | C | C | | |
| NC_011770 | 4004071 | C | T | 75 | 0 | 2 | 0 | NON_SYNONYMOU | MODERAT | MISSENSE | cGa/cAa | R3Q | pscJ | | | | | | | | | | | T | T |
| NC_011770 | 4002913 | G | T | 64 | 0 | 4 | 0 | NON_SYNONYMOU | MODERAT | MISSENSE | gaC/gaA | D137E | pscK | T | T | | | | | | | T | T | | |
| NC_011770 | 4002810 | C | T | 59 | 0 | 1 | 0 | NON_SYNONYMOU | MODERAT | MISSENSE | Gcc/Acc | A172T | pscK | | | | | | | | T | | | | |
| NC_011770 | 4002297 | G | A | 63 | 0 | 2 | 0 | NON_SYNONYMOU | MODERAT | MISSENSE | gCc/gTc | A143V | pscL | | | | | | | A | A | | | | |
| NC_011770 | 4022778 | A | G | 59 | 0 | 6 | 0 | NON_SYNONYMOU | MODERAT | MISSENSE | Acc/Gcc | T94A | pscO | G | G | | | | | G | G | | | G | G |
| NC_011770 | 4024048 | A | G | 75 | 0 | 6 | 0 | NON_SYNONYMOU | MODERAT | MISSENSE | Acg/Gcg | T4A | pscQ | | | G | G | G | G | G | G | | | | |
| NC_011770 | 4024076 | G | A | 71 | 0 | 2 | 0 | NON_SYNONYMOU | MODERAT | MISSENSE | cGc/cAc | R13H | pscQ | | | | | | | | | A | A | | |
| NC_011770 | 4024127 | T | C | 71 | 0 | 4 | 0 | NON_SYNONYMOU | MODERAT | MISSENSE | gTc/gCc | V30A | pscQ | | | | | | | | | C | C | C | C |
| NC_011770 | 4024177 | G | A | 75 | 0 | 2 | 0 | NON_SYNONYMOU | MODERAT | MISSENSE | Gtc/Atc | V47I | pscQ | | | | | | | | | | | A | A |
| NC_011770 | 4024250 | A | G | 77 | 0 | 6 | 0 | NON_SYNONYMOU | MODERAT | MISSENSE | cAc/cGc | H71R | pscQ | G | G | G | G | G | G | | | | | | |
| NC_011770 | 4024525 | C | A | 81 | 0 | 2 | 0 | NON_SYNONYMOU | MODERAT | MISSENSE | Ctt/Att | L163I | pscQ | | | | | | | A | A | | | | |
| NC_011770 | 4024551 | C | G | 77 | 0 | 2 | 0 | NON_SYNONYMOU | MODERAT | MISSENSE | gaC/gaG | D171E | pscQ | G | G | | | | | | | | | | |
| NC_011770 | 4024552 | A | G | 76 | 0 | 12 | 0 | NON_SYNONYMOU | MODERAT | MISSENSE | Acc/Gcc | T172A | pscQ | G | G | G | G | G | G | G | G | G | G | G | G |
| NC_011770 | 4024555 | G | A | 79 | 0 | 2 | 0 | NON_SYNONYMOU | MODERAT | MISSENSE | Ggc/Agc | G173S | pscQ | | | | | | | | | A | A | | |
| NC_011770 | 4024560 | G | T | 78 | 0 | 2 | 0 | NON_SYNONYMOU | MODERAT | MISSENSE | gaG/gaT | E174D | pscQ | | | | | | | T | T | | | | |
| NC_011770 | 4024706 | C | T | 81 | 0 | 2 | 0 | NON_SYNONYMOU | MODERAT | MISSENSE | aCc/aTc | T223I | pscQ | T | T | | | | | | | | | | |
| NC_011770 | 4024709 | C | A | 80 | 0 | 2 | 0 | NON_SYNONYMOU | MODERAT | MISSENSE | cCc/cAc | P224H | pscQ | | | | | | | A | A | | | | |
| NC_011770 | 4024723 | G | A | 77 | 0 | 4 | 0 | NON_SYNONYMOU | MODERAT | MISSENSE | Ggc/Agc | G229S | pscQ | A | A | | | | | | | | | A | A |
| NC_011770 | 4025028 | C | T | 76 | 0 | 12 | 0 | NON_SYNONYMOU | MODERAT | MISSENSE | Ctc/Ttc | L22F | pscR | T | T | T | T | T | T | T | T | T | T | T | T |
| NC_011770 | 4026077 | A | G | 70 | 0 | 2 | 0 | NON_SYNONYMOU | MODERAT | MISSENSE | gAc/gGc | D65G | pscT | G | G | | | | | | | | | | |
| NC_011770 | 4026220 | T | C | 74 | 0 | 2 | 0 | NON_SYNONYMOU | MODERAT | MISSENSE | Ttc/Ctc | F113L | pscT | C | C | | | | | | | | | | |
| NC_011770 | 4027161 | C | T | 86 | 0 | 2 | 0 | NON_SYNONYMOU | MODERAT | MISSENSE | Ctc/Ttc | L165F | pscU | T | T | | | | | | | | | | |
| NC_011770 | 4027515 | G | A | 81 | 0 | 2 | 0 | NON_SYNONYMOU | MODERAT | MISSENSE | Gtc/Atc | V283I | pscU | A | A | | | | | | | | | | |
| NC_011770 | 4027596 | A | G | 75 | 0 | 2 | 0 | NON_SYNONYMOU | MODERAT | MISSENSE | Att/Gtt | I310V | pscU | G | G | | | | | | | | | | |

| | | | | | | | | | | | | | | | | | | | | | | | | | |
|---|---|---|---|---|---|---|---|---|---|---|---|---|---|---|---|---|---|---|---|---|---|---|---|---|---|
| NC_011770 | 5894879 | T | C | 65 | 0 | 2 | 0 | NON_SYNONYMOU | MODERAT | MISSENSE | Ttc/Ctc | F43L | psd | C | C | | | | | | | | | | |
| NC_011770 | 5894937 | A | T | 66 | 0 | 2 | 0 | NON_SYNONYMOU | MODERAT | MISSENSE | tAt/tTt | Y62F | psd | | | | | | | T | T | | | | |
| NC_011770 | 5894967 | C | G | 65 | 0 | 2 | 0 | NON_SYNONYMOU | MODERAT | MISSENSE | gCc/gGc | A72G | psd | | | | | G | G | | | | | | |
| NC_011770 | 5895398 | G | A | 65 | 0 | 2 | 0 | NON_SYNONYMOU | MODERAT | MISSENSE | Gcc/Acc | A216T | psd | | | | | | | A | A | | | | |
| NC_011770 | 5895567 | G | A | 59 | 0 | 4 | 0 | NON_SYNONYMOU | MODERAT | MISSENSE | gGg/gAg | G272E | psd | | | A | A | A | A | | | | | | |
| NC_011770 | 3389404 | C | A | 82 | 2 | 2 | 0 | STOP_GAINED | HIGH | NONSENSE | Gag/Tag | E407* | pslA | . | . | A | A | | | | | | | | |
| NC_011770 | 3389710 | G | C | 69 | 2 | 4 | 0 | NON_SYNONYMOU | MODERAT | MISSENSE | Ctg/Gtg | L305V | pslA | . | . | C | C | C | C | | | | | | |
| NC_011770 | 3388834 | A | G | 81 | 2 | 2 | 0 | NON_SYNONYMOU | MODERAT | MISSENSE | aTc/aCc | I118T | pslB | . | . | | | | | G | G | | | | |
| NC_011770 | 3388916 | T | C | 74 | 2 | 2 | 0 | NON_SYNONYMOU | MODERAT | MISSENSE | Atg/Gtg | M91V | pslB | . | . | | | | | C | C | | | | |
| NC_011770 | 3387554 | T | G | 68 | 2 | 2 | 0 | NON_SYNONYMOU | MODERAT | MISSENSE | gAg/gCg | E56A | pslC | . | . | | | | | G | G | | | | |
| NC_011770 | 3387650 | G | A | 69 | 2 | 4 | 0 | NON_SYNONYMOU | MODERAT | MISSENSE | gCc/gTc | A24V | pslC | . | . | | | | | A | A | | | A | A |
| NC_011770 | 3384278 | C | T | 80 | 0 | 12 | 0 | NON_SYNONYMOU | MODERAT | MISSENSE | Gtc/Atc | V573I | pslE | T | T | T | T | T | T | T | T | T | T | T | T |
| NC_011770 | 3385652 | G | T | 62 | 2 | 2 | 0 | NON_SYNONYMOU | MODERAT | MISSENSE | Cgc/Agc | R115S | pslE | . | . | | | | | T | T | | | | |
| NC_011770 | 3382908 | C | T | 75 | 0 | 2 | 0 | NON_SYNONYMOU | MODERAT | MISSENSE | Gcg/Acg | A367T | pslF | T | T | | | | | | | | | | |
| NC_011770 | 3383831 | G | C | 63 | 0 | 10 | 0 | NON_SYNONYMOU | MODERAT | MISSENSE | gCc/gGc | A59G | pslF | C | C | C | C | C | C | | | C | C | C | C |
| NC_011770 | 3381646 | G | A | 80 | 0 | 1 | 0 | NON_SYNONYMOU | MODERAT | MISSENSE | tCg/tTg | S395L | pslG | | | | | | | A | | | | | |
| NC_011770 | 3381716 | C | T | 83 | 0 | 6 | 0 | NON_SYNONYMOU | MODERAT | MISSENSE | Gac/Aac | D372N | pslG | | | T | T | T | T | T | T | | | | |
| NC_011770 | 3382587 | G | C | 85 | 0 | 1 | 0 | NON_SYNONYMOU | MODERAT | MISSENSE | caC/caG | H81Q | pslG | | | | | | | C | | | | | |
| NC_011770 | 3382212 | C | T | 88 | 0 | 1 | 0 | NON_SYNONYMOU | MODERAT | MISSENSE | atG/atA | M206I | pslG | | | | T | | | | | | | | |
| NC_011770 | 3380550 | C | T | 76 | 0 | 1 | 0 | NON_SYNONYMOU | MODERAT | MISSENSE | Gcc/Acc | A315T | pslH | | | | | | | T | | | | | |
| NC_011770 | 3380732 | C | T | 86 | 0 | 2 | 0 | NON_SYNONYMOU | MODERAT | MISSENSE | tGc/tAc | C254Y | pslH | | | | | | | T | T | | | | |
| NC_011770 | 3380908 | C | T | 81 | 0 | 1 | 0 | NON_SYNONYMOU | MODERAT | MISSENSE | atG/atA | M195I | pslH | | | | | | | T | | | | | |
| NC_011770 | 3380924 | T | C | 77 | 0 | 12 | 0 | NON_SYNONYMOU | MODERAT | MISSENSE | cAg/cGg | Q190R | pslH | C | C | C | C | C | C | C | C | C | C | C | C |
| NC_011770 | 3381191 | G | T | 87 | 0 | 6 | 0 | NON_SYNONYMOU | MODERAT | MISSENSE | aCg/aAg | T101K | pslH | | | T | T | T | T | | | T | T | | |
| NC_011770 | 3379294 | C | T | 75 | 0 | 2 | 0 | NON_SYNONYMOU | MODERAT | MISSENSE | Gcg/Acg | A334T | pslI | T | T | | | | | | | | | | |
| NC_011770 | 3379354 | G | A | 73 | 0 | 2 | 0 | NON_SYNONYMOU | MODERAT | MISSENSE | Ccg/Tcg | P314S | pslI | | | | | | | | | A | A | | |
| NC_011770 | 3379630 | C | T | 66 | 0 | 8 | 0 | NON_SYNONYMOU | MODERAT | MISSENSE | Gcc/Acc | A222T | pslI | | | T | T | T | T | | | T | T | T | T |
| NC_011770 | 3379722 | T | A | 68 | 0 | 2 | 0 | NON_SYNONYMOU | MODERAT | MISSENSE | gAc/gTc | D191V | pslI | | | | | | | A | A | | | | |
| NC_011770 | 3379731 | C | T | 70 | 0 | 12 | 0 | NON_SYNONYMOU | MODERAT | MISSENSE | gGa/gAa | G188E | pslI | T | T | T | T | T | T | T | T | T | T | T | T |
| NC_011770 | 3379885 | C | T | 79 | 0 | 2 | 0 | NON_SYNONYMOU | MODERAT | MISSENSE | Gtc/Atc | V137I | pslI | | | | | | | T | T | | | | |
| NC_011770 | 3380049 | A | G | 78 | 0 | 12 | 0 | NON_SYNONYMOU | MODERAT | MISSENSE | gTc/gCc | V82A | pslI | G | G | G | G | G | G | G | G | G | G | G | G |
| NC_011770 | 3380076 | T | C | 73 | 0 | 12 | 0 | NON_SYNONYMOU | MODERAT | MISSENSE | cAc/cGc | H73R | pslI | C | C | C | C | C | C | C | C | C | C | C | C |
| NC_011770 | 3380190 | G | A | 66 | 0 | 2 | 0 | NON_SYNONYMOU | MODERAT | MISSENSE | aCg/aTg | T35M | pslI | | | | | | | | | | | A | A |
| NC_011770 | 3379045 | A | AAC | 82 | 0 | 1 | 0 | FRAME_SHIFT | HIGH | | ctg/cGTtg | L48R? | pslJ | | | | | | | AAC | | | | | |
| NC_011770 | 3378833 | C | T | 81 | 0 | 4 | 0 | NON_SYNONYMOU | MODERAT | MISSENSE | Gtg/Atg | V119M | pslJ | | | T | T | T | T | | | | | | |
| NC_011770 | 3377112 | G | A | 81 | 0 | 6 | 0 | NON_SYNONYMOU | MODERAT | MISSENSE | aCc/aTc | T213I | pslK | | | A | A | A | A | | | A | A | | |
| NC_011770 | 3377113 | T | C | 81 | 0 | 6 | 0 | NON_SYNONYMOU | MODERAT | MISSENSE | Acc/Gcc | T213A | pslK | | | C | C | C | C | | | C | C | | |
| NC_011770 | 3376995 | G | A | 75 | 0 | 1 | 0 | NON_SYNONYMOU | MODERAT | MISSENSE | aCc/aTc | T252I | pslK | | | | | | | | A | | | | |
| NC_011770 | 3375225 | T | C | 55 | 0 | 10 | 0 | NON_SYNONYMOU | MODERAT | MISSENSE | Acc/Gcc | T345A | pslL | C | C | C | C | C | C | C | C | C | C | | |
| NC_011770 | 3375365 | G | A | 67 | 0 | 8 | 0 | NON_SYNONYMOU | MODERAT | MISSENSE | gCc/gTc | A298V | pslL | A | A | | | | | A | A | A | A | A | A |
| NC_011770 | 3375987 | C | G | 91 | 0 | 2 | 0 | NON_SYNONYMOU | MODERAT | MISSENSE | Ggc/Cgc | G91R | pslL | G | G | | | | | | | | | | |
| NC_011770 | 3373574 | T | C | 69 | 0 | 2 | 0 | NON_SYNONYMOU | MODERAT | MISSENSE | tAt/tGt | Y477C | pslM | | | | | | | | | C | C | | |
| NC_011770 | 3373686 | A | G | 69 | 0 | 2 | 0 | NON_SYNONYMOU | MODERAT | MISSENSE | Tcc/Ccc | S440P | pslM | | | | | | | | | G | G | | |
| NC_011770 | 3373947 | G | C | 64 | 0 | 4 | 0 | NON_SYNONYMOU | MODERAT | MISSENSE | Ctg/Gtg | L353V | pslM | | | | | | | C | C | C | C | | |
| NC_011770 | 3374550 | G | A | 82 | 0 | 2 | 0 | NON_SYNONYMOU | MODERAT | MISSENSE | Ccg/Tcg | P152S | pslM | | | | | | | A | A | | | | |
| NC_011770 | 3372479 | C | T | 77 | 0 | 1 | 0 | NON_SYNONYMOU | MODERAT | MISSENSE | Gcc/Acc | A246T | pslN | | | | | | | T | | | | | |
| NC_011770 | 3372627 | A | C | 69 | 0 | 4 | 0 | NON_SYNONYMOU | MODERAT | MISSENSE | caT/caG | H196Q | pslN | | | C | C | C | C | | | | | | |
| NC_011770 | 3371909 | A | G | 87 | 0 | 10 | 0 | NON_SYNONYMOU | MODERAT | MISSENSE | Tat/Cat | Y93H | pslO | G | G | G | G | G | G | G | G | | | G | G |
| NC_011770 | 2227729 | A | T | 75 | 0 | 2 | 0 | NON_SYNONYMOU | MODERAT | MISSENSE | cAg/cTg | Q76L | psrA | | | | | | | | | T | T | | |
| NC_011770 | 2228175 | C | T | 75 | 0 | 4 | 0 | NON_SYNONYMOU | MODERAT | MISSENSE | Ccg/Tcg | P225S | psrA | | | T | T | T | T | | | | | | |
| NC_011770 | 6371514 | A | T | 61 | 0 | 12 | 0 | NON_SYNONYMOU | MODERAT | MISSENSE | Tct/Act | S57T | pstA | T | T | T | T | T | T | T | T | T | T | T | T |
| NC_011770 | 6372577 | C | T | 57 | 0 | 12 | 0 | NON_SYNONYMOU | MODERAT | MISSENSE | Gtc/Atc | V387I | pstC | T | T | T | T | T | T | T | T | T | T | T | T |
| NC_011770 | 6372787 | G | C | 74 | 0 | 2 | 0 | NON_SYNONYMOU | MODERAT | MISSENSE | Ctg/Gtg | L317V | pstC | | | | | | | | | C | C | | |
| NC_011770 | 6372834 | T | G | 71 | 0 | 2 | 0 | NON_SYNONYMOU | MODERAT | MISSENSE | gAc/gCc | D301A | pstC | G | G | | | | | | | | | | |
| NC_011770 | 6373338 | T | G | 97 | 0 | 2 | 0 | NON_SYNONYMOU | MODERAT | MISSENSE | gAc/gCc | D133A | pstC | | | | | | | G | G | | | | |
| NC_011770 | 4928213 | C | G | 90 | 0 | 2 | 0 | NON_SYNONYMOU | MODERAT | MISSENSE | Cag/Gag | Q454E | pta | | | | | | | | | | | G | G |
| NC_011770 | 2257595 | T | C | 74 | 0 | 4 | 0 | NON_SYNONYMOU | MODERAT | MISSENSE | gTc/gCc | V46A | ptpA | | | C | C | C | C | | | | | | |
| NC_011770 | 2257696 | T | C | 77 | 0 | 8 | 0 | NON_SYNONYMOU | MODERAT | MISSENSE | Tgc/Cgc | C80R | ptpA | C | C | C | C | C | C | | | | | C | C |
| NC_011770 | 2257615 | C | T | 72 | 0 | 1 | 0 | NON_SYNONYMOU | MODERAT | MISSENSE | Cgc/Tgc | R53C | ptpA | | | | | | | | T | | | | |
| NC_011770 | 3357816 | G | A | 75 | 0 | 1 | 0 | NON_SYNONYMOU | MODERAT | MISSENSE | Ggc/Agc | G274S | ptxR | | | | | | | A | | | | | |
| NC_011770 | 3357927 | G | A | 59 | 0 | 12 | 0 | NON_SYNONYMOU | MODERAT | MISSENSE | Ggc/Agc | G311S | ptxR | A | A | A | A | A | A | A | A | A | A | A | A |
| NC_011770 | 3355742 | T | C | 68 | 0 | 12 | 0 | NON_SYNONYMOU | MODERAT | MISSENSE | Aac/Gac | N232D | ptxS | C | C | C | C | C | C | C | C | C | C | C | C |

| | | | | | | | | | | | | | | | | | | | | | | | | | |
|---|---|---|---|---|---|---|---|---|---|---|---|---|---|---|---|---|---|---|---|---|---|---|---|---|---|
| NC_011770 | 3355786 | G | A | 69 | 0 | 1 | 0 | NON_SYNONYMOU | MODERAT | MISSENSE | cCg/cTg | P217L | ptxS | | | | | | | A | | | | | |
| NC_011770 | 5874574 | G | A | 83 | 0 | 4 | 0 | NON_SYNONYMOU | MODERAT | MISSENSE | cCc/cTc | P376L | purA | | | A | A | A | A | | | | | | |
| NC_011770 | 2664137 | T | C | 55 | 0 | 2 | 0 | NON_SYNONYMOU | MODERAT | MISSENSE | aAg/aGg | K454R | purB | C | C | | | | | | | | | | |
| NC_011770 | 2664899 | C | T | 59 | 0 | 2 | 0 | NON_SYNONYMOU | MODERAT | MISSENSE | gGc/gAc | G200D | purB | T | T | | | | | | | | | | |
| NC_011770 | 2665017 | A | G | 54 | 0 | 6 | 0 | NON_SYNONYMOU | MODERAT | MISSENSE | Ttc/Ctc | F161L | purB | | | | | | | G | G | G | G | G | G |
| NC_011770 | 2665047 | C | T | 57 | 0 | 2 | 0 | NON_SYNONYMOU | MODERAT | MISSENSE | Gcc/Acc | A151T | purB | | | | | | | | | T | T | | |
| NC_011770 | 2665273 | C | A | 57 | 0 | 2 | 0 | NON_SYNONYMOU | MODERAT | MISSENSE | caG/caT | Q75H | purB | | | | | | | | | | | A | A |
| NC_011770 | 2665350 | G | T | 54 | 0 | 2 | 0 | NON_SYNONYMOU | MODERAT | MISSENSE | Cac/Aac | H50N | purB | | | | | | | T | T | | | | |
| NC_011770 | 4750793 | G | A | 74 | 0 | 2 | 0 | NON_SYNONYMOU | MODERAT | MISSENSE | gCc/gTc | A169V | purC | | | | | | | A | A | | | | |
| NC_011770 | 4750853 | G | A | 81 | 0 | 2 | 0 | NON_SYNONYMOU | MODERAT | MISSENSE | gCg/gTg | A149V | purC | | | | | | | | | A | A | | |
| NC_011770 | 4751046 | C | T | 84 | 0 | 2 | 0 | NON_SYNONYMOU | MODERAT | MISSENSE | Gac/Aac | D85N | purC | | | | | | | | | T | T | | |
| NC_011770 | 5783355 | T | C | 32 | 0 | 1 | 0 | NON_SYNONYMOU | MODERAT | MISSENSE | gTc/gCc | V28A | purD | | | | | | | C | | | | | |
| NC_011770 | 5783676 | A | G | 20 | 0 | 6 | 0 | NON_SYNONYMOU | MODERAT | MISSENSE | cAc/cGc | H135R | purD | | | | | | | G | G | G | G | G | G |
| NC_011770 | 5782033 | G | A | 80 | 0 | 2 | 0 | NON_SYNONYMOU | MODERAT | MISSENSE | Gcc/Acc | A158T | purH | | | | | | | | | | | A | A |
| NC_011770 | 6441233 | G | T | 97 | 0 | 2 | 0 | NON_SYNONYMOU | MODERAT | MISSENSE | gCt/gAt | A311D | purK | | | | | | | | | | | T | T |
| NC_011770 | 6441515 | C | G | 95 | 0 | 8 | 0 | NON_SYNONYMOU | MODERAT | MISSENSE | gGg/gCg | G217A | purK | | | G | G | G | G | | | G | G | G | G |
| NC_011770 | 6441686 | C | T | 95 | 0 | 1 | 0 | NON_SYNONYMOU | MODERAT | MISSENSE | gGc/gAc | G160D | purK | | | | | | | T | | | | | |
| NC_011770 | 1320007 | G | A | 62 | 0 | 2 | 0 | NON_SYNONYMOU | MODERAT | MISSENSE | aGc/aAc | S648N | purL | | | | | | | | | A | A | | |
| NC_011770 | 1319956 | T | C | 64 | 0 | 1 | 0 | NON_SYNONYMOU | MODERAT | MISSENSE | cTg/cCg | L631P | **purL** | | C | | | | | | | | | | |
| NC_011770 | 1321935 | C | T | 70 | 0 | 1 | 0 | NON_SYNONYMOU | MODERAT | MISSENSE | Cgc/Tgc | R1291C | purL | | | | | | | | | | | T | |
| NC_011770 | 1320802 | C | T | 88 | 0 | 1 | 0 | NON_SYNONYMOU | MODERAT | MISSENSE | gCc/gTc | A913V | **purL** | | | | | | | | | | | | T |
| NC_011770 | 1318198 | C | T | 88 | 0 | 1 | 0 | NON_SYNONYMOU | MODERAT | MISSENSE | gCg/gTg | A45V | **purL** | | | | T | | | | | | | | |
| NC_011770 | 4804727 | G | A | 80 | 0 | 1 | 0 | NON_SYNONYMOU | MODERAT | MISSENSE | Ggc/Agc | G55S | purM | | | | | | | A | | | | | |
| NC_011770 | 4805093 | C | G | 86 | 0 | 4 | 0 | NON_SYNONYMOU | MODERAT | MISSENSE | Cag/Gag | Q177E | purM | | | G | G | G | G | | | | | | |
| NC_011770 | 4805099 | G | A | 85 | 0 | 1 | 0 | NON_SYNONYMOU | MODERAT | MISSENSE | Ggc/Agc | G179S | purM | | | | | | | A | | | | | |
| NC_011770 | 4805193 | A | C | 89 | 0 | 2 | 0 | NON_SYNONYMOU | MODERAT | MISSENSE | gAg/gCg | E210A | purM | | | | | | | | | C | C | | |
| NC_011770 | 4805794 | A | G | 79 | 0 | 2 | 0 | NON_SYNONYMOU | MODERAT | MISSENSE | Acc/Gcc | T57A | purN | | | | | | | G | G | | | | |
| NC_011770 | 4805842 | G | C | 79 | 0 | 8 | 0 | NON_SYNONYMOU | MODERAT | MISSENSE | Gag/Cag | E73Q | purN | C | C | C | C | C | C | | | | | C | C |
| NC_011770 | 1333942 | A | C | 78 | 0 | 2 | 0 | NON_SYNONYMOU | MODERAT | MISSENSE | Atc/Ctc | I315L | purT | | | | | | | C | C | | | | |
| NC_011770 | 5176588 | T | C | 90 | 0 | 1 | 0 | NON_SYNONYMOU | MODERAT | MISSENSE | gAg/gGg | E224G | purU | | | | | | | C | | | | | |
| NC_011770 | 5013536 | G | C | 84 | 0 | 2 | 0 | NON_SYNONYMOU | MODERAT | MISSENSE | gaC/gaG | D881E | putA | C | C | | | | | | | | | | |
| NC_011770 | 5014521 | T | C | 39 | 0 | 2 | 0 | NON_SYNONYMOU | MODERAT | MISSENSE | aAc/aGc | N553S | putA | C | C | | | | | | | | | | |
| NC_011770 | 5015695 | A | G | 89 | 0 | 12 | 0 | NON_SYNONYMOU | MODERAT | MISSENSE | Tgg/Cgg | W162R | putA | G | G | G | G | G | G | G | G | G | G | G | G |
| NC_011770 | 3360561 | G | T | 66 | 0 | 6 | 0 | NON_SYNONYMOU | MODERAT | MISSENSE | gCc/gAc | A227D | pvcB | | | | | | | T | T | T | T | T | T |
| NC_011770 | 3360643 | C | T | 70 | 0 | 8 | 0 | NON_SYNONYMOU | MODERAT | MISSENSE | Gtc/Atc | V200I | pvcB | T | T | T | T | T | T | G | G | | | | |
| NC_011770 | 3360666 | A | G | 73 | 0 | 6 | 0 | NON_SYNONYMOU | MODERAT | MISSENSE | aTc/aCc | I192T | pvcB | G | G | G | G | G | G | | | | | | |
| NC_011770 | 3360759 | T | C | 72 | 0 | 12 | 0 | NON_SYNONYMOU | MODERAT | MISSENSE | cAc/cGc | H161R | pvcB | C | C | C | C | C | C | C | C | C | C | C | C |
| NC_011770 | 3361209 | T | C | 81 | 0 | 12 | 0 | NON_SYNONYMOU | MODERAT | MISSENSE | cAt/cGt | H11R | pvcB | C | C | C | C | C | C | C | C | C | C | C | C |
| NC_011770 | 3359665 | G | C | 83 | 0 | 4 | 0 | NON_SYNONYMOU | MODERAT | MISSENSE | Ctg/Gtg | L217V | pvcC | C | C | | | | | | | | | C | C |
| NC_011770 | 3360041 | C | G | 76 | 0 | 2 | 0 | NON_SYNONYMOU | MODERAT | MISSENSE | gaG/gaC | E91D | pvcC | | | | | | | G | G | | | | |
| NC_011770 | 3358433 | C | T | 73 | 0 | 2 | 0 | NON_SYNONYMOU | MODERAT | MISSENSE | cGc/cAc | R129H | pvcD | | | | | | | | | | | T | T |
| NC_011770 | 3358649 | T | C | 59 | 0 | 12 | 0 | NON_SYNONYMOU | MODERAT | MISSENSE | cAg/cGg | Q57R | pvcD | C | C | C | C | C | C | C | C | C | C | C | C |
| NC_011770 | 3358803 | G | A | 75 | 0 | 4 | 0 | NON_SYNONYMOU | MODERAT | MISSENSE | Cgc/Tgc | R6C | pvcD | | | A | A | A | A | | | | | | |
| NC_011770 | 3205273 | T | TGGTC | 90 | 0 | 2 | 0 | FRAME_SHIFT | HIGH | | agc/GTCGa | S138VE? | pvdA | | | | | | | | | TGGTC | TGGTC | | |
| NC_011770 | 3205277 | CGCCG | C | 90 | 0 | 2 | 0 | FRAME_SHIFT | HIGH | | ggccag/ | GQ140 | pvdA | | | | | | | | | C | C | | |
| NC_011770 | 3204909 | G | A | 80 | 0 | 2 | 0 | NON_SYNONYMOU | MODERAT | MISSENSE | Gtc/Atc | V16I | pvdA | | | | | | | | | A | A | | |
| NC_011770 | 3204951 | C | G | 81 | 0 | 2 | 0 | NON_SYNONYMOU | MODERAT | MISSENSE | Cag/Gag | Q30E | pvdA | | | | | | | | | G | G | | |
| NC_011770 | 3204963 | C | A | 81 | 0 | 2 | 0 | NON_SYNONYMOU | MODERAT | MISSENSE | Cag/Aag | Q34K | pvdA | | | | | | | | | A | A | | |
| NC_011770 | 3204964 | A | G | 80 | 0 | 2 | 0 | NON_SYNONYMOU | MODERAT | MISSENSE | cAg/cGg | Q34R | pvdA | | | | | | | | | G | G | | |
| NC_011770 | 3204966 | G | A | 81 | 0 | 2 | 0 | NON_SYNONYMOU | MODERAT | MISSENSE | Gcg/Acg | A35T | pvdA | | | | | | | | | A | A | | |
| NC_011770 | 3204976 | C | A | 81 | 0 | 2 | 0 | NON_SYNONYMOU | MODERAT | MISSENSE | gCc/gAc | A38D | pvdA | | | | | | | | | A | A | | |
| NC_011770 | 3204981 | G | C | 84 | 0 | 2 | 0 | NON_SYNONYMOU | MODERAT | MISSENSE | Gaa/Caa | E40Q | pvdA | | | | | | | | | C | C | | |
| NC_011770 | 3205005 | G | A | 83 | 0 | 2 | 0 | NON_SYNONYMOU | MODERAT | MISSENSE | Ggc/Agc | G48S | pvdA | | | | | | | | | A | A | | |
| NC_011770 | 3205006 | G | C | 79 | 0 | 2 | 0 | NON_SYNONYMOU | MODERAT | MISSENSE | gGc/gCc | G48A | pvdA | | | | | | | | | C | C | | |
| NC_011770 | 3205038 | T | A | 84 | 0 | 2 | 0 | NON_SYNONYMOU | MODERAT | MISSENSE | Tcg/Acg | S59T | pvdA | | | | | | | | | A | A | | |
| NC_011770 | 3205039 | C | G | 84 | 0 | 2 | 0 | NON_SYNONYMOU | MODERAT | MISSENSE | tCg/tGg | S59W | pvdA | | | | | | | | | G | G | | |
| NC_011770 | 3205135 | A | G | 86 | 0 | 2 | 0 | NON_SYNONYMOU | MODERAT | MISSENSE | gAt/gGt | D91G | pvdA | | | | | | | | | G | G | | |
| NC_011770 | 3205136 | T | G | 86 | 0 | 2 | 0 | NON_SYNONYMOU | MODERAT | MISSENSE | gaT/gaG | D91E | pvdA | | | | | | | | | G | G | | |
| NC_011770 | 3205215 | A | G | 91 | 0 | 2 | 0 | NON_SYNONYMOU | MODERAT | MISSENSE | Agc/Ggc | S118G | pvdA | | | | | | | | | G | G | | |
| NC_011770 | 3205224 | C | G | 95 | 0 | 2 | 0 | NON_SYNONYMOU | MODERAT | MISSENSE | Cag/Gag | Q121E | pvdA | | | | | | | | | G | G | | |
| NC_011770 | 3205225 | A | C | 94 | 0 | 2 | 0 | NON_SYNONYMOU | MODERAT | MISSENSE | cAg/cCg | Q121P | pvdA | | | | | | | | | C | C | | |
| NC_011770 | 3205226 | G | C | 96 | 0 | 2 | 0 | NON_SYNONYMOU | MODERAT | MISSENSE | caG/caC | Q121H | pvdA | | | | | | | | | C | C | | |

| | | | | | | | | | | | | | | | | | | | | | | | | | |
|---|---|---|---|---|---|---|---|---|---|---|---|---|---|---|---|---|---|---|---|---|---|---|---|---|---|
| NC_011770 | 3205257 | C | T | 91 | 0 | 2 | 0 | NON_SYNONYMOU | MODERAT | MISSENSE | Cgc/Tgc | R132C | pvdA | | | | | | | | | T | T | | |
| NC_011770 | 3205258 | G | C | 90 | 0 | 2 | 0 | NON_SYNONYMOU | MODERAT | MISSENSE | cGc/cCc | R132P | pvdA | | | | | | | | | C | C | | |
| NC_011770 | 3205260 | A | G | 89 | 0 | 2 | 0 | NON_SYNONYMOU | MODERAT | MISSENSE | Atc/Gtc | I133V | pvdA | | | | | | | | | G | G | | |
| NC_011770 | 3205306 | T | C | 88 | 0 | 2 | 0 | NON_SYNONYMOU | MODERAT | MISSENSE | aTc/aCc | I148T | pvdA | | | | | | | | | C | C | | |
| NC_011770 | 3205322 | C | A | 88 | 0 | 2 | 0 | NON_SYNONYMOU | MODERAT | MISSENSE | gaC/gaA | D153E | pvdA | | | | | | | | | A | A | | |
| NC_011770 | 3205350 | G | T | 85 | 1 | 1 | 0 | NON_SYNONYMOU | MODERAT | MISSENSE | Gcc/Tcc | A163S | pvdA | | | | | | | | | . | T | | |
| NC_011770 | 3205353 | C | G | 87 | 1 | 1 | 0 | NON_SYNONYMOU | MODERAT | MISSENSE | Ctg/Gtg | L164V | pvdA | | | | | | | | | . | G | | |
| NC_011770 | 3205567 | C | T | 85 | 1 | 2 | 0 | NON_SYNONYMOU | MODERAT | MISSENSE | gCc/gTc | A235V | pvdA | | | . | | | | | | T | T | | |
| NC_011770 | 3205572 | A | C | 88 | 1 | 2 | 0 | NON_SYNONYMOU | MODERAT | MISSENSE | Atg/Ctg | M237L | pvdA | | | . | | | | | | C | C | | |
| NC_011770 | 3205585 | C | G | 84 | 1 | 2 | 0 | NON_SYNONYMOU | MODERAT | MISSENSE | gCc/gGc | A241G | pvdA | | | . | | | | | | G | G | | |
| NC_011770 | 3205641 | A | G | 82 | 1 | 2 | 0 | NON_SYNONYMOU | MODERAT | MISSENSE | Aag/Gag | K260E | pvdA | | | . | | | | | | G | G | | |
| NC_011770 | 3205663 | G | A | 82 | 1 | 2 | 0 | NON_SYNONYMOU | MODERAT | MISSENSE | aGc/aAc | S267N | pvdA | | | . | | | | | | A | A | | |
| NC_011770 | 3205666 | G | A | 78 | 1 | 2 | 0 | NON_SYNONYMOU | MODERAT | MISSENSE | cGc/cAc | R268H | pvdA | | | . | | | | | | A | A | | |
| NC_011770 | 3205671 | C | G | 80 | 1 | 2 | 0 | NON_SYNONYMOU | MODERAT | MISSENSE | Cat/Gat | H270D | pvdA | | | . | | | | | | G | G | | |
| NC_011770 | 3205672 | A | G | 78 | 1 | 2 | 0 | NON_SYNONYMOU | MODERAT | MISSENSE | cAt/cGt | H270R | pvdA | | | . | | | | | | G | G | | |
| NC_011770 | 3205692 | C | G | 79 | 1 | 2 | 0 | NON_SYNONYMOU | MODERAT | MISSENSE | Ctg/Gtg | L277V | pvdA | | | . | | | | | | G | G | | |
| NC_011770 | 3205732 | C | T | 88 | 1 | 2 | 0 | NON_SYNONYMOU | MODERAT | MISSENSE | aCc/aTc | T290I | pvdA | | | . | | | | | | T | T | | |
| NC_011770 | 3205758 | G | A | 87 | 1 | 2 | 0 | NON_SYNONYMOU | MODERAT | MISSENSE | Gtc/Atc | V299I | pvdA | | | . | | | | | | A | A | | |
| NC_011770 | 3205785 | A | G | 92 | 0 | 2 | 0 | NON_SYNONYMOU | MODERAT | MISSENSE | Atc/Gtc | I308V | pvdA | | | | | | | | | G | G | | |
| NC_011770 | 3205897 | C | T | 83 | 2 | 4 | 0 | NON_SYNONYMOU | MODERAT | MISSENSE | aCc/aTc | T345I | pvdA | | | T | T | T | T | | | . | . | | |
| NC_011770 | 3206055 | A | T | 79 | 0 | 2 | 0 | NON_SYNONYMOU | MODERAT | MISSENSE | Agc/Tgc | S398C | pvdA | | | | | | | | | T | T | | |
| NC_011770 | 3206154 | G | A | 76 | 0 | 4 | 0 | NON_SYNONYMOU | MODERAT | MISSENSE | Gct/Act | A431T | pvdA | A | A | | | | | | | | | A | A |
| NC_011770 | 3206157 | C | G | 77 | 0 | 4 | 0 | NON_SYNONYMOU | MODERAT | MISSENSE | Ccg/Gcg | P432A | pvdA | G | G | | | | | | | | | G | G |
| NC_011770 | 3205993 | G | A | 74 | 2 | 1 | 0 | NON_SYNONYMOU | MODERAT | MISSENSE | gGc/gAc | G377D | pvdA | | | | | | A | | | . | . | | |
| NC_011770 | 3205375 | C | T | 78 | 2 | 1 | 0 | NON_SYNONYMOU | MODERAT | MISSENSE | aCc/aTc | T171I | pvdA | | | T | | | | | | . | . | | |
| NC_011770 | 3205143 | G | A | 88 | 0 | 1 | 0 | NON_SYNONYMOU | MODERAT | MISSENSE | Gtc/Atc | V94I | pvdA | | | | | | | | | | | A | |
| NC_011770 | 3164359 | A | T | 51 | 4 | 4 | 0 | NON_SYNONYMOU | MODERAT | MISSENSE | Agt/Tgt | S294C | pvdD | T | T | | | | | . | . | . | . | T | T |
| NC_011770 | 3164360 | G | C | 52 | 4 | 4 | 0 | NON_SYNONYMOU | MODERAT | MISSENSE | aGt/aCt | S294T | pvdD | C | C | | | | | . | . | . | . | C | C |
| NC_011770 | 3164368 | A | G | 52 | 4 | 4 | 0 | NON_SYNONYMOU | MODERAT | MISSENSE | Acc/Gcc | T297A | pvdD | G | G | | | | | . | . | . | . | G | G |
| NC_011770 | 3164395 | C | G | 73 | 2 | 2 | 0 | NON_SYNONYMOU | MODERAT | MISSENSE | Ccg/Gcg | P306A | pvdD | . | . | | | | | G | T | . | . | . | . |
| NC_011770 | 3164400 | A | T | 74 | 2 | 6 | 0 | NON_SYNONYMOU | MODERAT | MISSENSE | gaA/gaT | E307D | pvdD | T | T | | | | | T | T | . | . | T | T |
| NC_011770 | 3164432 | C | A | 79 | 0 | 8 | 0 | NON_SYNONYMOU | MODERAT | MISSENSE | aCc/aAc | T318N | pvdD | A | A | | | | | A | A | A | A | A | A |
| NC_011770 | 3164444 | C | T | 82 | 0 | 2 | 0 | NON_SYNONYMOU | MODERAT | MISSENSE | aCc/aTc | T322I | pvdD | | | | | | | T | T | | | | |
| NC_011770 | 3164484 | G | C | 81 | 2 | 6 | 0 | NON_SYNONYMOU | MODERAT | MISSENSE | ttG/ttC | L335F | pvdD | C | C | | | | | C | C | . | . | C | C |
| NC_011770 | 3164608 | C | G | 74 | 0 | 8 | 0 | NON_SYNONYMOU | MODERAT | MISSENSE | Cag/Gag | Q377E | pvdD | G | G | | | | | G | G | G | G | G | G |
| NC_011770 | 3164627 | G | A | 71 | 0 | 2 | 0 | NON_SYNONYMOU | MODERAT | MISSENSE | aGc/aAc | S383N | pvdD | | | | | | | A | A | | | | |
| NC_011770 | 3164642 | T | C | 72 | 0 | 2 | 0 | NON_SYNONYMOU | MODERAT | MISSENSE | gTg/gCg | V388A | pvdD | | | | | | | | | C | C | | |
| NC_011770 | 3164644 | A | C | 64 | 0 | 4 | 0 | NON_SYNONYMOU | MODERAT | MISSENSE | Atg/Ctg | M389L | pvdD | C | C | | | | | C | C | | | | |
| NC_011770 | 3164648 | A | T | 64 | 0 | 6 | 0 | NON_SYNONYMOU | MODERAT | MISSENSE | tAc/tTc | Y390F | pvdD | T | T | | | | | T | T | | | T | T |
| NC_011770 | 3164653 | C | T | 62 | 0 | 4 | 0 | NON_SYNONYMOU | MODERAT | MISSENSE | Cac/Tac | H392Y | pvdD | T | T | | | | | T | T | | | | |
| NC_011770 | 3165082 | A | G | 37 | 8 | 4 | 0 | NON_SYNONYMOU | MODERAT | MISSENSE | Ata/Gta | I535V | pvdD | . | . | G | G | G | G | . | . | . | . | . | . |
| NC_011770 | 3165211 | A | C | 36 | 6 | 2 | 0 | NON_SYNONYMOU | MODERAT | MISSENSE | Atg/Ctg | M578L | pvdD | . | . | | | | | . | . | C | C | . | . |
| NC_011770 | 3166105 | A | C | 26 | 6 | 2 | 0 | NON_SYNONYMOU | MODERAT | MISSENSE | Atc/Ctc | I876L | pvdD | . | . | | | | | C | . | . | . | C | . |
| NC_011770 | 3166107 | C | G | 28 | 5 | 1 | 0 | NON_SYNONYMOU | MODERAT | MISSENSE | atC/atG | I876M | pvdD | . | . | | | | | G | . | . | . | | |
| NC_011770 | 3166186 | A | G | 29 | 3 | 2 | 0 | NON_SYNONYMOU | MODERAT | MISSENSE | Atc/Gtc | I903V | pvdD | . | . | | | | | G | . | | | G | . |
| NC_011770 | 3166242 | G | C | 24 | 7 | 1 | 0 | NON_SYNONYMOU | MODERAT | MISSENSE | gaG/gaC | E921D | pvdD | . | . | | | | | . | . | . | C | . | . |
| NC_011770 | 3166243 | T | A | 21 | 7 | 1 | 0 | NON_SYNONYMOU | MODERAT | MISSENSE | Tgc/Agc | C922S | pvdD | . | . | | | | | . | . | . | A | . | . |
| NC_011770 | 3166580 | G | A | 32 | 4 | 2 | 0 | NON_SYNONYMOU | MODERAT | MISSENSE | cGg/cAg | R1034Q | pvdD | . | . | | | | | . | . | A | A | | |
| NC_011770 | 3166591 | A | G | 32 | 4 | 2 | 0 | NON_SYNONYMOU | MODERAT | MISSENSE | Acc/Gcc | T1038A | pvdD | . | . | | | | | . | . | G | G | | |
| NC_011770 | 3166618 | G | C | 32 | 4 | 2 | 0 | NON_SYNONYMOU | MODERAT | MISSENSE | Gac/Cac | D1047H | pvdD | . | . | | | | | . | . | | | C | C |
| NC_011770 | 3167159 | T | C | 40 | 5 | 1 | 0 | NON_SYNONYMOU | MODERAT | MISSENSE | gTg/gCg | V1227A | pvdD | . | . | | | | | . | . | . | | . | C |
| NC_011770 | 3167239 | A | G | 37 | 8 | 4 | 0 | NON_SYNONYMOU | MODERAT | MISSENSE | Aac/Gac | N1254D | pvdD | . | . | G | G | G | G | . | . | . | . | . | . |
| NC_011770 | 3167660 | T | G | 36 | 8 | 4 | 0 | NON_SYNONYMOU | MODERAT | MISSENSE | cTg/cGg | L1394R | pvdD | . | . | G | G | G | G | . | . | . | . | . | . |
| NC_011770 | 3167945 | A | G | 35 | 6 | 2 | 0 | NON_SYNONYMOU | MODERAT | MISSENSE | gAc/gGc | D1489G | pvdD | . | . | | | | | . | . | . | . | G | G |
| NC_011770 | 3167962 | G | C | 35 | 7 | 1 | 0 | NON_SYNONYMOU | MODERAT | MISSENSE | Ggc/Cgc | G1495R | pvdD | . | . | | | | | . | . | . | . | . | C |
| NC_011770 | 3167963 | G | A | 35 | 7 | 1 | 0 | NON_SYNONYMOU | MODERAT | MISSENSE | gGc/gAc | G1495D | pvdD | . | . | | | | | . | . | . | . | . | A |
| NC_011770 | 3167969 | C | G | 35 | 6 | 2 | 0 | NON_SYNONYMOU | MODERAT | MISSENSE | aCc/aGc | T1497S | pvdD | . | . | | | | | . | . | . | . | G | G |
| NC_011770 | 3167983 | G | A | 36 | 6 | 2 | 0 | NON_SYNONYMOU | MODERAT | MISSENSE | Gtg/Atg | V1502M | pvdD | . | . | | | | | . | . | . | . | A | A |
| NC_011770 | 3168250 | G | C | 38 | 7 | 1 | 0 | NON_SYNONYMOU | MODERAT | MISSENSE | Gtg/Ctg | V1591L | pvdD | . | . | | | | | . | . | . | C | . | . |
| NC_011770 | 3168927 | A | T | 33 | 7 | 1 | 0 | NON_SYNONYMOU | MODERAT | MISSENSE | gaA/gaT | E1816D | pvdD | . | . | | | | | . | . | . | T | . | . |
| NC_011770 | 3168934 | T | G | 34 | 7 | 1 | 0 | NON_SYNONYMOU | MODERAT | MISSENSE | Tgc/Ggc | C1819G | pvdD | . | . | | | | | . | . | . | G | . | . |
| NC_011770 | 3168935 | G | T | 33 | 7 | 1 | 0 | NON_SYNONYMOU | MODERAT | MISSENSE | tGc/tTc | C1819F | pvdD | . | . | | | | | . | . | . | T | . | . |

| | | | | | | | | | | | | | | | | | | | | | | | | | |
|---|---|---|---|---|---|---|---|---|---|---|---|---|---|---|---|---|---|---|---|---|---|---|---|---|---|
| NC_011770 | 3168985 | C | G | 38 | 4 | 3 | 0 | NON_SYNONYMOU | MODERAT | MISSENSE | Cag/Gag | Q1836E | pvdD | . | G | | | | | . | . | G | G | . | . |
| NC_011770 | 3168986 | A | C | 37 | 4 | 1 | 0 | NON_SYNONYMOU | MODERAT | MISSENSE | cAg/cCg | Q1836P | pvdD | . | C | | | | | . | . | | | . | . |
| NC_011770 | 3169626 | G | C | 36 | 7 | 1 | 0 | NON_SYNONYMOU | MODERAT | MISSENSE | caG/caC | Q2049H | pvdD | . | . | | | | | . | . | . | C | . | . |
| NC_011770 | 3169633 | G | A | 39 | 7 | 1 | 0 | NON_SYNONYMOU | MODERAT | MISSENSE | Gtg/Atg | V2052M | pvdD | . | . | | | | | . | . | . | A | . | . |
| NC_011770 | 3169683 | TTCG | T | 39 | 6 | 2 | 0 | CODON_DELETION | MODERATE | | tcg/ | S2069 | pvdD | . | . | | | | | . | . | T | T | . | . |
| NC_011770 | 3169699 | G | A | 54 | 0 | 2 | 0 | NON_SYNONYMOU | MODERAT | MISSENSE | Gtg/Atg | V2074M | pvdD | | | | | | | | | A | A | | |
| NC_011770 | 3170011 | G | A | 42 | 0 | 6 | 0 | NON_SYNONYMOU | MODERAT | MISSENSE | Gtg/Atg | V2178M | pvdD | A | A | | | | | A | A | | | A | A |
| NC_011770 | 3170012 | T | C | 40 | 0 | 2 | 0 | NON_SYNONYMOU | MODERAT | MISSENSE | gTg/gCg | V2178A | pvdD | | | | | | | | | C | C | | |
| NC_011770 | 3170014 | T | G | 41 | 0 | 8 | 0 | NON_SYNONYMOU | MODERAT | MISSENSE | Tcg/Gcg | S2179A | pvdD | G | G | | | | | G | G | G | G | G | G |
| NC_011770 | 3170023 | C | G | 39 | 0 | 8 | 0 | NON_SYNONYMOU | MODERAT | MISSENSE | Cgt/Ggt | R2182G | pvdD | G | G | | | | | G | G | G | G | G | G |
| NC_011770 | 3170024 | G | T | 39 | 0 | 8 | 0 | NON_SYNONYMOU | MODERAT | MISSENSE | cGt/cTt | R2182L | pvdD | T | T | | | | | T | T | T | T | T | T |
| NC_011770 | 3170048 | C | T | 47 | 0 | 8 | 0 | NON_SYNONYMOU | MODERAT | MISSENSE | aCg/aTg | T2190M | pvdD | T | T | | | | | T | T | T | T | T | T |
| NC_011770 | 3170051 | G | C | 46 | 0 | 8 | 0 | NON_SYNONYMOU | MODERAT | MISSENSE | aGc/aCc | S2191T | pvdD | C | C | | | | | C | C | C | C | C | C |
| NC_011770 | 3170057 | C | A | 48 | 0 | 6 | 0 | NON_SYNONYMOU | MODERAT | MISSENSE | gCg/gAg | A2193E | pvdD | A | A | | | | | A | A | | | A | A |
| NC_011770 | 3170098 | G | T | 48 | 0 | 6 | 0 | NON_SYNONYMOU | MODERAT | MISSENSE | Gcc/Tcc | A2207S | pvdD | T | T | | | | | T | T | . | . | T | T |
| NC_011770 | 3170146 | G | A | 42 | 0 | 8 | 0 | NON_SYNONYMOU | MODERAT | MISSENSE | Gtg/Atg | V2223M | pvdD | A | A | | | | | A | A | A | A | A | A |
| NC_011770 | 3170161 | A | C | 33 | 0 | 8 | 0 | NON_SYNONYMOU | MODERAT | MISSENSE | Atc/Ctc | I2228L | pvdD | C | C | | | | | C | C | C | C | C | C |
| NC_011770 | 3170163 | C | G | 33 | 0 | 8 | 0 | NON_SYNONYMOU | MODERAT | MISSENSE | atC/atG | I2228M | pvdD | G | G | | | | | G | G | G | G | G | G |
| NC_011770 | 3170176 | G | C | 31 | 0 | 8 | 0 | NON_SYNONYMOU | MODERAT | MISSENSE | Gag/Cag | E2233Q | pvdD | C | C | | | | | C | C | C | C | C | C |
| NC_011770 | 3170177 | A | G | 31 | 0 | 6 | 0 | NON_SYNONYMOU | MODERAT | MISSENSE | gAg/gGg | E2233G | pvdD | G | G | | | | | G | G | | | G | G |
| NC_011770 | 3170188 | C | G | 31 | 0 | 6 | 0 | NON_SYNONYMOU | MODERAT | MISSENSE | Cag/Gag | Q2237E | pvdD | G | G | | | | | G | G | | | G | G |
| NC_011770 | 3170212 | A | G | 27 | 0 | 8 | 0 | NON_SYNONYMOU | MODERAT | MISSENSE | Atc/Gtc | I2245V | pvdD | G | G | | | | | G | G | G | G | G | G |
| NC_011770 | 3170214 | C | G | 27 | 0 | 6 | 0 | NON_SYNONYMOU | MODERAT | MISSENSE | atC/atG | I2245M | pvdD | G | G | | | | | G | G | | | G | G |
| NC_011770 | 3170216 | G | C | 28 | 0 | 2 | 0 | NON_SYNONYMOU | MODERAT | MISSENSE | gGg/gCg | G2246A | pvdD | | | | | | | | | C | C | | |
| NC_011770 | 3170317 | C | G | 29 | 0 | 6 | 0 | NON_SYNONYMOU | MODERAT | MISSENSE | Ccg/Gcg | P2280A | pvdD | G | G | | | | | G | G | | | G | G |
| NC_011770 | 3170334 | G | C | 11 | 6 | 2 | 0 | NON_SYNONYMOU | MODERAT | MISSENSE | gaG/gaC | E2285D | pvdD | . | . | | | | | . | . | C | C | . | . |
| NC_011770 | 3170360 | G | A | 6 | 8 | 4 | 0 | NON_SYNONYMOU | MODERAT | MISSENSE | gGc/gAc | G2294D | pvdD | . | . | A | A | A | A | . | . | . | . | . | . |
| NC_011770 | 3170521 | C | G | 18 | 7 | 1 | 0 | NON_SYNONYMOU | MODERAT | MISSENSE | Ctg/Gtg | L2348V | pvdD | . | . | | | | | . | . | . | G | . | . |
| NC_011770 | 3170527 | C | G | 19 | 7 | 1 | 0 | NON_SYNONYMOU | MODERAT | MISSENSE | Ctg/Gtg | L2350V | pvdD | . | . | | | | | . | . | . | G | . | . |
| NC_011770 | 3170580 | C | G | 30 | 0 | 8 | 0 | NON_SYNONYMOU | MODERAT | MISSENSE | gaC/gaG | D2367E | pvdD | G | G | | | | | G | G | G | G | G | G |
| NC_011770 | 3170666 | C | G | 33 | 3 | 5 | 0 | NON_SYNONYMOU | MODERAT | MISSENSE | gCa/gGa | A2396G | pvdD | G | . | | | | | G | G | . | . | G | G |
| NC_011770 | 3170680 | C | G | 36 | 2 | 6 | 0 | NON_SYNONYMOU | MODERAT | MISSENSE | Ctg/Gtg | L2401V | pvdD | G | G | | | | | G | G | . | . | G | G |
| NC_011770 | 3170944 | A | T | 42 | 2 | 6 | 0 | NON_SYNONYMOU | MODERAT | MISSENSE | Acc/Tcc | T2489S | pvdD | T | T | | | | | T | T | . | . | T | T |
| NC_011770 | 3170949 | G | C | 42 | 2 | 6 | 0 | NON_SYNONYMOU | MODERAT | MISSENSE | caG/caC | Q2490H | pvdD | C | C | | | | | C | C | . | . | C | C |
| NC_011770 | 3170950 | T | C | 40 | 2 | 6 | 0 | NON_SYNONYMOU | MODERAT | MISSENSE | Tgg/Cgg | W2491R | pvdD | C | C | | | | | C | C | . | . | C | C |
| NC_011770 | 3170951 | G | T | 41 | 2 | 6 | 0 | NON_SYNONYMOU | MODERAT | MISSENSE | tGg/tTg | W2491L | pvdD | T | T | | | | | T | T | . | . | T | T |
| NC_011770 | 3170953 | G | T | 40 | 2 | 6 | 0 | NON_SYNONYMOU | MODERAT | MISSENSE | Gtg/Ttg | V2492L | pvdD | T | T | | | | | T | T | . | . | T | T |
| NC_011770 | 3170965 | G | C | 39 | 1 | 6 | 0 | NON_SYNONYMOU | MODERAT | MISSENSE | Gcg/Ccg | A2496P | pvdD | C | C | | | | | C | C | . | | C | C |
| NC_011770 | 3170966 | C | G | 39 | 1 | 6 | 0 | NON_SYNONYMOU | MODERAT | MISSENSE | gCg/gGg | A2496G | pvdD | G | G | | | | | G | G | . | | G | G |
| NC_011770 | 3170968 | T | A | 39 | 1 | 7 | 0 | NON_SYNONYMOU | MODERAT | MISSENSE | Ttg/Atg | L2497M | pvdD | A | A | | | | | A | A | . | A | A | A |
| NC_011770 | 3171096 | T | C | 39 | 3 | 5 | 0 | NON_SYNONYMOU | MODERAT | MISSENSE | gaT/gaC | D2539E | pvdD | . | . | | | | | C | C | G | G | C | . |
| NC_011770 | 3171113 | G | A | 39 | 6 | 2 | 0 | NON_SYNONYMOU | MODERAT | MISSENSE | cGg/cAg | R2545Q | pvdD | . | . | | | | | . | . | A | A | . | . |
| NC_011770 | 3171121 | A | C | 38 | 6 | 2 | 0 | NON_SYNONYMOU | MODERAT | MISSENSE | Atg/Ctg | M2548L | pvdD | . | . | | | | | . | . | C | C | . | . |
| NC_011770 | 3171142 | C | A | 38 | 6 | 2 | 0 | NON_SYNONYMOU | MODERAT | MISSENSE | Ctc/Atc | L2555I | pvdD | . | . | | | | | . | . | A | A | . | . |
| NC_011770 | 3171172 | C | G | 37 | 6 | 2 | 0 | NON_SYNONYMOU | MODERAT | MISSENSE | Ctg/Gtg | L2565V | pvdD | . | . | | | | | . | . | G | G | . | . |
| NC_011770 | 3171682 | G | A | 41 | 5 | 3 | 0 | NON_SYNONYMOU | MODERAT | MISSENSE | Gta/Ata | V2735I | pvdD | . | A | | | | | . | . | A | A | . | . |
| NC_011770 | 3171695 | T | C | 41 | 3 | 1 | 0 | NON_SYNONYMOU | MODERAT | MISSENSE | gTg/gCg | V2739A | pvdD | . | C | | | | | . | . | | | . | . |
| NC_011770 | 3172122 | T | A | 39 | 6 | 2 | 0 | NON_SYNONYMOU | MODERAT | MISSENSE | gaT/gaA | D2881E | pvdD | . | . | | | | | . | . | A | A | . | . |
| NC_011770 | 3172130 | A | C | 38 | 6 | 2 | 0 | NON_SYNONYMOU | MODERAT | MISSENSE | gAg/gCg | E2884A | pvdD | . | . | | | | | . | . | C | C | . | . |
| NC_011770 | 3172131 | G | C | 38 | 6 | 2 | 0 | NON_SYNONYMOU | MODERAT | MISSENSE | gaG/gaC | E2884D | pvdD | . | . | | | | | . | . | C | C | . | . |
| NC_011770 | 3172134 | T | G | 40 | 6 | 2 | 0 | NON_SYNONYMOU | MODERAT | MISSENSE | gaT/gaG | D2885E | pvdD | . | . | | | | | . | . | G | G | . | . |
| NC_011770 | 3172145 | G | C | 41 | 6 | 2 | 0 | NON_SYNONYMOU | MODERAT | MISSENSE | aGc/aCc | S2889T | pvdD | . | . | | | | | . | . | C | C | . | . |
| NC_011770 | 3172146 | C | G | 40 | 6 | 2 | 0 | NON_SYNONYMOU | MODERAT | MISSENSE | agC/agG | S2889R | pvdD | . | . | | | | | . | . | G | G | . | . |
| NC_011770 | 3172252 | A | G | 38 | 7 | 1 | 0 | NON_SYNONYMOU | MODERAT | MISSENSE | Aac/Gac | N2925D | pvdD | . | . | | | | | . | . | . | . | . | G |
| NC_011770 | 3172261 | A | G | 39 | 7 | 1 | 0 | NON_SYNONYMOU | MODERAT | MISSENSE | Atc/Gtc | I2928V | pvdD | . | . | | | | | . | . | . | . | . | G |
| NC_011770 | 3173661 | A | C | 42 | 2 | 6 | 0 | NON_SYNONYMOU | MODERAT | MISSENSE | gaA/gaC | E3394D | pvdD | C | C | | | | | . | . | C | C | C | C |
| NC_011770 | 3173662 | G | A | 44 | 2 | 6 | 0 | NON_SYNONYMOU | MODERAT | MISSENSE | Gtg/Atg | V3395M | pvdD | A | A | | | | | . | . | A | A | A | A |
| NC_011770 | 3173682 | G | T | 42 | 2 | 4 | 0 | NON_SYNONYMOU | MODERAT | MISSENSE | caG/caT | Q3401H | pvdD | T | T | | | | | . | . | | | T | T |
| NC_011770 | 3173693 | G | C | 36 | 7 | 1 | 0 | NON_SYNONYMOU | MODERAT | MISSENSE | gGa/gCa | G3405A | pvdD | . | . | | | | | . | . | C | . | . | . |
| NC_011770 | 3175076 | A | G | 3 | 8 | 4 | 0 | NON_SYNONYMOU | MODERAT | MISSENSE | gAc/gGc | D3866G | pvdD | . | . | G | G | G | G | . | . | . | . | . | . |
| NC_011770 | 3175298 | G | A | 18 | 6 | 2 | 0 | NON_SYNONYMOU | MODERAT | MISSENSE | gGc/gAc | G3940D | pvdD | . | . | | | | | . | . | A | A | . | . |
| NC_011770 | 3175300 | C | A | 18 | 6 | 2 | 0 | NON_SYNONYMOU | MODERAT | MISSENSE | Ctc/Atc | L3941I | pvdD | . | . | | | | | . | . | A | A | . | . |

| | | | | | | | | | | | | | | | | | | | | | | | | | |
|---|---|---|---|---|---|---|---|---|---|---|---|---|---|---|---|---|---|---|---|---|---|---|---|---|---|
| NC_011770 | 3175301 | T | G | 18 | 6 | 2 | 0 | NON_SYNONYMOU | MODERAT | MISSENSE | cTc/cGc | L3941R | pvdD | . | . | | | | | . | . | G | G | . | . |
| NC_011770 | 3175556 | C | A | 28 | 8 | 4 | 0 | NON_SYNONYMOU | MODERAT | MISSENSE | gCg/gAg | A4026E | pvdD | . | . | A | A | A | A | . | . | . | . | . | . |
| NC_011770 | 3175887 | T | G | 34 | 8 | 4 | 0 | NON_SYNONYMOU | MODERAT | MISSENSE | ttT/ttG | F4136L | pvdD | . | . | G | G | G | G | . | . | . | . | . | . |
| NC_011770 | 3169975 | C | A | 55 | 0 | 5 | 0 | NON_SYNONYMOU | MODERAT | MISSENSE | Ctg/Atg | L2166M | pvdD | A | | | | | | A | A | | | A | A |
| NC_011770 | 3164419 | A | G | 75 | 2 | 1 | 0 | NON_SYNONYMOU | MODERAT | MISSENSE | Atc/Gtc | I314V | pvdD | | | | | | | | G | . | . | | |
| NC_011770 | 3179688 | A | C | 11 | 6 | 2 | 0 | NON_SYNONYMOU | MODERAT | MISSENSE | aTc/aGc | I530S | pvdE | . | . | | | | | . | . | C | C | . | . |
| NC_011770 | 3179773 | C | T | 12 | 6 | 2 | 0 | NON_SYNONYMOU | MODERAT | MISSENSE | Gtg/Atg | V502M | pvdE | . | . | | | | | . | . | T | T | . | . |
| NC_011770 | 3179785 | C | T | 12 | 6 | 2 | 0 | NON_SYNONYMOU | MODERAT | MISSENSE | Gca/Aca | A498T | pvdE | . | . | | | | | . | . | T | T | . | . |
| NC_011770 | 3179831 | G | C | 13 | 6 | 2 | 0 | NON_SYNONYMOU | MODERAT | MISSENSE | gaC/gaG | D482E | pvdE | . | . | | | | | . | . | C | C | . | . |
| NC_011770 | 3179833 | C | T | 13 | 6 | 2 | 0 | NON_SYNONYMOU | MODERAT | MISSENSE | Gac/Aac | D482N | pvdE | . | . | | | | | . | . | T | T | . | . |
| NC_011770 | 3179834 | G | C | 13 | 6 | 2 | 0 | NON_SYNONYMOU | MODERAT | MISSENSE | gaC/gaG | D481E | pvdE | . | . | | | | | . | . | C | C | . | . |
| NC_011770 | 3179839 | G | T | 13 | 6 | 2 | 0 | NON_SYNONYMOU | MODERAT | MISSENSE | Ctg/Atg | L480M | pvdE | . | . | | | | | . | . | T | T | . | . |
| NC_011770 | 3179851 | T | C | 13 | 6 | 2 | 0 | NON_SYNONYMOU | MODERAT | MISSENSE | Atc/Gtc | I476V | pvdE | . | . | | | | | . | . | C | C | . | . |
| NC_011770 | 3180166 | C | T | 12 | 3 | 5 | 0 | NON_SYNONYMOU | MODERAT | MISSENSE | Gtc/Atc | V371I | pvdE | . | . | | | | | T | T | T | . | T | T |
| NC_011770 | 3180519 | G | A | 20 | 3 | 5 | 0 | NON_SYNONYMOU | MODERAT | MISSENSE | gCg/gTg | A253V | pvdE | A | . | | | | | A | A | . | . | A | A |
| NC_011770 | 3184994 | C | T | 11 | 6 | 2 | 0 | NON_SYNONYMOU | MODERAT | MISSENSE | Gtg/Atg | V502M | pvdE2 | . | . | | | | | . | . | T | T | . | . |
| NC_011770 | 3185006 | C | T | 11 | 6 | 2 | 0 | NON_SYNONYMOU | MODERAT | MISSENSE | Gca/Aca | A498T | pvdE2 | . | . | | | | | . | . | T | T | . | . |
| NC_011770 | 3185052 | G | C | 12 | 6 | 2 | 0 | NON_SYNONYMOU | MODERAT | MISSENSE | gaC/gaG | D482E | pvdE2 | . | . | | | | | . | . | C | C | . | . |
| NC_011770 | 3185054 | C | T | 12 | 6 | 2 | 0 | NON_SYNONYMOU | MODERAT | MISSENSE | Gac/Aac | D482N | pvdE2 | . | . | | | | | . | . | T | T | . | . |
| NC_011770 | 3185055 | G | C | 12 | 6 | 2 | 0 | NON_SYNONYMOU | MODERAT | MISSENSE | gaC/gaG | D481E | pvdE2 | . | . | | | | | . | . | C | C | . | . |
| NC_011770 | 3185116 | G | C | 13 | 5 | 1 | 0 | NON_SYNONYMOU | MODERAT | MISSENSE | aCc/aGc | T461S | pvdE2 | . | . | | | | | . | C | | | . | . |
| NC_011770 | 3185387 | C | T | 11 | 3 | 5 | 0 | NON_SYNONYMOU | MODERAT | MISSENSE | Gtc/Atc | V371I | pvdE2 | . | . | | | | | T | T | . | T | T | T |
| NC_011770 | 3185740 | G | A | 20 | 3 | 5 | 0 | NON_SYNONYMOU | MODERAT | MISSENSE | gCg/gTg | A253V | pvdE2 | A | . | | | | | A | A | . | . | A | A |
| NC_011770 | 3190215 | C | T | 10 | 7 | 1 | 0 | NON_SYNONYMOU | MODERAT | MISSENSE | Gtg/Atg | V502M | pvdE3 | . | . | | | | | . | . | . | T | . | . |
| NC_011770 | 3190227 | C | T | 10 | 7 | 1 | 0 | NON_SYNONYMOU | MODERAT | MISSENSE | Gca/Aca | A498T | pvdE3 | . | . | | | | | . | . | . | T | . | . |
| NC_011770 | 3190273 | G | C | 9 | 7 | 1 | 0 | NON_SYNONYMOU | MODERAT | MISSENSE | gaC/gaG | D482E | pvdE3 | . | . | | | | | . | . | . | C | . | . |
| NC_011770 | 3190275 | C | T | 10 | 7 | 1 | 0 | NON_SYNONYMOU | MODERAT | MISSENSE | Gac/Aac | D482N | pvdE3 | . | . | | | | | . | . | . | T | . | . |
| NC_011770 | 3190276 | G | C | 10 | 7 | 1 | 0 | NON_SYNONYMOU | MODERAT | MISSENSE | gaC/gaG | D481E | pvdE3 | . | . | | | | | . | . | . | C | . | . |
| NC_011770 | 3190281 | G | T | 10 | 7 | 1 | 0 | NON_SYNONYMOU | MODERAT | MISSENSE | Ctg/Atg | L480M | pvdE3 | . | . | | | | | . | . | . | T | . | . |
| NC_011770 | 3190293 | T | C | 10 | 7 | 1 | 0 | NON_SYNONYMOU | MODERAT | MISSENSE | Atc/Gtc | I476V | pvdE3 | . | . | | | | | . | . | . | C | . | . |
| NC_011770 | 3190961 | G | A | 18 | 3 | 5 | 0 | NON_SYNONYMOU | MODERAT | MISSENSE | gCg/gTg | A253V | pvdE3 | A | . | | | | | A | A | . | . | A | A |
| NC_011770 | 3192651 | C | T | 97 | 2 | 6 | 0 | NON_SYNONYMOU | MODERAT | MISSENSE | Ctc/Ttc | L192F | pvdF | T | T | | | | | T | T | . | . | T | T |
| NC_011770 | 3192687 | T | G | 98 | 2 | 6 | 0 | NON_SYNONYMOU | MODERAT | MISSENSE | Tcg/Gcg | S204A | pvdF | G | G | | | | | G | G | . | . | G | G |
| NC_011770 | 3192693 | C | A | 99 | 2 | 6 | 0 | NON_SYNONYMOU | MODERAT | MISSENSE | Ctg/Atg | L206M | pvdF | A | A | | | | | A | A | . | . | A | A |
| NC_011770 | 3192798 | G | A | 87 | 0 | 2 | 0 | NON_SYNONYMOU | MODERAT | MISSENSE | Gag/Aag | E241K | pvdF | | | | | | | | | A | A | | |
| NC_011770 | 3192812 | G | C | 87 | 0 | 8 | 0 | NON_SYNONYMOU | MODERAT | MISSENSE | gaG/gaC | E245D | pvdF | C | C | | | | | C | C | C | C | C | C |
| NC_011770 | 3192849 | A | G | 90 | 0 | 2 | 0 | NON_SYNONYMOU | MODERAT | MISSENSE | Aac/Gac | N258D | pvdF | | | | | | | | | G | G | | |
| NC_011770 | 3112732 | G | A | 77 | 0 | 2 | 0 | NON_SYNONYMOU | MODERAT | MISSENSE | Ggc/Agc | G31S | pvdG | | | | | | | | | A | A | | |
| NC_011770 | 3112842 | A | C | 79 | 0 | 10 | 0 | NON_SYNONYMOU | MODERAT | MISSENSE | caA/caC | Q67H | pvdG | C | C | C | C | C | C | | | C | C | C | C |
| NC_011770 | 3113175 | C | A | 79 | 0 | 2 | 0 | NON_SYNONYMOU | MODERAT | MISSENSE | gaC/gaA | D178E | pvdG | A | A | | | | | | | | | | |
| NC_011770 | 3139398 | A | G | 63 | 0 | 12 | 0 | NON_SYNONYMOU | MODERAT | MISSENSE | gAc/gGc | D121G | pvdH | G | G | G | G | G | G | G | G | G | G | G | G |
| NC_011770 | 3140133 | T | G | 77 | 0 | 12 | 0 | NON_SYNONYMOU | MODERAT | MISSENSE | cTc/cGc | L366R | pvdH | G | G | G | G | G | G | G | G | G | G | G | G |
| NC_011770 | 3140300 | T | C | 63 | 0 | 2 | 0 | NON_SYNONYMOU | MODERAT | MISSENSE | Tgc/Cgc | C422R | pvdH | C | C | | | | | | | | | | |
| NC_011770 | 3140414 | G | A | 72 | 0 | 4 | 0 | NON_SYNONYMOU | MODERAT | MISSENSE | Gcc/Acc | A460T | pvdH | | | | | | | A | A | | | A | A |
| NC_011770 | 3140366 | C | T | 72 | 0 | 1 | 0 | NON_SYNONYMOU | MODERAT | MISSENSE | Cca/Tca | P444S | pvdH | | | | T | | | | | | | | |
| NC_011770 | 3150712 | T | TCG | 54 | 0 | 2 | 0 | FRAME_SHIFT | HIGH | | ata/atCGa | I554I? | pvdI | TCG | | | | | | | | . | . | | TCG |
| NC_011770 | 3159097 | GGGAAA | G | 80 | 5 | 1 | 0 | FRAME_SHIFT | HIGH | | gaaagc/ | ES3350 | pvdI | . | G | | | | | . | . | | | . | . |
| NC_011770 | 3149221 | G | A | 50 | 0 | 8 | 0 | NON_SYNONYMOU | MODERAT | MISSENSE | cGg/cAg | R57Q | pvdI | A | A | | | | | A | A | A | A | A | A |
| NC_011770 | 3149509 | G | A | 36 | 7 | 1 | 0 | NON_SYNONYMOU | MODERAT | MISSENSE | cGg/cAg | R153Q | pvdI | . | . | | | | | . | . | . | . | A | . |
| NC_011770 | 3149535 | C | T | 58 | 0 | 8 | 0 | NON_SYNONYMOU | MODERAT | MISSENSE | Ctt/Ttt | L162F | pvdI | T | T | | | | | T | T | T | T | T | T |
| NC_011770 | 3149572 | A | G | 61 | 0 | 8 | 0 | NON_SYNONYMOU | MODERAT | MISSENSE | cAg/cGg | Q174R | pvdI | G | G | | | | | G | G | G | G | G | G |
| NC_011770 | 3149994 | T | A | 61 | 0 | 8 | 0 | NON_SYNONYMOU | MODERAT | MISSENSE | Tct/Act | S315T | pvdI | A | A | | | | | A | A | A | A | A | A |
| NC_011770 | 3150142 | G | C | 42 | 3 | 5 | 0 | NON_SYNONYMOU | MODERAT | MISSENSE | aGc/aCc | S364T | pvdI | . | C | | | | | C | C | . | . | C | C |
| NC_011770 | 3150170 | G | T | 71 | 0 | 8 | 0 | NON_SYNONYMOU | MODERAT | MISSENSE | gaG/gaT | E373D | pvdI | T | T | | | | | T | T | T | T | T | T |
| NC_011770 | 3150199 | T | C | 75 | 0 | 2 | 0 | NON_SYNONYMOU | MODERAT | MISSENSE | tTc/tCc | F383S | pvdI | | | | | | | | | C | C | | |
| NC_011770 | 3150204 | A | G | 75 | 0 | 8 | 0 | NON_SYNONYMOU | MODERAT | MISSENSE | Atc/Gtc | I385V | pvdI | G | G | | | | | G | G | G | G | G | G |
| NC_011770 | 3150213 | A | C | 76 | 0 | 2 | 0 | NON_SYNONYMOU | MODERAT | MISSENSE | Agc/Cgc | S388R | pvdI | | | | | | | | | C | C | | |
| NC_011770 | 3150262 | A | T | 58 | 0 | 8 | 0 | NON_SYNONYMOU | MODERAT | MISSENSE | cAa/cTa | Q404L | pvdI | T | T | | | | | T | T | T | T | T | T |
| NC_011770 | 3150277 | G | A | 54 | 0 | 8 | 0 | NON_SYNONYMOU | MODERAT | MISSENSE | gGc/gAc | G409D | pvdI | A | A | | | | | A | A | A | A | A | A |
| NC_011770 | 3150279 | T | G | 54 | 0 | 8 | 0 | NON_SYNONYMOU | MODERAT | MISSENSE | Tcg/Gcg | S410A | pvdI | G | G | | | | | G | G | G | G | G | G |
| NC_011770 | 3150288 | G | A | 56 | 0 | 8 | 0 | NON_SYNONYMOU | MODERAT | MISSENSE | Gac/Aac | D413N | pvdI | A | A | | | | | A | A | A | A | A | A |
| NC_011770 | 3150289 | A | G | 55 | 0 | 8 | 0 | NON_SYNONYMOU | MODERAT | MISSENSE | gAc/gGc | D413G | pvdI | G | G | | | | | G | G | G | G | G | G |

| | | | | | | | | | | | | | | | | | | | | | | | | | |
|---|---|---|---|---|---|---|---|---|---|---|---|---|---|---|---|---|---|---|---|---|---|---|---|---|---|
| NC_011770 | 3150303 | C | A | 55 | 0 | 8 | 0 | NON_SYNONYMOU | MODERAT | MISSENSE | Cgt/Agt | R418S | pvdI | A | A | | | | | A | A | A | A | A | A |
| NC_011770 | 3150310 | C | G | 55 | 0 | 8 | 0 | NON_SYNONYMOU | MODERAT | MISSENSE | gCt/gGt | A420G | pvdI | G | G | | | | | G | G | G | G | G | G |
| NC_011770 | 3150313 | C | A | 53 | 0 | 8 | 0 | NON_SYNONYMOU | MODERAT | MISSENSE | cCt/cAt | P421H | pvdI | A | A | | | | | A | A | A | A | A | A |
| NC_011770 | 3150384 | C | T | 57 | 0 | 8 | 0 | NON_SYNONYMOU | MODERAT | MISSENSE | Cat/Tat | H445Y | pvdI | T | T | | | | | T | T | T | T | T | T |
| NC_011770 | 3150423 | G | C | 55 | 0 | 8 | 0 | NON_SYNONYMOU | MODERAT | MISSENSE | Gag/Cag | E458Q | pvdI | C | C | | | | | C | C | C | C | C | C |
| NC_011770 | 3150424 | A | G | 54 | 0 | 8 | 0 | NON_SYNONYMOU | MODERAT | MISSENSE | gAg/gGg | E458G | pvdI | G | G | | | | | G | G | G | G | G | G |
| NC_011770 | 3150429 | A | G | 55 | 0 | 8 | 0 | NON_SYNONYMOU | MODERAT | MISSENSE | Atc/Gtc | I460V | pvdI | G | G | | | | | G | G | G | G | G | G |
| NC_011770 | 3150790 | G | C | 41 | 3 | 5 | 0 | NON_SYNONYMOU | MODERAT | MISSENSE | gGa/gCa | G580A | pvdI | C | C | | | | | C | C | . | . | C | . |
| NC_011770 | 3150822 | T | G | 49 | 4 | 3 | 0 | NON_SYNONYMOU | MODERAT | MISSENSE | Ttg/Gtg | L591V | pvdI | G | . | | | | | . | . | G | G | . | . |
| NC_011770 | 3150840 | C | G | 47 | 4 | 4 | 0 | NON_SYNONYMOU | MODERAT | MISSENSE | Cag/Gag | Q597E | pvdI | G | G | | | | | . | . | G | G | . | . |
| NC_011770 | 3150850 | T | A | 46 | 4 | 4 | 0 | NON_SYNONYMOU | MODERAT | MISSENSE | cTt/cAt | L600H | pvdI | A | A | | | | | . | . | A | A | . | . |
| NC_011770 | 3150852 | T | G | 47 | 4 | 4 | 0 | NON_SYNONYMOU | MODERAT | MISSENSE | Tcc/Gcc | S601A | pvdI | G | G | | | | | . | . | G | G | . | . |
| NC_011770 | 3151057 | G | A | 41 | 4 | 4 | 0 | NON_SYNONYMOU | MODERAT | MISSENSE | cGa/cAa | R669Q | pvdI | . | . | | | | | A | A | . | . | A | A |
| NC_011770 | 3151093 | C | G | 63 | 0 | 8 | 0 | NON_SYNONYMOU | MODERAT | MISSENSE | aCc/aGc | T681S | pvdI | G | G | | | | | G | G | G | G | G | G |
| NC_011770 | 3151101 | C | T | 61 | 0 | 8 | 0 | NON_SYNONYMOU | MODERAT | MISSENSE | Ctt/Ttt | L684F | pvdI | T | T | | | | | T | T | T | T | T | T |
| NC_011770 | 3151116 | G | C | 40 | 6 | 2 | 0 | NON_SYNONYMOU | MODERAT | MISSENSE | Gat/Cat | D689H | pvdI | C | . | | | | | . | C | . | . | . | . |
| NC_011770 | 3151118 | T | G | 39 | 5 | 3 | 0 | NON_SYNONYMOU | MODERAT | MISSENSE | gaT/gaG | D689E | pvdI | G | . | | | | | . | G | . | . | G | . |
| NC_011770 | 3151600 | C | T | 48 | 1 | 7 | 0 | NON_SYNONYMOU | MODERAT | MISSENSE | gCg/gTg | A850V | pvdI | T | T | | | | | T | T | . | T | T | T |
| NC_011770 | 3151827 | A | G | 41 | 7 | 1 | 0 | NON_SYNONYMOU | MODERAT | MISSENSE | Acg/Gcg | T926A | pvdI | . | . | | | | | . | . | . | . | . | G |
| NC_011770 | 3151986 | A | G | 59 | 0 | 8 | 0 | NON_SYNONYMOU | MODERAT | MISSENSE | Atg/Gtg | M979V | pvdI | G | G | | | | | G | G | G | G | G | G |
| NC_011770 | 3151987 | T | C | 59 | 0 | 8 | 0 | NON_SYNONYMOU | MODERAT | MISSENSE | aTg/aCg | M979T | pvdI | C | C | | | | | C | C | C | C | C | C |
| NC_011770 | 3151995 | G | C | 59 | 0 | 8 | 0 | NON_SYNONYMOU | MODERAT | MISSENSE | Gtc/Ctc | V982L | pvdI | C | C | | | | | C | C | C | C | C | C |
| NC_011770 | 3152133 | G | C | 38 | 6 | 2 | 0 | NON_SYNONYMOU | MODERAT | MISSENSE | Gaa/Caa | E1028Q | pvdI | C | . | | | | | . | . | C | . | . | . |
| NC_011770 | 3152137 | C | A | 37 | 6 | 2 | 0 | NON_SYNONYMOU | MODERAT | MISSENSE | gCg/gAg | A1029E | pvdI | A | . | | | | | . | . | A | . | . | . |
| NC_011770 | 3152148 | A | G | 38 | 5 | 3 | 0 | NON_SYNONYMOU | MODERAT | MISSENSE | Atc/Gtc | I1033V | pvdI | G | . | | | | | . | . | G | . | G | . |
| NC_011770 | 3152181 | G | A | 36 | 5 | 2 | 0 | NON_SYNONYMOU | MODERAT | MISSENSE | Gcc/Acc | A1044T | pvdI | A | . | | | | | . | . | . | . | A | . |
| NC_011770 | 3152182 | C | G | 35 | 4 | 2 | 0 | NON_SYNONYMOU | MODERAT | MISSENSE | gCc/gGc | A1044G | pvdI | G | . | | | | | . | . | . | . | G | . |
| NC_011770 | 3156915 | C | G | 33 | 4 | 2 | 0 | NON_SYNONYMOU | MODERAT | MISSENSE | Caa/Gaa | Q2622E | pvdI | . | . | | | | | . | . | G | G | . | . |
| NC_011770 | 3156938 | A | C | 46 | 0 | 2 | 0 | NON_SYNONYMOU | MODERAT | MISSENSE | caA/caC | Q2629H | pvdI | | | | | | | . | | C | C | | |
| NC_011770 | 3156948 | G | C | 39 | 1 | 2 | 0 | NON_SYNONYMOU | MODERAT | MISSENSE | Gac/Cac | D2633H | pvdI | | | | | | | . | | C | C | | |
| NC_011770 | 3156950 | C | A | 39 | 1 | 1 | 0 | NON_SYNONYMOU | MODERAT | MISSENSE | gaC/gaA | D2633E | pvdI | . | . | | | | | . | A | | | . | . |
| NC_011770 | 3157491 | G | T | 5 | 5 | 1 | 0 | NON_SYNONYMOU | MODERAT | MISSENSE | Gtg/Ttg | V2814L | pvdI | | | | . | | . | . | . | . | T | | |
| NC_011770 | 3157507 | C | G | 1 | 10 | 1 | 0 | NON_SYNONYMOU | MODERAT | MISSENSE | aCc/aGc | T2819S | pvdI | . | . | | . | . | . | . | . | . | G | . | . |
| NC_011770 | 3157540 | C | T | 2 | 9 | 2 | 0 | NON_SYNONYMOU | MODERAT | MISSENSE | gCc/gTc | A2830V | pvdI | . | . | T | . | . | T | . | . | . | . | . | . |
| NC_011770 | 3157641 | A | T | 11 | 4 | 4 | 0 | NON_SYNONYMOU | MODERAT | MISSENSE | Agt/Tgt | S2864C | pvdI | T | T | | | | | . | . | . | . | T | T |
| NC_011770 | 3157642 | G | C | 11 | 4 | 4 | 0 | NON_SYNONYMOU | MODERAT | MISSENSE | aGt/aCt | S2864T | pvdI | C | C | | | | | . | . | . | . | C | C |
| NC_011770 | 3157643 | T | G | 11 | 4 | 2 | 0 | NON_SYNONYMOU | MODERAT | MISSENSE | agT/agG | S2864R | pvdI | . | . | | | | | . | . | . | . | G | G |
| NC_011770 | 3157650 | A | G | 12 | 4 | 4 | 0 | NON_SYNONYMOU | MODERAT | MISSENSE | Acc/Gcc | T2867A | pvdI | G | G | | | | | . | . | . | . | G | G |
| NC_011770 | 3157677 | C | G | 25 | 2 | 2 | 0 | NON_SYNONYMOU | MODERAT | MISSENSE | Ccg/Gcg | P2876A | pvdI | . | . | | | | | G | T | . | . | . | . |
| NC_011770 | 3157682 | A | T | 26 | 2 | 4 | 0 | NON_SYNONYMOU | MODERAT | MISSENSE | gaA/gaT | E2877D | pvdI | T | T | | | | | T | T | . | . | . | . |
| NC_011770 | 3157714 | C | A | 30 | 0 | 8 | 0 | NON_SYNONYMOU | MODERAT | MISSENSE | aCc/aAc | T2888N | pvdI | A | A | | | | | A | A | A | A | A | A |
| NC_011770 | 3157726 | C | T | 32 | 0 | 2 | 0 | NON_SYNONYMOU | MODERAT | MISSENSE | aCc/aTc | T2892I | pvdI | | | | | | | T | T | | | | |
| NC_011770 | 3157766 | G | C | 29 | 1 | 6 | 0 | NON_SYNONYMOU | MODERAT | MISSENSE | ttG/ttC | L2905F | pvdI | C | C | | | | | C | C | | . | C | C |
| NC_011770 | 3157801 | T | C | 9 | 8 | 1 | 0 | NON_SYNONYMOU | MODERAT | MISSENSE | gTc/gCc | V2917A | pvdI | . | . | . | C | | | . | . | . | . | . | . |
| NC_011770 | 3157877 | G | T | 24 | 0 | 2 | 0 | NON_SYNONYMOU | MODERAT | MISSENSE | gaG/gaT | E2942D | pvdI | | | | | | | | | T | T | | |
| NC_011770 | 3157890 | C | G | 18 | 4 | 4 | 0 | NON_SYNONYMOU | MODERAT | MISSENSE | Cag/Gag | Q2947E | pvdI | . | . | | | | | . | . | G | G | G | G |
| NC_011770 | 3157908 | A | G | 18 | 6 | 2 | 0 | NON_SYNONYMOU | MODERAT | MISSENSE | Agt/Ggt | S2953G | pvdI | . | . | | | | | . | . | G | G | . | . |
| NC_011770 | 3157909 | G | C | 17 | 6 | 2 | 0 | NON_SYNONYMOU | MODERAT | MISSENSE | aGt/aCt | S2953T | pvdI | . | . | | | | | . | . | C | C | . | . |
| NC_011770 | 3157910 | T | C | 17 | 6 | 6 | 0 | NON_SYNONYMOU | MODERAT | MISSENSE | agT/agG | S2953R | pvdI | . | . | C | C | C | C | . | . | G | G | . | . |
| NC_011770 | 3158381 | G | C | 37 | 7 | 1 | 0 | NON_SYNONYMOU | MODERAT | MISSENSE | caG/caC | Q3110H | pvdI | . | . | | | | | . | . | . | C | . | . |
| NC_011770 | 3158730 | C | A | 87 | 0 | 6 | 0 | NON_SYNONYMOU | MODERAT | MISSENSE | Ctg/Atg | L3227M | pvdI | A | A | | | | | A | A | | | A | A |
| NC_011770 | 3158766 | G | A | 72 | 0 | 6 | 0 | NON_SYNONYMOU | MODERAT | MISSENSE | Gtg/Atg | V3239M | pvdI | A | A | | | | | A | A | | | A | A |
| NC_011770 | 3158769 | T | G | 73 | 0 | 8 | 0 | NON_SYNONYMOU | MODERAT | MISSENSE | Tcg/Gcg | S3240A | pvdI | G | G | | | | | G | G | G | G | G | G |
| NC_011770 | 3158778 | C | G | 75 | 0 | 8 | 0 | NON_SYNONYMOU | MODERAT | MISSENSE | Cgt/Ggt | R3243G | pvdI | G | G | | | | | G | G | G | G | G | G |
| NC_011770 | 3158779 | G | T | 74 | 0 | 8 | 0 | NON_SYNONYMOU | MODERAT | MISSENSE | cGt/cTt | R3243L | pvdI | T | T | | | | | T | T | T | T | T | T |
| NC_011770 | 3158803 | C | T | 85 | 0 | 8 | 0 | NON_SYNONYMOU | MODERAT | MISSENSE | aCg/aTg | T3251M | pvdI | T | T | | | | | T | T | T | T | T | T |
| NC_011770 | 3158806 | G | C | 82 | 0 | 8 | 0 | NON_SYNONYMOU | MODERAT | MISSENSE | aGc/aCc | S3252T | pvdI | C | C | | | | | C | C | C | C | C | C |
| NC_011770 | 3158812 | C | A | 84 | 0 | 6 | 0 | NON_SYNONYMOU | MODERAT | MISSENSE | gCg/gAg | A3254E | pvdI | A | A | | | | | A | A | | | A | A |
| NC_011770 | 3158853 | G | T | 87 | 0 | 7 | 0 | NON_SYNONYMOU | MODERAT | MISSENSE | Gcc/Tcc | A3268S | pvdI | T | T | | | | | T | T | T | . | T | T |
| NC_011770 | 3158901 | G | A | 87 | 0 | 8 | 0 | NON_SYNONYMOU | MODERAT | MISSENSE | Gtg/Atg | V3284M | pvdI | A | A | | | | | A | A | A | A | A | A |
| NC_011770 | 3158916 | A | C | 79 | 0 | 8 | 0 | NON_SYNONYMOU | MODERAT | MISSENSE | Atc/Ctc | I3289L | pvdI | C | C | | | | | C | C | C | C | C | C |
| NC_011770 | 3158918 | C | G | 79 | 0 | 8 | 0 | NON_SYNONYMOU | MODERAT | MISSENSE | atC/atG | I3289M | pvdI | G | G | | | | | G | G | G | G | G | G |

| | | | | | | | | | | | | | | | | | | | | | | | | | |
|---|---|---|---|---|---|---|---|---|---|---|---|---|---|---|---|---|---|---|---|---|---|---|---|---|---|
| NC_011770 | 3158931 | G | C | 79 | 0 | 8 | 0 | NON_SYNONYMOU | MODERAT | MISSENSE | Gag/Cag | E3294Q | pvdI | C | C | | | | | C | C | C | C | C | C |
| NC_011770 | 3158932 | A | G | 77 | 0 | 6 | 0 | NON_SYNONYMOU | MODERAT | MISSENSE | gAg/gGg | E3294G | pvdI | G | G | | | | | G | G | | | G | G |
| NC_011770 | 3158943 | C | G | 80 | 0 | 6 | 0 | NON_SYNONYMOU | MODERAT | MISSENSE | Cag/Gag | Q3298E | pvdI | G | G | | | | | G | G | | | G | G |
| NC_011770 | 3158967 | A | G | 78 | 0 | 8 | 0 | NON_SYNONYMOU | MODERAT | MISSENSE | Atc/Gtc | I3306V | pvdI | G | G | | | | | G | G | G | G | G | G |
| NC_011770 | 3158969 | C | G | 78 | 0 | 6 | 0 | NON_SYNONYMOU | MODERAT | MISSENSE | atC/atG | I3306M | pvdI | G | G | | | | | G | G | | | G | G |
| NC_011770 | 3158971 | G | C | 79 | 0 | 2 | 0 | NON_SYNONYMOU | MODERAT | MISSENSE | gGg/gCg | G3307A | pvdI | | | | | | | | | C | C | | |
| NC_011770 | 3159072 | C | G | 96 | 0 | 6 | 0 | NON_SYNONYMOU | MODERAT | MISSENSE | Ccg/Gcg | P3341A | pvdI | G | G | | | | | G | G | | | G | G |
| NC_011770 | 3159089 | G | C | 79 | 5 | 2 | 0 | NON_SYNONYMOU | MODERAT | MISSENSE | gaG/gaC | E3346D | pvdI | . | . | | | | | . | . | C | C | . | . |
| NC_011770 | 3159099 | G | C | 76 | 6 | 2 | 0 | NON_SYNONYMOU | MODERAT | MISSENSE | Gaa/Caa | E3350Q | pvdI | . | . | | | | | . | . | C | C | . | . |
| NC_011770 | 3159100 | A | G | 76 | 6 | 2 | 0 | NON_SYNONYMOU | MODERAT | MISSENSE | gAa/gGa | E3350G | pvdI | . | . | | | | | . | . | G | G | . | . |
| NC_011770 | 3159101 | A | C | 76 | 7 | 1 | 0 | NON_SYNONYMOU | MODERAT | MISSENSE | gaA/gaC | E3350D | pvdI | . | . | | | | | . | . | . | C | . | . |
| NC_011770 | 3159102 | A | G | 75 | 6 | 2 | 0 | NON_SYNONYMOU | MODERAT | MISSENSE | Agc/Ggc | S3351G | pvdI | . | . | | | | | . | . | G | G | . | . |
| NC_011770 | 3159103 | G | C | 75 | 6 | 2 | 0 | NON_SYNONYMOU | MODERAT | MISSENSE | aGc/aCc | S3351T | pvdI | . | . | | | | | . | . | C | C | . | . |
| NC_011770 | 3159104 | C | G | 74 | 6 | 2 | 0 | NON_SYNONYMOU | MODERAT | MISSENSE | agC/agG | S3351R | pvdI | . | . | | | | | . | . | G | G | . | . |
| NC_011770 | 3159114 | G | C | 78 | 6 | 2 | 0 | NON_SYNONYMOU | MODERAT | MISSENSE | Gac/Cac | D3355H | pvdI | . | . | | | | | . | . | C | C | . | . |
| NC_011770 | 3159116 | C | A | 78 | 6 | 2 | 0 | NON_SYNONYMOU | MODERAT | MISSENSE | gaC/gaA | D3355E | pvdI | . | . | | | | | . | . | A | A | . | . |
| NC_011770 | 3159276 | C | G | 59 | 6 | 2 | 0 | NON_SYNONYMOU | MODERAT | MISSENSE | Ctg/Gtg | L3409V | pvdI | . | . | | | | | . | . | G | G | . | . |
| NC_011770 | 3159282 | C | G | 62 | 6 | 2 | 0 | NON_SYNONYMOU | MODERAT | MISSENSE | Ctg/Gtg | L3411V | pvdI | . | . | | | | | . | . | G | G | . | . |
| NC_011770 | 3159335 | C | G | 64 | 1 | 7 | 0 | NON_SYNONYMOU | MODERAT | MISSENSE | gaC/gaG | D3428E | pvdI | G | G | | | | | G | G | . | G | G | G |
| NC_011770 | 3159421 | C | G | 62 | 2 | 6 | 0 | NON_SYNONYMOU | MODERAT | MISSENSE | gCa/gGa | A3457G | pvdI | G | G | | | | | G | G | . | . | G | G |
| NC_011770 | 3159435 | C | G | 62 | 2 | 6 | 0 | NON_SYNONYMOU | MODERAT | MISSENSE | Ctg/Gtg | L3462V | pvdI | G | G | | | | | G | G | . | . | G | G |
| NC_011770 | 3159699 | A | T | 55 | 2 | 6 | 0 | NON_SYNONYMOU | MODERAT | MISSENSE | Aca/Tca | T3550S | pvdI | T | T | | | | | T | T | . | . | T | T |
| NC_011770 | 3159704 | G | C | 56 | 2 | 6 | 0 | NON_SYNONYMOU | MODERAT | MISSENSE | caG/caC | Q3551H | pvdI | C | C | | | | | C | C | . | . | C | C |
| NC_011770 | 3159705 | T | C | 55 | 2 | 6 | 0 | NON_SYNONYMOU | MODERAT | MISSENSE | Tgg/Cgg | W3552R | pvdI | C | C | | | | | C | C | . | . | C | C |
| NC_011770 | 3159706 | G | T | 55 | 2 | 6 | 0 | NON_SYNONYMOU | MODERAT | MISSENSE | tGg/tTg | W3552L | pvdI | T | T | | | | | T | T | . | . | T | T |
| NC_011770 | 3159708 | G | T | 56 | 2 | 6 | 0 | NON_SYNONYMOU | MODERAT | MISSENSE | Gtg/Ttg | V3553L | pvdI | T | T | | | | | T | T | . | . | T | T |
| NC_011770 | 3159720 | G | C | 54 | 2 | 6 | 0 | NON_SYNONYMOU | MODERAT | MISSENSE | Gcg/Ccg | A3557P | pvdI | C | C | | | | | C | C | . | . | C | C |
| NC_011770 | 3159721 | C | G | 53 | 2 | 6 | 0 | NON_SYNONYMOU | MODERAT | MISSENSE | gCg/gGg | A3557G | pvdI | G | G | | | | | G | G | . | . | G | G |
| NC_011770 | 3159723 | C | A | 53 | 1 | 6 | 0 | NON_SYNONYMOU | MODERAT | MISSENSE | Ctg/Atg | L3558M | pvdI | A | A | | | | | A | A | . | . | A | A |
| NC_011770 | 3157843 | C | G | 17 | 2 | 1 | 0 | NON_SYNONYMOU | MODERAT | MISSENSE | gCc/gGc | A2931G | pvdI | . | | | | | | . | | | **G** | | |
| NC_011770 | 3156944 | G | T | 45 | 0 | 1 | 0 | NON_SYNONYMOU | MODERAT | MISSENSE | gaG/gaT | E2631D | pvdI | | | | | | | | | T | | | |
| NC_011770 | 3150259 | C | T | 62 | 0 | 1 | 0 | NON_SYNONYMOU | MODERAT | MISSENSE | cCt/cTt | P403L | pvdI | | | | | | | | T | | | | |
| NC_011770 | 3157425 | A | C | 23 | 2 | 2 | 0 | NON_SYNONYMOU | MODERAT | MISSENSE | Atg/Ctg | M2792L | pvdI | | | | | . | . | | C | C | . | | |
| NC_011770 | 3157701 | A | G | 28 | 2 | 1 | 0 | NON_SYNONYMOU | MODERAT | MISSENSE | Atc/Gtc | I2884V | pvdI | | | | | | | | G | . | . | | |
| NC_011770 | 3160812 | G | GAC | 69 | 6 | 1 | 0 | FRAME_SHIFT | HIGH | | cgg/ACcgg | R240T? | pvdJ | . | . | | | | | . | . | . | . | . | GAC |
| NC_011770 | 3160816 | CAG | C | 69 | 6 | 2 | 0 | FRAME_SHIFT | HIGH | | cag/ | Q241 | pvdJ | . | . | | | | | . | . | . | . | C | C |
| NC_011770 | 3160264 | C | G | 45 | 1 | 7 | 0 | NON_SYNONYMOU | MODERAT | MISSENSE | Caa/Gaa | Q57E | pvdJ | . | G | | | | | G | G | G | G | G | G |
| NC_011770 | 3160279 | C | A | 90 | 0 | 1 | 0 | NON_SYNONYMOU | MODERAT | MISSENSE | Ctc/Atc | L62I | pvdJ | | | | | | | | | | A | | |
| NC_011770 | 3160287 | A | C | 91 | 0 | 2 | 0 | NON_SYNONYMOU | MODERAT | MISSENSE | caA/caC | Q64H | pvdJ | | | | | | | | | C | C | | |
| NC_011770 | 3160286 | A | G | 91 | 0 | 1 | 0 | NON_SYNONYMOU | MODERAT | MISSENSE | cAa/cGa | Q64R | pvdJ | | | | | | | | | | G | | |
| NC_011770 | 3160297 | G | C | 83 | 0 | 2 | 0 | NON_SYNONYMOU | MODERAT | MISSENSE | Gac/Cac | D68H | pvdJ | | | | | | | | | C | C | | |
| NC_011770 | 3160299 | C | A | 82 | 0 | 2 | 0 | NON_SYNONYMOU | MODERAT | MISSENSE | gaC/gaA | D68E | pvdJ | . | . | | | | | A | A | | | . | . |
| NC_011770 | 3160351 | A | G | 46 | 8 | 4 | 0 | NON_SYNONYMOU | MODERAT | MISSENSE | Atc/Gtc | I86V | pvdJ | . | . | G | G | G | G | . | . | . | . | . | . |
| NC_011770 | 3160790 | G | A | 73 | 3 | 3 | 0 | NON_SYNONYMOU | MODERAT | MISSENSE | cGt/cAt | R232H | pvdJ | . | A | | | | | . | . | | | A | A |
| NC_011770 | 3160820 | A | T | 68 | 6 | 2 | 0 | NON_SYNONYMOU | MODERAT | MISSENSE | cAg/cTg | Q242L | pvdJ | . | . | | | | | . | . | . | . | T | T |
| NC_011770 | 3160855 | A | T | 70 | 3 | 5 | 0 | NON_SYNONYMOU | MODERAT | MISSENSE | Acc/Tcc | T254S | pvdJ | T | T | | | | | T | . | . | . | T | T |
| NC_011770 | 3160856 | C | T | 71 | 3 | 5 | 0 | NON_SYNONYMOU | MODERAT | MISSENSE | aCc/aTc | T254I | pvdJ | T | T | | | | | T | . | . | . | T | T |
| NC_011770 | 3160990 | A | T | 55 | 7 | 1 | 0 | NON_SYNONYMOU | MODERAT | MISSENSE | Agt/Tgt | S299C | pvdJ | . | . | | | | | . | . | . | . | T | . |
| NC_011770 | 3160991 | G | C | 57 | 7 | 1 | 0 | NON_SYNONYMOU | MODERAT | MISSENSE | aGt/aCt | S299T | pvdJ | . | . | | | | | . | . | . | . | C | . |
| NC_011770 | 3160992 | T | G | 56 | 7 | 1 | 0 | NON_SYNONYMOU | MODERAT | MISSENSE | agT/agG | S299R | pvdJ | . | . | | | | | . | . | . | . | G | . |
| NC_011770 | 3161008 | C | T | 56 | 6 | 1 | 0 | NON_SYNONYMOU | MODERAT | MISSENSE | Cat/Tat | H305Y | pvdJ | . | . | | | | | . | . | | . | T | . |
| NC_011770 | 3161026 | C | T | 61 | 5 | 1 | 0 | NON_SYNONYMOU | MODERAT | MISSENSE | Ccg/Tcg | P311S | pvdJ | . | . | | | | | . | . | . | . | . | T |
| NC_011770 | 3161031 | A | C | 63 | 5 | 1 | 0 | NON_SYNONYMOU | MODERAT | MISSENSE | gaA/gaC | E312D | pvdJ | . | . | | | | | . | . | . | . | . | C |
| NC_011770 | 3161063 | C | A | 64 | 5 | 3 | 0 | NON_SYNONYMOU | MODERAT | MISSENSE | aCc/aAc | T323N | pvdJ | A | . | | | | | . | . | . | . | A | A |
| NC_011770 | 3161069 | T | C | 70 | 5 | 3 | 0 | NON_SYNONYMOU | MODERAT | MISSENSE | gTa/gCa | V325A | pvdJ | C | . | | | | | . | . | . | . | C | C |
| NC_011770 | 3161074 | A | G | 69 | 5 | 3 | 0 | NON_SYNONYMOU | MODERAT | MISSENSE | Acc/Gcc | T327A | pvdJ | G | . | | | | | . | . | . | . | G | G |
| NC_011770 | 3161075 | C | T | 68 | 5 | 3 | 0 | NON_SYNONYMOU | MODERAT | MISSENSE | aCc/aTc | T327I | pvdJ | T | . | | | | | . | . | . | . | T | T |
| NC_011770 | 3161192 | C | G | 61 | 4 | 2 | 0 | NON_SYNONYMOU | MODERAT | MISSENSE | gCc/gGc | A366G | pvdJ | . | . | | | | | . | . | G | G | | |
| NC_011770 | 3161239 | C | G | 64 | 2 | 6 | 0 | NON_SYNONYMOU | MODERAT | MISSENSE | Cag/Gag | Q382E | pvdJ | G | G | | | | | . | . | G | G | G | G |
| NC_011770 | 3161654 | A | C | 42 | 8 | 4 | 0 | NON_SYNONYMOU | MODERAT | MISSENSE | aAa/aCa | K520T | pvdJ | . | . | C | C | C | C | . | . | . | . | . | . |
| NC_011770 | 3161749 | C | T | 44 | 8 | 1 | 0 | NON_SYNONYMOU | MODERAT | MISSENSE | Ccg/Tcg | P552S | pvdJ | . | . | | | T | | . | . | . | . | . | . |
| NC_011770 | 3162647 | C | G | 11 | 6 | 1 | 0 | NON_SYNONYMOU | MODERAT | MISSENSE | gCc/gGc | A851G | pvdJ | . | . | | | | | . | . | . | G | . | . |

| | | | | | | | | | | | | | | | | | | | | | | | | | |
|---|---|---|---|---|---|---|---|---|---|---|---|---|---|---|---|---|---|---|---|---|---|---|---|---|---|
| NC_011770 | 3162653 | C | A | 11 | 6 | 1 | 0 | NON_SYNONYMOU | MODERAT | MISSENSE | gCc/gAc | A853D | pvdJ | . | . | | | | | . | . | . | A | . | . |
| NC_011770 | 3162736 | A | C | 18 | 3 | 3 | 0 | NON_SYNONYMOU | MODERAT | MISSENSE | Atc/Ctc | I881L | pvdJ | C | C | | | | | . | . | | . | . | C |
| NC_011770 | 3162799 | C | G | 19 | 7 | 1 | 0 | NON_SYNONYMOU | MODERAT | MISSENSE | Ctg/Gtg | L902V | pvdJ | . | . | | | | | . | G | . | . | . | . |
| NC_011770 | 3162853 | G | A | 28 | 0 | 2 | 0 | NON_SYNONYMOU | MODERAT | MISSENSE | Gcg/Acg | A920T | pvdJ | | | | | | | | | A | A | | |
| NC_011770 | 3162870 | G | T | 22 | 5 | 1 | 0 | NON_SYNONYMOU | MODERAT | MISSENSE | caG/caT | Q925H | pvdJ | . | . | | | | | . | . | | | T | . |
| NC_011770 | 3162871 | G | C | 22 | 5 | 1 | 0 | NON_SYNONYMOU | MODERAT | MISSENSE | Gag/Cag | E926Q | pvdJ | . | . | | | | | . | . | | | C | . |
| NC_011770 | 3162872 | A | C | 21 | 5 | 1 | 0 | NON_SYNONYMOU | MODERAT | MISSENSE | gAg/gCg | E926A | pvdJ | . | . | | | | | . | . | | | C | . |
| NC_011770 | 3162873 | G | C | 22 | 5 | 2 | 0 | NON_SYNONYMOU | MODERAT | MISSENSE | gaG/gaC | E926D | pvdJ | . | . | | | | | . | . | C | C | | . |
| NC_011770 | 3162874 | T | A | 20 | 5 | 2 | 0 | NON_SYNONYMOU | MODERAT | MISSENSE | Tgc/Agc | C927S | pvdJ | . | . | | | | | . | . | A | A | | . |
| NC_011770 | 3162876 | C | G | 21 | 5 | 1 | 0 | NON_SYNONYMOU | MODERAT | MISSENSE | tgC/tgG | C927W | pvdJ | . | . | | | | | . | . | | | G | . |
| NC_011770 | 3162890 | T | C | 21 | 5 | 1 | 0 | NON_SYNONYMOU | MODERAT | MISSENSE | gTg/gCg | V932A | pvdJ | . | . | | | | | . | . | | | C | . |
| NC_011770 | 3160293 | G | T | 91 | 0 | 1 | 0 | NON_SYNONYMOU | MODERAT | MISSENSE | gaG/gaT | E66D | pvdJ | | | | | | | | | | T | | |
| NC_011770 | 3160774 | A | C | 100 | 0 | 1 | 0 | NON_SYNONYMOU | MODERAT | MISSENSE | Atg/Ctg | M227L | pvdJ | | | | | | | | C | . | . | | |
| NC_011770 | 3113623 | C | T | 89 | 0 | 1 | 0 | NON_SYNONYMOU | MODERAT | MISSENSE | Cgc/Tgc | R49C | pvdL | | | | | | | T | | | | | |
| NC_011770 | 3113927 | A | T | 79 | 0 | 4 | 0 | NON_SYNONYMOU | MODERAT | MISSENSE | cAg/cTg | Q150L | pvdL | | | T | T | T | T | | | | | | |
| NC_011770 | 3114569 | C | A | 62 | 0 | 10 | 0 | NON_SYNONYMOU | MODERAT | MISSENSE | gCa/gAa | A364E | pvdL | A | A | A | A | A | A | A | A | | | A | A |
| NC_011770 | 3114635 | C | T | 66 | 0 | 2 | 0 | NON_SYNONYMOU | MODERAT | MISSENSE | gCg/gTg | A386V | pvdL | | | | | | | | | T | T | | |
| NC_011770 | 3114647 | T | C | 68 | 0 | 2 | 0 | NON_SYNONYMOU | MODERAT | MISSENSE | gTc/gCc | V390A | pvdL | | | | | | | | | C | C | | |
| NC_011770 | 3114719 | C | T | 74 | 0 | 2 | 0 | NON_SYNONYMOU | MODERAT | MISSENSE | cCg/cTg | P414L | pvdL | T | T | | | | | | | | | | |
| NC_011770 | 3115690 | G | A | 74 | 0 | 2 | 0 | NON_SYNONYMOU | MODERAT | MISSENSE | Gcg/Acg | A738T | pvdL | A | A | | | | | | | | | | |
| NC_011770 | 3115751 | C | T | 74 | 0 | 2 | 0 | NON_SYNONYMOU | MODERAT | MISSENSE | gCg/gTg | A758V | pvdL | | | | | | | | | T | T | | |
| NC_011770 | 3115790 | G | A | 74 | 0 | 2 | 0 | NON_SYNONYMOU | MODERAT | MISSENSE | cGc/cAc | R771H | pvdL | A | A | | | | | | | | | | |
| NC_011770 | 3115798 | G | T | 74 | 0 | 2 | 0 | NON_SYNONYMOU | MODERAT | MISSENSE | Gcc/Tcc | A774S | pvdL | | | | | | | | | T | T | | |
| NC_011770 | 3116089 | C | T | 70 | 0 | 4 | 0 | NON_SYNONYMOU | MODERAT | MISSENSE | Cgc/Tgc | R871C | pvdL | | | T | T | T | T | | | | | | |
| NC_011770 | 3116101 | G | A | 68 | 0 | 4 | 0 | NON_SYNONYMOU | MODERAT | MISSENSE | Gcg/Acg | A875T | pvdL | | | | | | | | | A | A | A | A |
| NC_011770 | 3116183 | T | C | 72 | 0 | 2 | 0 | NON_SYNONYMOU | MODERAT | MISSENSE | gTc/gCc | V902A | pvdL | | | | | | | | | C | C | | |
| NC_011770 | 3116725 | A | G | 72 | 0 | 12 | 0 | NON_SYNONYMOU | MODERAT | MISSENSE | Acc/Gcc | T1083A | pvdL | G | G | G | G | G | G | G | G | G | G | G | G |
| NC_011770 | 3116819 | G | A | 71 | 0 | 10 | 0 | NON_SYNONYMOU | MODERAT | MISSENSE | cGc/cAc | R1114H | pvdL | | | A | A | A | A | A | A | A | A | A | A |
| NC_011770 | 3116864 | G | T | 73 | 0 | 2 | 0 | NON_SYNONYMOU | MODERAT | MISSENSE | tGg/tTg | W1129L | pvdL | | | | | | | | | | | T | T |
| NC_011770 | 3116894 | C | T | 73 | 0 | 2 | 0 | NON_SYNONYMOU | MODERAT | MISSENSE | gCg/gTg | A1139V | pvdL | | | | | | | T | T | | | | |
| NC_011770 | 3116911 | G | A | 73 | 0 | 2 | 0 | NON_SYNONYMOU | MODERAT | MISSENSE | Gtc/Atc | V1145I | pvdL | A | A | | | | | | | | | | |
| NC_011770 | 3117061 | A | C | 78 | 0 | 12 | 0 | NON_SYNONYMOU | MODERAT | MISSENSE | Atg/Ctg | M1195L | pvdL | C | C | C | C | C | C | C | C | C | C | C | C |
| NC_011770 | 3117397 | G | A | 83 | 0 | 4 | 0 | NON_SYNONYMOU | MODERAT | MISSENSE | Gcc/Acc | A1307T | pvdL | A | A | | | | | | | A | A | | |
| NC_011770 | 3117560 | G | A | 75 | 0 | 6 | 0 | NON_SYNONYMOU | MODERAT | MISSENSE | gGg/gAg | G1361E | pvdL | | | A | A | A | A | | | | | A | A |
| NC_011770 | 3117974 | C | A | 66 | 0 | 8 | 0 | NON_SYNONYMOU | MODERAT | MISSENSE | gCt/gAt | A1499D | pvdL | | | A | A | A | A | A | A | | | A | A |
| NC_011770 | 3118088 | C | T | 77 | 0 | 2 | 0 | NON_SYNONYMOU | MODERAT | MISSENSE | cCg/cTg | P1537L | pvdL | | | | | | | | | | | T | T |
| NC_011770 | 3118199 | T | C | 80 | 0 | 2 | 0 | NON_SYNONYMOU | MODERAT | MISSENSE | gTc/gCc | V1574A | pvdL | | | | | | | C | C | | | | |
| NC_011770 | 3118223 | A | G | 82 | 0 | 1 | 0 | NON_SYNONYMOU | MODERAT | MISSENSE | cAg/cGg | Q1582R | pvdL | | | | | | | G | | | | | |
| NC_011770 | 3118279 | G | A | 77 | 0 | 2 | 0 | NON_SYNONYMOU | MODERAT | MISSENSE | Gcg/Acg | A1601T | pvdL | | | | | | | | | A | A | | |
| NC_011770 | 3118873 | G | A | 80 | 0 | 6 | 0 | NON_SYNONYMOU | MODERAT | MISSENSE | Ggc/Agc | G1799S | pvdL | | | A | A | A | A | A | A | | | | |
| NC_011770 | 3118934 | A | G | 86 | 0 | 2 | 0 | NON_SYNONYMOU | MODERAT | MISSENSE | cAg/cGg | Q1819R | pvdL | G | G | | | | | | | | | | |
| NC_011770 | 3119048 | T | C | 96 | 0 | 2 | 0 | NON_SYNONYMOU | MODERAT | MISSENSE | gTg/gCg | V1857A | pvdL | C | C | | | | | | | | | | |
| NC_011770 | 3119947 | C | T | 74 | 0 | 1 | 0 | NON_SYNONYMOU | MODERAT | MISSENSE | Cgg/Tgg | R2157W | pvdL | | | | | | | T | | | | | |
| NC_011770 | 3120346 | C | T | 83 | 0 | 2 | 0 | NON_SYNONYMOU | MODERAT | MISSENSE | Cac/Tac | H2290Y | pvdL | | | | | | | T | T | | | | |
| NC_011770 | 3120574 | T | C | 82 | 0 | 12 | 0 | NON_SYNONYMOU | MODERAT | MISSENSE | Tgc/Cgc | C2366R | pvdL | C | C | C | C | C | C | C | C | C | C | C | C |
| NC_011770 | 3120820 | G | A | 80 | 0 | 2 | 0 | NON_SYNONYMOU | MODERAT | MISSENSE | Gtg/Atg | V2448M | pvdL | A | A | | | | | | | | | | |
| NC_011770 | 3121597 | G | A | 74 | 0 | 4 | 0 | NON_SYNONYMOU | MODERAT | MISSENSE | Gtg/Atg | V2707M | pvdL | | | A | A | A | A | | | | | | |
| NC_011770 | 3121942 | C | A | 88 | 0 | 12 | 0 | NON_SYNONYMOU | MODERAT | MISSENSE | Ctc/Atc | L2822I | pvdL | A | A | A | A | A | A | A | A | A | A | A | A |
| NC_011770 | 3122032 | T | C | 87 | 0 | 2 | 0 | NON_SYNONYMOU | MODERAT | MISSENSE | Tgg/Cgg | W2852R | pvdL | C | C | | | | | | | | | | |
| NC_011770 | 3122057 | G | A | 90 | 0 | 4 | 0 | NON_SYNONYMOU | MODERAT | MISSENSE | cGc/cAc | R2860H | pvdL | A | A | | | | | A | A | | | | |
| NC_011770 | 3122228 | C | T | 139 | 0 | 6 | 0 | NON_SYNONYMOU | MODERAT | MISSENSE | gCg/gTg | A2917V | pvdL | | | T | T | T | T | T | T | | | | |
| NC_011770 | 3122287 | C | T | 91 | 0 | 8 | 0 | NON_SYNONYMOU | MODERAT | MISSENSE | Ctc/Ttc | L2937F | pvdL | | | T | T | T | T | T | T | T | T | | |
| NC_011770 | 3122291 | A | C | 89 | 0 | 10 | 0 | NON_SYNONYMOU | MODERAT | MISSENSE | gAc/gCc | D2938A | pvdL | C | C | C | C | C | C | C | C | C | C | | |
| NC_011770 | 3122357 | A | T | 86 | 0 | 2 | 0 | NON_SYNONYMOU | MODERAT | MISSENSE | cAg/cTg | Q2960L | pvdL | | | | | | | T | T | | | | |
| NC_011770 | 3122425 | T | C | 79 | 0 | 8 | 0 | NON_SYNONYMOU | MODERAT | MISSENSE | Tcg/Ccg | S2983P | pvdL | | | C | C | C | C | C | C | C | C | | |
| NC_011770 | 3122429 | C | T | 80 | 0 | 8 | 0 | NON_SYNONYMOU | MODERAT | MISSENSE | gCc/gTc | A2984V | pvdL | | | T | T | T | T | T | T | T | T | | |
| NC_011770 | 3122453 | A | G | 81 | 0 | 2 | 0 | NON_SYNONYMOU | MODERAT | MISSENSE | cAg/cGg | Q2992R | pvdL | G | G | | | | | | | | | | |
| NC_011770 | 3122626 | G | C | 93 | 0 | 2 | 0 | NON_SYNONYMOU | MODERAT | MISSENSE | Gtg/Ctg | V3050L | pvdL | | | | | | | C | C | | | | |
| NC_011770 | 3123596 | A | G | 84 | 0 | 10 | 0 | NON_SYNONYMOU | MODERAT | MISSENSE | gAc/gGc | D3373G | pvdL | G | G | G | G | G | G | G | G | | | G | G |
| NC_011770 | 3123767 | G | A | 89 | 0 | 4 | 0 | NON_SYNONYMOU | MODERAT | MISSENSE | cGc/cAc | R3430H | pvdL | A | A | | | | | | | | | A | A |
| NC_011770 | 3125198 | C | G | 72 | 0 | 4 | 0 | NON_SYNONYMOU | MODERAT | MISSENSE | gCc/gGc | A3907G | pvdL | | | G | G | G | G | | | | | | |
| NC_011770 | 3125536 | T | C | 69 | 0 | 2 | 0 | NON_SYNONYMOU | MODERAT | MISSENSE | Ttc/Ctc | F4020L | pvdL | | | | | | | C | C | | | | |

| | | | | | | | | | | | | | | | | | | | | | | | | | |
|---|---|---|---|---|---|---|---|---|---|---|---|---|---|---|---|---|---|---|---|---|---|---|---|---|---|
| NC_011770 | 3125818 | G | C | 101 | 0 | 4 | 0 | NON_SYNONYMOU | MODERAT | MISSENSE | Gtg/Ctg | V4114L | pvdL | | | C | C | C | C | | | | | | |
| NC_011770 | 3125926 | G | T | 84 | 0 | 4 | 0 | NON_SYNONYMOU | MODERAT | MISSENSE | Gcg/Tcg | A4150S | pvdL | | | T | T | T | T | | | | | | |
| NC_011770 | 3125996 | C | A | 82 | 0 | 10 | 0 | NON_SYNONYMOU | MODERAT | MISSENSE | cCg/cAg | P4173Q | pvdL | A | A | A | A | A | A | | | A | A | A | A |
| NC_011770 | 3126076 | T | G | 75 | 0 | 12 | 0 | NON_SYNONYMOU | MODERAT | MISSENSE | Tac/Gac | Y4200D | pvdL | G | G | G | G | G | G | G | G | G | G | G | G |
| NC_011770 | 3126077 | A | C | 74 | 0 | 12 | 0 | NON_SYNONYMOU | MODERAT | MISSENSE | tAc/tCc | Y4200S | pvdL | C | C | C | C | C | C | C | C | C | C | C | C |
| NC_011770 | 3126283 | G | A | 65 | 0 | 4 | 0 | NON_SYNONYMOU | MODERAT | MISSENSE | Gtg/Atg | V4269M | pvdL | | | A | A | A | A | | | | | | |
| NC_011770 | 3126430 | T | C | 57 | 0 | 2 | 0 | NON_SYNONYMOU | MODERAT | MISSENSE | Tcc/Ccc | S4318P | pvdL | | | | | | | | | | | C | C |
| NC_011770 | 3114190 | G | A | 74 | 0 | 1 | 0 | NON_SYNONYMOU | MODERAT | MISSENSE | Gtg/Atg | V238M | pvdL | | | | | | | | | | | | A |
| NC_011770 | 3117085 | G | A | 78 | 0 | 1 | 0 | NON_SYNONYMOU | MODERAT | MISSENSE | Ggc/Agc | G1203S | pvdL | | | | | | | | | | | | A |
| NC_011770 | 3120038 | C | T | 73 | 0 | 1 | 0 | NON_SYNONYMOU | MODERAT | MISSENSE | aCc/aTc | T2187I | pvdL | | | | | | | | | | | | T |
| NC_011770 | 3124688 | A | G | 53 | 0 | 1 | 0 | NON_SYNONYMOU | MODERAT | MISSENSE | cAg/cGg | Q3737R | pvdL | | | | | | | | G | | | | |
| NC_011770 | 3194053 | C | T | 69 | 0 | 6 | 0 | NON_SYNONYMOU | MODERAT | MISSENSE | aGc/aAc | S358N | pvdN | T | T | T | T | T | T | | | | | | |
| NC_011770 | 3194090 | C | T | 67 | 0 | 6 | 0 | NON_SYNONYMOU | MODERAT | MISSENSE | Ggc/Agc | G346S | pvdN | T | T | T | T | T | T | | | | | | |
| NC_011770 | 3194876 | T | C | 64 | 0 | 8 | 0 | NON_SYNONYMOU | MODERAT | MISSENSE | Acc/Gcc | T84A | pvdN | C | C | | | | | C | C | C | C | C | C |
| NC_011770 | 3195047 | T | G | 30 | 8 | 4 | 0 | NON_SYNONYMOU | MODERAT | MISSENSE | Acc/Ccc | T27P | pvdN | . | . | G | G | G | G | . | . | . | . | . | . |
| NC_011770 | 3195086 | T | G | 51 | 0 | 8 | 0 | NON_SYNONYMOU | MODERAT | MISSENSE | Atc/Ctc | I14L | pvdN | G | G | | | | | G | G | G | G | G | G |
| NC_011770 | 3193765 | CT | C | 62 | 0 | 1 | 0 | FRAME_SHIFT | HIGH | | cca/ | P18 | pvdO | | | | | | | C | | | | | |
| NC_011770 | 3193008 | G | T | 63 | 2 | 4 | 0 | NON_SYNONYMOU | MODERAT | MISSENSE | aCc/aAc | T271N | pvdO | T | T | | | | | T | T | | | . | . |
| NC_011770 | 3193009 | T | C | 62 | 2 | 6 | 0 | NON_SYNONYMOU | MODERAT | MISSENSE | Acc/Gcc | T271A | pvdO | C | C | | | | | C | C | C | C | . | . |
| NC_011770 | 3193011 | T | C | 64 | 2 | 4 | 0 | NON_SYNONYMOU | MODERAT | MISSENSE | gAc/gGc | D270G | pvdO | C | C | | | | | C | C | | | . | . |
| NC_011770 | 3193012 | C | T | 63 | 2 | 4 | 0 | NON_SYNONYMOU | MODERAT | MISSENSE | Gac/Aac | D270N | pvdO | T | T | | | | | T | T | | | . | . |
| NC_011770 | 3193017 | T | G | 63 | 2 | 4 | 0 | NON_SYNONYMOU | MODERAT | MISSENSE | tAc/tCc | Y268S | pvdO | G | G | | | | | G | G | | | . | . |
| NC_011770 | 3193018 | A | C | 63 | 2 | 4 | 0 | NON_SYNONYMOU | MODERAT | MISSENSE | Tac/Gac | Y268D | pvdO | C | C | | | | | C | C | | | . | . |
| NC_011770 | 3193057 | G | C | 85 | 0 | 4 | 0 | NON_SYNONYMOU | MODERAT | MISSENSE | Ccg/Gcg | P255A | pvdO | | | C | C | C | C | | | | | | |
| NC_011770 | 3193093 | C | T | 89 | 0 | 4 | 0 | NON_SYNONYMOU | MODERAT | MISSENSE | Gag/Aag | E243K | pvdO | | | T | T | T | T | | | | | | |
| NC_011770 | 3193471 | G | T | 68 | 0 | 4 | 0 | NON_SYNONYMOU | MODERAT | MISSENSE | Cag/Aag | Q117K | pvdO | | | T | T | T | T | | | | | | |
| NC_011770 | 3193774 | T | C | 60 | 0 | 4 | 0 | NON_SYNONYMOU | MODERAT | MISSENSE | Acc/Gcc | T16A | pvdO | | | C | C | C | C | | | | | | |
| NC_011770 | 3193807 | G | A | 62 | 0 | 12 | 0 | NON_SYNONYMOU | MODERAT | MISSENSE | Ccg/Tcg | P5S | pvdO | A | A | A | A | A | A | A | A | A | A | A | A |
| NC_011770 | 3196786 | G | C | 30 | 8 | 4 | 0 | NON_SYNONYMOU | MODERAT | MISSENSE | aGc/aCc | S18T | pvdP | . | . | C | C | C | C | . | . | . | . | . | . |
| NC_011770 | 3196959 | A | C | 49 | 2 | 6 | 0 | NON_SYNONYMOU | MODERAT | MISSENSE | Atg/Ctg | M76L | pvdP | C | C | | | | | C | C | . | . | C | C |
| NC_011770 | 3197395 | A | G | 64 | 0 | 8 | 0 | NON_SYNONYMOU | MODERAT | MISSENSE | cAa/cGa | Q221R | pvdP | G | G | | | | | G | G | G | G | G | G |
| NC_011770 | 3197483 | A | C | 57 | 2 | 6 | 0 | NON_SYNONYMOU | MODERAT | MISSENSE | agA/agC | R250S | pvdP | C | C | | | | | C | C | . | . | C | C |
| NC_011770 | 3197514 | C | T | 59 | 0 | 8 | 0 | NON_SYNONYMOU | MODERAT | MISSENSE | Ctt/Ttt | L261F | pvdP | T | T | | | | | T | T | T | T | T | T |
| NC_011770 | 3197538 | C | T | 66 | 0 | 4 | 0 | NON_SYNONYMOU | MODERAT | MISSENSE | Cat/Tat | H269Y | pvdP | | | T | T | T | T | | | | | | |
| NC_011770 | 3197574 | A | G | 58 | 2 | 6 | 0 | NON_SYNONYMOU | MODERAT | MISSENSE | Aag/Gag | K281E | pvdP | G | G | | | | | G | G | . | . | G | G |
| NC_011770 | 3197575 | A | G | 58 | 2 | 4 | 0 | NON_SYNONYMOU | MODERAT | MISSENSE | aAg/aGg | K281R | pvdP | | | G | G | G | G | | | . | . | | |
| NC_011770 | 3197593 | C | T | 55 | 2 | 6 | 0 | NON_SYNONYMOU | MODERAT | MISSENSE | gCg/gTg | A287V | pvdP | T | T | | | | | T | T | . | . | T | T |
| NC_011770 | 3197608 | T | A | 60 | 2 | 6 | 0 | NON_SYNONYMOU | MODERAT | MISSENSE | tTc/tAc | F292Y | pvdP | A | A | | | | | A | A | . | . | A | A |
| NC_011770 | 3197683 | G | A | 55 | 0 | 8 | 0 | NON_SYNONYMOU | MODERAT | MISSENSE | gGg/gAg | G317E | pvdP | A | A | | | | | A | A | A | A | A | A |
| NC_011770 | 3197698 | G | C | 61 | 0 | 2 | 0 | NON_SYNONYMOU | MODERAT | MISSENSE | aGc/aCc | S322T | pvdP | | | | | | | | | C | C | | |
| NC_011770 | 3197710 | G | A | 59 | 0 | 6 | 0 | NON_SYNONYMOU | MODERAT | MISSENSE | cGt/cAt | R326H | pvdP | A | A | | | | | A | A | | | A | A |
| NC_011770 | 3197731 | C | G | 56 | 0 | 2 | 0 | NON_SYNONYMOU | MODERAT | MISSENSE | gCg/gGg | A333G | pvdP | | | | | | | | | G | G | | |
| NC_011770 | 3197739 | T | G | 55 | 0 | 8 | 0 | NON_SYNONYMOU | MODERAT | MISSENSE | Tcc/Gcc | S336A | pvdP | G | G | | | | | G | G | G | G | G | G |
| NC_011770 | 3197752 | C | T | 56 | 0 | 8 | 0 | NON_SYNONYMOU | MODERAT | MISSENSE | gCc/gTc | A340V | pvdP | T | T | | | | | T | T | T | T | T | T |
| NC_011770 | 3197778 | G | A | 50 | 0 | 2 | 0 | NON_SYNONYMOU | MODERAT | MISSENSE | Gaa/Aaa | E349K | pvdP | | | | | | | | | | | A | A |
| NC_011770 | 3197779 | A | C | 50 | 0 | 6 | 0 | NON_SYNONYMOU | MODERAT | MISSENSE | gAa/gCa | E349A | pvdP | C | C | | | | | C | C | | | C | C |
| NC_011770 | 3197820 | A | C | 44 | 0 | 6 | 0 | NON_SYNONYMOU | MODERAT | MISSENSE | Atg/Ctg | M363L | pvdP | C | C | | | | | C | C | | | C | C |
| NC_011770 | 3197862 | A | T | 32 | 1 | 7 | 0 | NON_SYNONYMOU | MODERAT | MISSENSE | Acc/Tcc | T377S | pvdP | T | . | | | | | T | T | T | T | T | T |
| NC_011770 | 3197889 | T | G | 32 | 1 | 7 | 0 | NON_SYNONYMOU | MODERAT | MISSENSE | Tcg/Gcg | S386A | pvdP | G | . | | | | | G | G | G | G | G | G |
| NC_011770 | 3197901 | G | A | 34 | 1 | 7 | 0 | NON_SYNONYMOU | MODERAT | MISSENSE | Ggc/Agc | G390S | pvdP | A | . | | | | | A | A | A | A | A | A |
| NC_011770 | 3197925 | T | G | 33 | 1 | 5 | 0 | NON_SYNONYMOU | MODERAT | MISSENSE | Tcg/Gcg | S398A | pvdP | G | . | | | | | G | G | | | G | G |
| NC_011770 | 3198076 | G | A | 33 | 1 | 7 | 0 | NON_SYNONYMOU | MODERAT | MISSENSE | cGg/cAg | R448Q | pvdP | A | . | | | | | A | A | A | A | A | A |
| NC_011770 | 3198090 | C | G | 34 | 1 | 7 | 0 | NON_SYNONYMOU | MODERAT | MISSENSE | Cag/Gag | Q453E | pvdP | G | . | | | | | G | G | G | G | G | G |
| NC_011770 | 3198100 | C | A | 34 | 1 | 7 | 0 | NON_SYNONYMOU | MODERAT | MISSENSE | cCc/cAc | P456H | pvdP | A | . | | | | | A | A | A | A | A | A |
| NC_011770 | 3198111 | G | C | 32 | 1 | 7 | 0 | NON_SYNONYMOU | MODERAT | MISSENSE | Gct/Cct | A460P | pvdP | C | . | | | | | C | C | C | C | C | C |
| NC_011770 | 3198133 | A | G | 33 | 1 | 7 | 0 | NON_SYNONYMOU | MODERAT | MISSENSE | gAc/gGc | D467G | pvdP | G | . | | | | | G | G | G | G | G | G |
| NC_011770 | 3198180 | C | G | 61 | 0 | 8 | 0 | NON_SYNONYMOU | MODERAT | MISSENSE | Ctg/Gtg | L483V | pvdP | G | G | | | | | G | G | G | G | G | G |
| NC_011770 | 3198234 | G | T | 63 | 0 | 8 | 0 | NON_SYNONYMOU | MODERAT | MISSENSE | Gta/Tta | V501L | pvdP | T | T | | | | | T | T | T | T | T | T |
| NC_011770 | 3198243 | A | G | 66 | 0 | 2 | 0 | NON_SYNONYMOU | MODERAT | MISSENSE | Atc/Gtc | I504V | pvdP | | | | | | | G | G | | | | |
| NC_011770 | 3206627 | G | A | 84 | 0 | 2 | 0 | NON_SYNONYMOU | MODERAT | MISSENSE | Gcc/Acc | A104T | pvdQ | | | | | | | | | | | A | A |
| NC_011770 | 3206874 | C | T | 72 | 0 | 2 | 0 | NON_SYNONYMOU | MODERAT | MISSENSE | gCg/gTg | A186V | pvdQ | T | T | | | | | | | | | | |
| NC_011770 | 3206885 | G | A | 72 | 0 | 8 | 0 | NON_SYNONYMOU | MODERAT | MISSENSE | Gcg/Acg | A190T | pvdQ | A | A | A | A | A | A | | | A | A | | |

| | | | | | | | | | | | | | | | | | | | | | | | | | |
|---|---|---|---|---|---|---|---|---|---|---|---|---|---|---|---|---|---|---|---|---|---|---|---|---|---|
| NC_011770 | 3206915 | G | A | 71 | 0 | 2 | 0 | NON_SYNONYMOU | MODERAT | MISSENSE | Gcg/Acg | A200T | pvdQ | A | A | | | | | | | | | | |
| NC_011770 | 3207128 | G | C | 87 | 0 | 4 | 0 | NON_SYNONYMOU | MODERAT | MISSENSE | Gtg/Ctg | V271L | pvdQ | | | | | | | C | C | | | C | C |
| NC_011770 | 3207558 | C | T | 82 | 0 | 2 | 0 | NON_SYNONYMOU | MODERAT | MISSENSE | gCc/gTc | A414V | pvdQ | | | | | | | T | T | | | | |
| NC_011770 | 3207573 | T | A | 79 | 0 | 12 | 0 | NON_SYNONYMOU | MODERAT | MISSENSE | cTg/cAg | L419Q | pvdQ | A | A | A | A | A | A | A | A | A | A | A | A |
| NC_011770 | 3207618 | C | T | 81 | 0 | 2 | 0 | NON_SYNONYMOU | MODERAT | MISSENSE | gCc/gTc | A434V | pvdQ | | | | | | | | | T | T | | |
| NC_011770 | 3207686 | C | A | 78 | 0 | 2 | 0 | NON_SYNONYMOU | MODERAT | MISSENSE | Cgc/Agc | R457S | pvdQ | A | A | | | | | | | | | | |
| NC_011770 | 3208003 | T | G | 80 | 0 | 6 | 0 | NON_SYNONYMOU | MODERAT | MISSENSE | aaT/aaG | N562K | pvdQ | | | | | | | G | G | G | G | G | G |
| NC_011770 | 3208031 | G | C | 80 | 0 | 2 | 0 | NON_SYNONYMOU | MODERAT | MISSENSE | Gcg/Ccg | A572P | pvdQ | | | | | | | C | C | | | | |
| NC_011770 | 3208199 | A | G | 84 | 0 | 2 | 0 | NON_SYNONYMOU | MODERAT | MISSENSE | Acc/Gcc | T628A | pvdQ | | | | | | | | | | | G | G |
| NC_011770 | 3208598 | C | A | 70 | 0 | 2 | 0 | NON_SYNONYMOU | MODERAT | MISSENSE | Cgc/Agc | R761S | pvdQ | | | | | | | | | A | A | | |
| NC_011770 | 3111769 | T | G | 69 | 0 | 1 | 0 | NON_SYNONYMOU | MODERAT | MISSENSE | gAt/gCt | D77A | pvdS | | | | | | | G | | | | | |
| NC_011770 | 3111913 | G | A | 67 | 0 | 2 | 0 | NON_SYNONYMOU | MODERAT | MISSENSE | gCc/gTc | A29V | pvdS | | | | | | | | | | | A | A |
| NC_011770 | 3111536 | T | G | 76 | 0 | 1 | 0 | NON_SYNONYMOU | MODERAT | MISSENSE | Acc/Ccc | T155P | pvdS | | | | | | | | G | | | | |
| NC_011770 | 5191200 | G | A | 60 | 0 | 1 | 0 | NON_SYNONYMOU | MODERAT | MISSENSE | Gac/Aac | D179N | pykA | | | | | | | A | | | | | |
| NC_011770 | 4325655 | G | GC | 68 | 0 | 1 | 0 | FRAME_SHIFT | HIGH | | ggt/Cggt | G102R? | pykF | | | | | | | GC | | | | | |
| NC_011770 | 4325652 | G | A | 68 | 0 | 2 | 0 | NON_SYNONYMOU | MODERAT | MISSENSE | Gac/Aac | D99N | pykF | | | | | | | | | | | A | A |
| NC_011770 | 4326388 | T | C | 74 | 0 | 6 | 0 | NON_SYNONYMOU | MODERAT | MISSENSE | gTc/gCc | V344A | pykF | | | C | C | C | C | C | C | | | | |
| NC_011770 | 4326784 | A | G | 66 | 0 | 4 | 0 | NON_SYNONYMOU | MODERAT | MISSENSE | gAt/gGt | D476G | pykF | | | G | G | G | G | | | | | | |
| NC_011770 | 438739 | C | T | 94 | 0 | 1 | 0 | NON_SYNONYMOU | MODERAT | MISSENSE | Gtc/Atc | V232I | pyrB | | | T | | | | | | | | | |
| NC_011770 | 437542 | T | C | 61 | 0 | 4 | 0 | NON_SYNONYMOU | MODERAT | MISSENSE | aAc/aGc | N297S | pyrC | C | C | | | | | C | C | | | | |
| NC_011770 | 438385 | C | T | 92 | 0 | 8 | 0 | NON_SYNONYMOU | MODERAT | MISSENSE | gGc/gAc | G16D | pyrC | | | T | T | T | T | | | T | T | T | T |
| NC_011770 | 1633996 | T | G | 66 | 0 | 2 | 0 | NON_SYNONYMOU | MODERAT | MISSENSE | aAt/aCt | N284T | pyrC | G | G | | | | | | | | | | |
| NC_011770 | 2179733 | G | A | 72 | 0 | 2 | 0 | NON_SYNONYMOU | MODERAT | MISSENSE | aGg/aAg | R96K | pyrD | | | | | | | | | A | A | | |
| NC_011770 | 6337413 | A | G | 94 | 0 | 12 | 0 | NON_SYNONYMOU | MODERAT | MISSENSE | gTc/gCc | V192A | pyrE | G | G | G | G | G | G | G | G | G | G | G | G |
| NC_011770 | 6337644 | G | A | 84 | 0 | 2 | 0 | NON_SYNONYMOU | MODERAT | MISSENSE | cCg/cTg | P115L | pyrE | A | A | | | | | | | | | | |
| NC_011770 | 1508165 | A | G | 102 | 0 | 1 | 0 | NON_SYNONYMOU | MODERAT | MISSENSE | cAc/cGc | H117R | pyrG | | G | | | | | | | | | | |
| NC_011770 | 1487834 | C | T | 84 | 0 | 1 | 0 | NON_SYNONYMOU | MODERAT | MISSENSE | Ctt/Ttt | L165F | pyrH | | | | | | | | | | | T | |
| NC_011770 | 6570478 | A | G | 74 | 0 | 2 | 0 | NON_SYNONYMOU | MODERAT | MISSENSE | cAa/cGa | Q2R | pyrQ | | | | | | | | | G | G | | |
| NC_011770 | 6570832 | A | T | 73 | 0 | 6 | 0 | NON_SYNONYMOU | MODERAT | MISSENSE | tAc/tTc | Y120F | pyrQ | | | T | T | T | T | T | T | | | | |
| NC_011770 | 6571275 | G | A | 89 | 0 | 2 | 0 | NON_SYNONYMOU | MODERAT | MISSENSE | Gac/Aac | D268N | pyrQ | | | | | | | | | | | A | A |
| NC_011770 | 6571339 | A | G | 91 | 0 | 2 | 0 | NON_SYNONYMOU | MODERAT | MISSENSE | gAg/gGg | E289G | pyrQ | G | G | | | | | | | | | | |
| NC_011770 | 4602051 | G | A | 38 | 6 | 1 | 0 | STOP_GAINED | HIGH | NONSENSE | Caa/Taa | Q568* | pys2 | | | . | . | . | . | | | . | . | | A |
| NC_011770 | 4603574 | G | A | 50 | 6 | 1 | 0 | NON_SYNONYMOU | MODERAT | MISSENSE | aCg/aTg | T60M | pys2 | | A | . | . | . | . | | | . | . | | |
| NC_011770 | 4601896 | T | G | 36 | 6 | 4 | 0 | NON_SYNONYMOU | MODERAT | MISSENSE | aaA/aaC | K619N | pys2 | | | . | . | . | . | G | G | . | . | G | G |
| NC_011770 | 4602162 | C | T | 44 | 6 | 2 | 0 | NON_SYNONYMOU | MODERAT | MISSENSE | Gtg/Atg | V531M | pys2 | | | . | . | . | . | | | . | . | T | T |
| NC_011770 | 4602181 | C | G | 43 | 6 | 2 | 0 | NON_SYNONYMOU | MODERAT | MISSENSE | gaG/gaC | E524D | pys2 | | | . | . | . | . | | | . | . | G | G |
| NC_011770 | 4602182 | T | C | 43 | 6 | 2 | 0 | NON_SYNONYMOU | MODERAT | MISSENSE | gAg/gGg | E524G | pys2 | | | . | . | . | . | | | . | . | C | C |
| NC_011770 | 4602186 | T | C | 45 | 3 | 3 | 0 | NON_SYNONYMOU | MODERAT | MISSENSE | Acc/Gcc | T523A | pys2 | | | C | . | C | C | | | . | . | | |
| NC_011770 | 4602204 | T | C | 66 | 2 | 4 | 0 | NON_SYNONYMOU | MODERAT | MISSENSE | Acc/Gcc | T517A | pys2 | | | | | | | C | C | . | . | C | C |
| NC_011770 | 4602206 | A | G | 67 | 2 | 8 | 0 | NON_SYNONYMOU | MODERAT | MISSENSE | gTg/gCg | V516A | pys2 | | | G | G | G | G | G | G | . | . | G | G |
| NC_011770 | 4602237 | C | T | 74 | 2 | 2 | 0 | NON_SYNONYMOU | MODERAT | MISSENSE | Gta/Ata | V506I | pys2 | | | | | | | T | T | . | . | | |
| NC_011770 | 4602242 | A | G | 73 | 2 | 10 | 0 | NON_SYNONYMOU | MODERAT | MISSENSE | cTg/cCg | L504P | pys2 | G | G | G | G | G | G | G | G | . | . | G | G |
| NC_011770 | 4603113 | T | C | 41 | 6 | 2 | 0 | NON_SYNONYMOU | MODERAT | MISSENSE | Aat/Gat | N214D | pys2 | | | . | . | . | . | | | . | . | C | C |
| NC_011770 | 4603232 | T | G | 52 | 6 | 4 | 0 | NON_SYNONYMOU | MODERAT | MISSENSE | gAg/gCg | E174A | pys2 | | | . | . | . | . | G | G | . | . | G | G |
| NC_011770 | 4603607 | G | A | 52 | 6 | 2 | 0 | NON_SYNONYMOU | MODERAT | MISSENSE | aCt/aTt | T49I | pys2 | | | . | . | . | . | | | . | . | A | A |
| NC_011770 | 25202 | T | C | 80 | 0 | 12 | 0 | NON_SYNONYMOU | MODERAT | MISSENSE | Aac/Gac | N111D | qor | C | C | C | C | C | C | C | C | C | C | C | C |
| NC_011770 | 1249382 | G | C | 67 | 0 | 10 | 0 | NON_SYNONYMOU | MODERAT | MISSENSE | Gac/Cac | D89H | queA | C | C | C | C | C | C | | | C | C | C | C |
| NC_011770 | 1249431 | C | T | 71 | 0 | 2 | 0 | NON_SYNONYMOU | MODERAT | MISSENSE | aCg/aTg | T105M | queA | T | T | | | | | | | | | | |
| NC_011770 | 1249617 | A | G | 82 | 0 | 2 | 0 | NON_SYNONYMOU | MODERAT | MISSENSE | cAg/cGg | Q167R | queA | | | | | | | | | | | G | G |
| NC_011770 | 1249830 | T | C | 79 | 0 | 6 | 0 | NON_SYNONYMOU | MODERAT | MISSENSE | gTc/gCc | V238A | queA | C | C | C | C | C | C | | | | | | |
| NC_011770 | 1249913 | C | A | 83 | 0 | 2 | 0 | NON_SYNONYMOU | MODERAT | MISSENSE | Ctg/Atg | L266M | queA | | | | | | | | | | | A | A |
| NC_011770 | 1249950 | A | T | 84 | 0 | 2 | 0 | NON_SYNONYMOU | MODERAT | MISSENSE | tAc/tTc | Y278F | queA | T | T | | | | | | | | | | |
| NC_011770 | 2439135 | G | A | 75 | 0 | 6 | 0 | NON_SYNONYMOU | MODERAT | MISSENSE | cGg/cAg | R200Q | queF | A | A | | | | | | | A | A | A | A |
| NC_011770 | 4725877 | G | A | 63 | 0 | 2 | 0 | NON_SYNONYMOU | MODERAT | MISSENSE | gGc/gAc | G249D | quiP | A | A | | | | | | | | | | |
| NC_011770 | 4726596 | G | A | 75 | 0 | 4 | 0 | NON_SYNONYMOU | MODERAT | MISSENSE | Ggc/Agc | G489S | quiP | | | A | A | A | A | | | | | | |
| NC_011770 | 4726660 | A | G | 80 | 0 | 4 | 0 | NON_SYNONYMOU | MODERAT | MISSENSE | cAg/cGg | Q510R | quiP | | | G | G | G | G | | | | | | |
| NC_011770 | 4726878 | C | G | 83 | 0 | 4 | 0 | NON_SYNONYMOU | MODERAT | MISSENSE | Ctg/Gtg | L583V | quiP | G | G | | | | | | | | | G | G |
| NC_011770 | 4726879 | T | C | 80 | 0 | 4 | 0 | NON_SYNONYMOU | MODERAT | MISSENSE | cTg/cCg | L583P | quiP | C | C | | | | | | | | | C | C |
| NC_011770 | 4727247 | G | A | 79 | 0 | 1 | 0 | NON_SYNONYMOU | MODERAT | MISSENSE | Ggg/Agg | G706R | quiP | | | | | | | A | | | | | |
| NC_011770 | 4727299 | A | G | 69 | 0 | 4 | 0 | NON_SYNONYMOU | MODERAT | MISSENSE | cAg/cGg | Q723R | quiP | G | G | | | | | | | | | G | G |
| NC_011770 | 4727562 | G | A | 67 | 0 | 1 | 0 | NON_SYNONYMOU | MODERAT | MISSENSE | Gcc/Acc | A811T | quiP | | | | | | | A | | | | | |
| NC_011770 | 4727630 | T | G | 62 | 0 | 4 | 0 | NON_SYNONYMOU | MODERAT | MISSENSE | gaT/gaG | D833E | quiP | | | G | G | G | G | | | | | | |

| | | | | | | | | | | | | | | | | | | | | | | | | | |
|---|---|---|---|---|---|---|---|---|---|---|---|---|---|---|---|---|---|---|---|---|---|---|---|---|---|
| NC_011770 | 4727671 | G | A | 61 | 0 | 4 | 0 | NON_SYNONYMOU | MODERAT | MISSENSE | cGc/cAc | R847H | quiP | | | A | A | A | A | | | | | | |
| NC_011770 | 5503555 | T | C | 70 | 0 | 2 | 0 | NON_SYNONYMOU | MODERAT | MISSENSE | Acc/Gcc | T76A | radA | | | | | | | C | C | | | | |
| NC_011770 | 5503620 | T | G | 65 | 0 | 2 | 0 | NON_SYNONYMOU | MODERAT | MISSENSE | cAg/cCg | Q54P | radA | G | G | | | | | | | | | | |
| NC_011770 | 5503696 | C | T | 67 | 0 | 2 | 0 | NON_SYNONYMOU | MODERAT | MISSENSE | Gcc/Acc | A29T | radA | | | | | | | | | | | T | T |
| NC_011770 | 6325046 | G | A | 71 | 0 | 1 | 0 | NON_SYNONYMOU | MODERAT | MISSENSE | gCc/gTc | A77V | radC | | | | | | | | | | | | A |
| NC_011770 | 5660770 | CTCG | C | 90 | 0 | 3 | 0 | CODON_CHANGE_ | MODERATE | | gacgag/ga | DE126E | rbfA | | | | | | | | | C | | C | C |
| NC_011770 | 3733196 | A | G | 68 | 0 | 12 | 0 | NON_SYNONYMOU | MODERAT | MISSENSE | aTc/aCc | I256T | rbsA | G | G | G | G | G | G | G | G | G | G | G | G |
| NC_011770 | 3733206 | T | G | 67 | 0 | 12 | 0 | NON_SYNONYMOU | MODERAT | MISSENSE | Aca/Cca | T253P | rbsA | G | G | G | G | G | G | G | G | G | G | G | G |
| NC_011770 | 3733231 | C | T | 72 | 0 | 12 | 0 | NON_SYNONYMOU | MODERAT | MISSENSE | gaG/gaC | E244D | rbsA | T | T | T | T | T | T | G | G | T | T | T | T |
| NC_011770 | 3733251 | T | C | 74 | 0 | 2 | 0 | NON_SYNONYMOU | MODERAT | MISSENSE | Acc/Gcc | T238A | rbsA | | | | | | | C | C | | | | |
| NC_011770 | 3733742 | G | C | 57 | 0 | 2 | 0 | NON_SYNONYMOU | MODERAT | MISSENSE | gCt/gGt | A74G | rbsA | | | | | | | | | C | C | | |
| NC_011770 | 3733832 | G | A | 65 | 0 | 1 | 0 | NON_SYNONYMOU | MODERAT | MISSENSE | gCc/gTc | A44V | rbsA | | | | | | | A | | | | | |
| NC_011770 | 3733263 | C | T | 75 | 0 | 1 | 0 | NON_SYNONYMOU | MODERAT | MISSENSE | Gac/Aac | D234N | rbsA | | | | | | | | | | | T | |
| NC_011770 | 3732641 | G | A | 83 | 0 | 1 | 0 | NON_SYNONYMOU | MODERAT | MISSENSE | gCc/gTc | A441V | rbsA | | | | | | | | A | | | | |
| NC_011770 | 3734046 | T | C | 67 | 0 | 2 | 0 | NON_SYNONYMOU | MODERAT | MISSENSE | Acc/Gcc | T300A | rbsB | | | | | | | | | | | C | C |
| NC_011770 | 3734076 | G | C | 70 | 0 | 2 | 0 | NON_SYNONYMOU | MODERAT | MISSENSE | Cag/Gag | Q290E | rbsB | | | | | | | | | C | C | | |
| NC_011770 | 3734304 | C | T | 68 | 0 | 2 | 0 | NON_SYNONYMOU | MODERAT | MISSENSE | Gcg/Acg | A214T | rbsB | | | | | | | | | T | T | | |
| NC_011770 | 3734378 | T | C | 66 | 0 | 8 | 0 | NON_SYNONYMOU | MODERAT | MISSENSE | aAg/aGg | K189R | rbsB | C | C | C | C | C | C | C | C | | | | |
| NC_011770 | 3729526 | G | A | 48 | 0 | 1 | 0 | NON_SYNONYMOU | MODERAT | MISSENSE | gCg/gTg | A271V | rbsK | | | | | | | A | | | | | |
| NC_011770 | 3729677 | T | C | 51 | 0 | 10 | 0 | NON_SYNONYMOU | MODERAT | MISSENSE | Atc/Gtc | I221V | rbsK | C | C | C | C | C | C | C | C | | | C | C |
| NC_011770 | 3729799 | A | G | 61 | 0 | 4 | 0 | NON_SYNONYMOU | MODERAT | MISSENSE | gTg/gCg | V180A | rbsK | | | G | G | G | G | | | | | | |
| NC_011770 | 3729813 | G | C | 60 | 0 | 12 | 0 | NON_SYNONYMOU | MODERAT | MISSENSE | gaC/gaG | D175E | rbsK | C | C | C | C | C | C | C | C | C | C | C | C |
| NC_011770 | 3729953 | A | T | 69 | 0 | 4 | 0 | NON_SYNONYMOU | MODERAT | MISSENSE | Ttg/Atg | L129M | rbsK | | | T | T | T | T | | | | | | |
| NC_011770 | 3730037 | C | A | 66 | 0 | 2 | 0 | NON_SYNONYMOU | MODERAT | MISSENSE | Gac/Tac | D101Y | rbsK | | | | | | | A | A | | | | |
| NC_011770 | 3730174 | C | T | 69 | 0 | 10 | 0 | NON_SYNONYMOU | MODERAT | MISSENSE | gGg/gAg | G55E | rbsK | T | T | T | T | T | T | T | T | | | T | T |
| NC_011770 | 3730332 | T | A | 77 | 0 | 2 | 0 | NON_SYNONYMOU | MODERAT | MISSENSE | caA/caT | Q2H | rbsK | | | | | | | | | A | A | | |
| NC_011770 | 3730776 | C | T | 75 | 0 | 4 | 0 | NON_SYNONYMOU | MODERAT | MISSENSE | cGc/cAc | R210H | rbsR | | | T | T | T | T | | | | | | |
| NC_011770 | 5168973 | G | A | 59 | 0 | 4 | 0 | NON_SYNONYMOU | MODERAT | MISSENSE | gCg/gTg | A127V | rcpA | | | | | | | A | A | A | A | | |
| NC_011770 | 5168982 | G | A | 60 | 0 | 4 | 0 | NON_SYNONYMOU | MODERAT | MISSENSE | gCc/gTc | A124V | rcpA | | | | | | | A | A | A | A | | |
| NC_011770 | 5168991 | G | A | 62 | 0 | 2 | 0 | NON_SYNONYMOU | MODERAT | MISSENSE | gCg/gTg | A121V | rcpA | A | A | | | | | | | | | | |
| NC_011770 | 5169045 | C | T | 64 | 0 | 2 | 0 | NON_SYNONYMOU | MODERAT | MISSENSE | cGc/cAc | R103H | rcpA | | | | | | | | | | | T | T |
| NC_011770 | 5169373 | G | A | 64 | 0 | 6 | 0 | NON_SYNONYMOU | MODERAT | MISSENSE | gCg/gTg | A302V | rcpC | | | | | | | A | A | A | A | A | A |
| NC_011770 | 5169633 | G | T | 83 | 0 | 2 | 0 | NON_SYNONYMOU | MODERAT | MISSENSE | gaC/gaA | D215E | rcpC | | | | | | | T | T | | | | |
| NC_011770 | 5169937 | G | A | 71 | 0 | 2 | 0 | NON_SYNONYMOU | MODERAT | MISSENSE | aCc/aTc | T114I | rcpC | A | A | | | | | | | | | | |
| NC_011770 | 5170190 | C | T | 84 | 0 | 2 | 0 | NON_SYNONYMOU | MODERAT | MISSENSE | Gcg/Acg | A30T | rcpC | T | T | | | | | | | | | | |
| NC_011770 | 1951257 | C | A | 61 | 0 | 4 | 0 | NON_SYNONYMOU | MODERAT | MISSENSE | Gcg/Tcg | A171S | rdgC | | | A | A | A | A | | | | | | |
| NC_011770 | 1951680 | G | A | 23 | 0 | 2 | 0 | NON_SYNONYMOU | MODERAT | MISSENSE | Ccc/Tcc | P30S | rdgC | | | | | | | A | A | | | | |
| NC_011770 | 5142832 | C | T | 91 | 0 | 1 | 0 | NON_SYNONYMOU | MODERAT | MISSENSE | Gta/Ata | V695I | recB | | | | | | | | | | | T | |
| NC_011770 | 5141499 | T | A | 83 | 0 | 2 | 0 | NON_SYNONYMOU | MODERAT | MISSENSE | cAc/cTc | H1139L | recB | | | | | | | A | A | | | | |
| NC_011770 | 5141547 | G | A | 77 | 0 | 4 | 0 | NON_SYNONYMOU | MODERAT | MISSENSE | aCc/aTc | T1123I | recB | | | A | A | A | A | | | | | | |
| NC_011770 | 5141828 | C | A | 80 | 0 | 6 | 0 | NON_SYNONYMOU | MODERAT | MISSENSE | gaG/gaT | E1029D | recB | A | A | | | | | | | A | A | A | A |
| NC_011770 | 5141929 | C | T | 77 | 0 | 2 | 0 | NON_SYNONYMOU | MODERAT | MISSENSE | Gcc/Acc | A996T | recB | | | | | | | | | | | T | T |
| NC_011770 | 5142024 | G | A | 67 | 0 | 12 | 0 | NON_SYNONYMOU | MODERAT | MISSENSE | gCg/gTg | A964V | recB | A | A | A | A | A | A | A | A | A | A | A | A |
| NC_011770 | 5142123 | A | T | 67 | 0 | 12 | 0 | NON_SYNONYMOU | MODERAT | MISSENSE | cTg/cAg | L931Q | recB | T | T | T | T | T | T | T | T | T | T | T | T |
| NC_011770 | 5142973 | T | C | 92 | 0 | 4 | 0 | NON_SYNONYMOU | MODERAT | MISSENSE | Acg/Gcg | T648A | recB | C | C | | | | | | | | | C | C |
| NC_011770 | 5143020 | A | G | 91 | 0 | 4 | 0 | NON_SYNONYMOU | MODERAT | MISSENSE | aTc/aCc | I632T | recB | G | G | | | | | | | | | G | G |
| NC_011770 | 5143021 | T | C | 93 | 0 | 4 | 0 | NON_SYNONYMOU | MODERAT | MISSENSE | Atc/Gtc | I632V | recB | C | C | | | | | | | | | C | C |
| NC_011770 | 5144155 | C | T | 74 | 0 | 2 | 0 | NON_SYNONYMOU | MODERAT | MISSENSE | Gcg/Acg | A254T | recB | | | | | | | | | | | T | T |
| NC_011770 | 5144194 | C | G | 75 | 0 | 4 | 0 | NON_SYNONYMOU | MODERAT | MISSENSE | Gac/Cac | D241H | recB | | | G | G | G | G | | | | | | |
| NC_011770 | 5144236 | T | C | 78 | 0 | 4 | 0 | NON_SYNONYMOU | MODERAT | MISSENSE | Aca/Gca | T227A | recB | | | | | | | C | C | C | C | | |
| NC_011770 | 5144326 | T | C | 81 | 0 | 6 | 0 | NON_SYNONYMOU | MODERAT | MISSENSE | Acg/Gcg | T197A | recB | | | C | C | C | C | C | C | | | | |
| NC_011770 | 5144647 | C | A | 75 | 0 | 2 | 0 | NON_SYNONYMOU | MODERAT | MISSENSE | Gcc/Tcc | A90S | recB | | | | | | | | | | | A | A |
| NC_011770 | 5148211 | ACCG | A | 77 | 0 | 1 | 0 | CODON_CHANGE_ | MODERATE | | ggcggt/ggt | GG71G | recC | | | | A | | | | | | | | |
| NC_011770 | 5145165 | G | C | 74 | 0 | 10 | 0 | NON_SYNONYMOU | MODERAT | MISSENSE | Ctg/Gtg | L1088V | recC | C | C | C | C | C | C | | | C | C | C | C |
| NC_011770 | 5145360 | C | T | 72 | 0 | 2 | 0 | NON_SYNONYMOU | MODERAT | MISSENSE | Ggc/Agc | G1023S | recC | | | | | | | | | | | T | T |
| NC_011770 | 5145647 | G | A | 68 | 0 | 4 | 0 | NON_SYNONYMOU | MODERAT | MISSENSE | gCg/gTg | A927V | recC | | | | | | | A | A | | | A | A |
| NC_011770 | 5145666 | C | T | 77 | 0 | 2 | 0 | NON_SYNONYMOU | MODERAT | MISSENSE | Gat/Aat | D921N | recC | | | | | | | | | T | T | | |
| NC_011770 | 5145891 | G | A | 73 | 0 | 8 | 0 | NON_SYNONYMOU | MODERAT | MISSENSE | Ccc/Tcc | P846S | recC | A | A | A | A | A | A | A | A | | | | |
| NC_011770 | 5145911 | G | C | 75 | 0 | 2 | 0 | NON_SYNONYMOU | MODERAT | MISSENSE | gCc/gGc | A839G | recC | | | | | | | | | | | C | C |
| NC_011770 | 5145916 | G | C | 72 | 0 | 2 | 0 | NON_SYNONYMOU | MODERAT | MISSENSE | gaC/gaG | D837E | recC | | | | | | | | | | | C | C |
| NC_011770 | 5145927 | T | G | 75 | 0 | 2 | 0 | NON_SYNONYMOU | MODERAT | MISSENSE | Agc/Cgc | S834R | recC | | | | | | | | | | | G | G |
| NC_011770 | 5146043 | C | T | 69 | 0 | 2 | 0 | NON_SYNONYMOU | MODERAT | MISSENSE | cGt/cAt | R795H | recC | | | | | | | | | T | T | | |

| | | | | | | | | | | | | | | | | | | | | | | | | | |
|---|---|---|---|---|---|---|---|---|---|---|---|---|---|---|---|---|---|---|---|---|---|---|---|---|---|
| NC_011770 | 5139068 | T | C | 78 | 0 | 2 | 0 | NON_SYNONYMOU | MODERAT | MISSENSE | Agg/Ggg | R705G | recD | | | | | | | | | | | C | C |
| NC_011770 | 5139731 | T | C | 90 | 0 | 2 | 0 | NON_SYNONYMOU | MODERAT | MISSENSE | Acg/Gcg | T484A | recD | | | | | | | C | C | | | | |
| NC_011770 | 5139988 | G | T | 85 | 0 | 4 | 0 | NON_SYNONYMOU | MODERAT | MISSENSE | aCc/aAc | T398N | recD | | | T | T | T | T | | | | | | |
| NC_011770 | 5140028 | C | T | 87 | 0 | 2 | 0 | NON_SYNONYMOU | MODERAT | MISSENSE | Gcc/Acc | A385T | recD | | | | | | | | | T | T | | |
| NC_011770 | 5140035 | A | C | 93 | 0 | 12 | 0 | NON_SYNONYMOU | MODERAT | MISSENSE | tgT/tgG | C382W | recD | C | C | C | C | C | C | C | C | C | C | C | C |
| NC_011770 | 5140739 | T | C | 74 | 0 | 2 | 0 | NON_SYNONYMOU | MODERAT | MISSENSE | Acg/Gcg | T148A | recD | | | | | | | | | C | C | | |
| NC_011770 | 5141104 | A | G | 75 | 0 | 12 | 0 | NON_SYNONYMOU | MODERAT | MISSENSE | gTa/gCa | V26A | recD | G | G | G | G | G | G | G | G | G | G | G | G |
| NC_011770 | 3806 | T | C | 95 | 0 | 2 | 0 | NON_SYNONYMOU | MODERAT | MISSENSE | gTt/gCt | V213A | recF | | | | | | | | | | | C | C |
| NC_011770 | 3955 | T | G | 106 | 0 | 6 | 0 | NON_SYNONYMOU | MODERAT | MISSENSE | Tca/Gca | S263A | recF | G | G | | | | | G | G | | | G | G |
| NC_011770 | 6349124 | G | A | 68 | 0 | 2 | 0 | NON_SYNONYMOU | MODERAT | MISSENSE | Ggc/Agc | G68S | recG | A | A | | | | | | | | | | |
| NC_011770 | 6349133 | G | A | 68 | 0 | 2 | 0 | NON_SYNONYMOU | MODERAT | MISSENSE | Gtg/Atg | V71M | recG | | | A | A | | | | | | | | |
| NC_011770 | 6349541 | G | A | 80 | 0 | 2 | 0 | NON_SYNONYMOU | MODERAT | MISSENSE | Gat/Aat | D207N | recG | | | | | | | | | | | A | A |
| NC_011770 | 6350904 | A | G | 90 | 0 | 2 | 0 | NON_SYNONYMOU | MODERAT | MISSENSE | gAt/gGt | D661G | recG | | | | | | | G | G | | | | |
| NC_011770 | 6350975 | G | A | 93 | 0 | 2 | 0 | NON_SYNONYMOU | MODERAT | MISSENSE | Ggc/Agc | G685S | recG | | | | | | | | | | | A | A |
| NC_011770 | 1365158 | A | T | 35 | 0 | 2 | 0 | NON_SYNONYMOU | MODERAT | MISSENSE | tAc/tTc | Y47F | recJ | | | | | | | | | | | T | T |
| NC_011770 | 1365515 | C | A | 48 | 0 | 4 | 0 | NON_SYNONYMOU | MODERAT | MISSENSE | gCg/gAg | A166E | recJ | | | A | A | A | A | | | | | | |
| NC_011770 | 5027357 | C | T | 74 | 0 | 4 | 0 | NON_SYNONYMOU | MODERAT | MISSENSE | Gtt/Att | V75I | recO | | | | | | | | | T | T | T | T |
| NC_011770 | 1528753 | C | T | 71 | 0 | 12 | 0 | NON_SYNONYMOU | MODERAT | MISSENSE | aCc/aTc | T3I | recX | T | T | T | T | T | T | T | T | T | T | T | T |
| NC_011770 | 4813693 | C | T | 73 | 0 | 2 | 0 | NON_SYNONYMOU | MODERAT | MISSENSE | Gag/Aag | E549K | relA | | | | | | | | | | | T | T |
| NC_011770 | 4813339 | T | C | 75 | 0 | 1 | 0 | NON_SYNONYMOU | MODERAT | MISSENSE | Acc/Gcc | T667A | relA | | | | | | | | C | | | | |
| NC_011770 | 4815109 | C | T | 96 | 0 | 1 | 0 | NON_SYNONYMOU | MODERAT | MISSENSE | Gac/Aac | D77N | relA | | | | | | | | T | | | | |
| NC_011770 | 1856135 | T | G | 63 | 0 | 12 | 0 | NON_SYNONYMOU | MODERAT | MISSENSE | Tcc/Gcc | S124A | rfaD | G | G | G | G | G | G | G | G | G | G | G | G |
| NC_011770 | 1856255 | G | A | 65 | 0 | 2 | 0 | NON_SYNONYMOU | MODERAT | MISSENSE | Gtc/Atc | V164I | rfaD | A | A | | | | | | | | | | |
| NC_011770 | 5947913 | C | T | 91 | 0 | 1 | 0 | NON_SYNONYMOU | MODERAT | MISSENSE | gGc/gAc | G152D | rfaE | | | | | | | | | | | | T |
| NC_011770 | 5947647 | C | G | 81 | 0 | 12 | 0 | NON_SYNONYMOU | MODERAT | MISSENSE | Gat/Cat | D241H | rfaE | G | G | G | G | G | G | G | G | G | G | G | G |
| NC_011770 | 5947967 | A | G | 92 | 0 | 2 | 0 | NON_SYNONYMOU | MODERAT | MISSENSE | gTg/gCg | V134A | rfaE | | | | | | | | | | | G | G |
| NC_011770 | 5894259 | G | A | 70 | 0 | 2 | 0 | NON_SYNONYMOU | MODERAT | MISSENSE | cGc/cAc | R109H | rhdA | | | | | | | A | A | | | | |
| NC_011770 | 5894366 | A | G | 62 | 0 | 8 | 0 | NON_SYNONYMOU | MODERAT | MISSENSE | Aac/Gac | N145D | rhdA | G | G | | | | | G | G | G | G | G | G |
| NC_011770 | 1666279 | A | T | 92 | 0 | 2 | 0 | NON_SYNONYMOU | MODERAT | MISSENSE | Acc/Tcc | T23S | rhlB | | | T | T | | | | | | | | |
| NC_011770 | 1666396 | G | A | 99 | 0 | 12 | 0 | NON_SYNONYMOU | MODERAT | MISSENSE | Gcc/Acc | A62T | rhlB | A | A | A | A | A | A | A | A | A | A | A | A |
| NC_011770 | 1667358 | C | A | 87 | 0 | 2 | 0 | NON_SYNONYMOU | MODERAT | MISSENSE | gaC/gaA | D382E | rhlB | | | | | | | A | A | | | | |
| NC_011770 | 1208444 | T | C | 58 | 0 | 1 | 0 | NON_SYNONYMOU | MODERAT | MISSENSE | Acc/Gcc | T33A | rhlB helicase | | C | | | | | | | | | | |
| NC_011770 | 4624403 | G | A | 90 | 0 | 4 | 0 | NON_SYNONYMOU | MODERAT | MISSENSE | aGc/aAc | S44N | rhlC | | | A | A | A | A | | | | | | |
| NC_011770 | 4624504 | G | T | 85 | 0 | 4 | 0 | NON_SYNONYMOU | MODERAT | MISSENSE | Gcg/Tcg | A78S | rhlC | | | T | T | T | T | | | | | | |
| NC_011770 | 4625173 | C | A | 64 | 0 | 6 | 0 | NON_SYNONYMOU | MODERAT | MISSENSE | Cgc/Agc | R301S | rhlC | | | A | A | A | A | A | A | | | | |
| NC_011770 | 4625174 | G | C | 65 | 0 | 6 | 0 | NON_SYNONYMOU | MODERAT | MISSENSE | cGc/cCc | R301P | rhlC | | | C | C | C | C | C | C | | | | |
| NC_011770 | 4625221 | T | C | 59 | 0 | 6 | 0 | NON_SYNONYMOU | MODERAT | MISSENSE | Tcc/Ccc | S317P | rhlC | | | C | C | C | C | C | C | | | | |
| NC_011770 | 4625224 | G | A | 60 | 0 | 6 | 0 | NON_SYNONYMOU | MODERAT | MISSENSE | Gta/Ata | V318I | rhlC | | | A | A | A | A | A | A | | | | |
| NC_011770 | 4625225 | T | C | 60 | 0 | 6 | 0 | NON_SYNONYMOU | MODERAT | MISSENSE | gTa/gCa | V318A | rhlC | | | C | C | C | C | C | C | | | | |
| NC_011770 | 4625228 | C | T | 59 | 0 | 6 | 0 | NON_SYNONYMOU | MODERAT | MISSENSE | gCt/gTt | A319V | rhlC | | | T | T | T | T | T | T | | | | |
| NC_011770 | 4625230 | T | C | 59 | 0 | 6 | 0 | NON_SYNONYMOU | MODERAT | MISSENSE | Tcc/Ccc | S320P | rhlC | | | C | C | C | C | C | C | | | | |
| NC_011770 | 4624658 | G | A | 88 | 0 | 1 | 0 | NON_SYNONYMOU | MODERAT | MISSENSE | cGc/cAc | R129H | rhlC | | | | | | | | A | | | | |
| NC_011770 | 1800487 | G | C | 63 | 0 | 6 | 0 | NON_SYNONYMOU | MODERAT | MISSENSE | Cag/Gag | Q77E | rhlG | | | | | | | C | C | C | C | C | C |
| NC_011770 | 1800495 | C | A | 61 | 0 | 2 | 0 | NON_SYNONYMOU | MODERAT | MISSENSE | cGc/cTc | R74L | rhlG | | | | | | | | | | | A | A |
| NC_011770 | 1800685 | T | C | 60 | 0 | 12 | 0 | NON_SYNONYMOU | MODERAT | MISSENSE | Atc/Gtc | I11V | rhlG | C | C | C | C | C | C | C | C | C | C | C | C |
| NC_011770 | 1002006 | C | A | 88 | 0 | 2 | 0 | NON_SYNONYMOU | MODERAT | MISSENSE | gaC/gaA | D12E | ribB | | | | | | | | | A | A | | |
| NC_011770 | 999740 | A | G | 79 | 0 | 2 | 0 | NON_SYNONYMOU | MODERAT | MISSENSE | aAc/aGc | N2S | ribD | | | | | | | G | G | | | | |
| NC_011770 | 999901 | C | T | 82 | 0 | 2 | 0 | NON_SYNONYMOU | MODERAT | MISSENSE | Cat/Tat | H56Y | ribD | T | T | | | | | | | | | | |
| NC_011770 | 1000097 | G | C | 80 | 0 | 2 | 0 | NON_SYNONYMOU | MODERAT | MISSENSE | gGc/gCc | G121A | ribD | | | | | | | | | C | C | | |
| NC_011770 | 5445082 | G | T | 73 | 0 | 4 | 0 | NON_SYNONYMOU | MODERAT | MISSENSE | Cta/Ata | L266I | ribF | | | T | T | T | T | | | | | | |
| NC_011770 | 5582774 | A | G | 80 | 0 | 4 | 0 | NON_SYNONYMOU | MODERAT | MISSENSE | Ttc/Ctc | F149L | rimI | | | G | G | G | G | | | | | | |
| NC_011770 | 3758954 | G | C | 38 | 0 | 2 | 0 | NON_SYNONYMOU | MODERAT | MISSENSE | ttG/ttC | L11F | rimJ | | | | | | | C | C | | | | |
| NC_011770 | 3759134 | C | A | 47 | 0 | 12 | 0 | NON_SYNONYMOU | MODERAT | MISSENSE | gaC/gaA | D71E | rimJ | A | A | A | A | A | A | A | A | A | A | A | A |
| NC_011770 | 3759406 | A | G | 54 | 0 | 2 | 0 | NON_SYNONYMOU | MODERAT | MISSENSE | aAg/aGg | K162R | rimJ | G | G | | | | | | | | | | |
| NC_011770 | 6185374 | T | C | 78 | 0 | 1 | 0 | NON_SYNONYMOU | MODERAT | MISSENSE | Tcg/Ccg | S103P | rimK | | | | | | | C | | | | | |
| NC_011770 | 1341294 | A | G | 85 | 0 | 1 | 0 | NON_SYNONYMOU | MODERAT | MISSENSE | gAc/gGc | D161G | rimM | | | | | | | | G | | | | |
| NC_011770 | 4839139 | C | A | 83 | 0 | 2 | 0 | NON_SYNONYMOU | MODERAT | MISSENSE | gaC/gaA | D172E | rimO | A | A | | | | | | | | | | |
| NC_011770 | 2181912 | G | A | 84 | 0 | 2 | 0 | NON_SYNONYMOU | MODERAT | MISSENSE | aGg/aAg | R226K | rlmL | | | | | | | A | A | | | | |
| NC_011770 | 2182001 | A | G | 78 | 0 | 4 | 0 | NON_SYNONYMOU | MODERAT | MISSENSE | Acc/Gcc | T256A | rlmL | | | | | | | G | G | G | G | | |
| NC_011770 | 2182016 | C | T | 83 | 0 | 2 | 0 | NON_SYNONYMOU | MODERAT | MISSENSE | Ccg/Tcg | P261S | rlmL | | | | | | | | | T | T | | |
| NC_011770 | 2183252 | C | T | 68 | 0 | 2 | 0 | NON_SYNONYMOU | MODERAT | MISSENSE | Ccg/Tcg | P673S | rlmL | | | | | | | | | | | T | T |
| NC_011770 | 6480523 | C | T | 85 | 0 | 1 | 0 | NON_SYNONYMOU | MODERAT | MISSENSE | cGc/cAc | R128H | rmd | | | | | | | | | | | | T |

| | | | | | | | | | | | | | | | | | | | | | | | | | |
|---|---|---|---|---|---|---|---|---|---|---|---|---|---|---|---|---|---|---|---|---|---|---|---|---|---|
| NC_011770 | 6480625 | C | T | 84 | 0 | 2 | 0 | NON_SYNONYMOU | MODERAT | MISSENSE | cGg/cAg | R94Q | rmd | | | | | | | | | | | T | T |
| NC_011770 | 6146934 | A | C | 79 | 0 | 1 | 0 | NON_SYNONYMOU | MODERAT | MISSENSE | Acc/Ccc | T56P | rmlA | | | | | | | | C | | | | |
| NC_011770 | 6147187 | T | A | 66 | 0 | 2 | 0 | NON_SYNONYMOU | MODERAT | MISSENSE | cTg/cAg | L140Q | rmlA | | | | | | | | | A | A | | |
| NC_011770 | 6147330 | A | G | 73 | 0 | 6 | 0 | NON_SYNONYMOU | MODERAT | MISSENSE | Aac/Gac | N188D | rmlA | G | G | | | | | | | G | G | G | G |
| NC_011770 | 6147637 | A | G | 73 | 0 | 2 | 0 | NON_SYNONYMOU | MODERAT | MISSENSE | gAg/gGg | E290G | rmlA | G | G | | | | | | | | | | |
| NC_011770 | 6144953 | A | G | 59 | 0 | 12 | 0 | NON_SYNONYMOU | MODERAT | MISSENSE | Aac/Gac | N49D | rmlB | G | G | G | G | G | G | G | G | G | G | G | G |
| NC_011770 | 6145463 | G | T | 68 | 0 | 1 | 0 | NON_SYNONYMOU | MODERAT | MISSENSE | Ggc/Tgc | G219C | rmlB | | | | | | | T | | | | | |
| NC_011770 | 6145820 | G | A | 88 | 0 | 1 | 0 | NON_SYNONYMOU | MODERAT | MISSENSE | Gta/Ata | V338I | rmlB | | | | | | | A | | | | | |
| NC_011770 | 6148085 | G | C | 77 | 0 | 2 | 0 | NON_SYNONYMOU | MODERAT | MISSENSE | Gtc/Ctc | V146L | rmlC | | | | | | | | | | | C | C |
| NC_011770 | 6148166 | G | A | 80 | 0 | 2 | 0 | NON_SYNONYMOU | MODERAT | MISSENSE | Gca/Aca | A173T | rmlC | A | A | | | | | | | | | | |
| NC_011770 | 6148173 | C | T | 78 | 0 | 2 | 0 | NON_SYNONYMOU | MODERAT | MISSENSE | gCc/gTc | A175V | rmlC | T | T | | | | | | | | | | |
| NC_011770 | 6146135 | G | C | 83 | 0 | 8 | 0 | NON_SYNONYMOU | MODERAT | MISSENSE | gGc/gCc | G91A | rmlD | C | C | | | | | C | C | C | C | C | C |
| NC_011770 | 6146195 | T | C | 90 | 0 | 6 | 0 | NON_SYNONYMOU | MODERAT | MISSENSE | gTg/gCg | V111A | rmlD | C | C | C | C | C | C | | | | | | |
| NC_011770 | 6146281 | G | C | 96 | 0 | 2 | 0 | NON_SYNONYMOU | MODERAT | MISSENSE | Ggc/Cgc | G140R | rmlD | | | | | | | | | | | C | C |
| NC_011770 | 6146375 | A | C | 92 | 0 | 2 | 0 | NON_SYNONYMOU | MODERAT | MISSENSE | gAc/gCc | D171A | rmlD | | | | | | | | | | | C | C |
| NC_011770 | 6146474 | T | C | 91 | 0 | 8 | 0 | NON_SYNONYMOU | MODERAT | MISSENSE | gTc/gCc | V204A | rmlD | C | C | | | | | C | C | C | C | C | C |
| NC_011770 | 6146620 | G | A | 86 | 0 | 2 | 0 | NON_SYNONYMOU | MODERAT | MISSENSE | Gcc/Acc | A253T | rmlD | | | | | | | | | | | A | A |
| NC_011770 | 4297579 | C | T | 78 | 0 | 2 | 0 | STOP_GAINED | HIGH | NONSENSE | tGg/tAg | W247* | rnd | T | T | | | | | | | | | | |
| NC_011770 | 4297316 | T | C | 85 | 0 | 8 | 0 | NON_SYNONYMOU | MODERAT | MISSENSE | Atc/Gtc | I335V | rnd | C | C | C | C | C | C | | | C | C | | |
| NC_011770 | 4297463 | A | C | 79 | 0 | 8 | 0 | NON_SYNONYMOU | MODERAT | MISSENSE | Tcg/Gcg | S286A | rnd | C | C | C | C | C | C | | | C | C | | |
| NC_011770 | 4297808 | C | T | 70 | 0 | 6 | 0 | NON_SYNONYMOU | MODERAT | MISSENSE | Gcg/Acg | A171T | rnd | | | T | T | T | T | | | T | T | | |
| NC_011770 | 4297600 | A | G | 80 | 0 | 1 | 0 | NON_SYNONYMOU | MODERAT | MISSENSE | gTg/gCg | V240A | rnd | | | | | | | | G | | | | |
| NC_011770 | 2259333 | G | A | 48 | 0 | 2 | 0 | NON_SYNONYMOU | MODERAT | MISSENSE | gCt/gTt | A980V | rne | | | | | | | | | A | A | | |
| NC_011770 | 2259618 | G | A | 56 | 0 | 4 | 0 | NON_SYNONYMOU | MODERAT | MISSENSE | gCc/gTc | A885V | rne | | | A | A | A | A | | | | | | |
| NC_011770 | 2259645 | C | T | 54 | 0 | 10 | 0 | NON_SYNONYMOU | MODERAT | MISSENSE | gGc/gAc | G876D | rne | T | T | T | T | T | T | | | T | T | T | T |
| NC_011770 | 2259706 | T | C | 52 | 0 | 8 | 0 | NON_SYNONYMOU | MODERAT | MISSENSE | Acc/Gcc | T856A | rne | | | C | C | C | C | | | C | C | C | C |
| NC_011770 | 2259726 | C | T | 51 | 0 | 2 | 0 | NON_SYNONYMOU | MODERAT | MISSENSE | gGc/gAc | G849D | rne | | | | | | | T | T | | | | |
| NC_011770 | 2259754 | A | G | 47 | 0 | 8 | 0 | NON_SYNONYMOU | MODERAT | MISSENSE | Tcg/Ccg | S840P | rne | | | G | G | G | G | | | G | G | G | G |
| NC_011770 | 2259787 | C | T | 53 | 0 | 12 | 0 | NON_SYNONYMOU | MODERAT | MISSENSE | Ggc/Agc | G829S | rne | T | T | T | T | T | T | T | T | T | T | T | T |
| NC_011770 | 2259928 | G | C | 57 | 0 | 2 | 0 | NON_SYNONYMOU | MODERAT | MISSENSE | Ctg/Gtg | L782V | rne | C | C | | | | | | | | | | |
| NC_011770 | 2260126 | C | T | 52 | 0 | 2 | 0 | NON_SYNONYMOU | MODERAT | MISSENSE | Gcc/Acc | A716T | rne | T | T | | | | | | | | | | |
| NC_011770 | 2260137 | G | A | 51 | 0 | 2 | 0 | NON_SYNONYMOU | MODERAT | MISSENSE | gCc/gTc | A712V | rne | | | | | | | A | A | | | | |
| NC_011770 | 2260229 | ACCTTCGC( | A | 49 | 0 | 2 | 0 | CODON_CHANGE_ | MODERATE | | ggccgcgaa( | GREG678( | rne | | | | | A | A | | | | | | |
| NC_011770 | 2260596 | T | C | 64 | 0 | 2 | 0 | NON_SYNONYMOU | MODERAT | MISSENSE | aAg/aGg | K559R | rne | | | | | | | C | C | | | | |
| NC_011770 | 2260713 | C | T | 64 | 0 | 1 | 0 | NON_SYNONYMOU | MODERAT | MISSENSE | cGc/cAc | R520H | rne | | | | | | | T | | | | | |
| NC_011770 | 2261764 | T | C | 78 | 0 | 12 | 0 | NON_SYNONYMOU | MODERAT | MISSENSE | Agc/Ggc | S170G | rne | C | C | C | C | C | C | C | C | C | C | C | C |
| NC_011770 | 2261817 | C | G | 81 | 0 | 12 | 0 | NON_SYNONYMOU | MODERAT | MISSENSE | gGc/gCc | G152A | rne | G | G | G | G | G | G | G | G | G | G | G | G |
| NC_011770 | 3894817 | C | A | 72 | 0 | 2 | 0 | NON_SYNONYMOU | MODERAT | MISSENSE | Gcc/Tcc | A103S | rnhA | | | | | | | | | | | A | A |
| NC_011770 | 6601187 | G | A | 82 | 0 | 2 | 0 | NON_SYNONYMOU | MODERAT | MISSENSE | Cct/Tct | P127S | rnpA | A | A | | | | | | | | | | |
| NC_011770 | 6601489 | A | G | 63 | 0 | 12 | 0 | NON_SYNONYMOU | MODERAT | MISSENSE | aTc/aCc | I26T | rnpA | G | G | G | G | G | G | G | G | G | G | G | G |
| NC_011770 | 5871222 | T | TGCC | 72 | 0 | 1 | 0 | CODON_INSERTIOI | MODERATE | | ggc/ggcGC | G841GG | rnr | | | | | | | | TGCC | | | | |
| NC_011770 | 5871129 | G | A | 75 | 0 | 2 | 0 | NON_SYNONYMOU | MODERAT | MISSENSE | Ccg/Tcg | P873S | rnr | | | | | | | | | A | A | | |
| NC_011770 | 5871137 | A | G | 71 | 0 | 12 | 0 | NON_SYNONYMOU | MODERAT | MISSENSE | gTc/gCc | V870A | rnr | G | G | G | G | G | G | G | G | G | G | G | G |
| NC_011770 | 5871155 | T | C | 73 | 0 | 4 | 0 | NON_SYNONYMOU | MODERAT | MISSENSE | gAc/gGc | D864G | rnr | | | | | | | C | C | C | C | | |
| NC_011770 | 5871357 | C | T | 68 | 0 | 2 | 0 | NON_SYNONYMOU | MODERAT | MISSENSE | Gcc/Acc | A797T | rnr | | | | | | | T | T | | | | |
| NC_011770 | 5871465 | T | C | 80 | 0 | 10 | 0 | NON_SYNONYMOU | MODERAT | MISSENSE | Acg/Gcg | T761A | rnr | C | C | C | C | C | C | C | C | C | C | | |
| NC_011770 | 5872010 | G | A | 86 | 0 | 1 | 0 | NON_SYNONYMOU | MODERAT | MISSENSE | cCg/cTg | P579L | rnr | | | | | | | A | | | | | |
| NC_011770 | 5872953 | T | C | 81 | 0 | 2 | 0 | NON_SYNONYMOU | MODERAT | MISSENSE | Atc/Gtc | I265V | rnr | | | | | | | | | C | C | | |
| NC_011770 | 1107531 | G | A | 84 | 0 | 4 | 0 | NON_SYNONYMOU | MODERAT | MISSENSE | aCc/aTc | T118I | rocA1 | A | A | | | | | | | A | A | | |
| NC_011770 | 1107669 | C | T | 82 | 0 | 4 | 0 | NON_SYNONYMOU | MODERAT | MISSENSE | gGa/gAa | G72E | rocA1 | T | T | | | | | | | T | T | | |
| NC_011770 | 1107828 | A | G | 66 | 0 | 1 | 0 | NON_SYNONYMOU | MODERAT | MISSENSE | gTc/gCc | V19A | rocA1 | | | | | | | G | | | | | |
| NC_011770 | 6339397 | T | C | 79 | 0 | 2 | 0 | NON_SYNONYMOU | MODERAT | MISSENSE | aAc/aGc | N216S | rph | | | | | | | C | C | | | | |
| NC_011770 | 365316 | C | G | 71 | 0 | 2 | 0 | NON_SYNONYMOU | MODERAT | MISSENSE | Gaa/Caa | E89Q | rpiA | | | | | | | | | G | G | | |
| NC_011770 | 726099 | C | T | 84 | 0 | 2 | 0 | NON_SYNONYMOU | MODERAT | MISSENSE | gCt/gTt | A92V | rplC | | | | | | | | | T | T | | |
| NC_011770 | 726185 | A | G | 78 | 0 | 1 | 0 | NON_SYNONYMOU | MODERAT | MISSENSE | Acc/Gcc | T121A | rplC | | | | | | | G | | | | | |
| NC_011770 | 726490 | G | A | 82 | 0 | 12 | 0 | NON_SYNONYMOU | MODERAT | MISSENSE | aGc/aAc | S6N | rplD | A | A | A | A | A | A | A | A | A | A | A | A |
| NC_011770 | 711555 | A | G | 75 | 0 | 1 | 0 | NON_SYNONYMOU | MODERAT | MISSENSE | aAc/aGc | N86S | rplJ | | | | | | | | **G** | | | | |
| NC_011770 | 1544673 | C | T | 76 | 0 | 1 | 0 | NON_SYNONYMOU | MODERAT | MISSENSE | cCg/cTg | P43L | rpmJ | | | | | | | | | | | T | |
| NC_011770 | 715480 | T | C | 76 | 0 | 1 | 0 | NON_SYNONYMOU | MODERAT | MISSENSE | Tac/Cac | Y1005H | rpoB | | | | | | | C | | | | | |
| NC_011770 | 719171 | G | T | 88 | 0 | 4 | 0 | NON_SYNONYMOU | MODERAT | MISSENSE | gaG/gaT | E855D | rpoC | | | T | T | T | T | | | | | | |
| NC_011770 | 719805 | T | C | 80 | 0 | 12 | 0 | NON_SYNONYMOU | MODERAT | MISSENSE | Tgt/Cgt | C1067R | rpoC | C | C | C | C | C | C | C | C | C | C | C | C |
| NC_011770 | 719325 | C | T | 82 | 0 | 1 | 0 | NON_SYNONYMOU | MODERAT | MISSENSE | Cat/Tat | H907Y | rpoC | | | | | | | | **T** | | | | |

| | | | | | | | | | | | | | | | | | | | | | | | | | |
|---|---|---|---|---|---|---|---|---|---|---|---|---|---|---|---|---|---|---|---|---|---|---|---|---|---|
| NC_011770 | 625646 | C | T | 59 | 0 | 1 | 0 | NON_SYNONYMOU | MODERAT | MISSENSE | Gac/Aac | D570N | rpoD | | | | | | | T | | | | | |
| NC_011770 | 414674 | A | G | 89 | 0 | 1 | 0 | NON_SYNONYMOU | MODERAT | MISSENSE | aAc/aGc | N84S | rpoH | | | | | | | G | | | | | |
| NC_011770 | 415103 | A | T | 71 | 0 | 2 | 0 | NON_SYNONYMOU | MODERAT | MISSENSE | cAc/cTc | H227L | rpoH | | | | | | | | | | | T | T |
| NC_011770 | 5327291 | G | A | 78 | 0 | 1 | 0 | NON_SYNONYMOU | MODERAT | MISSENSE | cGc/cAc | R238H | rpoN | | A | | | | | | | | | | |
| NC_011770 | 5327595 | G | A | 76 | 0 | 1 | 0 | NON_SYNONYMOU | MODERAT | MISSENSE | atG/atA | M339I | rpoN | | | | | | | | | | | | A |
| NC_011770 | 5327414 | G | C | 74 | 0 | 1 | 0 | NON_SYNONYMOU | MODERAT | MISSENSE | cGg/cCg | R279P | rpoN | | | | | | | | C | | | | |
| NC_011770 | 5326810 | C | A | 71 | 0 | 2 | 0 | NON_SYNONYMOU | MODERAT | MISSENSE | Ctc/Atc | L78I | rpoN | | | | | | | | | | | A | A |
| NC_011770 | 5327360 | T | C | 75 | 0 | 1 | 0 | NON_SYNONYMOU | MODERAT | MISSENSE | cTc/cCc | L261P | rpoN | | | | | | | C | | | | | |
| NC_011770 | 6342059 | G | A | 89 | 0 | 1 | 0 | NON_SYNONYMOU | MODERAT | MISSENSE | Gac/Aac | D72N | rpoZ | | | | | | | A | | | | | |
| NC_011770 | 2053659 | C | T | 61 | 0 | 1 | 0 | NON_SYNONYMOU | MODERAT | MISSENSE | gCa/gTa | A207V | rpsA | | | | | | | | T | | | | |
| NC_011770 | 1485436 | G | A | 71 | 0 | 1 | 0 | NON_SYNONYMOU | MODERAT | MISSENSE | Gcc/Acc | A12T | rpsB | | | | | | | | | | | A | |
| NC_011770 | 4589886 | C | G | 66 | 0 | 2 | 0 | NON_SYNONYMOU | MODERAT | MISSENSE | gCc/gGc | A70G | rrmA | G | G | | | | | | | | | | |
| NC_011770 | 4589910 | C | T | 62 | 0 | 2 | 0 | NON_SYNONYMOU | MODERAT | MISSENSE | gCc/gTc | A78V | rrmA | | | | | | | T | T | | | | |
| NC_011770 | 4847240 | T | C | 73 | 0 | 2 | 0 | NON_SYNONYMOU | MODERAT | MISSENSE | gAa/gGa | E46G | rsmA | | | | | | | | | | | C | C |
| NC_011770 | 1522713 | T | C | 52 | 0 | 4 | 0 | INTRAGENIC | MODIFIER | | | | rsmZ | | | C | C | C | C | | | | | | |
| NC_011770 | 5470819 | G | A | 57 | 0 | 1 | 0 | NON_SYNONYMOU | MODERAT | MISSENSE | gGt/gAt | G333D | rtcA | | | | | | A | | | | | | |
| NC_011770 | 5470507 | T | A | 64 | 0 | 12 | 0 | NON_SYNONYMOU | MODERAT | MISSENSE | gTc/gAc | V229D | rtcA | A | A | A | A | A | A | A | A | A | A | A | A |
| NC_011770 | 5470668 | C | T | 60 | 0 | 2 | 0 | NON_SYNONYMOU | MODERAT | MISSENSE | Cac/Tac | H283Y | rtcA | | | | | | | | | T | T | | |
| NC_011770 | 5465369 | T | C | 61 | 0 | 1 | 0 | NON_SYNONYMOU | MODERAT | MISSENSE | cAg/cGg | Q178R | rtcR | C | | | | | | | | | | | |
| NC_011770 | 5465604 | C | T | 64 | 0 | 1 | 0 | NON_SYNONYMOU | MODERAT | MISSENSE | Gcc/Acc | A100T | rtcR | | | | | | | | T | | | | |
| NC_011770 | 5465106 | C | T | 65 | 0 | 2 | 0 | NON_SYNONYMOU | MODERAT | MISSENSE | Gtc/Atc | V266I | rtcR | | | | | | | | | T | T | | |
| NC_011770 | 5465582 | A | C | 63 | 0 | 10 | 0 | NON_SYNONYMOU | MODERAT | MISSENSE | cTc/cGc | L107R | rtcR | | | C | C | C | C | C | C | C | C | C | C |
| NC_011770 | 4815913 | T | C | 60 | 0 | 12 | 0 | NON_SYNONYMOU | MODERAT | MISSENSE | Atg/Gtg | M288V | rumA | C | C | C | C | C | C | C | C | C | C | C | C |
| NC_011770 | 4816045 | C | T | 77 | 0 | 2 | 0 | NON_SYNONYMOU | MODERAT | MISSENSE | Gcc/Acc | A244T | rumA | T | T | | | | | | | | | | |
| NC_011770 | 4816101 | G | A | 78 | 0 | 1 | 0 | NON_SYNONYMOU | MODERAT | MISSENSE | aCg/aTg | T225M | rumA | | | | | | | A | | | | | |
| NC_011770 | 4816582 | G | C | 74 | 0 | 2 | 0 | NON_SYNONYMOU | MODERAT | MISSENSE | Ctg/Gtg | L65V | rumA | C | C | | | | | | | | | | |
| NC_011770 | 4816725 | C | T | 73 | 0 | 4 | 0 | NON_SYNONYMOU | MODERAT | MISSENSE | cGc/cAc | R17H | rumA | | | T | T | T | T | | | | | | |
| NC_011770 | 6351127 | GC | G | 83 | 0 | 1 | 0 | FRAME_SHIFT | HIGH | | cag/ | Q16 | sadB | | | | | | | G | | | | | |
| NC_011770 | 6351757 | G | A | 78 | 0 | 1 | 0 | NON_SYNONYMOU | MODERAT | MISSENSE | Gaa/Aaa | E224K | sadB | | | | | | | A | | | | | |
| NC_011770 | 6352393 | C | A | 68 | 0 | 2 | 0 | NON_SYNONYMOU | MODERAT | MISSENSE | Ctg/Atg | L436M | sadB | | | | | | | A | A | | | | |
| NC_011770 | 5178264 | C | T | 57 | 0 | 4 | 0 | NON_SYNONYMOU | MODERAT | MISSENSE | Gcc/Acc | A386T | sbcB | | | T | T | T | T | | | | | | |
| NC_011770 | 5178443 | T | C | 60 | 0 | 8 | 0 | NON_SYNONYMOU | MODERAT | MISSENSE | cAa/cGa | Q326R | sbcB | | | C | C | C | C | C | C | C | C | | |
| NC_011770 | 5178454 | A | C | 62 | 0 | 8 | 0 | NON_SYNONYMOU | MODERAT | MISSENSE | gaT/gaG | D322E | sbcB | | | C | C | C | C | C | C | C | C | | |
| NC_011770 | 5178704 | G | A | 50 | 0 | 10 | 0 | NON_SYNONYMOU | MODERAT | MISSENSE | gCc/gTc | A239V | sbcB | | | A | A | A | A | A | A | A | A | A | A |
| NC_011770 | 5134456 | A | T | 81 | 0 | 2 | 0 | NON_SYNONYMOU | MODERAT | MISSENSE | Ata/Tta | I113L | sbcD | | | | | T | T | | | | | | |
| NC_011770 | 5134897 | A | G | 75 | 0 | 12 | 0 | NON_SYNONYMOU | MODERAT | MISSENSE | Atc/Gtc | I260V | sbcD | G | G | G | G | G | G | G | G | G | G | G | G |
| NC_011770 | 5135299 | A | T | 66 | 0 | 2 | 0 | NON_SYNONYMOU | MODERAT | MISSENSE | Agc/Tgc | S394C | sbcD | | | | | | | | | | | T | T |
| NC_011770 | 3091961 | G | A | 69 | 0 | 4 | 0 | NON_SYNONYMOU | MODERAT | MISSENSE | Gtc/Atc | V46I | sdaA | | | | | | | A | A | | | A | A |
| NC_011770 | 3092058 | C | A | 65 | 0 | 2 | 0 | NON_SYNONYMOU | MODERAT | MISSENSE | aCg/aAg | T78K | sdaA | | | | | | | | | A | A | | |
| NC_011770 | 3092379 | C | T | 65 | 0 | 6 | 0 | NON_SYNONYMOU | MODERAT | MISSENSE | gCc/gTc | A185V | sdaA | T | T | T | T | T | T | | | | | | |
| NC_011770 | 3092453 | A | G | 70 | 0 | 2 | 0 | NON_SYNONYMOU | MODERAT | MISSENSE | Acc/Gcc | T210A | sdaA | G | G | | | | | | | | | | |
| NC_011770 | 3092799 | C | T | 77 | 0 | 2 | 0 | NON_SYNONYMOU | MODERAT | MISSENSE | gCg/gTg | A325V | sdaA | | | | | | | | | | | T | T |
| NC_011770 | 3092061 | T | C | 66 | 0 | 1 | 0 | NON_SYNONYMOU | MODERAT | MISSENSE | gTg/gCg | V79A | sdaA | | | | | | | | C | | | | |
| NC_011770 | 6392915 | G | A | 102 | 0 | 2 | 0 | NON_SYNONYMOU | MODERAT | MISSENSE | Gct/Act | A32T | sdaB | | | | | | | A | A | | | | |
| NC_011770 | 6393098 | G | C | 106 | 0 | 4 | 0 | NON_SYNONYMOU | MODERAT | MISSENSE | Gat/Cat | D93H | sdaB | | | C | C | C | C | | | | | | |
| NC_011770 | 6393435 | C | T | 99 | 0 | 4 | 0 | NON_SYNONYMOU | MODERAT | MISSENSE | aCc/aTc | T205I | sdaB | | | T | T | T | T | | | | | | |
| NC_011770 | 4145425 | T | C | 93 | 0 | 2 | 0 | NON_SYNONYMOU | MODERAT | MISSENSE | gAa/gGa | E422G | sdhA | C | C | | | | | | | | | | |
| NC_011770 | 4145596 | T | C | 98 | 0 | 1 | 0 | NON_SYNONYMOU | MODERAT | MISSENSE | aAc/aGc | N365S | sdhA | | | | | | | C | | | | | |
| NC_011770 | 4145146 | G | A | 98 | 0 | 1 | 0 | NON_SYNONYMOU | MODERAT | MISSENSE | gCg/gTg | A515V | sdhA | | | | A | | | | | | | | |
| NC_011770 | 5059830 | C | T | 94 | 0 | 8 | 0 | NON_SYNONYMOU | MODERAT | MISSENSE | Gaa/Aaa | E444K | sdsA1 | T | T | T | T | T | T | | | T | T | | |
| NC_011770 | 5059874 | C | G | 88 | 0 | 2 | 0 | NON_SYNONYMOU | MODERAT | MISSENSE | aGc/aCc | S429T | sdsA1 | | | | | | | G | G | | | | |
| NC_011770 | 5060060 | A | T | 100 | 0 | 2 | 0 | NON_SYNONYMOU | MODERAT | MISSENSE | tTc/tAc | F367Y | sdsA1 | | | | | | | T | T | | | | |
| NC_011770 | 5060898 | G | C | 53 | 0 | 4 | 0 | NON_SYNONYMOU | MODERAT | MISSENSE | Cgc/Ggc | R88G | sdsA1 | | | C | C | C | C | | | | | | |
| NC_011770 | 5061001 | T | G | 56 | 0 | 8 | 0 | NON_SYNONYMOU | MODERAT | MISSENSE | gaA/gaC | E53D | sdsA1 | G | G | G | G | G | G | G | G | | | | |
| NC_011770 | 1251935 | G | A | 83 | 0 | 2 | 0 | NON_SYNONYMOU | MODERAT | MISSENSE | Gac/Aac | D68N | secD | A | A | | | | | | | | | | |
| NC_011770 | 1252313 | G | A | 76 | 0 | 2 | 0 | NON_SYNONYMOU | MODERAT | MISSENSE | Gcc/Acc | A194T | secD | | | | | | | | | A | A | | |
| NC_011770 | 5727737 | C | T | 88 | 0 | 2 | 0 | NON_SYNONYMOU | MODERAT | MISSENSE | aGg/aAg | R205K | selA | | | | | | | | | T | T | | |
| NC_011770 | 5727962 | G | A | 86 | 0 | 12 | 0 | NON_SYNONYMOU | MODERAT | MISSENSE | aCc/aTc | T130I | selA | A | A | A | A | A | A | A | A | A | A | A | A |
| NC_011770 | 5728205 | G | A | 96 | 0 | 2 | 0 | NON_SYNONYMOU | MODERAT | MISSENSE | gCc/gTc | A49V | selA | | | | | | | | | A | A | | |
| NC_011770 | 5725450 | G | A | 76 | 0 | 2 | 0 | NON_SYNONYMOU | MODERAT | MISSENSE | Cac/Tac | H500Y | selB | | | | | | | A | A | | | | |
| NC_011770 | 5725513 | G | C | 76 | 0 | 2 | 0 | NON_SYNONYMOU | MODERAT | MISSENSE | Ctc/Gtc | L479V | selB | | | | | | | | | C | C | | |
| NC_011770 | 5725675 | G | T | 64 | 0 | 4 | 0 | NON_SYNONYMOU | MODERAT | MISSENSE | Ccc/Acc | P425T | selB | | | T | T | T | T | | | | | | |

| | | | | | | | | | | | | | | | | | | | | | | | | | |
|---|---|---|---|---|---|---|---|---|---|---|---|---|---|---|---|---|---|---|---|---|---|---|---|---|---|
| NC_011770 | 4080351 | C | T | 90 | 0 | 2 | 0 | NON_SYNONYMOU | MODERAT | MISSENSE | Gag/Aag | E196K | selD | | | | | | | | | T | T | | |
| NC_011770 | 2683936 | C | T | 79 | 0 | 2 | 0 | NON_SYNONYMOU | MODERAT | MISSENSE | Cac/Tac | H146Y | serS | T | T | | | | | | | | | | |
| NC_011770 | 2683558 | A | G | 87 | 0 | 1 | 0 | NON_SYNONYMOU | MODERAT | MISSENSE | Acc/Gcc | T20A | serS | G | | | | | | | | | | | |
| NC_011770 | 3947073 | A | G | 82 | 0 | 1 | 0 | NON_SYNONYMOU | MODERAT | MISSENSE | Tac/Cac | Y148H | sigX | | | | | | | G | | | | | |
| NC_011770 | 3147978 | C | A | 32 | 8 | 2 | 0 | NON_SYNONYMOU | MODERAT | MISSENSE | tgG/tgT | W310C | sip | . | . | | | A | A | . | . | . | . | . | . |
| NC_011770 | 3148861 | A | T | 31 | 8 | 4 | 0 | NON_SYNONYMOU | MODERAT | MISSENSE | cTt/cAt | L16H | sip | . | . | T | T | T | T | . | . | . | . | . | . |
| NC_011770 | 2198519 | G | T | 57 | 0 | 12 | 0 | NON_SYNONYMOU | MODERAT | MISSENSE | gGc/gTc | G25V | snr1 | T | T | T | T | T | T | T | T | T | T | T | T |
| NC_011770 | 2198725 | A | G | 50 | 0 | 6 | 0 | NON_SYNONYMOU | MODERAT | MISSENSE | Atc/Gtc | I94V | snr1 | G | G | G | G | G | G | | | | | | |
| NC_011770 | 2198753 | G | A | 50 | 0 | 4 | 0 | NON_SYNONYMOU | MODERAT | MISSENSE | aGc/aAc | S103N | snr1 | | | A | A | A | A | | | | | | |
| NC_011770 | 2199803 | C | T | 59 | 0 | 2 | 0 | NON_SYNONYMOU | MODERAT | MISSENSE | tCc/tTc | S453F | snr1 | | | | | | | | | T | T | | |
| NC_011770 | 5331430 | G | GC | 69 | 0 | 1 | 0 | FRAME_SHIFT | HIGH | | ggc/ggGc | G110G? | sodM | | | | | | | | GC | | | | |
| NC_011770 | 5331550 | C | CG | 75 | 0 | 1 | 0 | FRAME_SHIFT | HIGH | | ccg/ccCg | P70P? | sodM | | | | | | | CG | | | | | |
| NC_011770 | 5331148 | C | T | 92 | 0 | 2 | 0 | SPLICE_SITE_REGIC | LOW | SILENT | taG/taA | *204 | sodM | | | | | | | | | T | T | | |
| NC_011770 | 5331434 | C | T | 62 | 0 | 2 | 0 | NON_SYNONYMOU | MODERAT | MISSENSE | gGg/gAg | G109E | sodM | | | | | | | | | | | T | T |
| NC_011770 | 6432968 | T | A | 78 | 0 | 4 | 0 | NON_SYNONYMOU | MODERAT | MISSENSE | Tcc/Acc | S8T | soxA | | | A | A | A | A | | | | | | |
| NC_011770 | 6433629 | C | T | 62 | 0 | 2 | 0 | NON_SYNONYMOU | MODERAT | MISSENSE | gCc/gTc | A228V | soxA | T | T | | | | | | | | | | |
| NC_011770 | 6434021 | G | C | 87 | 0 | 2 | 0 | NON_SYNONYMOU | MODERAT | MISSENSE | Gcc/Ccc | A359P | soxA | | | | | | | C | C | | | | |
| NC_011770 | 6434022 | C | T | 87 | 0 | 2 | 0 | NON_SYNONYMOU | MODERAT | MISSENSE | gCc/gTc | A359V | soxA | | | | | | | T | T | | | | |
| NC_011770 | 6434042 | A | G | 85 | 0 | 8 | 0 | NON_SYNONYMOU | MODERAT | MISSENSE | Atc/Gtc | I366V | soxA | G | G | G | G | G | G | | | | | G | G |
| NC_011770 | 6434351 | G | A | 73 | 0 | 2 | 0 | NON_SYNONYMOU | MODERAT | MISSENSE | Gag/Aag | E469K | soxA | A | A | | | | | | | | | | |
| NC_011770 | 6434414 | A | G | 75 | 0 | 6 | 0 | NON_SYNONYMOU | MODERAT | MISSENSE | Acc/Gcc | T490A | soxA | | | G | G | G | G | G | G | | | | |
| NC_011770 | 6434459 | T | C | 80 | 0 | 2 | 0 | NON_SYNONYMOU | MODERAT | MISSENSE | Tcc/Ccc | S505P | soxA | | | | | | | C | C | | | | |
| NC_011770 | 6435444 | T | C | 89 | 0 | 1 | 0 | NON_SYNONYMOU | MODERAT | MISSENSE | gTg/gCg | V833A | soxA | | | | | | | C | | | | | |
| NC_011770 | 6432294 | G | A | 83 | 0 | 1 | 0 | NON_SYNONYMOU | MODERAT | MISSENSE | Ggc/Agc | G337S | soxB | | | | | | | A | | | | | |
| NC_011770 | 6432322 | C | T | 78 | 0 | 1 | 0 | NON_SYNONYMOU | MODERAT | MISSENSE | gCc/gTc | A346V | soxB | | | | | | | T | | | | | |
| NC_011770 | 6432729 | G | A | 66 | 0 | 2 | 0 | NON_SYNONYMOU | MODERAT | MISSENSE | Gcc/Acc | A34T | soxD | | | | | | | | | | | A | A |
| NC_011770 | 6432924 | C | G | 75 | 0 | 6 | 0 | NON_SYNONYMOU | MODERAT | MISSENSE | Ccc/Gcc | P99A | soxD | | | G | G | G | G | | | | | G | G |
| NC_011770 | 6436235 | A | G | 71 | 0 | 12 | 0 | NON_SYNONYMOU | MODERAT | MISSENSE | Acc/Gcc | T60A | soxG | G | G | G | G | G | G | G | G | G | G | G | G |
| NC_011770 | 6436386 | G | A | 65 | 0 | 2 | 0 | NON_SYNONYMOU | MODERAT | MISSENSE | cGc/cAc | R110H | soxG | A | A | | | | | | | | | | |
| NC_011770 | 1377584 | C | T | 61 | 0 | 2 | 0 | NON_SYNONYMOU | MODERAT | MISSENSE | Gcc/Acc | A593T | spdH | | | | | | | | | | | T | T |
| NC_011770 | 1378162 | C | T | 73 | 0 | 4 | 0 | NON_SYNONYMOU | MODERAT | MISSENSE | cGc/cAc | R400H | spdH | | | | | | | T | T | T | T | | |
| NC_011770 | 1378169 | C | T | 72 | 0 | 2 | 0 | NON_SYNONYMOU | MODERAT | MISSENSE | Gcc/Acc | A398T | spdH | T | T | | | | | | | | | | |
| NC_011770 | 1378627 | C | T | 73 | 0 | 2 | 0 | NON_SYNONYMOU | MODERAT | MISSENSE | aGc/aAc | S245N | spdH | | | | | | | | | | | T | T |
| NC_011770 | 5765764 | C | T | 16 | 0 | 1 | 0 | NON_SYNONYMOU | MODERAT | MISSENSE | Ggc/Agc | G11S | speA | | | | | | | | | | | | T |
| NC_011770 | 5765652 | G | A | 31 | 0 | 1 | 0 | NON_SYNONYMOU | MODERAT | MISSENSE | gCc/gTc | A48V | speA | | | | | | | | A | | | | |
| NC_011770 | 5765493 | C | T | 39 | 0 | 2 | 0 | NON_SYNONYMOU | MODERAT | MISSENSE | aGc/aAc | S101N | speA | | | | | | | | | T | T | | |
| NC_011770 | 685656 | G | A | 73 | 0 | 4 | 0 | NON_SYNONYMOU | MODERAT | MISSENSE | Gcg/Acg | A239T | speD | A | A | | | | | A | A | | | | |
| NC_011770 | 4031113 | C | T | 75 | 0 | 2 | 0 | NON_SYNONYMOU | MODERAT | MISSENSE | cGc/cAc | R236H | speE | | | | | | | | | T | T | | |
| NC_011770 | 4031473 | T | G | 74 | 0 | 6 | 0 | NON_SYNONYMOU | MODERAT | MISSENSE | gAg/gCg | E116A | speE | | | G | G | G | G | | | | | G | G |
| NC_011770 | 6343534 | C | G | 87 | 0 | 1 | 0 | NON_SYNONYMOU | MODERAT | MISSENSE | Ccg/Gcg | P453A | spoT | | | | | | G | | | | | | |
| NC_011770 | 332416 | G | A | 71 | 0 | 1 | 0 | NON_SYNONYMOU | MODERAT | MISSENSE | Gtc/Atc | V338I | spuD | | | | **A** | | | | | | | | |
| NC_011770 | 332985 | GC | G | 69 | 0 | 1 | 0 | FRAME_SHIFT | HIGH | | ccc/ | P21 | spuE | | | | G | | | | | | | | |
| NC_011770 | 333776 | G | T | 92 | 0 | 4 | 0 | NON_SYNONYMOU | MODERAT | MISSENSE | Gcc/Tcc | A284S | spuE | | | | | | | | | T | T | T | T |
| NC_011770 | 333827 | G | T | 94 | 0 | 6 | 0 | NON_SYNONYMOU | MODERAT | MISSENSE | Gcc/Tcc | A301S | spuE | | | T | T | T | T | T | T | | | | |
| NC_011770 | 334310 | A | C | 103 | 0 | 12 | 0 | NON_SYNONYMOU | MODERAT | MISSENSE | Agg/Tgg | R78W | spuF | C | C | C | C | C | C | T | C | C | C | C | C |
| NC_011770 | 334571 | C | T | 109 | 0 | 1 | 0 | NON_SYNONYMOU | MODERAT | MISSENSE | Cgc/Tgc | R165C | spuF | | | | | | | | **T** | | | | |
| NC_011770 | 335264 | C | T | 115 | 0 | 1 | 0 | NON_SYNONYMOU | MODERAT | MISSENSE | cCc/cTc | P2L | spuG | | | | | | | T | | | | | |
| NC_011770 | 335277 | T | G | 117 | 0 | 2 | 0 | NON_SYNONYMOU | MODERAT | MISSENSE | caT/caG | H6Q | spuG | G | G | | | | | | | | | | |
| NC_011770 | 336810 | G | A | 88 | 0 | 2 | 0 | NON_SYNONYMOU | MODERAT | MISSENSE | gGc/gAc | G196D | spuH | A | A | | | | | | | | | | |
| NC_011770 | 5294960 | G | A | 77 | 0 | 4 | 0 | NON_SYNONYMOU | MODERAT | MISSENSE | Ccg/Tcg | P104S | sspB | | | A | A | A | A | | | | | | |
| NC_011770 | 1746187 | G | A | 72 | 0 | 2 | 0 | NON_SYNONYMOU | MODERAT | MISSENSE | cGc/cAc | R74H | ssuB | | | | | | | | | A | A | | |
| NC_011770 | 1746270 | C | T | 63 | 0 | 1 | 0 | NON_SYNONYMOU | MODERAT | MISSENSE | Ccg/Tcg | P102S | ssuB | | | | | | | T | | | | | |
| NC_011770 | 1746334 | T | A | 62 | 0 | 4 | 0 | NON_SYNONYMOU | MODERAT | MISSENSE | cTg/cAg | L123Q | ssuB | | | | | | | | | A | A | G | G |
| NC_011770 | 1745120 | G | A | 82 | 0 | 1 | 0 | NON_SYNONYMOU | MODERAT | MISSENSE | Gtg/Atg | V370M | ssuD | | | | | | | | A | | | | |
| NC_011770 | 2245024 | A | G | 74 | 0 | 1 | 0 | NON_SYNONYMOU | MODERAT | MISSENSE | aAg/aGg | K30R | sth | | | | | | | | G | | | | |
| NC_011770 | 4044280 | A | C | 91 | 0 | 12 | 0 | NON_SYNONYMOU | MODERAT | MISSENSE | gaT/gaG | D194E | stk1 | C | C | C | C | C | C | C | C | C | C | C | C |
| NC_011770 | 4044353 | A | G | 88 | 0 | 2 | 0 | NON_SYNONYMOU | MODERAT | MISSENSE | gTt/gCt | V170A | stk1 | | | | | | | | | | | G | G |
| NC_011770 | 4044591 | A | T | 76 | 0 | 2 | 0 | NON_SYNONYMOU | MODERAT | MISSENSE | Tcg/Acg | S91T | stk1 | | | | | | | | | | | T | T |
| NC_011770 | 4045480 | G | A | 68 | 0 | 2 | 0 | NON_SYNONYMOU | MODERAT | MISSENSE | gCg/gTg | A36V | stp1 | | | | | | | | | | | A | A |
| NC_011770 | 3849400 | G | A | 78 | 0 | 2 | 0 | NON_SYNONYMOU | MODERAT | MISSENSE | Gca/Aca | A38T | str | | | | | | | A | A | | | | |
| NC_011770 | 3849614 | G | A | 80 | 0 | 2 | 0 | NON_SYNONYMOU | MODERAT | MISSENSE | cGc/cAc | R109H | str | A | A | | | | | | | | | | |
| NC_011770 | 3849949 | A | G | 73 | 0 | 2 | 0 | NON_SYNONYMOU | MODERAT | MISSENSE | Atc/Gtc | I221V | str | | | | | | | | | | | G | G |

| | | | | | | | | | | | | | | | | | | | | | | | | | |
|---|---|---|---|---|---|---|---|---|---|---|---|---|---|---|---|---|---|---|---|---|---|---|---|---|---|
| NC_011770 | 3849986 | A | T | 79 | 0 | 2 | 0 | NON_SYNONYMOU | MODERAT | MISSENSE | cAa/cTa | Q233L | str | | | | | | | | | | | T | T |
| NC_011770 | 3850056 | A | C | 80 | 0 | 2 | 0 | NON_SYNONYMOU | MODERAT | MISSENSE | gaA/gaC | E256D | str | | | | | | | | | | | C | C |
| NC_011770 | 4140819 | AGCG | A | 54 | 0 | 4 | 0 | CODON_CHANGE_ | MODERATE | | gccgct/gct | AA82A | sucB | | | A | A | A | A | | | | | | |
| NC_011770 | 4136036 | T | C | 63 | 0 | 1 | 0 | NON_SYNONYMOU | MODERAT | MISSENSE | gAc/gGc | D268G | sucD | | | | | | | | C | | | | |
| NC_011770 | 644298 | C | T | 72 | 0 | 4 | 0 | NON_SYNONYMOU | MODERAT | MISSENSE | aGg/aAg | R202K | surA | T | T | | | | | | | | | T | T |
| NC_011770 | 644395 | T | A | 81 | 0 | 2 | 0 | NON_SYNONYMOU | MODERAT | MISSENSE | Atg/Ttg | M170L | surA | | | | | | | | | | | A | A |
| NC_011770 | 644473 | T | C | 86 | 0 | 2 | 0 | NON_SYNONYMOU | MODERAT | MISSENSE | Agc/Ggc | S144G | surA | | | | | | | C | C | | | | |
| NC_011770 | 1518964 | C | G | 77 | 0 | 4 | 0 | NON_SYNONYMOU | MODERAT | MISSENSE | aCc/aGc | T12S | surE | | | | | | | | | G | G | G | G |
| NC_011770 | 1519073 | A | C | 74 | 0 | 12 | 0 | NON_SYNONYMOU | MODERAT | MISSENSE | gaA/gaC | E48D | surE | C | C | C | C | C | C | C | C | C | C | C | C |
| NC_011770 | 5166880 | C | T | 83 | 0 | 2 | 0 | NON_SYNONYMOU | MODERAT | MISSENSE | gGc/gAc | G11D | tadA | | | | | | | | | | | T | T |
| NC_011770 | 5165376 | T | C | 86 | 0 | 2 | 0 | NON_SYNONYMOU | MODERAT | MISSENSE | Atc/Gtc | I92V | tadB | | | | | | | C | C | | | | |
| NC_011770 | 5165378 | G | A | 90 | 0 | 2 | 0 | NON_SYNONYMOU | MODERAT | MISSENSE | aCc/aTc | T91I | tadB | | | | | | | A | A | | | | |
| NC_011770 | 5165391 | C | G | 92 | 0 | 2 | 0 | NON_SYNONYMOU | MODERAT | MISSENSE | Ggc/Cgc | G87R | tadB | | | | | | | G | G | | | | |
| NC_011770 | 5163195 | T | C | 75 | 0 | 6 | 0 | NON_SYNONYMOU | MODERAT | MISSENSE | gAc/gGc | D218G | tadD | C | C | | | | | | | C | C | C | C |
| NC_011770 | 5163240 | G | A | 76 | 0 | 2 | 0 | NON_SYNONYMOU | MODERAT | MISSENSE | gCg/gTg | A203V | tadD | | | | | | | | | | | A | A |
| NC_011770 | 5163550 | G | C | 71 | 0 | 4 | 0 | NON_SYNONYMOU | MODERAT | MISSENSE | Ccc/Gcc | P100A | tadD | | | C | C | C | C | | | | | | |
| NC_011770 | 5162343 | TGCC | T | 75 | 0 | 5 | 0 | CODON_DELETION | MODERATE | | ggc/ | G145 | tadG | T | T | | | | | T | T | | | T | |
| NC_011770 | 5161329 | T | C | 64 | 0 | 4 | 0 | NON_SYNONYMOU | MODERAT | MISSENSE | Acc/Gcc | T484A | tadG | | | C | C | C | C | | | | | | |
| NC_011770 | 5161497 | C | T | 66 | 0 | 4 | 0 | NON_SYNONYMOU | MODERAT | MISSENSE | Gtc/Atc | V428I | tadG | | | T | T | T | T | | | | | | |
| NC_011770 | 5161739 | T | C | 71 | 0 | 2 | 0 | NON_SYNONYMOU | MODERAT | MISSENSE | cAg/cGg | Q347R | tadG | | | | | | | C | C | | | | |
| NC_011770 | 5161983 | T | C | 79 | 0 | 6 | 0 | NON_SYNONYMOU | MODERAT | MISSENSE | Acg/Gcg | T266A | tadG | C | C | | | | | | | C | C | C | C |
| NC_011770 | 5162112 | T | C | 81 | 0 | 2 | 0 | NON_SYNONYMOU | MODERAT | MISSENSE | Atc/Gtc | I223V | tadG | | | | | | | C | C | | | | |
| NC_011770 | 5162127 | C | T | 82 | 0 | 2 | 0 | NON_SYNONYMOU | MODERAT | MISSENSE | Gat/Aat | D218N | tadG | | | | | | | T | T | | | | |
| NC_011770 | 5162130 | C | G | 83 | 0 | 2 | 0 | NON_SYNONYMOU | MODERAT | MISSENSE | Gcg/Ccg | A217P | tadG | | | | | | | G | G | | | | |
| NC_011770 | 5162200 | C | G | 81 | 0 | 8 | 0 | NON_SYNONYMOU | MODERAT | MISSENSE | caG/caC | Q193H | tadG | G | G | | | | | G | G | G | G | G | G |
| NC_011770 | 5162334 | C | T | 79 | 0 | 2 | 0 | NON_SYNONYMOU | MODERAT | MISSENSE | Gtc/Atc | V149I | tadG | | | | | | | T | T | | | | |
| NC_011770 | 5162409 | T | C | 74 | 0 | 8 | 0 | NON_SYNONYMOU | MODERAT | MISSENSE | Aag/Gag | K124E | tadG | C | C | | | | | C | C | C | C | C | C |
| NC_011770 | 5162723 | G | A | 75 | 0 | 2 | 0 | NON_SYNONYMOU | MODERAT | MISSENSE | aCc/aTc | T19I | tadG | A | A | | | | | | | | | | |
| NC_011770 | 13615 | G | A | 67 | 0 | 2 | 0 | NON_SYNONYMOU | MODERAT | MISSENSE | Gcg/Acg | A30T | tag | A | A | | | | | | | | | | |
| NC_011770 | 13873 | G | A | 74 | 0 | 2 | 0 | NON_SYNONYMOU | MODERAT | MISSENSE | Gtc/Atc | V116I | tag | | | | | | | A | A | | | | |
| NC_011770 | 2445629 | T | C | 66 | 0 | 12 | 0 | NON_SYNONYMOU | MODERAT | MISSENSE | Act/Gct | T270A | tal | C | C | C | C | C | C | C | C | C | C | C | C |
| NC_011770 | 2445874 | A | G | 70 | 0 | 6 | 0 | NON_SYNONYMOU | MODERAT | MISSENSE | gTc/gCc | V188A | tal | | | G | G | G | G | | | | | G | G |
| NC_011770 | 2445962 | C | T | 65 | 0 | 2 | 0 | NON_SYNONYMOU | MODERAT | MISSENSE | Gcc/Acc | A159T | tal | | | | | | | | | T | T | | |
| NC_011770 | 2446024 | A | G | 62 | 0 | 2 | 0 | NON_SYNONYMOU | MODERAT | MISSENSE | aTt/aCt | I138T | tal | | | | | | | G | G | | | | |
| NC_011770 | 6041752 | T | C | 53 | 0 | 8 | 0 | NON_SYNONYMOU | MODERAT | MISSENSE | Tcg/Ccg | S88P | tatB | C | C | C | C | C | C | | | | | C | C |
| NC_011770 | 6041761 | C | A | 51 | 0 | 8 | 0 | NON_SYNONYMOU | MODERAT | MISSENSE | Ccc/Acc | P91T | tatB | A | A | A | A | A | A | | | | | A | A |
| NC_011770 | 6041765 | A | C | 51 | 0 | 8 | 0 | NON_SYNONYMOU | MODERAT | MISSENSE | cAg/cCg | Q92P | tatB | C | C | C | C | C | C | | | | | C | C |
| NC_011770 | 6041770 | C | T | 51 | 0 | 2 | 0 | NON_SYNONYMOU | MODERAT | MISSENSE | Ccg/Tcg | P94S | tatB | T | T | | | | | | | | | | |
| NC_011770 | 6041773 | A | G | 49 | 0 | 8 | 0 | NON_SYNONYMOU | MODERAT | MISSENSE | Acg/Gcg | T95A | tatB | G | G | G | G | G | G | | | | | G | G |
| NC_011770 | 6041777 | T | G | 51 | 0 | 8 | 0 | NON_SYNONYMOU | MODERAT | MISSENSE | gTc/gGc | V96G | tatB | G | G | C | C | C | C | | | | | C | C |
| NC_011770 | 6041783 | G | A | 51 | 0 | 2 | 0 | NON_SYNONYMOU | MODERAT | MISSENSE | aGc/aAc | S98N | tatB | A | A | | | | | | | | | | |
| NC_011770 | 6041803 | A | G | 48 | 0 | 2 | 0 | NON_SYNONYMOU | MODERAT | MISSENSE | Acg/Gcg | T105A | tatB | G | G | | | | | | | | | | |
| NC_011770 | 1122355 | G | A | 72 | 0 | 2 | 0 | NON_SYNONYMOU | MODERAT | MISSENSE | Ggc/Agc | G29S | tauD | | | | | A | A | | | | | | |
| NC_011770 | 2386305 | A | G | 46 | 0 | 8 | 0 | NON_SYNONYMOU | MODERAT | MISSENSE | aAg/aGg | K146R | tesA | G | G | G | G | G | G | | | G | G | | |
| NC_011770 | 1251279 | C | A | 79 | 0 | 4 | 0 | NON_SYNONYMOU | MODERAT | MISSENSE | cCg/cAg | P369Q | tgt | | | A | A | A | A | | | | | | |
| NC_011770 | 5914747 | T | A | 89 | 0 | 4 | 0 | NON_SYNONYMOU | MODERAT | MISSENSE | Acc/Tcc | T412S | thiC | | | A | A | A | A | | | | | | |
| NC_011770 | 5915580 | G | A | 58 | 0 | 1 | 0 | NON_SYNONYMOU | MODERAT | MISSENSE | cCg/cTg | P134L | thiC | | | | | | | A | | | | | |
| NC_011770 | 1079828 | C | T | 72 | 0 | 4 | 0 | NON_SYNONYMOU | MODERAT | MISSENSE | aGc/aAc | S194N | thiD | | | T | T | T | T | | | | | | |
| NC_011770 | 1078976 | C | T | 84 | 0 | 2 | 0 | NON_SYNONYMOU | MODERAT | MISSENSE | Gcc/Acc | A209T | thiE | | | | | | | | | | | T | T |
| NC_011770 | 1079137 | G | T | 78 | 0 | 12 | 0 | NON_SYNONYMOU | MODERAT | MISSENSE | cCc/cAc | P155H | thiE | T | T | T | T | T | T | T | T | T | T | T | T |
| NC_011770 | 1079153 | C | T | 77 | 0 | 2 | 0 | NON_SYNONYMOU | MODERAT | MISSENSE | Gcc/Acc | A150T | thiE | | | | | | | T | T | | | | |
| NC_011770 | 1079420 | A | G | 73 | 0 | 12 | 0 | NON_SYNONYMOU | MODERAT | MISSENSE | Tac/Cac | Y61H | thiE | G | G | G | G | G | G | G | G | G | G | G | G |
| NC_011770 | 417641 | A | G | 90 | 0 | 2 | 0 | NON_SYNONYMOU | MODERAT | MISSENSE | gAc/gGc | D61G | thiG | | | | | | | | | G | G | | |
| NC_011770 | 1004920 | G | A | 71 | 0 | 1 | 0 | NON_SYNONYMOU | MODERAT | MISSENSE | cGc/cAc | R255H | thiL | | | | | | | A | | | | | |
| NC_011770 | 1005010 | C | T | 73 | 0 | 4 | 0 | NON_SYNONYMOU | MODERAT | MISSENSE | aCg/aTg | T285M | thiL | T | T | | | | | | | T | T | | |
| NC_011770 | 6523538 | A | G | 55 | 0 | 4 | 0 | NON_SYNONYMOU | MODERAT | MISSENSE | Acc/Gcc | T233A | thrB | | | G | G | G | G | | | | | | |
| NC_011770 | 1350110 | G | T | 83 | 0 | 6 | 0 | NON_SYNONYMOU | MODERAT | MISSENSE | Gcg/Tcg | A266S | thrC | | | T | T | T | T | | | | | T | T |
| NC_011770 | 1350446 | T | G | 97 | 0 | 2 | 0 | NON_SYNONYMOU | MODERAT | MISSENSE | Tcc/Gcc | S378A | thrC | | | | | | | | | | | G | G |
| NC_011770 | 1350449 | T | A | 99 | 0 | 2 | 0 | NON_SYNONYMOU | MODERAT | MISSENSE | Tgt/Agt | C379S | thrC | | | | | | | A | A | | | | |
| NC_011770 | 3971100 | T | C | 57 | 0 | 2 | 0 | NON_SYNONYMOU | MODERAT | MISSENSE | gAc/gGc | D118G | thrH | | | | | | | | | C | C | | |
| NC_011770 | 599333 | T | A | 89 | 0 | 2 | 0 | NON_SYNONYMOU | MODERAT | MISSENSE | Tcc/Acc | S298T | tktA | | | | | | | | | | | A | A |
| NC_011770 | 599939 | G | A | 89 | 0 | 2 | 0 | NON_SYNONYMOU | MODERAT | MISSENSE | Gcg/Acg | A500T | tktA | | | | | | | A | A | | | | |

| | | | | | | | | | | | | | | | | | | | | | | | | | |
|---|---|---|---|---|---|---|---|---|---|---|---|---|---|---|---|---|---|---|---|---|---|---|---|---|---|
| NC_011770 | 599990 | A | G | 83 | 0 | 6 | 0 | NON_SYNONYMOU | MODERAT | MISSENSE | Atc/Gtc | I517V | tktA | | | G | G | G | G | | | G | G | | |
| NC_011770 | 2273708 | C | A | 75 | 0 | 2 | 0 | NON_SYNONYMOU | MODERAT | MISSENSE | Ctc/Atc | L78I | tmk | | | | | | | A | A | | | | |
| NC_011770 | 2273736 | C | T | 72 | 0 | 1 | 0 | NON_SYNONYMOU | MODERAT | MISSENSE | gCg/gTg | A87V | tmk | | | | | | | T | | | | | |
| NC_011770 | 4781516 | C | G | 92 | 0 | 2 | 0 | NON_SYNONYMOU | MODERAT | MISSENSE | aGc/aCc | S276T | tolB | | | | | | | | | | | G | G |
| NC_011770 | 4783979 | C | T | 85 | 0 | 1 | 0 | NON_SYNONYMOU | MODERAT | MISSENSE | Gcc/Acc | A189T | tolQ | | | T | | | | | | | | | |
| NC_011770 | 4783865 | G | A | 85 | 0 | 1 | 0 | NON_SYNONYMOU | MODERAT | MISSENSE | Cac/Tac | H227Y | tolQ | | | | A | | | | | | | | |
| NC_011770 | 6563047 | T | G | 20 | 0 | 2 | 0 | NON_SYNONYMOU | MODERAT | MISSENSE | gaA/gaC | E76D | tonB | | | | | | | G | G | | | | |
| NC_011770 | 226316 | G | A | 43 | 0 | 2 | 0 | NON_SYNONYMOU | MODERAT | MISSENSE | Gtt/Att | V36I | tonB2 | | | | | | | A | A | | | | |
| NC_011770 | 226361 | A | G | 37 | 0 | 2 | 0 | NON_SYNONYMOU | MODERAT | MISSENSE | Acg/Gcg | T51A | tonB2 | | | | | | | | | G | G | | |
| NC_011770 | 226565 | A | G | 17 | 0 | 12 | 0 | NON_SYNONYMOU | MODERAT | MISSENSE | Atc/Gtc | I119V | tonB2 | G | G | G | G | G | G | G | G | G | G | G | G |
| NC_011770 | 226720 | GCCGGCCG | G | 8 | 0 | 2 | 0 | CODON_CHANGE_ | MODERATE | | gccgccgccc | AAAPA172 | tonB2 | | | | | | | | | G | G | | |
| NC_011770 | 2224377 | T | C | 77 | 0 | 2 | 0 | NON_SYNONYMOU | MODERAT | MISSENSE | gTc/gCc | V637A | topA | C | C | | | | | | | | | | |
| NC_011770 | 2224820 | G | A | 76 | 0 | 1 | 0 | NON_SYNONYMOU | MODERAT | MISSENSE | Gcc/Acc | A785T | topA | | | | | | | | A | | | | |
| NC_011770 | 4605357 | C | T | 79 | 0 | 1 | 0 | NON_SYNONYMOU | MODERAT | MISSENSE | gCa/gTa | A171V | toxA | T | | | | | | | | | | | |
| NC_011770 | 4605029 | C | A | 50 | 0 | 12 | 0 | NON_SYNONYMOU | MODERAT | MISSENSE | Ccc/Acc | P62T | toxA | A | A | A | A | A | A | A | A | A | A | A | A |
| NC_011770 | 4605263 | G | T | 76 | 0 | 2 | 0 | NON_SYNONYMOU | MODERAT | MISSENSE | Gtg/Ttg | V140L | toxA | | | | | | | T | T | | | | |
| NC_011770 | 4605314 | A | C | 76 | 0 | 4 | 0 | NON_SYNONYMOU | MODERAT | MISSENSE | Atc/Ctc | I157L | toxA | | | C | C | C | C | | | | | | |
| NC_011770 | 4605356 | G | T | 77 | 0 | 4 | 0 | NON_SYNONYMOU | MODERAT | MISSENSE | Gca/Tca | A171S | toxA | | | T | T | T | T | | | | | | |
| NC_011770 | 4605455 | G | A | 82 | 0 | 2 | 0 | NON_SYNONYMOU | MODERAT | MISSENSE | Gcc/Acc | A204T | toxA | | | | | | | | | | | A | A |
| NC_011770 | 4605565 | G | C | 75 | 0 | 8 | 0 | NON_SYNONYMOU | MODERAT | MISSENSE | aaG/aaC | K240N | toxA | C | C | | | | | C | C | C | C | C | C |
| NC_011770 | 4606011 | G | A | 64 | 0 | 4 | 0 | NON_SYNONYMOU | MODERAT | MISSENSE | aGc/aAc | S389N | toxA | | | A | A | A | A | | | | | | |
| NC_011770 | 4606127 | A | G | 65 | 0 | 12 | 0 | NON_SYNONYMOU | MODERAT | MISSENSE | Aac/Gac | N428D | toxA | G | G | G | G | G | G | G | G | G | G | G | G |
| NC_011770 | 4606139 | G | A | 66 | 0 | 4 | 0 | NON_SYNONYMOU | MODERAT | MISSENSE | Gtc/Atc | V432I | toxA | A | A | | | | | | | A | A | | |
| NC_011770 | 4606347 | A | C | 65 | 0 | 12 | 0 | NON_SYNONYMOU | MODERAT | MISSENSE | gAg/gCg | E501A | toxA | C | C | C | C | C | C | C | C | C | C | C | C |
| NC_011770 | 4606488 | C | T | 49 | 0 | 1 | 0 | NON_SYNONYMOU | MODERAT | MISSENSE | gCg/gTg | A548V | toxA | | | | | | | T | | | | | |
| NC_011770 | 4606668 | A | C | 39 | 0 | 4 | 0 | NON_SYNONYMOU | MODERAT | MISSENSE | gAc/gCc | D608A | toxA | | | C | C | C | C | | | | | | |
| NC_011770 | 4606688 | A | C | 41 | 0 | 6 | 0 | NON_SYNONYMOU | MODERAT | MISSENSE | Aag/Cag | K615Q | toxA | C | C | C | C | C | C | | | | | | |
| NC_011770 | 4606709 | C | A | 45 | 0 | 2 | 0 | NON_SYNONYMOU | MODERAT | MISSENSE | Ctg/Atg | L622M | toxA | | | | | | | | | A | A | | |
| NC_011770 | 5074983 | A | G | 82 | 0 | 2 | 0 | NON_SYNONYMOU | MODERAT | MISSENSE | Atc/Gtc | I132V | toxR | G | G | | | | | | | | | | |
| NC_011770 | 5075079 | A | G | 84 | 0 | 12 | 0 | NON_SYNONYMOU | MODERAT | MISSENSE | Aat/Gat | N164D | toxR | G | G | G | G | G | G | G | G | G | G | G | G |
| NC_011770 | 5667077 | G | C | 79 | 0 | 2 | 0 | NON_SYNONYMOU | MODERAT | MISSENSE | gCg/gGg | A84G | tpiA | C | C | | | | | | | | | | |
| NC_011770 | 5667078 | C | T | 79 | 0 | 2 | 0 | NON_SYNONYMOU | MODERAT | MISSENSE | Gcg/Acg | A84T | tpiA | T | T | | | | | | | | | | |
| NC_011770 | 2409448 | A | G | 78 | 0 | 4 | 0 | NON_SYNONYMOU | MODERAT | MISSENSE | cAg/cGg | Q40R | tpm | | | | | | | G | G | G | G | | |
| NC_011770 | 2409574 | G | A | 71 | 0 | 2 | 0 | NON_SYNONYMOU | MODERAT | MISSENSE | cGc/cAc | R82H | tpm | A | A | | | | | | | | | | |
| NC_011770 | 2409595 | A | G | 73 | 0 | 2 | 0 | NON_SYNONYMOU | MODERAT | MISSENSE | gAt/gGt | D89G | tpm | | | | | | | | | G | G | | |
| NC_011770 | 2409954 | G | A | 60 | 0 | 2 | 0 | NON_SYNONYMOU | MODERAT | MISSENSE | Gac/Aac | D209N | tpm | | | | | | | | | A | A | | |
| NC_011770 | 2409910 | T | C | 59 | 0 | 1 | 0 | NON_SYNONYMOU | MODERAT | MISSENSE | cTc/cCc | L194P | tpm | | C | | | | | | | | | | |
| NC_011770 | 2968402 | C | A | 67 | 0 | 2 | 0 | NON_SYNONYMOU | MODERAT | MISSENSE | Ctc/Atc | L17I | tpx | | | | | | | | | | | A | A |
| NC_011770 | 2968480 | G | T | 72 | 0 | 2 | 0 | NON_SYNONYMOU | MODERAT | MISSENSE | Gcc/Tcc | A43S | tpx | | | | | | | | | | | T | T |
| NC_011770 | 3135144 | C | T | 75 | 0 | 4 | 0 | NON_SYNONYMOU | MODERAT | MISSENSE | gGc/gAc | G540D | treA | | | T | T | T | T | | | | | | |
| NC_011770 | 3135195 | T | C | 78 | 0 | 2 | 0 | NON_SYNONYMOU | MODERAT | MISSENSE | gAc/gGc | D523G | treA | | | | | | | | | C | C | | |
| NC_011770 | 3135215 | ACCG | A | 74 | 0 | 4 | 0 | CODON_CHANGE_ | MODERATE | | ggcggt/ggt | GG515G | treA | | | A | A | A | A | | | | | | |
| NC_011770 | 3135496 | T | C | 81 | 0 | 4 | 0 | NON_SYNONYMOU | MODERAT | MISSENSE | Acc/Gcc | T423A | treA | C | C | | | | | | | | | C | C |
| NC_011770 | 3135715 | A | G | 83 | 0 | 2 | 0 | NON_SYNONYMOU | MODERAT | MISSENSE | Tac/Cac | Y350H | treA | | | | | G | G | | | | | | |
| NC_011770 | 3135777 | T | C | 74 | 0 | 10 | 0 | NON_SYNONYMOU | MODERAT | MISSENSE | gAc/gGc | D329G | treA | | | C | C | C | C | C | C | C | C | C | C |
| NC_011770 | 3136617 | C | T | 36 | 0 | 12 | 0 | NON_SYNONYMOU | MODERAT | MISSENSE | gGc/gAc | G49D | treA | T | T | T | T | T | T | T | T | T | T | T | T |
| NC_011770 | 3136620 | G | A | 37 | 0 | 2 | 0 | NON_SYNONYMOU | MODERAT | MISSENSE | cCc/cTc | P48L | treA | | | | | | | | | A | A | | |
| NC_011770 | 3136661 | G | GGACG/ | 21 | 1 | 4 | 0 | CODON_CHANGE_ | MODERATE | | tcc/tcTTCG | S34SSS | treA | | . | | | | | GGACGAA | GGACGAA | | | A | A |
| NC_011770 | 3136659 | C | CAGGAC | 38 | 0 | 3 | 0 | CODON_INSERTIOI | MODERATE | | tgg/tCGTC | W35SSW | treA | | CAGGACG | CAGGACG | | | CAGGACG | | | | | | |
| NC_011770 | 17450 | C | T | 89 | 0 | 2 | 0 | NON_SYNONYMOU | MODERAT | MISSENSE | cGc/cAc | R426H | trkA | | | | | | | | | | | T | T |
| NC_011770 | 2003303 | C | G | 90 | 0 | 2 | 0 | NON_SYNONYMOU | MODERAT | MISSENSE | aGc/aCc | S197T | trkH | G | G | | | | | | | | | | |
| NC_011770 | 5633632 | A | G | 62 | 0 | 2 | 0 | NON_SYNONYMOU | MODERAT | MISSENSE | gAc/gGc | D208G | trmA | | | | | | | | | | | G | G |
| NC_011770 | 5633704 | T | C | 66 | 0 | 2 | 0 | NON_SYNONYMOU | MODERAT | MISSENSE | gTg/gCg | V232A | trmA | | | | | C | C | | | | | | |
| NC_011770 | 5633881 | G | A | 79 | 0 | 1 | 0 | NON_SYNONYMOU | MODERAT | MISSENSE | gGc/gAc | G291D | trmA | | | | | | | A | | | | | |
| NC_011770 | 5634094 | G | A | 83 | 0 | 2 | 0 | NON_SYNONYMOU | MODERAT | MISSENSE | cGc/cAc | R362H | trmA | | | | | | | A | A | | | | |
| NC_011770 | 418520 | T | C | 64 | 0 | 4 | 0 | NON_SYNONYMOU | MODERAT | MISSENSE | gTc/gCc | V37A | trmB | | | C | C | C | C | | | | | | |
| NC_011770 | 418871 | A | G | 77 | 0 | 2 | 0 | NON_SYNONYMOU | MODERAT | MISSENSE | aAg/aGg | K154R | trmB | G | G | | | | | | | | | | |
| NC_011770 | 1341402 | C | T | 86 | 0 | 4 | 0 | NON_SYNONYMOU | MODERAT | MISSENSE | gCg/gTg | A19V | trmD | | | T | T | T | T | | | | | | |
| NC_011770 | 1341458 | A | G | 87 | 0 | 4 | 0 | NON_SYNONYMOU | MODERAT | MISSENSE | Acc/Gcc | T38A | trmD | | | G | G | G | G | | | | | | |
| NC_011770 | 6597784 | G | C | 96 | 0 | 2 | 0 | NON_SYNONYMOU | MODERAT | MISSENSE | gaC/gaG | D442E | trmE | C | C | | | | | | | | | | |
| NC_011770 | 37439 | G | T | 72 | 0 | 2 | 0 | NON_SYNONYMOU | MODERAT | MISSENSE | Ctc/Atc | L148I | trpA | T | T | | | | | | | | | | |
| NC_011770 | 37460 | G | C | 75 | 0 | 2 | 0 | NON_SYNONYMOU | MODERAT | MISSENSE | Cac/Gac | H141D | trpA | | | | | | | | | | | C | C |

| | | | | | | | | | | | | | | | | | | | | | | | | | |
|---|---|---|---|---|---|---|---|---|---|---|---|---|---|---|---|---|---|---|---|---|---|---|---|---|---|
| NC_011770 | 37605 | C | G | 81 | 0 | 12 | 0 | NON_SYNONYMOU | MODERAT | MISSENSE | gaG/gaC | E92D | trpA | G | G | G | G | G | G | G | G | G | G | G | G |
| NC_011770 | 37662 | G | T | 83 | 0 | 2 | 0 | NON_SYNONYMOU | MODERAT | MISSENSE | gaC/gaA | D73E | trpA | | | | | | | | | | | T | T |
| NC_011770 | 37160 | C | T | 81 | 0 | 1 | 0 | NON_SYNONYMOU | MODERAT | MISSENSE | Gcc/Acc | A241T | trpA | | T | | | | | | | | | | |
| NC_011770 | 682467 | G | A | 83 | 0 | 2 | 0 | NON_SYNONYMOU | MODERAT | MISSENSE | Gcc/Acc | A28T | trpC | | | | | | | | | | | A | A |
| NC_011770 | 682746 | A | G | 80 | 0 | 2 | 0 | NON_SYNONYMOU | MODERAT | MISSENSE | Atc/Gtc | I121V | trpC | | | | | | | G | G | | | | |
| NC_011770 | 682956 | G | A | 80 | 0 | 2 | 0 | NON_SYNONYMOU | MODERAT | MISSENSE | Gtg/Atg | V191M | trpC | | | | | | | | | A | A | | |
| NC_011770 | 662270 | T | C | 80 | 0 | 2 | 0 | NON_SYNONYMOU | MODERAT | MISSENSE | gTg/gCg | V49A | trpE | C | C | | | | | | | | | | |
| NC_011770 | 662343 | C | G | 77 | 0 | 4 | 0 | NON_SYNONYMOU | MODERAT | MISSENSE | gaC/gaG | D73E | trpE | | | G | G | G | G | | | | | | |
| NC_011770 | 662371 | G | A | 75 | 0 | 10 | 0 | NON_SYNONYMOU | MODERAT | MISSENSE | Gtg/Atg | V83M | trpE | | | A | A | A | A | A | A | A | A | A | A |
| NC_011770 | 662719 | A | G | 76 | 0 | 4 | 0 | NON_SYNONYMOU | MODERAT | MISSENSE | Atc/Gtc | I199V | trpE | | | G | G | G | G | | | | | | |
| NC_011770 | 2101021 | G | A | 74 | 0 | 8 | 0 | NON_SYNONYMOU | MODERAT | MISSENSE | Gtg/Atg | V106M | trpF | A | A | A | A | A | A | | | | | A | A |
| NC_011770 | 681063 | A | G | 75 | 0 | 8 | 0 | NON_SYNONYMOU | MODERAT | MISSENSE | Aac/Gac | N111D | trpG | | | G | G | G | G | G | G | | | G | G |
| NC_011770 | 681159 | A | G | 80 | 0 | 2 | 0 | NON_SYNONYMOU | MODERAT | MISSENSE | Atc/Gtc | I143V | trpG | | | | | | | | | | | G | G |
| NC_011770 | 39532 | G | A | 72 | 0 | 12 | 0 | NON_SYNONYMOU | MODERAT | MISSENSE | cGg/cAg | R115Q | trpI | A | A | A | A | A | A | A | A | A | A | A | A |
| NC_011770 | 39763 | C | T | 84 | 0 | 2 | 0 | NON_SYNONYMOU | MODERAT | MISSENSE | cCg/cTg | P192L | trpI | T | T | | | | | | | | | | |
| NC_011770 | 39841 | G | A | 77 | 0 | 2 | 0 | NON_SYNONYMOU | MODERAT | MISSENSE | cGc/cAc | R218H | trpI | A | A | | | | | | | | | | |
| NC_011770 | 39921 | C | G | 74 | 0 | 4 | 0 | NON_SYNONYMOU | MODERAT | MISSENSE | Ccg/Gcg | P245A | trpI | | | G | G | G | G | | | | | | |
| NC_011770 | 5306236 | T | G | 91 | 0 | 6 | 0 | NON_SYNONYMOU | MODERAT | MISSENSE | gaA/gaC | E269D | trpS | G | G | | | | | | | G | G | G | G |
| NC_011770 | 5306277 | T | A | 88 | 0 | 4 | 0 | NON_SYNONYMOU | MODERAT | MISSENSE | Agc/Tgc | S256C | trpS | | | A | A | A | A | | | | | | |
| NC_011770 | 2100513 | A | G | 63 | 0 | 1 | 0 | NON_SYNONYMOU | MODERAT | MISSENSE | tAc/tGc | Y252C | truA | | | | | | | G | | | | | |
| NC_011770 | 5660181 | C | T | 105 | 0 | 2 | 0 | NON_SYNONYMOU | MODERAT | MISSENSE | gGc/gAc | G192D | truB | T | T | | | | | | | | | | |
| NC_011770 | 5660190 | T | C | 100 | 0 | 2 | 0 | NON_SYNONYMOU | MODERAT | MISSENSE | cAg/cGg | Q189R | truB | | | | | | | | | | | C | C |
| NC_011770 | 5660424 | G | A | 101 | 0 | 2 | 0 | NON_SYNONYMOU | MODERAT | MISSENSE | gCc/gTc | A111V | truB | A | A | | | | | | | | | | |
| NC_011770 | 1518226 | C | T | 75 | 0 | 6 | 0 | NON_SYNONYMOU | MODERAT | MISSENSE | Cgc/Tgc | R118C | truD | T | T | T | T | T | T | | | | | | |
| NC_011770 | 1518316 | C | G | 74 | 0 | 6 | 0 | NON_SYNONYMOU | MODERAT | MISSENSE | Ctt/Gtt | L148V | truD | | | | | | | G | G | G | G | G | G |
| NC_011770 | 1518724 | A | C | 72 | 0 | 4 | 0 | NON_SYNONYMOU | MODERAT | MISSENSE | Agc/Cgc | S284R | truD | | | C | C | C | C | | | | | | |
| NC_011770 | 1486824 | A | G | 58 | 0 | 12 | 0 | NON_SYNONYMOU | MODERAT | MISSENSE | aAc/aGc | N184S | tsf | G | G | G | G | G | G | G | G | G | G | G | G |
| NC_011770 | 708129 | A | G | 26 | 0 | 4 | 0 | NON_SYNONYMOU | MODERAT | MISSENSE | aAt/aGt | N185S | tufB | | | G | G | G | G | | | | | | |
| NC_011770 | 1111124 | GGT | G | 82 | 0 | 1 | 0 | FRAME_SHIFT | HIGH | | gtg/ | V561 | two-compo | | | | | | | G | | | | | |
| NC_011770 | 1109523 | T | A | 77 | 0 | 4 | 0 | NON_SYNONYMOU | MODERAT | MISSENSE | Tcc/Acc | S27T | two-compo | | | A | A | A | A | | | | | | |
| NC_011770 | 1110640 | G | T | 87 | 0 | 6 | 0 | NON_SYNONYMOU | MODERAT | MISSENSE | aGc/aTc | S399I | two-compo | | | T | T | T | T | | | T | T | | |
| NC_011770 | 1110726 | T | A | 85 | 0 | 2 | 0 | NON_SYNONYMOU | MODERAT | MISSENSE | Tcg/Acg | S428T | two-compo | | | | | | | A | A | | | | |
| NC_011770 | 1110909 | G | A | 84 | 0 | 2 | 0 | NON_SYNONYMOU | MODERAT | MISSENSE | Ggc/Agc | G489S | two-compo | | | | | | | | | | | A | A |
| NC_011770 | 1111020 | A | G | 83 | 0 | 2 | 0 | NON_SYNONYMOU | MODERAT | MISSENSE | Acc/Gcc | T526A | two-compo | | | | | | | | | G | G | | |
| NC_011770 | 1111083 | C | T | 81 | 0 | 2 | 0 | NON_SYNONYMOU | MODERAT | MISSENSE | Ctc/Ttc | L547F | two-compo | | | | | | | T | T | | | | |
| NC_011770 | 1111126 | T | C | 72 | 1 | 1 | 0 | NON_SYNONYMOU | MODERAT | MISSENSE | gTg/gCg | V561A | two-compo | | | | | | | . | C | | | | |
| NC_011770 | 1111675 | T | C | 69 | 0 | 2 | 0 | NON_SYNONYMOU | MODERAT | MISSENSE | gTg/gCg | V744A | two-compo | | | | | | | | | C | C | | |
| NC_011770 | 1112275 | A | G | 71 | 0 | 4 | 0 | NON_SYNONYMOU | MODERAT | MISSENSE | aAc/aGc | N944S | two-compo | | | | | | | | | G | G | G | G |
| NC_011770 | 1112299 | G | A | 70 | 0 | 4 | 0 | NON_SYNONYMOU | MODERAT | MISSENSE | aGt/aAt | S952N | two-compo | | | | | | | | | A | A | A | A |
| NC_011770 | 6097583 | A | ACT | 65 | 0 | 2 | 0 | FRAME_SHIFT | HIGH | | cag/cagAG | Q522Q? | typA | | | | | | | | | ACT | ACT | | |
| NC_011770 | 6098866 | A | G | 72 | 0 | 1 | 0 | NON_SYNONYMOU | MODERAT | MISSENSE | gTg/gCg | V95A | typA | | | | | | | G | | | | | |
| NC_011770 | 907229 | G | A | 83 | 0 | 2 | 0 | NON_SYNONYMOU | MODERAT | MISSENSE | Gcg/Acg | A387T | tyrS | | | | | | | | | | | A | A |
| NC_011770 | 698566 | T | G | 64 | 0 | 4 | 0 | NON_SYNONYMOU | MODERAT | MISSENSE | gaT/gaG | D318E | tyrZ | G | G | | | | | | | | | G | G |
| NC_011770 | 698759 | A | G | 53 | 0 | 12 | 0 | NON_SYNONYMOU | MODERAT | MISSENSE | Atg/Gtg | M383V | tyrZ | G | G | G | G | G | G | G | G | G | G | G | G |
| NC_011770 | 6038769 | G | T | 92 | 0 | 2 | 0 | NON_SYNONYMOU | MODERAT | MISSENSE | Gtc/Ttc | V6F | ubiB | | | | | | | | | | | T | T |
| NC_011770 | 6039501 | A | C | 69 | 0 | 2 | 0 | NON_SYNONYMOU | MODERAT | MISSENSE | Atc/Ctc | I250L | ubiB | | | | | | | | | C | C | | |
| NC_011770 | 6040191 | A | G | 77 | 0 | 10 | 0 | NON_SYNONYMOU | MODERAT | MISSENSE | Acg/Gcg | T480A | ubiB | G | G | G | G | G | G | G | G | G | G | | |
| NC_011770 | 2040512 | C | T | 66 | 0 | 10 | 0 | NON_SYNONYMOU | MODERAT | MISSENSE | aGc/aAc | S71N | ubiG | | | T | T | T | T | T | T | T | T | T | T |
| NC_011770 | 2040323 | C | T | 64 | 0 | 1 | 0 | NON_SYNONYMOU | MODERAT | MISSENSE | cGc/cAc | R134H | ubiG | | T | | | | | | | | | | |
| NC_011770 | 6216893 | G | A | 96 | 0 | 1 | 0 | NON_SYNONYMOU | MODERAT | MISSENSE | Cgc/Tgc | R236C | ubiH | | | | | | | | | | | | A |
| NC_011770 | 6216647 | G | A | 91 | 0 | 2 | 0 | NON_SYNONYMOU | MODERAT | MISSENSE | Cac/Tac | H318Y | ubiH | | | | | | | | | A | A | | |
| NC_011770 | 6216770 | A | G | 90 | 0 | 12 | 0 | NON_SYNONYMOU | MODERAT | MISSENSE | Tgc/Cgc | C277R | ubiH | G | G | G | G | G | G | G | G | G | G | G | G |
| NC_011770 | 5049292 | TA | T | 68 | 0 | 2 | 0 | FRAME_SHIFT | HIGH | | cct/ | P223 | ung | | | | | | | | | | | T | T |
| NC_011770 | 5548071 | G | A | 42 | 0 | 1 | 0 | NON_SYNONYMOU | MODERAT | MISSENSE | Gcc/Acc | A32T | upp | | | | | | | | A | | | | |
| NC_011770 | 1489370 | C | G | 80 | 0 | 2 | 0 | NON_SYNONYMOU | MODERAT | MISSENSE | aCc/aGc | T239S | uppS | | | | | | | | | | | G | G |
| NC_011770 | 5549879 | G | C | 68 | 0 | 4 | 0 | NON_SYNONYMOU | MODERAT | MISSENSE | Gaa/Caa | E421Q | uraA | C | C | | | | | | | | | C | C |
| NC_011770 | 5795927 | G | A | 75 | 0 | 2 | 0 | NON_SYNONYMOU | MODERAT | MISSENSE | cGc/cAc | R6H | ureA | A | A | | | | | | | | | | |
| NC_011770 | 5796104 | A | G | 80 | 0 | 2 | 0 | NON_SYNONYMOU | MODERAT | MISSENSE | aAc/aGc | N65S | ureA | | | | | | | | | | | G | G |
| NC_011770 | 5797209 | A | C | 62 | 0 | 4 | 0 | NON_SYNONYMOU | MODERAT | MISSENSE | Atc/Ctc | I29L | ureC | | | | | | | | | C | C | C | C |
| NC_011770 | 5797641 | C | T | 65 | 0 | 4 | 0 | NON_SYNONYMOU | MODERAT | MISSENSE | Ccc/Tcc | P173S | ureC | | | T | T | T | T | | | | | | |
| NC_011770 | 5795175 | G | T | 31 | 0 | 6 | 0 | NON_SYNONYMOU | MODERAT | MISSENSE | Gcc/Tcc | A37S | ureD | | | T | T | T | T | | | T | T | | |
| NC_011770 | 5795364 | T | G | 48 | 0 | 4 | 0 | NON_SYNONYMOU | MODERAT | MISSENSE | Tcg/Gcg | S100A | ureD | | | | | | | G | G | | | G | G |

| | | | | | | | | | | | | | | | | | | | | | | | | | |
|---|---|---|---|---|---|---|---|---|---|---|---|---|---|---|---|---|---|---|---|---|---|---|---|---|---|
| NC_011770 | 5795548 | C | T | 66 | 0 | 4 | 0 | NON_SYNONYMOU | MODERAT | MISSENSE | gCc/gTc | A161V | ureD | | | T | T | T | T | | | | | | |
| NC_011770 | 5821809 | G | A | 70 | 0 | 1 | 0 | NON_SYNONYMOU | MODERAT | MISSENSE | Gcg/Acg | A153T | ureJ | | | | | | | | A | | | | |
| NC_011770 | 5821689 | G | C | 60 | 0 | 12 | 0 | NON_SYNONYMOU | MODERAT | MISSENSE | Gcg/Ccg | A113P | ureJ | C | C | C | C | C | C | C | C | C | C | C | C |
| NC_011770 | 5821759 | A | C | 68 | 0 | 6 | 0 | NON_SYNONYMOU | MODERAT | MISSENSE | gAc/gCc | D136A | ureJ | C | C | | | | | C | C | C | C | | |
| NC_011770 | 741548 | G | A | 75 | 0 | 2 | 0 | NON_SYNONYMOU | MODERAT | MISSENSE | Ccc/Tcc | P930S | uvrA | | | | | | | | | | | A | A |
| NC_011770 | 742744 | C | G | 59 | 0 | 2 | 0 | NON_SYNONYMOU | MODERAT | MISSENSE | gGc/gCc | G531A | uvrA | | | | | | | | | | | G | G |
| NC_011770 | 743328 | T | G | 56 | 0 | 4 | 0 | NON_SYNONYMOU | MODERAT | MISSENSE | gaA/gaC | E336D | uvrA | | | G | G | G | G | | | | | | |
| NC_011770 | 742475 | G | A | 64 | 0 | 1 | 0 | NON_SYNONYMOU | MODERAT | MISSENSE | Cgc/Tgc | R621C | uvrA | | | | | | | | | | | A | |
| NC_011770 | 743604 | ATCCTCG | A | 72 | 0 | 9 | 0 | CODON_CHANGE_ | MODERATE | | gacgaggat, | DED242D | uvrA | A | A | | A | A | A | A | | A | A | A | |
| NC_011770 | 2073963 | T | G | 81 | 0 | 2 | 0 | NON_SYNONYMOU | MODERAT | MISSENSE | gTg/gGg | V447G | uvrB | | | | | G | G | | | | | | |
| NC_011770 | 2794677 | C | T | 76 | 0 | 1 | 0 | NON_SYNONYMOU | MODERAT | MISSENSE | cCc/cTc | P178L | uvrC | | | | | | | T | | | | | |
| NC_011770 | 2794749 | G | A | 65 | 0 | 2 | 0 | NON_SYNONYMOU | MODERAT | MISSENSE | aGg/aAg | R202K | uvrC | A | A | | | | | | | | | | |
| NC_011770 | 2795288 | C | T | 69 | 0 | 2 | 0 | NON_SYNONYMOU | MODERAT | MISSENSE | Ctt/Ttt | L382F | uvrC | T | T | | | | | | | | | | |
| NC_011770 | 2795528 | G | A | 79 | 0 | 2 | 0 | NON_SYNONYMOU | MODERAT | MISSENSE | Gac/Aac | D462N | uvrC | A | A | | | | | | | | | | |
| NC_011770 | 6468032 | A | G | 96 | 0 | 2 | 0 | NON_SYNONYMOU | MODERAT | MISSENSE | aAg/aGg | K344R | uvrD | | | | | | | G | G | | | | |
| NC_011770 | 6468986 | G | A | 74 | 0 | 12 | 0 | NON_SYNONYMOU | MODERAT | MISSENSE | aGt/aAt | S662N | uvrD | A | A | A | A | A | A | A | A | A | A | A | A |
| NC_011770 | 6468998 | A | G | 77 | 0 | 12 | 0 | NON_SYNONYMOU | MODERAT | MISSENSE | aAc/aGc | N666S | uvrD | G | G | G | G | G | G | G | G | G | G | G | G |
| NC_011770 | 1975561 | T | C | 68 | 0 | 2 | 0 | NON_SYNONYMOU | MODERAT | MISSENSE | aAg/aGg | K233R | vacJ | C | C | | | | | | | | | | |
| NC_011770 | 1236509 | A | G | 84 | 0 | 1 | 0 | NON_SYNONYMOU | MODERAT | MISSENSE | aTc/aCc | I705T | valS | | | | | | | G | | | | | |
| NC_011770 | 1236857 | T | G | 82 | 0 | 2 | 0 | NON_SYNONYMOU | MODERAT | MISSENSE | gAg/gCg | E589A | valS | | | | | | | | | G | G | | |
| NC_011770 | 5835605 | C | G | 67 | 0 | 2 | 0 | NON_SYNONYMOU | MODERAT | MISSENSE | gaC/gaG | D76E | vanA | | | | | | | | | | | G | G |
| NC_011770 | 5836255 | A | G | 53 | 0 | 2 | 0 | NON_SYNONYMOU | MODERAT | MISSENSE | aAg/aGg | K293R | vanA | | | | | | | | | | | G | G |
| NC_011770 | 5836716 | G | A | 68 | 0 | 8 | 0 | NON_SYNONYMOU | MODERAT | MISSENSE | cGc/cAc | R90H | vanB | | | A | A | A | A | A | A | A | A | | |
| NC_011770 | 5836826 | G | A | 71 | 0 | 4 | 0 | NON_SYNONYMOU | MODERAT | MISSENSE | Gcc/Acc | A127T | vanB | A | A | | | | | | | | | A | A |
| NC_011770 | 5837127 | A | G | 69 | 0 | 12 | 0 | NON_SYNONYMOU | MODERAT | MISSENSE | aAt/aGt | N227S | vanB | G | G | G | G | G | G | G | G | G | G | G | G |
| NC_011770 | 5837129 | G | A | 69 | 0 | 4 | 0 | NON_SYNONYMOU | MODERAT | MISSENSE | Gcg/Acg | A228T | vanB | | | A | A | A | A | | | | | | |
| NC_011770 | 683425 | C | T | 72 | 0 | 1 | 0 | NON_SYNONYMOU | MODERAT | MISSENSE | atG/atA | M168I | vfr | | | | | | | T | | | | | |
| NC_011770 | 5936169 | C | T | 59 | 0 | 2 | 0 | NON_SYNONYMOU | MODERAT | MISSENSE | gGg/gAg | G422E | waaA | T | T | | | | | | | | | | |
| NC_011770 | 5936286 | G | A | 69 | 0 | 2 | 0 | NON_SYNONYMOU | MODERAT | MISSENSE | gCg/gTg | A383V | waaA | | | | | | | A | A | | | | |
| NC_011770 | 5936287 | C | T | 71 | 0 | 2 | 0 | NON_SYNONYMOU | MODERAT | MISSENSE | Gcg/Acg | A383T | waaA | T | T | | | | | | | | | | |
| NC_011770 | 5936896 | T | C | 92 | 0 | 2 | 0 | NON_SYNONYMOU | MODERAT | MISSENSE | Atc/Gtc | I180V | waaA | | | | | | | | | | | C | C |
| NC_011770 | 5937354 | T | C | 69 | 0 | 2 | 0 | NON_SYNONYMOU | MODERAT | MISSENSE | cAg/cGg | Q27R | waaA | | | | | | | | | C | C | | |
| NC_011770 | 5964963 | T | A | 85 | 0 | 6 | 0 | NON_SYNONYMOU | MODERAT | MISSENSE | gAc/gTc | D189V | waaC | | | A | A | A | A | A | A | | | | |
| NC_011770 | 5965171 | G | A | 75 | 0 | 1 | 0 | NON_SYNONYMOU | MODERAT | MISSENSE | Cgc/Tgc | R120C | waaC | | | | | | | A | | | | | |
| NC_011770 | 5965183 | G | C | 76 | 0 | 2 | 0 | NON_SYNONYMOU | MODERAT | MISSENSE | Cgc/Ggc | R116G | waaC | | | | | | | C | C | | | | |
| NC_011770 | 5966253 | G | A | 81 | 0 | 1 | 0 | NON_SYNONYMOU | MODERAT | MISSENSE | Cgc/Tgc | R104C | waaF | | | | | | | | A | | | | |
| NC_011770 | 5964235 | T | G | 74 | 0 | 2 | 0 | NON_SYNONYMOU | MODERAT | MISSENSE | gAc/gCc | D77A | waaG | | | | | | | | | | | G | G |
| NC_011770 | 5951529 | C | T | 34 | 4 | 2 | 0 | NON_SYNONYMOU | MODERAT | MISSENSE | gCc/gTc | A182V | waaL | | | | | | | T | T | . | . | . | . |
| NC_011770 | 5951789 | G | A | 51 | 0 | 6 | 0 | NON_SYNONYMOU | MODERAT | MISSENSE | Gcc/Acc | A269T | waaL | | | | | | | A | A | A | A | A | A |
| NC_011770 | 5951790 | C | G | 50 | 0 | 6 | 0 | NON_SYNONYMOU | MODERAT | MISSENSE | gCc/gGc | A269G | waaL | | | | | | | G | G | G | G | G | G |
| NC_011770 | 5951794 | C | A | 50 | 0 | 6 | 0 | NON_SYNONYMOU | MODERAT | MISSENSE | gaC/gaA | D270E | waaL | | | | | | | A | A | A | A | A | A |
| NC_011770 | 5951795 | G | T | 50 | 0 | 6 | 0 | NON_SYNONYMOU | MODERAT | MISSENSE | Gcc/Tcc | A271S | waaL | | | | | | | T | T | T | T | T | T |
| NC_011770 | 5951810 | A | G | 51 | 0 | 6 | 0 | NON_SYNONYMOU | MODERAT | MISSENSE | Agc/Ggc | S276G | waaL | | | | | | | G | G | G | G | G | G |
| NC_011770 | 5951825 | C | T | 52 | 0 | 4 | 0 | NON_SYNONYMOU | MODERAT | MISSENSE | Ctc/Ttc | L281F | waaL | | | T | T | T | T | | | | | | |
| NC_011770 | 5952044 | G | T | 70 | 0 | 6 | 0 | NON_SYNONYMOU | MODERAT | MISSENSE | Gcc/Tcc | A354S | waaL | | | | | | | T | T | T | T | T | T |
| NC_011770 | 5952071 | C | A | 74 | 0 | 6 | 0 | NON_SYNONYMOU | MODERAT | MISSENSE | Ctg/Atg | L363M | waaL | | | | | | | A | A | A | A | A | A |
| NC_011770 | 5952074 | C | T | 73 | 0 | 6 | 0 | NON_SYNONYMOU | MODERAT | MISSENSE | Ccc/Tcc | P364S | waaL | | | | | | | T | T | T | T | T | T |
| NC_011770 | 5952113 | A | T | 54 | 1 | 5 | 0 | NON_SYNONYMOU | MODERAT | MISSENSE | Atg/Ttg | M377L | waaL | | | | | | | T | T | . | T | T | T |
| NC_011770 | 5952137 | A | G | 49 | 5 | 1 | 0 | NON_SYNONYMOU | MODERAT | MISSENSE | Atc/Gtc | I385V | waaL | | | | | | | . | . | . | . | . | G |
| NC_011770 | 2069571 | CTG | C | 92 | 0 | 1 | 0 | FRAME_SHIFT | HIGH | | ctgttg/ | LL356 | wbpM | | | | | | | C | | | | | |
| NC_011770 | 2068812 | C | G | 45 | 0 | 4 | 0 | NON_SYNONYMOU | MODERAT | MISSENSE | Ctg/Gtg | L103V | wbpM | | | G | G | G | G | | | | | | |
| NC_011770 | 2068830 | T | G | 45 | 0 | 4 | 0 | NON_SYNONYMOU | MODERAT | MISSENSE | Tcc/Gcc | S109A | wbpM | | | G | G | G | G | | | | | | |
| NC_011770 | 2068831 | C | G | 46 | 0 | 4 | 0 | NON_SYNONYMOU | MODERAT | MISSENSE | tCc/tGc | S109C | wbpM | | | G | G | G | G | | | | | | |
| NC_011770 | 2068833 | C | G | 47 | 0 | 4 | 0 | NON_SYNONYMOU | MODERAT | MISSENSE | Ccg/Gcg | P110A | wbpM | | | G | G | G | G | | | | | | |
| NC_011770 | 2068842 | G | C | 46 | 0 | 4 | 0 | NON_SYNONYMOU | MODERAT | MISSENSE | Gtg/Ctg | V113L | wbpM | | | C | C | C | C | | | | | | |
| NC_011770 | 2068843 | T | C | 44 | 0 | 4 | 0 | NON_SYNONYMOU | MODERAT | MISSENSE | gTg/gCg | V113A | wbpM | | | C | C | C | C | | | | | | |
| NC_011770 | 2070085 | A | G | 67 | 0 | 2 | 0 | NON_SYNONYMOU | MODERAT | MISSENSE | cAg/cGg | Q527R | wbpM | G | G | | | | | | | | | | |
| NC_011770 | 2070189 | A | G | 76 | 0 | 1 | 0 | NON_SYNONYMOU | MODERAT | MISSENSE | Agc/Ggc | S562G | wbpM | | | | | | | G | | | | | |
| NC_011770 | 2070382 | C | T | 63 | 0 | 4 | 0 | NON_SYNONYMOU | MODERAT | MISSENSE | gCc/gTc | A626V | wbpM | | | T | T | T | T | | | | | | |
| NC_011770 | 6477873 | G | A | 75 | 0 | 4 | 0 | NON_SYNONYMOU | MODERAT | MISSENSE | Ccg/Tcg | P383S | wbpW | | | A | A | A | A | | | | | | |
| NC_011770 | 6478271 | G | A | 76 | 0 | 2 | 0 | NON_SYNONYMOU | MODERAT | MISSENSE | gCc/gTc | A250V | wbpW | | | | | | | A | A | | | | |
| NC_011770 | 6478376 | T | C | 81 | 0 | 6 | 0 | NON_SYNONYMOU | MODERAT | MISSENSE | cAg/cGg | Q215R | wbpW | | | C | C | C | C | C | C | | | | |

| | | | | | | | | | | | | | | | | | | | | | | | | | |
|---|---|---|---|---|---|---|---|---|---|---|---|---|---|---|---|---|---|---|---|---|---|---|---|---|---|
| NC_011770 | 6474539 | T | A | 90 | 0 | 12 | 0 | NON_SYNONYMOU | MODERAT | MISSENSE | cAg/cTg | Q328L | wbpX | A | A | A | A | A | A | A | A | A | A | A | A |
| NC_011770 | 6474614 | G | A | 88 | 0 | 2 | 0 | NON_SYNONYMOU | MODERAT | MISSENSE | gCg/gTg | A303V | wbpX | A | A | | | | | | | | | | |
| NC_011770 | 6474624 | C | T | 92 | 0 | 2 | 0 | NON_SYNONYMOU | MODERAT | MISSENSE | Ggt/Agt | G300S | wbpX | | | | | | | T | T | | | | |
| NC_011770 | 6474947 | G | C | 79 | 0 | 2 | 0 | NON_SYNONYMOU | MODERAT | MISSENSE | gCc/gGc | A192G | wbpX | | | | | | | | | | | C | C |
| NC_011770 | 6473749 | C | T | 85 | 0 | 2 | 0 | NON_SYNONYMOU | MODERAT | MISSENSE | gGg/gAg | G136E | wbpY | | | | | | | | | T | T | | |
| NC_011770 | 6474054 | G | T | 83 | 0 | 4 | 0 | NON_SYNONYMOU | MODERAT | MISSENSE | ttC/ttA | F34L | wbpY | | | T | T | T | T | | | | | | |
| NC_011770 | 6472487 | T | C | 99 | 0 | 10 | 0 | NON_SYNONYMOU | MODERAT | MISSENSE | Aca/Gca | T181A | wbpZ | C | C | C | C | C | C | C | C | | | C | C |
| NC_011770 | 6472505 | C | A | 99 | 0 | 6 | 0 | NON_SYNONYMOU | MODERAT | MISSENSE | Ggc/Tgc | G175C | wbpZ | A | A | | | | | A | A | | | A | A |
| NC_011770 | 6472507 | T | G | 100 | 0 | 6 | 0 | NON_SYNONYMOU | MODERAT | MISSENSE | gAc/gCc | D174A | wbpZ | G | G | | | | | G | G | | | G | G |
| NC_011770 | 6472594 | G | A | 79 | 0 | 6 | 0 | NON_SYNONYMOU | MODERAT | MISSENSE | gCa/gTa | A145V | wbpZ | A | A | | | | | A | A | | | A | A |
| NC_011770 | 6472597 | A | G | 78 | 0 | 6 | 0 | NON_SYNONYMOU | MODERAT | MISSENSE | gTc/gCc | V144A | wbpZ | G | G | | | | | G | G | | | G | G |
| NC_011770 | 6472626 | A | C | 80 | 0 | 6 | 0 | NON_SYNONYMOU | MODERAT | MISSENSE | agT/agG | S134R | wbpZ | C | C | | | | | C | C | | | C | C |
| NC_011770 | 6472628 | T | C | 79 | 0 | 6 | 0 | NON_SYNONYMOU | MODERAT | MISSENSE | Agt/Ggt | S134G | wbpZ | C | C | | | | | C | C | | | C | C |
| NC_011770 | 6472937 | A | G | 86 | 0 | 6 | 0 | NON_SYNONYMOU | MODERAT | MISSENSE | Tcg/Ccg | S31P | wbpZ | | | G | G | G | G | | | | | G | G |
| NC_011770 | 4801375 | G | A | 86 | 0 | 2 | 0 | NON_SYNONYMOU | MODERAT | MISSENSE | Gcg/Acg | A56T | wrbA | A | A | | | | | | | | | | |
| NC_011770 | 4801516 | G | A | 85 | 0 | 4 | 0 | NON_SYNONYMOU | MODERAT | MISSENSE | Ggc/Agc | G103S | wrbA | | | | | | | A | A | | | A | A |
| NC_011770 | 1385963 | C | T | 76 | 0 | 1 | 0 | STOP_GAINED | HIGH | NONSENSE | Cag/Tag | Q268* | wspA | | | | | | | T | | | | | |
| NC_011770 | 1387896 | T | C | 75 | 0 | 12 | 0 | NON_SYNONYMOU | MODERAT | MISSENSE | gTa/gCa | V197A | wspC | C | C | C | C | C | C | C | C | C | C | C | C |
| NC_011770 | 1388153 | G | A | 74 | 0 | 4 | 0 | NON_SYNONYMOU | MODERAT | MISSENSE | Gcc/Acc | A283T | wspC | | | A | A | A | A | | | | | | |
| NC_011770 | 1388355 | G | A | 71 | 0 | 1 | 0 | NON_SYNONYMOU | MODERAT | MISSENSE | cGt/cAt | R350H | wspC | | | | | | | | A | | | | |
| NC_011770 | 1388624 | C | G | 76 | 0 | 2 | 0 | NON_SYNONYMOU | MODERAT | MISSENSE | aaC/aaG | N19K | wspD | | | G | G | | | | | | | | |
| NC_011770 | 1390311 | A | G | 77 | 0 | 10 | 0 | INTRAGENIC | MODIFIER | | | | wspE | G | G | G | G | G | G | | | G | G | G | G |
| NC_011770 | 1390463 | G | T | 79 | 0 | 4 | 0 | INTRAGENIC | MODIFIER | | | | wspE | | | T | T | T | T | | | | | | |
| NC_011770 | 1390614 | C | T | 91 | 0 | 2 | 0 | INTRAGENIC | MODIFIER | | | | wspE | T | T | | | | | | | | | | |
| NC_011770 | 1390869 | T | C | 92 | 0 | 12 | 0 | INTRAGENIC | MODIFIER | | | | wspE | C | C | C | C | C | C | C | C | C | C | C | C |
| NC_011770 | 1390890 | G | A | 90 | 0 | 2 | 0 | INTRAGENIC | MODIFIER | | | | wspE | | | | | | | A | A | | | | |
| NC_011770 | 1390953 | G | T | 85 | 0 | 4 | 0 | INTRAGENIC | MODIFIER | | | | wspE | | | T | T | T | T | | | | | | |
| NC_011770 | 1391019 | G | C | 85 | 0 | 2 | 0 | INTRAGENIC | MODIFIER | | | | wspE | C | C | | | | | | | | | | |
| NC_011770 | 1391064 | C | T | 85 | 0 | 2 | 0 | INTRAGENIC | MODIFIER | | | | wspE | | | | | | | T | T | | | | |
| NC_011770 | 1391355 | C | T | 97 | 0 | 2 | 0 | INTRAGENIC | MODIFIER | | | | wspE | T | T | | | | | | | | | | |
| NC_011770 | 1391430 | C | G | 98 | 0 | 2 | 0 | INTRAGENIC | MODIFIER | | | | wspE | | | | | | | | | G | G | | |
| NC_011770 | 1391505 | C | T | 85 | 0 | 6 | 0 | INTRAGENIC | MODIFIER | | | | wspE | | | T | T | T | T | T | T | | | | |
| NC_011770 | 1390726 | A | C | 91 | 0 | 1 | 0 | INTRAGENIC | MODIFIER | | | | wspE | | | | | | | | | | C | | |
| NC_011770 | 1391729 | G | C | 85 | 0 | 2 | 0 | INTRAGENIC | MODIFIER | | | | wspF | | | | | | | | | | | C | C |
| NC_011770 | 1391952 | T | C | 83 | 0 | 12 | 0 | INTRAGENIC | MODIFIER | | | | wspF | C | C | C | C | C | C | C | C | C | C | C | C |
| NC_011770 | 1392032 | C | T | 81 | 0 | 2 | 0 | INTRAGENIC | MODIFIER | | | | wspF | T | T | | | | | | | | | | |
| NC_011770 | 1392056 | A | C | 80 | 0 | 12 | 0 | INTRAGENIC | MODIFIER | | | | wspF | C | C | C | C | C | C | C | C | C | C | C | C |
| NC_011770 | 1392098 | G | A | 78 | 0 | 6 | 0 | INTRAGENIC | MODIFIER | | | | wspF | A | A | | | | | | | A | A | A | A |
| NC_011770 | 1392128 | C | A | 79 | 0 | 10 | 0 | INTRAGENIC | MODIFIER | | | | wspF | A | A | A | A | A | A | A | A | A | A | | |
| NC_011770 | 1392255 | C | T | 85 | 0 | 4 | 0 | INTRAGENIC | MODIFIER | | | | wspF | T | T | | | | | | | | | T | T |
| NC_011770 | 1392344 | T | C | 81 | 0 | 4 | 0 | INTRAGENIC | MODIFIER | | | | wspF | C | C | | | | | | | | | C | C |
| NC_011770 | 1392347 | T | C | 82 | 0 | 4 | 0 | INTRAGENIC | MODIFIER | | | | wspF | C | C | | | | | | | | | C | C |
| NC_011770 | 1392359 | T | C | 80 | 0 | 4 | 0 | INTRAGENIC | MODIFIER | | | | wspF | C | C | | | | | | | | | C | C |
| NC_011770 | 1392395 | C | T | 82 | 0 | 4 | 0 | INTRAGENIC | MODIFIER | | | | wspF | | | T | T | T | T | | | | | | |
| NC_011770 | 1392431 | T | C | 80 | 0 | 8 | 0 | INTRAGENIC | MODIFIER | | | | wspF | C | C | C | C | C | C | | | | | C | C |
| NC_011770 | 1392467 | A | G | 76 | 0 | 10 | 0 | INTRAGENIC | MODIFIER | | | | wspF | G | G | T | T | T | T | G | G | | | G | G |
| NC_011770 | 1392501 | G | A | 77 | 0 | 4 | 0 | INTRAGENIC | MODIFIER | | | | wspF | | | A | A | A | A | | | | | | |
| NC_011770 | 1392506 | A | C | 76 | 0 | 8 | 0 | INTRAGENIC | MODIFIER | | | | wspF | C | C | C | C | C | C | | | | | C | C |
| NC_011770 | 1392515 | G | A | 81 | 0 | 2 | 0 | INTRAGENIC | MODIFIER | | | | wspF | | | | | | | | | A | A | | |
| NC_011770 | 1392561 | G | A | 81 | 0 | 6 | 0 | INTRAGENIC | MODIFIER | | | | wspF | | | A | A | A | A | A | A | | | | |
| NC_011770 | 1392562 | A | G | 82 | 0 | 2 | 0 | INTRAGENIC | MODIFIER | | | | wspF | G | G | | | | | | | | | | |
| NC_011770 | 1392313 | CCATCGAC( | C | 83 | 0 | 1 | 0 | INTRAGENIC | MODIFIER | | | | wspF | | | | | | C | | | | | | |
| NC_011770 | 1393550 | C | T | 67 | 0 | 6 | 0 | NON_SYNONYMOU | MODERAT | MISSENSE | Cac/Tac | H293Y | wspR | T | T | T | T | T | T | | | | | | |
| NC_011770 | 2111114 | T | G | 52 | 0 | 2 | 0 | NON_SYNONYMOU | MODERAT | MISSENSE | aAg/aCg | K143T | xcpP | G | G | | | | | | | | | | |
| NC_011770 | 2109383 | A | C | 71 | 0 | 2 | 0 | NON_SYNONYMOU | MODERAT | MISSENSE | Tcg/Gcg | S483A | xcpQ | | | | | | | | | | | C | C |
| NC_011770 | 2109414 | C | A | 73 | 0 | 2 | 0 | NON_SYNONYMOU | MODERAT | MISSENSE | caG/caT | Q472H | xcpQ | | | | | | | | | | | A | A |
| NC_011770 | 2109416 | G | T | 74 | 0 | 2 | 0 | NON_SYNONYMOU | MODERAT | MISSENSE | Cag/Aag | Q472K | xcpQ | | | | | | | | | | | T | T |
| NC_011770 | 2112955 | T | C | 67 | 0 | 1 | 0 | NON_SYNONYMOU | MODERAT | MISSENSE | Tgc/Cgc | C399R | xcpR | | | | | | | | C | | | | |
| NC_011770 | 2113299 | C | A | 76 | 0 | 4 | 0 | NON_SYNONYMOU | MODERAT | MISSENSE | Ccc/Acc | P11T | xcpS | | | A | A | A | A | | | | | | |
| NC_011770 | 2114729 | C | T | 75 | 0 | 2 | 0 | NON_SYNONYMOU | MODERAT | MISSENSE | gCg/gTg | A80V | xcpT | | | | | | | T | T | | | | |
| NC_011770 | 2114845 | A | G | 82 | 0 | 4 | 0 | NON_SYNONYMOU | MODERAT | MISSENSE | Acc/Gcc | T119A | xcpT | | | G | G | G | G | | | | | | |
| NC_011770 | 2114947 | C | T | 86 | 0 | 1 | 0 | NON_SYNONYMOU | MODERAT | MISSENSE | Cgg/Tgg | R2W | xcpU | | | T | | | | | | | | | |

| | | | | | | | | | | | | | | | | | | | | | | | | | |
|---|---|---|---|---|---|---|---|---|---|---|---|---|---|---|---|---|---|---|---|---|---|---|---|---|---|
| NC_011770 | 2115642 | A | G | 81 | 0 | 2 | 0 | NON_SYNONYMOU | MODERAT | MISSENSE | Acg/Gcg | T62A | xcpV | | | | | | | | | | | G | G |
| NC_011770 | 2116118 | T | C | 78 | 0 | 2 | 0 | NON_SYNONYMOU | MODERAT | MISSENSE | Ttc/Ctc | F92L | xcpW | | | | | | | C | C | | | | |
| NC_011770 | 2116340 | G | A | 65 | 0 | 4 | 0 | NON_SYNONYMOU | MODERAT | MISSENSE | Gac/Aac | D166N | xcpW | | | A | A | A | A | | | | | | |
| NC_011770 | 2117306 | AG | A | 26 | 0 | 10 | 0 | FRAME_SHIFT | HIGH | | gag/ | E252 | xcpX | . | A | A | A | A | A | | A | A | A | A | A |
| NC_011770 | 2117310 | G | GCA | 26 | 0 | 12 | 0 | FRAME_SHIFT | HIGH | | aag/CAaag | K253Q? | xcpX | GCA | GCA | GCA | GCA | GCA | GCA | GCA | GCA | GCA | GCA | GCA | GCA |
| NC_011770 | 2117315 | CT | C | 25 | 0 | 12 | 0 | FRAME_SHIFT | HIGH | | act/ | T254 | xcpX | C | C | C | C | C | C | C | C | C | C | C | C |
| NC_011770 | 2116571 | A | G | 73 | 0 | 4 | 0 | NON_SYNONYMOU | MODERAT | MISSENSE | aAc/aGc | N6S | xcpX | | | G | G | G | G | | | | | | |
| NC_011770 | 2116820 | C | G | 79 | 0 | 2 | 0 | NON_SYNONYMOU | MODERAT | MISSENSE | aCg/aGg | T89R | xcpX | | | G | G | | | | | | | | |
| NC_011770 | 2117282 | C | A | 26 | 0 | 12 | 0 | NON_SYNONYMOU | MODERAT | MISSENSE | gCg/gAg | A243E | xcpX | A | A | A | A | A | A | A | A | A | A | A | A |
| NC_011770 | 2117319 | G | T | 25 | 0 | 12 | 0 | NON_SYNONYMOU | MODERAT | MISSENSE | gaG/gaT | E255D | xcpX | T | T | T | T | T | T | T | T | T | T | T | T |
| NC_011770 | 2117324 | G | C | 25 | 0 | 12 | 0 | NON_SYNONYMOU | MODERAT | MISSENSE | aGc/aCc | S257T | xcpX | C | C | C | C | C | C | C | C | C | C | C | C |
| NC_011770 | 2117336 | A | C | 25 | 0 | 12 | 0 | NON_SYNONYMOU | MODERAT | MISSENSE | aAc/aCc | N261T | xcpX | C | C | C | C | C | C | C | C | C | C | C | C |
| NC_011770 | 2117338 | G | C | 25 | 0 | 12 | 0 | NON_SYNONYMOU | MODERAT | MISSENSE | Gag/Cag | E262Q | xcpX | C | C | C | C | C | C | C | C | C | C | C | C |
| NC_011770 | 2117353 | G | A | 25 | 0 | 12 | 0 | NON_SYNONYMOU | MODERAT | MISSENSE | Gca/Aca | A267T | xcpX | A | A | A | A | A | A | A | A | A | A | A | A |
| NC_011770 | 2117356 | A | G | 25 | 0 | 12 | 0 | NON_SYNONYMOU | MODERAT | MISSENSE | Agt/Ggt | S268G | xcpX | G | G | G | G | G | G | G | G | G | G | G | G |
| NC_011770 | 2117358 | T | A | 25 | 0 | 12 | 0 | NON_SYNONYMOU | MODERAT | MISSENSE | agT/agA | S268R | xcpX | A | A | A | A | A | A | A | A | A | A | A | A |
| NC_011770 | 2117366 | A | G | 26 | 0 | 12 | 0 | NON_SYNONYMOU | MODERAT | MISSENSE | aAc/aGc | N271S | xcpX | G | G | G | G | G | G | G | G | G | G | G | G |
| NC_011770 | 2117369 | T | A | 25 | 0 | 12 | 0 | NON_SYNONYMOU | MODERAT | MISSENSE | tTt/tAt | F272Y | xcpX | A | A | A | A | A | A | A | A | A | A | A | A |
| NC_011770 | 2117425 | T | C | 73 | 0 | 12 | 0 | NON_SYNONYMOU | MODERAT | MISSENSE | Tgg/Cgg | W291R | xcpX | C | C | C | C | C | C | C | C | C | C | C | C |
| NC_011770 | 2117822 | G | A | 66 | 0 | 2 | 0 | NON_SYNONYMOU | MODERAT | MISSENSE | atG/atA | M90I | xcpY | | | | | | | A | A | | | | |
| NC_011770 | 2118117 | G | C | 64 | 0 | 12 | 0 | NON_SYNONYMOU | MODERAT | MISSENSE | Gcc/Ccc | A189P | xcpY | C | C | C | C | C | C | C | C | C | C | C | C |
| NC_011770 | 2118121 | C | T | 68 | 0 | 9 | 0 | NON_SYNONYMOU | MODERAT | MISSENSE | cCg/cTg | P190L | xcpY | T | . | T | T | T | T | | | T | T | T | T |
| NC_011770 | 2118141 | C | G | 69 | 0 | 10 | 0 | NON_SYNONYMOU | MODERAT | MISSENSE | Ccc/Gcc | P197A | xcpY | G | G | G | G | G | G | G | G | | | G | G |
| NC_011770 | 2118655 | G | A | 71 | 0 | 2 | 0 | NON_SYNONYMOU | MODERAT | MISSENSE | aGc/aAc | S368N | xcpY | | | | | | | | | | | A | A |
| NC_011770 | 2118838 | A | G | 77 | 0 | 8 | 0 | NON_SYNONYMOU | MODERAT | MISSENSE | Atg/Gtg | M46V | xcpZ | | | G | G | G | G | | | G | G | G | G |
| NC_011770 | 2118841 | C | T | 79 | 0 | 2 | 0 | NON_SYNONYMOU | MODERAT | MISSENSE | Ctc/Ttc | L47F | xcpZ | | | | | | | T | T | | | | |
| NC_011770 | 2118955 | C | G | 75 | 0 | 2 | 0 | NON_SYNONYMOU | MODERAT | MISSENSE | Cag/Gag | Q85E | xcpZ | | | | | | | | | | | G | G |
| NC_011770 | 4209532 | A | G | 57 | 0 | 2 | 0 | NON_SYNONYMOU | MODERAT | MISSENSE | aAg/aGg | K165R | xdhA | | | | | | | | | G | G | | |
| NC_011770 | 4209730 | T | C | 58 | 0 | 2 | 0 | NON_SYNONYMOU | MODERAT | MISSENSE | gTc/gCc | V231A | xdhA | | | | | | | | | | | C | C |
| NC_011770 | 4210134 | A | G | 64 | 0 | 2 | 0 | NON_SYNONYMOU | MODERAT | MISSENSE | Agc/Ggc | S366G | xdhA | G | G | | | | | | | | | | |
| NC_011770 | 4210234 | C | T | 65 | 0 | 2 | 0 | NON_SYNONYMOU | MODERAT | MISSENSE | gCc/gTc | A399V | xdhA | | | | | | | | | | | T | T |
| NC_011770 | 4210588 | GC | G | 66 | 0 | 1 | 0 | FRAME_SHIFT | HIGH | | ccc/ | P36 | xdhB | | | | | | | | G | | | | |
| NC_011770 | 4210657 | G | A | 68 | 0 | 1 | 0 | NON_SYNONYMOU | MODERAT | MISSENSE | Gtc/Atc | V58I | xdhB | | | | | | | A | | | | | |
| NC_011770 | 4210886 | A | T | 63 | 0 | 6 | 0 | NON_SYNONYMOU | MODERAT | MISSENSE | aAg/aTg | K134M | xdhB | T | T | | | | | T | T | T | T | | |
| NC_011770 | 4211044 | A | G | 68 | 0 | 4 | 0 | NON_SYNONYMOU | MODERAT | MISSENSE | Acg/Gcg | T187A | xdhB | | | G | G | G | G | | | | | | |
| NC_011770 | 4211692 | A | G | 83 | 0 | 4 | 0 | NON_SYNONYMOU | MODERAT | MISSENSE | Aac/Gac | N403D | xdhB | G | G | | | | | G | G | | | | |
| NC_011770 | 4211815 | G | A | 80 | 0 | 2 | 0 | NON_SYNONYMOU | MODERAT | MISSENSE | Gcc/Acc | A444T | xdhB | A | A | | | | | | | | | | |
| NC_011770 | 4212196 | G | A | 67 | 0 | 4 | 0 | NON_SYNONYMOU | MODERAT | MISSENSE | Gtc/Atc | V571I | xdhB | | | A | A | A | A | | | | | | |
| NC_011770 | 4212256 | G | A | 68 | 0 | 6 | 0 | NON_SYNONYMOU | MODERAT | MISSENSE | Gtc/Atc | V591I | xdhB | A | A | | | | | A | A | A | A | | |
| NC_011770 | 5218053 | G | A | 77 | 0 | 2 | 0 | NON_SYNONYMOU | MODERAT | MISSENSE | Gcc/Acc | A248T | xenB | | | | | | | | | A | A | | |
| NC_011770 | 5218111 | A | G | 77 | 0 | 2 | 0 | NON_SYNONYMOU | MODERAT | MISSENSE | gAc/gGc | D267G | xenB | | | | | | | G | G | | | | |
| NC_011770 | 5218170 | C | A | 69 | 0 | 4 | 0 | NON_SYNONYMOU | MODERAT | MISSENSE | Cag/Aag | Q287K | xenB | | | | | | | A | A | A | A | | |
| NC_011770 | 6280258 | G | A | 78 | 0 | 2 | 0 | NON_SYNONYMOU | MODERAT | MISSENSE | cGc/cAc | R2H | xerC | | | | | | | A | A | | | | |
| NC_011770 | 6280836 | G | A | 82 | 0 | 10 | 0 | NON_SYNONYMOU | MODERAT | MISSENSE | Gcc/Acc | A195T | xerC | A | A | A | A | A | A | | | A | A | A | A |
| NC_011770 | 6280878 | G | A | 84 | 0 | 10 | 0 | NON_SYNONYMOU | MODERAT | MISSENSE | Ggc/Agc | G209S | xerC | A | A | A | A | A | A | | | A | A | A | A |
| NC_011770 | 1346008 | C | A | 90 | 0 | 2 | 0 | NON_SYNONYMOU | MODERAT | MISSENSE | Cat/Aat | H6N | xerD | | | | | | | | | | | A | A |
| NC_011770 | 3836813 | T | C | 81 | 0 | 4 | 0 | NON_SYNONYMOU | MODERAT | MISSENSE | Att/Gtt | I768V | xqhA | | | C | C | C | C | | | | | | |
| NC_011770 | 3837073 | A | G | 86 | 0 | 6 | 0 | NON_SYNONYMOU | MODERAT | MISSENSE | cTg/cCg | L681P | xqhA | | | G | G | G | G | | | | | G | G |
| NC_011770 | 3837190 | A | C | 85 | 0 | 2 | 0 | NON_SYNONYMOU | MODERAT | MISSENSE | gTc/gGc | V642G | xqhA | | | | | | | C | C | | | | |
| NC_011770 | 3837259 | G | A | 83 | 0 | 4 | 0 | NON_SYNONYMOU | MODERAT | MISSENSE | gCc/gTc | A619V | xqhA | | | | | | | | | A | A | A | A |
| NC_011770 | 3837448 | T | A | 82 | 0 | 10 | 0 | NON_SYNONYMOU | MODERAT | MISSENSE | cAg/cTg | Q556L | xqhA | | | A | A | A | A | A | A | A | A | A | A |
| NC_011770 | 3838565 | A | C | 73 | 0 | 4 | 0 | NON_SYNONYMOU | MODERAT | MISSENSE | Tcc/Gcc | S184A | xqhA | | | | | | | | | C | C | C | C |
| NC_011770 | 3838726 | T | G | 78 | 0 | 2 | 0 | NON_SYNONYMOU | MODERAT | MISSENSE | aAg/aCg | K130T | xqhA | | | | | | | | | | | G | G |
| NC_011770 | 3838738 | T | G | 77 | 0 | 2 | 0 | NON_SYNONYMOU | MODERAT | MISSENSE | gAc/gCc | D126A | xqhA | | | | | | | | | G | G | | |
| NC_011770 | 3839062 | A | G | 82 | 0 | 1 | 0 | NON_SYNONYMOU | MODERAT | MISSENSE | gTc/gCc | V18A | xqhA | | | | | | | G | | | | | |
| NC_011770 | 1294393 | C | A | 82 | 0 | 2 | 0 | NON_SYNONYMOU | MODERAT | MISSENSE | caG/caT | Q354H | xseA | | | | | | | | | A | A | | |
| NC_011770 | 1294637 | C | T | 70 | 0 | 2 | 0 | NON_SYNONYMOU | MODERAT | MISSENSE | gGc/gAc | G273D | xseA | | | | | | | | | T | T | | |
| NC_011770 | 1294956 | G | T | 73 | 0 | 2 | 0 | NON_SYNONYMOU | MODERAT | MISSENSE | Cgc/Agc | R167S | xseA | T | T | | | | | | | | | | |
| NC_011770 | 1295036 | A | G | 73 | 0 | 10 | 0 | NON_SYNONYMOU | MODERAT | MISSENSE | cTa/cCa | L140P | xseA | G | G | G | G | G | G | G | G | | | G | G |
| NC_011770 | 1295156 | T | C | 73 | 0 | 2 | 0 | NON_SYNONYMOU | MODERAT | MISSENSE | gAc/gGc | D100G | xseA | C | C | | | | | | | | | | |
| NC_011770 | 1295207 | C | T | 74 | 0 | 1 | 0 | NON_SYNONYMOU | MODERAT | MISSENSE | gGg/gAg | G83E | xseA | | T | | | | | | | | | | |
| NC_011770 | 1012289 | C | T | 65 | 0 | 1 | 0 | NON_SYNONYMOU | MODERAT | MISSENSE | cGc/cAc | R22H | xseB | | | | | | | | | | | T | |

| | | | | | | | | | | | | | | | | | | | | | | | | | |
|---|---|---|---|---|---|---|---|---|---|---|---|---|---|---|---|---|---|---|---|---|---|---|---|---|---|
| NC_011770 | 2951403 | T | G | 54 | 0 | 2 | 0 | NON_SYNONYMOU | MODERAT | MISSENSE | aAc/aCc | N224T | xthA | | | | | | | | | G | G | | |
| NC_011770 | 2991975 | C | A | 78 | 0 | 4 | 0 | NON_SYNONYMOU | MODERAT | MISSENSE | Cta/Ata | L159I | xylS | | | A | A | A | A | | | | | | |
| NC_011770 | 2992330 | G | A | 78 | 0 | 2 | 0 | NON_SYNONYMOU | MODERAT | MISSENSE | gGc/gAc | G277D | xylS | | | | | | | | | | | A | A |
| NC_011770 | 2992745 | G | A | 69 | 0 | 4 | 0 | NON_SYNONYMOU | MODERAT | MISSENSE | Gtc/Atc | V58I | xylX | | | A | A | A | A | | | | | | |
| NC_011770 | 2993292 | C | A | 73 | 0 | 6 | 0 | NON_SYNONYMOU | MODERAT | MISSENSE | cCg/cAg | P240Q | xylX | | | A | A | A | A | | | A | A | | |
| NC_011770 | 2993897 | C | T | 86 | 0 | 2 | 0 | NON_SYNONYMOU | MODERAT | MISSENSE | Cgc/Tgc | R442C | xylX | | | | | | | | | | | T | T |
| NC_011770 | 2994257 | T | C | 91 | 0 | 8 | 0 | NON_SYNONYMOU | MODERAT | MISSENSE | gTg/gCg | V107A | xylY | | | C | C | C | C | C | C | C | C | | |
| NC_011770 | 2995402 | C | T | 79 | 0 | 2 | 0 | NON_SYNONYMOU | MODERAT | MISSENSE | Cgc/Tgc | R315C | xylZ | | | | | | | | | | | T | T |
| NC_011770 | 440510 | G | A | 81 | 0 | 2 | 0 | NON_SYNONYMOU | MODERAT | MISSENSE | gCc/gTc | A2V | yqgF | | | | | | | | | | | A | A |
| NC_011770 | 4202335 | T | C | 66 | 0 | 2 | 0 | NON_SYNONYMOU | MODERAT | MISSENSE | Atc/Gtc | I278V | zipA | C | C | | | | | | | | | | |
| NC_011770 | 4202815 | T | G | 64 | 0 | 4 | 0 | NON_SYNONYMOU | MODERAT | MISSENSE | Acg/Ccg | T118P | zipA | G | G | | | | | | | G | G | | |
| NC_011770 | 4202920 | C | T | 66 | 0 | 4 | 0 | NON_SYNONYMOU | MODERAT | MISSENSE | Gcc/Acc | A83T | zipA | T | T | | | | | | | T | T | | |
| NC_011770 | 6529674 | G | GC | 76 | 0 | 1 | 0 | FRAME_SHIFT | HIGH | | ctg/cCtg | L22P? | znuB | | | | | | | GC | | | | | |
| NC_011770 | 6529929 | A | G | 78 | 0 | 2 | 0 | NON_SYNONYMOU | MODERAT | MISSENSE | aAg/aGg | K105R | znuB | | | | | | | G | G | | | | |
| NC_011770 | 6529547 | C | G | 84 | 0 | 2 | 0 | NON_SYNONYMOU | MODERAT | MISSENSE | gCc/gGc | A245G | znuC | | | | | | | G | G | | | | |
| NC_011770 | 2029197 | C | G | 81 | 0 | 2 | 0 | NON_SYNONYMOU | MODERAT | MISSENSE | Ccc/Gcc | P54A | zwf | | | | | | | | | G | G | | |
| NC_011770 | 2029267 | T | C | 71 | 0 | 4 | 0 | NON_SYNONYMOU | MODERAT | MISSENSE | gTg/gCg | V77A | zwf | | | | | | | | | C | C | C | C |
| NC_011770 | 15 | C | T | 30 | 0 | 2 | 0 | | | | | | | | | | | | | | | T | T | | |
| NC_011770 | 3160 | A | T | 61 | 0 | 6 | 0 | | | | | | | T | T | | | | | T | T | | | T | T |
| NC_011770 | 3163 | T | C | 58 | 0 | 6 | 0 | | | | | | | C | C | | | | | C | C | | | C | C |
| NC_011770 | 6697 | G | A | 51 | 1 | 3 | 0 | | | | | | | | | A | A | . | A | | | | | | |
| NC_011770 | 6778 | T | C | 50 | 0 | 2 | 0 | | | | | | | C | C | | | | | | | | | | |
| NC_011770 | 6780 | G | A | 51 | 0 | 2 | 0 | | | | | | | A | A | | | | | | | | | | |
| NC_011770 | 6792 | C | T | 54 | 0 | 4 | 0 | | | | | | | | | T | T | T | T | | | | | | |
| NC_011770 | 6796 | G | GC | 53 | 0 | 4 | 0 | | | | | | | GC | GC | | | | | | | | | GC | GC |
| NC_011770 | 6812 | A | G | 54 | 0 | 4 | 0 | | | | | | | | | G | G | G | G | | | | | | |
| NC_011770 | 6825 | T | C | 55 | 0 | 4 | 0 | | | | | | | | | C | C | C | C | | | | | | |
| NC_011770 | 6828 | A | G | 55 | 0 | 4 | 0 | | | | | | | | | G | G | G | G | | | | | | |
| NC_011770 | 6832 | A | G | 54 | 0 | 4 | 0 | | | | | | | | | G | G | G | G | | | | | | |
| NC_011770 | 6849 | G | T | 57 | 0 | 4 | 0 | | | | | | | | | T | T | T | T | | | | | | |
| NC_011770 | 6858 | T | C | 57 | 0 | 4 | 0 | | | | | | | | | C | C | C | C | | | | | | |
| NC_011770 | 6872 | T | A | 54 | 0 | 4 | 0 | | | | | | | | | A | A | A | A | | | | | | |
| NC_011770 | 6882 | G | A | 56 | 0 | 4 | 0 | | | | | | | | | A | A | A | A | | | | | | |
| NC_011770 | 6897 | T | G | 57 | 0 | 4 | 0 | | | | | | | | | G | G | G | G | | | | | | |
| NC_011770 | 6901 | G | A | 56 | 0 | 4 | 0 | | | | | | | | | A | A | A | A | | | | | | |
| NC_011770 | 6907 | C | T | 57 | 0 | 2 | 0 | | | | | | | | | | | | | | | | | T | T |
| NC_011770 | 6983 | C | T | 64 | 0 | 2 | 0 | | | | | | | | | | | | | | | | | T | T |
| NC_011770 | 8404 | G | GC | 84 | 0 | 6 | 0 | | | | | | | | | GC | GC | GC | GC | GC | GC | | | | |
| NC_011770 | 8411 | C | T | 90 | 0 | 2 | 0 | | | | | | | | | | | | | | | T | T | | |
| NC_011770 | 8472 | T | C | 85 | 0 | 2 | 0 | | | | | | | | | | | | | | | | | C | C |
| NC_011770 | 8473 | C | T | 86 | 0 | 2 | 0 | | | | | | | | | | | | | | | | | T | T |
| NC_011770 | 8542 | T | C | 85 | 0 | 6 | 0 | | | | | | | | | C | C | C | C | C | C | | | | |
| NC_011770 | 13494 | T | C | 50 | 1 | 1 | 0 | | | | | | | | | | | | | | | | | C | . |
| NC_011770 | 14095 | C | T | 97 | 0 | 2 | 0 | | | | | | | | | | | | | T | T | | | | |
| NC_011770 | 14173 | G | GC | 96 | 0 | 2 | 0 | | | | | | | | | | | | | GC | GC | | | | |
| NC_011770 | 15158 | AGCTCT | A | 88 | 0 | 2 | 0 | | | | | | | | | | | | | | | A | A | | |
| NC_011770 | 15167 | G | A | 87 | 0 | 2 | 0 | | | | | | | | | | | | | | | A | A | | |
| NC_011770 | 15468 | A | G | 96 | 0 | 8 | 0 | | | | | | | G | G | G | G | G | G | | | G | G | | |
| NC_011770 | 15509 | A | C | 98 | 0 | 2 | 0 | | | | | | | C | C | | | | | | | | | | |
| NC_011770 | 15521 | C | T | 91 | 0 | 2 | 0 | | | | | | | | | | | | | T | T | | | | |
| NC_011770 | 15605 | C | T | 90 | 0 | 8 | 0 | | | | | | | | | T | T | T | T | T | T | T | T | | |
| NC_011770 | 16266 | T | C | 62 | 0 | 4 | 0 | | | | | | | C | C | | | | | C | C | | | | |
| NC_011770 | 16276 | A | G | 62 | 0 | 2 | 0 | | | | | | | G | G | | | | | | | | | | |
| NC_011770 | 16295 | C | CA | 58 | 0 | 2 | 0 | | | | | | | | | | | | | CA | CA | | | | |
| NC_011770 | 16300 | G | GTGA | 63 | 0 | 2 | 0 | | | | | | | | | | | | | GTGA | GTGA | | | | |
| NC_011770 | 16321 | A | G | 63 | 0 | 4 | 0 | | | | | | | G | G | | | | | G | G | | | | |
| NC_011770 | 16621 | T | C | 85 | 0 | 2 | 0 | | | | | | | C | C | | | | | | | | | | |
| NC_011770 | 16623 | A | G | 84 | 0 | 2 | 0 | | | | | | | G | G | | | | | | | | | | |
| NC_011770 | 16642 | T | G | 82 | 0 | 10 | 0 | | | | | | | G | G | G | G | G | G | G | G | G | G | | |
| NC_011770 | 16679 | T | C | 77 | 0 | 10 | 0 | | | | | | | C | C | C | C | C | C | C | C | C | C | | |
| NC_011770 | 16687 | C | CA | 80 | 0 | 12 | 0 | | | | | | | CA | CA | CA | CA | CA | CA | CA | CA | CA | CA | CA | CA |
| NC_011770 | 16695 | G | T | 78 | 0 | 4 | 0 | | | | | | | | | | | | | T | T | T | T | | |

| | | | | | | | | | | | | | | | | | | | |
|---|---|---|---|---|---|---|---|---|---|---|---|---|---|---|---|---|---|---|---|
| NC_011770 | 16710 | T | C | 77 | 0 | 2 | 0 | C | C | | | | | | | | | | |
| NC_011770 | 16797 | T | C | 82 | 0 | 6 | 0 | C | C | C | C | C | C | | | | | | |
| NC_011770 | 16807 | C | G | 81 | 0 | 2 | 0 | G | G | | | | | | | | | | |
| NC_011770 | 16814 | C | G | 81 | 0 | 2 | 0 | G | G | | | | | | | | | | |
| NC_011770 | 16845 | G | T | 80 | 0 | 2 | 0 | T | T | | | | | | | | | | |
| NC_011770 | 16846 | G | A | 80 | 0 | 2 | 0 | A | A | | | | | | | | | | |
| NC_011770 | 16854 | C | A | 80 | 0 | 2 | 0 | A | A | | | | | | | | | | |
| NC_011770 | 16863 | G | A | 78 | 0 | 6 | 0 | | | A | A | A | A | | | | | A | A |
| NC_011770 | 16870 | A | G | 74 | 1 | 4 | 0 | . | . | G | G | G | G | | | | | | |
| NC_011770 | 16876 | T | C | 82 | 0 | 4 | 0 | | | C | C | C | C | | | | | | |
| NC_011770 | 17268 | A | G | 105 | 0 | 6 | 0 | G | G | | | | | | | G | G | G | G |
| NC_011770 | 17272 | A | C | 99 | 0 | 2 | 0 | | | | | | | | | | | C | C |
| NC_011770 | 17280 | G | C | 99 | 0 | 2 | 0 | C | C | | | | | | | | | | |
| NC_011770 | 17306 | G | C | 101 | 0 | 2 | 0 | | | | | | | | | | | C | C |
| NC_011770 | 18732 | GA | G | 89 | 0 | 2 | 0 | | | | | | | | | | | G | G |
| NC_011770 | 22815 | C | T | 69 | 0 | 2 | 0 | | | | | | | T | T | | | | |
| NC_011770 | 22858 | A | G | 66 | 0 | 8 | 0 | G | G | G | G | G | G | | | G | G | | |
| NC_011770 | 25609 | C | T | 73 | 0 | 1 | 0 | | | | | | | T | | | | | |
| NC_011770 | 25696 | C | T | 70 | 0 | 2 | 0 | T | T | | | | | | | | | | |
| NC_011770 | 27534 | G | A | 70 | 0 | 2 | 0 | | | | | | | | | A | A | | |
| NC_011770 | 27535 | C | A | 71 | 0 | 2 | 0 | A | A | | | | | | | | | | |
| NC_011770 | 27556 | A | G | 73 | 0 | 2 | 0 | G | G | | | | | | | | | | |
| NC_011770 | 27565 | C | T | 72 | 0 | 2 | 0 | T | T | | | | | | | | | | |
| NC_011770 | 32974 | A | G | 79 | 0 | 8 | 0 | | | G | G | G | G | G | G | | | G | G |
| NC_011770 | 34605 | T | G | 71 | 0 | 6 | 0 | G | G | | | | | G | G | | | G | G |
| NC_011770 | 35757 | C | T | 86 | 0 | 4 | 0 | T | T | | | | | | | T | T | | |
| NC_011770 | 35871 | C | T | 91 | 0 | 2 | 0 | | | | | | | | | | | T | T |
| NC_011770 | 40423 | A | T | 82 | 0 | 6 | 0 | T | T | | | | | | | T | T | T | T |
| NC_011770 | 40433 | G | A | 83 | 0 | 6 | 0 | A | A | | | | | | | A | A | A | A |
| NC_011770 | 40444 | C | T | 80 | 0 | 4 | 0 | T | T | | | | | | | T | T | | |
| NC_011770 | 40883 | C | T | 69 | 0 | 4 | 0 | | | T | T | T | T | | | | | | |
| NC_011770 | 40884 | T | C | 69 | 0 | 2 | 0 | | | | | | | | | | | C | C |
| NC_011770 | 40885 | G | A | 70 | 0 | 4 | 0 | | | A | A | A | A | | | | | | |
| NC_011770 | 40969 | A | G | 67 | 0 | 4 | 0 | | | G | G | G | G | | | | | | |
| NC_011770 | 40994 | T | C | 70 | 0 | 4 | 0 | | | | | | | | | C | C | C | C |
| NC_011770 | 40995 | T | C | 72 | 0 | 4 | 0 | | | C | C | C | C | | | | | | |
| NC_011770 | 53534 | A | G | 66 | 6 | 6 | 0 | G | G | G | G | G | G | . | . | . | . | . | . |
| NC_011770 | 53552 | T | G | 68 | 6 | 2 | 0 | G | G | | | | | . | . | . | . | . | . |
| NC_011770 | 53589 | T | C | 68 | 6 | 2 | 0 | C | C | | | | | . | . | . | . | . | . |
| NC_011770 | 53852 | ATTTGAAT( | A | 40 | 4 | 2 | 0 | A | A | | | | | . | . | | | . | . |
| NC_011770 | 53883 | G | A | 34 | 6 | 6 | 0 | A | A | A | A | A | A | . | . | . | . | . | . |
| NC_011770 | 54013 | A | G | 13 | 8 | 4 | 0 | . | . | G | G | G | G | . | . | . | . | . | . |
| NC_011770 | 54030 | T | A | 13 | 8 | 4 | 0 | . | . | A | A | A | A | . | . | . | . | . | . |
| NC_011770 | 54065 | C | A | 13 | 8 | 4 | 0 | . | . | A | A | A | A | . | . | . | . | . | . |
| NC_011770 | 54262 | G | A | 120 | 2 | 2 | 0 | | | | | | | A | A | | | . | . |
| NC_011770 | 55123 | G | A | 16 | 8 | 4 | 0 | . | . | . | . | . | . | A | A | . | . | A | A |
| NC_011770 | 55741 | C | T | 28 | 6 | 2 | 0 | . | . | . | . | . | . | | | | | T | T |
| NC_011770 | 55748 | G | T | 28 | 6 | 2 | 0 | . | . | . | . | . | . | | | | | T | T |
| NC_011770 | 55775 | T | C | 29 | 6 | 2 | 0 | . | . | . | . | . | . | | | | | C | C |
| NC_011770 | 55803 | T | C | 29 | 6 | 2 | 0 | . | . | . | . | . | . | | | | | C | C |
| NC_011770 | 55817 | T | C | 28 | 6 | 4 | 0 | . | . | . | . | . | . | C | C | | | C | C |
| NC_011770 | 55818 | A | G | 28 | 6 | 2 | 0 | . | . | . | . | . | . | | | | | G | G |
| NC_011770 | 55834 | G | A | 24 | 6 | 2 | 0 | . | . | . | . | . | . | | | | | A | A |
| NC_011770 | 55934 | C | T | 65 | 0 | 8 | 0 | T | T | | | | | T | T | T | T | T | T |
| NC_011770 | 55945 | G | A | 66 | 0 | 8 | 0 | | | A | A | A | A | A | A | A | A | | |
| NC_011770 | 55999 | A | G | 76 | 0 | 6 | 0 | G | G | | | | | | | G | G | G | G |
| NC_011770 | 56009 | A | G | 75 | 0 | 2 | 0 | | | | | | | | | | | G | G |
| NC_011770 | 56012 | G | GA | 79 | 0 | 2 | 0 | GA | GA | | | | | | | | | | |
| NC_011770 | 56063 | G | A | 82 | 0 | 2 | 0 | | | | | | | | | | | A | A |
| NC_011770 | 56086 | T | C | 85 | 0 | 6 | 0 | | | | | | | C | C | C | C | C | C |
| NC_011770 | 56100 | T | G | 82 | 0 | 8 | 0 | G | G | | | | | C | C | C | C | C | C |
| NC_011770 | 56101 | A | T | 82 | 0 | 6 | 0 | | | | | | | T | T | T | T | T | T |

| | | | | | | | | | | | | | | | | | | | |
|---|---|---|---|---|---|---|---|---|---|---|---|---|---|---|---|---|---|---|---|
| NC_011770 | 56103 | G | A | 82 | 0 | 6 | 0 | | | | | | | A | A | A | A | A | A |
| NC_011770 | 56104 | G | A | 81 | 0 | 6 | 0 | | | | | | | A | A | A | A | A | A |
| NC_011770 | 56147 | T | C | 81 | 0 | 6 | 0 | | | | | | | C | C | C | C | C | C |
| NC_011770 | 56152 | A | G | 81 | 0 | 6 | 0 | | | | | | | G | G | G | G | G | G |
| NC_011770 | 56153 | A | G | 81 | 0 | 2 | 0 | | | | | | | G | G | | | | |
| NC_011770 | 56173 | C | G | 86 | 0 | 2 | 0 | G | G | | | | | | | | | | |
| NC_011770 | 56188 | A | G | 81 | 0 | 2 | 0 | | | | | | | | | | | G | G |
| NC_011770 | 56191 | C | T | 85 | 0 | 2 | 0 | | | | | | | | | | | T | T |
| NC_011770 | 56204 | C | CT | 87 | 0 | 2 | 0 | | | | | | | | | | | CT | CT |
| NC_011770 | 56294 | A | G | 86 | 0 | 4 | 0 | | | | | | | G | G | G | G | | |
| NC_011770 | 56356 | A | C | 88 | 0 | 6 | 0 | | | C | C | C | C | | | | | C | C |
| NC_011770 | 56765 | T | C | 95 | 0 | 2 | 0 | | | | | | | | | | | C | C |
| NC_011770 | 56783 | T | C | 96 | 0 | 8 | 0 | | | C | C | C | C | C | C | | | C | C |
| NC_011770 | 56799 | C | T | 99 | 0 | 2 | 0 | | | | | | | | | | | T | T |
| NC_011770 | 56818 | G | C | 91 | 0 | 10 | 0 | C | C | C | C | C | C | C | C | C | C | | |
| NC_011770 | 56840 | G | A | 90 | 0 | 2 | 0 | | | | | | | | | | | A | A |
| NC_011770 | 56976 | C | T | 82 | 0 | 2 | 0 | | | | | | | | | | | T | T |
| NC_011770 | 56996 | T | C | 80 | 0 | 4 | 0 | | | | | | | C | C | | | C | C |
| NC_011770 | 58452 | G | A | 71 | 0 | 2 | 0 | | | | | | | A | A | | | | |
| NC_011770 | 60011 | G | A | 64 | 0 | 4 | 0 | | | | | | | A | A | | | A | A |
| NC_011770 | 60051 | G | T | 63 | 0 | 8 | 0 | T | T | T | T | T | T | | | T | T | | |
| NC_011770 | 60057 | C | G | 62 | 0 | 8 | 0 | G | G | G | G | G | G | | | G | G | | |
| NC_011770 | 60073 | T | G | 62 | 0 | 8 | 0 | G | G | G | G | G | G | | | G | G | | |
| NC_011770 | 60251 | G | C | 42 | 0 | 6 | 0 | C | C | C | C | C | C | | | | | | |
| NC_011770 | 60318 | A | T | 42 | 0 | 12 | 0 | T | T | T | T | T | T | T | T | T | T | T | T |
| NC_011770 | 61614 | T | C | 87 | 0 | 12 | 0 | C | C | C | C | C | C | C | C | C | C | C | C |
| NC_011770 | 62756 | C | T | 69 | 0 | 4 | 0 | | | T | T | T | T | | | | | | |
| NC_011770 | 64614 | G | GGC | 26 | 0 | 2 | 0 | | | | | | | GGC | GGC | | | | |
| NC_011770 | 64627 | A | G | 26 | 0 | 2 | 0 | | | | | | | G | G | | | | |
| NC_011770 | 64664 | G | T | 22 | 0 | 4 | 0 | | | T | T | T | T | | | | | | |
| NC_011770 | 64668 | G | C | 23 | 0 | 2 | 0 | | | | | | | | | | | C | C |
| NC_011770 | 64694 | C | T | 17 | 0 | 2 | 0 | T | T | | | | | | | | | | |
| NC_011770 | 64703 | A | G | 19 | 0 | 6 | 0 | G | G | G | G | G | G | | | | | | |
| NC_011770 | 64730 | G | A | 21 | 0 | 4 | 0 | | | A | A | A | A | | | | | | |
| NC_011770 | 64752 | A | G | 24 | 0 | 4 | 0 | | | | | | | G | G | | | G | G |
| NC_011770 | 64801 | T | TG | 37 | 0 | 12 | 0 | TG | TG | TG | TG | TG | TG | TG | TG | TG | TG | TG | TG |
| NC_011770 | 65529 | G | T | 78 | 0 | 6 | 0 | | | T | T | T | T | | | | | T | T |
| NC_011770 | 65564 | C | A | 81 | 0 | 6 | 0 | | | A | A | A | A | | | | | A | A |
| NC_011770 | 65606 | C | G | 85 | 0 | 6 | 0 | | | G | G | G | G | | | | | G | G |
| NC_011770 | 65620 | T | C | 85 | 0 | 6 | 0 | | | C | C | C | C | | | | | C | C |
| NC_011770 | 65702 | C | T | 83 | 0 | 8 | 0 | | | T | T | T | T | | | T | T | T | T |
| NC_011770 | 65713 | A | C | 75 | 0 | 8 | 0 | | | C | C | C | C | | | C | C | C | C |
| NC_011770 | 65715 | C | G | 75 | 0 | 8 | 0 | | | G | G | G | G | | | G | G | G | G |
| NC_011770 | 65745 | C | T | 75 | 0 | 2 | 0 | | | | | | | | | | | T | T |
| NC_011770 | 65746 | T | C | 75 | 0 | 8 | 0 | | | C | C | C | C | | | C | C | C | C |
| NC_011770 | 65774 | C | T | 76 | 0 | 6 | 0 | | | T | T | T | T | | | | | T | T |
| NC_011770 | 65803 | T | C | 71 | 0 | 8 | 0 | | | C | C | C | C | | | A | A | C | C |
| NC_011770 | 65816 | C | T | 70 | 0 | 2 | 0 | | | | | | | | | | | T | T |
| NC_011770 | 65857 | GT | G | 70 | 0 | 4 | 0 | | | G | G | G | G | | | | | | |
| NC_011770 | 65960 | C | A | 75 | 0 | 2 | 0 | A | A | | | | | | | | | | |
| NC_011770 | 65963 | A | C | 76 | 0 | 8 | 0 | | | C | C | C | C | | | T | T | C | C |
| NC_011770 | 65965 | T | C | 76 | 0 | 2 | 0 | | | | | | | | | C | C | | |
| NC_011770 | 66026 | G | C | 75 | 0 | 2 | 0 | | | | | | | | | C | C | | |
| NC_011770 | 67960 | G | C | 72 | 0 | 2 | 0 | | | | | | | | | C | C | | |
| NC_011770 | 68553 | CT | C | 71 | 0 | 2 | 0 | | | | | | | C | C | | | | |
| NC_011770 | 69208 | G | T | 81 | 0 | 4 | 0 | | | | | | | | | T | T | T | T |
| NC_011770 | 69209 | C | T | 82 | 0 | 2 | 0 | | | | | | | | | | | T | T |
| NC_011770 | 70723 | C | T | 82 | 0 | 2 | 0 | | | | | | | | | T | T | | |
| NC_011770 | 70750 | A | G | 77 | 0 | 12 | 0 | G | G | G | G | G | G | G | G | G | G | G | G |
| NC_011770 | 71705 | G | A | 81 | 0 | 4 | 0 | | | A | A | A | A | | | | | | |
| NC_011770 | 71711 | C | T | 80 | 0 | 4 | 0 | | | T | T | T | T | | | | | | |
| NC_011770 | 72488 | A | G | 87 | 0 | 8 | 0 | | | G | G | G | G | G | G | | | G | G |

| | | | | | | | | | | | | | | | | | | | |
|---|---|---|---|---|---|---|---|---|---|---|---|---|---|---|---|---|---|---|---|
| NC_011770 | 73822 | C | T | 67 | 0 | 2 | 0 | | | | | | | T | T | | | | |
| NC_011770 | 73854 | T | C | 67 | 0 | 8 | 0 | | | C | C | C | C | C | C | C | C | | |
| NC_011770 | 76518 | A | G | 35 | 1 | 11 | 0 | G | . | G | G | G | G | G | G | G | G | G | G |
| NC_011770 | 80149 | C | T | 47 | 0 | 2 | 0 | | | | | | | | | T | T | | |
| NC_011770 | 80173 | C | G | 41 | 0 | 2 | 0 | | | | | | | | | G | G | | |
| NC_011770 | 80197 | C | T | 27 | 0 | 2 | 0 | T | T | | | | | | | | | | |
| NC_011770 | 81334 | T | C | 65 | 0 | 2 | 0 | | | | | | | | | | | C | C |
| NC_011770 | 82446 | G | A | 86 | 0 | 2 | 0 | | | | | | | | | | | A | A |
| NC_011770 | 82448 | G | A | 84 | 0 | 1 | 0 | | | | | | | A | | | | | |
| NC_011770 | 94021 | G | A | 94 | 0 | 12 | 0 | A | A | A | A | A | A | A | A | A | A | A | A |
| NC_011770 | 94121 | G | A | 87 | 0 | 2 | 0 | | | | | | | | | A | A | | |
| NC_011770 | 98969 | A | G | 74 | 0 | 4 | 0 | G | G | | | | | | | | | G | G |
| NC_011770 | 100240 | C | T | 74 | 0 | 2 | 0 | | | | | | | | | T | T | | |
| NC_011770 | 112006 | T | C | 93 | 0 | 2 | 0 | | | | | | | | | C | C | | |
| NC_011770 | 112016 | T | C | 98 | 0 | 2 | 0 | | | | | | | | | C | C | | |
| NC_011770 | 112021 | A | G | 97 | 0 | 2 | 0 | | | | | | | G | G | | | | |
| NC_011770 | 112072 | C | T | 97 | 0 | 4 | 0 | | | T | T | T | T | | | | | | |
| NC_011770 | 113691 | G | A | 57 | 0 | 6 | 0 | | | A | A | A | A | | | | | A | A |
| NC_011770 | 114252 | G | A | 80 | 0 | 4 | 0 | | | A | A | A | A | | | | | | |
| NC_011770 | 114255 | G | A | 79 | 0 | 2 | 0 | | | | | | | A | A | | | | |
| NC_011770 | 114300 | T | C | 82 | 0 | 4 | 0 | | | C | C | C | C | | | | | | |
| NC_011770 | 114302 | C | T | 82 | 0 | 4 | 0 | | | T | T | T | T | | | | | | |
| NC_011770 | 114333 | T | C | 83 | 0 | 6 | 0 | C | C | | | | | | | C | C | C | C |
| NC_011770 | 114347 | C | T | 82 | 0 | 2 | 0 | | | | | | | | | T | T | | |
| NC_011770 | 122498 | T | C | 73 | 0 | 2 | 0 | C | C | | | | | | | | | | |
| NC_011770 | 122552 | G | A | 74 | 0 | 2 | 0 | | | | | | | | | A | A | | |
| NC_011770 | 122629 | G | T | 69 | 0 | 10 | 0 | T | T | T | T | T | T | T | T | T | T | | |
| NC_011770 | 122638 | C | T | 74 | 0 | 10 | 0 | T | T | T | T | T | T | T | T | T | T | | |
| NC_011770 | 126055 | G | A | 86 | 0 | 2 | 0 | | | | | | | | | A | A | | |
| NC_011770 | 126083 | C | T | 83 | 0 | 2 | 0 | | | | | | | | | T | T | | |
| NC_011770 | 128975 | T | C | 87 | 0 | 2 | 0 | | | | | | | C | C | | | | |
| NC_011770 | 134764 | C | T | 68 | 0 | 4 | 0 | | | | | | | T | T | | | T | T |
| NC_011770 | 134768 | C | T | 66 | 0 | 4 | 0 | | | | | | | T | T | | | T | T |
| NC_011770 | 134787 | C | T | 67 | 0 | 4 | 0 | | | | | | | T | T | | | T | T |
| NC_011770 | 135264 | A | G | 83 | 0 | 2 | 0 | | | | | | | | | G | G | | |
| NC_011770 | 135269 | G | A | 85 | 0 | 10 | 0 | A | A | A | A | A | A | | | A | A | A | A |
| NC_011770 | 135348 | G | A | 97 | 0 | 2 | 0 | | | | | | | | | | | A | A |
| NC_011770 | 135867 | G | C | 90 | 0 | 4 | 0 | | | C | C | C | C | | | | | | |
| NC_011770 | 135870 | T | C | 88 | 0 | 12 | 0 | C | C | C | C | C | C | C | C | C | C | C | C |
| NC_011770 | 136030 | G | C | 81 | 0 | 6 | 0 | C | C | | | | | | | C | C | C | C |
| NC_011770 | 137451 | C | T | 54 | 0 | 6 | 0 | | | T | T | T | T | | | | | T | T |
| NC_011770 | 137497 | A | G | 56 | 0 | 4 | 0 | | | G | G | G | G | | | | | | |
| NC_011770 | 137510 | C | T | 60 | 0 | 8 | 0 | | | T | T | T | T | | | T | T | T | T |
| NC_011770 | 137511 | C | G | 57 | 0 | 8 | 0 | | | G | G | G | G | | | G | G | G | G |
| NC_011770 | 137523 | G | C | 62 | 0 | 4 | 0 | | | C | C | C | C | | | | | | |
| NC_011770 | 137530 | C | T | 61 | 0 | 2 | 0 | T | T | | | | | | | | | | |
| NC_011770 | 137532 | T | G | 60 | 0 | 2 | 0 | | | | | | | | | G | G | | |
| NC_011770 | 137533 | G | A | 60 | 0 | 2 | 0 | | | | | | | | | A | A | | |
| NC_011770 | 139781 | T | C | 81 | 0 | 2 | 0 | | | | | | | C | C | | | | |
| NC_011770 | 139792 | A | T | 77 | 0 | 4 | 0 | T | T | | | | | T | T | | | | |
| NC_011770 | 139824 | C | G | 78 | 0 | 2 | 0 | | | | | | | G | G | | | | |
| NC_011770 | 140700 | T | C | 84 | 0 | 12 | 0 | C | C | C | C | C | C | C | C | C | C | C | C |
| NC_011770 | 142216 | G | A | 80 | 0 | 10 | 0 | | | A | A | A | A | A | A | A | A | A | A |
| NC_011770 | 142261 | G | C | 82 | 0 | 2 | 0 | | | | | | | | | C | C | | |
| NC_011770 | 142273 | A | G | 83 | 0 | 2 | 0 | G | G | | | | | | | | | | |
| NC_011770 | 142293 | A | T | 78 | 0 | 8 | 0 | | | T | T | T | T | | | T | T | T | T |
| NC_011770 | 142298 | G | T | 76 | 0 | 2 | 0 | T | T | | | | | | | | | | |
| NC_011770 | 142303 | C | T | 75 | 0 | 6 | 0 | | | T | T | T | T | | | T | T | | |
| NC_011770 | 142311 | A | G | 71 | 0 | 12 | 0 | G | G | G | G | G | G | G | G | G | G | G | G |
| NC_011770 | 142312 | C | T | 71 | 0 | 6 | 0 | | | T | T | T | T | | | T | T | | |
| NC_011770 | 142313 | C | T | 71 | 0 | 2 | 0 | | | | | | | | | | | T | T |
| NC_011770 | 142314 | C | A | 70 | 0 | 6 | 0 | | | A | A | A | A | | | A | A | | |

| | | | | | | | | | | | | | | | | | | | |
|---|---|---|---|---|---|---|---|---|---|---|---|---|---|---|---|---|---|---|---|
| NC_011770 | 142315 | A | G | 69 | 0 | 12 | 0 | G | G | G | G | G | G | G | G | G | G | G | G |
| NC_011770 | 142350 | G | T | 75 | 0 | 4 | 0 | | | T | T | T | T | | | | | | |
| NC_011770 | 142362 | A | G | 74 | 0 | 4 | 0 | | | | | | | G | G | G | G | | |
| NC_011770 | 142364 | A | G | 74 | 0 | 4 | 0 | | | G | G | G | G | | | | | | |
| NC_011770 | 142367 | A | C | 74 | 0 | 12 | 0 | C | C | C | C | C | C | C | C | C | C | C | C |
| NC_011770 | 142369 | G | A | 74 | 0 | 12 | 0 | A | A | A | A | A | A | A | A | A | A | A | A |
| NC_011770 | 142386 | G | T | 77 | 0 | 4 | 0 | | | T | T | T | T | | | | | | |
| NC_011770 | 142420 | T | C | 82 | 0 | 2 | 0 | | | | | | | | | | | C | C |
| NC_011770 | 142945 | A | G | 69 | 0 | 12 | 0 | G | G | G | G | G | G | G | G | G | G | G | G |
| NC_011770 | 142946 | T | C | 70 | 0 | 12 | 0 | C | C | C | C | C | C | C | C | C | C | C | C |
| NC_011770 | 143037 | A | G | 76 | 0 | 2 | 0 | | | | | | | G | G | | | | |
| NC_011770 | 143800 | G | A | 97 | 0 | 2 | 0 | | | | | | | | | | | A | A |
| NC_011770 | 144340 | G | C | 81 | 0 | 2 | 0 | | | | | | | | | | | C | C |
| NC_011770 | 144809 | A | C | 67 | 0 | 8 | 0 | C | C | C | C | C | C | | | C | C | | |
| NC_011770 | 146370 | G | A | 71 | 0 | 2 | 0 | | | | | | | | | | | A | A |
| NC_011770 | 150688 | C | G | 71 | 0 | 8 | 0 | G | G | | | | | G | G | G | G | G | G |
| NC_011770 | 150689 | T | C | 70 | 0 | 8 | 0 | C | C | | | | | C | C | C | C | C | C |
| NC_011770 | 150696 | G | T | 73 | 0 | 2 | 0 | | | | | | | | | | | T | T |
| NC_011770 | 150701 | G | A | 72 | 0 | 4 | 0 | | | | | | | A | A | A | A | | |
| NC_011770 | 150718 | T | C | 70 | 0 | 4 | 0 | C | C | | | | | | | C | C | | |
| NC_011770 | 150723 | C | T | 73 | 0 | 2 | 0 | | | | | | | T | T | | | | |
| NC_011770 | 150730 | T | C | 72 | 0 | 4 | 0 | C | C | | | | | | | C | C | | |
| NC_011770 | 150732 | C | T | 72 | 0 | 4 | 0 | T | T | | | | | | | T | T | | |
| NC_011770 | 150745 | A | G | 70 | 0 | 4 | 0 | G | G | | | | | | | G | G | | |
| NC_011770 | 150795 | A | G | 74 | 0 | 4 | 0 | G | G | | | | | | | | | G | G |
| NC_011770 | 152187 | C | T | 86 | 0 | 10 | 0 | T | T | T | T | T | T | T | T | T | T | | |
| NC_011770 | 152308 | G | T | 76 | 0 | 2 | 0 | | | | | | | | | | | T | T |
| NC_011770 | 152313 | A | G | 75 | 0 | 2 | 0 | | | | | | | | | | | G | G |
| NC_011770 | 152352 | T | G | 74 | 0 | 2 | 0 | | | | | | | | | | | G | G |
| NC_011770 | 152370 | G | A | 72 | 0 | 2 | 0 | | | | | | | | | | | A | A |
| NC_011770 | 152391 | A | G | 75 | 0 | 2 | 0 | | | | | | | | | | | G | G |
| NC_011770 | 152464 | C | G | 73 | 0 | 4 | 0 | | | G | G | G | G | | | | | | |
| NC_011770 | 152468 | T | C | 72 | 0 | 2 | 0 | | | | | | | | | | | C | C |
| NC_011770 | 152511 | A | G | 69 | 0 | 2 | 0 | G | G | | | | | | | | | | |
| NC_011770 | 152530 | G | A | 68 | 0 | 2 | 0 | A | A | | | | | | | | | | |
| NC_011770 | 152544 | C | T | 65 | 0 | 4 | 0 | | | T | T | T | T | | | | | | |
| NC_011770 | 152553 | G | A | 68 | 0 | 2 | 0 | A | A | | | | | | | | | | |
| NC_011770 | 152555 | T | C | 68 | 0 | 2 | 0 | C | C | | | | | | | | | | |
| NC_011770 | 152702 | G | A | 77 | 0 | 2 | 0 | | | | | | | | | | | A | A |
| NC_011770 | 152706 | C | A | 74 | 0 | 10 | 0 | A | A | A | A | A | A | | | A | A | A | A |
| NC_011770 | 152711 | T | C | 77 | 0 | 4 | 0 | | | C | C | C | C | | | | | | |
| NC_011770 | 152818 | C | G | 73 | 0 | 2 | 0 | | | | | | | | | | | G | G |
| NC_011770 | 152821 | G | A | 76 | 0 | 2 | 0 | | | | | | | | | | | A | A |
| NC_011770 | 152839 | A | G | 72 | 0 | 2 | 0 | | | | | | | | | | | G | G |
| NC_011770 | 152844 | C | T | 74 | 0 | 2 | 0 | | | | | | | | | | | T | T |
| NC_011770 | 152851 | T | G | 73 | 0 | 2 | 0 | | | | | | | | | | | G | G |
| NC_011770 | 152853 | C | T | 73 | 0 | 2 | 0 | | | | | | | | | | | T | T |
| NC_011770 | 152883 | A | G | 78 | 0 | 2 | 0 | | | | | | | | | | | G | G |
| NC_011770 | 152908 | A | G | 75 | 0 | 8 | 0 | | | G | G | G | G | | | G | G | G | G |
| NC_011770 | 152928 | C | T | 71 | 0 | 2 | 0 | | | | | | | | | | | T | T |
| NC_011770 | 152931 | C | T | 74 | 0 | 2 | 0 | | | | | | | | | | | T | T |
| NC_011770 | 152936 | G | A | 75 | 0 | 2 | 0 | | | | | | | | | | | A | A |
| NC_011770 | 152964 | C | T | 73 | 0 | 6 | 0 | | | | | | | T | T | T | T | G | G |
| NC_011770 | 152969 | G | A | 74 | 0 | 4 | 0 | | | | | | | A | A | A | A | | |
| NC_011770 | 152983 | C | T | 73 | 0 | 2 | 0 | | | | | | | | | | | T | T |
| NC_011770 | 152993 | A | G | 74 | 0 | 2 | 0 | | | | | | | | | | | G | G |
| NC_011770 | 153075 | T | C | 82 | 0 | 2 | 0 | | | | | | | | | C | C | | |
| NC_011770 | 153095 | C | T | 82 | 0 | 2 | 0 | | | | | | | | | T | T | | |
| NC_011770 | 153136 | C | T | 83 | 0 | 2 | 0 | T | T | | | | | | | | | | |
| NC_011770 | 153146 | C | T | 82 | 0 | 2 | 0 | | | | | | | | | | | T | T |
| NC_011770 | 153154 | A | G | 82 | 0 | 12 | 0 | G | G | G | G | G | G | G | G | G | G | G | G |
| NC_011770 | 153171 | G | C | 84 | 0 | 4 | 0 | C | C | | | | | | | | | C | C |

| | | | | | | | | | | | | | | | | | | | |
|---|---|---|---|---|---|---|---|---|---|---|---|---|---|---|---|---|---|---|---|
| NC_011770 | 153179 | C | T | 85 | 0 | 4 | 0 | | | T | T | T | T | | | | | | |
| NC_011770 | 153193 | T | C | 85 | 0 | 4 | 0 | C | C | | | | | | | | | C | C |
| NC_011770 | 153249 | C | T | 93 | 0 | 2 | 0 | | | | | | | | | | | T | T |
| NC_011770 | 153250 | C | G | 93 | 0 | 4 | 0 | G | G | | | | | | | | | G | G |
| NC_011770 | 153269 | T | C | 89 | 0 | 4 | 0 | C | C | | | | | | | | | C | C |
| NC_011770 | 155948 | A | AG | 87 | 0 | 4 | 0 | AG | AG | | | | | | | | | AG | AG |
| NC_011770 | 157023 | AT | A | 77 | 0 | 2 | 0 | | | | | | | A | A | | | | |
| NC_011770 | 162118 | G | T | 44 | 0 | 8 | 0 | T | T | T | T | T | T | | | T | T | | |
| NC_011770 | 162124 | C | T | 43 | 0 | 8 | 0 | T | T | T | T | T | T | | | T | T | | |
| NC_011770 | 162125 | G | T | 44 | 0 | 8 | 0 | T | T | T | T | T | T | | | T | T | | |
| NC_011770 | 162129 | A | G | 44 | 0 | 8 | 0 | G | G | G | G | G | G | | | G | G | | |
| NC_011770 | 162130 | C | T | 44 | 0 | 8 | 0 | T | T | T | T | T | T | | | T | T | | |
| NC_011770 | 162131 | A | G | 44 | 0 | 2 | 0 | | | | | | | | | | | G | G |
| NC_011770 | 162133 | A | G | 45 | 0 | 8 | 0 | G | G | G | G | G | G | | | G | G | | |
| NC_011770 | 162137 | G | A | 44 | 0 | 8 | 0 | A | A | A | A | A | A | | | A | A | | |
| NC_011770 | 162143 | A | G | 44 | 0 | 8 | 0 | G | G | G | G | G | G | | | G | G | | |
| NC_011770 | 162144 | G | T | 44 | 0 | 8 | 0 | T | T | T | T | T | T | | | T | T | | |
| NC_011770 | 162176 | G | C | 48 | 0 | 2 | 0 | | | | | | | | | | | C | C |
| NC_011770 | 162182 | C | T | 48 | 0 | 2 | 0 | | | | | | | | | | | T | T |
| NC_011770 | 162188 | T | C | 49 | 0 | 2 | 0 | | | | | | | | | | | C | C |
| NC_011770 | 162198 | G | T | 47 | 0 | 2 | 0 | | | | | | | | | | | T | T |
| NC_011770 | 162199 | T | C | 47 | 0 | 2 | 0 | | | | | | | | | | | C | C |
| NC_011770 | 162205 | T | C | 45 | 0 | 2 | 0 | | | | | | | | | | | C | C |
| NC_011770 | 163286 | C | T | 65 | 2 | 2 | 0 | T | T | | | | | | | | | . | . |
| NC_011770 | 164609 | GC | G | 78 | 0 | 2 | 0 | | | | | | | | | | | G | G |
| NC_011770 | 164680 | A | G | 76 | 0 | 2 | 0 | | | | | | | | | | | G | G |
| NC_011770 | 164687 | G | A | 78 | 0 | 2 | 0 | | | | | | | | | | | A | A |
| NC_011770 | 164700 | A | G | 75 | 0 | 2 | 0 | | | | | | | | | | | G | G |
| NC_011770 | 165957 | A | G | 89 | 0 | 6 | 0 | | | G | G | G | G | | | C | C | | |
| NC_011770 | 165963 | C | T | 94 | 0 | 2 | 0 | | | | | | | T | T | | | | |
| NC_011770 | 166968 | T | C | 89 | 0 | 6 | 0 | C | C | | | | | C | C | C | C | | |
| NC_011770 | 168010 | A | G | 78 | 0 | 12 | 0 | G | G | G | G | G | G | G | G | G | G | G | G |
| NC_011770 | 168049 | T | C | 75 | 0 | 6 | 0 | | | C | C | C | C | C | C | | | | |
| NC_011770 | 168060 | T | C | 77 | 0 | 4 | 0 | C | C | | | | | | | C | C | | |
| NC_011770 | 168064 | T | G | 76 | 0 | 12 | 0 | G | G | G | G | G | G | G | G | G | G | G | G |
| NC_011770 | 168085 | A | C | 75 | 0 | 4 | 0 | | | C | C | C | C | | | | | | |
| NC_011770 | 168086 | A | G | 74 | 0 | 4 | 0 | | | G | G | G | G | | | | | | |
| NC_011770 | 168169 | T | C | 69 | 0 | 2 | 0 | | | | | | | | | | | C | C |
| NC_011770 | 169732 | T | C | 52 | 0 | 6 | 0 | | | C | C | C | C | | | C | C | | |
| NC_011770 | 169738 | C | T | 52 | 0 | 2 | 0 | | | | | | | | | | | T | T |
| NC_011770 | 169740 | G | C | 52 | 0 | 6 | 0 | | | C | C | C | C | | | C | C | | |
| NC_011770 | 169785 | C | G | 49 | 0 | 6 | 0 | | | G | G | G | G | | | G | G | | |
| NC_011770 | 169789 | C | G | 48 | 0 | 6 | 0 | | | G | G | G | G | | | G | G | | |
| NC_011770 | 169830 | G | A | 48 | 0 | 2 | 0 | | | | | | | | | | | A | A |
| NC_011770 | 172287 | T | C | 53 | 0 | 2 | 0 | | | | | | | | | | | C | C |
| NC_011770 | 172314 | G | GC | 49 | 2 | 6 | 0 | GC | GCC | | | | | GC | GC | GC | GC | . | . |
| NC_011770 | 172365 | C | T | 40 | 2 | 4 | 0 | | | | | | | T | T | T | T | . | . |
| NC_011770 | 172376 | TC | T | 38 | 2 | 4 | 0 | | | | | | | T | T | T | T | . | . |
| NC_011770 | 172494 | A | T | 44 | 0 | 12 | 0 | T | T | T | T | T | T | T | T | T | T | C | C |
| NC_011770 | 172502 | C | T | 43 | 0 | 2 | 0 | | | | | | | | | | | T | T |
| NC_011770 | 172527 | C | T | 47 | 0 | 2 | 0 | | | | | | | | | | | T | T |
| NC_011770 | 172538 | C | A | 47 | 0 | 6 | 0 | A | A | | | | | | | A | A | A | A |
| NC_011770 | 172544 | G | A | 49 | 0 | 2 | 0 | | | | | | | | | | | A | A |
| NC_011770 | 172635 | T | C | 54 | 0 | 2 | 0 | | | | | | | | | | | C | C |
| NC_011770 | 173468 | A | G | 95 | 0 | 2 | 0 | | | | | | | | | | | G | G |
| NC_011770 | 173493 | G | A | 88 | 0 | 2 | 0 | | | | | | | | | | | A | A |
| NC_011770 | 173523 | C | T | 82 | 0 | 2 | 0 | | | | | | | | | | | T | T |
| NC_011770 | 173532 | C | T | 82 | 0 | 2 | 0 | | | | | | | T | T | | | | |
| NC_011770 | 174938 | C | T | 72 | 0 | 4 | 0 | | | T | T | T | T | | | | | | |
| NC_011770 | 175086 | G | T | 68 | 0 | 4 | 0 | | | T | T | T | T | | | | | | |
| NC_011770 | 176097 | G | T | 80 | 0 | 6 | 0 | | | T | T | T | T | | | | | T | T |
| NC_011770 | 181403 | A | G | 39 | 0 | 10 | 0 | | | G | G | G | G | G | G | G | G | G | G |

| | | | | | | | | | | | | | | | | | | | |
|---|---|---|---|---|---|---|---|---|---|---|---|---|---|---|---|---|---|---|---|
| NC_011770 | 181406 | G | T | 38 | 0 | 10 | 0 | | | T | T | T | T | T | T | T | T | T | T |
| NC_011770 | 181416 | A | G | 32 | 0 | 10 | 0 | | | G | G | G | G | G | G | G | G | G | G |
| NC_011770 | 181501 | C | T | 36 | 0 | 4 | 0 | | | T | T | T | T | | | | | | |
| NC_011770 | 181546 | T | C | 48 | 0 | 4 | 0 | | | | | | | | | C | C | C | C |
| NC_011770 | 182932 | GT | G | 75 | 0 | 6 | 0 | | | | | | | G | G | G | G | G | G |
| NC_011770 | 183114 | G | A | 70 | 0 | 12 | 0 | A | A | A | A | A | A | A | A | A | A | A | A |
| NC_011770 | 183298 | C | T | 78 | 0 | 2 | 0 | | | | | | | | | T | T | | |
| NC_011770 | 183361 | A | G | 82 | 0 | 12 | 0 | G | G | G | G | G | G | G | G | G | G | G | G |
| NC_011770 | 187339 | G | A | 70 | 0 | 6 | 0 | | | A | A | A | A | | | | | A | A |
| NC_011770 | 187351 | G | A | 72 | 0 | 6 | 0 | A | A | | | | | A | A | | | A | A |
| NC_011770 | 187364 | T | A | 70 | 0 | 10 | 0 | A | A | A | A | A | A | A | A | | | A | A |
| NC_011770 | 187584 | G | A | 68 | 0 | 6 | 0 | | | A | A | A | A | | | | | A | A |
| NC_011770 | 188804 | G | A | 86 | 0 | 2 | 0 | | | | | | | | | | | A | A |
| NC_011770 | 188876 | G | T | 90 | 0 | 4 | 0 | | | T | T | T | T | | | | | | |
| NC_011770 | 188903 | T | C | 89 | 0 | 6 | 0 | | | C | C | C | C | | | C | C | | |
| NC_011770 | 188928 | C | T | 88 | 0 | 4 | 0 | | | T | T | T | T | | | | | | |
| NC_011770 | 188989 | T | A | 83 | 0 | 2 | 0 | | | | | | | | | | | A | A |
| NC_011770 | 189000 | T | C | 59 | 0 | 4 | 0 | | | C | C | C | C | | | | | | |
| NC_011770 | 189015 | T | C | 50 | 3 | 1 | 0 | | | . | . | C | . | | | | | | |
| NC_011770 | 189023 | G | GC | 78 | 0 | 4 | 0 | | | GC | GC | GC | GC | | | | | | |
| NC_011770 | 195642 | A | G | 35 | 0 | 2 | 0 | G | G | | | | | | | | | | |
| NC_011770 | 195770 | G | A | 5 | 3 | 2 | 0 | | . | . | | | | A | A | . | | | |
| NC_011770 | 195798 | CG | C | 4 | 2 | 1 | 0 | . | | | | | | C | . | | | | |
| NC_011770 | 198400 | C | T | 72 | 0 | 2 | 0 | | | | | | | T | T | | | | |
| NC_011770 | 202916 | A | C | 81 | 0 | 2 | 0 | | | | | | | | | | | C | C |
| NC_011770 | 204826 | T | G | 65 | 0 | 4 | 0 | | | G | G | G | G | | | | | | |
| NC_011770 | 204834 | G | A | 65 | 0 | 4 | 0 | | | A | A | A | A | | | | | | |
| NC_011770 | 205858 | A | G | 69 | 0 | 2 | 0 | | | | | | | G | G | | | | |
| NC_011770 | 206714 | C | T | 76 | 0 | 12 | 0 | T | T | T | T | T | T | T | T | T | T | T | T |
| NC_011770 | 208392 | A | G | 44 | 0 | 2 | 0 | | | | | | | | | G | G | | |
| NC_011770 | 208422 | A | AG | 40 | 0 | 11 | 0 | AG | AG | AG | AG | AG | AG | AG | AG | AG | | AG | AG |
| NC_011770 | 210923 | A | C | 56 | 0 | 6 | 0 | C | C | | | | | | | C | C | C | C |
| NC_011770 | 210964 | C | A | 57 | 0 | 6 | 0 | A | A | | | | | | | A | A | A | A |
| NC_011770 | 211037 | G | T | 57 | 0 | 2 | 0 | | | | | | | | | | | T | T |
| NC_011770 | 211051 | C | T | 60 | 0 | 2 | 0 | | | | | | | | | | | T | T |
| NC_011770 | 211054 | G | T | 60 | 0 | 2 | 0 | | | | | | | | | T | T | | |
| NC_011770 | 211156 | A | T | 64 | 0 | 2 | 0 | | | | | | | | | | | T | T |
| NC_011770 | 212255 | G | C | 62 | 0 | 2 | 0 | | | | | | | | | | | C | C |
| NC_011770 | 212630 | C | T | 57 | 0 | 2 | 0 | T | T | | | | | | | | | | |
| NC_011770 | 216722 | C | G | 43 | 2 | 6 | 0 | G | G | | | | | G | G | . | . | G | G |
| NC_011770 | 217764 | A | G | 61 | 0 | 12 | 0 | G | G | G | G | G | G | G | G | G | G | G | G |
| NC_011770 | 217778 | C | T | 71 | 0 | 2 | 0 | | | | | | | T | T | | | | |
| NC_011770 | 217826 | C | G | 61 | 0 | 4 | 0 | | | G | G | G | G | | | | | | |
| NC_011770 | 217874 | C | T | 65 | 0 | 4 | 0 | | | T | T | T | T | | | | | | |
| NC_011770 | 217908 | G | GC | 31 | 0 | 3 | 0 | | | . | GC | GCC | GCC | | | | | | |
| NC_011770 | 217978 | C | T | 61 | 0 | 4 | 0 | | | T | T | T | T | | | | | | |
| NC_011770 | 217987 | C | T | 61 | 0 | 4 | 0 | | | T | T | T | T | | | | | | |
| NC_011770 | 221357 | C | A | 93 | 0 | 8 | 0 | | | A | A | A | A | A | A | | | A | A |
| NC_011770 | 221375 | A | C | 90 | 0 | 2 | 0 | C | C | | | | | | | | | | |
| NC_011770 | 222288 | G | A | 78 | 0 | 2 | 0 | | | | | | | A | A | | | | |
| NC_011770 | 222426 | C | T | 72 | 0 | 2 | 0 | | | | | | | | | T | T | | |
| NC_011770 | 222495 | C | T | 70 | 0 | 6 | 0 | | | T | T | T | T | T | T | | | | |
| NC_011770 | 222519 | A | C | 70 | 0 | 12 | 0 | C | C | C | C | C | C | C | C | C | C | C | C |
| NC_011770 | 222584 | C | G | 70 | 0 | 10 | 0 | G | G | G | G | G | G | G | G | G | G | | |
| NC_011770 | 222621 | A | G | 71 | 0 | 10 | 0 | G | G | G | G | G | G | G | G | G | G | | |
| NC_011770 | 222681 | G | A | 67 | 0 | 2 | 0 | A | A | | | | | | | | | | |
| NC_011770 | 222712 | G | C | 73 | 0 | 2 | 0 | C | C | | | | | | | | | | |
| NC_011770 | 222726 | T | C | 70 | 0 | 10 | 0 | C | C | C | C | C | C | C | C | C | C | | |
| NC_011770 | 222733 | C | T | 70 | 0 | 2 | 0 | | | | | | | T | T | | | | |
| NC_011770 | 224285 | C | T | 102 | 0 | 4 | 0 | T | T | | | | | | | T | T | | |
| NC_011770 | 226056 | T | C | 53 | 0 | 10 | 0 | C | C | C | C | C | C | C | C | | | C | C |
| NC_011770 | 226077 | C | T | 53 | 0 | 2 | 0 | | | | | | | | | | | T | T |

| | | | | | | | | | | | | | | | | | | | |
|---|---|---|---|---|---|---|---|---|---|---|---|---|---|---|---|---|---|---|---|
| NC_011770 | 228281 | T | C | 85 | 0 | 8 | 0 | C | C | | | | | C | C | C | C | C | C |
| NC_011770 | 228284 | G | A | 84 | 0 | 4 | 0 | A | A | | | | | | | | | A | A |
| NC_011770 | 228301 | T | C | 80 | 0 | 2 | 0 | | | | | | | C | C | | | | |
| NC_011770 | 228335 | C | T | 81 | 0 | 2 | 0 | T | T | | | | | | | | | | |
| NC_011770 | 228352 | C | T | 82 | 0 | 2 | 0 | | | | | | | T | T | | | | |
| NC_011770 | 228594 | C | T | 84 | 0 | 2 | 0 | | | | | | | T | T | | | | |
| NC_011770 | 228608 | G | C | 84 | 0 | 2 | 0 | | | | | | | | | C | C | | |
| NC_011770 | 228681 | G | A | 83 | 0 | 8 | 0 | A | A | | | | | A | A | A | A | A | A |
| NC_011770 | 229456 | C | T | 89 | 0 | 2 | 0 | | | | | | | | | | | T | T |
| NC_011770 | 236907 | CCAGCGGT | C | 70 | 0 | 4 | 0 | | | | | | | C | C | | | C | C |
| NC_011770 | 236923 | T | C | 74 | 0 | 12 | 0 | C | C | C | C | C | C | C | C | C | C | C | C |
| NC_011770 | 237906 | C | T | 45 | 0 | 2 | 0 | | | | | | | T | T | | | | |
| NC_011770 | 238642 | A | G | 60 | 0 | 2 | 0 | | | | | | | | | | | G | G |
| NC_011770 | 238649 | G | A | 60 | 0 | 2 | 0 | | | | | | | | | | | A | A |
| NC_011770 | 241463 | C | A | 96 | 0 | 2 | 0 | A | A | | | | | | | | | | |
| NC_011770 | 244377 | C | A | 71 | 0 | 4 | 0 | A | A | | | | | | | | | A | A |
| NC_011770 | 246535 | T | C | 80 | 0 | 10 | 0 | C | C | C | C | C | C | | | C | C | C | C |
| NC_011770 | 248065 | G | A | 80 | 0 | 10 | 0 | A | A | A | A | A | A | | | A | A | A | A |
| NC_011770 | 251046 | G | T | 94 | 0 | 2 | 0 | T | T | | | | | | | | | | |
| NC_011770 | 251055 | G | A | 91 | 0 | 2 | 0 | | | | | | | | | | | A | A |
| NC_011770 | 251104 | T | G | 89 | 0 | 2 | 0 | | | | | | | | | | | G | G |
| NC_011770 | 254703 | G | A | 97 | 0 | 2 | 0 | | | | | | | | | | | A | A |
| NC_011770 | 255113 | T | C | 89 | 0 | 2 | 0 | | | | | | | | | | | C | C |
| NC_011770 | 255117 | C | CG | 89 | 0 | 2 | 0 | | | | | | | | | | | CG | CG |
| NC_011770 | 255123 | C | G | 86 | 0 | 2 | 0 | | | | | | | | | | | G | G |
| NC_011770 | 255128 | G | T | 87 | 0 | 4 | 0 | T | T | | | | | | | | | T | T |
| NC_011770 | 255207 | C | T | 53 | 4 | 4 | 0 | . | . | T | T | T | T | | | | | . | . |
| NC_011770 | 255212 | G | A | 59 | 0 | 6 | 0 | | | A | A | A | A | | | A | A | | |
| NC_011770 | 255234 | T | A | 76 | 0 | 6 | 0 | | | A | A | A | A | | | A | A | | |
| NC_011770 | 255282 | C | T | 78 | 0 | 2 | 0 | | | | | | | T | T | | | | |
| NC_011770 | 256269 | C | T | 93 | 0 | 2 | 0 | | | | | | | | | | | T | T |
| NC_011770 | 256270 | C | T | 91 | 0 | 2 | 0 | | | | | | | | | | | T | T |
| NC_011770 | 256273 | C | T | 91 | 0 | 2 | 0 | | | | | | | | | | | T | T |
| NC_011770 | 256297 | C | T | 93 | 0 | 2 | 0 | | | | | | | | | | | T | T |
| NC_011770 | 256367 | C | A | 91 | 0 | 2 | 0 | A | A | | | | | | | | | | |
| NC_011770 | 257342 | A | G | 45 | 0 | 4 | 0 | | | | | | | G | G | G | G | | |
| NC_011770 | 257367 | G | T | 41 | 0 | 4 | 0 | | | T | T | T | T | | | | | | |
| NC_011770 | 257370 | C | T | 40 | 0 | 2 | 0 | | | | | | | | | T | T | | |
| NC_011770 | 257384 | G | T | 34 | 2 | 4 | 0 | | | T | T | T | T | | | . | . | | |
| NC_011770 | 257389 | C | T | 33 | 2 | 4 | 0 | | | T | T | T | T | | | . | . | | |
| NC_011770 | 257393 | G | A | 32 | 2 | 4 | 0 | | | A | A | A | A | | | . | . | | |
| NC_011770 | 257408 | G | T | 24 | 2 | 2 | 0 | | | | | | | | | . | . | T | T |
| NC_011770 | 257443 | G | A | 16 | 2 | 2 | 0 | | | | | | | A | A | . | . | | |
| NC_011770 | 257449 | G | A | 16 | 2 | 1 | 0 | | | | | | | A | | . | . | | |
| NC_011770 | 257468 | T | G | 9 | 5 | 4 | 0 | . | G | G | . | . | G | | | . | . | A | . |
| NC_011770 | 257471 | C | T | 9 | 4 | 4 | 0 | . | T | T | T | . | T | | | . | . | | |
| NC_011770 | 257473 | T | C | 9 | 4 | 8 | 0 | . | C | C | C | . | C | C | C | . | . | C | C |
| NC_011770 | 257485 | G | T | 4 | 8 | 2 | 0 | . | . | . | . | . | . | T | T | . | . | | |
| NC_011770 | 257486 | C | A | 4 | 8 | 4 | 0 | . | . | . | . | . | . | A | A | . | . | A | A |
| NC_011770 | 257490 | C | T | 5 | 8 | 2 | 0 | . | . | . | . | . | . | T | T | . | . | | |
| NC_011770 | 257494 | G | A | 4 | 8 | 2 | 0 | . | . | . | . | . | . | A | A | . | . | | |
| NC_011770 | 257508 | T | C | 2 | 9 | 3 | 0 | . | . | . | . | . | . | C | . | . | . | C | C |
| NC_011770 | 257509 | G | T | 2 | 9 | 1 | 0 | . | . | . | . | . | . | | . | . | . | T | . |
| NC_011770 | 257513 | T | C | 2 | 10 | 1 | 0 | . | . | . | . | . | . | . | . | . | . | C | . |
| NC_011770 | 257660 | G | A | 15 | 8 | 2 | 0 | . | . | . | . | . | . | A | A | . | . | | |
| NC_011770 | 257670 | A | G | 17 | 8 | 2 | 0 | . | . | . | . | . | . | G | G | . | . | | |
| NC_011770 | 257687 | G | GCC | 22 | 3 | 5 | 0 | . | GCC | GCC | GCC | GCC | GCC | | | . | . | | |
| NC_011770 | 257696 | A | G | 26 | 3 | 7 | 0 | . | G | G | G | G | G | G | G | . | . | | |
| NC_011770 | 257699 | A | G | 28 | 3 | 5 | 0 | . | G | G | G | G | G | | | . | . | | |
| NC_011770 | 257705 | G | A | 32 | 2 | 4 | 0 | | | | | | | A | A | . | . | A | A |
| NC_011770 | 257752 | G | A | 46 | 2 | 1 | 0 | | | | | | | A | | . | . | | |
| NC_011770 | 257771 | T | G | 50 | 2 | 2 | 0 | G | G | | | | | | | . | . | | |

| | | | | | | | | | | | | | | | | | | | |
|---|---|---|---|---|---|---|---|---|---|---|---|---|---|---|---|---|---|---|---|
| NC_011770 | 257788 | G | T | 56 | 0 | 8 | 0 | | | T | T | T | T | T | T | GCC | GCC | | |
| NC_011770 | 257793 | C | T | 58 | 0 | 6 | 0 | | | T | T | T | T | T | T | | | | |
| NC_011770 | 257797 | G | A | 58 | 0 | 6 | 0 | | | A | A | A | A | A | A | | | | |
| NC_011770 | 257800 | A | G | 64 | 0 | 2 | 0 | | | | | | | | | G | G | | |
| NC_011770 | 257806 | A | G | 64 | 0 | 2 | 0 | | | | | | | | | G | G | | |
| NC_011770 | 257816 | C | T | 64 | 0 | 4 | 0 | | | T | T | T | T | | | | | | |
| NC_011770 | 259263 | GT | G | 71 | 0 | 2 | 0 | | | | | | | | | G | G | | |
| NC_011770 | 259282 | G | A | 67 | 0 | 2 | 0 | A | A | | | | | | | | | | |
| NC_011770 | 259455 | A | G | 74 | 0 | 8 | 0 | G | G | G | G | G | G | G | G | | | | |
| NC_011770 | 259523 | T | C | 72 | 0 | 8 | 0 | C | C | C | C | C | C | C | C | | | | |
| NC_011770 | 259540 | C | T | 72 | 0 | 8 | 0 | T | T | T | T | T | T | T | T | | | | |
| NC_011770 | 259543 | C | T | 73 | 0 | 8 | 0 | T | T | T | T | T | T | T | T | | | | |
| NC_011770 | 260354 | T | C | 83 | 0 | 12 | 0 | C | C | C | C | C | C | C | C | C | C | C | C |
| NC_011770 | 260362 | G | A | 80 | 0 | 10 | 0 | | | A | A | A | A | A | A | A | A | A | A |
| NC_011770 | 260393 | C | T | 83 | 0 | 10 | 0 | | | T | T | T | T | T | T | T | T | T | T |
| NC_011770 | 260396 | CG | C | 84 | 0 | 10 | 0 | | | C | C | C | C | C | C | C | C | C | C |
| NC_011770 | 260410 | A | G | 80 | 0 | 10 | 0 | | | G | G | G | G | G | G | G | G | G | G |
| NC_011770 | 260436 | G | T | 68 | 0 | 10 | 0 | | | T | T | T | T | T | T | T | T | T | T |
| NC_011770 | 260438 | T | C | 68 | 0 | 10 | 0 | | | C | C | C | C | C | C | C | C | C | C |
| NC_011770 | 260439 | C | G | 68 | 0 | 4 | 0 | | | | | | | G | G | G | G | | |
| NC_011770 | 260441 | T | C | 68 | 0 | 10 | 0 | | | C | C | C | C | C | C | C | C | C | C |
| NC_011770 | 260444 | T | C | 68 | 0 | 12 | 0 | C | C | C | C | C | C | C | C | C | C | C | C |
| NC_011770 | 260447 | C | T | 67 | 0 | 10 | 0 | | | T | T | T | T | T | T | T | T | T | T |
| NC_011770 | 260448 | G | A | 68 | 0 | 10 | 0 | | | A | A | A | A | A | A | A | A | A | A |
| NC_011770 | 260449 | C | G | 69 | 0 | 10 | 0 | | | G | G | G | G | G | G | G | G | G | G |
| NC_011770 | 260481 | A | G | 79 | 0 | 10 | 0 | | | G | G | G | G | G | G | G | G | G | G |
| NC_011770 | 260483 | T | C | 80 | 0 | 10 | 0 | | | C | C | C | C | C | C | C | C | C | C |
| NC_011770 | 260484 | C | A | 79 | 0 | 10 | 0 | | | A | A | A | A | A | A | A | A | A | A |
| NC_011770 | 260485 | C | T | 80 | 0 | 8 | 0 | | | T | T | T | T | T | T | T | T | | |
| NC_011770 | 260503 | T | C | 83 | 0 | 10 | 0 | | | C | C | C | C | C | C | A | A | C | C |
| NC_011770 | 260507 | T | C | 85 | 0 | 10 | 0 | | | C | C | C | C | C | C | C | C | C | C |
| NC_011770 | 260534 | C | T | 78 | 0 | 10 | 0 | | | T | T | T | T | A | A | A | A | T | T |
| NC_011770 | 260535 | T | G | 79 | 0 | 2 | 0 | G | G | | | | | | | | | | |
| NC_011770 | 260538 | G | A | 78 | 0 | 10 | 0 | | | A | A | A | A | A | A | A | A | A | A |
| NC_011770 | 260539 | A | G | 78 | 0 | 10 | 0 | | | G | G | G | G | G | G | G | G | G | G |
| NC_011770 | 260568 | C | T | 86 | 0 | 2 | 0 | T | T | | | | | | | | | | |
| NC_011770 | 263486 | C | G | 89 | 0 | 2 | 0 | | | | | | | | | G | G | | |
| NC_011770 | 264796 | C | T | 72 | 0 | 8 | 0 | | | T | T | T | T | T | T | | | T | T |
| NC_011770 | 264809 | T | C | 71 | 0 | 2 | 0 | C | C | | | | | | | | | | |
| NC_011770 | 266189 | T | C | 41 | 0 | 4 | 0 | C | C | | | | | | | C | C | | |
| NC_011770 | 266328 | T | C | 37 | 0 | 12 | 0 | C | C | C | C | C | C | C | C | C | C | C | C |
| NC_011770 | 266547 | G | T | 56 | 0 | 8 | 0 | | | T | T | T | T | | | T | T | T | T |
| NC_011770 | 269395 | T | C | 91 | 0 | 12 | 0 | C | C | C | C | C | C | C | C | C | C | C | C |
| NC_011770 | 269411 | C | T | 92 | 0 | 12 | 0 | T | T | T | T | T | T | T | T | T | T | T | T |
| NC_011770 | 270760 | A | G | 89 | 0 | 2 | 0 | | | | | | | | | G | G | | |
| NC_011770 | 270792 | G | A | 92 | 0 | 2 | 0 | | | | | | | | | A | A | | |
| NC_011770 | 272390 | A | C | 69 | 0 | 2 | 0 | | | | | | | C | C | | | | |
| NC_011770 | 272403 | C | G | 62 | 0 | 10 | 0 | G | G | G | G | G | G | | | G | G | G | G |
| NC_011770 | 272407 | G | A | 64 | 0 | 4 | 0 | | | | | | | | | A | A | A | A |
| NC_011770 | 272408 | A | T | 64 | 0 | 4 | 0 | | | | | | | | | T | T | T | T |
| NC_011770 | 272418 | T | A | 64 | 0 | 2 | 0 | | | | | | | | | A | A | | |
| NC_011770 | 272422 | C | T | 63 | 0 | 2 | 0 | | | | | | | | | T | T | | |
| NC_011770 | 272439 | T | C | 65 | 0 | 8 | 0 | C | C | C | C | C | C | | | | | C | C |
| NC_011770 | 272473 | A | C | 68 | 0 | 2 | 0 | | | | | | | | | C | C | | |
| NC_011770 | 273783 | A | G | 68 | 0 | 2 | 0 | | | | | | | | | G | G | | |
| NC_011770 | 273803 | G | T | 61 | 0 | 2 | 0 | T | T | | | | | | | | | | |
| NC_011770 | 273830 | G | C | 64 | 0 | 6 | 0 | | | C | C | C | C | | | C | C | | |
| NC_011770 | 275184 | A | G | 75 | 0 | 10 | 0 | | | G | G | G | G | G | G | G | G | G | G |
| NC_011770 | 275236 | T | C | 79 | 0 | 10 | 0 | | | C | C | C | C | C | C | C | C | C | C |
| NC_011770 | 275240 | T | G | 78 | 0 | 10 | 0 | | | G | G | G | G | G | G | G | G | G | G |
| NC_011770 | 275247 | T | C | 77 | 0 | 2 | 0 | | | | | | | | | | | C | C |
| NC_011770 | 275249 | T | C | 79 | 0 | 2 | 0 | | | | | | | | | | | C | C |

| | | | | | | | | | | | | | | | | | | | |
|---|---|---|---|---|---|---|---|---|---|---|---|---|---|---|---|---|---|---|---|
| NC_011770 | 275250 | A | C | 80 | 0 | 2 | 0 | | | | | | | | | | | C | C |
| NC_011770 | 275690 | G | A | 86 | 0 | 8 | 0 | A | A | A | A | A | A | A | A | | | | |
| NC_011770 | 275737 | G | A | 92 | 0 | 2 | 0 | | | | | | | | | | | A | A |
| NC_011770 | 275792 | G | A | 4 | 3 | 9 | 0 | A | A | A | A | A | A | A | A | A | . | . | . |
| NC_011770 | 275793 | C | T | 2 | 5 | 7 | 0 | . | . | T | T | T | T | T | T | T | . | . | . |
| NC_011770 | 275794 | CG | C | 3 | 5 | 6 | 0 | . | . | C | C | C | C | . | C | C | . | . | . |
| NC_011770 | 275800 | C | T | 73 | 0 | 12 | 0 | T | T | T | T | T | T | T | T | T | T | T | T |
| NC_011770 | 276500 | A | G | 22 | 0 | 12 | 0 | G | G | G | G | G | G | G | G | G | G | G | G |
| NC_011770 | 276521 | A | G | 10 | 1 | 11 | 0 | G | . | G | G | G | G | G | G | G | G | G | G |
| NC_011770 | 276534 | GGCA | G | 13 | 0 | 12 | 0 | G | G | G | G | G | G | G | G | G | G | G | G |
| NC_011770 | 276553 | T | G | 14 | 0 | 12 | 0 | G | G | G | G | G | G | G | G | G | G | G | G |
| NC_011770 | 276559 | G | T | 16 | 0 | 12 | 0 | T | T | T | T | T | T | T | T | T | T | T | T |
| NC_011770 | 276573 | A | G | 17 | 0 | 12 | 0 | G | G | G | G | G | G | G | G | G | G | G | G |
| NC_011770 | 276574 | T | A | 17 | 0 | 12 | 0 | A | A | A | A | A | A | A | A | A | A | A | A |
| NC_011770 | 276580 | C | G | 23 | 0 | 12 | 0 | G | G | G | G | G | G | G | G | G | G | G | G |
| NC_011770 | 276586 | C | T | 25 | 0 | 12 | 0 | T | T | T | T | T | T | T | T | T | T | T | T |
| NC_011770 | 276705 | C | T | 40 | 0 | 6 | 0 | | | T | T | T | T | T | T | | | | |
| NC_011770 | 277072 | AG | A | 65 | 0 | 2 | 0 | | | | | | | A | A | | | | |
| NC_011770 | 277116 | G | A | 73 | 0 | 2 | 0 | A | A | | | | | | | | | | |
| NC_011770 | 279181 | C | T | 84 | 0 | 6 | 0 | T | T | | | | | | | T | T | T | T |
| NC_011770 | 279187 | C | T | 66 | 5 | 1 | 0 | . | . | | | | | | | . | . | . | T |
| NC_011770 | 279220 | C | T | 64 | 6 | 6 | 0 | . | . | T | T | T | T | T | T | . | . | . | . |
| NC_011770 | 281167 | T | C | 64 | 0 | 6 | 0 | C | C | | | | | | | C | C | C | C |
| NC_011770 | 281176 | C | T | 62 | 0 | 2 | 0 | | | | | | | T | T | | | | |
| NC_011770 | 281177 | C | T | 61 | 0 | 4 | 0 | | | T | T | T | T | | | | | | |
| NC_011770 | 281182 | C | A | 64 | 0 | 2 | 0 | | | | | | | | | | | A | A |
| NC_011770 | 281186 | A | G | 63 | 0 | 4 | 0 | G | G | | | | | | | G | G | | |
| NC_011770 | 281226 | C | CT | 57 | 0 | 4 | 0 | CT | CT | | | | | | | CT | CT | | |
| NC_011770 | 281245 | C | T | 55 | 0 | 12 | 0 | T | T | T | T | T | T | T | T | T | T | T | T |
| NC_011770 | 281261 | T | C | 55 | 0 | 2 | 0 | | | | | | | C | C | | | | |
| NC_011770 | 281267 | C | T | 56 | 0 | 6 | 0 | T | T | | | | | | | T | T | T | T |
| NC_011770 | 281483 | T | A | 45 | 4 | 4 | 0 | A | A | | | | | . | . | A | A | . | . |
| NC_011770 | 281581 | C | A | 40 | 4 | 4 | 0 | A | A | | | | | . | . | A | A | . | . |
| NC_011770 | 281621 | G | T | 35 | 4 | 4 | 0 | | | T | T | T | T | . | . | | | . | . |
| NC_011770 | 281642 | G | A | 36 | 4 | 4 | 0 | A | A | | | | | . | . | A | A | . | . |
| NC_011770 | 281660 | A | G | 37 | 4 | 8 | 0 | G | G | G | G | G | G | . | . | G | G | . | . |
| NC_011770 | 281697 | A | G | 40 | 4 | 8 | 0 | G | G | G | G | G | G | . | . | G | G | . | . |
| NC_011770 | 281766 | C | T | 42 | 4 | 4 | 0 | | | T | T | T | T | . | . | | | . | . |
| NC_011770 | 281834 | T | G | 46 | 4 | 4 | 0 | | | G | G | G | G | . | . | | | . | . |
| NC_011770 | 281836 | A | G | 47 | 4 | 4 | 0 | G | G | | | | | . | . | G | G | . | . |
| NC_011770 | 281919 | C | T | 50 | 4 | 4 | 0 | T | T | | | | | . | . | T | T | . | . |
| NC_011770 | 282083 | G | A | 51 | 4 | 4 | 0 | A | A | | | | | . | . | A | A | . | . |
| NC_011770 | 282342 | C | T | 52 | 4 | 4 | 0 | | | T | T | T | T | . | . | | | . | . |
| NC_011770 | 282365 | C | T | 52 | 4 | 4 | 0 | T | T | | | | | . | . | T | T | . | . |
| NC_011770 | 282463 | G | T | 55 | 4 | 4 | 0 | | | T | T | T | T | . | . | | | . | . |
| NC_011770 | 283951 | C | T | 55 | 4 | 4 | 0 | | | T | T | T | T | . | . | | | . | . |
| NC_011770 | 283984 | G | A | 53 | 4 | 4 | 0 | | | A | A | A | A | . | . | | | . | . |
| NC_011770 | 283987 | C | T | 52 | 4 | 4 | 0 | | | T | T | T | T | . | . | | | . | . |
| NC_011770 | 283988 | C | G | 53 | 4 | 4 | 0 | | | G | G | G | G | . | . | | | . | . |
| NC_011770 | 284000 | G | A | 56 | 4 | 4 | 0 | | | A | A | A | A | . | . | | | . | . |
| NC_011770 | 289822 | T | C | 61 | 0 | 4 | 0 | C | C | | | | | C | C | | | | |
| NC_011770 | 289829 | T | G | 53 | 0 | 4 | 0 | G | G | | | | | G | G | | | | |
| NC_011770 | 289832 | C | T | 52 | 0 | 4 | 0 | T | T | | | | | T | T | | | | |
| NC_011770 | 289837 | G | A | 52 | 0 | 4 | 0 | A | A | | | | | A | A | | | | |
| NC_011770 | 289844 | C | T | 53 | 0 | 4 | 0 | T | T | | | | | T | T | | | | |
| NC_011770 | 289845 | C | T | 53 | 0 | 4 | 0 | T | T | | | | | T | T | | | | |
| NC_011770 | 289848 | C | A | 51 | 0 | 4 | 0 | A | A | | | | | A | A | | | | |
| NC_011770 | 289849 | A | C | 51 | 0 | 2 | 0 | | | | | | | | | | | C | C |
| NC_011770 | 289853 | G | A | 51 | 0 | 4 | 0 | A | A | | | | | A | A | | | | |
| NC_011770 | 289860 | A | G | 50 | 0 | 4 | 0 | G | G | | | | | G | G | | | | |
| NC_011770 | 289864 | G | A | 49 | 0 | 4 | 0 | A | A | | | | | A | A | | | | |
| NC_011770 | 289866 | GCC | G | 49 | 0 | 4 | 0 | G | G | | | | | G | G | | | | |

| | | | | | | | | | | | | | | | | | | | |
|---|---|---|---|---|---|---|---|---|---|---|---|---|---|---|---|---|---|---|---|
| NC_011770 | 289891 | C | G | 49 | 0 | 4 | 0 | G | G | | | | | G | G | | | | |
| NC_011770 | 289892 | T | C | 49 | 0 | 4 | 0 | C | C | | | | | C | C | | | | |
| NC_011770 | 289902 | C | CT | 54 | 0 | 4 | 0 | CT | CT | | | | | CT | CT | | | | |
| NC_011770 | 289910 | G | T | 53 | 0 | 4 | 0 | T | T | | | | | T | T | | | | |
| NC_011770 | 289916 | T | G | 54 | 0 | 4 | 0 | G | G | | | | | G | G | | | | |
| NC_011770 | 289929 | A | C | 54 | 0 | 4 | 0 | C | C | | | | | C | C | | | | |
| NC_011770 | 289963 | T | C | 61 | 0 | 2 | 0 | | | | | | | C | C | | | | |
| NC_011770 | 290564 | T | A | 61 | 0 | 2 | 0 | | | | | | | A | A | | | | |
| NC_011770 | 290637 | C | G | 54 | 1 | 2 | 0 | | | | | | | . | | | | G | G |
| NC_011770 | 290654 | G | A | 55 | 2 | 2 | 0 | | | | | | | . | . | A | A | | |
| NC_011770 | 290690 | C | T | 55 | 2 | 4 | 0 | | | T | T | T | T | . | . | | | | |
| NC_011770 | 290691 | G | T | 54 | 2 | 4 | 0 | | | T | T | T | T | . | . | | | | |
| NC_011770 | 290712 | C | T | 56 | 2 | 2 | 0 | | | | | | | . | . | T | T | | |
| NC_011770 | 290757 | G | A | 50 | 2 | 2 | 0 | | | | | | | . | . | | | A | A |
| NC_011770 | 290759 | T | G | 48 | 2 | 2 | 0 | | | | | | | . | . | | | G | G |
| NC_011770 | 290765 | C | A | 47 | 2 | 2 | 0 | | | | | | | . | . | | | A | A |
| NC_011770 | 290858 | G | C | 42 | 4 | 6 | 0 | | | C | C | C | C | . | . | C | C | . | . |
| NC_011770 | 290889 | C | A | 44 | 4 | 4 | 0 | | | A | A | A | A | . | . | | | . | . |
| NC_011770 | 290961 | G | C | 43 | 4 | 2 | 0 | | | | | | | . | . | C | C | . | . |
| NC_011770 | 290967 | C | T | 43 | 4 | 4 | 0 | | | T | T | T | T | . | . | | | . | . |
| NC_011770 | 291004 | G | T | 41 | 4 | 8 | 0 | T | T | T | T | T | T | . | . | T | T | . | . |
| NC_011770 | 291054 | A | G | 41 | 4 | 6 | 0 | | | G | G | G | G | . | . | G | G | . | . |
| NC_011770 | 291107 | C | G | 37 | 4 | 4 | 0 | | | G | G | G | G | . | . | | | . | . |
| NC_011770 | 291208 | G | C | 34 | 4 | 2 | 0 | | | | | | | . | . | C | C | . | . |
| NC_011770 | 291300 | T | C | 36 | 4 | 4 | 0 | | | C | C | C | C | . | . | | | . | . |
| NC_011770 | 291346 | C | T | 39 | 4 | 6 | 0 | | | T | T | T | T | . | . | T | T | . | . |
| NC_011770 | 291378 | TAACGTTG( | T | 39 | 4 | 2 | 0 | | | | | T | T | . | . | | | . | . |
| NC_011770 | 291466 | C | T | 38 | 4 | 2 | 0 | T | T | | | | | . | . | | | . | . |
| NC_011770 | 291538 | AG | A | 44 | 4 | 8 | 0 | A | A | A | A | A | A | . | . | A | A | . | . |
| NC_011770 | 291619 | C | A | 47 | 4 | 4 | 0 | A | A | | | | | . | . | A | A | . | . |
| NC_011770 | 291675 | G | A | 46 | 4 | 4 | 0 | | | A | A | A | A | . | . | | | . | . |
| NC_011770 | 291734 | G | A | 41 | 4 | 4 | 0 | | | A | A | A | A | . | . | | | . | . |
| NC_011770 | 291935 | T | C | 61 | 0 | 6 | 0 | C | C | | | | | C | C | C | C | | |
| NC_011770 | 291953 | G | A | 62 | 0 | 2 | 0 | | | | | | | | | A | A | | |
| NC_011770 | 293925 | C | T | 98 | 0 | 2 | 0 | | | | | | | T | T | | | | |
| NC_011770 | 293934 | G | A | 95 | 0 | 2 | 0 | | | | | | | A | A | | | | |
| NC_011770 | 294021 | A | T | 87 | 0 | 2 | 0 | | | | | | | T | T | | | | |
| NC_011770 | 294091 | C | T | 92 | 0 | 2 | 0 | | | | | | | T | T | | | | |
| NC_011770 | 294106 | A | G | 89 | 0 | 1 | 0 | | | | | | | G | | | | | |
| NC_011770 | 295452 | T | G | 50 | 0 | 2 | 0 | | | | | | | G | G | | | | |
| NC_011770 | 295578 | C | G | 53 | 0 | 6 | 0 | G | G | | | | | | | G | G | G | G |
| NC_011770 | 296985 | C | T | 78 | 0 | 2 | 0 | T | T | | | | | | | | | | |
| NC_011770 | 296990 | T | C | 77 | 0 | 8 | 0 | | | C | C | C | C | C | C | C | C | | |
| NC_011770 | 297013 | G | A | 85 | 0 | 6 | 0 | | | A | A | A | A | A | A | | | | |
| NC_011770 | 297030 | A | G | 79 | 0 | 8 | 0 | | | G | G | G | G | G | G | G | G | | |
| NC_011770 | 297034 | A | G | 82 | 0 | 12 | 0 | G | G | G | G | G | G | G | G | G | G | G | G |
| NC_011770 | 297052 | TG | T | 86 | 0 | 8 | 0 | | | T | T | T | T | T | T | T | T | | |
| NC_011770 | 297078 | G | T | 85 | 0 | 6 | 0 | | | T | T | T | T | | | T | T | | |
| NC_011770 | 297094 | A | T | 84 | 0 | 8 | 0 | | | T | T | T | T | T | T | T | T | | |
| NC_011770 | 297099 | CCT | C | 83 | 0 | 8 | 0 | | | C | C | C | C | C | C | C | C | | |
| NC_011770 | 297104 | G | A | 79 | 0 | 8 | 0 | | | A | A | A | A | A | A | A | A | | |
| NC_011770 | 297106 | G | A | 83 | 0 | 2 | 0 | | | | | | | A | A | | | | |
| NC_011770 | 297108 | C | T | 86 | 0 | 2 | 0 | T | T | | | | | | | | | | |
| NC_011770 | 297118 | C | T | 86 | 0 | 2 | 0 | | | | | | | T | T | | | | |
| NC_011770 | 297136 | A | G | 86 | 0 | 4 | 0 | | | | | | | G | G | G | G | | |
| NC_011770 | 297141 | C | A | 84 | 0 | 2 | 0 | | | | | | | | | A | A | | |
| NC_011770 | 298658 | T | C | 67 | 0 | 2 | 0 | | | | | | | C | C | | | | |
| NC_011770 | 299135 | A | G | 64 | 0 | 4 | 0 | | | G | G | G | G | | | | | | |
| NC_011770 | 299564 | C | T | 77 | 0 | 4 | 0 | | | T | T | T | T | | | | | | |
| NC_011770 | 300840 | T | C | 89 | 0 | 4 | 0 | | | C | C | C | C | | | | | | |
| NC_011770 | 302080 | A | G | 51 | 0 | 4 | 0 | | | G | G | G | G | | | | | | |
| NC_011770 | 302178 | A | G | 55 | 0 | 6 | 0 | | | G | G | G | G | G | G | | | | |

| | | | | | | | | | | | | | | | | | | | |
|---|---|---|---|---|---|---|---|---|---|---|---|---|---|---|---|---|---|---|---|
| NC_011770 | 302182 | T | G | 23 | 2 | 4 | 0 | | | . | G | G | G | G | . | | | | |
| NC_011770 | 302189 | C | G | 36 | 0 | 4 | 0 | | | G | G | G | G | | | | | | |
| NC_011770 | 303032 | A | G | 87 | 0 | 2 | 0 | | | | | | | G | G | | | | |
| NC_011770 | 304380 | A | G | 63 | 0 | 6 | 0 | | | G | G | G | G | | | | | G | G |
| NC_011770 | 304381 | A | C | 62 | 0 | 6 | 0 | | | C | C | C | C | | | | | C | C |
| NC_011770 | 304409 | G | T | 66 | 0 | 4 | 0 | T | T | | | | | | | T | T | | |
| NC_011770 | 304410 | T | C | 62 | 0 | 4 | 0 | C | C | | | | | | | C | C | | |
| NC_011770 | 304411 | G | C | 65 | 0 | 4 | 0 | C | C | | | | | | | C | C | | |
| NC_011770 | 305251 | GTC | G | 73 | 0 | 2 | 0 | | | | | | | G | G | | | | |
| NC_011770 | 305257 | C | T | 70 | 0 | 2 | 0 | | | | | | | T | T | | | | |
| NC_011770 | 305275 | A | G | 71 | 0 | 6 | 0 | G | G | G | G | G | G | | | | | | |
| NC_011770 | 306901 | C | T | 70 | 0 | 4 | 0 | | | T | T | T | T | | | | | | |
| NC_011770 | 313697 | C | T | 80 | 0 | 4 | 0 | | | | | | | | | T | T | T | T |
| NC_011770 | 314914 | GCC | G | 59 | 0 | 12 | 0 | G | G | G | G | G | G | G | G | G | G | G | G |
| NC_011770 | 315044 | T | C | 76 | 0 | 12 | 0 | C | C | C | C | C | C | C | C | C | C | C | C |
| NC_011770 | 315109 | G | T | 76 | 0 | 2 | 0 | T | T | | | | | | | | | | |
| NC_011770 | 318558 | G | A | 72 | 0 | 6 | 0 | | | | | | | A | A | A | A | A | A |
| NC_011770 | 319747 | A | G | 65 | 0 | 2 | 0 | | | | | | | | | G | G | | |
| NC_011770 | 319794 | G | A | 72 | 0 | 12 | 0 | A | A | A | A | A | A | A | A | A | A | A | A |
| NC_011770 | 319829 | T | A | 72 | 0 | 4 | 0 | | | | | | | | | A | A | A | A |
| NC_011770 | 320117 | C | T | 77 | 0 | 2 | 0 | T | T | | | | | | | | | | |
| NC_011770 | 321663 | T | G | 57 | 0 | 4 | 0 | | | G | G | G | G | | | | | | |
| NC_011770 | 321669 | T | A | 64 | 0 | 4 | 0 | | | A | A | A | A | | | | | | |
| NC_011770 | 321695 | T | C | 58 | 0 | 4 | 0 | | | C | C | C | C | | | | | | |
| NC_011770 | 322936 | C | T | 74 | 0 | 4 | 0 | | | T | T | T | T | | | | | | |
| NC_011770 | 323843 | A | G | 12 | 0 | 12 | 0 | G | G | G | G | G | G | G | G | G | G | G | G |
| NC_011770 | 324720 | G | A | 52 | 0 | 2 | 0 | | | | | | | A | A | | | | |
| NC_011770 | 325879 | C | T | 80 | 0 | 2 | 0 | T | T | | | | | | | | | | |
| NC_011770 | 326016 | G | A | 71 | 0 | 2 | 0 | | | | | | | | | A | A | | |
| NC_011770 | 327522 | C | T | 79 | 0 | 2 | 0 | | | | | | | | | | | T | T |
| NC_011770 | 329866 | G | A | 74 | 0 | 4 | 0 | | | | | | | | | A | A | A | A |
| NC_011770 | 331308 | G | A | 86 | 0 | 2 | 0 | | | | | | | A | A | | | | |
| NC_011770 | 332523 | G | A | 60 | 0 | 2 | 0 | | | | | | | A | A | | | | |
| NC_011770 | 332560 | A | T | 61 | 0 | 2 | 0 | | | | | | | T | T | | | | |
| NC_011770 | 332588 | C | T | 58 | 0 | 4 | 0 | | | | | | | | | T | T | T | T |
| NC_011770 | 332590 | A | G | 56 | 0 | 10 | 0 | G | G | G | G | G | G | | | G | G | G | G |
| NC_011770 | 337182 | A | G | 70 | 0 | 12 | 0 | G | G | G | G | G | G | G | G | G | G | G | G |
| NC_011770 | 337210 | C | G | 67 | 0 | 6 | 0 | | | | | | | G | G | G | G | G | G |
| NC_011770 | 337239 | C | G | 69 | 0 | 10 | 0 | | | G | G | G | G | G | G | G | G | G | G |
| NC_011770 | 337253 | A | G | 72 | 0 | 10 | 0 | | | G | G | G | G | G | G | G | G | G | G |
| NC_011770 | 341041 | C | T | 74 | 0 | 12 | 0 | T | T | T | T | T | T | T | T | T | T | T | T |
| NC_011770 | 341099 | A | G | 67 | 0 | 1 | 0 | | | | | | | G | | | | | |
| NC_011770 | 341236 | G | T | 48 | 0 | 12 | 0 | T | T | T | T | T | T | T | T | T | T | T | T |
| NC_011770 | 341256 | C | T | 44 | 0 | 2 | 0 | T | T | | | | | | | | | | |
| NC_011770 | 341344 | G | A | 46 | 0 | 12 | 0 | A | A | A | A | A | A | A | A | A | A | A | A |
| NC_011770 | 341369 | T | G | 46 | 0 | 4 | 0 | | | G | G | G | G | | | | | | |
| NC_011770 | 343053 | G | T | 80 | 0 | 6 | 0 | T | T | | | | | | | T | T | T | T |
| NC_011770 | 344577 | C | T | 78 | 0 | 2 | 0 | | | | | | | T | T | | | | |
| NC_011770 | 345297 | C | G | 48 | 0 | 2 | 0 | G | G | | | | | | | | | | |
| NC_011770 | 345310 | C | T | 50 | 0 | 2 | 0 | | | | | | | | | | | T | T |
| NC_011770 | 345318 | T | C | 49 | 0 | 6 | 0 | C | C | | | | | | | C | C | C | C |
| NC_011770 | 345352 | C | T | 52 | 0 | 2 | 0 | | | | | | | | | | | T | T |
| NC_011770 | 345360 | G | A | 55 | 0 | 2 | 0 | | | | | | | | | | | A | A |
| NC_011770 | 345893 | A | ACG | 80 | 0 | 2 | 0 | ACG | ACG | | | | | | | | | | |
| NC_011770 | 346657 | A | G | 73 | 0 | 8 | 0 | | | G | G | G | G | G | G | | | G | G |
| NC_011770 | 347681 | T | C | 82 | 0 | 2 | 0 | | | | | | | | | | | C | C |
| NC_011770 | 347689 | G | A | 82 | 0 | 2 | 0 | | | | | | | | | | | A | A |
| NC_011770 | 347693 | G | A | 81 | 0 | 2 | 0 | | | | | | | | | A | A | | |
| NC_011770 | 351050 | G | T | 76 | 0 | 4 | 0 | | | T | T | T | T | | | | | | |
| NC_011770 | 351810 | G | A | 67 | 0 | 2 | 0 | | | | | | | A | A | | | | |
| NC_011770 | 351816 | C | G | 67 | 0 | 4 | 0 | | | G | G | G | G | | | | | | |
| NC_011770 | 351820 | T | G | 61 | 2 | 4 | 0 | | | G | G | G | G | | | . | . | | |

| | | | | | | | | | | | | | | | | | | | |
|---|---|---|---|---|---|---|---|---|---|---|---|---|---|---|---|---|---|---|---|
| NC_011770 | 351828 | G | A | 62 | 0 | 6 | 0 | | | A | A | A | A | | | A | A | | |
| NC_011770 | 351830 | C | T | 63 | 0 | 6 | 0 | | | T | T | T | T | | | T | T | | |
| NC_011770 | 351852 | A | G | 67 | 0 | 6 | 0 | | | G | G | G | G | | | G | G | | |
| NC_011770 | 352960 | C | T | 69 | 0 | 2 | 0 | | | | | | | | | | | T | T |
| NC_011770 | 353353 | GAAGA | G | 50 | 0 | 4 | 0 | | | G | G | G | G | | | | | | |
| NC_011770 | 353386 | G | T | 50 | 0 | 4 | 0 | | | T | T | T | T | | | | | | |
| NC_011770 | 353399 | C | T | 51 | 0 | 4 | 0 | | | T | T | T | T | | | | | | |
| NC_011770 | 355894 | T | G | 62 | 0 | 4 | 0 | | | G | G | G | G | | | | | | |
| NC_011770 | 356209 | A | G | 49 | 0 | 2 | 0 | G | G | | | | | | | | | | |
| NC_011770 | 356218 | G | A | 49 | 0 | 2 | 0 | A | A | | | | | | | | | | |
| NC_011770 | 356223 | C | T | 48 | 0 | 2 | 0 | T | T | | | | | | | | | | |
| NC_011770 | 363480 | G | T | 87 | 0 | 2 | 0 | T | T | | | | | | | | | | |
| NC_011770 | 363482 | T | G | 86 | 0 | 2 | 0 | G | G | | | | | | | | | | |
| NC_011770 | 363483 | G | A | 86 | 0 | 2 | 0 | A | A | | | | | | | | | | |
| NC_011770 | 363491 | G | T | 89 | 0 | 2 | 0 | T | T | | | | | | | | | | |
| NC_011770 | 363502 | T | G | 86 | 0 | 2 | 0 | G | G | | | | | | | | | | |
| NC_011770 | 364431 | T | C | 65 | 2 | 4 | 0 | . | . | C | C | C | C | | | | | | |
| NC_011770 | 369273 | G | A | 98 | 0 | 4 | 0 | | | A | A | A | A | | | | | | |
| NC_011770 | 369282 | A | G | 101 | 0 | 6 | 0 | G | G | G | G | G | G | | | | | | |
| NC_011770 | 369312 | C | T | 101 | 0 | 4 | 0 | | | T | T | T | T | | | | | | |
| NC_011770 | 369351 | C | T | 97 | 0 | 4 | 0 | | | T | T | T | T | | | | | | |
| NC_011770 | 369384 | G | A | 93 | 0 | 4 | 0 | | | A | A | A | A | | | | | | |
| NC_011770 | 369405 | C | T | 89 | 0 | 4 | 0 | | | T | T | T | T | | | | | | |
| NC_011770 | 369406 | G | A | 87 | 0 | 4 | 0 | | | A | A | A | A | | | | | | |
| NC_011770 | 369498 | G | A | 90 | 0 | 2 | 0 | A | A | | | | | | | | | | |
| NC_011770 | 370947 | T | G | 62 | 0 | 8 | 0 | G | G | | | | | G | G | G | G | G | G |
| NC_011770 | 371682 | C | T | 84 | 0 | 4 | 0 | | | | | | | | | T | T | T | T |
| NC_011770 | 374641 | C | T | 88 | 0 | 2 | 0 | | | | | | | | | T | T | | |
| NC_011770 | 378297 | A | G | 83 | 0 | 12 | 0 | G | G | G | G | G | G | G | G | G | G | G | G |
| NC_011770 | 380102 | T | TG | 72 | 0 | 2 | 0 | TG | TG | | | | | | | | | | |
| NC_011770 | 383029 | G | A | 70 | 0 | 12 | 0 | A | A | A | A | A | A | A | A | A | A | A | A |
| NC_011770 | 383040 | C | T | 72 | 0 | 6 | 0 | | | T | T | T | T | T | T | | | | |
| NC_011770 | 383481 | T | C | 82 | 0 | 4 | 0 | | | | | | | | | C | C | C | C |
| NC_011770 | 384681 | G | T | 66 | 0 | 8 | 0 | T | T | T | T | T | T | T | T | | | | |
| NC_011770 | 388151 | A | G | 39 | 0 | 10 | 0 | | | G | G | G | G | G | G | G | G | G | G |
| NC_011770 | 388159 | A | G | 42 | 0 | 2 | 0 | | | | | | | | | G | G | | |
| NC_011770 | 388177 | G | C | 43 | 0 | 4 | 0 | | | C | C | C | C | | | | | | |
| NC_011770 | 389611 | G | A | 79 | 0 | 8 | 0 | | | A | A | A | A | A | A | | | A | A |
| NC_011770 | 389636 | C | T | 79 | 0 | 4 | 0 | T | T | | | | | | | T | T | | |
| NC_011770 | 389719 | G | A | 79 | 0 | 4 | 0 | | | | | | | A | A | | | A | A |
| NC_011770 | 391878 | G | A | 74 | 0 | 2 | 0 | | | | | | | A | A | | | | |
| NC_011770 | 393864 | T | C | 81 | 0 | 8 | 0 | C | C | C | C | C | C | C | C | | | | |
| NC_011770 | 393947 | C | T | 81 | 0 | 4 | 0 | | | T | T | T | T | | | | | | |
| NC_011770 | 396809 | G | A | 66 | 0 | 4 | 0 | | | | | | | | | A | A | A | A |
| NC_011770 | 396890 | C | T | 75 | 0 | 4 | 0 | | | | | | | | | T | T | T | T |
| NC_011770 | 396926 | G | C | 81 | 0 | 4 | 0 | | | C | C | C | C | | | | | | |
| NC_011770 | 398044 | C | T | 67 | 0 | 4 | 0 | T | T | | | | | T | T | | | | |
| NC_011770 | 398096 | A | G | 69 | 0 | 4 | 0 | | | | | | | | | G | G | G | G |
| NC_011770 | 398119 | G | A | 72 | 0 | 4 | 0 | | | A | A | A | A | | | | | | |
| NC_011770 | 399894 | A | G | 69 | 0 | 10 | 0 | G | G | G | G | G | G | | | G | G | G | G |
| NC_011770 | 400326 | A | G | 67 | 0 | 4 | 0 | | | G | G | G | G | | | | | | |
| NC_011770 | 400352 | C | T | 66 | 0 | 4 | 0 | | | T | T | T | T | | | | | | |
| NC_011770 | 400958 | C | G | 63 | 0 | 4 | 0 | | | G | G | G | G | | | | | | |
| NC_011770 | 400975 | CCGG | C | 61 | 0 | 4 | 0 | | | | | | | | | C | C | C | C |
| NC_011770 | 400976 | C | G | 46 | 4 | 4 | 0 | | | G | G | G | G | | | . | . | . | . |
| NC_011770 | 403219 | A | G | 79 | 0 | 4 | 0 | | | G | G | G | G | | | | | | |
| NC_011770 | 404854 | G | T | 81 | 0 | 2 | 0 | | | | | | | | | T | T | | |
| NC_011770 | 404856 | C | G | 83 | 0 | 2 | 0 | | | | | | | | | G | G | | |
| NC_011770 | 404889 | A | G | 85 | 0 | 4 | 0 | | | | | | | | | G | G | G | G |
| NC_011770 | 405616 | A | G | 79 | 0 | 2 | 0 | G | G | | | | | | | | | | |
| NC_011770 | 405870 | C | G | 76 | 0 | 2 | 0 | G | G | | | | | | | | | | |
| NC_011770 | 407602 | G | A | 55 | 0 | 2 | 0 | A | A | | | | | | | | | | |

| | | | | | | | | | | | | | | | | | | | |
|---|---|---|---|---|---|---|---|---|---|---|---|---|---|---|---|---|---|---|---|
| NC_011770 | 407612 | A | G | 55 | 0 | 12 | 0 | G | G | G | G | G | G | G | G | G | G | G | G |
| NC_011770 | 414318 | C | G | 85 | 0 | 4 | 0 | | | | | | | | | G | G | G | G |
| NC_011770 | 414320 | G | A | 85 | 0 | 2 | 0 | | | | | | | | | | | A | A |
| NC_011770 | 414334 | A | G | 79 | 0 | 2 | 0 | | | | | | | | | | | G | G |
| NC_011770 | 417103 | G | A | 78 | 0 | 4 | 0 | | | | | | | | | A | A | A | A |
| NC_011770 | 419123 | C | T | 79 | 0 | 2 | 0 | | | | | | | T | T | | | | |
| NC_011770 | 419130 | T | C | 82 | 0 | 4 | 0 | | | C | C | C | C | | | | | | |
| NC_011770 | 419165 | T | G | 78 | 0 | 4 | 0 | | | G | G | G | G | | | | | | |
| NC_011770 | 422513 | C | T | 4 | 3 | 1 | 0 | . | | | | | | | | . | | T | . |
| NC_011770 | 423613 | G | A | 79 | 0 | 8 | 0 | | | A | A | A | A | | | A | A | A | A |
| NC_011770 | 423658 | GC | G | 81 | 0 | 2 | 0 | | | | | | | G | G | | | | |
| NC_011770 | 423706 | A | AG | 76 | 0 | 2 | 0 | AG | AG | | | | | | | | | | |
| NC_011770 | 424367 | A | G | 80 | 0 | 2 | 0 | G | G | | | | | | | | | | |
| NC_011770 | 427629 | G | A | 14 | 0 | 2 | 0 | | | | | | | | | A | A | | |
| NC_011770 | 430235 | T | C | 95 | 0 | 10 | 0 | | | C | C | C | C | C | C | C | C | C | C |
| NC_011770 | 431355 | C | T | 76 | 0 | 2 | 0 | | | | | | | T | T | | | | |
| NC_011770 | 431357 | T | A | 73 | 0 | 10 | 0 | A | A | A | A | A | A | | | A | A | A | A |
| NC_011770 | 431473 | C | T | 77 | 0 | 2 | 0 | | | | | | | T | T | | | | |
| NC_011770 | 431499 | C | T | 75 | 0 | 2 | 0 | T | T | | | | | | | | | | |
| NC_011770 | 433695 | G | A | 76 | 0 | 10 | 0 | | | A | A | A | A | A | A | A | A | A | A |
| NC_011770 | 434137 | G | T | 67 | 0 | 2 | 0 | T | T | | | | | | | | | | |
| NC_011770 | 434150 | G | A | 65 | 0 | 4 | 0 | | | | | | | | | A | A | A | A |
| NC_011770 | 434151 | T | C | 64 | 0 | 4 | 0 | | | | | | | | | C | C | C | C |
| NC_011770 | 434190 | G | T | 62 | 0 | 6 | 0 | T | T | | | | | | | T | T | T | T |
| NC_011770 | 434195 | A | G | 62 | 0 | 4 | 0 | | | | | | | | | G | G | G | G |
| NC_011770 | 434222 | G | C | 62 | 0 | 4 | 0 | | | C | C | C | C | | | | | | |
| NC_011770 | 436965 | G | T | 61 | 0 | 2 | 0 | T | T | | | | | | | | | | |
| NC_011770 | 437001 | C | T | 57 | 0 | 4 | 0 | | | T | T | T | T | | | | | | |
| NC_011770 | 437004 | T | C | 58 | 0 | 12 | 0 | C | C | C | C | C | C | C | C | C | C | C | C |
| NC_011770 | 437058 | C | T | 58 | 0 | 4 | 0 | T | T | | | | | T | T | | | | |
| NC_011770 | 437147 | G | A | 52 | 0 | 2 | 0 | A | A | | | | | | | | | | |
| NC_011770 | 439446 | G | A | 95 | 0 | 4 | 0 | | | A | A | A | A | | | | | | |
| NC_011770 | 440055 | C | A | 75 | 0 | 10 | 0 | | | A | A | A | A | A | A | A | A | A | A |
| NC_011770 | 457637 | G | C | 77 | 0 | 6 | 0 | C | C | C | C | C | C | | | | | | |
| NC_011770 | 457640 | G | T | 76 | 0 | 6 | 0 | T | T | T | T | T | T | | | | | | |
| NC_011770 | 457641 | G | T | 76 | 0 | 6 | 0 | T | T | T | T | T | T | | | | | | |
| NC_011770 | 457642 | A | C | 75 | 0 | 6 | 0 | C | C | C | C | C | C | | | | | | |
| NC_011770 | 457645 | C | T | 76 | 0 | 6 | 0 | T | T | T | T | T | T | | | | | | |
| NC_011770 | 457667 | T | A | 80 | 0 | 6 | 0 | A | A | A | A | A | A | | | | | | |
| NC_011770 | 457671 | A | AC | 82 | 0 | 6 | 0 | AC | AC | AC | AC | AC | AC | | | | | | |
| NC_011770 | 458302 | A | C | 74 | 0 | 6 | 0 | | | C | C | C | C | | | C | C | | |
| NC_011770 | 459752 | A | C | 78 | 0 | 2 | 0 | C | C | | | | | | | | | | |
| NC_011770 | 459754 | G | A | 81 | 0 | 4 | 0 | | | A | A | A | A | | | | | | |
| NC_011770 | 460506 | G | A | 77 | 0 | 2 | 0 | | | | | | | | | A | A | | |
| NC_011770 | 460530 | G | A | 78 | 0 | 4 | 0 | | | A | A | A | A | | | | | | |
| NC_011770 | 460555 | A | G | 81 | 0 | 4 | 0 | | | G | G | G | G | | | | | | |
| NC_011770 | 460573 | G | T | 81 | 0 | 12 | 0 | T | T | T | T | T | T | T | T | T | T | T | T |
| NC_011770 | 463702 | G | A | 72 | 0 | 6 | 0 | | | A | A | A | A | | | | | A | A |
| NC_011770 | 463745 | T | G | 76 | 0 | 1 | 0 | | | | | | | G | | | | | |
| NC_011770 | 465495 | T | C | 71 | 0 | 4 | 0 | | | C | C | C | C | | | | | | |
| NC_011770 | 465538 | T | C | 72 | 0 | 4 | 0 | | | C | C | C | C | | | | | | |
| NC_011770 | 465548 | A | G | 73 | 0 | 2 | 0 | | | G | G | | | | | | | | |
| NC_011770 | 465757 | G | A | 66 | 0 | 12 | 0 | A | A | A | A | A | A | A | A | A | A | A | A |
| NC_011770 | 471539 | T | C | 62 | 0 | 12 | 0 | C | C | C | C | C | C | C | C | C | C | C | C |
| NC_011770 | 473691 | A | G | 17 | 0 | 2 | 0 | G | G | | | | | | | | | | |
| NC_011770 | 474893 | A | T | 97 | 0 | 6 | 0 | T | T | | | | | T | T | T | T | | |
| NC_011770 | 476440 | C | G | 35 | 0 | 12 | 0 | G | G | G | G | G | G | G | G | G | G | G | G |
| NC_011770 | 477918 | G | A | 77 | 0 | 10 | 0 | A | A | A | A | A | A | | | A | A | A | A |
| NC_011770 | 478084 | G | C | 72 | 0 | 12 | 0 | C | C | C | C | C | C | C | C | C | C | C | C |
| NC_011770 | 478086 | C | T | 72 | 0 | 4 | 0 | | | | | | | T | T | | | T | T |
| NC_011770 | 478088 | C | T | 70 | 0 | 12 | 0 | T | T | T | T | T | T | T | T | T | T | T | T |
| NC_011770 | 478095 | C | G | 73 | 0 | 12 | 0 | G | G | G | G | G | G | G | G | G | G | G | G |

| | | | | | | | | | | | | | | | | | | | |
|---|---|---|---|---|---|---|---|---|---|---|---|---|---|---|---|---|---|---|---|
| NC_011770 | 478662 | G | A | 32 | 0 | 2 | 0 | | | | | | | | | | | A | A |
| NC_011770 | 478672 | C | T | 33 | 0 | 2 | 0 | | | | | | | | | T | T | | |
| NC_011770 | 483121 | C | G | 82 | 0 | 4 | 0 | | | G | G | G | G | | | | | | |
| NC_011770 | 485804 | C | T | 66 | 0 | 6 | 0 | | | | | | | T | T | T | T | T | T |
| NC_011770 | 492154 | G | A | 70 | 0 | 2 | 0 | A | A | | | | | | | | | | |
| NC_011770 | 493491 | T | G | 50 | 0 | 6 | 0 | G | G | | | | | | | G | G | G | G |
| NC_011770 | 493492 | A | T | 51 | 0 | 6 | 0 | T | T | | | | | | | T | T | T | T |
| NC_011770 | 493496 | T | G | 49 | 0 | 4 | 0 | | | G | G | G | G | | | | | | |
| NC_011770 | 493506 | A | G | 42 | 0 | 2 | 0 | G | G | | | | | | | | | | |
| NC_011770 | 493512 | A | C | 41 | 0 | 4 | 0 | | | C | C | C | C | | | | | | |
| NC_011770 | 493519 | T | C | 40 | 0 | 6 | 0 | C | C | C | C | C | C | | | | | | |
| NC_011770 | 493533 | G | T | 38 | 0 | 8 | 0 | T | T | | | | | A | A | T | T | T | T |
| NC_011770 | 493541 | G | C | 38 | 0 | 10 | 0 | C | C | GC | GC | GC | GC | | | C | C | C | C |
| NC_011770 | 493551 | A | G | 38 | 0 | 2 | 0 | | | | | | | G | G | | | | |
| NC_011770 | 493578 | C | G | 30 | 0 | 2 | 0 | | | . | . | . | . | G | G | | | | |
| NC_011770 | 493580 | C | T | 30 | 0 | 2 | 0 | | | . | . | . | . | T | T | | | | |
| NC_011770 | 493583 | T | C | 30 | 0 | 2 | 0 | | | | | | | C | C | | | | |
| NC_011770 | 493584 | G | T | 29 | 0 | 2 | 0 | | | | | | | T | T | | | | |
| NC_011770 | 493585 | T | C | 29 | 0 | 6 | 0 | | | C | C | C | C | C | C | | | | |
| NC_011770 | 493609 | A | G | 21 | 0 | 10 | 0 | | | G | G | G | G | G | G | G | G | G | G |
| NC_011770 | 493614 | C | T | 21 | 0 | 4 | 0 | | | T | T | T | T | | | | | | |
| NC_011770 | 493634 | G | C | 20 | 0 | 2 | 0 | | | | | | | C | C | | | | |
| NC_011770 | 496317 | C | A | 77 | 0 | 12 | 0 | A | A | A | A | A | A | A | A | A | A | A | A |
| NC_011770 | 496385 | C | T | 81 | 0 | 2 | 0 | T | T | | | | | | | | | | |
| NC_011770 | 497336 | G | GC | 68 | 0 | 4 | 0 | | | GC | GC | GC | GC | | | | | | |
| NC_011770 | 499564 | A | T | 64 | 0 | 4 | 0 | | | T | T | T | T | | | | | | |
| NC_011770 | 499584 | G | T | 65 | 0 | 6 | 0 | T | T | T | T | T | T | | | | | | |
| NC_011770 | 499826 | G | C | 52 | 0 | 12 | 0 | C | C | C | C | C | C | C | C | C | C | C | C |
| NC_011770 | 502026 | C | T | 90 | 0 | 6 | 0 | T | T | T | T | T | T | | | | | | |
| NC_011770 | 502781 | A | C | 63 | 0 | 10 | 0 | C | C | C | C | C | C | C | C | C | C | | |
| NC_011770 | 502805 | A | C | 63 | 0 | 10 | 0 | C | C | C | C | C | C | C | C | C | C | | |
| NC_011770 | 505109 | G | T | 76 | 0 | 2 | 0 | | | | | | | | | | | T | T |
| NC_011770 | 505253 | C | T | 69 | 0 | 2 | 0 | T | T | | | | | | | | | | |
| NC_011770 | 505258 | A | G | 68 | 0 | 2 | 0 | G | G | | | | | | | | | | |
| NC_011770 | 507935 | A | C | 82 | 0 | 2 | 0 | | | | | | | | | C | C | | |
| NC_011770 | 507940 | A | G | 86 | 0 | 2 | 0 | | | | | | | | | G | G | | |
| NC_011770 | 507948 | A | G | 86 | 0 | 2 | 0 | | | | | | | | | G | G | | |
| NC_011770 | 507952 | C | G | 89 | 0 | 2 | 0 | | | | | | | | | G | G | | |
| NC_011770 | 507962 | C | G | 94 | 0 | 2 | 0 | | | | | | | | | G | G | | |
| NC_011770 | 507965 | A | G | 93 | 0 | 2 | 0 | | | | | | | | | G | G | | |
| NC_011770 | 507967 | A | G | 95 | 0 | 2 | 0 | | | | | | | | | G | G | | |
| NC_011770 | 507968 | A | G | 94 | 0 | 2 | 0 | | | | | | | | | G | G | | |
| NC_011770 | 507984 | A | C | 91 | 0 | 2 | 0 | | | | | | | | | C | C | | |
| NC_011770 | 507986 | C | T | 92 | 0 | 2 | 0 | | | | | | | | | T | T | | |
| NC_011770 | 507998 | T | A | 90 | 0 | 2 | 0 | A | A | | | | | | | | | | |
| NC_011770 | 508006 | T | C | 91 | 0 | 2 | 0 | | | | | | | | | C | C | | |
| NC_011770 | 508010 | T | C | 94 | 0 | 2 | 0 | | | | | | | | | C | C | | |
| NC_011770 | 508015 | A | T | 93 | 0 | 2 | 0 | | | | | | | | | T | T | | |
| NC_011770 | 508193 | G | C | 84 | 2 | 2 | 0 | C | C | | | | | | | . | . | | |
| NC_011770 | 508213 | G | T | 84 | 2 | 2 | 0 | | | | | | | | | . | . | T | T |
| NC_011770 | 508237 | G | A | 84 | 2 | 4 | 0 | | | A | A | A | A | | | . | . | | |
| NC_011770 | 509897 | A | G | 70 | 0 | 8 | 0 | G | G | | | | | G | G | G | G | G | G |
| NC_011770 | 509944 | T | C | 74 | 0 | 12 | 0 | C | C | C | C | C | C | C | C | C | C | C | C |
| NC_011770 | 512937 | T | C | 70 | 0 | 12 | 0 | C | C | C | C | C | C | C | C | C | C | C | C |
| NC_011770 | 513614 | C | T | 47 | 0 | 4 | 0 | | | T | T | T | T | | | | | | |
| NC_011770 | 513624 | T | C | 46 | 0 | 4 | 0 | | | C | C | C | C | | | | | | |
| NC_011770 | 513653 | A | G | 42 | 0 | 4 | 0 | | | G | G | G | G | | | | | | |
| NC_011770 | 513660 | T | C | 41 | 0 | 4 | 0 | | | C | C | C | C | | | | | | |
| NC_011770 | 513713 | C | T | 33 | 0 | 6 | 0 | T | T | | | | | T | T | T | T | | |
| NC_011770 | 513716 | T | G | 32 | 0 | 6 | 0 | G | G | | | | | G | G | G | G | | |
| NC_011770 | 513772 | A | G | 34 | 0 | 6 | 0 | G | G | | | | | | | G | G | G | G |
| NC_011770 | 513774 | C | G | 35 | 0 | 6 | 0 | G | G | | | | | | | G | G | G | G |

| | | | | | | | | | | | | | | | | | | | |
|---|---|---|---|---|---|---|---|---|---|---|---|---|---|---|---|---|---|---|---|
| NC_011770 | 513780 | G | A | 36 | 1 | 5 | 0 | A | A | | | | | | | . | A | A | A |
| NC_011770 | 513790 | C | CCCGCA | 35 | 0 | 2 | 0 | | | | | | | | | CCCGCA | CCCGCA | | |
| NC_011770 | 513795 | A | T | 34 | 2 | 2 | 0 | | | | | | | T | T | . | . | | |
| NC_011770 | 513834 | T | C | 24 | 0 | 1 | 0 | | | | | | | | | | | . | C |
| NC_011770 | 513837 | G | T | 24 | 0 | 2 | 0 | | | | | | | | | | | T | T |
| NC_011770 | 513876 | C | A | 28 | 0 | 2 | 0 | | | | | | | | | A | A | | |
| NC_011770 | 513893 | C | T | 34 | 0 | 2 | 0 | | | | | | | | | | | T | T |
| NC_011770 | 517649 | C | T | 82 | 0 | 4 | 0 | T | T | | | | | | | T | T | | |
| NC_011770 | 519112 | A | G | 66 | 0 | 2 | 0 | G | G | | | | | | | | | | |
| NC_011770 | 519459 | T | C | 73 | 0 | 12 | 0 | C | C | C | C | C | C | C | C | C | C | C | C |
| NC_011770 | 519564 | G | A | 71 | 0 | 2 | 0 | A | A | | | | | | | | | | |
| NC_011770 | 519567 | C | T | 70 | 0 | 2 | 0 | T | T | | | | | | | | | | |
| NC_011770 | 519597 | C | CG | 71 | 0 | 4 | 0 | | | CG | CG | CG | CG | | | | | | |
| NC_011770 | 521207 | C | T | 89 | 0 | 4 | 0 | | | | | | | | | T | T | T | T |
| NC_011770 | 521223 | T | G | 90 | 0 | 12 | 0 | G | G | G | G | G | G | G | G | G | G | G | G |
| NC_011770 | 522139 | T | C | 64 | 0 | 2 | 0 | C | C | | | | | | | | | | |
| NC_011770 | 522172 | A | G | 62 | 0 | 2 | 0 | G | G | | | | | | | | | | |
| NC_011770 | 522173 | T | A | 61 | 0 | 2 | 0 | A | A | | | | | | | | | | |
| NC_011770 | 522184 | G | A | 62 | 0 | 2 | 0 | A | A | | | | | | | | | | |
| NC_011770 | 522206 | T | A | 58 | 0 | 2 | 0 | A | A | | | | | | | | | | |
| NC_011770 | 522287 | A | G | 54 | 0 | 2 | 0 | G | G | | | | | | | | | | |
| NC_011770 | 524772 | G | A | 51 | 0 | 6 | 0 | | | A | A | A | A | A | A | | | | |
| NC_011770 | 524784 | G | A | 53 | 0 | 2 | 0 | | | | | | | | | | | A | A |
| NC_011770 | 527302 | G | A | 89 | 0 | 2 | 0 | A | A | | | | | | | | | | |
| NC_011770 | 527314 | C | T | 89 | 0 | 2 | 0 | | | | | | | T | T | | | | |
| NC_011770 | 527321 | C | T | 88 | 0 | 2 | 0 | | | | | | | | | | | T | T |
| NC_011770 | 527355 | A | G | 99 | 0 | 4 | 0 | G | G | | | | | | | | | G | G |
| NC_011770 | 528424 | T | C | 74 | 0 | 12 | 0 | C | C | C | C | C | C | C | C | C | C | C | C |
| NC_011770 | 530201 | G | A | 48 | 0 | 4 | 0 | | | A | A | A | A | | | | | | |
| NC_011770 | 530270 | C | A | 52 | 0 | 10 | 0 | A | A | A | A | A | A | A | A | | | A | A |
| NC_011770 | 530278 | T | C | 53 | 0 | 10 | 0 | C | C | C | C | C | C | C | C | | | C | C |
| NC_011770 | 530280 | T | C | 54 | 0 | 10 | 0 | C | C | C | C | C | C | C | C | | | C | C |
| NC_011770 | 530281 | T | C | 52 | 0 | 2 | 0 | | | | | | | | | | | C | C |
| NC_011770 | 530301 | A | G | 57 | 0 | 10 | 0 | G | G | G | G | G | G | G | G | | | G | G |
| NC_011770 | 530302 | C | G | 56 | 0 | 10 | 0 | G | G | G | G | G | G | G | G | | | G | G |
| NC_011770 | 530341 | C | T | 59 | 0 | 10 | 0 | T | T | T | T | T | T | T | T | | | T | T |
| NC_011770 | 530373 | G | A | 61 | 0 | 10 | 0 | A | A | A | A | A | A | A | A | | | A | A |
| NC_011770 | 530383 | G | A | 61 | 0 | 4 | 0 | | | A | A | A | A | | | | | | |
| NC_011770 | 530402 | G | A | 59 | 0 | 4 | 0 | | | A | A | A | A | | | | | | |
| NC_011770 | 530409 | A | G | 55 | 2 | 10 | 0 | . | . | G | G | G | G | G | G | G | G | G | G |
| NC_011770 | 530412 | T | G | 55 | 0 | 10 | 0 | G | G | G | G | G | G | G | G | | | G | G |
| NC_011770 | 530414 | G | A | 56 | 0 | 10 | 0 | A | A | A | A | A | A | A | A | | | A | A |
| NC_011770 | 530424 | A | G | 57 | 0 | 10 | 0 | G | G | G | G | G | G | G | G | | | G | G |
| NC_011770 | 530425 | G | C | 57 | 0 | 10 | 0 | C | C | T | T | T | T | C | C | | | C | C |
| NC_011770 | 530428 | T | C | 58 | 0 | 10 | 0 | C | C | C | C | C | C | C | C | | | C | C |
| NC_011770 | 530471 | C | T | 67 | 0 | 4 | 0 | | | T | T | T | T | | | | | | |
| NC_011770 | 531434 | C | A | 95 | 0 | 2 | 0 | | | | | | | | | A | A | | |
| NC_011770 | 531439 | A | G | 88 | 0 | 10 | 0 | G | G | G | G | G | G | G | G | | | G | G |
| NC_011770 | 531447 | G | T | 91 | 0 | 2 | 0 | | | | | | | | | | | T | T |
| NC_011770 | 532016 | C | T | 85 | 0 | 2 | 0 | T | T | | | | | | | | | | |
| NC_011770 | 533070 | G | A | 82 | 0 | 10 | 0 | | | A | A | A | A | A | A | A | A | A | A |
| NC_011770 | 533925 | G | C | 93 | 0 | 4 | 0 | C | C | | | | | | | | | C | C |
| NC_011770 | 533938 | G | C | 87 | 0 | 6 | 0 | C | C | | | | | C | C | | | C | C |
| NC_011770 | 536808 | C | T | 96 | 0 | 2 | 0 | T | T | | | | | | | | | | |
| NC_011770 | 536832 | C | T | 96 | 0 | 2 | 0 | | | | | | | T | T | | | | |
| NC_011770 | 536845 | A | G | 95 | 0 | 2 | 0 | G | G | | | | | | | | | | |
| NC_011770 | 536912 | T | TC | 91 | 0 | 6 | 0 | | | TC | TC | TC | TC | TC | TC | | | | |
| NC_011770 | 538048 | T | A | 99 | 0 | 2 | 0 | A | A | | | | | | | | | | |
| NC_011770 | 538112 | G | A | 86 | 0 | 2 | 0 | | | | | | | | | | | A | A |
| NC_011770 | 538130 | A | G | 87 | 0 | 2 | 0 | | | | | | | | | | | G | G |
| NC_011770 | 538208 | A | G | 87 | 0 | 2 | 0 | | | | | | | | | G | G | | |
| NC_011770 | 538249 | C | A | 85 | 0 | 2 | 0 | | | | | | | | | A | A | | |

| | | | | | | | | | | | | | | | | | | | |
|---|---|---|---|---|---|---|---|---|---|---|---|---|---|---|---|---|---|---|---|
| NC_011770 | 538254 | C | G | 88 | 0 | 4 | 0 | | | G | G | G | G | | | | | | |
| NC_011770 | 538260 | T | C | 88 | 0 | 2 | 0 | | | | | | | | | | | C | C |
| NC_011770 | 538264 | C | A | 86 | 0 | 2 | 0 | | | | | | | | | | | A | A |
| NC_011770 | 538279 | C | T | 87 | 0 | 2 | 0 | | | | | | | | | T | T | | |
| NC_011770 | 538301 | CT | C | 85 | 0 | 2 | 0 | | | | | | | C | C | | | | |
| NC_011770 | 538344 | A | C | 86 | 0 | 2 | 0 | | | | | | | | | | | C | C |
| NC_011770 | 538391 | T | G | 86 | 0 | 2 | 0 | | | | | | | | | | | G | G |
| NC_011770 | 538397 | C | T | 84 | 0 | 2 | 0 | | | | | | | | | T | T | | |
| NC_011770 | 538446 | C | A | 87 | 0 | 12 | 0 | A | A | A | A | A | A | A | A | A | A | A | A |
| NC_011770 | 538484 | C | A | 88 | 0 | 8 | 0 | A | A | A | A | A | A | A | A | | | | |
| NC_011770 | 539560 | C | T | 81 | 0 | 2 | 0 | | | | | | | | | T | T | | |
| NC_011770 | 539568 | C | A | 77 | 0 | 2 | 0 | | | | | | | A | A | | | | |
| NC_011770 | 539601 | G | A | 73 | 0 | 2 | 0 | | | | | | | A | A | | | | |
| NC_011770 | 539604 | A | G | 71 | 0 | 2 | 0 | | | | | | | G | G | | | | |
| NC_011770 | 539606 | C | G | 74 | 0 | 2 | 0 | | | | | | | G | G | | | | |
| NC_011770 | 539664 | C | T | 79 | 0 | 2 | 0 | | | | | | | | | T | T | | |
| NC_011770 | 539700 | C | T | 74 | 0 | 2 | 0 | | | | | | | T | T | | | | |
| NC_011770 | 540719 | C | G | 88 | 0 | 4 | 0 | | | G | G | G | G | | | | | | |
| NC_011770 | 541515 | G | A | 81 | 0 | 2 | 0 | | | | | | | A | A | | | | |
| NC_011770 | 541911 | T | C | 73 | 0 | 12 | 0 | C | C | C | C | C | C | C | C | C | C | C | C |
| NC_011770 | 541924 | A | C | 72 | 0 | 10 | 0 | C | C | C | C | C | C | C | C | | | C | C |
| NC_011770 | 541935 | C | T | 76 | 0 | 1 | 0 | | | | | | | T | | | | | |
| NC_011770 | 544142 | C | T | 84 | 0 | 4 | 0 | | | | | | | T | T | | | T | T |
| NC_011770 | 548527 | G | A | 71 | 0 | 2 | 0 | | | | | | | | | A | A | | |
| NC_011770 | 550660 | C | T | 87 | 0 | 1 | 0 | | | | | | | T | | | | | |
| NC_011770 | 551466 | G | A | 78 | 0 | 4 | 0 | | | | | | | | | A | A | A | A |
| NC_011770 | 551723 | G | A | 78 | 0 | 6 | 0 | | | A | A | A | A | A | A | | | | |
| NC_011770 | 551765 | G | A | 81 | 0 | 6 | 0 | | | A | A | A | A | A | A | | | | |
| NC_011770 | 551776 | T | C | 81 | 0 | 6 | 0 | | | C | C | C | C | C | C | | | | |
| NC_011770 | 551780 | G | T | 80 | 0 | 8 | 0 | T | T | A | A | A | A | A | A | | | | |
| NC_011770 | 551781 | G | A | 80 | 0 | 6 | 0 | | | A | A | A | A | A | A | | | | |
| NC_011770 | 551806 | C | T | 89 | 0 | 6 | 0 | | | T | T | T | T | T | T | | | | |
| NC_011770 | 551844 | A | G | 91 | 0 | 2 | 0 | G | G | | | | | | | | | | |
| NC_011770 | 553006 | T | C | 84 | 0 | 2 | 0 | | | | | | | C | C | | | | |
| NC_011770 | 556584 | C | T | 73 | 0 | 2 | 0 | | | | | | | | | T | T | | |
| NC_011770 | 556606 | G | A | 76 | 0 | 4 | 0 | A | A | | | | | | | A | A | | |
| NC_011770 | 556875 | C | T | 66 | 0 | 2 | 0 | T | T | | | | | | | | | | |
| NC_011770 | 556890 | C | T | 71 | 0 | 4 | 0 | | | | | | | T | T | | | T | T |
| NC_011770 | 556918 | G | A | 71 | 0 | 2 | 0 | | | | | | | | | | | A | A |
| NC_011770 | 559117 | C | T | 72 | 0 | 2 | 0 | | | | | | | | | T | T | | |
| NC_011770 | 559156 | C | T | 71 | 0 | 2 | 0 | | | | | | | T | T | | | | |
| NC_011770 | 561157 | C | A | 72 | 0 | 2 | 0 | | | | | | | A | A | | | | |
| NC_011770 | 561158 | C | A | 72 | 0 | 2 | 0 | | | | | | | | | | | A | A |
| NC_011770 | 561193 | G | T | 75 | 0 | 2 | 0 | T | T | | | | | | | | | | |
| NC_011770 | 561279 | T | TC | 74 | 0 | 4 | 0 | | | TC | TC | TC | TC | | | | | | |
| NC_011770 | 563122 | G | T | 9 | 2 | 2 | 0 | | | T | T | . | . | | | | | | |
| NC_011770 | 563147 | G | GGT | 3 | 5 | 2 | 0 | . | . | | | | . | | . | GGT | GGT | | . |
| NC_011770 | 563185 | A | T | 13 | 1 | 11 | 0 | T | . | T | T | T | T | T | T | T | T | T | T |
| NC_011770 | 563213 | T | G | 27 | 0 | 4 | 0 | | | G | G | G | G | | | | | | |
| NC_011770 | 563311 | G | A | 33 | 0 | 2 | 0 | | | | | | | | | A | A | | |
| NC_011770 | 563321 | T | C | 34 | 0 | 2 | 0 | | | | | | | | | | | C | C |
| NC_011770 | 572556 | A | G | 104 | 0 | 1 | 0 | | | | | | | G | | | | | |
| NC_011770 | 574257 | A | G | 76 | 0 | 8 | 0 | | | G | G | G | G | | | G | G | G | G |
| NC_011770 | 577980 | T | G | 84 | 0 | 4 | 0 | | | G | G | G | G | | | | | | |
| NC_011770 | 578200 | T | C | 76 | 0 | 10 | 0 | C | C | C | C | C | C | C | C | | | C | C |
| NC_011770 | 578216 | T | C | 76 | 0 | 6 | 0 | C | C | | | | | C | C | | | C | C |
| NC_011770 | 578256 | C | G | 75 | 0 | 4 | 0 | | | G | G | G | G | | | | | | |
| NC_011770 | 578261 | C | T | 75 | 0 | 10 | 0 | T | T | T | T | T | T | T | T | | | T | T |
| NC_011770 | 578958 | C | T | 75 | 0 | 1 | 0 | | | | | | | T | | | | | |
| NC_011770 | 578977 | C | G | 76 | 0 | 12 | 0 | G | G | G | G | G | G | G | G | G | G | G | G |
| NC_011770 | 578993 | C | A | 76 | 0 | 2 | 0 | A | A | | | | | | | | | | |
| NC_011770 | 579001 | G | A | 76 | 0 | 12 | 0 | A | A | A | A | A | A | A | A | A | A | A | A |

| | | | | | | | | | | | | | | | | | | | |
|---|---|---|---|---|---|---|---|---|---|---|---|---|---|---|---|---|---|---|---|
| NC_011770 | 579011 | T | C | 79 | 0 | 12 | 0 | C | C | C | C | C | C | C | C | C | C | C | C |
| NC_011770 | 579025 | G | C | 75 | 0 | 10 | 0 | | | C | C | C | C | C | C | C | C | C | C |
| NC_011770 | 579032 | G | C | 75 | 0 | 10 | 0 | | | C | C | C | C | C | C | C | C | C | C |
| NC_011770 | 579037 | C | T | 77 | 0 | 10 | 0 | | | T | T | T | T | T | T | T | T | T | T |
| NC_011770 | 579039 | C | T | 75 | 0 | 10 | 0 | | | T | T | T | T | T | T | T | T | T | T |
| NC_011770 | 579126 | T | C | 82 | 0 | 12 | 0 | C | C | C | C | C | C | C | C | C | C | C | C |
| NC_011770 | 579283 | C | A | 83 | 0 | 8 | 0 | A | A | | | | | A | A | A | A | A | A |
| NC_011770 | 579285 | A | G | 83 | 0 | 8 | 0 | G | G | | | | | G | G | G | G | G | G |
| NC_011770 | 580230 | G | A | 88 | 0 | 4 | 0 | A | A | | | | | | | | | A | A |
| NC_011770 | 580236 | C | A | 89 | 0 | 2 | 0 | | | | | | | | | A | A | | |
| NC_011770 | 581112 | A | T | 92 | 0 | 2 | 0 | T | T | | | | | | | | | | |
| NC_011770 | 581114 | C | A | 93 | 0 | 4 | 0 | | | A | A | A | A | | | | | | |
| NC_011770 | 581131 | G | A | 92 | 0 | 2 | 0 | A | A | | | | | | | | | | |
| NC_011770 | 581132 | C | G | 91 | 0 | 2 | 0 | G | G | | | | | | | | | | |
| NC_011770 | 581135 | C | T | 89 | 0 | 2 | 0 | T | T | | | | | | | | | | |
| NC_011770 | 581147 | A | G | 98 | 0 | 2 | 0 | G | G | | | | | | | | | | |
| NC_011770 | 581156 | G | A | 98 | 0 | 2 | 0 | A | A | | | | | | | | | | |
| NC_011770 | 581169 | T | C | 97 | 0 | 2 | 0 | C | C | | | | | | | | | | |
| NC_011770 | 583140 | C | T | 72 | 0 | 4 | 0 | | | T | T | T | T | | | | | | |
| NC_011770 | 585536 | C | T | 69 | 0 | 12 | 0 | T | T | T | T | T | T | T | T | T | T | T | T |
| NC_011770 | 586852 | C | T | 67 | 0 | 2 | 0 | T | T | | | | | | | | | | |
| NC_011770 | 587420 | A | G | 24 | 0 | 2 | 0 | | | | | | | | | | | G | G |
| NC_011770 | 587432 | G | T | 23 | 0 | 4 | 0 | | | | | | | T | T | | | T | T |
| NC_011770 | 589248 | T | G | 82 | 0 | 4 | 0 | | | | | | | | | G | G | G | G |
| NC_011770 | 589889 | C | T | 86 | 0 | 2 | 0 | T | T | | | | | | | | | | |
| NC_011770 | 589944 | C | T | 82 | 0 | 2 | 0 | T | T | | | | | | | | | | |
| NC_011770 | 589949 | G | A | 82 | 0 | 2 | 0 | A | A | | | | | | | | | | |
| NC_011770 | 589965 | C | T | 92 | 0 | 2 | 0 | T | T | | | | | | | | | | |
| NC_011770 | 589984 | C | A | 89 | 0 | 2 | 0 | A | A | | | | | | | | | | |
| NC_011770 | 589987 | G | A | 89 | 0 | 2 | 0 | A | A | | | | | | | | | | |
| NC_011770 | 590865 | A | T | 79 | 0 | 2 | 0 | | | | | | | | | T | T | | |
| NC_011770 | 590876 | T | C | 83 | 0 | 2 | 0 | C | C | | | | | | | | | | |
| NC_011770 | 591863 | C | T | 74 | 0 | 6 | 0 | | | | | | | T | T | T | T | T | T |
| NC_011770 | 591871 | C | T | 67 | 0 | 10 | 0 | | | T | T | T | T | G | G | G | G | G | G |
| NC_011770 | 591875 | C | G | 69 | 0 | 4 | 0 | | | G | G | G | G | | | | | | |
| NC_011770 | 591879 | A | G | 67 | 0 | 4 | 0 | | | G | G | G | G | | | | | | |
| NC_011770 | 591886 | G | T | 65 | 0 | 2 | 0 | T | T | | | | | | | | | | |
| NC_011770 | 591890 | G | C | 66 | 0 | 4 | 0 | | | C | C | C | C | | | | | | |
| NC_011770 | 591918 | T | C | 44 | 4 | 6 | 0 | C | C | . | . | . | . | C | C | | | C | C |
| NC_011770 | 591975 | G | A | 45 | 4 | 6 | 0 | | | . | . | . | . | A | A | A | A | A | A |
| NC_011770 | 591980 | G | A | 45 | 4 | 6 | 0 | | | . | . | . | . | A | A | A | A | A | A |
| NC_011770 | 591983 | G | A | 45 | 4 | 6 | 0 | | | . | . | . | . | A | A | A | A | A | A |
| NC_011770 | 592001 | A | G | 70 | 0 | 10 | 0 | | | G | G | G | G | G | G | G | G | G | G |
| NC_011770 | 592029 | A | C | 79 | 0 | 2 | 0 | | | | | | | | | C | C | | |
| NC_011770 | 592659 | A | G | 76 | 0 | 4 | 0 | | | G | G | G | G | | | | | | |
| NC_011770 | 592687 | A | G | 76 | 0 | 4 | 0 | | | G | G | G | G | | | | | | |
| NC_011770 | 594465 | G | A | 72 | 0 | 4 | 0 | A | A | | | | | | | | | A | A |
| NC_011770 | 595946 | G | GA | 74 | 0 | 2 | 0 | | | | | | | | | | | GA | GA |
| NC_011770 | 595960 | G | A | 75 | 0 | 2 | 0 | | | | | | | | | | | A | A |
| NC_011770 | 595974 | T | TC | 74 | 0 | 4 | 0 | TC | TC | | | | | | | | | TC | TC |
| NC_011770 | 598409 | C | T | 70 | 0 | 2 | 0 | | | | | | | | | | | T | T |
| NC_011770 | 602325 | G | A | 73 | 0 | 10 | 0 | A | A | A | A | A | A | | | A | A | A | A |
| NC_011770 | 602374 | T | A | 68 | 0 | 2 | 0 | | | | | | | A | A | | | | |
| NC_011770 | 602377 | A | G | 68 | 0 | 2 | 0 | | | | | | | G | G | | | | |
| NC_011770 | 602388 | C | T | 70 | 0 | 6 | 0 | T | T | | | | | | | T | T | T | T |
| NC_011770 | 602400 | C | T | 70 | 0 | 2 | 0 | | | | | | | T | T | | | | |
| NC_011770 | 605129 | C | T | 68 | 0 | 2 | 0 | T | T | | | | | | | | | | |
| NC_011770 | 606735 | C | T | 73 | 0 | 4 | 0 | | | | | | | T | T | T | T | | |
| NC_011770 | 609865 | A | C | 80 | 0 | 6 | 0 | C | C | | | | | C | C | C | C | | |
| NC_011770 | 611226 | A | C | 24 | 0 | 2 | 0 | | | | | | | | | | | C | C |
| NC_011770 | 611762 | A | C | 95 | 0 | 4 | 0 | | | | | | | C | C | C | C | | |
| NC_011770 | 612804 | C | A | 84 | 0 | 8 | 0 | | | A | A | A | A | | | A | A | A | A |

| | | | | | | | | | | | | | | | | | | | |
|---|---|---|---|---|---|---|---|---|---|---|---|---|---|---|---|---|---|---|---|
| NC_011770 | 616035 | A | G | 93 | 0 | 6 | 0 | | | | | | | G | G | G | G | G | G |
| NC_011770 | 616046 | A | C | 96 | 0 | 6 | 0 | | | | | | | C | C | C | C | C | C |
| NC_011770 | 616094 | T | C | 87 | 0 | 8 | 0 | C | C | | | | | C | C | C | C | C | C |
| NC_011770 | 616115 | A | G | 86 | 0 | 8 | 0 | G | G | | | | | G | G | G | G | G | G |
| NC_011770 | 618999 | C | T | 88 | 0 | 1 | 0 | | | | | | | T | | | | | |
| NC_011770 | 619085 | G | A | 94 | 0 | 2 | 0 | | | | | | | | | | | A | A |
| NC_011770 | 619099 | T | A | 91 | 0 | 6 | 0 | A | A | | | | | | | A | A | A | A |
| NC_011770 | 619101 | T | A | 90 | 0 | 2 | 0 | A | A | | | | | | | | | | |
| NC_011770 | 619185 | G | T | 85 | 0 | 6 | 0 | | | T | T | T | T | T | T | | | | |
| NC_011770 | 619208 | T | C | 90 | 0 | 6 | 0 | C | C | | | | | | | C | C | C | C |
| NC_011770 | 619225 | C | G | 86 | 0 | 10 | 0 | G | G | G | G | G | G | | | G | G | G | G |
| NC_011770 | 619373 | T | G | 94 | 0 | 10 | 0 | G | G | G | G | G | G | | | G | G | G | G |
| NC_011770 | 619839 | G | A | 95 | 0 | 10 | 0 | A | A | A | A | A | A | | | A | A | A | A |
| NC_011770 | 619873 | A | G | 94 | 0 | 2 | 0 | G | G | | | | | | | | | | |
| NC_011770 | 621073 | G | C | 84 | 0 | 4 | 0 | | | | | | | | | C | C | C | C |
| NC_011770 | 621076 | A | G | 84 | 0 | 4 | 0 | | | | | | | | | G | G | G | G |
| NC_011770 | 621372 | GT | G | 74 | 0 | 4 | 0 | | | G | G | G | G | | | | | | |
| NC_011770 | 621405 | C | A | 74 | 0 | 4 | 0 | | | A | A | A | A | | | | | | |
| NC_011770 | 621421 | A | G | 74 | 0 | 6 | 0 | G | G | G | G | G | G | | | | | | |
| NC_011770 | 621425 | C | T | 77 | 0 | 4 | 0 | | | T | T | T | T | | | | | | |
| NC_011770 | 621456 | A | G | 72 | 0 | 4 | 0 | | | G | G | G | G | | | | | | |
| NC_011770 | 621463 | T | G | 72 | 0 | 4 | 0 | | | G | G | G | G | | | | | | |
| NC_011770 | 621481 | C | T | 69 | 0 | 4 | 0 | | | T | T | T | T | | | | | | |
| NC_011770 | 621504 | T | C | 65 | 0 | 4 | 0 | | | C | C | C | C | | | | | | |
| NC_011770 | 621534 | C | A | 60 | 0 | 2 | 0 | | | | | | | A | A | | | | |
| NC_011770 | 625463 | GA | G | 57 | 0 | 2 | 0 | | | | | | | | | G | G | | |
| NC_011770 | 629441 | A | G | 78 | 0 | 2 | 0 | | | | | | | | | G | G | | |
| NC_011770 | 629984 | G | A | 73 | 0 | 2 | 0 | | | | | | | | | | | A | A |
| NC_011770 | 634887 | GC | G | 68 | 0 | 10 | 0 | | | G | G | G | G | G | G | G | G | G | G |
| NC_011770 | 634924 | C | T | 67 | 0 | 2 | 0 | T | T | | | | | | | | | | |
| NC_011770 | 641700 | T | C | 84 | 0 | 12 | 0 | C | C | C | C | C | C | C | C | C | C | C | C |
| NC_011770 | 651330 | C | T | 56 | 0 | 6 | 0 | | | T | T | T | T | | | T | T | | |
| NC_011770 | 651396 | C | T | 51 | 0 | 2 | 0 | T | T | | | | | | | | | | |
| NC_011770 | 651434 | G | A | 56 | 0 | 2 | 0 | | | | | | | | | | | A | A |
| NC_011770 | 655743 | C | T | 55 | 0 | 8 | 0 | T | T | | | | | T | T | T | T | T | T |
| NC_011770 | 655756 | C | T | 55 | 0 | 4 | 0 | T | T | | | | | | | T | T | | |
| NC_011770 | 655772 | C | T | 58 | 0 | 6 | 0 | T | T | | | | | T | T | | | T | T |
| NC_011770 | 655773 | T | C | 60 | 0 | 6 | 0 | C | C | | | | | C | C | | | C | C |
| NC_011770 | 655783 | C | CG | 61 | 0 | 2 | 0 | | | | | | | | | CG | CG | | |
| NC_011770 | 655787 | G | GT | 25 | 7 | 4 | 0 | . | . | GT | GT | GT | GT | . | . | . | . | . | . |
| NC_011770 | 658158 | A | T | 76 | 0 | 8 | 0 | | | T | T | T | T | | | T | T | T | T |
| NC_011770 | 658172 | G | A | 76 | 0 | 2 | 0 | | | | | | | | | | | A | A |
| NC_011770 | 658204 | T | C | 82 | 0 | 12 | 0 | C | C | C | C | C | C | C | C | C | C | C | C |
| NC_011770 | 658218 | GC | G | 79 | 0 | 2 | 0 | | | | | | | G | G | | | | |
| NC_011770 | 658219 | C | T | 64 | 2 | 8 | 0 | | | T | T | T | T | . | . | T | T | T | T |
| NC_011770 | 658220 | C | T | 65 | 2 | 8 | 0 | | | T | T | T | T | . | . | T | T | T | T |
| NC_011770 | 659503 | G | C | 84 | 0 | 2 | 0 | | | | | | | | | | | C | C |
| NC_011770 | 660475 | C | T | 76 | 0 | 2 | 0 | | | | | | | | | | | T | T |
| NC_011770 | 662090 | C | T | 77 | 0 | 4 | 0 | | | | | | | T | T | T | T | | |
| NC_011770 | 663646 | T | C | 56 | 0 | 6 | 0 | | | C | C | C | C | | | C | C | | |
| NC_011770 | 663654 | CT | C | 59 | 0 | 2 | 0 | | | | | | | | | C | C | | |
| NC_011770 | 663661 | A | G | 59 | 0 | 6 | 0 | G | G | | | | | G | G | | | G | G |
| NC_011770 | 663669 | C | CG | 61 | 0 | 2 | 0 | CG | CG | | | | | | | | | | |
| NC_011770 | 663677 | A | T | 61 | 0 | 6 | 0 | | | T | T | T | T | | | T | T | | |
| NC_011770 | 663690 | G | GC | 63 | 0 | 12 | 0 | GC | GC | GC | GC | GC | GC | GC | GC | GC | GC | GC | GC |
| NC_011770 | 663700 | C | A | 61 | 0 | 10 | 0 | | | A | A | A | A | A | A | A | A | A | A |
| NC_011770 | 663701 | A | G | 63 | 0 | 4 | 0 | | | | | | | G | G | | | G | G |
| NC_011770 | 663727 | G | A | 61 | 0 | 6 | 0 | | | A | A | A | A | | | A | A | | |
| NC_011770 | 663745 | G | A | 39 | 6 | 2 | 0 | A | A | . | . | . | . | | | . | . | | |
| NC_011770 | 663843 | A | AG | 40 | 6 | 4 | 0 | AG | AG | . | . | . | . | AG | AG | . | . | | |
| NC_011770 | 664255 | A | G | 80 | 0 | 2 | 0 | | | | | | | | | G | G | | |
| NC_011770 | 664264 | A | G | 78 | 0 | 2 | 0 | | | | | | | G | G | | | | |

| | | | | | | | | | | | | | | | | | | | |
|---|---|---|---|---|---|---|---|---|---|---|---|---|---|---|---|---|---|---|---|
| NC_011770 | 664265 | A | G | 79 | 0 | 10 | 0 | G | G | G | G | G | G | G | G | | | G | G |
| NC_011770 | 664268 | A | G | 79 | 0 | 6 | 0 | | | G | G | G | G | | | | | G | G |
| NC_011770 | 664289 | C | T | 81 | 0 | 6 | 0 | | | T | T | T | T | | | T | T | | |
| NC_011770 | 664290 | T | C | 80 | 0 | 2 | 0 | | | | | | | | | C | C | | |
| NC_011770 | 664294 | T | G | 81 | 0 | 2 | 0 | | | | | | | | | G | G | | |
| NC_011770 | 664304 | A | G | 83 | 0 | 2 | 0 | | | | | | | | | G | G | | |
| NC_011770 | 664313 | A | T | 81 | 0 | 2 | 0 | | | | | | | | | T | T | | |
| NC_011770 | 665105 | C | T | 77 | 0 | 2 | 0 | | | | | | | | | T | T | | |
| NC_011770 | 665109 | A | G | 74 | 0 | 12 | 0 | G | G | G | G | G | G | G | G | G | G | G | G |
| NC_011770 | 665131 | T | C | 80 | 0 | 2 | 0 | | | | | | | | | C | C | | |
| NC_011770 | 665203 | G | A | 80 | 0 | 4 | 0 | | | A | A | A | A | | | | | | |
| NC_011770 | 665213 | T | A | 79 | 0 | 12 | 0 | A | A | A | A | A | A | A | A | A | A | A | A |
| NC_011770 | 665274 | C | T | 78 | 0 | 2 | 0 | | | | | | | | | T | T | | |
| NC_011770 | 665288 | G | A | 81 | 0 | 2 | 0 | | | | | | | A | A | | | | |
| NC_011770 | 665314 | C | T | 79 | 0 | 2 | 0 | | | | | | | | | T | T | | |
| NC_011770 | 665370 | G | A | 76 | 0 | 2 | 0 | | | | | | | A | A | | | | |
| NC_011770 | 665378 | C | T | 76 | 0 | 2 | 0 | | | | | | | T | T | | | | |
| NC_011770 | 665418 | T | C | 77 | 0 | 12 | 0 | C | C | C | C | C | C | C | C | C | C | C | C |
| NC_011770 | 665469 | C | T | 80 | 0 | 2 | 0 | T | T | | | | | | | | | | |
| NC_011770 | 666184 | A | C | 78 | 0 | 2 | 0 | | | | | | | | | C | C | | |
| NC_011770 | 666187 | GT | G | 76 | 0 | 2 | 0 | | | | | | | | | G | G | | |
| NC_011770 | 666220 | A | G | 73 | 0 | 6 | 0 | | | G | G | G | G | | | G | G | | |
| NC_011770 | 666255 | G | A | 74 | 0 | 2 | 0 | | | | | | | | | | | A | A |
| NC_011770 | 666256 | C | T | 74 | 0 | 4 | 0 | T | T | | | | | | | | | T | T |
| NC_011770 | 666260 | C | T | 75 | 0 | 8 | 0 | T | T | T | T | T | T | | | | | T | T |
| NC_011770 | 666298 | A | G | 70 | 0 | 4 | 0 | G | G | | | | | | | G | G | | |
| NC_011770 | 666299 | T | G | 70 | 0 | 4 | 0 | G | G | | | | | | | G | G | | |
| NC_011770 | 666341 | C | CT | 72 | 0 | 2 | 0 | | | | | | | | | CT | CT | | |
| NC_011770 | 666348 | C | T | 70 | 0 | 6 | 0 | | | T | T | T | T | | | T | T | | |
| NC_011770 | 666361 | T | G | 70 | 0 | 2 | 0 | | | | | | | | | G | G | | |
| NC_011770 | 666364 | G | T | 69 | 0 | 4 | 0 | | | T | T | T | T | | | | | | |
| NC_011770 | 666399 | C | T | 71 | 0 | 4 | 0 | T | T | | | | | | | T | T | | |
| NC_011770 | 666484 | G | A | 67 | 0 | 2 | 0 | | | | | | | | | A | A | | |
| NC_011770 | 666982 | A | G | 54 | 6 | 1 | 0 | . | . | | | | | G | | . | . | . | . |
| NC_011770 | 668193 | C | T | 48 | 6 | 4 | 0 | . | . | T | T | T | T | | | . | . | . | . |
| NC_011770 | 668216 | G | A | 49 | 6 | 4 | 0 | . | . | A | A | A | A | | | . | . | . | . |
| NC_011770 | 668218 | A | G | 48 | 6 | 4 | 0 | . | . | G | G | G | G | | | . | . | . | . |
| NC_011770 | 669961 | C | T | 58 | 4 | 2 | 0 | T | T | | | | | | | . | . | . | . |
| NC_011770 | 672098 | G | T | 24 | 8 | 4 | 0 | . | . | T | T | T | T | . | . | . | . | . | . |
| NC_011770 | 672104 | C | T | 23 | 8 | 4 | 0 | . | . | T | T | T | T | . | . | . | . | . | . |
| NC_011770 | 672107 | T | C | 24 | 8 | 4 | 0 | . | . | C | C | C | C | . | . | . | . | . | . |
| NC_011770 | 672473 | G | A | 0 | 11 | 1 | 0 | . | . | . | . | . | A | . | . | . | . | . | . |
| NC_011770 | 672483 | C | CCG | 0 | 10 | 1 | 0 | . | . | . | . | . | CCG | . | . | . | . | . | . |
| NC_011770 | 672495 | C | G | 1 | 9 | 3 | 0 | . | . | G | . | G | G | . | . | . | . | . | . |
| NC_011770 | 672498 | T | C | 1 | 9 | 3 | 0 | . | . | C | . | C | C | . | . | . | . | . | . |
| NC_011770 | 672525 | T | C | 46 | 6 | 4 | 0 | . | . | C | C | C | C | | | . | . | . | . |
| NC_011770 | 674682 | C | T | 63 | 6 | 4 | 0 | . | . | T | T | T | T | | | . | . | . | . |
| NC_011770 | 678075 | C | A | 48 | 6 | 2 | 0 | . | . | | | | | A | A | . | . | . | . |
| NC_011770 | 678097 | T | G | 48 | 6 | 6 | 0 | . | . | G | G | G | G | G | G | . | . | . | . |
| NC_011770 | 678115 | A | G | 50 | 6 | 2 | 0 | . | . | | | | | G | G | . | . | . | . |
| NC_011770 | 678117 | C | T | 50 | 6 | 2 | 0 | . | . | | | | | T | T | . | . | . | . |
| NC_011770 | 679131 | G | GC | 61 | 5 | 1 | 0 | . | . | | | | | | | . | GC | . | . |
| NC_011770 | 680470 | G | A | 60 | 0 | 10 | 0 | | | A | A | A | A | A | A | A | A | A | A |
| NC_011770 | 680561 | G | A | 69 | 0 | 2 | 0 | A | A | | | | | | | | | | |
| NC_011770 | 680569 | C | T | 70 | 0 | 4 | 0 | | | | | | | T | T | | | T | T |
| NC_011770 | 680616 | C | T | 73 | 0 | 4 | 0 | | | T | T | T | T | | | | | | |
| NC_011770 | 680629 | G | A | 71 | 0 | 4 | 0 | | | A | A | A | A | | | | | | |
| NC_011770 | 680633 | A | T | 73 | 0 | 4 | 0 | | | | | | | T | T | | | T | T |
| NC_011770 | 680641 | C | T | 71 | 0 | 8 | 0 | | | T | T | T | T | T | T | | | T | T |
| NC_011770 | 680655 | A | T | 70 | 0 | 2 | 0 | | | | | | | | | | | T | T |
| NC_011770 | 680668 | C | A | 74 | 0 | 8 | 0 | | | A | A | A | A | A | A | | | A | A |
| NC_011770 | 680705 | A | G | 70 | 0 | 6 | 0 | | | G | G | G | G | G | G | | | | |

| | | | | | | | | | | | | | | | | | | | |
|---|---|---|---|---|---|---|---|---|---|---|---|---|---|---|---|---|---|---|---|
| NC_011770 | 683237 | T | C | 62 | 0 | 2 | 0 | | | | | | | C | C | | | | |
| NC_011770 | 683258 | C | T | 62 | 0 | 2 | 0 | | | | | | | T | T | | | | |
| NC_011770 | 683995 | G | A | 87 | 0 | 2 | 0 | A | A | | | | | | | | | | |
| NC_011770 | 684092 | G | A | 83 | 0 | 6 | 0 | | | A | A | A | A | A | A | | | | |
| NC_011770 | 684858 | T | C | 95 | 0 | 1 | 0 | | | | | | | C | | | | | |
| NC_011770 | 685740 | C | T | 71 | 0 | 2 | 0 | | | | | | | | | | | T | T |
| NC_011770 | 686537 | C | T | 89 | 0 | 8 | 0 | | | T | T | T | T | | | T | T | T | T |
| NC_011770 | 688439 | C | T | 74 | 0 | 4 | 0 | | | T | T | T | T | | | | | | |
| NC_011770 | 688440 | G | A | 74 | 0 | 4 | 0 | | | | | | | A | A | A | A | | |
| NC_011770 | 688446 | T | G | 72 | 0 | 4 | 0 | | | | | | | G | G | G | G | | |
| NC_011770 | 688452 | C | T | 71 | 0 | 4 | 0 | | | | | | | T | T | T | T | | |
| NC_011770 | 688454 | T | C | 72 | 0 | 4 | 0 | | | | | | | C | C | C | C | | |
| NC_011770 | 688455 | T | C | 71 | 0 | 4 | 0 | | | | | | | C | C | C | C | | |
| NC_011770 | 688475 | A | G | 71 | 0 | 4 | 0 | | | | | | | G | G | G | G | | |
| NC_011770 | 691967 | C | T | 79 | 0 | 4 | 0 | | | T | T | T | T | | | | | | |
| NC_011770 | 692102 | C | G | 77 | 0 | 12 | 0 | G | G | G | G | G | G | G | G | G | G | G | G |
| NC_011770 | 694445 | T | C | 61 | 0 | 6 | 0 | | | C | C | C | C | | | C | C | | |
| NC_011770 | 694849 | C | T | 51 | 0 | 8 | 0 | | | T | T | T | T | T | T | T | T | | |
| NC_011770 | 697588 | T | C | 74 | 0 | 4 | 0 | | | C | C | C | C | | | | | | |
| NC_011770 | 698859 | C | G | 72 | 0 | 12 | 0 | G | G | G | G | G | G | G | G | G | G | G | G |
| NC_011770 | 698967 | A | G | 110 | 0 | 12 | 0 | G | G | G | G | G | G | G | G | G | G | G | G |
| NC_011770 | 698981 | C | T | 114 | 0 | 6 | 0 | T | T | T | T | T | T | | | | | | |
| NC_011770 | 699319 | A | G | 120 | 0 | 4 | 0 | | | G | G | G | G | | | | | | |
| NC_011770 | 701056 | C | A | 185 | 0 | 2 | 0 | | | | | | | | | A | A | | |
| NC_011770 | 701135 | TA | T | 162 | 0 | 2 | 0 | T | T | | | | | | | | | | |
| NC_011770 | 701172 | G | A | 158 | 0 | 4 | 0 | | | | | | | A | A | | | A | A |
| NC_011770 | 701173 | A | G | 159 | 0 | 4 | 0 | | | | | | | G | G | | | G | G |
| NC_011770 | 704541 | G | A | 88 | 0 | 4 | 0 | | | A | A | A | A | | | | | | |
| NC_011770 | 704555 | T | C | 84 | 0 | 2 | 0 | | | | | | | C | C | | | | |
| NC_011770 | 707177 | A | G | 73 | 0 | 2 | 0 | G | G | | | | | | | | | | |
| NC_011770 | 707500 | G | A | 68 | 0 | 2 | 0 | | | | | | | A | A | | | | |
| NC_011770 | 708784 | C | A | 66 | 0 | 4 | 0 | | | A | A | A | A | | | | | | |
| NC_011770 | 708796 | C | T | 68 | 0 | 1 | 0 | | | | | | | T | | | | | |
| NC_011770 | 720840 | C | T | 92 | 0 | 1 | 0 | | | | | | | T | | | | | |
| NC_011770 | 720909 | T | C | 96 | 0 | 2 | 0 | | | | | | | C | C | | | | |
| NC_011770 | 732611 | A | G | 90 | 0 | 2 | 0 | G | G | | | | | | | | | | |
| NC_011770 | 739123 | C | T | 62 | 0 | 10 | 0 | T | T | T | T | T | T | T | T | | | T | T |
| NC_011770 | 739169 | C | A | 67 | 0 | 10 | 0 | A | A | A | A | A | A | A | A | | | A | A |
| NC_011770 | 739175 | G | T | 68 | 0 | 10 | 0 | T | T | T | T | T | T | T | T | | | T | T |
| NC_011770 | 739195 | T | C | 69 | 0 | 10 | 0 | C | C | C | C | C | C | C | C | | | C | C |
| NC_011770 | 740841 | C | T | 79 | 0 | 2 | 0 | | | | | | | | | T | T | | |
| NC_011770 | 740918 | C | G | 77 | 0 | 2 | 0 | | | | | | | | | G | G | | |
| NC_011770 | 741425 | AC | A | 71 | 0 | 4 | 0 | A | A | | | | | | | A | A | | |
| NC_011770 | 741461 | T | C | 72 | 0 | 4 | 0 | C | C | | | | | | | C | C | | |
| NC_011770 | 744498 | C | T | 80 | 0 | 2 | 0 | | | | | | | | | | | T | T |
| NC_011770 | 746466 | A | G | 58 | 0 | 12 | 0 | G | G | G | G | G | G | G | G | G | G | G | G |
| NC_011770 | 750700 | T | C | 37 | 0 | 10 | 0 | C | C | C | C | C | C | C | C | | | C | C |
| NC_011770 | 750726 | A | G | 36 | 0 | 12 | 0 | G | G | G | G | G | G | G | G | G | G | G | G |
| NC_011770 | 750734 | C | A | 37 | 0 | 4 | 0 | | | A | A | A | A | | | | | | |
| NC_011770 | 751825 | T | C | 65 | 0 | 10 | 0 | | | C | C | C | C | C | C | C | C | C | C |
| NC_011770 | 751948 | A | G | 52 | 0 | 2 | 0 | G | G | | | | | | | | | | |
| NC_011770 | 766232 | C | T | 65 | 0 | 8 | 0 | T | T | T | T | T | T | T | T | | | | |
| NC_011770 | 768796 | T | C | 64 | 0 | 6 | 0 | C | C | | | | | C | C | | | C | C |
| NC_011770 | 768966 | C | T | 60 | 0 | 4 | 0 | T | T | | | | | | | T | T | | |
| NC_011770 | 771479 | G | A | 20 | 0 | 2 | 0 | | | | | | | A | A | | | | |
| NC_011770 | 771524 | G | T | 25 | 0 | 5 | 0 | | | | | | | T | T | T | T | T | . |
| NC_011770 | 772859 | T | C | 59 | 0 | 2 | 0 | | | | | | | | | C | C | | |
| NC_011770 | 772864 | G | A | 59 | 0 | 2 | 0 | | | | | | | A | A | | | | |
| NC_011770 | 772890 | T | C | 59 | 0 | 2 | 0 | | | | | | | | | C | C | | |
| NC_011770 | 772903 | G | A | 60 | 0 | 2 | 0 | | | | | | | | | A | A | | |
| NC_011770 | 772904 | C | A | 58 | 0 | 2 | 0 | | | | | | | | | A | A | | |
| NC_011770 | 772908 | A | G | 56 | 0 | 2 | 0 | | | | | | | | | G | G | | |

| | | | | | | | | | | | | | | | | | | | |
|---|---|---|---|---|---|---|---|---|---|---|---|---|---|---|---|---|---|---|---|
| NC_011770 | 772914 | G | A | 58 | 0 | 2 | 0 | | | | | | | | | A | A | | |
| NC_011770 | 772923 | G | T | 59 | 0 | 2 | 0 | | | | | | | | | T | T | | |
| NC_011770 | 772944 | G | A | 42 | 0 | 6 | 0 | | | A | A | A | A | | | A | A | | |
| NC_011770 | 772946 | G | A | 43 | 0 | 6 | 0 | | | A | A | A | A | | | A | A | | |
| NC_011770 | 772951 | C | T | 45 | 0 | 2 | 0 | | | | | | | T | T | | | | |
| NC_011770 | 772953 | T | C | 44 | 0 | 8 | 0 | | | C | C | C | C | C | C | C | C | | |
| NC_011770 | 772995 | A | G | 65 | 0 | 2 | 0 | | | | | | | | | | | G | G |
| NC_011770 | 773003 | T | C | 64 | 0 | 6 | 0 | | | C | C | C | C | | | C | C | | |
| NC_011770 | 779322 | C | G | 81 | 0 | 2 | 0 | G | G | | | | | | | | | | |
| NC_011770 | 779381 | T | C | 75 | 0 | 4 | 0 | | | C | C | C | C | | | | | | |
| NC_011770 | 779471 | C | T | 74 | 0 | 2 | 0 | T | T | | | | | | | | | | |
| NC_011770 | 779528 | G | A | 74 | 0 | 12 | 0 | A | A | A | A | A | A | A | A | A | A | A | A |
| NC_011770 | 779636 | G | T | 76 | 0 | 2 | 0 | T | T | | | | | | | | | | |
| NC_011770 | 779816 | G | T | 75 | 0 | 4 | 0 | | | T | T | T | T | | | | | | |
| NC_011770 | 779965 | T | TA | 63 | 0 | 12 | 0 | TA | TA | TA | TA | TA | TA | TA | TA | TA | TA | TA | TA |
| NC_011770 | 781009 | T | C | 67 | 0 | 4 | 0 | | | C | C | C | C | | | | | | |
| NC_011770 | 787287 | G | A | 22 | 0 | 12 | 0 | A | A | A | A | A | A | A | A | A | A | A | A |
| NC_011770 | 787307 | A | G | 15 | 0 | 12 | 0 | G | G | G | G | G | G | G | G | G | G | G | G |
| NC_011770 | 787310 | T | G | 15 | 0 | 10 | 0 | G | G | G | G | G | G | G | G | | | A | A |
| NC_011770 | 787311 | C | A | 15 | 0 | 2 | 0 | | | | | | | | | A | A | | |
| NC_011770 | 787329 | C | T | 12 | 1 | 11 | 0 | . | T | T | T | T | T | T | T | T | T | T | T |
| NC_011770 | 787344 | A | G | 12 | 1 | 3 | 0 | . | G | | | | | | | | | G | G |
| NC_011770 | 787377 | A | G | 7 | 1 | 11 | 0 | G | G | G | G | G | G | G | G | . | G | G | G |
| NC_011770 | 787386 | G | C | 6 | 1 | 2 | 0 | | | | | | | C | C | . | | | |
| NC_011770 | 787687 | C | T | 42 | 0 | 12 | 0 | T | T | T | T | T | T | T | T | T | T | T | T |
| NC_011770 | 787708 | C | G | 42 | 0 | 12 | 0 | G | G | G | G | G | G | G | G | G | G | G | G |
| NC_011770 | 787720 | A | G | 47 | 0 | 2 | 0 | | | | | | | G | G | | | | |
| NC_011770 | 789905 | G | A | 61 | 0 | 4 | 0 | | | A | A | A | A | | | | | | |
| NC_011770 | 793183 | A | G | 70 | 0 | 12 | 0 | G | G | G | G | G | G | G | G | G | G | G | G |
| NC_011770 | 793237 | GA | G | 73 | 0 | 2 | 0 | | | | | | | G | G | | | | |
| NC_011770 | 793248 | C | T | 70 | 0 | 2 | 0 | | | | | | | T | T | | | | |
| NC_011770 | 795068 | G | T | 79 | 0 | 2 | 0 | | | | | | | | | | | T | T |
| NC_011770 | 795132 | G | T | 83 | 0 | 4 | 0 | | | T | T | T | T | | | | | | |
| NC_011770 | 797029 | A | G | 71 | 0 | 8 | 0 | G | G | G | G | G | G | | | G | G | | |
| NC_011770 | 797039 | C | T | 73 | 0 | 8 | 0 | T | T | T | T | T | T | | | T | T | | |
| NC_011770 | 799996 | A | G | 56 | 0 | 2 | 0 | G | G | | | | | | | | | | |
| NC_011770 | 800018 | G | T | 55 | 0 | 2 | 0 | T | T | | | | | | | | | | |
| NC_011770 | 808002 | G | T | 56 | 0 | 2 | 0 | | | | | | | T | T | | | | |
| NC_011770 | 809505 | T | C | 78 | 0 | 8 | 0 | | | C | C | C | C | | | C | C | C | C |
| NC_011770 | 809515 | A | C | 86 | 0 | 8 | 0 | | | C | C | C | C | | | C | C | C | C |
| NC_011770 | 811693 | T | G | 64 | 0 | 2 | 0 | | | | | | | G | G | | | | |
| NC_011770 | 811722 | G | A | 54 | 0 | 2 | 0 | | | | | | | A | A | | | | |
| NC_011770 | 811724 | T | C | 50 | 0 | 2 | 0 | | | | | | | C | C | | | | |
| NC_011770 | 811725 | C | T | 51 | 0 | 2 | 0 | T | T | | | | | | | | | | |
| NC_011770 | 811747 | G | GT | 42 | 0 | 1 | 0 | | | | | | | GT | | | | | |
| NC_011770 | 811760 | C | T | 39 | 0 | 2 | 0 | | | | | | | T | T | | | | |
| NC_011770 | 811765 | C | T | 40 | 0 | 2 | 0 | | | | | | | T | T | | | | |
| NC_011770 | 811798 | T | G | 37 | 0 | 2 | 0 | | | | | | | G | G | | | | |
| NC_011770 | 811802 | A | G | 37 | 0 | 12 | 0 | G | G | G | G | G | G | G | G | G | G | G | G |
| NC_011770 | 811805 | T | C | 36 | 0 | 2 | 0 | | | | | | | C | C | | | | |
| NC_011770 | 811807 | A | G | 37 | 0 | 2 | 0 | G | G | | | | | | | | | | |
| NC_011770 | 811840 | C | T | 31 | 0 | 2 | 0 | | | | | | | T | T | | | | |
| NC_011770 | 811925 | C | T | 16 | 0 | 2 | 0 | | | | | | | | | T | T | | |
| NC_011770 | 813489 | G | A | 60 | 0 | 12 | 0 | A | A | A | A | A | A | A | A | A | A | A | A |
| NC_011770 | 814152 | C | G | 66 | 0 | 2 | 0 | | | | | | | | | G | G | | |
| NC_011770 | 814155 | C | T | 66 | 0 | 2 | 0 | | | | | | | | | T | T | | |
| NC_011770 | 815035 | A | C | 39 | 0 | 12 | 0 | C | C | C | C | C | C | C | C | C | C | C | C |
| NC_011770 | 819152 | C | T | 80 | 0 | 4 | 0 | T | T | | | | | T | T | | | | |
| NC_011770 | 819188 | G | A | 83 | 0 | 4 | 0 | A | A | | | | | A | A | | | | |
| NC_011770 | 819213 | A | G | 80 | 0 | 4 | 0 | G | G | | | | | G | G | | | | |
| NC_011770 | 819219 | C | T | 81 | 0 | 4 | 0 | T | T | | | | | T | T | | | | |
| NC_011770 | 819645 | G | A | 75 | 0 | 2 | 0 | | | | | | | | | | | A | A |

| | | | | | | | | | | | | | | | | | | | |
|---|---|---|---|---|---|---|---|---|---|---|---|---|---|---|---|---|---|---|---|
| NC_011770 | 819656 | G | C | 74 | 0 | 6 | 0 | C | C | | | | | C | C | C | C | | |
| NC_011770 | 819686 | T | G | 73 | 0 | 2 | 0 | | | | | | | | | | | G | G |
| NC_011770 | 819692 | C | T | 74 | 0 | 2 | 0 | | | | | | | | | | | T | T |
| NC_011770 | 819703 | C | T | 72 | 0 | 2 | 0 | | | | | | | | | | | T | T |
| NC_011770 | 819712 | C | T | 73 | 0 | 2 | 0 | | | | | | | | | | | T | T |
| NC_011770 | 819718 | C | T | 70 | 0 | 2 | 0 | | | | | | | | | | | T | T |
| NC_011770 | 819732 | G | T | 70 | 0 | 2 | 0 | | | | | | | | | | | T | T |
| NC_011770 | 820069 | G | A | 67 | 0 | 8 | 0 | A | A | A | A | A | A | A | A | | | | |
| NC_011770 | 820107 | G | A | 59 | 0 | 8 | 0 | A | A | A | A | A | A | A | A | | | | |
| NC_011770 | 820131 | A | G | 51 | 0 | 8 | 0 | G | G | G | G | G | G | G | G | | | | |
| NC_011770 | 820136 | G | A | 51 | 0 | 8 | 0 | A | A | A | A | A | A | A | A | | | | |
| NC_011770 | 820139 | A | G | 51 | 0 | 8 | 0 | G | G | G | G | G | G | G | G | | | | |
| NC_011770 | 820189 | T | C | 12 | 1 | 9 | 0 | C | C | C | C | C | C | C | C | C | . | | |
| NC_011770 | 820193 | G | A | 12 | 1 | 9 | 0 | A | A | A | A | A | A | A | A | A | . | | |
| NC_011770 | 820223 | T | C | 36 | 0 | 10 | 0 | C | C | C | C | C | C | C | C | C | C | | |
| NC_011770 | 820249 | A | G | 20 | 0 | 4 | 0 | | | | | | | | | G | G | G | G |
| NC_011770 | 820252 | A | G | 22 | 0 | 8 | 0 | G | G | G | G | G | G | G | G | | | | |
| NC_011770 | 820264 | T | C | 23 | 0 | 12 | 0 | C | C | C | C | C | C | C | C | C | C | C | C |
| NC_011770 | 820273 | C | CGTTAT | 1 | 10 | 1 | 0 | . | . | . | . | . | . | . | . | . | CGTTATGC | . | . |
| NC_011770 | 820280 | T | C | 26 | 0 | 12 | 0 | C | C | C | C | C | C | C | C | C | C | C | C |
| NC_011770 | 820285 | AC | A | 25 | 0 | 10 | 0 | A | A | A | A | A | A | A | A | A | A | | |
| NC_011770 | 820290 | T | C | 26 | 0 | 10 | 0 | C | C | C | C | C | C | C | C | C | C | | |
| NC_011770 | 820294 | A | G | 38 | 0 | 2 | 0 | | | | | | | | | | | G | G |
| NC_011770 | 820300 | T | C | 39 | 0 | 12 | 0 | C | C | C | C | C | C | C | C | C | C | C | C |
| NC_011770 | 820307 | T | C | 39 | 0 | 12 | 0 | C | C | C | C | C | C | C | C | C | C | C | C |
| NC_011770 | 820310 | C | G | 40 | 0 | 10 | 0 | G | G | G | G | G | G | G | G | G | G | | |
| NC_011770 | 820312 | C | T | 39 | 0 | 10 | 0 | T | T | T | T | T | T | T | T | T | T | | |
| NC_011770 | 820316 | G | A | 41 | 0 | 12 | 0 | A | A | A | A | A | A | A | A | A | A | A | A |
| NC_011770 | 820321 | A | G | 42 | 0 | 12 | 0 | G | G | G | G | G | G | G | G | G | G | G | G |
| NC_011770 | 820324 | C | T | 47 | 0 | 2 | 0 | | | | | | | | | | | T | T |
| NC_011770 | 820327 | A | G | 47 | 0 | 2 | 0 | | | | | | | | | | | G | G |
| NC_011770 | 820353 | C | T | 58 | 0 | 2 | 0 | | | | | | | | | | | T | T |
| NC_011770 | 820358 | G | A | 58 | 0 | 2 | 0 | | | | | | | | | | | A | A |
| NC_011770 | 821909 | G | A | 68 | 0 | 10 | 0 | A | A | A | A | A | A | A | A | A | A | | |
| NC_011770 | 822043 | A | G | 70 | 0 | 10 | 0 | G | G | G | G | G | G | G | G | G | G | | |
| NC_011770 | 822062 | A | G | 67 | 0 | 10 | 0 | G | G | G | G | G | G | G | G | G | G | | |
| NC_011770 | 822160 | C | T | 72 | 0 | 1 | 0 | | | | | | | T | | | | | |
| NC_011770 | 822294 | C | T | 64 | 0 | 2 | 0 | | | | | | | | | | | T | T |
| NC_011770 | 823346 | C | CG | 80 | 0 | 12 | 0 | CG | CG | CG | CG | CG | CG | CG | CG | CG | CG | CG | CG |
| NC_011770 | 826258 | A | G | 74 | 0 | 2 | 0 | | | | | | | | | | | G | G |
| NC_011770 | 826259 | C | A | 74 | 0 | 2 | 0 | | | | | | | | | | | A | A |
| NC_011770 | 826260 | G | T | 75 | 0 | 2 | 0 | | | | | | | | | | | T | T |
| NC_011770 | 826264 | T | C | 77 | 0 | 2 | 0 | | | | | | | | | | | C | C |
| NC_011770 | 826300 | G | C | 81 | 0 | 2 | 0 | | | | | | | | | | | C | C |
| NC_011770 | 826311 | C | T | 82 | 0 | 2 | 0 | | | | | | | | | | | T | T |
| NC_011770 | 826312 | T | C | 83 | 0 | 2 | 0 | | | | | | | | | | | C | C |
| NC_011770 | 826319 | TCTC | T | 79 | 0 | 2 | 0 | | | | | | | | | | | T | T |
| NC_011770 | 826329 | T | C | 76 | 0 | 2 | 0 | | | | | | | | | | | C | C |
| NC_011770 | 826337 | C | T | 85 | 0 | 2 | 0 | | | | | | | | | | | T | T |
| NC_011770 | 826779 | G | A | 90 | 0 | 2 | 0 | | | | | | | | | | | A | A |
| NC_011770 | 829361 | G | T | 26 | 0 | 10 | 0 | T | T | T | T | T | T | T | T | T | T | | |
| NC_011770 | 829391 | A | G | 28 | 0 | 2 | 0 | | | | | | | | | | | G | G |
| NC_011770 | 829398 | G | T | 28 | 0 | 10 | 0 | T | T | T | T | T | T | T | T | T | T | | |
| NC_011770 | 829443 | T | C | 26 | 0 | 2 | 0 | | | | | | | | | | | C | C |
| NC_011770 | 829444 | T | C | 26 | 0 | 10 | 0 | C | C | C | C | C | C | C | C | C | C | | |
| NC_011770 | 829450 | G | C | 27 | 0 | 2 | 0 | | | | | | | | | | | C | C |
| NC_011770 | 830414 | A | G | 93 | 0 | 2 | 0 | | | | | | | | | | | G | G |
| NC_011770 | 830424 | CGGTGCAT | C | 78 | 1 | 1 | 0 | | | | | | | | | | | C | . |
| NC_011770 | 834888 | C | A | 88 | 0 | 12 | 0 | A | A | A | A | A | A | A | A | A | A | A | A |
| NC_011770 | 838699 | G | A | 79 | 0 | 2 | 0 | | | | | | | A | A | | | | |
| NC_011770 | 838703 | A | C | 76 | 0 | 10 | 0 | C | C | C | C | C | C | C | C | C | C | | |
| NC_011770 | 838740 | G | A | 81 | 0 | 10 | 0 | A | A | A | A | A | A | A | A | A | A | | |

| | | | | | | | | | | | | | | | | | | | |
|---|---|---|---|---|---|---|---|---|---|---|---|---|---|---|---|---|---|---|---|
| NC_011770 | 838745 | T | C | 81 | 0 | 12 | 0 | C | C | C | C | C | C | C | C | C | C | C | C |
| NC_011770 | 840543 | C | T | 73 | 0 | 8 | 0 | | | T | T | T | T | | | T | T | T | T |
| NC_011770 | 840553 | T | C | 69 | 0 | 6 | 0 | | | C | C | C | C | | | C | C | | |
| NC_011770 | 840555 | C | A | 70 | 0 | 2 | 0 | A | A | | | | | | | | | | |
| NC_011770 | 840576 | C | T | 70 | 0 | 12 | 0 | T | T | T | T | T | T | T | T | T | T | T | T |
| NC_011770 | 840583 | C | T | 70 | 0 | 2 | 0 | T | T | | | | | | | | | | |
| NC_011770 | 840595 | A | T | 70 | 0 | 8 | 0 | | | T | T | T | T | | | T | T | T | T |
| NC_011770 | 840612 | C | T | 68 | 0 | 2 | 0 | | | | | | | T | T | | | | |
| NC_011770 | 840620 | G | T | 69 | 0 | 6 | 0 | | | T | T | T | T | | | T | T | | |
| NC_011770 | 840647 | A | G | 65 | 0 | 2 | 0 | G | G | | | | | | | | | | |
| NC_011770 | 844172 | G | GA | 79 | 0 | 4 | 0 | GA | GA | | | | | | | | | GA | GA |
| NC_011770 | 844224 | G | A | 79 | 0 | 4 | 0 | A | A | | | | | | | | | A | A |
| NC_011770 | 854403 | C | G | 83 | 0 | 2 | 0 | | | | | | | | | | | G | G |
| NC_011770 | 855319 | G | A | 71 | 0 | 8 | 0 | A | A | A | A | A | A | | | A | A | | |
| NC_011770 | 860658 | T | C | 66 | 0 | 10 | 0 | C | C | C | C | C | C | C | C | C | C | | |
| NC_011770 | 860738 | C | A | 70 | 0 | 2 | 0 | | | | | | | | | | | A | A |
| NC_011770 | 863351 | C | T | 61 | 0 | 10 | 0 | T | T | T | T | T | T | | | T | T | T | T |
| NC_011770 | 863492 | C | T | 51 | 0 | 2 | 0 | | | | | | | | | | | T | T |
| NC_011770 | 863516 | G | A | 52 | 0 | 10 | 0 | A | A | A | A | A | A | | | A | A | A | A |
| NC_011770 | 863594 | T | G | 51 | 0 | 10 | 0 | G | G | G | G | G | G | | | G | G | G | G |
| NC_011770 | 863710 | G | T | 46 | 0 | 10 | 0 | T | T | T | T | T | T | | | T | T | T | T |
| NC_011770 | 863758 | T | G | 42 | 0 | 10 | 0 | G | G | G | G | G | G | | | G | G | G | G |
| NC_011770 | 866750 | G | A | 13 | 10 | 2 | 0 | . | . | . | . | A | A | . | . | . | . | . | . |
| NC_011770 | 866759 | T | A | 0 | 11 | 1 | 0 | . | . | . | . | . | A | . | . | . | . | . | . |
| NC_011770 | 871728 | G | A | 6 | 10 | 2 | 0 | . | . | . | . | A | A | . | . | . | . | . | . |
| NC_011770 | 871730 | T | C | 6 | 10 | 2 | 0 | . | . | . | . | C | C | . | . | . | . | . | . |
| NC_011770 | 871736 | G | T | 6 | 10 | 2 | 0 | . | . | . | . | T | T | . | . | . | . | . | . |
| NC_011770 | 872319 | A | C | 5 | 10 | 2 | 0 | . | . | . | . | C | C | . | . | . | . | . | . |
| NC_011770 | 875013 | G | A | 15 | 10 | 2 | 0 | . | . | . | . | A | A | . | . | . | . | . | . |
| NC_011770 | 875017 | C | T | 15 | 10 | 2 | 0 | . | . | . | . | T | T | . | . | . | . | . | . |
| NC_011770 | 875179 | T | C | 5 | 10 | 2 | 0 | . | . | . | . | C | C | . | . | . | . | . | . |
| NC_011770 | 875194 | T | C | 5 | 10 | 2 | 0 | . | . | . | . | C | C | . | . | . | . | . | . |
| NC_011770 | 879537 | T | C | 9 | 10 | 2 | 0 | . | . | . | . | C | C | . | . | . | . | . | . |
| NC_011770 | 879857 | T | C | 12 | 10 | 2 | 0 | . | . | . | . | C | C | . | . | . | . | . | . |
| NC_011770 | 879866 | A | G | 12 | 10 | 2 | 0 | . | . | . | . | G | G | . | . | . | . | . | . |
| NC_011770 | 879884 | C | A | 11 | 10 | 2 | 0 | . | . | . | . | A | A | . | . | . | . | . | . |
| NC_011770 | 879893 | G | A | 12 | 10 | 2 | 0 | . | . | . | . | A | A | . | . | . | . | . | . |
| NC_011770 | 879908 | G | A | 12 | 10 | 2 | 0 | . | . | . | . | A | A | . | . | . | . | . | . |
| NC_011770 | 880204 | C | T | 12 | 10 | 2 | 0 | . | . | . | . | T | T | . | . | . | . | . | . |
| NC_011770 | 880210 | A | G | 12 | 10 | 2 | 0 | . | . | . | . | G | G | . | . | . | . | . | . |
| NC_011770 | 880219 | A | G | 12 | 10 | 2 | 0 | . | . | . | . | G | G | . | . | . | . | . | . |
| NC_011770 | 880221 | G | A | 11 | 10 | 2 | 0 | . | . | . | . | A | A | . | . | . | . | . | . |
| NC_011770 | 880234 | C | A | 11 | 10 | 2 | 0 | . | . | . | . | A | A | . | . | . | . | . | . |
| NC_011770 | 880280 | GC | G | 12 | 10 | 2 | 0 | . | . | . | . | G | G | . | . | . | . | . | . |
| NC_011770 | 880308 | A | T | 10 | 10 | 2 | 0 | . | . | . | . | T | T | . | . | . | . | . | . |
| NC_011770 | 880310 | C | T | 10 | 10 | 2 | 0 | . | . | . | . | T | T | . | . | . | . | . | . |
| NC_011770 | 880312 | T | A | 10 | 10 | 2 | 0 | . | . | . | . | A | A | . | . | . | . | . | . |
| NC_011770 | 880315 | C | T | 10 | 10 | 2 | 0 | . | . | . | . | T | T | . | . | . | . | . | . |
| NC_011770 | 880497 | A | G | 10 | 10 | 2 | 0 | . | . | . | . | G | G | . | . | . | . | . | . |
| NC_011770 | 881329 | T | C | 14 | 6 | 6 | 0 | . | . | . | . | C | C | . | . | C | C | C | C |
| NC_011770 | 881336 | G | C | 14 | 6 | 2 | 0 | . | . | . | . | C | C | . | . | | | | |
| NC_011770 | 881345 | A | T | 14 | 6 | 2 | 0 | . | . | . | . | T | T | . | . | | | | |
| NC_011770 | 881346 | C | T | 14 | 6 | 4 | 0 | . | . | . | . | T | T | . | . | | | T | T |
| NC_011770 | 881347 | T | C | 14 | 6 | 4 | 0 | . | . | . | . | C | C | . | . | | | C | C |
| NC_011770 | 881353 | G | C | 14 | 6 | 2 | 0 | . | . | . | . | C | C | . | . | | | | |
| NC_011770 | 882323 | C | T | 28 | 8 | 4 | 0 | . | . | . | . | T | T | . | . | . | . | T | T |
| NC_011770 | 882326 | A | G | 28 | 8 | 4 | 0 | . | . | . | . | G | G | . | . | . | . | G | G |
| NC_011770 | 882329 | T | G | 28 | 8 | 4 | 0 | . | . | . | . | G | G | . | . | . | . | G | G |
| NC_011770 | 882333 | A | C | 28 | 8 | 4 | 0 | . | . | . | . | C | C | . | . | . | . | C | C |
| NC_011770 | 882334 | A | G | 28 | 8 | 4 | 0 | . | . | . | . | G | G | . | . | . | . | G | G |
| NC_011770 | 882335 | G | C | 28 | 8 | 4 | 0 | . | . | . | . | C | C | . | . | . | . | C | C |
| NC_011770 | 882341 | C | A | 29 | 8 | 2 | 0 | . | . | . | . | | | . | . | . | . | A | A |

| | | | | | | | | | | | | | | | | | | | |
|---|---|---|---|---|---|---|---|---|---|---|---|---|---|---|---|---|---|---|---|
| NC_011770 | 882343 | T | C | 30 | 8 | 4 | 0 | . | . | . | . | C | C | . | . | . | . | C | C |
| NC_011770 | 882353 | T | C | 32 | 8 | 4 | 0 | . | . | . | . | C | C | . | . | . | . | C | C |
| NC_011770 | 882371 | T | A | 34 | 8 | 2 | 0 | . | . | . | . | A | A | . | . | . | . | | |
| NC_011770 | 882383 | G | A | 33 | 8 | 2 | 0 | . | . | . | . | | | . | . | . | . | A | A |
| NC_011770 | 882407 | G | C | 33 | 8 | 2 | 0 | . | . | . | . | C | C | . | . | . | . | | |
| NC_011770 | 882437 | A | G | 32 | 8 | 4 | 0 | . | . | . | . | G | G | . | . | . | . | G | G |
| NC_011770 | 882438 | C | A | 32 | 8 | 2 | 0 | . | . | . | . | A | A | . | . | . | . | | |
| NC_011770 | 882440 | T | G | 32 | 8 | 2 | 0 | . | . | . | . | G | G | . | . | . | . | | |
| NC_011770 | 882449 | C | T | 33 | 8 | 2 | 0 | . | . | . | . | T | T | . | . | . | . | | |
| NC_011770 | 882464 | C | T | 34 | 8 | 2 | 0 | . | . | . | . | T | T | . | . | . | . | | |
| NC_011770 | 882471 | A | G | 33 | 8 | 2 | 0 | . | . | . | . | | | . | . | . | . | G | G |
| NC_011770 | 882499 | C | A | 33 | 8 | 4 | 0 | . | . | . | . | A | A | . | . | . | . | A | A |
| NC_011770 | 882508 | A | T | 33 | 8 | 4 | 0 | . | . | . | . | T | T | . | . | . | . | T | T |
| NC_011770 | 882533 | T | C | 32 | 8 | 4 | 0 | . | . | . | . | C | C | . | . | . | . | C | C |
| NC_011770 | 883747 | A | T | 6 | 10 | 2 | 0 | . | . | . | . | . | . | . | . | . | . | T | T |
| NC_011770 | 883955 | T | G | 3 | 10 | 2 | 0 | . | . | . | . | . | . | . | . | . | . | G | G |
| NC_011770 | 884032 | C | T | 2 | 10 | 2 | 0 | . | . | . | . | . | . | . | . | . | . | T | T |
| NC_011770 | 884086 | A | G | 0 | 11 | 1 | 0 | . | . | . | . | . | . | . | . | . | . | G | . |
| NC_011770 | 888565 | C | T | 16 | 10 | 2 | 0 | . | . | . | . | . | . | . | . | . | . | T | T |
| NC_011770 | 890340 | T | C | 8 | 10 | 2 | 0 | . | . | . | . | . | . | . | . | . | . | C | C |
| NC_011770 | 890703 | G | C | 8 | 8 | 2 | 0 | . | . | . | . | | | . | . | . | . | C | C |
| NC_011770 | 890706 | G | C | 9 | 8 | 2 | 0 | . | . | . | . | | | . | . | . | . | C | C |
| NC_011770 | 890742 | A | G | 11 | 8 | 4 | 0 | . | . | . | . | G | G | . | . | . | . | T | T |
| NC_011770 | 890792 | G | A | 14 | 8 | 2 | 0 | . | . | . | . | | | . | . | . | . | A | A |
| NC_011770 | 890793 | T | C | 14 | 8 | 4 | 0 | . | . | . | . | C | C | . | . | . | . | G | G |
| NC_011770 | 890796 | T | C | 14 | 8 | 4 | 0 | . | . | . | . | C | C | . | . | . | . | C | C |
| NC_011770 | 890807 | G | A | 16 | 8 | 4 | 0 | . | . | . | . | A | A | . | . | . | . | A | A |
| NC_011770 | 890812 | G | A | 15 | 8 | 4 | 0 | . | . | . | . | A | A | . | . | . | . | A | A |
| NC_011770 | 890838 | C | G | 19 | 8 | 2 | 0 | . | . | . | . | | | . | . | . | . | G | G |
| NC_011770 | 890840 | C | G | 19 | 8 | 2 | 0 | . | . | . | . | | | . | . | . | . | G | G |
| NC_011770 | 892481 | A | C | 20 | 8 | 2 | 0 | . | . | . | . | | | . | . | . | . | C | C |
| NC_011770 | 893168 | A | T | 17 | 8 | 2 | 0 | . | . | . | . | | | . | . | . | . | T | T |
| NC_011770 | 893172 | T | TTA | 17 | 8 | 1 | 0 | . | . | . | . | | | . | . | . | . | TTA | |
| NC_011770 | 893181 | A | G | 16 | 8 | 2 | 0 | . | . | . | . | | | . | . | . | . | G | G |
| NC_011770 | 893183 | G | C | 16 | 8 | 2 | 0 | . | . | . | . | | | . | . | . | . | C | C |
| NC_011770 | 893194 | T | C | 16 | 8 | 2 | 0 | . | . | . | . | | | . | . | . | . | C | C |
| NC_011770 | 893196 | T | C | 17 | 8 | 2 | 0 | . | . | . | . | | | . | . | . | . | C | C |
| NC_011770 | 893234 | A | G | 18 | 8 | 2 | 0 | . | . | . | . | | | . | . | . | . | G | G |
| NC_011770 | 893285 | G | A | 19 | 8 | 2 | 0 | . | . | . | . | | | . | . | . | . | A | A |
| NC_011770 | 893321 | G | A | 19 | 8 | 2 | 0 | . | . | . | . | | | . | . | . | . | A | A |
| NC_011770 | 893339 | C | T | 19 | 8 | 2 | 0 | . | . | . | . | | | . | . | . | . | T | T |
| NC_011770 | 893406 | G | A | 19 | 8 | 2 | 0 | . | . | . | . | | | . | . | . | . | A | A |
| NC_011770 | 893432 | T | C | 19 | 8 | 2 | 0 | . | . | . | . | C | C | . | . | . | . | | |
| NC_011770 | 893452 | G | T | 20 | 8 | 2 | 0 | . | . | . | . | T | T | . | . | . | . | | |
| NC_011770 | 893525 | T | A | 19 | 8 | 2 | 0 | . | . | . | . | | | . | . | . | . | A | A |
| NC_011770 | 893527 | G | A | 19 | 8 | 2 | 0 | . | . | . | . | | | . | . | . | . | A | A |
| NC_011770 | 893534 | T | C | 19 | 8 | 2 | 0 | . | . | . | . | C | C | . | . | . | . | | |
| NC_011770 | 893593 | C | A | 21 | 8 | 2 | 0 | . | . | . | . | A | A | . | . | . | . | | |
| NC_011770 | 896118 | A | G | 26 | 8 | 2 | 0 | . | . | . | . | G | G | . | . | . | . | | |
| NC_011770 | 896151 | T | G | 28 | 8 | 4 | 0 | . | . | . | . | G | G | . | . | . | . | G | G |
| NC_011770 | 896177 | T | C | 27 | 8 | 4 | 0 | . | . | . | . | C | C | . | . | . | . | C | C |
| NC_011770 | 896183 | C | A | 25 | 8 | 4 | 0 | . | . | . | . | A | A | . | . | . | . | A | A |
| NC_011770 | 896184 | G | A | 26 | 8 | 4 | 0 | . | . | . | . | A | A | . | . | . | . | A | A |
| NC_011770 | 896204 | C | T | 30 | 8 | 4 | 0 | . | . | . | . | T | T | . | . | . | . | T | T |
| NC_011770 | 896336 | C | T | 36 | 8 | 2 | 0 | . | . | . | . | | | . | . | . | . | T | T |
| NC_011770 | 896351 | A | G | 35 | 8 | 2 | 0 | . | . | . | . | | | . | . | . | . | G | G |
| NC_011770 | 896365 | A | G | 36 | 8 | 2 | 0 | . | . | . | . | | | . | . | . | . | G | G |
| NC_011770 | 896370 | T | C | 37 | 8 | 2 | 0 | . | . | . | . | | | . | . | . | . | C | C |
| NC_011770 | 896402 | T | C | 36 | 8 | 2 | 0 | . | . | . | . | | | . | . | . | . | C | C |
| NC_011770 | 896405 | C | G | 35 | 8 | 2 | 0 | . | . | . | . | | | . | . | . | . | G | G |
| NC_011770 | 896408 | C | T | 36 | 8 | 2 | 0 | . | . | . | . | | | . | . | . | . | T | T |
| NC_011770 | 896435 | A | C | 35 | 8 | 2 | 0 | . | . | . | . | | | . | . | . | . | C | C |

| | | | | | | | | | | | | | | | | | | | |
|---|---|---|---|---|---|---|---|---|---|---|---|---|---|---|---|---|---|---|---|
| NC_011770 | 896441 | T | G | 37 | 8 | 2 | 0 | . | . | . | . | | | . | . | . | . | G | G |
| NC_011770 | 896444 | A | C | 35 | 8 | 2 | 0 | . | . | . | . | | | . | . | . | . | C | C |
| NC_011770 | 896453 | C | G | 34 | 8 | 2 | 0 | . | . | . | . | | | . | . | . | . | G | G |
| NC_011770 | 896456 | C | G | 33 | 8 | 2 | 0 | . | . | . | . | | | . | . | . | . | G | G |
| NC_011770 | 896459 | T | C | 34 | 8 | 2 | 0 | . | . | . | . | | | . | . | . | . | C | C |
| NC_011770 | 896463 | A | G | 34 | 8 | 2 | 0 | . | . | . | . | | | . | . | . | . | G | G |
| NC_011770 | 896468 | G | A | 35 | 8 | 2 | 0 | . | . | . | . | | | . | . | . | . | A | A |
| NC_011770 | 896489 | C | T | 35 | 8 | 2 | 0 | . | . | . | . | | | . | . | . | . | T | T |
| NC_011770 | 896513 | G | C | 35 | 8 | 2 | 0 | . | . | . | . | | | . | . | . | . | C | C |
| NC_011770 | 896519 | C | T | 35 | 8 | 2 | 0 | . | . | . | . | | | . | . | . | . | T | T |
| NC_011770 | 896537 | C | T | 36 | 8 | 2 | 0 | . | . | . | . | | | . | . | . | . | T | T |
| NC_011770 | 896552 | A | G | 36 | 8 | 2 | 0 | . | . | . | . | | | . | . | . | . | G | G |
| NC_011770 | 896573 | C | A | 37 | 8 | 2 | 0 | . | . | . | . | | | . | . | . | . | A | A |
| NC_011770 | 896606 | C | G | 37 | 8 | 2 | 0 | . | . | . | . | | | . | . | . | . | G | G |
| NC_011770 | 896607 | A | G | 38 | 8 | 2 | 0 | . | . | . | . | | | . | . | . | . | G | G |
| NC_011770 | 896618 | G | A | 39 | 8 | 2 | 0 | . | . | . | . | | | . | . | . | . | A | A |
| NC_011770 | 896725 | C | A | 36 | 8 | 2 | 0 | . | . | . | . | | | . | . | . | . | A | A |
| NC_011770 | 896730 | C | T | 37 | 8 | 2 | 0 | . | . | . | . | | | . | . | . | . | T | T |
| NC_011770 | 896758 | T | A | 35 | 8 | 2 | 0 | . | . | . | . | | | . | . | . | . | A | A |
| NC_011770 | 896767 | T | A | 34 | 8 | 2 | 0 | . | . | . | . | | | . | . | . | . | A | A |
| NC_011770 | 896779 | C | A | 33 | 8 | 2 | 0 | . | . | . | . | | | . | . | . | . | A | A |
| NC_011770 | 896780 | C | G | 33 | 8 | 2 | 0 | . | . | . | . | | | . | . | . | . | G | G |
| NC_011770 | 896788 | G | A | 33 | 8 | 2 | 0 | . | . | . | . | A | A | . | . | . | . | | |
| NC_011770 | 896791 | C | T | 33 | 8 | 2 | 0 | . | . | . | . | | | . | . | . | . | T | T |
| NC_011770 | 896794 | C | T | 33 | 8 | 4 | 0 | . | . | . | . | T | T | . | . | . | . | T | T |
| NC_011770 | 896803 | C | T | 33 | 8 | 2 | 0 | . | . | . | . | | | . | . | . | . | T | T |
| NC_011770 | 896812 | G | A | 35 | 8 | 2 | 0 | . | . | . | . | | | . | . | . | . | A | A |
| NC_011770 | 896896 | G | A | 37 | 8 | 4 | 0 | . | . | . | . | A | A | . | . | . | . | C | C |
| NC_011770 | 896909 | A | G | 39 | 8 | 2 | 0 | . | . | . | . | G | G | . | . | . | . | | |
| NC_011770 | 896911 | G | A | 39 | 8 | 2 | 0 | . | . | . | . | | | . | . | . | . | A | A |
| NC_011770 | 896923 | T | C | 38 | 8 | 2 | 0 | . | . | . | . | | | . | . | . | . | C | C |
| NC_011770 | 896955 | C | T | 34 | 8 | 2 | 0 | . | . | . | . | T | T | . | . | . | . | | |
| NC_011770 | 896959 | G | C | 35 | 8 | 2 | 0 | . | . | . | . | | | . | . | . | . | C | C |
| NC_011770 | 896962 | T | G | 34 | 8 | 2 | 0 | . | . | . | . | | | . | . | . | . | G | G |
| NC_011770 | 896965 | A | G | 35 | 8 | 2 | 0 | . | . | . | . | | | . | . | . | . | G | G |
| NC_011770 | 896977 | G | T | 34 | 8 | 2 | 0 | . | . | . | . | | | . | . | . | . | T | T |
| NC_011770 | 896980 | A | G | 34 | 8 | 2 | 0 | . | . | . | . | | | . | . | . | . | G | G |
| NC_011770 | 896992 | G | A | 34 | 8 | 2 | 0 | . | . | . | . | | | . | . | . | . | A | A |
| NC_011770 | 896995 | A | G | 34 | 8 | 2 | 0 | . | . | . | . | | | . | . | . | . | G | G |
| NC_011770 | 897001 | A | T | 34 | 8 | 2 | 0 | . | . | . | . | | | . | . | . | . | T | T |
| NC_011770 | 897034 | G | T | 33 | 8 | 2 | 0 | . | . | . | . | | | . | . | . | . | T | T |
| NC_011770 | 897055 | A | G | 34 | 8 | 2 | 0 | . | . | . | . | | | . | . | . | . | G | G |
| NC_011770 | 901854 | G | A | 26 | 8 | 2 | 0 | . | . | . | . | | | . | . | . | . | A | A |
| NC_011770 | 901886 | T | C | 26 | 8 | 2 | 0 | . | . | . | . | C | C | . | . | . | . | | |
| NC_011770 | 901890 | T | A | 26 | 8 | 2 | 0 | . | . | . | . | A | A | . | . | . | . | | |
| NC_011770 | 901901 | T | G | 26 | 8 | 2 | 0 | . | . | . | . | G | G | . | . | . | . | | |
| NC_011770 | 901934 | C | A | 28 | 8 | 2 | 0 | . | . | . | . | A | A | . | . | . | . | | |
| NC_011770 | 901943 | A | G | 27 | 8 | 4 | 0 | . | . | . | . | G | G | . | . | . | . | G | G |
| NC_011770 | 901949 | C | A | 27 | 8 | 2 | 0 | . | . | . | . | A | A | . | . | . | . | | |
| NC_011770 | 902027 | T | A | 29 | 8 | 2 | 0 | . | . | . | . | A | A | . | . | . | . | | |
| NC_011770 | 902040 | A | C | 30 | 8 | 2 | 0 | . | . | . | . | C | C | . | . | . | . | | |
| NC_011770 | 902890 | A | C | 13 | 8 | 4 | 0 | . | . | . | . | C | C | . | . | . | . | C | C |
| NC_011770 | 902899 | G | A | 13 | 8 | 4 | 0 | . | . | . | . | A | A | . | . | . | . | A | A |
| NC_011770 | 902900 | T | C | 13 | 8 | 4 | 0 | . | . | . | . | C | C | . | . | . | . | C | C |
| NC_011770 | 902902 | G | C | 13 | 8 | 4 | 0 | . | . | . | . | C | C | . | . | . | . | C | C |
| NC_011770 | 902906 | A | G | 13 | 8 | 4 | 0 | . | . | . | . | G | G | . | . | . | . | G | G |
| NC_011770 | 902907 | A | C | 13 | 8 | 4 | 0 | . | . | . | . | C | C | . | . | . | . | C | C |
| NC_011770 | 902914 | A | C | 16 | 8 | 4 | 0 | . | . | . | . | C | C | . | . | . | . | T | T |
| NC_011770 | 902953 | A | C | 16 | 8 | 4 | 0 | . | . | . | . | C | C | . | . | . | . | C | C |
| NC_011770 | 902968 | G | C | 16 | 8 | 2 | 0 | . | . | . | . | | | . | . | . | . | C | C |
| NC_011770 | 902977 | G | C | 16 | 8 | 2 | 0 | . | . | . | . | C | C | . | . | . | . | | |
| NC_011770 | 902983 | T | C | 14 | 8 | 2 | 0 | . | . | . | . | C | C | . | . | . | . | | |

| | | | | | | | | | | | | | | | | | | | |
|---|---|---|---|---|---|---|---|---|---|---|---|---|---|---|---|---|---|---|---|
| NC_011770 | 902986 | G | T | 15 | 8 | 2 | 0 | . | . | . | . | T | T | . | . | . | . | | |
| NC_011770 | 902987 | C | G | 14 | 8 | 2 | 0 | . | . | . | . | G | G | . | . | . | . | | |
| NC_011770 | 902989 | G | A | 15 | 8 | 2 | 0 | . | . | . | . | A | A | . | . | . | . | | |
| NC_011770 | 902995 | T | C | 15 | 8 | 2 | 0 | . | . | . | . | C | C | . | . | . | . | | |
| NC_011770 | 903004 | T | G | 15 | 8 | 4 | 0 | . | . | . | . | G | G | . | . | . | . | G | G |
| NC_011770 | 903010 | T | G | 16 | 8 | 4 | 0 | . | . | . | . | G | G | . | . | . | . | A | A |
| NC_011770 | 903013 | T | C | 16 | 8 | 2 | 0 | . | . | . | . | C | C | . | . | . | . | | |
| NC_011770 | 903019 | T | G | 16 | 8 | 2 | 0 | . | . | . | . | G | G | . | . | . | . | | |
| NC_011770 | 903022 | G | T | 16 | 8 | 2 | 0 | . | . | . | . | T | T | . | . | . | . | | |
| NC_011770 | 903028 | C | T | 15 | 8 | 2 | 0 | . | . | . | . | T | T | . | . | . | . | | |
| NC_011770 | 903038 | C | A | 16 | 8 | 2 | 0 | . | . | . | . | A | A | . | . | . | . | | |
| NC_011770 | 903039 | GT | G | 16 | 8 | 2 | 0 | . | . | . | . | G | G | . | . | . | . | | |
| NC_011770 | 903043 | C | CT | 16 | 8 | 2 | 0 | . | . | . | . | CT | CT | . | . | . | . | | |
| NC_011770 | 903049 | T | G | 15 | 8 | 2 | 0 | . | . | . | . | G | G | . | . | . | . | | |
| NC_011770 | 903052 | A | T | 15 | 8 | 2 | 0 | . | . | . | . | T | T | . | . | . | . | | |
| NC_011770 | 903073 | G | T | 17 | 8 | 2 | 0 | . | . | . | . | T | T | . | . | . | . | | |
| NC_011770 | 903085 | C | T | 16 | 8 | 2 | 0 | . | . | . | . | T | T | . | . | . | . | | |
| NC_011770 | 903088 | C | G | 16 | 8 | 4 | 0 | . | . | . | . | G | G | . | . | . | . | T | T |
| NC_011770 | 903112 | G | C | 6 | 10 | 2 | 0 | . | . | . | . | C | C | . | . | . | . | . | . |
| NC_011770 | 903115 | G | A | 6 | 10 | 2 | 0 | . | . | . | . | A | A | . | . | . | . | . | . |
| NC_011770 | 903118 | A | G | 6 | 10 | 2 | 0 | . | . | . | . | G | G | . | . | . | . | . | . |
| NC_011770 | 903139 | G | T | 5 | 10 | 2 | 0 | . | . | . | . | T | T | . | . | . | . | . | . |
| NC_011770 | 903142 | T | C | 5 | 10 | 2 | 0 | . | . | . | . | C | C | . | . | . | . | . | . |
| NC_011770 | 903145 | T | C | 5 | 10 | 2 | 0 | . | . | . | . | C | C | . | . | . | . | . | . |
| NC_011770 | 903148 | T | C | 5 | 10 | 2 | 0 | . | . | . | . | C | C | . | . | . | . | . | . |
| NC_011770 | 903154 | C | T | 5 | 10 | 2 | 0 | . | . | . | . | T | T | . | . | . | . | . | . |
| NC_011770 | 903166 | G | T | 5 | 10 | 2 | 0 | . | . | . | . | T | T | . | . | . | . | . | . |
| NC_011770 | 903172 | C | T | 5 | 10 | 2 | 0 | . | . | . | . | T | T | . | . | . | . | . | . |
| NC_011770 | 903175 | T | A | 5 | 10 | 2 | 0 | . | . | . | . | A | A | . | . | . | . | . | . |
| NC_011770 | 903988 | C | T | 28 | 8 | 2 | 0 | . | . | . | . | | | . | . | T | T | . | . |
| NC_011770 | 903990 | A | G | 28 | 8 | 2 | 0 | . | . | . | . | | | . | . | G | G | . | . |
| NC_011770 | 903991 | G | C | 28 | 8 | 2 | 0 | . | . | . | . | | | . | . | C | C | . | . |
| NC_011770 | 903996 | G | T | 28 | 8 | 2 | 0 | . | . | . | . | | | . | . | T | T | . | . |
| NC_011770 | 904378 | G | C | 0 | 11 | 1 | 0 | . | . | . | . | . | . | . | . | . | C | . | . |
| NC_011770 | 908698 | T | C | 80 | 0 | 10 | 0 | C | C | C | C | C | C | | | C | C | C | C |
| NC_011770 | 908714 | T | C | 79 | 0 | 10 | 0 | C | C | C | C | C | C | | | C | C | C | C |
| NC_011770 | 908719 | A | C | 79 | 0 | 10 | 0 | C | C | C | C | C | C | | | C | C | C | C |
| NC_011770 | 908771 | T | A | 81 | 0 | 10 | 0 | A | A | A | A | A | A | | | A | A | A | A |
| NC_011770 | 908785 | C | T | 80 | 0 | 2 | 0 | | | | | | | T | T | | | | |
| NC_011770 | 916041 | C | T | 73 | 0 | 2 | 0 | | | | | | | | | | | T | T |
| NC_011770 | 918283 | A | C | 76 | 0 | 2 | 0 | | | | | | | | | C | C | | |
| NC_011770 | 921322 | T | C | 88 | 0 | 2 | 0 | | | | | | | C | C | | | | |
| NC_011770 | 925855 | T | A | 90 | 0 | 2 | 0 | | | | | | | | | | | A | A |
| NC_011770 | 927692 | G | A | 77 | 0 | 12 | 0 | A | A | A | A | A | A | A | A | A | A | A | A |
| NC_011770 | 927698 | A | T | 73 | 0 | 6 | 0 | | | T | T | T | T | T | T | | | | |
| NC_011770 | 927699 | C | G | 74 | 0 | 6 | 0 | | | G | G | G | G | G | G | | | | |
| NC_011770 | 927700 | T | C | 74 | 0 | 6 | 0 | | | C | C | C | C | C | C | | | | |
| NC_011770 | 927713 | C | T | 73 | 0 | 6 | 0 | | | T | T | T | T | T | T | | | | |
| NC_011770 | 927715 | G | T | 73 | 0 | 6 | 0 | | | T | T | T | T | T | T | | | | |
| NC_011770 | 927723 | A | G | 73 | 0 | 6 | 0 | | | G | G | G | G | G | G | | | | |
| NC_011770 | 927729 | A | G | 73 | 0 | 6 | 0 | | | G | G | G | G | G | G | | | | |
| NC_011770 | 927735 | A | G | 71 | 0 | 6 | 0 | | | G | G | G | G | G | G | | | | |
| NC_011770 | 927786 | C | T | 38 | 3 | 3 | 0 | | | T | T | . | T | . | . | | | | |
| NC_011770 | 927787 | G | A | 38 | 3 | 3 | 0 | | | A | A | . | A | . | . | | | | |
| NC_011770 | 931265 | T | A | 77 | 0 | 2 | 0 | | | | | | | | | | | A | A |
| NC_011770 | 932771 | T | C | 74 | 0 | 2 | 0 | | | | | | | C | C | | | | |
| NC_011770 | 932788 | G | A | 74 | 0 | 2 | 0 | | | | | | | A | A | | | | |
| NC_011770 | 932867 | G | A | 72 | 0 | 2 | 0 | | | | | | | A | A | | | | |
| NC_011770 | 932971 | T | C | 75 | 0 | 12 | 0 | C | C | C | C | C | C | C | C | C | C | C | C |
| NC_011770 | 933000 | G | A | 74 | 0 | 4 | 0 | A | A | | | | | A | A | | | | |
| NC_011770 | 933004 | A | G | 73 | 0 | 4 | 0 | G | G | | | | | G | G | | | | |
| NC_011770 | 933011 | G | A | 72 | 0 | 4 | 0 | A | A | | | | | A | A | | | | |

| | | | | | | | | | | | | | | | | | | | |
|---|---|---|---|---|---|---|---|---|---|---|---|---|---|---|---|---|---|---|---|
| NC_011770 | 933776 | C | T | 69 | 0 | 2 | 0 | | | | | | | | | T | T | | |
| NC_011770 | 935041 | C | T | 91 | 0 | 2 | 0 | | | | | | | | | | | T | T |
| NC_011770 | 935086 | T | G | 92 | 0 | 12 | 0 | G | G | G | G | G | G | G | G | G | G | G | G |
| NC_011770 | 939484 | G | A | 71 | 0 | 12 | 0 | A | A | A | A | A | A | A | A | A | A | A | A |
| NC_011770 | 939533 | C | G | 53 | 0 | 2 | 0 | | | | | | | | | G | G | | |
| NC_011770 | 939971 | A | C | 17 | 0 | 12 | 0 | C | C | C | C | C | C | C | C | C | C | C | C |
| NC_011770 | 940027 | C | T | 28 | 0 | 2 | 0 | T | T | | | | | | | | | | |
| NC_011770 | 941275 | G | A | 65 | 0 | 2 | 0 | | | | | | | A | A | | | | |
| NC_011770 | 941277 | T | C | 66 | 0 | 1 | 0 | | | | | | | C | | | | | |
| NC_011770 | 942590 | C | T | 38 | 4 | 2 | 0 | | | . | . | . | . | T | T | | | | |
| NC_011770 | 942600 | G | T | 37 | 4 | 4 | 0 | | | . | . | . | . | | | T | T | T | T |
| NC_011770 | 942603 | G | A | 36 | 4 | 4 | 0 | | | . | . | . | . | | | A | A | A | A |
| NC_011770 | 942619 | A | G | 36 | 4 | 2 | 0 | | | . | . | . | . | G | G | | | | |
| NC_011770 | 942635 | T | C | 38 | 4 | 2 | 0 | | | . | . | . | . | C | C | | | | |
| NC_011770 | 942644 | A | G | 39 | 4 | 2 | 0 | | | . | . | . | . | G | G | | | | |
| NC_011770 | 942651 | C | T | 40 | 4 | 2 | 0 | | | . | . | . | . | | | | | T | T |
| NC_011770 | 942689 | C | T | 35 | 4 | 2 | 0 | | | . | . | . | . | T | T | | | | |
| NC_011770 | 942729 | C | T | 67 | 0 | 2 | 0 | | | | | | | T | T | | | | |
| NC_011770 | 942740 | C | T | 67 | 0 | 4 | 0 | | | T | T | T | T | | | | | | |
| NC_011770 | 942749 | C | T | 69 | 0 | 2 | 0 | | | | | | | T | T | | | | |
| NC_011770 | 942756 | G | A | 69 | 0 | 10 | 0 | | | A | A | A | A | A | A | A | A | A | A |
| NC_011770 | 942800 | T | C | 71 | 0 | 2 | 0 | | | | | | | C | C | | | | |
| NC_011770 | 942804 | C | CG | 72 | 0 | 2 | 0 | | | | | | | CG | CG | | | | |
| NC_011770 | 942809 | G | C | 70 | 0 | 10 | 0 | | | C | C | C | C | C | C | C | C | C | C |
| NC_011770 | 944103 | C | A | 80 | 0 | 2 | 0 | A | A | | | | | | | | | | |
| NC_011770 | 944315 | T | C | 78 | 0 | 12 | 0 | C | C | C | C | C | C | C | C | C | C | C | C |
| NC_011770 | 944894 | T | C | 82 | 0 | 1 | 0 | | | | | | | C | | | | | |
| NC_011770 | 944916 | T | C | 80 | 0 | 2 | 0 | C | C | | | | | | | | | | |
| NC_011770 | 946553 | T | G | 33 | 0 | 12 | 0 | G | G | G | G | G | G | G | G | G | G | G | G |
| NC_011770 | 946564 | C | T | 30 | 0 | 12 | 0 | T | T | T | T | T | T | T | T | T | T | T | T |
| NC_011770 | 946574 | T | C | 29 | 0 | 12 | 0 | C | C | C | C | C | C | C | C | C | C | C | C |
| NC_011770 | 946586 | C | T | 0 | 11 | 1 | 0 | . | . | . | . | . | . | . | . | . | . | . | T |
| NC_011770 | 946651 | A | G | 25 | 0 | 12 | 0 | G | G | G | G | G | G | G | G | G | G | G | G |
| NC_011770 | 946715 | A | C | 43 | 0 | 4 | 0 | | | | | | | | | C | C | C | C |
| NC_011770 | 948056 | G | A | 96 | 0 | 2 | 0 | | | | | | | A | A | | | | |
| NC_011770 | 950366 | A | G | 82 | 0 | 4 | 0 | | | | | | | G | G | G | G | | |
| NC_011770 | 953869 | A | G | 80 | 0 | 1 | 0 | | | | | | | G | | | | | |
| NC_011770 | 957066 | G | A | 39 | 0 | 2 | 0 | | | | | | | | | | | A | A |
| NC_011770 | 957632 | G | A | 50 | 0 | 6 | 0 | A | A | | | | | A | A | A | A | | |
| NC_011770 | 957874 | T | C | 55 | 0 | 4 | 0 | | | C | C | C | C | | | | | | |
| NC_011770 | 962995 | C | CG | 82 | 0 | 6 | 0 | | | CG | CG | CG | CG | | | CG | CG | | |
| NC_011770 | 965160 | G | T | 66 | 0 | 2 | 0 | | | | | | | | | T | T | | |
| NC_011770 | 965198 | CG | C | 65 | 0 | 6 | 0 | | | C | C | C | C | C | C | | | | |
| NC_011770 | 965203 | A | G | 64 | 0 | 2 | 0 | | | | | | | | | G | G | | |
| NC_011770 | 968313 | T | C | 76 | 0 | 10 | 0 | | | C | C | C | C | C | C | C | C | C | C |
| NC_011770 | 968343 | A | G | 70 | 0 | 8 | 0 | | | G | G | G | G | G | G | | | G | G |
| NC_011770 | 968350 | C | T | 72 | 0 | 2 | 0 | | | | | | | | | T | T | | |
| NC_011770 | 968379 | T | TG | 65 | 0 | 10 | 0 | | | TG | TG | TG | TG | TG | G | G | G | TG | TG |
| NC_011770 | 968415 | T | C | 69 | 0 | 8 | 0 | | | C | C | C | C | C | C | | | C | C |
| NC_011770 | 968439 | A | G | 67 | 0 | 10 | 0 | | | G | G | G | G | G | G | G | G | G | G |
| NC_011770 | 968441 | C | T | 67 | 0 | 10 | 0 | | | T | T | T | T | T | T | T | T | T | T |
| NC_011770 | 968520 | C | T | 72 | 0 | 12 | 0 | T | T | T | T | T | T | T | T | T | T | T | T |
| NC_011770 | 968593 | C | T | 72 | 0 | 8 | 0 | | | T | T | T | T | T | T | | | T | T |
| NC_011770 | 968648 | C | T | 82 | 0 | 2 | 0 | T | T | | | | | | | | | | |
| NC_011770 | 970140 | T | C | 81 | 0 | 12 | 0 | C | C | C | C | C | C | C | C | C | C | C | C |
| NC_011770 | 970300 | G | GGGCG/ | 81 | 0 | 4 | 0 | | | | | GGGCGAA | | GGGCGAA | GGGCGAA | GGGCGAA | | | |
| NC_011770 | 970317 | AC | A | 75 | 0 | 2 | 0 | A | A | | | | | | | | | | |
| NC_011770 | 976788 | A | T | 22 | 4 | 2 | 0 | | | | | | | . | . | . | . | T | T |
| NC_011770 | 976801 | C | T | 21 | 4 | 2 | 0 | | | | | | | . | . | . | . | T | T |
| NC_011770 | 976803 | G | C | 24 | 1 | 2 | 0 | | | | | | | | | | . | C | C |
| NC_011770 | 976809 | C | A | 26 | 0 | 4 | 0 | | | | | | | A | A | A | A | | |
| NC_011770 | 976819 | T | C | 35 | 0 | 2 | 0 | | | | | | | | | | | C | C |

| | | | | | | | | | | | | | | | | | | | |
|---|---|---|---|---|---|---|---|---|---|---|---|---|---|---|---|---|---|---|---|
| NC_011770 | 976832 | C | T | 37 | 0 | 2 | 0 | | | | | | | | | | | T | T |
| NC_011770 | 978026 | C | T | 52 | 0 | 10 | 0 | T | T | T | T | T | T | T | T | | | T | T |
| NC_011770 | 978197 | G | A | 56 | 0 | 2 | 0 | | | | | | | | | A | A | | |
| NC_011770 | 978204 | A | G | 60 | 0 | 2 | 0 | | | | | | | G | G | | | | |
| NC_011770 | 978221 | G | A | 60 | 0 | 2 | 0 | A | A | | | | | | | | | | |
| NC_011770 | 978224 | A | G | 58 | 0 | 4 | 0 | | | G | G | G | G | | | | | | |
| NC_011770 | 978254 | C | G | 60 | 0 | 2 | 0 | | | | | | | | | | | G | G |
| NC_011770 | 978277 | C | A | 61 | 0 | 2 | 0 | A | A | | | | | | | | | | |
| NC_011770 | 978292 | A | C | 60 | 0 | 2 | 0 | | | | | | | | | | | C | C |
| NC_011770 | 978294 | G | T | 61 | 0 | 4 | 0 | | | | | | | T | T | | | T | T |
| NC_011770 | 978307 | G | C | 61 | 0 | 8 | 0 | C | C | | | | | C | C | C | C | C | C |
| NC_011770 | 978312 | G | C | 60 | 0 | 8 | 0 | C | C | | | | | C | C | C | C | C | C |
| NC_011770 | 978313 | G | A | 60 | 0 | 2 | 0 | A | A | | | | | | | | | | |
| NC_011770 | 978369 | A | C | 50 | 2 | 10 | 0 | . | . | C | C | C | C | C | C | C | C | C | C |
| NC_011770 | 978370 | A | G | 51 | 2 | 10 | 0 | . | . | G | G | G | G | G | G | G | G | G | G |
| NC_011770 | 978381 | G | A | 57 | 0 | 8 | 0 | | | A | A | A | A | A | A | | | A | A |
| NC_011770 | 978387 | G | A | 57 | 0 | 12 | 0 | A | A | A | A | A | A | A | A | A | A | A | A |
| NC_011770 | 978392 | G | T | 56 | 0 | 2 | 0 | T | T | | | | | | | | | | |
| NC_011770 | 978393 | G | A | 56 | 0 | 12 | 0 | A | A | A | A | A | A | A | A | A | A | A | A |
| NC_011770 | 978435 | A | G | 56 | 0 | 2 | 0 | G | G | | | | | | | | | | |
| NC_011770 | 978538 | G | A | 60 | 0 | 4 | 0 | | | | | | | A | A | | | A | A |
| NC_011770 | 980057 | CG | C | 87 | 0 | 6 | 0 | C | C | | | | | C | C | | | C | C |
| NC_011770 | 983080 | G | T | 29 | 0 | 2 | 0 | | | | | | | | | T | T | | |
| NC_011770 | 983242 | C | A | 44 | 0 | 4 | 0 | A | A | | | | | A | A | | | | |
| NC_011770 | 983850 | T | C | 52 | 0 | 4 | 0 | C | C | | | | | | | C | C | | |
| NC_011770 | 983856 | T | A | 50 | 0 | 12 | 0 | A | A | G | G | G | G | A | A | A | A | A | A |
| NC_011770 | 983866 | T | G | 47 | 0 | 8 | 0 | G | G | | | | | G | G | G | G | G | G |
| NC_011770 | 983879 | A | C | 45 | 0 | 8 | 0 | C | C | | | | | C | C | C | C | C | C |
| NC_011770 | 983883 | G | A | 45 | 0 | 4 | 0 | A | A | | | | | | | | | A | A |
| NC_011770 | 983885 | G | T | 44 | 0 | 8 | 0 | T | T | | | | | T | T | T | T | T | T |
| NC_011770 | 983889 | G | C | 45 | 0 | 8 | 0 | C | C | | | | | C | C | C | C | C | C |
| NC_011770 | 983892 | T | C | 45 | 0 | 8 | 0 | C | C | | | | | C | C | C | C | C | C |
| NC_011770 | 983916 | G | C | 5 | 8 | 4 | 0 | . | . | . | . | . | C | . | . | C | C | C | . |
| NC_011770 | 983917 | C | T | 5 | 8 | 1 | 0 | . | . | T | . | . | . | . | . | | | . | . |
| NC_011770 | 983922 | A | G | 37 | 0 | 12 | 0 | G | G | G | G | G | G | G | G | G | G | G | G |
| NC_011770 | 983935 | G | A | 42 | 0 | 12 | 0 | A | A | A | A | A | A | A | A | A | A | A | A |
| NC_011770 | 983937 | T | G | 42 | 0 | 12 | 0 | G | G | G | G | G | G | G | G | G | G | G | G |
| NC_011770 | 983962 | C | T | 41 | 0 | 12 | 0 | T | T | T | T | T | T | T | T | T | T | T | T |
| NC_011770 | 983974 | T | C | 39 | 0 | 12 | 0 | C | C | C | C | C | C | C | C | C | C | C | C |
| NC_011770 | 984945 | G | A | 83 | 0 | 4 | 0 | | | A | A | A | A | | | | | | |
| NC_011770 | 984952 | T | A | 83 | 0 | 4 | 0 | | | A | A | A | A | | | | | | |
| NC_011770 | 985014 | T | C | 87 | 0 | 4 | 0 | | | C | C | C | C | | | | | | |
| NC_011770 | 985027 | T | C | 93 | 0 | 6 | 0 | | | C | C | C | C | | | C | C | | |
| NC_011770 | 985091 | G | A | 89 | 0 | 2 | 0 | A | A | | | | | | | | | | |
| NC_011770 | 985131 | C | T | 85 | 0 | 6 | 0 | | | T | T | T | T | | | T | T | | |
| NC_011770 | 986660 | G | GA | 74 | 0 | 2 | 0 | | | | | | | | | GA | GA | | |
| NC_011770 | 986669 | G | T | 72 | 0 | 2 | 0 | | | | | | | | | T | T | | |
| NC_011770 | 986720 | G | A | 68 | 0 | 2 | 0 | | | | | | | A | A | | | | |
| NC_011770 | 986727 | C | T | 67 | 0 | 2 | 0 | | | | | | | | | T | T | | |
| NC_011770 | 988925 | A | G | 80 | 0 | 2 | 0 | G | G | | | | | | | | | | |
| NC_011770 | 988948 | T | A | 76 | 0 | 2 | 0 | | | | | | | | | A | A | | |
| NC_011770 | 990016 | A | T | 78 | 0 | 2 | 0 | | | | | | | T | T | | | | |
| NC_011770 | 992420 | A | G | 80 | 0 | 2 | 0 | | | | | | | G | G | | | | |
| NC_011770 | 999153 | G | T | 85 | 0 | 2 | 0 | | | | | | | | | | | T | T |
| NC_011770 | 999208 | A | G | 85 | 0 | 12 | 0 | G | G | G | G | G | G | G | G | G | G | G | G |
| NC_011770 | 1000914 | C | T | 65 | 0 | 2 | 0 | | | | | | | | | | | T | T |
| NC_011770 | 1001133 | A | G | 43 | 0 | 2 | 0 | | | | | | | | | | | G | G |
| NC_011770 | 1001242 | T | C | 34 | 0 | 12 | 0 | C | C | C | C | C | C | C | C | C | C | C | C |
| NC_011770 | 1001253 | C | T | 36 | 0 | 6 | 0 | T | T | T | T | T | T | | | | | | |
| NC_011770 | 1005641 | A | C | 76 | 0 | 12 | 0 | C | C | C | C | C | C | C | C | C | C | C | C |
| NC_011770 | 1007055 | C | T | 75 | 0 | 2 | 0 | | | | | | | T | T | | | | |
| NC_011770 | 1009004 | T | C | 58 | 0 | 10 | 0 | | | C | C | C | C | C | C | C | C | C | C |

| | | | | | | | | | | | | | | | | | | | |
|---|---|---|---|---|---|---|---|---|---|---|---|---|---|---|---|---|---|---|---|
| NC_011770 | 1009039 | C | T | 55 | 0 | 2 | 0 | | | | | | | T | T | | | | |
| NC_011770 | 1009046 | G | A | 56 | 0 | 2 | 0 | | | | | | | | | | | A | A |
| NC_011770 | 1009048 | T | C | 57 | 0 | 2 | 0 | | | | | | | | | | | C | C |
| NC_011770 | 1009056 | CG | C | 56 | 0 | 10 | 0 | | | C | C | C | C | C | C | C | C | C | C |
| NC_011770 | 1009076 | C | T | 54 | 0 | 2 | 0 | T | T | | | | | | | | | | |
| NC_011770 | 1009116 | C | T | 52 | 2 | 10 | 0 | T | T | T | T | T | T | T | T | T | T | . | . |
| NC_011770 | 1009153 | T | C | 63 | 0 | 2 | 0 | | | | | | | | | | | C | C |
| NC_011770 | 1009162 | T | C | 64 | 0 | 2 | 0 | | | | | | | | | | | C | C |
| NC_011770 | 1009177 | T | TG | 56 | 0 | 2 | 0 | TG | TG | | | | | | | | | | |
| NC_011770 | 1012428 | G | A | 58 | 0 | 6 | 0 | A | A | A | A | A | A | | | | | | |
| NC_011770 | 1012465 | C | A | 62 | 0 | 2 | 0 | | | | | | | | | | | A | A |
| NC_011770 | 1012482 | T | G | 65 | 0 | 12 | 0 | G | G | G | G | G | G | G | G | G | G | G | G |
| NC_011770 | 1016726 | A | G | 69 | 0 | 2 | 0 | | | | | | | | | G | G | | |
| NC_011770 | 1018420 | T | A | 75 | 0 | 2 | 0 | | | | | | | | | A | A | | |
| NC_011770 | 1018485 | A | G | 71 | 0 | 12 | 0 | G | G | G | G | G | G | G | G | G | G | G | G |
| NC_011770 | 1018486 | C | A | 71 | 0 | 2 | 0 | | | | | | | A | A | | | | |
| NC_011770 | 1018494 | A | G | 71 | 0 | 6 | 0 | G | G | | | | | G | G | | | G | G |
| NC_011770 | 1018532 | A | G | 70 | 0 | 4 | 0 | | | G | G | G | G | | | | | | |
| NC_011770 | 1018607 | T | C | 72 | 0 | 2 | 0 | | | | | | | C | C | | | | |
| NC_011770 | 1018624 | C | T | 57 | 2 | 2 | 0 | | | | | | | . | . | T | T | | |
| NC_011770 | 1018637 | G | A | 58 | 1 | 1 | 0 | | | | | | | . | A | | | | |
| NC_011770 | 1018642 | T | G | 68 | 0 | 2 | 0 | | | | | | | G | G | | | | |
| NC_011770 | 1018661 | C | T | 66 | 0 | 4 | 0 | | | T | T | T | T | | | | | | |
| NC_011770 | 1018678 | C | T | 68 | 0 | 2 | 0 | | | | | | | T | T | | | | |
| NC_011770 | 1018743 | A | G | 69 | 0 | 2 | 0 | | | | | | | G | G | | | | |
| NC_011770 | 1021110 | C | T | 91 | 0 | 6 | 0 | | | T | T | T | T | | | T | T | | |
| NC_011770 | 1023142 | T | A | 64 | 2 | 2 | 0 | A | A | | | | | . | . | | | | |
| NC_011770 | 1023625 | G | A | 76 | 0 | 12 | 0 | A | A | A | A | A | A | A | A | A | A | A | A |
| NC_011770 | 1024502 | A | C | 78 | 0 | 2 | 0 | | | | | | | C | C | | | | |
| NC_011770 | 1024507 | GGT | G | 81 | 0 | 2 | 0 | | | | | | | G | G | | | | |
| NC_011770 | 1026948 | T | C | 49 | 0 | 6 | 0 | C | C | | | | | | | C | C | C | C |
| NC_011770 | 1026950 | G | C | 50 | 0 | 6 | 0 | C | C | | | | | | | C | C | C | C |
| NC_011770 | 1027085 | C | A | 57 | 0 | 2 | 0 | | | | | | | A | A | | | | |
| NC_011770 | 1027172 | G | C | 60 | 0 | 10 | 0 | C | C | C | C | C | C | | | C | C | C | C |
| NC_011770 | 1027244 | A | G | 64 | 0 | 6 | 0 | | | G | G | G | G | G | G | | | | |
| NC_011770 | 1027397 | G | A | 63 | 0 | 2 | 0 | A | A | | | | | | | | | | |
| NC_011770 | 1027398 | T | G | 64 | 0 | 6 | 0 | | | G | G | G | G | G | G | | | | |
| NC_011770 | 1027416 | C | T | 62 | 0 | 6 | 0 | | | T | T | T | T | T | T | | | | |
| NC_011770 | 1027451 | G | A | 63 | 0 | 6 | 0 | | | A | A | A | A | A | A | | | | |
| NC_011770 | 1027460 | G | C | 64 | 0 | 6 | 0 | | | C | C | C | C | C | C | | | | |
| NC_011770 | 1027467 | G | A | 64 | 0 | 2 | 0 | | | | | | | A | A | | | | |
| NC_011770 | 1027559 | G | A | 59 | 0 | 2 | 0 | | | | | | | A | A | | | | |
| NC_011770 | 1027640 | A | G | 62 | 0 | 2 | 0 | | | | | | | G | G | | | | |
| NC_011770 | 1027928 | G | A | 68 | 0 | 2 | 0 | | | | | | | A | A | | | | |
| NC_011770 | 1027967 | C | T | 69 | 0 | 2 | 0 | | | | | | | T | T | | | | |
| NC_011770 | 1028150 | C | T | 65 | 0 | 2 | 0 | | | | | | | T | T | | | | |
| NC_011770 | 1028183 | T | C | 68 | 0 | 2 | 0 | | | | | | | C | C | | | | |
| NC_011770 | 1028204 | C | T | 70 | 0 | 2 | 0 | | | | | | | T | T | | | | |
| NC_011770 | 1028240 | C | G | 69 | 0 | 2 | 0 | | | | | | | G | G | | | | |
| NC_011770 | 1028279 | T | C | 68 | 0 | 2 | 0 | | | | | | | C | C | | | | |
| NC_011770 | 1028317 | C | G | 68 | 0 | 2 | 0 | | | | | | | | | G | G | | |
| NC_011770 | 1029289 | A | G | 85 | 0 | 4 | 0 | | | | | | | | | G | G | G | G |
| NC_011770 | 1029885 | T | C | 78 | 0 | 2 | 0 | | | | | | | | | C | C | | |
| NC_011770 | 1029899 | T | A | 72 | 0 | 2 | 0 | | | | | | | | | A | A | | |
| NC_011770 | 1032184 | AT | A | 60 | 0 | 2 | 0 | | | | | | | | | | | A | A |
| NC_011770 | 1033769 | T | C | 63 | 0 | 10 | 0 | C | C | C | C | C | C | | | C | C | C | C |
| NC_011770 | 1033847 | G | A | 62 | 0 | 12 | 0 | A | A | A | A | A | A | A | A | A | A | A | A |
| NC_011770 | 1035453 | A | T | 87 | 0 | 2 | 0 | | | | | | | T | T | | | | |
| NC_011770 | 1035586 | T | C | 89 | 0 | 10 | 0 | C | C | C | C | C | C | C | C | | | C | C |
| NC_011770 | 1035670 | C | G | 86 | 0 | 10 | 0 | G | G | G | G | G | G | G | G | | | G | G |
| NC_011770 | 1037676 | T | G | 66 | 1 | 11 | 0 | G | G | G | . | G | G | G | G | G | G | G | G |
| NC_011770 | 1037741 | C | T | 65 | 1 | 3 | 0 | | | T | . | T | T | | | | | | |

| | | | | | | | | | | | | | | | | | | | |
|---|---|---|---|---|---|---|---|---|---|---|---|---|---|---|---|---|---|---|---|
| NC_011770 | 1040097 | G | A | 75 | 0 | 6 | 0 | A | A | | | | | A | A | A | A | | |
| NC_011770 | 1042658 | C | CT | 33 | 0 | 2 | 0 | | | | | | | CT | CT | | | | |
| NC_011770 | 1042666 | AG | A | 32 | 0 | 4 | 0 | | | A | A | A | A | | | | | | |
| NC_011770 | 1049322 | T | C | 78 | 0 | 4 | 0 | | | C | C | C | C | | | | | | |
| NC_011770 | 1049335 | C | T | 78 | 0 | 4 | 0 | | | T | T | T | T | | | | | | |
| NC_011770 | 1049418 | C | G | 84 | 0 | 4 | 0 | | | G | G | G | G | | | | | | |
| NC_011770 | 1049436 | T | C | 80 | 0 | 4 | 0 | | | C | C | C | C | | | | | | |
| NC_011770 | 1049457 | G | C | 79 | 0 | 4 | 0 | | | C | C | C | C | | | | | | |
| NC_011770 | 1049458 | A | G | 76 | 0 | 4 | 0 | | | G | G | G | G | | | | | | |
| NC_011770 | 1049459 | T | C | 77 | 0 | 4 | 0 | | | C | C | C | C | | | | | | |
| NC_011770 | 1052321 | C | T | 69 | 0 | 2 | 0 | | | | | | | T | T | | | | |
| NC_011770 | 1060675 | G | T | 59 | 0 | 4 | 0 | | | | | | | | | T | T | T | T |
| NC_011770 | 1060691 | C | CT | 56 | 0 | 4 | 0 | | | | | | | | | CT | CT | CT | CT |
| NC_011770 | 1060731 | G | A | 54 | 0 | 4 | 0 | | | | | | | | | A | A | A | A |
| NC_011770 | 1060769 | T | C | 56 | 0 | 4 | 0 | | | | | | | | | C | C | C | C |
| NC_011770 | 1060804 | G | A | 56 | 0 | 4 | 0 | | | A | A | A | A | | | | | | |
| NC_011770 | 1060806 | A | G | 56 | 0 | 4 | 0 | | | G | G | G | G | | | | | | |
| NC_011770 | 1060815 | G | T | 54 | 0 | 12 | 0 | T | T | T | T | T | T | T | T | T | T | T | T |
| NC_011770 | 1061768 | G | C | 78 | 0 | 6 | 0 | C | C | | | | | | | C | C | C | C |
| NC_011770 | 1061795 | TCG | T | 73 | 0 | 12 | 0 | T | T | T | T | T | T | T | T | T | T | T | T |
| NC_011770 | 1062718 | A | G | 60 | 0 | 6 | 0 | G | G | | | | | | | G | G | G | G |
| NC_011770 | 1062740 | A | G | 63 | 0 | 2 | 0 | | | | | | | | | G | G | | |
| NC_011770 | 1062776 | G | A | 62 | 0 | 2 | 0 | | | | | | | A | A | | | | |
| NC_011770 | 1062813 | T | C | 61 | 0 | 4 | 0 | C | C | | | | | C | C | | | | |
| NC_011770 | 1064216 | G | A | 52 | 0 | 4 | 0 | | | A | A | A | A | | | | | | |
| NC_011770 | 1064261 | T | C | 50 | 0 | 12 | 0 | C | C | C | C | C | C | C | C | C | C | C | C |
| NC_011770 | 1064295 | C | T | 45 | 0 | 2 | 0 | | | | | | | T | T | | | | |
| NC_011770 | 1064566 | G | A | 22 | 0 | 2 | 0 | | | | | | | | | A | A | | |
| NC_011770 | 1064637 | T | C | 8 | 1 | 11 | 0 | . | C | C | C | C | C | C | C | C | C | C | C |
| NC_011770 | 1064931 | AGC | A | 33 | 0 | 2 | 0 | | | | | | | A | A | | | | |
| NC_011770 | 1066025 | A | G | 48 | 0 | 1 | 0 | | | | | | | G | | | | | |
| NC_011770 | 1066747 | T | C | 84 | 0 | 2 | 0 | | | | | | | | | | | C | C |
| NC_011770 | 1069497 | A | G | 81 | 0 | 12 | 0 | G | G | G | G | G | G | G | G | G | G | G | G |
| NC_011770 | 1069499 | G | C | 79 | 0 | 12 | 0 | C | C | C | C | C | C | C | C | C | C | C | C |
| NC_011770 | 1069979 | C | T | 75 | 0 | 10 | 0 | T | T | T | T | T | T | | | T | T | T | T |
| NC_011770 | 1069984 | G | C | 77 | 0 | 2 | 0 | | | | | | | C | C | | | | |
| NC_011770 | 1072566 | AG | A | 78 | 0 | 1 | 0 | | | | | | | A | | | | | |
| NC_011770 | 1075010 | T | C | 69 | 0 | 12 | 0 | C | C | C | C | C | C | C | C | C | C | C | C |
| NC_011770 | 1076435 | G | C | 80 | 0 | 12 | 0 | C | C | C | C | C | C | C | C | C | C | C | C |
| NC_011770 | 1077583 | C | A | 52 | 0 | 2 | 0 | | | | | | | | | | | A | A |
| NC_011770 | 1077613 | T | A | 46 | 0 | 10 | 0 | A | A | A | A | A | A | A | A | A | A | | |
| NC_011770 | 1077628 | G | A | 53 | 0 | 2 | 0 | | | | | | | | | | | A | A |
| NC_011770 | 1077649 | G | A | 54 | 0 | 10 | 0 | A | A | A | A | A | A | A | A | A | A | | |
| NC_011770 | 1078948 | C | CTCT | 85 | 0 | 1 | 0 | | | | | | | | | CTCT | | | |
| NC_011770 | 1080438 | C | T | 79 | 0 | 2 | 0 | | | | | | | | | | | T | T |
| NC_011770 | 1080470 | C | G | 76 | 0 | 2 | 0 | | | | | | | | | G | G | | |
| NC_011770 | 1083011 | G | T | 80 | 0 | 6 | 0 | | | T | T | T | T | T | T | | | | |
| NC_011770 | 1087414 | G | C | 54 | 0 | 4 | 0 | | | C | C | C | C | | | | | | |
| NC_011770 | 1087424 | T | C | 54 | 0 | 4 | 0 | | | C | C | C | C | | | | | | |
| NC_011770 | 1087434 | T | G | 51 | 0 | 12 | 0 | G | G | G | G | G | G | G | G | G | G | G | G |
| NC_011770 | 1087435 | C | G | 52 | 0 | 4 | 0 | | | G | G | G | G | | | | | | |
| NC_011770 | 1087453 | C | T | 56 | 0 | 6 | 0 | T | T | | | | | T | T | | | T | T |
| NC_011770 | 1087477 | T | C | 57 | 0 | 4 | 0 | | | C | C | C | C | | | | | | |
| NC_011770 | 1087509 | C | T | 59 | 0 | 4 | 0 | | | T | T | T | T | | | | | | |
| NC_011770 | 1087566 | T | C | 64 | 0 | 4 | 0 | | | C | C | C | C | | | | | | |
| NC_011770 | 1087601 | A | G | 66 | 0 | 12 | 0 | G | G | G | G | G | G | G | G | G | G | G | G |
| NC_011770 | 1087606 | T | C | 66 | 0 | 10 | 0 | C | C | C | C | C | C | C | C | | | C | C |
| NC_011770 | 1087615 | G | A | 68 | 0 | 6 | 0 | A | A | | | | | A | A | | | A | A |
| NC_011770 | 1087693 | G | T | 69 | 0 | 4 | 0 | | | T | T | T | T | | | | | | |
| NC_011770 | 1087705 | G | A | 71 | 0 | 4 | 0 | | | A | A | A | A | | | | | | |
| NC_011770 | 1087729 | T | C | 68 | 0 | 4 | 0 | | | C | C | C | C | | | | | | |
| NC_011770 | 1087753 | T | C | 64 | 0 | 4 | 0 | | | C | C | C | C | | | | | | |

| | | | | | | | | | | | | | | | | | | | |
|---|---|---|---|---|---|---|---|---|---|---|---|---|---|---|---|---|---|---|---|
| NC_011770 | 1087757 | T | C | 69 | 0 | 4 | 0 | | | C | C | C | C | | | | | | |
| NC_011770 | 1087933 | C | T | 69 | 0 | 6 | 0 | | | T | T | T | T | | | | | T | T |
| NC_011770 | 1087980 | C | G | 70 | 0 | 2 | 0 | | | | | | | | | | | G | G |
| NC_011770 | 1088056 | C | T | 72 | 0 | 12 | 0 | T | T | T | T | T | T | T | T | T | T | T | T |
| NC_011770 | 1089274 | C | T | 44 | 0 | 2 | 0 | | | | | | | T | T | | | | |
| NC_011770 | 1090368 | A | G | 74 | 0 | 10 | 0 | | | G | G | G | G | G | G | G | G | G | G |
| NC_011770 | 1090387 | G | A | 69 | 0 | 2 | 0 | A | A | | | | | | | | | | |
| NC_011770 | 1090752 | C | T | 74 | 0 | 8 | 0 | | | T | T | T | T | | | T | T | T | T |
| NC_011770 | 1091318 | C | T | 78 | 0 | 8 | 0 | | | T | T | T | T | | | T | T | T | T |
| NC_011770 | 1091477 | A | C | 74 | 0 | 2 | 0 | | | | | | | | | | | C | C |
| NC_011770 | 1093423 | G | A | 51 | 0 | 2 | 0 | | | | | | | A | A | | | | |
| NC_011770 | 1093843 | T | G | 65 | 0 | 12 | 0 | G | G | G | G | G | G | G | G | G | G | G | G |
| NC_011770 | 1096940 | G | A | 85 | 0 | 2 | 0 | | | | | | | | | A | A | | |
| NC_011770 | 1098910 | T | C | 77 | 0 | 6 | 0 | C | C | | | | | C | C | | | C | C |
| NC_011770 | 1099807 | T | C | 65 | 0 | 2 | 0 | C | C | | | | | | | | | | |
| NC_011770 | 1100253 | C | T | 71 | 0 | 2 | 0 | | | | | | | | | T | T | | |
| NC_011770 | 1102254 | C | A | 69 | 0 | 2 | 0 | | | | | | | | | | | A | A |
| NC_011770 | 1103121 | C | T | 96 | 0 | 4 | 0 | | | T | T | T | T | | | | | | |
| NC_011770 | 1103718 | C | T | 89 | 0 | 2 | 0 | | | | | | | | | | | T | T |
| NC_011770 | 1103719 | G | A | 92 | 0 | 2 | 0 | | | | | | | | | | | A | A |
| NC_011770 | 1106977 | A | G | 57 | 0 | 2 | 0 | | | G | G | | | | | | | | |
| NC_011770 | 1107044 | G | A | 65 | 0 | 2 | 0 | | | | | | | A | A | | | | |
| NC_011770 | 1107087 | C | G | 70 | 0 | 2 | 0 | G | G | | | | | | | | | | |
| NC_011770 | 1107088 | A | G | 70 | 0 | 2 | 0 | G | G | | | | | | | | | | |
| NC_011770 | 1107096 | G | A | 74 | 0 | 2 | 0 | A | A | | | | | | | | | | |
| NC_011770 | 1107110 | G | A | 72 | 0 | 2 | 0 | A | A | | | | | | | | | | |
| NC_011770 | 1107113 | A | G | 71 | 0 | 2 | 0 | G | G | | | | | | | | | | |
| NC_011770 | 1107121 | T | C | 73 | 0 | 2 | 0 | | | | | | | | | C | C | | |
| NC_011770 | 1107150 | C | T | 77 | 0 | 8 | 0 | T | T | T | T | T | T | | | T | T | | |
| NC_011770 | 1107171 | C | T | 73 | 0 | 2 | 0 | T | T | | | | | | | | | | |
| NC_011770 | 1107193 | G | A | 76 | 0 | 2 | 0 | A | A | | | | | | | | | | |
| NC_011770 | 1107213 | G | A | 74 | 0 | 2 | 0 | A | A | | | | | | | | | | |
| NC_011770 | 1107220 | CG | C | 73 | 0 | 2 | 0 | C | C | | | | | | | | | | |
| NC_011770 | 1107221 | G | C | 73 | 0 | 2 | 0 | . | . | | | | | C | C | | | | |
| NC_011770 | 1107230 | T | G | 72 | 0 | 2 | 0 | G | G | | | | | | | | | | |
| NC_011770 | 1107243 | C | G | 72 | 0 | 2 | 0 | G | G | | | | | | | | | | |
| NC_011770 | 1107911 | A | G | 62 | 0 | 4 | 0 | G | G | | | | | | | G | G | | |
| NC_011770 | 1107917 | T | G | 61 | 0 | 4 | 0 | G | G | | | | | | | G | G | | |
| NC_011770 | 1107939 | C | T | 59 | 0 | 4 | 0 | T | T | | | | | | | T | T | | |
| NC_011770 | 1109411 | C | T | 81 | 0 | 6 | 0 | | | T | T | T | T | | | T | T | | |
| NC_011770 | 1109413 | T | A | 81 | 0 | 6 | 0 | | | A | A | A | A | | | A | A | | |
| NC_011770 | 1113086 | C | T | 72 | 0 | 8 | 0 | T | T | T | T | T | T | | | | | T | T |
| NC_011770 | 1113155 | A | G | 74 | 0 | 12 | 0 | G | G | G | G | G | G | G | G | G | G | G | G |
| NC_011770 | 1113183 | T | G | 77 | 0 | 8 | 0 | G | G | | | | | G | G | G | G | G | G |
| NC_011770 | 1113197 | A | G | 76 | 0 | 8 | 0 | G | G | | | | | G | G | G | G | G | G |
| NC_011770 | 1113261 | G | A | 81 | 0 | 4 | 0 | | | A | A | A | A | | | | | | |
| NC_011770 | 1113271 | A | G | 79 | 0 | 4 | 0 | | | G | G | G | G | | | | | | |
| NC_011770 | 1117656 | A | G | 52 | 0 | 10 | 0 | G | G | G | G | G | G | | | G | G | G | G |
| NC_011770 | 1118150 | G | A | 72 | 0 | 2 | 0 | A | A | | | | | | | | | | |
| NC_011770 | 1118201 | AG | A | 72 | 0 | 2 | 0 | | | | | | | A | A | | | | |
| NC_011770 | 1118256 | G | A | 66 | 0 | 2 | 0 | A | A | | | | | | | | | | |
| NC_011770 | 1118261 | A | C | 64 | 0 | 2 | 0 | | | | | | | | | | | C | C |
| NC_011770 | 1118269 | G | A | 65 | 0 | 2 | 0 | A | A | | | | | | | | | | |
| NC_011770 | 1119286 | CG | C | 62 | 0 | 8 | 0 | | | C | C | C | C | C | C | | | CGG | CGG |
| NC_011770 | 1119319 | G | A | 56 | 0 | 2 | 0 | A | A | | | | | | | | | | |
| NC_011770 | 1119321 | G | T | 55 | 0 | 4 | 0 | | | | | | | T | T | | | T | T |
| NC_011770 | 1119326 | A | G | 54 | 0 | 2 | 0 | | | | | | | | | | | G | G |
| NC_011770 | 1119341 | C | A | 57 | 0 | 2 | 0 | | | | | A | A | | | | | | |
| NC_011770 | 1119367 | C | A | 53 | 0 | 8 | 0 | | | A | A | A | A | A | A | A | A | | |
| NC_011770 | 1119368 | G | T | 53 | 0 | 6 | 0 | | | T | T | T | T | | | | | T | T |
| NC_011770 | 1119411 | C | T | 60 | 0 | 2 | 0 | | | | | | | | | T | T | | |
| NC_011770 | 1120509 | C | A | 65 | 0 | 4 | 0 | | | | | | | A | A | A | A | | |

| | | | | | | | | | | | | | | | | | | | |
|---|---|---|---|---|---|---|---|---|---|---|---|---|---|---|---|---|---|---|---|
| NC_011770 | 1122203 | T | C | 65 | 0 | 4 | 0 | | | | | | | | | C | C | C | C |
| NC_011770 | 1123151 | C | G | 76 | 0 | 8 | 0 | G | G | | | | | G | G | G | G | G | G |
| NC_011770 | 1128581 | C | T | 46 | 0 | 6 | 0 | T | T | | | | | | | T | T | T | T |
| NC_011770 | 1128606 | CG | C | 47 | 0 | 1 | 0 | | | | | | | C | | | | | |
| NC_011770 | 1129520 | G | C | 51 | 0 | 2 | 0 | | | | | | | | | C | C | | |
| NC_011770 | 1129526 | T | C | 49 | 0 | 4 | 0 | | | C | C | C | C | | | | | | |
| NC_011770 | 1129527 | T | G | 50 | 0 | 4 | 0 | | | G | G | G | G | | | | | | |
| NC_011770 | 1129631 | T | C | 44 | 0 | 6 | 0 | | | C | C | C | C | | | | | C | C |
| NC_011770 | 1129634 | T | G | 45 | 0 | 4 | 0 | | | G | G | G | G | | | | | | |
| NC_011770 | 1129645 | A | C | 51 | 0 | 4 | 0 | | | C | C | C | C | | | | | | |
| NC_011770 | 1129649 | A | G | 50 | 0 | 4 | 0 | | | G | G | G | G | | | | | | |
| NC_011770 | 1129693 | T | A | 66 | 0 | 2 | 0 | | | | | | | | | | | A | A |
| NC_011770 | 1132689 | C | G | 92 | 0 | 10 | 0 | G | G | G | G | G | G | G | G | G | G | | |
| NC_011770 | 1132746 | A | G | 99 | 0 | 8 | 0 | G | G | G | G | G | G | | | G | G | | |
| NC_011770 | 1132752 | C | T | 98 | 0 | 2 | 0 | | | | | | | | | | | T | T |
| NC_011770 | 1132762 | C | G | 100 | 0 | 2 | 0 | | | | | | | G | G | | | | |
| NC_011770 | 1132838 | A | T | 88 | 0 | 10 | 0 | T | T | T | T | T | T | T | T | T | T | | |
| NC_011770 | 1132840 | T | C | 90 | 0 | 10 | 0 | C | C | C | C | C | C | C | C | C | C | | |
| NC_011770 | 1132846 | G | T | 93 | 0 | 10 | 0 | T | T | T | T | T | T | T | T | T | T | | |
| NC_011770 | 1133714 | C | G | 68 | 0 | 2 | 0 | G | G | | | | | | | | | | |
| NC_011770 | 1136216 | T | C | 85 | 0 | 2 | 0 | | | | | | | C | C | | | | |
| NC_011770 | 1136248 | C | T | 85 | 0 | 2 | 0 | | | | | T | T | | | | | | |
| NC_011770 | 1138057 | C | T | 83 | 0 | 2 | 0 | | | | | | | T | T | | | | |
| NC_011770 | 1138108 | C | G | 82 | 0 | 8 | 0 | G | G | G | G | G | G | | | | | G | G |
| NC_011770 | 1138160 | G | T | 81 | 0 | 2 | 0 | | | | | T | T | | | | | | |
| NC_011770 | 1138212 | C | T | 84 | 0 | 6 | 0 | | | T | T | T | T | | | | | T | T |
| NC_011770 | 1141774 | C | T | 72 | 0 | 4 | 0 | | | T | T | T | T | | | | | | |
| NC_011770 | 1141914 | C | T | 70 | 0 | 2 | 0 | | | | | | | | | T | T | | |
| NC_011770 | 1144667 | C | T | 80 | 0 | 2 | 0 | | | | | | | | | | | T | T |
| NC_011770 | 1147096 | C | T | 48 | 0 | 2 | 0 | | | | | | | | | T | T | | |
| NC_011770 | 1147220 | AGGAGCAC | A | 26 | 0 | 2 | 0 | | | | | | | | | A | A | | |
| NC_011770 | 1147223 | A | G | 23 | 2 | 6 | 0 | G | G | | | | | G | G | . | . | G | G |
| NC_011770 | 1147289 | G | A | 21 | 1 | 2 | 0 | | . | | | | | A | A | | | | |
| NC_011770 | 1148961 | G | A | 69 | 0 | 8 | 0 | | | A | A | A | A | A | A | | | A | A |
| NC_011770 | 1148993 | G | A | 67 | 0 | 4 | 0 | | | A | A | A | A | | | | | | |
| NC_011770 | 1150488 | AC | A | 74 | 0 | 4 | 0 | | | | | | | A | A | | | A | A |
| NC_011770 | 1154827 | A | G | 46 | 0 | 2 | 0 | | | | | | | | | G | G | | |
| NC_011770 | 1154842 | GC | G | 45 | 0 | 2 | 0 | | | | | | | | | G | G | | |
| NC_011770 | 1154878 | C | T | 50 | 0 | 2 | 0 | | | | | | | T | T | | | | |
| NC_011770 | 1154886 | A | G | 49 | 0 | 12 | 0 | G | G | G | G | G | G | G | G | G | G | G | G |
| NC_011770 | 1154899 | G | A | 49 | 0 | 2 | 0 | | | | | | | A | A | | | | |
| NC_011770 | 1154951 | T | C | 51 | 0 | 2 | 0 | | | | | | | C | C | | | | |
| NC_011770 | 1154988 | C | T | 56 | 0 | 2 | 0 | | | | | | | T | T | | | | |
| NC_011770 | 1156619 | G | T | 70 | 0 | 8 | 0 | T | T | T | T | T | T | | | | | T | T |
| NC_011770 | 1159038 | T | G | 53 | 0 | 4 | 0 | | | G | G | G | G | | | | | | |
| NC_011770 | 1159039 | C | A | 55 | 0 | 4 | 0 | | | A | A | A | A | | | | | | |
| NC_011770 | 1159062 | G | A | 55 | 0 | 4 | 0 | | | A | A | A | A | | | | | | |
| NC_011770 | 1159069 | T | TG | 56 | 0 | 2 | 0 | | | | | TG | TG | | | | | | |
| NC_011770 | 1159102 | G | A | 54 | 0 | 2 | 0 | | | | | | | | | A | A | | |
| NC_011770 | 1159105 | G | A | 53 | 0 | 2 | 0 | | | | | | | | | | | A | A |
| NC_011770 | 1159106 | T | C | 52 | 0 | 2 | 0 | | | | | | | | | C | C | | |
| NC_011770 | 1159109 | C | T | 52 | 0 | 4 | 0 | | | T | T | T | T | | | | | | |
| NC_011770 | 1159115 | C | A | 50 | 0 | 2 | 0 | | | | | | | | | A | A | | |
| NC_011770 | 1159116 | T | C | 49 | 0 | 2 | 0 | | | | | | | | | C | C | | |
| NC_011770 | 1159128 | G | A | 49 | 0 | 4 | 0 | | | A | A | A | A | | | | | | |
| NC_011770 | 1159137 | G | GT | 48 | 0 | 4 | 0 | | | GT | GT | GT | GT | | | | | | |
| NC_011770 | 1159141 | T | G | 47 | 0 | 2 | 0 | | | | | | | | | | | G | G |
| NC_011770 | 1159159 | C | G | 41 | 0 | 2 | 0 | | | | | | | G | G | | | | |
| NC_011770 | 1159182 | T | C | 20 | 5 | 1 | 0 | | | . | . | C | . | . | . | . | . | | |
| NC_011770 | 1159194 | G | A | 30 | 3 | 3 | 0 | | | A | . | A | A | . | . | | | | |
| NC_011770 | 1159196 | C | T | 31 | 3 | 3 | 0 | | | T | . | T | T | . | . | | | | |
| NC_011770 | 1159211 | A | G | 29 | 2 | 2 | 0 | | | | | | | . | . | | | G | G |

| | | | | | | | | | | | | | | | | | | | |
|---|---|---|---|---|---|---|---|---|---|---|---|---|---|---|---|---|---|---|---|
| NC_011770 | 1159216 | T | C | 29 | 2 | 2 | 0 | | | | | | | . | . | | | C | C |
| NC_011770 | 1159233 | C | T | 21 | 3 | 7 | 0 | | | T | . | T | T | . | . | T | T | T | T |
| NC_011770 | 1159235 | T | C | 20 | 3 | 7 | 0 | | | C | . | C | C | . | . | C | C | C | C |
| NC_011770 | 1159306 | TC | T | 13 | 5 | 1 | 0 | | | | | . | T | . | . | | | . | . |
| NC_011770 | 1159323 | G | A | 9 | 7 | 1 | 0 | | | . | . | . | A | . | . | | | . | . |
| NC_011770 | 1159325 | C | T | 10 | 7 | 1 | 0 | | | . | . | . | T | . | . | | | . | . |
| NC_011770 | 1159330 | G | A | 9 | 7 | 1 | 0 | | | . | . | . | A | . | . | | | . | . |
| NC_011770 | 1159373 | A | C | 9 | 6 | 2 | 0 | | | C | . | . | . | . | . | . | G | . | . |
| NC_011770 | 1159374 | T | C | 9 | 6 | 2 | 0 | | | | . | . | | . | . | C | C | . | . |
| NC_011770 | 1159386 | G | A | 11 | 5 | 3 | 0 | | | A | . | A | A | . | . | | | . | . |
| NC_011770 | 1159395 | G | GT | 12 | 3 | 3 | 0 | | | GT | . | GT | GT | . | . | | | | |
| NC_011770 | 1159402 | T | C | 11 | 4 | 6 | 0 | | | C | . | C | C | . | . | C | C | C | . |
| NC_011770 | 1159417 | G | C | 13 | 3 | 5 | 0 | | | C | . | C | C | . | . | | | C | C |
| NC_011770 | 1159418 | T | C | 12 | 3 | 7 | 0 | | | C | . | C | C | . | . | C | C | C | C |
| NC_011770 | 1159420 | A | C | 12 | 3 | 7 | 0 | | | C | . | C | C | . | . | C | C | C | C |
| NC_011770 | 1159435 | TG | T | 19 | 1 | 3 | 0 | | | T | . | T | T | | | | | | |
| NC_011770 | 1159436 | G | T | 18 | 1 | 6 | 0 | | | . | . | . | . | T | T | C | C | C | C |
| NC_011770 | 1159438 | T | A | 19 | 1 | 3 | 0 | | | | . | | | | | . | A | A | A |
| NC_011770 | 1159440 | C | T | 19 | 1 | 3 | 0 | | | | . | | | | | . | T | T | T |
| NC_011770 | 1159441 | G | A | 20 | 1 | 2 | 0 | | | | . | | | A | A | | | | |
| NC_011770 | 1159452 | G | A | 25 | 1 | 3 | 0 | | | A | . | A | A | | | | | | |
| NC_011770 | 1159454 | C | T | 25 | 1 | 3 | 0 | | | T | . | T | T | | | | | | |
| NC_011770 | 1159455 | TG | T | 25 | 1 | 2 | 0 | | | | . | | | T | T | | | | |
| NC_011770 | 1159459 | A | G | 27 | 1 | 1 | 0 | | | | . | | | | | . | G | | |
| NC_011770 | 1159469 | A | G | 37 | 0 | 2 | 0 | | | | | | | | | | | G | G |
| NC_011770 | 1159474 | C | T | 37 | 0 | 8 | 0 | | | T | T | T | T | T | T | T | T | | |
| NC_011770 | 1159480 | T | C | 39 | 0 | 1 | 0 | | | | | | | | | . | C | | |
| NC_011770 | 1159493 | C | T | 40 | 0 | 2 | 0 | | | | | | | T | T | | | | |
| NC_011770 | 1159496 | CG | C | 40 | 0 | 2 | 0 | | | | | | | | | | | C | C |
| NC_011770 | 1159502 | G | C | 42 | 0 | 2 | 0 | | | | | | | C | C | | | | |
| NC_011770 | 1159503 | C | T | 42 | 0 | 2 | 0 | | | | | | | T | T | | | | |
| NC_011770 | 1159512 | T | A | 43 | 0 | 10 | 0 | | | A | A | A | A | A | A | A | A | A | A |
| NC_011770 | 1159534 | C | T | 45 | 0 | 6 | 0 | | | T | T | T | T | | | | | T | T |
| NC_011770 | 1159543 | G | C | 44 | 0 | 6 | 0 | | | C | C | C | C | | | | | C | C |
| NC_011770 | 1159547 | CCCCACCA | C | 42 | 0 | 6 | 0 | | | C | C | C | C | | | | | C | C |
| NC_011770 | 1159551 | A | T | 19 | 6 | 2 | 0 | | | . | . | . | . | T | T | | | . | . |
| NC_011770 | 1162412 | G | T | 57 | 0 | 2 | 0 | | | | | | | | | T | T | | |
| NC_011770 | 1162528 | C | T | 61 | 0 | 2 | 0 | | | | | | | | | T | T | | |
| NC_011770 | 1162536 | G | A | 60 | 0 | 2 | 0 | | | | | | | | | | | A | A |
| NC_011770 | 1162566 | A | AG | 56 | 0 | 1 | 0 | | | | | | | | | AG | | | |
| NC_011770 | 1162601 | G | A | 55 | 0 | 6 | 0 | A | A | A | A | A | A | | | | | | |
| NC_011770 | 1162632 | A | G | 54 | 0 | 8 | 0 | G | G | G | G | G | G | | | G | G | | |
| NC_011770 | 1162646 | T | C | 51 | 0 | 6 | 0 | C | C | C | C | C | C | | | | | | |
| NC_011770 | 1162704 | G | C | 51 | 0 | 6 | 0 | | | C | C | C | C | | | C | C | | |
| NC_011770 | 1162713 | C | A | 53 | 0 | 4 | 0 | | | A | A | A | A | | | | | | |
| NC_011770 | 1162726 | C | T | 55 | 0 | 6 | 0 | | | T | T | T | T | | | T | T | | |
| NC_011770 | 1165029 | T | A | 73 | 0 | 12 | 0 | A | A | A | A | A | A | A | A | A | A | A | A |
| NC_011770 | 1165104 | C | T | 74 | 0 | 8 | 0 | T | T | T | T | T | T | | | T | T | | |
| NC_011770 | 1165105 | A | C | 74 | 0 | 8 | 0 | C | C | C | C | C | C | | | C | C | | |
| NC_011770 | 1170085 | A | C | 76 | 0 | 10 | 0 | C | C | C | C | C | C | | | C | C | C | C |
| NC_011770 | 1172094 | A | G | 55 | 0 | 2 | 0 | | | | | | | | | G | G | | |
| NC_011770 | 1172097 | G | T | 57 | 0 | 2 | 0 | | | | | | | | | T | T | | |
| NC_011770 | 1172099 | G | A | 57 | 0 | 2 | 0 | | | | | | | | | A | A | | |
| NC_011770 | 1173131 | T | C | 74 | 0 | 12 | 0 | C | C | C | C | C | C | C | C | C | C | C | C |
| NC_011770 | 1173223 | C | G | 74 | 0 | 2 | 0 | | | G | G | | | | | | | | |
| NC_011770 | 1177069 | G | C | 92 | 0 | 6 | 0 | | | C | C | C | C | | | | | C | C |
| NC_011770 | 1177091 | G | C | 97 | 0 | 4 | 0 | | | C | C | C | C | | | | | | |
| NC_011770 | 1177093 | T | C | 97 | 0 | 4 | 0 | | | C | C | C | C | | | | | | |
| NC_011770 | 1177176 | T | C | 108 | 0 | 2 | 0 | C | C | | | | | | | | | | |
| NC_011770 | 1178139 | G | A | 40 | 0 | 4 | 0 | | | A | A | A | A | | | | | | |
| NC_011770 | 1178144 | G | T | 42 | 0 | 4 | 0 | | | T | T | T | T | | | | | | |
| NC_011770 | 1181975 | C | T | 71 | 0 | 2 | 0 | | | | | | | | | T | T | | |

| | | | | | | | | | | | | | | | | | | | |
|---|---|---|---|---|---|---|---|---|---|---|---|---|---|---|---|---|---|---|---|
| NC_011770 | 1185499 | T | G | 76 | 0 | 8 | 0 | G | G | | | | | G | G | G | G | G | G |
| NC_011770 | 1186800 | C | T | 45 | 0 | 1 | 0 | | | | | | | T | | | | | |
| NC_011770 | 1186801 | G | A | 44 | 0 | 10 | 0 | A | A | A | A | A | A | | | A | A | A | A |
| NC_011770 | 1186871 | C | T | 51 | 0 | 6 | 0 | T | T | | | | | | | T | T | T | T |
| NC_011770 | 1188215 | G | A | 84 | 0 | 8 | 0 | | | A | A | A | A | A | A | A | A | | |
| NC_011770 | 1188230 | G | T | 82 | 0 | 8 | 0 | | | T | T | T | T | T | T | T | T | | |
| NC_011770 | 1188279 | C | G | 82 | 0 | 2 | 0 | | | | | | | G | G | | | | |
| NC_011770 | 1191296 | A | G | 90 | 0 | 10 | 0 | G | G | G | G | G | G | G | G | | | G | G |
| NC_011770 | 1194204 | C | T | 64 | 0 | 2 | 0 | | | | | | | | | | | T | T |
| NC_011770 | 1201903 | T | C | 84 | 0 | 12 | 0 | C | C | C | C | C | C | C | C | C | C | C | C |
| NC_011770 | 1201928 | C | A | 84 | 0 | 12 | 0 | A | A | A | A | A | A | A | A | A | A | A | A |
| NC_011770 | 1201930 | C | G | 84 | 0 | 12 | 0 | G | G | G | G | G | G | G | G | G | G | G | G |
| NC_011770 | 1201936 | T | C | 87 | 0 | 12 | 0 | C | C | A | A | A | A | A | A | A | A | A | A |
| NC_011770 | 1202950 | G | A | 86 | 0 | 2 | 0 | | | | | | | A | A | | | | |
| NC_011770 | 1202956 | C | T | 84 | 0 | 2 | 0 | | | | | | | T | T | | | | |
| NC_011770 | 1202957 | C | A | 84 | 0 | 2 | 0 | | | | | | | A | A | | | | |
| NC_011770 | 1202958 | T | G | 84 | 0 | 2 | 0 | | | | | | | G | G | | | | |
| NC_011770 | 1202977 | C | G | 84 | 0 | 2 | 0 | | | | | | | G | G | | | | |
| NC_011770 | 1203047 | G | GCC | 73 | 0 | 6 | 0 | | | GCC | GCC | GCC | GCC | A | A | | | | |
| NC_011770 | 1203052 | C | A | 43 | 0 | 2 | 0 | | | | | | | A | A | | | | |
| NC_011770 | 1203079 | A | T | 72 | 0 | 2 | 0 | | | | | | | T | T | | | | |
| NC_011770 | 1203083 | A | G | 73 | 0 | 2 | 0 | | | | | | | G | G | | | | |
| NC_011770 | 1203089 | T | C | 70 | 0 | 2 | 0 | | | | | | | C | C | | | | |
| NC_011770 | 1203098 | C | A | 72 | 0 | 2 | 0 | | | | | | | A | A | | | | |
| NC_011770 | 1203141 | A | G | 68 | 0 | 8 | 0 | G | G | G | G | G | G | | | G | G | | |
| NC_011770 | 1203144 | G | T | 70 | 0 | 8 | 0 | T | T | T | T | T | T | | | T | T | | |
| NC_011770 | 1203145 | G | A | 69 | 0 | 8 | 0 | A | A | A | A | A | A | | | A | A | | |
| NC_011770 | 1203160 | T | C | 77 | 0 | 2 | 0 | C | C | | | | | | | | | | |
| NC_011770 | 1203199 | G | A | 76 | 0 | 8 | 0 | A | A | A | A | A | A | | | A | A | | |
| NC_011770 | 1203244 | C | T | 79 | 0 | 8 | 0 | T | T | T | T | T | T | | | T | T | | |
| NC_011770 | 1203464 | C | T | 80 | 0 | 2 | 0 | | | | | | | T | T | | | | |
| NC_011770 | 1203476 | A | G | 83 | 0 | 2 | 0 | G | G | | | | | | | | | | |
| NC_011770 | 1203480 | A | G | 87 | 0 | 2 | 0 | G | G | | | | | | | | | | |
| NC_011770 | 1203495 | G | A | 82 | 0 | 2 | 0 | | | | | | | A | A | | | | |
| NC_011770 | 1203504 | A | G | 81 | 0 | 2 | 0 | | | | | | | | | G | G | | |
| NC_011770 | 1203514 | A | G | 88 | 0 | 2 | 0 | G | G | | | | | | | | | | |
| NC_011770 | 1203516 | G | A | 88 | 0 | 2 | 0 | A | A | | | | | | | | | | |
| NC_011770 | 1203518 | G | A | 85 | 0 | 2 | 0 | A | A | | | | | | | | | | |
| NC_011770 | 1203531 | T | G | 88 | 0 | 2 | 0 | | | | | | | G | G | | | | |
| NC_011770 | 1203536 | T | C | 86 | 0 | 2 | 0 | C | C | | | | | | | | | | |
| NC_011770 | 1205049 | A | G | 60 | 0 | 4 | 0 | G | G | | | | | G | G | | | | |
| NC_011770 | 1206233 | G | A | 98 | 0 | 4 | 0 | | | A | A | A | A | | | | | | |
| NC_011770 | 1207234 | G | T | 71 | 0 | 2 | 0 | | | | | | | T | T | | | | |
| NC_011770 | 1207241 | T | C | 65 | 0 | 8 | 0 | C | C | C | C | C | C | C | C | | | | |
| NC_011770 | 1207260 | T | C | 62 | 0 | 8 | 0 | C | C | C | C | C | C | C | C | | | | |
| NC_011770 | 1208552 | G | A | 53 | 0 | 2 | 0 | | | | | | | | | | | A | A |
| NC_011770 | 1208565 | C | T | 52 | 0 | 4 | 0 | | | T | T | T | T | | | | | | |
| NC_011770 | 1208632 | C | G | 45 | 0 | 10 | 0 | G | G | G | G | G | G | G | G | | | G | G |
| NC_011770 | 1208682 | C | T | 43 | 0 | 2 | 0 | T | T | | | | | | | | | | |
| NC_011770 | 1208711 | C | G | 41 | 1 | 9 | 0 | G | G | G | G | G | G | G | . | | | G | G |
| NC_011770 | 1208804 | G | A | 50 | 0 | 2 | 0 | | | | | | | | | | | A | A |
| NC_011770 | 1208956 | C | T | 56 | 0 | 4 | 0 | | | T | T | T | T | | | | | | |
| NC_011770 | 1208974 | G | T | 56 | 0 | 4 | 0 | | | T | T | T | T | | | | | | |
| NC_011770 | 1208986 | G | C | 56 | 0 | 4 | 0 | | | C | C | C | C | | | | | | |
| NC_011770 | 1209000 | T | G | 54 | 0 | 6 | 0 | G | G | | | | | G | G | | | G | G |
| NC_011770 | 1209014 | T | C | 58 | 0 | 4 | 0 | | | C | C | C | C | | | | | | |
| NC_011770 | 1209022 | A | G | 56 | 0 | 4 | 0 | | | G | G | G | G | | | | | | |
| NC_011770 | 1209023 | C | T | 55 | 0 | 4 | 0 | | | T | T | T | T | | | | | | |
| NC_011770 | 1209031 | C | T | 56 | 0 | 4 | 0 | | | T | T | T | T | | | | | | |
| NC_011770 | 1209040 | T | C | 55 | 0 | 10 | 0 | C | C | C | C | C | C | C | C | | | C | C |
| NC_011770 | 1209071 | A | G | 59 | 0 | 4 | 0 | | | G | G | G | G | | | | | | |
| NC_011770 | 1211745 | C | T | 81 | 0 | 2 | 0 | | | | | | | | | T | T | | |

| | | | | | | | | | | | | | | | | | | | |
|---|---|---|---|---|---|---|---|---|---|---|---|---|---|---|---|---|---|---|---|
| NC_011770 | 1211852 | A | G | 86 | 0 | 2 | 0 | G | G | | | | | | | | | | |
| NC_011770 | 1211864 | T | C | 83 | 0 | 2 | 0 | | | | | | | | | C | C | | |
| NC_011770 | 1211886 | C | G | 82 | 0 | 8 | 0 | | | G | G | G | G | G | G | | | G | G |
| NC_011770 | 1211905 | T | C | 80 | 0 | 2 | 0 | | | | | | | | | C | C | | |
| NC_011770 | 1213877 | A | G | 63 | 0 | 2 | 0 | | | | | | | | | G | G | | |
| NC_011770 | 1213893 | C | G | 63 | 0 | 2 | 0 | | | | | | | | | G | G | | |
| NC_011770 | 1213916 | A | C | 65 | 0 | 2 | 0 | | | | | | | | | C | C | | |
| NC_011770 | 1216216 | G | A | 61 | 0 | 4 | 0 | A | A | | | | | | | A | A | | |
| NC_011770 | 1216251 | C | T | 59 | 0 | 4 | 0 | T | T | | | | | | | T | T | | |
| NC_011770 | 1216281 | T | C | 65 | 0 | 2 | 0 | C | C | | | | | | | | | | |
| NC_011770 | 1218029 | C | T | 73 | 0 | 2 | 0 | | | | | | | T | T | | | | |
| NC_011770 | 1218039 | C | G | 75 | 0 | 2 | 0 | | | | | | | G | G | | | | |
| NC_011770 | 1218046 | T | G | 75 | 0 | 2 | 0 | | | | | | | | | | | G | G |
| NC_011770 | 1218087 | A | T | 80 | 0 | 2 | 0 | | | | | | | | | | | T | T |
| NC_011770 | 1222143 | A | G | 89 | 0 | 1 | 0 | | | | | | | G | | | | | |
| NC_011770 | 1223611 | G | A | 72 | 0 | 2 | 0 | | | | | | | A | A | | | | |
| NC_011770 | 1223612 | A | G | 71 | 0 | 2 | 0 | | | | | | | G | G | | | | |
| NC_011770 | 1223666 | T | C | 73 | 0 | 2 | 0 | | | | | | | C | C | | | | |
| NC_011770 | 1223675 | T | C | 73 | 0 | 2 | 0 | | | | | | | C | C | | | | |
| NC_011770 | 1223677 | G | A | 74 | 0 | 2 | 0 | | | | | | | A | A | | | | |
| NC_011770 | 1224340 | C | G | 62 | 0 | 2 | 0 | | | | | | | G | G | | | | |
| NC_011770 | 1224367 | C | T | 64 | 0 | 2 | 0 | | | | | | | | | T | T | | |
| NC_011770 | 1224369 | A | T | 63 | 0 | 2 | 0 | | | | | | | T | T | | | | |
| NC_011770 | 1224370 | T | C | 64 | 0 | 2 | 0 | | | | | | | | | | | C | C |
| NC_011770 | 1224399 | C | T | 65 | 0 | 6 | 0 | T | T | | | | | T | T | T | T | | |
| NC_011770 | 1224411 | G | T | 66 | 0 | 2 | 0 | | | | | | | | | T | T | | |
| NC_011770 | 1224465 | G | A | 66 | 0 | 4 | 0 | | | A | A | A | A | | | | | | |
| NC_011770 | 1224484 | G | A | 69 | 0 | 4 | 0 | | | A | A | A | A | | | | | | |
| NC_011770 | 1224490 | T | C | 71 | 0 | 4 | 0 | | | C | C | C | C | | | | | | |
| NC_011770 | 1224512 | G | C | 65 | 0 | 4 | 0 | | | C | C | C | C | | | | | | |
| NC_011770 | 1225309 | G | A | 63 | 0 | 8 | 0 | | | A | A | A | A | | | A | A | A | A |
| NC_011770 | 1225312 | C | T | 65 | 0 | 2 | 0 | | | | | | | T | T | | | | |
| NC_011770 | 1226843 | A | G | 71 | 0 | 4 | 0 | | | G | G | G | G | | | | | | |
| NC_011770 | 1229966 | C | A | 86 | 0 | 4 | 0 | | | A | A | A | A | | | | | | |
| NC_011770 | 1231693 | T | A | 66 | 0 | 12 | 0 | A | A | A | A | A | A | A | A | A | A | A | A |
| NC_011770 | 1231703 | C | T | 66 | 0 | 2 | 0 | | | | | | | | | | | T | T |
| NC_011770 | 1232782 | C | T | 29 | 0 | 2 | 0 | | | | | | | | | T | T | | |
| NC_011770 | 1232853 | C | T | 19 | 3 | 6 | 0 | T | T | . | . | | . | T | T | | | T | T |
| NC_011770 | 1232869 | GGGGC | G | 15 | 4 | 2 | 0 | G | G | . | . | | . | | | | . | | |
| NC_011770 | 1232892 | GC | G | 3 | 6 | 2 | 0 | G | G | | | | . | . | . | | . | . | . |
| NC_011770 | 1232916 | C | G | 3 | 7 | 2 | 0 | G | G | | | | . | . | . | . | . | . | . |
| NC_011770 | 1232995 | A | C | 3 | 6 | 2 | 0 | C | C | | | | . | . | . | | . | . | . |
| NC_011770 | 1233080 | GT | G | 2 | 7 | 1 | 0 | | | | | . | . | . | . | G | . | . | . |
| NC_011770 | 1233284 | G | A | 6 | 5 | 7 | 0 | A | A | A | A | A | A | . | . | A | . | . | . |
| NC_011770 | 1234909 | C | T | 26 | 4 | 8 | 0 | T | T | T | T | T | T | . | . | T | T | . | . |
| NC_011770 | 1234978 | T | C | 27 | 4 | 8 | 0 | C | C | C | C | C | C | . | . | C | C | . | . |
| NC_011770 | 1235250 | T | C | 33 | 4 | 4 | 0 | | | C | C | C | C | . | . | | | . | . |
| NC_011770 | 1235262 | G | T | 33 | 4 | 4 | 0 | | | T | T | T | T | . | . | | | . | . |
| NC_011770 | 1235273 | C | T | 33 | 4 | 4 | 0 | | | T | T | T | T | . | . | | | . | . |
| NC_011770 | 1235296 | A | G | 35 | 4 | 8 | 0 | G | G | G | G | G | G | . | . | G | G | . | . |
| NC_011770 | 1235315 | T | TA | 34 | 4 | 4 | 0 | | | TA | TA | TA | TA | . | . | | | . | . |
| NC_011770 | 1235321 | G | C | 32 | 4 | 4 | 0 | | | C | C | C | C | . | . | | | . | . |
| NC_011770 | 1235340 | T | C | 35 | 4 | 4 | 0 | | | C | C | C | C | . | . | | | . | . |
| NC_011770 | 1238694 | A | G | 44 | 0 | 8 | 0 | | | G | G | G | G | | | G | G | G | G |
| NC_011770 | 1238737 | T | C | 40 | 0 | 2 | 0 | | | | | | | C | C | | | | |
| NC_011770 | 1241217 | G | A | 91 | 0 | 2 | 0 | A | A | | | | | | | | | | |
| NC_011770 | 1242202 | A | G | 66 | 0 | 4 | 0 | | | G | G | G | G | | | | | | |
| NC_011770 | 1242251 | T | C | 69 | 0 | 6 | 0 | C | C | | | | | | | C | C | C | C |
| NC_011770 | 1243198 | A | G | 66 | 0 | 2 | 0 | | | | | | | | | | | G | G |
| NC_011770 | 1245597 | C | T | 72 | 0 | 4 | 0 | | | T | T | T | T | | | | | | |
| NC_011770 | 1245598 | A | T | 72 | 0 | 4 | 0 | | | T | T | T | T | | | | | | |
| NC_011770 | 1245611 | A | G | 74 | 0 | 12 | 0 | G | G | G | G | G | G | G | G | G | G | G | G |

| | | | | | | | | | | | | | | | | | | | |
|---|---|---|---|---|---|---|---|---|---|---|---|---|---|---|---|---|---|---|---|
| NC_011770 | 1245624 | G | A | 74 | 0 | 4 | 0 | | | A | A | A | A | | | | | | |
| NC_011770 | 1245639 | C | T | 75 | 0 | 4 | 0 | | | T | T | T | T | | | | | | |
| NC_011770 | 1245646 | C | T | 73 | 0 | 2 | 0 | | | | | | | | | T | T | | |
| NC_011770 | 1245656 | A | G | 76 | 0 | 4 | 0 | | | G | G | G | G | | | | | | |
| NC_011770 | 1245662 | C | G | 80 | 0 | 4 | 0 | | | G | G | G | G | | | | | | |
| NC_011770 | 1248885 | T | C | 56 | 0 | 2 | 0 | C | C | | | | | | | | | | |
| NC_011770 | 1248897 | T | G | 57 | 0 | 6 | 0 | G | G | C | C | C | C | | | | | | |
| NC_011770 | 1248900 | TC | T | 53 | 0 | 6 | 0 | T | T | T | T | T | T | | | | | | |
| NC_011770 | 1248905 | C | T | 24 | 3 | 2 | 0 | . | . | | | . | | T | T | | | | |
| NC_011770 | 1254580 | T | TTG | 84 | 0 | 2 | 0 | | | | | | | | | TTG | TTG | | |
| NC_011770 | 1255265 | A | G | 66 | 0 | 10 | 0 | G | G | G | G | G | G | G | G | | | G | G |
| NC_011770 | 1261080 | T | C | 83 | 0 | 12 | 0 | C | C | C | C | C | C | C | C | C | C | C | C |
| NC_011770 | 1263321 | GT | G | 89 | 0 | 4 | 0 | G | G | | | | | G | G | | | | |
| NC_011770 | 1263330 | C | T | 79 | 0 | 1 | 0 | | | | | | | T | | | | | |
| NC_011770 | 1264233 | A | G | 72 | 0 | 6 | 0 | | | G | G | G | G | G | G | | | | |
| NC_011770 | 1272992 | C | T | 68 | 0 | 6 | 0 | | | T | T | T | T | T | T | | | | |
| NC_011770 | 1273017 | G | A | 65 | 0 | 12 | 0 | A | A | A | A | A | A | A | A | A | A | A | A |
| NC_011770 | 1273020 | T | G | 69 | 0 | 4 | 0 | | | | | | | | | G | G | G | G |
| NC_011770 | 1273057 | G | T | 5 | 8 | 2 | 0 | . | . | . | | . | . | . | | T | T | . | . |
| NC_011770 | 1273060 | C | CCCGCG | 60 | 0 | 2 | 0 | | | | | | | | | CCCGCGG( | CCCGCGGGACGAA | | |
| NC_011770 | 1273078 | C | T | 70 | 0 | 2 | 0 | | | | | | | | | T | T | | |
| NC_011770 | 1273116 | C | T | 65 | 0 | 12 | 0 | T | T | T | T | T | T | T | T | T | T | T | T |
| NC_011770 | 1273121 | G | A | 69 | 0 | 12 | 0 | A | A | A | A | A | A | A | A | A | A | A | A |
| NC_011770 | 1273171 | C | T | 69 | 0 | 10 | 0 | T | T | T | T | T | T | T | T | | | T | T |
| NC_011770 | 1277043 | C | T | 52 | 0 | 2 | 0 | | | | | | | T | T | | | | |
| NC_011770 | 1277046 | G | T | 53 | 0 | 4 | 0 | | | T | T | T | T | | | | | | |
| NC_011770 | 1278012 | TG | T | 71 | 0 | 6 | 0 | T | T | | | | | | | T | T | T | T |
| NC_011770 | 1278043 | CT | C | 73 | 0 | 12 | 0 | C | C | C | C | C | C | C | C | C | C | C | C |
| NC_011770 | 1279959 | G | T | 67 | 0 | 2 | 0 | | | | | | | | | T | T | | |
| NC_011770 | 1279990 | T | G | 64 | 0 | 2 | 0 | | | | | | | | | | | G | G |
| NC_011770 | 1279995 | T | C | 66 | 0 | 2 | 0 | C | C | | | | | | | | | | |
| NC_011770 | 1280008 | T | C | 67 | 0 | 4 | 0 | | | C | C | C | C | | | | | | |
| NC_011770 | 1280071 | C | G | 66 | 0 | 2 | 0 | G | G | | | | | | | | | | |
| NC_011770 | 1280145 | A | T | 63 | 0 | 4 | 0 | | | T | T | T | T | | | | | | |
| NC_011770 | 1283008 | G | A | 79 | 0 | 2 | 0 | | | | | | | | | A | A | | |
| NC_011770 | 1283009 | T | C | 76 | 0 | 2 | 0 | | | | | | | | | C | C | | |
| NC_011770 | 1284928 | T | C | 44 | 0 | 4 | 0 | | | | | | | C | C | C | C | | |
| NC_011770 | 1285875 | G | A | 41 | 0 | 12 | 0 | A | A | A | A | A | A | A | A | A | A | A | A |
| NC_011770 | 1285895 | C | A | 46 | 0 | 4 | 0 | | | A | A | A | A | | | | | | |
| NC_011770 | 1285909 | A | G | 46 | 0 | 12 | 0 | G | G | G | G | G | G | G | G | G | G | G | G |
| NC_011770 | 1286456 | G | A | 73 | 0 | 2 | 0 | | | | | | | | | A | A | | |
| NC_011770 | 1287005 | C | T | 76 | 0 | 2 | 0 | | | | | | | | | | | T | T |
| NC_011770 | 1287608 | TG | T | 38 | 2 | 6 | 0 | T | T | . | . | AG | AG | T | T | | | | |
| NC_011770 | 1288282 | G | A | 79 | 0 | 12 | 0 | A | A | A | A | A | A | A | A | A | A | A | A |
| NC_011770 | 1288292 | C | T | 78 | 0 | 2 | 0 | T | T | | | | | | | | | | |
| NC_011770 | 1288373 | C | T | 78 | 0 | 2 | 0 | | | | | | | T | T | | | | |
| NC_011770 | 1289336 | A | G | 68 | 0 | 12 | 0 | G | G | G | G | G | G | G | G | G | G | G | G |
| NC_011770 | 1289338 | A | C | 69 | 0 | 6 | 0 | | | C | C | C | C | C | C | | | | |
| NC_011770 | 1289515 | G | A | 82 | 0 | 2 | 0 | | | | | | | | | | | A | A |
| NC_011770 | 1289521 | A | G | 81 | 0 | 12 | 0 | G | G | G | G | G | G | G | G | G | G | G | G |
| NC_011770 | 1289593 | A | C | 85 | 0 | 12 | 0 | C | C | C | C | C | C | C | C | C | C | C | C |
| NC_011770 | 1289649 | G | A | 87 | 0 | 2 | 0 | A | A | | | | | | | | | | |
| NC_011770 | 1289680 | C | T | 80 | 0 | 4 | 0 | | | T | T | T | T | | | | | | |
| NC_011770 | 1289813 | A | G | 77 | 0 | 1 | 0 | | | | | | | G | | | | | |
| NC_011770 | 1290037 | G | T | 64 | 0 | 4 | 0 | | | T | T | T | T | | | | | | |
| NC_011770 | 1290073 | G | A | 74 | 0 | 2 | 0 | A | A | | | | | | | | | | |
| NC_011770 | 1290090 | T | C | 73 | 0 | 6 | 0 | C | C | | | | | C | C | C | C | | |
| NC_011770 | 1291870 | T | C | 64 | 0 | 2 | 0 | | | | | | | C | C | | | | |
| NC_011770 | 1296409 | C | A | 75 | 0 | 2 | 0 | A | A | | | | | | | | | | |
| NC_011770 | 1296417 | C | T | 75 | 0 | 6 | 0 | | | T | T | T | T | T | T | | | | |
| NC_011770 | 1300604 | T | A | 50 | 0 | 6 | 0 | A | A | | | | | | | A | A | A | A |
| NC_011770 | 1300616 | T | G | 50 | 0 | 6 | 0 | G | G | | | | | | | G | G | G | G |

| | | | | | | | | | | | | | | | | | | | |
|---|---|---|---|---|---|---|---|---|---|---|---|---|---|---|---|---|---|---|---|
| NC_011770 | 1300639 | G | T | 57 | 0 | 2 | 0 | | | | | | | T | T | | | | |
| NC_011770 | 1301665 | T | TC | 80 | 0 | 12 | 0 | TC | TC | TC | TC | TC | TC | TC | TC | TC | TC | TC | TC |
| NC_011770 | 1301706 | G | T | 81 | 0 | 2 | 0 | | | | | | | T | T | | | | |
| NC_011770 | 1301763 | G | A | 74 | 0 | 2 | 0 | | | | | | | | | | | A | A |
| NC_011770 | 1301833 | G | C | 72 | 0 | 10 | 0 | C | C | C | C | C | C | C | C | C | C | | |
| NC_011770 | 1307117 | T | C | 0 | 11 | 1 | 0 | . | . | . | . | . | . | . | . | C | . | . | . |
| NC_011770 | 1311987 | A | AC | 34 | 0 | 11 | 0 | AC | AC | AC | AC | AC | AC | AGTAC | | AC | AC | AC | AC |
| NC_011770 | 1313988 | T | A | 81 | 0 | 6 | 0 | A | A | | | | | A | A | A | A | | |
| NC_011770 | 1314001 | C | A | 79 | 0 | 2 | 0 | | | | | | | A | A | | | | |
| NC_011770 | 1314115 | G | A | 81 | 0 | 2 | 0 | | | | | A | A | | | | | | |
| NC_011770 | 1315614 | T | C | 41 | 0 | 12 | 0 | C | C | C | C | C | C | C | C | C | C | C | C |
| NC_011770 | 1315629 | G | A | 44 | 0 | 2 | 0 | | | | | | | | | | | A | A |
| NC_011770 | 1315655 | C | T | 47 | 0 | 8 | 0 | T | T | T | T | T | T | T | T | | | | |
| NC_011770 | 1316297 | AG | A | 75 | 0 | 1 | 0 | | | | | | | A | | | | | |
| NC_011770 | 1316303 | C | T | 78 | 0 | 4 | 0 | | | T | T | T | T | | | | | | |
| NC_011770 | 1317814 | T | C | 93 | 0 | 1 | 0 | | | | | | | C | | | | | |
| NC_011770 | 1317912 | C | T | 93 | 0 | 2 | 0 | | | | | | | | | | | T | T |
| NC_011770 | 1317974 | G | T | 91 | 0 | 4 | 0 | | | T | T | T | T | | | | | | |
| NC_011770 | 1322247 | G | A | 64 | 0 | 2 | 0 | | | | | | | A | A | | | | |
| NC_011770 | 1322826 | A | G | 68 | 0 | 12 | 0 | G | G | G | G | G | G | G | G | G | G | G | G |
| NC_011770 | 1322874 | GA | G | 68 | 0 | 1 | 0 | | | | | | | G | | | | | |
| NC_011770 | 1330225 | T | C | 58 | 0 | 8 | 0 | C | C | | | | | C | C | C | C | C | C |
| NC_011770 | 1330301 | T | C | 63 | 0 | 8 | 0 | C | C | | | | | C | C | C | C | C | C |
| NC_011770 | 1331518 | T | C | 18 | 0 | 8 | 0 | C | C | C | C | C | C | C | C | | | | |
| NC_011770 | 1332244 | A | G | 72 | 0 | 10 | 0 | G | G | G | G | G | G | G | G | G | G | | |
| NC_011770 | 1338692 | A | T | 77 | 0 | 4 | 0 | T | T | | | | | T | T | | | | |
| NC_011770 | 1338769 | G | A | 75 | 0 | 4 | 0 | A | A | | | | | A | A | | | | |
| NC_011770 | 1338796 | C | A | 79 | 0 | 2 | 0 | A | A | | | | | | | | | | |
| NC_011770 | 1338800 | C | T | 78 | 0 | 2 | 0 | T | T | | | | | | | | | | |
| NC_011770 | 1338806 | A | G | 76 | 0 | 2 | 0 | G | G | | | | | | | | | | |
| NC_011770 | 1340364 | G | A | 72 | 0 | 8 | 0 | | | A | A | A | A | A | A | | | A | A |
| NC_011770 | 1340374 | T | G | 74 | 0 | 10 | 0 | G | G | G | G | G | G | G | G | | | G | G |
| NC_011770 | 1340384 | G | A | 75 | 0 | 4 | 0 | | | A | A | A | A | | | | | | |
| NC_011770 | 1342503 | C | T | 63 | 0 | 10 | 0 | T | T | T | T | T | T | | | T | T | T | T |
| NC_011770 | 1343075 | G | T | 47 | 0 | 4 | 0 | T | T | | | | | | | | | T | T |
| NC_011770 | 1343082 | G | A | 45 | 0 | 6 | 0 | A | A | | | | | | | A | A | A | A |
| NC_011770 | 1346973 | C | T | 76 | 0 | 4 | 0 | T | T | | | | | | | T | T | | |
| NC_011770 | 1346996 | C | T | 74 | 0 | 6 | 0 | | | T | T | T | T | | | | | T | T |
| NC_011770 | 1347897 | C | A | 68 | 0 | 12 | 0 | A | A | A | A | A | A | A | A | A | A | A | A |
| NC_011770 | 1349302 | A | G | 80 | 0 | 12 | 0 | G | G | G | G | G | G | G | G | G | G | G | G |
| NC_011770 | 1354044 | A | T | 67 | 0 | 2 | 0 | | | | | | | T | T | | | | |
| NC_011770 | 1354060 | G | T | 70 | 0 | 2 | 0 | | | | | | | T | T | | | | |
| NC_011770 | 1354071 | C | A | 74 | 0 | 2 | 0 | | | | | | | | | | | A | A |
| NC_011770 | 1354116 | A | G | 79 | 0 | 2 | 0 | | | | | | | G | G | | | | |
| NC_011770 | 1364387 | G | A | 79 | 0 | 2 | 0 | | | | | | | | | | | A | A |
| NC_011770 | 1368527 | C | T | 74 | 0 | 10 | 0 | T | T | T | T | T | T | T | T | T | T | | |
| NC_011770 | 1368534 | C | T | 76 | 0 | 6 | 0 | T | T | | | | | T | T | T | T | | |
| NC_011770 | 1369802 | A | G | 52 | 0 | 12 | 0 | G | G | G | G | G | G | G | G | G | G | G | G |
| NC_011770 | 1369890 | A | G | 53 | 0 | 4 | 0 | | | G | G | G | G | | | | | | |
| NC_011770 | 1369933 | T | C | 58 | 0 | 2 | 0 | | | | | | | | | | | C | C |
| NC_011770 | 1370332 | T | A | 63 | 0 | 4 | 0 | | | | | | | A | A | | | A | A |
| NC_011770 | 1370334 | G | T | 65 | 0 | 2 | 0 | | | | | | | T | T | | | | |
| NC_011770 | 1371790 | T | C | 69 | 0 | 12 | 0 | C | C | C | C | C | C | C | C | C | C | C | C |
| NC_011770 | 1371897 | T | C | 73 | 0 | 2 | 0 | | | | | | | | | | | C | C |
| NC_011770 | 1373200 | G | A | 54 | 0 | 2 | 0 | A | A | | | | | | | | | | |
| NC_011770 | 1373201 | G | A | 53 | 0 | 4 | 0 | A | A | | | | | | | | | A | A |
| NC_011770 | 1373205 | C | T | 53 | 0 | 4 | 0 | T | T | | | | | | | | | T | T |
| NC_011770 | 1373229 | TTGGTTCG( | T | 38 | 3 | 1 | 0 | T | . | | | | | | | | | . | . |
| NC_011770 | 1373246 | A | G | 43 | 2 | 8 | 0 | | | G | G | G | G | G | G | G | G | . | . |
| NC_011770 | 1373636 | C | T | 82 | 0 | 2 | 0 | | | | | | | | | T | T | | |
| NC_011770 | 1373663 | G | A | 79 | 0 | 2 | 0 | | | | | | | | | | | A | A |
| NC_011770 | 1373684 | A | C | 77 | 0 | 4 | 0 | C | C | | | | | | | | | G | G |

| | | | | | | | | | | | | | | | | | | | |
|---|---|---|---|---|---|---|---|---|---|---|---|---|---|---|---|---|---|---|---|
| NC_011770 | 1373689 | T | C | 79 | 0 | 2 | 0 | | | | | | | | | | | C | C |
| NC_011770 | 1373740 | C | T | 77 | 0 | 2 | 0 | | | | | | | T | T | | | | |
| NC_011770 | 1373742 | T | C | 76 | 0 | 2 | 0 | | | | | | | | | | | C | C |
| NC_011770 | 1375537 | C | T | 73 | 0 | 2 | 0 | | | | | | | | | | | T | T |
| NC_011770 | 1375539 | G | A | 71 | 0 | 8 | 0 | | | A | A | A | A | A | A | | | C | C |
| NC_011770 | 1375549 | T | C | 76 | 0 | 2 | 0 | | | | | | | | | | | C | C |
| NC_011770 | 1375592 | C | G | 65 | 0 | 2 | 0 | | | | | | | | | | | G | G |
| NC_011770 | 1375602 | T | C | 67 | 0 | 2 | 0 | | | | | | | | | | | C | C |
| NC_011770 | 1375605 | C | T | 70 | 0 | 2 | 0 | | | | | | | | | | | T | T |
| NC_011770 | 1376431 | C | T | 69 | 0 | 6 | 0 | T | T | T | T | T | T | | | | | | |
| NC_011770 | 1376476 | G | T | 71 | 0 | 2 | 0 | | | | | | | | | | | T | T |
| NC_011770 | 1377191 | G | T | 63 | 0 | 2 | 0 | T | T | | | | | | | | | | |
| NC_011770 | 1377198 | A | G | 63 | 0 | 4 | 0 | | | G | G | G | G | | | | | | |
| NC_011770 | 1377228 | G | A | 63 | 0 | 2 | 0 | | | | | | | | | | | A | A |
| NC_011770 | 1377317 | C | T | 60 | 0 | 2 | 0 | | | | | | | | | T | T | | |
| NC_011770 | 1377383 | T | G | 54 | 0 | 2 | 0 | | | | | | | | | | | G | G |
| NC_011770 | 1379467 | G | A | 58 | 0 | 2 | 0 | A | A | | | | | | | | | | |
| NC_011770 | 1380385 | G | A | 69 | 0 | 2 | 0 | | | | | | | | | | | A | A |
| NC_011770 | 1380388 | A | C | 66 | 0 | 12 | 0 | C | C | C | C | C | C | C | C | T | T | T | T |
| NC_011770 | 1381475 | T | C | 43 | 0 | 12 | 0 | C | C | C | C | C | C | C | C | C | C | C | C |
| NC_011770 | 1384921 | G | A | 88 | 0 | 2 | 0 | | | | | | | | | | | A | A |
| NC_011770 | 1384974 | T | A | 84 | 0 | 6 | 0 | A | A | | | | | A | A | A | A | | |
| NC_011770 | 1389427 | G | T | 88 | 0 | 2 | 0 | | | | | | | T | T | | | | |
| NC_011770 | 1389565 | T | C | 78 | 0 | 12 | 0 | C | C | C | C | C | C | C | C | C | C | C | C |
| NC_011770 | 1389640 | C | T | 75 | 0 | 6 | 0 | | | T | T | T | T | | | | | T | T |
| NC_011770 | 1389854 | C | A | 84 | 0 | 6 | 0 | | | A | A | A | A | | | | | A | A |
| NC_011770 | 1389896 | C | CG | 85 | 0 | 12 | 0 | CG | CG | CG | CG | CG | CG | CG | CG | CG | CG | CG | CG |
| NC_011770 | 1392572 | T | C | 76 | 0 | 8 | 0 | C | C | C | C | C | C | C | C | | | | |
| NC_011770 | 1392573 | T | G | 76 | 0 | 8 | 0 | G | G | G | G | G | G | G | G | | | | |
| NC_011770 | 1392577 | C | G | 77 | 0 | 8 | 0 | G | G | G | G | G | G | G | G | | | | |
| NC_011770 | 1392583 | C | T | 79 | 0 | 2 | 0 | T | T | | | | | | | | | | |
| NC_011770 | 1392611 | T | G | 75 | 0 | 12 | 0 | G | G | G | G | G | G | G | G | G | G | G | G |
| NC_011770 | 1392615 | C | T | 76 | 0 | 2 | 0 | T | T | | | | | | | | | | |
| NC_011770 | 1393776 | T | C | 71 | 0 | 6 | 0 | C | C | C | C | C | C | | | | | | |
| NC_011770 | 1393778 | T | C | 71 | 0 | 6 | 0 | C | C | C | C | C | C | | | | | | |
| NC_011770 | 1393779 | G | A | 70 | 0 | 6 | 0 | A | A | A | A | A | A | | | | | | |
| NC_011770 | 1393860 | C | T | 82 | 0 | 2 | 0 | | | | | | | T | T | | | | |
| NC_011770 | 1396709 | G | A | 71 | 0 | 2 | 0 | | | | | | | A | A | | | | |
| NC_011770 | 1396750 | T | C | 65 | 0 | 2 | 0 | | | | | | | | | | | C | C |
| NC_011770 | 1403485 | G | C | 69 | 0 | 2 | 0 | | | | | | | | | | | C | C |
| NC_011770 | 1406954 | T | TGG | 66 | 0 | 1 | 0 | | | | | | | TGG | | | | | |
| NC_011770 | 1410571 | T | C | 70 | 0 | 2 | 0 | | | | | | | | | | | C | C |
| NC_011770 | 1410594 | G | A | 65 | 0 | 2 | 0 | A | A | | | | | | | | | | |
| NC_011770 | 1411791 | C | G | 81 | 0 | 2 | 0 | | | | | | | G | G | | | | |
| NC_011770 | 1412680 | A | G | 66 | 0 | 10 | 0 | | | G | G | G | G | G | G | G | G | G | G |
| NC_011770 | 1414263 | T | C | 64 | 0 | 2 | 0 | | | | | | | C | C | | | | |
| NC_011770 | 1414269 | T | C | 67 | 0 | 2 | 0 | | | | | | | C | C | | | | |
| NC_011770 | 1414295 | C | T | 68 | 0 | 2 | 0 | | | | | | | | | T | T | | |
| NC_011770 | 1414306 | C | T | 69 | 0 | 2 | 0 | | | | | | | | | | | T | T |
| NC_011770 | 1416405 | G | A | 81 | 0 | 6 | 0 | | | A | A | A | A | | | | | A | A |
| NC_011770 | 1421369 | G | A | 78 | 0 | 6 | 0 | A | A | A | A | A | A | | | | | | |
| NC_011770 | 1422095 | G | C | 77 | 0 | 2 | 0 | | | | | | | C | C | | | | |
| NC_011770 | 1422173 | A | T | 70 | 0 | 2 | 0 | T | T | | | | | | | | | | |
| NC_011770 | 1422609 | C | CA | 59 | 0 | 2 | 0 | CA | CA | | | | | | | | | | |
| NC_011770 | 1422713 | G | A | 61 | 0 | 2 | 0 | A | A | | | | | | | | | | |
| NC_011770 | 1422728 | C | T | 62 | 0 | 2 | 0 | T | T | | | | | | | | | | |
| NC_011770 | 1422729 | T | C | 61 | 0 | 2 | 0 | C | C | | | | | | | | | | |
| NC_011770 | 1425375 | G | A | 69 | 0 | 4 | 0 | | | A | A | A | A | | | | | | |
| NC_011770 | 1431709 | A | G | 84 | 0 | 8 | 0 | G | G | G | G | G | G | | | G | G | | |
| NC_011770 | 1433439 | C | T | 58 | 0 | 4 | 0 | | | T | T | T | T | | | | | | |
| NC_011770 | 1433846 | T | C | 14 | 10 | 2 | 0 | . | . | . | . | . | . | . | . | . | . | C | C |
| NC_011770 | 1435198 | T | C | 21 | 8 | 2 | 0 | . | . | . | . | . | . | . | . | | | C | C |

| | | | | | | | | | | | | | | | | | | | |
|---|---|---|---|---|---|---|---|---|---|---|---|---|---|---|---|---|---|---|---|
| NC_011770 | 1435235 | A | C | 20 | 8 | 4 | 0 | . | . | . | . | . | . | . | . | C | C | C | C |
| NC_011770 | 1435245 | A | AT | 21 | 8 | 4 | 0 | . | . | . | . | . | . | . | . | AT | AT | AT | AT |
| NC_011770 | 1435255 | GT | G | 20 | 8 | 2 | 0 | . | . | . | . | . | . | . | . | | | G | G |
| NC_011770 | 1435259 | G | GA | 5 | 8 | 1 | 0 | . | . | . | . | . | . | . | . | | | . | GA |
| NC_011770 | 1435303 | T | G | 20 | 8 | 4 | 0 | . | . | . | . | . | . | . | . | G | G | G | G |
| NC_011770 | 1435306 | C | T | 20 | 8 | 4 | 0 | . | . | . | . | . | . | . | . | T | T | T | T |
| NC_011770 | 1435318 | G | A | 21 | 8 | 4 | 0 | . | . | . | . | . | . | . | . | A | A | A | A |
| NC_011770 | 1435342 | A | G | 20 | 8 | 4 | 0 | . | . | . | . | . | . | . | . | G | G | G | G |
| NC_011770 | 1435345 | T | G | 20 | 8 | 4 | 0 | . | . | . | . | . | . | . | . | G | G | G | G |
| NC_011770 | 1435352 | A | G | 19 | 8 | 4 | 0 | . | . | . | . | . | . | . | . | G | G | G | G |
| NC_011770 | 1435362 | T | G | 15 | 8 | 4 | 0 | . | . | . | . | . | . | . | . | G | G | G | G |
| NC_011770 | 1435363 | G | T | 15 | 8 | 4 | 0 | . | . | . | . | . | . | . | . | T | T | T | T |
| NC_011770 | 1435369 | G | A | 15 | 8 | 4 | 0 | . | . | . | . | . | . | . | . | A | A | A | A |
| NC_011770 | 1435371 | G | T | 15 | 8 | 4 | 0 | . | . | . | . | . | . | . | . | T | T | T | T |
| NC_011770 | 1435372 | A | G | 15 | 8 | 4 | 0 | . | . | . | . | . | . | . | . | G | G | G | G |
| NC_011770 | 1435383 | T | TG | 12 | 9 | 2 | 0 | . | . | . | . | . | . | . | . | | . | TG | TG |
| NC_011770 | 1435385 | G | GAA | 11 | 10 | 2 | 0 | . | . | . | . | . | . | . | . | . | . | GAA | GAA |
| NC_011770 | 1435715 | A | T | 3 | 10 | 2 | 0 | . | . | . | . | . | . | . | . | T | T | . | . |
| NC_011770 | 1435719 | G | A | 3 | 10 | 2 | 0 | . | . | . | . | . | . | . | . | A | A | . | . |
| NC_011770 | 1435720 | A | G | 3 | 10 | 2 | 0 | . | . | . | . | . | . | . | . | G | G | . | . |
| NC_011770 | 1435741 | T | C | 19 | 6 | 4 | 0 | C | C | . | . | . | . | . | . | | | C | C |
| NC_011770 | 1441298 | G | T | 6 | 9 | 1 | 0 | . | . | . | . | . | . | . | . | T | . | | |
| NC_011770 | 1441300 | T | C | 5 | 9 | 3 | 0 | . | . | . | . | . | . | . | . | C | . | A | A |
| NC_011770 | 1441308 | T | G | 7 | 8 | 4 | 0 | . | . | . | . | . | . | . | . | G | G | G | G |
| NC_011770 | 1441632 | C | A | 5 | 10 | 2 | 0 | . | . | . | . | . | . | . | . | . | . | A | A |
| NC_011770 | 1441634 | G | A | 5 | 10 | 2 | 0 | . | . | . | . | . | . | . | . | . | . | A | A |
| NC_011770 | 1441650 | A | G | 5 | 10 | 2 | 0 | . | . | . | . | . | . | . | . | . | . | G | G |
| NC_011770 | 1441654 | T | G | 7 | 8 | 2 | 0 | . | . | . | . | . | . | . | . | | | G | G |
| NC_011770 | 1441665 | G | A | 8 | 8 | 2 | 0 | . | . | . | . | . | . | . | . | | | A | A |
| NC_011770 | 1441673 | A | C | 8 | 8 | 2 | 0 | . | . | . | . | . | . | . | . | C | C | | |
| NC_011770 | 1441684 | A | G | 8 | 8 | 4 | 0 | . | . | . | . | . | . | . | . | G | G | G | G |
| NC_011770 | 1441686 | C | T | 8 | 8 | 2 | 0 | . | . | . | . | . | . | . | . | T | T | | |
| NC_011770 | 1441692 | A | C | 8 | 8 | 2 | 0 | . | . | . | . | . | . | . | . | C | C | | |
| NC_011770 | 1441708 | A | G | 8 | 8 | 2 | 0 | . | . | . | . | . | . | . | . | | | G | G |
| NC_011770 | 1441712 | C | G | 8 | 8 | 4 | 0 | . | . | . | . | . | . | . | . | G | G | A | A |
| NC_011770 | 1441719 | G | C | 8 | 8 | 4 | 0 | . | . | . | . | . | . | . | . | C | C | A | A |
| NC_011770 | 1441722 | G | T | 8 | 8 | 2 | 0 | . | . | . | . | . | . | . | . | T | T | | |
| NC_011770 | 1441727 | A | C | 8 | 8 | 2 | 0 | . | . | . | . | . | . | . | . | C | C | | |
| NC_011770 | 1441728 | C | T | 8 | 8 | 2 | 0 | . | . | . | . | . | . | . | . | | | T | T |
| NC_011770 | 1441731 | C | T | 8 | 8 | 2 | 0 | . | . | . | . | . | . | . | . | | | T | T |
| NC_011770 | 1441734 | C | G | 8 | 8 | 2 | 0 | . | . | . | . | . | . | . | . | G | G | | |
| NC_011770 | 1441738 | C | T | 8 | 8 | 2 | 0 | . | . | . | . | . | . | . | . | T | T | | |
| NC_011770 | 1441740 | T | A | 8 | 8 | 4 | 0 | . | . | . | . | . | . | . | . | A | A | G | G |
| NC_011770 | 1441742 | G | T | 8 | 8 | 2 | 0 | . | . | . | . | . | . | . | . | T | T | | |
| NC_011770 | 1441743 | T | C | 8 | 8 | 4 | 0 | . | . | . | . | . | . | . | . | C | C | A | A |
| NC_011770 | 1441749 | T | C | 8 | 8 | 2 | 0 | . | . | . | . | . | . | . | . | C | C | | |
| NC_011770 | 1441752 | C | T | 8 | 8 | 2 | 0 | . | . | . | . | . | . | . | . | T | T | | |
| NC_011770 | 1441755 | A | G | 8 | 8 | 4 | 0 | . | . | . | . | . | . | . | . | G | G | G | G |
| NC_011770 | 1441758 | A | T | 8 | 8 | 4 | 0 | . | . | . | . | . | . | . | . | T | T | T | T |
| NC_011770 | 1441761 | C | A | 8 | 8 | 4 | 0 | . | . | . | . | . | . | . | . | A | A | T | T |
| NC_011770 | 1441771 | G | A | 9 | 8 | 2 | 0 | . | . | . | . | . | . | . | . | A | A | | |
| NC_011770 | 1441779 | A | C | 9 | 8 | 4 | 0 | . | . | . | . | . | . | . | . | C | C | C | C |
| NC_011770 | 1441785 | G | A | 10 | 8 | 2 | 0 | . | . | . | . | . | . | . | . | | | A | A |
| NC_011770 | 1441808 | TGCTGCCG | T | 9 | 8 | 2 | 0 | . | . | . | . | . | . | . | . | T | T | | |
| NC_011770 | 1441811 | T | C | 9 | 8 | 2 | 0 | . | . | . | . | . | . | . | . | . | . | C | C |
| NC_011770 | 1441830 | A | G | 10 | 8 | 4 | 0 | . | . | . | . | . | . | . | . | G | G | G | G |
| NC_011770 | 1442088 | A | G | 12 | 8 | 2 | 0 | . | . | . | . | . | . | . | . | G | G | | |
| NC_011770 | 1442483 | T | TGG | 10 | 8 | 2 | 0 | . | . | . | . | . | . | . | . | TGG | TGG | | |
| NC_011770 | 1442485 | A | G | 9 | 10 | 2 | 0 | . | . | . | . | . | . | . | . | . | . | G | G |
| NC_011770 | 1442489 | A | G | 10 | 8 | 2 | 0 | . | . | . | . | . | . | . | . | G | G | | |
| NC_011770 | 1442500 | A | G | 10 | 8 | 4 | 0 | . | . | . | . | . | . | . | . | G | G | G | G |
| NC_011770 | 1442515 | C | CGGCGC | 9 | 8 | 2 | 0 | . | . | . | . | . | . | . | . | CGGCGGC | CGGCGGCGCGATTGTTTGCCGATG | | |

| | | | | | | | | | | | | | | | | | | | |
|---|---|---|---|---|---|---|---|---|---|---|---|---|---|---|---|---|---|---|---|
| NC_011770 | 1442519 | A | G | 11 | 8 | 2 | 0 | . | . | . | . | . | . | . | . | | | G | G |
| NC_011770 | 1442520 | T | G | 10 | 8 | 2 | 0 | . | . | . | . | . | . | . | . | | | G | G |
| NC_011770 | 1442521 | T | C | 10 | 8 | 2 | 0 | . | . | . | . | . | . | . | . | | | C | C |
| NC_011770 | 1442534 | C | G | 11 | 8 | 4 | 0 | . | . | . | . | . | . | . | . | G | G | G | G |
| NC_011770 | 1442542 | A | G | 11 | 8 | 4 | 0 | . | . | . | . | . | . | . | . | G | G | G | G |
| NC_011770 | 1442559 | A | G | 11 | 8 | 4 | 0 | . | . | . | . | . | . | . | . | G | G | G | G |
| NC_011770 | 1442560 | A | T | 11 | 8 | 4 | 0 | . | . | . | . | . | . | . | . | T | T | T | T |
| NC_011770 | 1442563 | G | A | 10 | 8 | 4 | 0 | . | . | . | . | . | . | . | . | A | A | A | A |
| NC_011770 | 1442572 | A | G | 12 | 8 | 4 | 0 | . | . | . | . | . | . | . | . | G | G | G | G |
| NC_011770 | 1442582 | C | A | 12 | 8 | 4 | 0 | . | . | . | . | . | . | . | . | A | A | A | A |
| NC_011770 | 1442583 | T | C | 12 | 8 | 4 | 0 | . | . | . | . | . | . | . | . | C | C | C | C |
| NC_011770 | 1443661 | G | A | 12 | 8 | 4 | 0 | . | . | . | . | . | . | . | . | A | A | A | A |
| NC_011770 | 1443671 | C | G | 12 | 8 | 2 | 0 | . | . | . | . | . | . | . | . | G | G | | |
| NC_011770 | 1443687 | G | A | 14 | 4 | 4 | 0 | | | . | . | | | . | . | A | A | A | A |
| NC_011770 | 1443688 | TG | T | 14 | 4 | 4 | 0 | | | . | . | | | . | . | T | T | T | T |
| NC_011770 | 1443704 | T | C | 14 | 4 | 2 | 0 | | | . | . | | | . | . | | | C | C |
| NC_011770 | 1443816 | G | A | 3 | 10 | 2 | 0 | . | . | . | . | . | . | . | . | . | . | A | A |
| NC_011770 | 1444094 | C | G | 4 | 10 | 2 | 0 | . | . | . | . | . | . | . | . | . | . | G | G |
| NC_011770 | 1444503 | A | C | 6 | 10 | 2 | 0 | . | . | . | . | . | . | . | . | C | C | . | . |
| NC_011770 | 1444553 | C | A | 4 | 10 | 2 | 0 | . | . | . | . | . | . | . | . | A | A | . | . |
| NC_011770 | 1444567 | G | A | 4 | 10 | 2 | 0 | . | . | . | . | . | . | . | . | A | A | . | . |
| NC_011770 | 1444575 | T | C | 4 | 10 | 2 | 0 | . | . | . | . | . | . | . | . | C | C | . | . |
| NC_011770 | 1444578 | A | T | 4 | 10 | 2 | 0 | . | . | . | . | . | . | . | . | T | T | . | . |
| NC_011770 | 1444590 | G | A | 3 | 10 | 2 | 0 | . | . | . | . | . | . | . | . | A | A | . | . |
| NC_011770 | 1444592 | T | G | 3 | 10 | 2 | 0 | . | . | . | . | . | . | . | . | G | G | . | . |
| NC_011770 | 1444604 | G | T | 3 | 10 | 2 | 0 | . | . | . | . | . | . | . | . | T | T | . | . |
| NC_011770 | 1444894 | A | G | 3 | 10 | 2 | 0 | . | . | . | . | . | . | . | . | G | G | . | . |
| NC_011770 | 1446444 | G | A | 7 | 10 | 2 | 0 | . | . | . | . | . | . | . | . | A | A | . | . |
| NC_011770 | 1446445 | G | T | 7 | 10 | 2 | 0 | . | . | . | . | . | . | . | . | T | T | . | . |
| NC_011770 | 1446458 | T | C | 7 | 10 | 2 | 0 | . | . | . | . | . | . | . | . | C | C | . | . |
| NC_011770 | 1446493 | T | A | 6 | 10 | 2 | 0 | . | . | . | . | . | . | . | . | A | A | . | . |
| NC_011770 | 1446494 | T | A | 6 | 10 | 2 | 0 | . | . | . | . | . | . | . | . | A | A | . | . |
| NC_011770 | 1446560 | A | G | 7 | 10 | 2 | 0 | . | . | . | . | . | . | . | . | G | G | . | . |
| NC_011770 | 1446604 | A | G | 7 | 10 | 2 | 0 | . | . | . | . | . | . | . | . | G | G | . | . |
| NC_011770 | 1446619 | A | G | 7 | 10 | 2 | 0 | . | . | . | . | . | . | . | . | G | G | . | . |
| NC_011770 | 1446652 | A | G | 7 | 10 | 2 | 0 | . | . | . | . | . | . | . | . | G | G | . | . |
| NC_011770 | 1446697 | T | C | 7 | 10 | 2 | 0 | . | . | . | . | . | . | . | . | C | C | . | . |
| NC_011770 | 1446712 | T | C | 6 | 10 | 2 | 0 | . | . | . | . | . | . | . | . | C | C | . | . |
| NC_011770 | 1446715 | C | T | 6 | 10 | 2 | 0 | . | . | . | . | . | . | . | . | T | T | . | . |
| NC_011770 | 1446717 | G | A | 6 | 10 | 2 | 0 | . | . | . | . | . | . | . | . | A | A | . | . |
| NC_011770 | 1446721 | C | T | 6 | 10 | 2 | 0 | . | . | . | . | . | . | . | . | T | T | . | . |
| NC_011770 | 1446730 | C | T | 6 | 10 | 2 | 0 | . | . | . | . | . | . | . | . | T | T | . | . |
| NC_011770 | 1446769 | G | A | 6 | 10 | 2 | 0 | . | . | . | . | . | . | . | . | A | A | . | . |
| NC_011770 | 1446781 | C | T | 5 | 10 | 2 | 0 | . | . | . | . | . | . | . | . | T | T | . | . |
| NC_011770 | 1446818 | A | AG | 5 | 10 | 2 | 0 | . | . | . | . | . | . | . | . | AG | AG | . | . |
| NC_011770 | 1446833 | T | G | 5 | 10 | 2 | 0 | . | . | . | . | . | . | . | . | G | G | . | . |
| NC_011770 | 1446868 | G | A | 5 | 10 | 2 | 0 | . | . | . | . | . | . | . | . | A | A | . | . |
| NC_011770 | 1446890 | G | C | 13 | 8 | 4 | 0 | . | . | . | . | . | . | . | . | C | C | C | C |
| NC_011770 | 1446921 | G | A | 13 | 8 | 4 | 0 | . | . | . | . | . | . | . | . | A | A | A | A |
| NC_011770 | 1446944 | A | G | 14 | 8 | 2 | 0 | . | . | . | . | . | . | . | . | G | G | | |
| NC_011770 | 1446946 | A | C | 13 | 8 | 2 | 0 | . | . | . | . | . | . | . | . | C | C | | |
| NC_011770 | 1447716 | C | A | 10 | 10 | 2 | 0 | . | . | . | . | . | . | . | . | . | . | A | A |
| NC_011770 | 1449373 | C | T | 16 | 10 | 2 | 0 | . | . | . | . | . | . | . | . | . | . | T | T |
| NC_011770 | 1449382 | C | A | 16 | 10 | 2 | 0 | . | . | . | . | . | . | . | . | . | . | A | A |
| NC_011770 | 1449397 | T | C | 16 | 10 | 2 | 0 | . | . | . | . | . | . | . | . | . | . | C | C |
| NC_011770 | 1449405 | A | T | 15 | 10 | 2 | 0 | . | . | . | . | . | . | . | . | . | . | T | T |
| NC_011770 | 1449407 | C | T | 15 | 10 | 2 | 0 | . | . | . | . | . | . | . | . | . | . | T | T |
| NC_011770 | 1449418 | G | C | 15 | 10 | 2 | 0 | . | . | . | . | . | . | . | . | . | . | C | C |
| NC_011770 | 1449421 | G | A | 15 | 10 | 2 | 0 | . | . | . | . | . | . | . | . | . | . | A | A |
| NC_011770 | 1449442 | G | T | 15 | 10 | 2 | 0 | . | . | . | . | . | . | . | . | . | . | T | T |
| NC_011770 | 1449448 | C | T | 15 | 10 | 2 | 0 | . | . | . | . | . | . | . | . | . | . | T | T |
| NC_011770 | 1449461 | T | G | 15 | 10 | 2 | 0 | . | . | . | . | . | . | . | . | . | . | G | G |

| | | | | | | | | | | | | | | | | | | | |
|---|---|---|---|---|---|---|---|---|---|---|---|---|---|---|---|---|---|---|---|
| NC_011770 | 1449466 | C | G | 15 | 10 | 2 | 0 | . | . | . | . | . | . | . | . | . | . | G | G |
| NC_011770 | 1449472 | G | C | 15 | 10 | 2 | 0 | . | . | . | . | . | . | . | . | . | . | C | C |
| NC_011770 | 1449476 | T | G | 15 | 10 | 2 | 0 | . | . | . | . | . | . | . | . | . | . | G | G |
| NC_011770 | 1449492 | C | A | 15 | 10 | 2 | 0 | . | . | . | . | . | . | . | . | . | . | A | A |
| NC_011770 | 1449494 | A | G | 15 | 10 | 2 | 0 | . | . | . | . | . | . | . | . | . | . | G | G |
| NC_011770 | 1449505 | T | G | 15 | 10 | 2 | 0 | . | . | . | . | . | . | . | . | . | . | G | G |
| NC_011770 | 1449511 | G | C | 15 | 10 | 2 | 0 | . | . | . | . | . | . | . | . | . | . | C | C |
| NC_011770 | 1449514 | G | A | 16 | 10 | 2 | 0 | . | . | . | . | . | . | . | . | . | . | A | A |
| NC_011770 | 1450148 | A | G | 11 | 10 | 2 | 0 | . | . | . | . | . | . | . | . | . | . | G | G |
| NC_011770 | 1450160 | C | G | 12 | 10 | 2 | 0 | . | . | . | . | . | . | . | . | . | . | G | G |
| NC_011770 | 1450184 | C | T | 12 | 10 | 2 | 0 | . | . | . | . | . | . | . | . | . | . | T | T |
| NC_011770 | 1450199 | A | G | 12 | 10 | 2 | 0 | . | . | . | . | . | . | . | . | . | . | G | G |
| NC_011770 | 1450235 | T | C | 10 | 10 | 2 | 0 | . | . | . | . | . | . | . | . | . | . | C | C |
| NC_011770 | 1450250 | T | C | 10 | 10 | 2 | 0 | . | . | . | . | . | . | . | . | . | . | C | C |
| NC_011770 | 1450258 | T | C | 13 | 8 | 2 | 0 | . | . | . | . | . | . | . | . | . | . | C | C |
| NC_011770 | 1450274 | C | G | 14 | 8 | 4 | 0 | . | . | . | . | . | . | . | . | G | G | G | G |
| NC_011770 | 1450277 | T | G | 14 | 8 | 4 | 0 | . | . | . | . | . | . | . | . | G | G | G | G |
| NC_011770 | 1450323 | G | C | 13 | 8 | 4 | 0 | . | . | . | . | . | . | . | . | C | C | C | C |
| NC_011770 | 1450352 | T | G | 13 | 8 | 4 | 0 | . | . | . | . | . | . | . | . | G | G | G | G |
| NC_011770 | 1450377 | C | A | 3 | 10 | 2 | 0 | . | . | . | . | . | . | . | . | A | A | . | . |
| NC_011770 | 1450790 | G | A | 3 | 10 | 2 | 0 | . | . | . | . | . | . | . | . | A | A | . | . |
| NC_011770 | 1450810 | A | G | 3 | 10 | 2 | 0 | . | . | . | . | . | . | . | . | G | G | . | . |
| NC_011770 | 1450829 | G | C | 3 | 10 | 2 | 0 | . | . | . | . | . | . | . | . | C | C | . | . |
| NC_011770 | 1450889 | A | T | 2 | 10 | 2 | 0 | . | . | . | . | . | . | . | . | T | T | . | . |
| NC_011770 | 1450895 | C | T | 2 | 10 | 2 | 0 | . | . | . | . | . | . | . | . | T | T | . | . |
| NC_011770 | 1450902 | T | G | 2 | 10 | 2 | 0 | . | . | . | . | . | . | . | . | G | G | . | . |
| NC_011770 | 1450912 | A | T | 1 | 10 | 2 | 0 | . | . | . | . | . | . | . | . | T | T | . | . |
| NC_011770 | 1450913 | C | T | 1 | 10 | 2 | 0 | . | . | . | . | . | . | . | . | T | T | . | . |
| NC_011770 | 1450928 | T | C | 1 | 10 | 2 | 0 | . | . | . | . | . | . | . | . | C | C | . | . |
| NC_011770 | 1452754 | C | G | 3 | 7 | 2 | 0 | . | . | . | . | | | . | . | G | G | . | |
| NC_011770 | 1452757 | C | G | 4 | 6 | 2 | 0 | . | . | . | . | | | . | . | G | G | | |
| NC_011770 | 1452758 | C | T | 4 | 6 | 2 | 0 | . | . | . | . | | | . | . | T | T | | |
| NC_011770 | 1452763 | A | G | 4 | 6 | 2 | 0 | . | . | . | . | | | . | . | G | G | | |
| NC_011770 | 1452766 | T | C | 4 | 6 | 6 | 0 | . | . | . | . | C | C | . | . | G | G | C | C |
| NC_011770 | 1452769 | G | T | 4 | 6 | 2 | 0 | . | . | . | . | | | . | . | T | T | | |
| NC_011770 | 1452772 | G | T | 5 | 6 | 2 | 0 | . | . | . | . | | | . | . | | | T | T |
| NC_011770 | 1452856 | T | C | 8 | 6 | 6 | 0 | . | . | . | . | C | C | . | . | C | C | C | C |
| NC_011770 | 1452880 | T | C | 8 | 6 | 6 | 0 | . | . | . | . | C | C | . | . | C | C | C | C |
| NC_011770 | 1452882 | G | T | 7 | 6 | 6 | 0 | . | . | . | . | T | T | . | . | T | T | A | A |
| NC_011770 | 1452886 | G | A | 8 | 6 | 2 | 0 | . | . | . | . | | | . | . | | | A | A |
| NC_011770 | 1452889 | A | G | 8 | 6 | 6 | 0 | . | . | . | . | G | G | . | . | G | G | G | G |
| NC_011770 | 1452894 | C | T | 7 | 6 | 2 | 0 | . | . | . | . | | | . | . | | | T | T |
| NC_011770 | 1452895 | T | C | 8 | 6 | 6 | 0 | . | . | . | . | C | C | . | . | C | C | C | C |
| NC_011770 | 1452899 | A | G | 8 | 6 | 4 | 0 | . | . | . | . | G | G | . | . | G | G | | |
| NC_011770 | 1452905 | G | T | 7 | 6 | 2 | 0 | . | . | . | . | | | . | . | T | T | | |
| NC_011770 | 1452907 | A | C | 8 | 6 | 6 | 0 | . | . | . | . | C | C | . | . | C | C | C | C |
| NC_011770 | 1452908 | A | C | 8 | 6 | 6 | 0 | . | . | . | . | C | C | . | . | C | C | C | C |
| NC_011770 | 1452916 | C | A | 10 | 6 | 6 | 0 | . | . | . | . | A | A | . | . | A | A | A | A |
| NC_011770 | 1452919 | A | C | 11 | 6 | 6 | 0 | . | . | . | . | C | C | . | . | C | C | C | C |
| NC_011770 | 1453173 | G | C | 16 | 8 | 2 | 0 | . | . | . | . | C | C | . | . | . | . | | |
| NC_011770 | 1453190 | C | A | 16 | 8 | 4 | 0 | . | . | . | . | A | A | . | . | . | . | A | A |
| NC_011770 | 1453940 | G | T | 19 | 8 | 2 | 0 | . | . | . | . | T | T | . | . | . | . | | |
| NC_011770 | 1453941 | A | G | 19 | 8 | 2 | 0 | . | . | . | . | G | G | . | . | . | . | | |
| NC_011770 | 1453942 | A | G | 19 | 8 | 4 | 0 | . | . | . | . | G | G | . | . | . | . | G | G |
| NC_011770 | 1453943 | T | C | 19 | 8 | 2 | 0 | . | . | . | . | C | C | . | . | . | . | | |
| NC_011770 | 1453961 | C | T | 21 | 8 | 2 | 0 | . | . | . | . | | | . | . | . | . | T | T |
| NC_011770 | 1453962 | G | C | 21 | 8 | 2 | 0 | . | . | . | . | | | . | . | . | . | C | C |
| NC_011770 | 1453963 | C | T | 21 | 8 | 2 | 0 | . | . | . | . | | | . | . | . | . | T | T |
| NC_011770 | 1453971 | T | C | 22 | 8 | 4 | 0 | . | . | . | . | C | C | . | . | . | . | C | C |
| NC_011770 | 1453977 | G | C | 21 | 8 | 2 | 0 | . | . | . | . | C | C | . | . | . | . | | |
| NC_011770 | 1460833 | G | C | 15 | 8 | 2 | 0 | . | . | . | . | | | . | . | . | . | C | C |
| NC_011770 | 1460836 | G | C | 14 | 8 | 2 | 0 | . | . | . | . | | | . | . | . | . | C | C |

| | | | | | | | | | | | | | | | | | | | |
|---|---|---|---|---|---|---|---|---|---|---|---|---|---|---|---|---|---|---|---|
| NC_011770 | 1460872 | G | T | 13 | 8 | 2 | 0 | . | . | . | . | | | . | . | . | . | T | T |
| NC_011770 | 1460882 | G | C | 13 | 8 | 2 | 0 | . | . | . | . | | | . | . | . | . | C | C |
| NC_011770 | 1460888 | C | T | 12 | 8 | 4 | 0 | . | . | . | . | T | T | . | . | . | . | T | T |
| NC_011770 | 1460922 | G | A | 10 | 8 | 2 | 0 | . | . | . | . | | | . | . | . | . | A | A |
| NC_011770 | 1460923 | C | G | 10 | 8 | 2 | 0 | . | . | . | . | | | . | . | . | . | G | G |
| NC_011770 | 1460947 | T | G | 9 | 8 | 4 | 0 | . | . | . | . | G | G | . | . | . | . | G | G |
| NC_011770 | 1460959 | T | C | 7 | 8 | 4 | 0 | . | . | . | . | C | C | . | . | . | . | C | C |
| NC_011770 | 1460968 | C | G | 7 | 8 | 2 | 0 | . | . | . | . | | | . | . | . | . | G | G |
| NC_011770 | 1460970 | G | C | 6 | 8 | 2 | 0 | . | . | . | . | C | C | . | . | . | . | | |
| NC_011770 | 1460971 | T | C | 6 | 8 | 4 | 0 | . | . | . | . | C | C | . | . | . | . | C | C |
| NC_011770 | 1460977 | C | G | 6 | 8 | 4 | 0 | . | . | . | . | G | G | . | . | . | . | G | G |
| NC_011770 | 1460980 | A | G | 6 | 8 | 4 | 0 | . | . | . | . | G | G | . | . | . | . | G | G |
| NC_011770 | 1461669 | G | T | 11 | 8 | 2 | 0 | . | . | . | . | | | . | . | . | . | T | T |
| NC_011770 | 1461670 | T | C | 11 | 8 | 2 | 0 | . | . | . | . | | | . | . | . | . | C | C |
| NC_011770 | 1461687 | G | A | 12 | 8 | 2 | 0 | . | . | . | . | | | . | . | . | . | A | A |
| NC_011770 | 1461690 | G | C | 12 | 8 | 4 | 0 | . | . | . | . | C | C | . | . | . | . | C | C |
| NC_011770 | 1461707 | T | C | 13 | 8 | 4 | 0 | . | . | . | . | C | C | . | . | . | . | C | C |
| NC_011770 | 1461795 | A | G | 17 | 9 | 1 | 0 | . | . | . | . | | | . | . | . | . | G | . |
| NC_011770 | 1463298 | A | T | 16 | 8 | 2 | 0 | . | . | . | . | | | . | . | . | . | T | T |
| NC_011770 | 1463302 | T | TTA | 16 | 8 | 2 | 0 | . | . | . | . | | | . | . | . | . | TTA | TTA |
| NC_011770 | 1463311 | A | G | 16 | 8 | 2 | 0 | . | . | . | . | | | . | . | . | . | G | G |
| NC_011770 | 1463313 | G | C | 16 | 8 | 2 | 0 | . | . | . | . | | | . | . | . | . | C | C |
| NC_011770 | 1463324 | T | C | 15 | 8 | 2 | 0 | . | . | . | . | | | . | . | . | . | C | C |
| NC_011770 | 1463326 | T | C | 15 | 8 | 2 | 0 | . | . | . | . | | | . | . | . | . | C | C |
| NC_011770 | 1463364 | A | G | 16 | 8 | 2 | 0 | . | . | . | . | | | . | . | . | . | G | G |
| NC_011770 | 1463415 | G | A | 16 | 8 | 2 | 0 | . | . | . | . | | | . | . | . | . | A | A |
| NC_011770 | 1463451 | G | A | 15 | 8 | 2 | 0 | . | . | . | . | | | . | . | . | . | A | A |
| NC_011770 | 1463469 | C | T | 15 | 8 | 2 | 0 | . | . | . | . | | | . | . | . | . | T | T |
| NC_011770 | 1463536 | G | A | 14 | 8 | 2 | 0 | . | . | . | . | | | . | . | . | . | A | A |
| NC_011770 | 1463562 | T | C | 13 | 8 | 2 | 0 | . | . | . | . | C | C | . | . | . | . | | |
| NC_011770 | 1463582 | G | T | 13 | 8 | 2 | 0 | . | . | . | . | T | T | . | . | . | . | | |
| NC_011770 | 1463655 | T | A | 14 | 8 | 2 | 0 | . | . | . | . | | | . | . | . | . | A | A |
| NC_011770 | 1463657 | G | A | 14 | 8 | 2 | 0 | . | . | . | . | | | . | . | . | . | A | A |
| NC_011770 | 1463664 | T | C | 14 | 8 | 2 | 0 | . | . | . | . | C | C | . | . | . | . | | |
| NC_011770 | 1463723 | C | A | 11 | 8 | 2 | 0 | . | . | . | . | A | A | . | . | . | . | | |
| NC_011770 | 1471536 | T | C | 28 | 8 | 4 | 0 | . | . | . | . | C | C | . | . | . | . | C | C |
| NC_011770 | 1471537 | C | T | 28 | 8 | 2 | 0 | . | . | . | . | | | . | . | . | . | T | T |
| NC_011770 | 1471538 | T | C | 29 | 8 | 2 | 0 | . | . | . | . | | | . | . | . | . | C | C |
| NC_011770 | 1471544 | T | A | 28 | 8 | 2 | 0 | . | . | . | . | | | . | . | . | . | A | A |
| NC_011770 | 1472160 | A | C | 1 | 10 | 2 | 0 | . | . | . | . | C | C | . | . | . | . | . | . |
| NC_011770 | 1472229 | C | A | 0 | 10 | 1 | 0 | . | . | . | . | . | A | . | . | . | . | . | . |
| NC_011770 | 1472244 | C | A | 0 | 11 | 1 | 0 | . | . | . | . | A | . | . | . | . | . | . | . |
| NC_011770 | 1472920 | A | G | 13 | 8 | 4 | 0 | . | . | . | . | G | G | . | . | . | . | G | G |
| NC_011770 | 1472940 | T | C | 13 | 8 | 4 | 0 | . | . | . | . | C | C | . | . | . | . | A | A |
| NC_011770 | 1472981 | A | G | 14 | 8 | 2 | 0 | . | . | . | . | G | G | . | . | . | . | | |
| NC_011770 | 1472986 | A | C | 15 | 8 | 2 | 0 | . | . | . | . | | | . | . | . | . | C | C |
| NC_011770 | 1473052 | C | A | 12 | 8 | 2 | 0 | . | . | . | . | | | . | . | . | . | A | A |
| NC_011770 | 1473061 | T | C | 12 | 8 | 2 | 0 | . | . | . | . | | | . | . | . | . | C | C |
| NC_011770 | 1473166 | G | T | 7 | 10 | 2 | 0 | . | . | . | . | T | T | . | . | . | . | . | . |
| NC_011770 | 1476279 | G | T | 0 | 11 | 1 | 0 | . | . | . | . | . | . | . | . | T | . | . | . |
| NC_011770 | 1476342 | T | C | 6 | 9 | 1 | 0 | . | . | . | . | . | . | . | . | C | . | | |
| NC_011770 | 1476350 | A | AAC | 6 | 9 | 1 | 0 | . | . | . | . | . | . | . | . | AAC | . | | |
| NC_011770 | 1476352 | AAG | A | 6 | 9 | 1 | 0 | . | . | . | . | . | . | . | . | A | . | | |
| NC_011770 | 1476371 | T | A | 6 | 9 | 3 | 0 | . | . | . | . | . | . | . | . | A | . | A | A |
| NC_011770 | 1476375 | C | A | 7 | 9 | 1 | 0 | . | . | . | . | . | . | . | . | A | . | | |
| NC_011770 | 1476387 | C | T | 7 | 9 | 1 | 0 | . | . | . | . | . | . | . | . | T | . | | |
| NC_011770 | 1476393 | A | G | 7 | 9 | 1 | 0 | . | . | . | . | . | . | . | . | G | . | | |
| NC_011770 | 1476454 | C | A | 7 | 10 | 2 | 0 | . | . | . | . | . | . | . | . | . | . | A | A |
| NC_011770 | 1476468 | C | A | 7 | 10 | 2 | 0 | . | . | . | . | . | . | . | . | . | . | A | A |
| NC_011770 | 1476490 | T | G | 7 | 10 | 2 | 0 | . | . | . | . | . | . | . | . | . | . | G | G |
| NC_011770 | 1476498 | A | G | 6 | 10 | 2 | 0 | . | . | . | . | . | . | . | . | . | . | G | G |
| NC_011770 | 1476515 | A | C | 7 | 10 | 2 | 0 | . | . | . | . | . | . | . | . | . | . | C | C |

| | | | | | | | | | | | | | | | | | | | |
|---|---|---|---|---|---|---|---|---|---|---|---|---|---|---|---|---|---|---|---|
| NC_011770 | 1476528 | G | A | 7 | 10 | 2 | 0 | . | . | . | . | . | . | . | . | . | . | A | A |
| NC_011770 | 1476620 | A | G | 34 | 0 | 2 | 0 | | | | | | | | | | | G | G |
| NC_011770 | 1476641 | C | T | 34 | 0 | 4 | 0 | T | T | | | | | | | | | T | T |
| NC_011770 | 1476807 | A | T | 37 | 0 | 6 | 0 | | | T | T | T | T | T | T | | | | |
| NC_011770 | 1476863 | C | T | 46 | 0 | 1 | 0 | | | | | | | T | | | | | |
| NC_011770 | 1477245 | C | A | 49 | 0 | 2 | 0 | | | | | | | | | | | A | A |
| NC_011770 | 1477666 | T | C | 59 | 0 | 8 | 0 | | | C | C | C | C | C | C | C | C | | |
| NC_011770 | 1478232 | T | A | 48 | 0 | 2 | 0 | | | | | | | | | | | A | A |
| NC_011770 | 1484211 | G | T | 71 | 0 | 2 | 0 | | | | | | | | | T | T | | |
| NC_011770 | 1484272 | C | T | 63 | 0 | 6 | 0 | T | T | | | | | | | T | T | T | T |
| NC_011770 | 1485299 | C | T | 74 | 0 | 2 | 0 | | | | | | | | | | | T | T |
| NC_011770 | 1486205 | C | T | 69 | 0 | 2 | 0 | | | | | | | | | T | T | | |
| NC_011770 | 1487206 | C | CA | 58 | 0 | 2 | 0 | | | | | | | CA | CA | | | | |
| NC_011770 | 1487260 | C | T | 58 | 0 | 4 | 0 | | | | | | | T | T | T | T | | |
| NC_011770 | 1499881 | T | C | 73 | 0 | 4 | 0 | | | C | C | C | C | | | | | | |
| NC_011770 | 1499883 | A | G | 74 | 0 | 4 | 0 | | | G | G | G | G | | | | | | |
| NC_011770 | 1499895 | T | C | 76 | 0 | 2 | 0 | C | C | | | | | | | | | | |
| NC_011770 | 1499917 | A | G | 74 | 0 | 2 | 0 | G | G | | | | | | | | | | |
| NC_011770 | 1499984 | G | A | 79 | 0 | 2 | 0 | A | A | | | | | | | | | | |
| NC_011770 | 1506248 | G | A | 68 | 0 | 2 | 0 | | | | | | | | | | | A | A |
| NC_011770 | 1506261 | A | C | 64 | 0 | 12 | 0 | C | C | C | C | C | C | C | C | C | C | C | C |
| NC_011770 | 1506297 | G | T | 63 | 0 | 4 | 0 | | | T | T | T | T | | | | | | |
| NC_011770 | 1512721 | C | A | 62 | 0 | 4 | 0 | | | | | | | A | A | A | A | | |
| NC_011770 | 1516455 | C | T | 76 | 0 | 4 | 0 | | | T | T | T | T | | | | | | |
| NC_011770 | 1516473 | C | CG | 81 | 0 | 4 | 0 | | | CG | CG | CG | CG | | | | | | |
| NC_011770 | 1517378 | T | C | 89 | 0 | 12 | 0 | C | C | C | C | C | C | C | C | C | C | C | C |
| NC_011770 | 1517397 | A | G | 91 | 0 | 12 | 0 | G | G | G | G | G | G | G | G | G | G | G | G |
| NC_011770 | 1519702 | G | A | 71 | 0 | 2 | 0 | | | | | | | | | A | A | | |
| NC_011770 | 1522404 | CT | C | 64 | 0 | 2 | 0 | | | | | | | | | C | C | | |
| NC_011770 | 1522549 | T | A | 55 | 0 | 4 | 0 | | | A | A | A | A | | | | | | |
| NC_011770 | 1522575 | G | A | 56 | 0 | 4 | 0 | | | A | A | A | A | | | | | | |
| NC_011770 | 1522618 | G | T | 57 | 0 | 2 | 0 | T | T | | | | | | | | | | |
| NC_011770 | 1522622 | T | A | 56 | 0 | 4 | 0 | | | A | A | A | A | | | | | | |
| NC_011770 | 1522623 | A | G | 57 | 0 | 10 | 0 | G | G | G | G | G | G | G | G | G | G | | |
| NC_011770 | 1522804 | T | A | 53 | 0 | 2 | 0 | A | A | | | | | | | | | | |
| NC_011770 | 1525821 | C | T | 46 | 0 | 4 | 0 | | | | | | | T | T | | | T | T |
| NC_011770 | 1525838 | A | G | 44 | 0 | 4 | 0 | | | | | | | G | G | | | G | G |
| NC_011770 | 1527571 | G | A | 84 | 0 | 2 | 0 | | | | | | | | | A | A | | |
| NC_011770 | 1530337 | C | T | 81 | 0 | 2 | 0 | | | | | | | T | T | | | | |
| NC_011770 | 1534463 | A | G | 35 | 0 | 6 | 0 | G | G | G | G | G | G | | | | | | |
| NC_011770 | 1535238 | C | T | 57 | 0 | 6 | 0 | | | T | T | T | T | T | T | | | | |
| NC_011770 | 1535258 | G | T | 57 | 0 | 6 | 0 | | | T | T | T | T | T | T | | | | |
| NC_011770 | 1535275 | C | CA | 58 | 0 | 2 | 0 | | | | | | | | | | | CA | CA |
| NC_011770 | 1536456 | A | C | 70 | 0 | 4 | 0 | C | C | | | | | C | C | | | | |
| NC_011770 | 1538133 | C | G | 84 | 0 | 6 | 0 | G | G | | | | | | | G | G | G | G |
| NC_011770 | 1539339 | A | G | 66 | 0 | 4 | 0 | | | | | | | G | G | | | G | G |
| NC_011770 | 1539363 | A | C | 65 | 0 | 10 | 0 | | | C | C | C | C | C | C | C | C | C | C |
| NC_011770 | 1539428 | T | A | 68 | 0 | 2 | 0 | | | | | | | | | A | A | | |
| NC_011770 | 1540202 | T | C | 81 | 0 | 4 | 0 | | | | | | | | | C | C | C | C |
| NC_011770 | 1546461 | G | A | 75 | 0 | 8 | 0 | A | A | | | | | A | A | A | A | A | A |
| NC_011770 | 1546551 | G | T | 85 | 0 | 8 | 0 | T | T | | | | | T | T | T | T | T | T |
| NC_011770 | 1546591 | C | T | 84 | 0 | 4 | 0 | | | T | T | T | T | | | | | | |
| NC_011770 | 1546601 | C | T | 82 | 0 | 4 | 0 | | | T | T | T | T | | | | | | |
| NC_011770 | 1547977 | G | GA | 81 | 0 | 12 | 0 | GA | GA | GA | GA | GA | GA | GA | GA | GA | GA | GA | GA |
| NC_011770 | 1551722 | G | C | 69 | 0 | 4 | 0 | C | C | | | | | C | C | | | | |
| NC_011770 | 1554724 | T | G | 80 | 0 | 2 | 0 | | | | | | | | | G | G | | |
| NC_011770 | 1557110 | C | G | 79 | 0 | 2 | 0 | G | G | | | | | | | | | | |
| NC_011770 | 1558368 | T | C | 52 | 0 | 12 | 0 | C | C | C | C | C | C | C | C | C | C | C | C |
| NC_011770 | 1558373 | T | C | 59 | 0 | 4 | 0 | | | | | | | C | C | | | C | C |
| NC_011770 | 1559636 | C | T | 89 | 0 | 4 | 0 | | | | | | | T | T | | | T | T |
| NC_011770 | 1559639 | T | C | 88 | 0 | 6 | 0 | | | | | | | C | C | C | C | C | C |
| NC_011770 | 1560650 | G | A | 85 | 0 | 2 | 0 | | | | | | | | | A | A | | |

| | | | | | | | | | | | | | | | | | | | |
|---|---|---|---|---|---|---|---|---|---|---|---|---|---|---|---|---|---|---|---|
| NC_011770 | 1560728 | A | G | 56 | 0 | 2 | 0 | | | | | | | | | G | G | | |
| NC_011770 | 1561849 | C | T | 82 | 0 | 2 | 0 | | | | | | | T | T | | | | |
| NC_011770 | 1562240 | T | C | 60 | 0 | 2 | 0 | C | C | | | | | | | | | | |
| NC_011770 | 1563896 | T | A | 64 | 0 | 2 | 0 | A | A | | | | | | | | | | |
| NC_011770 | 1563917 | C | T | 65 | 0 | 2 | 0 | | | | | | | | | | | T | T |
| NC_011770 | 1563951 | T | G | 61 | 0 | 4 | 0 | G | G | | | | | | | G | G | | |
| NC_011770 | 1563954 | C | T | 60 | 0 | 4 | 0 | T | T | | | | | | | T | T | | |
| NC_011770 | 1563985 | G | A | 57 | 0 | 2 | 0 | | | | | | | A | A | | | | |
| NC_011770 | 1564011 | G | T | 57 | 0 | 2 | 0 | | | | | | | | | | | T | T |
| NC_011770 | 1564017 | C | A | 48 | 0 | 8 | 0 | A | A | | | | | A | A | A | A | A | A |
| NC_011770 | 1564031 | A | G | 53 | 0 | 4 | 0 | | | | | | | | | G | G | G | G |
| NC_011770 | 1564047 | G | A | 52 | 0 | 2 | 0 | A | A | | | | | | | | | | |
| NC_011770 | 1564068 | A | G | 50 | 0 | 8 | 0 | G | G | | | | | G | G | G | G | G | G |
| NC_011770 | 1564069 | G | C | 51 | 0 | 8 | 0 | C | C | | | | | C | C | C | C | C | C |
| NC_011770 | 1564075 | G | T | 50 | 0 | 2 | 0 | T | T | | | | | | | | | | |
| NC_011770 | 1564086 | T | C | 50 | 0 | 8 | 0 | C | C | | | | | C | C | C | C | C | C |
| NC_011770 | 1564131 | GA | G | 53 | 0 | 4 | 0 | | | | | | | | | G | G | G | G |
| NC_011770 | 1564132 | A | G | 53 | 0 | 4 | 0 | G | G | | | | | G | G | . | . | . | . |
| NC_011770 | 1564954 | CG | CGG | 55 | 0 | 2 | 0 | | | | | | | CGG | C | | | | |
| NC_011770 | 1565052 | G | C | 65 | 0 | 8 | 0 | | | C | C | C | C | | | C | C | C | C |
| NC_011770 | 1565053 | C | G | 66 | 0 | 4 | 0 | | | G | G | G | G | | | | | | |
| NC_011770 | 1565063 | T | C | 67 | 0 | 12 | 0 | C | C | C | C | C | C | C | C | C | C | C | C |
| NC_011770 | 1565083 | G | T | 68 | 0 | 4 | 0 | | | | | | | T | T | | | T | T |
| NC_011770 | 1566651 | G | A | 85 | 0 | 6 | 0 | | | A | A | A | A | A | A | | | | |
| NC_011770 | 1567657 | C | G | 38 | 0 | 2 | 0 | | | | | | | | | | | G | G |
| NC_011770 | 1567730 | G | A | 41 | 0 | 4 | 0 | | | A | A | A | A | | | | | | |
| NC_011770 | 1568241 | A | ATC | 62 | 0 | 4 | 0 | | | ATC | ATC | ATC | ATC | | | | | | |
| NC_011770 | 1568251 | C | T | 64 | 0 | 2 | 0 | | | | | | | | | | | T | T |
| NC_011770 | 1569787 | G | A | 77 | 0 | 2 | 0 | | | | | | | | | | | A | A |
| NC_011770 | 1570762 | AG | A | 50 | 0 | 2 | 0 | | | | | | | | | A | A | | |
| NC_011770 | 1570766 | G | C | 45 | 2 | 2 | 0 | | | | | | | C | C | . | . | | |
| NC_011770 | 1570803 | C | T | 46 | 0 | 6 | 0 | T | T | T | T | T | T | | | | | | |
| NC_011770 | 1570813 | T | C | 42 | 0 | 10 | 0 | C | C | C | C | C | C | C | C | | | C | C |
| NC_011770 | 1570831 | G | A | 37 | 0 | 4 | 0 | | . | A | A | A | A | | | | | | |
| NC_011770 | 1570832 | G | A | 38 | 0 | 4 | 0 | | | A | A | A | A | | | | | | |
| NC_011770 | 1570838 | A | G | 35 | 0 | 4 | 0 | | | G | G | G | G | | | | | | |
| NC_011770 | 1570841 | C | T | 35 | 0 | 2 | 0 | | | | | | | | | | | T | T |
| NC_011770 | 1570845 | C | T | 33 | 0 | 8 | 0 | | | T | T | T | T | T | T | | | T | T |
| NC_011770 | 1570866 | C | CG | 21 | 4 | 2 | 0 | | | . | . | . | . | CG | CG | | | | |
| NC_011770 | 1570915 | A | G | 19 | 6 | 2 | 0 | . | . | . | . | . | . | G | G | | | | |
| NC_011770 | 1570921 | G | A | 17 | 6 | 2 | 0 | . | . | . | . | . | . | A | A | | | | |
| NC_011770 | 1570948 | T | C | 17 | 6 | 6 | 0 | . | . | . | . | . | . | C | C | C | C | C | C |
| NC_011770 | 1571010 | G | A | 16 | 6 | 2 | 0 | . | . | . | . | . | . | | | A | A | | |
| NC_011770 | 1571069 | G | A | 12 | 8 | 4 | 0 | . | . | . | . | . | . | . | . | A | A | A | A |
| NC_011770 | 1571116 | C | A | 12 | 7 | 2 | 0 | . | . | . | . | . | . | . | . | A | A | | |
| NC_011770 | 1571202 | A | G | 12 | 6 | 6 | 0 | . | . | . | . | . | . | G | G | G | G | G | G |
| NC_011770 | 1571208 | C | A | 13 | 6 | 4 | 0 | . | . | . | . | . | . | A | A | | | A | A |
| NC_011770 | 1571316 | A | G | 27 | 2 | 2 | 0 | | | G | . | G | . | | | | | | |
| NC_011770 | 1571403 | A | G | 19 | 6 | 4 | 0 | . | . | . | . | . | . | G | G | | | G | G |
| NC_011770 | 1571465 | C | T | 14 | 6 | 4 | 0 | . | . | . | . | . | . | T | T | | | T | T |
| NC_011770 | 1571499 | T | C | 11 | 6 | 4 | 0 | . | . | . | . | . | . | C | C | | | C | C |
| NC_011770 | 1571500 | T | G | 11 | 6 | 4 | 0 | . | . | . | . | . | . | G | G | | | G | G |
| NC_011770 | 1571506 | GC | G | 11 | 6 | 2 | 0 | . | . | . | . | . | . | G | G | | | | |
| NC_011770 | 1571508 | C | T | 10 | 8 | 2 | 0 | . | . | . | . | . | . | . | . | | | T | T |
| NC_011770 | 1571529 | T | C | 9 | 6 | 2 | 0 | . | . | . | . | . | . | C | C | | | | |
| NC_011770 | 1571538 | G | A | 9 | 6 | 4 | 0 | . | . | . | . | . | . | A | A | | | A | A |
| NC_011770 | 1571539 | A | C | 9 | 6 | 4 | 0 | . | . | . | . | . | . | C | C | | | C | C |
| NC_011770 | 1571541 | G | C | 9 | 6 | 2 | 0 | . | . | . | . | . | . | C | C | | | | |
| NC_011770 | 1571542 | C | T | 9 | 6 | 4 | 0 | . | . | . | . | . | . | T | T | | | T | T |
| NC_011770 | 1571544 | C | G | 9 | 6 | 4 | 0 | . | . | . | . | . | . | G | G | | | G | G |
| NC_011770 | 1571546 | C | T | 9 | 6 | 4 | 0 | . | . | . | . | . | . | T | T | | | T | T |
| NC_011770 | 1571548 | GC | G | 9 | 6 | 4 | 0 | . | . | . | . | . | . | G | G | | | G | G |

| | | | | | | | | | | | | | | | | | | | |
|---|---|---|---|---|---|---|---|---|---|---|---|---|---|---|---|---|---|---|---|
| NC_011770 | 1571672 | A | G | 34 | 6 | 4 | 0 | . | . | . | . | . | . | G | G | G | G | | |
| NC_011770 | 1571691 | C | T | 36 | 6 | 4 | 0 | . | . | . | . | . | . | T | T | T | T | | |
| NC_011770 | 1571739 | C | T | 34 | 6 | 6 | 0 | . | . | . | . | . | . | T | T | T | T | T | T |
| NC_011770 | 1572621 | C | T | 41 | 6 | 2 | 0 | . | . | . | . | . | . | T | T | | | | |
| NC_011770 | 1572703 | C | G | 60 | 0 | 8 | 0 | G | G | G | G | G | G | | | G | G | | |
| NC_011770 | 1572711 | T | G | 57 | 0 | 6 | 0 | G | G | G | G | G | G | | | | | | |
| NC_011770 | 1572716 | C | T | 58 | 0 | 8 | 0 | T | T | T | T | T | T | | | T | T | | |
| NC_011770 | 1572744 | C | T | 58 | 0 | 2 | 0 | | | | | | | | | T | T | | |
| NC_011770 | 1572884 | G | A | 35 | 6 | 6 | 0 | . | . | . | . | . | . | A | A | A | A | A | A |
| NC_011770 | 1572917 | T | A | 60 | 0 | 4 | 0 | | | A | A | A | A | | | | | | |
| NC_011770 | 1572971 | G | A | 73 | 0 | 2 | 0 | | | | | | | | | | | A | A |
| NC_011770 | 1572972 | C | T | 71 | 0 | 6 | 0 | T | T | T | T | T | T | | | | | | |
| NC_011770 | 1572986 | G | A | 66 | 0 | 2 | 0 | | | | | | | | | | | A | A |
| NC_011770 | 1572988 | A | G | 67 | 0 | 2 | 0 | | | | | | | | | | | G | G |
| NC_011770 | 1572999 | G | C | 65 | 0 | 8 | 0 | C | C | C | C | C | C | | | | | C | C |
| NC_011770 | 1573002 | G | C | 64 | 0 | 8 | 0 | C | C | C | C | C | C | | | | | C | C |
| NC_011770 | 1573012 | A | C | 62 | 0 | 8 | 0 | C | C | C | C | C | C | | | | | C | C |
| NC_011770 | 1573028 | C | A | 40 | 2 | 2 | 0 | | | . | . | A | A | | | | | | |
| NC_011770 | 1573070 | T | C | 72 | 0 | 6 | 0 | C | C | C | C | C | C | | | | | | |
| NC_011770 | 1573074 | G | C | 71 | 0 | 2 | 0 | C | C | | | | | | | | | | |
| NC_011770 | 1573127 | A | T | 76 | 0 | 4 | 0 | | | | | | | | | T | T | T | T |
| NC_011770 | 1573140 | G | A | 78 | 0 | 6 | 0 | A | A | A | A | A | A | | | | | | |
| NC_011770 | 1573142 | TG | T | 76 | 0 | 4 | 0 | | | | | | | . | | T | T | T | T |
| NC_011770 | 1573143 | G | A | 47 | 4 | 2 | 0 | | | | | | | A | A | . | . | . | . |
| NC_011770 | 1573160 | T | C | 79 | 0 | 12 | 0 | C | C | C | C | C | C | C | C | C | C | C | C |
| NC_011770 | 1573200 | G | A | 81 | 0 | 2 | 0 | A | A | | | | | | | | | | |
| NC_011770 | 1573210 | C | G | 79 | 0 | 8 | 0 | G | G | | | | | G | G | G | G | G | G |
| NC_011770 | 1573265 | C | T | 71 | 0 | 4 | 0 | | | | | | | | | T | T | T | T |
| NC_011770 | 1573288 | T | TG | 69 | 0 | 4 | 0 | | | TG | TG | TG | TG | | | | | | |
| NC_011770 | 1573355 | A | C | 69 | 0 | 4 | 0 | | | C | C | C | C | | | | | | |
| NC_011770 | 1573364 | C | G | 69 | 0 | 2 | 0 | | | | | | | | | | | G | G |
| NC_011770 | 1573370 | T | C | 66 | 0 | 6 | 0 | | | | | | | C | C | C | C | C | C |
| NC_011770 | 1573377 | C | T | 66 | 0 | 4 | 0 | | | T | T | T | T | | | | | | |
| NC_011770 | 1573829 | C | CG | 50 | 0 | 8 | 0 | | | CG | CG | CG | CG | CG | CG | | | CG | CG |
| NC_011770 | 1573839 | G | A | 54 | 0 | 6 | 0 | | | A | A | A | A | | | | | A | A |
| NC_011770 | 1573910 | G | A | 63 | 0 | 2 | 0 | A | A | | | | | | | | | | |
| NC_011770 | 1574769 | A | G | 98 | 0 | 2 | 0 | | | | | | | | | | | G | G |
| NC_011770 | 1575460 | C | G | 77 | 0 | 4 | 0 | | | G | G | G | G | | | | | | |
| NC_011770 | 1575561 | T | C | 72 | 0 | 6 | 0 | | | C | C | C | C | | | | | C | C |
| NC_011770 | 1575576 | T | A | 68 | 0 | 5 | 0 | | | A | A | A | A | C | | | | | |
| NC_011770 | 1575605 | C | T | 66 | 0 | 4 | 0 | | | T | T | T | T | | | | | | |
| NC_011770 | 1575664 | C | T | 70 | 0 | 2 | 0 | | | | | | | | | T | T | | |
| NC_011770 | 1575752 | CCG | C | 66 | 0 | 2 | 0 | | | | | | | | | | | C | C |
| NC_011770 | 1575768 | C | T | 68 | 0 | 2 | 0 | | | | | | | | | T | T | | |
| NC_011770 | 1577183 | G | A | 67 | 0 | 2 | 0 | | | | | | | | | | | A | A |
| NC_011770 | 1577290 | C | T | 66 | 0 | 2 | 0 | | | | | | | | | | | T | T |
| NC_011770 | 1577291 | G | C | 67 | 0 | 2 | 0 | | | | | | | | | | | C | C |
| NC_011770 | 1579850 | A | G | 61 | 0 | 2 | 0 | | | | | | | | | | | G | G |
| NC_011770 | 1579851 | C | T | 63 | 0 | 2 | 0 | | | | | | | | | T | T | | |
| NC_011770 | 1580757 | C | T | 86 | 0 | 4 | 0 | | | T | T | T | T | | | | | | |
| NC_011770 | 1580782 | T | C | 82 | 0 | 2 | 0 | | | | | | | | | | | C | C |
| NC_011770 | 1580790 | G | A | 82 | 0 | 2 | 0 | | | | | | | A | A | | | | |
| NC_011770 | 1582783 | C | T | 80 | 0 | 2 | 0 | | | | | | | | | | | T | T |
| NC_011770 | 1583817 | A | G | 43 | 0 | 2 | 0 | | | | | | | | | G | G | | |
| NC_011770 | 1583818 | A | G | 43 | 0 | 2 | 0 | | | | | | | | | G | G | | |
| NC_011770 | 1585270 | A | G | 73 | 2 | 10 | 0 | G | G | G | G | G | G | G | G | . | . | G | G |
| NC_011770 | 1587326 | G | GC | 70 | 0 | 12 | 0 | GC | GC | GC | GC | GC | GC | GC | GC | GC | GC | GC | GC |
| NC_011770 | 1587413 | A | T | 72 | 0 | 2 | 0 | T | T | | | | | | | | | | |
| NC_011770 | 1593147 | G | T | 83 | 0 | 4 | 0 | | | T | T | T | T | | | | | | |
| NC_011770 | 1593157 | A | G | 83 | 0 | 4 | 0 | G | G | | | | | | | | | G | G |
| NC_011770 | 1593232 | T | C | 52 | 0 | 6 | 0 | | | C | C | C | C | | | C | C | | |
| NC_011770 | 1593264 | C | T | 82 | 0 | 2 | 0 | | | | | | | | | T | T | | |

| | | | | | | | | | | | | | | | | | | | |
|---|---|---|---|---|---|---|---|---|---|---|---|---|---|---|---|---|---|---|---|
| NC_011770 | 1602240 | G | T | 64 | 0 | 4 | 0 | | | T | T | T | T | | | | | | |
| NC_011770 | 1602270 | T | A | 61 | 0 | 6 | 0 | | | A | A | A | A | A | A | | | | |
| NC_011770 | 1602310 | C | T | 57 | 0 | 4 | 0 | | | T | T | T | T | | | | | | |
| NC_011770 | 1602343 | T | C | 57 | 0 | 4 | 0 | | | C | C | C | C | | | | | | |
| NC_011770 | 1602435 | A | G | 52 | 0 | 8 | 0 | | | G | G | G | G | G | G | | | G | G |
| NC_011770 | 1604105 | C | T | 56 | 0 | 2 | 0 | | | | | | | | | T | T | | |
| NC_011770 | 1607775 | T | C | 68 | 0 | 2 | 0 | | | | | | | C | C | | | | |
| NC_011770 | 1617788 | G | C | 93 | 0 | 2 | 0 | C | C | | | | | | | | | | |
| NC_011770 | 1617793 | A | AGG | 91 | 0 | 2 | 0 | AGG | AG | | | | | | | | | | |
| NC_011770 | 1619210 | A | G | 87 | 0 | 2 | 0 | G | G | | | | | | | | | | |
| NC_011770 | 1619379 | C | A | 71 | 0 | 2 | 0 | | | | | | | A | A | | | | |
| NC_011770 | 1619426 | A | G | 66 | 0 | 8 | 0 | | | G | G | G | G | G | G | | | G | G |
| NC_011770 | 1619593 | G | T | 68 | 0 | 2 | 0 | | | | | | | | | | | T | T |
| NC_011770 | 1619718 | C | T | 68 | 0 | 2 | 0 | | | | | | | | | | | T | T |
| NC_011770 | 1619749 | A | G | 65 | 0 | 2 | 0 | G | G | | | | | | | | | | |
| NC_011770 | 1619752 | G | A | 65 | 0 | 2 | 0 | A | A | | | | | | | | | | |
| NC_011770 | 1619805 | T | C | 62 | 0 | 2 | 0 | | | | | | | | | | | C | C |
| NC_011770 | 1619814 | G | A | 60 | 0 | 2 | 0 | | | | | | | | | | | A | A |
| NC_011770 | 1619820 | T | C | 60 | 0 | 2 | 0 | | | | | | | | | | | C | C |
| NC_011770 | 1619821 | C | T | 61 | 0 | 6 | 0 | | | T | T | T | T | | | | | A | A |
| NC_011770 | 1619822 | T | G | 61 | 0 | 2 | 0 | | | | | | | | | | | G | G |
| NC_011770 | 1619823 | G | A | 61 | 0 | 2 | 0 | A | A | | | | | | | | | | |
| NC_011770 | 1619833 | C | T | 59 | 0 | 2 | 0 | | | | | | | | | | | T | T |
| NC_011770 | 1619858 | C | A | 62 | 0 | 2 | 0 | | | | | | | | | A | A | | |
| NC_011770 | 1619861 | G | A | 60 | 0 | 2 | 0 | A | A | | | | | | | | | | |
| NC_011770 | 1619921 | G | A | 57 | 0 | 2 | 0 | | | | | | | | | | | A | A |
| NC_011770 | 1619924 | A | G | 57 | 0 | 2 | 0 | | | | | | | | | | | G | G |
| NC_011770 | 1619939 | C | G | 57 | 0 | 4 | 0 | | | G | G | G | G | | | | | | |
| NC_011770 | 1619953 | G | C | 54 | 0 | 2 | 0 | | | | | | | | | | | C | C |
| NC_011770 | 1619954 | T | TCC | 54 | 0 | 2 | 0 | | | | | | | | | | | TCC | TCC |
| NC_011770 | 1620001 | A | G | 56 | 0 | 2 | 0 | | | | | | | | | | | G | G |
| NC_011770 | 1620010 | C | CG | 54 | 0 | 2 | 0 | | | | | | | | | | | CG | CG |
| NC_011770 | 1620017 | A | G | 48 | 0 | 2 | 0 | | | | | | | | | | | G | G |
| NC_011770 | 1623065 | G | A | 59 | 0 | 2 | 0 | | | | | | | | | A | A | | |
| NC_011770 | 1623102 | G | C | 53 | 0 | 8 | 0 | C | C | | | | | C | C | A | A | C | C |
| NC_011770 | 1623170 | C | T | 53 | 0 | 8 | 0 | T | T | T | T | T | T | | | T | T | | |
| NC_011770 | 1623181 | G | A | 54 | 0 | 2 | 0 | | | | | | | A | A | | | | |
| NC_011770 | 1623662 | C | A | 31 | 0 | 8 | 0 | A | A | A | A | A | A | A | A | | | | |
| NC_011770 | 1623694 | G | C | 26 | 0 | 8 | 0 | C | C | C | C | C | C | | | C | C | | |
| NC_011770 | 1623729 | A | G | 24 | 0 | 8 | 0 | G | G | G | G | G | G | | | G | G | | |
| NC_011770 | 1623736 | C | T | 23 | 0 | 2 | 0 | | | | | | | | | | | T | T |
| NC_011770 | 1627073 | C | T | 73 | 0 | 12 | 0 | T | T | T | T | T | T | T | T | T | T | T | T |
| NC_011770 | 1627101 | A | G | 71 | 0 | 12 | 0 | G | G | G | G | G | G | G | G | G | G | G | G |
| NC_011770 | 1627303 | G | A | 69 | 0 | 12 | 0 | A | A | A | A | A | A | A | A | A | A | A | A |
| NC_011770 | 1629895 | C | T | 61 | 0 | 2 | 0 | | | | | | | | | T | T | | |
| NC_011770 | 1631126 | C | CG | 54 | 0 | 2 | 0 | | | | | | | | | CG | CG | | |
| NC_011770 | 1631145 | G | A | 52 | 0 | 2 | 0 | | | | | | | | | A | A | | |
| NC_011770 | 1631155 | C | T | 52 | 0 | 2 | 0 | | | | | | | | | T | T | | |
| NC_011770 | 1631686 | G | T | 65 | 0 | 4 | 0 | T | T | | | | | T | T | | | | |
| NC_011770 | 1631698 | C | T | 64 | 0 | 10 | 0 | T | T | T | T | T | T | T | T | T | T | | |
| NC_011770 | 1631724 | A | G | 65 | 0 | 2 | 0 | | | G | G | | | | | | | | |
| NC_011770 | 1631819 | A | AG | 23 | 0 | 2 | 0 | AG | AG | | | | | | | | | | |
| NC_011770 | 1632187 | C | T | 63 | 0 | 2 | 0 | T | T | | | | | | | | | | |
| NC_011770 | 1632250 | T | C | 66 | 0 | 4 | 0 | | | C | C | C | C | | | | | | |
| NC_011770 | 1632251 | T | C | 65 | 0 | 4 | 0 | | | C | C | C | C | | | | | | |
| NC_011770 | 1632269 | G | T | 64 | 0 | 2 | 0 | T | T | | | | | | | | | | |
| NC_011770 | 1632273 | C | G | 66 | 0 | 2 | 0 | G | G | | | | | | | | | | |
| NC_011770 | 1632276 | C | A | 68 | 0 | 2 | 0 | | | | | | | | | A | A | | |
| NC_011770 | 1632289 | A | G | 68 | 0 | 2 | 0 | G | G | | | | | | | | | | |
| NC_011770 | 1632315 | G | T | 67 | 0 | 2 | 0 | | | | | | | | | T | T | | |
| NC_011770 | 1632974 | A | G | 58 | 2 | 2 | 0 | . | . | | | | | | | | | G | G |
| NC_011770 | 1633008 | T | C | 62 | 0 | 4 | 0 | | | C | C | C | C | | | | | | |

| | | | | | | | | | | | | | | | | | | | |
|---|---|---|---|---|---|---|---|---|---|---|---|---|---|---|---|---|---|---|---|
| NC_011770 | 1633014 | G | A | 61 | 0 | 4 | 0 | | | A | A | A | A | | | | | | |
| NC_011770 | 1633104 | G | A | 63 | 0 | 4 | 0 | | | | | | | A | A | A | A | | |
| NC_011770 | 1633126 | G | A | 61 | 0 | 2 | 0 | A | A | | | | | | | | | | |
| NC_011770 | 1634892 | AG | A | 2 | 6 | 1 | 0 | . | . | | | | . | A | . | | . | | . |
| NC_011770 | 1637280 | C | T | 53 | 0 | 4 | 0 | | | T | T | T | T | | | | | | |
| NC_011770 | 1637284 | C | T | 47 | 0 | 4 | 0 | | | T | T | T | T | | | | | | |
| NC_011770 | 1637286 | T | C | 47 | 0 | 12 | 0 | C | C | C | C | C | C | C | C | C | C | C | C |
| NC_011770 | 1637291 | C | T | 47 | 0 | 4 | 0 | T | T | | | | | T | T | | | | |
| NC_011770 | 1637297 | A | C | 49 | 0 | 8 | 0 | C | C | C | C | C | C | C | C | | | | |
| NC_011770 | 1637319 | C | T | 48 | 0 | 1 | 0 | | | | | | | T | | | | | |
| NC_011770 | 1637340 | C | A | 48 | 0 | 4 | 0 | | | A | A | A | A | | | | | | |
| NC_011770 | 1637342 | T | G | 47 | 0 | 6 | 0 | | | G | G | G | G | | | G | G | | |
| NC_011770 | 1637365 | C | G | 45 | 0 | 12 | 0 | G | G | G | G | G | G | G | G | G | G | G | G |
| NC_011770 | 1638026 | A | T | 44 | 0 | 12 | 0 | T | T | T | T | T | T | T | T | T | T | T | T |
| NC_011770 | 1638029 | G | GA | 44 | 0 | 12 | 0 | GA | GA | GA | GA | GA | GA | GA | GA | GA | GA | GA | GA |
| NC_011770 | 1638065 | T | C | 53 | 0 | 10 | 0 | C | C | C | C | C | C | | | C | C | C | C |
| NC_011770 | 1643923 | AGCCGCCG | A | 62 | 0 | 2 | 0 | A | A | | | | | | | | | | |
| NC_011770 | 1643966 | G | A | 68 | 0 | 4 | 0 | | | | | | | | | A | A | A | A |
| NC_011770 | 1644039 | G | A | 74 | 0 | 2 | 0 | A | A | | | | | | | | | | |
| NC_011770 | 1644143 | C | A | 76 | 0 | 2 | 0 | A | A | | | | | | | | | | |
| NC_011770 | 1644169 | T | C | 76 | 0 | 2 | 0 | | | | | | | C | C | | | | |
| NC_011770 | 1644182 | T | C | 74 | 0 | 2 | 0 | C | C | | | | | | | | | | |
| NC_011770 | 1644218 | T | C | 73 | 0 | 6 | 0 | | | C | C | C | C | C | C | | | | |
| NC_011770 | 1644224 | A | G | 73 | 0 | 2 | 0 | G | G | | | | | | | | | | |
| NC_011770 | 1644226 | C | T | 73 | 0 | 2 | 0 | T | T | | | | | | | | | | |
| NC_011770 | 1644229 | G | A | 72 | 0 | 8 | 0 | A | A | A | A | A | A | A | A | | | | |
| NC_011770 | 1644237 | T | C | 73 | 0 | 8 | 0 | C | C | C | C | C | C | C | C | | | | |
| NC_011770 | 1644293 | C | A | 71 | 0 | 6 | 0 | | | A | A | A | A | A | A | | | | |
| NC_011770 | 1644294 | T | C | 74 | 0 | 2 | 0 | C | C | | | | | | | | | | |
| NC_011770 | 1644298 | G | A | 73 | 0 | 8 | 0 | A | A | A | A | A | A | A | A | | | | |
| NC_011770 | 1644319 | C | G | 68 | 0 | 4 | 0 | | | | | | | | | G | G | G | G |
| NC_011770 | 1644335 | C | T | 70 | 0 | 8 | 0 | T | T | T | T | T | T | T | T | | | | |
| NC_011770 | 1644361 | T | A | 65 | 0 | 8 | 0 | A | A | A | A | A | A | A | A | | | | |
| NC_011770 | 1644362 | T | A | 66 | 0 | 8 | 0 | A | A | A | A | A | A | A | A | | | | |
| NC_011770 | 1644374 | C | T | 65 | 0 | 2 | 0 | T | T | | | | | | | | | | |
| NC_011770 | 1644383 | G | T | 68 | 0 | 8 | 0 | T | T | T | T | T | T | T | T | | | | |
| NC_011770 | 1644400 | A | C | 70 | 0 | 8 | 0 | C | C | C | C | C | C | C | C | | | | |
| NC_011770 | 1644763 | T | A | 74 | 0 | 6 | 0 | | | A | A | A | A | A | A | | | | |
| NC_011770 | 1644797 | T | C | 72 | 0 | 2 | 0 | C | C | | | | | | | | | | |
| NC_011770 | 1644801 | C | T | 72 | 0 | 2 | 0 | T | T | | | | | | | | | | |
| NC_011770 | 1644802 | G | T | 73 | 0 | 2 | 0 | T | T | | | | | | | | | | |
| NC_011770 | 1644843 | G | T | 72 | 0 | 2 | 0 | T | T | | | | | | | | | | |
| NC_011770 | 1644902 | A | C | 69 | 0 | 4 | 0 | C | C | | | | | C | C | | | | |
| NC_011770 | 1644936 | C | T | 69 | 0 | 4 | 0 | | | T | T | T | T | | | | | | |
| NC_011770 | 1644941 | C | A | 67 | 0 | 12 | 0 | A | A | A | A | A | A | A | A | A | A | A | A |
| NC_011770 | 1645030 | G | C | 68 | 0 | 2 | 0 | C | C | | | | | | | | | | |
| NC_011770 | 1650116 | C | T | 98 | 0 | 6 | 0 | T | T | T | T | T | T | | | | | | |
| NC_011770 | 1651397 | G | A | 60 | 0 | 6 | 0 | | | | | | | A | A | A | A | A | A |
| NC_011770 | 1651404 | T | TG | 60 | 0 | 6 | 0 | | | | | | | TG | TG | TG | TG | TG | TG |
| NC_011770 | 1658220 | CG | C | 39 | 0 | 2 | 0 | | | | | | | C | C | | | | |
| NC_011770 | 1658297 | C | T | 37 | 2 | 4 | 0 | | | T | T | T | T | . | . | | | | |
| NC_011770 | 1663069 | TC | T | 68 | 0 | 2 | 0 | | | | | | | | | | | T | T |
| NC_011770 | 1664174 | A | C | 66 | 0 | 10 | 0 | | | C | C | C | C | C | C | C | C | C | C |
| NC_011770 | 1664851 | G | T | 38 | 0 | 2 | 0 | | | | | | | | | T | T | | |
| NC_011770 | 1664902 | T | C | 36 | 0 | 8 | 0 | C | C | C | C | C | C | C | C | | | | |
| NC_011770 | 1664913 | C | T | 38 | 0 | 4 | 0 | | | T | T | T | T | | | | | | |
| NC_011770 | 1664933 | A | C | 38 | 0 | 4 | 0 | C | C | | | | | C | C | | | | |
| NC_011770 | 1664972 | A | G | 42 | 0 | 4 | 0 | G | G | | | | | G | G | | | | |
| NC_011770 | 1664975 | C | G | 44 | 0 | 4 | 0 | G | G | | | | | G | G | | | | |
| NC_011770 | 1665043 | C | T | 50 | 0 | 4 | 0 | | | T | T | T | T | | | | | | |
| NC_011770 | 1665133 | G | C | 64 | 0 | 2 | 0 | | | | | | | C | C | | | | |
| NC_011770 | 1665178 | C | T | 69 | 0 | 6 | 0 | | | T | T | T | T | T | T | | | | |

| | | | | | | | | | | | | | | | | | | | |
|---|---|---|---|---|---|---|---|---|---|---|---|---|---|---|---|---|---|---|---|
| NC_011770 | 1668344 | T | A | 71 | 0 | 2 | 0 | | | | | | | A | A | | | | |
| NC_011770 | 1668347 | T | G | 69 | 0 | 2 | 0 | | | | | | | G | G | | | | |
| NC_011770 | 1668348 | T | C | 69 | 0 | 2 | 0 | | | | | | | C | C | | | | |
| NC_011770 | 1668362 | T | C | 63 | 0 | 2 | 0 | | | | | | | C | C | | | | |
| NC_011770 | 1669140 | G | A | 74 | 0 | 10 | 0 | | | A | A | A | A | A | A | A | A | A | A |
| NC_011770 | 1669282 | C | T | 77 | 0 | 1 | 0 | | | | | | | T | | | | | |
| NC_011770 | 1670097 | C | G | 81 | 0 | 2 | 0 | G | G | | | | | | | | | | |
| NC_011770 | 1671952 | C | G | 66 | 0 | 2 | 0 | G | G | | | | | | | | | | |
| NC_011770 | 1672747 | C | A | 76 | 0 | 10 | 0 | | | A | A | A | A | A | A | A | A | A | A |
| NC_011770 | 1672830 | G | A | 75 | 0 | 6 | 0 | | | A | A | A | A | | | | | A | A |
| NC_011770 | 1672972 | T | G | 75 | 0 | 8 | 0 | G | G | G | G | G | G | | | | | G | G |
| NC_011770 | 1674712 | C | T | 83 | 0 | 2 | 0 | | | | | | | | | | | T | T |
| NC_011770 | 1674713 | A | T | 81 | 0 | 4 | 0 | | | | | | | T | T | T | T | | |
| NC_011770 | 1674730 | G | A | 82 | 0 | 2 | 0 | | | | | | | | | | | A | A |
| NC_011770 | 1674731 | G | A | 86 | 0 | 2 | 0 | | | | | | | | | | | A | A |
| NC_011770 | 1677201 | CG | C | 7 | 9 | 1 | 0 | . | . | | . | . | . | . | C | . | . | . | |
| NC_011770 | 1678824 | T | C | 60 | 0 | 12 | 0 | C | C | C | C | C | C | C | C | C | C | C | C |
| NC_011770 | 1678889 | T | C | 62 | 0 | 12 | 0 | C | C | C | C | C | C | C | C | C | C | C | C |
| NC_011770 | 1680237 | C | T | 57 | 0 | 4 | 0 | | | | | | | | | T | T | T | T |
| NC_011770 | 1680263 | C | T | 58 | 0 | 4 | 0 | | | | | | | | | T | T | T | T |
| NC_011770 | 1680335 | A | G | 67 | 0 | 12 | 0 | G | G | G | G | G | G | G | G | G | G | G | G |
| NC_011770 | 1680346 | A | T | 66 | 0 | 12 | 0 | T | T | T | T | T | T | T | T | T | T | T | T |
| NC_011770 | 1682207 | C | T | 79 | 0 | 4 | 0 | | | | | | | | | T | T | T | T |
| NC_011770 | 1683634 | C | T | 4 | 3 | 1 | 0 | | | . | | | | T | | | . | | . |
| NC_011770 | 1683669 | T | G | 7 | 1 | 1 | 0 | . | . | . | G | . | | | | . | . | . | |
| NC_011770 | 1683677 | T | A | 7 | 1 | 1 | 0 | | | . | | | | | | . | A | . | . |
| NC_011770 | 1683685 | G | T | 7 | 1 | 1 | 0 | | | . | | | | | | . | T | . | . |
| NC_011770 | 1683689 | C | T | 8 | 1 | 1 | 0 | | | . | | | | | | . | T | . | . |
| NC_011770 | 1683706 | G | A | 9 | 1 | 7 | 0 | | . | . | A | A | A | | | A | A | A | A |
| NC_011770 | 1683754 | G | A | 9 | 2 | 3 | 0 | | | . | A | A | A | | | . | | | |
| NC_011770 | 1683760 | T | G | 9 | 2 | 10 | 0 | G | G | . | G | G | G | G | G | . | G | G | G |
| NC_011770 | 1683792 | A | T | 9 | 2 | 2 | 0 | T | T | . | | | | | | . | . | | |
| NC_011770 | 1683831 | C | T | 10 | 1 | 11 | 0 | T | T | . | T | T | T | T | T | T | T | T | T |
| NC_011770 | 1683858 | G | A | 12 | 0 | 3 | 0 | | | A | . | A | A | | | | | | |
| NC_011770 | 1683861 | T | C | 12 | 0 | 5 | 0 | C | C | C | . | C | C | | | | | | |
| NC_011770 | 1683869 | A | G | 14 | 0 | 2 | 0 | G | G | | | | | | | | | | |
| NC_011770 | 1683892 | A | G | 17 | 0 | 2 | 0 | G | G | | | | | | | | | | |
| NC_011770 | 1683899 | T | A | 16 | 0 | 6 | 0 | A | A | G | G | G | G | | | | | | |
| NC_011770 | 1683907 | A | T | 16 | 0 | 6 | 0 | T | T | T | T | T | T | | | | | | |
| NC_011770 | 1684002 | G | T | 18 | 10 | 2 | 0 | . | . | . | . | T | T | . | . | . | . | . | . |
| NC_011770 | 1684011 | C | T | 18 | 10 | 2 | 0 | . | . | . | . | T | T | . | . | . | . | . | . |
| NC_011770 | 1684024 | G | A | 19 | 10 | 2 | 0 | . | . | . | . | A | A | . | . | . | . | . | . |
| NC_011770 | 1684044 | C | G | 13 | 10 | 2 | 0 | . | . | . | . | G | G | . | . | . | . | . | . |
| NC_011770 | 1684048 | C | G | 13 | 10 | 2 | 0 | . | . | . | . | G | G | . | . | . | . | . | . |
| NC_011770 | 1684049 | G | A | 13 | 10 | 2 | 0 | . | . | . | . | A | A | . | . | . | . | . | . |
| NC_011770 | 1684051 | T | C | 13 | 10 | 2 | 0 | . | . | . | . | C | C | . | . | . | . | . | . |
| NC_011770 | 1684055 | G | C | 13 | 10 | 2 | 0 | . | . | . | . | C | C | . | . | . | . | . | . |
| NC_011770 | 1684057 | A | G | 13 | 10 | 2 | 0 | . | . | . | . | G | G | . | . | . | . | . | . |
| NC_011770 | 1684058 | C | G | 13 | 10 | 2 | 0 | . | . | . | . | G | G | . | . | . | . | . | . |
| NC_011770 | 1684059 | A | T | 12 | 10 | 2 | 0 | . | . | . | . | T | T | . | . | . | . | . | . |
| NC_011770 | 1684070 | A | T | 16 | 10 | 2 | 0 | . | . | . | . | T | T | . | . | . | . | . | . |
| NC_011770 | 1684858 | G | A | 18 | 10 | 2 | 0 | . | . | . | . | A | A | . | . | . | . | . | . |
| NC_011770 | 1684864 | T | A | 19 | 10 | 2 | 0 | . | . | . | . | A | A | . | . | . | . | . | . |
| NC_011770 | 1684872 | C | T | 19 | 10 | 2 | 0 | . | . | . | . | T | T | . | . | . | . | . | . |
| NC_011770 | 1684879 | C | A | 19 | 10 | 2 | 0 | . | . | . | . | A | A | . | . | . | . | . | . |
| NC_011770 | 1684881 | T | G | 18 | 10 | 2 | 0 | . | . | . | . | G | G | . | . | . | . | . | . |
| NC_011770 | 1684887 | G | A | 18 | 10 | 2 | 0 | . | . | . | . | A | A | . | . | . | . | . | . |
| NC_011770 | 1684888 | A | G | 18 | 10 | 2 | 0 | . | . | . | . | G | G | . | . | . | . | . | . |
| NC_011770 | 1684972 | C | CA | 18 | 10 | 2 | 0 | . | . | . | . | CA | CA | . | . | . | . | . | . |
| NC_011770 | 1689524 | A | G | 19 | 10 | 2 | 0 | . | . | . | . | G | G | . | . | . | . | . | . |
| NC_011770 | 1689531 | C | T | 17 | 10 | 2 | 0 | . | . | . | . | T | T | . | . | . | . | . | . |
| NC_011770 | 1689545 | A | C | 13 | 10 | 2 | 0 | . | . | . | . | C | C | . | . | . | . | . | . |

| | | | | | | | | | | | | | | | | | | | |
|---|---|---|---|---|---|---|---|---|---|---|---|---|---|---|---|---|---|---|---|
| NC_011770 | 1689547 | G | A | 13 | 10 | 2 | 0 | . | . | . | . | A | A | . | . | . | . | . | . |
| NC_011770 | 1689549 | T | C | 13 | 10 | 2 | 0 | . | . | . | . | C | C | . | . | . | . | . | . |
| NC_011770 | 1689550 | T | G | 13 | 10 | 2 | 0 | . | . | . | . | G | G | . | . | . | . | . | . |
| NC_011770 | 1689551 | G | A | 13 | 10 | 2 | 0 | . | . | . | . | A | A | . | . | . | . | . | . |
| NC_011770 | 1689552 | G | T | 13 | 10 | 2 | 0 | . | . | . | . | T | T | . | . | . | . | . | . |
| NC_011770 | 1689556 | C | G | 13 | 10 | 2 | 0 | . | . | . | . | G | G | . | . | . | . | . | . |
| NC_011770 | 1689585 | A | G | 17 | 10 | 2 | 0 | . | . | . | . | G | G | . | . | . | . | . | . |
| NC_011770 | 1692008 | C | T | 18 | 10 | 2 | 0 | . | . | . | . | T | T | . | . | . | . | . | . |
| NC_011770 | 1692018 | A | C | 0 | 11 | 1 | 0 | . | . | . | . | . | C | . | . | . | . | . | . |
| NC_011770 | 1692254 | A | G | 0 | 11 | 1 | 0 | . | . | . | . | . | G | . | . | . | . | . | . |
| NC_011770 | 1693221 | C | A | 15 | 10 | 2 | 0 | . | . | . | . | A | A | . | . | . | . | . | . |
| NC_011770 | 1693371 | A | G | 12 | 10 | 2 | 0 | . | . | . | . | G | G | . | . | . | . | . | . |
| NC_011770 | 1693391 | G | A | 12 | 10 | 2 | 0 | . | . | . | . | A | A | . | . | . | . | . | . |
| NC_011770 | 1695076 | T | C | 20 | 10 | 2 | 0 | . | . | . | . | C | C | . | . | . | . | . | . |
| NC_011770 | 1695077 | T | C | 20 | 10 | 2 | 0 | . | . | . | . | C | C | . | . | . | . | . | . |
| NC_011770 | 1695111 | C | T | 22 | 10 | 2 | 0 | . | . | . | . | T | T | . | . | . | . | . | . |
| NC_011770 | 1695324 | T | C | 18 | 10 | 2 | 0 | . | . | . | . | C | C | . | . | . | . | . | . |
| NC_011770 | 1701242 | T | C | 25 | 10 | 2 | 0 | . | . | . | . | C | C | . | . | . | . | . | . |
| NC_011770 | 1701248 | C | T | 25 | 10 | 2 | 0 | . | . | . | . | T | T | . | . | . | . | . | . |
| NC_011770 | 1701398 | C | A | 23 | 10 | 2 | 0 | . | . | . | . | A | A | . | . | . | . | . | . |
| NC_011770 | 1701434 | T | C | 23 | 10 | 2 | 0 | . | . | . | . | C | C | . | . | . | . | . | . |
| NC_011770 | 1701442 | G | C | 22 | 10 | 2 | 0 | . | . | . | . | C | C | . | . | . | . | . | . |
| NC_011770 | 1701482 | A | G | 21 | 10 | 2 | 0 | . | . | . | . | G | G | . | . | . | . | . | . |
| NC_011770 | 1701485 | C | T | 21 | 10 | 2 | 0 | . | . | . | . | T | T | . | . | . | . | . | . |
| NC_011770 | 1701497 | A | G | 21 | 10 | 2 | 0 | . | . | . | . | G | G | . | . | . | . | . | . |
| NC_011770 | 1701503 | G | T | 21 | 10 | 2 | 0 | . | . | . | . | T | T | . | . | . | . | . | . |
| NC_011770 | 1701518 | G | C | 22 | 10 | 2 | 0 | . | . | . | . | C | C | . | . | . | . | . | . |
| NC_011770 | 1701578 | C | T | 15 | 10 | 2 | 0 | . | . | . | . | T | T | . | . | . | . | . | . |
| NC_011770 | 1701581 | G | A | 14 | 10 | 2 | 0 | . | . | . | . | A | A | . | . | . | . | . | . |
| NC_011770 | 1701582 | A | G | 15 | 10 | 2 | 0 | . | . | . | . | G | G | . | . | . | . | . | . |
| NC_011770 | 1701586 | T | C | 15 | 10 | 2 | 0 | . | . | . | . | C | C | . | . | . | . | . | . |
| NC_011770 | 1701590 | G | A | 14 | 10 | 2 | 0 | . | . | . | . | A | A | . | . | . | . | . | . |
| NC_011770 | 1701599 | T | G | 14 | 10 | 2 | 0 | . | . | . | . | G | G | . | . | . | . | . | . |
| NC_011770 | 1701602 | C | T | 13 | 10 | 2 | 0 | . | . | . | . | T | T | . | . | . | . | . | . |
| NC_011770 | 1701606 | A | G | 14 | 10 | 2 | 0 | . | . | . | . | G | G | . | . | . | . | . | . |
| NC_011770 | 1701609 | G | A | 14 | 10 | 2 | 0 | . | . | . | . | A | A | . | . | . | . | . | . |
| NC_011770 | 1701611 | G | A | 14 | 10 | 2 | 0 | . | . | . | . | A | A | . | . | . | . | . | . |
| NC_011770 | 1701614 | C | A | 14 | 10 | 2 | 0 | . | . | . | . | A | A | . | . | . | . | . | . |
| NC_011770 | 1701622 | T | C | 18 | 10 | 2 | 0 | . | . | . | . | C | C | . | . | . | . | . | . |
| NC_011770 | 1701645 | T | C | 19 | 10 | 2 | 0 | . | . | . | . | C | C | . | . | . | . | . | . |
| NC_011770 | 1701647 | A | C | 19 | 10 | 2 | 0 | . | . | . | . | C | C | . | . | . | . | . | . |
| NC_011770 | 1701669 | A | C | 20 | 10 | 2 | 0 | . | . | . | . | C | C | . | . | . | . | . | . |
| NC_011770 | 1701671 | G | A | 19 | 10 | 2 | 0 | . | . | . | . | A | A | . | . | . | . | . | . |
| NC_011770 | 1702883 | C | A | 21 | 10 | 2 | 0 | . | . | . | . | A | A | . | . | . | . | . | . |
| NC_011770 | 1702891 | G | A | 20 | 10 | 2 | 0 | . | . | . | . | A | A | . | . | . | . | . | . |
| NC_011770 | 1702899 | T | C | 19 | 10 | 2 | 0 | . | . | . | . | C | C | . | . | . | . | . | . |
| NC_011770 | 1702963 | AT | A | 19 | 10 | 2 | 0 | . | . | . | . | A | A | . | . | . | . | . | . |
| NC_011770 | 1712029 | A | G | 12 | 8 | 4 | 0 | . | . | . | . | G | G | . | . | G | G | . | . |
| NC_011770 | 1712030 | T | TC | 13 | 8 | 4 | 0 | . | . | . | . | TC | TC | . | . | TC | TC | . | . |
| NC_011770 | 1713934 | C | A | 49 | 6 | 2 | 0 | A | A | . | . | | | . | . | | | . | . |
| NC_011770 | 1713937 | G | A | 50 | 6 | 2 | 0 | A | A | . | . | | | . | . | | | . | . |
| NC_011770 | 1713943 | G | A | 48 | 6 | 2 | 0 | A | A | . | . | | | . | . | | | . | . |
| NC_011770 | 1713949 | T | G | 49 | 6 | 2 | 0 | G | G | . | . | | | . | . | | | . | . |
| NC_011770 | 1716430 | G | A | 28 | 8 | 2 | 0 | . | . | . | . | A | A | . | . | | | . | . |
| NC_011770 | 1716433 | G | GATAT | 28 | 8 | 2 | 0 | . | . | . | . | GATAT | GATAT | . | . | | | . | . |
| NC_011770 | 1716819 | A | G | 44 | 8 | 2 | 0 | . | . | . | . | G | G | . | . | | | . | . |
| NC_011770 | 1716858 | G | A | 42 | 8 | 2 | 0 | . | . | . | . | A | A | . | . | | | . | . |
| NC_011770 | 1716861 | C | T | 41 | 8 | 2 | 0 | . | . | . | . | T | T | . | . | | | . | . |
| NC_011770 | 1716865 | T | C | 41 | 8 | 2 | 0 | . | . | . | . | C | C | . | . | | | . | . |
| NC_011770 | 1716874 | G | A | 40 | 8 | 2 | 0 | . | . | . | . | A | A | . | . | | | . | . |
| NC_011770 | 1716876 | A | C | 40 | 8 | 2 | 0 | . | . | . | . | C | C | . | . | | | . | . |
| NC_011770 | 1716879 | C | G | 40 | 8 | 2 | 0 | . | . | . | . | G | G | . | . | | | . | . |

| | | | | | | | | | | | | | | | | | | | |
|---|---|---|---|---|---|---|---|---|---|---|---|---|---|---|---|---|---|---|---|
| NC_011770 | 1720580 | G | A | 7 | 10 | 2 | 0 | . | . | . | . | A | A | . | . | . | . | . | . |
| NC_011770 | 1720585 | G | C | 7 | 10 | 2 | 0 | . | . | . | . | C | C | . | . | . | . | . | . |
| NC_011770 | 1720600 | T | C | 3 | 10 | 2 | 0 | . | . | . | . | C | C | . | . | . | . | . | . |
| NC_011770 | 1720602 | G | T | 3 | 10 | 2 | 0 | . | . | . | . | T | T | . | . | . | . | . | . |
| NC_011770 | 1720604 | T | G | 3 | 10 | 2 | 0 | . | . | . | . | G | G | . | . | . | . | . | . |
| NC_011770 | 1720830 | T | C | 18 | 8 | 2 | 0 | . | . | . | . | | | . | . | C | C | . | . |
| NC_011770 | 1720839 | A | C | 18 | 8 | 2 | 0 | . | . | . | . | C | C | . | . | | | . | . |
| NC_011770 | 1720840 | T | G | 17 | 8 | 2 | 0 | . | . | . | . | G | G | . | . | | | . | . |
| NC_011770 | 1720852 | T | G | 16 | 8 | 2 | 0 | . | . | . | . | G | G | . | . | | | . | . |
| NC_011770 | 1720853 | G | C | 16 | 8 | 2 | 0 | . | . | . | . | C | C | . | . | | | . | . |
| NC_011770 | 1720854 | G | C | 16 | 8 | 2 | 0 | . | . | . | . | C | C | . | . | | | . | . |
| NC_011770 | 1720863 | T | C | 18 | 8 | 2 | 0 | . | . | . | . | C | C | . | . | | | . | . |
| NC_011770 | 1720868 | G | T | 17 | 8 | 2 | 0 | . | . | . | . | T | T | . | . | | | . | . |
| NC_011770 | 1720871 | C | T | 18 | 8 | 2 | 0 | . | . | . | . | T | T | . | . | | | . | . |
| NC_011770 | 1720881 | T | G | 17 | 8 | 2 | 0 | . | . | . | . | G | G | . | . | | | . | . |
| NC_011770 | 1720928 | A | C | 23 | 0 | 6 | 0 | | | | | | | C | C | C | C | C | C |
| NC_011770 | 1721177 | T | G | 58 | 0 | 6 | 0 | G | G | G | G | G | G | | | | | | |
| NC_011770 | 1724085 | G | A | 38 | 0 | 2 | 0 | | | | | | | A | A | | | | |
| NC_011770 | 1724106 | T | C | 34 | 0 | 8 | 0 | | | C | C | C | C | | | C | C | C | C |
| NC_011770 | 1729586 | C | T | 54 | 0 | 8 | 0 | | | T | T | T | T | | | T | T | T | T |
| NC_011770 | 1730403 | A | G | 63 | 0 | 2 | 0 | G | G | | | | | | | | | | |
| NC_011770 | 1732458 | C | G | 75 | 0 | 4 | 0 | | | | | | | | | G | G | G | G |
| NC_011770 | 1732459 | T | A | 74 | 0 | 12 | 0 | A | A | A | A | A | A | A | A | A | A | A | A |
| NC_011770 | 1734094 | T | C | 68 | 0 | 4 | 0 | C | C | | | | | C | C | | | | |
| NC_011770 | 1734215 | C | A | 57 | 0 | 8 | 0 | A | A | | | | | A | A | A | A | A | A |
| NC_011770 | 1734236 | T | A | 60 | 0 | 8 | 0 | A | A | | | | | A | A | A | A | A | A |
| NC_011770 | 1735430 | C | T | 50 | 0 | 4 | 0 | | | T | T | T | T | | | | | | |
| NC_011770 | 1735455 | T | C | 52 | 0 | 4 | 0 | | | C | C | C | C | | | | | | |
| NC_011770 | 1736226 | C | G | 52 | 0 | 4 | 0 | G | G | | | | | G | G | | | | |
| NC_011770 | 1736291 | GC | G | 51 | 0 | 4 | 0 | | | G | G | G | G | | | | | | |
| NC_011770 | 1736295 | G | A | 49 | 0 | 6 | 0 | A | A | | | | | | | A | A | A | A |
| NC_011770 | 1736325 | C | G | 50 | 0 | 12 | 0 | G | G | G | G | G | G | G | G | G | G | G | G |
| NC_011770 | 1736366 | C | T | 47 | 0 | 2 | 0 | T | T | | | | | | | | | | |
| NC_011770 | 1738060 | A | G | 69 | 0 | 2 | 0 | | | | | | | | | | | G | G |
| NC_011770 | 1738187 | G | A | 62 | 0 | 6 | 0 | | | A | A | A | A | A | A | | | | |
| NC_011770 | 1738203 | A | C | 63 | 0 | 4 | 0 | | | | | | | | | C | C | C | C |
| NC_011770 | 1738529 | G | A | 49 | 0 | 2 | 0 | | | | | | | A | A | | | | |
| NC_011770 | 1738548 | G | T | 47 | 0 | 2 | 0 | T | T | | | | | | | | | | |
| NC_011770 | 1738608 | T | C | 48 | 0 | 4 | 0 | | | C | C | C | C | | | | | | |
| NC_011770 | 1739404 | G | A | 43 | 0 | 2 | 0 | | | | | | | | | A | A | | |
| NC_011770 | 1742245 | T | TC | 59 | 0 | 5 | 0 | | | TC | TCC | TC | TC | | | TC | | | |
| NC_011770 | 1742908 | A | T | 60 | 0 | 6 | 0 | T | T | T | T | T | T | | | | | | |
| NC_011770 | 1743953 | A | G | 73 | 0 | 2 | 0 | | | | | | | | | | | G | G |
| NC_011770 | 1743997 | C | T | 76 | 0 | 2 | 0 | | | | | | | | | | | T | T |
| NC_011770 | 1746792 | C | G | 73 | 0 | 4 | 0 | G | G | | | | | G | G | | | | |
| NC_011770 | 1747071 | T | C | 55 | 0 | 12 | 0 | C | C | C | C | C | C | C | C | C | C | C | C |
| NC_011770 | 1747116 | C | CG | 52 | 0 | 2 | 0 | | | | | | | | | | | CG | CG |
| NC_011770 | 1747118 | C | T | 41 | 0 | 2 | 0 | | | | | | | | | | | T | T |
| NC_011770 | 1747135 | C | T | 51 | 0 | 2 | 0 | | | | | | | | | T | T | | |
| NC_011770 | 1747205 | C | T | 53 | 0 | 1 | 0 | | | | | | | T | | | | | |
| NC_011770 | 1747206 | T | C | 52 | 0 | 12 | 0 | C | C | C | C | C | C | C | C | C | C | C | C |
| NC_011770 | 1750405 | C | T | 64 | 0 | 4 | 0 | T | T | | | | | T | T | | | | |
| NC_011770 | 1750487 | G | C | 69 | 0 | 6 | 0 | C | C | | | | | | | C | C | C | C |
| NC_011770 | 1750548 | T | C | 69 | 0 | 8 | 0 | | | C | C | C | C | | | C | C | C | C |
| NC_011770 | 1750572 | A | G | 71 | 0 | 8 | 0 | | | G | G | G | G | | | G | G | G | G |
| NC_011770 | 1750609 | A | T | 71 | 0 | 4 | 0 | | | | | | | | | T | T | T | T |
| NC_011770 | 1751121 | C | G | 53 | 0 | 10 | 0 | | | G | G | G | G | G | G | G | G | G | G |
| NC_011770 | 1751123 | G | A | 52 | 0 | 10 | 0 | | | A | A | A | A | A | A | A | A | A | A |
| NC_011770 | 1751178 | G | A | 38 | 0 | 7 | 0 | A | . | A | A | A | A | A | A | | | | |
| NC_011770 | 1751179 | A | G | 39 | 0 | 6 | 0 | G | . | G | . | G | G | G | G | | | | |
| NC_011770 | 1751235 | C | T | 74 | 0 | 7 | 0 | T | T | T | T | T | T | T | | | | | |
| NC_011770 | 1752248 | T | C | 50 | 0 | 8 | 0 | | | C | C | C | C | | | C | C | C | C |

| | | | | | | | | | | | | | | | | | | | |
|---|---|---|---|---|---|---|---|---|---|---|---|---|---|---|---|---|---|---|---|
| NC_011770 | 1753385 | C | T | 71 | 0 | 2 | 0 | T | T | | | | | | | | | | |
| NC_011770 | 1753394 | A | G | 70 | 0 | 2 | 0 | | | | | | | | | | | G | G |
| NC_011770 | 1755150 | C | T | 76 | 0 | 2 | 0 | | | | | | | | | | | T | T |
| NC_011770 | 1755549 | C | T | 77 | 0 | 4 | 0 | T | T | | | | | | | T | T | | |
| NC_011770 | 1759241 | A | C | 80 | 0 | 2 | 0 | C | C | | | | | | | | | | |
| NC_011770 | 1759269 | A | G | 76 | 0 | 4 | 0 | | | | | | | | | G | G | G | G |
| NC_011770 | 1759282 | T | C | 75 | 0 | 12 | 0 | C | C | C | C | C | C | C | C | C | C | C | C |
| NC_011770 | 1760090 | G | A | 71 | 0 | 4 | 0 | | | | | | | | | A | A | A | A |
| NC_011770 | 1760110 | A | T | 74 | 0 | 12 | 0 | T | T | T | T | T | T | T | T | T | T | T | T |
| NC_011770 | 1760171 | C | T | 78 | 0 | 4 | 0 | | | | | | | | | T | T | T | T |
| NC_011770 | 1760249 | C | T | 87 | 0 | 4 | 0 | T | T | | | | | T | T | | | | |
| NC_011770 | 1763555 | C | T | 50 | 0 | 10 | 0 | T | T | T | T | T | T | T | T | | | T | T |
| NC_011770 | 1763557 | T | G | 50 | 0 | 8 | 0 | G | G | G | G | G | G | | | | | G | G |
| NC_011770 | 1763580 | G | T | 51 | 0 | 10 | 0 | T | T | T | T | T | T | T | T | | | T | T |
| NC_011770 | 1766169 | C | A | 69 | 0 | 6 | 0 | | | A | A | A | A | | | A | A | | |
| NC_011770 | 1767007 | G | A | 62 | 0 | 12 | 0 | A | A | A | A | A | A | A | A | A | A | A | A |
| NC_011770 | 1767120 | C | G | 63 | 0 | 8 | 0 | | | G | G | G | G | | | G | G | G | G |
| NC_011770 | 1767133 | T | G | 62 | 0 | 12 | 0 | G | G | G | G | G | G | G | G | G | G | G | G |
| NC_011770 | 1767198 | G | A | 61 | 0 | 8 | 0 | | | A | A | A | A | | | A | A | A | A |
| NC_011770 | 1768304 | G | A | 67 | 0 | 2 | 0 | | | | | | | | | | | A | A |
| NC_011770 | 1768326 | C | T | 63 | 0 | 4 | 0 | | | T | T | T | T | | | | | | |
| NC_011770 | 1770491 | G | A | 76 | 0 | 8 | 0 | A | A | | | | | A | A | A | A | A | A |
| NC_011770 | 1772020 | A | G | 5 | 6 | 6 | 0 | . | . | G | . | G | G | G | G | . | . | G | . |
| NC_011770 | 1773023 | G | C | 70 | 0 | 12 | 0 | C | C | C | C | C | C | C | C | C | C | C | C |
| NC_011770 | 1773045 | T | C | 73 | 0 | 12 | 0 | C | C | C | C | C | C | C | C | C | C | C | C |
| NC_011770 | 1773049 | A | C | 76 | 0 | 4 | 0 | | | C | C | C | C | | | | | | |
| NC_011770 | 1773093 | C | T | 76 | 0 | 2 | 0 | | | | | | | T | T | | | | |
| NC_011770 | 1773105 | G | A | 74 | 0 | 2 | 0 | A | A | | | | | | | | | | |
| NC_011770 | 1773477 | T | C | 63 | 0 | 12 | 0 | C | C | C | C | C | C | C | C | C | C | C | C |
| NC_011770 | 1773486 | C | T | 61 | 0 | 4 | 0 | | | T | T | T | T | | | | | | |
| NC_011770 | 1773489 | G | A | 62 | 0 | 2 | 0 | | | | | | | | | | | A | A |
| NC_011770 | 1773818 | G | T | 60 | 0 | 4 | 0 | | | | | | | | | T | T | T | T |
| NC_011770 | 1773871 | T | C | 64 | 0 | 10 | 0 | | | C | C | C | C | C | C | C | C | C | C |
| NC_011770 | 1774410 | A | T | 53 | 0 | 2 | 0 | T | T | | | | | | | | | | |
| NC_011770 | 1775538 | T | C | 64 | 0 | 12 | 0 | C | C | C | C | C | C | C | C | C | C | C | C |
| NC_011770 | 1779119 | G | A | 63 | 0 | 6 | 0 | | | A | A | A | A | | | A | A | | |
| NC_011770 | 1779247 | G | C | 87 | 0 | 12 | 0 | C | C | C | C | C | C | C | C | C | C | C | C |
| NC_011770 | 1783765 | T | G | 80 | 0 | 12 | 0 | G | G | G | G | G | G | G | G | G | G | G | G |
| NC_011770 | 1784058 | C | T | 72 | 0 | 2 | 0 | | | | | | | T | T | | | | |
| NC_011770 | 1784673 | T | C | 59 | 0 | 6 | 0 | | | C | C | C | C | | | C | C | | |
| NC_011770 | 1789412 | GC | G | 73 | 0 | 12 | 0 | G | G | G | G | G | G | G | G | G | G | G | G |
| NC_011770 | 1791968 | T | C | 54 | 0 | 8 | 0 | C | C | C | C | C | C | | | | | C | C |
| NC_011770 | 1792007 | C | T | 59 | 0 | 2 | 0 | T | T | | | | | | | | | | |
| NC_011770 | 1792012 | C | T | 59 | 0 | 2 | 0 | | | | | | | | | T | T | | |
| NC_011770 | 1792053 | T | C | 66 | 0 | 4 | 0 | | | | | | | | | C | C | C | C |
| NC_011770 | 1792071 | G | A | 67 | 0 | 2 | 0 | A | A | | | | | | | | | | |
| NC_011770 | 1792086 | T | C | 69 | 0 | 2 | 0 | | | | | | | | | | | C | C |
| NC_011770 | 1792112 | A | G | 69 | 0 | 2 | 0 | | | | | | | | | G | G | | |
| NC_011770 | 1792113 | T | C | 68 | 0 | 2 | 0 | | | | | | | | | C | C | | |
| NC_011770 | 1792115 | C | T | 69 | 0 | 2 | 0 | | | | | | | | | | | T | T |
| NC_011770 | 1792162 | A | G | 65 | 0 | 4 | 0 | | | | | | | | | G | G | G | G |
| NC_011770 | 1792178 | G | T | 64 | 0 | 2 | 0 | | | | | | | | | T | T | | |
| NC_011770 | 1792223 | G | A | 63 | 0 | 2 | 0 | | | | | | | | | | | A | A |
| NC_011770 | 1792256 | A | G | 65 | 0 | 4 | 0 | | | | | | | | | G | G | G | G |
| NC_011770 | 1792304 | G | A | 67 | 0 | 2 | 0 | | | | | | | | | | | A | A |
| NC_011770 | 1792362 | C | T | 62 | 0 | 4 | 0 | | | T | T | T | T | | | | | | |
| NC_011770 | 1792412 | C | T | 69 | 0 | 2 | 0 | | | | | | | | | T | T | | |
| NC_011770 | 1792601 | C | T | 76 | 0 | 2 | 0 | | | | | | | | | | | T | T |
| NC_011770 | 1792604 | C | T | 77 | 0 | 2 | 0 | | | | | | | | | | | T | T |
| NC_011770 | 1792635 | T | G | 77 | 0 | 2 | 0 | | | | | | | | | G | G | | |
| NC_011770 | 1792648 | G | A | 80 | 0 | 2 | 0 | | | | | | | | | | | A | A |
| NC_011770 | 1792694 | G | A | 79 | 0 | 2 | 0 | | | | | | | A | A | | | | |

| | | | | | | | | | | | | | | | | | | | |
|---|---|---|---|---|---|---|---|---|---|---|---|---|---|---|---|---|---|---|---|
| NC_011770 | 1792748 | C | T | 80 | 0 | 2 | 0 | T | T | | | | | | | | | | |
| NC_011770 | 1792783 | C | T | 79 | 0 | 2 | 0 | | | | | | | T | T | | | | |
| NC_011770 | 1796032 | G | GA | 94 | 0 | 1 | 0 | | | | | | | | | GA | | | |
| NC_011770 | 1798344 | A | G | 71 | 0 | 2 | 0 | G | G | | | | | | | | | | |
| NC_011770 | 1798346 | A | C | 69 | 0 | 2 | 0 | C | C | | | | | | | | | | |
| NC_011770 | 1798365 | C | G | 68 | 0 | 2 | 0 | G | G | | | | | | | | | | |
| NC_011770 | 1800721 | T | A | 60 | 0 | 2 | 0 | A | A | | | | | | | | | | |
| NC_011770 | 1800722 | G | C | 60 | 0 | 2 | 0 | C | C | | | | | | | | | | |
| NC_011770 | 1800737 | A | G | 59 | 0 | 2 | 0 | | | | | | | G | G | | | | |
| NC_011770 | 1800742 | A | T | 58 | 0 | 2 | 0 | T | T | | | | | | | | | | |
| NC_011770 | 1800779 | T | C | 62 | 0 | 2 | 0 | C | C | | | | | | | | | | |
| NC_011770 | 1801185 | C | T | 64 | 0 | 4 | 0 | | | T | T | T | T | | | | | | |
| NC_011770 | 1801843 | A | G | 75 | 0 | 12 | 0 | G | G | G | G | G | G | G | G | G | G | G | G |
| NC_011770 | 1801939 | G | T | 73 | 0 | 2 | 0 | T | T | | | | | | | | | | |
| NC_011770 | 1801955 | T | C | 78 | 0 | 2 | 0 | C | C | | | | | | | | | | |
| NC_011770 | 1802009 | A | AG | 78 | 0 | 2 | 0 | AG | AG | | | | | | | | | | |
| NC_011770 | 1802866 | A | G | 69 | 0 | 4 | 0 | | | G | G | G | G | | | | | | |
| NC_011770 | 1803943 | C | T | 65 | 0 | 2 | 0 | | | | | | | | | T | T | | |
| NC_011770 | 1809432 | G | A | 87 | 0 | 2 | 0 | | | | | | | | | A | A | | |
| NC_011770 | 1812875 | A | G | 79 | 0 | 8 | 0 | | | G | G | G | G | | | G | G | G | G |
| NC_011770 | 1813078 | CT | C | 77 | 0 | 2 | 0 | C | C | | | | | | | | | | |
| NC_011770 | 1813103 | A | G | 72 | 0 | 1 | 0 | | | | | | | G | | | | | |
| NC_011770 | 1813533 | A | C | 68 | 0 | 12 | 0 | C | C | C | C | C | C | C | C | C | C | C | C |
| NC_011770 | 1813557 | G | T | 77 | 0 | 2 | 0 | T | T | | | | | | | | | | |
| NC_011770 | 1813560 | C | A | 74 | 0 | 4 | 0 | A | A | | | | | T | T | | | | |
| NC_011770 | 1813561 | C | T | 75 | 0 | 2 | 0 | T | T | | | | | | | | | | |
| NC_011770 | 1813562 | T | C | 74 | 0 | 4 | 0 | C | C | | | | | | | | | C | C |
| NC_011770 | 1813563 | G | T | 74 | 0 | 2 | 0 | T | T | | | | | | | | | | |
| NC_011770 | 1813574 | A | G | 80 | 0 | 2 | 0 | G | G | | | | | | | | | | |
| NC_011770 | 1813591 | A | T | 77 | 0 | 2 | 0 | | | | | | | | | T | T | | |
| NC_011770 | 1813621 | A | G | 80 | 0 | 2 | 0 | | | | | | | | | G | G | | |
| NC_011770 | 1814583 | C | T | 29 | 8 | 2 | 0 | . | . | . | . | . | . | | | . | . | T | T |
| NC_011770 | 1814704 | T | C | 29 | 8 | 2 | 0 | . | . | . | . | . | . | | | . | . | C | C |
| NC_011770 | 1814806 | C | T | 30 | 8 | 1 | 0 | . | . | . | . | . | . | T | | . | . | | |
| NC_011770 | 1814831 | G | T | 18 | 8 | 2 | 0 | . | . | . | . | . | . | T | T | . | . | | |
| NC_011770 | 1814833 | GA | G | 15 | 8 | 2 | 0 | . | . | . | . | . | . | | | . | . | G | G |
| NC_011770 | 1814834 | A | G | 15 | 8 | 2 | 0 | . | . | . | . | . | . | G | G | . | . | . | . |
| NC_011770 | 1814838 | G | A | 12 | 9 | 3 | 0 | . | . | . | . | . | . | A | A | . | . | . | A |
| NC_011770 | 1814866 | A | T | 30 | 8 | 2 | 0 | . | . | . | . | . | . | | | . | . | T | T |
| NC_011770 | 1814891 | T | TA | 30 | 8 | 2 | 0 | . | . | . | . | . | . | TA | TA | . | . | | |
| NC_011770 | 1814905 | C | A | 28 | 8 | 2 | 0 | . | . | . | . | . | . | | | . | . | A | A |
| NC_011770 | 1814949 | GA | G | 27 | 8 | 2 | 0 | . | . | . | . | . | . | | | . | . | G | G |
| NC_011770 | 1815236 | C | CT | 29 | 8 | 2 | 0 | . | . | . | . | . | . | | | . | . | CT | CT |
| NC_011770 | 1815347 | T | C | 26 | 8 | 2 | 0 | . | . | . | . | . | . | | | . | . | C | C |
| NC_011770 | 1815364 | C | A | 26 | 8 | 2 | 0 | . | . | . | . | . | . | | | . | . | A | A |
| NC_011770 | 1815489 | C | T | 22 | 8 | 2 | 0 | . | . | . | . | . | . | | | . | . | T | T |
| NC_011770 | 1815610 | G | T | 13 | 10 | 2 | 0 | . | . | . | . | . | . | T | T | . | . | . | . |
| NC_011770 | 1815621 | C | T | 12 | 10 | 2 | 0 | . | . | . | . | . | . | T | T | . | . | . | . |
| NC_011770 | 1817093 | A | C | 23 | 4 | 2 | 0 | C | C | . | . | . | . | | | | | | |
| NC_011770 | 1817095 | C | A | 22 | 4 | 2 | 0 | | | . | . | . | . | | | | | A | A |
| NC_011770 | 1820245 | A | G | 8 | 10 | 2 | 0 | . | . | . | . | . | . | G | G | . | . | . | . |
| NC_011770 | 1820338 | A | G | 9 | 10 | 2 | 0 | . | . | . | . | . | . | G | G | . | . | . | . |
| NC_011770 | 1820349 | T | G | 9 | 10 | 2 | 0 | . | . | . | . | . | . | G | G | . | . | . | . |
| NC_011770 | 1820361 | G | T | 9 | 10 | 2 | 0 | . | . | . | . | . | . | T | T | . | . | . | . |
| NC_011770 | 1820401 | T | A | 8 | 10 | 2 | 0 | . | . | . | . | . | . | A | A | . | . | . | . |
| NC_011770 | 1820442 | T | C | 7 | 10 | 2 | 0 | . | . | . | . | . | . | C | C | . | . | . | . |
| NC_011770 | 1820481 | A | AG | 21 | 5 | 1 | 0 | . | . | | | | | AG | | . | . | . | |
| NC_011770 | 1820485 | G | GA | 14 | 6 | 2 | 0 | . | . | . | . | GA | GA | . | . | . | . | . | |
| NC_011770 | 1820509 | C | T | 23 | 5 | 7 | 0 | . | . | T | T | T | T | T | T | . | . | . | T |
| NC_011770 | 1820513 | G | A | 23 | 5 | 2 | 0 | . | . | | | | | A | A | . | . | . | |
| NC_011770 | 1820516 | G | A | 23 | 5 | 2 | 0 | . | . | | | | | A | A | . | . | . | |
| NC_011770 | 1820528 | G | T | 22 | 5 | 4 | 0 | . | . | . | T | T | T | | | . | . | . | T |

| | | | | | | | | | | | | | | | | | | | |
|---|---|---|---|---|---|---|---|---|---|---|---|---|---|---|---|---|---|---|---|
| NC_011770 | 1820534 | C | A | 18 | 5 | 5 | 0 | . | . | A | A | A | A | | | . | . | . | A |
| NC_011770 | 1820535 | T | C | 18 | 5 | 5 | 0 | . | . | C | C | C | C | | | . | . | . | C |
| NC_011770 | 1820536 | A | G | 18 | 5 | 5 | 0 | . | . | G | G | G | G | | | . | . | . | G |
| NC_011770 | 1820538 | C | T | 17 | 5 | 7 | 0 | . | . | T | T | T | T | T | T | . | . | . | T |
| NC_011770 | 1820551 | C | T | 17 | 5 | 5 | 0 | . | . | T | T | T | T | | | . | . | . | T |
| NC_011770 | 1820572 | G | A | 13 | 5 | 5 | 0 | . | . | A | A | A | A | | | . | . | . | A |
| NC_011770 | 1820600 | C | T | 11 | 5 | 2 | 0 | . | . | | | | | T | T | . | . | . | . |
| NC_011770 | 1820607 | T | G | 10 | 5 | 4 | 0 | . | . | G | G | G | G | | | . | . | . | . |
| NC_011770 | 1820655 | GA | G | 121 | 0 | 2 | 0 | G | G | | | | | | | | | | |
| NC_011770 | 1820662 | C | A | 121 | 0 | 2 | 0 | A | A | | | | | | | | | | |
| NC_011770 | 1820669 | T | A | 116 | 0 | 2 | 0 | A | A | | | | | | | | | | |
| NC_011770 | 1820675 | A | G | 115 | 0 | 2 | 0 | G | G | | | | | | | | | | |
| NC_011770 | 1820740 | A | G | 73 | 2 | 7 | 0 | . | . | G | G | G | G | G | | G | G | | |
| NC_011770 | 1820935 | A | T | 295 | 0 | 2 | 0 | T | T | | | | | | | | | | |
| NC_011770 | 1821830 | T | TGTCCC | 4 | 9 | 2 | 0 | . | . | . | . | . | . | | . | . | . | TGTCCCCT | TGTCCCCTTTATGCTC |
| NC_011770 | 1821849 | A | G | 6 | 10 | 2 | 0 | . | . | . | . | . | . | . | . | . | . | G | G |
| NC_011770 | 1822042 | AC | A | 4 | 10 | 2 | 0 | . | . | . | . | . | . | . | . | . | . | A | A |
| NC_011770 | 1822154 | T | C | 11 | 10 | 2 | 0 | . | . | . | . | . | . | . | . | . | . | C | C |
| NC_011770 | 1822157 | C | T | 11 | 10 | 2 | 0 | . | . | . | . | . | . | . | . | . | . | T | T |
| NC_011770 | 1822199 | A | T | 10 | 10 | 2 | 0 | . | . | . | . | . | . | . | . | . | . | T | T |
| NC_011770 | 1822245 | A | G | 9 | 10 | 2 | 0 | . | . | . | . | . | . | . | . | . | . | G | G |
| NC_011770 | 1822331 | A | G | 10 | 10 | 2 | 0 | . | . | . | . | . | . | . | . | . | . | G | G |
| NC_011770 | 1822342 | GA | G | 10 | 10 | 2 | 0 | . | . | . | . | . | . | . | . | . | . | G | G |
| NC_011770 | 1822352 | G | A | 10 | 10 | 2 | 0 | . | . | . | . | . | . | . | . | . | . | A | A |
| NC_011770 | 1822362 | T | G | 10 | 10 | 2 | 0 | . | . | . | . | . | . | . | . | . | . | G | G |
| NC_011770 | 1822399 | G | A | 11 | 10 | 2 | 0 | . | . | . | . | . | . | . | . | . | . | A | A |
| NC_011770 | 1822484 | C | G | 0 | 11 | 1 | 0 | . | . | . | . | . | . | G | . | . | . | . | . |
| NC_011770 | 1822497 | T | C | 65 | 4 | 8 | 0 | . | . | C | C | C | C | C | C | . | . | C | C |
| NC_011770 | 1822501 | C | T | 70 | 4 | 8 | 0 | . | . | T | T | T | T | T | T | . | . | T | T |
| NC_011770 | 1822545 | C | A | 98 | 4 | 4 | 0 | . | . | A | A | A | A | | | . | . | | |
| NC_011770 | 1822546 | T | C | 98 | 4 | 4 | 0 | . | . | C | C | C | C | | | . | . | | |
| NC_011770 | 1822562 | C | T | 95 | 4 | 4 | 0 | . | . | T | T | T | T | | | . | . | | |
| NC_011770 | 1822583 | G | A | 79 | 4 | 4 | 0 | . | . | A | A | A | A | | | . | . | | |
| NC_011770 | 1822599 | A | G | 58 | 4 | 4 | 0 | . | . | | | | | G | G | . | . | G | G |
| NC_011770 | 1822611 | C | T | 59 | 4 | 2 | 0 | . | . | | | | | T | T | . | . | | |
| NC_011770 | 1822617 | A | C | 58 | 4 | 8 | 0 | . | . | C | C | C | C | C | C | . | . | C | C |
| NC_011770 | 1822618 | A | G | 58 | 4 | 6 | 0 | . | . | G | G | G | G | T | T | . | . | . | . |
| NC_011770 | 1822625 | T | C | 58 | 4 | 6 | 0 | . | . | C | C | C | C | C | C | . | . | | |
| NC_011770 | 1822634 | T | TC | 62 | 4 | 4 | 0 | . | . | TC | TC | TC | TC | | | . | . | | |
| NC_011770 | 1822636 | G | GA | 58 | 4 | 4 | 0 | . | . | | | | | GA | GA | . | . | GA | GA |
| NC_011770 | 1825917 | G | A | 62 | 0 | 2 | 0 | A | A | | | | | | | | | | |
| NC_011770 | 1829938 | C | T | 89 | 0 | 4 | 0 | | | T | T | T | T | | | | | | |
| NC_011770 | 1830064 | C | G | 85 | 0 | 2 | 0 | | | | | | | G | G | | | | |
| NC_011770 | 1830081 | A | G | 86 | 0 | 12 | 0 | G | G | G | G | G | G | G | G | G | G | T | T |
| NC_011770 | 1830113 | T | C | 84 | 0 | 12 | 0 | C | C | C | C | C | C | C | C | C | C | C | C |
| NC_011770 | 1836312 | T | C | 75 | 0 | 12 | 0 | C | C | C | C | C | C | C | C | C | C | C | C |
| NC_011770 | 1836324 | CG | C | 77 | 0 | 12 | 0 | C | C | C | C | C | C | C | C | C | C | C | C |
| NC_011770 | 1836342 | T | G | 81 | 0 | 2 | 0 | | | | | | | | | | | G | G |
| NC_011770 | 1836366 | T | C | 78 | 0 | 4 | 0 | | | C | C | C | C | | | | | | |
| NC_011770 | 1839711 | G | C | 75 | 0 | 2 | 0 | | | | | | | C | C | | | | |
| NC_011770 | 1840923 | T | C | 82 | 0 | 2 | 0 | | | | | | | C | C | | | | |
| NC_011770 | 1841958 | C | T | 71 | 0 | 4 | 0 | | | T | T | T | T | | | | | | |
| NC_011770 | 1842884 | T | G | 44 | 1 | 1 | 0 | | | | | | | . | G | | | | |
| NC_011770 | 1849081 | T | C | 75 | 0 | 2 | 0 | | | | | | | | | | | C | C |
| NC_011770 | 1849082 | C | G | 75 | 0 | 2 | 0 | | | | | | | | | | | G | G |
| NC_011770 | 1850298 | A | G | 72 | 0 | 8 | 0 | G | G | G | G | G | G | | | G | G | | |
| NC_011770 | 1850977 | C | T | 88 | 0 | 12 | 0 | T | T | T | T | T | T | T | T | T | T | T | T |
| NC_011770 | 1850978 | C | T | 88 | 0 | 12 | 0 | T | T | T | T | T | T | T | T | T | T | T | T |
| NC_011770 | 1858029 | C | T | 65 | 0 | 6 | 0 | | | T | T | T | T | T | T | | | | |
| NC_011770 | 1858817 | G | A | 88 | 0 | 2 | 0 | | | | | | | | | A | A | | |
| NC_011770 | 1858877 | C | T | 84 | 0 | 2 | 0 | T | T | | | | | | | | | | |
| NC_011770 | 1872319 | C | T | 85 | 0 | 2 | 0 | | | | | | | T | T | | | | |

| | | | | | | | | | | | | | | | | | | | |
|---|---|---|---|---|---|---|---|---|---|---|---|---|---|---|---|---|---|---|---|
| NC_011770 | 1872367 | T | G | 86 | 0 | 2 | 0 | | | | | | | | | G | G | | |
| NC_011770 | 1872484 | A | G | 80 | 0 | 2 | 0 | | | | | | | | | G | G | | |
| NC_011770 | 1872485 | T | C | 79 | 0 | 2 | 0 | | | | | | | | | C | C | | |
| NC_011770 | 1872516 | T | A | 81 | 0 | 2 | 0 | | | | | | | | | A | A | | |
| NC_011770 | 1872572 | C | T | 78 | 0 | 2 | 0 | | | | | | | | | T | T | | |
| NC_011770 | 1872670 | A | G | 74 | 0 | 1 | 0 | | | | | | | G | | | | | |
| NC_011770 | 1872676 | G | A | 73 | 0 | 8 | 0 | A | A | | | | | A | A | A | A | A | A |
| NC_011770 | 1872688 | A | G | 78 | 0 | 2 | 0 | | | | | | | | | | | G | G |
| NC_011770 | 1872696 | GC | G | 69 | 0 | 2 | 0 | G | G | | | | | | | | | | |
| NC_011770 | 1872747 | C | T | 71 | 0 | 2 | 0 | | | | | | | | | | | T | T |
| NC_011770 | 1873375 | T | C | 50 | 0 | 2 | 0 | C | C | | | | | | | | | | |
| NC_011770 | 1873376 | T | A | 51 | 0 | 2 | 0 | A | A | | | | | | | | | | |
| NC_011770 | 1873379 | G | T | 49 | 0 | 2 | 0 | | | | | | | T | T | | | | |
| NC_011770 | 1873384 | C | T | 49 | 0 | 2 | 0 | T | T | | | | | | | | | | |
| NC_011770 | 1873389 | T | C | 53 | 0 | 2 | 0 | C | C | | | | | | | | | | |
| NC_011770 | 1873422 | C | T | 52 | 0 | 2 | 0 | T | T | | | | | | | | | | |
| NC_011770 | 1873426 | T | C | 50 | 0 | 4 | 0 | C | C | | | | | | | C | C | | |
| NC_011770 | 1873498 | GCA | GTCA | 50 | 0 | 6 | 0 | GTCA | GTCA | | | | | G | G | | | G | G |
| NC_011770 | 1873499 | CA | C | 44 | 2 | 2 | 0 | . | . | | | | | . | . | C | C | . | . |
| NC_011770 | 1873502 | T | TC | 51 | 0 | 4 | 0 | | | | | | | TC | TC | | | TC | TC |
| NC_011770 | 1873517 | GC | G | 51 | 0 | 2 | 0 | G | G | | | | | | | | | | |
| NC_011770 | 1873520 | G | A | 48 | 2 | 4 | 0 | . | . | | | | | A | A | | | A | A |
| NC_011770 | 1873546 | A | G | 52 | 0 | 2 | 0 | G | G | | | | | | | | | | |
| NC_011770 | 1873551 | G | A | 46 | 0 | 2 | 0 | A | A | | | | | | | | | | |
| NC_011770 | 1873585 | A | C | 47 | 0 | 2 | 0 | C | C | | | | | | | | | | |
| NC_011770 | 1873597 | C | T | 49 | 0 | 2 | 0 | | | | | | | | | T | T | | |
| NC_011770 | 1873616 | G | A | 49 | 0 | 2 | 0 | | | | | | | A | A | | | | |
| NC_011770 | 1873639 | C | A | 48 | 0 | 6 | 0 | A | A | | | | | A | A | | | A | A |
| NC_011770 | 1873640 | C | T | 48 | 0 | 2 | 0 | T | T | | | | | | | | | | |
| NC_011770 | 1873649 | A | AG | 52 | 0 | 2 | 0 | AG | AG | | | | | | | | | | |
| NC_011770 | 1877439 | G | T | 79 | 0 | 8 | 0 | T | T | | | | | T | T | T | T | T | T |
| NC_011770 | 1877446 | C | T | 76 | 0 | 8 | 0 | T | T | | | | | T | T | T | T | T | T |
| NC_011770 | 1877457 | G | A | 75 | 0 | 8 | 0 | A | A | | | | | A | A | A | A | A | A |
| NC_011770 | 1877504 | C | G | 75 | 0 | 6 | 0 | | | | | | | G | G | G | G | G | G |
| NC_011770 | 1877510 | C | T | 76 | 0 | 6 | 0 | | | | | | | T | T | T | T | T | T |
| NC_011770 | 1877515 | A | C | 76 | 0 | 6 | 0 | | | | | | | C | C | C | C | C | C |
| NC_011770 | 1877526 | A | G | 74 | 0 | 8 | 0 | G | G | | | | | G | G | G | G | G | G |
| NC_011770 | 1879358 | A | G | 77 | 0 | 2 | 0 | G | G | | | | | | | | | | |
| NC_011770 | 1879377 | G | C | 75 | 0 | 4 | 0 | | | | | | | C | C | C | C | | |
| NC_011770 | 1883516 | A | G | 59 | 0 | 8 | 0 | G | G | | | | | G | G | G | G | G | G |
| NC_011770 | 1883518 | T | TCC | 59 | 0 | 8 | 0 | TCC | TCC | | | | | TCC | TCC | TCC | TCC | TCC | TCC |
| NC_011770 | 1890385 | GGT | G | 42 | 0 | 1 | 0 | G | . | | | | | | | | | | |
| NC_011770 | 1890387 | T | G | 37 | 2 | 4 | 0 | . | . | | | | | | | G | G | G | G |
| NC_011770 | 1890390 | C | CG | 41 | 0 | 2 | 0 | CG | CG | | | | | | | | | | |
| NC_011770 | 1892084 | G | A | 65 | 0 | 2 | 0 | | | | | | | A | A | | | | |
| NC_011770 | 1892852 | G | A | 82 | 0 | 2 | 0 | | | | | | | | | A | A | | |
| NC_011770 | 1895850 | A | C | 65 | 0 | 2 | 0 | | | | | | | C | C | | | | |
| NC_011770 | 1895865 | T | G | 66 | 0 | 8 | 0 | G | G | G | G | G | G | G | G | | | | |
| NC_011770 | 1895937 | C | T | 62 | 0 | 2 | 0 | | | | | | | T | T | | | | |
| NC_011770 | 1896314 | A | G | 60 | 0 | 12 | 0 | G | G | G | G | G | G | G | G | G | G | G | G |
| NC_011770 | 1901929 | G | A | 73 | 0 | 4 | 0 | | | A | A | A | A | | | | | | |
| NC_011770 | 1901948 | A | G | 71 | 0 | 12 | 0 | G | G | G | G | G | G | G | G | G | G | G | G |
| NC_011770 | 1902026 | G | A | 69 | 0 | 8 | 0 | A | A | | | | | A | A | A | A | A | A |
| NC_011770 | 1902027 | T | C | 71 | 0 | 4 | 0 | | | C | C | C | C | | | | | | |
| NC_011770 | 1902034 | A | G | 74 | 0 | 4 | 0 | | | G | G | G | G | | | | | | |
| NC_011770 | 1902037 | C | T | 74 | 0 | 4 | 0 | | | T | T | T | T | | | | | | |
| NC_011770 | 1903483 | C | T | 74 | 0 | 2 | 0 | T | T | | | | | | | | | | |
| NC_011770 | 1905441 | C | T | 69 | 0 | 4 | 0 | | | T | T | T | T | | | | | | |
| NC_011770 | 1913346 | A | G | 86 | 0 | 2 | 0 | | | | | | | G | G | | | | |
| NC_011770 | 1913422 | C | A | 86 | 0 | 4 | 0 | | | A | A | A | A | | | | | | |
| NC_011770 | 1914037 | C | T | 92 | 0 | 2 | 0 | T | T | | | | | | | | | | |
| NC_011770 | 1914049 | G | A | 95 | 0 | 2 | 0 | A | A | | | | | | | | | | |

| | | | | | | | | | | | | | | | | | | | |
|---|---|---|---|---|---|---|---|---|---|---|---|---|---|---|---|---|---|---|---|
| NC_011770 | 1914097 | A | C | 86 | 0 | 6 | 0 | | | C | C | C | C | C | C | | | | |
| NC_011770 | 1914098 | C | G | 86 | 0 | 6 | 0 | | | G | G | G | G | G | G | | | | |
| NC_011770 | 1914102 | C | A | 89 | 0 | 6 | 0 | | | A | A | A | A | A | A | | | | |
| NC_011770 | 1914106 | T | A | 87 | 0 | 6 | 0 | | | A | A | A | A | A | A | | | | |
| NC_011770 | 1914114 | G | T | 90 | 0 | 6 | 0 | | | T | T | T | T | T | T | | | | |
| NC_011770 | 1914118 | C | G | 92 | 0 | 6 | 0 | | | G | G | G | G | G | G | | | | |
| NC_011770 | 1914158 | C | A | 93 | 0 | 6 | 0 | | | A | A | A | A | A | A | | | | |
| NC_011770 | 1917115 | C | T | 59 | 0 | 2 | 0 | | | | | | | | | | | T | T |
| NC_011770 | 1917118 | G | A | 58 | 0 | 2 | 0 | A | A | | | | | | | | | | |
| NC_011770 | 1917119 | G | A | 55 | 0 | 8 | 0 | | | A | A | A | A | | | A | A | A | A |
| NC_011770 | 1917130 | C | CA | 58 | 0 | 2 | 0 | | | | | | | CA | CA | | | | |
| NC_011770 | 1917156 | G | C | 58 | 0 | 2 | 0 | C | C | | | | | | | | | | |
| NC_011770 | 1917186 | T | C | 59 | 0 | 12 | 0 | C | C | C | C | C | C | C | C | C | C | C | C |
| NC_011770 | 1917191 | G | T | 63 | 0 | 2 | 0 | T | T | | | | | | | | | | |
| NC_011770 | 1917227 | A | G | 67 | 0 | 10 | 0 | | | G | G | G | G | G | G | G | G | G | G |
| NC_011770 | 1917411 | T | C | 71 | 0 | 10 | 0 | | | C | C | C | C | C | C | C | C | C | C |
| NC_011770 | 1917427 | A | G | 73 | 0 | 2 | 0 | | | | | | | G | G | | | | |
| NC_011770 | 1917480 | C | T | 74 | 0 | 8 | 0 | | | T | T | T | T | | | T | T | T | T |
| NC_011770 | 1917489 | C | T | 74 | 0 | 4 | 0 | | | T | T | T | T | | | | | | |
| NC_011770 | 1917491 | T | C | 72 | 0 | 8 | 0 | | | C | C | C | C | | | C | C | C | C |
| NC_011770 | 1917531 | T | C | 77 | 0 | 8 | 0 | | | C | C | C | C | | | C | C | C | C |
| NC_011770 | 1917607 | T | A | 79 | 0 | 8 | 0 | | | A | A | A | A | | | A | A | A | A |
| NC_011770 | 1917653 | C | T | 79 | 0 | 4 | 0 | | | | | | | | | T | T | T | T |
| NC_011770 | 1917663 | C | T | 80 | 0 | 4 | 0 | | | | | | | | | T | T | T | T |
| NC_011770 | 1917703 | C | A | 81 | 0 | 2 | 0 | | | | | | | A | A | | | | |
| NC_011770 | 1917709 | C | T | 78 | 0 | 4 | 0 | | | T | T | T | T | | | | | | |
| NC_011770 | 1917776 | G | T | 81 | 0 | 4 | 0 | | | | | | | | | T | T | T | T |
| NC_011770 | 1917815 | C | T | 78 | 0 | 8 | 0 | | | T | T | T | T | | | T | T | T | T |
| NC_011770 | 1917847 | A | G | 77 | 0 | 4 | 0 | | | G | G | G | G | | | | | | |
| NC_011770 | 1917849 | A | T | 77 | 0 | 8 | 0 | | | T | T | T | T | | | T | T | T | T |
| NC_011770 | 1917857 | C | T | 79 | 0 | 2 | 0 | | | | | | | T | T | | | | |
| NC_011770 | 1917872 | G | T | 81 | 0 | 2 | 0 | T | T | | | | | | | | | | |
| NC_011770 | 1917878 | C | T | 75 | 0 | 4 | 0 | | | | | | | | | T | T | T | T |
| NC_011770 | 1917898 | G | A | 78 | 0 | 2 | 0 | | | | | | | A | A | | | | |
| NC_011770 | 1917910 | C | T | 80 | 0 | 2 | 0 | | | | | | | T | T | | | | |
| NC_011770 | 1917927 | A | G | 77 | 0 | 10 | 0 | G | G | G | G | G | G | | | G | G | G | G |
| NC_011770 | 1917928 | G | C | 77 | 0 | 10 | 0 | C | C | C | C | C | C | | | C | C | C | C |
| NC_011770 | 1917952 | G | A | 76 | 0 | 2 | 0 | | | | | | | A | A | | | | |
| NC_011770 | 1917954 | C | T | 76 | 0 | 2 | 0 | | | | | | | T | T | | | | |
| NC_011770 | 1917964 | C | T | 80 | 0 | 6 | 0 | | | | | | | T | T | T | T | T | T |
| NC_011770 | 1917967 | C | T | 81 | 0 | 2 | 0 | | | | | | | T | T | | | | |
| NC_011770 | 1918002 | G | A | 78 | 0 | 10 | 0 | A | A | A | A | A | A | | | A | A | A | A |
| NC_011770 | 1918003 | C | A | 78 | 0 | 4 | 0 | | | | | | | | | A | A | A | A |
| NC_011770 | 1918005 | T | C | 78 | 0 | 12 | 0 | C | C | C | C | C | C | C | C | C | C | C | C |
| NC_011770 | 1918037 | C | T | 77 | 0 | 2 | 0 | | | | | | | T | T | | | | |
| NC_011770 | 1918074 | TA | T | 77 | 0 | 12 | 0 | T | T | T | T | T | T | T | T | T | T | T | T |
| NC_011770 | 1918083 | A | G | 75 | 0 | 6 | 0 | G | G | G | G | G | G | | | | | | |
| NC_011770 | 1918088 | G | T | 77 | 0 | 2 | 0 | | | | | | | T | T | | | | |
| NC_011770 | 1921323 | T | A | 66 | 0 | 12 | 0 | A | A | A | A | A | A | A | A | A | A | A | A |
| NC_011770 | 1922955 | T | G | 24 | 0 | 10 | 0 | G | G | G | G | G | G | G | G | | | G | G |
| NC_011770 | 1925117 | G | A | 62 | 0 | 4 | 0 | | | A | A | A | A | | | | | | |
| NC_011770 | 1925263 | G | A | 54 | 0 | 2 | 0 | | | | | | | | | | | A | A |
| NC_011770 | 1925405 | T | C | 48 | 0 | 4 | 0 | C | C | | | | | C | C | | | | |
| NC_011770 | 1925421 | A | G | 48 | 0 | 4 | 0 | G | G | | | | | G | G | | | | |
| NC_011770 | 1925434 | A | G | 50 | 0 | 4 | 0 | G | G | | | | | G | G | | | | |
| NC_011770 | 1925439 | T | C | 49 | 0 | 4 | 0 | C | C | | | | | C | C | | | | |
| NC_011770 | 1925445 | T | TC | 51 | 0 | 2 | 0 | | | | | | | | | | | TC | TC |
| NC_011770 | 1925447 | C | T | 42 | 2 | 4 | 0 | T | T | | | | | T | T | | | . | . |
| NC_011770 | 1925657 | G | T | 66 | 0 | 6 | 0 | T | T | | | | | T | T | T | T | | |
| NC_011770 | 1925674 | G | A | 67 | 0 | 2 | 0 | A | A | | | | | | | | | | |
| NC_011770 | 1928335 | A | C | 76 | 0 | 10 | 0 | C | C | C | C | C | C | C | C | C | C | | |
| NC_011770 | 1929937 | T | C | 80 | 0 | 2 | 0 | C | C | | | | | | | | | | |

| | | | | | | | | | | | | | | | | | | | |
|---|---|---|---|---|---|---|---|---|---|---|---|---|---|---|---|---|---|---|---|
| NC_011770 | 1932139 | A | G | 64 | 0 | 4 | 0 | | | G | G | G | G | | | | | | |
| NC_011770 | 1933996 | G | A | 54 | 0 | 12 | 0 | A | A | A | A | A | A | A | A | A | A | A | A |
| NC_011770 | 1934041 | G | A | 51 | 0 | 12 | 0 | A | A | A | A | A | A | A | A | A | A | A | A |
| NC_011770 | 1934961 | G | A | 69 | 0 | 4 | 0 | | | A | A | A | A | | | | | | |
| NC_011770 | 1942894 | C | T | 91 | 0 | 2 | 0 | | | | | | | | | T | T | | |
| NC_011770 | 1942946 | A | G | 85 | 0 | 12 | 0 | G | G | G | G | G | G | G | G | G | G | G | G |
| NC_011770 | 1951892 | C | G | 45 | 0 | 2 | 0 | | | | | | | | | G | G | | |
| NC_011770 | 1951894 | A | G | 45 | 0 | 6 | 0 | G | G | | | | | | | G | G | G | G |
| NC_011770 | 1952111 | C | T | 66 | 0 | 2 | 0 | T | T | | | | | | | | | | |
| NC_011770 | 1952112 | A | C | 64 | 0 | 2 | 0 | C | C | | | | | | | | | | |
| NC_011770 | 1952132 | C | A | 70 | 0 | 2 | 0 | | | | | | | A | A | | | | |
| NC_011770 | 1952153 | T | G | 66 | 0 | 4 | 0 | G | G | | | | | | | G | G | | |
| NC_011770 | 1952157 | T | C | 69 | 0 | 2 | 0 | | | | | | | | | C | C | | |
| NC_011770 | 1952174 | A | G | 68 | 0 | 4 | 0 | G | G | | | | | | | G | G | | |
| NC_011770 | 1952196 | A | G | 67 | 0 | 2 | 0 | G | G | | | | | | | | | | |
| NC_011770 | 1953121 | T | C | 69 | 0 | 4 | 0 | C | C | | | | | C | C | | | | |
| NC_011770 | 1953396 | C | T | 76 | 0 | 2 | 0 | T | T | | | | | | | | | | |
| NC_011770 | 1953574 | G | A | 81 | 0 | 2 | 0 | | | | | | | | | | | A | A |
| NC_011770 | 1953612 | G | A | 80 | 0 | 2 | 0 | | | | | | | | | | | A | A |
| NC_011770 | 1953739 | T | C | 63 | 0 | 2 | 0 | | | | | | | C | C | | | | |
| NC_011770 | 1953764 | C | T | 67 | 0 | 4 | 0 | | | | | | | T | T | T | T | | |
| NC_011770 | 1953787 | G | T | 72 | 0 | 2 | 0 | | | | | | | T | T | | | | |
| NC_011770 | 1953812 | G | T | 67 | 0 | 2 | 0 | | | | | | | T | T | | | | |
| NC_011770 | 1953841 | C | T | 68 | 0 | 2 | 0 | | | | | | | | | | | T | T |
| NC_011770 | 1953852 | G | C | 68 | 0 | 2 | 0 | | | | | | | C | C | | | | |
| NC_011770 | 1953865 | A | G | 69 | 0 | 2 | 0 | | | | | | | G | G | | | | |
| NC_011770 | 1955190 | G | T | 56 | 0 | 8 | 0 | | | T | T | T | T | T | T | | | T | T |
| NC_011770 | 1955211 | A | C | 60 | 0 | 6 | 0 | | | C | C | C | C | C | C | | | | |
| NC_011770 | 1960968 | C | G | 73 | 0 | 6 | 0 | G | G | G | G | G | G | | | | | | |
| NC_011770 | 1961002 | A | G | 76 | 0 | 4 | 0 | | | G | G | G | G | | | | | | |
| NC_011770 | 1965123 | G | C | 56 | 0 | 2 | 0 | C | C | | | | | | | | | | |
| NC_011770 | 1966350 | G | A | 79 | 0 | 2 | 0 | A | A | | | | | | | | | | |
| NC_011770 | 1966362 | G | A | 79 | 0 | 2 | 0 | | | | | | | | | | | A | A |
| NC_011770 | 1966451 | G | A | 85 | 0 | 10 | 0 | A | A | A | A | A | A | A | A | | | A | A |
| NC_011770 | 1966452 | C | G | 87 | 0 | 2 | 0 | | | | | | | | | | | G | G |
| NC_011770 | 1966465 | C | G | 82 | 0 | 12 | 0 | G | G | G | G | G | G | G | G | G | G | G | G |
| NC_011770 | 1966492 | G | A | 82 | 0 | 2 | 0 | | | | | | | | | A | A | | |
| NC_011770 | 1967934 | C | G | 51 | 0 | 2 | 0 | | | | | | | | | G | G | | |
| NC_011770 | 1969260 | C | T | 55 | 0 | 2 | 0 | | | | | | | | | | | T | T |
| NC_011770 | 1972140 | A | G | 68 | 0 | 4 | 0 | G | G | | | | | G | G | | | | |
| NC_011770 | 1974395 | C | T | 75 | 0 | 2 | 0 | | | | | | | | | | | T | T |
| NC_011770 | 1975356 | G | A | 70 | 0 | 2 | 0 | A | A | | | | | | | | | | |
| NC_011770 | 1975357 | T | C | 69 | 0 | 2 | 0 | C | C | | | | | | | | | | |
| NC_011770 | 1975359 | C | A | 70 | 0 | 2 | 0 | A | A | | | | | | | | | | |
| NC_011770 | 1975382 | G | A | 68 | 0 | 2 | 0 | A | A | | | | | | | | | | |
| NC_011770 | 1975398 | G | T | 68 | 0 | 2 | 0 | T | T | | | | | | | | | | |
| NC_011770 | 1975405 | G | A | 68 | 0 | 2 | 0 | A | A | | | | | | | | | | |
| NC_011770 | 1975408 | T | C | 67 | 0 | 2 | 0 | C | C | | | | | | | | | | |
| NC_011770 | 1975412 | T | C | 69 | 0 | 2 | 0 | C | C | | | | | | | | | | |
| NC_011770 | 1975417 | A | G | 68 | 0 | 2 | 0 | G | G | | | | | | | | | | |
| NC_011770 | 1975433 | AC | A | 66 | 0 | 2 | 0 | A | A | | | | | | | | | | |
| NC_011770 | 1977669 | G | A | 81 | 0 | 4 | 0 | | | A | A | A | A | | | | | | |
| NC_011770 | 1978164 | G | A | 90 | 0 | 2 | 0 | A | A | | | | | | | | | | |
| NC_011770 | 1978214 | G | A | 83 | 0 | 2 | 0 | | | | | | | | | A | A | | |
| NC_011770 | 1979159 | A | G | 54 | 0 | 4 | 0 | G | G | | | | | G | G | | | | |
| NC_011770 | 1979188 | GT | G | 57 | 0 | 2 | 0 | | | | | | | | | | | G | G |
| NC_011770 | 1979235 | G | A | 58 | 0 | 4 | 0 | | | A | A | A | A | | | | | | |
| NC_011770 | 1984119 | C | T | 57 | 0 | 2 | 0 | T | T | | | | | | | | | | |
| NC_011770 | 1984120 | G | GTGGTT | 53 | 0 | 1 | 0 | | | | | | | | | GTGGTTGTCCGCCGTCC | | | |
| NC_011770 | 1984253 | C | T | 60 | 0 | 2 | 0 | | | | | | | | | T | T | | |
| NC_011770 | 1984269 | T | A | 59 | 0 | 6 | 0 | | | | | | | A | A | A | A | A | A |
| NC_011770 | 1985489 | C | G | 84 | 0 | 2 | 0 | G | G | | | | | | | | | | |

| | | | | | | | | | | | | | | | | | | | |
|---|---|---|---|---|---|---|---|---|---|---|---|---|---|---|---|---|---|---|---|
| NC_011770 | 1985889 | G | A | 73 | 0 | 2 | 0 | A | A | | | | | | | | | | |
| NC_011770 | 1985989 | A | C | 68 | 0 | 2 | 0 | | | | | | | C | C | | | | |
| NC_011770 | 1987921 | G | A | 81 | 0 | 4 | 0 | | | A | A | A | A | | | | | | |
| NC_011770 | 1990380 | G | A | 81 | 0 | 2 | 0 | | | | | | | | | A | A | | |
| NC_011770 | 1990852 | T | C | 56 | 0 | 2 | 0 | | | | | | | | | C | C | | |
| NC_011770 | 1991632 | C | A | 66 | 0 | 2 | 0 | | | | | | | A | A | | | | |
| NC_011770 | 1991665 | C | T | 65 | 0 | 4 | 0 | | | T | T | T | T | | | | | | |
| NC_011770 | 1993806 | C | T | 74 | 0 | 6 | 0 | | | | | | | T | T | T | T | T | T |
| NC_011770 | 1993809 | G | C | 75 | 0 | 6 | 0 | | | | | | | C | C | C | C | C | C |
| NC_011770 | 1993861 | G | T | 78 | 0 | 10 | 0 | | | T | T | T | T | T | T | T | T | T | T |
| NC_011770 | 1994719 | C | T | 64 | 0 | 4 | 0 | | | | | | | | | T | T | T | T |
| NC_011770 | 1995244 | G | A | 77 | 0 | 4 | 0 | | | A | A | A | A | | | | | | |
| NC_011770 | 1995378 | C | T | 75 | 0 | 1 | 0 | | | | | | | T | | | | | |
| NC_011770 | 1995430 | G | T | 74 | 0 | 4 | 0 | | | T | T | T | T | | | | | | |
| NC_011770 | 1995490 | C | T | 79 | 0 | 2 | 0 | | | | | | | T | T | | | | |
| NC_011770 | 1995632 | T | C | 78 | 0 | 6 | 0 | | | | | | | C | C | C | C | C | C |
| NC_011770 | 1995643 | G | A | 81 | 0 | 6 | 0 | A | A | A | A | A | A | | | | | | |
| NC_011770 | 1995646 | G | A | 81 | 0 | 6 | 0 | A | A | A | A | A | A | | | | | | |
| NC_011770 | 1995682 | T | G | 78 | 0 | 6 | 0 | | | | | | | G | G | G | G | G | G |
| NC_011770 | 1995700 | G | A | 80 | 0 | 2 | 0 | | | | | | | A | A | | | | |
| NC_011770 | 1997717 | C | T | 75 | 0 | 4 | 0 | | | T | T | T | T | | | | | | |
| NC_011770 | 2003902 | G | T | 67 | 0 | 2 | 0 | T | T | | | | | | | | | | |
| NC_011770 | 2003925 | C | T | 62 | 0 | 2 | 0 | | | | | | | | | T | T | | |
| NC_011770 | 2004440 | T | C | 77 | 0 | 8 | 0 | | | C | C | C | C | | | C | C | C | C |
| NC_011770 | 2005521 | G | T | 79 | 0 | 2 | 0 | | | | | | | | | | | T | T |
| NC_011770 | 2006907 | C | T | 52 | 0 | 4 | 0 | | | | | | | T | T | T | T | | |
| NC_011770 | 2008247 | A | G | 70 | 0 | 6 | 0 | G | G | G | G | G | G | | | | | | |
| NC_011770 | 2008711 | G | A | 74 | 0 | 4 | 0 | | | A | A | A | A | | | | | | |
| NC_011770 | 2008715 | G | A | 74 | 0 | 2 | 0 | | | | | | | A | A | | | | |
| NC_011770 | 2009706 | A | G | 90 | 0 | 2 | 0 | G | G | | | | | | | | | | |
| NC_011770 | 2009802 | G | A | 86 | 0 | 2 | 0 | | | | | | | | | | | A | A |
| NC_011770 | 2009812 | A | C | 84 | 0 | 2 | 0 | C | C | | | | | | | | | | |
| NC_011770 | 2009846 | G | A | 85 | 0 | 2 | 0 | | | | | | | | | | | A | A |
| NC_011770 | 2011442 | G | C | 74 | 0 | 4 | 0 | | | C | C | C | C | | | | | | |
| NC_011770 | 2012356 | T | C | 74 | 0 | 4 | 0 | | | C | C | C | C | | | | | | |
| NC_011770 | 2012363 | C | T | 76 | 0 | 4 | 0 | | | | | | | T | T | T | T | | |
| NC_011770 | 2013554 | C | T | 21 | 2 | 5 | 0 | T | . | T | T | T | T | | | | | . | . |
| NC_011770 | 2013555 | CG | C | 21 | 1 | 6 | 0 | . | . | C | C | C | C | | | | | C | C |
| NC_011770 | 2014170 | T | A | 77 | 0 | 12 | 0 | A | A | A | A | A | A | A | A | A | A | A | A |
| NC_011770 | 2014189 | A | G | 80 | 0 | 12 | 0 | G | G | G | G | G | G | G | G | G | G | G | G |
| NC_011770 | 2015476 | C | T | 90 | 0 | 2 | 0 | | | | | | | | | | | T | T |
| NC_011770 | 2018483 | C | T | 77 | 0 | 4 | 0 | T | T | | | | | | | T | T | | |
| NC_011770 | 2020670 | T | C | 49 | 0 | 8 | 0 | C | C | C | C | C | C | | | C | C | | |
| NC_011770 | 2020673 | G | C | 49 | 0 | 8 | 0 | C | C | C | C | C | C | | | C | C | | |
| NC_011770 | 2020692 | G | A | 50 | 0 | 4 | 0 | | | A | A | A | A | | | | | | |
| NC_011770 | 2020695 | C | T | 50 | 0 | 4 | 0 | | | T | T | T | T | | | | | | |
| NC_011770 | 2020714 | C | T | 52 | 0 | 8 | 0 | T | T | T | T | T | T | | | T | T | | |
| NC_011770 | 2020821 | A | C | 55 | 0 | 2 | 0 | | | | | | | C | C | | | | |
| NC_011770 | 2020844 | G | A | 57 | 0 | 2 | 0 | A | A | | | | | | | | | | |
| NC_011770 | 2020957 | G | C | 61 | 0 | 2 | 0 | | | | | | | | | | | C | C |
| NC_011770 | 2021033 | A | G | 64 | 0 | 4 | 0 | G | G | | | | | | | G | G | | |
| NC_011770 | 2021046 | T | C | 64 | 0 | 10 | 0 | C | C | C | C | C | C | | | C | C | C | C |
| NC_011770 | 2021074 | A | G | 70 | 0 | 2 | 0 | | | | | | | | | | | G | G |
| NC_011770 | 2021134 | A | G | 71 | 0 | 2 | 0 | | | | | | | | | | | G | G |
| NC_011770 | 2024331 | A | AC | 69 | 0 | 2 | 0 | | | | | | | AC | AC | | | | |
| NC_011770 | 2026920 | C | T | 71 | 0 | 6 | 0 | T | T | | | | | | | T | T | T | T |
| NC_011770 | 2026953 | G | A | 69 | 0 | 6 | 0 | A | A | | | | | | | A | A | A | A |
| NC_011770 | 2026979 | GT | G | 69 | 0 | 2 | 0 | | | | | | | | | | | G | G |
| NC_011770 | 2028922 | T | C | 72 | 0 | 12 | 0 | C | C | G | G | G | G | G | G | C | C | C | C |
| NC_011770 | 2031956 | G | A | 62 | 0 | 12 | 0 | A | A | A | A | A | A | A | A | A | A | A | A |
| NC_011770 | 2031967 | A | G | 63 | 0 | 6 | 0 | G | G | G | G | G | G | | | | | | |
| NC_011770 | 2031976 | G | C | 62 | 0 | 4 | 0 | | | C | C | C | C | | | | | | |

| | | | | | | | | | | | | | | | | | | | |
|---|---|---|---|---|---|---|---|---|---|---|---|---|---|---|---|---|---|---|---|
| NC_011770 | 2033810 | T | A | 76 | 0 | 6 | 0 | A | A | A | A | A | A | | | | | | |
| NC_011770 | 2033827 | T | TC | 80 | 0 | 12 | 0 | TC | TC | TC | TC | TC | TC | TC | TC | TC | TC | TC | TC |
| NC_011770 | 2034195 | C | A | 78 | 0 | 12 | 0 | A | A | A | A | A | A | A | A | A | A | A | A |
| NC_011770 | 2034219 | A | G | 79 | 0 | 12 | 0 | G | G | G | G | G | G | G | G | G | G | G | G |
| NC_011770 | 2034267 | C | T | 77 | 0 | 4 | 0 | | | | | | | T | T | T | T | | |
| NC_011770 | 2034290 | T | C | 77 | 0 | 2 | 0 | C | C | | | | | | | | | | |
| NC_011770 | 2035389 | C | A | 73 | 0 | 10 | 0 | | | A | A | A | A | A | A | A | A | A | A |
| NC_011770 | 2040767 | G | C | 46 | 0 | 4 | 0 | | | C | C | C | C | | | | | | |
| NC_011770 | 2040805 | G | A | 48 | 0 | 4 | 0 | | | A | A | A | A | | | | | | |
| NC_011770 | 2042295 | G | C | 76 | 0 | 6 | 0 | C | C | C | C | C | C | | | | | | |
| NC_011770 | 2043412 | C | A | 70 | 0 | 2 | 0 | | | | | | | | | | | A | A |
| NC_011770 | 2043475 | C | T | 76 | 0 | 2 | 0 | | | T | T | | | | | | | | |
| NC_011770 | 2046421 | G | A | 69 | 0 | 6 | 0 | | | | | | | A | A | A | A | A | A |
| NC_011770 | 2048715 | C | T | 69 | 0 | 2 | 0 | T | T | | | | | | | | | | |
| NC_011770 | 2048717 | T | C | 68 | 0 | 1 | 0 | | | | | | | C | | | | | |
| NC_011770 | 2052798 | T | C | 68 | 0 | 8 | 0 | C | C | C | C | C | C | | | | | C | C |
| NC_011770 | 2052850 | T | C | 61 | 0 | 8 | 0 | C | C | C | C | C | C | | | | | C | C |
| NC_011770 | 2052852 | G | A | 62 | 0 | 4 | 0 | | | A | A | A | A | | | | | | |
| NC_011770 | 2052911 | T | C | 64 | 0 | 12 | 0 | C | C | C | C | C | C | C | C | C | C | C | C |
| NC_011770 | 2053011 | C | T | 61 | 0 | 4 | 0 | | | T | T | T | T | | | | | | |
| NC_011770 | 2054728 | T | C | 69 | 0 | 2 | 0 | | | | | | | | | C | C | | |
| NC_011770 | 2070552 | C | T | 66 | 0 | 4 | 0 | | | T | T | T | T | | | | | | |
| NC_011770 | 2070613 | T | G | 66 | 0 | 2 | 0 | | | | | | | | | G | G | | |
| NC_011770 | 2070679 | T | C | 78 | 0 | 1 | 0 | | | | | | | C | | | | | |
| NC_011770 | 2071180 | G | A | 69 | 0 | 2 | 0 | | | | | | | | | | | A | A |
| NC_011770 | 2071233 | A | T | 69 | 0 | 2 | 0 | | | | | | | | | T | T | | |
| NC_011770 | 2074637 | T | A | 88 | 0 | 2 | 0 | | | | | | | | | A | A | | |
| NC_011770 | 2077296 | C | T | 77 | 0 | 2 | 0 | T | T | | | | | | | | | | |
| NC_011770 | 2078252 | C | T | 82 | 0 | 2 | 0 | | | | | | | T | T | | | | |
| NC_011770 | 2079779 | T | C | 65 | 0 | 2 | 0 | | | | | | | C | C | | | | |
| NC_011770 | 2079781 | C | T | 63 | 0 | 4 | 0 | | | | | | | | | T | T | T | T |
| NC_011770 | 2079824 | G | T | 57 | 0 | 4 | 0 | | | | | | | | | T | T | T | T |
| NC_011770 | 2079906 | G | A | 60 | 0 | 4 | 0 | | | | | | | | | A | A | A | A |
| NC_011770 | 2080201 | A | G | 45 | 0 | 4 | 0 | | | | | | | | | G | G | G | G |
| NC_011770 | 2080210 | C | T | 42 | 0 | 2 | 0 | | | | | | | | | | | T | T |
| NC_011770 | 2080211 | G | A | 42 | 0 | 2 | 0 | | | | | | | | | A | A | | |
| NC_011770 | 2080339 | C | T | 59 | 0 | 2 | 0 | | | | | | | | | T | T | | |
| NC_011770 | 2080343 | C | T | 58 | 0 | 4 | 0 | | | T | T | T | T | | | | | | |
| NC_011770 | 2080461 | CT | C | 56 | 0 | 2 | 0 | | | | | | | | | | | C | C |
| NC_011770 | 2080517 | GT | G | 53 | 0 | 2 | 0 | | | | | | | | | | | G | G |
| NC_011770 | 2080550 | C | T | 51 | 0 | 4 | 0 | | | | | | | | | T | T | T | T |
| NC_011770 | 2080595 | T | C | 51 | 0 | 4 | 0 | | | | | | | | | C | C | C | C |
| NC_011770 | 2080596 | C | T | 51 | 0 | 4 | 0 | | | | | | | | | T | T | T | T |
| NC_011770 | 2081168 | A | G | 69 | 0 | 6 | 0 | G | G | | | | | G | G | G | G | | |
| NC_011770 | 2082050 | T | C | 45 | 0 | 4 | 0 | | | C | C | C | C | | | | | | |
| NC_011770 | 2082118 | A | G | 72 | 0 | 8 | 0 | | | G | G | G | G | | | G | G | G | G |
| NC_011770 | 2082136 | G | A | 76 | 0 | 8 | 0 | | | A | A | A | A | | | A | A | A | A |
| NC_011770 | 2084287 | C | G | 90 | 0 | 8 | 0 | G | G | G | G | G | G | G | G | | | | |
| NC_011770 | 2085049 | C | T | 90 | 0 | 2 | 0 | | | | | | | | | T | T | | |
| NC_011770 | 2085053 | A | G | 89 | 0 | 2 | 0 | G | G | | | | | | | | | | |
| NC_011770 | 2087840 | T | C | 75 | 0 | 4 | 0 | | | C | C | C | C | | | | | | |
| NC_011770 | 2087894 | G | A | 71 | 0 | 10 | 0 | | | A | A | A | A | A | A | A | A | A | A |
| NC_011770 | 2087900 | G | A | 73 | 0 | 10 | 0 | | | A | A | A | A | A | A | A | A | A | A |
| NC_011770 | 2087903 | G | A | 73 | 0 | 2 | 0 | | | | | | | | | A | A | | |
| NC_011770 | 2087905 | A | C | 73 | 0 | 2 | 0 | | | | | | | | | C | C | | |
| NC_011770 | 2087914 | C | T | 75 | 0 | 4 | 0 | | | T | T | T | T | | | | | | |
| NC_011770 | 2088866 | A | T | 65 | 0 | 2 | 0 | | | | | | | | | T | T | | |
| NC_011770 | 2088871 | T | C | 68 | 0 | 2 | 0 | | | | | | | | | C | C | | |
| NC_011770 | 2088872 | A | G | 68 | 0 | 2 | 0 | | | | | | | | | G | G | | |
| NC_011770 | 2088880 | A | C | 66 | 0 | 2 | 0 | | | | | | | | | C | C | | |
| NC_011770 | 2088889 | C | T | 65 | 0 | 2 | 0 | | | | | | | | | T | T | | |
| NC_011770 | 2088901 | G | C | 65 | 0 | 4 | 0 | C | C | | | | | C | C | | | | |

| | | | | | | | | | | | | | | | | | | | |
|---|---|---|---|---|---|---|---|---|---|---|---|---|---|---|---|---|---|---|---|
| NC_011770 | 2088904 | T | A | 66 | 0 | 2 | 0 | | | | | | | A | A | | | | |
| NC_011770 | 2088923 | C | T | 69 | 0 | 2 | 0 | | | | | | | | | T | T | | |
| NC_011770 | 2088924 | C | A | 68 | 0 | 2 | 0 | | | | | | | | | A | A | | |
| NC_011770 | 2088925 | A | C | 68 | 0 | 2 | 0 | | | | | | | | | C | C | | |
| NC_011770 | 2089368 | C | T | 72 | 0 | 2 | 0 | | | | | | | | | T | T | | |
| NC_011770 | 2089385 | AC | A | 77 | 0 | 2 | 0 | | | | | | | A | A | | | | |
| NC_011770 | 2089396 | C | T | 74 | 0 | 2 | 0 | | | | | | | | | | | T | T |
| NC_011770 | 2090447 | G | A | 90 | 0 | 2 | 0 | | | | | | | | | | | A | A |
| NC_011770 | 2092545 | C | G | 81 | 0 | 2 | 0 | G | G | | | | | | | | | | |
| NC_011770 | 2092601 | C | T | 77 | 0 | 2 | 0 | T | T | | | | | | | | | | |
| NC_011770 | 2092609 | G | A | 82 | 0 | 2 | 0 | A | A | | | | | | | | | | |
| NC_011770 | 2092612 | T | C | 80 | 0 | 12 | 0 | C | C | C | C | C | C | C | C | C | C | C | C |
| NC_011770 | 2092648 | C | T | 77 | 0 | 2 | 0 | | | | | | | | | T | T | | |
| NC_011770 | 2092661 | T | G | 79 | 0 | 4 | 0 | G | G | | | | | | | | | G | G |
| NC_011770 | 2092762 | C | T | 78 | 0 | 4 | 0 | T | T | | | | | | | | | T | T |
| NC_011770 | 2093385 | G | A | 71 | 0 | 6 | 0 | A | A | | | | | A | A | | | A | A |
| NC_011770 | 2095712 | T | C | 58 | 0 | 4 | 0 | | | | | | | C | C | | | C | C |
| NC_011770 | 2095723 | C | T | 60 | 0 | 2 | 0 | | | | | | | | | | | T | T |
| NC_011770 | 2099755 | C | T | 44 | 0 | 2 | 0 | | | | | | | | | T | T | | |
| NC_011770 | 2101485 | A | G | 73 | 0 | 4 | 0 | | | G | G | G | G | | | | | | |
| NC_011770 | 2101498 | C | T | 73 | 0 | 6 | 0 | | | T | T | T | T | T | T | | | | |
| NC_011770 | 2105159 | A | G | 63 | 0 | 8 | 0 | | | G | G | G | G | G | G | | | G | G |
| NC_011770 | 2108812 | GC | G | 71 | 0 | 4 | 0 | | | G | G | G | G | | | | | | |
| NC_011770 | 2119244 | C | G | 61 | 0 | 6 | 0 | | | G | G | G | G | | | | | G | G |
| NC_011770 | 2119265 | G | A | 60 | 0 | 2 | 0 | | | | | | | | | | | A | A |
| NC_011770 | 2119270 | T | C | 60 | 0 | 2 | 0 | | | | | | | | | | | C | C |
| NC_011770 | 2119272 | G | C | 61 | 0 | 2 | 0 | | | | | | | | | | | C | C |
| NC_011770 | 2119580 | A | G | 45 | 0 | 2 | 0 | | | | | | | G | G | | | | |
| NC_011770 | 2119582 | T | C | 45 | 0 | 4 | 0 | | | C | C | C | C | | | | | | |
| NC_011770 | 2119608 | G | A | 43 | 0 | 10 | 0 | A | A | A | A | A | A | | | A | A | A | A |
| NC_011770 | 2119623 | T | C | 44 | 0 | 2 | 0 | | | | | | | C | C | | | | |
| NC_011770 | 2119808 | C | T | 55 | 0 | 2 | 0 | | | | | | | T | T | | | | |
| NC_011770 | 2119815 | T | C | 54 | 0 | 2 | 0 | | | | | | | C | C | | | | |
| NC_011770 | 2119829 | G | T | 54 | 0 | 2 | 0 | | | | | | | T | T | | | | |
| NC_011770 | 2119976 | T | C | 53 | 0 | 2 | 0 | | | | | | | C | C | | | | |
| NC_011770 | 2119982 | A | G | 54 | 0 | 2 | 0 | | | | | | | G | G | | | | |
| NC_011770 | 2122253 | A | G | 94 | 0 | 1 | 0 | | | | | | | G | | | | | |
| NC_011770 | 2122314 | C | G | 90 | 0 | 4 | 0 | | | G | G | G | G | | | | | | |
| NC_011770 | 2124453 | A | G | 69 | 0 | 1 | 0 | | | | | | | G | | | | | |
| NC_011770 | 2124518 | T | TCGGCG | 63 | 0 | 2 | 0 | | | TCGGCG | TCGGCG | | | | | | | | |
| NC_011770 | 2124537 | C | T | 64 | 0 | 8 | 0 | T | T | | | | | T | T | T | T | T | T |
| NC_011770 | 2126198 | C | T | 73 | 0 | 4 | 0 | T | T | | | | | T | T | | | | |
| NC_011770 | 2128103 | A | T | 41 | 0 | 4 | 0 | | | | | | | | | T | T | T | T |
| NC_011770 | 2128141 | G | T | 42 | 0 | 4 | 0 | | | | | | | | | T | T | T | T |
| NC_011770 | 2134816 | G | A | 78 | 0 | 6 | 0 | A | A | | | | | | | A | A | A | A |
| NC_011770 | 2135140 | G | T | 70 | 0 | 4 | 0 | | | | | | | T | T | T | T | | |
| NC_011770 | 2137239 | C | T | 75 | 0 | 2 | 0 | | | | | | | T | T | | | | |
| NC_011770 | 2138647 | C | T | 76 | 0 | 6 | 0 | T | T | | | | | | | T | T | T | T |
| NC_011770 | 2138695 | T | TC | 76 | 0 | 6 | 0 | TC | TC | | | | | | | TC | TC | TC | TC |
| NC_011770 | 2139927 | C | T | 68 | 0 | 2 | 0 | | | | | | | | | T | T | | |
| NC_011770 | 2142317 | G | A | 77 | 0 | 2 | 0 | | | | | | | | | A | A | | |
| NC_011770 | 2142325 | C | T | 76 | 0 | 2 | 0 | | | | | | | | | T | T | | |
| NC_011770 | 2142343 | A | C | 75 | 0 | 2 | 0 | | | | | | | | | C | C | | |
| NC_011770 | 2142358 | T | TC | 56 | 0 | 2 | 0 | | | | | | | | | TC | TC | | |
| NC_011770 | 2142360 | T | C | 57 | 0 | 4 | 0 | C | . | | | | | | | C | C | . | C |
| NC_011770 | 2142375 | T | C | 77 | 0 | 6 | 0 | | | C | C | C | C | | | C | C | | |
| NC_011770 | 2152427 | C | G | 51 | 0 | 2 | 0 | | | | | | | G | G | | | | |
| NC_011770 | 2152464 | C | CG | 54 | 0 | 2 | 0 | | | | | | | CG | CG | | | | |
| NC_011770 | 2152604 | C | T | 52 | 0 | 4 | 0 | | | T | T | T | T | | | | | | |
| NC_011770 | 2152638 | T | G | 52 | 0 | 2 | 0 | | | | | | | G | G | | | | |
| NC_011770 | 2152641 | G | C | 53 | 0 | 2 | 0 | | | | | | | C | C | | | | |
| NC_011770 | 2152765 | T | C | 48 | 0 | 6 | 0 | | | C | C | C | C | C | C | | | | |

| | | | | | | | | | | | | | | | | | | | |
|---|---|---|---|---|---|---|---|---|---|---|---|---|---|---|---|---|---|---|---|
| NC_011770 | 2158611 | C | G | 66 | 0 | 2 | 0 | | | | | | | | | | | G | G |
| NC_011770 | 2158615 | A | C | 68 | 0 | 2 | 0 | | | | | | | | | | | C | C |
| NC_011770 | 2158693 | C | A | 69 | 0 | 2 | 0 | | | | | | | | | | | A | A |
| NC_011770 | 2160865 | G | A | 71 | 0 | 2 | 0 | | | | | | | | | A | A | | |
| NC_011770 | 2160876 | T | C | 70 | 0 | 12 | 0 | C | C | C | C | C | C | C | C | C | C | C | C |
| NC_011770 | 2160897 | T | C | 69 | 0 | 6 | 0 | C | C | | | | | C | C | | | C | C |
| NC_011770 | 2160914 | A | G | 66 | 0 | 6 | 0 | G | G | | | | | G | G | | | G | G |
| NC_011770 | 2160936 | G | C | 72 | 0 | 4 | 0 | | | | | | | C | C | | | C | C |
| NC_011770 | 2161025 | C | G | 72 | 0 | 6 | 0 | G | G | | | | | G | G | | | G | G |
| NC_011770 | 2161037 | G | A | 73 | 0 | 6 | 0 | A | A | | | | | A | A | | | A | A |
| NC_011770 | 2161050 | G | C | 74 | 0 | 4 | 0 | | | C | C | C | C | | | | | | |
| NC_011770 | 2161098 | G | A | 69 | 0 | 10 | 0 | A | A | A | A | A | A | | | A | A | A | A |
| NC_011770 | 2161101 | C | T | 69 | 0 | 8 | 0 | T | T | | | | | T | T | T | T | T | T |
| NC_011770 | 2167717 | C | T | 59 | 0 | 12 | 0 | T | T | T | T | T | T | T | T | T | T | T | T |
| NC_011770 | 2174948 | C | T | 63 | 0 | 2 | 0 | | | | | | | | | T | T | | |
| NC_011770 | 2174985 | T | A | 65 | 0 | 2 | 0 | A | A | | | | | | | | | | |
| NC_011770 | 2176914 | T | C | 73 | 0 | 4 | 0 | C | C | | | | | | | | | C | C |
| NC_011770 | 2176919 | A | G | 73 | 0 | 4 | 0 | | | G | G | G | G | | | | | | |
| NC_011770 | 2178175 | A | C | 75 | 0 | 2 | 0 | | | | | | | C | C | | | | |
| NC_011770 | 2178238 | T | C | 73 | 0 | 8 | 0 | | | C | C | C | C | | | C | C | C | C |
| NC_011770 | 2178253 | G | A | 75 | 0 | 6 | 0 | | | A | A | A | A | | | A | A | | |
| NC_011770 | 2178313 | A | C | 68 | 0 | 12 | 0 | C | C | C | C | C | C | C | C | C | C | C | C |
| NC_011770 | 2178330 | C | G | 68 | 0 | 6 | 0 | | | G | G | G | G | G | G | | | | |
| NC_011770 | 2178359 | T | C | 67 | 0 | 6 | 0 | | | C | C | C | C | C | C | | | | |
| NC_011770 | 2178406 | G | A | 70 | 0 | 2 | 0 | | | | | | | | | A | A | | |
| NC_011770 | 2178609 | C | T | 78 | 0 | 2 | 0 | | | | | | | | | T | T | | |
| NC_011770 | 2178676 | C | G | 74 | 0 | 8 | 0 | | | G | G | G | G | G | G | G | G | | |
| NC_011770 | 2178715 | C | G | 72 | 0 | 2 | 0 | G | G | | | | | | | | | | |
| NC_011770 | 2178814 | G | A | 61 | 0 | 2 | 0 | | | | | | | | | A | A | | |
| NC_011770 | 2178925 | C | G | 52 | 0 | 8 | 0 | | | G | G | G | G | | | G | G | G | G |
| NC_011770 | 2179063 | G | GT | 39 | 0 | 12 | 0 | GT | GT | GT | GT | GT | GT | GT | GT | GC | GC | GT | GT |
| NC_011770 | 2179396 | C | T | 54 | 0 | 2 | 0 | | | | | | | | | T | T | | |
| NC_011770 | 2180911 | G | A | 85 | 0 | 2 | 0 | | | | | | | | | | | A | A |
| NC_011770 | 2183423 | C | T | 69 | 0 | 4 | 0 | | | | | | | | | T | T | T | T |
| NC_011770 | 2183463 | CTT | C | 68 | 0 | 4 | 0 | | | | | | | | | C | C | C | C |
| NC_011770 | 2183476 | T | C | 68 | 0 | 2 | 0 | | | | | | | | | | | C | C |
| NC_011770 | 2184977 | GA | G | 39 | 0 | 2 | 0 | | | | | | | | | G | G | | |
| NC_011770 | 2185121 | T | C | 46 | 0 | 10 | 0 | C | C | C | C | C | C | C | C | C | C | | |
| NC_011770 | 2185124 | C | A | 47 | 0 | 2 | 0 | | | | | | | | | A | A | | |
| NC_011770 | 2185127 | G | GGA | 46 | 0 | 9 | 0 | GGA | . | GGA | GGA | GGA | GGA | GGA | GGA | GGA | GGA | | |
| NC_011770 | 2185143 | C | T | 46 | 0 | 2 | 0 | | | | | | | T | T | | | | |
| NC_011770 | 2185159 | C | T | 46 | 0 | 2 | 0 | | | | | | | | | T | T | | |
| NC_011770 | 2185615 | CT | C | 59 | 0 | 2 | 0 | | | | | | | | | | | C | C |
| NC_011770 | 2185620 | G | A | 60 | 0 | 4 | 0 | | | A | A | A | A | | | | | | |
| NC_011770 | 2185690 | G | GC | 62 | 0 | 2 | 0 | | | | | | | GC | GC | | | | |
| NC_011770 | 2185727 | C | T | 61 | 0 | 2 | 0 | | | | | | | T | T | | | | |
| NC_011770 | 2188855 | C | T | 81 | 0 | 1 | 0 | | | | | | | T | | | | | |
| NC_011770 | 2188862 | T | G | 81 | 0 | 6 | 0 | G | G | G | G | G | G | | | | | | |
| NC_011770 | 2190248 | T | C | 68 | 0 | 6 | 0 | C | C | C | C | C | C | | | | | | |
| NC_011770 | 2190307 | A | G | 57 | 0 | 2 | 0 | G | G | | | | | | | | | | |
| NC_011770 | 2191389 | GT | G | 73 | 0 | 4 | 0 | | | G | G | G | G | | | | | | |
| NC_011770 | 2191467 | C | T | 69 | 0 | 2 | 0 | | | | | | | | | T | T | | |
| NC_011770 | 2191571 | G | A | 66 | 0 | 12 | 0 | A | A | A | A | A | A | A | A | A | A | A | A |
| NC_011770 | 2193064 | T | C | 14 | 1 | 4 | 0 | | . | C | C | C | C | | | | | | |
| NC_011770 | 2193109 | C | T | 14 | 1 | 4 | 0 | | . | T | T | T | T | | | | | | |
| NC_011770 | 2197085 | C | T | 65 | 0 | 2 | 0 | | | | | | | T | T | | | | |
| NC_011770 | 2197097 | C | T | 61 | 0 | 4 | 0 | | | T | T | T | T | | | | | | |
| NC_011770 | 2197116 | G | A | 62 | 0 | 4 | 0 | | | A | A | A | A | | | | | | |
| NC_011770 | 2197135 | G | A | 59 | 0 | 6 | 0 | | | A | A | A | A | A | A | | | | |
| NC_011770 | 2197198 | C | G | 61 | 0 | 12 | 0 | G | G | G | G | G | G | G | G | G | G | G | G |
| NC_011770 | 2198125 | T | G | 75 | 0 | 4 | 0 | | | G | G | G | G | | | | | | |
| NC_011770 | 2198241 | T | C | 77 | 0 | 4 | 0 | | | C | C | C | C | | | | | | |

| | | | | | | | | | | | | | | | | | | | |
|---|---|---|---|---|---|---|---|---|---|---|---|---|---|---|---|---|---|---|---|
| NC_011770 | 2198254 | C | T | 73 | 0 | 4 | 0 | | | T | T | T | T | | | | | | |
| NC_011770 | 2199868 | C | T | 52 | 0 | 2 | 0 | | | | | | | | | | | T | T |
| NC_011770 | 2199894 | GT | G | 53 | 0 | 2 | 0 | | | | | | | | | | | G | G |
| NC_011770 | 2200111 | T | C | 56 | 0 | 2 | 0 | | | | | | | C | C | | | | |
| NC_011770 | 2202915 | C | T | 62 | 0 | 2 | 0 | | | | | | | T | T | | | | |
| NC_011770 | 2202921 | G | C | 62 | 0 | 2 | 0 | | | | | | | | | | | C | C |
| NC_011770 | 2210829 | A | C | 30 | 0 | 2 | 0 | | | | | | | | | | | C | C |
| NC_011770 | 2211251 | G | A | 53 | 0 | 2 | 0 | | | | | | | | | A | A | | |
| NC_011770 | 2213422 | A | T | 65 | 0 | 12 | 0 | T | T | T | T | T | T | T | T | T | T | T | T |
| NC_011770 | 2216223 | G | C | 61 | 0 | 2 | 0 | | | | | | | C | C | | | | |
| NC_011770 | 2216811 | G | C | 82 | 0 | 12 | 0 | C | C | C | C | C | C | C | C | C | C | C | C |
| NC_011770 | 2217319 | T | G | 94 | 0 | 12 | 0 | G | G | G | G | G | G | G | G | G | G | G | G |
| NC_011770 | 2218251 | T | C | 78 | 0 | 12 | 0 | C | C | C | C | C | C | C | C | C | C | C | C |
| NC_011770 | 2218252 | G | A | 77 | 0 | 4 | 0 | | | A | A | A | A | | | | | | |
| NC_011770 | 2218504 | G | A | 76 | 0 | 12 | 0 | A | A | A | A | A | A | A | A | A | A | A | A |
| NC_011770 | 2221929 | C | A | 69 | 0 | 2 | 0 | | | | | | | A | A | | | | |
| NC_011770 | 2221993 | T | C | 70 | 0 | 2 | 0 | | | | | | | C | C | | | | |
| NC_011770 | 2222403 | A | G | 96 | 0 | 1 | 0 | | | | | | | G | | | | | |
| NC_011770 | 2222428 | C | T | 90 | 0 | 2 | 0 | | | | | | | T | T | | | | |
| NC_011770 | 2225117 | G | A | 81 | 0 | 4 | 0 | | | A | A | A | A | | | | | | |
| NC_011770 | 2225731 | C | A | 81 | 0 | 2 | 0 | | | | | | | A | A | | | | |
| NC_011770 | 2225733 | T | C | 82 | 0 | 4 | 0 | | | C | C | C | C | | | | | | |
| NC_011770 | 2225825 | G | C | 82 | 0 | 2 | 0 | | | | | | | C | C | | | | |
| NC_011770 | 2226145 | TC | T | 62 | 0 | 2 | 0 | | | | | | | T | T | | | | |
| NC_011770 | 2227322 | A | G | 78 | 0 | 8 | 0 | G | G | | | | | G | G | G | G | G | G |
| NC_011770 | 2227492 | A | G | 74 | 0 | 12 | 0 | G | G | G | G | G | G | G | G | G | G | G | G |
| NC_011770 | 2228206 | C | T | 77 | 0 | 8 | 0 | | | T | T | T | T | T | T | | | T | T |
| NC_011770 | 2228220 | C | G | 79 | 0 | 2 | 0 | | | | | | | | | G | G | | |
| NC_011770 | 2228234 | C | T | 7 | 2 | 6 | 0 | . | | T | T | T | T | A | A | . | | | |
| NC_011770 | 2228240 | T | C | 6 | 1 | 2 | 0 | | | | | | | C | C | . | | | |
| NC_011770 | 2228262 | C | T | 75 | 0 | 4 | 0 | | | T | T | T | T | | | | | | |
| NC_011770 | 2228283 | G | A | 79 | 0 | 4 | 0 | A | A | | | | | | | | | A | A |
| NC_011770 | 2228313 | C | T | 83 | 0 | 2 | 0 | | | | | | | | | T | T | | |
| NC_011770 | 2228317 | C | T | 83 | 0 | 2 | 0 | | | | | | | | | T | T | | |
| NC_011770 | 2228322 | A | G | 81 | 0 | 2 | 0 | | | | | | | | | G | G | | |
| NC_011770 | 2230176 | G | GA | 40 | 0 | 2 | 0 | | | | | | | GA | GA | | | | |
| NC_011770 | 2237554 | T | G | 63 | 0 | 4 | 0 | | | G | G | G | G | | | | | | |
| NC_011770 | 2237643 | C | T | 71 | 0 | 2 | 0 | | | | | | | | | T | T | | |
| NC_011770 | 2244775 | A | G | 75 | 0 | 2 | 0 | | | | | | | G | G | | | | |
| NC_011770 | 2247322 | C | T | 70 | 0 | 2 | 0 | | | | | | | | | | | T | T |
| NC_011770 | 2251370 | A | T | 27 | 0 | 2 | 0 | | | | | | | T | T | | | | |
| NC_011770 | 2251376 | A | C | 27 | 0 | 4 | 0 | | | | | | | C | C | T | T | | |
| NC_011770 | 2251380 | C | T | 26 | 0 | 2 | 0 | | | | | | | T | T | | | | |
| NC_011770 | 2251398 | C | T | 22 | 0 | 4 | 0 | | | | | | | T | T | T | T | | |
| NC_011770 | 2256481 | C | G | 76 | 0 | 2 | 0 | G | G | | | | | | | | | | |
| NC_011770 | 2258972 | G | C | 47 | 0 | 2 | 0 | | | | | | | C | C | | | | |
| NC_011770 | 2265464 | G | A | 39 | 0 | 2 | 0 | A | A | | | | | | | | | | |
| NC_011770 | 2265466 | T | C | 37 | 0 | 2 | 0 | C | C | | | | | | | | | | |
| NC_011770 | 2265497 | C | T | 30 | 0 | 2 | 0 | T | T | | | | | | | | | | |
| NC_011770 | 2266118 | A | G | 73 | 0 | 2 | 0 | G | G | | | | | | | | | | |
| NC_011770 | 2266125 | G | A | 70 | 0 | 2 | 0 | | | | | | | | | A | A | | |
| NC_011770 | 2276274 | G | A | 78 | 0 | 2 | 0 | | | | | | | | | | | A | A |
| NC_011770 | 2276276 | G | A | 79 | 0 | 2 | 0 | | | | | | | | | | | A | A |
| NC_011770 | 2276277 | C | A | 78 | 0 | 2 | 0 | | | | | | | | | | | A | A |
| NC_011770 | 2276292 | C | A | 78 | 0 | 2 | 0 | | | | | | | | | | | A | A |
| NC_011770 | 2276492 | T | C | 90 | 0 | 2 | 0 | | | | | | | | | | | C | C |
| NC_011770 | 2278435 | G | C | 80 | 0 | 2 | 0 | | | | | | | | | | | C | C |
| NC_011770 | 2278445 | G | C | 82 | 0 | 2 | 0 | | | | | | | | | C | C | | |
| NC_011770 | 2280696 | C | T | 75 | 0 | 12 | 0 | T | T | T | T | T | T | T | T | T | T | T | T |
| NC_011770 | 2282514 | A | AG | 87 | 0 | 5 | 0 | AG | AG | | | | | AG | | | | AG | AG |
| NC_011770 | 2285748 | T | G | 62 | 0 | 4 | 0 | | | | | | | G | G | G | G | | |
| NC_011770 | 2285785 | C | A | 59 | 0 | 10 | 0 | A | A | A | A | A | A | | | A | A | A | A |

| | | | | | | | | | | | | | | | | | | | |
|---|---|---|---|---|---|---|---|---|---|---|---|---|---|---|---|---|---|---|---|
| NC_011770 | 2288073 | C | G | 68 | 0 | 2 | 0 | G | G | | | | | | | | | | |
| NC_011770 | 2288083 | A | G | 72 | 0 | 6 | 0 | G | G | G | G | G | G | | | | | | |
| NC_011770 | 2288096 | G | A | 73 | 0 | 2 | 0 | A | A | | | | | | | | | | |
| NC_011770 | 2288097 | CA | C | 73 | 0 | 4 | 0 | | | C | C | C | C | | | | | | |
| NC_011770 | 2288123 | T | G | 65 | 0 | 2 | 0 | | | | | | | G | G | | | | |
| NC_011770 | 2288139 | T | G | 66 | 0 | 12 | 0 | G | G | G | G | G | G | G | G | G | G | G | G |
| NC_011770 | 2289308 | C | T | 72 | 0 | 2 | 0 | | | | | | | | | T | T | | |
| NC_011770 | 2289387 | G | A | 77 | 0 | 1 | 0 | | | | | | | A | | | | | |
| NC_011770 | 2294558 | C | G | 62 | 0 | 4 | 0 | | | G | G | G | G | | | | | | |
| NC_011770 | 2294560 | A | G | 62 | 0 | 4 | 0 | | | G | G | G | G | | | | | | |
| NC_011770 | 2295781 | C | T | 61 | 0 | 8 | 0 | T | T | T | T | T | T | | | T | T | | |
| NC_011770 | 2296172 | C | A | 50 | 0 | 2 | 0 | A | A | | | | | | | | | | |
| NC_011770 | 2296197 | G | C | 47 | 0 | 4 | 0 | | | C | C | C | C | | | | | | |
| NC_011770 | 2299177 | G | GC | 7 | 2 | 4 | 0 | . | . | GC | . | . | GC | | . | GC | GC | | |
| NC_011770 | 2299180 | A | G | 19 | 1 | 7 | 0 | | | G | G | G | G | | | . | G | G | G |
| NC_011770 | 2301004 | A | G | 55 | 0 | 2 | 0 | | | | | | | | | G | G | | |
| NC_011770 | 2301058 | G | A | 59 | 0 | 6 | 0 | A | A | | | | | | | A | A | A | A |
| NC_011770 | 2301072 | T | C | 59 | 0 | 1 | 0 | | | | | | | C | | | | | |
| NC_011770 | 2301115 | CG | C | 57 | 0 | 2 | 0 | | | | | | | C | C | | | | |
| NC_011770 | 2301178 | G | GA | 60 | 0 | 2 | 0 | | | | | | | | | GA | GA | | |
| NC_011770 | 2302830 | C | G | 65 | 0 | 2 | 0 | | | | | | | | | G | G | | |
| NC_011770 | 2303975 | A | C | 74 | 0 | 4 | 0 | C | C | | | | | | | C | C | | |
| NC_011770 | 2306684 | G | A | 50 | 0 | 2 | 0 | | | | | | | | | | | A | A |
| NC_011770 | 2307848 | T | G | 81 | 0 | 12 | 0 | G | G | G | G | G | G | G | G | G | G | G | G |
| NC_011770 | 2307851 | C | T | 81 | 0 | 12 | 0 | T | T | T | T | T | T | T | T | T | T | T | T |
| NC_011770 | 2307882 | T | C | 84 | 0 | 2 | 0 | | | | | | | | | | | C | C |
| NC_011770 | 2307887 | A | G | 84 | 0 | 4 | 0 | | | G | G | G | G | | | | | | |
| NC_011770 | 2310347 | T | C | 68 | 0 | 12 | 0 | C | C | C | C | C | C | C | C | C | C | C | C |
| NC_011770 | 2310380 | C | G | 69 | 0 | 12 | 0 | G | G | G | G | G | G | G | G | G | G | G | G |
| NC_011770 | 2310465 | G | C | 75 | 0 | 2 | 0 | C | C | | | | | | | | | | |
| NC_011770 | 2310519 | G | A | 73 | 0 | 2 | 0 | | | | | | | | | | | A | A |
| NC_011770 | 2310593 | A | G | 72 | 0 | 2 | 0 | | | | | | | G | G | | | | |
| NC_011770 | 2310661 | A | C | 71 | 0 | 2 | 0 | | | | | | | | | C | C | | |
| NC_011770 | 2310849 | G | A | 64 | 0 | 4 | 0 | | | A | A | A | A | | | | | | |
| NC_011770 | 2310956 | C | T | 61 | 0 | 2 | 0 | | | | | | | T | T | | | | |
| NC_011770 | 2311169 | G | T | 48 | 0 | 10 | 0 | T | T | T | T | T | T | T | T | T | T | | |
| NC_011770 | 2311190 | C | T | 54 | 0 | 2 | 0 | | | | | | | | | T | T | | |
| NC_011770 | 2311209 | C | T | 53 | 0 | 2 | 0 | | | | | | | | | T | T | | |
| NC_011770 | 2311319 | T | C | 57 | 0 | 2 | 0 | | | | | | | | | C | C | | |
| NC_011770 | 2311469 | G | A | 55 | 0 | 4 | 0 | | | A | A | A | A | | | | | | |
| NC_011770 | 2311772 | G | A | 42 | 1 | 1 | 0 | A | . | | | | | | | | | | |
| NC_011770 | 2311788 | A | G | 42 | 1 | 3 | 0 | G | . | | | | | G | G | | | | |
| NC_011770 | 2311798 | T | G | 42 | 1 | 3 | 0 | G | . | | | | | G | G | | | | |
| NC_011770 | 2311809 | GGGC | G | 37 | 3 | 1 | 0 | . | . | | | | | G | . | | | | |
| NC_011770 | 2311958 | T | G | 52 | 0 | 2 | 0 | | | | | | | G | G | | | | |
| NC_011770 | 2312053 | G | A | 64 | 0 | 4 | 0 | A | A | | | | | A | A | | | | |
| NC_011770 | 2312134 | G | T | 69 | 0 | 2 | 0 | | | | | | | | | | | T | T |
| NC_011770 | 2312288 | G | A | 75 | 0 | 2 | 0 | | | | | | | | | | | A | A |
| NC_011770 | 2312347 | G | A | 76 | 0 | 2 | 0 | A | A | | | | | | | | | | |
| NC_011770 | 2312926 | G | A | 74 | 0 | 2 | 0 | | | | | | | | | A | A | | |
| NC_011770 | 2312944 | A | C | 73 | 0 | 2 | 0 | | | | | | | | | C | C | | |
| NC_011770 | 2312968 | C | T | 74 | 0 | 4 | 0 | | | | | | | | | T | T | T | T |
| NC_011770 | 2313055 | G | A | 73 | 0 | 2 | 0 | | | | | | | | | | | A | A |
| NC_011770 | 2313090 | G | T | 75 | 0 | 2 | 0 | | | | | | | | | | | T | T |
| NC_011770 | 2313482 | A | G | 70 | 0 | 2 | 0 | | | | | | | | | | | G | G |
| NC_011770 | 2317776 | G | C | 20 | 0 | 2 | 0 | | | | | | | | | | | C | C |
| NC_011770 | 2317811 | C | A | 20 | 0 | 2 | 0 | | | | | | | | | A | A | | |
| NC_011770 | 2320555 | GCGC | G | 67 | 0 | 2 | 0 | G | G | | | | | | | | | | |
| NC_011770 | 2321677 | C | T | 79 | 0 | 2 | 0 | | | | | | | | | | | T | T |
| NC_011770 | 2322701 | C | T | 76 | 0 | 2 | 0 | | | | | | | T | T | | | | |
| NC_011770 | 2322705 | T | C | 72 | 0 | 2 | 0 | | | | | | | C | C | | | | |
| NC_011770 | 2322725 | C | A | 79 | 0 | 2 | 0 | | | | | | | | | | | A | A |

| | | | | | | | | | | | | | | | | | | | |
|---|---|---|---|---|---|---|---|---|---|---|---|---|---|---|---|---|---|---|---|
| NC_011770 | 2323380 | C | G | 69 | 0 | 4 | 0 | | | G | G | G | G | | | | | | |
| NC_011770 | 2327138 | C | T | 69 | 0 | 2 | 0 | | | | | | | T | T | | | | |
| NC_011770 | 2327143 | G | C | 71 | 0 | 2 | 0 | | | | | | | C | C | | | | |
| NC_011770 | 2329320 | G | A | 52 | 0 | 10 | 0 | A | A | A | A | A | A | | | A | A | A | A |
| NC_011770 | 2329468 | T | C | 63 | 0 | 6 | 0 | C | C | | | | | | | C | C | C | C |
| NC_011770 | 2329535 | T | C | 62 | 0 | 12 | 0 | C | C | C | C | C | C | C | C | C | C | C | C |
| NC_011770 | 2329701 | A | T | 66 | 0 | 2 | 0 | | | | | | | T | T | | | | |
| NC_011770 | 2329886 | A | G | 73 | 0 | 12 | 0 | G | G | G | G | G | G | G | G | G | G | G | G |
| NC_011770 | 2330522 | T | C | 57 | 0 | 8 | 0 | | | C | C | C | C | | | C | C | C | C |
| NC_011770 | 2330541 | G | A | 55 | 0 | 6 | 0 | | | A | A | A | A | | | A | A | | |
| NC_011770 | 2333911 | C | A | 65 | 0 | 2 | 0 | | | | | | | A | A | | | | |
| NC_011770 | 2340554 | T | C | 69 | 0 | 6 | 0 | | | C | C | C | C | | | C | C | | |
| NC_011770 | 2340562 | T | C | 69 | 0 | 8 | 0 | | | C | C | C | C | C | C | C | C | | |
| NC_011770 | 2340579 | G | C | 73 | 0 | 4 | 0 | | | C | C | C | C | | | | | | |
| NC_011770 | 2340645 | C | A | 76 | 0 | 8 | 0 | | | A | A | A | A | A | A | A | A | | |
| NC_011770 | 2340690 | A | G | 76 | 0 | 2 | 0 | | | | | | | G | G | | | | |
| NC_011770 | 2341386 | C | CG | 62 | 0 | 1 | 0 | | | | | | | CG | | | | | |
| NC_011770 | 2341879 | G | A | 49 | 0 | 2 | 0 | | | | | | | A | A | | | | |
| NC_011770 | 2341926 | T | C | 46 | 0 | 6 | 0 | C | C | | | | | C | C | | | C | C |
| NC_011770 | 2341932 | G | A | 46 | 0 | 2 | 0 | | | | | | | A | A | | | | |
| NC_011770 | 2345227 | C | T | 67 | 0 | 8 | 0 | T | T | | | | | T | T | T | T | T | T |
| NC_011770 | 2345307 | A | T | 66 | 0 | 2 | 0 | | | | | | | | | T | T | | |
| NC_011770 | 2345496 | A | C | 53 | 0 | 12 | 0 | C | C | C | C | C | C | C | C | C | C | C | C |
| NC_011770 | 2352863 | A | G | 70 | 0 | 2 | 0 | G | G | | | | | | | | | | |
| NC_011770 | 2352938 | T | G | 69 | 0 | 4 | 0 | | | | | | | G | G | G | G | | |
| NC_011770 | 2357328 | G | A | 80 | 0 | 2 | 0 | | | | | | | A | A | | | | |
| NC_011770 | 2357343 | T | C | 78 | 0 | 12 | 0 | C | C | C | C | C | C | C | C | C | C | C | C |
| NC_011770 | 2357392 | G | C | 79 | 0 | 2 | 0 | | | | | | | C | C | | | | |
| NC_011770 | 2357396 | T | G | 77 | 0 | 2 | 0 | | | | | | | G | G | | | | |
| NC_011770 | 2357486 | G | C | 77 | 0 | 4 | 0 | | | C | C | C | C | | | | | | |
| NC_011770 | 2359022 | A | G | 66 | 0 | 4 | 0 | | | G | G | G | G | | | | | | |
| NC_011770 | 2359044 | C | T | 66 | 0 | 2 | 0 | | | | | | | | | | | T | T |
| NC_011770 | 2359081 | C | T | 62 | 0 | 4 | 0 | | | T | T | T | T | | | | | | |
| NC_011770 | 2359101 | GC | G | 62 | 0 | 2 | 0 | | | | | | | G | G | | | | |
| NC_011770 | 2359104 | C | A | 51 | 2 | 4 | 0 | | | A | A | A | A | . | . | | | | |
| NC_011770 | 2359111 | C | T | 61 | 0 | 4 | 0 | | | T | T | T | T | | | | | | |
| NC_011770 | 2359387 | A | G | 71 | 0 | 4 | 0 | | | G | G | G | G | | | | | | |
| NC_011770 | 2359411 | A | G | 71 | 0 | 4 | 0 | | | G | G | G | G | | | | | | |
| NC_011770 | 2359417 | C | T | 70 | 0 | 4 | 0 | | | T | T | T | T | | | | | | |
| NC_011770 | 2359430 | C | T | 70 | 0 | 4 | 0 | | | T | T | T | T | | | | | | |
| NC_011770 | 2359479 | A | G | 69 | 0 | 6 | 0 | | | G | G | G | G | G | G | | | | |
| NC_011770 | 2359492 | C | T | 68 | 0 | 4 | 0 | | | T | T | T | T | | | | | | |
| NC_011770 | 2365729 | AT | A | 66 | 0 | 2 | 0 | | | | | | | | | | | A | A |
| NC_011770 | 2370572 | C | CA | 79 | 0 | 4 | 0 | | | CA | CA | CA | CA | | | | | | |
| NC_011770 | 2373977 | G | A | 64 | 0 | 8 | 0 | A | A | | | | | A | A | A | A | A | A |
| NC_011770 | 2373980 | G | C | 65 | 0 | 8 | 0 | C | C | | | | | C | C | C | C | C | C |
| NC_011770 | 2374005 | G | A | 67 | 0 | 1 | 0 | | | | | | | A | | | | | |
| NC_011770 | 2375539 | AG | A | 58 | 0 | 2 | 0 | | | | | | | | | A | A | | |
| NC_011770 | 2375564 | C | A | 59 | 0 | 4 | 0 | | | | | | | A | A | A | A | | |
| NC_011770 | 2376495 | G | A | 72 | 0 | 10 | 0 | A | A | A | A | A | A | | | A | A | A | A |
| NC_011770 | 2376541 | G | A | 69 | 0 | 6 | 0 | A | A | | | | | | | A | A | A | A |
| NC_011770 | 2378214 | G | T | 86 | 0 | 2 | 0 | | | | | | | | | T | T | | |
| NC_011770 | 2378759 | CA | C | 60 | 0 | 6 | 0 | | | | | | | C | C | C | C | C | C |
| NC_011770 | 2378823 | CG | C | 58 | 0 | 2 | 0 | | | | | | | C | C | | | | |
| NC_011770 | 2378841 | T | C | 60 | 0 | 4 | 0 | | | | | | | | | C | C | C | C |
| NC_011770 | 2378854 | T | C | 57 | 0 | 4 | 0 | | | | | | | | | C | C | C | C |
| NC_011770 | 2379736 | C | T | 11 | 2 | 2 | 0 | | | | | | | | . | . | | T | T |
| NC_011770 | 2379756 | C | T | 10 | 4 | 2 | 0 | | | . | . | T | T | | . | . | | | |
| NC_011770 | 2379759 | T | A | 10 | 4 | 2 | 0 | | | . | . | A | A | | . | . | | | |
| NC_011770 | 2379775 | G | C | 9 | 5 | 4 | 0 | C | C | . | . | | | | . | . | . | C | C |
| NC_011770 | 2379800 | G | T | 9 | 5 | 4 | 0 | T | T | . | . | | | | . | . | . | T | T |
| NC_011770 | 2381007 | C | T | 49 | 0 | 2 | 0 | | | | | | | | | | | T | T |

| | | | | | | | | | | | | | | | | | | | |
|---|---|---|---|---|---|---|---|---|---|---|---|---|---|---|---|---|---|---|---|
| NC_011770 | 2382110 | G | A | 78 | 0 | 2 | 0 | | | | | | | | | A | A | | |
| NC_011770 | 2382114 | C | T | 81 | 0 | 10 | 0 | T | T | T | T | T | T | T | T | T | T | | |
| NC_011770 | 2382627 | C | A | 83 | 0 | 2 | 0 | | | | | | | A | A | | | | |
| NC_011770 | 2388449 | G | C | 49 | 0 | 2 | 0 | | | | | | | | | | | C | C |
| NC_011770 | 2388639 | C | T | 47 | 0 | 2 | 0 | | | | | | | | | T | T | | |
| NC_011770 | 2388832 | T | A | 64 | 0 | 4 | 0 | | | A | A | A | A | | | | | | |
| NC_011770 | 2391152 | G | A | 85 | 0 | 2 | 0 | | | | | | | A | A | | | | |
| NC_011770 | 2391162 | C | T | 84 | 0 | 6 | 0 | | | T | T | T | T | | | | | T | T |
| NC_011770 | 2391216 | T | G | 85 | 0 | 2 | 0 | | | | | | | | | | | G | G |
| NC_011770 | 2391219 | C | T | 84 | 0 | 10 | 0 | | | T | T | T | T | T | T | T | T | A | A |
| NC_011770 | 2391233 | G | T | 82 | 0 | 4 | 0 | | | T | T | T | T | | | | | | |
| NC_011770 | 2391246 | C | A | 85 | 0 | 2 | 0 | | | | | | | | | A | A | | |
| NC_011770 | 2391276 | G | A | 85 | 0 | 4 | 0 | A | A | | | | | A | A | | | | |
| NC_011770 | 2391773 | A | G | 90 | 0 | 2 | 0 | G | G | | | | | | | | | | |
| NC_011770 | 2391833 | C | T | 87 | 0 | 2 | 0 | T | T | | | | | | | | | | |
| NC_011770 | 2392872 | C | T | 75 | 0 | 2 | 0 | | | | | | | | | | | T | T |
| NC_011770 | 2399262 | CG | C | 48 | 0 | 1 | 0 | | | | | | | C | | | | | |
| NC_011770 | 2401322 | C | T | 71 | 0 | 2 | 0 | | | | | | | | | T | T | | |
| NC_011770 | 2401330 | G | A | 69 | 0 | 8 | 0 | A | A | A | A | A | A | | | A | A | | |
| NC_011770 | 2401360 | C | T | 70 | 0 | 10 | 0 | T | T | T | T | T | T | T | T | T | T | | |
| NC_011770 | 2401378 | G | A | 70 | 0 | 2 | 0 | | | | | | | A | A | | | | |
| NC_011770 | 2401379 | T | C | 69 | 0 | 2 | 0 | | | | | | | | | C | C | | |
| NC_011770 | 2401384 | T | C | 72 | 0 | 6 | 0 | | | C | C | C | C | | | C | C | | |
| NC_011770 | 2401398 | G | A | 71 | 0 | 4 | 0 | | | | | | | | | A | A | A | A |
| NC_011770 | 2401451 | T | C | 76 | 0 | 2 | 0 | | | | | | | | | | | C | C |
| NC_011770 | 2401459 | A | C | 72 | 0 | 12 | 0 | C | C | G | G | G | G | G | G | G | G | G | G |
| NC_011770 | 2401470 | G | A | 70 | 0 | 2 | 0 | | | | | | | | | | | A | A |
| NC_011770 | 2401491 | C | G | 76 | 0 | 2 | 0 | G | G | | | | | | | | | | |
| NC_011770 | 2401523 | T | C | 67 | 0 | 2 | 0 | | | | | | | | | | | C | C |
| NC_011770 | 2401527 | G | A | 66 | 0 | 2 | 0 | | | | | | | | | | | A | A |
| NC_011770 | 2401540 | G | A | 71 | 0 | 2 | 0 | | | | | | | | | | | A | A |
| NC_011770 | 2401599 | G | T | 69 | 0 | 2 | 0 | | | | | | | T | T | | | | |
| NC_011770 | 2401615 | C | A | 70 | 0 | 2 | 0 | | | | | | | | | | | A | A |
| NC_011770 | 2402409 | G | A | 82 | 0 | 6 | 0 | A | A | | | | | A | A | | | A | A |
| NC_011770 | 2402436 | T | G | 83 | 0 | 2 | 0 | | | | | | | G | G | | | | |
| NC_011770 | 2402481 | C | T | 95 | 0 | 6 | 0 | T | T | | | | | T | T | | | T | T |
| NC_011770 | 2402486 | G | T | 96 | 0 | 2 | 0 | | | | | | | T | T | | | | |
| NC_011770 | 2407640 | A | G | 42 | 0 | 10 | 0 | G | G | G | G | G | G | G | G | G | G | | |
| NC_011770 | 2408738 | G | T | 89 | 0 | 2 | 0 | T | T | | | | | | | | | | |
| NC_011770 | 2411194 | T | C | 55 | 0 | 2 | 0 | | | | | | | | | C | C | | |
| NC_011770 | 2412192 | A | G | 63 | 0 | 2 | 0 | | | | | G | G | | | | | | |
| NC_011770 | 2412221 | G | C | 60 | 0 | 2 | 0 | | | | | | | C | C | | | | |
| NC_011770 | 2414630 | A | C | 76 | 0 | 2 | 0 | | | | | | | | | C | C | | |
| NC_011770 | 2414640 | G | T | 77 | 0 | 12 | 0 | T | T | T | T | T | T | T | T | T | T | T | T |
| NC_011770 | 2414647 | G | A | 78 | 0 | 2 | 0 | | | | | | | | | | | A | A |
| NC_011770 | 2414670 | A | G | 75 | 0 | 12 | 0 | G | G | G | G | G | G | G | G | G | G | G | G |
| NC_011770 | 2420204 | G | A | 76 | 0 | 12 | 0 | A | A | A | A | A | A | A | A | A | A | A | A |
| NC_011770 | 2420230 | A | G | 76 | 0 | 10 | 0 | G | G | G | G | G | G | | | G | G | G | G |
| NC_011770 | 2421397 | G | A | 40 | 2 | 4 | 0 | A | A | | | | | | | A | A | . | . |
| NC_011770 | 2421410 | G | A | 38 | 2 | 2 | 0 | | | | | | | A | A | | | . | . |
| NC_011770 | 2421742 | G | T | 34 | 4 | 2 | 0 | | | . | . | . | . | | | | | T | T |
| NC_011770 | 2422116 | C | T | 23 | 4 | 8 | 0 | T | T | . | . | . | . | T | T | T | T | T | T |
| NC_011770 | 2423655 | T | C | 42 | 4 | 8 | 0 | C | C | . | . | . | . | C | C | C | C | C | C |
| NC_011770 | 2425845 | A | G | 32 | 4 | 2 | 0 | | | . | . | . | . | G | G | | | | |
| NC_011770 | 2425860 | C | T | 32 | 4 | 2 | 0 | | | . | . | . | . | T | T | | | | |
| NC_011770 | 2426797 | T | C | 33 | 4 | 8 | 0 | C | C | . | . | . | . | C | C | C | C | C | C |
| NC_011770 | 2426834 | T | C | 30 | 4 | 8 | 0 | C | C | . | . | . | . | C | C | C | C | C | C |
| NC_011770 | 2429445 | C | T | 54 | 0 | 2 | 0 | T | T | | | | | | | | | | |
| NC_011770 | 2432843 | C | T | 86 | 0 | 2 | 0 | | | | | | | | | | | T | T |
| NC_011770 | 2435265 | A | C | 71 | 0 | 2 | 0 | | | | | | | C | C | | | | |
| NC_011770 | 2435268 | C | T | 71 | 0 | 2 | 0 | | | | | | | T | T | | | | |
| NC_011770 | 2435274 | A | G | 60 | 0 | 2 | 0 | | | | | | | G | G | | | | |

| | | | | | | | | | | | | | | | | | | | |
|---|---|---|---|---|---|---|---|---|---|---|---|---|---|---|---|---|---|---|---|
| NC_011770 | 2435276 | A | AC | 58 | 0 | 2 | 0 | | | | | | | AC | AC | | | | |
| NC_011770 | 2437347 | A | G | 82 | 0 | 4 | 0 | | | G | G | G | G | | | | | | |
| NC_011770 | 2441146 | C | T | 53 | 0 | 2 | 0 | | | | | | | | | | | T | T |
| NC_011770 | 2443199 | A | G | 65 | 0 | 10 | 0 | G | G | G | G | G | G | G | G | | | G | G |
| NC_011770 | 2445426 | C | T | 83 | 0 | 2 | 0 | | | | | | | | | T | T | | |
| NC_011770 | 2445441 | C | T | 81 | 0 | 4 | 0 | | | | | | | | | T | T | T | T |
| NC_011770 | 2445446 | T | C | 80 | 0 | 4 | 0 | | | | | | | | | C | C | C | C |
| NC_011770 | 2445457 | TCC | T | 79 | 0 | 4 | 0 | | | | | | | | | T | T | T | T |
| NC_011770 | 2445467 | C | T | 75 | 0 | 4 | 0 | | | | | | | | | T | T | T | T |
| NC_011770 | 2447525 | G | T | 23 | 0 | 1 | 0 | T | . | | | | | | | | | | |
| NC_011770 | 2447526 | T | C | 23 | 0 | 4 | 0 | | | | | | | | | C | C | C | C |
| NC_011770 | 2447531 | G | A | 20 | 0 | 6 | 0 | A | A | | | | | | | A | A | A | A |
| NC_011770 | 2447649 | C | G | 26 | 6 | 4 | 0 | . | . | G | G | G | G | | | . | . | . | . |
| NC_011770 | 2447668 | T | A | 29 | 6 | 2 | 0 | . | . | | | | | A | A | . | . | . | . |
| NC_011770 | 2450341 | T | C | 42 | 6 | 6 | 0 | . | . | C | C | C | C | C | C | . | . | . | . |
| NC_011770 | 2450726 | A | G | 42 | 6 | 6 | 0 | . | . | G | G | G | G | G | G | . | . | . | . |
| NC_011770 | 2450886 | T | C | 54 | 0 | 8 | 0 | | | C | C | C | C | C | C | | | C | C |
| NC_011770 | 2450907 | A | T | 51 | 0 | 8 | 0 | | | T | T | T | T | T | T | | | T | T |
| NC_011770 | 2450911 | T | A | 52 | 0 | 8 | 0 | | | A | A | A | A | A | A | | | A | A |
| NC_011770 | 2450912 | G | T | 52 | 0 | 8 | 0 | | | T | T | T | T | T | T | | | T | T |
| NC_011770 | 2450933 | C | T | 57 | 0 | 8 | 0 | | | T | T | T | T | T | T | | | T | T |
| NC_011770 | 2450968 | G | A | 67 | 0 | 2 | 0 | | | | | | | | | A | A | | |
| NC_011770 | 2452018 | AGG | A | 83 | 0 | 4 | 0 | A | A | | | | | | | A | A | | |
| NC_011770 | 2452023 | G | C | 64 | 4 | 4 | 0 | . | . | | | | | C | C | . | . | C | C |
| NC_011770 | 2452034 | C | T | 79 | 0 | 2 | 0 | T | T | | | | | | | | | | |
| NC_011770 | 2452047 | CGATCGGC | C | 83 | 0 | 2 | 0 | C | C | | | | | | | | | | |
| NC_011770 | 2452063 | A | G | 84 | 0 | 2 | 0 | | | | | | | | | G | G | | |
| NC_011770 | 2452072 | C | CA | 84 | 0 | 2 | 0 | | | | | | | | | CA | CA | | |
| NC_011770 | 2452706 | G | A | 70 | 0 | 2 | 0 | | | | | | | | | A | A | | |
| NC_011770 | 2452733 | A | G | 67 | 0 | 4 | 0 | G | G | | | | | | | G | G | | |
| NC_011770 | 2452734 | G | A | 69 | 0 | 4 | 0 | A | A | | | | | | | A | A | | |
| NC_011770 | 2452761 | T | C | 63 | 0 | 2 | 0 | C | C | | | | | | | | | | |
| NC_011770 | 2452771 | C | A | 64 | 0 | 4 | 0 | A | A | | | | | | | A | A | | |
| NC_011770 | 2453143 | C | T | 7 | 8 | 4 | 0 | T | T | . | . | . | . | . | . | T | T | . | . |
| NC_011770 | 2453230 | A | C | 4 | 8 | 4 | 0 | C | C | . | . | . | . | . | . | C | C | . | . |
| NC_011770 | 2455362 | G | A | 13 | 8 | 2 | 0 | | | . | . | . | . | . | . | A | A | . | . |
| NC_011770 | 2455382 | T | C | 13 | 8 | 2 | 0 | | | . | . | . | . | . | . | C | C | . | . |
| NC_011770 | 2456102 | C | T | 65 | 0 | 2 | 0 | | | | | | | T | T | | | | |
| NC_011770 | 2456154 | C | T | 77 | 0 | 2 | 0 | | | | | | | T | T | | | | |
| NC_011770 | 2456277 | C | T | 76 | 0 | 2 | 0 | T | T | | | | | | | | | | |
| NC_011770 | 2456311 | C | T | 76 | 0 | 2 | 0 | | | | | | | T | T | | | | |
| NC_011770 | 2457529 | C | A | 38 | 0 | 2 | 0 | A | A | | | | | | | | | | |
| NC_011770 | 2457552 | T | A | 41 | 0 | 2 | 0 | | | | | | | A | A | | | | |
| NC_011770 | 2460519 | T | C | 40 | 0 | 10 | 0 | | | C | C | C | C | C | C | C | C | C | C |
| NC_011770 | 2461983 | T | C | 72 | 0 | 8 | 0 | | | C | C | C | C | | | C | C | C | C |
| NC_011770 | 2461984 | T | C | 72 | 0 | 8 | 0 | | | C | C | C | C | | | C | C | C | C |
| NC_011770 | 2461986 | G | A | 72 | 0 | 6 | 0 | | | A | A | A | A | | | A | A | | |
| NC_011770 | 2462031 | A | T | 71 | 0 | 2 | 0 | | | | | | | | | | | T | T |
| NC_011770 | 2462032 | T | G | 70 | 0 | 6 | 0 | | | G | G | G | G | | | A | A | | |
| NC_011770 | 2462046 | A | G | 72 | 0 | 8 | 0 | | | G | G | G | G | | | G | G | G | G |
| NC_011770 | 2462067 | T | C | 69 | 0 | 8 | 0 | C | C | TC | TCC | TC | TC | | | TC | TC | | |
| NC_011770 | 2462071 | C | CT | 37 | 2 | 2 | 0 | | | | | | | | | . | . | CT | CT |
| NC_011770 | 2462079 | C | T | 75 | 0 | 8 | 0 | | | T | T | T | T | T | T | T | T | | |
| NC_011770 | 2462082 | CG | C | 77 | 0 | 2 | 0 | | | | | | | | | | | C | C |
| NC_011770 | 2462090 | T | C | 77 | 0 | 10 | 0 | | | C | C | C | C | C | C | C | C | C | C |
| NC_011770 | 2464695 | A | G | 51 | 0 | 2 | 0 | | | | | | | | | G | G | | |
| NC_011770 | 2464942 | T | G | 64 | 0 | 4 | 0 | | | G | G | G | G | | | | | | |
| NC_011770 | 2464959 | G | C | 66 | 0 | 4 | 0 | | | | | | | | | C | C | C | C |
| NC_011770 | 2465027 | G | C | 73 | 0 | 8 | 0 | C | C | | | | | C | C | C | C | C | C |
| NC_011770 | 2465032 | T | C | 70 | 0 | 6 | 0 | | | | | | | C | C | C | C | C | C |
| NC_011770 | 2465048 | A | T | 73 | 0 | 6 | 0 | | | | | | | T | T | T | T | T | T |
| NC_011770 | 2466024 | T | C | 53 | 0 | 4 | 0 | | | C | C | C | C | | | | | | |

| | | | | | | | | | | | | | | | | | | | |
|---|---|---|---|---|---|---|---|---|---|---|---|---|---|---|---|---|---|---|---|
| NC_011770 | 2466033 | G | A | 55 | 0 | 2 | 0 | | | | | | | | | A | A | | |
| NC_011770 | 2466037 | G | A | 50 | 0 | 2 | 0 | | | | | | | | | A | A | | |
| NC_011770 | 2466043 | G | A | 49 | 0 | 4 | 0 | | | A | A | A | A | | | | | | |
| NC_011770 | 2466054 | T | C | 46 | 0 | 2 | 0 | C | C | | | | | | | | | | |
| NC_011770 | 2466091 | C | G | 49 | 0 | 2 | 0 | | | | | | | | | G | G | | |
| NC_011770 | 2466098 | G | GC | 51 | 0 | 12 | 0 | GC | GC | GC | GC | GC | GC | GC | GC | GC | GC | GC | GC |
| NC_011770 | 2466173 | G | A | 56 | 0 | 2 | 0 | | | | | | | A | A | | | | |
| NC_011770 | 2467576 | C | G | 57 | 0 | 2 | 0 | | | | | | | | | G | G | | |
| NC_011770 | 2467580 | C | T | 57 | 0 | 2 | 0 | | | | | | | | | | | T | T |
| NC_011770 | 2468610 | A | C | 64 | 0 | 8 | 0 | C | C | C | C | C | C | | | | | C | C |
| NC_011770 | 2470214 | G | A | 39 | 0 | 2 | 0 | | | | | | | A | A | | | | |
| NC_011770 | 2470257 | T | C | 40 | 0 | 6 | 0 | | | C | C | C | C | C | C | | | | |
| NC_011770 | 2470359 | A | G | 43 | 0 | 2 | 0 | | | | | | | | | G | G | | |
| NC_011770 | 2472222 | G | C | 71 | 0 | 2 | 0 | C | C | | | | | | | | | | |
| NC_011770 | 2474017 | CA | C | 34 | 0 | 4 | 0 | C | C | | | | | | | C | C | | |
| NC_011770 | 2474513 | C | T | 58 | 1 | 1 | 0 | | | | | | | | | . | T | | |
| NC_011770 | 2474531 | T | G | 68 | 0 | 2 | 0 | | | | | | | | | G | G | | |
| NC_011770 | 2474532 | C | A | 70 | 0 | 2 | 0 | | | | | | | | | A | A | | |
| NC_011770 | 2474564 | T | C | 76 | 0 | 6 | 0 | | | | | | | C | C | C | C | C | C |
| NC_011770 | 2475800 | CAA | C | 86 | 0 | 2 | 0 | | | | | | | C | C | | | | |
| NC_011770 | 2477423 | A | G | 80 | 0 | 10 | 0 | | | G | G | G | G | G | G | G | G | G | G |
| NC_011770 | 2477434 | C | A | 87 | 0 | 4 | 0 | | | | | | | A | A | | | A | A |
| NC_011770 | 2477439 | A | G | 86 | 0 | 2 | 0 | | | | | | | | | | | G | G |
| NC_011770 | 2478096 | C | T | 85 | 0 | 2 | 0 | T | T | | | | | | | | | | |
| NC_011770 | 2478159 | C | T | 80 | 0 | 2 | 0 | | | | | | | T | T | | | | |
| NC_011770 | 2478236 | G | A | 75 | 0 | 8 | 0 | A | A | A | A | A | A | | | | | A | A |
| NC_011770 | 2480073 | A | G | 59 | 0 | 4 | 0 | | | | | | | G | G | | | G | G |
| NC_011770 | 2480090 | A | G | 59 | 0 | 4 | 0 | | | | | | | G | G | | | G | G |
| NC_011770 | 2480140 | G | T | 53 | 0 | 4 | 0 | | | | | | | T | T | | | T | T |
| NC_011770 | 2480190 | A | T | 57 | 0 | 4 | 0 | | | | | | | T | T | | | T | T |
| NC_011770 | 2480198 | A | G | 57 | 0 | 4 | 0 | | | | | | | G | G | | | G | G |
| NC_011770 | 2480598 | T | C | 57 | 0 | 6 | 0 | | | C | C | C | C | | | | | G | G |
| NC_011770 | 2480799 | T | G | 69 | 0 | 2 | 0 | | | | | | | | | G | G | | |
| NC_011770 | 2481275 | C | T | 85 | 0 | 6 | 0 | | | T | T | T | T | | | T | T | | |
| NC_011770 | 2481278 | C | A | 84 | 0 | 6 | 0 | A | A | | | | | A | A | | | A | A |
| NC_011770 | 2481284 | C | T | 85 | 0 | 6 | 0 | T | T | | | | | T | T | | | T | T |
| NC_011770 | 2481306 | C | G | 87 | 0 | 12 | 0 | G | G | G | G | G | G | G | G | G | G | G | G |
| NC_011770 | 2481330 | G | A | 87 | 0 | 2 | 0 | | | | | | | | | A | A | | |
| NC_011770 | 2481749 | A | G | 92 | 0 | 12 | 0 | G | G | G | G | G | G | G | G | G | G | G | G |
| NC_011770 | 2482193 | T | C | 93 | 0 | 2 | 0 | C | C | | | | | | | | | | |
| NC_011770 | 2482198 | A | G | 96 | 0 | 12 | 0 | G | G | G | G | G | G | G | G | G | G | G | G |
| NC_011770 | 2482297 | C | T | 93 | 0 | 2 | 0 | T | T | | | | | | | | | | |
| NC_011770 | 2483659 | C | T | 81 | 0 | 1 | 0 | | | | | | | T | | | | | |
| NC_011770 | 2483770 | G | T | 84 | 0 | 2 | 0 | | | | | | | T | T | | | | |
| NC_011770 | 2483780 | A | C | 78 | 0 | 2 | 0 | C | C | | | | | | | | | | |
| NC_011770 | 2484427 | A | G | 82 | 0 | 2 | 0 | | | | | | | | | G | G | | |
| NC_011770 | 2484457 | G | C | 87 | 0 | 2 | 0 | C | C | | | | | | | | | | |
| NC_011770 | 2484463 | C | T | 84 | 0 | 2 | 0 | | | | | | | | | | | T | T |
| NC_011770 | 2485355 | G | A | 79 | 0 | 2 | 0 | | | | | | | | | | | A | A |
| NC_011770 | 2485372 | A | C | 78 | 0 | 12 | 0 | C | C | C | C | C | C | C | C | C | C | C | C |
| NC_011770 | 2485391 | C | T | 79 | 0 | 2 | 0 | | | | | | | T | T | | | | |
| NC_011770 | 2485404 | T | G | 77 | 0 | 12 | 0 | G | G | G | G | G | G | G | G | G | G | G | G |
| NC_011770 | 2485418 | C | T | 78 | 0 | 2 | 0 | | | | | | | T | T | | | | |
| NC_011770 | 2486474 | C | A | 40 | 0 | 2 | 0 | | | | | | | | | | | A | A |
| NC_011770 | 2486477 | C | A | 40 | 0 | 2 | 0 | | | | | | | | | | | A | A |
| NC_011770 | 2486486 | GA | G | 39 | 0 | 4 | 0 | | | G | G | G | G | | | | | | |
| NC_011770 | 2487040 | C | T | 51 | 0 | 2 | 0 | | | | | | | | | T | T | | |
| NC_011770 | 2487104 | T | C | 44 | 0 | 12 | 0 | C | C | C | C | C | C | C | C | C | C | C | C |
| NC_011770 | 2487183 | G | A | 41 | 0 | 2 | 0 | | | | | | | | | A | A | | |
| NC_011770 | 2487247 | G | A | 44 | 0 | 4 | 0 | | | | | | | A | A | A | A | | |
| NC_011770 | 2487338 | GT | G | 42 | 0 | 6 | 0 | | | | | | | G | G | G | G | G | G |
| NC_011770 | 2487592 | C | T | 38 | 0 | 8 | 0 | T | T | | | | | T | T | T | T | T | T |

| | | | | | | | | | | | | | | | | | | | |
|---|---|---|---|---|---|---|---|---|---|---|---|---|---|---|---|---|---|---|---|
| NC_011770 | 2487631 | C | T | 35 | 0 | 6 | 0 | T | T | T | T | T | T | | | | | | |
| NC_011770 | 2487692 | T | C | 42 | 0 | 8 | 0 | C | C | C | C | C | C | | | C | C | | |
| NC_011770 | 2487704 | C | A | 42 | 0 | 12 | 0 | A | A | A | A | A | A | A | A | A | A | A | A |
| NC_011770 | 2487708 | G | A | 43 | 0 | 2 | 0 | | | | | | | | | A | A | | |
| NC_011770 | 2487736 | G | T | 50 | 0 | 4 | 0 | | | T | T | T | T | | | | | | |
| NC_011770 | 2488007 | C | T | 58 | 0 | 2 | 0 | | | | | | | | | T | T | | |
| NC_011770 | 2488422 | A | C | 70 | 0 | 12 | 0 | C | C | C | C | C | C | C | C | C | C | C | C |
| NC_011770 | 2488431 | C | T | 68 | 0 | 2 | 0 | | | | | | | | | T | T | | |
| NC_011770 | 2488925 | C | T | 52 | 0 | 4 | 0 | | | T | T | T | T | | | | | | |
| NC_011770 | 2488936 | C | T | 52 | 0 | 8 | 0 | T | T | T | T | T | T | T | T | | | | |
| NC_011770 | 2489443 | G | T | 55 | 0 | 4 | 0 | | | | | | | | | T | T | T | T |
| NC_011770 | 2489449 | C | T | 56 | 0 | 4 | 0 | | | T | T | T | T | | | | | | |
| NC_011770 | 2489680 | C | T | 62 | 0 | 4 | 0 | T | T | | | | | T | T | | | | |
| NC_011770 | 2489706 | G | A | 59 | 0 | 12 | 0 | A | A | A | A | A | A | A | A | A | A | A | A |
| NC_011770 | 2489708 | G | A | 59 | 0 | 4 | 0 | A | A | | | | | A | A | | | | |
| NC_011770 | 2489775 | C | G | 68 | 0 | 2 | 0 | | | | | | | | | | | G | G |
| NC_011770 | 2490831 | C | T | 68 | 0 | 4 | 0 | | | T | T | T | T | | | | | | |
| NC_011770 | 2491241 | CG | C | 65 | 0 | 2 | 0 | C | C | | | | | | | | | | |
| NC_011770 | 2491386 | C | A | 67 | 0 | 2 | 0 | | | | | | | | | | | A | A |
| NC_011770 | 2491407 | C | T | 64 | 0 | 2 | 0 | | | | | | | T | T | | | | |
| NC_011770 | 2492824 | C | T | 82 | 0 | 2 | 0 | | | | | | | T | T | | | | |
| NC_011770 | 2492830 | C | T | 70 | 0 | 2 | 0 | | | | | | | T | T | | | | |
| NC_011770 | 2492868 | A | G | 86 | 0 | 12 | 0 | G | G | G | G | G | G | G | G | G | G | G | G |
| NC_011770 | 2492872 | G | A | 87 | 0 | 2 | 0 | | | | | | | | | A | A | | |
| NC_011770 | 2492877 | C | A | 88 | 0 | 2 | 0 | | | | | | | A | A | | | | |
| NC_011770 | 2493061 | C | T | 91 | 0 | 2 | 0 | | | | | | | T | T | | | | |
| NC_011770 | 2493105 | G | A | 86 | 0 | 12 | 0 | A | A | A | A | A | A | A | A | A | A | A | A |
| NC_011770 | 2493116 | A | T | 85 | 0 | 2 | 0 | | | | | | | T | T | | | | |
| NC_011770 | 2493929 | TGC | T | 77 | 0 | 2 | 0 | T | T | | | | | | | | | | |
| NC_011770 | 2493931 | C | A | 71 | 2 | 6 | 0 | . | . | A | A | A | A | A | A | | | | |
| NC_011770 | 2493956 | A | G | 77 | 0 | 12 | 0 | G | G | G | G | G | G | G | G | G | G | G | G |
| NC_011770 | 2494245 | G | A | 73 | 0 | 2 | 0 | | | | | | | | | | | A | A |
| NC_011770 | 2494288 | T | G | 69 | 0 | 12 | 0 | G | G | G | G | G | G | G | G | G | G | G | G |
| NC_011770 | 2494289 | T | A | 69 | 0 | 2 | 0 | | | | | | | A | A | | | | |
| NC_011770 | 2494771 | G | GC | 69 | 0 | 12 | 0 | GC | GC | GC | GC | GC | GC | GC | GC | GC | GC | GC | GC |
| NC_011770 | 2494791 | G | A | 73 | 0 | 2 | 0 | | | | | | | A | A | | | | |
| NC_011770 | 2494819 | C | T | 69 | 0 | 4 | 0 | | | T | T | T | T | | | | | | |
| NC_011770 | 2494827 | C | T | 70 | 0 | 2 | 0 | | | | | | | | | T | T | | |
| NC_011770 | 2494996 | A | G | 75 | 0 | 2 | 0 | G | G | | | | | | | | | | |
| NC_011770 | 2495260 | A | G | 79 | 0 | 2 | 0 | G | G | | | | | | | | | | |
| NC_011770 | 2495669 | G | A | 76 | 0 | 2 | 0 | A | A | | | | | | | | | | |
| NC_011770 | 2495726 | C | T | 81 | 0 | 4 | 0 | T | T | | | | | | | | | T | T |
| NC_011770 | 2495734 | C | G | 80 | 0 | 4 | 0 | | | G | G | G | G | | | | | | |
| NC_011770 | 2496704 | C | T | 84 | 0 | 2 | 0 | | | | | | | T | T | | | | |
| NC_011770 | 2500039 | C | A | 96 | 0 | 2 | 0 | | | | | | | | | | | A | A |
| NC_011770 | 2503861 | C | G | 69 | 0 | 2 | 0 | G | G | | | | | | | | | | |
| NC_011770 | 2504178 | G | A | 66 | 0 | 2 | 0 | | | | | | | A | A | | | | |
| NC_011770 | 2504244 | T | C | 51 | 0 | 12 | 0 | C | C | C | C | C | C | C | C | C | C | C | C |
| NC_011770 | 2504254 | T | C | 49 | 0 | 6 | 0 | | | C | C | C | C | C | C | | | | |
| NC_011770 | 2504347 | A | C | 35 | 1 | 4 | 0 | . | . | | | | | C | C | | | T | T |
| NC_011770 | 2504349 | C | T | 37 | 0 | 2 | 0 | | | | | | | | | T | T | | |
| NC_011770 | 2504350 | C | T | 37 | 0 | 1 | 0 | | | | | | | T | | | | | |
| NC_011770 | 2504351 | T | A | 41 | 0 | 1 | 0 | | | | | | | A | . | | | | |
| NC_011770 | 2504353 | A | G | 40 | 0 | 8 | 0 | G | G | G | G | | | | | G | G | G | G |
| NC_011770 | 2504362 | G | A | 41 | 0 | 4 | 0 | | | A | A | | | | | | | A | A |
| NC_011770 | 2504370 | A | G | 41 | 0 | 12 | 0 | G | G | G | G | G | G | G | G | G | G | G | G |
| NC_011770 | 2504372 | G | T | 42 | 0 | 2 | 0 | | | | | | | T | T | | | | |
| NC_011770 | 2504376 | G | A | 41 | 0 | 2 | 0 | | | | | | | | | A | A | | |
| NC_011770 | 2504377 | C | T | 41 | 0 | 4 | 0 | | | T | T | | | | | | | T | T |
| NC_011770 | 2504392 | C | T | 22 | 6 | 2 | 0 | | | . | . | T | T | | | . | . | . | . |
| NC_011770 | 2504397 | A | T | 15 | 8 | 2 | 0 | | | . | . | T | T | . | . | . | . | . | . |
| NC_011770 | 2504467 | C | T | 6 | 10 | 2 | 0 | T | T | . | . | . | . | . | . | . | . | . | . |

| | | | | | | | | | | | | | | | | | | | |
|---|---|---|---|---|---|---|---|---|---|---|---|---|---|---|---|---|---|---|---|
| NC_011770 | 2510837 | T | C | 7 | 10 | 2 | 0 | C | C | . | . | . | . | . | . | . | . | . | . |
| NC_011770 | 2511674 | A | G | 7 | 8 | 2 | 0 | G | G | . | . | | | . | . | . | . | . | . |
| NC_011770 | 2512812 | G | T | 2 | 10 | 2 | 0 | T | T | . | . | . | . | . | . | . | . | . | . |
| NC_011770 | 2513920 | G | A | 6 | 10 | 2 | 0 | A | A | . | . | . | . | . | . | . | . | . | . |
| NC_011770 | 2515157 | A | G | 6 | 10 | 2 | 0 | G | G | . | . | . | . | . | . | . | . | . | . |
| NC_011770 | 2521988 | C | T | 4 | 10 | 2 | 0 | T | T | . | . | . | . | . | . | . | . | . | . |
| NC_011770 | 2522001 | A | G | 4 | 10 | 2 | 0 | G | G | . | . | . | . | . | . | . | . | . | . |
| NC_011770 | 2522048 | G | A | 3 | 10 | 2 | 0 | A | A | . | . | . | . | . | . | . | . | . | . |
| NC_011770 | 2522079 | T | C | 4 | 10 | 2 | 0 | C | C | . | . | . | . | . | . | . | . | . | . |
| NC_011770 | 2522139 | G | A | 5 | 10 | 2 | 0 | A | A | . | . | . | . | . | . | . | . | . | . |
| NC_011770 | 2522147 | T | A | 5 | 10 | 2 | 0 | A | A | . | . | . | . | . | . | . | . | . | . |
| NC_011770 | 2523088 | G | C | 3 | 10 | 2 | 0 | C | C | . | . | . | . | . | . | . | . | . | . |
| NC_011770 | 2523090 | G | C | 3 | 10 | 2 | 0 | C | C | . | . | . | . | . | . | . | . | . | . |
| NC_011770 | 2523094 | T | C | 3 | 10 | 2 | 0 | C | C | . | . | . | . | . | . | . | . | . | . |
| NC_011770 | 2523101 | A | T | 3 | 10 | 2 | 0 | T | T | . | . | . | . | . | . | . | . | . | . |
| NC_011770 | 2523104 | G | A | 3 | 10 | 2 | 0 | A | A | . | . | . | . | . | . | . | . | . | . |
| NC_011770 | 2523108 | A | G | 3 | 10 | 2 | 0 | G | G | . | . | . | . | . | . | . | . | . | . |
| NC_011770 | 2523119 | A | G | 4 | 10 | 2 | 0 | G | G | . | . | . | . | . | . | . | . | . | . |
| NC_011770 | 2523122 | T | C | 4 | 10 | 2 | 0 | C | C | . | . | . | . | . | . | . | . | . | . |
| NC_011770 | 2523129 | A | T | 4 | 10 | 2 | 0 | T | T | . | . | . | . | . | . | . | . | . | . |
| NC_011770 | 2523137 | C | T | 4 | 10 | 2 | 0 | T | T | . | . | . | . | . | . | . | . | . | . |
| NC_011770 | 2523149 | C | G | 4 | 10 | 2 | 0 | G | G | . | . | . | . | . | . | . | . | . | . |
| NC_011770 | 2523171 | G | A | 4 | 10 | 2 | 0 | A | A | . | . | . | . | . | . | . | . | . | . |
| NC_011770 | 2523176 | T | A | 4 | 10 | 2 | 0 | A | A | . | . | . | . | . | . | . | . | . | . |
| NC_011770 | 2523203 | G | A | 5 | 10 | 2 | 0 | A | A | . | . | . | . | . | . | . | . | . | . |
| NC_011770 | 2523219 | T | C | 5 | 10 | 2 | 0 | C | C | . | . | . | . | . | . | . | . | . | . |
| NC_011770 | 2523224 | T | C | 5 | 10 | 2 | 0 | C | C | . | . | . | . | . | . | . | . | . | . |
| NC_011770 | 2523255 | C | G | 5 | 10 | 2 | 0 | G | G | . | . | . | . | . | . | . | . | . | . |
| NC_011770 | 2523258 | G | A | 5 | 10 | 2 | 0 | A | A | . | . | . | . | . | . | . | . | . | . |
| NC_011770 | 2523269 | C | A | 5 | 10 | 2 | 0 | A | A | . | . | . | . | . | . | . | . | . | . |
| NC_011770 | 2523280 | T | C | 5 | 10 | 2 | 0 | C | C | . | . | . | . | . | . | . | . | . | . |
| NC_011770 | 2523284 | A | T | 5 | 10 | 2 | 0 | T | T | . | . | . | . | . | . | . | . | . | . |
| NC_011770 | 2523290 | G | C | 5 | 10 | 2 | 0 | C | C | . | . | . | . | . | . | . | . | . | . |
| NC_011770 | 2523291 | C | T | 5 | 10 | 2 | 0 | T | T | . | . | . | . | . | . | . | . | . | . |
| NC_011770 | 2523298 | A | G | 5 | 10 | 2 | 0 | G | G | . | . | . | . | . | . | . | . | . | . |
| NC_011770 | 2523306 | C | T | 5 | 10 | 2 | 0 | T | T | . | . | . | . | . | . | . | . | . | . |
| NC_011770 | 2523311 | T | C | 5 | 10 | 2 | 0 | C | C | . | . | . | . | . | . | . | . | . | . |
| NC_011770 | 2523338 | A | G | 4 | 10 | 2 | 0 | G | G | . | . | . | . | . | . | . | . | . | . |
| NC_011770 | 2523350 | C | T | 4 | 10 | 2 | 0 | T | T | . | . | . | . | . | . | . | . | . | . |
| NC_011770 | 2523356 | C | A | 4 | 10 | 2 | 0 | A | A | . | . | . | . | . | . | . | . | . | . |
| NC_011770 | 2523362 | T | C | 5 | 10 | 2 | 0 | C | C | . | . | . | . | . | . | . | . | . | . |
| NC_011770 | 2523367 | T | C | 5 | 10 | 2 | 0 | C | C | . | . | . | . | . | . | . | . | . | . |
| NC_011770 | 2523392 | C | A | 4 | 10 | 2 | 0 | A | A | . | . | . | . | . | . | . | . | . | . |
| NC_011770 | 2523468 | C | G | 5 | 10 | 2 | 0 | G | G | . | . | . | . | . | . | . | . | . | . |
| NC_011770 | 2523483 | T | C | 5 | 10 | 2 | 0 | C | C | . | . | . | . | . | . | . | . | . | . |
| NC_011770 | 2523496 | C | A | 5 | 10 | 2 | 0 | A | A | . | . | . | . | . | . | . | . | . | . |
| NC_011770 | 2527146 | T | C | 3 | 10 | 2 | 0 | C | C | . | . | . | . | . | . | . | . | . | . |
| NC_011770 | 2527158 | T | C | 3 | 10 | 2 | 0 | C | C | . | . | . | . | . | . | . | . | . | . |
| NC_011770 | 2527181 | A | G | 3 | 10 | 2 | 0 | G | G | . | . | . | . | . | . | . | . | . | . |
| NC_011770 | 2527190 | G | C | 3 | 10 | 2 | 0 | C | C | . | . | . | . | . | . | . | . | . | . |
| NC_011770 | 2527202 | C | T | 3 | 10 | 2 | 0 | T | T | . | . | . | . | . | . | . | . | . | . |
| NC_011770 | 2527222 | C | G | 3 | 10 | 2 | 0 | G | G | . | . | . | . | . | . | . | . | . | . |
| NC_011770 | 2527224 | G | C | 3 | 10 | 2 | 0 | C | C | . | . | . | . | . | . | . | . | . | . |
| NC_011770 | 2527255 | G | A | 3 | 10 | 2 | 0 | A | A | . | . | . | . | . | . | . | . | . | . |
| NC_011770 | 2527279 | A | G | 3 | 10 | 2 | 0 | G | G | . | . | . | . | . | . | . | . | . | . |
| NC_011770 | 2527289 | G | A | 3 | 10 | 2 | 0 | A | A | . | . | . | . | . | . | . | . | . | . |
| NC_011770 | 2527291 | A | G | 3 | 10 | 2 | 0 | G | G | . | . | . | . | . | . | . | . | . | . |
| NC_011770 | 2527301 | T | A | 2 | 10 | 2 | 0 | A | A | . | . | . | . | . | . | . | . | . | . |
| NC_011770 | 2527302 | A | T | 2 | 10 | 2 | 0 | T | T | . | . | . | . | . | . | . | . | . | . |
| NC_011770 | 2527305 | T | C | 2 | 10 | 2 | 0 | C | C | . | . | . | . | . | . | . | . | . | . |
| NC_011770 | 2527314 | G | T | 3 | 10 | 2 | 0 | T | T | . | . | . | . | . | . | . | . | . | . |
| NC_011770 | 2527336 | A | T | 3 | 10 | 2 | 0 | T | T | . | . | . | . | . | . | . | . | . | . |

| | | | | | | | | | | | | | | | | | | | |
|---|---|---|---|---|---|---|---|---|---|---|---|---|---|---|---|---|---|---|---|
| NC_011770 | 2527365 | C | G | 4 | 10 | 2 | 0 | G | G | . | . | . | . | . | . | . | . | . | . |
| NC_011770 | 2527376 | C | T | 4 | 10 | 2 | 0 | T | T | . | . | . | . | . | . | . | . | . | . |
| NC_011770 | 2527379 | C | A | 4 | 10 | 2 | 0 | A | A | . | . | . | . | . | . | . | . | . | . |
| NC_011770 | 2527395 | C | T | 3 | 10 | 2 | 0 | T | T | . | . | . | . | . | . | . | . | . | . |
| NC_011770 | 2527399 | T | C | 4 | 10 | 2 | 0 | C | C | . | . | . | . | . | . | . | . | . | . |
| NC_011770 | 2527404 | C | T | 4 | 10 | 2 | 0 | T | T | . | . | . | . | . | . | . | . | . | . |
| NC_011770 | 2527419 | A | G | 3 | 10 | 2 | 0 | G | G | . | . | . | . | . | . | . | . | . | . |
| NC_011770 | 2527421 | G | A | 3 | 10 | 2 | 0 | A | A | . | . | . | . | . | . | . | . | . | . |
| NC_011770 | 2527423 | A | C | 3 | 10 | 2 | 0 | C | C | . | . | . | . | . | . | . | . | . | . |
| NC_011770 | 2527428 | T | C | 4 | 10 | 2 | 0 | C | C | . | . | . | . | . | . | . | . | . | . |
| NC_011770 | 2527438 | C | G | 4 | 10 | 2 | 0 | G | G | . | . | . | . | . | . | . | . | . | . |
| NC_011770 | 2527479 | G | A | 2 | 10 | 2 | 0 | A | A | . | . | . | . | . | . | . | . | . | . |
| NC_011770 | 2527483 | G | C | 2 | 10 | 2 | 0 | C | C | . | . | . | . | . | . | . | . | . | . |
| NC_011770 | 2527485 | G | A | 2 | 10 | 2 | 0 | A | A | . | . | . | . | . | . | . | . | . | . |
| NC_011770 | 2527506 | G | T | 0 | 10 | 2 | 0 | T | T | . | . | . | . | . | . | . | . | . | . |
| NC_011770 | 2527516 | T | G | 4 | 10 | 2 | 0 | G | G | . | . | . | . | . | . | . | . | . | . |
| NC_011770 | 2527522 | T | C | 4 | 10 | 2 | 0 | C | C | . | . | . | . | . | . | . | . | . | . |
| NC_011770 | 2527527 | C | G | 4 | 10 | 2 | 0 | G | G | . | . | . | . | . | . | . | . | . | . |
| NC_011770 | 2527530 | G | A | 4 | 10 | 2 | 0 | A | A | . | . | . | . | . | . | . | . | . | . |
| NC_011770 | 2527533 | G | A | 4 | 10 | 2 | 0 | A | A | . | . | . | . | . | . | . | . | . | . |
| NC_011770 | 2527581 | A | C | 5 | 10 | 2 | 0 | C | C | . | . | . | . | . | . | . | . | . | . |
| NC_011770 | 2527582 | G | A | 5 | 10 | 2 | 0 | A | A | . | . | . | . | . | . | . | . | . | . |
| NC_011770 | 2527593 | T | C | 5 | 10 | 2 | 0 | C | C | . | . | . | . | . | . | . | . | . | . |
| NC_011770 | 2527599 | T | C | 5 | 10 | 2 | 0 | C | C | . | . | . | . | . | . | . | . | . | . |
| NC_011770 | 2527605 | C | T | 5 | 10 | 2 | 0 | T | T | . | . | . | . | . | . | . | . | . | . |
| NC_011770 | 2527614 | G | A | 5 | 10 | 2 | 0 | A | A | . | . | . | . | . | . | . | . | . | . |
| NC_011770 | 2527626 | G | A | 5 | 10 | 2 | 0 | A | A | . | . | . | . | . | . | . | . | . | . |
| NC_011770 | 2527632 | G | C | 5 | 10 | 2 | 0 | C | C | . | . | . | . | . | . | . | . | . | . |
| NC_011770 | 2527635 | C | G | 5 | 10 | 2 | 0 | G | G | . | . | . | . | . | . | . | . | . | . |
| NC_011770 | 2527654 | T | G | 5 | 10 | 2 | 0 | G | G | . | . | . | . | . | . | . | . | . | . |
| NC_011770 | 2527690 | A | G | 6 | 10 | 2 | 0 | G | G | . | . | . | . | . | . | . | . | . | . |
| NC_011770 | 2527695 | T | C | 5 | 10 | 2 | 0 | C | C | . | . | . | . | . | . | . | . | . | . |
| NC_011770 | 2527703 | C | T | 5 | 10 | 2 | 0 | T | T | . | . | . | . | . | . | . | . | . | . |
| NC_011770 | 2527718 | T | C | 5 | 10 | 2 | 0 | C | C | . | . | . | . | . | . | . | . | . | . |
| NC_011770 | 2527737 | G | A | 5 | 10 | 2 | 0 | A | A | . | . | . | . | . | . | . | . | . | . |
| NC_011770 | 2527777 | A | C | 5 | 10 | 2 | 0 | C | C | . | . | . | . | . | . | . | . | . | . |
| NC_011770 | 2527778 | C | T | 5 | 10 | 2 | 0 | T | T | . | . | . | . | . | . | . | . | . | . |
| NC_011770 | 2527783 | A | C | 5 | 10 | 2 | 0 | C | C | . | . | . | . | . | . | . | . | . | . |
| NC_011770 | 2527794 | T | A | 6 | 10 | 2 | 0 | A | A | . | . | . | . | . | . | . | . | . | . |
| NC_011770 | 2527816 | T | G | 5 | 10 | 2 | 0 | G | G | . | . | . | . | . | . | . | . | . | . |
| NC_011770 | 2527821 | G | T | 5 | 10 | 2 | 0 | T | T | . | . | . | . | . | . | . | . | . | . |
| NC_011770 | 2527830 | A | G | 5 | 10 | 2 | 0 | G | G | . | . | . | . | . | . | . | . | . | . |
| NC_011770 | 2527833 | T | C | 5 | 10 | 2 | 0 | C | C | . | . | . | . | . | . | . | . | . | . |
| NC_011770 | 2527867 | A | G | 6 | 10 | 2 | 0 | G | G | . | . | . | . | . | . | . | . | . | . |
| NC_011770 | 2527872 | A | T | 6 | 10 | 2 | 0 | T | T | . | . | . | . | . | . | . | . | . | . |
| NC_011770 | 2527882 | G | A | 6 | 10 | 2 | 0 | A | A | . | . | . | . | . | . | . | . | . | . |
| NC_011770 | 2527914 | T | C | 6 | 10 | 2 | 0 | C | C | . | . | . | . | . | . | . | . | . | . |
| NC_011770 | 2527995 | A | G | 6 | 10 | 2 | 0 | G | G | . | . | . | . | . | . | . | . | . | . |
| NC_011770 | 2528004 | T | G | 6 | 10 | 2 | 0 | G | G | . | . | . | . | . | . | . | . | . | . |
| NC_011770 | 2528016 | T | C | 6 | 10 | 2 | 0 | C | C | . | . | . | . | . | . | . | . | . | . |
| NC_011770 | 2528330 | T | C | 6 | 10 | 2 | 0 | C | C | . | . | . | . | . | . | . | . | . | . |
| NC_011770 | 2528364 | G | T | 6 | 10 | 2 | 0 | T | T | . | . | . | . | . | . | . | . | . | . |
| NC_011770 | 2528392 | C | T | 5 | 10 | 2 | 0 | T | T | . | . | . | . | . | . | . | . | . | . |
| NC_011770 | 2528424 | C | T | 4 | 10 | 2 | 0 | T | T | . | . | . | . | . | . | . | . | . | . |
| NC_011770 | 2540009 | A | AGT | 6 | 10 | 2 | 0 | AGT | AGT | . | . | . | . | . | . | . | . | . | . |
| NC_011770 | 2540186 | T | A | 6 | 9 | 1 | 0 | | | . | . | . | A | . | . | . | . | . | . |
| NC_011770 | 2540187 | A | G | 6 | 9 | 1 | 0 | | | . | . | . | G | . | . | . | . | . | . |
| NC_011770 | 2540248 | T | C | 6 | 8 | 2 | 0 | | | . | . | C | C | . | . | . | . | . | . |
| NC_011770 | 2540249 | T | C | 6 | 8 | 2 | 0 | | | . | . | C | C | . | . | . | . | . | . |
| NC_011770 | 2545402 | G | A | 24 | 4 | 2 | 0 | | | . | . | A | A | | | | | . | . |
| NC_011770 | 2545412 | CG | C | 21 | 4 | 2 | 0 | | . | . | . | | | | | C | C | . | . |
| NC_011770 | 2545421 | A | T | 16 | 7 | 1 | 0 | | | . | . | . | . | T | . | | | . | . |

| | | | | | | | | | | | | | | | | | | | |
|---|---|---|---|---|---|---|---|---|---|---|---|---|---|---|---|---|---|---|---|
| NC_011770 | 2546171 | C | CA | 6 | 10 | 2 | 0 | CA | CA | . | . | . | . | . | . | . | . | . | . |
| NC_011770 | 2548438 | A | G | 102 | 0 | 6 | 0 | | | | | | | G | G | G | G | G | G |
| NC_011770 | 2548456 | G | A | 104 | 0 | 6 | 0 | | | | | | | A | A | A | A | A | A |
| NC_011770 | 2548468 | A | G | 104 | 0 | 5 | 0 | | | . | . | . | G | G | G | G | G | . | . |
| NC_011770 | 2548496 | C | T | 116 | 0 | 2 | 0 | | | | | | | | | T | T | | |
| NC_011770 | 2548555 | A | G | 107 | 0 | 4 | 0 | | | | | | | G | G | G | G | | |
| NC_011770 | 2548562 | C | G | 94 | 0 | 4 | 0 | | | | | | | G | G | G | G | | |
| NC_011770 | 2548565 | C | G | 94 | 0 | 4 | 0 | | | | | | | G | G | G | G | | |
| NC_011770 | 2548566 | A | C | 94 | 0 | 4 | 0 | | | | | | | C | C | C | C | | |
| NC_011770 | 2548574 | G | C | 95 | 0 | 4 | 0 | | | | | | | C | C | C | C | | |
| NC_011770 | 2548576 | T | G | 94 | 0 | 4 | 0 | | | . | . | . | . | G | G | G | G | | |
| NC_011770 | 2548577 | A | G | 93 | 0 | 10 | 0 | | | G | G | G | G | G | G | G | G | G | G |
| NC_011770 | 2548579 | C | T | 93 | 0 | 4 | 0 | | | | | | | T | T | T | T | | |
| NC_011770 | 2548598 | A | G | 85 | 0 | 10 | 0 | | | G | G | G | G | G | G | G | G | G | G |
| NC_011770 | 2548633 | A | G | 82 | 0 | 10 | 0 | | | G | G | G | G | G | G | G | G | G | G |
| NC_011770 | 2548636 | G | A | 82 | 0 | 4 | 0 | | | | | | | A | A | A | A | | |
| NC_011770 | 2548639 | T | C | 82 | 0 | 4 | 0 | | | | | | | C | C | C | C | | |
| NC_011770 | 2548652 | C | T | 81 | 0 | 4 | 0 | | | | | | | T | T | T | T | | |
| NC_011770 | 2548741 | G | A | 16 | 6 | 2 | 0 | | | . | . | | | . | . | . | . | A | A |
| NC_011770 | 2548749 | A | G | 13 | 7 | 3 | 0 | | | . | . | . | G | . | . | . | . | G | G |
| NC_011770 | 2548758 | A | G | 13 | 7 | 2 | 0 | | | . | . | . | | . | . | . | . | G | G |
| NC_011770 | 2559569 | G | A | 76 | 0 | 6 | 0 | | | A | A | A | A | | | A | A | | |
| NC_011770 | 2560174 | G | C | 34 | 0 | 12 | 0 | C | C | C | C | C | C | C | C | C | C | C | C |
| NC_011770 | 2560488 | T | C | 63 | 0 | 10 | 0 | | | C | C | C | C | C | C | C | C | C | C |
| NC_011770 | 2561416 | G | C | 76 | 0 | 4 | 0 | | | | | | | C | C | | | C | C |
| NC_011770 | 2561472 | C | T | 84 | 0 | 2 | 0 | | | | | | | | | T | T | | |
| NC_011770 | 2561542 | G | A | 81 | 0 | 4 | 0 | | | | | | | A | A | | | A | A |
| NC_011770 | 2562325 | C | T | 82 | 0 | 4 | 0 | | | T | T | T | T | | | | | | |
| NC_011770 | 2562346 | G | T | 86 | 0 | 8 | 0 | | | T | T | T | T | T | T | T | T | | |
| NC_011770 | 2562414 | T | C | 84 | 0 | 6 | 0 | | | C | C | C | C | | | C | C | | |
| NC_011770 | 2562446 | A | G | 79 | 0 | 6 | 0 | | | G | G | G | G | | | G | G | | |
| NC_011770 | 2562450 | C | CGT | 79 | 0 | 6 | 0 | | | CGT | CGT | CGT | CGT | | | CGT | CGT | | |
| NC_011770 | 2563771 | C | T | 84 | 0 | 2 | 0 | T | T | | | | | | | | | | |
| NC_011770 | 2563802 | T | A | 75 | 0 | 2 | 0 | | | | | | | A | A | | | | |
| NC_011770 | 2563846 | C | T | 76 | 0 | 2 | 0 | | | | | | | T | T | | | | |
| NC_011770 | 2563862 | G | A | 77 | 0 | 2 | 0 | | | | | | | A | A | | | | |
| NC_011770 | 2563905 | A | G | 65 | 1 | 3 | 0 | | | | | | | G | . | G | G | | |
| NC_011770 | 2563910 | C | A | 77 | 0 | 2 | 0 | | | | | | | A | A | | | | |
| NC_011770 | 2564836 | C | T | 68 | 0 | 2 | 0 | | | | | | | | | T | T | | |
| NC_011770 | 2564838 | G | A | 70 | 0 | 2 | 0 | | | | | | | | | A | A | | |
| NC_011770 | 2564842 | G | T | 67 | 0 | 8 | 0 | | | T | T | T | T | | | T | T | T | T |
| NC_011770 | 2566157 | G | A | 57 | 0 | 6 | 0 | A | A | A | A | A | A | | | | | | |
| NC_011770 | 2568949 | A | C | 78 | 0 | 2 | 0 | | | | | | | | | | | C | C |
| NC_011770 | 2568991 | T | C | 78 | 0 | 12 | 0 | C | C | C | C | C | C | C | C | C | C | C | C |
| NC_011770 | 2569019 | T | C | 80 | 0 | 4 | 0 | | | | | | | | | C | C | C | C |
| NC_011770 | 2569022 | C | T | 79 | 0 | 2 | 0 | | | | | | | | | T | T | | |
| NC_011770 | 2569024 | T | C | 78 | 0 | 12 | 0 | C | C | C | C | C | C | C | C | C | C | C | C |
| NC_011770 | 2569034 | C | T | 84 | 0 | 4 | 0 | | | | | | | | | T | T | T | T |
| NC_011770 | 2569054 | C | T | 79 | 0 | 2 | 0 | | | | | | | | | | | T | T |
| NC_011770 | 2569087 | G | A | 78 | 0 | 2 | 0 | | | | | | | | | A | A | | |
| NC_011770 | 2569152 | T | C | 75 | 0 | 12 | 0 | C | C | C | C | C | C | C | C | C | C | C | C |
| NC_011770 | 2569669 | C | T | 64 | 0 | 2 | 0 | | | | | | | | | | | T | T |
| NC_011770 | 2569676 | C | T | 64 | 0 | 4 | 0 | | | T | T | T | T | | | | | | |
| NC_011770 | 2569686 | T | A | 67 | 0 | 8 | 0 | A | A | A | A | A | A | A | A | | | | |
| NC_011770 | 2570637 | C | T | 26 | 0 | 4 | 0 | T | T | | | | | | | | | T | T |
| NC_011770 | 2570677 | A | G | 25 | 0 | 4 | 0 | G | G | | | | | | | | | G | G |
| NC_011770 | 2570679 | A | C | 24 | 0 | 4 | 0 | C | C | | | | | | | | | C | C |
| NC_011770 | 2570707 | A | G | 21 | 0 | 6 | 0 | | | G | G | G | G | | | G | G | | |
| NC_011770 | 2570755 | A | C | 18 | 0 | 4 | 0 | C | C | | | | | | | | | C | C |
| NC_011770 | 2570787 | T | C | 23 | 0 | 10 | 0 | C | C | C | C | C | C | | | C | C | C | C |
| NC_011770 | 2572146 | G | A | 64 | 0 | 4 | 0 | | | A | A | A | A | | | | | | |
| NC_011770 | 2572247 | T | G | 59 | 0 | 12 | 0 | G | G | G | G | G | G | G | G | G | G | G | G |

| | | | | | | | | | | | | | | | | | | | |
|---|---|---|---|---|---|---|---|---|---|---|---|---|---|---|---|---|---|---|---|
| NC_011770 | 2572248 | A | C | 59 | 0 | 12 | 0 | C | C | C | C | C | C | C | C | C | C | C | C |
| NC_011770 | 2572250 | G | C | 58 | 0 | 12 | 0 | C | C | C | C | C | C | C | C | C | C | C | C |
| NC_011770 | 2572261 | A | G | 62 | 0 | 8 | 0 | G | G | | | | | G | G | G | G | G | G |
| NC_011770 | 2572279 | T | C | 66 | 0 | 12 | 0 | C | C | C | C | C | C | C | C | C | C | C | C |
| NC_011770 | 2576167 | G | A | 74 | 0 | 6 | 0 | | | A | A | A | A | | | | | A | A |
| NC_011770 | 2576187 | C | T | 72 | 0 | 4 | 0 | | | T | T | T | T | | | | | | |
| NC_011770 | 2579036 | T | C | 66 | 0 | 4 | 0 | | | C | C | C | C | | | | | | |
| NC_011770 | 2579091 | G | T | 72 | 0 | 2 | 0 | T | T | | | | | | | | | | |
| NC_011770 | 2580042 | A | G | 54 | 0 | 12 | 0 | G | G | G | G | G | G | G | G | G | G | G | G |
| NC_011770 | 2580066 | C | G | 56 | 0 | 12 | 0 | G | G | G | G | G | G | G | G | G | G | G | G |
| NC_011770 | 2580086 | A | G | 57 | 0 | 12 | 0 | G | G | G | G | G | G | G | G | G | G | G | G |
| NC_011770 | 2580087 | T | G | 57 | 0 | 12 | 0 | G | G | G | G | G | G | G | G | G | G | G | G |
| NC_011770 | 2583032 | G | A | 40 | 0 | 2 | 0 | | | | | | | A | A | | | | |
| NC_011770 | 2585080 | T | G | 63 | 0 | 12 | 0 | G | G | G | G | G | G | G | G | G | G | G | G |
| NC_011770 | 2585098 | C | T | 63 | 0 | 2 | 0 | T | T | | | | | | | | | | |
| NC_011770 | 2585138 | A | G | 61 | 0 | 8 | 0 | G | G | G | G | G | G | G | G | | | | |
| NC_011770 | 2585187 | C | A | 62 | 0 | 4 | 0 | | | A | A | A | A | | | | | | |
| NC_011770 | 2585190 | A | G | 60 | 0 | 12 | 0 | G | G | G | G | G | G | G | G | G | G | G | G |
| NC_011770 | 2585364 | T | C | 60 | 0 | 12 | 0 | C | C | C | C | C | C | C | C | C | C | C | C |
| NC_011770 | 2585372 | A | G | 61 | 0 | 2 | 0 | | | | | | | G | G | | | | |
| NC_011770 | 2585379 | C | A | 57 | 0 | 4 | 0 | | | | | | | | | A | A | A | A |
| NC_011770 | 2585455 | C | A | 70 | 0 | 12 | 0 | A | A | A | A | A | A | A | A | A | A | A | A |
| NC_011770 | 2585469 | C | T | 70 | 0 | 2 | 0 | T | T | | | | | | | | | | |
| NC_011770 | 2586226 | T | C | 64 | 0 | 4 | 0 | | | | | | | | | C | C | C | C |
| NC_011770 | 2586235 | C | T | 66 | 0 | 2 | 0 | T | T | | | | | | | | | | |
| NC_011770 | 2586567 | C | G | 77 | 0 | 2 | 0 | | | | | | | | | G | G | | |
| NC_011770 | 2587683 | C | T | 83 | 0 | 2 | 0 | | | | | | | | | T | T | | |
| NC_011770 | 2590465 | T | C | 77 | 0 | 4 | 0 | C | C | | | | | | | | | C | C |
| NC_011770 | 2590500 | T | C | 77 | 0 | 4 | 0 | C | C | | | | | | | | | C | C |
| NC_011770 | 2590534 | A | C | 75 | 0 | 12 | 0 | C | C | C | C | C | C | C | C | C | C | C | C |
| NC_011770 | 2590641 | T | C | 70 | 0 | 2 | 0 | | | | | | | C | C | | | | |
| NC_011770 | 2590643 | T | G | 69 | 0 | 2 | 0 | | | | | | | G | G | | | | |
| NC_011770 | 2590649 | A | G | 68 | 0 | 2 | 0 | G | G | | | | | | | | | | |
| NC_011770 | 2590670 | A | G | 70 | 0 | 2 | 0 | | | | | | | | | G | G | | |
| NC_011770 | 2590695 | A | C | 64 | 0 | 12 | 0 | C | C | C | C | C | C | C | C | C | C | C | C |
| NC_011770 | 2590705 | C | T | 65 | 0 | 4 | 0 | | | T | T | T | T | | | | | | |
| NC_011770 | 2590715 | C | T | 64 | 0 | 1 | 0 | | | | | | | T | | | | | |
| NC_011770 | 2594346 | G | T | 76 | 0 | 2 | 0 | T | T | | | | | | | | | | |
| NC_011770 | 2596635 | A | G | 69 | 0 | 4 | 0 | | | | | | | G | G | G | G | | |
| NC_011770 | 2596645 | G | A | 71 | 0 | 2 | 0 | | | | | | | | | | | A | A |
| NC_011770 | 2601094 | G | A | 63 | 0 | 2 | 0 | | | | | | | | | A | A | | |
| NC_011770 | 2605140 | C | T | 78 | 0 | 8 | 0 | T | T | | | | | T | T | T | T | T | T |
| NC_011770 | 2605280 | G | A | 76 | 0 | 2 | 0 | A | A | | | | | | | | | | |
| NC_011770 | 2605291 | G | C | 76 | 0 | 4 | 0 | | | | | | | C | C | | | C | C |
| NC_011770 | 2605342 | C | A | 71 | 0 | 2 | 0 | A | A | | | | | | | | | | |
| NC_011770 | 2605400 | A | C | 66 | 0 | 8 | 0 | C | C | C | C | C | C | | | C | C | | |
| NC_011770 | 2605431 | G | A | 62 | 0 | 2 | 0 | | | | | | | A | A | | | | |
| NC_011770 | 2607725 | T | C | 27 | 0 | 2 | 0 | C | C | | | | | | | | | | |
| NC_011770 | 2607750 | T | G | 25 | 0 | 12 | 0 | G | G | G | G | G | G | G | G | G | G | G | G |
| NC_011770 | 2607797 | A | G | 20 | 0 | 12 | 0 | G | G | G | G | G | G | G | G | G | G | G | G |
| NC_011770 | 2607803 | C | A | 22 | 0 | 2 | 0 | A | A | | | | | | | | | | |
| NC_011770 | 2607813 | A | C | 22 | 0 | 4 | 0 | | | C | C | C | C | | | | | | |
| NC_011770 | 2609808 | C | T | 92 | 0 | 2 | 0 | T | T | | | | | | | | | | |
| NC_011770 | 2609929 | G | A | 90 | 0 | 4 | 0 | | | A | A | A | A | | | | | | |
| NC_011770 | 2610781 | C | T | 74 | 0 | 2 | 0 | | | | | | | | | | | T | T |
| NC_011770 | 2610855 | G | GCA | 72 | 0 | 4 | 0 | | | GCA | GCA | GCA | GCA | | | | | | |
| NC_011770 | 2610912 | C | G | 72 | 0 | 4 | 0 | | | G | G | G | G | | | | | | |
| NC_011770 | 2610917 | G | T | 67 | 0 | 8 | 0 | | | T | T | T | T | | | T | T | T | T |
| NC_011770 | 2611016 | T | C | 75 | 0 | 2 | 0 | | | | | | | C | C | | | | |
| NC_011770 | 2611049 | C | A | 74 | 0 | 6 | 0 | A | A | A | A | A | A | | | | | | |
| NC_011770 | 2611098 | T | A | 74 | 0 | 4 | 0 | | | A | A | A | A | | | | | | |
| NC_011770 | 2611110 | C | T | 74 | 0 | 4 | 0 | | | T | T | T | T | | | | | | |

| | | | | | | | | | | | | | | | | | | | |
|---|---|---|---|---|---|---|---|---|---|---|---|---|---|---|---|---|---|---|---|
| NC_011770 | 2611122 | T | C | 74 | 0 | 4 | 0 | | | C | C | C | C | | | | | | |
| NC_011770 | 2611210 | G | A | 72 | 0 | 6 | 0 | A | A | A | A | A | A | | | | | | |
| NC_011770 | 2611264 | C | A | 69 | 0 | 6 | 0 | | | A | A | A | A | | | | | A | A |
| NC_011770 | 2611270 | C | T | 70 | 0 | 2 | 0 | T | T | | | | | | | | | | |
| NC_011770 | 2618003 | T | C | 80 | 0 | 12 | 0 | C | C | C | C | C | C | C | C | C | C | C | C |
| NC_011770 | 2619632 | C | T | 69 | 0 | 2 | 0 | | | | | | | | | | | T | T |
| NC_011770 | 2619729 | C | T | 65 | 0 | 6 | 0 | T | T | | | | | T | T | | | T | T |
| NC_011770 | 2619739 | C | G | 69 | 0 | 2 | 0 | | | | | | | G | G | | | | |
| NC_011770 | 2619741 | T | C | 68 | 0 | 2 | 0 | | | | | | | | | | | C | C |
| NC_011770 | 2619774 | T | TG | 68 | 0 | 10 | 0 | | | TG | TG | TG | TG | TG | TG | TG | TG | TG | TG |
| NC_011770 | 2619780 | G | C | 67 | 0 | 2 | 0 | C | C | | | | | | | | | | |
| NC_011770 | 2619910 | C | T | 67 | 0 | 4 | 0 | | | | | | | T | T | | | T | T |
| NC_011770 | 2619920 | G | A | 68 | 0 | 4 | 0 | | | | | | | A | A | | | A | A |
| NC_011770 | 2620036 | C | T | 71 | 0 | 2 | 0 | | | | | | | T | T | | | | |
| NC_011770 | 2620170 | A | AG | 67 | 0 | 12 | 0 | AG | AG | AG | AG | AG | AG | AG | AG | AG | AG | AG | AG |
| NC_011770 | 2620187 | T | A | 68 | 0 | 12 | 0 | A | A | A | A | A | A | A | A | A | A | A | A |
| NC_011770 | 2620191 | G | A | 68 | 0 | 2 | 0 | | | | | | | A | A | | | | |
| NC_011770 | 2620643 | A | C | 67 | 0 | 10 | 0 | C | C | C | C | C | C | C | C | C | C | | |
| NC_011770 | 2620648 | C | T | 70 | 0 | 2 | 0 | | | | | | | | | | | T | T |
| NC_011770 | 2620708 | G | A | 64 | 0 | 2 | 0 | A | A | | | | | | | | | | |
| NC_011770 | 2620735 | A | G | 67 | 0 | 2 | 0 | | | | | | | G | G | | | | |
| NC_011770 | 2620845 | G | A | 72 | 0 | 2 | 0 | A | A | | | | | | | | | | |
| NC_011770 | 2621273 | G | A | 71 | 0 | 2 | 0 | | | | | | | A | A | | | | |
| NC_011770 | 2621331 | C | G | 71 | 0 | 4 | 0 | G | G | | | | | | | | | G | G |
| NC_011770 | 2621332 | T | C | 71 | 0 | 4 | 0 | C | C | | | | | | | | | C | C |
| NC_011770 | 2622966 | C | T | 77 | 0 | 4 | 0 | | | T | T | T | T | | | | | | |
| NC_011770 | 2622970 | C | T | 79 | 0 | 4 | 0 | | | T | T | T | T | | | | | | |
| NC_011770 | 2625780 | A | G | 66 | 0 | 12 | 0 | G | G | G | G | G | G | G | G | G | G | G | G |
| NC_011770 | 2625884 | G | A | 65 | 0 | 10 | 0 | A | A | A | A | A | A | | | A | A | A | A |
| NC_011770 | 2625964 | T | G | 65 | 0 | 4 | 0 | | | G | G | G | G | | | | | | |
| NC_011770 | 2626896 | G | T | 65 | 0 | 4 | 0 | | | T | T | T | T | | | | | | |
| NC_011770 | 2628025 | A | C | 59 | 0 | 2 | 0 | | | | | | | | | | | C | C |
| NC_011770 | 2633435 | C | A | 86 | 0 | 2 | 0 | | | | | | | | | | | A | A |
| NC_011770 | 2633455 | T | G | 89 | 0 | 2 | 0 | | | | | | | G | G | | | | |
| NC_011770 | 2637996 | A | G | 71 | 0 | 2 | 0 | | | | | | | | | | | G | G |
| NC_011770 | 2637997 | A | C | 71 | 0 | 2 | 0 | | | | | | | | | | | C | C |
| NC_011770 | 2639457 | G | A | 55 | 2 | 2 | 0 | . | . | | | | | A | A | | | | |
| NC_011770 | 2640400 | C | T | 13 | 0 | 4 | 0 | | | T | T | T | T | | | | | | |
| NC_011770 | 2640426 | C | G | 10 | 0 | 2 | 0 | | | | | | | G | G | | | | |
| NC_011770 | 2654296 | G | A | 60 | 0 | 4 | 0 | | | A | A | A | A | | | | | | |
| NC_011770 | 2655526 | G | A | 66 | 0 | 2 | 0 | | | | | A | A | | | | | | |
| NC_011770 | 2655617 | G | C | 65 | 0 | 10 | 0 | C | C | C | C | C | C | C | C | C | C | | |
| NC_011770 | 2655633 | C | A | 69 | 0 | 2 | 0 | | | | | | | | | A | A | | |
| NC_011770 | 2655635 | C | T | 68 | 0 | 10 | 0 | T | T | T | T | T | T | T | T | T | T | | |
| NC_011770 | 2655690 | G | T | 63 | 0 | 10 | 0 | T | T | T | T | T | T | T | T | T | T | | |
| NC_011770 | 2655708 | A | G | 61 | 0 | 10 | 0 | G | G | G | G | G | G | G | G | G | G | | |
| NC_011770 | 2655710 | C | T | 62 | 0 | 8 | 0 | T | T | T | T | T | T | T | T | | | | |
| NC_011770 | 2655814 | T | G | 64 | 0 | 2 | 0 | | | | | | | | | | | G | G |
| NC_011770 | 2655817 | C | A | 66 | 0 | 2 | 0 | | | | | | | | | | | A | A |
| NC_011770 | 2655818 | C | T | 64 | 0 | 2 | 0 | | | | | | | | | | | T | T |
| NC_011770 | 2658798 | G | A | 57 | 0 | 2 | 0 | | | | | | | | | A | A | | |
| NC_011770 | 2664119 | T | G | 52 | 0 | 2 | 0 | | | | | | | | | | | G | G |
| NC_011770 | 2664122 | C | G | 52 | 0 | 2 | 0 | G | G | | | | | | | | | | |
| NC_011770 | 2668782 | A | G | 52 | 0 | 12 | 0 | G | G | G | G | G | G | G | G | G | G | G | G |
| NC_011770 | 2668794 | TCG | T | 55 | 0 | 2 | 0 | | | | | | | | | | | T | T |
| NC_011770 | 2668809 | C | T | 56 | 0 | 4 | 0 | | | T | T | T | T | | | | | | |
| NC_011770 | 2673267 | C | T | 80 | 0 | 2 | 0 | T | T | | | | | | | | | | |
| NC_011770 | 2683464 | C | A | 85 | 0 | 2 | 0 | A | A | | | | | | | | | | |
| NC_011770 | 2687206 | T | C | 74 | 0 | 4 | 0 | | | | | | | C | C | C | C | | |
| NC_011770 | 2694222 | G | A | 6 | 10 | 2 | 0 | . | . | . | . | . | . | . | . | A | A | . | . |
| NC_011770 | 2694234 | C | T | 6 | 10 | 2 | 0 | . | . | . | . | . | . | . | . | T | T | . | . |
| NC_011770 | 2694267 | A | G | 5 | 10 | 2 | 0 | . | . | . | . | . | . | . | . | G | G | . | . |

| | | | | | | | | | | | | | | | | | | | |
|---|---|---|---|---|---|---|---|---|---|---|---|---|---|---|---|---|---|---|---|
| NC_011770 | 2696819 | T | A | 6 | 10 | 2 | 0 | . | . | . | . | . | . | . | . | . | . | A | A |
| NC_011770 | 2696827 | T | G | 8 | 8 | 4 | 0 | . | . | . | . | . | . | . | . | G | G | G | G |
| NC_011770 | 2697151 | C | A | 8 | 10 | 2 | 0 | . | . | . | . | . | . | . | . | . | . | A | A |
| NC_011770 | 2697153 | G | A | 8 | 10 | 2 | 0 | . | . | . | . | . | . | . | . | . | . | A | A |
| NC_011770 | 2697169 | A | G | 8 | 10 | 2 | 0 | . | . | . | . | . | . | . | . | . | . | G | G |
| NC_011770 | 2697173 | T | G | 10 | 8 | 2 | 0 | . | . | . | . | . | . | . | . | | | G | G |
| NC_011770 | 2697184 | G | A | 10 | 8 | 2 | 0 | . | . | . | . | . | . | . | . | | | A | A |
| NC_011770 | 2697203 | A | G | 10 | 8 | 4 | 0 | . | . | . | . | . | . | . | . | G | G | G | G |
| NC_011770 | 2697227 | A | G | 10 | 8 | 2 | 0 | . | . | . | . | . | . | . | . | | | G | G |
| NC_011770 | 2697231 | C | G | 9 | 8 | 4 | 0 | . | . | . | . | . | . | . | . | G | G | A | A |
| NC_011770 | 2697238 | G | C | 9 | 8 | 4 | 0 | . | . | . | . | . | . | . | . | C | C | A | A |
| NC_011770 | 2697247 | C | T | 8 | 8 | 2 | 0 | . | . | . | . | . | . | . | . | | | T | T |
| NC_011770 | 2697250 | C | T | 9 | 8 | 2 | 0 | . | . | . | . | . | . | . | . | | | T | T |
| NC_011770 | 2697259 | T | A | 8 | 8 | 4 | 0 | . | . | . | . | . | . | . | . | A | A | G | G |
| NC_011770 | 2697262 | T | C | 8 | 8 | 4 | 0 | . | . | . | . | . | . | . | . | C | C | A | A |
| NC_011770 | 2697274 | A | G | 8 | 8 | 4 | 0 | . | . | . | . | . | . | . | . | G | G | G | G |
| NC_011770 | 2697277 | A | T | 8 | 8 | 4 | 0 | . | . | . | . | . | . | . | . | T | T | T | T |
| NC_011770 | 2697280 | C | A | 8 | 8 | 4 | 0 | . | . | . | . | . | . | . | . | A | A | T | T |
| NC_011770 | 2697298 | A | C | 9 | 8 | 4 | 0 | . | . | . | . | . | . | . | . | C | C | C | C |
| NC_011770 | 2697304 | G | A | 9 | 8 | 2 | 0 | . | . | . | . | . | . | . | . | | | A | A |
| NC_011770 | 2697330 | T | C | 10 | 8 | 2 | 0 | . | . | . | . | . | . | . | . | . | . | C | C |
| NC_011770 | 2697349 | A | G | 10 | 8 | 4 | 0 | . | . | . | . | . | . | . | . | G | G | G | G |
| NC_011770 | 2697607 | A | G | 11 | 8 | 2 | 0 | . | . | . | . | . | . | . | . | G | G | | |
| NC_011770 | 2698004 | A | G | 7 | 10 | 2 | 0 | . | . | . | . | . | . | . | . | . | . | G | G |
| NC_011770 | 2698008 | A | G | 8 | 9 | 1 | 0 | . | . | . | . | . | . | . | . | G | . | | |
| NC_011770 | 2698019 | A | G | 8 | 9 | 3 | 0 | . | . | . | . | . | . | . | . | G | . | G | G |
| NC_011770 | 2698034 | C | CGGCGC | 8 | 9 | 1 | 0 | . | . | . | . | . | . | . | . | CGGCGGC | . | | |
| NC_011770 | 2698038 | A | G | 9 | 8 | 2 | 0 | . | . | . | . | . | . | . | . | | | G | G |
| NC_011770 | 2698039 | T | G | 9 | 8 | 2 | 0 | . | . | . | . | . | . | . | . | | | G | G |
| NC_011770 | 2698040 | T | C | 9 | 8 | 2 | 0 | . | . | . | . | . | . | . | . | | | C | C |
| NC_011770 | 2698053 | C | G | 10 | 8 | 4 | 0 | . | . | . | . | . | . | . | . | G | G | G | G |
| NC_011770 | 2698061 | A | G | 10 | 8 | 4 | 0 | . | . | . | . | . | . | . | . | G | G | G | G |
| NC_011770 | 2698078 | A | G | 10 | 8 | 4 | 0 | . | . | . | . | . | . | . | . | G | G | G | G |
| NC_011770 | 2698079 | A | T | 10 | 8 | 4 | 0 | . | . | . | . | . | . | . | . | T | T | T | T |
| NC_011770 | 2698082 | G | A | 10 | 8 | 4 | 0 | . | . | . | . | . | . | . | . | A | A | A | A |
| NC_011770 | 2698091 | A | G | 11 | 8 | 4 | 0 | . | . | . | . | . | . | . | . | G | G | G | G |
| NC_011770 | 2698101 | C | A | 10 | 8 | 4 | 0 | . | . | . | . | . | . | . | . | A | A | A | A |
| NC_011770 | 2698102 | T | C | 11 | 8 | 4 | 0 | . | . | . | . | . | . | . | . | C | C | C | C |
| NC_011770 | 2699180 | G | A | 10 | 8 | 4 | 0 | . | . | . | . | . | . | . | . | A | A | A | A |
| NC_011770 | 2699190 | C | G | 11 | 8 | 2 | 0 | . | . | . | . | . | . | . | . | G | G | | |
| NC_011770 | 2699206 | G | A | 12 | 6 | 4 | 0 | . | . | . | . | | | . | . | A | A | A | A |
| NC_011770 | 2699207 | TG | T | 12 | 6 | 4 | 0 | . | . | . | . | | | . | . | T | T | T | T |
| NC_011770 | 2699223 | T | C | 11 | 5 | 2 | 0 | | . | . | . | | | . | . | | | C | C |
| NC_011770 | 2699335 | G | A | 3 | 10 | 2 | 0 | . | . | . | . | . | . | . | . | . | . | A | A |
| NC_011770 | 2699613 | C | G | 4 | 10 | 2 | 0 | . | . | . | . | . | . | . | . | . | . | G | G |
| NC_011770 | 2699632 | G | A | 13 | 6 | 2 | 0 | A | A | . | . | | | . | . | . | . | | |
| NC_011770 | 2699801 | A | G | 18 | 6 | 2 | 0 | G | G | . | . | | | . | . | . | . | | |
| NC_011770 | 2699823 | G | T | 13 | 6 | 2 | 0 | | | . | . | | | . | . | . | . | T | T |
| NC_011770 | 2699830 | T | C | 4 | 10 | 2 | 0 | C | C | . | . | . | . | . | . | . | . | . | . |
| NC_011770 | 2699836 | A | G | 4 | 10 | 2 | 0 | G | G | . | . | . | . | . | . | . | . | . | . |
| NC_011770 | 2699842 | A | G | 4 | 10 | 2 | 0 | G | G | . | . | . | . | . | . | . | . | . | . |
| NC_011770 | 2699878 | A | G | 4 | 10 | 2 | 0 | G | G | . | . | . | . | . | . | . | . | . | . |
| NC_011770 | 2699908 | T | G | 3 | 10 | 2 | 0 | G | G | . | . | . | . | . | . | . | . | . | . |
| NC_011770 | 2700655 | T | A | 0 | 11 | 1 | 0 | A | . | . | . | . | . | . | . | . | . | . | . |
| NC_011770 | 2700663 | G | A | 0 | 11 | 1 | 0 | A | . | . | . | . | . | . | . | . | . | . | . |
| NC_011770 | 2700677 | C | T | 0 | 11 | 1 | 0 | T | . | . | . | . | . | . | . | . | . | . | . |
| NC_011770 | 2700678 | A | G | 0 | 11 | 1 | 0 | G | . | . | . | . | . | . | . | . | . | . | . |
| NC_011770 | 2700692 | A | C | 0 | 11 | 1 | 0 | C | . | . | . | . | . | . | . | . | . | . | . |
| NC_011770 | 2700700 | G | A | 0 | 11 | 1 | 0 | A | . | . | . | . | . | . | . | . | . | . | . |
| NC_011770 | 2700711 | T | G | 0 | 11 | 1 | 0 | G | . | . | . | . | . | . | . | . | . | . | . |
| NC_011770 | 2700713 | G | A | 0 | 11 | 1 | 0 | A | . | . | . | . | . | . | . | . | . | . | . |
| NC_011770 | 2700719 | ATCCT | A | 0 | 11 | 1 | 0 | A | . | . | . | . | . | . | . | . | . | . | . |

| | | | | | | | | | | | | | | | | | | | |
|---|---|---|---|---|---|---|---|---|---|---|---|---|---|---|---|---|---|---|---|
| NC_011770 | 2700752 | C | G | 1 | 9 | 2 | 0 | | . | . | . | G | G | . | . | . | . | . | . |
| NC_011770 | 2700757 | A | G | 1 | 9 | 2 | 0 | | . | . | . | G | G | . | . | . | . | . | . |
| NC_011770 | 2700760 | T | C | 1 | 9 | 2 | 0 | | . | . | . | C | C | . | . | . | . | . | . |
| NC_011770 | 2700765 | C | T | 1 | 9 | 2 | 0 | | . | . | . | T | T | . | . | . | . | . | . |
| NC_011770 | 2700775 | C | T | 1 | 9 | 2 | 0 | | . | . | . | T | T | . | . | . | . | . | . |
| NC_011770 | 2700785 | C | A | 1 | 9 | 1 | 0 | A | . | . | . | | | . | . | . | . | . | . |
| NC_011770 | 2700786 | C | A | 1 | 9 | 2 | 0 | | . | . | . | A | A | . | . | . | . | . | . |
| NC_011770 | 2700790 | A | G | 0 | 9 | 3 | 0 | G | . | . | . | G | G | . | . | . | . | . | . |
| NC_011770 | 2703669 | C | A | 9 | 10 | 2 | 0 | A | A | . | . | . | . | . | . | . | . | . | . |
| NC_011770 | 2703731 | C | G | 9 | 10 | 2 | 0 | G | G | . | . | . | . | . | . | . | . | . | . |
| NC_011770 | 2703738 | G | A | 10 | 10 | 2 | 0 | A | A | . | . | . | . | . | . | . | . | . | . |
| NC_011770 | 2703767 | C | T | 9 | 10 | 2 | 0 | T | T | . | . | . | . | . | . | . | . | . | . |
| NC_011770 | 2703843 | G | A | 9 | 10 | 2 | 0 | A | A | . | . | . | . | . | . | . | . | . | . |
| NC_011770 | 2703864 | C | T | 9 | 10 | 2 | 0 | T | T | . | . | . | . | . | . | . | . | . | . |
| NC_011770 | 2703979 | A | T | 7 | 10 | 2 | 0 | T | T | . | . | . | . | . | . | . | . | . | . |
| NC_011770 | 2704472 | G | A | 17 | 8 | 2 | 0 | A | A | . | . | | | . | . | . | . | . | . |
| NC_011770 | 2704527 | T | A | 17 | 8 | 2 | 0 | A | A | . | . | | | . | . | . | . | . | . |
| NC_011770 | 2704533 | G | T | 18 | 8 | 4 | 0 | T | T | . | . | T | T | . | . | . | . | . | . |
| NC_011770 | 2704575 | T | C | 20 | 8 | 4 | 0 | C | C | . | . | C | C | . | . | . | . | . | . |
| NC_011770 | 2704628 | G | A | 17 | 10 | 2 | 0 | . | . | . | . | A | A | . | . | . | . | . | . |
| NC_011770 | 2704702 | T | C | 18 | 10 | 2 | 0 | . | . | . | . | C | C | . | . | . | . | . | . |
| NC_011770 | 2704707 | G | A | 18 | 10 | 2 | 0 | . | . | . | . | A | A | . | . | . | . | . | . |
| NC_011770 | 2704717 | T | C | 20 | 10 | 2 | 0 | . | . | . | . | C | C | . | . | . | . | . | . |
| NC_011770 | 2704727 | G | A | 18 | 10 | 2 | 0 | . | . | . | . | A | A | . | . | . | . | . | . |
| NC_011770 | 2704732 | G | A | 18 | 10 | 2 | 0 | . | . | . | . | A | A | . | . | . | . | . | . |
| NC_011770 | 2704733 | A | T | 18 | 10 | 2 | 0 | . | . | . | . | T | T | . | . | . | . | . | . |
| NC_011770 | 2704737 | A | G | 18 | 10 | 2 | 0 | . | . | . | . | G | G | . | . | . | . | . | . |
| NC_011770 | 2704744 | T | C | 18 | 10 | 2 | 0 | . | . | . | . | C | C | . | . | . | . | . | . |
| NC_011770 | 2704746 | G | A | 18 | 10 | 2 | 0 | . | . | . | . | A | A | . | . | . | . | . | . |
| NC_011770 | 2704751 | T | G | 19 | 10 | 2 | 0 | . | . | . | . | G | G | . | . | . | . | . | . |
| NC_011770 | 2704790 | T | C | 23 | 10 | 2 | 0 | . | . | . | . | C | C | . | . | . | . | . | . |
| NC_011770 | 2708048 | T | G | 25 | 8 | 4 | 0 | G | G | . | . | C | C | . | . | . | . | . | . |
| NC_011770 | 2708050 | A | C | 25 | 8 | 2 | 0 | | | . | . | C | C | . | . | . | . | . | . |
| NC_011770 | 2708052 | A | C | 25 | 8 | 2 | 0 | | | . | . | C | C | . | . | . | . | . | . |
| NC_011770 | 2708084 | A | G | 26 | 8 | 2 | 0 | | | . | . | G | G | . | . | . | . | . | . |
| NC_011770 | 2708617 | C | T | 17 | 9 | 3 | 0 | . | T | . | . | T | T | . | . | . | . | . | . |
| NC_011770 | 2708626 | T | C | 21 | 8 | 2 | 0 | C | C | . | . | | | . | . | . | . | . | . |
| NC_011770 | 2708641 | T | A | 20 | 8 | 2 | 0 | | | . | . | A | A | . | . | . | . | . | . |
| NC_011770 | 2708653 | A | G | 21 | 8 | 2 | 0 | G | G | . | . | | | . | . | . | . | . | . |
| NC_011770 | 2708656 | C | A | 20 | 8 | 2 | 0 | | | . | . | A | A | . | . | . | . | . | . |
| NC_011770 | 2708694 | C | A | 22 | 8 | 2 | 0 | | | . | . | A | A | . | . | . | . | . | . |
| NC_011770 | 2708728 | G | A | 15 | 10 | 2 | 0 | . | . | . | . | A | A | . | . | . | . | . | . |
| NC_011770 | 2708758 | G | T | 14 | 10 | 2 | 0 | . | . | . | . | T | T | . | . | . | . | . | . |
| NC_011770 | 2711968 | G | C | 29 | 8 | 4 | 0 | C | C | . | . | C | C | . | . | . | . | . | . |
| NC_011770 | 2711980 | C | G | 29 | 8 | 2 | 0 | G | G | . | . | | | . | . | . | . | . | . |
| NC_011770 | 2711988 | T | C | 30 | 8 | 2 | 0 | C | C | . | . | | | . | . | . | . | . | . |
| NC_011770 | 2712009 | C | A | 30 | 8 | 2 | 0 | A | A | . | . | | | . | . | . | . | . | . |
| NC_011770 | 2712026 | C | T | 29 | 8 | 2 | 0 | T | T | . | . | | | . | . | . | . | . | . |
| NC_011770 | 2712039 | T | A | 28 | 8 | 2 | 0 | | | . | . | A | A | . | . | . | . | . | . |
| NC_011770 | 2712040 | C | A | 27 | 8 | 2 | 0 | | | . | . | A | A | . | . | . | . | . | . |
| NC_011770 | 2712133 | C | A | 17 | 10 | 2 | 0 | . | . | . | . | A | A | . | . | . | . | . | . |
| NC_011770 | 2712172 | A | T | 18 | 10 | 2 | 0 | . | . | . | . | T | T | . | . | . | . | . | . |
| NC_011770 | 2712188 | CT | C | 20 | 8 | 2 | 0 | C | C | . | . | | | . | . | . | . | . | . |
| NC_011770 | 2712242 | T | C | 17 | 10 | 2 | 0 | . | . | . | . | C | C | . | . | . | . | . | . |
| NC_011770 | 2712255 | T | C | 16 | 10 | 2 | 0 | . | . | . | . | C | C | . | . | . | . | . | . |
| NC_011770 | 2712341 | GCA | G | 1 | 10 | 2 | 0 | . | . | . | . | G | G | . | . | . | . | . | . |
| NC_011770 | 2712346 | GA | G | 1 | 10 | 2 | 0 | . | . | . | . | G | G | . | . | . | . | . | . |
| NC_011770 | 2712385 | A | C | 1 | 10 | 2 | 0 | . | . | . | . | C | C | . | . | . | . | . | . |
| NC_011770 | 2712392 | G | C | 1 | 10 | 2 | 0 | . | . | . | . | C | C | . | . | . | . | . | . |
| NC_011770 | 2712395 | G | T | 1 | 10 | 2 | 0 | . | . | . | . | T | T | . | . | . | . | . | . |
| NC_011770 | 2712397 | C | T | 0 | 10 | 2 | 0 | . | . | . | . | T | T | . | . | . | . | . | . |
| NC_011770 | 2712398 | AGCAGG | A | 0 | 10 | 2 | 0 | . | . | . | . | A | A | . | . | . | . | . | . |

| | | | | | | | | | | | | | | | | | | | |
|---|---|---|---|---|---|---|---|---|---|---|---|---|---|---|---|---|---|---|---|
| NC_011770 | 2712426 | A | C | 13 | 10 | 2 | 0 | . | . | . | . | C | C | . | . | . | . | . | . |
| NC_011770 | 2712482 | T | A | 18 | 8 | 2 | 0 | | | . | . | A | A | . | . | . | . | . | . |
| NC_011770 | 2712502 | C | A | 18 | 8 | 2 | 0 | | | . | . | A | A | . | . | . | . | . | . |
| NC_011770 | 2712807 | A | T | 18 | 8 | 2 | 0 | T | T | . | . | | | . | . | . | . | . | . |
| NC_011770 | 2712809 | A | T | 18 | 8 | 4 | 0 | T | T | . | . | T | T | . | . | . | . | . | . |
| NC_011770 | 2713748 | A | G | 21 | 8 | 2 | 0 | | | . | . | G | G | . | . | . | . | . | . |
| NC_011770 | 2713769 | G | A | 21 | 8 | 2 | 0 | | | . | . | A | A | . | . | . | . | . | . |
| NC_011770 | 2713771 | T | G | 21 | 8 | 4 | 0 | G | G | . | . | G | G | . | . | . | . | . | . |
| NC_011770 | 2713772 | T | C | 21 | 8 | 4 | 0 | C | C | . | . | A | A | . | . | . | . | . | . |
| NC_011770 | 2713781 | A | G | 22 | 8 | 4 | 0 | G | G | . | . | G | G | . | . | . | . | . | . |
| NC_011770 | 2713793 | T | G | 20 | 8 | 4 | 0 | G | G | . | . | C | C | . | . | . | . | . | . |
| NC_011770 | 2713796 | G | A | 20 | 8 | 2 | 0 | | | . | . | A | A | . | . | . | . | . | . |
| NC_011770 | 2713799 | C | T | 15 | 10 | 2 | 0 | . | . | . | . | T | T | . | . | . | . | . | . |
| NC_011770 | 2713802 | G | C | 15 | 10 | 2 | 0 | . | . | . | . | C | C | . | . | . | . | . | . |
| NC_011770 | 2713808 | A | G | 16 | 10 | 2 | 0 | . | . | . | . | G | G | . | . | . | . | . | . |
| NC_011770 | 2713815 | T | G | 16 | 10 | 2 | 0 | . | . | . | . | G | G | . | . | . | . | . | . |
| NC_011770 | 2713823 | T | C | 18 | 8 | 2 | 0 | | | . | . | C | C | . | . | . | . | . | . |
| NC_011770 | 2713824 | A | G | 17 | 8 | 2 | 0 | | | . | . | G | G | . | . | . | . | . | . |
| NC_011770 | 2713826 | G | T | 17 | 8 | 2 | 0 | T | T | . | . | | | . | . | . | . | . | . |
| NC_011770 | 2713847 | C | T | 0 | 11 | 1 | 0 | . | . | . | . | . | T | . | . | . | . | . | . |
| NC_011770 | 2714355 | G | A | 17 | 10 | 2 | 0 | . | . | . | . | A | A | . | . | . | . | . | . |
| NC_011770 | 2714361 | T | C | 18 | 10 | 2 | 0 | . | . | . | . | C | C | . | . | . | . | . | . |
| NC_011770 | 2714371 | G | A | 17 | 10 | 2 | 0 | . | . | . | . | A | A | . | . | . | . | . | . |
| NC_011770 | 2714385 | G | A | 19 | 10 | 2 | 0 | . | . | . | . | A | A | . | . | . | . | . | . |
| NC_011770 | 2714821 | T | A | 14 | 8 | 4 | 0 | A | A | . | . | G | G | . | . | . | . | . | . |
| NC_011770 | 2714824 | C | T | 14 | 8 | 4 | 0 | T | T | . | . | T | T | . | . | . | . | . | . |
| NC_011770 | 2714830 | C | T | 14 | 8 | 4 | 0 | T | T | . | . | T | T | . | . | . | . | . | . |
| NC_011770 | 2714831 | G | A | 14 | 8 | 4 | 0 | A | A | . | . | A | A | . | . | . | . | . | . |
| NC_011770 | 2714833 | C | T | 14 | 8 | 2 | 0 | T | T | . | . | | | . | . | . | . | . | . |
| NC_011770 | 2714834 | A | G | 14 | 8 | 2 | 0 | | | . | . | G | G | . | . | . | . | . | . |
| NC_011770 | 2714835 | C | T | 14 | 8 | 2 | 0 | | | . | . | T | T | . | . | . | . | . | . |
| NC_011770 | 2714836 | A | G | 14 | 8 | 2 | 0 | | | . | . | G | G | . | . | . | . | . | . |
| NC_011770 | 2714839 | G | A | 16 | 8 | 2 | 0 | A | A | . | . | | | . | . | . | . | . | . |
| NC_011770 | 2714849 | T | TA | 16 | 8 | 2 | 0 | | | . | . | TA | TA | . | . | . | . | . | . |
| NC_011770 | 2714852 | T | G | 16 | 8 | 2 | 0 | | | . | . | G | G | . | . | . | . | . | . |
| NC_011770 | 2714855 | A | C | 17 | 8 | 2 | 0 | C | C | . | . | | | . | . | . | . | . | . |
| NC_011770 | 2714863 | G | T | 17 | 8 | 2 | 0 | T | T | . | . | | | . | . | . | . | . | . |
| NC_011770 | 2714866 | T | C | 17 | 8 | 2 | 0 | | | . | . | C | C | . | . | . | . | . | . |
| NC_011770 | 2714867 | A | T | 17 | 8 | 2 | 0 | | | . | . | T | T | . | . | . | . | . | . |
| NC_011770 | 2714869 | T | C | 17 | 8 | 2 | 0 | | | . | . | C | C | . | . | . | . | . | . |
| NC_011770 | 2714870 | G | T | 17 | 8 | 2 | 0 | T | T | . | . | | | . | . | . | . | . | . |
| NC_011770 | 2714872 | T | C | 16 | 8 | 4 | 0 | C | C | . | . | C | C | . | . | . | . | . | . |
| NC_011770 | 2714876 | G | T | 18 | 8 | 2 | 0 | T | T | . | . | | | . | . | . | . | . | . |
| NC_011770 | 2714887 | T | G | 17 | 8 | 4 | 0 | G | G | . | . | G | G | . | . | . | . | . | . |
| NC_011770 | 2714888 | C | A | 18 | 8 | 4 | 0 | A | A | . | . | T | T | . | . | . | . | . | . |
| NC_011770 | 2714890 | G | T | 18 | 8 | 4 | 0 | T | T | . | . | T | T | . | . | . | . | . | . |
| NC_011770 | 2714897 | G | T | 18 | 8 | 2 | 0 | | | . | . | T | T | . | . | . | . | . | . |
| NC_011770 | 2714901 | G | T | 19 | 8 | 4 | 0 | T | T | . | . | T | T | . | . | . | . | . | . |
| NC_011770 | 2714916 | T | C | 18 | 8 | 2 | 0 | | | . | . | C | C | . | . | . | . | . | . |
| NC_011770 | 2714918 | A | T | 19 | 8 | 2 | 0 | | | . | . | T | T | . | . | . | . | . | . |
| NC_011770 | 2714930 | G | T | 18 | 8 | 2 | 0 | | | . | . | T | T | . | . | . | . | . | . |
| NC_011770 | 2714976 | C | T | 18 | 8 | 2 | 0 | | | . | . | T | T | . | . | . | . | . | . |
| NC_011770 | 2714991 | A | C | 17 | 8 | 4 | 0 | C | C | . | . | C | C | . | . | . | . | . | . |
| NC_011770 | 2714996 | G | A | 17 | 8 | 4 | 0 | A | A | . | . | A | A | . | . | . | . | . | . |
| NC_011770 | 2714997 | T | G | 17 | 8 | 4 | 0 | G | G | . | . | G | G | . | . | . | . | . | . |
| NC_011770 | 2718540 | C | T | 20 | 8 | 4 | 0 | T | T | . | . | T | T | . | . | . | . | . | . |
| NC_011770 | 2718552 | A | G | 20 | 8 | 4 | 0 | G | G | . | . | G | G | . | . | . | . | . | . |
| NC_011770 | 2718591 | T | C | 22 | 8 | 4 | 0 | C | C | . | . | C | C | . | . | . | . | . | . |
| NC_011770 | 2718600 | T | C | 22 | 8 | 4 | 0 | C | C | . | . | C | C | . | . | . | . | . | . |
| NC_011770 | 2718618 | T | C | 21 | 8 | 4 | 0 | C | C | . | . | C | C | . | . | . | . | . | . |
| NC_011770 | 2720727 | T | C | 22 | 8 | 2 | 0 | C | C | . | . | | | . | . | . | . | . | . |
| NC_011770 | 2720728 | T | C | 22 | 8 | 2 | 0 | | | . | . | C | C | . | . | . | . | . | . |

| | | | | | | | | | | | | | | | | | | | |
|---|---|---|---|---|---|---|---|---|---|---|---|---|---|---|---|---|---|---|---|
| NC_011770 | 2720741 | G | T | 19 | 8 | 4 | 0 | T | T | . | . | T | T | . | . | . | . | . | . |
| NC_011770 | 2720742 | A | C | 19 | 8 | 4 | 0 | C | C | . | . | C | C | . | . | . | . | . | . |
| NC_011770 | 2720743 | G | T | 20 | 8 | 4 | 0 | T | T | . | . | T | T | . | . | . | . | . | . |
| NC_011770 | 2720744 | A | C | 20 | 8 | 4 | 0 | C | C | . | . | C | C | . | . | . | . | . | . |
| NC_011770 | 2720769 | C | T | 23 | 8 | 2 | 0 | T | T | . | . | | | . | . | . | . | . | . |
| NC_011770 | 2720774 | C | T | 22 | 8 | 2 | 0 | T | T | . | . | | | . | . | . | . | . | . |
| NC_011770 | 2720794 | A | G | 22 | 8 | 2 | 0 | G | G | . | . | | | . | . | . | . | . | . |
| NC_011770 | 2720795 | A | G | 22 | 8 | 4 | 0 | G | G | . | . | G | G | . | . | . | . | . | . |
| NC_011770 | 2720800 | C | T | 22 | 8 | 4 | 0 | T | T | . | . | T | T | . | . | . | . | . | . |
| NC_011770 | 2720812 | C | T | 24 | 8 | 2 | 0 | T | T | . | . | | | . | . | . | . | . | . |
| NC_011770 | 2720821 | C | T | 24 | 8 | 2 | 0 | T | T | . | . | | | . | . | . | . | . | . |
| NC_011770 | 2720851 | T | C | 24 | 8 | 2 | 0 | C | C | . | . | | | . | . | . | . | . | . |
| NC_011770 | 2720863 | T | C | 23 | 8 | 2 | 0 | C | C | . | . | | | . | . | . | . | . | . |
| NC_011770 | 2720890 | T | A | 23 | 8 | 2 | 0 | A | A | . | . | | | . | . | . | . | . | . |
| NC_011770 | 2722149 | G | T | 4 | 10 | 2 | 0 | . | . | . | . | T | T | . | . | . | . | . | . |
| NC_011770 | 2722154 | C | G | 4 | 10 | 2 | 0 | . | . | . | . | G | G | . | . | . | . | . | . |
| NC_011770 | 2722167 | A | G | 4 | 10 | 2 | 0 | . | . | . | . | G | G | . | . | . | . | . | . |
| NC_011770 | 2722169 | C | T | 4 | 10 | 2 | 0 | . | . | . | . | T | T | . | . | . | . | . | . |
| NC_011770 | 2722170 | A | C | 4 | 10 | 2 | 0 | . | . | . | . | C | C | . | . | . | . | . | . |
| NC_011770 | 2722173 | T | C | 4 | 10 | 2 | 0 | . | . | . | . | C | C | . | . | . | . | . | . |
| NC_011770 | 2722179 | GC | G | 4 | 10 | 2 | 0 | . | . | . | . | G | G | . | . | . | . | . | . |
| NC_011770 | 2722184 | GAGTC | G | 4 | 10 | 2 | 0 | . | . | . | . | G | G | . | . | . | . | . | . |
| NC_011770 | 2722900 | A | C | 10 | 7 | 5 | 0 | C | . | . | . | T | T | . | . | T | T | . | . |
| NC_011770 | 2722901 | A | T | 10 | 7 | 5 | 0 | T | . | . | . | T | T | . | . | C | C | . | . |
| NC_011770 | 2722903 | T | C | 10 | 7 | 5 | 0 | C | . | . | . | A | A | . | . | C | C | . | . |
| NC_011770 | 2722914 | T | C | 10 | 7 | 1 | 0 | C | . | . | . | | | . | . | | | . | . |
| NC_011770 | 2724134 | T | A | 21 | 8 | 4 | 0 | A | A | . | . | C | C | . | . | . | . | . | . |
| NC_011770 | 2726252 | C | T | 6 | 10 | 2 | 0 | T | T | . | . | . | . | . | . | . | . | . | . |
| NC_011770 | 2726268 | T | G | 6 | 10 | 2 | 0 | G | G | . | . | . | . | . | . | . | . | . | . |
| NC_011770 | 2727479 | T | G | 3 | 10 | 2 | 0 | G | G | . | . | . | . | . | . | . | . | . | . |
| NC_011770 | 2727481 | A | G | 3 | 10 | 2 | 0 | G | G | . | . | . | . | . | . | . | . | . | . |
| NC_011770 | 2727505 | CG | C | 3 | 10 | 2 | 0 | C | C | . | . | . | . | . | . | . | . | . | . |
| NC_011770 | 2733985 | T | G | 0 | 11 | 1 | 0 | . | . | . | . | . | G | . | . | . | . | . | . |
| NC_011770 | 2739263 | G | A | 0 | 10 | 2 | 0 | . | . | . | . | A | A | . | . | . | . | . | . |
| NC_011770 | 2739287 | CCGG | C | 12 | 10 | 2 | 0 | . | . | . | . | C | C | . | . | . | . | . | . |
| NC_011770 | 2739299 | C | CT | 12 | 10 | 2 | 0 | . | . | . | . | CT | CT | . | . | . | . | . | . |
| NC_011770 | 2739323 | G | A | 16 | 10 | 2 | 0 | . | . | . | . | A | A | . | . | . | . | . | . |
| NC_011770 | 2739329 | G | T | 16 | 10 | 2 | 0 | . | . | . | . | T | T | . | . | . | . | . | . |
| NC_011770 | 2739332 | G | T | 16 | 10 | 2 | 0 | . | . | . | . | T | T | . | . | . | . | . | . |
| NC_011770 | 2739341 | T | G | 17 | 10 | 2 | 0 | . | . | . | . | G | G | . | . | . | . | . | . |
| NC_011770 | 2739359 | G | A | 16 | 10 | 2 | 0 | . | . | . | . | A | A | . | . | . | . | . | . |
| NC_011770 | 2739362 | G | A | 16 | 10 | 2 | 0 | . | . | . | . | A | A | . | . | . | . | . | . |
| NC_011770 | 2739371 | T | C | 16 | 10 | 2 | 0 | . | . | . | . | C | C | . | . | . | . | . | . |
| NC_011770 | 2739378 | A | G | 16 | 10 | 2 | 0 | . | . | . | . | G | G | . | . | . | . | . | . |
| NC_011770 | 2739380 | C | G | 16 | 10 | 2 | 0 | . | . | . | . | G | G | . | . | . | . | . | . |
| NC_011770 | 2739391 | G | A | 17 | 10 | 2 | 0 | . | . | . | . | A | A | . | . | . | . | . | . |
| NC_011770 | 2739396 | G | A | 16 | 10 | 2 | 0 | . | . | . | . | A | A | . | . | . | . | . | . |
| NC_011770 | 2739404 | T | C | 14 | 10 | 2 | 0 | . | . | . | . | C | C | . | . | . | . | . | . |
| NC_011770 | 2739405 | A | C | 14 | 10 | 2 | 0 | . | . | . | . | C | C | . | . | . | . | . | . |
| NC_011770 | 2739407 | C | A | 15 | 10 | 2 | 0 | . | . | . | . | A | A | . | . | . | . | . | . |
| NC_011770 | 2739426 | T | C | 16 | 10 | 2 | 0 | . | . | . | . | C | C | . | . | . | . | . | . |
| NC_011770 | 2739431 | C | G | 16 | 10 | 2 | 0 | . | . | . | . | G | G | . | . | . | . | . | . |
| NC_011770 | 2739437 | A | G | 14 | 10 | 2 | 0 | . | . | . | . | G | G | . | . | . | . | . | . |
| NC_011770 | 2739440 | C | G | 13 | 10 | 2 | 0 | . | . | . | . | G | G | . | . | . | . | . | . |
| NC_011770 | 2739441 | T | C | 13 | 10 | 2 | 0 | . | . | . | . | C | C | . | . | . | . | . | . |
| NC_011770 | 2739442 | T | G | 14 | 10 | 2 | 0 | . | . | . | . | G | G | . | . | . | . | . | . |
| NC_011770 | 2739447 | A | C | 14 | 10 | 2 | 0 | . | . | . | . | C | C | . | . | . | . | . | . |
| NC_011770 | 2739452 | C | T | 14 | 10 | 2 | 0 | . | . | . | . | T | T | . | . | . | . | . | . |
| NC_011770 | 2739455 | A | T | 14 | 10 | 2 | 0 | . | . | . | . | T | T | . | . | . | . | . | . |
| NC_011770 | 2739467 | G | A | 15 | 10 | 2 | 0 | . | . | . | . | A | A | . | . | . | . | . | . |
| NC_011770 | 2739495 | G | A | 15 | 10 | 2 | 0 | . | . | . | . | A | A | . | . | . | . | . | . |
| NC_011770 | 2739525 | GA | G | 1 | 10 | 2 | 0 | . | . | . | . | G | G | . | . | . | . | . | . |

| | | | | | | | | | | | | | | | | | | | |
|---|---|---|---|---|---|---|---|---|---|---|---|---|---|---|---|---|---|---|---|
| NC_011770 | 2740546 | A | T | 0 | 11 | 1 | 0 | . | . | . | . | . | . | T | . | . | . | . | . |
| NC_011770 | 2742994 | G | T | 32 | 0 | 4 | 0 | | | T | T | T | T | | | | | | |
| NC_011770 | 2744061 | G | A | 74 | 0 | 10 | 0 | A | A | A | A | A | A | A | A | A | A | | |
| NC_011770 | 2744080 | A | C | 72 | 0 | 10 | 0 | C | C | C | C | C | C | C | C | C | C | | |
| NC_011770 | 2744106 | G | T | 73 | 0 | 2 | 0 | | | | | | | | | | | T | T |
| NC_011770 | 2744151 | T | A | 75 | 0 | 2 | 0 | | | | | | | | | A | A | | |
| NC_011770 | 2744178 | A | T | 73 | 0 | 2 | 0 | | | | | | | T | T | | | | |
| NC_011770 | 2744206 | A | C | 69 | 0 | 12 | 0 | C | C | C | C | C | C | C | C | C | C | C | C |
| NC_011770 | 2748608 | AG | A | 94 | 0 | 4 | 0 | A | A | | | | | | | A | A | | |
| NC_011770 | 2748619 | G | A | 90 | 0 | 2 | 0 | | | | | | | A | A | | | | |
| NC_011770 | 2750613 | C | T | 55 | 0 | 2 | 0 | | | | | | | | | | | T | T |
| NC_011770 | 2750641 | T | C | 58 | 0 | 1 | 0 | | | | | | | C | | | | | |
| NC_011770 | 2750661 | A | C | 58 | 0 | 12 | 0 | C | C | C | C | C | C | C | C | C | C | C | C |
| NC_011770 | 2750673 | C | T | 62 | 0 | 2 | 0 | | | | | | | | | | | T | T |
| NC_011770 | 2752922 | A | G | 12 | 10 | 2 | 0 | . | . | . | . | . | . | . | . | . | . | G | G |
| NC_011770 | 2755697 | G | A | 20 | 10 | 2 | 0 | . | . | . | . | . | . | . | . | . | . | A | A |
| NC_011770 | 2755893 | G | A | 17 | 10 | 2 | 0 | . | . | . | . | . | . | . | . | . | . | A | A |
| NC_011770 | 2764259 | T | G | 9 | 10 | 2 | 0 | . | . | . | . | . | . | . | . | . | . | G | G |
| NC_011770 | 2764331 | A | G | 8 | 10 | 2 | 0 | . | . | . | . | . | . | . | . | . | . | G | G |
| NC_011770 | 2764419 | G | A | 9 | 10 | 2 | 0 | . | . | . | . | . | . | . | . | . | . | A | A |
| NC_011770 | 2764469 | T | G | 10 | 10 | 2 | 0 | . | . | . | . | . | . | . | . | . | . | G | G |
| NC_011770 | 2764727 | G | A | 14 | 10 | 2 | 0 | . | . | . | . | . | . | . | . | . | . | A | A |
| NC_011770 | 2784203 | C | T | 62 | 0 | 2 | 0 | | | | | | | | | T | T | | |
| NC_011770 | 2784265 | T | C | 66 | 0 | 12 | 0 | C | C | C | C | C | C | C | C | C | C | C | C |
| NC_011770 | 2784283 | G | A | 66 | 0 | 2 | 0 | | | | | | | | | | | A | A |
| NC_011770 | 2786369 | C | T | 71 | 0 | 2 | 0 | | | | | | | T | T | | | | |
| NC_011770 | 2791543 | C | T | 82 | 0 | 2 | 0 | | | | | | | | | | | T | T |
| NC_011770 | 2791553 | G | A | 78 | 0 | 4 | 0 | | | A | A | A | A | | | | | | |
| NC_011770 | 2791563 | G | A | 75 | 0 | 1 | 0 | | | | | | | A | | | | | |
| NC_011770 | 2791597 | T | C | 72 | 0 | 2 | 0 | | | | | | | | | | | C | C |
| NC_011770 | 2791600 | G | A | 72 | 0 | 2 | 0 | | | | | | | | | | | A | A |
| NC_011770 | 2791602 | A | G | 71 | 0 | 2 | 0 | | | | | | | | | | | G | G |
| NC_011770 | 2791609 | T | C | 71 | 0 | 2 | 0 | | | | | | | | | | | C | C |
| NC_011770 | 2791677 | T | C | 71 | 0 | 2 | 0 | | | | | | | | | | | C | C |
| NC_011770 | 2791679 | C | T | 71 | 0 | 4 | 0 | | | T | T | T | T | | | | | | |
| NC_011770 | 2791750 | G | T | 67 | 0 | 2 | 0 | | | | | | | | | | | T | T |
| NC_011770 | 2791751 | C | A | 66 | 0 | 2 | 0 | | | | | | | | | | | A | A |
| NC_011770 | 2791803 | A | T | 69 | 0 | 10 | 0 | T | T | T | T | T | T | T | T | T | T | | |
| NC_011770 | 2791830 | C | T | 71 | 0 | 2 | 0 | | | | | | | | | | | T | T |
| NC_011770 | 2793220 | G | A | 76 | 0 | 4 | 0 | | | | | | | A | A | | | A | A |
| NC_011770 | 2793282 | A | G | 68 | 0 | 8 | 0 | | | G | G | G | G | G | G | | | G | G |
| NC_011770 | 2793315 | T | C | 71 | 0 | 12 | 0 | C | C | C | C | C | C | C | C | C | C | C | C |
| NC_011770 | 2796710 | G | A | 27 | 4 | 8 | 0 | A | A | A | A | A | A | . | . | . | . | A | A |
| NC_011770 | 2796711 | A | T | 27 | 4 | 4 | 0 | T | T | T | T | | | . | . | . | . | | |
| NC_011770 | 2796713 | A | T | 27 | 5 | 6 | 0 | T | T | T | T | . | | . | . | . | . | T | T |
| NC_011770 | 2796714 | G | A | 27 | 5 | 7 | 0 | A | A | A | A | . | T | . | . | . | . | T | T |
| NC_011770 | 2796715 | T | A | 28 | 5 | 1 | 0 | | | | | . | A | . | . | . | . | | |
| NC_011770 | 2796720 | T | C | 29 | 5 | 2 | 0 | | | | | . | | . | . | . | . | C | C |
| NC_011770 | 2796730 | T | C | 31 | 6 | 4 | 0 | C | C | C | C | . | . | . | . | . | . | | |
| NC_011770 | 2796735 | A | G | 29 | 6 | 2 | 0 | | | | | . | . | . | . | . | . | G | G |
| NC_011770 | 2796736 | C | G | 29 | 6 | 2 | 0 | | | | | . | . | . | . | . | . | G | G |
| NC_011770 | 2796744 | A | G | 31 | 6 | 2 | 0 | | | | | . | . | . | . | . | . | G | G |
| NC_011770 | 2796748 | G | A | 30 | 6 | 6 | 0 | A | A | A | A | . | . | . | . | . | . | A | A |
| NC_011770 | 2796756 | G | A | 30 | 6 | 2 | 0 | | | | | . | . | . | . | . | . | A | A |
| NC_011770 | 2796760 | G | A | 30 | 6 | 6 | 0 | A | A | A | A | . | . | . | . | . | . | A | A |
| NC_011770 | 2796765 | T | C | 30 | 6 | 6 | 0 | C | C | C | C | . | . | . | . | . | . | C | C |
| NC_011770 | 2796769 | C | T | 30 | 6 | 6 | 0 | T | T | T | T | . | . | . | . | . | . | T | T |
| NC_011770 | 2796789 | G | A | 28 | 6 | 4 | 0 | A | A | A | A | . | . | . | . | . | . | | |
| NC_011770 | 2796790 | C | A | 28 | 6 | 6 | 0 | A | A | A | A | . | . | . | . | . | . | T | T |
| NC_011770 | 2796800 | A | G | 28 | 6 | 4 | 0 | G | G | G | G | . | . | . | . | . | . | | |
| NC_011770 | 2796806 | C | T | 12 | 8 | 2 | 0 | . | . | T | T | . | . | . | . | . | . | | |
| NC_011770 | 2796807 | C | G | 12 | 8 | 2 | 0 | . | . | G | G | . | . | . | . | . | . | | |

| | | | | | | | | | | | | | | | | | | | |
|---|---|---|---|---|---|---|---|---|---|---|---|---|---|---|---|---|---|---|---|
| NC_011770 | 2796813 | T | C | 13 | 8 | 1 | 0 | . | . | C | . | . | . | . | . | . | . | | |
| NC_011770 | 2796817 | G | A | 13 | 8 | 2 | 0 | . | . | A | A | . | . | . | . | . | . | | |
| NC_011770 | 2796833 | T | A | 12 | 10 | 2 | 0 | . | . | . | . | . | . | . | . | . | . | A | A |
| NC_011770 | 2796835 | G | A | 12 | 10 | 2 | 0 | . | . | . | . | . | . | . | . | . | . | A | A |
| NC_011770 | 2796852 | T | G | 11 | 10 | 2 | 0 | . | . | . | . | . | . | . | . | . | . | G | G |
| NC_011770 | 2796857 | A | T | 11 | 10 | 2 | 0 | . | . | . | . | . | . | . | . | . | . | T | T |
| NC_011770 | 2796890 | A | G | 11 | 10 | 2 | 0 | . | . | . | . | . | . | . | . | . | . | G | G |
| NC_011770 | 2796905 | T | A | 11 | 10 | 2 | 0 | . | . | . | . | . | . | . | . | . | . | A | A |
| NC_011770 | 2796906 | A | G | 12 | 10 | 2 | 0 | . | . | . | . | . | . | . | . | . | . | G | G |
| NC_011770 | 2796919 | A | T | 12 | 10 | 2 | 0 | . | . | . | . | . | . | . | . | . | . | T | T |
| NC_011770 | 2796929 | C | T | 13 | 10 | 2 | 0 | . | . | . | . | . | . | . | . | . | . | T | T |
| NC_011770 | 2799840 | C | A | 0 | 11 | 1 | 0 | A | . | . | . | . | . | . | . | . | . | . | . |
| NC_011770 | 2799844 | C | A | 0 | 11 | 1 | 0 | A | . | . | . | . | . | . | . | . | . | . | . |
| NC_011770 | 2799859 | G | A | 0 | 11 | 1 | 0 | A | . | . | . | . | . | . | . | . | . | . | . |
| NC_011770 | 2799862 | T | A | 0 | 11 | 1 | 0 | A | . | . | . | . | . | . | . | . | . | . | . |
| NC_011770 | 2799868 | C | T | 0 | 11 | 1 | 0 | T | . | . | . | . | . | . | . | . | . | . | . |
| NC_011770 | 2799871 | C | G | 0 | 11 | 1 | 0 | G | . | . | . | . | . | . | . | . | . | . | . |
| NC_011770 | 2799883 | G | C | 0 | 11 | 1 | 0 | C | . | . | . | . | . | . | . | . | . | . | . |
| NC_011770 | 2799886 | A | T | 0 | 11 | 1 | 0 | T | . | . | . | . | . | . | . | . | . | . | . |
| NC_011770 | 2840927 | T | C | 46 | 6 | 2 | 0 | C | C | | | . | . | . | . | . | . | | |
| NC_011770 | 2840928 | C | G | 46 | 6 | 6 | 0 | G | G | G | G | . | . | . | . | . | . | G | G |
| NC_011770 | 2840937 | G | C | 20 | 9 | 1 | 0 | . | . | C | . | . | . | . | . | . | . | | |
| NC_011770 | 2840938 | A | T | 19 | 9 | 1 | 0 | . | . | T | . | . | . | . | . | . | . | | |
| NC_011770 | 2840942 | C | G | 22 | 7 | 3 | 0 | G | . | G | G | . | . | . | . | . | . | | |
| NC_011770 | 2840950 | G | C | 42 | 6 | 6 | 0 | C | C | C | C | . | . | . | . | . | . | C | C |
| NC_011770 | 2840964 | G | C | 44 | 6 | 4 | 0 | C | C | C | C | . | . | . | . | . | . | | |
| NC_011770 | 2840973 | A | G | 44 | 6 | 6 | 0 | G | G | G | G | . | . | . | . | . | . | G | G |
| NC_011770 | 2841017 | A | G | 42 | 6 | 6 | 0 | G | G | G | G | . | . | . | . | . | . | G | G |
| NC_011770 | 2841053 | C | A | 40 | 6 | 4 | 0 | A | A | A | A | . | . | . | . | . | . | | |
| NC_011770 | 2841058 | C | G | 39 | 6 | 2 | 0 | G | G | . | . | . | . | . | . | . | . | . | . |
| NC_011770 | 2841078 | C | G | 38 | 6 | 4 | 0 | G | G | G | G | . | . | . | . | . | . | . | . |
| NC_011770 | 2841088 | A | T | 27 | 6 | 6 | 0 | T | T | T | T | . | . | . | . | . | . | T | T |
| NC_011770 | 2841089 | C | G | 27 | 6 | 4 | 0 | G | G | G | G | . | . | . | . | . | . | | |
| NC_011770 | 2841095 | G | C | 27 | 6 | 2 | 0 | | | | | . | . | . | . | . | . | C | C |
| NC_011770 | 2841099 | A | G | 13 | 6 | 6 | 0 | G | G | G | G | . | . | . | . | . | . | G | G |
| NC_011770 | 2841103 | G | A | 12 | 9 | 1 | 0 | . | . | A | . | . | . | . | . | . | . | | |
| NC_011770 | 2841110 | T | C | 12 | 10 | 2 | 0 | . | . | . | . | . | . | . | . | . | . | C | C |
| NC_011770 | 2841122 | C | T | 1 | 10 | 2 | 0 | . | . | . | . | . | . | . | . | . | . | T | T |
| NC_011770 | 2841124 | G | T | 6 | 10 | 2 | 0 | . | . | . | . | . | . | . | . | . | . | T | T |
| NC_011770 | 2841158 | G | GGTT | 6 | 10 | 2 | 0 | . | . | . | . | . | . | . | . | . | . | GGTT | GGTT |
| NC_011770 | 2841167 | C | T | 1 | 11 | 1 | 0 | . | . | . | . | . | . | . | . | . | . | T | . |
| NC_011770 | 2841169 | G | A | 1 | 11 | 1 | 0 | . | . | . | . | . | . | . | . | . | . | A | . |
| NC_011770 | 2841176 | T | A | 30 | 6 | 4 | 0 | A | A | A | A | . | . | . | . | . | . | | |
| NC_011770 | 2841186 | A | G | 33 | 6 | 2 | 0 | | | | | . | . | . | . | . | . | G | G |
| NC_011770 | 2841188 | T | C | 32 | 6 | 6 | 0 | C | C | C | C | . | . | . | . | . | . | C | C |
| NC_011770 | 2841196 | A | G | 34 | 6 | 4 | 0 | G | G | G | G | . | . | . | . | . | . | | |
| NC_011770 | 2841199 | A | C | 35 | 6 | 2 | 0 | | | | | . | . | . | . | . | . | C | C |
| NC_011770 | 2841219 | T | C | 39 | 6 | 2 | 0 | | | | | . | . | . | . | . | . | C | C |
| NC_011770 | 2841227 | A | G | 39 | 6 | 4 | 0 | G | G | G | G | . | . | . | . | . | . | | |
| NC_011770 | 2841229 | C | T | 39 | 6 | 2 | 0 | | | | | . | . | . | . | . | . | T | T |
| NC_011770 | 2841230 | T | C | 39 | 6 | 2 | 0 | | | | | . | . | . | . | . | . | C | C |
| NC_011770 | 2841236 | C | A | 40 | 6 | 6 | 0 | A | A | A | A | . | . | . | . | . | . | A | A |
| NC_011770 | 2841238 | G | C | 41 | 6 | 4 | 0 | C | C | C | C | . | . | . | . | . | . | | |
| NC_011770 | 2841869 | A | G | 48 | 6 | 4 | 0 | G | G | G | G | . | . | . | . | . | . | | |
| NC_011770 | 2841872 | G | A | 48 | 6 | 4 | 0 | A | A | A | A | . | . | . | . | . | . | | |
| NC_011770 | 2841876 | C | T | 47 | 6 | 2 | 0 | | | | | . | . | . | . | . | . | T | T |
| NC_011770 | 2841877 | C | T | 47 | 6 | 4 | 0 | T | T | T | T | . | . | . | . | . | . | | |
| NC_011770 | 2842524 | A | G | 44 | 6 | 4 | 0 | G | G | G | G | . | . | . | . | . | . | | |
| NC_011770 | 2842526 | A | G | 44 | 6 | 4 | 0 | G | G | G | G | . | . | . | . | . | . | | |
| NC_011770 | 2842539 | G | C | 44 | 6 | 6 | 0 | C | C | C | C | . | . | . | . | . | . | C | C |
| NC_011770 | 2842540 | C | G | 44 | 6 | 2 | 0 | | | | | . | . | . | . | . | . | G | G |
| NC_011770 | 2842545 | C | A | 43 | 6 | 6 | 0 | A | A | A | A | . | . | . | . | . | . | G | G |

| | | | | | | | | | | | | | | | | | | | |
|---|---|---|---|---|---|---|---|---|---|---|---|---|---|---|---|---|---|---|---|
| NC_011770 | 2842553 | G | A | 44 | 6 | 4 | 0 | A | A | A | A | . | . | . | . | . | . | | |
| NC_011770 | 2842558 | G | C | 44 | 6 | 4 | 0 | C | C | C | C | . | . | . | . | . | . | | |
| NC_011770 | 2842564 | T | A | 43 | 6 | 2 | 0 | | | | | . | . | . | . | . | . | A | A |
| NC_011770 | 2842581 | A | T | 35 | 6 | 4 | 0 | T | T | T | T | . | . | . | . | . | . | | |
| NC_011770 | 2842582 | A | G | 34 | 6 | 6 | 0 | G | G | G | G | . | . | . | . | . | . | G | G |
| NC_011770 | 2842583 | C | T | 35 | 6 | 6 | 0 | T | T | T | T | . | . | . | . | . | . | T | T |
| NC_011770 | 2842964 | C | T | 2 | 8 | 1 | 0 | | . | | | . | . | . | . | . | . | T | . |
| NC_011770 | 2842965 | C | G | 2 | 8 | 3 | 0 | G | . | G | G | . | . | . | . | . | . | | . |
| NC_011770 | 2842967 | G | A | 2 | 8 | 3 | 0 | A | . | A | A | . | . | . | . | . | . | | . |
| NC_011770 | 2842969 | C | T | 2 | 9 | 3 | 0 | T | . | T | T | . | . | . | . | . | . | . | . |
| NC_011770 | 2842970 | T | C | 2 | 9 | 3 | 0 | C | . | C | C | . | . | . | . | . | . | . | . |
| NC_011770 | 2842973 | C | G | 2 | 9 | 3 | 0 | G | . | G | G | . | . | . | . | . | . | . | . |
| NC_011770 | 2844499 | C | T | 35 | 6 | 2 | 0 | | | | | . | . | . | . | . | . | T | T |
| NC_011770 | 2844500 | C | T | 34 | 6 | 4 | 0 | T | T | T | T | . | . | . | . | . | . | | |
| NC_011770 | 2844501 | C | G | 34 | 6 | 2 | 0 | | | | | . | . | . | . | . | . | G | G |
| NC_011770 | 2844503 | ATC | A | 34 | 6 | 4 | 0 | A | A | A | A | . | . | . | . | . | . | | |
| NC_011770 | 2844504 | T | C | 12 | 10 | 2 | 0 | . | . | . | . | . | . | . | . | . | . | C | C |
| NC_011770 | 2847624 | G | A | 22 | 6 | 2 | 0 | | | | | . | . | . | . | . | . | A | A |
| NC_011770 | 2847626 | C | G | 22 | 6 | 2 | 0 | | | | | . | . | . | . | . | . | G | G |
| NC_011770 | 2847627 | C | A | 22 | 6 | 6 | 0 | A | A | A | A | . | . | . | . | . | . | A | A |
| NC_011770 | 2847628 | G | T | 22 | 6 | 4 | 0 | T | T | T | T | . | . | . | . | . | . | | |
| NC_011770 | 2847629 | G | A | 22 | 6 | 2 | 0 | | | | | . | . | . | . | . | . | A | A |
| NC_011770 | 2847632 | G | A | 22 | 6 | 6 | 0 | A | A | A | A | . | . | . | . | . | . | A | A |
| NC_011770 | 2847647 | C | T | 24 | 6 | 4 | 0 | T | T | T | T | . | . | . | . | . | . | | |
| NC_011770 | 2847649 | G | A | 23 | 6 | 2 | 0 | | | | | . | . | . | . | . | . | A | A |
| NC_011770 | 2847650 | C | T | 24 | 6 | 4 | 0 | T | T | T | T | . | . | . | . | . | . | | |
| NC_011770 | 2847657 | C | G | 24 | 6 | 2 | 0 | | | | | . | . | . | . | . | . | G | G |
| NC_011770 | 2847658 | T | G | 20 | 6 | 4 | 0 | G | G | G | G | . | . | . | . | . | . | | |
| NC_011770 | 2847661 | A | T | 20 | 6 | 4 | 0 | T | T | T | T | . | . | . | . | . | . | | |
| NC_011770 | 2847664 | G | A | 20 | 6 | 6 | 0 | A | A | A | A | . | . | . | . | . | . | A | A |
| NC_011770 | 2847665 | G | A | 20 | 6 | 4 | 0 | A | A | A | A | . | . | . | . | . | . | | |
| NC_011770 | 2847667 | A | C | 20 | 6 | 4 | 0 | C | C | C | C | . | . | . | . | . | . | | |
| NC_011770 | 2847671 | T | C | 20 | 6 | 2 | 0 | | | | | . | . | . | . | . | . | C | C |
| NC_011770 | 2847673 | T | G | 20 | 6 | 6 | 0 | G | G | G | G | . | . | . | . | . | . | G | G |
| NC_011770 | 2847680 | G | A | 20 | 6 | 4 | 0 | A | A | A | A | . | . | . | . | . | . | | |
| NC_011770 | 2847681 | C | T | 20 | 6 | 4 | 0 | T | T | T | T | . | . | . | . | . | . | | |
| NC_011770 | 2847688 | T | C | 19 | 6 | 6 | 0 | C | C | C | C | . | . | . | . | . | . | C | C |
| NC_011770 | 2847689 | T | G | 19 | 6 | 6 | 0 | G | G | G | G | . | . | . | . | . | . | G | G |
| NC_011770 | 2847690 | G | A | 19 | 6 | 2 | 0 | | | | | . | . | . | . | . | . | A | A |
| NC_011770 | 2847702 | A | G | 20 | 6 | 6 | 0 | G | G | G | G | . | . | . | . | . | . | G | G |
| NC_011770 | 2847705 | A | C | 20 | 6 | 6 | 0 | C | C | C | C | . | . | . | . | . | . | C | C |
| NC_011770 | 2847706 | G | A | 20 | 6 | 6 | 0 | A | A | A | A | . | . | . | . | . | . | A | A |
| NC_011770 | 2847716 | G | GC | 21 | 6 | 4 | 0 | GC | GC | GC | GC | . | . | . | . | . | . | | |
| NC_011770 | 2847721 | G | C | 20 | 6 | 6 | 0 | C | C | C | C | . | . | . | . | . | . | C | C |
| NC_011770 | 2847724 | G | C | 20 | 6 | 4 | 0 | C | C | C | C | . | . | . | . | . | . | | |
| NC_011770 | 2847726 | C | G | 20 | 6 | 6 | 0 | G | G | G | G | . | . | . | . | . | . | T | T |
| NC_011770 | 2847729 | G | C | 26 | 6 | 2 | 0 | | | | | . | . | . | . | . | . | C | C |
| NC_011770 | 2847730 | G | A | 26 | 6 | 2 | 0 | | | | | . | . | . | . | . | . | A | A |
| NC_011770 | 2847754 | G | GGC | 28 | 6 | 4 | 0 | GGC | GGC | GGC | GGC | . | . | . | . | . | . | | |
| NC_011770 | 2847755 | C | G | 27 | 6 | 2 | 0 | | | | | . | . | . | . | . | . | G | G |
| NC_011770 | 2847756 | A | G | 27 | 6 | 2 | 0 | | | | | . | . | . | . | . | . | G | G |
| NC_011770 | 2847757 | CTG | C | 27 | 6 | 4 | 0 | C | C | C | C | . | . | . | . | . | . | | |
| NC_011770 | 2847778 | T | C | 14 | 10 | 2 | 0 | . | . | . | . | . | . | . | . | . | . | C | C |
| NC_011770 | 2847780 | GC | G | 14 | 10 | 2 | 0 | . | . | . | . | . | . | . | . | . | . | G | G |
| NC_011770 | 2847788 | G | A | 14 | 10 | 2 | 0 | . | . | . | . | . | . | . | . | . | . | A | A |
| NC_011770 | 2847794 | G | A | 29 | 6 | 2 | 0 | | | | | . | . | . | . | . | . | A | A |
| NC_011770 | 2847815 | T | A | 15 | 10 | 2 | 0 | . | . | . | . | . | . | . | . | . | . | A | A |
| NC_011770 | 2848338 | G | A | 14 | 8 | 4 | 0 | A | A | A | A | . | . | . | . | . | . | . | . |
| NC_011770 | 2848348 | A | C | 16 | 8 | 4 | 0 | C | C | C | C | . | . | . | . | . | . | . | . |
| NC_011770 | 2848349 | A | G | 16 | 8 | 4 | 0 | G | G | G | G | . | . | . | . | . | . | . | . |
| NC_011770 | 2848354 | G | GAT | 15 | 8 | 4 | 0 | GAT | GAT | GAT | GAT | . | . | . | . | . | . | . | . |
| NC_011770 | 2848364 | C | CG | 18 | 8 | 4 | 0 | CG | CG | CG | CG | . | . | . | . | . | . | . | . |

| | | | | | | | | | | | | | | | | | | | |
|---|---|---|---|---|---|---|---|---|---|---|---|---|---|---|---|---|---|---|---|
| NC_011770 | 2848366 | A | G | 16 | 8 | 4 | 0 | G | G | G | G | . | . | . | . | . | . | . | . |
| NC_011770 | 2848427 | C | CG | 20 | 8 | 4 | 0 | CG | CG | CG | CG | . | . | . | . | . | . | . | . |
| NC_011770 | 2848430 | G | A | 2 | 9 | 3 | 0 | A | . | A | A | . | . | . | . | . | . | . | . |
| NC_011770 | 2848438 | T | C | 21 | 7 | 1 | 0 | | | | | . | . | . | . | . | . | . | C |
| NC_011770 | 2848439 | T | C | 21 | 7 | 5 | 0 | C | C | C | C | . | . | . | . | . | . | . | C |
| NC_011770 | 2848456 | CG | C | 29 | 6 | 2 | 0 | | | | | . | . | . | . | . | . | C | C |
| NC_011770 | 2848457 | GA | G | 21 | 7 | 4 | 0 | G | G | G | G | . | . | . | . | . | . | . | . |
| NC_011770 | 2848460 | A | G | 0 | 11 | 1 | 0 | . | . | . | . | . | . | . | . | . | . | G | . |
| NC_011770 | 2848468 | G | A | 28 | 6 | 2 | 0 | | | | | . | . | . | . | . | . | A | A |
| NC_011770 | 2848471 | A | T | 28 | 6 | 6 | 0 | T | T | T | T | . | . | . | . | . | . | C | C |
| NC_011770 | 2848472 | G | T | 28 | 6 | 4 | 0 | T | T | T | T | . | . | . | . | . | . | | |
| NC_011770 | 2848483 | G | A | 24 | 6 | 6 | 0 | A | A | A | A | . | . | . | . | . | . | A | A |
| NC_011770 | 2848485 | C | T | 24 | 6 | 6 | 0 | T | T | T | T | . | . | . | . | . | . | A | A |
| NC_011770 | 2848487 | A | G | 25 | 6 | 6 | 0 | G | G | G | G | . | . | . | . | . | . | G | G |
| NC_011770 | 2848488 | G | A | 25 | 6 | 6 | 0 | A | A | A | A | . | . | . | . | . | . | A | A |
| NC_011770 | 2848490 | C | T | 24 | 6 | 6 | 0 | T | T | T | T | . | . | . | . | . | . | T | T |
| NC_011770 | 2848504 | C | G | 33 | 6 | 6 | 0 | G | G | G | G | . | . | . | . | . | . | G | G |
| NC_011770 | 2848511 | GA | G | 32 | 6 | 6 | 0 | G | G | G | G | . | . | . | . | . | . | G | G |
| NC_011770 | 2849313 | G | T | 42 | 6 | 2 | 0 | | | | | . | . | . | . | . | . | T | T |
| NC_011770 | 2849314 | T | C | 41 | 6 | 2 | 0 | | | | | . | . | . | . | . | . | C | C |
| NC_011770 | 2852213 | A | G | 43 | 5 | 5 | 0 | G | G | G | G | . | . | . | G | . | . | | |
| NC_011770 | 2852214 | C | T | 44 | 5 | 5 | 0 | T | T | T | T | . | . | . | T | . | . | | |
| NC_011770 | 2852215 | G | C | 44 | 5 | 4 | 0 | C | C | C | C | . | . | . | | . | . | | |
| NC_011770 | 2852217 | G | T | 42 | 5 | 4 | 0 | T | T | T | T | . | . | . | | . | . | | |
| NC_011770 | 2852224 | C | A | 46 | 4 | 4 | 0 | A | A | A | A | | . | . | | . | . | | |
| NC_011770 | 2852230 | C | G | 47 | 4 | 4 | 0 | | | | | G | . | . | G | . | . | G | G |
| NC_011770 | 2852239 | G | C | 40 | 4 | 6 | 0 | C | C | C | C | | . | . | | . | . | C | C |
| NC_011770 | 2852241 | T | C | 40 | 5 | 7 | 0 | C | C | C | C | C | . | . | . | . | . | C | C |
| NC_011770 | 2852242 | G | T | 40 | 5 | 3 | 0 | | | | | T | . | . | . | . | . | A | A |
| NC_011770 | 2852251 | G | C | 36 | 5 | 6 | 0 | C | C | C | C | | . | . | . | . | . | C | C |
| NC_011770 | 2852253 | G | A | 36 | 5 | 5 | 0 | A | A | A | A | A | . | . | . | . | . | | |
| NC_011770 | 2852254 | C | G | 37 | 5 | 7 | 0 | G | G | G | G | G | . | . | . | . | . | G | G |
| NC_011770 | 2852257 | G | A | 36 | 5 | 2 | 0 | | | | | | . | . | . | . | . | A | A |
| NC_011770 | 2852260 | G | A | 40 | 5 | 5 | 0 | A | A | A | A | C | . | . | . | . | . | | |
| NC_011770 | 2852263 | G | C | 39 | 5 | 6 | 0 | C | C | C | C | | . | . | . | . | . | C | C |
| NC_011770 | 2852266 | C | G | 43 | 5 | 1 | 0 | | | | | G | . | . | . | . | . | | |
| NC_011770 | 2852275 | G | C | 41 | 6 | 6 | 0 | C | C | C | C | . | . | . | . | . | . | C | C |
| NC_011770 | 2852278 | C | A | 41 | 6 | 2 | 0 | | | | | . | . | . | . | . | . | A | A |
| NC_011770 | 2852281 | C | G | 42 | 6 | 4 | 0 | G | G | G | G | . | . | . | . | . | . | | |
| NC_011770 | 2852284 | C | A | 42 | 6 | 4 | 0 | A | A | A | A | . | . | . | . | . | . | | |
| NC_011770 | 2852290 | A | G | 41 | 6 | 4 | 0 | G | G | G | G | . | . | . | . | . | . | | |
| NC_011770 | 2852299 | G | C | 44 | 6 | 2 | 0 | | | | | . | . | . | . | . | . | C | C |
| NC_011770 | 2852305 | A | G | 43 | 6 | 6 | 0 | G | G | G | G | . | . | . | . | . | . | G | G |
| NC_011770 | 2852317 | G | A | 45 | 6 | 2 | 0 | | | | | . | . | . | . | . | . | A | A |
| NC_011770 | 2852323 | C | G | 43 | 6 | 2 | 0 | | | | | . | . | . | . | . | . | G | G |
| NC_011770 | 2852326 | G | A | 44 | 6 | 2 | 0 | | | | | . | . | . | . | . | . | A | A |
| NC_011770 | 2852335 | A | G | 42 | 6 | 2 | 0 | | | | | . | . | . | . | . | . | G | G |
| NC_011770 | 2852338 | G | C | 43 | 6 | 2 | 0 | | | | | . | . | . | . | . | . | C | C |
| NC_011770 | 2852344 | G | A | 42 | 6 | 4 | 0 | A | A | A | A | . | . | . | . | . | . | | |
| NC_011770 | 2852347 | C | G | 42 | 6 | 2 | 0 | | | | | . | . | . | . | . | . | G | G |
| NC_011770 | 2852359 | G | A | 42 | 6 | 2 | 0 | | | | | . | . | . | . | . | . | A | A |
| NC_011770 | 2852361 | G | A | 43 | 6 | 2 | 0 | | | | | . | . | . | . | . | . | A | A |
| NC_011770 | 2852368 | A | G | 43 | 6 | 2 | 0 | | | | | . | . | . | . | . | . | G | G |
| NC_011770 | 2852377 | G | C | 43 | 6 | 2 | 0 | | | | | . | . | . | . | . | . | C | C |
| NC_011770 | 2852401 | G | A | 41 | 6 | 4 | 0 | A | A | A | A | . | . | . | . | . | . | | |
| NC_011770 | 2852402 | A | G | 41 | 6 | 2 | 0 | | | | | . | . | . | . | . | . | G | G |
| NC_011770 | 2852403 | C | T | 42 | 6 | 2 | 0 | | | | | . | . | . | . | . | . | T | T |
| NC_011770 | 2852404 | A | G | 40 | 6 | 2 | 0 | | | | | . | . | . | . | . | . | G | G |
| NC_011770 | 2852407 | T | C | 39 | 6 | 2 | 0 | | | | | . | . | . | . | . | . | C | C |
| NC_011770 | 2852415 | G | C | 40 | 6 | 2 | 0 | | | | | . | . | . | . | . | . | C | C |
| NC_011770 | 2852425 | G | C | 39 | 6 | 2 | 0 | | | | | . | . | . | . | . | . | C | C |
| NC_011770 | 2852434 | G | C | 38 | 6 | 2 | 0 | | | | | . | . | . | . | . | . | C | C |

| | | | | | | | | | | | | | | | | | | | |
|---|---|---|---|---|---|---|---|---|---|---|---|---|---|---|---|---|---|---|---|
| NC_011770 | 2852436 | G | T | 38 | 6 | 2 | 0 | | | | | . | . | . | . | . | . | T | T |
| NC_011770 | 2852440 | A | G | 37 | 6 | 2 | 0 | | | | | . | . | . | . | . | . | G | G |
| NC_011770 | 2852455 | C | T | 35 | 6 | 2 | 0 | | | | | . | . | . | . | . | . | T | T |
| NC_011770 | 2852485 | T | C | 21 | 8 | 4 | 0 | C | C | C | C | . | . | . | . | . | . | . | . |
| NC_011770 | 2852500 | G | C | 3 | 7 | 2 | 0 | | . | | | . | . | . | . | . | . | C | C |
| NC_011770 | 2852506 | G | A | 24 | 6 | 2 | 0 | | | | | . | . | . | . | . | . | A | A |
| NC_011770 | 2852512 | C | T | 24 | 6 | 6 | 0 | T | T | T | T | . | . | . | . | . | . | G | G |
| NC_011770 | 2852514 | C | T | 25 | 6 | 2 | 0 | | | | | . | . | . | . | . | . | T | T |
| NC_011770 | 2852541 | G | T | 34 | 6 | 2 | 0 | | | | | . | . | . | . | . | . | T | T |
| NC_011770 | 2852545 | A | G | 34 | 6 | 2 | 0 | | | | | . | . | . | . | . | . | G | G |
| NC_011770 | 2852553 | T | G | 34 | 6 | 2 | 0 | | | | | . | . | . | . | . | . | G | G |
| NC_011770 | 2852554 | C | G | 35 | 6 | 2 | 0 | | | | | . | . | . | . | . | . | G | G |
| NC_011770 | 2852578 | G | A | 34 | 6 | 2 | 0 | | | | | . | . | . | . | . | . | A | A |
| NC_011770 | 2852587 | G | A | 34 | 6 | 4 | 0 | A | A | A | A | . | . | . | . | . | . | | |
| NC_011770 | 2852611 | C | T | 14 | 7 | 3 | 0 | T | . | T | T | . | . | . | . | . | . | | |
| NC_011770 | 2852612 | A | G | 15 | 7 | 5 | 0 | G | . | G | G | . | . | . | . | . | . | G | G |
| NC_011770 | 2852618 | C | G | 14 | 7 | 2 | 0 | | . | | | . | . | . | . | . | . | G | G |
| NC_011770 | 2854045 | G | A | 25 | 6 | 2 | 0 | | | | | . | . | . | . | . | . | A | A |
| NC_011770 | 2854049 | C | T | 25 | 6 | 4 | 0 | T | T | T | T | . | . | . | . | . | . | | |
| NC_011770 | 2854074 | C | T | 25 | 6 | 2 | 0 | | | | | . | . | . | . | . | . | T | T |
| NC_011770 | 2854080 | C | G | 25 | 6 | 2 | 0 | | | | | . | . | . | . | . | . | G | G |
| NC_011770 | 2854092 | A | C | 26 | 6 | 2 | 0 | | | | | . | . | . | . | . | . | C | C |
| NC_011770 | 2854106 | G | T | 26 | 6 | 2 | 0 | | | | | . | . | . | . | . | . | T | T |
| NC_011770 | 2854107 | A | G | 26 | 6 | 2 | 0 | | | | | . | . | . | . | . | . | G | G |
| NC_011770 | 2854109 | T | C | 25 | 6 | 2 | 0 | | | | | . | . | . | . | . | . | C | C |
| NC_011770 | 2854113 | C | T | 26 | 6 | 2 | 0 | | | | | . | . | . | . | . | . | T | T |
| NC_011770 | 2854116 | C | T | 26 | 6 | 2 | 0 | | | | | . | . | . | . | . | . | T | T |
| NC_011770 | 2854125 | C | T | 26 | 6 | 2 | 0 | | | | | . | . | . | . | . | . | T | T |
| NC_011770 | 2854131 | G | A | 27 | 6 | 2 | 0 | | | | | . | . | . | . | . | . | A | A |
| NC_011770 | 2854137 | G | A | 27 | 6 | 2 | 0 | | | | | . | . | . | . | . | . | A | A |
| NC_011770 | 2854140 | G | C | 27 | 6 | 2 | 0 | | | | | . | . | . | . | . | . | C | C |
| NC_011770 | 2854145 | C | G | 27 | 6 | 6 | 0 | G | G | G | G | . | . | . | . | . | . | G | G |
| NC_011770 | 2854149 | G | C | 29 | 6 | 2 | 0 | | | | | . | . | . | . | . | . | C | C |
| NC_011770 | 2854158 | G | C | 32 | 6 | 6 | 0 | C | C | C | C | . | . | . | . | . | . | C | C |
| NC_011770 | 2854161 | G | A | 32 | 6 | 2 | 0 | | | | | . | . | . | . | . | . | A | A |
| NC_011770 | 2854182 | G | C | 32 | 6 | 2 | 0 | | | | | . | . | . | . | . | . | C | C |
| NC_011770 | 2854194 | C | A | 33 | 6 | 2 | 0 | | | | | . | . | . | . | . | . | A | A |
| NC_011770 | 2854206 | C | T | 35 | 6 | 2 | 0 | | | | | . | . | . | . | . | . | T | T |
| NC_011770 | 2854251 | G | C | 28 | 6 | 2 | 0 | | | | | . | . | . | . | . | . | C | C |
| NC_011770 | 2854254 | C | G | 27 | 6 | 2 | 0 | | | | | . | . | . | . | . | . | G | G |
| NC_011770 | 2854256 | C | T | 25 | 6 | 6 | 0 | T | T | T | T | . | . | . | . | . | . | T | T |
| NC_011770 | 2854259 | C | T | 25 | 6 | 6 | 0 | T | T | T | T | . | . | . | . | . | . | T | T |
| NC_011770 | 2854264 | T | C | 25 | 6 | 6 | 0 | C | C | C | C | . | . | . | . | . | . | C | C |
| NC_011770 | 2854267 | G | C | 24 | 6 | 6 | 0 | C | C | C | C | . | . | . | . | . | . | C | C |
| NC_011770 | 2854268 | C | T | 24 | 6 | 2 | 0 | | | | | . | . | . | . | . | . | T | T |
| NC_011770 | 2854271 | C | T | 23 | 6 | 6 | 0 | T | T | T | T | . | . | . | . | . | . | T | T |
| NC_011770 | 2854273 | G | C | 24 | 6 | 6 | 0 | C | C | C | C | . | . | . | . | . | . | C | C |
| NC_011770 | 2854274 | T | C | 27 | 6 | 4 | 0 | C | C | C | C | . | . | . | . | . | . | | |
| NC_011770 | 2854281 | A | C | 30 | 6 | 6 | 0 | C | C | C | C | . | . | . | . | . | . | C | C |
| NC_011770 | 2854293 | A | G | 31 | 6 | 2 | 0 | | | | | . | . | . | . | . | . | G | G |
| NC_011770 | 2854296 | C | G | 31 | 6 | 4 | 0 | G | G | G | G | . | . | . | . | . | . | | |
| NC_011770 | 2854302 | T | C | 31 | 6 | 6 | 0 | C | C | C | C | . | . | . | . | . | . | C | C |
| NC_011770 | 2854308 | T | C | 31 | 6 | 4 | 0 | C | C | C | C | . | . | . | . | . | . | | |
| NC_011770 | 2854314 | G | C | 30 | 6 | 6 | 0 | C | C | C | C | . | . | . | . | . | . | A | A |
| NC_011770 | 2854320 | C | T | 31 | 6 | 2 | 0 | | | | | . | . | . | . | . | . | T | T |
| NC_011770 | 2854344 | C | G | 32 | 6 | 6 | 0 | G | G | G | G | . | . | . | . | . | . | G | G |
| NC_011770 | 2854353 | T | C | 31 | 6 | 2 | 0 | | | | | . | . | . | . | . | . | C | C |
| NC_011770 | 2854362 | T | C | 31 | 6 | 2 | 0 | | | | | . | . | . | . | . | . | C | C |
| NC_011770 | 2854366 | T | C | 31 | 6 | 2 | 0 | | | | | . | . | . | . | . | . | C | C |
| NC_011770 | 2854368 | C | G | 31 | 6 | 2 | 0 | | | | | . | . | . | . | . | . | G | G |
| NC_011770 | 2854380 | G | A | 31 | 6 | 2 | 0 | | | | | . | . | . | . | . | . | A | A |
| NC_011770 | 2854384 | G | C | 31 | 6 | 2 | 0 | | | | | . | . | . | . | . | . | C | C |

| | | | | | | | | | | | | | | | | | | | |
|---|---|---|---|---|---|---|---|---|---|---|---|---|---|---|---|---|---|---|---|
| NC_011770 | 2854392 | G | C | 32 | 6 | 2 | 0 | | | | | . | . | . | . | . | . | C | C |
| NC_011770 | 2854406 | C | T | 31 | 6 | 2 | 0 | | | | | . | . | . | . | . | . | T | T |
| NC_011770 | 2854410 | A | C | 31 | 6 | 6 | 0 | C | C | C | C | . | . | . | . | . | . | C | C |
| NC_011770 | 2854419 | G | C | 31 | 6 | 6 | 0 | C | C | C | C | . | . | . | . | . | . | C | C |
| NC_011770 | 2854428 | C | T | 31 | 6 | 2 | 0 | | | | | . | . | . | . | . | . | T | T |
| NC_011770 | 2854430 | G | A | 31 | 6 | 4 | 0 | A | A | A | A | . | . | . | . | . | . | | |
| NC_011770 | 2854434 | T | C | 29 | 6 | 6 | 0 | C | C | C | C | . | . | . | . | . | . | C | C |
| NC_011770 | 2854445 | T | G | 29 | 6 | 4 | 0 | G | G | G | G | . | . | . | . | . | . | | |
| NC_011770 | 2854449 | C | T | 29 | 6 | 2 | 0 | | | | | . | . | . | . | . | . | T | T |
| NC_011770 | 2854455 | G | A | 28 | 6 | 2 | 0 | | | | | . | . | . | . | . | . | A | A |
| NC_011770 | 2854467 | C | G | 20 | 8 | 4 | 0 | G | G | G | G | . | . | . | . | . | . | . | . |
| NC_011770 | 2854482 | A | G | 20 | 8 | 4 | 0 | G | G | G | G | . | . | . | . | . | . | . | . |
| NC_011770 | 2854581 | A | G | 21 | 7 | 1 | 0 | | | | | . | . | . | . | . | . | . | G |
| NC_011770 | 2854599 | C | T | 31 | 6 | 2 | 0 | | | | | . | . | . | . | . | . | T | T |
| NC_011770 | 2854602 | C | T | 31 | 6 | 2 | 0 | | | | | . | . | . | . | . | . | T | T |
| NC_011770 | 2854617 | C | G | 31 | 6 | 2 | 0 | | | | | . | . | . | . | . | . | G | G |
| NC_011770 | 2854628 | G | C | 32 | 6 | 2 | 0 | | | | | . | . | . | . | . | . | C | C |
| NC_011770 | 2854629 | C | T | 31 | 6 | 6 | 0 | T | T | T | T | . | . | . | . | . | . | T | T |
| NC_011770 | 2854632 | T | C | 32 | 6 | 2 | 0 | | | | | . | . | . | . | . | . | C | C |
| NC_011770 | 2854635 | G | C | 31 | 6 | 2 | 0 | | | | | . | . | . | . | . | . | C | C |
| NC_011770 | 2854638 | T | C | 32 | 6 | 2 | 0 | | | | | . | . | . | . | . | . | C | C |
| NC_011770 | 2854641 | G | A | 32 | 6 | 2 | 0 | | | | | . | . | . | . | . | . | A | A |
| NC_011770 | 2854682 | G | A | 36 | 6 | 2 | 0 | | | | | . | . | . | . | . | . | A | A |
| NC_011770 | 2854688 | A | G | 35 | 6 | 2 | 0 | | | | | . | . | . | . | . | . | G | G |
| NC_011770 | 2854734 | A | G | 35 | 6 | 6 | 0 | G | G | G | G | . | . | . | . | . | . | G | G |
| NC_011770 | 2854740 | A | G | 35 | 6 | 2 | 0 | | | | | . | . | . | . | . | . | G | G |
| NC_011770 | 2854749 | G | C | 37 | 6 | 2 | 0 | | | | | . | . | . | . | . | . | C | C |
| NC_011770 | 2854752 | C | T | 36 | 6 | 2 | 0 | | | | | . | . | . | . | . | . | T | T |
| NC_011770 | 2854758 | T | C | 36 | 6 | 2 | 0 | | | | | . | . | . | . | . | . | C | C |
| NC_011770 | 2854764 | C | G | 36 | 6 | 2 | 0 | | | | | . | . | . | . | . | . | G | G |
| NC_011770 | 2854773 | G | A | 37 | 6 | 2 | 0 | | | | | . | . | . | . | . | . | A | A |
| NC_011770 | 2854776 | C | T | 37 | 6 | 2 | 0 | | | | | . | . | . | . | . | . | T | T |
| NC_011770 | 2854788 | C | G | 39 | 6 | 2 | 0 | | | | | . | . | . | . | . | . | G | G |
| NC_011770 | 2854794 | C | G | 40 | 6 | 2 | 0 | | | | | . | . | . | . | . | . | G | G |
| NC_011770 | 2854800 | G | A | 40 | 6 | 4 | 0 | A | A | A | A | . | . | . | . | . | . | | |
| NC_011770 | 2854815 | T | C | 42 | 6 | 2 | 0 | | | | | . | . | . | . | . | . | C | C |
| NC_011770 | 2854818 | C | G | 42 | 6 | 2 | 0 | | | | | . | . | . | . | . | . | G | G |
| NC_011770 | 2854821 | A | G | 41 | 6 | 2 | 0 | | | | | . | . | . | . | . | . | G | G |
| NC_011770 | 2854839 | T | C | 41 | 6 | 2 | 0 | | | | | . | . | . | . | . | . | C | C |
| NC_011770 | 2854844 | G | A | 41 | 6 | 2 | 0 | | | | | . | . | . | . | . | . | A | A |
| NC_011770 | 2854845 | T | C | 42 | 6 | 2 | 0 | | | | | . | . | . | . | . | . | C | C |
| NC_011770 | 2854851 | G | A | 42 | 6 | 2 | 0 | | | | | . | . | . | . | . | . | A | A |
| NC_011770 | 2854860 | A | G | 42 | 6 | 6 | 0 | G | G | G | G | . | . | . | . | . | . | G | G |
| NC_011770 | 2854875 | C | T | 42 | 6 | 6 | 0 | T | T | T | T | . | . | . | . | . | . | T | T |
| NC_011770 | 2854878 | C | A | 42 | 6 | 4 | 0 | A | A | A | A | . | . | . | . | . | . | | |
| NC_011770 | 2854882 | G | A | 41 | 6 | 2 | 0 | | | | | . | . | . | . | . | . | A | A |
| NC_011770 | 2854884 | G | A | 41 | 6 | 2 | 0 | | | | | . | . | . | . | . | . | A | A |
| NC_011770 | 2854885 | A | G | 40 | 6 | 2 | 0 | | | | | . | . | . | . | . | . | G | G |
| NC_011770 | 2854886 | C | T | 40 | 6 | 4 | 0 | T | T | T | T | . | . | . | . | . | . | | |
| NC_011770 | 2854887 | G | A | 41 | 6 | 2 | 0 | | | | | . | . | . | . | . | . | A | A |
| NC_011770 | 2854890 | T | A | 40 | 6 | 2 | 0 | | | | | . | . | . | . | . | . | A | A |
| NC_011770 | 2854893 | C | G | 40 | 6 | 2 | 0 | | | | | . | . | . | . | . | . | G | G |
| NC_011770 | 2854907 | G | A | 44 | 6 | 2 | 0 | | | | | . | . | . | . | . | . | A | A |
| NC_011770 | 2854908 | T | C | 44 | 6 | 2 | 0 | | | | | . | . | . | . | . | . | C | C |
| NC_011770 | 2854922 | C | T | 44 | 6 | 2 | 0 | | | | | . | . | . | . | . | . | T | T |
| NC_011770 | 2854925 | G | A | 45 | 6 | 4 | 0 | A | A | A | A | . | . | . | . | . | . | | |
| NC_011770 | 2855959 | G | C | 37 | 6 | 4 | 0 | C | C | C | C | . | . | . | . | . | . | | |
| NC_011770 | 2855965 | A | G | 35 | 6 | 6 | 0 | G | G | G | G | . | . | . | . | . | . | G | G |
| NC_011770 | 2855968 | G | A | 35 | 6 | 4 | 0 | A | A | A | A | . | . | . | . | . | . | | |
| NC_011770 | 2855983 | A | G | 35 | 6 | 6 | 0 | G | G | G | G | . | . | . | . | . | . | G | G |
| NC_011770 | 2855995 | C | G | 37 | 6 | 4 | 0 | G | G | G | G | . | . | . | . | . | . | | |
| NC_011770 | 2855998 | G | C | 36 | 6 | 2 | 0 | | | | | . | . | . | . | . | . | C | C |

| | | | | | | | | | | | | | | | | | | | |
|---|---|---|---|---|---|---|---|---|---|---|---|---|---|---|---|---|---|---|---|
| NC_011770 | 2856001 | C | G | 37 | 6 | 2 | 0 | | | | | . | . | . | . | . | . | G | G |
| NC_011770 | 2856004 | C | T | 36 | 6 | 4 | 0 | T | T | T | T | . | . | . | . | . | . | | |
| NC_011770 | 2856031 | G | A | 38 | 6 | 2 | 0 | | | | | . | . | . | . | . | . | A | A |
| NC_011770 | 2856046 | GC | G | 39 | 6 | 4 | 0 | G | G | G | G | . | . | . | . | . | . | | |
| NC_011770 | 2856413 | G | A | 38 | 6 | 2 | 0 | | | | | . | . | . | . | . | . | A | A |
| NC_011770 | 2856414 | G | T | 38 | 6 | 2 | 0 | | | | | . | . | . | . | . | . | T | T |
| NC_011770 | 2857052 | T | G | 31 | 6 | 2 | 0 | | | G | G | . | . | . | . | . | . | | |
| NC_011770 | 2857067 | G | C | 31 | 6 | 2 | 0 | | | | | . | . | . | . | . | . | C | C |
| NC_011770 | 2857068 | G | T | 32 | 6 | 2 | 0 | | | | | . | . | . | . | . | . | T | T |
| NC_011770 | 2857069 | T | A | 30 | 6 | 2 | 0 | | | | | . | . | . | . | . | . | A | A |
| NC_011770 | 2857093 | T | C | 30 | 6 | 2 | 0 | | | | | . | . | . | . | . | . | C | C |
| NC_011770 | 2857114 | T | C | 28 | 6 | 2 | 0 | | | | | . | . | . | . | . | . | C | C |
| NC_011770 | 2857115 | G | T | 27 | 6 | 2 | 0 | | | | | . | . | . | . | . | . | T | T |
| NC_011770 | 2857121 | C | T | 26 | 6 | 2 | 0 | | | | | . | . | . | . | . | . | T | T |
| NC_011770 | 2857166 | G | T | 19 | 8 | 4 | 0 | T | T | T | T | . | . | . | . | . | . | . | . |
| NC_011770 | 2859456 | T | G | 27 | 8 | 4 | 0 | G | G | G | G | . | . | . | . | . | . | . | . |
| NC_011770 | 2859471 | A | G | 28 | 8 | 2 | 0 | G | G | | | . | . | . | . | . | . | . | . |
| NC_011770 | 2859629 | G | C | 29 | 8 | 2 | 0 | C | C | | | . | . | . | . | . | . | . | . |
| NC_011770 | 2864574 | A | T | 34 | 6 | 4 | 0 | T | T | T | T | . | . | . | . | . | . | | |
| NC_011770 | 2864613 | C | A | 32 | 6 | 6 | 0 | A | A | A | A | . | . | . | . | . | . | A | A |
| NC_011770 | 2864623 | T | A | 31 | 6 | 4 | 0 | A | A | A | A | . | . | . | . | . | . | | |
| NC_011770 | 2864746 | G | A | 37 | 6 | 6 | 0 | A | A | A | A | . | . | . | . | . | . | A | A |
| NC_011770 | 2864761 | T | A | 35 | 6 | 2 | 0 | | | | | . | . | . | . | . | . | A | A |
| NC_011770 | 2864764 | T | C | 35 | 6 | 6 | 0 | C | C | C | C | . | . | . | . | . | . | A | A |
| NC_011770 | 2864767 | A | C | 35 | 6 | 6 | 0 | C | C | C | C | . | . | . | . | . | . | C | C |
| NC_011770 | 2864772 | G | A | 36 | 6 | 2 | 0 | | | | | . | . | . | . | . | . | A | A |
| NC_011770 | 2864779 | C | T | 36 | 6 | 2 | 0 | | | | | . | . | . | . | . | . | T | T |
| NC_011770 | 2864791 | T | C | 36 | 6 | 6 | 0 | C | C | C | C | . | . | . | . | . | . | C | C |
| NC_011770 | 2864812 | C | T | 36 | 6 | 2 | 0 | | | | | . | . | . | . | . | . | T | T |
| NC_011770 | 2864821 | G | C | 35 | 6 | 2 | 0 | | | | | . | . | . | . | . | . | C | C |
| NC_011770 | 2864839 | A | G | 35 | 6 | 2 | 0 | | | | | . | . | . | . | . | . | G | G |
| NC_011770 | 2864854 | T | C | 36 | 6 | 2 | 0 | | | | | . | . | . | . | . | . | C | C |
| NC_011770 | 2864870 | A | G | 38 | 6 | 2 | 0 | | | | | . | . | . | . | . | . | G | G |
| NC_011770 | 2864872 | T | C | 36 | 6 | 4 | 0 | C | C | C | C | . | . | . | . | . | . | | |
| NC_011770 | 2864899 | G | C | 38 | 6 | 2 | 0 | | | | | . | . | . | . | . | . | C | C |
| NC_011770 | 2864900 | C | G | 38 | 6 | 2 | 0 | | | | | . | . | . | . | . | . | G | G |
| NC_011770 | 2864901 | T | C | 38 | 6 | 2 | 0 | | | | | . | . | . | . | . | . | C | C |
| NC_011770 | 2864906 | C | T | 38 | 6 | 2 | 0 | | | | | . | . | . | . | . | . | T | T |
| NC_011770 | 2864916 | C | G | 38 | 6 | 6 | 0 | G | G | G | G | . | . | . | . | . | . | G | G |
| NC_011770 | 2864924 | G | A | 37 | 6 | 6 | 0 | A | A | A | A | . | . | . | . | . | . | A | A |
| NC_011770 | 2864951 | T | G | 39 | 6 | 2 | 0 | | | | | . | . | . | . | . | . | G | G |
| NC_011770 | 2864958 | A | G | 39 | 6 | 2 | 0 | | | | | . | . | . | . | . | . | G | G |
| NC_011770 | 2864962 | C | G | 38 | 6 | 2 | 0 | | | | | . | . | . | . | . | . | G | G |
| NC_011770 | 2864968 | C | A | 39 | 6 | 2 | 0 | | | | | . | . | . | . | . | . | A | A |
| NC_011770 | 2864971 | G | A | 40 | 6 | 2 | 0 | | | | | . | . | . | . | . | . | A | A |
| NC_011770 | 2864980 | C | T | 41 | 6 | 2 | 0 | | | | | . | . | . | . | . | . | T | T |
| NC_011770 | 2864986 | C | G | 41 | 6 | 2 | 0 | | | | | . | . | . | . | . | . | G | G |
| NC_011770 | 2864989 | G | T | 40 | 6 | 2 | 0 | | | | | . | . | . | . | . | . | T | T |
| NC_011770 | 2865000 | G | A | 43 | 6 | 2 | 0 | | | | | . | . | . | . | . | . | A | A |
| NC_011770 | 2865019 | G | C | 43 | 6 | 2 | 0 | | | | | . | . | . | . | . | . | C | C |
| NC_011770 | 2865031 | C | G | 43 | 6 | 2 | 0 | | | | | . | . | . | . | . | . | G | G |
| NC_011770 | 2865061 | G | A | 43 | 6 | 2 | 0 | | | | | . | . | . | . | . | . | A | A |
| NC_011770 | 2865067 | G | A | 43 | 6 | 2 | 0 | | | | | . | . | . | . | . | . | A | A |
| NC_011770 | 2865070 | T | C | 43 | 6 | 2 | 0 | | | | | . | . | . | . | . | . | C | C |
| NC_011770 | 2865081 | G | A | 44 | 6 | 2 | 0 | | | | | . | . | . | . | . | . | A | A |
| NC_011770 | 2865094 | G | A | 43 | 6 | 2 | 0 | | | | | . | . | . | . | . | . | A | A |
| NC_011770 | 2865169 | G | A | 39 | 6 | 4 | 0 | A | A | A | A | . | . | . | . | . | . | | |
| NC_011770 | 2865172 | G | C | 38 | 6 | 2 | 0 | | | | | . | . | . | . | . | . | C | C |
| NC_011770 | 2865178 | C | T | 37 | 6 | 2 | 0 | | | | | . | . | . | . | . | . | T | T |
| NC_011770 | 2865181 | G | A | 37 | 6 | 2 | 0 | | | | | . | . | . | . | . | . | A | A |
| NC_011770 | 2865186 | G | A | 37 | 6 | 2 | 0 | | | | | . | . | . | . | . | . | A | A |
| NC_011770 | 2865187 | C | G | 36 | 6 | 2 | 0 | | | | | . | . | . | . | . | . | G | G |

| | | | | | | | | | | | | | | | | | | | |
|---|---|---|---|---|---|---|---|---|---|---|---|---|---|---|---|---|---|---|---|
| NC_011770 | 2865190 | G | A | 37 | 6 | 2 | 0 | | | | | . | . | . | . | . | . | A | A |
| NC_011770 | 2865196 | C | T | 38 | 6 | 4 | 0 | T | T | T | T | . | . | . | . | . | . | | |
| NC_011770 | 2865219 | G | A | 36 | 6 | 4 | 0 | A | A | A | A | . | . | . | . | . | . | | |
| NC_011770 | 2865220 | C | G | 36 | 6 | 6 | 0 | G | G | G | G | . | . | . | . | . | . | G | G |
| NC_011770 | 2865223 | C | T | 35 | 6 | 4 | 0 | T | T | T | T | . | . | . | . | . | . | | |
| NC_011770 | 2865229 | C | T | 37 | 6 | 4 | 0 | T | T | T | T | . | . | . | . | . | . | | |
| NC_011770 | 2865232 | G | A | 38 | 6 | 2 | 0 | | | | | . | . | . | . | . | . | A | A |
| NC_011770 | 2865258 | C | T | 36 | 6 | 4 | 0 | T | T | T | T | . | . | . | . | . | . | | |
| NC_011770 | 2865262 | A | G | 36 | 6 | 2 | 0 | | | | | . | . | . | . | . | . | G | G |
| NC_011770 | 2867965 | G | A | 10 | 6 | 2 | 0 | | | | | . | . | . | . | . | . | A | A |
| NC_011770 | 2867974 | CGTGAA | C | 10 | 6 | 2 | 0 | | | | | . | . | . | . | . | . | C | C |
| NC_011770 | 2867982 | CGAGA | C | 9 | 6 | 2 | 0 | | | | | . | . | . | . | . | . | C | C |
| NC_011770 | 2867993 | G | A | 8 | 7 | 1 | 0 | | | | | . | . | . | . | . | . | . | A |
| NC_011770 | 2868000 | G | A | 13 | 6 | 2 | 0 | | | | | . | . | . | . | . | . | A | A |
| NC_011770 | 2868004 | C | T | 12 | 6 | 2 | 0 | | | | | . | . | . | . | . | . | T | T |
| NC_011770 | 2869316 | G | GTC | 28 | 6 | 2 | 0 | | | | | . | . | . | . | . | . | GTC | GTC |
| NC_011770 | 2869318 | G | T | 26 | 6 | 2 | 0 | | | | | . | . | . | . | . | . | T | T |
| NC_011770 | 2869327 | G | A | 28 | 6 | 2 | 0 | | | | | . | . | . | . | . | . | A | A |
| NC_011770 | 2870438 | G | T | 19 | 6 | 6 | 0 | T | T | T | T | . | . | . | . | . | . | T | T |
| NC_011770 | 2870442 | T | A | 19 | 6 | 2 | 0 | | | | | . | . | . | . | . | . | A | A |
| NC_011770 | 2870445 | T | C | 18 | 6 | 6 | 0 | C | C | C | C | . | . | . | . | . | . | C | C |
| NC_011770 | 2870446 | G | A | 17 | 6 | 6 | 0 | A | A | A | A | . | . | . | . | . | . | A | A |
| NC_011770 | 2870452 | C | T | 17 | 6 | 6 | 0 | T | T | T | T | . | . | . | . | . | . | T | T |
| NC_011770 | 2870456 | A | T | 17 | 6 | 2 | 0 | | | | | . | . | . | . | . | . | T | T |
| NC_011770 | 2870457 | C | G | 17 | 6 | 6 | 0 | G | G | G | G | . | . | . | . | . | . | G | G |
| NC_011770 | 2870464 | A | C | 19 | 6 | 4 | 0 | C | C | C | C | . | . | . | . | . | . | | |
| NC_011770 | 2870467 | C | G | 19 | 6 | 4 | 0 | G | G | G | G | . | . | . | . | . | . | | |
| NC_011770 | 2870470 | C | G | 18 | 6 | 6 | 0 | G | G | G | G | . | . | . | . | . | . | T | T |
| NC_011770 | 2870482 | A | G | 18 | 6 | 2 | 0 | | | | | . | . | . | . | . | . | G | G |
| NC_011770 | 2870485 | G | A | 8 | 10 | 2 | 0 | . | . | . | . | . | . | . | . | . | . | A | A |
| NC_011770 | 2870491 | C | T | 8 | 10 | 2 | 0 | . | . | . | . | . | . | . | . | . | . | T | T |
| NC_011770 | 2870500 | C | A | 8 | 10 | 2 | 0 | . | . | . | . | . | . | . | . | . | . | A | A |
| NC_011770 | 2870520 | C | T | 8 | 10 | 2 | 0 | . | . | . | . | . | . | . | . | . | . | T | T |
| NC_011770 | 2870524 | A | C | 8 | 10 | 2 | 0 | . | . | . | . | . | . | . | . | . | . | C | C |
| NC_011770 | 2870541 | A | G | 8 | 10 | 2 | 0 | . | . | . | . | . | . | . | . | . | . | G | G |
| NC_011770 | 2870551 | C | G | 8 | 10 | 2 | 0 | . | . | . | . | . | . | . | . | . | . | G | G |
| NC_011770 | 2870590 | C | G | 7 | 10 | 2 | 0 | . | . | . | . | . | . | . | . | . | . | G | G |
| NC_011770 | 2870610 | C | G | 7 | 10 | 2 | 0 | . | . | . | . | . | . | . | . | . | . | G | G |
| NC_011770 | 2870611 | A | G | 7 | 10 | 2 | 0 | . | . | . | . | . | . | . | . | . | . | G | G |
| NC_011770 | 2870614 | C | T | 7 | 10 | 2 | 0 | . | . | . | . | . | . | . | . | . | . | T | T |
| NC_011770 | 2870617 | C | T | 7 | 10 | 2 | 0 | . | . | . | . | . | . | . | . | . | . | T | T |
| NC_011770 | 2870619 | A | C | 6 | 10 | 2 | 0 | . | . | . | . | . | . | . | . | . | . | C | C |
| NC_011770 | 2873006 | AT | A | 49 | 6 | 2 | 0 | | | | | . | . | . | . | . | . | A | A |
| NC_011770 | 2873035 | A | G | 53 | 6 | 2 | 0 | | | | | . | . | . | . | . | . | G | G |
| NC_011770 | 2873056 | A | G | 51 | 6 | 4 | 0 | G | G | G | G | . | . | . | . | . | . | | |
| NC_011770 | 2873058 | A | T | 50 | 6 | 2 | 0 | | | | | . | . | . | . | . | . | T | T |
| NC_011770 | 2873059 | G | A | 51 | 6 | 2 | 0 | | | | | . | . | . | . | . | . | A | A |
| NC_011770 | 2873065 | A | G | 49 | 6 | 4 | 0 | G | G | G | G | . | . | . | . | . | . | | |
| NC_011770 | 2873074 | T | C | 48 | 6 | 6 | 0 | C | C | C | C | . | . | . | . | . | . | C | C |
| NC_011770 | 2873077 | G | A | 47 | 6 | 6 | 0 | A | A | A | A | . | . | . | . | . | . | C | C |
| NC_011770 | 2873089 | G | A | 48 | 6 | 4 | 0 | A | A | A | A | . | . | . | . | . | . | | |
| NC_011770 | 2873098 | T | C | 46 | 6 | 2 | 0 | | | C | C | . | . | . | . | . | . | | |
| NC_011770 | 2873102 | C | T | 46 | 6 | 6 | 0 | T | T | T | T | . | . | . | . | . | . | T | T |
| NC_011770 | 2873107 | C | A | 45 | 6 | 2 | 0 | | | | | . | . | . | . | . | . | A | A |
| NC_011770 | 2873110 | A | G | 46 | 6 | 6 | 0 | G | G | G | G | . | . | . | . | . | . | G | G |
| NC_011770 | 2873112 | G | C | 47 | 6 | 2 | 0 | | | | | . | . | . | . | . | . | C | C |
| NC_011770 | 2873445 | A | G | 47 | 6 | 4 | 0 | G | G | G | G | . | . | . | . | . | . | | |
| NC_011770 | 2873446 | G | T | 47 | 6 | 4 | 0 | T | T | T | T | . | . | . | . | . | . | | |
| NC_011770 | 2873453 | A | G | 47 | 6 | 4 | 0 | G | G | G | G | . | . | . | . | . | . | | |
| NC_011770 | 2873493 | T | A | 49 | 6 | 2 | 0 | | | | | . | . | . | . | . | . | A | A |
| NC_011770 | 2873495 | G | A | 50 | 6 | 2 | 0 | | | | | . | . | . | . | . | . | A | A |
| NC_011770 | 2873500 | G | A | 50 | 6 | 4 | 0 | A | A | A | A | . | . | . | . | . | . | | |

| | | | | | | | | | | | | | | | | | | | |
|---|---|---|---|---|---|---|---|---|---|---|---|---|---|---|---|---|---|---|---|
| NC_011770 | 2873515 | G | GA | 49 | 6 | 2 | 0 | | | | | . | . | . | . | . | . | GA | GA |
| NC_011770 | 2873517 | C | G | 30 | 6 | 1 | 0 | | | | | . | . | . | . | . | . | . | G |
| NC_011770 | 2873855 | G | T | 48 | 6 | 2 | 0 | | | | | . | . | . | . | . | . | T | T |
| NC_011770 | 2873856 | T | G | 47 | 6 | 2 | 0 | | | | | . | . | . | . | . | . | G | G |
| NC_011770 | 2875026 | A | T | 38 | 6 | 4 | 0 | T | T | T | T | . | . | . | . | . | . | | |
| NC_011770 | 2875027 | C | G | 39 | 6 | 6 | 0 | G | G | G | G | . | . | . | . | . | . | G | G |
| NC_011770 | 2875035 | T | C | 39 | 6 | 6 | 0 | C | C | C | C | . | . | . | . | . | . | C | C |
| NC_011770 | 2875041 | A | G | 40 | 6 | 2 | 0 | | | | | . | . | . | . | . | . | G | G |
| NC_011770 | 2875044 | G | A | 39 | 6 | 2 | 0 | | | | | . | . | . | . | . | . | A | A |
| NC_011770 | 2875057 | GGT | G | 39 | 6 | 2 | 0 | | | | | . | . | . | . | . | . | G | G |
| NC_011770 | 2875729 | A | G | 16 | 10 | 2 | 0 | . | . | . | . | . | . | . | . | . | . | G | G |
| NC_011770 | 2875748 | C | T | 15 | 10 | 2 | 0 | . | . | . | . | . | . | . | . | . | . | T | T |
| NC_011770 | 2875749 | T | C | 15 | 10 | 2 | 0 | . | . | . | . | . | . | . | . | . | . | C | C |
| NC_011770 | 2875756 | A | G | 16 | 10 | 2 | 0 | . | . | . | . | . | . | . | . | . | . | G | G |
| NC_011770 | 2875766 | C | CGG | 15 | 10 | 2 | 0 | . | . | . | . | . | . | . | . | . | . | CGG | CGG |
| NC_011770 | 2876050 | C | T | 19 | 10 | 2 | 0 | . | . | . | . | . | . | . | . | . | . | T | T |
| NC_011770 | 2876484 | C | G | 13 | 10 | 2 | 0 | . | . | . | . | . | . | . | . | . | . | G | G |
| NC_011770 | 2876485 | G | A | 13 | 10 | 2 | 0 | . | . | . | . | . | . | . | . | . | . | A | A |
| NC_011770 | 2876487 | A | G | 13 | 10 | 2 | 0 | . | . | . | . | . | . | . | . | . | . | G | G |
| NC_011770 | 2876492 | T | A | 13 | 10 | 2 | 0 | . | . | . | . | . | . | . | . | . | . | A | A |
| NC_011770 | 2876497 | T | C | 13 | 10 | 2 | 0 | . | . | . | . | . | . | . | . | . | . | C | C |
| NC_011770 | 2876498 | G | A | 13 | 10 | 2 | 0 | . | . | . | . | . | . | . | . | . | . | A | A |
| NC_011770 | 2876499 | A | G | 13 | 10 | 2 | 0 | . | . | . | . | . | . | . | . | . | . | G | G |
| NC_011770 | 2876546 | A | G | 13 | 10 | 2 | 0 | . | . | . | . | . | . | . | . | . | . | G | G |
| NC_011770 | 2876548 | C | G | 12 | 10 | 2 | 0 | . | . | . | . | . | . | . | . | . | . | G | G |
| NC_011770 | 2876557 | A | T | 0 | 11 | 1 | 0 | . | . | . | . | . | . | . | . | . | . | T | . |
| NC_011770 | 2876935 | A | G | 26 | 6 | 6 | 0 | G | G | G | G | . | . | . | . | . | . | G | G |
| NC_011770 | 2876937 | T | A | 26 | 6 | 6 | 0 | A | A | A | A | . | . | . | . | . | . | A | A |
| NC_011770 | 2876938 | G | A | 26 | 6 | 6 | 0 | A | A | A | A | . | . | . | . | . | . | A | A |
| NC_011770 | 2876942 | A | G | 26 | 6 | 6 | 0 | G | G | G | G | . | . | . | . | . | . | G | G |
| NC_011770 | 2876943 | G | A | 26 | 6 | 2 | 0 | | | | | . | . | . | . | . | . | A | A |
| NC_011770 | 2876946 | C | G | 26 | 6 | 4 | 0 | G | G | G | G | . | . | . | . | . | . | | |
| NC_011770 | 2876948 | T | C | 26 | 6 | 6 | 0 | C | C | C | C | . | . | . | . | . | . | A | A |
| NC_011770 | 2876949 | C | T | 27 | 6 | 4 | 0 | T | T | T | T | . | . | . | . | . | . | | |
| NC_011770 | 2876958 | G | A | 29 | 6 | 4 | 0 | A | A | A | A | . | . | . | . | . | . | | |
| NC_011770 | 2876961 | A | G | 29 | 6 | 6 | 0 | G | G | G | G | . | . | . | . | . | . | G | G |
| NC_011770 | 2876964 | G | A | 29 | 6 | 4 | 0 | A | A | A | A | . | . | . | . | . | . | | |
| NC_011770 | 2876967 | A | G | 29 | 6 | 4 | 0 | G | G | G | G | . | . | . | . | . | . | | |
| NC_011770 | 2876972 | G | A | 34 | 6 | 2 | 0 | | | | | . | . | . | . | . | . | A | A |
| NC_011770 | 2876996 | C | G | 33 | 6 | 2 | 0 | | | | | . | . | . | . | . | . | G | G |
| NC_011770 | 2877003 | C | T | 33 | 6 | 6 | 0 | T | T | T | T | . | . | . | . | . | . | T | T |
| NC_011770 | 2877010 | G | A | 35 | 6 | 2 | 0 | | | | | . | . | . | . | . | . | A | A |
| NC_011770 | 2878756 | A | G | 21 | 6 | 6 | 0 | G | G | G | G | . | . | . | . | . | . | G | G |
| NC_011770 | 2879778 | A | G | 29 | 6 | 4 | 0 | G | G | G | G | . | . | . | . | . | . | | |
| NC_011770 | 2879780 | G | A | 29 | 6 | 6 | 0 | A | A | A | A | . | . | . | . | . | . | A | A |
| NC_011770 | 2879799 | A | T | 29 | 6 | 6 | 0 | T | T | T | T | . | . | . | . | . | . | T | T |
| NC_011770 | 2879834 | G | A | 29 | 6 | 4 | 0 | A | A | A | A | . | . | . | . | . | . | | |
| NC_011770 | 2879842 | C | A | 26 | 6 | 2 | 0 | | | | | . | . | . | . | . | . | A | A |
| NC_011770 | 2879851 | C | T | 27 | 6 | 2 | 0 | | | | | . | . | . | . | . | . | T | T |
| NC_011770 | 2879868 | C | T | 26 | 6 | 2 | 0 | | | | | . | . | . | . | . | . | T | T |
| NC_011770 | 2879884 | G | A | 26 | 6 | 6 | 0 | A | A | A | A | . | . | . | . | . | . | A | A |
| NC_011770 | 2879885 | A | G | 26 | 6 | 4 | 0 | G | G | G | G | . | . | . | . | . | . | | |
| NC_011770 | 2879903 | C | T | 24 | 6 | 6 | 0 | T | T | T | T | . | . | . | . | . | . | T | T |
| NC_011770 | 2879913 | G | C | 23 | 6 | 6 | 0 | C | C | C | C | . | . | . | . | . | . | C | C |
| NC_011770 | 2879930 | G | A | 23 | 6 | 6 | 0 | A | A | A | A | . | . | . | . | . | . | A | A |
| NC_011770 | 2879942 | C | T | 22 | 6 | 4 | 0 | T | T | T | T | . | . | . | . | . | . | | |
| NC_011770 | 2879947 | C | T | 21 | 6 | 6 | 0 | T | T | T | T | . | . | . | . | . | . | T | T |
| NC_011770 | 2879951 | G | C | 21 | 6 | 4 | 0 | C | C | C | C | . | . | . | . | . | . | | |
| NC_011770 | 2879955 | C | T | 21 | 6 | 4 | 0 | T | T | T | T | . | . | . | . | . | . | | |
| NC_011770 | 2879968 | G | A | 1 | 9 | 1 | 0 | . | . | A | . | . | . | | | . | . | . | . |
| NC_011770 | 2879972 | C | A | 1 | 9 | 1 | 0 | . | . | A | . | . | . | | | . | . | . | . |
| NC_011770 | 2879973 | A | G | 1 | 9 | 1 | 0 | . | . | G | . | . | . | | | . | . | . | . |

| | | | | | | | | | | | | | | | | | | | |
|---|---|---|---|---|---|---|---|---|---|---|---|---|---|---|---|---|---|---|---|
| NC_011770 | 2880844 | A | G | 20 | 6 | 6 | 0 | G | G | G | G | . | . | . | . | . | . | G | G |
| NC_011770 | 2880851 | A | G | 18 | 6 | 2 | 0 | | | | | . | . | . | . | . | . | G | G |
| NC_011770 | 2885844 | A | T | 11 | 6 | 6 | 0 | T | T | T | T | . | . | . | . | . | . | T | T |
| NC_011770 | 2885856 | A | AG | 52 | 6 | 4 | 0 | AG | AG | AG | AG | . | . | . | . | . | . | | |
| NC_011770 | 2885857 | A | G | 49 | 6 | 2 | 0 | | | | | . | . | . | . | . | . | G | G |
| NC_011770 | 2885903 | C | T | 47 | 6 | 4 | 0 | T | T | T | T | . | . | . | . | . | . | | |
| NC_011770 | 2885907 | G | A | 49 | 6 | 2 | 0 | | | | | . | . | . | . | . | . | A | A |
| NC_011770 | 2885909 | T | C | 49 | 6 | 2 | 0 | | | | | . | . | . | . | . | . | C | C |
| NC_011770 | 2885915 | C | G | 49 | 6 | 2 | 0 | | | | | . | . | . | . | . | . | G | G |
| NC_011770 | 2885920 | G | C | 49 | 6 | 6 | 0 | C | C | C | C | . | . | . | . | . | . | C | C |
| NC_011770 | 2885925 | G | A | 48 | 6 | 2 | 0 | | | | | . | . | . | . | . | . | A | A |
| NC_011770 | 2885927 | A | G | 46 | 6 | 4 | 0 | G | G | G | G | . | . | . | . | . | . | | |
| NC_011770 | 2885929 | G | A | 46 | 6 | 4 | 0 | A | A | A | A | . | . | . | . | . | . | | |
| NC_011770 | 2885932 | G | A | 45 | 6 | 4 | 0 | A | A | A | A | . | . | . | . | . | . | | |
| NC_011770 | 2885933 | A | C | 45 | 6 | 4 | 0 | C | C | C | C | . | . | . | . | . | . | | |
| NC_011770 | 2885935 | A | T | 45 | 6 | 4 | 0 | T | T | T | T | . | . | . | . | . | . | | |
| NC_011770 | 2885948 | T | C | 48 | 6 | 4 | 0 | C | C | C | C | . | . | . | . | . | . | | |
| NC_011770 | 2885955 | C | T | 47 | 6 | 4 | 0 | T | T | T | T | . | . | . | . | . | . | | |
| NC_011770 | 2885956 | G | A | 47 | 6 | 2 | 0 | | | | | . | . | . | . | . | . | A | A |
| NC_011770 | 2885967 | C | T | 48 | 6 | 6 | 0 | T | T | T | T | . | . | . | . | . | . | T | T |
| NC_011770 | 2885976 | C | T | 47 | 6 | 4 | 0 | T | T | T | T | . | . | . | . | . | . | | |
| NC_011770 | 2885978 | A | G | 46 | 6 | 6 | 0 | G | G | G | G | . | . | . | . | . | . | G | G |
| NC_011770 | 2885984 | T | A | 47 | 6 | 2 | 0 | | | | | . | . | . | . | . | . | A | A |
| NC_011770 | 2885994 | C | T | 46 | 6 | 6 | 0 | T | T | T | T | . | . | . | . | . | . | T | T |
| NC_011770 | 2886016 | G | A | 43 | 6 | 2 | 0 | | | | | . | . | . | . | . | . | A | A |
| NC_011770 | 2886022 | C | A | 41 | 6 | 6 | 0 | A | A | A | A | . | . | . | . | . | . | G | G |
| NC_011770 | 2886028 | C | T | 29 | 6 | 2 | 0 | | | | | . | . | . | . | . | . | T | T |
| NC_011770 | 2886031 | G | GGAAT | 15 | 6 | 2 | 0 | | | | | . | . | . | . | . | . | GGAAT | GGAAT |
| NC_011770 | 2886036 | GGTCT | G | 18 | 6 | 6 | 0 | G | G | G | G | . | . | . | . | . | . | G | G |
| NC_011770 | 2886052 | G | A | 26 | 6 | 2 | 0 | | | | | . | . | . | . | . | . | A | A |
| NC_011770 | 2886057 | C | T | 26 | 6 | 2 | 0 | | | | | . | . | . | . | . | . | T | T |
| NC_011770 | 2886058 | G | C | 26 | 6 | 2 | 0 | | | | | . | . | . | . | . | . | C | C |
| NC_011770 | 2886063 | C | A | 25 | 6 | 2 | 0 | | | | | . | . | . | . | . | . | A | A |
| NC_011770 | 2886064 | G | C | 26 | 6 | 4 | 0 | C | C | C | C | . | . | . | . | . | . | | |
| NC_011770 | 2886066 | A | G | 25 | 6 | 2 | 0 | | | | | . | . | . | . | . | . | G | G |
| NC_011770 | 2886067 | C | T | 26 | 6 | 6 | 0 | T | T | T | T | . | . | . | . | . | . | T | T |
| NC_011770 | 2886069 | A | G | 26 | 6 | 6 | 0 | G | G | G | G | . | . | . | . | . | . | G | G |
| NC_011770 | 2886072 | A | C | 25 | 6 | 6 | 0 | C | C | C | C | . | . | . | . | . | . | C | C |
| NC_011770 | 2886076 | T | C | 26 | 6 | 6 | 0 | C | C | C | C | . | . | . | . | . | . | C | C |
| NC_011770 | 2886082 | T | A | 33 | 6 | 4 | 0 | A | A | A | A | . | . | . | . | . | . | | |
| NC_011770 | 2886084 | T | C | 33 | 6 | 6 | 0 | C | C | C | C | . | . | . | . | . | . | C | C |
| NC_011770 | 2886092 | G | A | 37 | 6 | 2 | 0 | | | | | . | . | . | . | . | . | A | A |
| NC_011770 | 2886094 | A | G | 38 | 6 | 4 | 0 | G | G | G | G | . | . | . | . | . | . | | |
| NC_011770 | 2886109 | G | GA | 40 | 6 | 2 | 0 | | | | | . | . | . | . | . | . | GA | GA |
| NC_011770 | 2886117 | C | CT | 39 | 6 | 2 | 0 | | | | | . | . | . | . | . | . | CT | CT |
| NC_011770 | 2886121 | A | G | 2 | 6 | 5 | 0 | G | G | . | G | . | . | . | . | . | . | T | T |
| NC_011770 | 2886122 | G | T | 20 | 6 | 2 | 0 | | | | | . | . | . | . | . | . | T | T |
| NC_011770 | 2886123 | GAA | G | 20 | 6 | 4 | 0 | G | G | G | G | . | . | . | . | . | . | | |
| NC_011770 | 2886124 | A | G | 23 | 6 | 2 | 0 | . | . | . | . | . | . | . | . | . | . | G | G |
| NC_011770 | 2886125 | A | G | 23 | 6 | 2 | 0 | . | . | . | . | . | . | . | . | . | . | G | G |
| NC_011770 | 2887242 | G | A | 27 | 6 | 4 | 0 | A | A | A | A | . | . | . | . | . | . | | |
| NC_011770 | 2887244 | T | A | 16 | 7 | 1 | 0 | | | | | . | . | . | . | . | . | A | . |
| NC_011770 | 2887270 | A | C | 13 | 8 | 4 | 0 | C | C | C | C | . | . | . | . | . | . | . | . |
| NC_011770 | 2887274 | C | T | 13 | 8 | 4 | 0 | T | T | T | T | . | . | . | . | . | . | . | . |
| NC_011770 | 2887291 | G | GA | 13 | 8 | 4 | 0 | GA | GA | GA | GA | . | . | . | . | . | . | . | . |
| NC_011770 | 2887304 | G | T | 12 | 8 | 4 | 0 | T | T | T | T | . | . | . | . | . | . | . | . |
| NC_011770 | 2888011 | A | C | 14 | 10 | 2 | 0 | . | . | . | . | . | . | . | . | . | . | C | C |
| NC_011770 | 2888012 | A | T | 13 | 10 | 2 | 0 | . | . | . | . | . | . | . | . | . | . | T | T |
| NC_011770 | 2888028 | AT | A | 12 | 10 | 2 | 0 | . | . | . | . | . | . | . | . | . | . | A | A |
| NC_011770 | 2888032 | A | G | 12 | 10 | 2 | 0 | . | . | . | . | . | . | . | . | . | . | G | G |
| NC_011770 | 2888045 | T | A | 11 | 10 | 2 | 0 | . | . | . | . | . | . | . | . | . | . | A | A |
| NC_011770 | 2888059 | A | G | 11 | 10 | 2 | 0 | . | . | . | . | . | . | . | . | . | . | G | G |

| | | | | | | | | | | | | | | | | | | | |
|---|---|---|---|---|---|---|---|---|---|---|---|---|---|---|---|---|---|---|---|
| NC_011770 | 2888182 | A | G | 0 | 10 | 2 | 0 | . | . | . | . | . | . | . | . | . | . | G | G |
| NC_011770 | 2888184 | G | A | 0 | 10 | 2 | 0 | . | . | . | . | . | . | . | . | . | . | A | A |
| NC_011770 | 2888591 | G | A | 41 | 6 | 2 | 0 | | | | | . | . | . | . | . | . | A | A |
| NC_011770 | 2888594 | C | T | 40 | 6 | 2 | 0 | | | | | . | . | . | . | . | . | T | T |
| NC_011770 | 2889024 | G | A | 37 | 6 | 2 | 0 | | | | | . | . | . | . | . | . | A | A |
| NC_011770 | 2889034 | A | G | 26 | 7 | 1 | 0 | | | | | . | . | . | . | . | . | . | G |
| NC_011770 | 2893795 | A | G | 22 | 8 | 4 | 0 | G | G | G | G | . | . | . | . | . | . | . | . |
| NC_011770 | 2895845 | A | G | 27 | 6 | 2 | 0 | | | | | . | . | . | . | . | . | G | G |
| NC_011770 | 2895866 | T | A | 28 | 6 | 2 | 0 | | | | | . | . | . | . | . | . | A | A |
| NC_011770 | 2895891 | C | G | 30 | 6 | 2 | 0 | | | | | . | . | . | . | . | . | G | G |
| NC_011770 | 2895901 | T | C | 29 | 6 | 2 | 0 | | | | | . | . | . | . | . | . | C | C |
| NC_011770 | 2895907 | C | T | 29 | 6 | 2 | 0 | | | | | . | . | . | . | . | . | T | T |
| NC_011770 | 2895923 | A | G | 28 | 6 | 2 | 0 | | | | | . | . | . | . | . | . | G | G |
| NC_011770 | 2895936 | AG | A | 27 | 6 | 2 | 0 | | | | | . | . | . | . | . | . | A | A |
| NC_011770 | 2895943 | A | G | 27 | 6 | 2 | 0 | | | | | . | . | . | . | . | . | G | G |
| NC_011770 | 2895949 | C | T | 28 | 6 | 2 | 0 | | | | | . | . | . | . | . | . | T | T |
| NC_011770 | 2895958 | A | G | 28 | 6 | 2 | 0 | | | | | . | . | . | . | . | . | G | G |
| NC_011770 | 2895969 | G | A | 25 | 6 | 2 | 0 | | | | | . | . | . | . | . | . | A | A |
| NC_011770 | 2895970 | T | G | 25 | 6 | 2 | 0 | | | | | . | . | . | . | . | . | G | G |
| NC_011770 | 2895972 | G | A | 25 | 6 | 2 | 0 | | | | | . | . | . | . | . | . | A | A |
| NC_011770 | 2895973 | T | G | 25 | 6 | 2 | 0 | | | | | . | . | . | . | . | . | G | G |
| NC_011770 | 2895978 | C | T | 24 | 6 | 2 | 0 | | | | | . | . | . | . | . | . | T | T |
| NC_011770 | 2895983 | A | G | 25 | 6 | 2 | 0 | | | | | . | . | . | . | . | . | G | G |
| NC_011770 | 2896016 | C | T | 24 | 6 | 2 | 0 | | | | | . | . | . | . | . | . | T | T |
| NC_011770 | 2896018 | G | A | 24 | 6 | 2 | 0 | | | | | . | . | . | . | . | . | A | A |
| NC_011770 | 2896020 | T | C | 24 | 6 | 2 | 0 | | | | | . | . | . | . | . | . | C | C |
| NC_011770 | 2896021 | G | A | 24 | 6 | 2 | 0 | | | | | . | . | . | . | . | . | A | A |
| NC_011770 | 2896022 | G | A | 24 | 6 | 2 | 0 | | | | | . | . | . | . | . | . | A | A |
| NC_011770 | 2896040 | AT | A | 26 | 6 | 2 | 0 | | | | | . | . | . | . | . | . | A | A |
| NC_011770 | 2896077 | A | G | 13 | 6 | 2 | 0 | | | | | . | . | . | . | . | . | G | G |
| NC_011770 | 2896093 | G | A | 13 | 6 | 2 | 0 | | | | | . | . | . | . | . | . | A | A |
| NC_011770 | 2896097 | G | C | 11 | 6 | 2 | 0 | | | | | . | . | . | . | . | . | C | C |
| NC_011770 | 2896098 | G | A | 11 | 6 | 2 | 0 | | | | | . | . | . | . | . | . | A | A |
| NC_011770 | 2896103 | A | G | 21 | 6 | 2 | 0 | | | | | . | . | . | . | . | . | G | G |
| NC_011770 | 2896113 | A | T | 23 | 6 | 2 | 0 | | | | | . | . | . | . | . | . | T | T |
| NC_011770 | 2896125 | C | T | 24 | 6 | 2 | 0 | | | | | . | . | . | . | . | . | T | T |
| NC_011770 | 2896141 | G | T | 24 | 6 | 2 | 0 | | | | | . | . | . | . | . | . | T | T |
| NC_011770 | 2898176 | A | G | 36 | 6 | 2 | 0 | | | | | . | . | . | . | . | . | G | G |
| NC_011770 | 2898195 | T | C | 35 | 6 | 2 | 0 | | | | | . | . | . | . | . | . | C | C |
| NC_011770 | 2898216 | G | A | 35 | 6 | 2 | 0 | | | | | . | . | . | . | . | . | A | A |
| NC_011770 | 2898234 | CAG | C | 21 | 6 | 1 | 0 | | | | | . | . | . | . | . | . | . | C |
| NC_011770 | 2898241 | T | TGG | 34 | 6 | 2 | 0 | | | | | . | . | . | . | . | . | TGG | TGG |
| NC_011770 | 2898244 | C | T | 20 | 6 | 2 | 0 | | | | | . | . | . | . | . | . | T | T |
| NC_011770 | 2898267 | G | C | 33 | 6 | 2 | 0 | | | | | . | . | . | . | . | . | C | C |
| NC_011770 | 2898269 | A | G | 33 | 6 | 2 | 0 | | | | | . | . | . | . | . | . | G | G |
| NC_011770 | 2898291 | A | G | 33 | 6 | 2 | 0 | | | | | . | . | . | . | . | . | G | G |
| NC_011770 | 2898298 | G | T | 29 | 6 | 2 | 0 | | | | | . | . | . | . | . | . | T | T |
| NC_011770 | 2898299 | A | C | 28 | 6 | 2 | 0 | | | | | . | . | . | . | . | . | C | C |
| NC_011770 | 2898301 | T | G | 28 | 6 | 2 | 0 | | | | | . | . | . | . | . | . | G | G |
| NC_011770 | 2898309 | A | T | 28 | 6 | 2 | 0 | | | | | . | . | . | . | . | . | T | T |
| NC_011770 | 2898312 | T | C | 28 | 6 | 2 | 0 | | | | | . | . | . | . | . | . | C | C |
| NC_011770 | 2898322 | G | C | 29 | 6 | 2 | 0 | | | | | . | . | . | . | . | . | C | C |
| NC_011770 | 2898324 | A | G | 29 | 6 | 2 | 0 | | | | | . | . | . | . | . | . | G | G |
| NC_011770 | 2898333 | G | A | 28 | 6 | 2 | 0 | | | | | . | . | . | . | . | . | A | A |
| NC_011770 | 2898336 | G | A | 28 | 6 | 2 | 0 | | | | | . | . | . | . | . | . | A | A |
| NC_011770 | 2898337 | A | G | 28 | 6 | 2 | 0 | | | | | . | . | . | . | . | . | G | G |
| NC_011770 | 2898338 | G | T | 28 | 6 | 2 | 0 | | | | | . | . | . | . | . | . | T | T |
| NC_011770 | 2898350 | C | G | 29 | 6 | 2 | 0 | | | | | . | . | . | . | . | . | G | G |
| NC_011770 | 2898354 | G | A | 29 | 6 | 2 | 0 | | | A | A | . | . | . | . | . | . | | |
| NC_011770 | 2898355 | AGC | A | 28 | 6 | 2 | 0 | | | | | . | . | . | . | . | . | A | A |
| NC_011770 | 2898368 | C | G | 29 | 6 | 2 | 0 | | | | | . | . | . | . | . | . | G | G |
| NC_011770 | 2898372 | G | A | 29 | 6 | 2 | 0 | | | | | . | . | . | . | . | . | A | A |

| | | | | | | | | | | | | | | | | | | | |
|---|---|---|---|---|---|---|---|---|---|---|---|---|---|---|---|---|---|---|---|
| NC_011770 | 2898382 | G | A | 29 | 6 | 2 | 0 | | | | | . | . | . | . | . | . | A | A |
| NC_011770 | 2898435 | G | A | 29 | 6 | 2 | 0 | | | | | . | . | . | . | . | . | A | A |
| NC_011770 | 2898440 | A | G | 27 | 6 | 2 | 0 | | | | | . | . | . | . | . | . | G | G |
| NC_011770 | 2898456 | C | G | 29 | 6 | 2 | 0 | | | | | . | . | . | . | . | . | G | G |
| NC_011770 | 2898458 | A | T | 29 | 6 | 2 | 0 | | | | | . | . | . | . | . | . | T | T |
| NC_011770 | 2898901 | T | G | 37 | 6 | 2 | 0 | | | | | . | . | . | . | . | . | G | G |
| NC_011770 | 2898904 | G | T | 36 | 6 | 2 | 0 | | | | | . | . | . | . | . | . | T | T |
| NC_011770 | 2898905 | A | G | 36 | 6 | 2 | 0 | | | | | . | . | . | . | . | . | G | G |
| NC_011770 | 2898939 | T | C | 37 | 6 | 2 | 0 | | | | | . | . | . | . | . | . | C | C |
| NC_011770 | 2898942 | G | C | 37 | 6 | 2 | 0 | | | | | . | . | . | . | . | . | C | C |
| NC_011770 | 2898943 | C | G | 37 | 6 | 2 | 0 | | | | | . | . | . | . | . | . | G | G |
| NC_011770 | 2898944 | T | C | 37 | 6 | 2 | 0 | | | | | . | . | . | . | . | . | C | C |
| NC_011770 | 2898957 | C | G | 39 | 6 | 2 | 0 | | | | | . | . | . | . | . | . | G | G |
| NC_011770 | 2898963 | G | C | 39 | 6 | 2 | 0 | | | | | . | . | . | . | . | . | C | C |
| NC_011770 | 2898965 | A | C | 39 | 6 | 2 | 0 | | | | | . | . | . | . | . | . | C | C |
| NC_011770 | 2900291 | T | C | 23 | 6 | 2 | 0 | | | C | C | . | . | . | . | . | . | | |
| NC_011770 | 2900293 | A | T | 23 | 6 | 4 | 0 | | | T | T | . | . | . | . | . | . | T | T |
| NC_011770 | 2900294 | T | C | 23 | 6 | 4 | 0 | | | C | C | . | . | . | . | . | . | C | C |
| NC_011770 | 2900301 | C | T | 23 | 6 | 2 | 0 | | | | | . | . | . | . | . | . | T | T |
| NC_011770 | 2900303 | G | T | 17 | 7 | 1 | 0 | | | . | T | . | . | . | . | . | . | | |
| NC_011770 | 2900304 | GT | G | 17 | 6 | 4 | 0 | | | G | G | . | . | . | . | . | . | TT | TT |
| NC_011770 | 2900309 | C | A | 17 | 6 | 4 | 0 | | | A | A | . | . | . | . | . | . | A | A |
| NC_011770 | 2900311 | G | A | 17 | 6 | 4 | 0 | | | A | A | . | . | . | . | . | . | A | A |
| NC_011770 | 2900317 | C | G | 23 | 6 | 4 | 0 | | | G | G | . | . | . | . | . | . | G | G |
| NC_011770 | 2900319 | C | T | 23 | 6 | 2 | 0 | | | T | T | . | . | . | . | . | . | | |
| NC_011770 | 2900323 | C | G | 24 | 6 | 2 | 0 | | | G | G | . | . | . | . | . | . | | |
| NC_011770 | 2900324 | A | T | 23 | 6 | 2 | 0 | | | | | . | . | . | . | . | . | T | T |
| NC_011770 | 2900327 | C | G | 24 | 6 | 4 | 0 | | | G | G | . | . | . | . | . | . | G | G |
| NC_011770 | 2900331 | T | C | 24 | 6 | 4 | 0 | | | C | C | . | . | . | . | . | . | G | G |
| NC_011770 | 2900333 | C | A | 24 | 6 | 2 | 0 | | | | | . | . | . | . | . | . | A | A |
| NC_011770 | 2900335 | G | A | 28 | 6 | 2 | 0 | | | A | A | . | . | . | . | . | . | | |
| NC_011770 | 2901958 | G | A | 30 | 6 | 2 | 0 | | | A | A | . | . | . | . | . | . | | |
| NC_011770 | 2902522 | C | CG | 19 | 6 | 4 | 0 | | | CG | CG | . | . | . | . | . | . | CT | CT |
| NC_011770 | 2904222 | C | T | 20 | 8 | 2 | 0 | | | T | T | . | . | . | . | . | . | . | . |
| NC_011770 | 2904245 | C | G | 9 | 8 | 2 | 0 | | | G | G | . | . | . | . | . | . | . | . |
| NC_011770 | 2904251 | GC | G | 17 | 8 | 2 | 0 | | | G | G | . | . | . | . | . | . | . | . |
| NC_011770 | 2904267 | T | G | 29 | 6 | 4 | 0 | | | G | G | . | . | . | . | . | . | G | G |
| NC_011770 | 2904276 | G | T | 30 | 6 | 2 | 0 | | | T | T | . | . | . | . | . | . | | |
| NC_011770 | 2904285 | C | T | 31 | 6 | 2 | 0 | | | | | . | . | . | . | . | . | T | T |
| NC_011770 | 2904297 | G | A | 25 | 6 | 4 | 0 | | | A | A | . | . | . | . | . | . | A | A |
| NC_011770 | 2904300 | C | G | 27 | 6 | 2 | 0 | | | G | G | . | . | . | . | . | . | | |
| NC_011770 | 2904302 | A | G | 22 | 7 | 3 | 0 | | | G | G | . | . | . | . | . | . | G | . |
| NC_011770 | 2904303 | T | G | 21 | 7 | 3 | 0 | | | G | G | . | . | . | . | . | . | G | . |
| NC_011770 | 2904304 | T | G | 21 | 7 | 1 | 0 | | | | | . | . | . | . | . | . | G | . |
| NC_011770 | 2904305 | C | T | 21 | 7 | 1 | 0 | | | | | . | . | . | . | . | . | T | . |
| NC_011770 | 2904306 | G | T | 21 | 7 | 3 | 0 | | | T | T | . | . | . | . | . | . | C | . |
| NC_011770 | 2904314 | G | A | 28 | 6 | 2 | 0 | | | A | A | . | . | . | . | . | . | | |
| NC_011770 | 2904315 | G | A | 28 | 6 | 2 | 0 | | | A | A | . | . | . | . | . | . | | |
| NC_011770 | 2904318 | A | G | 28 | 6 | 4 | 0 | | | G | G | . | . | . | . | . | . | G | G |
| NC_011770 | 2904321 | C | T | 30 | 6 | 2 | 0 | | | | | . | . | . | . | . | . | T | T |
| NC_011770 | 2904324 | T | C | 31 | 6 | 2 | 0 | | | | | . | . | . | . | . | . | C | C |
| NC_011770 | 2904330 | A | G | 30 | 6 | 4 | 0 | | | G | G | . | . | . | . | . | . | G | G |
| NC_011770 | 2904333 | C | G | 31 | 6 | 4 | 0 | | | G | G | . | . | . | . | . | . | G | G |
| NC_011770 | 2904342 | C | G | 34 | 6 | 2 | 0 | | | G | G | . | . | . | . | . | . | | |
| NC_011770 | 2904345 | C | G | 33 | 6 | 4 | 0 | | | G | G | . | . | . | . | . | . | G | G |
| NC_011770 | 2904354 | G | A | 34 | 6 | 2 | 0 | | | | | . | . | . | . | . | . | A | A |
| NC_011770 | 2904356 | C | T | 34 | 6 | 4 | 0 | | | T | T | . | . | . | . | . | . | G | G |
| NC_011770 | 2904366 | C | T | 34 | 6 | 2 | 0 | | | T | T | . | . | . | . | . | . | | |
| NC_011770 | 2904375 | C | T | 32 | 6 | 4 | 0 | | | T | T | . | . | . | . | . | . | G | G |
| NC_011770 | 2904378 | G | A | 32 | 6 | 2 | 0 | | | A | A | . | . | . | . | . | . | | |
| NC_011770 | 2904381 | C | A | 33 | 6 | 2 | 0 | | | | | . | . | . | . | . | . | A | A |
| NC_011770 | 2904382 | G | C | 33 | 6 | 2 | 0 | | | C | C | . | . | . | . | . | . | | |

| | | | | | | | | | | | | | | | | | | |
|---|---|---|---|---|---|---|---|---|---|---|---|---|---|---|---|---|---|---|
| NC_011770 | 2904384 | C | A | 33 | 6 | 2 | 0 | | A | A | . | . | . | . | . | . | | |
| NC_011770 | 2904387 | G | A | 33 | 6 | 2 | 0 | | A | A | . | . | . | . | . | . | | |
| NC_011770 | 2904390 | C | T | 33 | 6 | 4 | 0 | | T | T | . | . | . | . | . | . | T | T |
| NC_011770 | 2904402 | G | A | 35 | 6 | 2 | 0 | | | | . | . | . | . | . | . | A | A |
| NC_011770 | 2904405 | C | T | 34 | 6 | 4 | 0 | | T | T | . | . | . | . | . | . | T | T |
| NC_011770 | 2904408 | A | G | 34 | 6 | 4 | 0 | | G | G | . | . | . | . | . | . | G | G |
| NC_011770 | 2904410 | C | T | 34 | 6 | 2 | 0 | | T | T | . | . | . | . | . | . | | |
| NC_011770 | 2904411 | G | C | 34 | 6 | 4 | 0 | | C | C | . | . | . | . | . | . | A | A |
| NC_011770 | 2904417 | G | A | 35 | 6 | 2 | 0 | | A | A | . | . | . | . | . | . | | |
| NC_011770 | 2904435 | C | A | 34 | 6 | 2 | 0 | | | | . | . | . | . | . | . | A | A |
| NC_011770 | 2904438 | C | T | 35 | 6 | 2 | 0 | | T | T | . | . | . | . | . | . | | |
| NC_011770 | 2904450 | G | A | 34 | 6 | 4 | 0 | | A | A | . | . | . | . | . | . | T | T |
| NC_011770 | 2904455 | C | A | 34 | 6 | 2 | 0 | | A | A | . | . | . | . | . | . | | |
| NC_011770 | 2904456 | C | T | 34 | 6 | 2 | 0 | | | | . | . | . | . | . | . | T | T |
| NC_011770 | 2904459 | C | T | 35 | 6 | 2 | 0 | | T | T | . | . | . | . | . | . | | |
| NC_011770 | 2905340 | C | T | 45 | 6 | 2 | 0 | | | | . | . | . | . | . | . | T | T |
| NC_011770 | 2905347 | G | T | 40 | 6 | 4 | 0 | | T | T | . | . | . | . | . | . | T | T |
| NC_011770 | 2905348 | C | T | 40 | 6 | 4 | 0 | | T | T | . | . | . | . | . | . | G | G |
| NC_011770 | 2905351 | G | A | 40 | 6 | 2 | 0 | | | | . | . | . | . | . | . | A | A |
| NC_011770 | 2905352 | G | C | 41 | 6 | 2 | 0 | | C | C | . | . | . | . | . | . | | |
| NC_011770 | 2905353 | A | G | 40 | 6 | 4 | 0 | | G | G | . | . | . | . | . | . | G | G |
| NC_011770 | 2905366 | C | A | 42 | 6 | 2 | 0 | | A | A | . | . | . | . | . | . | | |
| NC_011770 | 2905367 | G | T | 43 | 6 | 2 | 0 | | | | . | . | . | . | . | . | T | T |
| NC_011770 | 2905370 | A | G | 43 | 6 | 2 | 0 | | G | G | . | . | . | . | . | . | | |
| NC_011770 | 2905373 | G | A | 43 | 6 | 2 | 0 | | | | . | . | . | . | . | . | A | A |
| NC_011770 | 2905376 | G | A | 43 | 6 | 2 | 0 | | A | A | . | . | . | . | . | . | | |
| NC_011770 | 2905612 | G | A | 47 | 6 | 4 | 0 | | A | A | . | . | . | . | . | . | A | A |
| NC_011770 | 2905636 | T | C | 51 | 6 | 2 | 0 | | | | . | . | . | . | . | . | C | C |
| NC_011770 | 2905648 | C | A | 50 | 6 | 4 | 0 | | A | A | . | . | . | . | . | . | A | A |
| NC_011770 | 2905652 | C | A | 49 | 6 | 2 | 0 | | | | . | . | . | . | . | . | A | A |
| NC_011770 | 2905655 | C | G | 47 | 6 | 2 | 0 | | | | . | . | . | . | . | . | G | G |
| NC_011770 | 2905656 | G | A | 48 | 6 | 2 | 0 | | A | A | . | . | . | . | . | . | | |
| NC_011770 | 2905657 | G | A | 48 | 6 | 4 | 0 | | A | A | . | . | . | . | . | . | A | A |
| NC_011770 | 2905683 | C | A | 45 | 6 | 2 | 0 | | A | A | . | . | . | . | . | . | | |
| NC_011770 | 2905686 | T | C | 45 | 6 | 2 | 0 | | C | C | . | . | . | . | . | . | | |
| NC_011770 | 2905699 | C | A | 38 | 6 | 4 | 0 | | A | A | . | . | . | . | . | . | A | A |
| NC_011770 | 2905704 | T | A | 38 | 6 | 4 | 0 | | A | A | . | . | . | . | . | . | A | A |
| NC_011770 | 2905705 | A | G | 37 | 6 | 2 | 0 | | G | G | . | . | . | . | . | . | | |
| NC_011770 | 2905707 | C | A | 38 | 6 | 2 | 0 | | A | A | . | . | . | . | . | . | | |
| NC_011770 | 2905708 | C | T | 38 | 6 | 2 | 0 | | T | T | . | . | . | . | . | . | | |
| NC_011770 | 2906458 | C | T | 23 | 8 | 2 | 0 | | T | T | . | . | . | . | . | . | . | . |
| NC_011770 | 2906474 | G | A | 22 | 8 | 2 | 0 | | A | A | . | . | . | . | . | . | . | . |
| NC_011770 | 2906475 | T | A | 22 | 8 | 2 | 0 | | A | A | . | . | . | . | . | . | . | . |
| NC_011770 | 2906500 | T | C | 8 | 8 | 2 | 0 | | C | C | . | . | . | . | . | . | . | . |
| NC_011770 | 2906626 | T | C | 12 | 8 | 2 | 0 | | C | C | . | . | . | . | . | . | . | . |
| NC_011770 | 2906640 | A | G | 12 | 8 | 2 | 0 | | G | G | . | . | . | . | . | . | . | . |
| NC_011770 | 2906722 | T | C | 12 | 8 | 2 | 0 | | C | C | . | . | . | . | . | . | . | . |
| NC_011770 | 2906724 | G | C | 12 | 8 | 2 | 0 | | C | C | . | . | . | . | . | . | . | . |
| NC_011770 | 2907673 | G | A | 14 | 8 | 2 | 0 | | A | A | . | . | . | . | . | . | . | . |
| NC_011770 | 2907681 | G | T | 14 | 8 | 2 | 0 | | T | T | . | . | . | . | . | . | . | . |
| NC_011770 | 2907691 | A | G | 14 | 8 | 2 | 0 | | G | G | . | . | . | . | . | . | . | . |
| NC_011770 | 2907692 | C | T | 14 | 8 | 2 | 0 | | T | T | . | . | . | . | . | . | . | . |
| NC_011770 | 2907711 | A | G | 19 | 6 | 2 | 0 | | G | G | . | . | . | . | . | . | | |
| NC_011770 | 2907727 | G | C | 15 | 6 | 2 | 0 | | C | C | . | . | . | . | . | . | | |
| NC_011770 | 2907730 | T | G | 15 | 6 | 4 | 0 | | G | G | . | . | . | . | . | . | G | G |
| NC_011770 | 2907731 | T | C | 15 | 6 | 2 | 0 | | C | C | . | . | . | . | . | . | | |
| NC_011770 | 2907740 | A | C | 13 | 6 | 4 | 0 | | C | C | . | . | . | . | . | . | G | G |
| NC_011770 | 2907745 | C | G | 14 | 6 | 2 | 0 | | G | G | . | . | . | . | . | . | | |
| NC_011770 | 2907746 | A | G | 14 | 6 | 4 | 0 | | G | G | . | . | . | . | . | . | C | C |
| NC_011770 | 2907748 | A | G | 13 | 6 | 2 | 0 | | G | G | . | . | . | . | . | . | | |
| NC_011770 | 2907750 | T | C | 13 | 6 | 4 | 0 | | C | C | . | . | . | . | . | . | C | C |
| NC_011770 | 2907752 | G | A | 13 | 6 | 4 | 0 | | A | A | . | . | . | . | . | . | A | A |

| | | | | | | | | | | | | | | | | | | | |
|---|---|---|---|---|---|---|---|---|---|---|---|---|---|---|---|---|---|---|---|
| NC_011770 | 2907757 | A | T | 16 | 6 | 2 | 0 | | | T | T | . | . | . | . | . | . | | |
| NC_011770 | 2907772 | G | A | 17 | 6 | 4 | 0 | | | A | A | . | . | . | . | . | . | A | A |
| NC_011770 | 2907773 | A | G | 16 | 6 | 4 | 0 | | | G | G | . | . | . | . | . | . | C | C |
| NC_011770 | 2907776 | C | T | 17 | 6 | 2 | 0 | | | T | T | . | . | . | . | . | . | | |
| NC_011770 | 2907784 | T | C | 17 | 6 | 4 | 0 | | | C | C | . | . | . | . | . | . | C | C |
| NC_011770 | 2907792 | A | C | 17 | 6 | 2 | 0 | | | C | C | . | . | . | . | . | . | | |
| NC_011770 | 2907795 | A | T | 17 | 6 | 4 | 0 | | | T | T | . | . | . | . | . | . | T | T |
| NC_011770 | 2907801 | T | C | 42 | 0 | 2 | 0 | | | C | C | | | | | | | | |
| NC_011770 | 2907973 | C | T | 54 | 0 | 2 | 0 | | | | | | | T | T | | | | |
| NC_011770 | 2908055 | A | AG | 57 | 0 | 2 | 0 | | | | | | | | | | | AG | AG |
| NC_011770 | 2908182 | C | T | 64 | 0 | 4 | 0 | | | T | T | T | T | | | | | | |
| NC_011770 | 2908211 | A | G | 67 | 0 | 2 | 0 | | | | | | | | | G | G | | |
| NC_011770 | 2908242 | G | A | 71 | 0 | 2 | 0 | | | | | | | | | A | A | | |
| NC_011770 | 2908260 | AG | A | 76 | 0 | 2 | 0 | | | | | | | | | A | A | | |
| NC_011770 | 2908322 | G | T | 74 | 0 | 2 | 0 | | | | | | | | | T | T | | |
| NC_011770 | 2908409 | T | C | 71 | 0 | 8 | 0 | C | C | | | | | C | C | C | C | C | C |
| NC_011770 | 2908428 | G | A | 69 | 0 | 4 | 0 | | | A | A | A | A | | | | | | |
| NC_011770 | 2908489 | G | A | 67 | 0 | 8 | 0 | A | A | | | | | A | A | A | A | A | A |
| NC_011770 | 2908501 | T | C | 65 | 0 | 8 | 0 | C | C | | | | | C | C | C | C | C | C |
| NC_011770 | 2908504 | G | GT | 63 | 0 | 8 | 0 | GT | GT | | | | | GT | GT | GT | GT | GT | GT |
| NC_011770 | 2908514 | C | G | 62 | 0 | 8 | 0 | G | G | | | | | G | G | G | G | G | G |
| NC_011770 | 2908515 | A | C | 62 | 0 | 8 | 0 | C | C | | | | | C | C | C | C | C | C |
| NC_011770 | 2908519 | C | T | 63 | 0 | 8 | 0 | T | T | | | | | T | T | T | T | T | T |
| NC_011770 | 2908520 | C | T | 63 | 0 | 8 | 0 | T | T | | | | | T | T | T | T | T | T |
| NC_011770 | 2908524 | A | G | 63 | 0 | 8 | 0 | G | G | | | | | G | G | G | G | G | G |
| NC_011770 | 2912070 | T | C | 60 | 0 | 8 | 0 | | | C | C | C | C | | | C | C | C | C |
| NC_011770 | 2912071 | A | G | 60 | 0 | 8 | 0 | | | G | G | G | G | | | G | G | G | G |
| NC_011770 | 2912073 | CG | C | 61 | 0 | 2 | 0 | C | C | | | | | | | | | | |
| NC_011770 | 2912097 | A | G | 55 | 0 | 8 | 0 | | | G | G | G | G | | | G | G | G | G |
| NC_011770 | 2912103 | C | T | 50 | 0 | 8 | 0 | | | T | T | T | T | | | T | T | T | T |
| NC_011770 | 2912104 | T | C | 53 | 0 | 8 | 0 | | | C | C | C | C | | | C | C | C | C |
| NC_011770 | 2912219 | C | T | 62 | 0 | 10 | 0 | T | T | T | T | T | T | | | T | T | T | T |
| NC_011770 | 2912277 | T | G | 62 | 0 | 4 | 0 | | | G | G | G | G | | | | | | |
| NC_011770 | 2912302 | G | A | 62 | 0 | 4 | 0 | | | A | A | A | A | | | | | | |
| NC_011770 | 2912308 | C | T | 61 | 0 | 8 | 0 | T | T | | | | | T | T | T | T | T | T |
| NC_011770 | 2912318 | G | A | 62 | 0 | 4 | 0 | | | A | A | A | A | | | | | | |
| NC_011770 | 2912448 | G | A | 63 | 0 | 6 | 0 | | | A | A | A | A | | | | | A | A |
| NC_011770 | 2912453 | G | A | 64 | 0 | 4 | 0 | | | A | A | A | A | | | | | | |
| NC_011770 | 2912454 | T | C | 64 | 0 | 4 | 0 | | | C | C | C | C | | | | | | |
| NC_011770 | 2912472 | TC | T | 64 | 0 | 2 | 0 | | | | | | | | | | | T | T |
| NC_011770 | 2912559 | T | C | 72 | 0 | 4 | 0 | | | C | C | C | C | | | | | | |
| NC_011770 | 2912626 | G | T | 75 | 0 | 2 | 0 | | | | | | | | | T | T | | |
| NC_011770 | 2912648 | C | T | 82 | 0 | 2 | 0 | | | | | | | | | T | T | | |
| NC_011770 | 2912668 | C | T | 78 | 0 | 2 | 0 | | | | | | | | | T | T | | |
| NC_011770 | 2912709 | T | G | 74 | 0 | 2 | 0 | | | | | | | G | G | | | | |
| NC_011770 | 2912745 | T | C | 74 | 0 | 2 | 0 | | | | | | | C | C | | | | |
| NC_011770 | 2912807 | C | T | 75 | 0 | 2 | 0 | | | | | | | | | T | T | | |
| NC_011770 | 2914001 | G | A | 88 | 0 | 1 | 0 | | | | | | | A | | | | | |
| NC_011770 | 2914188 | A | T | 73 | 0 | 4 | 0 | | | | | | | T | T | T | T | | |
| NC_011770 | 2914202 | T | G | 77 | 0 | 2 | 0 | G | G | | | | | | | | | | |
| NC_011770 | 2914889 | A | G | 63 | 0 | 8 | 0 | | | G | G | G | G | G | G | G | G | | |
| NC_011770 | 2914922 | G | A | 61 | 0 | 4 | 0 | | | A | A | A | A | | | | | | |
| NC_011770 | 2914954 | T | C | 59 | 0 | 4 | 0 | | | C | C | C | C | | | | | | |
| NC_011770 | 2914982 | C | G | 57 | 0 | 4 | 0 | | | G | G | G | G | | | | | | |
| NC_011770 | 2915026 | G | A | 61 | 0 | 6 | 0 | A | A | | | | | | | A | A | A | A |
| NC_011770 | 2915058 | C | G | 66 | 0 | 4 | 0 | | | G | G | G | G | | | | | | |
| NC_011770 | 2915113 | C | CT | 68 | 0 | 2 | 0 | | | | | | | | | CT | CT | | |
| NC_011770 | 2916081 | C | T | 73 | 0 | 6 | 0 | | | T | T | T | T | | | | | T | T |
| NC_011770 | 2917126 | GT | G | 67 | 0 | 2 | 0 | | | | | | | | | | | G | G |
| NC_011770 | 2918811 | G | T | 75 | 0 | 2 | 0 | | | | | | | T | T | | | | |
| NC_011770 | 2918847 | A | G | 73 | 0 | 8 | 0 | | | G | G | G | G | | | G | G | G | G |
| NC_011770 | 2918870 | G | A | 68 | 0 | 4 | 0 | | | A | A | A | A | | | | | | |

| | | | | | | | | | | | | | | | | | | | |
|---|---|---|---|---|---|---|---|---|---|---|---|---|---|---|---|---|---|---|---|
| NC_011770 | 2918919 | A | C | 64 | 0 | 8 | 0 | | | C | C | C | C | | | T | T | C | C |
| NC_011770 | 2918956 | G | A | 65 | 0 | 4 | 0 | A | A | | | | | A | A | | | | |
| NC_011770 | 2918972 | A | C | 66 | 0 | 6 | 0 | | | C | C | C | C | | | | | C | C |
| NC_011770 | 2919031 | G | C | 69 | 0 | 2 | 0 | | | | | | | | | C | C | | |
| NC_011770 | 2919050 | C | G | 69 | 0 | 4 | 0 | G | G | | | | | G | G | | | | |
| NC_011770 | 2919054 | G | T | 69 | 0 | 4 | 0 | | | T | T | T | T | | | | | | |
| NC_011770 | 2919062 | C | G | 67 | 0 | 6 | 0 | | | G | G | G | G | | | G | G | | |
| NC_011770 | 2919151 | G | A | 68 | 0 | 2 | 0 | | | | | | | | | A | A | | |
| NC_011770 | 2920354 | A | G | 64 | 0 | 2 | 0 | | | | | | | | | G | G | | |
| NC_011770 | 2920365 | C | T | 64 | 0 | 2 | 0 | | | | | | | | | T | T | | |
| NC_011770 | 2923453 | G | A | 71 | 0 | 4 | 0 | | | A | A | A | A | | | | | | |
| NC_011770 | 2924884 | G | A | 14 | 0 | 4 | 0 | | | A | A | A | A | | | | | | |
| NC_011770 | 2925246 | G | T | 49 | 0 | 10 | 0 | T | T | T | T | T | T | T | T | T | T | | |
| NC_011770 | 2925304 | T | C | 49 | 0 | 12 | 0 | C | C | C | C | C | C | C | C | C | C | C | C |
| NC_011770 | 2925364 | G | A | 50 | 0 | 2 | 0 | | | | | | | | | | | A | A |
| NC_011770 | 2925860 | C | G | 32 | 1 | 1 | 0 | G | . | | | | | | | | | | |
| NC_011770 | 2925862 | C | CG | 33 | 0 | 2 | 0 | CG | CG | | | | | | | | | | |
| NC_011770 | 2925864 | G | GA | 32 | 2 | 5 | 0 | . | . | GA | GA | . | GA | | | GA | GA | | |
| NC_011770 | 2925895 | T | A | 22 | 4 | 8 | 0 | A | A | A | A | A | A | . | . | . | . | A | A |
| NC_011770 | 2925899 | G | A | 32 | 0 | 10 | 0 | A | A | A | A | A | A | A | A | A | A | | |
| NC_011770 | 2925977 | A | G | 29 | 1 | 9 | 0 | G | . | G | G | G | G | G | G | G | G | | |
| NC_011770 | 2925993 | A | G | 28 | 1 | 9 | 0 | G | . | G | G | G | G | G | G | G | G | | |
| NC_011770 | 2926023 | T | G | 24 | 5 | 1 | 0 | G | . | . | . | . | . | . | . | . | . | | |
| NC_011770 | 2926481 | C | T | 71 | 0 | 2 | 0 | | | | | | | | | T | T | | |
| NC_011770 | 2926500 | T | C | 74 | 0 | 2 | 0 | | | | | | | | | C | C | | |
| NC_011770 | 2926524 | T | A | 69 | 0 | 4 | 0 | | | A | A | A | A | | | | | | |
| NC_011770 | 2926540 | C | A | 68 | 0 | 2 | 0 | | | | | | | | | A | A | | |
| NC_011770 | 2926541 | G | T | 68 | 0 | 2 | 0 | | | | | | | | | T | T | | |
| NC_011770 | 2926542 | C | T | 68 | 0 | 2 | 0 | | | | | | | | | | | T | T |
| NC_011770 | 2926561 | C | CGTT | 70 | 0 | 2 | 0 | | | | | | | | | | | CGTT | CGTT |
| NC_011770 | 2927001 | T | C | 86 | 0 | 10 | 0 | C | C | C | C | C | C | | | C | C | C | C |
| NC_011770 | 2927024 | G | C | 88 | 0 | 10 | 0 | C | C | C | C | C | C | | | C | C | C | C |
| NC_011770 | 2927031 | T | C | 88 | 0 | 4 | 0 | | | C | C | C | C | | | | | | |
| NC_011770 | 2927037 | G | T | 87 | 0 | 4 | 0 | | | T | T | T | T | | | | | | |
| NC_011770 | 2927045 | A | G | 82 | 0 | 12 | 0 | G | G | G | G | G | G | G | G | G | G | G | G |
| NC_011770 | 2928887 | G | A | 52 | 0 | 4 | 0 | | | A | A | A | A | | | | | | |
| NC_011770 | 2928923 | G | A | 52 | 0 | 4 | 0 | | | | | | | A | A | A | A | | |
| NC_011770 | 2928994 | G | C | 47 | 0 | 4 | 0 | | | | | | | C | C | C | C | | |
| NC_011770 | 2928995 | T | C | 48 | 0 | 4 | 0 | | | | | | | C | C | C | C | | |
| NC_011770 | 2928998 | A | T | 47 | 0 | 4 | 0 | | | | | | | T | T | T | T | | |
| NC_011770 | 2929004 | T | G | 47 | 0 | 4 | 0 | | | | | | | G | G | G | G | | |
| NC_011770 | 2929005 | C | T | 50 | 0 | 4 | 0 | | | | | | | T | T | T | T | | |
| NC_011770 | 2929017 | G | A | 44 | 0 | 4 | 0 | | | | | | | A | A | A | A | | |
| NC_011770 | 2929020 | A | G | 44 | 0 | 4 | 0 | | | | | | | G | G | G | G | | |
| NC_011770 | 2929024 | T | C | 45 | 0 | 4 | 0 | | | | | | | C | C | C | C | | |
| NC_011770 | 2929325 | G | GC | 18 | 10 | 2 | 0 | . | . | . | . | . | . | . | . | . | . | GC | GC |
| NC_011770 | 2931218 | T | G | 73 | 0 | 4 | 0 | | | G | G | G | G | | | | | | |
| NC_011770 | 2931269 | C | G | 76 | 0 | 6 | 0 | G | G | G | G | G | G | | | | | | |
| NC_011770 | 2931936 | C | A | 55 | 1 | 10 | 0 | A | A | A | A | A | A | | . | A | A | A | A |
| NC_011770 | 2933734 | T | A | 86 | 0 | 8 | 0 | A | A | A | A | A | A | | | | | A | A |
| NC_011770 | 2933804 | A | G | 87 | 0 | 12 | 0 | G | G | G | G | G | G | G | G | G | G | G | G |
| NC_011770 | 2934209 | T | G | 71 | 0 | 2 | 0 | | | | | | | | | G | G | | |
| NC_011770 | 2934773 | T | TG | 67 | 0 | 2 | 0 | | | | | | | | | TG | TG | | |
| NC_011770 | 2935374 | A | C | 86 | 0 | 10 | 0 | C | C | C | C | C | C | C | C | | | C | C |
| NC_011770 | 2935659 | C | T | 88 | 0 | 2 | 0 | | | | | | | | | T | T | | |
| NC_011770 | 2935662 | T | A | 87 | 0 | 2 | 0 | | | | | | | | | | | A | A |
| NC_011770 | 2936368 | TG | T | 92 | 0 | 2 | 0 | | | | | | | | | | | T | T |
| NC_011770 | 2936369 | GA | G | 91 | 0 | 6 | 0 | | | G | G | G | G | G | G | | | . | . |
| NC_011770 | 2936378 | A | G | 90 | 0 | 12 | 0 | G | G | G | G | G | G | G | G | G | G | G | G |
| NC_011770 | 2936412 | G | A | 92 | 0 | 2 | 0 | | | | | | | | | | | A | A |
| NC_011770 | 2936432 | T | C | 95 | 0 | 2 | 0 | | | | | | | | | | | C | C |
| NC_011770 | 2938290 | C | T | 62 | 0 | 2 | 0 | T | T | | | | | | | | | | |

| | | | | | | | | | | | | | | | | | | | |
|---|---|---|---|---|---|---|---|---|---|---|---|---|---|---|---|---|---|---|---|
| NC_011770 | 2938309 | A | C | 60 | 0 | 12 | 0 | C | C | C | C | C | C | C | C | C | C | C | C |
| NC_011770 | 2939524 | C | T | 71 | 0 | 2 | 0 | T | T | | | | | | | | | | |
| NC_011770 | 2939547 | G | A | 69 | 0 | 2 | 0 | | | | | | | | | A | A | | |
| NC_011770 | 2939551 | T | C | 69 | 0 | 8 | 0 | C | C | C | C | C | C | | | | | C | C |
| NC_011770 | 2944513 | T | G | 89 | 0 | 4 | 0 | | | G | G | G | G | | | | | | |
| NC_011770 | 2946887 | C | G | 76 | 0 | 8 | 0 | G | G | | | | | G | G | G | G | G | G |
| NC_011770 | 2946898 | C | A | 67 | 0 | 4 | 0 | | | A | A | A | A | | | | | | |
| NC_011770 | 2946903 | A | G | 64 | 0 | 2 | 0 | | | | | | | | | G | G | | |
| NC_011770 | 2946904 | T | G | 65 | 0 | 4 | 0 | | | G | G | G | G | | | | | | |
| NC_011770 | 2946928 | C | G | 65 | 0 | 4 | 0 | | | | | | | G | G | G | G | | |
| NC_011770 | 2946931 | C | T | 66 | 0 | 2 | 0 | | | | | | | | | T | T | | |
| NC_011770 | 2947033 | G | A | 67 | 0 | 6 | 0 | A | A | | | | | A | A | A | A | | |
| NC_011770 | 2949666 | G | A | 71 | 0 | 2 | 0 | | | | | | | | | A | A | | |
| NC_011770 | 2949668 | A | G | 69 | 0 | 2 | 0 | | | | | | | | | G | G | | |
| NC_011770 | 2949671 | G | A | 71 | 0 | 2 | 0 | | | | | | | | | A | A | | |
| NC_011770 | 2950619 | G | C | 66 | 0 | 2 | 0 | | | | | | | | | C | C | | |
| NC_011770 | 2950659 | C | G | 64 | 0 | 2 | 0 | | | | | | | | | G | G | | |
| NC_011770 | 2950673 | A | G | 60 | 0 | 2 | 0 | | | | | | | | | G | G | | |
| NC_011770 | 2950693 | T | C | 58 | 0 | 4 | 0 | | | | | | | C | C | C | C | | |
| NC_011770 | 2950709 | C | T | 57 | 0 | 2 | 0 | | | | | | | | | T | T | | |
| NC_011770 | 2950711 | A | T | 57 | 0 | 2 | 0 | | | | | | | | | T | T | | |
| NC_011770 | 2950713 | A | G | 56 | 0 | 12 | 0 | G | G | G | G | G | G | G | G | G | G | G | G |
| NC_011770 | 2951160 | T | C | 37 | 0 | 10 | 0 | C | C | C | C | C | C | C | C | | | C | C |
| NC_011770 | 2951162 | G | C | 38 | 0 | 10 | 0 | C | C | C | C | C | C | C | C | | | C | C |
| NC_011770 | 2951163 | C | T | 37 | 0 | 2 | 0 | | | | | | | T | T | | | | |
| NC_011770 | 2951176 | G | C | 38 | 0 | 2 | 0 | | | | | | | | | C | C | | |
| NC_011770 | 2951180 | A | C | 36 | 0 | 12 | 0 | C | C | C | C | C | C | C | C | T | T | C | C |
| NC_011770 | 2951200 | G | A | 39 | 0 | 2 | 0 | | | | | | | | | A | A | | |
| NC_011770 | 2951202 | A | G | 40 | 0 | 2 | 0 | | | | | | | | | G | G | | |
| NC_011770 | 2951208 | T | C | 43 | 0 | 2 | 0 | | | | | | | | | C | C | | |
| NC_011770 | 2951214 | T | C | 41 | 0 | 2 | 0 | | | | | | | | | C | C | | |
| NC_011770 | 2951251 | G | A | 48 | 0 | 2 | 0 | | | | | | | | | A | A | | |
| NC_011770 | 2951259 | C | T | 48 | 0 | 2 | 0 | | | | | | | | | T | T | | |
| NC_011770 | 2952086 | A | G | 72 | 0 | 12 | 0 | G | G | G | G | G | G | G | G | G | G | G | G |
| NC_011770 | 2952178 | C | T | 72 | 0 | 2 | 0 | | | | | | | | | T | T | | |
| NC_011770 | 2952937 | C | T | 76 | 0 | 2 | 0 | | | | | | | T | T | | | | |
| NC_011770 | 2961109 | A | G | 60 | 0 | 2 | 0 | G | G | | | | | | | | | | |
| NC_011770 | 2964579 | T | C | 67 | 0 | 2 | 0 | | | | | | | C | C | | | | |
| NC_011770 | 2964646 | G | A | 68 | 0 | 12 | 0 | A | A | A | A | A | A | A | A | A | A | A | A |
| NC_011770 | 2964674 | C | T | 70 | 0 | 2 | 0 | | | | | | | T | T | | | | |
| NC_011770 | 2965751 | T | C | 78 | 0 | 12 | 0 | C | C | C | C | C | C | C | C | C | C | C | C |
| NC_011770 | 2966732 | T | C | 60 | 0 | 12 | 0 | C | C | C | C | C | C | C | C | C | C | C | C |
| NC_011770 | 2968883 | T | C | 76 | 0 | 10 | 0 | C | C | C | C | C | C | | | C | C | C | C |
| NC_011770 | 2968935 | T | C | 73 | 0 | 10 | 0 | C | C | C | C | C | C | | | C | C | C | C |
| NC_011770 | 2968959 | A | T | 74 | 0 | 2 | 0 | | | | | | | | | | | T | T |
| NC_011770 | 2968964 | G | A | 71 | 0 | 2 | 0 | | | | | | | | | A | A | | |
| NC_011770 | 2970198 | T | G | 61 | 0 | 4 | 0 | | | G | G | G | G | | | | | | |
| NC_011770 | 2970212 | T | C | 56 | 0 | 4 | 0 | | | C | C | C | C | | | | | | |
| NC_011770 | 2970245 | G | A | 59 | 0 | 4 | 0 | | | A | A | A | A | | | | | | |
| NC_011770 | 2970246 | G | A | 59 | 0 | 4 | 0 | A | A | | | | | A | A | | | | |
| NC_011770 | 2970258 | T | G | 57 | 0 | 6 | 0 | | | G | G | G | G | | | | | G | G |
| NC_011770 | 2970259 | C | T | 55 | 0 | 6 | 0 | | | T | T | T | T | | | | | G | G |
| NC_011770 | 2970261 | T | A | 55 | 0 | 2 | 0 | | | | | | | | | | | A | A |
| NC_011770 | 2970278 | G | C | 55 | 0 | 2 | 0 | | | | | | | | | | | C | C |
| NC_011770 | 2970279 | C | A | 53 | 0 | 2 | 0 | | | | | | | | | | | A | A |
| NC_011770 | 2970280 | G | A | 52 | 0 | 4 | 0 | A | A | | | | | A | A | | | | |
| NC_011770 | 2970285 | T | C | 51 | 0 | 2 | 0 | | | | | | | | | | | C | C |
| NC_011770 | 2970286 | C | T | 54 | 0 | 4 | 0 | T | T | | | | | T | T | | | | |
| NC_011770 | 2970289 | G | C | 54 | 0 | 2 | 0 | | | | | | | | | | | C | C |
| NC_011770 | 2970296 | G | T | 53 | 0 | 2 | 0 | | | | | | | | | | | T | T |
| NC_011770 | 2970297 | A | C | 51 | 0 | 2 | 0 | | | | | | | | | | | C | C |
| NC_011770 | 2970298 | A | G | 53 | 0 | 2 | 0 | | | | | | | | | | | G | G |

| | | | | | | | | | | | | | | | | | | | |
|---|---|---|---|---|---|---|---|---|---|---|---|---|---|---|---|---|---|---|---|
| NC_011770 | 2970304 | G | C | 56 | 0 | 2 | 0 | | | | | | | | | | | C | C |
| NC_011770 | 2970315 | T | C | 51 | 0 | 6 | 0 | | | C | C | C | C | | | | | C | C |
| NC_011770 | 2970317 | T | C | 52 | 0 | 6 | 0 | | | C | C | C | C | | | | | C | C |
| NC_011770 | 2970320 | A | G | 53 | 0 | 6 | 0 | | | G | G | G | G | | | | | G | G |
| NC_011770 | 2970321 | T | C | 52 | 0 | 6 | 0 | | | C | C | C | C | | | | | C | C |
| NC_011770 | 2973142 | C | T | 80 | 0 | 12 | 0 | T | T | T | T | T | T | T | T | T | T | T | T |
| NC_011770 | 2973194 | G | A | 76 | 0 | 4 | 0 | | | | | | | A | A | A | A | | |
| NC_011770 | 2973278 | T | C | 76 | 0 | 4 | 0 | C | C | | | | | | | | | C | C |
| NC_011770 | 2984901 | A | G | 77 | 0 | 6 | 0 | | | G | G | G | G | G | G | | | | |
| NC_011770 | 2985029 | A | G | 74 | 0 | 4 | 0 | G | G | | | | | | | G | G | | |
| NC_011770 | 2985057 | CT | C | 75 | 0 | 2 | 0 | C | C | | | | | | | | | | |
| NC_011770 | 2986635 | GC | G | 75 | 0 | 1 | 0 | | | | | | | G | | | | | |
| NC_011770 | 2991432 | A | G | 72 | 0 | 2 | 0 | | | | | | | | | | | G | G |
| NC_011770 | 2994439 | C | T | 106 | 0 | 2 | 0 | | | | | | | | | T | T | | |
| NC_011770 | 2995481 | CG | C | 76 | 0 | 8 | 0 | C | C | | | | | C | C | C | C | C | C |
| NC_011770 | 2995494 | T | C | 75 | 0 | 8 | 0 | | | C | C | C | C | C | C | C | C | | |
| NC_011770 | 2999210 | T | G | 72 | 0 | 2 | 0 | | | | | | | | | | | G | G |
| NC_011770 | 2999225 | G | C | 71 | 0 | 12 | 0 | C | C | C | C | C | C | C | C | C | C | C | C |
| NC_011770 | 2999239 | A | G | 68 | 0 | 12 | 0 | G | G | G | G | G | G | G | G | G | G | G | G |
| NC_011770 | 2999244 | T | C | 67 | 0 | 12 | 0 | C | C | C | C | C | C | C | C | C | C | C | C |
| NC_011770 | 2999289 | G | C | 68 | 0 | 12 | 0 | C | C | C | C | C | C | C | C | C | C | C | C |
| NC_011770 | 2999290 | T | C | 68 | 0 | 12 | 0 | C | C | C | C | C | C | C | C | C | C | C | C |
| NC_011770 | 2999293 | G | A | 66 | 0 | 10 | 0 | A | A | A | A | A | A | A | A | | | A | A |
| NC_011770 | 2999296 | C | T | 68 | 0 | 2 | 0 | | | | | | | | | | | T | T |
| NC_011770 | 2999312 | A | G | 67 | 0 | 12 | 0 | G | G | G | G | G | G | G | G | G | G | G | G |
| NC_011770 | 2999316 | C | G | 66 | 0 | 12 | 0 | G | G | G | G | G | G | G | G | G | G | G | G |
| NC_011770 | 2999332 | T | G | 61 | 0 | 12 | 0 | G | G | G | G | G | G | G | G | G | G | G | G |
| NC_011770 | 2999333 | T | C | 61 | 0 | 12 | 0 | C | C | C | C | C | C | C | C | C | C | C | C |
| NC_011770 | 2999335 | C | G | 61 | 0 | 12 | 0 | G | G | G | G | G | G | G | G | G | G | G | G |
| NC_011770 | 2999336 | G | A | 60 | 0 | 12 | 0 | A | A | A | A | A | A | A | A | A | A | A | A |
| NC_011770 | 2999492 | T | C | 72 | 0 | 2 | 0 | C | C | | | | | | | | | | |
| NC_011770 | 3000514 | G | A | 84 | 0 | 2 | 0 | | | | | | | | | A | A | | |
| NC_011770 | 3000515 | T | C | 83 | 0 | 2 | 0 | | | | | | | | | C | C | | |
| NC_011770 | 3001416 | A | G | 61 | 0 | 4 | 0 | | | G | G | G | G | | | | | | |
| NC_011770 | 3001459 | C | T | 56 | 0 | 2 | 0 | | | | | | | | | | | T | T |
| NC_011770 | 3001460 | C | T | 56 | 0 | 2 | 0 | | | | | | | | | | | T | T |
| NC_011770 | 3001463 | A | G | 54 | 0 | 4 | 0 | | | | | | | | | G | G | G | G |
| NC_011770 | 3001466 | G | A | 56 | 0 | 4 | 0 | | | | | | | | | A | A | A | A |
| NC_011770 | 3001504 | A | G | 55 | 0 | 8 | 0 | | | G | G | G | G | | | G | G | G | G |
| NC_011770 | 3001524 | A | G | 54 | 0 | 2 | 0 | G | G | | | | | | | | | | |
| NC_011770 | 3003973 | T | C | 59 | 0 | 4 | 0 | C | C | | | | | | | | | C | C |
| NC_011770 | 3006064 | C | T | 69 | 0 | 2 | 0 | T | T | | | | | | | | | | |
| NC_011770 | 3009645 | T | C | 79 | 0 | 2 | 0 | | | | | | | | | C | C | | |
| NC_011770 | 3009650 | A | G | 81 | 0 | 2 | 0 | | | | | | | | | G | G | | |
| NC_011770 | 3009679 | T | C | 82 | 0 | 4 | 0 | | | C | C | C | C | | | | | | |
| NC_011770 | 3012324 | A | G | 88 | 0 | 12 | 0 | G | G | G | G | G | G | G | G | G | G | G | G |
| NC_011770 | 3013875 | G | C | 61 | 0 | 2 | 0 | | | | | | | | | C | C | | |
| NC_011770 | 3013880 | C | CG | 59 | 0 | 12 | 0 | CG | CG | CG | CG | CG | CG | CG | CG | CG | CG | CG | CG |
| NC_011770 | 3017774 | T | C | 88 | 0 | 12 | 0 | C | C | C | C | C | C | C | C | C | C | C | C |
| NC_011770 | 3019960 | T | C | 75 | 0 | 8 | 0 | C | C | | | | | C | C | C | C | C | C |
| NC_011770 | 3022271 | G | A | 41 | 0 | 2 | 0 | | | | | | | A | A | | | | |
| NC_011770 | 3022277 | C | T | 39 | 0 | 2 | 0 | T | T | | | | | | | | | | |
| NC_011770 | 3022300 | T | G | 38 | 0 | 4 | 0 | | | | | | | | | G | G | G | G |
| NC_011770 | 3022317 | A | G | 11 | 7 | 3 | 0 | | | . | . | . | . | G | . | G | G | . | . |
| NC_011770 | 3024487 | T | C | 77 | 0 | 12 | 0 | C | C | C | C | C | C | C | C | C | C | C | C |
| NC_011770 | 3025930 | G | T | 90 | 0 | 2 | 0 | T | T | | | | | | | | | | |
| NC_011770 | 3025936 | A | G | 92 | 0 | 4 | 0 | G | G | | | | | G | G | | | | |
| NC_011770 | 3026038 | T | C | 77 | 0 | 8 | 0 | | | C | C | C | C | | | C | C | C | C |
| NC_011770 | 3026048 | C | T | 75 | 0 | 2 | 0 | T | T | | | | | | | | | | |
| NC_011770 | 3026065 | C | T | 75 | 0 | 8 | 0 | | | T | T | T | T | | | T | T | T | T |
| NC_011770 | 3027077 | G | T | 59 | 0 | 2 | 0 | | | | | | | T | T | | | | |
| NC_011770 | 3027083 | AG | A | 59 | 0 | 2 | 0 | | | | | | | A | A | | | | |

| | | | | | | | | | | | | | | | | | | | |
|---|---|---|---|---|---|---|---|---|---|---|---|---|---|---|---|---|---|---|---|
| NC_011770 | 3027091 | T | C | 58 | 0 | 2 | 0 | | | | | | | C | C | | | | |
| NC_011770 | 3027099 | A | T | 62 | 0 | 2 | 0 | | | | | | | T | T | | | | |
| NC_011770 | 3027124 | C | T | 58 | 0 | 4 | 0 | | | T | T | T | T | | | | | | |
| NC_011770 | 3028738 | A | G | 51 | 0 | 12 | 0 | G | G | G | G | G | G | G | G | G | G | G | G |
| NC_011770 | 3028743 | G | C | 49 | 0 | 12 | 0 | C | C | C | C | C | C | C | C | C | C | C | C |
| NC_011770 | 3028769 | C | G | 45 | 0 | 12 | 0 | G | G | G | G | G | G | G | G | G | G | G | G |
| NC_011770 | 3028770 | T | G | 45 | 0 | 12 | 0 | G | G | G | G | G | G | G | G | G | G | G | G |
| NC_011770 | 3028773 | G | T | 45 | 0 | 2 | 0 | | | | | | | T | T | | | | |
| NC_011770 | 3028791 | G | A | 43 | 0 | 12 | 0 | A | A | A | A | A | A | A | A | A | A | A | A |
| NC_011770 | 3028819 | G | C | 43 | 0 | 12 | 0 | C | C | C | C | C | C | C | C | C | C | C | C |
| NC_011770 | 3028837 | CG | C | 44 | 0 | 10 | 0 | C | C | C | C | C | C | C | C | | | C | C |
| NC_011770 | 3028851 | G | A | 47 | 0 | 2 | 0 | | | | | | | | | A | A | | |
| NC_011770 | 3028861 | A | G | 42 | 0 | 8 | 0 | G | G | G | G | G | G | | | | | G | G |
| NC_011770 | 3028863 | T | C | 41 | 0 | 8 | 0 | C | C | C | C | C | C | | | | | C | C |
| NC_011770 | 3028864 | T | C | 42 | 0 | 12 | 0 | C | C | C | C | C | C | C | C | C | C | C | C |
| NC_011770 | 3028894 | T | C | 5 | 10 | 2 | 0 | . | . | . | . | . | . | . | . | C | C | . | . |
| NC_011770 | 3031055 | G | T | 77 | 0 | 2 | 0 | T | T | | | | | | | | | | |
| NC_011770 | 3037417 | G | A | 72 | 0 | 10 | 0 | A | A | A | A | A | A | A | A | | | A | A |
| NC_011770 | 3038796 | C | T | 57 | 0 | 6 | 0 | T | T | | | | | | | T | T | T | T |
| NC_011770 | 3038845 | A | G | 50 | 0 | 6 | 0 | G | G | | | | | | | G | G | G | G |
| NC_011770 | 3043296 | T | C | 62 | 0 | 2 | 0 | | | | | | | | | C | C | | |
| NC_011770 | 3043299 | T | C | 62 | 0 | 2 | 0 | | | | | | | | | C | C | | |
| NC_011770 | 3043320 | A | G | 62 | 0 | 2 | 0 | G | G | | | | | | | | | | |
| NC_011770 | 3043401 | C | T | 64 | 0 | 2 | 0 | | | | | | | | | T | T | | |
| NC_011770 | 3045887 | C | G | 69 | 0 | 2 | 0 | | | | | | | | | | | G | G |
| NC_011770 | 3045906 | C | T | 70 | 0 | 2 | 0 | | | | | | | | | T | T | | |
| NC_011770 | 3045928 | A | G | 66 | 0 | 12 | 0 | G | G | G | G | G | G | G | G | G | G | G | G |
| NC_011770 | 3045982 | GC | G | 67 | 0 | 2 | 0 | G | G | | | | | | | | | | |
| NC_011770 | 3047173 | G | A | 72 | 0 | 2 | 0 | A | A | | | | | | | | | | |
| NC_011770 | 3047871 | C | T | 36 | 6 | 6 | 0 | T | T | T | T | T | T | . | . | . | . | . | . |
| NC_011770 | 3066973 | C | T | 37 | 8 | 4 | 0 | . | . | T | T | T | T | . | . | . | . | . | . |
| NC_011770 | 3066995 | G | A | 37 | 8 | 4 | 0 | . | . | A | A | A | A | . | . | . | . | . | . |
| NC_011770 | 3067249 | T | G | 35 | 8 | 4 | 0 | . | . | G | G | G | G | . | . | . | . | . | . |
| NC_011770 | 3067295 | C | T | 47 | 6 | 2 | 0 | . | . | | | | | . | . | T | T | . | . |
| NC_011770 | 3067298 | G | A | 46 | 6 | 2 | 0 | . | . | | | | | . | . | A | A | . | . |
| NC_011770 | 3067336 | A | G | 38 | 6 | 2 | 0 | . | . | . | . | . | . | . | . | G | G | . | . |
| NC_011770 | 3067359 | C | T | 34 | 8 | 4 | 0 | . | . | T | T | T | T | . | . | . | . | . | . |
| NC_011770 | 3067374 | T | C | 32 | 8 | 4 | 0 | . | . | C | C | C | C | . | . | . | . | . | . |
| NC_011770 | 3067469 | C | T | 14 | 8 | 4 | 0 | . | . | T | T | T | T | . | . | . | . | . | . |
| NC_011770 | 3068260 | C | T | 14 | 8 | 2 | 0 | . | . | T | T | | | . | . | . | . | . | . |
| NC_011770 | 3068263 | G | A | 12 | 8 | 2 | 0 | . | . | A | A | | | . | . | . | . | . | . |
| NC_011770 | 3068266 | T | TC | 12 | 8 | 2 | 0 | . | . | TC | TC | | | . | . | . | . | . | . |
| NC_011770 | 3068367 | G | T | 30 | 8 | 4 | 0 | . | . | T | T | T | T | . | . | . | . | . | . |
| NC_011770 | 3068383 | G | A | 31 | 8 | 4 | 0 | . | . | A | A | A | A | . | . | . | . | . | . |
| NC_011770 | 3068634 | T | C | 12 | 10 | 2 | 0 | . | . | . | . | C | C | . | . | . | . | . | . |
| NC_011770 | 3068637 | T | C | 11 | 10 | 2 | 0 | . | . | . | . | C | C | . | . | . | . | . | . |
| NC_011770 | 3068649 | C | T | 12 | 10 | 2 | 0 | . | . | . | . | T | T | . | . | . | . | . | . |
| NC_011770 | 3068673 | T | C | 11 | 10 | 2 | 0 | . | . | . | . | C | C | . | . | . | . | . | . |
| NC_011770 | 3068719 | G | A | 9 | 10 | 2 | 0 | . | . | . | . | A | A | . | . | . | . | . | . |
| NC_011770 | 3068738 | C | T | 8 | 10 | 2 | 0 | . | . | . | . | T | T | . | . | . | . | . | . |
| NC_011770 | 3068775 | G | A | 8 | 10 | 2 | 0 | . | . | . | . | A | A | . | . | . | . | . | . |
| NC_011770 | 3068800 | A | G | 8 | 10 | 2 | 0 | . | . | . | . | G | G | . | . | . | . | . | . |
| NC_011770 | 3068829 | A | G | 7 | 10 | 2 | 0 | . | . | . | . | G | G | . | . | . | . | . | . |
| NC_011770 | 3068832 | C | G | 7 | 10 | 2 | 0 | . | . | . | . | G | G | . | . | . | . | . | . |
| NC_011770 | 3068885 | G | C | 9 | 6 | 6 | 0 | C | C | C | C | C | C | . | . | . | . | . | . |
| NC_011770 | 3068930 | G | A | 9 | 6 | 2 | 0 | A | A | | | | | . | . | . | . | . | . |
| NC_011770 | 3068961 | C | T | 4 | 10 | 2 | 0 | T | T | . | . | . | . | . | . | . | . | . | . |
| NC_011770 | 3068993 | G | A | 21 | 6 | 2 | 0 | A | A | | | | | . | . | . | . | . | . |
| NC_011770 | 3068999 | C | T | 19 | 6 | 6 | 0 | T | T | T | T | T | T | . | . | . | . | . | . |
| NC_011770 | 3069024 | T | C | 20 | 6 | 6 | 0 | C | C | C | C | C | C | . | . | . | . | . | . |
| NC_011770 | 3069051 | A | G | 19 | 6 | 2 | 0 | | | G | G | | | . | . | . | . | . | . |
| NC_011770 | 3069063 | T | TGTGCA | 15 | 6 | 2 | 0 | | | TGTGCAA | TGTGCAA | | | . | . | . | . | . | . |

| | | | | | | | | | | | | | | | | | | | |
|---|---|---|---|---|---|---|---|---|---|---|---|---|---|---|---|---|---|---|---|
| NC_011770 | 3069073 | T | G | 39 | 6 | 2 | 0 | | | G | G | | | . | . | . | . | . | . |
| NC_011770 | 3069078 | T | C | 40 | 6 | 2 | 0 | | | C | C | | | . | . | . | . | . | . |
| NC_011770 | 3069081 | T | A | 41 | 6 | 2 | 0 | | | A | A | | | . | . | . | . | . | . |
| NC_011770 | 3069100 | T | C | 79 | 6 | 2 | 0 | | | C | C | | | . | . | . | . | . | . |
| NC_011770 | 3069162 | A | AG | 12 | 8 | 2 | 0 | | | . | . | AG | AG | . | . | . | . | . | . |
| NC_011770 | 3069174 | C | T | 41 | 6 | 2 | 0 | | | T | T | | | . | . | . | . | . | . |
| NC_011770 | 3069178 | G | A | 42 | 6 | 2 | 0 | | | A | A | | | . | . | . | . | . | . |
| NC_011770 | 3069182 | T | C | 41 | 6 | 2 | 0 | C | C | | | | | . | . | . | . | . | . |
| NC_011770 | 3069197 | A | G | 40 | 6 | 2 | 0 | G | G | | | | | . | . | . | . | . | . |
| NC_011770 | 3069205 | G | A | 41 | 6 | 2 | 0 | A | A | | | | | . | . | . | . | . | . |
| NC_011770 | 3069254 | T | TG | 39 | 6 | 6 | 0 | TG | TG | TG | TG | TG | TG | . | . | . | . | . | . |
| NC_011770 | 3069300 | G | C | 34 | 6 | 4 | 0 | | | C | C | C | C | . | . | . | . | . | . |
| NC_011770 | 3069522 | T | A | 11 | 6 | 2 | 0 | | | A | A | | | . | . | . | . | . | . |
| NC_011770 | 3069532 | C | T | 11 | 6 | 6 | 0 | T | T | T | T | T | T | . | . | . | . | . | . |
| NC_011770 | 3069543 | A | G | 12 | 6 | 1 | 0 | G | . | | | | | . | . | . | . | . | . |
| NC_011770 | 3069554 | G | A | 12 | 7 | 3 | 0 | A | . | A | A | | | . | . | . | . | . | . |
| NC_011770 | 3069765 | T | G | 2 | 7 | 1 | 0 | G | . | | | | | . | . | . | . | . | . |
| NC_011770 | 3069790 | T | C | 0 | 10 | 2 | 0 | . | . | C | . | C | . | . | . | . | . | . | . |
| NC_011770 | 3069792 | C | T | 1 | 9 | 1 | 0 | | . | | . | T | . | . | . | . | . | . | . |
| NC_011770 | 3069807 | C | T | 0 | 10 | 1 | 0 | T | . | . | . | . | . | . | . | . | . | . | . |
| NC_011770 | 3070098 | C | T | 3 | 10 | 2 | 0 | T | T | . | . | . | . | . | . | . | . | . | . |
| NC_011770 | 3070243 | G | A | 5 | 10 | 2 | 0 | A | A | . | . | . | . | . | . | . | . | . | . |
| NC_011770 | 3070256 | G | A | 5 | 10 | 2 | 0 | A | A | . | . | . | . | . | . | . | . | . | . |
| NC_011770 | 3070318 | C | T | 5 | 10 | 2 | 0 | T | T | . | . | . | . | . | . | . | . | . | . |
| NC_011770 | 3070402 | A | G | 4 | 10 | 2 | 0 | G | G | . | . | . | . | . | . | . | . | . | . |
| NC_011770 | 3070439 | A | G | 3 | 10 | 2 | 0 | G | G | . | . | . | . | . | . | . | . | . | . |
| NC_011770 | 3070466 | G | A | 4 | 10 | 2 | 0 | A | A | . | . | . | . | . | . | . | . | . | . |
| NC_011770 | 3070486 | C | T | 3 | 10 | 2 | 0 | T | T | . | . | . | . | . | . | . | . | . | . |
| NC_011770 | 3070489 | C | T | 3 | 10 | 2 | 0 | T | T | . | . | . | . | . | . | . | . | . | . |
| NC_011770 | 3070498 | A | G | 18 | 6 | 2 | 0 | G | G | | | | | . | . | . | . | . | . |
| NC_011770 | 3070515 | G | T | 40 | 0 | 2 | 0 | T | T | | | | | | | | | | |
| NC_011770 | 3070529 | A | G | 39 | 0 | 2 | 0 | G | G | | | | | | | | | | |
| NC_011770 | 3070538 | T | C | 37 | 0 | 4 | 0 | C | C | | | | | | | C | C | | |
| NC_011770 | 3070545 | C | T | 36 | 0 | 2 | 0 | | | | | | | | | T | T | | |
| NC_011770 | 3070548 | G | A | 35 | 0 | 2 | 0 | | | | | | | | | A | A | | |
| NC_011770 | 3070567 | C | T | 32 | 2 | 2 | 0 | . | . | | | | | | | | | T | T |
| NC_011770 | 3070574 | G | T | 31 | 2 | 1 | 0 | | . | | | | | | | | | . | T |
| NC_011770 | 3070577 | GC | G | 31 | 2 | 1 | 0 | | . | | | | | | . | | | . | G |
| NC_011770 | 3070580 | T | C | 30 | 2 | 7 | 0 | C | . | C | C | C | C | | | C | C | . | |
| NC_011770 | 3070582 | G | A | 31 | 1 | 1 | 0 | | | | | | | | . | | | . | A |
| NC_011770 | 3070586 | G | T | 33 | 1 | 3 | 0 | | | | | | | T | T | | | . | T |
| NC_011770 | 3070589 | C | T | 35 | 1 | 1 | 0 | | | | | | | | . | | | . | T |
| NC_011770 | 3070597 | A | G | 35 | 1 | 3 | 0 | | | | | | | G | G | | | . | G |
| NC_011770 | 3070598 | G | A | 36 | 1 | 3 | 0 | | | | | | | A | A | | | . | A |
| NC_011770 | 3070604 | T | C | 34 | 1 | 3 | 0 | | | | | | | C | C | | | . | C |
| NC_011770 | 3070606 | GT | G | 36 | 1 | 3 | 0 | | | | | | | G | G | | | . | G |
| NC_011770 | 3070613 | T | C | 37 | 1 | 3 | 0 | | | | | | | C | C | | | . | C |
| NC_011770 | 3070620 | T | C | 37 | 1 | 3 | 0 | | | | | | | C | C | | | . | C |
| NC_011770 | 3070635 | T | G | 36 | 2 | 2 | 0 | | | | | | | G | . | | | . | G |
| NC_011770 | 3070636 | C | A | 36 | 2 | 2 | 0 | | | | | | | A | . | | | . | A |
| NC_011770 | 3070644 | T | G | 33 | 3 | 1 | 0 | | | | | | | G | . | | | . | . |
| NC_011770 | 3070677 | A | G | 29 | 4 | 2 | 0 | G | G | | | | | . | . | | | . | . |
| NC_011770 | 3070706 | G | C | 35 | 0 | 3 | 0 | | | | | | | . | C | | | C | C |
| NC_011770 | 3070730 | T | TCG | 15 | 2 | 1 | 0 | . | . | | | | | | | . | TCG | | |
| NC_011770 | 3070743 | A | AGAG | 5 | 10 | 2 | 0 | . | . | . | . | . | . | . | . | AGAG | AGAG | . | . |
| NC_011770 | 3070765 | C | T | 5 | 10 | 2 | 0 | . | . | . | . | . | . | . | . | T | T | . | . |
| NC_011770 | 3070890 | C | A | 5 | 10 | 2 | 0 | . | . | . | . | . | . | . | . | A | A | . | . |
| NC_011770 | 3070931 | G | A | 5 | 10 | 2 | 0 | . | . | . | . | . | . | . | . | A | A | . | . |
| NC_011770 | 3071072 | G | A | 5 | 10 | 2 | 0 | . | . | . | . | . | . | . | . | A | A | . | . |
| NC_011770 | 3071277 | C | A | 5 | 10 | 2 | 0 | . | . | . | . | . | . | . | . | A | A | . | . |
| NC_011770 | 3071308 | G | A | 4 | 10 | 2 | 0 | . | . | . | . | . | . | . | . | A | A | . | . |
| NC_011770 | 3071409 | A | G | 24 | 3 | 6 | 0 | . | . | . | | | | G | G | G | G | G | G |

| | | | | | | | | | | | | | | | | | | | |
|---|---|---|---|---|---|---|---|---|---|---|---|---|---|---|---|---|---|---|---|
| NC_011770 | 3071417 | T | G | 19 | 5 | 2 | 0 | . | . | . | | | | | | G | G | . | . |
| NC_011770 | 3071427 | G | A | 9 | 8 | 1 | 0 | . | . | . | . | | | . | A | . | . | . | . |
| NC_011770 | 3071428 | A | G | 12 | 8 | 2 | 0 | . | . | . | . | | | G | G | . | . | . | . |
| NC_011770 | 3071433 | C | T | 12 | 7 | 1 | 0 | . | . | . | . | | | | | T | . | . | . |
| NC_011770 | 3071435 | CCAGTG | C | 12 | 7 | 1 | 0 | . | . | . | . | | | | | C | . | . | . |
| NC_011770 | 3071439 | T | C | 10 | 8 | 2 | 0 | . | . | . | . | | | C | C | . | . | . | . |
| NC_011770 | 3071446 | A | G | 16 | 6 | 3 | 0 | . | . | . | . | | | G | . | G | G | . | . |
| NC_011770 | 3071451 | G | A | 17 | 6 | 4 | 0 | . | . | . | . | | | A | A | A | A | . | . |
| NC_011770 | 3071455 | A | G | 17 | 6 | 4 | 0 | . | . | . | . | | | G | G | G | G | . | . |
| NC_011770 | 3071457 | G | A | 17 | 6 | 4 | 0 | . | . | . | . | | | A | A | A | A | . | . |
| NC_011770 | 3071467 | GC | G | 21 | 6 | 4 | 0 | . | . | . | . | | | G | G | G | G | . | . |
| NC_011770 | 3071496 | G | A | 27 | 4 | 2 | 0 | . | . | . | . | | | | | | | A | A |
| NC_011770 | 3071506 | T | C | 28 | 4 | 2 | 0 | . | . | . | . | | | | | | | C | C |
| NC_011770 | 3071510 | G | A | 28 | 4 | 2 | 0 | . | . | . | . | | | | | A | A | | |
| NC_011770 | 3071517 | G | A | 30 | 4 | 6 | 0 | . | . | . | . | | | A | A | A | A | A | A |
| NC_011770 | 3071523 | G | A | 29 | 4 | 2 | 0 | . | . | . | . | | | | | A | A | | |
| NC_011770 | 3071526 | G | A | 29 | 4 | 2 | 0 | . | . | . | . | | | | | | | A | A |
| NC_011770 | 3071529 | T | C | 29 | 4 | 2 | 0 | . | . | . | . | | | | | | | C | C |
| NC_011770 | 3071535 | A | G | 31 | 2 | 2 | 0 | . | . | | | | | | | G | G | | |
| NC_011770 | 3071540 | T | G | 32 | 2 | 2 | 0 | . | . | G | G | | | | | | | | |
| NC_011770 | 3071541 | A | G | 31 | 2 | 2 | 0 | . | . | | | | | | | G | G | | |
| NC_011770 | 3071560 | T | C | 29 | 1 | 1 | 0 | . | C | | | | | | | | | | |
| NC_011770 | 3071566 | T | C | 18 | 5 | 1 | 0 | . | C | . | . | | | | | | . | | . |
| NC_011770 | 3074639 | G | T | 10 | 6 | 2 | 0 | . | . | . | . | . | . | | . | | | T | T |
| NC_011770 | 3075792 | C | T | 19 | 8 | 2 | 0 | . | . | . | . | . | . | | | . | . | T | T |
| NC_011770 | 3075803 | G | A | 18 | 8 | 2 | 0 | . | . | . | . | . | . | | | . | . | A | A |
| NC_011770 | 3075820 | G | A | 18 | 8 | 2 | 0 | . | . | . | . | . | . | | | . | . | A | A |
| NC_011770 | 3075829 | G | A | 18 | 8 | 2 | 0 | . | . | . | . | . | . | | | . | . | A | A |
| NC_011770 | 3075863 | A | G | 18 | 8 | 4 | 0 | . | . | . | . | . | . | G | G | . | . | G | G |
| NC_011770 | 3075890 | T | G | 21 | 8 | 4 | 0 | . | . | . | . | . | . | G | G | . | . | G | G |
| NC_011770 | 3076063 | A | G | 0 | 10 | 2 | 0 | . | . | . | . | . | . | . | . | . | . | G | G |
| NC_011770 | 3076072 | G | A | 0 | 10 | 2 | 0 | . | . | . | . | . | . | . | . | . | . | A | A |
| NC_011770 | 3076073 | C | T | 0 | 11 | 1 | 0 | . | . | . | . | . | . | . | . | . | . | . | T |
| NC_011770 | 3076075 | A | G | 0 | 10 | 2 | 0 | . | . | . | . | . | . | . | . | . | . | G | G |
| NC_011770 | 3076086 | A | G | 0 | 10 | 2 | 0 | . | . | . | . | . | . | . | . | . | . | G | G |
| NC_011770 | 3076089 | C | G | 0 | 10 | 2 | 0 | . | . | . | . | . | . | . | . | . | . | G | G |
| NC_011770 | 3076090 | G | A | 0 | 10 | 2 | 0 | . | . | . | . | . | . | . | . | . | . | A | A |
| NC_011770 | 3076099 | C | T | 0 | 10 | 2 | 0 | . | . | . | . | . | . | . | . | . | . | T | T |
| NC_011770 | 3076101 | C | G | 0 | 10 | 2 | 0 | . | . | . | . | . | . | . | . | . | . | G | G |
| NC_011770 | 3076102 | A | G | 0 | 10 | 2 | 0 | . | . | . | . | . | . | . | . | . | . | G | G |
| NC_011770 | 3076169 | T | A | 31 | 8 | 4 | 0 | . | . | . | . | . | . | A | A | . | . | A | A |
| NC_011770 | 3076225 | A | G | 35 | 8 | 4 | 0 | . | . | . | . | . | . | G | G | . | . | G | G |
| NC_011770 | 3076254 | C | A | 34 | 8 | 4 | 0 | . | . | . | . | . | . | A | A | . | . | A | A |
| NC_011770 | 3077465 | G | A | 24 | 8 | 2 | 0 | . | . | . | . | . | . | | | . | . | A | A |
| NC_011770 | 3077503 | A | G | 22 | 8 | 4 | 0 | . | . | . | . | . | . | G | G | . | . | G | G |
| NC_011770 | 3077556 | G | T | 20 | 8 | 4 | 0 | . | . | . | . | . | . | T | T | . | . | T | T |
| NC_011770 | 3077562 | T | G | 20 | 8 | 4 | 0 | . | . | . | . | . | . | G | G | . | . | G | G |
| NC_011770 | 3077626 | G | A | 19 | 8 | 2 | 0 | . | . | . | . | . | . | | | . | . | A | A |
| NC_011770 | 3077779 | G | A | 22 | 8 | 4 | 0 | . | . | . | . | . | . | A | A | . | . | A | A |
| NC_011770 | 3077797 | A | G | 23 | 8 | 2 | 0 | . | . | . | . | . | . | G | G | . | . | | |
| NC_011770 | 3077840 | A | G | 23 | 8 | 2 | 0 | . | . | . | . | . | . | | | . | . | G | G |
| NC_011770 | 3077868 | C | T | 23 | 8 | 1 | 0 | . | . | . | . | . | . | T | | . | . | | |
| NC_011770 | 3079326 | C | T | 71 | 0 | 4 | 0 | | | | | | | | | T | T | T | T |
| NC_011770 | 3079334 | C | T | 69 | 0 | 2 | 0 | T | T | | | | | | | | | | |
| NC_011770 | 3079673 | T | C | 40 | 0 | 2 | 0 | | | | | | | C | C | | | | |
| NC_011770 | 3079706 | T | G | 37 | 0 | 2 | 0 | | | | | | | G | G | | | | |
| NC_011770 | 3079717 | A | C | 36 | 0 | 6 | 0 | C | C | | | | | | | C | C | C | C |
| NC_011770 | 3081362 | T | G | 65 | 0 | 2 | 0 | | | | | | | G | G | | | | |
| NC_011770 | 3081373 | G | A | 72 | 0 | 6 | 0 | | | A | A | A | A | | | | | A | A |
| NC_011770 | 3081393 | T | C | 68 | 0 | 12 | 0 | C | C | C | C | C | C | C | C | C | C | C | C |
| NC_011770 | 3082454 | G | A | 83 | 0 | 4 | 0 | | | A | A | A | A | | | | | | |
| NC_011770 | 3085890 | A | G | 51 | 0 | 4 | 0 | G | G | | | | | G | G | | | | |

| | | | | | | | | | | | | | | | | | | | |
|---|---|---|---|---|---|---|---|---|---|---|---|---|---|---|---|---|---|---|---|
| NC_011770 | 3086993 | G | T | 65 | 0 | 2 | 0 | | | | | | | T | T | | | | |
| NC_011770 | 3087016 | T | C | 63 | 0 | 12 | 0 | C | C | C | C | C | C | C | C | C | C | C | C |
| NC_011770 | 3087049 | A | T | 66 | 0 | 2 | 0 | | | | | | | T | T | | | | |
| NC_011770 | 3087070 | C | T | 71 | 0 | 2 | 0 | | | | | | | | | | | T | T |
| NC_011770 | 3087498 | C | T | 61 | 0 | 10 | 0 | T | T | T | T | T | T | T | T | T | T | | |
| NC_011770 | 3090429 | C | A | 58 | 0 | 10 | 0 | A | A | A | A | A | A | A | A | A | A | | |
| NC_011770 | 3090486 | A | G | 55 | 0 | 2 | 0 | | | | | | | | | | | G | G |
| NC_011770 | 3091785 | A | G | 72 | 0 | 8 | 0 | | | G | G | G | G | G | G | | | G | G |
| NC_011770 | 3091793 | G | A | 73 | 0 | 8 | 0 | | | A | A | A | A | A | A | | | A | A |
| NC_011770 | 3093210 | CGCTG | C | 71 | 0 | 2 | 0 | | | | | | | | | C | C | | |
| NC_011770 | 3093231 | T | C | 74 | 0 | 8 | 0 | | | C | C | C | C | | | C | C | C | C |
| NC_011770 | 3093267 | C | A | 74 | 0 | 2 | 0 | | | | | | | A | A | | | | |
| NC_011770 | 3096579 | C | T | 78 | 0 | 4 | 0 | | | T | T | T | T | | | | | | |
| NC_011770 | 3096941 | T | C | 73 | 0 | 2 | 0 | | | | | | | | | C | C | | |
| NC_011770 | 3101071 | T | C | 32 | 0 | 12 | 0 | C | C | C | C | C | C | C | C | C | C | C | C |
| NC_011770 | 3101657 | C | T | 80 | 0 | 2 | 0 | | | | | | | T | T | | | | |
| NC_011770 | 3104225 | G | A | 47 | 0 | 2 | 0 | | | | | | | | | A | A | | |
| NC_011770 | 3104267 | G | A | 45 | 0 | 2 | 0 | A | A | | | | | | | | | | |
| NC_011770 | 3104276 | C | T | 44 | 0 | 6 | 0 | T | T | | | | | T | T | T | T | | |
| NC_011770 | 3104286 | T | G | 45 | 0 | 2 | 0 | | | | | | | G | G | | | | |
| NC_011770 | 3104290 | A | G | 42 | 0 | 10 | 0 | G | G | G | G | G | G | G | G | G | G | | |
| NC_011770 | 3104700 | G | C | 67 | 0 | 2 | 0 | | | | | | | | | C | C | | |
| NC_011770 | 3105740 | T | C | 68 | 0 | 4 | 0 | C | C | | | | | | | C | C | | |
| NC_011770 | 3105773 | G | T | 63 | 0 | 2 | 0 | | | | | | | | | T | T | | |
| NC_011770 | 3108214 | G | A | 81 | 0 | 2 | 0 | A | A | | | | | | | | | | |
| NC_011770 | 3108250 | C | T | 84 | 0 | 2 | 0 | | | | | | | T | T | | | | |
| NC_011770 | 3108289 | C | T | 87 | 0 | 6 | 0 | T | T | T | T | T | T | | | | | | |
| NC_011770 | 3108301 | C | T | 83 | 0 | 10 | 0 | T | T | T | T | T | T | T | T | | | T | T |
| NC_011770 | 3108390 | A | G | 78 | 0 | 6 | 0 | G | G | G | G | G | G | | | | | | |
| NC_011770 | 3109892 | C | T | 72 | 0 | 6 | 0 | | | T | T | T | T | | | A | A | | |
| NC_011770 | 3112138 | C | A | 66 | 0 | 2 | 0 | | | | | | | A | A | | | | |
| NC_011770 | 3112141 | C | T | 66 | 0 | 2 | 0 | | | | | | | T | T | | | | |
| NC_011770 | 3112282 | G | T | 69 | 0 | 2 | 0 | T | T | | | | | | | | | | |
| NC_011770 | 3112289 | A | C | 69 | 0 | 2 | 0 | C | C | | | | | | | | | | |
| NC_011770 | 3112290 | T | C | 69 | 0 | 2 | 0 | C | C | | | | | | | | | | |
| NC_011770 | 3112298 | C | G | 65 | 0 | 4 | 0 | | | | | | | G | G | G | G | | |
| NC_011770 | 3112309 | G | A | 64 | 0 | 4 | 0 | | | | | | | A | A | A | A | | |
| NC_011770 | 3112331 | T | G | 64 | 0 | 8 | 0 | | | G | G | G | G | G | G | G | G | | |
| NC_011770 | 3112351 | T | C | 61 | 0 | 2 | 0 | | | | | | | | | C | C | | |
| NC_011770 | 3112382 | G | A | 64 | 0 | 2 | 0 | | | | | | | | | | | A | A |
| NC_011770 | 3112435 | G | C | 67 | 0 | 2 | 0 | | | | | | | | | C | C | | |
| NC_011770 | 3126536 | G | A | 36 | 4 | 2 | 0 | | | . | . | . | . | A | A | | | | |
| NC_011770 | 3126556 | A | G | 37 | 4 | 2 | 0 | | | . | . | . | . | G | G | | | | |
| NC_011770 | 3126572 | C | T | 40 | 4 | 2 | 0 | | | . | . | . | . | T | T | | | | |
| NC_011770 | 3126578 | T | C | 36 | 4 | 2 | 0 | | | . | . | . | . | C | C | | | | |
| NC_011770 | 3126598 | G | A | 39 | 4 | 2 | 0 | | | . | . | . | . | A | A | | | | |
| NC_011770 | 3126599 | T | G | 38 | 4 | 2 | 0 | | | . | . | . | . | G | G | | | | |
| NC_011770 | 3126604 | A | ATT | 39 | 4 | 2 | 0 | | | . | . | . | . | ATT | ATT | | | | |
| NC_011770 | 3126612 | A | G | 39 | 4 | 2 | 0 | | | . | . | . | . | G | G | | | | |
| NC_011770 | 3126622 | A | G | 43 | 4 | 2 | 0 | | | . | . | . | . | G | G | | | | |
| NC_011770 | 3126627 | G | T | 40 | 4 | 6 | 0 | T | T | . | . | . | . | T | T | T | T | | |
| NC_011770 | 3127479 | T | C | 38 | 4 | 2 | 0 | | | . | . | . | . | | | | | C | C |
| NC_011770 | 3127522 | A | G | 39 | 4 | 8 | 0 | G | G | . | . | . | . | G | G | G | G | G | G |
| NC_011770 | 3127560 | G | C | 38 | 4 | 4 | 0 | | | . | . | . | . | | | C | C | C | C |
| NC_011770 | 3127565 | G | A | 38 | 4 | 8 | 0 | A | A | . | . | . | . | A | A | A | A | A | A |
| NC_011770 | 3127614 | C | T | 35 | 4 | 4 | 0 | T | T | . | . | . | . | T | T | | | | |
| NC_011770 | 3127623 | C | T | 21 | 6 | 2 | 0 | T | T | . | . | . | . | . | . | | | | |
| NC_011770 | 3127629 | T | C | 21 | 4 | 4 | 0 | C | C | . | . | . | . | C | C | | | | |
| NC_011770 | 3127636 | T | G | 18 | 8 | 2 | 0 | . | . | . | . | . | . | . | . | | | G | G |
| NC_011770 | 3127642 | T | G | 19 | 6 | 6 | 0 | . | G | . | . | . | . | G | . | G | G | G | G |
| NC_011770 | 3127664 | C | T | 30 | 4 | 4 | 0 | T | T | . | . | . | . | T | T | | | | |
| NC_011770 | 3127665 | T | C | 30 | 4 | 4 | 0 | C | C | . | . | . | . | C | C | | | | |

| | | | | | | | | | | | | | | | | | | | |
|---|---|---|---|---|---|---|---|---|---|---|---|---|---|---|---|---|---|---|---|
| NC_011770 | 3127666 | T | C | 30 | 4 | 4 | 0 | C | C | . | . | . | . | C | C | | | | |
| NC_011770 | 3127674 | G | A | 29 | 4 | 4 | 0 | A | A | . | . | . | . | A | A | | | | |
| NC_011770 | 3127675 | A | G | 29 | 4 | 4 | 0 | G | G | . | . | . | . | G | G | | | | |
| NC_011770 | 3127676 | T | G | 30 | 4 | 4 | 0 | G | G | . | . | . | . | G | G | | | | |
| NC_011770 | 3127677 | A | T | 30 | 4 | 4 | 0 | T | T | . | . | . | . | T | T | | | | |
| NC_011770 | 3127682 | C | G | 31 | 4 | 4 | 0 | G | G | . | . | . | . | G | G | | | | |
| NC_011770 | 3127687 | T | C | 32 | 4 | 4 | 0 | C | C | . | . | . | . | C | C | | | | |
| NC_011770 | 3127698 | C | T | 39 | 4 | 2 | 0 | | | . | . | . | . | T | T | | | | |
| NC_011770 | 3127702 | G | A | 37 | 4 | 4 | 0 | A | A | . | . | . | . | A | A | | | | |
| NC_011770 | 3127745 | A | G | 38 | 4 | 6 | 0 | | | . | . | . | . | G | G | G | G | G | G |
| NC_011770 | 3127754 | C | T | 14 | 4 | 6 | 0 | | | . | . | . | . | T | T | T | T | T | T |
| NC_011770 | 3127814 | T | A | 37 | 4 | 6 | 0 | | | . | . | . | . | A | A | A | A | A | A |
| NC_011770 | 3127815 | C | T | 37 | 4 | 6 | 0 | | | . | . | . | . | T | T | T | T | T | T |
| NC_011770 | 3127818 | C | T | 38 | 4 | 2 | 0 | | | . | . | . | . | T | T | | | | |
| NC_011770 | 3127827 | C | T | 41 | 4 | 2 | 0 | | | . | . | . | . | T | T | | | | |
| NC_011770 | 3127842 | A | T | 40 | 4 | 2 | 0 | | | . | . | . | . | T | T | | | | |
| NC_011770 | 3127917 | ACCTGTTG/ | A | 31 | 4 | 1 | 0 | | | . | . | . | . | A | | | | | |
| NC_011770 | 3127969 | T | C | 43 | 4 | 8 | 0 | C | C | . | . | . | . | C | C | C | C | C | C |
| NC_011770 | 3127971 | T | G | 43 | 4 | 6 | 0 | | | . | . | . | . | G | G | G | G | G | G |
| NC_011770 | 3128893 | A | G | 46 | 0 | 12 | 0 | G | G | G | G | G | G | G | G | G | G | G | G |
| NC_011770 | 3128908 | G | A | 49 | 0 | 2 | 0 | | | | | | | | | A | A | | |
| NC_011770 | 3129015 | A | G | 46 | 0 | 6 | 0 | | | G | G | G | G | | | G | G | | |
| NC_011770 | 3129077 | A | G | 56 | 0 | 2 | 0 | | | | | | | | | G | G | | |
| NC_011770 | 3129406 | T | G | 37 | 0 | 2 | 0 | G | G | | | | | | | | | | |
| NC_011770 | 3129417 | G | A | 39 | 0 | 6 | 0 | | | A | A | A | A | | | A | A | | |
| NC_011770 | 3129527 | T | C | 48 | 0 | 2 | 0 | C | C | | | | | | | | | | |
| NC_011770 | 3129643 | A | G | 54 | 0 | 12 | 0 | G | G | G | G | G | G | G | G | G | G | G | G |
| NC_011770 | 3129662 | G | T | 55 | 0 | 4 | 0 | | | T | T | T | T | | | | | | |
| NC_011770 | 3129675 | C | T | 53 | 0 | 2 | 0 | | | | | | | T | T | | | | |
| NC_011770 | 3129677 | C | G | 54 | 0 | 2 | 0 | | | | | | | G | G | | | | |
| NC_011770 | 3129710 | T | C | 59 | 0 | 4 | 0 | | | C | C | C | C | | | | | | |
| NC_011770 | 3129728 | A | G | 57 | 0 | 2 | 0 | G | G | | | | | | | | | | |
| NC_011770 | 3129729 | C | A | 57 | 0 | 2 | 0 | A | A | | | | | | | | | | |
| NC_011770 | 3129739 | C | T | 53 | 0 | 2 | 0 | | | | | | | | | | | T | T |
| NC_011770 | 3129740 | G | A | 56 | 0 | 2 | 0 | | | | | | | | | | | A | A |
| NC_011770 | 3129744 | C | T | 56 | 0 | 2 | 0 | | | | | | | | | | | T | T |
| NC_011770 | 3129778 | C | T | 50 | 0 | 6 | 0 | T | T | | | | | | | T | T | T | T |
| NC_011770 | 3129796 | T | A | 47 | 0 | 6 | 0 | . | . | | | | | A | A | A | A | A | A |
| NC_011770 | 3129826 | C | T | 48 | 0 | 2 | 0 | | | | | | | | | T | T | | |
| NC_011770 | 3129837 | C | G | 20 | 2 | 2 | 0 | | | | | | | G | G | | | . | . |
| NC_011770 | 3129838 | C | T | 18 | 2 | 2 | 0 | | | | | | | T | T | | | . | . |
| NC_011770 | 3129839 | T | TG | 12 | 6 | 2 | 0 | | | . | . | . | . | TG | TG | | | . | . |
| NC_011770 | 3129847 | T | G | 32 | 0 | 8 | 0 | | | G | G | G | G | G | G | | | G | G |
| NC_011770 | 3129853 | T | C | 35 | 0 | 6 | 0 | | | C | C | C | C | C | C | | | | |
| NC_011770 | 3129854 | G | A | 36 | 0 | 2 | 0 | | | | | | | | | | | A | A |
| NC_011770 | 3129858 | T | C | 37 | 0 | 6 | 0 | | | C | C | C | C | C | C | | | | |
| NC_011770 | 3129866 | T | C | 40 | 0 | 6 | 0 | | | C | C | C | C | C | C | | | | |
| NC_011770 | 3129881 | A | G | 45 | 0 | 12 | 0 | G | G | G | G | G | G | G | G | G | G | G | G |
| NC_011770 | 3129892 | C | T | 45 | 0 | 6 | 0 | T | T | | | | | | | T | T | T | T |
| NC_011770 | 3129910 | T | A | 44 | 0 | 8 | 0 | A | A | | | | | A | A | A | A | A | A |
| NC_011770 | 3129948 | A | G | 49 | 0 | 12 | 0 | G | G | G | G | G | G | G | G | G | G | G | G |
| NC_011770 | 3129951 | C | G | 48 | 0 | 8 | 0 | G | G | | | | | G | G | G | G | A | A |
| NC_011770 | 3129952 | C | T | 48 | 0 | 2 | 0 | | | | | | | T | T | | | | |
| NC_011770 | 3129953 | C | T | 49 | 0 | 8 | 0 | T | T | | | | | T | T | T | T | T | T |
| NC_011770 | 3129977 | A | G | 57 | 0 | 12 | 0 | G | G | G | G | G | G | G | G | G | G | G | G |
| NC_011770 | 3129978 | C | T | 57 | 0 | 2 | 0 | | | | | | | | | | | T | T |
| NC_011770 | 3129991 | T | C | 59 | 0 | 2 | 0 | | | | | | | C | C | | | | |
| NC_011770 | 3130918 | G | A | 34 | 0 | 10 | 0 | A | A | A | A | A | A | A | A | A | A | | |
| NC_011770 | 3130919 | A | G | 34 | 0 | 10 | 0 | G | G | T | T | T | T | G | G | G | G | | |
| NC_011770 | 3132387 | A | G | 75 | 2 | 10 | 0 | G | G | G | G | G | G | G | G | . | . | G | G |
| NC_011770 | 3133159 | A | G | 69 | 0 | 2 | 0 | | | | | | | G | G | | | | |
| NC_011770 | 3133177 | C | T | 67 | 0 | 2 | 0 | | | | | | | T | T | | | | |

| | | | | | | | | | | | | | | | | | | | |
|---|---|---|---|---|---|---|---|---|---|---|---|---|---|---|---|---|---|---|---|
| NC_011770 | 3133182 | C | T | 68 | 0 | 2 | 0 | | | | | | | | | T | T | | |
| NC_011770 | 3133196 | G | A | 66 | 0 | 2 | 0 | | | | | | | A | A | | | | |
| NC_011770 | 3133210 | A | AG | 69 | 0 | 2 | 0 | | . | | | | | | | AG | AG | | |
| NC_011770 | 3133213 | G | GGC | 48 | 2 | 2 | 0 | | . | | | | | GGC | GGC | . | | | |
| NC_011770 | 3136827 | T | G | 32 | 0 | 12 | 0 | G | G | G | G | G | G | G | G | G | G | G | G |
| NC_011770 | 3136830 | C | G | 32 | 0 | 2 | 0 | G | G | | | | | | | | | | |
| NC_011770 | 3136844 | G | A | 32 | 0 | 2 | 0 | | | | | | | | | A | A | | |
| NC_011770 | 3136848 | G | A | 33 | 0 | 2 | 0 | A | A | | | | | | | | | | |
| NC_011770 | 3136858 | G | T | 33 | 0 | 8 | 0 | T | T | T | T | T | T | | | | | T | T |
| NC_011770 | 3136863 | A | C | 35 | 0 | 2 | 0 | C | C | | | | | | | | | | |
| NC_011770 | 3136880 | T | C | 35 | 0 | 2 | 0 | C | C | | | | | | | | | | |
| NC_011770 | 3136885 | G | C | 36 | 0 | 2 | 0 | C | C | | | | | | | | | | |
| NC_011770 | 3136903 | G | C | 40 | 0 | 2 | 0 | C | C | | | | | | | | | | |
| NC_011770 | 3136945 | A | G | 46 | 0 | 2 | 0 | G | G | | | | | | | | | | |
| NC_011770 | 3136992 | GC | G | 46 | 0 | 2 | 0 | G | G | | | | | | | | | | |
| NC_011770 | 3137030 | C | T | 51 | 0 | 2 | 0 | T | T | | | | | | | | | | |
| NC_011770 | 3137050 | T | C | 53 | 0 | 2 | 0 | | | | | | | C | C | | | | |
| NC_011770 | 3137096 | C | T | 58 | 0 | 2 | 0 | T | T | | | | | | | | | | |
| NC_011770 | 3138864 | C | T | 69 | 0 | 2 | 0 | | | | | | | | | T | T | | |
| NC_011770 | 3138902 | ACGGTGCT | A | 66 | 0 | 6 | 0 | A | A | | | | | A | A | TCGGTGCT | TCGGTGCTGCGG | | |
| NC_011770 | 3138914 | T | C | 45 | 0 | 4 | 0 | | | C | C | C | C | | | | | | |
| NC_011770 | 3138936 | T | G | 66 | 0 | 4 | 0 | | | G | G | G | G | | | | | | |
| NC_011770 | 3138960 | C | A | 69 | 0 | 4 | 0 | | | A | A | A | A | | | | | | |
| NC_011770 | 3138973 | T | C | 67 | 0 | 8 | 0 | C | C | | | | | C | C | C | C | C | C |
| NC_011770 | 3138978 | T | C | 67 | 0 | 8 | 0 | C | C | | | | | C | C | C | C | C | C |
| NC_011770 | 3138987 | C | A | 69 | 0 | 8 | 0 | A | A | A | A | A | A | A | A | | | | |
| NC_011770 | 3138995 | T | C | 70 | 0 | 4 | 0 | | | | | | | | | C | C | C | C |
| NC_011770 | 3138996 | C | G | 70 | 0 | 4 | 0 | | | | | | | | | G | G | G | G |
| NC_011770 | 3139012 | T | C | 71 | 0 | 8 | 0 | C | C | | | | | C | C | C | C | C | C |
| NC_011770 | 3139019 | G | A | 72 | 0 | 2 | 0 | | | | | | | | | | | A | A |
| NC_011770 | 3141553 | GA | G | 50 | 0 | 12 | 0 | G | G | G | G | G | G | G | G | G | G | G | G |
| NC_011770 | 3141568 | T | G | 52 | 0 | 6 | 0 | G | G | | | | | | | G | G | G | G |
| NC_011770 | 3141574 | G | T | 52 | 0 | 2 | 0 | T | T | | | | | | | | | | |
| NC_011770 | 3146674 | C | T | 62 | 0 | 1 | 0 | | | | | | | T | | | | | |
| NC_011770 | 3146682 | T | A | 65 | 0 | 2 | 0 | | | | | | | | | A | A | | |
| NC_011770 | 3146779 | G | A | 70 | 0 | 8 | 0 | | | A | A | A | A | A | A | | | A | A |
| NC_011770 | 3179220 | C | T | 10 | 8 | 4 | 0 | . | . | T | T | T | T | . | . | . | . | . | . |
| NC_011770 | 3179247 | C | A | 10 | 8 | 4 | 0 | . | . | A | A | A | A | . | . | . | . | . | . |
| NC_011770 | 3179384 | C | A | 11 | 8 | 4 | 0 | . | . | A | A | A | A | . | . | . | . | . | . |
| NC_011770 | 3181448 | G | A | 9 | 8 | 4 | 0 | . | . | A | A | A | A | . | . | . | . | . | . |
| NC_011770 | 3184441 | C | T | 10 | 8 | 4 | 0 | . | . | T | T | T | T | . | . | . | . | . | . |
| NC_011770 | 3184468 | C | A | 10 | 8 | 4 | 0 | . | . | A | A | A | A | . | . | . | . | . | . |
| NC_011770 | 3184605 | C | A | 12 | 8 | 4 | 0 | . | . | A | A | A | A | . | . | . | . | . | . |
| NC_011770 | 3186669 | G | A | 10 | 8 | 4 | 0 | . | . | A | A | A | A | . | . | . | . | . | . |
| NC_011770 | 3189662 | C | T | 13 | 8 | 4 | 0 | . | . | T | T | T | T | . | . | . | . | . | . |
| NC_011770 | 3189689 | C | A | 13 | 8 | 4 | 0 | . | . | A | A | A | A | . | . | . | . | . | . |
| NC_011770 | 3189826 | C | A | 11 | 8 | 4 | 0 | . | . | A | A | A | A | . | . | . | . | . | . |
| NC_011770 | 3191890 | G | A | 24 | 8 | 4 | 0 | . | . | A | A | A | A | . | . | . | . | . | . |
| NC_011770 | 3192942 | C | T | 41 | 8 | 4 | 0 | . | . | T | T | T | T | . | . | . | . | . | . |
| NC_011770 | 3192957 | C | CT | 43 | 8 | 4 | 0 | . | . | CT | CT | CT | CT | . | . | . | . | . | . |
| NC_011770 | 3196500 | A | G | 36 | 2 | 6 | 0 | G | G | | | | | G | G | . | . | G | G |
| NC_011770 | 3196503 | T | C | 36 | 2 | 6 | 0 | C | C | | | | | C | C | . | . | C | C |
| NC_011770 | 3196512 | G | T | 36 | 2 | 6 | 0 | T | T | | | | | T | T | . | . | T | T |
| NC_011770 | 3196602 | T | C | 28 | 8 | 4 | 0 | . | . | C | C | C | C | . | . | . | . | . | . |
| NC_011770 | 3196654 | C | T | 55 | 0 | 2 | 0 | | | | | | | | | T | T | | |
| NC_011770 | 3196658 | C | A | 55 | 0 | 6 | 0 | A | A | | | | | A | A | | | A | A |
| NC_011770 | 3196683 | G | A | 56 | 0 | 8 | 0 | A | A | | | | | A | A | A | A | A | A |
| NC_011770 | 3196689 | G | A | 56 | 0 | 8 | 0 | A | A | | | | | A | A | A | A | A | A |
| NC_011770 | 3196690 | C | T | 56 | 0 | 2 | 0 | | | | | | | | | T | T | | |
| NC_011770 | 3196693 | T | C | 57 | 0 | 2 | 0 | | | | | | | | | C | C | | |
| NC_011770 | 3196696 | AC | A | 57 | 0 | 2 | 0 | | | | | | | | | A | A | | |
| NC_011770 | 3196697 | C | T | 56 | 0 | 6 | 0 | T | T | | | | | T | T | . | . | T | T |

| | | | | | | | | | | | | | | | | | | | |
|---|---|---|---|---|---|---|---|---|---|---|---|---|---|---|---|---|---|---|---|
| NC_011770 | 3196727 | A | C | 55 | 0 | 8 | 0 | C | C | | | | | C | C | C | C | C | C |
| NC_011770 | 3196730 | T | A | 55 | 0 | 8 | 0 | A | A | | | | | A | A | A | A | A | A |
| NC_011770 | 3198348 | A | G | 60 | 0 | 2 | 0 | | | | | | | | | | | G | G |
| NC_011770 | 3199821 | A | G | 69 | 0 | 8 | 0 | | | G | G | G | G | G | G | G | G | | |
| NC_011770 | 3204146 | T | G | 69 | 0 | 2 | 0 | | | | | | | | | G | G | | |
| NC_011770 | 3204248 | C | T | 71 | 0 | 2 | 0 | | | | | | | | | T | T | | |
| NC_011770 | 3204766 | A | G | 78 | 0 | 2 | 0 | | | | | | | | | G | G | | |
| NC_011770 | 3204773 | T | G | 79 | 0 | 2 | 0 | | | | | | | | | G | G | | |
| NC_011770 | 3204856 | A | AC | 77 | 0 | 2 | 0 | | | | | | | | | AC | AC | | |
| NC_011770 | 3206196 | T | C | 76 | 0 | 2 | 0 | C | C | | | | | | | | | | |
| NC_011770 | 3206203 | C | T | 77 | 0 | 2 | 0 | | | | | | | | | T | T | | |
| NC_011770 | 3211329 | A | G | 55 | 0 | 4 | 0 | | | G | G | G | G | | | | | | |
| NC_011770 | 3211335 | T | G | 18 | 3 | 5 | 0 | | | G | G | . | G | | | G | . | . | G |
| NC_011770 | 3211674 | T | C | 55 | 0 | 12 | 0 | C | C | C | C | C | C | C | C | C | C | C | C |
| NC_011770 | 3211754 | A | G | 59 | 0 | 8 | 0 | | | G | G | G | G | | | G | G | G | G |
| NC_011770 | 3211770 | C | T | 68 | 0 | 2 | 0 | | | | | | | T | T | | | | |
| NC_011770 | 3211841 | G | T | 66 | 0 | 4 | 0 | | | T | T | T | T | | | | | | |
| NC_011770 | 3211875 | A | G | 65 | 0 | 4 | 0 | | | | | | | | | G | G | G | G |
| NC_011770 | 3211884 | C | T | 67 | 0 | 2 | 0 | | | | | | | T | T | | | | |
| NC_011770 | 3212173 | T | G | 68 | 0 | 4 | 0 | | | G | G | G | G | | | | | | |
| NC_011770 | 3212178 | T | A | 65 | 0 | 4 | 0 | | | | | | | | | A | A | A | A |
| NC_011770 | 3212179 | G | C | 68 | 0 | 4 | 0 | | | | | | | | | C | C | C | C |
| NC_011770 | 3212180 | G | T | 67 | 0 | 4 | 0 | | | | | | | | | T | T | T | T |
| NC_011770 | 3212183 | G | T | 65 | 0 | 4 | 0 | | | | | | | | | T | T | T | T |
| NC_011770 | 3212184 | A | C | 62 | 0 | 4 | 0 | | | | | | | | | C | C | C | C |
| NC_011770 | 3212186 | C | T | 65 | 0 | 4 | 0 | | | | | | | | | T | T | T | T |
| NC_011770 | 3212193 | C | CT | 68 | 0 | 4 | 0 | | | CT | CT | CT | CT | | | | | | |
| NC_011770 | 3212229 | G | T | 71 | 0 | 10 | 0 | | | T | T | T | T | T | T | T | T | T | T |
| NC_011770 | 3212240 | C | T | 71 | 0 | 2 | 0 | T | T | | | | | | | | | | |
| NC_011770 | 3212291 | G | A | 75 | 0 | 4 | 0 | | | A | A | A | A | | | | | | |
| NC_011770 | 3212311 | G | C | 75 | 0 | 6 | 0 | | | C | C | C | C | C | C | | | | |
| NC_011770 | 3212347 | C | T | 78 | 0 | 4 | 0 | | | T | T | T | T | | | | | | |
| NC_011770 | 3212366 | G | A | 75 | 0 | 4 | 0 | | | A | A | A | A | | | | | | |
| NC_011770 | 3212371 | TG | T | 80 | 0 | 4 | 0 | T | T | | | | | T | T | | | | |
| NC_011770 | 3212412 | C | A | 82 | 0 | 4 | 0 | | | A | A | A | A | | | | | | |
| NC_011770 | 3215219 | T | A | 64 | 0 | 4 | 0 | | | A | A | A | A | | | | | | |
| NC_011770 | 3215220 | C | G | 64 | 0 | 4 | 0 | | | G | G | G | G | | | | | | |
| NC_011770 | 3215227 | G | A | 67 | 0 | 4 | 0 | | | A | A | A | A | | | | | | |
| NC_011770 | 3215229 | C | G | 67 | 0 | 2 | 0 | | | | | | | G | G | | | | |
| NC_011770 | 3215234 | C | T | 65 | 0 | 2 | 0 | | | | | | | | | T | T | | |
| NC_011770 | 3215314 | T | C | 65 | 0 | 4 | 0 | | | C | C | C | C | | | | | | |
| NC_011770 | 3215320 | T | C | 67 | 0 | 4 | 0 | | | C | C | C | C | | | | | | |
| NC_011770 | 3215338 | T | C | 66 | 0 | 4 | 0 | | | C | C | C | C | | | | | | |
| NC_011770 | 3215353 | G | C | 65 | 0 | 4 | 0 | | | C | C | C | C | | | | | | |
| NC_011770 | 3215362 | G | T | 66 | 0 | 4 | 0 | | | T | T | T | T | | | | | | |
| NC_011770 | 3215386 | C | T | 64 | 0 | 4 | 0 | | | T | T | T | T | | | | | | |
| NC_011770 | 3215394 | C | T | 63 | 0 | 4 | 0 | | | T | T | T | T | | | | | | |
| NC_011770 | 3215437 | C | T | 64 | 0 | 4 | 0 | | | T | T | T | T | | | | | | |
| NC_011770 | 3215471 | C | T | 56 | 0 | 6 | 0 | | | T | T | T | T | A | A | | | | |
| NC_011770 | 3215477 | A | G | 57 | 0 | 2 | 0 | | | | | | | G | G | | | | |
| NC_011770 | 3215485 | G | A | 60 | 0 | 2 | 0 | | | | | | | | | A | A | | |
| NC_011770 | 3215493 | C | A | 60 | 0 | 2 | 0 | | | | | | | | | A | A | | |
| NC_011770 | 3215497 | C | T | 58 | 0 | 6 | 0 | | | T | T | T | T | | | T | T | | |
| NC_011770 | 3215516 | G | T | 61 | 0 | 2 | 0 | | | | | | | | | T | T | | |
| NC_011770 | 3215524 | A | G | 61 | 0 | 4 | 0 | | | G | G | G | G | | | | | | |
| NC_011770 | 3215555 | G | A | 64 | 0 | 2 | 0 | | | | | | | | | A | A | | |
| NC_011770 | 3215571 | A | T | 65 | 0 | 6 | 0 | | | T | T | T | T | T | T | | | | |
| NC_011770 | 3216922 | C | T | 73 | 0 | 2 | 0 | | | | | | | | | T | T | | |
| NC_011770 | 3216926 | A | C | 69 | 0 | 2 | 0 | | | | | | | | | C | C | | |
| NC_011770 | 3217027 | G | A | 80 | 0 | 2 | 0 | | | | | | | | | A | A | | |
| NC_011770 | 3217030 | A | G | 80 | 0 | 2 | 0 | | | | | | | | | G | G | | |
| NC_011770 | 3217031 | C | T | 80 | 0 | 2 | 0 | | | | | | | | | T | T | | |

| | | | | | | | | | | | | | | | | | | | |
|---|---|---|---|---|---|---|---|---|---|---|---|---|---|---|---|---|---|---|---|
| NC_011770 | 3217035 | A | C | 79 | 0 | 12 | 0 | C | C | C | C | C | C | C | C | C | C | C | C |
| NC_011770 | 3217691 | A | T | 69 | 0 | 10 | 0 | | | T | T | T | T | T | T | T | T | T | T |
| NC_011770 | 3217708 | A | G | 71 | 0 | 2 | 0 | | | | | | | | | G | G | | |
| NC_011770 | 3217710 | T | G | 71 | 0 | 10 | 0 | | | G | G | G | G | G | G | G | G | G | G |
| NC_011770 | 3217735 | G | A | 69 | 0 | 2 | 0 | | | | | | | | | A | A | | |
| NC_011770 | 3217808 | G | A | 66 | 0 | 4 | 0 | | | A | A | A | A | | | | | | |
| NC_011770 | 3217823 | C | G | 69 | 0 | 2 | 0 | | | | | | | | | G | G | | |
| NC_011770 | 3228989 | G | A | 60 | 0 | 2 | 0 | | | | | | | A | A | | | | |
| NC_011770 | 3228990 | A | G | 59 | 0 | 2 | 0 | | | | | | | G | G | | | | |
| NC_011770 | 3231087 | GC | G | 78 | 0 | 6 | 0 | | | G | G | G | G | G | G | | | | |
| NC_011770 | 3231088 | C | G | 75 | 0 | 6 | 0 | G | G | . | . | . | . | . | . | G | G | G | G |
| NC_011770 | 3231732 | G | A | 92 | 0 | 6 | 0 | | | A | A | A | A | A | A | | | | |
| NC_011770 | 3231758 | A | G | 95 | 0 | 2 | 0 | | | | | | | | | | | G | G |
| NC_011770 | 3238806 | T | G | 44 | 0 | 8 | 0 | G | G | G | G | G | G | | | | | G | G |
| NC_011770 | 3238807 | G | A | 45 | 0 | 6 | 0 | A | A | A | A | A | A | | | | | | |
| NC_011770 | 3238808 | A | G | 40 | 0 | 6 | 0 | G | G | G | G | G | G | | | | | | |
| NC_011770 | 3238810 | C | G | 40 | 0 | 2 | 0 | | | | | | | G | G | | | | |
| NC_011770 | 3238812 | G | T | 41 | 0 | 6 | 0 | T | T | T | T | T | T | | | | | | |
| NC_011770 | 3238815 | T | C | 41 | 0 | 6 | 0 | C | C | C | C | C | C | | | | | | |
| NC_011770 | 3238821 | T | C | 45 | 0 | 6 | 0 | C | C | C | C | C | C | | | | | | |
| NC_011770 | 3238869 | A | G | 45 | 0 | 12 | 0 | G | G | G | G | G | G | G | G | G | G | G | G |
| NC_011770 | 3240004 | T | G | 71 | 0 | 2 | 0 | | | | | | | G | G | | | | |
| NC_011770 | 3240007 | T | C | 74 | 0 | 6 | 0 | | | | | | | C | C | C | C | C | C |
| NC_011770 | 3240031 | C | T | 73 | 0 | 2 | 0 | | | | | | | | | | | T | T |
| NC_011770 | 3240076 | A | C | 67 | 0 | 2 | 0 | | | | | | | | | | | C | C |
| NC_011770 | 3240589 | C | T | 64 | 0 | 2 | 0 | | | | | | | | | | | T | T |
| NC_011770 | 3242388 | C | T | 43 | 0 | 2 | 0 | T | T | | | | | | | | | | |
| NC_011770 | 3253907 | C | T | 38 | 0 | 2 | 0 | | | | | | | | | T | T | | |
| NC_011770 | 3253916 | G | T | 40 | 0 | 2 | 0 | | | | | | | | | T | T | | |
| NC_011770 | 3254006 | T | C | 42 | 0 | 2 | 0 | | | | | | | | | C | C | | |
| NC_011770 | 3254015 | T | C | 43 | 0 | 10 | 0 | C | C | C | C | C | C | | | C | C | C | C |
| NC_011770 | 3254069 | G | T | 51 | 0 | 4 | 0 | | | T | T | T | T | | | | | | |
| NC_011770 | 3254080 | G | A | 51 | 0 | 2 | 0 | A | A | | | | | | | | | | |
| NC_011770 | 3254082 | T | G | 51 | 0 | 8 | 0 | | | G | G | G | G | | | G | G | G | G |
| NC_011770 | 3254189 | G | A | 54 | 0 | 8 | 0 | A | A | A | A | A | A | | | | | A | A |
| NC_011770 | 3254230 | C | T | 56 | 0 | 2 | 0 | T | T | | | | | | | | | | |
| NC_011770 | 3254231 | C | T | 56 | 0 | 2 | 0 | T | T | | | | | | | | | | |
| NC_011770 | 3254241 | T | C | 54 | 0 | 8 | 0 | C | C | C | C | C | C | | | | | C | C |
| NC_011770 | 3256516 | G | GC | 83 | 0 | 4 | 0 | | | GC | GC | GC | GC | | | | | | |
| NC_011770 | 3256519 | A | G | 51 | 0 | 4 | 0 | | | G | G | G | G | | | | | | |
| NC_011770 | 3259506 | G | A | 58 | 0 | 1 | 0 | | | | | | | A | | | | | |
| NC_011770 | 3263990 | C | G | 57 | 0 | 2 | 0 | | | | | | | | | G | G | | |
| NC_011770 | 3263994 | A | C | 56 | 0 | 2 | 0 | | | | | | | | | C | C | | |
| NC_011770 | 3265035 | T | A | 35 | 0 | 2 | 0 | | | | | | | | | A | A | | |
| NC_011770 | 3269894 | C | T | 88 | 0 | 4 | 0 | | | T | T | T | T | | | | | | |
| NC_011770 | 3277040 | C | T | 24 | 0 | 10 | 0 | | | T | T | T | T | T | T | T | T | T | T |
| NC_011770 | 3277063 | G | T | 7 | 6 | 4 | 0 | | | T | T | T | T | . | . | . | . | . | . |
| NC_011770 | 3277064 | T | C | 18 | 0 | 4 | 0 | | | C | C | C | C | | | | | | |
| NC_011770 | 3277071 | G | GA | 20 | 0 | 5 | 0 | | | | | | | GA | GA | GA | | GA | GA |
| NC_011770 | 3277097 | T | C | 15 | 1 | 11 | 0 | C | C | C | C | C | C | C | C | . | C | C | C |
| NC_011770 | 3277120 | A | T | 14 | 0 | 2 | 0 | | | | | | | T | T | | | | |
| NC_011770 | 3277302 | C | T | 16 | 2 | 1 | 0 | . | | | | | | | . | | | . | T |
| NC_011770 | 3277312 | C | T | 16 | 2 | 1 | 0 | . | | | | | | | . | | | . | T |
| NC_011770 | 3277349 | A | G | 9 | 3 | 7 | 0 | . | G | G | G | G | G | . | . | . | G | G | . |
| NC_011770 | 3277420 | C | G | 2 | 6 | 2 | 0 | . | . | . | . | G | G | | | | . | . | . |
| NC_011770 | 3277421 | C | T | 2 | 6 | 3 | 0 | . | . | T | . | T | T | | | | . | . | . |
| NC_011770 | 3277427 | T | A | 2 | 7 | 3 | 0 | . | . | A | . | A | A | | . | | . | . | . |
| NC_011770 | 3277436 | C | T | 1 | 7 | 2 | 0 | . | . | T | . | . | T | | | | . | . | . |
| NC_011770 | 3277459 | A | AT | 1 | 9 | 2 | 0 | . | . | . | . | . | . | AT | AT | | . | . | . |
| NC_011770 | 3277467 | ACGGGATT | A | 1 | 9 | 2 | 0 | . | . | . | . | . | . | A | . | A | . | . | . |
| NC_011770 | 3279073 | C | CG | 71 | 0 | 2 | 0 | | | | | | | | | CG | CG | | |
| NC_011770 | 3279084 | A | G | 71 | 0 | 8 | 0 | G | G | G | G | G | G | | | G | G | | |

| | | | | | | | | | | | | | | | | | | | |
|---|---|---|---|---|---|---|---|---|---|---|---|---|---|---|---|---|---|---|---|
| NC_011770 | 3279118 | C | G | 75 | 0 | 2 | 0 | | | | | | | | | | | G | G |
| NC_011770 | 3281649 | A | G | 60 | 0 | 2 | 0 | G | G | | | | | | | | | | |
| NC_011770 | 3281735 | C | A | 68 | 0 | 2 | 0 | | | | | | | | | | | A | A |
| NC_011770 | 3281816 | G | T | 74 | 0 | 2 | 0 | T | T | | | | | | | | | | |
| NC_011770 | 3283545 | T | A | 45 | 0 | 4 | 0 | | | A | A | A | A | | | | | | |
| NC_011770 | 3283615 | TG | T | 44 | 0 | 2 | 0 | | | | | | | T | T | | | | |
| NC_011770 | 3283801 | A | G | 58 | 0 | 4 | 0 | | | | | | | G | G | | | G | G |
| NC_011770 | 3285227 | T | C | 68 | 0 | 2 | 0 | | | | | | | C | C | | | | |
| NC_011770 | 3285255 | A | T | 65 | 0 | 2 | 0 | T | T | | | | | | | | | | |
| NC_011770 | 3285818 | G | A | 77 | 0 | 2 | 0 | A | A | | | | | | | | | | |
| NC_011770 | 3285930 | G | A | 74 | 0 | 2 | 0 | | | | | | | | | A | A | | |
| NC_011770 | 3287061 | C | T | 45 | 0 | 12 | 0 | T | T | T | T | T | T | T | T | T | T | T | T |
| NC_011770 | 3287065 | T | C | 44 | 0 | 10 | 0 | C | C | C | C | C | C | C | C | C | C | | |
| NC_011770 | 3287066 | T | A | 43 | 0 | 10 | 0 | A | A | A | A | A | A | A | A | A | A | | |
| NC_011770 | 3287072 | T | G | 44 | 0 | 10 | 0 | G | G | G | G | G | G | G | G | G | G | | |
| NC_011770 | 3287080 | T | G | 44 | 0 | 2 | 0 | G | G | | | | | | | | | | |
| NC_011770 | 3287083 | C | A | 43 | 0 | 2 | 0 | A | A | | | | | | | | | | |
| NC_011770 | 3287088 | C | T | 41 | 0 | 8 | 0 | | | T | T | T | T | T | T | T | T | | |
| NC_011770 | 3287102 | G | A | 37 | 0 | 6 | 0 | A | A | A | A | A | A | | | | | | |
| NC_011770 | 3287108 | G | GC | 34 | 0 | 8 | 0 | GC | GC | | | GC | GC | GC | GC | GC | GC | | |
| NC_011770 | 3287113 | G | A | 33 | 0 | 4 | 0 | | | A | A | A | A | | | | | | |
| NC_011770 | 3287136 | A | G | 30 | 0 | 4 | 0 | | | | | | | G | G | G | G | | |
| NC_011770 | 3287179 | G | A | 25 | 2 | 2 | 0 | A | . | | | | | | | . | A | | |
| NC_011770 | 3287181 | G | A | 25 | 0 | 4 | 0 | A | A | | | | | | | A | A | | |
| NC_011770 | 3287211 | T | G | 16 | 0 | 2 | 0 | G | G | | | | | | | . | | | |
| NC_011770 | 3287221 | T | C | 15 | 1 | 8 | 0 | C | C | C | C | C | C | C | C | . | . | | |
| NC_011770 | 3287233 | A | G | 17 | 1 | 2 | 0 | | | | | | | G | G | . | | | |
| NC_011770 | 3287245 | G | A | 17 | 1 | 4 | 0 | | | A | A | A | A | | | . | | | |
| NC_011770 | 3287268 | G | A | 19 | 0 | 3 | 0 | | | A | A | . | A | | | | | | |
| NC_011770 | 3287311 | G | A | 19 | 0 | 3 | 0 | . | A | | | | | | | A | A | | |
| NC_011770 | 3287313 | G | A | 19 | 0 | 4 | 0 | A | A | | | | | | | A | A | | |
| NC_011770 | 3287328 | T | C | 20 | 0 | 8 | 0 | | | C | C | C | C | C | C | C | C | | |
| NC_011770 | 3287355 | G | GA | 8 | 6 | 1 | 0 | | | . | . | . | . | . | . | . | GA | | |
| NC_011770 | 3287399 | A | G | 16 | 6 | 2 | 0 | | | . | . | . | . | . | . | G | G | | |
| NC_011770 | 3287454 | T | C | 18 | 0 | 3 | 0 | | | . | C | C | . | . | C | | | | |
| NC_011770 | 3287466 | T | G | 16 | 0 | 6 | 0 | | | G | G | G | G | G | G | | | | |
| NC_011770 | 3287474 | G | T | 18 | 0 | 8 | 0 | | | T | T | T | T | T | T | | | T | T |
| NC_011770 | 3287482 | T | C | 18 | 0 | 1 | 0 | | | | | | | | | | | C | . |
| NC_011770 | 3287487 | A | G | 18 | 0 | 8 | 0 | | | G | G | G | G | G | G | | | G | G |
| NC_011770 | 3287496 | A | G | 21 | 0 | 3 | 0 | | | | | | | G | G | | | G | . |
| NC_011770 | 3287507 | G | A | 20 | 0 | 10 | 0 | | | A | A | A | A | A | A | A | A | A | A |
| NC_011770 | 3287508 | G | A | 22 | 0 | 4 | 0 | | | A | A | A | A | | | | | | |
| NC_011770 | 3287531 | G | A | 26 | 0 | 5 | 0 | | | A | A | A | A | | | | | A | . |
| NC_011770 | 3287586 | C | T | 15 | 0 | 6 | 0 | | | T | . | T | T | T | T | | | . | T |
| NC_011770 | 3287587 | T | TG | 14 | 0 | 7 | 0 | | | TG | TG | TG | TG | TG | . | | | TG | TG |
| NC_011770 | 3287597 | T | G | 32 | 0 | 8 | 0 | | | G | G | G | G | G | G | | | G | G |
| NC_011770 | 3287601 | G | GCGCAC | 10 | 6 | 1 | 0 | | | . | . | . | . | . | . | | | GCGCACG | . |
| NC_011770 | 3287608 | T | C | 35 | 0 | 8 | 0 | | | C | C | C | C | C | C | | | C | C |
| NC_011770 | 3287619 | T | C | 35 | 0 | 8 | 0 | | | C | C | C | C | C | C | | | C | C |
| NC_011770 | 3287620 | A | T | 35 | 0 | 8 | 0 | | | T | T | T | T | T | T | | | T | T |
| NC_011770 | 3287622 | G | T | 35 | 0 | 8 | 0 | | | T | T | T | T | T | T | | | T | T |
| NC_011770 | 3287623 | C | T | 35 | 0 | 8 | 0 | | | T | T | T | T | T | T | | | T | T |
| NC_011770 | 3290347 | A | G | 63 | 0 | 8 | 0 | | | G | G | G | G | G | G | | | G | G |
| NC_011770 | 3290349 | C | T | 62 | 0 | 4 | 0 | | | T | T | T | T | | | | | | |
| NC_011770 | 3292875 | T | A | 92 | 0 | 2 | 0 | A | A | | | | | | | | | | |
| NC_011770 | 3292925 | A | T | 93 | 0 | 10 | 0 | | | T | T | T | T | T | T | T | T | T | T |
| NC_011770 | 3292927 | G | A | 98 | 0 | 2 | 0 | A | A | | | | | | | | | | |
| NC_011770 | 3292977 | C | T | 92 | 0 | 4 | 0 | | | | | | | | | T | T | T | T |
| NC_011770 | 3293001 | A | G | 88 | 0 | 10 | 0 | | | G | G | G | G | G | G | G | G | G | G |
| NC_011770 | 3293007 | G | T | 90 | 0 | 2 | 0 | | | | | | | T | T | | | | |
| NC_011770 | 3293737 | G | A | 59 | 0 | 6 | 0 | A | A | | | | | A | A | A | A | | |
| NC_011770 | 3294367 | G | A | 75 | 0 | 2 | 0 | | | | | | | A | A | | | | |

| | | | | | | | | | | | | | | | | | | | |
|---|---|---|---|---|---|---|---|---|---|---|---|---|---|---|---|---|---|---|---|
| NC_011770 | 3294547 | T | C | 70 | 0 | 2 | 0 | | | | | | | C | C | | | | |
| NC_011770 | 3294591 | C | T | 66 | 0 | 2 | 0 | | | | | | | | | | | T | T |
| NC_011770 | 3294644 | C | CA | 63 | 0 | 4 | 0 | | | | | | | CA | CA | | | CA | CA |
| NC_011770 | 3294649 | A | G | 41 | 0 | 8 | 0 | | | G | G | G | G | G | G | | | G | G |
| NC_011770 | 3294691 | G | A | 58 | 0 | 8 | 0 | A | A | A | A | A | A | | | A | A | | |
| NC_011770 | 3294794 | G | C | 69 | 0 | 4 | 0 | | | C | C | C | C | | | | | | |
| NC_011770 | 3295737 | C | T | 93 | 0 | 2 | 0 | | | | | | | | | | | T | T |
| NC_011770 | 3295739 | T | C | 90 | 0 | 2 | 0 | | | | | | | | | | | C | C |
| NC_011770 | 3297640 | G | A | 70 | 0 | 2 | 0 | A | A | | | | | | | | | | |
| NC_011770 | 3297645 | A | G | 68 | 0 | 2 | 0 | G | G | | | | | | | | | | |
| NC_011770 | 3297646 | T | C | 68 | 0 | 4 | 0 | C | C | | | | | C | C | | | | |
| NC_011770 | 3311639 | A | G | 51 | 0 | 8 | 0 | G | G | G | G | G | G | G | G | | | | |
| NC_011770 | 3312300 | C | T | 66 | 0 | 2 | 0 | | | | | | | T | T | | | | |
| NC_011770 | 3313912 | G | A | 68 | 0 | 6 | 0 | | | A | A | A | A | A | A | | | | |
| NC_011770 | 3313924 | T | C | 68 | 0 | 4 | 0 | C | C | | | | | | | C | C | | |
| NC_011770 | 3313931 | C | T | 69 | 0 | 4 | 0 | T | T | | | | | | | T | T | | |
| NC_011770 | 3313934 | T | C | 68 | 0 | 4 | 0 | C | C | | | | | | | C | C | | |
| NC_011770 | 3313951 | G | A | 68 | 0 | 4 | 0 | A | A | | | | | | | A | A | | |
| NC_011770 | 3313954 | A | T | 69 | 0 | 4 | 0 | T | T | | | | | | | T | T | | |
| NC_011770 | 3313963 | T | C | 53 | 1 | 5 | 0 | C | . | | | | | G | G | C | C | | |
| NC_011770 | 3313968 | C | G | 64 | 0 | 4 | 0 | G | G | | | | | | | G | G | | |
| NC_011770 | 3313970 | A | C | 63 | 0 | 4 | 0 | C | C | | | | | | | C | C | | |
| NC_011770 | 3314026 | A | G | 75 | 0 | 6 | 0 | | | G | G | G | G | | | | | G | G |
| NC_011770 | 3314028 | C | T | 76 | 0 | 6 | 0 | | | T | T | T | T | | | | | T | T |
| NC_011770 | 3314047 | T | C | 76 | 0 | 2 | 0 | | | | | | | | | | | C | C |
| NC_011770 | 3314048 | A | G | 77 | 0 | 6 | 0 | G | G | | | | | G | G | G | G | | |
| NC_011770 | 3314050 | C | T | 76 | 0 | 6 | 0 | | | T | T | T | T | | | | | T | T |
| NC_011770 | 3314052 | C | G | 77 | 0 | 8 | 0 | G | G | | | | | G | G | G | G | G | G |
| NC_011770 | 3314062 | T | G | 78 | 0 | 6 | 0 | G | G | | | | | G | G | G | G | | |
| NC_011770 | 3314063 | G | A | 76 | 0 | 6 | 0 | A | A | | | | | A | A | A | A | | |
| NC_011770 | 3316558 | G | GC | 68 | 0 | 12 | 0 | GC | GC | GC | GC | GC | GC | GC | GC | GC | GC | GC | GC |
| NC_011770 | 3316936 | C | T | 78 | 0 | 6 | 0 | | | | | | | T | T | T | T | T | T |
| NC_011770 | 3316942 | G | A | 77 | 0 | 8 | 0 | A | A | | | | | A | A | A | A | A | A |
| NC_011770 | 3316964 | A | G | 78 | 0 | 8 | 0 | G | G | | | | | G | G | G | G | G | G |
| NC_011770 | 3321372 | C | T | 70 | 0 | 2 | 0 | | | | | | | | | | | T | T |
| NC_011770 | 3321398 | A | C | 73 | 0 | 6 | 0 | C | C | | | | | C | C | C | C | | |
| NC_011770 | 3321450 | A | G | 71 | 0 | 2 | 0 | | | | | | | | | | | G | G |
| NC_011770 | 3321457 | A | G | 69 | 0 | 12 | 0 | G | G | G | G | G | G | G | G | G | G | G | G |
| NC_011770 | 3321470 | A | G | 72 | 0 | 12 | 0 | G | G | G | G | G | G | G | G | G | G | G | G |
| NC_011770 | 3321599 | T | C | 70 | 0 | 8 | 0 | C | C | | | | | C | C | C | C | C | C |
| NC_011770 | 3321650 | C | T | 68 | 0 | 2 | 0 | | | | | | | | | | | T | T |
| NC_011770 | 3327933 | A | T | 77 | 0 | 4 | 0 | | | | | | | T | T | | | G | G |
| NC_011770 | 3333821 | C | A | 56 | 0 | 2 | 0 | | | | | | | A | A | | | | |
| NC_011770 | 3333927 | A | T | 62 | 0 | 6 | 0 | | | T | T | T | T | T | T | | | | |
| NC_011770 | 3338657 | T | C | 71 | 0 | 2 | 0 | | | | | | | | | | | C | C |
| NC_011770 | 3338728 | T | C | 69 | 0 | 8 | 0 | | | C | C | C | C | | | C | C | C | C |
| NC_011770 | 3338841 | A | C | 71 | 0 | 2 | 0 | | | | | | | C | C | | | | |
| NC_011770 | 3343854 | T | C | 78 | 0 | 4 | 0 | | | C | C | C | C | | | | | | |
| NC_011770 | 3343894 | T | G | 88 | 0 | 8 | 0 | G | G | G | G | G | G | G | G | | | | |
| NC_011770 | 3346265 | C | T | 84 | 0 | 4 | 0 | T | T | | | | | | | T | T | | |
| NC_011770 | 3347181 | T | G | 55 | 0 | 4 | 0 | G | G | | | | | | | G | G | | |
| NC_011770 | 3351197 | A | T | 61 | 0 | 2 | 0 | | | | | | | T | T | | | | |
| NC_011770 | 3356504 | T | G | 71 | 0 | 12 | 0 | G | G | G | G | G | G | G | G | G | G | G | G |
| NC_011770 | 3356552 | G | GA | 73 | 0 | 12 | 0 | GA | GA | GA | GA | GA | GA | GA | GA | GA | GA | GA | GA |
| NC_011770 | 3357939 | T | C | 61 | 0 | 12 | 0 | C | C | C | C | C | C | C | C | C | C | C | C |
| NC_011770 | 3357952 | T | TG | 48 | 0 | 2 | 0 | | | | | | | | | TG | TG | | |
| NC_011770 | 3357980 | A | C | 52 | 0 | 10 | 0 | C | C | C | C | C | C | C | C | | | C | C |
| NC_011770 | 3357996 | T | C | 48 | 0 | 12 | 0 | C | C | C | C | C | C | C | C | C | C | C | C |
| NC_011770 | 3357998 | G | A | 47 | 0 | 12 | 0 | A | A | A | A | A | A | A | A | A | A | A | A |
| NC_011770 | 3358003 | G | A | 50 | 0 | 12 | 0 | A | A | A | A | A | A | A | A | A | A | A | A |
| NC_011770 | 3358026 | A | G | 50 | 0 | 12 | 0 | G | G | G | G | G | G | G | G | G | G | G | G |
| NC_011770 | 3358043 | ACGG | A | 50 | 0 | 2 | 0 | | | | | | | | | A | A | | |

| | | | | | | | | | | | | | | | | | | | |
|---|---|---|---|---|---|---|---|---|---|---|---|---|---|---|---|---|---|---|---|
| NC_011770 | 3358070 | T | C | 40 | 4 | 1 | 0 | . | . | | | | | C | | | | . | . |
| NC_011770 | 3358110 | A | C | 47 | 0 | 4 | 0 | C | C | | | | | | | | | C | C |
| NC_011770 | 3358129 | C | T | 48 | 0 | 2 | 0 | | | | | | | T | T | | | | |
| NC_011770 | 3358131 | T | C | 48 | 0 | 2 | 0 | | | | | | | C | C | | | | |
| NC_011770 | 3358133 | A | G | 49 | 0 | 4 | 0 | | | G | G | G | G | | | | | | |
| NC_011770 | 3358134 | T | C | 49 | 0 | 10 | 0 | C | C | C | C | C | C | G | G | | | C | C |
| NC_011770 | 3362372 | G | T | 36 | 0 | 6 | 0 | T | T | T | T | T | T | | | | | | |
| NC_011770 | 3362397 | T | C | 35 | 0 | 10 | 0 | C | C | C | C | C | C | C | C | C | C | | |
| NC_011770 | 3362398 | GT | G | 36 | 0 | 2 | 0 | | | | | | | | | G | G | | |
| NC_011770 | 3362456 | C | T | 33 | 0 | 2 | 0 | | | | | | | | | T | T | | |
| NC_011770 | 3363540 | G | A | 59 | 0 | 2 | 0 | | | | | | | | | | | A | A |
| NC_011770 | 3363542 | A | G | 59 | 0 | 12 | 0 | G | G | G | G | G | G | G | G | G | G | G | G |
| NC_011770 | 3365131 | A | G | 85 | 0 | 12 | 0 | G | G | G | G | G | G | G | G | G | G | G | G |
| NC_011770 | 3366005 | G | T | 46 | 0 | 2 | 0 | | | | | | | | | | | T | T |
| NC_011770 | 3366054 | A | T | 44 | 0 | 4 | 0 | | | T | T | T | T | | | | | | |
| NC_011770 | 3371223 | G | A | 12 | 1 | 2 | 0 | | | | | | | A | A | . | | | |
| NC_011770 | 3371225 | C | G | 13 | 1 | 2 | 0 | | | | | | | G | G | . | | | |
| NC_011770 | 3371862 | T | C | 84 | 0 | 2 | 0 | | | | | | | C | C | | | | |
| NC_011770 | 3371867 | G | A | 81 | 0 | 4 | 0 | A | A | | | | | | | | | A | A |
| NC_011770 | 3371868 | C | T | 80 | 0 | 4 | 0 | | | T | T | T | T | | | | | | |
| NC_011770 | 3373252 | C | T | 67 | 0 | 2 | 0 | | | | | | | | | T | T | | |
| NC_011770 | 3375022 | T | G | 59 | 0 | 8 | 0 | G | G | | | | | G | G | G | G | G | G |
| NC_011770 | 3375079 | A | G | 57 | 0 | 6 | 0 | | | G | G | G | G | | | | | G | G |
| NC_011770 | 3375132 | T | C | 54 | 0 | 6 | 0 | | | C | C | C | C | | | | | C | C |
| NC_011770 | 3375162 | C | T | 53 | 0 | 1 | 0 | | | | | | | T | | | | | |
| NC_011770 | 3390931 | G | A | 55 | 2 | 2 | 0 | . | . | | | | | | | A | A | | |
| NC_011770 | 3390933 | T | C | 54 | 2 | 6 | 0 | . | . | C | C | C | C | | | | | C | C |
| NC_011770 | 3390957 | G | A | 57 | 2 | 2 | 0 | . | . | | | | | | | | | A | A |
| NC_011770 | 3390958 | T | C | 57 | 2 | 8 | 0 | . | . | C | C | C | C | | | C | C | C | C |
| NC_011770 | 3390967 | T | C | 55 | 2 | 8 | 0 | . | . | C | C | C | C | C | C | C | C | | |
| NC_011770 | 3390982 | C | A | 57 | 2 | 2 | 0 | . | . | | | | | A | A | | | | |
| NC_011770 | 3390994 | C | T | 60 | 2 | 2 | 0 | . | . | | | | | | | | | T | T |
| NC_011770 | 3391002 | G | A | 58 | 2 | 2 | 0 | . | . | | | | | | | | | A | A |
| NC_011770 | 3391006 | C | A | 58 | 2 | 2 | 0 | . | . | | | | | | | | | A | A |
| NC_011770 | 3391013 | C | G | 59 | 2 | 6 | 0 | . | . | G | G | G | G | | | | | G | G |
| NC_011770 | 3391095 | TTGTCCG | T | 60 | 2 | 4 | 0 | . | . | T | T | T | T | | | | | | |
| NC_011770 | 3391123 | T | G | 61 | 2 | 4 | 0 | . | . | G | G | G | G | | | | | | |
| NC_011770 | 3391135 | C | T | 60 | 2 | 2 | 0 | . | . | | | | | | | T | T | | |
| NC_011770 | 3391152 | GC | G | 56 | 3 | 1 | 0 | . | . | | | | | | | G | . | | |
| NC_011770 | 3391849 | C | CG | 59 | 2 | 2 | 0 | . | . | | | | | | | CG | CG | | |
| NC_011770 | 3391850 | G | GA | 54 | 4 | 4 | 0 | . | . | GA | GA | GA | GA | | | . | . | | |
| NC_011770 | 3392614 | G | A | 8 | 2 | 4 | 0 | . | . | A | A | A | A | | | | | | |
| NC_011770 | 3392627 | T | C | 14 | 2 | 4 | 0 | . | . | C | C | C | C | | | | | | |
| NC_011770 | 3394022 | A | G | 29 | 6 | 6 | 0 | . | . | . | . | . | . | G | G | G | G | G | G |
| NC_011770 | 3394091 | A | G | 33 | 6 | 6 | 0 | . | . | . | . | . | . | G | G | G | G | G | G |
| NC_011770 | 3394188 | T | C | 31 | 6 | 2 | 0 | . | . | . | . | . | . | C | C | | | | |
| NC_011770 | 3394257 | C | T | 37 | 6 | 2 | 0 | . | . | . | . | . | . | T | T | | | | |
| NC_011770 | 3394279 | T | C | 32 | 6 | 4 | 0 | . | . | . | . | . | . | | | C | C | C | C |
| NC_011770 | 3395489 | G | T | 22 | 4 | 2 | 0 | | | . | . | . | . | | | | | T | T |
| NC_011770 | 3396030 | T | C | 30 | 6 | 6 | 0 | C | C | C | C | C | C | . | . | . | . | . | . |
| NC_011770 | 3396032 | G | A | 30 | 6 | 6 | 0 | A | A | A | A | A | A | . | . | . | . | . | . |
| NC_011770 | 3396042 | C | G | 31 | 6 | 6 | 0 | G | G | G | G | G | G | . | . | . | . | . | . |
| NC_011770 | 3397987 | CA | C | 32 | 6 | 2 | 0 | C | C | | | | | . | . | . | . | . | . |
| NC_011770 | 3399001 | C | T | 35 | 6 | 4 | 0 | | | T | T | T | T | . | . | . | . | . | . |
| NC_011770 | 3404129 | G | A | 42 | 6 | 4 | 0 | | | A | A | A | A | . | . | . | . | . | . |
| NC_011770 | 3405608 | G | A | 49 | 6 | 2 | 0 | A | A | | | | | . | . | . | . | . | . |
| NC_011770 | 3405704 | T | A | 49 | 6 | 6 | 0 | A | A | A | A | A | A | . | . | . | . | . | . |
| NC_011770 | 3405828 | G | A | 47 | 6 | 6 | 0 | A | A | A | A | A | A | . | . | . | . | . | . |
| NC_011770 | 3405878 | A | G | 44 | 6 | 4 | 0 | | | G | G | G | G | . | . | . | . | . | . |
| NC_011770 | 3405895 | G | A | 44 | 6 | 4 | 0 | | | A | A | A | A | . | . | . | . | . | . |
| NC_011770 | 3405905 | A | T | 44 | 6 | 2 | 0 | T | T | | | | | . | . | . | . | . | . |
| NC_011770 | 3405990 | G | A | 47 | 6 | 6 | 0 | A | A | A | A | A | A | . | . | . | . | . | . |

| | | | | | | | | | | | | | | | | | | | |
|---|---|---|---|---|---|---|---|---|---|---|---|---|---|---|---|---|---|---|---|
| NC_011770 | 3405997 | CA | C | 45 | 6 | 4 | 0 | | | C | C | C | C | . | . | . | . | . | . |
| NC_011770 | 3406078 | A | G | 49 | 6 | 6 | 0 | G | G | G | G | G | G | . | . | . | . | . | . |
| NC_011770 | 3406083 | G | A | 48 | 6 | 6 | 0 | A | A | A | A | A | A | . | . | . | . | . | . |
| NC_011770 | 3406782 | A | G | 48 | 6 | 6 | 0 | G | G | G | G | G | G | . | . | . | . | . | . |
| NC_011770 | 3406812 | C | T | 48 | 6 | 4 | 0 | | | T | T | T | T | . | . | . | . | . | . |
| NC_011770 | 3406838 | T | C | 48 | 6 | 2 | 0 | C | C | | | | | . | . | . | . | . | . |
| NC_011770 | 3406848 | A | T | 47 | 6 | 6 | 0 | T | T | T | T | T | T | . | . | . | . | . | . |
| NC_011770 | 3409311 | G | A | 56 | 6 | 4 | 0 | | | A | A | A | A | . | . | . | . | . | . |
| NC_011770 | 3409323 | T | C | 56 | 6 | 6 | 0 | C | C | C | C | C | C | . | . | . | . | . | . |
| NC_011770 | 3409375 | G | A | 54 | 6 | 4 | 0 | | | A | A | A | A | . | . | . | . | . | . |
| NC_011770 | 3409392 | A | T | 54 | 6 | 6 | 0 | T | T | T | T | T | T | . | . | . | . | . | . |
| NC_011770 | 3409462 | C | T | 47 | 6 | 4 | 0 | | | T | T | T | T | . | . | . | . | . | . |
| NC_011770 | 3409530 | T | A | 47 | 6 | 2 | 0 | A | A | | | | | . | . | . | . | . | . |
| NC_011770 | 3410221 | C | T | 53 | 6 | 2 | 0 | T | T | | | | | . | . | . | . | . | . |
| NC_011770 | 3410535 | T | C | 53 | 6 | 6 | 0 | C | C | C | C | C | C | . | . | . | . | . | . |
| NC_011770 | 3413008 | A | G | 31 | 6 | 6 | 0 | G | G | G | G | G | G | . | . | . | . | . | . |
| NC_011770 | 3414398 | A | C | 37 | 6 | 4 | 0 | | | C | C | C | C | . | . | . | . | . | . |
| NC_011770 | 3415734 | T | C | 46 | 6 | 4 | 0 | | | C | C | C | C | . | . | . | . | . | . |
| NC_011770 | 3419569 | G | A | 51 | 6 | 2 | 0 | A | A | | | | | . | . | . | . | . | . |
| NC_011770 | 3419570 | T | C | 50 | 6 | 2 | 0 | C | C | | | | | . | . | . | . | . | . |
| NC_011770 | 3419797 | A | G | 47 | 6 | 4 | 0 | | | G | G | G | G | . | . | . | . | . | . |
| NC_011770 | 3419837 | C | T | 46 | 6 | 6 | 0 | T | T | T | T | T | T | . | . | . | . | . | . |
| NC_011770 | 3421863 | G | A | 37 | 6 | 4 | 0 | | | A | A | A | A | . | . | . | . | . | . |
| NC_011770 | 3423616 | T | C | 14 | 6 | 4 | 0 | | | C | C | C | C | . | . | . | . | . | . |
| NC_011770 | 3423634 | G | A | 11 | 6 | 4 | 0 | | | A | A | A | A | . | . | . | . | . | . |
| NC_011770 | 3427259 | T | C | 42 | 6 | 2 | 0 | C | C | | | | | . | . | . | . | . | . |
| NC_011770 | 3427287 | C | T | 42 | 6 | 2 | 0 | T | T | | | | | . | . | . | . | . | . |
| NC_011770 | 3427288 | G | T | 42 | 6 | 2 | 0 | T | T | | | | | . | . | . | . | . | . |
| NC_011770 | 3432964 | A | G | 37 | 6 | 6 | 0 | G | G | G | G | G | G | . | . | . | . | . | . |
| NC_011770 | 3433099 | C | T | 38 | 6 | 6 | 0 | T | T | T | T | T | T | . | . | . | . | . | . |
| NC_011770 | 3433400 | A | C | 25 | 6 | 2 | 0 | C | C | | | | | . | . | . | . | . | . |
| NC_011770 | 3433405 | C | T | 25 | 6 | 2 | 0 | T | T | | | | | . | . | . | . | . | . |
| NC_011770 | 3435083 | C | A | 69 | 0 | 4 | 0 | | | A | A | A | A | | | | | | |
| NC_011770 | 3435098 | G | A | 70 | 0 | 2 | 0 | | | | | | | A | A | | | | |
| NC_011770 | 3435179 | G | A | 73 | 0 | 2 | 0 | A | A | | | | | | | | | | |
| NC_011770 | 3435182 | C | G | 71 | 0 | 2 | 0 | G | G | | | | | | | | | | |
| NC_011770 | 3435185 | C | T | 72 | 0 | 2 | 0 | T | T | | | | | | | | | | |
| NC_011770 | 3435186 | G | C | 75 | 0 | 2 | 0 | C | C | | | | | | | | | | |
| NC_011770 | 3435190 | T | C | 75 | 0 | 2 | 0 | C | C | | | | | | | | | | |
| NC_011770 | 3435202 | T | C | 72 | 0 | 2 | 0 | C | C | | | | | | | | | | |
| NC_011770 | 3435205 | G | A | 74 | 0 | 2 | 0 | A | A | | | | | | | | | | |
| NC_011770 | 3435206 | G | A | 74 | 0 | 2 | 0 | A | A | | | | | | | | | | |
| NC_011770 | 3435207 | CT | C | 74 | 0 | 2 | 0 | C | C | | | | | | | | | | |
| NC_011770 | 3438797 | C | T | 89 | 0 | 2 | 0 | T | T | | | | | | | | | | |
| NC_011770 | 3438808 | C | T | 86 | 0 | 1 | 0 | | | | | | | T | | | | | |
| NC_011770 | 3438860 | G | A | 89 | 0 | 2 | 0 | | | | | | | | | | | A | A |
| NC_011770 | 3438894 | A | C | 90 | 0 | 6 | 0 | C | C | | | | | C | C | | | C | C |
| NC_011770 | 3438897 | C | A | 88 | 0 | 2 | 0 | | | | | | | | | | | A | A |
| NC_011770 | 3438934 | A | G | 80 | 0 | 6 | 0 | G | G | | | | | G | G | | | G | G |
| NC_011770 | 3440353 | C | T | 75 | 0 | 4 | 0 | | | | | | | | | T | T | T | T |
| NC_011770 | 3443794 | A | G | 73 | 0 | 12 | 0 | G | G | G | G | G | G | G | G | G | G | G | G |
| NC_011770 | 3443810 | T | C | 73 | 0 | 4 | 0 | | | C | C | C | C | | | | | | |
| NC_011770 | 3444951 | C | T | 78 | 0 | 2 | 0 | T | T | | | | | | | | | | |
| NC_011770 | 3448597 | G | A | 68 | 0 | 2 | 0 | | | | | | | | | A | A | | |
| NC_011770 | 3448629 | A | G | 64 | 0 | 8 | 0 | | | G | G | G | G | G | G | G | G | | |
| NC_011770 | 3448694 | C | A | 66 | 0 | 2 | 0 | A | A | | | | | | | | | | |
| NC_011770 | 3448790 | C | CG | 53 | 0 | 2 | 0 | | | | | CG | CG | | | | | | |
| NC_011770 | 3451169 | T | G | 68 | 0 | 12 | 0 | G | G | G | G | G | G | G | G | G | G | G | G |
| NC_011770 | 3451172 | G | A | 68 | 0 | 2 | 0 | A | A | | | | | | | | | | |
| NC_011770 | 3452186 | C | T | 80 | 0 | 8 | 0 | | | T | T | T | T | | | T | T | T | T |
| NC_011770 | 3453845 | G | A | 68 | 0 | 2 | 0 | A | A | | | | | | | | | | |
| NC_011770 | 3456157 | C | T | 75 | 0 | 12 | 0 | T | T | T | T | T | T | T | T | T | T | T | T |

| | | | | | | | | | | | | | | | | | | | |
|---|---|---|---|---|---|---|---|---|---|---|---|---|---|---|---|---|---|---|---|
| NC_011770 | 3456188 | A | G | 76 | 0 | 6 | 0 | | | | | | | G | G | G | G | G | G |
| NC_011770 | 3456224 | C | T | 71 | 0 | 10 | 0 | T | T | T | T | T | T | T | T | | | T | T |
| NC_011770 | 3456259 | A | T | 71 | 0 | 10 | 0 | T | T | T | T | T | T | T | T | | | T | T |
| NC_011770 | 3456858 | G | T | 62 | 0 | 10 | 0 | T | T | T | T | T | T | T | T | | | T | T |
| NC_011770 | 3456949 | G | A | 56 | 0 | 10 | 0 | A | A | A | A | A | A | A | A | | | A | A |
| NC_011770 | 3460012 | G | A | 64 | 0 | 2 | 0 | A | A | | | | | | | | | | |
| NC_011770 | 3460076 | G | A | 64 | 0 | 4 | 0 | | | A | A | A | A | | | | | | |
| NC_011770 | 3460177 | G | A | 68 | 0 | 2 | 0 | | | | | | | A | A | | | | |
| NC_011770 | 3460243 | C | T | 65 | 0 | 6 | 0 | T | T | T | T | T | T | | | | | | |
| NC_011770 | 3460253 | G | C | 66 | 0 | 2 | 0 | | | C | C | | | | | | | | |
| NC_011770 | 3460269 | C | T | 65 | 0 | 8 | 0 | T | T | | | | | T | T | T | T | T | T |
| NC_011770 | 3460867 | G | A | 56 | 0 | 10 | 0 | A | A | A | A | A | A | A | A | | | A | A |
| NC_011770 | 3460868 | T | C | 56 | 0 | 10 | 0 | C | C | C | C | C | C | C | C | | | C | C |
| NC_011770 | 3460901 | C | T | 63 | 0 | 4 | 0 | | | | | | | | | T | T | T | T |
| NC_011770 | 3460922 | G | A | 62 | 0 | 2 | 0 | | | | | | | | | | | A | A |
| NC_011770 | 3460950 | T | C | 61 | 0 | 6 | 0 | C | C | C | C | C | C | | | | | | |
| NC_011770 | 3462185 | G | A | 59 | 0 | 10 | 0 | A | A | A | A | A | A | A | A | A | A | | |
| NC_011770 | 3462232 | G | A | 58 | 0 | 6 | 0 | | | A | A | A | A | | | A | A | | |
| NC_011770 | 3462745 | G | T | 63 | 0 | 2 | 0 | | | | | | | | | | | T | T |
| NC_011770 | 3463442 | A | C | 80 | 0 | 2 | 0 | C | C | | | | | | | | | | |
| NC_011770 | 3463443 | T | G | 80 | 0 | 2 | 0 | G | G | | | | | | | | | | |
| NC_011770 | 3463445 | CT | GT | 80 | 0 | 6 | 0 | GT | GT | C | C | C | C | | | | | | |
| NC_011770 | 3463552 | GC | G | 75 | 0 | 4 | 0 | G | G | | | | | G | G | | | | |
| NC_011770 | 3463567 | C | CG | 78 | 0 | 1 | 0 | | | | | | | CG | | | | | |
| NC_011770 | 3464835 | A | G | 80 | 0 | 10 | 0 | G | G | G | G | G | G | | | G | G | G | G |
| NC_011770 | 3465373 | C | T | 53 | 0 | 6 | 0 | | | T | T | T | T | | | T | T | | |
| NC_011770 | 3465614 | G | A | 59 | 0 | 4 | 0 | | | A | A | A | A | | | | | | |
| NC_011770 | 3467594 | C | T | 72 | 0 | 4 | 0 | T | T | | | | | | | T | T | | |
| NC_011770 | 3467701 | C | G | 75 | 0 | 4 | 0 | | | G | G | G | G | | | | | | |
| NC_011770 | 3467780 | G | C | 79 | 0 | 6 | 0 | | | C | C | C | C | | | | | C | C |
| NC_011770 | 3467802 | C | G | 79 | 0 | 10 | 0 | G | G | G | G | G | G | G | G | | | G | G |
| NC_011770 | 3468124 | G | C | 76 | 0 | 10 | 0 | C | C | C | C | C | C | C | C | C | C | | |
| NC_011770 | 3468162 | C | G | 74 | 0 | 2 | 0 | | | | | | | G | G | | | | |
| NC_011770 | 3469376 | T | A | 47 | 0 | 12 | 0 | A | A | A | A | A | A | A | A | A | A | A | A |
| NC_011770 | 3469401 | T | C | 43 | 0 | 12 | 0 | C | C | C | C | C | C | C | C | C | C | C | C |
| NC_011770 | 3469404 | C | T | 44 | 0 | 12 | 0 | T | T | T | T | T | T | T | T | T | T | T | T |
| NC_011770 | 3469457 | A | G | 43 | 0 | 1 | 0 | | | | | | | G | | | | | |
| NC_011770 | 3469474 | C | T | 44 | 0 | 4 | 0 | | | T | T | T | T | | | | | | |
| NC_011770 | 3471869 | TCCAGGCT( | T | 73 | 0 | 4 | 0 | T | T | | | | | | | T | T | | |
| NC_011770 | 3471953 | T | C | 69 | 0 | 4 | 0 | C | C | | | | | | | C | C | | |
| NC_011770 | 3471991 | T | C | 65 | 0 | 12 | 0 | C | C | C | C | C | C | C | C | C | C | C | C |
| NC_011770 | 3472068 | T | TG | 69 | 0 | 2 | 0 | TG | TG | | | | | | | | | | |
| NC_011770 | 3472119 | C | T | 66 | 0 | 2 | 0 | T | T | | | | | | | | | | |
| NC_011770 | 3472183 | G | C | 72 | 0 | 10 | 0 | C | C | C | C | C | C | C | C | | | C | C |
| NC_011770 | 3475984 | A | G | 56 | 0 | 10 | 0 | G | G | G | G | G | G | G | G | | | G | G |
| NC_011770 | 3476016 | T | C | 58 | 0 | 8 | 0 | C | C | C | C | C | C | | | | | C | C |
| NC_011770 | 3476193 | T | C | 51 | 0 | 2 | 0 | | | | | | | C | C | | | | |
| NC_011770 | 3477241 | T | G | 79 | 0 | 6 | 0 | G | G | | | | | G | G | | | G | G |
| NC_011770 | 3478840 | G | A | 47 | 0 | 2 | 0 | | | | | | | | | | | A | A |
| NC_011770 | 3479606 | G | A | 65 | 0 | 4 | 0 | | | A | A | A | A | | | | | | |
| NC_011770 | 3479654 | C | T | 69 | 0 | 2 | 0 | | | | | | | T | T | | | | |
| NC_011770 | 3479655 | A | G | 68 | 0 | 2 | 0 | | | | | | | G | G | | | | |
| NC_011770 | 3481584 | A | G | 65 | 0 | 4 | 0 | | | G | G | G | G | | | | | | |
| NC_011770 | 3481588 | G | C | 64 | 0 | 4 | 0 | | | C | C | C | C | | | | | | |
| NC_011770 | 3481591 | C | T | 65 | 0 | 2 | 0 | | | | | | | | | T | T | | |
| NC_011770 | 3481592 | A | G | 65 | 0 | 4 | 0 | | | G | G | G | G | | | | | | |
| NC_011770 | 3481861 | A | T | 69 | 0 | 2 | 0 | | | | | | | | | | | T | T |
| NC_011770 | 3482342 | A | G | 62 | 0 | 2 | 0 | | | | | | | G | G | | | | |
| NC_011770 | 3482462 | T | G | 59 | 0 | 2 | 0 | G | G | | | | | | | | | | |
| NC_011770 | 3482498 | A | G | 56 | 0 | 6 | 0 | | | | | | | G | G | G | G | G | G |
| NC_011770 | 3482585 | T | C | 58 | 0 | 6 | 0 | | | | | | | C | C | C | C | C | C |
| NC_011770 | 3493272 | C | T | 75 | 0 | 2 | 0 | | | | | | | | | | | T | T |

| | | | | | | | | | | | | | | | | | | | |
|---|---|---|---|---|---|---|---|---|---|---|---|---|---|---|---|---|---|---|---|
| NC_011770 | 3493747 | A | G | 69 | 0 | 10 | 0 | | | G | G | G | G | G | G | G | G | G | G |
| NC_011770 | 3493749 | T | A | 69 | 0 | 2 | 0 | | | | | | | A | A | | | | |
| NC_011770 | 3493767 | C | T | 70 | 0 | 2 | 0 | | | | | | | | | T | T | | |
| NC_011770 | 3506468 | A | G | 63 | 0 | 4 | 0 | | | | | | | G | G | | | G | G |
| NC_011770 | 3508213 | T | C | 81 | 0 | 6 | 0 | C | C | C | C | C | C | | | | | | |
| NC_011770 | 3508247 | G | A | 79 | 0 | 2 | 0 | | | | | | | | | A | A | | |
| NC_011770 | 3508383 | G | A | 89 | 0 | 2 | 0 | A | A | | | | | | | | | | |
| NC_011770 | 3510930 | C | T | 72 | 0 | 8 | 0 | T | T | T | T | T | T | | | T | T | | |
| NC_011770 | 3510973 | T | C | 71 | 0 | 12 | 0 | C | C | C | C | C | C | C | C | C | C | C | C |
| NC_011770 | 3511036 | A | C | 74 | 0 | 2 | 0 | | | | | | | | | | | C | C |
| NC_011770 | 3511238 | C | T | 73 | 0 | 8 | 0 | T | T | T | T | T | T | | | T | T | | |
| NC_011770 | 3511239 | C | T | 75 | 0 | 2 | 0 | | | | | | | | | T | T | | |
| NC_011770 | 3511257 | C | T | 77 | 0 | 2 | 0 | | | | | | | T | T | | | | |
| NC_011770 | 3511275 | G | T | 77 | 0 | 2 | 0 | | | | | | | | | T | T | | |
| NC_011770 | 3514589 | G | T | 52 | 0 | 4 | 0 | | | | | | | T | T | T | T | | |
| NC_011770 | 3514598 | C | A | 51 | 0 | 12 | 0 | A | A | A | A | A | A | A | A | A | A | A | A |
| NC_011770 | 3514628 | G | T | 54 | 0 | 4 | 0 | | | T | T | T | T | | | | | | |
| NC_011770 | 3514925 | C | T | 59 | 0 | 2 | 0 | T | T | | | | | | | | | | |
| NC_011770 | 3516598 | A | G | 80 | 0 | 6 | 0 | G | G | | | | | G | G | G | G | | |
| NC_011770 | 3516872 | G | A | 85 | 0 | 2 | 0 | A | A | | | | | | | | | | |
| NC_011770 | 3519428 | T | C | 77 | 0 | 2 | 0 | | | | | | | | | C | C | | |
| NC_011770 | 3519442 | G | C | 78 | 0 | 4 | 0 | C | C | | | | | | | | | C | C |
| NC_011770 | 3520640 | A | G | 78 | 0 | 4 | 0 | | | | | | | G | G | | | G | G |
| NC_011770 | 3520817 | A | G | 85 | 0 | 2 | 0 | | | | | | | G | G | | | | |
| NC_011770 | 3520897 | C | G | 76 | 0 | 2 | 0 | | | | | | | G | G | | | | |
| NC_011770 | 3523064 | T | G | 70 | 0 | 12 | 0 | G | G | G | G | G | G | G | G | G | G | G | G |
| NC_011770 | 3530205 | T | C | 68 | 0 | 2 | 0 | | | | | | | | | | | C | C |
| NC_011770 | 3530238 | C | T | 68 | 0 | 6 | 0 | | | T | T | T | T | | | T | T | | |
| NC_011770 | 3530248 | T | C | 69 | 0 | 10 | 0 | C | C | C | C | C | C | C | C | C | C | | |
| NC_011770 | 3530420 | T | C | 79 | 0 | 2 | 0 | | | | | | | | | | | C | C |
| NC_011770 | 3530490 | T | C | 80 | 0 | 12 | 0 | C | C | C | C | C | C | C | C | C | C | C | C |
| NC_011770 | 3530774 | G | GA | 75 | 0 | 9 | 0 | GA | GA | GA | GA | GA | GA | | | GA | | GA | GA |
| NC_011770 | 3532156 | C | T | 85 | 0 | 4 | 0 | | | T | T | T | T | | | | | | |
| NC_011770 | 3537359 | G | A | 77 | 0 | 1 | 0 | | | | | | | A | | | | | |
| NC_011770 | 3538559 | T | C | 69 | 0 | 12 | 0 | C | C | C | C | C | C | C | C | C | C | C | C |
| NC_011770 | 3539547 | A | G | 70 | 0 | 2 | 0 | | | | | | | | | | | G | G |
| NC_011770 | 3539548 | T | C | 71 | 0 | 2 | 0 | | | | | | | | | | | C | C |
| NC_011770 | 3539560 | G | C | 61 | 0 | 2 | 0 | | | | | | | | | | | C | C |
| NC_011770 | 3539609 | A | G | 57 | 0 | 2 | 0 | | | | | | | | | | | G | G |
| NC_011770 | 3539625 | C | G | 54 | 0 | 2 | 0 | | | | | | | | | | | G | G |
| NC_011770 | 3539634 | A | T | 53 | 0 | 2 | 0 | | | | | | | | | | | T | T |
| NC_011770 | 3539635 | A | G | 53 | 0 | 2 | 0 | | | | | | | | | | | G | G |
| NC_011770 | 3539640 | C | T | 54 | 0 | 2 | 0 | | | | | | | | | | | T | T |
| NC_011770 | 3539644 | A | C | 55 | 0 | 2 | 0 | | | | | | | | | | | C | C |
| NC_011770 | 3540262 | G | C | 17 | 6 | 2 | 0 | C | C | . | . | . | . | | | . | . | | |
| NC_011770 | 3540326 | A | C | 17 | 6 | 2 | 0 | | | . | . | . | . | C | C | . | . | | |
| NC_011770 | 3540410 | C | T | 22 | 6 | 4 | 0 | T | T | . | . | . | . | | | . | . | T | T |
| NC_011770 | 3540435 | A | AC | 23 | 6 | 4 | 0 | AC | AC | . | . | . | . | AC | AC | . | . | | |
| NC_011770 | 3541620 | T | G | 54 | 0 | 2 | 0 | | | | | | | G | G | | | | |
| NC_011770 | 3541659 | A | G | 52 | 0 | 8 | 0 | | | G | G | G | G | G | G | G | G | | |
| NC_011770 | 3541669 | A | T | 58 | 0 | 8 | 0 | | | T | T | T | T | T | T | T | T | | |
| NC_011770 | 3541918 | A | AC | 77 | 0 | 6 | 0 | AC | AC | | | | | AC | AC | | | AC | AC |
| NC_011770 | 3541919 | C | CG | 36 | 4 | 6 | 0 | . | . | CG | CG | CG | CG | . | . | CG | CG | | . |
| NC_011770 | 3541922 | GA | G | 46 | 4 | 2 | 0 | | | G | . | . | G | | | . | . | | |
| NC_011770 | 3542082 | T | C | 90 | 0 | 2 | 0 | | | | | | | C | C | | | | |
| NC_011770 | 3543206 | A | G | 70 | 0 | 2 | 0 | | | | | | | G | G | | | | |
| NC_011770 | 3543248 | A | G | 70 | 0 | 2 | 0 | | | | | | | G | G | | | | |
| NC_011770 | 3543254 | A | G | 71 | 0 | 2 | 0 | | | | | | | G | G | | | | |
| NC_011770 | 3543257 | T | C | 70 | 0 | 2 | 0 | | | | | | | C | C | | | | |
| NC_011770 | 3543259 | C | T | 69 | 0 | 2 | 0 | | | | | | | T | T | | | | |
| NC_011770 | 3543274 | T | C | 71 | 0 | 2 | 0 | | | | | | | C | C | | | | |
| NC_011770 | 3543279 | A | G | 73 | 0 | 2 | 0 | | | | | | | G | G | | | | |

| | | | | | | | | | | | | | | | | | | | |
|---|---|---|---|---|---|---|---|---|---|---|---|---|---|---|---|---|---|---|---|
| NC_011770 | 3543286 | T | C | 71 | 0 | 2 | 0 | | | | | | | C | C | | | | |
| NC_011770 | 3543308 | G | C | 71 | 0 | 6 | 0 | | | C | C | C | C | | | C | C | | |
| NC_011770 | 3543311 | G | T | 73 | 0 | 2 | 0 | | | | | | | T | T | | | | |
| NC_011770 | 3543343 | T | C | 75 | 0 | 2 | 0 | | | | | | | C | C | | | | |
| NC_011770 | 3545128 | G | A | 70 | 0 | 2 | 0 | A | A | | | | | | | | | | |
| NC_011770 | 3545133 | C | G | 69 | 0 | 2 | 0 | G | G | | | | | | | | | | |
| NC_011770 | 3545143 | A | G | 69 | 0 | 2 | 0 | G | G | | | | | | | | | | |
| NC_011770 | 3545192 | A | G | 64 | 0 | 12 | 0 | G | G | G | G | G | G | G | G | G | G | G | G |
| NC_011770 | 3546267 | C | T | 52 | 0 | 2 | 0 | | | | | | | | | | | T | T |
| NC_011770 | 3546404 | G | GC | 61 | 0 | 4 | 0 | | | GC | GC | GC | GC | | | | | | |
| NC_011770 | 3547710 | A | G | 68 | 0 | 2 | 0 | | | | | | | | | G | G | | |
| NC_011770 | 3548977 | T | A | 95 | 0 | 2 | 0 | | | | | | | A | A | | | | |
| NC_011770 | 3553752 | G | A | 63 | 0 | 2 | 0 | | | | | | | A | A | | | | |
| NC_011770 | 3554296 | C | T | 41 | 4 | 4 | 0 | T | T | . | . | . | . | T | T | | | | |
| NC_011770 | 3557902 | A | G | 55 | 0 | 4 | 0 | | | G | G | G | G | | | | | | |
| NC_011770 | 3567801 | T | G | 70 | 0 | 6 | 0 | | | G | G | G | G | | | | | G | G |
| NC_011770 | 3567803 | A | G | 70 | 0 | 12 | 0 | G | G | G | G | G | G | G | G | G | G | G | G |
| NC_011770 | 3567813 | G | A | 71 | 0 | 4 | 0 | | | A | A | A | A | | | | | | |
| NC_011770 | 3570795 | G | C | 77 | 0 | 10 | 0 | C | C | C | C | C | C | C | C | | | C | C |
| NC_011770 | 3570796 | G | T | 76 | 0 | 10 | 0 | T | T | T | T | T | T | T | T | | | T | T |
| NC_011770 | 3574127 | T | G | 68 | 0 | 12 | 0 | G | G | G | G | G | G | G | G | G | G | G | G |
| NC_011770 | 3574149 | A | G | 69 | 0 | 4 | 0 | | | G | G | G | G | | | | | | |
| NC_011770 | 3574163 | C | G | 71 | 0 | 4 | 0 | | | G | G | G | G | | | | | | |
| NC_011770 | 3574168 | G | T | 70 | 0 | 4 | 0 | | | T | T | T | T | | | | | | |
| NC_011770 | 3574196 | G | C | 71 | 0 | 2 | 0 | | | | | | | | | | | C | C |
| NC_011770 | 3574688 | G | A | 71 | 0 | 6 | 0 | A | A | | | | | | | A | A | A | A |
| NC_011770 | 3574689 | C | T | 68 | 0 | 12 | 0 | T | T | T | T | T | T | T | T | T | T | T | T |
| NC_011770 | 3574702 | G | T | 70 | 0 | 2 | 0 | | | | | | | | | T | T | | |
| NC_011770 | 3574730 | C | T | 66 | 0 | 4 | 0 | | | T | T | T | T | | | | | | |
| NC_011770 | 3574743 | G | T | 66 | 0 | 4 | 0 | | | T | T | T | T | | | | | | |
| NC_011770 | 3574799 | C | T | 69 | 0 | 8 | 0 | T | T | T | T | T | T | T | T | | | | |
| NC_011770 | 3576712 | C | T | 49 | 0 | 10 | 0 | | | T | T | T | T | T | T | T | T | T | T |
| NC_011770 | 3576718 | T | C | 48 | 0 | 10 | 0 | | | C | C | C | C | C | C | C | C | C | C |
| NC_011770 | 3576726 | G | A | 48 | 0 | 10 | 0 | | | A | A | A | A | A | A | A | A | A | A |
| NC_011770 | 3576738 | T | C | 50 | 0 | 2 | 0 | | | | | | | | | | | C | C |
| NC_011770 | 3576757 | A | G | 48 | 0 | 10 | 0 | | | G | G | G | G | G | G | G | G | G | G |
| NC_011770 | 3578303 | C | G | 77 | 0 | 4 | 0 | | | G | G | G | G | | | | | | |
| NC_011770 | 3578411 | AG | A | 79 | 0 | 4 | 0 | | | A | A | A | A | | | | | | |
| NC_011770 | 3582384 | T | C | 50 | 0 | 12 | 0 | C | C | C | C | C | C | C | C | C | C | C | C |
| NC_011770 | 3582395 | C | T | 50 | 0 | 2 | 0 | | | | | | | | | | | T | T |
| NC_011770 | 3582408 | C | G | 49 | 0 | 12 | 0 | G | G | G | G | G | G | G | G | G | G | G | G |
| NC_011770 | 3583328 | A | G | 66 | 0 | 2 | 0 | G | G | | | | | | | | | | |
| NC_011770 | 3583398 | A | T | 60 | 0 | 4 | 0 | | | T | T | T | T | | | | | | |
| NC_011770 | 3583408 | A | G | 64 | 0 | 2 | 0 | G | G | | | | | | | | | | |
| NC_011770 | 3583436 | G | A | 59 | 0 | 2 | 0 | A | A | | | | | | | | | | |
| NC_011770 | 3583448 | C | A | 60 | 0 | 2 | 0 | A | A | | | | | | | | | | |
| NC_011770 | 3583453 | T | G | 61 | 0 | 2 | 0 | G | G | | | | | | | | | | |
| NC_011770 | 3583455 | A | T | 62 | 0 | 2 | 0 | T | T | | | | | | | | | | |
| NC_011770 | 3583590 | G | A | 60 | 0 | 2 | 0 | | | | | | | A | A | | | | |
| NC_011770 | 3583672 | CG | C | 51 | 0 | 12 | 0 | C | C | C | C | C | C | C | C | C | C | C | C |
| NC_011770 | 3583748 | T | C | 51 | 0 | 10 | 0 | C | C | C | C | C | C | C | C | | | C | C |
| NC_011770 | 3584173 | C | T | 61 | 0 | 2 | 0 | | | | | | | | | T | T | | |
| NC_011770 | 3585916 | C | T | 46 | 0 | 2 | 0 | | | | | | | | | T | T | | |
| NC_011770 | 3585940 | G | A | 41 | 0 | 2 | 0 | | | | | | | | | | | A | A |
| NC_011770 | 3585942 | T | C | 38 | 2 | 8 | 0 | C | C | C | C | C | C | C | C | . | . | | |
| NC_011770 | 3585946 | T | C | 38 | 2 | 10 | 0 | C | C | C | C | C | C | C | C | . | . | C | C |
| NC_011770 | 3585948 | C | T | 37 | 2 | 2 | 0 | | | | | | | | | . | . | T | T |
| NC_011770 | 3588300 | T | C | 18 | 10 | 2 | 0 | . | . | . | . | . | . | . | . | . | . | C | C |
| NC_011770 | 3588380 | C | G | 59 | 0 | 6 | 0 | G | G | G | G | G | G | | | | | | |
| NC_011770 | 3588387 | T | C | 58 | 0 | 6 | 0 | C | C | C | C | C | C | | | | | | |
| NC_011770 | 3588443 | G | T | 52 | 0 | 6 | 0 | T | T | T | T | T | T | | | | | | |
| NC_011770 | 3588445 | C | T | 52 | 0 | 6 | 0 | T | T | T | T | T | T | | | | | | |

| | | | | | | | | | | | | | | | | | | | |
|---|---|---|---|---|---|---|---|---|---|---|---|---|---|---|---|---|---|---|---|
| NC_011770 | 3591153 | C | T | 62 | 0 | 2 | 0 | | | | | | | | | T | T | | |
| NC_011770 | 3593410 | G | A | 78 | 0 | 2 | 0 | | | | | | | | | | | A | A |
| NC_011770 | 3593464 | C | T | 73 | 0 | 2 | 0 | T | T | | | | | | | | | | |
| NC_011770 | 3593480 | C | T | 73 | 0 | 2 | 0 | | | | | | | | | | | T | T |
| NC_011770 | 3593490 | G | A | 72 | 0 | 2 | 0 | | | | | | | | | | | A | A |
| NC_011770 | 3593547 | C | T | 66 | 0 | 2 | 0 | | | | | | | | | | | T | T |
| NC_011770 | 3600713 | C | T | 74 | 0 | 6 | 0 | | | T | T | T | T | | | T | T | | |
| NC_011770 | 3600736 | C | T | 78 | 0 | 2 | 0 | T | T | | | | | | | | | | |
| NC_011770 | 3602666 | C | T | 86 | 0 | 4 | 0 | | | T | T | T | T | | | | | | |
| NC_011770 | 3602765 | C | G | 93 | 0 | 4 | 0 | | | | | | | G | G | | | G | G |
| NC_011770 | 3602795 | T | C | 85 | 0 | 1 | 0 | | | | | | | C | | | | | |
| NC_011770 | 3602889 | A | G | 80 | 0 | 4 | 0 | | | G | G | G | G | | | | | | |
| NC_011770 | 3603294 | G | A | 88 | 0 | 4 | 0 | | | | | | | A | A | | | A | A |
| NC_011770 | 3603511 | A | G | 89 | 0 | 1 | 0 | | | | | | | G | | | | | |
| NC_011770 | 3603575 | C | T | 88 | 0 | 2 | 0 | T | T | | | | | | | | | | |
| NC_011770 | 3605084 | G | A | 63 | 0 | 4 | 0 | | | | | | | A | A | | | A | A |
| NC_011770 | 3605152 | C | CG | 6 | 3 | 4 | 0 | . | . | . | CG | . | CG | CG | . | . | . | CG | . |
| NC_011770 | 3606360 | G | A | 77 | 0 | 4 | 0 | | | A | A | A | A | | | | | | |
| NC_011770 | 3606365 | A | C | 75 | 0 | 4 | 0 | | | C | C | C | C | | | | | | |
| NC_011770 | 3606469 | T | A | 76 | 0 | 8 | 0 | A | A | | | | | A | A | A | A | A | A |
| NC_011770 | 3614662 | G | A | 63 | 0 | 2 | 0 | | | | | | | | | | | A | A |
| NC_011770 | 3614701 | C | G | 60 | 0 | 2 | 0 | G | G | | | | | | | | | | |
| NC_011770 | 3614780 | T | C | 56 | 0 | 4 | 0 | | | C | C | C | C | | | | | | |
| NC_011770 | 3614792 | C | T | 59 | 0 | 4 | 0 | | | T | T | T | T | | | | | | |
| NC_011770 | 3614880 | G | A | 57 | 0 | 2 | 0 | A | A | | | | | | | | | | |
| NC_011770 | 3614888 | C | G | 56 | 0 | 10 | 0 | G | G | G | G | G | G | | | G | G | G | G |
| NC_011770 | 3614889 | A | T | 55 | 0 | 10 | 0 | T | T | T | T | T | T | | | T | T | T | T |
| NC_011770 | 3614890 | G | A | 55 | 0 | 10 | 0 | A | A | A | A | A | A | | | A | A | A | A |
| NC_011770 | 3614979 | A | G | 52 | 0 | 2 | 0 | G | G | | | | | | | | | | |
| NC_011770 | 3615057 | G | A | 49 | 0 | 2 | 0 | A | A | | | | | | | | | | |
| NC_011770 | 3615061 | G | A | 45 | 0 | 2 | 0 | A | A | | | | | | | | | | |
| NC_011770 | 3615070 | G | A | 47 | 0 | 4 | 0 | | | | | | | | | A | A | A | A |
| NC_011770 | 3615073 | A | G | 45 | 0 | 6 | 0 | G | G | | | | | | | G | G | G | G |
| NC_011770 | 3615075 | A | G | 44 | 0 | 2 | 0 | | | | | | | G | G | | | | |
| NC_011770 | 3615076 | G | T | 44 | 0 | 6 | 0 | T | T | | | | | | | T | T | T | T |
| NC_011770 | 3615082 | G | A | 45 | 0 | 4 | 0 | | | | | | | | | A | A | A | A |
| NC_011770 | 3615099 | T | C | 44 | 0 | 2 | 0 | C | C | | | | | | | | | | |
| NC_011770 | 3615108 | T | C | 40 | 0 | 6 | 0 | C | C | C | C | C | C | | | | | | |
| NC_011770 | 3615113 | C | G | 41 | 0 | 8 | 0 | G | G | G | G | G | G | | | G | G | | |
| NC_011770 | 3615115 | A | C | 41 | 0 | 8 | 0 | C | C | C | C | C | C | | | C | C | | |
| NC_011770 | 3615120 | A | T | 34 | 1 | 6 | 0 | T | . | T | T | T | T | | | T | . | | |
| NC_011770 | 3615121 | C | T | 34 | 4 | 4 | 0 | . | . | T | T | T | T | | | . | . | | |
| NC_011770 | 3615126 | A | C | 34 | 4 | 4 | 0 | . | . | C | C | C | C | | | . | . | | |
| NC_011770 | 3615132 | C | G | 45 | 0 | 4 | 0 | | | | | | | G | G | | | G | G |
| NC_011770 | 3615133 | G | C | 46 | 0 | 2 | 0 | | | | | | | | | C | C | | |
| NC_011770 | 3615137 | A | C | 46 | 0 | 12 | 0 | C | C | C | C | C | C | C | C | C | C | C | C |
| NC_011770 | 3615151 | C | T | 47 | 0 | 2 | 0 | | | | | | | T | T | | | | |
| NC_011770 | 3615155 | T | G | 46 | 0 | 12 | 0 | G | G | G | G | G | G | G | G | G | G | G | G |
| NC_011770 | 3615163 | G | A | 47 | 0 | 12 | 0 | A | A | A | A | A | A | A | A | A | A | A | A |
| NC_011770 | 3616129 | A | G | 71 | 0 | 2 | 0 | G | G | | | | | | | | | | |
| NC_011770 | 3616130 | T | G | 72 | 0 | 2 | 0 | G | G | | | | | | | | | | |
| NC_011770 | 3616232 | G | A | 73 | 0 | 2 | 0 | | | | | | | | | | | A | A |
| NC_011770 | 3618563 | A | G | 84 | 0 | 2 | 0 | G | G | | | | | | | | | | |
| NC_011770 | 3618575 | A | G | 76 | 0 | 2 | 0 | G | G | | | | | | | | | | |
| NC_011770 | 3618622 | C | T | 74 | 0 | 2 | 0 | T | T | | | | | | | | | | |
| NC_011770 | 3618628 | T | C | 73 | 0 | 4 | 0 | C | C | | | | | | | C | C | | |
| NC_011770 | 3619329 | T | C | 57 | 0 | 10 | 0 | C | C | C | C | C | C | C | C | | | C | C |
| NC_011770 | 3619359 | T | C | 59 | 0 | 2 | 0 | C | C | | | | | | | | | | |
| NC_011770 | 3621487 | C | T | 50 | 0 | 4 | 0 | | | | | | | | | T | T | T | T |
| NC_011770 | 3621600 | C | G | 40 | 0 | 2 | 0 | G | G | | | | | | | | | | |
| NC_011770 | 3621824 | G | A | 49 | 0 | 2 | 0 | A | A | | | | | | | | | | |
| NC_011770 | 3621949 | C | G | 49 | 0 | 2 | 0 | G | G | | | | | | | | | | |

| | | | | | | | | | | | | | | | | | | | |
|---|---|---|---|---|---|---|---|---|---|---|---|---|---|---|---|---|---|---|---|
| NC_011770 | 3621970 | T | C | 44 | 0 | 2 | 0 | C | C | | | | | | | | | | |
| NC_011770 | 3621991 | A | G | 43 | 0 | 8 | 0 | G | G | G | G | G | G | G | G | | | | |
| NC_011770 | 3622000 | G | T | 50 | 0 | 4 | 0 | | | T | T | T | T | | | | | | |
| NC_011770 | 3622090 | A | G | 59 | 0 | 2 | 0 | G | G | | | | | | | | | | |
| NC_011770 | 3622105 | A | G | 63 | 0 | 12 | 0 | G | G | G | G | G | G | G | G | G | G | G | G |
| NC_011770 | 3623648 | G | A | 73 | 0 | 4 | 0 | | | | | | | | | A | A | A | A |
| NC_011770 | 3623685 | C | A | 68 | 0 | 2 | 0 | | | | | | | A | A | | | | |
| NC_011770 | 3623686 | A | G | 67 | 0 | 2 | 0 | | | | | | | G | G | | | | |
| NC_011770 | 3624325 | T | G | 62 | 0 | 2 | 0 | G | G | | | | | | | | | | |
| NC_011770 | 3624328 | G | A | 61 | 0 | 2 | 0 | A | A | | | | | | | | | | |
| NC_011770 | 3624434 | C | T | 53 | 0 | 8 | 0 | T | T | T | T | T | T | T | T | | | | |
| NC_011770 | 3624466 | G | C | 58 | 0 | 2 | 0 | | | | | | | C | C | | | | |
| NC_011770 | 3624475 | C | T | 60 | 0 | 8 | 0 | T | T | T | T | T | T | T | T | | | | |
| NC_011770 | 3624540 | C | A | 61 | 0 | 4 | 0 | A | A | | | | | A | A | | | | |
| NC_011770 | 3625603 | T | C | 42 | 0 | 10 | 0 | C | C | C | C | C | C | | | C | C | C | C |
| NC_011770 | 3625932 | G | A | 76 | 0 | 2 | 0 | | | | | | | A | A | | | | |
| NC_011770 | 3626305 | C | G | 82 | 0 | 4 | 0 | | | G | G | G | G | | | | | | |
| NC_011770 | 3626364 | C | A | 75 | 0 | 4 | 0 | | | A | A | A | A | | | | | | |
| NC_011770 | 3626427 | C | T | 72 | 0 | 4 | 0 | | | T | T | T | T | | | | | | |
| NC_011770 | 3626514 | T | C | 76 | 0 | 4 | 0 | | | | | | | | | C | C | C | C |
| NC_011770 | 3627313 | C | T | 49 | 0 | 8 | 0 | | | T | T | T | T | | | T | T | T | T |
| NC_011770 | 3627399 | T | C | 50 | 0 | 10 | 0 | C | C | C | C | C | C | | | C | C | C | C |
| NC_011770 | 3629383 | C | T | 76 | 0 | 2 | 0 | T | T | | | | | | | | | | |
| NC_011770 | 3629402 | G | GC | 77 | 0 | 2 | 0 | GC | GC | | | | | | | | | | |
| NC_011770 | 3629432 | A | G | 75 | 0 | 2 | 0 | | | | | | | | | | | G | G |
| NC_011770 | 3630465 | A | T | 74 | 0 | 4 | 0 | | | T | T | T | T | | | | | | |
| NC_011770 | 3630492 | C | T | 74 | 0 | 2 | 0 | | | | | | | T | T | | | | |
| NC_011770 | 3630511 | GA | G | 73 | 0 | 2 | 0 | G | G | | | | | | | | | | |
| NC_011770 | 3631870 | A | G | 58 | 0 | 8 | 0 | G | G | | | | | G | G | G | G | G | G |
| NC_011770 | 3631971 | TG | T | 58 | 0 | 2 | 0 | | | | | | | | | | | T | T |
| NC_011770 | 3632028 | G | A | 52 | 0 | 2 | 0 | A | A | | | | | | | | | | |
| NC_011770 | 3632041 | C | T | 51 | 0 | 8 | 0 | T | T | | | | | T | T | T | T | T | T |
| NC_011770 | 3632058 | G | A | 53 | 0 | 4 | 0 | | | A | A | A | A | | | | | | |
| NC_011770 | 3633657 | G | A | 60 | 0 | 2 | 0 | | | | | | | A | A | | | | |
| NC_011770 | 3633689 | T | A | 59 | 0 | 2 | 0 | | | | | | | | | | | A | A |
| NC_011770 | 3635239 | A | C | 66 | 0 | 6 | 0 | | | C | C | C | C | | | C | C | | |
| NC_011770 | 3635249 | A | G | 67 | 0 | 10 | 0 | G | G | G | G | G | G | | | G | G | G | G |
| NC_011770 | 3635256 | G | A | 72 | 0 | 4 | 0 | | | A | A | A | A | | | | | | |
| NC_011770 | 3635271 | C | A | 67 | 0 | 4 | 0 | | | | | | | A | A | | | A | A |
| NC_011770 | 3635277 | G | A | 67 | 0 | 4 | 0 | | | | | | | A | A | | | A | A |
| NC_011770 | 3635278 | C | T | 67 | 0 | 8 | 0 | | | T | T | T | T | A | A | | | A | A |
| NC_011770 | 3635492 | C | T | 62 | 0 | 2 | 0 | | | | | | | | | | | T | T |
| NC_011770 | 3635519 | C | T | 61 | 0 | 2 | 0 | T | T | | | | | | | | | | |
| NC_011770 | 3635530 | C | T | 60 | 0 | 2 | 0 | | | | | | | T | T | | | | |
| NC_011770 | 3635553 | C | G | 57 | 0 | 2 | 0 | | | | | G | G | | | | | | |
| NC_011770 | 3635575 | T | G | 56 | 0 | 12 | 0 | G | G | G | G | G | G | G | G | G | G | G | G |
| NC_011770 | 3638394 | T | C | 9 | 10 | 2 | 0 | . | . | . | . | . | . | C | C | . | . | . | . |
| NC_011770 | 3638402 | G | A | 10 | 10 | 2 | 0 | . | . | . | . | . | . | A | A | . | . | . | . |
| NC_011770 | 3638413 | A | G | 10 | 10 | 2 | 0 | . | . | . | . | . | . | G | G | . | . | . | . |
| NC_011770 | 3638414 | C | G | 10 | 10 | 2 | 0 | . | . | . | . | . | . | G | G | . | . | . | . |
| NC_011770 | 3638419 | A | C | 10 | 10 | 2 | 0 | . | . | . | . | . | . | C | C | . | . | . | . |
| NC_011770 | 3638421 | A | C | 10 | 10 | 2 | 0 | . | . | . | . | . | . | C | C | . | . | . | . |
| NC_011770 | 3638975 | C | T | 17 | 8 | 2 | 0 | | | . | . | . | . | T | T | . | . | . | . |
| NC_011770 | 3639050 | C | T | 18 | 8 | 2 | 0 | | | . | . | . | . | T | T | . | . | . | . |
| NC_011770 | 3639055 | A | G | 19 | 8 | 2 | 0 | G | G | . | . | . | . | | | . | . | . | . |
| NC_011770 | 3639098 | C | T | 19 | 8 | 2 | 0 | | | . | . | . | . | T | T | . | . | . | . |
| NC_011770 | 3639197 | T | TA | 18 | 8 | 2 | 0 | TA | TA | . | . | . | . | | | . | . | . | . |
| NC_011770 | 3639232 | G | A | 19 | 8 | 2 | 0 | A | A | . | . | . | . | | | . | . | . | . |
| NC_011770 | 3639240 | G | GC | 17 | 8 | 2 | 0 | GC | GC | . | . | . | . | | | . | . | . | . |
| NC_011770 | 3639252 | A | G | 17 | 8 | 2 | 0 | G | G | . | . | . | . | | | . | . | . | . |
| NC_011770 | 3639257 | AC | A | 14 | 8 | 2 | 0 | A | A | . | . | . | . | | | . | . | . | . |
| NC_011770 | 3639274 | A | G | 17 | 8 | 2 | 0 | G | G | . | . | . | . | | | . | . | . | . |

| | | | | | | | | | | | | | | | | | | | |
|---|---|---|---|---|---|---|---|---|---|---|---|---|---|---|---|---|---|---|---|
| NC_011770 | 3639280 | G | A | 17 | 8 | 2 | 0 | A | A | . | . | . | . | | | . | . | . | . |
| NC_011770 | 3639290 | A | C | 17 | 8 | 2 | 0 | C | C | . | . | . | . | | | . | . | . | . |
| NC_011770 | 3639297 | T | C | 17 | 8 | 2 | 0 | C | C | . | . | . | . | | | . | . | . | . |
| NC_011770 | 3639321 | A | C | 18 | 8 | 2 | 0 | C | C | . | . | . | . | | | . | . | . | . |
| NC_011770 | 3639324 | T | C | 18 | 8 | 2 | 0 | C | C | . | . | . | . | | | . | . | . | . |
| NC_011770 | 3639331 | C | T | 17 | 8 | 2 | 0 | T | T | . | . | . | . | | | . | . | . | . |
| NC_011770 | 3639341 | C | A | 17 | 8 | 2 | 0 | A | A | . | . | . | . | | | . | . | . | . |
| NC_011770 | 3639346 | A | G | 17 | 8 | 2 | 0 | G | G | . | . | . | . | | | . | . | . | . |
| NC_011770 | 3639354 | T | G | 15 | 8 | 2 | 0 | G | G | . | . | . | . | | | . | . | . | . |
| NC_011770 | 3639356 | C | A | 16 | 8 | 2 | 0 | A | A | . | . | . | . | | | . | . | . | . |
| NC_011770 | 3639357 | T | G | 15 | 8 | 2 | 0 | G | G | . | . | . | . | | | . | . | . | . |
| NC_011770 | 3639498 | C | T | 16 | 8 | 2 | 0 | T | T | . | . | . | . | | | . | . | . | . |
| NC_011770 | 3639519 | T | G | 16 | 8 | 2 | 0 | G | G | . | . | . | . | | | . | . | . | . |
| NC_011770 | 3639527 | T | C | 16 | 8 | 2 | 0 | C | C | . | . | . | . | | | . | . | . | . |
| NC_011770 | 3639529 | G | T | 16 | 8 | 2 | 0 | T | T | . | . | . | . | | | . | . | . | . |
| NC_011770 | 3639533 | T | A | 16 | 8 | 4 | 0 | A | A | . | . | . | . | A | A | . | . | . | . |
| NC_011770 | 3639558 | T | C | 14 | 8 | 2 | 0 | C | C | . | . | . | . | | | . | . | . | . |
| NC_011770 | 3639562 | C | G | 15 | 8 | 2 | 0 | G | G | . | . | . | . | | | . | . | . | . |
| NC_011770 | 3639564 | A | G | 15 | 8 | 2 | 0 | G | G | . | . | . | . | | | . | . | . | . |
| NC_011770 | 3639565 | C | T | 14 | 8 | 2 | 0 | T | T | . | . | . | . | | | . | . | . | . |
| NC_011770 | 3639569 | G | A | 16 | 8 | 2 | 0 | | | . | . | . | . | A | A | . | . | . | . |
| NC_011770 | 3639626 | G | A | 48 | 0 | 2 | 0 | A | A | | | | | | | | | | |
| NC_011770 | 3639636 | G | A | 50 | 0 | 2 | 0 | A | A | | | | | | | | | | |
| NC_011770 | 3639650 | G | C | 50 | 0 | 2 | 0 | C | C | | | | | | | | | | |
| NC_011770 | 3642958 | T | C | 78 | 0 | 8 | 0 | C | C | C | C | C | C | | | C | C | | |
| NC_011770 | 3642961 | A | C | 76 | 0 | 8 | 0 | C | C | C | C | C | C | | | C | C | | |
| NC_011770 | 3642978 | G | A | 78 | 0 | 8 | 0 | A | A | A | A | A | A | | | A | A | | |
| NC_011770 | 3642983 | G | A | 79 | 0 | 2 | 0 | | | | | | | | | | | A | A |
| NC_011770 | 3642985 | A | G | 78 | 0 | 2 | 0 | | | | | | | | | G | G | | |
| NC_011770 | 3643026 | C | T | 84 | 0 | 8 | 0 | T | T | T | T | T | T | | | T | T | | |
| NC_011770 | 3643058 | T | C | 87 | 0 | 8 | 0 | C | C | C | C | C | C | | | C | C | | |
| NC_011770 | 3645230 | G | A | 31 | 4 | 2 | 0 | | | . | . | . | . | | | A | A | | |
| NC_011770 | 3645564 | G | A | 65 | 0 | 2 | 0 | A | A | | | | | | | | | | |
| NC_011770 | 3646084 | G | A | 74 | 0 | 2 | 0 | | | | | | | A | A | | | | |
| NC_011770 | 3646115 | G | A | 68 | 0 | 2 | 0 | | | | | | | A | A | | | | |
| NC_011770 | 3646674 | G | A | 84 | 0 | 12 | 0 | A | A | A | A | A | A | A | A | A | A | A | A |
| NC_011770 | 3647751 | A | C | 56 | 0 | 2 | 0 | C | C | | | | | | | | | | |
| NC_011770 | 3647752 | G | T | 57 | 0 | 2 | 0 | T | T | | | | | | | | | | |
| NC_011770 | 3647774 | C | T | 58 | 0 | 2 | 0 | | | | | | | T | T | | | | |
| NC_011770 | 3649708 | A | G | 45 | 0 | 12 | 0 | G | G | G | G | G | G | G | G | G | G | G | G |
| NC_011770 | 3649714 | G | A | 44 | 0 | 2 | 0 | | | | | | | | | A | A | | |
| NC_011770 | 3649728 | A | G | 47 | 0 | 4 | 0 | G | G | | | | | | | G | G | | |
| NC_011770 | 3649762 | A | G | 47 | 0 | 2 | 0 | | | | | | | G | G | | | | |
| NC_011770 | 3649819 | C | T | 48 | 0 | 4 | 0 | | | T | T | T | T | | | | | | |
| NC_011770 | 3652133 | G | A | 73 | 0 | 4 | 0 | A | A | | | | | | | | | A | A |
| NC_011770 | 3652163 | G | A | 76 | 0 | 2 | 0 | A | A | | | | | | | | | | |
| NC_011770 | 3652242 | C | T | 75 | 0 | 4 | 0 | | | T | T | T | T | | | | | | |
| NC_011770 | 3652246 | A | G | 74 | 0 | 12 | 0 | G | G | G | G | G | G | G | G | G | G | G | G |
| NC_011770 | 3652537 | G | GGGA | 69 | 0 | 2 | 0 | | | | | | | | | | | GGGA | GGGA |
| NC_011770 | 3652558 | G | A | 66 | 0 | 2 | 0 | | | | | | | | | | | A | A |
| NC_011770 | 3652561 | G | T | 63 | 0 | 4 | 0 | | | T | T | T | T | | | | | | |
| NC_011770 | 3652570 | G | A | 65 | 0 | 4 | 0 | | | A | A | A | A | | | | | | |
| NC_011770 | 3652594 | G | A | 72 | 0 | 2 | 0 | A | A | | | | | | | | | | |
| NC_011770 | 3652616 | GT | G | 76 | 0 | 2 | 0 | | | G | G | | | | | | | | |
| NC_011770 | 3653167 | G | A | 67 | 0 | 2 | 0 | | | | | | | | | | | A | A |
| NC_011770 | 3657702 | T | C | 41 | 0 | 1 | 0 | | | | | | | C | | | | | |
| NC_011770 | 3657716 | T | C | 43 | 0 | 2 | 0 | | | | | | | | | | | C | C |
| NC_011770 | 3657818 | C | T | 48 | 0 | 2 | 0 | | | | | | | | | | | T | T |
| NC_011770 | 3658879 | C | T | 70 | 0 | 2 | 0 | | | | | | | | | | | T | T |
| NC_011770 | 3658952 | A | G | 73 | 0 | 12 | 0 | G | G | G | G | G | G | G | G | G | G | G | G |
| NC_011770 | 3658971 | C | T | 69 | 0 | 2 | 0 | | | | | | | | | | | T | T |
| NC_011770 | 3658976 | C | A | 71 | 0 | 4 | 0 | | | A | A | A | A | | | | | | |

| | | | | | | | | | | | | | | | | | | | |
|---|---|---|---|---|---|---|---|---|---|---|---|---|---|---|---|---|---|---|---|
| NC_011770 | 3659011 | A | T | 68 | 0 | 2 | 0 | | | | | | | | | | | T | T |
| NC_011770 | 3659015 | T | C | 67 | 0 | 12 | 0 | C | C | C | C | C | C | C | C | C | C | C | C |
| NC_011770 | 3659425 | C | A | 70 | 0 | 2 | 0 | | | | | | | | | | | A | A |
| NC_011770 | 3659430 | A | G | 70 | 0 | 2 | 0 | | | | | | | | | | | G | G |
| NC_011770 | 3660645 | C | T | 77 | 0 | 2 | 0 | T | T | | | | | | | | | | |
| NC_011770 | 3660656 | T | G | 76 | 0 | 6 | 0 | G | G | G | G | G | G | | | | | | |
| NC_011770 | 3662367 | C | T | 68 | 0 | 4 | 0 | T | T | | | | | | | T | T | | |
| NC_011770 | 3666068 | C | T | 64 | 0 | 2 | 0 | T | T | | | | | | | | | | |
| NC_011770 | 3666080 | G | A | 62 | 0 | 2 | 0 | | | | | | | | | | | A | A |
| NC_011770 | 3666091 | G | A | 65 | 0 | 2 | 0 | A | A | | | | | | | | | | |
| NC_011770 | 3666099 | T | C | 67 | 0 | 2 | 0 | | | | | | | | | | | C | C |
| NC_011770 | 3666103 | A | G | 66 | 0 | 2 | 0 | | | | | | | | | | | G | G |
| NC_011770 | 3666106 | A | G | 64 | 0 | 2 | 0 | | | | | | | | | | | G | G |
| NC_011770 | 3666108 | C | G | 65 | 0 | 2 | 0 | | | | | | | | | | | G | G |
| NC_011770 | 3666110 | T | C | 64 | 0 | 2 | 0 | | | | | | | | | | | C | C |
| NC_011770 | 3666127 | A | G | 68 | 0 | 2 | 0 | | | | | | | | | | | G | G |
| NC_011770 | 3666979 | C | T | 52 | 0 | 8 | 0 | | | T | T | T | T | T | T | T | T | | |
| NC_011770 | 3670378 | G | A | 65 | 0 | 2 | 0 | | | | | | | | | A | A | | |
| NC_011770 | 3673295 | G | T | 68 | 0 | 2 | 0 | T | T | | | | | | | | | | |
| NC_011770 | 3673370 | G | A | 70 | 0 | 8 | 0 | | | A | A | A | A | A | A | | | A | A |
| NC_011770 | 3673383 | T | C | 70 | 0 | 2 | 0 | | | | | | | | | C | C | | |
| NC_011770 | 3675748 | A | T | 60 | 0 | 2 | 0 | | | | | | | | | | | T | T |
| NC_011770 | 3675766 | C | A | 58 | 0 | 2 | 0 | | | | | | | | | | | A | A |
| NC_011770 | 3675831 | C | T | 54 | 0 | 2 | 0 | | | | | | | | | T | T | | |
| NC_011770 | 3675847 | A | T | 50 | 0 | 4 | 0 | | | T | T | T | T | | | | | | |
| NC_011770 | 3675870 | A | G | 54 | 0 | 4 | 0 | G | G | | | | | | | G | G | | |
| NC_011770 | 3675876 | T | G | 54 | 0 | 2 | 0 | G | G | | | | | | | | | | |
| NC_011770 | 3675877 | C | A | 54 | 0 | 2 | 0 | | | | | | | | | | | A | A |
| NC_011770 | 3675882 | A | C | 55 | 0 | 2 | 0 | | | | | | | | | | | C | C |
| NC_011770 | 3675893 | T | G | 55 | 0 | 2 | 0 | G | G | | | | | | | | | | |
| NC_011770 | 3675894 | C | A | 56 | 0 | 2 | 0 | A | A | | | | | | | | | | |
| NC_011770 | 3675915 | GCCAGGTC | G | 51 | 1 | 1 | 0 | G | . | | | | | | | | | | |
| NC_011770 | 3675933 | T | C | 53 | 0 | 4 | 0 | | | C | C | C | C | | | | | | |
| NC_011770 | 3678812 | G | A | 27 | 0 | 6 | 0 | A | A | A | A | A | A | | | | | | |
| NC_011770 | 3678824 | G | C | 23 | 0 | 8 | 0 | C | C | C | C | C | C | | | | | C | C |
| NC_011770 | 3680373 | G | T | 69 | 0 | 6 | 0 | T | T | T | T | T | T | | | | | | |
| NC_011770 | 3681389 | GAAC | G | 77 | 0 | 2 | 0 | | | | | | | | | G | G | | |
| NC_011770 | 3684712 | T | C | 70 | 0 | 4 | 0 | | | C | C | C | C | | | | | | |
| NC_011770 | 3686004 | T | C | 79 | 0 | 6 | 0 | C | C | | | | | C | C | C | C | | |
| NC_011770 | 3694108 | GC | G | 56 | 0 | 4 | 0 | G | G | | | | | G | G | | | | |
| NC_011770 | 3694131 | AG | A | 60 | 0 | 4 | 0 | | | A | A | A | A | | | | | | |
| NC_011770 | 3694189 | A | G | 61 | 0 | 12 | 0 | G | G | G | G | G | G | G | G | G | G | G | G |
| NC_011770 | 3694232 | G | A | 61 | 0 | 2 | 0 | A | A | | | | | | | | | | |
| NC_011770 | 3694310 | C | A | 61 | 0 | 4 | 0 | | | A | A | A | A | | | | | | |
| NC_011770 | 3695916 | C | T | 69 | 0 | 2 | 0 | T | T | | | | | | | | | | |
| NC_011770 | 3696402 | G | A | 66 | 0 | 2 | 0 | | | | | | | | | | | A | A |
| NC_011770 | 3696533 | A | C | 62 | 0 | 12 | 0 | C | C | C | C | C | C | C | C | C | C | C | C |
| NC_011770 | 3696535 | T | G | 61 | 0 | 4 | 0 | | | | | | | G | G | | | G | G |
| NC_011770 | 3696579 | T | C | 61 | 0 | 4 | 0 | | | | | | | C | C | | | C | C |
| NC_011770 | 3696586 | G | A | 61 | 0 | 2 | 0 | | | | | | | A | A | | | | |
| NC_011770 | 3699261 | GC | G | 74 | 0 | 4 | 0 | | | G | G | G | G | | | | | | |
| NC_011770 | 3699263 | C | G | 42 | 4 | 4 | 0 | | | . | . | . | . | G | G | | | G | G |
| NC_011770 | 3700821 | T | C | 60 | 0 | 1 | 0 | | | | | | | C | | | | | |
| NC_011770 | 3701066 | C | T | 47 | 0 | 6 | 0 | T | T | T | T | T | T | | | | | | |
| NC_011770 | 3701110 | T | C | 44 | 0 | 4 | 0 | | | C | C | C | C | | | | | | |
| NC_011770 | 3701117 | C | G | 44 | 0 | 4 | 0 | | | G | G | G | G | | | | | | |
| NC_011770 | 3701118 | G | C | 45 | 0 | 4 | 0 | | | C | C | C | C | | | | | | |
| NC_011770 | 3701153 | G | A | 46 | 0 | 2 | 0 | | | | | A | A | | | | | | |
| NC_011770 | 3701157 | A | G | 45 | 0 | 2 | 0 | | | | | | | | | G | G | | |
| NC_011770 | 3706625 | G | C | 31 | 0 | 2 | 0 | | | | | | | | | C | C | | |
| NC_011770 | 3707916 | T | G | 59 | 0 | 6 | 0 | | | G | G | G | G | | | | | G | G |
| NC_011770 | 3707963 | G | A | 65 | 0 | 2 | 0 | | | | | | | | | A | A | | |

| | | | | | | | | | | | | | | | | | | | |
|---|---|---|---|---|---|---|---|---|---|---|---|---|---|---|---|---|---|---|---|
| NC_011770 | 3708072 | A | G | 61 | 0 | 10 | 0 | G | G | G | G | G | G | G | G | | | G | G |
| NC_011770 | 3708103 | A | G | 59 | 0 | 12 | 0 | G | G | G | G | G | G | G | G | G | G | G | G |
| NC_011770 | 3710510 | A | G | 87 | 0 | 10 | 0 | | | G | G | G | G | G | G | G | G | G | G |
| NC_011770 | 3712226 | C | T | 54 | 0 | 2 | 0 | | | | | | | | | | | T | T |
| NC_011770 | 3712305 | G | A | 50 | 0 | 8 | 0 | | | A | A | A | A | A | A | A | A | | |
| NC_011770 | 3712351 | T | C | 51 | 0 | 2 | 0 | C | C | | | | | | | | | | |
| NC_011770 | 3712353 | G | A | 52 | 0 | 2 | 0 | | | | | | | | | | | A | A |
| NC_011770 | 3712354 | G | A | 52 | 0 | 2 | 0 | A | A | | | | | | | | | | |
| NC_011770 | 3712362 | G | T | 50 | 0 | 2 | 0 | | | | | | | | | | | T | T |
| NC_011770 | 3712365 | G | A | 50 | 0 | 2 | 0 | A | A | | | | | | | | | | |
| NC_011770 | 3712373 | G | A | 51 | 0 | 6 | 0 | A | A | | | | | A | A | A | A | | |
| NC_011770 | 3712379 | T | G | 48 | 0 | 12 | 0 | G | G | G | G | G | G | G | G | G | G | G | G |
| NC_011770 | 3712380 | G | T | 48 | 0 | 6 | 0 | | | T | T | T | T | T | T | | | | |
| NC_011770 | 3712396 | G | T | 49 | 0 | 4 | 0 | | | | | | | T | T | T | T | | |
| NC_011770 | 3712409 | G | A | 49 | 0 | 2 | 0 | A | A | | | | | | | | | | |
| NC_011770 | 3712424 | C | A | 47 | 0 | 6 | 0 | A | A | A | A | A | A | | | | | | |
| NC_011770 | 3712426 | T | C | 49 | 0 | 6 | 0 | C | C | C | C | C | C | | | | | | |
| NC_011770 | 3712428 | T | C | 48 | 0 | 6 | 0 | C | C | C | C | C | C | | | | | | |
| NC_011770 | 3712445 | T | TA | 49 | 0 | 4 | 0 | | | | | | | | | TA | TA | TA | TA |
| NC_011770 | 3712446 | C | A | 50 | 0 | 2 | 0 | | | | | | | A | A | | | | |
| NC_011770 | 3712453 | C | T | 53 | 0 | 4 | 0 | | | | | | | T | T | T | T | | |
| NC_011770 | 3712466 | A | G | 51 | 0 | 2 | 0 | | | | | | | G | G | | | | |
| NC_011770 | 3712467 | C | T | 50 | 0 | 4 | 0 | | | T | T | T | T | | | | | | |
| NC_011770 | 3712470 | A | G | 50 | 0 | 4 | 0 | | | G | G | G | G | | | | | | |
| NC_011770 | 3712496 | A | G | 50 | 0 | 2 | 0 | | | | | | | | | | | G | G |
| NC_011770 | 3712497 | C | T | 51 | 0 | 4 | 0 | | | | | | | T | T | T | T | | |
| NC_011770 | 3712504 | C | G | 50 | 0 | 2 | 0 | | | | | | | | | | | G | G |
| NC_011770 | 3712515 | T | C | 49 | 0 | 10 | 0 | C | C | C | C | C | C | C | C | C | C | | |
| NC_011770 | 3712516 | G | A | 49 | 0 | 6 | 0 | A | A | A | A | A | A | | | | | | |
| NC_011770 | 3712518 | T | C | 50 | 0 | 2 | 0 | | | | | | | | | | | C | C |
| NC_011770 | 3712592 | C | A | 40 | 0 | 2 | 0 | A | A | | | | | | | | | | |
| NC_011770 | 3712617 | T | C | 40 | 0 | 10 | 0 | C | C | C | C | C | C | C | C | C | C | | |
| NC_011770 | 3712620 | C | A | 40 | 0 | 10 | 0 | A | A | A | A | A | A | A | A | A | A | | |
| NC_011770 | 3712629 | C | T | 40 | 0 | 10 | 0 | T | T | T | T | T | T | G | G | T | T | | |
| NC_011770 | 3712642 | A | G | 38 | 0 | 10 | 0 | G | G | G | G | G | G | G | G | G | G | | |
| NC_011770 | 3714064 | G | A | 56 | 0 | 1 | 0 | | | | | | | A | | | | | |
| NC_011770 | 3714082 | T | C | 58 | 0 | 6 | 0 | | | C | C | C | C | | | | | C | C |
| NC_011770 | 3714132 | T | G | 58 | 0 | 8 | 0 | | | G | G | G | G | | | G | G | G | G |
| NC_011770 | 3714138 | A | G | 59 | 0 | 6 | 0 | | | G | G | G | G | | | | | G | G |
| NC_011770 | 3714218 | A | G | 62 | 0 | 6 | 0 | | | G | G | G | G | | | | | G | G |
| NC_011770 | 3715297 | T | C | 61 | 0 | 12 | 0 | C | C | C | C | C | C | C | C | C | C | C | C |
| NC_011770 | 3715317 | C | A | 60 | 0 | 4 | 0 | | | A | A | A | A | | | | | | |
| NC_011770 | 3715374 | G | A | 61 | 0 | 4 | 0 | | | A | A | A | A | | | | | | |
| NC_011770 | 3715427 | C | T | 60 | 0 | 2 | 0 | | | | | | | | | T | T | | |
| NC_011770 | 3715450 | A | G | 62 | 0 | 4 | 0 | | | G | G | G | G | | | | | | |
| NC_011770 | 3716467 | T | G | 58 | 0 | 2 | 0 | | | | | | | | | | | G | G |
| NC_011770 | 3716495 | G | A | 61 | 0 | 6 | 0 | A | A | | | | | A | A | A | A | | |
| NC_011770 | 3717000 | G | A | 62 | 0 | 4 | 0 | | | A | A | A | A | | | | | | |
| NC_011770 | 3717067 | C | G | 71 | 0 | 2 | 0 | | | | | | | | | | | G | G |
| NC_011770 | 3717215 | C | T | 81 | 0 | 8 | 0 | | | T | T | T | T | T | T | T | T | | |
| NC_011770 | 3718863 | T | C | 62 | 0 | 2 | 0 | | | | | | | | | C | C | | |
| NC_011770 | 3718880 | A | G | 63 | 0 | 6 | 0 | G | G | | | | | G | G | | | G | G |
| NC_011770 | 3719243 | G | A | 64 | 0 | 4 | 0 | A | A | | | | | | | | | A | A |
| NC_011770 | 3719298 | G | A | 63 | 0 | 4 | 0 | | | A | A | | | | | | | A | A |
| NC_011770 | 3721761 | G | A | 44 | 0 | 12 | 0 | A | A | A | A | A | A | A | A | A | A | A | A |
| NC_011770 | 3721770 | A | G | 43 | 0 | 12 | 0 | G | G | G | G | G | G | G | G | G | G | G | G |
| NC_011770 | 3721791 | T | C | 48 | 0 | 12 | 0 | C | C | C | C | C | C | C | C | C | C | C | C |
| NC_011770 | 3721797 | TG | T | 46 | 0 | 12 | 0 | T | T | T | T | T | T | T | T | T | T | T | T |
| NC_011770 | 3722799 | A | C | 47 | 0 | 12 | 0 | C | C | C | C | C | C | C | C | C | C | C | C |
| NC_011770 | 3722801 | A | G | 47 | 0 | 12 | 0 | G | G | G | G | G | G | G | G | G | G | G | G |
| NC_011770 | 3722815 | G | C | 46 | 0 | 12 | 0 | C | C | C | C | C | C | C | C | C | C | C | C |
| NC_011770 | 3722821 | G | A | 46 | 0 | 12 | 0 | A | A | A | A | A | A | A | A | A | A | A | A |

| | | | | | | | | | | | | | | | | | | | |
|---|---|---|---|---|---|---|---|---|---|---|---|---|---|---|---|---|---|---|---|
| NC_011770 | 3722831 | C | T | 45 | 0 | 12 | 0 | T | T | T | T | T | T | T | T | T | T | T | T |
| NC_011770 | 3722836 | G | A | 44 | 0 | 12 | 0 | A | A | A | A | A | A | A | A | A | A | A | A |
| NC_011770 | 3722841 | C | T | 42 | 0 | 12 | 0 | T | T | T | T | T | T | T | T | T | T | T | T |
| NC_011770 | 3722863 | A | T | 12 | 1 | 11 | 0 | T | T | T | T | T | T | T | T | . | T | T | T |
| NC_011770 | 3722866 | C | T | 15 | 1 | 11 | 0 | T | T | T | T | T | T | T | T | . | T | T | T |
| NC_011770 | 3724025 | C | A | 71 | 0 | 8 | 0 | | | A | A | A | A | A | A | A | A | | |
| NC_011770 | 3724045 | G | A | 71 | 0 | 8 | 0 | | | A | A | A | A | A | A | A | A | | |
| NC_011770 | 3724046 | C | G | 71 | 0 | 8 | 0 | | | G | G | G | G | G | G | G | G | | |
| NC_011770 | 3724062 | G | A | 73 | 0 | 8 | 0 | | | A | A | A | A | A | A | A | A | | |
| NC_011770 | 3724064 | G | A | 72 | 0 | 8 | 0 | | | A | A | A | A | A | A | A | A | | |
| NC_011770 | 3724071 | G | C | 76 | 0 | 8 | 0 | | | C | C | C | C | C | C | C | C | | |
| NC_011770 | 3724134 | A | G | 76 | 0 | 8 | 0 | | | G | G | G | G | G | G | G | G | | |
| NC_011770 | 3724140 | G | T | 73 | 0 | 8 | 0 | | | T | T | T | T | T | T | T | T | | |
| NC_011770 | 3724155 | C | G | 74 | 0 | 8 | 0 | | | G | G | G | G | G | G | G | G | | |
| NC_011770 | 3724156 | G | A | 73 | 0 | 8 | 0 | | | A | A | A | A | A | A | A | A | | |
| NC_011770 | 3724161 | C | A | 75 | 0 | 8 | 0 | | | A | A | A | A | A | A | A | A | | |
| NC_011770 | 3724166 | T | C | 75 | 0 | 8 | 0 | | | C | C | C | C | C | C | C | C | | |
| NC_011770 | 3724208 | T | G | 76 | 0 | 2 | 0 | | | | | | | G | G | | | | |
| NC_011770 | 3724239 | G | T | 76 | 0 | 6 | 0 | | | T | T | T | T | | | T | T | | |
| NC_011770 | 3726503 | T | C | 47 | 0 | 2 | 0 | | | | | | | | | C | C | | |
| NC_011770 | 3726511 | A | G | 50 | 0 | 4 | 0 | G | G | | | | | | | G | G | | |
| NC_011770 | 3727978 | T | C | 80 | 0 | 2 | 0 | C | C | | | | | | | | | | |
| NC_011770 | 3729301 | A | ATCGTG | 64 | 0 | 2 | 0 | ATCGTG | ATCGTG | | | | | | | | | | |
| NC_011770 | 3729337 | G | A | 65 | 0 | 2 | 0 | A | A | | | | | | | | | | |
| NC_011770 | 3729400 | C | T | 61 | 0 | 2 | 0 | | | | | | | | | | | T | T |
| NC_011770 | 3729408 | C | T | 63 | 0 | 2 | 0 | T | T | | | | | | | | | | |
| NC_011770 | 3730380 | T | C | 72 | 0 | 12 | 0 | C | C | C | C | C | C | C | C | C | C | C | C |
| NC_011770 | 3732428 | A | G | 84 | 0 | 12 | 0 | G | G | G | G | G | G | G | G | G | G | G | G |
| NC_011770 | 3733983 | C | A | 69 | 0 | 2 | 0 | | | | | | | | | | | A | A |
| NC_011770 | 3734992 | G | A | 64 | 0 | 2 | 0 | | | | | | | | | A | A | | |
| NC_011770 | 3735020 | G | A | 64 | 0 | 2 | 0 | | | | | | | | | A | A | | |
| NC_011770 | 3735060 | C | T | 65 | 0 | 2 | 0 | | | | | | | | | | | T | T |
| NC_011770 | 3735088 | C | G | 64 | 0 | 2 | 0 | G | G | | | | | | | | | | |
| NC_011770 | 3735145 | G | C | 63 | 0 | 2 | 0 | | | | | | | | | | | C | C |
| NC_011770 | 3735160 | A | G | 63 | 0 | 4 | 0 | G | G | | | | | | | | | G | G |
| NC_011770 | 3738127 | GT | G | 81 | 0 | 4 | 0 | | | G | G | G | G | | | | | | |
| NC_011770 | 3738128 | T | G | 81 | 0 | 2 | 0 | | | . | . | . | . | | | G | G | | |
| NC_011770 | 3739439 | G | A | 70 | 0 | 2 | 0 | | | | | | | | | A | A | | |
| NC_011770 | 3739787 | G | A | 61 | 0 | 10 | 0 | A | A | A | A | A | A | A | A | | | A | A |
| NC_011770 | 3739789 | T | C | 62 | 0 | 10 | 0 | C | C | C | C | C | C | C | C | | | C | C |
| NC_011770 | 3739806 | G | A | 63 | 0 | 2 | 0 | | | | | | | | | A | A | | |
| NC_011770 | 3739812 | A | C | 62 | 0 | 10 | 0 | C | C | C | C | C | C | C | C | | | C | C |
| NC_011770 | 3739814 | A | C | 60 | 0 | 4 | 0 | | | C | C | C | C | | | | | | |
| NC_011770 | 3739825 | TC | T | 60 | 0 | 10 | 0 | T | T | T | T | T | T | T | T | | | T | T |
| NC_011770 | 3739829 | G | A | 33 | 3 | 1 | 0 | | | . | . | . | A | | | | | | |
| NC_011770 | 3739862 | T | C | 64 | 0 | 12 | 0 | C | C | C | C | C | C | G | G | C | C | G | G |
| NC_011770 | 3739978 | C | A | 68 | 0 | 12 | 0 | A | A | A | A | A | A | A | A | A | A | A | A |
| NC_011770 | 3741985 | A | G | 14 | 0 | 4 | 0 | | | | | | | G | G | G | G | | |
| NC_011770 | 3743219 | A | G | 3 | 8 | 4 | 0 | . | G | . | . | . | . | G | G | . | G | . | . |
| NC_011770 | 3743221 | A | G | 3 | 8 | 4 | 0 | . | G | . | . | . | . | G | G | . | G | . | . |
| NC_011770 | 3743225 | G | GGGAA | 3 | 8 | 3 | 0 | . | . | . | . | . | . | GGGAA | GGGAA | . | GGGAA | . | . |
| NC_011770 | 3743233 | G | T | 4 | 8 | 1 | 0 | . | | . | . | . | . | | | . | T | . | . |
| NC_011770 | 3743241 | C | T | 4 | 7 | 2 | 0 | | | . | . | . | . | T | T | . | | . | . |
| NC_011770 | 3743248 | C | A | 4 | 8 | 1 | 0 | . | A | . | . | . | . | | | . | | . | . |
| NC_011770 | 3743250 | T | G | 4 | 8 | 1 | 0 | . | G | . | . | . | . | | | . | | . | . |
| NC_011770 | 3743252 | T | G | 3 | 8 | 1 | 0 | | . | . | . | . | . | | | . | G | . | . |
| NC_011770 | 3743254 | T | C | 3 | 8 | 2 | 0 | | . | . | . | . | . | C | C | . | | . | . |
| NC_011770 | 3743255 | C | CA | 4 | 6 | 4 | 0 | CA | CA | . | . | . | . | | | CA | CA | . | . |
| NC_011770 | 3743263 | T | G | 4 | 7 | 1 | 0 | | | . | . | . | . | | | . | G | . | . |
| NC_011770 | 3743270 | A | G | 4 | 7 | 3 | 0 | | | . | . | . | . | G | G | . | G | . | . |
| NC_011770 | 3743274 | A | G | 4 | 7 | 1 | 0 | | | . | . | . | . | | | . | G | . | . |
| NC_011770 | 3743276 | T | C | 4 | 7 | 1 | 0 | | | . | . | . | . | | | . | C | . | . |

| | | | | | | | | | | | | | | | | | | | |
|---|---|---|---|---|---|---|---|---|---|---|---|---|---|---|---|---|---|---|---|
| NC_011770 | 3743284 | T | G | 4 | 7 | 3 | 0 | G | G | . | . | . | . | | | . | G | . | . |
| NC_011770 | 3744266 | C | T | 8 | 10 | 2 | 0 | . | . | . | . | . | . | . | . | . | . | T | T |
| NC_011770 | 3744269 | A | G | 8 | 10 | 2 | 0 | . | . | . | . | . | . | . | . | . | . | G | G |
| NC_011770 | 3744270 | A | C | 8 | 10 | 2 | 0 | . | . | . | . | . | . | . | . | . | . | C | C |
| NC_011770 | 3744272 | C | T | 8 | 10 | 2 | 0 | . | . | . | . | . | . | . | . | . | . | T | T |
| NC_011770 | 3744275 | G | A | 8 | 10 | 2 | 0 | . | . | . | . | . | . | . | . | . | . | A | A |
| NC_011770 | 3744279 | G | C | 8 | 10 | 2 | 0 | . | . | . | . | . | . | . | . | . | . | C | C |
| NC_011770 | 3744280 | A | G | 8 | 10 | 2 | 0 | . | . | . | . | . | . | . | . | . | . | G | G |
| NC_011770 | 3744291 | T | C | 11 | 10 | 2 | 0 | . | . | . | . | . | . | . | . | . | . | C | C |
| NC_011770 | 3744297 | C | T | 11 | 10 | 2 | 0 | . | . | . | . | . | . | . | . | . | . | T | T |
| NC_011770 | 3744326 | A | T | 11 | 10 | 2 | 0 | . | . | . | . | . | . | . | . | . | . | T | T |
| NC_011770 | 3744338 | T | C | 0 | 11 | 1 | 0 | . | . | . | . | . | . | . | . | . | . | C | . |
| NC_011770 | 3744341 | G | A | 0 | 11 | 1 | 0 | . | . | . | . | . | . | . | . | . | . | A | . |
| NC_011770 | 3744416 | C | G | 2 | 10 | 2 | 0 | . | . | . | . | . | . | . | . | . | . | G | G |
| NC_011770 | 3744417 | C | A | 2 | 10 | 2 | 0 | . | . | . | . | . | . | . | . | . | . | A | A |
| NC_011770 | 3744422 | T | A | 2 | 10 | 2 | 0 | . | . | . | . | . | . | . | . | . | . | A | A |
| NC_011770 | 3744437 | G | A | 2 | 10 | 2 | 0 | . | . | . | . | . | . | . | . | . | . | A | A |
| NC_011770 | 3744443 | A | T | 2 | 10 | 2 | 0 | . | . | . | . | . | . | . | . | . | . | T | T |
| NC_011770 | 3744551 | G | T | 0 | 11 | 1 | 0 | . | . | . | . | . | . | . | . | . | . | T | . |
| NC_011770 | 3744560 | C | T | 9 | 10 | 2 | 0 | . | . | . | . | . | . | . | . | . | . | T | T |
| NC_011770 | 3744563 | A | T | 10 | 8 | 2 | 0 | . | . | . | . | . | . | . | . | | | T | T |
| NC_011770 | 3744574 | C | G | 10 | 8 | 2 | 0 | . | . | . | . | . | . | . | . | | | G | G |
| NC_011770 | 3744575 | T | G | 10 | 8 | 4 | 0 | . | . | . | . | . | . | . | . | G | G | C | C |
| NC_011770 | 3744581 | A | G | 10 | 8 | 2 | 0 | . | . | . | . | . | . | . | . | | | G | G |
| NC_011770 | 3744582 | A | C | 10 | 8 | 2 | 0 | . | . | . | . | . | . | . | . | | | C | C |
| NC_011770 | 3744587 | G | C | 10 | 8 | 4 | 0 | . | . | . | . | . | . | . | . | C | C | C | C |
| NC_011770 | 3744602 | A | G | 12 | 8 | 2 | 0 | . | . | . | . | . | . | . | . | | | G | G |
| NC_011770 | 3744617 | C | T | 13 | 8 | 2 | 0 | . | . | . | . | . | . | . | . | | | T | T |
| NC_011770 | 3744618 | CA | C | 13 | 8 | 1 | 0 | . | . | . | . | . | . | . | . | . | C | | |
| NC_011770 | 3744623 | G | A | 12 | 8 | 2 | 0 | . | . | . | . | . | . | . | . | | | A | A |
| NC_011770 | 3744635 | G | A | 13 | 8 | 2 | 0 | . | . | . | . | . | . | . | . | | | A | A |
| NC_011770 | 3744644 | A | G | 13 | 8 | 2 | 0 | . | . | . | . | . | . | . | . | G | G | | |
| NC_011770 | 3744659 | A | G | 12 | 8 | 4 | 0 | . | . | . | . | . | . | . | . | G | G | G | G |
| NC_011770 | 3744662 | A | C | 11 | 8 | 2 | 0 | . | . | . | . | . | . | . | . | C | C | | |
| NC_011770 | 3744669 | C | T | 10 | 8 | 2 | 0 | . | . | . | . | . | . | . | . | | | T | T |
| NC_011770 | 3744671 | A | G | 10 | 8 | 4 | 0 | . | . | . | . | . | . | . | . | G | G | G | G |
| NC_011770 | 3744672 | A | C | 10 | 8 | 2 | 0 | . | . | . | . | . | . | . | . | | | C | C |
| NC_011770 | 3744674 | A | G | 10 | 8 | 2 | 0 | . | . | . | . | . | . | . | . | | | G | G |
| NC_011770 | 3744677 | A | C | 10 | 8 | 2 | 0 | . | . | . | . | . | . | . | . | | | C | C |
| NC_011770 | 3744680 | G | T | 10 | 8 | 4 | 0 | . | . | . | . | . | . | . | . | T | T | T | T |
| NC_011770 | 3744683 | A | C | 11 | 8 | 2 | 0 | . | . | . | . | . | . | . | . | C | C | | |
| NC_011770 | 3744686 | C | T | 11 | 8 | 2 | 0 | . | . | . | . | . | . | . | . | | | T | T |
| NC_011770 | 3744692 | C | T | 11 | 8 | 2 | 0 | . | . | . | . | . | . | . | . | | | T | T |
| NC_011770 | 3744696 | C | T | 11 | 8 | 2 | 0 | . | . | . | . | . | . | . | . | T | T | | |
| NC_011770 | 3744698 | T | C | 11 | 8 | 2 | 0 | . | . | . | . | . | . | . | . | C | C | | |
| NC_011770 | 3744707 | T | G | 11 | 8 | 2 | 0 | . | . | . | . | . | . | . | . | | | G | G |
| NC_011770 | 3744710 | G | A | 11 | 8 | 2 | 0 | . | . | . | . | . | . | . | . | A | A | | |
| NC_011770 | 3744716 | G | C | 10 | 8 | 2 | 0 | . | . | . | . | . | . | . | . | | | C | C |
| NC_011770 | 3744719 | C | G | 10 | 8 | 4 | 0 | . | . | . | . | . | . | . | . | G | G | T | T |
| NC_011770 | 3744725 | C | T | 10 | 8 | 2 | 0 | . | . | . | . | . | . | . | . | | | T | T |
| NC_011770 | 3744728 | A | C | 10 | 8 | 4 | 0 | . | . | . | . | . | . | . | . | C | C | T | T |
| NC_011770 | 3744731 | G | C | 10 | 8 | 2 | 0 | . | . | . | . | . | . | . | . | C | C | | |
| NC_011770 | 3744743 | T | C | 9 | 8 | 2 | 0 | . | . | . | . | . | . | . | . | | | C | C |
| NC_011770 | 3744749 | A | C | 9 | 8 | 2 | 0 | . | . | . | . | . | . | . | . | | | C | C |
| NC_011770 | 3744755 | G | A | 9 | 8 | 2 | 0 | . | . | . | . | . | . | . | . | | | A | A |
| NC_011770 | 3744761 | G | A | 8 | 8 | 2 | 0 | . | . | . | . | . | . | . | . | | | A | A |
| NC_011770 | 3744767 | G | A | 8 | 8 | 2 | 0 | . | . | . | . | . | . | . | . | A | A | | |
| NC_011770 | 3744770 | T | G | 8 | 8 | 4 | 0 | . | . | . | . | . | . | . | . | G | G | C | C |
| NC_011770 | 3744776 | C | T | 1 | 8 | 2 | 0 | . | . | . | . | . | . | . | . | | | T | T |
| NC_011770 | 3751791 | C | A | 26 | 2 | 6 | 0 | A | A | A | A | A | A | | | . | . | | |
| NC_011770 | 3751843 | C | T | 31 | 2 | 4 | 0 | | | T | T | T | T | | | . | . | | |
| NC_011770 | 3751856 | C | G | 30 | 2 | 8 | 0 | G | G | G | G | G | G | G | G | . | . | | |

| | | | | | | | | | | | | | | | | | | | |
|---|---|---|---|---|---|---|---|---|---|---|---|---|---|---|---|---|---|---|---|
| NC_011770 | 3751865 | T | C | 29 | 2 | 4 | 0 | | | C | C | C | C | | | . | . | | |
| NC_011770 | 3751867 | G | A | 29 | 2 | 4 | 0 | | | A | A | A | A | | | . | . | | |
| NC_011770 | 3751868 | T | A | 29 | 2 | 4 | 0 | | | A | A | A | A | | | . | . | | |
| NC_011770 | 3751874 | A | G | 29 | 2 | 4 | 0 | | | G | G | G | G | | | . | . | | |
| NC_011770 | 3751878 | G | C | 29 | 2 | 4 | 0 | | | C | C | C | C | | | . | . | | |
| NC_011770 | 3751912 | G | C | 34 | 0 | 4 | 0 | | | C | C | C | C | | | | | | |
| NC_011770 | 3751914 | A | G | 33 | 0 | 2 | 0 | | | | | | | G | G | | | | |
| NC_011770 | 3751916 | G | A | 33 | 0 | 8 | 0 | A | A | | | | | A | A | A | A | A | A |
| NC_011770 | 3751925 | G | A | 35 | 0 | 3 | 0 | A | . | | | | | A | A | | | | |
| NC_011770 | 3751962 | G | A | 33 | 0 | 2 | 0 | | | | | | | A | A | | | | |
| NC_011770 | 3751976 | G | A | 36 | 0 | 4 | 0 | | | | | | | A | A | A | A | | |
| NC_011770 | 3752000 | C | G | 30 | 0 | 2 | 0 | | | | | | | G | G | | | | |
| NC_011770 | 3752011 | A | C | 35 | 0 | 6 | 0 | | | C | C | C | C | T | T | | | | |
| NC_011770 | 3752013 | A | T | 36 | 0 | 6 | 0 | | | T | T | T | T | T | T | | | | |
| NC_011770 | 3752017 | GA | G | 39 | 0 | 2 | 0 | | | | | | | | | G | G | | |
| NC_011770 | 3752019 | A | G | 34 | 2 | 4 | 0 | | | G | G | G | G | | | . | . | | |
| NC_011770 | 3752024 | GC | G | 39 | 0 | 2 | 0 | | | | | | | G | G | | | | |
| NC_011770 | 3752043 | A | C | 41 | 0 | 2 | 0 | C | C | | | | | | | | | | |
| NC_011770 | 3752044 | G | A | 41 | 0 | 2 | 0 | A | A | | | | | | | | | | |
| NC_011770 | 3752049 | G | A | 40 | 0 | 4 | 0 | | | | | | | A | A | A | A | | |
| NC_011770 | 3752131 | T | A | 45 | 0 | 2 | 0 | A | A | | | | | | | | | | |
| NC_011770 | 3752134 | C | T | 46 | 0 | 2 | 0 | | | | | | | | | | | T | T |
| NC_011770 | 3752161 | G | C | 43 | 0 | 10 | 0 | C | C | C | C | C | C | C | C | | | C | C |
| NC_011770 | 3752162 | T | C | 44 | 0 | 10 | 0 | C | C | C | C | C | C | C | C | | | C | C |
| NC_011770 | 3752607 | T | G | 57 | 0 | 12 | 0 | G | G | G | G | G | G | G | G | G | G | G | G |
| NC_011770 | 3752635 | G | C | 57 | 0 | 2 | 0 | C | C | | | | | | | | | | |
| NC_011770 | 3752636 | C | T | 57 | 0 | 8 | 0 | T | T | T | T | T | T | | | T | T | | |
| NC_011770 | 3752721 | G | A | 54 | 0 | 4 | 0 | | | A | A | A | A | | | | | | |
| NC_011770 | 3752722 | C | T | 56 | 0 | 2 | 0 | | | | | | | T | T | | | | |
| NC_011770 | 3752730 | T | C | 55 | 0 | 2 | 0 | | | | | | | | | | | C | C |
| NC_011770 | 3752733 | G | C | 53 | 0 | 4 | 0 | | | C | C | C | C | | | | | | |
| NC_011770 | 3752734 | G | A | 54 | 0 | 4 | 0 | | | A | A | A | A | | | | | | |
| NC_011770 | 3752735 | C | A | 53 | 0 | 4 | 0 | | | A | A | A | A | | | | | | |
| NC_011770 | 3752744 | A | G | 53 | 0 | 6 | 0 | | | G | G | G | G | | | G | G | | |
| NC_011770 | 3752762 | C | T | 53 | 0 | 6 | 0 | | | T | T | T | T | | | T | T | | |
| NC_011770 | 3752765 | C | T | 52 | 0 | 6 | 0 | T | T | | | | | T | T | | | T | T |
| NC_011770 | 3752766 | C | G | 53 | 0 | 4 | 0 | | | G | G | G | G | | | | | | |
| NC_011770 | 3752784 | A | G | 37 | 3 | 3 | 0 | G | . | | | | | G | . | | | G | . |
| NC_011770 | 3752785 | C | A | 38 | 3 | 3 | 0 | A | . | | | | | A | . | | | A | . |
| NC_011770 | 3752791 | G | A | 36 | 4 | 2 | 0 | A | . | | | | | . | . | | | A | . |
| NC_011770 | 3752793 | A | G | 34 | 4 | 2 | 0 | G | . | | | | | . | . | | | G | . |
| NC_011770 | 3752813 | C | T | 34 | 6 | 2 | 0 | . | . | | | | | . | . | T | T | . | . |
| NC_011770 | 3752910 | G | A | 28 | 6 | 2 | 0 | . | . | | | | | . | . | A | A | . | . |
| NC_011770 | 3756818 | C | T | 35 | 6 | 4 | 0 | . | . | T | T | T | T | . | . | | | . | . |
| NC_011770 | 3758229 | G | A | 60 | 0 | 2 | 0 | | | | | | | A | A | | | | |
| NC_011770 | 3758318 | C | T | 59 | 0 | 6 | 0 | T | T | | | | | T | T | T | T | | |
| NC_011770 | 3758352 | A | G | 58 | 0 | 4 | 0 | G | G | | | | | G | G | | | | |
| NC_011770 | 3758379 | T | C | 58 | 0 | 10 | 0 | C | C | C | C | C | C | C | C | | | C | C |
| NC_011770 | 3758459 | C | T | 54 | 0 | 6 | 0 | | | T | T | T | T | | | T | T | | |
| NC_011770 | 3758513 | A | G | 53 | 0 | 2 | 0 | | | | | | | | | G | G | | |
| NC_011770 | 3758519 | C | A | 54 | 0 | 2 | 0 | | | | | | | | | A | A | | |
| NC_011770 | 3758549 | A | C | 53 | 0 | 2 | 0 | C | C | | | | | | | | | | |
| NC_011770 | 3758550 | T | C | 53 | 0 | 2 | 0 | C | C | | | | | | | | | | |
| NC_011770 | 3758557 | A | C | 49 | 0 | 2 | 0 | C | C | | | | | | | | | | |
| NC_011770 | 3758563 | T | C | 53 | 0 | 2 | 0 | C | C | | | | | | | | | | |
| NC_011770 | 3758885 | T | C | 32 | 0 | 6 | 0 | | | C | C | C | C | | | C | C | | |
| NC_011770 | 3758908 | G | A | 32 | 0 | 6 | 0 | | | A | A | A | A | | | A | A | | |
| NC_011770 | 3758920 | T | G | 33 | 0 | 8 | 0 | G | G | G | G | G | G | | | G | G | | |
| NC_011770 | 3759509 | G | C | 49 | 0 | 4 | 0 | C | C | | | | | | | C | C | | |
| NC_011770 | 3759533 | C | G | 53 | 0 | 2 | 0 | G | G | | | | | | | | | | |
| NC_011770 | 3761886 | G | A | 75 | 0 | 4 | 0 | A | A | | | | | | | | | A | A |
| NC_011770 | 3761894 | T | G | 73 | 0 | 8 | 0 | | | G | G | G | G | G | G | G | G | | |

| | | | | | | | | | | | | | | | | | | | |
|---|---|---|---|---|---|---|---|---|---|---|---|---|---|---|---|---|---|---|---|
| NC_011770 | 3761898 | G | A | 72 | 0 | 8 | 0 | | | A | A | A | A | A | A | A | A | | |
| NC_011770 | 3761965 | G | A | 77 | 0 | 2 | 0 | | | | | | | | | A | A | | |
| NC_011770 | 3763894 | C | T | 73 | 0 | 6 | 0 | T | T | | | | | T | T | T | T | | |
| NC_011770 | 3763924 | G | A | 76 | 0 | 10 | 0 | A | A | A | A | A | A | A | A | A | A | | |
| NC_011770 | 3764047 | A | G | 69 | 0 | 10 | 0 | G | G | G | G | G | G | G | G | G | G | | |
| NC_011770 | 3764049 | G | T | 69 | 0 | 4 | 0 | | | T | T | T | T | | | | | | |
| NC_011770 | 3764173 | A | G | 52 | 0 | 10 | 0 | G | G | G | G | G | G | G | G | G | G | | |
| NC_011770 | 3770982 | C | T | 58 | 0 | 2 | 0 | | | | | | | | | | | T | T |
| NC_011770 | 3770992 | C | A | 56 | 0 | 4 | 0 | | | | | | | A | A | T | T | | |
| NC_011770 | 3770993 | T | A | 56 | 0 | 4 | 0 | | | | | | | A | A | C | C | | |
| NC_011770 | 3771025 | A | G | 60 | 0 | 4 | 0 | | | | | | | | | G | G | G | G |
| NC_011770 | 3771100 | T | A | 63 | 0 | 8 | 0 | | | A | A | A | A | A | A | A | A | | |
| NC_011770 | 3771937 | A | G | 76 | 0 | 4 | 0 | | | G | G | G | G | | | | | | |
| NC_011770 | 3771993 | C | G | 79 | 0 | 4 | 0 | | | G | G | G | G | | | | | | |
| NC_011770 | 3772010 | T | C | 77 | 0 | 2 | 0 | | | | | | | | | | | C | C |
| NC_011770 | 3772029 | C | A | 83 | 0 | 2 | 0 | | | | | | | | | | | A | A |
| NC_011770 | 3772070 | A | G | 80 | 0 | 4 | 0 | | | G | G | G | G | | | | | | |
| NC_011770 | 3772071 | G | A | 80 | 0 | 4 | 0 | | | A | A | A | A | | | | | | |
| NC_011770 | 3774173 | A | G | 74 | 0 | 8 | 0 | | | G | G | G | G | | | G | G | G | G |
| NC_011770 | 3775062 | T | C | 45 | 0 | 2 | 0 | | | | | | | | | | | C | C |
| NC_011770 | 3775099 | C | T | 32 | 0 | 4 | 0 | | | | | | | T | T | | | T | T |
| NC_011770 | 3775141 | C | T | 23 | 0 | 12 | 0 | T | T | T | T | T | T | T | T | T | T | T | T |
| NC_011770 | 3776589 | T | C | 62 | 0 | 12 | 0 | C | C | C | C | C | C | C | C | C | C | C | C |
| NC_011770 | 3778259 | G | A | 82 | 0 | 2 | 0 | A | A | | | | | | | | | | |
| NC_011770 | 3779839 | GC | G | 44 | 0 | 10 | 0 | | | G | G | G | G | G | G | G | G | G | G |
| NC_011770 | 3779903 | T | C | 42 | 0 | 2 | 0 | | | | | | | C | C | | | | |
| NC_011770 | 3779909 | C | G | 43 | 0 | 2 | 0 | | | | | | | G | G | | | | |
| NC_011770 | 3780019 | G | A | 18 | 6 | 1 | 0 | | | . | . | . | . | A | | . | . | | |
| NC_011770 | 3780093 | G | C | 13 | 6 | 2 | 0 | | | . | . | . | . | C | C | . | . | | |
| NC_011770 | 3780137 | A | G | 7 | 6 | 6 | 0 | G | G | . | . | . | . | G | G | . | . | G | G |
| NC_011770 | 3781666 | A | G | 64 | 0 | 12 | 0 | G | G | G | G | G | G | G | G | G | G | G | G |
| NC_011770 | 3782465 | A | G | 87 | 0 | 2 | 0 | | | G | G | | | | | | | | |
| NC_011770 | 3782497 | T | C | 78 | 0 | 2 | 0 | | | | | | | C | C | | | | |
| NC_011770 | 3783985 | C | G | 33 | 0 | 2 | 0 | G | G | | | | | | | | | | |
| NC_011770 | 3783987 | GC | G | 33 | 0 | 2 | 0 | G | G | | | | | | | | | | |
| NC_011770 | 3783990 | C | T | 31 | 2 | 2 | 0 | . | . | | | | | T | T | | | | |
| NC_011770 | 3784020 | G | C | 35 | 0 | 12 | 0 | C | C | C | C | C | C | C | C | C | C | C | C |
| NC_011770 | 3784036 | G | A | 35 | 0 | 4 | 0 | | | | | | | | | A | A | A | A |
| NC_011770 | 3784056 | T | G | 36 | 0 | 4 | 0 | | | G | G | G | G | | | | | | |
| NC_011770 | 3784081 | G | A | 36 | 0 | 2 | 0 | A | A | | | | | | | | | | |
| NC_011770 | 3787952 | C | T | 30 | 0 | 2 | 0 | T | T | | | | | | | | | | |
| NC_011770 | 3791406 | C | T | 55 | 0 | 2 | 0 | T | T | | | | | | | | | | |
| NC_011770 | 3791408 | T | C | 55 | 0 | 2 | 0 | C | C | | | | | | | | | | |
| NC_011770 | 3791424 | C | G | 53 | 0 | 2 | 0 | | | | | | | | | G | G | | |
| NC_011770 | 3791447 | A | G | 54 | 0 | 6 | 0 | | | G | G | G | G | | | | | G | G |
| NC_011770 | 3791457 | C | G | 52 | 0 | 8 | 0 | G | G | G | G | G | G | | | | | G | G |
| NC_011770 | 3791492 | C | T | 52 | 0 | 6 | 0 | T | T | T | T | T | T | | | | | | |
| NC_011770 | 3791510 | C | A | 52 | 0 | 2 | 0 | | | | | | | A | A | | | | |
| NC_011770 | 3791545 | A | G | 45 | 2 | 6 | 0 | G | G | | | | | G | G | G | G | . | . |
| NC_011770 | 3791552 | C | T | 50 | 0 | 8 | 0 | T | T | | | | | T | T | T | T | T | T |
| NC_011770 | 3791568 | G | C | 47 | 0 | 12 | 0 | C | C | C | C | C | C | C | C | C | C | C | C |
| NC_011770 | 3791591 | G | A | 45 | 1 | 2 | 0 | . | | | | | | A | A | | | | |
| NC_011770 | 3791598 | G | A | 44 | 2 | 4 | 0 | . | . | | | | | A | A | | | A | A |
| NC_011770 | 3791601 | G | C | 47 | 0 | 2 | 0 | | | | | | | C | C | | | | |
| NC_011770 | 3791636 | C | T | 42 | 0 | 8 | 0 | T | T | T | T | T | T | | | T | T | | |
| NC_011770 | 3791647 | C | G | 42 | 0 | 2 | 0 | | | | | | | | | G | G | | |
| NC_011770 | 3791654 | C | A | 43 | 0 | 2 | 0 | | | | | | | A | A | | | | |
| NC_011770 | 3791663 | T | C | 33 | 0 | 12 | 0 | C | C | C | C | C | C | C | C | C | C | C | C |
| NC_011770 | 3791668 | C | G | 34 | 0 | 2 | 0 | | | | | | | | | G | G | | |
| NC_011770 | 3791680 | G | A | 34 | 0 | 8 | 0 | A | A | A | A | A | A | | | A | A | | |
| NC_011770 | 3791688 | C | T | 35 | 0 | 4 | 0 | | | | | | | T | T | T | T | | |
| NC_011770 | 290673 | C | T | 54 | 2 | 5 | 0 | | | T | T | T | | . | . | T | T | | |

| | | | | | | | | | | | | | | | | | | | |
|---|---|---|---|---|---|---|---|---|---|---|---|---|---|---|---|---|---|---|---|
| NC_011770 | 515465 | GGCGCGAT | G | 85 | 0 | 1 | 0 | | | | | G | | | | | | | |
| NC_011770 | 3798020 | C | T | 65 | 0 | 4 | 0 | | | T | T | T | T | | | | | | |
| NC_011770 | 3798065 | A | C | 63 | 0 | 12 | 0 | C | C | C | C | C | C | C | C | C | C | C | C |
| NC_011770 | 3798085 | A | G | 65 | 0 | 4 | 0 | | | | | | | | | G | G | G | G |
| NC_011770 | 3798105 | G | A | 63 | 0 | 4 | 0 | | | | | | | | | A | A | A | A |
| NC_011770 | 3798249 | T | A | 53 | 0 | 8 | 0 | A | A | A | A | A | A | | | | | A | A |
| NC_011770 | 3799310 | A | G | 69 | 0 | 2 | 0 | G | G | | | | | | | | | | |
| NC_011770 | 3799342 | G | T | 71 | 0 | 2 | 0 | | | | | | | | | T | T | | |
| NC_011770 | 3799478 | G | A | 73 | 0 | 2 | 0 | | | | | | | | | A | A | | |
| NC_011770 | 3799641 | T | A | 62 | 0 | 4 | 0 | | | A | A | A | A | | | | | | |
| NC_011770 | 3799708 | A | T | 65 | 0 | 2 | 0 | | | | | | | T | T | | | | |
| NC_011770 | 3799935 | A | G | 75 | 0 | 2 | 0 | G | G | | | | | | | | | | |
| NC_011770 | 3807413 | T | C | 70 | 0 | 8 | 0 | | | C | C | C | C | C | C | C | C | | |
| NC_011770 | 3807458 | C | T | 75 | 0 | 4 | 0 | | | T | T | T | T | | | | | | |
| NC_011770 | 3809004 | C | T | 29 | 2 | 2 | 0 | | | | | | | . | . | T | T | | |
| NC_011770 | 3809188 | G | A | 12 | 3 | 1 | 0 | A | . | | | | | . | . | | | | |
| NC_011770 | 3813130 | T | G | 68 | 0 | 2 | 0 | | | | | | | | | | | G | G |
| NC_011770 | 3814140 | A | G | 41 | 0 | 4 | 0 | | | G | G | G | G | | | | | | |
| NC_011770 | 3814146 | G | A | 40 | 0 | 4 | 0 | | | A | A | A | A | | | | | | |
| NC_011770 | 3814480 | G | A | 65 | 0 | 2 | 0 | | | | | | | A | A | | | | |
| NC_011770 | 3814606 | T | A | 68 | 0 | 4 | 0 | | | A | A | A | A | | | | | | |
| NC_011770 | 3814609 | C | T | 68 | 0 | 4 | 0 | | | T | T | T | T | | | | | | |
| NC_011770 | 3814618 | G | A | 69 | 0 | 4 | 0 | | | A | A | A | A | | | | | | |
| NC_011770 | 3814621 | T | A | 67 | 0 | 4 | 0 | | | A | A | A | A | | | | | | |
| NC_011770 | 3814658 | C | T | 72 | 0 | 6 | 0 | | | | | | | T | T | T | T | T | T |
| NC_011770 | 3814667 | A | G | 71 | 0 | 12 | 0 | G | G | G | G | G | G | G | G | G | G | G | G |
| NC_011770 | 3814711 | A | G | 77 | 0 | 12 | 0 | G | G | G | G | G | G | G | G | G | G | G | G |
| NC_011770 | 3814752 | C | T | 78 | 0 | 2 | 0 | | | | | | | T | T | | | | |
| NC_011770 | 3817525 | A | T | 49 | 0 | 2 | 0 | | | | | | | | | | | T | T |
| NC_011770 | 3817545 | T | TCC | 50 | 0 | 1 | 0 | | | | | | | TCC | | | | | |
| NC_011770 | 3818187 | G | A | 77 | 0 | 6 | 0 | | | A | A | A | A | | | A | A | | |
| NC_011770 | 3818855 | G | A | 76 | 0 | 2 | 0 | | | | | | | | | A | A | | |
| NC_011770 | 513755 | C | A | 37 | 0 | 1 | 0 | | | | | | | | | A | | | |
| NC_011770 | 539592 | AG | A | 76 | 0 | 2 | 0 | | | | | A | | AGG | | | | | |
| NC_011770 | 778258 | G | A | 55 | 0 | 1 | 0 | | | | | A | | | | | | | |
| NC_011770 | 778265 | C | CA | 53 | 0 | 1 | 0 | | | | | CA | | | | | | | |
| NC_011770 | 3831752 | A | G | 40 | 0 | 4 | 0 | | | | | | | | | G | G | G | G |
| NC_011770 | 3831816 | T | C | 38 | 0 | 12 | 0 | C | C | C | C | C | C | C | C | C | C | C | C |
| NC_011770 | 3831900 | C | A | 44 | 0 | 12 | 0 | A | A | A | A | A | A | A | A | A | A | A | A |
| NC_011770 | 3831924 | T | G | 51 | 0 | 12 | 0 | G | G | G | G | G | G | G | G | G | G | G | G |
| NC_011770 | 3831946 | TA | T | 52 | 0 | 4 | 0 | | | | | | | | | T | T | T | T |
| NC_011770 | 3831973 | C | T | 56 | 0 | 2 | 0 | T | T | | | | | | | | | | |
| NC_011770 | 3832019 | C | A | 57 | 0 | 4 | 0 | A | A | | | | | A | A | | | | |
| NC_011770 | 3832036 | G | A | 60 | 0 | 4 | 0 | | | A | A | A | A | | | | | | |
| NC_011770 | 3832076 | T | C | 59 | 0 | 4 | 0 | | | | | | | | | C | C | C | C |
| NC_011770 | 3832099 | C | T | 59 | 0 | 4 | 0 | | | | | | | | | T | T | T | T |
| NC_011770 | 3832114 | T | C | 59 | 0 | 4 | 0 | | | | | | | | | C | C | C | C |
| NC_011770 | 3832136 | G | A | 56 | 0 | 2 | 0 | | | | | | | | | | | A | A |
| NC_011770 | 3832139 | A | C | 59 | 0 | 2 | 0 | | | | | | | | | | | C | C |
| NC_011770 | 3835582 | T | G | 76 | 0 | 4 | 0 | | | G | G | G | G | | | | | | |
| NC_011770 | 3835662 | T | TGC | 77 | 0 | 4 | 0 | | | TGC | TGC | TGC | TGC | | | | | | |
| NC_011770 | 3836096 | A | G | 60 | 0 | 4 | 0 | | | G | G | G | G | | | | | | |
| NC_011770 | 3836115 | G | A | 59 | 0 | 2 | 0 | | | | | | | | | | | A | A |
| NC_011770 | 3836121 | A | G | 56 | 0 | 4 | 0 | | | G | G | G | G | | | | | | |
| NC_011770 | 3836122 | C | G | 57 | 0 | 2 | 0 | | | | | | | G | G | | | | |
| NC_011770 | 3836125 | G | A | 55 | 0 | 4 | 0 | | | A | A | A | A | | | | | | |
| NC_011770 | 3836129 | G | C | 58 | 0 | 4 | 0 | | | C | C | C | C | | | | | | |
| NC_011770 | 3836137 | A | G | 53 | 0 | 4 | 0 | | | G | G | G | G | | | | | | |
| NC_011770 | 3836139 | G | A | 53 | 0 | 4 | 0 | | | A | A | A | A | | | | | | |
| NC_011770 | 3836140 | C | A | 53 | 0 | 4 | 0 | | | A | A | A | A | | | | | | |
| NC_011770 | 3836141 | T | C | 52 | 0 | 4 | 0 | | | C | C | C | C | | | | | | |
| NC_011770 | 3836146 | T | C | 55 | 0 | 4 | 0 | | | C | C | C | C | | | | | | |

| | | | | | | | | | | | | | | | | | | | |
|---|---|---|---|---|---|---|---|---|---|---|---|---|---|---|---|---|---|---|---|
| NC_011770 | 3836150 | T | C | 54 | 0 | 4 | 0 | | | C | C | C | C | | | | | | |
| NC_011770 | 3836170 | G | C | 57 | 0 | 4 | 0 | | | C | C | C | C | | | | | | |
| NC_011770 | 3836202 | C | T | 58 | 0 | 4 | 0 | | | T | T | T | T | | | | | | |
| NC_011770 | 3836210 | A | T | 59 | 0 | 4 | 0 | | | T | T | T | T | | | | | | |
| NC_011770 | 3836212 | T | C | 56 | 0 | 4 | 0 | | | C | C | C | C | | | | | | |
| NC_011770 | 3836218 | C | T | 58 | 0 | 4 | 0 | | | T | T | T | T | | | | | | |
| NC_011770 | 3836715 | T | C | 73 | 0 | 4 | 0 | | | C | C | C | C | | | | | | |
| NC_011770 | 3836742 | A | T | 72 | 0 | 4 | 0 | | | T | T | T | T | | | | | | |
| NC_011770 | 3836759 | G | C | 74 | 0 | 4 | 0 | | | C | C | C | C | | | | | | |
| NC_011770 | 3836765 | C | T | 76 | 0 | 2 | 0 | | | | | | | | | T | T | | |
| NC_011770 | 3840029 | C | T | 69 | 0 | 12 | 0 | T | T | T | T | T | T | T | T | T | T | T | T |
| NC_011770 | 3844099 | A | G | 73 | 0 | 10 | 0 | G | G | G | G | G | G | | | G | G | G | G |
| NC_011770 | 3844100 | T | C | 72 | 0 | 10 | 0 | C | C | C | C | C | C | | | C | C | C | C |
| NC_011770 | 3844897 | GC | G | 36 | 0 | 8 | 0 | | | G | G | G | G | | | G | G | G | G |
| NC_011770 | 3850146 | G | A | 79 | 0 | 4 | 0 | | | A | A | A | A | | | | | | |
| NC_011770 | 3851170 | G | A | 59 | 0 | 2 | 0 | | | | | | | A | A | | | | |
| NC_011770 | 3851175 | G | A | 60 | 0 | 2 | 0 | | | | | | | | | | | A | A |
| NC_011770 | 3851237 | C | T | 58 | 0 | 1 | 0 | | | | | | | T | | | | | |
| NC_011770 | 3851247 | CT | C | 58 | 0 | 2 | 0 | | | | | | | | | | | C | C |
| NC_011770 | 3851255 | T | C | 57 | 0 | 2 | 0 | | | | | | | | | | | C | C |
| NC_011770 | 3851309 | A | C | 66 | 0 | 4 | 0 | | | C | C | C | C | | | | | | |
| NC_011770 | 3851311 | A | G | 65 | 0 | 2 | 0 | | | | | | | | | G | G | | |
| NC_011770 | 3851329 | G | T | 64 | 0 | 2 | 0 | | | | | | | | | | | T | T |
| NC_011770 | 3851419 | A | G | 74 | 0 | 2 | 0 | G | G | | | | | | | | | | |
| NC_011770 | 3851475 | A | G | 71 | 0 | 2 | 0 | | | | | | | | | G | G | | |
| NC_011770 | 3853217 | A | G | 67 | 0 | 2 | 0 | | | | | | | | | | | G | G |
| NC_011770 | 3855350 | C | G | 75 | 0 | 4 | 0 | | | G | G | G | G | | | | | | |
| NC_011770 | 3855359 | G | A | 78 | 0 | 6 | 0 | | | A | A | A | A | A | A | | | | |
| NC_011770 | 3855360 | T | A | 77 | 0 | 6 | 0 | | | A | A | A | A | A | A | | | | |
| NC_011770 | 3855384 | G | A | 80 | 0 | 2 | 0 | | | | | | | A | A | | | | |
| NC_011770 | 3855399 | T | C | 77 | 0 | 6 | 0 | | | C | C | C | C | C | C | | | | |
| NC_011770 | 3855400 | A | T | 78 | 0 | 6 | 0 | | | T | T | T | T | T | T | | | | |
| NC_011770 | 3855513 | C | G | 81 | 0 | 2 | 0 | | | | | | | G | G | | | | |
| NC_011770 | 3855833 | C | T | 90 | 0 | 6 | 0 | T | T | | | | | | | T | T | T | T |
| NC_011770 | 3855856 | C | T | 90 | 0 | 2 | 0 | | | | | | | T | T | | | | |
| NC_011770 | 3855863 | CT | C | 90 | 0 | 2 | 0 | | | | | | | C | C | | | | |
| NC_011770 | 3857114 | G | A | 76 | 0 | 12 | 0 | A | A | A | A | A | A | A | A | A | A | A | A |
| NC_011770 | 3857148 | A | G | 77 | 0 | 12 | 0 | G | G | G | G | G | G | G | G | G | G | G | G |
| NC_011770 | 3859820 | C | T | 79 | 0 | 8 | 0 | | | T | T | T | T | T | T | T | T | | |
| NC_011770 | 3862879 | A | G | 63 | 0 | 8 | 0 | | | G | G | G | G | | | G | G | G | G |
| NC_011770 | 3862898 | G | A | 59 | 0 | 2 | 0 | | | | | | | | | A | A | | |
| NC_011770 | 3862917 | A | G | 59 | 0 | 4 | 0 | | | | | | | | | G | G | G | G |
| NC_011770 | 3862918 | C | T | 61 | 0 | 4 | 0 | | | T | T | T | T | | | | | | |
| NC_011770 | 3862951 | T | A | 57 | 0 | 2 | 0 | A | A | | | | | | | | | | |
| NC_011770 | 3863060 | G | C | 61 | 0 | 2 | 0 | | | | | | | | | | | C | C |
| NC_011770 | 3863110 | C | T | 62 | 0 | 12 | 0 | T | T | T | T | T | T | T | T | T | T | T | T |
| NC_011770 | 3864260 | T | TC | 50 | 0 | 12 | 0 | TC | TC | TC | TC | TC | TC | TC | TC | TC | TC | TC | TC |
| NC_011770 | 3864286 | C | G | 50 | 0 | 2 | 0 | | | | | | | G | G | | | | |
| NC_011770 | 3864290 | C | T | 49 | 0 | 12 | 0 | T | T | T | T | T | T | T | T | T | T | T | T |
| NC_011770 | 3864303 | C | T | 51 | 0 | 2 | 0 | | | | | | | | | T | T | | |
| NC_011770 | 3864339 | G | A | 53 | 0 | 10 | 0 | A | A | A | A | A | A | A | A | A | A | | |
| NC_011770 | 3864362 | G | A | 51 | 0 | 12 | 0 | A | A | A | A | A | A | A | A | A | A | A | A |
| NC_011770 | 3864363 | C | T | 52 | 0 | 12 | 0 | T | T | T | T | T | T | T | T | T | T | T | T |
| NC_011770 | 3864378 | G | A | 53 | 0 | 12 | 0 | A | A | A | A | A | A | A | A | A | A | A | A |
| NC_011770 | 3864388 | T | C | 54 | 0 | 12 | 0 | C | C | C | C | C | C | C | C | C | C | C | C |
| NC_011770 | 3864415 | A | G | 52 | 0 | 12 | 0 | G | G | G | G | G | G | G | G | G | G | G | G |
| NC_011770 | 3864416 | G | A | 52 | 0 | 12 | 0 | A | A | A | A | A | A | A | A | A | A | A | A |
| NC_011770 | 3864427 | T | C | 52 | 0 | 12 | 0 | C | C | C | C | C | C | C | C | C | C | C | C |
| NC_011770 | 3868585 | T | C | 67 | 0 | 4 | 0 | | | | | | | | | C | C | C | C |
| NC_011770 | 3868593 | C | A | 70 | 0 | 2 | 0 | | | | | | | A | A | | | | |
| NC_011770 | 3871058 | T | C | 83 | 0 | 4 | 0 | | | C | C | C | C | | | | | | |
| NC_011770 | 3873416 | C | A | 38 | 1 | 2 | 0 | . | | | | | | A | A | | | | |

| | | | | | | | | | | | | | | | | | | | |
|---|---|---|---|---|---|---|---|---|---|---|---|---|---|---|---|---|---|---|---|
| NC_011770 | 3873420 | C | G | 38 | 1 | 2 | 0 | . | | | | | | G | G | | | | |
| NC_011770 | 3873430 | G | A | 40 | 0 | 6 | 0 | A | A | | | | | A | A | | | A | A |
| NC_011770 | 3873459 | A | G | 39 | 0 | 2 | 0 | | | | | | | G | G | | | | |
| NC_011770 | 3873469 | T | C | 40 | 0 | 4 | 0 | | | C | C | C | C | | | | | | |
| NC_011770 | 3874076 | T | A | 49 | 0 | 2 | 0 | A | A | | | | | | | | | | |
| NC_011770 | 3874095 | C | T | 50 | 0 | 2 | 0 | | | | | | | T | T | | | | |
| NC_011770 | 3874173 | C | T | 64 | 0 | 2 | 0 | T | T | | | | | | | | | | |
| NC_011770 | 3875534 | C | A | 62 | 0 | 8 | 0 | A | A | | | | | A | A | A | A | A | A |
| NC_011770 | 3877745 | A | G | 76 | 0 | 2 | 0 | | | | | | | | | | | G | G |
| NC_011770 | 3878974 | C | T | 71 | 0 | 2 | 0 | | | | | | | | | | | T | T |
| NC_011770 | 3879102 | C | T | 63 | 0 | 8 | 0 | T | T | T | T | T | T | T | T | | | | |
| NC_011770 | 3881171 | G | A | 67 | 0 | 2 | 0 | A | A | | | | | | | | | | |
| NC_011770 | 3881172 | GGACA | G | 68 | 0 | 2 | 0 | | | | | | | | | G | G | | |
| NC_011770 | 3882106 | A | G | 93 | 0 | 2 | 0 | | | | | | | G | G | | | | |
| NC_011770 | 3887069 | T | C | 84 | 0 | 4 | 0 | | | C | C | C | C | | | | | | |
| NC_011770 | 3888117 | C | A | 72 | 0 | 2 | 0 | | | | | | | | | A | A | | |
| NC_011770 | 3893402 | G | T | 68 | 0 | 2 | 0 | | | | | | | | | T | T | | |
| NC_011770 | 3893412 | G | A | 68 | 0 | 2 | 0 | | | | | | | | | A | A | | |
| NC_011770 | 3894645 | C | T | 74 | 0 | 8 | 0 | T | T | T | T | T | T | | | | | T | T |
| NC_011770 | 3894656 | C | T | 77 | 0 | 2 | 0 | | | | | | | | | | | T | T |
| NC_011770 | 3898526 | A | G | 71 | 0 | 12 | 0 | G | G | G | G | G | G | G | G | G | G | G | G |
| NC_011770 | 3898533 | A | G | 74 | 0 | 2 | 0 | G | G | | | | | | | | | | |
| NC_011770 | 3898540 | C | A | 69 | 0 | 10 | 0 | A | A | A | A | A | A | | | A | A | A | A |
| NC_011770 | 3898620 | G | A | 72 | 0 | 1 | 0 | | | | | | | A | | | | | |
| NC_011770 | 3906054 | G | A | 61 | 0 | 2 | 0 | | | | | | | | | A | A | | |
| NC_011770 | 3906057 | C | A | 58 | 0 | 10 | 0 | A | A | A | A | A | A | A | A | A | A | | |
| NC_011770 | 3906892 | CA | C | 32 | 0 | 4 | 0 | | | | | | | C | C | | | C | C |
| NC_011770 | 3906905 | A | C | 32 | 0 | 4 | 0 | | | | | | | C | C | | | C | C |
| NC_011770 | 3909033 | A | T | 66 | 0 | 8 | 0 | T | T | | | | | T | T | T | T | T | T |
| NC_011770 | 3909041 | C | A | 64 | 0 | 8 | 0 | A | A | | | | | A | A | A | A | A | A |
| NC_011770 | 3914062 | G | A | 90 | 0 | 4 | 0 | | | A | A | A | A | | | | | | |
| NC_011770 | 3917671 | A | T | 61 | 0 | 4 | 0 | | | T | T | T | T | | | | | | |
| NC_011770 | 3917693 | T | C | 60 | 0 | 8 | 0 | | | C | C | C | C | C | C | C | C | | |
| NC_011770 | 3917743 | A | G | 62 | 0 | 8 | 0 | | | G | G | G | G | G | G | G | G | | |
| NC_011770 | 3919584 | G | A | 74 | 0 | 2 | 0 | A | A | | | | | | | | | | |
| NC_011770 | 3919665 | C | T | 73 | 0 | 2 | 0 | T | T | | | | | | | | | | |
| NC_011770 | 3919667 | A | G | 70 | 0 | 2 | 0 | G | G | | | | | | | | | | |
| NC_011770 | 3919671 | C | T | 74 | 0 | 2 | 0 | T | T | | | | | | | | | | |
| NC_011770 | 3919697 | A | G | 74 | 0 | 4 | 0 | G | G | | | | | | | G | G | | |
| NC_011770 | 3919827 | T | C | 73 | 0 | 2 | 0 | C | C | | | | | | | | | | |
| NC_011770 | 3919831 | A | C | 75 | 0 | 2 | 0 | C | C | | | | | | | | | | |
| NC_011770 | 3920132 | A | G | 88 | 0 | 4 | 0 | G | G | | | | | | | G | G | | |
| NC_011770 | 3920154 | C | G | 85 | 0 | 2 | 0 | G | G | | | | | | | | | | |
| NC_011770 | 3920156 | C | T | 89 | 0 | 2 | 0 | T | T | | | | | | | | | | |
| NC_011770 | 3920180 | CG | C | 88 | 0 | 2 | 0 | C | C | | | | | | | | | | |
| NC_011770 | 3920193 | G | A | 88 | 0 | 2 | 0 | | | | | | | | | | | A | A |
| NC_011770 | 3920195 | A | G | 85 | 0 | 2 | 0 | G | G | | | | | | | | | | |
| NC_011770 | 3920202 | A | G | 89 | 0 | 2 | 0 | G | G | | | | | | | | | | |
| NC_011770 | 3920230 | T | A | 85 | 0 | 2 | 0 | A | A | | | | | | | | | | |
| NC_011770 | 820209 | C | G | 37 | 0 | 1 | 0 | | | | | | | | | G | | | |
| NC_011770 | 3920835 | T | A | 68 | 0 | 2 | 0 | | | | | | | | | | | A | A |
| NC_011770 | 3920957 | G | A | 53 | 0 | 2 | 0 | A | A | | | | | | | | | | |
| NC_011770 | 3921922 | T | A | 39 | 0 | 4 | 0 | | | A | A | A | A | | | | | | |
| NC_011770 | 3921923 | T | A | 40 | 0 | 4 | 0 | | | A | A | A | A | | | | | | |
| NC_011770 | 3923339 | T | C | 83 | 0 | 6 | 0 | | | C | C | C | C | | | | | C | C |
| NC_011770 | 3926393 | AGC | A | 37 | 0 | 2 | 0 | | | | | | | | | | | A | A |
| NC_011770 | 3926399 | G | A | 36 | 0 | 2 | 0 | | | | | | | | | | | A | A |
| NC_011770 | 3926424 | G | A | 36 | 0 | 2 | 0 | A | A | | | | | | | | | | |
| NC_011770 | 3926441 | C | T | 36 | 0 | 2 | 0 | | | | | | | | | | | T | T |
| NC_011770 | 3926469 | TC | T | 34 | 0 | 2 | 0 | | | | | | | | | | | T | T |
| NC_011770 | 3927868 | T | C | 63 | 0 | 2 | 0 | | | | | | | C | C | | | | |
| NC_011770 | 3927874 | G | A | 63 | 0 | 4 | 0 | A | A | | | | | | | | | A | A |

| | | | | | | | | | | | | | | | | | | | |
|---|---|---|---|---|---|---|---|---|---|---|---|---|---|---|---|---|---|---|---|
| NC_011770 | 3928522 | C | T | 81 | 0 | 10 | 0 | T | T | T | T | T | T | T | T | | | T | T |
| NC_011770 | 3928570 | T | C | 77 | 0 | 10 | 0 | C | C | C | C | C | C | C | C | | | C | C |
| NC_011770 | 3929547 | GA | G | 56 | 0 | 2 | 0 | | | | | | | | | | | G | G |
| NC_011770 | 3929548 | A | G | 47 | 2 | 4 | 0 | | | G | G | G | G | | | | | . | . |
| NC_011770 | 3929552 | A | C | 46 | 1 | 2 | 0 | | | | | | | C | C | | | . | |
| NC_011770 | 3930114 | C | T | 53 | 0 | 2 | 0 | | | | | | | | | T | T | | |
| NC_011770 | 3933172 | T | G | 46 | 0 | 2 | 0 | G | G | | | | | | | | | | |
| NC_011770 | 3933193 | G | A | 47 | 0 | 2 | 0 | A | A | | | | | | | | | | |
| NC_011770 | 3933204 | T | C | 46 | 0 | 8 | 0 | C | C | C | C | C | C | C | C | | | | |
| NC_011770 | 3933215 | C | T | 51 | 0 | 4 | 0 | | | T | T | T | T | | | | | | |
| NC_011770 | 3935966 | A | G | 68 | 0 | 12 | 0 | G | G | G | G | G | G | G | G | G | G | G | G |
| NC_011770 | 3936105 | T | A | 73 | 0 | 4 | 0 | | | A | A | A | A | | | | | | |
| NC_011770 | 3936165 | C | T | 74 | 0 | 4 | 0 | | | T | T | T | T | | | | | | |
| NC_011770 | 3937541 | C | T | 73 | 0 | 4 | 0 | | | | | | | T | T | T | T | | |
| NC_011770 | 3937583 | C | T | 73 | 0 | 4 | 0 | | | | | | | T | T | T | T | | |
| NC_011770 | 3939190 | A | T | 49 | 0 | 8 | 0 | | | T | T | T | T | T | T | | | T | T |
| NC_011770 | 3939449 | A | C | 68 | 0 | 4 | 0 | | | | | | | | | C | C | C | C |
| NC_011770 | 3941915 | C | T | 58 | 0 | 6 | 0 | | | T | T | T | T | | | | | T | T |
| NC_011770 | 3941923 | G | C | 60 | 0 | 2 | 0 | | | | | | | C | C | | | | |
| NC_011770 | 3941943 | T | C | 58 | 0 | 12 | 0 | C | C | C | C | C | C | C | C | C | C | C | C |
| NC_011770 | 3945826 | A | G | 44 | 1 | 4 | 0 | G | G | . | | | | G | G | | | | |
| NC_011770 | 3945841 | T | C | 53 | 0 | 2 | 0 | C | C | | | | | | | | | | |
| NC_011770 | 3948766 | G | A | 76 | 0 | 4 | 0 | | | A | A | A | A | | | | | | |
| NC_011770 | 3948787 | A | C | 64 | 0 | 4 | 0 | C | C | | | | | | | C | C | | |
| NC_011770 | 3948798 | T | C | 65 | 0 | 10 | 0 | C | C | C | C | C | C | C | C | | | C | C |
| NC_011770 | 3949886 | C | T | 68 | 0 | 2 | 0 | | | | | | | T | T | | | | |
| NC_011770 | 3949912 | G | A | 70 | 0 | 2 | 0 | | | | | | | A | A | | | | |
| NC_011770 | 3949920 | A | G | 66 | 0 | 2 | 0 | | | | | | | G | G | | | | |
| NC_011770 | 3949923 | C | T | 67 | 0 | 2 | 0 | | | | | | | T | T | | | | |
| NC_011770 | 3950484 | A | G | 65 | 0 | 12 | 0 | G | G | G | G | G | G | G | G | G | G | G | G |
| NC_011770 | 3950531 | A | T | 65 | 0 | 2 | 0 | | | | | | | | | | | T | T |
| NC_011770 | 3955072 | T | C | 46 | 0 | 12 | 0 | C | C | C | C | C | C | C | C | C | C | C | C |
| NC_011770 | 3955104 | T | A | 49 | 0 | 6 | 0 | A | A | | | | | | | A | A | A | A |
| NC_011770 | 3955112 | A | G | 50 | 0 | 4 | 0 | G | G | | | | | | | G | G | | |
| NC_011770 | 3958395 | G | A | 65 | 0 | 2 | 0 | | | | | | | | | A | A | | |
| NC_011770 | 3958396 | T | C | 66 | 0 | 2 | 0 | C | C | | | | | | | | | | |
| NC_011770 | 3958425 | C | T | 62 | 0 | 12 | 0 | T | T | T | T | T | T | T | T | T | T | T | T |
| NC_011770 | 3963720 | T | C | 72 | 0 | 4 | 0 | | | | | | | C | C | | | C | C |
| NC_011770 | 3971547 | G | A | 46 | 0 | 4 | 0 | | | A | A | A | A | | | | | | |
| NC_011770 | 3972428 | A | G | 69 | 0 | 8 | 0 | G | G | G | G | G | G | G | G | | | | |
| NC_011770 | 3973958 | C | T | 74 | 0 | 2 | 0 | | | | | | | T | T | | | | |
| NC_011770 | 3974496 | A | G | 81 | 0 | 2 | 0 | | | | | | | | | | | G | G |
| NC_011770 | 3975958 | T | G | 62 | 0 | 10 | 0 | G | G | G | G | G | G | | | G | G | G | G |
| NC_011770 | 3975999 | G | A | 60 | 0 | 4 | 0 | | | A | A | A | A | | | | | | |
| NC_011770 | 3976005 | A | C | 59 | 0 | 2 | 0 | | | | | | | C | C | | | | |
| NC_011770 | 3976008 | C | G | 59 | 0 | 2 | 0 | | | | | | | G | G | | | | |
| NC_011770 | 3976060 | A | G | 54 | 0 | 2 | 0 | | | | | | | G | G | | | | |
| NC_011770 | 3977395 | G | A | 37 | 0 | 6 | 0 | A | A | | | | | A | A | | | A | A |
| NC_011770 | 3977844 | G | C | 67 | 0 | 2 | 0 | | | | | | | C | C | | | | |
| NC_011770 | 3977847 | A | G | 68 | 0 | 2 | 0 | | | | | | | G | G | | | | |
| NC_011770 | 3977867 | C | T | 69 | 0 | 2 | 0 | | | | | | | | | | | T | T |
| NC_011770 | 3977905 | C | T | 71 | 0 | 2 | 0 | | | | | | | T | T | | | | |
| NC_011770 | 3977927 | C | G | 70 | 0 | 2 | 0 | | | | | | | G | G | | | | |
| NC_011770 | 3977946 | A | G | 70 | 0 | 2 | 0 | | | | | | | G | G | | | | |
| NC_011770 | 3977958 | C | G | 72 | 0 | 2 | 0 | | | | | | | G | G | | | | |
| NC_011770 | 3977959 | T | C | 71 | 0 | 2 | 0 | | | | | | | C | C | | | | |
| NC_011770 | 3977966 | GC | G | 70 | 0 | 2 | 0 | | | | | | | G | G | | | | |
| NC_011770 | 3977974 | A | G | 71 | 0 | 2 | 0 | | | | | | | | | | | G | G |
| NC_011770 | 3977978 | A | G | 71 | 0 | 2 | 0 | | | | | | | G | G | | | | |
| NC_011770 | 3978025 | A | G | 70 | 0 | 2 | 0 | | | | | | | G | G | | | | |
| NC_011770 | 3978028 | C | G | 71 | 0 | 2 | 0 | | | | | | | G | G | | | | |
| NC_011770 | 3978769 | T | G | 53 | 0 | 12 | 0 | G | G | G | G | G | G | G | G | G | G | G | G |

| | | | | | | | | | | | | | | | | | | | |
|---|---|---|---|---|---|---|---|---|---|---|---|---|---|---|---|---|---|---|---|
| NC_011770 | 3978770 | C | G | 52 | 0 | 4 | 0 | | | G | G | G | G | | | | | | |
| NC_011770 | 3978788 | A | G | 56 | 0 | 10 | 0 | G | G | G | G | G | G | | | G | G | G | G |
| NC_011770 | 3978841 | A | C | 53 | 0 | 10 | 0 | C | C | C | C | C | C | | | C | C | C | C |
| NC_011770 | 3978842 | G | T | 53 | 0 | 10 | 0 | T | T | T | T | T | T | | | T | T | T | T |
| NC_011770 | 3978870 | A | G | 54 | 0 | 10 | 0 | G | G | G | G | G | G | | | G | G | G | G |
| NC_011770 | 3978879 | G | A | 57 | 0 | 2 | 0 | A | A | | | | | | | | | | |
| NC_011770 | 3978880 | A | G | 57 | 0 | 10 | 0 | G | G | G | G | G | G | | | G | G | G | G |
| NC_011770 | 3978900 | G | C | 56 | 0 | 12 | 0 | C | C | C | C | C | C | C | C | C | C | C | C |
| NC_011770 | 3979166 | T | C | 47 | 0 | 10 | 0 | C | C | C | C | C | C | | | C | C | C | C |
| NC_011770 | 3979775 | G | A | 75 | 0 | 2 | 0 | A | A | | | | | | | | | | |
| NC_011770 | 3979776 | A | G | 73 | 0 | 2 | 0 | G | G | | | | | | | | | | |
| NC_011770 | 3979793 | G | C | 75 | 0 | 2 | 0 | | | | | | | | | | | C | C |
| NC_011770 | 3979875 | T | A | 82 | 0 | 2 | 0 | | | | | | | A | A | | | | |
| NC_011770 | 3980395 | G | A | 59 | 0 | 2 | 0 | A | A | | | | | | | | | | |
| NC_011770 | 3980468 | G | T | 59 | 0 | 2 | 0 | | | | | | | | | T | T | | |
| NC_011770 | 3980470 | C | T | 58 | 0 | 8 | 0 | T | T | A | A | A | A | | | A | A | | |
| NC_011770 | 3981077 | A | G | 62 | 0 | 6 | 0 | G | G | | | | | G | G | | | G | G |
| NC_011770 | 3984441 | A | G | 60 | 0 | 2 | 0 | | | | | | | | | G | G | | |
| NC_011770 | 3984467 | G | GC | 57 | 0 | 6 | 0 | GC | GC | | | | | GC | GC | GC | GC | | |
| NC_011770 | 3984468 | C | T | 39 | 6 | 6 | 0 | . | . | T | T | T | T | . | . | . | . | T | T |
| NC_011770 | 3984549 | C | T | 55 | 0 | 2 | 0 | | | | | | | | | T | T | | |
| NC_011770 | 3984557 | G | A | 55 | 0 | 2 | 0 | A | A | | | | | | | | | | |
| NC_011770 | 3985585 | TCCGCCGA | T | 43 | 2 | 2 | 0 | T | T | | | | | | | . | . | | |
| NC_011770 | 3985594 | C | T | 36 | 4 | 2 | 0 | . | . | | | | | | | . | . | T | T |
| NC_011770 | 3985600 | T | C | 43 | 2 | 2 | 0 | | | | | | | C | C | . | . | | |
| NC_011770 | 3985619 | T | C | 39 | 2 | 2 | 0 | | | | | | | C | C | . | . | | |
| NC_011770 | 3985620 | G | A | 39 | 2 | 2 | 0 | | | | | | | A | A | . | . | | |
| NC_011770 | 3985640 | A | C | 35 | 3 | 1 | 0 | | | | | | | C | . | . | . | | |
| NC_011770 | 3985641 | A | G | 35 | 3 | 1 | 0 | | | | | | | G | . | . | . | | |
| NC_011770 | 3985645 | C | A | 34 | 3 | 2 | 0 | | | | | | | | . | . | . | A | A |
| NC_011770 | 3985667 | C | T | 36 | 2 | 2 | 0 | | | | | | | T | T | . | . | | |
| NC_011770 | 3985670 | C | A | 34 | 2 | 2 | 0 | | | | | | | A | A | . | . | | |
| NC_011770 | 3985674 | G | A | 33 | 2 | 2 | 0 | | | | | | | A | A | . | . | | |
| NC_011770 | 3985687 | C | G | 36 | 2 | 8 | 0 | | | G | G | G | G | G | G | . | . | G | G |
| NC_011770 | 3985741 | A | G | 38 | 2 | 4 | 0 | | | G | G | G | G | | | . | . | | |
| NC_011770 | 3985823 | G | A | 52 | 0 | 8 | 0 | A | A | A | A | A | A | | | A | A | | |
| NC_011770 | 3989428 | G | C | 83 | 0 | 2 | 0 | | | | | | | C | C | | | | |
| NC_011770 | 3989473 | C | T | 66 | 0 | 4 | 0 | | | T | T | T | T | | | | | | |
| NC_011770 | 3989476 | G | C | 64 | 0 | 4 | 0 | | | | | | | C | C | C | C | | |
| NC_011770 | 3989773 | AC | A | 57 | 0 | 4 | 0 | | | | | | | | | A | A | A | A |
| NC_011770 | 3989803 | C | T | 58 | 0 | 2 | 0 | T | T | | | | | | | | | | |
| NC_011770 | 3990778 | C | T | 81 | 0 | 4 | 0 | T | T | | | | | T | T | | | | |
| NC_011770 | 3995745 | G | A | 78 | 0 | 2 | 0 | | | | | | | | | A | A | | |
| NC_011770 | 3995856 | G | A | 75 | 0 | 2 | 0 | | | | | | | | | | | A | A |
| NC_011770 | 3995872 | A | C | 77 | 0 | 2 | 0 | C | C | | | | | | | | | | |
| NC_011770 | 3995899 | T | C | 73 | 0 | 4 | 0 | C | C | | | | | C | C | | | | |
| NC_011770 | 3995921 | G | A | 74 | 0 | 2 | 0 | | | | | | | | | | | A | A |
| NC_011770 | 3995942 | G | C | 77 | 0 | 2 | 0 | | | | | | | C | C | | | | |
| NC_011770 | 3997128 | CAG | C | 55 | 0 | 1 | 0 | | | | | | | C | | | | | |
| NC_011770 | 3997142 | A | G | 53 | 0 | 2 | 0 | | | | | | | | | G | G | | |
| NC_011770 | 3997174 | T | C | 53 | 0 | 2 | 0 | | | | | | | | | | | C | C |
| NC_011770 | 3997182 | GT | G | 55 | 0 | 2 | 0 | G | G | | | | | | | | | | |
| NC_011770 | 3997409 | G | T | 64 | 0 | 2 | 0 | T | T | | | | | | | | | | |
| NC_011770 | 3997423 | G | C | 60 | 0 | 12 | 0 | C | C | C | C | C | C | C | C | C | C | C | C |
| NC_011770 | 3999528 | T | C | 71 | 0 | 12 | 0 | C | C | C | C | C | C | C | C | C | C | C | C |
| NC_011770 | 3999535 | G | T | 74 | 0 | 2 | 0 | | | | | | | T | T | | | | |
| NC_011770 | 3999548 | C | G | 74 | 0 | 2 | 0 | | | | | | | G | G | | | | |
| NC_011770 | 4002026 | T | C | 47 | 0 | 2 | 0 | C | C | | | | | | | | | | |
| NC_011770 | 4002030 | A | C | 50 | 0 | 4 | 0 | | | C | C | C | C | | | | | | |
| NC_011770 | 4002058 | C | CCGGGC | 52 | 0 | 2 | 0 | | | | | | | CCGGGGG | CCGGGGG | | | | |
| NC_011770 | 4002067 | G | T | 41 | 0 | 4 | 0 | | | T | T | T | T | | | | | | |
| NC_011770 | 4010983 | A | G | 58 | 0 | 10 | 0 | | | G | G | G | G | G | G | G | G | G | G |

| | | | | | | | | | | | | | | | | | | | |
|---|---|---|---|---|---|---|---|---|---|---|---|---|---|---|---|---|---|---|---|
| NC_011770 | 4010986 | G | T | 58 | 0 | 6 | 0 | | | | | | | T | T | T | T | T | T |
| NC_011770 | 4011041 | G | T | 57 | 0 | 2 | 0 | T | T | | | | | | | | | | |
| NC_011770 | 4011067 | A | G | 58 | 0 | 2 | 0 | | | | | | | G | G | | | | |
| NC_011770 | 4011120 | C | T | 55 | 0 | 2 | 0 | | | | | | | | | | | T | T |
| NC_011770 | 4011149 | T | C | 54 | 0 | 4 | 0 | | | C | C | C | C | | | | | | |
| NC_011770 | 4011174 | A | C | 56 | 0 | 4 | 0 | | | | | | | | | C | C | C | C |
| NC_011770 | 4011211 | C | T | 58 | 0 | 2 | 0 | | | | | | | T | T | | | | |
| NC_011770 | 4011225 | A | G | 56 | 0 | 12 | 0 | G | G | G | G | G | G | G | G | G | G | G | G |
| NC_011770 | 4011701 | A | G | 63 | 0 | 6 | 0 | | | G | G | G | G | | | | | G | G |
| NC_011770 | 4011713 | A | G | 62 | 0 | 6 | 0 | | | G | G | G | G | | | | | G | G |
| NC_011770 | 4011723 | G | A | 67 | 0 | 4 | 0 | | | | | | | A | A | A | A | | |
| NC_011770 | 4011731 | A | G | 65 | 0 | 6 | 0 | | | G | G | G | G | | | | | G | G |
| NC_011770 | 4011734 | A | G | 65 | 0 | 6 | 0 | | | G | G | G | G | | | | | G | G |
| NC_011770 | 4011797 | C | G | 63 | 0 | 8 | 0 | G | G | G | G | G | G | | | | | G | G |
| NC_011770 | 4011841 | GC | G | 67 | 0 | 12 | 0 | G | G | G | G | G | G | G | G | G | G | G | G |
| NC_011770 | 4011851 | C | T | 65 | 0 | 8 | 0 | T | T | T | T | T | T | | | | | T | T |
| NC_011770 | 4011861 | A | T | 66 | 0 | 8 | 0 | T | T | T | T | T | T | | | | | T | T |
| NC_011770 | 4011912 | G | A | 67 | 0 | 8 | 0 | A | A | A | A | A | A | | | | | A | A |
| NC_011770 | 4014552 | T | C | 62 | 0 | 2 | 0 | C | C | | | | | | | | | | |
| NC_011770 | 4014772 | C | T | 84 | 0 | 2 | 0 | T | T | | | | | | | | | | |
| NC_011770 | 4014808 | G | A | 80 | 0 | 6 | 0 | | | A | A | A | A | | | A | A | | |
| NC_011770 | 4014927 | G | A | 98 | 0 | 2 | 0 | | | | | | | | | A | A | | |
| NC_011770 | 4014990 | G | A | 90 | 0 | 4 | 0 | | | A | A | A | A | | | | | | |
| NC_011770 | 4015006 | G | T | 90 | 0 | 2 | 0 | | | | | | | | | T | T | | |
| NC_011770 | 4021001 | T | C | 61 | 0 | 4 | 0 | | | | | | | | | C | C | C | C |
| NC_011770 | 4021038 | T | C | 56 | 0 | 10 | 0 | C | C | C | C | C | C | | | C | C | C | C |
| NC_011770 | 4021041 | A | G | 57 | 0 | 12 | 0 | G | G | G | G | G | G | G | G | G | G | G | G |
| NC_011770 | 4021043 | T | C | 56 | 0 | 4 | 0 | | | C | C | C | C | | | | | | |
| NC_011770 | 4021044 | G | A | 57 | 0 | 2 | 0 | | | | | | | A | A | | | | |
| NC_011770 | 4021045 | C | G | 58 | 0 | 4 | 0 | | | G | G | G | G | | | | | | |
| NC_011770 | 4021174 | C | T | 63 | 0 | 6 | 0 | T | T | T | T | T | T | | | | | | |
| NC_011770 | 4023015 | C | T | 47 | 0 | 2 | 0 | | | | | | | | | T | T | | |
| NC_011770 | 4023084 | C | T | 42 | 0 | 2 | 0 | T | T | | | | | | | | | | |
| NC_011770 | 4023124 | G | A | 35 | 0 | 6 | 0 | A | A | A | A | A | A | | | | | | |
| NC_011770 | 4023210 | T | C | 38 | 0 | 8 | 0 | | | C | C | C | C | C | C | | | C | C |
| NC_011770 | 4023224 | A | G | 35 | 0 | 6 | 0 | | | G | G | G | G | | | | | G | G |
| NC_011770 | 4023322 | A | G | 36 | 0 | 6 | 0 | | | G | G | G | G | G | G | | | | |
| NC_011770 | 4023324 | T | C | 37 | 0 | 2 | 0 | | | | | | | | | C | C | | |
| NC_011770 | 4023326 | A | T | 36 | 0 | 6 | 0 | | | T | T | T | T | T | T | | | | |
| NC_011770 | 4023354 | G | C | 35 | 0 | 6 | 0 | | | C | C | C | C | C | C | | | | |
| NC_011770 | 4023372 | C | T | 35 | 0 | 10 | 0 | | | T | T | T | T | T | T | T | T | T | T |
| NC_011770 | 4023421 | C | A | 37 | 0 | 2 | 0 | | | | | | | A | A | | | | |
| NC_011770 | 4023429 | A | T | 35 | 0 | 12 | 0 | T | T | T | T | T | T | T | T | T | T | T | T |
| NC_011770 | 4023431 | A | G | 37 | 0 | 12 | 0 | G | G | G | G | G | G | G | G | G | G | G | G |
| NC_011770 | 4023454 | A | G | 38 | 0 | 6 | 0 | | | G | G | G | G | G | G | | | | |
| NC_011770 | 4023482 | A | G | 37 | 0 | 6 | 0 | | | G | G | G | G | G | G | | | | |
| NC_011770 | 4023500 | T | C | 36 | 0 | 6 | 0 | | | C | C | C | C | C | C | | | | |
| NC_011770 | 4023519 | T | C | 38 | 0 | 6 | 0 | | | C | C | C | C | C | C | | | | |
| NC_011770 | 4023606 | A | T | 37 | 0 | 12 | 0 | T | T | T | T | T | T | T | T | T | T | T | T |
| NC_011770 | 4023631 | G | A | 45 | 0 | 2 | 0 | | | | | | | | | A | A | | |
| NC_011770 | 4023640 | C | T | 43 | 0 | 12 | 0 | T | T | T | T | T | T | T | T | T | T | T | T |
| NC_011770 | 4023676 | T | C | 48 | 0 | 12 | 0 | C | C | C | C | C | C | C | C | C | C | C | C |
| NC_011770 | 4023747 | C | G | 54 | 0 | 12 | 0 | G | G | G | G | G | G | G | G | G | G | G | G |
| NC_011770 | 4023751 | T | C | 53 | 0 | 12 | 0 | C | C | C | C | C | C | C | C | C | C | C | C |
| NC_011770 | 4023801 | G | A | 56 | 0 | 6 | 0 | | | A | A | A | A | A | A | | | | |
| NC_011770 | 4023860 | G | A | 65 | 0 | 6 | 0 | | | A | A | A | A | A | A | | | | |
| NC_011770 | 4023895 | T | G | 62 | 0 | 6 | 0 | | | G | G | G | G | G | G | | | | |
| NC_011770 | 4023917 | T | C | 64 | 0 | 2 | 0 | | | | | | | C | C | | | | |
| NC_011770 | 4023973 | C | T | 71 | 0 | 6 | 0 | | | T | T | T | T | T | T | | | | |
| NC_011770 | 4023991 | C | T | 69 | 0 | 6 | 0 | | | T | T | T | T | T | T | | | | |
| NC_011770 | 4027719 | T | A | 73 | 0 | 4 | 0 | A | A | | | | | A | A | | | | |
| NC_011770 | 4027735 | C | T | 73 | 0 | 4 | 0 | T | T | | | | | T | T | | | | |

| | | | | | | | | | | | | | | | | | | | |
|---|---|---|---|---|---|---|---|---|---|---|---|---|---|---|---|---|---|---|---|
| NC_011770 | 4027736 | A | G | 73 | 0 | 12 | 0 | G | G | G | G | G | G | G | G | G | G | G | G |
| NC_011770 | 4027743 | G | T | 73 | 0 | 8 | 0 | T | T | T | T | T | T | T | T | | | | |
| NC_011770 | 4027754 | G | A | 76 | 0 | 4 | 0 | A | A | | | | | A | A | | | | |
| NC_011770 | 4027758 | A | C | 77 | 0 | 4 | 0 | C | C | | | | | C | C | | | | |
| NC_011770 | 4027808 | A | T | 79 | 0 | 4 | 0 | T | T | | | | | T | T | | | | |
| NC_011770 | 4029927 | G | T | 91 | 0 | 2 | 0 | T | T | | | | | | | | | | |
| NC_011770 | 4030844 | GA | G | 70 | 0 | 2 | 0 | | | | | | | G | G | | | | |
| NC_011770 | 4031951 | C | A | 63 | 0 | 10 | 0 | | | A | A | A | A | A | A | A | A | A | A |
| NC_011770 | 4033861 | A | C | 73 | 0 | 12 | 0 | C | C | C | C | C | C | C | C | C | C | C | C |
| NC_011770 | 4036198 | T | C | 89 | 0 | 6 | 0 | | | C | C | C | C | | | C | C | | |
| NC_011770 | 4036202 | T | G | 89 | 0 | 6 | 0 | | | G | G | G | G | | | G | G | | |
| NC_011770 | 4037342 | A | G | 71 | 0 | 2 | 0 | | | | | | | | | G | G | | |
| NC_011770 | 4037343 | T | G | 71 | 0 | 4 | 0 | | | G | G | G | G | | | | | | |
| NC_011770 | 4041049 | GCACGGTT | G | 18 | 0 | 3 | 0 | | | G | G | . | G | | | | | | |
| NC_011770 | 4041466 | A | G | 55 | 0 | 12 | 0 | G | G | G | G | G | G | G | G | G | G | G | G |
| NC_011770 | 4042128 | T | C | 92 | 0 | 2 | 0 | | | C | C | | | | | | | | |
| NC_011770 | 4042145 | C | T | 90 | 0 | 4 | 0 | | | T | T | T | T | | | | | | |
| NC_011770 | 4042754 | C | T | 80 | 0 | 2 | 0 | | | | | | | | | T | T | | |
| NC_011770 | 778272 | CGG | C | 53 | 0 | 1 | 0 | | | | | C | | | | | | | |
| NC_011770 | 4043357 | C | T | 74 | 0 | 4 | 0 | | | T | T | T | T | | | | | | |
| NC_011770 | 4043373 | C | T | 76 | 0 | 2 | 0 | T | T | | | | | | | | | | |
| NC_011770 | 4043390 | T | C | 74 | 0 | 4 | 0 | | | C | C | C | C | | | | | | |
| NC_011770 | 4051839 | G | A | 86 | 0 | 2 | 0 | A | A | | | | | | | | | | |
| NC_011770 | 4053247 | T | A | 73 | 0 | 4 | 0 | A | A | | | | | | | | | A | A |
| NC_011770 | 4053255 | T | A | 71 | 0 | 4 | 0 | A | A | | | | | | | | | A | A |
| NC_011770 | 4060001 | G | A | 61 | 0 | 2 | 0 | A | A | | | | | | | | | | |
| NC_011770 | 4060163 | T | C | 61 | 0 | 6 | 0 | | | C | C | C | C | | | C | C | | |
| NC_011770 | 4060375 | CAGTACA | C | 57 | 0 | 2 | 0 | C | C | | | | | | | | | | |
| NC_011770 | 4060376 | A | G | 57 | 0 | 10 | 0 | . | . | G | G | G | G | G | G | G | G | G | G |
| NC_011770 | 4060452 | C | A | 53 | 0 | 2 | 0 | | | | | | | | | | | A | A |
| NC_011770 | 4060460 | A | G | 54 | 0 | 2 | 0 | | | | | | | | | | | G | G |
| NC_011770 | 4060474 | T | C | 54 | 0 | 12 | 0 | C | C | C | C | C | C | C | C | C | C | C | C |
| NC_011770 | 4060493 | G | T | 54 | 0 | 6 | 0 | | | T | T | T | T | | | T | T | | |
| NC_011770 | 4062457 | G | A | 25 | 2 | 2 | 0 | . | . | | | | | | | A | A | | |
| NC_011770 | 4064638 | A | G | 66 | 0 | 2 | 0 | | | | | | | | | | | G | G |
| NC_011770 | 4064656 | G | A | 69 | 0 | 2 | 0 | | | | | | | | | | | A | A |
| NC_011770 | 4064891 | A | C | 65 | 0 | 2 | 0 | C | C | | | | | | | | | | |
| NC_011770 | 4064902 | C | A | 64 | 0 | 2 | 0 | | | | | | | | | | | A | A |
| NC_011770 | 4064911 | G | A | 65 | 0 | 4 | 0 | | | A | A | A | A | | | | | | |
| NC_011770 | 4064962 | T | C | 69 | 0 | 2 | 0 | | | | | | | | | | | C | C |
| NC_011770 | 4065014 | C | A | 70 | 0 | 2 | 0 | | | | | | | A | A | | | | |
| NC_011770 | 4065044 | A | G | 69 | 0 | 2 | 0 | | | | | | | | | | | G | G |
| NC_011770 | 4065070 | A | G | 63 | 0 | 8 | 0 | | | G | G | G | G | G | G | | | G | G |
| NC_011770 | 4065074 | T | C | 62 | 0 | 2 | 0 | | | | | | | | | C | C | | |
| NC_011770 | 4065094 | T | A | 60 | 0 | 2 | 0 | | | | | | | | | A | A | | |
| NC_011770 | 4065105 | T | A | 60 | 0 | 8 | 0 | | | A | A | A | A | A | A | | | A | A |
| NC_011770 | 4065137 | C | A | 61 | 0 | 2 | 0 | | | | | | | A | A | | | | |
| NC_011770 | 4065146 | G | A | 59 | 0 | 8 | 0 | | | A | A | A | A | A | A | | | T | T |
| NC_011770 | 4065169 | T | C | 58 | 0 | 2 | 0 | | | | | | | C | C | | | | |
| NC_011770 | 4065170 | G | GA | 58 | 0 | 8 | 0 | | | GA | GA | GA | GA | GA | GA | | | GA | GA |
| NC_011770 | 4065193 | A | C | 58 | 0 | 2 | 0 | | | | | | | | | | | C | C |
| NC_011770 | 4065195 | G | A | 58 | 0 | 6 | 0 | | | A | A | A | A | | | | | A | A |
| NC_011770 | 4065258 | C | T | 57 | 0 | 4 | 0 | | | T | T | T | T | | | | | | |
| NC_011770 | 4065289 | G | A | 52 | 0 | 8 | 0 | | | A | A | A | A | A | A | | | A | A |
| NC_011770 | 4065470 | C | T | 37 | 0 | 2 | 0 | | | | | | | | | | | T | T |
| NC_011770 | 4065680 | C | T | 26 | 0 | 2 | 0 | | | | | | | | | | | T | T |
| NC_011770 | 4066355 | G | A | 68 | 0 | 2 | 0 | | | | | | | | | | | A | A |
| NC_011770 | 4066369 | T | C | 65 | 0 | 6 | 0 | C | C | | | | | C | C | C | C | | |
| NC_011770 | 4066370 | C | G | 68 | 0 | 6 | 0 | G | G | | | | | G | G | G | G | | |
| NC_011770 | 4066372 | G | A | 69 | 0 | 2 | 0 | | | | | | | A | A | | | | |
| NC_011770 | 4067582 | G | A | 66 | 0 | 2 | 0 | | | | | | | | | | | A | A |
| NC_011770 | 4067623 | A | G | 65 | 0 | 2 | 0 | | | | | | | | | | | G | G |

| | | | | | | | | | | | | | | | | | | | |
|---|---|---|---|---|---|---|---|---|---|---|---|---|---|---|---|---|---|---|---|
| NC_011770 | 4068886 | G | A | 78 | 0 | 1 | 0 | | | | | | | A | | | | | |
| NC_011770 | 4068907 | G | C | 76 | 0 | 4 | 0 | | | C | C | C | C | | | | | | |
| NC_011770 | 4068983 | G | A | 76 | 0 | 2 | 0 | | | | | | | | | | | A | A |
| NC_011770 | 4071172 | G | A | 35 | 0 | 2 | 0 | | | | | | | | | A | A | | |
| NC_011770 | 4071224 | C | T | 28 | 0 | 8 | 0 | | | T | T | T | T | | | T | T | T | T |
| NC_011770 | 4071270 | G | A | 22 | 0 | 2 | 0 | A | A | | | | | | | | | | |
| NC_011770 | 4073393 | C | A | 51 | 0 | 2 | 0 | | | | | | | A | A | | | | |
| NC_011770 | 4077160 | G | A | 72 | 0 | 4 | 0 | | | A | A | A | A | | | | | | |
| NC_011770 | 4077259 | G | A | 74 | 0 | 4 | 0 | | | A | A | A | A | | | | | | |
| NC_011770 | 4077720 | G | A | 65 | 0 | 2 | 0 | | | | | | | A | A | | | | |
| NC_011770 | 4078406 | A | G | 69 | 0 | 2 | 0 | | | | | | | G | G | | | | |
| NC_011770 | 4078428 | G | A | 71 | 0 | 6 | 0 | A | A | A | A | A | A | | | | | | |
| NC_011770 | 4078655 | C | T | 84 | 0 | 2 | 0 | | | | | | | T | T | | | | |
| NC_011770 | 4078715 | C | T | 88 | 0 | 8 | 0 | T | T | T | T | T | T | | | T | T | | |
| NC_011770 | 4078724 | C | T | 86 | 0 | 2 | 0 | T | T | | | | | | | | | | |
| NC_011770 | 4078774 | G | A | 83 | 0 | 6 | 0 | | | A | A | A | A | | | A | A | | |
| NC_011770 | 4084091 | C | G | 75 | 0 | 12 | 0 | G | G | G | G | G | G | G | G | G | G | G | G |
| NC_011770 | 4084105 | G | A | 75 | 0 | 12 | 0 | A | A | A | A | A | A | A | A | A | A | A | A |
| NC_011770 | 4084112 | C | CG | 77 | 0 | 12 | 0 | CG | CG | CG | CG | CG | CG | CG | CG | CG | CG | CG | CG |
| NC_011770 | 4084161 | G | A | 72 | 0 | 12 | 0 | A | A | A | A | A | A | A | A | A | A | A | A |
| NC_011770 | 4084968 | A | G | 82 | 0 | 4 | 0 | | | | | | | G | G | | | G | G |
| NC_011770 | 4084989 | G | A | 82 | 0 | 4 | 0 | | | | | | | A | A | | | A | A |
| NC_011770 | 4088331 | C | T | 74 | 0 | 2 | 0 | T | T | | | | | | | | | | |
| NC_011770 | 4090437 | G | A | 74 | 0 | 2 | 0 | | | | | | | A | A | | | | |
| NC_011770 | 4092261 | A | G | 95 | 0 | 12 | 0 | G | G | G | G | G | G | G | G | G | G | G | G |
| NC_011770 | 4092317 | C | A | 100 | 0 | 12 | 0 | A | A | A | A | A | A | A | A | A | A | A | A |
| NC_011770 | 4092324 | C | T | 93 | 0 | 6 | 0 | | | | | | | T | T | T | T | T | T |
| NC_011770 | 4092348 | A | G | 98 | 0 | 2 | 0 | | | | | G | G | | | | | | |
| NC_011770 | 4092375 | G | C | 89 | 0 | 8 | 0 | | | C | C | C | C | C | C | | | C | C |
| NC_011770 | 4092388 | G | C | 92 | 0 | 2 | 0 | | | | | | | C | C | | | | |
| NC_011770 | 4093600 | C | A | 87 | 0 | 4 | 0 | A | A | | | | | | | A | A | | |
| NC_011770 | 4093601 | T | C | 87 | 0 | 12 | 0 | C | C | C | C | C | C | C | C | C | C | C | C |
| NC_011770 | 4094636 | G | A | 68 | 0 | 10 | 0 | A | A | A | A | A | A | A | A | A | A | | |
| NC_011770 | 4094642 | A | G | 67 | 0 | 10 | 0 | G | G | G | G | G | G | G | G | G | G | | |
| NC_011770 | 4094668 | C | A | 68 | 0 | 12 | 0 | A | A | A | A | A | A | A | A | A | A | A | A |
| NC_011770 | 4094705 | T | C | 69 | 0 | 12 | 0 | C | C | C | C | C | C | C | C | C | C | C | C |
| NC_011770 | 4097082 | A | G | 66 | 0 | 12 | 0 | G | G | G | G | G | G | G | G | G | G | G | G |
| NC_011770 | 4097090 | C | T | 67 | 0 | 12 | 0 | T | T | T | T | T | T | T | T | T | T | T | T |
| NC_011770 | 4097150 | C | T | 68 | 0 | 2 | 0 | | | | | | | | | | | T | T |
| NC_011770 | 4097157 | G | T | 69 | 0 | 6 | 0 | | | | | | | T | T | T | T | T | T |
| NC_011770 | 4103689 | C | G | 32 | 0 | 2 | 0 | | | | | | | | | G | G | | |
| NC_011770 | 4109548 | G | T | 52 | 0 | 4 | 0 | | | T | T | T | T | | | | | | |
| NC_011770 | 4111789 | C | T | 85 | 0 | 4 | 0 | | | T | T | T | T | | | | | | |
| NC_011770 | 4116518 | A | C | 50 | 0 | 10 | 0 | C | C | C | C | C | C | C | C | | | C | C |
| NC_011770 | 4116536 | T | C | 49 | 0 | 10 | 0 | C | C | C | C | C | C | C | C | | | C | C |
| NC_011770 | 4118340 | G | A | 49 | 0 | 10 | 0 | A | A | A | A | A | A | | | A | A | A | A |
| NC_011770 | 4118387 | G | GC | 48 | 0 | 4 | 0 | | | GC | GC | GC | GC | | | | | | |
| NC_011770 | 4118848 | G | A | 60 | 0 | 2 | 0 | A | A | | | | | | | | | | |
| NC_011770 | 4118855 | G | A | 59 | 0 | 2 | 0 | A | A | | | | | | | | | | |
| NC_011770 | 4118856 | A | T | 55 | 0 | 10 | 0 | T | T | T | T | T | T | T | T | | | T | T |
| NC_011770 | 4118866 | T | C | 58 | 0 | 2 | 0 | C | C | | | | | | | | | | |
| NC_011770 | 4118867 | T | G | 58 | 0 | 2 | 0 | G | G | | | | | | | | | | |
| NC_011770 | 4118875 | C | T | 47 | 0 | 4 | 0 | | | T | T | T | T | | | | | | |
| NC_011770 | 4118879 | C | A | 47 | 0 | 2 | 0 | A | A | | | | | | | | | | |
| NC_011770 | 4118898 | G | A | 43 | 2 | 2 | 0 | A | A | | | | | | | | | . | . |
| NC_011770 | 4118903 | T | C | 44 | 2 | 2 | 0 | | | | | | | C | C | | | . | . |
| NC_011770 | 4118907 | T | C | 43 | 2 | 4 | 0 | | | C | C | C | C | | | | | . | . |
| NC_011770 | 4118914 | A | G | 58 | 0 | 4 | 0 | G | G | | | | | G | G | | | | |
| NC_011770 | 4118972 | C | G | 64 | 0 | 2 | 0 | G | G | | | | | | | | | | |
| NC_011770 | 4121501 | A | G | 62 | 0 | 2 | 0 | G | G | | | | | | | | | | |
| NC_011770 | 4121505 | C | T | 63 | 0 | 2 | 0 | T | T | | | | | | | | | | |
| NC_011770 | 4121523 | T | C | 65 | 0 | 2 | 0 | C | C | | | | | | | | | | |

| | | | | | | | | | | | | | | | | | | | |
|---|---|---|---|---|---|---|---|---|---|---|---|---|---|---|---|---|---|---|---|
| NC_011770 | 4121983 | G | A | 63 | 0 | 2 | 0 | A | A | | | | | | | | | | |
| NC_011770 | 4121989 | C | T | 68 | 0 | 2 | 0 | T | T | | | | | | | | | | |
| NC_011770 | 4122034 | G | A | 71 | 0 | 4 | 0 | | | | | | | A | A | | | A | A |
| NC_011770 | 4122043 | G | A | 72 | 0 | 2 | 0 | A | A | | | | | | | | | | |
| NC_011770 | 4122045 | T | C | 72 | 0 | 2 | 0 | C | C | | | | | | | | | | |
| NC_011770 | 4122079 | T | A | 74 | 0 | 2 | 0 | A | A | | | | | | | | | | |
| NC_011770 | 4122084 | A | G | 74 | 0 | 12 | 0 | G | G | G | G | G | G | G | G | G | G | G | G |
| NC_011770 | 4122087 | T | C | 75 | 0 | 2 | 0 | C | C | | | | | | | | | | |
| NC_011770 | 4122089 | T | C | 75 | 0 | 12 | 0 | C | C | C | C | C | C | C | C | C | C | C | C |
| NC_011770 | 4122103 | T | C | 75 | 0 | 2 | 0 | C | C | | | | | | | | | | |
| NC_011770 | 4122111 | T | C | 76 | 0 | 2 | 0 | C | C | | | | | | | | | | |
| NC_011770 | 4122136 | G | C | 74 | 0 | 2 | 0 | C | C | | | | | | | | | | |
| NC_011770 | 4124937 | G | T | 69 | 0 | 2 | 0 | T | T | | | | | | | | | | |
| NC_011770 | 4127066 | A | G | 78 | 0 | 8 | 0 | G | G | | | | | G | G | G | G | G | G |
| NC_011770 | 4127089 | C | T | 84 | 0 | 2 | 0 | | | | | | | T | T | | | | |
| NC_011770 | 4130700 | T | G | 71 | 0 | 12 | 0 | G | G | G | G | G | G | G | G | G | G | G | G |
| NC_011770 | 4133093 | C | T | 50 | 0 | 2 | 0 | | | | | | | | | | | T | T |
| NC_011770 | 4135707 | A | T | 70 | 0 | 2 | 0 | | | | | | | T | T | | | | |
| NC_011770 | 4135769 | T | C | 64 | 0 | 2 | 0 | | | | | | | C | C | | | | |
| NC_011770 | 4135785 | A | G | 61 | 0 | 2 | 0 | | | | | | | G | G | | | | |
| NC_011770 | 4135843 | C | T | 60 | 0 | 2 | 0 | | | T | T | | | | | | | | |
| NC_011770 | 4135904 | G | A | 53 | 0 | 2 | 0 | | | | | | | | | A | A | | |
| NC_011770 | 4138060 | A | G | 59 | 0 | 12 | 0 | G | G | G | G | G | G | G | G | G | G | G | G |
| NC_011770 | 4138082 | C | T | 58 | 0 | 12 | 0 | T | T | T | T | T | T | T | T | T | T | T | T |
| NC_011770 | 4138197 | T | G | 60 | 0 | 12 | 0 | G | G | G | G | G | G | G | G | G | G | G | G |
| NC_011770 | 4138313 | G | T | 63 | 0 | 4 | 0 | | | T | T | T | T | | | | | | |
| NC_011770 | 4144088 | A | AT | 81 | 0 | 2 | 0 | AT | AT | | | | | | | | | | |
| NC_011770 | 4149088 | C | T | 52 | 0 | 2 | 0 | | | | | | | | | T | T | | |
| NC_011770 | 4149179 | A | G | 49 | 0 | 2 | 0 | | | | | | | | | G | G | | |
| NC_011770 | 4149833 | T | C | 58 | 0 | 1 | 0 | | | | | | | C | | | | | |
| NC_011770 | 4150270 | G | A | 54 | 0 | 2 | 0 | | | | | | | A | A | | | | |
| NC_011770 | 4153437 | G | C | 52 | 0 | 2 | 0 | | | | | | | | | C | C | | |
| NC_011770 | 4153456 | T | C | 52 | 0 | 8 | 0 | | | C | C | C | C | C | C | | | C | C |
| NC_011770 | 4153468 | T | G | 53 | 0 | 8 | 0 | | | G | G | G | G | G | G | | | G | G |
| NC_011770 | 4153507 | A | G | 54 | 0 | 8 | 0 | | | G | G | G | G | G | G | | | G | G |
| NC_011770 | 4155368 | A | G | 65 | 0 | 2 | 0 | | | | | | | G | G | | | | |
| NC_011770 | 4155457 | A | C | 57 | 0 | 10 | 0 | | | C | C | C | C | C | C | C | C | C | C |
| NC_011770 | 4155668 | C | T | 37 | 0 | 2 | 0 | | | | | | | T | T | | | | |
| NC_011770 | 4155707 | CT | C | 35 | 0 | 4 | 0 | | | C | C | C | C | | | | | | |
| NC_011770 | 4155720 | G | C | 34 | 0 | 4 | 0 | | | C | C | C | C | | | | | | |
| NC_011770 | 4156629 | T | C | 64 | 0 | 2 | 0 | | | | | | | C | C | | | | |
| NC_011770 | 4156718 | T | C | 66 | 0 | 2 | 0 | | | | | | | C | C | | | | |
| NC_011770 | 4156838 | A | T | 64 | 0 | 2 | 0 | | | | | | | T | T | | | | |
| NC_011770 | 4156844 | G | T | 63 | 0 | 2 | 0 | | | | | | | T | T | | | | |
| NC_011770 | 4156883 | A | T | 64 | 0 | 2 | 0 | | | | | | | T | T | | | | |
| NC_011770 | 4156894 | G | A | 65 | 0 | 2 | 0 | | | | | | | A | A | | | | |
| NC_011770 | 4156897 | A | G | 63 | 0 | 4 | 0 | G | G | | | | | | | | | G | G |
| NC_011770 | 4156991 | T | A | 74 | 0 | 2 | 0 | | | | | | | A | A | | | | |
| NC_011770 | 4162010 | C | T | 69 | 0 | 2 | 0 | | | | | | | | | | | T | T |
| NC_011770 | 4162048 | G | T | 67 | 0 | 2 | 0 | | | | | | | T | T | | | | |
| NC_011770 | 4163402 | G | A | 76 | 0 | 2 | 0 | | | | | | | A | A | | | | |
| NC_011770 | 4163701 | G | A | 70 | 0 | 2 | 0 | | | | | | | | | | | A | A |
| NC_011770 | 4163729 | C | A | 71 | 0 | 2 | 0 | A | A | | | | | | | | | | |
| NC_011770 | 4164919 | A | C | 79 | 0 | 2 | 0 | | | | | | | | | C | C | | |
| NC_011770 | 4167921 | C | G | 56 | 0 | 2 | 0 | | | | | | | | | | | G | G |
| NC_011770 | 4167925 | A | G | 53 | 0 | 4 | 0 | | | | | | | G | G | | | G | G |
| NC_011770 | 4167944 | G | A | 56 | 0 | 2 | 0 | | | | | | | | | | | A | A |
| NC_011770 | 4167952 | G | T | 56 | 0 | 2 | 0 | | | | | | | | | | | T | T |
| NC_011770 | 4167985 | G | A | 56 | 0 | 2 | 0 | | | | | | | | | | | A | A |
| NC_011770 | 4167987 | T | C | 56 | 0 | 4 | 0 | | | C | C | C | C | | | | | | |
| NC_011770 | 4168011 | G | A | 57 | 0 | 2 | 0 | | | | | | | | | | | A | A |
| NC_011770 | 4168038 | C | G | 60 | 0 | 2 | 0 | | | | | | | | | | | G | G |

| | | | | | | | | | | | | | | | | | | | |
|---|---|---|---|---|---|---|---|---|---|---|---|---|---|---|---|---|---|---|---|
| NC_011770 | 4168044 | C | T | 59 | 0 | 2 | 0 | | | | | | | | | | | T | T |
| NC_011770 | 4168062 | C | T | 61 | 0 | 2 | 0 | | | | | | | | | | | T | T |
| NC_011770 | 4168079 | G | A | 59 | 0 | 4 | 0 | | | A | A | A | A | | | | | | |
| NC_011770 | 4169824 | G | A | 79 | 0 | 4 | 0 | | | A | A | A | A | | | | | | |
| NC_011770 | 4171796 | C | T | 88 | 0 | 2 | 0 | | | | | | | | | | | T | T |
| NC_011770 | 4175256 | C | T | 30 | 0 | 6 | 0 | T | T | | | | | T | T | T | T | | |
| NC_011770 | 4178812 | GC | GCCC | 66 | 0 | 2 | 0 | | | | | | | GCCC | G | | | | |
| NC_011770 | 4184124 | T | C | 85 | 0 | 12 | 0 | C | C | C | C | C | C | C | C | C | C | C | C |
| NC_011770 | 4184171 | T | C | 83 | 0 | 2 | 0 | | | | | | | | | C | C | | |
| NC_011770 | 4185636 | A | G | 51 | 0 | 2 | 0 | | | | | | | | | G | G | | |
| NC_011770 | 4185645 | C | A | 53 | 0 | 12 | 0 | A | A | A | A | A | A | A | A | A | A | A | A |
| NC_011770 | 4186189 | GC | G | 70 | 0 | 4 | 0 | | | G | G | G | G | | | | | | |
| NC_011770 | 4186235 | T | C | 73 | 0 | 4 | 0 | | | C | C | C | C | | | | | | |
| NC_011770 | 4186289 | T | C | 72 | 0 | 2 | 0 | | | | | | | | | C | C | | |
| NC_011770 | 4187124 | A | G | 64 | 0 | 2 | 0 | | | | | | | | | | | G | G |
| NC_011770 | 4187139 | G | A | 65 | 0 | 4 | 0 | | | A | A | A | A | | | | | | |
| NC_011770 | 4187751 | G | C | 75 | 0 | 2 | 0 | C | C | | | | | | | | | | |
| NC_011770 | 4187770 | C | T | 74 | 0 | 2 | 0 | T | T | | | | | | | | | | |
| NC_011770 | 4187774 | C | T | 77 | 0 | 2 | 0 | T | T | | | | | | | | | | |
| NC_011770 | 4187776 | C | T | 78 | 0 | 2 | 0 | T | T | | | | | | | | | | |
| NC_011770 | 4187839 | G | A | 79 | 0 | 2 | 0 | A | A | | | | | | | | | | |
| NC_011770 | 4187913 | A | G | 76 | 0 | 4 | 0 | | | G | G | G | G | | | | | | |
| NC_011770 | 4187942 | T | C | 80 | 0 | 4 | 0 | | | C | C | C | C | | | | | | |
| NC_011770 | 4188828 | G | A | 43 | 0 | 2 | 0 | | | | | | | A | A | | | | |
| NC_011770 | 4188841 | C | T | 43 | 0 | 2 | 0 | T | T | | | | | | | | | | |
| NC_011770 | 4188861 | G | A | 45 | 0 | 2 | 0 | | | | | | | | | A | A | | |
| NC_011770 | 4188883 | G | C | 45 | 0 | 2 | 0 | | | | | | | | | C | C | | |
| NC_011770 | 4188989 | G | A | 42 | 0 | 12 | 0 | A | A | A | A | A | A | A | A | A | A | A | A |
| NC_011770 | 4189030 | T | C | 38 | 0 | 2 | 0 | C | C | | | | | | | | | | |
| NC_011770 | 4189111 | A | C | 38 | 0 | 2 | 0 | | | | | | | | | C | C | | |
| NC_011770 | 4189194 | C | T | 42 | 0 | 2 | 0 | | | | | | | | | | | T | T |
| NC_011770 | 4190720 | T | G | 40 | 0 | 12 | 0 | G | G | G | G | G | G | G | G | G | G | G | G |
| NC_011770 | 4193263 | C | T | 84 | 0 | 8 | 0 | T | T | | | | | T | T | T | T | T | T |
| NC_011770 | 4195075 | T | C | 85 | 0 | 12 | 0 | C | C | C | C | C | C | C | C | C | C | C | C |
| NC_011770 | 4195079 | T | C | 83 | 0 | 2 | 0 | C | C | | | | | | | | | | |
| NC_011770 | 4195094 | G | A | 87 | 0 | 2 | 0 | A | A | | | | | | | | | | |
| NC_011770 | 4195767 | G | A | 77 | 0 | 2 | 0 | A | A | | | | | | | | | | |
| NC_011770 | 4195769 | G | A | 79 | 0 | 2 | 0 | A | A | | | | | | | | | | |
| NC_011770 | 4199599 | G | T | 72 | 0 | 2 | 0 | | | T | T | | | | | | | | |
| NC_011770 | 4199617 | T | C | 71 | 0 | 1 | 0 | | | | | | | C | | | | | |
| NC_011770 | 4199695 | G | A | 72 | 0 | 2 | 0 | | | | | | | | | A | A | | |
| NC_011770 | 4202265 | C | CCT | 59 | 0 | 2 | 0 | CCT | CCT | | | | | | | | | | |
| NC_011770 | 4202272 | T | C | 62 | 0 | 2 | 0 | C | C | | | | | | | | | | |
| NC_011770 | 4203199 | T | TA | 74 | 0 | 4 | 0 | TA | TA | | | | | | | TA | TA | | |
| NC_011770 | 4207552 | G | A | 68 | 0 | 2 | 0 | A | A | | | | | | | | | | |
| NC_011770 | 4207632 | T | C | 65 | 0 | 12 | 0 | C | C | C | C | C | C | C | C | C | C | C | C |
| NC_011770 | 4213755 | C | G | 33 | 0 | 12 | 0 | G | G | G | G | G | G | G | G | G | G | G | G |
| NC_011770 | 4213802 | A | AC | 34 | 0 | 2 | 0 | | | | | | | | | AC | AC | | |
| NC_011770 | 4215932 | A | G | 72 | 0 | 10 | 0 | G | G | G | G | G | G | | | G | G | G | G |
| NC_011770 | 4217692 | A | C | 18 | 0 | 8 | 0 | C | C | | | | | C | C | C | C | C | C |
| NC_011770 | 4217697 | C | T | 24 | 0 | 4 | 0 | | | T | T | T | T | | | | | | |
| NC_011770 | 4217698 | C | T | 24 | 0 | 2 | 0 | | | | | | | | | T | T | | |
| NC_011770 | 4217701 | C | A | 23 | 0 | 12 | 0 | A | A | A | A | A | A | A | A | A | A | A | A |
| NC_011770 | 4217703 | A | G | 24 | 0 | 12 | 0 | G | G | G | G | G | G | G | G | G | G | G | G |
| NC_011770 | 4217704 | T | C | 31 | 0 | 8 | 0 | C | C | | | | | C | C | C | C | C | C |
| NC_011770 | 4218932 | C | G | 84 | 0 | 2 | 0 | | | | | | | | | G | G | | |
| NC_011770 | 4218935 | C | T | 85 | 0 | 2 | 0 | | | | | | | | | T | T | | |
| NC_011770 | 4219028 | T | C | 84 | 0 | 2 | 0 | | | | | | | | | C | C | | |
| NC_011770 | 4219047 | G | T | 83 | 0 | 2 | 0 | | | | | | | | | T | T | | |
| NC_011770 | 4219050 | G | A | 85 | 0 | 2 | 0 | | | | | | | | | | | A | A |
| NC_011770 | 4221057 | T | C | 6 | 10 | 2 | 0 | C | C | . | . | . | . | . | . | . | . | . | . |
| NC_011770 | 4221221 | G | T | 4 | 10 | 2 | 0 | T | T | . | . | . | . | . | . | . | . | . | . |

| | | | | | | | | | | | | | | | | | | | |
|---|---|---|---|---|---|---|---|---|---|---|---|---|---|---|---|---|---|---|---|
| NC_011770 | 4221231 | C | T | 4 | 10 | 2 | 0 | T | T | . | . | . | . | . | . | . | . | . | . |
| NC_011770 | 4221263 | C | T | 4 | 10 | 2 | 0 | T | T | . | . | . | . | . | . | . | . | . | . |
| NC_011770 | 4221286 | T | C | 4 | 10 | 2 | 0 | C | C | . | . | . | . | . | . | . | . | . | . |
| NC_011770 | 4221409 | G | A | 4 | 10 | 2 | 0 | A | A | . | . | . | . | . | . | . | . | . | . |
| NC_011770 | 4221493 | C | T | 5 | 10 | 2 | 0 | T | T | . | . | . | . | . | . | . | . | . | . |
| NC_011770 | 4221501 | T | C | 5 | 10 | 2 | 0 | C | C | . | . | . | . | . | . | . | . | . | . |
| NC_011770 | 4221529 | G | A | 6 | 10 | 2 | 0 | A | A | . | . | . | . | . | . | . | . | . | . |
| NC_011770 | 4221652 | G | A | 5 | 10 | 2 | 0 | A | A | . | . | . | . | . | . | . | . | . | . |
| NC_011770 | 4221908 | A | G | 4 | 10 | 2 | 0 | G | G | . | . | . | . | . | . | . | . | . | . |
| NC_011770 | 4222483 | T | C | 3 | 10 | 2 | 0 | C | C | . | . | . | . | . | . | . | . | . | . |
| NC_011770 | 4222574 | G | GC | 2 | 10 | 2 | 0 | GC | GC | . | . | . | . | . | . | . | . | . | . |
| NC_011770 | 4222619 | C | CT | 4 | 8 | 2 | 0 | | | . | . | . | . | CT | CT | . | . | . | . |
| NC_011770 | 4222639 | C | A | 14 | 1 | 2 | 0 | A | A | | | | | | | . | | | |
| NC_011770 | 4222640 | C | A | 14 | 1 | 2 | 0 | A | A | | | | | | | . | | | |
| NC_011770 | 4223505 | C | CA | 22 | 8 | 4 | 0 | . | . | . | . | . | . | CA | CA | CA | CA | . | . |
| NC_011770 | 4223644 | A | G | 27 | 8 | 2 | 0 | . | . | . | . | . | . | | | G | G | . | . |
| NC_011770 | 4223749 | A | G | 27 | 8 | 2 | 0 | . | . | . | . | . | . | | | G | G | . | . |
| NC_011770 | 4224682 | G | C | 8 | 8 | 2 | 0 | . | . | . | . | . | . | | | C | C | . | . |
| NC_011770 | 4224728 | T | G | 5 | 8 | 2 | 0 | . | . | . | . | . | . | | | G | G | . | . |
| NC_011770 | 4224868 | G | T | 14 | 0 | 2 | 0 | | | | | | | | | T | T | | |
| NC_011770 | 4229322 | C | CTA | 41 | 0 | 1 | 0 | | | | | | | CTA | | | | | |
| NC_011770 | 4229348 | G | A | 40 | 0 | 2 | 0 | A | A | | | | | | | | | | |
| NC_011770 | 4229375 | C | T | 34 | 0 | 4 | 0 | | | T | T | T | T | | | | | | |
| NC_011770 | 1159276 | C | T | 20 | 2 | 3 | 0 | | | | | T | | . | . | T | T | . | |
| NC_011770 | 4229485 | C | T | 32 | 0 | 2 | 0 | | | | | | | | | | | T | T |
| NC_011770 | 4229540 | G | C | 26 | 0 | 10 | 0 | C | C | C | C | C | C | C | C | | | C | C |
| NC_011770 | 4229541 | T | C | 26 | 0 | 10 | 0 | C | C | C | C | C | C | C | C | | | C | C |
| NC_011770 | 4229592 | G | A | 24 | 0 | 4 | 0 | | | A | A | A | A | | | | | | |
| NC_011770 | 4229612 | T | C | 22 | 0 | 2 | 0 | | | | | | | | | C | C | | |
| NC_011770 | 4229640 | C | T | 18 | 0 | 2 | 0 | | | | | | | | | T | T | | |
| NC_011770 | 4229657 | T | C | 18 | 0 | 12 | 0 | C | C | C | C | C | C | C | C | C | C | C | C |
| NC_011770 | 4229664 | T | C | 18 | 0 | 1 | 0 | | | | | | | C | | | | | |
| NC_011770 | 4233652 | G | A | 76 | 0 | 4 | 0 | | | | | | | | | A | A | A | A |
| NC_011770 | 4233670 | G | A | 81 | 0 | 2 | 0 | | | | | | | A | A | | | | |
| NC_011770 | 4234793 | T | C | 87 | 0 | 10 | 0 | | | C | C | C | C | C | C | C | C | C | C |
| NC_011770 | 4235204 | G | A | 78 | 0 | 2 | 0 | | | | | | | A | A | | | | |
| NC_011770 | 4235263 | T | C | 82 | 0 | 4 | 0 | | | C | C | C | C | | | | | | |
| NC_011770 | 4236277 | G | A | 43 | 0 | 2 | 0 | A | A | | | | | | | | | | |
| NC_011770 | 4236331 | C | T | 48 | 0 | 2 | 0 | T | T | | | | | | | | | | |
| NC_011770 | 4239692 | G | A | 57 | 0 | 6 | 0 | | | A | A | A | A | | | A | A | | |
| NC_011770 | 4239703 | C | G | 62 | 0 | 2 | 0 | G | G | | | | | | | | | | |
| NC_011770 | 4240185 | T | C | 72 | 0 | 6 | 0 | C | C | | | | | C | C | C | C | | |
| NC_011770 | 4240188 | CG | C | 71 | 0 | 1 | 0 | | | | | | | C | | | | | |
| NC_011770 | 4240256 | A | T | 58 | 0 | 2 | 0 | | | | | | | T | T | | | | |
| NC_011770 | 4240273 | A | G | 58 | 0 | 6 | 0 | G | G | | | | | G | G | G | G | | |
| NC_011770 | 4241771 | C | T | 76 | 0 | 2 | 0 | T | T | | | | | | | | | | |
| NC_011770 | 4243392 | G | A | 81 | 0 | 2 | 0 | | | | | | | | | | | A | A |
| NC_011770 | 4245077 | A | G | 69 | 0 | 4 | 0 | | | | | | | | | G | G | G | G |
| NC_011770 | 4245132 | C | A | 74 | 0 | 2 | 0 | | | | | | | A | A | | | | |
| NC_011770 | 4246053 | T | C | 59 | 0 | 10 | 0 | | | C | C | C | C | C | C | C | C | C | C |
| NC_011770 | 4246056 | T | C | 60 | 0 | 2 | 0 | | | | | | | C | C | | | | |
| NC_011770 | 4246057 | T | C | 60 | 0 | 2 | 0 | | | | | | | C | C | | | | |
| NC_011770 | 4246059 | C | T | 61 | 0 | 2 | 0 | | | | | | | T | T | | | | |
| NC_011770 | 4246060 | T | G | 60 | 0 | 2 | 0 | | | | | | | G | G | | | | |
| NC_011770 | 4246525 | T | G | 44 | 0 | 2 | 0 | | | | | | | G | G | | | | |
| NC_011770 | 4246527 | G | T | 45 | 0 | 2 | 0 | | | | | | | | | T | T | | |
| NC_011770 | 4246550 | G | T | 46 | 0 | 2 | 0 | | | | | | | T | T | | | | |
| NC_011770 | 4246661 | C | A | 50 | 0 | 2 | 0 | | | | | | | A | A | | | | |
| NC_011770 | 4246685 | C | G | 51 | 0 | 2 | 0 | | | | | | | | | | | G | G |
| NC_011770 | 4246797 | C | T | 47 | 0 | 2 | 0 | | | | | | | T | T | | | | |
| NC_011770 | 4246798 | G | A | 48 | 0 | 2 | 0 | | | | | | | A | A | | | | |
| NC_011770 | 4246799 | G | T | 47 | 0 | 2 | 0 | | | | | | | T | T | | | | |

| | | | | | | | | | | | | | | | | | | | |
|---|---|---|---|---|---|---|---|---|---|---|---|---|---|---|---|---|---|---|---|
| NC_011770 | 4246814 | A | G | 43 | 0 | 2 | 0 | | | | | | | G | G | | | | |
| NC_011770 | 4247889 | GC | G | 28 | 0 | 4 | 0 | | | G | G | G | G | | | | | | |
| NC_011770 | 4247928 | GC | G | 23 | 0 | 2 | 0 | | | | | | | | | | | G | G |
| NC_011770 | 4247941 | G | A | 22 | 0 | 10 | 0 | A | A | A | A | A | A | A | A | | | A | A |
| NC_011770 | 4248028 | C | T | 21 | 0 | 4 | 0 | | | T | T | T | T | | | | | | |
| NC_011770 | 4248061 | G | A | 26 | 0 | 2 | 0 | | | | | | | | | | | A | A |
| NC_011770 | 4253158 | A | C | 41 | 0 | 2 | 0 | | | | | | | C | C | | | | |
| NC_011770 | 4253242 | G | T | 38 | 0 | 2 | 0 | | | | | | | T | T | | | | |
| NC_011770 | 4255176 | C | G | 57 | 0 | 2 | 0 | | | | | | | | | G | G | | |
| NC_011770 | 4256477 | GA | G | 64 | 0 | 6 | 0 | G | G | G | G | G | G | | | | | | |
| NC_011770 | 4256480 | C | G | 30 | 5 | 1 | 0 | . | . | G | . | . | . | | | | | | |
| NC_011770 | 1287607 | GT | G | 62 | 0 | 3 | 0 | | | G | G | G | | | | | | | |
| NC_011770 | 4256507 | T | C | 6 | 9 | 1 | 0 | . | . | . | . | . | C | . | . | | | . | . |
| NC_011770 | 4256508 | G | A | 6 | 9 | 1 | 0 | . | . | . | . | . | A | . | . | | | . | . |
| NC_011770 | 4256518 | A | G | 61 | 0 | 10 | 0 | G | G | G | G | G | G | G | G | | | G | G |
| NC_011770 | 4256535 | G | A | 64 | 0 | 10 | 0 | A | A | A | A | A | A | A | A | | | A | A |
| NC_011770 | 4256555 | A | T | 65 | 0 | 2 | 0 | | | | | | | | | T | T | | |
| NC_011770 | 4256566 | C | T | 66 | 0 | 2 | 0 | | | | | | | | | T | T | | |
| NC_011770 | 4256572 | A | G | 64 | 0 | 4 | 0 | | | G | G | G | G | | | | | | |
| NC_011770 | 4256577 | A | G | 62 | 0 | 12 | 0 | G | G | G | G | G | G | G | G | G | G | G | G |
| NC_011770 | 4256588 | T | C | 60 | 0 | 6 | 0 | C | C | C | C | C | C | | | | | | |
| NC_011770 | 4256626 | C | A | 61 | 0 | 2 | 0 | | | | | | | | | A | A | | |
| NC_011770 | 4256666 | T | G | 60 | 0 | 8 | 0 | | | G | G | G | G | G | G | G | G | | |
| NC_011770 | 4256701 | C | T | 61 | 0 | 2 | 0 | | | | | | | | | T | T | | |
| NC_011770 | 4256715 | T | C | 60 | 0 | 2 | 0 | | | | | | | | | C | C | | |
| NC_011770 | 4256716 | A | G | 60 | 0 | 2 | 0 | | | | | | | | | G | G | | |
| NC_011770 | 4256726 | G | A | 59 | 0 | 2 | 0 | | | | | | | | | A | A | | |
| NC_011770 | 4257870 | C | A | 65 | 0 | 8 | 0 | | | A | A | A | A | A | A | A | A | | |
| NC_011770 | 4257934 | T | G | 62 | 0 | 4 | 0 | | | G | G | G | G | | | | | | |
| NC_011770 | 4258302 | T | C | 70 | 0 | 12 | 0 | C | C | C | C | C | C | C | C | C | C | C | C |
| NC_011770 | 4258380 | C | T | 69 | 0 | 6 | 0 | T | T | T | T | T | T | | | | | | |
| NC_011770 | 4259002 | G | C | 37 | 0 | 2 | 0 | | | | | C | C | | | | | | |
| NC_011770 | 4259170 | G | A | 29 | 0 | 4 | 0 | | | A | A | A | A | | | | | | |
| NC_011770 | 4260802 | T | C | 64 | 0 | 6 | 0 | | | | | | | C | C | C | C | C | C |
| NC_011770 | 4262057 | G | A | 65 | 0 | 2 | 0 | A | A | | | | | | | | | | |
| NC_011770 | 4267185 | C | G | 65 | 0 | 2 | 0 | | | | | | | | | | | G | G |
| NC_011770 | 4267216 | T | C | 60 | 0 | 10 | 0 | | | C | C | C | C | C | C | C | C | C | C |
| NC_011770 | 4267840 | G | A | 82 | 0 | 10 | 0 | A | A | A | A | A | A | | | A | A | A | A |
| NC_011770 | 4267846 | C | T | 80 | 0 | 8 | 0 | T | T | T | T | T | T | | | T | T | | |
| NC_011770 | 4268282 | G | A | 77 | 0 | 4 | 0 | | | A | A | A | A | | | | | | |
| NC_011770 | 4270542 | G | A | 68 | 0 | 2 | 0 | | | | | | | A | A | | | | |
| NC_011770 | 4270567 | T | C | 62 | 0 | 8 | 0 | C | C | | | | | C | C | C | C | C | C |
| NC_011770 | 4270638 | G | C | 68 | 0 | 2 | 0 | | | | | | | C | C | | | | |
| NC_011770 | 4270654 | C | T | 70 | 0 | 2 | 0 | | | | | | | T | T | | | | |
| NC_011770 | 4270670 | A | T | 65 | 0 | 4 | 0 | | | T | T | T | T | | | | | | |
| NC_011770 | 4270726 | CT | C | 70 | 0 | 6 | 0 | C | C | | | | | | | C | C | C | C |
| NC_011770 | 4276125 | A | G | 69 | 0 | 2 | 0 | | | | | | | G | G | | | | |
| NC_011770 | 4276194 | C | T | 61 | 0 | 4 | 0 | | | T | T | T | T | | | | | | |
| NC_011770 | 4276198 | A | G | 61 | 0 | 10 | 0 | G | G | G | G | G | G | | | G | G | G | G |
| NC_011770 | 4276200 | G | A | 62 | 0 | 4 | 0 | | | A | A | A | A | | | | | | |
| NC_011770 | 4277837 | C | A | 88 | 0 | 4 | 0 | | | A | A | A | A | | | | | | |
| NC_011770 | 4278535 | G | A | 87 | 0 | 2 | 0 | | | | | | | | | A | A | | |
| NC_011770 | 4280364 | G | A | 79 | 0 | 4 | 0 | | | A | A | A | A | | | | | | |
| NC_011770 | 4280382 | C | T | 76 | 0 | 4 | 0 | | | T | T | T | T | | | | | | |
| NC_011770 | 4280399 | A | T | 73 | 0 | 12 | 0 | T | T | T | T | T | T | T | T | T | T | T | T |
| NC_011770 | 4281338 | A | G | 32 | 0 | 8 | 0 | G | G | | | | | G | G | G | G | G | G |
| NC_011770 | 4286795 | G | A | 32 | 0 | 1 | 0 | | | | | | | A | | | | | |
| NC_011770 | 4288953 | AG | A | 77 | 0 | 6 | 0 | | | A | A | A | A | | | A | A | | |
| NC_011770 | 4288959 | C | A | 71 | 0 | 1 | 0 | | | | | | | | | . | A | | |
| NC_011770 | 4289528 | T | C | 88 | 0 | 12 | 0 | C | C | C | C | C | C | C | C | C | C | C | C |
| NC_011770 | 4294330 | G | A | 53 | 0 | 4 | 0 | A | A | | | | | | | | | A | A |
| NC_011770 | 4294355 | T | C | 58 | 0 | 1 | 0 | | | | | | | C | | | | | |

| | | | | | | | | | | | | | | | | | | | |
|---|---|---|---|---|---|---|---|---|---|---|---|---|---|---|---|---|---|---|---|
| NC_011770 | 4296832 | A | G | 75 | 0 | 12 | 0 | G | G | G | G | G | G | G | G | G | G | G | G |
| NC_011770 | 4298342 | G | A | 82 | 0 | 2 | 0 | A | A | | | | | | | | | | |
| NC_011770 | 4298417 | T | C | 101 | 0 | 10 | 0 | C | C | C | C | C | C | C | C | C | C | | |
| NC_011770 | 4299629 | GA | G | 41 | 0 | 4 | 0 | | | | | | | | | G | G | G | G |
| NC_011770 | 4299656 | C | T | 44 | 0 | 3 | 0 | T | . | | | | | | | . | . | T | T |
| NC_011770 | 4299657 | A | C | 43 | 0 | 6 | 0 | C | C | | | | | | | C | C | C | C |
| NC_011770 | 4300498 | A | G | 83 | 0 | 2 | 0 | | | | | | | | | | | G | G |
| NC_011770 | 4300528 | G | A | 83 | 0 | 12 | 0 | A | A | A | A | A | A | A | A | A | A | A | A |
| NC_011770 | 4300562 | A | G | 88 | 0 | 2 | 0 | | | | | | | | | | | G | G |
| NC_011770 | 4300614 | G | T | 92 | 0 | 2 | 0 | | | | | | | | | | | T | T |
| NC_011770 | 4300650 | C | A | 88 | 0 | 2 | 0 | | | | | | | | | | | A | A |
| NC_011770 | 4302181 | A | G | 30 | 0 | 4 | 0 | | | G | G | G | G | | | | | | |
| NC_011770 | 4302219 | G | A | 29 | 0 | 4 | 0 | | | A | A | A | A | | | | | | |
| NC_011770 | 4302246 | G | A | 24 | 0 | 4 | 0 | | | A | A | A | A | | | | | | |
| NC_011770 | 4302251 | T | C | 25 | 0 | 6 | 0 | | | C | C | C | C | | | | | C | C |
| NC_011770 | 4302258 | C | G | 22 | 1 | 5 | 0 | | | G | G | G | G | | | | | G | . |
| NC_011770 | 4302259 | C | T | 22 | 1 | 5 | 0 | | | T | T | T | T | | | | | T | . |
| NC_011770 | 4302269 | G | A | 24 | 1 | 4 | 0 | | | A | A | A | A | | | | | | . |
| NC_011770 | 4302270 | T | C | 24 | 1 | 5 | 0 | | | C | C | C | C | | | | | C | . |
| NC_011770 | 4302274 | C | T | 23 | 2 | 4 | 0 | | | T | T | T | T | | | | | . | . |
| NC_011770 | 4302290 | T | G | 15 | 5 | 1 | 0 | | | . | G | . | . | | | | | . | . |
| NC_011770 | 4302331 | A | C | 24 | 0 | 6 | 0 | | | C | C | C | C | | | | | C | C |
| NC_011770 | 4302332 | C | T | 24 | 0 | 6 | 0 | | | T | T | T | T | | | | | T | T |
| NC_011770 | 4302351 | A | G | 25 | 2 | 2 | 0 | | | G | G | | | | | | | . | . |
| NC_011770 | 4302362 | T | C | 28 | 0 | 2 | 0 | | | C | C | | | | | | | | |
| NC_011770 | 4302372 | G | A | 26 | 0 | 2 | 0 | | | | | | | | | A | A | | |
| NC_011770 | 4302385 | G | A | 27 | 0 | 6 | 0 | | | A | A | A | A | | | | | A | A |
| NC_011770 | 4302387 | A | G | 26 | 0 | 6 | 0 | | | G | G | G | G | | | | | G | G |
| NC_011770 | 4302394 | G | A | 28 | 0 | 6 | 0 | | | A | A | A | A | | | | | A | A |
| NC_011770 | 4302898 | T | C | 43 | 0 | 8 | 0 | C | C | | | | | C | C | C | C | C | C |
| NC_011770 | 4302929 | A | G | 42 | 0 | 4 | 0 | | | G | G | G | G | | | | | | |
| NC_011770 | 4302988 | G | A | 42 | 0 | 4 | 0 | | | A | A | A | A | | | | | | |
| NC_011770 | 4303006 | C | A | 42 | 0 | 2 | 0 | | | | | | | | | A | A | | |
| NC_011770 | 4303017 | G | A | 40 | 0 | 4 | 0 | | | A | A | A | A | | | | | | |
| NC_011770 | 4305651 | C | T | 71 | 0 | 2 | 0 | T | T | | | | | | | | | | |
| NC_011770 | 4307526 | T | G | 72 | 0 | 2 | 0 | | | | | | | | | G | G | | |
| NC_011770 | 4307667 | T | C | 70 | 0 | 2 | 0 | | | | | | | | | C | C | | |
| NC_011770 | 4307785 | T | C | 70 | 0 | 6 | 0 | | | C | C | C | C | | | | | C | C |
| NC_011770 | 4307825 | G | A | 67 | 0 | 2 | 0 | | | | | | | | | | | A | A |
| NC_011770 | 4307903 | A | G | 65 | 0 | 4 | 0 | | | G | G | G | G | | | | | | |
| NC_011770 | 4307908 | T | C | 64 | 0 | 4 | 0 | C | C | | | | | | | | | C | C |
| NC_011770 | 4307929 | G | A | 63 | 0 | 4 | 0 | A | A | | | | | | | | | A | A |
| NC_011770 | 4307939 | C | T | 61 | 0 | 12 | 0 | T | T | T | T | T | T | T | T | T | T | T | T |
| NC_011770 | 4307953 | C | T | 59 | 0 | 4 | 0 | T | T | | | | | | | | | T | T |
| NC_011770 | 4308002 | G | C | 60 | 0 | 2 | 0 | | | | | | | | | | | C | C |
| NC_011770 | 4308075 | A | T | 61 | 0 | 2 | 0 | | | | | | | T | T | | | | |
| NC_011770 | 4308676 | A | G | 62 | 0 | 2 | 0 | | | | | | | G | G | | | | |
| NC_011770 | 4308710 | T | G | 54 | 0 | 4 | 0 | G | G | | | | | G | G | | | | |
| NC_011770 | 4308723 | A | C | 55 | 0 | 4 | 0 | C | C | | | | | C | C | | | | |
| NC_011770 | 4308729 | C | A | 53 | 0 | 4 | 0 | A | A | | | | | A | A | | | | |
| NC_011770 | 4308734 | C | CA | 55 | 0 | 4 | 0 | CA | CA | | | | | CA | CA | | | | |
| NC_011770 | 4308747 | A | G | 52 | 0 | 4 | 0 | G | G | | | | | G | G | | | | |
| NC_011770 | 4308748 | G | C | 53 | 0 | 4 | 0 | C | C | | | | | C | C | | | | |
| NC_011770 | 4308775 | C | T | 51 | 0 | 4 | 0 | T | T | | | | | T | T | | | | |
| NC_011770 | 4308779 | T | C | 50 | 0 | 4 | 0 | C | C | | | | | C | C | | | | |
| NC_011770 | 4308786 | C | T | 51 | 0 | 4 | 0 | T | T | | | | | T | T | | | | |
| NC_011770 | 4308790 | T | G | 51 | 0 | 2 | 0 | | | | | | | | | | | G | G |
| NC_011770 | 4308791 | G | T | 50 | 0 | 4 | 0 | T | T | | | | | T | T | | | | |
| NC_011770 | 4308794 | G | A | 52 | 0 | 4 | 0 | A | A | | | | | A | A | | | | |
| NC_011770 | 4308795 | G | A | 52 | 0 | 4 | 0 | A | A | | | | | A | A | | | | |
| NC_011770 | 4308802 | C | T | 52 | 0 | 4 | 0 | T | T | | | | | T | T | | | | |
| NC_011770 | 4308807 | G | A | 53 | 0 | 4 | 0 | A | A | | | | | A | A | | | | |

| | | | | | | | | | | | | | | | | | | | |
|---|---|---|---|---|---|---|---|---|---|---|---|---|---|---|---|---|---|---|---|
| NC_011770 | 4308810 | A | C | 53 | 0 | 4 | 0 | C | C | | | | | C | C | | | | |
| NC_011770 | 4308817 | A | G | 60 | 0 | 4 | 0 | G | G | | | | | G | G | | | | |
| NC_011770 | 983912 | A | AT | 46 | 0 | 2 | 0 | | | | | | | AT | | AT | | | |
| NC_011770 | 1631813 | T | TA | 57 | 0 | 7 | 0 | | | TA | TA | TA | TA | TA | TA | TA | | | |
| NC_011770 | 4314206 | A | G | 81 | 0 | 4 | 0 | | | | | | | G | G | G | G | | |
| NC_011770 | 4314501 | A | C | 70 | 0 | 10 | 0 | C | C | C | C | C | C | C | C | C | C | | |
| NC_011770 | 4314566 | T | C | 71 | 0 | 2 | 0 | | | | | | | | | | | C | C |
| NC_011770 | 4314572 | C | A | 70 | 0 | 10 | 0 | A | A | A | A | A | A | A | A | A | A | | |
| NC_011770 | 4314583 | A | C | 73 | 0 | 4 | 0 | C | C | | | | | | | | | C | C |
| NC_011770 | 1480210 | GA | G | 60 | 0 | 9 | 0 | G | G | G | G | G | | G | G | | | G | G |
| NC_011770 | 4314596 | C | G | 39 | 1 | 1 | 0 | | | G | . | . | . | | | | | | |
| NC_011770 | 4314629 | C | T | 70 | 0 | 4 | 0 | | | T | T | T | T | | | | | | |
| NC_011770 | 4314696 | G | A | 74 | 0 | 2 | 0 | | | | | | | | | A | A | | |
| NC_011770 | 4314721 | G | A | 79 | 0 | 2 | 0 | | | | | | | | | | | A | A |
| NC_011770 | 4314758 | A | C | 79 | 0 | 4 | 0 | | | C | C | C | C | | | | | | |
| NC_011770 | 4314804 | G | A | 75 | 0 | 2 | 0 | | | | | | | | | | | A | A |
| NC_011770 | 4314805 | G | T | 76 | 0 | 2 | 0 | T | T | | | | | | | | | | |
| NC_011770 | 4314811 | T | C | 76 | 0 | 2 | 0 | | | | | | | | | | | C | C |
| NC_011770 | 4314814 | C | T | 75 | 0 | 8 | 0 | T | T | T | T | T | T | | | | | T | T |
| NC_011770 | 4314826 | G | T | 74 | 0 | 2 | 0 | | | | | | | | | | | T | T |
| NC_011770 | 4316975 | C | T | 66 | 0 | 6 | 0 | T | T | | | | | T | T | T | T | | |
| NC_011770 | 4316984 | G | T | 66 | 0 | 6 | 0 | | | | | | | T | T | T | T | T | T |
| NC_011770 | 4317048 | T | C | 69 | 0 | 2 | 0 | | | | | | | | | | | C | C |
| NC_011770 | 4317053 | G | A | 69 | 0 | 2 | 0 | | | | | | | | | | | A | A |
| NC_011770 | 4317063 | A | G | 68 | 0 | 6 | 0 | | | G | G | G | G | | | | | G | G |
| NC_011770 | 4317070 | T | A | 68 | 0 | 2 | 0 | | | | | | | | | | | A | A |
| NC_011770 | 4317079 | G | A | 71 | 0 | 4 | 0 | | | A | A | A | A | | | | | | |
| NC_011770 | 4317114 | T | C | 71 | 0 | 8 | 0 | | | C | C | C | C | C | C | | | C | C |
| NC_011770 | 4318143 | CG | C | 73 | 0 | 6 | 0 | C | C | | | | | C | C | C | C | | |
| NC_011770 | 4318218 | G | C | 67 | 0 | 2 | 0 | | | | | | | | | C | C | | |
| NC_011770 | 4318220 | G | A | 68 | 0 | 6 | 0 | A | A | | | | | A | A | A | A | | |
| NC_011770 | 4319051 | A | T | 62 | 0 | 2 | 0 | | | | | | | | | | | T | T |
| NC_011770 | 4319060 | T | C | 68 | 0 | 2 | 0 | | | | | | | | | | | C | C |
| NC_011770 | 4319095 | C | T | 69 | 0 | 2 | 0 | | | | | | | T | T | | | | |
| NC_011770 | 4319335 | G | A | 69 | 0 | 4 | 0 | | | A | A | A | A | | | | | | |
| NC_011770 | 4319342 | A | G | 64 | 0 | 10 | 0 | G | G | G | G | G | G | G | G | G | G | | |
| NC_011770 | 4319343 | G | A | 63 | 0 | 12 | 0 | A | A | A | A | A | A | A | A | A | A | A | A |
| NC_011770 | 4319344 | T | G | 66 | 0 | 10 | 0 | G | G | G | G | G | G | G | G | G | G | | |
| NC_011770 | 4319367 | C | T | 70 | 0 | 8 | 0 | T | T | T | T | T | T | T | T | | | | |
| NC_011770 | 4319369 | T | C | 71 | 0 | 10 | 0 | C | C | C | C | C | C | C | C | C | C | | |
| NC_011770 | 4319403 | A | G | 70 | 0 | 10 | 0 | G | G | G | G | G | G | G | G | G | G | | |
| NC_011770 | 4319405 | G | A | 70 | 0 | 2 | 0 | | | | | | | | | A | A | | |
| NC_011770 | 4319896 | T | G | 68 | 0 | 2 | 0 | | | | | | | | | | | G | G |
| NC_011770 | 4319961 | C | A | 70 | 0 | 2 | 0 | | | | | | | | | A | A | | |
| NC_011770 | 4319971 | A | G | 66 | 0 | 2 | 0 | | | | | | | | | | | G | G |
| NC_011770 | 4319974 | A | G | 65 | 0 | 2 | 0 | | | | | | | G | G | | | | |
| NC_011770 | 4319995 | T | C | 63 | 0 | 2 | 0 | | | | | | | C | C | | | | |
| NC_011770 | 4320028 | A | G | 58 | 0 | 6 | 0 | G | G | | | | | G | G | G | G | | |
| NC_011770 | 4320029 | A | C | 59 | 0 | 6 | 0 | C | C | | | | | C | C | C | C | | |
| NC_011770 | 4320032 | G | T | 58 | 0 | 10 | 0 | T | T | T | T | T | T | T | T | T | T | | |
| NC_011770 | 4320033 | G | A | 59 | 0 | 8 | 0 | A | A | | | | | A | A | A | A | T | T |
| NC_011770 | 4320034 | T | C | 59 | 0 | 6 | 0 | C | C | | | | | C | C | C | C | | |
| NC_011770 | 4320037 | T | C | 58 | 0 | 6 | 0 | C | C | | | | | C | C | C | C | | |
| NC_011770 | 4320039 | C | A | 58 | 0 | 6 | 0 | A | A | | | | | A | A | A | A | | |
| NC_011770 | 4320053 | A | T | 61 | 0 | 10 | 0 | T | T | T | T | T | T | T | T | T | T | | |
| NC_011770 | 4320055 | G | A | 61 | 0 | 6 | 0 | A | A | | | | | A | A | A | A | | |
| NC_011770 | 4320062 | T | G | 61 | 0 | 6 | 0 | G | G | | | | | G | G | G | G | | |
| NC_011770 | 4320070 | G | T | 62 | 0 | 2 | 0 | T | T | | | | | | | | | | |
| NC_011770 | 4320071 | C | T | 62 | 0 | 2 | 0 | | | | | | | | | | | T | T |
| NC_011770 | 4320081 | C | A | 63 | 0 | 6 | 0 | A | A | | | | | A | A | A | A | | |
| NC_011770 | 4320131 | T | C | 72 | 0 | 2 | 0 | C | C | | | | | | | | | | |
| NC_011770 | 4322858 | C | T | 69 | 0 | 4 | 0 | | | T | T | T | T | | | | | | |

| | | | | | | | | | | | | | | | | | | | |
|---|---|---|---|---|---|---|---|---|---|---|---|---|---|---|---|---|---|---|---|
| NC_011770 | 4324051 | T | C | 66 | 0 | 4 | 0 | | | C | C | C | C | | | | | | |
| NC_011770 | 4326820 | T | TC | 58 | 0 | 1 | 0 | | | | | | | TC | | | | | |
| NC_011770 | 4326836 | G | A | 64 | 0 | 4 | 0 | | | A | A | A | A | | | | | | |
| NC_011770 | 4326928 | T | C | 60 | 0 | 1 | 0 | | | | | | | C | | | | | |
| NC_011770 | 4327067 | T | C | 40 | 0 | 2 | 0 | | | | | | | | | | | C | C |
| NC_011770 | 4327075 | A | C | 41 | 0 | 4 | 0 | | | C | C | C | C | | | | | | |
| NC_011770 | 4327077 | A | C | 43 | 0 | 4 | 0 | | | C | C | C | C | | | | | | |
| NC_011770 | 4327091 | G | A | 38 | 0 | 8 | 0 | | | A | A | A | A | A | A | | | A | A |
| NC_011770 | 4327138 | C | T | 46 | 0 | 6 | 0 | | | T | T | T | T | | | | | T | T |
| NC_011770 | 4329626 | G | A | 62 | 0 | 2 | 0 | | | | | | | | | | | A | A |
| NC_011770 | 4329649 | C | T | 65 | 0 | 2 | 0 | | | | | | | | | | | T | T |
| NC_011770 | 4329650 | G | A | 63 | 0 | 4 | 0 | | | A | A | A | A | | | | | | |
| NC_011770 | 4329660 | C | T | 69 | 0 | 4 | 0 | | | T | T | T | T | | | | | | |
| NC_011770 | 4329675 | G | A | 70 | 0 | 2 | 0 | | | | | | | | | | | A | A |
| NC_011770 | 4329702 | C | T | 70 | 0 | 4 | 0 | | | T | T | T | T | | | | | | |
| NC_011770 | 4331454 | G | T | 77 | 0 | 2 | 0 | T | T | | | | | | | | | | |
| NC_011770 | 4331480 | C | T | 72 | 0 | 2 | 0 | T | T | | | | | | | | | | |
| NC_011770 | 4332521 | T | C | 47 | 0 | 2 | 0 | | | | | | | | | | | C | C |
| NC_011770 | 4332586 | C | A | 50 | 0 | 2 | 0 | | | | | | | | | A | A | | |
| NC_011770 | 4332962 | G | GA | 71 | 0 | 2 | 0 | | | | | | | | | | | GA | GA |
| NC_011770 | 4332999 | G | A | 66 | 0 | 2 | 0 | | | | | | | | | | | A | A |
| NC_011770 | 4334372 | C | T | 56 | 0 | 2 | 0 | | | | | | | | | T | T | | |
| NC_011770 | 4334383 | G | T | 56 | 0 | 2 | 0 | | | | | | | | | | | T | T |
| NC_011770 | 4334387 | G | A | 55 | 0 | 2 | 0 | | | | | | | A | A | | | | |
| NC_011770 | 4334391 | G | T | 54 | 0 | 2 | 0 | | | T | T | | | | | | | | |
| NC_011770 | 4335376 | C | G | 47 | 0 | 2 | 0 | G | G | | | | | | | | | | |
| NC_011770 | 4335389 | A | C | 50 | 0 | 2 | 0 | | | | | | | | | | | C | C |
| NC_011770 | 4335393 | G | A | 56 | 0 | 2 | 0 | A | A | | | | | | | | | | |
| NC_011770 | 4335416 | C | T | 55 | 0 | 2 | 0 | T | T | | | | | | | | | | |
| NC_011770 | 4335480 | G | A | 62 | 0 | 2 | 0 | A | A | | | | | | | | | | |
| NC_011770 | 4335516 | G | T | 63 | 0 | 2 | 0 | | | | | | | | | T | T | | |
| NC_011770 | 4335517 | C | A | 63 | 0 | 2 | 0 | | | | | | | | | A | A | | |
| NC_011770 | 4335653 | G | A | 67 | 0 | 4 | 0 | | | A | A | A | A | | | | | | |
| NC_011770 | 4342136 | T | G | 76 | 0 | 2 | 0 | G | G | | | | | | | | | | |
| NC_011770 | 4342213 | G | A | 73 | 0 | 2 | 0 | | | | | | | | | | | A | A |
| NC_011770 | 4349237 | C | A | 59 | 0 | 4 | 0 | | | A | A | A | A | | | | | | |
| NC_011770 | 4350884 | A | G | 17 | 1 | 9 | 0 | G | G | G | G | G | G | | | . | G | G | G |
| NC_011770 | 4350885 | T | C | 18 | 1 | 9 | 0 | C | C | C | C | C | C | | | . | C | C | C |
| NC_011770 | 4352771 | GC | G | 57 | 0 | 4 | 0 | | | | | | | G | G | G | G | | |
| NC_011770 | 4354693 | T | A | 80 | 0 | 2 | 0 | | | | | | | A | A | | | | |
| NC_011770 | 4354737 | G | A | 71 | 0 | 4 | 0 | | | A | A | A | A | | | | | | |
| NC_011770 | 4354739 | G | A | 71 | 0 | 4 | 0 | | | A | A | A | A | | | | | | |
| NC_011770 | 4354756 | G | C | 72 | 0 | 4 | 0 | | | C | C | C | C | | | | | | |
| NC_011770 | 4355590 | C | T | 69 | 0 | 10 | 0 | T | T | T | T | T | T | T | T | T | T | | |
| NC_011770 | 4355596 | A | G | 71 | 0 | 4 | 0 | | | G | G | G | G | | | | | | |
| NC_011770 | 4357092 | C | T | 68 | 0 | 2 | 0 | | | | | T | T | | | | | | |
| NC_011770 | 4357093 | G | A | 67 | 0 | 10 | 0 | A | A | A | A | A | A | A | A | A | A | | |
| NC_011770 | 4357127 | C | A | 73 | 0 | 2 | 0 | | | | | | | A | A | | | | |
| NC_011770 | 4357130 | T | C | 73 | 0 | 10 | 0 | C | C | C | C | C | C | C | C | C | C | | |
| NC_011770 | 4357149 | G | A | 74 | 0 | 2 | 0 | | | | | | | A | A | | | | |
| NC_011770 | 4362413 | C | G | 83 | 0 | 2 | 0 | | | | | | | G | G | | | | |
| NC_011770 | 4365981 | G | A | 60 | 0 | 2 | 0 | | | | | | | | | A | A | | |
| NC_011770 | 4372673 | T | C | 63 | 0 | 4 | 0 | | | C | C | C | C | | | | | | |
| NC_011770 | 4372683 | C | T | 62 | 0 | 2 | 0 | | | | | | | | | | | T | T |
| NC_011770 | 4372709 | G | GA | 64 | 0 | 2 | 0 | | | | | | | | | | | GA | GA |
| NC_011770 | 4375381 | G | A | 78 | 0 | 6 | 0 | A | A | A | A | A | A | | | | | | |
| NC_011770 | 4375386 | G | A | 80 | 0 | 2 | 0 | A | A | | | | | | | | | | |
| NC_011770 | 4375418 | T | C | 82 | 0 | 2 | 0 | | | | | | | | | | | C | C |
| NC_011770 | 4380967 | T | C | 73 | 0 | 2 | 0 | | | | | | | C | C | | | | |
| NC_011770 | 4380974 | A | G | 75 | 0 | 2 | 0 | | | | | | | G | G | | | | |
| NC_011770 | 4380982 | C | T | 76 | 0 | 2 | 0 | T | T | | | | | | | | | | |
| NC_011770 | 4380983 | A | G | 73 | 0 | 10 | 0 | G | G | G | G | G | G | G | G | | | G | G |

| | | | | | | | | | | | | | | | | | | | |
|---|---|---|---|---|---|---|---|---|---|---|---|---|---|---|---|---|---|---|---|
| NC_011770 | 4381060 | T | C | 71 | 0 | 2 | 0 | | | | | | | C | C | | | | |
| NC_011770 | 4383602 | G | T | 79 | 0 | 10 | 0 | T | T | T | T | T | T | | | T | T | T | T |
| NC_011770 | 4390107 | G | C | 11 | 1 | 2 | 0 | | | | | | | C | C | | . | | |
| NC_011770 | 4390109 | G | A | 11 | 1 | 4 | 0 | | | A | A | A | A | | | | . | | |
| NC_011770 | 4393037 | A | G | 82 | 0 | 12 | 0 | G | G | G | G | G | G | G | G | G | G | G | G |
| NC_011770 | 4393699 | A | T | 66 | 0 | 12 | 0 | T | T | T | T | T | T | T | T | T | T | T | T |
| NC_011770 | 1803928 | G | A | 65 | 0 | 1 | 0 | | | | | A | | | | | | | |
| NC_011770 | 4394905 | G | A | 94 | 0 | 2 | 0 | | | | | | | | | | | A | A |
| NC_011770 | 4394971 | T | G | 96 | 0 | 4 | 0 | | | G | G | G | G | | | | | | |
| NC_011770 | 4395051 | G | A | 98 | 0 | 4 | 0 | | | A | A | A | A | | | | | | |
| NC_011770 | 4395070 | G | A | 94 | 0 | 1 | 0 | | | | | | | A | | | | | |
| NC_011770 | 4395116 | G | A | 96 | 0 | 4 | 0 | A | A | | | | | | | A | A | | |
| NC_011770 | 4395140 | C | G | 89 | 0 | 12 | 0 | G | G | G | G | G | G | G | G | G | G | G | G |
| NC_011770 | 4395154 | G | A | 89 | 0 | 4 | 0 | A | A | | | | | | | A | A | | |
| NC_011770 | 4395157 | G | A | 88 | 0 | 12 | 0 | A | A | A | A | A | A | A | A | A | A | A | A |
| NC_011770 | 4395177 | G | T | 89 | 0 | 4 | 0 | T | T | | | | | | | T | T | | |
| NC_011770 | 4398620 | A | G | 55 | 0 | 8 | 0 | | | G | G | G | G | G | G | | | G | G |
| NC_011770 | 4398639 | A | G | 55 | 0 | 6 | 0 | | | G | G | G | G | G | G | | | | |
| NC_011770 | 4398905 | A | C | 50 | 2 | 2 | 0 | C | C | | | | | . | . | | | | |
| NC_011770 | 4399062 | C | T | 52 | 2 | 2 | 0 | | | | | | | . | . | T | T | | |
| NC_011770 | 4399107 | G | A | 53 | 2 | 4 | 0 | | | A | A | A | A | . | . | | | | |
| NC_011770 | 4399206 | G | A | 52 | 2 | 4 | 0 | | | A | A | A | A | . | . | | | | |
| NC_011770 | 4399353 | A | C | 48 | 2 | 8 | 0 | C | C | C | C | C | C | . | . | | | C | C |
| NC_011770 | 4399419 | C | A | 49 | 2 | 4 | 0 | A | A | | | | | . | . | | | A | A |
| NC_011770 | 4399545 | C | G | 53 | 2 | 2 | 0 | | | | | | | . | . | G | G | | |
| NC_011770 | 4399654 | G | A | 50 | 2 | 4 | 0 | | | A | A | A | A | . | . | | | | |
| NC_011770 | 4399742 | A | G | 53 | 2 | 10 | 0 | G | G | G | G | G | G | . | . | G | G | G | G |
| NC_011770 | 4401975 | C | T | 66 | 0 | 4 | 0 | | | T | T | T | T | | | | | | |
| NC_011770 | 4402251 | G | A | 73 | 0 | 2 | 0 | A | A | | | | | | | | | | |
| NC_011770 | 4403102 | A | G | 18 | 0 | 2 | 0 | | | | | | | | | G | G | | |
| NC_011770 | 4403104 | G | T | 17 | 0 | 2 | 0 | | | | | | | | | | | T | T |
| NC_011770 | 4405481 | C | CG | 55 | 0 | 2 | 0 | | | | | | | | | | | CG | CG |
| NC_011770 | 4405496 | G | A | 54 | 0 | 2 | 0 | | | | | | | | | | | A | A |
| NC_011770 | 4405499 | A | T | 53 | 0 | 4 | 0 | | | | | | | | | T | T | T | T |
| NC_011770 | 4405502 | A | G | 52 | 0 | 4 | 0 | | | | | | | | | G | G | G | G |
| NC_011770 | 4405526 | G | A | 54 | 0 | 2 | 0 | | | | | | | | | A | A | | |
| NC_011770 | 4405630 | G | A | 44 | 0 | 2 | 0 | | | | | | | | | A | A | | |
| NC_011770 | 4406632 | G | A | 79 | 0 | 2 | 0 | | | | | | | | | | | A | A |
| NC_011770 | 4408603 | G | T | 83 | 0 | 12 | 0 | T | T | T | T | T | T | T | T | T | T | T | T |
| NC_011770 | 4413165 | T | C | 87 | 0 | 2 | 0 | | | | | | | | | | | C | C |
| NC_011770 | 4413927 | G | A | 57 | 0 | 12 | 0 | A | A | A | A | A | A | A | A | A | A | A | A |
| NC_011770 | 4415149 | G | A | 65 | 0 | 2 | 0 | | | | | | | | | A | A | | |
| NC_011770 | 4415203 | G | T | 61 | 0 | 6 | 0 | T | T | T | T | T | T | | | | | | |
| NC_011770 | 4417330 | G | A | 80 | 0 | 1 | 0 | | | | | | | A | | | | | |
| NC_011770 | 4417333 | G | A | 81 | 0 | 4 | 0 | | | A | A | A | A | | | | | | |
| NC_011770 | 4417431 | C | T | 78 | 0 | 4 | 0 | | | T | T | T | T | | | | | | |
| NC_011770 | 4417436 | G | C | 77 | 0 | 4 | 0 | | | C | C | C | C | | | | | | |
| NC_011770 | 4417437 | G | A | 76 | 0 | 4 | 0 | | | A | A | A | A | | | | | | |
| NC_011770 | 4417442 | G | A | 77 | 0 | 4 | 0 | | | A | A | A | A | | | | | | |
| NC_011770 | 4417452 | T | C | 77 | 0 | 4 | 0 | | | C | C | C | C | | | | | | |
| NC_011770 | 4417456 | T | C | 77 | 0 | 4 | 0 | | | C | C | C | C | | | | | | |
| NC_011770 | 4417483 | C | G | 83 | 0 | 4 | 0 | | | G | G | G | G | | | | | | |
| NC_011770 | 4417504 | TGGCCGC | T | 82 | 0 | 4 | 0 | | | T | T | T | T | | | | | | |
| NC_011770 | 4419785 | C | A | 65 | 0 | 2 | 0 | | | | | | | | | A | A | | |
| NC_011770 | 4419816 | C | T | 67 | 0 | 2 | 0 | | | | | | | | | T | T | | |
| NC_011770 | 2259052 | CA | C | 48 | 0 | 7 | 0 | C | C | C | C | C | | | | C | C | | |
| NC_011770 | 4422300 | A | G | 71 | 0 | 12 | 0 | G | G | G | G | G | G | G | G | G | G | G | G |
| NC_011770 | 4422310 | G | T | 73 | 0 | 4 | 0 | | | | | | | | | T | T | T | T |
| NC_011770 | 4423320 | T | C | 52 | 0 | 2 | 0 | | | | | | | | | | | C | C |
| NC_011770 | 4424106 | G | A | 69 | 0 | 2 | 0 | | | | | | | | | | | A | A |
| NC_011770 | 4425740 | G | A | 73 | 0 | 12 | 0 | A | A | A | A | A | A | A | A | A | A | A | A |
| NC_011770 | 4425750 | G | A | 73 | 0 | 12 | 0 | A | A | A | A | A | A | A | A | A | A | A | A |

| | | | | | | | | | | | | | | | | | | | |
|---|---|---|---|---|---|---|---|---|---|---|---|---|---|---|---|---|---|---|---|
| NC_011770 | 4425778 | C | T | 69 | 0 | 10 | 0 | T | T | T | T | T | T | T | T | | | T | T |
| NC_011770 | 4425838 | C | G | 69 | 0 | 2 | 0 | | | | | | | | | G | G | | |
| NC_011770 | 4425840 | C | T | 68 | 0 | 10 | 0 | T | T | T | T | T | T | T | T | | | T | T |
| NC_011770 | 4426060 | A | T | 69 | 0 | 2 | 0 | | | | | | | | | T | T | | |
| NC_011770 | 4426125 | C | G | 67 | 0 | 2 | 0 | G | G | | | | | | | | | | |
| NC_011770 | 4426472 | G | A | 56 | 0 | 8 | 0 | A | A | | | | | A | A | A | A | A | A |
| NC_011770 | 4426484 | T | TC | 56 | 0 | 12 | 0 | TC | TC | TC | TC | TC | TC | TC | TC | TC | TC | TC | TC |
| NC_011770 | 4426493 | C | A | 57 | 0 | 8 | 0 | A | A | | | | | A | A | A | A | A | A |
| NC_011770 | 4426494 | AC | A | 56 | 0 | 4 | 0 | | | A | A | A | A | | | | | | |
| NC_011770 | 4426517 | C | A | 57 | 0 | 10 | 0 | A | A | A | A | A | A | | | A | A | A | A |
| NC_011770 | 4426521 | C | G | 59 | 0 | 4 | 0 | | | G | G | G | G | | | | | | |
| NC_011770 | 4426528 | C | G | 60 | 0 | 2 | 0 | G | G | | | | | | | | | | |
| NC_011770 | 4426532 | T | C | 59 | 0 | 4 | 0 | | | C | C | C | C | | | | | | |
| NC_011770 | 4426536 | TG | T | 59 | 0 | 6 | 0 | | | | | | | T | T | T | T | T | T |
| NC_011770 | 4426575 | A | C | 58 | 0 | 4 | 0 | | | | | | | C | C | C | C | | |
| NC_011770 | 4426577 | G | A | 59 | 0 | 2 | 0 | | | | | | | | | | | A | A |
| NC_011770 | 4426581 | C | T | 61 | 0 | 4 | 0 | T | T | | | | | | | | | T | T |
| NC_011770 | 4427249 | T | G | 74 | 0 | 4 | 0 | G | G | | | | | | | | | G | G |
| NC_011770 | 4428021 | T | C | 88 | 0 | 2 | 0 | | | | | | | C | C | | | | |
| NC_011770 | 4431819 | G | A | 73 | 0 | 2 | 0 | | | | | | | A | A | | | | |
| NC_011770 | 4431844 | C | A | 73 | 0 | 2 | 0 | A | A | | | | | | | | | | |
| NC_011770 | 4431900 | A | C | 72 | 0 | 2 | 0 | | | | | | | | | | | C | C |
| NC_011770 | 4431901 | C | A | 72 | 0 | 2 | 0 | | | | | | | A | A | | | | |
| NC_011770 | 4431912 | C | T | 78 | 0 | 2 | 0 | T | T | | | | | | | | | | |
| NC_011770 | 4431917 | A | G | 77 | 0 | 8 | 0 | G | G | | | | | G | G | G | G | G | G |
| NC_011770 | 4431927 | A | G | 71 | 0 | 2 | 0 | G | G | | | | | | | | | | |
| NC_011770 | 4431938 | T | G | 70 | 0 | 2 | 0 | | | | | | | | | | | G | G |
| NC_011770 | 4431968 | G | A | 70 | 0 | 2 | 0 | | | | | | | | | | | A | A |
| NC_011770 | 4432916 | A | T | 82 | 0 | 4 | 0 | | | T | T | T | T | | | | | | |
| NC_011770 | 4433285 | T | A | 71 | 0 | 2 | 0 | | | | | | | | | | | A | A |
| NC_011770 | 4433314 | G | A | 69 | 0 | 4 | 0 | A | A | | | | | | | A | A | | |
| NC_011770 | 4433319 | C | T | 60 | 2 | 2 | 0 | | | | | | | T | T | | | . | . |
| NC_011770 | 4434096 | A | C | 87 | 0 | 12 | 0 | C | C | C | C | C | C | C | C | C | C | C | C |
| NC_011770 | 4435741 | G | A | 89 | 0 | 10 | 0 | A | A | A | A | A | A | A | A | A | A | | |
| NC_011770 | 4435748 | A | C | 87 | 0 | 4 | 0 | C | C | | | | | | | C | C | | |
| NC_011770 | 4435756 | G | A | 88 | 0 | 4 | 0 | A | A | | | | | | | A | A | | |
| NC_011770 | 4435759 | C | T | 89 | 0 | 2 | 0 | | | | | | | T | T | | | | |
| NC_011770 | 4435760 | G | T | 88 | 0 | 6 | 0 | | | T | T | T | T | T | T | | | | |
| NC_011770 | 4436673 | C | T | 40 | 6 | 2 | 0 | . | . | | | | | . | . | . | . | T | T |
| NC_011770 | 4436681 | A | G | 40 | 6 | 2 | 0 | . | . | | | | | . | . | . | . | G | G |
| NC_011770 | 4436721 | A | T | 39 | 6 | 2 | 0 | . | . | | | | | . | . | . | . | T | T |
| NC_011770 | 4437003 | T | C | 49 | 0 | 12 | 0 | C | C | C | C | C | C | C | C | C | C | C | C |
| NC_011770 | 4437082 | C | T | 54 | 0 | 10 | 0 | T | T | T | T | T | T | T | T | | | T | T |
| NC_011770 | 4438843 | T | G | 88 | 0 | 2 | 0 | | | | | | | G | G | | | | |
| NC_011770 | 4438845 | C | T | 88 | 0 | 6 | 0 | | | | | | | T | T | T | T | T | T |
| NC_011770 | 4438945 | G | A | 90 | 0 | 4 | 0 | | | | | | | | | A | A | A | A |
| NC_011770 | 4439575 | G | A | 82 | 0 | 2 | 0 | | | | | | | A | A | | | | |
| NC_011770 | 4439595 | G | A | 82 | 0 | 4 | 0 | | | | | | | | | A | A | A | A |
| NC_011770 | 4439606 | C | T | 84 | 0 | 4 | 0 | | | | | | | | | T | T | T | T |
| NC_011770 | 4439607 | T | C | 82 | 0 | 6 | 0 | | | | | | | C | C | C | C | C | C |
| NC_011770 | 4439651 | T | C | 83 | 0 | 2 | 0 | | | | | | | | | | | C | C |
| NC_011770 | 4439654 | C | T | 84 | 0 | 2 | 0 | | | | | | | | | | | T | T |
| NC_011770 | 4440321 | C | T | 88 | 0 | 2 | 0 | T | T | | | | | | | | | | |
| NC_011770 | 4441727 | A | G | 73 | 0 | 10 | 0 | G | G | G | G | G | G | G | G | G | G | | |
| NC_011770 | 4444251 | T | C | 90 | 0 | 2 | 0 | | | | | | | C | C | | | | |
| NC_011770 | 4446189 | AGC | A | 61 | 0 | 2 | 0 | | | | | | | | | A | A | | |
| NC_011770 | 4446819 | C | T | 73 | 0 | 4 | 0 | T | T | | | | | | | T | T | | |
| NC_011770 | 4446825 | C | T | 76 | 0 | 2 | 0 | | | | | | | | | T | T | | |
| NC_011770 | 4446860 | T | A | 77 | 0 | 4 | 0 | A | A | | | | | | | A | A | | |
| NC_011770 | 4449094 | G | A | 87 | 0 | 6 | 0 | | | A | A | A | A | A | A | | | | |
| NC_011770 | 4456622 | C | G | 70 | 0 | 4 | 0 | G | G | | | | | G | G | | | | |
| NC_011770 | 4457268 | T | G | 80 | 0 | 10 | 0 | G | G | G | G | G | G | G | G | | | G | G |

| | | | | | | | | | | | | | | | | | | | |
|---|---|---|---|---|---|---|---|---|---|---|---|---|---|---|---|---|---|---|---|
| NC_011770 | 4457300 | C | A | 82 | 0 | 2 | 0 | | | | | | | | | | | A | A |
| NC_011770 | 4459158 | G | A | 90 | 0 | 2 | 0 | | | | | | | A | A | | | | |
| NC_011770 | 4459227 | A | G | 89 | 0 | 4 | 0 | | | G | G | G | G | | | | | | |
| NC_011770 | 4459326 | T | G | 92 | 0 | 12 | 0 | G | G | G | G | G | G | G | G | G | G | G | G |
| NC_011770 | 4461610 | T | C | 70 | 0 | 2 | 0 | C | C | | | | | | | | | | |
| NC_011770 | 4461614 | T | C | 73 | 0 | 12 | 0 | C | C | C | C | C | C | C | C | C | C | C | C |
| NC_011770 | 4461649 | A | G | 53 | 0 | 4 | 0 | | | G | G | G | G | | | | | | |
| NC_011770 | 4461666 | T | C | 4 | 10 | 2 | 0 | . | . | . | . | . | . | . | . | C | C | . | . |
| NC_011770 | 4461726 | A | G | 64 | 0 | 8 | 0 | G | G | | | | | G | G | G | G | G | G |
| NC_011770 | 4461731 | A | G | 65 | 0 | 8 | 0 | | | G | G | G | G | G | G | | | G | G |
| NC_011770 | 4461761 | C | T | 70 | 0 | 2 | 0 | | | | | | | T | T | | | | |
| NC_011770 | 4461772 | G | C | 68 | 0 | 6 | 0 | C | C | | | | | C | C | | | C | C |
| NC_011770 | 4461773 | C | G | 68 | 0 | 6 | 0 | G | G | | | | | G | G | | | G | G |
| NC_011770 | 4461778 | G | A | 69 | 0 | 6 | 0 | A | A | | | | | A | A | | | A | A |
| NC_011770 | 4461780 | A | G | 68 | 0 | 10 | 0 | G | G | G | G | G | G | G | G | | | G | G |
| NC_011770 | 4461817 | C | T | 81 | 0 | 6 | 0 | T | T | | | | | T | T | | | T | T |
| NC_011770 | 4461826 | C | T | 82 | 0 | 6 | 0 | T | T | | | | | T | T | | | T | T |
| NC_011770 | 4462525 | C | A | 69 | 0 | 8 | 0 | A | A | | | | | A | A | A | A | A | A |
| NC_011770 | 4462545 | C | T | 63 | 0 | 8 | 0 | T | T | | | | | T | T | T | T | T | T |
| NC_011770 | 4462546 | T | C | 63 | 0 | 8 | 0 | C | C | | | | | C | C | C | C | C | C |
| NC_011770 | 4462575 | G | A | 59 | 0 | 4 | 0 | | | A | A | A | A | | | | | | |
| NC_011770 | 4462576 | C | T | 58 | 0 | 8 | 0 | T | T | | | | | T | T | T | T | T | T |
| NC_011770 | 4462610 | G | A | 55 | 0 | 2 | 0 | A | A | | | | | | | | | | |
| NC_011770 | 4464767 | G | A | 60 | 0 | 2 | 0 | | | | | | | | | A | A | | |
| NC_011770 | 4468084 | C | T | 63 | 0 | 12 | 0 | T | T | T | T | T | T | T | T | T | T | T | T |
| NC_011770 | 4468095 | G | T | 66 | 0 | 2 | 0 | T | T | | | | | | | | | | |
| NC_011770 | 4468098 | G | A | 66 | 0 | 2 | 0 | A | A | | | | | | | | | | |
| NC_011770 | 4468136 | G | A | 67 | 0 | 2 | 0 | | | | | | | | | | | A | A |
| NC_011770 | 4468193 | G | A | 66 | 0 | 6 | 0 | | | A | A | A | A | | | A | A | | |
| NC_011770 | 4468213 | C | G | 64 | 0 | 12 | 0 | G | G | G | G | G | G | G | G | G | G | G | G |
| NC_011770 | 4468270 | A | G | 66 | 0 | 12 | 0 | G | G | G | G | G | G | G | G | G | G | G | G |
| NC_011770 | 4468276 | C | G | 66 | 0 | 12 | 0 | G | G | G | G | G | G | G | G | G | G | G | G |
| NC_011770 | 4468295 | T | A | 66 | 0 | 2 | 0 | | | | | | | | | | | A | A |
| NC_011770 | 4468329 | C | A | 68 | 0 | 2 | 0 | | | | | | | | | | | A | A |
| NC_011770 | 4469264 | T | C | 57 | 0 | 12 | 0 | C | C | C | C | C | C | C | C | C | C | C | C |
| NC_011770 | 4469270 | C | T | 56 | 0 | 2 | 0 | | | | | | | T | T | | | | |
| NC_011770 | 4469288 | T | C | 58 | 0 | 4 | 0 | | | C | C | C | C | | | | | | |
| NC_011770 | 4469292 | T | C | 59 | 0 | 2 | 0 | | | | | | | C | C | | | | |
| NC_011770 | 4470759 | G | A | 24 | 0 | 2 | 0 | | | | | | | | | | | A | A |
| NC_011770 | 4470816 | G | A | 15 | 0 | 4 | 0 | | | A | A | A | A | | | | | | |
| NC_011770 | 4471686 | A | C | 62 | 0 | 6 | 0 | C | C | | | | | C | C | | | C | C |
| NC_011770 | 3028877 | GGCGCGCG | G | 40 | 0 | 6 | 0 | . | G | G | G | G | | | | | | G | G |
| NC_011770 | 4471742 | C | G | 60 | 0 | 4 | 0 | | | G | G | G | G | | | | | | |
| NC_011770 | 4471778 | TA | T | 69 | 0 | 8 | 0 | T | T | T | T | T | T | | | T | T | | |
| NC_011770 | 4471799 | C | T | 64 | 0 | 4 | 0 | | | T | T | T | T | | | | | | |
| NC_011770 | 4472695 | C | T | 78 | 0 | 2 | 0 | T | T | | | | | | | | | | |
| NC_011770 | 4476514 | G | A | 58 | 0 | 4 | 0 | | | A | A | A | A | | | | | | |
| NC_011770 | 4477638 | C | T | 44 | 0 | 2 | 0 | T | T | | | | | | | | | | |
| NC_011770 | 4477683 | C | T | 44 | 0 | 2 | 0 | | | T | T | | | | | | | | |
| NC_011770 | 4477687 | T | C | 45 | 0 | 6 | 0 | C | C | C | C | C | C | | | | | | |
| NC_011770 | 4477711 | A | G | 45 | 0 | 8 | 0 | G | G | G | G | G | G | C | C | | | | |
| NC_011770 | 4478645 | A | G | 51 | 0 | 10 | 0 | G | G | G | G | G | G | G | G | | | G | G |
| NC_011770 | 4478650 | C | G | 50 | 0 | 6 | 0 | G | G | | | | | G | G | | | G | G |
| NC_011770 | 4478673 | G | A | 53 | 0 | 4 | 0 | A | A | | | | | | | | | A | A |
| NC_011770 | 4478690 | G | A | 51 | 0 | 4 | 0 | A | A | | | | | | | | | A | A |
| NC_011770 | 4480325 | G | A | 75 | 0 | 2 | 0 | | | | | | | | | | | A | A |
| NC_011770 | 4481439 | G | A | 59 | 0 | 4 | 0 | | | A | A | A | A | | | | | | |
| NC_011770 | 4481444 | A | G | 58 | 0 | 2 | 0 | | | | | | | | | G | G | | |
| NC_011770 | 4481466 | A | G | 54 | 0 | 4 | 0 | G | G | | | | | | | | | G | G |
| NC_011770 | 4481555 | A | G | 53 | 0 | 4 | 0 | | | G | G | G | G | | | | | | |
| NC_011770 | 4481579 | C | A | 45 | 0 | 4 | 0 | A | A | | | | | | | | | A | A |
| NC_011770 | 4481582 | C | A | 47 | 0 | 4 | 0 | A | A | | | | | | | | | A | A |

| | | | | | | | | | | | | | | | | | | | |
|---|---|---|---|---|---|---|---|---|---|---|---|---|---|---|---|---|---|---|---|
| NC_011770 | 4481594 | T | C | 47 | 0 | 4 | 0 | | | C | C | C | C | | | | | | |
| NC_011770 | 4481613 | A | G | 55 | 0 | 4 | 0 | | | G | G | G | G | | | | | | |
| NC_011770 | 4483283 | A | G | 64 | 0 | 8 | 0 | G | G | G | G | G | G | | | | | G | G |
| NC_011770 | 4483921 | T | C | 69 | 0 | 12 | 0 | C | C | C | C | C | C | C | C | C | C | C | C |
| NC_011770 | 4483922 | G | T | 68 | 0 | 12 | 0 | T | T | T | T | T | T | T | T | T | T | T | T |
| NC_011770 | 4485500 | C | T | 79 | 0 | 12 | 0 | T | T | T | T | T | T | T | T | T | T | T | T |
| NC_011770 | 1784708 | TCCGCCCG | T | 61 | 0 | 7 | 0 | | | T | T | T | T | T | T | T | | | |
| NC_011770 | 4485577 | T | C | 73 | 0 | 6 | 0 | | | C | C | C | C | | | C | C | | |
| NC_011770 | 4485632 | A | G | 75 | 0 | 2 | 0 | | | | | | | | | G | G | | |
| NC_011770 | 4485645 | A | G | 78 | 0 | 2 | 0 | | | | | | | | | G | G | | |
| NC_011770 | 4485649 | G | A | 78 | 0 | 2 | 0 | | | | | | | | | A | A | | |
| NC_011770 | 4485661 | C | T | 79 | 0 | 2 | 0 | | | | | | | | | T | T | | |
| NC_011770 | 4485711 | T | C | 81 | 0 | 10 | 0 | C | C | C | C | C | C | C | C | | | C | C |
| NC_011770 | 4491761 | GC | CC | 60 | 0 | 3 | 0 | | | | | | | CC | | | | G | G |
| NC_011770 | 4491847 | A | G | 52 | 0 | 12 | 0 | G | G | G | G | G | G | G | G | G | G | G | G |
| NC_011770 | 4491850 | A | T | 52 | 0 | 2 | 0 | T | T | | | | | | | | | | |
| NC_011770 | 4491906 | A | G | 49 | 0 | 4 | 0 | | | G | G | G | G | | | | | | |
| NC_011770 | 4492024 | G | A | 48 | 0 | 12 | 0 | A | A | A | A | A | A | A | A | A | A | A | A |
| NC_011770 | 4492095 | C | A | 43 | 0 | 4 | 0 | | | | | | | A | A | | | A | A |
| NC_011770 | 4492096 | A | C | 43 | 0 | 2 | 0 | C | C | | | | | | | | | | |
| NC_011770 | 4492104 | C | A | 42 | 0 | 2 | 0 | A | A | | | | | | | | | | |
| NC_011770 | 4492194 | A | G | 35 | 0 | 12 | 0 | G | G | G | G | G | G | G | G | G | G | G | G |
| NC_011770 | 4492239 | G | A | 30 | 0 | 2 | 0 | | | | | | | | | | | A | A |
| NC_011770 | 4492265 | A | C | 26 | 0 | 4 | 0 | | | | | | | C | C | | | C | C |
| NC_011770 | 4492945 | G | T | 36 | 0 | 2 | 0 | | | | | | | | | T | T | | |
| NC_011770 | 4492946 | A | G | 36 | 0 | 8 | 0 | G | G | G | G | G | G | | | G | G | | |
| NC_011770 | 4492982 | T | G | 36 | 0 | 6 | 0 | G | G | | | | | G | G | | | G | G |
| NC_011770 | 4492986 | C | T | 36 | 0 | 6 | 0 | T | T | | | | | T | T | | | T | T |
| NC_011770 | 4492989 | A | G | 36 | 0 | 6 | 0 | G | G | | | | | G | G | | | G | G |
| NC_011770 | 4493026 | A | G | 36 | 0 | 6 | 0 | G | G | | | | | G | G | | | G | G |
| NC_011770 | 4493034 | A | G | 36 | 0 | 6 | 0 | G | G | | | | | G | G | | | G | G |
| NC_011770 | 4493046 | A | G | 33 | 0 | 2 | 0 | | | | | | | | | G | G | | |
| NC_011770 | 4493049 | GCACCGAG | G | 28 | 0 | 5 | 0 | G | G | | | | | G | G | | | G | . |
| NC_011770 | 4493051 | A | G | 33 | 0 | 6 | 0 | . | . | G | G | G | G | . | . | G | G | . | . |
| NC_011770 | 4493060 | CTG | C | 34 | 0 | 6 | 0 | C | C | | | | | C | C | | | C | C |
| NC_011770 | 4493072 | G | C | 34 | 0 | 6 | 0 | C | C | | | | | C | C | | | C | C |
| NC_011770 | 4493084 | C | CG | 34 | 0 | 11 | 0 | CG | . | CG | CG | CG | CG | CG | CG | CG | CG | CG | CG |
| NC_011770 | 4493126 | A | G | 47 | 0 | 6 | 0 | G | G | | | | | G | G | | | G | G |
| NC_011770 | 4493171 | C | T | 53 | 0 | 2 | 0 | T | T | | | | | | | | | | |
| NC_011770 | 4495753 | C | G | 76 | 0 | 2 | 0 | | | | | | | G | G | | | | |
| NC_011770 | 4495759 | C | A | 76 | 0 | 2 | 0 | | | | | | | | | | | A | A |
| NC_011770 | 4497559 | T | G | 67 | 0 | 6 | 0 | G | G | | | | | | | G | G | G | G |
| NC_011770 | 4497583 | G | A | 67 | 0 | 8 | 0 | | | A | A | A | A | A | A | A | A | | |
| NC_011770 | 4497680 | A | G | 61 | 0 | 2 | 0 | | | | | | | G | G | | | | |
| NC_011770 | 4497765 | G | A | 65 | 0 | 8 | 0 | A | A | | | | | A | A | A | A | A | A |
| NC_011770 | 4497835 | T | C | 66 | 0 | 8 | 0 | C | C | | | | | C | C | C | C | C | C |
| NC_011770 | 4502905 | A | G | 34 | 0 | 8 | 0 | G | G | | | | | G | G | G | G | G | G |
| NC_011770 | 4502906 | C | A | 35 | 0 | 2 | 0 | | | | | | | | | A | A | | |
| NC_011770 | 4502939 | T | C | 38 | 0 | 10 | 0 | | | C | C | C | C | C | C | C | C | C | C |
| NC_011770 | 4503471 | G | T | 77 | 0 | 8 | 0 | T | T | | | | | T | T | T | T | T | T |
| NC_011770 | 4506940 | C | G | 73 | 0 | 6 | 0 | | | | | | | G | G | G | G | G | G |
| NC_011770 | 4506961 | A | G | 73 | 0 | 6 | 0 | G | G | | | | | G | G | | | G | G |
| NC_011770 | 4507012 | AC | A | 71 | 0 | 6 | 0 | A | A | | | | | A | A | | | A | A |
| NC_011770 | 4507017 | C | A | 73 | 1 | 1 | 0 | A | . | | | | | | | | | | |
| NC_011770 | 4507838 | G | A | 57 | 0 | 2 | 0 | | | | | | | A | A | | | | |
| NC_011770 | 4507876 | T | TC | 54 | 0 | 2 | 0 | | | | | | | TC | TC | | | | |
| NC_011770 | 4508352 | A | C | 61 | 0 | 2 | 0 | | | | | | | C | C | | | | |
| NC_011770 | 4508355 | A | G | 59 | 0 | 2 | 0 | | | G | G | | | | | | | | |
| NC_011770 | 4509878 | T | C | 77 | 0 | 2 | 0 | | | | | | | | | C | C | | |
| NC_011770 | 4509887 | C | T | 76 | 0 | 2 | 0 | | | | | | | T | T | | | | |
| NC_011770 | 4509898 | C | T | 74 | 0 | 12 | 0 | T | T | T | T | T | T | T | T | T | T | T | T |
| NC_011770 | 4509906 | A | G | 74 | 0 | 6 | 0 | | | | | | | G | G | G | G | G | G |

| | | | | | | | | | | | | | | | | | | | |
|---|---|---|---|---|---|---|---|---|---|---|---|---|---|---|---|---|---|---|---|
| NC_011770 | 4509907 | G | A | 75 | 0 | 4 | 0 | A | A | | | | | | | | | A | A |
| NC_011770 | 4509910 | G | A | 72 | 0 | 6 | 0 | | | | | | | A | A | A | A | A | A |
| NC_011770 | 4509937 | G | GC | 73 | 0 | 1 | 0 | | | | | | | GC | | | | | |
| NC_011770 | 4511391 | G | A | 72 | 2 | 4 | 0 | . | . | A | A | A | A | | | | | | |
| NC_011770 | 4511428 | C | T | 73 | 2 | 2 | 0 | . | . | | | | | | | | | T | T |
| NC_011770 | 4511429 | G | T | 73 | 2 | 4 | 0 | . | . | T | T | T | T | | | | | | |
| NC_011770 | 4512443 | C | T | 60 | 0 | 4 | 0 | | | T | T | T | T | | | | | | |
| NC_011770 | 4513635 | T | G | 55 | 0 | 2 | 0 | | | | | | | | | | | G | G |
| NC_011770 | 4513691 | C | A | 51 | 0 | 4 | 0 | A | A | | | | | | | | | A | A |
| NC_011770 | 4513762 | T | C | 52 | 0 | 4 | 0 | C | C | | | | | | | | | C | C |
| NC_011770 | 4513825 | C | A | 54 | 0 | 2 | 0 | | | | | | | | | A | A | | |
| NC_011770 | 4513866 | C | A | 52 | 0 | 4 | 0 | | | A | A | A | A | | | | | | |
| NC_011770 | 4513897 | A | G | 55 | 0 | 6 | 0 | | | G | G | G | G | | | G | G | | |
| NC_011770 | 4513916 | G | A | 59 | 0 | 6 | 0 | | | A | A | A | A | | | A | A | | |
| NC_011770 | 4514025 | T | A | 61 | 0 | 12 | 0 | A | A | C | C | C | C | C | C | A | A | A | A |
| NC_011770 | 4514100 | T | A | 62 | 0 | 6 | 0 | A | A | | | | | | | A | A | A | A |
| NC_011770 | 4514157 | T | G | 63 | 0 | 6 | 0 | G | G | | | | | | | G | G | G | G |
| NC_011770 | 4514229 | T | A | 58 | 0 | 6 | 0 | A | A | | | | | | | A | A | A | A |
| NC_011770 | 4514230 | T | G | 57 | 0 | 6 | 0 | G | G | | | | | | | G | G | G | G |
| NC_011770 | 4514246 | C | T | 57 | 0 | 6 | 0 | T | T | | | | | | | T | T | T | T |
| NC_011770 | 4514254 | C | T | 57 | 0 | 6 | 0 | T | T | | | | | | | T | T | T | T |
| NC_011770 | 4514256 | T | C | 57 | 0 | 6 | 0 | C | C | | | | | | | C | C | C | C |
| NC_011770 | 4514257 | A | G | 56 | 0 | 6 | 0 | G | G | | | | | | | G | G | G | G |
| NC_011770 | 4514272 | C | T | 56 | 0 | 6 | 0 | T | T | | | | | | | T | T | T | T |
| NC_011770 | 4514293 | C | T | 57 | 0 | 4 | 0 | | | T | T | T | T | | | | | | |
| NC_011770 | 4526915 | G | T | 62 | 0 | 6 | 0 | T | T | T | T | T | T | | | | | | |
| NC_011770 | 4526976 | GC | G | 60 | 0 | 9 | 0 | G | G | G | G | G | G | GCC | | | | G | G |
| NC_011770 | 4527726 | G | A | 59 | 0 | 6 | 0 | A | A | A | A | A | A | | | | | | |
| NC_011770 | 4527842 | T | C | 61 | 0 | 4 | 0 | | | C | C | C | C | | | | | | |
| NC_011770 | 4527867 | C | G | 62 | 0 | 6 | 0 | | | | | | | G | G | G | G | G | G |
| NC_011770 | 4532917 | G | A | 66 | 0 | 6 | 0 | A | A | A | A | A | A | | | | | | |
| NC_011770 | 4532928 | A | G | 70 | 0 | 6 | 0 | G | G | G | G | G | G | | | | | | |
| NC_011770 | 4534532 | T | A | 88 | 0 | 6 | 0 | A | A | A | A | A | A | | | | | | |
| NC_011770 | 4534537 | C | T | 85 | 0 | 6 | 0 | T | T | T | T | T | T | | | | | | |
| NC_011770 | 4534541 | C | CG | 85 | 0 | 6 | 0 | CG | CG | CG | CG | CG | CG | | | | | | |
| NC_011770 | 4535733 | G | T | 73 | 0 | 2 | 0 | T | T | | | | | | | | | | |
| NC_011770 | 4535774 | A | T | 75 | 0 | 12 | 0 | T | T | T | T | T | T | T | T | T | T | T | T |
| NC_011770 | 4535783 | A | T | 74 | 0 | 2 | 0 | T | T | | | | | | | | | | |
| NC_011770 | 4535837 | G | A | 76 | 0 | 2 | 0 | A | A | | | | | | | | | | |
| NC_011770 | 4535853 | T | C | 75 | 0 | 2 | 0 | C | C | | | | | | | | | | |
| NC_011770 | 4535880 | C | A | 80 | 0 | 4 | 0 | | | | | | | | | A | A | A | A |
| NC_011770 | 4535881 | G | C | 79 | 0 | 2 | 0 | C | C | | | | | | | | | | |
| NC_011770 | 4535890 | A | G | 77 | 0 | 12 | 0 | G | G | G | G | G | G | T | T | G | G | G | G |
| NC_011770 | 4535927 | T | C | 81 | 0 | 2 | 0 | C | C | | | | | | | | | | |
| NC_011770 | 4535929 | T | G | 79 | 0 | 2 | 0 | G | G | | | | | | | | | | |
| NC_011770 | 4535934 | C | T | 79 | 0 | 2 | 0 | T | T | | | | | | | | | | |
| NC_011770 | 4535936 | C | T | 78 | 0 | 10 | 0 | | | T | T | T | T | T | T | T | T | T | T |
| NC_011770 | 4535939 | G | GT | 75 | 0 | 2 | 0 | GT | GT | | | | | | | | | | |
| NC_011770 | 4535940 | T | TG | 70 | 2 | 10 | 0 | . | . | TG | TG | TG | TG | TG | TG | TG | TG | TG | TG |
| NC_011770 | 4535950 | C | T | 77 | 0 | 2 | 0 | T | T | | | | | | | | | | |
| NC_011770 | 4535951 | A | G | 75 | 0 | 2 | 0 | G | G | | | | | | | | | | |
| NC_011770 | 4539697 | C | A | 93 | 0 | 4 | 0 | | | A | A | A | A | | | | | | |
| NC_011770 | 4541147 | T | A | 59 | 0 | 6 | 0 | | | | | | | A | A | A | A | A | A |
| NC_011770 | 4541280 | A | G | 69 | 0 | 4 | 0 | | | | | | | | | G | G | G | G |
| NC_011770 | 4542146 | T | G | 42 | 0 | 6 | 0 | | | G | G | G | G | G | G | | | | |
| NC_011770 | 4542149 | T | C | 43 | 0 | 12 | 0 | C | C | C | C | C | C | C | C | C | C | C | C |
| NC_011770 | 4544309 | C | T | 87 | 0 | 2 | 0 | T | T | | | | | | | | | | |
| NC_011770 | 4544318 | G | A | 85 | 0 | 6 | 0 | | | | | | | A | A | A | A | A | A |
| NC_011770 | 4547609 | T | C | 30 | 4 | 2 | 0 | . | . | | | . | . | | | C | C | | |
| NC_011770 | 4547780 | G | A | 18 | 10 | 2 | 0 | . | . | . | . | . | . | A | A | . | . | . | . |
| NC_011770 | 4547793 | T | C | 18 | 10 | 2 | 0 | . | . | . | . | . | . | C | C | . | . | . | . |
| NC_011770 | 4549080 | A | G | 23 | 10 | 2 | 0 | . | . | . | . | . | . | G | G | . | . | . | . |

| | | | | | | | | | | | | | | | | | | | |
|---|---|---|---|---|---|---|---|---|---|---|---|---|---|---|---|---|---|---|---|
| NC_011770 | 4550569 | G | A | 17 | 10 | 2 | 0 | . | . | . | . | . | . | A | A | . | . | . | . |
| NC_011770 | 4550591 | T | G | 18 | 10 | 2 | 0 | . | . | . | . | . | . | G | G | . | . | . | . |
| NC_011770 | 4551193 | G | A | 18 | 10 | 2 | 0 | . | . | . | . | . | . | A | A | . | . | . | . |
| NC_011770 | 4551305 | C | G | 15 | 4 | 2 | 0 | . | . | | | . | . | | | G | G | | |
| NC_011770 | 4551783 | A | G | 31 | 4 | 8 | 0 | . | . | G | G | . | . | G | G | G | G | G | G |
| NC_011770 | 4551784 | G | T | 32 | 4 | 8 | 0 | . | . | T | T | . | . | T | T | T | T | T | T |
| NC_011770 | 4551788 | G | A | 31 | 4 | 2 | 0 | . | . | | | . | . | | | | | A | A |
| NC_011770 | 4552279 | G | A | 53 | 4 | 2 | 0 | . | . | | | . | . | | | A | A | | |
| NC_011770 | 4552280 | G | A | 53 | 4 | 2 | 0 | . | . | | | . | . | | | A | A | | |
| NC_011770 | 4552281 | G | T | 53 | 4 | 2 | 0 | . | . | T | T | . | . | | | | | | |
| NC_011770 | 4555785 | C | G | 23 | 0 | 4 | 0 | | | | | | | | | G | G | G | G |
| NC_011770 | 4555803 | T | C | 28 | 0 | 2 | 0 | C | C | | | | | | | | | | |
| NC_011770 | 4556523 | T | TGG | 91 | 0 | 1 | 0 | | | | | | | TGG | | | | | |
| NC_011770 | 4557244 | C | T | 65 | 0 | 1 | 0 | | | | | | | T | | | | | |
| NC_011770 | 4559064 | A | G | 82 | 0 | 6 | 0 | | | | | | | G | G | G | G | G | G |
| NC_011770 | 4560282 | G | A | 64 | 0 | 4 | 0 | | | A | A | A | A | | | | | | |
| NC_011770 | 4562129 | C | T | 71 | 0 | 2 | 0 | T | T | | | | | | | | | | |
| NC_011770 | 4564607 | C | A | 72 | 0 | 4 | 0 | | | A | A | A | A | | | | | | |
| NC_011770 | 4564610 | T | C | 71 | 0 | 2 | 0 | | | | | | | | | | | C | C |
| NC_011770 | 4564613 | G | A | 69 | 0 | 8 | 0 | A | A | A | A | A | A | A | A | | | | |
| NC_011770 | 4564614 | T | TA | 71 | 0 | 8 | 0 | TA | TA | TA | TA | TA | TA | TA | TA | | | | |
| NC_011770 | 4564644 | A | T | 74 | 0 | 6 | 0 | | | T | T | T | T | T | T | | | | |
| NC_011770 | 4564672 | A | G | 74 | 0 | 12 | 0 | G | G | G | G | G | G | G | G | G | G | G | G |
| NC_011770 | 4572055 | G | A | 77 | 0 | 2 | 0 | A | A | | | | | | | | | | |
| NC_011770 | 4572137 | G | A | 84 | 0 | 4 | 0 | | | A | A | A | A | | | | | | |
| NC_011770 | 4576800 | A | G | 60 | 0 | 2 | 0 | | | | | | | G | G | | | | |
| NC_011770 | 4578066 | T | C | 48 | 0 | 4 | 0 | | | | | | | C | C | | | C | C |
| NC_011770 | 4578203 | A | G | 51 | 0 | 4 | 0 | G | G | | | | | | | | | G | G |
| NC_011770 | 4578204 | T | C | 51 | 0 | 4 | 0 | C | C | | | | | | | | | C | C |
| NC_011770 | 4578234 | G | A | 50 | 0 | 4 | 0 | | | A | A | A | A | | | | | | |
| NC_011770 | 4578314 | G | T | 52 | 0 | 2 | 0 | | | | | | | T | T | | | | |
| NC_011770 | 4579205 | A | G | 63 | 0 | 10 | 0 | G | G | G | G | G | G | G | G | | | G | G |
| NC_011770 | 4581338 | C | T | 71 | 0 | 4 | 0 | | | T | T | T | T | | | | | | |
| NC_011770 | 4581356 | A | G | 73 | 0 | 2 | 0 | | | | | | | G | G | | | | |
| NC_011770 | 4581688 | G | T | 62 | 0 | 8 | 0 | T | T | | | | | T | T | T | T | T | T |
| NC_011770 | 4581702 | G | A | 63 | 0 | 8 | 0 | A | A | | | | | A | A | A | A | A | A |
| NC_011770 | 4581708 | A | G | 61 | 0 | 8 | 0 | G | G | | | | | G | G | G | G | G | G |
| NC_011770 | 4581712 | G | T | 61 | 0 | 8 | 0 | T | T | | | | | T | T | T | T | T | T |
| NC_011770 | 4581713 | A | G | 61 | 0 | 8 | 0 | G | G | | | | | G | G | G | G | G | G |
| NC_011770 | 4581715 | C | T | 60 | 0 | 8 | 0 | T | T | | | | | T | T | T | T | T | T |
| NC_011770 | 4581755 | T | C | 66 | 0 | 4 | 0 | C | C | | | | | C | C | | | | |
| NC_011770 | 4581759 | T | C | 64 | 0 | 8 | 0 | C | C | | | | | C | C | C | C | C | C |
| NC_011770 | 4581799 | G | A | 65 | 0 | 2 | 0 | A | A | | | | | | | | | | |
| NC_011770 | 4581800 | A | G | 64 | 0 | 4 | 0 | | | G | G | G | G | | | | | | |
| NC_011770 | 4581805 | T | C | 63 | 0 | 8 | 0 | C | C | | | | | C | C | C | C | C | C |
| NC_011770 | 4581849 | A | G | 66 | 0 | 8 | 0 | G | G | | | | | G | G | G | G | G | G |
| NC_011770 | 4581864 | A | C | 63 | 0 | 2 | 0 | | | | | | | C | C | | | | |
| NC_011770 | 4581882 | A | G | 61 | 0 | 8 | 0 | G | G | | | | | G | G | G | G | G | G |
| NC_011770 | 4581909 | C | T | 34 | 2 | 6 | 0 | T | T | | | | | . | . | T | T | T | T |
| NC_011770 | 4581912 | A | C | 34 | 2 | 6 | 0 | C | C | | | | | . | . | C | C | C | C |
| NC_011770 | 4581915 | A | G | 35 | 2 | 6 | 0 | G | G | | | | | . | . | G | G | G | G |
| NC_011770 | 4581922 | T | G | 34 | 2 | 6 | 0 | G | G | | | | | . | . | G | G | G | G |
| NC_011770 | 4581925 | C | T | 34 | 3 | 5 | 0 | . | T | | | | | . | . | T | T | T | T |
| NC_011770 | 4581926 | C | G | 34 | 3 | 5 | 0 | . | G | | | | | . | . | G | G | G | G |
| NC_011770 | 4581927 | G | A | 34 | 3 | 5 | 0 | . | A | | | | | . | . | A | A | A | A |
| NC_011770 | 4581932 | T | C | 35 | 3 | 5 | 0 | . | C | | | | | . | . | C | C | C | C |
| NC_011770 | 4581937 | C | A | 34 | 3 | 5 | 0 | . | A | | | | | . | . | A | A | A | A |
| NC_011770 | 4581944 | C | G | 34 | 3 | 5 | 0 | . | G | | | | | . | . | G | G | G | G |
| NC_011770 | 4581947 | C | A | 34 | 3 | 5 | 0 | . | A | | | | | . | . | A | A | A | A |
| NC_011770 | 4581948 | G | T | 35 | 3 | 5 | 0 | . | T | | | | | . | . | T | T | T | T |
| NC_011770 | 4581952 | C | A | 34 | 3 | 5 | 0 | . | A | | | | | . | . | A | A | A | A |
| NC_011770 | 4581955 | T | C | 35 | 3 | 5 | 0 | . | C | | | | | . | . | C | C | C | C |

| | | | | | | | | | | | | | | | | | | | |
|---|---|---|---|---|---|---|---|---|---|---|---|---|---|---|---|---|---|---|---|
| NC_011770 | 4581958 | A | G | 36 | 3 | 9 | 0 | . | G | G | G | G | G | . | . | G | G | G | G |
| NC_011770 | 4581968 | C | A | 43 | 2 | 6 | 0 | A | A | | | | | . | . | A | A | A | A |
| NC_011770 | 4581972 | C | G | 42 | 2 | 10 | 0 | G | G | G | G | G | G | . | . | G | G | G | G |
| NC_011770 | 4581975 | T | C | 43 | 2 | 6 | 0 | C | C | | | | | . | . | C | C | C | C |
| NC_011770 | 4581982 | G | A | 45 | 2 | 6 | 0 | A | A | | | | | . | . | A | A | A | A |
| NC_011770 | 4581988 | G | C | 45 | 2 | 6 | 0 | C | C | | | | | . | . | C | C | C | C |
| NC_011770 | 4581989 | G | A | 45 | 1 | 7 | 0 | A | A | | | | | A | . | A | A | A | A |
| NC_011770 | 4582002 | G | A | 51 | 0 | 4 | 0 | A | A | | | | | A | A | | | | |
| NC_011770 | 4582007 | C | A | 52 | 0 | 8 | 0 | A | A | | | | | A | A | A | A | A | A |
| NC_011770 | 4582018 | A | G | 50 | 0 | 8 | 0 | G | G | | | | | G | G | G | G | G | G |
| NC_011770 | 4582019 | T | C | 50 | 0 | 4 | 0 | | | | | | | | | C | C | C | C |
| NC_011770 | 4582021 | T | C | 52 | 0 | 8 | 0 | C | C | | | | | C | C | C | C | C | C |
| NC_011770 | 4582023 | A | C | 52 | 0 | 8 | 0 | C | C | | | | | C | C | C | C | C | C |
| NC_011770 | 4582039 | A | G | 50 | 0 | 12 | 0 | G | G | G | G | G | G | G | G | G | G | G | G |
| NC_011770 | 4582044 | A | G | 51 | 0 | 8 | 0 | G | G | | | | | G | G | G | G | G | G |
| NC_011770 | 4582045 | G | C | 52 | 0 | 8 | 0 | C | C | | | | | C | C | C | C | C | C |
| NC_011770 | 4582047 | G | A | 51 | 0 | 8 | 0 | A | A | | | | | A | A | A | A | A | A |
| NC_011770 | 4582053 | T | C | 52 | 0 | 8 | 0 | C | C | | | | | C | C | C | C | C | C |
| NC_011770 | 4582055 | A | C | 51 | 0 | 8 | 0 | C | C | | | | | C | C | C | C | C | C |
| NC_011770 | 4582057 | A | G | 51 | 0 | 8 | 0 | G | G | | | | | G | G | G | G | G | G |
| NC_011770 | 4582064 | C | T | 58 | 0 | 12 | 0 | T | T | T | T | T | T | T | T | T | T | T | T |
| NC_011770 | 4582069 | C | T | 60 | 0 | 12 | 0 | T | T | T | T | T | T | T | T | T | T | T | T |
| NC_011770 | 4582076 | G | A | 61 | 0 | 8 | 0 | A | A | | | | | A | A | A | A | A | A |
| NC_011770 | 4582092 | C | T | 59 | 0 | 8 | 0 | T | T | | | | | T | T | T | T | T | T |
| NC_011770 | 4582096 | C | G | 58 | 0 | 12 | 0 | G | G | T | T | T | T | G | G | G | G | G | G |
| NC_011770 | 4582100 | C | A | 59 | 0 | 8 | 0 | A | A | | | | | A | A | A | A | A | A |
| NC_011770 | 4582106 | C | T | 59 | 0 | 8 | 0 | T | T | | | | | T | T | T | T | T | T |
| NC_011770 | 4582108 | T | C | 59 | 0 | 12 | 0 | C | C | C | C | C | C | C | C | C | C | C | C |
| NC_011770 | 4582109 | C | T | 59 | 0 | 8 | 0 | T | T | | | | | T | T | T | T | T | T |
| NC_011770 | 4582123 | T | C | 59 | 0 | 12 | 0 | C | C | C | C | C | C | C | C | C | C | C | C |
| NC_011770 | 4582140 | A | G | 59 | 0 | 8 | 0 | G | G | | | | | G | G | G | G | G | G |
| NC_011770 | 4582143 | A | G | 58 | 0 | 8 | 0 | G | G | | | | | G | G | G | G | G | G |
| NC_011770 | 4582151 | G | A | 57 | 0 | 12 | 0 | A | A | A | A | A | A | A | A | A | A | A | A |
| NC_011770 | 4582186 | C | T | 59 | 0 | 2 | 0 | | | | | | | | | T | T | | |
| NC_011770 | 4582917 | G | C | 74 | 0 | 2 | 0 | | | | | | | C | C | | | | |
| NC_011770 | 4583014 | C | T | 77 | 0 | 2 | 0 | | | | | | | T | T | | | | |
| NC_011770 | 4583827 | A | T | 65 | 0 | 12 | 0 | T | T | T | T | T | T | T | T | T | T | T | T |
| NC_011770 | 4583933 | T | A | 73 | 0 | 12 | 0 | A | A | A | A | A | A | A | A | A | A | A | A |
| NC_011770 | 4583968 | T | C | 71 | 0 | 12 | 0 | C | C | C | C | C | C | C | C | C | C | C | C |
| NC_011770 | 4584016 | A | G | 70 | 0 | 12 | 0 | G | G | G | G | G | G | G | G | G | G | G | G |
| NC_011770 | 4584039 | A | G | 68 | 0 | 12 | 0 | G | G | G | G | G | G | G | G | G | G | G | G |
| NC_011770 | 4585723 | C | T | 77 | 0 | 6 | 0 | | | | | | | T | T | T | T | T | T |
| NC_011770 | 4588414 | G | A | 50 | 0 | 2 | 0 | A | A | | | | | | | | | | |
| NC_011770 | 4588418 | A | G | 45 | 0 | 2 | 0 | | | | | | | G | G | | | | |
| NC_011770 | 4588441 | T | C | 45 | 0 | 2 | 0 | | | | | | | C | C | | | | |
| NC_011770 | 4588471 | G | A | 43 | 0 | 4 | 0 | A | A | | | | | A | A | | | | |
| NC_011770 | 4588520 | A | C | 43 | 0 | 12 | 0 | C | C | C | C | C | C | C | C | C | C | C | C |
| NC_011770 | 4590912 | A | G | 49 | 0 | 6 | 0 | | | G | G | G | G | | | | | G | G |
| NC_011770 | 4590930 | G | A | 50 | 0 | 1 | 0 | | | | | | | A | | | | | |
| NC_011770 | 4592895 | T | C | 90 | 0 | 12 | 0 | C | C | C | C | C | C | C | C | C | C | C | C |
| NC_011770 | 4592912 | A | C | 89 | 0 | 12 | 0 | C | C | C | C | C | C | C | C | C | C | C | C |
| NC_011770 | 4593011 | C | T | 86 | 0 | 2 | 0 | | | | | | | T | T | | | | |
| NC_011770 | 4593962 | T | C | 88 | 0 | 8 | 0 | C | C | | | | | C | C | C | C | C | C |
| NC_011770 | 4594077 | G | A | 87 | 0 | 6 | 0 | | | A | A | A | A | | | A | A | | |
| NC_011770 | 4594107 | G | T | 82 | 0 | 2 | 0 | | | | | | | | | T | T | | |
| NC_011770 | 4594380 | C | T | 87 | 0 | 2 | 0 | | | | | | | | | T | T | | |
| NC_011770 | 4594392 | C | T | 86 | 0 | 4 | 0 | | | T | T | T | T | | | | | | |
| NC_011770 | 4594414 | T | A | 82 | 0 | 4 | 0 | A | A | | | | | A | A | | | | |
| NC_011770 | 4594525 | T | G | 85 | 0 | 4 | 0 | G | G | | | | | G | G | | | | |
| NC_011770 | 4597589 | C | T | 77 | 0 | 4 | 0 | | | T | T | T | T | | | | | | |
| NC_011770 | 4597602 | T | C | 75 | 0 | 4 | 0 | | | | | | | | | C | C | C | C |
| NC_011770 | 4597604 | G | C | 74 | 0 | 8 | 0 | | | C | C | C | C | | | C | C | C | C |

| | | | | | | | | | | | | | | | | | | | |
|---|---|---|---|---|---|---|---|---|---|---|---|---|---|---|---|---|---|---|---|
| NC_011770 | 4597605 | C | T | 73 | 0 | 4 | 0 | | | | | | | | | T | T | T | T |
| NC_011770 | 4597609 | G | A | 75 | 0 | 6 | 0 | | | A | A | A | A | | | A | A | | |
| NC_011770 | 4597612 | T | C | 75 | 0 | 6 | 0 | | | C | C | C | C | | | C | C | | |
| NC_011770 | 4597613 | CG | C | 77 | 0 | 2 | 0 | C | C | | | | | | | | | | |
| NC_011770 | 4597617 | C | T | 76 | 0 | 4 | 0 | | | T | T | T | T | | | | | | |
| NC_011770 | 4597620 | C | T | 75 | 0 | 2 | 0 | | | | | | | | | T | T | | |
| NC_011770 | 4599084 | C | T | 58 | 2 | 4 | 0 | | | T | T | T | T | . | . | | | | |
| NC_011770 | 4599093 | A | G | 56 | 2 | 8 | 0 | | | G | G | G | G | . | . | G | G | G | G |
| NC_011770 | 4599132 | A | G | 51 | 4 | 6 | 0 | | | G | G | G | G | . | . | . | . | G | G |
| NC_011770 | 4599344 | A | G | 35 | 6 | 4 | 0 | | | G | G | G | G | . | . | . | . | . | . |
| NC_011770 | 4599366 | A | G | 35 | 6 | 4 | 0 | | | G | G | G | G | . | . | . | . | . | . |
| NC_011770 | 4599375 | A | C | 36 | 6 | 4 | 0 | | | C | C | C | C | . | . | . | . | . | . |
| NC_011770 | 4599956 | C | T | 30 | 6 | 4 | 0 | | | T | T | T | T | . | . | . | . | . | . |
| NC_011770 | 4599981 | C | T | 31 | 6 | 4 | 0 | | | T | T | T | T | . | . | . | . | . | . |
| NC_011770 | 4599986 | T | C | 31 | 6 | 4 | 0 | | | C | C | C | C | . | . | . | . | . | . |
| NC_011770 | 4600014 | G | A | 30 | 6 | 4 | 0 | | | A | A | A | A | . | . | . | . | . | . |
| NC_011770 | 4600031 | A | G | 29 | 6 | 4 | 0 | | | G | G | G | G | . | . | . | . | . | . |
| NC_011770 | 4600071 | A | G | 28 | 6 | 4 | 0 | | | G | G | G | G | . | . | . | . | . | . |
| NC_011770 | 4600121 | G | A | 32 | 2 | 4 | 0 | | | A | A | A | A | | | . | . | | |
| NC_011770 | 4600174 | A | G | 27 | 6 | 4 | 0 | | | G | G | G | G | . | . | . | . | . | . |
| NC_011770 | 4600263 | G | A | 27 | 6 | 4 | 0 | | | A | A | A | A | . | . | . | . | . | . |
| NC_011770 | 4600432 | C | T | 20 | 6 | 4 | 0 | | | T | T | T | T | . | . | . | . | . | . |
| NC_011770 | 4600438 | T | A | 19 | 6 | 4 | 0 | | | A | A | A | A | . | . | . | . | . | . |
| NC_011770 | 4600538 | A | G | 31 | 2 | 8 | 0 | | | G | G | G | G | G | G | . | . | G | G |
| NC_011770 | 4600551 | G | A | 33 | 2 | 2 | 0 | | | | | | | A | A | . | . | | |
| NC_011770 | 4600565 | A | G | 32 | 2 | 10 | 0 | G | G | G | G | G | G | G | G | . | . | G | G |
| NC_011770 | 4601043 | C | T | 31 | 2 | 4 | 0 | | | T | T | T | T | | | . | . | | |
| NC_011770 | 4601046 | C | T | 31 | 2 | 4 | 0 | | | T | T | T | T | | | . | . | | |
| NC_011770 | 4601048 | T | C | 31 | 2 | 4 | 0 | | | C | C | C | C | | | . | . | | |
| NC_011770 | 4601170 | G | A | 25 | 6 | 2 | 0 | | | . | . | . | . | A | A | . | . | | |
| NC_011770 | 4601199 | G | A | 26 | 6 | 4 | 0 | | | . | . | . | . | A | A | . | . | A | A |
| NC_011770 | 4601224 | A | C | 29 | 6 | 4 | 0 | | | . | . | . | . | C | C | . | . | C | C |
| NC_011770 | 4601301 | G | A | 30 | 6 | 4 | 0 | | | . | . | . | . | A | A | . | . | A | A |
| NC_011770 | 4601342 | T | C | 33 | 6 | 4 | 0 | | | . | . | . | . | C | C | . | . | C | C |
| NC_011770 | 4601397 | T | C | 32 | 6 | 4 | 0 | | | . | . | . | . | C | C | . | . | C | C |
| NC_011770 | 4601470 | G | A | 33 | 6 | 4 | 0 | | | . | . | . | . | A | A | . | . | A | A |
| NC_011770 | 4601502 | G | A | 31 | 6 | 4 | 0 | | | . | . | . | . | A | A | . | . | A | A |
| NC_011770 | 4601516 | A | G | 29 | 6 | 4 | 0 | | | . | . | . | . | G | G | . | . | G | G |
| NC_011770 | 4601526 | A | C | 29 | 6 | 4 | 0 | | | . | . | . | . | C | C | . | . | C | C |
| NC_011770 | 4601538 | T | C | 31 | 6 | 4 | 0 | | | . | . | . | . | C | C | . | . | C | C |
| NC_011770 | 4601577 | C | T | 32 | 6 | 4 | 0 | | | . | . | . | . | T | T | . | . | T | T |
| NC_011770 | 4601731 | T | C | 33 | 6 | 4 | 0 | | | . | . | . | . | C | C | . | . | C | C |
| NC_011770 | 4603805 | C | A | 40 | 6 | 4 | 0 | | | . | . | . | . | A | A | . | . | A | A |
| NC_011770 | 4603830 | C | T | 41 | 6 | 2 | 0 | | | . | . | . | . | | | . | . | T | T |
| NC_011770 | 4603905 | A | G | 40 | 6 | 4 | 0 | | | . | . | . | . | G | G | . | . | T | T |
| NC_011770 | 4603944 | T | G | 41 | 6 | 2 | 0 | | | . | . | . | . | G | G | . | . | | |
| NC_011770 | 4603993 | G | A | 50 | 4 | 2 | 0 | | | | | . | . | A | A | . | . | | |
| NC_011770 | 4604014 | T | C | 50 | 2 | 2 | 0 | | | C | C | . | . | | | | | | |
| NC_011770 | 4604064 | G | C | 46 | 2 | 4 | 0 | | | C | C | . | . | | | C | C | | |
| NC_011770 | 4604105 | C | G | 43 | 2 | 2 | 0 | | | | | . | . | | | G | G | | |
| NC_011770 | 4604106 | T | C | 42 | 2 | 4 | 0 | | | C | C | . | . | | | C | C | | |
| NC_011770 | 4604117 | G | A | 40 | 2 | 2 | 0 | | | A | A | . | . | | | | | | |
| NC_011770 | 4604147 | C | A | 39 | 2 | 4 | 0 | | | A | A | . | . | A | A | | | | |
| NC_011770 | 4604218 | G | A | 36 | 2 | 2 | 0 | | | | | . | . | | | | | A | A |
| NC_011770 | 4604220 | A | C | 35 | 2 | 2 | 0 | | | | | . | . | | | | | C | C |
| NC_011770 | 4604736 | T | C | 6 | 3 | 1 | 0 | C | . | | | | | . | . | | | | |
| NC_011770 | 4609797 | T | C | 81 | 0 | 8 | 0 | C | C | | | | | C | C | C | C | C | C |
| NC_011770 | 4610853 | C | G | 68 | 0 | 2 | 0 | | | | | | | G | G | | | | |
| NC_011770 | 4610855 | C | A | 68 | 0 | 4 | 0 | A | A | | | | | A | A | | | | |
| NC_011770 | 4610881 | T | G | 70 | 0 | 12 | 0 | G | G | G | G | G | G | G | G | G | G | G | G |
| NC_011770 | 4610900 | A | C | 68 | 0 | 12 | 0 | C | C | C | C | C | C | C | C | C | C | C | C |
| NC_011770 | 4610927 | T | C | 69 | 0 | 6 | 0 | C | C | | | | | C | C | C | C | | |

| | | | | | | | | | | | | | | | | | | | |
|---|---|---|---|---|---|---|---|---|---|---|---|---|---|---|---|---|---|---|---|
| NC_011770 | 4610929 | C | CT | 72 | 0 | 2 | 0 | CT | CT | | | | | | | | | | |
| NC_011770 | 4610931 | T | C | 67 | 2 | 2 | 0 | . | . | | | | | | | C | C | | |
| NC_011770 | 4610945 | T | C | 75 | 0 | 4 | 0 | | | C | C | C | C | | | | | | |
| NC_011770 | 4610951 | T | C | 74 | 0 | 2 | 0 | C | C | | | | | | | | | | |
| NC_011770 | 4610956 | A | G | 73 | 0 | 10 | 0 | G | G | G | G | G | G | G | G | G | G | | |
| NC_011770 | 4611066 | G | A | 82 | 0 | 2 | 0 | A | A | | | | | | | | | | |
| NC_011770 | 4611087 | T | C | 82 | 0 | 12 | 0 | C | C | C | C | C | C | C | C | C | C | C | C |
| NC_011770 | 4611125 | G | A | 83 | 0 | 2 | 0 | | | | | | | A | A | | | | |
| NC_011770 | 4612563 | T | C | 77 | 0 | 2 | 0 | C | C | | | | | | | | | | |
| NC_011770 | 4613301 | G | GC | 52 | 0 | 2 | 0 | . | GC | | | | | | | | | GC | . |
| NC_011770 | 4614108 | C | T | 58 | 2 | 2 | 0 | | | | | | | | | . | . | T | T |
| NC_011770 | 4614137 | C | T | 57 | 2 | 10 | 0 | T | T | T | T | T | T | T | T | . | . | T | T |
| NC_011770 | 4615151 | C | T | 60 | 0 | 2 | 0 | T | T | | | | | | | | | | |
| NC_011770 | 4615153 | C | A | 62 | 0 | 2 | 0 | | | | | | | | | A | A | | |
| NC_011770 | 4615199 | G | A | 63 | 0 | 2 | 0 | | | | | | | | | A | A | | |
| NC_011770 | 4615220 | C | A | 60 | 0 | 10 | 0 | A | A | A | A | A | A | A | A | | | A | A |
| NC_011770 | 4616162 | T | G | 65 | 0 | 10 | 0 | G | G | G | G | G | G | G | G | | | G | G |
| NC_011770 | 4618073 | A | C | 68 | 0 | 8 | 0 | C | C | | | | | C | C | C | C | C | C |
| NC_011770 | 4618117 | G | A | 66 | 0 | 2 | 0 | | | | | | | | | | | A | A |
| NC_011770 | 4618126 | C | T | 64 | 0 | 6 | 0 | | | | | | | T | T | T | T | T | T |
| NC_011770 | 4619875 | A | C | 61 | 0 | 6 | 0 | C | C | C | C | C | C | | | | | | |
| NC_011770 | 4619880 | A | G | 59 | 0 | 6 | 0 | G | G | G | G | G | G | | | | | | |
| NC_011770 | 4619882 | G | A | 59 | 0 | 6 | 0 | A | A | A | A | A | A | | | | | | |
| NC_011770 | 4619895 | C | T | 58 | 0 | 10 | 0 | | | T | T | T | T | T | T | T | T | T | T |
| NC_011770 | 4619954 | T | C | 56 | 0 | 2 | 0 | C | C | | | | | | | | | | |
| NC_011770 | 4619957 | T | C | 60 | 0 | 2 | 0 | C | C | | | | | | | | | | |
| NC_011770 | 4619960 | G | A | 61 | 0 | 2 | 0 | A | A | | | | | | | | | | |
| NC_011770 | 4619970 | T | C | 61 | 0 | 2 | 0 | | | | | | | C | C | | | | |
| NC_011770 | 4620004 | C | G | 64 | 0 | 2 | 0 | G | G | | | | | | | | | | |
| NC_011770 | 4620013 | G | A | 62 | 0 | 10 | 0 | | | A | A | A | A | A | A | A | A | A | A |
| NC_011770 | 4621043 | A | G | 53 | 0 | 2 | 0 | | | | | | | | | | | G | G |
| NC_011770 | 4621050 | C | T | 54 | 0 | 4 | 0 | | | T | T | T | T | | | | | | |
| NC_011770 | 4621100 | A | C | 55 | 0 | 4 | 0 | | | C | C | C | C | | | | | | |
| NC_011770 | 4621642 | A | C | 79 | 0 | 4 | 0 | | | C | C | C | C | | | | | | |
| NC_011770 | 4622746 | C | A | 65 | 0 | 4 | 0 | | | A | A | A | A | | | | | | |
| NC_011770 | 4622901 | G | A | 71 | 0 | 4 | 0 | A | A | | | | | | | | | A | A |
| NC_011770 | 4622903 | C | A | 72 | 0 | 2 | 0 | A | A | | | | | | | | | | |
| NC_011770 | 4622939 | G | A | 72 | 0 | 2 | 0 | | | | | | | | | A | A | | |
| NC_011770 | 4622964 | G | C | 70 | 0 | 6 | 0 | | | C | C | C | C | | | C | C | | |
| NC_011770 | 4622984 | G | A | 73 | 0 | 6 | 0 | | | A | A | A | A | | | A | A | | |
| NC_011770 | 4623009 | G | A | 71 | 0 | 6 | 0 | | | A | A | A | A | | | A | A | | |
| NC_011770 | 4625680 | G | A | 71 | 0 | 4 | 0 | A | A | | | | | A | A | | | | |
| NC_011770 | 4625681 | T | G | 71 | 0 | 12 | 0 | G | G | G | G | G | G | G | G | G | G | G | G |
| NC_011770 | 4626605 | A | C | 76 | 0 | 4 | 0 | | | | | | | C | C | C | C | | |
| NC_011770 | 4626691 | A | G | 74 | 0 | 6 | 0 | G | G | | | | | G | G | G | G | | |
| NC_011770 | 4626706 | A | G | 75 | 0 | 4 | 0 | | | | | | | G | G | G | G | | |
| NC_011770 | 4626707 | G | C | 77 | 0 | 4 | 0 | | | | | | | C | C | C | C | | |
| NC_011770 | 4627743 | C | CA | 74 | 0 | 12 | 0 | CA | CA | CA | CA | CA | CA | CA | CA | CA | CA | CA | CA |
| NC_011770 | 4627776 | G | A | 80 | 0 | 2 | 0 | | | | | | | | | | | A | A |
| NC_011770 | 4627810 | T | G | 77 | 0 | 12 | 0 | G | G | G | G | G | G | G | G | G | G | G | G |
| NC_011770 | 4630778 | G | C | 60 | 0 | 4 | 0 | C | C | | | | | | | C | C | | |
| NC_011770 | 4630832 | C | T | 59 | 0 | 4 | 0 | | | T | T | T | T | | | | | | |
| NC_011770 | 4631457 | A | G | 56 | 0 | 8 | 0 | G | G | | | | | G | G | G | G | G | G |
| NC_011770 | 4631462 | T | C | 56 | 0 | 8 | 0 | C | C | | | | | C | C | C | C | C | C |
| NC_011770 | 4631466 | G | A | 56 | 0 | 4 | 0 | A | A | | | | | | | A | A | | |
| NC_011770 | 4631473 | A | C | 57 | 0 | 2 | 0 | | | | | | | | | | | C | C |
| NC_011770 | 4631479 | T | C | 55 | 0 | 4 | 0 | | | | | | | C | C | | | C | C |
| NC_011770 | 4636837 | T | G | 81 | 0 | 4 | 0 | | | | | | | G | G | G | G | | |
| NC_011770 | 4636841 | G | A | 81 | 0 | 2 | 0 | | | | | | | | | A | A | | |
| NC_011770 | 4636890 | A | G | 79 | 0 | 2 | 0 | | | | | | | G | G | | | | |
| NC_011770 | 3189851 | TTGGGCGA | T | 10 | 8 | 1 | 0 | . | . | | | T | | . | . | . | . | . | . |
| NC_011770 | 4641593 | G | A | 74 | 0 | 2 | 0 | A | A | | | | | | | | | | |

| | | | | | | | | | | | | | | | | | | | |
|---|---|---|---|---|---|---|---|---|---|---|---|---|---|---|---|---|---|---|---|
| NC_011770 | 4641713 | C | T | 68 | 0 | 10 | 0 | T | T | T | T | T | T | | | T | T | T | T |
| NC_011770 | 4641732 | T | C | 72 | 0 | 2 | 0 | | | | | | | | | | | C | C |
| NC_011770 | 4641736 | A | T | 71 | 0 | 2 | 0 | | | | | | | | | | | T | T |
| NC_011770 | 4641742 | G | C | 68 | 0 | 2 | 0 | | | | | | | | | | | C | C |
| NC_011770 | 4641760 | CA | C | 67 | 0 | 2 | 0 | | | | | | | | | | | C | C |
| NC_011770 | 4641766 | C | T | 68 | 0 | 2 | 0 | | | | | | | | | | | T | T |
| NC_011770 | 4641769 | A | G | 68 | 0 | 2 | 0 | | | | | | | | | | | G | G |
| NC_011770 | 4641803 | AG | A | 69 | 0 | 2 | 0 | | | | | | | | | | | A | A |
| NC_011770 | 4641842 | A | G | 68 | 0 | 8 | 0 | | | G | G | G | G | | | G | G | G | G |
| NC_011770 | 4641850 | A | G | 68 | 0 | 2 | 0 | | | | | | | | | | | G | G |
| NC_011770 | 4641881 | G | A | 68 | 0 | 2 | 0 | | | | | | | | | A | A | | |
| NC_011770 | 4641889 | A | C | 66 | 0 | 2 | 0 | | | | | | | | | | | C | C |
| NC_011770 | 4642241 | A | T | 71 | 0 | 2 | 0 | | | | | | | | | | | T | T |
| NC_011770 | 4642288 | G | T | 67 | 0 | 4 | 0 | | | T | T | T | T | | | | | | |
| NC_011770 | 4643775 | G | A | 79 | 0 | 4 | 0 | | | A | A | A | A | | | | | | |
| NC_011770 | 4646091 | G | A | 67 | 0 | 2 | 0 | | | | | | | | | A | A | | |
| NC_011770 | 4646092 | G | T | 66 | 0 | 4 | 0 | T | T | | | | | | | | | T | T |
| NC_011770 | 4646095 | G | A | 69 | 0 | 2 | 0 | | | | | | | | | A | A | | |
| NC_011770 | 4647307 | G | A | 79 | 0 | 6 | 0 | | | A | A | A | A | A | A | | | | |
| NC_011770 | 4647385 | T | C | 75 | 0 | 6 | 0 | | | C | C | C | C | | | C | C | | |
| NC_011770 | 4649965 | C | G | 84 | 0 | 2 | 0 | | | | | | | G | G | | | | |
| NC_011770 | 4655375 | A | C | 77 | 0 | 2 | 0 | | | | | | | | | | | C | C |
| NC_011770 | 4655421 | T | A | 81 | 0 | 2 | 0 | | | | | | | | | | | A | A |
| NC_011770 | 4655539 | A | C | 80 | 0 | 4 | 0 | C | C | | | | | | | | | C | C |
| NC_011770 | 4658269 | T | A | 66 | 0 | 10 | 0 | A | A | A | A | A | A | | | A | A | A | A |
| NC_011770 | 4659898 | C | T | 41 | 0 | 10 | 0 | T | T | T | T | T | T | | | T | T | T | T |
| NC_011770 | 4659967 | A | G | 47 | 0 | 10 | 0 | G | G | G | G | G | G | | | G | G | G | G |
| NC_011770 | 4660813 | AG | A | 38 | 6 | 4 | 0 | . | . | A | A | A | A | . | . | | | . | . |
| NC_011770 | 4662816 | C | A | 42 | 6 | 4 | 0 | . | . | A | A | A | A | . | . | | | . | . |
| NC_011770 | 4664491 | A | G | 32 | 6 | 6 | 0 | . | . | G | G | G | G | . | . | G | G | . | . |
| NC_011770 | 4664499 | C | T | 30 | 6 | 2 | 0 | . | . | | | | | . | . | T | T | . | . |
| NC_011770 | 4664510 | C | T | 30 | 6 | 4 | 0 | . | . | T | T | T | T | . | . | | | . | . |
| NC_011770 | 4664546 | A | C | 29 | 6 | 2 | 0 | . | . | | | C | C | . | . | | | . | . |
| NC_011770 | 4664582 | A | G | 32 | 6 | 2 | 0 | . | . | | | | | . | . | G | G | . | . |
| NC_011770 | 4664585 | G | A | 32 | 6 | 2 | 0 | . | . | | | | | . | . | A | A | . | . |
| NC_011770 | 4664587 | ACGC | A | 32 | 6 | 2 | 0 | . | . | | | | | . | . | A | A | . | . |
| NC_011770 | 4673109 | A | G | 49 | 0 | 6 | 0 | G | G | | | | | G | G | | | G | G |
| NC_011770 | 4673110 | C | T | 50 | 0 | 6 | 0 | T | T | | | | | T | T | | | T | T |
| NC_011770 | 4673112 | G | A | 47 | 0 | 2 | 0 | | | | | | | | | A | A | | |
| NC_011770 | 4677530 | A | G | 78 | 0 | 2 | 0 | | | | | | | | | G | G | | |
| NC_011770 | 4679104 | T | A | 80 | 0 | 12 | 0 | A | A | A | A | A | A | A | A | A | A | A | A |
| NC_011770 | 4679910 | C | T | 62 | 0 | 2 | 0 | | | | | | | | | | | T | T |
| NC_011770 | 4679965 | T | C | 55 | 0 | 2 | 0 | | | | | | | C | C | | | | |
| NC_011770 | 4680018 | T | G | 56 | 0 | 10 | 0 | | | G | G | G | G | G | G | G | G | G | G |
| NC_011770 | 4680049 | G | A | 56 | 0 | 2 | 0 | | | | | | | A | A | | | | |
| NC_011770 | 4683114 | C | T | 59 | 0 | 4 | 0 | | | T | T | T | T | | | | | | |
| NC_011770 | 4683128 | G | A | 56 | 0 | 2 | 0 | | | | | | | A | A | | | | |
| NC_011770 | 4683165 | G | A | 54 | 0 | 2 | 0 | | | | | | | A | A | | | | |
| NC_011770 | 4683167 | G | A | 56 | 0 | 4 | 0 | | | A | A | A | A | | | | | | |
| NC_011770 | 4683296 | A | C | 59 | 0 | 10 | 0 | | | C | C | C | C | C | C | C | C | C | C |
| NC_011770 | 4683317 | G | A | 58 | 0 | 2 | 0 | | | | | | | A | A | | | | |
| NC_011770 | 4685583 | C | T | 64 | 0 | 2 | 0 | | | | | | | | | | | T | T |
| NC_011770 | 4685609 | TGGACG | T | 65 | 0 | 2 | 0 | T | T | | | | | | | | | | |
| NC_011770 | 4689399 | G | A | 63 | 0 | 2 | 0 | | | | | | | | | | | A | A |
| NC_011770 | 4689424 | A | C | 63 | 0 | 2 | 0 | | | | | | | | | | | C | C |
| NC_011770 | 4689425 | A | G | 63 | 0 | 2 | 0 | | | | | | | | | | | G | G |
| NC_011770 | 4689438 | A | G | 60 | 0 | 4 | 0 | | | G | G | G | G | | | | | | |
| NC_011770 | 4689445 | GT | G | 62 | 0 | 4 | 0 | | | G | G | G | G | | | | | | |
| NC_011770 | 4689461 | G | A | 64 | 0 | 4 | 0 | | | A | A | A | A | | | | | | |
| NC_011770 | 4693737 | G | A | 89 | 0 | 10 | 0 | A | A | A | A | A | A | A | A | A | A | | |
| NC_011770 | 4693772 | T | G | 90 | 0 | 10 | 0 | G | G | G | G | G | G | G | G | G | G | | |
| NC_011770 | 4694747 | T | C | 83 | 0 | 4 | 0 | | | | | C | C | | | C | C | | |

| | | | | | | | | | | | | | | | | | | | |
|---|---|---|---|---|---|---|---|---|---|---|---|---|---|---|---|---|---|---|---|
| NC_011770 | 4696611 | C | CGCA | 74 | 0 | 2 | 0 | | | | | CGCA | | CGCA | | | | | |
| NC_011770 | 4696615 | A | G | 23 | 2 | 6 | 0 | | | G | . | G | G | G | G | G | . | | |
| NC_011770 | 4696621 | CT | C | 71 | 0 | 8 | 0 | | | C | C | C | C | C | C | C | C | | |
| NC_011770 | 4697402 | C | T | 78 | 0 | 2 | 0 | | | | | | | | | T | T | | |
| NC_011770 | 4699443 | GC | G | 70 | 0 | 6 | 0 | G | G | | | | | G | G | | | G | G |
| NC_011770 | 4699444 | C | T | 34 | 6 | 2 | 0 | . | . | | | | | . | . | T | T | . | . |
| NC_011770 | 4699446 | G | A | 70 | 0 | 2 | 0 | | | | | | | | | A | A | | |
| NC_011770 | 4699483 | G | A | 74 | 0 | 2 | 0 | | | | | | | | | A | A | | |
| NC_011770 | 4699489 | T | C | 73 | 0 | 2 | 0 | | | | | | | | | C | C | | |
| NC_011770 | 4699530 | G | T | 77 | 0 | 2 | 0 | | | | | | | | | T | T | | |
| NC_011770 | 4705603 | T | C | 62 | 0 | 2 | 0 | | | | | | | | | C | C | | |
| NC_011770 | 4706469 | G | A | 82 | 0 | 1 | 0 | | | | | | | A | | | | | |
| NC_011770 | 4707646 | A | G | 84 | 0 | 12 | 0 | G | G | G | G | G | G | G | G | G | G | G | G |
| NC_011770 | 4709108 | G | A | 31 | 0 | 2 | 0 | | | | | | | | | A | A | | |
| NC_011770 | 4709109 | C | A | 31 | 0 | 6 | 0 | A | A | | | | | A | A | | | A | A |
| NC_011770 | 4709168 | T | C | 43 | 0 | 2 | 0 | | | | | | | | | C | C | | |
| NC_011770 | 4709187 | T | TC | 43 | 0 | 2 | 0 | | | | | | | | | TC | TC | | |
| NC_011770 | 4710313 | C | T | 56 | 0 | 6 | 0 | T | T | T | T | T | T | | | | | | |
| NC_011770 | 4711053 | G | A | 68 | 0 | 2 | 0 | | | | | | | | | A | A | | |
| NC_011770 | 4712111 | G | C | 73 | 0 | 4 | 0 | | | C | C | C | C | | | | | | |
| NC_011770 | 4715682 | C | A | 87 | 0 | 8 | 0 | A | A | A | A | A | A | | | A | A | | |
| NC_011770 | 4715686 | G | A | 87 | 0 | 8 | 0 | A | A | A | A | A | A | | | A | A | | |
| NC_011770 | 4715704 | T | C | 45 | 0 | 8 | 0 | C | C | C | C | C | C | | | C | C | | |
| NC_011770 | 4715724 | G | A | 92 | 0 | 2 | 0 | | | | | | | | | A | A | | |
| NC_011770 | 4715730 | C | A | 88 | 0 | 4 | 0 | A | A | | | | | | | A | A | | |
| NC_011770 | 4715735 | TC | T | 85 | 0 | 6 | 0 | | | T | T | T | T | | | T | T | | |
| NC_011770 | 4715745 | G | A | 86 | 0 | 8 | 0 | A | A | A | A | A | A | | | A | A | | |
| NC_011770 | 4715746 | C | T | 86 | 0 | 8 | 0 | T | T | T | T | T | T | | | T | T | | |
| NC_011770 | 4717929 | A | G | 70 | 0 | 4 | 0 | | | G | G | G | G | | | | | | |
| NC_011770 | 4718501 | T | G | 55 | 0 | 2 | 0 | | | | | | | | | | | G | G |
| NC_011770 | 4718520 | C | T | 56 | 0 | 2 | 0 | T | T | | | | | | | | | | |
| NC_011770 | 4719711 | G | A | 62 | 0 | 2 | 0 | | | | | | | A | A | | | | |
| NC_011770 | 4719737 | C | T | 68 | 0 | 2 | 0 | | | | | | | | | T | T | | |
| NC_011770 | 4719744 | T | A | 67 | 0 | 2 | 0 | | | | | | | | | A | A | | |
| NC_011770 | 4720494 | C | T | 80 | 0 | 2 | 0 | | | | | | | T | T | | | | |
| NC_011770 | 4722505 | A | G | 60 | 0 | 2 | 0 | | | | | | | | | G | G | | |
| NC_011770 | 4722516 | GC | G | 58 | 0 | 2 | 0 | | | | | | | | | G | G | | |
| NC_011770 | 4722525 | C | T | 59 | 0 | 2 | 0 | | | | | | | | | T | T | | |
| NC_011770 | 4722546 | T | G | 53 | 0 | 2 | 0 | | | | | | | | | G | G | | |
| NC_011770 | 4722557 | C | T | 58 | 0 | 2 | 0 | T | T | | | | | | | | | | |
| NC_011770 | 4722558 | G | A | 56 | 0 | 2 | 0 | A | A | | | | | | | | | | |
| NC_011770 | 4722569 | CT | C | 56 | 0 | 2 | 0 | | | | | | | | | C | C | | |
| NC_011770 | 4723210 | A | C | 72 | 0 | 12 | 0 | C | C | C | C | C | C | C | C | C | C | C | C |
| NC_011770 | 4723240 | G | A | 75 | 0 | 2 | 0 | | | | | | | | | A | A | | |
| NC_011770 | 4723290 | C | T | 73 | 0 | 2 | 0 | | | | | | | | | T | T | | |
| NC_011770 | 4723390 | T | C | 76 | 0 | 2 | 0 | | | | | | | | | C | C | | |
| NC_011770 | 4724312 | G | C | 64 | 0 | 4 | 0 | C | C | | | | | C | C | | | | |
| NC_011770 | 4725012 | A | G | 45 | 0 | 2 | 0 | | | | | | | | | | | G | G |
| NC_011770 | 4725048 | C | T | 44 | 0 | 2 | 0 | | | | | | | | | | | T | T |
| NC_011770 | 4727691 | C | CT | 60 | 0 | 4 | 0 | | | CT | CT | CT | CT | | | | | | |
| NC_011770 | 4727825 | C | T | 62 | 0 | 2 | 0 | T | T | | | | | | | | | | |
| NC_011770 | 4729192 | G | C | 70 | 0 | 8 | 0 | | | C | C | C | C | C | C | C | C | | |
| NC_011770 | 4729195 | T | C | 70 | 0 | 8 | 0 | | | C | C | C | C | C | C | C | C | | |
| NC_011770 | 4729252 | G | T | 75 | 0 | 4 | 0 | | | T | T | T | T | | | | | | |
| NC_011770 | 4729393 | G | A | 76 | 0 | 4 | 0 | | | A | A | A | A | | | | | | |
| NC_011770 | 4729469 | A | G | 72 | 0 | 4 | 0 | | | G | G | G | G | | | | | | |
| NC_011770 | 4729478 | A | G | 69 | 0 | 10 | 0 | G | G | G | G | G | G | G | G | G | G | | |
| NC_011770 | 4729504 | G | A | 70 | 0 | 2 | 0 | | | | | | | | | A | A | | |
| NC_011770 | 4729517 | T | C | 69 | 0 | 2 | 0 | | | | | | | | | C | C | | |
| NC_011770 | 4729582 | C | T | 62 | 0 | 2 | 0 | | | | | | | T | T | | | | |
| NC_011770 | 4729875 | G | C | 70 | 0 | 4 | 0 | | | | | | | C | C | C | C | | |
| NC_011770 | 4729901 | G | A | 72 | 0 | 4 | 0 | | | A | A | A | A | | | | | | |

| | | | | | | | | | | | | | | | | | | | |
|---|---|---|---|---|---|---|---|---|---|---|---|---|---|---|---|---|---|---|---|
| NC_011770 | 4729908 | C | G | 70 | 0 | 12 | 0 | G | G | G | G | G | G | G | G | G | G | G | G |
| NC_011770 | 4729948 | G | GC | 72 | 0 | 8 | 0 | | | GC | GC | GC | GC | GC | GC | GC | GC | | |
| NC_011770 | 4730784 | G | T | 101 | 0 | 4 | 0 | T | T | | | | | | | T | T | | |
| NC_011770 | 4730825 | GC | G | 105 | 0 | 2 | 0 | | | | | | | | | | | G | G |
| NC_011770 | 4730847 | C | T | 100 | 0 | 6 | 0 | T | T | | | | | | | T | T | T | T |
| NC_011770 | 4731255 | T | C | 90 | 0 | 2 | 0 | | | | | | | C | C | | | | |
| NC_011770 | 4731279 | T | C | 89 | 0 | 4 | 0 | | | C | C | C | C | | | | | | |
| NC_011770 | 4731288 | A | G | 92 | 0 | 6 | 0 | G | G | | | | | | | G | G | G | G |
| NC_011770 | 4732598 | A | C | 69 | 0 | 12 | 0 | C | C | C | C | C | C | C | C | C | C | C | C |
| NC_011770 | 4734251 | G | C | 70 | 0 | 8 | 0 | C | C | C | C | C | C | | | C | C | | |
| NC_011770 | 4734299 | C | T | 73 | 0 | 4 | 0 | | | T | T | T | T | | | | | | |
| NC_011770 | 4735086 | C | T | 71 | 0 | 12 | 0 | T | T | T | T | T | T | T | T | T | T | T | T |
| NC_011770 | 4735101 | G | A | 76 | 0 | 2 | 0 | | | | | | | | | | | A | A |
| NC_011770 | 4735150 | G | A | 77 | 0 | 6 | 0 | A | A | A | A | A | A | | | | | | |
| NC_011770 | 4735152 | G | T | 77 | 0 | 12 | 0 | T | T | T | T | T | T | T | T | T | T | T | T |
| NC_011770 | 4736447 | A | G | 77 | 0 | 6 | 0 | G | G | | | | | G | G | | | G | G |
| NC_011770 | 4736448 | C | G | 77 | 0 | 4 | 0 | | | G | G | G | G | | | | | | |
| NC_011770 | 4741466 | A | C | 77 | 0 | 2 | 0 | C | C | | | | | | | | | | |
| NC_011770 | 4741977 | T | G | 79 | 0 | 2 | 0 | | | | | | | | | | | G | G |
| NC_011770 | 4743383 | G | A | 72 | 0 | 2 | 0 | | | | | | | | | A | A | | |
| NC_011770 | 4743393 | G | A | 71 | 0 | 2 | 0 | | | | | | | | | | | A | A |
| NC_011770 | 4743410 | A | G | 75 | 0 | 10 | 0 | G | G | G | G | G | G | | | G | G | G | G |
| NC_011770 | 4743419 | C | A | 78 | 0 | 4 | 0 | | | A | A | A | A | | | | | | |
| NC_011770 | 4746431 | G | C | 76 | 0 | 4 | 0 | | | C | C | C | C | | | | | | |
| NC_011770 | 4746513 | G | A | 76 | 0 | 6 | 0 | A | A | A | A | A | A | | | | | | |
| NC_011770 | 4747726 | A | G | 79 | 0 | 4 | 0 | G | G | | | | | G | G | | | | |
| NC_011770 | 4747776 | G | C | 81 | 0 | 2 | 0 | C | C | | | | | | | | | | |
| NC_011770 | 4748141 | C | T | 71 | 0 | 4 | 0 | T | T | | | | | T | T | | | | |
| NC_011770 | 4748202 | A | G | 82 | 0 | 2 | 0 | G | G | | | | | | | | | | |
| NC_011770 | 4749044 | C | A | 76 | 0 | 6 | 0 | A | A | | | | | A | A | | | A | A |
| NC_011770 | 4749054 | G | A | 71 | 0 | 2 | 0 | A | A | | | | | | | | | | |
| NC_011770 | 4749057 | C | A | 74 | 0 | 2 | 0 | A | A | | | | | | | | | | |
| NC_011770 | 4750097 | C | A | 65 | 0 | 4 | 0 | A | A | | | | | A | A | | | | |
| NC_011770 | 4750118 | G | A | 64 | 0 | 2 | 0 | | | | | | | | | | | A | A |
| NC_011770 | 4750135 | T | C | 61 | 0 | 4 | 0 | C | C | | | | | C | C | | | | |
| NC_011770 | 4750158 | A | G | 65 | 0 | 4 | 0 | G | G | | | | | G | G | | | | |
| NC_011770 | 4750159 | A | G | 65 | 0 | 4 | 0 | G | G | | | | | G | G | | | | |
| NC_011770 | 4750197 | A | G | 65 | 0 | 4 | 0 | G | G | | | | | G | G | | | | |
| NC_011770 | 4750213 | G | A | 64 | 0 | 12 | 0 | A | A | A | A | A | A | A | A | A | A | A | A |
| NC_011770 | 4750232 | A | G | 65 | 0 | 4 | 0 | G | G | | | | | G | G | | | | |
| NC_011770 | 4750238 | C | T | 66 | 0 | 2 | 0 | | | | | | | | | | | T | T |
| NC_011770 | 4750267 | G | A | 64 | 0 | 4 | 0 | A | A | | | | | A | A | | | | |
| NC_011770 | 4750269 | A | G | 63 | 0 | 4 | 0 | G | G | | | | | G | G | | | | |
| NC_011770 | 4750285 | A | G | 65 | 0 | 4 | 0 | G | G | | | | | G | G | | | | |
| NC_011770 | 4750286 | A | G | 65 | 0 | 4 | 0 | G | G | | | | | G | G | | | | |
| NC_011770 | 4750287 | T | C | 64 | 0 | 4 | 0 | C | C | | | | | C | C | | | | |
| NC_011770 | 4750311 | C | T | 66 | 0 | 4 | 0 | T | T | | | | | T | T | | | | |
| NC_011770 | 4750322 | C | G | 66 | 0 | 4 | 0 | G | G | | | | | G | G | | | | |
| NC_011770 | 4750333 | C | T | 65 | 0 | 4 | 0 | T | T | | | | | T | T | | | | |
| NC_011770 | 4750366 | T | C | 67 | 0 | 4 | 0 | C | C | | | | | C | C | | | | |
| NC_011770 | 4750567 | A | G | 70 | 0 | 2 | 0 | | | | | | | | | G | G | | |
| NC_011770 | 4750579 | A | G | 66 | 0 | 2 | 0 | | | | | | | G | G | | | | |
| NC_011770 | 4750582 | T | A | 69 | 0 | 2 | 0 | | | | | | | A | A | | | | |
| NC_011770 | 4750583 | T | A | 67 | 0 | 6 | 0 | | | A | A | A | A | G | G | | | | |
| NC_011770 | 4755347 | G | GA | 54 | 0 | 5 | 0 | | | GA | GA | GA | GA | GA | | | | | |
| NC_011770 | 4755349 | A | C | 26 | 2 | 2 | 0 | | | | | . | | . | | | | C | C |
| NC_011770 | 4756840 | C | CG | 79 | 0 | 2 | 0 | | | | | | | | | CG | CG | | |
| NC_011770 | 4758351 | G | A | 74 | 0 | 2 | 0 | A | A | | | | | | | | | | |
| NC_011770 | 4758356 | A | G | 74 | 0 | 2 | 0 | G | G | | | | | | | | | | |
| NC_011770 | 4758358 | A | G | 76 | 0 | 2 | 0 | | | | | | | | | G | G | | |
| NC_011770 | 4758372 | A | G | 74 | 0 | 4 | 0 | G | G | | | | | | | G | G | | |
| NC_011770 | 4758377 | G | A | 73 | 0 | 2 | 0 | | | | | | | | | A | A | | |

| | | | | | | | | | | | | | | | | | | | |
|---|---|---|---|---|---|---|---|---|---|---|---|---|---|---|---|---|---|---|---|
| NC_011770 | 4758381 | A | T | 73 | 0 | 4 | 0 | T | T | | | | | | | T | T | | |
| NC_011770 | 4758386 | A | G | 74 | 0 | 4 | 0 | G | G | | | | | | | G | G | | |
| NC_011770 | 4758429 | G | A | 68 | 0 | 2 | 0 | | | | | | | | | A | A | | |
| NC_011770 | 4758435 | C | G | 69 | 0 | 2 | 0 | | | | | | | | | G | G | | |
| NC_011770 | 4758443 | G | C | 65 | 0 | 2 | 0 | C | C | | | | | | | | | | |
| NC_011770 | 4759584 | A | C | 84 | 0 | 2 | 0 | | | | | | | | | C | C | | |
| NC_011770 | 4759589 | A | G | 87 | 0 | 2 | 0 | | | | | | | | | G | G | | |
| NC_011770 | 4759622 | T | C | 81 | 0 | 2 | 0 | | | | | | | | | C | C | | |
| NC_011770 | 4759624 | TC | T | 78 | 0 | 10 | 0 | | | T | T | T | T | T | T | AC | AC | T | T |
| NC_011770 | 4759626 | C | A | 14 | 4 | 2 | 0 | | | | | . | . | | . | A | A | | . |
| NC_011770 | 4759631 | T | C | 79 | 0 | 2 | 0 | | | | | | | | | C | C | | |
| NC_011770 | 4759636 | A | G | 80 | 0 | 2 | 0 | | | | | | | | | G | G | | |
| NC_011770 | 4759639 | C | G | 80 | 0 | 2 | 0 | | | | | | | | | G | G | | |
| NC_011770 | 4759641 | C | T | 79 | 0 | 2 | 0 | | | | | | | | | T | T | | |
| NC_011770 | 4759646 | G | GT | 78 | 0 | 2 | 0 | | | | | | | | | GT | GT | | |
| NC_011770 | 4759652 | T | C | 79 | 0 | 12 | 0 | C | C | C | C | C | C | C | C | C | C | C | C |
| NC_011770 | 4759704 | A | G | 82 | 0 | 2 | 0 | | | | | | | | | G | G | | |
| NC_011770 | 4759800 | C | T | 73 | 0 | 4 | 0 | | | T | T | T | T | | | | | | |
| NC_011770 | 4759801 | CT | C | 73 | 0 | 6 | 0 | C | C | AT | AT | AT | AT | | | | | | |
| NC_011770 | 4759802 | T | G | 72 | 0 | 2 | 0 | . | . | | | | | | | G | G | | |
| NC_011770 | 4759803 | G | A | 71 | 0 | 6 | 0 | | | A | A | A | A | | | A | A | | |
| NC_011770 | 4759831 | A | ACATGC | 71 | 0 | 1 | 0 | | | | | | | | | | | ACATGCCGG | |
| NC_011770 | 4759837 | C | A | 59 | 0 | 2 | 0 | A | A | | | | | | | | | | |
| NC_011770 | 4759881 | A | G | 70 | 0 | 4 | 0 | | | | | | | | | G | G | G | G |
| NC_011770 | 4763442 | A | C | 63 | 0 | 2 | 0 | C | C | | | | | | | | | | |
| NC_011770 | 188996 | C | CG | 53 | 0 | 2 | 0 | | | CG | . | | CG | | | | | | |
| NC_011770 | 4769078 | T | A | 70 | 0 | 2 | 0 | | | | | | | | | A | A | | |
| NC_011770 | 4769168 | G | A | 63 | 0 | 12 | 0 | A | A | A | A | A | A | A | A | A | A | A | A |
| NC_011770 | 4769301 | G | A | 74 | 0 | 2 | 0 | | | | | | | | | A | A | | |
| NC_011770 | 4769313 | AG | A | 75 | 0 | 2 | 0 | | | | | | | | | A | A | | |
| NC_011770 | 4770043 | G | A | 77 | 0 | 2 | 0 | | | | | | | | | A | A | | |
| NC_011770 | 4770050 | T | G | 79 | 0 | 6 | 0 | | | G | G | G | G | | | | | G | G |
| NC_011770 | 4772608 | A | C | 78 | 0 | 2 | 0 | | | | | | | C | C | | | | |
| NC_011770 | 4772659 | A | G | 75 | 0 | 2 | 0 | | | | | | | G | G | | | | |
| NC_011770 | 4773398 | T | A | 86 | 0 | 2 | 0 | | | | | | | | | | | A | A |
| NC_011770 | 4773406 | C | T | 89 | 0 | 2 | 0 | | | | | | | T | T | | | | |
| NC_011770 | 4773437 | C | T | 87 | 0 | 2 | 0 | | | | | | | T | T | | | | |
| NC_011770 | 4774093 | G | A | 75 | 0 | 2 | 0 | | | | | | | | | A | A | | |
| NC_011770 | 4774096 | T | C | 76 | 0 | 2 | 0 | | | | | | | C | C | | | | |
| NC_011770 | 4774215 | G | A | 65 | 0 | 2 | 0 | | | | | | | A | A | | | | |
| NC_011770 | 4774216 | A | C | 63 | 0 | 2 | 0 | | | | | | | C | C | | | | |
| NC_011770 | 4774238 | C | T | 65 | 0 | 2 | 0 | | | | | | | | | | | T | T |
| NC_011770 | 4774250 | T | C | 69 | 0 | 2 | 0 | | | | | | | C | C | | | | |
| NC_011770 | 4774255 | C | T | 66 | 0 | 4 | 0 | | | | | | | T | T | T | T | | |
| NC_011770 | 4774260 | A | G | 68 | 0 | 2 | 0 | | | | | | | G | G | | | | |
| NC_011770 | 4774265 | T | C | 68 | 0 | 2 | 0 | | | | | | | C | C | | | | |
| NC_011770 | 4774268 | T | C | 68 | 0 | 2 | 0 | | | | | | | C | C | | | | |
| NC_011770 | 4774272 | C | CA | 65 | 0 | 4 | 0 | | | CA | CA | CA | CA | | | | | | |
| NC_011770 | 4774276 | C | A | 38 | 1 | 5 | 0 | | | A | A | . | A | G | G | | | | |
| NC_011770 | 4774278 | A | C | 40 | 1 | 2 | 0 | | | | | . | | C | C | | | | |
| NC_011770 | 4774300 | G | GC | 65 | 0 | 2 | 0 | | | | | | | GC | GC | | | | |
| NC_011770 | 4774315 | G | A | 66 | 0 | 6 | 0 | A | A | A | A | A | A | | | | | | |
| NC_011770 | 4774328 | A | G | 67 | 0 | 2 | 0 | | | | | | | G | G | | | | |
| NC_011770 | 4774332 | A | G | 65 | 0 | 2 | 0 | | | | | | | G | G | | | | |
| NC_011770 | 4774342 | C | A | 65 | 0 | 2 | 0 | | | | | | | A | A | | | | |
| NC_011770 | 4774352 | C | T | 64 | 0 | 2 | 0 | | | | | | | T | T | | | | |
| NC_011770 | 4774369 | G | A | 57 | 1 | 1 | 0 | | | | | | | . | A | | | | |
| NC_011770 | 4774371 | C | T | 57 | 1 | 1 | 0 | | | | | | | . | T | | | | |
| NC_011770 | 4774372 | T | A | 57 | 1 | 1 | 0 | | | | | | | . | A | | | | |
| NC_011770 | 4774381 | T | C | 59 | 1 | 1 | 0 | | | | | | | . | C | | | | |
| NC_011770 | 4774387 | T | TG | 59 | 1 | 1 | 0 | | | | | | | . | TG | | | | |
| NC_011770 | 4774399 | G | A | 63 | 0 | 2 | 0 | | | | | | | A | A | | | | |

| | | | | | | | | | | | | | | | | | | | |
|---|---|---|---|---|---|---|---|---|---|---|---|---|---|---|---|---|---|---|---|
| NC_011770 | 4774400 | G | A | 62 | 0 | 2 | 0 | | | | | | | A | A | | | | |
| NC_011770 | 4774412 | C | T | 65 | 0 | 2 | 0 | | | | | | | T | T | | | | |
| NC_011770 | 4774416 | A | T | 63 | 0 | 2 | 0 | | | | | | | T | T | | | | |
| NC_011770 | 4774417 | A | T | 64 | 0 | 2 | 0 | | | | | | | T | T | | | | |
| NC_011770 | 4774420 | G | A | 64 | 0 | 2 | 0 | | | | | | | A | A | | | | |
| NC_011770 | 4774435 | T | A | 67 | 0 | 8 | 0 | A | A | A | A | A | A | A | A | | | | |
| NC_011770 | 4774443 | T | G | 67 | 0 | 2 | 0 | | | | | | | G | G | | | | |
| NC_011770 | 4774483 | C | G | 71 | 0 | 2 | 0 | | | | | | | G | G | | | | |
| NC_011770 | 4774484 | C | T | 70 | 0 | 2 | 0 | | | | | | | T | T | | | | |
| NC_011770 | 4774491 | T | C | 70 | 0 | 2 | 0 | | | | | | | C | C | | | | |
| NC_011770 | 4774494 | T | C | 69 | 0 | 2 | 0 | | | | | | | C | C | | | | |
| NC_011770 | 4774499 | T | A | 68 | 0 | 6 | 0 | A | A | A | A | A | A | | | | | | |
| NC_011770 | 4774927 | A | G | 77 | 0 | 12 | 0 | G | G | G | G | G | G | G | G | G | G | G | G |
| NC_011770 | 4774937 | C | CG | 80 | 0 | 12 | 0 | CG | CG | CG | CG | CG | CG | CG | CG | CG | CG | CG | CG |
| NC_011770 | 4774943 | C | T | 85 | 0 | 2 | 0 | | | | | | | T | T | | | | |
| NC_011770 | 4775078 | A | T | 87 | 0 | 2 | 0 | | | | | | | T | T | | | | |
| NC_011770 | 4775382 | G | C | 81 | 0 | 2 | 0 | | | | | | | C | C | | | | |
| NC_011770 | 4775435 | C | A | 81 | 0 | 2 | 0 | A | A | | | | | | | | | | |
| NC_011770 | 4776251 | G | A | 61 | 0 | 2 | 0 | | | | | | | | | | | A | A |
| NC_011770 | 4776297 | G | A | 65 | 0 | 4 | 0 | | | A | A | A | A | | | | | | |
| NC_011770 | 4776307 | G | GC | 64 | 0 | 1 | 0 | | | | | | | GC | | | | | |
| NC_011770 | 4776309 | A | C | 58 | 1 | 1 | 0 | | | | | | | . | C | | | | |
| NC_011770 | 4776330 | C | T | 65 | 0 | 4 | 0 | | | T | T | T | T | | | | | | |
| NC_011770 | 4776394 | T | C | 73 | 0 | 6 | 0 | C | C | C | C | C | C | | | | | | |
| NC_011770 | 4777055 | G | T | 83 | 0 | 2 | 0 | T | T | | | | | | | | | | |
| NC_011770 | 4777700 | G | A | 57 | 0 | 2 | 0 | A | A | | | | | | | | | | |
| NC_011770 | 4777840 | T | C | 49 | 0 | 12 | 0 | C | C | C | C | C | C | C | C | C | C | C | C |
| NC_011770 | 4777917 | T | G | 47 | 0 | 12 | 0 | G | G | G | G | G | G | G | G | G | G | G | G |
| NC_011770 | 4777921 | C | T | 47 | 0 | 8 | 0 | | | T | T | T | T | T | T | | | T | T |
| NC_011770 | 4777936 | G | T | 46 | 0 | 8 | 0 | | | T | T | T | T | T | T | | | T | T |
| NC_011770 | 4778079 | C | T | 62 | 0 | 4 | 0 | | | T | T | T | T | | | | | | |
| NC_011770 | 4778084 | T | C | 62 | 0 | 4 | 0 | | | C | C | C | C | | | | | | |
| NC_011770 | 4778773 | A | G | 90 | 0 | 4 | 0 | | | G | G | G | G | | | | | | |
| NC_011770 | 4779588 | C | T | 74 | 0 | 2 | 0 | | | | | | | T | T | | | | |
| NC_011770 | 4779597 | C | A | 73 | 0 | 4 | 0 | | | A | A | A | A | | | | | | |
| NC_011770 | 4779650 | A | G | 76 | 0 | 4 | 0 | | | G | G | G | G | | | | | | |
| NC_011770 | 4786106 | G | A | 76 | 0 | 4 | 0 | | | A | A | A | A | | | | | | |
| NC_011770 | 4787284 | T | C | 66 | 0 | 12 | 0 | C | C | C | C | C | C | C | C | C | C | C | C |
| NC_011770 | 4787290 | T | TC | 65 | 0 | 12 | 0 | TC | TC | TC | TC | TC | TC | TC | TC | TC | TC | TC | TC |
| NC_011770 | 4788138 | T | A | 55 | 0 | 4 | 0 | | | A | A | A | A | | | | | | |
| NC_011770 | 4790103 | C | T | 63 | 0 | 4 | 0 | | | T | T | T | T | | | | | | |
| NC_011770 | 4790104 | T | C | 63 | 0 | 10 | 0 | C | C | C | C | C | C | C | C | C | C | | |
| NC_011770 | 4790213 | T | G | 63 | 0 | 6 | 0 | | | G | G | G | G | TC | TC | | | | |
| NC_011770 | 4790215 | C | A | 54 | 1 | 4 | 0 | | | A | A | A | A | . | | | | | |
| NC_011770 | 4790863 | G | A | 24 | 0 | 2 | 0 | | | | | | | | | | | A | A |
| NC_011770 | 4790881 | T | C | 23 | 0 | 4 | 0 | | | C | C | C | C | | | | | | |
| NC_011770 | 4790889 | TCC | T | 20 | 1 | 6 | 0 | T | T | T | T | T | T | | . | | | | |
| NC_011770 | 4790950 | C | G | 29 | 0 | 2 | 0 | | | | | | | | | | | G | G |
| NC_011770 | 4790961 | G | A | 31 | 0 | 4 | 0 | | | A | A | A | A | | | | | | |
| NC_011770 | 4790976 | G | T | 32 | 0 | 2 | 0 | T | T | | | | | | | | | | |
| NC_011770 | 4791395 | A | G | 37 | 0 | 4 | 0 | | | G | G | G | G | | | | | | |
| NC_011770 | 4791404 | A | G | 37 | 0 | 2 | 0 | G | G | | | | | | | | | | |
| NC_011770 | 4791415 | G | A | 37 | 0 | 4 | 0 | | | A | A | A | A | | | | | | |
| NC_011770 | 4791432 | G | A | 36 | 0 | 4 | 0 | | | A | A | A | A | | | | | | |
| NC_011770 | 4791434 | A | G | 37 | 0 | 4 | 0 | | | G | G | G | G | | | | | | |
| NC_011770 | 4792637 | T | A | 76 | 0 | 4 | 0 | | | A | A | A | A | | | | | | |
| NC_011770 | 4792648 | A | G | 74 | 0 | 4 | 0 | | | G | G | G | G | | | | | | |
| NC_011770 | 4792653 | A | G | 74 | 0 | 4 | 0 | | | G | G | G | G | | | | | | |
| NC_011770 | 4792680 | C | T | 75 | 0 | 4 | 0 | | | T | T | T | T | | | | | | |
| NC_011770 | 4792721 | A | G | 72 | 0 | 4 | 0 | | | G | G | G | G | | | | | | |
| NC_011770 | 4792807 | G | A | 75 | 0 | 4 | 0 | | | A | A | A | A | | | | | | |
| NC_011770 | 4792821 | A | G | 76 | 0 | 6 | 0 | G | G | G | G | G | G | | | | | | |

| | | | | | | | | | | | | | | | | | | | |
|---|---|---|---|---|---|---|---|---|---|---|---|---|---|---|---|---|---|---|---|
| NC_011770 | 4792861 | C | G | 81 | 0 | 4 | 0 | | | G | G | G | G | | | | | | |
| NC_011770 | 4792912 | G | A | 84 | 0 | 4 | 0 | | | A | A | A | A | | | | | | |
| NC_011770 | 4792933 | G | A | 82 | 0 | 4 | 0 | | | A | A | A | A | | | | | | |
| NC_011770 | 4792935 | A | G | 81 | 0 | 4 | 0 | | | G | G | G | G | | | | | | |
| NC_011770 | 4792937 | G | C | 81 | 0 | 4 | 0 | | | C | C | C | C | | | | | | |
| NC_011770 | 4794452 | T | C | 68 | 0 | 4 | 0 | | | C | C | C | C | | | | | | |
| NC_011770 | 4798559 | C | T | 72 | 0 | 4 | 0 | | | T | T | T | T | | | | | | |
| NC_011770 | 4798580 | A | G | 71 | 0 | 4 | 0 | | | G | G | G | G | | | | | | |
| NC_011770 | 4799446 | T | C | 74 | 0 | 1 | 0 | | | | | | | C | | | | | |
| NC_011770 | 4800769 | G | T | 78 | 0 | 10 | 0 | T | T | T | T | T | T | T | T | | | T | T |
| NC_011770 | 4802245 | C | A | 91 | 0 | 2 | 0 | A | A | | | | | | | | | | |
| NC_011770 | 4802293 | G | A | 77 | 0 | 10 | 0 | A | A | A | A | A | A | A | A | | | A | A |
| NC_011770 | 4802294 | A | G | 77 | 0 | 10 | 0 | G | G | G | G | G | G | G | G | | | G | G |
| NC_011770 | 4802297 | C | T | 76 | 0 | 10 | 0 | T | T | T | T | T | T | T | T | | | T | T |
| NC_011770 | 4802345 | C | T | 84 | 0 | 2 | 0 | | | | | | | T | T | | | | |
| NC_011770 | 4803187 | G | A | 75 | 0 | 4 | 0 | | | A | A | A | A | | | | | | |
| NC_011770 | 4803213 | C | T | 76 | 0 | 2 | 0 | | | | | | | T | T | | | | |
| NC_011770 | 4803294 | A | G | 73 | 0 | 6 | 0 | | | G | G | G | G | G | G | | | | |
| NC_011770 | 4803311 | G | A | 72 | 0 | 4 | 0 | | | A | A | A | A | | | | | | |
| NC_011770 | 4807004 | T | C | 69 | 0 | 2 | 0 | | | | | | | | | | | C | C |
| NC_011770 | 4807633 | CA | C | 94 | 0 | 2 | 0 | | | | | | | | | | | C | C |
| NC_011770 | 4807635 | A | T | 81 | 0 | 2 | 0 | | | | | | | | | T | T | | |
| NC_011770 | 4807673 | G | A | 85 | 0 | 2 | 0 | | | | | | | A | A | | | | |
| NC_011770 | 4807675 | G | C | 85 | 0 | 6 | 0 | | | C | C | C | C | C | C | | | | |
| NC_011770 | 4807676 | C | T | 85 | 0 | 4 | 0 | | | T | T | T | T | | | | | | |
| NC_011770 | 4807689 | GA | G | 91 | 0 | 4 | 0 | | | G | G | G | G | | | | | | |
| NC_011770 | 4807713 | A | C | 92 | 0 | 2 | 0 | C | C | | | | | | | | | | |
| NC_011770 | 4807718 | C | T | 90 | 0 | 6 | 0 | | | | | | | T | T | T | T | T | T |
| NC_011770 | 4808587 | G | A | 94 | 0 | 2 | 0 | | | | | | | | | A | A | | |
| NC_011770 | 4808598 | A | T | 93 | 0 | 4 | 0 | | | T | T | T | T | | | | | | |
| NC_011770 | 4808861 | C | T | 74 | 0 | 2 | 0 | | | | | | | T | T | | | | |
| NC_011770 | 4808879 | C | A | 76 | 0 | 2 | 0 | | | | | | | | | A | A | | |
| NC_011770 | 4810269 | A | G | 74 | 0 | 4 | 0 | | | G | G | G | G | | | | | | |
| NC_011770 | 4810278 | T | A | 74 | 0 | 6 | 0 | | | A | A | A | A | G | G | | | | |
| NC_011770 | 4810292 | G | A | 73 | 0 | 2 | 0 | | | | | | | A | A | | | | |
| NC_011770 | 4810403 | G | A | 73 | 0 | 6 | 0 | | | A | A | A | A | A | A | | | | |
| NC_011770 | 4810954 | G | A | 52 | 0 | 2 | 0 | | | | | | | | | A | A | | |
| NC_011770 | 4812154 | T | TC | 59 | 0 | 2 | 0 | | | | | | | | | TC | TC | | |
| NC_011770 | 4813086 | A | G | 68 | 0 | 8 | 0 | | | G | G | G | G | | | G | G | G | G |
| NC_011770 | 4817820 | C | T | 88 | 0 | 2 | 0 | | | | | | | T | T | | | | |
| NC_011770 | 4817822 | A | G | 88 | 0 | 1 | 0 | | | | | | | G | | | | | |
| NC_011770 | 4817916 | A | G | 83 | 0 | 6 | 0 | | | G | G | G | G | | | | | G | G |
| NC_011770 | 4820296 | G | A | 85 | 0 | 12 | 0 | A | A | A | A | A | A | A | A | A | A | A | A |
| NC_011770 | 4820319 | G | A | 86 | 0 | 2 | 0 | | | | | | | A | A | | | | |
| NC_011770 | 4820320 | A | C | 84 | 0 | 12 | 0 | C | C | C | C | C | C | C | C | C | C | C | C |
| NC_011770 | 4820323 | G | A | 86 | 0 | 2 | 0 | | | | | | | | | | | A | A |
| NC_011770 | 4820335 | C | T | 83 | 0 | 12 | 0 | T | T | T | T | T | T | T | T | T | T | T | T |
| NC_011770 | 4820337 | C | A | 84 | 0 | 4 | 0 | | | | | | | A | A | A | A | | |
| NC_011770 | 4820343 | A | G | 88 | 0 | 2 | 0 | | | | | | | G | G | | | | |
| NC_011770 | 4820364 | T | C | 87 | 0 | 4 | 0 | | | C | C | C | C | | | | | | |
| NC_011770 | 4820366 | C | T | 86 | 0 | 8 | 0 | T | T | T | T | T | T | | | | | T | T |
| NC_011770 | 4820369 | G | A | 86 | 0 | 4 | 0 | | | A | A | A | A | | | | | | |
| NC_011770 | 4822458 | C | A | 45 | 0 | 2 | 0 | | | | | | | A | A | | | | |
| NC_011770 | 4822494 | C | T | 51 | 0 | 2 | 0 | | | | | | | | | | | T | T |
| NC_011770 | 4822579 | A | G | 55 | 0 | 12 | 0 | G | G | G | G | G | G | G | G | G | G | G | G |
| NC_011770 | 4826878 | C | T | 77 | 0 | 8 | 0 | T | T | T | T | T | T | | | T | T | | |
| NC_011770 | 4830564 | T | C | 50 | 2 | 2 | 0 | C | C | | | | | | | | | . | . |
| NC_011770 | 4830714 | G | A | 49 | 2 | 4 | 0 | | | A | A | A | A | | | | | . | . |
| NC_011770 | 4830724 | T | C | 54 | 2 | 4 | 0 | | | C | C | C | C | | | | | . | . |
| NC_011770 | 4830727 | C | T | 53 | 2 | 4 | 0 | | | | | | | T | T | T | T | . | . |
| NC_011770 | 4831163 | T | C | 68 | 0 | 4 | 0 | C | C | | | | | | | | | C | C |
| NC_011770 | 4831540 | T | TC | 51 | 0 | 2 | 0 | TC | TC | | | | | | | | | | |

| | | | | | | | | | | | | | | | | | | | |
|---|---|---|---|---|---|---|---|---|---|---|---|---|---|---|---|---|---|---|---|
| NC_011770 | 4836425 | C | T | 49 | 0 | 2 | 0 | | | | | | | T | T | | | | |
| NC_011770 | 4836443 | A | G | 50 | 0 | 2 | 0 | | | | | | | G | G | | | | |
| NC_011770 | 4836453 | C | CT | 52 | 0 | 2 | 0 | | | | | | | CT | CT | | | | |
| NC_011770 | 4836468 | G | A | 52 | 0 | 2 | 0 | | | | | | | A | A | | | | |
| NC_011770 | 4838471 | G | T | 74 | 0 | 4 | 0 | | | T | T | T | T | | | | | | |
| NC_011770 | 4838594 | A | G | 71 | 0 | 2 | 0 | G | G | | | | | | | | | | |
| NC_011770 | 4839950 | T | C | 67 | 2 | 2 | 0 | C | C | | | | | | | | | . | . |
| NC_011770 | 4839976 | C | CT | 64 | 0 | 4 | 0 | CT | CT | | | | | | | | | CT | CT |
| NC_011770 | 4842362 | A | AC | 81 | 0 | 4 | 0 | | | AC | AC | AC | AC | | | | | | |
| NC_011770 | 4842455 | G | T | 84 | 0 | 2 | 0 | | | | | | | T | T | | | | |
| NC_011770 | 4844248 | G | T | 66 | 0 | 4 | 0 | T | T | | | | | T | T | | | | |
| NC_011770 | 4845048 | C | T | 31 | 0 | 2 | 0 | | | | | | | T | T | | | | |
| NC_011770 | 4845122 | C | A | 42 | 0 | 2 | 0 | | | | | | | | | A | A | | |
| NC_011770 | 4845130 | C | T | 42 | 0 | 2 | 0 | | | | | | | T | T | | | | |
| NC_011770 | 4845131 | A | C | 42 | 0 | 2 | 0 | | | | | | | C | C | | | | |
| NC_011770 | 4845134 | C | T | 41 | 0 | 2 | 0 | | | | | | | T | T | | | | |
| NC_011770 | 4845761 | G | C | 79 | 0 | 6 | 0 | | | | | | | C | C | C | C | C | C |
| NC_011770 | 4845763 | T | C | 78 | 0 | 6 | 0 | | | | | | | C | C | C | C | C | C |
| NC_011770 | 4845806 | G | C | 85 | 0 | 6 | 0 | | | | | | | C | C | C | C | C | C |
| NC_011770 | 4845839 | G | T | 91 | 0 | 6 | 0 | | | | | | | T | T | T | T | T | T |
| NC_011770 | 4845842 | C | A | 92 | 0 | 6 | 0 | | | | | | | A | A | A | A | A | A |
| NC_011770 | 4846611 | C | T | 67 | 0 | 2 | 0 | | | | | | | | | | | T | T |
| NC_011770 | 4846638 | G | A | 68 | 0 | 4 | 0 | | | | | | | A | A | | | A | A |
| NC_011770 | 4846812 | C | G | 52 | 0 | 2 | 0 | G | G | | | | | | | | | | |
| NC_011770 | 4846816 | G | GC | 61 | 0 | 2 | 0 | | | | | | | | | | | GC | GC |
| NC_011770 | 4847506 | G | A | 78 | 0 | 8 | 0 | A | A | | | | | A | A | A | A | A | A |
| NC_011770 | 4847555 | C | T | 75 | 0 | 6 | 0 | | | | | | | T | T | T | T | T | T |
| NC_011770 | 3796943 | G | A | 54 | 0 | 1 | 0 | | | | | A | | | | | | | |
| NC_011770 | 4848919 | A | G | 70 | 0 | 6 | 0 | | | | | | | G | G | G | G | G | G |
| NC_011770 | 4848932 | G | A | 72 | 0 | 2 | 0 | | | | | | | | | | | A | A |
| NC_011770 | 4854150 | C | T | 73 | 0 | 4 | 0 | | | | | | | T | T | | | T | T |
| NC_011770 | 4859214 | G | T | 67 | 0 | 2 | 0 | | | | | | | | | | | T | T |
| NC_011770 | 4860641 | G | A | 84 | 0 | 2 | 0 | | | | | | | A | A | | | | |
| NC_011770 | 4865437 | G | A | 98 | 0 | 4 | 0 | A | A | | | | | A | A | | | | |
| NC_011770 | 4866299 | A | G | 50 | 0 | 12 | 0 | G | G | G | G | G | G | G | G | G | G | G | G |
| NC_011770 | 4866556 | C | A | 58 | 0 | 2 | 0 | | | | | | | A | A | | | | |
| NC_011770 | 4866559 | G | A | 58 | 0 | 2 | 0 | | | | | | | | | A | A | | |
| NC_011770 | 4866565 | A | G | 59 | 0 | 2 | 0 | | | | | | | | | | | G | G |
| NC_011770 | 4866689 | T | C | 54 | 0 | 1 | 0 | | | | | | | C | | | | | |
| NC_011770 | 4868828 | C | A | 78 | 0 | 12 | 0 | A | A | A | A | A | A | A | A | A | A | A | A |
| NC_011770 | 4868881 | C | T | 77 | 0 | 2 | 0 | | | | | | | | | T | T | | |
| NC_011770 | 4872810 | C | T | 70 | 0 | 2 | 0 | T | T | | | | | | | | | | |
| NC_011770 | 4874048 | G | A | 83 | 0 | 2 | 0 | | | | | | | | | A | A | | |
| NC_011770 | 4874059 | A | G | 79 | 0 | 12 | 0 | G | G | G | G | G | G | G | G | G | G | G | G |
| NC_011770 | 4875432 | T | C | 79 | 0 | 12 | 0 | C | C | C | C | C | C | C | C | C | C | C | C |
| NC_011770 | 4875459 | A | G | 77 | 0 | 8 | 0 | | | G | G | G | G | G | G | G | G | | |
| NC_011770 | 4875848 | C | T | 85 | 0 | 2 | 0 | | | | | | | T | T | | | | |
| NC_011770 | 4877058 | A | G | 69 | 0 | 12 | 0 | G | G | G | G | G | G | G | G | G | G | G | G |
| NC_011770 | 4878928 | A | G | 58 | 0 | 6 | 0 | | | G | G | G | G | | | G | G | | |
| NC_011770 | 4878972 | A | AT | 59 | 0 | 10 | 0 | AT | AT | AT | AT | AT | AT | | | AT | AT | AT | AT |
| NC_011770 | 4882425 | G | C | 85 | 0 | 2 | 0 | | | C | C | | | | | | | | |
| NC_011770 | 4882966 | G | C | 72 | 0 | 2 | 0 | | | | | | | | | | | C | C |
| NC_011770 | 4883045 | T | C | 73 | 0 | 4 | 0 | C | C | | | | | | | C | C | | |
| NC_011770 | 4884817 | A | G | 68 | 0 | 2 | 0 | | | | | | | | | | | G | G |
| NC_011770 | 4885776 | A | T | 71 | 0 | 2 | 0 | | | | | | | T | T | | | | |
| NC_011770 | 4885784 | G | A | 71 | 0 | 2 | 0 | | | | | | | A | A | | | | |
| NC_011770 | 4885827 | C | T | 71 | 0 | 2 | 0 | | | | | | | T | T | | | | |
| NC_011770 | 4885828 | A | C | 71 | 0 | 2 | 0 | | | | | | | C | C | | | | |
| NC_011770 | 4885842 | G | C | 72 | 0 | 2 | 0 | | | | | | | C | C | | | | |
| NC_011770 | 4887490 | G | A | 68 | 0 | 2 | 0 | | | | | | | | | | | A | A |
| NC_011770 | 4887551 | T | A | 70 | 0 | 2 | 0 | | | | | | | | | | | A | A |
| NC_011770 | 4887554 | T | C | 71 | 0 | 2 | 0 | | | | | | | | | | | C | C |

| | | | | | | | | | | | | | | | | | | | |
|---|---|---|---|---|---|---|---|---|---|---|---|---|---|---|---|---|---|---|---|
| NC_011770 | 4887571 | A | C | 69 | 0 | 4 | 0 | | | C | C | C | C | | | | | | |
| NC_011770 | 4887601 | T | C | 69 | 0 | 4 | 0 | | | C | C | C | C | | | | | | |
| NC_011770 | 4889119 | G | T | 69 | 0 | 2 | 0 | | | | | | | | | | | T | T |
| NC_011770 | 4889171 | C | A | 64 | 0 | 2 | 0 | A | A | | | | | | | | | | |
| NC_011770 | 4889658 | T | A | 67 | 0 | 2 | 0 | | | | | | | A | A | | | | |
| NC_011770 | 4889715 | A | G | 65 | 0 | 12 | 0 | G | G | G | G | G | G | G | G | G | G | G | G |
| NC_011770 | 4889718 | A | G | 66 | 0 | 4 | 0 | | | G | G | G | G | | | | | | |
| NC_011770 | 4891187 | A | G | 49 | 0 | 6 | 0 | | | G | G | G | G | | | | | G | G |
| NC_011770 | 4891307 | G | A | 42 | 0 | 2 | 0 | | | | | | | | | | | A | A |
| NC_011770 | 4891311 | G | A | 42 | 0 | 2 | 0 | | | | | | | | | | | A | A |
| NC_011770 | 4891411 | G | A | 43 | 0 | 2 | 0 | A | A | | | | | | | | | | |
| NC_011770 | 4891462 | G | C | 46 | 0 | 4 | 0 | | | | | | | C | C | | | C | C |
| NC_011770 | 4892607 | G | A | 50 | 0 | 10 | 0 | A | A | A | A | A | A | A | A | A | A | | |
| NC_011770 | 4892754 | A | G | 34 | 0 | 10 | 0 | G | G | G | G | G | G | G | G | G | G | | |
| NC_011770 | 4892767 | G | T | 31 | 0 | 12 | 0 | T | T | T | T | T | T | T | T | T | T | T | T |
| NC_011770 | 4892832 | C | T | 24 | 0 | 4 | 0 | T | T | | | | | T | T | | | | |
| NC_011770 | 4893669 | T | C | 76 | 0 | 4 | 0 | | | | | | | | | C | C | A | A |
| NC_011770 | 4893679 | C | T | 77 | 0 | 2 | 0 | | | | | | | | | T | T | | |
| NC_011770 | 4893680 | G | T | 76 | 0 | 4 | 0 | | | T | T | T | T | | | | | | |
| NC_011770 | 4893718 | C | T | 77 | 0 | 2 | 0 | | | | | | | | | T | T | | |
| NC_011770 | 4893719 | A | G | 76 | 0 | 2 | 0 | | | | | | | | | G | G | | |
| NC_011770 | 4893732 | G | A | 77 | 0 | 1 | 0 | | | | | | | A | | | | | |
| NC_011770 | 4894926 | A | G | 40 | 0 | 10 | 0 | G | G | G | G | G | G | G | G | G | G | | |
| NC_011770 | 4894929 | G | A | 40 | 0 | 12 | 0 | A | A | A | A | A | A | A | A | A | A | A | A |
| NC_011770 | 4894932 | C | T | 40 | 0 | 6 | 0 | | | T | T | T | T | | | T | T | | |
| NC_011770 | 4894948 | C | T | 41 | 0 | 6 | 0 | | | T | T | T | T | | | T | T | | |
| NC_011770 | 4894956 | A | G | 47 | 0 | 6 | 0 | | | G | G | G | G | | | G | G | | |
| NC_011770 | 4895756 | T | TA | 84 | 0 | 4 | 0 | | | TA | TA | TA | TA | | | | | | |
| NC_011770 | 4895761 | G | C | 79 | 0 | 2 | 0 | | | | | | | | | | | C | C |
| NC_011770 | 4895803 | C | T | 80 | 0 | 2 | 0 | | | | | | | | | | | T | T |
| NC_011770 | 4898307 | C | T | 75 | 0 | 2 | 0 | T | T | | | | | | | | | | |
| NC_011770 | 4898320 | G | A | 74 | 0 | 2 | 0 | A | A | | | | | | | | | | |
| NC_011770 | 4898418 | T | C | 71 | 0 | 2 | 0 | C | C | | | | | | | | | | |
| NC_011770 | 4898499 | A | G | 65 | 0 | 12 | 0 | G | G | G | G | G | G | G | G | G | G | G | G |
| NC_011770 | 4898552 | A | G | 67 | 0 | 2 | 0 | | | | | | | | | G | G | | |
| NC_011770 | 4902080 | G | A | 85 | 0 | 2 | 0 | | | | | | | | | | | A | A |
| NC_011770 | 4903091 | G | T | 66 | 0 | 2 | 0 | | | | | | | T | T | | | | |
| NC_011770 | 4904148 | G | C | 61 | 0 | 2 | 0 | | | | | | | | | | | C | C |
| NC_011770 | 4904214 | C | G | 59 | 0 | 4 | 0 | | | | | | | G | G | G | G | | |
| NC_011770 | 4906361 | C | G | 28 | 0 | 2 | 0 | | | | | | | | | | | G | G |
| NC_011770 | 4909326 | G | C | 88 | 0 | 2 | 0 | | | | | | | | | | | C | C |
| NC_011770 | 4909419 | C | T | 83 | 0 | 2 | 0 | | | | | | | | | | | T | T |
| NC_011770 | 4909440 | A | C | 87 | 0 | 2 | 0 | | | | | | | | | | | C | C |
| NC_011770 | 4909484 | C | T | 84 | 0 | 2 | 0 | | | | | | | | | | | T | T |
| NC_011770 | 4909531 | C | T | 85 | 0 | 1 | 0 | | | | | | | T | | | | | |
| NC_011770 | 4914253 | G | A | 37 | 0 | 6 | 0 | | | | | | | A | A | A | A | A | A |
| NC_011770 | 4914273 | G | A | 36 | 0 | 2 | 0 | | | | | A | A | | | | | | |
| NC_011770 | 4914463 | GC | G | 18 | 1 | 2 | 0 | | | | | | | G | G | | . | | |
| NC_011770 | 4914481 | A | C | 23 | 1 | 1 | 0 | | | | | | | | | C | . | | |
| NC_011770 | 4914510 | A | G | 32 | 0 | 4 | 0 | | | G | G | G | G | | | | | | |
| NC_011770 | 4916615 | G | T | 79 | 0 | 2 | 0 | | | | | | | T | T | | | | |
| NC_011770 | 4916624 | G | A | 78 | 0 | 4 | 0 | | | A | A | A | A | | | | | | |
| NC_011770 | 4916671 | TA | T | 75 | 0 | 8 | 0 | T | T | T | T | T | T | | | | | T | T |
| NC_011770 | 4916717 | G | T | 75 | 0 | 2 | 0 | | | | | | | | | T | T | | |
| NC_011770 | 4916723 | C | T | 74 | 0 | 2 | 0 | | | | | | | T | T | | | | |
| NC_011770 | 4916813 | T | C | 70 | 0 | 2 | 0 | | | | | | | | | | | C | C |
| NC_011770 | 4919790 | A | C | 61 | 0 | 2 | 0 | | | | | | | | | | | C | C |
| NC_011770 | 4922101 | C | G | 59 | 0 | 8 | 0 | | | G | G | G | G | G | G | | | G | G |
| NC_011770 | 4922126 | C | T | 59 | 0 | 4 | 0 | | | | | | | T | T | | | T | T |
| NC_011770 | 4922131 | C | G | 60 | 0 | 4 | 0 | | | | | | | G | G | | | G | G |
| NC_011770 | 4922133 | G | A | 60 | 0 | 4 | 0 | | | A | A | A | A | | | | | | |
| NC_011770 | 4922188 | C | T | 60 | 0 | 4 | 0 | | | | | | | T | T | | | T | T |

| | | | | | | | | | | | | | | | | | | | |
|---|---|---|---|---|---|---|---|---|---|---|---|---|---|---|---|---|---|---|---|
| NC_011770 | 4922190 | G | C | 60 | 0 | 4 | 0 | | | | | | | C | C | | | C | C |
| NC_011770 | 4922202 | G | A | 57 | 0 | 4 | 0 | | | | | | | A | A | | | A | A |
| NC_011770 | 4922210 | G | A | 60 | 0 | 4 | 0 | | | | | | | A | A | | | A | A |
| NC_011770 | 4922217 | C | T | 60 | 0 | 4 | 0 | | | | | | | T | T | | | T | T |
| NC_011770 | 4924031 | G | A | 73 | 0 | 2 | 0 | | | | | | | A | A | | | | |
| NC_011770 | 4924036 | A | G | 72 | 0 | 8 | 0 | G | G | | | | | G | G | G | G | G | G |
| NC_011770 | 4924052 | G | A | 73 | 0 | 2 | 0 | | | | | | | A | A | | | | |
| NC_011770 | 4924074 | G | C | 70 | 0 | 2 | 0 | | | | | | | C | C | | | | |
| NC_011770 | 4924120 | C | CGG | 67 | 0 | 2 | 0 | | | | | | | CGG | CGG | | | | |
| NC_011770 | 4924658 | G | A | 75 | 0 | 4 | 0 | | | A | A | A | A | | | | | | |
| NC_011770 | 4925353 | A | G | 79 | 0 | 2 | 0 | | | | | | | | | | | G | G |
| NC_011770 | 4930049 | A | G | 46 | 0 | 12 | 0 | G | G | G | G | G | G | G | G | G | G | G | G |
| NC_011770 | 4930884 | A | C | 68 | 0 | 12 | 0 | C | C | C | C | C | C | C | C | C | C | C | C |
| NC_011770 | 4931543 | A | G | 28 | 8 | 4 | 0 | . | . | G | G | G | G | . | . | . | . | . | . |
| NC_011770 | 4931546 | T | C | 28 | 8 | 4 | 0 | . | . | C | C | C | C | . | . | . | . | . | . |
| NC_011770 | 4931580 | A | G | 19 | 8 | 4 | 0 | . | . | G | G | G | G | . | . | . | . | . | . |
| NC_011770 | 4931582 | T | C | 19 | 8 | 4 | 0 | . | . | C | C | C | C | . | . | . | . | . | . |
| NC_011770 | 4931587 | T | A | 18 | 8 | 4 | 0 | . | . | A | A | A | A | . | . | . | . | . | . |
| NC_011770 | 4931589 | A | T | 19 | 8 | 4 | 0 | . | . | T | T | T | T | . | . | . | . | . | . |
| NC_011770 | 4931590 | G | C | 18 | 8 | 4 | 0 | . | . | C | C | C | C | . | . | . | . | . | . |
| NC_011770 | 4931591 | A | T | 19 | 8 | 4 | 0 | . | . | T | T | T | T | . | . | . | . | . | . |
| NC_011770 | 3796950 | C | CA | 51 | 0 | 1 | 0 | | | | | CA | | | | | | | |
| NC_011770 | 4949582 | A | G | 25 | 5 | 1 | 0 | | | | | | | . | . | G | . | . | . |
| NC_011770 | 4960937 | G | A | 37 | 0 | 12 | 0 | A | A | A | A | A | A | A | A | A | A | A | A |
| NC_011770 | 4960940 | G | C | 36 | 0 | 12 | 0 | C | C | C | C | C | C | C | C | C | C | C | C |
| NC_011770 | 4960949 | G | A | 37 | 0 | 12 | 0 | A | A | A | A | A | A | A | A | A | A | A | A |
| NC_011770 | 4961026 | A | G | 40 | 0 | 4 | 0 | | | G | G | G | G | | | | | | |
| NC_011770 | 4961072 | T | C | 39 | 0 | 4 | 0 | | | C | C | C | C | | | | | | |
| NC_011770 | 4963085 | A | G | 69 | 0 | 12 | 0 | G | G | G | G | G | G | G | G | G | G | G | G |
| NC_011770 | 4963099 | A | T | 70 | 0 | 2 | 0 | | | | | | | | | T | T | | |
| NC_011770 | 4963107 | T | C | 62 | 1 | 10 | 0 | C | C | C | C | C | C | C | C | . | . | C | C |
| NC_011770 | 4963135 | C | T | 68 | 0 | 12 | 0 | T | T | T | T | T | T | T | T | T | T | T | T |
| NC_011770 | 4963188 | C | A | 70 | 0 | 10 | 0 | A | A | A | A | A | A | A | A | | | A | A |
| NC_011770 | 4964193 | A | G | 78 | 0 | 8 | 0 | | | G | G | G | G | G | G | G | G | | |
| NC_011770 | 4964197 | A | G | 76 | 0 | 8 | 0 | | | G | G | G | G | G | G | G | G | | |
| NC_011770 | 4964237 | C | T | 80 | 0 | 4 | 0 | | | T | T | T | T | | | | | | |
| NC_011770 | 4964991 | C | T | 75 | 0 | 2 | 0 | | | | | | | | | T | T | | |
| NC_011770 | 4965939 | T | C | 61 | 0 | 12 | 0 | C | C | C | C | C | C | C | C | C | C | C | C |
| NC_011770 | 4965949 | T | A | 64 | 0 | 2 | 0 | | | | | | | | | | | A | A |
| NC_011770 | 4966314 | C | T | 37 | 0 | 4 | 0 | T | T | | | | | T | T | | | | |
| NC_011770 | 4966315 | G | C | 37 | 0 | 4 | 0 | C | C | | | | | T | T | | | | |
| NC_011770 | 4966448 | G | A | 41 | 0 | 6 | 0 | | | | | | | A | A | A | A | A | A |
| NC_011770 | 4966474 | C | T | 41 | 0 | 6 | 0 | | | | | | | T | T | T | T | T | T |
| NC_011770 | 4966493 | T | C | 43 | 0 | 6 | 0 | | | | | | | C | C | C | C | C | C |
| NC_011770 | 4966540 | G | T | 50 | 0 | 2 | 0 | | | | | | | T | T | | | | |
| NC_011770 | 4966550 | T | C | 51 | 0 | 6 | 0 | | | | | | | C | C | C | C | C | C |
| NC_011770 | 4966581 | A | C | 52 | 0 | 4 | 0 | | | | | | | | | C | C | C | C |
| NC_011770 | 4966602 | C | T | 51 | 0 | 6 | 0 | | | | | | | T | T | T | T | T | T |
| NC_011770 | 4966665 | G | T | 53 | 0 | 4 | 0 | | | T | T | T | T | | | | | | |
| NC_011770 | 4966683 | T | C | 61 | 0 | 6 | 0 | | | | | | | C | C | C | C | C | C |
| NC_011770 | 4967476 | T | C | 58 | 2 | 6 | 0 | | | C | C | C | C | C | C | . | . | | |
| NC_011770 | 4967494 | T | G | 56 | 2 | 6 | 0 | | | G | G | G | G | G | G | . | . | | |
| NC_011770 | 4967495 | G | C | 56 | 2 | 6 | 0 | | | C | C | C | C | C | C | . | . | | |
| NC_011770 | 4967499 | T | C | 56 | 2 | 6 | 0 | | | C | C | C | C | C | C | . | . | | |
| NC_011770 | 4967503 | C | A | 56 | 2 | 6 | 0 | | | A | A | A | A | A | A | . | . | | |
| NC_011770 | 4967509 | T | C | 60 | 2 | 4 | 0 | | | C | C | C | C | | | . | . | | |
| NC_011770 | 4967518 | C | T | 60 | 2 | 2 | 0 | | | | | | | T | T | . | . | | |
| NC_011770 | 4967552 | C | T | 58 | 2 | 6 | 0 | | | T | T | T | T | T | T | . | . | | |
| NC_011770 | 4967553 | C | T | 57 | 2 | 6 | 0 | | | T | T | T | T | T | T | . | . | | |
| NC_011770 | 4967579 | C | T | 61 | 2 | 6 | 0 | | | T | T | T | T | T | T | . | . | | |
| NC_011770 | 4967594 | C | T | 62 | 2 | 6 | 0 | | | T | T | T | T | T | T | . | . | | |
| NC_011770 | 4967614 | G | C | 64 | 2 | 2 | 0 | | | | | | | C | C | . | . | | |

| | | | | | | | | | | | | | | | | | | | |
|---|---|---|---|---|---|---|---|---|---|---|---|---|---|---|---|---|---|---|---|
| NC_011770 | 4967636 | G | A | 66 | 2 | 6 | 0 | | | A | A | A | A | A | A | . | . | | |
| NC_011770 | 4968043 | T | C | 51 | 2 | 8 | 0 | | | C | C | C | C | C | C | . | . | C | C |
| NC_011770 | 4968045 | A | G | 51 | 2 | 6 | 0 | | | G | G | G | G | G | G | . | . | | |
| NC_011770 | 4968114 | T | C | 45 | 2 | 6 | 0 | | | C | C | C | C | C | C | . | . | | |
| NC_011770 | 4968138 | G | A | 43 | 2 | 2 | 0 | | | | | | | | | . | . | A | A |
| NC_011770 | 4968144 | A | G | 46 | 2 | 2 | 0 | | | | | | | G | G | . | . | | |
| NC_011770 | 4968829 | C | T | 50 | 4 | 6 | 0 | | | T | T | T | T | T | T | . | . | . | . |
| NC_011770 | 4968842 | G | A | 51 | 4 | 4 | 0 | | | A | A | A | A | | | . | . | . | . |
| NC_011770 | 4968884 | G | A | 50 | 4 | 4 | 0 | | | A | A | A | A | | | . | . | . | . |
| NC_011770 | 4970528 | T | C | 54 | 4 | 2 | 0 | | | | | | | C | C | . | . | . | . |
| NC_011770 | 4970530 | A | G | 54 | 4 | 2 | 0 | | | | | | | G | G | . | . | . | . |
| NC_011770 | 4970537 | A | C | 55 | 4 | 2 | 0 | | | | | | | C | C | . | . | . | . |
| NC_011770 | 4970669 | C | T | 56 | 4 | 4 | 0 | | | T | T | T | T | | | . | . | . | . |
| NC_011770 | 4970675 | C | T | 56 | 4 | 2 | 0 | | | | | | | T | T | . | . | . | . |
| NC_011770 | 4970728 | C | T | 61 | 4 | 2 | 0 | | | | | | | T | T | . | . | . | . |
| NC_011770 | 4970745 | C | T | 60 | 4 | 4 | 0 | | | T | T | T | T | | | . | . | . | . |
| NC_011770 | 4970821 | G | T | 64 | 4 | 2 | 0 | | | | | | | T | T | . | . | . | . |
| NC_011770 | 4971237 | A | G | 70 | 4 | 2 | 0 | | | | | | | G | G | . | . | . | . |
| NC_011770 | 4971305 | A | G | 66 | 4 | 2 | 0 | | | | | | | G | G | . | . | . | . |
| NC_011770 | 4971395 | T | G | 59 | 4 | 2 | 0 | | | | | | | G | G | . | . | . | . |
| NC_011770 | 4971533 | T | TAC | 58 | 4 | 1 | 0 | | | | | TAC | | | | . | . | . | . |
| NC_011770 | 4971566 | C | T | 52 | 4 | 2 | 0 | T | T | | | | | | | . | . | . | . |
| NC_011770 | 4971569 | A | G | 52 | 4 | 8 | 0 | G | G | G | G | G | G | G | G | . | . | . | . |
| NC_011770 | 4971592 | T | C | 52 | 4 | 2 | 0 | | | | | | | C | C | . | . | . | . |
| NC_011770 | 4971594 | C | T | 53 | 4 | 2 | 0 | T | T | | | | | | | . | . | . | . |
| NC_011770 | 4971596 | T | C | 53 | 4 | 4 | 0 | C | C | | | | | C | C | . | . | . | . |
| NC_011770 | 4971604 | C | G | 51 | 4 | 4 | 0 | G | G | | | | | G | G | . | . | . | . |
| NC_011770 | 4971623 | A | C | 51 | 4 | 4 | 0 | C | C | | | | | C | C | . | . | . | . |
| NC_011770 | 4971626 | A | G | 51 | 4 | 4 | 0 | G | G | | | | | G | G | . | . | . | . |
| NC_011770 | 4971630 | A | T | 50 | 4 | 2 | 0 | | | | | | | T | T | . | . | . | . |
| NC_011770 | 4971632 | G | A | 51 | 4 | 2 | 0 | A | A | | | | | | | . | . | . | . |
| NC_011770 | 4971653 | C | T | 46 | 4 | 4 | 0 | T | T | | | | | T | T | . | . | . | . |
| NC_011770 | 4971654 | G | C | 48 | 4 | 4 | 0 | C | C | | | | | C | C | . | . | . | . |
| NC_011770 | 4971658 | G | C | 48 | 4 | 2 | 0 | C | C | | | | | | | . | . | . | . |
| NC_011770 | 4971660 | C | T | 48 | 4 | 4 | 0 | T | T | | | | | T | T | . | . | . | . |
| NC_011770 | 4971666 | A | C | 48 | 4 | 4 | 0 | C | C | | | | | C | C | . | . | . | . |
| NC_011770 | 4971676 | A | C | 45 | 4 | 6 | 0 | | | C | C | C | C | C | C | . | . | . | . |
| NC_011770 | 4971683 | C | T | 46 | 4 | 2 | 0 | | | | | | | T | T | . | . | . | . |
| NC_011770 | 4971699 | G | A | 44 | 4 | 2 | 0 | A | A | | | | | | | . | . | . | . |
| NC_011770 | 4971758 | G | GTC | 36 | 4 | 4 | 0 | | | GTC | GTC | GTC | GTC | | | . | . | . | . |
| NC_011770 | 4971765 | G | T | 27 | 4 | 2 | 0 | | | | | | | T | T | . | . | . | . |
| NC_011770 | 4971766 | A | C | 27 | 4 | 2 | 0 | | | | | | | C | C | . | . | . | . |
| NC_011770 | 4971795 | T | C | 30 | 4 | 4 | 0 | C | C | | | | | C | C | . | . | . | . |
| NC_011770 | 4971831 | C | T | 37 | 1 | 2 | 0 | T | T | | | | | | | | | | . |
| NC_011770 | 4971849 | G | T | 38 | 2 | 1 | 0 | | | | | | | | | . | T | . | . |
| NC_011770 | 4971903 | A | G | 37 | 2 | 10 | 0 | G | G | G | G | G | G | G | G | . | . | G | G |
| NC_011770 | 4971904 | CA | C | 37 | 2 | 4 | 0 | | | | | | | C | C | . | . | C | C |
| NC_011770 | 4971915 | A | G | 37 | 2 | 4 | 0 | | | | | | | G | G | . | . | G | G |
| NC_011770 | 4971919 | GGT | G | 36 | 2 | 4 | 0 | | | | | | | G | G | . | . | G | G |
| NC_011770 | 4971978 | A | C | 13 | 7 | 3 | 0 | . | . | . | . | . | . | | | C | . | C | C |
| NC_011770 | 4971982 | G | C | 11 | 7 | 5 | 0 | . | . | . | . | . | . | C | C | C | . | C | C |
| NC_011770 | 4971983 | T | G | 11 | 7 | 5 | 0 | . | . | . | . | . | . | G | G | G | . | G | G |
| NC_011770 | 4972002 | A | G | 13 | 7 | 2 | 0 | . | . | . | . | . | . | G | G | . | . | | |
| NC_011770 | 4972119 | G | A | 40 | 0 | 4 | 0 | | | A | A | A | A | | | | | | |
| NC_011770 | 4972124 | G | T | 38 | 0 | 4 | 0 | | | | | | | | | T | T | T | T |
| NC_011770 | 4972125 | A | C | 38 | 0 | 4 | 0 | | | C | C | C | C | | | | | | |
| NC_011770 | 4972126 | C | T | 39 | 0 | 2 | 0 | | | | | | | | | T | T | | |
| NC_011770 | 4972129 | G | A | 39 | 0 | 2 | 0 | A | A | | | | | | | | | | |
| NC_011770 | 4972131 | T | C | 38 | 0 | 6 | 0 | C | C | C | C | C | C | | | | | | |
| NC_011770 | 4972137 | T | C | 38 | 0 | 2 | 0 | C | C | | | | | | | | | | |
| NC_011770 | 4972139 | G | C | 39 | 0 | 4 | 0 | | | | | | | | | C | C | C | C |
| NC_011770 | 4972147 | T | C | 39 | 0 | 4 | 0 | | | C | C | C | C | | | | | | |

| | | | | | | | | | | | | | | | | | | | |
|---|---|---|---|---|---|---|---|---|---|---|---|---|---|---|---|---|---|---|---|
| NC_011770 | 4972149 | C | T | 39 | 0 | 4 | 0 | | | T | T | T | T | | | | | | |
| NC_011770 | 4972252 | T | C | 33 | 0 | 1 | 0 | | | | | | | C | | | | | |
| NC_011770 | 4972255 | T | C | 33 | 0 | 12 | 0 | C | C | C | C | C | C | C | C | C | C | C | C |
| NC_011770 | 4972291 | T | C | 32 | 0 | 8 | 0 | | | C | C | C | C | | | C | C | C | C |
| NC_011770 | 4972567 | T | C | 1 | 10 | 2 | 0 | . | . | . | . | C | C | . | . | . | . | . | . |
| NC_011770 | 4972572 | C | T | 45 | 0 | 4 | 0 | | | T | T | T | T | | | | | | |
| NC_011770 | 4973013 | C | T | 64 | 0 | 2 | 0 | T | T | | | | | | | | | | |
| NC_011770 | 4973090 | C | T | 65 | 0 | 4 | 0 | | | T | T | T | T | | | | | | |
| NC_011770 | 4975053 | G | C | 60 | 0 | 4 | 0 | C | C | | | | | C | C | | | | |
| NC_011770 | 4976841 | G | A | 83 | 0 | 2 | 0 | | | | | | | | | A | A | | |
| NC_011770 | 4976905 | G | A | 77 | 0 | 2 | 0 | | | | | | | | | A | A | | |
| NC_011770 | 4977056 | C | G | 76 | 0 | 8 | 0 | | | G | G | G | G | G | G | G | G | | |
| NC_011770 | 4980389 | C | A | 68 | 0 | 2 | 0 | | | | | | | | | | | A | A |
| NC_011770 | 4980396 | GC | G | 65 | 0 | 2 | 0 | | | | | | | | | | | G | G |
| NC_011770 | 4982355 | G | A | 30 | 0 | 4 | 0 | A | A | | | | | | | | | A | A |
| NC_011770 | 4982384 | C | T | 28 | 0 | 4 | 0 | T | T | | | | | | | | | T | T |
| NC_011770 | 4982387 | T | C | 30 | 0 | 4 | 0 | C | C | | | | | | | | | C | C |
| NC_011770 | 4982390 | A | G | 31 | 0 | 4 | 0 | G | G | | | | | | | | | G | G |
| NC_011770 | 4983270 | G | A | 67 | 0 | 8 | 0 | A | A | A | A | A | A | A | A | | | | |
| NC_011770 | 4983315 | G | A | 67 | 0 | 2 | 0 | A | A | | | | | | | | | | |
| NC_011770 | 4983364 | TG | T | 58 | 0 | 4 | 0 | | | | | | | | | T | T | T | T |
| NC_011770 | 4983371 | A | G | 60 | 0 | 4 | 0 | G | G | | | | | G | G | | | | |
| NC_011770 | 4983396 | A | C | 57 | 0 | 12 | 0 | C | C | C | C | C | C | C | C | C | C | C | C |
| NC_011770 | 4983415 | C | G | 56 | 0 | 4 | 0 | | | G | G | G | G | | | | | | |
| NC_011770 | 4983416 | C | T | 55 | 0 | 4 | 0 | | | T | T | T | T | | | | | | |
| NC_011770 | 4983478 | G | C | 62 | 0 | 4 | 0 | C | C | | | | | C | C | | | | |
| NC_011770 | 4983515 | G | A | 64 | 0 | 4 | 0 | A | A | | | | | A | A | | | | |
| NC_011770 | 4983690 | C | T | 61 | 0 | 4 | 0 | T | T | | | | | T | T | | | | |
| NC_011770 | 4983692 | G | A | 61 | 0 | 4 | 0 | A | A | | | | | A | A | | | | |
| NC_011770 | 4984199 | C | G | 44 | 0 | 2 | 0 | | | | | | | | | G | G | | |
| NC_011770 | 4984262 | G | A | 48 | 0 | 2 | 0 | | | | | | | A | A | | | | |
| NC_011770 | 4984270 | T | C | 47 | 0 | 6 | 0 | | | C | C | C | C | C | C | | | | |
| NC_011770 | 4984308 | G | A | 50 | 0 | 6 | 0 | | | A | A | A | A | A | A | | | | |
| NC_011770 | 4984310 | G | A | 51 | 0 | 4 | 0 | | | A | A | A | A | | | | | | |
| NC_011770 | 4984318 | C | T | 51 | 0 | 2 | 0 | | | | | | | T | T | | | | |
| NC_011770 | 4984357 | G | C | 52 | 0 | 6 | 0 | C | C | | | | | C | C | C | C | | |
| NC_011770 | 4984378 | A | C | 50 | 0 | 2 | 0 | | | | | | | C | C | | | | |
| NC_011770 | 4984399 | A | AC | 52 | 0 | 2 | 0 | | | | | | | | | AC | AC | | |
| NC_011770 | 4984405 | C | A | 52 | 0 | 2 | 0 | A | A | | | | | | | | | | |
| NC_011770 | 4984470 | C | A | 54 | 0 | 2 | 0 | | | | | | | | | A | A | | |
| NC_011770 | 4984499 | C | A | 55 | 0 | 10 | 0 | A | A | A | A | A | A | T | T | T | T | | |
| NC_011770 | 4984522 | C | G | 55 | 0 | 4 | 0 | | | G | G | G | G | | | | | | |
| NC_011770 | 4984574 | T | C | 60 | 0 | 12 | 0 | C | C | C | C | C | C | C | C | C | C | C | C |
| NC_011770 | 4984711 | C | T | 66 | 0 | 2 | 0 | | | | | | | | | | | T | T |
| NC_011770 | 4985045 | A | T | 69 | 0 | 2 | 0 | | | | | | | T | T | | | | |
| NC_011770 | 4986527 | A | G | 55 | 0 | 2 | 0 | | | | | | | | | | | G | G |
| NC_011770 | 4988961 | C | T | 72 | 0 | 2 | 0 | | | | | | | | | T | T | | |
| NC_011770 | 4988963 | C | T | 71 | 0 | 2 | 0 | | | | | | | | | | | T | T |
| NC_011770 | 4989004 | T | C | 73 | 0 | 2 | 0 | | | | | | | | | | | C | C |
| NC_011770 | 4989047 | A | T | 78 | 0 | 2 | 0 | T | T | | | | | | | | | | |
| NC_011770 | 4989166 | C | T | 75 | 0 | 10 | 0 | T | T | T | T | T | T | T | T | T | T | | |
| NC_011770 | 4989176 | G | A | 79 | 0 | 1 | 0 | | | | | | | A | | | | | |
| NC_011770 | 4989340 | G | A | 77 | 0 | 2 | 0 | | | | | | | | | | | A | A |
| NC_011770 | 4991346 | G | C | 80 | 0 | 2 | 0 | | | | | | | | | C | C | | |
| NC_011770 | 4992054 | T | G | 62 | 0 | 12 | 0 | G | G | G | G | G | G | G | G | G | G | G | G |
| NC_011770 | 4992065 | T | C | 60 | 0 | 12 | 0 | C | C | C | C | C | C | C | C | C | C | C | C |
| NC_011770 | 4992066 | G | A | 61 | 0 | 4 | 0 | | | A | A | A | A | | | | | | |
| NC_011770 | 4992082 | C | A | 60 | 0 | 4 | 0 | | | A | A | A | A | | | | | | |
| NC_011770 | 4992111 | C | T | 59 | 0 | 4 | 0 | | | T | T | T | T | | | | | | |
| NC_011770 | 4992112 | G | A | 59 | 0 | 2 | 0 | A | A | | | | | | | | | | |
| NC_011770 | 4992120 | A | T | 63 | 0 | 2 | 0 | | | | | | | | | | | T | T |
| NC_011770 | 4993756 | A | G | 95 | 0 | 4 | 0 | | | G | G | G | G | | | | | | |

| | | | | | | | | | | | | | | | | | | | |
|---|---|---|---|---|---|---|---|---|---|---|---|---|---|---|---|---|---|---|---|
| NC_011770 | 4994980 | T | C | 56 | 0 | 8 | 0 | | | C | C | C | C | | | C | C | C | C |
| NC_011770 | 4997743 | A | G | 10 | 0 | 2 | 0 | | | | | | | | | G | G | | |
| NC_011770 | 4999035 | C | T | 74 | 0 | 4 | 0 | | | T | T | T | T | | | | | | |
| NC_011770 | 5001414 | C | T | 98 | 0 | 2 | 0 | T | T | | | | | | | | | | |
| NC_011770 | 5001425 | C | T | 95 | 0 | 2 | 0 | | | | | | | | | | | T | T |
| NC_011770 | 5001426 | G | A | 96 | 0 | 4 | 0 | | | | | | | A | A | A | A | | |
| NC_011770 | 5001462 | C | T | 92 | 0 | 4 | 0 | | | T | T | T | T | | | | | | |
| NC_011770 | 5002378 | A | G | 74 | 0 | 4 | 0 | | | G | G | G | G | | | | | | |
| NC_011770 | 5003944 | A | G | 80 | 0 | 2 | 0 | | | | | | | | | | | G | G |
| NC_011770 | 5004047 | A | G | 77 | 0 | 6 | 0 | | | G | G | G | G | G | G | | | | |
| NC_011770 | 5004149 | C | T | 81 | 0 | 2 | 0 | | | | | | | | | T | T | | |
| NC_011770 | 5004198 | G | A | 76 | 0 | 10 | 0 | A | A | A | A | A | A | A | A | | | A | A |
| NC_011770 | 5004284 | T | G | 64 | 0 | 12 | 0 | G | G | G | G | G | G | G | G | G | G | G | G |
| NC_011770 | 5004589 | G | A | 74 | 0 | 2 | 0 | | | | | | | | | | | A | A |
| NC_011770 | 5004593 | G | A | 74 | 0 | 2 | 0 | | | | | | | A | A | | | | |
| NC_011770 | 5011034 | C | T | 77 | 0 | 2 | 0 | | | | | | | | | T | T | | |
| NC_011770 | 5011043 | C | G | 74 | 0 | 4 | 0 | | | G | G | G | G | | | | | | |
| NC_011770 | 5011053 | G | A | 78 | 0 | 2 | 0 | | | | | A | A | | | | | | |
| NC_011770 | 5011071 | A | C | 69 | 0 | 12 | 0 | C | C | C | C | C | C | C | C | C | C | C | C |
| NC_011770 | 5011072 | G | C | 68 | 0 | 12 | 0 | C | C | C | C | C | C | C | C | C | C | C | C |
| NC_011770 | 5011077 | G | A | 72 | 0 | 2 | 0 | | | | | | | | | A | A | | |
| NC_011770 | 5011144 | T | C | 75 | 0 | 2 | 0 | | | | | | | | | C | C | | |
| NC_011770 | 5012821 | C | A | 84 | 0 | 8 | 0 | | | A | A | A | A | | | A | A | A | A |
| NC_011770 | 5012933 | C | T | 68 | 0 | 4 | 0 | T | T | | | | | T | T | | | | |
| NC_011770 | 5016256 | C | G | 69 | 0 | 6 | 0 | | | G | G | G | G | | | | | G | G |
| NC_011770 | 5016264 | G | A | 69 | 0 | 6 | 0 | | | A | A | A | A | | | | | A | A |
| NC_011770 | 5016280 | G | A | 72 | 0 | 6 | 0 | | | A | A | A | A | | | | | A | A |
| NC_011770 | 5016585 | G | A | 66 | 0 | 12 | 0 | A | A | A | A | A | A | A | A | A | A | A | A |
| NC_011770 | 5016599 | C | T | 66 | 0 | 2 | 0 | | | | | | | T | T | | | | |
| NC_011770 | 5018736 | G | A | 76 | 0 | 2 | 0 | A | A | | | | | | | | | | |
| NC_011770 | 5018743 | G | C | 75 | 0 | 4 | 0 | | | C | C | C | C | | | | | | |
| NC_011770 | 5018820 | T | C | 79 | 0 | 4 | 0 | | | C | C | C | C | | | | | | |
| NC_011770 | 5019668 | G | A | 46 | 0 | 4 | 0 | | | A | A | A | A | | | | | | |
| NC_011770 | 5022199 | G | A | 48 | 0 | 4 | 0 | | | A | A | A | A | | | | | | |
| NC_011770 | 5022207 | C | T | 47 | 0 | 4 | 0 | | | T | T | T | T | | | | | | |
| NC_011770 | 5022220 | A | G | 48 | 0 | 4 | 0 | | | G | G | G | G | | | | | | |
| NC_011770 | 5022231 | A | G | 48 | 0 | 4 | 0 | | | G | G | G | G | | | | | | |
| NC_011770 | 5022235 | G | A | 50 | 0 | 4 | 0 | | | | | | | A | A | A | A | | |
| NC_011770 | 5022694 | C | T | 51 | 0 | 4 | 0 | | | T | T | T | T | | | | | | |
| NC_011770 | 5022695 | G | C | 52 | 0 | 4 | 0 | | | C | C | C | C | | | | | | |
| NC_011770 | 5022722 | G | A | 53 | 0 | 4 | 0 | | | A | A | A | A | | | | | | |
| NC_011770 | 5022729 | T | C | 54 | 0 | 2 | 0 | C | C | | | | | | | | | | |
| NC_011770 | 5022772 | C | T | 57 | 0 | 6 | 0 | | | | | | | T | T | T | T | T | T |
| NC_011770 | 5024288 | A | G | 78 | 0 | 4 | 0 | | | G | G | G | G | | | | | | |
| NC_011770 | 5026054 | G | A | 67 | 0 | 2 | 0 | | | | | | | A | A | | | | |
| NC_011770 | 5026055 | A | G | 65 | 0 | 4 | 0 | | | | | | | | | G | G | G | G |
| NC_011770 | 5026061 | T | A | 70 | 0 | 6 | 0 | | | A | A | A | A | A | A | | | | |
| NC_011770 | 5026088 | TTCA | T | 70 | 0 | 2 | 0 | | | | | | | T | T | | | | |
| NC_011770 | 5026091 | A | G | 55 | 2 | 4 | 0 | | | | | | | . | . | G | G | G | G |
| NC_011770 | 5026094 | C | G | 70 | 0 | 4 | 0 | | | G | G | G | G | | | | | | |
| NC_011770 | 5026095 | T | C | 70 | 0 | 4 | 0 | | | C | C | C | C | | | | | | |
| NC_011770 | 5026104 | C | A | 72 | 0 | 2 | 0 | | | | | | | | | A | A | | |
| NC_011770 | 5026109 | C | A | 71 | 0 | 4 | 0 | | | A | A | A | A | | | | | | |
| NC_011770 | 5026120 | C | G | 72 | 0 | 4 | 0 | | | | | | | | | G | G | G | G |
| NC_011770 | 5027604 | A | G | 70 | 0 | 2 | 0 | G | G | | | | | | | | | | |
| NC_011770 | 5027608 | T | G | 70 | 0 | 6 | 0 | G | G | | | | | | | G | G | G | G |
| NC_011770 | 5027637 | T | C | 71 | 0 | 12 | 0 | C | C | C | C | C | C | C | C | C | C | C | C |
| NC_011770 | 5029672 | G | C | 68 | 0 | 2 | 0 | | | | | | | C | C | | | | |
| NC_011770 | 5032443 | G | A | 76 | 0 | 1 | 0 | | | | | | | A | | | | | |
| NC_011770 | 5032482 | G | A | 72 | 0 | 2 | 0 | | | | | | | | | A | A | | |
| NC_011770 | 2228232 | TTCCTTTC | T | 48 | 0 | 3 | 0 | T | T | | | | | | | T | | | |
| NC_011770 | 772955 | G | A | 45 | 0 | 5 | 0 | | | A | A | | A | | | A | A | | |

| | | | | | | | | | | | | | | | | | | | |
|---|---|---|---|---|---|---|---|---|---|---|---|---|---|---|---|---|---|---|---|
| NC_011770 | 5038756 | G | A | 86 | 0 | 6 | 0 | | | | | | | A | A | A | A | A | A |
| NC_011770 | 5038809 | A | G | 82 | 0 | 4 | 0 | | | | | | | | | G | G | G | G |
| NC_011770 | 5038893 | C | A | 84 | 0 | 4 | 0 | | | | | | | | | A | A | A | A |
| NC_011770 | 5038974 | G | A | 85 | 0 | 2 | 0 | | | | | | | | | | | A | A |
| NC_011770 | 5038999 | C | T | 85 | 0 | 2 | 0 | | | | | | | | | | | T | T |
| NC_011770 | 5039055 | G | A | 89 | 0 | 2 | 0 | | | | | | | A | A | | | | |
| NC_011770 | 5039426 | C | T | 82 | 0 | 2 | 0 | | | | | | | T | T | | | | |
| NC_011770 | 5039526 | C | T | 78 | 0 | 8 | 0 | | | T | T | T | T | T | T | T | T | | |
| NC_011770 | 5041502 | G | A | 17 | 4 | 5 | 0 | A | . | . | . | A | A | | | . | . | A | A |
| NC_011770 | 5043638 | C | T | 78 | 0 | 2 | 0 | T | T | | | | | | | | | | |
| NC_011770 | 5043672 | C | T | 78 | 0 | 8 | 0 | | | T | T | T | T | T | T | | | T | T |
| NC_011770 | 5043718 | T | C | 74 | 0 | 10 | 0 | | | C | C | C | C | C | C | C | C | C | C |
| NC_011770 | 5049263 | C | T | 64 | 0 | 4 | 0 | | | T | T | T | T | | | | | | |
| NC_011770 | 5051862 | C | T | 67 | 0 | 4 | 0 | | | T | T | T | T | | | | | | |
| NC_011770 | 5055564 | G | A | 57 | 0 | 4 | 0 | | | | | | | A | A | | | A | A |
| NC_011770 | 5056727 | G | C | 93 | 0 | 2 | 0 | | | | | | | | | C | C | | |
| NC_011770 | 5056789 | G | T | 89 | 0 | 2 | 0 | | | | | | | | | | | T | T |
| NC_011770 | 5057905 | C | T | 96 | 0 | 12 | 0 | T | T | T | T | T | T | T | T | T | T | T | T |
| NC_011770 | 5057948 | A | G | 101 | 0 | 12 | 0 | G | G | G | G | G | G | G | G | G | G | G | G |
| NC_011770 | 5058403 | G | A | 80 | 0 | 2 | 0 | | | | | | | | | A | A | | |
| NC_011770 | 5058423 | T | G | 79 | 0 | 12 | 0 | G | G | G | G | G | G | G | G | G | G | G | G |
| NC_011770 | 5058447 | A | G | 82 | 0 | 6 | 0 | | | G | G | G | G | | | G | G | | |
| NC_011770 | 5061215 | G | A | 73 | 0 | 4 | 0 | | | | | | | | | A | A | A | A |
| NC_011770 | 5062457 | G | A | 83 | 0 | 6 | 0 | | | | | | | A | A | A | A | A | A |
| NC_011770 | 5062803 | C | T | 77 | 0 | 2 | 0 | | | | | | | T | T | | | | |
| NC_011770 | 5062830 | G | A | 73 | 0 | 6 | 0 | | | | | | | A | A | A | A | A | A |
| NC_011770 | 5063670 | T | G | 70 | 0 | 2 | 0 | G | G | | | | | | | | | | |
| NC_011770 | 5063724 | G | A | 72 | 0 | 2 | 0 | | | | | | | A | A | | | | |
| NC_011770 | 5065725 | C | T | 73 | 0 | 12 | 0 | T | T | T | T | T | T | T | T | T | T | T | T |
| NC_011770 | 5065731 | G | A | 72 | 0 | 12 | 0 | A | A | A | A | A | A | A | A | A | A | A | A |
| NC_011770 | 5066744 | G | T | 55 | 0 | 4 | 0 | | | | | | | T | T | T | T | | |
| NC_011770 | 5066769 | G | A | 51 | 0 | 12 | 0 | A | A | A | A | A | A | A | A | A | A | A | A |
| NC_011770 | 5066786 | A | G | 53 | 0 | 4 | 0 | | | G | G | G | G | | | | | | |
| NC_011770 | 5066833 | A | G | 53 | 0 | 4 | 0 | | | G | G | G | G | | | | | | |
| NC_011770 | 5068695 | A | G | 62 | 0 | 1 | 0 | | | | | | | G | | | | | |
| NC_011770 | 5068713 | G | A | 62 | 0 | 2 | 0 | | | | | | | | | A | A | | |
| NC_011770 | 5068719 | T | C | 63 | 0 | 4 | 0 | | | C | C | C | C | | | | | | |
| NC_011770 | 5068724 | T | C | 65 | 0 | 4 | 0 | | | C | C | C | C | | | | | | |
| NC_011770 | 5068736 | A | C | 61 | 0 | 4 | 0 | | | C | C | C | C | | | | | | |
| NC_011770 | 5068737 | T | C | 61 | 0 | 4 | 0 | | | C | C | C | C | | | | | | |
| NC_011770 | 5068749 | A | G | 63 | 0 | 4 | 0 | | | G | G | G | G | | | | | | |
| NC_011770 | 5068757 | T | C | 65 | 0 | 4 | 0 | | | C | C | C | C | | | | | | |
| NC_011770 | 5068758 | A | G | 64 | 0 | 4 | 0 | | | G | G | G | G | | | | | | |
| NC_011770 | 5068765 | T | C | 60 | 0 | 4 | 0 | | | C | C | C | C | | | | | | |
| NC_011770 | 5068766 | C | G | 61 | 0 | 4 | 0 | | | G | G | G | G | | | | | | |
| NC_011770 | 5068787 | A | T | 60 | 0 | 4 | 0 | | | T | T | T | T | | | | | | |
| NC_011770 | 5068788 | GC | TC | 60 | 0 | 5 | 0 | | | TC | TC | TC | TC | G | | | | | |
| NC_011770 | 5068792 | C | T | 54 | 0 | 4 | 0 | | | T | T | T | T | | | | | | |
| NC_011770 | 5068793 | C | T | 54 | 0 | 4 | 0 | | | T | T | T | T | | | | | | |
| NC_011770 | 5068807 | A | C | 63 | 0 | 4 | 0 | | | C | C | C | C | | | | | | |
| NC_011770 | 5068828 | T | C | 63 | 0 | 4 | 0 | | | C | C | C | C | | | | | | |
| NC_011770 | 5068868 | T | C | 62 | 0 | 4 | 0 | | | C | C | C | C | | | | | | |
| NC_011770 | 5068871 | C | T | 61 | 0 | 4 | 0 | | | T | T | T | T | | | | | | |
| NC_011770 | 5068873 | A | G | 61 | 0 | 4 | 0 | | | G | G | G | G | | | | | | |
| NC_011770 | 5068928 | C | T | 63 | 0 | 2 | 0 | | | | | | | | | T | T | | |
| NC_011770 | 5068952 | A | T | 62 | 0 | 8 | 0 | T | T | | | | | T | T | T | T | T | T |
| NC_011770 | 5068960 | G | A | 63 | 0 | 8 | 0 | A | A | | | | | A | A | A | A | A | A |
| NC_011770 | 5068962 | C | T | 63 | 0 | 8 | 0 | T | T | | | | | T | T | T | T | T | T |
| NC_011770 | 5068967 | A | G | 63 | 0 | 8 | 0 | G | G | | | | | G | G | G | G | G | G |
| NC_011770 | 5068971 | G | A | 64 | 0 | 8 | 0 | A | A | | | | | A | A | A | A | A | A |
| NC_011770 | 5068981 | A | C | 63 | 0 | 8 | 0 | C | C | | | | | C | C | C | C | C | C |
| NC_011770 | 5068982 | A | T | 63 | 0 | 8 | 0 | T | T | | | | | T | T | T | T | T | T |

| | | | | | | | | | | | | | | | | | | | |
|---|---|---|---|---|---|---|---|---|---|---|---|---|---|---|---|---|---|---|---|
| NC_011770 | 5068994 | T | C | 66 | 0 | 4 | 0 | | | C | C | C | C | | | | | | |
| NC_011770 | 5069011 | T | C | 65 | 0 | 8 | 0 | C | C | | | | | C | C | C | C | C | C |
| NC_011770 | 5069022 | T | C | 68 | 0 | 8 | 0 | C | C | | | | | C | C | C | C | C | C |
| NC_011770 | 5069121 | A | T | 70 | 0 | 12 | 0 | T | T | T | T | T | T | T | T | T | T | T | T |
| NC_011770 | 5070173 | A | G | 19 | 0 | 8 | 0 | G | G | | | | | G | G | G | G | G | G |
| NC_011770 | 5070267 | A | G | 33 | 0 | 4 | 0 | | | | | | | | | G | G | G | G |
| NC_011770 | 5070313 | T | TC | 38 | 0 | 4 | 0 | | | | | | | | | TC | TC | TC | TC |
| NC_011770 | 5070319 | G | C | 31 | 0 | 3 | 0 | | | | | | | | | C | . | C | C |
| NC_011770 | 5070339 | G | C | 41 | 0 | 4 | 0 | | | | | | | | | C | C | C | C |
| NC_011770 | 5070344 | GC | G | 41 | 0 | 8 | 0 | G | G | G | G | G | G | G | G | | | | |
| NC_011770 | 5070349 | CA | C | 12 | 5 | 4 | 0 | . | . | . | . | . | | | | C | C | C | C |
| NC_011770 | 5070361 | C | T | 41 | 0 | 4 | 0 | T | T | | | | | T | T | | | | |
| NC_011770 | 5070366 | A | G | 41 | 0 | 8 | 0 | G | G | | | | | G | G | G | G | G | G |
| NC_011770 | 5070380 | T | C | 43 | 0 | 4 | 0 | | | | | | | | | C | C | C | C |
| NC_011770 | 5070389 | T | C | 43 | 0 | 12 | 0 | C | C | C | C | C | C | C | C | C | C | C | C |
| NC_011770 | 5071024 | T | G | 69 | 0 | 8 | 0 | G | G | | | | | G | G | G | G | G | G |
| NC_011770 | 5071054 | A | G | 70 | 0 | 4 | 0 | G | G | | | | | G | G | | | | |
| NC_011770 | 5071405 | T | C | 93 | 0 | 2 | 0 | C | C | | | | | | | | | | |
| NC_011770 | 5071462 | C | G | 87 | 0 | 12 | 0 | G | G | G | G | G | G | G | G | G | G | G | G |
| NC_011770 | 5071548 | G | A | 89 | 0 | 2 | 0 | A | A | | | | | | | | | | |
| NC_011770 | 5071562 | G | T | 87 | 0 | 4 | 0 | | | | | | | | | T | T | T | T |
| NC_011770 | 5071584 | C | T | 91 | 0 | 4 | 0 | | | | | | | | | T | T | T | T |
| NC_011770 | 5071600 | G | A | 92 | 0 | 8 | 0 | | | A | A | A | A | | | A | A | A | A |
| NC_011770 | 5071625 | A | G | 89 | 0 | 4 | 0 | | | G | G | G | G | | | | | | |
| NC_011770 | 5071628 | C | T | 91 | 0 | 4 | 0 | | | T | T | T | T | | | | | | |
| NC_011770 | 5071673 | G | A | 94 | 0 | 4 | 0 | | | | | | | | | A | A | A | A |
| NC_011770 | 5071687 | C | A | 95 | 0 | 6 | 0 | | | | | | | A | A | A | A | A | A |
| NC_011770 | 5071705 | C | T | 102 | 0 | 4 | 0 | T | T | | | | | T | T | | | | |
| NC_011770 | 5072596 | G | A | 90 | 0 | 4 | 0 | | | | | | | | | A | A | A | A |
| NC_011770 | 5072999 | T | C | 86 | 0 | 2 | 0 | C | C | | | | | | | | | | |
| NC_011770 | 5073010 | A | G | 85 | 0 | 6 | 0 | G | G | | | | | | | G | G | G | G |
| NC_011770 | 5073397 | C | T | 87 | 0 | 2 | 0 | T | T | | | | | | | | | | |
| NC_011770 | 5073421 | C | T | 85 | 0 | 2 | 0 | T | T | | | | | | | | | | |
| NC_011770 | 5073427 | A | G | 83 | 0 | 4 | 0 | | | | | | | | | G | G | G | G |
| NC_011770 | 5073428 | T | G | 82 | 0 | 4 | 0 | | | | | | | | | G | G | G | G |
| NC_011770 | 5073430 | G | A | 83 | 0 | 4 | 0 | | | | | | | | | A | A | A | A |
| NC_011770 | 5073447 | G | T | 89 | 0 | 4 | 0 | | | | | | | | | T | T | T | T |
| NC_011770 | 5074311 | T | C | 67 | 0 | 6 | 0 | C | C | | | | | | | C | C | C | C |
| NC_011770 | 5074314 | T | A | 67 | 0 | 2 | 0 | A | A | | | | | | | | | | |
| NC_011770 | 5074438 | G | A | 66 | 0 | 2 | 0 | | | | | | | A | A | | | | |
| NC_011770 | 5074542 | T | C | 64 | 0 | 12 | 0 | C | C | C | C | C | C | C | C | C | C | C | C |
| NC_011770 | 5075372 | G | A | 84 | 0 | 4 | 0 | | | A | A | A | A | | | | | | |
| NC_011770 | 5075379 | C | T | 83 | 0 | 4 | 0 | | | T | T | T | T | | | | | | |
| NC_011770 | 5075417 | G | A | 84 | 0 | 2 | 0 | | | | | | | A | A | | | | |
| NC_011770 | 5075456 | A | G | 78 | 0 | 4 | 0 | | | | | | | | | G | G | G | G |
| NC_011770 | 5076096 | G | A | 86 | 0 | 2 | 0 | | | | | | | A | A | | | | |
| NC_011770 | 5076097 | T | A | 85 | 0 | 10 | 0 | A | A | A | A | A | A | | | A | A | A | A |
| NC_011770 | 5076144 | A | G | 85 | 0 | 8 | 0 | G | G | | | | | G | G | G | G | G | G |
| NC_011770 | 5076150 | G | A | 91 | 0 | 8 | 0 | A | A | | | | | A | A | A | A | A | A |
| NC_011770 | 5076209 | C | G | 92 | 0 | 2 | 0 | | | | | | | G | G | | | | |
| NC_011770 | 5076217 | A | G | 90 | 0 | 8 | 0 | G | G | | | | | G | G | G | G | G | G |
| NC_011770 | 5076236 | C | A | 92 | 0 | 8 | 0 | A | A | | | | | A | A | A | A | A | A |
| NC_011770 | 5076244 | T | C | 89 | 0 | 2 | 0 | | | | | | | C | C | | | | |
| NC_011770 | 5076253 | A | C | 89 | 0 | 8 | 0 | C | C | | | | | C | C | C | C | C | C |
| NC_011770 | 5076288 | G | A | 83 | 0 | 8 | 0 | A | A | | | | | A | A | A | A | A | A |
| NC_011770 | 5076294 | T | C | 76 | 0 | 8 | 0 | C | C | | | | | C | C | C | C | C | C |
| NC_011770 | 5076301 | A | G | 88 | 0 | 6 | 0 | G | G | | | | | | | G | G | G | G |
| NC_011770 | 5077301 | C | T | 65 | 0 | 4 | 0 | T | T | | | | | | | T | T | | |
| NC_011770 | 5077312 | A | G | 63 | 0 | 2 | 0 | | | | | | | | | | | G | G |
| NC_011770 | 5077326 | C | T | 65 | 0 | 2 | 0 | | | | | | | | | | | T | T |
| NC_011770 | 5078766 | G | A | 39 | 0 | 2 | 0 | | | | | | | | | | | A | A |
| NC_011770 | 5080169 | G | A | 80 | 0 | 2 | 0 | | | | | | | A | A | | | | |

| | | | | | | | | | | | | | | | | | | | |
|---|---|---|---|---|---|---|---|---|---|---|---|---|---|---|---|---|---|---|---|
| NC_011770 | 5080179 | T | C | 79 | 0 | 4 | 0 | | | C | C | C | C | | | | | | |
| NC_011770 | 5080186 | A | G | 78 | 0 | 2 | 0 | | | | | | | G | G | | | | |
| NC_011770 | 5081138 | A | G | 72 | 0 | 12 | 0 | G | G | G | G | G | G | G | G | G | G | G | G |
| NC_011770 | 5081202 | G | A | 67 | 0 | 6 | 0 | A | A | | | | | | | A | A | A | A |
| NC_011770 | 5081264 | A | G | 67 | 0 | 12 | 0 | G | G | G | G | G | G | G | G | G | G | G | G |
| NC_011770 | 5081294 | A | G | 60 | 0 | 12 | 0 | G | G | G | G | G | G | G | G | G | G | G | G |
| NC_011770 | 5081295 | C | T | 62 | 0 | 2 | 0 | T | T | | | | | | | | | | |
| NC_011770 | 5081342 | G | A | 60 | 0 | 4 | 0 | | | | | | | | | A | A | A | A |
| NC_011770 | 5081512 | C | T | 65 | 0 | 4 | 0 | | | T | T | T | T | | | | | | |
| NC_011770 | 5081574 | G | T | 66 | 0 | 6 | 0 | T | T | | | | | | | T | T | T | T |
| NC_011770 | 5081582 | G | A | 68 | 0 | 6 | 0 | A | A | | | | | | | A | A | A | A |
| NC_011770 | 5081634 | A | C | 71 | 0 | 4 | 0 | | | | | | | | | C | C | C | C |
| NC_011770 | 5081688 | C | A | 74 | 0 | 4 | 0 | | | A | A | A | A | | | | | | |
| NC_011770 | 5081828 | A | G | 73 | 0 | 4 | 0 | | | | | | | | | G | G | G | G |
| NC_011770 | 5084699 | G | A | 84 | 0 | 6 | 0 | | | | | | | A | A | A | A | A | A |
| NC_011770 | 5084719 | G | A | 83 | 0 | 4 | 0 | | | A | A | A | A | | | | | | |
| NC_011770 | 5085414 | C | T | 95 | 0 | 2 | 0 | T | T | | | | | | | | | | |
| NC_011770 | 5090236 | G | A | 73 | 0 | 4 | 0 | A | A | | | | | A | A | | | | |
| NC_011770 | 5091956 | G | A | 92 | 0 | 4 | 0 | A | A | | | | | A | A | | | | |
| NC_011770 | 5091977 | G | A | 84 | 0 | 4 | 0 | A | A | | | | | A | A | | | | |
| NC_011770 | 5091978 | C | T | 84 | 0 | 4 | 0 | T | T | | | | | T | T | | | | |
| NC_011770 | 5092004 | G | A | 81 | 0 | 4 | 0 | A | A | | | | | A | A | | | | |
| NC_011770 | 5092007 | G | GC | 80 | 0 | 4 | 0 | GC | GC | | | | | GC | GC | | | | |
| NC_011770 | 5092010 | C | T | 62 | 4 | 4 | 0 | . | . | | | | | . | . | T | T | T | T |
| NC_011770 | 5092024 | C | T | 82 | 0 | 4 | 0 | T | T | | | | | T | T | | | | |
| NC_011770 | 5092050 | T | C | 85 | 0 | 4 | 0 | C | C | | | | | C | C | | | | |
| NC_011770 | 5092687 | A | T | 83 | 0 | 2 | 0 | T | T | | | | | | | | | | |
| NC_011770 | 5092743 | T | G | 78 | 0 | 10 | 0 | G | G | G | G | G | G | | | G | G | G | G |
| NC_011770 | 5092829 | G | A | 72 | 0 | 6 | 0 | A | A | A | A | A | A | | | | | | |
| NC_011770 | 5092868 | A | G | 72 | 0 | 2 | 0 | G | G | | | | | | | | | | |
| NC_011770 | 5092912 | G | A | 70 | 0 | 6 | 0 | A | A | A | A | A | A | | | | | | |
| NC_011770 | 5092919 | A | G | 71 | 0 | 6 | 0 | G | G | G | G | G | G | | | | | | |
| NC_011770 | 5092930 | A | G | 68 | 0 | 10 | 0 | G | G | G | G | G | G | | | G | G | G | G |
| NC_011770 | 5092933 | G | A | 70 | 0 | 1 | 0 | | | | | | | A | | | | | |
| NC_011770 | 5092985 | G | GC | 68 | 0 | 2 | 0 | GC | GC | | | | | | | | | | |
| NC_011770 | 5092986 | C | CCG | 59 | 2 | 8 | 0 | . | . | CCG | CCG | CCG | CCG | | | CCG | CCG | CCG | CCG |
| NC_011770 | 5092993 | AG | A | 61 | 0 | 8 | 0 | | | A | A | A | A | | | A | A | A | A |
| NC_011770 | 5092994 | G | A | 16 | 9 | 1 | 0 | A | . | . | . | . | . | | | . | . | . | . |
| NC_011770 | 5105663 | G | T | 42 | 4 | 2 | 0 | T | T | . | . | . | . | | | | | | |
| NC_011770 | 5105666 | T | C | 42 | 4 | 2 | 0 | C | C | . | . | . | . | | | | | | |
| NC_011770 | 5105676 | A | T | 60 | 0 | 4 | 0 | | | T | T | T | T | | | | | | |
| NC_011770 | 5105684 | T | C | 61 | 0 | 4 | 0 | | | C | C | C | C | | | | | | |
| NC_011770 | 5105717 | G | T | 38 | 6 | 2 | 0 | . | . | . | . | . | . | | | | | T | T |
| NC_011770 | 5105761 | A | C | 40 | 6 | 4 | 0 | . | . | . | . | . | . | C | C | | | C | C |
| NC_011770 | 5106956 | T | C | 32 | 4 | 2 | 0 | C | C | . | . | . | . | | | | | | |
| NC_011770 | 5108139 | T | C | 62 | 0 | 2 | 0 | C | C | | | | | | | | | | |
| NC_011770 | 5108248 | C | G | 65 | 0 | 2 | 0 | G | G | | | | | | | | | | |
| NC_011770 | 5108250 | C | G | 66 | 0 | 2 | 0 | G | G | | | | | | | | | | |
| NC_011770 | 5108251 | A | G | 65 | 0 | 2 | 0 | G | G | | | | | | | | | | |
| NC_011770 | 5108252 | G | C | 66 | 0 | 2 | 0 | C | C | | | | | | | | | | |
| NC_011770 | 5108255 | T | A | 66 | 0 | 2 | 0 | A | A | | | | | | | | | | |
| NC_011770 | 5108400 | A | G | 68 | 0 | 2 | 0 | G | G | | | | | | | | | | |
| NC_011770 | 5108435 | T | C | 71 | 0 | 2 | 0 | C | C | | | | | | | | | | |
| NC_011770 | 5108456 | T | C | 67 | 0 | 2 | 0 | C | C | | | | | | | | | | |
| NC_011770 | 5108461 | C | T | 64 | 1 | 1 | 0 | T | . | | | | | | | | | | |
| NC_011770 | 5108463 | G | T | 66 | 1 | 1 | 0 | T | . | | | | | | | | | | |
| NC_011770 | 5108464 | A | C | 64 | 1 | 1 | 0 | C | . | | | | | | | | | | |
| NC_011770 | 5108465 | T | C | 65 | 1 | 1 | 0 | C | . | | | | | | | | | | |
| NC_011770 | 5108466 | T | A | 66 | 1 | 1 | 0 | A | . | | | | | | | | | | |
| NC_011770 | 5108468 | C | T | 65 | 1 | 1 | 0 | T | . | | | | | | | | | | |
| NC_011770 | 5108502 | G | A | 70 | 1 | 4 | 0 | | . | A | A | A | A | | | | | | |
| NC_011770 | 5108504 | A | G | 69 | 1 | 1 | 0 | G | . | | | | | | | | | | |

| | | | | | | | | | | | | | | | | | | | |
|---|---|---|---|---|---|---|---|---|---|---|---|---|---|---|---|---|---|---|---|
| NC_011770 | 5108505 | T | G | 67 | 1 | 11 | 0 | G | . | G | G | G | G | G | G | G | G | G | G |
| NC_011770 | 5108508 | T | A | 70 | 1 | 1 | 0 | A | . | | | | | | | | | | |
| NC_011770 | 5108510 | C | CGA | 72 | 1 | 1 | 0 | CGA | . | | | | | | | | | | |
| NC_011770 | 5108528 | ACGGCGGA | A | 69 | 1 | 1 | 0 | A | . | | | | | | | | | | |
| NC_011770 | 5108550 | T | G | 71 | 1 | 1 | 0 | G | . | | | | | | | | | | |
| NC_011770 | 5108553 | T | C | 70 | 1 | 5 | 0 | C | . | C | C | C | C | | | | | | |
| NC_011770 | 5108556 | C | T | 71 | 1 | 1 | 0 | T | . | | | | | | | | | | |
| NC_011770 | 5108559 | A | T | 70 | 1 | 1 | 0 | T | . | | | | | | | | | | |
| NC_011770 | 5108569 | G | T | 79 | 0 | 2 | 0 | T | T | | | | | | | | | | |
| NC_011770 | 5108579 | A | G | 79 | 0 | 2 | 0 | G | G | | | | | | | | | | |
| NC_011770 | 5108582 | T | C | 82 | 0 | 4 | 0 | C | C | | | | | | | C | C | | |
| NC_011770 | 5108586 | G | A | 80 | 0 | 4 | 0 | | | A | A | A | A | | | | | | |
| NC_011770 | 5114385 | G | C | 63 | 0 | 2 | 0 | | | | | | | | | C | C | | |
| NC_011770 | 5114424 | T | C | 57 | 0 | 8 | 0 | C | C | | | | | C | C | C | C | C | C |
| NC_011770 | 5114475 | C | T | 62 | 0 | 6 | 0 | T | T | | | | | T | T | T | T | | |
| NC_011770 | 5114482 | A | G | 63 | 0 | 6 | 0 | G | G | | | | | G | G | G | G | | |
| NC_011770 | 5114623 | C | T | 66 | 0 | 4 | 0 | T | T | | | | | T | T | | | | |
| NC_011770 | 5114652 | G | A | 65 | 0 | 8 | 0 | A | A | | | | | A | A | A | A | A | A |
| NC_011770 | 5114690 | A | T | 73 | 0 | 2 | 0 | | | | | | | | | | | T | T |
| NC_011770 | 5114763 | G | A | 67 | 0 | 2 | 0 | | | | | | | | | | | A | A |
| NC_011770 | 5114794 | A | G | 71 | 0 | 8 | 0 | G | G | | | | | G | G | G | G | G | G |
| NC_011770 | 5114795 | T | C | 72 | 0 | 8 | 0 | C | C | | | | | C | C | C | C | C | C |
| NC_011770 | 5114832 | A | T | 71 | 0 | 8 | 0 | T | T | | | | | T | T | T | T | T | T |
| NC_011770 | 5114853 | C | T | 78 | 0 | 2 | 0 | | | | | | | | | | | T | T |
| NC_011770 | 5114905 | A | G | 82 | 0 | 2 | 0 | | | | | | | | | | | G | G |
| NC_011770 | 5114965 | G | C | 77 | 0 | 2 | 0 | | | | | | | | | | | C | C |
| NC_011770 | 5114994 | C | A | 75 | 0 | 2 | 0 | | | | | | | | | A | A | | |
| NC_011770 | 5115033 | T | C | 78 | 0 | 2 | 0 | | | | | | | | | | | C | C |
| NC_011770 | 5115071 | G | A | 76 | 0 | 2 | 0 | | | | | | | | | | | A | A |
| NC_011770 | 5115108 | G | A | 76 | 0 | 2 | 0 | | | | | | | A | A | | | | |
| NC_011770 | 5115141 | A | G | 77 | 0 | 6 | 0 | G | G | | | | | G | G | | | G | G |
| NC_011770 | 5115162 | G | A | 75 | 0 | 2 | 0 | | | | | | | A | A | | | | |
| NC_011770 | 5115180 | A | G | 76 | 0 | 4 | 0 | G | G | | | | | | | | | G | G |
| NC_011770 | 5115270 | C | T | 83 | 0 | 4 | 0 | T | T | | | | | | | | | T | T |
| NC_011770 | 5115295 | A | G | 80 | 0 | 4 | 0 | G | G | | | | | | | | | G | G |
| NC_011770 | 5115338 | A | G | 76 | 0 | 4 | 0 | G | G | | | | | | | | | G | G |
| NC_011770 | 5115339 | A | G | 76 | 0 | 4 | 0 | G | G | | | | | | | | | G | G |
| NC_011770 | 5115343 | T | G | 78 | 0 | 2 | 0 | G | G | | | | | | | | | | |
| NC_011770 | 5115398 | C | T | 74 | 0 | 4 | 0 | T | T | | | | | | | | | T | T |
| NC_011770 | 5115402 | C | T | 76 | 0 | 4 | 0 | T | T | | | | | | | | | T | T |
| NC_011770 | 5116916 | T | G | 76 | 0 | 2 | 0 | G | G | | | | | | | | | | |
| NC_011770 | 5117871 | C | CG | 59 | 0 | 6 | 0 | | | CG | CG | CG | CG | | | | | CG | CG |
| NC_011770 | 5117905 | A | G | 62 | 0 | 12 | 0 | G | G | G | G | G | G | G | G | G | G | G | G |
| NC_011770 | 5117958 | C | A | 67 | 0 | 2 | 0 | | | | | | | | | | | A | A |
| NC_011770 | 5120095 | T | G | 78 | 0 | 4 | 0 | | | | | | | G | G | G | G | | |
| NC_011770 | 5121876 | C | T | 79 | 0 | 6 | 0 | T | T | | | | | T | T | T | T | | |
| NC_011770 | 5121892 | T | C | 77 | 0 | 6 | 0 | C | C | | | | | C | C | C | C | | |
| NC_011770 | 5121915 | A | G | 77 | 0 | 12 | 0 | G | G | G | G | G | G | G | G | G | G | G | G |
| NC_011770 | 5121935 | C | A | 76 | 0 | 2 | 0 | | | | | | | | | A | A | | |
| NC_011770 | 5122632 | A | G | 78 | 0 | 12 | 0 | G | G | G | G | G | G | G | G | G | G | G | G |
| NC_011770 | 5122633 | T | C | 78 | 0 | 8 | 0 | C | C | | | | | C | C | C | C | C | C |
| NC_011770 | 5122640 | G | A | 82 | 0 | 4 | 0 | | | A | A | A | A | | | | | | |
| NC_011770 | 5122681 | A | G | 79 | 0 | 8 | 0 | G | G | | | | | G | G | G | G | G | G |
| NC_011770 | 3796957 | CGG | C | 51 | 0 | 1 | 0 | | | | | C | | | | | | | |
| NC_011770 | 5128058 | T | C | 75 | 0 | 4 | 0 | | | C | C | C | C | | | | | | |
| NC_011770 | 5128068 | T | C | 77 | 0 | 6 | 0 | | | | | | | C | C | C | C | C | C |
| NC_011770 | 5128073 | T | C | 79 | 0 | 6 | 0 | | | | | | | C | C | C | C | C | C |
| NC_011770 | 5128096 | T | C | 78 | 0 | 4 | 0 | | | | | | | C | C | | | C | C |
| NC_011770 | 5128116 | C | A | 80 | 0 | 4 | 0 | | | | | | | A | A | | | A | A |
| NC_011770 | 5128125 | A | G | 74 | 0 | 12 | 0 | G | G | G | G | G | G | G | G | G | G | G | G |
| NC_011770 | 5128129 | G | C | 76 | 0 | 2 | 0 | | | | | | | C | C | | | | |
| NC_011770 | 5128135 | G | A | 77 | 0 | 4 | 0 | | | | | | | A | A | | | A | A |

| | | | | | | | | | | | | | | | | | | | |
|---|---|---|---|---|---|---|---|---|---|---|---|---|---|---|---|---|---|---|---|
| NC_011770 | 5128142 | C | A | 77 | 0 | 2 | 0 | | | | | | | A | A | | | | |
| NC_011770 | 5128144 | C | T | 75 | 0 | 4 | 0 | | | T | T | T | T | | | | | | |
| NC_011770 | 5128145 | C | T | 75 | 0 | 2 | 0 | | | | | | | T | T | | | | |
| NC_011770 | 5128146 | A | C | 75 | 0 | 2 | 0 | | | | | | | | | | | C | C |
| NC_011770 | 5128156 | G | C | 77 | 0 | 2 | 0 | | | | | | | | | C | C | | |
| NC_011770 | 5128188 | G | A | 81 | 0 | 2 | 0 | | | | | | | | | | | A | A |
| NC_011770 | 5128204 | G | T | 78 | 0 | 6 | 0 | | | | | | | T | T | T | T | T | T |
| NC_011770 | 5128239 | G | A | 75 | 0 | 2 | 0 | | | | | | | A | A | | | | |
| NC_011770 | 5128241 | G | A | 74 | 0 | 4 | 0 | | | A | A | A | A | | | | | | |
| NC_011770 | 5128475 | G | GT | 87 | 0 | 2 | 0 | GT | GT | | | | | | | | | | |
| NC_011770 | 5128530 | G | A | 84 | 0 | 2 | 0 | | | | | | | | | | | A | A |
| NC_011770 | 5131627 | T | C | 78 | 0 | 4 | 0 | | | | | | | C | C | | | C | C |
| NC_011770 | 5131628 | C | T | 77 | 0 | 4 | 0 | | | | | | | T | T | | | T | T |
| NC_011770 | 5131660 | GT | G | 80 | 0 | 2 | 0 | G | G | | | | | | | | | | |
| NC_011770 | 5131744 | G | T | 62 | 0 | 2 | 0 | | | | | | | | | T | T | | |
| NC_011770 | 5133481 | T | C | 83 | 0 | 4 | 0 | | | C | C | C | C | | | | | | |
| NC_011770 | 5133819 | G | A | 40 | 0 | 8 | 0 | A | A | A | A | A | A | | | | | T | T |
| NC_011770 | 5133831 | G | A | 39 | 0 | 12 | 0 | A | A | A | A | A | A | A | A | A | A | A | A |
| NC_011770 | 5133833 | T | C | 39 | 0 | 12 | 0 | C | C | C | C | C | C | C | C | C | C | C | C |
| NC_011770 | 5133848 | A | G | 43 | 0 | 10 | 0 | | | G | G | G | G | G | G | G | G | G | G |
| NC_011770 | 5133874 | C | T | 51 | 0 | 2 | 0 | | | | | | | | | | | T | T |
| NC_011770 | 5133878 | G | T | 51 | 0 | 4 | 0 | | | T | T | T | T | | | | | | |
| NC_011770 | 5133895 | T | C | 55 | 0 | 4 | 0 | | | C | C | C | C | | | | | | |
| NC_011770 | 5133896 | C | A | 55 | 0 | 4 | 0 | | | A | A | A | A | | | | | | |
| NC_011770 | 5133925 | G | A | 59 | 0 | 2 | 0 | A | A | | | | | | | | | | |
| NC_011770 | 5133941 | G | C | 63 | 0 | 12 | 0 | C | T | C | C | C | C | C | C | C | C | C | C |
| NC_011770 | 5134053 | T | C | 77 | 0 | 12 | 0 | C | C | C | C | C | C | C | C | C | C | C | C |
| NC_011770 | 5149184 | T | C | 80 | 0 | 2 | 0 | C | C | | | | | | | | | | |
| NC_011770 | 5150184 | A | G | 87 | 0 | 10 | 0 | G | G | G | G | G | G | | | G | G | G | G |
| NC_011770 | 5150185 | G | C | 87 | 0 | 10 | 0 | C | C | C | C | C | C | | | C | C | C | C |
| NC_011770 | 5150188 | G | C | 87 | 0 | 10 | 0 | C | C | C | C | C | C | | | C | C | C | C |
| NC_011770 | 5150225 | T | C | 90 | 0 | 2 | 0 | C | C | | | | | | | | | | |
| NC_011770 | 5154707 | G | A | 64 | 0 | 2 | 0 | A | A | | | | | | | | | | |
| NC_011770 | 5156244 | GA | G | 55 | 0 | 8 | 0 | | | G | G | G | G | G | G | | | G | G |
| NC_011770 | 5156271 | GT | G | 57 | 0 | 6 | 0 | | | G | G | G | G | | | | | G | G |
| NC_011770 | 5156291 | A | G | 56 | 0 | 2 | 0 | | | | | | | G | G | | | | |
| NC_011770 | 5159604 | T | C | 73 | 0 | 2 | 0 | | | | | | | | | | | C | C |
| NC_011770 | 5160133 | T | C | 68 | 0 | 2 | 0 | | | | | | | | | C | C | | |
| NC_011770 | 5160146 | G | A | 65 | 0 | 2 | 0 | | | | | | | | | A | A | | |
| NC_011770 | 5160203 | A | G | 79 | 0 | 2 | 0 | | | | | | | | | G | G | | |
| NC_011770 | 2920318 | C | CT | 64 | 0 | 1 | 0 | | | | | | | | | CT | | | |
| NC_011770 | 5161100 | A | G | 76 | 0 | 10 | 0 | G | G | G | G | G | G | | | G | G | G | G |
| NC_011770 | 5170503 | C | T | 65 | 0 | 12 | 0 | T | T | T | T | T | T | T | T | T | T | T | T |
| NC_011770 | 5170553 | C | T | 57 | 0 | 12 | 0 | T | T | T | T | T | T | T | T | T | T | T | T |
| NC_011770 | 5170901 | GTCCTCTTC | G | 23 | 0 | 12 | 0 | G | G | G | G | G | G | G | G | G | G | G | G |
| NC_011770 | 5170917 | A | C | 23 | 0 | 12 | 0 | C | C | C | C | C | C | C | C | C | C | C | C |
| NC_011770 | 5170964 | A | G | 52 | 0 | 12 | 0 | G | G | G | G | G | G | G | G | G | G | G | G |
| NC_011770 | 5170966 | G | A | 51 | 0 | 12 | 0 | A | A | A | A | A | A | A | A | A | A | A | A |
| NC_011770 | 5170967 | C | A | 52 | 0 | 12 | 0 | A | A | A | A | A | A | A | A | A | A | A | A |
| NC_011770 | 5170988 | G | T | 63 | 0 | 12 | 0 | T | T | T | T | T | T | T | T | T | T | T | T |
| NC_011770 | 5171015 | T | TC | 63 | 0 | 12 | 0 | TC | TC | TC | TC | TC | TC | TC | TC | TC | TC | TC | TC |
| NC_011770 | 5173006 | G | A | 27 | 0 | 2 | 0 | | | | | | | | | | | A | A |
| NC_011770 | 5173050 | C | G | 27 | 0 | 2 | 0 | G | G | | | | | | | | | | |
| NC_011770 | 5173124 | G | C | 39 | 0 | 12 | 0 | C | C | C | C | C | C | C | C | C | C | C | C |
| NC_011770 | 5173169 | C | G | 43 | 0 | 12 | 0 | G | G | G | G | G | G | G | G | G | G | G | G |
| NC_011770 | 5173171 | G | A | 42 | 0 | 12 | 0 | A | A | A | A | A | A | A | A | A | A | A | A |
| NC_011770 | 5173173 | A | G | 42 | 0 | 12 | 0 | G | G | G | G | G | G | G | G | G | G | G | G |
| NC_011770 | 5173216 | T | C | 49 | 0 | 12 | 0 | C | C | C | C | C | C | C | C | C | C | C | C |
| NC_011770 | 5173218 | G | A | 46 | 0 | 4 | 0 | | | A | A | A | A | | | | | | |
| NC_011770 | 5176191 | T | C | 73 | 0 | 4 | 0 | | | C | C | C | C | | | | | | |
| NC_011770 | 5176223 | C | T | 77 | 0 | 4 | 0 | | | T | T | T | T | | | | | | |
| NC_011770 | 5176271 | G | T | 86 | 0 | 4 | 0 | | | T | T | T | T | | | | | | |

| | | | | | | | | | | | | | | | | | | | |
|---|---|---|---|---|---|---|---|---|---|---|---|---|---|---|---|---|---|---|---|
| NC_011770 | 5176281 | A | G | 84 | 0 | 4 | 0 | | | | | | | G | G | G | G | | |
| NC_011770 | 5176307 | G | A | 83 | 0 | 4 | 0 | | | | | | | A | A | A | A | | |
| NC_011770 | 5177326 | G | A | 78 | 0 | 4 | 0 | | | A | A | A | A | | | | | | |
| NC_011770 | 5177363 | A | G | 78 | 0 | 4 | 0 | | | G | G | G | G | | | | | | |
| NC_011770 | 5177952 | T | C | 66 | 0 | 4 | 0 | | | C | C | C | C | | | | | | |
| NC_011770 | 5177956 | T | C | 66 | 0 | 4 | 0 | | | C | C | C | C | | | | | | |
| NC_011770 | 5177959 | G | A | 64 | 0 | 4 | 0 | | | A | A | A | A | | | | | | |
| NC_011770 | 5177974 | C | T | 66 | 0 | 4 | 0 | | | T | T | T | T | | | | | | |
| NC_011770 | 5179586 | T | C | 74 | 0 | 8 | 0 | C | C | C | C | C | C | | | C | C | | |
| NC_011770 | 5188240 | A | C | 87 | 0 | 4 | 0 | | | C | C | C | C | | | | | | |
| NC_011770 | 5188247 | T | G | 86 | 0 | 4 | 0 | | | G | G | G | G | | | | | | |
| NC_011770 | 5188732 | C | T | 95 | 0 | 4 | 0 | T | T | | | | | T | T | | | | |
| NC_011770 | 5192142 | T | C | 207 | 0 | 2 | 0 | C | C | | | | | | | | | | |
| NC_011770 | 5192143 | A | G | 207 | 0 | 2 | 0 | G | G | | | | | | | | | | |
| NC_011770 | 5193025 | C | CG | 98 | 0 | 2 | 0 | CG | CG | | | | | | | | | | |
| NC_011770 | 5198244 | G | A | 78 | 0 | 2 | 0 | | | | | | | | | A | A | | |
| NC_011770 | 5198288 | T | A | 77 | 0 | 12 | 0 | A | A | A | A | A | A | A | A | A | A | A | A |
| NC_011770 | 5198367 | G | C | 77 | 0 | 2 | 0 | | | | | | | | | C | C | | |
| NC_011770 | 5198688 | T | G | 76 | 0 | 2 | 0 | | | | | | | | | G | G | | |
| NC_011770 | 5198695 | G | T | 85 | 0 | 2 | 0 | T | T | | | | | | | | | | |
| NC_011770 | 5199349 | G | A | 83 | 0 | 2 | 0 | A | A | | | | | | | | | | |
| NC_011770 | 5199382 | C | T | 86 | 0 | 2 | 0 | | | | | | | | | | | T | T |
| NC_011770 | 5201377 | C | T | 81 | 0 | 4 | 0 | | | T | T | T | T | | | | | | |
| NC_011770 | 5208256 | G | A | 75 | 0 | 2 | 0 | | | | | | | | | A | A | | |
| NC_011770 | 5208984 | G | A | 67 | 0 | 4 | 0 | | | A | A | A | A | | | | | | |
| NC_011770 | 5210503 | C | T | 59 | 0 | 2 | 0 | | | | | | | | | T | T | | |
| NC_011770 | 5215006 | A | G | 78 | 0 | 8 | 0 | G | G | | | | | G | G | G | G | G | G |
| NC_011770 | 5215634 | C | G | 79 | 0 | 8 | 0 | G | G | | | | | G | G | G | G | G | G |
| NC_011770 | 5218378 | C | G | 60 | 0 | 6 | 0 | G | G | | | | | G | G | | | G | G |
| NC_011770 | 5218379 | AC | A | 60 | 0 | 4 | 0 | | | | | | | A | A | | | A | A |
| NC_011770 | 5218398 | C | T | 59 | 0 | 2 | 0 | | | | | | | | | | | T | T |
| NC_011770 | 5221248 | T | C | 55 | 0 | 12 | 0 | C | C | C | C | C | C | C | C | C | C | C | C |
| NC_011770 | 5221315 | C | T | 51 | 0 | 12 | 0 | T | T | T | T | T | T | T | T | T | T | T | T |
| NC_011770 | 5221405 | CG | C | 49 | 0 | 12 | 0 | C | C | C | C | C | C | C | C | C | C | C | C |
| NC_011770 | 5222283 | G | A | 68 | 0 | 2 | 0 | | | | | | | A | A | | | | |
| NC_011770 | 5222324 | G | A | 67 | 0 | 4 | 0 | | | A | A | A | A | | | | | | |
| NC_011770 | 5222430 | C | T | 71 | 0 | 10 | 0 | | | T | T | T | T | T | T | T | T | T | T |
| NC_011770 | 5222436 | G | C | 74 | 0 | 4 | 0 | | | | | | | C | C | C | C | | |
| NC_011770 | 5222473 | T | C | 73 | 0 | 12 | 0 | C | C | C | C | C | C | C | C | C | C | C | C |
| NC_011770 | 5222476 | C | T | 73 | 0 | 2 | 0 | T | T | | | | | | | | | | |
| NC_011770 | 5222495 | G | A | 74 | 0 | 2 | 0 | | | | | | | | | A | A | | |
| NC_011770 | 5222524 | T | A | 74 | 0 | 12 | 0 | A | A | A | A | A | A | A | A | A | A | A | A |
| NC_011770 | 5223827 | C | T | 82 | 0 | 12 | 0 | T | T | T | T | T | T | T | T | T | T | T | T |
| NC_011770 | 5226033 | C | T | 76 | 0 | 2 | 0 | | | | | | | | | T | T | | |
| NC_011770 | 5226039 | G | A | 75 | 0 | 2 | 0 | | | | | | | | | | | A | A |
| NC_011770 | 5227068 | T | C | 73 | 0 | 6 | 0 | C | C | | | | | C | C | | | C | C |
| NC_011770 | 5227073 | T | A | 72 | 0 | 4 | 0 | A | A | | | | | A | A | | | | |
| NC_011770 | 5227094 | A | G | 69 | 0 | 10 | 0 | G | G | G | G | G | G | G | G | | | G | G |
| NC_011770 | 5227109 | G | A | 72 | 0 | 4 | 0 | A | A | | | | | A | A | | | | |
| NC_011770 | 5227116 | T | C | 69 | 0 | 6 | 0 | C | C | | | | | C | C | | | C | C |
| NC_011770 | 5227254 | G | A | 70 | 0 | 4 | 0 | | | A | A | A | A | | | | | | |
| NC_011770 | 5228049 | G | C | 51 | 0 | 10 | 0 | C | C | C | C | C | C | C | C | | | C | C |
| NC_011770 | 814913 | GGGCATTC | G | 19 | 0 | 7 | 0 | . | G | G | G | | G | | | G | . | G | G |
| NC_011770 | 5231723 | A | C | 67 | 0 | 10 | 0 | C | C | C | C | C | C | C | C | | | C | C |
| NC_011770 | 5231727 | T | G | 67 | 0 | 10 | 0 | G | G | G | G | G | G | G | G | | | G | G |
| NC_011770 | 5231730 | G | A | 68 | 0 | 8 | 0 | A | A | A | A | A | A | | | | | A | A |
| NC_011770 | 5231821 | A | G | 59 | 0 | 10 | 0 | G | G | G | G | G | G | G | G | | | G | G |
| NC_011770 | 5231859 | A | G | 56 | 0 | 2 | 0 | | | | | | | G | G | | | | |
| NC_011770 | 5233252 | T | C | 61 | 0 | 4 | 0 | | | C | C | C | C | | | | | | |
| NC_011770 | 5233307 | T | A | 59 | 0 | 4 | 0 | | | A | A | A | A | | | | | | |
| NC_011770 | 5233353 | C | T | 63 | 0 | 4 | 0 | | | T | T | T | T | | | | | | |
| NC_011770 | 5233388 | T | TC | 64 | 0 | 2 | 0 | | | | | | | | | TC | TC | | |

| | | | | | | | | | | | | | | | | | | | |
|---|---|---|---|---|---|---|---|---|---|---|---|---|---|---|---|---|---|---|---|
| NC_011770 | 5237009 | G | A | 80 | 0 | 2 | 0 | | | | | | | | | | | A | A |
| NC_011770 | 5237014 | G | A | 82 | 0 | 6 | 0 | A | A | A | A | A | A | | | | | | |
| NC_011770 | 5237103 | C | T | 80 | 0 | 4 | 0 | | | | | | | T | T | T | T | | |
| NC_011770 | 5237127 | G | A | 74 | 0 | 2 | 0 | A | A | | | | | | | | | | |
| NC_011770 | 5238306 | T | G | 78 | 0 | 4 | 0 | | | G | G | G | G | | | | | | |
| NC_011770 | 5238319 | C | T | 81 | 0 | 4 | 0 | | | T | T | T | T | | | | | | |
| NC_011770 | 5238320 | G | A | 80 | 0 | 6 | 0 | | | A | A | A | A | | | A | A | | |
| NC_011770 | 5238326 | T | A | 78 | 0 | 2 | 0 | A | A | | | | | | | | | | |
| NC_011770 | 5238329 | G | C | 79 | 0 | 12 | 0 | C | C | C | C | C | C | C | C | C | C | C | C |
| NC_011770 | 5241529 | A | T | 71 | 0 | 2 | 0 | | | | | | | | | | | T | T |
| NC_011770 | 5242762 | T | C | 64 | 0 | 8 | 0 | C | C | C | C | C | C | | | | | C | C |
| NC_011770 | 5242769 | A | G | 65 | 0 | 4 | 0 | | | | | | | G | G | G | G | | |
| NC_011770 | 1235223 | GT | G | 10 | 5 | 1 | 0 | | | | | . | | . | . | | G | . | . |
| NC_011770 | 5246560 | C | T | 95 | 0 | 6 | 0 | T | T | T | T | T | T | | | | | | |
| NC_011770 | 5247581 | G | GC | 80 | 0 | 12 | 0 | GC | GC | GC | GC | GC | GC | GC | GC | GC | GC | GC | GC |
| NC_011770 | 5247785 | A | G | 79 | 0 | 12 | 0 | G | G | G | G | G | G | G | G | G | G | G | G |
| NC_011770 | 5248262 | G | A | 63 | 0 | 4 | 0 | | | | | | | | | A | A | A | A |
| NC_011770 | 5248422 | C | T | 62 | 0 | 4 | 0 | | | | | | | | | T | T | T | T |
| NC_011770 | 5248440 | GC | G | 58 | 0 | 2 | 0 | | | | | | | | | | | G | G |
| NC_011770 | 5249163 | T | C | 69 | 0 | 8 | 0 | C | C | C | C | C | C | | | | | C | C |
| NC_011770 | 5249177 | G | A | 67 | 0 | 8 | 0 | A | A | A | A | A | A | | | | | A | A |
| NC_011770 | 5251363 | A | AT | 75 | 0 | 4 | 0 | AT | AT | AT | AT | | | | | | | | |
| NC_011770 | 5252693 | G | A | 73 | 0 | 10 | 0 | | | A | A | A | A | A | A | A | A | A | A |
| NC_011770 | 5257903 | G | A | 70 | 0 | 2 | 0 | A | A | | | | | | | | | | |
| NC_011770 | 5257923 | T | C | 67 | 0 | 2 | 0 | C | C | | | | | | | | | | |
| NC_011770 | 5257947 | C | T | 67 | 0 | 2 | 0 | T | T | | | | | | | | | | |
| NC_011770 | 5258810 | GA | G | 62 | 0 | 2 | 0 | | | | | | | G | G | | | | |
| NC_011770 | 5258915 | A | G | 61 | 0 | 12 | 0 | G | G | G | G | G | G | G | G | G | G | G | G |
| NC_011770 | 5264413 | GA | G | 54 | 0 | 2 | 0 | | | | | | | | | | | G | G |
| NC_011770 | 5265437 | CCGG | C | 71 | 0 | 8 | 0 | C | C | C | C | C | C | | | | | C | C |
| NC_011770 | 5267472 | T | C | 78 | 0 | 6 | 0 | C | C | | | | | | | C | C | C | C |
| NC_011770 | 5270349 | T | C | 82 | 0 | 1 | 0 | | | | | | | C | | | | | |
| NC_011770 | 5271397 | A | T | 54 | 0 | 2 | 0 | | | | | | | T | T | | | | |
| NC_011770 | 1683699 | C | T | 8 | 1 | 8 | 0 | T | T | . | . | | T | T | T | T | T | T | . |
| NC_011770 | 5289787 | T | C | 84 | 0 | 1 | 0 | | | | | | | C | | | | | |
| NC_011770 | 5289928 | A | G | 79 | 0 | 4 | 0 | | | G | G | G | G | | | | | | |
| NC_011770 | 5289965 | C | A | 78 | 0 | 2 | 0 | A | A | | | | | | | | | | |
| NC_011770 | 5291313 | T | A | 46 | 0 | 4 | 0 | | | A | A | A | A | | | | | | |
| NC_011770 | 5298718 | G | C | 82 | 0 | 2 | 0 | | | | | | | C | C | | | | |
| NC_011770 | 5298816 | C | T | 85 | 0 | 2 | 0 | | | | | | | T | T | | | | |
| NC_011770 | 5300940 | C | G | 51 | 0 | 6 | 0 | G | G | | | | | G | G | | | G | G |
| NC_011770 | 5301007 | G | C | 58 | 0 | 2 | 0 | | | | | | | | | | | C | C |
| NC_011770 | 5301022 | C | T | 58 | 0 | 2 | 0 | T | T | | | | | | | | | | |
| NC_011770 | 58508 | T | TA | 69 | 0 | 1 | 0 | | TA | | | | | | | | | | |
| NC_011770 | 5302232 | C | T | 75 | 0 | 8 | 0 | T | T | T | T | T | T | | | T | T | | |
| NC_011770 | 5303325 | G | T | 78 | 0 | 2 | 0 | | | | | | | | | | | T | T |
| NC_011770 | 5303329 | A | T | 77 | 0 | 2 | 0 | | | | | | | T | T | | | | |
| NC_011770 | 5303369 | C | T | 82 | 0 | 2 | 0 | | | | | | | T | T | | | | |
| NC_011770 | 5303422 | T | C | 72 | 0 | 2 | 0 | | | | | | | | | | | C | C |
| NC_011770 | 5304454 | C | T | 75 | 0 | 2 | 0 | | | | | | | | | | | T | T |
| NC_011770 | 5304471 | G | T | 74 | 0 | 4 | 0 | T | T | | | | | | | | | T | T |
| NC_011770 | 5304506 | A | G | 76 | 0 | 2 | 0 | | | | | | | | | G | G | | |
| NC_011770 | 5310314 | T | C | 75 | 0 | 2 | 0 | | | | | | | | | | | C | C |
| NC_011770 | 183279 | CT | C | 79 | 0 | 11 | 0 | C | C | C | | C | C | C | C | C | C | C | C |
| NC_011770 | 5314718 | G | A | 57 | 0 | 2 | 0 | | | | | | | | | | | A | A |
| NC_011770 | 5314719 | C | T | 56 | 0 | 4 | 0 | | | T | T | T | T | | | | | | |
| NC_011770 | 5322642 | GC | G | 98 | 0 | 4 | 0 | G | G | | | | | G | G | | | | |
| NC_011770 | 5322666 | C | G | 103 | 0 | 2 | 0 | | | | | | | G | G | | | | |
| NC_011770 | 5322966 | G | A | 81 | 0 | 2 | 0 | | | | | | | A | A | | | | |
| NC_011770 | 5326358 | A | G | 81 | 0 | 2 | 0 | | | | | | | | | G | G | | |
| NC_011770 | 5326383 | G | A | 83 | 0 | 2 | 0 | A | A | | | | | | | | | | |
| NC_011770 | 5326412 | C | T | 83 | 0 | 4 | 0 | | | T | T | T | T | | | | | | |

| | | | | | | | | | | | | | | | | | | | |
|---|---|---|---|---|---|---|---|---|---|---|---|---|---|---|---|---|---|---|---|
| NC_011770 | 150741 | TA | T | 77 | 0 | 3 | 0 | | T | | | | | | | T | T | | |
| NC_011770 | 5328118 | C | T | 86 | 0 | 1 | 0 | | | | | | | T | | | | | |
| NC_011770 | 5329806 | C | T | 61 | 0 | 2 | 0 | | | | | | | | | T | T | | |
| NC_011770 | 5330101 | G | C | 60 | 0 | 4 | 0 | | | | | | | C | C | C | C | | |
| NC_011770 | 5330112 | C | A | 60 | 0 | 4 | 0 | | | | | | | A | A | A | A | | |
| NC_011770 | 5330117 | A | G | 58 | 0 | 6 | 0 | G | G | G | G | G | G | | | | | | |
| NC_011770 | 5330159 | G | A | 58 | 0 | 2 | 0 | | | | | | | A | A | | | | |
| NC_011770 | 5330179 | T | G | 56 | 0 | 12 | 0 | G | G | G | G | G | G | G | G | G | G | G | G |
| NC_011770 | 5330182 | G | C | 57 | 0 | 6 | 0 | C | C | C | C | C | C | | | | | | |
| NC_011770 | 5330192 | A | G | 59 | 0 | 4 | 0 | | | | | | | G | G | G | G | | |
| NC_011770 | 5331145 | A | G | 88 | 0 | 2 | 0 | | | | | | | | | G | G | | |
| NC_011770 | 245583 | C | CAA | 66 | 0 | 1 | 0 | | CAA | | | | | | | | | | |
| NC_011770 | 5332222 | G | T | 71 | 0 | 2 | 0 | | | T | T | | | | | | | | |
| NC_011770 | 5334057 | C | T | 74 | 0 | 4 | 0 | | | T | T | T | T | | | | | | |
| NC_011770 | 5334061 | G | A | 73 | 0 | 2 | 0 | | | | | | | A | A | | | | |
| NC_011770 | 5334089 | G | A | 71 | 0 | 2 | 0 | | | | | | | A | A | | | | |
| NC_011770 | 5334098 | T | C | 69 | 0 | 2 | 0 | | | | | | | C | C | | | | |
| NC_011770 | 5334139 | T | A | 67 | 0 | 2 | 0 | | | | | | | | | | | A | A |
| NC_011770 | 5334141 | T | C | 67 | 0 | 2 | 0 | | | | | | | C | C | | | | |
| NC_011770 | 5334144 | T | C | 69 | 0 | 2 | 0 | | | | | | | C | C | | | | |
| NC_011770 | 2548478 | C | T | 113 | 0 | 5 | 0 | | | | | | T | T | T | T | T | | |
| NC_011770 | 5335532 | T | TGGCCG | 80 | 0 | 2 | 0 | | | | | TGGCCGCCCACGC | | | | TGGCCGCCCACGC | | | |
| NC_011770 | 5335571 | G | A | 84 | 0 | 2 | 0 | | | | | | | A | A | | | | |
| NC_011770 | 5335582 | T | C | 87 | 0 | 12 | 0 | C | C | C | C | C | C | C | C | C | C | C | C |
| NC_011770 | 5336128 | C | T | 71 | 0 | 2 | 0 | | | | | | | T | T | | | | |
| NC_011770 | 5338457 | T | C | 63 | 0 | 4 | 0 | | | | | | | C | C | | | C | C |
| NC_011770 | 5338473 | G | A | 63 | 0 | 2 | 0 | A | A | | | | | | | | | | |
| NC_011770 | 5338482 | CA | C | 63 | 0 | 4 | 0 | | | | | | | C | C | | | C | C |
| NC_011770 | 5344446 | C | CG | 90 | 0 | 4 | 0 | | | CG | CG | CG | CG | | | | | | |
| NC_011770 | 5344494 | GA | G | 93 | 0 | 10 | 0 | G | G | G | G | G | G | G | G | G | G | | |
| NC_011770 | 5346074 | A | G | 80 | 0 | 12 | 0 | G | G | G | G | G | G | G | G | G | G | G | G |
| NC_011770 | 266311 | G | A | 38 | 0 | 1 | 0 | | A | | | | | | | | | | |
| NC_011770 | 5351547 | T | C | 85 | 0 | 2 | 0 | | | | | | | C | C | | | | |
| NC_011770 | 5351635 | C | A | 87 | 0 | 2 | 0 | | | | | | | | | | | A | A |
| NC_011770 | 5351639 | T | C | 94 | 0 | 2 | 0 | | | | | | | C | C | | | | |
| NC_011770 | 5351646 | A | G | 86 | 0 | 12 | 0 | G | G | G | G | G | G | G | G | G | G | G | G |
| NC_011770 | 5351654 | G | C | 84 | 0 | 2 | 0 | | | | | | | | | | | C | C |
| NC_011770 | 5362051 | C | T | 73 | 0 | 12 | 0 | T | T | T | T | T | T | T | T | T | T | T | T |
| NC_011770 | 5362053 | C | T | 74 | 0 | 12 | 0 | T | T | T | T | T | T | T | T | T | T | T | T |
| NC_011770 | 5364163 | G | A | 90 | 0 | 8 | 0 | A | A | | | | | A | A | A | A | A | A |
| NC_011770 | 5364192 | A | G | 87 | 0 | 12 | 0 | G | G | G | G | G | G | G | G | G | G | G | G |
| NC_011770 | 5365060 | G | A | 89 | 0 | 2 | 0 | | | | | | | | | | | A | A |
| NC_011770 | 5365099 | G | A | 89 | 0 | 4 | 0 | | | A | A | A | A | | | | | | |
| NC_011770 | 5366874 | G | A | 64 | 0 | 2 | 0 | | | | | | | | | | | A | A |
| NC_011770 | 5368546 | T | C | 64 | 0 | 12 | 0 | C | C | C | C | C | C | C | C | C | C | C | C |
| NC_011770 | 183274 | G | A | 80 | 0 | 1 | 0 | | | | | | | | | | | A | |
| NC_011770 | 5370557 | G | T | 63 | 0 | 6 | 0 | T | T | T | T | T | T | | | | | | |
| NC_011770 | 5372395 | T | C | 83 | 0 | 2 | 0 | C | C | | | | | | | | | | |
| NC_011770 | 5375719 | C | T | 83 | 0 | 2 | 0 | | | | | | | T | T | | | | |
| NC_011770 | 5377733 | C | T | 81 | 0 | 8 | 0 | | | T | T | T | T | T | T | T | T | | |
| NC_011770 | 5380489 | T | C | 85 | 0 | 2 | 0 | | | | | | | C | C | | | | |
| NC_011770 | 5384371 | C | T | 68 | 0 | 2 | 0 | | | | | | | T | T | | | | |
| NC_011770 | 5384382 | G | C | 67 | 0 | 4 | 0 | C | C | | | | | C | C | | | | |
| NC_011770 | 5384408 | G | GACGCC | 85 | 0 | 2 | 0 | . | | | | | | GACGCCA | GACGCCA | . | | | |
| NC_011770 | 5384431 | C | A | 94 | 0 | 4 | 0 | | | A | A | A | A | | | | | | |
| NC_011770 | 5384473 | A | G | 64 | 0 | 4 | 0 | | | G | G | G | G | | | | | | |
| NC_011770 | 5384476 | C | A | 64 | 0 | 4 | 0 | | | A | A | A | A | | | | | | |
| NC_011770 | 5384479 | C | G | 64 | 0 | 4 | 0 | | | G | G | G | G | | | | | | |
| NC_011770 | 5384487 | A | G | 67 | 0 | 2 | 0 | | | | | | | | | G | G | | |
| NC_011770 | 5384496 | G | A | 69 | 0 | 2 | 0 | | | | | | | | | A | A | | |
| NC_011770 | 5387275 | T | C | 82 | 0 | 4 | 0 | | | | | | | C | C | C | C | | |
| NC_011770 | 5387278 | A | G | 80 | 0 | 4 | 0 | | | | | | | G | G | G | G | | |

| | | | | | | | | | | | | | | | | | | | |
|---|---|---|---|---|---|---|---|---|---|---|---|---|---|---|---|---|---|---|---|
| NC_011770 | 5387312 | T | A | 79 | 0 | 4 | 0 | | | | | | | A | A | A | A | | |
| NC_011770 | 5393261 | G | A | 76 | 0 | 2 | 0 | | | | | | | | | A | A | | |
| NC_011770 | 5393283 | G | A | 73 | 0 | 2 | 0 | | | | | | | | | A | A | | |
| NC_011770 | 5393773 | A | G | 75 | 0 | 2 | 0 | | | | | | | G | G | | | | |
| NC_011770 | 5393810 | A | G | 74 | 0 | 2 | 0 | | | | | | | | | G | G | | |
| NC_011770 | 5393837 | G | A | 80 | 0 | 2 | 0 | | | | | | | A | A | | | | |
| NC_011770 | 5395439 | C | T | 75 | 0 | 2 | 0 | | | | | | | | | T | T | | |
| NC_011770 | 5397538 | G | A | 102 | 0 | 4 | 0 | A | A | | | | | | | A | A | | |
| NC_011770 | 5397566 | T | C | 100 | 0 | 2 | 0 | C | C | | | | | | | | | | |
| NC_011770 | 5397644 | C | T | 98 | 0 | 4 | 0 | | | T | T | T | T | | | | | | |
| NC_011770 | 5397662 | T | C | 95 | 0 | 8 | 0 | | | C | C | C | C | C | C | | | C | C |
| NC_011770 | 5397674 | G | A | 97 | 0 | 12 | 0 | A | A | A | A | A | A | A | A | A | A | A | A |
| NC_011770 | 5399130 | C | T | 66 | 0 | 2 | 0 | | | | | | | | | | | T | T |
| NC_011770 | 5399163 | G | T | 64 | 0 | 2 | 0 | | | | | | | | | | | T | T |
| NC_011770 | 5399190 | A | C | 61 | 0 | 6 | 0 | | | C | C | C | C | | | | | C | C |
| NC_011770 | 5399197 | A | G | 60 | 0 | 2 | 0 | | | | | | | | | G | G | | |
| NC_011770 | 5399198 | C | T | 61 | 0 | 2 | 0 | | | | | | | | | T | T | | |
| NC_011770 | 281858 | A | AGC | 47 | 4 | 7 | 0 | | AGC | AGC | AGC | AGC | AGC | . | . | AGC | AGC | . | . |
| NC_011770 | 5401668 | G | A | 89 | 0 | 2 | 0 | | | | | | | | | A | A | | |
| NC_011770 | 5401691 | C | T | 86 | 0 | 4 | 0 | | | T | T | T | T | | | | | | |
| NC_011770 | 5401728 | C | T | 80 | 0 | 4 | 0 | T | T | | | | | T | T | | | | |
| NC_011770 | 5402630 | T | C | 54 | 0 | 8 | 0 | C | C | | | | | C | C | C | C | C | C |
| NC_011770 | 5402631 | A | G | 55 | 0 | 2 | 0 | | | | | | | | | | | G | G |
| NC_011770 | 5402633 | T | A | 56 | 0 | 2 | 0 | | | | | | | | | | | A | A |
| NC_011770 | 5402662 | AT | TT | 54 | 0 | 4 | 0 | | | | | | | TT | TT | | | A | A |
| NC_011770 | 5402663 | T | A | 44 | 2 | 2 | 0 | | | | | | | A | A | | | . | . |
| NC_011770 | 5402770 | A | G | 47 | 4 | 4 | 0 | G | G | | | | | G | G | . | . | . | . |
| NC_011770 | 5404229 | C | T | 6 | 10 | 2 | 0 | T | T | . | . | . | . | . | . | . | . | . | . |
| NC_011770 | 5404236 | C | T | 6 | 10 | 2 | 0 | T | T | . | . | . | . | . | . | . | . | . | . |
| NC_011770 | 5404240 | A | G | 5 | 10 | 2 | 0 | G | G | . | . | . | . | . | . | . | . | . | . |
| NC_011770 | 5404242 | A | C | 6 | 10 | 2 | 0 | C | C | . | . | . | . | . | . | . | . | . | . |
| NC_011770 | 5404249 | A | G | 6 | 10 | 2 | 0 | G | G | . | . | . | . | . | . | . | . | . | . |
| NC_011770 | 5404254 | A | G | 10 | 8 | 2 | 0 | G | G | . | . | . | . | | | . | . | . | . |
| NC_011770 | 5404725 | A | T | 67 | 0 | 2 | 0 | | | | | | | | | T | T | | |
| NC_011770 | 5404728 | A | C | 65 | 0 | 2 | 0 | | | | | | | | | C | C | | |
| NC_011770 | 5404735 | TG | T | 67 | 0 | 2 | 0 | | | | | | | | | | | T | T |
| NC_011770 | 5404752 | A | T | 66 | 2 | 2 | 0 | | | | | | | | | . | . | T | T |
| NC_011770 | 5404767 | T | C | 60 | 4 | 8 | 0 | C | C | C | C | C | C | C | C | . | . | . | . |
| NC_011770 | 257422 | C | T | 21 | 2 | 1 | 0 | | | | | | | | | . | . | T | |
| NC_011770 | 5409559 | A | G | 70 | 0 | 2 | 0 | | | | | | | G | G | | | | |
| NC_011770 | 5409839 | A | G | 72 | 0 | 4 | 0 | G | G | | | | | G | G | | | | |
| NC_011770 | 5409854 | A | G | 73 | 0 | 2 | 0 | | | | | | | | | G | G | | |
| NC_011770 | 5409873 | C | T | 74 | 0 | 4 | 0 | T | T | | | | | T | T | | | | |
| NC_011770 | 5411506 | A | G | 67 | 0 | 12 | 0 | G | G | G | G | G | G | G | G | G | G | G | G |
| NC_011770 | 5411539 | G | A | 69 | 0 | 2 | 0 | | | | | | | A | A | | | | |
| NC_011770 | 5411561 | G | A | 69 | 0 | 8 | 0 | A | A | A | A | A | A | | | | | A | A |
| NC_011770 | 5411568 | G | A | 69 | 0 | 8 | 0 | A | A | A | A | A | A | | | | | A | A |
| NC_011770 | 5411575 | T | C | 70 | 0 | 8 | 0 | C | C | C | C | C | C | | | | | C | C |
| NC_011770 | 5414265 | A | G | 80 | 0 | 4 | 0 | | | | | | | G | G | G | G | | |
| NC_011770 | 5415392 | G | T | 70 | 0 | 10 | 0 | T | T | T | T | T | T | | | T | T | T | T |
| NC_011770 | 5415417 | A | C | 69 | 0 | 10 | 0 | C | C | C | C | C | C | | | C | C | C | C |
| NC_011770 | 5415420 | G | T | 69 | 0 | 10 | 0 | T | T | T | T | T | T | | | T | T | T | T |
| NC_011770 | 5415429 | G | GA | 66 | 0 | 10 | 0 | GA | GA | GA | GA | GA | GA | | | GA | GA | GA | GA |
| NC_011770 | 5415434 | A | C | 65 | 0 | 10 | 0 | C | C | C | C | C | C | | | C | C | C | C |
| NC_011770 | 5415474 | T | C | 65 | 0 | 10 | 0 | C | C | C | C | C | C | | | C | C | C | C |
| NC_011770 | 5415477 | A | C | 62 | 0 | 4 | 0 | | | C | C | C | C | | | | | | |
| NC_011770 | 5415553 | CT | C | 63 | 0 | 6 | 0 | C | C | C | C | C | C | | | | | | |
| NC_011770 | 5415583 | A | T | 61 | 0 | 4 | 0 | | | | | | | | | T | T | T | T |
| NC_011770 | 5415593 | G | A | 62 | 0 | 10 | 0 | A | A | A | A | A | A | | | A | A | A | A |
| NC_011770 | 5415665 | C | A | 63 | 0 | 2 | 0 | | | | | | | | | A | A | | |
| NC_011770 | 5415703 | C | T | 69 | 0 | 4 | 0 | | | | | | | | | T | T | T | T |
| NC_011770 | 5415706 | A | G | 68 | 0 | 12 | 0 | G | G | G | G | G | G | G | G | G | G | G | G |

| | | | | | | | | | | | | | | | | | | | |
|---|---|---|---|---|---|---|---|---|---|---|---|---|---|---|---|---|---|---|---|
| NC_011770 | 5415750 | CG | C | 76 | 0 | 10 | 0 | C | C | C | C | C | C | | | C | C | C | C |
| NC_011770 | 5415771 | T | C | 73 | 0 | 10 | 0 | C | C | C | C | C | C | | | C | C | C | C |
| NC_011770 | 5415779 | C | T | 73 | 0 | 6 | 0 | T | T | T | T | T | T | | | | | | |
| NC_011770 | 5415780 | C | T | 74 | 0 | 2 | 0 | | | | | | | | | T | T | | |
| NC_011770 | 5415856 | G | A | 69 | 0 | 10 | 0 | A | A | A | A | A | A | | | A | A | A | A |
| NC_011770 | 5415859 | G | T | 71 | 0 | 10 | 0 | T | T | T | T | T | T | | | T | T | T | T |
| NC_011770 | 5415865 | C | T | 70 | 0 | 10 | 0 | T | T | T | T | T | T | | | T | T | T | T |
| NC_011770 | 5415871 | G | A | 72 | 0 | 10 | 0 | A | A | A | A | A | A | | | A | A | A | A |
| NC_011770 | 5415898 | T | G | 74 | 0 | 6 | 0 | G | G | G | G | G | G | | | | | | |
| NC_011770 | 5415900 | C | A | 77 | 0 | 2 | 0 | | | | | | | A | A | | | | |
| NC_011770 | 5415905 | C | T | 78 | 0 | 2 | 0 | | | | | | | T | T | | | | |
| NC_011770 | 5415920 | C | T | 76 | 0 | 2 | 0 | | | | | | | T | T | | | | |
| NC_011770 | 968558 | GCGT | G | 72 | 0 | 11 | 0 | G | | G | G | G | G | G | G | G | G | G | G |
| NC_011770 | 5421908 | A | AT | 49 | 0 | 2 | 0 | | | | | | | AT | AT | | | | |
| NC_011770 | 5421909 | C | T | 40 | 2 | 10 | 0 | T | T | T | T | T | T | . | . | T | T | T | T |
| NC_011770 | 5421936 | C | CG | 51 | 0 | 2 | 0 | | | | | | | CG | CG | | | | |
| NC_011770 | 5421945 | A | T | 51 | 0 | 2 | 0 | | | | | | | T | T | | | | |
| NC_011770 | 5421961 | C | A | 49 | 0 | 2 | 0 | | | | | | | A | A | | | | |
| NC_011770 | 5421964 | C | A | 52 | 0 | 2 | 0 | | | | | | | A | A | | | | |
| NC_011770 | 5421977 | C | T | 52 | 0 | 2 | 0 | T | T | | | | | | | | | | |
| NC_011770 | 5421995 | A | G | 50 | 0 | 2 | 0 | | | | | | | G | G | | | | |
| NC_011770 | 5421999 | G | T | 51 | 0 | 2 | 0 | | | | | | | T | T | | | | |
| NC_011770 | 5422002 | T | C | 50 | 0 | 2 | 0 | | | | | | | C | C | | | | |
| NC_011770 | 5422026 | C | T | 49 | 0 | 2 | 0 | | | | | | | T | T | | | | |
| NC_011770 | 5422032 | T | C | 49 | 0 | 2 | 0 | C | C | | | | | | | | | | |
| NC_011770 | 5422034 | G | A | 49 | 0 | 6 | 0 | | | A | A | A | A | | | A | A | | |
| NC_011770 | 5422376 | A | G | 61 | 0 | 10 | 0 | | | G | G | G | G | G | G | G | G | G | G |
| NC_011770 | 5422468 | A | G | 59 | 0 | 12 | 0 | G | G | G | G | G | G | G | G | G | G | G | G |
| NC_011770 | 5422473 | C | T | 62 | 0 | 10 | 0 | | | T | T | T | T | T | T | T | T | T | T |
| NC_011770 | 5428175 | C | T | 69 | 0 | 6 | 0 | | | T | T | T | T | | | | | T | T |
| NC_011770 | 5428419 | C | A | 65 | 0 | 4 | 0 | | | | | | | A | A | A | A | | |
| NC_011770 | 257458 | G | A | 15 | 2 | 9 | 0 | A | A | A | A | A | A | A | A | . | . | A | |
| NC_011770 | 5431459 | G | C | 81 | 0 | 2 | 0 | | | | | | | | | | | C | C |
| NC_011770 | 5432648 | A | T | 61 | 2 | 2 | 0 | T | T | | | | | | | . | . | | |
| NC_011770 | 5432714 | A | G | 62 | 2 | 2 | 0 | G | G | | | | | | | . | . | | |
| NC_011770 | 5433321 | C | A | 79 | 2 | 10 | 0 | A | A | A | A | A | A | A | A | . | . | A | A |
| NC_011770 | 4229402 | AGATG | A | 36 | 0 | 3 | 0 | | | A | A | A | | | | | | | |
| NC_011770 | 5440092 | A | G | 47 | 2 | 8 | 0 | | | G | G | G | G | G | G | . | . | G | G |
| NC_011770 | 2548504 | G | C | 117 | 0 | 9 | 0 | | | C | C | | C | C | C | C | C | C | C |
| NC_011770 | 257464 | T | C | 14 | 2 | 1 | 0 | | | | | | | | | . | . | C | |
| NC_011770 | 431445 | G | A | 78 | 0 | 1 | 0 | | | | | | | | | | | A | |
| NC_011770 | 5447705 | C | T | 66 | 0 | 8 | 0 | T | T | T | T | T | T | | | | | T | T |
| NC_011770 | 5451029 | C | T | 72 | 0 | 6 | 0 | | | T | T | T | T | | | | | T | T |
| NC_011770 | 5451159 | T | C | 79 | 0 | 2 | 0 | | | | | | | | | C | C | | |
| NC_011770 | 5451840 | G | A | 72 | 0 | 1 | 0 | | | | | | | A | | | | | |
| NC_011770 | 418382 | C | T | 42 | 0 | 1 | 0 | | | | T | | | | | | | | |
| NC_011770 | 5451945 | A | C | 59 | 0 | 11 | 0 | C | C | C | C | . | C | C | C | C | C | C | C |
| NC_011770 | 541937 | C | T | 75 | 0 | 1 | 0 | | | | | | | | | | | T | |
| NC_011770 | 5453501 | G | A | 75 | 0 | 2 | 0 | | | | | | | A | A | | | | |
| NC_011770 | 5453506 | C | T | 76 | 0 | 2 | 0 | | | | | | | T | T | | | | |
| NC_011770 | 5453510 | A | C | 76 | 0 | 12 | 0 | C | C | C | C | C | C | C | C | C | C | C | C |
| NC_011770 | 5453518 | T | C | 79 | 0 | 2 | 0 | | | | | | | C | C | | | | |
| NC_011770 | 561027 | A | G | 61 | 0 | 1 | 0 | | | | | | | | | | | G | |
| NC_011770 | 5453580 | T | C | 83 | 0 | 2 | 0 | | | | | | | | | | | C | C |
| NC_011770 | 5453594 | T | C | 82 | 0 | 10 | 0 | C | C | C | C | C | C | | | C | C | C | C |
| NC_011770 | 5453648 | C | G | 93 | 0 | 12 | 0 | G | G | G | G | G | G | G | G | G | G | G | G |
| NC_011770 | 5455705 | A | C | 53 | 0 | 12 | 0 | C | C | C | C | C | C | C | C | C | C | C | C |
| NC_011770 | 5455749 | G | T | 56 | 0 | 12 | 0 | T | T | T | T | T | T | T | T | T | T | T | T |
| NC_011770 | 5456453 | G | A | 78 | 0 | 4 | 0 | A | A | | | | | | | | | A | A |
| NC_011770 | 5456857 | T | C | 71 | 0 | 6 | 0 | | | C | C | C | C | | | C | C | | |
| NC_011770 | 5456877 | G | C | 73 | 0 | 2 | 0 | | | | | | | C | C | | | | |
| NC_011770 | 5456984 | T | C | 76 | 0 | 12 | 0 | C | C | C | C | C | C | C | C | C | C | C | C |

| | | | | | | | | | | | | | | | | | | | |
|---|---|---|---|---|---|---|---|---|---|---|---|---|---|---|---|---|---|---|---|
| NC_011770 | 5458138 | CG | C | 56 | 0 | 2 | 0 | | | | | | | | | | | C | C |
| NC_011770 | 5458148 | C | T | 60 | 0 | 2 | 0 | T | T | | | | | | | | | | |
| NC_011770 | 5458162 | A | G | 58 | 0 | 2 | 0 | G | G | | | | | | | | | | |
| NC_011770 | 5458172 | A | G | 62 | 0 | 2 | 0 | G | G | | | | | | | | | | |
| NC_011770 | 5458175 | G | A | 60 | 0 | 2 | 0 | A | A | | | | | | | | | | |
| NC_011770 | 5458182 | A | G | 60 | 0 | 2 | 0 | G | G | | | | | | | | | | |
| NC_011770 | 5458184 | G | A | 60 | 0 | 2 | 0 | A | A | | | | | | | | | | |
| NC_011770 | 1873531 | C | CT | 51 | 0 | 1 | 0 | CT | | | | | | | | | | | |
| NC_011770 | 5465971 | A | G | 40 | 0 | 2 | 0 | | | | | | | G | G | | | | |
| NC_011770 | 5465986 | G | A | 41 | 0 | 4 | 0 | | | A | A | A | A | | | | | | |
| NC_011770 | 5465999 | T | C | 42 | 0 | 2 | 0 | | | | | | | C | C | | | | |
| NC_011770 | 5466003 | C | A | 41 | 0 | 2 | 0 | A | A | | | | | | | | | | |
| NC_011770 | 5466019 | C | T | 40 | 0 | 4 | 0 | T | T | | | | | T | T | | | | |
| NC_011770 | 5466026 | G | T | 42 | 0 | 2 | 0 | T | T | | | | | | | | | | |
| NC_011770 | 5466369 | C | G | 51 | 0 | 6 | 0 | | | G | G | G | G | | | | | G | G |
| NC_011770 | 5466459 | T | C | 60 | 0 | 2 | 0 | | | | | | | | | C | C | | |
| NC_011770 | 5466504 | A | G | 60 | 0 | 2 | 0 | | | | | | | | | G | G | | |
| NC_011770 | 5466537 | C | T | 63 | 0 | 2 | 0 | T | T | | | | | | | | | | |
| NC_011770 | 5467733 | A | G | 77 | 0 | 2 | 0 | | | | | | | | | G | G | | |
| NC_011770 | 5467734 | A | G | 76 | 0 | 2 | 0 | | | | | | | | | G | G | | |
| NC_011770 | 5467739 | G | A | 76 | 0 | 2 | 0 | | | | | | | | | A | A | | |
| NC_011770 | 3069600 | T | A | 9 | 8 | 3 | 0 | . | . | A | A | | A | . | . | . | . | . | . |
| NC_011770 | 5470909 | T | C | 59 | 0 | 2 | 0 | | | | | | | | | C | C | | |
| NC_011770 | 5470918 | GC | G | 60 | 0 | 2 | 0 | | | | | | | | | G | G | | |
| NC_011770 | 5470931 | G | A | 60 | 0 | 2 | 0 | A | A | | | | | | | | | | |
| NC_011770 | 5470945 | G | A | 64 | 0 | 2 | 0 | | | | | | | | | A | A | | |
| NC_011770 | 5471387 | G | A | 78 | 0 | 10 | 0 | A | A | A | A | A | A | A | A | | | A | A |
| NC_011770 | 5472527 | G | GTT | 69 | 0 | 3 | 0 | | | | | | | | | GTT | GTT | GTT | |
| NC_011770 | 5472537 | T | G | 69 | 0 | 4 | 0 | | | | | | | | | G | G | G | G |
| NC_011770 | 5472539 | TG | T | 69 | 0 | 2 | 0 | | | | | | | | | | | T | T |
| NC_011770 | 1873635 | C | CG | 49 | 0 | 1 | 0 | CG | | | | | | | | | | | |
| NC_011770 | 5474143 | C | G | 52 | 0 | 4 | 0 | | | G | G | G | G | | | | | | |
| NC_011770 | 5474177 | T | C | 60 | 0 | 12 | 0 | C | C | C | C | C | C | C | C | C | C | C | C |
| NC_011770 | 5475635 | GA | G | 59 | 0 | 4 | 0 | | | G | G | G | G | | | | | | |
| NC_011770 | 5475643 | G | A | 60 | 0 | 2 | 0 | | | | | | | | | A | A | | |
| NC_011770 | 5475700 | A | G | 54 | 0 | 4 | 0 | | | G | G | G | G | | | | | | |
| NC_011770 | 5475723 | A | G | 53 | 0 | 8 | 0 | | | G | G | G | G | | | G | G | G | G |
| NC_011770 | 5475770 | G | C | 53 | 0 | 2 | 0 | | | | | C | C | | | | | | |
| NC_011770 | 5475812 | A | G | 49 | 0 | 8 | 0 | G | G | | | | | G | G | G | G | G | G |
| NC_011770 | 5476392 | G | A | 52 | 0 | 6 | 0 | A | A | | | | | A | A | A | A | | |
| NC_011770 | 5476432 | A | T | 51 | 0 | 4 | 0 | | | T | T | T | T | | | | | | |
| NC_011770 | 5476467 | G | A | 55 | 0 | 4 | 0 | | | A | A | A | A | | | | | | |
| NC_011770 | 5476482 | A | AT | 52 | 0 | 6 | 0 | | | AT | AT | AT | AT | | | | | AT | AT |
| NC_011770 | 5476491 | G | A | 54 | 0 | 4 | 0 | | | A | A | A | A | | | | | | |
| NC_011770 | 5476577 | AG | A | 58 | 0 | 4 | 0 | | | A | A | A | A | | | | | | |
| NC_011770 | 5476597 | G | A | 56 | 0 | 4 | 0 | | | A | A | A | A | | | | | | |
| NC_011770 | 5476602 | G | A | 58 | 0 | 2 | 0 | | | | | | | | | | | A | A |
| NC_011770 | 5476603 | A | G | 58 | 0 | 4 | 0 | | | G | G | G | G | | | | | | |
| NC_011770 | 5476604 | G | A | 57 | 0 | 4 | 0 | | | A | A | A | A | | | | | | |
| NC_011770 | 5476611 | A | T | 59 | 0 | 8 | 0 | T | T | G | G | G | G | T | T | | | | |
| NC_011770 | 5476654 | T | A | 60 | 0 | 4 | 0 | | | A | A | A | A | | | | | | |
| NC_011770 | 5476659 | G | A | 59 | 0 | 8 | 0 | A | A | | | | | A | A | A | A | A | A |
| NC_011770 | 5476663 | A | G | 59 | 0 | 4 | 0 | | | G | G | G | G | | | | | | |
| NC_011770 | 5476690 | A | G | 60 | 0 | 12 | 0 | G | G | G | G | G | G | G | G | G | G | G | G |
| NC_011770 | 5476708 | G | A | 63 | 0 | 6 | 0 | A | A | | | | | A | A | A | A | | |
| NC_011770 | 5476744 | C | A | 66 | 0 | 8 | 0 | A | A | | | | | A | A | A | A | A | A |
| NC_011770 | 5476755 | G | A | 68 | 0 | 4 | 0 | | | A | A | A | A | | | | | | |
| NC_011770 | 5476759 | G | A | 67 | 0 | 4 | 0 | | | A | A | A | A | | | | | | |
| NC_011770 | 5476768 | T | C | 67 | 0 | 4 | 0 | | | C | C | C | C | | | | | | |
| NC_011770 | 5476769 | G | A | 68 | 0 | 4 | 0 | | | A | A | A | A | | | | | | |
| NC_011770 | 5476782 | G | A | 68 | 0 | 4 | 0 | | | A | A | A | A | | | | | | |
| NC_011770 | 5476792 | C | T | 67 | 0 | 6 | 0 | T | T | | | | | T | T | | | T | T |

| | | | | | | | | | | | | | | | | | | | |
|---|---|---|---|---|---|---|---|---|---|---|---|---|---|---|---|---|---|---|---|
| NC_011770 | 5476858 | G | A | 72 | 0 | 4 | 0 | | | A | A | A | A | | | | | | |
| NC_011770 | 5476873 | G | T | 73 | 0 | 4 | 0 | | | T | T | T | T | | | | | | |
| NC_011770 | 5476874 | G | T | 72 | 0 | 2 | 0 | | | | | | | T | T | | | | |
| NC_011770 | 5476888 | C | G | 71 | 0 | 8 | 0 | | | G | G | G | G | | | G | G | G | G |
| NC_011770 | 563142 | TG | T | 10 | 4 | 1 | 0 | | . | | T | . | . | | . | . | | | |
| NC_011770 | 5481538 | A | G | 45 | 0 | 2 | 0 | | | | | | | | | G | G | | |
| NC_011770 | 5481784 | A | C | 61 | 0 | 12 | 0 | C | C | C | C | C | C | C | C | C | C | C | C |
| NC_011770 | 5481853 | A | G | 60 | 0 | 6 | 0 | G | G | | | | | G | G | G | G | | |
| NC_011770 | 5481913 | T | G | 61 | 0 | 4 | 0 | G | G | | | | | | | TC | TC | | |
| NC_011770 | 5482037 | T | C | 70 | 0 | 10 | 0 | C | C | C | C | C | C | C | C | C | C | | |
| NC_011770 | 5483808 | C | T | 64 | 0 | 6 | 0 | | | T | T | T | T | T | T | | | | |
| NC_011770 | 5484065 | C | T | 69 | 0 | 4 | 0 | | | T | T | T | T | | | | | | |
| NC_011770 | 5484170 | A | C | 70 | 0 | 2 | 0 | | | | | | | | | C | C | | |
| NC_011770 | 5484187 | C | T | 72 | 0 | 2 | 0 | | | | | | | T | T | | | | |
| NC_011770 | 627395 | C | T | 89 | 0 | 1 | 0 | | | | | | | | | | | T | |
| NC_011770 | 964631 | CG | C | 44 | 0 | 1 | 0 | | | | C | | | | | | | | |
| NC_011770 | 5490615 | T | C | 78 | 2 | 2 | 0 | C | C | | | | | | | . | . | | |
| NC_011770 | 5490620 | G | A | 76 | 2 | 10 | 0 | A | A | A | A | A | A | A | A | . | . | A | A |
| NC_011770 | 5490621 | C | A | 78 | 2 | 2 | 0 | | | | | | | | | . | . | A | A |
| NC_011770 | 5490679 | A | C | 79 | 2 | 10 | 0 | C | C | C | C | C | C | C | C | . | . | C | C |
| NC_011770 | 257412 | C | T | 22 | 2 | 3 | 0 | | | T | | T | T | | | . | . | | |
| NC_011770 | 5491264 | T | C | 62 | 2 | 10 | 0 | C | C | C | C | C | C | C | C | . | . | C | C |
| NC_011770 | 5491285 | C | A | 65 | 2 | 2 | 0 | | | | | | | A | A | . | . | | |
| NC_011770 | 5491374 | G | C | 64 | 2 | 10 | 0 | C | C | C | C | C | C | C | C | . | . | C | C |
| NC_011770 | 5491377 | C | T | 63 | 2 | 10 | 0 | T | T | T | T | T | T | T | T | . | . | T | T |
| NC_011770 | 1917133 | G | GC | 48 | 2 | 9 | 0 | GC | | GC | GC | GC | GC | . | . | GC | GC | GC | GC |
| NC_011770 | 5495781 | A | G | 76 | 0 | 4 | 0 | | | | | | | G | G | G | G | | |
| NC_011770 | 5497147 | A | C | 74 | 0 | 2 | 0 | | | | | | | | | | | C | C |
| NC_011770 | 5497161 | C | CG | 78 | 0 | 12 | 0 | CG | CG | CG | CG | CG | CG | CG | CG | CG | CG | CG | CG |
| NC_011770 | 5497588 | T | C | 94 | 0 | 12 | 0 | C | C | C | C | C | C | C | C | C | C | C | C |
| NC_011770 | 5497598 | A | G | 92 | 0 | 12 | 0 | G | G | G | G | G | G | G | G | G | G | G | G |
| NC_011770 | 1564018 | G | GC | 49 | 0 | 1 | 0 | | | | | | | | | | | GC | |
| NC_011770 | 5501177 | A | G | 88 | 0 | 2 | 0 | | | | | | | | | | | G | G |
| NC_011770 | 5501203 | C | T | 85 | 0 | 2 | 0 | | | | | | | T | T | | | | |
| NC_011770 | 5501274 | C | A | 80 | 0 | 2 | 0 | | | | | | | | | | | A | A |
| NC_011770 | 5501322 | A | G | 81 | 0 | 10 | 0 | | | G | G | G | G | G | G | G | G | G | G |
| NC_011770 | 5501981 | C | A | 81 | 0 | 4 | 0 | | | A | A | A | A | | | | | | |
| NC_011770 | 5504657 | C | G | 82 | 0 | 2 | 0 | | | | | | | | | | | G | G |
| NC_011770 | 5505272 | A | G | 75 | 0 | 2 | 0 | | | | | | | G | G | | | | |
| NC_011770 | 579117 | T | TC | 82 | 0 | 1 | 0 | | TC | | | | | | | | | | |
| NC_011770 | 5507052 | T | C | 60 | 0 | 4 | 0 | | | | | | | C | C | | | C | C |
| NC_011770 | 5507079 | CG | C | 63 | 0 | 2 | 0 | | | | | | | | | | | C | C |
| NC_011770 | 5507115 | C | T | 63 | 0 | 2 | 0 | | | | | | | | | | | T | T |
| NC_011770 | 2926022 | GT | G | 24 | 5 | 3 | 0 | | . | . | . | . | . | G | G | G | | | |
| NC_011770 | 5507874 | A | G | 55 | 0 | 6 | 0 | | | G | G | G | G | G | G | | | | |
| NC_011770 | 5507878 | T | C | 56 | 0 | 6 | 0 | | | C | C | C | C | C | C | | | | |
| NC_011770 | 5507902 | C | A | 62 | 0 | 6 | 0 | | | A | A | A | A | A | A | | | | |
| NC_011770 | 1587421 | C | T | 76 | 0 | 1 | 0 | | | | | | | | | | | T | |
| NC_011770 | 5509802 | C | A | 73 | 0 | 2 | 0 | | | | | | | A | A | | | | |
| NC_011770 | 5509849 | G | A | 72 | 0 | 4 | 0 | | | A | A | A | A | | | | | | |
| NC_011770 | 5509862 | T | A | 71 | 0 | 4 | 0 | | | A | A | A | A | | | | | | |
| NC_011770 | 5509870 | A | G | 70 | 0 | 4 | 0 | | | G | G | G | G | | | | | | |
| NC_011770 | 5511261 | T | C | 71 | 0 | 1 | 0 | | | | | | | C | | | | | |
| NC_011770 | 5511298 | C | T | 77 | 0 | 2 | 0 | T | T | | | | | | | | | | |
| NC_011770 | 5511306 | A | G | 78 | 0 | 2 | 0 | G | G | | | | | | | | | | |
| NC_011770 | 5511514 | G | C | 75 | 0 | 6 | 0 | | | C | C | C | C | C | C | | | | |
| NC_011770 | 5511531 | T | G | 73 | 0 | 2 | 0 | | | | | | | | | | | G | G |
| NC_011770 | 5511728 | G | T | 59 | 0 | 4 | 0 | T | T | | | | | | | | | T | T |
| NC_011770 | 5511806 | G | T | 59 | 0 | 2 | 0 | | | | | | | T | T | | | | |
| NC_011770 | 589978 | AT | A | 93 | 0 | 1 | 0 | | A | | | | | | | | | | |
| NC_011770 | 5517568 | A | C | 72 | 0 | 12 | 0 | C | C | C | C | C | C | C | C | C | C | C | C |
| NC_011770 | 5517712 | A | C | 67 | 0 | 6 | 0 | | | C | C | C | C | C | C | | | | |

| | | | | | | | | | | | | | | | | | | | |
|---|---|---|---|---|---|---|---|---|---|---|---|---|---|---|---|---|---|---|---|
| NC_011770 | 5518945 | T | C | 69 | 0 | 6 | 0 | | | C | C | C | C | | | C | C | | |
| NC_011770 | 5519075 | G | T | 73 | 0 | 6 | 0 | | | T | T | T | T | | | T | T | | |
| NC_011770 | 5519080 | G | C | 73 | 0 | 6 | 0 | | | C | C | C | C | | | C | C | | |
| NC_011770 | 5519583 | T | G | 77 | 0 | 8 | 0 | | | G | G | G | G | G | G | G | G | | |
| NC_011770 | 5519595 | C | T | 81 | 0 | 8 | 0 | | | T | T | T | T | T | T | T | T | | |
| NC_011770 | 5519600 | C | G | 78 | 0 | 8 | 0 | | | G | G | G | G | G | G | G | G | | |
| NC_011770 | 5519631 | A | G | 72 | 0 | 12 | 0 | G | G | G | G | G | G | G | G | G | G | G | G |
| NC_011770 | 5519635 | C | T | 74 | 0 | 8 | 0 | | | T | T | T | T | T | T | T | T | | |
| NC_011770 | 5519637 | A | G | 75 | 0 | 8 | 0 | | | G | G | G | G | G | G | G | G | | |
| NC_011770 | 5519663 | G | A | 74 | 0 | 12 | 0 | A | A | A | A | A | A | A | A | A | A | A | A |
| NC_011770 | 5519690 | A | G | 72 | 0 | 2 | 0 | | | | | | | G | G | | | | |
| NC_011770 | 5519710 | G | A | 70 | 0 | 4 | 0 | A | A | | | | | | | | | A | A |
| NC_011770 | 5519711 | G | A | 71 | 0 | 4 | 0 | | | | | | | A | A | A | A | | |
| NC_011770 | 793263 | GCGACAAC | G | 71 | 0 | 3 | 0 | | G | | | | | | | G | G | | |
| NC_011770 | 3071367 | AGCGAATA | A | 4 | 10 | 1 | 0 | . | . | . | . | . | . | . | . | A | | . | . |
| NC_011770 | 3277059 | CG | C | 21 | 0 | 1 | 0 | | | | | | | | | C | | | |
| NC_011770 | 1984302 | CT | C | 57 | 0 | 5 | 0 | | | | | | | C | C | | C | C | C |
| NC_011770 | 1088036 | G | A | 72 | 0 | 1 | 0 | | | | A | | | | | | | | |
| NC_011770 | 3287323 | C | T | 20 | 0 | 7 | 0 | | | T | T | T | T | T | T | T | | | |
| NC_011770 | 792197 | C | CCCGCC | 63 | 0 | 1 | 0 | | | CCCGCCGCCGGGGCGGG | | | | | | | | | |
| NC_011770 | 1159245 | T | C | 22 | 2 | 7 | 0 | | | C | | C | C | . | . | C | C | C | C |
| NC_011770 | 1273056 | A | AGCCCG | 55 | 0 | 9 | 0 | AGCCCGC( | . | AGCCCGCGGGAC | | AGCCCGC( | AGCCCGC( | AGCCCGCGGGAC | | G | G | AGCCCGC( | AGCCCGCGGGAC |
| NC_011770 | 3649800 | A | ACCC | 47 | 0 | 7 | 0 | | | ACCC | ACCC | | ACCC | | | AC | AC | AC | AC |
| NC_011770 | 1658231 | C | T | 39 | 0 | 1 | 0 | | | | | | | | | | | T | |
| NC_011770 | 1822539 | G | T | 102 | 4 | 1 | 0 | . | . | T | | | | | | . | . | | |
| NC_011770 | 1991593 | AC | A | 62 | 0 | 1 | 0 | | | | | | | | | | A | | |
| NC_011770 | 54226 | C | T | 159 | 0 | 1 | 0 | | | | | | | | | | | | T |
| NC_011770 | 3920537 | C | T | 61 | 0 | 3 | 0 | | | | | | | | | T | | T | T |
| NC_011770 | 4023218 | C | G | 31 | 2 | 7 | 0 | . | . | G | G | | G | G | G | | | G | G |
| NC_011770 | 4355546 | T | TGC | 78 | 0 | 4 | 0 | | | TGC | TGC | | TGC | TGC | | | | | |
| NC_011770 | 1159238 | C | T | 22 | 2 | 1 | 0 | | | | T | | | . | . | | | | |
| NC_011770 | 2009847 | C | T | 87 | 0 | 1 | 0 | T | | | | | | | | | | | |
| NC_011770 | 1668483 | C | T | 71 | 0 | 1 | 0 | | | | | | | | | | | T | |
| NC_011770 | 5528197 | C | A | 71 | 0 | 12 | 0 | A | A | A | A | A | A | A | A | A | A | A | A |
| NC_011770 | 5528241 | G | T | 70 | 0 | 12 | 0 | T | T | T | T | T | T | T | T | T | T | T | T |
| NC_011770 | 5528310 | G | A | 68 | 0 | 2 | 0 | | | | | | | A | A | | | | |
| NC_011770 | 5528331 | T | G | 70 | 0 | 12 | 0 | G | G | G | G | G | G | G | G | G | G | G | G |
| NC_011770 | 5532955 | G | A | 82 | 0 | 6 | 0 | A | A | | | | | A | A | | | A | A |
| NC_011770 | 81447 | C | T | 70 | 0 | 1 | 0 | | | | | | | | | | | | T |
| NC_011770 | 1159257 | G | A | 23 | 2 | 1 | 0 | | | | A | | | . | . | | | | |
| NC_011770 | 4314593 | CA | C | 42 | 0 | 3 | 0 | | | C | C | C | | | | | | | |
| NC_011770 | 5538666 | A | G | 91 | 0 | 4 | 0 | G | G | | | | | | | | | G | G |
| NC_011770 | 5538703 | AC | A | 93 | 0 | 2 | 0 | | | | | | | | | A | A | | |
| NC_011770 | 5538762 | A | T | 94 | 0 | 6 | 0 | | | T | T | T | T | | | T | T | | |
| NC_011770 | 5538765 | G | A | 95 | 0 | 4 | 0 | A | A | | | | | | | | | A | A |
| NC_011770 | 5538796 | C | T | 94 | 0 | 2 | 0 | | | | | | | | | | | T | T |
| NC_011770 | 5542817 | A | G | 50 | 0 | 4 | 0 | G | G | | | | | | | | | G | G |
| NC_011770 | 5542868 | G | A | 43 | 0 | 2 | 0 | A | A | | | | | | | | | | |
| NC_011770 | 5542921 | A | G | 34 | 0 | 12 | 0 | G | G | G | G | G | G | G | G | G | G | G | G |
| NC_011770 | 5543555 | C | T | 60 | 0 | 2 | 0 | | | | | | | | | | | T | T |
| NC_011770 | 1747114 | G | A | 53 | 0 | 1 | 0 | | | | | | | | | | | A | |
| NC_011770 | 5545128 | C | CCG | 59 | 0 | 1 | 0 | | | | | | | | | CCG | | | |
| NC_011770 | 5545136 | C | T | 60 | 0 | 6 | 0 | | | T | T | T | T | | | T | T | | |
| NC_011770 | 5545587 | TG | T | 73 | 0 | 12 | 0 | T | T | T | T | T | T | T | T | T | T | T | T |
| NC_011770 | 5545598 | G | A | 71 | 0 | 2 | 0 | | | | | | | | | A | A | | |
| NC_011770 | 5545632 | A | G | 69 | 0 | 12 | 0 | G | G | G | G | G | G | G | G | G | G | G | G |
| NC_011770 | 5545735 | T | A | 64 | 0 | 2 | 0 | | | | | | | A | A | | | | |
| NC_011770 | 5545771 | C | T | 65 | 0 | 12 | 0 | T | T | T | T | T | T | T | T | T | T | T | T |
| NC_011770 | 5545799 | G | A | 67 | 0 | 2 | 0 | | | | | | | A | A | | | | |
| NC_011770 | 5545821 | C | A | 66 | 0 | 2 | 0 | | | | | | | | | A | A | | |
| NC_011770 | 1159266 | G | GT | 22 | 2 | 3 | 0 | | | | GT | | | . | . | GT | GT | | |
| NC_011770 | 5546754 | G | A | 83 | 0 | 4 | 0 | | | A | A | A | A | | | | | | |

| | | | | | | | | | | | | | | | | | | | |
|---|---|---|---|---|---|---|---|---|---|---|---|---|---|---|---|---|---|---|---|
| NC_011770 | 294020 | G | A | 90 | 0 | 1 | 0 | | | | | | | | | | | | A |
| NC_011770 | 2033753 | C | CG | 74 | 0 | 11 | 0 | CG | CG | CG | | CG | CG | CG | CG | CG | CG | CG | CG |
| NC_011770 | 5547955 | G | GC | 29 | 0 | 2 | 0 | | | | | | | GC | GC | | | | |
| NC_011770 | 5549905 | A | C | 67 | 0 | 4 | 0 | C | C | | | | | | | | | C | C |
| NC_011770 | 5549912 | A | G | 67 | 0 | 4 | 0 | G | G | | | | | | | | | G | G |
| NC_011770 | 5549936 | A | G | 66 | 0 | 4 | 0 | G | G | | | | | | | | | G | G |
| NC_011770 | 5549938 | T | A | 66 | 0 | 4 | 0 | A | A | | | | | | | | | A | A |
| NC_011770 | 5549950 | A | T | 67 | 0 | 4 | 0 | T | T | | | | | | | | | T | T |
| NC_011770 | 5549960 | CA | C | 67 | 0 | 4 | 0 | C | C | | | | | | | | | C | C |
| NC_011770 | 5549973 | C | T | 69 | 0 | 4 | 0 | T | T | | | | | | | | | T | T |
| NC_011770 | 5550005 | C | T | 65 | 0 | 4 | 0 | T | T | | | | | | | | | T | T |
| NC_011770 | 5550008 | T | C | 65 | 0 | 4 | 0 | C | C | | | | | | | | | C | C |
| NC_011770 | 5550013 | T | C | 63 | 0 | 4 | 0 | C | C | | | | | | | | | C | C |
| NC_011770 | 5550022 | T | C | 63 | 0 | 4 | 0 | C | C | | | | | | | | | C | C |
| NC_011770 | 5550027 | T | C | 65 | 0 | 4 | 0 | C | C | | | | | | | | | C | C |
| NC_011770 | 5550044 | G | A | 65 | 0 | 4 | 0 | A | A | | | | | | | | | A | A |
| NC_011770 | 5550045 | C | T | 65 | 0 | 4 | 0 | T | T | | | | | | | | | T | T |
| NC_011770 | 5550046 | T | C | 65 | 0 | 4 | 0 | C | C | | | | | | | | | C | C |
| NC_011770 | 5550049 | T | C | 63 | 0 | 4 | 0 | C | C | | | | | | | | | C | C |
| NC_011770 | 5550086 | AG | A | 59 | 0 | 4 | 0 | A | A | | | | | | | | | A | A |
| NC_011770 | 5550096 | C | T | 58 | 0 | 4 | 0 | T | T | | | | | | | | | T | T |
| NC_011770 | 5550118 | G | T | 60 | 0 | 4 | 0 | T | T | | | | | | | | | T | T |
| NC_011770 | 5550129 | C | A | 61 | 0 | 4 | 0 | A | A | | | | | | | | | A | A |
| NC_011770 | 5550136 | G | T | 61 | 0 | 4 | 0 | T | T | | | | | | | | | T | T |
| NC_011770 | 5550178 | A | C | 59 | 0 | 4 | 0 | C | C | | | | | | | | | C | C |
| NC_011770 | 5550183 | T | C | 57 | 0 | 2 | 0 | | | | | | | C | C | | | | |
| NC_011770 | 5550215 | G | A | 56 | 0 | 4 | 0 | A | A | | | | | | | | | A | A |
| NC_011770 | 1128676 | A | G | 47 | 0 | 1 | 0 | | G | | | | | | | | | | |
| NC_011770 | 403205 | G | A | 77 | 0 | 1 | 0 | | | | | | | | | | | | A |
| NC_011770 | 1829942 | C | T | 90 | 0 | 1 | 0 | | | | | | | | | | | T | |
| NC_011770 | 2119737 | C | T | 56 | 0 | 1 | 0 | | | | | | | | | | | T | |
| NC_011770 | 5557447 | A | AG | 67 | 0 | 4 | 0 | AG | AG | | | | | | | | | AG | AG |
| NC_011770 | 5557497 | A | G | 60 | 0 | 4 | 0 | G | G | | | | | | | | | G | G |
| NC_011770 | 5557652 | T | C | 63 | 0 | 2 | 0 | | | | | | | | | | | C | C |
| NC_011770 | 2178909 | G | A | 51 | 0 | 1 | 0 | | | | | | | | | | | A | |
| NC_011770 | 5557659 | C | A | 61 | 0 | 2 | 0 | | | | | | | | | | | A | A |
| NC_011770 | 5557671 | T | TC | 63 | 0 | 2 | 0 | | | | | | | | | | | TC | TC |
| NC_011770 | 5559699 | C | T | 72 | 0 | 10 | 0 | | | T | T | T | T | T | T | T | T | T | T |
| NC_011770 | 5559724 | A | G | 83 | 0 | 12 | 0 | G | G | G | G | G | G | G | G | G | G | G | G |
| NC_011770 | 5559810 | C | T | 83 | 0 | 12 | 0 | T | T | T | T | T | T | T | T | T | T | T | T |
| NC_011770 | 5559913 | A | G | 79 | 0 | 8 | 0 | | | G | G | G | G | G | G | G | G | | |
| NC_011770 | 5559931 | C | T | 82 | 0 | 12 | 0 | T | T | T | T | T | T | T | T | T | T | T | T |
| NC_011770 | 5559959 | C | T | 81 | 0 | 12 | 0 | T | T | T | T | T | T | T | T | T | T | T | T |
| NC_011770 | 433722 | A | AG | 79 | 0 | 1 | 0 | | | | | | | | | | | | AG |
| NC_011770 | 5561123 | G | A | 95 | 0 | 2 | 0 | | | | | | | | | A | A | | |
| NC_011770 | 5561156 | T | C | 94 | 0 | 6 | 0 | | | C | C | C | C | | | C | C | | |
| NC_011770 | 4481520 | G | GCTCGA | 50 | 0 | 7 | 0 | GCTCGAG( | GCTCGAGCACAT | | | | GCTCGAGCACAT | | | GCTCGAG( | GCTCGAG( | GCTCGAG( | GCTCGAGCACAT |
| NC_011770 | 1167521 | GA | G | 83 | 0 | 1 | 0 | | | | G | | | | | | | | |
| NC_011770 | 1427178 | A | G | 74 | 0 | 1 | 0 | | G | | | | | | | | | | |
| NC_011770 | 5564535 | T | C | 60 | 0 | 4 | 0 | C | C | | | | | C | C | | | | |
| NC_011770 | 5564542 | G | T | 60 | 0 | 10 | 0 | T | T | T | T | T | T | T | T | T | T | | |
| NC_011770 | 5564559 | G | C | 63 | 0 | 4 | 0 | C | C | | | | | C | C | | | | |
| NC_011770 | 5565240 | G | A | 63 | 0 | 10 | 0 | A | A | A | A | A | A | A | A | A | A | | |
| NC_011770 | 1368655 | C | CA | 46 | 0 | 1 | 0 | | | | CA | . | . | | | | | | |
| NC_011770 | 2247397 | G | A | 68 | 0 | 1 | 0 | | | | | | | | | | | A | |
| NC_011770 | 5571917 | C | T | 96 | 0 | 2 | 0 | T | T | | | | | | | | | | |
| NC_011770 | 5574507 | G | A | 78 | 0 | 2 | 0 | | | | | | | | | A | A | | |
| NC_011770 | 5577008 | A | G | 13 | 0 | 2 | 0 | | | | | | | | | | | G | G |
| NC_011770 | 5577034 | CT | C | 11 | 0 | 1 | 0 | | | | | | | | | . | . | C | . |
| NC_011770 | 5577042 | T | C | 10 | 0 | 2 | 0 | | | | | | | | | C | C | | |
| NC_011770 | 5577046 | C | T | 10 | 0 | 4 | 0 | | | | | | | | | T | T | T | T |
| NC_011770 | 5577047 | G | A | 10 | 0 | 2 | 0 | | | | | | | | | A | A | | |

| | | | | | | | | | | | | | | | | | | | |
|---|---|---|---|---|---|---|---|---|---|---|---|---|---|---|---|---|---|---|---|
| NC_011770 | 5577048 | T | C | 10 | 0 | 4 | 0 | | | | | | | | | C | C | C | C |
| NC_011770 | 5577058 | C | T | 11 | 0 | 4 | 0 | | | | | | | | | T | T | T | T |
| NC_011770 | 5577064 | G | T | 11 | 0 | 4 | 0 | | | | | | | | | T | T | T | T |
| NC_011770 | 5577123 | C | G | 17 | 0 | 2 | 0 | | | | | | | | | | | G | G |
| NC_011770 | 5577145 | C | T | 18 | 0 | 2 | 0 | | | | | | | | | | | T | T |
| NC_011770 | 5577280 | C | CT | 55 | 0 | 1 | 0 | | | | | | | CT | | | | | |
| NC_011770 | 5577298 | AGACTACA | A | 46 | 0 | 1 | 0 | | | | | | | A | | | | | |
| NC_011770 | 5577355 | T | TA | 49 | 0 | 12 | 0 | TA | TA | TA | TA | TA | TA | TA | TA | TA | TA | TA | TA |
| NC_011770 | 5577396 | T | C | 54 | 0 | 2 | 0 | | | | | | | C | C | | | | |
| NC_011770 | 5577801 | C | T | 82 | 0 | 12 | 0 | T | T | T | T | T | T | T | T | T | T | T | T |
| NC_011770 | 5577874 | C | T | 81 | 0 | 2 | 0 | | | | | | | T | T | | | | |
| NC_011770 | 5577875 | G | C | 81 | 0 | 8 | 0 | C | C | C | C | C | C | | | C | C | | |
| NC_011770 | 5577976 | G | A | 84 | 0 | 2 | 0 | | | | | | | | | | | A | A |
| NC_011770 | 5578021 | C | T | 84 | 0 | 8 | 0 | T | T | T | T | T | T | | | T | T | | |
| NC_011770 | 5578033 | T | C | 87 | 0 | 8 | 0 | C | C | C | C | C | C | | | C | C | | |
| NC_011770 | 4763448 | C | CT | 64 | 0 | 5 | 0 | . | | CT | CT | | CT | | | CT | CT | | |
| NC_011770 | 5578150 | GC | G | 88 | 0 | 2 | 0 | | | | | G | G | | | | | | |
| NC_011770 | 5578151 | C | G | 48 | 2 | 2 | 0 | | | G | G | . | . | | | | | | |
| NC_011770 | 5580595 | T | G | 97 | 0 | 4 | 0 | | | G | G | G | G | | | | | | |
| NC_011770 | 5580598 | T | C | 96 | 0 | 4 | 0 | | | C | C | C | C | | | | | | |
| NC_011770 | 5580599 | G | A | 96 | 0 | 4 | 0 | | | A | A | A | A | | | | | | |
| NC_011770 | 5580602 | A | G | 97 | 0 | 4 | 0 | | | G | G | G | G | | | | | | |
| NC_011770 | 5582660 | G | A | 83 | 0 | 2 | 0 | | | | | | | | | | | A | A |
| NC_011770 | 5582707 | A | G | 81 | 0 | 12 | 0 | G | G | G | G | G | G | G | G | G | G | G | G |
| NC_011770 | 5582759 | G | A | 80 | 0 | 2 | 0 | | | | | | | | | A | A | | |
| NC_011770 | 5583929 | C | T | 63 | 0 | 2 | 0 | | | | | | | | | | | T | T |
| NC_011770 | 5583962 | G | A | 65 | 0 | 6 | 0 | | | A | A | A | A | A | A | | | | |
| NC_011770 | 508185 | C | T | 83 | 2 | 1 | 0 | | | | | | | | | . | . | | T |
| NC_011770 | 5584732 | C | G | 73 | 0 | 4 | 0 | | | G | G | G | G | | | | | | |
| NC_011770 | 5584734 | C | G | 72 | 0 | 12 | 0 | G | G | G | G | G | G | G | G | G | G | G | G |
| NC_011770 | 5586919 | C | G | 78 | 0 | 8 | 0 | G | G | G | G | G | G | | | | | G | G |
| NC_011770 | 5587648 | A | T | 70 | 0 | 4 | 0 | T | T | | | | | T | T | | | | |
| NC_011770 | 5587680 | G | A | 69 | 0 | 2 | 0 | | | | | | | | | | | A | A |
| NC_011770 | 5587756 | G | A | 75 | 0 | 2 | 0 | | | | | | | | | A | A | | |
| NC_011770 | 5587773 | T | C | 72 | 0 | 4 | 0 | C | C | | | | | C | C | | | | |
| NC_011770 | 5587782 | G | A | 73 | 0 | 12 | 0 | A | A | A | A | A | A | A | A | A | A | A | A |
| NC_011770 | 5587855 | C | T | 75 | 0 | 12 | 0 | T | T | T | T | T | T | T | T | T | T | T | T |
| NC_011770 | 5592783 | G | A | 69 | 0 | 2 | 0 | | | | | | | A | A | | | | |
| NC_011770 | 5592790 | G | A | 70 | 0 | 2 | 0 | | | | | | | A | A | | | | |
| NC_011770 | 5596577 | G | A | 82 | 0 | 2 | 0 | | | | | | | | | | | A | A |
| NC_011770 | 5596704 | G | A | 90 | 0 | 2 | 0 | | | | | | | A | A | | | | |
| NC_011770 | 5596727 | T | C | 91 | 0 | 10 | 0 | C | C | C | C | C | C | | | C | C | C | C |
| NC_011770 | 5596760 | C | T | 87 | 0 | 10 | 0 | T | T | T | T | T | T | | | T | T | T | T |
| NC_011770 | 5596900 | A | G | 77 | 0 | 2 | 0 | | | | | | | | | | | G | G |
| NC_011770 | 5596922 | G | A | 78 | 0 | 4 | 0 | A | A | | | | | A | A | | | | |
| NC_011770 | 5596985 | C | A | 74 | 0 | 6 | 0 | A | A | | | | | T | T | A | A | | |
| NC_011770 | 5597000 | G | A | 71 | 0 | 4 | 0 | | | | | | | | | A | A | A | A |
| NC_011770 | 5597011 | G | A | 74 | 0 | 2 | 0 | | | | | | | | | | | A | A |
| NC_011770 | 5597063 | G | A | 83 | 0 | 2 | 0 | | | | | | | | | A | A | | |
| NC_011770 | 5597878 | G | A | 89 | 0 | 2 | 0 | A | A | | | | | | | | | | |
| NC_011770 | 5274199 | C | T | 93 | 0 | 1 | 0 | | | | | | T | | | | | | |
| NC_011770 | 5597896 | G | A | 91 | 0 | 2 | 0 | A | A | | | | | | | | | | |
| NC_011770 | 5598110 | G | GT | 97 | 0 | 2 | 0 | GT | GT | | | | | | | | | | |
| NC_011770 | 5598165 | G | A | 93 | 0 | 2 | 0 | | | | | | | | | | | A | A |
| NC_011770 | 5601262 | T | C | 83 | 0 | 4 | 0 | | | | | | | C | C | | | C | C |
| NC_011770 | 5601263 | C | T | 83 | 0 | 4 | 0 | | | | | | | T | T | | | T | T |
| NC_011770 | 5601295 | GT | G | 85 | 0 | 2 | 0 | G | G | | | | | | | | | | |
| NC_011770 | 5601379 | G | T | 66 | 0 | 2 | 0 | | | | | | | | | T | T | | |
| NC_011770 | 5603116 | T | C | 57 | 0 | 4 | 0 | | | C | C | C | C | | | | | | |
| NC_011770 | 5603276 | C | T | 38 | 0 | 12 | 0 | T | T | T | T | T | T | T | T | T | T | T | T |
| NC_011770 | 5603312 | G | T | 57 | 0 | 1 | 0 | | | | | | | T | | | | | |
| NC_011770 | 2012336 | GCGCGC | G | 76 | 0 | 5 | 0 | | | | | | | G | G | | G | G | G |

| | | | | | | | | | | | | | | | | | | | |
|---|---|---|---|---|---|---|---|---|---|---|---|---|---|---|---|---|---|---|---|
| NC_011770 | 2401394 | TC | T | 71 | 0 | 3 | 0 | | | | | | | | | | T | T | T |
| NC_011770 | 5603454 | A | G | 92 | 0 | 3 | 0 | | | | | | | G | | . | . | G | G |
| NC_011770 | 5603495 | A | G | 96 | 0 | 10 | 0 | G | G | G | G | G | G | G | G | | | G | G |
| NC_011770 | 2380957 | C | T | 53 | 0 | 1 | 0 | | | | | | | | | | | T | |
| NC_011770 | 2473952 | G | A | 29 | 0 | 1 | 0 | | | | | | | | | | | A | |
| NC_011770 | 2660572 | A | AG | 65 | 0 | 1 | 0 | | | | | | | | | | | AG | |
| NC_011770 | 2660834 | G | A | 69 | 0 | 1 | 0 | | | | | | | | | | | A | |
| NC_011770 | 2690227 | C | T | 45 | 0 | 1 | 0 | | | | | | | | | | | T | |
| NC_011770 | 5606163 | A | G | 73 | 0 | 2 | 0 | G | G | | | | | | | | | | |
| NC_011770 | 5606168 | A | T | 73 | 0 | 6 | 0 | | | T | T | T | T | | | | | T | T |
| NC_011770 | 5606256 | C | T | 71 | 0 | 2 | 0 | T | T | | | | | | | | | | |
| NC_011770 | 1632973 | G | GA | 63 | 0 | 1 | 0 | | GA | | | | | | | | | | |
| NC_011770 | 5609895 | C | G | 41 | 0 | 2 | 0 | | | | | | | | | G | G | | |
| NC_011770 | 5609940 | G | T | 38 | 0 | 2 | 0 | | | | | | | T | T | | | | |
| NC_011770 | 5610059 | G | A | 37 | 0 | 2 | 0 | | | | | | | A | A | | | | |
| NC_011770 | 5610584 | A | T | 24 | 0 | 6 | 0 | | | T | T | T | T | T | T | | | | |
| NC_011770 | 5610585 | T | G | 24 | 0 | 6 | 0 | | | G | G | G | G | G | G | | | | |
| NC_011770 | 5610622 | G | GA | 25 | 0 | 2 | 0 | | | | | | | | | | | GA | GA |
| NC_011770 | 2699195 | GTGAT | G | 10 | 8 | 1 | 0 | . | . | . | . | . | . | . | . | | G | | |
| NC_011770 | 1616158 | C | T | 78 | 0 | 1 | 0 | | | | T | | | | | | | | |
| NC_011770 | 1162548 | AGCGATGC | A | 59 | 0 | 2 | 0 | | | | A | | A | | | | | | |
| NC_011770 | 5614199 | A | G | 33 | 0 | 6 | 0 | G | G | | | | | | | G | G | G | G |
| NC_011770 | 5615902 | T | C | 54 | 0 | 2 | 0 | | | | | | | C | C | | | | |
| NC_011770 | 5615930 | G | A | 53 | 0 | 2 | 0 | | | | | | | | | A | A | | |
| NC_011770 | 5615968 | G | A | 60 | 0 | 10 | 0 | A | A | A | A | A | A | | | A | A | A | A |
| NC_011770 | 5616856 | T | TC | 62 | 0 | 2 | 0 | | | | | | | | | | | TC | TC |
| NC_011770 | 630311 | G | A | 79 | 0 | 1 | 0 | | | | | | | | | | | | A |
| NC_011770 | 2938279 | G | T | 64 | 0 | 1 | 0 | | | T | | | | | | | | | |
| NC_011770 | 5618167 | A | G | 80 | 0 | 2 | 0 | G | G | | | | | | | | | | |
| NC_011770 | 720877 | C | T | 99 | 0 | 1 | 0 | | | | | | | | | | | | T |
| NC_011770 | 5620878 | G | A | 60 | 0 | 6 | 0 | | | A | A | A | A | A | A | | | | |
| NC_011770 | 5622901 | C | CA | 39 | 0 | 5 | 0 | CA | . | | | | | | | CA | CA | CA | CA |
| NC_011770 | 5622936 | A | G | 33 | 0 | 12 | 0 | G | G | G | G | G | G | G | G | G | G | G | G |
| NC_011770 | 3419801 | CCTGCCATC | C | 46 | 6 | 1 | 0 | | | C | | | | . | . | . | . | . | . |
| NC_011770 | 3751895 | C | T | 24 | 2 | 1 | 0 | | | T | | . | | | | . | . | | |
| NC_011770 | 5623004 | G | A | 21 | 0 | 1 | 0 | | | | | | | A | | | | | |
| NC_011770 | 4471732 | G | GT | 62 | 0 | 11 | 0 | GT | GT | GT | GT | GT | | GT | GT | GT | GT | GT | GT |
| NC_011770 | 4848865 | T | A | 64 | 0 | 1 | 0 | | | | | A | | | | | | | |
| NC_011770 | 5626852 | GC | G | 76 | 0 | 4 | 0 | G | G | | | | | | | G | G | | |
| NC_011770 | 5626904 | A | G | 79 | 0 | 2 | 0 | | | | | | | G | G | | | | |
| NC_011770 | 5627029 | C | T | 63 | 0 | 2 | 0 | | | | | | | | | | | T | T |
| NC_011770 | 2888024 | AT | A | 13 | 10 | 1 | 0 | . | . | . | . | . | . | . | . | . | . | A | |
| NC_011770 | 5627551 | G | A | 62 | 0 | 12 | 0 | A | A | A | A | A | A | A | A | A | A | A | A |
| NC_011770 | 5627577 | G | A | 61 | 0 | 12 | 0 | A | A | A | A | A | A | A | A | A | A | A | A |
| NC_011770 | 5628870 | G | A | 67 | 0 | 12 | 0 | A | A | A | A | A | A | A | A | A | A | A | A |
| NC_011770 | 5628895 | C | A | 66 | 0 | 2 | 0 | | | | | | | | | A | A | | |
| NC_011770 | 5628917 | C | T | 64 | 0 | 2 | 0 | | | | | | | T | T | | | | |
| NC_011770 | 957659 | C | T | 54 | 0 | 1 | 0 | | | | | | | | | | | | T |
| NC_011770 | 5630823 | A | G | 62 | 0 | 4 | 0 | G | G | | | | | | | | | G | G |
| NC_011770 | 5631370 | A | C | 69 | 0 | 2 | 0 | C | C | | | | | | | | | | |
| NC_011770 | 5634101 | A | AGC | 84 | 0 | 6 | 0 | AGC | AGC | | | | | AGC | AGC | | | AGC | AGC |
| NC_011770 | 5634127 | T | C | 83 | 0 | 6 | 0 | C | C | | | | | C | C | | | C | C |
| NC_011770 | 1751170 | C | CG | 224 | 0 | 5 | 0 | | CG | | | | | CG | CG | CG | CG | | |
| NC_011770 | 5636712 | C | T | 56 | 0 | 2 | 0 | | | | | | | | | T | T | | |
| NC_011770 | 5642355 | A | C | 58 | 0 | 8 | 0 | C | C | C | C | C | C | | | C | C | | |
| NC_011770 | 5642404 | T | C | 54 | 0 | 2 | 0 | | | | | | | | | C | C | | |
| NC_011770 | 5642538 | T | C | 49 | 0 | 12 | 0 | C | C | C | C | C | C | C | C | C | C | C | C |
| NC_011770 | 5642785 | A | AC | 43 | 0 | 1 | 0 | | | | | | | | | | | AC | |
| NC_011770 | 5642797 | C | T | 43 | 0 | 2 | 0 | | | | | | | | | | | T | T |
| NC_011770 | 5642803 | C | T | 42 | 0 | 2 | 0 | | | | | | | | | | | T | T |
| NC_011770 | 5642807 | G | C | 42 | 0 | 2 | 0 | | | | | | | C | C | | | | |
| NC_011770 | 5642910 | C | T | 52 | 0 | 2 | 0 | | | | | | | | | | | T | T |

| | | | | | | | | | | | | | | | | | | | |
|---|---|---|---|---|---|---|---|---|---|---|---|---|---|---|---|---|---|---|---|
| NC_011770 | 2895625 | CAGGCCAA | C | 26 | 6 | 1 | 0 | | | | | . | . | . | . | . | . | C | |
| NC_011770 | 5642986 | G | A | 59 | 0 | 2 | 0 | | | | | | | | | | | A | A |
| NC_011770 | 5643010 | G | A | 60 | 0 | 2 | 0 | | | | | | | | | | | A | A |
| NC_011770 | 983203 | AG | A | 48 | 0 | 1 | 0 | | | | | | | | | | | | A |
| NC_011770 | 5645079 | A | AC | 87 | 0 | 12 | 0 | AC | AC | AC | AC | AC | AC | AC | AC | AC | AC | AC | AC |
| NC_011770 | 5645145 | G | A | 78 | 0 | 6 | 0 | A | A | A | A | A | A | | | | | | |
| NC_011770 | 5645267 | A | C | 89 | 0 | 6 | 0 | C | C | C | C | C | C | | | | | | |
| NC_011770 | 5646999 | G | C | 70 | 1 | 11 | 0 | C | C | C | C | C | C | . | C | C | C | C | C |
| NC_011770 | 5649206 | C | T | 55 | 0 | 12 | 0 | T | T | T | T | T | T | T | T | T | T | T | T |
| NC_011770 | 5649209 | T | G | 55 | 0 | 12 | 0 | G | G | G | G | G | G | G | G | G | G | G | G |
| NC_011770 | 5649262 | A | T | 57 | 0 | 8 | 0 | T | T | | | | | T | T | T | T | T | T |
| NC_011770 | 5649292 | A | G | 66 | 0 | 8 | 0 | G | G | | | | | G | G | G | G | G | G |
| NC_011770 | 5651306 | G | GC | 59 | 0 | 12 | 0 | GC | GC | GC | GC | GC | GC | GC | GC | GC | GC | GC | GC |
| NC_011770 | 5651335 | G | T | 57 | 0 | 12 | 0 | T | T | T | T | T | T | T | T | T | T | T | T |
| NC_011770 | 5651383 | T | C | 64 | 0 | 12 | 0 | C | C | C | C | C | C | C | C | C | C | C | C |
| NC_011770 | 1923267 | C | T | 55 | 0 | 1 | 0 | | T | | | | | | | | | | |
| NC_011770 | 5656025 | C | T | 67 | 0 | 2 | 0 | | | | | | | | | | | T | T |
| NC_011770 | 5656329 | C | G | 50 | 0 | 2 | 0 | | | | | | | | | G | G | | |
| NC_011770 | 5656332 | T | C | 49 | 0 | 2 | 0 | | | | | | | | | C | C | | |
| NC_011770 | 5656572 | C | T | 57 | 0 | 2 | 0 | T | T | | | | | | | | | | |
| NC_011770 | 5656998 | C | G | 55 | 0 | 2 | 0 | | | | | | | | | G | G | | |
| NC_011770 | 5656999 | T | C | 56 | 0 | 2 | 0 | | | | | | | | | C | C | | |
| NC_011770 | 5657041 | G | T | 59 | 0 | 2 | 0 | | | | | | | T | T | | | | |
| NC_011770 | 5657092 | T | C | 58 | 0 | 6 | 0 | | | C | C | C | C | C | C | | | | |
| NC_011770 | 5657094 | G | A | 58 | 0 | 2 | 0 | | | | | | | | | | | A | A |
| NC_011770 | 5657120 | A | G | 58 | 0 | 8 | 0 | | | G | G | G | G | G | G | | | G | G |
| NC_011770 | 5657174 | G | A | 62 | 0 | 2 | 0 | A | A | | | | | | | | | | |
| NC_011770 | 5659447 | C | T | 61 | 1 | 7 | 0 | | | . | T | T | T | T | T | | | T | T |
| NC_011770 | 4485572 | T | TA | 80 | 0 | 1 | 0 | | | | | | | | | TA | | | |
| NC_011770 | 5661179 | T | C | 83 | 0 | 2 | 0 | | | | | | | | | | | C | C |
| NC_011770 | 993222 | G | A | 80 | 0 | 1 | 0 | | | | | | | | | | | | A |
| NC_011770 | 5666020 | C | G | 72 | 0 | 2 | 0 | | | | | | | G | G | | | | |
| NC_011770 | 5666024 | C | T | 75 | 0 | 2 | 0 | | | | | | | | | T | T | | |
| NC_011770 | 5666047 | G | A | 74 | 0 | 2 | 0 | | | | | | | | | A | A | | |
| NC_011770 | 5672761 | T | C | 74 | 0 | 8 | 0 | C | C | | | | | C | C | C | C | C | C |
| NC_011770 | 5673210 | G | A | 74 | 0 | 2 | 0 | A | A | | | | | | | | | | |
| NC_011770 | 2912150 | C | T | 63 | 0 | 1 | 0 | | | | | | | | | | | T | |
| NC_011770 | 5678910 | T | C | 71 | 0 | 8 | 0 | | | C | C | C | C | C | C | | | C | C |
| NC_011770 | 5679750 | T | G | 75 | 0 | 4 | 0 | | | G | G | G | G | | | | | | |
| NC_011770 | 5680919 | T | C | 59 | 0 | 1 | 0 | | | | | | | C | | | | | |
| NC_011770 | 5680960 | A | G | 59 | 0 | 12 | 0 | G | G | G | G | G | G | G | G | G | G | G | G |
| NC_011770 | 5680997 | G | A | 57 | 0 | 4 | 0 | | | A | A | A | A | | | | | | |
| NC_011770 | 5681019 | G | A | 56 | 0 | 4 | 0 | | | A | A | A | A | | | | | | |
| NC_011770 | 5685462 | A | C | 65 | 0 | 2 | 0 | | | | | | | | | C | C | | |
| NC_011770 | 5128001 | AG | A | 75 | 0 | 3 | 0 | | | A | A | A | | | | | | | |
| NC_011770 | 5686592 | A | C | 35 | 0 | 2 | 0 | C | C | | | | | | | | | | |
| NC_011770 | 5686597 | T | A | 33 | 0 | 8 | 0 | A | A | G | G | G | G | G | G | | | | |
| NC_011770 | 404 | A | G | 65 | 0 | 1 | 0 | | | | | | | | G | | | | |
| NC_011770 | 5687415 | G | A | 78 | 0 | 4 | 0 | | | A | A | A | A | | | | | | |
| NC_011770 | 5688089 | G | A | 78 | 0 | 8 | 0 | A | A | | | | | A | A | A | A | A | A |
| NC_011770 | 5688098 | G | T | 80 | 0 | 12 | 0 | T | T | T | T | T | T | T | T | T | T | T | T |
| NC_011770 | 2388283 | G | T | 62 | 0 | 1 | 0 | | | | T | | | | | | | | |
| NC_011770 | 5688936 | A | G | 73 | 0 | 6 | 0 | | | | | | | G | G | G | G | G | G |
| NC_011770 | 5688988 | G | C | 79 | 0 | 4 | 0 | | | C | C | C | C | | | | | | |
| NC_011770 | 5689163 | A | G | 73 | 0 | 8 | 0 | | | G | G | G | G | G | G | | | G | G |
| NC_011770 | 5690941 | G | GT | 96 | 0 | 4 | 0 | | | GT | GT | GT | GT | | | | | | |
| NC_011770 | 5690942 | C | T | 56 | 1 | 1 | 0 | | | . | T | . | . | | | | | | |
| NC_011770 | 5690951 | G | A | 91 | 0 | 4 | 0 | | | A | A | A | A | | | | | | |
| NC_011770 | 5690992 | G | A | 99 | 0 | 4 | 0 | | | A | A | A | A | | | | | | |
| NC_011770 | 2363267 | A | AG | 60 | 0 | 1 | 0 | AG | | | | | | | | | | | |
| NC_011770 | 2540152 | C | CT | 6 | 10 | 1 | 0 | CT | | . | . | . | . | . | . | . | . | . | . |
| NC_011770 | 5695090 | GC | G | 18 | 8 | 2 | 0 | . | . | . | . | . | . | G | G | . | . | | |

| | | | | | | | | | | | | | | | | | | | |
|---|---|---|---|---|---|---|---|---|---|---|---|---|---|---|---|---|---|---|---|
| NC_011770 | 5695103 | T | C | 66 | 0 | 10 | 0 | C | C | C | C | C | C | C | C | C | C | | |
| NC_011770 | 5695126 | G | A | 64 | 0 | 2 | 0 | | | | | | | | | | | A | A |
| NC_011770 | 5695128 | G | A | 63 | 0 | 6 | 0 | | | A | A | A | A | | | A | A | | |
| NC_011770 | 5695131 | G | A | 63 | 0 | 2 | 0 | A | A | | | | | | | | | | |
| NC_011770 | 5695161 | G | C | 58 | 0 | 6 | 0 | | | C | C | C | C | | | C | C | | |
| NC_011770 | 5695165 | C | A | 56 | 0 | 2 | 0 | | | | | | | | | | | A | A |
| NC_011770 | 1262966 | C | T | 92 | 0 | 1 | 0 | | | | | | | | | | | | T |
| NC_011770 | 5695186 | G | A | 43 | 0 | 12 | 0 | A | A | A | A | A | A | A | A | A | A | A | A |
| NC_011770 | 5695189 | G | A | 45 | 0 | 12 | 0 | A | A | A | A | A | A | A | A | A | A | A | A |
| NC_011770 | 5695192 | GC | G | 44 | 0 | 10 | 0 | G | G | G | G | G | G | G | G | G | G | | |
| NC_011770 | 5695202 | T | C | 56 | 0 | 3 | 0 | C | C | | | | | C | | | | | |
| NC_011770 | 5695225 | A | G | 58 | 0 | 8 | 0 | | | G | G | G | G | | | G | G | G | G |
| NC_011770 | 5695228 | G | A | 58 | 0 | 2 | 0 | | | | | | | | | | | A | A |
| NC_011770 | 5695229 | C | A | 56 | 0 | 2 | 0 | A | A | | | | | | | | | | |
| NC_011770 | 5695263 | C | G | 59 | 0 | 4 | 0 | G | G | | | | | | | | | G | G |
| NC_011770 | 5695267 | C | A | 59 | 0 | 2 | 0 | | | | | | | | | | | A | A |
| NC_011770 | 5695271 | G | T | 58 | 0 | 10 | 0 | T | T | T | T | T | T | . | . | T | T | T | T |
| NC_011770 | 5695273 | C | T | 62 | 0 | 4 | 0 | T | T | | | | | | | | | T | T |
| NC_011770 | 5695294 | G | GC | 67 | 0 | 2 | 0 | | | | | | | | | | | GC | GC |
| NC_011770 | 5695353 | G | A | 73 | 0 | 4 | 0 | | | | | | | A | A | A | A | | |
| NC_011770 | 5695412 | C | CA | 77 | 0 | 12 | 0 | CA | CA | CA | CA | CA | CA | CA | CA | CA | CA | CA | CA |
| NC_011770 | 5695462 | TC | T | 85 | 0 | 2 | 0 | | | T | T | | | | | | | | |
| NC_011770 | 1928476 | A | G | 75 | 0 | 1 | 0 | | G | | | | | | | | | | |
| NC_011770 | 5695998 | G | A | 74 | 0 | 2 | 0 | | | | | | | A | A | | | | |
| NC_011770 | 5698678 | C | T | 62 | 0 | 4 | 0 | | | T | T | T | T | | | | | | |
| NC_011770 | 54261 | C | T | 120 | 2 | 1 | 0 | | | | | | | | T | | | . | . |
| NC_011770 | 5700675 | C | A | 95 | 0 | 2 | 0 | | | | | | | | | | | A | A |
| NC_011770 | 5700697 | C | T | 93 | 0 | 2 | 0 | | | | | | | | | | | T | T |
| NC_011770 | 5700756 | T | C | 95 | 0 | 12 | 0 | C | C | C | C | C | C | C | C | C | C | C | C |
| NC_011770 | 5700853 | A | G | 100 | 0 | 8 | 0 | G | G | G | G | G | G | | | G | G | | |
| NC_011770 | 5700863 | G | A | 97 | 0 | 2 | 0 | | | | | | | A | A | | | | |
| NC_011770 | 5701082 | A | T | 83 | 0 | 12 | 0 | T | T | T | T | T | T | T | T | T | T | T | T |
| NC_011770 | 1571516 | A | G | 10 | 6 | 1 | 0 | . | . | . | . | . | . | | | | | | G |
| NC_011770 | 5705587 | A | T | 51 | 0 | 2 | 0 | | | | | | | T | T | | | | |
| NC_011770 | 5705716 | C | T | 58 | 0 | 6 | 0 | | | T | T | T | T | | | | | T | T |
| NC_011770 | 5706255 | C | G | 50 | 0 | 8 | 0 | G | G | | | | | G | G | G | G | G | G |
| NC_011770 | 5706286 | T | G | 47 | 0 | 8 | 0 | G | G | | | | | G | G | G | G | G | G |
| NC_011770 | 5706325 | A | C | 45 | 0 | 4 | 0 | | | C | C | C | C | | | | | | |
| NC_011770 | 5706354 | G | A | 46 | 0 | 4 | 0 | | | A | A | A | A | | | | | | |
| NC_011770 | 2560198 | C | T | 37 | 0 | 1 | 0 | T | | | | | | | | | | | |
| NC_011770 | 5708154 | A | G | 74 | 0 | 10 | 0 | G | G | G | G | G | G | G | G | G | G | | |
| NC_011770 | 5708157 | G | C | 75 | 0 | 6 | 0 | C | C | | | | | C | C | C | C | | |
| NC_011770 | 5708206 | A | G | 76 | 0 | 12 | 0 | G | G | G | G | G | G | G | G | G | G | G | G |
| NC_011770 | 5708238 | C | T | 76 | 0 | 8 | 0 | T | T | T | T | T | T | | | | | T | T |
| NC_011770 | 5708259 | C | G | 83 | 0 | 12 | 0 | G | G | G | G | G | G | G | G | G | G | G | G |
| NC_011770 | 5708292 | G | A | 80 | 0 | 8 | 0 | A | A | A | A | A | A | | | | | A | A |
| NC_011770 | 5708321 | T | G | 84 | 0 | 2 | 0 | G | G | | | | | | | | | | |
| NC_011770 | 5708325 | C | T | 82 | 0 | 8 | 0 | T | T | T | T | T | T | | | | | T | T |
| NC_011770 | 5708402 | C | A | 89 | 0 | 4 | 0 | | | | | | | A | A | A | A | | |
| NC_011770 | 5708403 | A | G | 89 | 0 | 4 | 0 | | | | | | | G | G | G | G | | |
| NC_011770 | 5708429 | GCC | G | 87 | 0 | 12 | 0 | G | G | G | G | G | G | G | G | G | G | G | G |
| NC_011770 | 5709411 | GA | G | 46 | 0 | 6 | 0 | | | G | G | G | G | | | | | G | G |
| NC_011770 | 5709412 | A | C | 18 | 5 | 2 | 0 | C | C | . | . | . | . | | | | | . | . |
| NC_011770 | 5709414 | T | C | 16 | 6 | 2 | 0 | C | C | . | . | . | . | | | | | . | . |
| NC_011770 | 5709422 | G | A | 44 | 0 | 4 | 0 | | | | | | | A | A | A | A | | |
| NC_011770 | 56083 | G | A | 86 | 0 | 5 | 0 | | | | | | | | A | A | A | A | A |
| NC_011770 | 5710619 | C | T | 21 | 0 | 8 | 0 | T | T | T | T | T | T | | | | | T | T |
| NC_011770 | 5710628 | G | A | 20 | 1 | 4 | 0 | A | A | | | | | T | T | | . | | |
| NC_011770 | 5711595 | G | A | 62 | 0 | 2 | 0 | A | A | | | | | | | | | | |
| NC_011770 | 1571524 | C | T | 10 | 6 | 1 | 0 | . | . | . | . | . | . | | | | | | T |
| NC_011770 | 5714424 | G | A | 59 | 0 | 2 | 0 | | | | | | | A | A | | | | |
| NC_011770 | 5716337 | A | G | 40 | 0 | 8 | 0 | G | G | | | | | G | G | G | G | G | G |

| | | | | | | | | | | | | | | | | | | | |
|---|---|---|---|---|---|---|---|---|---|---|---|---|---|---|---|---|---|---|---|
| NC_011770 | 5716338 | G | A | 40 | 0 | 2 | 0 | A | A | | | | | | | | | | |
| NC_011770 | 5716340 | T | C | 41 | 0 | 2 | 0 | | | | | | | C | C | | | | |
| NC_011770 | 2842572 | GCATT | G | 42 | 6 | 1 | 0 | G | | | | . | . | . | . | . | . | | |
| NC_011770 | 5718777 | T | C | 54 | 0 | 12 | 0 | C | C | C | C | C | C | C | C | C | C | C | C |
| NC_011770 | 5721180 | G | C | 91 | 0 | 4 | 0 | | | C | C | C | C | | | | | | |
| NC_011770 | 3211343 | A | G | 16 | 3 | 1 | 0 | . | . | | | . | . | | | | . | G | |
| NC_011770 | 3069030 | A | T | 21 | 6 | 1 | 0 | T | | | | | | . | . | . | . | . | . |
| NC_011770 | 5722624 | T | G | 74 | 0 | 12 | 0 | G | G | G | G | G | G | G | G | G | G | G | G |
| NC_011770 | 2488964 | CG | C | 50 | 0 | 3 | 0 | | | | C | C | C | | | | | | |
| NC_011770 | 1942927 | C | T | 86 | 0 | 1 | 0 | | T | | | | | | | | | | |
| NC_011770 | 5735494 | G | A | 80 | 0 | 6 | 0 | A | A | A | A | A | A | | | | | | |
| NC_011770 | 5735557 | C | T | 74 | 0 | 2 | 0 | T | T | | | | | | | | | | |
| NC_011770 | 2072511 | C | CG | 64 | 0 | 1 | 0 | | CG | | | | | | | | | | |
| NC_011770 | 5737883 | A | G | 39 | 0 | 8 | 0 | G | G | | | | | G | G | G | G | G | G |
| NC_011770 | 5737897 | CG | C | 36 | 0 | 2 | 0 | | | | | | | | | C | C | | |
| NC_011770 | 5737922 | G | A | 42 | 0 | 2 | 0 | | | | | | | | | | | A | A |
| NC_011770 | 5737939 | T | C | 37 | 0 | 2 | 0 | | | | | | | | | C | C | | |
| NC_011770 | 5738510 | C | T | 88 | 0 | 2 | 0 | | | | | | | | | T | T | | |
| NC_011770 | 5738600 | C | T | 91 | 0 | 2 | 0 | | | | | | | | | | | T | T |
| NC_011770 | 5738607 | A | C | 91 | 0 | 10 | 0 | C | C | C | C | C | C | | | C | C | C | C |
| NC_011770 | 1571527 | G | C | 9 | 6 | 1 | 0 | . | . | . | . | . | . | | | | | | C |
| NC_011770 | 56316 | C | T | 87 | 0 | 1 | 0 | | | | | | | | T | | | | |
| NC_011770 | 5744759 | A | G | 40 | 0 | 2 | 0 | G | G | | | | | | | | | | |
| NC_011770 | 5744776 | C | T | 48 | 0 | 4 | 0 | | | | | | | T | T | | | T | T |
| NC_011770 | 5744798 | A | G | 46 | 0 | 4 | 0 | | | | | | | G | G | | | G | G |
| NC_011770 | 5744815 | G | C | 49 | 0 | 8 | 0 | C | C | | | | | C | C | C | C | C | C |
| NC_011770 | 5744831 | C | A | 49 | 0 | 6 | 0 | A | A | | | | | A | A | | | A | A |
| NC_011770 | 5744839 | G | A | 50 | 0 | 2 | 0 | A | A | | | | | | | | | | |
| NC_011770 | 5744844 | G | C | 49 | 0 | 2 | 0 | | | | | | | | | C | C | | |
| NC_011770 | 5744882 | G | A | 52 | 0 | 2 | 0 | | | | | | | | | A | A | | |
| NC_011770 | 5744952 | A | C | 56 | 0 | 4 | 0 | | | | | | | C | C | | | C | C |
| NC_011770 | 4715699 | A | AT | 88 | 0 | 7 | 0 | AT | AT | AT | AT | AT | AT | | | | AT | | |
| NC_011770 | 5747026 | C | T | 82 | 0 | 2 | 0 | T | T | | | | | | | | | | |
| NC_011770 | 5747901 | C | G | 76 | 0 | 12 | 0 | G | G | G | G | G | G | G | G | G | G | G | G |
| NC_011770 | 1587374 | G | A | 78 | 0 | 1 | 0 | | | | | | | | | | | | A |
| NC_011770 | 5750834 | G | A | 85 | 0 | 10 | 0 | | | A | A | A | A | A | A | A | A | A | A |
| NC_011770 | 5750847 | C | G | 83 | 0 | 2 | 0 | | | | | | | | | G | G | | |
| NC_011770 | 5751222 | A | C | 87 | 0 | 2 | 0 | | | | | | | | | C | C | | |
| NC_011770 | 5751264 | C | T | 77 | 0 | 2 | 0 | | | | | | | T | T | | | | |
| NC_011770 | 5751276 | C | CG | 74 | 0 | 4 | 0 | | | CG | CG | CG | CG | | | | | | |
| NC_011770 | 5751384 | C | T | 70 | 0 | 4 | 0 | | | T | T | T | T | | | | | | |
| NC_011770 | 212244 | A | G | 62 | 0 | 1 | 0 | | | | | | | | G | | | | |
| NC_011770 | 5752290 | A | C | 50 | 0 | 12 | 0 | C | C | C | C | C | C | C | C | C | C | C | C |
| NC_011770 | 5752295 | G | A | 47 | 0 | 2 | 0 | A | A | | | | | | | | | | |
| NC_011770 | 5752301 | T | C | 49 | 0 | 4 | 0 | | | C | C | C | C | | | | | | |
| NC_011770 | 302179 | C | CGTGGC | 54 | 0 | 2 | 0 | | | | | CGTGGCGA | | | CGTGGCGA | | | | |
| NC_011770 | 1801254 | C | T | 65 | 0 | 1 | 0 | | | | | | | | | | | | T |
| NC_011770 | 5754295 | CGTAGA | C | 75 | 0 | 2 | 0 | | | | | | | | | | | C | C |
| NC_011770 | 5756409 | T | G | 82 | 0 | 10 | 0 | G | G | | | G | G | G | G | G | G | G | G |
| NC_011770 | 5756435 | A | G | 79 | 0 | 2 | 0 | | | | | | | | | | | G | G |
| NC_011770 | 5756436 | C | A | 80 | 0 | 2 | 0 | A | A | | | | | | | | | | |
| NC_011770 | 5756516 | A | G | 78 | 0 | 4 | 0 | | | G | G | G | G | | | | | | |
| NC_011770 | 5756524 | A | C | 79 | 0 | 2 | 0 | | | | | | | | | | | C | C |
| NC_011770 | 5756574 | C | A | 84 | 0 | 2 | 0 | A | A | | | | | | | | | | |
| NC_011770 | 5756575 | C | T | 83 | 0 | 6 | 0 | | | | | | | T | T | T | T | T | T |
| NC_011770 | 3228976 | G | A | 58 | 0 | 1 | 0 | | | | | | | | | | | A | |
| NC_011770 | 2185951 | AGAATTTTT | A | 71 | 0 | 1 | 0 | | | | | | | | | | | | A |
| NC_011770 | 5762378 | G | A | 88 | 0 | 2 | 0 | A | A | | | | | | | | | | |
| NC_011770 | 5762428 | G | T | 85 | 0 | 4 | 0 | T | T | | | | | | | T | T | | |
| NC_011770 | 5762443 | C | T | 89 | 0 | 12 | 0 | T | T | T | T | T | T | T | T | T | T | T | T |
| NC_011770 | 5762480 | C | A | 94 | 0 | 2 | 0 | | | | | | | | | | | A | A |
| NC_011770 | 3277113 | A | G | 14 | 1 | 6 | 0 | G | G | | | | | C | C | . | G | G | |

| | | | | | | | | | | | | | | | | | | | |
|---|---|---|---|---|---|---|---|---|---|---|---|---|---|---|---|---|---|---|---|
| NC_011770 | 5763691 | C | A | 48 | 0 | 4 | 0 | | | A | A | A | A | | | | | | |
| NC_011770 | 5763706 | C | A | 52 | 0 | 6 | 0 | | | A | A | A | A | | | A | A | | |
| NC_011770 | 5763707 | T | C | 51 | 0 | 4 | 0 | | | C | C | C | C | | | | | | |
| NC_011770 | 5763770 | T | C | 48 | 0 | 4 | 0 | | | C | C | C | C | | | | | | |
| NC_011770 | 5763771 | A | G | 48 | 0 | 4 | 0 | | | G | G | G | G | | | | | | |
| NC_011770 | 5763779 | A | AG | 49 | 0 | 1 | 0 | | | | | AG | | | | | | | |
| NC_011770 | 5763793 | C | T | 44 | 0 | 4 | 0 | | | T | T | T | T | | | | | | |
| NC_011770 | 5763799 | G | C | 38 | 0 | 4 | 0 | | | C | C | C | C | | | | | | |
| NC_011770 | 5763801 | C | T | 38 | 0 | 4 | 0 | | | T | T | T | T | | | | | | |
| NC_011770 | 5763815 | C | T | 47 | 0 | 4 | 0 | | | T | T | T | T | | | | | | |
| NC_011770 | 5763817 | G | C | 47 | 0 | 2 | 0 | | | | | | | | | | | C | C |
| NC_011770 | 5763837 | C | A | 45 | 0 | 4 | 0 | | | A | A | A | A | | | | | | |
| NC_011770 | 2441119 | G | A | 61 | 0 | 1 | 0 | | | | | | | | | | | | A |
| NC_011770 | 303067 | C | T | 89 | 0 | 1 | 0 | | | | | | | | T | | | | |
| NC_011770 | 2443566 | G | A | 66 | 0 | 1 | 0 | | | | | | | | | | | | A |
| NC_011770 | 5765833 | C | T | 10 | 0 | 2 | 0 | T | T | | | | | | | | | | |
| NC_011770 | 4556453 | A | G | 101 | 0 | 1 | 0 | | | G | | | | | | | | | |
| NC_011770 | 5766523 | G | A | 83 | 0 | 4 | 0 | | | A | A | A | A | | | | | | |
| NC_011770 | 5768519 | G | A | 95 | 0 | 2 | 0 | | | | | | | | | A | A | | |
| NC_011770 | 340919 | C | T | 85 | 0 | 1 | 0 | | | | | | | | T | | | | |
| NC_011770 | 5770228 | G | T | 91 | 0 | 2 | 0 | | | | | T | T | | | | | | |
| NC_011770 | 5770229 | C | T | 92 | 0 | 2 | 0 | | | | | | | | | T | T | | |
| NC_011770 | 386981 | G | GGCCCG | 68 | 0 | 10 | 0 | GGCCCGC | GGCCCGC | GGCCCGC | GGCCCGC | GGCCCGC | GGCCCGCACAGTT | | GGCCCGC | GGCCCGCACAGTT | | GGCCCGC | GGCCCGCACAGTT |
| NC_011770 | 5772395 | C | T | 76 | 0 | 12 | 0 | T | T | T | T | T | T | T | T | T | T | T | T |
| NC_011770 | 396779 | GA | G | 64 | 0 | 1 | 0 | | | | | | | | G | | | | |
| NC_011770 | 5160219 | T | TA | 78 | 0 | 1 | 0 | | | | | | | | | TA | | | |
| NC_011770 | 2411238 | TA | T | 52 | 0 | 1 | 0 | | T | | | | | | | | | | |
| NC_011770 | 2484283 | G | GC | 74 | 0 | 2 | 0 | | GC | | | | | GC | | | | | |
| NC_011770 | 513872 | T | C | 28 | 0 | 1 | 0 | | | | | | | | C | | | | |
| NC_011770 | 5778594 | A | G | 71 | 0 | 8 | 0 | G | G | | | | | G | G | G | G | G | G |
| NC_011770 | 5778631 | T | C | 74 | 0 | 4 | 0 | | | C | C | C | C | | | | | | |
| NC_011770 | 5778667 | T | C | 73 | 0 | 8 | 0 | C | C | | | | | C | C | C | C | C | C |
| NC_011770 | 2885837 | TG | T | 51 | 6 | 1 | 0 | | | | | . | . | . | . | . | . | | T |
| NC_011770 | 5780002 | C | A | 84 | 0 | 4 | 0 | | | A | A | A | A | | | | | | |
| NC_011770 | 5780023 | G | A | 82 | 0 | 2 | 0 | | | | | | | | | | | A | A |
| NC_011770 | 2898238 | C | A | 20 | 6 | 1 | 0 | | | | | . | . | . | . | . | . | | A |
| NC_011770 | 625452 | C | T | 56 | 0 | 1 | 0 | | | | | | | | T | | | | |
| NC_011770 | 5783197 | T | C | 40 | 0 | 6 | 0 | | | C | C | C | C | C | C | | | | |
| NC_011770 | 941274 | C | T | 65 | 0 | 1 | 0 | | | | | | | | T | | | | |
| NC_011770 | 5784571 | G | A | 73 | 0 | 2 | 0 | | | | | | | | | | | A | A |
| NC_011770 | 5787583 | G | C | 204 | 0 | 2 | 0 | C | C | | | | | | | | | | |
| NC_011770 | 5787626 | T | C | 87 | 0 | 10 | 0 | C | C | C | C | C | C | C | C | C | C | | |
| NC_011770 | 5787676 | G | A | 90 | 0 | 1 | 0 | | | | | | | A | | | | | |
| NC_011770 | 5787685 | A | G | 87 | 0 | 6 | 0 | G | G | | | | | | | G | G | G | G |
| NC_011770 | 5787717 | C | A | 85 | 0 | 12 | 0 | A | A | A | A | A | A | A | A | A | A | A | A |
| NC_011770 | 5788350 | G | A | 86 | 0 | 2 | 0 | A | A | | | | | | | | | | |
| NC_011770 | 5788385 | A | G | 85 | 0 | 12 | 0 | G | G | G | G | G | G | G | G | G | G | G | G |
| NC_011770 | 5788409 | T | A | 83 | 0 | 6 | 0 | | | A | A | A | A | A | A | | | | |
| NC_011770 | 3287442 | A | G | 16 | 0 | 1 | 0 | | | | | | | | | | | G | |
| NC_011770 | 5790042 | GC | GCC | 41 | 0 | 2 | 0 | | | | | | | GCC | G | | | | |
| NC_011770 | 5790064 | C | A | 38 | 0 | 4 | 0 | | | A | A | A | A | | | | | | |
| NC_011770 | 5790115 | C | T | 28 | 0 | 10 | 0 | | | T | T | T | T | T | T | T | T | T | T |
| NC_011770 | 5790136 | G | A | 22 | 0 | 2 | 0 | | | | | | | | | A | A | | |
| NC_011770 | 5790200 | T | G | 19 | 0 | 4 | 0 | | | | | | | G | G | | | G | G |
| NC_011770 | 5790565 | T | A | 52 | 0 | 10 | 0 | | | A | A | A | A | A | A | A | A | A | A |
| NC_011770 | 5790800 | A | G | 66 | 0 | 2 | 0 | | | | | | | G | G | | | | |
| NC_011770 | 5796219 | G | C | 75 | 0 | 2 | 0 | | | | | | | | | | | C | C |
| NC_011770 | 5796751 | G | GC | 58 | 0 | 1 | 0 | | | | | | | GC | | | | | |
| NC_011770 | 5797080 | T | C | 55 | 0 | 4 | 0 | | | | | | | | | C | C | C | C |
| NC_011770 | 5797084 | C | G | 56 | 0 | 4 | 0 | | | | | | | | | G | G | G | G |
| NC_011770 | 5797085 | C | G | 56 | 0 | 4 | 0 | | | | | | | | | G | G | G | G |
| NC_011770 | 5799006 | G | A | 68 | 0 | 10 | 0 | A | A | A | A | A | A | A | A | A | A | | |

| | | | | | | | | | | | | | | | | | | | |
|---|---|---|---|---|---|---|---|---|---|---|---|---|---|---|---|---|---|---|---|
| NC_011770 | 2898396 | T | A | 28 | 6 | 1 | 0 | | | | | . | . | . | . | . | . | | A |
| NC_011770 | 5800692 | C | A | 100 | 0 | 2 | 0 | | | | | | | A | A | | | | |
| NC_011770 | 5800701 | G | A | 104 | 0 | 2 | 0 | | | | | | | A | A | | | | |
| NC_011770 | 5801328 | G | A | 99 | 0 | 8 | 0 | | | A | A | A | A | A | A | | | A | A |
| NC_011770 | 5801333 | A | C | 99 | 0 | 2 | 0 | | | | | | | | | C | C | | |
| NC_011770 | 5801341 | C | A | 97 | 0 | 2 | 0 | | | | | | | | | A | A | | |
| NC_011770 | 5801369 | T | C | 89 | 0 | 8 | 0 | | | C | C | C | C | C | C | | | C | C |
| NC_011770 | 5801401 | G | A | 101 | 0 | 2 | 0 | A | A | | | | | | | | | | |
| NC_011770 | 5801413 | A | G | 94 | 0 | 8 | 0 | | | G | G | G | G | G | G | | | G | G |
| NC_011770 | 5801414 | A | G | 96 | 0 | 8 | 0 | | | G | G | G | G | G | G | | | G | G |
| NC_011770 | 5802285 | G | A | 61 | 0 | 4 | 0 | | | A | A | A | A | | | | | | |
| NC_011770 | 5802348 | A | G | 89 | 0 | 2 | 0 | | | | | | | | | G | G | | |
| NC_011770 | 5802436 | T | C | 87 | 0 | 6 | 0 | | | C | C | C | C | | | | | C | C |
| NC_011770 | 5802437 | A | G | 86 | 0 | 6 | 0 | | | G | G | G | G | | | | | G | G |
| NC_011770 | 5803882 | G | A | 73 | 0 | 2 | 0 | | | | | | | | | A | A | | |
| NC_011770 | 5803894 | T | C | 74 | 0 | 2 | 0 | | | | | | | | | C | C | | |
| NC_011770 | 5803920 | G | A | 79 | 0 | 6 | 0 | | | | | | | A | A | T | T | A | A |
| NC_011770 | 5803925 | T | C | 80 | 0 | 2 | 0 | | | | | | | | | C | C | | |
| NC_011770 | 5803929 | G | A | 79 | 0 | 2 | 0 | | | | | | | | | A | A | | |
| NC_011770 | 1147170 | C | T | 36 | 0 | 1 | 0 | | | | | | | | T | | | | |
| NC_011770 | 5804482 | C | T | 85 | 0 | 10 | 0 | T | T | T | T | T | T | | | T | T | T | T |
| NC_011770 | 5804498 | T | C | 88 | 0 | 2 | 0 | | | | | | | | | C | C | | |
| NC_011770 | 5804651 | C | T | 82 | 0 | 2 | 0 | | | | | | | | | | | T | T |
| NC_011770 | 5804697 | C | T | 85 | 0 | 6 | 0 | T | T | T | T | T | T | | | | | | |
| NC_011770 | 5805995 | G | C | 79 | 0 | 2 | 0 | C | C | | | | | | | | | | |
| NC_011770 | 5806049 | G | C | 80 | 0 | 4 | 0 | | | C | C | C | C | | | | | | |
| NC_011770 | 5806885 | A | C | 80 | 0 | 2 | 0 | | | | | | | | | | | C | C |
| NC_011770 | 5809169 | T | C | 68 | 0 | 8 | 0 | | | C | C | C | C | C | C | C | C | | |
| NC_011770 | 5809786 | T | G | 56 | 0 | 4 | 0 | | | G | G | G | G | | | | | | |
| NC_011770 | 5809854 | A | G | 51 | 0 | 2 | 0 | | | | | G | G | | | | | | |
| NC_011770 | 5809892 | C | A | 49 | 0 | 2 | 0 | A | A | | | | | | | | | | |
| NC_011770 | 5809937 | A | G | 50 | 0 | 8 | 0 | | | G | G | G | G | G | G | G | G | | |
| NC_011770 | 5809949 | T | C | 51 | 0 | 10 | 0 | | | C | C | C | C | C | C | C | C | C | C |
| NC_011770 | 5809950 | A | G | 52 | 0 | 2 | 0 | G | G | | | | | | | | | | |
| NC_011770 | 5809964 | G | A | 56 | 0 | 2 | 0 | A | A | | | | | | | | | | |
| NC_011770 | 5809991 | T | C | 62 | 0 | 2 | 0 | | | | | | | | | | | C | C |
| NC_011770 | 1184797 | C | T | 61 | 0 | 1 | 0 | | | | | | | | T | | | | |
| NC_011770 | 1280165 | A | G | 62 | 0 | 1 | 0 | | | | | | | | G | | | | |
| NC_011770 | 5810415 | C | T | 62 | 0 | 2 | 0 | | | | | | | | | T | T | | |
| NC_011770 | 5810499 | A | G | 65 | 0 | 2 | 0 | | | | | | | G | G | | | | |
| NC_011770 | 5810502 | CA | C | 66 | 0 | 4 | 0 | | | C | C | C | C | | | | | | |
| NC_011770 | 5812664 | G | A | 84 | 0 | 6 | 0 | A | A | | | | | | | A | A | A | A |
| NC_011770 | 5812694 | T | C | 83 | 0 | 12 | 0 | C | C | C | C | C | C | C | C | C | C | C | C |
| NC_011770 | 1286424 | T | C | 68 | 0 | 1 | 0 | | | | | | | | C | | | | |
| NC_011770 | 1375548 | G | A | 76 | 0 | 1 | 0 | | | | | | | | A | | | | |
| NC_011770 | 5816130 | T | A | 76 | 0 | 2 | 0 | | | | | | | | | A | A | | |
| NC_011770 | 5816134 | A | G | 74 | 0 | 2 | 0 | | | | | | | | | G | G | | |
| NC_011770 | 5816206 | A | G | 70 | 0 | 4 | 0 | | | | | | | | | G | G | G | G |
| NC_011770 | 5816233 | T | C | 73 | 0 | 2 | 0 | | | | | | | | | C | C | | |
| NC_011770 | 1478317 | A | G | 41 | 0 | 1 | 0 | | | | | | | | G | | | | |
| NC_011770 | 5818510 | G | C | 65 | 0 | 2 | 0 | | | | | | | C | C | | | | |
| NC_011770 | 5818513 | A | G | 65 | 0 | 2 | 0 | | | | | | | G | G | | | | |
| NC_011770 | 5818627 | A | G | 60 | 0 | 2 | 0 | | | | | | | G | G | | | | |
| NC_011770 | 5819302 | G | A | 59 | 0 | 2 | 0 | | | | | | | A | A | | | | |
| NC_011770 | 5819312 | A | T | 58 | 0 | 4 | 0 | | | T | T | T | T | | | | | | |
| NC_011770 | 5819336 | T | C | 58 | 0 | 2 | 0 | | | | | | | C | C | | | | |
| NC_011770 | 5819429 | G | A | 70 | 0 | 2 | 0 | | | | | | | | | A | A | | |
| NC_011770 | 5820705 | A | C | 64 | 0 | 10 | 0 | C | C | C | C | C | C | | | C | C | C | C |
| NC_011770 | 5821330 | C | T | 60 | 0 | 12 | 0 | T | T | T | T | T | T | T | T | T | T | T | T |
| NC_011770 | 5821349 | G | A | 61 | 0 | 6 | 0 | A | A | | | | | | | A | A | A | A |
| NC_011770 | 1486189 | G | A | 68 | 0 | 1 | 0 | | | | | | | | A | | | | |
| NC_011770 | 5821937 | TC | T | 66 | 0 | 2 | 0 | | | | | | | | | | | T | T |

| | | | | | | | | | | | | | | | | | | | |
|---|---|---|---|---|---|---|---|---|---|---|---|---|---|---|---|---|---|---|---|
| NC_011770 | 3287244 | A | G | 17 | 0 | 1 | 0 | G | | | | | | | | | | | |
| NC_011770 | 5826696 | A | G | 80 | 0 | 4 | 0 | | | | | | | G | G | G | G | | |
| NC_011770 | 5826712 | C | T | 80 | 0 | 2 | 0 | | | | | | | | | | | T | T |
| NC_011770 | 5828073 | A | G | 73 | 0 | 2 | 0 | | | | | | | | | G | G | | |
| NC_011770 | 5832671 | C | T | 56 | 0 | 12 | 0 | T | T | T | T | T | T | T | T | T | T | T | T |
| NC_011770 | 5507842 | C | CT | 57 | 0 | 1 | 0 | | | | | | | | | CT | | | |
| NC_011770 | 3287444 | A | G | 17 | 0 | 1 | 0 | | | | | | | | | | | G | |
| NC_011770 | 5838137 | T | A | 81 | 0 | 4 | 0 | | | | | | | | | A | A | A | A |
| NC_011770 | 3371171 | C | T | 11 | 0 | 1 | 0 | | | | | | | | | | | T | |
| NC_011770 | 5839086 | C | A | 78 | 0 | 2 | 0 | A | A | | | | | | | | | | |
| NC_011770 | 5839231 | A | G | 76 | 0 | 4 | 0 | G | G | | | | | | | G | G | | |
| NC_011770 | 5839232 | G | T | 76 | 0 | 2 | 0 | | | | | | | | | T | T | | |
| NC_011770 | 5839236 | T | C | 74 | 0 | 4 | 0 | | | C | C | C | C | | | | | | |
| NC_011770 | 5840293 | C | T | 61 | 0 | 6 | 0 | T | T | | | | | G | G | T | T | | |
| NC_011770 | 2626880 | A | G | 64 | 0 | 1 | 0 | | G | | | | | | | | | | |
| NC_011770 | 3530643 | G | A | 77 | 0 | 1 | 0 | | | | | | | | | | | A | |
| NC_011770 | 5622975 | C | T | 26 | 0 | 1 | 0 | | | T | | | | | | | | | |
| NC_011770 | 5845676 | T | C | 75 | 0 | 2 | 0 | | | | | C | C | | | | | | |
| NC_011770 | 5845817 | C | G | 67 | 0 | 12 | 0 | G | G | G | G | G | G | T | T | T | T | G | G |
| NC_011770 | 5845830 | G | T | 69 | 0 | 4 | 0 | | | T | T | T | T | | | | | | |
| NC_011770 | 5845966 | G | C | 65 | 0 | 8 | 0 | C | C | C | C | C | C | | | | | C | C |
| NC_011770 | 5845969 | G | C | 65 | 0 | 2 | 0 | | | | | | | | | | | C | C |
| NC_011770 | 5845996 | C | T | 67 | 0 | 8 | 0 | T | T | T | T | T | T | | | | | T | T |
| NC_011770 | 5847003 | C | T | 79 | 0 | 2 | 0 | | | | | | | | | | | T | T |
| NC_011770 | 5847011 | C | T | 78 | 0 | 4 | 0 | T | T | | | | | | | | | T | T |
| NC_011770 | 5850075 | G | A | 79 | 0 | 2 | 0 | | | | | | | A | A | | | | |
| NC_011770 | 5852956 | G | A | 61 | 0 | 1 | 0 | | | | | | | A | | | | | |
| NC_011770 | 5852979 | T | A | 67 | 0 | 8 | 0 | A | A | A | A | A | A | | | | | A | A |
| NC_011770 | 5853971 | T | G | 60 | 0 | 2 | 0 | G | G | | | | | | | | | | |
| NC_011770 | 5854498 | T | C | 86 | 0 | 12 | 0 | C | C | C | C | C | C | C | C | C | C | C | C |
| NC_011770 | 5854588 | G | A | 88 | 0 | 2 | 0 | | | | | | | | | | | A | A |
| NC_011770 | 5854637 | C | T | 86 | 0 | 8 | 0 | T | T | T | T | T | T | | | T | T | | |
| NC_011770 | 5856083 | G | A | 81 | 0 | 2 | 0 | | | | | | | | | | | A | A |
| NC_011770 | 5856178 | G | A | 82 | 0 | 4 | 0 | A | A | | | | | | | | | A | A |
| NC_011770 | 5857082 | A | G | 62 | 0 | 12 | 0 | G | G | G | G | G | G | G | G | G | G | G | G |
| NC_011770 | 5857138 | G | A | 57 | 0 | 2 | 0 | | | | | | | | | | | A | A |
| NC_011770 | 5857238 | A | G | 58 | 0 | 8 | 0 | G | G | | | | | G | G | G | G | G | G |
| NC_011770 | 5857260 | G | A | 60 | 0 | 4 | 0 | A | A | | | | | | | | | A | A |
| NC_011770 | 5857279 | G | A | 61 | 0 | 4 | 0 | | | | | | | A | A | A | A | | |
| NC_011770 | 5857280 | A | G | 61 | 0 | 8 | 0 | G | G | | | | | G | G | G | G | G | G |
| NC_011770 | 5857287 | C | T | 60 | 0 | 2 | 0 | | | | | T | T | | | | | | |
| NC_011770 | 5603438 | T | C | 91 | 0 | 1 | 0 | | | | | | | | | | C | | |
| NC_011770 | 5860778 | G | A | 75 | 0 | 4 | 0 | A | A | | | | | | | | | A | A |
| NC_011770 | 5860783 | A | C | 75 | 0 | 4 | 0 | C | C | | | | | | | | | C | C |
| NC_011770 | 5860822 | C | G | 76 | 0 | 2 | 0 | | | | | | | | | | | G | G |
| NC_011770 | 5860823 | G | C | 76 | 0 | 2 | 0 | | | | | | | | | | | C | C |
| NC_011770 | 5860824 | A | G | 74 | 0 | 2 | 0 | | | | | | | | | | | G | G |
| NC_011770 | 5860832 | T | C | 78 | 0 | 2 | 0 | | | | | | | | | | | C | C |
| NC_011770 | 5860835 | T | C | 79 | 0 | 2 | 0 | | | | | | | | | | | C | C |
| NC_011770 | 5860846 | A | T | 78 | 0 | 2 | 0 | | | | | | | | | | | T | T |
| NC_011770 | 5860876 | G | A | 79 | 0 | 2 | 0 | | | A | A | | | | | | | | |
| NC_011770 | 5860880 | T | C | 79 | 0 | 2 | 0 | | | | | | | | | | | C | C |
| NC_011770 | 5860905 | T | C | 75 | 0 | 6 | 0 | | | C | C | C | C | | | | | C | C |
| NC_011770 | 5860909 | G | A | 76 | 0 | 6 | 0 | | | A | A | A | A | | | | | A | A |
| NC_011770 | 5860914 | A | G | 77 | 0 | 6 | 0 | | | G | G | G | G | | | | | G | G |
| NC_011770 | 5860921 | T | C | 78 | 0 | 6 | 0 | | | C | C | C | C | | | | | C | C |
| NC_011770 | 5860940 | A | T | 71 | 0 | 6 | 0 | | | T | T | T | T | | | | | T | T |
| NC_011770 | 5860942 | G | A | 72 | 0 | 2 | 0 | | | | | | | | | | | A | A |
| NC_011770 | 5860944 | A | C | 70 | 0 | 4 | 0 | | | C | C | C | C | | | | | | |
| NC_011770 | 5860945 | A | G | 72 | 0 | 6 | 0 | | | G | G | G | G | | | | | G | G |
| NC_011770 | 5860956 | G | A | 74 | 0 | 6 | 0 | | | A | A | A | A | | | | | A | A |
| NC_011770 | 5860962 | G | C | 73 | 0 | 6 | 0 | | | C | C | C | C | | | | | C | C |

| | | | | | | | | | | | | | | | | | | | |
|---|---|---|---|---|---|---|---|---|---|---|---|---|---|---|---|---|---|---|---|
| NC_011770 | 1683810 | C | T | 9 | 2 | 8 | 0 | . | T | . | T | T | T | | T | . | T | T | T |
| NC_011770 | 5863265 | G | A | 63 | 0 | 2 | 0 | A | A | | | | | | | | | | |
| NC_011770 | 5863282 | CG | C | 61 | 0 | 4 | 0 | C | C | | | | | | | | | C | C |
| NC_011770 | 5863343 | T | C | 64 | 0 | 4 | 0 | C | C | | | | | | | | | C | C |
| NC_011770 | 5863360 | A | G | 64 | 0 | 2 | 0 | | | | | | | | | | | G | G |
| NC_011770 | 5863376 | A | C | 66 | 0 | 4 | 0 | C | C | | | | | | | | | C | C |
| NC_011770 | 5863381 | T | C | 65 | 0 | 4 | 0 | C | C | | | | | | | | | C | C |
| NC_011770 | 5863382 | C | T | 65 | 0 | 2 | 0 | | | | | | | | | | | T | T |
| NC_011770 | 5863417 | G | A | 70 | 0 | 2 | 0 | | | | | | | | | | | A | A |
| NC_011770 | 5863453 | G | A | 74 | 0 | 4 | 0 | | | A | A | A | A | | | | | | |
| NC_011770 | 5865194 | C | A | 61 | 0 | 2 | 0 | A | A | | | | | | | | | | |
| NC_011770 | 5865236 | G | A | 62 | 0 | 2 | 0 | | | | | | | | | | | A | A |
| NC_011770 | 5865324 | G | A | 55 | 0 | 2 | 0 | | | | | | | | | | | A | A |
| NC_011770 | 5866456 | T | G | 79 | 0 | 2 | 0 | | | | | | | | | | | G | G |
| NC_011770 | 2840936 | TG | T | 25 | 6 | 2 | 0 | T | . | | T | . | . | . | . | . | . | | |
| NC_011770 | 5870147 | T | C | 73 | 0 | 4 | 0 | | | | | | | C | C | | | C | C |
| NC_011770 | 5870172 | G | A | 67 | 0 | 2 | 0 | | | | | | | | | A | A | | |
| NC_011770 | 3516828 | CG | C | 89 | 0 | 1 | 0 | C | | | | | | | | | | | |
| NC_011770 | 1892114 | T | TCC | 62 | 0 | 1 | 0 | | | | | | | | TCC | | | | |
| NC_011770 | 5874055 | T | A | 50 | 1 | 2 | 0 | | | . | | | | | | A | A | | |
| NC_011770 | 5874084 | A | G | 56 | 1 | 2 | 0 | | | . | | | | | | | | G | G |
| NC_011770 | 5874260 | A | G | 65 | 0 | 10 | 0 | G | G | G | G | G | G | G | G | G | G | | |
| NC_011770 | 5874300 | A | AT | 61 | 0 | 4 | 0 | | | | | | | AT | AT | AT | AT | | |
| NC_011770 | 5874324 | C | T | 67 | 0 | 4 | 0 | | | | | | | T | T | T | T | | |
| NC_011770 | 5874325 | T | C | 64 | 0 | 2 | 0 | | | | | | | | | C | C | | |
| NC_011770 | 5874335 | A | G | 63 | 0 | 4 | 0 | | | | | | | G | G | G | G | | |
| NC_011770 | 2908138 | CCAGCGCT | C | 60 | 0 | 1 | 0 | | | | | | | | | | | | C |
| NC_011770 | 3130925 | C | CT | 34 | 0 | 8 | 0 | CT | | CT | CT | CT | CT | CT | CT | | CT | | |
| NC_011770 | 2913872 | C | T | 84 | 0 | 1 | 0 | | | | | | | | | | | | T |
| NC_011770 | 2964492 | G | A | 68 | 0 | 1 | 0 | | | | | | | | | | | | A |
| NC_011770 | 5881052 | A | T | 83 | 0 | 2 | 0 | | | | | | | | | | | T | T |
| NC_011770 | 5881053 | G | C | 82 | 0 | 2 | 0 | | | | | | | | | | | C | C |
| NC_011770 | 5881057 | T | C | 83 | 0 | 2 | 0 | | | | | | | | | | | C | C |
| NC_011770 | 5882063 | C | A | 76 | 0 | 4 | 0 | A | A | | | | | | | A | A | | |
| NC_011770 | 5882098 | G | T | 74 | 0 | 2 | 0 | | | | | | | T | T | | | | |
| NC_011770 | 5882099 | C | T | 73 | 0 | 2 | 0 | | | | | | | T | T | | | | |
| NC_011770 | 1991620 | GT | G | 63 | 0 | 5 | 0 | | | G | G | G | G | | G | | | | |
| NC_011770 | 5888601 | C | T | 68 | 0 | 2 | 0 | | | | | | | | | | | T | T |
| NC_011770 | 3045890 | C | T | 68 | 0 | 1 | 0 | | | | | | | | | | | | T |
| NC_011770 | 5891348 | G | A | 89 | 0 | 4 | 0 | | | | | | | A | A | | | A | A |
| NC_011770 | 2178478 | G | A | 76 | 0 | 1 | 0 | | | | | | | | A | | | | |
| NC_011770 | 5893866 | T | C | 70 | 0 | 2 | 0 | | | | | | | | | | | C | C |
| NC_011770 | 5895650 | C | T | 52 | 0 | 4 | 0 | | | | | | | T | T | T | T | | |
| NC_011770 | 5895660 | G | A | 53 | 0 | 4 | 0 | | | | | | | A | A | A | A | | |
| NC_011770 | 5895678 | G | A | 54 | 0 | 4 | 0 | A | A | | | | | | | | | A | A |
| NC_011770 | 5895809 | G | A | 47 | 0 | 2 | 0 | | | | | | | A | A | | | | |
| NC_011770 | 5578132 | GCAT | G | 86 | 0 | 3 | 0 | | | G | G | | G | | | | | | |
| NC_011770 | 5895932 | A | T | 40 | 0 | 2 | 0 | | | | | | | | | T | T | | |
| NC_011770 | 5897801 | T | C | 82 | 0 | 6 | 0 | C | C | | | | | C | C | | | C | C |
| NC_011770 | 820184 | AC | A | 35 | 0 | 6 | 0 | | A | A | | A | A | A | | | A | | |
| NC_011770 | 3472171 | AC | A | 69 | 0 | 3 | 0 | | | | A | | | | | | | ACC | ACC |
| NC_011770 | 5903538 | G | A | 58 | 0 | 10 | 0 | A | A | A | A | A | A | A | A | | | A | A |
| NC_011770 | 5904471 | T | C | 71 | 0 | 1 | 0 | | | | | | | C | | | | | |
| NC_011770 | 5904488 | C | A | 71 | 0 | 2 | 0 | | | A | A | | | | | | | | |
| NC_011770 | 3071544 | A | G | 31 | 2 | 1 | 0 | . | . | | | | | | | | | | G |
| NC_011770 | 2254378 | CT | C | 72 | 0 | 1 | 0 | | | | | | | | C | | | | |
| NC_011770 | 5910912 | G | A | 68 | 0 | 2 | 0 | | | | | | | | | | | A | A |
| NC_011770 | 5910933 | A | C | 71 | 0 | 2 | 0 | | | | | | | C | C | | | | |
| NC_011770 | 5910986 | T | C | 68 | 0 | 2 | 0 | | | | | | | C | C | | | | |
| NC_011770 | 5910995 | T | C | 68 | 0 | 2 | 0 | | | | | | | C | C | | | | |
| NC_011770 | 5911004 | A | T | 66 | 0 | 2 | 0 | | | | | | | | | | | T | T |
| NC_011770 | 5911006 | C | CG | 67 | 0 | 2 | 0 | | | | | | | CG | CG | | | | |

| | | | | | | | | | | | | | | | | | | | |
|---|---|---|---|---|---|---|---|---|---|---|---|---|---|---|---|---|---|---|---|
| NC_011770 | 3277114 | C | T | 14 | 0 | 3 | 0 | T | T | | | | | | | . | | | T |
| NC_011770 | 5913140 | G | C | 69 | 0 | 4 | 0 | | | C | C | C | C | | | | | | |
| NC_011770 | 5913151 | G | C | 68 | 0 | 4 | 0 | | | C | C | C | C | | | | | | |
| NC_011770 | 5916255 | A | G | 58 | 0 | 12 | 0 | G | G | G | G | G | G | G | G | G | G | G | G |
| NC_011770 | 5917892 | C | G | 63 | 0 | 2 | 0 | | | | | | | | | | | G | G |
| NC_011770 | 5917919 | C | G | 63 | 0 | 2 | 0 | | | | | | | G | G | | | | |
| NC_011770 | 5917927 | T | C | 61 | 0 | 1 | 0 | | | | | | | C | | | | | |
| NC_011770 | 5918703 | G | T | 90 | 0 | 4 | 0 | | | T | T | T | T | | | | | | |
| NC_011770 | 5918704 | T | G | 86 | 0 | 2 | 0 | | | | | | | | | G | G | | |
| NC_011770 | 3873415 | AC | A | 38 | 0 | 1 | 0 | A | | | | | | | | | | | |
| NC_011770 | 5921618 | T | TC | 83 | 0 | 4 | 0 | . | | | | | | | | TC | TC | TC | TC |
| NC_011770 | 5925828 | C | T | 75 | 0 | 4 | 0 | | | | | | | | | T | T | T | T |
| NC_011770 | 5925916 | A | G | 72 | 0 | 2 | 0 | G | G | | | | | | | | | | |
| NC_011770 | 5925964 | G | A | 69 | 0 | 4 | 0 | | | A | A | A | A | | | | | | |
| NC_011770 | 5925966 | T | A | 69 | 0 | 12 | 0 | A | A | A | A | A | A | A | A | A | A | A | A |
| NC_011770 | 2284400 | GT | G | 61 | 0 | 1 | 0 | | | | | | | | G | | | | |
| NC_011770 | 2289404 | A | G | 81 | 0 | 1 | 0 | | | | | | | | G | | | | |
| NC_011770 | 5927445 | C | T | 79 | 0 | 12 | 0 | T | T | T | T | T | T | T | T | T | T | T | T |
| NC_011770 | 5927456 | G | A | 82 | 0 | 12 | 0 | A | A | A | A | A | A | A | A | A | A | A | A |
| NC_011770 | 3735034 | G | A | 60 | 0 | 1 | 0 | | | | | | | | | | | | A |
| NC_011770 | 5931397 | T | G | 61 | 0 | 10 | 0 | G | G | G | G | G | G | G | G | G | G | | |
| NC_011770 | 5931442 | G | C | 59 | 0 | 4 | 0 | | | C | C | C | C | | | | | | |
| NC_011770 | 5932209 | G | A | 79 | 0 | 2 | 0 | | | | | | | | | | | A | A |
| NC_011770 | 2311808 | GGGGC | G | 41 | 2 | 1 | 0 | . | . | | | | | | G | | | | |
| NC_011770 | 2341440 | GA | G | 62 | 0 | 1 | 0 | | | | | | | | G | | | | |
| NC_011770 | 4072238 | C | CCGCCG | 68 | 0 | 1 | 0 | CCGCCGG | | | | | | | | | | | |
| NC_011770 | 5936112 | C | G | 58 | 0 | 4 | 0 | G | G | | | | | G | G | | | | |
| NC_011770 | 5936139 | G | C | 57 | 0 | 2 | 0 | C | C | | | | | | | | | | |
| NC_011770 | 5936152 | A | G | 57 | 0 | 2 | 0 | G | G | | | | | | | | | | |
| NC_011770 | 5936155 | T | G | 58 | 0 | 6 | 0 | G | G | | | | | G | G | G | G | | |
| NC_011770 | 5938369 | A | C | 74 | 0 | 6 | 0 | | | C | C | C | C | | | C | C | | |
| NC_011770 | 3758245 | G | A | 62 | 0 | 1 | 0 | | | | | | | | | | | A | |
| NC_011770 | 5940926 | T | C | 45 | 0 | 1 | 0 | | | | | | | C | | | | | |
| NC_011770 | 5941905 | G | A | 67 | 0 | 12 | 0 | A | A | A | A | A | A | A | A | A | A | A | A |
| NC_011770 | 5943142 | A | G | 91 | 0 | 2 | 0 | | | | | | | | | | | G | G |
| NC_011770 | 5943167 | A | G | 86 | 0 | 2 | 0 | | | | | | | | | | | G | G |
| NC_011770 | 5943174 | A | C | 86 | 0 | 2 | 0 | | | | | | | | | | | C | C |
| NC_011770 | 3790846 | G | GC | 55 | 0 | 3 | 0 | | | | | | | GC | GC | | | | GC |
| NC_011770 | 4062453 | AG | A | 38 | 0 | 3 | 0 | A | A | | | | | | | | | | A |
| NC_011770 | 5950274 | T | C | 95 | 0 | 10 | 0 | C | C | C | C | C | C | C | C | C | C | | |
| NC_011770 | 5953095 | A | G | 83 | 0 | 2 | 0 | | | | | | | | | | | G | G |
| NC_011770 | 5953107 | C | T | 84 | 0 | 2 | 0 | | | | | | | | | | | T | T |
| NC_011770 | 5597894 | T | C | 90 | 0 | 5 | 0 | | | C | C | | C | | | | | C | C |
| NC_011770 | 5954232 | G | T | 81 | 0 | 2 | 0 | | | | | | | | | T | T | | |
| NC_011770 | 2357578 | A | G | 81 | 0 | 1 | 0 | | | | | | | | G | | | | |
| NC_011770 | 2359709 | C | T | 60 | 0 | 1 | 0 | | | | | | | | T | | | | |
| NC_011770 | 5970594 | G | C | 72 | 0 | 2 | 0 | | | | | | | | | C | C | | |
| NC_011770 | 5970742 | G | A | 66 | 0 | 12 | 0 | A | A | A | A | A | A | A | A | A | A | A | A |
| NC_011770 | 5975287 | G | T | 77 | 0 | 4 | 0 | | | T | T | T | T | | | | | | |
| NC_011770 | 5975301 | C | T | 77 | 0 | 4 | 0 | | | T | T | T | T | | | | | | |
| NC_011770 | 4084937 | G | A | 84 | 0 | 1 | 0 | | | | | | | | | | | A | |
| NC_011770 | 5978469 | A | T | 73 | 0 | 2 | 0 | T | T | | | | | | | | | | |
| NC_011770 | 5979992 | G | A | 63 | 0 | 4 | 0 | | | A | A | A | A | | | | | | |
| NC_011770 | 4099138 | G | GC | 68 | 0 | 11 | 0 | GC | GC | GC | GC | GC | GC | GC | GC | GC | GC | | GC |
| NC_011770 | 2887249 | CCAA | C | 15 | 7 | 1 | 0 | | C | | | . | . | . | . | . | . | | . |
| NC_011770 | 3129752 | C | T | 55 | 0 | 3 | 0 | | T | | | | | | | | | T | T |
| NC_011770 | 3211340 | G | GGGAA/ | 23 | 1 | 1 | 0 | | GGGAAA | | | | | | | | . | | |
| NC_011770 | 5987558 | C | A | 61 | 0 | 2 | 0 | | | | | | | | | | | A | A |
| NC_011770 | 5987574 | C | T | 60 | 0 | 4 | 0 | | | T | T | T | T | | | | | | |
| NC_011770 | 5989086 | G | A | 88 | 0 | 2 | 0 | | | | | | | | | A | A | | |
| NC_011770 | 5989106 | G | A | 88 | 0 | 2 | 0 | | | | | | | | | A | A | | |
| NC_011770 | 5989107 | A | G | 88 | 0 | 4 | 0 | G | G | | | | | | | T | T | | |

| | | | | | | | | | | | | | | | | | | | |
|---|---|---|---|---|---|---|---|---|---|---|---|---|---|---|---|---|---|---|---|
| NC_011770 | 5989116 | A | C | 87 | 0 | 12 | 0 | C | C | C | C | C | C | C | C | C | C | C | C |
| NC_011770 | 5989118 | A | G | 87 | 0 | 2 | 0 | | | | | | | | | G | G | | |
| NC_011770 | 5989152 | C | T | 86 | 0 | 2 | 0 | | | | | | | | | | | T | T |
| NC_011770 | 5989432 | A | G | 81 | 0 | 8 | 0 | | | G | G | G | G | G | G | | | G | G |
| NC_011770 | 5989469 | G | A | 76 | 0 | 8 | 0 | | | A | A | A | A | A | A | | | A | A |
| NC_011770 | 5990228 | A | G | 52 | 0 | 10 | 0 | | | G | G | G | G | G | G | G | G | G | G |
| NC_011770 | 5990229 | G | A | 53 | 0 | 6 | 0 | | | A | A | A | A | | | | | A | A |
| NC_011770 | 2412791 | CG | CGG | 38 | 0 | 3 | 0 | | | | | | | | CGG | C | C | | |
| NC_011770 | 5991655 | C | T | 76 | 0 | 4 | 0 | | | T | T | T | T | | | | | | |
| NC_011770 | 5992165 | A | C | 73 | 0 | 8 | 0 | | | C | C | C | C | C | C | C | C | | |
| NC_011770 | 5992225 | A | G | 75 | 0 | 4 | 0 | G | G | | | | | | | G | G | | |
| NC_011770 | 3287335 | T | G | 13 | 6 | 1 | 0 | | G | . | . | . | . | . | . | | | | |
| NC_011770 | 4246567 | G | A | 46 | 0 | 1 | 0 | | | | | | | | | | | A | |
| NC_011770 | 2426805 | A | AG | 31 | 4 | 7 | 0 | AG | AG | . | . | . | . | | AGG | AG | AG | AG | AG |
| NC_011770 | 5994870 | A | T | 85 | 0 | 8 | 0 | | | T | T | T | T | | | T | T | T | T |
| NC_011770 | 5994884 | A | C | 82 | 0 | 2 | 0 | C | C | | | | | | | | | | |
| NC_011770 | 5995018 | A | T | 92 | 0 | 12 | 0 | T | T | T | T | T | T | C | T | T | T | T | T |
| NC_011770 | 2473938 | C | T | 26 | 0 | 1 | 0 | | | | | | | | T | | | | |
| NC_011770 | 5996357 | T | G | 66 | 0 | 6 | 0 | G | G | | | | | G | G | G | G | | |
| NC_011770 | 5996359 | C | T | 66 | 0 | 6 | 0 | T | T | | | | | T | T | T | T | | |
| NC_011770 | 5997308 | T | C | 76 | 0 | 12 | 0 | C | C | C | C | C | C | C | C | C | C | C | C |
| NC_011770 | 5997345 | C | T | 82 | 0 | 2 | 0 | | | | | | | | | | | T | T |
| NC_011770 | 5997351 | A | G | 79 | 0 | 2 | 0 | G | G | | | | | | | | | | |
| NC_011770 | 5997379 | G | A | 75 | 0 | 2 | 0 | A | A | | | | | | | | | | |
| NC_011770 | 5997389 | G | A | 77 | 0 | 2 | 0 | A | A | | | | | | | | | | |
| NC_011770 | 5998495 | G | A | 73 | 0 | 2 | 0 | A | A | | | | | | | | | | |
| NC_011770 | 5998497 | G | A | 75 | 0 | 2 | 0 | | | | | | | | | | | A | A |
| NC_011770 | 5998499 | C | T | 75 | 0 | 2 | 0 | T | T | | | | | | | | | | |
| NC_011770 | 5998509 | T | C | 84 | 0 | 4 | 0 | | | | | | | C | C | C | C | | |
| NC_011770 | 5998521 | C | T | 76 | 0 | 2 | 0 | | | | | | | T | T | | | | |
| NC_011770 | 5999556 | G | A | 31 | 0 | 2 | 0 | A | A | | | | | | | | | | |
| NC_011770 | 5999559 | G | T | 35 | 0 | 2 | 0 | T | T | | | | | | | | | | |
| NC_011770 | 5999574 | A | G | 33 | 0 | 12 | 0 | G | G | G | G | G | G | G | G | G | G | G | G |
| NC_011770 | 5999586 | A | C | 32 | 0 | 2 | 0 | | | | | | | | | C | C | | |
| NC_011770 | 5999595 | G | A | 35 | 0 | 2 | 0 | A | A | | | | | | | | | | |
| NC_011770 | 2492826 | A | AG | 82 | 0 | 5 | 0 | | | | | | | | AG | AG | AG | AG | AG |
| NC_011770 | 6000785 | G | A | 71 | 0 | 2 | 0 | A | A | | | | | | | | | | |
| NC_011770 | 6000911 | A | G | 62 | 0 | 6 | 0 | G | G | G | G | G | G | | | | | | |
| NC_011770 | 6002364 | CTTCGG | C | 73 | 0 | 12 | 0 | C | C | C | C | C | C | C | C | C | C | C | C |
| NC_011770 | 2583065 | C | T | 35 | 0 | 1 | 0 | | | | | | | | T | | | | |
| NC_011770 | 2585116 | C | T | 63 | 0 | 1 | 0 | | | | | | | | T | | | | |
| NC_011770 | 2585229 | A | G | 58 | 0 | 1 | 0 | | | | | | | | G | | | | |
| NC_011770 | 4636975 | C | T | 77 | 0 | 1 | 0 | | | | | | | | | | | T | |
| NC_011770 | 6021774 | C | T | 80 | 0 | 2 | 0 | | | | | | | T | T | | | | |
| NC_011770 | 6022086 | T | C | 78 | 0 | 2 | 0 | | | | | | | | | | | C | C |
| NC_011770 | 6022133 | CG | C | 78 | 0 | 2 | 0 | | | | | | | | | | | C | C |
| NC_011770 | 6024451 | C | T | 63 | 0 | 2 | 0 | | | | | | | T | T | | | | |
| NC_011770 | 6024634 | GA | G | 55 | 0 | 2 | 0 | G | G | | | | | | | | | | |
| NC_011770 | 6029733 | G | A | 67 | 0 | 6 | 0 | A | A | | | | | | | A | A | A | A |
| NC_011770 | 6029779 | G | A | 68 | 0 | 2 | 0 | | | | | | | | | | | A | A |
| NC_011770 | 6029908 | G | A | 64 | 0 | 1 | 0 | | | | | | | A | | | | | |
| NC_011770 | 4160296 | G | A | 57 | 0 | 1 | 0 | | | | | | | | | | | | A |
| NC_011770 | 6031745 | A | T | 50 | 0 | 6 | 0 | | | T | T | T | T | | | | | T | T |
| NC_011770 | 6031809 | T | C | 48 | 0 | 6 | 0 | | | C | C | C | C | | | | | C | C |
| NC_011770 | 6031854 | C | T | 48 | 0 | 10 | 0 | T | T | T | T | T | T | | | T | T | T | T |
| NC_011770 | 4203308 | C | T | 84 | 0 | 1 | 0 | | | | | | | | | | | | T |
| NC_011770 | 6032794 | T | C | 117 | 0 | 2 | 0 | | | | | | | | | C | C | | |
| NC_011770 | 6032875 | C | T | 116 | 0 | 4 | 0 | | | T | T | T | T | | | | | | |
| NC_011770 | 6034724 | T | C | 83 | 0 | 4 | 0 | | | C | C | C | C | | | | | | |
| NC_011770 | 6034727 | GGCCC | G | 82 | 0 | 2 | 0 | | | | | | | | | | | G | G |
| NC_011770 | 6034740 | A | G | 82 | 0 | 12 | 0 | G | G | G | G | G | G | G | G | G | G | G | G |
| NC_011770 | 6034755 | C | G | 82 | 0 | 4 | 0 | | | G | G | G | G | | | | | | |

| | | | | | | | | | | | | | | | | | | | |
|---|---|---|---|---|---|---|---|---|---|---|---|---|---|---|---|---|---|---|---|
| NC_011770 | 3727992 | T | C | 81 | 0 | 1 | 0 | | | | C | | | | | | | | |
| NC_011770 | 6036931 | C | CG | 67 | 0 | 10 | 0 | CG | CG | CG | CG | CG | CG | | | CG | CG | CG | CG |
| NC_011770 | 6038120 | A | G | 68 | 0 | 2 | 0 | | | | | | | | | | | G | G |
| NC_011770 | 4826920 | G | A | 77 | 0 | 1 | 0 | | | | | | | | | | | A | |
| NC_011770 | 4831758 | C | T | 60 | 0 | 1 | 0 | | | | | | | | | | | T | |
| NC_011770 | 5248270 | G | A | 64 | 0 | 1 | 0 | | | | | | | | | | | A | |
| NC_011770 | 6048928 | C | T | 81 | 0 | 4 | 0 | T | T | | | | | | | T | T | | |
| NC_011770 | 4394976 | G | A | 100 | 0 | 1 | 0 | | | | | | | | | | | | A |
| NC_011770 | 4493042 | T | TCGC | 36 | 0 | 1 | 0 | | | | | | | | | | | | TCGC |
| NC_011770 | 4851616 | G | A | 77 | 0 | 1 | 0 | | | | | | | | | | | | A |
| NC_011770 | 6053292 | T | C | 64 | 0 | 10 | 0 | C | C | C | C | C | C | C | C | | | C | C |
| NC_011770 | 6053394 | C | G | 59 | 0 | 2 | 0 | G | G | | | | | | | | | | |
| NC_011770 | 6053413 | C | G | 60 | 0 | 2 | 0 | | | | | | | G | G | | | | |
| NC_011770 | 6055070 | A | G | 76 | 0 | 2 | 0 | G | G | | | | | | | | | | |
| NC_011770 | 6055097 | G | C | 83 | 0 | 2 | 0 | | | | | | | | | | | C | C |
| NC_011770 | 6055098 | G | A | 83 | 0 | 2 | 0 | | | | | | | | | | | A | A |
| NC_011770 | 3540177 | C | T | 13 | 6 | 1 | 0 | | T | . | . | . | . | | | . | . | | |
| NC_011770 | 3751819 | T | TC | 30 | 2 | 2 | 0 | | | | TC | TC | | | | . | . | | |
| NC_011770 | 3881223 | A | C | 60 | 0 | 3 | 0 | | | | C | | | | | C | C | | |
| NC_011770 | 5453572 | C | T | 86 | 0 | 1 | 0 | | | | | | | | | | | T | |
| NC_011770 | 6058238 | T | C | 91 | 0 | 12 | 0 | C | C | C | C | C | C | C | C | C | C | C | C |
| NC_011770 | 6059921 | G | C | 112 | 0 | 2 | 0 | C | C | | | | | | | | | | |
| NC_011770 | 6059933 | G | C | 99 | 0 | 2 | 0 | C | C | | | | | | | | | | |
| NC_011770 | 6059934 | T | C | 99 | 0 | 12 | 0 | C | C | C | C | C | C | C | C | C | C | C | C |
| NC_011770 | 6059936 | G | A | 98 | 0 | 2 | 0 | A | A | | | | | | | | | | |
| NC_011770 | 6059942 | C | A | 58 | 0 | 6 | 0 | A | A | A | A | A | A | | | | | | |
| NC_011770 | 6059982 | C | T | 31 | 6 | 1 | 0 | . | . | . | . | . | . | T | | | | | |
| NC_011770 | 6060040 | A | G | 30 | 6 | 2 | 0 | . | . | . | . | . | . | | | G | G | | |
| NC_011770 | 6060059 | G | A | 31 | 6 | 2 | 0 | . | . | . | . | . | . | | | | | A | A |
| NC_011770 | 6060072 | A | G | 31 | 6 | 2 | 0 | . | . | . | . | . | . | | | G | G | | |
| NC_011770 | 6060083 | C | T | 32 | 6 | 2 | 0 | . | . | . | . | . | . | | | T | T | | |
| NC_011770 | 6060085 | A | G | 32 | 6 | 2 | 0 | . | . | . | . | . | . | | | G | G | | |
| NC_011770 | 6060129 | A | G | 33 | 6 | 2 | 0 | . | . | . | . | . | . | | | G | G | | |
| NC_011770 | 6060991 | T | C | 56 | 2 | 2 | 0 | . | . | | | | | | | C | C | | |
| NC_011770 | 6066584 | C | G | 73 | 0 | 4 | 0 | | | G | G | G | G | | | | | | |
| NC_011770 | 6066587 | C | A | 71 | 0 | 8 | 0 | A | A | | | | | A | A | A | A | A | A |
| NC_011770 | 6066656 | C | A | 67 | 0 | 4 | 0 | | | A | A | A | A | | | | | | |
| NC_011770 | 6066719 | A | G | 67 | 0 | 8 | 0 | G | G | | | | | G | G | G | G | G | G |
| NC_011770 | 6066744 | A | C | 70 | 0 | 2 | 0 | | | | | | | | | | | C | C |
| NC_011770 | 6066772 | T | C | 68 | 0 | 4 | 0 | | | C | C | C | C | | | | | | |
| NC_011770 | 6066781 | T | TC | 67 | 0 | 12 | 0 | TC | TC | TC | TC | TC | TC | TC | TC | TC | TC | TC | TC |
| NC_011770 | 6066787 | G | A | 69 | 0 | 8 | 0 | A | A | | | | | A | A | A | A | A | A |
| NC_011770 | 6073067 | G | A | 52 | 0 | 2 | 0 | | | | | | | | | A | A | | |
| NC_011770 | 6073086 | T | C | 49 | 0 | 2 | 0 | | | | | | | | | | | C | C |
| NC_011770 | 6074524 | C | G | 60 | 0 | 10 | 0 | G | G | G | G | G | G | G | G | | | G | G |
| NC_011770 | 6076154 | GT | G | 67 | 0 | 2 | 0 | | | | | | | G | G | | | | |
| NC_011770 | 3939416 | CT | C | 63 | 0 | 1 | 0 | | | | C | | | | | | | | |
| NC_011770 | 6077710 | GC | G | 75 | 0 | 2 | 0 | G | G | | | | | | | | | | |
| NC_011770 | 6077718 | G | A | 78 | 0 | 2 | 0 | | | | | | | | | A | A | | |
| NC_011770 | 6077742 | T | C | 79 | 0 | 2 | 0 | | | | | | | C | C | | | | |
| NC_011770 | 2925887 | CT | C | 33 | 0 | 2 | 0 | | | | | | | | C | | C | | |
| NC_011770 | 2931935 | G | GC | 62 | 0 | 1 | 0 | | | | | | | | GC | | | | |
| NC_011770 | 6079466 | C | G | 73 | 0 | 10 | 0 | G | G | G | G | G | G | G | G | G | G | | |
| NC_011770 | 6079579 | G | A | 78 | 0 | 2 | 0 | | | | | | | | | | | A | A |
| NC_011770 | 6079624 | G | T | 77 | 0 | 2 | 0 | | | | | | | | | T | T | | |
| NC_011770 | 6079644 | G | T | 79 | 0 | 2 | 0 | | | | | | | | | T | T | | |
| NC_011770 | 6079731 | T | G | 80 | 0 | 2 | 0 | G | G | | | | | | | | | | |
| NC_011770 | 6082620 | A | G | 71 | 0 | 2 | 0 | | | | | | | | | | | G | G |
| NC_011770 | 6082627 | G | T | 72 | 0 | 2 | 0 | | | | | | | | | | | T | T |
| NC_011770 | 6082646 | C | T | 74 | 0 | 2 | 0 | | | | | | | | | | | T | T |
| NC_011770 | 6082669 | T | C | 71 | 0 | 10 | 0 | C | C | C | C | C | C | | | C | C | C | C |
| NC_011770 | 6082684 | G | A | 72 | 0 | 2 | 0 | | | | | | | | | | | A | A |

| | | | | | | | | | | | | | | | | | | | |
|---|---|---|---|---|---|---|---|---|---|---|---|---|---|---|---|---|---|---|---|
| NC_011770 | 6084106 | C | T | 86 | 0 | 6 | 0 | T | T | T | T | T | T | | | | | | |
| NC_011770 | 6084109 | A | G | 84 | 0 | 2 | 0 | | | | | | | | | | | G | G |
| NC_011770 | 3627391 | GC | G | 53 | 0 | 1 | 0 | | G | | | | | | | | | | |
| NC_011770 | 3043382 | T | C | 71 | 0 | 1 | 0 | | | | | | | | C | | | | |
| NC_011770 | 6088262 | A | G | 67 | 0 | 12 | 0 | G | G | G | G | G | G | G | G | G | G | G | G |
| NC_011770 | 6088938 | A | T | 67 | 0 | 2 | 0 | | | | | | | T | T | | | | |
| NC_011770 | 4906383 | C | T | 30 | 0 | 1 | 0 | | | | | | | | | | | | T |
| NC_011770 | 4023128 | G | GCCGAA | 31 | 0 | 2 | 0 | | | | GCCGAAGAGC | | GCCGAAGAGC | | | | | | |
| NC_011770 | 3204854 | G | A | 81 | 0 | 1 | 0 | | | | | | | | A | | | | |
| NC_011770 | 3293741 | C | CG | 60 | 0 | 3 | 0 | | . | | | | | | CG | CG | CG | | |
| NC_011770 | 6096092 | G | C | 57 | 0 | 6 | 0 | C | C | | | | | C | C | | | C | C |
| NC_011770 | 6096095 | G | A | 58 | 0 | 2 | 0 | | | | | | | | | | | A | A |
| NC_011770 | 6097261 | C | T | 51 | 0 | 2 | 0 | | | | | | | | | T | T | | |
| NC_011770 | 6097313 | G | A | 53 | 0 | 2 | 0 | | | | | | | A | A | | | | |
| NC_011770 | 5501175 | C | T | 85 | 0 | 1 | 0 | | | | | | | | | | | T | |
| NC_011770 | 4030925 | GA | G | 70 | 0 | 1 | 0 | | | | G | | | | | | | | |
| NC_011770 | 6100966 | A | G | 76 | 0 | 2 | 0 | | | | | | | | | | | G | G |
| NC_011770 | 6100997 | G | A | 79 | 0 | 12 | 0 | A | A | A | A | A | A | A | A | A | A | A | A |
| NC_011770 | 3311632 | C | T | 51 | 0 | 1 | 0 | | | | | | | | T | | | | |
| NC_011770 | 6102571 | C | A | 100 | 0 | 4 | 0 | | | | | | | | | A | A | A | A |
| NC_011770 | 6102573 | C | G | 100 | 0 | 4 | 0 | | | | | | | | | G | G | G | G |
| NC_011770 | 6102636 | C | A | 95 | 0 | 6 | 0 | A | A | A | A | A | A | | | | | | |
| NC_011770 | 6102648 | C | A | 98 | 0 | 2 | 0 | | | | | | | | | | | A | A |
| NC_011770 | 6102709 | C | T | 104 | 0 | 2 | 0 | | | | | | | T | T | | | | |
| NC_011770 | 6105475 | C | T | 74 | 0 | 2 | 0 | T | T | | | | | | | | | | |
| NC_011770 | 6105595 | C | A | 78 | 0 | 12 | 0 | A | A | A | A | A | A | A | A | A | A | A | A |
| NC_011770 | 6106723 | A | G | 85 | 0 | 6 | 0 | | | G | G | G | G | G | G | | | | |
| NC_011770 | 6106725 | G | C | 85 | 0 | 6 | 0 | | | C | C | C | C | C | C | | | | |
| NC_011770 | 6106729 | G | C | 83 | 0 | 2 | 0 | | | | | | | | | C | C | | |
| NC_011770 | 6106747 | T | A | 85 | 0 | 2 | 0 | | | | | | | | | | | A | A |
| NC_011770 | 6106756 | C | A | 85 | 0 | 8 | 0 | A | A | A | A | A | A | | | | | T | T |
| NC_011770 | 6106785 | C | T | 88 | 0 | 2 | 0 | | | | | | | T | T | | | | |
| NC_011770 | 6106796 | G | A | 89 | 0 | 2 | 0 | | | A | A | | | | | | | | |
| NC_011770 | 3476585 | TC | T | 8 | 2 | 2 | 0 | T | . | | | | | | T | . | . | | |
| NC_011770 | 6106862 | C | A | 74 | 0 | 12 | 0 | A | A | A | A | A | A | A | A | A | A | A | A |
| NC_011770 | 6109553 | A | G | 69 | 0 | 12 | 0 | G | G | G | G | G | G | G | G | G | G | G | G |
| NC_011770 | 6109573 | A | G | 88 | 0 | 4 | 0 | G | G | | | | | G | G | | | | |
| NC_011770 | 6109597 | A | C | 87 | 0 | 6 | 0 | C | C | | | | | C | C | | | C | C |
| NC_011770 | 6109601 | A | G | 87 | 0 | 4 | 0 | G | G | | | | | | | | | G | G |
| NC_011770 | 6109602 | C | T | 89 | 0 | 4 | 0 | T | T | | | | | | | | | T | T |
| NC_011770 | 6109605 | G | A | 88 | 0 | 4 | 0 | A | A | | | | | | | | | A | A |
| NC_011770 | 6109611 | T | C | 91 | 0 | 4 | 0 | C | C | | | | | | | | | C | C |
| NC_011770 | 6109615 | A | G | 90 | 0 | 4 | 0 | G | G | | | | | | | | | G | G |
| NC_011770 | 6109655 | C | G | 65 | 0 | 4 | 0 | | | G | G | G | G | | | | | | |
| NC_011770 | 6109656 | G | T | 64 | 0 | 4 | 0 | | | T | T | T | T | | | | | | |
| NC_011770 | 6109662 | G | A | 61 | 0 | 2 | 0 | | | | | | | A | A | | | | |
| NC_011770 | 6109664 | A | T | 58 | 0 | 2 | 0 | T | T | | | | | | | | | | |
| NC_011770 | 6109754 | C | T | 63 | 0 | 2 | 0 | | | | | | | | | T | T | | |
| NC_011770 | 6109848 | T | C | 63 | 0 | 4 | 0 | C | C | | | | | C | C | | | | |
| NC_011770 | 6109870 | A | C | 57 | 0 | 2 | 0 | | | | | | | | | | | C | C |
| NC_011770 | 6109917 | A | G | 62 | 0 | 12 | 0 | G | G | G | G | G | G | G | G | G | G | G | G |
| NC_011770 | 6110012 | G | A | 63 | 0 | 4 | 0 | | | | | | | | | A | A | A | A |
| NC_011770 | 6110052 | A | C | 64 | 0 | 4 | 0 | | | C | C | C | C | | | | | | |
| NC_011770 | 6110085 | C | T | 67 | 0 | 4 | 0 | T | T | | | | | T | T | | | | |
| NC_011770 | 6112503 | G | C | 73 | 0 | 2 | 0 | | | | | | | | | C | C | | |
| NC_011770 | 6112602 | C | A | 77 | 0 | 2 | 0 | | | | | | | | | A | A | | |
| NC_011770 | 6114282 | A | G | 45 | 0 | 2 | 0 | | | | | | | | | G | G | | |
| NC_011770 | 6114435 | T | TCC | 47 | 0 | 1 | 0 | | | | | | | TCC | | | | | |
| NC_011770 | 6115350 | G | A | 58 | 0 | 2 | 0 | | | | | | | A | A | | | | |
| NC_011770 | 6115477 | G | A | 47 | 0 | 2 | 0 | A | A | | | | | | | | | | |
| NC_011770 | 4962199 | C | T | 90 | 0 | 1 | 0 | | | | | | | | | | | | T |
| NC_011770 | 3624114 | T | C | 58 | 0 | 1 | 0 | | | | | | | | C | | | | |

| | | | | | | | | | | | | | | | | | | | |
|---|---|---|---|---|---|---|---|---|---|---|---|---|---|---|---|---|---|---|---|
| NC_011770 | 6120453 | A | G | 87 | 0 | 12 | 0 | G | G | G | G | G | G | G | G | G | G | G | G |
| NC_011770 | 6124971 | G | A | 75 | 0 | 10 | 0 | A | A | A | A | A | A | A | A | A | A | | |
| NC_011770 | 6126303 | G | A | 83 | 0 | 2 | 0 | A | A | | | | | | | | | | |
| NC_011770 | 6126345 | A | T | 83 | 0 | 4 | 0 | T | T | | | | | | | T | T | | |
| NC_011770 | 6126773 | T | C | 81 | 0 | 2 | 0 | | | | | | | C | C | | | | |
| NC_011770 | 6126835 | T | C | 82 | 0 | 2 | 0 | | | | | | | | | | | C | C |
| NC_011770 | 4293745 | GC | G | 72 | 0 | 3 | 0 | G | G | | G | | | | | | | | |
| NC_011770 | 6126847 | G | A | 81 | 0 | 4 | 0 | | | | | | | A | A | | | A | A |
| NC_011770 | 6126877 | G | A | 85 | 0 | 4 | 0 | | | | | | | A | A | | | A | A |
| NC_011770 | 6126889 | T | G | 85 | 0 | 8 | 0 | G | G | | | | | G | G | G | G | G | G |
| NC_011770 | 6126895 | C | T | 81 | 0 | 4 | 0 | | | | | | | T | T | | | T | T |
| NC_011770 | 6126914 | T | A | 80 | 0 | 2 | 0 | | | | | | | | | | | A | A |
| NC_011770 | 6126919 | C | T | 81 | 0 | 4 | 0 | | | | | | | T | T | | | T | T |
| NC_011770 | 6126925 | G | A | 82 | 0 | 4 | 0 | | | | | | | A | A | | | A | A |
| NC_011770 | 6126940 | G | A | 83 | 0 | 4 | 0 | | | | | | | A | A | | | A | A |
| NC_011770 | 6127001 | G | A | 83 | 0 | 2 | 0 | | | | | | | | | A | A | | |
| NC_011770 | 6127021 | A | G | 79 | 0 | 2 | 0 | | | | | | | G | G | | | | |
| NC_011770 | 6128217 | C | T | 55 | 0 | 2 | 0 | T | T | | | | | | | | | | |
| NC_011770 | 4357879 | A | AAGCGC | 73 | 0 | 3 | 0 | | | | AAGCGCCAGCGCC | | | | | AAGCGCC | AAGCGCC | | |
| NC_011770 | 3625776 | C | T | 70 | 0 | 1 | 0 | | | | | | | | T | | | | |
| NC_011770 | 6132992 | A | G | 67 | 0 | 10 | 0 | G | G | G | G | G | G | | | G | G | G | G |
| NC_011770 | 6133042 | A | G | 64 | 0 | 10 | 0 | G | G | G | G | G | G | | | G | G | G | G |
| NC_011770 | 5543559 | C | T | 62 | 0 | 1 | 0 | | | | | | | | | | | T | |
| NC_011770 | 5622980 | C | T | 26 | 0 | 3 | 0 | | | T | | | | T | T | | | | |
| NC_011770 | 6134817 | A | G | 74 | 0 | 2 | 0 | | | | | | | | | | | G | G |
| NC_011770 | 6134970 | C | A | 74 | 0 | 4 | 0 | | | | | | | A | A | | | A | A |
| NC_011770 | 3694170 | GA | G | 61 | 0 | 1 | 0 | | | | | | | | G | | | | |
| NC_011770 | 6138044 | G | C | 75 | 0 | 4 | 0 | | | C | C | C | C | | | | | | |
| NC_011770 | 6138104 | C | T | 74 | 0 | 4 | 0 | | | T | T | T | T | | | | | | |
| NC_011770 | 6138128 | T | C | 70 | 0 | 2 | 0 | C | C | | | | | | | | | | |
| NC_011770 | 6138230 | C | A | 67 | 0 | 2 | 0 | A | A | | | | | | | | | | |
| NC_011770 | 6138251 | G | A | 65 | 0 | 6 | 0 | | | | | | | A | A | A | A | A | A |
| NC_011770 | 6138313 | G | A | 68 | 0 | 2 | 0 | A | A | | | | | | | | | | |
| NC_011770 | 6139402 | A | C | 67 | 0 | 12 | 0 | C | C | C | C | C | C | C | C | C | C | C | C |
| NC_011770 | 6139424 | G | A | 66 | 0 | 6 | 0 | A | A | A | A | A | A | | | | | | |
| NC_011770 | 6139426 | C | T | 67 | 0 | 6 | 0 | | | | | | | T | T | T | T | T | T |
| NC_011770 | 6139476 | T | A | 65 | 0 | 4 | 0 | | | A | A | A | A | | | | | | |
| NC_011770 | 6139483 | T | A | 66 | 0 | 12 | 0 | A | A | A | A | A | A | A | A | A | A | A | A |
| NC_011770 | 6139494 | A | G | 67 | 0 | 6 | 0 | | | | | | | G | G | G | G | G | G |
| NC_011770 | 6139501 | G | A | 64 | 0 | 6 | 0 | A | A | A | A | A | A | | | | | | |
| NC_011770 | 6139508 | C | T | 67 | 0 | 6 | 0 | | | | | | | T | T | T | T | T | T |
| NC_011770 | 6139593 | G | A | 75 | 0 | 8 | 0 | A | A | A | A | A | A | A | A | | | | |
| NC_011770 | 6139605 | A | G | 76 | 0 | 6 | 0 | G | G | | | | | | | G | G | G | G |
| NC_011770 | 6139616 | C | T | 78 | 0 | 8 | 0 | T | T | T | T | T | T | T | T | | | | |
| NC_011770 | 6139645 | T | C | 81 | 0 | 6 | 0 | C | C | C | C | C | C | | | | | | |
| NC_011770 | 6139661 | G | C | 80 | 0 | 6 | 0 | C | C | C | C | C | C | | | | | | |
| NC_011770 | 6139683 | A | G | 79 | 0 | 12 | 0 | G | G | G | G | G | G | G | G | G | G | G | G |
| NC_011770 | 6139722 | ATCCCC | A | 80 | 0 | 6 | 0 | A | A | A | A | A | A | | | | | | |
| NC_011770 | 6139731 | C | A | 40 | 2 | 6 | 0 | . | . | | | | | A | A | A | A | A | A |
| NC_011770 | 5557423 | C | T | 67 | 0 | 1 | 0 | | | | | | | | | | | T | |
| NC_011770 | 3720056 | T | TC | 71 | 0 | 11 | 0 | TC | TC | TC | TC | TC | TC | | TC | TC | TC | TC | TC |
| NC_011770 | 6144622 | G | C | 52 | 1 | 1 | 0 | C | . | | | | | | | | | | |
| NC_011770 | 6144637 | A | G | 55 | 0 | 2 | 0 | G | G | | | | | | | | | | |
| NC_011770 | 6144652 | A | C | 57 | 0 | 2 | 0 | C | C | | | | | | | | | | |
| NC_011770 | 6144660 | G | A | 58 | 0 | 2 | 0 | | | | | | | | | | | A | A |
| NC_011770 | 6144663 | G | A | 57 | 0 | 2 | 0 | | | | | | | | | | | A | A |
| NC_011770 | 5557657 | C | T | 62 | 0 | 1 | 0 | | | | | | | | | | | T | |
| NC_011770 | 4729246 | G | A | 75 | 0 | 1 | 0 | A | | | | | | | | | | | |
| NC_011770 | 513663 | CCCGCA | C | 39 | 0 | 2 | 0 | | | | | C | | | C | | | | |
| NC_011770 | 6151674 | C | T | 63 | 0 | 2 | 0 | T | T | | | | | | | | | | |
| NC_011770 | 6151676 | C | T | 63 | 0 | 2 | 0 | T | T | | | | | | | | | | |
| NC_011770 | 6154929 | C | T | 43 | 0 | 6 | 0 | T | T | T | T | T | T | | | | | | |

| | | | | | | | | | | | | | | | | | | | |
|---|---|---|---|---|---|---|---|---|---|---|---|---|---|---|---|---|---|---|---|
| NC_011770 | 6155094 | C | T | 44 | 0 | 2 | 0 | | | | | | | | | T | T | | |
| NC_011770 | 6155188 | C | T | 55 | 0 | 4 | 0 | | | T | T | T | T | | | | | | |
| NC_011770 | 6155193 | G | C | 54 | 0 | 10 | 0 | C | C | C | C | C | C | | | C | C | C | C |
| NC_011770 | 6155245 | T | A | 57 | 0 | 6 | 0 | A | A | | | | | | | A | A | A | A |
| NC_011770 | 6155255 | A | T | 59 | 0 | 6 | 0 | T | T | | | | | | | T | T | T | T |
| NC_011770 | 6155272 | C | T | 60 | 0 | 4 | 0 | | | T | T | T | T | | | | | | |
| NC_011770 | 6155278 | C | G | 61 | 0 | 2 | 0 | G | G | | | | | | | | | | |
| NC_011770 | 6155319 | G | A | 58 | 0 | 6 | 0 | A | A | | | | | | | A | A | A | A |
| NC_011770 | 6155406 | T | G | 57 | 0 | 6 | 0 | G | G | | | | | | | G | G | G | G |
| NC_011770 | 3836225 | CA | C | 60 | 0 | 1 | 0 | | | | | | | | C | | | | |
| NC_011770 | 6158217 | G | GC | 83 | 0 | 2 | 0 | GC | GC | | | | | | | | | | |
| NC_011770 | 6159282 | G | C | 74 | 0 | 4 | 0 | | | C | C | C | C | | | | | | |
| NC_011770 | 6160260 | C | T | 65 | 0 | 2 | 0 | | | | | | | | | T | T | | |
| NC_011770 | 6160313 | A | G | 62 | 0 | 8 | 0 | G | G | | | | | G | G | G | G | G | G |
| NC_011770 | 6160316 | T | C | 62 | 0 | 8 | 0 | C | C | | | | | C | C | C | C | C | C |
| NC_011770 | 6160339 | G | T | 66 | 0 | 2 | 0 | | | | | | | | | | | T | T |
| NC_011770 | 6160359 | C | A | 70 | 0 | 2 | 0 | | | | | | | A | A | | | | |
| NC_011770 | 6160372 | C | T | 69 | 0 | 6 | 0 | T | T | | | | | T | T | | | T | T |
| NC_011770 | 6160414 | C | T | 68 | 0 | 6 | 0 | | | T | T | T | T | | | T | T | | |
| NC_011770 | 6160545 | G | A | 67 | 0 | 1 | 0 | | | | | | | A | | | | | |
| NC_011770 | 6160561 | C | T | 66 | 0 | 6 | 0 | T | T | | | | | T | T | | | T | T |
| NC_011770 | 6162576 | A | G | 51 | 0 | 10 | 0 | G | G | G | G | G | G | G | G | | | G | G |
| NC_011770 | 4774388 | A | AC | 55 | 2 | 7 | 0 | AC | AC | | AC | AC | AC | . | . | AC | | AC | |
| NC_011770 | 6162800 | G | A | 13 | 10 | 2 | 0 | . | . | . | . | . | . | . | . | . | . | A | A |
| NC_011770 | 6162803 | A | G | 13 | 10 | 2 | 0 | . | . | . | . | . | . | . | . | . | . | G | G |
| NC_011770 | 6162840 | C | T | 13 | 10 | 2 | 0 | . | . | . | . | . | . | . | . | . | . | T | T |
| NC_011770 | 6162874 | C | T | 14 | 10 | 2 | 0 | . | . | . | . | . | . | . | . | . | . | T | T |
| NC_011770 | 6163116 | T | G | 16 | 10 | 2 | 0 | . | . | . | . | . | . | . | . | . | . | G | G |
| NC_011770 | 6163149 | C | T | 15 | 10 | 2 | 0 | . | . | . | . | . | . | . | . | . | . | T | T |
| NC_011770 | 6163153 | A | G | 15 | 10 | 2 | 0 | . | . | . | . | . | . | . | . | . | . | G | G |
| NC_011770 | 6163162 | A | G | 14 | 10 | 2 | 0 | . | . | . | . | . | . | . | . | . | . | G | G |
| NC_011770 | 6163170 | C | T | 14 | 10 | 2 | 0 | . | . | . | . | . | . | . | . | . | . | T | T |
| NC_011770 | 6163179 | C | A | 14 | 10 | 2 | 0 | . | . | . | . | . | . | . | . | . | . | A | A |
| NC_011770 | 6163193 | T | C | 15 | 10 | 2 | 0 | . | . | . | . | . | . | . | . | . | . | C | C |
| NC_011770 | 6163228 | T | C | 16 | 10 | 2 | 0 | . | . | . | . | . | . | . | . | . | . | C | C |
| NC_011770 | 6163333 | A | G | 15 | 10 | 2 | 0 | . | . | . | . | . | . | . | . | . | . | G | G |
| NC_011770 | 6163410 | A | G | 15 | 10 | 2 | 0 | . | . | . | . | . | . | . | . | . | . | G | G |
| NC_011770 | 6163418 | T | C | 15 | 10 | 2 | 0 | . | . | . | . | . | . | . | . | . | . | C | C |
| NC_011770 | 6163496 | G | T | 14 | 10 | 2 | 0 | . | . | . | . | . | . | . | . | . | . | T | T |
| NC_011770 | 6163498 | G | A | 14 | 10 | 2 | 0 | . | . | . | . | . | . | . | . | . | . | A | A |
| NC_011770 | 6163504 | C | T | 15 | 10 | 2 | 0 | . | . | . | . | . | . | . | . | . | . | T | T |
| NC_011770 | 6163516 | A | G | 15 | 10 | 2 | 0 | . | . | . | . | . | . | . | . | . | . | G | G |
| NC_011770 | 6163521 | G | A | 15 | 10 | 2 | 0 | . | . | . | . | . | . | . | . | . | . | A | A |
| NC_011770 | 6163601 | G | A | 14 | 10 | 2 | 0 | . | . | . | . | . | . | . | . | . | . | A | A |
| NC_011770 | 6163611 | C | T | 12 | 10 | 2 | 0 | . | . | . | . | . | . | . | . | . | . | T | T |
| NC_011770 | 5603499 | G | A | 96 | 0 | 1 | 0 | | | | | | | | | | | A | |
| NC_011770 | 6166774 | C | T | 66 | 0 | 4 | 0 | | | T | T | T | T | | | | | | |
| NC_011770 | 6166777 | G | C | 66 | 0 | 4 | 0 | | | C | C | C | C | | | | | | |
| NC_011770 | 6166786 | T | C | 65 | 0 | 4 | 0 | | | C | C | C | C | | | | | | |
| NC_011770 | 6166848 | C | T | 70 | 0 | 2 | 0 | | | | | | | | | | | T | T |
| NC_011770 | 6167736 | T | C | 80 | 0 | 4 | 0 | | | C | C | C | C | | | | | | |
| NC_011770 | 6167738 | A | C | 78 | 0 | 4 | 0 | | | C | C | C | C | | | | | | |
| NC_011770 | 6167739 | C | T | 80 | 0 | 4 | 0 | | | T | T | T | T | | | | | | |
| NC_011770 | 6167749 | C | G | 84 | 0 | 4 | 0 | | | G | G | G | G | | | | | | |
| NC_011770 | 6167751 | C | T | 84 | 0 | 4 | 0 | | | T | T | T | T | | | | | | |
| NC_011770 | 6167774 | T | C | 84 | 0 | 4 | 0 | | | C | C | C | C | | | | | | |
| NC_011770 | 6167777 | C | T | 82 | 0 | 4 | 0 | | | T | T | T | T | | | | | | |
| NC_011770 | 6167783 | T | A | 85 | 0 | 4 | 0 | | | A | A | A | A | | | | | | |
| NC_011770 | 6167798 | T | C | 84 | 0 | 4 | 0 | | | C | C | C | C | | | | | | |
| NC_011770 | 6167799 | G | T | 83 | 0 | 4 | 0 | | | T | T | T | T | | | | | | |
| NC_011770 | 6168673 | C | T | 65 | 0 | 4 | 0 | | | T | T | T | T | | | | | | |
| NC_011770 | 6171003 | T | C | 49 | 0 | 10 | 0 | | | C | C | C | C | C | C | C | C | C | C |

| | | | | | | | | | | | | | | | | | | | |
|---|---|---|---|---|---|---|---|---|---|---|---|---|---|---|---|---|---|---|---|
| NC_011770 | 6171904 | C | T | 82 | 0 | 6 | 0 | | | T | T | T | T | | | T | T | | |
| NC_011770 | 6172370 | G | A | 71 | 0 | 2 | 0 | | | | | | | A | A | | | | |
| NC_011770 | 6172685 | C | A | 82 | 0 | 2 | 0 | | | | | | | | | | | A | A |
| NC_011770 | 6175015 | T | C | 83 | 0 | 2 | 0 | | | | | | | | | | | C | C |
| NC_011770 | 4963149 | T | TA | 71 | 0 | 1 | 0 | | | | | | | | | | | | TA |
| NC_011770 | 6178077 | A | G | 85 | 0 | 12 | 0 | G | G | G | G | G | G | G | G | G | G | G | G |
| NC_011770 | 4811011 | T | C | 46 | 0 | 1 | 0 | | | | C | | | | | | | | |
| NC_011770 | 6179726 | T | TG | 88 | 0 | 8 | 0 | | | TG | TG | TG | TG | TG | TG | TG | TG | | |
| NC_011770 | 6179749 | C | A | 90 | 0 | 4 | 0 | | | A | A | A | A | | | | | | |
| NC_011770 | 6179840 | A | G | 77 | 0 | 6 | 0 | | | G | G | G | G | | | G | G | | |
| NC_011770 | 6180354 | A | C | 64 | 0 | 12 | 0 | C | C | C | C | C | C | C | C | C | C | C | C |
| NC_011770 | 5491483 | T | C | 57 | 2 | 1 | 0 | C | | | | | | | | . | . | | |
| NC_011770 | 3851480 | G | A | 75 | 0 | 1 | 0 | | | | | | | | A | | | | |
| NC_011770 | 4134275 | G | A | 34 | 0 | 1 | 0 | | | | | | | | A | | | | |
| NC_011770 | 4147749 | TG | T | 73 | 0 | 1 | 0 | | | | | | | | T | | | | |
| NC_011770 | 6183887 | C | T | 75 | 0 | 2 | 0 | | | | | | | | | T | T | | |
| NC_011770 | 6184308 | T | C | 84 | 0 | 2 | 0 | | | | | | | | | C | C | | |
| NC_011770 | 6184319 | C | T | 84 | 0 | 2 | 0 | | | | | | | | | T | T | | |
| NC_011770 | 4977085 | G | A | 82 | 0 | 1 | 0 | | | | | | | | | | | | A |
| NC_011770 | 6184354 | G | A | 81 | 0 | 2 | 0 | A | A | | | | | | | | | | |
| NC_011770 | 6184377 | G | A | 79 | 0 | 2 | 0 | A | A | | | | | | | | | | |
| NC_011770 | 5171051 | C | T | 67 | 0 | 1 | 0 | | | | | | | | | | | | T |
| NC_011770 | 4077292 | G | GA | 75 | 0 | 9 | 0 | | GA | GA | GA | GA | GA | GA | GA | | | GA | GA |
| NC_011770 | 6185056 | C | T | 73 | 0 | 2 | 0 | | | T | T | | | | | | | | |
| NC_011770 | 6186003 | G | A | 78 | 0 | 2 | 0 | | | | | | | | | A | A | | |
| NC_011770 | 6186006 | C | G | 77 | 0 | 4 | 0 | | | | | | | | | G | G | G | G |
| NC_011770 | 6186022 | A | G | 80 | 0 | 4 | 0 | | | G | G | G | G | | | | | | |
| NC_011770 | 6186113 | T | C | 78 | 0 | 2 | 0 | | | | | | | | | C | C | | |
| NC_011770 | 6186119 | C | A | 77 | 0 | 6 | 0 | | | A | A | A | A | A | A | | | | |
| NC_011770 | 5603508 | C | A | 99 | 0 | 1 | 0 | | | | | | | | | | | A | |
| NC_011770 | 6187103 | T | C | 41 | 0 | 2 | 0 | | | | | | | | | C | C | | |
| NC_011770 | 6187125 | G | T | 40 | 0 | 2 | 0 | | | | | | | | | T | T | | |
| NC_011770 | 6187126 | C | T | 40 | 0 | 2 | 0 | | | | | | | | | T | T | | |
| NC_011770 | 6192276 | G | A | 71 | 0 | 12 | 0 | A | A | A | A | A | A | A | A | A | A | A | A |
| NC_011770 | 6192329 | C | T | 68 | 0 | 2 | 0 | | | | | | | | | | | T | T |
| NC_011770 | 6192337 | G | T | 66 | 0 | 10 | 0 | T | T | T | T | T | T | T | T | T | T | | |
| NC_011770 | 6192354 | C | A | 68 | 0 | 10 | 0 | A | A | A | A | A | A | A | A | A | A | | |
| NC_011770 | 6194043 | C | G | 70 | 0 | 6 | 0 | | | G | G | G | G | G | G | | | | |
| NC_011770 | 6195384 | T | TG | 83 | 0 | 2 | 0 | | | | | | | TG | TG | | | | |
| NC_011770 | 6195518 | T | C | 67 | 0 | 10 | 0 | | | C | C | C | C | C | C | C | C | C | C |
| NC_011770 | 4302344 | GA | G | 24 | 0 | 3 | 0 | | | | | | | | G | | | G | G |
| NC_011770 | 6195546 | C | G | 56 | 1 | 8 | 0 | | | G | G | G | G | | . | G | G | G | G |
| NC_011770 | 6195548 | G | A | 62 | 0 | 8 | 0 | | | A | A | A | A | | | A | A | A | A |
| NC_011770 | 6195551 | C | G | 62 | 0 | 8 | 0 | | | G | G | G | G | | | G | G | G | G |
| NC_011770 | 6195573 | G | A | 67 | 0 | 1 | 0 | | | | | | | A | | | | | |
| NC_011770 | 6195577 | C | T | 62 | 0 | 6 | 0 | | | T | T | T | T | | | | | T | T |
| NC_011770 | 6195582 | G | A | 61 | 0 | 2 | 0 | | | | | | | A | A | | | | |
| NC_011770 | 6195588 | A | C | 59 | 0 | 8 | 0 | | | C | C | C | C | C | C | | | C | C |
| NC_011770 | 6195610 | T | C | 57 | 0 | 6 | 0 | | | C | C | C | C | | | | | C | C |
| NC_011770 | 6195646 | A | G | 59 | 0 | 6 | 0 | | | G | G | G | G | | | | | G | G |
| NC_011770 | 6195696 | A | C | 64 | 0 | 8 | 0 | | | C | C | C | C | C | C | | | C | C |
| NC_011770 | 6197552 | C | T | 72 | 0 | 12 | 0 | T | T | T | T | T | T | T | T | T | T | T | T |
| NC_011770 | 6197555 | C | G | 76 | 0 | 2 | 0 | | | | | | | | | | | G | G |
| NC_011770 | 6197591 | G | A | 73 | 0 | 12 | 0 | A | A | A | A | A | A | A | A | A | A | A | A |
| NC_011770 | 6197640 | C | T | 74 | 0 | 10 | 0 | | | T | T | T | T | T | T | T | T | T | T |
| NC_011770 | 6197643 | A | C | 76 | 0 | 12 | 0 | C | C | C | C | C | C | C | C | C | C | C | C |
| NC_011770 | 4097193 | T | C | 69 | 0 | 1 | 0 | | C | | | | | | | | | | |
| NC_011770 | 6198929 | A | G | 47 | 0 | 10 | 0 | G | G | G | G | G | G | G | G | G | G | | |
| NC_011770 | 6198933 | G | A | 47 | 0 | 10 | 0 | A | A | A | A | A | A | A | A | A | A | | |
| NC_011770 | 6198938 | GAA | G | 47 | 0 | 10 | 0 | G | G | G | G | G | G | G | G | G | G | | |
| NC_011770 | 6198942 | G | GT | 45 | 0 | 10 | 0 | GT | GT | GT | GT | GT | GT | GT | GT | GT | GT | | |
| NC_011770 | 6198952 | A | G | 59 | 0 | 10 | 0 | G | G | G | G | G | G | G | G | G | G | | |

| | | | | | | | | | | | | | | | | | | | |
|---|---|---|---|---|---|---|---|---|---|---|---|---|---|---|---|---|---|---|---|
| NC_011770 | 6198953 | C | A | 59 | 0 | 10 | 0 | A | A | A | A | A | A | A | A | A | A | | |
| NC_011770 | 6203068 | T | C | 42 | 0 | 12 | 0 | C | C | C | C | C | C | C | C | C | C | C | C |
| NC_011770 | 6203071 | C | T | 40 | 0 | 2 | 0 | | | | | | | | | | | T | T |
| NC_011770 | 6203114 | T | C | 44 | 0 | 2 | 0 | | | | | | | | | | | C | C |
| NC_011770 | 6203119 | G | A | 47 | 0 | 2 | 0 | | | | | | | | | | | A | A |
| NC_011770 | 4121969 | C | CT | 70 | 0 | 1 | 0 | | CT | | | | | | | | | | |
| NC_011770 | 6203647 | A | G | 53 | 0 | 10 | 0 | G | G | G | G | G | G | | | G | G | G | G |
| NC_011770 | 4023217 | GC | G | 35 | 0 | 6 | 0 | G | G | | AC | | AC | | | | | AC | AC |
| NC_011770 | 6204077 | C | A | 63 | 0 | 4 | 0 | A | A | | | | | | | A | A | | |
| NC_011770 | 6208787 | A | G | 74 | 0 | 12 | 0 | G | G | G | G | G | G | G | G | G | G | G | G |
| NC_011770 | 5200007 | C | T | 80 | 0 | 1 | 0 | | | | | | | | | | | | T |
| NC_011770 | 6211520 | A | G | 82 | 0 | 1 | 0 | | | | | | | G | | | | | |
| NC_011770 | 6211577 | C | A | 80 | 0 | 4 | 0 | | | A | A | A | A | | | | | | |
| NC_011770 | 4292206 | T | C | 70 | 0 | 1 | 0 | | C | | | | | | | | | | |
| NC_011770 | 5070387 | C | T | 46 | 0 | 9 | 0 | T | T | | T | T | T | | | T | T | T | T |
| NC_011770 | 4547695 | G | C | 20 | 10 | 1 | 0 | . | . | . | . | . | . | | C | . | . | . | . |
| NC_011770 | 5300945 | AG | A | 53 | 0 | 3 | 0 | | | | | | | A | A | | | | A |
| NC_011770 | 5322728 | C | A | 96 | 0 | 3 | 0 | | | A | A | | | | | | | | T |
| NC_011770 | 6218944 | G | A | 91 | 0 | 2 | 0 | | | | | | | A | A | | | | |
| NC_011770 | 6219556 | G | A | 99 | 0 | 2 | 0 | | | | | | | A | A | | | | |
| NC_011770 | 6220221 | G | A | 71 | 0 | 2 | 0 | | | | | | | A | A | | | | |
| NC_011770 | 6220294 | T | C | 67 | 0 | 1 | 0 | | | | | | | C | | | | | |
| NC_011770 | 6221127 | A | G | 47 | 0 | 8 | 0 | | | G | G | G | G | | | G | G | G | G |
| NC_011770 | 6226854 | G | A | 80 | 0 | 2 | 0 | | | | | | | | | A | A | | |
| NC_011770 | 6226866 | G | A | 80 | 0 | 2 | 0 | | | | | | | | | | | A | A |
| NC_011770 | 6226879 | G | T | 83 | 0 | 2 | 0 | | | | | | | T | T | | | | |
| NC_011770 | 6227562 | C | T | 70 | 0 | 2 | 0 | | | | | | | T | T | | | | |
| NC_011770 | 6228620 | G | C | 62 | 0 | 8 | 0 | | | C | C | C | C | C | C | C | C | | |
| NC_011770 | 6228757 | C | T | 68 | 0 | 2 | 0 | T | T | | | | | | | | | | |
| NC_011770 | 6228788 | G | T | 74 | 0 | 2 | 0 | | | | | | | T | T | | | | |
| NC_011770 | 5362054 | A | AG | 73 | 0 | 1 | 0 | | | | | | | | | | | | AG |
| NC_011770 | 6234725 | T | C | 30 | 5 | 1 | 0 | . | . | | | | | . | C | | | . | . |
| NC_011770 | 6234726 | T | G | 31 | 5 | 1 | 0 | . | . | | | | | . | G | | | . | . |
| NC_011770 | 6234728 | T | C | 30 | 5 | 1 | 0 | . | . | | | | | . | C | | | . | . |
| NC_011770 | 6234729 | T | C | 30 | 5 | 1 | 0 | . | . | | | | | . | C | | | . | . |
| NC_011770 | 6234752 | T | C | 27 | 5 | 1 | 0 | . | . | | | | | . | C | | | . | . |
| NC_011770 | 6234758 | CA | C | 30 | 5 | 1 | 0 | . | . | | | | | . | C | | | . | . |
| NC_011770 | 6234771 | C | CT | 30 | 5 | 1 | 0 | . | . | | | | | . | CT | | | . | . |
| NC_011770 | 6234786 | G | C | 32 | 3 | 3 | 0 | . | . | | | | | C | . | | | C | C |
| NC_011770 | 6236202 | A | G | 70 | 0 | 2 | 0 | | | | | | | | | | | G | G |
| NC_011770 | 2852492 | G | GCGCCA | 20 | 8 | 2 | 0 | | GCGCCAC | GCGCCAC | | . | . | . | . | . | . | . | . |
| NC_011770 | 6236667 | G | A | 80 | 0 | 4 | 0 | | | A | A | A | A | | | | | | |
| NC_011770 | 5451944 | T | C | 67 | 0 | 3 | 0 | | | | C | TC | | C | | | | | |
| NC_011770 | 5534616 | GC | G | 76 | 0 | 1 | 0 | | | | G | | | | | | | | |
| NC_011770 | 6241556 | GT | G | 83 | 0 | 12 | 0 | G | G | G | G | G | G | G | G | G | G | G | G |
| NC_011770 | 4534971 | TG | T | 65 | 0 | 9 | 0 | | T | T | T | T | T | T | T | | | T | T |
| NC_011770 | 6242400 | G | A | 81 | 0 | 2 | 0 | A | A | | | | | | | | | | |
| NC_011770 | 6242426 | T | C | 86 | 0 | 6 | 0 | | | | | | | C | C | C | C | C | C |
| NC_011770 | 6246313 | A | G | 85 | 0 | 2 | 0 | | | | | | | G | G | | | | |
| NC_011770 | 6246316 | A | G | 84 | 0 | 2 | 0 | | | | | | | G | G | | | | |
| NC_011770 | 6247130 | C | T | 81 | 0 | 2 | 0 | | | | | | | | | T | T | | |
| NC_011770 | 6247148 | G | A | 81 | 0 | 10 | 0 | A | A | A | A | A | A | | | A | A | A | A |
| NC_011770 | 4564519 | A | AC | 75 | 0 | 1 | 0 | | | | | | | | AC | | | | |
| NC_011770 | 4636796 | TC | T | 80 | 0 | 1 | 0 | | T | | | | | | | | | | |
| NC_011770 | 4593056 | G | A | 83 | 0 | 1 | 0 | | | | | | | | A | | | | |
| NC_011770 | 6250538 | G | A | 75 | 0 | 4 | 0 | | | A | A | A | A | | | | | | |
| NC_011770 | 6250632 | G | T | 74 | 0 | 4 | 0 | | | T | T | T | T | | | | | | |
| NC_011770 | 6250677 | T | C | 63 | 0 | 4 | 0 | | | C | C | C | C | | | | | | |
| NC_011770 | 6250749 | C | T | 64 | 0 | 4 | 0 | | | T | T | T | T | | | | | | |
| NC_011770 | 6250860 | A | G | 66 | 0 | 4 | 0 | | | G | G | G | G | | | | | | |
| NC_011770 | 6250869 | C | T | 67 | 0 | 4 | 0 | | | T | T | T | T | | | | | | |
| NC_011770 | 6250871 | G | A | 66 | 0 | 4 | 0 | | | A | A | A | A | | | | | | |

| | | | | | | | | | | | | | | | | | | | |
|---|---|---|---|---|---|---|---|---|---|---|---|---|---|---|---|---|---|---|---|
| NC_011770 | 189013 | A | AC | 81 | 0 | 2 | 0 | | | | AC | AC | | | | | | | |
| NC_011770 | 513691 | GC | G | 38 | 0 | 7 | 0 | . | G | G | | G | G | G | G | | | | G |
| NC_011770 | 1159244 | C | G | 22 | 2 | 6 | 0 | | | G | | G | | . | . | G | G | G | G |
| NC_011770 | 1822547 | A | G | 98 | 4 | 2 | 0 | . | . | G | | G | | | | . | . | | |
| NC_011770 | 2276283 | TA | T | 81 | 0 | 2 | 0 | | | | T | T | | | | | | | |
| NC_011770 | 4685601 | A | G | 62 | 0 | 1 | 0 | | | | | | | | G | | | | |
| NC_011770 | 6252841 | G | A | 79 | 0 | 4 | 0 | | | A | A | A | A | | | | | | |
| NC_011770 | 4719665 | A | G | 65 | 0 | 1 | 0 | | | | | | | | G | | | | |
| NC_011770 | 6253421 | G | A | 86 | 0 | 2 | 0 | A | A | | | | | | | | | | |
| NC_011770 | 6253450 | T | C | 86 | 0 | 6 | 0 | | | C | C | C | C | | | | | C | C |
| NC_011770 | 6253472 | G | C | 83 | 0 | 4 | 0 | | | C | C | C | C | | | | | | |
| NC_011770 | 6253570 | C | T | 87 | 0 | 4 | 0 | | | T | T | T | T | | | | | | |
| NC_011770 | 6253590 | A | G | 91 | 0 | 2 | 0 | | | | | | | | | | | G | G |
| NC_011770 | 6253596 | C | A | 88 | 0 | 6 | 0 | | | A | A | A | A | | | | | G | G |
| NC_011770 | 6253607 | A | G | 89 | 0 | 2 | 0 | | | | | | | | | | | G | G |
| NC_011770 | 6254173 | T | G | 68 | 0 | 10 | 0 | G | G | G | G | G | G | | | G | G | G | G |
| NC_011770 | 6254175 | C | A | 68 | 0 | 10 | 0 | A | A | A | A | A | A | | | A | A | A | A |
| NC_011770 | 6254189 | A | G | 69 | 0 | 12 | 0 | G | G | G | G | G | G | G | G | G | G | G | G |
| NC_011770 | 6254263 | G | GT | 69 | 0 | 10 | 0 | GT | GT | GT | GT | GT | GT | | | GT | GT | GT | GT |
| NC_011770 | 4723486 | C | CG | 77 | 0 | 1 | 0 | | | | | | | | CG | | | | |
| NC_011770 | 5603513 | G | C | 103 | 0 | 1 | 0 | | | . | . | . | . | | | | | C | |
| NC_011770 | 1291917 | CGGGG | CG | 56 | 0 | 6 | 0 | CG | CG | C | | | | C | C | | | | CG |
| NC_011770 | 3970792 | G | GA | 57 | 0 | 5 | 0 | | | GA | | GA | | GA | GA | | | GA | |
| NC_011770 | 6260385 | C | CA | 65 | 0 | 2 | 0 | | | | | | | CA | CA | | | | |
| NC_011770 | 6260483 | A | G | 70 | 0 | 12 | 0 | G | G | G | G | G | G | G | G | G | G | G | G |
| NC_011770 | 6261942 | T | C | 39 | 6 | 4 | 0 | . | . | . | . | . | . | | | C | C | C | C |
| NC_011770 | 6261969 | A | G | 40 | 6 | 4 | 0 | . | . | . | . | . | . | | | G | G | G | G |
| NC_011770 | 6261994 | T | G | 43 | 6 | 2 | 0 | . | . | . | . | . | . | | | G | G | | |
| NC_011770 | 6261997 | A | G | 41 | 6 | 4 | 0 | . | . | . | . | . | . | | | G | G | G | G |
| NC_011770 | 6263051 | C | T | 38 | 6 | 4 | 0 | . | . | . | . | . | . | | | T | T | T | T |
| NC_011770 | 6263081 | C | T | 37 | 6 | 4 | 0 | . | . | . | . | . | . | | | T | T | T | T |
| NC_011770 | 6263130 | A | G | 40 | 6 | 2 | 0 | . | . | . | . | . | . | | | G | G | | |
| NC_011770 | 6263262 | A | G | 42 | 6 | 4 | 0 | . | . | . | . | . | . | | | G | G | G | G |
| NC_011770 | 4776101 | GC | G | 58 | 0 | 1 | 0 | | . | | | | | | G | | | | |
| NC_011770 | 6268749 | G | A | 77 | 0 | 8 | 0 | A | A | A | A | A | A | A | A | | | | |
| NC_011770 | 5041501 | G | GA | 40 | 0 | 1 | 0 | | GA | | | | | | | | | | |
| NC_011770 | 5895896 | TC | T | 41 | 0 | 6 | 0 | T | T | | | | TCC | | | | TCC | T | T |
| NC_011770 | 6269423 | C | T | 76 | 0 | 6 | 0 | T | T | | | | | | | T | T | T | T |
| NC_011770 | 6269453 | G | A | 75 | 0 | 6 | 0 | A | A | | | | | | | A | A | A | A |
| NC_011770 | 6269510 | G | A | 80 | 0 | 4 | 0 | | | A | A | A | A | | | | | | |
| NC_011770 | 6269521 | C | G | 82 | 0 | 2 | 0 | G | G | | | | | | | | | | |
| NC_011770 | 6269538 | C | T | 83 | 0 | 12 | 0 | T | T | T | T | T | T | T | T | T | T | T | T |
| NC_011770 | 6269542 | A | G | 83 | 0 | 12 | 0 | G | G | G | G | G | G | G | G | G | G | G | G |
| NC_011770 | 6269553 | G | A | 82 | 0 | 12 | 0 | A | A | A | A | A | A | A | A | A | A | A | A |
| NC_011770 | 6269558 | T | G | 83 | 0 | 12 | 0 | G | G | G | G | G | G | G | G | G | G | G | G |
| NC_011770 | 6270988 | A | G | 62 | 0 | 4 | 0 | | | G | G | G | G | | | | | | |
| NC_011770 | 6272194 | G | A | 90 | 0 | 4 | 0 | | | | | | | | | A | A | A | A |
| NC_011770 | 6272196 | C | G | 91 | 0 | 2 | 0 | | | | | | | G | G | | | | |
| NC_011770 | 6272213 | C | T | 91 | 0 | 4 | 0 | | | | | | | | | T | T | T | T |
| NC_011770 | 1381343 | AGGG | A | 36 | 0 | 4 | 0 | | A | | A | | | | | | | A | A |
| NC_011770 | 5049195 | CT | C | 67 | 0 | 4 | 0 | | | | C | | C | C | C | | | | |
| NC_011770 | 6277036 | G | GA | 78 | 0 | 12 | 0 | GA | GA | GA | GA | GA | GA | GA | GA | GA | GA | GA | GA |
| NC_011770 | 6277165 | C | T | 69 | 0 | 2 | 0 | | | | | | | T | T | | | | |
| NC_011770 | 6281888 | GC | G | 72 | 0 | 6 | 0 | | | | | | | G | G | G | G | G | G |
| NC_011770 | 4788145 | G | A | 58 | 0 | 1 | 0 | | | | | | | | A | | | | |
| NC_011770 | 6283191 | T | C | 83 | 0 | 4 | 0 | | | C | C | C | C | | | | | | |
| NC_011770 | 6283197 | T | C | 77 | 0 | 12 | 0 | C | C | C | C | C | C | C | C | C | C | C | C |
| NC_011770 | 6283198 | G | C | 78 | 0 | 12 | 0 | C | C | C | C | C | C | C | C | C | C | C | C |
| NC_011770 | 6283203 | A | G | 78 | 0 | 12 | 0 | G | G | G | G | G | G | G | G | G | G | G | G |
| NC_011770 | 6283207 | G | A | 79 | 0 | 12 | 0 | A | A | A | A | A | A | A | A | A | A | A | A |
| NC_011770 | 6283221 | A | G | 84 | 0 | 12 | 0 | G | G | G | G | G | G | G | G | G | G | G | G |
| NC_011770 | 6285608 | A | C | 70 | 0 | 4 | 0 | | | C | C | C | C | | | | | | |

| | | | | | | | | | | | | | | | | | | | |
|---|---|---|---|---|---|---|---|---|---|---|---|---|---|---|---|---|---|---|---|
| NC_011770 | 6285876 | G | GGGGC( | 41 | 0 | 2 | 0 | | | GGGGCGA | GGGGCGAGAT | | | | | | | | |
| NC_011770 | 6285880 | T | C | 40 | 0 | 4 | 0 | | | C | C | C | C | | | | | | |
| NC_011770 | 5036931 | T | C | 73 | 0 | 1 | 0 | | | | | | | | C | | | | |
| NC_011770 | 6286923 | T | G | 64 | 0 | 12 | 0 | G | G | G | G | G | G | G | G | G | G | G | G |
| NC_011770 | 6236244 | C | CA | 63 | 0 | 3 | 0 | | | CA | | CA | CA | | | | | | |
| NC_011770 | 6288883 | T | C | 71 | 0 | 2 | 0 | C | C | | | | | | | | | | |
| NC_011770 | 5603525 | G | A | 103 | 0 | 1 | 0 | | | | | | | | | | | A | |
| NC_011770 | 6289451 | T | G | 73 | 0 | 2 | 0 | G | G | | | | | | | | | | |
| NC_011770 | 6291012 | C | T | 72 | 0 | 8 | 0 | T | T | T | T | T | T | | | | | T | T |
| NC_011770 | 1573278 | TG | T | 68 | 0 | 2 | 0 | | T | | T | | | | | | | | |
| NC_011770 | 6295359 | G | T | 81 | 0 | 2 | 0 | T | T | | | | | | | | | | |
| NC_011770 | 5603540 | T | A | 102 | 0 | 1 | 0 | | | | | | | | | | | A | |
| NC_011770 | 6298601 | C | T | 75 | 0 | 4 | 0 | | | T | T | T | T | | | | | | |
| NC_011770 | 5062821 | G | A | 72 | 0 | 1 | 0 | | | | | | | | A | | | | |
| NC_011770 | 6302529 | C | T | 74 | 0 | 2 | 0 | | | | | | | | | T | T | | |
| NC_011770 | 5128259 | T | C | 76 | 0 | 3 | 0 | | C | | | | | | | | | C | C |
| NC_011770 | 6305296 | T | G | 68 | 0 | 6 | 0 | | | | | | | G | G | G | G | G | G |
| NC_011770 | 6305299 | C | T | 72 | 0 | 6 | 0 | | | | | | | T | T | T | T | T | T |
| NC_011770 | 6305898 | T | A | 71 | 0 | 2 | 0 | A | A | | | | | | | | | | |
| NC_011770 | 6309511 | A | G | 69 | 0 | 12 | 0 | G | G | G | G | G | G | G | G | G | G | G | G |
| NC_011770 | 6309512 | G | C | 68 | 0 | 12 | 0 | C | C | C | C | C | C | C | C | C | C | C | C |
| NC_011770 | 6309563 | C | T | 76 | 0 | 12 | 0 | T | T | T | T | T | T | G | G | T | T | T | T |
| NC_011770 | 6309641 | T | C | 75 | 0 | 8 | 0 | | | C | C | C | C | C | C | | | C | C |
| NC_011770 | 6309661 | G | A | 77 | 0 | 2 | 0 | | | | | | | | | | | A | A |
| NC_011770 | 6309739 | G | A | 77 | 0 | 2 | 0 | | | | | | | | | A | A | | |
| NC_011770 | 6313570 | A | T | 59 | 0 | 8 | 0 | T | T | T | T | T | T | | | | | T | T |
| NC_011770 | 6313572 | G | A | 57 | 0 | 8 | 0 | A | A | A | A | A | A | | | | | A | A |
| NC_011770 | 6313574 | A | G | 58 | 0 | 8 | 0 | G | G | G | G | G | G | | | | | G | G |
| NC_011770 | 6313576 | T | G | 57 | 0 | 8 | 0 | G | G | G | G | G | G | | | | | G | G |
| NC_011770 | 6313577 | T | C | 58 | 0 | 8 | 0 | C | C | C | C | C | C | | | | | C | C |
| NC_011770 | 6313655 | GT | G | 60 | 0 | 8 | 0 | | | G | G | G | G | G | G | | | G | G |
| NC_011770 | 6315081 | T | C | 73 | 0 | 2 | 0 | | | | | | | C | C | | | | |
| NC_011770 | 6321136 | GC | G | 72 | 0 | 8 | 0 | G | G | G | G | G | G | G | G | | | | |
| NC_011770 | 6321548 | T | G | 62 | 0 | 8 | 0 | G | G | G | G | G | G | G | G | | | | |
| NC_011770 | 6321606 | C | CG | 61 | 0 | 4 | 0 | | | CG | CG | CG | CG | | | | | | |
| NC_011770 | 6321642 | G | GT | 59 | 0 | 4 | 0 | GT | GT | | | | | GT | GT | | | | |
| NC_011770 | 6322174 | C | T | 73 | 0 | 2 | 0 | | | | | | | | | T | T | | |
| NC_011770 | 5546769 | C | CCGAAG( | 82 | 0 | 9 | 0 | CCGAAG | | CCGAAG | CCGAAG | | CCGAAG | CCGAAG | CCGAAG | CCGAAG | | CCGAAG | CCGAAG |
| NC_011770 | 2142351 | G | GT | 77 | 0 | 2 | 0 | GT | | | | | | | | | | | GT |
| NC_011770 | 6327102 | A | G | 77 | 0 | 12 | 0 | G | G | G | G | G | G | G | G | G | G | G | G |
| NC_011770 | 6327213 | C | G | 63 | 0 | 2 | 0 | | | | | | | G | G | | | | |
| NC_011770 | 6327257 | G | A | 59 | 0 | 8 | 0 | | | A | A | A | A | | | A | A | A | A |
| NC_011770 | 6327272 | C | T | 56 | 0 | 4 | 0 | T | T | | | | | T | T | | | | |
| NC_011770 | 6327294 | G | T | 57 | 0 | 2 | 0 | | | T | T | | | | | | | | |
| NC_011770 | 6327557 | C | T | 68 | 0 | 2 | 0 | | | | | | | T | T | | | | |
| NC_011770 | 6327654 | C | T | 73 | 0 | 8 | 0 | | | T | T | T | T | | | T | T | T | T |
| NC_011770 | 6327805 | C | T | 66 | 0 | 2 | 0 | T | T | | | | | | | | | | |
| NC_011770 | 6327965 | G | C | 62 | 0 | 2 | 0 | | | | | | | | | C | C | | |
| NC_011770 | 6328004 | A | G | 65 | 0 | 12 | 0 | G | G | G | G | G | G | G | G | G | G | G | G |
| NC_011770 | 6328190 | A | AGCGCC( | 71 | 0 | 5 | 0 | | | AGCGCCG | AGCGCCG | AGCGCCG | AGCGCCGGTG | | | | AGCGCCGGTG | | |
| NC_011770 | 5076291 | G | GC | 84 | 0 | 1 | 0 | | | | | | | | GC | | | | |
| NC_011770 | 5628921 | C | T | 67 | 0 | 1 | 0 | | | | | | | | | | | | T |
| NC_011770 | 6330643 | TC | T | 78 | 0 | 6 | 0 | | | | | | | T | T | T | T | T | T |
| NC_011770 | 6330657 | T | C | 77 | 0 | 12 | 0 | C | C | C | C | C | C | C | C | C | C | C | C |
| NC_011770 | 6331760 | G | A | 81 | 0 | 2 | 0 | | | | | | | | | A | A | | |
| NC_011770 | 6332984 | G | T | 61 | 0 | 4 | 0 | | | T | T | T | T | | | | | | |
| NC_011770 | 6335662 | A | G | 88 | 0 | 12 | 0 | G | G | G | G | G | G | G | G | G | G | G | G |
| NC_011770 | 6336190 | T | G | 90 | 0 | 2 | 0 | G | G | | | | | | | | | | |
| NC_011770 | 6338861 | C | T | 72 | 0 | 2 | 0 | | | | | | | T | T | | | | |
| NC_011770 | 6338895 | T | C | 71 | 0 | 2 | 0 | | | | | | | C | C | | | | |
| NC_011770 | 6338897 | C | A | 72 | 0 | 2 | 0 | | | | | | | A | A | | | | |
| NC_011770 | 6340045 | G | A | 91 | 0 | 2 | 0 | | | | | | | A | A | | | | |

| | | | | | | | | | | | | | | | | | | | |
|---|---|---|---|---|---|---|---|---|---|---|---|---|---|---|---|---|---|---|---|
| NC_011770 | 6340098 | C | T | 88 | 0 | 4 | 0 | | | T | T | T | T | | | | | | |
| NC_011770 | 6340126 | G | A | 84 | 0 | 2 | 0 | | | | | | | | | | | A | A |
| NC_011770 | 6340180 | T | A | 87 | 0 | 2 | 0 | A | A | | | | | | | | | | |
| NC_011770 | 6340196 | C | A | 88 | 0 | 2 | 0 | A | A | | | | | | | | | | |
| NC_011770 | 5706423 | C | CA | 41 | 0 | 9 | 0 | CA | | CA | CA | | CA | CA | CA | . | CA | CA | CA |
| NC_011770 | 6345512 | C | G | 86 | 0 | 12 | 0 | G | G | G | G | G | G | G | G | G | G | G | G |
| NC_011770 | 6346179 | A | G | 81 | 0 | 6 | 0 | | | | | | | G | G | G | G | G | G |
| NC_011770 | 5349147 | T | C | 61 | 0 | 1 | 0 | | C | | | | | | | | | | |
| NC_011770 | 5133802 | A | G | 47 | 0 | 3 | 0 | | | | | | | | G | G | G | | |
| NC_011770 | 6360624 | G | A | 92 | 0 | 4 | 0 | | | A | A | A | A | | | | | | |
| NC_011770 | 5627539 | G | GT | 62 | 0 | 11 | 0 | GT | GT | GT | GT | GT | GT | GT | GT | GT | GT | GT | |
| NC_011770 | 6362161 | A | G | 83 | 0 | 4 | 0 | | | | | | | G | G | | | G | G |
| NC_011770 | 6362164 | A | G | 86 | 0 | 4 | 0 | | | | | | | G | G | | | G | G |
| NC_011770 | 6362625 | C | T | 81 | 0 | 6 | 0 | | | T | T | T | T | T | T | | | | |
| NC_011770 | 6363434 | A | G | 84 | 0 | 6 | 0 | | | G | G | G | G | G | G | | | | |
| NC_011770 | 6363462 | T | TC | 86 | 0 | 2 | 0 | | | | | | | | | | | TC | TC |
| NC_011770 | 5695171 | T | C | 53 | 0 | 11 | 0 | C | C | C | C | C | C | C | C | C | C | | C |
| NC_011770 | 6364820 | G | A | 86 | 0 | 6 | 0 | A | A | A | A | A | A | | | | | | |
| NC_011770 | 6364830 | G | A | 85 | 0 | 6 | 0 | A | A | A | A | A | A | | | | | | |
| NC_011770 | 6364841 | G | A | 84 | 0 | 4 | 0 | | | A | A | A | A | | | | | | |
| NC_011770 | 6364843 | C | T | 85 | 0 | 2 | 0 | | | | | | | | | T | T | | |
| NC_011770 | 6367301 | G | GA | 75 | 0 | 2 | 0 | GA | GA | | | | | | | | | | |
| NC_011770 | 6369105 | A | G | 86 | 0 | 2 | 0 | | | | | | | | | G | G | | |
| NC_011770 | 6369132 | T | C | 84 | 0 | 1 | 0 | | | | | | | C | | | | | |
| NC_011770 | 6373787 | G | C | 88 | 0 | 2 | 0 | | | | | | | | | | | C | C |
| NC_011770 | 6373829 | T | C | 83 | 0 | 4 | 0 | C | C | | | | | C | C | | | | |
| NC_011770 | 6374086 | A | G | 82 | 0 | 2 | 0 | G | G | | | | | | | | | | |
| NC_011770 | 6374109 | C | T | 88 | 0 | 2 | 0 | | | | | | | | | | | T | T |
| NC_011770 | 5642960 | G | A | 61 | 0 | 1 | 0 | | | | | | | | | | | A | |
| NC_011770 | 6375330 | C | G | 57 | 2 | 10 | 0 | G | G | G | G | G | G | G | G | G | G | . | . |
| NC_011770 | 5703035 | G | A | 88 | 0 | 1 | 0 | | | | | | | | | | | | A |
| NC_011770 | 6375716 | G | GT | 110 | 0 | 2 | 0 | | | GT | GT | | | | | | | | |
| NC_011770 | 2387932 | T | TA | 64 | 0 | 7 | 0 | TA | TA | TA | TA | | TA | | TA | | TA | | |
| NC_011770 | 5447774 | GT | G | 62 | 0 | 1 | 0 | | | | | | | | G | | | | |
| NC_011770 | 6378868 | T | C | 81 | 0 | 4 | 0 | | | | | | | C | C | | | C | C |
| NC_011770 | 6378869 | C | T | 81 | 0 | 4 | 0 | | | | | | | T | T | | | T | T |
| NC_011770 | 6378901 | GT | G | 84 | 0 | 2 | 0 | G | G | | | | | | | | | | |
| NC_011770 | 6378985 | G | T | 66 | 0 | 2 | 0 | | | | | | | | | T | T | | |
| NC_011770 | 6380722 | T | C | 83 | 0 | 4 | 0 | | | C | C | C | C | | | | | | |
| NC_011770 | 5456442 | T | C | 75 | 0 | 1 | 0 | | | | | | | | C | | | | |
| NC_011770 | 6381024 | C | T | 57 | 0 | 10 | 0 | T | T | T | T | T | T | | | T | T | T | T |
| NC_011770 | 4260780 | GCGGCCGC | G | 67 | 0 | 2 | 0 | | | G | | G | | | | | | | |
| NC_011770 | 6381060 | A | T | 57 | 0 | 2 | 0 | | | | | | | | | T | T | . | . |
| NC_011770 | 6381061 | T | C | 57 | 0 | 6 | 0 | C | C | | | | | | | C | C | C | C |
| NC_011770 | 6381068 | C | T | 58 | 0 | 2 | 0 | | | | | | | T | T | | | | |
| NC_011770 | 6381085 | A | G | 55 | 0 | 10 | 0 | G | G | G | G | G | G | | | G | G | G | G |
| NC_011770 | 6381043 | A | G | 59 | 0 | 1 | 0 | | | | | G | | | | | | | |
| NC_011770 | 6381115 | T | C | 54 | 0 | 8 | 0 | C | C | C | C | C | C | C | C | | | | |
| NC_011770 | 6381119 | G | C | 57 | 0 | 4 | 0 | | | C | C | C | C | | | | | | |
| NC_011770 | 6381166 | G | A | 60 | 0 | 2 | 0 | A | A | | | | | | | | | | |
| NC_011770 | 3070641 | T | TG | 34 | 2 | 3 | 0 | TG | | | TG | | TG | | . | | | . | . |
| NC_011770 | 6382603 | G | A | 70 | 0 | 2 | 0 | | | | | | | | | A | A | | |
| NC_011770 | 6382627 | G | C | 68 | 0 | 2 | 0 | | | | | | | | | C | C | | |
| NC_011770 | 6382725 | T | C | 63 | 0 | 2 | 0 | | | | | | | | | | | C | C |
| NC_011770 | 5565253 | G | GAACGA | 64 | 0 | 1 | 0 | | GAACGACTA | | | | | | | | | | |
| NC_011770 | 6387549 | G | A | 87 | 0 | 2 | 0 | | | | | | | | | A | A | | |
| NC_011770 | 6389247 | G | A | 58 | 0 | 2 | 0 | A | A | | | | | | | | | | |
| NC_011770 | 6389248 | G | A | 57 | 0 | 4 | 0 | | | | | | | | | A | A | A | A |
| NC_011770 | 6389254 | G | A | 56 | 0 | 10 | 0 | | | A | A | A | A | A | A | A | A | A | A |
| NC_011770 | 6389259 | T | C | 55 | 0 | 6 | 0 | | | C | C | C | C | C | C | | | | |
| NC_011770 | 6389263 | G | C | 57 | 0 | 6 | 0 | | | C | C | C | C | C | C | | | | |
| NC_011770 | 6389266 | C | A | 56 | 0 | 6 | 0 | | | A | A | A | A | A | A | | | | |

| | | | | | | | | | | | | | | | | | | | |
|---|---|---|---|---|---|---|---|---|---|---|---|---|---|---|---|---|---|---|---|
| NC_011770 | 6389274 | G | A | 57 | 0 | 6 | 0 | | | A | A | A | A | A | A | | | | |
| NC_011770 | 6389278 | G | A | 56 | 0 | 6 | 0 | | | A | A | A | A | A | A | | | | |
| NC_011770 | 6389296 | G | A | 56 | 0 | 6 | 0 | | | A | A | A | A | A | A | | | | |
| NC_011770 | 6389330 | C | T | 55 | 0 | 4 | 0 | T | T | | | | | T | T | | | | |
| NC_011770 | 5507292 | CG | C | 65 | 0 | 1 | 0 | | | | | | | | C | | | | |
| NC_011770 | 6389350 | A | T | 56 | 0 | 2 | 0 | T | T | | | | | | | | | | |
| NC_011770 | 6389354 | G | T | 61 | 0 | 2 | 0 | | | | | | | | | T | T | | |
| NC_011770 | 6394254 | G | A | 83 | 0 | 1 | 0 | | | | | | | A | | | | | |
| NC_011770 | 6394267 | T | C | 86 | 0 | 4 | 0 | | | C | C | C | C | | | | | | |
| NC_011770 | 6394285 | A | G | 85 | 0 | 4 | 0 | | | G | G | G | G | | | | | | |
| NC_011770 | 6394333 | C | G | 83 | 0 | 2 | 0 | G | G | | | | | | | | | | |
| NC_011770 | 6394336 | G | C | 82 | 0 | 8 | 0 | | | C | C | C | C | C | C | C | C | | |
| NC_011770 | 6394374 | C | T | 84 | 0 | 4 | 0 | | | T | T | T | T | | | | | | |
| NC_011770 | 6394377 | G | A | 82 | 0 | 4 | 0 | | | A | A | A | A | | | | | | |
| NC_011770 | 6394397 | C | T | 82 | 0 | 4 | 0 | | | T | T | T | T | | | | | | |
| NC_011770 | 6394425 | T | C | 83 | 0 | 4 | 0 | | | C | C | C | C | | | | | | |
| NC_011770 | 6394427 | C | A | 85 | 0 | 4 | 0 | | | A | A | A | A | | | | | | |
| NC_011770 | 6394431 | T | G | 86 | 0 | 2 | 0 | | | | | | | | | G | G | | |
| NC_011770 | 6394438 | A | G | 84 | 0 | 4 | 0 | | | G | G | G | G | | | | | | |
| NC_011770 | 6394527 | C | T | 76 | 0 | 2 | 0 | | | | | | | T | T | | | | |
| NC_011770 | 3211331 | CGAATG | C | 56 | 0 | 3 | 0 | | | | | | C | | | | C | C | |
| NC_011770 | 6395723 | G | C | 63 | 0 | 4 | 0 | | | C | C | C | C | | | | | | |
| NC_011770 | 6395743 | G | A | 65 | 0 | 2 | 0 | A | A | | | | | | | | | | |
| NC_011770 | 6395751 | GA | G | 66 | 0 | 4 | 0 | | | G | G | G | G | | | | | | |
| NC_011770 | 6396150 | G | A | 62 | 0 | 4 | 0 | | | A | A | A | A | | | | | | |
| NC_011770 | 6396267 | T | C | 63 | 0 | 4 | 0 | | | C | C | C | C | | | | | | |
| NC_011770 | 6396278 | A | G | 64 | 0 | 4 | 0 | | | G | G | G | G | | | | | | |
| NC_011770 | 6396283 | A | G | 64 | 0 | 6 | 0 | G | G | G | G | G | G | | | | | | |
| NC_011770 | 6397239 | G | T | 69 | 0 | 2 | 0 | | | | | | | | | T | T | | |
| NC_011770 | 6397240 | C | A | 70 | 0 | 2 | 0 | | | A | A | | | | | | | | |
| NC_011770 | 6399787 | A | C | 93 | 0 | 6 | 0 | | | C | C | C | C | C | C | | | | |
| NC_011770 | 6399798 | G | T | 93 | 0 | 4 | 0 | | | T | T | T | T | | | | | | |
| NC_011770 | 6400783 | G | A | 68 | 0 | 2 | 0 | A | A | | | | | | | | | | |
| NC_011770 | 6400795 | A | G | 67 | 0 | 4 | 0 | | | | | | | | | G | G | G | G |
| NC_011770 | 5762377 | C | T | 89 | 0 | 2 | 0 | | | | | | | T | | | | | T |
| NC_011770 | 6402742 | G | A | 98 | 0 | 2 | 0 | | | | | | | | | | | A | A |
| NC_011770 | 6402835 | A | G | 96 | 0 | 12 | 0 | G | G | G | G | G | G | G | G | G | G | G | G |
| NC_011770 | 6402873 | T | C | 95 | 0 | 12 | 0 | C | C | C | C | C | C | C | C | C | C | C | C |
| NC_011770 | 5603437 | A | G | 91 | 0 | 3 | 0 | | | | | | | G | | | G | G | |
| NC_011770 | 5867991 | T | TG | 71 | 0 | 1 | 0 | | | | TG | | | | | | | | |
| NC_011770 | 6407684 | G | T | 50 | 0 | 4 | 0 | T | T | | | | | T | T | | | | |
| NC_011770 | 6407724 | A | T | 47 | 0 | 4 | 0 | T | T | | | | | T | T | | | | |
| NC_011770 | 6407755 | A | G | 46 | 0 | 4 | 0 | G | G | | | | | G | G | | | | |
| NC_011770 | 6409362 | AG | A | 56 | 0 | 6 | 0 | A | A | A | A | A | A | | | | | | |
| NC_011770 | 6409368 | G | A | 29 | 0 | 2 | 0 | | | | | | | | | A | A | | |
| NC_011770 | 6409969 | A | T | 61 | 0 | 12 | 0 | T | T | T | T | T | T | T | T | T | T | T | T |
| NC_011770 | 6409978 | C | G | 69 | 0 | 4 | 0 | | | G | G | G | G | | | | | | |
| NC_011770 | 6410022 | T | C | 68 | 0 | 6 | 0 | C | C | C | C | C | C | | | | | | |
| NC_011770 | 6410088 | C | T | 71 | 0 | 6 | 0 | T | T | T | T | T | T | | | | | | |
| NC_011770 | 5623003 | G | A | 22 | 0 | 1 | 0 | | | | | | | | A | | | | |
| NC_011770 | 6411177 | C | A | 86 | 0 | 6 | 0 | | | | | | | A | A | A | A | A | A |
| NC_011770 | 6411184 | G | C | 81 | 0 | 6 | 0 | | | | | | | C | C | C | C | C | C |
| NC_011770 | 6413908 | G | GC | 81 | 0 | 1 | 0 | | | | | | | GC | | | | | |
| NC_011770 | 6415962 | G | A | 85 | 0 | 4 | 0 | | | A | A | A | A | | | | | | |
| NC_011770 | 6415965 | C | T | 85 | 0 | 6 | 0 | T | T | T | T | T | T | | | | | | |
| NC_011770 | 6415974 | G | A | 88 | 0 | 4 | 0 | | | | | | | | | A | A | A | A |
| NC_011770 | 6415987 | C | T | 86 | 0 | 4 | 0 | | | T | T | T | T | | | | | | |
| NC_011770 | 6416022 | A | G | 83 | 0 | 4 | 0 | | | G | G | G | G | | | | | | |
| NC_011770 | 6416066 | T | C | 83 | 0 | 4 | 0 | | | C | C | C | C | | | | | | |
| NC_011770 | 6416138 | G | C | 82 | 0 | 4 | 0 | | | C | C | C | C | | | | | | |
| NC_011770 | 6416184 | A | G | 76 | 0 | 4 | 0 | | | G | G | G | G | | | | | | |
| NC_011770 | 6416214 | C | G | 73 | 0 | 6 | 0 | G | G | G | G | G | G | | | | | | |

| | | | | | | | | | | | | | | | | | | | |
|---|---|---|---|---|---|---|---|---|---|---|---|---|---|---|---|---|---|---|---|
| NC_011770 | 6416272 | C | T | 71 | 0 | 2 | 0 | T | T | | | | | | | | | | |
| NC_011770 | 6416276 | C | G | 72 | 0 | 4 | 0 | | | G | G | G | G | | | | | | |
| NC_011770 | 6416287 | T | C | 73 | 0 | 4 | 0 | | | C | C | C | C | | | | | | |
| NC_011770 | 6416290 | C | T | 73 | 0 | 4 | 0 | | | T | T | T | T | | | | | | |
| NC_011770 | 6418799 | T | C | 101 | 0 | 2 | 0 | | | | | | | | | C | C | | |
| NC_011770 | 6419444 | T | TCCCCA | 75 | 0 | 2 | 0 | | | | | | | | | TCCCCATC | TCCCCATCCAC | | |
| NC_011770 | 6421060 | C | G | 52 | 0 | 2 | 0 | | | | | | | | | G | G | | |
| NC_011770 | 6421086 | G | A | 47 | 0 | 4 | 0 | | | A | A | A | A | | | | | | |
| NC_011770 | 6421152 | T | C | 36 | 0 | 12 | 0 | C | C | C | C | C | C | C | C | C | C | C | C |
| NC_011770 | 6421263 | G | A | 45 | 0 | 10 | 0 | A | A | A | A | A | A | A | A | | | A | A |
| NC_011770 | 6422910 | T | C | 57 | 0 | 6 | 0 | | | C | C | C | C | C | C | | | | |
| NC_011770 | 6424177 | A | G | 84 | 0 | 12 | 0 | G | G | G | G | G | G | G | G | G | G | G | G |
| NC_011770 | 6424217 | C | A | 86 | 0 | 2 | 0 | A | A | | | | | | | | | | |
| NC_011770 | 6424400 | T | C | 89 | 0 | 2 | 0 | | | | | | | C | C | | | | |
| NC_011770 | 6424406 | G | C | 89 | 0 | 2 | 0 | | | | | | | C | C | | | | |
| NC_011770 | 6424567 | T | C | 94 | 0 | 1 | 0 | | | | | | | C | | | | | |
| NC_011770 | 6424651 | A | G | 79 | 0 | 12 | 0 | G | G | G | G | G | G | G | G | G | G | G | G |
| NC_011770 | 6424652 | C | T | 78 | 0 | 4 | 0 | T | T | | | | | | | | | T | T |
| NC_011770 | 6424660 | C | T | 82 | 0 | 2 | 0 | | | | | | | T | T | | | | |
| NC_011770 | 6424930 | T | C | 71 | 0 | 12 | 0 | C | C | C | C | C | C | C | C | C | C | C | C |
| NC_011770 | 6425108 | C | A | 81 | 0 | 2 | 0 | | | | | | | | | | | A | A |
| NC_011770 | 6425213 | G | C | 79 | 0 | 10 | 0 | C | C | C | C | C | C | | | C | C | C | C |
| NC_011770 | 6425249 | G | A | 80 | 0 | 4 | 0 | | | | | | | A | A | | | A | A |
| NC_011770 | 6425465 | G | A | 75 | 0 | 2 | 0 | | | | | | | | | | | A | A |
| NC_011770 | 6425494 | C | T | 82 | 0 | 10 | 0 | T | T | T | T | T | T | T | T | | | T | T |
| NC_011770 | 6425519 | T | A | 79 | 0 | 4 | 0 | | | | | | | A | A | | | A | A |
| NC_011770 | 6425550 | C | T | 76 | 0 | 8 | 0 | T | T | T | T | T | T | | | T | T | | |
| NC_011770 | 6425573 | C | A | 71 | 0 | 12 | 0 | A | A | A | A | A | A | A | A | A | A | A | A |
| NC_011770 | 6425576 | C | G | 73 | 0 | 12 | 0 | G | G | G | G | G | G | G | G | G | G | G | G |
| NC_011770 | 6425601 | C | T | 72 | 0 | 4 | 0 | T | T | | | | | | | T | T | | |
| NC_011770 | 6425670 | G | T | 74 | 0 | 8 | 0 | T | T | T | T | T | T | | | T | T | | |
| NC_011770 | 6425743 | C | T | 80 | 0 | 4 | 0 | | | | | | | T | T | | | T | T |
| NC_011770 | 6425765 | T | C | 78 | 0 | 4 | 0 | | | | | | | C | C | | | C | C |
| NC_011770 | 5780125 | C | T | 87 | 0 | 1 | 0 | | | | | | | | | | | | T |
| NC_011770 | 6426000 | A | G | 79 | 0 | 12 | 0 | G | G | G | G | G | G | G | G | G | G | G | G |
| NC_011770 | 6426136 | C | G | 78 | 0 | 4 | 0 | | | G | G | G | G | | | | | | |
| NC_011770 | 6426152 | C | T | 77 | 0 | 4 | 0 | | | T | T | T | T | | | | | | |
| NC_011770 | 6426176 | T | G | 76 | 0 | 4 | 0 | | | G | G | G | G | | | | | | |
| NC_011770 | 6426188 | G | C | 73 | 0 | 4 | 0 | | | C | C | C | C | | | | | | |
| NC_011770 | 6426196 | G | T | 72 | 0 | 4 | 0 | | | T | T | T | T | | | | | | |
| NC_011770 | 6426197 | G | T | 73 | 0 | 4 | 0 | | | T | T | T | T | | | | | | |
| NC_011770 | 6426201 | A | G | 75 | 0 | 4 | 0 | G | G | | | | | G | G | | | | |
| NC_011770 | 6426212 | T | C | 72 | 0 | 4 | 0 | | | C | C | C | C | | | | | | |
| NC_011770 | 6426239 | G | A | 70 | 0 | 4 | 0 | | | A | A | A | A | | | | | | |
| NC_011770 | 6426242 | G | A | 69 | 0 | 4 | 0 | | | A | A | A | A | | | | | | |
| NC_011770 | 6426251 | G | A | 70 | 0 | 4 | 0 | | | A | A | A | A | | | | | | |
| NC_011770 | 6426254 | A | G | 70 | 0 | 12 | 0 | G | G | G | G | G | G | G | G | G | G | G | G |
| NC_011770 | 6426262 | A | G | 73 | 0 | 4 | 0 | | | G | G | G | G | | | | | | |
| NC_011770 | 6426398 | G | A | 74 | 0 | 2 | 0 | | | | | | | | | | | A | A |
| NC_011770 | 6426427 | C | T | 75 | 0 | 4 | 0 | | | T | T | T | T | | | | | | |
| NC_011770 | 6426438 | A | G | 79 | 0 | 4 | 0 | | | G | G | G | G | | | | | | |
| NC_011770 | 6426467 | A | C | 80 | 0 | 4 | 0 | | | C | C | C | C | | | | | | |
| NC_011770 | 6426518 | G | A | 79 | 0 | 2 | 0 | | | | | | | | | A | A | | |
| NC_011770 | 6426616 | G | A | 71 | 0 | 4 | 0 | A | A | | | | | | | A | A | | |
| NC_011770 | 6426633 | A | C | 66 | 0 | 10 | 0 | C | C | T | T | T | T | C | C | C | C | | |
| NC_011770 | 6426634 | G | C | 66 | 0 | 4 | 0 | | | C | C | C | C | | | | | | |
| NC_011770 | 6426638 | A | C | 65 | 0 | 4 | 0 | | | C | C | C | C | | | | | | |
| NC_011770 | 6426640 | G | GCT | 66 | 0 | 4 | 0 | | | GCT | GCT | GCT | GCT | | | | | | |
| NC_011770 | 6426644 | A | T | 66 | 0 | 4 | 0 | | | T | T | T | T | | | | | | |
| NC_011770 | 6426657 | G | T | 74 | 0 | 4 | 0 | | | T | T | T | T | | | | | | |
| NC_011770 | 6426715 | T | C | 24 | 6 | 2 | 0 | | | . | . | . | . | | | C | C | . | . |
| NC_011770 | 6426734 | C | T | 24 | 6 | 2 | 0 | T | T | . | . | . | . | | | | | . | . |

| | | | | | | | | | | | | | | | | | | | |
|---|---|---|---|---|---|---|---|---|---|---|---|---|---|---|---|---|---|---|---|
| NC_011770 | 6426807 | T | C | 26 | 6 | 4 | 0 | C | C | . | . | . | . | | | C | C | . | . |
| NC_011770 | 6426832 | G | GC | 26 | 6 | 2 | 0 | GC | GC | . | . | . | . | | | | | . | . |
| NC_011770 | 6426840 | C | G | 44 | 0 | 6 | 0 | | | G | G | G | G | | | | | G | G |
| NC_011770 | 6426844 | C | G | 44 | 0 | 6 | 0 | | | G | G | G | G | | | | | G | G |
| NC_011770 | 6426886 | G | C | 24 | 6 | 2 | 0 | C | C | . | . | . | . | | | | | . | . |
| NC_011770 | 6426892 | T | C | 27 | 6 | 2 | 0 | | | . | . | . | . | | | C | C | . | . |
| NC_011770 | 6426912 | G | A | 23 | 6 | 4 | 0 | A | A | . | . | . | . | A | A | | | . | . |
| NC_011770 | 6426913 | C | A | 24 | 6 | 4 | 0 | A | A | . | . | . | . | A | A | | | . | . |
| NC_011770 | 6426957 | A | C | 26 | 5 | 1 | 0 | | | . | . | . | . | | | | | . | C |
| NC_011770 | 6426971 | T | A | 24 | 5 | 1 | 0 | | | . | . | . | . | | | | | . | A |
| NC_011770 | 6426972 | T | C | 25 | 5 | 3 | 0 | | | . | . | . | . | | | C | C | . | C |
| NC_011770 | 6426975 | T | C | 25 | 5 | 1 | 0 | | | . | . | . | . | | | | | . | C |
| NC_011770 | 6426990 | A | G | 24 | 5 | 1 | 0 | | | . | . | . | . | | | | | . | G |
| NC_011770 | 6426995 | G | T | 24 | 5 | 1 | 0 | | | . | . | . | . | | | | | . | T |
| NC_011770 | 6426998 | G | A | 23 | 6 | 2 | 0 | | | . | . | . | . | | | A | A | . | . |
| NC_011770 | 6426999 | CA | C | 23 | 5 | 1 | 0 | | | . | . | . | . | | | | | . | C |
| NC_011770 | 6427007 | G | C | 24 | 5 | 1 | 0 | | | . | . | . | . | | | | | . | C |
| NC_011770 | 6427008 | C | G | 25 | 5 | 1 | 0 | | | . | . | . | . | | | | | . | G |
| NC_011770 | 6427011 | G | A | 25 | 5 | 1 | 0 | | | . | . | . | . | | | | | . | A |
| NC_011770 | 6427013 | C | G | 25 | 5 | 1 | 0 | | | . | . | . | . | | | | | . | G |
| NC_011770 | 6427018 | C | G | 26 | 1 | 5 | 0 | | | . | G | G | G | | | | | G | G |
| NC_011770 | 6427043 | A | C | 65 | 0 | 8 | 0 | | | C | C | C | C | G | G | | | C | C |
| NC_011770 | 6427062 | A | C | 66 | 0 | 6 | 0 | | | C | C | C | C | | | | | C | C |
| NC_011770 | 5645071 | G | A | 85 | 0 | 1 | 0 | | | | | | | | A | | | | |
| NC_011770 | 6427065 | G | GC | 59 | 0 | 1 | 0 | | | | | | | GC | . | | | | |
| NC_011770 | 6427068 | C | G | 56 | 1 | 6 | 0 | | | G | G | G | G | . | | | | G | G |
| NC_011770 | 6427089 | C | T | 67 | 0 | 6 | 0 | | | T | T | T | T | | | | | T | T |
| NC_011770 | 6427111 | T | C | 60 | 0 | 6 | 0 | | | C | C | C | C | | | | | C | C |
| NC_011770 | 6427115 | G | T | 59 | 0 | 6 | 0 | | | T | T | T | T | | | | | T | T |
| NC_011770 | 6427125 | A | C | 57 | 0 | 6 | 0 | | | C | C | C | C | | | | | C | C |
| NC_011770 | 6427128 | A | G | 56 | 0 | 6 | 0 | | | G | G | G | G | | | | | G | G |
| NC_011770 | 6427143 | C | G | 55 | 0 | 6 | 0 | | | G | G | G | G | | | | | G | G |
| NC_011770 | 6427233 | A | G | 28 | 6 | 2 | 0 | | | . | . | . | . | | | G | G | . | . |
| NC_011770 | 6427272 | A | G | 79 | 0 | 6 | 0 | | | G | G | G | G | | | | | G | G |
| NC_011770 | 6427291 | G | A | 78 | 0 | 6 | 0 | | | A | A | A | A | | | | | A | A |
| NC_011770 | 6427302 | A | G | 76 | 0 | 6 | 0 | | | G | G | G | G | | | | | G | G |
| NC_011770 | 6427306 | A | G | 77 | 0 | 6 | 0 | | | G | G | G | G | | | | | G | G |
| NC_011770 | 6427308 | A | G | 77 | 0 | 6 | 0 | | | G | G | G | G | | | | | G | G |
| NC_011770 | 6427313 | A | T | 76 | 0 | 6 | 0 | | | T | T | T | T | | | | | T | T |
| NC_011770 | 6427316 | T | C | 75 | 0 | 6 | 0 | | | C | C | C | C | | | | | C | C |
| NC_011770 | 6427331 | C | G | 74 | 0 | 6 | 0 | | | G | G | G | G | | | | | G | G |
| NC_011770 | 6427334 | C | A | 74 | 0 | 6 | 0 | | | A | A | A | A | | | | | A | A |
| NC_011770 | 6427335 | A | G | 73 | 0 | 6 | 0 | | | G | G | G | G | | | | | G | G |
| NC_011770 | 6427349 | C | A | 74 | 0 | 6 | 0 | | | A | A | A | A | | | | | A | A |
| NC_011770 | 6427355 | G | C | 72 | 0 | 6 | 0 | | | C | C | C | C | | | | | C | C |
| NC_011770 | 6427367 | T | C | 56 | 0 | 8 | 0 | | | C | C | C | C | | | G | G | C | C |
| NC_011770 | 6427372 | T | A | 55 | 0 | 6 | 0 | | | A | A | A | A | | | | | A | A |
| NC_011770 | 6427373 | C | G | 55 | 0 | 6 | 0 | | | G | G | G | G | | | | | G | G |
| NC_011770 | 6427378 | A | G | 55 | 0 | 6 | 0 | | | G | G | G | G | | | | | G | G |
| NC_011770 | 6427384 | T | G | 52 | 0 | 6 | 0 | | | G | G | G | G | | | | | G | G |
| NC_011770 | 6427387 | C | G | 52 | 0 | 6 | 0 | | | G | G | G | G | | | | | G | G |
| NC_011770 | 6427388 | C | G | 52 | 0 | 6 | 0 | | | G | G | G | G | | | | | G | G |
| NC_011770 | 6427391 | A | C | 53 | 0 | 6 | 0 | | | C | C | C | C | | | | | C | C |
| NC_011770 | 6427392 | C | T | 53 | 0 | 6 | 0 | | | T | T | T | T | | | | | T | T |
| NC_011770 | 6427394 | G | T | 53 | 0 | 6 | 0 | | | T | T | T | T | | | | | T | T |
| NC_011770 | 6427400 | T | C | 53 | 0 | 6 | 0 | | | C | C | C | C | | | | | C | C |
| NC_011770 | 6427402 | C | A | 53 | 0 | 6 | 0 | | | A | A | A | A | | | | | A | A |
| NC_011770 | 6427406 | C | T | 55 | 0 | 6 | 0 | | | T | T | T | T | | | | | T | T |
| NC_011770 | 6427423 | T | G | 73 | 0 | 6 | 0 | | | G | G | G | G | | | | | G | G |
| NC_011770 | 6427428 | T | C | 73 | 0 | 6 | 0 | | | C | C | C | C | | | | | C | C |
| NC_011770 | 6427429 | C | T | 73 | 0 | 6 | 0 | | | T | T | T | T | | | | | T | T |
| NC_011770 | 6427459 | C | G | 51 | 0 | 6 | 0 | | | G | G | G | G | | | | | G | G |

| | | | | | | | | | | | | | | | | | | | |
|---|---|---|---|---|---|---|---|---|---|---|---|---|---|---|---|---|---|---|---|
| NC_011770 | 6427460 | T | G | 50 | 0 | 6 | 0 | | | G | G | G | G | | | | | G | G |
| NC_011770 | 6427465 | G | C | 50 | 0 | 4 | 0 | | | C | C | C | C | | | | | | |
| NC_011770 | 6427466 | A | G | 49 | 0 | 6 | 0 | | | G | G | G | G | | | | | G | G |
| NC_011770 | 6427467 | T | C | 49 | 0 | 6 | 0 | | | C | C | C | C | | | | | C | C |
| NC_011770 | 6427468 | T | C | 50 | 0 | 6 | 0 | | | C | C | C | C | | | | | C | C |
| NC_011770 | 6427471 | T | A | 50 | 0 | 6 | 0 | | | A | A | A | A | | | | | A | A |
| NC_011770 | 6427472 | T | A | 50 | 0 | 6 | 0 | | | A | A | A | A | | | | | A | A |
| NC_011770 | 6427475 | A | G | 50 | 0 | 6 | 0 | | | G | G | G | G | | | | | G | G |
| NC_011770 | 6427476 | T | C | 51 | 0 | 6 | 0 | | | C | C | C | C | | | | | C | C |
| NC_011770 | 6427480 | C | T | 51 | 0 | 6 | 0 | | | T | T | T | T | | | | | T | T |
| NC_011770 | 6427487 | A | C | 51 | 0 | 6 | 0 | | | C | C | C | C | | | | | C | C |
| NC_011770 | 6427489 | T | A | 51 | 0 | 6 | 0 | | | A | A | A | A | | | | | A | A |
| NC_011770 | 6427495 | A | G | 52 | 0 | 6 | 0 | | | G | G | G | G | | | | | G | G |
| NC_011770 | 6427499 | A | G | 53 | 0 | 6 | 0 | | | G | G | G | G | | | | | G | G |
| NC_011770 | 6427542 | T | G | 67 | 0 | 6 | 0 | | | G | G | G | G | | | | | G | G |
| NC_011770 | 6427547 | T | G | 67 | 0 | 6 | 0 | | | G | G | G | G | | | | | G | G |
| NC_011770 | 6427550 | A | G | 65 | 0 | 6 | 0 | | | G | G | G | G | | | | | G | G |
| NC_011770 | 6427552 | T | A | 66 | 0 | 6 | 0 | | | A | A | A | A | | | | | A | A |
| NC_011770 | 6427558 | G | C | 65 | 0 | 6 | 0 | | | C | C | C | C | | | | | C | C |
| NC_011770 | 6427572 | C | T | 26 | 5 | 1 | 0 | | | . | . | . | T | | | | | . | . |
| NC_011770 | 6427574 | GT | G | 25 | 5 | 1 | 0 | | | . | . | . | G | | | | | . | . |
| NC_011770 | 6427593 | T | C | 57 | 0 | 6 | 0 | | | C | C | C | C | | | | | C | C |
| NC_011770 | 6427596 | C | T | 57 | 0 | 6 | 0 | | | T | T | T | T | | | | | T | T |
| NC_011770 | 6427597 | A | C | 57 | 0 | 6 | 0 | | | C | C | C | C | | | | | C | C |
| NC_011770 | 6427602 | C | T | 54 | 0 | 6 | 0 | | | T | T | T | T | | | | | T | T |
| NC_011770 | 6427605 | G | C | 56 | 0 | 6 | 0 | | | C | C | C | C | | | | | C | C |
| NC_011770 | 6427607 | A | C | 56 | 0 | 6 | 0 | | | C | C | C | C | | | | | C | C |
| NC_011770 | 6427608 | G | A | 55 | 0 | 6 | 0 | | | A | A | A | A | | | | | A | A |
| NC_011770 | 6427617 | A | G | 76 | 0 | 6 | 0 | | | G | G | G | G | | | | | G | G |
| NC_011770 | 6427640 | G | T | 77 | 0 | 6 | 0 | | | T | T | T | T | | | | | T | T |
| NC_011770 | 6427648 | C | T | 77 | 0 | 6 | 0 | | | T | T | T | T | | | | | T | T |
| NC_011770 | 6427654 | G | T | 79 | 0 | 6 | 0 | | | T | T | T | T | | | | | T | T |
| NC_011770 | 6427660 | C | G | 74 | 0 | 6 | 0 | | | G | G | G | G | | | | | G | G |
| NC_011770 | 6427663 | T | C | 74 | 0 | 6 | 0 | | | C | C | C | C | | | | | C | C |
| NC_011770 | 6427664 | G | T | 74 | 0 | 6 | 0 | | | T | T | T | T | | | | | T | T |
| NC_011770 | 6427700 | G | A | 81 | 0 | 6 | 0 | | | A | A | A | A | | | | | A | A |
| NC_011770 | 6427707 | G | A | 81 | 0 | 6 | 0 | | | A | A | A | A | | | | | A | A |
| NC_011770 | 6427718 | G | A | 81 | 0 | 6 | 0 | | | A | A | A | A | | | | | A | A |
| NC_011770 | 6427725 | T | C | 81 | 0 | 6 | 0 | | | C | C | C | C | | | | | C | C |
| NC_011770 | 6427728 | C | G | 82 | 0 | 6 | 0 | | | G | G | G | G | | | | | G | G |
| NC_011770 | 6427762 | G | A | 85 | 0 | 6 | 0 | | | A | A | A | A | | | | | A | A |
| NC_011770 | 6427763 | T | C | 85 | 0 | 6 | 0 | | | C | C | C | C | | | | | C | C |
| NC_011770 | 6427785 | A | C | 83 | 0 | 6 | 0 | | | C | C | C | C | | | | | C | C |
| NC_011770 | 6427792 | A | C | 75 | 0 | 6 | 0 | | | C | C | C | C | | | | | C | C |
| NC_011770 | 6427796 | C | T | 75 | 0 | 6 | 0 | | | T | T | T | T | | | | | T | T |
| NC_011770 | 6427798 | C | G | 75 | 0 | 6 | 0 | | | G | G | G | G | | | | | G | G |
| NC_011770 | 6427801 | C | T | 75 | 0 | 6 | 0 | | | T | T | T | T | | | | | T | T |
| NC_011770 | 6427802 | T | G | 75 | 0 | 6 | 0 | | | G | G | G | G | | | | | G | G |
| NC_011770 | 6427811 | G | A | 76 | 0 | 6 | 0 | | | A | A | A | A | | | | | A | A |
| NC_011770 | 6427832 | G | A | 56 | 0 | 6 | 0 | | | A | A | A | A | | | | | A | A |
| NC_011770 | 6427833 | G | A | 54 | 0 | 6 | 0 | | | A | A | A | A | | | | | A | A |
| NC_011770 | 6427835 | G | A | 53 | 0 | 6 | 0 | | | A | A | A | A | | | | | A | A |
| NC_011770 | 6427836 | T | C | 54 | 0 | 6 | 0 | | | C | C | C | C | | | | | C | C |
| NC_011770 | 6427838 | T | C | 55 | 0 | 6 | 0 | | | C | C | C | C | | | | | C | C |
| NC_011770 | 6427840 | G | C | 55 | 0 | 6 | 0 | | | C | C | C | C | | | | | C | C |
| NC_011770 | 6427845 | G | A | 55 | 0 | 2 | 0 | | | | | | | | | | | A | A |
| NC_011770 | 6427849 | C | T | 56 | 0 | 6 | 0 | | | T | T | T | T | | | | | T | T |
| NC_011770 | 6427853 | A | G | 56 | 0 | 6 | 0 | | | G | G | G | G | | | | | G | G |
| NC_011770 | 6427857 | A | G | 56 | 0 | 6 | 0 | | | G | G | G | G | | | | | G | G |
| NC_011770 | 6427858 | C | T | 55 | 0 | 6 | 0 | | | T | T | T | T | | | | | T | T |
| NC_011770 | 6427862 | C | G | 55 | 0 | 6 | 0 | | | G | G | G | G | | | | | G | G |
| NC_011770 | 6427864 | A | C | 55 | 0 | 6 | 0 | | | C | C | C | C | | | | | C | C |

| | | | | | | | | | | | | | | | | | | | |
|---|---|---|---|---|---|---|---|---|---|---|---|---|---|---|---|---|---|---|---|
| NC_011770 | 6427867 | G | A | 55 | 0 | 2 | 0 | | | | | | | | | A | A | | |
| NC_011770 | 6427868 | A | G | 54 | 0 | 6 | 0 | | | G | G | G | G | | | | | G | G |
| NC_011770 | 6427919 | T | G | 89 | 0 | 6 | 0 | | | G | G | G | G | | | | | G | G |
| NC_011770 | 6427920 | T | A | 89 | 0 | 6 | 0 | | | A | A | A | A | | | | | A | A |
| NC_011770 | 6427923 | G | A | 90 | 0 | 8 | 0 | | | A | A | A | A | | | A | A | A | A |
| NC_011770 | 6427943 | C | G | 85 | 0 | 6 | 0 | | | G | G | G | G | | | | | G | G |
| NC_011770 | 6427944 | G | T | 84 | 0 | 6 | 0 | | | T | T | T | T | | | | | T | T |
| NC_011770 | 6427945 | C | T | 85 | 0 | 6 | 0 | | | T | T | T | T | | | | | T | T |
| NC_011770 | 6427949 | G | A | 88 | 0 | 6 | 0 | | | A | A | A | A | | | | | A | A |
| NC_011770 | 6427951 | G | A | 87 | 0 | 6 | 0 | | | A | A | A | A | | | | | A | A |
| NC_011770 | 6427956 | A | G | 86 | 0 | 8 | 0 | G | G | G | G | G | G | | | | | G | G |
| NC_011770 | 5944545 | C | CGAA | 42 | 0 | 1 | 0 | | | | | | | | | | | | CGAA |
| NC_011770 | 6429018 | C | CGGA | 75 | 0 | 2 | 0 | CGGA | CGGA | | | | | | | | | | |
| NC_011770 | 6429050 | A | G | 71 | 0 | 2 | 0 | G | G | | | | | | | | | | |
| NC_011770 | 6429059 | T | C | 68 | 0 | 2 | 0 | C | C | | | | | | | | | | |
| NC_011770 | 6429065 | C | T | 69 | 0 | 2 | 0 | T | T | | | | | | | | | | |
| NC_011770 | 6429070 | G | A | 71 | 0 | 4 | 0 | | | A | A | A | A | | | | | | |
| NC_011770 | 6429073 | C | G | 71 | 0 | 2 | 0 | G | G | | | | | | | | | | |
| NC_011770 | 6429744 | G | A | 87 | 0 | 2 | 0 | | | | | | | A | A | | | | |
| NC_011770 | 6429799 | T | A | 85 | 0 | 2 | 0 | | | | | | | A | A | | | | |
| NC_011770 | 6432577 | G | T | 69 | 0 | 10 | 0 | T | T | T | T | T | T | T | T | | | T | T |
| NC_011770 | 6432583 | A | G | 69 | 0 | 10 | 0 | G | G | G | G | G | G | G | G | | | G | G |
| NC_011770 | 2228407 | CT | C | 91 | 0 | 9 | 0 | C | C | C | C | C | C | | C | | C | | C |
| NC_011770 | 6053116 | C | T | 65 | 0 | 1 | 0 | | | | | | | | | | | | T |
| NC_011770 | 6435975 | A | G | 68 | 0 | 6 | 0 | | | G | G | G | G | G | G | | | | |
| NC_011770 | 6435976 | T | C | 68 | 0 | 4 | 0 | | | C | C | C | C | | | | | | |
| NC_011770 | 6435977 | G | T | 67 | 0 | 6 | 0 | | | T | T | T | T | T | T | | | | |
| NC_011770 | 6435981 | T | C | 67 | 0 | 6 | 0 | | | C | C | C | C | C | C | | | | |
| NC_011770 | 6435999 | T | C | 71 | 0 | 6 | 0 | | | C | C | C | C | C | C | | | | |
| NC_011770 | 6437672 | A | T | 88 | 0 | 4 | 0 | | | | | | | | | T | T | T | T |
| NC_011770 | 6439010 | T | A | 67 | 0 | 8 | 0 | | | A | A | A | A | | | A | A | A | A |
| NC_011770 | 6439032 | C | T | 70 | 0 | 6 | 0 | | | T | T | T | T | | | T | T | | |
| NC_011770 | 6439045 | G | T | 69 | 0 | 8 | 0 | | | T | T | T | T | | | T | T | T | T |
| NC_011770 | 6439056 | C | T | 67 | 0 | 10 | 0 | | | T | T | T | T | T | T | T | T | T | T |
| NC_011770 | 6441051 | G | A | 102 | 0 | 2 | 0 | | | | | | | | | | | A | A |
| NC_011770 | 6441071 | A | G | 101 | 0 | 6 | 0 | G | G | G | G | G | G | | | | | | |
| NC_011770 | 5784565 | G | A | 73 | 0 | 1 | 0 | | | | | | | | A | | | | |
| NC_011770 | 6115483 | G | A | 46 | 0 | 1 | 0 | | | | | | | | | | | | A |
| NC_011770 | 6446642 | A | G | 79 | 0 | 4 | 0 | G | G | | | | | | | | | G | G |
| NC_011770 | 6446959 | G | A | 75 | 0 | 2 | 0 | | | | | | | A | A | | | | |
| NC_011770 | 6449715 | G | A | 79 | 0 | 2 | 0 | | | | | | | A | A | | | | |
| NC_011770 | 6449743 | G | A | 79 | 0 | 2 | 0 | | | | | | | | | | | A | A |
| NC_011770 | 6450969 | G | A | 79 | 0 | 2 | 0 | | | | | | | A | A | | | | |
| NC_011770 | 5901370 | C | CG | 74 | 0 | 1 | 0 | | | | CG | | | | | | | | |
| NC_011770 | 6452340 | G | A | 73 | 0 | 2 | 0 | | | | | | | | | | | A | A |
| NC_011770 | 5845532 | GT | G | 79 | 0 | 1 | 0 | | G | | | | | | | | | | |
| NC_011770 | 2082047 | G | GT | 71 | 0 | 9 | 0 | GT | GT | | GT | | GT | | GT | GT | GT | GT | GT |
| NC_011770 | 6455860 | G | T | 77 | 0 | 2 | 0 | T | T | | | | | | | | | | |
| NC_011770 | 6456839 | A | C | 82 | 0 | 12 | 0 | C | C | C | C | C | C | C | C | C | C | C | C |
| NC_011770 | 6456902 | G | A | 83 | 0 | 2 | 0 | | | | | | | A | A | | | | |
| NC_011770 | 6457057 | A | G | 83 | 0 | 4 | 0 | | | | | | | G | G | | | G | G |
| NC_011770 | 6457087 | G | A | 81 | 0 | 4 | 0 | | | A | A | A | A | | | | | | |
| NC_011770 | 5921617 | GT | G | 85 | 0 | 1 | 0 | G | | | | | | | | | | | |
| NC_011770 | 6457995 | C | T | 90 | 0 | 2 | 0 | T | T | | | | | | | | | | |
| NC_011770 | 6184321 | G | A | 81 | 0 | 1 | 0 | | | | | | | | | | | | A |
| NC_011770 | 6461099 | C | T | 86 | 0 | 8 | 0 | T | T | | | | | T | T | T | T | T | T |
| NC_011770 | 6461126 | A | G | 82 | 0 | 12 | 0 | G | G | G | G | G | G | G | G | G | G | G | G |
| NC_011770 | 6461150 | A | G | 80 | 0 | 8 | 0 | G | G | | | | | G | G | G | G | G | G |
| NC_011770 | 6461180 | A | G | 85 | 0 | 8 | 0 | G | G | | | | | G | G | G | G | G | G |
| NC_011770 | 6461462 | G | A | 91 | 0 | 8 | 0 | A | A | | | | | A | A | A | A | A | A |
| NC_011770 | 6461477 | T | C | 91 | 0 | 8 | 0 | C | C | | | | | C | C | C | C | C | C |
| NC_011770 | 3313959 | C | CG | 69 | 0 | 3 | 0 | | CG | | | | | | CG | | CG | | |

| | | | | | | | | | | | | | | | | | | | |
|---|---|---|---|---|---|---|---|---|---|---|---|---|---|---|---|---|---|---|---|
| NC_011770 | 6466920 | G | A | 71 | 0 | 12 | 0 | A | A | A | A | A | A | A | A | A | A | A | A |
| NC_011770 | 6466952 | A | C | 72 | 0 | 2 | 0 | C | C | | | | | | | | | | |
| NC_011770 | 6469199 | C | G | 85 | 0 | 4 | 0 | G | G | | | | | | | G | G | | |
| NC_011770 | 6288795 | G | C | 87 | 0 | 1 | 0 | C | | | | | | | | | | | |
| NC_011770 | 6469658 | A | C | 88 | 0 | 12 | 0 | C | C | C | C | C | C | C | C | C | C | C | C |
| NC_011770 | 6471273 | T | C | 83 | 0 | 2 | 0 | | | | | | | | | C | C | | |
| NC_011770 | 6471345 | G | C | 78 | 0 | 2 | 0 | | | | | | | | | | | C | C |
| NC_011770 | 6471362 | C | T | 80 | 0 | 2 | 0 | T | T | | | | | | | | | | |
| NC_011770 | 6471379 | G | C | 81 | 0 | 2 | 0 | | | | | | | C | C | | | | |
| NC_011770 | 6471382 | T | C | 82 | 0 | 12 | 0 | C | C | C | C | C | C | C | C | C | C | C | C |
| NC_011770 | 6471391 | C | T | 82 | 0 | 2 | 0 | | | | | | | T | T | | | | |
| NC_011770 | 6471416 | T | C | 87 | 0 | 2 | 0 | | | | | | | | | C | C | | |
| NC_011770 | 6471841 | C | T | 94 | 0 | 4 | 0 | | | | | | | T | T | | | T | T |
| NC_011770 | 6479566 | C | G | 93 | 0 | 2 | 0 | | | | | | | G | G | | | | |
| NC_011770 | 6479583 | A | G | 88 | 0 | 12 | 0 | G | G | G | G | G | G | G | G | G | G | G | G |
| NC_011770 | 6479728 | G | A | 92 | 0 | 6 | 0 | A | A | | | | | | | A | A | A | A |
| NC_011770 | 6479857 | C | T | 89 | 0 | 12 | 0 | T | T | T | T | T | T | T | T | T | T | T | T |
| NC_011770 | 6184391 | C | T | 83 | 0 | 1 | 0 | | | | | | | | | | | | T |
| NC_011770 | 6375500 | C | T | 65 | 0 | 3 | 0 | T | T | | | | | | | | | | T |
| NC_011770 | 6480937 | G | A | 85 | 0 | 2 | 0 | | | | | | | | | A | A | | |
| NC_011770 | 6480967 | C | A | 94 | 0 | 2 | 0 | | | | | | | A | A | | | | |
| NC_011770 | 6481165 | G | A | 83 | 0 | 4 | 0 | A | A | | | | | | | A | A | | |
| NC_011770 | 6481175 | C | G | 81 | 0 | 2 | 0 | | | | | | | G | G | | | | |
| NC_011770 | 6100856 | AG | A | 70 | 0 | 1 | 0 | | | | A | | | | | | | | |
| NC_011770 | 5999628 | GA | G | 36 | 0 | 5 | 0 | | G | | | | | G | G | G | G | | |
| NC_011770 | 6106846 | A | G | 75 | 0 | 1 | 0 | | | | | | | | G | | | | |
| NC_011770 | 6487073 | A | G | 41 | 0 | 4 | 0 | | | G | G | G | G | | | | | | |
| NC_011770 | 6487884 | G | A | 61 | 0 | 4 | 0 | | | A | A | A | A | | | | | | |
| NC_011770 | 6487904 | T | C | 63 | 0 | 2 | 0 | | | | | | | | | | | C | C |
| NC_011770 | 6487976 | C | T | 74 | 0 | 2 | 0 | | | | | | | | | | | T | T |
| NC_011770 | 6490616 | CATG | C | 81 | 0 | 2 | 0 | | | | | | | | | | | C | C |
| NC_011770 | 6490622 | G | C | 77 | 0 | 12 | 0 | C | C | C | C | C | C | C | C | C | C | C | C |
| NC_011770 | 6490626 | G | A | 79 | 0 | 10 | 0 | A | A | A | A | A | A | A | A | A | A | | |
| NC_011770 | 6490635 | C | T | 81 | 0 | 10 | 0 | T | T | T | T | T | T | T | T | T | T | | |
| NC_011770 | 6490690 | C | G | 79 | 0 | 2 | 0 | | | | | | | | | | | G | G |
| NC_011770 | 6195541 | G | GC | 63 | 0 | 1 | 0 | | | | | | | | GC | | | | |
| NC_011770 | 1570776 | A | AG | 50 | 0 | 4 | 0 | | | | | | | | AG | AG | AG | AG | |
| NC_011770 | 6126844 | G | A | 87 | 0 | 1 | 0 | | | | A | | | | | | | | |
| NC_011770 | 6492502 | T | C | 50 | 0 | 6 | 0 | | | | | | | C | C | C | C | C | C |
| NC_011770 | 6492732 | A | G | 50 | 0 | 12 | 0 | G | G | G | G | G | G | G | G | G | G | G | G |
| NC_011770 | 6496806 | G | A | 86 | 0 | 4 | 0 | | | | | | | A | A | A | A | | |
| NC_011770 | 6496899 | A | G | 94 | 0 | 4 | 0 | | | | | | | G | G | G | G | | |
| NC_011770 | 6496923 | C | T | 93 | 0 | 2 | 0 | | | | | | | | | | | T | T |
| NC_011770 | 6496941 | A | G | 91 | 0 | 4 | 0 | | | | | | | G | G | G | G | | |
| NC_011770 | 6496964 | T | C | 90 | 0 | 4 | 0 | | | | | | | C | C | C | C | | |
| NC_011770 | 6496969 | A | G | 90 | 0 | 4 | 0 | | | | | | | G | G | G | G | | |
| NC_011770 | 6381089 | G | A | 56 | 0 | 3 | 0 | A | A | | | A | | | | | | | |
| NC_011770 | 6503438 | C | G | 65 | 0 | 2 | 0 | G | G | | | | | | | | | | |
| NC_011770 | 6503492 | G | A | 64 | 0 | 2 | 0 | | | | | | | | | | | A | A |
| NC_011770 | 6503496 | T | C | 63 | 0 | 2 | 0 | | | | | | | C | C | | | | |
| NC_011770 | 6503533 | C | T | 62 | 0 | 2 | 0 | T | T | | | | | | | | | | |
| NC_011770 | 6503548 | A | C | 59 | 0 | 2 | 0 | | | | | | | C | C | | | | |
| NC_011770 | 6503553 | C | CG | 55 | 0 | 2 | 0 | | | | | | | CG | CG | | | | |
| NC_011770 | 6503554 | A | C | 52 | 0 | 2 | 0 | | | | | | | C | C | | | | |
| NC_011770 | 6503598 | A | G | 57 | 0 | 2 | 0 | | | | | | | G | G | | | | |
| NC_011770 | 6503604 | C | T | 59 | 0 | 2 | 0 | | | | | | | T | T | | | | |
| NC_011770 | 6214672 | C | G | 64 | 0 | 1 | 0 | | | | G | | | | | | | | |
| NC_011770 | 6507575 | G | A | 91 | 0 | 2 | 0 | | | | | | | | | A | A | | |
| NC_011770 | 6507607 | A | G | 88 | 0 | 4 | 0 | | | G | G | G | G | | | | | | |
| NC_011770 | 6185049 | A | AC | 67 | 0 | 1 | 0 | | AC | | | | | | | | | | |
| NC_011770 | 6507904 | G | C | 70 | 0 | 8 | 0 | | | C | C | C | C | | | C | C | C | C |
| NC_011770 | 6507938 | G | A | 71 | 0 | 6 | 0 | | | A | A | A | A | | | | | A | A |

| | | | | | | | | | | | | | | | | | | | |
|---|---|---|---|---|---|---|---|---|---|---|---|---|---|---|---|---|---|---|---|
| NC_011770 | 6507985 | A | T | 59 | 0 | 10 | 0 | T | T | T | T | T | T | | | T | T | T | T |
| NC_011770 | 6508107 | T | G | 68 | 0 | 2 | 0 | | | | | | | | | | | G | G |
| NC_011770 | 6508843 | A | G | 79 | 0 | 4 | 0 | | | | | | | | | G | G | G | G |
| NC_011770 | 6508876 | A | AT | 77 | 0 | 4 | 0 | | | AT | AT | AT | AT | | | | | | |
| NC_011770 | 6508917 | G | C | 74 | 0 | 4 | 0 | | | | | | | | | C | C | C | C |
| NC_011770 | 6508945 | C | T | 71 | 0 | 4 | 0 | | | | | | | | | T | T | T | T |
| NC_011770 | 6508980 | T | C | 76 | 0 | 1 | 0 | | | | | | | C | | | | | |
| NC_011770 | 6425865 | G | A | 79 | 0 | 1 | 0 | | | | | | | | | | | | A |
| NC_011770 | 6509146 | A | G | 73 | 0 | 8 | 0 | | | G | G | G | G | | | G | G | G | G |
| NC_011770 | 5839062 | G | A | 77 | 0 | 1 | 0 | | | | | | | | | | | A | |
| NC_011770 | 6241569 | G | A | 83 | 0 | 1 | 0 | | A | | | | | | | | | | |
| NC_011770 | 6513222 | G | GAA | 89 | 0 | 1 | 0 | | | | | | | | | | GAA | | |
| NC_011770 | 6513237 | T | C | 87 | 0 | 2 | 0 | | | | | | | | | C | C | | |
| NC_011770 | 6513913 | G | A | 85 | 0 | 2 | 0 | A | A | | | | | | | | | | |
| NC_011770 | 6458001 | G | A | 92 | 0 | 1 | 0 | | | | | | | | | | | | A |
| NC_011770 | 6518618 | A | AG | 77 | 0 | 2 | 0 | | | | | | | | | | | AG | AG |
| NC_011770 | 6512422 | C | CTTT | 92 | 0 | 9 | 0 | | CTTT | CTTT | CTTT | CTTT | CTTT | | | CTTT | CTTT | CTTT | CTTT |
| NC_011770 | 6283084 | C | T | 84 | 0 | 1 | 0 | | | | | | | | T | | | | |
| NC_011770 | 6522815 | T | C | 78 | 0 | 4 | 0 | | | C | C | C | C | | | | | | |
| NC_011770 | 6522824 | T | G | 77 | 0 | 2 | 0 | | | | | | | G | G | | | | |
| NC_011770 | 6523812 | G | A | 65 | 0 | 2 | 0 | | | | | | | | | A | A | | |
| NC_011770 | 6523861 | T | C | 58 | 0 | 4 | 0 | | | | | | | C | C | | | C | C |
| NC_011770 | 6523894 | T | C | 47 | 0 | 2 | 0 | | | | | | | | | | | C | C |
| NC_011770 | 6523908 | C | A | 46 | 0 | 2 | 0 | | | | | | | | | | | A | A |
| NC_011770 | 6523943 | A | C | 36 | 0 | 2 | 0 | | | | | | | | | C | C | | |
| NC_011770 | 6523944 | T | C | 36 | 0 | 2 | 0 | | | | | | | | | C | C | | |
| NC_011770 | 6523946 | G | T | 35 | 0 | 2 | 0 | | | | | | | | | T | T | | |
| NC_011770 | 6523947 | A | C | 36 | 0 | 2 | 0 | | | | | | | | | C | C | | |
| NC_011770 | 6523953 | C | G | 34 | 0 | 12 | 0 | G | G | G | G | G | G | G | G | G | G | G | G |
| NC_011770 | 6523963 | C | T | 29 | 2 | 4 | 0 | T | T | | | | | | | T | T | . | . |
| NC_011770 | 6523971 | T | G | 27 | 2 | 2 | 0 | | | | | | | | | G | G | . | . |
| NC_011770 | 6523976 | G | A | 25 | 2 | 4 | 0 | | | A | A | A | A | | | | | . | . |
| NC_011770 | 6523978 | G | A | 25 | 2 | 2 | 0 | | | | | | | | | A | A | . | . |
| NC_011770 | 6523986 | C | T | 22 | 2 | 4 | 0 | | | T | T | T | T | | | | | . | . |
| NC_011770 | 6523998 | G | A | 21 | 2 | 2 | 0 | A | A | | | | | | | | | . | . |
| NC_011770 | 6523999 | A | G | 20 | 2 | 2 | 0 | G | G | | | | | | | | | . | . |
| NC_011770 | 6524003 | T | C | 19 | 2 | 2 | 0 | C | C | | | | | | | | | . | . |
| NC_011770 | 6524018 | A | AT | 16 | 2 | 2 | 0 | AT | AT | . | | . | . | | | | | . | . |
| NC_011770 | 6524060 | C | A | 16 | 6 | 1 | 0 | . | . | . | . | . | A | | | | | . | . |
| NC_011770 | 6524061 | G | C | 15 | 6 | 2 | 0 | . | . | . | C | . | C | | | | | . | . |
| NC_011770 | 6524081 | G | GA | 18 | 3 | 5 | 0 | . | GA | GA | GA | GA | GA | | | | | . | . |
| NC_011770 | 6524111 | C | T | 21 | 2 | 5 | 0 | . | T | T | T | T | T | | | | | . | . |
| NC_011770 | 6524127 | C | T | 28 | 2 | 2 | 0 | | | | | | | | | T | T | . | . |
| NC_011770 | 6524146 | GGC | G | 31 | 1 | 1 | 0 | | | | | | | | | | | G | . |
| NC_011770 | 6524152 | T | C | 29 | 1 | 1 | 0 | | | | | | | | | | | C | . |
| NC_011770 | 6524160 | C | A | 35 | 0 | 6 | 0 | A | A | A | A | A | A | | | | | | |
| NC_011770 | 6524161 | C | T | 35 | 0 | 6 | 0 | T | T | T | T | T | T | | | | | | |
| NC_011770 | 6524163 | T | G | 36 | 0 | 6 | 0 | G | G | G | G | G | G | | | | | | |
| NC_011770 | 6524164 | C | A | 36 | 0 | 6 | 0 | A | A | A | A | A | A | | | | | | |
| NC_011770 | 6524169 | C | A | 38 | 0 | 2 | 0 | | | | | | | | | | | A | A |
| NC_011770 | 6524170 | G | C | 37 | 0 | 2 | 0 | | | | | | | | | | | C | C |
| NC_011770 | 6524180 | C | T | 35 | 0 | 2 | 0 | T | T | | | | | | | | | | |
| NC_011770 | 6524188 | T | C | 38 | 0 | 2 | 0 | | | | | | | | | | | C | C |
| NC_011770 | 6524190 | G | GA | 40 | 0 | 2 | 0 | | | | | | | | | | | GA | GA |
| NC_011770 | 6524193 | A | G | 39 | 0 | 6 | 0 | G | G | | | | | | | G | G | G | G |
| NC_011770 | 6524203 | T | C | 44 | 0 | 6 | 0 | C | C | | | | | | | C | C | C | C |
| NC_011770 | 6524215 | G | A | 49 | 0 | 2 | 0 | A | A | | | | | | | | | | |
| NC_011770 | 6524216 | A | G | 48 | 0 | 2 | 0 | G | G | | | | | | | | | | |
| NC_011770 | 6524220 | T | C | 47 | 0 | 6 | 0 | C | C | | | | | | | C | C | C | C |
| NC_011770 | 6524234 | C | A | 51 | 0 | 2 | 0 | | | | | | | | | | | A | A |
| NC_011770 | 6524236 | C | T | 54 | 0 | 4 | 0 | T | T | | | | | | | T | T | | |
| NC_011770 | 6524241 | A | G | 48 | 0 | 12 | 0 | G | G | G | G | G | G | G | G | G | G | G | G |

| | | | | | | | | | | | | | | | | | | | |
|---|---|---|---|---|---|---|---|---|---|---|---|---|---|---|---|---|---|---|---|
| NC_011770 | 6524267 | G | GCC | 62 | 0 | 2 | 0 | | | | | | | | | GCC | GCC | | |
| NC_011770 | 6524271 | TG | T | 62 | 0 | 2 | 0 | | | | | | | | | T | T | | |
| NC_011770 | 6524290 | T | G | 63 | 0 | 4 | 0 | | | | | | | G | G | G | G | | |
| NC_011770 | 6524291 | C | A | 63 | 0 | 4 | 0 | | | | | | | A | A | A | A | | |
| NC_011770 | 6524293 | A | C | 63 | 0 | 4 | 0 | | | | | | | C | C | C | C | | |
| NC_011770 | 6524295 | T | G | 64 | 0 | 4 | 0 | | | | | | | G | G | G | G | | |
| NC_011770 | 6166682 | G | A | 71 | 0 | 1 | 0 | | | | | | | | | | | A | |
| NC_011770 | 6527229 | G | A | 90 | 0 | 4 | 0 | | | A | A | A | A | | | | | | |
| NC_011770 | 6527261 | C | T | 96 | 0 | 2 | 0 | T | T | | | | | | | | | | |
| NC_011770 | 6527306 | A | G | 91 | 0 | 8 | 0 | G | G | G | G | G | G | | | | | G | G |
| NC_011770 | 6375327 | GC | G | 64 | 0 | 1 | 0 | | | | | | | | | | | G | |
| NC_011770 | 6531236 | A | G | 74 | 0 | 6 | 0 | | | G | G | G | G | | | | | G | G |
| NC_011770 | 6531238 | C | T | 75 | 0 | 2 | 0 | | | | | | | | | | | T | T |
| NC_011770 | 6531244 | T | A | 78 | 0 | 4 | 0 | | | A | A | A | A | | | | | | |
| NC_011770 | 6531247 | C | T | 78 | 0 | 2 | 0 | | | | | | | | | | | T | T |
| NC_011770 | 6531280 | C | A | 70 | 0 | 4 | 0 | | | A | A | A | A | | | | | | |
| NC_011770 | 6531281 | A | G | 72 | 0 | 4 | 0 | | | G | G | G | G | | | | | | |
| NC_011770 | 6531288 | G | A | 73 | 0 | 4 | 0 | | | A | A | A | A | | | | | | |
| NC_011770 | 6531306 | A | T | 70 | 0 | 4 | 0 | | | T | T | T | T | | | | | | |
| NC_011770 | 6531307 | T | C | 71 | 0 | 4 | 0 | | | | | | | C | C | C | C | | |
| NC_011770 | 6380918 | G | T | 72 | 0 | 1 | 0 | | | | | | | | T | | | | |
| NC_011770 | 6534039 | T | G | 53 | 0 | 8 | 0 | G | G | G | G | G | G | | | G | G | | |
| NC_011770 | 6534050 | G | C | 54 | 0 | 6 | 0 | | | C | C | C | C | | | C | C | | |
| NC_011770 | 6534071 | G | A | 56 | 0 | 6 | 0 | | | A | A | A | A | | | A | A | | |
| NC_011770 | 6534076 | G | A | 58 | 0 | 6 | 0 | | | A | A | A | A | | | A | A | | |
| NC_011770 | 6534125 | C | A | 58 | 0 | 2 | 0 | | | | | | | | | | | A | A |
| NC_011770 | 6537780 | A | G | 82 | 0 | 10 | 0 | | | G | G | G | G | G | G | G | G | G | G |
| NC_011770 | 6537795 | T | C | 84 | 0 | 6 | 0 | C | C | | | | | C | C | C | C | | |
| NC_011770 | 6537828 | C | T | 83 | 0 | 2 | 0 | | | | | | | | | | | T | T |
| NC_011770 | 6539230 | G | A | 87 | 0 | 2 | 0 | | | | | | | A | A | | | | |
| NC_011770 | 6539234 | C | G | 90 | 0 | 10 | 0 | G | G | G | G | G | G | G | G | G | G | | |
| NC_011770 | 6542419 | C | T | 84 | 0 | 8 | 0 | | | T | T | T | T | T | T | | | T | T |
| NC_011770 | 6543316 | G | A | 88 | 0 | 2 | 0 | | | | | | | A | A | | | | |
| NC_011770 | 6543334 | C | T | 89 | 0 | 2 | 0 | T | T | | | | | | | | | | |
| NC_011770 | 6543352 | A | G | 87 | 0 | 2 | 0 | | | | | | | G | G | | | | |
| NC_011770 | 6543355 | A | AC | 83 | 0 | 8 | 0 | | | AC | AC | AC | AC | AC | AC | | | AC | AC |
| NC_011770 | 6543357 | C | T | 18 | 4 | 2 | 0 | | | | . | | | . | | T | T | . | . |
| NC_011770 | 6389343 | C | CG | 58 | 0 | 1 | 0 | CG | | | | | | | | | | | |
| NC_011770 | 6545671 | T | G | 42 | 8 | 4 | 0 | . | . | G | G | G | G | . | . | . | . | . | . |
| NC_011770 | 6545692 | C | T | 41 | 8 | 4 | 0 | . | . | T | T | T | T | . | . | . | . | . | . |
| NC_011770 | 6545693 | A | C | 42 | 8 | 4 | 0 | . | . | C | C | C | C | . | . | . | . | . | . |
| NC_011770 | 6545764 | G | C | 44 | 6 | 2 | 0 | . | . | | | | | . | . | C | C | . | . |
| NC_011770 | 6545809 | T | C | 44 | 8 | 4 | 0 | . | . | C | C | C | C | . | . | . | . | . | . |
| NC_011770 | 6545935 | C | T | 45 | 8 | 4 | 0 | . | . | T | T | T | T | . | . | . | . | . | . |
| NC_011770 | 6545974 | T | C | 46 | 8 | 4 | 0 | . | . | C | C | C | C | . | . | . | . | . | . |
| NC_011770 | 6546052 | A | G | 41 | 8 | 4 | 0 | . | . | G | G | G | G | . | . | . | . | . | . |
| NC_011770 | 6546083 | A | G | 40 | 8 | 4 | 0 | . | . | G | G | G | G | . | . | . | . | . | . |
| NC_011770 | 6546268 | A | G | 36 | 8 | 4 | 0 | . | . | G | G | G | G | . | . | . | . | . | . |
| NC_011770 | 6546341 | T | G | 36 | 8 | 4 | 0 | . | . | G | G | G | G | . | . | . | . | . | . |
| NC_011770 | 6546394 | C | T | 37 | 8 | 4 | 0 | . | . | T | T | T | T | . | . | . | . | . | . |
| NC_011770 | 6546679 | T | C | 37 | 8 | 4 | 0 | . | . | C | C | C | C | . | . | . | . | . | . |
| NC_011770 | 6546697 | A | G | 40 | 8 | 4 | 0 | . | . | G | G | G | G | . | . | . | . | . | . |
| NC_011770 | 6546701 | C | A | 38 | 8 | 4 | 0 | . | . | A | A | A | A | . | . | . | . | . | . |
| NC_011770 | 6479958 | G | A | 92 | 0 | 1 | 0 | | | | | | | | | | | | A |
| NC_011770 | 6549209 | C | CAAGG( | 70 | 0 | 2 | 0 | CAAGGCG( | CAAGGCGGA | | | | | | | | | | |
| NC_011770 | 6549220 | G | A | 64 | 1 | 4 | 0 | . | | A | A | A | A | | | | | | |
| NC_011770 | 6549222 | C | T | 65 | 0 | 4 | 0 | | | T | T | T | T | | | | | | |
| NC_011770 | 6549310 | G | T | 73 | 0 | 4 | 0 | | | T | T | T | T | | | | | | |
| NC_011770 | 6549350 | C | A | 70 | 0 | 10 | 0 | | | A | A | A | A | A | A | A | A | A | A |
| NC_011770 | 6549388 | G | T | 67 | 0 | 4 | 0 | | | T | T | T | T | | | | | | |
| NC_011770 | 6549408 | C | T | 68 | 0 | 4 | 0 | | | T | T | T | T | | | | | | |
| NC_011770 | 6549453 | G | T | 69 | 0 | 4 | 0 | | | T | T | T | T | | | | | | |

| | | | | | | | | | | | | | | | | | | | |
|---|---|---|---|---|---|---|---|---|---|---|---|---|---|---|---|---|---|---|---|
| NC_011770 | 6549482 | C | T | 72 | 0 | 2 | 0 | | | | | | | | | T | T | | |
| NC_011770 | 6549493 | C | T | 72 | 0 | 6 | 0 | | | T | T | T | T | | | | | T | T |
| NC_011770 | 6550133 | T | G | 76 | 0 | 6 | 0 | | | G | G | G | G | | | | | G | G |
| NC_011770 | 6508988 | G | A | 73 | 0 | 1 | 0 | | | | | | | | | | | | A |
| NC_011770 | 6554636 | C | T | 67 | 0 | 4 | 0 | | | T | T | T | T | | | | | | |
| NC_011770 | 6554639 | G | A | 67 | 0 | 2 | 0 | | | | | | | | | | | A | A |
| NC_011770 | 6555458 | A | G | 80 | 0 | 6 | 0 | G | G | | | | | G | G | G | G | | |
| NC_011770 | 6556595 | G | C | 80 | 0 | 2 | 0 | | | | | | | C | C | | | | |
| NC_011770 | 6557121 | A | G | 63 | 0 | 2 | 0 | | | | | | | | | | | G | G |
| NC_011770 | 6557195 | T | G | 66 | 0 | 2 | 0 | | | | | | | G | G | | | | |
| NC_011770 | 5695089 | CG | C | 34 | 0 | 4 | 0 | C | | C | | C | | | | . | C | | |
| NC_011770 | 6560276 | T | C | 76 | 0 | 10 | 0 | C | C | C | C | C | C | | | C | C | C | C |
| NC_011770 | 6560334 | T | A | 76 | 0 | 4 | 0 | | | A | A | A | A | | | | | | |
| NC_011770 | 6560361 | C | T | 81 | 0 | 2 | 0 | | | | | | | | | | | T | T |
| NC_011770 | 6560412 | G | A | 79 | 0 | 2 | 0 | A | A | | | | | | | | | | |
| NC_011770 | 6561845 | T | C | 65 | 0 | 4 | 0 | C | C | | | | | | | C | C | | |
| NC_011770 | 6561874 | C | A | 61 | 0 | 2 | 0 | | | | | | | | | | | A | A |
| NC_011770 | 6561972 | C | T | 55 | 0 | 6 | 0 | T | T | | | | | T | T | T | T | | |
| NC_011770 | 6561975 | G | A | 54 | 0 | 6 | 0 | A | A | | | | | A | A | A | A | | |
| NC_011770 | 6561976 | A | C | 54 | 0 | 6 | 0 | C | C | | | | | C | C | C | C | | |
| NC_011770 | 6561993 | C | T | 55 | 0 | 6 | 0 | T | T | | | | | T | T | T | T | | |
| NC_011770 | 6562007 | C | T | 56 | 0 | 6 | 0 | T | T | | | | | T | T | T | T | | |
| NC_011770 | 6562009 | C | T | 56 | 0 | 4 | 0 | T | T | | | | | | | T | T | | |
| NC_011770 | 6562025 | T | C | 54 | 0 | 12 | 0 | C | C | C | C | C | C | C | C | C | C | C | C |
| NC_011770 | 6562092 | T | C | 52 | 0 | 4 | 0 | C | C | | | | | | | C | C | | |
| NC_011770 | 6562126 | A | G | 52 | 0 | 12 | 0 | G | G | G | G | G | G | G | G | G | G | G | G |
| NC_011770 | 4256502 | TGGCGTGG | T | 63 | 0 | 7 | 0 | T | T | T | T | T | | | T | | | T | |
| NC_011770 | 6562219 | T | C | 47 | 0 | 2 | 0 | | | | | | | | | C | C | | |
| NC_011770 | 6427064 | GGC | G | 60 | 0 | 1 | 0 | | | | | | | | G | | | | |
| NC_011770 | 6564394 | A | C | 84 | 0 | 2 | 0 | | | | | | | | | C | C | | |
| NC_011770 | 6564843 | A | G | 89 | 0 | 12 | 0 | G | G | G | G | G | G | G | G | G | G | G | G |
| NC_011770 | 6567219 | G | A | 58 | 0 | 4 | 0 | | | | | | | A | A | A | A | | |
| NC_011770 | 6568933 | A | T | 75 | 0 | 2 | 0 | T | T | | | | | | | | | | |
| NC_011770 | 6568947 | C | T | 82 | 0 | 4 | 0 | | | T | T | T | T | | | | | | |
| NC_011770 | 6568976 | C | A | 81 | 0 | 4 | 0 | | | A | A | A | A | | | | | | |
| NC_011770 | 6568999 | A | G | 84 | 0 | 8 | 0 | G | G | G | G | G | G | | | | | G | G |
| NC_011770 | 6442715 | G | A | 75 | 0 | 1 | 0 | | | | | | | | A | | | | |
| NC_011770 | 6571814 | T | A | 91 | 0 | 2 | 0 | | | | | | | | | A | A | | |
| NC_011770 | 6571819 | T | C | 93 | 0 | 2 | 0 | C | C | | | | | | | | | | |
| NC_011770 | 6571843 | T | C | 89 | 0 | 4 | 0 | C | C | | | | | | | C | C | | |
| NC_011770 | 6571845 | C | A | 89 | 0 | 6 | 0 | | | A | A | A | A | A | A | | | | |
| NC_011770 | 6571853 | G | A | 91 | 0 | 2 | 0 | | | | | | | | | | | A | A |
| NC_011770 | 6573166 | G | T | 56 | 0 | 8 | 0 | | | T | T | T | T | T | T | T | T | | |
| NC_011770 | 6573170 | T | G | 53 | 0 | 10 | 0 | | | G | G | G | G | G | G | G | G | G | G |
| NC_011770 | 6519444 | CA | C | 39 | 0 | 2 | 0 | | C | | | | | | C | | | | |
| NC_011770 | 6578000 | TG | T | 97 | 0 | 2 | 0 | T | T | | | | | | | | | | |
| NC_011770 | 6578177 | T | C | 99 | 0 | 2 | 0 | | | | | | | | | | | C | C |
| NC_011770 | 6578186 | C | G | 105 | 0 | 2 | 0 | | | | | | | | | | | G | G |
| NC_011770 | 6578187 | A | T | 105 | 0 | 2 | 0 | | | | | | | | | | | T | T |
| NC_011770 | 6578190 | C | G | 104 | 0 | 2 | 0 | | | | | | | | | | | G | G |
| NC_011770 | 6578212 | C | T | 102 | 0 | 2 | 0 | | | | | | | | | | | T | T |
| NC_011770 | 6578852 | G | T | 58 | 0 | 2 | 0 | | | | | | | | | | | T | T |
| NC_011770 | 6578861 | A | G | 49 | 2 | 2 | 0 | | | | | | | . | . | | | G | G |
| NC_011770 | 6578862 | C | T | 49 | 2 | 2 | 0 | | | | | | | . | . | | | T | T |
| NC_011770 | 6578871 | A | G | 50 | 2 | 2 | 0 | | | | | | | . | . | | | G | G |
| NC_011770 | 6578901 | C | T | 42 | 2 | 2 | 0 | | | | | | | . | . | | | T | T |
| NC_011770 | 6578907 | C | G | 43 | 2 | 2 | 0 | | | | | | | . | . | | | G | G |
| NC_011770 | 6578922 | T | A | 44 | 2 | 2 | 0 | | | | | | | . | . | | | A | A |
| NC_011770 | 6578926 | A | G | 44 | 2 | 2 | 0 | | | | | | | . | . | | | G | G |
| NC_011770 | 6578945 | G | A | 42 | 2 | 2 | 0 | | | | | | | . | . | | | A | A |
| NC_011770 | 6578950 | A | G | 41 | 2 | 2 | 0 | | | | | | | . | . | | | G | G |
| NC_011770 | 6578961 | A | T | 38 | 2 | 2 | 0 | | | | | | | . | . | | | T | T |

| | | | | | | | | | | | | | | | | | | | |
|---|---|---|---|---|---|---|---|---|---|---|---|---|---|---|---|---|---|---|---|
| NC_011770 | 6578964 | A | G | 37 | 2 | 2 | 0 | | | | | | | . | . | | | G | G |
| NC_011770 | 6578970 | G | A | 38 | 2 | 2 | 0 | | | | | | | . | . | | | A | A |
| NC_011770 | 6579003 | A | C | 41 | 2 | 2 | 0 | | | | | | | . | . | | | C | C |
| NC_011770 | 6579012 | G | T | 39 | 2 | 2 | 0 | | | | | | | . | . | | | T | T |
| NC_011770 | 6579025 | T | C | 40 | 2 | 4 | 0 | | | C | C | C | C | . | . | | | | |
| NC_011770 | 6579026 | C | CGAT | 41 | 2 | 2 | 0 | | | | | | | . | . | | | CGAT | CGAT |
| NC_011770 | 6579043 | C | G | 39 | 2 | 2 | 0 | | | | | | | . | . | | | G | G |
| NC_011770 | 6579047 | A | G | 39 | 2 | 2 | 0 | | | | | | | . | . | | | G | G |
| NC_011770 | 6579055 | C | G | 39 | 2 | 2 | 0 | | | | | | | . | . | | | G | G |
| NC_011770 | 6579058 | C | T | 39 | 2 | 2 | 0 | | | | | | | . | . | | | T | T |
| NC_011770 | 6579064 | T | C | 40 | 2 | 2 | 0 | | | | | | | . | . | | | C | C |
| NC_011770 | 6579087 | T | C | 37 | 2 | 2 | 0 | | | | | | | . | . | | | C | C |
| NC_011770 | 6579091 | T | C | 37 | 2 | 2 | 0 | | | | | | | . | . | | | C | C |
| NC_011770 | 6579094 | A | G | 36 | 2 | 2 | 0 | | | | | | | . | . | | | G | G |
| NC_011770 | 6579098 | A | C | 36 | 2 | 2 | 0 | | | | | | | . | . | | | C | C |
| NC_011770 | 6579102 | G | A | 36 | 2 | 2 | 0 | | | | | | | . | . | | | A | A |
| NC_011770 | 6579107 | G | C | 37 | 2 | 2 | 0 | | | | | | | . | . | | | C | C |
| NC_011770 | 6579110 | C | T | 38 | 2 | 2 | 0 | | | | | | | . | . | | | T | T |
| NC_011770 | 6579137 | A | G | 42 | 0 | 2 | 0 | | | | | | | | | | | G | G |
| NC_011770 | 6579154 | A | G | 42 | 0 | 2 | 0 | | | | | | | | | | | G | G |
| NC_011770 | 6579168 | C | A | 44 | 0 | 2 | 0 | | | | | | | | | A | A | | |
| NC_011770 | 6562193 | T | TA | 47 | 0 | 10 | 0 | TA | TA | TA | TA | TA | | TA | TA | TA | TA | TA | |
| NC_011770 | 6580414 | GGCGAGC | G | 95 | 0 | 6 | 0 | G | G | | | | | G | G | G | G | | |
| NC_011770 | 6580428 | G | T | 58 | 5 | 6 | 0 | . | . | T | T | T | T | | . | . | . | T | T |
| NC_011770 | 6580456 | TC | T | 90 | 0 | 4 | 0 | | | T | T | T | T | | | | | | |
| NC_011770 | 6580463 | C | T | 91 | 0 | 6 | 0 | | | T | T | T | T | | | | | T | T |
| NC_011770 | 6562240 | G | A | 49 | 0 | 1 | 0 | | | | | | | | A | | | | |
| NC_011770 | 6583150 | A | C | 75 | 0 | 10 | 0 | C | C | C | C | C | C | C | C | C | C | | |
| NC_011770 | 6583154 | C | T | 75 | 0 | 4 | 0 | | | T | T | T | T | | | | | | |
| NC_011770 | 6587932 | G | A | 71 | 0 | 2 | 0 | | | | | | | | | A | A | | |
| NC_011770 | 6596564 | G | A | 81 | 0 | 12 | 0 | A | A | A | A | A | A | A | A | A | A | A | A |
| NC_011770 | 6596602 | G | A | 84 | 0 | 1 | 0 | | | | | | | | | | | A | |
| NC_011770 | 6596666 | G | A | 82 | 0 | 2 | 0 | | | | | | | | | | | A | A |
| NC_011770 | 6596700 | A | AAAAAT | 83 | 0 | 1 | 0 | | | | | | | | | | AAAAAT | | |
| NC_011770 | 6596715 | G | A | 83 | 0 | 2 | 0 | | | | | | | A | A | | | | |
| NC_011770 | 6596754 | G | A | 85 | 0 | 12 | 0 | A | A | A | A | A | A | A | A | A | A | A | A |
| NC_011770 | 6596781 | T | G | 81 | 0 | 12 | 0 | G | G | G | G | G | G | G | G | G | G | G | G |
| NC_011770 | 6596804 | C | T | 87 | 0 | 2 | 0 | | | | | | | | | | | T | T |
| NC_011770 | 6596815 | G | A | 88 | 0 | 6 | 0 | | | A | A | A | A | A | A | | | | |
| NC_011770 | 6596840 | G | A | 87 | 0 | 2 | 0 | A | A | | | | | | | | | | |
| NC_011770 | 6596882 | C | T | 87 | 0 | 12 | 0 | T | T | T | T | T | T | T | T | T | T | T | T |
| NC_011770 | 6596934 | T | G | 95 | 0 | 4 | 0 | | | | | | | G | G | G | G | | |
| NC_011770 | 6597015 | G | A | 96 | 0 | 10 | 0 | A | A | A | A | A | A | A | A | | | A | A |
| NC_011770 | 6597262 | T | C | 96 | 0 | 10 | 0 | C | C | C | C | C | C | C | C | | | C | C |
| NC_011770 | 6597284 | C | G | 89 | 0 | 10 | 0 | G | G | G | G | G | G | G | G | | | G | G |
| NC_011770 | 6597316 | G | C | 92 | 0 | 10 | 0 | C | C | C | C | C | C | C | C | | | C | C |
| NC_011770 | 6597388 | A | G | 101 | 0 | 2 | 0 | | | | | | | | | G | G | | |
| NC_011770 | 6597389 | T | TA | 101 | 0 | 12 | 0 | TA | TA | TA | TA | TA | TA | TA | TA | TA | TA | TA | TA |
| NC_011770 | 6597392 | C | T | 94 | 1 | 1 | 0 | | | | | | | | | T | . | | |
| NC_011770 | 6597407 | T | A | 104 | 0 | 10 | 0 | A | A | A | A | A | A | A | A | | | A | A |
| NC_011770 | 6597442 | G | A | 103 | 0 | 12 | 0 | A | A | A | A | A | A | A | A | A | A | A | A |
| NC_011770 | 6597443 | G | A | 105 | 0 | 2 | 0 | | | | | | | | | | | A | A |
| NC_011770 | 6597453 | G | A | 103 | 0 | 10 | 0 | A | A | A | A | A | A | A | A | | | A | A |
| NC_011770 | 6597516 | T | G | 98 | 0 | 10 | 0 | G | G | G | G | G | G | G | G | | | G | G |
| NC_011770 | 6597586 | C | A | 98 | 0 | 2 | 0 | | | | | | | | | | | A | A |
| NC_011770 | 6597605 | G | C | 99 | 0 | 2 | 0 | | | | | | | | | C | C | | |
| NC_011770 | 6597661 | G | A | 98 | 0 | 6 | 0 | | | A | A | A | A | A | A | | | | |
| NC_011770 | 6597665 | C | T | 99 | 0 | 2 | 0 | | | | | | | | | | | T | T |
| NC_011770 | 6597674 | A | G | 103 | 0 | 6 | 0 | | | G | G | G | G | G | G | | | | |
| NC_011770 | 6597685 | CAGAACCG | C | 97 | 0 | 2 | 0 | | | | | | | | | | | C | C |
| NC_011770 | 5763677 | TCCCCCC | TC | 55 | 0 | 6 | 0 | TC | | | | | T | | TC | TC | TC | | TC |
| NC_011770 | 6599142 | AG | A | 84 | 0 | 10 | 0 | A | A | A | A | A | | A | A | | A | A | A |
